# Genomic analysis of the recent monkeypox outbreak

**DOI:** 10.1101/2022.06.01.494368

**Authors:** Federico M. Giorgi, Daniele Pozzobon, Antonio Di Meglio, Daniele Mercatelli

## Abstract

The Monkeypox virus is the etiological cause of a recent multi-country outbreak, with nearly one hundred distinct cases detected outside the endemic areas of Africa in May 2022. In this article, we analyze the sequences of two full genomes of Monkeypox virus from Portugal and Belgium, and compare them with all available Monkeypox sequences, annotated by year and geographic origin, as well as related Cowpox and Variola (smallpox) virus sequences. Our results show that the recent outbreak is most likely originating from the West African clade of Monkeypox, with >99% sequence identity with sequences derived from historical and recent cases, dating from 1971 to 2017. We analyze specific mutations occurring in viral proteins A42R (for which a crystal structure is available) and H3L (an important epitope in host immune recognition), highlighting specific amino acids varying between the current outbreak, previous Monkeypox and Cowpox sequences and the historical Variola virus. Genome-wide sequence analysis of the recent outbreak and other monkeypox/cowpox/variola viruses shows a very high conservation, with 97.9% (protein-based) and 97.8% (nucleotide-based) sequence identity.

## Introduction

Monkeypox is an infectious viral disease affecting humans with symptoms similar to smallpox, including fever, muscle pains and blisters. Monkeypox and smallpox are caused by the Monkeypox virus and the Variola virus, respectively, both members of the Orthopoxvirus family of doublestranded DNA viruses. However, while the Variola virus has been eradicated worldwide in 1980 [1], the Monkeypox virus is still endemic in sub-Saharan Africa [2].

In May 2022, nearly a hundred independent cases of monkeypox have been reported in 12 non-African countries, namely Australia, Belgium Canada, France, Germany, Italy, Netherlands, Portugal, Spain, Sweden, United Kingdom and United States of America [3,4]. While the main source of contagion in the endemic areas is believed to be zoonotic, i.e. deriving from the contact with infected animals in the wild [5], human-to-human transmission has been confirmed in cases from the United Kingdom in 2018 [6] and also in the recent multi-country outbreak [7]. Previous monkeypox cases in countries outside the African continent have been reported since 2003 [5], but never with case numbers as high as the recent outbreak [8,9].

From the genetic point of view, there are two distinct clades of the Monkeypox virus, namely the West African clade and the Central African (or Congolese) clade [10], with similar clinical and pathological features [11]. The Monkeypox virus is generally considered closely related to the Variola and Cowpox viruses, both in etiological aspects and genetic measurements [12–14]. This similarity is reflected also in immunological contexts, for example it has been reported that smallpox vaccination offers 85% protection against monkeypox in humans [15]. The reference Monkeypox virus, available on NCBI with accession code NC_003010.1 and derived from the sample Zaire-96-I-16 [14], carries a 196,858 nucleotide-long double-stranded DNA linear genome, containing 191 non-overlapping genes [16] **(Figure 1 A**), and is a representative of the Central African clade.

**Figure 1.**
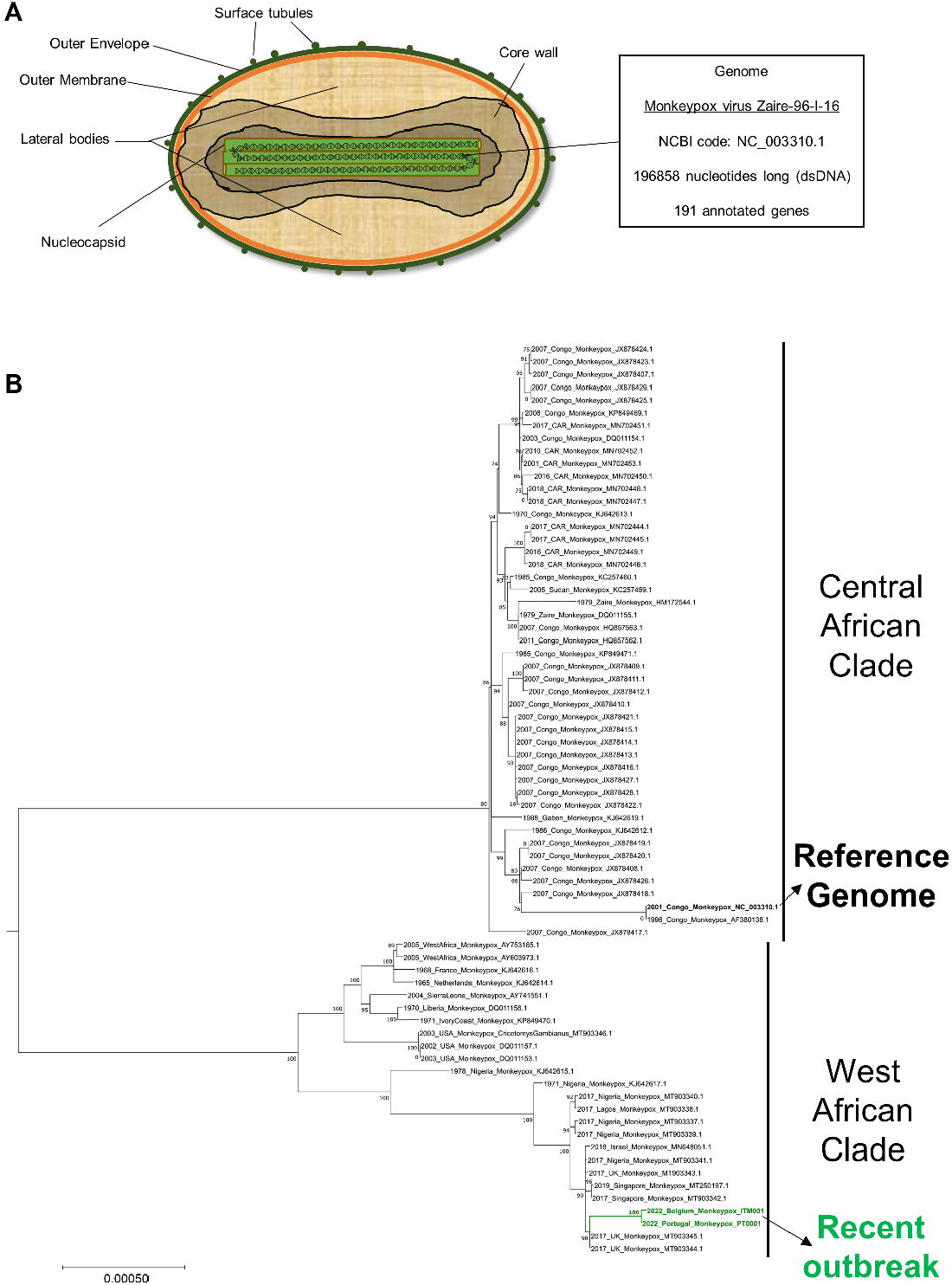
A) Schematic representation of the structure of the Monkeypox virus. Reported genome information is based on the reference sequence NC_003310.1, associated with the Central African Monkeypox virus sample Zaire-96-I-96. B) Phylogenetic tree of Monkeypox viruses. Phylogenetic analysis was based on a core alignment including all available Monkeypox viral sequences from 1971 to May 2022. The percentage of bootstraps supporting each branch is reported. The ruler in the phylogenetic tree reports the number of nucleotide substitutions per site. A likely origin from the West African clade is supported for the recent Belgium and Portugal Monkeypox sequences (highlighted in green).

As shown by the recent worldwide pandemic caused by the SARS-CoV-2[17], a strict monitoring of the genomic features of the Monkeypox virus is of paramount importance, as well as the need for continued genomic surveillance on the evolution of this potential threat. In this article, we will analyze the feature of the Monkeypox virus genome, including two full sequences from the recent outbreak of May 2022, and establish its relationship with historical sequences from Monkeypox viruses, as well as Cowpox viruses and the Variola virus.

## Results

We retrieved two draft full-genome Monkeypox virus sequences from recent (May 2022) cases of monkeypox in Lisbon, Portugal [18] and Antwerpen, Belgium [19], named respectively PT0001 and ITM001 by the original authors. Using BLAST [20] with the PT0001 Monkeypox genome sequence as query, we retrieved the 100 most similar sequences according to BLAST bitscore. These were formed by 72 Monkeypox sequences and 28 Cowpox sequences, dating back to samples retrieved from 1971. We added to this selection the two reference sequences of Central African Monkeypox virus NC_003310.1 [14] and the Indian Variola virus NC_001611.1 [21]. The total of analyzed genomes is 104 and is available, in FASTA format, as **Supplementary File 1**.

Phylogenetic analysis of Monkeypox virus genome sequences shows a clear distinction between Central African sequences (including NC_003310.1) and West African sequences. The two sequences from the recent cases in Portugal and Belgium appear highly related to each other, and robustly located within the West African clade **(Figure 1 B**).

Beyond the Monkeypox sequences, Cowpox genomes constitute a clearly different genetic group, highlighted by both phylogenetic analysis **(Supplementary File 2**) and by clustering based on sequence similarity **(Figure 2 A**). The Variola virus appears genetically distinct from both Monkeypox and Cowpox viruses **(Figure 2 A**), and several proteins have a low conservation within the variola/monkeypox/cowpox proteomes **(Figure 2 B**). Nevertheless, the Variola virus genome is highly similar, in terms of genome-wide sequence identity, to the Monkeypox virus **(Figure 3 A**), with 96.6% DNA identity with the Central African Monkeypox (sequence NC_003310.1), 96.2% with the West African Monkeypox (representative sequence from Nigeria MT903340.1), 96.8% with the recent Monkeypox case from Belgium (sequence ITM001), and 95.7% with the recent Monkeypox case from Portugal (sequence PT0001). Overall, all West African Monkeypox, Central African Monkeypox, Cowpox and Variola virus genome sequences analyzed possess a sequence identity above 95% (on average, 97.8%) **(Figure 3 A**). On the proteomic side, the 191 monkeypox proteins also possess a high conservation in the cowpox/monkeypox/variola clades, with an average protein sequence identity of 97.9% (**Figure 2 B**). The only proteins with sequence identities within the pox virus groups that are below 90% are D16L (82.9%) and D14L (79.5%). Both are currently poorly characterized and annotated on both the Uniprot and NCBI databases as hypothetical proteins, with domain homology with complement-binding proteins (D14L) and KELCH-motif containing proteins (D16L) [22].

**Figure 2.**
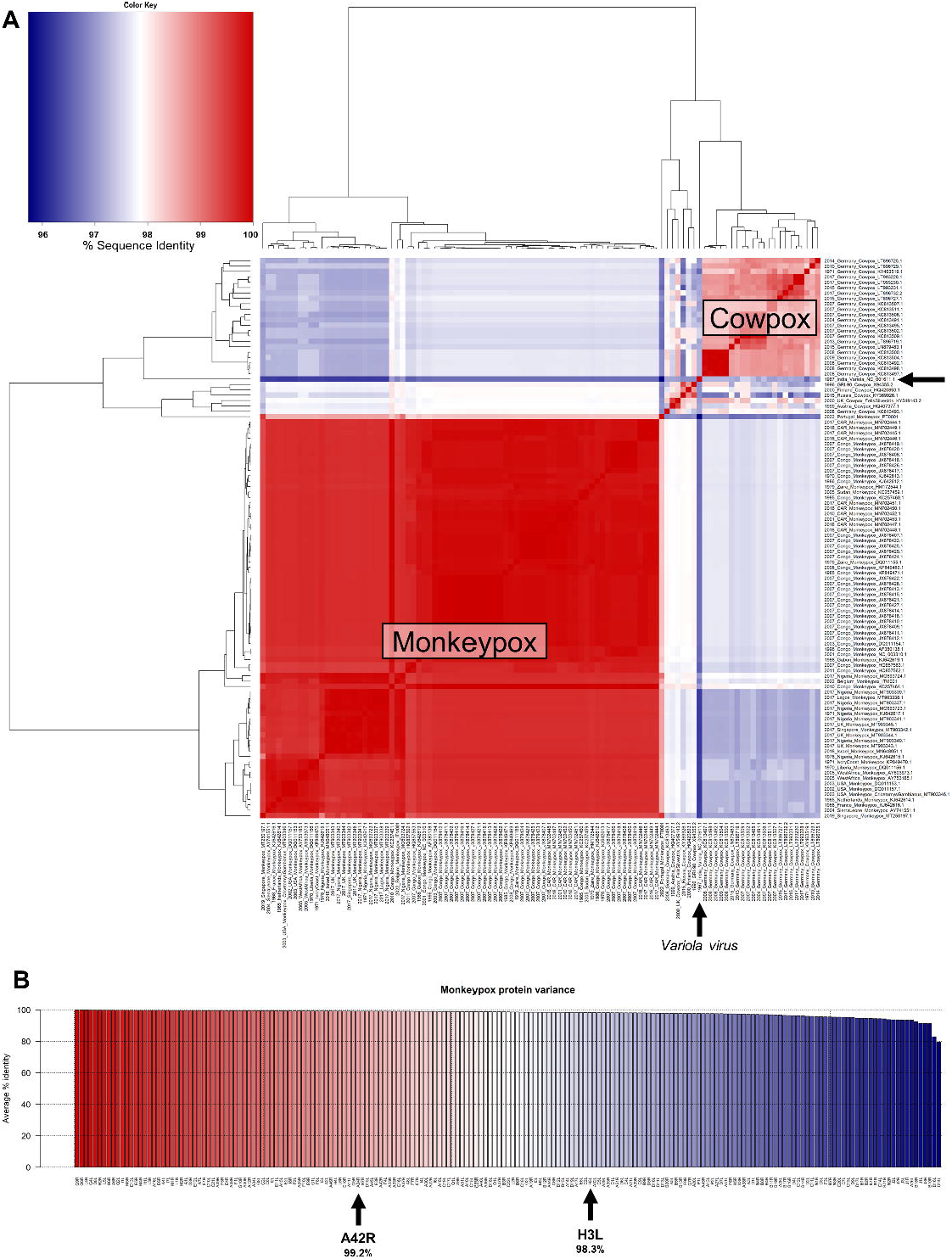
A) Sequence similarity heat map of Monkeypox, Cowpox, and Variola viruses, based on sequence identity. The color intensity in the heatmap reflects the percentage of identity, calculated via the BLASTN algorithm, between genomes in a scale ranging from blue (95% identity) to red (100% identity). The position of the Variola virus is indicated by a black arrow. B) Average protein-based sequence identity for all 191 monkeypox protein-coding genes. The average for each protein is calculated using the NC_003310.1 Central African monkeypox reference vs. all sequences reported in Figure 1 A.

**Figure 3.**
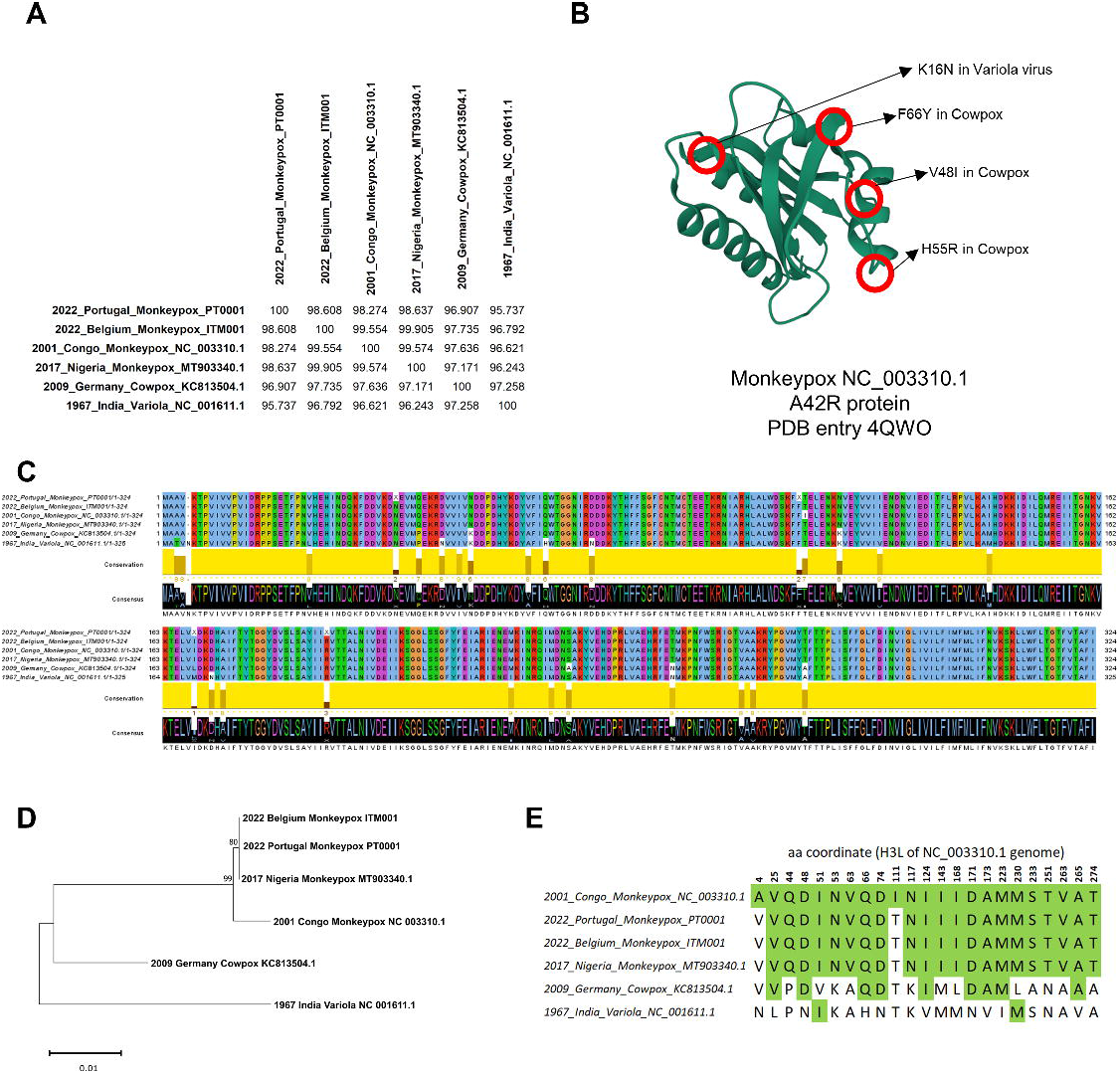
A) Percentage sequence identity highlight between the Variola virus genome and representative Monkeypox or Cowpox virus genomes. A sequence identity >95% between all highlighted genomes was detected, confirming high sequence similarity between these viruses. B) 3D representation of the A42R Profilin-like Protein from Monkeypox Virus Zaire-96-I-16 (PDB entry 4QWO). The location of the 4 amino acids differing between Monkeypox and other closely related Orthopoxvirus A42R sequences are indicated by red circles. C) Multiple sequence alignment of the H3L protein encoded by poxvirus. The two Monkeypox virus sequences from recent cases of monkeypox (PT0001 and ITM001) were compared with representative sequences of West African (MT903340.1) and Central African Monkeypox (NC_003310.1), Cowpox (KC813504.1), and Variola (NC_001611.1) viruses. D) Phylogenetic tree based on H3L protein sequences in the previously indicated representative genomes. The percentage of bootstraps supporting each branch is reported, from a total of 1000 bootstraps. The ruler in the phylogenetic tree reports the number of amino acid substitutions per site. E) Coordinates of amino acids differing in the H3L protein sequence between Monkeypox reference NC_003310.1 and other selected Orthopoxviruses. Amino acids corresponding to the reference NC_003310.1 H3L sequence are highlighted in green.

We proceeded then to analyze specific sequences within the genomes of Monkeypox and related viruses, focusing on proteins of relevance and representative of the average conservation shown in our analysis within the pox proteomes analyzed: A42R (99.2% sequence identity) and H3L (98.3% sequence identity) **(Figure 2 B**). A42R is a short (~130 aminoacid long) protein homologous to eukaryote profilins, actin-binding proteins involved in cytoskeletal structure and function, such as membrane trafficking [23]. A search in the Protein Data Bank (PDB) [24] for protein structures found that A42R is currently the only entry for a Monkeypox protein: a crystallography-inferred model is available in PDB entry 4QWO, based on A42R the protein from from Central African Monkeypox strain Zaire-96-I-16, represented in our genome analysis by reference sequence NC_003310.1 [25]. Our analysis showed a 100% identity between Monkeypox sequences from the recent outbreak PT0001 and ITM001 and the other Monkeypox A42R proteins; only 4 amino acids differ between Monkeypox A42R sequences and other closely related Orthopoxvirus, namely Lysin 16, which is replaced by an Asparagin in Variola virus **(Figure 3 B**), and Valine 48, Histidine 55 and Phenylalanine 66, which are Isoleucine, Arginine and Tyrosine in Cowpox virus **(Figure 3 B**). All these varying amino acids are located either outside or on the border of structural alpha-helices and beta-sheets and are therefore unlikely to cause a major structural rearrangement of this protein.

We then focused our analysis on H3L, an Orthopoxvirus 324 amino acid long glycosiltransferase located on the viral envelope, which plays a role in the attachment to the human target cell and in the general poxvirus entry [26]. We considered H3L due to its pivotal immunological importance, since it offers one of the main epitopes recognized by the host immune system [27]; studies showed how antibodies elicited by smallpox vaccines are binding H3 proteins [28], with specific recognition of H3L as an antigen by immune CD8+ T cells [28]. As with A42R, we extracted H3L proteins between the two recent Monkeypox genome sequences (PT0001 and ITM001) and compared them with representative database sequences for West African Monkeypox (MT903340.1), Central African Monkeypox (NC_003310.1), Cowpox (KC813504.1) and Variola (NC_001611.1) viruses. Multiple sequence alignment shows a high similarity between H3L sequences, however with a selection of amino acids diverging between major clades **(Figure 3 C**). Phylogenetic analysis based on this single protein confirms what previously inferred from genome-wide analysis, specifically a high similarity between the recent Belgium and Portugal Monkeypox sequences, and their likely origin from the West African clade **(Figure 3 D**). H3L proteins from Central African Monkeypox, Cowpox and Variola viruses are more distantly related to the recent outbreak. 21 amino acids differ between the reference Monkeypox and the reference Smallpox (Variola) virus in H3L, suggesting a higher variance for this protein, with respect to the core protein A42R **(Figure 3 E**). The sequence of this important protein epitope has indeed remained identical in the available sequences form the recent 2022 outbreak (PT0001 and ITM001) when compared to the West African sequences, represented here by the MT903340.1 sequences, retrieved from a 2017 Nigerian sample [8].

## Discussion

Our preliminary analysis shows that sequences retrieved from the recent multi-country Monkeypox outbreak, started in May 2022, are likely derived from the West African Monkeypox clade, both from genome-wide **(Figure 1 B**) and protein-specific **(Figure 3 D**) phylogenetic analyses. Our analyses on specific Monkeypox proteins, A42R (selected due to the availability of a crystal structure) and H3L (selected due to its centrality in immune recognition) show in fact complete identity between the recent outbreak and West African Monkeypox sequences **(Figure 3 B** and **Figure 3 E**). Genome-wide sequence identity between all Monkeypox viruses is well above 98% (**Figure 3 A**) and above 95% when considering also the more distantly related Cowpox and Variola viruses. A higher variability (21 amino acids over 324 amino acids, 6.5% of the whole protein sequence) is found between the Variola and the Monkeypox virus H3L protein **(Figure 3**). All these results suggest a low variability within the Orthopoxvirus family, but at the same time the differences between the Variola and Monkeypox viruses in H3L, a host recognition protein, may suggest host affinity and adaption of these viruses. While both pathogens are able to infect different species, Variola virus is an ancient actor in human history and it is considered highly specific to human hosts [29], while Monkeypox has been diagnosed in humans only since 1970 [30].

The roles of H3 in virus/host recognition and in the viral entry process, as well as its high immunogenic potential, allow to observe functional analogies with the Spike protein in coronavirus infections [31]. In the case of Monkeypox H3 can therefore, similarly to Spike, constitute an ideal target for vaccine development, resembling the recent success of Spike mRNA-based vaccines against COVID-19 [32]. Overall, the high conservation observed between the proteome of the recent outbreak Monkeypox and other Orthopoxviruses, including the Variola virus **(Figure 2 B**) are one of the likely molecular causes for the efficacy of smallpox vaccines in protecting against monkeypox [15]. Alongside protein epitopes, it is very likely that also molecular mechanisms of disease transmission are conserved, allowing to epidemiologically predict the course of this monkeypox outbreak based on historical data [33].

While the multi-country monkeypox case number of May 2022 remains the largest ever occurred for this disease outside the endemic regions in Africa [9], it must be noted that thankfully no death related to this outbreak has been reported, and that it is unlikely that this disease will reach pandemic status [34]. Moreover, as shown in our work here, there has been virtually no genetic divergence from previously known endemic Monkeypox viruses **(Figure 1 B**). However, a close and continuous inspection on the evolution of this protein is of paramount importance to monitor the progression of monkeypox, during this outbreak and in the future years, in order to maintain the capability of developing efficacious vaccines.

## Methods

Pairwise sequence analysis, including sequence identity calculation, was performed using BLASTN version 2.9.0+ [20]. Sequences were retrieved from the NCBI database [35] and, for the recent outbreak, from the virological.org database [18,19]. For pairwise protein identity, we used BLASTP, keeping the highest bitscore pairwise match in the calculation of the average percentage sequence identity for each protein vs. protein protein. This last precaution was taken in case multiple suboptimal local alignments were generated by BLASTP for the same protein pair.

Genome-wide multiple sequence alignment was performed using Parsnp version 1.2 [36] with default parameters, using the RefSeq Monkeypox sequence NC_003310.1 as reference. Protein multiple sequence alignment was performed using Muscle with default parameters [37]. Multiple sequence alignment visualization was performed using Jalview version 2.11.2.0 [38].

Phylogenetic trees were constructed using the Neighbor-Joining method [39]. Tree evaluation was performed with 1000 bootstrap replicates [40]. Sequence distances in phylogenetic trees were computed using the Poisson method [41]. All phylogenetic analysis was performed using the MEGA tool, version 11 [42].

Protein structure visualization was performed using the PDB web tools [24]. The structure of Monkeypox protein A42R was obtained from PDB entry 4QWO [25]. All reported amino acid numbers are based on the reference sequence of Central African Monkeypox NC_003310.1 [14].

All sequence preprocessing and the plots in **Figure 2** have been generated using the R statistical software [43].

## Supplementary files

**Supplementary** 1: FASTA files. Top 100 BLAST hits. Date reported is collection date. When not specified (e.g., HQ857562.1), the date of submission or publication is indicated.

**Supplementary 2:** phylogenetic tree including cowpox and monkeypox sequences (built using the same methods described for the tree in Figure 1 A).

## Acknowledgments

We sincerely thank the teams of Philippe Selhorts (https://virological.org/t/belgian-case-of-monkeypox-virus-linked-to-outbreak-in-portugal/801) and Joao Paulo Gomes (https://virological.org/t/first-draft-genome-sequence-of-monkeypox-virus-associated-with-the-suspected-multi-country-outbreak-may-2022-confirmed-case-in-portugal/799) for providing the draft genome sequences of the two recent cases of Monkeypox virus infection, available on the aggregator site virological.org. We also thank Rossana Bartone, Angela Tomasini and Giulia Minarini for their administrative support.

