## Supplementary File S1 for "Genomic analysis of the recent monkeypox outbreak"

>1967_India_Variola_NC_001611.1

CTCGAGAGTATATGTTGTTGAACGTTATTGTTTGAGAAATAGTTGATGCATCAGAATGGTTTGCATTTATTGTAGACGTAATAATGCACTACATTGTTTCCTCGTCTATCTATATGTTTTGATAAGTTATAACACGTTTCAATCTCTAGTTTTATTTTTTTGTACGTCACATCTTCATCCAGTAGACGACATAGAATACATGTGCAATCTTTGCCACAATCCATAGCTATTTTGGTGCTAATTATTCCTATATTATATTTCACAAAAAATGATGAAGGCAATCATTCCTCATAAGATAATAAAAAGTGTATTGAGAGTGAGTAAGCATGAAAAAGATTTAGTATTTAGCAGTGCGGATATGATCCAAGAGGGTGAAATAGTCGTTCTCGTTCAGAATCTTTTGCAGCATAAGTAGTATGTCGATATACTTATCGTTGAAGACTCTTCCAGAGACGATAGCTAATTGAGTACAAAGTCCAATGATTGCACAAAGTTCTTTGGCGGTTTTCATGGAGTCATTTCTGATGAAACATTTAATGATCTCCACGCAATTGTCTATATTGTCCCACGGAAGTGAATCCGAGAACTCCTTCAACTCGCTACCAAAGAGCTCCGTTGCATCAGTTCTGAAAGAGATGAGAAGCCTGTAAAGAGACCCTGCGCTTTCTCTATGGGTCCATCTATGAGAAACCCACAGGATGTATTCAGTCAGACAATGTCTGACGTCTGCCACGGTATTCAGGGAGTCCTTAGTAGCATGGCAATGACAGGGACTGAACTGGGCACAAGAAGAGGCCATTGTGAAGGTAGACGAAGGTTAACCCGATGGTTAACCGTAGCCGTCTGTGCTAATAGACGGCATTAATTTCCATTTTTTAATGGGGTTGTGGATGAGGAATGAGAGTGTCTTATCTCGTCCTTGGTTTACATGGATCAGTGTGAGAAAAAATATCTTGTATATTATTAACTAAACTAACTCCTTGGTTTCAATTCATGTTTAAAATGATCTATATGTTTTATTAATTTATTATACTGTTAAATGTACTGTATATAATAGTTTCTAACGTGGAGTACAGAATTTTCGGTTAATGAGTTCATGGTAAAGGGCGAATGAATGCCTGCCTGTATTATAATATACATAAGTTAATAGTTTTTATCATATTTTTCTAATACTATATAAAAATATCATTATGTATAATCATTACTGTCGCTATCATTATTGTGTTTGTGTAGTTCTGTTCTATCATCTACATCACTGTCACTGTCACTCTCCCTATATCTTCTAAAATTACAAACAACTGGATATTCGATAACAGCATTTGTGTAGTTTTTGTCTTTTATAGTATATACGTTATTGTCAAAATCTAAACAAATATTGGCATAATACCTATCTATAAGATTAGGATCCATGTTCGAGCATACTAGCATATTTGTAACTTCGTCGTAAAGCGTTAAATCAATAGAATAAACCATTTTGTGGTGCAACTTTTTACGAACTAGTTGTATGAGTTTATCGTTTACATAAGCCATTAACAGCTTTAACAGATGATCTGAGTAATATCCTCTATTATACATTTAATGTTCACGGTCTTAGTATTTTTAGATATCAATTGTAATTTACACCCATATTCTACTCCCTTGTGTAACGTTAGACATTCTAAATCTATAGTATTATCTATTACAGCGTAAAACACATTCAATATTGTATTGTTATATTATTAGTTTATATTACTGAATTAATAATATAAAATTCCCAATCTTGTTATAAACACACACTGAGAAACAGCATAAACACAAAATCCATCAAAAATGTCGATGAAATATCTGATGTTGTTGTTCGCTGCTATGATAATTAGATCATTCGCCAATAGTGGTAACGCTATCGAAACGACATTGTCAGAAATTACAAACACTACAACAGATATTCCAGCTATCAGATTATGCGGTCCAGAGGGAGATAGGTATTGTTTTCACGGTATCTGTATCCACGCTAGAGATATTGACGGTATGTATTGTAGATGCTCTCATGGTTATACAGGCATTAGATGTCAGCATGTAGTATTAGTAGACTATCAACGTTCAGAAAAACCAAACACTACAACGTCATATATCCCATCTCCCGGTATTGTGCTTGTATTATTAGTAAGCATTATTGTGTGTTGTCTATTATTTGTTTATAGGTTCACTCGAAGAACTAATAAATTACCTTTACAAGATATGGTTGTGCCATAATTTTTATAAATTTTTTTATGAGTATTTTTATGAGTATTTTTCAAAAAATGTATAAAGTGTATGTCTTATGTATATTTATAAAAATGCTAAATGTGATGTATCTATGTTATTTGTATTTATCTAAACAATACCTCTACCTCTAGATATTATACAAAAATTTTTTATTTCAGTATATTAAAGTAAAATCTAGTTACCTTGAAAATAAATACAGTGGGTGGTTCTGTATCACCAGTAAGAACATAATGGTCGAACACAGTATCCGATTGAGATTTTGCATACAATACTAGTCTAGAAAGAAATTTGTAATCATCTTCTGTGACGGGAGTCCATATATCTGTATCATCGTCTAGTTTATCAGTGTCCCATGCTATATTCCTGTTATCATCATTAGTTAATGAAAATAACTCTCGTGCTTCAGAAAAGTCAAATATTGTATCCATACATACATCTCCAAAACTATCACTTATACGTTTATCTTTAACGATACCTATACCTAGATGGTTATTTACTAACAGACATTTTCCAGATCTATTGACTATAACTCCTATAGTTTCCACATTAACCAAGTAATGATCATCTATTGTTATATAACAATAACATAACTCTTTTCCATTTTTATCAGTATGTATATCTATATCAACGTCGCCGTAGTGAATAGTAGTCATTGATCTATTATATGAAACGGATATGTCTAGAACGGCAATTGTCTTACGTCCAGTTAACACTTTCTTTGATTTAAAGTCTAGAGTCTTTGCAAACATAATATCCTTATCCGACTTTATATTTCCTGTAGGGTGGTATAATTTTATTTTGCCTCCACATATCGGTGTTTCCAAATATATTACTAGACAATATTCCATATAGTTATTAGTTAAGGGTACCCAATTAGAACACGTACGCTTATTATCATCATTCGGATCGTATTTCATAAAAGTTATTGTACTATCGATGTCAACACATTCTACATTTTTTAATCGTCTATATAGTATTTTTCTGATATTTTCTATAATATCAGAATTGTCTTCCATCGGAAGTTGTATACTATCGGAATCAGTTACATGTTTAAATAATTCTCTGATGTCATTCCTTATACAATCAAATTCATTATTAAACAGTTTAATAGTCTGTAGACCTTTATCGTTGTAAATATCCATTGTCTTATTAGTTACGCTTATTTTTATGTGTTTTATGTTGATTTATTATATTTATAAGAATGATTGTTTGACAAATCACGAGAACTATTAAGACATATATTATTAGAGGTATATATTATAAAAAAGTTTTTGATTACGATGTTATAAGAGGAAAAAGGACACATTAACATCATACATCAATTAACATTCTTATAACATCGTAATCAAAAGAATTGCAATTTTGATGTATAACAACTGTCAATGAATTATGGAATTGTATATTACACATTATACGGTATGTTGGCAACGACAAATACCGATTGGTAATTGTTTACCGGTGTAATAGAATTTATATATCTATTACACCGGCCAAGTATGCATAATAATAAGTTGTGGGTAGTATCATCCACATATTTATAATTTAGGACTTTGTATTCAGTATTTTTGGAATCATAAAAAAAAGTTTTACCATTTTAAAATTTAAAAAGTATTTACATTTTTTTCACTGTTTAGTCGCGGATATGGAATTTGATCCTACCAAAATCAATATATCATCTATAGATCATGTAACAATATTACAATACATAGATGAACCAAATGATATAAGACTAACAGTATGCATTATCCAAAATATTAATAACATTACATATTATATCAATATCACAAAAATAAATCCACATTTGGCTAATCAATTTCGGGCTTGGAAAAAACGTATCGCTGGAAGGGACTATATGACTAACTTATCTAGAGATACAGGAATACAACAATCAAACCTTACTGAAACTATACGTAACTGTCAAAAAAATAGAAACATATATGGTCTATATATACACTACAATTTAGTTATTAATGTGGTTATTGATTGGATAACCGATGTGATTGTTCAATCAATATTAAGAGGGTTGGTAAATTGGTACATAGATAATAATACCTATACTCCAAATACACCCAATAATACAACAACCATTTCTGAGTTGGATATCATCAAAATACTGGATAAATACGAGGACGTATATAAAGTAAGTAAAGAAAAAGAATGTGGAATTTGCTATGAAGTTGTCTACTCAAAACGATTAGAAAACGATCGATACTTTGGTTTATTGGATTCGTGTAATCATATATTTTGCATAACATGTATCAATATATGGCATAGAACACGAAGAGAAACTGGTGCGTCGGATAATTGTCCTATATGTCGTACCCGTTTTAGAAACATAACAATGAGTAAGTTCTATAAGCTAGTTAACTAATAAATAAAAAGTTTAATTTGTTGACGACATATATCGTTATTTTTCTCGTATGAAAGATTAATTTGATTCTAATATAATCTTTAGTATTAGATGAACTTTAATTCAAATTTCCATTAGATTATATCATAAATAAAAATAGTAGCACGTGCTACTTTAGCCAAATATTCTTTTTTGAAACGCCATTTAGCGTAGTGAGGACACAAGTGAACCTATAATGAGCAAATTTATTAGTATCGGTTACATGAAGGACTTTACGTAGAGTGGTGATTCCACCATCTGTGGTACGAATGGTTTCATCTTCTTTGATGCCATCACCCAGATGTTCTATAAACTTGGCATACTCGTCCGATTTCATATCCTTTGCCAACCAATACATATAGCTAAACTTAGGCATATGTTCCACACATCCTGAACAATAAAATTCTCCAGATGATGTTACAATGTCTAGATTTGGACATTTGGTTTCAACCGCGTTAACATATGAGTGAACACAGCCATACATGAAAGCGATGAGAAATAGGATTCTCATCTCGCCAAAATATCACTAGAAAAAATTTATTTATCAATTTTAAAGGTATAAAATACTTATTGTTGTCCTCGTTTGTCAATCATGTTAATGTCTTTAAGATAAGGCAGGCAAATGTTTATAGTATTAAGAATTGGGCAAGCATAAGACATGACACAAAGACCCTTTTTGTATATATAAGTGTAGAAATTATAACATCCATAGTTGGATTTACATAGGTGTCCAATTGGGATCTCACCATCATCGAGATAATTGATGACATCTCCCCCTTTCTTTTTTAGTAGATATTTCATCGTGTAAGAATCAATATTAATATTTCTAAAGTATCCGTGTATAGCCTCTTTATTTACCACAGCTCCATATGCCAACATGCATTCCACTAGATGGATATCGATATCGTCGAATGTCATATACTCAATTAGTATATGTTGGAGGATATTCGAGTTCATTGTTTTCAATATCAAAGAGATGGTTTCCTTATCATTTCTCCATAGTGGTACAATACTACACATTATTCCGTGTGGCTTTCCATTTTCCAAAAACAATTTGACCAAATCTAAATTCACATCTTCATTGTATCTATAATCACTATTCAGATAATCAGCCATAATTACTCGAGTGCAACGTGTTAGATCGTCTATATATGAATAAGCAGTGTTATCTATTCCTTTCATTAACAATTTAACGATGTCTATATCTATATGAGATGACTTAATATAATATTGAAGAGCTGTACAATAGTTTTTATCTATAGAAGACGGCTTGATTCCGTGATTAATTAGACATTTAACAACTTCCGGACGCACATATGCTCTCGTATCCGACTCTGAATACAGATGAGAGATAATATACAAATGCAATACGGTACCGCAATTTTGTGGTTGATAATCATCATACGTGTATCCATACTCGTCATCCTCATAAAGAACATTGCAGCCATTTTCTATGAACAAATCAATAATTTCAGGAACAGGATCATCTGTCATTACATAATTTTCTATAACTGAACGATGGTTTTCACATTTAACACTCAAGTCAAATCCATGTTCTACCAACACCTTTATTAAGTCAACGTTTACATTTTTGGATTTCATATAGTTGAATATATTAAAGTTATTTATGTTGCTAAATCCAGTGGCTTCTAGTAGAGCCATCGCTATATCCTTATTAACTTTAACATGTCTACTATTTGTGTATTCTTCTAATGGGGTAAAATGTCTCCAATTTTTGTGCAATGGATTAGTGCCACTGTCTAGTAGTAGTTTGACGACCTCGACATTATTACAATGCTCATTAAAAAGGTATGCATGTAAAGCATTATTCTTGAATTGGTTCCTGGTATCATTAGGATCTCTGTCTTTCAACATCTGTTTAAGTTCATCGAGAGCTACCTCCTCATTTTCCAGATAGTTAAACATTTTGACTGAATAGAAGTGAATGAGCTACTGTGAACTCTGTACACCCACACAACTAATGTTATTAAATATCATTTTTTGAATATATTTATACCATGTCAAAACTTGTTCAATTATTAATAAAAATAATTTAGTGTTTAAATTTTACCAGTTCCAGATTTCACACCTCCATTAACCCCACTTTTTACACCACAGGACGATCCTCCCCACATTTCACCGCCACCAGATGTATAAGTTTTAGATCCTTTATTACTACCATCATGTCCATGGATAAAGACACTCCACATGCCGCCACTACCTCCTTTAGAAGACATATTAATAAGACTTAAGACAAGTTTAACAATAAAATTAATCACGAGTACCCTACTACCAACCTACACTATTATATGATTATAGTTTCTATTTTTACAGTACCTTGACTAAAGTCTTTAGTCACACAATACTACCAACCTACACTATTATATGATTATAGTTTCTATTTTTATAGGAACACGTACGAGAAAATCAAATGTCTAAGTTCTAACGGTAGTGTTGATAAACGATTGTTATCCACGGATACCTCATTTATCATGTTGTCTATTTTCTTACTTTGTTCTATTAACTTATTAGCGTTGTATATTATTTGATTATAAAACTTATATTGCTGTTTAGTCCACTCTATAAATATCGGATTATTCACATATCGTTTCTTTGTAGGTTTATTTAACTTGTACATCACTGTAAGCATGTCCGTACCATTTATTTTAATTTGACACATATCAGCAATTTCTTTTTTGCATTCGGTTCTACCAACGTAATAATGAAGTGTGGTATATGATCTATAAGTATAATATAGTATCTCTTTTTTCGTAAGATATTAAACAGTTTATCACATCTAGAGATAATTTATCTCGTGATAATATAATAATAGAGTGCGGTTTTACCTATTTTATCATCTATATAATAATAATATAATACTAATAGTAATCGTTATCATTGTACCCACTTGAGAAGTTAGTAACTTTTTTTCTATTATAATCTTGTATCCGTAAGATACATTACTACACATAGGAATTCCCTGATCGAGCAATAGTTTAAATACATCTACATTTGGATTTGATGTAGTTGCGTATTTCTCTACAATATTAATACCATTTTTGCAACTATTTATTTCTAGACCTTTTGTGATTAGTAATCTCAATAATTCTACGTCAATATTATCAGATTCTATATATTCGAATATATCAAAGTCATTGATATTTTTATAATTGGTAGAAGACAATAATGACACCACAACATCAGTTTTGATATTCTTATTTTTCTTGGTAACGTATACATTTAATGAATTTTCATTACGTTCTACCAATGATTGTGCACTGCAGGCATCAAAAGTTTTACAACTATCATAAAGCATACTATCCTATCCAACCATTATGGTATTGATGCTATAGTATATTTTAATTCATTTCATTTATGTGTATTTGGTACGAGCAGACAGCATGAGATCGTATGATTTCAGTACCGTAGTGACAATACTATTAGTTTCAATAAGATCAATCTCTAGATCTATAGAATCAAAACATGATAGGTTAGAAGATAATGAATATCTGTAGGCTTCTTGTTGCACTGTAACTTCTCGTTTTGTTAGATGTTTGTATCATGCTTTAACATTAATGGTACAAATTTTATCCTCGCTTTGTGTATCATATTCGTCCCTACTATAAAATTGTATATTCAGATTATCATGAGATGTGTTTACACTAACTGTATCAATAAACGGCACACCATTTAGTCATAACCGTAATCCAAAAATTTTTAAAGTATATCTTAACGAAAGAAGTTGTATCATCGTTAGGATTTGGTAAATTATTATCTACAGTGTATGGTACTAAATCCTCATAAGTGTATATATCTAGAGTAATGTTTAATTTATCAAATGGTTCATAATATGGATCCTCATGACAATTTCCGAAGACGAAAATGAGATATAGACATGGTTACTCCTTAAAAAAATACGAATAATCGCCTTGGCTATTTAGTAAGTGTCATTTAACACTATACTTATATTAATCCATGGACTCATAATCTCTATACGGGATTAACGGATGTTCTATATACGGGGATGAGTAGTTCTCTTCTTTAACTTTATACTTTTTACTAATCATATTTAGACTAATGTATGGGTAATAGTGTTTGAAGAGCTCGTTCTCATCATCAGAATAAATCAATATCTCTGTTTTTTTGTTATACAGATGTATTACAGCCTCATATATTACGTAATAGAACGTGTCATCTACCTTATTAACTTTCACCGCATAGTTGTTTGCAAATACGGTTAATCCTTTGACCTCGTCGATTTCCGACCAATCTGGGCGTATAATGAATCTGAACTTTAATTTCTTGTAATCATTCGAAATAATTTTTAGTTTGCATCCGTAGTTATCTCCTCTATGCAACTGTAAATTTCTCAACGCGATATCTCCATTAATAATGATGTCAAATTCGTGTTGTATACCCATACTGAATTGATGAACGAACGAATACCGTCGGCGTTAATAGTAATTTACTTTTTCATCTTTACATACTCGGTACTAGTTTTACTATCATAAGTTTATAAATTCCACAAGCTACTATGGAATATACCAACCATCTTAGTATAGAACACATGTCTTAAAGTTATTAATTAATTACATATTGTTTTATATATCGCTACGAATTTAAACAGAGAAATCAGTTAGGAAGAAAAAATTATCTGTCTTCTTCATCATCGTCTATTGGATAACGTCTCTGTATTCTACGATAGAGTGCTACTTTAAGATGAGACATATCCGTGTTATCAAAAATATACTCCATTAAAATGATTATTCCGGCAGCGAACTTGATATTGGATATATCACAACCTTTGTTAATATCTACGACAATAGACAGCAGTCCCATGGTTCCATAAACAGTGAGTTTATCTTTCTTTGAAGAGATATTTTGTAGAGATCTTATAAAACTGTCAAATGACATCGCATTTATATCTTTAGCTAAATCGTATATGTTACCATCGTAATATCTAACCGCGTCTATCTTAAACGTTTCCATCGCTTTAAAGACGTTTCCGATAGATGGTCTCATTTCATCAGTCATACTAAGCCAACAAATATAATCGTGTATAACATCTTTGATAGAATCAGACTCTAAAGAAAACGAATCGGCTTTATTATACACATTCATGATAAACTTAATGAAAAATGTTTTCGTTGTTTAAGTTGGATGAATAGTATGTCTTAATAATTGTTATTATTTCACTAATTAATATTTAGTAATTAGTGCACTCTATAAAAACGAGAATGACATAACTAATCATAACTAGTTATCAAAGTGTCTAGGATGCGTAATTTTTTATATGGTATAGATCCTGTAAGCGTTGTCTGTATTCTGGAGCTATTTTCTCTATCGCATTAGTGAGTTCAGAATATGTTATAAATTTAAATCGAATAACGAACATAACTTTAGTAAAGTTGTCTATATTAACTCTTTTATTTTCCAGCCATCGTAATACCCATGTTTAAGATAGTATATTCTCTAGTTACACTATGGTCTCATCGTTGTTTAGAATATCACATACCGAATCTACATCAAATTTTAGAAATTGGTCTGTGTTACGTATCTCTTCTATATTATTGTTGATGTATTGTCGTAGAAAACTATTACGTAGACCATTTTCTTTATAAAACGAATATATAGTATACCAATTATCTTTTCCGATAATATATTTACACACATAATCCATTCTCTCAATCACTACATCTTTAAGTTTTTCGTTGTTAAGATATTTGGCTAAATTATATAATTCTATTAGATCATTAACAGAATCAGTATATATTTTTCTAGATCCAAAGACGAACTCTTTGGCGTCCTCTATAATATTCCCAGAAAAGATATTTTCGTGTTTTAGTTTATCGAGATCTAATCTGTTCATATACGTATCCATGATTGCGCGGTACGTTATGATAGCCGCATAAAATAAAATCCATTTTCATTTTTAACCAATACTATCTCCTCTATTCATAATTGAGATTGATGTAATACTTTGTTACTTTGAACGTAAAAACAGTACACGGATCCGTATCTCCAACAAGCATGTAGTAATCAAATTTGGTGTTGTTAAACTTCGCAATATTCATTAATTTAGATAGAAATTTATACTCATCATCTGTTTTAGGAATCCATGTATTATTACCACTTTCCAACTTATCATTATCCCAGGCTATGTTTCGTCCATCATCATTGCGCAGAGTGAATAATTCTTCTGTATCCGGCAGTTCAAATATATGATCCATGCATAGATCGGCAAAGCTATCGTAGATGTGATTTTTCCTAAATCGAGTATAAAACTCGTTTACTAGCAAACACTTTCCTGATTTATTGACCAAGACACATATGGTTTCTAAGTCTATCAAGTGGTGGGGATCCATAGTTATGACGCAGTAACATAGATTATTACCTTCTTGACTGTCGCTAATATCTATATATTTATTGTTATCGTATTGGATTCTGCATATAGATGGCTTGTATGTCAAAGATATAGAACACATAACCAATTTATAGTCGCGCTTTACATTCTCGAATCTAAAGTTAAGAGATTTAGAAAACATTATATCCTCGGATGATGTTATCACTGTTTCTGGAGTAGGATATATTAAAGTCTTTACAGATTTCATCCGATTCAAATAAATCACTAAATAATATCCCACATTGTCATCTGTTAGAGTAGTATCATTAAATCTATTATATTTTATGAAAGATATATTACTGCTCACCTCTATATTTCGTACATTTTTAAACTGTTTGTATAATATCTCTCTGATACAATTAGATATATCTATTGTGTCGGTAGACGATACCGTTACATTTGAATTAATGGTGTTCCATTTTACAACTTTTAACAAGTTGACCAATTTATTTCTAATAGTATCAAACTCTCCATGATTAAATATTTTAATAGTATCCATTTTATATCAACGGACACAAAGTAGCTAACATAAACCATTGTATAATTTTTATGTTTTATGTTTATTAGCGTACACATTTTGGAAGTTCCGGTTGCCATGTATTTCCTGGAGAGCAAGTAGATGATGAGGAACCAGATAGTTTATATCCGTACTTGCACGTAAAGTCTACATTGTCATTATACGAATATGATCTTTTAAACCCGCTAGACAAGTATCCGTTTAATATTGTAGGATGTGGACATTTAACAATCTGACACGTGGGTGGATTAGACCATTCTCCTCCTGAACACAGGACACCAGAGTTACCAATCAACGAATATCCACTATTGCAACTATAAGTTACAACGCTTCCATCGGTATAAAAATCATTATATCCGTTATGTCTTCCGTTGGATATAGATGGAGGTAATTGGCATTTAACAGATTCGCAAATAGGTGCCTTGGGATTCCATACCATAGATCCAGTAGATCCTAATTTACAATACGATTTATATTCACCAATCAAATAATATCCGCTATTACAAGAGTACGTTATACTAGAGCCAAAGTCTACTCCGCCAATATCAAGATGGCCATTATCGATATCTCGAGGCGACGGGCATCTCCGTTTAATACATTGATTAAAGAGTGTCCATCCAGTACCTGTACATTTAGCATATATAGGTCCCATTTTTTGCTTTCTGTATCCAGGTAGACATAGATATTCTATAGTGTCTCCTATGTTGTAATTAGCATTAGCATCAGTCTCCACACTATTCTTAAATGTCATATTAATGGGTCGTGACGGAATAGTACAGCATGATAGAACGCATCCTATTCCCAACAATGTCAGGAACGTCACTCTCTCCACCTTCATATTTATTTATCCGTAAAAATGTTATCCTGGACATTGTGCCAATAATAAAAAGCCCATATATGTTCGCTATTGTAGAAATTGTTTTTCATAGTTGCTCAAAAACGATGGCGGTGACTTATGAGTCACGTTACACTTTGGAGTCTCATCTTTAGTAAACATATCATAATATTCGATATTACGAGTTGACACATTGAACAAATCCAAGTATTTGATTTTGGATAATATTCGTATTTTGCATCTGCTATAATTAAGATATAATCGCCACAAAAACACGCGAACATCTTTCCTACATGGTTAAAGTACATGTACAATTCTATCCATTTGTCTTCCTTAACTATATATTTGTATAGATAATTACGAGTCTCGTGTGTAATTCCAGTAATTGCATAGATGTTACCATCGTACTCTACAGCATAAACTATACTATGATGTCTAGGCATGGGAGACTTTTTTATCCAACAATTTTTAGTGAAACATTCCACATCGTTTAATATTACATATTTTTCATACGTGGTATAAACTCCACCCATTACATATATATCATCATTTACGAATACCGACGCGTCTGAATATCTAGGAGTAATTAAGTTTGGATGTCTTATCCATTTCGAAGTGTAGCGTTTTAAATATTTTGCTACACCTGTTGAAATAGAAAATTTTAATTCTCCTATTACATATAACTTTCCATCATTAACACACGTACTAACTTTTGATTTTAACGACGACATATTAGTAACTGTTTTCCATTTTTTTTTTTTTTTTTTTTTGTTTTAAGATCTACCCACGGATACGAAATAAACATGTCTATTGTTAATCATGCCACCAATAATGTATAGACAATTATATAAAACATTTGCATTATAGAATTGTCTATCTATCTGTATTACCGACGATTGTCCAATATTCTGTTCTAGGAGAGTAATGGGTTATTGTGGATATATAATCAGAGTTTTTAATGACTACTATATTATGTTTTATACCATTTCGTGTCACAGGCTTTGTAGATTTGGATATAGCATTGCACATAGTATTAGTCATAAACTTGGGATGTAAACTGTTGATGATATCTACATCGTTTGGATTTTTATGTATCCACTTTAATAATATCATAGCTGTAACATCCTCGTGATTTACGTTGACGTTTTCGTGGGATAAGATAGTTGTTCATCCTTTGATAATTTTCCAAATTCTGGATTGGATGTCACTGCAGTAATATTGTTAATTATTTCTAACATTGACGCATTATATAGTTTTTACACATAGATCTACAACAGGTGGAATATCAATTAATACGGCAGCAACTAGTATCATCTCTACATTGTTTACGGTGATGTTTATCTTCTTCCAGCGTATATAGTCTAATAGCGATTCAAACGCGTGATAGTTTATACCATTCAATATAATCGCTTCATCATTTATATGGTGTTTCTAAATTCGTTTAAAAAATTATACGGAGACGCCGTAATAATTTCCCCATTGATAGAAAATATCACGCTTTCCATTCTTGAAGTACTATAAGTAATTATAGTATAATGTAACGTTTATATATTCAATATTTTTTATAAAAAAATCATTTCGACATTAATTCCTTTTTAAATTTCCGTCTATCATCTATAGAAACATATTCTATGAATTTATAAAATGCTTTTACGTGTCCTATCGTAGGCGATAGAACCGCTAAAAAGCCTATCGAATTTCTACAAAAGAATCTATTATATGGTATAGGGAGAGTATAAAACATTAAATGTCCGTACTTATTAAAGTATTCAGTAGCCAATCCTAACTCTTTCGAATACTTATTAATGGCTCTTATTCTGTACGAATCTATTTTTTTGAACAACGGACCTAGTGGTATATCTTGTTCTATGTATCTAAAATAATGTCTGACAAGATCCGTTAGTTTAATATCCTTAGTCATCTTGTCTAGAATGGCGTATCTAACTGCGGGTTTAGGCTTTAGTTTAGTTTCTATATCTATATCTACATCTATGTCTTTATCCAACACTAAAAATATAATAGCTAATATTTTATTACAATCATCCGGATATTCTTCTACGATCTCACTAACTAATGTTTCTTTGGTTATACTAGTATAGTCACGATCAGACAAATAAAGAAAATCAGATGATTGATGAATAATACATTTAAATTTATCATCTGTAAGATTTTTGAGATATCTCATTAAAATATTATTAGGGTCAGTACTCATTATCATATATTGGCAGCAGCTATTACTTATTTTATTTTATTTTTCTTTATTTTATTATTTTTCACCATATAGATCAATCATTAGATCATCAAAATATGTTTCAATCATCCTAAAGAGTATGGTAAATGACTCTTCCCATCTAATTTCTGGACGTTCACCAATGTCTCTAGCCACTTTGGCACTAATAGCGATCATTCGCTTAACATCTTCTACATTATTAACTGGTTGATTTAATCTATCTAGCAATGGACCGTCGGACAGTGTCATTCTCATGTTCTTAATCAATGTACATACATCACTGTCATCTACCAATTCATCCAACAACATAAGCTTTTCAAAATCATCATTATAATATGATGGATCGCCATCATTTCTCCAAAGAATATATCTAATAAGTAGAGTCCTCATGCTTAGTAATTTAACTATTTTAGTTAACAACTATTTTTTATGTTAAATCGAGTAGAACCACTATGTTAATACTTATTCATATTTTAGTTTTTAGGATTGATAATCAACATAAAAATTAATACATCATCAATTTTGAAAATACTTAGTTTCCACGTAGTCAATGAAACATTTGAGCTCATCGTACAGAACGTTCTCGTACAGGACGTAACTATAAACTGGTTTATATTTGTTCAAGATAGATACAAATCCGATAACTTTTTTGACAAATTCTACGGGATCCACTTTAAAAGTGTCATACCGGGTTCTTTTTATTTTTTTAAACAGATCAATGGTGTGATGTTGATTAGGTCTTTTACGAATTTGATATAAAATAGCGTTCACATATCCTCCATAATGGTCAATCGCCATTTGTTCGTATGTCATAAATTCTTTAATTATATGGCACTGTGTATTATTTAGTTCATCCTTGTTCATCATTAGGAATCTATCCAATATGGCAATTATACTAGAACTATAAGTGTGTTGTAAACACATATTGATGTGTCTGTTTATACAATTAATGATATTTGGATCCATGCTACTACCTTCGGGTAAAATTGGAGAATCGTATACCATTTCTAGTACTTTAGGTTCATTGTTATCCATTGTAGAGGATGACATGCTCGAATCATACAAAAATATATTATTTTTATGTTATTTTGTTAAAAATAATCATCGAATACTTTGTAAGATACTCCTTCATGAACATAATCAGTTACAAAACGTTTATATGAAGTAAAGTATCTACGATTTTTACAAAAGTCCGGATGCATAAGTACAAAGTACGCGATAAACGGAATAATAATAGATTTATCTAGTTTATCTTTTTCTATCGCTTTCATAGTTAGATACATAGTCTCGGAAGTCGGATTATGTAACATCAACTTCGATAAAATTACTGGGTTATTTAGTCTTACGCATTCACTCATACATGTATTACCGTTAACTACAGAGTCTACACTAAAATGATTAAACAATAGATAGTCTACCATTGTTTCGTATTCAGATAGTACAGCGTAGTACATGGCATCTTCACAAATTATATCATTGTCTAATAGATATTTGATGCATCTTATGGATCCCACTTCGACAGCCATCTTAAAATCGGTAGAATCATATTGCTTTCCTTTATCATTAATAATTTCTAGAATATCATCTCTATCATAAAAGATACAAATATTAACTGTTTGATCCGTAATAACATTACTAGTTGATAACAATTTGTTAATAAGATGTGCTGGGCTCAATGTCTTAATAAGAAGTGTAAGAGGACTATCTCCAAATTTGTTTTGTTTATTAACATCCGTTGATGGAAGTAAAAGATTTATAATGTCTACATTCTTGACTGTTTTAGAGCATACAATATGGAGAGGTGTATTTCCATCATGATCTGGTTTTGAGGGGCTAATTCCTAGTTTCATCATCCATGAGATTGTAGAAGCTTTGGGATTGTCTGACATAAGATGTCTATGAATATGATTTTTGCCAAATTTATCTACTATCCTGGCTTCAAATCCAATGGACATTATTTTTTTAAACACTCTTTCTGAAGGATCGGTAGACGCCAACAACGGACCACATCCTTCTTCATCAACCGAGTTGTTAATCTTGGCTCCATACTGTACCAATAAATTTATTCTCTCTATGACTTCATCATCTGTTCCCGAAAGATAATATAGAGGCGTTTTATGCTGTTTATCGCACGCGTTTGGATCTGCGCCGTGCGTCAATAGCATCGCAACTATTCTATTATTATTAATTTTAGAAGCTATATGTAATGGATAATTTCCATCATCATCCGTCTCATTTGGAGAGTATCCTCTATGAAGAAGTTCTTCGACAAATCGTTCATCTAGTCCTTTAATTCCACAATACGCATGTAGAATGTGATAATTTCCAGAAGGTTCAATAGCTTGTAGCATATTCCTAAATACATCTAAATTTTTACTATTATATTTGGTATAAAGAGATAGATAATACTCGGCCGACATAATGTTGTCCATTGTAGTATAAAAATTAATATTTCTATTTCTATTTCTACTATAATTTACTCTCTATAACAAATGTCATAACTTAGTTCTTTTATGTCAAGAAGGCACTGGTTTAGTTCATCTATGAATGTCACACCATAACTACCACGCATACCATACTTAGAGTTATGATAAAGATATTTATCCTTAGGGTGCAGGTAATGGGGATTAATCTTTGTTGGATCAGTCTCTAAGTTAACACATGTCACACAGGATCCATTTATAGTTATATCACATGATGATTTATGGTGAATTGATTCCGGAAGATCGCTATTGTATTTTGTGGTTCCACAATTCATTTCCATACATGTTATTATCACACTGATATTATGATGAACTTTATCCAGCCGCTGGGTTGTAAACAACATAACAGATAGTTTATTATCTTTACCAACTCCCTCTGCCGCCGCAACGAATCTCTGATCCGTATCCATGATGGTCATGTTTATTTCTAGCCCGTATCCAGTCAACACTATGTTAGCATTTCTATCGATATAGCTTTCACTCATATAACACTCACCAATAATAGTAGAATTAATGTCGTAATTTACACCAATAGTAAGTTCGGCGGCAAAGTACCAATACCTGTAATCTTGTCGAGGAGGACATATAGTATTCTTGTATTCTACCGAATACCCAAGAGATGCAATACAAAAGAGTAAGACTAATTTGTAAACCATCTTACTTAAAATATGTAACAATAGTACGATGAATGAGTAATAAGAAATATGAAATCTATCTTATATACACATAATTATTCTATCAATTTTACCAATTAGTTAGTGTTTAATAAAAATATGGGAATCTAATTAATTTTTCCTTACACAATCGACGTACATGAGTCTGAGTTCCTTGTTTTTGCTAATTATTTCATCCAATTTATTATTTTTGACGATGTCAAGATCTTTTGTATAGGAGTCAGACTTGTATTCAACATGCTTTTCTATAATCATTTTTAGCTATTTCGGCATCATCCAATAGTACATTTTCCAGATTAGCAGAATAGATATTAATGTCGTATTTGAACAGAGCCTGTAACACCTCAATGTCTTTATTATCTATAGCCAATTTAATGTCCGGAATGAAGAGAAGGAAATTATTGGTGTTTGACGGCGTCATATAGTCGAGCAAAAGAATCATCATATCCACGTGTCCATTTTTATAGTGGTGTGAATACAACTAAGGAGAATAGCCAGATCAAAAGTAGATGGTATTTCTGAAAGAAAGTATGATATAATACTTACATCATTAAGCATGACGGTATGATAAAATGAAGTTTTCCATCCAGTTTTCCCATAGAACATAAGTCTCCAATTTTTCTTAAAACAGTTTTACCGTTTGTATGTTACCACTATCAACCACATAATACAATACAGTGTTCCCTTGTCATTAAATTGTGAATCATCCAGTCCACTGAATAGCAAAATTTTTACTATTTTGGTATCTTCCAATGTGGCTGCCTAATGTAATGGAAATTCATTCTCTAGAAGATTTTTTAATGCTCCAGCGTTCAACAACGTACATACTAGACGCATGTTATTATCAGCTATTGCATAATACAAGGCACTATGTCCATGGACATCCGCCTTAAATGTATCTTTACTAGAGAGAAAGCTTTTCAGCTGCTTAGACTTCCGAGTATTAATTCGTGACAGATCCATGTCTGAAACGAGACGCTAATTAGTGTATAATTTTGTCATACTGCTCCAGAATTAATAATATCTCTAATAGATTTGATTAGTAGATATATGGCTATCGCAAAACAACATATACACATTTAATAAAAATAATATTCATTAAGGAGATTCAGATTCCGCCGTATCCATAAACAATATTAAGGAGATTCTACCTTACCCATAAACAATATAAATCCAGTAATATCATGTCTAATGATGAACACAAATGGTGTATTAAATTCCAGTTCTTCAGGAGATGATCTCACCGTAGCTACCATGATAGTAGATGCCTCCGCTACAGTTCCTTGTTCGTTGACATCTATCTTTGCATTCTGAAACATTTTATAAATATATAATGGGTCCCTAGTCATATGTTTAAACGACGCATTATCTGGATTAAACATACTAGGAGCCATCATTTCGGCTATCGACTTAATATCCCTCTTATTTTCGATAGAAAATCTAGGGAGTTTAAGATTGTACACTTTATTCCCTAATTGAGACGACCAATAGTCTAATTTTGCAGCCATGATAGAATCTGTGAAATGGGTCATATTATCACCTATTGCCAGATACATACTAATATTAGCATCCTTATACTGAAGGCGCACCATATCATATTCTTCGTCATCGATTGTGATTGTATTTCCTTGTAATTTAGTAACTACGCTCATCATGGGAACCGTTTTCGTGCCGTACTTATTAGTAAAACTAGTATTGTGTGTTTTAGTGATATCAAATGGATATTGCCACGTACCTTTAAAATATATAGTATTAATGATTGCCCATAGAGTATTATTGTCGAGCATAGTAGAATCTACTACATTAGACATACCTGATCTACGTTCTACTATGGAATTAATTTTATTAACCGCATCTCGTCTAAAGTTTAATCTATATAGGCCGAATCTATGATATTGTTGATAATACGACGGTTTAATACACACAGTATTATCTACGAAACTTTGATAAGTTAGATCAGTGTACGTATATTTAGATGTTTTTGGCTTAGCTAATCCTGATATTAATTCTGTAAATGCTGGACCCAGATCTATTTTTCTCAAATCCATAGTCTTCAATAATTCTACTTTAGTATTACCTGATGCAGGCAATAGTGACATAAACATAGAAAACGAATAGCCAAACGGCGAGAAGACAATATTATCATCCTCATTCCCATCTTGAATATTTTTATACGCTACTATACCGGCATTGGTAAATCCTTGCAGGCGATAGGTGAACGCTGTACACGCTAACGATAGTATCAATAACACAATCATGATTTTATGGTATTAATATTTTACCTTATTTTTTATATTTGGTATAAAAATTATTGATGTCTACACATCCTTTTGTAATTGACATCTATATATCCTTTTGTATAATCAATTCTAATCACTTTAACTTTTACAGTTTTCCCTACCAGTTTATCCCTATATTTAAAATATCTATTCATATGCATCTGAGTACTCTCTACCAAGATAGCTTCAGAGTGAGGATAGTCAAAAAGATCAACATATAGAACATAACCATTCTCGTATACTTTGCCCTTTAATACATCACCCACATTGGGCAACGAATAACAAAATACAAGCATCTTGTTAACGGGCTCGTAAATTGGGATAAAAATTATGTTTTTATTATTTTATATTCATTTAATATTCATTCAATATTTACAATATATGTAATCATTATCCAGATTGAACATACAGTTTGCACTTGCCATGATTCGTTATATAACTATTAATATTAACAGTTCGTTTAACGATCATATTATTTTTATGTTTTATTAATAATTGTAAAACATACAATTAAATCAATATAGAGGAAGGGACGGATACTGTCTTTTGTGAGATATAGTTATGGCGACTAAATCAGATTATGAGGATGCTGTTTTTTACTTTGTGGATGATGATGAAATATGTAGTCGTGACTCCATCATCGATCTAATAGATGAATATATCACGTGGAGAAATCATGTTATAGTGTTTAACAAAGATATTACCAGTTGTGGAAGACTCTACAAGGAATTGATGAAGTTCGATGATGCTGCTATACGATACTATGGTATTGATAAAATTAATGAGATTGTCGAGGCTATGAGTGAAGGAGACCACTACATCAATCTTACAGAAGTTCATGATCAGGAAAGTCTATTCGCTACCATAGGAATATGTGCTAAAATCACTGAACATTGGGGATACAAAAAGATTTCAGAATCTAGATTCCAATCATTGGGAAACATTACAGATCTAATGACCGACGATAATATAAACATCTTGATACTTTTTCTAGAAAAAAAATTGAATTGATGATATAGGGTTCTTTATAATGCATTCGTATTCTATATCCGGTGTTAAAAAAATTATCCTATCATGTATTTGAGAGTCTTATATGTAGCAAACATGATAGCTGTGATACCAATAAGTTTTAGATATTCACGCGTGCTAATGTTGGGGATGGTATTATCTGGCGGTGAAATGTCCGTTATATAATCTATAAAACAATCATCGCATATAGTATGCGATAGTAGAGTAAACATTTTTATAGTTTCTACTGGATTCATACATCGTTTACCCAATTCGGTTATGAATGAAATTGTCGCCAATCTTACACCCAACCCCTTGTTATCCATTAGTATAGTATTAACTTCGTTATTTATGTCATAAACTGTAAATGATTCTGTAGATGCCATATCATACATGATATTCATGTCCCTATTATAATCATTATTAACTTTATCACAATACGTGTTGATAATATCTATATATGATCTAGTTTTTGTGGGCAATTGTCTATACAAGTCGTCTAAACGTTGTTTACTCATATAGTATTGAACAGCCATCATTACATGGTCCCGTTCCGTTGATAGATAATCGAGTATGTTAGTGGACTTGTTAAATCTATATACCATATTTTCTGGAAGTGGATATACATAGTTGTGATCATTATTGCTAGCCTTATCTTCTATATCATCTATATCATCTATATCATCTATATCATCTATATCATCTATATCATCTATATCATCTATATCATCTATATCATCTATATCATCTATATCATCTATATCATCTACGATATTATTACACATAAACATCGGCAACATACTATTGTACATTATCTAAGTCCGGTTGATCCAAACCCTTGATCTCCTCTATCTGTACTATCTAGAGATTGTACTTCTTTCAGTTCTGGATAATATATACGTTGATAGATTAGCTGAGCTATTCTATCCCCCGTATTTACATTAAATGTATATTTTCCATTATTAATAAGAATGACTCCTATATTTCCCCTATAATCTTCGTCTATTACACCGCCTCCTATATCAATGCCTTTTAGGGACAGACCAGACCTAGGAGCTATTCTACCATAGCAGAACTTAGGCATGGACATACTAATATCTGTCTTAATTAATTGTCTTTCTCCTGGAGGGATAGTATAATCATAAGCGCTATATAAATCATATCCGGCGGCGTACGGTGATTGTCTAGTAGGAGATTTAGCTCTGTTAGTTTCCTTAACAAATCTAACTGGTGAGTTAATATTCATGTTGAACATAAAACTAATATTTTATTTCAAAATTATTTACCATCCCATATATTCCATGAATAAGTGCGATGATTGTACACTTCTATAGTATCTATTACGAACCACGATAAAATCCTCCTATCAATAGCAGTTTATTATCCACTATTATCAATTCTGGATTATCCCTTTGATAAATAGTATCATCTATCAGAGACCATGTATTGCTGTATTTGTAATAAAATTTAGCATTTCTACCAACCAAGAATAACCTTCTACCGAACACTAACGCGCATGATTTATAATGTGGATAATAAGTTGACGGTCCAAACTGCCACTGATCATGATTGGGTAGCAAATATTCTGTAGTTGTATCCGTTTCAGAATGTCCTCCTATTACGTACATAACATTGTTTATGGATGCCATTGCTGGATTACATCTAGGTTTCAAAAGACTCGGCATATTAACCCAAGCAGCATTCCCGTGGAACCAACGCTCAACAGATGTGGGATTTGGTAGACCTCCTACTACGTATAATTTATTGTTAGCGGGTATCCCGCTAGCATACAGTTTGGGGCTATTCATCGGAGGAATTGGAATCCAATTGTTTGATATATAATTTACCGCTATAGCATTGTTATGTATTTCATTGTTCATCCATCACCGATGAGATATACTACTTCTCCAACATGTGTACTTGTACACATATGGAATATATCTATAATTTGATCCATGTTCATAGGATACTTTATGAATGGATACCTGTATTATTTGCGTGGTTGTTTATCACAATGAAATATTTTGGTACAGTCTAATATCCATTTTACATTATTTATACCTCTGGGTGAAAGATAATTTGACCTGATTACATTTTTGATAAGGAGTAGCAAATTTCCTAATTTATTTCTTCTCTTCGTATACCACTTAATGACAAAATCAACTACATAATCTTCGTCTGGAACATTTAACTCATTGCTTTCTAGAATAAGTTTCAAGATAAATAATCAAAATTGTCTATGATATCCTCTTCCAGTTCCAAAAAGTGTTTGGCAATAAAGTCTTTAGTATGACATAAGAGATTGGATAGTCCGTATTCTATACCCATTATGTAACACTCGACACAATATTCCTTTCTAAAATCTCGTAGTATAAAGTTTATACAAGTGTAGATGATAAATTCTACAGACGTTAATATAGACACGCGTAATAAATTGATGACGTCATGTCTATTATGTATACCTTTCCAGTATATGAGTAAATAACTATAGAAGTTAAACTGTGAATGTCAAGGTCTAGGCAAACCCTCGTAACTTGATCTTTATTTTTTGTGTATTTTTGACGTAAATGTGTGCGAAAATACGGAGATAACTTTTTCAATATTGTAGAATTGACTATTATATTGCCTCCATAGCATTAATAATTGTTTTGAATTTCTTAGTCATAGACAATGCTAATATATTCTTACAGTACATAGTATTGACAAATATCGGCATTTATGTTTCTTTAAAAGTCAACATCTAAAGAAAAATGATTGTCTTCTTGAGACATAACTCCCATTTTTTGGTATTCACCCACACGTTTTTCGAAAAAATTAGTTTTACCTTCTAATGATATATTTTCCATGAAATCGGATGGATTGGTAACATTATAGATTCTTTTAAATCCCAATTCAGAAATCAATCTATCTGCGACAAACTCCATATATGTTTTCATCATTTCACAATTCATTCCTATAAGTTTAACTGGAAGAGCCACAGTAAGAAATTCTTGTTCAATAGAGACCGCGTCTGTTATAATAGATCTTACTGTTTCTTCACTCGGTGGATACAATAAATGTTTAAACATCAAACATGCGAAATCGCAGTGTAGACCCTCGTCTCTACTAATTAATTCGTTGGAAAACGTGAGTCCGGGCATTAGGCCACGCTTTTTAAGCCAAAATATGGACGCAAATGATCCAGAAAAGAAAATTCCTTCTACTGCCGCAAAGGCAATAAGTCTTTCTCCATAACCGGCGCTGTCATGTATCCACTTTTGAGCCCAATCGGCCTTCTTTTTTACACAAGGCATTGTCTCTATAGCATTAAAGAGATAGTTTTTTTCATTACTATCTTTAACATATGTATCAATCAAAAGACTATACATTTCCGAATGAATGTTTTCAATGGCCATTTGAAATCCGTAGAAACATCTAGCCTCGGTAATCTGTACTTCTATACAAAATCGTTCTGCTAAATTTTCATTCACTATTCCGTCACTGGCTGCAAAAAACGCCAATACATGTTTTATAAAATATTTTTCGTCTGGTGTTAGTTTATTCCAATCATTGATATCTTTAGATATATCCACTTCTTCCACTGTCCAAAATGATGCCTCTGCCTTTTTATACATATTCCAGATGTCATGATATTGAATTGGAAAAATAACAAATCTATTTGGATTTTTTGCAAGGATTGGTTCCATAACTAAATTAACAATAGTAGTAATTTTTTTTTCAGTTATCATCTATATGACTGTATTTGGATCTTTTGTACATCGCTATTATCGCAATCACTACAATAATTATAATTATTATTGATAGCATTGTTATTAGCACTATCATAATTAAATTATTGATATTCATGGGTGCTGAATAATCGTTATTATCATCATTATCATTTTGTAATTGTGACATCATACTAGATAAATCGTTTGCGCGATTGTTGTGGGAAACGGGCATGGAGGATGAATTATCGTTATTATTATTTAAAGCCTCCCATTCGGATTCACAAATATGACGCGCGTTCAACATTTTATGGAAACTATAATTTTGTGAAAACAGGTAACAAGAAAACTCGTCATCGTTATTTTTAACGATAGTAAACCGATTAAACGTCGCGCTAATTTCTAACGCTAGCGACTCTATTGGATATGGATTTCCAGATATATATCTTTTCAGCTCCCCTACGTATCTATAATCATCTGTGGGAAATGGAAGGTATTTCCATCTATCTAGTGTTCCTAATATCATATGTAGTGGTGTAATAGAACCATTAAGCGCGAAAGATTTTATTTCACATCGTATTTTAACTTCGCAATAATTTCTGGTTAGATAACACACTCTACCAGTTAAGTCAATAATATTAGCCTTTACAGATATATTCATAGTAGTTGTGACAATGACTCCATCTTTTAGATGCGATACTCCTTTATATGAGCCAGAATCTTCGTACCTCAAACTCGATATATTCAAACAAGTTAATGAGATATTAACGCGTTTTATGAACGATGATATGTATCCAGAAGTTCTATCCTCCGTGGCTAACGCTATAACCTTATTATTATAGTACCAAAAAGTGTGATTAATGTGTGACACATTAGCGCATGTACATATTTGTACATTATTGTCTACGTCGTATTCGACACATCCACATACAGCCAACAAATATAAAATGATAAATGCTCTAATGGTGTTCGTACCCATCTTAATGCGGCTTAATAAATGTTTCGATTTCAATTTATTGTAAAAAAGAATCGGTTTTATACTGTTCGAGATTCTCATTGATTATATTCTCATCTATCATCTCCACACAGTCAAATCCGTGGTTAGCATATACCTCATTAACCGATAATAGACTATCAGACTCTTCTATCGTTATAACTCGAGAATATTTAATTTGGTCATTCTTAATCAAGTCAATTATCTTATTTTTAACAAATGTGAGTATTTTACTCATTTTTTATAAAAAACTTTTAGAAATATACAGACTCTATCGCGTGTCTATATCTTCTTTTTATATCCAATGTATTTATATCTGATTTTTCTTCATTTATTATATATAATGGTCCAAATTCTATACGTGTTGCTTCATCCAGATCATTAAGGTTCTTATAATCGCAATATCCTTCTCCTTCCTCTTCTATATCGTCCTTCTTAAACTTATTCTTAGCGTCACAGAATCGACCACAGCAGGATCCCATGACAAGCGTCATATTAAACTAATTCATTTTCAATAATATACTAATTAGTAACGACTATTAAAATAAATATTCTTCATAACCGGTAAGAATAGTAAAAAGACTAAATCTTCACATTGTAACTATTCAGTAGTGTTTACATAAATGAGATGAGAAGATATTCTCTATTTTGGTGGAGGATTATATGATATAATTCGTGGATAATCATTCTTAAGACACATTTCTTCATTCGTAAATCTTTTCACGTTAAATGAGTGTCCATATTTTGCAATTTCTTCATAGGATGGCGGTGTACGTGGACGGTGCTGTTCCTGTTCTTGTTGTGGTCTCCGACTTTCGTGTTTGCGTTTAGATCCCTCCATTATCGCGATTGCATAGATGAGGTACTATTTTATACCTTGTAATTAAATTTTTTTATTAATTAATTAATTAAACGTATAAAAACGTTCCGTATCTGTATTTAAGAGCCAGATTTCGTCTAATAGAACAAATAGCTACAGTAAAAATAACTAGAATAATCGCTACACCCACTAGAAACCACGGATCGTAATACGGCAATCGGTTTTCGATAATAGGTGGAACGTATATTTTATTTAAGGACTTAAGAATTGTCTGTAAACCACAATTTGCTCCCGCCGATCCTGTATTAACTATCTGTAAAAGCATATGTTGGCCGGGCGGAGCCGAACATTCTCCGATATTCAATTTTTGTATATCTATAATGTTATTAACCTCCGCATACGCATTACAGTTTTTTTCTAGCTTGGATACGGCACTAGGTACATCATTTAGATCTATTCCTATTTCCTCAGCGATAGCTCTTCTATCCTTTTCCGGAAGCAATGAAATCACTTCAATAAATGATTCAACCATGAGTGTGAAACTAAGTCTAGTATTACTCATGCATTTGTTAGTTATTCGGAGCGCGCAATTTTTAAACTGTCCTATAACCTCTCCTATATGAATAGCACAAGTGACATTAGTAGGGATAGAATGTTGAGCTAATTTTTGTAAATAACTATCTATAAAAAGATTATACAAAGTTTTAAACTCTTTAGTTTCTGCCATTTATCCAGTCTGAGAAAATGTCCCTCATAATAAATTTTTCCAAGAAACTAATTGGGTGAAGAATGGAAACCTTTAATCTATATTTATCACAGTCTGTTTTGGTACACATGATGAATTCTTCTAATGCTGTACTAAATTCAATATCTTTTTCGATTTCTGGATATGTTTTTAATAAAGTATGAACAAAGAAATGGAAATCGTAATACCAGTTATGTTCAACTTTGAAATTGTTTTTTATTTTCTTGTTAATGATTCCAGCCACTTGGGAAAAGTCAAAGTCGTTTAATGCCGATTTAATACGTTCATTAAAAACAAACTTTTTATCCTTTAGATGAATTATTATTGGTTCATTGGAATCAAAAAGTAAGATATTATCGGGTTTAAGATCTGCGTGTAAAAAGTTGTCGCAACGGGGTAGTTCGTAGATTTTAATGTATAACAGAGCCATCTGTAAAAAGATAAACTTTATGTATTGTACCAAAGATTTAAATCCTAATTTAATAGCTAACTCGGTATCTACTTTATCTGCCGAATACAGTGCTAGGGGAAAAATTATAATATTTCCTCTTTCGTATTCGTAGTTAGTTCGCTTTTCATGTTCGAAAAAGTGAAACATGCGGTTAAAATAGTTTATAACATTAATATTACTGTTAATAACTGCCGGATAAAAGTGGGATAGTAATTTCACGAATTTGATACTGTCCTTTCTCTCGTTAAACGCCTTTAGAAAAACTTTAGAAGAATATCTCAATGAGAGTTCCTGACCGTCCATAGTTTGTATCAATAATAGTAACATATTAAGAACCCGTTTATACAGTGTATGTAAAAATGTTAATTTATAGTTTAATCCCATGGCCAATGCACACACGATTAATTTTTTTTCGTCTCCCTTTAGATTGTTGTATAGAAATTTTGGTACCGTGAACTCCGCCGTAGTTTCCATGGGACTATATAATTTTGTGGCCTCGAATACAAATTTTACTACATAGTTATCTATCTTAAAGACTATACCATATCCTCCTGTAGATATGTGATAAAAATCGTCGTTTATAGGATAAAATCGTTTATCTTTTTGTTGGAAAAATGATGAATTAATGTAATCATTCTCTTCTATCTTTAGTAGTGTTTCCCTATTAAAATTCTTAAAATAATTTAACAATCTAACTGACGGAGCCCAATTTTGGTGTAAATCTAATTGTGACATTATGTTGTTAAAATATAAACAGTCTCCTAATATAACAGTATCAGATAATCTATGTGGAGACATCCATTGATATTCAGGGGATGAATCATTGGCAACACCCATTTATTGTACAAAAAGCCCCAATTTACAAATAAAAGTCCAGGTTTGATAGAGACAAACTATTAACTATCGTGTCTCTGTTTTTAACACCTCCACAGTTTTTAATTTCTTTGGTAATAAAATTATTCACTATATCAGTATCTTCTTTATCTACCAGAGATTTTACTAACTTGATAACCTTGGCTGTCTCATTCAATAGGGTAGTGATATTTGTATGAGTGATATTGATATCTTTTTGAATTGTTTCTTTTAGAAGTGATTCTTTGATGGTGTTAGCATACAAATTACAATAATGCAGAAACTCAGTTAACATACAGGAATTATAGTAAGCCAATTCCAATTGCTGTCCGTATTGTATTAGAGTATCGATATGAGTAATGGTGTCCTTGCGTTTTTCTGATAGAATGCGAGCAGCGATTTTGGCGTTATCATTTGATGATATGTTTGGAATGACGAATCCTGTTTCTACTAACTTTTTGGTAGGACAAAGTGAAACAATCAAGAAGATAGCTTCTCCTCCTATTTGTAGAAGAAATTGAACTCCTCTAGATGATCTACTGACGATAGTATCTCCTTGACAGATATTGGTCCGAATTACAGAAGTACCTGGAATGTAAAGCCCTGAAACCCCCTCATTTTTTAAGCAGATTGTTGCCGTAAATCCTGCACTATGTCCAAGATAGAGAGCTCCTTTGGTGAATCCATCTCTATGTTTCAGTTTAACCAAGAAACAGTCAGCTGGTCTAAAATTTCCATCTCTATCTAATACAGCATCCAACTTGATGTCAGGAACTATGACCGGTTTAATGTTATATGTAACATTAAGTAAATCCTTAAGTTCATAATCATCACTGTCATCAGTTATGTACGATCCAAACAATGTTTCTACCGACATAGTGGATACGAAGATACTATCCATCAGAATGTTTCCCTGATTAGTATTTTCTATATAGCTATTCTTCTTTAAACGATTTTCCAAATCAGTAACTATGTTCATTTTTTTAGGAGTATGACGCCTAGCCAGCATGGAAGAGAATTTTCTAGATCCTCTCTTCAACATCTTTAATCTCAATGGGATGCAAAACCCCATGGTGTAACAACTAACGATAAAAATAATATTGTTTTTCACTTTTTATAATTTTACCATTTGACTCATGACTTCATTGATATCTTTATACGAGCTACGAATATATAATTCTTTATAACTGAACTGAGATATATACACTGGATCTATGGTTTCCATAATTGAGTAAATAAATGCTCTGCAATAACTAATGGCAAATGTATAAAACAACAAAATTATACTAGAGTTGTTAAAGTTAATATTTTCTATTAGTTGTTCCAATAAATTATTTGTTGTGACTGTGTTCAAGTCATAAATCATTTTGATACTATCCAATAAACAGTCTTTAAGTTCTGGAATATTATCATCCCATTGTAAAGCCCCTAATTCGACTATCGAATACCCTGCTCTGGTAGCCGTTTCAATATCGACGGACGTCAATACTGTAATAAAGGTGGTAGTATTGTCATCATCATGATACACAACAGGAATATGGTCGTTAGTAGGTACGGTGACTTTACACAACGCGATATATAACTTTCCTTTTGTGCCATTTTTAACGTAGTTGGGACGTCCTGCAGGGTATTGTTTTGAAGAAATGATATCGAGAACAGATTTGATACGATATTTGTTGGATTTCTGATTATTCACTATAATATAATCTAGACAGATAGATGATTCGATAAATAAAGAAGATATATCGTTGGTAGGATAATACATCCCCATTCCAGTATTCTTGGATACTCTATTGATGACACTAGTTAAGAACATGTCTTCAATTCTAGAAAACGAAAACATCCTACACGGACTCATTAAAACTTCTAGCGCTCCTGATTGTGTCTCAAATGCCTCATACAATGATTTCAAGGATGTCATAGATTCTTTAACCCACGATTTAATATTGCGTTTAGCATCTATTTTTTTTATTAAATCGAATGGTCGGCTCTCTGGTTTGCTACCCCAATGATAACAATAGTCTTGTAAAGATAAACCGCAAGAAAATTTATACGCATCCATCCAAATAACTCTTGCACCATCAGATGATATCAGTATATTATTATAGATTTTCCATCCACAATTATTGGGCCAGTATACTGTTAGCAACGGTATATCGAATAGATTACTCATGTAACCTACTAGAATGATAGTTCGTGTACTAGTCATAATATCTTTAATCCAATCTAAGAAATTTAAAATTAGATCTTTTACACTGTTAAAGTTAACAAAGGTATTACCCGGGTACGTGGATATCATATATGGCATTGGTCCATTATCAGTAATAGCTCCATAAACTGATACGGTGATGGTTTTTATATGTGTTTGATCTAACGAGGAAGAAATTTGCGCCCATAATTCATCTCTAGATATGTATTTAATATCAAACGGTAACACATCAATTTCGGGACGCGTATATGTTTCTAAATTTTTAATCCAAATATAATGATGACCTATATGCCCTATTATCATACTGTCAACTATAGTACACTTAGAGAACTTACGATACATCTGTTTCCTATAATCGTTAAATTTTACAAATCTATAACATGCTAAACCTTTTGACGACAACCATTCATTAATTTCTGATATGGAATCTGTATTCTCGATACAGTATCGTTTTAAAGCCAGTGCTATATCTCCCTGTTCGTGAGAACGCTTTCGTATAATATCAATCAACGGATAATCTGAGGTTTTTGGAGAATAATATGACTCATGATCTATTTCGTCCATAAACAATCTAGACATAGGAATTGGAGGCGATGATCTTAATTTTGTGCAATGAGTAGTTAATCCTATAACTTCTAATCTTGTAATATTCATCATCGACATAATACTATCTATGTTATCATCGTATATTAGTATACCATGACCTTCTTCATTTCGTGCCAAAATGATATACAGTCTTAAATAGTTACGCAATATCTCAATAGTTTCATAATTGTTAGCTGTTTTCATTAAGATTTGTACCCTGTTTAACATGATGGTGTTCTATACGTCTTTATTTTCTATTTTTAATTTTTTAAATTTTTAACGATTTACTGTGGCTAGATACCCAATCTCTCTCAAATATTTTTTTAGCCTCGCTTACAAGCTGTTTATCTATACTATTAAAACTAACAAATCCGTGATTTTGGTAATGGGTTCCGTCAAAATTTGCCGACGTGATATGAACATATTCATCGTCGACTATCAACAATTTTGTATTATTCTGAATAGTGAAAACCTTCACAGATAGATCATTTTGAACACACAACGCATCTAGACTTTCGGCGGTTGCCATAGAATATACGTCGTTCTTATCCCAATTACCAACTAAAAGTCTGATCTTAACTCCTCTATTAATGGCTGCTTCTATAATGGAGTTGTAAATGTCGGGCCAATAGTAGCTATTACCATCGACACGTGTAGTGGGAACTATTGCCAAATGTTCAATATCTATACTAGTTTTAGCTGACCTAAGTTTATCAATAACTACATCGGTATCTAGATCTCTAGAATATCCCAATAGGTGTTCCGGAGAATCAGTAAAGAACACTCCACCTATAGGATTCTTAATATGATACGCAGTGCTAACTGGCAGACAACAAGCCGAAGAGTATAAATTCAACCATGAATTTTTTACGCTATTAAAGGCTTTAAAAGTATCAAATCTTCTACGAAGATCTGTGGCCAGTGGTGGATAATCAGAATATACACCTAACGTTTTAATTGTATGTATAGATCCTCCAGTAAATGACGCGTTTCCTACATAACATCTTTCATCATCTGATACCCAAAAACATCCAAGTAGTAGTCCTACGTTATTTTTTTTATCAATATTAACGGTTATAAAATTTATATCCGGGCAGTGACTTTGTAGCTCTCCCAGATTTCTTTTCCCTCGTTCATCTAGCAAAACTATTATTTTAATCCCTTTTTCAGATGCCTCTTTTAGTTTATCAAAAATAAGCGCTCCCCTAGTCGTACTCAGAGGATTACAACAAAAAGATGCTATGTATATATATTTCTTAGCTAGAGTGATAATTTCGTTAAAACATTCAAATGTTGTCAAATGATCGGATCTAAAATCCATATTTTCTGGTAGTGTTTCTACCAGTCTACATTTTGCTCCCGCAGGTGCTGATGTAAATGGCCACATTTAGTTAACAAAAAAAATTTACATATCCTGTTCCATCAACGATTCTAGAACGTCATCGGTTATATCGCTAAAATCTTCATCAAAGTCGACATCACAACCTAGCTCAGTCAATATATTAAGAAGTTCCGTGATATTATCTTCATCGATTTCTATATCCATATCGGATGTAGATAATTGACTGATTATTGAGTTAAAATCAGTACTAATTTCCAACCCTTCAGAATATACTTTGTGTTTCATGGTAATATCGGTAGCGTATTTAAGTTGGTAGATTTTTCAATTATGCATCAATATAGCAACAGTAGTTCTTGCTCCTCCTTAATTCTAGCATCTTCTTCATTATTTTCTTCTACATACATAAGCATCTCCAATACGTTAGACAGCACACCGATGATGGCGGCCGCCACAGACACGAATATGACTAGACTGATGACCATTTAAAAAAACACTCTAGCTTTCACTTAAACTGTATCGATCATTCTTTTAGCACATGTATAATATAAAAACATTATTCTATTTCGAATTTAGGCTTCCAAAAATTTTTCATCCGTAAACCGATAATAATATATATAGACTTGTTAATAGTTGGAATAAATAGATTAATGCTTAAACTATCATCATCTCCACGATTAGAGATACAATATTTACATTCTTTTTGCTGTTTCGAAACTTTATCAATACACGTTAATACAAACCCAGGAAGGAGATATTGAAACTGAGGCTGTTGAAAATGAAACGGTGAATACAATAGTTCAGATAATGTAAAATCATGATTCCGTATTCTGATGATATTAGAACTACTAATGGATGTCAATGGTATGTATCTAGGAGTATCTATTTTAACAAAGCATCTATTTGCTAATATACAATTATCCTTTTGATTAATTGTTATTTTATTCATATTCTTAAAAGGTTTCATATTTATCAATTCTTCTACATTAAAAATTTCCATTTTTAATTTATGTAGCCCGCAATACTCCTCATTACGTTTCATTTTTTGTCTATAATATCCATTTTGTTTATCTCAGTACATAGATTATCCAATTGAGAAGCACATTTAGTAGTTTTGTACATTTTAAGTTTATTGACGAATCGTCGAAAACTAGTTATAGTTAACATTTTATTATTTGATACCCTGATATTAATACCCCTACCGTTACTATTATTTATAACTGATGTAACCCACGTAACATTAGAATTAATTATCGATAGTAATGCATTCACGCTTCCAAAATTGTCTATTATAAACTCACCAATAATTTTTTTATTGCATGTTTTCATATTCATTAGGATTATCAAATCTTTAATCTTATTACGATTGTATGCGTTAATATTACAAGACGTCATTCTAAAAGACGGAGGATCTCCATCAAATGCCAGACAATCACGTACAAAGAACATGGAAATAGGTTTTGTTCTATTGCGCATCATAGATTCATATAGAACACCCGTAGAAATACTAATTTGTTTTACTCTATAAAATACTAATGCATCTATTTCATCGTTTTGTATAACGTCTTTCCAAGTGGCGAATTCCAAATTTTTTTCATTGATAGTACCAAATTCTTCTATCTCTTTAACTATTTGCATAGATAGGTAATTACAGTGATGTCTACATGCCGTTTTTTGAAACTGAATAGATGCATCTAAAAGCGATGCTACGCTAGTCACGATCACCACTTTCATATTTAGAATATATGTATGTAAAAATATAGTAGAATTTCATTTTGTTTTTTTCTATGCTATAAATGAATTCTCATTTTGCATCTGCCCATACTCCGTTTTATATCAATACCAAAGAAGGAAGATATCTGGTTCTAAAAGCTGTTAAAGTATGCGATGTTAGAACTGTAGAATTCGAAGGAAGTAAAGCTTCCTGCGTACTCAAAGTAGATAAACCCTCATCGCCCGCGAGCGAGAGAAGACCTTCGTCACCGTCCAGATGCGAGAGAATGAATAACCCAGGAAAACAAGTTCCGTTTATGAGGACGGACATGCTACAAAATATGTTCGCGGCTAATCGCGATAATGTAGCTTCTAGACTTTTGTCCTAAAATACTATTATATCTTTTTCGATATTAATAAATCCGTGTCGCCCATGTTTTTTATCTCTTTCAGTATGTGAATAGATAGGTATTTTATCTCTATTCATCATCGAATTTAAGAGATCCGATAAACATTGTTTGTATGCTCCAGATGTCAGCATCTGATACAACAATATATGTGCACATAAACCTCTGGCACTTATTTCGTGTACCTTCCCCTTATCACTAAGGAGAATAGTATTTGAGAAATATGTATACATAATATTATCATGAATTAGATATACAGAATTTGTAACACTCTCGAAATCACACGATGTGTCTGCGTTAAGATCTAATATATCACTCGATAACACATTTTCATCTAGATACACTAGACATTTTTTAAAGCTAAAATAGTCTTTAGTAGTAACAGTAACTACGCGATTATTTTCATCGATGATGCATTTCATCGGCATATTATTACGCTTACCATCAAAGACTATACCATGTGTATATCTAACGTATTCTAGCATAGTTGCCATACGCGCATTAAACTTTTCAGGATCTTTGGATAGATCTTCCAATCTATCTATTTGAGAAAACATTTTTATCATGTTCAATAGTTGAAACGTCGGGTCCACTATATAGATATTATCTATAAAGATTTTAGGAACTACGTTCATGGTATCCTGGCGAATATTAAAACTATCAATGATATGATTATCGTTTTCATCTTTTATCACCATATAGTTTCTAAGATATGGGATTTTACTTAATATAATATTATTTCCCGTAATAAATTTTATTAGAAATGCCAAATCTATAAGAAAAGTCCTAGAATTAGTCTGAAGAATATCTATATCACCGTACCGTATATTTGGATTAATTAGATATAGAGAATATGATCCGTAACATATACAACTTTTATTATGGCGTCTAAGATATTCTTCCATCAACTTATTAACATTTTTGACTAGGGAAGATACATTATGACGTCCCATCACTTTTGCCTTGTCTATTACTGCAACGTTCATAGAATTTAGCATATCTCTTGCCAATTCTTCCATTGATGTTACATTATAAGAAATTTTAGATGAAATTACATTTGGAGCTTTAATAGTAAGAACTCCTAATATGTTCGTGTATGTGGTCACTAATACAGATTGTAGTTCTATAATCGTAAATAATTTACCTATATTATATGTTTGAGTCTGTTTAGAAAAGTAGCTAAGTATACGATCTTTTATTTCTGATGCAGATGTATCAACATCGGAAAAAAATCTTTTTTTATTCTTTTTTACTAAAGATACAAATATGTCTTTGTTAAAAACAGTTATTTTCTGAATATTTCTAGTTTGTAATTTTAACATATGATATTCATTCACACTAGGTAATCTACCTAAATAGGTTTCTATAATCTTTAATGTAATATTAGGAAAAGTATTATGATCAGGATTCCTATTCATTTTGAGGATTTAAAACTCTGATTATTGTCTAATATGGTCTCAACGCAAACTTTTTCACAGAGCGATAGAGTTTTTGATAACTCGTTTTTCTTAAGAAATATAAAACTACTGTCTCCAGAGCTTGCTCTATCTTTTATTTTATCTAATTCGATACAAACTCCTGATACTGGTTCAGAAAGTAATTCATTAATTTTCAGTCCTTTATAGAAGATATTTAATATAGATAATACAAAATTTTCAGTTTTTGATATCGATCTGATTGATCCTAGAACTAGATATATTAATAACGTGCTCATTAGGCAGTTTATGGCAGCTTGATAATTAGATATAGTATATTCCAGTTCATATTTATTAGATACCGCATTGCCCAGATTTTGATATTCTATGAATTCCTCTGAAAATAAATCCAAAATAACTAGACATTCTATTTTTTGTGGATTAGTGTACTCTCTTCCCTCTATCATGTTCACAACTGGTGTCCACGATGATAAATATCTAGAGGGAATATAATATAGTCCATAGGATGCCAATCTAGCAATGTCGAATAACTGTAATTTTATTCTTCGCTCTTCATTATGAATTGATTCTTGAGGTATAAACCTAACACAAATTATATTATTAGACTTTTCGTATGTAATGTCTTTCATGTTATAAGTTTTTAATCCTGGAATAGAATCTATTTTAATGAGGCTTTTAAACGCAGAGTTCTCCAACAAGTCAAAACATAATACTCTGTTGGTTTTCTTATATACAATATTACGATTTTCTTCTTTGAATGGAATAGGTTTTTGAATTAGTTTATAATTACAACATAATAAATAAGGAAGTGTGCAAATAGTACGCGGAAAAAACATAATAGCTCCCCTGTTTTCATCCATGGTTTTAAGTAAATGATCACTGGCTTCTTTAGTCAATGGATATTCGAACATTAACCGTTTCATCATCATTGGACAAAATCCATATTTCTTAATGTAAAGAGTGATCAAATCATTATGTTTATTGTACCATCTTGTTGTAAATGTGTATTCGGTTATCGGATCTGCTCCTTTTTCTATTAAAGTATCGATGTCGATCTCGTCTAAGAATTCAACTATATCGACATATTTCATTTGTATACACATAACCATTACTAACGTAGAATGTATAGGAAGAGATGTAACGGGAACAGGGTTTGTTGATTCGCAAACTATTCTAATACATAATTCTTCTGTTAATACGTCTTGCACGTAATCTATTATAGATGCCAAGATATCTATATAATTATTTTGTAAGATGATGTTAACTATGTGATCTATATAAGTAGTGTAATAATTTATGTATTTCGATATATGTTCCAACTCTGTCTTTGTGATGTCTAGTTTTGTAATATCTATAGCATCCTCAAAAAATATATTCGCATATATTCCCAAGTCTTCAGTTCTATCTTCTAAAAAATCTTCAACGTATGGAATATAATAATCTATTTTACCTCTTCTGATATCATTAATGATATAGTTTTTGACACTATCTTCTGTCAATTGATTCTTATTCACTATATCTAAGAAACGGATAGCGTCCCTAGGACGAACTACTGCCATTAATATCTCTATTATAGCTTCTGGATATAATTCATCTATTATACCAGAATTAATAGGAACTATTCCGTATTTATCTAACATAGTTTTAAGAAAGTCAGAATCTAAGACCTGATGTTCATATATTGGTTCATACATGAAATGATCTCTATTGATGATAGTGACTATTTCGTTCTCTGAAAATTGGTAACTCATTCTATATATGCTTTCCTTGTTGATGAAGGATAGGATATACTCAATAGAATTTGTACCAACAAACTGTTCTCTTATGAATCGTATATCATCATCTGAAATAATCATGTAAGGCATACATTTAACAATAAGAGACTTGTCTCCTGTTATCAATATACTATTCTTGTGATAATTTATGTGTGAGGCAAATTTGTCCACGTTCTTTAATTTTGTTATAGTAGATATCAAATCCAATGGAGCTACAGTTCTTGGCTTAAACAGATATAGTTTTTCTGGAACGAATTCTACAACATTATTATAAAGGACTTTGGGTAGATAAGTGGGATGAAATCCTATTTTAATTAATGCAATAGCCTTGTCCTCGTGCAGATATCCAAACGCTTTTGTGATAGTATGGCATTCATTGTCTAAAAACGCTCTACGAATATCTGTGACAGATATCATCTTTAGAGAATACTAGTCGCGTTAATAGTACTAAAATTTGTATTTTTAATCTATCTCAATAAAAAATTAATATTTATGATTTAATGTATAACTTTACTAACTGTTATTGATAACTAGAATCAGAATCTAATGATGACGTAACCAAGAAGTTTATCTACAGCCAATTTAGCTGCATTATTTTTAGCATCTCGTTTAGATTTTCCATCGGCCTTATCGAATACTCTCCCGTCGATGTCTACACAGGCATAAAATGTAGGAGAGTTACTAGGCCCCACTGATTCAATACGAAAAGACCAATCTCTCTTAGTTATTTGGCAGTACTCATTAATAATGGTGACAGGGTTAGCATTTTTCCAATCAATAATTTTTTTAGCCGGAATAACATCATCAAAAGACTTATGATCCTCTCTCATTGATTTTTCGCGGGATACATCATCTATTATGACGTCAGCCATAGTCATAGCATCCGGCTTATCCGCCTCAGTTGTCATAAACCAACGAGGAGGAATATCGTCGGAGCTGTACACCATAGCACTACGTTGAAGATCGTACAGAGCTTTATTAACTTCTCGCTTCTCCATATTAAGTTGTCTAGTTAGTTGTACAGCAGTAACTCCTTCAAGTCCAATGTTTTTAATAGCCTCACATACAATCTCTGCGTCAGAACGCTCGTCAATATAGATCTTAGACATTTTTAGAGAGAACTACCGCAACTAGTAATAAAACTAATCTATTTTATCATTTTTTTATTCATCATCTTCTGGTGGTTCATCGTTCCTATCGAATGTGACTCTGATTAACCCATCATCTATAGGTGATGCTGGTTCTGGAGATTCTGGAGGAGATGGATTATTATCTGGAAGAATCTCTGTTATTTCCTTGTTTTTATGTATCGATTGCGTTGTAACATTAAGATTGCGAAATGCTCTAAATTTGGGAGGCTTGAAGTGTTGTTTGCAATCTCTACACGCGTGTCTAACTAATGGAGGTTCATCAGCGGCTCTAGTTTGAATCATCATCGGTGTAGTATTCCTACTTTTACAGTTAGGACACGGTGTATTGTATTTCTCGTTGAGAACGTTAAAGTAATCATTGTAACTCACATCCTTTATTTTATCTATATTGTATTCTACTCCTTTCTTAATGCATTTTATACCGAATAAGAGATAGCGAAGGAATTCTTTTTCGGTACCGCTAGTACCCTTAATCATGTCACATAGTGTTTTATATTCCAAATTTGTGGCAATGAACGGTTTATTTCTATACGATAGTTTGTTTCTAGAATCCTTTGAGTATTCTATACCAATATTATTCTTTGATTCGAATTTAGTTTCTTCGATATTGGATTTTGTATTACCTATATTCTTGATGTAGTACTTTGATGATTTTTCCATGGCCCATTCTATTAAGTCTTCCAAGTTGGCATCATCCACATATTGTGATAGTAATTCTCGGATATTAGTAGCGGCTACCGCCATTGATGTTTGTTCATTGGATGAGTAACTACTAATGTATACATTTTCCATTTATAACACGTATGTATTAACTTTGTTCATTTATATTTTTTCATTATTATGTTGATATTAACAAAAGTGAATATATATATGTTAATAATTGTATTGTGGTTATATGGCTACAATTTCATAATGAGCGGAAGTCAGTGTCCGATGATTAATGACGATAGATTTACTCTGAAAAGAAAGTATCAAATCGATAGTGTAGAGTCGACAATGAAAATGGATAAGAAGAGGACAAAGTTTCAAAATAGAGCCAAAATGGTAAAAGAAATAAATCAGACAATAAGAGCAGCACAAACTCATTACGAGACATTGAAACTAGGATACATAAAATTTAAGAAAATGATTAGGACTACTACTTTAGAAGATATAACAACATCTATTCCAAATATTCAGAAAATTTATAAACTATTCTCGGACATTTCAGCCATTGGCAAAGTATCACAGAATCCGAGTAAAATGGCATATGCTTTACTGCTTTACATGTTCCCCAATTTGTTTGGAGATGACCATAGATTCATTCTTTATAGAATGTTTCCAATGAGTAAAATCAAACACAAGATCTTCTCTCCTTTCAAACTTAATCTTATTAGAATATTAGTGGAAGAAAGATTCTATAATAATGAATGCAGATCTAATAAATGGAGAATAATTGGAACACAAGTTGATAAAATGTTGATAGCTGAATCTGATAAATATACAATAGATGCAAGGTATCGCCTAAGACCTATATATAGAATCAAGGGAAAATCTGAAGAAGATACCCTCTTCATCAAACAGATGGTAGACCAATGTGTGACATCCCAGGAATTGGTGGAAAAAGTGTTGAAGATACTGTTTAGAGATTTGTTCAAGAGTGGAGAATACAAAGCGTACAGATACGATGATGACGTAGAAAATGGATTCATTGGATTGGATAAACTAAAATTAAACATTGTTCATGATATAGTTGAACCATGTATGCCTGTTCGTAGGCCAGTGGCTAAGATACTGTGTAAAGAAATGGTAAATAAATACTTTGAGAATCCGCTGCATATTATTGGTAAGAATCTTCAAGAGTGCATTGACTTTGTTAGTGAATAGGCATTTCATCTTTCTCCAATACTAATTCAAATTGTTAAATTAATAATGGATATAGTATAAATAGTTATTAGAATAAGAGTGTAGTATCATAGATAAGTATTTACTTTCTTCTATAAAATGGATTTTATTCGTAGAAAGTATCTTATATACACAGTAGAAAATAATATAGATTTTTTAAAGGATGATACATTAAGTAAAGTAAACAATTTTACCCTCAATCATGTACTAGCTCTCAAGTATCTAGTTAGCAATTTTCCTCAACACGTTATTACTAAGGATGTATTAGCTAATACCAATTTTTTTGTTTTCATACATATGGTACGATGTTGTAAAGTGTACGAAGCGGTTTTACGACACGCATTTGATGCACCCACGTTGTACGTTAAAGCATTGACTAAGAATTATTTATCGTTTAGTAACGCAATACAATCGTACAAGGAAACAGTGCATAAACTAACACAAGATGAAAAATTTTTAGAGGTAGCCGAATACATGGAAGAATTAGGAGAACTTATAGGGGTAAATTATGACTTAGTTCTTAATCCATTATTTCACGGAGGGGAACCCATCAAAGATATGGAAATCATTTTTTTAAAACTGTTTAAGAAAACAGACTTTAAAGTTGTTAAAAAATTAAGTGTTATAAGATTACTTATTTGGGCTTACCTAAGCAAGAAAGATACAGGTATAGAGTTTGCGGATAATGATAGACAAGATATATATACTCTATTTCAACACACTGGTAGAATAGTCCATAGCAATCTAACAGAAACGTTTAGAGATTATATCTTTCCCGGAGATAAGACTAGCTATTGGGTGTGGTTAAACGAAAGTATAGCTAATGATGCAGATATCGTTCTTAATAGACCCGCCATTACCATGTATGATAAAATTCTTAGTTATATATACTCTGAGATAAAACAGGGACGTGTTAATAAAAACATGCTTAAGTTAGTTTATATCTTTGAGCCTGAAAAAGATATCAGAGAACTTCTGCTAGAAATCATATATGATATTCCTGGAGATATCCTATCTATTATTGATGCAAAAAACGACGATTGGAAAAAATATTTTATTAGTTTTTACAAAGCTAATTTTATTAACGGTAATACATTTATTAGCGATAGAACGTTTAACGAGGATTTATTCAGAGTTGTTGTTCAAATAGATCCTGAATATTTTGATAATGAACGAATTATGTCTTTATTCTCTACGAGTGCTGTGGAAATTAAACGATTTGATGAGTTAGATATTAATAACAGTTATATATCTAATATAATTTATGAGGTGAACGATATCACATTAGATACAATGGATGATATGAAGAAGTGCCAAATCTTTAACGAGGATACATCGTATTATGTTAAGGAATACAATACATACCTGTTTTTGAACGAGACGGATCCCATGGTCATAGAGAACGGAATACTAAAGAAACTGTCATCTATAAAATCCAAGAGTAGACGACTGAACTTGTTTAGCAAAAACATTTTAAAATATTATTTAGACGGACAATTGGCTCGTCTAGGTCTTGTGTTAGATGATTATAAAGGAGACTTGTTAGTTAAAATGATAAACCATCTCAAGTTTGTGGAGGATGTATCAGCATTCGTTAGATTTTCTACAGATAAAAACCCTAGTGTTCTTCCATCGCTAATCAACACTATTTTAGCTAGTTATAATATTTCCATCATTGTCCTATTTCAAAAGTTTTTAAGAGATAATCTATATCATGTAGAAAAATTCTTGGATAAAAGCATCCATCTAACCAAGACGGATAAGAAATATATACTTCAATTGATAAGGCACGGCAGATCATAGAACAAACCAAATATATTATTAATAATTTGTATATACATAGATATAATTATCACATATTAAAATTCACACATTTTTGATAAATGGGAACTACTGCAACAATTTAGACTCCCACCAAATTAATGAATAAAGAGAATGCAGAAATGATTTTGGAAAAATTGTTGATCATATAGTTATGTATATTAGTGACGAATCAAGAGATGAAAATAATCCTGAATATATTGATTTTCGTAACAGATACGGAGACTATAGATCTTTCATTATAAAAAGTGATCACGGGTTTGTAAAGATCATGCAGAGAAAAGTTCTCCAAAAACACAACAAATGATAATCAAACACATCTACGAACAATATCTTATTTCGGTATCTGAAGTACTTATTAAAACCTATGATATCCATGGGTGACATAATTACATATAACGGATGTAAAGACAATAAATGGATGCTAGAACAACTCTCTACCCTAAACTTTAACAATCTCCACGTATGGAACTCATGTAGCATAGGTAATGTAACGCGTATATTTTATACATTTTTTAGTTATCTGATGAAAGATAAACTAGATATATAAATAATCCCATTCTAATACTTTAACATGTGTATTACATGCATCTTATTAGAATATTAACCTAACTAAAAGACATAATATAAAAACTTATTACATAGTTGATAAAAAGCGGTAGGCTATAAATATTATGGCAGCAGTCGTTCCGCGTTTTGATGACGTGTACAAAAATGCACAGCGAAGAATTCTAGATCAAGAAACATTTTTTAGTAGAGGTCTAGGTAGACCGTTAATGAAGAACACATATTTGTTTGATAATTACGCGTATGGATGGATACCAGAAACGGCAATTTGGAGTAGTCGATACGCAAACCTAGATGCAAGTGACTATTATCCCATTTCGTTGGGATTACTTAAAAAGTTTAAGTTTCTCATGTCTCTATATAAAGGTCCTATTCCAGTATACGAAGAAAAAGTAAATACTGAATTCATAGCCAATGGATCGTTCTCTGGTAGATACGTATCATATCTTCGAAAGTTTTCTGCTCTTCCAACAAACGAGTTTATTAGTTTTTTGTTACTGACTTCCATTCCAATCTATAATATCTTGTTCTGGTTTAAAAATACTCAGTTTGATATTACTAAACACACATTATTCAGATACGTCTATACAGATAATGCCAAACACCTGGCATTGGCTAGGTATATGCATCAAACAGGAGACTATAAGCCTTTGTTTAGTCGTCTCAAAGAGAATTATATATTTACCGGTCCCGTTCCAATAGGTATCAAAGATATAGATCACCCTAATCTTAGTAGAGCAAGAAGTCCATCCGACTATGAGACATTAGCTAATATTAGTACTATATTGTACTTTACCAAGTATGATCCGGTATTAATGTTTTTATTGTTTTACGTACCTGGGTATTCAATTACTACAAAAATTACTCCAGCAGTAGAGTATCTAATGGATAAACTGAAACTAACAAAGAACGATGTACAACTGTTGTGAATATTTTATGCTTCGTAAAATGTAGGTCTTGAACCAAACATTCTTTCAAAGAATGAGATACATAAAACTTTATTATCCAATAGATTGACTATTTCGGACGTCAATCGTTTAAAGTAAACTTCGTAAAATATTCTTTGATCACTACCGAGTTTAAAACTTCTATCGATAATTGTTTCATATGTTTTAATATTTACAAGTTTTTTGGTCCATGGTACATTAGCCGGACAAATATATGCAAAATAATATCGTTCTCCAAGTTCTATAGTCTCTGGATTATTTTTATTATATTCAGTAACCAAATACATATTAGGGTTATCTGCGGATTTATAATTTAAGTGATGCATTCGACTCAACATAAATAATTCTAGAGGAGACGATCTACTATCAAATTCGGATCGTAAATCTGTTTCTAAAGAACGGAGAATATCTATACATACCTGATTAGAATTCATCCGTCCTTCAGACAACATCTCAGACAGTCTGGTCTTGTATATCTTAATCATATTCTTATGAAACTTGGAAACATCTCTTCTAGTTTCACTAGTACCTTTATTAATTCTCTCAGGTACAGATTTTGAATTCGATGATGCCGAGTATTTCATCGTTGTATATTTCTTCTTCGATTGCATAATCAGATTCTTATATACCGCCTCAAACTCTATTTTAAAATTATTAAACAATACTCTACTATTAATCAGTCGTTCTAATTCCTTTGCTATTTCTATGGACTTATCTACATCTTGACTGTCTATCTCTGTAAACACGGAGTCGGTATCTCCGTACACGCTACGAAAACGAAATCTGTAATCTATAGGCAACGATGTTTTCACAATCGGATTAATATCTCTATCGTCCATATAAAATGGATTACTTAATGGATTGGCAAACCGTAACATACCGTTAGATAACTCTGCTCCATTTAGTACCGATTCTAGATACAAGATCATTCTACGTCCTATGGATGTGCAACTCTTAGCCGAAGCGTATGAGTATAGAGCACTATTTCTAAATCCCATCAGACCATATACTGAGTTGGCTATGATCTTGTACGTATACTGCATGGAATCATAGATGGCCTTTTCAGTTGAACTGGTAGCCTGTTTTAGCATCTTTTTATATCTGGCTCTCTCTGCCAAAAATGTTCTTAATAGTCTAGGAATGGTTCCTTCTATTGATCTATCGAAAATTGCTATTTCAGAGATGAGGTTAGGTAGTCTAGGTTCACAATGAACCGTAATATATCTAGGCGGTGGATATTTCTGAAGCAAGAGCTGATTATTTATTTCTTCTTCCAATCTATTGCTACTAACAACGACACCGACTAATGTTTCCGGAGATAGATTTCCAAAGATACACACATTAGGATACAGACTGTTATAATCAAAGATTAATACATTATTACTAAACATTTTTTGTTTTGGAGCAAATACCTTACCGCCTTCATAAGGAAACTTTTGTTTTGTTTCTGATCTAACTAAGATAGTTTTAGTTTCCAACAATAGCTTTAACAGTGGACCCTTGATGACTGTACTCGCTTTATACCCGAATACCATGGATTGAGGAAGCACATATGTTGACGCACCCGCGTCTGTTTTTGTTTCTACTCCATAATACTCCCACAAATACTGACACAAGCAAGCATCATGAATACAGTATCTAGCCATATCTAAAGCTATGTTTAGATTATAATCCTTATACATCTGTGCTAGATCAACGTCATCCTTTCCGAACGATAATTTATACGTATCATTTGTTAAAGTGGGACATGATAGTACAACTTTGAATCCATTTTCCCAAATATCTTTATGAATTACTTTACATATAATATCATCAACCGTCACATAATTACCTGTGGTTAAAACCTTTGCAAATACAGCGGCTTTGCCTTTAGCGTCCGTAGTATCGTCACCGATGAACGTCATTTCTCTAACTCCTCTATTTAATACTTTACCCATGCAACTAAACGCGTTCTTGGATATAGAATCCAATTTGTACGAATCCAATTTTTCAGATTTTTGAATGAATGAATACAGATCAAAAAATATAGTTCCATTATTGTTATTAACGTGAAACGTAGTATTGGCCATGCCACCTACCCCCTTGTGACTAGACTGATTTCTCTCATAAATACAGAGATGTACAGCTTCCTTTTTGTCTGGAGATCTAAAGATAATCTTCTCTCCGGTTAATAATTCTAGACGATTAGTAATATATCTCAGATCAAAGTTATGTCCGTTAAAGGTAACAATATAGTCGAACGTTAGTTCCAACAATTGTTTAGCTATTTGTAGCAAAACTATTTCAGAACACAGAACTAGTTCTCTCTCATAATCCATTTCCATCAGTGACTGTATCCTCAAACATCCTCTATCGACGGCTTCTTGTATTTCCTGTTCCGTTAACATCTCTTCATTAATAAGCGTAAACAATAATCGTTTACCACTTAAATCGATATAACAGTAACTTGTATGAGAGATTGGGTTAATAAATACAGAAGGAAACTTCTTATCGAAGTGACACTCTATATCTAGAAATAAGTACGATCTTGGGATATCGAATCTAGGTATTTCTTTAGCGAAACAGTTACGTGGATCGCCACAATGATAACATCCATTGTTAATCTTTGTTAAATATTGATCGTCCAACGAATAACATCCGTCTGGAGATATCCCGTTAGAAATATAAAACCAACTAATATTGAGAAATTCATCCATAGTGGCGTTTTGTATGTTGCGTTTCTTTGGCTCTTCTATTAACCACATATCTGCGACGGAGCATTTTCTATCTTTAATATCTAGATTATAACTTATTGTCTCGTCAATGTCTATAGTTCTCATCTTTCCCATTGGCCTCGCATTAAATGGAGGAGGAGCCAATGACTGATATATTTCGTCTGTCACTACGTAATAAAAGTAATGAGGAAATCGTATAAATACTGTCTCGCCATTTCGACATCTGGATTTCAGATATAAAAATCTGTTTTCACCATGACTTTCGAACCAATTAATACACCGAACATCCATTTCTAGAATTTAGAAATATATTTTCATTTAAATGAATCCCAAACATTGGGGAAGAGCCGCTTGGACCATTATTTTTATAGTACTTTCACAAGCGGGTTTAGACGGCAACATAGAAGCGTGTAAACGAAAACTATATACTATAGTCAGCACTCTTCCATGTCCTGCCTGCAGACGACACGCAACTATCGCTATAGAGGACAATAATATTATGTCTAGCAATGATCTAAATTATATTTATTATTTTTTCATCAGATTATTTAACAATTTGGCATCTGATCCCAAATACGCAATCGATGTGTCAAAGGTTAAACCTTTATAAACTTAACCCATTATAAAACTTATGATTAGTTACGACTGAAATAACCGCATGATTATTTTTTGGTATAATTCTACACGGCATAGTTTCTGTAACTATGAATTCTACCCCCGTTACATTTGTATAATCTTTAACAAACAGTAAGGGCTCGTCAAATACGTAAAACTCATTACTTACGATAGAAATAGATCCCTTATCACACTTAAAATAAAAAATATCCTTATCCTTTACCACCAAATAAAATTCTGATTGGTCAATGTGAATGTATTCACTTAACAATTCCACAAATTTATTTATTAACTCCGAGGCACATACATCGTCGGTATTTTTTATGACAAACTTTACTCTTCCAGCATCCGTTTCTAAAAAAATATTAACGAGTTCCATTTATATCATCCAATATTATTGAAATGACGTTGATGGACAGATGATGCAAATAAGAAGGTACAATACCTTTGTCTACCATCTCCTCCAATTCATGCTCTATTTTGTCATTAACTTTAATGTGTGAAAACAGTACGCCACATGCTTCCATAACAGTGTGTAACACTTTTAATACGAAATGTCTGATGGCGACATAATTGTTCAAAACTGCCAATCTATAATAGATGGTGGCCACAATATATTCTATGATGGTATTGAAGAAGATAACAACCTTGGCATATTGATCATTTGATACGGACATGGTATCAACAGATAGCTTGAATGAAAGAGACTCAGTAATCGGAATAAGTGTTTTCTCAATGGAGTGTCCGTATACCAACATGTCTGATATTTTGATGTACACCATCAAATTGTTTAGTTTTCTCTTTTTATCCTCATTAAACAGCATTTCTGTCAATGGACCCCAACATCTTTGACCGATTAAGTTTTGATTGATTCTTCCATGTAAGGCGTATCTAGTCAGATCGTATAATCTAGCCAACAATCCATCGTCGGTGTTTAGGTCACATCGTATACTTTTTAATTCTCTATAGAAGAGAGACACACACCTGGAACAATTACAGACAGTAATTTCTTTATTATCTACAGATGTAAGATACTTGAAGACATTCCTATGATGATGTAGAATTTTTGATAACACTGTATTGATGGTATCTGTTACCATAATACCTTTTATGGCGGACAGTGTCAGAGCACAGGATTTCCAATCTTTTGCAATTTTCAGTACCATTATCTTTGTTTTGATATCTATATTAGATAGCATGGTGCGTCTGACAACACAGGGATTAAGACGAAAAGATGAAATGATTCTATCTACATCTTCAATAGATACCTTGCTATTTTTTCTGGCATTATCTATATGTGAGAGAATATCCTCTAGAGAATCAGTATCCTTTTTGATAATAGTGGATCTCAATGACATGGGACGTCTAAACCTTCTTATTTTATCACCTGTTTGCATGGTGATTTGTCTGCTTTCTTTTAACATGATGTAATCTCTAAATTCATTGGCAAATTGTCTATATCTAAAATCATAATATGAGATGTTTACCTCTACAAATATCTGTTCGTCCAATGTTAGAGTATCTACATCAGTCTTGTATTCCAAATTAAACATGGCAACGGATTTAATTTTATATTCCTCTATTAGGTCCTTGTCGACAATAACAGAGTGCAGATAATCATTTAATCCGTCGTACATGGTTGGAAGATGCGCATTAACAAAATCTTTAATTGTCTTGATGAAGGTGGGACTATATCTAACATCTTGATTAATAAAATTTATAACATTTTCCATAGGATACTTTGTAACTAGCTTTATACACATCTCTTCATCTGTAAGTTTAGACAGAATATCGTGAACAGGTGGTATATTATATTCATCAGATATACGAAGAATAATGTTCAAATCTAGATTGTTTAATATATTATATAGATGTAGTGTAGCTCCTACAGGAATATCTTTAACTAAGTCAATTATTTCATCAACCGTTAGATCTATTTTAAAGTTAATCATATAGGCATTGATTTTTAAAAGGTATGTAGCCTTGATTACATTCTCATTAATTAACCATTCCAAGTCACTATGTGTAAGAAGATTATATTCTATCATAAGCTTGACTACATTTGGTCCCGATACCATTAAAGAATTCTTATGATATAAGGAAACAGCTTTTAGGTACTCATCTACTCTACAAGAATTTTGGAGAGCCTTAACGATATCAGTGACGTTTATTATTTCAGGAGGAAAAAACCTAACATTGAGAATGTCGGAGTTAATAGCTTCCAGATACAGTGATTTTGGCAATAGTCCGTGTAATCCATAATCCAGTAACACGAGCTGGTGCTTGCTAGACACCTTTTCAATGTTTAATTTTTTTGAAATAAGCTTTGATAAAGCCTTCCTTGCAAATTCCGGATACATGAACATGTCGCCGACATGAATGAGTATTTTTTTTCAGTATTTTATATTTTTCTTAACAAGTTCTCAATACACCAATAGATGATAGAATATCACCCAATGCGTCCATGTTGTCTATTTCCAACAGGTCGCTATATCCACCAATAGAAGTTTTCCCAAAAAATATTCTAGGAACAGTTCTACCACCAGTAATTTGTTCAAAATAGTCGTGCAATTTATTTTCGGGTTTAAATTCTTTAATATCGACAATTTCATACGCTCCTCTTTTGAAACTAAACTTATTTAGAATATCCAGTGCGTTTCTACAAAAAGGACATGTAAACTTGACAAAAATTGTGACTTTGTTATTGGTCAACCTTTGTTGTACAAATTCCTCGGCCATTTTAATATTTAAGTGATATAAAACATTCTCGACTTATTTAACTCTTTAGTCGAGATATATGGACGCAGATAGCTATATGATAACCAACTACAGAAGGCGAACGCTATAAAAAACATAATTACGACGAGCATATTCATAAATATTTTTATTCAGCATTACTTGATATAGTAATATTAGGCACAGTCAAACATTCAACCACTCTCGATACATCAACTCTCTCATTTTCTTCAACAAATTCTGCAATATCTTCGTAAAAAGATTCTTGAAACTTTTCAGAATATCTATCGACTTCAGATGAAATAGCGTTCGTCAACATACTATGTTTTGTATACATAAAGGCGCCCATTTCAACAGTTTCTAGTGACAAAATGCTAGCGATCCTAGGATCCTTCAGAATCACATAGATTGACGATTCGTCTCTCTTAGTAACTCTAGTAAAATAATCATACAATCTAGTACTCGAAATAATATTATCCTTGACTTGAGGAGATCTAAACGATCTAGTTTTGAGAACATCGATAAGTTCATCGGGAATGACATACATACTATCTTTAATAGAACTCTTTTCATCCAGTTGAATGGACTCGTCCTTAACCAACTGATTAATAAGATCTTCTATTTTATCATTTTCCAGATGATATGTATGTCCATTAAAGTTAAATTGTGTAGCGCTTCTTTTTAGTCTAGCAGCCAATACTTTAACATCACTAATATCAATATACAAAGGAGATGATTTATCGACGGTATTAAGAATTCGTTTTTCAACATCCGTCAAAACCAATTCCTTTTTGCCTGTATCATCCAGTTTGCCATTCTTTGTAAAGAAATTATTTTCTACTAGACTATTAATAAGACTGATAAGGATTCCTCCATAATTGCACAATCCAAACTTTTTTACAAAACTAGACTTTACGAGATCTACAGGAATGCGTACTTCAGGTTTCTTAGCTTGTGATTTTTTCTTTTGTGGACATTTTCTAGTGACCAACTCATCTACCATTTCATTGATTTTAGCAGTGAAATAAGCTTTCAATGCACGGGCACTGATACTATTAAAAACGAGTTGATCTTCAAATTCCACCATTTAAGTTCACCAAACAACTTTTAAATACAAATATATCAATAGTAGTAGAATAAGAACTATAAAAAAAATAATAATTAACCAATACCAACCCCAACAACCGGTATTATTAGTTGATGTGACAGTTTTCTCATCACTTAGAACAGATTTAACAATTTCTATAAAGTCTGTCAAATCATCTTCCGGAGAACCCATAAATACACCAAATATAGCGGCGTACAACTTATCCATTTATACATTGAATATTGGCTTTTCTTTATCACTATCTTCATCATATTCATCATCGATATCAACAAGTCCCAAATTACGAGCCAAATCTTCTTCTACATTTTCAGTCATTGATACACGTTCACTATCTCCAGAGAGTCCGATAACGTTAGCCACCACTTCTCTATCAATGATTAGTTTCTTAAGTGCAAATGTAATTTTTGTTTCAGTTCCGGATCTATAGAAAACTACAGGTGTGATAATTGCCTTGGCTAATTGTCTTTCTCTTTTACTGAGTGATTCTAGTTCACCTTCTATAGATCTGAGAATGGATAATTCTCCAGTCGAAACATATTCTACCATGGCTCCGTTTAATTTGTTAATGAAGATGGATTCATCCTTAAATGTTTTCTCTGTAATAGTTTCCACCAAAAGACTATGCAAAGAATTTGGAATGCGTTCCTTGTGCTTAATGTTTCCATAGACAGCTTCTAGAAGTTGATACAACATAGGACTAGCCGCGGTAACTTTTATTTTTAGAAAGTATCCATCACTTCTATCTTGTTTAGATTTATTTTTATAAAGTTTAGTCTCTCCTTCCAACATAATAAAAGTGGAAGTCATTTGACTAGATAAACTATCAGTAAGTTTTATAGAGATAGATGAACAATTAGCGTATTGAGAAGCATTTAGTGTGACGTATTCGATACATTTTGCATTAGATTTACTAATCGATTTTGCATACTCTATAACACCCGCACAAGTCTGTAGAGAATCGCTAGATGCTGTAGGTCTTGGTGAAGTTTCAACTCTCTTTTTGATTACCTTACTCATGATTAAACCTAAATAATTATACTTTGTAATATAATGATATATATTTTCACTTTATCTCATTTGAAAATAAAAATGTTTTTGTTTAACCGCTGCATGATGTACAGATTTCGGAATCACAAACCACCAGTGGTTTTATTTTATCCTTGTCCAATGTGAATTGAATGGGAGCGGATGCGGGTTTCGTACGTAGATAGTACATTCCCGTTTTTAGACCGAGACTCCATCCGTAAAAATGCATACTTGTTAGTTTGGAATAACTCGGATCTGCTATATGGATATTCATAGATTGACTTTGATCGATGAAGGCTCCCCTGTCTGCAGCCATTTTTATGATCGTCTTTTGTGGAATTTCCCAAATAGTTTTATAAACTCGCTTAATATCTTCTGGAAGGTTTGTATTCTGAATAGATCCACCATCTACCATAATCCTATTCTTGATCTCATCATTCCATAATTTTCTCTCGGTTAAAACTCTAAGGAGATGCGGATTAACTACTTGAAATTCTCCAGACAATACTCTCCGAGTGTAAATATTACTGGTATACGGTTCCACAGACTCATTATTTCCCAAAATTTGAGCAGTTGATGCAGTCGGCATAGGTGCCACCAATAAACTATTTCTAAGACCGTATGTTCTGATTTTATCTTTTAGAGGTTCCCAATTCCAAAGATCCGACGGTACAACATTCCAAAGATCATATTGTAGAATACCGTTACTGGCGTACGATCCTACATATGTATCGTATGGTCCTTCCTTCTCAGCTAGTTCACAACTCGCCTCTAATGCACCGTAATAAATGGTTTCGAAGATCTTCTTATTTAGATCTTGTGCTTCCAGGCTATCAAATGGATAATTTAAGAGAATAAACGCGTCTGCTAATCCTTGAACACCAATACCGATAGGTCTATGTCTCTTATTAGAGATTTCAGCTTCTGGAATAGGATAATAATTAATATCTATAATTTTATTGAGGTTTCTGACAATTACTTTGACCACATCCTTCAGTTTGAGAAAATCAAATTGTCCATCTATTACAAACATGTTCAATGCAATAGATGCCAGATTACAAACGGCTACCTCATTAGCATCCGCATATTGTATTATCTCAGTGCAAAGATTACTACACTTGATGGTTCCTAAATTTTGTTGATTACTCTTTTTGTTACACGCATCCTTATAAAGAATGAATGGCGTACCAGTTTCAATCTGGGATTCTATAATCGCTTTCCAGACGACACGAGCCTTTATTATAGATTTGTATCTCTTTTCTCTTTCGTATAGTGTATACAATCGTTCAAACTCGTCTCCCCAAACATTGTCCAATCCAGGACATTCATCCGGACACATCAACGACCACTCTCCGTCATCCTTCACTCGTTTCATAAAGAGATCAGGAATCCAAAGAGCTATAAATAGATCTCTGGTTCTATGTTCCTCGTTTCCTGTATTCTTTTTAAGATCGAGGAACGCCATAATATCAGAATGCCACGGTTCCAAGTATATGGTCATAACTCCAGGCCGTTTGTTTCCTCCCTGATCTATGTATCTAGCGGTGTTATTATAAACTCTCAACATTGGAATAATACCATTGGATGCACCATTGGTCCCGGAGATATAGCTTCCACTGGCACGAATATTACTAATTGATAGACCTATCCCCCCTGCCATTTTAGAGATTAATGCGCATCGTTTTAACGTATCATAGATGCCCTCTATACTATCATCCATCATGTTAAGTAGAAAACAGCTAGACATTTGGTGACGACTAGTTCCCGCATTAAATAAGGTAGGAGAAGCGTGCGTAAACCATTTTTCAGAAAGTAGATTGTACGTCTCGATAGCTGAGTCTATATCCCATTGATGAATTCCTACTGCGACACGCATTAACATGTGCTGAGGTCTTTCAACGATCTTGTTGTTTATTTTCAGCAAGTAGGATTTTTCCAAAGTTTTAAAACCAAAATAGTTGTATGAAAAGTCTCGTTCGTAAATAATAACTGAGTTGAGTTTATCCTTATATTTGTTAACTACATCCATGGTAATACTTGAAATAATCGGAGAATGTTTCCCATTTTTAGGATTGACATAGTTAAATAAATCCTTCATCACTTCACTAAATAGTTTTTTTGTTTCTTTGTGTAGATTTGATACGGCTATTCTAGCGGCTAGAATGGCATAATCCGGATGTTGTGTAGTACAAGTGGCTGCTATTTCGGCTGTCAGAGTGTCCAATTCTACCGTTGTTACTCCATTATATATTCCTTGGATAACCTTCATAGCTATTTTAATAGGATCTATATGATCAGTGTTTAAGCCATAACATAATTTTCTAATACGAGATGTGATTTTATCAAACATGACATTTTCCTTGTATCCATTTCGTTTAATGACAAACATTTTTGTTGGTGTAATAAAAAAAATTATTTAACTTTTCATTAATAGGGATTTGATGTACGTAGCGTACAAAATTATCGTTCCTGGTATATAGATAAAGAGTCCTATATATTTGAAAATCGTTACGGCTCGATTAAACTTTAATGATTGCATTGTGAATATATCATTAGGATTTAACTCCTTGACTATCATGGCGGTGCCAGAAATTACCATCAAAAGCATTAATACAGTTATACCGATCGCAGTTAGAACGGTTATAGCATCCGCCATTTATATCTAAAAATTAGATCAAAGAATATGTGACAAAGTCCTAGTTGTATACTGAGAATTAACGAAACAATGTTTCTTACATATTTTTTTCTTATTAGTAACTGACTTAATAGTAGGAACTGGAAAGCTAGACTTGATTATTCTATAAGTATAAATACCCTTCCAAATAATATTCTCTTTGATAAAAGTTCCAGAAAATGTAGAATTTTTTAAAAAGTTATCTTTTGCTATTACCAAGATTGTGTTTAGACGCTTATTATTAATATGAGTGATAAAATCCACACCGCCTCTAGATATAGCTTTTATTTCCACATTAGATGGTAAATCCAATAGTGAAACTATCTTTTTAGGAATGTATGGACTCGCGTTTAGAGGAGTGAACGTCTTAGGCGTCGGAAAGGATGATTCATCAAACGAATAAACAATTTCACAAATGGATGTTAATGTATTAGTAGGAAATTTTTTGACGCTAGTGGAATTGAAAATTCTAATGGATGATGTTCTACCTATTTCATCAGATAACATGTTAATTTCCGACATCAACGGTTTTAATATTTCGATAATATACGGTAATCTCTCTTTCGGACTTATATAGCTTATTCCACAATACGAGTCATTATATACTCCAAAAAACAAAATAACTAGTATAAAATCTGTATCGAATGGGAAAAACGAAATTATCGACATAGGTATAGAATCCGGAACATTGAACGTATTAATACTTAATTCTTTTTCAGTGGTAAGTACCGATAGGTTATTGACATTGTATGGTTTTAAATATTCTATAACTTGAGACTTGATAGATATTAATGACGAATTGAAAATTATTTTTATCACCACGTGTGTTTCAGGATCGTCATCGATACCCGTCAACCAACCAAACGGAGTAAAATAAATATCATTAATATATGCTCTAGATATTAGTATTTTTATTAATCCTTTGATTATCATCTTCTCGTACGCGAATGATTCCATGATCAAGAGTGATTTGAGAACATCCTCCGGAGTATTAATGGGCTTAGTAAACAGTCCATCGTTGCAATAATAAAAGTTATCCAAGTTAAAGGATATTATGCATTCGTTTAAAGATATTACCTCATCTGACGGAGACAATTTTTTGGTAGGTTTTAGAGACTTTGAAGCTACTTGTTTAACAAAGTTATTCATCGTCGTCTACTATTCTATTTAATTTTGTAGTTAATTTATCACATATCACATTAATTGACTTTTTGGTCCACTTTTCCATACGTTTATATTCTTTTAATCCTGCGTTATCCGTTTCCGTTATATCCAGGGATAGATCTTGCAAGTTAAATAGAATGCTCTTAAATAATGTCATTTTTTTATCCGCTAAAAATTTAAAGAATGTATAAACCTTTTTTAGAGATTTGAAACTCTTAGGTGGTGTCCTAGTACACAATATCATAAACAAACTAATAAACATTCCACATTCAGATTCCAACAGTTGATTAACTTCCACATTAATACAGCCTATTTTCGCTCCAAATATACATTCGAAAAATCTGAATAAAACATCAATGTCGCAATTTGTATTATCCAATACAGAATGTCTGTGATTCGTGTTAAAACCATCGGAGAAGGAATAGAAATAAAAATTATTATAGTGGTGGAATTCAGTTGGAATATTGCCTCCGGAGTCATAAAAGGATACTAAACATTGTTTTTTATCATAAATTACACATTTCCAATGAGACAAATAACAAAATCCAAACATTACAAATCTAGAGGTAGAACTTTTAATTTTGTCTTTAAGTATATACGATAAGATATGTTTATTCATAAACGCGTCAAATTTTTCATGAATCGCTAAGGAGTTTAAGAATCTCATGTCAAATTGTCCTATATAATCCACTTCGGATCCATAAGCAAACTGAGAGACTAAGTTCTTAATACTTCGATTGCTCATCCAGGCTCCTCTCTCAGGCTCTATTTTCATCTTGACGACCTTTGGATTTTCACCAGTATGTATTCCTTTACGTGATAAATCATCGATTTTCAAATCCATTTGTGAGAAGTCTATTGCCTTAGATACTTTTTCCCGTAGTCGAGGTTTAAAGAAATACGCTAACGGTATACTAGTAGGTAACTCAAAGACATCATATATAGAATGATAACGCGTCTTTAACTCATCGGTTAACTCTTTCTTTTGATCGAGTTCGTCGCTACTATTGGGTCTGCTCAGGTGCCTCGACTCTACTAGTTCCAACATCATACCAATAGGAATACAAGACACTTTGCCAGCGGTTGTAGATTTATCATATTTCTCCACTACATATCCGTTACAATTTGTTAAGAATTTAGATACATCTATATTGCTACATAATCCAGCTAGTGAATATATATGACATAATAAATTGGTAAATCCTAGTTCTGGTATTTTACTAATTACTAAATCTGTATATCTTTCCATTTATCATGGAAAAGAATTTACCAGATATCTTCTTTTTTCCAAACTGCGTTAATGTATTCTCTTACAAATATTCACAAGATGAATTTAGTAATATGAGTAACATGGAACGTGATAGTTTCTCTTTGGCGGTGTTTCCAGTTATAAAACATAGATGGCATAACGCACACGTTGTAAAACATAAAGGAATATACAAAGTTAGTACAGAAGCACATGGAAAAAAAGTATCTCCTCCATCACTAGGAAAACCCTCACACATAAACCTAACCGCGAAGCAATATATATACAGTGAACACACAATAAGCTTTGAATGTTATAGTTTTCTAAAATGTATAACAAATGCAGAAATTAATTCGTTCGATGAGTATATATTAAGAGGACTATTAGAAGCTGGTAATAGTTTACAGATATTTTCCAATTCCGTAGGTAAACGAACAGATACTATAGGTGTACTAGGGAATAAGTATCCATTTAGCAAAATTCCATTGGCCTCATTAACTCCTAAAGCACAACGAGAGATATTTTCAGCATGGATTTCTCATAGACCCGTAGTTTTAACTGGAGGAACCGGAGTGGGTAAGACGTCACAGGTACCCAAGTTATTGCTTTGGTTTAATTATTTATTTGGTGGATTCTCTACTCTAGATAAAATCACTGACTTTCACGAAAGACCAGTCATTCTATCTCTTCCTAGGATAGCTTTAGTTAGATTGCATAGCAATACCATTTTAAAATCATTGGGATTTAAGGTACTAGATGGATCTCCTATTTCTTTACGGTACGGATCTATACCGGAAGAATTAATAAACAAACAACCAAAAAAATATGGAATTGTATTTTCTACCCATAAGTTATCTCTAACAAAACTATTTAGTTATGGCACTCTTATTATAGACGAAGTTCATGAGCATGATCAAATAGGAGATATTATTATAGCAGTAGCGAGAAAGCATCATACGAAAATAGATTCTATGTTTTTAATGACTGCCACGTTAGAGGATGACAGGGAACGGCTAAAAGTATTTTTACCTAATCCCGCATTTATACATATTCCTGGAGATACACTGTTTAAAATTAGCGAGGTATTTATTCATAATAAGATAAATCCATCTTCCAGAATGGCATACATAGAAGAAGAAAAGAGAAATTTAGTTACTGCTATACAGATGTATACTCCTCCTGATGGATCATCTGGTATAGTCTTTGTGGCATCCGTTGCACAGTGTCACGAATATAAATCATATTTAGAAAAAAGATTACCGTATGATATGTATATTATTCATGGTAAGGTCTTAGAAATAGACAAAATATTAGAAAAAGTGTATTCATCGCCTAATGTATCGATAATTATTTCTGCTCCTTATTTGGAATCCAGCGTTACTATACACAATGTTACTCACATTTATGATATGGGTAGAGTTTTTGTTCCCGCTCCTTTTGGAGGATCACAACAATTTATTTCTAAATCTATGAGAGATCAACGAAAAGGAAGAGTAGGAAGAGTTAATCCTGGTACATACGTCTATTTCTATGATCTGTCTTATATGAAGTCTATACAGCGAATAAATTCAGAATTTCTACATAATTATATATTGTACGCTAATAAGTTTAATTTAACACTTCCCGAAGATTTGTTTATAATCCCTACAAATTTGGATATTCTATGGCGTACAAAGGAATATATAGACTCGTTCGATATTAGTACAGAAACATGGAATAAATTATTATCCAATTATTATATGAAGATGATAGAGTATGCTAAACTTTATGTACTAAGTCCTATTCTCGCTGAGGAGTTGGATAACTTTGAGAGGACGGGAGAATTAACTAGTATTGTACAAGAAGCCATTTTATCTCTAAATTTACGAATTAAGATTTTAAATTTTAAACATAAAGATAATGATACGTATATACACTTTTGTAAAATATTATTTGGTGTCTATAACGGAACAAACGCTACTATATATTATCATAGACCTCTAACGGGATATATGAATATGATTTCAGATACTATATTTGTTCCTGTAGATAATAACTAAAAATCAAACTCTAATGACCACATCTTTTTTTAGAGATGAAAAATTTTCCACATCTCCTTTTGTAGATATGACTAAACATTTTGCAGAAAAAAGTTTATTAGTGTTTAGATAATCGTATACTTCATCAGTGTAGATAGTAAATGTGAACAGATAAAAGGTATTCTTGCTCAATAGATTGGTAAATTCCATAGAATATATTAATCCTTTTTTCTTGAGATCCCACATCATTTCAACCAGAGAAGTTTTATCCAATAATTTACCTCGTACTATACCACATACAAAACTAGATTTTGTAGTGACGTCGTACCTGGCATTCCTACCAAACAAAATTTTACTTTTAGTTTTTTTAGAAAATTCTAAGGTAGAATCTCTATTTGTCAATATGTCATCTATGGAATTACCACTAGCAAAAAATGATAGAAATATATATTGATACATCGCAGCTGGTTTTGATCTACTATATTTTAAAAACGAATCAGATTCCATAATTGCCTGTATATCATCAGCTGAAAAACTATGTTTTACACGTATTCCTTCGGCATTTCTTTTTAATGATATATCTTGTTTAGACAATGATAAAGTTATCATGTCCATGAGAGACGCGTCTCCGTATCGTATAAATATTTCATTAGATGTTAGACGCTTCATTAGGGGTATACTTCTATAAGGTTTCTTGATTAGCCCATCATTTGTTGCGTCAAGAACTACTATCGGATGTTGTTGGGTATCTCTAGTGTTACACATGGCCTTACTAAAGTTTGGGTAAATAACTATGATATCTCTATTAATTATAGATGTATATATTTCATTCGTCAAGGATATTAGTATCGACTTGCTATCGTCATTAATACGTGTAATGTAATCATATAAATCATGCGATAGCCAAGGAAAATTCAAATAGATGTTCATCATATAATCATCGCTATAATTCATATTAATACGTTGACATTGACTAATTTGTAATATAGCCTCGCCACGAAGAAAGCTCTCGTATTCAGTTTCATCAATAAAGGATACCGTTAAATATAACTGGTTGCCGATTGTCTCATAGTCTATTAAGTGGTAAGTTTCGTACAAATACAGAATCCCTAAAATATTATCTAATGTTGGATTAATCTTTACCATAACTGTATAAAATGGAGACGGAGTCATAACTATTTTACCGTTTGTACTTACTGGAATAGATGAAGGAATAATCTCCGGACATGCTGGTAAAGACCCAAATGTCTGGTTGAAGAAATCCAATGTTCCGGGTCCTAATCTCTTGACAAAAATTACGATATTCGATCCCGATATCCTTTGCATTCTATTTACCAGCATATCACGAACTATATTAAGATTATCTATCATGTCTATTCTCCCACCGTTATATAAATCGCCTCCGCTAAGAAACGTTAGTATATCCATACAATGGAATACTTCATTTCTAAAATAGTATTCGTTTTCTAATTCTTTAATGTGAAATCGTATACTAGAAAGGGAAAAATTATCTTTAAGTTTTCCGTTAGAAAAGAACCACGAAACTAATGTTCTGATTGCGTCTGACTCCGTCGCTGAATTAATGGATTTACACCAAAAACTCATATAACTTCTAGATGTAGAAGCATTAGCTAAAAAAATAGTAGAATCAAAGGATATAAGTAGATGTTCCAACAAGTGAGCAATTCCCAAAATTTCATCTATATCATTCTCAAATCCGAAATTAGAAATTCCCAAGTAGATATCTTTTTTCATCCGATCATTAATAAAAATACGAACTTTATTCGGTAAGACGATCATTTACTAAGGAGTAAAATAGGAAGTAATGTTCGTATATCGTTATCATCGTATAAATTAAAGGTGTGTTTTTTACCATTAAGTGACATTATAATTTTACCAATATTGGAATTATAATATAGGTATATTTGCGCACTTGCGACAGTTGATGCATCGGTAAATATAGCTGTATCTAATGTTCTAGTCGGTATTTCATCATTTCGCTGTATAATAATAGCGTTTTCTCTATCTGTTTCCATTACAGCTGCCTGAAGTTTATTGGTCGGATAATATGTAAAATAATAAGAAATACATACGAATAACAAAAATAAAATAAAATATAATAAAGATGCCATTTAGAGATCTAATTTTGTTCAACTTGTCCAAATTCCTACTTACAGAAGATAAGGAATCGTTGGAGATAGTGTCTTCCTTATGTAGAGGATTTGAAATATCTTATGATGACTTGATAACTTACTTTCCAGATAGAAAATATCATAAATATATTTATAAAGTATTTGAACATGTAGATTTATCGGAGGAATTAAGTATGGAATTCCATGATACAACTCTGCGAGATTTAGTCTATCTTAAATTGTATAAGTATTCCAAGTGTATACGACCGTGTTATAAATTAGGAGATAATCTAAAAGGCATAGTTGTTATAAAGGACAGAAATATATATATTAGAGAAGCAAATGATGACTTGATAGAATATCTCCTCAAGGAATACACTCCTCAGATTTATACATATTCTAATGAGCACGTTCCCATAGCTGGTTCAAAATTAATTCTTTGCGGATTTTCTCAAGTTACATTTATGGCGTATACAACGTCTCATATAACAACAAATAAAAAGGTAGATGTTCTCGTTTCCAAAAAATGTATAGATAAACTAGTCGATCCAATAAATTATCAAATACTTCAAAATTTATTTGATAAAGGAAGCGGAACAATAAACAAAATACTCAGGAAGATATTTTATTCGGTAACCGGTGGCCAAACTCCATAATTTGCTTTTTCTATTTCGGATTTTAGAATTTCCAAATTCACCAGCGATTTATCTGTTTTGGTGAAATCCAAGGATTTATTAATGTCCACAAATGCCATTTGTTTTGTCTGTGGATTGTATTTGAAAATGGAAACGATGTAGTTAGATAGATGCGCTGCAAAGTTTCCTATTAGGGTTCCGCGCTTTACGTCACCTAGCATACTTGAATCACCATCCTTTAAAAAAAATGATAAGATATCAACATGGAGTATATCATACTCTGATTTTAATTCTTCTACTGCCTCACTGACATTTTCACAAATACTACAATACGGTTTACCGAAAATAATCAGTACGTTCTTCATTTATGGGTATCAAAAACTTAAAATCGTTACTGCTGGAAAATAAATCACTGACGATATTAGATGATAATTTATACAAAGTATACAATGGAATATTTGTGGATACAATGAGTATTTATATAGCCGTCGCCAATTGTGTCCGAAACTTAGAAGAGTTAACTACGGTATTCATAAAATACGTAAACGGATGGGTAAAAAAGGGAGGGCATGTAACCCTTTTTATCGATAGAGGAAGTATAAAAATTAAACAAAACGTTAGAGACAAAAGACGTAAATATTCTAAATCAACCAAGGACAGAAAAATGTTAGAATTAGAAAAGTGTACATCCAAAATACAAAATGTTACCGGATTTATGGAAGAAGAAATAAAGGCAGAAATACAATTAAAAATCGATAAACTCACATTTCAAATATATTTATCTGATTCTGATAACATAAAAATATCATTGAATGAGATACTAACACATTTCAACAATAATGAGAATGTTACATTATTTTATTGTGATGAACGAGATGCAGAATTTGTTATGTGTCTAGAGGCTAAAACATATTTCTTTACCACAGGAGAATGGCCGTTGATAATAAGTACCGATCAGGATACTATGCTATTTGCGTCTGTTGATAATCATCCTAAGATGATAAAAAACTTAACTCAACTGTTTAAATTTGTTCCATCTGCAGAGGATAACTATTTAGCAAAATTAACGGCGTTAGTGAATGGATGTGATTTCTTTCCTGGACTCTATGGGGCATCTATAACACCCACCAACTTAAACAAAATACAATTGTTTAGTGATTTTACAATCAATAATATAGTCACTAGTTTGGCAATTAAAAATTATTATAGAAAGACTAACTCTACCGTAGACGTGCGTAATATTGTTACGTTTATAAACGATTACGCTAATTTAGACGATGTCTACTCGTATATTCCTCCTTGTCAATGCACTGTTCAAGAATTTATATTTTCCGCATTAGATGAAAAATGGAACGATTTTAAATCATCTTATTTAGAGACCGTTCCGTTACCCTGTCAATTAATGTACGCGTTAGAACCACGTAAGGAGATTGATGTTTCAGAAGTTAAAACTTTATCATCTTATATAGATTTCGAAAATACTAAATCAGATATCGATGTTATAAAATCTATATCCTCGATCTTTGGATATTCTAACGAAAACTGTAACACGATAGTGTTTGGCATCTATAAGGATAATTTACTACTGAGTATAAATAGTTCATTTTACTTTAACAATAGTCTGTTAATAACCAATACTAAAAGTGATAATATAATAAATATAGGTTACTAGATTAAAAATGGTTTTCCAAGTAGTGTGCTCTACATGCGGTAAAGATATTTCTCACGAACGATATAAATTGATTATACGAAAAAAATCATTAAAGGATGTACTCGTTAGTGTAAAGAACAAATGTTGTAGGTTAAAATTATCTACACAAATAGAACCTCAACGTAACTTAACAGTGCAACCTCTATTGGATATAAACTAATATGGATCCGGTTAATTTTATCAAGACATATGCACCTAGAGGTTCTATTATTTTTATTAATTATACTATGTCATTAACAAGTCATTTAAATCCATCGATAGAAAAACATGTGGGTATTTATTATGGTACGTTATTATCGGAACACTTGGTAGTTGAATCAACATATAGAAAAGGAGTTCAAATAGTCCCATTGGATAGTTTTTTTGAAGGATATCTTAGTGCAAAAGTATACATGTTAGAGAATATTCAAGTTATGAAAATAGCAGCTGATACGTCATTAACTTTACTGGGTATTCCATATGGATTTGGTCATAATAGAATGTATTGTTTTAAATTGGTAGCTGAATGTTATAAAAATGCCGGTATTAATACATCATCTAAACGAATATTGGGCAAAGATATTTTTCTAAGCCAAAACTTCACAGATGATAATAGATGGATAAAGATATATGATTCTAATAATTTAACATTTTGGCAAATTGATTACCTTAAAGGGTGAGTTAATATGCATAACTACTCCTCCGTTGTTTTTTCCCTCGTTCTTTTTCTTAACGTTGTTTGCCATCACTCTCATAATGTAAAGATATTCTAAAATGGTAAACTTTTGCATATCGGACGCAGAAATTGGTATAAATGTTGTAATTGTATTATTTCCCGTCAATGGACTAGTCACAGCTCCATCAGTTTTATATCCTTTAGAGTATTTCTCACTAGTGTCTAGCATTTTAGAGCATTCCATGATCTGTTTATCATTGATATTGGCCGGAAAGATAGATTTTTTATTTTTTATTATATTACTATTGGCAATTGTAGATATAACTTCTGGTAAATATTTTTCTACCTTTTCAATCTCTTCTATTTTCAAGCCGGCTATATATTCTGCTATATTATTGCTAGTATCAATACCTTTTCTGGCTAAGAAGTCATATGTGGTATTAACTATATCAGTTTTAACTGGTAGTTCCATTAGCCTTTCCACTTCTGCAGAATAATTAGAAATTGGTTCTTTACCAGAAAATCCAGCTACTATAATAGGCTCACCGATGATCATTGGCAAAATCCTATATTGTACCAAATTAATGAGAGCATATTTCATTTCCAATAATTCTGCTAGTTCTTGAGACATTGATTTATTTGATGAATCTAGTTGGTTCTCTAGATACTCTACCATTTCTGCCGCATACAATAACTTGTTAGATAAAATCAGGGTTATCAAAGTGTTTAGCGTGGCTAGAATAGTGGGCTTACATGTATTAAAGAATGCGGTAGTATGAGTAAACCGTTTTAACGAATTATATAGTCTCCAGAAATCTGTGGCGTTGCATACATGAGCTGAATGACATCGAAGATTGTCCAATATTTTTAATAGCTGCTCTTTGTCCATTATTTCTATATTTGACTCGCAACAATTGTAAATACCATTAATCACTGATTCCTTTTTTGATGCCGGACAATAGCACAATTGTTTAGCTTTGGACTCTATGTATTCAGAATTAATAGATATATCTCTCAATACAGATTGTACTATACATTTTGAAACTATGTCAAAAATTGTAGAACGACGCTGTTCTGCAGCCATTTAACTTTAAATAATTTACAAAAATTTAAAATGAGTATCCGTATAAAAATCGATAAACTGCGCCAAATTGTGGCATATTTTTCAGAGTTCAGTGAAGAAGTGTCTATAAATGTAGACTCAACGGATGAGTTAATGTATATTTTTGCCGCCTTGGGCGGATCTGTAAACATTTGGGCCATTATACCTCTTAGTGCATCAGTGTTCTACCGCGGAGCCGAAAATATTGTGTTTAATCTTCCGGTGTCCAAGGTAAAATCGTGTTTGTGTAGTTTTCACAATGATGCCATCATAGATATAGAACCTGATCTGGAAAATAATCTAGTAAAACTTTCTAGTTATCATGTAGTAAGTGTCGATTGTAACAAGGAACTGATGCCTATTAGGACAGATACTACTATTTGTCTAAGTATAGATCAAAAGAAATCTTACGTGTTTAATTTTCACAAGTATGAAGAAAAATGTTGTGGTAGAACCGTCATTCATCTAGAATGGTTGTTGGGCTTTATTAAGTGTATTAGTCAGCATCAGCATTTGACTATTATGTTTAAAGATGACAATATTATTATGAAGACTCCTGGTAATACTGATGCTTTTTCCAGGGAATATTCTATGACTGAATGTTCTCAAGAACTACAAAAGTTTTCTTTCAAAATAGCTATCTCGTCTCTCAACAAACTACGAGGATTCAAAAAGAGAGTCAATGTTTTTGAAACCAGAATCGTAATGGATAATGACGATAACATTCTAGGAATGTTGTTTTCGGATAGAGTTCAATCCTTTAAGATCAACATCTTTATGGCGTTTTTAGATTAATACTTTCAATGAGATAAATATGGGTGGCGGAGTAAGTGTTGAGCTCCCTAAACGGGATCCGCCTCCGGGAGTACCCACTGATGAGATGTTATTAAATGTTGATAAAATGCATGACGTGATAGCTCCCGCTAAGCTTTTAGAATATGTGCATATAGGACCACTAGCAAAAGATAAAGAGGATAAAGTAAAGAAAAGATATCCAGAGTTTAGATTAGTTAATACAGGACCCGGTGGTCTTTCGGCATTATTAAGACAATCGTATAATGGAACCGCACCCAACTGCTGTCACACTTTTAATCGTACTCATTATTGGAAGAAGGATGGAAAGATATCAGATAAGTATGAAGAGGGTGCAGTATTAGAATCGTGTTGGCCAGACGTTCACGATACCGGAAAATGCGATGTTAATTTATTTGACTGGTGTCAGGGGGATACGTTCGATAGAAACATATGCCATCAGTGGATCGGTTCAGCATTTAATAGGAGTGATAGAACTGTAGAGGGTCAACAATCGTTAATAAATCTGTATAATAAGATGCAAACATTATGTAGTAAAGATGCTAGTGTACCAATATGCGAATCATTTTTGCATCATTTACGCGCACACAATACAGAAGATAGCAAAGAGATGATCGATTATATTCTAAGACAACAGTCTGCGAACTTTAAACAGAAATATATGAGATGTAGTTATCCCACTAGAGATAAGTTAGAAGAGTCATTAAAATATGCGGAACCTCGAGAATGTTGGGATCCAGAGTGTTCGAATGCCAATGTTAATTTCTTACTAACACGTAATTATAATAATTTAGGACTTTGTAATATTGTACGATGTAATACCAGCGTGAACAACTTACAGATGGATAAAACTTCCTCATTAAGATTGTCATGTGGATTAAGCAATAGTGATAGATTTTCTACTGTTCCCGTCAATAGAGCAAAAGTAGTTCAACATAATATTAAACACTCGTTCGATCTAAAATTGCATTTGATTAGTTTATTATCTCTCTTGGTAATATGGATACTAATTGTAGCTATTTAAATGGGTGCCGCGGCAAGCATACAGACGACGGTGAATACACTCAGCGAACGTATCTCGTCTAAATTAGAACAAGAAGCGAATGCTAGTGCTCAAACAAAATGTGATATAGAAATCGGAAATTTTTATATCCGACAAAACCATGGATGTAACCTCACTGTTAAAAATATGTGCTCTGCAGATGCGGATGCTCAGTTGGATGCTGTGTTATCAGCCGCTACAGAAACATATAGTGGATTAACACCGGAACAAAAAGCATACGTACCAGCTATGTTTACGGCTGCGTTAAACATTCAGACGAGTGTAAACACTGTTGTTAGAGATTTTGAAAATTATGTGAAACAGACTTGTAATTCTAGCGCGGTCGTCGATAACAAATTAAAGATACAAAACGTAATTATAGATGAATGTTACGGAGCCCCAGGATCTCCAACAAATTTGGAATTTATTAATACAGGATCTAGCAAAGGAAATTGTGCCATTAAGGCGTTGATGCAATTGACGACTAAGGCCACTACTCAAATAGCACCTAGACAAGTTGCTGGTACAGGAGTTCAGTTTTATATGATTGTTATCGGTGTTATAATATTGGCAGCGTTGTTTATGTACTATGCCAAGCGTATGCTGTTCACGTCCACCAATGATAAAATCAAACTTATTTTAGCCAATAAGGAAAACGTCCATTGGACTACTTACATGGACACATTCTTTAGAACTTCTCCAATGGTTATTGCTACCACGGATATACAAAACTGAAAATATATTGATAATATTTTAATAGATTAACATGGAAGTTATCGCTGATCGTCTAGATGATATAGTGAAACAAAATATAGCGGATGAAAAATTTGTAGATTTTGTTATACACGGTCTAGAGCATCAATGTCCTGCTATACTTCGGCCATTAATTAGGTTGTTTATTGATATACTATTATTTGTTATAGTAATTTATATTTTTACGGTACGTCTAGTAAGTAGAAATTATCAAATGTTGTTGGCGTTATTGGCGCTAGTCATCTCATTAACTATTTTTTATTACTTTATACTATAATAGTACTAGACTGACTTCTAACAAACATCTCACCTGCCATAAATAAATGCTTGATATTAAAGTCTTCTATTTCTAACACTATTCCATCTGTGGAAAATAATACTCTGACATTATCGCTAATTGACACATCGGTGAGTGATATGCCTATAAAGTAATAATCTTCTTTGGGCACATATACCAGTGTACCAGGTTCTAACAACCTATTTACTGGTGCTCCTGTAGCATACTTTTTCTTTACCTTAAGAATATCCATCGTTTGCTTGGTCAATAGCGACAGGTGATTTTTTATCAACCATTCAAAAAAGTAATTGGAGTGTTCATATCCTCTACAGGCTATTGTCTCATGACCGTGTATGAAATTTAAGTAACACGACTGTGGTAGATTTGTTCTATAGAGCCGGTTGCCGCAAATAGATAGAACTACCAATATGTCTGTACAAATGTTAAACATTAATTGATTAACAGAAAAAACAATGTTCGTTCTGGGAATAGAAACCAAATCAAAACAAAATTCATTAGAATATATGTCACGTTTATACATCGAATATAAAATAACTACAGTTTGAAAAATAACAGTATCATTTAAACATTTAACTTGCGGGGTTAATCTCACAACTTTACTGTTTTTGAACTGTTCAAAATATAGCATAGATCCGTGAGAAATACGTTTAGCCGCCTTTAATAGAGGAAATCCCACCGCCTTTTTGGATCTTACCAACGACGATAGTTCTGACCAGCAACTCATTTCTTCATCATCCACCTGTTTTAACATATAATAGGCAGGAGATAGATATCCGTCATTGCAATATTCCTTCTCGTAGGCACACAATCGAATATTGATAAAATCTCCATTCTCTTCTCTGCATTTATTATCTTGTCTTGGTGGTGGGCTAGGCTGTAGTCTTGGTCTATTATTACCATCGTTGTTGAATCTATTTCTGTCATTAAATCTTTCATTTCTTCCTGGTATATTTTTATTACCTCGCTTGGTTGGATTTTTGTCTATATTATCGTTTGTAACATCGGTACGGGTATTCATTTATCACAAAAAAAACTTCTCTAAATGAGTCTACTGCTAGAAAACCTCATCGAAGAAGATACCATATTTTTTGCAGGAAGTATATCTGAGTATGATGATTTACAAATGGTTATTGCCGGTGCAAAATCCAAATTTCCAAGATCTATGCTTTCTATTTTTAATATAGTACCTAGAACAATGTCAAAATATGAGTTGGAGTTAATTCATAACGAGAATATCACAGGGGCAATGTTTACCACAATGTATAATATAAGAAACAATTTGGGTTTAGGTGATGATAAACTAACTATTGAAGCCATTGAAAACTATTTCTTGGATCCTAACAATGAGGTTATGCCTCTTATTATTAATAATACGGATATGACGGCCGTCATTCCTAAAAAAAGTGGTAGGAGAAAGAATAAGAACATGGTTATCTTCCGTCAAGGATCATCACCTATCTTGTGTATTTTCGAAACTCGTAAAAAGATTAATATTTATAAAGAAAATATGGAATCCGCATCGACTGAGTATACACCTATCGGAGACAACAAGGCTTTAATATCTAAATATGCGGGAATTAATGTCCTGAATGTGTATTCTCCTTCCACATCCATGAGATTGAATGCCATTTACGGATTCACCAATAAAAATAAACTAGAAAAACTTAGTACTAATAAGGAACTAGAATTGTATAGTTCTAGCCCTCTTCAAGAACCCATTAGGTTAAATGATTTTCTGGGACTATTGGAATGTGTTAAAAAGAATATTCCTCTAACAGATATTCCGACAAAGGATTGATTACTATAAATGGAGAATGTTCCTAATGTATACTTTAATCCTGTGTTTATAGAGCCCACGTTTAAACATTCTTTATTAAGTGTTTATAAACACAGATTAATAGTTTTATTTGAAGTATTTGTTGTATTCATTCTAATATATGTATTTTTTAGATCTGAATTAAATATGTTCTTCATGCATAAACGAAAAATACCCGATCCTATTGATAGATTACGACGTGCTAATCTAGCGTGTGAAGACGATAAATTAATGATCTATGGATTACCATGGATAACAACTCAAACATCTGCATTATCAATAAATAGTAAACCGATAGTGTATAAAGATTGTGCAAAGCTTTTGCGATCAATAAATGGATCACAACCAGTATCTCTTAACGATGTTCTTCGCAGATGATGATTCATTTTTTAAGTATTTGGCTAGTCAAGATGATGAATCTTCATTATCTGATATATTGCAAATCACTCAATATCTAGACTTTCTGTTATTATTATTGATCCAATCAAAAAATAAATTAGAAGCTGTTGGTCATTGTTATGAATCTCTTTCAGAGGAATACAGACAATTGACAAAATTCACAGACTCTCAAGATTTTAAAAAACTGTTTAACAAGGTCCCTATTGTTACAGATGGAAGGGTCAAACTTAATAAAGGATATTTGTTCGACTTTGTGATTAGTTTGATGCGATTCAAAAAAGAATCAGCTCTAGCTACCACCGCAATAGATCCTGTTAGATACATAGATCCTCGTCGCGATATCGCATTTTCTAACGTGATGGATATATTAAAGTCAAATAAAGCGAAAAATAATTATTCTTTATTGTCATCATGAACGGCGGGCATATTCAGTTGATAATTGGCCCCATGTTTTCAGGTAAAAGTACAGAATTAATTAGACGAGTTAGACGTTATCAAATAGCTCAATATAAATGCGTGACTATAAAATATTCTAACGATAATAGATACGGAACGGGACTATGGACGCATGATAAGAATAATTTTGAAGCATTGGAAGCAACTAAACTATGCGATGTCTTGGAAGCAATTACAGATTTCTCCGTGATAGGTATTGATGAAGGACAGTTCTTTCCAGACATTGTTGAATTCTGTGAGCGTATGGCAAACGAAGGAAAAATAGTTATAGTAGCCGCACTCGATGGGACATTTCAACGTAAACCGTTTAATAATATCTTGGATCTTATTCCATTATCTGAAATGGTGGTAAAACTAACTGCAGTGTGTATGAAATGCTTTAAGGAGGCGTCCTTTTCTAAACGATTAGGTACAGAAACCAAGATAGAAATAATAGGAGGAATTGATATGTATCAATCGGTGTGTAGAAAGTGTTACATCGACTCATAATATTATATTTTTTATCTAAAAAACTAAAAATAAACATTGATTAAATTTTAATATAATACTTAAAAATGGATGTTGTGTCGTTAGATAAACCGTTTATGTATTTTGAGGAAATTGATAATGAGTTAGATTACGAACCAGAAAGTGCAAATGAGGTAGCAAAAAAACTGCCGTATCAAGGACAGTTAAAACTATTACTAGGAGAATTATTTTTTCTTAGTAAGTTACAGCGACACGGTATATTAGATGGTGCCACCGTAGTGTATATAGGATCTGCTCCCGGTACACATATACGTTATTTGAGAGATCATTTCTATAATTTAGGAGTGATCATCAAATGGATGCTAATTGACGGCCGTCATCATGATCCTATTCTAAATGGATTACGTGATGTGACTCTAGTAACTCGGTTCGTTGATGAGGAATATCTACGATCCATCAAAAAACAACTGCATCCTTCTAAGATTATTTTAATTTCTGATGTGCGATCCAAACGAGGAGGAAATGAACCTAGTACTGAGGATTTACTAAGTAATTACGCTCTACAAAATGTCATGATTAGTATTTTAAACCCCGTAGCGTCTAGTCTTAAATGGAGATGCCCGTTTCCAGATCAATGGATTAAGGACTTTTATATCCCACACGGTAATAAAATGTCACAACCTTTTGCTCCTTCATATTCAGCTGAAATGAGATTATTAAGTATTTATACCGGTGAGAACATGAGACTTACTCGAGTTACCAAATCAGATGCTGTAAATTATGAAAAAAAGATGTATTACCTTAATAAGATAGTTCGCAACAAAGTAGTTGTTAACTTTGATTATCCTAATCAGGAATATGACTATTTTCACATGTACTTTATGTTGAGGACCGTATACTGCAATAAAACATTTCCTACTACTAAAGCAAAGGTACTATTTCTACAACAATCTATATTTCGTTTCTTAAATATTCCAACAACATCAACTGAAAAAGTTAGTCATGAACCAATACAATGTAAAGTATCTAGCAAAGATTCTGTGTATAAAAACAGAAATAGCAAGAGATCCGTACGCGGTAATAAATAGAAATGTGCTACTAAGATATACTACCGATATACAGTATAATGATTTAGTTACTTTAATAACCGTTAGACATAAAATTGATTCTATGAAAACTGTGTTTCAGGTATTTAACGAATCATCCATAAATTATACTCCGGTCGATGATGATTACGGAGAACCAATCATTATAACATCGTATCTTCAAAAAGGTCATAACAAGTTTCCTGTAAATTTTCTATACATAGATGTAGTAATATCTGATTTATTTCCTAGCTTTGTTAGACTAAATACTACAGAAACTAATATAGTTAATAGTGTACTACAAACAGGCGATGGTAAAAAGACTCTTCGTCTTCCCAAAATGTTAGAGACGGAAATAGTTGTCAAGATTCTCTATCGTCCTAATATACCATTAAAAATTGTTAGATTTTTCCGCAATAACATGATAACTGGTGTAGAGATAGCCGATAGATCTGTTATTTCAGTCGCTGATTAATCAATTAGTAGAGATGAGATAAGAACATAATAATATATCTTATATCTCGTTTAGAAAAATGCTAATATTAAAATAGCTAACGCTAGTAATCCAATCGGAAGCCATTTGATATCTATAATAGGGTATCTAATTTCCTGATTCAGATAGCGGACAGCTATATTCTCTGTAGCTACTCGTTTGGAATCACAAACATTATTTACATCTAATTTACTATCTGTAATGGAAACGTTTCCCAATGAAATGGTACAATCAGATACATTACATCTTGATATATTTTTTTTTAAAGAGGCTGGTAACAACGCATCGCTTCGTTTACATGGCTCGTACCAACAGTAATAGGGTAATCTTGTATCTATTCCTATCCGTACTATGCTTTTATCAGGATAAATACATTTACATCGTATATCGTCTTTGTTAGCATCACAGAATGCATAAATTTGTTCGTCTGTCATGATAAAAATTTAAAGTGTAAATATAACTATTATTTTTATAGTTGTAATAAAAAGGGAAATTTAATTGTATACCTTTGGTTCTTTAAAAGAAACTGACTTGATAAAAATGGCTGTAATCTCTAAGGTTACGTATAGTCTATATGATCAAAAAGAGATTAATGCTACAGATATTATCATTAGTCATGTTAAAAATGACGACGATATCGGTACCGTTAAAGATGGTAGACTAGGTGCTATGGATGGGGCATTATGTAAGACTTGTGGGAAAACGGAATTGGAATGTTTCGGTCACTGGGGTAAAGTAAGTATCTATAAAACTCATATAGTTAAGCCTGAATTTATTTCAGAAATTATTCGTTTACTGAATCATATATGTATTCACTGCGGATTATTGCGTTCACGAGAACCGTATTCCAACGATATTAACCTAAAAGAGTTATCGGGACACGCTCTTAGGAGATTAAAGGATAAAATATTATCCAAGAAAAAGTCATGTTGGAATAGCGAATGTATGCAACCGTATCAAAAAATTTCTTTTTCAAAGAAAAAGGTTTGTTTCGTCAACAAGTTGGATGATATTAACGTTCCTAATTCTCTCATTTATCAAAAGTTAATTTCTATTCATGAAAAGTTTTGGCCATTATTAGAAATTTATCAATATCCAGCTAACTTATTTTATACAGACTACTTTCCTATCCCTCCGTTGATTATTAGACCGGCTATTAGTTTTTGGATAGATAGTATACCCAAAGAGACCAATGAATTAACTTACTTATTAGGTATGATCGTTAAGAATTGTAACTTGAATGCTGATGAACAGGTTATCCAAAAGGCGGTAATAGAATACGATGATATTAAAATTATTTCTAATAACACTACCAGTATCAATTTATCATATATCACATCCGGCAAAAATAATATGATTAGAAGTTATATCGTCGCCCGGCGAAAAGATCAGACCGCTAGATCTGTAATTGGTCCCAGTACATCTATCACAGTTAATGAGGTAGGAATGCCCGCATATATTAGAAATACACTTACAGAAAAGATATTTGTTAATGCCTTTACAGTGAATAAAGTTAAACAACTATTAGCGTCAAACCAAGTTAAATTTTACTTTAATAAACGATTAAACCAATTAACAAGAATACGTCAAGGAAAGTTTATCAAAAATAAAATACATTTATTGCCTGGTGATTGGGTAGAAGTAGCTGTTCAAGAATATACAAGTATTATTTTTGGAAGACAGCCGTCGCTACATAGATACAACGTCATCGCTTCATCTATCAGAGCTACCGAAGGAGATACTATCAAAATATCTCCCGGAATTGCCAACTCTCAAAATGCTGATTTCGACGGAGATGAAGAATGGATGATATTGGAGCAAAATCCTAAAGCCGTAGTTGAACAGAGTATTCTTATGTATCCGACAACGTTACTCAAACACGATATTCATGGAGCCCCCGTTTATGGATCTATTCAAGATGAAATCGTAGCAGCATATTCATTGTTTAGGATACAAGATCTTTGTTTAGATGAAGTATTGAACATCTTGGGGAAATATGGAAGAGAGTTCGATCCTAAAGGTAAATGTAAATTCAGTGGTAAAGATATCTATACTTACTTAATAGGTGAAAAGATTAATTATCCGGGTCTCTTAAAGGATGGCGAAATTATTGCAAACGATGTAGATAGTAATTTTGTTGTGGCTATGAGGCATCTGTCATTGGCTGGACTCTTATCCGATCATAAGTCGAACGTGGAAGGTATCAACTTTATTATCAAGTCATCTTATGTTTTTAAGAGATATCTATCTATTTACGGTTTTGGGGTGACATTCAAAGATTTGAGACCAAATTCGACGTTCACTAATAAATTGGAGGCCATCAACGTAGAAAAAATAGAACTTATCAAAGAAGCATACGCCAAATATCTCAAAGATGTAAGAGACGGAAAAATAGTTCCATTATCTAAAGCTTTAGAGGCGGACTATGTGGAATCCATGTTATCCAACTTAACAAATCTTAATATCAGAGAGATAGAAGAACATATGAGACAAACGCTGATAGATAATCCAGATAATAACCTCCTGAAAATGGCCAAAGCGGGTTATAAAGTAAATCCCACAGAACTAATGTATATTTTAGGTACTTATGGACAACAGAGGATAGATGGCGAACCAGCAGAGACTCGAGTATTGGGTAGAGTCTTACCTTACTATCTTCCAGACTCTAAGGATCCAGAAGGAAGAGGTTATATTCTTAATTCTTTAACAAAAGGATTAACGGGTTCTCAATATTACTTTTCAATGCTGGTTGCAAGATCTCAATCTACTGATATTGTCTGTGAAACATCGCGTACCGGAACACTGGCTAGAAAAATCATTAAAAAGATGGAGGATATGGTGGTCGACGGATACGGACAAGTAGTTATAGGTAATACGCTCATCAAGTACGCCGCCAATTATACCAAAATTCTAGGCTCAGTATGTAAACCTGTAGATCTTATCTATCCAGATGAGTCCATGACTTGGTATTTGGAAATTAGTGCTCTGTGGAATAAAATAAAACAAGGATTCGTTTACTCTCAGAAACAGAAACTTGCAAAGAAGACATTGGCGCCGTTTAATTTCCTAGTATTTGTCAAACCCACCACTGAGGATAATGCTATTAAGGTTAAGGATCTGTACGATATGATTCATAACGTCATTGATGATGTGAGAGAGAAATACTTCTTTACGGTATCTAATATAGATTTTATGGAGTATATATTCTTGACGCATCTTAATCCTTCTAGAATTAGAATTACAAAAGAAACGGCTATCACTATCTTTGAAAAGTTCTATGAAAAACTCAATTATACTCTAGGTGGTGGAACTCCTATTGGAATTATTTCTGCACAGGTATTGTCTGAGAAGTTTACACAACAAGCCCTGTCCAGTTTTCACACTACTGAAAAAAGCGGTGCTGTCAAACAAAAACTTGGTTTCAATGAGTTTAATAACTTGACTAATTTGAGTAAGAATAAGACCGAAATTATCACTCTGGTATCCGATGATATCTCTAAACTTCAATCTGTTAAGATTAATTTTGAATTTGTATGTTTGGGAGAATTAAATCCAAACATCACTCTTCGAAAAGAAACAGATAGATATGTAGTAGACATAATAGTCAATAGATTATACATCAAGAGAGCAGAAATAACCGAATTAGTCGTCGAATATATGATTGAACGATTTATCTCCTTTAGCGTCATTGTAAAGGAATGGGGTATGGAGACATTCATTGAGGACGAGGATAATATTAGATTTACTGTTTATCTAAATTTCGTTGAACCAGAGGAATTGAATCTTAGTAAGTTTATGATGGTTCTTCCGGGTGCCGCCAACAAGGGCAAGATTAGTAAATTCAAGATTCCTATCTCTGATTATACAGGTTATGACGATTTCAATCAAACAAAAAAGCTCAATAAGATGACTGTAGAACTCATGAATCTAAAAGAATTGGGTTCTTTCGATTTGGAGAACGTTAACGTGTATCCTGGAGTATGGAATACATACGATATCTTCGGTATCGAGGCCGCTCGTGGATACTTATGCGAAGCCATGTTAAACACCTATGGAGAAGGTTTCGATTATCTGTATCAGCCTTGTGATCTTCTCGCTAGTTTACTATGTGCTAGTTACGAACCAGAATCAGTTAATAAATTCAAGTTCGGCGCAGCTAGTACTCTTAAGAGAGCTACGTTTGGAGACAATAAAGCATTGTTAAACGCGGCTCTTCATAAAAAGTCAGAACCTATTAACGATAATAGTAGCTGCCACTTTTTTAGCAAGGTCCCTAATATAGGAACTGGATATTACAAATACTTTATCGACTTGGGTCTTCTCATGAGAATGGAAAGGAAACTATCTGATAAGATATCTTCTCAAAAGATTAAGGAGATGGAAGAAACAGAAGACTTTTAATTCTTATCAATAACATATTTTTCTATGATCTGTCTTTTAAACGATGGATTTTCCACAAATGCGCCTCTCAAGTCCCTCATAGAATGATACACGTATAAAAAATATAACATAGGTGATGACTCCTTATTTTTAGACATTAGATATGCCAAAATCATAGCCCCGCTTCTATTTACTCCCGCAACACAATGAACCAACACGGGCTCGTTTCGTTGATCACATTTAGATAAGAAGGCGGTTACGTCGTCAAAATATTTACTAATATCGGTAGTTGTATCATCTACCAACGGTATATGAATAATATTAATATTAGAGTTAGGTAATGTATATTTATCCATCGTCAAATTTAAAACATATTTGAACTTAACTTCAGATGATGGTGCATTCATAGCATTTTTATAATTTCCCAAATACACATTATTGGTTACTCTTGTCATTATAGTGGGAGATTTGGCTCTGCGCATATCTCCAGTTGAACGTAGTAGTAAGTATTTATACAAACTTTTCTTATCCATTTATAACGTACAAATGGATAAAACTACTTTATCAGTAAACGCGTGTAATTTAGAATACGTTAGAGAAAAGGCTATAGTAGGCGTACAAGCAGCCAAGACATCAACACTTATATTCTTTGTTATTATATTGGCAATTAGTGCGCTATTACTCTGGTTTCAGACGTCTGATAATCCAGTCTTTAATGAATTAACGAGATATATGCGAATTAAAAATACGGTTAACGATTGGAAATCATTAACGGATAGTAAAACAAAATTAGAAAGTGATAGAGGTAGACTTCTAGCCGCTGGTAAGGATGATATATTCGAATTCAAATGTGTGGATTTTGGTGCCTATTTTATAGCTATGCGATTGGATAAGAAAACATATCTACCGCAAGCTATTAGGCGAGGTACTGGAGACGCGTGGATGGTTAAAAAGGCGGCAAAGGTTGATCCATCTGCTCAACAATTTTGTCAGTATTTGATAAAACACAAGTCTAATAATGTTATTACTTGTGGTAATGAGATGTTAAATGAATTAGGTTATAGCGGTTATTTTATGTCGCCACATTGGTGTTCCGATCTTAGTAATATGGAATAGTGTTAGATAAATGCGGTAACAAATGTTCCTGTAAGGAACCATAACAGCTTAGATTTAACGTTAAAGATGAGCATAAACATAATAAACAAAATTACAATCAAACCTATAACATTAATATCAAACAATCCAAAAAATGAAATCAGTGGAGTAGTAAATGCGTACATAACTCCTGGATAACGTTTAACAGCTGCCGTTCCTATTCTAGACCAAAAATTTGGTTTCATGTTTTCAAAGCGGTGTTCTGCAACAAGACGAGGATCGTGTTCTACGTATTTGGCAGAGTTATCCATTATTTGCCTGTTAATCTTTATTTCGTTTTCGATTCTGGCTATTTCAAAATAAAATCCCGATGATAGACCTCCAGACTTTATAATTTCATCTACAATGTTCAGCGCCGTAGTAACTCTAATAATATAGGCTGACAAGCTAACATCATACCCTCCTGTATATGTGAATATGACATGATTTTTGTCCATTACTAGCTCGGTTTTAACTTTATTGCCTGTAATAATTTCTCTCATCTGTAGGATATCTATTTTCTTGTCATGCATTGCCTTTAAGACTGGACGAAGAAACGTAATATCCTCAATAACGTTATCATTTTCTACAATAACTACATATTCTACCTTTTTATTTTCTAACTCGGTAAAAAATTTAGAATCCCATAGGGCTAAATGTCTAGCGATATTTCTTTTCGTTTCCTCTGTACACATGGTGTTACAAAACCCTGAAAAGAAGTGAGTATACTTGTCATCATTTCTAATGTTTCCTCCAGTCCAGTGTATAAACGCATAATCCTTGTAATGATCTGGATCATCCTTGACTATCACAACATTTCTTTTTTCTGGCATAACTTCGTTGTCCTTCACATCATCGAACTTCTGATCATTAATATGCTCATGAAGATTAGGAAATGTTTCTGATGGAGGTCTATCAATAACTGGCACAACAATAACAGGAGTTTTATTCACAGTCGCCATTTAGTTATTGAAATTAATCATATACAACTCTCTAATACGAGTTATATTTTCGTCTATCCATTGTTTCACATTGACATATTTCGACAAAAAGATATAAAATGCGTATTCCAATGCTTCTCTGTTTAATGAATTACTAAAATATACAAACACGTCACTGTCTGGCAATAAATGATATCTTAGAATATTGTAACAATTTATTTTGTATTGCGCATGTTCGTGATCTATGAGTTCTTCTTCGAATGGCATAGGATCTCCGAATCTGAAAATGTATAAATAGGAGTTAGAATAATAATATTTGAGAGTATTGGTAATGTACAAACTCTTTAGCGGTATAATTAGTTTTTTTCTCTCGATTTCTATTTTTAGATGTGATGGAAAAATGACTAATTTTGTAGCATTAGTATCATGAACTCTAATCAAAATCTTAATATCTTCGTCACACGTTAGCTCTTTGAACTGTTTAAGAGATGCATCAGTTGGTTCCACGGATGGAGTAGGTGCAACAATTTTTTGTTCGACATATGTATGTACTGGAGCCATTGTCTTAACTATAATGGTGCTTGTATCGAAAAACTTTAATGTAGATAGCGGAAGCTCTTCGCCGCGACTTTCTACGTCGTAATTAGGTTCTAATGCCGATCTTTGAATGGATACTAGTTTTCTAAGTTCTAATGTGATTCTCTGAAAATGTAAATCCAATTCCTCCGGCATTATAGATGTGTATACATCGGTAAATAAAACTATAGTATCCAACGATCCCTTCTCGCAAATTCTAGTCTTAACCAAGAAATCGTATATAACTACGGAAATGGCGTATTTAAGAGTGGATTCTTCTACCGTTTTGTTCTTGGATTTCATATAAGAAACTATAAAGTCCGCACTACTGTTAAGAATGATTACTAACGCAACTATATAGTTTAAATTAAGCATCTTGGAAACATAAAATAACTCTGTAGACGATACTTGACTTTCGAATAAGTTTGCAGACAAACGAAGAAAGAACAGACCTCTCTTAATTTCAGAAGAAAACTTTTTTTCGTATTCCTGACGTCTAGAGTTTATATCAATAAAAAAGTTAAGAATTAGTCGGTTAATGTTGTATTTCATTACCCAAGTTTGAGATTTCATAATATTATCAAAAGACATGATAATATTAAAGATAAAGCGCTGACTATGCACGAAATAGCTATATGGTTCGCTCAAGAATATAGTCTTGTTAAACGTGGAAACGATAACTGTATTTTTAATTACGTCAGCTGCATCTAAATTAAATATAGGTATATTTATTCCACACACTCTACAATATGCCACACCATCTTCATAATAAATAAATTCATTAGCAAAGTTATTAATTTTAGTGAAATAGTTAGCGTCAACTTTCATAGCTTCCTTCAATCTAATTTGATGCTCACACGGTGCGAATTCCACTTTAACATCTCTTTTCCATGCCTCAGGTTCATCGATCTCTATAATATCTAGTTTCTTGCGTTTAATAAACACGGGTTCGTCTTTCGCAATGAGGTCTGTATAGTAACTATGTAAATGATAACTAGATAGAAAGATGTAGCTATATAGATGACGATCCTTTAAGAGAGGTATGATAACTTTACCCCAATCAGATAAACTATTGTTATGGTCTTCGGAAAAAGAATTTTTATAAATTTTTCCAGTATTTTCCAAATATACGTACTTGACATCTAAAAAATCCTTAATGATAATAGGAATGGATAATCCGTCTATTTTATAAAGAAATACATATCGCACATTATACTTTTTTTTGGAAATGGGAATACCGATGTGTCTACATAAATATGCAAAGTCTAAATATTTTTTAGAGAATCTTAATTGGTCCAAATTCTTTTCCAAGTACGGTAATAGATTTTTCATATTGAACGGTATCTTCTTGATTTCTGGTTCTAATTCCGCATTAAATGATGAAACTAAGTCGCTATTTTTATAACTAACGATTACATCACCTCTAACATCATCATTTACCAGGATACTGATCTTCTTTTGTCGTAAATACATGTCTAATGTGTTAAAAAAAAGATCATACAAGTTATACGTCATTTCATCTGTAGTATTCTTGTCATTGAGTGATAAACTTGTGCTAATCTCTTCTTTAACAGTCTGTTCAAATTTATATCCTATATACGAAAAAATAGCAACCAATGTTTGATCATTCGCGTCAATATTCTGTTCTATTGTAGTGTATAACAATCTTATATCTTCTTCCGTGATAGCTGATACGTTATAAAGGTTGATAACGAAAATATTTTTATTTCGTGAGATAAAGTCATCGTAGGATTTTGGACTTATATTCGCGTCTAGTAGATATGCTTTTATTTTTGGAATAATCTCAATTAGAATAGTCTCTTTAGAGTCCATTTAAAGTTACAAACAACTAGGAAATTGGTTTATAATGTATAATTTTTTTAGTTTTTATAGATTCTTTATTCTATACTTAAAAAATGAAAATAAATACAAAGGTTCTTGAGGGTTGTGTTAAATTGAAAGCGAGAAATAATCATAAATTATTTCATTATCGCGATATCCGTTAAGTTTGTATTGTAATGGCGTGGTCAATTACGAATAAAGCGGATACTAGTAGTTTCACAAAGATGGCTGAAATTAGAGCTCATCTAAGAAATAGCGCTGAAAATAAAGATAAAAACGATGATATTTTCCCGGAAGATGTAATAATTCCATCTACTAAGCCCAAAACCAAACGAGCCACTACTCCTCGTAAACCAGCGGCTACTAAACGATCAACCAAAAAGGATAAAGAAAAGGAGGAAGTGGAAGAAGAAGAAGTAGTTATAGAGGAATATCATCAAACAACTGAAGAAAATTCTCCACCTCCGTCATCATCTCCTGGAGTCGGCAACATTGTAGAAAGCGTGACCGCTGTAGAGCTTGATGATAGCAACGGGGATGATGATAATGATAATGATAATGACGATAATGAACCTATGGTACAAGTTGAAGCTGGTAAAGTAAATCATAGTGCTAGAAGCGATCTCTCTGACCTAAAGGTGGCTACCGACAATATTGTTAAAGATCTTAAGAAAATTATTACTAGAATCTCTGCAGTATCGACTGTTCTAGAGGATGTTCAAGCAGCTGGTATCTCTAGACAATTTACTTCTATGACTAAATCTATTACAACACTATCTGATCTAGTCACCGAGGGAAAATCTAAAGTTGTTCGTAAAAAAGTTAAAACTTGTAAGAAGTAAATGCGTGCACTTTTTTATAAAGATGGTAAACTGTTTACCGATAATAATTTTTTAAATCCTGTATCAGACAATAATCCAGCGTATGAGGTTTTGCAACATGTTAAAATTCCTACTCATTTAACAGATGTAGTAGTATATGGACAAACGTGGGAGGAGGCGTTAACTAGATTAATTTTTGTGGGAAGTGATTCAAAAGGACGTAGACAATACTTTTACGGAAAAATGCATGTACAGAATCGCAACGCTAAAAGAGATCGTATTTTTGTTAGAGTATATAACGTTATGAAACGAATTAATTGTTTTATAAACAAAAATATAAAGAAATCATCCACAGATTCCAATTATCAGTTGGCAGTTTTTATGTTAATGGAAACTATGTTTTTTATTAGATTTGGTAAAATGAAATATCTTAAGGAGAATGAAACAGTAGGGTTATTAACACTAAAAAATAAACACATAGAAATAAGTCCCGATAAAATAGTTATCAAGTTTGTAGGAAAGGACAAAGTTTCACATGAATTTGTTGTTCATAAGTCTAATAGACTATATAAACCGCTATTGAAACTGACGGATGATTCTAGTCCCGAAGAATTTCTGTTCAACAAACTAAGTGAACGAAAGGTATACGAATGTATCAAACAGTTTGGTATTAGAATCAAGGATCTCCGAACGTATGGAGTCAATTATACGTTTTTATATAATTTTTGGACAAATGTAAAGTCCATATCTCCTCTTCCATCACCAAAAAAGTTAATAGCGTTAACTATCAAACAAACTGCTGAAGTGGTAGGTCATACTCCATCAATTTCAAAAAGAGCTTATATGGCAACGACTATTTTAGAAATGGTAAAGGATAAAAATTTTTTAGATGTAGTATCTAAAACTACGTTCGATGAATTCCTATCTATAGTCGTAGATCACGTTAAATCATCTACGGATGGATGATATAGATCCTTACACAAATAATTACAAAACCGATAAATGGAAATGGATAAGCGTATGAAATCTCTCGCAATGACCGCTTTCTTTGGAGAGCTAACCACATTAGATATTATGGCATTAATAATGTCTATATTTAAACGCCATCCAAACAATACCATTTTTTCAGTGGATAAGGATGGTCAGTTTATGATTGATTTCGAATACGATACTTATAAGGCTTCTCAATATTTGGATTTGCCCCTTACTCCGATATCTGGAGATGAATGCAAGACTCACGCATCGAGTATAGCCAAACAATTGGCGTGTGTGGACATTATTAAAGAGGATATTAGCGAATATATCAAAACTACACCCCGTCTTAAACGATTTATAAAAAAATACCGCAATAGATCAGATACTCGTATCAGTCAAGATACAGAAAAGCTTAAAATAGCTCTAGCTAAAGGCATAGATTACGAATATATAAAAGACGCTTGTTAATAAGTAAATGAAAAAAAACTAGTCGTTTATAATAAAACACGATATGGATGCCAACGTAGTATCATCTTCCACTATCGCGACATATATAGACGCTTTAGCAAAGAATGCTTCGGAATTAGAACAGGGTTCTACCGCATACGAAATAAATAATGAATTGGAACTAGTATTTATTAAGCCGCCATTGATTACTTTGACAAATGTAGTAAATATCTCCACGATTCAGGAATCGTTTATCCGATTTACCGTTACTAATAAGGAAGGCGTCAAAATTAGAACTAAGATTCCATTATCTAAGGTACATGGTCTAGATGTAAAAAATGTACAGTTAGTAGATGCTATAGATAACATAGTTTGGGAAAAGAAATCATTAGTGACGGAAAATCGTCTTCATAAAGCATGCTTGTTGAGACTATCAACAGAGGAACGCCATATATTTTTGGATTACAAGAAATATGGATCCTCTATCCGACTAGAATTAGTTAATCTTATTCAAGCAAAAACAAAAAACTTTACGATAGACTTTAAGCTAAAATATTTTCTAGGATCCGGTGCGCAGTCTAAAAGTTCTTTATTGCACGCTATTAATCATCCAAAGTCAAGGCCTAATACATCTCTGGAAATAGAATTCACACCTAGAGACAATGAAACAGTTCCATATGATGAACTAATAAAGGAATTGACGACTTTCTCGCGTCATATATTTATGGCTTCTCCAGAGAATGTAATTCTTTCTCCACCTATTAACGCACCTATAAAGACTTTTATGTTGCCTAAACAAGATATAGTAGGTATGGATCTGGAAAATCTATATGCCGTAACTAAGACTGACGGCATTCCTATAACTATCAGAGTTACATCAAAGGGGTTGTATTGTTATTTTACACATCTTGGTTATATTATTAGATATCCAGTTAAGAGAATAATAGATTCTGAAGTAGTAGTCTTTGGTGAGGCAGTTAAGGATAAGAACTGGACTGTATATCTCATTAAGCTAATAGAGCCCGTAAATGCAATCAGTGATAGACTAGAAGAAAGTAAGTATGTCGAATCTAAACTAGTGGATATTTGTGATCGGATAGTATTCAAGTCAAAGAAATACGAAGGTCCTTTTACTACAACTAGTGAAGTCGTTGATATGTTATCTACATATTTACCAAAGCAACCAGAAGGTGTTATTTTGTTTTATTCAAAGGGACCTAAATCTAACATTGATTTTAAAATCAAAAAGGAGAATACTATAGACCAAACTGTAAATGTAGTATTTAGGTACATGTCCAGTGAACCAATTATCTTTGGAGAATCATCTATCTTTATAGAGTATAAGAAATTTACCAACGATAAAGGCTTTCCTAAAGAATATGGTTCTGGTAAAATTGTGTTATACAACGGCGTTAATTATCTAAATAATATCTATTGTTTGGAATATATTAATACACATAATGAAGTGGGTATTAAGTCCGTGGTTGTACCTATTAAGTTTATAGCAGAATTCTTAGTCAATGGAGAAATACTTAAACCTAGAATCGATAAAACCATGAAATATATTAACTCAGAAGACTATTATGGAAATCAACATAATGTCATAGTTGAACATTTAAGAGATCAAAGCATCAAAATAGGAGATGTCTTTAACGAGGACAAACTATCTGATGTAGGACATCAATATGCCAATAATGATAAATTTAGATTAAATCCAGAAGTTAGTTATTTTACGAATAAAAGAACTAGAGGACCGTTGGGAATTTTATCGAACTACGTCAAGACTCTTCTTATTTCTATGTATTGTTCCAAAACATTTTTAGACGATTCCAACAAACGAAAGGTATTGGCGATTGATTTTGAAAACGGTGCTGACCTGGAAAAATACTTTTATGGAGAGATTGCGTTATTGGTAGCGACGGATCCAGATGCTGATGCTATAGCTAGAGGAAATGAAAGATACAACAAATTAAATTCTGGAATTAAAACCAAGTACTACAAATTTGACTACATTCAGGAAACTATTCGATCCAATACATTTGTATCTAGTGTCAGAGAAGTATTTTATTTTGGAAAGTTTAATATCATCGACTGGCAGTTTGCTATCCATTATTCTTTCCATCCGAGACATTATGCTACCATCATGAATAACTTATCCGAACTAACTGCTTCTGGAGGCAAGGTATTAATTACTACCATGGACGGAGACAAATTATCAAAATTAACCGATAAAAAGACTTTTATAATTCATAAGAATCTACCTAGTAGCGAAAACTATATGTCTGTAGAAAAAATAGCTGATGATAGAATAGTGGTATATAATCCATCAACAATGTCTACTCCAATGACTGAATACATTATCAAAAAGAACGATATAGTTAGAGTGTTTAACGAATACGGATTTGTTCTTGTAGATAATGTTGATTTCGCTACAATTATAGAACGAAGTAAAAAGTTTATTAATGGTGCATCTACAATGGAAGATAGACCGTCTACAAGAAACTTTTTCGAACTAAATAGAGGAGCCATTAAATGTGAAGGTTTAGATGTCGAAGACTTACTTAGTTACTATGTTGTTTATGTCTTTTCTAAGCGGTAAATAATAATATGGTATGGGTTCTGATATCCCCGTTTTAAATGCATTAAATAATTCCAATAGAGCGATTTTTGTTCCTATAGGACCTTCCAACTGTGGATACTCTGTATTGTTAATAGATATATTAATACTTTTGTCGGGTAACAAAGGTTCTACGTCTTCTAAAAATAAAAGTTTGATAACATCTGGCCTGTTCATAAATAAAAACTTGGCGATTCTATATATACTCTTATTATCAAATCTAGCCATTGTCTTATAGATGTGAGCTACTGTAGGTGTACCATTTGATTTTCTTTCTAATACTATATATTTCTCTCGAAGAAGTTCTTGCACATCATCTGGGAATAAAATACTACTGTTGAGTAAATCAGTTATTTTTTTTATATCGATATTGATGGACATTTTTATAGTTAAGGATAATAAGTATCCCAAAGTAGATAACGACGATAACGAAGTATTTATACTTTTAGGAAATCACAATGACTTTATCAGATCAAAATTAACAAAATTAAAGGAGCATGTATTTTTTTCTAAATATATTGTAACTCCAGATACATATGGATCTTTATGTGTAGAATTAAATGGATCTAGTTTTCAGCACGGTGGTAGATATATAGAGGTGGAGGAATTTATAGATGATGGAAGACAAGTTAGATGGTGTTCTACATCCAATCATATATCTGAAGATATACCCGAAGATATACACACTAATAAATTTATTATTTATGATATTTATACGTTTGATTCGTTCAAAAATAAACGATTGGTATTTGTACAGGTACCTACATCATTAGGAGATGATAGCTATTTAACTAACCCGTTATTGTCACCGTATTATTGTAATTCAGTAGCTAGACAAATGGTCAATGATATGATTTTTAATCAAGATTCATTTTTAAAATATTTATTAGAACATCTGATTAGAAGCCACTATAGAGTTTCTAAACATATAACAATAGTTAGATACAAAGATACCGAAGAATTAAATCTAACGAGAATATGTTATAATAGAGATAAGTTTAAGGCGTTTGCATTCGCTTGGTTTAACGGCGTTTTGGAAAATGAAAAAGTACTAGATACGTATAAAAAGGTATCTGATTTGATATAATGAATTCAGTGACTGTATCACACGCACCATATACTATTACTTATCACGATGATTGGGAACCAGTCATGAATCAATTAGTAGAGTTTTATAACGAAGTAGCCAGTTGGCTGCTACGAGACGAGACGTCGCCTATTCCTGATAAGTTCTTTATACAGTTGAAACAACCGCTTAGAAATAAGCGCGTATGTGTGTGTGGTATAGATCCGTATCCGAAAGATGGAACTGGTGTACCGTTCGAGTCACCAAATTTTACAAAAAAATCAATTAAGGAGATAGCATCTTCTATATCTAGATTAACCGGAGTAATTGATTATAAAGGTTATAACCTTAATATAATAGACGGGGTTATACCCTGGAATTATTACTTAAGTTGTAAATTAGGAGAAACAAAAAGTCACGCGATTTACTGGGATAAGATTTCTAAGTTACTGCTGCATCATATAACTAAACACGTTAGGTTTCTTTATTGTTTGGGTAAAACAGATTTCTCGAATATACGGGCCAAGTTAGAATCTCCAGTAACTACCATAGTGGGATATCATCCAGCAGCTAGAGACCGTCAATTCGAGAAAGATAGATCATTTGAAATTATCAACGTTTTACTGGAATTAGACAACAAGGCACCTATAAATTGGGCTCAAGGGTTTATTTATTAATGCTTTAGTGAAATTTTAACTTGTGTTCTAAATGGATGCGGTTATTAGAGGTAATGATGTTATCTTTGTTCTTAAGACTATAGGTGTCCCGTCAGTATGCAGACAAAATGAAGATCCAAGATTTGTAGAAGCATTTAAATGCGACGAGTTAGAAAGATATATTAAGAATAATCCAGAATGTACACTATTCGAAAGTCTTAGGGATGAGGAAGCATATTCTATAGTCAGAATTTTCATGGATGTAGATTTAGACGCGTGTCTAGACGAAATAGATTATTTAACAGCTATTCAAGATTTTATTATCGAGGTGTCAAACTGTGTAGCTAGATTCGCGTTTACAGAATGCGGTGCCATTCATGAAAATGTAATAAAATCCATGAGATCTAATTTTTCATTAACTAAGTCTACAAATAGAGATAAAACAAGTTTTCATATCATCTTTTTAGATACGTATACCACTATGGATACCTTGATAGCCATGAAACGAACACTATTAGAATTAAGTAGATCATCTGAAAATCCACTAACAAGATCGATAGACACTGCCGTATATAGGAAAAAAACAACTCTTCGGGTTGTAGGTACTAGGAAAAATCCAAATTGCGACACTATTCATGTAATGCAACCACCGCATGATAATATAGAAGATTACCTATTCACTTACGTGGATATGAACAACAATAGTTATTACTTTTCTCTACAACGACGATTGGAGGATTTAGTTCCTGATAAGTTATGGGAACCAGGGTTTATTTCATTCGAAGACGCTATAAAAAGAGTTTCAAAAATATTCATTAATTCTATAATAAACTTTAATGATCTCGATGAAAATAATTTTACGACGGTACCACTGGTCATAGATTACGTAACACCTTGTGCATTATGTAAAAAACGATCGCATAAACATCCGCATCAACTATCGTTGGAAAATGATGCTATTAGAATTTACAAAACTGGTAATCCACATAGTTGTAAAGTTAAAATTGTTCCGTTGGATGGTAATAAACTGTTTAATATCGCACAAAGAATTTTAGACACTAACTCTGTTTTATTAACCGAACGAGGAGACCATATAGTTTGGATTAATAATTCATGGAAATTTAACAGCGAAGAACCCTTAATAACAAAACTAATTCTGTCAATAAGACATCAACTACCTAAGGAATATTCAAGCGAATTACTCTGTCCGAGGAAACGAAAGACTGTAGAAGCTAACATACGAGACATGTTAGTAGATTCAGTAGAGACCGATACCTATCCGGATAAACTTCCGTTTAAAAATGGTGTATTGGACCTGGTAGACGGAATGTTTTACTCTGGAGATGATGCTAAAAAATATACGTGTACTGTATCAACCGGATTTAAATTTGACGATACAAAGTTCGTCGAAGATAGTCCAGAAATGGAAGAGTTAATAAATATCATTAACGATATCCAACCATTAACGGATGAAAATAAGAAAAATAGAGAGTTGTACGAAAAAACTTTATCTAGTTGTTTATGCGGTGCTACCAAAGGATGTTTAACATTCTTTTTTGGAGAAACGGCAACTGGAAAGTCGACAACCAAACGTTTGTTAAAGTCTGCTATCAGTGATCTGTTTGTTGAGACGGGTCAAACAATTTTAACAGATGTATTGGATAAAGGACCTAATCCATTTATCGCTAACATGCATTTGAAAAGATCTGTGTTCTGTAGCGAACTACCTGATTTTGCATGTAGTGGGACAAAGAAAATCAGATCTGATAATATTAAAAAGTTGACAGAACCTTGTGTCATTGGAAGACCGTGTTTCTCCAATAAAATTAATAATAGAAACCATGCTACAATCATTATCGATACTAATTACAAACCTGTCTTTGATAGGATAGATAACGCATTAATGAGAAGAATCGCCGTTGTGCGATTCAGAACACACTTTTCTCAACCTTCTGGTAGAGAGGCTGCTGAAAATAATGACGCGTACGATAAAGTCAAACTATTAGACGAGGGGTTAGATGGTAAAATACAAAATAATAGATATAGATTCGCATTTCTATACTTGTTGGTGAAATGGTACAGAAAATATCATGTTCCTATTATGAAACTATATCCTACACCGGAAGAGATTCCTGACTTTGCATTCTATCTCAAAATAGGTACTCTGTTAGTATCTAGCTCTGTAAAGCATATTCCATTAATGACGGACCTCTCCAAAAAGGGATATATATTGCACGATAATGTGGTTACTCTTCCGTTGACTACTTTCCAACAGAAAATATCCAAGTATTTTAATTCTAGACTATTTGGACACGACATAGAGAGCTTCATCAATAGACATAAGAAATTTGCCAATGTTAGTGATGAATATCTACAATATATATTCATAGAGGATATTTCATCTCCGTAAATATATGCTCATATATTTATAGAAGATATTACATATCTAAATGAATACCGGAATCATAGATTTATTTGATAATCATGTTGATAGTATACCAACTATATTACCTCATCAGTTAGCTACTCTAGATTATCTAGTCAGAACTATCATAGATGAGAACAGAAGCGTGTTATTGTTCCATATTATGGGATCAGGTAAAACAATAATCGCTTTGTTGTTCGCCTTGATAGCTTCCAGATTTAAAAAGGTTCACATTCTAGTGCCGAACATTAACATTTTGAAAATTTTTAATTATAATATGGGTGTAGCTATGAACTTGTTTAATGATGAATTCATAGCTGAGAATATCTTTATTCATTCAACAACAAGTTTTTATTCTCTTAATTATAACGATAACGTCATTAATTATAACGGATTATCTCGCTACAATAACTCTATTTTTATCGTTGATGAGGCACATAATATCTTTGGGAATAATACTGGAGAACTTATGACCGTAATAAAAAATAAAAACAAGATTCCTTTTCTACTATTGTCTGGATCTCCCATTACTAACACACCTAATACTCTGGGTCATATTATAGATTTAATGTCCGAAGAGACGATAGATTTTGGTGAGATTATTAGTCGTGGTAAGAAAGTAATTCAGACACTTCTTAACGAACGCGGTGTGAATGTACTCAAGGATTTGCTTAAAGGAAGAATATCATATTACGAAATGCCTGATAAAGATTTACCAACGATAAGATATCACGGACGTAAGTTTCTAGATACTAGAGTAGTATATTGTCACATGTCTAAACTTCAAGAGAGAGATTATATGATTACTAGACGACAGCTATGTTATCATGAAATGTTTGATAAAAATATGTATAACGTGTCAATGGCAGTATTGGGACAACTTAATCTGATGAATAATTTAGATACTTTATTTCAGGAACAGGATAAGGAATTGTACCCAAATCTGAAAATAAATAATGGTGTGTTATACGGAGAAGAATTGGTAACGTTAAACATTAGTTCCAAATTTAAATACTTTATCAATCGGATACAGACACTCAACGGAAAACATTTTATATACTTTTCTAATTCTACATATGGTGGATTAGTAATTAAATATATCATGCTCAGTAATGGATATTCTGAATATAATGGTTCTCAGGGAACTAATCCACATATGATAAACGGCAAACCAAAAACATTTGCTATCGTTACTAGTAAAATGAAATCGTCTTTAGAGGATCTATTAGATGTGTATAATTCTCCTGAAAACGATGATGGTAGTCAATTGATGTTTTTGTTTTCGTCAAACATTATGTCCGAATCCTATACTCTGAAAGAGGTAAGGCATATTTGGTTTATGACTATCCCGGATACTTTTTCTCAATACAACCAAATTCTTGGACGATCTATTAGAAAATTCTCTTACGTCGATATTTCTGAACCCGTTAATGTATATCTTTTAGCAGCCGTATATTCCGATTTCAATGACGAAGTGACGTCATTAAACGATTACACGCAGGATGAATTGATTAATGTTTTACCCTTTGACATTAAAAAGCTGTTGTATCTAAAATTTAAGACTAAAGAAACGAATAGAATATACTCTATTCTTCAAGAGATGTCTGAAACGTATTCTCTTCCACCACATCCATCAATTGTAAAAGTTTTATTGGGAGAATTGGTCAGACAATTTTTTTATAATAATTCTCGTATTAAGTATAACGATGCCAAGTTACTTAAAATGGTTACATCAGTTATAAAAAATAAAGAAGACGCTAGGAATTACATAGATGATATTGTAAACGGTCACTTCTTTGTATCGAATAAAGTATTTGATAAATCTCTTTTATACAAATACGAAAACGATATTATTACAGTACCGTTTAGACTTTCCTACGAACCGTTTGTTTGGGGAGTTAACTTTCGTAAGGAATATAACGTGGTATCTTCTCCATAAAACTGATAAAATATATAAAGAAATAAATGTCGAGCTTTGTTACCAATGGATACCTTCCAGTTACATTGGAACCACATGAGCTGACGTTAGACATAAAAACTAATATTAGGAATGCCGTATATAAGACGTATCTCCATAAAGAAATTAGTGGTAAAATGGCCAAGAAAATAGAAATTTGTAAAGACGTGGAATTACCTCTCGGCGAAATAGTTAATAATTCTGTAGTTATAAACGTTCCGTGTGTAATAACCTACGCGTATTATCACGTTGGGGATATAGTTAGAGGAACATTAAACATCGAAGATGAATCAAATGTAACTATTCAATGCGGAGATTTAATCTGTAAACTAAGTAGAGATTCGGGTACTGTATCATTTAGCGATTCAAAGTACTGCTTTTTTCGAAATGGTAATGCGTATGATAACGGCAGCGAAGTCTCCGCCGTTCTAATGGAGGCTCAACAAGGTACCGAATCTAGTTTTGTTTTTCTCGCGAATATCGTCGACTCATAAGAAAGAGAATAGCGGTAAGTATATATACGAATACTATGGCAATAATTGCGAATGTTTTATTCCCTTCGATATATTTTTGATAATATGAAAAACATGTCTCTCTCAAATCAGACAACCATCTCATAAAATAGTTCTCGCGTGCTGGAGAGGTAGTTGCTGCTCGTATAATCTCCCCAGAATAATATACTTCTGTGTCATCGTTCAATTTATACGGATTTCTATAGTTCTCTGTTATATAATGCGGTTTGCCCTCATGATTAGACAACGACAATAGTGTTCTGAATTTAGATAGTTGATCAGAATGAATGTTTATTGGCGTTGGAAAAATTATCCATACAGCGTCTGCAGAGTGGTTGATAGTTGTTCCTAGATATTTAAAATAATCCAACTTACTAGGCAGCAAATTGTCTAGATAAAATACTGAATCAAACGGTGCAGACGTATTGGCGGTTCTAATGGAATCCAATTGATTGACTATCTTTTGAAAATATACATTTTTATGATCCGATACTTGTAAGAATATAGAAATAATGATAAGTCCATCATCGTGTTTTTTTGCCTCTTCATAAGAACTATATTTTTTCTTATTCCAATGAACAAGATTAATCTCTCCAGAGTATTTGTACACATCTATCAAGTGATTGGATCCATAATCGTCTTCCTTTCCCCAATATATATGTAGTGATGATAACACATATTCATTGGGAAGAAACCCTCCACTTAAATATCCTCCTTTAAAATTAATCCTTACTAGTTTTCCAGTATTCTGGATAGTGGTTGGTTTCGACTCATTATAATGTATGTTTAACGGCTTTAATCGCGCGTTAGAAATTGCTTTTTTAGTTTCTATATTAATAGGAGATAGTTGTTGTGACATAGTAAAAATGAAATGATAACTGTTTAGAAATAGCTCTTAGTATGGGAATTACAATGGATGAGGAAGTGATATTTGAAACTCCTAGAGAATTAATATCTATTAAACGAATAAAAGATATTCCAAGATCAAAAGACACGCACGTGTTTGCTGCGTGTATAACAAGTGACGGATATCCGCTAATAGGAGCTAGAAGAACTTCATTCGCATTCCAGGCGATATTATCTCAACAAAATTCAGATTCTATCTTTAGAGTATCCACTAAACTATTACGGTTTATGTATTACAATGAACTAAGAGAAATCTTTAGACGGTTGAGAAAAGGTTCTATCAACAATATCGATCCTCACTTCGAAGAGTTAATATTATTGGGTGGTAAACTAGATAAAAAGGAATCTATTAAAGATTGTTTAAGAAGAGAATTAAAAGAGGAAAGTGATGAACGTATAACAGTAAAAGAATTCGGAAATGTAATTCTAAAACTTACAACACAGGATAAATTATTTAATAAAGTGTATATAGGTTATTGCATGTCGTGTTTTATTAATCAATCATTAGAGGATTTATCACATACTAGTATTTACAATGTAGAAATTAGAAAGATTAAATCATTAAATGATTGTATTAACGATGATAAATACGAATATCTGTCTTATATTTATAATATGCTAGTTAATAGTAAATGAACTTTTACAGATCTAGTATAATTAGTCAGATTATTAAGTATAATAGACGACTAGCTAAGTCTATTATTTGCGAGGATGACTCTCAAATTATTACACTTACGGCATTCGTTAACCAATGCCTATGGTGTCATAAACGAGTATCCGTGTCCGCTATTTTATTAACTACGGATAACAAAATATTAGTATGTAACAGACGAGATAGTTTTCTCTATTCTGAAATAATTAGAACTAGAAACATGTCTAGAAAGAAACGATTATTTCTAAATTATTCCAATTATTTGAACAAACAGGAAAGAAGTATACTATCGTCATTTTTTTCTCTAGATCCAGCTACGATTGATAATGATAGAATAGACGCTATTTATCCGGGCGGTATACTCAAAAGGGGTGAGAACGTTCCAGAGTGTCTATCCAGGGAAATCAAAGAAGAAGTTAATATAGACAATTCTTTTGTATTCATAGACACTCGGTTTTTTATTCATGGTATCATAGAAGATACCATTATTAACAAATTTTTTGAAGTAATTTTCTTTGTTGGAAGAATATCTCTAACGAGTGATCAAATTATTGATACCTTTAAAAGTAATCATGAAATTAAGGATCTAATATTTTTAGATCCGAATTCAGGTAATGGACTCCAATACGAAATTGCAAAATATGCTCTAGATACTGCAAAACTTAAATGTTATGGCCATAGAGGATGTTATTATGAATCATTAAAAAAATTAACTGAGGATGATTGATTAGAAAATATAAATTAATTTACCATCGTGTATTTTTATAACGGGATTGTCCGGCATATCATGTAGATAGTTACCATCTACATCGTATACTCGACCATCTACGCCTTTAAATCCTCTATTTATTGACATTAATCTATTAGAATTGGAATACCAAATATTAGTACCCTCAATTAGTTTATTGGTAATATTTTTTTTAGACGATAGATCGATGGCTCTTGAAACCAAGGTTTTCCAACCGGACTCGTTATCGATCGGTGAAAAGTCTTTTTCATTAGCATGAATCCATTCTAATGATGTATGTTTAAACACTCTAAACAATTGGACAAATTCTTTTGATTTGCTTTGAATAATTTCAAATAGGTCTTCGTCTACAGTAGGCATACCATTAGATAATCTAGCCATTATAAAGTGCACGTTTACATATCTACGTTCTGGAGGAGTAAGAACGTGACTATTAAGACGAATGGCTCTTCCTACTATTTGACGAAGAGATGCCTCGTTCCAAGTCATATCTAGAATGAAGATATCATTAATTGAGAAAAAACTAATACCCTCGCCTCCACTAGAAGAGAATACACATGTTTTAATGCATTCTCCGTTAGTGTTTGATTCTTGGTTAAACTCAGCCACCGCCTTGATTCTAGTATCTTTTGTTCTAGATGAGAACTCTATATTAGAGATACCAAAGACTTTGAAATATAGTAATAAAATTTCTATTCCTGACTGATTAACAAATGGTTCAAAGACTAGACATTTACCATGTGATGCTAATATTCCCAAACATACATCTATGAATTTGGCGCTTTTCTCTTTTAATTCAGTAAATAGAGAGATATCAGCCGCACTAGCGTCCCCTCCCAATAGTTCTCCCTTTTTAAAGGTATCTAATGCGGATTTAGAAAACTCTCTATCTCTTAATGAGCGTTTAAAGTCATTATATAGGGTTGCTATCTCTTGCGCGTATTCACCCGGATCACGATTTTGTCTTTCAGGAAAGCTATCGAACGTAAACGTAGTAGCCATACGTCGCAGAATTCTAAATGATGATATACCTGTTTTTATTTCAGTAAGTTTAGCCTTTTGATAAATCTCTTCTTGCTTTTTTGACATATTAACGTATCGCATTAATACTGTTTTCTTAGCGAATGATGCAGACCCTTCTACGTCATCAAAAATAGAAAACTCGTTATTAACTATGTACGAACATAGGCCTCCTAGTTTGGAGACTAATTCTTTTTCATTAACTAGACGTTTATTCTCAAATAGCGATTGGTGTTGTAAGGATCCTGGTCGTAGTAAGTTAACCAACATGGTGAATTCTTGCACACTATTGACGATAGGTGTAGCCGATAAACAAATCATCTTATGGTTTTTTAACGCAATGGTCTTAGATAAAAAATTATATACTGAACGAGTAGGACGGATCTTACCATCTTCTTTGATTAATGATTTAGAAATGAAGTTATGACATTCATCAATGATGACGCATATTCTACTCTTGGAATTAATAGTTTTGATATTAGTAAAAAATTTATTTCTAAAATTTTGATCATCGTAATTAATAAAAATACAATCCTTCGTTATCTCTGGAGCGTATCTGAGTATAGTGTTCATCCAAGGATCTTCTATCAAAGCTTTTTTCACTAATAAGATAATAGCCCAATTCGTATAAATATCCTTAAGATGTTTGAGAATATATACAGTAGTCATTGTTTTACCGACACCTGTTTCATGGAACAATAAAAGAGAATGCATACTGTCTAATCCTAAGAAAACTCTTGCTACAAAATGTTGATAATCCTTGAGGCGTACTACGTCTGTTCCCATCATTTCAACAGGCATATTAGTAGTTCTGCGCAAGGCATAATCGATATAGGCCGCATGTGATTTACTCATTTATAGTGATAAGTAATAACTATGTTTTAAAAATCATAACAGTAGTTTAACTAGTCTTCTCTGATGTTTGTTTTCGATACTTTTCGAATCAGAAGTCATACTAGAATAAAGCAACGAGTGAACGTAATAGAGAGCTTCGTATACTCTATTCGAAAACTCTAAGAACTTATTAATGAATTCCGTATCCACTGGATCGTTTAGAATACTAAATTGAACACTGTTCACATCCTTCCAAGAAGAAGACTTAGTAACGGACTTAACATGAGACATAAATAAATCCAAATTTTTTTTACAAACATTACTAGCCACCATAATGGCGCTATCTTTTAACCAGCTATCGCTTACACATTTCAGCAATCTAACATTTTTAAAGAGACTACAATATATTCTCATGGTATCGATTACACCTCTACCGAATAGAGTAGGAAGTTTAATAATACAATATTTTTCGTTTACAAAATCAAATAATGGTCGAAACACGTCAAAGGTTAACATCTTATAATCGCTAATGTATAGATTGTTTTCAGTAAGATGATTATTAGATTTAATAGCATCTCGTTCGCGTTTAAACAGTTTATTGCGTGCGCTAAGATCGGCAACTACTGCGTCCGCTTTAGTACTTCTCCCATAATACTTTACGCTATTAATCTTTAAAATTTCATAGACTTTATCTAGATCGCTTTCTGGTAACATGATATCATGTGTAAAAAGTTTTAACATGTCGGTTGGCATTCTATTTAGATCATTAACTCTAGAAATCTGAAGAAAGTAATTAGCTCCGTATTCCAAACTAGGTAATGGGCTTTTACCTAGAGACAGATTAAGTTCTGGCAATGTTTCATAAAATGGAAGAAGGACATGCGTCCCCTCCCGGATATTTTTTACAATTTCATCCATTTACAACTCTATAGTTTGTTTTCATTATTATTAGTTATTATCTCCCATTATCTTGGTAATACTTACCCCTTGATCGTAAGATACCTTATACAGGTCATTACATACAACTACCAATTGTTTTTGTACATAATAGATTGGATGGTTGACATCCATGGTGGAATAAACTACTCGAACAGATAGTTTATCTTTCCCCCTAGATACATTGGCCGTAATAGTTGTCGGTCTAAAGAATATCTTTGGTGTAAAGTTAAAAGTTAGGGTTCTTGTTCCATTATTGCTTTTTGTCAGTAGTTCATTATAAATTCTCGAGATGGGCCCGTTCTCTGAATATAGAACATCATTTCCAAATCTAACTTCTAGTCTAGAAATAATATCGGTCTTATTCTTAAAATCTATTCCCTTGATGAAGGGATCGTTAATGAACAAATCCTTGGCCTTTGATTCGGCTGATCTATTATCTCCGTTATAGACGTTACGTTGACTAGTCCAAAGACTTACAGGAATAGATGTATCGATGATGTTGATACTATGTGATATGTGTGCAAAGACTGTTCTCTTGGTGGCGTCGCTATATGTTCCAGTAATGGCGGAAAACTTTTTAGAAATGTTATATATAAAAGAATTTTTTCGGGTTCCAAACATTAACAGATTAGTATGAAGATAAACACTCATATTATCAGGAACATTATCAATTTTTACATACACATCAGCATCTTGAATAGAAACGATACCATCTTCTGGAACCTCAACAATCTCGGCAGACTCCGGATAACCAGTTGGTGGGCCATCACTAACAATAACTAGATCATCCAACAATCTACTCACATATGCATCTATATAATCTTTTTCATCTTGTGAGTACCCTGGATACGAAATAAATTTATTATCCGTATTTCCATAATAAGGTTTAGTATAAACAGAGAGCGATGTTGCCGCATGAACTTCAGTTACAGTCGCCGTTGGTTGGTTTATTTGACCTATTACTCTCCTAGGTTTCTCTATAAACGATGGTTTAATTTGTACATTCTTAACCATATATCCAATAAAGCTCAATTCAGGAACATAAACAAATTCTTTGTTGAACGTTTCAAAGTCGAACGAAGAGTCGCGAATAACGATATCGGATACTGGATTGAAGGTCACCGTTACGGTAATTTTTGAATCGGATAGTTTAAGACTGCTAAATGTATCTTCCACATCAAACGGAGTTTTAATATAAACGTATACTGTAGATGGTTCTTTAATAGTGTCATTAGGAGTTAAGCCAATAGAAATATCATTAAGTTCACTAGAATATCCAGAATGTTTCAAAGCAATTGTATTATTGATACAATTATTATATAATTCTTCGCCCTCAATTTCCCAAATAACACCATTACACGAAGAGACCGATACATGATTAATACATTTATATCCAACATATGGTACGTAACCGAATCTTCCCATACCTTTAACTTCTGGAAGTTCCAAACTTAGAACCAAATGATTAAGCGCAGTAATATACTGATCCCTAATTTCGAAGCTAGCGATAGTCTGATTGTCGGGACCATTGTTTGTCATAACTCCGGATAGAGAAATATATTGCGGCATATATAAAGTTGGAATTTGACTATCGACTGCGAAGACATTAGACCGTTTAATAGAGTCATCCCCACCAATCAACGAATTAATGATAGTATTATTCATTTTCTATTTAAAATGGAAAAAGCTTACAATAAACTCCGTAGAGAAATATCTATAATTTGTGAGTTTTCCTTAAAGTAACAACTTCCGTAAACGCCGTCTTTATCTCTTAGTAAGTTTATTGTATTTATGACCTTTTCCTTATCTTCATAAAATACTAAAGGCAACAAAGAAATTTTTGGTTCTTCTCTAAGAGCTACGTGAGACTTAACCATAGACGCCAACGAATCCCTACATATTTTAGAACAGAAATACCCAACTTCGCCACCCTTGAATGTCTCAATACTAATAGGTCTAAAAACCAAATCTTGATTACAAAACCAACACTTATCAATTACACTATTTGTCTTAATAGACATATCTGCCATAGATTTATAATACTTTGGTAGTATACAAGCGAGTGCTTCTTCTTTAGCGGGCTTAAAGACTGCTTTAGGTGCTGAAATAACCACATCTGGAAGGCTTACTCGCTTAGCCATTTAATTACGGAACTATTTTTTTATACTTCTAATGAGCAAGTAGAAAACCTCTCATCTACAAAAACATACTCGTGTCCATAATCCTCTACCATAGTAACACGTTTTTTAGATCTCATATGTGCTAAAAAGTTTTCCCATACTAACTGATTACTATTATTTTTCGTATAATTTTTAACTGTTTGAGGTTTTAGATTTTTAGTTACAGAAGTAATATCGAATATTTTATCCAAAAAGAATGAGTAATTAATTGTTTTAGAAGGAGTGTTTTCTTGGCAAAAGAATACCAAGTGCTTAAATATTTCTACTACTTCATTAATCTTTTCGGTACTCAGATTCAGTTTCTCATCTTTTACTTGATTGATTATTTCAAAGACTAACTTATAATCCTTTTTATTTATCCTCTCGTTAGCCTTAAGAAAACTAGATACAAAATTTGCATCTACATCATCCGTGGATATTTGATTTTTTTCCATGATATCCAAAAGTTCCGAGATAATTTCTCCAGAACATTGATGAGACAATAATCTCCGCAATACATTTCTCAAATGAATAAGTTTATTAGACACGTGGAAGTTTGACTTTTTTTGTACCTTTGTACATTTTTGAAATACAGACTCGCAAAAAATACAATATTCATATCCTTGTTCAGATACTATACCGTTGTGTCTACAACCGCTACATAATCGTAGATTCATGTTAACACTCTACGTATCTCGTCGTCCAATATTTTATATAAAAACATTTTATTTCTAGACGTTGCCAAAAAATCCTGTAATACTTTTAGTTTTTGGGGTTGTGAATAAAGTATCGCCCTAATATTATTACCATCTTCCGCCAATATAGTAGTTAGATTATCAGCACATGAAGAAAAACACCTCTTAGGTGGATTCAGTACAATGTTATATTTTTCGTACCAACTCATTTAAATATCATAATCTAAAATAGTTCTGTAATATGTCTAGCGCTAATATATTGATCATAATCCTGTGCATAAATTAAAATACAACAATGTCTCGAAATCATTGACATGGCTTCTTCCATAGTTAGAAGATCGTCGTCAAAGTTAGCAACGTGATTCATCAACATTTGCTGTTTTGAGGCAGCAAATACTGAACCGTCACCATTCAACCATTCATAAAAACCATCGTCTGAATCCATTGATAATTTCTTGTACTGGTTTTTGAGAGCTCGCATCAATCTAGCATTTCTAGCTCCCGGATTGAAAACAGAAAGAGGATCGTACATCCAGGGTCCATTTTCTGTAAATAGAATAGTATAATGTCCCTTTAAGAAGATATCAGACGATCCACAATCAAAAAATTGGTCTCCGAGTTTGTAACAGACTGCGGACTTTAACCTATACATGATACCGTTTAGCATGATTTCTGGTGATACGTCAATTGGAGTATCATCTATTAGAGATCTAAAGCCGGTGTAACATTCTCCACCAAACATATTCTTATTCTGACGTCGTTCTACATAAAACATCATTGCTCCATTAACGATAACAGGGGAATGAACAGCACTACCCATCACATTAGTTCCCAATGAATCAATGTGTGTAACTCCAGAACATCTTCCATAGCCTATGTTAGGAGGAGCGAACACTACTCTTCCACTATTGCCATCGAATGCCATAGAATAAATATCCTTGGAATTGATAGAAATCGGACTGTCGGATGTTGTGATCATCTTCATAGGATTAACAACAATGTATGGTGCCGCCTGAAGTTTCATATCGTAACTGATGCCGTTTATAGGTCTAGCCACAGAAACCAATGTAGGTCTAAATCCAACTATAGACAAAATAGAAGCCAATATCTGTTCCTCATCTGTCATAACTTGAGAGCATCCAGTATGAATAATCTTCATTAGATGGGGATCTACCGCATCATCATCGTTACAATAAAAAATTCCCATTCTAATGTTCATAATTGCTTTTCTAATCATGGTATGCATGTTTGCCCTTTGAATCTCTGTGGAAATTAGATCTGATACACCTGTAATCACTATCGGATTATCCTCCGTAAGACGATTAACCAACAACATATAATTATAAGACTTTACTCTTCTAAATTCATAAAGTTGCTGGATTAGACTATATGTGTCTCCATGTACATACGCGTTCTCAAGCGCGGGAAGTTTAATGCCAAATAGTGCCATCAGAATAGGATGAATGTAGTAATTAGTTTCTGGTTTTCTATAAATAAAAGACAAATCTTGCGAGCTAGACATATCGGTAAAATGCATGGATTGGAATCGTGTAGTCGACAGAAGAATATGATGATTAGACGGAGAGTATATTTTATCTAACTCTTTGAGTTGGTCACCAATTCTAGGACTAGCTCGAGAATGAATAAGTACTAAAGGATGAGTACATTTCACAGAAACACTGGCGTTGTTCAATGTGCTCTTTACATGGGTAAGGAGTTGAAATAGCTCGTTTCTATTTGTTCTGACAATATTTAGTTTATTCATAATGTTAAGCATATCCTGAATAGTAAAATTAGATGTGTCATACTTGTTAGTAGTTAGATATTTAGCAATTGCATTCCCATCATTTCTCAATCTCGTACTCCAATCATGTGTGGATGCTACTTCATCTATAGAAACCATACAATCCTTTTTGGTAGTCTGTTGAGCTTGATCATTTCCTGCACGTTTAGGTTTGGTACGTTGATTTCTAGCCCCTGCGGATATAAAGTCATCGTCTACAATTTGGGACAATGAATTGCATACACTACATGACAAAGATTTATCAGAAGTGTGAATATGATCTTCATCTACCAAAGAAAGAGTTTGATTAGTATAACTAGATTTTAGTCCTGTGTTAGATGTTAAAAAAACATCGCTATTGACCACGGCTTCCATTATTTATATTCGTAGTTTTTACTCGAAAGCGTGATTTTAATATCCAATCTTATTACTTTTGGAATCGTTCAAAACCTTTGACTAGTTGTAGAATTTGATCTATTGCCCTACGCGTATACTCCCTTGCATCATATACGTTCGTCACCAGATCGTTTGTTTCGGCCTGAAGTTGGCGCATATCTTTTTCAACACTCGACATGAGATCCTTAAGGGCCATATTGTCTAGATTTTGTTGAGATGTTGCTCCTGGATTTGGATTTTGTTGTGCTGTTGTACATACTGTACCACCAGTAGGTGTAGGAGTACATACAGTGGCCACAATAGGAGGTTGAGGAGGTGTAACTGTCGGAGTAGTACAAGAAATACTTCCATCCGATTGTTGTGTACATGTAGTTGTTGGTAACGTCTGAGAAGGTTGGGTAGATGGCGTCGTCGTCGTCTTTTGATCTTTATTAAATTTAGAGATAATATCCTGAACAGCATTGATCGGCGTCAACGCTGGAAGGAGTGAACTCGGTTGTGTATTATCATCTGCCGACAACCAATCAAAAAGATTAGACATATCCGATGATGTATTAGTTTGTTGTTGTCGTGGTTTTGGTGTAGGAGCCGGTGTAGGTACCTGCAGTGGAGTTATATGAATAGTTGGTTGTAGCGGTTGGATAGGCTGTCTGCTGGCGGCCATCATATTATCTCTAGCTAGTTGTTCTCGCAACTGTCTTTGATAATACGACTCTTGAGACTTTAGTCCTATTTCAATCGCTTCATCCTTTTTTATATCCAGATCCTTTTCTTCAGAATAATAGATTGACGACTTTGGTGTAGAGGATTCTGCCAACCCCTGTGAGAACTTGTTAAAGAAGTCCATTTAAGGCTTTAAAATTGAATTGCGATTATAAGATTAAATGGCAGACACAGACGATATTATCGACTATGAATCCGATGATCTCACTGAATACGAGGATGATGATGAAGAGGAAGAAGATGGAGAGTCACTAGAAACTAGTGATATAGATCCCAAATCTTCTTATAAGATTGTAGAATCAGCATCCACTCATATAGAAGATGCACATTCCAATCTTAAACATATAGGGAATCATATATCTGCTCTTAAACGACGCTATACTAGACGTATAAGTCTATTTGAAATAGCGGGTATAATAGCAGAAAGCTATAACTTGCTTCAACGAGGAAGATTACCTCTAGTTTCAGAATTTTCTAACGAAACGATGAAGCAAAATATGCTACATGTAATTATACAAGAGATAGAGGAGGGTTCTTGTCCTATAGTCATCGAAAAGAACGGAGAATTGTTGTCGGTAAACGATTTTGACAAAGATGGTCTGAAATTCCATCTAGACTATATTATCAAAATTTGGAAACTTCAAAAACGATATTAGAATTTATACGAATATCGTTCTCTAAATGTCACAATCAAGTCTCTCATATTTAGCAGGTTATTGTCGTACTTTATATCGTGTTCATTAACGATATTTTGCAAAATAGTAATGATTCTATCTTCCTTCGATAGATATTCTTCAGAGATTATTGTCTTATATTCTTTCTTGTTATCCGATATAAATTTAATAAGACTTTGAACATTATTAATACCCGTTTGTTTAAATTTTTCTACAGATATTTTAGTTTTGGTAGATTCTATTGTGTCTGTTAATAGACACCCAACATCGACATTCGACGTTAATTGTCTATAAATTAGAGTATAAATTTTAGAAATAACATTAGCAAATTGTTGTGCGTTGATGTCGTTATTCTGAAACAGTATGATTTTAGGTAGCATTTTCTTAACAAAGAGAACGTATTTATTGTTACTCAGTTGAACAGATGATATATCCAGATTACTAACGCATCTGATTCCATATACCAAACTTTCAGAAGAAATGGTGTACAATTGTTTGTATTCATTCAATGTCTCCTTTTCAGAAATTAGTTTAGAGTCAAATACTGCAATAATTTTCAAAAGATAGTTTTCATCAGATAAGATTTTATTTAGTGTAGATATGATAAAACTATTGTTTTGTTGGAGAACTTGATACGCCGCATTCTCTGTAGTCGACGCTCTCAAATGGGAAACAATCTCCATTATTTTTTTGGAATCGGATACTATATCTTCGGTATCTTGACGCAGTCTAGTATACATAGAGTTAAGAGAAATTAGAGTTTGTACATTAAGCAACATGTCTTTAAATGTGGCTACAAACTTTTCCTTTTCCGCATCATCTAGTTTATTATATACCGATTTTACAACGGCGCCAGATTTAAGAAACCAGAATGAAAAACTCTGATAACTACAATATTTCATCATAGTTACAATTTTATCATCTTCTATAGTTGGTGTGATAACACATACCTTTTTCTCCAAGACTGGAACCAACGTCATAAAAATGTTTAAATCAAAATCCATATCAACATCTGATGCGCTAAGACCAGTCTCGCGTTCAAGATTATCTTTACTAATGGTGACGAACTCATCGTATAGAACTCTAAGTTTGTCCATTATTTATTTACAGATTTAGTTGTTTAATTTATTTGTACTCTTCCAAAGTTGGGATAGTATTTTTCTAACGTTGGTATTATATTATTAGGATCTACGTTCATATGTATCATAATATTAATCATCCACGTTTTGATAAATCTATCTTTAGCTTCTGAAATAACGTATTTAAACAAAGGAGAAAAATATTTAGCTACGGCATCAGACGTAATAACATTTTTTGTAAATGTAACGTATTTAGACGAGAGATCTTCGTTAAAAAGTTTTCCATCTATGTAGAATCCATCGGTTGTTAACACCATTCCCGCGTCAGATTGAATAGGAGTTTGAATAGTTTGTTTTGGAAATAGATCCTTCAATAACTTATAGTTGGGTGGGAAAAAATCGATTTTATCACTAGACTCTTTCTTTTTTACTATCATTACCTCATGAACTATTTCTTGAATGAGTATATGTATTTTCTTTCCTATATCGGACGCGTTCATTGGAAAATATACCATGTCGTTAACTATAAGAATATTTTTATCCTCGTTTACAAACTGAATAATATTAGACGTAGTTCGTAAACGAACTATATCATTACCAGTACAACATCTAACTATATGATATCCACTAGTTTCCTTTAGCCGTTTATTATCTTGTTCCATATTAGCAGTCATTCCATCATTTAAGAAGGCGTCAAAGATAATAGGGAGAAATGACATTTTGGATTCTGTTACTACTTTACCAAAATTAAGGATATACGGACTTACTATCTTTTTCTCAACGTCGATTTGATGAACACACGATGAAAATGTGCTTCTATGAGATTGATCATGTAGAAAACAACAAGGGATACAATATTTCCGCATATCATGAAATATATTAAGAAATCCCACCTTATTATATTTCCCCAAAGGATCCATGCACGTAAACATTATATCGTTATCATTAATAAAGACTTCTTTCTCATCGGATCTGTAAAAGTTGTTACTGATTTTTTTCATTCCAGGATCTAGATAATTAATAATGACGGGTTTTCTATTTTTATTCTTTGTATTTTGGCATATCCTAGACCAGTAAACAGTTTCCACTTTGGTAAAATCAGCAGACTTTTGAACGCTATTAAACATGGCATTAATGGCAATAACTAAAAATGTAAAATATTTTTCTATGTTAGGAATATGGTTTTTCACTTTAATAGATATATGGTTTTTGGCCAAAATGATAGATATTTTTTTATCCGAGGATAGTAAAATATTATTAGTCGCCGTCTCTATAAAAATGAAGCTAGTCTCGATATCCAATTTTATTCTAGAATTGATAGGAGTCGCCAAATGTACCTTATACGTTATATCTCCCTTGATGCGTTCCATTTGTGTATCTATATCTGACACAAGATCGGTAAATAGTTTTACGTTATTAATCATCACGGTATCACCGTCGCTAGATAATGCTAATGTTCCATCCAAGTCCCAAATGGAGAGATTTAACTGTTCATCGTTTAGAATAAAATGATTACCGGTCATATTAATAAAGTGTTCATCGTATCTAGATAACAACGACTTATAATTAATGTCCAAGTCTTGAACTCGCTGAATGATCTTTTTTAACCCAGTTAGTTTTAGATTGGTACGAAATATATTGTTAAACTTTGATTCTACAGTAATGTCCAAATCTAGTTGTGGAAATACGTCCATCAGCATTGTTTCAAACTTGATAATATTATTATCCACATCTTCATATGATCCAAATTCTGGAATAGATGTATCACACGCTCTGGCTACCCAGATAACCAAAAAGTCGCACGCTCCAGGATATACATTGTATAAAAAGCTATCGTTTTTAAGTAGTGTTTTTTTTTGCGTATATACGAAGGGATTGAAAATAGTATTATCGACGTAACTATATTCCAAATTATTCTTATGCGAATAGATAATAATATCGTCCTTAATATCTAACAAATTTCCTAAATATCCCTTTAATTGAGTCATTCGAAGCGTCAATAGAATATGTCTCTTAACTATTTCCGGCTGTTGTATATTTAAATGACTTCGTAAGAAATAATATATGGGCGACTTTTCATCTATGTAATCATATGGAGTGAGATATAGGGCTCGTTCTACCTCCTGCCCTTTACCCACCTGTAATACCAATTGTGGACTCACTATATATCGCATATTTATATCGTGGGGTAAAGTGAAAATCTACTACCGATGATGTAAGTCTTACAATGTTTGAACCAGTACCAGATCTTAATTTGGAGGCCTCCGTAGAACTAGGGGAGGTAAATATAGATCAAACAACACCTATGATAAAGGAGAATATCGGTTTTATATCCCGCAGTAGACGTCTATTCGCCCATAGATCTAAGGATGATGAGAGAAAACTAGCACTACGATTCTTTTTACAAAGACTTTATTTTTTAGATCATAGAGAGATTCATTATTTGTTCAGATGCGTTGACGCTGTAAAAGACGTCACTATTACCAAAAAAAATAACATTATTGTGGCGCCTTATATAGCACTTTTAACTATTGCATCAAAAGGATGCAAACTTACAGAAACAATGATTGAAGCATTCTTTCCAGAACTATATAATGAACATAGTAAGAAATTCAAATTCAACTCTCAAGTATCCATCATCCAAGAAAAACTTGGATATCAGTCTGGAAACTATCACGTTTATGATTTTGAACCGTATTACTCTACAGTAGCTTTGGCTATTCGAGATGAACATTCATCTGGCATTTTTAATATCCGTCAAGAGAGTTATCTGGTAAGTTCATTATCTGAAATAACATATAGATTTTATCTAATTAATCTAAAATCTGATCTTGTTCAATGGAGTGCTAGTACGGGCGCTGTAATTAATCAAATGGTAAATACTGTATTGATTACAGTGTATGAAAAGTTACAACTGGTCATAGAAAATGATTCACAATTTATATGTTCATTGGCTGTGGAATCAGAACTTCCAATAAAATTACTTAAAGATAGAAATGAATTATTTACAAAATTCATCAACGAGTTAAAAAAGACCAGTTCATTCAAGATAAGCAAACGTGATAAGGATACGTTACTAAAATATTTTACTTAGGAGTTAGAATTTATAGACGACTCATTTAGTTTATTACTATTACTAGCATTATTGGTATTCTTCTTGTCATCTTGTTCAGAAATATATAACAATGCTATACCTAATACTAAATACATTATCATGCTTGCAATGGCTCTAACAACGACGAACCAAAATGAATTTGGTCGTAGCTTTTGTTCACAAAAATACATAAAGAAATGTCTACATAAATCTATGGCGCCATTGGCATCTTGAAATAGCGCCAGTCCTCCTACAGATTTTAATATAGCTGTATAACATGACATTTATTCATCATCAAAAGAGACAGAGTCACCATCTGTCATATTTAGATTTTTTTTCATGTGTTCAAAGTATCCTCTACTCATTTCATTATAATAGTTTATCATACTTAGAATTTTAGGACGGATCAATGAGTAAGACTTGACTAAATCGTCAGTAGTAATTTGCGCATCGTCTATTCTACATCCGCTTCGTCGAATAATGTATAGCATTGCTTTGAGATTCTCCATAGCTATCAAGTCTTTATACAATGACATGGAAATATCTGTGAACACTTTATACTTCTCCAACATCGATGCCTTAACATCATCACCTACTTTAGCATTGAAAATACGTTCTATTGTGTAGATGGATGTAGCAAGATTTTTAAACAACAATGCCATCTTACATGATGATTGCCTTAAGTCTCCAATCTTTTGTTTAGAATGATTAGCTACAGAGTCCAACGCTTGGCTGACTAGCATATTATTATCTTTAGAAATTGTATTCTTCAATGAGGCGTTTATCATATCTGTAATTTCGTTAGTCATATTACAGTCTGACTGGGTTGTAATGTTATCCAACATATCACCTATGGATACGGTACACGTACCAGCATTTGTAATAATCCTATCTAAGATGTTGTATGGCATTGCGCAGAAAATATCTTCTCCTGTAATATCTCCACTCTCGATAAATCTACTTAGATTATTCTTAAATGCCTTATTCTCTGGAGAAAAGATATCAGTGTCCATCATTTCATTAATAGTATACGCAGAAAAGATACCACGAGTATCAATTCTATCCAAGATACTTATTGGTTCCGAGTCACAAATAATGGTTTCCTCTCCTTCAGGAGATCCTGCATAGAAATATCTAGGACAATAGTTTCTATACTGTCTGTAACTCTGATAATCTCTAAAGTCACTAACTGATACCATGAAATTGAGAAGATCAAACGCTGAAGTAATCAATTTTTCTGCCTCGTTTTTACTACAACTAGTTTTCATCAATGTAGTGACGATGTATTGTTTAGTTACTCTTGGTCTAATACTGATGATAGAGATATTATTACTTCCCATAATGGATCTTCTAGTAGTCACCTTAAAGCCCATTGATGCGAATAGCAGATAGATAAAGTCTTGGTATGACTCCTTTCTAATATAGTACGGACTACCTTTGTCACCCAACTTTATACCCACATAAGCCATAACAACCTCTTTAATAGCCGTTTCATGAGGTTTATCAGCCATGAGCCTAAGTAGTTGGAAGAATCTCATGAATCCCGTCTCAGAAAGTCCTATATGCATGATAGAAGATTTATCTTTCTTGGGAAACTCTCGTATAGTCATAGATGAAATACTCTTTAAAGTTTCTGAAATAAGATCAGTAACAGTCTTACCTCCGACTACTCTGGGTAACAAACATACTCTAATAGGTGTTTTCTCTGCGGAGATAATATCAGAAAGGATAGAGCAATAAGTAGTATTATTGTGATTATAAAGACCGAATACATAACAGGTAGAATTTATAAACATCATGTCCTGAAGGTTTTTAGACTTGTATTCCTCGTAATCCATACCGTCCCAAAACATGGATTTGGTAACTTTGATAGCCGTAGATCTTTGTTCCTTCGCCAACAGGTTAAAGAAATTAATAAAGAATTTGTTGTTTCTATTTATGTCCACAAATTGCATATTTGGAAGTCCCACCGTTACGTTCACTGCAGCATTTTGAGGATCTCGAGTATGAAGTACGATGTTATTGTTTACTGGTATATCTGGAAAGAATTCTACCAGTCTAGGAATAAGAGATTGATATCGCATAGAAATACAAAAGTTAATAATCTCATCATCGAAGATTACTCGGTTACCATTGTAATAAATTGGTACTCTATCATAATCGTCGACAAAGTACTGTTCATACATGATGAGATGTTTATATGTTGGCATAGTAGTGAGATCGACGTTTGGTAATGGCAATGTATTAAGATTAACTCCATAATGTCTAGCAGCATCCGCGATGGCATAAGTGTTGTCAAAGCGGGGTTGATCTTGTGCTGTTATATATTGTCTAACACCTATAAGATTATCAAAATCTTGTCTGCTTAATACACCGTTAACAATTTTCGCCTTGAATTCTTTTATTGGTGCATTAATAACATCCTTATAGAGGATGTTAAACAAATAAGTATTATCAAAGTTAAGATCTGGGTATTTCTTTTCTGCTAGAACATCCATTGAGTCGGAGCCATCTGGTTTAATATAACCACCGATAAATCTAGCTCTGTATTCTGTATCCGTCAATCTAATATTAAGAAGGTGTTGAGTGAAAGGTGGAAGATCGTAAAAGCTGTGAGTATTAATGATAGGGTTAGTTTCCGAACTAATGTTAATTGGGGTATTAATAATATCGATATTTCCAGCGTTAAGTGTAACATTAAACAGTTTTAATTCACGTGACGTGGTATCAATTAAATAATTAATGCCCAATTTGGATATAGCAGCCTGAAGCTCATCTTGTTTAGTTACGGATCCTAATGAGTTATTAAGAAATACATCAAACGGATGAACGAAGGTTGTTTTAAGTTGGTCACATACTTTGTAATCTAGACATAGATGCGGAAGAACGGTAGAAACTATACGAAATAGATATTCAGAGTCCTCTAATTGATCAAGAGTAACTATTGACTTAATAGGCATCATTTATTTAGTATTAAATGACGACCGTACCAGTGACAGATATACAAAACGACTTAATTACAGAGTTTTCAGAAGATAATTATCCATCTAACAAAAATTATGAAATAACTCTTCGTCAAATGTCTATTCTAACTCACGTTAATAACGTGGTAGATAGAGAACATAATGCCGCCGTAGTGTCATCTCCAGAGGAAATATCATCACAACTTAATGAAGATTTATTTCCAGACGATGATTCTCCGGCCACTATTATTGAAAGAGTACAACAACCGCATACTACTATTATTGACGATACGCCACCTCCTACGTTTCGTAGAGAGTTATTGATATCGGAACAACGTCAGCAACGAGAAAAAAGATTTAATATTACGGTATCAAAAAATGCCGAAGCGATAATGGAATCTAGATCGATGATAACTTCTATGCCAACACAAACACCATCCTTGGGAGTAGTTTATGATAAAGATAAAAGAATTCAGATGCTAGAGGATGAAGTGGTTAATCTTAGAAATCAACAATCTAATACAAAATCATCTAATAATTTGGATAATTTTACCAGAATACTATTTGGTAAGACTCCGTATAAATCAACAGAAGTTAATAAGCGTATAGCCATCGTTAATTATGCAAATTTGAATGGGTCCCCCTTATCAGTAGAGGACTTGGATGTTTGTTCAGAGGATGAAATAGATAGAATCTATAAAACGATTAAACAATATCACGAAAGTAGAAAACGAAAAATTATCGTTACTAACGTGATTATTATTGTCATAAACATTATTGAACAGGCATTGCTAAAACTTGGATTTGACGAAATCAAAGGACTGAGTACCGACATCACTTCAGAAATTATCGATGTGGAGATCGGAGACGACTGTGATGCTGTAGCATCAAAACTAGGAATCGGCAACAGTCCGGTTCTTAATATTGTATTGTTTATACTCAAGATATTTGTTAAACGAATTAAAATTATTTAATTTAATACATTCCCATATCCAGACAACAATCGTTTGGATTAATCTGTTCCTGTCGTCTCATACCGGACGACATATTAATCTTTTTATTAGTGGGCATCTTTTTAGATGGTTTCTTTTTCCCAGCATTAACTGATTCGATACCTAGAAGATCGTGATTGATCTCTCCGACCATTCCACGAACTTCTAATTGACCGTCTCTAACGGTACCATAAACTATTTTACCAGCATTAGTAACAGCTTGGACAATCTGACCATCCATTGCGTTGAATGATGTAGTTGCTGTTGTTCTACGTCTAGGAGCACCAGAAGTATTTTTAGAGCTCTTGGATGTTGATGTAGAAGAAGACGAGGATTTTGATTTACATGTGATACATTTTGAACTCTTTGATTTTGTGTCACATGCGCCGGCAGTCACATCTGTTTGAGAATTAAGATTATTGTTGCCTCCTTTGACGGCTGCATCTCCACCGATCTGTGCTAGTAGATTTTTAAGCTGTGGTGTAATCTTATTAACTGTTTCGATATAATCATCGTAACTGCTTCTAACGGCTAAATTTTTTTTATCCGCCATTTAGAAGCTAAAAATATTTTTATTTATACAGAAGATTTAACTAGATTATACAATGAACTAATATGATCCTTTTCCAGATTATTTACAAACCTGGTATTTCTTGTTTCTGGAGGAGGCAAGTTTACATTCGGACTTGGATTTTGTGAGTTCTTGATCTTATTATACATCGCGTATAGGATGGCGACGGTAACTGCTACACAAATACCAATCAAAAGAAGAATACCAATCATTTATTGATAATAACTTCACTATTGATCAAGTATACAATATATCATCTTTTCACTAAATAAGTAGTAATAATGATTCAACAATGTCGAGATATATGAACGATAATAACTTAGTTTACGGAAATATCGCTATGATTGGTATGAATGACTCCACTAACTCTGTGGGGTGCGCAGTGCTTTCCCCACATAGAATAAATTAGCATTCCGACTGTGATAATAATACCAAGTATAAACGACATAATACTCAATACTTTCCATGTACGAGTGGGACTGGTAGACTTACTAAAGTCAATAAAGGCAAAGATACACGAAAGAATCAAAAGAATGATTCCAGCAATTAACACTCCGGAAAAATAATTTCCAATCATAAGCATCATGTCCATTTAACTAATAAAAATTTTAAATCGCCGAATGAACAAAGTGGAATATAAACCATATAAAAACAATAGTTTGTACTGCAAAAATAATATCTATTTTTGTTTTCGAAGATATGGTAAAATTAAATAGTAGTACACAACATGTTATAACTAACAGCAGCAACGGTTCGTAATTACTTATCATTTACTAGACGAAAAGGTGGCGGGATATTTTCTTGCTCAAATAATACGAATATATTACCCATCCATTTTATGCGATGTTTATATACTCTAATCTTTAATAGATCTATAGACGACGGGTTTACCAACAATATAGATTTTATCGATTCATCTAATTTAAACCCTTCCTTAAACGTGAATGATCTATTATCTGGCATAACGATGACCCTACCTGATGAATCAGACAATGTACTGGGCCATGTAGAATAAATTATCAACGAATCATCGTCTACGAACATTTATATCATTTGTTTTAATTTTAGGACGCGAATAAATAGATATAAAATAGAAAATAACAGATATTACAACCAGTGTTATGGCCGCACCCAACCAGGTAGGCAGTTTTATTTTATCTTTTACTACAGGTTCTCCTGGATGTATGTCACCAACTGTAGACGTAGTTCTAGTACAATTAGACGTAAGTTCCGCTTGGGAATTTTTTAACGCTAAAGAGTTAACGTTAATCGTACACCCAACGTATTTACATCTAGTTCTTTGAACATCTTGATTATAATATAACCATTTTCTATCTCTAGATTCGTCAGTGCACTCATGTAACCAACATACCCTAGGTCCTAAATATTTATCTCCGGAATTAGATTTTGGATAATTCGCACACCAACAATTTCTATTTCCTTTATGATCGTTACAAAAGACGTATAATGCCGTATCCCCAAAAGTAAAATAATCAGGACGAATAATTCTAATAAACTCAGAACAATATCTCGCATCCATATGTTTGGAGCAAATATCGGAATAAGTAGACATAGCCGGTTTCCGTTTTACACGTAACCATTCTAAACAATTGGGGTTTCCAGGATCGTTTCTACAAAATCCAGTCATAAAATCGTCACAATGTTCTGTCTTGTAATTATTATTAAATATTTTTGGACAGTGTTTGGTATTTGTCTTTGAACAACATTTTGCTACGCTATCACTATCTCCCAGGAGATAATCCTTTTTTATAAAATGACATCGTTGCCCGGATGCTATATAATCAGTAGCGTGTTTTAAATCCTTAATATATTCAGGAGTTACCTCGTTCTGATAATAGATTAATGTTCCAGGACGAAATTTGAAAGAACTACATGGTTCTCCATGAATTAATACATATTGTTTAGCAAACTCAGGAACTATAAAACTACTACAATGATCTATCGACATACCATCTATCAAACAAAATTTGGGTTTAATTTCTCCTGGAGACGTTTCATAATAATACATATAACTTTCTTCGGCAAACTTAACAGTTCTATTATATTTAGGATAATTAAAACCTAATTCCATATATTTGTCTCGTATATCTGCTATTCCTGATGCTATGTTGATTCTATTAAGAGTAACGGCTGCCCCCATTCTTAATAATCGTCAGTATTTAAACTGTTAAATGTTGGTATATCAACATCTACCTTATTTCCCGCAGTATAAGGTTTGTTGCAGGTATACTGTTCAGGAATGGTTACATTTATACTTCTTCTATAGTCCTGTCTTTCGATGTTCATCACATATGCAAAGAACAAAATAAACAAAATAATGTAAGAAATAATATTAAATATCTGTGAATTCGTAAATACATTAATTGCCATAATAATTACAGCAGCTACAATACACACAATAGACATTCCCACAGTGTTGCCATTACCTCCACGATACATTTGAGTTACTAAGCAATAGGTAATAACTAAGCTAGTAAGAGGCAATAGAAAAGATGAGATAAATATCATCAATATAGAGATTAGAGGAGGGCTATATAGAGCCAAGACGAACAAAATCAAACCGAGTAACGTTCTAACATCATTATTTTTGAAGATTCCCAAATAATCATTCATTCCTCCATAATCGTTTTGCATCATACCTCCATCTTTAGGCATAAACGATTGCTGCTGTTCCTCTGTAAATAAATCTTTATCAAGCACTCCAGCACCCGCAGAGAAGTCGTCAAGCATATTGTAATATCTTAAATAACTCATTTATATATTAAAAAATGTCACTATTAAAGATGGAGTATAATCTTTATGCCGAACTAAAAAAAATAACTTGTGGTCAATCCCTAAGTCTTTTTAACGAAGACGGGGATTTCGTAGAAGTTGAACCGGGATCATTATTTAAGTTTTTGATACCTAAGGGATTTTACTCCTCTCCTTCCGTAAAGACGAGTTTAGTATTCGAGACATTAACAACGACCGATAATAAAATTACTAGTATCAATCCAACAAATGCACCAAAGTTATATCCTCTTCAACACAAAGTCGTATCTGAAGTAGTTTCTAATATGAGGAAAATGATTAAATTAAAACGTCCTCTATACATCACTCTTCACTTGGCGTGTGGATTTGGTAAGACTATTACCACGTGTTATCTTATGGCTACACACGGAAGAAAAACCGTCATTTGTGTACCCAATAAAATGTTAATACATCAATGGAAGACACAGGTAGAGGCAGTCGGATTGGAACATAAGATATCTATAGATGGAGTAAGTAGTTTATTAAAGGAACTAAAGACTCAAAGTCCGGATGTATTAATAGTAGTTAGTAGACATCTGACAAACGATGCCTTTTGTAAATATATCAATAAGCATTATGATTTGTTCATCTTGGATGAATCACATACGTATAATCTGATGAACAATACAGCAGTTACAAGATTTTTAGCGTATTATCCTCCGATGATGTGTTATTTTTTAACTGCTACACCTAGACCATCTAACAGAATTTATTGTAACAGTATTATTAATATTGCCAAGTTATCCGATCTAAAAAAAACTATCTATGCTGTAGATAGTTTTTTTGAGCCATATTCCACAGATAATATTAGACATATGATAAAACGACTAGATGGACCATCTAATAAATATCATATATATACTGAAAAGTTATTATCTGTAGACGAGCCTAGAAATCAACTTATTCTTAATACCCTGGTAGAAGAATTCAAGTCAGGAACTATTAATCGCGTTTTAGTTATTACTAAACTACGTGAACATATGGTATTATTCTACAAACGATTATTAGATCTTTTCGGACCAGAGGTTGTATTTATAGGAGACGCTCAAAATAGACGTACTCCAGATATGGTTAAATCAATCAAGGAACTAAATAGATTTATATTCGTATCCACCTTATTTTATTCCGGTACTGGTTTAGATATTCCTAGTTTGGATTCGTTGTTCATTTGCTCGGCAGTAATCAACAATATGCAAATAGAGCAATTACTAGGGAGGGTATGTCGAGAAACAGAACTATTAGATAGGACGGTATATGTATTTCCTAACACATCCGTCAAAGAAATAAAGTACATGATAGGAAATTTTGTGCAACGAATTATTAGTCTGTCTGTAGATAAACTAGGATTTAAACAAGAAAGTTATCGGAAACATCAAGAATCCGATCCCACTTCTGTATGTACAGCATCCTCCAGAGAAGAACGTGTATTAAATAGAATATTTAACTCGCAAAATCGTTAAGAAGTTTAAGCGACGATCCACATGCTGTACAGGCCAGTGTATTACCCCTCATAGTATTAATATAATCCAATGATACTTTTGTGATGTCGGAAATCTTAACCAATTTAGACTGACAGACAGAACACGTCATGCAATCATCCTCATCGATAACTGTAGTCTTAGGCTTCTTTTTGCGACTCTTCATTCCGGAACGCGCATTGGTGCTATCCATTTAGGTAGTAAAAAATAAGTCAGAATATGCCCTATAGCACGATCGTGTAAAACCTGGTATATCGTCTCTATCTTTATCACAGTATAGTGTATCGACATCTTTATTATTATTGACCTCGTTTATCTCGGAACATGGAATGGGAACATTTTTGTTATCAACGGCCACCTTTGCCTTAATTCCAGATGTTGTAAAATTATAACTAAACAGTCTATCATCGACACAAATGAAATTTTTGTTTAAACGTTTGTAGTTTACGTATGCGGCTCGTTCTCGTCTCATTTTTTCAGATATTGCAGGTACTATAATATTAAAAATAAGAATGAAATAACATAGGATTAAAAATAAAGTTATCATGACTTCTAGCGCTGATTTAACTAACTTAAAAGAATTACTTAGTCTGTACAAAAGTTTGAGATTTTCAGATTCTGCGGCTATAGAGAAGTATAATTCTTTGGTAGAATGGGGAACATCTACTTACTGGAAAATAGGCGTACAAAAGGTAGCTAATGTCGAGACGTCCATATCTGATTATTATGATGAGGTAAAAAATAAACCGTTTAATATTGATCCAGGGTATTATATTTTCTTACCGGTATATTTTGGAAGCGTCTTTATTTATTCGAAGGGTAAAAATATGGTAGAACTTGGATCTGGAAACTCTTTTCAAATACCAGATGATATGCGAAGTGTGTGTAACAAAGTATTAGACGGCGATAACGGAATAGACTTTCTGAGATTTGTTTTGTTAAACAATAGATGGATAATGGAAGACGCTATATCAAAATATCAGTCACCAGTTAATATATTTAAACTAGCGAGTGAGTACGGATTAAACATACCCAACTATTTAGAAATTGAAATAGAGGAAGACACATTATTTGACGACGAGTTATACTCTATTATAGAACGCTCTTTCGATGATAATTTTCCAAAAATATCCATATCGTATATTAAGTTGGGAGAACTTAGGCGGCAAGTTGTAGACTTTTTCAAATTCTCATTCATGTATATTGAGTCCATCAAGGTAGATCGTATAGGGGATAATATTTTTATTCCTAGCGTTATAACAAAATCAGGAAAAAAGATATTAGTAAAAGATGTAGACCATTTAATACGATCCAAGGTTAGAGAACATACATTTGTAAAAGTAAAAAAGAAAAACACATTTTCCATTTTATACGACTATGATGGGAACGGAACAGAAACTAGAGGAGAAGTAATAAAACGAATTATAGACACTATAGGACGAGACTATTATGTTAACGGAAAGTATTTCTCTAAGGTTGGTAGTGCGGGCTTAAAGCAATTGACTAATAAATTAAATATTAATGAGTGCACAACTGTCGATGAGTTAGTTGATGAGATTAATAAATCCGGAACTGTAAAACGAAAAATAAAAACCCAATCAGCATTTGATTTAAGTAGAGAATGTTTGGGATATCCAGAAGCGGATTTTATAACGTTAGTTAATAACATGCGGTTCAAAATAGAAAATTGTAAGGTTGTAAATTTTAATATTGAAAATACTAATTGTTTAAATAACCCAAGTATTGAAACTATATATGGAAACTTTAACCAGTTCGTCTCAATCTTTAATATCGTCACCGATGTCAAAAAAAGATTATTCGAGTGAAATAATATGTGCCTTTGATATAGGTGCAAAAAATCCTGCCAGAACTGTTTTAGAAGTCAAGGATAACTCCGTTAGGGTATTGGATATATCAAAATTAGACTGGAGTTCTGATTGGGAAAGGCGCATAGCTAAAGATTTGTCACAATATGAATACACTACAGTTCTTCTAGAACGTCAGCCTAGAAGGTCGCCATACGTCAAATTTATCTATTTTATTAAAGGCTTTTTATATCATACATCGGCTACCAAAGTTATTTGTGTCTCGCCTGTCATGTCTGGTAATTCATATAGAGATCGAAAAAAGAGATCTGTCGAAGCATTTTTTGATTGGATGGACATATTCGGATTGCGAGACTCCGTTCCAGATAGACGCAAATTAGACGATGTAGCGGATAGTTTCAATTTGGCTATGAGATACGTATTAGATAAATGGAATACTAATTATACACATTATAATAGGTGTAAATCTAGAAATTACATAAAAAAAATGTAATAACGTTAGTAACGCCATTATGGATAATCTATTTACCTTTCTACATGAAATAGAAGATAGATATACCAGAACTATTTTTAACTTTCATCTAATAAGTTGCGATGAAATAGGAGATATATATGGTCTTATGAAAGAACGCATTTCCTCAGAGGATATGTTTGATAATATAGTATATAATAAAGATATACATCCTGCCATTAAGAAACTAGTTTATTGTGACATTCAACTTACTAAACACATTATTAATCAGAATACGTATCCGGTATTTAACGATTCTTCACAAGTGAAATGTTGTCATTATTTCGATATAAACTCAGATAATAGCAATATTAGCTCTCGTACAGTAGAGATATTTGAGAGGGAAAAGTCATCTCTTGTATCATATATTAAAACTACCAATAAGAAGAGAAAAGTCAATTATGGGGAAATAAAGAAAACTGTACATGGAGGCACTAATGCAAATTACTTTTCCGGTAAAAAATCTGATGAGTATCTGAGCACTACAGTCAGGTCCAATATTAATCAACCTTGGATCAAAACCATCTCTAAGAGGATGAGAGTCAATATCATTAATCACTCTATAGTAACGCGTGGAAAAAGCTCCATATTACAAACTATAGAAATTATTTTTACTAATAGAACATGTGTGAAAATATTCAAGGATTCTACTATGCACATTATTCTATCCAAGGACAAGGATGAAAAGGGGTGTATACACATGATTGACAAATTATTCTATGTCTATTATAATTTATTTCTGTTGTTCGAGGATATCATCCAAAACGAGTACTTTAAAGAAGTAGCTAATGTTGTGAACCATGTACTCACGGCTACGGCATTAGATGAGAAATTATTCCTAATTAAGAAAATGGCTAAACACGATGTTTATGGAGTTAGCAATTTCAAAATAGGGATGTTTAACTTGACATTTATTAAGTCATTGGATCATACCGTTTTCCCCTCTCTGTTAGATGAGGATAGTAAAATAAAGTTTTTTAAGGGGAAAAAGCTCAATATTGTAGCATTACGCTCTCTGGAGGATTGTATAAATTACGTGACTAAATCCGAGAATATGATAGAAATGATGAAGGAAAGATCGACTATTTTAAATAGCATAGATATAGAAACGGAATCGGTAGATCGTCTAAAAGATTTGCTTCTAAAATGAAAAAAAACACTGATTCAGAAATGGATCAACGACTCGGGTATAAGTTTTTGGTACCTGATCCTAAAGCCGGAGTTTTTTATAGACCGTTACATTTCCAATATGTATCGTATTCTAATTTTATATTGCATCGATTGCATGGAATCTTGACCGTCAAGCGGCCACTTTTATCGTTTAAGAATAATACAGAACGAATTATGATAGAAATTAGCAATGTTAAAGTGACTCCTCCAGATTACTCACCTATAATCGCGAGTATTAAAGGTAAGAGTTATGACGCATTAGCCACGTTCACTGTAAATATCTTTAAAGAGGTAATGGCCAAAGAGGGTATATCCATCACTAAAATAAGTAGCTATGAGGGAAAAGATTCTCATTTGATAAAAATTCCGCTACTAATAGGATACGGGAATAAAAATCCACTTGATACAGCCAAGTATCTTGTCCCTAACGTCATAGGTGGGGTATTTATCAATAAACAATCTGTCGAAAAAGTAGGAATTAATCTAGTAGAAAAGATTACAACATGGCCAAAATTTAGGGTTGTTAAGCCAAACTCATTCACGTTCTCGTTTTCCTCCGTATCCCCTCCTAATATATTACCGACAAGATATCGCCATTACAAGATATCTCTGGATATATCACAATTGGAAGCGTCGAATATATCATCGACAAAGACATTTATAACGGTCAATATTGTTTTACTGTCTCAATATTTATCTAGAGTGAGTTTAGGATTCATTAGACGTAGTTTATCATACGATATGCCTCCAGAAGTTGTCTATTTAGTAAACGCGATAATAGATAGTGCTAAACGACTTACCGAATCTATTACTGACTTTAATATTGATACGTATATTAATGACCTGGTGGAAGCTGAACACGTTAAACAAAAATCTCAGTTAACGATTAACGAGTTCAAATATGAAATGCTACATAACTTTTTACCTCATATGAACTATACACCCGATCAACTAAAGGGATTTTATATGATATCTTTACTAAGAAAGTTTCTTTACTGTATCTACTACACTTCTAGATATCCAGATAGAGATTCAATGGTTTGTCATCGCATCCTAACATACGGCAAATATTTTGAGACGTTGGCACATGATGAATTAGAGAATTACATAGGCAACATCAGAAACGACATCATGAACAATCACAAGAATAGAGGCACTTACGCGGTAAACATTCATGTACTAACAACTCCCGGACTTAATCACGCGTTTTCTAGCTTATTGAGTGGAAAGTTCAAAAAGTCAGACGGTAGTTATCGAACACATCCTCACTATTCATGGATGCAGAATATTTCTATTCCTAGAAGTGTTGGGTTTTATCCGGATCAAGTAAAGATTTCAAAGATGTTTTCTGTCAGAAAATACCATCCAAGTCAATATCTTTACTTTTGTTCATCAGATGTTCCGGAAAGAGGTCCTCAGGTAGGTTTAGTATCTCAATTGTCTGTCTTAAGTTCCATTACAAATATACTAACGTCTGAGTATTTGGATTTGGAAAAGAAAATTTGTGAGTATATCAGATCATATTATAAAGATGATATAAGTTACTTTGAAACAGGATTTCCAATCACTATAGAAAATGCTCTAGTCGCATCTCTTAATCCAAATATGATATGTGATTTTGTAACTGACTTTAGACGTAGAAAACGGATGGGATTCTTCGGTAACTTGGAGGTAGGTATTACTTTAGTTAGGGATCACATGAATGAAATTCGTATTAATATTGGAGCAGGAAGATTAGTCAGACCATTCTTGGTGGTGGATAACGGAGAGCTCATGATGGATGTGTGTCCGGAGTTAGAAAGCAGATTAGATGACATGACATTCTCTGACATTCAGAAAGAGTTCCCACATGTCATCGAAATGGTAGATATAGAACAATTTACTTTTAGTAACGTATGTGAATCTGTTCAAAAATTTAGAATGATGTCAAAGGATGAAAGAAAGCAATACGATTTATGTGACTTTCCTGCCGAATTTAGAGATGGATATGTAGCATCTTCACTAGTGGGAATCAATCACAATTCTGGTCCAAGAGCTATTCTTGGATGTGCTCAAGCTAAACAAGCTATCTCTTGTCTGAGTTCGGATATACGAAATAAAATAGACAATGGAATTCATTTGATGTATCCAGAGAGGCCAATTGTCATTAGTAAGGCTTTAGAAACTTCAAAGATTGCGGCTAATTGCTTCGGACAACATGTTACTATAGCATTAATGTCGTACAAAGGTATCAATCAAGAGGATGGAATTATCATCAAAAAACAATTTATTCAGAGAGGCGGTCTTGATATTGTTACAGCCAAGAAACACCAAGTAGAAATTCCATTGGAAAACTTTAATAACAAAGAAAGAGATAGGTCTAACGCCTATTCGAAATTAGAAAGTAATGGATTAGTTAGACTGAATGCTTTCTTGGAATCCGGAGACGCTATGGCACGAAATATCTCATCAAGAACTCTTGAAGATGATTTTGCTAGAGATAATCAGATTAGCTTTGATGTTTCCGAGAAATATACCGATATGTACAAATCTCGCGTTGAACGAGTGCAAGTAGAACTTACTGACAAAGTTAAGGTGCGAGTATTAACCATGAAAGAAAGAAGACCCATTCTAGGAGATAAATTTACCACTAGAACGAGTCAAAAGGGAACAGTCGCGTATATCGCAGATGAAACGGAACTTCCATACGACGAAAATGGTATCACACCAGATGTCATTATTAATTCTACATCCATCTTCTCTAGAAAAACTATATCTATGTTGATAGAAGTTATTTTAACAGCCGCATATTCCGCTAAGCCGTACAACAATAAGGGAGAAAACCGACCTGTCTGTTTTCCTAGTAGTAACGAAACATCCATCGATACATATATGCAATTCGCTAAACAATGTTATGAGCATTTAAATCCGAAATTGACCGAGAAAGAATTATCGGATAAAATCTTTTGTGAAAAGATTCTTTATGATCCTGAAACGGATAAGCCTTATGCATCCAAAGTATTTTTTGGACCAATTTATTACTTGCGTCTGAGACATTTAACTCAGGACAAGGCAACCGTTAGATGTAGAGGTAAAAAGACGAAGCTAATTAGACAAGCGAATGAGGGACGAAAACGTGGAGGAGGTATCAAGTTTGGAGAAATGGAGAGAGACTGTTTAATAGCGCATGGCGCAGCCAATACTATTACAGAAGTTTTAAAAGACTCAGAAGAGGATTATCAAGATGTGTATATTTGTGAAAATTGTGGAGACATAGCAGCACAAATCAAAAGTATTAATACATGTCTTAGATGTTCAAAACTTAATCTCTCTCCTCTCTTAACAAAAATTGATACCACGCACGTATCTAAAGTATTTCTTACTCAAATGAACGCCAGAGGCGTAAAAGTCAAATTAGATTTCGAACGAAGACCTCCTTCGTTTTATAAACCATTAGATAAAGTTGATCTCAAGCCGTCTTTTCTGAAGTAGGTTTCTTATTTAGTAGTTGTTGTAGATAATCGTTAATATTATCAAGATAATCCAAATTTAATTCTTCAAATGAAGTAGGATCTTCTATAACTACTTTATCATAGTCTAAAGATGACTGATATTCAGGTTTTGTTGGCGGTTGAGGTGTAACAGCTACTTGAGATACCAAAGTCTGATATTCAGAAAGCTGTGGATGTTTGGGTTCAACATCCACTGATGGTGTTACACCAATCGGTTCAGTGACGTCTGTGGATGGAGGTGCTACTTTTACAGAACCTGTAGCCTTAGTTGTCGACGGAGATACATCTTCAATGCGAGGAAATGTATAATTTGGTAATGTTTCTTATGTGGATCTGAAGAAGAGGTAAGATATCTACTAGAAAGATACCGATCACGTTCTAGTTCTCTTTTGTAGAACTTAACTTTTTCTTTCTCAGCATCTAGTTGATATTCCAACCTCTTCACGTTACTACGTTCATATTCCAATTCACGTTCGCATGGGTTACCTTCGCAGTTTTTACGAGCGATTTCACGTTCCAGATTACGTTTAGCCTTCATGTGTCTCTCACTCTCTCTATCGAGTTTATCAGAGCAGTCTTTCTAAAGGCGATCGAACTCCATAAATTTCTCCAACGATTTGATTGTTTCCATAGATTTCCGAAGTTCAGCTTCTAGGACGGCGATTCTTTTTCTTTCGAATTCACAGCTGGATGTACAACCGTTTCCTCCGTTTCCATTACCATCTCTAAGTTTCTTTCCTAGATCGGCAATCTTTCTCAACATTTCATCCCCATACATTTTACATTCCTCGAGTCTACCGTCGTCGAAATATCTTTCCAACTCCTTTTCGACCTCAATAACTTTAACACGTTGTCTTTCAAGCTCTCGTTTGTAGCAACTATCATTTTATCTGATTCCCTGGCGCGTTTAAGATCTTCATGCAATTGAGTCAGCTCTTAACGCAATCTCTTGCTTCTTCGTCATAGTACTCACAATCACTGTGAGATCCATTGCTACCACGTCTACACTTGGTGAGCTCGCGTTTAAGAGATTCAATTTCCCGTTTGTATTGGTCCATGTCTCCATTGCCGCAACCATTAGATTTACAGGCTGTTAGCTGTCGTTCAAGATCAGAAATACGGTTTTTCTTGGAATTGATTTCGTCAATGTACTTAACATGATTAACATCGAAACACTTATTAAGTTCTTTTTCCAATTCTACGATTTTATTTCTTTCGCGAGTCAATTCCCTCCTGTAGTAGCTATCGGTTTTGTCAGATTCACGCTCTCTACGTAGACTTTCTTGCAAGTTACTAATTTGTTCCCTGACACGTCCAAGTTCAGTTTTATATGCCGAATAGAGTTCTGATTTATCCTTTAAGCAGATCTCTAGCGATTGTTTAAGATCACTAATTCTAGTCTTTAGCCTATTTACCTCCTCAGAAGATGTTCCGTTACCGTTGCGTTTACACTCGTTAAGCTGTCTATCAAGATCCATGATTCTATCTCTAAGACGTTGTATCTCTTTTTCCATATCAGCATTGCTTTCATTATTACGTCTGTAGTCACTTAACTATCTTTCAAGATTTGAAATTCTATCTCTAAGACGTCTCATTTCTATCTGTTTCGGTATTGGTCTTATTATTATGTCTACAGTCGTTCAACTGTCTTTCAAGATCTAATATTCTAGATTGGAGTCTACTAATCTCTGTAAGATTTCCTCCTCCGCTCTCAATGCAGTCGGTCAACTTATTCTCTAGTTCGCTAATACGCGAACGCAGTGCATCAACTTCTTGCGTGTCTTTCTGGTTGTGTGTACATTCATCGAGTCTAGATTCGAGATCTCTAACGCGTCGTCGTTCTTCCTCAAGTTCTCCGCGTACTGCAGAAAGCGTGTCCCTATCTTGTTGATATTTAGCAATTTCTGATTCTAGAGTACTGATTCTGCTCACGTAGTTACTAATAGTTGTCTTAGCCTTATCAAGTTCCTCCTTGTATTTGTCGCATTCCTTGATATCCCTACGAAGTTTGGACAGTTCCCATTCGACATTACGACGTTTATCGATTTCAGCTCGGAGATCGTCATCGCGTTGTTTTAGCCACATACGACTAAGTTCAAGTCTTCCTTGACAAGATCCATCTACTTTTCCATCCCTAATAGTATCCAGTTCCTTTTCTAGTTCTGAACGCATTTCTCGTTCCCTATCAAGCGATTCTTTCAATTCTCGGATAGTCTTCTTATCAATTTCTGATGAATCTGAACCATCATCTGTCCCATTTTGAATATCCCTAAGTTCTTTGATCTCTTTTGTAAGTCGGTCGATTCTTTCGGTTTTATAAACAGAATCCCTTTCCAAAGTCCTAATCTTACTGAGTTTATCACTAAGTTCTGCATTCAATTCGGTGAGTTTTCTCTTGGCTTCTTCCAACTCTGTTTTAAACTCTCCACCATTTCCGCATTCTTCCTCGCATTTATCTAACCATTCAATTAGTTTATTAATAACTAGTTGGTAATCAGCGATTCCTATAGCCGTTCTTGTAATTGTGGGAACATAATTAGGATCTTCTAATGGATTGTATGACTTAATAGCATCATCTTTATCATTATTAGGGGGATGGACAACCTTAATTGGTTGGTCCTCCTTATCTCCTCCAGTAGCATGTGGTTCTTCAATACCAGTGTTAGTAATAGGCTTAGGCAAATGCTTGTCGTACGCGGGCACTTCCTCATCCATTAAGTATTTATAATCGGGTTCTGCTTCAGAATATTCTTTTCTAAGAGACGCGACTTCAGGAGTTAGTAGAAGAACTCTGTTTCTGTATCTATCAACGCTGGAATCAATACTCAAGTTAAGGATAGCGAATACCTCATCGTCATCATCCGTATCTTCTGAAACACCATCATATGACATTTCATGAAGTCTAACGTATTGATAAATAGAATCAGATTTAGTATTAAACAGATCCTTAACCTTTTTAGTAAATGCATATATATATATTAGATCTCCAGATTTCATAATATGATCACATGCCTTAAATGTCAGTGCTTCCATAATATAGTCTGGAACACTAATGGGTGACGAAAAAGATACCGCACCATATGCTACGTTAATAAATAAATCTGAACCACTAAGTAGATAATGATTAATATTAAGGAAGAGGAAATATTCAGTATATAGATATGCCTTAGCATCATATCTTGTACTAAACACACTAAACAGTTTATTGATGTGATCAATTTCCAACAAAACAATTAGAGCAGCAGGAATACCAACAAACATATTACCACAGCCGTATTTTTTATGAATATCACATATCATGTTAAAAAATCTTGATAGAAGAGCAAATATCTCATCTGACTTAATGAGACGTAGTTCAGCAGCAGCATAAGTCATAACTGTAAATAGAACATACTTTCCTGTAGTATTGATTCTAGACTCCGCATCAACACCATTATTAAAAATAGTTTTATATACATCTTTAATCTGCTCACCGTTAATCGTCGAACGTTCTAGTATACGGAAACACTTTGATTTCTTATCTGTAGTTAATGACTTAGTGATATCACGAAGAATATTACGAATTACATTTCTTGTTTTTCTTGAGAGACATGATTCAGAACTCAACTCATCGTTCCATAGTTTTTCTACCTCAGTGGCGAAATCTTTGGAGTGCTTGGTACATTTTTCAATAAGGTTCGTGACCTCCATTTATTATAAAAAATTTTATTCAAAACTTAACTACAATCGGGTAATTATAAGATCGTAGATCTCCCATGTGGTGGAATACTACCATCTATCGCATGTGGATGGACAGTGGGTAATAGCCATGGGAACAGTAATGATTGCATATTTATCTTTCTTGCTAGTATTACTGCATATTGTCCCAATGTTTCGATGTGGTGTTCTAACCTATCAACTGCCGCTGTATCACAACAATAGTGTCCGATGGAATTAAGATTATGATCCAATGTGTTTAATATATGATTATCAAGTCTTATACGATCCGCGTCTTTTTTGACAGGATCAGGTTCTTCTACATGAAGAAGTTTCGGCCTCTTATAATATTCATGTCTGGGAAACGGTGGTCTAGGGTAAGGCTCCGGTATCGGAGTGGGTTTTGGATTATAATCATCATCGTCTACATCATCATCTTTGATATTTATTTTGCTATCTTGATGATGTCCTGTATCAGTTGCATTTTCAACACTCGACTGAATATTAGCACATTCATTGTCTATTATTACCATATTTCTAAACCCAAAATGTATGTGTTGAACATCAGTACTATTGTTGATGAGTCTTATAGCATGAATTCGCTTATCGTTATCGGGTTTAGTTTCTGTAACCTTAACAATTCCTTTTTTATTAAACTCTACATAATCATAACCATTTCTATTGTTTGTTTTAATATAAACGAGTATAGCATCATTGCTAAATTTTTCAATAGTATCGAAAACAGAATATCCCAAACCATATAATATATATTCAGGGACACTCAAACTAAATGTCCAGGATTCTCCTAAATACGTAAACTTTAATAGTGCGAAATCATTCAAAAATCTATCACTTATAGATAGATAGTACATGAATGCGTATAGTAGTCTACCGATCTCTTTATTATGAAAACCGACATTACCATCATATATTTCGTGATATACATGTGATCCGTTTACGTTAAACCATAAATACATGGGTGACCCTATAAACATGAATTTATTTCTAATTCTCAGAGCCATAGTTAATTGACCGTGTAATATTTGCTTACATGCATACTTGATACGATCATTAATAAGATTTTTATTATTGCTCGTTATTTCAGAATCGTATATATAAGGAGTACCATTGTGATTCTTACCAGATATTATACAAAATACTATATATAAAATATATTGACCCACGTTAGTAATCATGTAAATGTTTAATGTTTTAAATTTTGTATTTAATGATTCATTATCATACGCTAGCATGGTCTTGTAATATTCATTCTTTAAAATATAATATTGTGTTAGCCATTGCATTGGAGCTCCTAATGGAGATTTTCTATTCTCGTCCATTTTAGGATATGCTTTCATAAAGTCCCTAATAACTTCGTGAATAATGTTTCTATGTTTTCTACTGATGCATGTATTTGCTTCGATTTTTTTATCCCATGTTTCATCTATCATAGATTTAAACGCAGTAATGCTTGCAACATTAACATCTTGAACCATTGGTACAATTCCGTTCCATAAATTTATAATGTTCACCATTTATATAACTCATTTTTTGAATATACTTTTAATTGAACAAAAGAGTTAAGTTACTCATATGGGCGCCGACCAGTCTGAACATCAATCTTTTTAGCCAGAGATATCATAGCCGCTCTTAGAGTTTCAGCGTGATTTTCCAACCTAAATAGAACGTCATCATTGCGTTTACAACACTTTTCTATTTGTTCAAACTTTGTTGTTACATTAGTAATTTTTTTTTCCAAATTAGTTAGCCGTTGTTTGAGAGTTTCCTCATTATTGTCTCCATCGGCTTTAACAATTGCTTCGCGTTTAGCCTCTGGCTTTTTAGCAGCCTTTGTAGAGAAAAATTCAGTTGCTGGAATTGCAAGATCATCATCTCCAGGGAAAAGAGTTCCGTCCATTTAAAGTACAGATTTTAAAAACTGACACTCTACGTTATTTATATTTGGTGCAACACATGGATTATAAATATCGATGTTAATAACATCAGAAAATGTAAAGTCTATACATTGCGCCATCGTGTTAAATTTTCTAATGGATCTAGTATTATTGGGTCCAACTTCAGCCTGAAATCCAAATATGGAAGCGGATACAAAACCGTTTCCTGGATAAGCCACACATCTCCACTTTTGCTTTACATCAGAAATTGTGTCATTGACATCTTGAACTCTCCTATCTAATGCTGGTGTACCACCTATAGATTTTGAATATTCGAATGCTGCATGAGTAGCATTAAATTCCTTAATATTGCCATAATTCTCATATATTGAGTAACCCTGGATAAAAAGTAAACACACTGCAGCCGTCGCTACCACAATAAAAAAAATTGATAGAGAGTTCATTTATAATCTATTAGAAGCTAACAAAATTTTTTTACACGCATCAGACAATGCTTTAATAAATAGTTCAACATCTACTTTTGTTATATCGAACCGATGATATGATTCTAACCTAGAATTACATCCGAAAAAGTTGACTATGTTCATTGTCATTAAGTCATTAACGAACAATATGCCAGACTCTGGATTATAAGACGATACTGTTTCATCACAATCACCTACCTTAATCATGTGATTATGAATATTGGCTATTAGAGCACCTTCTAAGAAATCTATAATATCTTTGAAACACGATTTAAAATCAAACCACGAATATACTTCTACGAAGAAAGTTAGTTTACCCATAGGAGAGATAACTATAAATGGAGATCTAGATACAAAATCCGGATTCATGATAGTTTTAACATTATTATATTCTCTATTAAATACCTCCACATCTAAAAATGTTAATTTTGAAACTATGTCTTCGTTTATTACCGTACCTGAACTAAACGCTATAAGCTCTATTGTTTGTGAACTCTTTAAACGATATTCTTGAAATACATGTAACAAAGTTTCCTTTAACTCGGTCGGTTTATCTACCATAGTTACAGAATTTGTATCCTTATCTATAATATAATAATCAAAATCGTATAAAGTTATATAATTATCGCGTTCAGATTGTGATCTTTTCAAATAGACTAAAAACCCCATTTCTCTAGTAAGTATCTTATGTATATGTTTGTAAAATATCTTCATGGTGGGAATATGCTCTACCACAGTTAGCCATTCCTCATTGATAGCGGTAGATGTATTAGACAAAACTATTCCAATGTTTAACAAGGGCCATTTTACAAGATTATTAAATCCTTGTTTGATAAATGTAGCTAATGAGGGTTCGAGTTCAACGACGATTGAATTCTCTTCCCGCGGATGCCGCATGATGAACGACTGGATGTTGTTCGATTGATTTGGAATTCTTTTTCGACTTTTTGTTTATATTAAATATTTTAAAATTTATAGCTGATAGCAATTCATGTACTACGGATAATGTAGACGCGTATTGTGCATCGATATCTTTATTATTAGATAAATTTATCAATAAATGTGAGAAGTTTGCCTCGTTAAGGTCTTCCATTTAAATATTATATAAACATTTGTGTTTGTATCTTATTCGTCTTTTATGGAATAGTTTTTAACTAGTAAAGCTGTAATTACATACTTTGTCCGTAAAACATAAATATAAACACCCGCTTTTATCAAACGTTCCAAAAAGTCGTTAGTAGACATTTTTAACATGGTATCTATTTTAAATACACTTAGGTTTTTAGAAAAAACATCATTTTATAATTGTAACGATTCAATAACTAAAGAAAAGATTAAGATTAAACATAAGGGAATGTCATTTGTATTTTATAAGCCAAAGCATTCTACCGTTGTTAAATACTTATCTGGAGGATGTATATATCATGATGATTTGGTTGTATTGGGGAAGGTAACAATTAATGATCTAAAGATGATGCTATTTTACATGGATTTATCATATCATGGAGTGACAAGTAGTGGAGTAATTTACAAATTGGGATCATCCATAGATAGACTTTCTCTAAATAGGACTATTGTTACAAAAGTTAATAACAATTATAACAATTATAACAATTATAACAATTATTATAATTGTTATAATTATGATGATACATTTTTTGACGATGATGATTGATCGCTATTACACAATTTTGTTTTTGTACTTTCTAATATAGTGTTTAGGTTCTTTTTCATATGAGAATATTGATTTACTAAAATATCTATGTTTAACTTTTGTTCTATAACGTCCTTATCGGCGGTATCGGTACATATACGTAATTCACCTTCACAAAATACGGAGTCTTCGATAATAATAGCCAATCGATTATTGGATCTAACCGTCTGTATCATATTCAACATGTTTAATATATCCTTTCGTTTACCCTTTACAGGCATCGATCGTAGCATATTTTCCGCGTCTGAGATGGAAATGTTAAAACTACAAAAATGCGTAATGTTAGCCCGTCCTAATATTGGTACATGCCTATAAGTTTGGCATAGTAGAATAATAGACGTGTTCAAATGCCTTCCAAAGTTTAAGAATTCTATTAGAGTATTGCATTTTGATAGTTTATCACCTACATCATCAAAAATAAGTAAAAAGTGTGCTGATTTTTTATGATTTTGTGCGACAGTAATACATTTTTCTATGTTACTTTTAGTTCGTATTAGATTATATTCTAGAGATTCCTGACTACTAACGAAATTAATATGATTTGGCCAAATGTACCCATCATAATCTGGGTTATAAACGGGTGTAAACAAGAATATATGTTTATATTTTTTAACTAGTGTAGAAAACAGAGATAGTAAATAGATAGTTTTTCCAGATCCAGATCCTCCCGTTAAAACCATTCTAAACGGCATTTTTAATAAATTTTCTCTTAAAAATTGTTTTTCTTGGAAACAATTCATAATTATATTTACAGTTACTAAATTAATTTGATAATAAATCAAAATATGGAAAACTAAGGTCGTTAGTAGGGAGGAGAACAAAGAAGGCACATCGTGATATAAATAATATTTGTTATCATGATGACACCAGAAAACGACGAAGAGCAAACATCTGTGTTCTCCGCCACTGTTTACGGAGACAAAATTCAGGGAAAAAATAAACGCAAACGCGTAATTGGTATATGTATTAGAATATCTATGGTTATTTCACTACTATCTATGATTACCATGTCCGCGTTTCTCATAGTGCGTCTAAATCAATGCATGTCTGCTAACGAGGCTGCTATTACTGACGCCACTGCAGTTGCTGCTGCATTATCTACTCATAGAAAGGTTGCGTCTAGCACTACACAATATAAACACCAAGAAAGCTGTAATGGTTTATATTACCAGGGTTCTTGTTATATATTCCATTCAGACTACCAGTTATTCTCGGATGCTAAAGCAAATTGCGCCACAGAATCATCAACACTACCCAATAAATCTGATGTCTTGACTACCTGGCTCATTGATTATGTTGAGGATACATGGGGATCTGATGGTAATCCAATTACAAAAACTACAACCGATTATCAAGATTCCGATGTATCACAAGAAGTTAGAAAGTATTTTTGTGTTAAAACAATGAACTAATATTTATTTTTGTACATTAATAAATGAAATCGCTTAATAGACAAACTGTAAGTAGGTTTAAGAAGTTGTCGGTGCCGGCCGCTATAATGATGATACTCTCAACCATTATTAGCGGCATAGGAACATTTCTGCATTACAAAGAAGAACTGATGCCTAGTGCTTGCGCCAATGGATGGATACAATACGATAAACATTGTTATTTAGATACTAACATTAAAATGTCTACAGATAATACGGTTTATCAGTGTCGTAAATTACGAGCCAGATTGCCTAGACCGGATACTAGACATCTAAGAGTATTGTTTAGTATTTTTTATAAAGATTATTGGGTAAGTTTAAAAAAGACCAATAATAAATGGTTAGATATTAATAATGATAAAGATATAGATATTAGTAAATTAACAAATTTTAAACAACTAAACAGTACAACGGATGCTGAAGCGTGTTATATATACAAGTCTGGAAAACTGGTTAAAACAGTATGTAAAAGTACTCAATCTGTACTATGTGTTAAAAGATTCTACAAGTAACAACAAAAAATAAAATAATAATAAGTCCTTAACGAACGTTGATGGACACCACGTTTGTTATTACTCCAATGGGTATGCTGACTATAACAGATACATTATATGATGATCTCGATATCTCAATAATGGACTTTATAGGACCATACATTATAGGTAACATAAAAACTGTCCAAATAGATGTACGGGATATAAAATATTCCGACATGCAAAAATGCTACTTTAGCTAAGGGTAAAATAGTTCCTTAGGATTCTAATGATTTGGTTAGATTCAACATTTATAGCATTTGTACCGCATACAGATCAAAAATACCATCATCATAGCATGCGACTATGATACCATGTTAGATATAGAAGGTAAACATCAGCCATTTTATCTATTTACATCTATTGATGTTTTTAACGCTACAATCATAGAAGCGTATAACCTGTATACAGCTGGAGATTATCATCTGATCATCAATCCTTCAGATAATCTGAAAATGAAATTGTAGTTTAATTCTTCATTCTGCATATCAAACGGCAATGGATGGATCATAATTGATGGGAAATGCAATAGTAATTTTTTATCATAAAAGTTGTAAAGTAAATAATAAAACAATAAATATTGAACTAGTAGTACGTATATTGAGCAATCAGAAATGATACTGGTACCTCTTATCACAGTGACCGTAGTTGCGGGAACAATATTAGTATGTTATATATTATATATTTGTAGGAAAAAGATACGTACTGTCTATAATGACAATAAAATTATTATGACAAAATTAAAAAAGATAAAGAGTTCTAATTCCAGCAAATCTAGTAAATCAACTGATAACGAATCAGACTGGGAGGATCACTGTAGTGCTATGGAACAAAATAATGACGTGGATAATATTTCTAGGAATGAGATATTGGATGATGATAGCTTCGCTGGTAGTTTAATATGGGATAACGAATCCAATGTCATAGCGCCTAGCACAGAACACATTTACGATAGTGTTGCTGGAAGCACGCGGCTAATAAATAATGATTGTAATGAACAAACTATTTATCAGAACACTACAGTAATTAATGAGACAGAGACTATTGAAGTACTTAATGAAGATACCAAACAGAATCCTAGCTATTCGTCCAATCCTTTCGTAAATTATAATAAAACCAGTATTTGTAGCAAGTCAAATCCGTTCATTACAGAACTCAACAATAAATTTAGTGAGAATAATCCGTTTAGGAGAGCACATAGCGATGATTACCTAAATAAGCAAGAACATGATGATATAGAATCATCTGTTGTATCATTAGTCTGATTAGTTTCCTTTTTATAAAATTGAAGTAATATTTAGTATTAATTGCTACCGTTACATTGTACAAATGGAGATATTCCCTGTATTTGGCATTTCTAAAATTAGCAATTTTATTGCTAATAATGACTGTAGATATTATATAGATACAGAACATCATTAAAAAATTATACCTAATGAGATCAATAGACAGATGGATGAAATGGTACTTCTTACCAACATCTTAAGCGTAGAAGTTGTAAATAACAATGAGATGTATCATCTTATTCCCCATAGACTATCGATGATTATACTCTGTATTAGTTCTATTGGAAGATGTGTTATCTCTATAGATAATGACGTTAATAACAAAAATATTCTAACCTTTCCCATTGATCATGCTGTAATCATATCCCACTAAGTAAATGTGTCGTAGTTAGTAAGGGTCCTACAACCATATTGGTTGTTAAAGCGGATATACCCAGCAAACGACATACTATATGTAAATAATCTGTCACTGATTAATTATTTGCCGTTGTCTGTATTCATTATTAGACGAGTTACAAACTATTTGGATAGACACATATGCGATCAGATATTTGCTAATAATAAGTGGTATTCCATTATAACCATCGACGATAAGCAGTTTCCTATTCCATCAAACTGTATAGGTATGTCCTCTACCAAGTACATAAATTCTAGCATCGAGCAAGATACTTTAATCCATGTTTGTAACCTCGAGCATCCATTCGACTCAGTATACAAAAATGCAGTCGTACAAATTCTGTACCTATCAAGGAACAAATATTGTACGGTAGAATTGATAATATAAATATGAGCATTAGTATTTCTGTGGATTAATAGATTTCTAGTATGGGATCATTAATCATTTCTAATCTCTAAATACCTCATAAAACAAAAAAGCTATTATCAAATACTGTACGGAATGGATTCATTCTCTTCTCTTTTTATGAAACTCTGTTGTATATCTACTGATAAAACTGGAAGCAAAAAATCTGATAGAAAGAATAAGAATAAGATCAAGGATTATATGGAACACGATTATTATAAAATAACAATAGTTCCTGGTTCCTCTTCCACGTCTACTAGCTCGTGGTATTATACGCATGCCTAGTAATAGTCTCTTTGCGTTGATGGAAAGCAGACTAGAAATAACAGGCTAAAATGTTCAGACACCATAATAGTTCCCAACCCAGATAATAACAGAGTTCCATCAACACATTCCTTTAGACTCAATCCCAATCCCAAAACCGTTAAAATGTATCCGGCCAATTGATAGTAAATAATGAGGTGTACAGCACATGATAATTTACACAGTAACCAAAATGAAAACACTTTAGTAATTATAAGAAATATAGACGGTAATGTCATCATTAACAATCCAATAATATGCCTGAGAGTAAACATTGACGGATAAAACAAAAATGCCCCGCATAACTCTATCATGGCAATAATACAACCAAACACTTGTAAAATTCCTAAATTAGTAGAAAATACAACTGATATCGATGTATAAGCGATTTCGAGGAATAATAAGAACAAAGTAATGCCTGTAAAAATAAACATCAACATTGTATGATGGTCATTAAACCAATTAGTATGACGTTGAACTAATTTCACAGTAGATTTTATTCCGGTGTTATCCTCGCATGTATAAGTACCGGGTAATATATCTTTATACTTTATAATTAATGAAACATCGTTATCAGATAACGAATGGAGTTTGGCACTAGTATGCCATTTACTTAATATGATCGTCTTGGAAGTTTTATTATAAGTTAAAATATTATGGTTGTCCAATTTCCATCTAATATACTTTGTTGGATTATCTATAGTACACGGAATAATGATGGTATCATTGCACGCCGTATACTCTATAGTCTTTGTAGATGTTATAACCACAAAAGTGCAGAGGTATATCAACAATATTCTAACTCTTAACATTTTTATTTATTTAAAATGATACCTTTGTTATTTATTTTATTCTATTTTGCGAACGGTATCGAATGGCATAAGTTTGAAACGAGTGAAAAAATAATTTCTACTTACTTAATAGATGACGTATTATACACGGGTGTTAATGGGGCAGTATATACATTTTCAAATAATAAACTAAACAAAACTGGTTTAGCTAATACTAATTATATCACAACATCTATAAAAGTAGAGGATGTGATACATTAGTATGCGGAACCAATAACGGAAATCCCAAATGTTGGAAAATAGACGGTTCCAAAGATCCAAAACATAGAGGTAGAGGATACGCTCCTTATCAAAATAGTAAAGTAACGATAATCAGTCATAACAAATGTATACTATCTAACATAAACATATCAAAAGAAGGAATTAAACGATGGAGAAGATTTGACGGACCATGTGGTTATGATTTATTTATACACGGCGGATAACGTAATTCCAAAAGATGGTTTACAAGGAGCATTTGTCGATAAAGACGGTACTTATGACAAAGTTTACATTCTTTTCACTGTTACTATCGGCTCAAAGAGAATTGTTAAAATTCCGTATATAGCACAAATGTGCTTAAACGACGAATGTGGTCCATCATCATTGTCTAGTCATAGATGGTCGACGTTGCTCAAAGTCGAATTAGAATGTGACATCGACGGAAGAAGTTATAGTCAAATTAATCATTCTAAAACTATAAAACAGATAATGATACGATACTATATGTATTCTTTGATAGTCCTTTTCCAAGTCCGCATTATGTACCTATTCTATGAATACCATTAAACAATCTTTTTCTACGTCAAATTGGGAGGATATACAAAGCAATTACTGTCTCCAGCTTTTGGTATATGTCTACCAGCTGGAAAAAGTTGTTCCACATAACACGTTTGACGTTATAGAACAATATAATGTACTAGATAATATTATAAAGCCTTTATTTAACCAACCTATCTTCAAAGGACCGTCTGATGTTAAATGGTTCGATATAAAGGAGAAGGAAAATGAACATCGGAAATATAGAATATACTTCATAAAAGAAAATACTATATATTCGTTCAATACAAAATCTAAACAAACTCGTAGCTCGCAAGTCGATGCGCAACTATTTTCAGTAATGGTAACTTCGAAACCGTTATTTATAGCAGATATAGGGATAGAAGTAGGAATGCCACGAATAAAAAATACTTAAAATGTAATCTTAATCGAGTACGCCATATGACAATGAACAAACCTAAGACAAATTATGCTGGTTATGCTTGCTGCGTAATATGCGGTCTAATTGTTGGAATTATTTTTACAGCGACACTATTAAAAGCTGTAGAACGTAAATTAATTCATACACCATTAATAGATAAAACGATAAAAGATGCATATATTAGAGAAGATTGTCCTACTGACTGAATAAGCTATATAATAAATGTATACATTTATCGACTAATCAAAAAACCTGGGAGGAAGGACGTAATGCATGCAAAGCTCTAAATCCAAATTCGGATCTAATTAATATAGAGACTCTAAACGAGTTAAGTTTTTTAAGAAGCCTTAGAAGAGGCTATTGGGTAGGAGAATTCAAAATATTAAACCAGACAACCACGTATAATTTTATAGCTAAAAATGTCACGAAGAATGGAACTAAAAAACGAAATATATTTGTAGTACAACGAATACTCCCAAACTGCATTCATGTTACACTATATAACAATTACACTACATTTTTATCATAACACTACTTTGGTTAGATGTTTTAGAAAAAAAATATCGCCGTACCGTTCTTGTTTTTATAAAAATAACAATTAACAATTATCAAATTTTTTCTTTAATATTTTACGTGGTTGACCATTCGTGGTGGTAAAATAATCTCTTAGTGTTGGAATGGAATGCTGTTTAATGTTTCCACACTCATCGTATATTTTGACGTATGCAGTTACATCGTTTACGCAATAGTCAGACTGTAGTTCTATTATGCTTCCTACATTAGGAGGAACAGTTTTAAAGTCTCTTGGTTTTAATCTATTACCGTTAGTTTTTATGAAATCCTTTGTTTTATCCACTTCACATTTTAAATAAATGTCCACTATACATTCTTCTGTTAATTTTACTAGATCGTCATATGTCATAGAATTTATAGGTTCCATAGTCCATGGATCCAAACTAGTAAACTTCGCGTATACAGTATCGCGATTAGTGTATACACCAACTGTATGAAAATTAAGAAAACAGTTTAATAGATCAACAGAAATATTTAATCCTCCGTTTGATACAGATGCACCATATTTATGGATTTCGGATTCACACGTTGTTTGTCTGAGGGGTTCGTCTAGCGTTGCTTCTACATAAACTTCTATTCCCATATATTCTTTATTATCAGAATCGCATACCGATTTATCATCATACACTGTTTGAAAACTAAATGGTATACACATCAAAATAACAAATACTAACGAGTACATTCTGCAATATTGTTATCGTAATTGGAAAATTAGTGTTCGGGTGAGTTGGATTATGTGAGTACTGGATTGCATATTATATTTTATATTTTATATTTTATATTTTGTAATAAGAATAGAATGCTAATGTCAAGTTTATTCCAATAAATGACTTATTAAAAAACATATATAATAAATAACAATGGCTGAATGGCATAAAATTATCGAGGATATTTCAAAAAATAATAATTTCGAGGATGCCGCCATCGTTGATTACAAGACTACAAAGAATGTTCTAGCGGCTATTCCTAACAGAACATTTGCAAAGATTAATCCGGGCGAAGTTATTCCGCTCATCACTAATCATAATATTCTAAAACCTCTTATTGGTCAGAAGTTTTGTATTGTATATACTAACTCTCTAATGGATGAGAACACGTATGCTATGGAGTTGCTTACTGGGTACGCCCCTGTATCTCCGATCGTTATAGCGAGAACTCATACCGCACTTATATTTTTGATGGGTAAGCCAACAACATCTAGACGTGATGTGTATAGAACGTGTAGAGATCACGCTACCCGTGTACGTGCAACTGGTAATTAAAATAAAAAGTAATATTCATATGTAGTGTTAATTTTAAATGATGATAAAATGGATAATATCCATATTGACGATGTCAATAATGCCGGTATTGGTATACAGCTCATCGATTTTTAGATTTCGTTCAGAGGATGTGGAATTATGTTATGGGAATTTGTATTTTGATAGGATCTATAATAATGTAGTAAATATAAAATATATTCCTGAGCATATTCCATATAAATATAATTTTATTAATCGTACGTTCTCCGTAGATGAACTAGACAATAATGTCTTTTTTACACATGGTTATTTTTTAAAACACAAATATGGTTCACTTAATCCTAGTTTGATTGTCTCATTATCAGGAAACTTAAAATATAATGATATACAATGCTCAGTAAATGTATCGTGCCTCATTAAAAATTTGGCAACGAGTATATCTACTATATTAACATCTAAACATAAGACGTATTCTCTACATCGGTCCAAGTGTATTACTATAATAGGATATGATTCTATTATATGGTATAAAGATATAAATGACAAGTATAATGATATCTATGATTTTACCGCAATATGTATGCTAATAGCGTCTACATTGATAGTGACCATATACGTGTTTAAAAAAATAAAAATGAACTCTTAATTATGCTATTAGAAATGGATAAAATCAAAATTACGGTTGATTCAAAAATTGGTAATGTTGTTACCATATCGTATAACTTGGAAAAAATAACTATTAACGACACACCAAAAAAGAAAAGGATGTATTATTAGCGCAATCAGTTGCTGTTGAAGAGGCAAAAGATGTCAAGGTGGAAGAAAAAAATATCGATATTGAAAATGACAATGACGATGATATGGATGTAGAAAGCGTGTAATACGATCTATAAAAAATAAGTATATAAATACTTTTTATTTACGGTACTCTTGTAGTGGTAATACCCCTACTCGATTATTTTTTTTAAAAAAAAATACTTATTCTGATTATTCTAGCCATTTCCGTGTTCGTTCAAATGCCACATCAAAGATATGGGAGTAGTTGAAATCTAGTTCTGCATTGTTGGCGCGCCTCAAATGTAGTGTTGGATATCTTCAACGTATAGTTGTTGAGTAGTGATGGTTTTCTAAATAGAATTCTCTTCATATCATTCTTGCACGTGTACATTTTTAGCATCCATCTTGGAATCCTAGATCCTTGTTCTATTCCCAATGGTTTCATCAATAGAAGATTAAACATATCGTACGAACACGATGGAGAGTAACCGTAGCAAAAGTAAGCATTTCCTTTAATCTCAGATCCCGGATACTGGATATATTTTACAGCCAACACATGCATCCATGCAACATTTCCTACATATACCCGGCTATGTACCGCGTCATCATCGACTGTACGATACATAATGTTACCGTGTTGCGTACATTGCTCGTAAAAGACTTTCGTTAATTTGTCTCCTCCTCCGTAAATTCCAGTGGGTCTTAGGCAACAAGTATACAATTTTGCACCATTCATAATTACAGAATTATTGGCTTTCGTGACCAGTTGCTCGACCATACGTTTACTTTTTGCGTATACATGTCCGGGTGATATATCGTACAGGGTATGCTCATGACCAATGAATGGATTACCGTGTTTATTTGGTCCTATTGCTTCCATGCTACCTAATATAAATCAAATACTTGATTCCTAGGTCTACAGAAGCTGCCAATATAGTCTGTGTTACATAATAGTTTACTTTCATGATTTCATTATCGGTGTATTTTCCAAATACATCCACTAGAGCAACCGTATGAATAATCAGATTTACCCCATCTAGTGCTTCTTTCACCTTATTAAGTCGTTTATATCACATTGTATATAGTTTATAACTTTAACCTTCGATACGAGAGGTTGTGGATCTTCTACGACATTGATAACTCTGATTTCTTGAACATCATCTGCGCTAATTAAAAGTTTTACTATATACCTGCCTAGAAATTCGGCACCACCAGTAACCGCGTACACGGTCATTGCTGCTGTCACTCATAAATATCGGACTACTTATTCTATTTTACAAATAATGGCTGTTTGTATAATAGACCACGATAATATCAGAGGAGTTATTTACTTTGAACCAGTCCATGGAAAAGATAAAGTTTTAGGATCAGTTATTGGATTAAAATCCGGAACGTATAATTTGATAATTCATCGTTACGGGGATATTAGTCGAGGATGTAATTCCATAGGCAGTCCAGAAATATTTATCGGTAACATCTTTGTAAACAGATATGGTGTAGCATATGTTTATTTAGATACAGATGTAAATATATCTACAATTATTGGAAAGGCGTTATCTATTTCAAAAAATGATCAGAGATTAGCGTGTGGAGTTATTGGTATTTCTTACATAAATGAAAAGATAATACATTTTCTTACAATTAACGAGAATGGCGTTTGATATATCAGTTAATGCATCTAAAACAATAAATGCATTAGTTTACTTTTCTACTCAGCAAAATAAATTAGTCATACGTAATGAAGTTAATGATACACACTATACTGTCGAATTTGATAGGGACAAAGTAGTTGACACGTTTATTTCGTATAATAGACATAATGACTCCATAGAGATAAGAGGGGTGCTTCCAGAGGAAACTAATATCGGTTGCACGGTTAATACGCCGGTTAGTATGACTTACTTGTATAATAAGTATAGTTTTAAACTAATTTTAGCAGAATATATAAGACACAGAAATACTGTATCCGGCAATATTTATTCGGCATTGATGACACTAGATGATTTGGTTATTAAACAGTATGGCGACATTGATCTATTATTTAATGAGAAACTTAAAGTGGACTCCGATTCGGGATTATTTGACTTTGTCAACTTTGTAAAGGATATTATATGTTGTGATTCTAGAATAGTAGTAGCTCTATCTAGTCTAGTATCTAAACATTGGGAATTGACAAATAAAAAGTATAGGTGTATGGCATTAGCCGAACATATAGCTGATAGTATTCCAATATCTGAGCTATCTAGACCACGATACAATCTATGTAAGTATTTACGAGGACACACTGAGAGCATAGAGGATGAATTTGATTATTTTGAAGACGATGATTCGTCGACATGTTCTGTCGTAACCGACAGGGAAACGGATGTATAATTTTTTATAGTGTGCAGGATATGATAAAAATATAATTGTTGTATCCATTCCCATTCTAATCACATTATATGATTCTGTAAAAAATTATACTGTAACACAATGAAGTAGTTGCATAGATGTATATAGGTCAGATACTGGTTTGATAAACTTTTTATTCCACATGAGTATGTTTGACTTTATGGTTAGACCCGCATACTTTAACAAATCACTGAAAATTGGAGTTAGGTATTGACATCTCAGAATCAGTTGCCGTTCTGGAACATTAAATGTATTTTTTATGATATATTCCAACGCATTTATGTGGGTATACAACAAGTCATTACTAATAGAGTATTCCAAGAGTTTTAATTGGCTAGTATTTAACAAGAGAAGAGATTTCAACAGACTGTTTATGAACTCGAATGCCGCCTTATTGTCGCTTATATTGATGATGTCGAATTCTCCCAATATCATCACTGATGAGTAGCTCATCTTGTTATCAGGATCCAAGCTTTCTAAAGACGTCATTAAACCCTCGATCATGAATGGATTTATCATCATCGTTTTTATGTTGGACATAAGCTTAGTCCGTTTGTCCACATCTATAGACGATGATTTCTGAATTATTGCATATATCTCTCTCTTTAACTCCAGGAACTTGTCAGGATGGTCCACTTTAACATGTTCTCGCCTAAGAGATAAAAATCTTTGGATGGTTGCATGTGACTTTTCTCTAAATGATGATGTTGCCCAAGATCCTCTCTTAAATGAATCCATCCTATCCTTGTACAAGATGGACAGTCTATTTTCCTTAGATGGTTTAATATTTTTGTTACCCATGATCTATAAAGGTAGACCTAATCGTCTTGGATGACCATATATTTATTTCCAGTTTTATTATACGCATAAATTGTAAAAATATGTTAGGTTTACTAAAATGTCTCGCGGGGCATTAATCGTTTTTGAAGGATTGGACAAATCTGGAAAAACAACACAATGTATGAACATCATGGAATCTATACCGACAAACACGATAAAATATCTTAACTTTCCGCAGAGATCCACTGTCACTGGAAAGATGATAGATGACTATCTAACTCGTAAAAAAACCTATAATGATCATATAGTTAATCTATTATTTTGTGCAAATAGATGGGAGTTTGCATCTTTTATACAAGAACAATTAGAACAGGGAATTACTTTAATAGTTGATAGATACGCGTTCTCTGGAGTAGCATATGCCACCGCTAAAGGCGCGTCAATGACTCTCAGTAAGAGTTATGAATCTGGATTGCCTAAACCCGACTTAGTTATATTCTTGGAATCTGGTAGCAAAGAAATTAATAGAAACGTCGGCGAGGAAATTTATGAAGATGTAGCATTCCAACAAAAGGTATTACAAGAATATAAAAAAATGATTGAAGAAGGAGAAGATATTCATTGGCAAATTATTTCTTCTGAATTCGAGGAAGATGTAAAGAAGGAGTTGATTAAGAATATAGTTATAGAGGCTATACATACGGTTACTGGACCAGTGGGGCAACTGTGGATGTAATAAAGTGAAATTACATTTTTATAAATAGATGTTAGTGCAGTGTTAAAAAATGGATGAAGGATATTACTCTGGCAACTTGGAATCAGTTCTCGGATACGTATCTGATATGCATACTAAACTCGCATCAATAACTCAATTAGTTATTGCCAAGATAGAAACTATAGATAATGATATATTAAACAACGACATTGTAAATTTCATTATGTGTAGATCAAACTTAAATAATCCATTTATCTCTTTCCTAGATACTGTATATACTATTATAGATCAAGAGATCTATCAGAACGAGTTGATTAATTCATTAGACGACAATAAAATTATCGATTGTATAGTTAATAAGTTTATGAGCTTTTATAAGGATAACCTAGAAAATATAGTAGATGCTATCATTACTCTAAAATATATAATGAATAATCCAGATTTTAAAACTACGTATGCAGAAGTACTCGGTTCCAGAATAGCGGATATAGATATTAAACAAGTAATACGTGAGAATATACTACAATTGTCTAATGATATCCGCGAACGATATTTGTGAAAATATTAAAAAAAAATACTTTTTTTATTAAATGACGTCTCTTCGTGAATTTAGAAAATTATGCTGTGCTATATATCACGCATCAGGATATAAAGAAAAATCTAAATTAATTAGAGACTTTATAACAGATAGGGATGATAAATATTTAATCATTAAGCTATTGCTTCCCGGATTAGACGATAGAATTTATAACATGAACGATAAACAAATTATAAAAATATATAGTATAATATTTAAACAATCTCAGAAAGATATGCTACAAGATTTAGGATACGGATATATAGGAGACACTATTAGTACATTCTTCAAAGAGAACACAGAAATCCGTCCAAGAAATAAAAGCATTTTAACTTTAGAAGACGTGGATAGTTTCTTAACTACATTATCATCCATAACTAAAGAATCGCATCAAATAAAATTATTGACTGACATCGCATCCGTTTGTACATGTAATGATTTAAAATGTGTAGTCATGCTTATTGATAAAGATCTAAAAATTAAAGCGGGTCCTCGGTACGTACTTAACGCTATCAGTCCTCATGCCTATGATGTTTTTAGAAAATCTAATAACTTGAAAGAGATAATAGAAAATGAATCTAAACAAAATCTAGACTCTATATCTGTTTCTGTTATGACTCCAATTAATCCCATGTTAGCGGAATCGTGTGATTCTGTCAATAAGGCATTTAAAAAATTTCCATCAGGAATGTTTGCTGAAGTCAAATACGATGGCGAGAGAGTACAAGTTCATAAAAATAATAACGAGTTTGCCTTTTTTAGTAGAAACATGAAACCAGTACTCTCTTATAAAGTGGATTATCTCAAAGAATACATACCGAAAGCATTTAAAAAAGCTACGTCTATCGTATTGGATTCTGAAATTGTTCTTGTAGACGAACATAATGTACAGCTCCCGTTTGGAAGTTTAGGTATACACAAAAAGAAAGAATATAAAAACTCTAACATGTGTTTGTTTGTGTTTGACTGTTTGTACTTTGATGGATTCGATATGACGGACATTCCATTGTACAAACGAAGATCTTTTCTCAAAGATGTTATGGTCGAGATACCCAATAGAATAGTATTCTCAGAGTTGACGAATATTAGTAACGAGTCTCAGTTAACTGACGTATTGGATGATGCACTAACAAGAAAATTAGAAGGATTGGTCTTAAAAGATATTAATGGAGTATACGAGCCGGGAAAAAGAAGATGGTTAAAAATAAAGCGAGACTATTTGAACGAGGGTTCCATGGCAGATTCTGCCGATTTAGTAGTACTAGGTGCTTACTATGGTAAAGGAGCAAAGGGTGGTATTATGGCAGTCTTTCTAATGGGTTGTTACGATGATGAATCCGGTAAATGGAAGACGGTAACTAAATGTTCAGGACATGATGATAATACGTTAAGGGTTTTGCAAGACCAATTAACGATGGTTAAAATTAACAAGGATCCCAAAAAAATTCCAGAGTGGTTGGTAGTTAATAAAATCTATATTCCCGATTTTGTAGTAGAGGATCCGAAACAATCTCAAATATGGGAAATTTCAGGAGCAGAGTTTACATCTTCCAAGTCACATACAGCGAATGGAATATCGATTAGATTTCCTAGATTTACTAGGATTAGAGAAGATAAAACGTGGAAAGAATCTACTCATCTAAACGATTTAGTAAATTTGACTAAATCTTAATAGTTACATACAAACTGAAAATTAAAATAACACTATTTAGTTGGTGATCGCCATGGATGGTGTTATTGTATACTGTCTAAACGCGTTAGTAAAACATGGTGAGGAAATAAATCATATAAAAAATGATTTCATGATTAAACCATGTTGTGAAAGAGTTTGTGAAAAAGTCAAGAACGTACACATCGACGGACAATCTAAAAACAATACAGTGATTGCAGATTTGCCATATCTGGATAATGCTGTATTGGATGTATGCAAATCAGTATATAAAAAGAATGTATCAAGAATATCCAGATTTGCTAATTTGATAAAGATAGATGATGATGACAAGACTCCTACCGGTGTATATAATTATTTTAAACCTAAAGATGCTATTTCTGTTATTATATCCATAGGAAAGGATAAAGATGTCTGTGAACTATTAATCGCATCCGATAAAGCGTGTGCGTGTATAGAGTTAAATTCATATAAAGTAGCCATTCTTCCCATGAATGTTTCCTTCTTTACCAAAGGAAATGCGTCATTGATTATTCTCCTGTTTGACTTCTCTATCAATGCGGCACCTCTCTTAAGAAGTGTAACCGATAATAATGTTGTTATATCTAGACACAAACGTCTTCATGGCGAGATTCCGAGTTCCAATTGGTTCAAGTTTTATATAAGTATAAAGTCCAACTATTGTTCTATATTATATATGGTAGTTGATGGATCTGTGATGTATGCAATAGCTGATAATAAAACTCACACAATTATTAGCAAAAATATATTAGACAATACTACAATTAACGATGAGTGCAGATGCTGTTATTTTGAACCACAGATTAAGATTCTCGATAGAGATGAGATGCTTAATGGATCATCGTGTGATATGAACAGACATTGTATTATGATGAATTTACCAGATATAGGAGAATTCGGTTCCAGTATATTGGGGAAATATGAACCTGACATGATTAAGATTGCTCTTTCAGTGGCTGGTAATTTAATAAGAAATCAAGACTACATTCCCGGGAGACGCGGCTATAGCTACTACGTTTACGGTATAGCCTCTAGATAATTTTTTAAGCACAAAATAAAAAACATAATTTTAAACTAGTCTATTTCATACTATTTTGTGTGATCACCATGGACATAAAGATAGATATTAGTATTTTTGGTGATAAATTTACGGTGACTACTAGGAGGGAAAATGAAGAAAGAAAAAATATCTACCTCTCCAAAAAGAAAAATTTACTACTGATGTTATCAAACCTAATTATCTTGAGCACGATAACTTATTAGATAGAGATGAGATGTCTACTATTCTAGAGGAATATTTTATGTACAGAGGTCTATTAGGCCTCAGAATAAAATATGGACGACTTTTTAACGAAATTAGAAAATTCGACAATGATGCGGAAGAACAATTCGGTACTATAGAAGAACTCAAACAGAAACTTAGATTAAATGCTGAAGAGGGAGCCGATAACTTTATAGATTATATAAAGGTACAAACAGGATATCTATGATAGGATTGTGTGCGTGTGTGGTAGATGTTTGGAGAAAGGAGAAACTGTTTTCTAGATGGAAATATTGTTTACGAGCTATTAAACTGTTTATTGATGATCCCATACTTGATAAGATAAAATCTATACTGCAGAATAGACTAGTGTATGTGGAAATGTTATAGAAAATTAAAAGTTAATGAGAGCAAAATATAATGTTGTATTCTAATCCCATATTTATTATTTTCACGGAGATATAGGTGTAGTTTTGCAGTGACCGTCCTGATAATATTATCTATATGATGGGTGGATATGATCAGTACCCGTATAGAAGTTCAAAGGTTATAGTGTACAATACATGTACAAATTCTTGGATATATGATATACCAGAGCTAAAATATCCTCGTTCTAATTGTGGAGGAGTTGCTGATGATGAATACATTTATTGCATAGGCGATCAGGATTCATCGTTGATATCTAGTATTGATAGATGGAAGCCATCAAAACCATATTGATAAACGTATGCTAAAATGCGAGAGACAAAATGTGATATTGGTGTAGCGATGTTAAACGGATTAATATATGTCATAGGCGGAGTTGTTAAAGGTGACACATGTACCGACACACTTGAGAGTTTATCAGAAGATGGATGGATGATGCATCAACGTCTTCCAATAAATGTCCAATATGTCGACGATTGTTCATACCGGCAAAATTTATATATCAGGAGGCTACACAATAGTAGTGTAGTTAATGGAATATCAAATCTAGTCCTTAGCTATAATCCGATATATGATGAATGGACCAAATTATCATCATTAAATATTCCTAGAATTAATCCTGCTCTATGGTCAGTGCATAATAAAGTATATGTAGGAGGAATATCTGATGATATTCAAACTAATACATCTGAAACATACAACAAAGAAAAAGATCGTTGGACATTGGATAATAGTCACGTGTTACCACGCAATTATATAATGTATAAATGCGAACCGATTAAACATAAATATCCATTGGAAAAACACAGTACACGAATGATTTTCTAAAGTACTTGGAAAGTTTTATATGTAGTTGATAGAACAAAATACATATTTTTTGTAAAAATAAATCACTTTTTATACTAATATGACACGATTGTCAATACTTTTGTTACTAATATCATTAGTATACTCTACACCTTATCCTCAGACACAGATATCTAAAAAAATAGGTGATGATGCAACTCTATCATGTAGTAGAAATAATATAAATGATTATGTTGTTATGAGTGCTTGGTATAAGGAGCCCAATTCCATTATTCTTTTAGCTGCCAAAAGTGACGTCTTGTATTTTGATAATTATACCAAGGATAAAATATCATACGATTCTCCATACGATGATCTAGTTACAACTATTACAATTAAATCATTGACTGCTAAAGATGCCGGTACTTATGTATGTGCATTCTTTATGACATCAACTACAAATGATACTGATAAAGTAGATTATGAAGAATACTCTACAGAGTTGATTGTAAACACAGATAGTGAATCGACTATAGACATAATACTATCTGGATCTTCACATTCACCGGAAACTAGTTCTGAGAAACCTGATTATATAAATAATTTTAATTGCTCGTTGGTATTTGAAATCGCGACTCCGGGACCAATTACTGATAATGTAGAAAATCATACAGACACTGTCACATACACTAGTGATATCATTAATACAGTAAGTACATCATCTAGAGAATCCACAACAGTCAAGACGTCGGGACCAATTACTAATAAAGAAGATCATACAGTCACAGACACTGTCTCATACACTACAGTAAGTACATCATCTGAAATTGTCACTACTAAATCAACCGCCAATGATGCGCACAATGATAATGAACCATCTACTGTGTCACCAACAACTGTAAAAAACATCACGAAATCTATAGGTAAGTATAGTACTAAAGACTATGTCAAAGTATTTGGTATTGCAGCATTAATTATATTGTCGGCCGTGGCAATTTTCTGTATTACGTATTATATATGTAATAAACGTTCACGTAAATACAAAACAGAAAACAAAGTCTAGATTTTGACTTACATAAATGTCTGGAATAGTAAAATCTATCATTTTGAGCGGGTCATCTGGTTTAGGAAAGACAGCCATAGCCAAAAGACTATGGGAATATATTTGGATTTGTGGTGTCCCATACCACTAGATTTCCTCGTCCTATGGAACGAGAAGGTGTTGATTACCATTACGTTAACAGAGAGGCAATCTGGAAGGGAATAGCCACCGGAAACTTTCTAGAACATACTGAGTTTTTAGGAAATATTTACGGAACTTCTAAAACAGCTGTGAATACAGCGGCTATTAATAATCGTATTTGTGTGATGGATTTAAACATCGACGGTGTTAGAAGTCTTAAAAATACTTACCTAATGCCTTACTCGGTGTATATAAAACCTACCTCTCTTAAAATGGTTGAGACCAAGCTTCGTCGTAGAAACACTGAAGCGGACGATAAGATTCATCGTCGCGCGATGTTGGCAAAAACGGATATGGATGAGGCCAACGAAGCAGGTCTATTCAACACTACTATTATTGAAGATAATGTGAATTTAGCATATAGTAAGTTAATTCAGATACTACAGGACCGTATTAGAATGTATTTTAACACTAATTAAGATATAATACTTGACTGACTTAAAACTTGATAATTAATAATATAATTCGTTTTTATATGTGACTATTTTAACGTCTAAATGTATTAGTTAAATATTAAACTTACCACGTAAAACTTAAAATTTAAAATGGTATTTCATTGACGGATGATCATACATTATGAACTTTCAAGGACTTGTCTTAACTGACAATTGCAAAAATCAATGGGTCGTTGGACCATTAATAGGAAAAGGTGGATTCGGTAGTATTTATACTACTAATGACAATAATTATGTAGTAAAAATAGAGCCCAAAGCTAACGGATCATTATTTACTGAACAGGCATTTTATACTAGAGTACTTAAACCATCCGTTATCGAAGAATGGAAAAAATCTCACCATATAAGCCACGTAGGAGTTATCACATGCAAGGCATTTGGTCTATACAAATCCATTAATACGGAATATAGATTCTTGGTAATTAATAGATTGGGTGTAGATCTAGATGCGGTGATCAGGGCTAACAATAATAGACTACCGAAAAGATCGGTGATGTTAGTAGGAATAGAAATCTTGAATACCATACAATTTATGCACGAGCAAGGATATTCTCATGGAAATATTAAAGCGAGCAATATAGTTTTGGATCAAATGGATAAGAATAAATTATATCTAGTGGATTACGGATTGGTTTCTAAATTCATGTCTAACGGCGAACATGTTCCATTTATAAGAAATCCAAATAAAATGGATAATGGTACTCTAGAATTTACACCTATAGATTCACATAAAGGATACGTTGTATCGAGACGTGGAGATCTAGAAACACTTGGATATTGTATGATTAGATGGTTGGGAGGTATCTTGCCATGGACTAAGATAGCTGAAACAAAGAATTGCGCATTAGTAAGTGCTACAAAACAGAAATATGTGAACAATACTACGACTTTGTTAATGACCAGTTTGCAATATGCGCCTAGAGAATTGCTGCAATATATTACCATGGTAAACTCTTTGACATATTTTGAGGAACCCAATTACGATAAGTTTCGGCACATATTAATGCAGGGTGCATATTATTAAGTGTGAGGTTTTGGTTTGATGTAAAATTTTTGTTGATAAAAATTAAAAAATAACTTAATTTATTATTGATCTCGTGTGTACAACCGAAATCATGGCGATTTTTACACACACGTCTTGGCGGGTACGACGAGAATCTTTATACCTTTCCTGGAATATCATTAACTGTTGCCAATGATGTTAGGAAATATTCTGTTGTGTCAGTTTATAATAACAAGTATGACATTGTAAAAGAAATATATGTGGTGTTACAGTTAGGTGAACAAGAGATATATTGGAGCATTGCTGCATATGTTTGAGTGCAATGAATATCTACAAATTGGAGATCCAATCCATGATCAAGAAGGAAATCAAATCTCTATCGTCACATATCGCTACAAAACTACTATGCTCTAAGCGGAATCGGGTACAAAAGTCTAGACTTGTGTTTGGAAGGAGTAGGGATTCATCATCACGTACTTGAAACAGGAAACGCTGTATATGGAAAAGTTCAACATGATTATTCTACTATCAAAGAGAAGGCCAAAGAAATGAGTGCACTTAGTCCAGGACCTATAATTGATTACCACGTCTGGATAGGAGATTGTATTGGTAAAGTTACTGCTGTAGACGTACATGGAAAGGAAATTATGAGAATGTGATTCAAAAAGGGTGCGGTACTACAGATACCAAATCTGGTAAAAGTTAAACTTGGGGATAATGATACAGAAAATCTTTCCACTACCATATCAGCTCTCCTGAATTCCGGCGGCGGTACCATCAAGGTAACCTCTTAGGAAGATCGCGTTAATCATGTACTCATAAAACGATTGGAATCTATACGCCATATGTGGTCTGTCTATGATCGTTTTAATATTGTGAATGGTAAAGAATGCTGTTATATACATCTTCATTCATCTAATCAGAATCTTATGCCTAGTACTGTAAAAACCAATTTGTATATGTGATCTATGACATCATGCATCCAAATGGATCTCATAACTGCTCTAGATTATCTTAGTGAACTTAAGGAATCAAGTGGATAAAGTCCCAAACCAGAATTGCCCAAATTCGATTACCCAGATGGAGTGCAAGACAATGGATCAATTGAGAGATTTGCAGAAGAGTTCTTCAATAGATCAGAACTTCAGGCGGGTGAATCAGTCAAATTCGGTAATTCCGTTAATATTAAAAATACATCTGTTTCAGCTAAGCAACTAAGAACACGTATACGGCAACAGCTTCCCTCTATACTCTCATCTTTTGCCAACACAGAGGGTGGATATTTGTTCATTGGATTGGATAATAATACACACAAAGTAATTGGATTCATGGCGGGACACGACTGCATTAAACTGATAGAGAATGAGATAGAAAAGTGTATTAGACGACTCATGTTGTACATTTCTGTGAGAAAAAGAGGATATCAAATACGCATATAGATTCATCAAGGTATATAAACCTGGGGAGGAGACTACCTCGACATACGTGTGCGCTATCAAAGTGGAAAGATGTTGCTGTGCTGTGTTTACAGATTGGCCAGAATCATGGTATATGGATACTAGTGGTATCAAGAAGTATTCTCCAGATGAATGGGTGTCATCTATAAAATTTTAATTAAGAACTATAGAGAACAAATAATAGGTTGTAATACATATAGACAATAACTAACAATTAATTAGTAACTGTTATCTCTTTTTAACTAACTCTTATATACCTATTAATACATCGTATTATAGTTCTTAATCATTGATTCACTTATTGTTTTAATAAACTAACATTGTTAATTGAAAAGGGATAACATGTTATAGAATATAAATTATATATGGATTTTTTTAAAAAGGAAATACTTGACTGGAGTATATATTTATCTCTTCATTATATAGCACATGCGTGTTCCAATTCTTCCACATCCCATATAATACAGGAATATAATCTTATTCGAACATACAAAAAAGTGGATAAAACAATAGTTGATTTTTTATCTAGGTGGCCAAATTTATTCCATATTTTAGAATATGGGGAAAATATTCTACATATTTATTCTATGGATGATGCTAATACGAATATTATAATTTTTTTTCTAGATAACGTATTAAATATTAATAAGAACGGGTCATTTATACACAATCTCGGGTTATCATCATCCATTAATATAAAAGAATATGTATATCAATTAGTTAATAATGATCATCTAGATAATGGGATAAGACTAATGCTTGAAAATGGACGTAGAACAAGACATTTTTTATCCTATATATTAGATACAGTTAATATCTATATATGTATTTTAATAAATCATGGATTTTACATAGATGCTGTGGACAGTTACGGTTGTACATTATTACATAGATGTATATATCACTATAAGAAATCAGAATCAGAATCATACAATGAATTAATTAAAATATTGTTAAATAATGGATCAGATGTAGATAAAAAAGATACGCATGGAAACACGCCATTTATCCTATTATGTAAACACGATATAGACAACGTAGAATTGTTTGAGATATGTTTAGAGAATGCTAATATAGACTCTGTAGACTTTAATGGATATACACCTCTTCATTATGTCTCATGTCGCAATAAATATGATTTTGTAAAGTCATTAATTTCTAAAGGAGCAAATGTTAATACGCGTAATAGATTCGGAACTACTCCATTTTATTGTGGAATTATACACGGTATCTCGCTTATAAAACTATATTTGGAATCAGACACAGAGTTAGAAATAGATAATGAACATATAGTTCGTCATTTAATAATTTTTGATGCCGTTGAATCTTTAGATTATCTATTATTTAGAGGAGTTATTGATATTAACTATCGTACTATATACAACGAAACATCTATTTACGACGCTGTTAGTTATAATGCGTATAATATGTTGGTCTATCTATTAAACAGAAATGGTGATTTTGAGACGATTACTACTAGTGGATGTACATGTATTTCGAAAGCAGTTGCGAACAACAACAAAATAATAATGGAAGTACTATTGTCTAAACAACCATCTTTGAAAATTATGATACTGTCTATAATAGCAATTACTAAACATAAACAACATAATACAAATTTATTAAAAATGTGTATAAAATATACTGCGTGTATGACAGATTATGATACTCTTATAGATGTACAGTCGTTACAGCAATATAAATGGTATATTTTAAAATGTTTCGATGAAATAGATATCATGAAGAGATGTTATATAAAAAATAAAACTGTATTCCAATTAGTTTTTTGTACTAAAGACATTAATACTTTAATGAGATACGGCAGACATCCTTCTTTCGTTAAATATACTAGTCTCGACGTATACGGAAGTCGTGTACGTAATATCATAGCATCTATTAGATATCGTCAGAGATTAATTAGTCTATTATCCAAGAAGCTGGATGTGGGAGATAAATGGGCATGTTTTCCTAACGAAATAAAATATAAAATATTGGAAAACTTTAACGATAACGAACTATCGACATATCTAAAAATCTTATAAACACTATTAAAATATAAAACCTAAGTAGGATAAAACACATTACATCATTGTTTCCTTTTATTGCTTAACAGTGTATACTATTTTTAACACTCATAAATAAAAATGAAAACGATTTCCGTTGTTACGTTGTTATGCGTACTACCTGCGGTTGTTTATTCAACATGTACTGTACCCACTATGAATAACGCTAAATTAACGTCTACCGAAACATCGTTTAATGATAAACAAAAAGTTACATTTACATGTGATTCGGGATATTATTCTTTGGATCCAAATGCTGTCTGTGAAACAGATAAATGGAAATACGAAAATCCATGTAAAAAAATGTGTACAGTTTCTGATTATGTCTCTGAACTATATAATAAACCGCTATACGAAGTAAATGCCATCATAACACTAATTTGTAAAGACGAAACAAAATATTTTCGTTGTGAAGAAAAAAATGGAAATACTTCTTGGAATGATACTGTTACGTGTCCTAATGCGGAATGTCAATCTCTTCAATTAGATCACGGATCGTGTCAACCAGTTAAAGAAAAATACTCATTTGGGGAACATATAACTATCAACTGTGATGTTGGATATGAGGTTATTGGTGCTTCGTACATAACTTGTACAGCTAATTCTTGGAATGTTATTCCATCATGTCAACAAAAATGTGATATACCGTCTCTATCTAATGGATTAATTTCCGGATCTACATTTTCTATCGGTGGCGTTATACATCTTAGTTGTAAAAGTGGTTTTATACTAACGGGATCTCCATCATCCACATGTATCGACGGTAAATGGAATCCCGTACTCCCAATATGTATACGATCTAACGAAGAATTTGATCCAGTGGAGGATGGTCCCGATGATGAGACAGATCTGAGCAAACTCTCAAAAGACGTTGTACAATATGAACAAGAAATAGAATCGTTAGAAGCAACTTATCATATAATCATAGTAGCGTTGACAATTATGGGCGTCATATTTTTAATCTCCGTTATAGTATTAGTTTGTTCCTGTAACAAAAATAATGACCAATATAAGTTCCATAAATTGTTGCTGTGAATATAAATCCGTTAAAATAATAAATAATTAATAACGAACAAGTATCAAAAGATTAAAGAATTATAGCTAGAATTAATTGAGATGTCTTCTTCAGTGGATGTTGATATCTACGATGCTGCCAGAGTATTTTTACTTAGGCACTATTATAACAAGAGATTTATTGTGTATGGAAGAAGTAACCATATTACATAATATATACAGGCTATTTACAAGATGCGCCGTACCGTTCGATGATATAGTACGTACTATGCCAAATGAATCACGTGTTTAACAATGGGTGATAGATACACTTAATGGCATAATGATGAATGAACACAATGTTGCTGTGTAGGTACCGGACTACTGTTCATGGAAATGTTTTTCGATTATAATAAAAATAATTCCAAAAATAGCATCATCAACTAATGTATGACATAATTAATAGCGTATCCATAATTCTAGCTAATGAGAGATATAGAAGCGCGTTTAACGACAATGGAATATACATCCGTAGAACTATGATTGACAAGTTGTACGGATACGCATCTCTAACTACTATTGCACGATCGCTGGAGGTGTTTGTTATTATCTGTTGATGCATCTAGTTAGTTTGTATAAATAATTATTTCAATATACTAGTTAAAATTTTAAGATTTTAAATGTATAAAAACTAATAACGTTTTTATTTGTAATAGGTGCAATAGCATTCTATTCGAATAATGAGTATACTCCGTTTAATAAACTTAGTGTAAAACTATATATAGATGGAGTAGATAATATAGAAAATTCATATACTGATAATAATGAATTAGTGTTAATTTTTAAAGAGTACACAGTTTCTATTATTACAGAGTCATACGACGTCGGATTTGATTCCATAGATATAGATGTTATAAACGACTATAAAATTATTGATATGTATACTATTGACTCGTCTACTATTTAATGCAAAGGACACACATGTAGAATATCTACCAAATTATCATGCCATTATGATAAACACCCTTATATTCACAAATATAAGGGTGATAAGCGACAATATTCTATTACTACAAAGGGAAAATGCTATAAAGGAATAAAATATGAAATAAGTACAATGAACGATAATATTCTATTGAGACAACATACTCTTAAAATTGGATCTACTAATATATTCGATAGTTATGGACATAGTAATACATATTATTCAAAATATGGTCTTTAAAAATTTAAAATATATTATCACTTCAACGACAATAGTCAAATAACAGCCAATATGAGATCTGTAATGCTAACAGTTTTGCTAATTAATAGCATAAATGCTACGATAACTAGCTATAAATTCGAATCCGTCAATTTTGATTCCAAAATTGAATGGACTGGCGATGGTCTATACAATATATCCCTTAAAAATTATGGGATAAAGACGTGGCAAACAATGTATACAAATGTACCAGAAGGAACATACGACATATCCGGATTTCCAAAGAATGATTTCGTATCTTTCTGGGTTAAATTTGAACAAGGTGACTATAAAGTGGAAGAGTATTGTACAGGACTATGCATCGAAGTAAAAATCGGACCACCAACTGTAACATTGACTGAATACGACGGCCATATCAATTTGTACATCGAGCATCCGTATGCCACTAGAGGTAGCAAGAAGATTCCTATTTACAAACGTAATGACATGTGTGATATCTACTTGTTGTATACGGCTAACTTCACATTCGGAGATTCTGAAGAACCAGTAACATATAATATTGATGACTACGATTGTACGTCTACAGGTTGCAGCATAGACTTTGCCACAACAGAAAAAGTGTGCGTGACGGCGCAGGGAGCCACAGAAGGATTTCTAGAAAAAATTACTCCATGGAGTTCGGAAGTATGTCTGACACCTAAGAAGAATGTATTTACATGCGCTATTAGATCCAAGGAAGATGTATCCAATTTCAAGGACAAAATGACCAGAGTTATCAAGAGAAAATTTAATAAACAGTCTCAAAATTATATGACCAAATTTCTCGGTACTACGGCTAATGATGTTACTACCGTGATTAGCATGCTAGATTAAATATTTTGGGCAAATCTAAATATTTTTTATAAAAATTAAAAAATCACTTTCGTTATCGATAACTCGAGAACATGGGTTACATATCAACATATTAATGTTGGTGGCATTGTCTAAAACAGTCACGATATCTCTTAAATGCTACAGAAAGGTAGGATTTTATAATCTTTATGGATCTAATGATAGCGAGCAGGAAAATTATTCTCCTGTATCTGTTAAAACTACGATTATCTATTAAAGACAGAGATACTAAATATAGACAATAACATACCATGGAAAAATATTACAGATTTTATTAAACTCTTTTATAGAGAAATGTAATAATTGCTCAAATAATACTATATACAATATCACTATAAAGTCTTGGACTAATTATACTTTGGTAAATGTTACAACTGACAATAGTGTCTACTTAAATGTATCTAGTGATATCGTGATCAATAAAACAGGAAATATTACTACATTCATTAATAATACAGTTAATGGTAATTTATCGATTAGTCTTAATGATACACTAAGTATATACACAGAATCAGAACCTGTGTTTATAGTGGATCCATTCATTACTCTTGATAAATTGTACAACTGTTCGTTCTCCAAACCAATCAATTATGTAAATATCTTTAATACAGAGTTTAATACTAATACTAATACTAATACTAATACTAATACTAATACTAATACTACTACTTGTAAGAATGAGACTGACTACACACCAATGGACATTTCTTATGTTATTAATGACAATATAACATTCATTAGTAATATCACATCAAGTACAAAATATATCAACTCATGTGTAAATTCGTCACTGAGTTTACAGATAGCGGTTGTCAGAATACCAGTATTGTTTAATAAAACAATGGAACAATTAGGTATTACTCTAGGATCTGATTTGGAAGCGTATTTCATGTGTAAGTTACAGGAACTTAATTGTGACGCGAGTATTCTCTTGAACAAGGCATCAAAAAATGTAATGGATAGACAATTATCAAAAACTCCAACGAGATAAAAGAGATCCACTGAGATGACTGGAAATCACTTCTGTCTACATAAGAATCAAGCTAGGCTGGAAACTATTTACGAAACTATCAAAATGCAGGGTACAACAAACAATAATGACGGAAGTAAACGAACGTCTCCAATATATGATAACGTTGGCAATACTAAGGATTCATTGACATATGTAAATATTAATAAAGTATATATGAGATACACGTTTTGATAGTCATCCTAATCGTATCGGAGTACGGAGATCATCAAGTTCATCATCCGATATTATTGTAACCACAATGCATTGGCATCAAAAGAAGATAGATGAACTGACATATAGCGAATATAAGGAGATCCAACACTATTAGGAGAAAGCAGGTCATATGGAAGATGTTCAAGCTGTTAATATCAATCTCGATAAACGTGAACAGGCTGTCGGAGCCACAGTTTCGAGACGAGGAGATTTAGAAATGTTGGGATTATTGCATGATAGAATGGTTCAGTGGCAAACTTCCATGGAAAAACAAAAGTAGTATAGCAATACAACAGAAAAGAGAATATAAAAAATTTATAGCTACTTTCTTTGAGGACTGTTTTCCGGAAGGAAATAAACCTTTGGAATTAGTATACACGTTAGATTATTCTCAAACTCCTAATTATGAAAGACTACGTAGACTAGACTGTTTATACAAGATTGAAATTATATTCTTTTTTATAGAGCGTGTGATGTTACGGATATCTAATAATATTAGACTATCTCTGACACTACACGACCAAATCGATTACTATGGATATCTTCAGGGAAATCGCATCTTCTACGAAAGGAGAGAATGTATTCATTTCTCCAGCGACAATCTCGTCAGTATTGACAATACTGTATTATGGAGCTAATGGATCCACTGCTGAACAGTTATCAAAATATGTAGAAAAGGAGGAGACCATGGATAAAGTTAGCGCTCAGAATATCTCATTCAAATCCATGAATAAAGTATATGGGCGATATTCTGCCGTGTTTAAAAATTCCTTTTTGGGAAAAATTGGCGATAATTTCCAAACTGTTGACTTCACTGATTGTCGCACTATAGATGCTATCAATAAGTGTGTTGATGTCTTTACTGAGGGGAAAATCAATCCACTATTGACTGAACAATTGTCTCCCAATACCTGTCTCCTAGCAATTAGTGCCGTATACTTTAAAGCAAAATGGTTGATACCATTCAAAAAGGAATTTACCAGTGATTATCCATTTTACGTATCACCAACGGAAATGGTAGACGTAAGTATGATGTCTATGTATGGTGAGTCATTTAATTACGCATCTGTAAAAGAATCATTCGGTAACTTTTCAATCATAGAACTGCCATATGTTGGAAATACTAGTATGATGGTCATTCTTCCAGACAAGATTGATGGATTAGAATCCATAAAACAAAATCTAACAGATACAAATTTTAAGAAATGGTGTAACTCTCTGGAAGCTACGTTTATCGATGTACACATTCCCAAGTTTAAGGTAACAGGTTCGTATAATCTGGTGGATACTCTAGTAAAGTTGGGACTGACAGATGTGTTCTATTCAACTGGCGATTATAGCAATATGTGTAATTCAGATGTGAGTGTTGACGCTATGATTCACAAAACGTATATAGATGTCAATGAAGAGTACACAGAAGCAGCTGCAGCAACTTCTGTACTAGTGGCAGACTGTGCATCAACAGTTACAAATGAGTTCTGTGCAGATCATCCGTTCATCTATGTGATTAGGCATGTCGATGGCAAAATTCTTTTCGTTGGTAGATATTGCTCTCCAACAACTAATTAAGCATATTCTTAATATTAGAATATTATATAGTTAAGATTTTTTTAAAAAATAGAAAAAACATGTGGTATTAGTGCAGGTCGTTATTTCTCCCATTGCAATTGGTAAGATGACGGCCAACTTTAGTACCCACGTCTTTTCACCACAGCACTGTGGATGTGACAGACTGACCAGTATTGATGACGTCAGACAATGTTTGACTGAATATATTTATTGGTCGTCCTATGCATACCGCAACAGGCAATGCGCTGGACAGTTGTATTCCACACTCCTCTCTTTTAGAGATGATGCGGAATCAGTGTTCATCGATGTTCGCGAGCTGGTGAAAAATATGCCGTGGGATGATGTCAAAGATTGTGTAGAAATCATCCGTTGTTATATACCGGATGAGCAAAAAACCATCAGAGAGATTTCGGCCATCATCGGACTTTGTGCATATGCTGCTACTTACTGGGGAGGCGAAGACCATCCCACTAGTAACAGTCTGAACGCATTGTTTGTGATGCTTAAAATGCTCAATTACGTGGATTATAACATCATATTCCGGCGTATGAATTGACATTTCTCCCTTTCCCAGAAATAAACTTTTTTACCCACTATAAAATAAAAGAGTATACTACCTATTATATTTCTTCCTATATTTTTTATTCTTCATTAGTTACGACTTTTAACACACCTAAATGTGTCGACAAAGGGCAATAGTTTGCGTCATTCATGGAGTTAGAAAACGAGCCAGTAATCTTACCATGTCCTCAAATAAATACTCTATCGTTTGGATATAATATATTAGATATTTTATGGGAAAACGAGAAGCGGATAATGATAGAATTATACCAATAGATAATTGTAACAATATGCTAATTCTGAACCCGACACAATCAGACTCTGGTATTTATATGTGCATTACCAAGAACGAGACCTACAGTGATATGATGTCGTTAAATTTGACAATTGTGTCTGTCTCAGAATCAAATATAGATCTTATTTAGTATCCACAAATAGTAAATGAGAGATCTACTGGTGAAATGTGACCCAGTAATGCATTTATTGCTAGTAACATAAACACAGATATTATATGGAGCGGACATCAACGCCTTAGAAATAAGAGACTTAAACAACGGACACCTGGAATTATTACCATAGAAGATGTTAGAAAAATGATGCTGGTTATTACACATGTGTTTTAATATATATATATATATATATATATATATATGGGAGTAAAGCATATAACGTAACCAGAATTGTAAAATTAGAGGTATGGGATAAAATAATACCTTCTACTATGCAATTACCAAAAGGCCTTGTAACTTCACTAGGTAGTAATTTGACTATTGTGTGTAAAGTATCGTTGAGACCTCCCACAACGGATGCAGATGTCTTTTGGATAAGTAATGGTATGTATTACGAAGAAGACAATGAGGACGGAGACGGTAGAATAAGTATAGCAAATAAAATCTATACGACCGATAAGAGACGTGTTATTACATTGTGATTAAACATTAATCCTGTCAAGGAAAAAGATGCTACAGCGTTTACTATTCCTAGCATCAGCAAAACAGTTACTATTAGTATAATGTGATTGTATGTTGTTACATTTCCATGTCAAATAGAGTTTATAAAAATTTTTTATACATTATCTTCCAACAAGCAATTGACGAACGTATTACTATGATTAACTCCCACGATACTATGCATATTATTAATCATTAACTTGCAGACTATACCTAGTGCTATTTTGACATACTTGTGTTCTTGTGTAATTGCGGTATCTATATTATTAAAGTACGTAAATCTAGCTATAGTTTTATTATTTAATTTTAGATAATATACCATCTCCTTATTTTTAAAAATTACCACATCCTTTATTAAATCATGGATGGGAATTTCTATGTCATCGTTAGTATATTGTGAACAACAAGAGCAGATATCTATAGGAAAGGGTGGAATGCGATACATTGATCTATGTAGTTTTAAAACACACGAGAACTTTGAAGAATTTATATAAATTATTCCATCGATACATCCTTCTATGTTAATATGTATATATCCAGGAATTCTTTTATGAATATCAGGAAATGTATAAACTAAAACATTGCCCGGATACGGTGCTTCTATTGGCATTATATCCGTTCTTAACTTGCAAAATGTAACCAATACCTTTGCATGGCTTGTTTTGTTCGGTAACGTTAGTTTAAACTTGACGAATGGATTAATTACAATAGCATGATCTGCGCATCTATTAAGTTTTTTTACTTTAACGCCCTTGTATGTTTTTACAGAGACTTTATCTAAATTTCTAGTGCTTGTATGTGTTATAAATATAACTGGATATAGAACTGAATCACCTACCTTAGATACCCAATTACATTTTATCAGATCCAGATAATAAACAAATTTTGTCGCCCTAACTAATTCTATATTGTTATATATTTTACAATTGGTTATGATATCATGTAATAACTTGGAATCTAACGCGCATCGTCGTACGTTTACACAATTGTGATTTAGTATAGTATATCTACACATATATTTTTCCGCGCTATAGTATTCTGGACTAGCGATAAAACTATCGTTATATCTGTCTTCGATGAACTCATCAAGATATTGCTCTCTGTCATATTCATACACCTGCATAAACTTTCTAGACATTGTACAATATGTGTTATTTTAGGATCATATTTACATATTTACGGTATATCAAAGATGTTAGATATAGTTAATAGGACACATATAATAATGAATATTAAACACTAGTATAGCAGAAGTTTATACTCACAAAACATCATAAAATGAGTCGTCGTCTGATTTATGTTTTAAATATCAACCGCAAATCAACTCATAAAATACAAGAGAATGAAATATATACATATTTTAGTTATTGCAATATAGACCATACTTCTACAGAACTTGATTTTGTAGTTAAAAACTATGATCTAAACAGACGACAGCCTGTAACTGGGTATACCGCACTACACTGCTATTTGTATAATAATTACTTTACAAACGATGTACTGAAGGTATTATTAAATCATGGAGTGGATGTAACGATAAAACCCAGTAGCGGACATATGCCTATTTATATATTGCTTACTAGATGTTGCAATATTTCACACAATGTAGTGATAGATATGATAAACAAAGATAAAACCCACTTATCGCATAAAGACTATTCCAACCTACTACTAGAGTATATAAAATCTCGTTACATGCTATTGAAGGAAGAGGATATCGATGAGAACATAGTATCAACTTTATTAGATAAGGGAATCGATCCTAACTTTAAACAAGACGGATATACAGCATTACATTATTATTATTTGTGTCTTGCACACGTTTATAAACCAGGCGAGTGTAGAAAACCGATAACGATAAAAAAGGCCAAGCGAATTATTTCTTTGTTTATACAACATGGAGCTAATCTAAACGCGTTAGATAATTGTGGTAATACACCATTCCATTTGTATCTTAGTATTGAAATGTGTAATAATATCCATATGACTAAAATGCTGTTGACTTTTAATCCGAATTTCGAAATATGTAATAATCATGGATTAACGCCTATACTATGTTATATAACTTCCGACTACATACAACACGATATTCTTGTCATGTTGATACATCACTATGAAACAAATGTTGGAGAAATGCCTATAGATGAGCGTCGTATGATCGTATTCGAGTTTATCAAAACATATTCTACACGTCCGTTAGATTCGATAACTTATTTGATGAATAGGTTTAAAAATATAGATATTCATACCCGCTATGAAGGAAAGACATTATTACACATAGCATGTGAATATAATAATACACACGTAATAGATTATCTTATACGTATTAACGGAGATATAAATGCGTTAACCGACAATAACAAACACGCTATCCAACTCATTATAGATAACAAAGAAAATTCCCAGTATACCATCGATTGTTTATTGTATATACTTAGATATATTGTAGATAAGAATGTGATAAGATCGTTGGTGGATCAACTTCCATATCTACCTATCTTCGATATAAAATCATTTGAGAAATTCATATCATACTGTATACTTTTAGATGACACATTTTACGATAGACACGTTCAGAATCGCGATTCTAAAACTTATCGATACACGTTTTCAAAATACATATCATTCGATAAATACGATAGTATAATAACTAAATGCTATGAAGAAACAATATTGCTCAAACTATCCACTGTTCTAGACACAACACTATATTCTGTTTTAAGATGTCATAATTCGAGAAAGTTAAAAAGATACCTCAGCGTGTTAAAAAAATATAATAATGATAAGTCCTTCAAAATATATTCTAATATTATGAATGAGAGATACCTTAATGTCTATTATAAAGATATGTATGTGTCAAAAGTATACGATAAACTATTTCCTGTTTTCACAGATAAAAAATGTCTACTAACATTACTACCTTCAGAAATTATATATGAAATATTATACATGTTAACAATTTACGATCTTTATAATATATCGTATCCACCTACCAAAGTATAGTTGTATTTGTAATTTTTCTCATACGATGTGTGTAAAAACTGATATTATAAATTATCTTAGTGTCGTATGATGAAGATGACGATGAAAATGATGGTACATATATATTTTGTATCATTATTGTTATTGCTATTCCACAGTTACGCCATAGACATTGAAAATGAAATCACAGATTTCTTCAATAAAATGAAAGATACTCTACCAGCTAAAGACTCTAAATGGTTGAATCCAACATGTATATTTGGAGGCACAATGAATAATATGGCTGCTATAGGAGAGCCATTCAGTGCAAAGTGTCCTCCTATTGAAGACAGTCTTTTATCGCGTAGATATATAAATAAAGACAACGTAGTTAACTGGGAGAAAATAGGAAAAACTAGACGACCTCTTAATAGACGTGTTAAAAACGGTGACTTATGGATAGCCAACTATACATCTAACGACAGTCATCGTATGTATTTGTGTACAGTGATTACAAAGAATGGCGACTGTATTCAAGGTATAGTTAGATCCCATGTACGGAAACCTTCTTCATGCATTCCAGAAATCTATGAACTAGGTACTCATGATAAGTATGGAATAGACTTATACTGTGGAATTATTTACGCAAAACATTATAATAATATAACTTGGTATAAAGATAATAAGGAAATTAATATCGATGATATTAAGTATTCACAAACGGGAAAGGAATTAATTATTCATAATCCAGCATTAGAAGATAGCGGAAGATATGACTGTTATGTTCATTACGACGACGTTAGAATCAAGAATGATATCGTAGTATCAAGATGTAAAATACTTACGGTTATACCGTCACAAGACCACAGGTTTAAACTAATACTAGATTCGAAAATCAACGTAATTATAGGAGAACCTGCCAATATAACATGCACTGCTGTATCAACGTCATTATTGTTCGATGATGTACTGATTGAATGGGAAAATCCATCCGGATGGCTTATAGGATTCGATTTTGATGTATACTCTGTTTTAACTAGTAGAGGCGGTATTACTGAGGCGACCTTGTATTTTAAAAATGTTACTGAAGAATATATAGGTAATACATATAAATGTCGTGGACACAACTATTATTTTGAAAAAACCCTTACAACTACAGTAGTATTGGAGTAAATACACAATGCATTTTTATATACATTACTGAATTATTATTATTAATTATATCGTATTTGTGCTATAGAATGGATGAAGATACGAAACTATCTAGGTATTTGTATCTCACCGATAGAGAACATATAAATGTAGACTCTATTAAACAGTTGTGTAAAATATCAAATCCTAATTCATGTTATAGATGTGGATGTACGGCTTTACATGAGTACTTTTATAATTATAGATCAGTCAACGGAAAATACAAGTATAGATACAACGGTTACTATCAATATTATTCATTTAGCGATTATGAAAATTATAATGAATATTATTATGATGATTATGATAGAACTGGTATGAATAGCGAGAGTGAGAGTGATAATATATCAATCAAAACAGAATATGAGAATGAATACGAATTCTATGATGAAACACAAGATCAAAGTACACAACTAGTAGGTTACGACATTAAACTCAAAACCAATGAGGATGATTTTGTGGATGAATTCTATGGCTATGATAGATCAGTGGGTGTCCATGATTATATAGATAAATCAATTAATAAAGTAGTATATGGAAGAGAGTCTTACGTAAGATGGCGGGATATATGGCAAGAACATAATGATGGTGTATACAGTATAGGAAAGGAGTGCATAGATAATATATACGAAGACAGACATACAGTAGACGAATTCTACAAGATAGACAGCGTATCAAATGTAAATGACGCGGAACACATATCTCAGATAACTAATGATGTATCTACACAAACATGGGAAAAGAAATCAGAGTTAGATAGATACATGGAAATGTATCCTCGTCATAGATATAGTAAGCATTCTATCTTTAAGGGATTTTCTGACAAAGTTAGAAAAAATGATTTAGACATGAATGTGGTAAAAGAATTACTTTCTAACGGCGCATCTCTAACAATCAAGGATAGCAGTAATAAGGATCCAATTGCTGTTTATTTTAGAAGAACGATAATGAATTTAGAAATGATTGATATCATTAACAAACATACAACTATCTATGAACGCCGGTATATAGTACACTCCTATCTAAAAAATTATAGAAATTTTGATTATCCATTTTTCAGGAAGTTAGTTTTGACTAATAAACATTGTCTCGATAATTATTATAATACAGGTGACCGCAGACATGGAACGCCGCTACATATATTGGCGACTAATAAAAAATTAATAACACCTAATTACATGAAGTTATTAGTGTATAACGGAAATGATATAAACGCACGCGGCGAAAATACACAAATGCAAACTCCATTACACAAATATTTGTGTAAATTTGTATATCATAATATTGAATATGGTATCCGATACTATAATGAAAAGATTATAGACGCATTTATAGAGTTAGGAGCCGATCTAACTATTCCAAATGATGATGAAATGATACCAGTAATTTACTGCATACACGTAAATGCTGAATATGGTTATAACAATATTACTAACATAAGGATAATACGTAAACTACTTAATCTTAGCAGACATGCTCCACATAATCTATTTAGAGATCGAGTCATGCACGATTATATAAGTAATACATATATTGATATAGAGTGTTTAGATATTATTAGATCACTGGATGGGTTCGATATTAATTGTTACTTTGAAGGACGTACACCTCTTCATTGCGCTATACAGTATAACTTCACTCATATTGCGGAATACTTATTAGATCGAGGAGCTGATATATCATTAAAGACAGGCGATGGAAAATCTGTATTCGATTTATCGTTATGTAGCTACATACCTCTTAAATGGACTAGTTTTTTGATTAGTCGTCTACCGCCTAAAAGTGTCATGAGCTCACTGACTAACCATATAATAGATTATGTTCTTACAGACAATAAAAATATTATTTGGCAAAGTCAAATGATTAATAAGTATGTGTTGTTACTTGATCCATCCTTTTATTCTAGATTCAGAAATGCTATTAAAAGCAAATTAAATCAATACGATTATCGTTGCTTCAATTACAATAGATATCGTGTTAATAAAAAGTATCGCAAAGTTTTACATGACATTGATATATATATTAAGGATGTACAAGTATTAAAATCTATTCTCATCGCCAATAATGTAACACTATACGACACTATTGTAAATAATACATCAGAGTTTCCTGTACATCGTCTAAACAACAAACAATTAATTAATACAGTAAATTCCAATATATATCATGATCTTATCGAAAAAGTTATTAAAAAATCATTAGAGAAATGTACTCTAACTAATGCCGTCCTCGAGTATATGATCGCATCTCGATCTCAATCATCTTATTTAAGTCGTATTCCTAATGAGATATTACTCAAAATACTATATAACCTCAGCGTCTACGATTTACGCAAACTATATGCAAGATACATGCTAGAGAATGAGACAGATTATCATATAGAGTCAAGGTCAATTTCTACGCAGACTGAATAATGTTATATACACTACATATTTTTATGTCATTATAACTACTTGTAATATTGAAATCGTAATTTGATATATAGTGAAAAATGGATTTTGATGATATTAAACATAATAGACGAGTAGTAAGTAACATTAGCAATCTTTTAGATGATGATATATATTATGCGATGTAATCATAACTATTGGTGATGGAGAAGAAATTAAAGCACATAAAACTATCTTGGCTGCTGGATCTAAATATTTTAGAACACTGTTTACAACACCTATGATAATAAGAGATCTAGTAACTAGAGTAAAAATCTACAGATGATCAATAAAGATTCCGTCAAAAATATTGTACAGTACTTATACAATAGGTATATAAGTTCTATGAATATAATAGATGTATTAAAATGCGCCAACTATCTTCTAATCGATTATCTAGTCACTGACTGCGAATCCTATATTAAAGATTATACTAATCATGATACCTGCATATGTATGTACCATAGGTTAATGAGATGGTTCATATTCCAATAGTCAAATATATTAAACGTATGCTGATGAGTAATATATCAACATTGATAACTACGGACGCATTTAAAAAAACAGTATTTGAAATATTATTTGATATTCTTTCTACTAACGACAATGTATATTTATACAGAGAAGGCTATAAAGTAACGATACTATTGAAGTGGCTAGAATACAACTACATCACAAAAGAACAGCTTTTGTATATACTGTCCTGTATAGATATACAATCTAGATAGAAGTATAGACTACTACTCTATTCAAATAAAACTATAAATATGCATCCCTCATGTATACAATTTCTATTAGATAATAAACAGAATAGAAACATTATACCCCAACCAGTTATGTTTAGCGTGTCATGATACTAAGTATAATGTATGTAACCCGTGCATATTAGTATATAACATTAATACTATGGAATATAGTGTTATTTCCACCATTCCTAATCACATAATTAATTACACATCCGCGATTGTAGATAATGAAATAATAATAGCTGGAGGATATAACTTTAATAATCCATCGTTAAACAAGGTATATAAAATAAACATTGAGAACAAAATACATGTCGAACTCCCACCCATGATAATGAATAGATGTCGTTTTTCTTTGGCTGTAATCGATAACACAATTTATGCTATAGGTGGTCAAAAACTGAAGTTTAACATTATTTATTTATGATATATATTATATATCGGTATCGTTATTGTTTGGTCTATACCATGGATATCTTTAAAGAACTAATCTTAAAATACCCTGATGAAAATGTTTTGATTTCTCCAGTTTCCATTTTATCTACTTTATCTATTCTGAATCATGGAGCAGCTGGTTCTACAGCTGAACAACTATCAAAATATATAGAGAATGTGAATGAAAATACACCCGATGATAAGAAGGATGACAATAATGACATGGACGTAGATGTTCCGTATTGTGCGACACTAGCTATTGCAAATAAAATATACTGTAGCGATAGTATCGAGTTCCACGCCTCCTTCCTACAAAAAATAAAAGATGATTTTCAAACTGTAAACTTTAATAATGCTAACCAAACAAAGGAACTAATCAATGAATGGGTTAAGACGATGACAAATGGTAAAATTAATTCCTTATTGACTAGTCCACTGCCCATTAATACTCGCATGACAGTTGTTAGCGCCGTCCATTTTAAAGCAATGTGGAAATATCCATTTTCTAAACATCTTACATATACAGACAAGTTTTATATTTCTAAAAATATAGTTACCAGCGTTGATATGATGGTGAGCACTGAAAACGACTTACAATATGTACATATTAATGAATTATTTGGAGGATTCTCTATTATCGATATTCCATACGAGGGAAACTCTAGTATGGTAATTATACTACCAGACGACATAGAAGGTTTATATAACATAGAAAAACACATAACTGAAGAAAATTTTAAAAAATGGTGTGGTAAGTTATACACTAAAAGTATAGACTTGTATATGCCAAAGTTTAAACTGAAAATGACAGAATCGTATAATCTGGTACCTATTCTAGAAAATTTAGGACTTACTAATATATTCGGATATTATGCAGATTTTAGCAAGATGTGTAATGAAACTATCACTGTAGAAAAATTTCTACACAAGACGTTTATAGATGTTAATGAGGAGTATACAGAAGCATCGGCTATTACAGGAGTATTTATGACTAACTTTTCGATGGTATATCGTACGAAGGTCTACATAAACCATCCATTCATATACATGATTAAAGACAACACAGGACGTATACTCTTTATAGGGAAATACTGTTATCCACAATAAATATAAACAAATAGACTTTTTATCTCATGTCTAAGAATTACAAAGTAATATAGTTAGTATAACTAAAGTTAATACTACATTAAAACTTAAAAAAAATAATAATATCACTATTACAATTAATAAGTATAAACTATAAACTAAAAATTAAACAATATCGTTATTATATAAGTAATACCAAAATGATGATATACGATTAATAGCCTGTCTTATATTCGTGACTTTATCCATTGCTAGTCACTTTACATTCCCGTTATTCCTCCCATTACGGAAGATAAATCGTTTAATAGTGTAGAGGTATTAGTTTCCTTGTTTAGAGATGATCAAAAAGACTATACGGTAACTTCTCAGTTCAATAACTACACATCGACACCAACGACTGGACTATAAATGTACTATCCGATGGTCTGGATATACCATTGACTAATATAACTTATTGGTCACGGTTTACTATAGGTCGTGTATTGTTCAAATCAGAGTCTGAGGATATTTTCCAAAAGAACATGAGTATTCTAGGTGTCTCTATTGAATGTAAGAAGCCGTCGGTATTACTTACTTTTTGACCGTGTGTAAAATGACTCGAGTATTTGACTTCCAGATATGGATTATTATCGAGGAGACTGTCTAGAAGTCGTTTATATAACAATGACTTATAAAAATACTAAAACTGAAGAGACTGATTACGCATACCTCTCTAATGTGCGGATTCCTGAATACTATCGATTGATGAGTAATGGGGTTGATGGTTGATTATTGATTAGTTTTCCTTATTCTTTTTATTCACATAAAAGAATATTTTTTATAAACATGAAACCACTGTCTAAATGTAATTATGATCTTGATTTATAGATGAAAATCAGCTGCTTTAGATGATTTTGGTTAGTATGTATAATATAAAAAAATAAATATAACTTATTTTAAGATTAAGTGCTATTGTACTTAATTTTTTGTTCTATAAACTGAATATATAGCTACAATTATTGACGGGTTTATTTATGACCGGCAACCATGAATTTACAGAGATTATCTCTGGCTATATATCTTACTGTGACATGTTCGTGGTGTTATGAAACATGTATGAGAAAAACTGCGTTGTTTCATGACAATCAATTGGGGCATGCGGAAGACAATCAAGATAGTGTAGCGTCGCTACCGTACAAGTATCTACAAGTAGTCAATAAAAGAGAACGTAGTAGATTGCTGGCTACGTTTAATTGGACAAGTATAGCTGAGGGTGTTAAAAATGATTTCATTAGAATATGTGATATCAACGGAACATATCTATATAATTATACTATTGCTGCTAGTATGATTATTGATTCCATGGAAGAACTACCAACAATTACAACATATGAACCGTCTACCTATAATTATACTTTCGATAATAGCACTGTTAGTACTACTGAAGAACTAAAAGTGACTCCATCACCAACTACATACGCGACTGTAACAACTCCTCTTCCAACATCATCAGTTCCATATGATCAACGATCTAATAACAATGTAGGTACTATATCTATTCAGATACTGAGTAAAATATTGGGAGTCAATGAAACAGAATTAACTAATTATCTTATTACGCATAAAAATGCCACGGTTGACAATAACACATTGTATGGTAATATAGGATTTTTGGAAATAAATAATTGCTACAATATTTCTGTATCAAATGCTAGTTTTAGAATAACATTAGTAAACAATACTTCTGAAGAAATTGTTATAATGCTAACAGGGACTAGTTCATCCGACACCTTCATATCTTCCACCAATACCACTGAATGTTTGAAAACATTAATCAATAATACGTCGAATATTAGTGATGTAAGTATAACACAAAATATGAATGTAACATCTAATTGTGATAAATGTTCAATGAATTTGATGACATCCGTTATCCCTGCCGTTAAAGAATTTAACAATACGTTGAAAAAAATTGGTGTAAAAGATGATAAAAACAATACAGTATATAACTATTATAATTGTAAACTAACTACAAATTCTACATGTGACGAGTTAATCAATTTAGATGAAGTCATTAACAATATAACTTTGACAAATATTATAAGTAGTAGTGTTTCTACAACCAACAGCAGAAAAAGACGAGATCTGAATGGTGAGTTTGAATTTTCTACATCCGAGGAATTAGATTGTCTTTACAAATCATATGGTGTAAGTGATGATGTAAGTCATTGTTTTTCATCACCTAGACGTAGACGATCTGACGACAAACAGGAGTACACAGAAATGAAATTACTCGACCACGCGAAAAAAGATTTAAGAATAGACAGTGTTATTCCTAGAGGTACAACTCATTTCCAAGTAGGTGCATCTGGAGCAAGTGGTGGTGTTGTAGGTGATAGTTCCCCATTTCAAAATGTTAAATCACGGGCCAGTCTATTGGCAGAAAAAATAATGCCTAGAGTACCCACTACTGCTACCGAAGAGCAGCTATATGCAACTATAAATAGACAAACAAAGTTGCCTGCAGGTGTTAAAAGTACTCCGTTTACAGAGGCGCTTGTGTCTACGATAAACCAAAAGCTTTCTAGTGTTAAAGAGGTAACTTATGCTTCGTCCAATCTGCCAGGATCAAGTGGTTATGTTCATAGACCATCTGATTCTGTTATTTACAGCACTATAAGACGGACACGTTTACCTAGTGATACCGATAGTGATTTTGAGGATATACAAACTGTTGTTAAGGAATATAATGAAAGATATGGTAGACGAGTTAGTAGAACACAGTCATCAAGTAGTGATTTTGAAGATATAGATGAGGTGGTTGCAGAATATAGACAAAAATATGGTGGAGCTAGCCGTGGTAGAACATCGTCATCAAGTAGTAGTGATTTTGAAGATATAGATGAGGTGGTTGCAGAATATAGACAAAAATATGGCAACGCTATGACAAAAGGACGTGGTTCTTCTAAACCGGATCCGTTATATAGTACTGTTAAGAAAACACCTAAAAGTATAGCATCTGGAGTAGACATAGTTTCAAAACAAACAGACTATTCTTTATTACCCGGCGTTAATACTGGCAGTTCTATCGTAACACCTCTCACCAGAAGAGGAGCTACTAGACGACCTAAACGCCCATCTACACCTCCACGCGAAGATCTACCACCGCTTCCTCTAAATCCTCCTTATCGACAACTTTCTCGTGGTGGTGATCATTCTCTACAACAAGTTCCTCAACGTGATTATTCTCCACCACATAGACCTCCTCCACCACTTCCTCCTAAACCAGTTCCAGCTATTCCACCTAGAGATAGTCAACCAAATAATAAAGGATTTAGTAAGTTTGTATCTCCTAGACGGTGTAGAAGATCAACCTCTGGAGTCGTATGTGGTATGATACAATCAAGACCAAACGATGATACATATTCACTTCTTCAACTACCAAAAATTGAACCAGAATATGCGGAGGTAGGTAATGGCTTACCCAAGAACAATGTTCCTGTGATAGGCAATAAACATAGTAAAAAATATACATCGTCGATGTCAAAAATATCAACAAAATTTGATAAATCTATGGCATTTGGAACAGCAATGTTACTAACTGGTCAACAGGCTATTAACCAACAGGATAGATCAACTGCGTTGATTAAAAAAGATCAAATGAGCAAGGACGAAAAAATATTCGAAGCAGTTACAATGACTCTATCAACTATAGGTTCAACGTTGACAACTGCAGGTATGATAGCTCCACCACTAATGATTGCTGGAATAGGTATATCGCTTATATCAGGTATAATAGATACGGCAAAAGATATATATTACCTATTTTCAGGACAGGAGAAGCCAGTAGATCCAGTTATTAAGTTTTTTAACACATACGCTGGATTAGTATCCGATAGTAGTAAAATGGGTGTAAGGAAATGTTTGACGCCTGGAGAAGACACACTTATTTACATCGCATACAAAAACGATTCCAGTTTTAAACAAAATACGGAGGCAATGGCTTTATATTTCTTAGATGTTATCAACTCAGAGATCCTATATCTAAACACATCAAATTTAGTTTTAGAGTATCATCTAAAGGTGGCTTGTCCCATAGGAACATTAAGATCTGTAGATGTGGACATAACTGCGTATACAATATTATATGATACAGCGGATAATATTAAAAAATACAAGTTTATCAGAATGGCAACGCTACTATCCAAACATCCAGTTATTAGATTGACATGTGGTTTAGCAGCAACATTGGTAATTAAACCGTACGAGGTACCCATCAGTGATATGCAACTACTAAAAATGGCAACGCATGGTGAACCAGAATCCACTAAATCAATACCATCTGATGTCTGTGATAGGTATCCTCTAAAGAAATTCTATCTTTTGGCTGGTGGTTGTCCATATGATACATCTCAAACTTTTATTGTACATACTACTTGCAGTATTTTACTAAGAACAGCTACATGGGATCAGTTTAGAAACAGATGGGTGTTACAAAATCCATTTAGGCAAGAAGGGACATATAAGCAACTGTTTACCTTTAGCAAATACGATTTTAACGACACCATAATAGATCCTAATGGTGTGGCGGGTCATGCTAGCTTTTGTACCAATAGAAGCAGCAACCAATGTTTCTGGTCAGAACCTATGATATTGGAAGATGTATCATCGTGTAGTTCTAGAACTAGAAAAATATACGTAAAACTGGGAATATTTAATACTGAAGGGTTTAATAGTTTTGTACTAAACTGTCCAACTGGGTCTACGCCAACATACATCAAAGATAAAAATACGGACAGTAACAATGTTATCATAGAGCTACCTGTAGGTGATTACGGCACTGCCAAATTGTATTCGGTAACAAAACCATCGAGAATAGCTGTATTCTGCACACATAACTATGATAAACGATTCAAATCAGATATTATAGTTCTAATATTTAATAGTATCAGTGGTATTCCATTTTCGAGCATATACACAGGAAGTGTGAACGGTAGAAATAGACTGTTCACCACATTGTCTAAAGGAATGCCATATAGATCAATGTATTGCGATAACAGACGACCAGGTTGTTATTATGCAGGAATACCATTTAATGAAAATAGTGTAGAATCAGATCTGCATTATGGACCAGAAATAATGCTTAAGGAAACATATGACACAAACAGTATTGATCCACAAGTTATAACAAAGTCAAAAACACACTTTCCTACTCCAATAAGTGTAAAATTTACGGTTGACAATTTAGGAAATGGGTATAACAAACCCGAAAATTTTTGGAAAGACGCTAAAAGTAAGAAAAGGACATATAGCGCAATGACAATAAAAATCCTACCATGTACAGTGAGAAATAAAAATGTAGACTTTGGATATAACTATGGACATATTATTTCCAATATGGTTTATGCACAATCTACTAGTCAGGATTATGGAGATGGTACCAACTATACATTTAAATCTGTAAATAGATCCGATCATGAGTGTGAATCTATTTTAGATTTGAAGGCTAAGGAAGTAACTGTGATGTGTCCCGCGTTTAGTATACCAAGAAATATATCAGCGTATGAAGGTCTATGCTTTAGTGTCACTACATCTAAAGATCATTGTGCCTCAAATAAGGAATGGTTAAAATCTTATGGTTATGGGAATACAGATGCTACTAAACAACGTGTCTGTTTTCATCATTGGAATTATGTCACAACGTCGTTGGATTATTACTGTTCATACGAAGATATTTGGAAAAGCGACTGGCCTGACTATGATCCATGTAAGTCATATATTTATATAGAGTATAGAGACATATGGATAGAATCTAAAGTGTTGCAGCAACCTCCTTATACATTCGAATTCACACATGACGATTCTAATGAATACGTGAATAAAGAAATTAGTAACAAATTGAATGATCTGTACAATGAATACAAGAACATTATGGAATATAGCGACGGATCATTGCCTGCTTCTATAAACAGATTAGCAAAATCATTAACTTCAGAGGGTAGAGAAATAGCAAGTGTTAATATAGATGGTAATCTGTTAGATATTGCATATCAAGCAGATAAGGAAAAGATGGCTGATATACAGAACAAAATAAATGACATCACAAGAGATTTATTCATTCACACTTTATCAAACAAAGATATAAAAGACATTATAGAATCCGAAGAAGGTAAGAGATGTTGTATAATAGATGTTAAGAACAATCGTGTTAAAAAGTACTATCCTATTGATAATTATCTATGTGGTACTTTAGATGATTATATATACACTTCTGTAGAATATAACAAATCCTATGTGTTAATAAACGACACTTACATGAGCTATGACTATCTTGAATCATCAGGTGTAGTTGTTTTATCATGTTATGAAATGACTATAATCTCCTTGGATACAAAAGACGCCAAAGATGCTATAGAAGATGAGATAGTAGCAAGTGCTGTAGCCGAAGCATTGAATGACATGTTTAAGGAATTTGATAAAAACGTAAGTGTTATTATAATAAAAGAAGAAGATAATTATCTAAACAGTTCTCCGAATATATACCATATAATATACATCATAGGCGGCACTATTCTGATACTGTTAGTCATTATTTTAATATTGGTAATTTATATAGCATGCAATAAATACAGAACCAGGAAATATAAAATAATGAAAGATGATACTATGAGCATTAAATCTGAGCATCATAATAGTCTTGAAACAGTATCTATGGAAATTATGGATAATCGGTACTAATAAAAAATAGTTTAACTCTTTTTAGAACCAGTTTGGTACTGGAATTTCAGTTCATTCGTTGAGAATATTGATGATTTTTTTTAAGATGATATTTACTTTTTATATGCTTGCATTACAGAATAATATTCACAGGTATTATTAAAATGAGTATCGGAGCTACATTACCATATCATCATCCATGCTCATGTGGATGGAAAATCCATCCATTATATAATTAATGATACGTGCATGTTTGATATATTCTTTAAATGGAACAGACATCACAGACTGAGATACGCAAAGAATCCTACATTCCTAAACTTTGTATCTGACATAAGAACAGTGAGAGGATCGACTCCATAGTGGACAACATTAATACAAACGATAACTTGATTTCAAAATTATCAATGGAGATACAACGTGAAATACTTTATTACTCCATTAAATAATGTATCACTTACGATAATGTCTCTATAAACGACCGGTTTATCATTAATTAATACAATGTCTAAATTCCATAGCATATTACAAAATCACCGTCTAAGATGATAAAAACACGTTTATTTTATATGTCAACCATTACTAAAAAATATATTGTTCTGTTTTTGTTTTCTTTTTCTTACACACATCGCTAATTATGAAAAAGTAAATGATGAAATGTATGAGATGGTGAGATGGACGAGATTGTGAACATTGTTCGTGATAGTATGTGGTTTATACCTAACGTATTTATGGACAACGGTGAGAATGACGGCCACGTTTCTGTCAACAATGTCTGTCATATGTATTTCGCGTTCTTTGATGTTGATACATCGTCTCATCTGTTTAAGCTAGTTATTAAACACTGCGATCTGAATAAACAACTAAAATGTGGTATGTCTCCATTACATTGCTATGTGATGAATACACGATTTAAGCCATCTGTATTAAAGATATTGTTACACAACGGTGTGAATAACTTTGATAATAAGGATAACAAAGGACATATTCCTCTACACCACTATCTGATTTATTCACTATCAATCGATAACAAGGTCTTTGATATACTAACTGACCCCATTGATGACTTTAGTAAATCATCCGATCTATTGCTGTGTTATCTTAGATATAAATTCAATGGGAGATTAAACTACTACGTTCTGTACAAATTATTGACTAAAGGATCTGACCCTAATTGCGTCGACGAGGATGGACTCACTTCTCTTCATTACTACTGTAAACACATATCCGCATTCCACGAAAGTAATTATTACAAGTCAAAGAGTTACACTAAGATGCGAGCTGAGAAGCGATTCATCTATACGATAATAAATCATGGGGCAAACATTAACGCGGTTACGAAAATCGGAAATACGCCGTTACACACTTACCTTCAACAGTATACCAAACATAGTCCTCGTGTGGTGTATGCTCTTTTATCTCGAGGAGCCGATACGAGAATACGTAATAATTTTGATTGTACACCCATCATGGAATACATAAAGAACGATTGTGTAACAGGTCATATTCTCATAATGTTACTCAATTGGCACGAACAAAAATACGGGAAATTACAAAAGGAAGAAGGACACCATTTACTTTATCTATTCATAAAACATAATCAAGGATACGGAAGTCACGCTTTCAATATATTACGGTATCTACTAGATAGGTTCGACATTCAGAAAGACGAATATTATAATACCATGACTCCTCTTCATACAGCCTTCCAGAATTGTAACAACAATGTTGCCTCATACCTTGTATACATCGGATACGACATCAACCTTCCTACTAAAGACGATAAGACAGTATTCGACTTGGTGTTTGAAAACAGAAACATCTTATTCAATGCGGGTGTTATTCACAACATCATCCACCACAGACTGAAAGTATCTCTACCTATGATTAAATCGTTGTTCTATAAGATGTTAGAATTCTCTCCCTACGACGATTACTACGTAAAGAAGATAATAGCCTACTGCATATTAAGGGACGAGTCATTCACGGAACTACATAGTAAATTCTGTTTAAACGAGGACTATAAAAGTGTATTTATGAAAAATATATCATTCGATAAGATAGATTCCATCATCAAAAAATGCAACTGGGACATAAGTCGCCTCAAAGATATTCAAATCTCTGACACCAACCTGTATACTGTATTAAGAACAGAAGACATCCGGTATCGTACCTATCTTAAAGCCATACATTTAGATAGTCACATTTCATTTCCCATGTACGACGATCTTATAGAACAGTGTCATCTATCGATGGAGCGTAAAAGTAAACTCGTCGACAAAGTACTCAATAAATTAAAGTCTACCATCGATGGTCAATCTAGACTATCGTATTTGCCTCCAGAAATTATACGCAACATCATAACCAAGCTAAGCGACTATCATCTAAACAGTATGTTGTACGGAAAGAACCATTACAAATATTATACATGATAGGAAGAAAATATTTAAAAAATATCTATATGATTGGAGAAGTAGGAAACAGGAACACGGTGATTACTACATTATTAAATCATGAAGTCCGTATTATACTTGTATATATTGTTTCTCTCATGTATAATAATAAACGGAAGAGATGCAGCACCGTATACACCACCCAATGGAAAGTGTAAAGACACCGAATACAAACGCCATAATCTGTGTTGTTTATCGTGTCCTCCGGGAACATACGCTTCCAGATTATGTGATAGCAAGACTAACACACAATGTACACCGTGTGGTTCGGGTACCTTTACATCTCGCAATAATCATTTACCCGCTTGTCTAAGTTGTAACGGAAGATGCAATAGTAATCAGGTAGAGACGCGATCGTGTAACACGACTCACAATAGAATCTGTGAATGCTCTCCCGGATATTATTGTCTTCTTAAAGGATCATCCGGATGCAAGGCATGTGTTTCCCAAACAAAATGTGGAATAGGATACGGAGTATCCGGACACACGTCTGTTGGAGACGTCATCTGTTCTCCGTGTGGTTTCGGAACATATTCTCACACCGTCTCTTCCGCAGATAAATGCGAACCCGTACCCAACAATACATTTAACTATATCGATGTGGAAATTACACTGTATCCAGTTAACGACACATCGTGTACTCGGACGACCACTACCGGTCTCAGCGAATCCATCTTAACGTCGGAACTAACTATTACTATGAATCATACAGATTGCAATCCCGTATTTCGTGAGGAATACTTCTCTGTCCTTAATAAGGTAGCAACTTCAGGATTTTTTACAGGAGAAAATAGATATCAAAATATTTCAAAGGTGTGTACTTTAAATTTTGAGATTAAATGTAATAACAAAGGTTCTTCCTTCAAACAGCTAACGAAAGCAAAGAATGATGACGGTATGATGTCGCATTCGGAGACGGTAACTCTAGCGGGTGACTGTCTATCTAGCGTCGACATCTATATACTATATAGTAATACCAATGCTCAAGACTACGAAACTGATACAATCTCTTATCGTGTGGGTAATGTTCTCGATGATGATAGCCATATGCCCGGTAGTTGCAATATACATAAACCGATCACTAATTCCAAACCCACCCGCTTTTTATAGTAAGTTTTTCACCCATAAATAGTAAATACAATAATTAATTTCTCGTAAAAGTAGAAAATATATTCTAATTTATTGCACGGTAAGGAAGTAGAATCATAAAGAACAGTACTCAATCAATAGCAATCATGAAACAATATATCGTCCTGGCATGCATGTGCCTGGCGGCAGCTGCTATGCCTGCCAGTCTTCAGCAATCCTCATCCTCGTGTACTGAAGAAGAAAACAAACATTATATGGGAATCGATGTTATTATCAAAGTCACAAAGCAAGACCAAACACCGACCAATGATAAGATTTGCCAATCCGTAACGGAAATTACAGAGTCCGAGTCAGATCCCGAGGTGGAATCAGAAGATGATTCCACATCAGTCGAGGATGTAGATCCTCCTACCACTTATTACTCCATTATTGGCGGAGGTCTGAGAATGAACTTTGGATTCACCAAATGTCCTCAGATTAAATCCATCTCAGAATCCGCTAATGGAAACGCAGTGAATGCTAGATTGTCCAGCGTGCCCCTAGGACAAGGTAAGGACTCTCCCGCTATCACTCGTGCAGAAGCTCTGGCTATGATCAAAGACTGTGAGCTGTCTATCGACATTAGATGTAGCGAAGAAGAGAAAGACAGCGACATCCAGACCCATCCAGTACTCGAGTCTAACATCTCTCATAAGAAAGTGAGTTACGAAGATATCATCGGTTCAACGATCGTTGATACAAAATGCGTCAAGAATCTAGAGTTTAGTGTTCGTATCGGAGACATGTGCAAGGAATCGTCTGATCTTGAGGTCAAGGATGGATTTAAGTATGTCGACGGATCGGTATCTGAAGGTGTAACCGATGATACTTCACTCATCGATTCAACAAAACTCAAATCGTGTGTCTGAATAAATAACTCTATTCATCCGAGATTCGGGTAAGTAGGGTTAATCGAACGATTTAGTCACACTACGAATTAAAAAAGTGTACCTGACACTATACTCCGGTTTACAAAACAAAAATGTTCTTAACTACATTAACAAAAAGTTATCTCTCGCGACTTCTTCTTTTTCTGTCTCAGTAGTGTGTACGATATGATCACTGTTCATATTCCTATTCTCTCTTCGATGGTCTCACAAAAAAGTTTTACAAAAATATTTTTATTCTCTTTCTCTCTTCGATGGTCTCACAAAAAATATTAAACCTCTTTCTGATGGAGTCGTAAAAAGTTTTATCTCTTTCTCTCTTCGATGGTCTCACAAAAAATATTAAACCTCTTTCTGATGGAGTCGTAAAAAGTTTTATCTCTTTCTCTCTTCGATGGTCTCACAAAAAATATTAAACCTCTTTCTGATGGAGTCGTAAAAAGTTTTATCTCTTTCTCTCTTCGATGGTCTCACAAAAAATATTAAACCTCTTTCTGATGGAGTCGTAAAAAGTTTTATCTCTTTCTCTCTTCGATGGTCTCACAAAAAATATTAAACCTCTTTCTGATGGAGTCGTAAAAAGTTTTATCTCTTTCTCTCTTCGATGGTCTCACAAAAAATATTAAACCTCTTTCTGATGGAGTCGTAAAAAGTTTTATCTCTTTCTCTCTTCGATGGTCTCACAAAAAATATTAAACCTCTTTCTGATGGAGTCGTAAAAAGTTTTATCTCTTTCTCTCTTCGATGGTCTCACAAAAAATATTAAACCTCTTTCTGATGGAGTCGTAAAAAGTTTTATCTCTTTCTCTCTTCGATGGTCTCACAAAAAATATTAAACCTCTTTCTGATGGAGTCGTAAAAAGTTTTATCTCTTTCTCTCTTCGATGGTCTCACAAAAAATATTAAACCTCTTTCTGATGGAGTCGTAAAAAGTTTTATCTCTTTCTCTCTTCGATGGTCTCACAAAAAATATTAAACCTCTTTCTGATGGAGTCACTAAAATATTTTTATTCTCATTCTGATGCATCAACTATTTCTTAAACAATAACGTTCAACAACATATACTCTCGAG

>2001_Congo_Monkeypox_NC_003310.1

GTTAGTAAATTATATACATAATTTTATAATTAATTTAATTTTACTATTTTATTTAGTGTCTAGAAAAAAATGTGTGACCCACGACCGTAGGAAACTCTAGAGGGTAAGAAAAATCAATCGTTTATAGAGACCATCAGAAAGAGGTTTAATATTTTTGTGAGACCCATCGAAGAGAGAAAGGATAAAAACTTTTTACGACTCCATCAGAAAGAGGTTTAATATTTTTGTGAGACCCATCGACGAGAGAAAGAGATGGTTAGTCAAGATATTTTTCTTAGTACAAAAGTCAATGTTTTAAAATATATGGACGAGAATTAATTTGTCTGTATAAAAACTTGTGTGAAATTATGTACTAGAGAAAAAACGTGAGCAGTGTCCCCTACATGGATTTTACAGATCATTTATATTCCAAAAATATTAACTATATACGTTTATTATATGATGTTAACGTGTAAATTATAAACATTATTTTATGATGCAATTGTCTGACAACCTAGATTGGCATAAGGATATTGATAAGCTCTACGAGAATATATTGTTGGACGTTATCGTTTACGAAATAGTTGAGACATCAGAAAGAGGTTTAATATTTTTGTGAGACCATCGAAGAGAGAAAGAGAATAAAAATATTTTTTTGTAAAACTTTTTATGAGACAAGAGAGAAAGAGAATACGAATAGTGATCATATCGTATCACATATTGAAACAGAAAGAAGAAGTAACGAGAGGTAACTTTTTGTGAATGTAGTTAAACATTTTTGTTTTGCAAACCGGAATATAGTACCCGGTACACTTTTTTAATTCGTGGTGCGGTGTCTGAATCGTTCGATTAACCCAACTCATCCATTTTCAGATGAATAGAGTTATCGATTCAGACACATGCTTTGAGTTTTGTTGAATCGATGAGTGAAGTATCATCGGTTGCACCTTCAGATGCCGATCCGTCGACATACTTGAATCCATCCTTGACTTCAAGTTCAGATGATTCCTCACACATGTCTCCGATACGTACGCTAAACTCTAGGTTCTTAACACATTTTGTATCAACGATCGTTGAACCGATGATATCTTTGTAACTCACTTTCTTATGTGAGATGTTAGACCCAAGTACTGGATGGGTCTTGATGTCGCTGTCTTTCTCTTCTTCGCTACATCTGATGTCGATAGACATCTCACAGTCTTTGATCATAGCCAGAGCTTCTTCACGCGTGATCGCGGGAGAGTCCTTACCTTGTCCTGGTGACACGCTGGACAATCTAGTATTCACAGTGTTTCCATCAGAGGATTCGGAGATGGATGAAATCTTTGGGCATTTGGTGAATCCAAAGTTCATGTTAAGACCCGCGCCGACGATAGTGTAATAAGTGGTGGGATCTCCTTTTACAACTTCTTCGGATACCTCATCATCTTCGGTCTCTGTAACTTCCGTTACGGATTGACAAATCTTATCATTGGTCGGTGTTTGGTCTTGCTTTGTGACTTTGATAATAACATCGATTCCCATATGATGTTTGTTTTCTTCTTCAGTACACGAGGATGAGGATTGTTGAAGACTAGTAGGCATAGCAGCTGCCACTAGGCACATGCATGCCAGGACAATATATTGTTTCATGATTGCTATTGATTGATTACTGTTCTAGATGATTCTACTTTCTTACCATATAATAAATTAGAATATATTTTCTACTTTTACGAGAAATTAATTATTGTATTTATTATTTATGGGTAAAAAAAAACTTACTATAAGTGGGTGGGATTCTGGGAATTAGTGATCAGTTTATGTATATCGCAACTAGCGGGCATATGGCTATTGACATCGAGAACATTACCCATATGATAAGAGATTGTATCCGTTTCGTAGTCTTGAGTATTGGTATTACTATATAGTATGTAGATGTCGACGCTAGATAGACAGTCGCCCACTAGAGTTACCGTCTCTGAATGCGGCATGATAGTATCATTCTTTGTTTTCGTTAACTGTTTGGAAGATGAATCTTTGTTGTTACATTTAATCTCGAAATTCAGAGTACATATCTTTGAAGTATTCTGATATCTATTTTCTCCTGTAAAGAATCCTGAAGTTGCTACATTATTAAGGACAGAGAAGTATTCTGCACGAAAGACGGGATCACAATCTTTATGATTCATGGTAATAGTTAGTTCCGACGTTGAGATGGATTCGCTGAGACCGGTAGTGGTCGTCCGAGTACACGATGTGTCGTTGACTGGATACAGGTTAATTTCCACATCGATATAGTTAAATGTATTGCTGGTTACGGGTTCGCATTTATCTGTGGAAGAGACGGTGTGAGAATATGTTCCGGGACCACACGGAGAACAGATGACGTCTCCGGTAGACGTGTATCCGGATACTCCGTATCCTATTCCACACTTTGTTTTAGAAATACATGTTCTACACCCTGATGATCCTTTGAGAAGACAATAATATCCTGGAGAGCATTCACAGATTCTATTGTGAGTCGTGTTACACGATCGCGTCTCTACCTGATTACTATCACATCTTCCGTTACAACTTAGACAAGCCTGTAAATGATTATTGTGAGATGTAAAGGTATCCGAACCACACGGCGTACATTGTGTATTAGTCTTGCTATCACATAATCTGGAAGCGTAAGTTCCCGGAGGACACGATAGACAACATAGATTACGGCTTCTGTATTCGTTGTCTTTACATTTTCCATTGGATGGTGCATGTGGTGCTAAATCTCTTCCGTTTATTATTATACATGAGAGAAACAATATATACGAGTATAATACGGACCTCATGATTTAATAATAGTAGTAATCGTCGTCTTGTTACTGTTTGTTTCCTACTTCTCCAATCATATAGATTATTTTTTAAATATTTTCTTTCTATCATGGATAATATTTGTAATGGTTCTTTCCGTACAACATACTGTTTAGATGGTAGTCGCTTAGCTTGGTTATGATATTGCGCATAATTTCCGGAGGCAAATACGATAGTCTAGATTGACTATCGATGGTAGACTCTAATTTATTGAGTGCTTTGTCGACGAGTTTACTTTTATGCTCCATCGATAGATGACACTGTTCTATGAGATCGTCGTACATGGGAAATGAAATGCGTTTGTCTGAATGTATGGCTTCGAGATAGGTGTGATACCGGATGTCTTCTGTTCTCAATACCGTATACAAGTTGGTGTCTGAGATTCGAATCTCTTTGAGGAGACTTATGTCACGACTACATTTTTCGATGATGGAATCTATCTTATCGAATGATATATTTTTCATAAATACACTTTTATAGTCCTCGTTTAAACAGAATTTAGTATGTAGTTCCGCAAATGACTCGTCCCTTAATAGGCAGTAGGCTATTATCTTCTTTACGTAGTGATCGTCGTAGGGAGAGAACTCCGACATCTTGTAGAACAACGATTTAATCATAGGTAGAGATACTTTCAGTCTGTGGTGGATGATGTCATTAACGACATCCGCCTTGTATATGATGTTTCTGTTTTCAAACACCAAGTCGAATACTGTCTTATCGTCTTTAGTCGGAAGGTTGATGTCGTATCCGATGTATACGAGGTATGAGGCAACATTGTTATTGCAATTCTGGAAGGCGGTATGAAGAGGAGTCATTGTATTATAGTATTCGTCTTTCTGAATGTCGAATCTATCTAGTAGATACCGTAGTATATTGAGAGAGCGACTTCCATATCCTTGATTATGTTTTATGAATAGATAAAGTAGATGTTGTCCTTCTTCCTTTTGTAATTTCCCGTATTTTTGTTCGTGCCAATTGAGTAACATTATGAGAATATGACCTGTTGCACAATCGTTCTTTATGTATTCCATGATGGGTGTACAATCAAGATTATTACGTATCCTCGTATCGGCTCCTCGAGATAAAAGAGCATACACCACACGAGGACTATGTTTGGTATACTGTTGAAGGTAAGTGTGTAACGGCGTATTTCCGATTTTCGTAACCGCGTTAATGTTTGCTCCATGATCTATTATCGCGTAGATGAATCGCTTCTCAGCTCGCATCTTAGTGTGACTCTTTGACTTGTAATAATTGCTTTCGTGGAACACGGATATGTGTTTACAGTAGTAATGAAGAGAAGTGAGTCCATCCTCATCGACGCAATTAGGGTCAGATCCTTTAGTCAATAATTTGTACAGAACGTAATAGTTTAAGCTCCCATTGAATTTATATCTAAGATAACACAGCAATAGATCGGATGATTTACTAAAGTCATCAATGGGGTCCGTTAGTATATCAAAGATCTTGTTATCGATTGATAGTGAATGAATCAGATAGTGGTGTAGAGGAATATGTCCTTTTTTATCCTTGCTATCAAAGTTACGCATGCCGTGGCGTAACAATATCTTTAATACAGATGGATTAAATCGTGTATTCATCGTATAGCAATGTAATGGAGAGTTACCACATTTTAGTCGTTTATTCAGATCGCAGTGTTTAATAACTAATTTAAACAGATGAGATGATATATCCACATCAAAGAATGCGAGATACATATGACAGACATTATTGACAGAAATGTGACCTTCATTATCACCGTCGTCCATAAATGCGTTAGGTACGTACCACATACTATCGTTAACGATGCGCACAATCTCGTCCATTTTCATCCATCTTCATAATGATTTACTTTTTCATAATTAGAGAAAAAGATCAAGGTATAAAAATTAGAAGTGTTAGACTATAAACTAACTTATAAAACTAACTTATGACTTAACTAACTTATGACTATTAACTCATTTCAAGAAAGGTGGGTGGAGAGAACTCTATATGACAGCTTGTGAAACAATTAGATCCCTAATTTCTAATGGAAGTTTTGATAGGAGATTGTCATCAGTTGATACATTGTTTATTATCTCATCTATTAGAGCACGTCTGTTTAGAGCTTTAGTGACCTGCTCGGTTACTTCTGTGTAAATCTTGAATCCTTTAGTGATACACTGTGTCAAAACTGGATGTTTAGAATACCTATGTAGAATATGGGAAGCATGCTTGTTTTTGTCTCTATTATAGATTAACTCATACATGGTTGTATTATGAATTTTCATCTGCCTAATGTACTCCAATTCTTGTTTACAATCAATTATATAATCAAAGAGTGATGATGCATACACATTACAAAGTGAATAATCTACCATCATAAAATACTTGATACAGAGCTTTATCACATCATGGTTTTCAATTGTATTATTAAGTATAGCTAATTTTATACAGTCAATAGACAATGGTTCTCTAAGCAATATTTCTAATATTTTAAGATGTGTTTCCCTACGGGCGATGACAGATCCCCTATCCACGGCCACGTCAAGACATGTATATCCATTACTCATTACTGCGTTGACATTTGCTCCATTTTCTAATAGCCATGATACTAAATCTATATAACCTGCATAGATAGCGCGATAAAGCAAGGTCCTTCCACCAGCATCTAGTTGATTGATATCTTCAATATATGGGATACAAAGCTTATAAATTTCTAATACTGTGGGTTCATCTACAAGGAATCCCCTAGTATACTGAATTATTTTATATAGATCTAATTTAACATCATTTTCATCTGGGATACCACAATTCAAAATAAACTCAACAACACTACTTTCCTTTTTACATATTCCCCTAAAATAGGCATTCAAGCATTCTATTTTATATATTACAGCCCCATGATCTACCATAAAATCAACAATGTCTATTTCTACATATGCATTAGATAGATAGTAAAGTAAGAGATCTTGTACAGAATTACAATTCTTAATAATTATAGAGAAAATATCTTCCATATAATTCTTTGACACTAATGCAGATATAATATCTTTATATGTAATATATGCAAACAGTCTATCTACTATATACTGATCAATATTATCTCTATGAATCCTAAAATAATCATACAGAACATCTACAGGATCACAAATTGGTTCAAGGAGAAATCTATCAAATATTTTCCTGTCAACAACTGGTTCTAGAACATAACAGTCAACACCTAATCCATGTTTTTTATAGTCATCTACCAAAGATAATGACCAAAGATCGAGGTCGTCGTGAAACTGCTCATCGACAGCCATGAAATCTGCCGACTCCATGGCGCGAATCGCACTGTCTTATTCGCCATTGATTTTCATTTTTTATAATTATGTACATGTTTTCCTTCTATTCTCAAGAGTCTACAAAAATATATTTTTTTTCGATATCTAAGTACTAAGTTTTTTTACTGTTTTTGTTACTGTCTTCCATTCTTCTAACTAAAGATCTGAGATAAATTATACAATCTTCGCTATCGAACCATTTTTGTAGTCTAAAGCCTGAAGTAATTAACCAACTGTTTTTATTAGTGGCTTTTTTCGATCTATCCTCGTCCTCATCATCCTTATATTATTATCATTATTATCATAGTCTATTAAACACAAATCATCTACGTTTATAACAACATTCTCATTATTAATTAGTTCTGTAGTAATATCTTTAATAATTTGGCTATACATCTGTTCAATACTATCTATTGATGATTTCTTTTTTAAGACTTAAACTAGTTATGGTAATGACGATGAAATCGAGTAGTAACTTCTAATAAAGACTTGATATCATTATCATATGTTTGATCGTCATAGTTAATAGTGTGGCTAAATGGTACTGTTAATAAGTTTATAGACAATATCATAGTATTTTCTTTCCAGAATTAGATTATTTTTTTAAATACTGATCCTCACAATTCCGTGATGTAGCAGTAGTTGGTGCATGGTCTATATCGTTAAAATGTATCATATATAATAGTTTTCTGACGTGGAGTACAGAATTTTCGATTAATGAGTTCATGGTAAGGAAGGGCAAATGTCTGTATATAATATACATAAGTTAATAGTTTTTTATCATATTTTCTAATACCATAATAAAAATTATCATTATGTATAATCATCACTGTCGCTATCATTATTGCGTTTGTGTAGTTCTGCCCTATCATCTACATCACTGTCACTCTCACTATATCTTCTAAAATTACAAACAACTGGATATTCGATAACAGCATTTGTGTAGTTTTTGTCTTTTACAGTATATACGTTATTGTCAAAATCTAAACAAATATTAGCATAATACATCTATAAGATCAGGATCCATGTTCGAGCATACTAGCCATGTATATTTGTAACTTCGTCATACAGCGTTAGATCAATAGAATAAACAATCGTGTGACGCAACTTTTTTACGATCTAGTTGTATGAGTTTATCGTTTACATAAGCAATTAACGGCTTTAACAGATGATCTGAGTAATAATATACCTCTGTTATACGTTTAATGTTCACGGTCTTAGTATTTTTAGATATCAATTGTGATTTACACCATATTCGACTCCCTGTGTGCAACGTTAGAAATTCTAAATCTATAGTATTATCTATTACAGCGTAAAACACATTCAATATTGTATTGTTATTTTTATATTATTTACACAATTAACAATGTATTATTAGTTTATATTACTGAATTAATAATATAAAATTCCCAATCTTGTCATAAACACAAAATCCATTAAAAATGTCGATAAAATATCTGATGTTGTTGTTCGCTGCTATGATAATCAGATCATTAGCCGATAGTGGTAACGCTATCGAAACAACATCGCCAGAAATTACAAACGCTACAACAGATATTCCAGCTATCAGATTATGTGGTCCAGAGGGAGATGGATATTGTTTACACGGTGACTGTATCCACGCTAGAGATATCGACGGTATGTATTGTAGATGCTCTCATGGTTATACAGGCATTAGATGTCAGCATGTAGTATTAGTAGACTATCAACGTTCAGGAAAACCAGACACTACAACATCATATATCCCATCTCTAGGTATTGTGCTTGTATTAGTAGGCATTATTATTACGTGTTGTCTATTATCTGTTTATAGGTTCACTCGAAGAACTAAACTACCTATACAAGATATGGTTGTGTTATATTTTTTATAAAATTTTTTTATGAGTATTTTTACAAAAATGTATATGTATAAAAAAATACTAAGTATGCGATGTATCCTGTATTATTTGTATTTATCTAAACAATACTTCTGCCTCTAGATGGGATACAAAAATTTTTTATTTCAGCATATTAAAGTAAAATCTAGTTACCTTGAAAATGAATACAGTGGGTGGTTCCGTATCACCAGTAAGAACATAATAGTCGAATACAGTATCCGATTGAGATTTTGCATACAATACTAGTCTAGAAAGAAATTTATAATCATCTTCTGTGACAGGAGTCCATATATCTGTATCATCGTCTAGTTTATCAGTGTCCTATGATATATTCCTGTTATCATCATTAGTTAATGAAAATAACTCTCGTGCTTCAGAAAAGTCAAATATTGTATCCATACATCTCCAAAACTATCACTTATACGTTTATCTTTAACGAACATATACCTGATGGTTATTTACTAACAGACATTTTTCAAGATCTATTGACAATAACTCCTATAGTTTCCACATCAACCAAGTAATGATCATCTATTGTTATATAACAATAACATAACTCTTTTCCATTTTTATCAGTATATATATATATATATATATATCAACGTCGTTGTAGTGAATAGTAGTCATTGATCTATTATATGAAACGGATATGTCTAGTTAATATTTTCTTTGATTTAAAGTCTAGAGTCTTTACAAACATAATATCCTTATCCGACTTTATATTTCCTGTAGGGTGGCATAATTTTATTCTGCCTCCACAATCAGTGTTTCCAAATATATTACTAGACAATATTCCATATGGTTATTAGTTAAGGGACCCGATTAGAACACGTACGCGCTTATTCATCATTTGGATCGTATTTCATAAAAGTTATTATGTTATCGATGTCAACACATTCTACATTTTTTAATCGTCTATATAGTATTTTTCTGATATTTTCTATAATATCAGAATTGTCTTCCATAGGAAGTTGTATACTATCGGAATCAGTTACATGTTTAAATAATTCTCTGATGTCATTCCTTATACAATCAAATTCATTATTAAACAGTTTAATAGTCTGTAGACCTTTATCGTCGTACATATCCATTGTCTTATTAGTTACGCTTATTTTTATGGGTTTTACATTGCTTTATTATATTTTATAAGAATGATTGTTTGACAATGTCGTAGTATAGATATATTATTAGAGGAGGTATAATTATAAAAAGTTTCTGAGTACGATGTTATAAGAGGAGAGGACACATTAACAATCATACATCAATTAACTCATTCTTATAACATTGTAATCAAAAGAATTGCAATTTTGATGTATAACAACTGTCAATGGAATTGTATATTACAAATTACGGTATGTTGTAACGACAAATACCGATCGGTAATTGTCTCTGTCGCTGTAATAGAATTAATTATATATCTATTACACCGACCTTGTATCATAATAAAGTTGTGGTAGTATGATCTCCATATTTATAATTTAGTACTTTGTATTTAGTATTTTTGGAATCATAAAAAAGTTTTACTAATTTAAAATTTAAAAAGTATTTACATTTTTTTCACTGTTTAGTCGCGGATATGGAATTCGATCCTGTCAAAATCAATACATCATCTATAGATCATGTAACAATATTACAATACATAGATGAACCAAATGATATAAGACTAACAGTATGCATTATCCGAAATATTAATAACATTACATATTATATCAATATCACAAAAATAAATCCACATTTGGCTAATCGATTTCGGGCTTGGAAAAAACGTATCGCCGGAAGGGACTATATGACTAACTTATCTAGAGATACAGGAATACAACAATCAAAACTTACTGAAACTATACGTAACTGTCAAAAAAATAAAAACATATATGGTCTATATATACACTACAATTTAGTTATTAATGTGGTTATTGATTGGATAACCGATGTGATTGTTCAATCAATATTAAGAGGGTTGGTAAATTGGTACATAGCTAATAATACATATACTCCAAATACACCCAATAATACTACAACCATTTCTGAGTTGGATATCATCAAAATACTGGATAAATACGAGGACATGTATAGAGTAAGTAAAGAAAAAGAATGTGGAATTTGCTATGAAGTTGTTTACTCAAAACGATTAGAAAACGATAGATACTTTGGTTTATTGGATTCGTGTAATCATATATTTTGCATAACATGTATCAATATATGGCATAGAACACGAAGAGAAACCGGTGCGTTAGATAATTGTCCTATATGCCGTACCCGTTTTAGAAAAATAACAATGAGCAAGTTCTATAAGCTAGTTAACTAATAAATAAAAAGTTTAATTATCGACGATATATGTCGTTATTTTTCTCTCATATGAAAGATTAATTTGATTCTAATATAATCTTCAGTATTGGATGAATCTCAATTCAAATTAATTCCATTAGATTAGATTAGATTAGATTAGATTAGATTAGATTAGATTAGATTAGATTAGATTAGATTAGATTAGATTAGATTAGATTAGATCATAAATAAAAATAGTAGCACGCACTACTTCAGCCAAATATTCTTTTTTGAAACGCCATCTAGCGTAATGAGAACACAAGTGAACCTATAATGAGCAAATTTATTAGTATCGGTTACATGAAGGACTTTACGTAGAGTGGTGATTCCTCCATCTGTGGTACGAACGGTTTCATCTTCTTTGATGCCATCACCCAGATGTTCTATAAACTTGGTATCCTCGTCCGATTTCATATCATTTGCCAACCAATACATATAGCTAAACCCAGGCATACGTTCCACACATCCGGAACAATGAAATTCTCCAGAAGATGTTACAATGGCTAGATTTGGACATTTGGTTTCAACCGCGTTAACATATGAGTGAACACACCCATACATGAAAGCGATGAGAAATAGGATTCTCATCTTGCCAAAATATCACTATAAAAAATTTATTTATCAATTTTAAAGGTATAAAAAAATACTTATTGTTGCTCGAATATTTTGTATTTGATGGTATACGGAAGATTAGAAATGTAGGTATTATCATCAACTGATTCTATGATGGTTTTATGAATTTTATTATGCTTCACTATTGCATCGGAAATAATATCATATGCTTCCACGTATATTTTATTTTGTTTTGACTCATAATACGCACGTATTTCTGGATTATTGGCATATCGATGAATAATTTTAGCTCCATGCTCAGTAAATATTAATGAGAACATAGTGTTGCCTCCTACCATTATTTTTTTCATCTCATTCAATTCTTGATTGCAGAGATCTATATAATCATTATAGCGTTGACTTATGGACTCTGGAATCTTAGACGATGTACAGTCATCTATAATCATGGTATATTTAATACATTGTTTTATAGCATAGGCATTATCTACGATATTAGATACTTCACTCAATGAATCAATCACACAATCTAATGTAGGTTTATGACATAATAGCATTTTCAGCAGTTCAATGTTTCTAGATTCGTTGATGGCAATGGCTATACATGTATATCCGTTATTTGATCTAATGTTGACATCTGAACCGGATTCTAGCAGTAAAGATACTAGAGATTGTTTATTATATCTAACAGCCTTGTGAAGAAGTGTTTCTCCTCGTTTGTCAATCATGTTAATGTCTTTAAGATAAGGTAGGCAAATGTTTATAGTACTAAGAATTGGGCAAGTATAAGACATGTCACAAAGACCCTTTTTGTATGTATAAGTGTAGAAATTATAACATCCATAGTTGGATTCACATAGGTGTCCAATCGGGATCTCTCCATCATCGAGATGATTGACGGCATCTCCCCCTTCCTTTTTTAGTAGATATTTCATCGTGTAAGAATCAATATTAATATTTCTAAAGTATCTGTGTATAGCCTCTTTATTTACCACAGCTCCATATTCCAACATGCATTCCACTAGAGGGATATCGATATCGCCGAATGTCATATACTCAATTAGTATATGTTGGAGGACATCCGAGTTCATTGTTTTCAATATCAAAGAGATGGTTTCCTTATCATTTCTCCATAGTGGTACAATACTACGCATTATTCCGTGCGGCTTTCCATTCTCCAAAAACAATTTTACCAAATCTAAATCTACATCTTTATTGTATCTATAATCACTATTTAGATAATCAGCCATAATTCCTCGAGTGCAACATGTTAGATCGTCTATATATAAATAAGCCGTGTTATCTATTCCTTTCATTAACAATTTAACGATGTCTATATCTATATGAGATGACTTAATATAATATTGAAGAGCTGTACAATAGTTTTTATCTATAGAAGACGGCTTGATTCCGTGATTAATTAGACATTTAACAACTTCCGGACGCACATATGCTCTCGTATCCGACTCTGAATACAGATGAGAGATGATATACAGATGCAATACGGTACCGCAATTTCGTGGTTGATAATCATCATACGCGTATCCGTACTCGTCATCCTCATAAAGAACACTGCAGCCATTTTCTATGAACAAATCAATAATTTCAGGAACAGGATCATCTGTCATTACATAATTTTCTATAACTGAACGATGGTTTTCACATTTAACACTCAAGTCAAATCCATGTTCTACCAACACCTTTATCAAGTCAACGTCTACATTTTTTGATTTAATATAGCTGAATATATTAAAGTCATTTATGTTGCTATATCCAGTAGCTTCTAGTAGAGCCATCGCTATATCCTTATTGACTTTAACATGTCTACTATTTGTGTATTCTTCTATTGGGGTAAACTGTCTCCAATTTTTGTGTAATGGATTAGTGCCACTGTCTAGTAGTAGTTTGACGACCTCAACATTATTACAATGCTCATTGAAAAGGTATGCGTGTAAAGCATTATTCTTGAATTGGTTCCTGGTATCATTAGGATCTCTGTCTCTCAACATCTGTTTAAGTTCATCGAGAACCACCTCCTCATTTTCCAGATAGTCAAACATTTTGACTGAATAGAAGTGAATGAGCTACTGTGAACTCTATACACCCGCACAACTAATGTCATTAAATATCATTTTTGAATGTATTTATACCATGTCAAAAACTTGTACAATTATTAATAAAAATAATTAGTGTTTAAATTTTACCAGTTCCAGATTTTACACCTCCGTTAACACCTCCATTAACCCCACTTTTTACACCACTGGACGATCCTCCTCCCCACATTCCACTGCCACCAGATGTATAAGTTTTAGATCCTTTATTACTACCATCATGTCCATGGATAAAGACACTCCACATGCCGCCACTACTACCCCCTTTAGAAGACATATTAATAAGACAAGTTTAACAATAAAATTAATCACGAGTACCCTACTCCAACCACTATTATATGATTATAGTTTCTATTTTTACAGTACCTTGACTAAAGTCTCTAGTCACAAGATCAATACTACCAACCTACACTATATATGATTATAGTTTCTATTTTTATAGGAACGCGTACGAGAAAATCAAATGTCTAAGTTCTAACGGTAGTGTTGATAAACGATTGTTATCCGCGGATACCTCATCTATCATGTTGTCTATTTTCTTACTTTGTTCTATTAACCTATTAGCATTATATATTATTTGATTATAAAACTTATATTGCTTATTAGCCCAATCTGTAAATATCGGATTATTAACATATCGTTTCTTTGTAGGTTTATTTAACTTGTACATCACTGTAAGCATGTCCGTACCATTTATTTTAATTTGACACATATCAGCAATTTCTTTTTCGCAGTCGGTTATATATTCTATATAAGATGGATACGTATCACATATGTACTTATAGTCTACTAATATGAAGTACTTAATACATATTTTCAGTAACGATTTAGCCTTATTACCTATTAATAAGTGCCTGTCGTTGGATAGGTAATCAACTGTTTTCTTAATACATTCGATGGTTGGTAATTTACTCAAAATAATTTCCAATATCTTAATATATATTTCTGCTATTTCTGGTATACATGCATGTGCCATTATAACACAAATACCAATACATGTAGACCCATATGTTGTTGTTATATTAATATCTGCGCCATTATCTATTAACCATTCTACTAGTGCAACACTATGCGACTCGATACAATAATAAAGTATACTACGTCCATGTTTATCTATTTTGTTTATATCATCGATATACGGCTTACAAATTTTTAGTATCGATAACACTTCTGACTCGTGAATAAATAAGGTAGGGAATAACGGCATAATATTTATTATGTTATCATCATTAACAACTACGTTTCCATTTTTTAAAATATACTCTACAACTTTAGGATCCCTATTGTCAAATCTTTTAAAATATTTATTTATATGCTTAAATCTATATAATATAGCTCCTTCCCTAATCATACATTTGATAACATTGATGTACACTGTATGATAAGATACATATTCTGACAATAGATCTTGTATAGAATCTGTATATCTTTTAAGAATTGAGGATATTATGACATTATTACGTAAACTATTACACAATTCTAAAATATAAAACGTATCATGGGCAGATAATAGTTTATCCACTATATAATTATCTATTTTATGATTTTTCTTCCTATATTGTTTACGTAAATAGATAGATAGAATATGCATTAGTTCATTACCGCTATAGTTACTATCGAATAACACGTCAAATATTTCCCGTTTAATATCGCATTTGTCAACATAATAATAGAGTATGGTACGTTCACGATAAGTATAATGACACATTTCGTTTTCGTGCGAAATTAAATAGTTTATCACGTCCAAAGATGTCACATAACCATCTTGTGACCTAGTAATAATATAATAATAGAGAACTGTTTTACCCATTCTATTATCATAATCAGTGGTGTAGTCATAATCTAAATAATCAAACTCGTCATCCCAATTAAAATAAATATAATCAGTACATTGAATGGGTATGATATTGTACCCATACTGTATGTTGCTACATGTAGGTATTCCTTTATCCAATAATAGTTTAAATACATCTATATTAGGATTTGATGTTGTCGCGTATTTCTCTACAATATTAATACCATTTTTGATACTATTTATTTCTATACCTTTCGAAATTAGTAATTTCAATAAGTCTATATCGATGTTATCAGAACATAGATATTCAAATATATCAAAATCATTGATATTTTTATAGTCGACTGACGACAATAACAAAATCACAACATCGTTTTTGATATTATTATTTTTTTTGGTAACGTATGCCTTTAATGGAGTTTCACCATCATACTCATATAATGGATTTGCACCACTTTCTATTAATGATTGTGCAATACTGGCATCGATGTTAAATGTTTTACAACTATCATAGAGTATCTTATCGTTAACCATGATTGGTTGTTGACGTTATCACATTTTTTGGTTTCTTTCATTTCAGTTATGTATGGATTTAGCACGTTTGGGAAGCATGAGCTCATATGATTTCAGTACTGTAGTGTCAGTACTATTAGTTTCGATCAGATCAATGTCTAGATCTATAGAATCAAAACACGATAGGTCAGAAGATAATGAATATCTGTACGCTTCTTCTTGTACTGTAACTTCTGGTTTTGTTAGATGGTTGCATCGTGCTTTAACGTCAATGGTACAAATTTTATCCTCGCTTTGTGTATCATATTCGTCTCTAGTATAAAATTCTATATTCAAATTATCATGCGATGTGTATACGCTAACGGTATCAATAAACGGAGCACAGCATTTAGTCAACAGTAATCCAAATTTTTTTAAAGTATATCTTAACGAAAGAAGTTGTCATCGTTAGAGTGTGGTAAATCATTGTCTACGGTACTAGATCCTCATAAGTGTATATATCTAGAGTAATATTTAATTTATCAAATGGTTGATAATATGGATGTCGTGGCAATTTCCTAATACGGAAATAAGACATAAACACGCAATAAATCTAATTGCGGACATGTTACACTCCTTAAAAATACGAATAAAAACTTTGGCTTTTAGTAAGTGTCATTTAACACTATACTCATATTAATCCATGGACTCATAATCTCTATACGGGATTAACGGATGTTCTATATACGGGGATGAGTAGTTCTCTTCTTTAATTTTATACTTTTTACTAATCATATTTAGACTGATGTATGGGTAATAGTGTTTGAAGAGCTCGTTCTCATCATCAGAATAAATCAATATCTCTGTTTTTTTATTATACAGATGTATTACAGCCTCATATATTACGTAATATAACGTGTAATCTACCTTATTAACTTTCACCGCATAGTTGTTTGCAAATACGGTTAATCCTTTGACCTCGTCGATTTCCGACCAATCTGGGCGTATAACGAATCTTAACTTTAATTTCTTGTAATCATTCGAAATAATTTTTAGTTTGCATCCGTAGTTATCTCCTCTATGTAACTGTAAATTTCTCAACGCGATATCTCCATTAATAATGATGTCGAATTCGTGTTGTATACCCATACTGAATTGATGAACGAATACCGACGGTGTGTGTTAATAGTAATTTACTTTTCATCTTTACATACTTGGTAATAGTTTTACTATCATAAGTTTATAAATTCCACAAGCTACTATGGAATATACCAACCATCTTAGTATAGAACACATGTCTTAAAGTTATTAATTAATTACATGTTGTTTTATATATCGCTACGAATTTAAACAGAGAAATCAGTTAGGAAGAAAAAATTATCTATCATCATCTATTGGATAACGTCTCTGTATTCTACGATAGAGTGCTATTTTAAGATGTGACAGATCCGTGTCATCAAATATATACTCCATTAAAATGATTATTCCGGCAGCGAACTTGATATTGGATACATCACGACCTTTGTTAATATCCACGACAATAGACAGCAATCCCATGGTTCCATAAACAGTGAGTTTATCTTTCTTTGAAGTGATATTTTGTAGAGATCTTATAAAACTGTCGAATGACATCGTATTTATATCTTTAGCTAAATCATATATGTTACCATCATAATATCTAACAGCATCTATCTTAAACGTTTCCATCGCTGTAAAGACGTTTCCGATAGATGGTCTCGTTTCATCAGTCATACTGAGCCAACAAATGTAATCGTGTATAACATCTTTGATAGAATCAGACTCTAAAGAAAAGGAATCGGCTTTATTATACACATTCATGATAAACTTAATGAAAAATGTTTTTCGTTGTTTAAGTTGGATGAATAGTATGTCTTAATAATTGTTATTATTTCACTAATTAATATTTAGTAACGAGTACACTCTATAAAAACGAGAATGACATAACTAATCATAACTAGTTATCAAAGAATGTCTAGGACGCGTAATTTTTTATGGTATAGATCCTGTAAGCGTTGTCTGTATTCTGGAGCTATTTTCTCTATCGCATTAGTGAGTTCAGAATATGTTATAAATTTAAATCGAATAACGAACATAACTTTAGTAAAGTCGTCTATATTAACTCTTTTATTTTCTAGCCATCGTAATACCATGTTTAAGATAGTATATTCTCTAGTTACTACGATCTCATCGTTGTCTAGAATATCGCATACTGAATCTACATCCAATTTTAGAAATTGGTCTGTGTTACATATCTCTTCTATATTATTGTTGATATATTGTCGTAGAAAACTATTACGTAGACCATTTTCTTTATAAAACGAATATATAGTACTCCAATTATCTTTACCGATATATTTGCATACATAATCCATTCTCTCAATCACTACATCTTTAAGAGTTTGGTTGTTAAGATATTTGGCTAAACTATATAATTCTATTAGATCATCAACAGAATCAGTATATATTTTTCTAGATCCAAAGATGAACTCTTTGGCATCCTCTATAATATTATCAGAAAAGATATTTTCGTGTTTTAGTTTATCAAGATCTAACCTGTTCATATCCATGATTAACGACGTCATATAACCACATAAAATAAAAATCCATTTTCATTTTTAGCACAATACTATTCATAATTGATATTGATGTAATATTTTGTTACTTTGAACGTAAAGACAGTACACGGGTCCGTATCTCCAACAAGCACGTAGTAATCAAATTTGGTGTTGTTAAACTTCGCAATATTCATCAATTTAGATAGAAACTTATACTCATCATCTGTTTTAGGAATCCATGTATTATTACTTTCCAACTTATCATTATCCCAGGCTATGTTTCGCCCATCATCGTTGTACAGAGTGAATAATTCTTTTGTATTCGGTAGTTCAAATATATGATCCATGCATATATCGACAAAGCTATTGTAGATGTGATTTTTCCTAAATCTAATATAAAACTCGTTTACTAGCAAACATTTTCCTGATTTATCGACCAAGACACACATGGTTTCTAAATCTATCAAGTGGTGGGGATCCATAGTTATAACGCAGTAACATAGATTATTACCTTCTTGACTGTCGCTAATATCTATATACTTATTGTTATCGTATTGGATTCTACATATAGATGGCTTGTATATCAAAGATATAGAACACATAACCAATTTATATTCTCGCTTTGTATTTTCGAATCTAAAGTTAAGAGATTTAGAAAACATTATATCCTCGGATGATGATATCACTGTTTCCAGAGTAGGATATATTAAAGTCTTTAAAGATTTTGTCCGATTCAAATAAATCACTAAATAATATCCCATATTATCATCTGTTATAGTCATGTCATTAAATCTATTATATTTTATGAAAGATATATCACTGCTCACCTCTATATTTCGTACATTTTTAAACTGTTTGTATAATATCTCTCTAATACAATCAGATATATCTATTGTGTCGGTAGACGATACCGTTACATTTGAATTAATGGTGTTCCATTTTACAACTTTTAACAAGTTGACCAATTCATTTCTAATAGTATCAAACTCTCCATGATTAAATATTTTAATAGTATCCATTTTATATCACTACGGACATAAACCATTGTATAATTTTTATGTTTATTAGTGTACACATTTTGGAAGTAAGTTCCGGCTGCCATGTATTTCCTGGAGAGCAAGTAGATGATGAGGAACCAGATAGTTTATATCCATACTTGCACTTAAAGTCTACATTGTAGTTGTATGAGAGTATGATCTTTTAAGCCGCTAGAAGTTTTCCGTTTGATATAGGATGTGGACATTTAACAATCTGACACGTGGGTGGATTGGACCATTCTCCTCCTGAACACATGACACCAGAGTTACCAATCAACGAATATCCACTATTGCAACTATAAGTTACAATGCTCCCATCGATATAAAAATCCTCGTATCCGTTATGTCTTCCGTTGGATATAGATGGAGGTGATTGGCATTTAACAGATTCGCAAATAGGTGCCTCAGGATTCCATACCATAGATCCAGTAGATCCTAATTCACAATACGATTTAGATTCACCGATCAAATGATATCCGCTATTACAAGAGTACGTTATACTAGAGCCAAAGTCTACTCCGCCAATATCAAGTTGGCCATTATCGATATCTCGAGGCGATGGGCATCTCCGTTTAATACATTGATTAAAGAGTGTCCATCCGGTACCGGTACATTTAGCATATATGGGTCCCATTTTTTGCTTTCTGTATCCAGGTAGACATAGATATTCTATAGTGTCTCCTATGTTGTAATTAGCATCAGTCTCTACACTATTCTTAAATTTCATATTAATGGGGCGTGACGGAATAGTACAGTATGATAGAACACATCCTATTCCCAACAATGTCAGGAACGTCACGCTCTCCACCTTCATATTTATTTATCCGTAAAATGTTATCCTGGACATCGTACAAATAATAAAAAGCCCATATATATGTTCGCTATTGTAGAAATTGTTTTTCACAGTTGCTCAAAAACAATGGCAGTGACTTATGAGTTAGTTACACTTTGGAGTCTCATCTTTAGTAAACATATCATAATATTCGATATTACGAGTTGACATATCGAACAAATTCCAAGTATTTGATTTTGGATAATATTCGTATTTTGCATCTGCTATAATTAAGATATAATCACCACAAGAACACACGAACGTCTTTCCTACATGGTTAAAGTACATGTACAATTCTATCCATTTGTCTTCCTTAACTATATATTTGTATAGATAATTACGAGTCTCATGAGTAATTCCAGTAATTGCATAGATGTCACCATCGTATTCTACAGCATAAACTATACTATGACGTCTAGGCATGGGAGACTTTTTTATCCAACGATTTTTAGTGAAACATTCCACATCGTTTAATACTACATATTTCTCATAGTGGTATAAACTCCACCCATTACATATATATCATCGTTTACGAATACTGATGCGCCTGAATATCTAGGAGTGATTAAGTTTGGAAGTCTTTTCCATTTCGAAGTGCCGTGTTTCAAATATTCTGCTATACCCGTTGAAATAGAAAATTCTAATCCTCCTATTACATATAACTTTCCATCGTTAACACAAGTACTAACTTCTGATTTTAACGACGACATATTAGTAACCGTTTTCCATTTTTTTTGTTTTAAGATCTACCCGCGATACGGAATAAACATGTCTATTGTTAATCATGCCGCCAATAATGTATAGACAATTATGTAAAACATTTGCATCATAGAATTGTCTATCTGTATTACCGACTATCGTCCAATATTCTGTTCTAGGAGAGTAATGGGTTATTGTGGATATATAATCAGAGTTTTTAATGACTACTATATTATGTTTTATACCATTTCGTGTCACAGCTTTGTAGATTTGGATATAGTTAATCCCAACAATGCTATAGCATTGCATATAGCATTAGTCATAAACTTGGGATGTAAAATGTTGATGATATCTACATCGTTTGGATTTTTATGTATCCACTTTAATAATATTATAGCGTAACATCCTCATGATTTACGTTAACGTTTTCGTGTGATAAGATAGTGGTCAGTTCATCCTTTGATAATTTTCCAAATTCTGGATCGGATGTCACCGCAGTAATATTGTTGATTATTTCTGACATCGACGCATTATATAGTTTTTTAATTCCATATCTTTTAGAAAAGTTAAACATCCTTATACAATTTGTGGAATTAATATTATGAATCATGGTTTTTACACATAGATCTACTACAGGCGGAACATCAATTATTATGGCAGCGACTAGTATCATTTCTACATTGTTTATGGTGATGTTTATCTTCTTCCAGCGCATATAGTCTAATATCGATTCAAACGCGTGATAGTTTATACCATTCAATATAATCGCTTCATCCTTTAGATGGTGATCCTGAATGTGTTTAAAAAATTATACGGAGACGCCGTAATAATTTCCCCATTGATAGAAAATATCACGCGTTCCATTCTCTTGAAGTACTATAAGTAATTATAATATAATGTAAAGGTTTATATATTCAATATTTTTTTTATAAAAAAAATCATTTCGACATTAATTCCTTTTTAAATTTCCGTCTATCATCTATAGAAACATATTCTATGAATTTATAAAATGCTTTTACGTATCCTATCGTAGGCGATAGAACCGCTAAAAAGCCTATCGAATTTCTACAAAAGAATCTGTTATATGGTATAGGGAGAGTATAAAACATTAAATGTCCGTACTTATTAAAGTATTCAGTAGCCAATCCTAACTCTTTCGAATAATTATTAATGGCTCTTATTCTGTACGAATCTATTTTTTTGAACAATGGACCTAGTGGTATATCTTGTTCTATGTATCTAAAATAATGTCTGACTAGATCCGTTAGTTTAATATCCGCAGTCATCTTGTCTAGAATGGCAAATCTAACTGCGGGTTTAGGTGTAGGCGTTAGTTTAGTTTCTATATCTACATCTATGTCTTTATCTAACACCAAAAATATAATAGCTAATATTTTATTACAATCATCCGGATATTCTTCTACGATCTCACTAACTAATGTTTCTTTGGTTATACTAGTATAGTCACGATCAGACAAATAAAGAAAATCAGATGATCGATGAATAATACATTTAAATTCATCATCTGTAAGATTTTTGAGATGTCTCATTAAAATATTATTAGTGTCAGTTCTCATTATCATATATTGACAGCAGCTATTACACTTATTTTATTTTTCTGTATTTTATTACTTTTCACCATATAGATCAGTCATTAGATCATCAAAATACTTTTCAATCATCCTAAAGAGTATGGTGAACGAATCTTCCCATCTAATTTCTGAACGTCTACCAATGTCTCTAGCCACTTTGGCACTAATAGCGATCATTCGCTTAACATCTTCTACATTATTAACTGGTTGATTCAATCTATCTAGCAATGGACCGTCGGATAGCGTCATTCTCATGTTCTTAATCAATGTACATACATCGTCATCATCTACCAATTCATCAAACAATATAAGCTTTTTAAAATCATCATTATAATAGGATGGATCGCCGTCATTTCTCCAAAGAATATATCTAATAAGTAGAGTCCTCATGCTTAGTAATTTAACTATTTTAGTTAACAACTATTTTTTATGTTAAATCAATTAGTAACACCGCTATGTTTAATACTTATTCATATTTTAGTTTTAGGATCGAGAATCAATACAAAAATTAATACATCAATTTTGGAAATACTTAGTTTCCACGTAGTCAATGAAACATTTGAGCTCATCGTAAAGGACGTTCTCGTACAGGACGTAACTATAAATTGGTTTATATTTGTTCAAGATAGATACAAATCCGATAACTTTTTTGACGAATTCTACGGGATTCACTTTAAAAGTGTCATACCGGGTTCTTTTTATTCTTTTAAACAGATCGATTGTGTGATGTTGATTAGGTCTTTTACGAATTTGATACAGAATAGCGTTTACATATCCACCATAGTAATCAATAGCCATTTGTTCGTATGTCATAAATTCTTTAATTATATGACACTGTGTATTATTTAGTTCGTCCTTGTTCATCATTAGGAATCTATCCAATATGGCAATTATATTAGAACTATAACTGCGTTGTATGCGCATGTTGATGTGTCTGTTTATACAATCAATTATACTAGGATCCATACCACTACATTCGGGTAAAATTGTAGCATCATATACCATTTCTAGTACTTTAGGTTCATTGTTATCCATTGCAGAGGACGTCATGATCGAATCCAAAAAAATATATTATTTTTATGTTATTTTGTTAAAAATAATCATCAAATACTTCGTAAGATACTCCTTCATGAACATAATCAGTTACAAAACGTTTATATGAAGTAAAGTATCTACGATTTTTACAAAAGTCAGGATGCATAAGTACAAAGTACGCGATAAACGGAATAATAATAGATTTATCTAGTTTATCTTTTTCTATCTCTTTCATAGTTATATACATGGTCTCAGAAGTCGGATTATGTAACATCAGCTTCGATAAAATGACTGGGTTATTTAGTCTTACACATTCGCTAATACATGTATGACCGTTAACTATAGAGTCTACACTAAAATGATTGAATAATAGATAGTCTACCATTGTTTCGTATTCAGATAGTACAGCGTAGTACATGGCATCTTCACAAATTATATCATTATCTAATAGATATTTGACGCATCTTATGGATCCCACTTCAACAGCCATCTTAAAATCGGTAGAATCATATTGCTTTCCTTTATCGTTAATAATTTCTAGAACATCATCTCTATCATAAAAGATACAAATATTAACTGTTTGATCAGTAATAACATTGCTAGTCGATATCAATTTGTTAATAAGATGCGCTGGGCTCAATGTCTTAATAAGAAGTGTAAGAGGACTATCTCCGAATTTGTTTTGTTTATTAACATCCGTTGATGGAAGTAAAAGATTTATAATGTCTACATACTTGACTGTTTTAGAGCATACAATATGGAGAGGCGTATTTCCATCATGATCTGGTTTTGAGGGACTAATTCCTAGTTTCATCATCCATGAGATTGTAGAAGCTTTTGGATTGTCTGACATAAGATGTCTATGAATATGATTTTTGCCAAATTTATCCACTATCCTGGCTTCGAATCCGATAGACATTATTTTTTTAAACACTCTTTCTGAAGGATCTGTATACGCCAACAACGGACCACATCCTTCTTCATCAACCGAGTTGTTAATCTTGGCTCCATACTGTACCAATAAATTTATTCTCTCTATGACTTCATCATCTGTTCCCGAGAGATAATATAGAGGTGTTTTATTATGTTTATCACATGCGTTTGGATCTGCGCCGTGCACCAGCAGCATCGCGACTATTCTATTATTATTAATTTTAGAAGCTATATGCAATGGATAATTTCCATCATCATCCGTCTCATTTGGAGAGTATCCTCTATGAAGAAGTTCTTCTATAAATCGTTCATCTAGTCCTTTAATGCCACAATACGCATGTAGAATGTGATAATTTCCAGAGGGTTCGATAACTTGTAGCATATTCCTAAATACATCTAAATTTTTACTATTATATTTGGCATAAAGAGATAGATAATACTCGACCGACATAATGTTGTGTTGTCCATTATAGTATAAAAATTAATATTTCTATTTCTATTTCTATATATTTGCAACAATTTACTCTCTATAACAAATATCATAACTTAGTTCTTTTATGTCAAGAAGGCACTGGTTTAATTCATCTATAAATGTCACGCCATAACTACCACGCATACTATACTCAGAATTATGATAAAGATATTTATTCTTGGGGTGTAAGTAATGGGGATTAATCTTTGTTGGATCAGTCTCTAAGTTAACACATGTCACACATGATCCATTTATAGTTATATCACACGATGATGATTTATGAATTGATTCCGGAAGATCGCTATTGTATTTTGTAGTTCCACAATTCATTTCCATACATGTTATTGTCACACTAATATTATGATGAACTTTATCTAGCCGCTGAGTGGTAAACAACAGAACAGATAGTTTATTATCTTTACCAACACCCTCAGCCGCTGCCACAAATCTCTGATCCGTATCCATGATGGTCATGTTTATTTTTAGTCCGTATCCAGTCAACACTATGTTAGCATTTCTGTCGATATAGCTTTCACTCATATGACACTCACCAATAATTGTAGAATTAATGTCGTAATTTACACCAATAGTGAGTTCGGCGACAAAGTACCAGTACCGGTAATCTTGTCGAGGAGGACATATAGTATTCTTGTATTCTACCGAATACCCGAGAGATGCGATACAAAAGAGTAAGACTAATTTGTAAACCATCTTACTCAAAATATGCGACAATAGTACGATGCAATGAGTAAGACAATAGGAAATCTATCTTATACACATAATTATTCTATCAATTTTACCAATTAGTTAGTGTAATGTTAACAAAAATGTGGGATAATTTAATAGTTTTTCCTTACATAATTGACATACATGAGTCTGAGTTCCTCGTTTTTGCTAATTATTTCGTCCAATTTATTATTCTTGACATCGTCAAGATCTTTTGTATAGGAGTCAGACTTGTATTCAACATGTTTTTCTATAATCATCTTAGCTATTTCGGCATCATCCAATAGTACATTTTCCAGATTAACAGAATAGATATTAATGTCGTATTTGAACAGAGCCTGTAACATCTCAATGTCTTTATTATCTATAGCCAATTTGATGTCCGGAATGAAGAGAAGGGAATTGGTGTTTGTCGACGTCATATAGTCGAGCAAGAGAATCATCATATCCACGTGTCCATTTTTTATAGTGGTGTGAATACAACTAAGGAGAATAGCTAGATCAAAAGGAGATGGTATCTCTGAAAGAAAGTAGGAAACAATACTTACATCATTAAGCATGACAGCATGATAAAATGAAGTTTTCCATCCAGTTTTCCCATAGAACATCAGTCTCCAATTTTTCTTAACAAACAGTTTTACCGTTTGCATGTTACCACTATCAACCGCATAATACAATGCGGTGTTTCCTTTGTCATCAAATTGTGAATCATCCATTCCACTGAATAGCAAAATCTTTACTATTTTGGTATCTTCTAATGTGGCTGCCTGATGTAATGGAAATTCATTATCTAGAAGATTTTTCAATGCTCCAGCGTTCAACAACGTACATACTAGACGCACGTTATTATCAGCTATTGCATAATACAAGACACTATGACCGTTGATATCCGCCTTAAATGCATCTTTGCTAGAGAGAAAGCTTTTCAGTTGCTTAGACTTCCAAGTATTAATTCGTGACAGATCCATGTCTGAAACGAGACGCTAATTAGTGTATATTTTTTCATTTTTTATAATTTTGTCATATTGCACCAGAATTAATAATATCTCTAATAGATCTGATTAGTAGATACATGGCTATCGCAAAACAACATATACACATTTAATAAAAATAATATTCATTAAGAAGATTCAGATTCCACTGTACCCATCAATATAAATAAAATAATTATTCCTTACATCGTACCATAAACAATATATTAAGTAGATTCCACCTTACCCATAAACAATATAAATCCAGTTATATCATGTCTAATGATGAACACAAATGGTGTATTAAATTCCAGTTCTTCAGGAGATGATCTCGCCGTAGCTACCATGATAGTAGATGCCTCCGCTACAGTTCCTTGTTCGTCTACATCTATCTTTACATTCTGAAACATTTTATAAATATATAATGGGTCCCTAGTCATATGTTTAAACGACGCCTTATCTGGATTAAACATACTAGGAGCCATCATTTCGGCTATCGACTTAATATCCCTCTTGTTTTCGATAGAAAATCTAGGGAGTTTAAGATTGTACATTTTATTCCCTAATTGAGATGACCAATATTCTAATTTTGCAGCCGTGATAGAATCTGTGAAATGGGTCATATTATCACCTATTGCCAGGTACATACTAATATTAGCATCCTTATACAGAAGGCGCACCATATCATATTCTTCGTCATCGATTGTGATTGTATTTCCTTGCAATTTAGTAACTACGTTCATCATGGGAACCGTTTTCGTACCGTACTTATTAGTAAAACTAGCATTGTGTGTTTTAGTGATATCAAACGGATATTGCCATGTACCTTTAAAATATATAGTATTAATGATTGCCCATAGAGTATTATCGTCGAGCATAGTAGAATCAACTACATTAGACATACCAGATCTACGTTCTACTATAGAATTAATTTTATTAACCGCATCTCGTCTAAAGTTTAATCTATATAGGCCGAATCTATGATATTGTTGATAATACGACGGTTTAATACACACAGTACTATCGACGAAACTTTGATACGTTAGATCGGTGTACGTATATTTAGATGTTTTCATCTTAGCTAATCCTGATATTAATTCTGTAAATGCTGGACCCAGATCTCTTTTTCTCAAATTCATAGTATTCAATAATTCTACTCTAGTATTACCTGATGCAGACAATAGCGACATAAACATAGAAAACGAATACCCAAACGGTGAGAAGACAATATTATCATTATCATCCTCATCCCCATTTTGAATATTTTTATACGCTAATATACCGGCATTGATAAATCCCTGCAGACGATATGCGGATACTGAACACGCTAATGATAGTATCAATAACGCAATCATGATTTTTATGGTATTAATAATTAACCTTATTTTTATGTTTGGTATAAAATTTATTGATGTCTACACATCCTTTTGTATAATCAACTCTAATCACTTTAACTTTTACAGTTTTCCCTACAAGTTTATCCCTATATTCAACATATCTATCCATATGCATCTCTTAACACTCTGCCAAGATAGCTTCAGAGTGAGGATAGTCAAAAAGATAAATATATAGAGTATAATCATTCTCGTATACTCTGCCCTTTATTACATCGCCCGCATTGGGCAACGAATAACAAAATGCAAGCATCTTGTTAACAGGCTCGTAAATTGGGATAAAATTATGTTTTTATTGTTTATCTATTTTATTCAAGAGAATATTCAGGAAGTTCCTTTTCCGGTTGTATCTCGTCGCAGTATATATCATTTGTACATTGTTTCATATTTTTTAATAGTCTACACCTTTTAGTAGGACTAGTATCGTACAATTCATAGCTGTATTTTGAATTCCAATCACGTATAAAAATATCTTCCAATTGTTGACGAAGACCTAATCCATCATCCGGTGTAATATTAATAGATGCTCCACATATATCCGTAAAGTAATTTCCTGTCCAATTTGATGTACCTATATACGCCGTTTTATCGGTTACCATATATTTTGCATGGTTTACCCTAGAATACGGAATGGGAGGATCAGCATCTGGTACAATAAATAGCTTTACTTCTATATCTATGTTTTTAGATTTTAGCATAGCTATAGATCTTAAAAAGTTTCTCATGATAAACGAAGATCGTTGCCAGCAACTAATCAATAGCTTAACGGATACTTGTCTGTCTATAGCGGATCTTCTTAATTCATCTTCTATATAAGGCCAAAACAAAATTTTACCCGCCTTTGAATAAATAATAGGAATAAAGTTCATAACAGATACATAAACGAATTTACTCGCATTTCTGATACATGACAATAAAGCGGTTAAATCATTGGTTCTTTCCATAGTACATAATTGTTGTGGTGCAGAAGCAATAAATACAGAGTGTGGAACACCGCTTACGTTAATACTAAGAGGATGATCTGTATTATAATACGACGGATAAAAGTTTTTCCAATTATATGGTAGATTGTTAACTCCAAGATACCAGTATACCTCAAAAATTTGAGTGAGATCCGCTGCCAAGTTCCTATTATTGAAGATCGCAATACCCAATTCCTTGACCTGAGTTAGTGATCTCCAATCCATGTTAGCGCTTCCTAAATAAATATGTGTATTATCAGATATCCAAAATTTTGTATGAAGAACTCCTCCTAGGATATTTGTAATATCTATGTATCGTACTTCAACTCCGGCCATTTGTAGTCTTTCAACATCCTTTAATGGTTTGTTGGATTTATTGACGGCTACTCTAACTCTTACTCCTCTTTTGGGTAATTGTACAATCTCGTTTAATATTACCGTGCCGAAATTCGTACCCACTTCATCCGATAAACTCCAATAAAAAGATGATATATCTAGTGTTTTTATGGTATTGGATAGAATTTCCCTCCACATGTTAAATGTAGTCAAATATACTTTATCAAATTGCATACCTATAGGAATAGTCTCTGTAATCACTGCGATTGTATTATCCGGATTCATTTTATTTGTTAAAAAATAATCCTATATCACTTCACTCTATTAAAAATCCAAGTTTCTATTTCTTTCATGACTGATTTTTTAACTTCATCCGTTTCCTTATGAAGATGATGTTTGGCACCTTCATAAATTTTTATTTCCCTATTACAATTTGCATGTTGCATGAAATAATATGCACCTGAAACATCGCTAATCTCATTGTTTGTTCCCTGGAGTATGAGAGTCGGGGTGTCAATCTTGGGAATTATTTTTCTAACCTTGTTGGTAGCCTTCAAGACCTGACTAGCAAATCCAGCCTTAATTTTTTCATGATTGACTAATGGATCGTATTGGTATTTATAAACTTCATCCATATCTCTAGATACTGATTCTGGACATAGCTTTCCGACTGACGCATTTGGTGTAATGGTTCCCATAAGTTTTGCAGCTAGCAGATTCAGTCTTGGAACAGCGTCTGCATTAACTAGAGGAGACATTAGAATCATTGCTGTAAACAAGTTTGGATTATCGCAAGCAGCCAGTATAGAAATTGTTGCTCCCATGGAATGACCCAATAAGAAGACTGGAACTCCTGGATAAGTAGATTTAATAGTCACTACGTGCTGTACCACATCTCTAACATACTTACCAAAGTCATCAATCATCATTTTTTCACCATTACTTCTTCCATGGCCAATATGATCATGTGAGAATACTAAAATTCCTAACGATGATATGTTTTCAGCTAGTTCGTCATAACGTCCAGAATGTTCACCAGCTCCATGACTTATGAATACTAATGCCTTAGGATATGTAATAGGTTTCCAATATTTACAATATATGTAATCATTGTCCAGATTGAACATACAGTTTGCACTCATGATTCACTATATAACTATCAATATTAACAGTTCGTTTAATGATCATATTATTTTTATGTTTTATTGATAATTGTAAAAATATACAATTAAATCAATATAGAGGAAGGAGACGGTACTGTATTTTGTGAGATAGTCATGGAGACTAAATCAGATTATGAGGATGCTGTTTTTTACTTTGTGGATGATGATGAAATATGTAGTCGCGACTCCATCATTGATCTAATAGATGAATATATCACGTGGAGAAATCATGTTATAGTGTTTAATAAAGATATTACCAGTTGTGGAAGACTGTACAAGGAATTGATAAAGTTCGATGATGCCGCTATACGGTACTATGGTATTGATAAAATTAATGAGATTGTCGAGGCTATGAGTGAAGGAGACCACTACATCAATCTTACAGAAGTCCATGATCAGGAAAGTCTATTCGCTACCATAGGAATATGTGCTAAAATCGCTGAACATTGGGGATACAAAAAGATTTCAGAATCTAAATTCCAATCATTGGGAAACATTACAGATCTGATGACCGACGATAATATAAACATCTTGATACTTTTTCTAGAAAAAAAATTGAATTGATGATATAGGTGTCTTCATAACGCATTATTACGTTAGCATTCTATTATCCTATCATGTATTTGAGAGTCTTATATGTAGCAAACATGATAACTGCAATACCCATAATCTTTAGATATTCACGCGTGCTATGGATGGCATTATCCCGCGGTGTGGAAATGTACGTTATATAATCTACAAAATAATCATCGCATATAGTATGAGATAGTAGAGTAAACATTTTTATCGTTTCTACTGGGTTCATACATCGTCTACCCAATTCGGTAATGAATGAAATTGTCGCCAATCTTACACCCAAACCCTTGTTGTTCATTAGTATAGTATTAACTTCATTATTTATGTCATAAACTGTAAATGATTCTGTAGATGCCATATCACACATGATATTCATGTCACTATTATAATCATTATTAACTTTATCACAATACGTGTTGATAATATCTACATATGATCTAGTTTTTGTGGGTAATTGTCCATACAAGTCGTCTAAACGTTGTTTACTCATATAGTATTGAACAGCCATCATTACATGGTCCCGTTCCGTTGATAGATAATCGAGTATGTTAGTAGACTTGTCAAATCTATATACCATATTTTCTGGAAGCGGATATACATAGTCGCGATCATCATTATCACTAGCCTCATCCTCTATATCATGTACATGTACATAATCTATGATATTATTATACATAAACATCGACAACATACTATTGTCTATTATCTAAGTCCTGTTGATCCAAACCCTTGATCTCCTCTATCTGTACTATCTAGAGATTGTACTTCTTCAAGTTCTGGATAATATATACGTTGATAGATTAGCTGAGCTATTCTATCTCCAGTATTTACATTAAACGTACATTTTCCATTATTAATAAGAATGACTCCTATGTTTCCCCTATAATCTTCGTCTATTACACCGCCTCCTATATCAATGCCTTTTAGGGACAGACCAGACCTAGGAGCTATTCTACCATAGCAGAACTTAGGCATGGACATACTAATATCTGTCTTAATTAACTGTCGTTCTCCAGGAGGGATAGTATAATCGTAAGCGCTATACAAATCATATCCGGCAGCACCCGGCGATTGCCTAGTAGGCGATTTAGCTCTGTTAGTTTCCTTAACAAATCTAACTGGTGAGTTAATATTCATGTTGAACATAAAAAATATCATTTTATTTCAAAATTATTTACCATCCCATCCCATCCCATCCCATCCCATTCCATATATTCCATGAATAAGTGCGATTATTGTACACTTCTATAGTATCTATATACGATCCACGATAAAATCCTCCTATCAATAGCAGTTTATTATCCACTATGATCAATTCTGGATTATCCCTCGGATAAATAGGATCATCTATCAGAGTCCATGTATTACTGGATTCACAATAAAATTCCGCATTTCTACCAACCAAGAATAACCTTCTACCAAACACTAACGCACATGATTTATAATGAGGATAATAAGTGGATGGTCCAAACTGCCACTGATCATGATTGGGTAGCAAATATTCTGTAGTTGTATCAGTTTCAGAATGTCCTCCCATTACGTATATAACATTGTTTATGGATGCCACTGCTGGATTACATCTAGGTTTCAGAAGACTCGGCATATTAACCCAAGCAGCATCCCCGTGGAACCAACGCTCAACAGATGTGGGATTTGGTAGACCTCCTACTACGTATAATTTATTGTTAGCGGGTATCCCGCTAGCATACAGTCTGGGGCTATTCATCGGAGGAATTGGAATCCAATTGTTTGATATATAATTTACCGCTATAGCATTGTTATGTATTTCATTGTTCATCCATCCACCGATGAGATATACTACTTCTCCAACATGAGTACTTGTACACATATGGAATATATCTATAATTTGATCCATGTTCATAGGATACTCTATGAATGGATACTTGTATGATTTGCGTGGTTGTTTATCACAATGAAATATTTTGTTACAGTCTAGTATCCATTTTACATTATGTATACCTCTGGGAGAAAGATAATTTGACCTGATTACATTTTTGATAAGAAGTAGCAGATTTCCTAATCTATTTCTTCGCCTCATATACCACTTAATGACAAAATCAACTACATAATCCTCATCTGGAACATTTAGTTCGTCGCTTTCTAGAATAAGTTTCATAGATAGATAATCAAAATTGTCTATGATGTCATCTTCTAGTTCCAAAAAGAGTTTGGTAATAAAGTCTTTAGTATGACATAAGAGATTGGATAGTCCGTATTCTATACCCATCATGTAACACACGATACAATATTCCTTTCTAAAATCTCGTAGGATAAAGTATATACAAGTGTAGATGATAAATTCTACAGATGTTAATATAGAAGCACGTAATAAATTGACGACGTTATGACTATCTATATATACCTTTCCAGTATATGAGTAAATAACTATAGAAGTTAGACTGTGAATGTCAAGGTCTAGACAAACCCTCGTAACTGGATCTTTATTTTTTGTGTATTTTTGACGTAAATGTGTGCGAAAGTATGGAGATAACTTTTTCAATATTGTAGAATTGACTATTATATTGCCTCCTATAGCTTCAATAATTGTTTTGAATTTCTTAGTCGTATACAATGCTAATATATTCTTACAGTACACAGTATTGACAAATATCGGCATTTATGTTTCTTTAAAAGTCAACATCTAAAGAAAAATGATTGTCTTCTTGAGACATAACTCCCATTTTTTGGTATTCACCCACACGTTTTTCGAAAAAATTAGTTTTTCCTTCCAATGATATATTTTCCATGAAATCAAACGGATTGGTAACATTGTAAATTTTTTTAAATCCCAATTCAGAAATCAATCTATCTGCGACGAATTCTATATATGTTTTCATCATTTCACAATTCATTCCTATGAGTTTAACTGGAAGAGCCACAGTAAGAAATTCTTGTTCAATGGATACCGCATTTGTTATAATAAATCTAACGGTTTCTTCACTCGGTGGATGTAATAAATGTTTAAACATCAAACATGCGAAATCGCAGTGCAGACCCTCGTCTCTACTAATTAATTCGTTAGAAAACGTGAGTCCGGGCATTAGGCCACGCTTTTTAAGCCAAAATATGGAAGCGAATGATCCGGAAAAGAAGATTCCTTCTACTGCAGCAAAGGCAATAAGTCTCTCTCCATAACCGGCGCTGTCATGTATCCACTTTTGAGCCCAATCGGCCTTCTTTTTTACACAAGGCATCGTTTCTATGGCATTAAAGAGATAGTTTTTTTCATTACTATCTTTAACATAAGTATCGATCAAAAGACTATACATTTCCGAATGAATGTTTTCAATGGCCATCTGAAATCCGTAGAAACATCTAGCCTCGGTAATCTGCACTTCTGTACAAAATCGTTCTGCTAAATTTTCATTCACTATTCCATCACTGGCTGCAAAAAACGCCAATACATGTTTTATAAAATATTTTTCGTCTGGTGTTAGTTTATTCCAATCATTGATATCTTTAGATATATCCACTTCTTCCACTGTCCAAAATGATGCCTCTGCCTTTTTATACATATTCCAGATGTCATGATATTGGATTGGGAAAATAACAAATCTATTTGGATTTGGTGCAAGGATAGGTTCCATAACTAAATTAACAATAGTAGTAATTTTTTTTTCAGTTATCTGTATGACTGTACTTGGATCTTTTGTATATCGCTATCGCCGCAATCACTACAATAATTACAAGTATTATTGATAGCATTGTTATTACTACTATCATAATTAAATTATCGACATTCGTGGGTGCTGAATAATCGTTATCATCATTTTGTAATTGTGACATCATACTAGATAAATCATTTGTGAGATTGTTGTGGGAAGCGGGCACGGAAGATGCATTATCATTATTATTTAACGCCTCCCATTCGGATTCACAAATGTTACGCACGTTCAACGTTTTATGGAAACTATAATTTTGTGAAAACAGATAACAAGAAAACTCGTCATCGTTCAAATTTTTAACGATAGTAAACCGATTAAACGTCGAGCTAATTTCTAACGCTAGCGACTCTGTTGGATATGGGTTTCCAGATATATATCTTTTCAGTTCCCCTACGTATCTATAATCATCTGTAGGAAATGGAAGATATTTCCATTTATCTACTGTTCCTAATATCATATGCGGTGGTGTAGAACCATTAAGCGCGAAAGATGTTATTTCGCATCGTATTTTAACTTCGCAATAATTTCTGGTTAGATAACGCACTCTACCAGTCAAGTCAATGATATTAGCCTTTACAGATATATTCATAGTAGTCGTAACGATGACTCCATCTTTTAGATGCGATACTCCTTTGTATGTACCAGAATCTTCGTACCTCAAACTCGATATATTTAAACAAGTTAATGATATATTAACGCGTTTTATGAATGATGATATATAACCAGAAGTTTTATCCTCTGTGGCTAGCGCTATAACCTTATCATTATAATACCAACTAGTGTAATTAATATGTGACACGACAGTGTGGGTACAAATATGTACATTATCGTCTACGTCGTATTTGATACATCCGCATACAGCCAACAAATATAAAATTACAAAAACTCTAACGACGTTCGTACACATCTTGATGCGGTTTAATAAATGTTTTGATTTCAATTTATTGTAAAAAAGATTCGGTTTTATACTGTTCGATATTCTCATTGCTTATATTCTCATCTATCATCTCCACACAGTCAAATCCATGGTTAACATGTACCTCATCAACCGGTAAAAGACTATCGGATTCTTCTATCATCATAACTCGAGAATATTTAATTTGGTGGTCATTATTAATCAAGTCAATTATCTTATTTTTAACAAACGTAAGTATTTTACTCATTTTTTATAAAAACTTTTAGAAATATACAGACTCTATCGTGTGTCTATATCTTCTTTTTATATCCAATGTATTTATGTCTGATTTTTCTTCATTTATCATATATAATGGTCCAAATTCTACACGTGCTTCGGATTCATCCAGATCATTAAGGTTCTTATAATCGCAACATCCTTCTCTTCCATCTTCTACATCTTCCTTCTTATTCTTAGCGTCACAGAATCTACCACAGCAGGATCCCATGACGAGCGTCACATTAAACTAATTCATTTTCAATTATAATATACTGATTAGTAATGACCATTAAAATAAAAATATTCTTCATAACCGGTAAGAAAGTAAAAAGTTCACATTGAAACTATGTCAGTAGTTATACATCATGAGATGATATACTCTATTTTGGTGGAGGATTATATGATATAATTCGTGGATAATCATTCTTAAGACACATTTCTTCATTCGTAAATCTTTTCACATTAAATGAGTGTCCATATTTTGCAATTTCTTCATATGATGGCGGTGTACGTGGACGAGGCTGCTCCTGTTCTTGTAGTCGCCGACTGTCGTGTTTGCGTTTAGATCCCTCCATTATCGCGATTGCGTAGTGAGTACTATTTATACCTTGTAATTAAATTTTTTTATTAATTAAACGTATAAAAACGTTCCGTATCTGTATTTAAGAGCCAGATTTCGTCTAATAGAACAAATAGCTACAGTAAAAATAACTAGAATAATCGCTACACCCACTAGAAACCACGGATCGTAATACGGCAATCGGTTTTCGATAATAGGTGGAACGTATATTTTATTTAAGGACTTAACAATTGTCTGTAAACCACAATTTGCTTCCGCCGATCCTGTATTAACTATCTGTAAAAGCATATGTTGGCCGGGCGGAGCCGAACATTCTCCGATATTCAATTTTTGTATATCTATAATGTTATTAACCTCCGCATACGCATTACAGTTCTTTTCTAGCTTGGATACTACACTAGGTACATCATCTAAATCTATTCCTATTTCCTCAGCGATAGCTCTTCTATCCTTTTCCGGAAGTAATGAAATCACTTCAATAAATGATTCAACCATGAGTGTGAAACTAAGTCGAGAATTACTCATGCATTTGTTAGTTATTCGGAGCGCGCAATTTTTAAACTGTCCTATAACCTCTCCTATATGAATAGCACAAGTGACATTAGTAGGGATAGAATGTTGAGCTAATTTTTGTAAATAACTATCTATAAAAAGATTATACAAAGTTTTAAACTCTTTAGTTTCCGCCATTTATCCAGTCTGAGAAAATGTCTCTCATAATAAATTTTTCCAAGAAACTAATTGGGTGAAGAATGGAAACCTTTAATCTATATTTATCACAGTCTGTTTTGGTACACATAATGAATTCTTCTAATGCTGTACTAAATTCGATATCTTTTTCGATTTCTGGATATGTTTTTAATAAAGTATGAACAAAGAAATGGAAATCGTAATACCAGTTATGTTCAACTTTGAAATTGTTTTTTATTTTCTTGTTAATGATACCAGCCACTTGGGAAAAGTCAAAGTCGTTTAATGCCGATTTAATACGTTCATTAAAAACAAACTTTTTATTCTTTAGATGAATTATTATTGGTTCATTGGAATCAAAAAGTAAGATATTATCGGGTTTAAGATCTGCATGTAAAAAGTTGTCACAACAGGGTAGTTCGTAGATTTTAATGTATAACAGAGACATCTGTAAAAAGATAAACTTTATGTATTGTACCAAAGATTTAAATCCTAATTTGATAGCTAACTCGGTATCTACTTTATCTGCCGAATACAGTACTAGGGGAAAAATTATAATATTTCCTCTTTCGTATTCGTAATTAGTTCTCTTTTCATGTTCGAAAAAGTGAAACATGCGGTTAAAATAGTTTATAACATTAATATTACTGTTAATAACTGCAGGATAAAAGTGGGATAGTAATTTCACGAATTTGATACTGTCCTTTCTCTCGTTAAACGCCTTTAGAAAAACTTTAGAAGAATATCTCAATGAGAGTTCCTGACCATCCATAGTTTGTATCAATAATAGCAACATATGAAGAACCCGCTTATACAGAGTATGTAAAAATGTTAATTTATAGTTTAATCCCATGGCCCACGCACACACGATTAATTTTTTTTCATCTCCCTTTAGATTGTTGTATAGAAATTTGGGTACTGTGAACTCCGCCGTAGTTTCCATGGGACTATATAATTTTGTGGCCTCGAATACAAATTTTACTACATAGTTATCTATCTTAAAGACTATACCATATCCTCCTGTAGATATGTGATAAAAATCGTCGTTTATAGGATAAAATCGTTTATCTTTTTGTTGGAAAAAAGATGAATTAATGTAATCATTCTCTTCTATCTTTAGTAGTGTTTCCTTATTAAAATTCTTAAAATAATTTAACAATCTAACTGACGGAGCCCAATTTTGGTGTAAATCTAATTGGGACATTATGTTGTTAAAATATAAACAGTCTCCTAATATAACAGTATCTGATAATCTATGGGGAGACATCCATTGATATTCAGGGGATGAATCATTGGCAACACCCATTTATTGTACAAAAAGCCCCAATTTACAAACGAAAGTCCAGGTTTGATAGAGACAAACTATTAACTATTTTGTCTCTGTTTTTAACACCTCCACAGTTTTTAATTTCTTTGGTAATGAAATTATTCACAATATCAGTATCTTCTTTATCTACCAGAGATTTTACTAACTTGATAACCTTGGCTGTCTCATTCAATAGGGTAGTGATATTTGTATGTATGATATTGATATCTTTTTGAATTGTTTCTTTTAGAAGTGATTCTTTGATGGTATCAGCATACGAATTACAATAATGCAGAAACTCAGTTAACATGCAGGAATTATAGTAAGCCAATTCCAATTGTTGCCTGTATTGTATTAGAGTATTAATATGCGCAATGATGTCCTTGCGTTTCTCTGATAGAATGCGAGCAGCGATTTTGGCGTTATCATTTGACGATATTTCTGGAATAACGAATCCTGTTTCTACTAACTTCTTGGTAGGACAAAGTGAAACAATCAAGAAAATAGCTTCTCCTCCTATTTGTGGAAGAAATTGAACTCCTCTAGATGATCTACTGACGATAGTATCTCCTTGACAGATATTGGACCGAACTACGGAAGTACCTGGAATGTAAAGCCCTGAAACCCCCTCATTTTTTAAGCAGATTGTTGCCGTAAATCCTGCACTATGCCCAAGATAGAGAGCTCCTTTGGTGAATCCATCACTATGTTTCAGTTTAACCAAGAAACAGTCAGCTGGTCTAAAATTTCCATCTCTATCTAATACAGAATCCAACTTGATGTCAGGGACTATGACCGGTTTAATGTTATATGTAACATTGAGTAAATCCTTAAGTTCATAATCATCGTTGTCATCAGTTATGTACGATCCAAACAATGTTTCTACCGGCATGGTGGATACGAAGATGCTATCCATCAGAATGTTTCCCTGATTAGTATTTTCTATATAGCTATTCTTCTTTAAACGATTTTCCGAATCAGTAACTATGTTCATTTTTTTAGGAGTAGGACGTCTAGCCAGTATGGAAGAGGATTTTCTAGATACTCTCTTCAACATCTTTGATCTCAATGGAATGCAAAACCCCATGGTGTAACAACCAACGATAAAAATAATATTGTTTTTTCACTTTTTATAATTTTACCATCTGACTCATGGATTCATTAATATCTTTATAAGAGCTACTAACGTATAATTCTTTATAACTGAACTGAGATATATACACCGGATCTATGGTTTCCATAATTGAGTAAATGAATGCTCGGCAATAACTAATGGCAAATGTATAGAACAACGAAATTATACTAGAGTTGTTAAAGTTAATATTTTCTATGAGTTGTTCCAATAAATTATTTGTTGTGACTGCGTTCAAGTCATAAATTATCTTGATACTATCCAGTAAACAGTCTTTAAGTTCTGGAATATTATCATCCCATTGTAAAGCCCCTAGTTCGACTATCGAATATCCTGCTCTGATAGCAGTTTCAATATCGACGGACGTCAATACTGTAATAAAGGTGGTAGTATTGTCATCATCGTGATAAACTACGGGAATATGGTCGTTAGTAGGTACCGTGACTTTACACAACGCGATATATAACTTTCCTTTTGTACCATTTTTAACGTAGTTGGGACGTCCTGCAGGGTATTGTTTTGAAGAAATGATATCGAGAACAGATTTGATACGATATTTGTTGGATTCCTGATTATTCACTATAATATAATCTAGACAGATAGATGATTCGATAAATAGAGAAGGTATATCGTTGGTAGGATAATACATCCCCATTCCAGTATTCTCGGATACTCTATTGATGACACTAGTTAAGAACATGTCTTCTATTCTAGAAAACGAAAACATCCTACATGGACTCATTAAAACTTCTAACGCTCCTGATTGTGTTTCGAATGCCTCGTACAAGGATTTCAAGGATGCCATAGATTCTTTGACCAACGATTTAGAATTGCGTTTAGCATCTGATTTTTTTATTAAATCAAATGGTCGGCTCTCTGGTTTACTACCCCAATGATAACAATAGTCTTGTAAAGATAAACCGCAAGAAAATTTATACGCATCCATCCAAATAACCCTAGCACCGTCGGATGATATTAATGTATTATTATAGATTTTCCATCCACAGTTATTGGGCCAGTATACTGTTAGCAACGGTATATCGAATAGATTACTCATGTAACCTACTAGAATGATAGTTCGTGTACTAGTCATAATATCTTTAATCCAATCTAAGAAATCTAAAATTAGATCTTTTACACTATTAAAGTTAACAAAGGTATTACCCGGGTACGTGGATATCATATATGGTATTGGTCCATTATCAGTAATGGCTCCATAAACTGATACGGCGATGGTTTTTATATGTGTTTGATCTAATGAGGACGAAATTCGCGCCCACAATTCATCTCTAGATATGTATTTAATATCGAACGGTAACACATCAATCTCGGGACGCGTATATGTTTCTAAATTCTTAATCCAAATATAATGATGACCTATATGCCCTATTATCATACTGTCAACTATAGTATACCTAGAGAACTTTCGATACATCTGCTGTTTCCTGTAATCGTTAAATTTTACAAATCTATAACATGCTAAACCTTTTGACGACAGCCATTCATTAATTTCTGATATGGAATCTGTAATCTCAATACCGTATCGTTCTAAAGCCAGTGCTATATCTCCCTGTTCGTGGGAACGCTTTCGTATAATATCGATCAATGGATAATATGAAGTTTTTGGAGAATAATATGATTCATGATCTATTTCGTCCATAAACAATCTAGACATAGGAATTGGAGGCGATGATCTTAATTTTGTGCAATGGGTCAATCCTATAACTTCTAATATTGTAATATTCATCATCGACATAACACTATCTATGTTATCATCGTATATTAGTATACCACGACCTTCTTCATTTCGTGCCAAAATGATATACAGTCTTAAATAATTACGCAATATCTCAATAGTTTCATAATTGTTAGCTGTTTTCATCAAGGTTTGTATCCTGTTTAACATGATGGCGTTCTATAACGTCTCTATTTTCTATTTTTTAATTTTTTAAATTTTTAACGATTTACTGTGGCTAGATACCCAATCTCTCTCAAATATTTTTTTAGCCTCGCTTACAAGCTGTTTATCTATACTATTAAAACTGACGAATCCGTGATTTTGGTAATGGGTTCCGTCGAAATTTGCCGAAGTGATATGAACATATTCGTCGTCGACTATCAACAATTTTGTATTATTCTGAATAGTGAAAACCTTCACAGATAGATCATTTTGAACACACAACGCATCTAGACTTCTGGCGGTTGCCATAGAATATACGTCGTTCTTATCCCAATTACCAACTAGAAGTCTGATCTTAACTCCTCTATTAATGGCTGCTTCTATAATGGAGTTGTAAATGTCAGGCCAATAGTAGCTACTACCGTCGACACGTGTAGTGGGAACTATGGCCAAATGTTCAATATCTATACTAGTCTTAGCCGACTTGAGTTTATCAATAACTACATCAGTGTCTAGATCTCTAGAATATCCCAATAGGTGTTCTGGAGAATCAGTAAAGAACACTCCACCTATAGGATTCTTAATATGATACGCAGTGCTAACTGGCAGACAACAAGCCGCAGAGCATAAATTCAACCATGAATTTTTTGCGCTATTAAAGGCTTTAAAAGTATCAAATCTTCTACGAAGATCTGTGGCCAGCGGAGGATAATCAGAATATACGCCTAACGTTTTAATCGTATGTATAGATCCTCCAGTAAATGACGCGTTTCCTACATAACATCTTTCATCATCAGACACCCAAAAACAACCGAGTAGTAGTCCCACATTATTTTTTTTATCTATATTAACGGTTATAAAATTTATATCCGGGGAGTGACTTTGTAGCTCTCCCAGATTTCTTTTCCCTCGTTCATCTAGCAAAACTATTATTTTAATCCCTTTTTCAGATACCTCTTTTAGTTTATCAAAAATAAGCGCTCCCCTAGTAGTACTCAGAGGATTACAACAAAAAGATGCTATGTATATATATTTCTTAGCTAGAGTGATAATTTCGTTAAAACATTCAAATGTTGTCAAATGATCGGATCTAAAATCCATATTTTCTGGTAGTGTTTCTACCAGCCTACATTTTGCTCCCGCAGGTACCGATACAAATGGCCACATTTAGTTAACATAAAAACTTATATATCCTGTTCTATCAACGATTCTAGAATATCATCGGCTATATCGCTAAAATTTTCATCAAAGTCGACATCACAACCTAACTCAGTCAATATATTAAGAAGTTCCATGATGTCATCTTCGTCTATTTCTATATCCGTATCCATTGTAGATTGTTGACCGATTATCGAGTTTAAATCATTACTAATACTCAATCCTTCAGAATACAATCTGTGTTTCATTGTAAATTTATAGGCGGTGTATTTAAGTTGGTAGATTTTCAATTATGTATCATTATAGCAACAGTAATTCTTGCTCCTCCTTGATTTTAGCATCCTCTTCATTATTTTCTTCTACGTACATAAGCATGTCCAATACGTTAGACAACACACCGACGATGGTGGCCGCCACAGACACGAATATGACTAGACCGATGACCATTTAAAAAATACTCTCTAGCTTTAACTTAAACTGTATCGATCATTCTTTTAGCACATGTATAATATAAAAACATTATTCTATTTCGAATTTAGGCTTCCAAAAATTTTTCATCCGTAAACCGATAATAATATATATAGACTTGTTAATAGTCGGAATAAATATATTAATGCTTAAACTATCATCATCTCCACGATTAGAGATACAATATTTACATTCTTTTTGCTGTTTCGAAACTTTATCAATACACGTTAATACAAACCCAGGAAGGAGATATTGAAACTGAGGCTGTTGAAAATGAAACGGTGAATACAATAATTCAGATAATGTAAAATCATGATTCCGTATTCTGATGATATTAGAACTGCTAATGGATGTCGATGGTATGTATCTAGGAGTATCTATTTTAACAAAGCATCGATTTGCTAATATACAATTATCCTTTTGATTAATTGTTATTTTATTCATATTCTTAAAAGGTTTCATATTTATCAATTCTTCTACATTAAAAATTTCCATTTTTAATTTATCTAGCCCCGCAATACTCCTCATTACGTTTCATTTTTTGTCTAGAATGCCCATTTTGTTCATCTTGGTACATAGATTATCCAATTGAGAAGCGCATTTAGTAGTTTTGTACATTTTAAGTTTATTAACGAATCGTCGAAAACTAGTTATAGTTAACATTTTATTATTTGATACCCTGATATTAATACCCCTGCCGTTACTATTATTTATAACTGATGTAACCCACGTAACATTGGAATTAATTATCGATAGTAATGCATCGACACTTCCAAAATTGTCTATTATAAACTCACCGATAATTTTTTTATTGCATGTTTTCATATTCATTAGGATTATCAAATCTTTAATCTTATTACGATTGTATGCGTTGATATTACAAGACGTCATTCTAAAAGACGGAGGATTTCCATCAAATGCCAGACAATCACGTACAAAGTACATGGAAATAGGTTTTGTTCTATTACGCATCATAGATTCATATAAAACACCCGTAGAAATACTAATTTGTTTTACTCTATAAAATACTATTGCATCTATTTCATCGTTTTGTATAACGTCTTTCCAAGTGTCAAATTCTAATTTTTTTTCATTGATAGTACCAAATTCTTCTATCTCTTTAACTACTTGCATAGATAGGTAATTACAGTGATGCCTACATGCCGTTTTTTGAAACTGAATAGATGCATCTAGAAGCGATGCTACGCTAGTCACGATCACCACTTTCATATTTAGAATATATGTATGTAAAAATATAGTAGAATTTCATTTTGTTTTTTCTATGCTATAAATGAATTCTCATTTTGTATCTGCACATACTCCGTTTTATATCAATACCAAAGAAGGAAGATATCTGGTTCTAAAAGCCGTTAAAGTATGCGATGTTAGAACTGTAGAATGCGAAGGAAGTAAAGCTTCCTGCGTACTCAAAGTAGATAAACCCTCATCACCCACGTGTGAGAGAAGACCTTCGTCCCCGTCCAGATGCGAGAGAATGAATAACCCTGGAAAACAAGTCCCGTTTATGAGAACGGACATGTTACAAAATATGTTTGCTGCTAATCGCGACAACGTAACGTCAAGACTTTTGAACTAAAATACAATTATATCTTTTTCGATATTAATAAATCCGTGTCTCCCGGGTTTTTTATCTCTTTCAGTATGTGAATAGATAGGTATTTTATCTCTATTCATCATCGAATTTAAGAGATCCGATAAACATTGTTTGTATTCTCCAGATGTCAGCATCTGATACAACAATATATGTGCACATAAACCTCTGGCACTTATTTCATGTACCTTCCCCTTATCACTAAGGAGAATAGTATTTGAGAAATATGTATACATGATATTATCATGAATTAGATATACAGAATTTGTAACACTCTCGAAATCACACGATGTGTCGGCGTTAAGATCTAATATATCACTCGATAACACATTTTCATCTAGATACACTAGACATTTTTTAAAGCTAAAATAGTCTTTAGTAGTAACAGTAACTATGCGATTATTTTCATCGATGATACATTTCATCGGCATATTATTACGCGTACCATCAAAGACTATACCATGTGTATATCTAACGTATTCTAGCATAGTTGCCATACGTGCATTAAACTTTTCAGGATCTTTGGATAGATCTTCCAATCTATCTATTTGAGAAAACATTTTTATCATGTTCAATAGTTGAAACGTCGGATCCACTATATAGATATTATCTATAAAGATTTTAGGAACTATGTTCATGGTATCCTGGCGAATATTAAAACTATCAATGATATGATTATCGTTTTCATCTTTTATCACCATATAGTTTCTAAGATATGGGATTTTACTTAATATAATATTATTTCCCGTAATAAATTTTATTAGAAATGCCAAATCTATAAGAAAAGTCCTAGAATTAGTCTGAAGAATATCTATATCACCGTACCGTATATTTGGATTAATTAGATATAGAGAATATGATCCGTAACATATACAACTTTTATTATGACGTCTAAGATATTCTTCCATCAACTTATTAACATTTTTGACTAGGGAAGATACATTATGACGTCCCATTACTTTTGCCTTGTCTATTACAGCGACGTTCATAGAATTTAGCATATCTCTTGCCAATTCTTCCATTGATGTTACATTATAAGAAATTTTAGATGAAATTACATTTGGAGCTTTAATAGTAAGAACTCCTAATATATCCGTGTATGTGGTCACTAATACAGATTGTAGTTCTATAATCGTAAATAATTTACCTATATTATATGTTTGAGTTTGTTTAGAAAAGTAGCTAAGTATACGATCTTTTATTTCTGATGCCGATGTATCAACATCGAAAAAAAATCTTTTTTTATTCTTTTTTACTAACGATACGAATATGTCTTTGTTAAAAACAGTTATTTTCTGAATATTTCTAGCTTGTAATTTTAACATATGATATTCGTTCACACTAGGTACTCTGCCTAAATAGGTTTCTATAATCTTTAATGTAATATTAGGAAGAGTATTCTGATCAGGATTCCTATTCATTTTGAGGATTTAAAACTCTGATTATTGTCTAATATGGTCTCAACACAAACTTTTTCACAGAGCGATAGAGTTTTTGATAACTCGTTTTTCTTAAGAAATATAAAACTACTGTCTCCAGAGCTCGCTCTATCTTTTATTTTATCTAATTCGATACAAACTCCTGATACTGGTTCAGAAAGTAATTCATTAATTTTCAGTCCTTTATAGAAGATATTTAATATAGATAATACAAAATCTTCAGTTCTTGATATCGATCTGATTGATCCTAGAACTAGATATATTAATAACGTGCTCATTAGGCAGTTTATGGCAGCTTGATAATTAGATATAGTATATTCCAGTTCATATTTATTAGATACCGCATTGCCCAGATTTTGATATTCTATGAATTCCTCTGAAAATAAATCCAAAATAACTAGACATTCTATTTTTTGTGGATTAGTGTACTCTCTTCCCTCTATCATGTTCACTACTGGTGTCCACAATGATAAATATCTAGAGGGAATATAATATAGTCCATATGATGCCAATCTAGCAATGTCGAATAACTGTAATTTTATTCTTCGCTCTTCATTATGAATTGAATCTTGAGGTATAAACCTAACACAAATTATATCATTAGACTTTTCGTATGTAATGTCTTTCATGTTATAAGTTTTTAATCCTGGAATAGAATCTATTTTAATGAGGCTTTTAAATGCAGCGTTCTCCAACGAGTCAAAGCATAATACTCTGTTGGTTTTCTTATATTCAATATTACGATTTTCTTCTTTGAATGGAATAGGTTTTTGAATTAGTTTATAATTACAACATAATAGATAAGGAAGTGTGTAAATAGTACGCGGAAAAAACATAATAGCTCCCCTGTTTTCATCCATGGTTTTAAGTAAATGATCACTGGCTTCTTTAGTCAATGGATATTCGAACATTAACCGTTTCATCATCATTGGACAGAATCCATATTTCTTAATGTAAAGAGTGATCAAATCATTGTGTTTATTGTACCATCTTGTTGTAAATGTGTATTCGGTTATCGGATCTGCTCCTTTTTCTATTAAAGTATCGATATCGATCTCGTCTAAGAATTCAACTATATCGACATATTTCATTTGTATACACATAACCATTACTAACGTAGAATGTATAGGAAGAGATGTAACGGGAACAGGGTTTGTTGATTCGCAAACTATTCTAATACATAATTCTTCTGTTAATACGTCTTGCACGTAATCTATTATAGATGCCAAGATATCTATATAATTATTTTGTAAGATGATATTAACTATGTGATCTATATAAGTAGTGTAATAATTCATGTATTTCGATATATGTTCCAACTCTGTCTTTGTGATGTCTAGTTTCGTAATATCTATAGCGTCCTCAAAAAATATATTCGCATATATTCCCAAGTCTTCAGTTCTATCTTCTAAAAAATCTTCAACGTATGGAATATAATAATCTATTTTACCTCTTCTGATGTCATTAATGATATAGTTTTTGACACTATTTTCCGTCAATTGATTCTTATTCACTATGTCTAAAAACCGGATAGCGTCCCTAGGACGAACTACTGCCATTAATATCTCTATTATAGCTTCTGGACATAAATCATCTATTATACCAGAATTAATGGGAACTATTCCGTATCTATCTAACATAGTTTTAAGAAAGTCAGAATCTAAGACCTGATGTTCATATATTGGTTCATACATGAAATGATCTCTATTGATGATAGTGACTATTTCATTCTCTGAAAATTGGTAACTCATTCTATACACGCTTTCCTTGTTGATAAAGGATAGTATATACTCGATGGAATTTGTACCAACAAACTGTTCTCTTATGAATCGTATATCATCATCTGAAATGATCATGTAAGGCATACATTTAACAATGAGAGACTTGTCTCCTGTTATCAATATACTATTCTTGTGATAATTTATGTGTGAGGCAAATTTGTCCACGTTCTTTAATTTTGTTATAGTAGATATCAAATCCAATGGAGATACAGTTCTTGGCTTAAACAGATATAGTTTTTCTGGAACGAATTCTACAACATTATTATAAAGGACTTTGGGTATATAAGTGGGATGAAATCCTATTTTAATTAATGCGATAGCCTTGTCCTCGTGCAGATATCCAAACGCTTTTGTGATAGTATGGCATTCATTGTCTAGAAACGCTCTACGAATATCTGTAACAGATATCATCTTTAGAGAATACTAGTCGCGTTAATAGTACTAAAATTTGTATTTTTTAATCTATCTCAATAAAAAATTAATATGTATGATTCAATGTATAACTAAACTACTAACTGTTATTGATAACTAGAATCAGAATCTAATGATGACATAACTAAGAAGTTTATCTACAGCCAATTTAGCTGCATTATTTTTAGCATCTCGTTTAGATTTTCCATCTGCCTTATCGAATACTCTTCCGTCAATGTCTACACAGGCATAAAATGTAGGAGAGTTACTAGGCCCCACTGATTCAATACGAAAAGACCAATCTCTCCTAGTTATTTGGCAGTACTCATTAATAACGGTGACAGGGTTAACACCTTTCCAATAAATAATTTTTTTAACCGGAATAACATCATCAAAAGACTTATTATCCTCTCTCATTGATTTTTCGCGGGATACATCATCTATTATAGCATCAGCATCAGAATCTGTAGGCCGTGTATCAGCATCCATTGTCGTAGACCAACGAGGAGGAGTATCGTCGGAACTGTACACCATAGTACTACGTTGAAGATCATACAGAGCTTTATTAACTTCTCGCTTCTCCATATTAAGTTGTTTAGTTAGTTGTGCAGTAGCTCCTTAGTCCAATGTTTTTAATAACCGCACACAATCTCTGTGTCAGAACGCTCGTCAATATAGATCGTAGAAATTTTTTAGAGAGAACTAACACAACTAGCAATAAAACTGATCTTATTTTATCATTTTTTTATTCATCATCCTCTGGTGGTTCGTCGTTCCTATCGAATGTAGCTCTGATTAACCCGTCATCTATAGGTGATGCTGGTTCTGGAGATTCTGGAGGAGATGGATTATTATCCGGAAGAATCTCTGTTATTTCCTTGTTTTCATGTATCGATTGCGTTGTAACATTAAGATTGCGAAATGCTCTAAATTTGGGAGGCTTAAAGTGTTGTTTACAATCTCTACACGCGTGTCTAACTAATGGAGGTTCGTCAGCGGCTCTAGTTTGAATCATCATCGGTGTAGTATTCCTACTTTTACAGTTAGGACACGGTGTATTGTATTTCTCGTCGAGAACGTTAAAATAATCGTTGTAACTCACATCCTTTATTTTATCTATATTGTATTCTACTCCTTTCTTAATGCATTTTATACCGAACAAGAGATAGCGAAGGAATTCTTTTTCGGTACCGCTAGTACCCTTAATCATATCACATAGTGTTTTATATTCTAAATTTGTGGCAATGGACGGTTTATTTCTATACGATAGTTTGTTTCTGGAATCCTTTGAGTATTCTATACCAATATTATTCTTTGATTCGAATTTAGTTTCTTCGATATTAGATTTTGTATTACCTATATTCTTGATGTAGTACTTTGATGATTTTTCCATGGCCCATTCTATTAAGTTTTCCAAGTTGGCATCATCCACATATTGTGATAGTAATTCTCGGATATCAGTAGTGACTACCGCCATTGATATTTGTTCATTTGATGAGTAACTACTAATGTATACATTTTCCATTTATAACACTTATGTATTAACTTTGTTTATTTATATTTTTTCATTATTATGTTGATATTAATAATCGTATTGTGGTTATATGGCTACAATTTCATAATGAGTTGAAGTCAGTGTCCTATGATCAATGACGATAGCTTTACTCTGAAAGAAAGTATCAAATCGATAGTGCAGAGTCAACAATGAAAATGGATAAGACGATGACAAAGTTTCAGAATAGAGTCAAAATGGTAAAAGAAATAAATCAGACGATAAGAGCAGCACAAACTCATTGAGACATTGAAACTAGGATATATAAAATTTAAGTGAATGATTAGGACAGGACTACTACTCTAGAAGATATAGCACCATCTATTATTCCAAATAATCAGAAAACTTATAAACTATTCTCGGACATTTCAGTCATTGGCAAAGCATCACAGAATCCGAGTAAGATGATATATGCTCGCTGCTTTACATGTTTCCCAATTTGTTTGGAGATGACCATAGATTCATTTGTTATAGAATGCATCCAATGAGTAAAATCAAACACAAAATCTTCGTTCAAACTTAATCTTATTAGAATATTAGTGGAAGAAAGATTCTATAATAATGAATGCAGAGATTATAAATGGAGAATAATTGGAACACAAGTTGATAAAATATTGATAGCTAAATATACAATAGATGCAATGTATCGCATAAGACCGATATATATAATACAAAGCAGTACAGATACAATGATGATGTAGAAAATGGATTCATTGGATTGGATAAACTAAAATTAAACATTGTTCATGATATAGTTGAATCATGTATACCTGTTCGTATGCCTGTGGCTAAGATACTGTGTAAAGAAATGGTAAATAAATACTTTGAGAATCTTTAAGAGTGTATTGACTTTGTTAGTGAATATGCATTCCATCTTTCTCCAATACTAATTCAAATTGTTAAATTAATAATGGAATAGTATAAATAGTTATTAGTGATAAGATAGTAAAAATAATTATTAGAATAGTGTAGTATCATAGATAACTCTCTTCTATAAAAAATGGATTTTATTCGTAGAAAGTATCTTATATACACAGTAGAAAATAATATAGATTTTTTAAAGGATGATACATTAAGTAAAGTAAACAATTTTACCCTCAATCATGTACTAGCTCTCAAGTATCTAGTTAGCAATTTTCCTCAACATGTTATTACTAAGGATGTATTAGCTAATACCAATTTTTTTGTTTTTATACATATGGTACGATGCTGTAAAGTATACGAAGCGGTTTTACGACACGCATTTGATGCACCCACGTTGTACGTTAAAGCATTGACTAAGAATTATTTATCGTTTAGTAACACAATACAGTCGTACAAGGAAACCGTGCATAAACTAACACAAGATGAAAAATTTTTAGAGGTTGCCGAATACATGGACGAATTAGGAGAACTTATAGGCGTAAATTATGACTTAGTTCTTAATCCATTATTTCACGGAGGGGAACCCATCAAAGATATGGAAATCATTTTTTTAAAACTGTTTAAGAAAACAGACTTCAAAGTTGTTAAAAAATTAAGTGTTATAAGATTACTTATTTGGGCATACCTAAGCAAGAAAGATACAGGCATAGAGTTTGCGGATAATGATAGACAAGATATATATACTCTATTTCAACAAACTGGTAGAATAGTCCATAGCAATCTAACAGAAACGTTTAGGGATTATATCTTTCCCGGAGATAAGACTAGCTATTGGGTGTGGTTAAACGAAAGTATAGCTAATGATGCGGATATCGTTATTAATAGATCCGCCATTACCATGTATGATAAAATTCTTAGTTATATATACTCTGAGATAAAACAGGGACGCGTTAATAAAAACATGCTTAAGTTAGTTTATATCTTTGAGCCTGAAAAAGATATCAGAGAACTTCTGCTAGAAATCATATATGATATTCCTGGAGATATCCTATCTATTATTGATGCAAAAAACGATGATTGGAAAAAATATTTTATTAGTTTTTACAAAGCTAATTTTATTAACGGTAATACATTTATTAGTGATAGAACGTTTAACGAGGACTTATTCAGAGTTGTTGTTCAAATAGATCCCGAATATTTCGATAATGAACGAATTATGTCTTTATTCTATACGAGTGCTGCGGACATTAAACGATTTGATGAGTTAGATATTAATAACAGTTATATATCTAATATAATTTATGAGGTGAACGATATCACATTAGATACAATGGATGATATGAAGAAGTGTCAAATCTTTAACGAGGATACGTTGTATTATGTTAAGGAATACAATACATACCTGTTTTTGCACGAGTCGGATCCCATGGTCATAGAGAACGGAATACTAAAGAAACTGTCATCTATAAAATCCAAGAGTAGACGGCTGAACTTGTTTAGCAAAAACATTTTAAAATATTATTTAGACGGACAATTGGCTCGTCTAGGTCTTGTGTTAGATGATTATAAAGGAGACTTATTAGTTAAAATGATAAACCATCTCAAATCTGTGGAGGATGTATCCGCATTCGTTAGATTTTCTACAGATAAAAACCCTAGTATTCTTCCATCGCTAATCAAAACTATTTTAGCTAGTTATAATATTTCCATCATCGTCTTATTTCAAAGGTTTTTAAGAGATAATCTATATCATGTAGAAGAATTCTTGGATAAAAGCATCCATCTAACCAAGACTGATAAGAAATATATACTTCAATTGATAAGGCACGGTAGATCATAGAACAAACCAAATATATTATTAATAATTTGTATATACATAGATATAATTATCATATATTAAAAAATAACACATTTTTGATAAATGGAAACTGTTGCAACAATTCAGACTCCCACCAAATTAATGAATAAAGAAAATGCAGAAATGATTTTGGAAAAAATTGTTAATCATATAGCTATGTATATTAGTGACGAATCAATATATTCAGAAAATAATCCTGAATATATTGATTTTCGTAACAGATACGGAGACTATAGATCTCTCATTATAAAAAGTGATCACGAGTTTGTAAAGCTATGTAAAGATCATGCAGAGAAAAGTTCTCCAGAAACGCAACAAATGATTATCAAACACATATACGAACAATATCTTATTCCAGTATCTGAAGTACTGTTAAAACCTATAATGTCCATGGGTGACATATTTACATATAACGGATGTAAAGACAATGAATGGATGCTAGAACAACTCTCTACCCTAAACTTTAACAATCTCTACACATGGAACTCATGTAGCATAGGCAATGTAACGCGTCTGTTTTATACATTTTTTAGTTATCTGATGAAAGATAAACTAAATATATAAGTATAATCCCATTAATACTTTAACCTGATGTATTATTACCTGCATCTTATTAGAATATTAACCTAACTAAAAGACATAAAAAGCGGGGATATAAATATTATGGCAGCAACCGTTCCGCGTTTTGACGATGTGTACAAAAATGCACAAAGAAGAATTCTAGATCAAGAAACATTTTTTAGTAGAGGTCTAAGTAGACCGTTAATGAAAAACACATATCTATTTGATAATTACGCGTATGGATGGATACCAGAAACTGCAATTTGGAGTAGTAGATACGCAAACCTAGATGCTAGTGACTATTATCCCATTTCGTTGGGATTACTTAAAAAGTTTGAATTTCTCATGTCTCTATATAAAGGTCCTATTCCCGTATATGAAGAAAAAGTAAATACTGAATTCATTGCTAATGGATCTTTCTCCGGTAGATACGTATCATATCTTAGAAAGTTTTCTGCCCTTCCAACAAACGAGTTTATTAGTTTTTTATTATTGACCTCCATCCCTATCTATAATATCTTATTCTGGTTTAAAAACACACAGTTTGATATTACTAAACACACATTATTCAGATACGTCTATACAGATAATACCAAACACCTTGCGTTGGCTAGGTATATACATCAAACAGGAGACTATAAGCCTTTGTTTAGTCGTCTCAAAGAGAATTATATATTTACCGGTCCCGTTCCAATAGGTATCAAAGATATAGATTACCCTAATCTTAGTAGAGCAAGAAGTCCATCCGATTATGAGACATTAGCTAATATTAGTACTATATTGTACTTTACCAAGTATGATCCAGTATTAATGTTTTTATTGTTTTACGTACCTGGGTATTCAATTACTACAAAAATTACTCCAGCCGTAGAATATCTAATGGATAAACTGAATCTAACAAAGAGCGACGTACAACTGTTGTAAATTATTTTTATGCTTCGTAAAATGTAGGTCTTGAACCAAACATTCTTTGAAAAAATGAGATGCATAAAACTTTATTATCCAATAGATTAACTATTTCAGACGTCAATCGTTTAAAGTAAACTTCGTAAAATATTCTTTGATTGCTGCCGAGTTTAAAACTTCTATCGATAATTGTTTCATATGTTTTAATATTTACAAGTTTTTTGGTCCATGGTACATTAGCTGGACAGATATATGCAAAATAATATCGTTCTCCAAGTTCTATAGTCTCTGGATTGTTTTTATTATATTCAGTAACCAAATACATATTAGGGTTATCTGCGGATTTATAATTTGAGTGATGCATTCGACTCAACATAAATAATTCTAGAGGAGACGATCTACTATCAAATTCGGATCGTAAATCTGTTTCTAAAGAACGGAGAATATCTATACATACCTGATTAGAATTCATCCGTCCTTCAGACAACATCTCAGACAGTCTGGTCTTGTATGTCTTAATCATATTCTTATGAAACTTGGAAACATCTCTTCTAGTTTCACTAGTACCTTTATTAATTCTCTCAGGTACAGATTTTGAATTCGACGATGCCGAGTATTTCATCGTTGTATATTTCTTCTTCGATTGCATAATCAAATTCTTATATACCGCCTCAAACTCTATTTTAAAATTATCAAACAATACTCTACTATTAATCAGTCGTTCTAACTCCTTTGCTATTTCTATGGACTTATCTACATCTTGACTGTCTATCTCTGTAAACACGGAGTCGGTATCTCCATACACGCTACGAAAACGAAATCTATAATCTATAGGCAACGATGTTTTCACAATCGGATTAATATCTCTATCGTCCATATAAAATGGATTACTTAATGTATTGGCAAACCGTAACATACCGTTGGATAACTCTGCTCCATTTAGTACCGATTCTAGATACAAGATCATTCTACGTCCTATGGATGTGCAACTCTTAGCCGAAGCGTATGAGTATAGAGCACTATTTCTAAATCCCATCAGACCATATACTGAGTTGGCTACTATCTTGTACGTATATTGCATGGAATCATAGATGGCCTTTTCAGTTGAACTGGTAGCCTGTTTTAACATCTTTTTATATCTGGCTCTCTCTGCCAAAAATGTTCTTAATAGTCTAGGAATGGTTCCTTCTATTGATCTATCGAAAATTGCTATTTCAGAGATGAGGTTCGGTAGTCTAGGTTCACAATGAACCGTAATATATCTAGGAGGTGGATATTTCTGAAGCAAGAGCTGATTATTTATTTCTTCTTCCAATCTATTGGTACTAACAACGACACCGACTAATGTTTCCGGAGATAGATTTCCAAAGATACACACATTAGGATACAGACTGTTATAATCAAAGATTAATACATTATTACTAAACATTTTTTGTTTTGGAGCAAATACCTTACCGCCTTCATAAGGAAACTTTTGTTTTGTTTCTGATCTGACTAAGATAGTTTTAGTTTCCAACAATAGCTTTAACAGTGGACCCTTGATGACTGTACTCGCTCTATATTCGAATACCATGGATTGAGGAAGCACATATGTTGACGCACCAGCGTCTGTTTTTGTTTCTACTCCATAATACTCCCACAAATACTGACACAAACAAGCATCATGAATACAGTATCTAGCCATATCTAAAGCTATGTTTAGATTATAATCCTTATACATCTGAGCTAAATCAATGTCATCCTTTCCGAAAGATAATTTATATGTATCATTAGGTAAAGTAGGACATGATAGTACGACTTTAAATCCATTTTCCCAAATATCTTTACGAATTACTTTACATATAATATCCTCATCAACAGTCACATAATTACCTGTTGTTAAAACCTTTGCAAATGTATCGGCTTTGCCTTTCGCGTCCGTAGTATCGTCACCGATGAACGTCATTTCTCTAACTCCTCTATTTAATACTTTACCCATGCAACTGAACGCGTTCTTGGATATAGAATCCAATTTGTACGAATCCAATTTTTCAGATTTTTGAATGAATGAATATAGATCGAAAAATATAGTTCCATTATTGTTATTAACGTGAAACGTAGTATTGGCCATGCCGCATACTCCCTTATGACTAGACTGATTTCTCTCATAAATACAGAGATGTACAGCTTCCTTTTTGTCTGGAGATCTAAAGATAATCTTCTCTCCTGTTAATAACTCTAGACGATTAGTAATATATCTCAGATCAAAGTTATGTCCGTTAAAGGTAACGACGTAGTCGAACGTTAGTTCCAACAATTGTTTAGCTATTCGTAACAAAACTATTTCAGAACATAGAACTAGTTCTCGTTCGTAATCCATTTCCATTAGCGACTGTATCCTCAAACATCCTCTATCGACGGCTTCTTGTATTTCCTGTTCCGTTAACATCTCTTCATTAATGAGCGTAAACAGTAATCGTTTACCACTTAAATCGATATAACAGTAACTTGTATGCGAGATTGGGTTAATAAATACAGAAGGAAACTTCTTATCGAAGTGACACTCTATATCTAGAAATAAGTACGATCTTGGGATATCGAATCTAGGTATTTCTTTAGCGAAACAGTTACGTGGATCGTCACAATGATAACATCCATTGTTAATCTTTGTCAAATATTGCTCGTCCAACGAGTAACATCCGTCTGGAGATATCCCGTTAGAAATATAAAACCAACTAATATTGAGAAATTCATCCATGGTGGCATTTTGTATGCTGCGTTTCTTTGGCTCTTCTATCAACCACATATCTGCGACGGAGCATTTTCTATCTTTAATATCTAGATTATAACTTATTGTCTCGTCAATGTCTATAGTTCTCATCTTTCCCATCGGCCTCGCATTAAATGGAGGAGGAGATAATGACTGATATATTTCGTCCGTCACTACGTAATAAAAGTAATGAGGAAATCGTATAAATACTGTCTCGCCATTTCGACATCTGGATTTCAGATATAAAAATCTGTTTTCACCGTGACTTTCAAACCAATTAATACACCTAACATCCATTTCTAGAATTTAGAAATATATTTTCATTTAAATGAATCCCAAACATTGGGGAAGAGCCATATGGACCATTATTTTTATAGTACTTTCGCAAGCGGGTTTAGACGGCAACATAGAAGCGTGTAAACGAAAAATATATACTATAGTCAGCACTCTTCCATGTCCTGCATGTAGACGACACGCGACTATCGCTATAGAGGACAATAATGTCATGTCTAGCGATGATCTGAATTATATTTATTATTTTTTCATCAGATTATTTAACAATTTGGCATTTGATCCCAAATACGCAATCGATGTGTCAAAGGTTAAACCTTTATAAACTTAACCCATTATAAAACTTATGATTAGTCACGACTGAAATAACCGCGTGATTATTTTTTGGTATAATTCTACACGGCATGGTTTCTGTGACTATGAATTCAACACCTGTTATCTTAGTGAAATCTTTAACAAACAGCAAGGGTTCGTCAAAGACATAAAACTCATTGTTTACGATCGAAATAGACCCCCTATCACACTTAAAATAAAAAATATCCTTATCCTTTACCACCAAATAAAATTCTGATTGGTCAATGTGAATGTATTCACTTAACAGTTCCACAAATTTATTTATTAACTCCGAGGCACATACATCGTCGGTATTTTTTATGACAAACTTTACTCTTCCAGCATCCGTTTCTAAAAAAATATTAACGAGTTCCATTTATATCATCCAATATTATTGAAATGACGTTGATGGACAGATGATACAAATAAGAAGGTACGGTACCTTTGTCCACCATCTCCTCCAATTCATACTCTATTTTGTCATTAACTTTAATGTGTGAAAACAGTACGCCACATGCTTCCATGACAGTGTGTAACACTTTGGATACAAAATGTTTGACATTAGTATAATTGTCCAAGACTGTCAATCTATAATAGATAGTAGCTATAATATATTCTATGATGGTATTGAAGAAGATGACAACCTTGGCATATTGATCATTTAACACAGACATGGTATCAACAAATAGCTTAAATGAAAGAGAATCAGTAATTGGAATAAGCGTCTTCTCGATGTAGTGTCCGTATACCAACATGTCTGATATTTTGATGTATTCCATTAAATTATTTAGTTTTTTCTTTTTATTCTCGTTAAACAGAATTTCTGTCAATGGACCCCAACATCGTTGACCTATTAAGTTTTGATTGATTTTTCCGTGTAAGGCGTATCTAGTCAGATCGTATAGCCTATCCAATAATCCATCGTCTGTGCGTAGATCACATCGTACACTTTTTAATTTTCTATAGAAGAGTGACAGACATCTGGAGCAATTACAGACAGCAATTTCTTTATTCTCTACAGATGTAAGATACTTGAAGACATTCCTATGATGATGCAGAATTTTGGATAACACGGTATTGATGGTATCTGTTACCATAATTCCTTTGACTGATAGTGTCAAAGTACAAGATTTCCAATCTTTTGCAATTTTCAGTACCATTATCTTTGTTTTGATATCTATATCAGACAGCATGGTACGTCTGACAACACAGGGATTAAGACGGAAAGATGAAATGATTCTCTCAACATCTTCAATAGATACCTTGCTATTTTTTTTGGCATTATCTATATGTGAGAGAATATCCTCTAGAGAATCAGTATCCTTTTTGATGATAGTGGATCTCAATGACATGGGACGTCTAAACCTTCTTATTCTATCACCAGATTGCATGGTGATTTGTCTTCTTTCTTTTATCATGATGTAATCTCTAAATTCATCGGCAAATTGTCTATATCTAAAATCATAATATGAGATGTTTACCTCTACAAATATCTGTTCGTCCAATGTTAGAGTATCTATATCAGTTTTGTATTCCAAATTAAACATGGCAACGGATTTAATTTTATATTCCTCTATTAAGTCCTCGTCGATAATAACAGAATGTAGATAATCATTTAATCCATCGTACATTGTTGGAAGATGCTCGTTGACAAAATCTTTAATTGTCTTGATGAAGGTGGGACTATATCTAACATCTTGATTAATAAAATTTATAACATTGTCCATAGGATACTTTGTAACTAGTTTTATACACATCTCTTCATCGGTAAGTTTAGACAGAATATCGTGAACAGGTGGTATATTATATTCATCAGATATACGAAGAATAATGTCCAAATCTATATTGTTTAATATATTATATAGATGTAGTGTAGCTCCTACAGGAATATCTTTAACTAAGTCAATGATTTCATCAACAGTTAGATCTATTTTAAAGTTAATCATATAGGCATTGATTTTTAAAAGGTATGTAGCCTTGACTACATTCTCATTAATTAACCATTCCAAGTCACTGTGTGTAAGAAGATTATATTCTATCATAAGCTTGACTACATTTGGTCCCGATACCATTAAAGAATTCTTATGATATAAGGAAACAGCTTTTAGGTACTCATCTACTCTACAAGAATTTTGGAGAGCCTTAACGATATCAGTGACGTTTATTATTTCAGGAGGAAAGAACCTAACATTGAGAATATCTGAATTAATAGCTTCCAGATACAGTGATTTTGGCAATAGTCCGTGTAATCCATAATCCAGTAACACGAGCTGGTGCTTGCTAGACACCTTTTCAATGTTTAATTTTTTTGAAATAAGCTTTGATAAAGCCTTCCTCGCAAATTCCGGATACATGAACATGTCGCCAACATGATTAAGTATTGTTTTTCATTATTTTTATATTTTCTCAACAAGTTCTCAATACCCCAATAGATAATAGAATATCACCCAATGCGTCCATGTTGTCTATTTCCAACAGGTCGCTATATCCACCAATAGAAGTTTTCCCAAAAAAGATTCTAGGAACAGTTCTACCACCAGTAATTTGTTCAAAATAGTCACGCAATTCATTTTCGGGTTTAAATTCTTTAATATCTACAATTTCATACGCTCCTCTTTTGAAACTAAACTTATTTAGAATATCCAGTGCGTTTCTACAAAAAGGACACGTAAACTTGACAAAAATTGTCACTTTGTTATTGGCCAACCTTTGTTGTACAAATTCCTCGGCCATTTTTAATATTTAAGTGATACAAAACTATCTCGACTTATTTAACTCTTTAGTCGAGATATATGGACACAGATAGTTATATGATAACCAACTACAGAAGACAAACGCTATAAAAAACATAATTACAACGAGCATATTTATAAATATTTTTATTCAGTATTACTTGATATAGTAATATTAGGTACAGTCAAACATTCAACCACTCTAGATACATTAACTCTCTCATTTTCTTTAACAAATTCTGCAATATCTTCGTAAAAAGATTCTTGAAACTTTTTAGAATATCTATCGACTCTAGATGAAATAGCGTTCGTCAACATACTATGTTTTGTATACATAAAGGCGCCCATTTTAACAGTTTCTAGTGACAAAATGCTAGCGATCCTAGGATCCTTTAGAATCACATAGATTGACGATTCGTCTCTCTTAGTAACTCTAGTAAAATAATCATACAATCTAGTACGCGAAATAATATTATCCTTGACTTGAGGAGATCTAAACAATCTAGTTTTGAGAACATCGATAAGTTCATCGGGAATGACATACATACTATCTTTAATAGAACTCTTTTCATCCAGTTGAATGGATTCGTCCTTAACCAACTGATTAATGAGATCTTCTATTTTATCATTTTCCAGATGATATGTATGTCCATTAAAGTTAAATTGTGTAGCGCTTCTTTTTAGCCTAGCAGCCAATACTTTAACATCACTAATATCGATATACAAAGGAGATGATTTATCGATGGTATTAAGAATTCGTTTTTCGACATCCGTCAAAACCAATTCCTTTTTGCCTGTATCATCCAGTTTGCCATTCTTTGTAAAGAAATTATTTTCTACTAGACTATTAATAAGACTGATAAGGATTCCTCCATAATTGCACAATCCAAACTTTTTCACAAAACTAGACTTTACGAGATCTACAGGAATGCGTACTTCAGGTTTCTTAGCTTGTGATTTTTTCTTTTGCGGACATTTTCTAGTGACCAACTCATCTACCATTTCATTGATTTTAGCAGTGAAATAAGCTTTCAATGCACGGGCACTGATACTATTGAAAACGAGTTGATCTTCAAATTCCGCCATTTAAGTTCACCAAACAACTTTTAAATACAAATATATCAATAGTAGTAGAATAAGAACTATAAAAAAAATAATAATTAACCAATACCAACCCCAACAACCTGTATTATTAGTTGATGTGACAGTTTTCTCATCACTTAGAACAGATTTAACAATTTCTATAAAGTCTGTCAAATCATCTTCCTGAGAACCCATAAATACACCAAATATAGCAGCGTACAACTTATCCATTTATACATTGAATATTGGCTTTTCTTTATCGCTATCTTCATCATATTCATCATCAATATCGACAAGTCCCAGATTACGAACCAGATCTTCTTCTACATTTTCAGTCATTGATACGCGTTCACTATCTCCAGAGAGTCCGATAACGTTAGCCACCACTTCTCTATCAATGATTAGTTTCTTGAGCGCGAATGTAATTTTTGTTTCCGTTCCGGATCTATAGAAAACTACAGGTGTGATAATTGCCTTGGCTAATTGTCTTTCTCTTTTACTGAGTGATTCTAGTTCACCTTCTATAGATCTGAGAATGGATGATTCTCCAGTCGAAACATATTCTACCATGGCTCCGTTTAATTTGTTGATGAAGATGGATTCATCCTTAAATGTTTTCTCTGTAATAGTTTCCGCCGAAAGACTATGCAAAGAATTTGGAATGCGTTCCTTGTGCGTAATGTTTCCATAGACAGCTTCTAGAAGTTGATACAACATAGGACTAGCCGCGGTAACTTTTATTTTTAGAAAGTATCCATCGCTTCTATCTTGTTTAGATTTATTTTTATAAAGTTTAGTCTCTCCTTCCAACATAATAAAAGTGGAAGTCATCTGACTAGATAAACTATCAGTAAGTTTTATAGAGATAGATGAACAATTAGCGTATTGAGAAGCATTTAGTGTAACGTATTCGATACATTTTGCATTAGATTTACTAATCGATTTTGCATACTCTATAACACCCGCACAAGTCTGTAGAGAATCGCTAGATGCTGTAGGTCTTGGTGAAGTTTCAACTCTCTTCTTGATTACCTTACTCATGATTAAACCTAAATAATTGTACTTTGCAATATAATGATATATATTTTCACTTTATCTCATTTGAGAATAAAAATGTTTTTGTTAACCGCTGCATGATGTACAGATTTCGGAATCGCAAACCACTTGTGGTTTTATTTTATCCTTGTCCAATGTGAATTGAATGGGAGCGGATGCGGGTTTCGTACGTAGATAGTACATTCCCGTTTTTAGACCGAGACTCCATCCGTAAAAATGCATACTCGTTAGTTTGGAATAACTCGGATCTGCTATATGGATATTCATAGATTGACTTTGATCGATGAAGGCTCCCCTGTCTGCAGCCATTTTTATGATCGTCTTTTGTGCAATTTCCCAAATAGTTTTATAAACTCGCTTAATATCTTCTGGAAGGTTTGTATTCTGAATGGATCCACCATCTACCATAATCCTATTCTTGATCTCATCATTCCATAATTTTCTCTCGGTTAAAACTCTAAGGAGATGCGGGTTAACTACTTGGAATTCTCCAGACAATACTCTCCGAGTGTAAATATTACTGGTATACGGTTCCACCGACTCATTATTTCCCAAAATTTGAGCAGTTGATGCAGTCGGCATAGGTGCCACCAATAAACTATTTCTAAGACCGTATGTTCTGATTTTATCTTTTAGATGTTCCCAATTCCAAAGATCCGACGGTACAACATTCCAAAGATCATATTGTAGAATACCGTTACTGGCGTACGATCCTACATATGTATCATATGGTCCTTCCTTCTCAGCTAGTTTACAACTCGCCTCTAATGCACCGTAATAAATGGTTTCAAAGATCTTCTTATTTAGATCTTGTGCTTCCGGGCTATCAAATGGATAATTTAAGAGAATAAACGCGTCCGCTAATCCTTGAACACCAATACCGATAGGTCTATGTCTCTTATTAGAGATTTCAGCTTCTGGAATAGGATAATAATTAATATCTATAATTTTATTGAGATTTCTGACAATTACTTTGACCACATCCTTCAGTTTGAGAAAATCAAATCGCCCATCTATTACAAACATGTTCAATGCAACAGATGCCAGATTACACACGGCTACCTCATTAGCATCCGCATATTGTATTATCTCAGTGCAAAGATTACTACACTTGATAGTTCCTAAATTTTGTTGATTACTCTTTTTGTTACACGCATCCTTATAAAGAATGAATGGAGTACCAGTTTCAATCTGAGATTCTATAATCGCTTTCCAGACGACTCGAGCCTTTATTATAGATTTGTATCTCCTTTCTCTTTCGTATAGAGTATACAATCGTTCGAACTCGTCTCCCCAAACATTGTCCAATCTAGGACATTCATCCGGACACATCAACGACCACTCTCCGTCATCCTTCACTCGTTTCATAAAGAGATCAGGAATCCAAAGAGCTATAAATAGATCTCTTGTTCTATGTTCCTCGTTTCCTGTATTCTTTTTAAGATCGAGGAACGCCATAATATCAGAATGCCACGGTTCCAAGTATATGGCCATAACTCCAGGCCGTTTGTTTCCTCCCTGATCTATGTATCTAGCGGTGTTATTATAAACTCTCAACATTGGAATAATACCGTTTGATATACCATTGGTACCGGAGATATAGCCTCCACTGGCACGAATATTACTAATTGATAGACCTATTCCCCCTGCCATTTTAGAGATTAATGCGCATCGTTTTAACGTGTCATAGATGCCTTCTATGCTATCATCGATCATGTTAAGTAGAAAACAGCTAGACATTTGGTGACGAGTAGTTCCCGCATTAAATAAGGTAGGAGAAGCGTGCGTAAACCATTTTTCAGAAAGTAGATTGTACGTCTCAATAGCTGAGTCTATATCCCATTGATGAATTCCTACTGCGACACGCATTAACATGTGCTGAGGTCTTTCAACAATTTTGTTGTTTATTTTCAACAAGTAGGATTTTTCCAAAGTTTTAAAACCAAAATAGTTGTATGAAAAGTCTCGTTCGTAAATAATAACCGAATTGAGCTTATCCTTATATTTGTTAACTATATCCATGGTAATACTTGAAATAATCGGAGAATGTTTCCCATTTTTAGGATTAACATAGTTGAATAAATCCTCCATCACTTCACTAAATAGTTTTTTTGTTTCCTTGTGTAGATTTGATATGGCTATTCTGGCGGCTAGAATGGCATAATCCGGATGTTGTGTAGTACAAGTGGCTGCTATTTCGGCTGCCAGAGTGTCCAATTCTACCGTTGTTACTCCATTATATATTCCTTGAATAACCTTCATAGCTATTTTAATAGGATCTATATGATCCGTGTTTAAGCCATAGCACAATTTTCTAATACGAGACGTGATTTTATCAAACATGACATTTTCCTTGTATCCATTTCGTTTAATGACAAACATTTTTGTTGGTGTAATAAAAAAAAATTATTTAATTTTTCATTAATAGGGATTTGACGTATGTAGCGTACAAAATTATCGTTCCTGGTATATAGATAAAGAGTCCTATATATTTGAAAATCGTTACGGTTCGATTAAACTTTAATGATTGCATAGTGAATATATCATTAGGATTTAACTCCTTGACTATCATGGCGGTGCCAGAAATTACCATCAAAAGCATTAATACAGTTATACAGATCGCAGTTAGAACGGTTATAGCATCCACCATTTATATCTAAAAATTAGATCAAAGAATATGTGACAACGTCCTAGTTGTATACTGAGAATTGACGAAACAATGTTTCTTACATATTTTTTTCTTATTAGTAACCGACTTAATAGTAGGAACTGGAAAACTAGACTTGATTATTCTATAAGTATAGATACCCTTCCAAATAATGTTCTCTTTGATAAAAGTTCCAGAAAATGTAGAATTTTTTAAAAAGTTATCTTTTGCTATTACTAATATCGTGGTTAGACGCTTATTATTAATATGAGTGATGAAATCCACACCGCTTCTAGATATCGCTTTTATTTCCACATTAGATGGTAAATCCAATAGTGAAACTATCTTTTTAGGAATGTATGGACTTGCGTTTAGAGGAGTGAACGTCTTCGGAGTAGTAAAGGATGATTCGTCAAATGAATAAACAATTTCACAAATGGATGTTAATGTATTAGTAGGAAATTTTTTGACGCTAGTGGAATTGAAGATTCTAATGGATGATGTTCTACCTATTTCATCCGATAACATGTTAATTTCCAACACCAACGGTTTTAATATTTCGATGATATACGGTAGTCTCTCTTTCGGACTTATATAGCTTATTCCACAATACGAGTCATTATATACTCCAAAAAACAAAATAACTAGTATAAAATCTTTATCGAATGGGAAAAACGAAATTATCGATATAGGTATAGAATCCGGAACATTGAACGTATTAATACTTAATTCTTTTTCAGTGGTAAGAACCGATAGGTTATTGACATTGTATGGTTTTAAATATTCTATAACTTGAGACTTGATAGATATTAATGACGAATTGAAAATTATTTTTATCACCACGTGTGTTTCAGGATCATCGTCGACGCCAGTCAACCAACCGAATGGAGTAAAATAAATATCATTAATATATGCTCTAGATATTAGTATTTTTATTAATCCTTTGATTATCATCTTCTCGTACGCGAATGATTCCATGATCAAGAGTGATTTGAGAACATCCTCCGGAGTATTAATGGGCTTAGTAAACAGTCCATCGTTGCAATAATAAAAGTTGTCCAAGTTAAAGGATATTATGCATTCGTTTAAAGATATCACCTCATCTAACGGAGACAATTTTTTGGTAGGTTTTAGAGACTTTGAAGCTACTTGTTTAACAAAGTTATTCATCGTCGTCTACTATTCTATTTAATTTTGTAGTTAATTTATCACATATCACATTAATTGACTTTTTGGTCCACTTTTCCATACGTTTATATTCTTTTAATCCTGCGTTATCCGTTTCCGTTATATACAGGGATAGATCTTGCAAGTTAAATAGAATGCTCTTAAATAATGTCATTTTTTTATCCGCTAAAAATTTAAAGAATGTATAAACTTTTTTCAAAGATTTAAAACTTTTAGGTGGAGTTCTGGTACACAATATCATAAACAAACTAATAAACATCCCACATTCAGATTCCAACAATTGATTAACTTCCACATTAATACAGCCTATTTTCGCTCCAAATGTACATTCGAAAAATCTGAATAAAACATCAATATCGCAATTTGTATTATCCAATACAGAATGTCTGTGATTCGTGTTAAAACCATCGGAAAAAGAATAGAAATAAAAATTATTATAATGGTGGAATTCAGTTGGAATATTGCCTCCGGAGTCATAAAAGGATACTAAACATTGTTTTTTATCATAAATTACACATTTCCAATGAGACAAATAACAAAATCCAAACATTACAAATCTAGAGGTAGAACTTTTAATTTTGTCTTTAAGTATATACGATAAGATATGTTTATTCATAAACGCGTCAAATTTTTCATGAATAGCTAAGGAGTTTAAGAATCTCATGTCAAATTGTCCTATATAATCCACTTCGGATCCATAAGCAAACTGAGAGACTAAGTTCTTAATACTTCGATTGCTCATCCAGGCTCCTCTCTCAGGCTCTATTTTCATCTTGACGACCTTTGGATTTTCACCAGTATGTATTCCTTTACGTGATAAATCATCGATTTTCAAATCCATTTGTGAGAAGTCTATCGCCTTAGATACTTTTTCCCGTAGTTGAGGTTTAAAGAAATACGCTAACGGTATACTAGTAGGTAACTCAAAGACATCATATATAGAATGGTAACGCGTCGTTAACTCGTCGGTTAACTCTTTCTTTTGATCGAGTTCGTCGCTACTATTGGGTCTGCTCAGGTGCCCCGACTCTACTAGTTCCAACATCATACCGATAGGAATACAAGACACTTTGCCAGCGGTTGTAGATTTATCATATTTCTCCACCACATATCCGTTACAATTTGTTAAGAATTTAGATACATCTATATTGCTACATAATCCAGCTAGTGAATATATATGACATAATAAATTGGTAAATCCTAGTTCTGGTATTTTACTAATTACTAAATCTGTATATCTTTCCATTTATCATGGAAAAGAATTTACCAGATATCTTCTTTTTTCCAAACTGCGTTAATGTATTCTCTTACAAATATTCACAAGATGAATTCAGTAATATGAGTAAAACGGAACGTGATAATTTCTCATTGGCTGTGTTTCCAGTGATAAAACATAGATGGCATAACGCACACGTTGTAAAACATAAAGGAATATACAAAGTTAGTACAGAAGCACGTGGAAAAAAAGTATCTCCTCCATCACTAGGAAAACCCGCACATATAAACCTAATGTCGAAGCAATATATATATAGTGAGTATGCAATAAGCTTTGAATGTTATAGTTTTCTAAAATGTATAACAAATACAGAAATCAATTCGTTCGATGAGTATATATTAAGAGGACTATTAGAAGCTGGTAATAGTTTACAGATATTTTCCAATTCCGTAGGCAAACGAATAGATACTATAGGTGTACTAGGGAATAAGTATCCATTTAGCAAAATTCCATTGGCCTCATTAACTCCTAAAGCACAACGAGAGATATTTTTAGCGTGGATTTCTCATAGACCTGTAGTTTTAACTGGAGGAACCGGAGTGGGTAAGACGTCACAGGTACCCAAGTTATTGCTTTGGTTTAATTATTTATTTGGTGGATTCTCTACTCTAGATAAAATCACTGACTTTCACGAAAGACCAGTCATTCTATCTCTTCCTAGGATAGCTTTAGTTAGATTGCATAGCAATACCATTTTAAAATCATTGGGATTTAAGGTACTAGATGGATCTCCTATCTCTTTACGGTACGGATCTATACCGGAAGAATTAATAAACAAACAACCAAAAAAATATGGAATTGTATTTTCTACCCATAAGTTATCTCTAACAAAACTATTTAGTTATGGCACTATTATTATAGACGAAGTTCATGAGCATGATCAAATAGGAGATATTATTATAGCAGTAGCGAGAAAGCATCATACGAAAATAGATTCTATGTTTTTAATGACTGCCACGTTAGAGGATGACAGGGAACGTCTAAAAATATTTTTACCTAATCCCGCATTTATACATATTCCTGGAGATACACTGTTTAAAATTAGCGAGGTATTTATTCATAATAAGATAAATCCATCTTCCAGAATGGCATATATAGAAGAAGAAAAGAGAAATTTAGTTACCGCTATACAGATGTATACTCCTCCTGATGGATCATCCGGTATAGTCTTTGTGGCATCCGTTGCACAGTGTCACGAATATAAATCATATTTAGAAAAAAGATTACCGTATGATATGTATATTATTCATGGTAAGGTCTTAGATATAGACGAAATATTAGAAAAAGTGTATTCATCACCTAATGTATCGATAATTATTTCTACTCCTTATTTGGAATCCAGCGTTACTATACGCAATGTTACACACATTTATGATATGGGTAGAGTTTTTGTCCCCGCTCCTTTTGGAGGATCACAACAATTTATTTCTAAATCTATGAGAGATCAACGAAAAGGAAGAGTAGGAAGAGTTAATCCTGGAACATACGTATATTTCTATGATCTGTCTTATATGAAGTCTATACAGCGAATAGATTCAGAATTTCTACATAATTATATATTGTACGCTAATAAGTTTAATCTAACACTCCCCGAAGATTTGTTTATAATCCCTACAAATTTGGATATTCTATGGCGTACAAAGGAATATATAGACTCGTTCGATATTAGTACAGAAACATGGAATAAATTATTATCCAATTATTATATGAAGATGATAGAGTATGCTAAACTTTATGTACTAAGTCCTATTCTCGCTGAGGAGTTGGATAACTTTGAGAGGACGGGAGAATTAACTAGTATTGTACAAGAAGCCATTTTATCTCTAAATTTACGAATTAAGATTTTAAAATTTAAACATAAAGATGATGATACGTATATACACTTTTGTAGAATATTATTCGGCGTCTATAACGGAACAAACGCTACTATATATTATCATAGACCTCTAACGGGATATATGAATATGATTTCAGATACTATATTTGTTCCTGTAGATAATAACTAAAAATCAAACTCTAATGACCACATCTTTTTTTAGAGATGAAAAATTTTCCACATCTCCTTTTGTAGACACGACTAAACATTTTGCAGAAAAAAGTTTATTATTATTTAGATAATCGTATACTTCATCAGTGTAGATAGTAAATGTGAACAGATAAAAGGTATTCTTGCTCAATAGATTGGTAAATTCCATAGAATATATTAATCCTTTCTTCTTGAGATCCCACATCATTTCAACCAAAGACGTTTTATCCAATGATTTACCTCGTACTATACCACATACAAAACTAGATTTTGCAGTGATGTCGTACCTGGTATTCCTACCAAACAAAATTTTACTTTTAGTTCTTTTAGAAAATTCTAAGGTAGAATCTCTATTTGTCAATATGTCATCTATGGAATTACCACTAGCAAAAAATGATAGAAATATATATTGATACATCGCAGCTGGTTTTGATCTACTATACTTTAAAAACGAATCAGATTCCATAATTGCCTGTATATCATCAGCTGAAAAACTATGTTTTACACGTATTCCTTCGGCATTTCTTTTTAATGATATATCTTGTTTAGACAATGATAAAGTTATCATGTCCATGGGAGATGCGTCTCCGTATCGTATAAATATTTCATTAGATGTTAGACGCTTCATTAGGGGTATACTTCTATAAGGTTTCTTAATTAGTCCATCATTGGTTGCGTCAAGAACTACTATCTGATGTTGTTGGGTATCTCTAGTGTTACACATGGCCTTACTAAAGTTTGGGTAAATAACTATGATATCTCTATTAATTATAGATGCATATATTTCATTCGTCAAGGATATTAATATCGACTTACTATCGTCATTAATACGTGTAATATAATCATATAAATCATGCGATAGCCAAGGAAAATTCAAATAGATGTTCATCATATAATCGTCGCTATAATTCATATTAATACTTTGACATTGACTAATTTGTAATATAGCCTCGCCACGAAGAAAGCTCTCGTATTCAGTTTCATCGATAAAGGATACCGTTAAATATAACTGGTTGCCGATAGTCTCATAGTCTATTAAGTGGTAAGTTTCGTACAAATACAGAATCCCTAAAATATTATCTAATGTGGGATTAATCCTTACCATAACTGTATAAAATGGAGCCGGAGTCATAACTATTTTACCGTTTGTACTTACTGGAATAGATGAAGGAATAATCTCCGGACATGATGGTAAAGACCCAAATGTCTGTTTGAAGAAATCCAATGTTCCAGGTCCTAATCTCTTGACAAAAATTACGATATTCGATCCCGATATCCTTTGCATTCTATTTACCAGCATATCACGAACTATATTAAGATTATCTATCATGTCTATTCTCCCACCGTTATATAAATCGCCTCCGCTAAGAAACGTTAGTATATCCATACAATGGAATACTTCATTTCTAAAATAGTATTCGTTTTCTAATTCTTTAATGTGAAATCGTATACTAGAAAGGGAAAAATTATCTTTGAGTTTTCCATTAGAAAAGAACCACGAAACTAATGTTCTGATTGCGTCTGTCTCCGTCGCTGAATTAATAGATTTACACCAAAAACTCATATAACTTCTAGATGTAGAAGCATTCGCTAAAAAATTAGTAGAATCAAAGGATATAAGTAGATGTTCCAACAAGTGAGCAATTCCCAAGATTTCATCTATATCATTCTCGAATCCGAAATTAGAAATTCCCAAGTAGATATCCTTTTTCATCCGATCATTGATGAAAATACGAACTTTATTCGGTAAGACGATCATTTACTAAGGAGTAAAATAGGAAGTAACGTTCGTATATCGTTATCGTCGTATAAATTAAAGGTGTGTTTTTTGCCATTAAGAGACATTATAATTTTACCAATATTGGAATTATAATATAGGTGTATTTGAGCACTAGAAACGGTCGATGCATCGGTAAATATAGCTGTATCTAATGTTCTAGTCGGTATTTCTTCATTTCGCTGTCTAATGATAGCGTTTTCTCTATCTGTTTCCATTACAGCTGCCTGAAGTTTATTGGTCGGGTAATATGTAAAATAATAAGAAATACATACGAATAACAAAAATAAAATAAGATATAATAAAGATGCCATTTAGAGATCTAATTTTGTTCAACTTGTCCAAATTCCTACTTACAGAAGATGAGGAATCGTTGGAGATAGTATCTTCCTTATGTAGAGGATTTGAAATATCTTACGATGACTTAATATCGTACTTTCCAGATAGGAAATACCATAAATATATTTCTAAGGTATTTGAACATGTAGATTTATCGGAGGAATTAAGTATGGAATTCCATGATACAACTCTGAGAGATTTAGTATATCTTAGATTGTACAAGTATTCCAAGTATATACGGCCGTGTTATAAATTAGGAGATAATCTAAAAGGTATAGTTGTTATAAAGGACAGAAATATATATATTAGAGAAGCAAATGATGACTTGATAGAATATCTCCTCAAGGAATACACTCCTCAGATTTATACATATTCTAATGAGCGAGTTCCCATAGCTGGTTCAAAATTAATTCTTTGTGGATTTTCTCAAGTTACATTTATGGCGTATACAACGTCGCATATAACAACAAATAAAAAGGTAGATGTTCTCGTTTCCAAAAAATGTATAGATGAACTAGTCGATCCAATAAATTATCAAATACTTCAAAATTTATTTGATAAAGGAAGCGGAACAATAAACAAAATACTCAGGAAGATATTTTATTCGGTAACAGGTGGCCAAACTCCATAGGTAGCTTTTTCTATTTCGGATTTTAGAATTTCCAAATTCACCAGCGATTTATCGGTTTTGGTGAAATCCAAGGATTTATTAATGTCCACAAATGCCATTTGTTTTGTCTGTGGATTGTATTTGAAAATGGAAACGATGTAGTTAGATAGATGCGCTGCGAAGTTTCCTATTAGGGTTCCGCGCTTCACGTCACCCAACATACTTGAATCACCATCCTTTAAAAAAAATGATAAGATATCAACATGGAGTATATCATACTCGGATTTTAATTCTTCTACTGCCTCACTGACATTTTCACAAATACTACAATACGGTTTACCGAAAATAATCAGTACGTTCTTCATTTATGGGTATCAAAAACTTAAAATCGTTACTGTTGGAAAATAAATCACTGACGATATTAGATGATAATTTATACAAAGTATACAATGGAATATTTGTGGATACAATGAGTATTTATATAGCCGTCGCCAATTGTGTCAGAAACTTAGAAGAGTTAACTACGGTATTCATAAAATACGTAAACGGATGGGTAAAAAAGGGAGGACATGTAACCCTTTTTATCGATAGAGGAAGTATAAAAATTAAACAAGACGTTAGAGACAAGAGACGTAAATATTCTAAATTAACCAAGGACAGAAAAATGTTAGAATTAGAAAAGTGTACATCCGAAATACAAAATGTTACCGGATTTATGGAAGAAGAAATAAAGGCAGAAATGCAATTAAAAATCGATAAACTCACATTTCAAATATATTTATCTGATTATGATAACATAAAAATATCATTGAATGAGATACTAACACATTTCAACAATAATGAGAATGTTACATTATTTTATTGTGATGAACGAGACGCAGAATTCGTTATGTGTCTAGAGGCTAAAACACAGTTCTCTACCACAGGAGAATGGCCGTTAATAATAAGTACCGATCAGGATACTATGCTATTCGCGTCTGCTGATAATCATCCTAAGATGATAATAAACTTAACTCAACTGTTTAAATTTGTTCCCTCGGCAGAGGATAACTATTTAGCAAAATTAACTGCATTAGTGAATGGATGTGATTTCTTTCCTGGACTCTATGGGGCATCTATAACACCCAACAACTTAAACAAAATACAATTGTTTAGTGATTTTACAATCGATAATATAGTCACTAGTTTGGCAATTAAAAATTATTATAGAAAGACTAACTCTACCGTAGACGTGCGTAATATTGTTACGTTTATAAACGATTACGCTAATTTAGAAGATGTCTACTCGTATATTCCTCCTTGTCAATGCACTGTTCAAGAATTTATATTCTCCGCATTAGATGAAAAATGGAATGAATTTAAATCATCTTATTTAGAGAGCGTGCCGTTACCCTGCCAATTAATGTACGCATTAGAACCACGTAAGGAGATTGATGTTTCAGAAGTTAAAACTTTATCATCTTATATAGATTTCGAAAATACTAAATCAGATATCGATGTTATAAAATCTATATCCTCGATTTTTGGATATTCTAATGAAAACTGTAACACCATAGTGTTCGGCATCTATAAGGATAATTTACTACTGAGTATAAATAATTCATTTTACTTTAACGATAGTCTGTTAATAACCAATACTAAAAGTGATAATATAATAAATATAGGTTACTAGATTAAAAATGGTGTTCCAGCTCGTGTGTTCTACGTGCGGCAAAGATATTTCTCACGAACGATATAAATTGATTATACGAAAAAAATCATTAAAGGATGTACTAGTCAGTGTAAAGAACGAATGTTGTAGGTTAAAATTATCTACACAAATAGAACCTCAACGTAACTTAACAGTGCAACCTCTATTGGATATAAACTAATGGATCCGGTTAATTTTATCAAGACATATGCGCCTAGAGGTTCTATTATTTCTATTAATTATGCCATGTCATTAACTAGTCATTTGAATCCATCGATAGAAAAACATGTGGGTATTTATTATGGTACGTTATTATCGGAACACTTGGTAGTTGAATCTACCTATAGAAAAGGAGTTAGAATAATCCCATTGGATAGATTTTTTGAAGGATATCTTAATGCAAAAGTATACATGTTAGAGAATATTCAAGTTATGAAAATAGCAGCTGATATGTCGTTAACTTTACTAGGTATTCCATATGGATTTGGTCATGATAGAATGTATTGTTTTAAATTGGTAGCTGAATGTTATAAAAATGCCGGTATTGATACATCGTCTAAACGAATATTAGGTAAAGATATTTTTCTGAGCCAAAACTTTACAGATGATAATAGATGGATAAAGATATATGATTCTAATAATTTAACATTTTGGCAAATTGATTACCTTAAAGGGTGAGTTAATATGCATAACTACTCCTCCGTTGTTTTTTCCCTCGTTCTTTTTCTTAACGTTGTTTGCCATCACTCTCATAATGTAAAGATATTCTAAAATGGTAAACTTTTGCATATCGGATGCAGAAATTGGTATAAATGTTGTAATTGTATTATTTCCCGTCAATGGACTAGTCACAGCTCCATCAGTTTTATATCCTTTAGAGTATTTCTCACTCGTGTCTAGCATTCTAGAGCATTCCATGATTTGTTTATCGTTGATATTGGCCGGAAAGATAGATTTTTTATTTTTTATTATATTACTATTGGCAATTGTAGATATAACTTCTGGTAAATATTTTTCTACCTTTTCATTCTCTTCTATTTTTAAGCCGGCTATATATTCTGCTATATTGTTGCTAGTATCAATACCTTTTCTGGCTAAGAAGTCATATGTGGTATTCACTATATCAGTTTTAACTGGTAGTTCCATTAGCCTTTCCACTTCTGCAGAATAATCAGAAATTGGTTCTTTACCAGAAAATCCAGCTACTATAATAGGCTCACCGATGATCATTGGCAAAATCCTATATTGTACCAGATTAATGAGAGCATATTTCATTTCCAATAATTCTGCTAGTTCTTGAGACATTGATTTATTTGATGAATCTATTTGGTTCTCTAGATACTCTACCATTTCTGCCGCATACAATAACTTGTTAGATAAAATCAGGGTTATCAAAGTGTTTAGTGTGGCTAGAATAGTGGGCTTGCACGTATTAAAGAATGCTGTAGTATGAGTAAACCGTTTTAACGAATTATATAGTCTCCAGAAATCTGTGGCGTTGCATACATGAACTGAATGACATCGAAGATTGTCCAATATTTTTAATAGCTGCTCTTTGTCCATTATTTCTATATTTGACTCGCAACAATTGTAGATACCATTAATCACTGATTCCTTTTTCGATGCCGGACAATAGCACAATTGTTTAGCTTTGGACTCTATGTATTCAGAATTAATAGATATATCTCTCAATACAGATTGCACTATACATTTTGAAACTATGTCAAAAATTGTAGAACGACGCTGTTCTGTAGCCATTTAACTTTAAATAATTTACAAAAATTTAAAATGAGCATCCGTATAAAAATCGATAAATTGCGCCAAATTGTGGCATATTTTTCAGAGTTCAGCGAAGAAGTGTCTATAAATGTAGACTCGACGGATGAGTTAATGTATATTTTTGCCGCCTTGGGCGGATCTGTAAACATTTGGGCCATTATACCTCTCAGTGCATCAGTGTTCTACCGCGGAGCCGAAAATATTGTGTTTAACCTTCCAGTGTCCAAGGTAAAATCGTGTTTGTGTAGTTTTCACAATGATGCTATCATAGATATAGAACCTGATCTGGAAAATAATCTAGTAAAACTTTCTAGTTATCATGTAGTAAGTGTCGATTGTAACAAGGAACTGATGCCTATTAGGACAGATACTACTATTTGTTTAAGTATAGATCAAAAGAAATCTTACGTATTTAATTTTCACAAGTATGAAGAAAAATGTTGTGGTAGAACCGTCATTCATCTAGAATGGTTGTTGGGCTTTATCAAGTGTATTAGTCAGCATCAGCATTTGGCTATTATGTTTAAAGATGACAATATTATTATGAAGACTCCTGGTAATACTGATGCGTTTTCCAGGGAATATTCTATGACTGAATGTTCTCAAGAACTACAAAAGTTTTCTTTCAAAATAGCTATCTCGTCTCTCAACAAACTACGAGGATTCAAAAAGAGAGTCAATGTTTTTGAAACTAGAATCGTAATGGATAATGACGATAACATTCTAGGAATGTTGTTTTCGGATAGAGTTCAATCCTTTAAGATTAACATCTTTATGGCGTTTTTAGACTAATACTTTCAATGAGATATATATGGGTGGCGGAGTAAGTGTTGAGCTCCCTAAACGGGATCCGCCTCCGGGAGTACCCACTGATGAGATGTTATTAAACGTGGATAAAATGCATGACGTGATAGCTCCCGCTAAGCTTTTAGAATATGTGCATATAGGACCACTAACAAAAGATAAAGAGGATAAAGTAAAGAAAAGATATCCAGAGTTTAGATTAGTCAACACAGGACCCGGTGGTCTTTCGGCATTATTAAGACAATCATATAATGGAACCGCACCCAATTGCTGTCGCACTTTTAATCGTACTCATTATTGGAAGAAGGATGGAAAGATATCAGATAAGTATGAAGAGGGCGCAGTATTAGAATCGTGTTGGCCCGACGTCCACGACACTGGAAAATGCGATGTTGATTTATTCGACTGGTGTCAGGGGGATACGTTCGATATGAACATATGCCATCAGTGGATCGGTTCAGCCTTTAATAGGAGTGATAGAACTGTAGAGGGTCGACAATCGTTAATAAATCTGTATAATAAGATGCAAACATTATGTAGTAAAGATGCTAGTGTACCAATATGTGAATTATTTTTGCATCATTTACGCGCACACAATACAGAAGATAGCAAAGAGATGATCGATTATATTCTAAGACAACAGTCGGCGGACTTTAAACAGAAATATATGAGATGTAGTTATCCCACTAGAGATAAGTTAGAAGAGTCATTAAAATATGCGGAACCTCGAGAATGTTGGGATCCAGAGTGTTCGAATGCCAATGTTAATTTCTTACTAACACGTAATTATAATAATTTAGGACTTTGCAATATTGTACGATGTAATACGAGCGTGAATAACTTACAGATGGATAAAACTTCCTCATTAAGATTATCATGTGGATTAAGCAATAGTGATAGATTTTCTACTGTTCCCGTCAATAGAGCAAAAGTAGTTCAACATAATATTAAACATTCGTTCGACCTAAAATTGCATTTGATCAGTTTATTATCTCTCTTGGTAATATGGATACTAATTGTAGCTATTTAAATGGGTGCCGCAGCAAGCATACAGACGACTGTGAATACACTCAGTGAACGTATCTCGTCTAAATTAGAACAAGAAGCGAACGCTAGTGCTCAAACAAAATGTGATATAGAAATCGGAAATTTTTATATCCGACAAAACCATGGATGTAACATCACTGTTAAAAATATGTGCTCTGCGGACGCGGATGCTCAGTTGGATGCTGTGTTATCAGCCGCTACAGAAACATATAGTGGATTAACACCGGAACAAAAAGCATACGTACCAGCTATGTTTACTGCTGCGTTAAACATTCAGACGAGTGTAAACACTGTTGTTAGAGATTTTGAAAATTATGTGAAACAGACTTGTAATTCTAGCGCTGTTGTCGATAACAAATTAAAGATACAAAACGTAATTATAGATGAATGTTACGGAGCCCCAGGATCTCCAACAAATTTGGAATTTATTAATACAGGATCTAGCAAAGGAAATTGTGCCATTAAGGCGTTGATGCAATTGACTACTAAGGCCACTACTCAAATAGCACCTAGACAAGTTGCTGGTACAGGTGTTCAGTTTTATATGATTGTTATCGGTGTTATAATATTGGCAGCGTTGTTTATGTACTATGCCAAGCGTATGCTGTTCACATCCACCAATGATAAAATCAAACTTATTTTAGCCAATAAGGAAAACGTCCATTGGACTACTTACATGGACACATTCTTTAGAACTTCTCCGATGATTATTGCTACCACGGATATACAAAACTGAAAATATATTGATAATATTTTAATAGATTAACATGGAAGTTATCGCTGATCGTCTAGACGATATAGTGAAACAAAATATAGCGGATGAAAAATTTGTAGATTTTGTTATACACGGTCTAGAGCATCAATGTCCTGCTATACTTCGACCATTAATTAGGTTGTTTATTGATATACTATTATTTGTTATAGTAATTTATATTTTTACGGTACGTCTAGTAAGTAGAAATTATCAAATGTTGTTGGTGTTGGTGGCGCTAGTCATCACATTAACTATTTTTTTATTACTTTATACTATAATAGTACTAGACTGACTTCTAACAAACATCTCACCTGCCATAAATAAATGCTTGATATTAAAGTCTTCTATTTCTAACACTATTCCATCTGTGGAAAATAATACTCTGACATTATCGCTAATTGATACATCGGTAAGTGATATGCCTATAAAGTAATAATCTTCTTTGGGTACATATACCAGTGTACCAGGTTCTAACAACCTATTTACTGGTGCTCCTGTAGCATACTTTTTTTTTACCTTGAGAATATCCATTGTTTGCTTGGTCAATAGCGATATGTGATTTTTTATCAACCACTCAAAAAAGTAATTGGAGTGTTCATATCCTCTACGGGCTATTGTCTCATGGCCGTGTATGAAATTTAAGTAACACGACTGTGGTAGATTTGTTCTATAGAGCCGGTTGCCGCAAATAGATAGAACTACCAATATGTTTGTACAAATGTTAAACATTAATTGATTAACAGAAAAAACAATGTTCGTTCTGGGAATAGAAACCAGATTAAAACAAAATTCATTAGAATATATGCCACGTTTATACATGGAATATAAAATAACTACAGTTTGAAAAATAACAGTATCATTTAAACATTTAACTTGCGGGGTTAATCTCACAACTTTACTGTTTTTGAACTGTTCAAAATATAGCATAGATCCATGAGAAATACGTTTAGCCGCCTTTAATAGAGGAAATCCAACCGCCTTTCTGGATCTCACCAACGACGATAGTTCTGACCAGCAACTCATTTCTTCATCATCCACCTGTTTTAACATATAATAGGCAGGAGATAGATATCCGTCATTGCAATATTCCTTCTCGTAGGCACACAATCTAATATTGATAAAATCTCCATTCTCTTCTCTGTATTTATTATCTTGTCTCGGTGGCTGATTAGGCTGTGGTCTATCGTTGTTGAATCTATTTTGGTCATTAAATCTTTCATTTCTTCCTGGTATATTTCTATCACCTCGTTTGGTTGGATTTTTGTCTATATTATCGTTTGTAACATCGGTACGGGTATTCATTTATCACAAAAAAAACTTCTCTAAATGAGTCTACTGCTAGAAAACCTCATCGAAGAAGATACCATATTTTTTGCAGGAAGTATATCTGAGTATGATGATTTACAAATGGTTATTGCTGGTGCAAAATCCAAATTTCCAAGATCTATGCTTTCTATTTTTAATATAGTACCTAGAACGATGTCAAAATATGAGTTGGAGTTGATTCATAACGAGAATATCACAGGGGCAATGTTTACCACAATGTATAATATAAGAAACAATTTGGGTCTAGGCGATGATAAACTAACTATTGAAGCCATTGAAAACTATTTCTTGGATCCTAACAATGAGGTTATGCCTCTTATCATTAATAATACGGATATGACTACCGTCATTCCTAAAAAAAGTGGTAGGAGAAAGAATAAGAACATGGTTATCTTCCGTCAAGGATCATCACCTATCTTGTGTATTTTCGAAACTCGTAAAAAGATTAATATTTATAAAGAAAATATGGAATCCGTATCGACTGAGTATACACCTATCGGAGACAACAAGGCTTTGATATCTAAATATGCGGGAATTAATATCCTGAATGTGTATTCTCCTTCCACGTCCATGAGATTGAATGCCATTTACGGATTCACCAATAAAAATAAACTAGAGAAACTTAGTACTAATAAGGAACTAGAATCGTATAGTTCTAGCCCTCTTCAAGAACCCATTAGGTTAAATGATTTTCTGGGACTATTGGAATGTGTTAAAAAGAATATTCCTCTAACAGATATTCCGACAAAGGATTGATTACTATAAATGGAGAATGTTCCTAATGTATACTTTAATCCTGTGTTTATAGAGCCCACGTTTAAACATTCTTTATTAAGTGTTTATAAACACAGATTAATAGTTTTATTTGAAGTATTCGTTGTATTCATTCTAATATATGTATTTTTTAGATCTGAATTAAATATGTTCTTCATGCCTAAACGAAAAATACCCGATCCTATTGATAGATTACGACGTGCTAATCTAGCGTGTGAAGACGATAAATTAATGATCTATGGATTACCATGGATAACAACTCAAACATCTGCGTTATCAATAAATAGTAAACCGATAGTGTATAAAGATTGTGCAAAGCTTTTGCGATCAATAAATGGATCACAACCAGTATCTCTTAACGATGTTCTTCGCAGATGATGATTCATTTTTTAAGTATTTTGCTAGTCAAGATGATGAATCTTCATTATCTGATATATTGCAAATCACTCAATATCTAGACTTTCTGTTATTATTATTGATCCAATCAAAAAATAAATTAGAAGCTGTGGGTCATTGTTATGAATCTCTTTCAGAGGAATACAGACAATTGACAAAATTCACAGACTCTCAAGATTTTAAAAAACTGTTTAACAAGGTCCCTATTGTTACAGATGGAAGGGTCAAACTTAATAAAGGATATTTGTTCGACTTTGTGATTAGTTTGATGCGATTCAAAAAAGAATCAGCTCTAGCTACCACCGCAATAGATCCTGTTAGATACATAGATCCTCGTCGTGATATCGCATTTTCTAACGTGATGGATATATTAAAGTCGAATAAAGTTGAACAATAATTAATTCTTTATTGTTATCATGAACGGCGGACATATTCAGTTGATAATCGGCCCCATGCTTTCAGGTAAAAGTACAGAATTAATTAGACGAGTTAGACGTTATCAAATAGCTCAATATAAATGTGTGACTATAAAATATTCTAACGATAATAGATACGGAACGGGACTATGGACGCATGATAAGAATAATTTTGCAGCATTGGAAGTAACTAAACTATGTGATGTCTTGGAAGCAATTACAGATTTCTCCGTGATAGGTATCGATGAAGGACAGTTCTTTCCAGACATTGTTGAATTCTGTGAGCGTATGGCAAACGAAGGAAAAATAGTTATAGTAGCCGCGCTCGATGGGACATTTCAACGTAGACCGTTTAATAATATTTTGAATCTTATTCCATTATCTGAAATGGTGGTAAAACTAACTGCAGTGTGTATGAAATGCTTTAAGGAGGCTTCCTTTTCTAAACGATTAGGTGCAGAAACCGAGATAGAAATAATAGGAGGTAATGATATGTATCAATCTGTGTGTAGAAAGTGTTACATCGACTCATAATATTATATTTTTTATCTAAAAAACTAAAAATAAACATTGATTAAATTTTAATATAATACTTAAAAATGGATGTTGTGTCGTTAGATAAACCGTTTATGTATTTTGAGGAAATTGATAATGAGTTAGATTACGAACCAGAAAGTGCAAATGAGGTCGCAAAAAAACTGCCGTATCAAGGACAGTTAAAACTATTACTAGGAGAATTATTTTTTCTTAGTAAGTTACAGCGACACGGTATATTAGATGGTGCCACCGTAGTGTATATAGGATCTGCTCCAGGTACACATATACGTTATTTGAGAGATCATTTCTATAATTTAGGAGTGATCATCAAATGGATGCTAATTGACGGCCGCCATCATGATCCTATTCTAAATGGATTGCGTGATGTGACTCTAGTGACTCGGTTTGTTGATGAGGAATATCTACGATCCATCAAAAAACAACTACATCCTTCTAAGATTATTTTAATTTCTGATGTGAGATCCAAACGAGGAGGAAATGAACCTAGTACTGCGGATTTACTAAGTAATTATGCTCTACAAAATGTCATGATTAGTATTTTAAACCCCGTGGCGTCTAGTCTTAAATGGAGATGCCCGTTTCCAGATCAATGGATCAAGGACTTTTATATCCCACACGGTAATAAAATGTTACAACCTTTTGCTCCTTCATATTCAGCTGAAATGAGATTATTAAGTATTTATACCGGTGAGAATATGAGACTGACTCGAGTTACCAAATCAGACGCTGTAAATTATGAAAAAAAGATGTATTACCTTAATAAGATAGTCCGCAACAAAGTAGTTGTTAACTTTGATTATCCTAATCAGGAATATGACTATTTTCACATGTACTTTATGTTGAGGACCGTATACTGCAATAAAACATTTCCTACTACTAAAGCAAAGATACTATTTCTACAACAATCTATATTTCGTTTCTTAAATATTCCAACGACATCAACTGAAAAAGTTAGTCATGAACCAATACAACGTAAAATATCTAGCAAAGATTCTATGTCTAAAAACAGAAATAGCAAGAGATCCGTACGCGGTAATAAATAGAAACGTACTACTGAGATATACTACCGATATAGAGTATAATGATTTAGTTACTTTAATAACCGTTAGACATAAAATTGATTCTATGAAAACTGTGTTTCAGGTATTTAACGAATCATCCATAAATTATACTCCGGTTGATGATGATTATGGAGAACCAATCATTATAACATCGTATCTTCAAAAAGGTCATAACAAGTTTCCTGTAAATTTTCTATACATAGATGTGGTAATATCTGACTTATTTCCTAGCTTTGTTAGACTAGATACTACAGAAACTAATATAGTTAATAGTGTACTACAAACAGGCGATGGTAAAAAGACTCTTCGTCTTCCTAAAATGTTAGAGACGGAAATAGTTGTCAAGATTCTCTATCGTCCTAATATACCATTAAAAATTGTTAGATTTTTCCGCAATAACATGGTAACTGGAGTAGAGATAGCCGATAGATCTGTTATTTCAGTCGCTGATTAATCAATTAGTAGAGATGAGATAAGAACATTATAATAATCAATAATATATCTTATATCTGTTTAGAAAAATGCTAATATTAAAATAGCTAACGCTAGTAATCCAATCGGAAGCCATTTGATATCTATAATAGGGTATCTAATTTCCTGATTCAGATAGCGTACGGCTATATTCTCGGTAGCTACTCGTTTGGAATCACAAACATTATTTACATCTAATTTACTATCTGTAATGGAAACGTTTCCCAATGAAATGGTACAATCAGATACATTACATCTTGATATATTTTTTTTTAAAGAGGCTGGTAACAACGCATCGCTTCGTTTACATGGCTCGTACCAACAATAATAGGGTAATCTTGTATCTATTCCTATCCGTACTATACTTTTATCAGGATAAATACATTTACATCGTATATCGTCTTTGTTAGCATCACAGAATGCATAAATTTGTTCGTCCGTCATGATAAAAATTTAAAGTGTAAATATAACTATTATTTTTATAGTTGTAATAAAAAGGGAAATTTGATTGTATACCTTCGGTTCTTTAAAAGAAACTGACTTGATAAAAATGGCTGTAATCTCTAAGGTTACGTATAGTCTATACGATCAAAAAGAGATTAATGCTACAGATATTATCATTAGTCATATTAAAAATGACGACGATATCGGTACCGTTAAAGATGGTAGACTAGGTGCTATGGATGGGGCATTATGTAAGACTTGTGGGAAAACGGAATTGGAATGTTTCGGTCACTGGGGTAAAGTAAGTATTTATAAAACTCATATAGTTAAGCCTGAATTTATTTCAGAAATTATTCGTTTACTGAATCATATATGTATTCATTGCGGATTATTGCGTTCACGAGAACCGTATTCCGACGATATTAACCTAAAAGAGTTATCGGGACACGCTCTTAGGAGATTAAAGGATAAAATATTATCCAAGAAAAAGTCATGTTGGAACAGCGAATGTATGCAACCGTATCAAAAAATTACTTTTTCAAAGAAAAAGGTTTGTTTCGTCAACAAGTTGGATGATATTAACGTTCCTAATTCTCTCATCTATCAAAAGTTAATTTCTATTCATGAAAAGTTTTGGCCATTATTAGAAATTCATCAATATCCAGCTAACTTATTTTATACAGACTACTTTCCCATCCCTCCGTTGATTATTAGACCGGCTATTAGTTTTTGGATAGATAGTATACCCAAAGAGACAAATGAATTAACTTACTTATTAGGTATGATCGTTAAGAATTGTAACTTGAATGCTGATGAACAGGTTATCCAGAAGGCGGTAATAGAATACGATGATATTAAAATTATTTCTAATAACACTACCAGTATCAATTTATCATATATCACATCCGGCAAAAATAATATGATTAGAAGTTATATCGTCGCTCGGCGAAAGATCAGACCAGCTAGATCTGTAATTGGTCCCAGTACATCTATCACCGTTAATGAGGTAGGAATGCCCACATATATTAGAAATACACTTACAGAAAAGATATTTGTTAATGCCTTTACAGTGGATAAAGTTAAACAACTATTAGCATCAAACCAAGTTAAATTTTACTTTAATAAACGATTAAACCAATTAACAAGAATACGTCAAGGAAAGTTTATCAAAAATAAAATACATTTATTGCCTGGTGATTGGGTAGAAGTAGCTGTTCAAGAATATACAAGTATTATTTTTGGAAGACAACCGTCTCTACATAGATACAACGTCATCGCTTCATCTATCAGAGCCACCGAAGGAGATACTATCAAAATATCTCCCGGAATTGCCAACTCTCAAAATGCTGATTTCGACGGAGATGAAGAATGGATGATATTGGAGCAAAATCCTAAAGTCGTAGTTGAACAAAGTATTCTTATGTATCCGACGACGTTACTCAAACACGATATTCATGGAGCCCCCGTTTATGGATCTATTCAAGATGAAATCGTAGCAGCGTATTCATTGTTTAGGATACAAGATCTTCGTTTAGATGAAGTATTGAACATCTTGGGGAAATATGGAAGAGAGTTCGATCCTAAAGGTAAATGTAAATTCAGCGGTAAAGATATCTATACTGACTTGATAGGTGAAAAGATTAATTATCCGGGTCTCTTAAAGGATGGCGAAATTATTGCAAACGACGTAGATAGTAATTTTGTTGTAGCTATGAGGCATCTGTCATTGGCTGGACTCTTATCCGATCATAAATCGAACGTGGAAGGTATCAACTTTATTATCAAGTCATCTTATGTTTTTAAGAGATATCTATCTATTTACGGTTTTGGGGTGACATTCAAAGATCTGAGACCAAATTCGACGTTCACTAATAAATTGGAGGCTATCAACGTAGAAAAAATAGAACTTATCAAAGAAGCATACGCCAAATATCTCAAAGATGTAAGAGACGGGAAAATAGTTCCATTATCTAAAGCTTTAGAGGCGGACTACTTGGAATCCATGTTATCCAACTTGACAAATCTTAATATCAGAGAGATAGAAGAACATATGAGACAAACGCTGATAGATGATCCAGATAATAACCTCCTGAAAATGGCCAAAGCGGGTTATAAAGTAAATCCCACAGAACTAATGTATATTCTAGGTACTTATGGACAACAGAGGATAGATGGCGAACCAGCAGAGACTCGAGTATTGGGTAGAGTCTTACCTTACTATCTTCCAGACTCTAAGGATCCAGAAGGAAGAGGTTATATTCTTAATTCTTTAACAAAAGGATTAACGGGTTCTCAATATTACTTTTCGATGCTGGTTGCAAGATCTCAATCTACTGATATCGTCTGTGAAACATCACGTACCGGAACACTGGCTAGAAAAATCATTAAAAAGATGGAGGATATGGTGGTCGACGGATACGGACAAGTAGTTATAGGTAATACGCTCATCAAGTACGCAGCCAATTATACCAAAATTCTAGGCTCAGTATGTAAACCTGTAGATCTTATCTATCCAGATGAGTCCATGACTTGGTATTTGGAAATTAGTGCTTTGTGGAATAAAATAAAACAGGGATTCGTTTACTCTCAGAAACAGAAACTTGCAAAGAAGACATTGGCGCCGTTTAATTTCCTAGTATTCGTCAAACCCACCACTGAGGATAATGCTATTAAGGTTAAGGATCTGTACGATATGATTCATAACGTCATTGATGATGTGAGAGAGAAATACTTCTTTACGGTATCTAATATAGATTTTATGGAGTATATATTCTTGACGCATCTTAATCCTTCTAGAATTAGAATTACAAAAGAAACGGCTATCACTATCTTTGAAAAGTTCTATGAAAAACTCAATTATACTCTAGGTGGTGGAACTCCTATTGGAATTATTTCTGCACAGGTATTGTCTGAGAAGTTTACACAACAAGCCCTGTTCAGTTTTCACACTACTGAAAAGAGTGGTGCTGTAAAACAAAAACTTGGTTTCAACGAGTTTAATAACTTGACTAATTTGAGTAAGAATAAGACCGAAATTATCACTCTGGTATCCGATGATATCTCTAAACTTCAATCTGTTAAGATTAATTTCGAATTTGTATGTTTGGGAGAATTAAATCCAGACATCACTCTTCGAAAAGAAACAGATAGATATGTAGTAGACATAATAGTCAATAGATTATACATCAAGAGAGCAGAAATAACCGAATTAGTCGTCGAATATATGATTGAACGATTTATCTCCTTTAGCGTCATTGTAAAGGAATGGGGCATGGAGACATTCATTGAGGACGAGGATAATATTAGATTTACTGTCTACCTAAATTTCGTTGAACCGGAGGAATTGAATCTTAGTAAGTTTATGATGGTTCTTCCGGGTGCCGCCAACAAGGGCAAGATTAGTAAATTCAAGATTCCTATCTCTGACTATACGGGATATAACGACTTCAATCAAACAAAAAAGCTCAATAAGATGACTGTAGAACTCATGAATCTAAAAGAATTGGGTTCTTTCGATTTGGAGAACGTCAACGTGTATCCTGGAGTATGGAATACATACGATATCTTCGGTATTGAGGCCGCTCGTGGATACTTGTGCGAAGCCATGTTAAACACCTATGGAGAAGGTTTCGATTATCTGTACCAGCCTTGTGATCTTCTCGCTAGTTTACTATGTGCTAGTTACGAACCAGAATCAGTTAATAAATTCAAGTTCGGTGCAGCTAGTACTCTTAAGAGAGCTACGTTCGGAGATAATAAAGCATTGTTAAACGCGGCTCTTCATAAAAAGTCAGAACCTATTAACGATAATAGTAGCTGCCACTTTTTTAGCAAGGTCCCTAATATAGGAACTGGATATTACAAATACTTTATCGACTTGGGTCTTCTCATGAGAATGGAAAGGAAATTATCTGATAAGATATCTTCTCAAAAGATCAAGGAGATAGAAGAAACAGAAGACTTTTAATTCTTATCAATAACATATTTTTCTATGATCTGTCTTTTAAACGATGGATTTTCCACAAATGCGCCTCTCAAGTCCCTCATAGAATGATACACGTATAAAAAATATAGCATAGGTGATGACTCCTTATTTTTAGACATTAGATATGCCAAAATCATAGCCCCGCTTCTATTTACTCCCGCAACACAATGAACCAACACGGGCTCGTTTCGTTGATCACATTTAGATAAGAAGGCGGTCACGTCGTCAAAATATTTACTAATATCAGTAGTTGTATCATCTACCAACGGTATATGAATAATATTAATATTAGAGTTAGGTAATGTATATTTATCCATCGTCAAATTTAAAACATATTTGAACTTAACTTCAGATGATGGTGCATCCATAGCATTTTTATAATTTCCCAAATACACATTATTTGTTACTCTTGTCATTATAGTGGGAGATTTGGCTCTGTGCATATCTCCAGTTGAACGTAGTAGTAAGTATTTATACAAACTTTTCTTATCCATTTATAACGTACAAATGGATAAAACTACTTTATCAGTAAACGCATGCAATTTAGAATACGTTAGAGAAAAGGCTATAGTAGGCGTACAAGCAGCCAAGACATCAACACTTATATTTTTTGTTATTATATTGGCAATTAGTGCGCTATTACTCTGGTTTCAGACGTCTGATAATCCAGTCTTTAATGAATTAACGAGATATATGCGAATTAAAAATACGGTTAACGATTGGAAATCATTAACGGATAGCAAAACAAAATTAGAAAGTGATAGAGGTAGACTTCTAGCCGCTGGTAAGGATGATATATTCGAATTCAAATGTGTGGATTTCGGCGCCTATTTTATAGCTATGCGATTGGATAAGAAAACATATCTGCCGCAAGCTATTAGGCGAGGTACTGGAGACGCGTGGATGGTTAAAAAGGCGGCAAAAGTCGATCCATCTGCTCAACAATTTTGTCAGTATTTGATAAAACACAAGTCTAATAATGTTATTACTTGTGGTAATGAGATGTTAAATGAATTAGGTTATAGCGGTTATTTTATGTCACCGCATTGGTGTTCCGATCTTAGTAATATGGAATAAGTGTTAGATAAATGCGGTAACAAATGTTCCTGTAAGGAACCATAACAGTTTAGATTTAACATTAAAGATGAGCATAAACATAATAAACAAAATTACAATCAAACCTATAACATTAATATCAAACAATCCAAAAAATGAAATCAGTGGAGTAGTAAACGTGTACATGACTCCTGGATAACGTTTAGCAGCTACCGTTCCTATTCTAGACCAAAAATTTGGTTTCATGGTTTCGAAGCGGTGTTCTGCAACAAGACGAGGATCGTGTTCTACATATTTGGCAGAGTTATCCATTATTTGCCTGTTAATCTTCATTTCGTTTTCGATTCTGGCTATTTCAAAATAAAATCCCGATGATAGACCTCCAGACTTTATAATTTCATCTACGATGTTCAGCGCCGTAGTAACTCTAATAATATAGGCTGATAAGCTAACATCATACCCTCCTGTATATGTGAATATGGCATGATCTTTGTCTATTACAAGCTCGGTTTTAACTTTATTTCCTGTAATAATTTCTCTCATCTGTAGGATATCTATTTTCTTGTCATGTATTGCCTTCAAGACGGGACGAAGAAACGTAATATCCTCAATAACGTTATCGTTTTCTATAATAACTACATATTCTACATTTTTATTTTCTAGCTCGATAAAAAATTTAGAATCCCATAGGGCTAAATGTCTAGCGATATTTCTTTTCGTTTCCTCTGTACACATAGTGTTACAAAACCCTGAAAAGAAGTGAGTATACTTGTCATCATCTCTAATATTTCCTCCAGTCCATTGTATAAACACATAATCCTTGTAATGATCTGGATCATCATTGACTATCACAACATCTCTTTTTTCTTGCATAACTTCATTGTCCTTCACATCATCGAACTTCTGATCATTAATATGCTCATGAACATTAGGAAATGTTTCTGATGGAGGTCTATCAATAACTGGCACAACAATAACAGGAGTTTTCGCCGCCGCCATTTAGTTATTGAAATTAATCATATACAACTCTCTAATACGAGTTATATTTTCGTCTATCCATTGTTTCACATTGACATATTTCGACAAAAAGATATAAAATGCGTATTCCAATGCTTCTCTGTTTAATGAATTACTAAAATATACAAACACGTCACTGTCTGGTAATAAATGATATCTTAGAATATTGTAACAATTTATTTTGTATTGTACATGTTCGTGATCTATGAGTTCTTCTTCGAATGGCATAGGATCTCCGAATCTGAAAACGTATAAATAGGAGTTAGAATAATAATATTTGAGAGTATTGGTAATGTATAAACTCTTTAGCGGTATAATTAGTTTTTTTCTCTCGATTTCTATTTTTAGATGTGATGGAAAAATGACTAATTTTGTAGCATTAGTATCATGAACTCTAATCAAAATCTTAATATCTTCGTCACACGTTAGCTCTTTGAAGTTTTTAAGAGATGCATCAGTTGGTTTTACAGATGGAGTAGGTGTAACAATTTTTTGTTTAATGCATGTATGTATTGGAGCCATTGTCTTAACTATAATGGTGCTTGTATCGAAAAACTTTAATGCGGATAACGGAAGCTCTTCGCCGCGACTTTCTACGTCGTAATTGGGTTCTAATGCCGATCTCTGAATGGATACTAGTTTTCTAAGTTCTAATGTGATTCTCTGAAAATGTAAATCCAATTCCTCCGGCATTATAGATGTGTATACATCGGTAAATAAAACTATAGTATCCAACGATCCCTTCTCGCAAATTCTAGTCTTAACCAAGAAATCGTATATAACTACGGAGATGGCGTATTTAAGAGTGGATTCTTCTACCGTTTTGTTCTTGGATTTCATATAAGAAACTATAAAGTCCGCACTACTGTTAAGAATGATCACTAACGCAACTATATAGTTCAAATTAAGCATCTTGGAAACATAAAATAACTCTGTAGATGATACTTGACTTTCGAATAAGTTTGCAGACAAACGAAGAAAGAACAGACCTCTCTTAATTTCAGAAGAAAACTTTTTTTCGTATTCCTGACGTCTAGAGTTTATATCAATAAGAAAGTTAAGAATTAGTCGGTTAATGTTGTATTTCATTACCCAAGTTTGAGATTTCATAATATTGTCAAAAGACATGATAATATTAAAGATAAAGCGCTGACTATGAACGAAATAGCTATATGGTTCGCTCAAGAATATAGTCTTGTTAAACGTGGAAACGATAACTGTATTTTTAATCACGTCAGCGACATCTAAATTAAATATAGGTATATTTATTCCACACACACTACAATATGCCACACCATCTTCATAATAAATAAATTCGTTAGCAAAATTATTAATTTTAGTGAAATAGTTAGCGTCAACTTTCATAGCTTCCTTCAATCTAATTTGATGCTCACATGGCGCGAATTCTACTCTAACATCCCTTTTCCATGCCTCAGGTTCATCGATCTCTATAATATCTAGTTTCTTGCGTTTCACAAACACAGGCTCGTCTCTCGCGATGAGATCTGTATAGTAACTATGTAAATGATAACTAGATAGAAAGATGTAGCTATATAGATGACGATCCTTTAAGAGAGGTATAATAACTTTACCCCAATCAGATAGACTGTTGTTATGGTCTTCGGAAAAAGAATTTTTATAAATTTTTCCAGTATTTTCTAAATATACATACTTGATATCTAAGAAATCCTTAATGATAATAGGAATGGATAATCCGTCTATTTTATAAAGAAATACATATCGCATATTATACTTTTTTTTGGAAATTGGAATACCGATGTGTCTACATAAATACGCAAAGTCTAAATATTTTTTAGAGAATCTTAGTTGGTCCAAATTCTTTTCCAAGTACGGTAATAGATTTTTCATATTGAACGGTATCTTCTTGATCTCTGGTTCTAATTCCGCATTAAATGATGAAACTAAGTCACTATTTTTATAACTAACGATTACATCACCTCTAACATCATCATTTACCAGGATACTGATCTTCTTTTGTCGTAAATACATGTCTAATGTGTTAAAAAAAAGATCATACAAGTTATACGTCATTTCATCTGTAGTATTCTTGTCATTGAAGGATAAACTCGTACTAATCTCTTCTTTAACAGTCTGTTCAAATTTATATCCTATATACGAAAAAATAGCAACCAGTGTTTGATCATCCGCGTCAATATTCTGTTCTATCGTAGTGTATAGCAATCTTATATCTTCTTCTGTGATAGTCGATACGTTATAAAGGTTGATAACGAAAATATTTTTATTTCGTGAAATAAAGTCATTGTAGGATTTTGGACTTATATTCGTGTCTAGTAGATATGATTTTATTTTTGGAATGATCTCAATTAAAATAGTCTCTTTAGAGTCCATTTAAAGTTACAAACAACTAGGAAATTGGTTTATGATGTATAATTTTTTTAGTTTTTATAGATTCTTTATTCTATACTTAAAAAATGAAAATAAATACAAAGGTTCTTGAGGGTTGTGTTAATTGAAAGCGAGAAATAATCATAAATTATTTCATTATCGCGATATCCGTTAAGTTTGTATCGTAATGGCGTGGTCAATTACGAATAAAGCGGATACTAGTAGTTTCACAAAGATGGCTGAAATCAGAGCTCATCTAAGAAATAGCGCTGAAAATAAAGATAAAAACGAGGATATTTTCCCGGAAGATGTAATAATTCCATCTACTAAGCCCAAAACCAAACGAACCACTACTCCTCGTAAACCAGCGGCTACTAAAAGATCAACCAAAAAGGATAAAGAAAAGGAGGAAGTGGAAGAAGTAGAAGAAGTAGTTATAGAGGAATATCATCAAACAACTGAAGAAAATTCTCCACCTCCGTCATCATCTCCTGGAGTCGGCGACATTGTAGAAAGCGTGGCCGCTGTAGAGCTCGATGATAGCGACGGGGATGATGAACCTATGGTACAAGTTGAAGCTGGTAAAGTAAATCATAGTGCTAGAAGCGATCTCTCTGACCTAAAGGTGGCTACCGACAATATCGTTAAAGATCTTAAGAAAATTATTACTAGAATCTCTGCAGTATCGACTGTTCTAGAGGATGTTCAAGCAGCTGGTATCTCTAGACAATTTACTTCTATGACTAAAGCTATTACAACACTATCTGATCTAGTCACCGAGGGAAAATCTAAAGTTGTTCGTAAAAAAGTTAAAACTTGTAAGAAGTAAATGCGTGCACTTTTTTATAAAGATGGTAAACTGTTTACCGATAATAATTTTTTAAATCCTGTATCAGACGATAATCCAGCGTATGAGGTTTTGCAACATGTTAAAATTCCTACTCATTTAACAGATGTAGTAGTATATGAACAAACGTGGGAAGAGGCATTAACTAGATTAATTTTTGTGGGAAGTGATTCAAAAGGACGTAGACAATACTTTTACGGAAAAATGCATGTACAGAATCGCAATGCTAAAAGAGATCGTATTTTTGTTAGAGTATATAACGTTATGAAACGAATTAATTGTTTTATAAACAAAAATATAAAGAAATCGTCCACAGATTCCAATTATCAGTTGGCGGTTTTTATGTTAATGGAAACTATGTTTTTTATTAGATTTGGTAAAATGAAATATCTTAAGGAGAATGAAACAGTAGGGTTATTAACACTAAAAAATAAACACATAGAAATAAGTCCCGATGAAATAGTTATCAAGTTTGTAGGAAAGGACAAAGTTTCACATGAATTTGTTGTTCATAAGTCTAATAGACTATATAAACCGTTATTGAAACTGACTGATGATTCTAGTCCCGAAGAATTTCTGTTCAACAAACTAAGTGAACGAAAGGTATACGAATGTATCAAACAGTTTGGTATTAGAATCAAGGATCTCCGAACGTATGGAGTCAATTATACGTTTTTATATAATTTTTGGACAAATGTAAAGTCCGTATCTCCTCTTCCATCACCAAAAAAGTTGATAGCGTTAACTATCAAACAAACTGCTGAAGTGGTAGGTCATACTCCATCAATTTCAAAAAGAGCTTATATGGCAACGACTATTTTAGAAATGGTAAAGGATAAAAATTTTTTAGACGTAGTATCTAAAACTACGTTCGATGAATTCCTATCTATAGTCGTAGATCACGTTAAATCATCTACGGATGGATGATAATAGATCTTTACACAAATAATTACAAGACCGATAAATGGAAATGGATAAACGGATGAAATCTCTCGCTATGACAGCTTTCTTCGGAGAGCTAAACACGTTAGATATTATGGCATTGATAATGTCTATATTTAAACGCCATCCAAACAATACCATTTTTTCAGTGGATAAGGATGGTCAATTTATGATTGATTTCGAATACGATAATTATAAGGCTTCTCAATATTTGGATCTGACCCTCACTCCGATATCTGGAAATGAATGCAAGACTCACGCATCGAGTATAGCCGAACAATTGGCGTGTGTGGATATTATTAAAGAGGATATTAGCGAATATATCAAAACTACTCCCCGTCTTAAACGATTTATAAAAAAATACCGCAATAGATCATATACTCGTATCAGTCGAGATACAGAAAAGCTTAAAATAGCTCTAGCTAAAGGCATAGATTACGAATATATAAAAGACGCTTGTTAATAAGTAAATGAAAAAAAACTAGTCGTTTATAATAAAACACGATATGGATGCCAACATAGTATCATCTTCTACTATTGCGACGTATATAGACGCTTTAGCGAAGAATGCTTCAGAATTAGAACAGAGGTCTACCGCATACGAAATAAATAATGAATTGGAACTAGTATTTATTAAACCGCCATTGATTACGTTGACAAATGTAGTAAATATCTCCACGATTCAGGAATCGTTTATTCGATTTACCGTTACTAATAAGGAAGGTATCAAAATTAGAACTAAGATTCCATTATCTAAGGTACATGGTCTAGATGTAAAAAATGTACAGTTGGTAGATGCTATAGATAACATAGTTTGGGAAAAGAAATCATTAGTGACGGAAAATCGTCTTCACAAAGAATGCTTGTTGAGACTATCGACAGAGGAACGTCATATATTTTTGGATTACAAGAAATATGGATCCTCTATCCGACTAGAATTAGTCAATCTTATTCAAGCAAAAACAAAAAACTTTACGATAGACTTTAAGCTAAAATATTTTCTAGGATCTGGTGCTCAATCTAAAAGTTCTTTATTGCACGCTATTAATCATCCAAAGTCAAGGCCTAATACATCTCTGGAAATAGAATTTACACCTAGAGACAATGAAACAGTTCCATATGATGAACTAATAAAGGAATTGACGACTCTCTCGCGTCATATATTTATGGCTTCTCCAGAGAATGTAATTCTTTCTCCACCTATTAACGCACCTATAAAGACTTTTATGTTGCCTAAACAAGATATAGTAGGTCTGGATCTGGAAAATCTATATGCCGTAACTAAGACTGATGGCATTCCTATAACTATCAGAGTTACATCAAAAGGGTTGTATTGTTATTTTACACATCTTGGTTATATTATTAGATATCCAGTTAAGAGAACAATAGATTCCGAAGTAGTAGTCTTTGGTGAGGCAGTTAAGGATAAGAACTGGACCGTATATCTCATTAAGCTAATAGAGCCCGTAAATGCAATCAGTGATAGACTAGAAGAAAGTAAGTATGTTGAATCTAAACTAGTGGATATTTGTGATCGGATAGTATTCAAGTCAAAGAAATACGAAGGTCCGTTTACTACAACTAGTGAAGTCGTCGATATGTTATCTACATATTTACCAAAGCAACCAGAAGGTGTTATTCTGTTCTATTCAAAGGGACCTAAATCTAACATTGATTTTAAAATCAAAAAGGAGAATACTATAGACCAAACTGCAAATGTAGTATTTAGGTACATGTCCAGTGAACCAATTATCTTTGGAGAGTCGTCTATCTTTATAGAGTATAAGAAATTTACCAACGATAAAGGCTTTCCTAAAGAATATGGTTCTGGTAAGATTGTGTTATATAACGGCGTTAATTATCTAAATAATATCTATTGTTTGGAATATATTAATACACATAATGAAGTGGGTATTAAGTCCGTTGTTGTACCTATTAAGTTTATAGCAGAATTCTTAGTCAATGGAGAAATACTTAAACCTAGAATCGATAAAACCATGAAATATATTAACTCAGAAGACTATTATGGAAATCAACATAATATCATAGTCGAACATTTAAGAGATCAAAGCATCAAAATAGGAGATGTCTTTAACGAGGATAAACTATCGGATATTGGACATCAATACGCCGCCAACAACGATAAATTTAGATTAAATCCAGAAGTTAGTTATTTTACTAATAAACGAACTAGAGGGCCGTTGGGAATTTTATCAAACTACGTCAAGACTCTTCTTATTTCTATGTATTGTTCCAAAACATTTTTAGACGATTCCAACAAACGAAAGGTATTAGCGATTGATTTTGGAAACGGTGCTGACCTGGAAAAATACTTTTATGGAGAGATTGCGTTATTGGTAGCGACGGATCCGGATGCTGATGCTATAGCTAGAGGAAATGAAAGATACAACAAATTAAATTCTGGAATTAAAACCAAGTACTACAAATTTGACTACATTCAGGAAACTATTCGATCCGATACATTTGTCTCTAGTGTCAGAGAAGTATTCTATTTTGGAAAGTTTAATATCATTGACTGGCAGTTCGCTATTCATTATTCTTTTCATCCAAGACATTATGCTACAGTCATGAATAACTTATCCGAACTAACTGCTTCTGGAGGCAAGGTATTAATTACTACCATGGATGGAGACAAATTATCAAAATTAACCGATAAAAAGACTTTTATAATTCATAAGAATCTACCTAGTAGCGAAAACTATATGTCTGTAGAAAAAATAGCTGATGATAGAATAGTGGTATATAATCCATCAACAATGTCTACTCCAATGACTGAATACATTATCAAAAAGAACGATATAGTCAGAGTGTTTAATGAATACGGATTTGTTCTTGTAGATAATGTTGATTTCGCTACAATTATAGAACGAAGTAAAAAGTTTATTAATGGCGCATCTACAATGGAAGATAGACCGTCTACAAGAAACTTTTTCGAACTAAATAGAGGAGCCATTAAATGTGAAGGTTTAGATGTCGAAGACTTACTTAGTTACTATGTTGTTTATGTCTTTTCTAAGCGGTAAATAATAATATGGTATGGGTTCTGATATCCCCGTTCTAAATGCATTAAATAATTCCAATAGAGCGATTTTTGTTCCTATAGGACCTTCCAACTGTGGATACTCTGTATTATTAATAGATATATTAATACTTTTGTAGGGTAACAGAGGTTCTACGTCTTCTAAAAATAAAAGTTTTATAACATCTGGCCTGTTCATAAATAAAAACTTGGCGATTCTATATATACTCTTATTATCAAATCTAGCCATTGTCTTATAGATGTGAGCTACTGTAGGTGTACCATTTGATTTTCTTTCTAATACTATATATTTCTCTCGAAGAAGTTCTTGCAGATCATCTGGGAATAAAATACTACTATTGAGTAAATCAGTTATTTTTTTTATATCGATATTGATGGACATTTTTATAGTTAAGGATAATAAGTATCCCAAAGTAGATAACGACGATAACGAAGTATTTATACTTTTAGGAAATCACAATGACTTTATCAGATCAAAATTAACAAAATTAAAGGAGCATGTATTTTTTTCTGAATATATTGTGACTCCAGATACATATGGATCTTTATGCGTCGAATTAAATGGGTCTAGTTTTCAGCACGGTGGTAGATATATAGAGGTGGAGGAATTTATAGATGCTGGAAGACAAGTTAGATGGTGTTCTACATCCAATCATATATCTGAAGATATACACACTGATAAATTTGTCATTTATGATATTTATACGTTTGATTCGTTCAAGAATAAACGATTGGTATTTGTACAGGTGCCTCCATCATTAGGAGATGATAGCTATTTAACTAATCCGTTATTGTCTCCGTATTATCGTAATTCAGTAGCCAGACAAATGGTCAATGATATGATTTTTAATCAAGATTCATTTTTAAAATATTTATTAGAACATCTGATTAGAAGCCACTATAGAGTTTCTAAACATATAACAATAGTTAGATACAAGGATACCGAAGAATTAAATCTAACAAGAATATGTTATAATAGAGATAAGTTTAAGGCATTTGTATTCGCTTGGTTTAACGGCGTTTCGGAAAATGAAAAGGTACTAGATACGTATAAAAAGGTATCTGATTTGATATAATGAATTCAGTGACTATATCACACGCACCATATACTATTACTTATCACGATGATTGGGAACCAGTAATGAGTCAATTGGTAGAGTTTTATAACGAAGTAGCCAGTTGGCTGCTACGAGACGAGACGTCGCCTATTCCTGATAAGTTCTTTATACAATTGAAACAGCCGCTTAGAAATAAACGAGTATGTGTGTGTGGTATAGATCCGTATCCAAAAGATGGAACTGGTGTACCGTTCGAATCACCAAATTTTACAAAAAAATCAATTAAGGAGATAGCTTCATCTATATCTAGATTAACCGGAGTAATTGATTATAAAGGTTATAACCTTAATATAATAGACGGGGTTATACCCTGGAATTATTACTTAAGTTGTAAATTAGGAGAAACAAAAAGTCACGCGATTTACTGGGATAAGATTTCCAAGTTACTGCTGCAGCATATAACTAAACACGTTAGTGTTCTTTATTGTTTGGGTAAAACAGATTTCTCGAATATACGGGCAAAGTTAGAATCCCCGGTAACTACCATAGTGGGATATCATCCAGCGGCCAGAGACCACCAATTCGAGAAAGATCGATCATTTGAAATTATCAACGTTTTACTGGAATTAGACAACAAGACACCTATAAATTGGGCTCAAGGGTTTATTTATTAATGCTTTAGTGAAATTTTAACTTGTGTTCTAAATGGATGCGGCTATTAGAGGTAATGATGTTATCTTTGTTCTTAAGACTATAGGTGTCCCGTCAGCATGCAGACAAAATGAAGATCCAAGATTCGTAGAAGCATTTAAATGCGACGAGTTAGAAAGATATATTGATAATAATCCAGAATGTACACTATTCGAAAGTCTTAGGGATGAGGAAGCATACTCTATAGTCAGAATTTTCATGGATGTAGATTTAGACGCGTGTCTAGACGAAATAGATTATTTAACGGCTATTCAAGATTTTATTATCGAGGTGTCAAACTGTGTAGCTAGATTCGCATTTACAGAATGCGGTGCCATTCATGAAAATGTAATAAAATCCATGAGATCTAATTTTTCATTGACTAAGTCTACAAATAGAGATAAAACAAGTTTTCATATTATCTTTTTAGACACGTATACCACTATGGATACATTGATAGCTATGAAACGAACACTATTAGAATTAAGTAGATCATCTGAAAATCCACTAACAAGATCGATAGACACTGCCGTATATAGGAGAAAAACAACTCTTCGGGTTGTAGGTACTAGGAAAAATCCAAATTGCGACACTATTCATGTAATGCAACCACCTCATGATAATATAGAAGATTACCTATTCACTTACGTGGATATGAACAACAATAGTTATTACTTTTCTCTACAACGACGATTGGAGGATTTAGTTCCTGATAAGTTATGGGAACCAGGGTTTATTTCGTTCGAAGACGCTATAAAAAGAGTTTCAAAAATATTCATTAATTCTATAATAAACTTTAATGATCTCGATGAAAATAATTTTACAACGGTACCACTGGTCATAGATTATGTAACACCTTGTGCATTATGTAAAAAACGATCGCATAAACATCCGCATCAACTATCGTTGGAAAATGGTGCTATTAGAATTTACAAAACTGGTAATCCACATAGTTGTAAAGTTAAAATTGTTCCGTTGGATGGTAATAAACTGTTTAATATTGCACAAAGAATTTTAGACACTAACTCTGTTTTATTAACCGAACGAGGAGACCATATAGTTTGGATTAATAATTCATGGAAATTTAACAGCGAAGAACCATTGATAACAAAACTAATTCTATCAATAAGACATCAACTACCTAAGGAATATTCAAGCGAATTACTCTGTCCGAGGAAACGAAAGACTGTAGAAGCTAACATACGAGACATGTTAGTAGATTCAGTAGAGACCGATACCTATCCGGATAAACTTCCGTTTAAAAATGGTGTATTGGACCTGGTAGACGGAATGTTTTACTCTGGAGATGATGCTAAAAAATATACGTGTACTGTATCGACCGGATTTAAATTTGACGATACAAAATTCGTCGAAAACAGTCCAGAAATGGAAGAGTTAATGAATATCATTAACGATATCCAACCATTAACGGATGAAAATAAGAAAAATAGAGAGCTGTATGAAAAAACATTATCTAGTTGTTTATGTGGTGCTACCAAAGGATGTTTAACATTCTTTTTTGGAGAAACCGCAACTGGAAAGTCGACAACCAAACGTTTGTTAAAGTCTGCTATCGGTGACCTGTTTGTCGAGACGGGTCAAACAATTTTAACAGATGTATTGGATAAAGGACCTAATCCATTTATCGCTAATATGCATTTAAAAAGATCTGTATTCTGTAGCGAACTACCTGATTTTGCATGTAGTGGATCAAAGAAAATTAGATCTGATAATATTAAAAAGTTGACAGAACCTTGTGTCATTGGAAGACCGTGTTTCTCCAATAAAATTAATAATAGAAACCATGCGACAATCATTATCGATACTAATTACAAACCTGTCTTTGATAGGATAGATAACGCATTAATGAGAAGAATTGCCGTCGTGCGATTCAGAACACACTTTTCTCAACCTTCTGGTAGAGAGGCTGCTGAAAATAATGACGCGTACGATAAAGTCAAACTATTAGACGAGGGATTAGATGGTAAAATACAGAATAATAGATATAGATTCGCATTTCTATACTTGTTGGTTAAATGGTACAAAAAATATCATATTCCTATTATGAAACTATATCCTACACCGGAAGAGATTCCGGACTTTGCATTCTATCTCAAAATAGGTACTCTGTTGGTATCTAGCTCTGTAAAGCATATTCCATTAATGACAGACCTCTCCAAAAAGGGATATATATTGTACGATAATGTGGTTACTCTTCCGTTGACTACTTTCCAACAGAAAATATCCAAGTATTTTAATTCTAGACTATTTGGACACGATATAGAGAGCTTCATCAATAGACATAAGAAATTTGCCAATGTTAGTGATGAATATCTGCAATATATATTCATAGAGGATATTTCATCTCCGTAAATATATGCCATATATTTATAGAAGATATCACATATCTAAATGAATACCGGAATCATAGATTTATTTGATAATCATGTTGATAGTATACCAACTATATTACCTCATCAGTTAGCTACTTTAGATTATCTAGTTAGAACTATCATAGATGAGAACAGAAGCGTGTTATTGTTCCATATTATGGGATCGGGTAAAACAATAATCGCTTTGTTGTTCGCCTTGGTAGCTTCCAGATTTAAAAAGGTTTACATTTTAGTACCGAACATCAACATCTTAAAAATTTTCAATTATAATATGGGTGTAGCTATGAACTTGTTTAATGACGAATTCATAGCTGAGAATATCTTTATTCATTCCACAACAAGTTTTTATTCTCTTAATTATAACGATAACGTCATTAATTATAACGGATTAAGTCGCTACAATAACTCTATTTTTATCGTTGATGAGGCACATAATATTTTTGGGAATAATACTGGAGAACTTATGACCGTGATAAAAAATAAAAACAAGATTCCTTTTCTACTATTGTCTGGATCTCCCATTACTAACACACCTAATACGCTGGGTCATATTATAGATTTAATGTCCGAAGAGACGATAGATTTTGGTGAGATTATTAGTCGTGGTAAGAAAGTAATTCAGACACTTCTTAACGAACGCGGAGTGAATGTACTCAAGGATTTGCTTAAAGGAAGAATATCATATTACGAAATGCCGGACAAAGATCTACCAACAATAAGATATCACGGACGTAAATTTCTAGATACTCGAGTAGTATATTGTCACATGTCTAAACTTCAAGAGAGAGATTATATGATTACTAGACGACAGCTATGTTATCATGAAATGTTTGATAAAAATATGTATAACGTGTCAATGGCAGTATTGGGACAACTTAATCTGATGAATAATTTAGATACGTTATTTCAGGAACAGGATAAGGAATTGTACCCAAATCTGAAAATAAATAATGGAGTGTTATACGGTGAAGAATTGGTAACGTTAAACATTAGTTCCAAATTTAAGTACTTTATCAATCGGATACAGACACTCAAGGGAAAACACTTTATATACTTCTCTAATTCTACATATGGTGGATTGGTAATTAAATATATCATGCTCAGTAATGGATATTCTGAATATAATGGTTCTCAGGGAACTAATCCACATATGATAAACGGCAAACCAAAAACATTTGCTATCGTTACTAGTAAAATGAAATCGTCTTTAGAGGATCTATTAGATGTGTATAATTCTCCTGAAAACGATGATGGCAGTCAATTGATGTTTTTGTTTTCGTCAAACATTATGTCTGAATCCTATACTCTGAAAGAGGTAAGGCATATTTGGTTTATGACTATCCCGGATACTTTTTCTCAATACAACCAAATTCTTGGACGATCTATTAGAAAATTCTCTTACGCCGATATTTCTGAACCCGTTAATGTATATCTTTTAGCAGCCGTATATTCAGATTTCAATGACGAAGTGACGTCATTAAACGATTATACACAGGATGAATTGATTAATGTTTTACCCTTTGACATCAAAAAGCTGTTGTATCTAAAATTTAAGACTAAAGAAACGAATAGAATATACTCTATTCTTCAAGAGATGTCTGAAACGTATTCTCTTCCACCACATCCATCAATTGTAAAAGTTTTATTGGGAGAATTGGTCAGACAATTTTTTTATAATAATTCTCGTATTAAGTATAACGACTCCAAGTTACTTAAAATGGTTACATCAGTTATAAAAAATAAAGAAGACGCTAGGAATTACATAGATGATATTGTAAACGGTCACTTCTTTGTATCGAATAAAGTATTTGATAAATCTCTTTTATACAAATACGAAAACGATATTATTACAGTACCGTTTAGACTTTCCTACGAACCATTTGTTTGGGGAGTTAACTTTCGTAAAGAATATAACGTGGTATCTTCTCCATAAAACTGATGAGATATATAAAGAAATAAATGTCGAGCTTTGTTACCAATGGATATCTTCCAGTTACATTGGAACCACATGAGTTGACATTAGACATAAAAACTAATATTAGGAATGCCGTATATAAGGCGTATCTCCATAGAGAAATTAGTGGTAAAATGGCCAAGAAAATAGAAATTCGTGAAGACGTGGAATTACCTCTCGGCGAAATAGTTAATAATTCTGTAGTTATAAACGTTCCGTGTGTAATAACCTACGCATATTATCACGTTGGGGATATAGTCAGAGGAACATTAAACATCGAAGATGAATCAAATGTAACTATTCAATGTGGAGATTTAATCTGTAAACTAAGTAGAGATTCGGGTACTGTATCATTTAGCGATTCAAAGTACTGCTTTTTTCGAAATGGTAATGCGTATGATAACGGCATCGAAGTCTCCGCCGTTCTAATGGAGGCTCAACAAGGTACCGAATCTAGTTTTGTTTTTCTCGCGAATATCGTTGACTCATAAGAAAGAGAATAGCGGTGAGTATAAATACGAATACTATGGCAATAATTGCGAATGTTTTATTCCCTTCGATATATTTTTGATAATATGAAAAACATGCCTCTCTCAAATCAGACAACCATTTCATAAAATAGTTCTCTCGCACTGGTGAGGTGGTTGCAGCTCGTATAATCTCCCCAGAATAATATACTTGCGTGTCGTCGTTCAATTTATACGGATTTCTATAATTCTCTGTTATATAATGAGGTTTACCCTCATGATTAGACGACGACAATAGTGTTCTGAATTTAGATAGTTGATCAGAATGAATGTTTATTGGTGTTGGAAAAATTATCCATGCTGCGTCTGCAGAGTGGTTGATAGTTGTTCCTAGATATGTAAAATAATCCAACGTACTAGGTAGCAAATTGTCTAGATAAAATACTGAATCAAATGGCGCAGACATATTAGCGGATCTAATGGAATCCAATTGATTGACTATCTTTTGAAAATATACATTTTTATGATCTGATACTTGTAAGAATATAGCAATAATGATAATTCCATCATCGTGTTTTTTTGCCTCTTCATAAGAACTATATTTTTTCTTATTCCAATGAACCAGATTAATCTCTCCAGAGTATTTGTATACATCTATCAAGTGATTGGATCCATAATCGTCTTCCTTCCCCCAATATATATGTATTGTTGATAACACATATTCATTGGGGAGAAACCCTCCACTTATATATCCTCCTTTAAAATTAATCCTTACTAGTTTTCCAGTATTCTGGATAGTGGTTGGTTTCGACTCATTATAATGTATGTCTAACGTCTTCAATCGCGTGTCAGAAATTGCTTTTTTAGTTTCTATATTAATAGGAGATAGTTGTTGAGGCATAGTAAAAATGAAATGATAACTGTCTAGAAATAGCTCTTAGTATGGGATTTACAATGGATGAGGAAGTGATATTTGAAACTCCTAGAGAATTAATATCTATTAAACGAATAAAAGATATTCCAAGATCAAAAGACACGCACGTGTTTGCTGCGTGTATAACAAGTGACGGATATCCGTTAATAGGAGCTAGAAGAACTTCATTCGCATTCCAGGCGATATTATCTCAACAAAATTCAGATTCTATCTTTAGAGTATCCACTAAACTATTACGGTTTATGTACTACAATGAACTAAGAGAAATCTTTAGACGGTTGAGAAAAGGTTCTATCAACAATATCGATCCTCACTTCGAAGAGTTAATATTATTGGGTGGTAAACTAGATAAAAAGGAATCTATTAAAGATTGTTTAAGAAGAGAATTAAAAGAGGAAAGTGATGAACATATAACAGTAAAAGAATTCGGAAATGTAATTCTAAAACTTACAACGAGTGATAAATTATTTAATAAAGTATATATAGGTTATTGCATGGCATGTTTTATTAATCAATCGTTGGAGGATTTATCACATACTAGTATTTACAATGTAGAAATTAGAAAGATTAAATCGTTAAATGATTGTATTAACGACGATAAATACGAATATCTGTCTTATATTTATAATATACTAATTAATAGTAAATGAGCTTTTACAGATCTAGTATAATTAGTCAGATTATTAAGTATAATAGACGACTAGCTAAGTCTATTATTTGCGAGGATGACTCTCAAATTATTACACTCACGGCATTCGTTAACCAATGCCTATGGTGTCATAAACGAGTATCCGTGTCCGCTATTTTATTAACTACTGATAACAAAATATTAGTATGTAACAGACGAGATAGTTTTCTCTATTCTGAAATAATTAGAACTAGAAACATGTATAGAAAGAAACGATTATTTCTGAATTATTCCAATTATTTGAACAAACAGGAAAGAAGTATACTATCGTCATTTTTTTCTCTAGATCCAGCTACTGCTGATAATGATAGAATAAACGCTATTTATCCGGGTGGTATACCCAAAAGGGGTGAGAACGTTCCAGAGTGTTTATCCAGGGAAATCAAAGAAGAAGTTAATATAGACAATTCTTTTGTATTCATAGACACTCGTTTTTTTATTCATGGTATCATAGAAGATACCATTATTAACAAATTTTTTGAGGTAATTTTCTTTGTTGGAAGAATATCTCTAACGAGTGATCAAATTATTGATACATTTAAAAGTAATCATGAAATAAAGGATCTAATATTTTTAGATCCGAATTCAGGTAATGGACTCCAATACGAAATTGCAAAATATGCTCTAGATACTGCAAAACTTAAATGTTACGGTCATAGAGGATGTTATTATGAATCATTAAAAAAATTAACTGAGGATGATTGATTAGAAAATATAAATTAATTTACCATCGTGTATTTTTATAACGGGATTGTCTGGCATATCATGTAGATAGTTACCGTCTACATCGTATACTCTACCATCTACGCCTTTAAATCCTCTATTTATTGATATTAATCTATTAGAATTGGAATACCAAATATTAGTACCCTCAATTAGTTTATTGGTAATATTTTTTTTAGACGATAGATCGATGGCTCTTGAAACCAAGGTTTTCCAACCGGACTCATTGTCGATCGGTGAGAAGTCTTTTTCATTAGCATGAATCCATTCTAATGATGTATGTTTAAACACTCTAAACAATTGTACAAATTCTTTTGATTTGTTTTGAATGATTTCAAATAGGTCTTCGTCTACAGTAGGCATACCATTAGATAATCTAGCCATTATAAAGTGCACGTTTACATATCTACGTTCTGGAGGAGTAAGAACGTGACTATTGAGACGAATGGCTCTTCCTACTATCTGACGAAGAGACGCCTCGTTCCATGTCATATCTAAAATGAAGATATCATTGATTGAGAAGAAACTAATACCCTCGCCTCCGCTAGAAGAGAATACGCATGTTTTAATGTATTCTCCGTTAGTGTTTGATTCTTGGTTAAACTCAGCCACAGCCTTGATTCTAGTATCTTTTGTTCTAGATGAGAACTCTATATTAGAGATACCAAAGACTTTGAAATATAGTAATAAGATTTCTATTCCTGACTGATTAACAAATGGTTCAAAGACTAGACATTTACCATGGGATGCTAATATTCCCAAACATACATCTATAAATTTGACGCTTTTCTCTTTTAATTCAGTAAATAGAGAGATATCAGCCGCAATAGCATCCCCTCCCAATAGTTCTCCCTTTTTAAAGGTGTCTAATGCGGATTTAGAAAATTCTCTATCTCTTAATGAATTTTTAAAATCATTATATAGGGTTGCTATCTCTTGTGTGTATTCTCCCGGATCACGATTTTGTCTTTCAGGAAAGCTATCGAATGTAAACGTAGTAGCCATACGTCTCAGAATTCTAAATGATGATATACCAGTTTTTATTTCTGCGAGTTTAGCCTTTTGATAAATCTCTTCTTGCTTTTTTGACATATTAACGTATCGCATTAATACTGTTTTCTTAGCGAATGATGCAGACCCTTCCACATCATCAAAAATAGAAAACTCGTTATTAACTATGTACGAACATAGGCCTCCTAGTTTGGAGACTAATTCTTTTTCATCGACTAGACGTTTATTCTCAAATAGCGATTGGTGTTGTAAGGATCCTGGTCGCAGTAAGTTAACCAACATGGTGAATTCTTGCACACTATTAACGATAGGTGTAGCCGATAAACAAATCATCTTATGGTTTTTTAACGCAGTGGTCTTAGATAAAAAATTATATACTGACCGAGTAGGACGGATCTTACCATCTTCTTTGATTAATGATTTAGAAATGAAGTTATGACATTCATCAATGATGACGCATATTCTACTCTTGGAATTAATAGTTTTGATATTAGTAAAAAATTTATTTCTAAAATTTTGATCATCGTAATTAATAAAAATACAATCCTTCGTTATCTCTGGAGCGTATCTGAGTATAGTGTTTATCCAAGGATCTTCTATCAAAGCCTTTTTTACCAATAAGATAATTGCCCAATTTGTATAAATATCCTTAAGATGTTTGAGAATATATACAGTAGTCATTGTTTTACCGACACCTGTTTCATGGAACAATAAAAGAGAATGCATACTGTCTAATCCTAAGAAAACTCTTGCTACAAAATGTTGATAATCCTTGAGGCGTACTACGTCTGACCCCATCATTTCAACGGGCATATTAGTAGTTCTGCGTAAGGCATAATCGATATAGGCCGCGTGTGATTTACTCATTTATGAGTGATAAGTAATAACTATGTTTTAAAAATCACAGCAGTAGTTTAACTAGTCTTCTCTGATGTTTGTTTTCGATACTTTTTGAATCAGAAGTCATACTAGAATAAAGCAGCGAGTGAACGTAATAGAGAGCTTCGTATACTCTATTCGAAAACTCTAAGAACTTATTAATGAATTCCGTATCCACTGGATCGTTTAAAATACTAAATTGAACAGTGTTCACATCCTTCCAAGACGAAGACTTAGTGACGGACTTAACATGAGACATAAATAAATCCAAATTTTTTTTATAAACATCACTAGCCACCATAATGGCGCTATCTTTCAACCAACTATCGCTTACGCATTTTAACAGTCTAACATTTTTAAAGAGACTACAATATATTCTCATAGTATCGATTACACCTCTACCGAATAGAGTGGGAAGTTTAATAATACAATATTTTTCGTTTACAAAATCAAATAATGGTCGAAACACGTCGAAGGTTAACATCTTATAATCGCTAATGTATAGATTGTTTTCAGTGAGATGATTATTAGATTTAATAGCATCTCGTTCACGTTTGAACAGTTTATTGCGTGCGCTGAGGTCGGCAACTACGGCATCCGCTCTAGTACTCCTCCCATAATACTTTACGCTATTAATCTTTAAAATTTCATAGACTTTATCTAGATCGCTTTCTGGTAACATGATATCATGTGTAAAAAGTTTTAACATGTCGGTCGGCATTCTATTTAGATCATTAACTCTAGAAATCTGAAGAAAGTAATTAGCTCCATATTCCAGACTAGGTAATGGGCTTTTACCTAAAGACAAGTTAAGTTCTGGCAATGTTTCATAAAATGGAAGAAGGACATGTGTCCCCTCCCGGATATTTTTTACAATTTCATCCATTTACAACTCTATAGTTTGTTTTCATTATTATTAGTTATTATCTCCCATAATCTTGGTAATACTTACACCTTGATCATAAGATACCTTATACAGGTCATTACATACAACTACCAATTGTTTTTGTACATAATAGATTGGATGATTGACATCCATGGTGGAATAAACTACTCGAACAGATAGTTTATCTTTCCCCCTAGATACATTGGCCGTAATAGTTGTCGGCCTAAAGAATATCTTTGGTGTAAAGTTAAAAGTTAGGGTTCTTGTTCCATTATTGCTTTTTGTCAGTAGTTCGTTATAAATTCTCGAGATGGGCCCGTTCTCTGAATATAGAACATCATTTCCAAATCTAACTTCTAGTCTAGAAATAATATCGGTCTTATTTTTAAAATCTATTCCCTTGATGAATGGATCGTTAATAAACAAATCCTTGGCCTTTGATTCGGCTGATCTATTATCTCCGTTATAGACGTTACGTTGACTAGTCCAAAGACTTACAGGAATAGATGTATCGATGATGTTGATAGTATGTGATATGTGAGCAAAGACTGTTCTCTTGGTGGCGTCGCTATATGTTCCAGTAATAGCGGAAAACTTTTTAGAAATGTTATATATAAAAGAATTTTTTCGGGTTCCAAACATTAACAGATTAGTATGAAGATAAACACTCATATTATCAGGAACATTATCAATTTTTACATAAACATCGGCATCTTGAATAGAAACAACACCATCTTCTGGAACCTCTACGATCTCGGCAGATTCCGGATAACCAGTCGGTGGACCATCACTAACAATAACTAGATCATCCAACAATCTACTCACATATGCGTCTATATAATCTTTTTCATCTTGTGAGTACCCTGGATACGAAATAAATTTGTTATCCGTATTTCCATAATAAGGTTTAGTATAAACAGAGAGAGATGTTGCTGCATGAACTTCGGTTACTGTCGCCGTTGGTTGGTTTATTTGACCTATTACTCTCCTAGGTTTCTCTATAAATGATGGTTTAATTTGTACATTCTTAACCATATATCCAATAAAGCTCAATTCAGGAACATAAACAAATTCTTTGTTGAACGTTTCAAAGTCGAACGAAGAGTCACGAATAACGATATCGGATACTGGATTGAAGGTCACCGTTACCGTAATTTTTGAATCGGATAGTTTAAGACTACTGAATGTATCTTCCACATCAAACGGAGTTTTAATATAAACGTATACTGTAGATGGTTCTTTAATAGTGTCATTAGGAGTTAGACCAATAGAAATATCATTAAGTTCACTAGAATATCCAGAATGTTTCAAAGCAATTGTATTATTGATACAATTATTATATAATTCTTCGCCATCAATTTCCCAAATAACACCGTTACACGAAGAGACAGATACATGATTAATACATTTATATCCAACATATGGTACGTAACCGAATCTTCCCATACCTTTAACTTCTGGAAGTTCCAAACTCAGAACCAAATGATTAAGCGCAGTAATATACTGATCCCTAATTTCGAAGCTAGCGATAGCCTGATTGTCGGGCCCATCGTTTGTCATAACTCCGGATAGAGAAATATATTGCGGCATATATAAAGTTGGAATTTGACTATCAACTGCGAAGACATTAGACCGTTTAATAAAGTCATCCCCACCGATCAAAGAATTAATGATAGTATTATTCATTTTCTATTTAAAATGGAAAAAAGCTTACAATAAACTCCGTAGAGAAATATCTATAATTTGTGAGTTTTCCTTAAAGTAACAGCTTCCGTAAACACCGTCTTTATCTCTTAATAAGTTTATTGTATTTATGACCTTTTCCTTATCTTCATAGAATACTAAAGGCAATAAAGAAATTTTTGGTTCTTCTCTAAGAGCTACGTGAGACTTAACCATAGACGCCAACGAATCCCTACATATTTTAGAACAGAAATACCCAACTTCACCACCCTTGAATGTCTCAATACTAATAGGTCTAAAAACCAAATCTTGATTACAAAACCAACACTTATCAATTACACTATTTGTCTTAATAGACATATCTGCCATAGATTTATAATACTTTGGTAGTATACAAGCGAGTGCTTCTTCTTTAGCGGGCTTAAAGACTGCTTTAGGTGCTGAAATAACCACATCTGGAAGACTTACTCGCTTAGCCATTTAATTACGGAACTATTTTTTTATACTTCTAATGAACAAGTAGAAAACCTCTCATCTACAAAAACATACTCGTGTCCATAATCCTCTACCATAGTAACACGTTTTTTAGATCTCATATGTGCTAAAAAGTTTTCCCATACTAATTGGTTACTATTATTTTTCGTATAATTTCTAACAGTTTGAGGTTTTAGATTTTTAGTTACAGAAGTGATATCGAATATTTTATCCAAAAAGAATGAGTAATTAATTGTCTTAGAAGGAGTGTTTTCTTGGCAAAAGAATACCAAGTGCTTAAATATTTCTACTACTTCATTAATCTTTTCTGTACTCAGATTCAGTTTCTCATCTTTTACTTGATTGATTATTTCAAAGACTAACTTATAATCCTTTTTATTTATTCTCTCGTTAGCCTTAAGAAAACTAGATACAAAATTTGCATCTACATCATCCGTGGATATTTGATTTTTTTCCATGATATCCAATAGTTCCGAGATAATTTCTCCAGAACATTGATGAGACAATAATCTCCGCAATACATTTCTCAAATGAATAAGTTTATTAGACACGTGGAAGTTTGACTTTTTTTGTACCTTTGTACATTTTTGAAATACAGACTCGCAAAAAATACAATATTCATATCCTTGTTCAGATACTATACCGTTGTGTCTACAACAGCTACATAATCGTAGATTCATGTTAACACTCTACGTATCTCGTCGTCCAATATTTTATATAAAAACATTTTATTTCTAGACGTTGTCAGAAAATCCTGTAATATTTTTAGTTTTTTTGGTTGTGAATAAAGTATCGCCCTAATAATATTGGTACCGTCTTCCGACAATATAGTAGTTAAATTATCCGAGCATGTAGAAGAACACCGCTTAGGCGGATTCAGTACAATGTTATATTTTTCGTACCAACTCATTTAAATATCATAATCTAAAATAGTTCTGTAATATGTCTAGCGCTAATATATTGATCATAATCCTGTGCATAAATTAAGATACAACAATGTCTTGAAATCATCGACATGGCTTCTTCCATAGTTAGAAGATCATCGTCAAAGTTAGCAACGTGATTCATCAACATTTGCTGTTTTGAGGCAGCAAATACTGAACCATCACCATTCAACCATTCATAAAAACCATCGTCTGAATCCATTGATAATTTCTTGTACTGGTTTTTGAGAGCTCGCATCAATCTAGCATTTCTAGCTCCCGGATTGAAAACAGAAAGAGGATCGTACATCCAGGGTCCATTTTCTGTAAATAGAATCGTATAATGTCCCTTCAAGAAGATATCAGACGATCCACAATCAAAGAATTGGTCTCCGAGTTTGTAACAGACTGCGGACTTTAACCTATACATGATACCGTTTAGCATGATTTCTGGTGATACGTCAATAGGAGTATCATCTATTAGAGATCTAAAGCCGGTGTAACATTCTCCACCAAACATATTCTTATTCTGACGTCGTTCTACATAAAACATCATTGCTCCATTAACGATAACAGGTGAATGAACAGCACTACCCATCACATTAGTTCCCAATGGATCAATGTGTGTAACTCCAGAACATCTTCCATAGCCTATGTTAGGAGGAGCGAACACCACTCTTCCACTATTGCCATCGAATGCCATAGAATAAATATCCTTGGAATTGATAGAAATCGGACTGTCGGATGTTGTTATCATCTTCATAGGATTAACAACGATGTATGGTGCAGCCTGAAGTTTCATATCATAACTGATGCCGTTCATAGGTCTAGCCACAGAAACCAACGTAGGTCTAAATCCAACTATAGACAAAATAGAAGCTAATATCTGTTCCTCATCTGTCATAACTTGAGAGCATCCAGTATGAATAATCTTCATTAGATGGGGATCTACCGCATCATCATCGTTACAATAAAAAATTCCCATTCTAATGTTCATAATTGCTTTTCTAATCATGGTATGAATGTTTGCTCTCTGAATCTCTGTGGAAATTAGATCTGATACACCTGTAATCACTATCGGATTATCCTCCGTAAGACGATTAACCAACAACATATAATTATAAGACTTTACTCTTCTAAATTCATAAAGTTGCTGGATTAGACTATATGTGTCTCCATGTACATACGCGTTCTCGAGCGCAGGAAGTTTAATACCGAATAGTGCCATCAGAATAGGATGAATGTAGTAATTAGTTTCTGGTTTTCTATAAATAAAAGACAAATCTTGTGAACTAGACATATCGGTAAAATGCATGGATTGGAATCGTGTAGTCGACAGAAGAATATGATGATTAGATGGAGAGTATATTTTATCTAACTCTTTGAGTTGGTCACCGATTCTAGGACTAGCTCGAGAATGAATAAGTACTAAGGGATGAGTACATTTCACAGAAACACTGGCGTTGTTCAACGTACTCTTTACATGGGAAAGGAGTTGAAATAGCTCGTTTCTATTTGTCCTGACAATATTTAGTTTATTCATAATATTAAGCATATCCTGAATAGTAAAGTTAGATGTGTCATACTTGTTAGTAGTTAGATATTTAGCAATTGCATTCCCATCATTTCTCAATCTCGTACTCCAATCATGTGTGGATGCTACTTCGTCGATGGAAACCATACAATCCTTTTTGATAGGCTGTTGAGATTGATCATTTCCTGTACGTTTAGGTTTGGTACGTTGATTTCTAGCCCCTGCGGATATAAAGTCATCGTCTACAATTTGGGATAATGAATTACATACACTACAAGACAAAGATTTATCAGAAGTGTGAATATGATCTTCATCTACCAAAGAAAGAGTTTGATTAGTATAACTAGATTTTAGTCCCGCGTTAGATGTTAAAAAAACATCGCTATTGACCACGGCTTCCATTATTTATATTCGTAGTTTTTACTCGAAAGCGTGATTTTAATATCCAATCTTATTACTTTTGGAATCGTTCAAAACCTTTGACTAGTTGTATAATTTGATCTATTGCCCTACGCGTATACTCCCTTGCATCATATACGTTCGTCACCAGATCGTTTGTTTCGGCCTGAAGTTGACGCATATCTTTTTCAACACTCGACATGAGATCCTTAAGGGTCATATCGTCTAGATTTTGTTGAGATGCTGCTCCTGGATTTGGATTTTGTTGTGCTGTTGTACATACTGTACCACCAGTAGGTGTGGGAGTACATACAGTGGCCACAATAGGAGGTTGAAGAGGTGTAACCGTTGGAGTAGTACAAGAAATACTTCCATCCGATTGTTGTGTACATGTGGTTGTTGGTAACGTCTGAGAAGGTTGGGTAGATGGCGGTGTCGTCATCTTTTGATCTTTATTAAATTTAGAGATAATATCCTGAACAGTATTGCTCGGCGTCAACGCTGGAAGGAGTGTACTCGCCGGCGCATCAGTATCTGCAGACAGCCAATCAAAAAGATTAGACATATCAGATGATGTATTAGTTTGTTGTCGTGGTTTTAGTACAGGAGCAGTACTACTAGGTAGAAGAATAGGAGCCGGTGTAGGTGTCGGAACCGGCTGTGGAGTTATATGAATAGTTGGTTGTAGCGGTTGGGTAGGCTGTCTGCTGGCGACCATCATATTATCTCTAGCTAGTTGTTCTCGCAACTGTCTTTGATAATACGACTCTTGAGACTTTAGTCCTATTTCAATCGCTTCATCCTTTTTCGTATCCGGATCCTTTTCTTCAGAATAATAGATTGACGACTTTGGTGTAGAGGATTCTGCCAGCCCCTGTGAGAACTTGTTAAAGAAGTCCATTTAAGGCTTTAAAATTGAATTGCGATTATAAGATTAAATGGCAGACACGGACGATATTATCGACTATGAATCCGATGATCTCACCGAATACGAGGATGATGAAGAAGATGGAGAGTCACTAGAAACTAGTGATATAGATCCCAAATCTTCTTATAAGATTGTAGAATCAACATCCACTCATATAGAAGATGCGCATTCCAATCTTAAACATATAGGGAATCATATATCTGCTCTTAAACGACGCTATACTAGACGTATAAGTCTATTTGAAATAGCGGGTATAATAGCAGAAAGCTATAACTTACTTCAACGAGGAAGGTTACCTCTAGTTTCAGAATTTTCTGACGAAACGATGAAGCAAAATATGCTACATGTAATTATACAAGAGATAGAGGAGGGTTCTTGTCCTATAGTCATCGAAAAGAACGGAGAATTGTTGTCGGTAAACGATTTTGATAAAGATGGTCTAAAATTCCATCTAGACTATATTATCAAAATTTGGAAACTTCAAAAACGATATTAGAATTTATACGAATATCGTTCTCTAAATGTTACAATCAAGTCTCTCATATTCAGCAGTTTATTGTCGTACTTTATATCGTGTTCATTAACGATATTTTGCAAAATAGTAATGATTCTATCTTCCTTCGATAGATATTCTTCAGAGATTATTGTCTTATATTCTTTCTTGTTATCCGATATGAATTTGATAAGACTTTGAACATTATTAATACCCGTCTGTTTAATTTTTTCTATAGATATTTTAGTTTTGGTAGATTCTATGGTGTCTGTTAATAGGCATCCAACATCGACATTCGACGTCAATTGTCTATAAATCAGAGTATAAATTTTAGAAATAACATTAGCAAATTGTTGTGCGTTGATGTCGTTATTCTGAAACAGTATGATTTTAGGTAGCATTTTCTTAACAAAGAGAACGTATTTATTGTTACTCAGTTGAACAGATGATATATCCAGATTACTAACGCATCTGATTCCATATACCAAACTTTCAGAAGAAATGGTGTACAATTGTTTGTATTCATTCAATGTCTCCTTTTCAGAAATTAGTTTAGAGTCGAATACTGCAATAATTTTCAAGAGATAGTTTTCATCAGATAAGATTTTATTTAGTGTAGATATGATAAAACTATTGTTTTGTTGGAGAACTTGATACGCCGCATTCTCTGTAGTCGACGCTCTCAAATGGGAAACAATCTCTATTATTTTTTTGGAATCGGATACTATATCTTCGGTATCTTGACGCAGTCTAGTATACATAGAGTTAAGAGAAATTAGAGTTTGTACATTAAGCAACATGTCTCTAAATGTGGCTACAAACTTTTCTTTTTCCACATCATCTAGTTTATTATATACCGATTTCACAACGGCACCAGATTTAAGGAACCAGAATGAAAAACTCTGATAACTACAATATTTCATCATAGTTACGATTTTATCATCTTCTATAGTTGGTGTGATAACACATACCTTTTTCTCCAAGACTGGAACCAACGTCATAAAAATGTTTAAATCAAAATCCATATCAACATCTGATGCGCTAAGACCAGTCTCGCGTTCAAGATTATCTTTACTAATGGTGACGAACTCATCGTATAGAACTCTAAGTTTGTCCATTATTTATTTACAGATTTAGTTGTTTAATTTATTTGTGCTCTTCCAGAGTTGGGATAGTATTTTTCTAACGTCGGTATTATATTATTAGGATCTACGTTCATATGTATCATAATATTAATCATCCACGTTTTGATAAATCTATCTTTAGCTTCTGAAATAACGTATTTAAACAAAGGAGAAAAATATTTAGTTACGGCATCAGACGCAATAACATTTTTTGTAAATGTAACGTATTTAGACGACAGATCTTCGTTAAAAAGTTTTCCATCTATGTAGAATCCATCGGTTGTTAACACCATTCCCGCGTCAGAGTGAATAGGAGTTTGAATAGTTTGTTTTGGAAATAGATCCTTCAATAACTTATAGTTGGGTGGGGAAAAATCAATTTTATCACTAGACTCTTTCTTTTTTACTATCATTACCTCATGAACTATTTCTTGAATGAGTATATGTATTTTCTTTCCTATATCGGTCGCGTTCATTGGAAAATATATCATGTCGTTAACTATAAGAATATTTTTATCCTCGTTTACAAACTGAATAATATCAGATATAGTTCGTAAACGAACTATATCATCACCAGCACAACATCTAACTATATGATATCCACTAGTTTCCTTTAGCCGTTTATTATCTTGTTCCATATTAGCAGTCATTCCATCATTTAAGAAGGCGTCAAAGATAATAGGGAGAAATGACATTTTGGATTCTGTTACGACTTTACCAAAATTAAGGATATACGGACTTACTATCTTTTTCTCAACGTCGATTTGATGAACACACGATGAAAATGTACTTCGATGAGATTGATCATGTAGAAAACAACAAGGGATACAATATTTCCGCATATCATGAAATATATTAAGAAATCCCACTTTATTATATTTCCCCAAAGGATCAATGCATGTAAACATTATGCCGTTATCATTAATAAAGACTTCTTTCTCATCGGATCTGTAAAAGTTGTTACTGATTTTTTTCATTCCAGGATCTAGATAATTAATAATAATGGGTTTTCTATTCTTATTCTTTGTATTTTGACATATCCTAGACCAGTAAACAGTTTCCACTTTGGTAAAATCAGAAGACTTTTGAACGCTATTAAACATGGCATTAATGGCAATAACTAAAAATGTAAAATATTTTTCTATGTTAGGAATATGGTTTTTCACTTTAATAGATATATGGTTTTTTGCCAAAATGATAGATATTTTTTTATCCGATGATAGCAAAATATTATTAGTCGCCGTCTCTATAAAAATGAAGCTAGTCTCGATATCCAATTTTATTCTAGAATTGATAGGAGTCGCCAAATGTACCTTATACGTTATATCTCCCTTGATGCGTTCCATTTGTGTATCTATATCGGACACAAGATCTGTAAATAGTTTTACGTTATTAATCATCACGGTATCGCCATCGCTAGATAATGCTAATGTACTATCCAAGTCCCAAATGGAGAGATTTAACTGTTCATCGTTTAGAATAAAATGATTACCTGTCATATTAATAAAGTGTTCATCGTATCTAGATAACAACGACTTATAATTAATGTCCAAGTCTTGAACTCGCTGAATGATCTTTTTTAACCCAGTTAGTTTTAGATTGGTACGAAATATATTGTTAAACTTTGATTCTACAGTAATGTCCAAATCTAGTTGTGGAAATACTTCCATCAACATTGTTTCAAACTTGATAATATTATTATCTACATCTTCGTACGATCCAAATTCCGGAATAGATGTATCGCACGCTCTGGCCACCCAGATAACCAAAAAGTCACACGCTCCAGAATATACATTGTATAAAAAGCTATCGTTTTTTAGTAGTGTTTTTTTCTGAGTATATACGAAAGGATTAAAAATAGTATTATCAACGTAACTATATTCCAAATTATTCTTATGAGAATAGATAATAATATCGTCCTTAATATCTAACAAATTTCCTAAATATCCCTTTAATTGAGTCATTCGAAGCGTTAATAAAATATGTCTCTTAACTATTTCCGGCCGTTGTATATTTAAATGACTTCGTAAGAAATAATATATAGGCGACTTCTCATCTATGTAATCATATGGAGTGAGATATAGGGCTCGTTCTACCTCCTGCCCCTTACCCACCTGTAATACCAATTGTGGACTCACTATATATCGCATATTTATATCGTGGGGTAAAGTGAAAATCTACTACCGATGATGTAAGTCTTACAATGTTCGAACCAGTACCAGATCTTAATTTGGAGGCCTCCGTAGAGCTAGGGGAGGTAAATATAGATCAAACAACACCTATGATAAAGGAGAATAGCGGTTTTATATCCCGTAGTAGACGTCTATTCGCCCATAGATCTAAGGATGATGAGAGAAAACTAGCACTACGATTCTTTTTACAAAGACTTTATTTTTTAGATCATAGAGAGATTCATTATTTGTTCAGATGCGTCGACGCTGTAAAAGACGTCACTATTACCAAAAAAAATAACATTATCGTGGCGCCTTATATAGCACTTTTAACTATCGCATCAAAAGGATGCAAACTTACAGAAACAATGATTGAAGCATTCTTTCCAGAACTATATAATGAACATAGTAAGAAATTCAAATTCAACTCTCAAGTATCCATCATCCAAGAAAAACTCGGATACCAGTCTGGAAACTATCACGTTTATGATTTTGAACCGTATTACTCTACAGTAGCTCTGGCTATTCGAGATGAACATTCATCTGGCATTTTTAATATCCGTCAAGAGAGTTATCTTGTAAGTTCATTATCTGAAATAACATATAGATTTTATCTAATTAATCTAAAATCTGATCTTGTTCAATGGAGTGCTAGTACGGGCGCTGTAATTAATCAAATGGTAAATACTGTATTGATTACAGTGTATGACAAATTACAACTGGCCATAGAAAATGATTCACAATTTACATGTTCATTGGCTGTGGAATCAGAACTTCCAATAAAATTACTTAAAGATAGAAATGAATTATTTACAAAATTCATTAACGAGTTAAAAAAGACCAGTTCATTCAAGATAAGCAAACGCGATAAGGATACGCTATTAAAACATTTTACTTATGACTGGAGTTAGAATTTATAGACGACTCATTTCGTTTATCATTATTACTATTACTATCATTATTAGTATTCTTCTTGTCATCTTGTTCAGAAATATACAGCAATGCTATACCTAATACCAAATACATTATCATGCTTGCAATGGCTCTAACAACAACGAACCAAAATGAATTTGGTCGTAGCTTTTGTTCACAAAAATACATAAAGAAATGTCTACATAAATCTATGGCGCCATTGGCTACTTGAAATAGCGCCAGTCCTCCTACAGATTTTAATATAGCTGTATAACATGACATTTATTCATCATCAAAAGAGACAGAGTCACCATCTGTCATATTTAGATTTTTTTTCATGTGTTCAAAGTATCCTCTACTCATTTCATTATAATAGTTTATCATACTTAGAATTTTAGGACGGATCAATGAGTAAGATTTGACTAGATCGTCAGTAGTAATTTGTGCATCATCTATTCTGCATCCGCTTCGTCGAATAATGTATAGCATCGCTTTGAGATTCTCCATAGCTATCAAGTCTTTATATAATGACATGGAAATATCTGTGAATGCTTTATACTTCTCCAACATCGATGCCTTAACATCATCACATACTTTAGCATTGAAAATACGTTCTATTGTGTAGATGGATGTAGCAAGATTTTTAAACAACAATGCCATCTTACATGATGATTGTCTCAAGTCTCCAATCGTTTGTTTAGAACGATTAGCTACAGAGTCCAATGCTTGGCTAACTAGCATATTATTATCTTTAGAAATTGTATTCTTCAATGAGGCGTTTATCATATCTGTGATTTCGTTAGTCATATTACAGTCTGACTGGGTTGTAATGTTATCCAACATATCACCTATGGATACGGTACACGTACCAGCATTTGTAATAATCCTATCTAAGATGTTGTATGGCATTGCGCAGAAAATATCTTCTCCTGTAATATCTCCACTCTCGATAAATCTACTCAGATTATTCTTAAATGCCTTATTCTCTGGAGAAAAGATATCAGTGTCCATCATTTCATTAATAGTATACGCAGAAAAGATACCACGAGTATCAATTCTATCCAAGATACTTATCGGTTCCGAGTCACAGATAATGGTTTCCTCTCCTTCGGGAGATCCTGCATAGAAATATCTAGGACAATAGTTTCTATACTGTCTGTAACTCTGATAATCTCTAAAGTCACTAACTGATACCATGAAATTGAGAAGATCAAACGCTGAAGTAATCAATTTTTCTGCCTCGTTTTTACTACAACTAGTTTTCATCAATGTAGTGACGATGTATTGTTTAGTTACTCTTGGTCTAATACTGATGATAGAGATATTATTGCTTCCCATAATGGATCTTCTAGTAGTCACCTTAAAGCCCATTGATGCGAATAGCAGATAGATAAAGTCTTGGTATGACTCCTTTCTAATATAGTACGGACTACCTTTGTCACCCAACTTTATACCCACATAAGCCATAACAACCTCTTTAATAGCCGTTTCATGAGGTTTATCAGCCATGAGCCTGAGTAGTTGAAAGAATCGCATGAATCCCGTCTCAGAAAGTCCTATATGCATGATAGATTTATCTTTCCTGGGAAACTCTCGTATAGTTATAGATGAAATACTCTTCAAAGTTTCTGAAATAAGATTAGTAACAGTCTTACCTCCGACTACTCTGGGTAACAAACATACTCTAATAGGTGTTTTCTCTGCGGAGATAATATCAGAAAGGATAGAGCAATAAGTAGTATTATTGTGATTATAAAGACCGAATACATAACAGGTAGAATTTATAAACATCATGTCCTGAAGGGTTTTAGACTTGTATTCCTCGTAATCTATACCGTCCCAAAACATGGATTTGGTAACTTTGATAGCCGTAGATCTTTGTTCCTTCGCCAACAGGTTAAAGAAATTAATAAAGAATTTGTTGTTTCTATTTATGTCCACAAATTGCACGTTTGGAAGCGCCACGGTTACATTCACTGCAGCATTTTGAGGATCGCGAGTATGAAGTACAATGTTATTGTTTACTGGTATATCTGGAAAGAAATCTACCAGTCTAGGAATAAGAGATTGATATCGCATAGAAATAGTAAAGTTTATAATCTCATCATTGTAGATTACTCTGTTACCATTGTAATAAATTGGTACTCTATCATAATCATTGACAAAGTACTGTTCATACATGATGAGATGTTTATATGTTGGCATAGTAGTGAGATCGACGTTTGGTAATGGCAATGTATTAAGATTAACTCCATAATGTCTAGCAGCATCTGTGATGTTATAAGTGATGTCAAAGCGGGGTTGATCTTGTGCTGTTATATATTGTCTAACACCTATAAGATTATCAAAATCTTGTCTGCTTAATACACCGTTAACAATTTTTGCCTTGAATTCTTTTATTGGTGCATTAATAACATCCTTATAGAGGATGTTAAACAAATAAGTATTATCAAAGTTAAGATCTGGGTATTTCTTTTCTGCTAGAACATCCATTGAGTCGGAGCCATCTGGTTTAATATAACCACCGATAAATCTAGCTCTGTATTCTGTATCCGTCAATCTAATATTAAGAAGGTGTTGAGTGAAAGGTGGAAGATCGTAAAAGCTGTGAGTATTAATAATAGGGTTAGTTTCCGAACTAATGTTAATTGGATGATTAATAATATCTATATTTCCAGCGTTAAGTGTAACATTAAACAGTTTTAATTCACGTGACGTGGTATCAATTAAATAATTAATGCCCAATTTGGATATAGTAGCCTGAAGCTCATCTTGTTTAGTTACGGATCCTAATGAGTTATTAAGAAATACATCGAACGGATGAACGAAGGTTGTTTTAAGTTGATCACATACTTTGTAATCTAGACATAGATGTGGAAGAACGGTAGAAACTATACGAAATAGATATTCAGAGTCCTCTAATTGATCAAGAGTAACTATTGACTTAATAGGCATCATTTATTTAGTATTAAATGACGACCGTACCAGTGACAGATATACAAAACGACTTAATTACAGAGTTTTCAGAAGATAATTATCCATCTAACAAAAATTATGAAATAACTCTTCGCCAAATGTCTATTCTAACTCACGTTAACAACGTGGTAGATAGAGAACATAATGCCGCCGTAGTGTCATCTCCAGAGGAAATATCATCACAACTTAATGAAGATCTATTTCCAGATGATGATTCACCGGCCACTATTATCGAACGAGTACAACCTCATACTACTATTATTGACGATACGCCACCTCCTACTTTTCGTAGAGAGTTATTGATATCGGAACAACGTCAACAACGAGAAAAAAGATTTAATATTACAGTATCAAAAAATTCTGAAGCAATAATGGAATCTAGATCTATGATAACTTCTATGCCAACACAAACACCATCCTTGGGAGTAGTTTATGATAAAGATAAAAGAATTCAGATGCTAGAGGATGAAGTGGTTAATCTTAGAAATCAACGATCTAATACAAAATCATCTGATAATTTAGATAATTTTACCAGAATACTATTTGGTAAGACTCCGTATAAATCAACAGAAGTTAATAAGCGTATAGCCATCGTTAATTATGCAAATTTGAACGGGTCCCCCTTATCAGTCGAGGACTTGGATGTTTGTTCGGAGGATGAAATAGATAGAATCTATAAAACGATTAAACAATATCACGAAAGTAGAAAACGAAAAATTATCGTCACTAACGTGATTATTATTGTCATAAACATTATCGAGCAGGCATTGCTAAAACTCGGATTTGAAGAAATCAAAGGACTGAGTACCGATATCACTTCAGAAATTATCGATGTGGAGATCGGAGATGACTGCGATGCTGTAGCATCAAAACTAGGAATCGGTAACAGTCCGGTTCTTAATATTGTATTGTTTATACTCAAGATATTCGTTAAACGAATTAAAATTATTTAATTTAATACATTCCCATATCCAGACAACAATCGTCTGGATTAATCTGTTCCTGTCGTCTCATACCGGACGACATATTAATCTTTTTATTAGTGGGCATCTTTTTAGATGGTTTCTTTTTCCCAGCATTAACTGATTCGATACCTAGAAGATCGTGATTGATTTCTCCGACCATTCCACGAACTTCTAATTGGCCGTCTCTAACGGTACCATAAACTATTTTACCAGCATTAGTAACAGCTTGGACAATCTGACCATCCATTGCGTTGAATGATGTAGTTGCTGTTGTTCTACGTCTAGGAGCACCAGAGGTATTTTTAGAGCTCTTGGATGTTGATGTAGAAGACGAGGATTTTGATTTTGGTTTACATGTAATACATTTTGAACTCTTTGATTTTGTATCACATGCGCCGGCAGTCACATCTGTTTGAGAATTAAGATTATTGTTGCCTCCTTTGACGGCTGCATCTCCACCGATCTGCGCTAGTAGATTTTTAAGCTGTGGTGTAATCTTATTAACTGTTTCGATATAATCATCGTAACTACTTCTAACGGCTAAATTTTTTTTATCCGCCATTTAGAAGCTAAAAATATTTTTATTTATGCAGAAGATTTAACTAGATTATACAATGAACTAATATGATCCTTTTCTAGATTATTTACGAACTTGGTATTTCTTGTTTCTGGAGGAGGAGAATTTAAATTCGGACTTGGATTCGGATTTTGTGGGTTCTTGATCTTATTATACAGCGTGTATAGGATGGCGACGGTAACTGCTACACAAATACCGATCAACAGAAGAATACCAATCATTTATTGACAATAACTTCACTATGATCAAGTATGTAATAATCATCTTTTCACTAAGTAAGTAGTAATAATGATTCAACAATGACAAGATATATGGACGATAATAATTTAGTTCATGGAAATATCGCTATGATTGGTGTGAATGACTCCGCTAACTCTGTGGGGTGCACAGTGCTTTCCCCACATAGAATAAATTAGCATTCCGACTGTGATAATAATACCAAGTATAAACGCCATAATACTCAATACTTTCCATGTACGAGTGGGACTGGTAGACTTACTAAAGTCAATAAAGGCGAAGATACACGAAAGAATCAAAAGAATGATTCCAGCGATTAGCACGCCGGAAAAATAATTTCCAATCATAAGCATCATGTCCATTTAACTAATAAAAATTTTAAATCGCCGAATAAACAAAGTGGAATATAAACCATATAAAAACAATAGTTTGTACTGCAAAAATAATATCTATTTTTGTTTTCGAAGATATGGTAAAATTAAATAGTAGTACACAGCATGTTATAACTAACAGCAGCAACGGCTCGTAATTACTTATCATTTACTAGACGAAAAGGTGGTGGGATATTTTCTTGCTCAAATAATACGAATATATCACCCATCCATTTTATACGATGTTTATATACTCTAATCTTTAATAGATCTATAGATGACGGGTTTACCAACAATATAGATTTTATCGATTCATCTAATTTAAACCCTTCCTTAAACGTGAATGATCTATTATCTGGCATAATGATGACCCTACCTGATGAATCTGACAATGTACTGGGCCATGTAGAATAAATTATCAACGAATTATCGTCTACGAACATTTATATCATTTGTTTTAATTTTAGGACGTGAATAAATAGATATAAAATAGAAAATAACAGATATTACAACCAGTGTTATGGACGCACCCAACCATGTAGGCAGTTTTATTTTATCGTTTACTACAGGTTCTCCTGGATGTACGTCACCAACTACAGACGTAGTTCTAGTACAATTAGACGTAAGTTCCGCTTGGGAATTTTTTAACACTAAAGAGTTAACGTTAATCGTACACCCAACGTATTTACATCTAGCTCTTTGAACATCTTGATTATAATATAACCATTTTCTATCTCTAGATTCGTCAGTGCACTCATGTAACCAACATACCCTAGGTCCTAAATATTTATCTCCGGAATTAGATTTTGGATAATTCGCGCACCAACAATTTCTATTTCCTTTATGGTCGTTACAAAAGACGTATAATGCCGTATCCCCAAAAGTAAAATAATCAGGACGAATAATTCTAATAAACTCAGAACAATATCTCGCATCCATATGTTTGGAGCAAATATCGGAATAAGTAGACATAGCCGGTTTCCGTTTTACACGTAACCATTCTAAACAATTGGGGTTTCCAGGATCGTTTCTACAAAAACCAGTCATGAAATCGTCACAATGTTCTGTCTTGTAATTATTATTAAATATTTTTGGACAGTGTTTGGTATTTGTCTTAGAACAACATTTTGCCACGCTATCACTATCACCCAGGAGATAATCCTTTTTTATAAAATGACACCGTTGCCCGGATGCTATATAATCAGTAGCATATTTTAAATCCTTAATATACTCAGGAGTTACCTCGTTCTGATAATAGATTAATGATCCAGGACGAAATTTGAAAGAACTACATGGTTCTCCATGAATTAATACATATTGTTTAGCAAATTCAGGAACTATAAAACTACTACAATGATCTATCGACATACCATCTATCAAACAAAATTTGGGTTTAATTTCTCCTGGAGACGTTTCATAATAATACATATAACTTTCTTCGGCAAACCTAACAGCTCTATTATATTCAGGATAATTAAAATCTAATACCATATATTTGTCTCGTATATCTGCTATTCCTGTCTCTATTTTGATTCTATTAAGAGTAACAGCTGCCCCCATTCTTAATAATCATCAGTATTTAAACTGTTAAATGTTGGTATATCAACATCTATCTTATTTCCCGCAATATAAGGTTTGTTGCAGGTATACTGTTCAGGAATGGGTACATTTATACTTCTTTTATAGTCCTGTCTTTCGATGTTCATCACAAATGCAAAGAACAGAATAAACAAAATAATGTAAGAAATAATATTAAATATCTGTGAATTCGTAAATACATTGATTGCCATAATAATTACAGCAGCTACAATACACACAATAGACATTCCCACAGTGTTGCCATTACCTCCACGATACATTTGAGTTACTAAGCAATAGGTAATAACTAAGCTAGTAAGAGGCAATAGAAAAGATGAGATAAATATCATCAATATAGAGATTAGAGGAGGGCTATATAGAGCCAAGACGAACAAAATCAAACCGAGTAACGTTCTAACATCATTATTTTTGAAGATTCCCAAATAATCATTCATTATTCCTCCATAATCGTTTTGCATCATACCCCCATCTTTAGGCATAAACGATTGCTGCTGTTCCTCTGTAAATAAATCTTTATCAAGCACTCCAGCACCCGCAGAGAAGTCATCAAGCATATTGTAATATCTTAAATAACTCATTTATATATTAAAAAATGTCACTATTAAAGATGGAGTATAATCTTTATGCCGAACTAAAAAAAATGACTTGTGGTCAGACCATAAGTCTTTTTAATGAAGACGGCGATTTCGTAGAAGTTGAACCAGGATCATCCTTTAAGTTTCTAATACCTAAGGGATTTTACTCCTCTCCTTGTGTAAAGACGAGTCTAGTATTCGAGACATTAACAACGACCGATAATAAAATTACTAGTATCAATCCAACAAATGCGCCAAAGTTATATCCTCTTCAACGCAAAGTCGTATCTGAAGTAGTTTCTAATATGAGGAAAATGATCGAATTAAAACGTCCTCTATACATCACTCTTCACTTGGCATGTGGATTTGGTAAGACTATTACCACGTGTTATCTTATGACCACACACGGCAGAAAAACCATCATTTGCGTACCCAATAAAATGTTAATACATCAATGGAAGACACAGGTAGAGGCAGTCGGATTGGAACATAAGATATCTATAGATGGAGTTAGTAGTCTATTAAAGGAACTAAAGACTCAAAGTCCGGATGTATTAATCGTAGTCAGTAGACATCTGACAAACGATGCATTTTGTAAATATATCAATAAGCATTATGATTTGTTTATCTTGGATGAATCACATACGTATAATCTGATGAACAATACAGCAGTTACAAGATTTTTAGCGTATTATCCTCCGATGATGTGTTATTTTTTAACTGCTACACCTAGACCAGCTAACCGAATTTATTGTAACAGTGTTATTAATATTGCCAAGTTATCAGATCTAAAAAAAACTATCTATACAGTAGATAGTTTTTTTGAGCCATATTCCACAGACAATATTAGAAATATGGTAAAACGACTAGATGGACCATCTAATAAATATCATATATATACCGAGAAGTTATTATCTGTAGACGAGCCTAGAAACCAACTTATTCTTGATACCCTGGTAGAAGAATTCAAGTCAGGAACTATTAATAGAATTTTAGTTATTACTAAACTACGTGAACATATGGTATTCTTCTACAAACGATTATTAGATCCTTTCGGACCAGAGGTTGTATTTATAGGAGACGCCCAAAATAGACGTACTCCAGATATGGTCAAATCAATTAAGGAACTAAATAGATTTATATTCGTATCCACCTTATTTTATTCCGGCACTGGTTTAGATATTCCGAGTTTGGATTCTTTGTTCATTTGCTCGGCAGTAATCAACAATATGCAAATAGAGCAATTACTAGGGAGGGTATGTCGAGAAACAGAACTATTAGATAGGACGGTATATGTATTTCCTAACACATCCATCAAAGAAATAAAGTACATGATAGGAAATTTCGTGCAACGAATTATTAGTCTGTCTGTAGATAAACTCGGATTTAAACAAGAAAGTTATCGGAAACATCAGGAATCTGAACCCGCTTCCGTGCCAACATCCTCCAGAGAAGAACGTGTATTAAATAGAATATTTAACTCGCAAAATCGTTAAGAAGTTTAAGAGACGATCCACATGCTGAGCAGGCCAGTGTATTACCCCTCATAGTATTAATATAATCCAATGATACTTTTGTGATGTCGGAAATCTTAACCAATTTAGACTGACAGGCAGAACACGTCATGCAATCATCACCGTCATCGATAACTGTAGTCTTGGGCTTCTTTTTGCGACTCTTCATTCCGGAACGCATATTGGTGCTATCCATTTAGGTAGTAAAAAATAAGTCAGAATATGCCCTATAACACGATCGTGCAAAACCTGGTATATCGTCTCTATCTTTATCACAATATAGTGTATCAACATCTTTATTATTGACCTCGTTTATCTTGGAACATGGAATGGGAACATTTTTGTTAACGGCCACCTTTGCCTTAATTCCAGATGTTGTAAAATTATAACCAAACAGTCTATCATCGACACAAATGAAATTCTTGTTTAGACGTTTGTAGTTTACGTATGCGGCTCGTTCTCGTCTCATTTTTTCAGATATTGCAGGTACTATAATATTAAAAATAAGAATGAAATAACATAGGATTAAAAATAAAGTTATCATGACTTCTAGTGCTGATTTAACTAACTTAAAAGAATTACTTAGTCTGTACAAAAGTTTGAGATTTTCAGATTCTGTGGCTATAGAGAAGTATAATTCTTTGGTAGAATGGGGAACATCTACTTACTGGAAAATAGGCGTACAAAAGGTAACTAATGTCGAGACGTCCATATCTGATTATTATGATGAGGTAAAAAATAAACCGTTCAATATTGATCCGGGGTATTATATTTTCTTACCAGTATACTTTGGAAGCGTCTTTATTTATTCGAAGGGTAAAAATATGGTAGAACTTGGATCTGGAAACTCTTTTCAAATACCGGATGAGATTCGAAGTGCGTGTAACAAAGTATTAGATAGTGATAACGGAATAGACTTTCTGAGATTTGTTTTGTTAAACAATAGATGGATAATGGAAGACGCTATATCAAAATACCAGTCTCCAGTTAATATATTTAAACTAGCTAGTGAGTACGGATTAAACATACCCAACTATTTAGAAATTGAAATAGAGGAAGACACATTATTTGACGATGAGTTATACTCTATTATGGAACGCTCTTTCGATGATACATTTCCAAAAATATCTATATCGTATATTAAGTTGGGAGAACTTAAACGGCAAGTTGTAGACTTTTTCAAATTCTCATTCATGTATATTGAGTCAATCAAGGTAGATCGTATAGGAGATAATATTTTTATTCCTAGCGTTATAACAAAATCAGGAAAAAAGATATTAGTAAAAGATGTAGACCATTTAATACGATCCAAGGTTAGAGAACATACATTTGTAAAAGTAAAAAAGAAAAACACATTTTCCATTTTATACGACTATGATGGGAACGGAACAGAAACTAGAGGAGAAGTAATAAAACGAATTATAGACACTATAGGACGAGACTATTATGTTAATGGAAAGTATTTCTCTAAGGTTGGTAGTGCAGGCTTAAAGCAATTGACTAATAAATTAGATATTAACGAGTGCGCAACTGTCGATGAGTTAGTTGATGAGATTAATAAATCCGGAACTGTAAAACGAAAAATAAAAAACCAATCAGTATTTGATTTAAGCAGAGAATGTTTGGGATATCCAGAAGCGGATTTTATAACGTTAGTTAATAACATGCGGTTCAAAATAGAAAATTGTAAGGTCGTAAATTTCAATATTGAAAATACTAATTGTTTAAATAACCCGAGTATTGAAACTATATATGGAAACTTCAACCAGTTCGTCTCAATCTTTAATACCGTTACCGATGTCAAAAAAAGATTATTCGAGTGAAATAATATGCGCCTTTGATATAGGTGCAAAAAATCCTGCCAGAACTGTTTTAGAAGTCAAGGATAACTCCGTTAGGGTATTGGATATATCAAAATTAGACTGGAGTTCTGATTGGGAAAGGCGCATAGCTCAAGATTTGTCACAATATGAATACACTACAGTTCTTCTAGAACGTCAGCCTAGAAGGTCACCGTACGTCAAATTTATCTATTTTATTAAAGGCTTTTTATATCATACATCTGCTGCCAAAGTTATTTGCGTCTCACCTGTCATGTCTGGTAATTCATATAGAGATCGAAAAAAGAGATCTGTTGAAGCATTTCTTGATTGGATGGACACATTCGGATTGCGAGACTCCGTTCCGGATAGACGCAAATTAGACGATGTAGCGGATAGTTTCAATTTGGCTATGAGATACGTATTAGATAAATGGAATACTAATTATACACCTTATAATAGGTGTAAATATAGAAATTACATAAAAAAAATGTAATAACGTTAGTAACGCCATTATGGATAATCTATTTACCTTTCTACATGAAATAGAAGATAGATATGCCAGAACTATTTTTAACTTTCATCTAATAAGTTGCGATGAAATAGGAGATATATATGGTCTTATGAAAGAACGCATTTCCTCAGAGGATATGTTTGACAATATAGTATATAATAAAGATATACATCCTGCCATTAAGAAACTAGTTTATTGCGACATCCAACTTACTAAACATATTATTAATCAGAATACGTATCCGGTATTTAACGATTCTTCACAAGTGAAATGTTGTCATTATTTCGATATAAACTCAGATAATAGCAATATTAGCTCTCGTACAGTAGAGATATTTGAGAGTGAAAAGTCATCTCTTGTATCATATATTAAAACTACCAATAAGAAGAGAAAGGTCAATTACGGCGAAATAAAGAAAACTGTACATGGAGGCACTAATGCAAATTACTTTTCCGGTAAAAAGTCTGATGAGTATCTGAGCACTACAGTCAGGTCCAACATTAATCAACCTTGGATCAAAACCATTTCTAAGAGAATGAGAGTAGATATCATTAATCACTCTATAGTAACGCGTGGAAAAAGCTCTATATTACAAACTATAGAAATTATTTTTACTAATAGAACATGTGTGAAAATATTCAAGGATTCTACTATGCACATTATTCTATCCAAGGACAAGGATGAAAAGGGATGTATAAACATGATTGATAAATTATTCTATGTATATTATAATTTATTTCTGTTGTTCGAGGATATCATCCAAAACGAGTACTTTAAAGAAGTAGCTAATGTTGTAAACCATGTACTCATGGCTACGGCATTAGATGAGAAATTATTCCTAATTAAGAAAATGGCTGAACACGATGTTTATGGAGTTAGCAATTTCAAAATAGGGATGTTTAACCTGACATTTATTAAGTCGTTGGATCATACCGTTTTCCCCTCTCTGTTAGATGAGGATAGCAAAATAAAGTTTTTTAAGGGGAAAAAGCTCAATATTGTAGCATTACGATCTCTGGAGGATTGTACAAATTACGTGACTAAATCCGAGAATATGATAGAAATGATGAAGGAAAGATCGACTATTTTAAATAGCATAGATATAGAAACGGAATCGGTAGATCGTCTAAAAGAATTGCTTCTAAAATGAAAAAAAACACTGTTTCAGAAATGGATCAACGACTCGGGTATAAGTTTTTGGTGCCTGATCCTAAAGCCGGAGTTTTTTATAGACCGTTACATTTCCAATATGTATCGTATTCTAATTTTATATTGCATCGATTGCATGAAATCTTGACCGTCAAGCGGCCACTCTTATCGTTTAAGAATAATACAGAACGAATTATGATAGAAATTAGCAATGTTAAAGTGACTCCTCCAGATTACTCACCTATAATTGCGAGTATTAAAGGTAAGAGTTATGACGCATTAGCCACGTTCACTGTAAATATCTTTAAAGAGGTAATGACCAAAGAGGGTATATCCATCACTAAAATAAGTAGTTATGAGGGAAAAGATTCTCATTTGATAAAAATTCCGCTACTAATAGGATATGGGAATAAAAATCCACTTGATACAGCCAAGTATCTTGTTCCTAATGTCATAGGTGGAGTCTTTATCAATAAACAATCTGTCGAAAAAGTAGGAATTAATCTAGTAGAAAAGATTACAACATGGCCAAAATTTAGGGTTGTTAAGCCAAACTCATTCACTTTCTCGTTTTCCTCCGTATCCCCTCCTAATGTATTACCGACAAGATATCGCCATTACAAGATATCTCTGGATATATCACAATTGGAAGCGTCTAATATATCATCGACAAAGACATTTATAACGGTCAATATTGTTTTGCTGTCTCAATATTTATCTAGAGTGAGTCTAGAATTCATTAGACGTAGTTTATCATACGATATGCCTCCAGAAGTTGTCTATCTAGTAAACGCGATAATAGATAGTGCTAAACGACTTACCGAATCTATTACTGACTTTGATATTGATACATACATTAATGACCTGGTGGAAGCTGAACACATTAAACAAAAATCTCAGTTAACGATTAACGAGTTTAAATATGAAATGCTGCATAACTTTTTACCTCATATGAACTATACACCCGATCAACTAAAGGGATTTTATATGATATCTTTACTAAGAAAGTTTCTCTACTGTATCTACCACACTTCTAGATATCCAGATAGAGATTCGATGGTTTGTCATCGCATCCTAACGTACGGCAAATATTTTGAGACGTTGGCACATGATGAATTAGAGAATTACATAGGTAACATCCGAAACGATATCATGAACAATCACAAGAACAGAGGCACTTACGCAGTAAACATTCATGTACTAACAACTCCTGGACTTAATCATGCATTTTCTAGTCTATTGAGTGGAAAGTTCAAAAAGTCAGACGGTAGTTATCGAACACATCCTCACTATTCATGGATGCAGAATATTTCTATTCCTAGAAGTGTTGGATTTTATCCGGATCAAGTAAAGATTTCAAAGATGTTTTCTGTCAGAAAATACCATCCAAGCCAATATCTTTACTTTTGTTCATCAGACGTTCCGGAAAGAGGTCCTCAGGTAGGTTTAGTATCTCAATTGTCTGTCTTGAGTTCCATTACAAATATACTAACGTCTGAGTATTTGGATTTGGAAAAGAAAATTTGTGAGTATATCAGATCATATTATAAAGATGATATAAGTTACTTTGAAACAGGATTTCCAATCACTATAGAAAATGCTCTAGTCGCATCTCTTAATCCAAATATGATATGTGATTTTGTAACTGACTTTAGACGTAGAAAACGGATGGGATTCTTCGGTAACTTGGAGGTAGGTATTACTTTAGTTAGGGATCACATGAATGAAATTCGCATTAATATTGGAGCAGGAAGATTAGTCAGACCATTCTTGGTTGTGGATAACGGAGAGCTCATGACGGATGTGTGTCCGGAGTTAGAAAGCAGATTAGACGACATGACATTCTCTGACATTCAGAAAGAGTTTCCACATGTCATCGAAATGGTAGATATAGACCAATTTACTTTTAGTAACGTATGTGAATCGGTTCAAAAATTTAGAATGATGTCAAAGGATGAAAGAAAGCAATACGATTTATGTGACTTTCCTGCCGAATTTAGAGATGGATATGTAGCATCTTCACTAGTGGGAATCAATCACAATTCTGGACCCAGAGCTATTCTTGGATGTGCTCAAGCTAAACAAGCTATCTCTTGTCTGAGTTCGGATATACGAAATAAAATAGACAATGGAATTCATTTGATGTATCCAGAGAGGCCAATTGTGATTAGTAAGGCTTTAGAAACTTCAAAGATTGCGGCTAATTGCTTCGGACAACATGTTACTATAGCATTAATGTCGTACAAAGGTATCAATCAAGAGGATGGAATTATCATCAAAAAACAATTTATTCAGAGAGGCGGTCTCGATATTGTTACAGCCAAGAAACATCAAGTAGAAATTCCATTGGAAAACTTTAATAACAAAGAAAGAGATAGGTCTAACGCCTATTCGAAATTAGAAAGTAATGGATTAGTTAGACTGAATGCTTTCTTGGAATCCGGAGACGCTATGGCAAGAAATATCTCATCAAGAACTCTTGAAGATGATTTTGCTAGAGATAATCAGATTAGCTTTGATGTTTCCGAGAAATATACAGATATGTACAAATCTCGCGTTGAACGAGTACAAGTAGAACTTACTGACAAAGTTAAGGTGCGAGTATTAACCATGAAAGAAAGAAGACCCATTCTAGGAGACAAATTTACTACTAGAACGAGTCAAAAGGGAACAGTCGCGTATATCGCAGATGAAACGGAACTTCCGTACGACGAAAATGGTATCACACCAGATGTCATTATTAATTCTACATCCATCTTCTCTAGAAAAACTATATCTATGTTGATAGAAGTTATTTTAACAGCCGCATATTCTACTAAGCCGTACAACAATAAGGGAGAAAACCGACCTGTCTGTTTTCCTAGTAGTAACGAAACATCCATTGATGCATATATGCAATTCGCTAAACAATGTTATGAGTATTCAAATCCGAAATTGTCCGAGGAAGAATTATCGGATAAAATCTTTTGTGAAAAGATTCTCTATGATCCTGAAACGGATAAGCCTTATGAATCCAAAGTATTTTTTGGACCAATTTATTACTTGCGTCTGAGACATTTAACTCAGGACAAGGCAACCGTTAGATGTAGAGGTAAAAAGACGAAGCTCATTAGACAAGCGAATGAGGGACGAAAACGTGGAGGAGGTATCAAGTTTGGAGAAATGGAGAGAGACTGTTTAATAGCACATGGCGCAGCCAATACTATTACAGAAGTTTTAAAAGACTCAGAAGAGGATTATCAAGATGTGTATATTTGTGAAAATTGTGGAGACATAGCAGCACAAATCAAAAGTATTAATACATGTCTTAGATGTTCAAAACTTAATCTCTCTCCTCTCTTAACAAAAATTGATACCACACACGTATCTAAAGTATTTCTTACTCAAATGAACGCCAGAGGCGTAAAAGTTAAATTAGATTTCGAACGAAGGCCTCCTTCGTTTTATAAACCATTAGATAAAGTTGATCTTAAACCGTCTTTTCTGGTATAATATTGTTTAGTAGATACTCATCAAGATTATCAAGATAAGCTAATTCACTAAACATATTATCGGATTCGGTATTGTTACTCGAGAATAGAGTTCGTTATGCTCCTGATATTCGGAAATCTGTGGAGTTTCAGGTTTTGGTGGAAGTGTAACTGCTACTTGGTGGGATACTGAAGGATATTTCAGAGAGTTGTGGATGTTCGGGTTCGACATCCACCGATGGTGTCACGCCACTAATCGGTTCGGTAACGTCTGTGGATGGAGGTGCTACTTCTACAGAACCTGTAGCCTCAGTTGTCAACGGAGATACATCTTCAATGCGCGGAAATGTATAATTTGGTAATGGTTTCTCATGTGGATCTTAAGAAGAAGAGGTAAGATATCTACGAAAGATACCGATCACGTTCTAGTTCTCTTTTGTAGAACTTTAACTTTTTCTTTCTCAGCATCTAGTTGATATTCCGACCTCTTCACGTTTCACATGGGTTACCTCCGCAGTTTTTACAAGCGATTTCACGTTCCAGATCACGTTCAGCCTTCATACGTCTCTCCCTCTCTCTATCGAGTTTATCAGAGCAGTCTTTCTGAAGGCGATCGAACTCCATAAATTTCTCCAACGCTTTGATTGTTTCCATAGATTTCCGAAGTTTAGCTTCTAGGACGGCGATTCTTTTTTCTTTCGAATTCACGGGGTACAACCGTTTCCATTACCACCATCTCTACGTTTCTTTTCTAGATCGGCAATCTTTCTCAACATTTCATCCCCATGCCTTTTCATTCCTCGAGTCTATCGTCGTCGAAATATCGTTCCAGCTCCTTTTCGACCTCAATAACTTTAGCACGTTGTCTCATCAAGCTCTCTCTTGTAGTACTATCATTTTTATCTGATTCCCTGGCACGTTTAAGATCTTCATGTAATTGAGTCAGCTCTTGACACAATCTCTTAACTAACTTCCTCTCTTGCTTCTTCGTCATAGTACTTACAATCACTATGGGATCCATTGTTACCACGTCTGTACTCGGCGAGCTCACGTTTAAGAGATTCAATTTCCAGTTTGTACATTGATTTCATTATTACGTCCGCAGTCGTTCAACTGTATTTCAAGATCTGAGATTCTAGATTGTAATCTCTGTAGCATTTCCACGGCATTCACTCAGTTGTCTTTCAAGATCTGAGATTCTAGATTGGAGTCTGCTAATCTCTGTAAGATTTCCTCCTCCGCTCTCGATGCAGTCGGTCAACTTATTCTCTAGTTCTCTAATACGTGAACGCAGTGCATCAACTTCTTGTGTGTCTTCTTGATTGCGTGTGCATTCATCGAGTCTAGATTCGAGATCTCTAACGTGTCGTCGTTCTTCCTCAAGTTCTCTGTGTACTACAGAAAGCGTGTCCCTATCTTGTTGATATTTAGCAATTTCTGATTCTAGAGTACTGATTCTGCTCACGTATGTACTAATAGTTGTCTTATCCTTATCAAGATCCTCCTTGTATTTGTCACATTCCTTGATATCCATACGAAGTCTGGACAGTTCCCATTCGACATTACGACGTTTATCGATTTCAGCTCGGAGATCGTCGTCGCGTTGTTTTAGCCACATACGACTGAGTTCAAGTTCTCGTTGACAAGATCCATCTACTTTTCCATCCCTAATAGTATCCAGTTCCTTTTCTAGTTCTGACCGCATTTCTCGTTCCATATCAAGAGATTCTCTCAATTCTCGTATAGTCTTCTTATCAATTTCTGATGAATCTGAACCATCATCTGTCCCATTTTGTTGCATATCCCTGAGTTCTTTGATCTCTGTTGTAAGTCTGTCGATTCTTTCGGTTTTATAAACAGAATCCCTTTCCAAAGTCCTAATCTTACTGAGTTTATCACTAAGTTCTTCATTCAATTCAGTGAGTTTTCTCTTGGCTTCTTCCAAGTCTGTTTTAAACTCTCCATCATTTCCGCATTCTTCCTCGCATTTATCTAACCATTCAATTAGTTTATTAATGACTAGTTGGTAATCAGCGATTCCTATAACCGTTCTTGTATTTGTGGGAACATAATTAGGATCTTCTAATGGATTGTATGGCTTGATAGCATCATCTTTATCATTATTAGGTGGGGGATGGACAACCTTAATTGGTTGGTCCTCCTTATCTCCTCCAGTAGCATGTGGTTCTTCAATACCAGTATTAGTAATAGGCTTAGACAAATGCTTGTCGTACGCGGGCACTTCCTCATCCATCAAGTATTTATAATCGGGTTCTGTTTCAGAATATTCTTTTCTAAGAGACGCGACTTCAGGAGTTAGTAGAAGAACTCTGTTTCTGTATCTATCAACGCTGGAATCGATACTCAAGTTAAGGATAGCGAATACCTCATCGTCATCATCCGTATCTTCTGAAACGCCATCATATGACATTTCATGAAGTCTAACGTATTGATAAACAGAATCAGATTTAGTATTAAACAGATCCTTGACCTTTTTAGTAAATGCATATGTATATTTTAGATCTCCAGATTTCATAATATGATCGCATGCCTTAAATGTCAATGCTTCCATGATATAGTCTGGAACACTAATGGGTGACGAAAAAGATACAGCACCATATGCTACGTTGATAAATAGATCTGAACCACTAAGTAGATAATGATTAATGTTAAGGAAGAGGAAATATTCAGTATATAGATATGCCTTAGCATCATATCTTGTACTAAACACGCTAAACAGTTTATTGATGTGATCAATTTCCAACAGAACAATTAGAGCGGCAGGAATACCAACAAACATATTACCACATCCGTATTTTCTATGAATATCACATATCATGTTAAAAAATCTTGATAGAAGAGCGAATATCTCGTCTGACTTAATGAGATGTAGTTCAGCAGCATAAGTCATAACTGTAAATAGAACATACTTTCCTGTAGTGTTGATTCTAGACTCCACATCAACACCATTATTAAAAATAGTTTTATATACATCTTTAATCTGCTCTCCGTTAATCGTCGAACGTTCTAGTATACGGAAACACTTTGATTTCTTATCTGTAGTTAATGACTTAGTGATATCACGAAGAATATTACGAATTACATTTCTTGTTTTTCTTGTGAGACCTGATTCAGAACTCAACTCATCATTCCATAGTTTTTCTACCTCAGTGGCGAAATCTTTGGAGTGTTTGGTACATTTTTTAATAAGGTTCGTGACCTCCATTTATTATAAAAAATTTTTATTCAAAACTTAACTACAATCGGGTAATTATAAGATCGTAGATCTCCCATGTGGTGGAATACTACCATCTATCGCATGTGGATGGACAGTAGGTAATGGCCATGGGAACAGTAATGTTTGCATATTTATCTTTCTTGCTAGTATTACTGTATATTGTCCCAATGTTTCAATGTGATGTTCTAACCTATCAACTGCCACTGTATCACAACAATAATGTCCGATGGAATTAAGATTATGATCCAATGTGTTTAATATATGATTATCAAGTCTTATACGATCAGCGTCTTTTTTGACAGGATCAGGCTCTTCTACAGGAAGAAGTTTCGGCCTCTTATGATAGTCATGTCTGGGAAATGGTGGTCTAGGATGAGGATCCGGTATCGGAGTGGGTTTTGGATTATAATCATCATCATCATCATCATCATCATCATCATCATCATCATCATCATCATCATCATCATCATCATCATCTATGATATCATCATCATCATCTTCGATATTTATTTTGCTATCTTGATGATGTCCTGTATCAGTTGCATTTTCAGCACTCGACTGAATACTAGTACATTCATTGTCTATTATTAACGTATTTCTAAACCCAAAATGTATGTGTTGAACATCACTACTATAGTTGATGAGTCTTATAGCATGAATTCGCTTATCGTTATCGGGTTTATCTTCTGTCACCTTAACAATTCCTTTTTTATTAAACTCTGCATAATCATAACCATTTCTATTGTTTGTTCTAATATAAACGAGTATAGCATCATTGCTAAATTTTTCAATCGTATCGAAAACAGAATATCCTAAACCATATAATATATATTCAGGAACACTCAAACTAAATGTCCAGGATTCTCCTAAATACGTAAACTTTAATAGTGCTAAATCATTCAAAAATCTACCGCTTATAGATAGATAGTACATGAATGCGTATAGTAGTCTACCGATCTCTTTATTATGAAAACCGACATTACGATCATATATTTCGTGATATACATGTGACCCGTTTACGTTAAACCATAAATACATGGGTGATCCTATAAACATGAATTTATTTCTAATTCTCAGAGCCATAGTTAATTGACCGTGTAATATTTGTTTACATGCATACTTGATACGATCATTAATAAGATTTTTATCATTGCTCGTTATTTCAGAATCGTATATATAAGGAGTACCATCATGATTCTTACCAGATATTATACAAAATACTATATATAAAATATATTGACCCACGTTAGTAATCATGTAAATGTTTAATGTTTTAAATTTTGTATTTAATGATCCATCATCATACGCTAGCATGGTCTTGTGATATTCATTCTTTAAAATATAATATTGTGTTAGCCATTGCATTGGAGCTCCTAATGGAGATTTTCTATTCTCGTCCATTTTAGGATATGCTTTCATAAAGTCCCTAATAACTTCGTGAATAATGTTTCTATGTTTTCTACTGATGCATGTATTTGCTTCGATTTTTTTATCCCATGTTTCATCTATCATAGATTTAAACGCAGTAATGCTCGCAACATTAACATCTTGAACCGTTGGTACAATTCCGTTCCATAAATTTATAATGTTCGCCATTTATATAACTCATTTTTTGAATATACTTTTAATTGAACAAAAGAGTTAAGTTACTCATATGGACGCCGTCCAGTCTGTACATCAATCTTTTTAGCCAGAGATATCATAGCCGCTCTTAGAGTTTCAGCGTGATTTTCCAACCTAAATAGAACTTCATCGTTGTGTTTACAACACTTTTCTATTTGTTCAAACTTTGTTGTTATATTAGTAATCTTTTTTTCCAAATTAGTTAGCCGTTGTTTGAGAGTTTCCTCATTGTCGTCTCCATAGGCTTTAACAATTGCTTCGCGTTTAGTCTCTGGATTTTTAGCAGCCTTTGTAGAGAAAAATTCAGTTGCTGGAATTGCAAGATCGTCATCTCCGGGGAAAAGAGTTCCGTCCATTTAAAGTACAGATTTTAGAAACTGACACTCTGTGTTATTTATATTTGGCGCAATACATGGATTATAAATATCGATGTTAATAACATCAGAAAATGTAAAGTCTATACATTGTCGCATCGTGTTAAATTTTCTAATGGATCTAGTATTATTGGGTCCAACTTCTGCCTGAAATCCAAATATGGAAGCGGATACAAAACCGTTTCCTGGATAAACCACACATCTCCACTTTTGCTTTACATCAGAAATTGTGTCATTGACATCTTGAACTCTCCTATCTAATGCCGGTGTTCCACCTATAGATTTTGAATACTCGAATGCTGCATGAGTAGCATTGAATTCCTTAATATTGCCATAATTTTCATATATTGAGTAACTCTGGATAAAAAGTAAACACACCGCAGCCGTCGCTACTACAATAAAAAAAATTGATAGAGAGTTCATTTATAATCTATTAGAAGCTGACAAAATTTTTTTACACGCGTCAGACAATGCTTTAATAAATAGTTCAACATCTACTTTTGTCATATCGAACCGATGGTATGATTCTAACCTAGAATTACATCCGAAAAAGTTGACCATGTTCATAGTCATTAAGTCATTAACAAACAACATTCCAGACTCTGGATTATAAGACGATACTGTTTCGTCACAATCACCCACCTTAATCATGTGATTATGAATATTGGCTATTAGAGCACCTTCTAAGAAATCTATAATATCTTTGAAACACGATTTAAAATCAAACCACGAATATACTTCTACGAAGAAAGTTAGTTTACCCATAGGAGATATAACTATAAATGGAGATCTAGATACAAAATCCGGATCTATGATAGTTTTAACATTATTATATTCTCTATTAAATACCTCCACATCTAAAAATGTTAATTTTGAAACCATGTCTTCGTTTATTACCGTACCTGAACTAAACGCTATAAGCTCTATTGTTTGAGAACTCTTTAAACGATATTCTTGAAATACATGTAACAAAGTTTCCTTTAACTCGGTCGGTTTATCTACCATAGTTACAGAATTTGTATCCTTATCTATAATATAATAATCAAAATCGTATAAAGTTATATAATTATCGTGTTCAGATTGTGATCTTTTCAAATAGACTAAAAACCCCATTTCTCTAGTAAGTATCTTATGTATATGTTTGTAAAATATCTTCATGGTGGGAATATGCTCTACAGCAGTTAGCCATTCCTCATTGACAGCTGTAGATGTATTAGACAAAACTACTCCAATGTTTAACAAGGGCCATTTTACGAGATTATTAAATCCTTGTTTGATAAATGTAGCCAATGCGGGTTCGAGTTCAACGACGATTGAATTCTCTTCCCGTGGATGCTGCATGATGAACGACGGGATGTTGTTCTATTGATTTGGAATTCTTTTTCGACTTTTTGTTTATATTAAATATTTTAAAATTTATGGCTGATAGTAATTCATGTACTACGGATAATGTAGACGTGTATTGCACATCGATATCTTTATTATTAGATAAATTTATCAATAAATGTGAGAAGTTTGCCTCGTTAAGGTCTTCCATTTAAATATTATATAAATATTTGTGTTTGTATTTTATTCGTCTTTTATGGGATAGTTTTTAACTAGTAAAGCTGTAATTACATACTTTGTCCGTAAAACATAAATATAAATACCCGCTTTTATCAAACGTTCCAAAAAGTCGGCAGCCGACATTTTTAACATGGCATCTATTTTAAATACACTTAGGTTTTTAGAAAAAACATCATTTTATAATTGTAACGATTCAATAACTAAAGAAAAGATTAAGATTAAACATAAGGGAATGTCATTTGTATTTTATAAGCCAAAGCATTCTACCGTTGTTAAATACTTGTCTGGAGGAGGTATATATCATGATGATTTGGTTGTATTGGGGAAGGTAACAATTAATGATCTAAAGATGATGCTATTTTACATGGATTTATCATATCATGGAGTGACAAGTAGTGGAGCAATTTACAAATTGGGATCGTCTATCGATAGACTTTCTCTAAATAGGACTATTGTTACAAAAGTTAATAACAATTATAACAATTATAACAATTATAATTGTTATAATAATTATAATTGTTATAATTATGATGATACATTTTTTGACGATGATGATTGATCACTATTACACAATTTTGTTTTTGTACTTTCTAATATAGTGTTTAGGTTCTTTTTCATATGAGAATATTGACTTACTAAAATATCTATGTTTAACTTTTGTTCTATAACGTCCTTATCGGCGGTATCGGTACATATACGTAATTCACCTTCACAAAATACGGAGTCTTCGATAATAATAGCCAATCGATTATTGGATCTAGCTGTCTGTATCATATTCAACATGTTTAATATATCCTTTCGTTTCCCCTTTACAGGCATCGATCGTAGCATATTTTCCGCGTCTGAGATGGAAATGTTAAAACTGCAAAAATGCGTAATGTTAGCCCGTCCTAATATTGGTACGTGTCTATAAGTTTGGCATAGTAGAATAATAGACGTGTTTAAATGCCTTCCAAAGTTTAAGAATTCTATTAGAGTATTACATTTTGATAGTTTATCACCTACATCATCAAAAATAAGTAAAAAGTGTGCTGATTTTTTATGATTTTGTGCGACAGCAATACATTTTTCTATGTTACTTTTAGTTCGTATCAGATTATATTCTAGAGCTTCCTGACTACTAACGAAATTAATATGATTTGGCCAAATGTATCCATCATAATCTGGGTTATAAACGGGTGTAAACAAGAATATATGTTTATATTTTTTAACTAGTGTAGAAAACAGAGATAGTAAATAGATAGTTCTTCCAGATCCAGATCCTCCTGTTAAAACCATTCTAAACGGCATTTTTAATAAATTTTCTCTTGAAAATTGTTTTTCTTGAAAACAATTCATAATTATATTTACAGTTACTAAATTAATTTGATAATAAATCAAAATATGGAAAACTAAGGTCGTTAGTAGGGAGGAGAACAACGAAGACATATCGTGATATAAATAACATTTATTATCATGATGACACCAGAAAACGACGAAGAGCAGACATCTGTGTTCTCCGCTACTGTTTACGGAGACAAAATTCAGGGAAAGAATAAACGCAAACGCGTGATTGGTCTATGTATTAGAATATCTATGGTTATTTCACTACTATCTATGATTACCATGTCCGCGTTTCTCATAGTGCGCCAAAATCAATGCATGTCTGCTAACGAGGCTGCTATTACTGACTCCGCTGTTGCCGTTGCTGCGGCATCATCTACTCATAGAAAGGTTGCGTCTAGCACTACACAATATGATCACAAAGAAAGCTGTAATGGTTTATATTACCAGGGTTCTTGTTATATATTACATTCAGACTATAAGTCATTCGAGGATGCTAAAGCAAACTGCGCTGCGGAATCATCAACGCTACCCAATAAATCCGATGTCTTGACTACCTGGCTCATTGATTATGTTGAGGATACATGGGGATCTGATGGTAATCCAATTACAAAAACTACATCCGATTATCAAGATTCTGATGTATCACAAGAAGTTAGAAAGTATTTTTGTACATAAATAAATGAAATCGCTTAATAGACAAACTGTAAGTAGGTTTAGGAAGTTGTCGGTGCCGGCCGCTATAATGATGTTACTCTCAACCATTATTAGCGGCATAGGAACATTTCTACATTACAGAGAAGAACTGATGCCTAGTGCTTGCGCCAATGGATGGATACAATACGATAAACATTGTTATCTGGATACCAACATTAAAATGTCTACGGATAATGCAGTTTATCAGTGTCGCAAATTACGAGCTAGATTGCCTGGACCTGATACTAGACATCTGAGAGTATTGTTTAGTATTTTTTATAAAGATTATTGGGTAAGTTTAAAAAAGACCAATGATAAATGGTTAGATATTAATAATGATAAAGATATAGATATTAGTAAATTAACAAATTTTAAGCAACTAAACAGCACAACGGATTCTGAAGCGTGTTATATATACAAGTCTGGAAAACTGGTTAAAACAGTATGTAAAAGTACTCAATCTGTACTATGCGTTAAAAGATTCTACAAGTGATAACAAAAAATGAATTAATAGTAAGTCGTTAACGTACGCCGCCATGGACGCCGCGTTTGTTATTACTCCAATGGGTGTGTTGACTATAACAGATACATTGTATGATGATCTCGATATCTCAATCATGGACTTTATAGGACCATACATTATAGGTAACATAAAAATTGTCCAAATAGATGCACGGGATATAAAATATTCCGACATGCAAAAATGCTACTTTAGCTATAAGGGTAAAATAGTTCCTCAGGATTCTAATGATTTGGCTAGATTCAACATTTATAGTATTTGTACCGCATACAGATCAAAAAATACCATCATCATAGCATGCGACTATGATATCATGTTAGATATAGAAGGTAAACATCAACCATTTTATCTATTCCCATCTATTGATGTTTTTAACGCTACAATCATAGAAGCGTATAATCTGTATACAGCTGGAGATTATCATCTGATCATCAATCCTTCAGATAATCTGAAAATGAAATTGTCGTTTAATTCTTCATTTTGTATATCAGACGGCAATGGATGGATTATAATTGATGGGAAATGTAATAGTAATTTTTTATCATAAAAGTTGTAAAGTAAATAATAAACAATAAATATTGAACTAGTAGTATGTTGTATATTGAGCAATCAGAGATGATGCTGGTACCTCTTATCACGGTGACCGTAGTTGCGGGAACAATATTAGTATGTTATATATTATATATTTGTAGGAAAAAGATACGTACTGTCTATAATGACAATAAAATTATCATGACAAAATTAAAAAAGATAAAGAGTCCTAATTCCAGCAAATCTAGTAAATCAACTGATAGCGAATCAGACTGGGAGGATCACTGTAGTGCTATGGAACAAAACTATGACGTAGATAATATTTCTAGAAATGAGATATTGAACGATGATAGCTTCGCTGGTAGTTTAATATGGGATAACGAATCCAATGTCATGGCGCCTAGCACAAAACACATTTACGATAGTGTTGCTGGAAGCACGCTGCTAATAAATAATGATCGTAATGAACTGACTATTTATCAGAATACTACAGTAGTAATTAATGATACAGAGACTGTTGAAATACTTAATGAAGATACCAAACAGATTCCTAGCTATTCTTCCAATCCTTTCGTAAATTATAATAAAACCAGTATTTGTAGCAAGTCAAATCCGTTCATTGCAGAACTCAACAATAAATTTAGTGATAATAATCCGTTTAGGAGAGCACATAGCGACGATTATCTTAATAAGCAATAACAATAACAATAACAAGATCATGAACACGATGATATAGAATCATCGGTTGTATCATTGTCTGATTAGTTTCCTTTTTATAAAATTGAAGTAATATTTAGTATTAATTACCGCCGATGCATTATACAAATATGGAGATATTCCCTGTATTCGGCATTTCTAAAATTAGCAATTTTATTGCTAATAATGACTGTAGATATTATATAGATGTAGAGCATCAAAAAATTATATCTGATGAGATCAATAGACAGATGGATGAAACGGTACTTCTTACCAACATCTTAAGCGTAGAAGTTGTAAATGGCAATGAGATGTACCATCTTATTCCCCATAGACTATCGACGATTATACTCTGTATTAGTTCTGTTGGAGGATGTGTTATCTCTATAGATAATGACGTCAATGATAAAAATATTCTAACATTTCCCATTAATCATGCTGTAATCATATCCCCACTGAGTAAATGTGTCGTAGTTAGCAAGGGCCCTACAACCACACTGGTTGTTAAAGCGGATATACCCAGCAAACGATTGGTAACATCGTTTACAAACGACATACTGTATGTAAACAATCTATCACTGATTAATTATTTACCGTTGTCTGTATTCATTATTAGACGAGTCACTGACTATTTGGATAGACACATATGCGATCAGATATTTGCTAATAATAAGTGGTATTCCATTATAACTATCGACGATAAGCAATATCCTATTCCATCAAATTGTATAGGTATGTCTTCTGCCAAGTACATAAATTCGAGCATCGAGCAAGATATTTTGATCCATGTTTGTAACCTCGAGCATCCATTCGACTCAGTCTACAAAAAAATGCAGTCGTACAATTCTCTACCTATCAAGGAACAAATATTGTATGGTAGAATTGATAATATAAATATGAGCATTAGTATTTCGGTGGATTAATAGATTTCTCTAGTATGGGATCATTAATCATCTCTAAATACATCATAAAAAAGCTATTATCAAATACTGTACTGAATGGATTCATTCTTTTCTCTTTTTATGAAACTCTGTTGTATATCTACGGATAAAACTAGAAGCAAAAAATCTGATAGGAAGAATAATGATTATATGGAGGAACACGATTATTATAAAATAACAATAGTTCCTGGTTCCTCTTCCACGTCTACTAGCTCGTGGTATTATACACATGCCTAGTAATAGTCTCTTTGCGTTGACGGAAAGCAGACTAGAAATAACAGGCCAAAATGTTCAGACACCATAATAGTTCCCAACCCAGATAATAACAGAGTTCCATCAACACATTCCTTTAAACTCAATCCCAAACCCAAAACCGTTAAAATGTATCCAGCCAATTGATAATAGATAATGAGGTGTACAGCACATGATAATTTACACAGTAACCAAAATGAAAACACTTTAGTAATTATAAGAAATATAGACGGTAATGTCATCATCAACAATCCAATAATATGCCTGAGAGTAAACATTGACGGATAAAACAAAAATGCCCCGCATAACTCTATCATGGCAATAACGCAACCAAACACTTGTAAAATTCCTAAATTAGTAGAAAATACAACTGATATCGATGTATAAGCGATTTCGAGGAATAATAAGAACAAAGTAATTCCCGTAAAGATAAACATCAACATTGTTTGGTAATCATTAAACCAATTAGTATGACGTTGAACTAATTTCACAGTAGATTTTATTCCAGTATTATCCCCGCATGTATACGTACCTGGTAAGATATCTTTATATTCCATAATCAATGAGACATCACTATCCGATAACGAATGAAGTCTAGCACTAGTATGCCATTTACTTAATATGGTCGTCTTGGAAGTTTTATTATAAGTTAAAATATCATGATTGTCCAATTTCCATCTAATATACTTTGTCGGATTATCTATAATACATGGAATAATGATGGTATCATTACATGCTGTATACTCTATAGTCTTTGTAGATGTTATAACCACAAAAGTACAGAGGTATATCAACAATATTCTAACTCTTAACATTTTTATTTATTTAAAATGATACCTTTGTTATTTATTTTATTCTTATTTTGCTAACGGTATCGAATGGCATAAGTTTGAAACGAGTGAAGAAATAATTTCTACTTACTTAATAGATGATGTGGTAACGGGTGTTATTAATGGGGCTGTATATACATTTTCAAATAATGAACTAAACAAAACTGGGTTAACTAATAACAATAATTATATCACAACATCTATAAAAGTAGAGGATAATGATACATTAGTAGTATGCGGAACCAATAACGGAAATCCCAAATGTTGGAAAATAGACGGTTCATACTACCCAAAACATATAGGTAGAGGATACGATCATCAAAATAGCAAAGTAACGATAATCAGTCACAATGAATGTGTACTATCCGACATAAACATATCAAAAGAAGGAATTAAACGATGGAGAAGATTTGACGGACCATGTGGTTATGATTAAACGAGTTAAGTTTTTTAAGAAGCCTTAGAAGAGGGCTATTGGGTATGAAAATCCGAAATATTAAACCAGACAACCCCATATAATTTTATAGCTAAGAATACCGCGAAGAATGGAACTAAAAACGGAAATATTTGTAGCACAACGAATAACTCCCAAACTGCATTCATGTTACACTATATAACAATTACAATACATTTTTATCATAACACTACTTCGGTTAGATGTTTTAGAAAAAAATAAATATCACTACCGTTCTTGTTTTATAAAAATAACAATTAACAATTATCAATTTTTTTCTTTAATATTTTACGTGGTTGACCATTCTTGGTGGTAAAATAATCTCTTAGTGTTGGAATGGAATGCTGTTTAATGTTTCCGCACTCATCGTATATTTTGACGTATGCAGTCACATCGTTTACGCAATAGTCAGACTGTAGTTCTATCATGCTTCCTACGTTAGAAGGAGGAACAGTTTTAAAGTCTCTTGGTTTTAATCTATTGTCATTAGTTTTCATGAAATCCTTTGTTTTATCCACTTCACATTTTAAATAAATGTCAACTATACATTCTTCTGTTAATTTTACTAGATCATCATAGGTCATAGAATTTATAGGTTCCGTAGTCCATGGATCCAAACTAGCAAACTTCGCGTATACAGTATCGCGATTAGTGTATACACCAACTGTATGAAAATTAAGAAAACAGTTTAATAAATCTACAGAAATATTTAATCCTCCGTTTGATACAGATGCGCCATATTTATGGATTTCGGATTCACACGTTGTTTGTCTAAGTGGTTCGTCTAGTGTTGCTTCTACATAGACTTCGATTCCCATATATTCTTTATTGTCAGAATCACATACCGATTTATCATACGCTGGTTCACTTGTTTGAAAACTAAATGGTAGTAGATACATCAAAATAATAAATAATAAGTACATTCTGCAATATTGTTATCGTAATTGGAAAATTGGTATTCAAGTGAGCTGGATTATGTGAGTATTGGATTGTATATTTTATTTTATTTTATATTTTATATTTTATATTTTATTTTATATTTTGTAGTAAGAATAGAATGCTAAATGTCAAGTTTATTCGAATAGATGTCTTATTAAAAACATATATAATAAATAACAATGGCTGAATGGCATAAAATTATCGAGGATATCTCAAAAAATAATAAGTTCGAGGATGCCGCCATCGTTGATTACAAGACTACAAAGAATGTTCTAGCGGCTATTCCTAACAGAACATTTGCAAAGATTAATCCGGGTGAAGTTATTCCCCTCATCACTAATCATAATATTCTAAAACCTCTTATTGGTCAGAAATTTTGTATTGTATATACTAACTCTCTAATGGATGAGAACACGTATGCTATGGAGTTGCTTACTGGGTACGCCCCTGTATCTCCGATCGTTATAGCGAGAACTCATACCGCACTTATATTTTTGATGGGTAAGCCAACAACATCCAGACGTGATGTGTATAGAACATGTAGAGATCACGCTACCCGTGTACGTGCAACTGGTAATTAAAATAAAAAGTAATATTCATATGTAGTGTCAATTTTAAATGATGATGATGAAATGGATAATATCCATATTGACGATGTCAATAATGCCGGTATTGACATACAGTTCATCGATTTTTAGATTTCATTCAGAGGATATTGAATTATGTTATGGGAATTTGTATTTTGATAGGATCTATAATAATGTAGTAAATATAAAATATATTCCTGAGCATATTCCATATAGATATAATTTTATTAATCGTACGTTCTCCGTAGATGAACTAGATGATAATGTCTTTTTTACACATGGTTATTTTTTAAAACACAAATATGGTTGTTCACTTAATCCTAGTTTGATTGTCTCATTATCAGGAAACTTAAAATATAATGATATACAATGCTCAGTAAATGTATCATGTCTCATTAAAAATTTGGTAACGAGTACATCTACTATATTAACATCTAAACATAAGACTTATTCTCTATATCGGTCCACGTGTATTGCTATAATAGGATACGATTCTATTATATGGTATAAAGATATAAATGACAGGTATAATGACATCTATGATTTTACTGCAATATGTATGCTAATAGCGTCTACATTGATAGTGATCATATACGTGTTTAAAAAAATAAAAATGAACTCTTAATTATGTTATACTATTAGAAATGGATAAAATCAAAATTACGATTGATTCAAAAATTGGTAATGTTGTTACCATATCGTATAACTTGGAAAAGATAACTATTGATGTCACACCAAAAAAGAAAAAAGAAAAGGATGTATTATTAGCGCAATCAGTTGCTGTCGAAGAGGCAAAAGATGTCAAGGTGGAAGAAAAAAATATTATCGATATTGAAGATGACGATGATATGGATATAGAAAACACGTAATACGATCTATAAAAATAAGGTATTAAATACTTTTTATTTACGGTACTCTTGTAGTGGTGATACCACTAATCGATTTTTTTTTAAAAAAAATACTTATTCTGATTCTTCTAGCCATTTCCGTGTTCGTTCGAATGCCACATCGACGTCAAAGATAGGGGAGTAGTTGAAATCTAGTTCTGCATTGTTGGTACGCACCTCAAATGTAGTGTTGGATATCTTCAACGTATAGTTGTTGAGTATTGATGGTTTTCTAAATAGAATTCTCTTCATATCATTCTTGCACGCGTACATTTTTAGCATCCATCTTGGAATCCTAGATCCTTGTTCTATTCCCAATGGTTTCATCAATAGAAGATTAAACATATCGTAAGAACACGATGGAGAGTAATCGTAGCAAAAGTAAGCATTTCCTTTAATCGCAGATCCCGGATACTGGATATATTTTGCAGCCAACACGTGCATCCATGCAACATTTCCTACATATACCCGGCTATGCACAGCGTCATCATCGACTGTACGATACATAATGTTACCGTGTTGCTTACATTGCTCGTAAAAGACTTTCGTCAATTTGTCTCCTTCTCCGTAAATTCCAGTGGGTCTTAGGCAACAAGTATACAATTTTGCGCCATTCATGATTACGGAATTATTGGCTTTCATAACCAGTTGCTCGGCCATACGTTTACTTTTTGCGTATACATGTCCTGGTGATATATCATAAAGGGTATGCTCATGACCGATGAATGGATTACCGTGTTTATTTGGTCCTATTGCTTCCATGCTACTAGTATAGATCAAATACTTGATTCCTAGGTCCACACAAGCTGCCAATATAGTCTGTGTTCCATAATAGTTTACTTTCATGATTTCATTATCAGTGTATTTTCCAAATACATCCACTAGAGCAGCCGTATGAATAATCAGATTTACCCCATCTAGCGCTTCTCTCACCTTATCAAAGTCGTTTATATCACATTGTATATAGTTTATAACCTTAACTTTCGAGGTTATTGGTTGTGGATCTTCTACAATATCTATGACTCTTATTTCTTGAACATCATCTGCGCTAATTAAAAGTTTTACTATATACCTGCCTAGAAATCCGGCACCGCCAGTAACCGCGTACACGGCCATTGCTGCCACTCATAATATCAGACTACTTATTCTATTTTACTAAATAATGGCTGTTTGTATAATAGACCACGATAATATCAGAGGAGTTATTTACGTTGAACAAGTCCATGGAAAAGATAAAGTTTTAGGATCAGTTATTGGATTAAAATCCGGAACGTATAGTTTGATAATTCATCGTTACGGAGATATTAGTCGAGGATGTGATTCCATAGGCAGTCCAGAAATATTTATCGGTAACATCTTTGTAAACAGATATGGTGTAGCATATGTTTATTTAGATACAGATGTAAATATATCTACAATTATTGGAAAGGCGTTATCTATTTCAAAAAATGATCAGAGATTAGCGTGTGGAGTTATTGGTATTTCTTACATAAATGAAAAGATAATACATTTTCTTACAATTAACGAGAATGGCGTTTGATATATCAGTTAATGCGTCTAAAACAATAAATGCATTAGTTTACTTTTCTACTCAGCAAAATAAATTAGTCATACGTAATGAAGTTAATGATATACACTACACTGTCGAATTTGATAGGGACAAAGTAGTTGATACGTTTATTTCATATAATAGACATAATGACTCCATAGAGATAAGAGGGGTGCTTCCAGAGGAAACTAATATTGGTCGCGTGGTTAATACGCCGGTTAGTATGACTTACTTGTATAATAAGTATAGTTTTAAACCGATTTTAGCAGAATATATAAGACACAGAAATACTATATCCGGCAATATTTATTCGGCATTGATGACACTGGATGATTTGGTTATTAAACAGTATGGAGACATTGATCTATTATTTAATGAGAAACTTAAAGTAGACTCCGATTCGGGACTATTTGACTTTGTCAACTTTGTAAAGGATATGATATGTTGTGATTCTAGAATAGTAGTAGCTCTATCTAGTCTAGTATCTAAACATTGGGAATTGACAAATAAAAAGTATAGGTGTATGGCATTAGCCGAACATATAGCTGATAGTATTCCAATATCTGAGCTATCTAGACTACGATACAATCTATGTAAGTATCTACGCGGACACACCGATAGCATAGAGGATGAATTTGATCATTTTGAAGACGATGATTCGTCTACATGTTCTGCCGTAACCGACAGGGAAACGGATGTATAATTTTTTTATAGTGTGATGGATATGATGGATATGATATATGATGGATATTATGATGGATATGATGGATATTATGATGGATATGATGGATATGATGGATATGATGGATATGATAAATATGATAAAAATATAATTGTTGTATATGATGGATATGATGGATATGATGGATATGATGGATATGATGGATATGATGGATATGATAAATATGATAAAAATATAATTGTTGTATCCATTCCCATTCAAATCACCTTATATGATTCTGTAACACAATGAAGGAGTCTCATAGATATATAGAGGTCAGATACTGGTTTGATAAACTTTTTATTCCACATGAGTATGTTTGACTTATGGTTAGACACGCATACTTTAACAAATCACTGAAAATTGGAGTTAGGTATTCCTCTCAGAATCAGTTGCCGTTCTGGAACATTAAATGTATTTTTTATGATATACTCCAACGCATTTATGTGGGTATACAACAAGTCATTAATAATGAGTATTTCCAAGAGTTTTAGCTGTCTAGTATTTAACAAGAGAAGAGATTTCATCAGACTGTTTATGAACTCGAATACCGCCTCATTGTCGCTTATATTGATGACATCGAATTCCCAATATCATCTCATCAGTGATGAGTAGCTCAATCTTGTTATCGGGATCCAATTTCTAAAGATGTCATTAAACCCTCGATCGTGAATGGATTTATCATCATCGTTTTTATGTTGGACATGAGCTTAGTCCGTTTGTCCACATCTATATACGATGATTTCTGAATTATTTCATATATCTCTCGTTAACTCCAGGAACTTGTCAGGGATCTAACTTTAATATGTTCTCGTCTAAGAGATGAAAATCTTTGGATGGTGGCATGTGACTTTTCTCTAAAGGATGATGTTGCCCGATCCTCTCTTAAATGAATCCATCTTATCCTTGGACAAGATGGACAGTCTATTTTCCTTAGATGGTTTAATATTTTTTACCCATGATCTATAAAGGTAGACAGACCTAATCGTCTCGGATGACCATATATTATTTTCAGTTTTATTATACGCATAAATTGTAAAAAATATGTTAGGTTTACGAAAATGTCTCGTGGGGCATTAATCGTTTTTGAAGGATTGGACAAATCTGGAAAAACAACACAATGTATGAACATCATGGAATCTATACCGGCAAACACGATAAAATATCTTAACTTTCCTCAGCGATCCACAGTCACTGGAAAGATGATAGATGACTATCTAACTCGTAAAAAAACCTATAATGATCATATAGTTAATCTATTATTTTGTGCAAATAGATGGGAGTTTGCATCTTTTATACAAGAACAACTAGAACAGGGAATTACTTTAATAGTTGACAGATACGCGTTCTCTGGAGTAGCGTATGCCACCGCTAAAGGCGCGTCAATGACTCTCAGTAAGAGTTATGAATCTGGATTGCCTAAACCCGACTTAGTTATATTCTTGGAATCTGGTAGCAAAGAAATTAATAGAAACGTCGGCGAGGAAATTTATGAAGATGTTGAATTCCAACAAAAGGTATTACAAGAATATAAAAAAATGATTGAAGAAGGAGATATTCATTGGCAAATTATTTCTTCTGAATTCGAGGAAGATGTAAAGAAGGAGTTGATTAAGAATATAGTTATAGAGGCTATACACACGGTTACTGGACCAGTGGGGCAACTGTGGATGTAATAAAATGAAATTACATTTTTATAAATAGATGTTAGTACAGTGTTATAAATGGATGAAGCATATTACTCTGGCAACTTGGAATCAGTACTCGGGGATACGTGTCCGATATGCATACCGAACTCGCATCAATATCTCAATTAGTTGCCAAGATAGAAACTATAGATAATGATTATTAAACAAGGACATTGTAAATTTTATCATATGTAGATCAAACTTGGATAATCCATTTATCTCTTTCCTAGATACTGCATATACTATCATAGATCAAGAGATCTATCAGAACGAGTTGATTAATTCATTAGACGATAATGAAATTATCGATTGTATAGTTAACAAGTTTATGAGCTTTTATAAGGATAACCTAGAAAATATGGTAGATGCTATCATTACTCTAAAATATTATAATTAATAATCCAGATTTTAAAACTACGTATGTGGAAGTACTCGGTTCCAGAATAGCTGATATAGATATTAAACAAGTGATACGTAAGAATATAATACAATTGTCTAATGATCCGCGAACGATATTTGTGAAAATATTAAAAAAAAATACTTTTTTTATTAAATGACGTCTCTTCGCGAATTTAGAAAATTATGCTGTGATATATATCACGCATCAGGATATAAAGAAAAATCTAAATTAATTAGAGACTTTATAACAGATAGAGATGATACCGATACATATTTGATCATTAAGCTATTGCTTCCCGGATTAGACGATAGAATGTATAACATGAACGATAAACAAATTATAAAATTATATAGTATAATATTTAAACAATCTCAGGAAGATATGCTACAAGATTTAGGATACGGATATATAGGAGACACTATTAGGACATTCTTCAAAGAGAACACGGAAATCCGTCCACGAGATAAAAGCATTTTAACTTTAGAAGAAGTGGATAGTTTTTTAACTACGTTATCATCAGTAACTAAAGAATCACATCAAATAAAATTATTGACTGATATAGCATCTGTTTGTACATGTAATGATTTAAAATGTGTAGTCATGCTTATTGATAAAGATCTAAAAATTAAAGCGGGTCCTCGGTACGTGCTTAACGCTATTAGTCCTCATGCCTATGATGTTTTTAGAAAATCTAATAACTTGAAAGAGATAATAGAAAATGCAGCTAAACAAAATCTAGACTCTATATCTATTTCTGTTATGACTCCAATTAATCCCATGTTAGCGGAATCATGTGATTCTGTCAATAAGGCGTTTAAAAAATTTCCATCAGGAATGTTTGCGGAAGTCAAATACGATGGTGAAAGAGTACAAGTTCATAAAAAAAATAACGAGTTTGCATTCTTTAGTAGAAACATGAAACCAGTACTCTCTCATAAAGTGGATTATCTCAAAGAATACATACCGAAAGCATTTAAAAAAGCTACGTCTATCGTATTGGATTCTGAAATTGTTCTTGTAGACGAACATAATGTACCGCTACCGTTTGGAAGTTTAGGTATACACAAAAAGAAAGAATATAAAAACTCTAACATGTGTTTGTTCGTGTTTGACTGTTTATACTTTGATGGATTCGATATGACAGACATTCCATTGTATGAACGAAGATCTTTTCTCAAAGATGTTATGGTCGAAATACCCAATAGAATAGTATTCTCAGAGTTGACGAATATTAGTAACGAGTCTCAGTTAACTGATGTATTAGATGATGCACTAACGAGAAAATTAGAAGGATTGGTCTTAAAAGATATTAATGGCGTATACGAACCGGGAAAGAGAAGATGGTTAAAAATAAAGCGAGACTATTTGAACGAGGGTTCCATGGCAGATTCTGCCGATTTAGTAGTACTAGGTGCCTACTATGGTAAAGGAGGAAAGGGTGGTATCATGGCAGTCTTTCTAATGGGTTGTTACGACGATGAATCCGGTAAATGGAAGACGGTAACTAAATGTTCCGGTCACGATGATAATACGTTAAGGGTTTTGCAAGACCAATTAACGATGGTTAAAATTAACAAGGATCCCAAAAAAATTCCAGAGTGGTTGGTAGTTAATAAAATCTATATTCCCGATTTTGTAGTAGATGATCCGAAACAATCTCAGATATGGGAAATTTCAGGAGCAGAGTTTACATCTTCCAAGTCACATACAGCGAATGGAATATCAATTAGATTTCCTAGATTTACTAGGATTAGAGAAGATAAAACGTGGAAAGAATCTACTCATCTAAACGATTTAGTAAACTTGACTAAATCTTAATAGTTACATATAAACTGAAAAATAAAATAATACTATTTTAGTTGGTGGTCGCCATGGATGGTGTTATCGTATACTGTCTAAATGCGTTAGTAAAACATGGCGAGGAAATAAATCATATAAAAAATGATTTCATGATTAAACCATGTTGTGAAAGAGTTTGTGAAAAAGTCAAGAACGTTCACATCGGCGGACAATCTAAAAACAATACAGTGATTGCAGATTTGCCATATCTGGATAATGCTGTATCAGATGTATGCAAATCAATATATAAAAAGAATGTATCAAGAATATCCAGATTTGCTAATATGATAAAAATAGATGACGATGACAAGACTCCTACCGGTGTATATAATTATTTTAAACCTAAAGATACTATTCCTGTTATTATATCCATAGGAAAGGATAAAGATGTCTGTGAACTATTAATCTCATATGATAAAGCGTGTGCGTGTATAAAGTTAAATTTATATAAAGTAGCCATTCTTCCCATGGATGTTTCCTTTTTTACCAAAGGAAATGCATCATTGATTATTCTCCTGTTTGATTTCTCTATCGATGCGGCACCTCTCTTAAGAAGTGTAACCGATAATAATGTTATTATATCTAGACACCAGCGTCTACATGACGAGCTTCCGAGTTCCAATTGGTTCAAGTTTTACATAAGTATAAAGTCCGACTATTGTTCTATATTATATATGGTTGTTGATGGATCTATGATGTATGCGATAGCTGATAATAGAACTCACGCAATTATTAGCAAAAATATATTAGACAATACTACGATTAACGATGAGTGTAGATGCTGTTATTCTGAACCACAGATTAGGATTCTTGATAGAGATGAGATGCTCAATGGATCATCGTGTTATATGAACAGACATTGTATTATGATGAATTTACCTGATGTAGGCGAATTTGGATCTAGTATGTTGGGGAAATATGAACCTGACATGATTAAGATTGCTCTTTCGGTGGCTGGTAATTTAATAAGAAATCGAGACTACATTCCCGGGAGACGAGGCTATAGCTACTACGTTTACGGTATAGCCTCTAGATAATTTTTTTAAGCACGAAATAAAAACATAATTTTAAACAATCTATTTCATACTATTTTGTGTGATCAACATGAACATAAAGATAGATGATATGATATATTAGTATTTCTGGTGATAAATTTACGGCGACTGCTAGGAGGGAAAATGAAGAAAGAAAAAATATCTACCTCTCCAAAAAGAAAAACTACTGATGTTATCAAACCTGATTATCTTGAGTACAATGACTTGTTAGATAGAGATGAGATGTCTACTATTCTAGAGGAATATTAGGCCTTAGAATAAAATATGGACGACTCTTAACGAAATTAGAAAATTCGATAATGATGTTGAAGAACAATTCGGTACTATAGAAGAACTCAAGCAGAAGCTTAGATTAAATTCTGAAGAGGGAGCAGATAATTTTTATAGATTATATAAAGGTACAAAAACAGGATATCGTCAAACTTACTGTATACGATTGCATATATCTATGATAGGATTGTATGCGTGCGTGGTAGATGTTTGGAGAAATGAGAAACTGTTTTCTAGATGGAAATATTGGTTACAAGCGATTAAACTGTTTATTGATGATCACATGCTTGATAAGATAAAATCTATACTGTAGAATAGACTAGTGTATGTGGAAATGTCATAGAAAGTTAAAAGTTAATGAGAGCAAAAATATATAAGGTTGTATTCCATATTTGTTATTTTTTTCTGTAATAGTTAGAAAATACATTCGATGGTCTATCTACCAGATTATTATGTGTTATAAGGTACTTTTCTCATAATAAACTAGAGTATGAGTAAGATAGTGTTTTTCAAAACATATAAATCTAAAATTGATGGATGAGATATACAGCTATTAATTTCGAAAATATATTTTAATCTGATAACTTTAAACATGGATTTTTGATGGTGGTTTAAGTTTAAAAAAGATTTTGTTATTGTAGTATGATAATATCAAAAAGATGGATATAAAGAATTGGTCAGTGTATAATAAATTATATGTAGGAGGAGGAATATCTGATGATGTTCAAACTAATACATCTGAAACATACGATAAAGAAAAAGATTGTTGGACATTGGATAATGGTCACTTGGTACCACATAATTATATAATGTATAAATGCGAACCGGTTAAACATAGATATCCATTGGAAAAAAACACAGTACACGAATGATTTTCTAAAGTATTTGGAAAGTTTTATAGGTAGTTGATAGAACAAAATACATAATTTTGTAAAAATAAATCACTTTTTATACTAATATGACACAATTACCAATACTTTTGTTACTAATATCATTAGTATACGTTACACCTTCTCCTCAGACATCTAAAAAAATAGGTGATGATGCAACTATATCATGTAGTCGAAATAATACAAATTACTACGTTGTTATGAGTGCTTGGTATAAGGAGCCCAATTCCATTATTCTCTTAGCTGCCAAAAGCGACGTCTTGTATTTTGATAATTATACCAAGGATAAAATATCTTACGACTCTCCATACGATGATCTAGTTACAACTATCACAATTAAATCATTGACTGCTGGAGATGCCGGTACTTATATATGTGCATTCTTTATGACATCGACTACAAATGATACTGATAAAGTAGATTATGAAGAATACTCCATAGAGTTGATTGTAAATACAGATAGTGAATCGACTATAGACATAATACTATCTGGATCTACACCGGAAACTATTTCTGAGAAACCAGAGGATATAGATAATTCTAATTGCTCGTCTGTATTCGAAATCGCGACTCCGGAACCAATTACTGATAATGTAGAAGACCATACAGACACCGTCACATACACTAGTGATAGCATTAATACAGTAAATGCATCATCTGGAGAATCCACAACAGACGAGACTCCGGAACCAATTACTGATAAAGAAGAAGATCATACAGTAACAGACACTGTCTCATACACTACAGTAAGTACATCATCTGGAATTGTCACTACTAAATCAACCACCGATGATGCGGATCTTTATGATACATACAATGATAATGATACAGTACCGCCAACTACTGTAGGTGGTAGTACAACCTCTATTAGCAATTATAAAACCAAGGACTTTGTAGAAATATTTGGTATTACCACATTAATTATATTGTCGGCAGTGGCGATTTTCTGTATTACGTATTATATATGTAATAAACACCCACGTAAATACAAAACAGAGAACAAAGTCTAGATTTTTGACTTACATAAATATCTGGGATAATAAAATCTATCATATTGAGAGGACCATCTGGTTCAGGAAAGACAGCCATAACCAAAAGACTGTTAAAAGACTATGGGAATATATTTGGATTTGTGGTGTCCCATACCACTAGATTTCCTCGTCCTATGGAACGACGAGAAGGTGTTGATTACCATTACTTAACAGAGAGGCAATCTGGAAGGGAATAGCCGCCGGAAACTTTCTAGAACATACTGAGTTTTTAGGAAATATTTACGGAACTTCTAAAACAGCTGTAAATACAGCGGCTATTAATAATCGTATTTGCGCGATGGATTTAAACATCAACGGTGTTAGAAGTCTTAAAAATACTTACCTAATGCATTACTTGGGTATATAAGACCTACCTCTCTTAAAATGGTTGAGGCCAATCTTCGTCGTAGAAACACTGAAGCGGACGACGAATCTCATCGTCGCGTGATGTTGGCAAAAAACGGATATGGATGAGGTCAACGAAGCAGGTCTATTCGACACTATTATTATTGAAGATGATGTGAATTTAGCATATAGTAAGTGTTAATTCAGATACTACAGGACCGTATTAGAATGTATTTTAACACTAATTAGAGACTTAAGATTTGACTTAAAACTTGATAATTAATAATATAACTCGTTTTTATATGTGGCTATTTCAACGTCTAATGTATTAGTTAAATATTAAAACTTACCACGTAAAACTTAAAATTTAAAATGGTATTTCATTGACAGATCATACATTATGAAGTTTCAAGGACTTGTGTTAATTGACAATTGCAAAAATCAATGGGTCGTTGGACCATTAATAGGAAAAGGTGGATTCGGTAGTATTTATACTACTAATGACAATAATTATGTAGTAAAAATAGAGCCCAAAGCTAACGGATCATTATTTACCGAACAGGCATTTTATACTAGAGTACTTAAACCATCCGTTATCGAAGAATGGAAAAAATCTCACAATATAAAGCACGTAGGTCTTATCACGTGCAAGGCATTTGGTTTATACAAATCCATTAATGTGGAATATCGATTCTTGGTAATTAATAGATTAGGTGCAGATCTAGATGCGGTGATCAGAGCCAATAATAATAGACTACCAGAAAGGTCGGTGATGTTGATCGGAATCGAAATCTTAAATACCATACAATTTATGCACGAGCAAGGATATTCTCACGGAGATATTAAAGCGAGTAATATAGTCTTGGATCAAATAGATAAGAATAAATTATATCTAGTGGATTACGGATTGGTTTCTAAATTCATGTCTAACGGCGAACATGTTCCATTTATAAGAAATCCAAATAAAATGGATAACGGTACTCTAGAATTTACACCTATAGATTCGCATAAAGGATACGTTGTATCTAGACGTGGTGATCTAGAAACACTTGGATATTGTATGATTAGATGGTTGGGAGGTATCTTGCCATGGACTAAGATATCTGAAACAAAGAATTCTGCATTAGTAAGTGCTGCAAAACAGAAATATGTTAACAATACTGCGACTTTGTTAATGACCAGTTTGCAATATGCACCTAGAGAATTGCTGCAATATATTACCATGGTAAACTCTTTGACATATTTTGAGGAACCCAATTACGACGAGTTTCGTCGAGTATTAATGAATGGAGTTATGTAAAATTTTTGTTGATAAAAAAATTAAAAAAATAACTTAGTTATTATCACTCTCGCGAGTGCAATAGAAACACGGCGATGTTTTACGCACACGCTTTCGGTGGGTACGACGAGAACCTTCATGCATTTCCTAGAATATCATCGACGGTTGCCAATGATGTCAGGAAATATTCTGTTGTGTCAGTTTATAATAAAAAGTATAACATTGTAAAAAACAAATATATGTGGTGTAACAGTCAAGTGAACAAGAGATATATTGGAGCACTACTGCCTATGTTTGAGTGCAATGAATATCTACAAATTGGAGATCCAATCCATGATCTAGAAGGAAATCAAATCTCTATTGTCACATATCGCCACAAAAACTACTATGCTCTAAGTGGAATTGGGTACGAGAGTCTAGACTTGTGTTTGGAAGGAGTAGGGATTCATCATCACGTACTTGAAACAGGAAACGCGGTATATGGAAAAGTTCAACATGAGTATTCTACTATCAAAGAGAAGGCCAAAGAAATGAATGCACTCAAACCAGGACCTATCATCGATTACCACGTCTGGATAGGAGATTGTGTCTGCCAAGTTACTACTGTAGACGTGCATGGAAAGGAAATTATGAGAATGAGATTCAAAAGGGGTGCGGTGCTTCCGATTCCAAATCTGGTAAAAGTTAAAGTTGGGGAGGAAAATGATACAATAAATCTTTCCACTTCCATATCAGCTCTCCTGAATTCCGGTGGCGGCACCATCGAGGTAACATCTAAGGAAGAACGTGTGGATTATGTACTCATGAAACGTTTGGAATCTATACATCATCTGTGGTCTGTAGTGTATGATCATCTTAATGTTGTGAATGGCGAAGAACGATGCTATATACATATGCATTCATCTCATCAAAGTCCTATGCTGAGTACTGTAAAAACAAATTTGTATATGAAGACTATGGGAGCATGTCTTCAAATGGACTCCATGGAAGCTCTAGAGTATCTTAGTGAACTGAAGGAATCAGGTGGGCGGAGTCCCAGACCAGAATTGCAGAAATTTGAATATCCAGATGGAGTGAAAGACACTGAATCAATTGAGAGATTGGCAGAGGAGTTCTTCAATAGATCAGAACTTCAGGCTGGTGAATCAGTCAAATTTGGTAATTCTATTAATGTTAAACATACATCTGTTTCAGCTAAGCAACTAAGAACACGTATACGGCAGCAGCTTCCTTCTATACTCTCATCTTTTGCCAACACAAAGGGTGGATATTTGTTCATTGGAGTTGATAATAATACACACAAAGTAATTGGATTCACGGTGGGTCACGACTACCTCAAACTGGTAGAGAGAGATATAGAAAAGTATATCCAAAAACTTCCTGTTGTGCATTTCTGCAAGAAAAAAGAGGACATCAAGTACGCATGTAGATTCATCAAGGTGTATAAACCTGGTGATGAGACTACCTCGACATATGTGTGCGCAATCAAAGTGGAAAGATGCTGCTGTGCTGTGTTTGCGGATTGGCCAGAATCATGGTACATGGATACTAGTGGTAGTATGAAGAAGTATTCTCCAGATGAATGGGTGTCACATATAAAATTTTAATTAGGGTAAGGTAAAACTATATATAATAACTAACAATTTGTGTATCATATAGACAATTAATTAGGTAACTGTTATCTCTTTTTAACTAACTAACTAACTCTTATATACTATTAATAATACATCTATTAATAATACATCTATTAATAATACATCTATTAATCATTGATTAGTTTATTGCTTTAATTGTTTTTGTAAACTAACACTGTTCATTGAAAAGGGATAACATGTTACAGAATATAAATTATATATGGATTTTTTTAAAAAGGAAATACTTGACTGGAGTATATATTTATTTCTTCATTACATAACACGTCTGTGTTCTAATTCTTCCAATTCTTCCACATCTCATATAATACAGGAATATAATCTTGTTCGAAAATACGAGAAAGTGGATAAAACAATAGTTGATTTTTTATCTAGGTGGCCAAATTTATTCCATATTTTAGAATATGGGGAAAATATTCTACATATTTATTTTATAGATGCTGCTAATACGAATATTATGATTTTTTTTCTAGATAGAGTATTAAATATTAATAAGAACCGTGGGTCATTTATACATAATCTCGGGTTATCATCCATTAATATAAAAGAATATGTATATCAATTAGTTAATAATGATCATCTAGATAATAGTATAAGACTAATGCTTGAAAATGGACGTAGAACAAGACATTTTTTGTCTTATATATTGGATACAGTTAATATCTATATAAGTATTTTAATAAATCATAGATTTTATATAGATGCCGAAGACAGTTACGGTTGTACATTATTACATAGATGTATATATAACTATAGGAAATCAGAATCAGAATCATATAATGAATTAATTAAGATATTGTTAAATAATGGATCAGATGTAGATAAAAAAGATACGTACGGAAACACACCGTTTATCCTATTATGTAAACACGATATCGACAACGCGGAATTGTTTGAGATATGTTTAGAGAATGCTAATATAGACTCTGTAGACTTTAATGGATATACACCTCTTCATTATGTCTCATGTCGTAATAAATATGATTTTGTAAAGTTATTAATTTCTAAAGGAGCAAATGTTAATGCACGTAATAGATTCGGAACTACTCCATTTTATTGTGGAATTATACACGGTATCTCGCTTATAAAACTATATTTGGAATCAGACACAGAGTTAGAAATAGATAATGAACATATAGTTCGTCATTTAATAATTTTTGATGCTGTTGAATCTTTAGATTATCTATTGTCCAGAGGAGTTATTGATATTAACTATCGTACTATATACAACGAAACATCTATTTACGACGCTGTCAGTTATAATGCGTATAATACGTTAGTCTATCTATTAAACAGAAATGGTGATTTTGAGACGATTACTACTAGTGGATGTACATGTATTTCGGAAGCAGTCGCGAACAACAACAAAATAATAATGGATATACTATTGTCTAAACGACCATCTTTGAAAATTATGATACCATCTATGATAGCAATTACTAAACATAAACAACATAATGCAGATTTATTGAAAATGTGTATAAAATATACTGCGTGTATGACCGATTATGATACTCTTATAGATGTACAATCGCTACATCAATATAAATGGTATATTTTAAAATGTTTTGATGAAATAGATATCATGAAGAGATGTTATATAAAAAATAAAACTGTATTCCAATTAGTTTTTTGTATCAAAGACATTAATACTTTAATGAGATACGGTAGACATCCTTCTTTCGTGAAATGTAATATTCTCGACGTATACGGAAGTCATGTACGTAATATCATAGCATCTATTAGATATCGTCAGAGATTAATTAGTCTATTATCCAAGAAGCTGGATGCTGGAGATAAATGGTCGTGTTTTCCTAACGAAATAAAATATAAAATATTGGAAAACTTTAACGATAACGAACTGACCACATATCTAAAAATCTTATAAACACTATTAAAATATAAAATCTAAGTAGGATAAAAACACTACATCATTGTTTCCTTTTAGTGCTCGACAGTGTATGCTATTTTTAACACTCATAAATAAAAATGAAAACGATTTCCGTTGTTACGTTGTTATGCGTACTACCTGCTGTTGTTTATTCAACATGTACTGTACCCACTATGAATAACGCTAAATTAACGTCTACCGAAACATCGTTTAATGATAAACAGAAAGTTACGTTTACATGTGATTCAGGATATCATTCTTTGGATCCAAATGCTGTCTGCGAAACAGATAAATGGAAATACGAAAATCCATGCAAGAAAATGTGCACAGTTTCTGATTATGTCTCTGAACTATATGATAAGCCATTATACGAAGTGAATTCCACCATGACACTAAGTTGCAACGGTGAAACAAAATATTTTCGTTGTGAAGAAAAAAATGGAAATACTTCTTGGAATGATACTGTCACGTGTCCTAATGCGGAATGTCAACCTCTTCAATTAGAACACGGATCGTGTCAACCAGTTAAAGAAAAATACTCATTTGGGGAATATATGACTATCAACTGTGATGTTGGATATGAGGTTATTGGTGTTTCGTATATAAGTTGTACGGCTAATTCTTGGAATGTTATTCCATCATGTCAACAAAAATGTGATATACCGTCTCTATCTAATGGATTAATTTCCGGATCTACATTTTCTATCGGTGGCGTTATACATCTTAGTTGTAAAAGTGGTTTTACACTAACGGGGTCTCCATCATCCACATGTATCGACGGTAAATGGAATCCCATACTCCCAACATGTGTACGATCTAACGAAGAATTTGATCCAGTGGATGATGGTCCCGACGATGAGACAGATCTGAGCAAACTCTCGAAAGACGTTGTACAATATGAACAAGAAATAGAATCGTTAGAAGCAACTTATCATATAATCATAATGGCGTTGACAATTATGGGTGTCATATTTCTAATCTCTATTATAGTATTAGTTTGTTCCTGTGACAAAAATAATGACCAATATAAGTTCCATAAATTGCTACCGTGAATATAAATCCGTTAAAATAATTAATAATTAATAATTAATAATTAATAACGAACAAGTATCAAAAGATTAAAGAATTAGCTAGAATCAATTAGATGTCTTCTTCAGTGGATGTTGATATCTACGATGCTGTTAGAGCATTTTTACTCAGGCACTATTATGACAAGAGATTTATTGTGTATGGAAGAAGTAACACCATATTACATAATATATACAGGCTATTTACAAGATGCACCGTTATACAGTTCGATGATATAGTACGTACTATGCCAAATGAATCACGTGTTAAACAATGGGTGATGGATACACTTAATGGTATAATGATGAATGAATGCGATACTGTATGTGTGGGTACCGGACTACGATTCATGGAAATGTTTTTCGATTACAATAAAAATAATCCCAAAAATAGCATCAACAATCAAATAATGTATGATATAATTAATAGCGTAACCATAATTCTAGCTAATGAGAGATATAGAAGCGCGTTTAACGACGATAGAATATACATCCGTAGAACTATGATGGACAAATTGTACGAATACGCATCTCTAACTACTATTGGTACGATCACTGGAGGTGTTTGTTATTATCTGTTGATGCATCTAGTTAGTTTGTATAAATAATTATTTCGATATACTAGTAAAATTTTAAGATTTTAAATGTATAAAAAACTAATAACGTTTTTATTTGTAATAGGTGCAGTTGCATCCTATTCGAATAATGAGTACACTCCGTTTAATAAACTGAGTGTAAAACTCTATATAGATGGAGTAGATAATATAGAAAATTCATATACTGATGATAATAATGAATTGGTGTTAAATTTTAAAGAGTACACAATTTCTATTATTACAGAGTCATGCGACGTCGGATTTGATTCCATAGATATAGATGTTATAAACGACTATAAAATTATTGATATGTATACCATTGACTCGTCTACTATTCAACGCAGAGAACATACGTGTAGAATATCTACCAAATTATCATGCTATTATGATAAGTACCCTTATATCCACAAATATGAGGGTGATGAACGACAATATTCTATTACTGCAGAGGGAAAATGCTATAAAGGAATAAAATATGAAATAAGTATGATGAACGATGATACTCTATTGAGAAAACATACTCTTAAAATTGGATTTACTTATATATTCGATCGTCATGGGCATAGTAATACATATTATTCAAAATATGATTTTTAAAAATTTAAAATATATTATCACTTCAGTGACAGTAGTCAAATAACAAACAACACCATGAGATATATTATAATTCTCGCAGTTTTGTTCATTAATAGTATACATGCTAAAATAACTAGTTATAAGTTTGAATCCGTCAATTTTGATTCCAAAATTGAATGGACTGGGGATGGTCTATACAATATATCCCTTAAAAATTATGGCATCAAGACGTGGCAAACAATGTATACAAATGTACCAGAAGGAACATACGACATATCCGGATTTCCAAAGAATGATTTCGTATCTTTCTGGGTTAAATTTGAACAAGGCGACTATAAAGTGGAAGAGTATTGTACGGGACTATGTGTCGAAGTAAAAATTGGACCACCAACTGTAAGATTGACTGAATATGACGATCATATCAATTTGTTCATCGAGCATCCGTATGCTACTAGAGGTAGCAAGAAGATTCCTATTTACAAACGCGGTGACATGTGTGATATCTACTTGTTGTATACGGCTAACTTCACATTCGGAGATTCTGAAGAACCAGTAACATATGATATCGATGACTACGATTGCACGTCTACAGGTTGCAGCATAGACTTTGCCACAACAGAAAAAGTGTGTGTGACAGCACAGGGAGCCACAGAAGGGTTTCTCGAAAAAATTACTCCATGGAGTTCGGAAGTATGTCTGACACCTAAAAAGAATGTATATACGTGCGCAATTAGATCTAAAGAAGATGTTCCCAATTTCAAGGACAAAATAGCCAGAGTTATCACGAGAAAATTTAATAAACAGTCTCAATCTTATTTGACTAAATTTCTCGGTAGCACATCGAATGATGTTACAACTTTTCTTAGCATTCTTGACTAAATATTCATAACTAATTTTTATTAATGATACAAAAATGAAATAAAACTGCATATTATACACTGGTTAACGCCCTTGGCCTAACCATTTTTAAGATGAGGTCCCTGATTATAGTCCTTCTGTTCCCCTCTATCATCTACTCCATGTCTATTAGACGATGCGAGAAGACTGAAGAGGAAACATGGGGATTAAAAATAGGGTTGTGTATAATTGCCAAAGATTTCTATCCCGAAAGAACTGATTGCAGTGTTCATCGCCCAACTGCAAGTGGAGGATTGATAACTGAAGGCAATGGATTCAGAGTAGTTATATATGATCAATGTACAGAACCACATGACTTTATTATCACCGATACTCAACAAACACGTCTTGGATCATCTCATACATATATTAAATTCAGTAACATGAATACAGGTGTCCCATCTAGTATTCCAAAATGTTCCAGAACTCTCTCTATTTCTGTATATTGTGATCAAGAGGCGGGAGACATAAAATTTGAGGAGTATACTCAAGAATCAAGTGATATCAGTATTAGAGTTAAGTATGATTCATCATGTATTGATTATCTGGGTATTAATCAAAGTTTCATGAATGAATGTATTCGAAGAATTACAACATGGGATAGAGAATCATGTGTCAGAATTGATACACAGACTATAAATAAATATCTTAAGTCTTGCACCAACACAAAATTCGACCGTAATGTCTACAAAAGGTACATACTGAAGAGTAAAGCACTCCATGCTAAAACAGAGTTGTAATAGATATAAAATACTTTTTATAATAATTAGGCTAGAAAAATCTCACTCACATGTAATCTTAAAAAATGATATGATAGTTCTTACAAGTAGAGATTGAGTTTTAAATGGATTCTATTAATTACCGGGGACTTAACAATTCGTTCTGATCTACAGACATTGGTTAATAAATCATCTTATTTTGCCGATATATTAAAATGTGGAAACTCCACTAATAATATTACATTGTGTGACTTTCAAGATGATGCGATATATAGGGTTATACAGTTTAACAATTATATAATAGAGATAGAAAGTACAAAAGATGTAGAATCAATGATATGGCACGCTAAACAGTTGGGTGTGGAATCATTGCTAAAAGAATGTCAAAATTATTTGCTTAGAATATTACGTATATAATTGTTTAGAAATTTATAGAATAACTAATATTAATACATTATCGTATATCTACAACGATGTAAGAAACTTCATATTGGATAATATTACTATTAATATATAAGGATCCAGATTTTATATATTTGCCTAAATACATTATTATAGATTTACTAGGACAATCACCTAAATGTTTTTAACGAAGATAATGTGGTAAAGATTATATACACTTATATATCTTCCGATATCTACAAGGATATTTCCATATCATCATTGTGTAAACTAAATAACGTTTTCTATGGCATTTAATAAGGCATTGGATATGTGGAAAAGATGATGGTGTATGGAAGTTAGTACATCATTATCAACTTCTCCTTATTGATTGAAAATGAAAATATAAATAGTTTTTATATATAGCGGTATCTACCCTATAGTTTTATTGCTTACTACTAACATGGATACAGATACAGATACAGATACAGATACAGATATAGATACAGATACAGATGTTAGATAGAGAGAAAGAAGAAATACTAAAAAATGTAGAAATTGAAAATAATAAAAACATTAACAAGAATCATCCCAGTGAATATATTAGAGAAGCACTCGTTATTAATACCAGTAGTAATAGTGATTCCATTGATAAAGAAGTTATAGAATATATCAGTCACGATGTAGGAATATAGATCATATCTACTAATTTTTATAATCGATACAAAACATAAAAAACAACTCGTTATTACATAGCAGGCATGGAATCCTTCAAGTATTGTTTTGATAACGATGGTAAGGAATGGATTATCGGAAATACTTTATATTCTGGTAATTCAATACTCTATAAGGTCAGAAAAAATTTCACTAGTTCGTTCTACAATTACGTAATGAAGATAGATCATAAATCACACAAGCCATTGTTGTCCGAAATACGATTCTATATATCTGTATTGGATCCTTTGACTATCGACAACTGGACACGGGAACGTGGTATAAAGTATTTGGCTATTCCAGATCTGTATGGAATTGGAGAAACCGATGATTATATGTTCTTCGTTATAAAGAATTTGGGAAGAGTATTCGCCCCAAAGGATAGTGAATCAGTTTTCGAAGCATGTGTCACTATGATAAACACGTTAGAGTTTATACACTCTCGAGGATTTACTCATGGAAAAATAGAACCGATGAATATACTGATTAGAAATAAACGTATTTCACTAATTGACTATTCTAGAACTAACAAACTATACAAAAGTGGAACACATATAGATTACAACGAGGACATGATAACTTCAGGAAATATCAATTATATGTGTGTAGACAATCATCTTGGAGCAACAGTTTCAAGACGAGGAGATTTAGAAATGTTGGGATATTGCATGATAGAATGGTTCGGTGGTAAACTTCCATGGAAAAACGAAAGTAGTATAAAAGTAATAAAACAAAAAAAAGAATATAAACAATTTATAGCTACTTTTTTTGAGGACTGTTTTCCTGAAGGAAATGAACCTCTGGAATTAGTTAGATATATAGAATTAGTATACATGTTAGATTATTCTCAAACTCCTAATTATGACAGACTACGTAGACTGTTTATACAAGATTGAAATTATATTCTTTTTTTTATAGAGTGTGGGGTAGTGTTACGGATATCTGATATCTAATATCTAATATTAATATTAGACTATCTCTATCGCGCTACACGACCAATATCGATTACTATGGATATCTTTAGGGAAATCGCATCTTCTATGAAAGGAAAGAATGTATTCATTTCTCCAGCGTCAATCTCGTCAGTATTGACAATACTGTATTATGGAGCTAATGGATCCACTGCTGAACAGCTATCAAAATATGTAGAAAAGGAGGAGAACATGGATAAGGTTAGCGCTCAGAATATCTCATTCAAATCCATGAATAAAGTATATGGGCGATATTCTGCCGTGTTTAAAGATTCCTTTTTGGGAAAAATTGGCGATAAGTTTCAAACTGTTGACTTCACTGATTGTCGCACTATAGATGCAATCAATAAGTGTGTAGATATCTTTACTGAGGGAAAAATCAATCCACTATTGGATGAACCATTGTCTCCTGATACCTGTCTCCTAGCAATTAGTGCCGTATACTTTAAAGCAAAATGGTTGATGCCATTCGAAAAGGAATTTACCAGTGATTATCCCTTTTACGTATCTCCAACGGAAATGGTAGATGTAAGTATGATGTCTATTTACGGCGAGCCATTTAATCACGCATCTGTAAAAGAATCATTCGGTAACTTTTCAATCATAGAACTGTCATATGTTGGAGATACTAGTATGATGGTCATTCTTCCAAACAAGATTGATGGATTAGAATCCATAGAACAAAATCTAACAGATACAAATTTTAAGAAATGGTGTAACTCTCTGGAAGCTACGTTTATCGATGTGCACATTCCTAAGTTTAAGGTAATAGGTTCGTATAATCTTGTGGATACGCTAATAAAGTTGGGACTGACAGATGTGTTCTATTCAACTGGTGATTATATCAATATGTGTAATTCAGATGTGAGTGTTGACGCTATGATTCACAAAACGTATATAGATGTCAATGAAGAGTATACAGAAGCAGCTGCAGCAACTTCTGTACTAGTGGCAGACTGTGCATCAACAGTTACAAATGAGTTCTGTGCAGATCATCCGTTCATCTATGTGATTAGACATGTCGATGGTAAAATTCTTTTCGTTGGTAGATATTGCTCTCCAACAACTAATTAAGCACATTCTTAATATTAGAATATTATATAGTTAAGATTTTACTAACAGGTTAACATTTTTTTTAAAAAATAGAAAAAACATGTGGTATTAGTGCAGGTCGTTATTCTTCCAATTGCAATTGGTAAGATGACGGCCAACTTTAGTACCCACGTCTTTTCACCACAGCACTGTGGATGTGACAGACTGACCAGTATTGATGACGTCAGACAATGTTTGACTGAATATATTTATTGGTCGTCGTATGCATACCGCAACAGGCAATGCGCTGGACAACTGTATGGCACACTCCTCTCTTTTAAAGATGATGCGGAATCAGTGTTCATCGACGTTCGTGAGCTGGTAAAAAATATGCCGTGGGATAATGTTAAGGATTGTACAGAGATCATCCGTTGTTATATACCGGATGAGCAAAAAACCATCAGAGAGATTTCGGCCATCATTGGACTTTGTGCATATGCTGCTACTTACTGGGGAGGTGAAGACCATCCCACTAGTAACAGTCTGAACGCATTGTTTGTGATGCTTGGGATGCTCAATTACATGGATTATACCATCATATTCTGGCGTATGAATTGATGAGTTACAGCTTGACATTTCTTCTTTCCTCCCTCTTCTTCTACTTTTCCCAGAAACAAACTTTTTTTACCTACTATAAAATAAAATGAGTATACTACCTGTTATATTTCTTCCTATATTTTTTTATTCTCCATTCGTTCAGACTTTTAACGTGCCTGAATGTATCGACAAAGGGCAATATTTTGCATCATTCATGGAGTTAGAAAACGAGCCAGTAATCTTACCATGTCCTCAAATAAATACGCTATCATCCGGATATAATATATTAGATATTTTATGGGAAAAACGAGGAGCGGATAATGATAGAATTATACAGATAGATAATGGTAGCAATATGCTAATTCTGAACCCGACACAATCAGACTCTGGTATTTATATATGCATTACCACGAACGAAACCTACTGTGACATGATGTCGTTAAATTTGACAATCGTGTCTGTCTCAGAATCAAATATAGATCTTATCTCGTATCCACAAATAGTAAATGAGAGATCTACTGGTGAAATGGTATGTCCCAATATTAATGCATTTATTTCTAGTAACGTAAACGCAGATATTATATGGAGCGGACATCGACGCCTTAGAAATAAGAGACTTAAACAACGGACACCTGGAATTATTACCATAGAAGATGTTAGAAAAAATGATGCTGGTTATTATACATGTGTTTTAGAATATATATATATGGGCAAAACATATAACGTAACCAGAATTATAAAATTAGAGGTACGTGATAGAATAATACCTCCTACTATGAAATTACCAGAAGGAGTAGTAACTTCAATAGGTAGTAATTTGACTATTACATGCAGAGTATCGTTGAGACTTCCCACAACGGACGCTGACGTCTTTTGGATAAGTAATGGTATGTATTACGAAGAAGAAGACGAGGACGGAGACGGTAGAATAAGTGTAGCAAATAAAATCTATATGACTGATAAGAGACGTGTTATTACATCCTGGTTAAACATTAATCCTGTCAAGGAAGAAGATGCTACAACGTTTACGTGTATGGCGTTTACTATTCCTAGCATCAGCAAAACAGTTACTGTTAGTATAACGTGAATGTATGTTGTTACATTTCCATATCAATTGAGTTTATAAGAATTTTTATACATTATCTTCCAACAAACAATTGACGAACGTATTGCTATGATTAACTCCCACAATACTATGCATATTATTAATCATTAACTTGCAGACTATACCTAGTAGTGCTATTTTGACATACTCATGTTCTTGTGTAATTGCAGTATCTATATTATTAAAGTACGTAAATCTAGCTATAGTTTTATTATTTAATTTTAGATAATATACTGTCTCCGTATTTTTAAAAAATTGCCACATCCTTTATTAAATCATGAATGGGAATTTCTGTGTCATCGTTAGTATATTGTGAACAACAAGAGCTGATATCTATAGGAAAGGGTGGAATGCGATACATTGATCTATGTAGTTTTAAAACATAAGCGAACTTTGAAGAATTTATATAAATCATCTCACGAGATATTGTTCTCTGTCATATTCATACACCTGCATAAACTTTCTATACATCTTACAATGTGTTATTTTATGATCATATTTACATATTTACTGGTATATCAAAGATGTTAGATTAGTTAATGGGAATCGTCTATAATAATGAATATTAAACAATTATAGGAGGAATTTATACCTACAAAAACATCATAAAAATGAGTCATCGTCCGATTTATGTTTTAAATATACTAACATTACTACCTTCAGAAATTATATACGAAATATTATACATGCTGACAATTAACGATCTTTATAATATATAGTATCCACCTACCAAAGTATAATTGTATTTTTTCTCATGTGATGTGTGTAAAAAACTGATATTATATAATTATCTTAGTACCTATGATGAAGATGAAGATGATGGTCCGTATATATTTTGTATCATTATCGTTATTGCTATTCCATAGTTACGCCATAGACATCGAAAATGAAATCACCGAATTCTTCAATAAAATGAGAGATACTCTACCAGCTAAAGACTCTAAATGGTTGAATCCAGTATGTATGTTTGGAGGCACAATGAATGATATGGCCGCTCTAGGAGAGCCATTCAGTGCAAAGTGTCCTCCTATTGAAGACAGTCTTTTATCGCATAGATATAAAGACTATGTGGTTAAATGGGAAAGGCTAGAAAAGAATAGACGGCGACAGGTTTCTAATAAACGTGTCAAACATGGTGATTTATGGATAGCCAACTATACATCTAAATTCAGTAACCGTAGGTATTTATGTACCGTAACCACAAAGAATGGTGACTGTGTTCAGGGTGTAGTTAGATCTCATGTGTGGAAACCTTCTTCATGCATTCCAAAAACATATGAACTAGGTACTTATGATAAGTATGGCATAGACTTATACTGTGGAATTCTTTATGCGAAACATTATAATAATATAACTTGGTATAAAGATAATAAGGAAATTAATATCGACGATTTTAAGTATTCACAAGCGGGAAAGGAATTAATTATTCATAATCCAGAGTTAGAAGATAGTGGAAGATACGACTGTTACGTTCATTACGACGACGTTAGAATCAAGAATGATATCGTAGTATCAAGATGTAAAATACTTACGGTTATACCGTCACAAGACCACAGGTTTAAACTAATACTAGATCCGAAAATCAACGTAACGATAGGAGAACCTGCCAATATAACATGCAGTGCTGTGTCAACGTCATTATTTGTCGACGATGTACTGATTGAATGGGAAAATCCATCCGGATGGATTATAGGATTAGATTTTGGTGTATACTCTATTTTAACTAGTAGAGGCGGTATCACCGAGGCGACTTTGTATTTTGAAAATGTTACTGAAGAATATATAGGCAATACATATACATGTCGTGGACACAACTATTATTTTGATAAAACTCTTACAACTACAGTAGTATTGGAGTAAATACACAATGCATTTTTATATACATTACTGAATTATTATTATTAATTATATCGTATTTGTGCTATAGAATGGATGAAGATACGCGACTATCTAGGTATTTGTATCTCACCGATAGAGAACATATAAATGTAGACTCTATTAAACAGTTGTGTAAAATATCAGATCCTAATGCATGTTATAGATGTGGATGTACGGCTTTACATGAGTACTTTTATAATTATAGATCAGTCAACGGAAAATACAAGTATAGATACAACGGTTACTATCAATATTATTCATCTAGCGATTATGAAAATTATAATGAATATTATTATGATGATTATGATAGAACTGGTATGAACAGTGAGAGTGATAATATATCAATCAAAACAGAATATGAGAATGAATATGAATTCTATGATGAAACACAAGATCAAAGTACACAACTAGTAGATTACGACATTAAACTCAAAACCAATGAGGATGATTTTGTTGATGAATTCTATGGTTATGATAGATCAGTGGGTGTCCATGATTATATAGATGTATCAATTAATAAAGTAGTATATGGAAGAGAGTCTCACGTAAGATGGCGGGATATATGGCAAGAACATAATGATGGCGTATACAGTATAGGAAAGGAGTGCATAGATAATATATACGAAGACAGACATACCGTAGACGAATTCTACAAGATAGACAGCGTATCAGATGTAGATGACGCAGAACATATATCTCAGATAACTAATGATGTATCTACACAAACATGGGAAAAGAAATCAGAGTTAGATAGATACATGGAAATGTATCCTCGTCATAGATATGGTAAGCATTCTGTCTTTAAGGGATTTTCTGACAAAGTTAGAAAAAATGATTTAGACATGAATGTGGTAAAAGAATTACTTTCTAACGGTGCATCTCTAACAATCAAGGATAGCAGTAATAAGGATCCAATTGCTGTTTATTTTAGAAGAACAATAATGAATTTAGAAATGATTGATATCATTAACAAACATACAACTATCTATGAACGCAGGTATATAGTACACTCCTATCTAAAAAATTATAGAAATTTCGATTATCCATTTTTCAGAAAGTTAGTTTTGACTAATAAACATTGTCTCAACAATTATTATAATATAAGCGACAGCAAATATGGAACACCACTACATATATTAGCATCTAATAAAAAAATAATAACTCCTAATTACATGAAGTTATTAGTGTATAACGGAAATGATATAAACGCACGAGGTGAAGATACACAAATGCGAACTCCATTACACAAATATTTGTGTAAATTTGTATATCATAATATTGAATATGGTATCCGATACTATAATGAAAAGATTATAGACGCATTTATAGAGTTAGGAGCCGATCTAACTATTCCAAATGACGATGGAATGATACCAGTAGTTTACTGTATACACTCAAATGCCGAATATGGTTATAACAATATTACTAACATAAAGATAATACGTAAACTACTTAATCTTAGTAGACATGCGTCACATAATCTATTTAGAGATCGAGTCATGCACGATTATATAAGTAATACATATATTGATCTTGAGTGTTTAGATATCATTAGATCACTGGATGGGTACGATATTAATTGTTACTTTGAAGGACGTACACCACTTCATTGCGCTATACAATATAACTTCACTCAGATTGCTGAGTACTTATTAGATCGAGGAGCTGATATATCATTAAAGACAGACGATGGTAAAACTGTATTTGATTTATCGTTATGTAGTTACATTCCTCTTAAATGGACTAGCTTTTTGATTAGTCGTCTACCGCCTAAAAGTGTCATATGCTCACTGACTAACCATATAATAGATTATGTTCTTACGAACAATAGACGTATTATTTGGCAGAGTCAAATGATTAATAAGTACGTACTGTTACTGGACCCATCCTTTTATTATAGATTCAGAAATGCTATCGAAAACAAATTAGACCAATACAATAATCGTTATAATATGTTCGAACACGATAGGGACGTTAATGAAAAGTATGGCAAAGTCTTACATGACCTCGATACATATATCAAGGATGTACAAGTATTAAAATCTACTTCCATCACTAATAATATAACACTATACGACACTATTATAAATAATAAGTCAGAGTTTCCTATACGTCGTGCAAACGACAAACAATTAATTAATCTCATAAAATCCAATACATATCATAATCTTATCGAAAAAGTTATTAAAAATACATTAGAGAAATATACTTTAACTAATATAGTCCTCGAGTATATGATCTCATCTCAATCTCAATCTCAATCTCAATCTCAATCATCTTATTTGAGTCGTATTCCTAATGAGATATTACTCGAAATATTATATAAACTCGACATGTACGATTTACGTAATCTATATACAAGATATATGAGAGAGAATGATATCACAGAGTATCATATAGAGAATACGAGGTCTGTTTCTACACAGACATGAATAATGAATACACATACAACGTTTTTTTTTTAATCTTAGATATAACACTAATTACATCAAGATTATATATTGAAATCGTAATTTGAGTTGTCTGATCATCATGGATATCGAAAATGATATACGTAACATTAGCAATCTTTTTTAGATGATTGATATATTATTATGCGATGTAATCATAACTATCGGAGATGTAGAAATTAAAGCGCATAAAACTATTTTGGCTGCCGGATCTACGTATTTTAAAACAATGTTCACAACACCTATGATAGCGAGAGATCTAGCAACTAGAGTAAATCTACAGATGTTCGATAAGATGCCGTCAAAAATATTGTACAGTACTTATACAATAGGTATATAAGTTCTATGAATGTGATAGACGTATTAAAATGCACCGACTAAGAACGTAGAACGAACTATAGAATGTTATACAATGGGTGATGATAAGTAGAAGATGTTACCCGATATACCCATAGCATTATCTAGTTATGGCATGTGTGTATTAGATCAATACATATACATTATAGGCGGTCGTACCCAACACTGATTATACATCGGTACATACAGTAAATAGCATAGATATGGAGGAGGATACAAATATTTCAAATAAAGTTATGAGATACGCACTGTCAATAATATATGGAAGACATTACCTAACTTCTGAACTGGAACTATAAATCCAGGCTCTTGCATAAAGATGAATATATATGTTGTATGCGACATCAAAGATGAAAAAATGTTAAGACTTATATATTTAGATATAACACGAATATGTATAACGGATGGGAATTGGTAACGATGACAGAAAGCAGATTGTCAGCTCTGCATACTATTCTTCATGACAATACCATAATGATGTTACATTGTTATGAAACGTATATGTTACAAGATACATTTAATGTGCTTACGGAACATATATTTAGAAACATCTACTAACGATTTTTTATGCTTGTATTATTAATGGTATGTAATATGATTTAATTGATTGTGTACACGATACCAATTTGTCAAGTATGAATACGGAGTACAAACATAAACTGAAGTTTAACATTATTTATTTATGATATACATTATATACATTATATACATTATATACATTATATACATTATATACATTATATACATTATATACATTATATACATTATATACATTATATACATTATATACATTATATACATTATATACATTATATACATTATATACATTATATATCGTTATATATCGTTATATATCGTTATTGTTTGGTCTATGCCATGGATATCTTTAAAGAACTAATCTTAAAACATACGGATGAAAATGTTTTGATTTCTCCAGTTTCCATTTTATCTACTTTATCTATTCTGAATCATGGAGTAGCTGGTTCTACAGCTGAACAACTATCAAAATATATAGAGAATATGAATGAGAATACACCCGATGATAAGAAGGATGACAATAATGACATGGACGTAGATATTCCGTATTGTGCGACACTAGCTACCGCAAATAAAATATACGGTAGTGATAGTATCGAGTTCCATGCCTCCTTCCTACAAAAAATAAAAGACGATTTTCAAACTGTAAACTTTAATAATGCTAACCAAACAAAGGAACTAATCAACGAATGGGTTAAGACAATGACAAATGGTAAAATTAATTCCTTATTGACTAGTCCGCTATCCATTAATACTCGTATGACAGTTATTAGCGCCGTCCATTTTAAAGCAATGTGGAAATATCCATTTTCTAAACATCTTACATATACAGACAAGTTTTATATTTCTAAGAATATAGTTACCAGTGTTGATATGATGGTGGGTACCGAGAATGACTTGCAATATGTACATATTAATGAATTATTCGGAGGATTCTCTATTATCGATATTCCATACGAGGGAAACTCTAGTATGGTGATTATACTGCCGGACGACATAGAAGGTATATATAACATAGAAAAAAATATAACAGATGAAAAATTTAAAAAATGGTGTGGTATGTTATCTACTAAAAGTATAGACTTGTATATGCCAAAGTTTAAAGTGGAAATGACGGAACCGTATAATCTGGTACCGATTCTAGAAAATTTAGGACTTACTAATATATTTGGATATTATGCAGATTTTAGTAAGATGTGTAATGAAACTATCACTGTAGAAAAATTTCTACATACGGCGTTTATAGATGTTAATGAGGAGTATACAGAAGCATCGGCCGTTACAGGAGTATTCATGACTAACTTTTCGATGGTATATCGTATGAAGGTCTACATAAACCATCCATTCATATACATGATTAAAGATAACACCGGACATACACTTTTTATAGGGAAATACTGCTATCCGCAATAAATATAAACAATAGACTTTTATCACGTTATCTCATGTATAAAATATTACAAATAGTATAGCATAAACTAAAGTCGATACATACATTAAAACTTAAAAAATAATGTAATTTACAATTAATAGTATAAACTAAAAAAATTAAAAAATTAAAAACAATATCATTATTATAAGTAATATCAAAATGACGATATACGGATTAATAGCGTATCTTGTATTCGTGACTTCATCCATCGCTAGTCCATTTTACATTCCCGTTATTCCGCCCATTTCGGAAGATAAATCGTTCAATAGTGTAGAGGTATTAGTTTCTTTGTTTCCCGATGACCAAAAAGACTATACAGTAACTTCTCAGTTCAATAACTACACTATCGGTACCAAAGACTGGACTATCAACGTACTATCCACACCTGATGGTCTGGACATACCATTGACTAATATAACTTATTGGTCACGGTTTACTATAGGTCGTGCATTGTTCAAATCAGAGTCTGAGGATATTTTCCAAAAGAAAATGAGTATTCTAGGTGTTTCTATAGAATGTAAGAAGCCGTCGACATTACTTACTTTTTTAACCGTGCGTAAAATGACTCGAGTATTTAATAGATTTCCAGATATGGCTTATTATCGAGGAGACTGTCTAGAAGCCGTTTATGTAACAATGACTTATAAAAATACTAAAACTGGAGAGACTGATTACACGTACCTCTCTAATGGGGGGTTGCCTGCATACTATCGTAATGGGGTCGATGGTTGATTATTGATTAGTATATTCCTTATTCTTTTTATTCACACAAAAAGAACATTTTTATAAACATGAAACCACTGTCTAAATGTAATTATGATCTTGATTTATAGATGATGATCAGCCTTCAGAGGATTTTGACCAGCATGTTTAATATGAAAAAAAACATAACTATTAAGCGCTATTGCGCTATTGTGCTTAATTATTTTGCTCTATAAACTGAATATATATAGCCACAATTATTGACGGGCTTGTTTGTGACCGGTAATCATGAATTTACAGAAATTATCTCTGGCTATATATCTTACGGTGACATGTTCGTGGTGTTATGAAACATGTATGAGAAAAACTGCGTTGTATCATGACATTCAATTGGAGCATGTAGAAGACAATAAAGATAGTGTAGCATCGCTACCGTACAAGTATCTACAAGTAGTCAAACAAAGAGAACGTAGTAGATTGTTGGCTACATTTAATTGGACGGATATAGCTGAGGGTGTTAGAAATGAGTTCATTAAAATATGTGATATCAACGGAACATATCTATATAATTATACTATTGATGTTAGTATAATTATTGATTCCACGGAAGAACTACCAACAGTTACTCCAATTACAACATATGAACCTTCTATATATAATTATACTATCGATTATAGCACTGTTATTACTACTGAAGAACTACAAGTGACTCCAACATATGCGCCTGTAACAACTCCTCTTCCAACATCAGCAGTTCCTTATGATCAACGATCGAATAACAATGTAAGTACTATATCTATTCAGATACTGAGTAAAATATTGGGAGTCAATGAAACAGAATTAACTAATTATCTTATTATGCATAAAAATGACACTGTTGACAATAACACCATGGTTGATGATGAGACATCTGACAATAACACATTGCATGGTAATATAGGATTTTTGGAAATAAATAATTGTTACAATGTTTCTGTGTCAGATGCTAGTTTTAGAATAACATTAGTAAACGATACTTCTGAAGAAATTTTGCTAATGCTAACAGGAACTAGTTCATCCGACACCTTCATATCTTCCACCAATATCACTGAATGTTTGAAAACATTAATCAATAATGTGTCGATTAATGATGTACTTATAACACAAAATATGAATGTAACATCTAATTGTGATAAATGCTCAATGAATTTGATGGCATCCGTTATTCCTGCAGTTAATGAATTTAACAATACGTTGATGAAAATTGGTGTAAAAGATGATGAAAACAATACGGTATATAACTATTATATTTGTAAACTAACTACAAATTCTACATGTGATGAGTTAATCAATTTAGATGAAGTCATTAACAACATAACTCTGACAAATATTATACGCAATAGTGTTTCGACAACTAACAGCAGAAAAAGACGAGATCTGAATGGTGAGTTTGAATTTTCCACTTCCAAGGAATTAGATTGTCTTTACGAATCATATGGTGTAAACGATGATATAAGTCATTGTTTTGCATCACCTAGACGTAGACGATCTGACGACAAAAAGGAGTACATGGACATGAAATTATTCGACCACGCGAAAAAAGATTTAGGAATAGACAGTGTTATTCCTAGAGGTACAACCCATTTCCAAGTAGGTGCATCTGGTGCAAGTGGTGGTGTTGTAGGAGATAGTTTCCCATTTCAAAATGTTAAATCGCGTGCCAGTCTATTGGCGGAAAAAATAATGCCTAGAGTACCTATTACTGCTACCGAAGCTGATCTATATGCAACTGTAAATAGACAACCCAAGTTACCAGCAGGTGTTAAAAGTACTCCGTTTACAGAGGCGCTTGCGTCTACGATAAACCAAAAGCTTTCTAATGTTAGAGAGGTAACTTATGCTTCGTCCAATCTGCCAGGATCAAGTGGCTATGTTCATAGACCATCTGATTCTGTTATTTACAGCAGTATAAGACGGTCACGTTTACCTAGTGATAGCGATAGTGATTATGAGGATATACAAACTGTTGTTAAGGAATATAATGAAAGATATGGTAGATCAGTCAGTAGAACACAGTCATCAAGTAGTGAAAGCGATTTTGAAGATATAGATACTGTTGTTAGGGAATATAGACAAAAATATGGCAATGCAATGGCAAAAGGACGTAGTAGTTCCCCTAAACCTGATCCATTATATAGTACTGTTAAGAAAACAACTAAAAGTCTATCTACTGGTGTAGACATAGTTACAAAACAATCAGACTATTCTCTATTACCTGACGTTAATACTGGCAGTTCTATTGTGTCACCTCTCACCAGAAAAGGAGCTACTAGACGACGACCTAGACGCCCTACAAATGATGGTCTACAGAGTCCAAATCCTCCTCTCCGTAATCCACTTCCTCAACATGATGATTATTCTCCTCCACAAGTACACAGACCTCCGACACTTCCTCCTAAACCAGTCCAAAATCCGACACAACTTCCCCCTAGACCAGTAGGTCAATTACCTCCTCCTATAGATCAACCAGATAAAGGATTTAGTAAGTTTGTATCACCTAGACGGTGTAGAAGAGCAAGCTCTGGAGTCATATGTGGTATGATACAATCAAAACCAAACGATGATACCTATTCACTTCTTCAACGGCCAAAAATTGAACCAGAATATGCGGAGGTTGGTAATGGTATACCCAAGAACAATGTTCCTGTAATAGGTAATAAACATAGTAAAAAATATACATCGACGATGTCAAAAATATCAACAAAATTTGATAAATCTACGGCATTTGGAGCAGCAATGTTACTAACTGGTCAGCAGGCCATTAGCCAACAGACTAGATCAACTACGTTGAGTAGAAAAGATCAGATGAGCAAGGAAGAAAAGATATTCGAAGCAGTTACAATGAGTCTATCAACTATAGGTTCAACGTTGACGTCTGCAGGTATGACGGGTGGTCCAAAACTAATGATTGCAGGAATGGCTATAACGGCTATAACTGGTATAATAGATACGATAAAAGATATATATTACATGTTTTCAGGACAGGAGAGGCCAGTAGATCCTGTTATTAAATTATTTAATAAGTACGCTGGCTTAATGTCCGATAATAATAAAATGGGTGTAAGAAAATGTTTGACACCCGGTGACGACACACTTATTTATATCGCATACAGAAACGATACCAGTTTTAAACAGAATACGGATGCGATGGCTTTGTATTTCTTAGATGTTATCGACTCAGAGATCCTATATCTAAACACATCAAATTTAGTTCTAGAGTATCAACTAAAGGTGGCTTGCCCCATAGGAACATTAAGATCTGTAGATGTGGACATAACTGCGTATACAATATTATATGATACAGCGGATAATATTAAGAAATACAAGTTTATCAGAATGGCAACGCTACTATCCAAACATCCAGTTATTAGATTGACATGTGGTTTAGCAGCAACATTGGTGATTAAACCGTACGAGGTACCCATCAGTGATATGCAACTACTAAAAATGGCGACGCCTGGTGAACCAGAATCCACTAAATCTATACCATCCGATGTCTGTGATAGGTATCCTCTAAAGAAATTCTATCTTTTAGCTGGTGGTTGTCCCTATGATACATCTCAAACTTTTATTGTACATACTACTTGCAGTATTCTACTAAGAACAGCTACACGGGATCAGTTTAGAAACAGATGGGTGTTACAAAATCCATTTAGACAAGAAGGGACATATAAGCAACTGTTTACCTTTAGCAAATACGATTTTAACGACACCATAATCGATCCTAATGGTGTGGTGGGTCATGCTAGCTTTTGTACCAATAGAAGCAGCAACCAATGTTTCTGGTCCGAACCTATGATATTGGAAGATGTATCATCGTGTAGTTCTAGAACTAGAAAAATATACGTAAAACTGGGAATATTTAATGCTGAAGGTTTTAATAGTTTTGTACTAAATTGTCCAACTGGGTCTACACCTACATACATCAAACATAAAAATGCGGACAGTAACAATGTTATCATAGAGCTACCTGTAGGTGATTACGGCACAGCCAAATTGTATTCAGCAACAAAACCATCGAGGATAGCTGTGTTCTGCACACATAACTATGATAAACGATTCAAATCAGATATTATAGTTCTAATGTTTAATAAAAACAGCGGTATTCCATTTTGGAGCATGTACACAGGAAGTGTAACTAGTAAAAATAGAATGTTTACCACATTGGCTAGAGGAATGCCGTTTAGATCAACGTATTGCGATAACAGACGACGATCAGGTTGTTATTATGCAGGAATACCATTTCATGAAGATAGTGTAGAAGCAGATATACATTATGGACCAGAAATAATGTTAAAGGAAACATATGACATAAACAGTATTGACCCACGAGTTATAACAAAGTCAAAGACCCATTTTCCTACTCCATTGAGTGTAAAATTCATGGTTGACAATTTAGGAAATGGATATGACAACCCTAATTCATTTTGGGAAGATGCTAAAACTAAGAAACGGACATATAGTGCAATGACGATAAAAGTCCTACCATGTACAGTGAGAAATAAAAATATAGACTTTGGATATAACTATGGAGATATTATTTCTAATATGGTTTATCTACAATCTACTAGTCAGGATTATGGAGATGGTACCAAATATACATTTAAATCCGTAACTAGATCAGATCATGAGTGTGAATCTAGCTTAGATCTGACGTCTAAGGAAGTAACTGTGACATGTCCTGCGTTTAGTATACCAAGAAATATATCAACATATGAAGGTCTATGCTTTAGTGTTACTACATCTAAAGATCATTGTGCTACAGGTATTGGTTGGTTAAAATCTAGTGGCTATGGGAAGGAAGATGCTGATAAACCACGTGCTTGTTTTCATCATTGGAATTATTACACACTGTCGTTGGATTATTACTGTTCATACGAAGATATTTGGAGAAGCACCTGGCCTGACTATGATCCATGTAAGTCATATATCCATATAGAGTATAGAGATACATGGATAGAATCTAACGTGTTACAGCAACCTCCTTACACATTCGAATTCATTCATGACAATTCTAACGAATATGTGGATAAAGAAATTAGTAACAAATTAAATGATCTGTACAATGAATACAAGAAGATTATGGAATATAGCGACGGATCATTGCCGGCGTCTATAAACAGATTAGCAAAGGCATTGACTTCAGAGGGTAGAGAAATAGCAAGTGTTAATATAGATGGTAATCTGTTAGATATCGCATATCAAGCAGATAAGGAAAAGATGGCCGACATACAGACAAGAATAAATGATATTATTAGAGATTTGTTTATACACACTCTATCAGACAAAGATATAAAAGACATTATAGAATCCGAAGAAGGTAAGAGATGTTGTATAATAGATGTTAAGAACAATCGTGTTAAAAAGTACTATTCTATTGATAATTATCTATGCGGTACTTTAGATGATTATATATACACCGTTGTAGAATATAACAAATCCTATGTGTTAGTAAACGATACTTATATGAGCTATGACTATCTTGAATCATCAGGTGTAGTTGTTCTATCATGTTATGAAATGACTATAATCTCCTTGGATACAAAAGACGCCAAAGATGCTATAGAAGATGTGATAGTAGCAAGTGCGGTAGCCGAAGCATTGAATGACATGTTTAAGGAATTTGATAAAAACGTAAGTGCTATTATAATAAAAGAAGAAGATAATTATCTAAACAGTTCGCCCGATATCTACCATATAATATATATCATAGGTGGCACTATTCTGCTATTGTTAGTCATTATTTTAATATTGGCAATTTATATAGCGCGCAATAAATACAGAACCAGGAAATATGAAATAATGAAATATGACAATATGAGCATTAAATCTGATCATCATGATAGTCTTGAAACAGTGTCTATGGAAATTATTGATAATCGGTACTAATAAAATAGTTTAACTCTTTTAGAACCAGTTTGGTACTGTAATTTCAGTTCATTACTCGTTGAGAATATTGATGATTTTTTTTAAAATGAGTATCGGTAGTTACATATTACCATATCATCCATTATATAATCGATGATGCATGTATTAAAATACTTTCCGAATAAGTCTTCTAAATATTGTATTAATTATGAAAAACTATGCTATGTGAGATGATTCAAAGATGTTTAATGATACGATACTAGATTTTATCTCTAGCGAGATTGTTTAGAATCATTTATCATAACTATGTTTAATAAATTCATCAACGAATATCGATAAAGACCTCTTGTAATTCGAGTATAGGAAGCAGTATTACCATATCAACTTCCGAGTTAACAATTACTCTAAAACATGAGGATTGTACTCCTGTCTTTATTGGAGATTACTATTCAGTCGTTGATAAACTAGTAACCTCAGGTTTCTTTACAAACGATAAAGTACAACATCAAGACCTCACAACGCAGTGCAAGATTAATCTAGAAATCAAATGTAATTATGGAGGAGAATCTAGACAACTAACACCCACGGCGAAGTATACTTTATGCCTCATTCAGAAACGGTAACTGTAGTAGGAGACTGTCTCTCTAATCTCGATGTATATATAATATATACCAATACGGACGCGATATATTCCGACATGAATGGCGTCGCTTATCATATGTTATATCCTAAATGTTGATCATATTCCACAAATGATTGTGAACGAGATTAAATCATCTAACAAATAATTAGTTTTTATGACATTAACATATAATAAATAAATTAATCATTATTGACTTAACGATGACGAAAGTTATCATCATCTTAGGATTCTTGATTATTAATACAAATTCGTTGTGTCTATGAAATGTGAACAATGTGTCTCATATTATAATACACAAGAATTAAAGTGTTGTAAACTATCTAAGCCAGGAACATATTCAGATCATCGATGTGATAAATACAGCGATACCATCTGTGGACATTGTCCAAGTGACACATTCACGTCAATATATAATCGTTCTCCTCGGTGTCATAGTTGTAGAGGTCACACCTTGTACACCTACCACAAATAGAATCTGTCATTGTGACTCGAATAGTTATCGTCTCCTTAAAGCTTCTGATGGTAACTGTGTTACATGTGCTCCTAAAACAAAATATGGTCGTGTGTACGGAAAGAAAGGAGAAAATGATATGGAATACCATTTGTAAGAAATGTCGGAAGGGTACTTATTCAGATATTGTATCTGACTCTGATCAATGTAAACCTATGACAAGATAAGACTTACTCGCATCTACTGGATAGACATAAATATCCTCCTCGTAATAATGAAATATAATATAATATACACTAATTATTAATATCAATCGAGTATTAACATATAAGTTATTTTTAAACCCCTTTTGGGTTCCGTCCTAAACGGCGTTTCGGTCTGTGTCGCCACCATGGTCACACCGAGCCTCTGCGTGCTCCTCCATCGAGGACGACTTCAACTATGACAGCTCGGTGGCGTCTGCCAGCGTGCACATACGAATGGCATTTCTAAGAAAAGTCTACGGTATCCTTGTCTACAATTTCCTTTAACAACGGCAACAGCTGCAGTATTTTTATACTTTGAATGCATTCGGACATTTATACAAGGGAGTCCTGTTCTAATATTGGCATCAATGTTCGGATCTATAGGCTTGATTTTCGCATTGACTTTACACAGACATAAACATCCCCTGAATCTGTACCTACTTTGTGGATTTACACTGTTAGAATCTCTAACGCTGGCCTCTGTTGTTACTTTCTATGATGCGCGTATCGTTATACAAGCTTTCATGTTGACTACTGCAGTGTTTCTTGCTCTGACTACATGTACTCTACAATCAAAGAGAGATTTCAGTAAACTTGTAACAGGATTGTTTGCTGCTTTCTGGATTTTAATTTTGTCAGGAGTCTTGAGGATAAAGTTTAAAATAGAATTAATAAAGAACATATAGGTCATTTTTTAAACATGGATAGAAACCAAGGTTGTTAGTTAATAATATACAAGATATTTTTTCTCACTCTGATCCATGTAAACCAAGGACGAGAGACACTCTCATTCCTCATTCACGACACCATTAAAAATGGAAATTAAAGCCCTCTATTAAGCACAGACGGCTACAGGTCTACCATCAGGTTAACCTTCGTCTACCTTCACAATGGCCTTTCCTTGTGCCCAGTTCAGTCCCTGTCATTGCCACGCTACTAAGGACTCCCTGAATACCGTGACTGACGTCAGACATTGTCTGACTGAATACATCCTGTGGGTTTCTCATAGATGGACCCATAGAGAAAGCGCAGGGCCTCTCTACAGGCTTCTCATCTCTTTCAGAATTGATGCAATGGAGCTATTTGGTAGCGAGTTGAAGGAGTTCTCGGATTCACTTCCGTGGGACAATATCGACAATTGCGTGGAGATCATTAAATGTTTCATCAGAAATGACTCCATGAAAACCGCCAAAGAACTTTGTGCAATAATTGGACTTTGTACTCAATCAGCTATTGTCACTGGAAGAGTCTTCAATGATAAGTATATCGACATACTACTTATGCTGCGAAAGATTCTGAATGAGAACGACTATCTCACCCTCTTGGATCATATCCTCACTGCTAAATACTAAATCTCCTTCATGCTCTCTCACTAATACTCTTACTCACTACACTTTTTATCATCTTATGATGAATGATTGCCTTCATCATTTTTCGTGGAATATAATATAGGAATAATTAGCACCAGAATAGCTATGGATATCTCGTTAAGAATATTCTCGATAAGAGACATAATGTAGACATAGTTATTATATCCTTCTTAGATAAGTGTTACGCTACTGGAAAGTTTCCATCGTTATTATTACGTGAAGATGATATCATTAAACCAACATTGAGATTGGCTCTTATGTTAGCTGGATTGAATTACTGTAATAAATGCATCGAGTATAGAGGGATATAGCAATTCTCGATAATAGTCATGCAATATTTGAATGAGACTGATAATTTAGGTAATACAGTACTACACACACATATCTTTCTAGATTATATATCGTTAAAAATCTGTAAGAGGTATATTTCTCATAAGTATCCACTGTGTAATATTATTAATGGATATATAGATAACACAATAGGGACTAATAGTATTGTAAAAGATATAATCGACTATTTGTACATATCCAGATATCTATATTCCTACTAGTTTGCTGCGTAGTTGCATCATTGATATGCATGATTTATCAGGATTCAGAGATGAATTACTAAGTAAACTACAATCCCACAATAAGTAAGAATCAAATATCAAAAACTCACTTTTGATTTTTCTAGTCTTAAGTAATACATATATTTATTAATAGACCTATGAAATAAAAAAGTTAACAATGGATTCGCGTATAGCTATTTACGTATTAGTATCGGCATCTCTTTTGTATCTTGTTAATTGTCACAAACTAGTACATTACTTCAATCTGAAAATAAATGGAAGTGATATAACTAATACAGCAGATATATTGCTGGACAATTATCCAATTATGACCTTTGATGGAAAGGATATTTATCCATCTATCGCGTTCATGGTCGGTAATAAACTTTTCCTAGATCTTTATAAAAATATCTTTGTAGAATTTTTCAGACTATTTCGAGTATCTGTAAGTAGTCAATACGAGGAATTAGAATATTATTATTCATGTGATTATACTAACAACCGTCCTACAATTAAACAACATTACTTTTATAACGGCGAAGAATATACTGAAATTGATAGATCGAAAAAAGCCACTAATAAAAACAGTTGGTTAATTACTTCAGGCTTTAGACTACAAAAATGGTTCGATAGCGAAGATTGTATAATTTATCTCAGATCTTTAGTTAGAAGAATGGAAGACAGTAACAAAAACAGTAAAAAAACTTAGTACTTAGATATCGAAAAAAAATATATTTTTGTAGACTCTTGAGAATAGAAGGAAAACATGTACATAATTATAAAAAATGAAAATCAATGGCGAATAAGACAGTGCGATTCGCGCCATGGAGTCGGCAGATTTCATGGCTGTCGATGAGCAGTTTCACGACGACCTCGATCTTTGGTCATTATCTTTGGTAGATGACTATAAAAAACATGGATTAGGTGTTGACTGTTATGTTCTAGAACCAGTTGTTGACAGGAAAATATTTGATAGATTTCTCCTTGAACCAATTTGTGATCCTGTAGATGTTCTGTATGATTATTTTAGGATTCATAGAGATAATATTGATCAGTATATAGTAGATAGACTGTTTGCATATATTACATATAAAGATATTATATCTGCATTAGTGTCAAAGAATTATATGGAAGATATTTTCTCTATAATTATTAAGAATTGTAATTCTGTACAAGATCTCTTACTTTACTATCTATCTAATGCATATGTAGAAATAGACATTGTTGATTTTATGGTAGATCATGGGGCTGTAATATATAAAATAGAATGCTTGAATGCCTATTTTAGGGGAATATGTAAAAAGGAAAGTAGTGTTGTTGAGTTTATTTTGAATTGTGGTATCCCAGATGAAAATGATGTTAAATTAGATCTATATAAAATAATTCAGTATACTAGGGGATTCCTTGTAGATGAACCCACAGTATTAGAAATTTATAAGCTTTGTATCCCATATATTGAAGATATCAATCAACTAGATGCTGGTGGAAGGACCTTGCTTTATCGCGCTATCTATGCAGGTTATATAGATTTAGTATCATGGCTATTAGAAAATGGAGCAAATGTCAACGCAGTAATGAGTAATGGATATACATGTCTTGACGTGGCCGTGGATAGGGGATCTGTCATCGCCCGTAGGGAAACACATCTTAAAATATTAGAAATATTGCTTAGAGAACCATTGTCTATTGACTGTATAAAATTAGCTATACTTAATAATACAATTGAAAACCATGATGTGATAAAGCTCTGTATCAAGTATTTTATGATGGTAGATTATTCACTTTGTAATGTGTATGCATCATCACTCTTTGATTATATAATTGATTGTAAACAAGAATTGGAGTACATTAGGCAGATGAAAATTCATAATACAACCATGTATGAGTTAATCTATAATAGAGACAAAAACAAGCATGCTTCCCATATTCTACATAGGTATTCTAAACATCCAGTTTTGACACAGTGTATCACTAAAGGATTCAAGATTTACACAGAAGTAACCGAGCAGGTCACTAAAGCTCTAAACAGACGTGCTCTAATAGATGAGATAATAAACAATGTATCAACTGATGACAATCTCCTATCAAAACTTCCATTAGAAATTAGGGATCTAATTGTTTCACAAGCTGTCATATAGAGTTCTCTCCACCCACCTTTCTTGAAATGAGTTAATAGTCATAAGTTAGTTAAGTCATAAGTTAGTTTTATAAGTTAGTTTATAGTCTAACACTTCTAATTTTTATACCTTGATCTTTTTCTCTAATTATGAAAAAGTAAATCATTATGAAGATGGATGAAAATGGACGAGATTGTGCGCATCGTTAACGATAGTATGTGGTACGTACCTAACGCATTTATGGACGACGGTGATAATGAAGGTCACATTTCTGTCAATAATGTCTGTCATATGTATCTCGCATTCTTTGATGTGGATATATCATCTCATCTGTTTAAATTAGTTATTAAACACTGCGATCTGAATAAACGACTAAAATGTGGTAACTCTCCATTACATTGCTATACGATGAATACACGATTTAATCCATCTGTATTAAAGATATTGTTACGCCACGGCATGCGTAACTTTGATAGCAAGGATAAAAAAGGACATATTCCTCTACACCACTATCTGATTCATTCACTATCAATCGATAACAAGATCTTTGATATACTAACGGACCCCATTGATGACTTTAGTAAATCATCCGATCTATTGCTGTGTTATCTTAGATATAAATTCAATGGGAGCTTAAACTATTACGTTCTGTACAAATTATTGACTAAAGGATCTGACCCTAATTGCGTCGATGAGGATGGACTCACTTCTCTTCATTACTACTGTAAACACATATCCGTGTTCCACGAAAGCAATTATTACAAGTCAAAGAGTCACACTAAGATGCGAGCTGAGAAGCGATTCATCTACGCGATAATAGATCATGGAGCAAACATTAACGCGGTTACGAAAATCGGAAATACGCCGTTACACACTTACCTTCAACAGTATACCAAACATAGTCCTCGTGTGGTGTATGCTCTTTTATCTCGAGGAGCCGATACGAGGATACGTAATAATCTTGATTGTACACCCATCATGGAATACATAAAGAACGATTGTGCAACAGGTCATATTCTCATAATGTTACTCAATTGGCACGAACAAAAATACGGGAAATTACAAAAGGAAGAAGGACAACATCTACTTTATCTATTCATAAAACATAATCAAGGATATGGAAGTCGCTCTCTCAATATACTACGGTATCTACTAGATAGATTCGACATTCAGAAAGACGAATACTATAATACAATGACTCCTCTTCATACCGCCTTCCAGAATTGCAATAACAATGTTGCCTCATACCTCGTATACATCGGATACGACATCAACCTTCCGACTAAAGACGATAAGACAGTATTCGACTTGGTGTTTGAAAACAGAAACATCATATACAAGGCGGATGTCGTTAATGACATCATCCACCACAGACTGAAAGTATCTCTACCTATGATTAAATCGTTGTTCTACAAGATGTCGGAGTTCTCTCCCTACGACGATCACTACGTAAAGAAGATAATAGCCTACTGCCTATTAAGGGACGAGTCATTTGCGGAACTACATACTAAATTCTGTTTAAACGAGGACTATAAAAGTGTATTTATGAAAAATATATCATTCGATAAGATAGATTCCATCATCGAAAAATGTAGTCGTGACATAAGTCTCCTCAAAGAGATTCGAATCTCAGACACCAACTTGTATACGGTATTGAGAACAGAAGACATCCGGTATCACACCTATCTCGAAGCCATACATTCAGACAAACGCATTTCATTTCCCATGTACGACGATCTCATAGAACAGTGTCATCTATCGATGGAGCATAAAAGTAAACTCGTCGACAAAGCACTCAATAAATTAGAGTCTACCATCGATAGTCAATCTAGACTATCGTATTTGCCTCCGGAAATTATGCGCAATATCATAACCAAGCTAAGCGACTACCATCTAAACAGTATGTTGTACGGAAAGAACCATTACAAATATTATCCATGATAGAAAGAAAATATTTAAAAAATAATCTATATGATTGGAGAAGTAGGAAACAAACAGTAACAAGACGACGATTACTACTATTATTAAATCATGAGGTCCGTATTATACTCGTATATATTGTTTCTCTCATGTATAATAATAAACGGAAGAGATTTAGCACCACATGCACCATCCAATGGAAAATGTAAAGACAACGAATACAGAAGCCGTAATCTATGTTGTCTATCGTGTCCTCCGGGAACTTACGCTTCCAGATTATGTGATAGCAAGACTAATACACAATGTACGCCGTGTGGTTCGGATACCTTTACATCTCACAATAATCATTTACAGGCTTGTCTAAGTTGTAACGGAAGATGTGATAGTAATCAGGTAGAGACGCGATCGTGTAACACGACTCACAATAGAATCTGTGAATGCTCTCCAGGATATTATTGTCTTCTCAAAGGATCATCAGGGTGTAGAACATGTATTTCTAAAACAAAGTGTGGAATAGGATACGGAGTATCCGGATACACGTCTACCGGAGACGTCATCTGTTCTCCGTGTGGTCCCGGAACATATTCTCACACCGTCTCTTCCACAGATAAATGCGAACCCGTAACCAGCAATACATTTAACTATATCGATGTGGAAATTAACCTGTATCCAGTCAACGACACATCGTGTACTCGGACGACCACTACCGGTCTCAGCGAATCCATCTCAACGTCGGAACTAACTATTACCATGAATCATAAAGATTGTGATCCCGTCTTTCGTGCAGAATACTTCTCTGTCCTTAATAATGTAGCAACTTCAGGATTCTTTACAGGAGAAAATAGATATCAGAATACTTCAAAGATATGTACTCTGAATTTCGAGATTAAATGTAACAACAAAGATTCATCTTCCAAACAGTTAACGAAAACAAAGAATGATACTATCATGCCGCATTCAGAGACGGTAACTCTAGTGGGCGACTGTCTATCTAGCGTCGACATCTACATACTATATAGTAATACCAATACTCAAGACTACGAAACGGATACAATCTCTTATCATATGGGTAATGTTCTCGATGTCAATAGCCATATGCCCGCTAGTTGCGATATACATAAACTGATCACTAATTCCCAGAATCCCACCCACTTATAGTAAGTTTTTTTTTACCCATAAATAATAAATACAATAATTAATTTCTCGTAAAAGTAGAAAATATATTCTAATTTATTATATGGTAAGAAAGTAGAATCATCTAGAACAGTAATCAATCAATAGCAATCATGAAACAATATATTGTCCTGGCATGCATGTGCCTAGTGGCAGCTGCTATGCCTACTAGTCTTCAACAATCCTCATCCTCGTGTACTGAAGAAGAAAACAAACATCATATGGGAATCGATGTTATTATCAAAGTCACAAAGCAAGACCAAACACCGACCAATGATAAGATTTGTCAATCCGTAACGGAAGTTACAGAGACCGAAGATGATGAGGTATCCGAAGAAGTTGTAAAAGGAGATCCCACCACTTATTACACTATCGTCGGCGCGGGTCTTAACATGAACTTTGGATTCACCAAATGCCCAAAGATTTCATCCATCTCCGAATCCTCTGATGGAAACACTGTGAATACTAGATTGTCCAGCGTGTCACCAGGACAAGGTAAGGACTCTCCCGCGATCACGCGTGAAGAAGCTCTGGCTATGATCAAAGACTGTGAGATGTCTATCGACATCAGATGTAGCGAAGAAGAGAAAGACAGCGACATCAAGACCCATCCAGTACTTGGGTCTAACATCTCACATAAGAAAGTGAGTTACAAAGATATCATCGGTTCAACGATCGTTGATACAAAATGTGTTAAGAACCTAGAGTTTAGCGTACGTATCGGAGACATGTGTGAGGAATCATCTGAACTTGAAGTCAAGGATGGATTCAAGTATGTCGACGGATCGGCATCTGAAGGTGCAACCGATGATACTTCACTCATCGATTCAACAAAACTCAAAGCATGTGTCTGAATCGATAACTCTATTCATCTGAAAATGGATGAGTTGGGTTAATCGAACGATTCAGACACCGCACCACGAATTAAAAAAGTGTACCGGGTACTATATTCCGGTTTGCAAAACAAAAATGTTTAACTACATTCACAAAAAGTTACCTCTCGTTACTTCTTCTTTCTGTTTCAATATGTGATACGATATGATCACTATTCGTATTCTCTTTCTCTCTTGTCTCATAAAAAGTTTTACAAAAAAATATTTTTATTCTCTTTCTCTCTTCGATGGTCTCACAAAAATATTAAACCTCTTTCTGATGTCTCAACTATTTCGTAAACGATAACGTCCAACAATATATTCTCGTAGAGCTTATCAATATCCTTATGCCAATCTAGGTTGTCAGACAATTGCATCATAAAATAATGTTTATAATTTACACGTTAACATCATATAATAAACGTATATAGTTAATATTTTTGGAATATAAATGATCTGTAAAATCCATGTAGGGGACACTGCTCACGTTTTTTCTCTAGTACATAATTTCACACAAGTTTTTATACAGACAAATTAATTCTCGTCCATATATTTTAAAACATTGACTTTTGTACTAAGAAAAATATCTTGACTAACCATCTCTTTCTCTCGTCGATGGGTCTCACAAAAATATTAAACCTCTTTCTGATGGAGTCGTAAAAAGTTTTTATCCTTTCTCTCTTCGATGGGTCTCACAAAAATATTAAACCTCTTTCTGATGGTCTCTATAAACGATTGATTTTTCTTACCCTCTAGAGTTTCCTACGGTCGTGGGTCACACATTTTTTTCTAGACACTAAATAAAATAGTAAAATTAAATTAATTATAAAATTATGTATATAATTTACTAAC

>2022_Portugal_Monkeypox_PT0001

CGAAGAGAGAAAGAGATGGTTAGTCAAGATATTTTTCTTAGTACAAAAGTCAATGTTTTAAAATATATGGACGAGAATTAATTTGTCTGTATAAAAACTTGTGTGAAATTATGTACTAGAGAAAAAACGTGAGCAGTGTCCCCTACATGGATTTTACAGATCATTTATATTCCAAAAATATTAACTATATACGTTTATTATATGATGTTAACGTGTAAATTATAAACATTATTTTATGATGCAATTGTCTGACAACCTAGATTGGTATAAGGATGTTGATAAGCTCTACGAGAATATATTGTTGGACGTTATCGTTTACGAAATAGTTGAGACATCAGNAAGAGGTTTAATATTTTTGTGAGACCATCGAAGAGAGAAAGAGAATAAAAATATTTTTTTTTTTTTTGTAAAACNTTTTTATGAGACCAAGAGAATACGAATAGTGATCATATCGTATCACATATTGAAACAGAAAGAAGAAGTAACGAGAGGTAACTTTTTGTGAATGTAGTTAAATATTTTTGTTTTGCAAACCGGAATATAGTGCCCGGTCTTTTTTAATTCGTGGTGCGGTGTCTGAATCGTTCGATTAACCCAACTCATCCATTTTCAGATGAATAGAGTTATCGATTCAGACACATGCTTTGAGTTTTGTTGAATCGATGAGTGAAGTATCATCGGTTGCACCTTCAGATGCCGATCCGTCGACATACTTGAATCCATCCTTGACTTCAAGTTCAGATGATTCCTCACACATGTCTCCGATACGTACGCTAAACTCTAGGTTCTTGACACATTTTGTATCAACGATCGTTGAACCGATGATATCTTTGTAACTCACTTTCTTATGTGAGATGTTAGACCCAAGTACTGGATGGGTCTTGATGTCACTGTCTTTCTCTTCTTCGCTACATCTGATGTCGATAGACATCTCACAGTCTTTGATCATAGCCAGAGCTTCTTCACGCGTGATCGCGGAGAGTCCTTACCTTGTCCCGGTGACACGCTGGACAATCTAGTATTCACAGTGTTTCCATCAGAGGATTCGGAGATGGATAAAATCTTTGGGCATTTGGTGAATCCAAAGTTCATGTTAAGACCCGCACCGACGATAGTGTAATAAGTGGTGGGATCTCCTTTTACAACTTCTTCGGATACCTCATCATCTTCGGTCTCTGTAACTTCCGTTACGGATTGACAAATCTTATCATTGGTCGGTGTTTGGTCTTGCTTTGTGACTTTGATAATAACATCGATTCCCATATGATGTTTGTTTTCTTCTTCAGTACACGAGGATGAAGATTGTTGAAGACTAGTAGGCATAGCAGCTGCCACTAGGCACATGCATGCCAGGACAATATATTGTTTCATGATTGCTATTGATTGATTACTGTTCTAGATGATTCTACTTTCTTACCATATAATAAATTAGAATATATTTTCTACTTTTACGAGAAATTAATTATTGTATTTATTATTTATAGGTNAAAAAACTTACTATAAGTGGGTGGGATTCTGGGAATTAGTGATCAGTTTATGTATATCGCAACTAGCGGGCATATGGCTATTGACATCGAGAACATTACCCATATGATAAGAGATTGTATCATTTTCGTAGTCTTGAGTATTGGTATTACTATATAGTATGTAGATGTCGACGCTAGATAGACAGTCGCCCACTAGAGTTACNGTCTCTGAATGCGGCATGATAGTATCATTCTTTGTTTTCGTTAACTGTTTGGAAGATGAATCTTTGTTGTTACATTTAATCTCGAAATTCAGAGTACATATCTTTGAAGTATTCTGATATCTATTTTCTCCTGTAAAGAATCCTGAAGTTGCTACATTATTAAGGACAGAGAAGTATTCTGCACGAAAGACTGGATCACAATCTTTATGATTCATGGTAATAGTTAGTTCCGACGTTGAGATGGATTCGCTGAGACCGGTAGTGGTCGTCCGAGTACACGATGTGTCGTTGACTGGATACAGGTTAATTTCCACATCGATATAGTTAAATGTATTGCTGGTTACGACGGGTTCGCATTTATCTGTGGAAGAGACGGTGTGAGAATATGTTCCGGGACCACACGGAGAACAGATGACGTCTCCGGTAGACGTGTATCCGGATACTCCGTATCCTATTCCACACTTTGTTTTAGAAATACATGTTCTACACCCTGATGCTCCNTTTGAGAAGACAATAATATCCTNGAGAGCATTCACAGATTCTATTGTGAGTCGTGTTACACGATCGCGTCTCTACCTGATTACTATCACATCTTCCGTTACAACTTAGACAAGCCTGTAAATGATTATTGTGAGATGTAAAGGTATCCGAACCACACGGTGTACATTGTGTATTAGTCTTGCTATCACATAATCTGAAAGCGTAAGTTCCCGGAGGACACGATAGACAACATAGATTACGGCTTCTGTATTCGTTGTCTTTACACTTTCCATTGGATGGTGCATGTGGTGCTATATCTCTTCCGTTTATTATTATACATGAGAGAAACAATATATACGAGTATAATACGGACCTCATGATTTAATAATGTAGTAATCGTCGTCTTGTTACTGTTTGTTTCCTACTTCTCCAATCATATAGATTATTTTTTAAATATTTTCTTTCTATCATGGATAATATTTGTAATGGTTCTTTCCGTACAACATACTGTTTAGATGGTAGTCGCTTAGCTTGGTTATGATATTGCGCATAATTTCCGGAGGCAAATACGATAGTCTAGATTGACTATCGATGGTAGACTCTAATTTATTGAGTGCTTTGTCGACGAGTTTACTTTTATGCTCCATCGATAGATGACACTGTTCTATGAGATCGTCGTACATGGGAAATGAAATGCGTTTGTCTGAATGTATGGCTTCGAGATATGTGTGATACCGAATGTCTTCTGTTCTCAATACCGTATACAAGTCGGTGTCTGAGATTCGAATCTCTTTGAGGAGACTTATGTCACGACTACATTTTTCGATGATGGAATCTATCTTATCGAATGATATATTTTTCATAAATACACTTTTATAGTCCTCGTTTAAACAGAATTTAGTATGTAGTTCCGCAAATGACTCGTCCCTTAATAGGCAGTAGGCTATTATCTTCTTTACGTAGTGATCGTCGTANGGAGAGAACTCCGACATCTTGTAGAACAACGATTTAATCATAGGTAGAGATACTTTCAGTCTGTGGTGGATAATGTCATTAACGACATCCGCCTTGTATATAATGTTTCTGTTTTCAAACACCAAGTCGAATACTGTCTTATCGTCTTTAGTCGGAAGGTTGATGTCGTATCCAATGTATACGAGGTATGAGGCAACATTGTTATTGCAATTCTGGAAGGCGGTATGAAGAGGAGTCATTGTATTATAGTATTCGTCTTTCTGAATGTCGAATCTATCTAGTAGATACNGTAGTATATTGAGAGAGCGACTTCCATATCCTTGATTATGTTTTATGAATAGATAAAGTAGATGTTGTCCTTCTTCCTTTTGTAATTTCCCGTATTTTTGTTCGTGCCAATTGAGTAACATTATGAGAATATGACCTGTTGCACAATCGTTCTTTATGTATTCCATGATGGGTGTACAATTNAGATTATTACGTATCCTCGTATCGGCTCCTCGAGATAAAAGAGCATACACCACACGAGGACTATGTTTGGTATACTGTTGAAGGTAANNGTGTAACGGCGTATTTCCGATTTTCGTAACCGCGTTAATGTTTGCTCCATGATCTATTATCGCGTAGATGAATCGCTTCTCAGCTCGCATCTTAGTGTGACTCTTTGACTTGTAATAATTGCTTTCGTGGAACGCGGATATGTGTTTACAGTAGTAATGAAGAGAAGTGAGTCCATCCTCATCGACGCAATTANGGTCAGATCCTTTAGTCAATAATTTGTACAGAACGTAATAGTTTAAGCTCCCATTGAATTTATATCTAAGATAACACAGCAATAGATCGGATGATTTACTAAAGTCATCAATGGGGTCCGTTAGTATATCAAAGATCTTGTTATCGATTGATAGTGAATGAATCAGATAGTGGTGTAGAGGAATATGTCCNNNTTTATCCTTGCTATCAAAGTTACGCATGCCGTGGCGTAACAATATCTTTAATACAGATGGATTAAATCGTGTATTCATCGTATAGCAATGTAATGGAGAGTTACCACATTTTAGTCGTTTATTCAGATCGCAGTGTTTAATAACTAATTTAAACAGATGAGATGATATATCCACATCAAAGAATGCGAGATACATATGACAGACATTATTGACAGAAATGTGACCTTCATTATCACCGTCGTCCATAAATGCGTTAGGTACGTANCACATACTATCGTTAACGATGCGCACAATCTCGTCCATTTCATCCATCTTCATAATGATTTACTTTTTCATAATTAGAGAAAAAGNNNNNNNNNNNNNNNNNNNNNNNNNNNNNNNNNNNNNNNNNNNNNNNNNNNNNNNNNNNNNNNNNNNNNNNNNNNNNNNNNNNNNNNNNNNNNNNNNNNNNNNNNNNNNNNNNNNNNNNNNNNNNNNNNNNNNNNNNNNNNNNNNNNNNNNNNNNNNNNNNNNNNNNNNNNNNNNNNNNNNNNNNNNNNNNNNNNNNNNNNNNNNNNNNNNNNNNNNNNNNNNNNNNNNNNNNNNNNNNNNNNNNNNNNNNNNNNNNNNNNNNNNNNNNNNNNNNNNNNNNNNNNNNNNNNNNNNNNNNNNNNNNNNNNNNNNNNNNNNNNNNNNNNNNNNNNNNNNNNNNNNNNNNNNNNNNNNNNNNNNNNNNNNNNNNNNNNNNNNNNNNNNNNNNNNNNNNNNNNNNNNNNNNNNNNNNNNNNNNNNNNTAACTAACTTATGANTTAACTAACTTATGACTTAACTAACTTATGACTTAACTAACTTATGACTTAACTAACTTATGACTTAACTAACTTATGACTTAACTAACTTATGACTTAACTAACTTATGACTTAACTAACTTATGACTTAACTAACTTATGACTATTAACTCATTTCAAGAAAGGTGGGTGGATAGAACTCTATATGACAGCTTGTGAAACAATTAGATCCCTAATTTCTAATGGAAGTTTTGATAGGAGATTGTCATCAGTTGATACATTGTTTATTATCTCATCTATTAGAGCACGTCTGTTTAGAGCTTTAGTGACCTGCTCGGTTACTTCTGTGTAAATCTTGAATCCTTTAGTGATACACTGTGTCAAAACTGGATGTTTAGAATACCTATGTAGAATATGGGAAGCATGCTTGTTTTTGTCTCTATTATAGATTAACTCATACATGGTTGTATTATGAATTTTCATCTGCCTAATGTACTCCAATTCTTGTTTACAATCAATTATATAATCAAAGAGTGATGATGCATACACATTACAAAGTGAATAATCTACCATCATAAAATACTTGATACAGAGCTTTATCACATCATGGTTTTCAATTGTATTATTAAGTATAGCTAATTTTATACAGTCAATAGACAATGGTTCTCTAAGCAATATTTCTAATATTTTAAGATGTGCTTCCCTACGGGCGATGACAGATCCCCTATCCACGGCCACGTCAANACATGTATATCCATTACTCATTACTGCGTTGACATTTGCTCCATTTTCTAATAGCCATGATACTAAATCTATATAACCTGCATAGATAGCGCGATAAAGCAAGGTCCTTCCACCAGCATCTAGTTGATTGATATCTTCAATATATGGGATACAAAGCTTATAAATTTCTAATACTGTGGGTTCATCTACAAGGAATCCCCTAGTATACTGAATTATTTTATATAGATCTAATTTAACATCATTTTCATCTGGGATACCACAATTCAAAATAAACTCAACAACACTACTTTCCTTTTTACATATTCCCCTAAAATAGGCATTCAAGCATTCTATTTTATATATTACAGCCCCATGATCTACCATAAGATCAACAATGTCTATTTCTACATATGCATTAGATAGATAGTAAAGTAAGAGATCTTGCACAGAATTACAATTCTTAATAATTATAGAGAAAATATCTTCCATATAATTCTTTGACACTAATGCAGATATAATATCTTTATATGTAATATATGCAAACAGTCTATCTACTATATACTGATCAATATTATCTCTATGAATCCTAAAATAATCATACAGAACATCTACAGGATCACAAATTGGTTCAAGGAGAAATCTATCAAATATTTTCCTGTCAACAACTGGTTCTAGAACATAACAGTCAACACCTAATCCATGTTTTTTATAATCATCTACCAAAGATAATGACCAAAGATCGAGGTCGTCGTGAAACTGCTCATCGACAGCCATGAAATCTACCGACTCCATGGTGCGAATCGCACTGTCTTATTCGCCATTGATTTTCATTTTTTATAATTATGTACATGTTTTCCTTCTATTCTCAAGAGTCTACAAAAATATANTTTTTCGATATCTAAGTACTAAGTTTTTTTACTGTTTTTGTTACTGTCTTCCATTCTTCTAACTAAAGATCTGAGATAAATTATACAATCTTCGCTATCGAACCATTTTTGTAGTCTAAAGCCTGAAGTAATTAACCAACTGTTTTTATTAGTGGCTTTTTTCGATCTATCCTCGTCCTCTCATCATCCTTATATTATTATCATTATTATCATAGTCTATTAAACACAAATCATCTACGTTTATAACAACATTCTCATTATTAATTAGTTCTGTAGAATATCTTTAATAATTTGGCTATACATCTGTTCAATACTATCTATTGATGATTTCTTTTTTAAGACTTAAACTAGTTATGGTAATGACGATGAAATCGAGTAGTAACTTCTAATAAAGACTTGATATCATTATCATATGTTTGATCGTCATAGTTAATAGTGTGGCTAAATGGTACTGTTAATAAGTTTATAGACAATATCATAGTATTTTCTTTCCAGAATTAGATTATTTTTTTAAATACTGATCCTCACAATTCCGTGATGTAGCAGTAGTTGGTGCATGGTCTATATCGTTAAAATGTATCATATATAATAGTTTTCTGACGTGGAGTACAGAATTTTCGATTAATGAGTTCATGGTAAGGAAGGGCAAATGCCTGTATATAATATACATAAGTTAATAGTTTTTTATCATATTTTCTAATACCATAATAAAAATTATCATTATTGCGTTTGGTAGTTCTGCCCTATCATCTATATCACTGTCACTCTCGCTCTCACTATATCTTCTAAAATTACAAACAACTGGATATTCGATAACAGCATTTGTGTAGTTTTTGTCTTTTACAGTATATACGTTATTGTCAAAATCTAAACAAATATTAGCATAATACATCTATCTATAAGATCAGGATCCATGTTCGAGCATACTAGCCATGTATATTTGTAACTTCGTCGTACAGCGTTAGATCAATAGAATAAACAATCGTGTGACGCAACTTTTTTACGATCTAGTTGTATGAGTTTATCGTTTACATAAGCAATTAACGGCTTTAACAGATGATCTGAGTAATAATATACCTCTGTTATACGTTTAATGTTCACGGTCTTAGTATTTTTAGATATCAATTGTGATTTACACCATATTCGACTCCCTTGTGTGTAACGTTAGAAATTCTAAATCTATATTATCTATTACAGCGTAAAACACATTCAATATTGTATTGTTATTTTTATATTATTTACACAATTAACAATGTATTATTAGTTTATATTACTGAATTAATAATATAAAATTCCCAATCTTGTCATAAACACAAAATCCATTAAAAATGTCGATAAAATATCTGATGTTGTTGTTCGCTGCTATGATAATCAGATCATTAGCCGATAGTGGTAATGCTATCGAAACAACATCGCCAGAAATTACAAACGCTACAACAGATATTCCAGCTATCAGATTATGTGGTCCAGAGGGAGATGGATATTGTTTACACGGTGACTGTATCCACGCTAGAGATATTGACGGTATGTATTGTAGATGCTCTCATGGTTATACAGGTATTAGATGTCAGCATGTAGTATTAGTAGACTATCAACGTTCAGGAAAACCAGACACTACAACATCATATATCCCATCTCTAGGTATTGTGCTTGTATTAGTAGGCATTATTATTACGTGTTGTCTATTATCTGTTTATATGTTCACTCGAAGAACTAAACTACCTATACAAGATATGGTTGTGTTATATTTTTTATAAAATTTTTTTATGAGTATTTTTACAAAAATGTATATGTATNAAAAAAATACTAAGTATACGATGTATCCTGTATTATTTGTATTTATCTAAACAATACTTCTGCCTCTAGATGGGATACAAAAATTTTTTATTTCAGCATATTAAAGTAAAATCTAGTTACCTTGAAAATGAATACAGTGGGTGGTTCCGTATCACCAGTAAGAACATAATAGTCGAATACAGTATCCGATTGAGATTTTGCATACAATACTAGTCTAGAAAGAAATTTGTAATCATCCTCTGTGACAGGAGTCCATATATCTGTATCATCGTCTAGTTTTTATCAGTGTCCTATGCTATATTCCTGTTATCATCATTAGTTAATGAAAATAACTCTCGTGCTTCAGAAAAGTCAAATATTGTATCCATACATATATCTCCAAAACTATCACTTATACGTTTATCTTTAACGAACATATACCTAGATGGTTATTTACTAACAGACATTTTTTCAAGATCTATTGACAATAACTCCTATAGTTTCCACATCAACCAAGTAATGATCATCTATTGTTATATAACAATAACATAACTCTTTTCCATTTTTATCAGTATCTATATCAACGTCGTTGTAGTGAATAGTAGTCATTGATCTATTATATGAAACGGATATGTCTAGTTAATATTTTCTTTGATTTAAAGTCTATAGTCTTTACAAACATAATATCCTTATCCGACTTTATATTTCCTGTAGGGTGGCATAATTTTATTCTGCCTCCACAATCAGTGTTTCCAAATATATTACTAGACAATATTCCATATAGTTATTAGTTAAGGGACCCAATTAGAACACGTACGCGCTTATTCATCATTTGGATCGTATTTCATAAAAGTTATTATGTTATAGATGTCAACACATTCTACATTTTTTTAATTGTCTATATAGTATTTTTCTGATATTTTCTATAATATCAGAATTGTCTTCCATAGGAAGTTGTATACTATTCGGAATCAGTTACATGTTTAAATAATTCTCTGATGTCATTCCTTATACAATCAAATTCATTATTAAACAGTTTAATAGTCTGTAGACCTTTATCGTCGTACATATCCATTGTCTTATTAGTTACGCTTATTTTTATGGGTTTTACGTTGCTTTATTATATTTTATAATAATGATTGTTTGACAATGTCGTAGTATAGATATATTATTAGAGGAGGTATAATTATAAAAAGTTTCTGAGTACGATGTTATAAGAGGAGAGGACACATTAACATCATACATCAATTAACTACATTCTTATAACATTGTAATCAAAAGAATTGCAATTTTGATGTATAACAACTGTCAATGGAATTGTATATTACAAATTACGGTATGTTGTAACAACAAATACCGATCGGTAATTGTCTCTGCCGCTGTAATAGAATTAATTATATATCTATTACACCGGCCTTGTATCATAATAAAGTTGTGGTAGTATGATCTCCATATTTATAATTTAGTACTTTGTATTTAGTTTTTTTGGAATCATNAAAAAAGTTTTACTAATTTAAAATTTAAAAAGTATTTACATTTTTCACTGTTTAGTCGCGGATATGGAATTCGATCCTGTCAAAATCAATACATCATCTATAGATCATGTAACAATATTACAATACATAGATGAACCAAATGATATAAGACTAACAGTATGCATTATNCGAAATGTTAATAACATTACATATTATATCAATATCACAAAAATAAATCCACATTTGGCTAATCGATTTCGGGCTTGGAAAAAACGTATCGCCGGAAGGGACTATATGACTAACTTATCTAGAGATACAGGAATACAACAATCAAAACTTACTGAAACTATACGTAACTGTCNAAAAAATAAAAACATATATGGTCTATATATACACTACAATTTAGTTATTAATGTGGTTATTGATTGGATAACCGATGTGATTGTTCAATCAATATTAAGAGGGTTGGTAAATTGGTACATAGCTAATAATACATATACTCCAAATACACCCAATAATACTACAACCATTTCTGAGTTGGATATCATCAAAATACTGGATAAATACGAGGACATGTATAGAGTAAGTAAAGAAAAAGAATGTGGAATTTGCTATGAAGTTGTTTACTCAAAACGATTAGAAAACGATAGATACTTTGGTTTATTGGATTCGTGTAATCATATATTTTGCATAACATGTATCAATATATGGCATAGAACACGAAGAGAAACCGGTGCGTCAGATAATTGTCCTATATGCCGTACACGTTTTNNAAAAATAACAATGAGCAAGTTCTATAAGCTAGTTAACTAATAAATNAAAAGTTTAATTATCGACGATATATGTCGTTATTTTTCTCTCATATGAAAGATTAATTTGATTCTAATATAATCTTCAGTATTGGATGAATCTCAATTCAAATTAATTCCATTAGATTAGATTAGATTAGATCATAAATAAAAATAGTAGCACGCACTACTTCAGCCAAATATTCNTTTTTGAAACGCCATCTAGCGTAATGAGAACACAAGTGAACCTATAATGAGCAAATTTATTAGTATCGGTTACATGAAGGACTTTACGTAGAGTGGTGATTCCTCCATCTGTGGTACGAACGGTTTCATCTTCTTTGATGCCACCACCCAGATGTTCTATAAACTTGGTATCCTCGTCCGATTTCATATCATTTGCCAACCAATACATATAGCTAAACCCAGGCATACGTTCCACACATCCGGAACAATGAAATTCTCCAGAAGATGTTACAATGGCTAGATTTGGACATTTGGTTTCAACCGCGTTAACATATGAGTGAACACACCCATACATGAAAGCGATGAGAAATAGGATTCTCATCTTGCCAAAATATCACTATAAAAAATTTATTTATCAATTTTAAAGGTATNAAAAAATACTTATTGTTGCTCGAATATTTTGTATTTGATGGTATACGGAAGATTAGAAATGTAGGTATTATCATCAACTGATTTTATGATGGTTTTATGAATTTTATTATGCTTCACTATTGCATCGGAAATAATATCATATGCTTCCACGTATATTTTATTTTGTTTTGACTCATAATACGCACGTAATTCTGGATTATTGGCATATCGATGAATAATTTTAGCTCCATGCTCAGTAAATATTAATGAGAACATAGTGTTGCCTCCTACCATTANTTTTTTCATCTCATTCAATTCTTGATTGCAGAGATCTATATAATCATTATAGCGTTGACTTATGGACTCTGGAATCTTAGACGATGTACAGTCATCTATAATCATGGTATATTTAATACATTGTTTTATAGCATAGGCATTATCTACGATATTAGATACTTCACTCAATGAATCAATCACACAATCTAATGTAGGTTTATGACATAATAGCATTTTCAGCAGTTCAATGTTTCTAGATTCGTTGATGGCAATGGTTATACATGTATATCCGTTATTTGATCTAATGTTGACATCTGAACCGGATTCTAGCAGTAAAGATACTAGAGATTGTTTATTATATCTAACAGCCTTGTGAAGAAGTGTTTCTCCTCGTTTGTCAATCATGTTAATGTCTTTAAGATAAGGTAGACAAATGTTTATAGTACTAAGAATTGGGCAAACATAAGACATGTCACAAAGACCCNTTTTGTATGTATAAGTGTAGAAATTATAACATCCATAGTTGGATTCACATAGGTGTCCAATCGGGATCTCTCCATCATCGAGATGATTGACGGCATCTCCCCCTTCCNNTTTTAGTAGATATTTCATCGTGTAAGAATCAATATTAATATTTCTAAAGTATCTGTGTATAGCCTCTTTATTTACCACAGCTCCATATTCCAACATGCATTCCACTAGAGGGATATCGATATCGCNGAATGTCATATACTCAATTAGTATATGTTGGAGGACATCCGAGTTCATTGTTTTCAATATCAAAGAGATGGTTTCCTTATCATTTCTCCATAGTGGTACAATACTACACATTATTCCGTACGGCTTTCCATTCTCCAAAAACAATTTTACCAAATCTAAATCTACATCTTTATTGTATCTATAATCACTATTTAGATAATCAGCCATAATTCCTCGAGTGCAACATGTTAGATCGTCTATATATGAATAAGCCGTNNTATCTATTCCTTTCATTAACAATTTAACGATGTCTATATCTATATGAGATGACTTAATATAATATTGAAGAGCTGTACAATAGTTTTTATCTATAGAAGACGGCTTGATTCCGTGATTAATTAGACATTTAACAACTTCCGGACGCACATATGCTCTCGTATCCGACTCTGAATACAGATGAGCGATGATATACAGATGCAATACGGTACCACAATTTCGTGGTTGATAATCATCATACACGTATCCGTACTNGTCATCCTCATAAAGAACACTGCATCCATTTTCTATGAACAAATCAATAATTTCAGGAACAGGATCATCTGTCATTACATAATTTTCTATAACTGAACGATGGTTTTCACATTTAACACTCAAGTCAAATCCATGTTCTACCAACACCTTTATCAAGTCAACGTCTACATTTTTTGATTTCATATAGCTGAATATATTAAAGTCATTTATGTTGCTATATCCAGTAGCTTCTAGTAGAGCCATCGCTATATCCTTATTGACTTTAACATGTCTACTATTTGTGTATTCTTCTATTGGGGTAAACTGTCTCCAATTTTTATGTAATGGATTAGTGCCACTATCTAGTAGTAGTTTGACGACCTCAACATTATTACAATGCTCATTGAAAAGGTATGCGTGTAAAGCATTATTCTTGAATTGGTTCCTGGTATCATTAGGATCTCTGTCTCTCAACATCTGTTTAAGTTCATCGAGAACCACCTCCTCATTTTCCAGATAGTCAAACATTTTGACTGAATAGAAGTGAATGAGCTACTGTGAACTCTATACACCCGCACAACTAATGTCATTAAATATCATTTTTGAATGTATTTATACCATGTCAAAAACTTGTACAATTATTAATAAAAATAATTAGTGTTTAAATTTTACCAGTTCCAGATTTTACACCTCCGTTAACACCTCCATTAACCCCACTTTTTACACCACTGGACGATCCTCCTCCCCACATTCCACTGCCACTAGATGTATAAGTTTTAGATCCTTTATTACTACCATCATGTCCATGGATAAAGACACTCCACATGCCGCCACTACTACCCCCTTTAGACGACATATTAATAAGACAAGTTTAACAATAAAATTAATCACGAGTACCCTACTACCAACCACTATTATATGATTACAGTACCTTGACTAAAGTCTCTAGTCACAAGATCAATACTACCAACCTACGCTATTATATGATTATAGTTTCTATTTTTATAGGAACGCGTACGAGAAAATCAAATGTCTAAGTTCTAACGGTAGTGTTGATAAACGATTGTTATCCGCGGATACCTCATCTATCATGTTGTCTATTTTCTTACTTTGTTCTATTAACCTATTAGCATTATATATTATTTGATTATAAAACTTATATTGCTTATTAGCCCAATCTGTAAATATCGGATTATTAACATATCGTTTCTTTGTAGGTTTATTTAACTTGTACATCACTGTAAGCATGTCCGTACCATTTATTTTAATTTGACACATATCAGCAATTTCTTTTTCGCAGTCGGTTATATATTCTATATAAGATGGATACGTATCACATATGTACTTATAGTCTACTAATATGAAGTACTTAATACATATTTTCAGTAACGATTTAGCCTTATTACCTATTAATAAGTGCCTGTCGTTGGATAGGTAATCAACTGTTTTCTTAATACATTCGATGGTTGGTAATTTACTCAAAATAATTTCCAATATCTTAATATATATTTCTGCTATTTCTGGTATACATGCATGTGCCATTATAACACAAATACCAATACATGTAGACCCATATGTTGTTGTTATATTAATATCTGCGCCATTATCTATTAACCATTCTATTAGGTCAACACTATGCGACTCGATACAATAATAAAGTATACTACGTCCATGTTTATCTATTTTGTTTATATCATCGATATACGGCTTACAAATTTCTAGTATCGATAACACTTCTGACTCGTGAATAAATAAGGTAGGGAATAACGGCATAATATTTATTATGTTATCATCATTAACAACTACGTTTCCANTTTTTAAAATATACTCTACAACTTTAGGATCCCTATTGTCAAATCTTTTAAAATATTTATTTATATGCTTAAATCTATATAATATAGCTCCTTCCCTAATCATACATTTGATAACATTGATGTACACTGTATGATAAGATACATATTCTGACAATAGATCTTGTATAGAATCTGTATATCTTTTAAGAATTGAGGATATTATGACATTATTACGTAAACTATTACACAATTCTAAAATATAAAATGTATCATGGGCAGATAATAGTTTATCCACTATATAATTATCTATTTTATGATTTTTCTTCCTATATTGTTTACGTAAATAGATAGATAGAATATGCATTAGTTCATTACCGCTATAGTTACTATCGAATAACACGTCAAATATTTCCCGTTTAATATCGCATTTGTCAACATAATAATAGAGTATGGTACGTTCACGATAAGTATAATGACACATCTCGTTTTCGTGCGAAATTAAATAGTTTATCACGTCCAAAGATGTCACATAACCATCTTGTGACCTAGTAATAATATAATAATAGAGAACTGTTTTACCCATTCTATTATCATAATCAGTGGTGTAGTCATAATCTAAATAATCAAACTCGTCATCCCAATTAAAATAAATATAATCAGTACATTGAATGGGTATGATATTGTACCCATACTGTATGTTGCTACATGTAGGTATTCCTTTATCCAATAATAGTTTNAATACATCTATATTANGATTTGATGTTGTCGCGTANTNNTCTACAATATTAATACCANTTTTGATACTATTTATTTCTATACCTTTCGAAATTAGTAATTTNAATAAGTCTATATCGATGTTATCAGAACATAGATATTCAAATATATCAAANNNATTNATATTTTTATAGTCGACTGACGACAATAACAAAATNACNACATCNTTTTTGANNNNNNNNNNNTTTNNNNTAACGTATGCCTTTNNTGGANNNTCACCATCATACNCNNATAATGGATTTGCACCACTTTCTATTAATGATTNTNNACTACTGGCANCGATGTTAAATGNNTTACAACTATCATAGNGTATNNTATCGTTAACCATGATTGGTTGTTGATGTTATCACANNTTTTTGGTNNNTTTCATTTCAGNTANNTATGGATTTAGCACGTTTNGGAAGCANGNGCTCNNATGATTTCAGTACTGTAGTGTCAGTNNTATTAGTTTCGNTCAGATCAATGTCTAGATCTATAGAATCNAAACACAATAGGTCAGAAGATAATGAATATCTGTACGCTTCNTCNNNNNNNNNNNNNNNNNNNNNNNNNNNNNNNNNNNNNNNNNNNNNNNNNNNNNNNNNNNNNNNNNNNNNNNNNNNNNNNNNNNNNNNNNNNNNNNNCTAGTATAAAATTCTATATTCAAATTATCATGCGANNTGTGTACGCTAACGGTATCAATAAACGGAGCACAGCATTTAGTCAACAGTAATCCAAAATTTTTTAAAGTATATCTTAACGAANNAAGTTGTCATCGTTAGAGTGTGGTAAATCATTGTCTACGGTACTAGATCCTCATAAGTGTATATATCTAGAGTAATATTTAATTTATCAAATGGTTGATAATATGGATGTTGTGGCAATTTCCTAATACGGAAATAAGACATAAACACGCAATAAATCTAATTGCGGACATGTTACACTCCTTAAAAAATACGAATAAACACTTTGGCTTTTAGTAAGTGTCATTTAACACTATACTCATATTAATCCATGGACTCATAATCTCTATACGGGATTAACGGATGTTCTATATACGGGGATGAGTAGTTCTCTTCTTTAACTTTATACTTTTTACTAATCATATTTAGACTGATGTATGGGTAATAGTGTTTGAAGAGCTCGTTCTCATCATCAGAATAAATCAATATCTCTGNTTTTTTGTTATACAGATGTATTACAGCCTCATATATTACGTAATAGAACGTGTAATCTACCTTATTAACTTTCACCGCATAGTTGTTTGCAAATACGGTTAATCCTTTGACCTCGTCGATTTCCGACCAATCTGGGCGTATAATGAATCTTAACTTTAATTTCNTGTAATCATTCGAAATAANTTTTAGTTTGCATCCGTAGTTATCTCCTCTATGTAACTGTAAATTTCTCAACGCGATATCTCCATTAATAATGATGTCGAATTCGTGTTGTATACCCATACTGAATTGATGAACTAATACCGACGGTATTAATAGTAATTTACTTTTCATCTTTACATACTTGGTAATAGTTTTACTATCATAAGTTTATAAATTCCACAAGCTACTATGGAATATACCAACCATCTTAGTATAGAACACATGTCTTAAAGTTATTAATTAATTACATGTTGTTTTATATATCGCTACGAATTTAAACAGNGGAATCAGTTANGAANAAAAAATTATCTGTCATCATCATCATCTATTNGATAACATCTCTGTATTCTACGATAGAGTGCTATTTTAAGATGTGACAGATCCGTGTCATCAAATATATACTCCATTAAAATGATTATTCCGGCAGCGAACTTGATATTNGATACATCACGACCNTTGTTAATATCCACGACAATAGACAGCAATCCCATTGTTCCATAAACAGTGAGTTTATCTTTCNTTGAAGTGATATTTTGTAGAGATCTTATAAAACTGTCGAATGACATCGTATTTATATCTTTAGCTAAATCATATATGTTACCATCATAATATCTAACAGCATCTATCTTAAACGTTTCCATCGCTGTAAAGACGTTTCCGATAGATGGTCTCGTTTCATCAGTCATACTGAGCCAACAAATGTAATCGTGTATAACATCTTTGATAGAATCAGACTCTAAAGAAAAGGAATCGGCTTTATTATACACATTCATGATAAACTTAATGAAAAATGTTTTTCGTTGTTTAAGTTGGATGAATAGTATGTCTTAATAATTGTTATTATTTCACTAATTAATATTTAGTAACGAGTACACTCTATAAAAACGAGAATGACATAACTAATCATAACTAGTTATCAAAGAATGTCTAGGACGCGTAANTTTTTATGGTATAGATCCNNTAAGCGTTGTCTGTATTCTGGAGCTATTTTCTCTATCGCATTAGTGAGTTCAGAATATGTTATAAATTTAAATCGAATAACGAACATAACTTTAGTAAAGTCGTCTATATTANCTCTTTTATTTTCTAGCCATCGTAATACCATGTTTAAGATAGTATATTCTCTAGTTACTACGATCTCATCGTTGTCTAGAATATCGCATACTGAATCTACATCCAATTTTAGAAATTGGTCTGTGCTACATATCTCTTCTATATTATTGTTGATGTATTGTCGTAGAAAACTATTACGTAGACCATTTTCTTTATAAAACGAATATATAGTACTCCAATTATCTTTACCGATATATTTGCATACATAATCCATTCTCTCAATCACTACATCTTTAAGAGTTTGGTTGTTAAGATATTTGGCTAAACTATATAATTCTATTAGATCATCAACAGAATCAGTATATATTTTTCTAGATCCAAAGATGAACTCTTTGGCATCCTCTATAATATTATCAGAAAAGATATTTTCGTGTTTTAGTTTATCAAGATTTAACCTGTTCATATCCATGATTAACGACGTCATATAACCACATAAAATAAAAATCCATTTTCATTTTTAGCACAATACTATTCATAATTGATATTGATGTAATATTTTGTTACTTTGAACGTAAAGACAGTACACGGGTCCGTATCTCCAACAAGCACGTAGTAATCAAATTTGGTGTTGTTAAACTTCGCAATATTCATCAATTTAGATAGAAACATATACTCATCATCTGTTTTAGGAATCCATGTATTATTACCACTTTCCAACTTATCATTATCCCAGGCTATGTTTCGCCCATCATCGTTGCACAGAGTGAATAATTCNTTTGTATTCGGTAGTTCAAATATATGATCCATGCATATATCGACAAAGCTATTGTAGATGTGATTTTTCCTAAATCTAATATAAAACTCGTTTACTAGCAAACATTTTCCTGATTTATCGACCAAGACACACATGGTTTCTAAATCTATCAAGTGGTGGGGATCCATAGTTATAACGCAGTAACATAAATTATTACCTTCTTGACTGTCGCTAATATCTATATACTTATTGTTATCGTATTNGATTCTACATATAGATGGCTTGTATATCAAAGATATAGAACACATAACCAATTTATATTCTCGCTTTGTATTTTCGAATCTAAAGTTAAGAGATTTAGAAAACATTATATCATCGGATGATGATATCACTGTTTCCAGAGTAGGATATATTAAAGTCTTTAAAGATTTTGTCCGATTCAAATAAATCACTAAATAATATCCCATATTATCATCTGTTATAGTCGTGTCATTAAATCTATTATATTTTATGAAAGATATATCACTGCTCACCTCTATATTTCGTACATTTTTAAACTGTTTGTATAATATCTCTCTAATACAATCAGATATATCTATTGTGTCGGTAGACGATACCGTTACATTTGAATTAATGGTGTTCCATTTTACAACTTTTAACAAGTTGACCAATTCATTTCTAATAGTATCAAACTCTCCATGATTAAATATTTTAATAGTATCCATTTTATATCACTACGGACATAAACCATTGTATATTTTTTATGTTTATTAGTGTACACATTTTGGAAGTAAGTTCCTGGATCGGATGTCACCGCAGTAATATTGTTGATTATTTCTGACATCGACGTATTATATAGTTTTTTAATTCCATATCTTTTAGAAAAGTTAAACATCCTTATACAATTTGTGGAATTAATATTATGAATCATGGTTTTTACACATAGATCTATTACAGGCGGAACATCAATTATTATGGCAGCAACTAGTATCATTTCTACATTGTTTATGGTGATGTTTATCTTCTTCCAGCGCATATAGTCTAATATCGATTCAAACGCGTGATAGTTTATACCATTCAATATAATCGCTTCATCCTTTAGATGGTGATCCTGAATGTGTTTAAAAAATTATACGGAGACGCCGTAATAATTTCCCCATTGATAGAAAATATCACGCTTTCCATTTTCTTGAAGTACTAAAAGTAATTATAATATAATGTAAAGGTTTATATATTCAATATTTTTTATNAAAAAAATCATTTTGACATTAATTCCTTTTTAAATTTCAGTCTATCATCTATAGAAACATATTCTATGAATTTATAAAATGCTTTTACGTATCCTATCGTAGGTGATAGAACCGCTAAAAAACCTATCGAATTTCTACAAAAGAATCTGTTATATGGTATAGGGAGAGTATAAAACATTAAATGTCCGTACTTATTAAAGTATTCAGTAGCCAATCCTAACTCTTTCGAATAATTATTAATGGCTCTTATTCTGTACGAATCTATTTTTTTGAACAATGGACCTAGTGGTATATCTTGTTCTATGTATCTAAAATAATGTCTGACTAGATCCGTTAGTTTAATATCCGCAGTCATCTTGTCTAGAATGGCAAATCTAACTGCGGGTTTAGGCGTAGGCGTTAGTTTAGTTTCTATATCTACATCTATGTCTTTATCTAACACCAAAAATATAATAGCTAATATTTTATTACAATCATCCGGATATTCTTCTACGATCTCACTAACTAATGTTTCTTTGGTTATACTAGTATAGTCACGATCAGACAAATAAAGAAAATCAGATGATCGATGAATAATACATTTAAATTCATCATCTGTAAGATTTTTGAGATGTCTCATTAAAATATTATTAGTGTCAGTTCTCATTATCATATATTGACAGCAGCTATTACACTTATTTTATTTTTCTGTATTTTATTACTTTTCACCATATAGATCAGTCATTAGATCATCAAAATACTTTTCAATCATCCTAAAGAGTATGGTGAACGAATCTTCCCATCTAATTTCTGAACGTCTACCAATGTCTCTAGCCACTTTGGCACTAATAGCGATCATTCGCTTAACATCTTCTACATTATTAACTGGTTGATTCAATCTATCTAGCAATGGACCGTCGGATAGCGTCATTCTCATGTTCTTAATCAATGTACATACATCGTCATCATCTACCAATTCATCAAACAATATAAGCTTTTTAAAATCATCATTATAATAGGATGGATCGCCGTCATTTCTCCAAAGAATATATCTAATAAGTAGAGTCCTCATGCTTAGTAATTTAACTATTTTAGTTAACAACTATTTTTTATGTTAAATCAATTAGTAACACCGCTATGTTTAATACTTATTCATATTTTAGTTTTAGGATCGAGAATCAATACAAAAATTAATACATCAATTTTGGAAATACTTAGTTTCCACGTAGTCAATGAAACATTTGAGCTCATCGTAAAGGACGTTCTCGTACAAGACGTAACTATAAATTGGTTTATATTTGTTCAAGATAGATACAAATCCGATAACTTTTTTGACGAATTCTACGGGATTCACTTTAAAAGTGTCATACCGGGTTCTTTTTATTCTTTTAAACAGATCGATTGTGTGATGTTGATTAGGTCTTTTACGAATTTGATACAGAATAGCGTTTACATATCCACCATAGTAATCAATAGCCATTTGTTCGTATGTCATAAATTCTTTAATTATATGACACTGTGTATTATTTAGTTCGTCCTTGTTCATCATTANGAATCTATCCAATATGGCAATTATATTAGAACTATAACTGCGTTGTATGCGCATGTTGATGTGTCTGTTTATACAATCAATTATACTANGATCCATACCACTACAATCGGGTAAAATTGTAGCATCATATACCATTTCTAGTACTTTAGATTCATTGTTATCCATTGCAGAGGACGTCATGATCGAATCCNAAAAAATATATTATTTTTATGTTATTTTGTTAAAAATAATCATCNAATACTTNGTAAGATACTCCTTCATGAACATAATCAGTTACAAAACGTTTATATGAAGTAAAGTATCTACAATTTTTACAAAAGTCAGGATGCATAAGTACAAAGTACGCGATAAACGGAATAATAATAGATTTATCTAGTTTATCTTTTTCTATCTCTTTCATAGTTATATACATGGTCTCAGAAGTCGGATTATGTAACATCAGCTTCGATAAAATGACTGGGTTATTTAGTCTTACACATTCGCTCATACATGTATGACCGTTAACTATAGAGTCTACACTAAAATGATTGAATAATAGATAGTCTACCATTGTTTCGTATTCAGATAGTACAGCGTAGTACATGGCATCTTCACAAATTATATCATTATCTAATAGATATTTGACGCATCTTATGGATCCCACTTCAACAGCCATCTTAAAATCGGTAGAATCATATTGCTTTCCTTTATCGTTAATAATTTCTAGAACATCATCTCTATCATAAAAGATACAAATATTAACTGTTTGATCAGTAATAACATTGCTAGTCGATATCAATTTGTTAATAAGATGCGCTGGGCTCAATGTCTTAATAAGAAGTGTAAGAGGACTATCTCCAAATTTGTTTTGTTTATTAACATCCGTTGATGGAANNAAAAGATTTATAATGTCTACATACTTGACTGTTTTAGAGCATACAATATGGAGAGGCGTATTTCCATCATGATCTGGTTTTGAGGGACTAATTCCTAGTTTCATCATCCATGAGATTGTAGAAGCTTTTGGATTGTCTGACATAAGATGTCTATGAATATGATTTTTGCCAAATTTATCCACTATCCTGGCTTCGAANCNGATAGACATTATTTTTTTAAACACTCTTTCTGAAGGATCTGTACACGCCAACAACGGACCACATCCTTCTTCATCAACCGAGTTGTTAATCTTGGCTCCATACTGTACCAATAAATTTATTCTCTCTATGACTTCATCATCTGTTCCCGAGAGATAATATAGAGGTGTTTTATTATGTTTATCACATGCGTTTNGATCTGCGCNGTGCACCAGCAGCATCGCGACTATTCTATTATTATTAATTTTAGAAGCTATATGCAATGGATAATTTCCATCATCATCCGTCTCATTTGGAGAGTATCCTCTATGAAGAAGTTCTTCTATAAATCGTTCATCTAGTCCTTTAATGCCACAATACGCATGTAGAATGTGATAATTTCCAGAGGGTTCGATAACTTGTAGCATATTCCTAAATACATCTAAATTTTTACTATTATATTTGGCATAAAGAGATAGATAATACTCGACCGACATAATGTTGTGTTGTCCATTATAGTATAAAAATTAATATTTCTATTTCTATATATTTGCAACAATTTACTCTCTATAACAAATATCATAACTTAGTTCTTTTATGTCAAGAAGGCACTGGTTTAATTCATCTATAAATGTCACTCCATAACTACCACGCATACTATACTCAGAATTATGATAAAGATATTTATTCTTGGGGTGTAAGTAATGGGGATTAATCTTTGTTGGATCAGTCTCTAAGTTAACACATGTCACACATGATCCATTTATAGTTATATCACACGATGATGATTTATGAATTGATTCCGGAAGATCGCTATTGTATTTTGTAGTTCCACAATTCATTTCCATACATGTTATTGTCACACTAATATTATGATGAACTTTATCTAGCCGCTGAGTGGTAAACAACAGAACAGATAGTTTATTATCTTTACCAACACCCTCAGCCGCTGCCACAAATCTCTGATCCGTATCCATGATGGTCATGTTTACTTTTAGTCCGTATCCAGTCAACACTATGTTAGCATTTCTGTCGATATAGCTTTCACTCATATGACACTCACCAATAATTGTAGAATTAATGTCGTAATTTACACCAATAGTGAGTTCGGCGACAAAGTACCAGTACCGGTAATCTTGTCGATGAGGACATATAGTATTCTTGTATTCTACCGAATACCCGAGAGATGCGATACAAAAGAGTAAGACTAATTTGTAAACCATCTTACTCAAAAATATGCGACAATAGGAAATCTATCTTATACACATAATTATTCTATCAATTTTACCAATTAGTTAGTGTAATGTTAACAAAAATGTGGGATAATTTAATAGTTTTTCCTTACACAATTGACATACATGAGTCTGAGTTCCTCGTTTTTGCTAATTATTTCGTCCAATTTATTATTCTTGACATCGTCAAGATCTTTTGTATAGGAGTCAGACTTGTATTCAACATGTTTTTCTATAATCATCTTAGCTATTTCGGCATCATCCAATAGTACATTTTCCAGATTAACAGAATAGATATTAATGTCGTATTTGAACAGAGCCTGTAACATCTCAATGTCTTTATTATCTATAGCCAATTTGATGTCCGGAATGAAGAGAAGGGAATTGGTGTTTGTTGACGTCATATAGTCGAGCAAGAGAATCATCATATCCACGTGTCCATTTTTTATAGTGGTGTGAATACAACTAAGGAGAATAGCCAGATCAAAAGGAGATGGTATCTCTGAAAGAAAGTAGGAAACAATACTTACATCATTAAGCATGACAGCATGATAAAATGAAGTTTTCCATCCAGTTTTCCCATAGAACATCAGTCTCCAATTTTTCTTAACAAACAGTTTTACCGTTTGCATGTTACCACTATCAACCGCATAATACAATGCGGTGTTTCCTTTGTCATCAAATTGTGAATCATCCATTCCACTGAATAGCAAAATCTTTACTATTTTGGTATCTTCTAATGTGGCTGCCTGATGTAATGGAAATTCATTATCTAGAAGATTTTTCAATGCTCCAGCGTTCAACAACGTACATACTAGACGCACGTTATTATCAGCTATTGCATAATACAAGACACTATGACCATTGATATCCGCCTTAAATGCATCTTTGCTAGAGAGAAAGCTTTTCAGTTGCTTAGACTTCCAAGTATTAATTCGTGACAGATCCATGTCTGAAACGAGACGCTAATTAGTGTATATTTTTTCATTTTTTATAATTTTGTCATATTGTACCAGAATTAATAATATCTCTAATAGATCTGATTAGTAGATATATGGCTATCGCNAAACAACATATACACATTTAATNAAAATAATATTCATTAAGAAGATTCAGATTCCACTGTACCCATCAATATAAAATAAAATAATTATTCCTTACATCGTACCCATAAACAATATATTAAGTATATTCCACCTTACCCATAAACAATATAAATCCAGTAATATCATGTCTAATGATGAACACAAATGGTGTATTAAATTCCAGTTCTTCAGGAGATGATCTCGCCGTAGCTACCATGATAGTAGATGCCTCCGCTACAGTTCCTTGTTCGTCTACATCTATCTTTACATTCTGAAACATTTTATAAATATATAATGGGTCCCTAGTCATATGTTTAAACGACGCCTTATCTGGATTAAACATACTANGAGCCATCATTTCGGCTATCGACTTAATATNCCTCTTGTTTTCGATAGAAAATCTANNGAGNTTAAGATTGTACATTTTATTCCCTAATTGAGATGACCAATATTCTAATTTTGCAGGCGNGATAGAATCTGTGAAATGGGTCATATTATCACCTATTGCCAGGTACATACTAATATTAGCATCCTTATACAGAAGGCGCACCATATCATATTCTTCGTCATCGATTGTGATTGTATTTCNTTGCAATTTAGTAACTACGTTCATCATGGGAACCGTTTTCGTACCGTACTTATTAGTAAAACTAGCATTGTGTGTTTTAGTGATATCAAACGGATATTGCCACGTACCTTTAAAATATATAGTATTAATGATTGCCCATAGAGTATTATCGTCGAGCATAGTAGAATCAACTACATTAGACATACCAGATCTACGTTCTACTATAGAATTAATTTTATTAACCGCATCTCGTCTAAAGTTTAATCTATATAGGCCGAATCTATGATATTGTTGATAATACGACGGTTTAATACACACAGTACTATCGACGAAACTTTGATACGTTAGATCTGTGTACGTATATTTAGATGTTTTCATCTTAGCTAATCCGGATATTAATTCTGTAAATGCTGGACCCAGATCTCTTTTTCTCAAATTCATAGTATTCAATAATTCTACTCTAGTATTACCTGATGCAGACAATAGCGACATAAACATAGAAAACGAATACCCAAACGGTGAGAAGACAATATTATCATTATCATCCTCATCCCCATTTTGAATATTTTTATACGCTAATATACCAGCATTGATAAATCCCTGCAGACGATATGCGGATACTGAACACGCTAATGATAGTATCAATAACGCAATCATGATTTTTATGGTATTAATAATTAACCTTATTTTTATGTTTGGTATAAAAATTATTGATGTCTACACATCCTTTATAATCAACTCTAATCACTTTAACTTTTACAGTTTTCCCTACAAGTTTATCCCTATATTCAACATATCTATCCATATGCATCTCTTAACACTCTGCCAAGATAGCTTCAAAGTGAGGATAGTCAAAAAGATAAATATATAGAGTATAATCATTCTCGTATACTCTGCCCTTTATTACATCGCCCGCATTGGGCAACGAATAACAAAATGCAAGCATCGTGTTAACGGGCTCGTAAATTGGGATAAAATTATGTTTTTATTGTTTATCTATTTTATTCAAGAGAATATTCAGGAAGTTCCTTTTCTGGTTGTATCTCGTCGCAGTATATATCATTTGTACATTGTTTCATATTTTTTAATAGTCTACACCTTTTAGTAGGACTAGTATCGTACAATTCATAGCTGTATTTTGAATTCCAATCACGTATAAAAATATCTTCCAATTGTTGACGAAGACCTAATCCATCATCCGGTGTAATATTAATAGATGCTCCACATATATCCGTAAAGTAATTTCCTGTCCAATTTGATGTACCTATATACGCCGTTTTATCGGTTACCATATATTTTGCATGGTTTACCCTAGAATACGGAATGGGAGGATCAGCATCTGGTACAATAAATAGCTTTACTTCTATATCTATGTTTTTAGATTTTAGCATAGCTATAGATCTTAAAAAGTTTCTCATGATAAACGAAGATCGTTGCCAGCAACTAATCAATAGCTTAACGGATACTTGTCTGTCTATAGCGGATCTTCTTAATTCATCTTCTATATAAGGCCAAAACAAAATTTTACCCGCCTTTGAATAAATAATAGGAATAAAGTTCATAACAGATACATAAACGAATTTACTCGCATTTCCGATACATGACAATAAAGCGGTTAAATCATTGGTTCTTTCCATAGTACATAATTGTTGTGGTGCAGAAGCAATAAATACAGAGTGTGGAACACCGCTTACGTTAATACTAAGAGGATGATCTGTATTATAATACGACGGATAAAAGTTTTTCCAATTATATGGTAGATTGTTAACTCCAAGATACCAGTATACCTCAAAAATTTGAGTGAGATCCGCTGCCAAGTTCCTATTATTGAAGATCGCAATACCCAATTCCTTGACCTGAGTTAGTGATCTCCAATCCATGTTAGCGCTTCCTAAATAAATATGTGTATTATCAGATATCCAAAATTTTGTATGAAGAACTCCTCCTAGGATATTTGTAATATCTATGTATCGTACTTCAACTCCGGCCATTTGTAGTCTTTCAACATCCTTTAATGGTTTGTTGGATTTATTGACGGCTACTCTAACTCTTACTCCTCTTTTGGGTAATTGTACAATCTCGTTTAATATTACCGTGCCGAAATTCGTACCCACTTCATCCGATAAACTCCAATAAAAAGATGATATATCTAGTGTTTTTATGGTATTGGATAGAATTTCCCTCCACATGTTAAATGTAGTCAAATATACTTTATCAAATTGCATACCTATAGGAATAGTCTCTGTAATCACTGCGATTGTATTATCCGGATTCATTTTATTTGTTAAAAAAATAATCCTATATCACTTCACTCTATTAAAAATCCAAGTTTCTATTTCTTTCATGACTGATTTTTTAACTTCATCCGTTTCCTTATGAAGATGATGTTTGGCGCCTTCATAAATTTTTATTTCCCTATTACAATTTGCATGTTGCATGAAATAATATGCACCTGAAACATCGCTAATCTCATTGTTTGTTCCCTGGAGTATGAGAGTCGGGGTGTTAATCTTGGGAATTATTTTTCTAACCTTGTTGGTAGCCTTCAAGACCTGACTAGCAAATCCAGCCTTAATTTTTTCATGATTGACTAATGGATCGTATTGGTATTTATAAACTTCATCCATATCTCTAGATACTGATTCTGGACATAGCTTTCCGACTGACGCATTTGGTGTAATGGTTCCCATAAGTTTTGCAGCTAGCAGATTCAGTCTTGGAACAGCGTCTGCATTAACTAGAGGAGACATTAGAATCATTGCTGTAAACAAGTTTGGATTATCGCAAGCAGCTAGTATAGAAATTGTTGCTCCCATGGAATGACCCAATAAGAAGACTGGAACTCCTGGATAAGTAGATTTAATAGTCACCACGTGCTGTACCACATCTCTAACATACTTACCAAAGTCATCAATCATCATTTTTTCACCATTACTTCTTCCATGGCCAATATGATCATGTGAGAATACTAAAATTCCTAACGATGATATGTTTTCAGCTAGTTCGTCATAACGTCCAGAATGTTCACCAGCTCCATGACTTATGAATACTAATGCCTTAGGATATGTAATAGGTTTCCAATATTTACAATATATGTAATCATTGTCCAGATTGAACATACAGTTTGTACTCATGATTCACTATATAACTATCAATATTAACAGTTCGTTTAATGATCATATTATTTTTATGTTTTATTGATAATTGTAAAAATATACAATTAAATCAATATAGAGGAAGGAGACGGTACTGTATTTTGTGAGATAGTAATGGAGACTAAATCAGATTATGAGGATGCTGTTTTTTACTTTGTGGATGATGATGAAATATGTAGTCGCGACTCCATCATTGATCTAATAGATGAATATATCACGTGGAGAAATCATGTTATAGTGTTTAATAAAGATATTACCAGTTGTGGAAGACTGTACAAGGAATTGATAAAGTTCGATGATGTCGCTATACGGTACTATGGTATTGATAAAATTAATGAGATTGTCGAGGCTATGAGTGAAGGAGACCACTACATCAATCTTACAGAAGTCCATGATCAGGAAAGTCTATTCGCTACCATAGGAATATGTGCTAAAATCACTGAACATTGGGGATACAAAAAGATTTCAGAATCTAAATTCCAATCATTGGGAAACATTACAGATCTGATGACCGACGATAATATAAACATCTTGATACTTTTTCTAGNAAAAAAAATGAATTGATGATATAAGTGTCTTCATAACGCATTATTACGTTAGCATTCTATTATCCAGTGTTNAAAAAATTATCCTATCATGTATTTGAGAGTCTTATATGTAGCAAACATGATAACTGCAATACCCATAATCTTTAGATATTCACGCGTGCTATGGATGGCATTATNCCGCGGTGCGGAAATGTACGTTATATAATCTACAAAATAATCATCGCATATAGTATGAGATAGTAGAGTAAACATTTTTATCGTTTCTACTGGGTTCATACATCGTCTACCCAATTCGGTAATGAATGAAATTGTCGCCAATCTTACACCCAAACCCTTGTTGTTCATTAGTATAGTATTAACTTCATTATTTATGTCATAAACTGTAAATGATTCTGTAGATGCCATATCACACATGATATTCATGTCACTATTATAATCATTATTAACTTTATCACAATACGTGTTGATAATATCTACATATGATCTAGTTTTTGTGGGTAATTGTCTATACAAGTCGTCTAAACGTTGTTTACTCATATAGTATTGAACAGCCATCATTACATGGTCCCGTTCCGTTGATAGATAATCGAGTATGTTAGTAGACTTGTCAAATCTATATACCATATTTTCTGGAAGCGGATATACATAGTCGCGATCATCATTATCACTAGCCTCATCCTCTATATCATGTACATGTACATAATCTATGATATTATTATACATAAACATCGACAACATACTATTGTCTATTATCTAAGTCCTGTTGATCCAAACCCTTGATCTCCTCTATCTGTACTATCTAGAGATTGTACTTCTTCAAGTTCTGGATAATATATACGTTGATAGATTAGCTGAGCTATTCTATCTCCAGTATTTACATTAAACGTACATTTTCCATTATTAATAAGAATGACTCCTATGCTTCCCCTATAATCTTCGTCTATTACACCGCCTCCTATATCAATGCCNTTTAGGGACAGACCAGACCTAGGAGCTATTCTACCATAGCAGAACTTAGGCATGGACATACTAATATCTGTCTTAATTAACTGTCGTTCTCCAGGAGGGATAGTATAATCGTAAGCGCTATACAAATCATATCCGGCAGCACCCGGCGATTGCCTAGTAGGTGATTTAGCTCTGTTAGTTTCCTTAACAAATCTAACTGGTGAGTTAATATTCATGTTGAACATAAAAAATATCATTTTATTTCAAAATTATTTACCATTCCATTCCATCCCATATATTCCATGAATAAGTGCGATTATTGTACACTTCTATAGTATCTATATACGATCCACGATAAAATCCTCCTATCAATAGCAGTTTATTATCCACTATGATCAATTCTGGATTATCCCTCGGATAAATAGGATCATCTATCAGAGTCCATGTATTGCTGGATTCACAATAAAATTCCGCATTTCTACCAACCAAGAATAACCTTCTACCAAACACTAACGCACATGATTTATAATGAGGATAATAAGTGGATGGTCCAAACTGCCACTGATCATGATTGGGTAGCAAATATTCTGTAGTTGTATCAGTTTCAGAATGTCCTCCCATTACGTATATAACATTGTTTATGGATGCCACTGCTGGATTACATCTAGGTTTCAGAAGACTCGGCATATTAACCCAAGCAGCATCCCCGTGGAACCAACGCTCAACAGATGTGGGATTTGGTAGACCTCCTACTACGTATAATTTATTGTTAGCGGGTATCCCGCTAGCATACAGTCTGGGGCTATTCATCGGAGGAATTGGAATCCAATTGTTTGATATATAATTTACCGCTATAGCATTGTTATGTATTTCATTGTTCATCCATCCACCGATAAGATATACTACTTCTCCAACATGAGTACTTGTACACATATGGAATATATCTATAATTTGATCCATGTTCATAGGATACTCTATGAATGGATACTTGTATGATTTGCGTGGTTGTTTATCACAATGAAATATTATGTTACAGTCTAGTATCCATTTTACATTATGTATACCTCTGGGAGAAAGATAATTTGACCTGATTACATTTTTGATAAGAAGTAGCAGATTTCCTAATCTATTTCTTCGCCTCATATACCACTTAATGACAAAATCAACTACATAATCCTCATCTGGAACATTTAGTTCGTCGCTTTCTAGAATAAGTTTCATAGATAGATAATCAAAATTGTCTATGATGTCATCTTCCAGTTCCAAAAAGTGTTTGGTAATAAAGTCTTTAGTATGACATAAGAGATTGGATAGTCCGTATTCTATACCCATCATGTAACACTCGATACAATATTCCTTTCTAAAATCTCGTAGGATAAAGTTTATACAAGTGTAGATGATAAATTCTACAGATGTTAATATAGAAGCACGTAATAAATTGACGACGTTATGACTATCTATATATACCTTTCCAGTATATGAGTAAATAACTATAGAAGTTAGACTGTGAATGTCAAGGTCTAGACAAACCCTCGTAACTGGATCTTTATTTTTTGTGTATTTTTGGCGTAAATGTGTGCAAAAGTATGGAGATAACTTTTTCAATATCGTAGAATTGACTATTATATTACCTCCTATAGCTTCAATAATTGTTTTGAATTTCTTAGTCGTGTACAATGCTAATATATTCTTACAGTACACAGTATTGACAAATATCGGCATTTATGTTTCTTTAAAAGTCAACATCTAAAGAAAAATGATTGTCTTCTTGAGACATAACTCCCATTTTTTGGTATTCACCCACACGTTTTTCGNAAAAATTAGTTTTTCCTTCCAATGATATATTTTCCATGAAATCAAACGGATTGGTAACATTGTAAANTTTTTTAAATCCCAATTCAGAAATCAATCTATCTGCGACGAATTCTATATATGTTTTCATCATTTCACAATTCATTCCTATGAGTTTAACTGGAAGAGCCACAGTAAGAAATTCTTGTTCAATGGATACCGCATTTGTTATAATAAATCTAACGGTTTCTTCACTCGGTGGATGCAATAAATGTTTAAACATCAAACATGCGAAATCGCAGTGCAGACCCTCGTCTCTACTAATTAATTCGTTAGAAAACGTGAGTCCGGGCATTAGGCCACGCNTTTTAAGCCAAAATATGGAAGCGAATGATCCGGNAAAGAAGATTCCTTCTACTGCAGCAAAGGCAATAAGTCTCTCTCCATAACCGGCGCTGTCATGTATCCACTTTTGAGCCCAATCGGCCTTCNTTTTTACACAAGGCATCGTTTCTATGGCATTAAAGAGGTAGTTTTTTTCATTACTATCTTTAACATAAGTATCGATCAAAAGACTATACATTTCCGAATGAATGTTTTCAATGGCCATCTGAAATCCGTAGAAACATCTAGCCTCGGTAATCTGCACTTCTGTACAAAATCGTTCTGCTAAATTTTCATTCACTATTCCGTCACTGGCTGCAAAAAACGCCAATACATGTTTTATAAAATATTTTTCGTCTGGTGTTAGTTTATTCCAGTCATTGATATCTTTAGATATATCCACTNNTTCCACTGTCCAAAATGATGNCTCTGCCNTTTTATACATATTCCAGATGTCATGATATTGGATTGGGAAAATAACAAATCTATTTGGATTTGGTGCAAGGATAGGTTCCATAACTAAATTAACAATAGTAGTAANTTTTTTTCAGTTATCTGTATGACGACTGTACTTGGATCTTTTGTATATCGCTATCGCCGCAATCACTACAATAATTACAAGTATTATTGATAGCATTGTTATTACTACTATCATAATTAAATTATCGACATTCATGGGTGTTGAATAATCGTTATCATCATTTTGTAATTGTGACGTCATACTAGATAAATCATTTGTGAGATTGTTGTGGGAAGCGGGCACGGAAGATGCATTATCATTATTATTTAACGCCTCCCATTCGGATTCACAAATGTTACGCACGTTCAAAGTTTTATGGAAACTATAATTTTGTGAAAACAGATAACAAGAAAACTCGTCATCGTTCAAATTTTTAACGATAGTAAACCGATTAAACGTCGAGCTAATTTCTAACGCTAGCGACTCTGTTGGATATGGGTTTCCAGATATATATCTTTTCAGTTNCCCTACGTATCTATAATCATCTGTAGGAAATGGAAGATATTTCCATTTATCTACTGTTCCTAATATCATATGCGGTGGTGTAGAACCATTAAGCGCGAAAGATGTTATTTCGCATCGTATTTTAACTTCGCAATAATTTCTGGTTAGATAACGCACTCTACCAGTCAAGTCAATGATATTAGCCTTTACAGATATATTCATAGTAGTCGTAACGATGACTCCATCTTTTAGATGTGATACTCCTTTGTATGTACCAGAATCTTCGTACCTCAAACTCGATATATTTAAACAAGTTAATGATATATTAACGCGTTTTATGAATGATGATATATAACCAGAAGTTTTATCCTCTGTGGCTAGCGCTATAACCTTATCATTATAATACCAACTAGTGTAATTAATATGTGACATGACAGTGTGGGTACAAATATGTACATTATCGTCTACGTCGTATTTGATACATCCGCATTCAGCCAACAAATATAAAATTACAAAAACTCTAACGACGTTTGTACACATCTTGATGCGGTTTAATAAATGTTTTGATTTCAATTTATTGTNNAAAAAGATTCGGTTTTATACTGTTCGATATTCTCATTGCTTATATTCTCATCTATCATCTCCACACAGTCAAATCCATGGTTAACATGTACCTCATCAACCGGTAAAAGACTATCGGATTCTTCTATCATCATAACTCGAGAATATTTAATTTGGTGGTCATTATTAATCAAGTCAATTATCTTATTTTTAACAAACGTAAGTATTTTACTCATTTTTTTATAAAAACTTTTAGAAATATACAGACTCTATCGTGTGTCTATATCTTCTTTTTATATCCAATGTATTTATGTCTGATTTTTCTTCATTTATCATATATAATGGTCCAAATTCTACACGTGCTTCGGATTCATCCAGATCATTAAGGTTCTTATAATCGCAACATCCTTCTCTTCCATCTTCTACATCTTCCTTCTTATTCTTAGCGTCACAGAATCTACCACAGCAGGATCCCATGACGAGAGTCACATTAAACTAATTCATTTTCAATTATAATATACTGATTAGTAATGACAATTAAAATAAAAATATTCTTCATAACCGGTAAGAAAGTAAAAAGTTCACATTGAAACTATGTCAGTAGTTATACATCATGAGATGATATACTCTATTTTGGTGGAGGATTATATGATATAATTCGTGGATAATCATTCTTAAGACACATTTCTTCATTCGTAAATCTTTTCACATTAAATGAGTGTCCATATTTTGCAATTTCTTCATATGATGGCGGTGTACGTGGACGAGGCTGCTCCTGTTCTTGTAGTCGTCGACTGTCGTGTTTGCGTTTAGATCCCTCCATTATCGCGATCGCGTAGTGAGTACTATTTATACCTTGTAATTAAATTTTTTTATTAATTAAACGTATAAAAACGTTCCGTATCTGTATTTAAGAGCCAGATTTCGTCTAATAGAACAAATAGCTACAGTAAAAATAACTAGAATAATCGCTACACCCACTAGAAACCACGGATCGTAATACGGCAATCGGTTTTCGATAATAGGTGGAACGTATATTTTATTTAAGGACTTAACAATTGTCTGTAAACCACAATTTGCTTCCGCCGATCCTGTATTAACTATCTGTAAAAGCATATGTTGACCGGGAGGAGCCGAACATTCTCCGATATTCAATTTTTGTATATCTATAATGTTATTAACCTCCGCATACGCATTACAGTTCTTTTCTAGCTTGGATACTACACTAGGTACATCATCTAAATCTATTCCTATTTCCTCAGCGATAGCTCTTCTATCCTTTTCCGAAAGTAATGAAATCACTTCAATAAATGATTCAACCATGAGTGTGAAACTAAGTCGAGAATTACTCATGCATTTGTTAGTTATTCGGAGCGCGCAATTTTTAAACTGTCCTATAACCTCNCCTATATGAATAGCACAAGTGACATTAGTAGGGATAGAATGTTGAGCTAANTTTTGTAAATAACTATCTATAAAAAGATTATACAAAGTTTTAAACTCTTTAGTTTCTGCCATTTATCCAGTCTGAGAAAATGTCTCTCATAATAAATTTTTCCAAGAAACTAATTGGGTGAAGAATGGAAACCTTTAATCTATATTTATCACAGTCTGTTTTGGTACACATGATGAATTCTTCTAATGCTGTACTAAATTCGATATCTTTTTCGATTTCTGGATATGNTTTTAATAAAGTATGAACAAAGAAATGGAAATCGTAATACCAGTTATGTTCAACTTTGAAATTGNTTTTTATTTTCTTGTTAATGATTCCAGCCACTTGGGAAAAGTCAAAGTCGTTTAATGCCGATTTAATACGTTCATTNAAAACAAACTTTTTATTCTTTAGATGAATTATTATTGGTTCATTGGAATCNAAAAGTAAGATATTATCGGGTTTAAGATCTGCATGTNAAAAGTTGTCACAACAGGGTNGTTCGTAGATTTTANNGTATAACNGAGACATNNNTNNAAAGATAAACTTTNTGTATTGTACCAAAGATTTAAATCCTAATTTGATAGCTNNNTCGGTATCTACTTTATCTGCCGAATACAGTGCTNNNGNNAAAATTATAATNTTTNNNCTTTCGTATTCGTAATTNGTTCNCTTTTCATGTTCGNAAAAGTGAAACNTGCGGTTAAAATNGTTTATAACATTAATATTACTGTTNNNNNCTGNCGGATNAAAGTGGGATAGTAATTTCACGAATTTGATACTGTCCTTNNNNTCGTTAAACGCCTTTAGNAAAACTNNNGAAGAATATCTCAATGNGAGTTCCTGACCATCCNNNNNNNNNNNNNNNNNNNNNNNNNNNNNNNNNNNNNNNNNNNNNNNNNNNNNNNNNNNNNNNNNNNNNNNNNNNNNNNNNNNNNNNNNNNNNNNNNNNNNNNNNNTTTTTTCATCTNCCTTTAGATTGTTGTATAGAAATTTGGGTACTGTAAACTCCNNCGTAGTTTCCATGGGACTATANNNNNNTGTNNCCTNGAATNNNNNNTTTACTACATAGTTATCTATCTTAAAGACTATACCATATCNNCCTGTAGATATGTGATNAAAATCGTCGTTTATAGGATAAANNCGTTTATCNNTTTGTTGNNNAAAGGATGAATTAATGTAATCATTCNNNTCTATCTTTAGTAGTGNTTCCNTATTAAAATTNTTAAAATAATTTAACAATCTAACTGACGGAGCCCNNNNTTNGTGTNAATCTAATTGGGACATTANNTTGTTAAAATATAAANNNTCTCCTAATATAACAGTATCTGATAANCTATNGGGAGNCATCCATTGATATTCANGGGATGAATCATTNGNAACACCCATTTATTGTACAAAAAGCCCCAATTTACAAACGAAAGTCCAGGTTTGATAGAGATAAACTATTAACTATTTTGTCTCTGTTTTTAACACCTCCACAGTTTTTAATTTCTTTGGTAATGAAATTATTCACAATATCAGTATCTTCTTTATCTACCAGAGATTTTACTAACTTGATAACCTTGGCTGTCTCATTCAATAGGGTAGTGATATTTGTATGTATGATATTGATATCNTTTTGAATTGTTTCTTTTAGAAGTGATTCTTTGATGGTATCAGCATACGAATTACAATAATGCAGAAACTCAGTTAACATGCAGGAATTATAGTAAGCCAATTNCAATTGTTGCCTGTATTGTATTANAGTATTAATATGCGCAATGATGTCCTTGCGTTTCTCTGATAGAATGCGAGCAGCGATTTTGGCGTTATCATTTGACGATATTTCTGGAATGACGAATCCTGTTTCTACTAACTTCTTGGTAGGACAAAGTGAAACAATCAAGAAAATAGCTTCTCCTCCTATTTGTGGAAGAAATTGAACTCCTCTAGATGATCTACTGACGATAGTATCTCCTTGACAGATATTGGACCGAACTANGGAAGTACCTGGAATGTAAAGCCCTGAAACCCCCTCANTTTTTAAGCAGATTGTTGCCGTAAATCCTGCACTATGCCCAAGATAGAGAGCTCCTTTGGTGAATCCATCACTATGTTTCAGTTTAANCANNAAACAGTCAGCTGGTCTAAAATTTCCATCTCTATCTAATACAGAATCCAACTTGATGTCAGGGACNATGNNCGGNTTAATGTNNTATGTAACATTGAGTAAATNCTTAAGTTCATAATCATCGTTGTCATCAGTTATGTACGATNCAAACAATGTTTCTANCGGCNNNNNNNNNNNNNNNNNNNNNNNNNNNNNNNNNNNNNNNNNNNNNNNNNNNNNNNNNNNNNNNNNNNNNNNNNNNNNNNNTNNCGAATCAGTAACTATGTTCANNTTTTTANNNGTANGACGTCTAGCCAGTATNGANNAGGATTTTCNAGATANNNNNNNNNNNNNNNNNNNNNNNNNNNNNNNNNNNNNNNNNNNNNNNNNNNNNNNNNNNNNNNNNNNNNNNNNNNNNNNNNNNNNNNNNNNNNNNNNNNNNNNNNNNNNNNNNNNNNNNNNNNNNNNNNNNNNNNNNNNNNNNNNNNNNNNNNNNNNNNNNNNNNNNNNNNNNNNNNNNNNNNNNNNNNNNNNNNNNNNNNNNNNNNNNNNNNNNNNNNNNNNNNNNNNNNNNNNNNNNNNNNNNNNNNNNNNNNNNNNNNNNNNNNNNNNNNNNNNNNNNTTCTATGAGTTGTTNCAATAAATTATTTGTTGTGACTGCGTTCAAGNCATAAATTANNTTGATACTATCCAGTAAACAGTCTTTAAGTTCTGGAATATTATCATCCCATTGTAAAGCCCCTAGTTCGACTATCGAATATCCTGCNCTGATAGCAGTTTNAATATCGACGGACGTCAATACTGTAATAAAGGTGGTAGTATTGTCATCATCGTGATAAACTACGGGAATATGGTCGTTAGTAGGTACCGTGACTTTACACAACGCGATATATAACTTTCCNTTTGTACCANTTTTAACGTAGTTGGGACGTCCTGCAGGGTATTGTTTTGAAGAAATGATATCGAGAACAGATTTGATACGATATTTGTTGGATTCCTGATTATTCACTATAATATAATCTAGACAGATAGATGATTCGATAAATAGAGAAGGTATATCGTTGGTAGGATAATACATCCCCATTCCAGTATTCTCGGATACTCTATTGATGACACTAGTTNAGAACATGTCTTCTATTCTNGAAAACGAAAACATCCTNNATNGACTCATTAAAACTTNTAACGNNCCNGNTTGTGTTTCNAATGCCTNNNNCAAGGANNTCAAGGATGCCATAGATNNTTTGACCAACGATTTAGTATTGCGTTTAGCATCTGANTTTTTTATTAAATCAAATNGTCGGCTCTCTGGTTTACTACCCCAANGATAACAATAGTCTTGTAAAGATANNNNNNNNNNNNNNNNNNNNNNNNCCATCCAAATAANCCTANNANCGTCGGATGATATTAATGTATTATTATAGATTTTCCATCCACAGTTATTGGGCCAGTATACTGTTAGCAACGGTATATCGAATAGATTACTCATGTAACCTACTAGAATGATAGTTCGTGTACTAGTCATAATATCTTTAATCCAATCTAAGAAATCTAAAATTAGATCTTTTACACTATTAAAGTTAACAAAGGTATTACCCGGGTACGTGGATATCATATATGGTATTGGTCCATTATCAGTAATGGCTCCATAAACTGATACGGCAATGGTTTTTATATGTGTTTGATCTAATGAGGACGAAATTCGCGCCCACAATTCATCTCTAGATATGCATTTAATATCGAACGGTAACACATCAATCTCGGGACGCGTATATGTTTCTAAATTCTTAATCCAAATATAATGATGACCTATATGCCCTATTATCATACTGTCAACTATAGTATACCTAGAGAACTTTCGATACATCTGCTGTTTCCTGTAATCGTTAAATTTTACAAATCTATAACATGCTAAACCNTTTGACGACAGCCATTCATTAATTTCTGATATGGAATCTGTATTCTTAATACCGTATCGTTCTAAAGCCAGTGCTATATCTCCCTGTTCGTGGGAACGCTTTCGTATAATATCGATCAATGGATAATATGAANTTTTTGGAGAATAATATGATTCATGATCTATTTCGTCCATAAACAATCTAGACATAGGAATTGGAGGCGATGATCTTAATTTTGTGCAATGGGTCAATCCTATAACTTCTAATATTGTAATATTCATCATCGACATAACACTATCTATGTTATCATCGTATATTAGTATACCACGACCTTCTTCATTTCGTGCCAAAATGATATACAGTCTTAAATAATTACGCAATATCTCAATAGTTTCATAATTGTTAGCTGTTTTCATCAAGGTTTGTATCCTGTTTAACATGATGGCGTTCTATAACGTCTCTATTTTCTATTTTTAANTTTTTTAAATTTTTAACGATTTACTGTGGCTAGATACCCAATCTCTTTCAAATANTTTTTTAGCCTTGCTTACAAGCTGTTTATCTATACTATTAAAACTGACGAATCCGTGATTTTGGTAATGGGTTCCGTCGAAATTTGCCGAAGTGATATGAACATATTCGTCGTCGACTATTAACAATTTTGTATTATTCTGAATAGTGAAAACCTTCACAGATAGATCATTTTGAACACACAACGCATCTAGACTTCTGGCGGTTGCCATAGAATATACGTCGTTCTTATCCCAATTACCAACTAGAAGTCTGATCTTAACTCCTCTATTAATGGCTGCTTCTATAATGGAGTTGTAAATGTCAGGCCAATAGTAGCTATTACCGTCGACACGTGTAGTGGGAACTATGGCCAAATGTTCAATATCTATACTAGTCTTAGCCGACTTGAGTTTATCAATAACTACATCAGTGTCTAGATCTCTAGAATATCCCAATAGGTGTTCTGGAGAATCAGTAAAGAACACTCCACCTATAGGATTCTTAATATGATACGCAGTGCTAACTGGCAGACAACAAGCCGCAGAGCATAAATTCAACCATGAATTTTTTGCGCTATTAAAGGCTTTNAAAGTATCAAATCTTCTACGAAGATCTGTGGCCAGCGGAGGATAATCAGAATATACGCCTAACGTTTTAATCGTATGTATAGATCCTCCAGTAAATGACGCGTTTCCTACATAACATCTTTCATCATCAGACACCCAAAAACAACCGAGTAGTAGTCCCACATTATTTTTTTTATCTATATTAACGGTTATAAAATTTATATCCGGGGAGTGACTTTGTAGCTCTCCCAGATTTCTTTTCCCTCGTTCATCTAGCAAAACTATTATTTTAATCCCTTTTTCAGATACCTCTTTTAGTTTATCAAAAATAAGCGCTCCCCTAGTAGTACTCAGAGGATTACAACAAAAAGATGCTATGTATATATATTTCTTAGCTAGAGTGATAATTTCGTTAAAACATTCAAATGTTGTCAAATGATCGGATCTAAAATCCATATTTTCTGGTAGTGTTTCTACCAGCCTACANTTTGCTCCCGCAGGTACCGATGCAAATGGCCACATTTAGTTAACATAAAAACTTATATATCCTGTTCTATCAACGATTCTAGAATATCATCGGCTATATCGCTAAAATTTTCATCAAAGTCGACATCACAACCTAACTCAGTCAATATATTAAGAAGTTCCATGATGTCATCTTCGTCTATTTCTATATCCGTATCCATTGTAGATTGTTGACCGATTATCGAGTTTAAATCATTACTAATACTCAATCCTTCAGAATACAATCTGTGTTTCATTGTAAATTTATAGGCGGTGTATTTAAGTTGGTAGATTTTCAATTATGTATCAATATAGCAACAGTAGTTCTTGCTCCTCCTTGATTTTAGCATCCTCTTCATTATTTTCTTCTACGTACATAATCATGTCTAATACGTTAGACAACACACCGACGATGGTGGCCGCCACAGACACGAATATGACTAGACCGATGACCATTTAAAAAATACTCTCTAGCTTTAACTTAAACTGTATCGATCATTCTTTTAGCACATGTATAATATAAAAACATTATTCTATTTCGAATTTAGGCTTCCAAAAATTTTTCATCCGTAAACCGATAATAATATATATAGACTTGTTAATAGTCGGAATAAATATATTAATGCTTAAACTATCATCATCTCCACGATTAGAGATACAATATTTACATTCTTTTTGCTGTTTCGAAACTTTATCAATACACGTTAATACAAACCCAGGAAGGAGATATTGAAACTGAGGCTGTTGAAAATGAAACGGCGAATACAATAATTCAGATAATGTAAAATCATGATTCCGTATTCTGATGATATTAGAACTGCTAATGGATGTCGATGGTATGTATCTAGGAGTATCTATTTTAACAAAGCATCGATTTGCTAATATACAATTATCCTTTTGATTAATTGTTATTTTATTCATATTCTTAAAAGGTTTCATATTTATCAATTCTTCTACATTAAAAATTTCCATTTTTAATTTATCTAGCCCCGCAATACTCCTCATTACGTTTCATTTTTTGTCTAGAATACCCATTTTGTTCATCTTGGTACATAGATTATCCAATTGAGAAGCGCATTTAGTAGTTTTGTACATTTTAAGTTTATTAACGAATCGTCGAAAACTAGTTATAGTTAACATTTTATTATTTGATACCCTGATATTAATACCCCTGCCGTTACTATTATTTATAACTGATGTAACCCACGTAACATTGGAATTAATTATCGATAGTAATGCATCGACACTTCCAAAATTGTCTATTATAAACTCACCGATAATTTTTTTATTGCATGTTTTCATATTCATTAGGATTATCAAATCTTTAATCTTATTACGATTGTATGCGTTGATATTACAAGACGTCATTCTAAAAGACGGAGGATTTCCATCAAATGCCAGACAATCACGTACAAAGTACATGGAAATAGGTTTTGTTCTATTACGCATCATAGATTCATATAAAACACCCGTAGAAATACTAATTTGTTTTACTCTATAAAATACTATTGCATCTATTTCATCGTTTTGTATAACGTCTTTCCAAGTGTCAAATTCCAATTTTTTTTCATTGATAGTACCAAATTCTTCTATCTCTTTAACTACTTGCATAGATAGGTAATTACAGTGATGCCTACATGCCGTTTTTTGAAACTGAATAGATGCATCTAGAAGCGATGCTACACTAGTCACGATCACCACTTTCATATTTAGAATATATGTATGTAAAAATATAGTAGAATTTCATTTTGTTTTTTTCTATGCTATAAATGAATTCTCATTTTGTATCCGCACATACTCCGTTTTATATCAATACCAAAGAAGGAAGATATCTGGTTCTAAAAGCCGTTAAAGTATGCGATGTTAGAACTGTAGAATGCGAAGGAAGTAAAGCTTCCTGCGTACTCAAAGTAGATAAACCCTCATCACCCACGTGTGAGAGAAGACCTTCGTCCCCGTCCAGATGCGAGAGAATGAATAACCCTGGAAAACAAGTCCCGTTTATGAGAACGGACATGTTACAAAATATGTTTGCTGCTAATCGCGACAACGTAACGTCAAGACTTTTGAACTAAAATACAATTATATCTTTTTCGATATTAATAAATCCGTGTCTCCCGGGTTTTTTATCTCTTTCAGTATGTGAATAGATAGGTATTTTATCTCTATTCATCATCGAATTTAAGAGATCCGATAAACATTGTTTGTATTCTCCAGATGTCAGCATCTGATACAACAATATATGTGCACATAAACCTCTGGCACTTATTTCATGTACCTTCCCCTTATCACTAAGGAGAATAGTATTTGAGAAATATGTATACATGATATTATCATGTATTAGATATACAGAATTTGTAACACTCTCGAAATCACACGATGTGTCGGCGTTAAGATCTAATATATCACTCGATAACACATTTTCATCTAGATACACTAGACATTTTTTAAAGCTAAAATAGTCTTTAGTAGTAACAGTAACTATGCGATTATTTTCATCGATGATACATTTCATCGGCATATTATTACGCGTACCATCAAAGACTATACCATGTGTATATCTAACGTATTCTAGCATAGTTGCCATACGTACATTAAACTTTTCAGGATCTTTGGATAGATCTTCCAATCTATCTATTTGAGAAAACATTTTTATCATGTTCAATAGTTGAAACGTCGGATCCACTATATAGATATTATCTATAAAGATTTTAGGAACTATGTTCATGGTATCCTGGCGAATATTAAAACTATCAATGATATGATTATCGTTTTCATCTTTTATCACCATATAGTTTCTAAGATATGGGATTTTACTTAATATAATATTATTTCCCGTAATAAATTTTATTAGAAATGCCAAATCTATAAGAAAAGTCCTAGAATTAGTCTGAAGAATATCTATATCACCGTACCGTATATTTGGATTAATTAGATATAGAGAATATGATCCGTAACATATACAACNTTTATTATGACGTCTAAGATATTCTTCCATCAACTTATTAACATTTTTGACTANGGAAGATACATTATGACGTCCCATTACTTTTGCCTTGTCTATTACAGCGACGTTCATAGAATTTAGNATATCTCTTGCCAATTCTTCCATTGATGTTACATTATAAGAAATTTTAGATGAAATTACATTTGGAGCTTTAATAGTAAGAACTNCTAATATATCCGTGTATGTGGTCACTAATACAGATTGTAGTTCTATAATCGTAAATAATTTACCTATATTATATGTTTGAGTTTGTTTAGAAAAGTAGCTAAGTATACGATCTTTTATTTCTGATGCCGATGTATCAACATCNNNAAAAAATCNTTTTTNATTNNTTTTTACTAACGATACGAATATGTCTTTGTTAAAAACAGTTATTTTCTGAATATTTCTAGCTTGTAATTTTAACATATGATATTCGTTCACACTAGGTACTCTGCCTAAATAGGTTTCTATAATCTTTAATGTAATATTAGGAAGAGTATTCTGATCAGGATTCCTATTCATTTTGAGGATTTAAAACTCTGATTATTGTCTAATATGGTCTCAACACAAACTTTTTCACAGAGTGATAGAGTTTTTGATAACTCNTTTTTCTTAAGAAATATAAAACTACTGTCTCCAGAGCTCGCTCTATCTTTTATTTTATCTAATTCGATACAAACTCCTGATACTGGTTCAGAAAGTAATTCATTAATTTTCAGTCCTTTATAGAAGATATTTAATATAGATAATACAAAATCTTCAGTTCTTGATATCGATCTGATTGATCCTAGAACTAGATATATTAATAACGTGCTCATTAGGCAGTTTATGGCAGCTTGATAATTAGATATAGTATATTCCAGTTCATATTTATTAGATACCGCATTGCCCAGATTTTGATATTCTATGAATTCCTCTGAAAATAAATCCAAAATAACTAGACATTCTATTTTTTGTGGATTAGTGTACTCTCTTCCCTCTATCATGTTCACTACTGGTGTCCACAATGATAAATATCTAGAGGGAATATAATATAGTCCATATGATGCCAATCTAGCAATGTCGAATAACTGTAATTTTATTCTTCGCTCTTCATTATGAATTGAATCTTGAGGTATAAACCTAACACAAATTATATCATTAGACTTTTCGTATGTAATGTCTTTCATGTTATAAGTTTTTAATCCTGGAATAGAATCTATTTTAATGAGGCTTTTAAACGCAGCGTTCTCCAACGAGTCAAAGCATAATACTCTGTTGGTTTTCTTATATTCAATATTACGATTTTCTTCTTTGAATGGAATAGGTTTTTGAATTAGTTTATAATTACAACATAATAGATAAGGAAGTGTGTAAATAGTACGCGGAAAAAACATAATAGCTCCCCTGTTTTCATCCATGGTTTTAAGTAAATGATCACTGGCTTCTTTAGTCAATGGATATTCGAACATTAACCGTTTCATCATCATTGGACAGAATCCATATTTCTTAATGTAAAGAGTGATCAAATCATTGTGTTTATTGTACCATCTTGTTGTAAATGTGTATTCGGTTATCGGATCTGCTCCTTTTTCTATTAAAGTATCGATATCGATCTCGTCTAAGAATTCAACTATATCGACATATTTCATTTGTATACACATAACCATTACTAACGTAGAATGTATAGGAAGAGATGTAACGGGAACAGGGTTTGTTGATTCGCAAACTATTCTAATACATAATTCTTCTGTTAATACGTCTTGCACGTAATCTATTATAGATGCCAAGATATCTATATAATTATTTTGTAAGATGATGTTAACAATGTGATCTATATAAGTAGTGTAATAATTCATGTATTTCGATATATGTTCCAACTCTGTCTTTGTGATGTCTAGTTTCGTAATATCTATAGCGTCCTCAAAAAATATATTCGCATATATTCCCAAGTCTTCAGTTCTATCTTCTAAAAAATCTTCAACGTATGGAATATAATAATCTATTTTACCTCTTCTGATGTCATTAATGATATAGTTTTTGACACTATTTTCCGTCAATTGATTCTTATTCACTATGTCTAAAAACCGGATAGCGTCCCTAGGACGAACTACTGCCATTAATATCTCTATTATAGCTTCTGGACATAAATCATCTATTATACCAGAATTAATGGGAACTATTCCGTATCTATCTAACATAGTTTTAAGAAAGTCAGAATCTAAGACCTGATGTTCATATATTGGTTCATACATGAAATGATCTCTATTGATGATAGTGACTATTTCATTCTCTGAAAATTGGTAACTCATTCTATACACGCTTTCCTTGTTGATAAAGGATAGTATATACTCAATGGAATTTGTACCAACAAACTGTTCTCTTATGAATCGTATATCATCATCTGAAATGATCATGTAAGGCATACATTTAACAATAAGAGACTTGTCTCCTGTTATCAATATACTATTCTTGTGATAATTTATGTGTGCGGCAAATTTGTCCACGTTCTTTAATTTTGTTATAGTAGATATCAAATCCAATGGAGATACAGTTCTTGGCTTAAACAGATATAGTTTTTCTGGAACGAATTCTACAACATTATTATAAAGGACTTTGGGTATATAAGTGGGATGAAATCCTATTTTAATTAATGCGATAGCCTTGTCCTCGTGCAGATATCCAAACGCTTTTGTGATAGTATGGCATTCATTGTCTAGAAACGCTCTACGAATATCTGTAACAGATATCATCTTTAGAGAATACTAGTCGCGTTAATAGTACTAAAATTTGTATTTTTTAATCTATCTCAATAAAAAATTAATATGTATGATTCAATGTATAACTAAACTACTAACTGTTATTGATAACTAGAATCAGAATCTAATGATGACATAACTAAGAAGTTTATCTACAGCCAATTTAGCTGCATTATTTTTAGCATCTCGTTTAGATTTTCCATCTGCCTTATCGAATACTCTTCCGTCAATGTCTACACAGGCATAAAATGTAGGAGAGTTACTAGGCCCCACTGATTCAATACGAAAAGACCAATCTCTCCTAGTTATTTGACAGTACTCATTAATAACGGTGACAGGGTTAACACCTTTCCAATAAATAATTTTTTTAACCGGAATAACATCATCAAAAGACTTATGATCCTCTCTCATTGATTTTTCGCGGGATACATCATCTATTATAGCATCAGCATCAGAATCTGTAGGCCGTGTATCAGCATCCATTGTCGTAGACCAACGAGGAGGAGTATCGTTGGAGCTGTAAACCATAGCACTACGTTGAAGATCATACAGAGCTTTATTAACTTCTCGCTTCTCCATATTAAGTTGTTTAGTTAGTTGTACAGCAGTAGCTCCTTAGTCCAATGTTTTTAATAACCGCACACAATCTCTGTGTCAGAACGCTCGTCAATATAGATCTTAGAAATTTTTTTAGAGAGAACTAACGCAACTAGCAATAAAACTGATCTTATTTTATCATTTTTTTTATTCATCATCCTCTGGTGGTTCGTCGTTCCTATCGAATGTAGCTCTGATTAACCCGTCATCTATAGGTGATGCTGGTTCTGGAGATTCTGGAGGAGATGGATTATTATCTGGAAGAATCTCTGTTATTTCCTTGTTTTCATGTATCGATTGCGTTGTAACATTAAGATTGCGAAATGCTCTAAATTTGGGAGGCTTAAAGTGTTGTTTACAATCTCTACACGCGTGTCTAACTAATGGAGGTTCATCAGCGGCTCTAGTTTGAATCATCATCGGTGTAGTATTCCTACTTTTACAGTTAGGACACGGTGTATTGTATTTCTCGTCGAGAACGTTAAAATAATCGTTGTAACTCACATCCTTTATTTTATCTATATTGTATTCTACTCCTTTCTTAATGCATTTTATACCGAACAAGAGATAGCGAAGGAATTCTTTTTCGGTACCGCTAGTACCCTTAATCATATCACATAGTGTTTTATATTCTAAATGTGTGGCAATGGACGGTTTATTTCTATACGATAGTTTGTTTTTGGAATCCTTTGAGTATTCTATACCAATATTATTCTTTGATTCGAATTTAGTTTCTTCGATATTAGATTTTGTATTACCTATATTCTTGATGTAGTACTTTGATGATTTTTCCATGGCCCATTCTATTAAGTTTTCCAAGTTGGCATCATCCACATATTGTGATAGTAATTCTCGGATATCAGTAGTGACTACCGCCATTGATATTTGTTCATTTGATGAGTAACTACTAATGTATACATTTTCCATTTATAACACTTATGTATTAACTTTGTTTATTTATATTTTTTCATTATTATGTTGATATTAATAATCGTATTGTGGTTATATGGCTACAATTTCATAATGAGTTGAAGTCAGTGTCCTATGATCAATGACGATAGCTTTACTCTGAAAAGAAAGTATCAAATCGATAGTGCAGAGTCAACAATGAAAATGGATAAGACGATGACAAAGTTTCAGAATAGAGTCAAAATGGTAAAAGAAATAAATCAGACGATAAGAGCAGCACAAACTCATTACGAGACATTGAAACTAGGATATATAAAATTTAAGGGAATGATTAGGACTACTACTCTAGAAGATATAGCACCATCTATTCCAAATAATCAGAAAACTTATAAACTATTCTCGGACATTTCAGTCATTGGCAAAGCATCACAGAATCCGAGTAAGATGATATATGCTCGCTGCTTTACATGTTTCCCAATTTGTTTGGAGATGACCATAGATTCATTTGTTATAGAATGCATCCAACATTGTTCATGATATAGTTGAATCATGTATGCCTGTTCGTATGCCTGTGGCTAAGATACTGTGTAAAGAAATGGTAAATAAATACTTTGAGAATCTTTAAGAGTGCATTGACTTTGTTAGTGAATAGGCATTCCATCTTTCTCCAATACTAATTCAAATTGTTAAATTAATAATGGAATAGTATAAATAGTTATTAGTGATAGGATAGTAAACATAATTATTAGAATAGTAGTGTAGTATCATAGATAACTCTCTTCTATAAAAAATGGATTTTATTCGTAGAAAGTATCTTATATACACAGTAGAAAATAATATAGATTTTTTAAAGGATGATACATTAAGTAAAGTAAACAATTTTACCCTCAATCATGTACTAGCTCTCAAGTATCTAGTTAGCAATTTTCCTCAACATGTTATTACTAAGGATGTATTAGCTAATACCAATTTTTTTGTTTTCATACATATGGTACGATGCTGTAAAGTATACGAAGCGGTTTTACGACACGCATTTGATGCACCCACGTTGTACGTTAAAGCATTGACTAAGAATTATTTATCGTTTAGTAACACAATACAGTCGTACAAGGAAACAGTGCATAAACTAACACAAGATGAAAAATTTTTAGAGGTTGCCGAATACATGGACGAATTAGGAGAACTTATAGGCGTAAATTATGACTTAGTTCTTAATCCATTATTTCACGGAGGGGAACCCATCAAAGATATGGAAATCATTTTTTTAAAACTGTTTAAGAAAACAGACTTCAAAGTTGTTAAAAAATTAAGTGTTATAAGATTACTTATTTGGGCATACCTAAGCAAGAAAGATACAGGCATAGAGTTTGCGGATAATGATAGACAAGATATATATACTCTATTTCAACAAACTGGTAGAATAGTCCATAGCAATCTAACAGAAACGTTTAGGGATTATATCTTTCCCGGAGATAAGACTAGCTATTGGGTGTGGTTAAACGAAAGTATAGCTAATGATGCGGATATCGTTATTAATAGACCCGCCATTACCATGTATGATAAAATTCTTAGTTATATATACTCTGAGATAAAACAGGGACGCGTTAATAAAAACATGCTTAAGTTAGTTTATATCTTTGAGCCTGAAAAAGATATCAGAGAACTTCTGCTAGAAATCATATATGATATTCCTGGAGATATCCTATCTATTATTGATGCAAAAAACGACGATTGGAAAAAATATTTTATTAGTTTTTACAAAGCTAATTTTATTAACGGTAATACATTTATTAGTGATAGAACGTTTAACGATGACNTATTCAGAGTTGTTGTTCAAATAGATNCCGAATATTTCGATAATGAACGAATTATGTCTTTATTCTATACGAGNNNTGCGGACATTAAACGATTTGATGAGTTAGATATTAATNACNNNNATATATCTAATATAATTTATGAGGTGAACGATNNNACATTAGATACAATGGATGATATNAAGAAGTGTCAAATCNTTAACGAGGATACGNNGTATTATGTTAAGGAATACAATACATACCTGNTTTTGCACNAGTCNGATCCCATGGTCNNAGAGAACGGAATACTAAAGAAACTGTCATCNATAAAATCCAAGAGTAGACGGCTNNACTTGNTTAGCNAAAACATTNNNAANNNNTATTTNNACGGACAATTNGCTCGTCTAGGTCNNNTGTTNGATGATTATAAAGGAGACTTATTAGTTAAAATGATAAACCATCTCAAATCTGNGGAGGATGTATCCGCATTCGTTAGATTTTCTACAGATAAAAANCCTAGTATTCTTCCATCGCTAATCAAAACTATTTTAGNTAGTTATAATATTTCCATCATCGTCTTATTTCAAAGGTTTTNNAGAGATAATCTATATCATGTAGAAGAATTCTTNGATNAAAGCATCCATCTAACCAAGACGGATAAGAAATATATACTTCAATTGATAAGGCACNNNNNNNNNNNNNNNNNNNNNNNNNNNNNNNNNNNNNNNNNNNNNNNNNNNNNNNNNNNNNNNNNNNNNNNNNNNNNNNNNNNNNTTTGATAAATGNNNACCGTTGCNACANNTCAGACTCCCACCAAATTAATGAATAAAGAAAATGCAGAAATGATTTTGGAAAAAATTGTTAATCATATAGCTATGTATATTAGTGACGAATCAATATATTCAGAAAATAATCCTGAATATATTGATTTTCGTAACAGATACGGAGACTATAGATCTCTCATTATAAAAAGTGATCACGAGTTTGTAAAGCTATGTAAAGATCATGCAGAGAAAAGTTCTCCAGAAACGCAACAAATGATTATCAAACACATATACGAACAATATCTTATTCCAGTATCTGAAGTACTATTAAAACCTATAATGTCCATGGGTGACATATTTACATATAACGGATGTAAAGACAATGAATGGATGCTAGAACAACTCTCTACCCTAAACTTTAACAATCTCTACACATGGAACTCATGTAGCATAGGCAATGTAACGCGTCTGTTTTATACATTTTTTAGTTATCTGATGAAAGATAAACTAAATATATAAGTATAATCCCATTCTAATANTTTAACCTGATGTNNTATTACCTGCATCTTATTAGAATATTAACCTAACTAAAAGACATAAAAANNNGTAGGATATNAATATTATGGCCGCAACCGTTCCGCGNTTTGACGANNTGTANAAAAATGCACAAAGAAGAATTNTAGATCNAGNAACANNTTTNNGTAGAGGTCTNNNTAGACCGTTAATGNAAAACACATATCTATTTGATAATTACGCGTATGGATGGATACCAGAAACTGCAATTTGGAGTAGTAGATACGCAAACCTAGATGCTAGTGACTATTATCCCATTTCGTTGGGATTACTTAAAAAGTTTGAATTTCTCATGTCTCNNTATAAANGTCCTATTNNCGTATATGAAGNAAAAGTAAATACTGAATTCATTGCTAATGGATCTTTCTNCGGTAGATACGTATCATATCTTNNAAAGTTTTCNNNCCTTCCAACAAACGAGTTTATTANNNNNTTANTATTGACCTCCATCCCTATCTNNNATATCNTATTCTNGTTTNNAANNACACAGTTTGATATTACTAAACACACATTATTCAGATACGTCTATACAGATAATACCAAACACCNTGCGTTGGCTANGTATATACATCAAACAGGAGACTATNNGNCTTTGTTTAGTCGTCTCAAAGAGAATTATATATTTACTNGTCCCGTTCCAATAGGTATCNAAGATATAGATCACCCTAATCNTAGTAGAGCANNAAGTCCATCCGATTATGAGACATTAGCTAATATTAGTACTATATTGTACTTTACCAAGTATGATCCAGTATTAATGNTTTTATTGNTTTACGTACCTGGGTATTCAATTACTACNNNNNNTACTCCAGCCGTAGAATATCTAATGGATAAACTGAATCTAACNAAGAGCGACGTACAANTNTNNTAAATTATTTTATGCTTCNNAAAATGTAGGTCTTGAACCAAACATTCTTTGNAAAAATGAGATGCATAAAACTTTATTATCCAATAGATTAACTATTTCAGACGTCAATNGTTTNNAGNAAACTNNNTAAAATATTCTTTGATTGCTGCCGAGNTTAAAACTTNTATCGATAATTGTTTCNNNNGTTTTAATATTTNNNNNNNNNNNNNNNNNNNNNNNNNNNNNNNNNNNNNNNNNNNNNNNNNNNNNNNNNNNNNNNNNNNNNNNNNNNNNNNNNNNNNNNNNNNNNNNNNNNNNAACCAAATACATATTANGGTTATCTGCGGATTTNNNNNNNNNNNNNNNNNNNNNNNNNNNNNNNNNNNNNNNNNNNNNNNNNNNNNNNNNNNNNNNNNNNNNNNNNNNNNNNNNNNNNNNNNNNNNNNNNNNNNNNNNNNNNNNNNNNNNNNNNNNNNNNNNNNNNNNNNNNNNNNNNNNNNNNNNNNNNNNNNNNNNNNNNNNNNNNNNNNNNNNNNNNNNNNNNNNNNNNNNNNNNNNNNNNNNNNNNNNNNNNNNNNNNNNNNNNNNNNNNNNNNNNNNNNNNNNNNNNNNNNNNNNNNNNNNNNNNNNNNNNNNNNNNNNNNNNNNNNNNNNNNNNNNNNNNNNNNNNNNNNNNNNNNNNNNNNNNNNNNNNNNNNNNNNNNNNNNNNNNCTATTAATCNGTCGTTCNAACTCCTTTGNTATTTCTATGGACTTATCTACATCTTGACTGTCTATCTCNGTNAACACNGAGTCGGTANNTCCATACACGCTACGAAAACGAAATCTATAATCTATAGGCAACGATGTTTTCACAATCGGATTAATATNTCTATCGTCCATATAAAATGGATTACTTAATGTATTNGCNAACCGTAACATACCGTTNGATNNCTCNGNTCCATTTAGTACCNANNCNAGATACAATATCATTCTACGTCCTATGGANNTGCAACTCTTAGCCNAAGNGTATGAGTATAGAGCACTATTTCTAAATCCCATCAGACCATATACTGAGTTGGCTACTATCTTGTACGTATATTGCATGGAATCATAGATGGCCTTTTCAGTTGAACTGGTANNNTGNTTTAANNNNNTTTTATATNNNNNNNNNTNNNCCNAAAATGTTCTTAATAGTCTAGGAATGGTTCCTTCTATTGATCTATCGAAAATTGCTATTTCAGAGATGAGGTTCGGTAGTCTAGGTTCACAATGAACCGTAATATATCTAGGAGGTGGATATTNNTGAAGCAAGAGTTGATTATTTATNNNTTCTTNCNATCTNTTGGTACTAACAACGACACCGACTAATGTTTCCGGAGATNNNNNNNNNNNNNNNNNNNNNNNNNGATACAGACNNNTATAATCAAAGATTAATACATTATTACTAAACATTTTTTGTTTTGGAGCAAATACCTTACCGCCTTCATAAGGAAACTTTTGTTTTGTTTCTGATCTAACTAAGATAGTTTTAGTTTCCAACAATAGCTTTAACAGTGGACCCTTGATGATTGTACTCGCTCTATATTCGAATACCATGGATTGAGGAAGCACATATGTTGCCGCACCAGCGTCTGNTTTTGTTTCTACTCCATAATACTCCCACAAATACTGACACAAACAAGCATCATGAATACAGTATCTAGCCATATCTAAAGCTATGTTTAGATTATAATCCTTATACATCTGAGCTAAATCAATGTCATCCTTTCCGAAAGATAATTTATATATATCATTAGGTAAAGTAGGACATGATAGTACGACTTTAAATCCATTTTCCAAAATATCTTTACGAATTACTTTACATATAATATCCTCATCAACAGTCACGTAATTACCTGTGGTTAAAACCTTTGCAAATGTATCGGCTTTGCCTTTCGCGTCCNNAGTATNGTCACCGATGAACGTCATTNNNNTAACTCCTCTATTTAATACTTTACCCATGCAACTGAACGCGTTCTTNGATATAGAATCCAATTTGTACGAATCCAATTTTTCAGANTTTTGAATGAATGAATATAGATCGAAAAATATAGTTCCATNATTGTTATTAACGTGAAACGTAGTATTGGCCATGCCGCATACTCCCTTATGACTAGACTGATTTCTCTCATAAATACAGAGATGTACAGCTTCCNTTTTGTCTGGAGATCTAAAGATAATCTTCTCTCCTGTTAATAACTCTAGACGATTAGTAATATATCTCAGATCAAAGTTATGTCCGTTAAAGGTAACGACGTAGTCGAACGTTAGTTCCAACAATTGTTTAGCTATTCGTAACAAAACTATTTCAGAACATAGAACTAGTTCTCGTTCGTAATCCATTTCCATTAGCGACTGTATCCTCAAACATCCTCTATCGACGGCTTCTTGTATTTCCTGTTCCGTTAACANNNNNTCATTAATGANNGTAAACNNNAATCGTTTACCACTTAAATCGATATAACAGTAACTTGTATGCGAGATTGGGTTAATAAATACAGAAGGAAACTTCTTATCGAAGTGACACTCTATATCTAGAAATAAGTACGATCTTGGGATATCGAATCTAGGTATTTCTTTAGCGAAACAGTTACGTGGATCGTCACAATGATAACATCCATTGTTAATCTTTGTCAAATATTGCTCGTCCAACGAGTAACATCCGTCTGGAGATATCCCGTTAGAAATATAAAACCAACTAATATTGAAAAATTCATCCATGGTGGCATTTTGTATGCTGCGTTTCTTTGGCTCTTCTATCAACCACATATCTGCGACGGAGCATTTTCTATCTTTAATATCTAGATTATAACTTATTGTCTCGTCAATGTCTATAGTTCTCATCTTTCCCATCGGCCTCGCATTAAATGGAGGAGGAGATAATGACTGATATATTTCGTCCGTCACTACGTAATAAAAGTAATGAGGAAATCGTATAAATACGGTCTCGCCATTTCGACATCTGGATTTCAGATATAAAAATCTGTTTTCACCGTGACTTTCAAACCAATTAATACACCTAACATCCATTTCTAGAATTTAGAAATATATTTTCATTTAAATGAATCCCAAACATTGGGGAAGAGCCGTATGGACCATTATTTTTATAGTACTTTCGCAAGCGGGTTTAGACGGCAACATAGAAGCGTGTAAACGAAAACTATATACTATAGTCAGCACTCTTCCATGTCCTGCATGTAGACGACACGCGACTATCGCTATAGAGAACAATAATGTCATGTCTAGCGATGATCTGAATTATATTTATTATTTTTTCATCAGATTATTTAACAATTTGGCATTTGATCCCAAATACGCAATCGATGTGTCAAAGGTTAAACCTTTATAAACTTAACCCATTATAAAACTTATGATTAGTCACGACTGAAATAACCGCGTGATTATTTTTTGGTATAATTCTACACGGCATGGTTTCTGTGACTATGAATTCAACACCTGTTATCTTAGTGAAATCTTTAACAAACAGCAAGGGTTCGTCAAAGACATAAAACTCATTGTTTACGATCGAAATAGANCCCCTATCACACTTAAAATAAAAAATATCCTTATCCTTTACCACCAAATAAAATTCTGATTGGTCAATGTGAATGTATTCACTTAACAGTTCCACAAATTTATTTATTAACTCCGAGGCACATACATCGTCGGTATTTTTTATGACAAACTTTACTCTTCCAGCATCCGTTTCTNAAAAAATATTAACGAGTTCCATTTATATCATCCAATATTATTGAAATGACGTTGATGGACAGATGATATAAATAAGAAGGTACAGTACCTTTGTCCACCATCTCCTCCAATTCATACTCTATTTTGTCATTAACTTTAATGTGTGAAAACAGTACGCCACATGCTTCCATGACAGTGTGTAACACTTTGGATACAAAATGTTTGACATTAGTATAATTGTCCAAGACTGTCAATCTATAATAGATAGTAGCTATAATATATTCTATGATGGTATTGAAGAAGATGACAACCTTGGCATATTGATCATTTAACACAGACATGGTATCAACAAATAGCTTAAATGAAAGAGAATCAGTAATTGGAATAAGCGTCTTCTCGATGTAGTGTCCGTATACCAACATGTCTGATATTTTGATGTATTCCATTAAATTATTTAGNTTTTTCTTTTTATTCTCGTTAAACAGAATTTCTGTCAATGGACCCCAACATCGTTGACCTATTAAGTTTTGATTGATTTTTCCGTGTAAGGCGTATCTAGTCAGATCGTATAGCCTATCCAATAATCCATCGTCTGTGCGTAGATCACATCGTACACTTTTTAATTTTCTATAGAAGAGTGACAGACATCTGGAGCAATTACAGACAGCAATTTCTTTATTCTCTACAGATGTAAGATACTTGAAGATATTCCTATGATGATGCAGAATTTTGGATAACACGGTATTGATGGTATCTGTTACCATAATTCCTTTGACTGATAGTGTCAAAGTACAAGATTTCCAATNTTTTGCAATTTTCAGTACCATTATCTTTGTTTTGATATCTATATCAGACAGCATGGTACGTCTGACAACACAGGGATTAAGACGGAAAGATGAAATGATTCTCTCAACATCTTCAATAGATACCTTGCTANTTTTTTTGGCATTATCTATATGTGAGAGAATATCCTCTAGAGAATCAGTATCCTTTTTGATGATAGTGGATCTCAATGACATGGGACGTCTAAACCTTCTTATTCTATCACCAGATTGCATGGTGATTTGTCTTCTTTCTTTTATCATGATGTAATCTCTAAATTCATCGGCAAATTGTCTATATCTAAAATCATAATATGAGATGTTTACCTCTACAAATATCTGTTCGTCCAATGTTAGAGTATCTATATCAGTTTTGTATTCCAAATTAAACATGGCAACGGATTTAATTTTATATTCCTCTATTAAGTCCTCGTCGATAATAACAGAATGTAGATAATCATTTAATCCATCGTACATGGTTGGAAGATGCTCGTTGACAAAATCTTTAATTGTCTTGATGAAGGTGGGACTATATCTAACATCTTGATTAATAAAATTTATAACATTGTCCATAGGATACTTTGTAACTAGTTTTATACACATCTCTTCATTGGTAAGTTTAGACAGAATATCGTGAACAGGTGGTATATTATATTCATCAGATATACGAAGAATAATGTCCAAATCTATATTGTTTAATATATTATATAGATGTAGTGTAGCTCCTACAGGAATATCTTTAACTAAGTCAATGATTTCATCAACAGTTAGATCTATTTTAAAGTTAATCATATAGGCATTGATTTTTAAAAGGTATGTAGCCTTGACTACATTCTCATTAATTAACCATTCCAAGTCACTGTGTGTAAGAAGATTATATTCTATCATAAGCTTGACTACATTTGGTCCCGATACCATTAAAGAATTCTTATGATATAAGGAAACAGCTTTTAGGTACTCATCTACTCTACAAGAATTTTGGAGAGCCTTAACGATATCAGTGACGTTTATTATTTCAGGAGGAAAGAACCTAACATTGAGAATATCTGAATTAATAGCTTCCAGATACAGTGATTTTGGCAATAGTCCGTGTAATCCATAATCCAGTAACACGAGCTGGTGCTTGCTAGACACCTTTTCAATGTTTAATTTTTTTGAAATAAGCTTTGATAAAGCCTTCCTCGCAAATTCCGGATACATGAACATGTCGCCAACATGATTAAGTATTGTTTTTCATTATTTTTATATTTTCTCAACAAGTTCTCAATACCCCAATAGATAATAGAATATCACCCAATGCGTCCATGTTGTCTATTTCCAACAGGTCGCTATATCCACCAATAGAAGTTTTCCCNAAAAAGATTCTAGGAACAGTTCTACCACCAGTAATTTGTTCAAAATAGTCACGCAATTCATTTTCGGGTTTAAATTCTTTAATATCTACAATTTCATACGCTCCTCTTTTGAAACTAAACTTATTTAGAATATCCAGTGCGTTTCTACAAAAAGGACATGTAAACTTGACAAAAATTGTCACTTTGTTATTGGCCAACCTTTGTTGTACAAATTCCTCGGCCATTTTTAATATTTAAGTGATACAAAACTATCTCGACTTATTTAACTCTTTAGTCGAGATATATGGACACAGATAGCTATATGATAACCAACTACAGAAGACAAACGCTATAAAAAACATAATTACGACGAGCATATTTATAAATATTTTTATTCAGTATTACTTGATATAGTAATATTAGGCACAGTCAAACATTCAACCACTCTAGATACATTAACTCTCTCATTTTCTTTAACAAATTCTGCAATATCTTCGTAAAAAGATTCTTGAAACTTTTTAGAATATCTATCGACTCTAGATGAAATAGCGTTCGTCAACATACTATGTTTTGTATACATAAAGGCGCCCATTTTAACAGTTTCTAGTGACAAAATGCTAGCGATCCTAGGATCCTTTAGAATCACATAGATTGACGATTCGTCTCTCTTAGTAACTCTAGTAAAATAATCATACAATCTAGTACGCGAAATAATATTATCCTTGACTTGAGGAGATCTAAACAATCTAGTTTTGAGAACATCGATAAGTTCATCGGGAATTACATACATACTATCTTTAATAGAACTCTTTTCATCCAGTTGAATGGATTCGTCCTTAACCAACTGATTAATGAGATCTTCTATTTTATCATTTTCTAGATGATATGTATGTCCATTAAAGTTAAATTGTGTAGCGCTTCTTTTTAGCCTAGCAGCCAATACTTTAACATCACTAATATCGATATACAAAGGAGATGATTTATCGATGGTATTAAGAATTCGTTTTTCGACATCCGTCAAAACCAATTCCTTTTTGCCTGTATCATCCAGTTTGCCATTCTTTGTAAAGAAATTATTTTCTACTAGACTATTAATAAGACTGATAAGGATTCCTCCATAATTGCACAATCCAAACTTTTTCACAAAACTAGACTTTACGAGATCTACAGGAATGCGTACTTCAGGTTTCTTAGCTTGTGATTTTTTCTTTTGTGGACATTTTCTAGTGACCAACTCATCTACCATTTCATTGATTTTAGCAGTGAAATAAGCTTTCAATGCACGGGCACTGATACTATTGAAAACGAGTTGATCTTCAAATTCCGCCATTTAAGTTCACCAAACAACTTTTAAATACAAATATATCAATAGTAGTAGAATAAGAACTATAAAAAAAATAATAATTAACCAATACCAACCCCAACAACCTGTATTATTAGTTGATGTGACAGTTTTCTCATCACTTAGAACAGATTTAACAATTTCTATAAAGTCTGTCAAATCATCTTCCTGAGAACCCATAAATACACCAAATATAGCAGCGTACAACTTATCCATTTATACATTGAATATTGGCTTTTCTTTATCGCTATCTTCATCATATTCATCATCAATATCAACAAGTCCCAGATTACGAACCAGATCTTCTTCTACATTTTCAGTCATTGATACGCGTTCACTATCTCCAGAGAGTCCGATAACGTTAGCCACTACTTCTCTATCAATGATTAGTTTCTTGAGCGCGAATGTAATTTTTGTTTCCGTTCCGGATCTATAGAAAACTACAGGTGTAATAATTGCCTTGGCTAATTGTCTTTCTCTTTTACTGAGTGATTCTAGTTCACCTTCTATAGATCTGAGAATGGATGATTCTCCAGTCGAAACATATTCTACCATGGCTCCGTTTAATTTGTTGATGAAGATGGATTCATCCTTAAATGTTTTCTCTGTAATAGTTTCCGCCGAAAGACTATGCAAAGAATTTGGAATGCGTTCCTTGTGTGTAATGTTTCCATAGACAGCTTCTAGAAGTTGATACAACATAGGACTAGCCGCGGTAACTTTTATTTTTNGAAAGTATCCATCGCTTCTATCTTGTTTAGATTTATTTTTATAAAGTTTAGTCTCTCCTTCCAACATAATAAAAGTGGAAGTCATCTGACTAGATAAACTATCAGTAAGTTTTATAGAGATAGATGAACAATTAGCGTATTGAGAAGCATTTAGTGTAACGCATTCGATACANTTTGCATTAGATTTACTAATCGATTTTGCATACTCTATAACACCCGCACAAGTCTGTAGAGAATCGCTAGATGCTGTAGGTCTTGGTGAAGTTTCAACTCTCTTCTTGATTACCTTACTCATGATTAAACCTAAATAATTGTACTTTGNAATATAATGATATATATTTTCACTTTATCTCATTTGAGAATAAAAATNTTTTTGTTAACCACTGCATGATGTACAGATTTCGGAATCGCAAACCACTTGTGGTTTTATTTTATCCTTGTCCAATGTGAATTGAATGGGAGCGGATGCGGGTTTCGTACGTAGATAGTACATTCCCGTTTTTAGACCGAGACTCCATCCGTAAAAATGCATACTCGTTAGTTTGGAATAACTCGGATCTGCTATATGGATATTCATAGATTGACTTTGATCGATGAAGGCTNCCCTGTCTGCAGCCATTTTTANGNTCGTCTTTTGTGGAATTTCCCAAATAGTTTTATAAACTCGCTTAATATCTTCTGGAAGGTTTGTATTCTGAATGGATCCACCATCTACCATAATCCTATTCTTGATCTCATCATTCCATAATTTTCTCTCGGTTAAAACTCTAAGGAGATGCGGGTTAACTACTTGGAATTCTCCAGACAATACTCTCCGAGTGTAAATATTACTGGTATACGGTTCCACCGACTCATTATTTCCCAAAATTTGAGCAGTTGATGCAGTCGGCATAGGTGCCACCAATAAACTATTTCTAAGACCGTATGTTCTGATTTTATCTTTTAGAGGTTCCCAATTCCAAAGATCCGACGGTACAACATTCCAAAGATCATATTGTAGAATACCGTTACTGGCGTACGATCCTACATATGTATCATATGGTCCTTCCTTCTCAGCTAGTTTACAACTCGCCTCTAATGCACCGTAATAAATGGTTTCAAAGATCTTCTTATTTAGATCTTGTGCTTCCAGGCTATCAAATGGATAATTTAAGAGAATAAACGCGTCCGCTAATCCTTGAACACCAATACCGATAGGTCTATGTCTCTTATTAGAGATTTCAGCTTCTGGAATAGGATAATAATTAATATCTATAATTTTATTGAGATTTCTGACAATTACTTTGACCACATCCTTCAGTTTGAGAAAATCAAATCGCCCATCTATTACAAACATGTTCAATGCAACAGATGCCAGATTACACACGGCTACCTCATTAGCATCCNCATATTGTATTATCTCAGTGCAAAGATTACTACACTTGATGGTTCCTAAATTTTGTTGATTACTCTTTTTGTTACACGCATCCTTATAAAGAATGAATGGAGTACCAGTTTCAATCTGAGATTCTATAATCGCTTTCCAGACGACTCGAGCCTTTATTATACATTTGTATCTCCTTTCTCTTTCGTATAGTGTATACAATCGTTCGAACTCGTCTCCCCAAACATTGTCCAATNCAGGACATTCATCCGGACACATCAACGACCACTCTCCGTCATCCTTCACTCGTTTCATAAAGAGATCAGGAATCCAAAGAGCTATAAATAGATCTCTTGTTCTATGTTCATCGTTTCCTGTATTCNTTTTAAGATCGAGGAACGCCATAATATCAGAATGCCACGGTTCCAAGTATATGGCCATAACTCCAGGCCGTTTGTTTCCTNCCTGATCTATGTATCTAGCGGTGTTATTATAAACTCNCAACATTNGAATAATACCGTTTGATATACCATTNGTACCNGAGATATAGCTTCCACTNGCACGAATATNNNTAATTGATAGACCTATTNCCCCTNCCATTTTAGAGATTAATGCGCATCGTTTTAACGTGTCATAGATGCCTTCNATGCTATCATCGATCATGTTAAGTNGAAAACAGCTAGACATTTGGTGACGNGTNGTTNCCGCATTNAANAAGGTAGGAGAAGCGTGCGTAAACCATTTTTCAGAAAGTAGATTGTACGTCTCAATAGCTGAGTCTATATCCCATTGATGAATTCCTACTGCGACACGCATTAACATGTGCTGAGGTCTTTCAACAATTTTGTTGTTTATTTTCAACAAGTAGGATTTTTCCAAANTTTTAAAACCAAAATAGTTGTANGAAAAGTCTCGTTCGTAAATAATAACCGAATTGAGCTTATCCTTATATTTGTTAACTATATCCATGGTAATACTTGAAATAATCNGAGAATGTTTCCCATTTTTAGGATTAACATAGTTGAATAAATCCTCCATCACTTCACTAAATAGTTTTTTTGTTTCCTTGTGTAGATTTGATATGGCTATTCTGGCGGCTAGAATGGCATAATCCGGATGTTGTGTAGTACAAGTGGCTGCTATTTCGGCTGCCAGAGTGTCCAATTCTACCGTTGTTACTCCATTATATATTCCTTGAATAACCTTCATAGCTATTTTAATAGGATCTATATGATCAGTGTTTAAGCCATAGCACAATTTTCTAATACGAGACGTGATTTTATCAAACATGACATTTTCCTTGTATCCATTTCGTTTAATGACAAACATTTTTGTTGGTGTAATAAAAAAAATTATTTAATTTTTCATTAATAGGGATTTGACGTATGTAGCGTACAAAATTATCGTTCCTGGTATATAGATAAAGAGTCCTATATATTTGAAAATCGTTACGGTTCGATTAAACTTTAATGATTGCATTGTGAATATATCATTAGGATTTAACTCCTTGACTANNATGGCGGTGCCAGAAATTACCATCNAAAGCATTAATACAGTTATGCAGATCGCAGTTAGAACGGTTATAGCATCCACCATTTATATCTAAAAATTAGATCAAAGAATATGTGACAACGTCCTAGTTGTATACTGAGAATTGACGAAACAATGTNNNTTACATATTTTTTTCTTATTAGTAACCGACTTAATAGTAGGAACTGGAAAACTAGACTTGATTATTCTATAAGTATAGATACCCTTCCAAATAATGTNNTCTTTGATAAAAGTTCCAGAAAATGTAGAATTTTTTAAAAAGTTATCNTTTGCTATTACTAATATCGTGGTTAGACGCTTATTATTAATATGAGTGATGAAATCCACACCGCTTCTAGATATCGCTTTTATTTCCACATTAGATGGTAAATCCAATAGTGAAACTATCNTTTTAGGAATGTATGGACTCGCGTTTAGAGGAGTGAACGTCTTCNGAGTAGTAAAGGATGATTCGTCAAATGAATAAACAATTTCACAAATGGATGTTAATGTATTAGTAGGAAATTTTTTGACGCTAGTGGAATTGAAGATTCTAATGGATGATGTTCTACCTATTTNANNNNNNNNNNNNNNNNNNNNNNNNNNNNNNNNNNNNAATATTTCGATGATATACGGTAGTCTCTCNTTCNGACTTNTATAGCNTATTNCACAATACGAGTCATTATATACNCCNNAAAANNNAATNNNTAGTATNAAATCNGTATCGAATGGNNAAAACGAAATTATCGATATAGGNNTAGAANCNNGAACATTGAACGTATTAATACTTAATNNTTTTTCANNNNNAAGAACCGATAGGTTATTGACATTGTATGGTTTTAAATATTCTATAACTTNNGACTTGATAGATATTAANGACGAATTGNAAATTANTTTTATNACCACNNGTGNNTCAGGANCNTNGTCGANNNCAGTTAACCAACCGAATGGAGTANNNNAAATATCATTAATATATGCNCTAGATATTAGTANNNNNNTTAANNCTTTGATTATCATCTNNTCGTACGCGAATGATTCCATGATCAAGAGTGATTTGAGNNCATCCTCCNGAGTATTAATNGGTTTAGTAAACAGTCCANCGTTGCNATAATAAAAGTTGTCCAAGTTAAAGGATATTATGCNNNCGNTTAAAGATNNNACCTCATCTAACGGAGACAANNTTNNNGTAGGNTTTNNAGACTTTGAAGCTACTTGTTTAACAAAGTTATTCATCGTCGTCTACTATTCTATTTAATTTTGTAGTTAATTTATCACATATCACATTAATTGACNTTTTGGTCCACTTTTCCATACGTTTATATTCTTTTAATCCTGCGTTATCCGTTTCCGTTATATACAGGGATAGATCTTGCAAGTTAAATAGAATGCTCTTAAATAATGTCANTTTTTTATCCGCTAAAAATTTAAAGAATGTATAAACTTTTTTCAAAGATTTAAAACTTTTAGGTGGAGTTCTGGTACACAATATCATAAACAAACTAATAAACATCCCACATTCAGATTCCAACAATTGATTAACTTCCACATTAATACAGCCTATTTTCGCTCCAAATGTACATTCGAAAAATCTGAATAAAACATCAATATCGCAATTTGTATTATCCAATACAGAATGTCTGTGATTCGTGTTAAAACCATCGGNAAAAGAATAGAAATNAAAATTATTATAATGGTGGAATTCAGTTNGAATATTGNCTCCNGAGTCATNAAAGGATACTAAACATTGTTTTTTATCGTAAATTACACATTTNCAATGAGACAAATAACAAAATCCAAACATTACAAATCTAGAGGTAGAACNTTTAATTTTGTCTTTAAGTATATACGATAAGATATGTTTATTCATAAACGCGTCAAATTTTTCATGAATAGCTAAGGAGTTTAAGAATCTCATGTCAAATTGTCCTATATAATCCACTTCGGATCCATAAGCAAACTGAGAGACTAAGTTCTTAATACTTCGATTGCTCATCCAGGCTCCTCTCTCAGGCTCTATTTTCATCTTGACGACCTTTGGATTTTCACCAGTATGTATTCCTTTACGTGATAAATCATCAATTTTCAAATCCATTTGTGAGAAGTCTATCGCCTTAGATACTTTTTCCCGTAGTTGAGGTTTAAAGAAATACGCTAACGGTATACTAGTAGGTAACTCAAAGACATCATATATAGAATGGTAACGCGTCGTTAACTCGTCGGTTAACTCTTTCTTTTGATCGAGTTCATCGCTACTATTGGGTCTGCTCAGGTGCCCCGACTCTACTAGTTCCAACATCATACCGATAGGAATACAAGACACTTTGCCAGCGGTTGTAGATTTATCATATTTCTCCACCACATATCCGTTACAATTTGTTAAGAATTTAGATACATCTATATTGCTACATAATCCAGCTAGTGAATATATATGACATAATAAATTGGTAAATCCTAGTTCTGGTATTTTACTAATTACTAAATCTGTATATCTTTCCATTTATCATGGAAAAGAATTTACCAGATATCTTCNTTTTTCCAAACTGCGTTAATGTATTCTCTTACAAATATTCACAAGATGAATTCAGTAATATGAGTAAAACGGAACGNGATAATTTCTCATTGGCTGTGTTTCCAGTGATAAAACATAGATGGCATAACGCACACGTTGTAAAACATAAAGGAATATACAAAGTTAGTACAGAAGCACGTGGNAAAAAAGTATCTCCTCCATCACTAGGAAAACCCGCACATATAAACCTAATGTCGAAGCAATATATATATAGTGAGTATGCAATAAGCTTTGAATGTTATAGTTTTCTAAAATGTATAACAAATACAGAAATCAATTCGTTCGATGAGTATATATTAAGAGGACTATTAGAAGCTGGTAATAGTTTACAGATATTTTCCAATTCCGTAGGTAAACGAATAGATACTATAGGTGTACTANGGAATAAGTATCCATTTAGCAAAATTCCATTGGCCTCATTAACTCCTAAAGCACAACGAGAGATATTTTTAGCGTGGATTTCTCATAGACCTGTAGTTTTAACTGGAGGAACCGGAGTGGGTAAGACGTCACAGGTACCCAAGTTATTGCTTTGGTTTAATTATTTATTTGGTGGATTCTCTTCTCTAGATAAAATCACTGACTTTCACGAAAGACCAGTCATTCTATCTCTTCCTAGGATAGCTTTAGTTAGATTGCATAGCAATACCATTTTAAAATCATTGGGATTTAAGGTACTAGATGGATCTCCTATCTCTTTACGGTACGGATCTATACCGGAAGAATTAATAAACAAACAACCNAAAAAATATGGAATTGTATTTTCTACCCATAAGTTATCTCTAACAAAACTATTTAGTTATGGCACTATTATTATAGACGAAGTTCATGAGCATGATCAAATAGGAGATATTATTATAGCAGTAGCGAGAAAACATCATACGAAAATAGATTCTATGTTTTTAATGACTGCCACGTTAGAGGATGACAGGGAACGTCNAAAAATATTTTTACCTAATCCCGCATTTATACATATTCCTGGAGATACACTGTTTAAAATTAGCGAGGTATTTATTCATAATAAGATAAATCCATCTTCCAGAATGGCATATATAGAAGAAGAAAAGAGAAATTTAGTTACTGCTATACAGATGTATACTCCTCCTGATGGATCATCCGGTATAGTCTTTGTGGCATCCGTTGCACAGTGTCACGAATATAAATCATATTTAGNAAAAAGATTACCGTATGATATGTATATTATTCATGGTAAGGTCTTAGATATAGACGAAATATTAGAAAAAGTGTATTCATCACCTAATGTATCGATAATTATTTCTACTCCTTATTTGGAATCCAGCGTTANTATACGCAATGTTACACACATTTATGATATGGGTAGAGTTTTTGTCCCCGCTCNTTTTGGAGGATCACAACAATTTATTTCTAAATCTATGAGAGATCAACGAAAAGGAAGAGTAGGAAGAGTTAATCCTGGAACATACGTATATTTCTATGATCTGTCTTATATGAAATCTATACAGCGAATAGATTCAGAATTTCTACATAATTATATATTGTACGCTAATAAGTTTAATCTAACACTCCCCGAAGATTTGTTTATAATCCCTACAAATTTGGATATTCTATGGCGTACAAAGGAATATATAGACTCGTTCGATATTAGTACAGAAACATGGAATAAATTATTATCCAATTATTATATGAAGATGATAGAGTATGCTAAACTTTATGTACTAAGTCCTATTCTCGCTGAGGAGTTGGATAACTTTGAGAGGACGGGAGAATTAACTAGTATTGTACAAGAAGCCATTTTATCTCTAAATTTACAAATTAAGATTTTAAAATTTAAACATAAAGATGATGATACGTATATACACNTTTGTAGAATATTATTCNGTGTCTATAACGGAACAAACGCTACTATATATTATCATAGACCTCTAACGGGATATATGAATATGATTTCAGATACTATATTTGTTCCTGTAGATAATAACTNAAAATCAAAATCTAATGACCACATCNNTTTTTAGAGATGNAAAATTTTCCACATCTCCTTTTGTAGACACGACTAAACATTTTGCAGNAAAAANTTTATTATTATTTAGATAATCGTATACTTCATCAGTGTAGATAGTAAATGTGAACAGATAAAAGGTATTCTTGCTCAATAGATTGGTAAATTCCATAGAATATATTAATCCTTTCTTCTTGAGATCCCACATCATTTCAACCAAAGACGTTTTATCCAATGATTTACCTCGTACTATACCACATACAAAACTAGATTTTGCAGTGATGTCGTACCTGGTATTCCTACCAAACAAAATTTTACTTTTAGTTCTTTTAGAAAATTCTAAGGTAGAATCTCTATTTGTCAATATGTCATCNATGGAATTACCACTAGCAAAAAATGATAGAAATATATATTGATACATCGCAGCTGGNTTTGATCTACTATACTTTNAAAACGAATCAGATTCCATAATTGCTTGTATATCATCAGCTGAAAAACTATGTTTTACACGTATTNCTTCGGCATTTCNNTTTAATGATATATCTTGTTTAGACAATGATAAAGTTATCATGTCCATGAGAGANNCGTCTNNGTATCGTATAAATATTTCATTAGATGTTAGACGCTTCATTANGGGTATACTTCNATAAGGTTTNTTAATTAGTCCATCATTGGTTGCGTCAAGAACTACTATCTGATGTTGTTNGGTATCTCTAGTGTTACACATGGCCTTACTAAAGTTTGGGTAAATAACTATGATATCTCTATTAATTATAGATGTATATATTTCATTCGTCAAGGATATTAATATCGACTTACTATCGTCATTAATACGNGTAATGTAATCATATAAATCATGCGATAGCCAAGGAAAATTCAAATAGATGTTCATCATATAATCGTCGCTATAATTCATATTAATACTTTGACATTGACTAATTTGTAATATAGCCTCGCCACGAAGAAAGCTCTCGTATTCAGTTTCATCGATAAAGGATACCGTTAAATATAACTGGTTGCCGATAGTCTCATAGTCTATTANNTGGTAAGTTTCGTATAAATACAGAATCCCTAAAATATTATCTAATGTGGGATTAATCCTTACCATAACTGTANAAAATGGAGCCGGAGTCATAACTATTTTACCGTTTGTACTTACTGGAATAGATGAAGGAATAATCTCCGGACATGATGGTAAAGACCCAAATGTCTGTTTGAAGAAATCCAATGTTCCAGGTCCTAATCTNTTGACAAAAATTACGATATTCGATCCCGATATCCTTTGCATTCTATTTACCAGCATATCACGAACTATATTAAGATTATCTATCATGTCTATTCTCCCACCGTTATATAAATCGCCTCCGCTAAGAAACGTTAGTATATCCATACAATGGAATACTTCATTTCNAAAATAGTATTCGTTTTCTAATTCTTTAATGTGAAATCGTATACTAGAAAGGGAAAAATTATCTTTGAGTTTTCCATTAGAAAAGAACCACGAAACTAATGTTCTGATTGCGTCTGACTCCGTCGCTGAATTAATAGATTTACACCAAAAACTCATATAACTTCTAGATGTAGAAGCATTCGCTAAAAAATTAGTAGAATCAAAGGATATAAGTAGATGTTCCAACAAGTGAGCAATTCCCAAGATTTCATCTATATCATTCTCGAATCCGAAATTAGAAATTCCCAAGTAGATATCCNTTTTCATCCGATCATTGATGAAAATACGAACTTTATTCGGTAAGACGATCATTTACTAAGGAGTAAAATAGGAAGTAACGTTCGTATATCGTTATCGTCGTATAAATTAAAGGTGTGTTTTTTGCCATTAAGAGACATTATAATTTTACCAATATTGGAATTATAATATAGGTGTATTTGAGCACTAGAAACGGTCGATGCATCGGTAAATATAGCTGTATCTAATGTTCTAGTCGGTATTTCTTCATTTCGCTGTCTAATGATAGCGTTTTCTCTATCTGTTTCCATTACAGCTGCCTGAAGTTTATTGGTCGGATAATATGTAAAATAATAAGAAATACATACGAATAACAAAAATAAAATAAGATATAATAAAGATGCCATTTAGAGATCTAATTTTGTTCAACTTGTCCAAATTCCTACTTACAGAAGATGAGGAATCGTTGGAGATAGTATCTTCCTTATGTAGAGGATTTGAAATATCTTACGATGACTTAATATCGTACTTTCCAGATAGGAAATACCATAAATATATTTCTAAGGTATTTGAACATGTAGATTTATCGGAGGAATTAAGTATGGAATTCCATGATACAACTCTGAGAGATTTAGTATATCTTAGATTGTACAAGTATTCCAAGTATATACGGCCGTGTTATAAATTAGGAGATAATCTAAAAGGTATAGTTGTTATAAAGGACAGAAATATATATATTAGAGAAGCAAATGATGACTTGATAGAATATCTCCTCAAGGAATACACTCCTCAGATTTATACATATTCTAATGAGCGAGTTCCCATAGCTGGTTCAAAATTAATTCTTTGTGGATTTTCTCAAGTTACATTTATGGCGTATACAACGTCGCATATAACAACAAATAAAAAGGTAGATGTTCTCGTTTCCAAAAAATGTATAGATGAACTAGTCGATCCAATAAATTATCAAATACTTCAAAATTTATTTGATAAAGGAAGCGGAACAATAAACAAAATACTCAGGAAGATATTTTATTCGGTAACAGGTGGCCAAACTCCATAGGTAGCTTTTTCTATTTCGGATTTTAGAATTTCCAAATTCACCAGCGATTTATCGGTTTTGGTGAAATCCAAGGATTTATTAATGTCCACAAATGCCATTTGTTTTGTCTGTGGATTGTATTTGAAAATGGAAACGATGTAGTTAGATAGATGCGCGGCGAAGTTTCCTATTAGGGTTCCGCGCTTCACGTCACCCAACATACTTGAATCACCATCCTTTNAAAAAAATGATAAGATATCAACATGGAGTATATCATACTCGGATTTTAATTCTTCTACTGCCTCACTGACATTTTCACAAATACTACAATACGGTTTACCGAAAATAATCAGTACGTTCTTCATTTATGGGTATCAAAAACTTAAAATCGTTACTGCTGGAAAATAAATCACTGACGATATTAGATGATAATTTATACAAAGTATACAATGGAATATTTGTGGATACAATGAGTATTTATATAGCCGTCGCCAATTGTGTCAGAAACTTAGAAGAGTTAACTACGGTATTCATAAAATACGTAAACGGATGGGTAAAAAAGGGAGGACATGTAACCCTTTTTATCGATAGAGGAAGTATAAAAATTAAACAAGACGTTAGAGACAAGAGACGTAAATATTCTAAATTAACCAAGGACAGAAAAATGTTAGAATTAGAAAAGTGTACATCCGAAATACAAAATGTTACCGGATTTATGGAAGAAGAAATAAAGGCAGAAATGCAATTAAAAATCGATAAACTCACATTTCAAATATATTTATCTGATTATGATAACATAAAAATATCATTGAATGAGATACTAACACATTTCAACAATAATGAGAATGTTACATTATTTTATTGTGATGAACGAGACGCAGAATTCGTTATGTGTCTAGAGGCTAAAACACAGTTCTCTACCACAGGAGAATGGCCGTTAATAATAAGTACCGATCAGGATACTATGCTATTCGCGTCTGCTGATAATCATCCTAAGATGATAAAAAACTTAACTCAACTGTTTAAATTTGTTCCCTCGGCAGAGGATAACTATTTAGCAAAATTAACTGCATTAGTGAATGGATGTGATTTCTTTCCTGGACTCTATGGGGCATCTATAACACCCAACAACTTAAACAAAATACAATTGTTTAGTGATTTTACAATCGATAATATAGTCACTAGTTTGGCAATTAAAAATTATTATAGAAAGACTAACTCTACCGTAGACGTGCGTAATATTGTTACGTTTATAAACGATTACGCTAATTTAGACGATGTCTACTCGTATATTCCTCCTTGTCAATGCACTGTTCAAGAATTTATATTCTCCGCATTAGATGNAAAATGGAATGAATTTAAATCATCTTATTTAGAGAGCNNGCCGTTACCCTGCCAATTAATGTACGCATTAGAACCACGTAAGGAGATTGATGTTTCNGAAGTTAAAACTTTATCATCTNNTATAGATTTCGAAAATACTAAATCAGATATCGATGTTATAAAATCTATATCCTCGATTTTTGGATATTCTAACGAAAACTGTAACACCATAGTGTTCGGCATCTATAAGGATAATTTACTACTGAGTATAAATAATTCATTTTACTTTAACGATAGTCTGTTAATAACCAATACTAAAAGTGATAATATAATAAATATAGGTTACTAGATTAAAAAATGGTGTTCCAGCTCGTGTGTTCTACATGCGGCAAAGATATTTCTCACGAACGATATAAATTGATTATACGNAAAAAATCATTAAAGGATGTACTAGTCAGTGTAAAGAACGAATGTTGTAGGTTAAAATTATCTACACAAATAGAACCTCAACGTAACTTAACAGTGCAACCTCTATTNGATATAAACTAATGGATCCGGTTAATTTTATCAAGACATATGCGCCTAGAGGTTCTATTATTTTTATTAATTATGCCATGTCATTAACTAGTNATTTGAATCCATCGATAGAAAAACANNTGGGTATTTATTATGGTACGTTATTATCGGAACACTTGGTAGTTGAATCTACCTATAGAAAAGGAGTTAGAATAGTCCCATTNGATAGATTTTTTNNANGATATCTTAGTGCNAAAGTATACATGTTAGAGAATATTCNAGTTATGAAAATAGNAGCTGATATGTCGTTAACTTTACTAGGTATTCCATATGGATTTGGTCATGATAGAATGTATTGTTTTAAATTGGTAGCTGAATGTTATNAAAATGCCGGTATTGATACATCGTCTAAACGAATATTAGGTAAAGATATTTTTCTGAGCCAAAACTTTACAGATGATAATAGATGGATAAAGATATATGATTCTAATAATTTAACATTTTGGCAAATTGATTACCTTAAAGGGTGAGTTAATATGCATAACTACTCCTCCGTTGTTTTTTCCCTCGTTCTTTTTCTTAACGTTGTTTGCCATCACTCTCATAATGTAAAGATATTCTAAAATGGTAAACTTTTGCATATCGGATGCAGAAATTGGTATAAATGTTGTAATTGTATTATTTCCCGTCAATGGACTAGTCACAGCTCCATCAGTTTTATATCCTTTAGNGTATTTNTCACTCGTGTCTAGCATTCTAGAGCATTCCATGATCTGTTTATCGTTGATATTGGCCGGAAAGATAGATTTTTTANTTTTTATTATATTACTATTGGCAATTGTAGATATAACTTCTGGTAAATATTTTTCTACCTTTTCAATCTCTTCTATTTTCAAGCCGGCTATATATTCTGCTATATTGTTACTAGTATCAATACCTTTTCTGGCTAAGAAGTCATATGTGGTATTCACTATATCAGTTTTAACTGGTAGTTCCATTAGCCTTTCCACTTCTGCAGAATAATTAGAAATTGGTTCTTTACCAGAAAATCCAGCTACTATAATAGGCTCACCGATGATCATTGGCAAAATCCTATATTGTACCAGATTAATGAGAGCATATTTCATTTCCAATAATTCTGCTAGTTCTTGAGACATTGATTTATTTGATGAATCTATTTGGTTCTCTAGATACTCTACCATTTCTGCCGCATACAATAACTTGTTAGATAAAATCAGGGTTATCAAAGTGTTTAGTGTGGCTAGAATAGTGGGCTTGCACGTATTAAAGAATGCTGTAGTATGAGTAAACCGTTTTAACGAATTATATAGTNTCCAGAAATCTGTGGCGTTGCATACATGAACTGAATGACATCGAAGATTGTCCAATANTTTTAATAGCTGCTCTTTGTCCATTATTTCTATATTTGACTNGCAACAATTGTAGATACCATTAATCACTGATTCCNNTTTCGATNNNGGACAATAGCACAATTGTTTAGCNNTNGACTCTANNTATNNAGAATTAATAGATATATCTCTCAATACAGATTGCACTATACATTTTGAAACTATGTCAAAAATTGTAGAACGACGCTGTTCTGTAGCCATTTAACTTTAAATAATTTACAAAAATTTAAAATGAGCATCCGTATNAAAATCGATAAATTGCGCCAAATTGTGGCATATTTTTCAGAGTTCAGCGAAGAAGTGTCTATAAATGTAGACTTGACGGATGAATTAATGTATATTTTTGCCGCCTTGGGCGGATCTNTAAACATTTGGGCCATTATACCTCTCAGTGCATCAGTGTTCTACCGCGGAGCCGAAAATATTGTGTTTAACCTTCCAGTGTCCAAGGTAAAATCGTGTTTGTGTAGTTTTCACAATGATGCTATCATAAATATAGAACCTGATCTGGAAAATAATCTAGTAAAACTTTCTAGTTATCATGTAGTAAGTGTCGATTGTAACAAGGAACTGATGCCTATTAGGACAGATACTACTATTTGTCTAAGTATAGATCAAAAGAAATCTTACGTATTTAATTTTCACAAGTATGAAGAAAAATGTTGTGGTAGAACCGTCATTCATCTAGAATGGTTGTTGGGCNTTATCAAGTGTATTAGTCAGCATCAGCATTTGGCTATTATGTTTAAAGATGACAATATTATTATGAAGACTNCTGGTAATACTGATGCGTTTTCCAGGGAATATTCTATGACTGAATGTTCTCAAGAACTACAAAAGTTTTCTTTCAAAATAGCTATCTCGTCTCTCAACAAACTACGAGGATTCAAAAAGAGAGTCAATGTTTTTGAAACTAGAATCGTAATGGATAATGACGATAACATTCTAGGAATGTTGTTTTCGGATAGAGTTCAANCCTTTAAGATTAACATCTTTATGGCGTTTTTAGACTAATACTTTCAATGAGATAAATATGGGTGGCGGAGTAAGTGTTGAGCTCCCTAAACGGGATCCACCTCCGGGAGTACCCACTGATGAGATGTTATTAAACGTGGATAAAATGCATGACGTGATAGCTCCCGCTAAGCTTTTAGAATATGTGCATATAGGACCACTAACAAAAGATAAAGAGGATAAAGTAAAGAAAAGATATCCAGAGTTTAGATTAGTCAACACAGGACCCGGTGGTCTTTCGGCATTATTAAGACAATCATATAATGGAACNGCACCCAATTGCTGTCGCACNTTTAATCGTACTCATTATTGGAAGAAGGATGGAAAGATATCAGATAAGTATGAAGAGGGTGCAGTATTAGAATCGTGTTGGCCCGACGTCCACGACACTGGAAAATGCGATGNTGATTTATTCGACTGGTGTCANGGGGATACGTTCGATATAAACATATGCCATCAGTGGATCGGTTCAGCCTTTAATAGGAGTGATAGAACTGTAGAGGGTCGACAATCGTTAATAAATCTGTATAATAAGATGCAAAGATTATGTAGTAAAGATGCTAGTGTACCAATATGTGAATTATTTTTGCATCATTTACGCGCACACAATACAGAAGATAGTAAAGAGATGATCGATTATATTCTAAGACAACAGTCGGCGGACTTTAAACAGAAATATATGAGATGTAGTTATCCCACTAGAGATAAGTTAGAAGAGTCATTAAAATATGCGGAACCTCGAGAATGTTGGGATCCAGAGTGTTCGAATGCCAATGTTAATTTCTTACTAACACGTAATTATAATAATTTAGGACTTTGCAATATTGTACGATGTAATACGAGCGTGAATAACTTACAGATGGATAAAACTTCCTCATTAAGATTATCATGTGGATTAAGCAATAGTGATAGATTTTCTACTGTTCCCGTCAATAGAGCAAAAGTAGTTCAACATAATATTAAACATTCGTTCGACCTAAAATTGCATTTGATCAGTTTATTANNTCTCTTNNTNATATGGATACTAATTGTAGCTATTTAAATGNGTGCCGCANCAAGCATACAGACGACTGTGAATACACTCAGTGAACGTATCTCGTCTAAATTAGAACAAGAAGCGAACGCTAGTGCTCAAACAAAATGTGATATAGAAATCGGAAATTTTTATATCCGACAAAACCATGGATGTAACATCACTGTTAAAAATATGTGCTCTGCGGACGCGGATGCTCAGTTGGATGCTGTGTTATCAGCCGCTACAGAAACATATAGTGGATTAACACCGGAACAAAAAGCATACGTACCAGCTATGTTTACTGCTGCGTTAAACATTCAGACGAGTGTAAACACTGTTGTTAGAGATTTTNAAAATTATGTGAAACAGACTTGTAATTCTAGCGCTGTTGTCGATAACAAATTAAAGATACAAAACGTAATTATAGATGAATGTTACGGAGCCCCAGGATCTCCAACAAATTTGGAATTTATTAATACAGGATCTAGCAAAGGAAATTGTGCCATTAAGGCGTTGATGCAATTGACTACTAAGGCCACTACTCAAATAGCACCTAGACAAGTTGCTGGTACAGGAGTTCAGTTTTATATGATTGTTATCGGTGTTATAATATTGGCAGCGTTGTTTATGTACTATGCCAAGCGTATGCTGTTCACATCCACCAATGATAAAATCAAACTTATTTTAGNNAATAAGGAAAACGTCCATTGGACTACTTACATGGACACATTCTTTAGAACTTCTCCGATGATTATTGCTACCACGGATATACAAAACTGAAAATATATTGATAATATTTTAATAGATTAACATGGAAGTTATCGCTGATCGTCTAGACGATATAGTGAAACAAAATATAGCGGATGAAAAATTTGTAGATTTTGTTATACACGGTCTAGAGCATCAATGTCCTGCTATACTTCGACCATTAATTAGGTTGTTTATTGATATACTATTATTTGTTATAGTAATTTATATTTTTACGGTACGTCTAGTAAGTAGAAATTATCAAATATTGTTGGTGTTGGTGGCGCTAGTCATCACATTAACTANNTTTTTATTACTTTATACTATAATAGTACTAGACTGACTTCTAACAAACATCTCACCNGCCATAAATAAATGCTTGATATTAAAGTCTTCTATTTCTAACACTATTCCATCTGTGGAAAATAATACTCTGACATTATCGCTAATTGATACATCGGTAAGTGATATGCCTATAAAGTANTAATCTTCTTTGGGCACATATACCAGTNTACCAGGTTCTNNNAACCTANTTACTGGTGCTCCTGTAGNATACNNNNNTTTTACCTTGAGAATATCCATTGTTTGCTTGGTCAATAGTGATATGTGATTTTTTATCAACCACTCAAAAAAGTAATTGGAGTGTTCATATCCTCTACGGGCTATTGTCTCATGACCGTGTATGAAATTTAAGTANNACGACTGTGGTAGATTTGTTCTATAGAGCCGGTTGCCGCAAATAGATAGAACTACCAATATGTCTGTACAAATGTTAAACATTAATTGATTAACAGNAAAAACAATGTTCGTTCTGGGAATAGAAACCAGATTAAAACAAAATTCATTAGAATATATGCCACGTTTATACATGGAATATAAAATAACTACAGTTTGAAAAATAACAGTATCATTTAAACATTTAACTTGCNGGGTTAATCTCACAACTTTACTGNTTTTGAACTGTTCAAAATATAGCATAGATCCATGAGAAATACGTTTAGCCGCCTTTAATAGAGGAAATNCAACCGCCTTTCTGGATCTCACCAACGACGATAGTTCNGACCAGCAACTCATTTCTTCATCATCCACCTGTTTTAACATATAATAGGCAGGAGATAGATATCCATCATTGCAATATTCCTTCTCGTAGGCACACAATCTAATATTGATAAAATCTCCATTCTCTTCTCTGTATTTATTATCTTGTCTCGGTGGCTGATTAGGCTGTGGTCTATCGTTGTTGAATCTATTTTGGTCATTAAATCTTTCATTTCTTCCTGGTATATTTCTATCACCTCGTTTGGTTGGATTTTTGTCTATATTATCGTTTGTAACATCGGTACGGGTATTCATTTATCACAAAAAAAACTTCTCTAAATGAGTCTACTACTAGAAAACCTCATCGAAGAAGATACCATATTTTTTGCAGGAAGTATATCTGAGTATGATGATTTACAAATGGTTATTGCTGGTGCAAAATCCAAATTTCCAAGATCTATGCTTTCTATTTTTAATATAGTACCTAGAACGATGTCAAAATATGAGTTGGAGTTGATTCATAACGAGAATATCACAGGGGCAATGTTTACCACAATGTATAATATAAGAAACAATTTGGGTCTAGGCGATGATAAACTAACTATTGAAGCCATTGAAAACTATTTCTTGGATCCTAACAATGAGGTTATGCCTCTTATCATTAATAATACGGATATGACTACCGTCATTCCTAAAAAAAGTGGTAGGAGAAAGAATAAGAACATGGTTATCTTCCGTCAAGGATCATCACCTATCTTGTGTATTTTCGAAACTCGTAAAAAGATTAATATTTATAAAGAAAATATGGAATCCGTATCGACTAAGTATACACCTATCGGAGACAACAAGGCTTTGATATCTAAATATGCGGGAATTAATATCCTGAATGTGTATTCTCCTTCCACGTCCATGAGATTGAATGCCATTTACGGATTCACCAATAAAAATAAACTAGAGAAACTTAGTACTAATAAGGAACTAGAATCGTATAGTTCTAGCCCTCTTCAAGAACCCATTAGGTTAAATGATTTTCTGGGACTATTGGAATGTGTTAAAAAGAATATTCCTCTAACAGATATTCCGACAAAGGATTGATTACTATAAATGGAGAATGTTCCTAATGTATACTTTAATCCTGTGTTTATAGAGCCCACGTTTAAACATTCTTTATTAAGTGTTTATAAACACAGATTAATAGTTTTATTTGAAGTATTCGTTGTATTCATTCTAATATATGTATTTTTTAGATCTGAATTAAATATGTTCTTCATGCCTAAACGAAAAATACCCGATCCTATTGATAGATTACGACGTGCTAATCTAGCGTGTGAAGACGATAAATTAATGATCTATGGATTACCATGGATAACAACTCAAACATCTGCGTTATCAATAAATAGTAAACCGATAGTGTATAAAGATTGTGCAAAGCTTTTGCGATCAATAAATGGATCACAACCAGTATCTCTTAACGATGTTCTTCGCAGATGATGATTCATTTTTTAAGTATTTTGCTAGTCAAGATGATGAATCTTCATTATCTGATATATTGCAAATCACTCAATATCTAGACTTTCTGTTATTATTATTGATCCAATCAAAAAATAAATTAGAAGCTGTGGGTCATTGTTATGAATCTCTTTCAGAGGAATACAGACAATTGACAAAATTCACAGACTCTCAAGATTTTAAAAAACTGTTTAACAAGGTCCCTATTGTTACAGATGGAAGGGTCAAACTTAATAAAGGATATTTGTTCGACTTTGTGATTAGTTTGATGCGATTCAAAAAAGAATCAGCTCTAGCTACCACCGCAATAGATCCTGTTAGATACATAGATCCTCGTCGTGATATCGCATTTTCTAACGTGATGGATATATTAAAGTCGAATAAAGTTGAAAAATAATTAATTCTTTATTGTTATCATGAACGGCGGACATATTCAGTTGATAATCGGCCCCATGTTTTCAGGTAAAAGTACAGAATTAATTAGACGAGTTAGACGTTATCAAATAGCTCAATATAAATGTGTGACTATAAAATATTCTAACGATAATAGATACGGAACGGGACTATGGACACATGATAAGAATAATTTTGCAGCATTGGAAGTAACTAAACTATGTGATGTCTTGGAAGCAATTACAGATTTCTCCGTGATAGGTATAGATGAAGGACAGTTCTTTCCAGACATTGTTGAATTCTGTGAGCGTATGGCAAACGAAGGAAAAATAGTTATAGTAGCCGCGCTCGATGGGACATTTCAACGTAGACCGTTTAATAATATTTTGAATCTTATTCCATTATCTGAAATGGTGGTAAAACTAACTGCAGTGTGTATGAAATGCTTTAAGGAGGCTTCCTTTTCTAAACGATTAGGTACAGAAACCGAGATAGAAATAATAGGAGGTAATGATATGTATCAATCTGTGTGTAGAAAGTGTTACATCGACTCATAATATTATATTTTTTATCTAAAAAACTAAAAATAAACATTGATTAAATTTTAATATAATACTTAAAAATGGATGTTGTGTCGTTAGATAAACCGTTTATGTATTTTGAGGAAATTGATAATGAGTTAGATTACGAACCAGAAAGTGCAAATGAGGTCGCNAAAAAACTGCCGTATCAAGGACAGTTAAAACTATTACTAGGAGAATTATTTTTTCTTAGTAAGTTACAGCGACACGGTATATTAGATGGCGCCACCGTAGTGTATATAGGATCTGCTCCAGGTACACATATACGTTATTTGAGAGATCATTTCTATAATTTAGGAGTGATCATCAAATGGATGCTAATTGACGGCCGCCATCATGATCCTATTCTAAATGGATTGCGTGATGTGACTCTAGTGACTCGGTTTGTTGATGAGGAATATCTACGATCCATCAAAAAACAACTACATCCTTCTAAGATTATTTTAATTTCTGATGTGCGATCCAAACGAGGAGGAAATGAACCTAGTACTGCGGATTTACTAAGTAATTATGCTCTACAAAATGTCATGATTAGTATTTTAAACCCCGTGGCGTCTAGTCTTAAATGGAGATGCCCGTTTCCAGATCAATGGATCAAGGACTTTTATATCCCACACGGTAATAAAATGTTACAACCTTTTGCTCCTTCATATTCAGCTGAAATGAGATTATTAAGTATTTATACCGGTGAGAATATGAGACTGACTCGAGTTACCAAATCAGACGCTGTAAATTATGAAAAAAAGATGTATTACCTTAATAAGATAGTCCGCAACAAAGTAGTTATTAACTTTGATTATCCTAATCAGGAATATGACTATTTTCACATGTACTTTATGTTGAGGACCGTATACTGCAATAAAACATTTCCTACTACTAAAGCAAAGATACTATTTCTACAACAATCTATATTTCGTTTCTTAAATATTCCAACGACATCAACTNAAAAAGTTAGTCATGAACCAATACAACGTAAAATATCTAGNAAAGATTCTATGTCTAAAAACAGAAATAGCAAGAGATCCGTACGCGGTAATAAATAGAAACGTACTACTGAGATATACTACCGATATAGAGTATAATGATTTAGTTACTTTAATAACCGTTAGACATAAAATTGATTCTATGAAAACTGTGTTTCAGGTATTTAACGAATCATCCATAAATTATACTCCGGTTGATGATGATTATGGAGAACCAATCATTATAACATCGTATCTTCAAAAAGGTCATAACAAGTTTCCTGTAAATTTTCTATACATAGATGTGGTAATATCTGACTTATTTCCTAGCTTTGTTAGACTAGATACTACAGAAACTAATATAGTTAATAGTGTACTACAAACAGGCGATGGTAAAAAGACTCTTCGTCTTCCTAAAATGTTAGAGACGGAAATAGTTGTCAAGATTCTCTATCGTCCTAATATACCATTAAAAATTGTTAGATTTTTCCGCAATAACATGGTAACTGGAGTAGAGATAGCCGATAGATCTGTTATTTCAGTCGCTGATTAATCAATTAGTAGAGATGAGATAAGAACATTATAATAATCAATAATATATCTTATATCTGTTTAGAAAAATGCTAATATTAAAATAGCTAACGCTAGTAATCCAATCGGAAGCCATTTGATATCTATAATAGGGTATCTAATTTCCTGATTCAGATAGCGTACGGCTATATTCTCGGTAGCTACTCGTTTGGAATCACAGACATTATTTACATCTAATTTACTATCTGTAATGGAAACGTTTCCCAATGAAATGGTACAATCAGATACATTACATCTTGATATANTTTTTTTTAAAGAGGCTGGTAACAACGCATCGCTTCGTTTACATGGCTCGTACCAACAATAATAGGGTAATCTTGTATCTATTCCTATCCGTACTATACTTTTATCAGGATAAATACATTTACATCGTATATCGTCTTTGTTAGTATCACAGAATGCATAAATTTGTTCGTCCGTCATGATAAAAATTTAAAGTGTAAATATAACTATTATTTTTATAGTTATAATAAAAAGGGAAATTTGATTGTATACCTTCGGTTCTTTAAAAGAAACTGACTTGATAAAAATGGCTGTAATCTCTAAGGTTACGTATAGTCTATACGATCAAAAAGAGATTAATGCCACAGATATTATCATTAGTCATATTAAAAATGACGACGATATCGGTACCGTTAAAGATGGTAGACTAGGTGCTATGGATGGGGCATTATGTAAAACTTGTGGGAAAACGGAATTGGAATGTTTCGGTCACTGGGGTAAAGTAAGTATTTATAAAACTCATATAGTTAAGCCTGAATTTATTTCAGAAATTATTCGTTTACTGAATCATATATGTATTCATTGCGGATTATTGCGTTCACGAGAACCGTATTCCGACGATATTAACCTAAAAGAGTTATCGGTACACGCTCTTAGGAGATTAAAGGATAAAATATTATCCAAGNAAAAGTCATGTTGGAACAGCGAATGTATGCAACCGTATCAAAAAATTACTTTTTCAAAGAAAAAGGTTTGTTTCGTCAACAAGTTGGATGATATTAACGTTCCTAATTCTCTCATCTATCAAAAGTTAATTTCTATTCATGAAAAGTTTTGGCCATTATTAGAAATTCATCAATATCCAGCTAACTTATTTTATACAGACTACTTTCCCATCCCTCCGTTGATTATTAGACCGGCTATTAGTTTTTGGATAGATAGTATACCCAAAGAGACAAATGAATTAACTTACTTATTAGGTATGATCGTTAAGAATTGTAACTTGAATGCTGATGAACAGGTTATCCAGAAGGCGGTAATAGAATACGATGATATTAAAATTATTTCTAATAACACTACCAGTATCAATTTATCATATATCACATCCGGCAAAAATAATATGATTAGAAGTTATATCGTCGCTCGGCGAAAAGATCAGACCGCTAGATCCGTAATTGGTCCCAGTACATCTATCACCGTTAATGAGGTAGGAATGCCCACATATATTAGAAATACACTTACAGAAAAGATATTTGTTAATGCCTTTACAGTGGATAAAGTTAAACAACTATTAGCATCAAACCAAGTTAAATTTTACTTTAATAAACGATTAAACCAATTAACAAGAATACGTCAAGGAAAGTTTATCAAAAATAAAATACATTTATTGCCTGGTGATTGGGTAGAAGTAGCTGTTCAAGAATATACAAGTATTATTTTTGGAAGACAACCGTCTCTACATAGATACAACGTCATCGCTTCATCTATCAGAGCTACCGAAGGAGATACTATCAAAATATCTCCCGGAATTGCCAACTCTCAAAATGCTGATTTTGACGGAGATGAAGAATGGATGATATTGGAGCAAAATCCTAAAGCCGTAGTTGAACAAAGTATTCTTATGTATCCGACAACGTTACTCAAACACGATATTCATGGAGCCCCCGTTTATGGATCTATTCAAGATGAAATCGTAGCAGCGTATTCATTGTTTAGGATACAAGATCTTTGTTTAGATGAAGTATTGAACATCTTGGGGAAATATGGAAGAGAGTTCGATCCTAAAGGTAAATGTAAATTCAGCGGTAAAGATATCTATACTTACTTGATAGGTGAAAAGATTAATTATCCGGGTCTCTTAAAGGATGGTGAAATTATTGCAAACGACGTAGATAGTAATTTTGTTGTAGCTATGAGGCATCTGTCATTGGCTGGACTCTTATCCGATCATAAATCGAACGTGGAAGGTATCAACTTTATTATCAAGTCATCTTATGTTTTTAAGAGATATCTATCTATATACGGTTTTGGGGTGACATTCAAAGATCTGAGACCAAATTCGACGTTCACTAATAAATTGGAGGCTATCAACGTAGAAAAAATAGAACTTATCAAAGAAGCATACGCCAAATATCTCAAAGATGTAAGAGACGGGAAAATAGTTCCATTATCTAAAGCTTTAGAGGCGGACTACTTGGAATCCATGTTATCCAACTTGACAAATCTTAATATCAGAGAGATAGAAGAACATATGAGACAAACGCTGATAGATGATCCAGATAATAACCTCCTGAAAATGGCCAAAGCGGGTTATAAAGTAAATNCCACAGAACTAATGTATATTCTAGGTACTTNTGGACAACNNAGGATAGATNNNGAANCAGCAGAGNNNNNNNNNNNNNNNNNNNNNNNNNNNNNNNNNNNNNNNNNNNNNNNNNNNNNNNNNNNNNNNNNNNNNNNNNNNNNNNNNNNNNNNNNNNNNNNNNNNNNNNNNNNNNNNNNNNNNNNNNNNNNNNNNNNNNNNNNNNNNNNNNNNNNNNNNNNNNNNNNNNNNNNNNNNNNNNNNNNNNNNNNNNNNNNNNNNNNNNNNNNNNNNNNNNNNNNNNNNNNNNNNNNNNNNNNNNNNNNNNNNNNNNNNNNNNNNNNNNNNNNNNNNNNNNNNNNNNNNNNNNNNNNNNNNNNNNNNNNNNNNNNNNNNNNNNNNNNNNNNNNNNNNNNNNNNNNNNNNNNNNNNNNNNNNNNNNNNNNNNNNNNNNNNNNNNNNNNNNNNNNNNNNNNNNNNNNNNNNNNNNNNNNNNNNNNNNNNNNNNNNNNNNNNNNNNNNNNNNNNNNNNNNNNNNNNNNNNNNNNNNNNNNNNNNNNNNNNNNNNNNNNNNNNNNNNNNNNNNNNNNNNNNNNNNNNNNNNNNNNNNNNNNNNNNNNNNNNNNNNNNNNNNNNNNNNNNNNNNNNNNNNNNNNNNNNNNNNNNNNNNNNNNNNNNNNNNNNNNNNNNNNNNNNNNNNNNNNNNNNNNNNNNNNNNNNNNNNNNNNNNNNNNNNNNNNNNNNNNNNNNNNNNNNNNNNNNNNNNNNNNNNNNNNNNNNNNNNNNNNNNNNNNNNNNNNNNNNNNNNNNNNNNNNNNNNNNNNNNNNNNNNNNNNNNNNNNNNNNNNNNNNNNNNNNNNNNNNNNNNNNNNNNNNNNNNNNNNNNNNNNNNNNNNNNNNNNNNNNNNNNNNNNNNNNNNNNNNNNNNNNNNNNNNNNNNNNNNNNNNNNNNNNNNNNNNNNNNNNNNNNNNNNNNNNNNNNNNNNNNNNNNNNNNNNNNNNNNNNNNNNNNNNNNNNNNNNNNNNNNNNNNNNNNNNNNNNNNNNNNNNNNNNNNNNNNNNNNNNNNNNNNNNNNNNNNNNNNNNNNNNNNNNNNNNNNNNNNNNNNNNNNNNNNNNNNNNNNNNNNNNNNNNNNNNNNNNNNNNNNNNNNNNNNNNNNNNNNNNNNNNNNNNNNNNNNNNNNNNNNNNNNNNNNNNNNNNNNNNNNNNNNNNNNNNNNNNNNNNNNNNNNNNNNNNNNNNNNNNNNNNNNNNNNNNNNNNNATGGTTCTTCCAGGTGNCGCCAACANGGGNNANNTTAGTAAATTCNAGATTCCTATCTCTGACTATACGGGATATAACGACNTCAATCAAACNAAANNNNNNNNNNNNNNNNNNNNNNNNNNNNNNNNNNNNNNNNNNNNNNNNNNNNNNNNNNNNNNNNNNNNNNNNNNNNNNNNNNNNNNNNNNNNNNNNNNNNNNNNNNNNNNNNNNNNNNNNNNNNNNNNNNNNNNNNNNNNNNNNNNNNNNNNNNNNNNNNNNNNNNNNNNNNNNNNNNNNNNNNNNNNNNNNNNNNNNNNNNNNNNNNNNNNNNNNNCTAGTTACGANNCAGAATCANTTAATAAATTCAAGTTCGGTGCAGCTAGTACTCTTANNAGAGNNNCGTTCGGAGATAATAAAGCATTGTTAAACGCGGCTCTTCNTAAAAAGTCAGAACCTANNNNNNNNNNNNNNNNNNNNNNNNNNNNNNNNNNNNNNNNNNNNNNNNNNNNNNNNNNNNNNNNNNNNNNNNNNNNNNNNNNNNNNNNNNNNNNNNNNNNAANGAAACTATCTGATAAGATATCTTCTCNAAAGATCNAGGAGATAGNAGAAACNGNAGACNTTTAATTCTTATCNNNAACATANTTTTCNATGATCTGTCTTTTAAACGATNGATTTTNCACAAATGCGCCTCTCNANTCCCTCATAGAATGATNNACGNATNNANNNNATAGCATAGGTNATGANNCCTTATTTTTAGACATTAGATATNNNAAAATCATAGCCCNNCTTCTATTTACTNCTGCAACACAATGAACCAACACNGGCTCGNTTNGTTGATCACATNNNGANAAGAAGGCGGTCNCNTCGTCAAAANNNTTACTAATATCNGTAGTTGTATCATCTACCAACGGTATATGAATAATATTAATATTAGAGTTAGGTAATGTATATTTATCCATCGTCAAATTTAAAACATATTTGAACTTAACTTCAGATGATGGTGCATCCATAGCATTTTTATAATTTCCCAAATNNACATTATTTGTTACTCTTGTCATTATAGTGGGAGATTTGGCTCTGTGCATATCTCCAGTTGAACGTAGTAGTAAGTATTTATACAAACTTTTCTTATCCATTTATAACGTACAAATGGATAAAACTACTTTATCAGTAAACGCATGCAATTTAGAATACGTTAGAGAAAAGGCTATAGTAGGCGTACAAGCAGNCAAGACATCAACACTTATANTTTTTGTTATTATATTGGCAATTAGTNNGCTATTACTCTGGTTTCAGACGTCTGATAATCCAGTCTTTAATGAATTAACNAGATATATGCGAATNAAAAATACNGTTAACGATTNNAAATCATTAACGGATAGCAAAACAAAATTAGAAAGCGATAGAGGTAGACTTCTAGCCGCTGGTAAGGATGATATATTCGAATTCAAATGTGTGGATTTCGGCGCCTATTTTATAGCTATGCGATTGGATAAGAAAACATATCTGCCGCAAGCTATTAGGCGAGGTACTGGAGACGCGTGGATGGTTNAAAAGGCGGCNAAAGTCGATCCATCTGCTCAACAATTTTGTCAGTATTTGATAAAACACAAGTCTAATAATGTTATTACTTGTGGTAATGAGATGTTAAATGAATTAGGTTATAGCGGTTATTTTATGTCACCGCATTGGTGTTCCGATCTTAGTAATATGGAATAAGTGTTAGATAAATGCGGTAACAAATGTTCCTGTAAGGAACCATAACAGTTTAGATTTAACATTAAAGATGAGCATAAACATAATAAACAAAATTACAATCAAACCTATAACATTAATATCAAACAATCCAAAAAATGAAATCAATGGAGTAGTAAACGTGTACATAACTCCTGGATAACGTTTAGCAGCTACCGTTCCTATTCTAGACCAAAAATTTGGTTTCATGGTTTCGAAGCGGTGTTCTGCAACAAGACGAGGATCGTGTTCTACATATTTGGCAGAGTTATCCATTATTTGCCTGTTAATCTTCATTTCGTTTTCGATTCTGGCTATTTCAAAATAAAATCCCGATGATAGACCTCCAGACTTTATAATTTCATCTACGATGTTCAGCGCCGTAGTAACTCNAATAATATAGGCGGATAAGCTAACATCATACCCTCCTGTATATGTAAATATGGCATGATCTTTGTCNANTACAAGCTCGGTTTTAACTTTATTTCCTGTAATAATTTCTCTCATCTGTAGGATATCTATTTTCTTGTCATGTATTGCCTTCAAGACGGGACGAAGAAACGTAATATCCTCAATAACGTTATCGTTTTCTATAATAACTACATATTCTACATTTTTATTTTCTAGCTCGGTNAAAAATTTAGAATCCCATAGGGCTAAATGTCTAGCGATATTTCTTTTCGTTTCCTCTGTACACATAGTGTTACAAAACCCTGAAAAGAAGTGAGTATACTTGTCATCATCTCTAATATTTCCTCCAGTCCATTGTATAAACACATAATCCTTGTAATGATCTGGATCATCATTGACTATCACAACATCTCTTTTTTCTTGCATAACTTCNTTGTCCTTCACATCATCGAACTTCTGATCATTAATATGCTCATGAACATTAGGAAATGTTTCTGATGGAGGTCTATCAATAACTGGCACAACAATAACAGGAGTTTTCACCGCCGCCATTTAGTTATTGAAATTAATCATATACAACTCTCTAATACGAGTTATATTTTCGTCTATCCATTGTTTCACATTGACATATTTCGACAAAAAGATATAAAATGCGTATTCCAATGCTTCTCTGTTTAATGAATTACTAAAATATACAAACACGTCACTGTCTGGTAATAAATAATATCTTAGAATATTGTAACAATTTATTTTGTATTGCACATGTTCGTGATCTATGAGTTCTTCTTCAAATGGCATAGGATCTCCGAATCTGAAAACGTATAAATAGGAGTTAGAATAATAATATTTGAGAGTATTGGTAATGTATAAACTCTTTAGCGGTATAATTAGTTTTTTTCTCTCGATTTCTATTTTTAGATGTGATGGAAAAATGACTAATTTTGTAGCATTAGTATCATGAACTCTAATCAAAATCTTAATATCTTCGTCACATGTTAGCTCTTTGAAGTTTTTAAGAGATGCATCAGTTGGTTTTACAGATGGAGTAGGTGCAACAATTTTTTGTTTAATGCATGCATGTATTGGAGCCATTGTCTTAACTATAATGGTGCTTGTATCGAAAAACTTTAATGCGGATAACGGAAGCTCTTCGCCGCGACTTTCTACGTCGTAATTGGGTTCTAATGCCGATCTCTGAATGGATACTAGTTTTCTAAGTTCTAATGTAATTCTCTGAAAATGTAAATCCAATTCCTCCGGCATTATAGATGTGTATACATCGGTAAATAAAACTATAGTATCCAACGATNCCTTCTCGCAAATTCTAGTCTTAACCAAGAAATCGTATATAACTACGGAGATGGCGTATTTNAGAGTGGATTCTTCTACCGTTTTGTTCTTGGATTTCATATAAGAAACTATAAAGTCCGCACTACTGTTAAGAATGATCACTAACGCAACTATATAGTTCAAATTAAGCATCTTGGAAACATAAAATAACTCTGTAGATGATACTTGACTTTCGAATAAGTTTGCAGACAAACGAAGAAAGAACAGACCTCTCTTAATTTCAGAAGAAAACNTTTTTTCGTATTCCTGACGTCTAGAGTTTATATCAATAAGAAAGTTAAGAATTAGTCGGTTAATGTTGTATTTCATTACCCAAGTTTGAGATTTCATAATATTGTCAAAAGACATGATAATATTNAAGATAAAGCGCTGACTATGAACGAAATAGCTATATGGTTCGCTCAAGAATATAGTCTTGTTAAACGTGGAAACGATAACTGTATTTTTAATCACGTCAGCGGCATCTAAATTAAATATAGGTATATTTATTCCACACACTCTACAATATGCCACACCATCTTCATAATAAATAAATTCGTTAGCAAAATTATTAATTTTAGTGAAATAGTTAGCGTCAACTTTCATAGCTTCCTTCAATCTAATTTGATGCTCACATGGCGCGAATTCTACTCTAACATCCCTTTTCCATGCCTCAGGTTCATCGATCTCTATAATATCTAGTTTCTTGCGTTTCACAAACACAGGCTCGTCTCTCGCGATGAGATCTGTATAGTAACTATGTAAATGATAACTAGATAGAAAGATGTAGCTATATAGATGACGATCCTTTAAGAGAGGTATAATAACTTTACCCCAATCAGATAGACTGTTGTTATGGTCTTCGGAAAAAGAATTTTTATAAANTTTTCCAGTATTTTCTAAATATACGTACTTGATATCTAAGAAATCCTTAATAATAATAGGAATGGATAATCCGTCTATTTTATAAAGAAATACATATCGCATATTATACNTTTTTTTGGAAATTGGAATACCGATGTGTCTACATAAATACGCAAAGTCTAAATATTTTTTAGAGAATCTTAGTTGGTCCAAATTCTTTTCCAAGTACGGTAATAGATTTTTCATATTGAACGGTATCTTCTTGATCTCTGGTTCTAATTCCGCATTAAATGATGAAACTAAGTCACTATTTTTATAACTAACGATTACATCACCTCTAACATCATCATTTACCAGGATACTGATCTTCTTTTGTCGTAAATACATGTCTAATGTGTTNNNNAAAAGATCNTNCAAGTTATACGTCATTTCATCTGTAGTATTCTTGTCATTGAAGGATNAACTCGTACTAATCTCTTCTTTAACAGTCTGTTCAAATTTATATNCTATATATGNAAAAATAGCAACCNGTGTTTGATCATNCGCGTCAATATTCTGTTCTATCGTANTNTANAACAATCTTATATCNTCTTCTGTGATAGTCGATACGTTATAAAGGTTGATAACGAAAATANTTTTATTTCGTGAAATAAAGTCATTGTAGGATTTTNGACTTATATTCGTGTCTAGTAGATATGATTTTATTTTTGGAATGATCTCAATTAAAATAGTCTCTTTAGAGTCCATTTAAAGTTACAAACAACTAGGAAATTGGTTTATGATGTATAANTTTTTTAGTTTTTATAGATTCTTTATTCTATACTTAAAAAATGAAAATAAATACNAAGGTTCTTGAGGGTTGTGTTAATTGAAAGCGATAAATAATCATAAATTATTTCATTATCGCGATATCCGTTAAGTTTGTATCGTAATGGCGTGGTCAATTACGAATAAAGCGGATACTAGTAGTTTCACAAAGATGGCTGAAATCAGAGCTCATCTAAGAAATAGCGCTGAAAATAAAGATAAAAACGAGGATATTTTCCCGGAAGATGTAATAATTCCATCTACTAAGCCCAAAACCAAACGAACCACTACTCCTCGTAAACCAGCGGCTACTAAAAGATCAACCAAAAAGGATAAAGAAAAGGAGGAAGTGGAAGAAGTAGTTATAGAGGAATATCATCAAACAACTGAAGAAAATTCTCCACCTCCGTCATCATCTCCTGGAGTCGGCGACATTGTAGAAAGCGTGGCCGCTGTAGAGCTCGATGATAGCGACGGGGATGATGAACCTATGGTACAAGTTGAAGCTGGTAAAGTAAATCATAGTGCTAGAAGCGATCTCTCTGACCTAAAGGTGGCTACCGACAATATCGTTAAAGATCTTAAGAAAATTATTACTAGAATCTCTGCAGTATCGACTGTTCTAGAGGATGTTCAAGCAGCTGGTATCTCTAGACAATTTACTTCTATGACTAAAGCTATTACAACACTATCTGATCTAGTCACCGAGGGAAAATCTAAAGTTGTTCGTAAAAAAGTTAAAACTTGTAAGAAGTAAATGCGTGCACTTTTTTATAAAGATGGTAAACTGTTTACCGATAATAANTTTTTAAATCCTGTATCAGACGATAATCCAGCGTATGAGGTTTTGCAACATGTTAAAATTCCTACTCATTTAACAGATGTAGTAGTATATGAACAAACGTGGGAAGAGGCATTAACTAGATTAATTTTTGTGGGAAGTGATTCNAAAGGACGTAGACAATACTTTTACGGAAAAATGCATATACAGAATCGCAATGCTAAAAGAGATCGTATTTTTGTTAGAGTATATAACGTTATGAAACGAATTAATTGTTTTATAAACAAAAATATAAAGAAATCGTCCACAGATTCCAATTATCAGTTGGCGGTTTTTATGTTAATGGAAACTATGTTTTTTATTAGATTTGGTAAAATGAAATATCTTAAGGAGAATGAAACAGTAGGGTTATTAACACTAAAAAATAAACACATAGAAATAAGTCCCGATGAAATAGTTATCAAGTTTGTAGGAAAGGACAAAGTTTCACATGAATTTGTTGTTCATAAGTCTAATAGACTATATAAACCGCTATTGAAACTGACTGATGATTCTAGTCCCGAAGAATTTCTGTTCAACAAACTAAGTGAACGAAAGGTATATGAATGTATCAAACAGTTTGGTATTAGAATCAAGGATCTCCGAACGTATGGAGTCAATTATACGTTTTTATATAATTTTTGGACAAATGTAAAGTCCGTATCTCCTCTTCCATCACCAAAAAAGTTGATAGCATTAACTATCAAACAAACTGCTGAAGTGGTAGGTCATACTCCATCAATTTCAAAAAGAGCTTATATGGCAACGACTATTTTAGAAATGGTAAAGGATAAAAATTTTTTAGACGTAGTATCTAAAACTACGTTCGATGAATTCCTATCTATAGTCGTAGATCACGTTAAATCATCTACGGATGGATGATAATAGATCTTTACACAAATAATTACAAGACCGATAAATGGAAATGGATAAACGGATGAAATCTCTCGCTATGACAGCTTTCTTCGGAGAGCTAAACACGTTAGATATTATGGCATTGATAATGTCTATATTTAAACACCATCCAAACAATACCATTTTTTCAGTGGATAAGGATGGTCAATTTATGATTGATTTCGAATACGATAATTATAAGGCTTCTCAATATTTGGATCTGACCCTCACTCCGATATCTGGAAATGAATGCAAGACTCACGCATCTAGTATAGCCGAACAATTGGCGTGTGTGGATATTATTAAAGAGGATATTAGCGAATATATCAAAACTACTCCCCGTCTTAAACGATTTATAAAAAAATACCGCAATAGATCATATACTCGTATCAGTCGAGATACAGAAAAGCTTAAAATAGCTCTAGCTAAAGGCATAGATTACGAATATATAAAAGACGCTTGTTAATAAGTAAATGAAAAAAAACTAGTCGTTTATAATAAAACACAATATGGATGCCAACATAGTATCATCTTCTACTATTGCGACGTATATAGACGCTTTAGCGAAGAATGCTTCAGAATTAGAACAGAGGTCTACCGCATACGAAATAAATAATGAATTNGAACTAGTATTTATTAAACCGCCATTGATTACGTTGACAAATGTAGTNAATATCTCCACGATTCAGGAATCGTTTATTCGATTTACCGTTACTAATAAGGAAGGTATCAAAATTAGAACTNAGNTTNNATTATCTNNNGTACATGGTCTNNNNNNNNNNNNNNNNNNNNNNNNNNNNNNNNNNGATANNATAGTNNNNGNNAAGAAATCATTAGTGACGGAAAATCGTCTTCACNAAGAATGCTTGTTGAGACNATCAACAGAGGAACGTCATATNTTTTTGGATTACAAGAAATATGGATCCTCTATCCGACTAGAATTAGTCAATCTTATTCAAGCAAAAACAAAAAANTTTACGATAGACTTTAAGCTAAAATATTTTCTANGATCTGGCGCTCAATCTAAAAGTTCTTTATTGCNCGCTATTAATCATCCNAAGTCAAGGCCTAATANATCTCTNGAAATAGAATTTACACCTAGAGACAATGAAACAGTTCCATATGATGAACTAATAAAGGAATTGACGACTCTCTCGCGTCATATATTTATGGCTTCTCCAGAGAATGTAATTNTTTCTCCACCTATTAACGCACCTATAAAGACNTTTATGTTGCCTAAACAAGATATAGTAGGTCTNGATCTGGAAAATCTATATGCCGTAACTAAGACTGACGGCATTCCTATAACTATCAGAGNTACATCAAAAGGGTTGNATTGTTANTTTANACATCTTGGTTATATTATTAGATATCCAGTTAAGAGAACAATAGATTCCGAAGTAGTAGTCTTTGGTGAGGCAGTTNAGGATAAGAACTGGACCGTATATCTCATTAAGCTAATAGAGCCCGTAAATGCAATCAGTGATAGACTAGAAGAAAGTAAGTATGTTGAATCTAAACTAGTGGATATTTGTGATCGGATAGTATTCAAGTCAAAGAAATACGAAGGTCCGTTTACTACAACTAGTGAAGTCGTCGATATGTTATCTACATATTTACCAAAGCAACCAGAAGGTGTTATTCTGTTCTATTCAAAGGGACCTAAATCTAACATTGATTTTAAAATCAAAAAGGAGAATACTATAGACCAAACTGCAAATGTAGTATTTAGGTACATGTCCAGTGAACCAATTATCTTTNGAGAGTCGTCTATCTTTATAGAGTATANNAAATTTACCAACGATANNNGCTTTNNNAAAGAATATGGTTCTNGTNNGATTGTGTNNTATAACGGCGTTAATTATCTAAATAATATCTATTGTTTGGAATATATTAATACACATAATGAAGTGGNNATTNAGTNNGTTGTNGTACCTATTAAGTTTATAGCAGAATTCTTAGTCAATGGAGAAATACTTAAACCTAGAATCGATAAAACCATGAAATATATTAACTCAGAAGACTATTATGGAAATCAACATAATATCATAGTCGAACATTTAAGAGATCNAAGCATCAAAATAGGAGATGTCTTTANCGAGGATAAACTATCGGATGTTGGACATCAATACGCTGCCAACAACGATAAATTTAGATTAAATCCAGAAGTTAGTTATTTTACTAATAAACGAACTAGAGGGCCGTTGGGAATTTTATCAAACTACGTCAAGACTCTTCTTATTTCTATGTATTGTTCCAAAACATTTTTAGACGATTCCAACAAACGAAAGGTATTAGCGATTGATTTTGGAAACGGTGCTGACCTGNAAAAATACTTTTATGGAGAGATTGCGTTATTGGTAGCGACGGATCCGGATGCTGATGCNNTAGCTAGAGGAAATGAAAGATACAACAAATTAAATTCTGGAATTAAAACCAAGTACTACAAATTTGACTACATTCAGGAAACTATTCGATCCGATACATTTGTCTCTAGTGTCNNNGAAGTATTCTATTTTGGAAAGTTTAATATCATTGACTGGCAGTTCGCTATTCATTATTCTTTTCATCCAAGACATTATGCTACAGTCATGAATAACTTATCCGAACTAACTGCTTCTGGAGGCAAGGTATTAATTACTACCATGGATGGAGACAAATTATCAAAATTAACCGATAAAAAGACTTTTATAATTCATAAGAATCTACCTAGTAGCGAAAACTATATGTCTGTAGAAAAAATAGCTGATGATAGAATAGTGGTATATAATCCATCAACAATGTCTACTCCAATGACTGAATACATTATCNANNNNNNNNNNNNNNNNNNNNNNNNNNNNNNNNNNNNNNNNNNNNNNNNNNNNNNNNNNNNNNNNNNNNNNNNNNNNGAACNAAGTNNAAAGNTTATTANNNGCGCATCTACAATGGAAGATAGACCGTCTACAANAAACTTTTTCGAACTAAATAGAGGAGCCATTAAANNTGAAGGTTTAGATGTCGAAGACTTACTTAGTTACTATGTTGTTTATGTCNTTTCTNNNNNGTNAANAATANNATGGTATNGGTTCTGNTATNCCCGTTCTAAATGCATTAAATAATTCCAATAGAGCGATTTTTGTTCCTATAGGACCTTCCAACTGTGGATACTCTGTATTATTAATAGATATATTAATACTTTTGTAGGGTAACAGAGGTTCTACGTCTTCTAAAAATNAAAGTTTTATAACATCTGGCCTGTTCATAAATAAAAACTTGGCGATTCTATATATACTCTTATTATCAAATCTAGNNATTGTCTTATAGATGTGAGCTACTGTAGGTGTACCATTTGATTTTCTTTCNAATACTATATATTTCTCTCGAAGAAGTTCTTGCAGATCATCTGGGAATAAAATACTACTGTTGAGTAAATCAGTTANTTTTTTTATATCGATATTGATGGACATTTTTATAGTTAAGGATAATAAGTATCCCAAAGTAGATAACGACGATAACGAAGTATTTATACTTTTNNGAAATCACAATGACTTTATCAGATCNAAATTAACAAAATTAAANGAGCATGTANNTTTTTCTGAATATATTGTGACTCCAGATACATATGGATCTTTATGCGTCGAATTAAATGGGTCTAGTTTTCAGCACGGTGGTAGATATATAGAGGTGGAGGAATTTATAGATGCTGGAAGACAAGTTAGATGGTGTTCTACATCCAATCATATATCTGAAGATATACACACTGATAAATTTGTCATTTATGATATTTATACGTTTGATTCGTTNAAGAATAAACGATTGGTATTTGTACAGGTACCTCCATCATTAGGAGATGATAGCTATTTAACTAATCCGTTATTGTCTCCGTATTATCGTAATTCAGTAGCCAGACAAATGGTCAATGATATGATTTTTAATCAAGATTCATTTTTAAAATATTTATTAGAACATCTGATTNGAAGCCACTATAGAGTTTCTAAACATATAACAATAGTTAGATACAAGGATACCGAAGAATTAAATCTAACAAGAATATGTTATAATAGAGATAAGTTTAAGGCGTTTGTATTCGCTTGGTTTAACGGCGTTTCGGAAAATGAAAAGGTACTAGATACGTATAAAAAGGTATCTGATTTGATATAATGAATTCAGTGACTATATCACACGCACCATATACTATTACTTATCACGATGATTGGGAACCAGTAATGAGTCAATTGGTAGAGTTTTATAACGAAGTAGCCAGTTGGTTGCTACGCGACGAGACGTCGCCTATTCCTGATAAGTTCTTTATACAATNGAAACAGCCGCTTAGAAATAAACGAGTANNTGTGTGTGGTATAGATCCGTATCCNAAAGATGGAACTGGTGTACCGTTCGAATCACCAAATTTTACNAAAAAATCAATTAAGGAGATAGCTTCATCTATATCTAGATTAACCGGAGTAATTGATTATAAAGGTTATAACCTTAATATAATAGACGGGGTTATACCCTGGAATTATTACTTAAGTTGTAAATTANGAGAAACAAAAAGTCACGCGATTTACTNGGATAAGATTTCCAAGTTACTGCTACAGCATATAACTAAACACGTTAGTGTTCTTTATTGTTTGGGTAAAACAGATTTCTCGAATATACGGGCAAAGTTAGAATCCCCGGTAACTACCATAGTGGGATATCATCCAGCGGCCAGAGACCACCAATTCGAGAAAGATCGATCATTTGAAATTATCAACGTTTTACTGGAATTAGACAACAAGACACCTATAAATTGGGCTCAAGGGTTTATTTATTAATGCTTTAGTGAAATTTTAACTTGTGTTCTAAATGGATGCGGCTATTAGAGGTAATGATGTTATCTTTGTTCTTAAGACTATANGTGTCCCGTCAGCATGCAGACAAAATGAAGATCCAAGATTCGTAGAAGCATTTAAATGCGACGAGTTAGAAAGATATATTGATAATAATCCAGAATGTACACTATTCGAAAGTCTTAGGGATGAGGAAGCATACTCTATAGTCAGAATTTTCATGGATGTAGATTTAGACGCGTGTCTAGACGAAATAGATTATTTAACGGCTATTCAAGATTTTATTATCGAGGTGTCAAACTGTGTAGCTAGATTCGCATTTACAGAATGCGGTGCCATTCATGAAAATGTAATAAAATCCATGAGATCTAATTTTTCATTGACTAAGTCTACAAATAGAGATAAAACAAGTTTTCATATTATCTTTTTAGACACGTATACCACTATGGATACATTGATAGCTATGAAACGAACACTATTAGAATTAAGTAGATCATCTGAAAATCCACTAACAAGATCGATAGACACTGCCGTATATAGGAGAAAAACAACTCTTCGGGTTGTAGGTACTAGNAAAAATCCAAATTGCGACACTATTCATGTAATGCAACCACCTCACGATAATATAGAAGATTACCTATTCACTTACGTGGATATGAACAACAATAGTTATTACTTTTCTCTACAACGACGATTGGAGGATTTAGTTCCTGATAAGTTATGGGAACCAGGGTTTATTTCGTTCGAAGACGCTATAAAAAGAGTTTCAAAAATATTCATTAATTCTATAATAAACTTTAATGATCTCGATGAAAATAATTTTACAACGGTACCACTGGTCATAGATTATGTAACACCTTGTGCATTATGTAAAAAACGATCGCATAAACATCCGCATCAACTATCGTTGGAAAATGGTGCTATTAGAATTTACAAAACTGGTAATCCACATAGTTGTAAAGTTAAAATTGTTCCGTTNGATGGTAATAAACTGTTTAATATTGCACAAAGAATTTTAGACACTAACTCTGTTTTATTAACCGAACGAGGAGACCATATAGTTTGGATTAATAATTCATGGAAATTTAACAGCGAAGAANCCTTGATAACNAAACTAATTCTATCAATAAGACATCAACTACCTAAGGAATATTCAAGCGAATTACTCTGTCCGAGGAAACGAAAGACTGTAGAAGCTAACATACGAGACATGTTAGTAGATTCAGTAGAGACCGATACCTATCCGGATAAACTTCCGTTTAAAAATGGTGTATTGGACCTGGTAGACGGAATGTTTTACTCTGGAGATGATGCTNAAAAATATACNNGTACTGTATCGACCGGATTTAAATTTGACGATACAAAATTCGTCGAAGACAGTCCAGAAATGGAAGAGTTAATGAATATCATTAACGATATCCAACCATTAACGGATGAAAATAAGNAAAATAGAGAGCTGTATGNNAAAACATTATCTAGTTGNTTATGTGGTGCTACNNNANGATGTTTAACATTCNNNNTTGGAGAAACCGCAANNNNGAAGTCGACAACCAAACGTTTGTTAAAGTNNNCTATCGGTGACCTGTTTGTCGAGACGGGTCAAACAANTTTAACAGATGTATTNGANAAAGGACCTAATCCATTTATCGCTAATATGCATTTNNAAAGATCTGTANTCNGTAGNNNACNACCTGNTTTTGCATGTANNGGATCNAAGNNNNTTANATCNNNNNNNNNTAAAAAGTTGACAGAACCTTGTGTCATTGGAAGACCGTGTTTCTNCAATAAAATTAATAATANAAACCATGCGACAATCATTATCGATACTAATTACAAACCTGTCTTTGATAGGATAGATAACGCATTAATNAGAAGAATTGCCGTCGTGCGATTCAGAACACACNTTTCTCAACCTTCTGGTNNAGAGGCTGCTGAAAATAATGACGCGTACGATAAAGTCAAACTATTAGACGAGGGATTAGATGGTAAAATACAGAATAATAGATATAGATTCGCATTTCTATACTTGTTGGTTAAATGGTACNAAAAATATCATATTNCTATTATGAAACTATATCCTACACCGGAAGAGATTNCGGACTTTGCNTTCTATCTCAAAATAGGTACTCTGTTGGTATCTAGCTCTGTAAAGCATATTNCNTTAATGACGGACCTCTNNNAAAAGGGATATATATTGTACGATAATGTGGTTANTCTTCCGTTGANTNCTNNNCAACAGAAAATATCCAAGTATTTTAATTNNNNNCTATTTGGNNACGATANNNANAGCTNNATCAATAGACATANGAANNNTNCCAATGTTAGTGATGAATATCTGCANNATATATTCATANNNGATANNTNATCTCCGTAAATATATGCCATATATTTATAGAATATATCACANATCTAAATGAATACCNGAATCATAGATTTATTTGATAATCATGTTGATAGTATACCAACTATATTACCTCATCAGTTAGCTACTTTAGNTTATCTAGTTNGAANTATCATAGATGAGAACNGAAGCGTGTTATTGTTCCNNATTATNNGATCGGGTAAAACAATAATCGCTTTGTTGTTNNNCTTGGTAGCTTNCAGATTTNNAAAGGTTTACANTTTAGTANCGAACATCAACATCTTAAAAATTTTCAATTATAATATNGGTGTAGCTATGAACNTGTTTAATGACGAATTCATAGCTGNNNATATNNTTATTCATTCCACAACAAGNTTTTATTCTCTTAATTATAACGATAACGTNATTAATTATAACNGATTNNNNCNCTACAATAACTCTANTTTTATCGTTGATGAGGCACATNATATTTTTNGGAATAANNNNNNNGNACTTATGACCGTGATNNNAAATAAAAACAAGANNNNNNNNNNNNNNNNNNNNNNNNNNNNNNNNNNNNNNNNNNNNNNNNNNNNNNNNNNNNNNNNNNNNNNNNNNNNNNNNNNNNNNNNNNNNNNNNGTGAGATTATTAGTCGTGGTNNNNAAGNAATTCAGACACTTCTTAACGAACNCNGAGNGANNNTACNCNNNNATTTGNNTANNNNAAGAATATCATATTACGAAATGCCGGACAAAGATCTACCAACAATAAGATATCACGGACGTAAATTTCTAGATACNNNAGTAGTATATTGTCACATGTCTAAACNNCANNAGAAAGATTANNNGATTACTAGACGGCAGCTATGTTATCATGAAATGTTTGATNAAANNNNNNNNNNNNNNNNNNNNNNNNNNNNNNNNNNNNNNNNNNNNNNNNNNNNNNNNNNNNNNNNNNNNNNNNNNNNNNNNNNNNNNNNNNNNNNNNNNNNNNNNNNNNNNNNNNNNNNNNNNNNNNNNNNNNNNNNNNNNNNNNNNNNNNNNNNNNNNNNNNNNNNNNNNNNNNNNNNNNNNNNNNNNNNNNNNNNNNNNNNNCACTTTNNNNNCTTCTNTAATTCNANATATGGTGGATTGGTAATTAAATATATCATGCTCNNNAATGGATATTCTGAATATAATGGTTCTCANGGNNCTAATCCACATATGATAAACGGCAAACCNAAAACATTTGCTATCGTTACTAGTAAAATGAAATCGTCTTTAGAGGATCTATTAGATGTGTATAATTCTCCTGAAAACGATGATGGCAATCAATTGATGTTTTTGTTTTCGTCAAACATTATGTCTGAATCCTATACTCTGAAAGAGGTAAGGCATATTTGGTTTATGACTATCCCGGATACTTTTTCTCAATACAACCAAATTNTTGGACGATCTATTAGAAAATTCTCTTACGTCGATATTTCTGAACCCGTTAATGTATATCNTTTAGCAGCCGTATATTCAGATTTCAATGACGAAGTGACGTCATTAAACGATTATACACAGGATGAATTGATTAATGTTTTACCCNTTGACATCNAAAANCTGTTGTATCTAAAATTTAAGACTAAAGAAACGAATAGAATATACTCTATTCTTCAAGAGATGTCTGAAACGTATTCTCTTCCACCACATCCATCAATNNNNNNNNNNNNNNNNNNNNNNNNNNNNNNNNNNNNNNNNNNNNATAATTNTCGTATTAAGTATAACGACTNCAAGTTACTTNAAATGGTTACATCAGTTATAAAAAATAAANAAGACGCTANGAATTACATNNNNGATATTGTAAACGGTCACTTCTTTGTATCGAATNAAGTATTTGATAANNNNCTTTTATACAAATACGAAAACGATATTANTACAGTACCGTTTAGACTTTCCTACGAACCATTTGTTTNGGGAGTTAANTNNNGTAAAGAATATAATGTGGTATCTTCTCCATAAAACTGATGAGNNATATAAAGAAATAAATGTCGAGCNTTGTTACCAATGGATATCTTCCAGTTACATTNGAACNNCATGAGNTGACGTTAGACATNAAAACTAATATTANGAATGCCGTATATAAGGCGTATNTCCATAGNNNAATNNGTGGTNAAATGGCCAAGNAAATAGAAATTCGTGAAGACGTGGAATTANNNNTCGGTGAAATAGTTAATAATTCTGTAGTTATAAACGTTCCNNNTGNANTAACCTACGCATATTATCACGTTNGGGATATAGTCAGAGGAACANNAAACATCGAAGATGAATCNAATGTAACTATTCANNNNNNAGATTTAANCTGTAAACTAAGTAGAGATTCNGGNACTGTATCATTTAGCGATTCNNAGTACNGNNTTTTTCGAAATGNNAANGCGTATGATAACGGCATCGAAGTCTCCGCCGTTCTAATGGAGGCTCAACAAGGTACCGAATCTAGTTTTGTTTTTNTCGCGAATATCGTTGACTNANAAGAAAGAGAATAGCGGTGAGTATAAATACGAATACTATGGCAATAATTGCGAATGTTTTATTNCCTTCGATATATTTTTGATAATATGNAAAACATGCNNCTCTCAAATCAGACAACCATTTCATAAAATAGTTCTCTCGCACTGGTGAGGTGGTTGCAGCTCNNATAATCTNCCCAGAATAATATACTTGCGTGTCGTCGTTCAATTTATACGGATTTCTATAATTNTCTGTTATATAATGAGGTTTACCCTCATGATTAGACGACGACAATAGTGTTCNGAATTTAGATAGTTGATCAGAATGAATGTTTATTGGTGTTNGNAAAANTATCCATGCTGCGTCTGCAGAGTGGTTGATAGTTGTTCCTAGATATGTAAAATAATCCAACGTACTAGGTAGCAAATTGTCTAGATAAAATACTGAATCAAATGGCGCAGACATATTAGCGGATCTAATGGAATCCAATTGATTGACTATCTTTTGAAAATATACANTTTTATGATCTGATACTTGTAAGAATATAGCAATAATGATAATTCCATCATCGTGNTTTTTTGCCTCTTCATAAGAACTATATTTTTTCTTATTCCAATGAACCAGATTAATCTCTCCAGAGTATTTGTATACATCTATCAAGTGATTGGATCCATAATCGTCTTCCTTTCCCCAATATATATGTATTGTTGATAACACATATTCATTGGGGAGAAANCCTCCACTTATATATCCTCCTTTNAAATTAATCCTTACTAGTTTTCCAGTATTCTNGATAGTGGTTGGTTTCGACTCATTATAATGTATGTCTAACGTCTTCAATCGCGCGTCAGAAATTGCTTTTTTAGTTTCTATATTAATAGGAGATAGTTGTTGAGGCANAGTAAAAATGAAATGATAACTGTCTAGAAATAGCTCTTAGTATGGGATTTACAATGGATGNGGAAGTGATATTTGAAACTCCTAGAGAATTAATATCTATTAAACGAATAAAAGATATTCCAAGATCAAAAGACACGCACGTGTTTGCTGCGTGTATAACAAGTGACGGATATCCGTTAATAGGAGCTAGAAGAACTTCATTCGCATTCCAGGCGATATTATCTCAACAAAATTCAGATTCTATCTTTAGAGTATCCACTAAACTATTACGGTTTATGTACTACAATGAACTAAGAGAAATCTTTAGACGGTTGAGAAAAGGTTCTATCAACAATATCGATCCTCACTTCGAAGAGTTAATATTATTGGGTGGTAAACTAGATAAAAAGGAATCTATTAAAGATTGTTTAAGAAGAGAATTAAAAGAGGAAAGTGATGAACATATAACAGTAAAAGAATTCGGAAATGTAATTCTAAAACTTACAACGAGTGATAAATTATTTAATAAAGTATATATAGGTTATTGCATGGCATGTTTTATTAATCAATCGTTGGAGGATTTATCACATACTAGTATTTACAATGTAGAAATTAGAAAGATTAAATCGTTAAATGATTGTATTAACGACGATAAATACGAATATCTGTCTTATATTTATAATATACTAATTAATAGTAAATGAGCTTTTACAGATCTAGTATAATTAGTCAGATTATTAAGTATAATAGACGACTAGCTAAGTCTATTANTTNCNAGGATGACTNTCAAATTATTACNCNCACGGCANNNGNTAACCAATGCCTATNGTNTCANAAACNANTATCCGTGTCCGCTATTTTATTAACTACTGATAACAAAATATTAGTATGNAACAGACGAGATAGNTTTNTCTATTNNNAAATAATTAGAACTNNAAACATGTATNGAAAGNAACGATTATTTCTGAATTATTNCAATTATTTGAACAAACAGNNNNGAAGTATACTATCGTCANNTTTTTCTCNNGNTNCANNTACTGCTGATAATGATAGAATAAACGCNATTTATCCNNGTGGTATACCCAAAANNGGTGAGAACGTTCCAGAGTGTTTATCCAGGGAAATCAAAGAAGAAGNNAATATAGACANNNNNNNNNNNNNNNNNNNNNNNNNNNNNNNNNNNNNNNNNNNNNNNGAAGATACCATTATTAACAAANTTTTTGAGGTAATTTTCTTTGTTNGACGAATATNTCTAACGAGTGATCAAATTATTGATACCTTTAAAAGTAATCATGAAATAAAGGATCTAATATTTTTAGATCCAANNTCAGGTAATGGACTCCAATACGAAATTGCAAAATATGCTCTAGATACTGCAAAACTTAAATGTTACGGTCATAGAGGATGTTATTATGAATCATTNAAAAAATTAACTGAGGATGATTGATTAGAAAATATAAATTAATTTACCATCGTGTATTTTTATAACGGGATTGTCTGGCATATCATGTAGATAGTTACCGTCTACATCGTATACTCTACCATCTACGCCTTTAAATCCTCTATTTATTGATATTAATCTATTAGAATTGGAATACCAAATATTAGTACCCTCAATTAGTTTATTGGTAATANTTTTTTTAGACGATAGATCGATGGCTCTTGAAACCAAGGTTTTCCAACCGGACTCATTGTCTATCGGTGAGAAGTCNTTTTCATTAGCATGAATCCATTCTAATGATGTATGTTTAAACACTCTAAACAATTGTACAAATTCTTTTGATTTGTTTTGAATGATTTCAAATAGGTCTTCGTCTACAGTAGGCATACCATTAGATAATCTAGCCATTATAAAGTGCACGTTTACATATCTACGTTCTGGAGGAGTAAGAACGTGACTATTGAGACGAATGGCTCTTCCTACTATCTGACGAAGAGACGCCTCGTTCCATGTCATATCTAAAATGAAGATATCATTGATTGAGAAGAAACTAATACCCTCGCCTCCGCTAGAAGAGAATACGCATGTTTTAATGCATTCTCCGTTAGTGTTTGATTCTTGGTTAAACTCAGCCACCGCCTTGATTCTAGTATCTTTTGTTCTAGATGAGAACTCTATATTAGAGATACCAAAGACTTTGAAATATAGTAATAAGATTTCTATTCCTGACTGATTAACAAATGGTTCAAAGACTAGACATTTACCATGGGATGCTAATATTCCCAAACATACATCTATAAATTTGACGCTTTTCTCTTTTAATTCAGTAAATAGAGAGATATCAGCCGCAATAGCATCCCCTCCCAATAGTTCTCCCNTTTTAAAGGTGTCTAATGCGGATTTAGAAAATTCTCTATCTCTTAATGAATTTTTAAAATCATTATATAGGGTTGCTATCTCTTGTGCGTATTCTCCCGGATCACGATTTTGTCTTTCAGGAAAGCTATCGAATGTAAACGTAGTAGCCATACGTCTCAGAATTCTAAATGATGATATACCAGTTTTTATTTCNGCGAGTTTAGCCNTTTGATAAATCTCTTCTTGCNTTTTTGACATATTAACGTATCGCATTAATACTGTTTTCTTAGCGAATGATGCAGACCCTTCCACATCATCAAAAATAGAAAACTCGTTATTAACTATGTACGAACATAGGCCTNCTAGTTTGGAGACTAATTCTTTTTCATCGACTAGACGTTTATTCTCAAATAGCGATTGGTGTTGTAAGGATCCTGGTCGCAGTAAGTTAACCAACATGGTGAATTCTTGCACACTATTAACGATAGGTGTAGCCGATAAACAAATCATCTTATGGNTTTTTAACGCAATGGTCTTAGATNAAAAATTATATACTGACCGAGTAGGACGGATCTTACCATCTTCTTTGATTAATGATTTAGAAATGAAGTTATGACATTCATCAATGATGACGCATATTCTACTCTTGGAATTAATAGTTTTGATATTAGTNAAAAATTTATTTCTAAAATTTTGATCATCGTAATTAATAAAAATACAATCCTTCGTTATCTCTGGAGCGTATCTGAGTATAGTGTTTATCCNNGGATCTTNTATCAAAGCCNTTTTTACCAATAAGATAATTGCCCAATTCGTATAAATATCCTTAAGATGTTTGNNNNNATATACAGTAGTCATTGNTTTACCGACACCTGNTTCATGGAACAATNAAAGAGAATGCATACTGTCTAATNCTNNGAAAACTCNNNNNNNNNNNNNNNNNNNNNNNNNNNNNNNNNNNNNNNNNNNNNNNNNNNTTTNNACGGGCATATTAGTAGTTCTGCGTAAGGCATAANCGATATAGGCNNNNNGTGATTTACTCATTTATNNGNGNTNAGTAATAACTATGTTTTNAAAATCNNNNCAGTAGTTTNNCTAGNCTTNTCTNATGTNNNNTTTCGATACNTTTTGAATNNNAAGTCATACTAGAATAAAGCAGCGAGTGAACGTAATAGAGAGCTTNGTATACTCTATTCGAAAACTCTAAGAACTTATTAATNAATTNCGTATCCACTGGATCGTTTNAAATNNTAAATTGAACAGTGTTCACATCCTTNCAAGACGAAGACTTAGTGNNGGACTTAACATGAGACATAAATNAATNCAAANTTTTTTTATAAACATCACTAGCCACCATAATGGCGCTATCNTTCNNCCAACTANCGCNTACGCATTTTAACAGTCNAANANTTTTAAAGAGACTACAATATANTCTCNTNGTATCGATTNNACCTCTACCGAATAGAGTNGGAAGTTTAATAATACAATATTTTTCGTTTACAAAATCAAATAATGGTCGAAACACGTCGAANGTTAACATCTTATAATCGCTAATGTATNGATTGNTTTCAGTGAGATGATTATTAGATTTAATAGCATCTCGTTCACGTTTGAACAGTTTATTGCGTGCGCTGAGGTCGGCAACTACGGCATCCGCTCTAGTACTCCTCCCATAATACTTTACGCTATTAATCTTTAAAATTTCATAGACTTTATCTAGATCGCTTTCTGGTAACATGATATCATGTGTAAAAAGTTTTAACATGTCGGTCGGCATTCTATTTAGATCATTAACTCTAGAAATCTGAAGAAAGTAATTAGCTCCATATTCCAGACTAGGTAATGGGCTTTTACCTAAAGACAAGTTAAGTTCTGGCAATGTTTCATAAAATGGAAGAAGGACATGTGTNCCCTCCCGGATATTTTTTACAATTTCATCCATTTACAACTCTATAGTTTGTTTTCATTATTATTAGTTATTATCTCCCATAATCTTGGTAATACTTACACCTTGATCATAAGATACCTTATACAGGTCATTACATACAACTACCAATTGTTTTTGTACATAATAGATTGGATGATTGATATCCATGGTGGAATAAACTACTCGAACAGATAGTTTATCTTTCCCCCTAGATACATTGGCCGTAATAGTTGTCGGCCTAAAGAATATCTTTGGTGTAAAGTTAAAAGTTAGGGTTCTTGTTCCATTATTGCTTTTTGTCAGTAGTTCGTTATAAATTCTCGAGATGGGCCCGTTCTCTGAATATAGAACATCATTTCCAAATCTAACTTCTAGTCTAGAAATAATATCGGTCTTATTTTTAAAATCTATTCCCTTGATGAATGGATCGTTAATAAACAAATCCTTGGCCTTTGATTCGGCTGATCTATTATCTCCGTTATAGACGTTACGTTGACTAGTCCAAAGACTTACAGGAATAGATGTATCGATGATGTTGATAGTATGTGATATGTGAGCAAAGACTGTTCTCTTGGTGGCGTCGCTATATGTTCCAGTAATGGCGGAAAACTTTTTAGAAATGTTATATATAAAAGAATTTTTTCGGGTTCCAAACATTAACAGATTAGTATGAAGATAAACACTCATATTATCAGGAACATTATCAATTTTTACATAAACATCGGCATCTTGAATAGAAACAACACCATCTTCTGGAACCTCTACGATCTCGGCAGATTCCGGATAACCAGTCGGTGGACCATCACTAACAATAACTAGATCATCCAACAATCTACTCACATATGCGTCTATATAATCTTTTTCATCTTGTGAGTACCCTGGATACGAAATAAATTTGTTATCAGTATTTCCATAATAAGGTTTAGTATAAACNGAGAGAGATGTTGCTGCATGAACTTCGGTTACTGTCGCCGTTGGTTGGTTTATTTGACCTATTACTCTCCTAGGTTTCTCTATAAATGATGGTTTAATTTGTACATTCTTAACCATATATCCAATAAAGCTCAATTCAGGAACATAAACAAATTCTTTGTTGAACGTTTCAAAGTCGAACGAAGAGTCACGAATAACGATATCGGATACTGGATTGAAGGTCACCGTTACGGTAATTTTTGAATCGGATAGTTTAAGACTACTGAATGTATCTTCCACATCAAACGGAGTTTTAATATAAACGTATACTGTAGATGGTTCTTTAATAGTGTCATTAGGAGTTAGGCCAATAGAAATATCATTAAGTTCACTAGAATATCCAGAATGTTTCAAAGCAATTGTATTATTGATACAATTATTATATAATTCTTCGCCCTCAATTTCCCAAATAACACCGTTACACGAAGAGACAGATACATGATTAATACATTTATATCCAACATATGGCACGTAACCGAATCTTCCCATACCTTTAACTTCTGGAAGTTCCAAACTCAGAACCAAATGATTAAGCGCAGTAATATACTGATCCCTAATTTCGAAGCTAGCGATAGCCTGATTGTCTGGCCCATCGTTTGTCATAACTCCGGATAGAGAAATATATTGCGGCATATATAAAGTTGGAATTTGACTATCAACTGCGAAGACATTAGACCGTTTAATAAAGTCATCCCCACCGATCAAAGAATTAATGATAGTATTATTCATTTTCTATTTAAAATGGAAAAAGCTTACAATAAACTCCGTAGAGAAATATCTATAATTTGTGAGTTTTCCTTAAAGTAACAGCTTCCGTAAACANCGTCTTTATCTCTTAGTAAGTTTATTGTATTTATGACCTTTTCCTTATCTTCATAGAATACTAAAGGCAATAAAGAAATTTTTGGTTCTTCTCTAAGAGCTACGTGAGACTTAACCATAGACGCCAACGAATCCCTACATATTTTAGAACAGAAATACCCAACTTCACCACCCTTGAATGTCTCAATACTAATAGGTCTNAAAACCAAATCTTGATTACAAAACCAACACTTATCAATTACACTATTTGTCTTAATAGACATATCTGCCATAGATTTATAATACTTTGGTAGTATACAAGNGAGTGCTTCTTCTTTANCGGGCTTAAAGACTGCTTTNNGTGCTGAAATAACCACATCTGGANNACTTACTNGCNNAGCCATTTAATTACGGAACTANTTTTTTATACTTCTAATGAACAAGTAGAAAACCTCTCATCTACNAAAACATACTCGTGTCCATAATCCTCTACCATAGTAACACGNTTTTTAGATCTCATATGTGCTNAAAAGTTTTCCCATACTAATTNGTTACTATTATTTTTCGTATANTTTTTAACAGTTTGAGGNTTTAGANTTTTAGTTACNGAAGTGATATCGAATATTTTATCCNAAAAGAATGAGTAATTAATTGTCTTAGAAGGAGTGTTTTCTTGGCNAAAGAATACCAAGTGCTTAAATATTTCTACTACTTCATTAATCNTTTCTGTACTCNGATTCAGNTTNTCATCNTTTANNTGATTGATTATTTNAAAGACTAACTTATAANNCTTTTTATTTATTCTCTCGTTAGCCTTAAGAAAACTAGATACAAAATTTGCATCTACATCATCCGTGGATATTTGANNTTTTTCCATGATATCCAATAGTTCCGAGATAATTTCTNCAGAACATTGATGAGACAATAATCTCCGCAATACATTNNTCAAATGAATAAGTTTATTAGACACNNGGAAGNTTGNNNNNTTTTGTACCTTTGTACATTTTTGAAATACNGACTNNCNAAAAATACAATATTCATATCCTTGTTCAGATACNNTACCNNNATGTCTANAACAGCTACATAATCGTAGATTCATGTTAACACTCTACGTATCTCGTCGTNCAATATTTTATATNAAAACATTTTATTTCTAGACGTTGTCAGAAAATCCTGTAATANTTTTANTTTTTTTGGTTGTGAATAAAGTATCGCCCTAATAATATTGGTACCGTCTTCCGACAATATAGTAGTTAAATTATCCGAGCATGTAGAAGAACACCGCTTAGGCGGATTCAGTACAATGTTATNTTTTTCGTACCAACTCNTTTAAATATCATAATCTAAAATAGTTCTGTAATATGTCTAGCGCTAATATATTGATCATAATCCTGTGCATAAATTAAGATACAACAATGTCTTGAAATCATCGACATGGCTTCTTCCATAGTTANAAGATCATCGTCAAAGTTAGCAACGTGATTCATCAACATTTGCTGTTTTGAGGCAGCAAATACTGAACCATCACCATTCAACCATTCATAAAAACCATCGTCTGAATCCATTGATAATTTCTTGTACTGGTTTTTGAGAGCTCGCATCAATCTAGCATTTCTANNTCNCGGATTGNAAACAGAAAGAGGATCGTACATCCAAGGTCCNTTTTCTGTAAATAGAATCGTATAATGTNCCTTCAAGAAGATATCNGACGATCCACAATCAAAGAATTGGTCTCCGAGTTTGTAACAGACTGCGGACTTTANCCTATACATGATACCGTTTAGCNTGATTTCTGGTGATNCGTCAATCGGAGTATCATCTATNAGAGATCTAAAGCCGGTGTAACATTCTCCGCCAAACATANNCNTATTCTGANNTNNTTCTACATNAAACATCATTGCTCCATTAACGATAACANNNGAATGAACAGCNCTACCCATCACNNTAGTTCCCAATGGATCAANNTGTGTAACTNCAGAACATCTTCCATAGCCTATGTTANGAGGAGCGAACACCACTCTTCCACTATTGCCATCGAATGCCATAGAATAAATATCCTTGGAATTGATAGAAATCGGACTGTCGGATGTTGTTATCATCTTCATAGGATTAACAACGATGTATGGTGCAGCCTGAAGTTTCATATCGTAACTGATGCCGTTCATAGGTCTAGCCACAGAAACCAACGTAGGTCTAAATCCAACTATAGACAAAATAGAAGCTAATATCTGTTCCTCATCTGTCATAACTTGAGAGCATCCAGTATGAATAATCTTCATTAGATGGGGATCTACCGCATCATCATCGTTACAATAAAAAATTCCCATTCTAATGTTCATAATTGCTTTTCTAATCATGGTATGAATGTTTGCTCTCTGAATCTCTGTGGAAATTAGATCTGATACACCTGTAATCACTATCGGATTATCCTCCGTAAGACGATTAACCAACAACATATAATTATAAGACTTTACTCTTCTAAATTCATAAAGTTGCTGGATTAGACTATATGTGTCTCCATGTACATACGCGTTCTCGAGCGCAGGAAGTTTAATACCGAATAGTGCCATCAGAATNGGATGAATGTAGTAATTAGTTTCTGGTTTTCTATAAATAAAAGACAAATCTTGTGAACTAGACATATCGGTAAAATGCATGGATTGGAATCGTGTAGTCGACAGAAGAATATGATGATTAGATGGAGAGTATATTTTATCTAACTCTTTGAGTTGGTCACCGATTCTAGGACTAGCTCGAGAATGAATAAGTACTAAGGGATGNGTNCATTTCACAGAAACACTGGCGTTGTTCAACGTACTCTTTACATNGGAAAGGAGTNGAAATNNCTCGNTTCTATTTGTCCTGACAATATTTAGTTTATTCATAATATTAAGCATATNCTGAATAGTAAAGNTAGATNNNTCATACTTGTTAGTAGTTAGATATTNNGCAATTGNANTCCNNNNNNNNNNNNNNNNNNNNNNNNNNNNNNNNNNNNNNNNNNNNNNNNNNNNNNNNNNNNNNNNNNNNNNNNNNNNNNNNNNNNNNNNNNNNNNNNNNNNNGTACGTTTAGGTTTGGTACGTTGATTTCNANNCCCTGCTGATATNAAGTCATCGTCTACAATTTNGGATAATGAATTACATACACTACAAGACAAAGATTTATCAGAANNGTGAATATGATCTTCATCTACCAAAGAAAGAGTTTGATTAGTATAACTAGATTTTAGTCCCGCGTTAGATGTTNAAAAAACATCGCTATTGACCACGGCTTCCATTATTTATATTCGTAGTTTTTACTCGAAAGCGTGANTTTAATATCCAATCTTATTACTTTTGGAATCGTTCAAAACCTTTGACTAGTTGTAGAATTTGATCTATTGCCCTACGCGTATACTCCCTTGCATCATATACGTTCGTCACCAGATCGTTTGTTTCNGNCTGAAGTTGACGCATATCNTTTTCAACACTNGACATGAGATCCNTAAGGGTCATATCGTCTAGATTTTGTTGAGATGCTGCTCCTNGATTTGGANTTTGTTGTGCTGTTGTACATACTGTACCACCAGTAGGTGTAGGAGNNCATACAGTGGCCACAATANGAGGTTGAAGAGGTGTAACCGTTNGAGTAGTACAAGAAATACTTCCATCCGATTGTTGTGTACATGTGGTTGTTGGTAACGTCTGAGAAGGTTGGGTAGATGGCGGTGTCGTCATCNTTTGATCTTTATTAAATTTAGAGATAATATCCTGAACAGTATTGCTCGGCGTCAACGCTGGAAGGAGTGTACTNGCCGGCGCATCAGTATCTGTAGACAACCAATCAAAAAGATTAGACATATCAGATGATGTATTAGTTTGTTGACGTGGTTTTAGTACAGGAGCAGTACTACTAGGTAGAAGAATAGGAGCCGGTGTAGGTGTCGGAACCGGCTGTGGAGTTATATGAATAGTTGGTTGTAGCGGTTGGGTAGGCTGTCTGCTGGCGGTCATCATATTATCTCTAGCTAGTTGTTCTCGCAACTGTCTTTGATAATACGACTCTTGAGACTTTAGTCCTATTTCAATCGCTTCATCCTTTTTCGTATCCGGATCCTTTTCTTCAGAATAATAGATTGACGACTTTGGTGTAGAGGATTCTGCCAGCCCCTGTGAGAACTTGTTAAAGAAGTCCATTTAAGGCTTTAAAATTGAATTGCGATTATAAGATTAAATGGCAGACACGGACGATATTATCGACTATGAATCCGATGATCTCACCGAATACGAGGATGATGAAGAAGATGGAGAGTCACTAGAAACTAGTGATATAGATCCCAAATCTTCTTATAAGATTGTAGAATCAACATCCACTCATATAGAAGATGCGCATTCCAATCTTAAACATATANGGAATCATATATCTGCTCTTAAACGACGCTATACTAGACGTATAAGTCTATTTGAAATAGCGGGTATAATAGCAGAAAGCTATAACTTGCTTCAACGAGGAAGATTACCTCTAGTTTCAGAATTTTCTGACGAAACGATGAAGCAAAATATGCTNCATGTAATTATACAAGAGATAGAGGAGGGTTCTTGTCCTATAGTCATCGAAAAGAACGGAGAATTGTTGTCGGTAAACGATTTTGATAAAGATGGTCTAAAATTCCATCTAGACTATATTATCAAAATTTGGAAACTTCAAAAACGATATTAGAATTTATACGAATATCGTTCTCTAAATGTCACAATCAAGTCTCTCATATTCAGCAGTTTATTGTCGTACTTTATATCGTGTTCATTAACGATATTTTGCAAAATAGTAATGATTCTATCTTCCTTCGATAGATATTCTTCAGAGATTATTGTCTTATATTCNTTCTTGTTATCCGATATGAATTTGATAAGACTTTGAACATTATTAATACCCGTCTGTTTAATTTTTTCTATAGATATTTTAGTTTTGGTAGATTCTATGGTGTCTGTTAATAGGCATCCAACATCGACATTCGACGTCAATTGTCTATAAATCAGAGTATAAATTTTAGAAATAACATTAGCAAATTGTTGTGCGTTGATGTCGTTATTCTGAAACAGTATGATTTTAGGTAGCATTTTCTTAACAAAGAGAACGTATTTATTGTTACTCAGTTGAACAGATGATATATCCAGATTACTAACGCATCTGATTCCATATACCAAACTTTCAGAAGAAATGGTGTACAATTGTTTGTATTCATTCAATGTCTCCTTTTCAGAAATTAGTTTAGAGTCGAATACTGCAATAATTTTCAAGAGATAGTTTTCATCAGATAAGATTTTATTTAGTGTAGATATGATAAAACTATTGTTTTGTTGGAGAACTTGATACGCCGCATTCTCTGTAGTCGACGCTCTCAAATGGGAAACAATCTCTATTATTTTTTTGGAATCGGATACTATATCTTCGGTATCTTGACGCAGTCTAGTATACATAGAGTTAAGAGAAATTAGAGTTTGTACATTAAGCAACATGTCTCTAAATGTGGCTACAAACTTTTCTTTTTCCACATCATCTAGTTTATTATATACCGATTTCACAACGGCACCAGATTTAAGGAACCAGAATGAAAAACTCTGATAACTACAATATTTCATCATAGTTACGATTTTATCATCTTCTATAGTTGGTGTGATAACACATACCTTTTTCTCCAAGACTGGAACCAACGTCATAAAAATGTTTAAATCAAAATCCATATCAACATCTGATGCGCTAAGACCAGTCTCGCGTTCAAGATTATCTTTACTAATGGTGACGAACTCATCGTATAGAACTCTAAGTTTGTCCATTATTTATTTACAGATTTAGTTGTTTAATTTATTTGTGCTCTTCCAGAGTTGGGATAGTATTTTTCTAACGTCGGTATTATATTATTAGGATCTACGTTCANNNNNNNNNNNNNNNNNNNCATCCACGNTTTGATAAATCTATCNTTANCTNNTGAAATAACGNNNNNNNNNNNNNNNNNNNNNNNNNNNNNNNNNNNNNNNNNNNNGATNACANNTTTTGTAAATGTAANGTANTTAGACGANAGATCTTCGTTNNAAAGTTTTNCATNTATNNNNAATNCATCGGTTGTTAACACCATTCCCGCGTCAGANTGAATANGAGTTTGAATNNNNNNNNNNNNNNNNNNNNNNNNNNNNNNNNNNNNNNNNNNNNNNNNNNNNNNNNNNNNNNNNNNNNNNNNNNNNNNNNNNNNNNNNNNNNNNNNNNNNNNNNNNNNNNNNNNNNNNNNNNNNNNNNNNNNNNNNNNNNNNNNNNNNNNNNNNNNNNNNNNNNNNNNNNNNNNNNNNNNNNNNNNNNNNNNNNNNNNNNNNNNNNNNNNNNNNNNNNNNNNNNNNNNNNNNNNNNNNNNNNNNNNNNNNNNNNNNNNNNNNNNNNNNNNNNNNNNNNNNNNNNNNNNNNNNNNNNNNNNNNNNNNNNNNNNNNNNNNNNNNNNNNNNNNNNNNNNNNNNNNNNNNNNNNNNNNNNNNNNNNNNNNNNNNNNNNNNNNNNNNNNNNNNNNNNNNNNNNNNNNNNNNNNNNNNNNNNNNNNNNNNNNNNNNNNNNNNNNNNNNNNNNNNNNNNNNNNNNNNNNNNNNNNNNNNNNNNNNNNNNNNNNNNNNNNNNNNNNNNNNNNNNNNNNNNNNNNNNNNNNNNNNNNNNNNNNNNNNNNNNNNNNNNNNNNNNNNNNNNNNNNNNNNNNNNNNNNNNNNNNNNNNNNNNNNNNNNNNNNNNNNNNNNNNNNNNNNNNNNNNNNNNNNNNNNNNNNNNNNNNNNNNNNNNNNNNNNNNNNNNNNNNNNNNNNNNNNNNNNNNNNNNNNNNNNNNNNNNNNNNNNNNNNNNNNNNNNNNNNNNNNNNNNNNNNNNNNNNNNNNNNNNNNNNNNNNNNNNNNNNNNNNNNNNNNNNNNNNNNNNNNNNNNNNNNNNNNNNNNNNNNNNNNNNNNNNNNNNNNNNNNNNNNNNNNNNNNNNNNNNNNNNNNNNNNNNNNNNNNNNNNNNNNNNNNNNNNNNNNNNNNNNNNNNNNNNNNNNNNNNNNNNNNNNNNNNNNNNNNNNNNNNNNNNNNNNNNNNNNNNNNNNNNNNNNNNNNNNNNNNNNNNNNNNNNNNNNNNNNNNNNNNNNNNNNNNNNNNNNNNNNNNNNNNNNNNNNNNNNNNNNNNNNNNNNNNNNNNNNNNNNNNNNNNNNNNNNNNNNNNNNNNNNNNNNNNNNNNNNNNNNNNNNNNNNNNNNNNNNNNNNNNNNNNNNNNNNNNNNNNNNNNNNNNNNNNNNNNNNNNNNNNNNNNNNNNNNNNNNNNNNNNNNNNNNNNNNNNNNNNNNNNNNNNNNNNNNNNNNNNNNNNNNNNNNNNNNNNNNNNNNNNNNNNNNNNNNNNNNNNNNNNNNNNNNNNNNNNNNNNNNNNNNNNNNNNNNNNNNNNNNNNNNNNNNNNNNNNNNNNNNNNNNNNNNNNNNNNNNNNNNNNNNNNNNNNNNNNNNNNNNNNNNNNNNNNNNNNNNNNNNNNNNNNNNNNNNNNNNNNNNNNNNNNNNNNNNNNNNNNNNNNNNNNNNNNNNNNNNNNNNNNNNNNNNNNNNNNNNNNNNNNNNNNNNNNNNNNNNNNNNNNNNNNNNNNNNNNNNNNNNNNNNNNNNNNNNNNNNNNNNNNNNNNNNNNNNNNNNNNNNNNNNNNNNNNNNNNNNNNNNNNNNNNNNNNNNNNNNNNNNNNNNNNNNNNNNNNNNNNNNNNNNNNNNNNNNNNNNNNNNNNNNNNNNNNNNNNNNNNNNNNNNNNNNNNNNNNNNNNNNNNNNNNNNNNNNNNNNNNNNNNNNNNNNNNNNNNNNNNNNNNNNNNNNNNNNNNNNNNNNNNNNNNNNNNNNNNNNNNNNNNNNNNNNNNNNNNNNNNNNNNNNNNNNNNNNNNNNNNNNNNNNNNNNNNNNNNNNNNNNNNNNNNNNNNNNNNNNNNNNNNNNNNNNNNNNNNNNNNNNNNNNNNNNNNNNNNNNNNNNNNNNNNNNNNNNNNNNNNNNNGATGNAAGTCTTACAATGTTCGAACCAGTACCAGATCTTAATTTNGAGGCCTCCGTAGAACTANGGGANGTAAATANAGATCNAACAACACCTATGATAAAGGAGAATANCGGNTTTATATCCCGTAGTNGNCGTCTATTNNCCCATANNNNTNNGGATNNNNNGANAAAACNNNCANNNNNNNNNNNNNNNNNNNNNNNNNNNNNNNNNNNNNNNNNNNNNNNNNNNNNNNNNNNNNNNNNNNNNNNNNNNNNNNNNNNNNNNNNNNNNNNNNNNNNNNNNNNNNNNNNNNNNNNNNNNNNNNNNNNNNNNNNNNNNNNNNNNNNNNNNNNNNNNNNNNNNNNNNNNNNNNNNNNNNNNNNNNNNNNNNNNNNNNNNNNNNNNNNNNNNNNNNNNNNNNNNNAANTCAACTCNNNAGTATCCATCATCCAAGNAAAACNCGGATACCAGTCTNGNNNNNNNNNNNNNNNNNNNNNNNNNNNNNNNNNNNNNNNNNNNNNNNNNNNNNNNNNNNNNNNNNNNNNNNNNNNNNNNNNNNNNNNNNNNNNNNNNNNNNNNNNNNNNAGTTCANTATCTGAAATAACNNATAGATTTTATNNNNTTAATCNAAAATCTGATCTTGTTCAATGGNGTGCTAGTACGGGCGCTGTAATTAATCAAATGGTAAATACTGTATTGATTACAGTGTATGAAAAATTACAACTGGCCATAGNAAATGATTCACNNNNNNNNNNNNNNNNNNNNNNNNNNNNNNNNNNNNNNNNNNNNNNNNNNNNNNNNNNNNNNNNNNNNNNNNNNNNNNNNNNNNNNNNNNNNNNNNNNNNNNNNNNNNNNNNNNNAAGCAAACGTGATNAGGATACGCTATTAAAACANTTTACTTATGACTGGNGTTAGAATTTATAGACGNNACATTTCGTTTATCATTGTTACTATTACTATCATTATTAGTATTCTTCTTGTCATCTTGTTCAGAAATATACAGCAATGCTATACCTAATACTAAATACATTATCATGCTTGCAATGGCTCTAACAACAACGAACCNAAATGAATTTGGTCGTAGCTTTTGTTCACAAAAATACATAAAGAAATGTCTACATAAATCTATGGCGCCATTGGCTACTTGAAATAGCGCCAGTCCTCCTACAGATTTTAATATAGCTGTATAACATGACATTTATTCATCATCAAAAGAGACAGAGTCACCATCTGTCATATTTAGANTTTTTTTCATGTGTTCAAAGTATCCTCTACTCATTTCATTATAATAGTTTATCATGCTTAGAATTTTAGGACGGATCAATGAGTAAGACTTGACTAGATCGTCAGTAGTAATTTGTGCATCATCTATTCTGCATCCGCTTCGTCGAATAATGTATAGCATCGCTTTGAGATTCTCCATAGCTATCAAGTCTTTATATAATGACATGGAAATATCTGTGAATGCTTTATNCTTCTNCAACATCGATGCCTTAACATCATCACATACTTTAGCATTGAAAATACGTTCTATTGTGTAGATGGATGTAGCAAGATTTTTAAACAACAATGCCATCTTACNNNATGATTGTCTCAAGTCTNCAATCGTTTGTTTNGNNCGATTAGCTNCAGAGTNCAATGCTTGGCTAACTAGCATATTATTATCTTTNGAAATTGTATTCTTCAATGAGGCGTTTNNNATATCNNTGATTTCGTTAGTNATATTACAGTCTGANTNGGTTGTAATGTTATNCAACATATCNNNNNTGGATACGGTNCACGTACCAGCATTTGTAATAATCCTATCTAAGATGTTGTATGGCATTGCGCAGAAAATATCTTCTCCTGTAATATCTCCACTCTCGATAAATCTACTCAGATTATTCTTAAATGCCTTATTCTCTGGAGAAAAGATATCAGTGTCCATCATTTCATTAATAGTATACGCAGAAAAGATACCACNAGTATCAATTCTATCCAAGATACTTATCGGTTCCGAGTCACAGATAATTGTTTCCTCTCCTTCGGGAGATCCTGCATANAAATATCTAGGACAATAGTTTCTATACTGTCTGTAACTCTGATAATCTCTAAAGTCACTAACTGATACCATGAAATTGAGAAGATCAAANNCTNAANTAATCAANTTTTNNNNNNNGTTTTTACTACAACTAGTTTTCATCANNGTAGTGACGATGNNTTGTTTAGTTNNNNNNNNNNNNNNNNNNNNNNNNNNNNNNNNNNNNNNNNNNNNNNNNNNNNNNNNNNNNNNNNNNNNNNNNNNNNNNNNNNNNNNNNNNNNNNNNNNNNNNNNNNNNNNNNNNNNNNNNNNNNNNNNNNNNNNNNNNNNNNNNNNNNNNNNNNNNNNNNNNNNNNNNNNNNNNNNNNNNNNNNNNNNNNNNNNNNNNNNNNNNNNNNNNNNNNNNNNNNNNNNNNNNNNNNNNNNNNNNNNNNNNNNNNNNNNNNNNNNNNNNNNNNNNNNNNNNNNNNNNNNNNNNNNNNNNNNNNNNNNNNNNNNNNNNNNNNNNNNNNNNNNNNNNNNNNNNNNNNNNNNNNNNNNNNNNNNNNNNNNNNNNNNNNNNNNNNNNNNNNNNNNNNNNNNNNNNNNNNNNNNNNNNNNNNNNNNNNNNNNNNNNNNNNNNNNNNNNNNNNNNNNNNNNNNNNNNNNNNNNNNNNNNNNNNNNNNNNNNNNNNNNNNNNNNNNNNNNNNNNNNNNNNNNNNNNNNNNNNNNNNNNNNNNNNNNNNNNNNNNNNNNNNNNNNNNNNNNTTGGTAACNTTGATAGCCGTAGATCTTTGTTCCNTCGCTAANNNGTTNANNAAATTAATAAAGAATTTGTTGTTTNNATTTATGTCNNNNNNNNNNNNNNNNNNNNNNNNNNNNNNNNNNNNNNNNNNNNNNNNNNNNNNNNNNNNNNNNNNNNNNNNNNNNNNNNNNNNNNNNNNNNNNNNNNNNNNNNNNNNNNNNNNNNNNNNNNNNNNNNNNNNNNNNNNNNNNNNNNNNNNNNNGNTTATAATCTNATCATTGANNNNTACTCTGTTACCATTGTAATAAATTGGTACTCTATCNNNATCANCGACNNAGTACTGTTCATANATGATGAGATGTTTATATGTTGGCATANNNNNNNNNNNGACGTTTNGTAATGGCAANNNNNNNNNATTAACNNCATAATGTCTAGCAGCANCTGCGATGTTATAAGTGATGTCAAAGCNNGGTTNATCTTNNNNNNTTATATATTGTCTAACACCTATAANATTATCAAAATCTTGTCTGCTTAATACACCGTTAACAANTTTTGCCTTGAATTCTTTTATTGGTGCATTAATAACATCCTTATAGAGGATGTTAAACAAATAAGTATTATCAAAGTTAAGATCTGGGTATTTCTTTTCTGCTAGAACATCCATTGAGTCGGAGCCATCTGGTTTAATATAACCACCGATAAATCNAGCTCTGTATTCTGTATCCGTCAATCTAATATTAAGAAGGTGTTGAGTGAAAGGTGGAAGATCGNAAAAGCTGTGAGTATTAATAATANGGTTAGNTTNCGAACTAATGNNNNTTNGATGATNNNTAATATTTATATTTCCAGCGTTAAGTGTAACATTAAACANTTTTAATTCACGTGACGTGGTATCAATTAAATAATTAATGCCCAATTTGGATATAGTAGCCTGAAGCTCATCTTGTTTAGTTACGGATCCTAATGAGTTATTAAGAAATACATCGAACGGATGAACGAAGGTTGTTTTAAGTTGGTCACATACTTTGTAATCTAGACATAGATGTGGAAGAACGGTAGAAACTATACGAAATAGATATTCAGAGTCCTCTAATTGATCAAGAGTAACTATTGACTTAATAGGCATCATTTATTTAGTATTAAATGACGACCGTACCAGTGACAGATATACAAAACGACTTAATTACAGAGTTTTCAGAAGATAATTATCCATCTAACNAAAATTATGAAATAACTCTTCGCCAAATGTCTATTCTAACTCACGTTAACAACGTGGTAGATAGAGAACATAATGCCGCCGTAGTGTCATCTCNNNAGGAAATATCATCACAACNNAATGAAGATCTNTTTCCAGNNNNNNNTTCACCGGCCACTATTATCGAACGAGTACAACCTCATACTACTATTATTGACGATACGCCACCTCCTACTTTTCGTNGAGAGTTATTGATATCGGAACAACGTCAACAACNAGNAAAAAGATTTAATATTACAGTATCAAAAAATTCTGAAGCAATAATGGAATCTAGATCTATGATAACTTCTATGCCAACACAAACACCATCCTTGGGAGTAGTTTNTGATAAAGATAAAAGAATTCAGATGCTAGAGGATGAAGTGGTTAATCTTAGAAATCAACGATCTAATACAAAATCATCTGATAATTTAGATAATTTTACCAGAATACTATTTGGTAAGACTCCGTATAAATCAACCGAAGTTAATAAGCGTATAGCCATCGTTAATTATGCAAATTTGAACGGGTCCCCCTTATCAGTCGAGGACTTGGATGTCTGTTCGGAGGATGAAATAGATAGAATCTATAAAACGATTAAACAATATCACGAAAGTAGAAAACGNAAAATTATCGTCACTAACGTGATTATTATTGTCATAAACATTATTGAGCAGGCATTGCTAAAACTCGGATTTGAAGAAATCNAAGGACTGAGTACCGATATCACTTCAGAAATTATCGATGTGGAGATCGGAGATGACTGCGATGCTGTAGCATCNAAACTAGGAATCGGTAACAGTCCGGTTCTTAATATTGTATTGTTTATACTCAAGATATTCGTTAAACGAATTAAAATTATTTAATTTAATACATTCCCATATCCAGACAACAATCGTCTGGATTAATCTGTTCCTGTCGTCTCATACCGGACGACATATTAATCNTTTTATTAGTGGGCATCTTTTTAGATGGTTTCNTTTTCCCAGCATTAACTGATTCGATACCTAGAAGATCGTGATTGATTTCTCCGACCATTCCACGAACTTCTAATTGGCCGTCTCTAACGGTACCATAAACTATTTTACCAGCATTAGTAACAGCTTGGACAATCTGACCATCCATTGCGTTGAATGATGTAGTTGCTGTTGTTCTACGTCTAGGAGCACCAGAGGTATTTTTAGAGCTCTTGGATGTTGATGTAGAAGACGAGGATTTTGATTTTGGTTTACATGTAATACATTTTGAACTCTTTGATTTTGTATCACATGCACCGGCAGTCACATCTGTTTGAGAATTAAGATTATTGTTGCCTCCNTTGACGGCTGCATCTCCACCGATCTGCGCTAGTAGNTTTTTAAGCTGTGGTGTAANNNNNNNNNNNNNNNNNNNNNNNNNNNNNNNNNNNNNNNNNNNNNNNNNNNNNNNNNNNNNNNNNNNNNNNNNNNNNNNNNNNNNNNNNNNNNNNNNNNNNNNNNNNNNNNNNNNNNNNNNNNNNNNNNNNNNNNNNNNNNNNNNNNNNNNNNNNNNNNNNNNNNNNNNNNNNNNNNNNNNNNNNNNNNNNNNNNNNNNNNNNNNNNNNNNNNNNNNNNNNNNNNNNNNNNNNNNNNNNNNNNNNNNNNNNNNNNNNNNNNNNNNNNNNNNNNNNNNNNNNNNNNNNNNNNNNNNNNNNNNNNNNNNNNNNNNNNNNNNNNNNNNNNNNNNNNNNNNNNNNNNNNNNNNNNNNNNNNNNNNNNNNNNNNNNNNNNNNNNNNNNNNNNNNNNNNNNNNNNNNNNNNNNNNNNNNNNNNNNNNNNNNNNNNNNNNNNNNNNNNNNNNNNNNNNNNNNNNNNNNNNNNNNNNNNGTGCTTNNNCCACATAGAATAAANTAGCATTCCGACTGTGNNNATNNTACCNNGTATAAACGCCATAATACTNAATACTTNNCATGNNCGAGTNGGACTGGTAGACTTACTNNAGTCANNAAAGGCGANNATACACGNAAGAATCNNNAGAATGATTCCAGCGATTAGCACGCCAGAAAAATAATTTCCAATCATAAGCATCATGTNCATTTAACTAATNAAAATTTTAAATCGCCGNNNNNNNNNNNNNNNNNNNNNNNNNNNNNNNNNNNNNNNNNNNNNNNNNNNNNNTAATATCTANTTTTGTTTTCGAAGATATGGTAAAATTAAATAGTAGTNNNCNGCATNTTATAACTAACAGCAGCAACGGCTCGTAANNNNNTATCATTTACTAGACGAAAAGGTGGTNGGANNNNNNNNNNNNNNNNNNNNNNNNNNNNNNNNNNNNNNNNNNNNNNNNNNNNNNNNNNNNNNNNNNNNNNNNNNNNNNNNNNNNNNNNNNNNNNNNNNNNNNNNNNNNNNNNNNNNNNNNNNNNNNNNNNNNNNNNNNNNNNNNNNNNNNNNNNNNNNNNNNNNNNNNNNNNNNNNNNNNNNNNNNNNNNNNNNNNNNNNNNNNNNNNNNNNNNNNNNNNNNNNNNNNNNNNNNNNNNNNNNNNNNNNNNNNNNNNNNNNNNNNNNNNNNNNNNNNNNNNNNNNNNNNNNNNNNNNNNNNNNNNNNNNNNNNNNNNNNNNNNNNNNNNNNNNNNNNNNNNNNNNNNNNNNNNNNNNNNNNNNNNNNNNNNNNNNNNNNNNNNNNNNNNNNNNNNNNNNNNNNNNNNNNNNNNNNNNNNNNNNNNNNNNNNNNNNNNNNNNNNNNNNNNNNNNNNNNNNNNNNNNNNNNNNNNNNNNNNNNNNNNNNNNNNNNNNNNNNNNNNNNNNNNNNNNNNNNNNNNNNNNNNNNNNNNNNNNNNNNNNNNNNNNNNNNNNNNNNNNNNNNNNNNNNNNNNNNNNNNNNNNNNNNNNNNNNNNNNNNNNNNNNNNNNNNNNNNNNNNNNNNNNNNNNNNNNNNNNNNNNNNNNNNNNNNNNNNNNNNNNNNNNACGTATAATGCNNNATCCCCAAAAGTAAAATAATCAGGACGAATAATTCTAATAAACTCAGAACAATATCTCGCATCCATATGTTTGGAGCNAATATCGGAATAAGTAGACATAGCCGGTTTCCGTTTTACACGTAACCATTCTAAACAATTGGGGTTTCCAGGATCGTTTCTACAAAAACCAGTCATGAAATCGTCACAATGTTCTGTCTTGTAATTATTATTAAATATTTTTGGACAGTGTTTGGTATTTGTCTTAGAACANNATTTTGCCACGCTATCACTATCACCCNGGAGATAATCNTTTTTTATAAAATGACATCGTTGCCCGGATGCTATATAATCANNAGNATATTTTAAATNCTTAATATATTCANGAGTTACCTCGTTCTGATAATAGATTAATGATCCAGGACGAAATTTGNAAGAACTACATGGTTCTCCANGAATTAATACATATTGTTTAGCAAATTCAGGAACTATNAAACTACTACAATGATCTATCGACATACCATCTATCAAACAAAATTTGGGTTTAATTTCTCCTGGAGACGTTTCATAATAATACATATAACTTTCTTCGGCAAACCTAACAGCTCTATTATATTCAGGATAATTAAAATCTAATACCATATATTTGTCTCGTATATCTGCTATTCCTGTCTCTATTTTGATTCTATTAAGAGTAACAGCTGCCCCCATTCTTAATAATCATCAGTATTTAAACTGTTAAATGTTGGTATATCAACATCTATCTTATTTNCCGCAGTATAAGGTTTGTTGCAGGTATACTGTTCAGGAATGGGTACATTTATACTTCTTTTATAGTCCTGTCTTTCGATGTTCATCACAAATGCAAAGAACAGAATAAACAAAATAATGTAAGAAATAATATTAAATATCTGTGAATTCGTAAATACATTGATTGCCATAATAATTACAGCAGCTACAATACACACAATAGACATTCCCACAGTGTTGCCATTACCTCCACGATACATTTGAGTTACTAAGCAATAGGTAATAACTAAGCTAGTAAGAGGCAATAGAAAAGATGAGATAAATATCATCAATATAGAGATTAGAGGAGGGCTATATAGAGCCAAGACGAACAAAATCAAACCGAGTAACGTTCTAACATCATTATTTTTGAAGATTCCCAAATAATCATTCATTATTCCTCCATAATCGTTTTGCATCATACCCCCATCTTTAGGCATAAACGATTGCTGCTGTTCCTCTGTAAATAAATCTTTATCAAGCACTCCAGCACCCGCAGAGAAGTCATCAAGCATATTGTAATATCTTAAATAACTCATTTATATATTAAAAAATGTCACTATTAAAGATGGAGTATAATCTTTATGCCGAACTNAAAAAAATGACTTGTGGTCAGACCATAAGTCNTTTTAATGAAGACGGCGATTTCGTAGAAGTTGAACCAGGATCATCCTTTAAGTTTCTAATACCTAAGGGATTTTACTCCTCTCCTTGTGTAAAGACGAGTCTAGTATTCAAGACATTAACAACGACCGATAATNAAATTACTAGTATCAATCCAACAAATGCGCCAAAGTTATATCCTCTTCAACGCAAAGTCGTATCTGAAGTAGTTTCTAATATGAGGAAAATGATCGAATTAAAACGTCCTCTATACATCACTCTTCACTTGGCATGTGGATTTGGTAAGACTATTACCACGTGTTATCTTATGACCACACACGGCAGAAAAACCATCATTTGCGTACCCAATAAAATGTTAATACATCAATGGAAGACACAGGTAGAGGCAGTCGGATTGGAACATAAGATATCTATAGATGGAGTTAGTAGTCTATTAAAGGAACTAAAGACTCAAAGTCCGGATGTATTAATCGTAGTCAGTAGACATCTGACAAACGATGCATTTTGTAAATATATCAATAAGCATTATGATTTGTTTATCTTGGATGAATCACATACGTATAATCTGATGAACAATACAGCAGTTACAAGATTTTTAGCGTATTATCCTCCGATGATGTGTTANTTTTTAACTGCTACACCTAGACCAGCTAACCAAATTTATTGTAATANTATTATTAATATTGCCAAGTTATCCGATCNNNAAAAAACTATCTATATAGTAGATAGTTTTTTTGAGCCATATTCCACAGACAATATTAGAAATATGGTAAAACGACTAGATGGACCATCTAATAAATATCATATATATACCGAGAAGTTATTATCTGTAGACGAGCCTAGAAACCAACTTATTCTTGATACCCTGGTAGAAGAATTCAAGTCAGGAACTATTAATAGAATTTTAGTTATTACTAAACTACGTGAACATATGGTATTCTTCTACAAACGATTATTAGATCTTTTCNGAGCAGAGGTTGTATTTATAGGAGACGCCCAAAATAGACGTACTCCAGATATGGTCAAATCGATTAAGGAACTAAATAGATTTATATTCGTATCCACCTTATTTTATTCCGGCACTGGTTTAGATATTCCGAGTTTGGATTCTTTGTTCATTTGCTCGGCAGTAATCAACAATATGCAAATAGAGCAATTACTAGGGAGGGTATGTCGAGAAACAGAACNATTAGATAGGACGGTATATGTATTTCCTAACACATCCATCNAAAAAATAAAGTACATGATANGAAATTTCGTGCAACGAATTATTAGTCTGTCTGTNGATAAACTCGGATTTAAACAAGAAAGTTATCAGAAACATCAGGAATCTGAACCCGCTTCCGTACCAACATCCTCCAGAGAAGAACNNGTATTAAATAGAATATTTAANNNNCAAAATCNNNNNNAAGNTTNNNAGACGATCNNCATGCTGNGCNNGCCAGTGTATTANCCCTCATAGTATTAATATAATCCAATGATACNTTTGTGATGTNNGAAATNTTAACCAATTTAGACTGACAGGCANNACACGTCATACANNNATNATCGTCATCGATAANTNTAGTCNTGGGCTTNNNNNNNNNNNNNNNNNNNNNNNNNNNNNNNNNNNNNNNNNNNNNNNNNNNNNNNNNNNNNNNNNNNNNNNNNNNNNNNNNNNNNNNNNNNNNNNNNCCTGGTATANNGTCTCTATCTTTATNNCAATATAGTGTATCAACATCTTNNTTATTGACCTCNTTTATCTTNGAACATGGAATNGGNACATTTTTGTTAACNGCCACNNNTNCCNTAATTCCAGATGTTGTNAAATTATAACTAAACAGTCTATCATCGACACAAATGAAATTNTTGTTTAGACGTTTGTAGTTTACGTATGCGGCTCGTTCTCGTCTCATTTTTTCAGATATTGCAGGTACTATAATATTAAAAATAAGAATGAAATAACATAGGATTNAAAATAAAGTTATCATGACTTCTAGTGCTGATTTAACTAACTTAAAAGAANNNNTTAGTCTGTACNAAAGTTTNNNNNNNNNNNNNNNNNNNNNNNNNNNNNNNNNNNNNNNNNNNNNNNNNNNNNNNNNNNNNNNNNNNNNNNNNNNNNNNNNNNNNNNGGTAACTAATGTCNNGACGTCCATATCTGATTATTATGATGAGGTAAAAAANAAACCGTTTAATATTGATCCGGGGTATTATATTTTCTTACCAGTATATTTTGGAAGCGTCTTTATTTATTCAAANGGTAAAAATATGGTAGAACTTGGATCTGGAAACTNTTTTCAAATACCGGATNAGATTCNAAGTGCGTGTAACAAAGTATTAGATAGTGATAACGGAATAGACTTTCTGAGATTTGTTTTGTTAAACAATAGATGGATAATGGAAGACGCTATATCAAAATACCAGTCTCCAGTTAATATATTTAAACTAGCTAGTGAGTACGGATTAAACATACCCAACTATTTAGAAATTGAAATAGANGAAGACACATTATTTGACGATGAGTTATACTCTATTATGGAACGCTCTTTCGATGATACATTTCCAAAAATATCTATATCGTATATTAAGTTGGGAGAACTTAAGCGGCAAGTTGTAGACTTTTTCAAATTCTCATTCATGTATATTGAGTCAATCAAGGTAGATNGTATAGGAGATAATNTTTTTATTCCTAGCGTTATAACAAAATCAGGNAAAAAGATATTAGTAAAAGATGTAGACCATTTAATACGATCCAAGGTTAGAGAACATACATTTGTAAAAGTAAAAAAGAAAAACACATTTTCCATTTTATACGACTATGATNGGAACGGAACAGAAACTAGAGGAGAAGTAATAAAACGAATTATNGACACTANNNGACGAGACTANTATGNNNNTGNNNANTANNNNNNNNNNGTTGGTATTGCAGGCNNNAAGCAATTGACTAATAAATTAGATATTAATGAGTGTGCAANNNTCGATGAGTTAGTTGATGAGATTAATAAATCNNNAACTNTNAAACGAAAAANNAAAAACCAANCAGTATTTGATTTNAGNAGAGAATGNTTGGGATATCCNNNAGCNGANTTTATAACNNNAGTNNNTAACATNNNGTTCNNANNAGNAAATNNTAAGGTTGTAAATTTCAATATTGAAAATACTAATTGTTTAAATAACCCGAGTATTGAAANTATATANGGAAACNNCAACCANTTCGTCTCAATCTTTAATACCGTTACCGATGTCNAAAAAAGATTATTCGAGTGAAATAATATGCGCCTTTGATATAGGTGCNAAAAATCCTGCCAGAACTGTTTTAGAAGTCAAGGATAACTCCGTTAGGGTATTGGATATATCAAAATTAGACTGGAGTTCTGATTGGGAAAGGCGCATAGCTCAAGATTTGTCACAATATGAATACACTACAGTTCTTCTAGAACGTCAGCCTAGNAGGTCACCGTANNTCAAATTTATCTATTTTATTNAAGGCTTTTTATATCATACATCTGCTGCCAAAGTTNTTTGCGTCTCACCTGTCATGTCTGGTAATTCATATAGAGATCGNAAAAAGAGATCTGTTGAAGCATTTCTTGATTGGATNNNNNCATTCGGATTNCGAGANTNCGTTNCNGATAGACNNNNATTAGACGATGTAGCGGATAGTTTCAATTTGGCTATGAGATACGTATTAGATAAATGGAATACTAATTATACACCTTATAATAGGTGTAAATATAGAAATTACATNAAAAAAATGTAATAACGTTAGTAACGCCATTATGGATAATCTATTTACCTTTCTACATGAAATAGAAGATAGATATGCCAGAACTATTTTTAACTTTCATCTAATAAGTTGTGATGAAATAGGAGATATATATGGTCTTATGAAAGAACGCATTTCCTCAGAGGATATGTTTGACAATATAGTATATAATAAAGATATACATCCTGCCATTAAGAAACTAGTTTATTGCGACATCCAACTTACTAAACATATTATTAATCAGAATACGTATCCGGTATTTAACGATTCTTCACAAGTGAAATGTTGTCATTATTTCGATATAAACTCAAATAATAGCAATATTAGCTCTCGTACAGTAGAGATATTTGAGAGTGAAAAGTCATCTCTTGTATCATATATTAAAACTACCAATAAGAAGAGAAAGGTCAATTACGGCGAAATAAAGAAAACTGTACATGGAGGCACTAATGCAAATTACTTTTCCGGTNAAAAGTCTGATGAGTATCTGAGCACTACAGTCAGGTCCAACATTAATCAACCTTGGATCAAAACCATTTCTAAGAGAATGAGAGTAGATATCATTAATCACTCTATAGTAACGCGTGGAAAAAGCTCTATATTACAAACTATAGAAATTATTTTTACTAATAGAACATGTGTGAAAATATTCAAGGATTCTACTATGCACATTATTCTATCCAAGGACAAGGATGAAAAGGGATGTATAAACATGATTGATAAATTATTCTATGTATATTATAATTTATTTCTGTTGTTCGAGGATATCATCCAAAACGATTACTTTAAAGAAGTAGCTAATGTTGTAAACCATGTACTCATGGCTACGGCATTAGATGAGAAATTATTCCTAATTAAGAAAATGGCTGAACACGATGTTTATGGAGTTAGCAATTTCAAAATAGGGATGTTTAACCTGACATTTATTAAGTTGTTGGATCATACCGNTTTNCCCTCTCTGTTAGATGAGGATAGCAAAATAAAGTTTTTTAAGGGNAAAAAGCTCAATATTGTAGCATTACGATCTCTGGAGGATTGTACAAATTACGTGACTAAATCCGAGAATATGATAGAAATGATGAAGGAAAGATCGACTATTTTAAATAGCATAGATATAGAAACGGAATCGGTAGATCGTCTAAAAGAATTGCTTCTAAAATGNAAAAAAACACTGATTCAGAAATGGATCAACGACTCGGGTATAAGNTTTTGGTGCCTGATCCTAAAGCCGGAGTTTTTTATAGANNNNTANATTTNCAATATGTATNNNNNNNNNNNNNNNNNNNNNNNNGATTGCANGAAATNTTGANNNNCAANCGGCCNCTCNTATCGTTTNANAATAATACAGAANNNNNNNNNNNNNNNNNNNNNNNNNNNNNNNNNNNNNNNNNNNNNNNNNNNNNNNNNNNNNNNNNNNNNNNNNNNNNNNNNNNNNACGCATTANCCACGTTCACNNNNNNNNNNNNNNNNNNNNNNNNNNNNNNNNNNNNNNNNNNNNNNNNNNNNNNNNNNNNNNNNNNNNNNNNNNNNNNNNNNNNNNNNNNNNNNNNNNNNNNNNNNNNNNNNNNNNNNNNNNNNNNNNNNNNNNNNNNNNNNNNTCNNGTTCNNNANNTCATAGGTGGAGNCTTTATCNNNNNACAATCNNNNNNAAAAGTANNNATTAATCTANTAGAAAAGATTACAACATGGCCAAAATTTANGGTTGTTNANNNNNNNNNNNNNNNNNNNNNNNNNNNNNNNNNNNNNNNNNNNNNNNNNNNNNNNNNNNNNNNNNNNNNNNNNNNNNNNNNNNNNNNNNNNNNNNNNNNNNNNNNNNNNNNNNNNNNNNNNNNNNNNNNNNNNNNNNNNNNNNNNNNNNNNNNNNNNNNNNNNNNNNNNNNNNNNNNNNNNNNNNNNTTCATTNGACGTAGTTTATCATACGNTATGCCTCNNNAAGTTGTCTATCTAGTAAANGCGATAATAGANANNGCTAANNNACTTANCGAATCTATTACTGACTTTNATATTGNNNNNTACATTAATGACCTGGTGNNAGCTGAACACNNNAAACNAAAATCTCAGTTAACGATTAACGAGTTTAAATATGAAATGCTGCATAACNTTTTACCTCNNATGAACTATACACCCGATCAACTAAAGGGATTTTATATGATATCTTTACTAAGAAAGTTTCTCTACTGTATCTACCACACTTCTAGATATCCAGATAGAGATTCGATGGTTTGTCATCGCATCCTAACNTACGGCAAATATTTTGAGACGTTAGCACATGATGAATTAGAGAATTACATAGGTAACATNCGAAACGATATCATGAACAATCACAAGAACAGAGGCACTTACGCAGTAAACATTCATGTACTAACAACTCCTGGACTTAATCATGCATTTTCTAGTCTATTGNGTGGAAAGTTCNNAAAGTCAGACGGTAGTTATCGAACACATCCTCACTATTCATGGATGCAGAATATTTCTATTCCTAGAAGTGTTGGATTTTATCCGGATCAAGTAAAGATTTCAAAGATGTTTTCTGTCAGAAAATACCATCCAAGCCAATATCTTTACTTTTGTTCATCAGACGTTNCGGAAAGAGGTCCTCAGGTAGGTTTAGTATCTCAATTGTCTGTCTTGAGTTCCATTACAAATATACTAACGTCTGAGTATTTGGATTTGGAAAAGAAAATTTGTGAGTATATCAGATCATATTATAAAGATGATATAAGTTACTTTGAAACAGGATTTCCAATCACTATAGAAAATGCTCTAGTCGCATCTCTTAATCCAAATATGATATGTGATTTTGTAACTGACTTTAGACGTAGAAAACGGATGGGATTCTTCGGTAACTTGGAGGTAGGTATTACTTTAGTTAGGGATCACATGAATGAAATTCGCATTAATATTGGAGCAGGAAGATTAGTCAGACCATTCTTGGTTGTGGATAACGGAGAGCTCATGATGGATGTGTGTCCGGAGTTAGAAAGCAGATTAGACGACATGACATTCTCTGACATTCAGNAAGAGTTTCCACATGTCATCGAAATGGTAGATATAGAACAATTTACTTTTAGTAACGTATGTGAATCGGTTCNAAAATTTAGAATGATGTCAAAGGATGAAAGAAAGCAATNNGATTTATGTGACTTTCCTGCCGAATTTAGAGATGGATATGTAGCATCTTCACTAGTGGGAATCAATCACAATTCTGGACCCAGAGCTATTCTTGGATGTGCTCAAGCTAAACAAGCTATCTCTTGTCTGAGTTCGGATATACGAAATAAAATAGACAATGGAATTCATTTGATGTATCCAGAGAGGCCAATTGTGATTAGTAAGGCTTTAGAAACTTCAAAGATTGCGGCTAATTGCTTCGGACAACATGTTACTATAGCATTAATGTCGTACAAAGGTATCAATCAAGAGGATGGAATTATCATCAAAAAACAATTTATTCAGAGAGGCGGTCTCGATATTGTTACAGCCAAGAAACATCAAGTAGAAATTCCATTGGAAAACTTTAATAACAAAGAAAGAGATAGGTCTAACGCCTATTCGAAATTAGAAAGTAATGGATTAGTTAGACTGAATGCTTTCTTGGAATCCGGAGACGCTATGGCAAGAAATATCTCATCAAGAACTCTTGAAGATGATTTTGCTAGAGATAATCAGATTAGCTTTGATGTTTCCGAGAAATATACAGATATGTACAAATCTCGCGTTGAACGAGTACAAGTAGAACTTACTGACAAAGTTAAGGTGCNAGTATTAACCATGAAAGAAAGAAGACCCATTCTAGGAGACAAATTTACTACTAGAACGAGTCAAAAGGGAACAGTCGCGTATATCGCAGATGAAACGGAACTTCCGTACGACGAAAATGGTATCACACCAGATGTCATTATTAATTCTACATCCATCTTCTCTAGAAAAACTATATCTATGTTGATAGAAGTTATTTTAACAGCCGCATATTCTACTAAGCCGTACAACAATAAGGGAGAAAACCGACCTGTCTGTTTTCCTAGTAGTAACGAAACATCTATCGATGCATATATGCAATTCGCTAAACAATGTTATGAGTATTCAAATCCGAAATTGTCCGAGGAAGAATTATCGGATAAAATCTTTTGTNAAAAGATTCTCTATGATCCTGAAACGGATAAGCCTTATGAATCCAAAGTATTTTTTGGACCAATTTATTACTTGCGTCTGAGACATTTAACTCAGGACAAGGCAACCGTTAGATGTAGAGGTNNAAAGACGAAGCTCATTAGACAAGCGAATGAGGGACGNNNACGTGGAGGAGGTATCAAGTTTGNAGAAATGGAGAGAGACTGTTTAATAGCACATGGTGCAGCCAATACTATTACAGAAGTTTTAAAAGACTCAGAAGAGGATTATCAAGATGTGTATATTTGTGAAAATTGTGGAGACATAGCAGCACAAATCAAAAGTATTAATACATGTCTTAGATGTTCAAAACTTAATCTCTCTCNNCTCTTAACAAAAATTGATACCACGCACGTATCTAAAGTATTTCTTACTCAAATGAACGCCAGAGGCGTAAAAGTTAAATTAGATTTCGAACGAAGGCCTCCTTCGTTTTATAAACCATTAGATAAAGTTGATCTTAAACCGTCTTTTCTGGTATAATATTGTTTAGTAGATACTCATCAAGATAAGCTAATTCACTAAACATATTATCGGATTCGGTATTGTTACTCGAGAATAGAGTTCGTTATGCTCCTGATATTCGGAAATCTGTGGAGTTTCAGGTTTTGGTGGAAGTGTAACTGCTACTTGGTGGGATACTGAAGGATATTTCAGAGAGTTGTGGATGTTCGGGTTCGACATCCACCGATGGTGTCACGCCACTAATCGGTTCGGTAACGTCTGTGGATGGAGGTGCTACTTCTACAGAACCTGTAGCCTCAGTTGTCAACGGAGATACATATTCAATGCGCGGAAATGTATAATTTGGTAATGGTTTCTCATGTGGATCTTAAGAAGAAGAGGTAAGATATCTACGAAAGATACCGATCACGTTTCTAGTTCTCTTTTGTAGAACTTTAACTTTTTCTTTCTCAGCATCTAGTTGATATTCCGACCTCTTCACGTTTCGCATGGGTTACCTCCGCAGTTTTTACAAGCGATTTCACGTTCCAGATCACGTTCAGCCTTCATACGTCTCTCCCTCTCTCTATCGAGTTTATCAGAGCAGTCTTTCTGAAGGCGATCGAACTCCATAAATTTCTCCAACGCTTTGATTGTTTCCATAGATTTCCGAAGTTTAGCTTCTAGGACGGCGATTCNTTTTTTTTTTTTTTTTTTCGAATTCACGGGGTACAACCGTTTCCATTACCACCATCTCTATGTTTCTTTTCTAGATCGGCAANNNNNNNCAATCTTTCTCAACATTTCATCCCCATACCTTTTCATTCCTCGAGTCTATTGTCGTCGAAATATCGTTCCAGCTCCTTTTCGACCTCAATAACTTTAGCACGTTGTTTCATCAAGCTCTCTCTTGTAGTACTATCATTTTTATCTGATTCCCTGACACGTTTAAGATCTTCATGTAATTGAGTCAGCTCTTGACGCAATCTCTTAACTAACTTCCTCTCTTGCTTCTTCGTCATAGTACTTACAATCACTATGGGATCCATTGTTACCACGTCTGTACTCGACGAGCTCACGTTTAAGAGATTCAATTTCCAGTTTGTATCGGTCCATGTCTCCATTGCTACACCACCATTAGATTTACAGGCTGCTAGTTGTCGTTCGAGATCAGAAATACGTGTTTTCTTGGAATGGATTTCGTCGATGTACTTGTCATGATTGGCATCGAAACACTTATTAAGTTCNTTTTTTCAATTCTACGATTTTATTTCTTTCGCGAGTCAATTCCCTCCTGTAGTAACTATCAGTTTTGTCAGATTCACGCTCTCTACGTAGACTTTCTTGTAAGTTACTAATTTGTTCCCTGGCATTACCGAGTTCAGTTTTATATGCCGAATAGAGTTCTGATTCATCCTTTGAGAAGATCTCTAGCGATCGTTCAAGATCCCTGATTCTAGTCTTTAGCCTATTTACCTCCTCAGAAGATGCTCCGTTACCGTTTTTACAATCGTTAAGATGTCTATCAAGATCCATGATTCTATCTCTTTTCCATATCAGCATTGATTTCATTATTACGTTCGCAGTCGTTCAACTGTATTTCAAGATCTGAGATTCTAGATTGTAATCTCTGTAGCATTTCCACGGCATTCACTCAGTTGTCTTTCAAGATCTGAGATTCTAGATTGGAGTCTGCTAATCTCTGTAAGATTTCCTCCTCCGCTCTCGATGCAGTCGGTCAACTTATTCTCTAGTTCTCTAATACGCGAACGCAGTGCATCAACTTCTTGTGTGTCTTCTTGATTGCGTGTGCATTCATCGAGTCTAGATTCGAGATCTCTAACGTGACGTCGTTCTTCCTCAAGTTCTCTGTGTACTACAGAAAGCGTGTCCCTATCTTGTTGATATTTAGCAATTTCTGATTCTAGAGTACTGATTCTACTCACGTATGTACTAATAGTTGTCTTAGCCTTATCAAGATCCTCCTTGTATTTGTCACATTCCTTGATATCCATACGAAGTCTGGACAGTTCCCATTCGACATTACGACGTTTATCGATTTCAGCTCGGAGATCGTCGTCGCGTTGTTTTAGCCACATACGACTAAGTTCAAGTTCTCGTTGACAAGATCCATCTACTTTTCCATCCCTAATAGTATCCAGTTCCTTTTCTAGTTCTGACCGCATTTCTCGTTCCATATCAAGAGATTCTCTCAATTCTCGTATAGTCTTCTTATCAATTTCTGATGAATCTGAACCATCATCTGTCCCATTTTGTTGCATATCCCTGAGTTCTTTGATCTCTGTTGTAAGTCTGTCGATTCTTTCGGTTTTATAAACAGAATCCCTTTCCAAAGTCCTAATCTTACTGAGTTTATCATTAAGTTCTTCATTCAATTCAGTGAGTTTTCTCTTGGCTTCTTCCAAGTCTGTTTTAAACTCTCCATCATTTCCGCATTCTTCCTCGCATTTATCTAACCATTCAATTAGTTTATTAATAACTAGTTGGTAATCAGCGATTCCTATAGCCGTTCTTGTATTTGTGGGAACATAATTAGGATCTTCTAATGGATTGTATGGCTTGATAGCATCATCTTTATCATTATTAGGTGGGGGATGGACAACCTTAATTGGTTGGTCCTCCTTATCTCCTCCAGTAGCATGTGGTTCTTCAATACCAGTATTAGTAATAGGCTTAGACAAATGCTTGTCGTACGCGGGCACTTCCTCATCCATCAAGTATTTATAATCGGGTTCTGTTTCAGAATATTCTTTTCTAAGAGACGCGACTTCAGGAGTTAGTAGAAGAACTCTGTTTCTGTATCTATCAACGCTGGAATCGATACTCAAGTTAAGGATAGCGAATACCTCATCGTCATCATCCNNATCTTCTGAAACGCCATCATATGACATTTCATGAAGTCTAACGTATTGATAAACAGAATCAGATTTAGTATTAAACAGATCCTTGACCNTTTTAGTAAATGCATATGTATATTTTAGATCTCCAGATTTCATAATATGATCGCATGCCTTAAATGTCNATGCTTCCNNGNNATAGTCTGGAACACTAATGGGTGACGNAAAAGATACAGCACCATATGCTACGTTGATAAATAGATCTGAACCACTAAGTAGATAATGATTAATGTTAAGGAAGAGGAAATATTCAGTATATAGATATGCCTTAGCATCATATCTTGTACTAAACACGCTAAACAGTTTATTGATGTGATCAATTTCCAACAGAACAATTAGNGCGGCNGGAATACCAACAAACATATTACCACATCCGTATTTTCTATGAATATNNCATATCATATTNNAAAATCTTGATNGAAGNNCGAATATCTCGTCTGACNNAATGAGATGTAGTTCAGCAGCATAAGTCATAACTGTAAATAGAACATACTTTNCTGTAGTGTTGATTCTAGACTCCACATCAACACCATTATTAAAAATAGTTTTATATACATCTTTAATCTGCTCTCCGTTAATCGTCGAACGTTCTAGTATACGGAAACACTTTGATTTCTTATCTGTAGTTAATGACTTAGTGATATCACGAAGAATATTACGAATTACATTTCTTGNTTTTCTTGAGAGACCTGATTCAGAACTCAACTCATCGTTCCATAGNTTTTCTACCTCAGTGGCGAAATCTTTGGNGTGTTTGGTACANTTTTTAATAAGGTTCGTGACCTCCATTTATTATAAAAAATTTTTATTCAAAACTTAACTACAATCGGGTAATTATAAGATCGTAGATCTCCCATGTGGTGGAATACTACCATCTATCGCATGTTGATGGACAGTAGGTAATGGCCATGGGAACAGTAATGNTTGCATATTTATCTTTCTTGCTAGTATTACTGTATATTGTCCCAATGTTTCAATGTGATGTTCTAACCTATCAACTGCCACTGTATCACAACAATAATGTCCGATGGAATTAAGATTATGATCCAATGTGTTTAATATATGATTATCAAGTCTTATACGATCCGCGTCTTTTTTGACAGGATCAGGCTCTTCTACAGGAAGAAGTTTCGGCCTCTTATGATAGTCATGTCTGGGAAATGGTGGTCTAGGATGAGGATCAGGTATCGGAGTAGGTTTTGGATTATAATCATCATCATCATCATCATCATCATCATCATCATCATCATCTATGATATCATCATCTTCGATATTTATTTTGCTATCTTGATAATGTCCTATATCAGTTGCATTTTCAGCACTCGACTGAATATTAGTACATTCATTGTCTATTATTAACGTATTTCTAAACCCAAAATGTATATGTTGAACATCACTACTATAGTTGATGAGTCTTATAGCATGAATTCGCTTATCGTTATCGGGTTTATCTTCTGTCACCTTAACAATTCCNTTTTTATTAAACTCTGCATAATCATAACCATTTCTATTGTTTGTTCTAATATAAACGAGTATAGCATCATTGCTAAATTTTTCAATAGTATCNAAAACAGAATATCCTAAACCATATAATATATATTCAGGAACACTCAAACTAAATGTCCAGGATTCTCCTAAATACGTAAACTTTAATAGTGCTAAATCATTCAAAAATCTACCGCTTATAGATAGATAGTACATGAATGCGTATAGTAGTCTACCTATCTCTTTATTATGAAAACCGACATTACGATCATATATTTCGTGATATACATGTGACCCGTTTACGTTAAACCATAAATACATGGGTGATCCTATAAACATGAATTTATTTCTAATTCTCAGAGCCATAGTTAATTGACCGTGTAATATTTGTTTACATGCATACTTGATACGATCATTAATAAGATTTTTATCATTGCTCGTTATTTCAGAATCGTATATATAAGGAGTACCATCATGATTCTTACCAGATATTATACAAAATACTATATATAAAATATATTGACCCACGTTAGTAATCATGTAAATGTTTAATGTTTTAAATTTTGTATTTAATGATCCATCATCATATGCTAGCATGGTCTTGTGATATTCATTCTTTAAAATATAATATTGTGTTAGCCATTGCATTGGAGCTCCTAATGGAGATTTTCTATTCTCGTCCATTTTAGGATATGCTTTCATAAAGTCCCTAATAACTTCGTGAATAATGTTTCTATGTTTTCTACTGATGCATGTATTTGCTTCGATTTTTTTATCCCATGTTTCATCTATCATAGATTTAAACGCAGTAATGCTCGCAACATTAACATCTTGAACCATTGGTACAATTCCGTTCCATAAATTTATAATGTTCGCCATTTATATAACTCATTTTTTGAATATACTTTTAATTGAACAAAAGAGTTAAGTTACTCATATGGATGCCGTCCAGTCTGTACATCAATCTTTTTAGCCAGAGATATCATAGCCGCTCTTAGAGTTTCAGCGTGATTTTCCAACCTAAATAGAACTTCATCGTTGCGTTTACAACACTTTTCTATTTGTTCAAACTTTGTTGTTATATTAGTAATCNTTTTTTCCAAATTAGTTAGCCGTTGTTTGAGAGTTTCCTCATTATCGTCTCCATAGGCTTTAACAATTGCTTCGCGTTTAGTCTCTGGATTTTTAGCAGCCTTTGTAGAGAAAAATTCAGTTGCTGGAATTGCAAGATCGTCATCTCCGGGGAAAAGAGTTCCGTCCATTTAAAGTACAGATTTTAGAAACTGACACTCTGTGTTATTTATATTTGGCGCAATACATGGATTATAAATATCGATGTTAATAACATCAGAAAATGTAAAGTCTATACATTGTCGCATCGTGTTAAATTTTCTAATGGATCTAGTATTATTGGGTCCAACTTCTGCCTGAAATCCAAATATGGAAGCGGATACAAAACCGTTTCCTGGATAAACCACACATCTCCACTTTTGCTTTACATCAGAAATTGTGTCATTGACATCTTGAACTCTCCTATCTAATGCCGGTGTTCCACCTATAGATTTTGAATACTCGAATGCTGCATGAGTAGCATTGAATTCCTTAATATTGCCATAATTTTCATATATTGAGTAACTCTGGATAAAAAGTAAACACACCGCAGCCGTCGCTACTACAANAAAAAAAATTGATAGAGAGTTCATTTATAATCTATTAGAAGCTGATAAAATTTTTTTACACGCGTCAGACAATGCTTTAATAAATAGTTCAACATCTACTTTTGTCATATCGAACCGATGGTATGATTCTAACCTAGAATTACATCCGAAAAAGTTGACTATGTTCATAGTCATTAAGTCATTAACGAACAACATTCCAGACTCTGGATTATAAGACGATACTGTTTCGTCACAATCACCCACCTTAATCATGTGATTATGAATATTGGCTATTAGAGTACCTTCTAAGAAATCTATAATATCTTTGAAACACGATTTAAAATCAAACCACGAATATACTTCTACGAAGAAAGTTAGTTTACCCATAGGAGAGATAACTATAAATGGAGATCTAGATACAAAATCCGGATCTATGATAGTTTTAACATTATTATATTCTCTATTAAATACCTCCACATCTAAAAATGTTAATTTTGAAACTATGTCTTCGTTTATTACCGTACCTGAACTAAACGCTATAAGCTCTATTGTTTGAGAACTCTTTAAACGATATTCTTGAAATACATGTAACAAAGTTTCCTTTAACTCGGTCGGTTTATCTACCATAGTTACAGAATTTGTATCCTTATCTATAATATAATAATCAAAATCGTATAAAGTTATATAATTATCGTGTTCAGATTGTGATCTTTTCAAATAGACTAAAAACCCCATTTCTCTAGTAAGTATCTTATGTATATGTTTGTAAAATATCTTCATGGTGGGAATATGCTCTACAGCAGTTAGCCATTCCTCATTGACAGCTGTAGATGTATTATACAAAACTACTCCAATGTTTAACAAGGGCCATTTTACGAGATTATTAAATCCTTGTTTGATAAATGTAGCCAATGCGGGTTCGAGTTCAACGACGATTGAATTCTCTTCCCGTGGATGCTGCATGATGAACGACGGGATGTTGTTGTTCTATTGATTTGGAATTCTTTTTCGACTTTTTGTTTATATTAAATATTTTAAAATTTATGGCTGATAGTAATTCATGTACTACGGATAATGTAGACGTGTATTGCATATCGATATCTTTATTATTAGATAAATTTATCAATAAATGTGAGAAGTTTGCCTCGTTAAGGTCTTCCATTTAAATATTATATAAATATTTGTGTTTGTATTTTATTCGTCTTTTATGGGATAGTTTTTAACTAGTAAAGCTGTAATTACATACTTTGTCCGTAAAACATAAATATAAATACCCGCTTTTATCAAACGTTCCAAAAAGTCGGCAGCTGACATTTTTAACATGACATCTATTTTAAATACACTTAGGTTTTTAGAAAAAACATCATTTTATAATTGTAACGATTCAATAACTAAAGAAAAGATTAAGATTAAACATAAGGGAATGTTATTTGTATTTTATAAGCCAAAGCATTCTACCGTTGTTAAATACTTGTCTGGAGGAGGTATATATCATGATGATTTGGTTGTATTGGGGAAGGTAACAATTAATGATCTAAAGATGATGCTATTTTACATGGATTTATCATATCATGGAGTGACAAGTAGTGGAGCAATTTACAAATTGGGATCGTCTATCGATAGACTTTCTCTAAATAGGACTATTGTTACAAAAGTTAATAACAATTATAACAATTATAACAATTATAACAATTATAATTGTTATAATAATTATAATTGTTATAATTATGATGATACATTTTTTGACGATGATGATTGATCACTATTACACAATTTTGTTTTTGTACTTTCTAATATAGTGTTTAGGTTCTTTTTCATATGAGAATATTGACTTACTAAAATATCTATGTTTAACTTTTGTTCTATAACGTCCTTATCGGCGGTATCGGTACATATACGTAATTCACCTTCACAAAATACGGAGTCTTCGATAATAATAGCCAATCGATTATTGGATCTAGCTGTCTGTATCATATTCAACATGTTTAATATATCCTTTCGTTTCCCCTTTACAGGCATCGATCGTAGCATATTTTCCGCGTCTGAGATGGAAATGTTAAAACTGCAAAAATGCGTAATGTTAGCCCGTCCTAATATTGGTACGTGTCTATAAGTTTGGCATAGTAGAATAATAGACGTGTTTAAATGCCTTCCAAAGTTTAAGAATTCTATTAGAGTATTACATTTTGATAGTTTATCACCTACATCATCAAAAATAAGTAAAAAGTGTGCTGATTTTTTATGATTTTGTGCGACAGCAATACATTTTTCTATGTTACTTTTAGTTCGTATCAGATTATATTCTAGAGCTTCCTGACTACTAACGAAATTAATATGATTTGGCCAAATGTATCCATCATAATCTGGGTTATAAACGGGTGTAAACAAGAATATATGTTTATATTTTTTAACTAGTGTAGAAAACAGAGATAGTAAATAGATAGTTTTTCCAGATCCAGATCCTCCTGTTAAAACCATTCTAAACGGCATTTTTAATAAATTTTCTCTTGAAAATTGTTTTTCTTGAAAACAATTCATAATTATATTTACAGTTACTAAATTAATTTGATAATAAATCAAAATATGGAAAACTAAGGTCGTTAGTAGGGAGGAGAACAACGAAGGCATATCGTGATATAAATAACATTTATTATCATGATGACACCAGAAAACGACGAAGAGCAGACATCTGTGTTCTCCGCTACTGTTTACGGAGACAAAATTCAGGGAAAGAATAAACGCAAACGCGTGATTGGTCTATGTATTAGAATATCTATGGTTATTTCACTACTATCTATGATTACCATGTCCGCGTTTCTCATAGTGCGCCTAAATCAATGCATGTCTGCTAACAAGGCTGCTATTACTGACTCCGCTGTTGCCGTTGCTGCGGCATCATCTACTCATAGAAAGGTTGTGTCTAGCACTACACAATATGATCACAAAGAAAGCTGTAATGGTTTATATTACCAGGGTTCTTGTTATATATTACATTCAGACTATAAGTCATTCGAGGATGCTAAAGCAAACTGCGCTGCGGAATCATCAACACTACCCAATAAATCCGATGTCTTGACTACCTGGCTCATTGATTATGTTGAGGATACATGGGGATCTGATGGTAATCCAATTACAAAAACTACATCCGATTATCAAGATTCTGATGTATCACAAGAAGTTAGAAAGTATTTTTGTACATAAATAAATGAAATCGCTTAATAGACAAACTGTAAGTAGGTTTAGGAAGTTGTCGGTGCCGGCCGCTATAATGATGTTACTCTCAACCATTATTAGCGGCATAGGAACATTTCTGCATTACAGAGAAGAACTGATGCCTAGTGCTTGCGCCAATGGATGGATACAATACGATAAACATTGTTATCTGGATACCAACATTAAAATGTCTACGGATAATGCAGTTTATCAGTGTCGCAAATTACGAGCTAGATTGCCTAGACCTGATACTAGACATCTGAGAGTATTGTTTAGTATTTTTTATAAAGATTATTGGGTAAGTTTNAAAAAGACCAATGATAAATGGTTAGATATTAATAATGATAAAGATATAGATATTAGTAAATTAACAAATTTTAAGCAACTAAACAGCACGACGGATTCTGAGGCGTGTTATATATACAAGTCTGGAAAACTGGTTAAAACAGTATGTAAAAGTACTCAATCTGTACTATGCGTTAAAAGATTCTACAAGTGACAACAAAAAATGAATTAATAGTAAGTCGTTAACGTACGCCGCCATGGACGCCGCGTTTGTTATTACTCCAATGGGTGTGTTGACTATAACAGATACATTGTATGATGATCTCGATATCTCAATCATGGACTTTATAGGACCATACATTATAGGTAACATNAAAATTGTCCAAATAGATGTACGGGATATAAAATATTCCGACATGCAAAAATGCTACTTTAGCTATAAGGGTAAAATAGTTCCTCAGGATTCTAATGATTTGGCTAGATTCAACATTTATAGTATTTGTACAGCATACAGATCNAAAAATACCATCATCATAGCATGCGACTATGATATCATGTTAGATATAGAAGGTAAACATCAACCATTTTATCTATTCCCATCTATTGATGTTTTTAACGCTACAATCATAGAAGCGTATAATCTGTATACAGCTGGAGATTATCATCTGATCATCAATCCTTCAGATAATCTGAAAATGAAATTGTCGTTTAATTCTTCATTTTGTATATCAGACGGCAATGGATGGATTATAATTGATNGGAAATGTAATAGTAANTTTTTATCATAAAAGTTGTAAAGTAAATAANAAAACAATAAATATTGAACTAGTAGTATGTTGTATATTGAGCAATCAGAGATGATGCTGGTACCTCTTATCACGGTGACCGTAGTTGCNGGAACAATATTAGTATGTTATATATTATATATTTGTAGGAAAAAGATACGTACTGTCTATAATGACAATAAAATTATCATGACAAAATTAAAAAAGATAAAGAGTCCTAATTCCAGCAAATCTAGTAAATCAACTGATAGCGAATCAGACTGGGAGGATCACTGTAGTGCTATGGAACAAAACAATGACGTAGATAATATTTCTAGAAATGAGATATTGAACGATGATAGCTTCGCTGGTAGTTTAATATGGGATAACGAATCCAATATCATGGCGCCTAGCACAGAACACATTTACGATAGTGTTGCTGGAAGCACGCTGCTAATAAATAATGATCGTAATGAACAGACTATTTATCAGAATACTACAGTAGTAATTAATGATACAGAGACTGTTNNAATACTTAATGAAGATACCAAACAGATTCCTAGCTATTCTTCCAATCCTTTCGTAAATTATAATAAAACCAGTATTTGTAGCAAGTCAAATCCGTTCATTGCAGAACTCAACAATAAATTTAGTGATAATAATCCGTTTAGGAGAGCACATAGTGACGATTATCTTAATAAGCAACAAGATCATGAATACGATGATATAGAATCATCGGTTGTATCATTGGTCTGATTAGTTTCCTTTTTATAAAATTGAAGTAATATTTAGTATTAATTACCGCCGATGCATTATACAAATATGGAGATATTCCCTGTATTCGGCATTTCTAAAATTAGCAATTTTATTGCTAATAATGACTGTAGATATTATATAGATGTAGAGCATCAAAAAATTATATCTGATGAGATCAATAGACAGATGGATGAAACGGTACTTNTTACCAACATCTTAAGCGTAGAAGTTGTAAATGACAATGAGATGTACCATCTTATTCCCCATAGACTATCGANTATTATACTCTGTATTAGTTCNGTTGGAGGATGTGTTATCTCTATAGATAATGACGTCAATGACAAAAATATTCTAACATTTCCCATTGATCATGCTGTAATCATATCCCCACTGAGTAAATGTGTCGTAGTTAGNAAGGGCCCTACAACCATACTGGTTGTTAAAGCGGATATACCCAGCAAACGATTGGTAACATCGTTTACAAACGACATACTGTATGTAAACAATCTATCACTGATTAATTATTTACCGTCGTCTGTATTCATTATTAGACGAGTCACCGACTATTTGGATAGACACATATGTGATCAGATATTTGCTAATAATAAGTGGTATTCCATTATAACTATCGACGATANNCAATATCCTATTCCATCAAATTGTATAGGTATGTCTTCTGCCAAGTACATAAATTCGAGCATCGAGCAAGATATTTTGATCCATGTTTGTAACCTCGAGCATCCATTCGACTCAGTCTACAAAAAAATGCAGTCGTACAATTCTCTACCTATCAAGGAACAAATATTGTATGGTAGAATTGATAATATAAATATGAGCATTAGTATTTCGGTGGATTAATAGATTTCTCTAGTATGGGATCATTAATCATCTCTAAATACATCATNAAAAAGCTATTATCAAATACTGTACTGAATGGATTCATTCTTTTCTCTTTTTATGAAACTCTGTTGTATATCTACGGATAAAACTAGAAGCAAAAAATCTGATAGGAAGAATAATGATTATATGGAGGAACACGATTATTATAAAATAACAATAGTTCCTGGTTCCTCTTCCACGTCTACTAGCTCGTGGTATTATACACATGCCTAGTAATAGTCTCTTTGCGTTGACGGAAAGCAGACTAGAAATAACAGGCCAAAATGTTCAGACACCATAATAGTTCCCAACCCAGATAATAACAGAGTTCCATCAACACATTCCTTTAAACTCAATCCCAAACCCAAAACCGTTAAAATGTATCCAGCCAATTGATAGTAGATAATGAGGTGTACAGCACATGATAATTTACACAGTAACCAAAATGAAAACACTTTAGTAATTATAAGAAATATAGACGGTAATGTCATCATCAACAATCCAATAATATGCCTGAGAGTAAACATTGACGGATAAAACAAAAATGCCCCGCATAACTCTATCATGGCAATAACGCAACCAAACACTTGTAAAATTCCTAAATTAGTAGAAAATACAACTGATATCGATGTATAAGCGATTTCGAGGAATAATAAGAACAAAGTAATTCCCGTAAAGATAAACATCAACATTGTTTGGTAATCATTAAACCAATTAGTATGACGTTGAATTAATTTCACAGTATATTTTATTCCAGTATTATCCCCGCATGTATACGTACCTGGTAAGATATCTTTATATTCCATAATCAATGAGACATCACTATCCGATAACGAATGAAGTCTAGCACTAGTATGCCATTTACTTAATATGGTCGTCTTGGAAGTTTTATTATAAGTTAAAATATCATGATTGTCCAATTTCCATCTAATATACTTTGTCGGATTATCTATAATACACGGAATAATGATGGTATCATTACATGCTGTATATTCTATAGTCTTTGTAGATGTTATAACCACAAAAGTACAGAGGTATATCAACAATATTCTAACTCTTAACATTTTTATTTATTTAAAATGATACCTTTGTTATTTATTTTATTCTTATTTTGCTAACGGTATCGAATGGCATAAGTTTGAAACGAGTGAAGAAATAATTTCTACTTACTTAATAGATTATGTGGTAACGGGTGTTATTAATGGGGATGTATATACATTTTCAAATAATGAACTAAACAAAACTGGGTTAACTAATAACAATAATTATATCACAACATCTATAAAAGTAGAGGATAAGGATACATTAGTAGTATGCGGAACCAATAACGGAAATCCCAAATGTTGGAAAATAGACGGTTCATACTACCCAAAACATATAGGTAGAGGATACGATCATCAAAATAGCAAAGTAACGATAATCAGTCACAATGAATGTGTACTATCCGACATAAACATATAAAAAGAAGGAATTAAACGATGGAGAAGATTTGACGGACCATGTGGTTATGATTAAACGAGTTAAGTTTTTTAAGAAGCCTTAGAAGAGAGGCTATTGGGTATGAGAATCCGAAATATTAAACCAGACAACCCCATATAATTTTATAGCTAAGAATGCCGCGAAGAATGGAACTAATAAAAACGGAAATATTTGTAGCACAACGAATAACTCCCAAACTGCATTCATGTTACACTATATAACACTACTTCGGTTAGATGTTTTAGAAAAAATAAATATCACCGTACCGTTTTGTTGTATAAAAATAACAATTAACAATTATCAATTTTTTTCTTTAATATTTTACGTGGTTGACCATTCTTGGTGGTAAAATAATCTCTTAGTGTTGGAATGGAATGCTGTTTAATGTTTCCACACTCATCGTATATTTTGACGTATGCAGTCACATCGTTTACGCAATAGTCAGACTGTAGTTCTATCATGCTTCCTACGTTAGAAGGAGGAACAGTTTTAAAGTCTCTTGGTTTTAATCTATTGTCATTAGTTTTCATGAAATCCTTTGTTTTATCCACTTCACATTTTAAATAAATGTCCACTATACATTCTTCTGTTAATTTTACTAGATCATCATGAGTCATAGAATTCATAGGTTCCGTAGTCCATGGATCCAAACTAGCAAACTTCGCGTATACGGTATCGCGATTAGTGTATACACCAACTGTATGAAAATTAAGAAAACAGTTTAATAAATCTACAGAAATATTTAATCCTCCGTTTGATACAGATGCGCCATATTTATGGATTTCGGATTCACACGTTGTTTGTCTAAGGGGTTCGTCTAGTGTTGCTTCTACATAGACTTCGATTCCCATATATTCTTTATTGCCAGAATCACATACCGATTTATCATACGCTGGTTCACTTGTTTGAAAACTAAATGGTAGTAGATACATCAAAATAATAAATAATAAGTACATTCTGCAATATTGTTATCGTAATTGGAAAATTGGTATTCAAGTGAGCTGGATTATGTGAGTATTGGATTGTATATTTTATTTTATATTTTATATTTTATATTTTATTTTATATTTTATATTTTATTTTATATTTTGTAGTAAGAATAGAATGCTAATGTCAAGTTTATTCGAATAGATGTCTTATTAAAAAACATATATAATAAATAACAATGGCTGAATGGCATAAAATTATCGAGGATATCTCAAAAAATAATAAGTTCGAGGATGCCGCCATCGTTGATTACAAGACTACAAAGAATGTTCTAGCGGCTATTCCTAACAGAACATTTGCAAAGATTAATCCGGGTGAAGTTATTCCCCTCATCACTAATCATAATATTCTAAAACCTCTTATTGGTCAGAANTTTTGTATTGTATATACTAACTCTCTAATGGATGAGAACACGTATGCTATGGAGTTGCTTACTGGGTACGCCCCTGTATCTCCGATCGTTATAGCGAGAACTCATACCGCACTTATATTTTTGATGGGTAAGCCAACAACATCCAGACGTGATGTGTATAGAACATGTAGAGATCACGCTACCCGTGTACGTGCAACTGGTAATTAAAATAAAAAGTAATATTCATATGTAGTGTCAATTTTAAATGATGATGAAATGGATAATATCCATATTGACGATGTCAATAATGCCGGTATTGACATACAGCTCATCGATTTTTAGATTTCATTCAGAGGATATTGAATTATGTTATGGGAATTTGTATTTTGATAGGATCTATAATAATGTAGTAAATATAAAATATATTCCTGAGCATATTCCATATAGATATAATTTTATTAATCGTACGTTCTCCGTAGATGAACTAGATGATAATGTCNTTTTTACACATGGTTATTTTTTAAAACACAAATATGGTTGTTCACTTAATCCTAGTTTGATTGTCTCATTATCAGGAAACTTAAAATATAATGATATACAATGCTCAGTAAATGTATCGTGTCTCATTAAAAATTTGGCAACGAGTACATCTACTATATTAACATCTAAACATAAGACTTATTCTCTATATCGGTCCATGTGTATTGCTATAATAGGATACGATTCTATTATATGGTATAAATATATAAATGACAGGTATAATGACATCTATGATTTTACTGCAATATGTATGCTAATAGCGTCTACATTGATAGTGATCATATACGTGTTTAAAAAAATAAAAATGAACTCTTAATTATGTTATACTATTAGAAATGGATAAAATCAAAATTACGATTGATTCAAAAATTGGTAATGTTGTTACCATATCGTATAACTTGGAAAAGATAACTATTGATGTCACGCCAAAAAAGAAAAAAGAAAAGGATGTATTATTAGCGCAATCAGTTGCTGTCGAAGAGGCAAAAGATGTCAAGGTGGAAGAAAAAAATATTATCGATATTGAAGATGACGATGATATGGATATAGAAAACACGTAATACGATCTATAAAAATAAGTATTAAATACTTTTTATTTACGGTACTCTTGTAGTGGTGATACCACTAATCGATTATTTTTTTTNAAAAAATACTTATTCTGATTCTTCTAGCCATTTCCGTGTTCGTTCGAATGCCACATCGACGTCAAAAATAGGGGAGTAGTTGAAATCTAGTTCTGCATTGTTGGTACGCACCTCAAATGTAGTGTTGGATATCTTCAACGTATAGTTGTTGAGTATTGATGGTTTTCTAAATAGAATTCTCTTCATATCATTCTTGCACGCGTACATTTTTAGCATCCATCTTGGAATCCTAGATCCTTGTTCTATTCCCAATGGTTTCATCAATAGAAGATTAAACATATCGTAAGAACACGATGGAGAGTAATCGTAGCAAAAGTAAGCATTTCCTTTAATCGCAGATCCCGGATACTGGATATATTTTGCAGCCAACACGTGCATCCATGCAACATTTCCTACATATACCCGGCTATGCACAGCGTCATCATCGACTGTACGATACATAATGTTACCGTGTTGCTTACATTGCTCGTAAAAGACTTTCGTCAATTTGTCTCCTTCTCCGTAAATTCCAGTGGGTCTTAGGCAACAAGTATACAATTTTGCGCCATTCATGATTACGGAATTATTGGCTTTCATAACCAGTTGCTCGGCCATACGTTTACTTTTTGCGTATACATGTCCTGGTGATATATCATAAAGGGTATGCTCATGACCGATGAATGGATTACCGTGTTTATTTGGTCCTATTGCTTCCATGCTACTAGTATAGATCAAATACTTGATTCCTAGGTCCACACAAGCTGCCAATATAGTCTGTGTTCCATAATAGTTTACTTTCATGATTTCATTATCAGTGTATTTTCCAAATACATCCACTAGAGCAGCCGTATGAATAATCAGATTTACCCCATCTAGCGCTTCTCTCACCTTATCAAAGTCGTTTATATCACATTGTATATAGTTTATAACCTTAACTTTCGAGGTTATTGGTTGTGGATCTTCTACAATATCTATGACTCTTATTTCTTGAACATCATCTGCGCTAATTAAAAGTTTTACTATATACCTGCCTAGAAATCCGGCACCGCCAGTAACCGCGTACACGGCCATTGCTGCCACTCNTAATATCAGACTACTTATTCTATTTTACTAAATAATGGCTGTTTGTATAATAGACCACGATAATATCAGAGNAGTTANTTACGTTGAACAAGTNCATGGNAAAGATAAAGTTTTAGGATCAGTTATTGGATTAAAATCCGGAACGTATAGTTTGATAATTCATCGTTACGGAGATATTAGTNGAGGATGTGATTCCATAGGCAGTCNNNAAATATTTATCGGTAACATCTTTGTAAACAGATATGGTGTAGCATATGTTTATTTAGATACAGATGTAAATATATCTACAATTATTGGAAAGGCGTTATCTATTTCAAAAAATGATCAGAGATTAGCGTGTGGAGTTATTGGTATTTCGTACATAAATGAAAAGATAATACANTTTCTTNNNATTAACGAGAATGGCGTTTNATATATCAGTTAATGCGTCTAAAACAATAAATGCATTAGTTTACTTTTCTACTCAGCAAGATAAATTAGTCATACGTAATGAAGTTAATGATATACACTACACTGTCGAATTTGATAGGGACAAAGTAGTTGATACGTTTATTTCATATAATAGACATAATGACTCCATAGAGATAAGAGGGGTGCTTCCAGANGAAACTAATATTGGTCGCGTGGTTAATACGCCGGTTAGTATGACTTACTTGTATAATAAGTATAGTTTTAAACCGATTTTAGCAGAATATATAAGACACNGAAATACTATATCNNNCAACATTTATTCGGCATTGATGACGCTAGATGATTTGGTTATTAAACAGTATGGANACATTGATCTATTATTTAATGAGAAACTTAAAGTAGACTCCGATTCGGGACTATTTGACTTTGTCAACTTTGTAAAGGATATGATATGTTGTGATTCTAGAATAGTAGTAGCTCTATCTAGTCTAGTATCTAAACATTGGGAATTGACAAATNAAAAGTATAGGTGTATGGCATTAGCCGAACATATAGCTGATAGTATTCCAATATCTGAGCTATCTAGACTACGATACAATCTATGTAAGTATCTACGCGGACACACTGATAGCATAGAGGATGAATTTGATTATTTTGAAGACGATGATTTGTCTACATGTTCTGCCGTAACCGATAGGGAAACGGATGTATAANNTTTTTTATAGTATGAAGGNNNNGATGNNNNNNNNNATATGATGGATATGATGATATGATGGATATGATGGATATGATGGATATGATGGATATGATNAAAAAATATAATTGTTGTATCCATTCCCATTCAAATCACCTTATATGATTCTGTAACACAATGAAGGAGTCTCATAGATATATAGAGGTCAGATACTGGTTTGATAAACTTTTTATTCCACATGAGCATGTTTGACTTATGGTTAGACACACATACTTTAACAAATCACTGAAAATTGGAGTTAGGTATTCCTCTCAGAATCAGTTGCCGTTCTGGAACATTAAATGTANTTTTTATGATATACTCCAACGCATTTATGTGGGTATACAACAAGTCATTAATAATGAGTATTTCCAAGAGTTTTAGTTGTCTAGTATTTAACAAGAGAAGAGATTTCAACAGACTGTTTATGAACTCGAATACCGCCTCATTGTCGCTTATATTGATGACATGACATCGAATTCCCAATATCAATCTCATCAGTGATGAGTAGCTCAATCTTGTTATCGGGATCCAATTTCTAAAGATGTCATTAAACCCTCGATCGTGAATGGATTTATCATCATCGTTTTTATGTTGGACATGAGCTTAGTCCGTTTGTCCACATCTATATACGATGATTTCTGAATTATTTCATATATCTCTCGTTAACTCCAGGAACTTGTCAGGGATCTAACTTTAATATGTTCTCGTCTAAGAGATGAAAATCTTTGGATGGTTGCATGTGACTTTTCTCTAAAGGATGATGTTACCCGATCCTCTCTTAAATGACTCCATCTTATCCTTGGACAAGATGGACAGTCTATTTTCCTTAGATGGTTTAATATTTTTTACCCATGATCTATAAAGGTAGACAGACCTAATCGTCTCGGATGACCATATATTATTTTCCGTTTTATTATACGCATAAATTGTAAAAAATATGTTAGGTTTACGAAAATGTCTCGTGGGGCATTAATCGTTTTTGAAGGATTGGACAAATCTGGAAAAACAACACAATGTATGAACATCATGGAATCTATACCGGCAAACACGATAAAATATCTTAACTTTCCTCAGCGATCCACAGTCACTGGAAAGATGATAGATGACTATCTAACTCGTNAAAAAACCTATAATGATCATATAGTTAATCTATTATTTTGTGCAAATAGATGGGAGTTTGCATCTTTTATACAAGAACAACTAGAACAGGGAATTACTTTAATAGTTGACAGATACGCGTTCTCTGGAGTAGCGTATGCCACCGCTAAAGGCGCGTCAATGACTCTCAGTAAGAGTTATGAATCTGGATTGCCTAAACCCGACTTAGTTATATTCTTGGAATCTGGTAGCAAAGAAATTAATAGAAACATCGGCGAGGAAATTTATGAAGATGTTGAATTCCAACAAAAGGTATTACAAGAATATNAAAAAATGATTGAAGAAGGAGATATTCATTGGCAAATTATTTCTTCTGAATTCGAGGAAGATGTAAAGAAGGAGTTGATTAAGAATATAGTTATAGAGGCTATACACACGGTTACTGGACCAGTGGGGCAACTGTGGATGTAATAAAATGAAATTACATTTTTATAAATAGATGTTAGTACAGTGTTATAAATGGATGAAGCATATTACTCTGGCAACTTGGAATCAGTACTCGGGGATACGTGTCCGATATGCATACCGAACTCGCATCAATATCTCAATTAGTTATTGCCAAGATAGAAACTATAGATAATGATTATTAAACAAGGACATTGTAAATTTTATCATGTGTAGATCAAACTTNGATAATCCATTTATNTCTTTCCTAGATACTGCATATACTATCATAGATCAAGAGATCTATCAGAACGAGTTGATTAATTCATTANNNGATAATGAAATTATCGATTGTATAGTTAACAAGTTTATGAGCTTTTATAAGGATAACCTAGAAAATATGGTAGATGCTATCATTACTCTAAAATATTATAATTAATAATCCAGATTTTAAAACTACGTATGTGGAAGTACTCGGTTCCAGAATAGCTGATATAGATATTAAACAAGTGATACGTAAGAATATAATACAATTGTCTAATGATCCGCGAACGATATTTGTGAAAATATTNNNNNNAAATACNTTTTTTATTAAATGACGTCTCTTCGCGAATTTAGAAAATTATGCTGTGATATATATCACGCATCAGGATATAAAGAAAAATCTAAATTAATTAGAGACTTTATAACAGATAGAGATGATACCGATACATATTTGATCATTAAGCTATTGCTTCCCGGATTAGACGATAGAATGTATAACATGAACGATAAACAAATTATAAAATTATATAGTATAATATTTAAACAATCTCAGGAAGATATGCTACAAGATTTAGGATACGGATATATANGAGACACTATTAGGACTTTCTTCAAAGAGAACACGGAAATCCGTCCACGAGATAAAAGCATTTTAACTTTAGAAGAAGTGGATAGTTTTTTAACTACGTTATCATCAGTAACTAAAGAATCACATCAAATAAAATTATTGACTGATATAGCATCTGTTTGTACATGTAATGATTTAAAATGTGTAGTCATGCTTATTGATAAAGATCTAAAAATTAAAGCGGGTCCTCGGTACGTGCTTAACGCTATTAGTCCTCATGCCTATGATGTTTTTAGAAAATCTAATAACTTGAAAGAGATAATAGAAAATGCAGCTAAACAAAATCTAGACTCTATATCTATTTCTGTTATGACTCCAATTAATCCCATGTTAGCGGAATCATGTGATTCTGTCAATAAGGCGTTTNAAAAATTTCCATCAGGAATGTTTGCGGAAGTCAAATACGATGGTGAAAGAGTACAAGTTCATAAAAAAAATAACGAGTTTGCATTCTTTAGTAGAAACATGAAACCAGTACTCTCTCATAAAGTGGATTATCTCAAAGAATACATACCGAAAGCATTTAAAAAAGCTACGTCTATCGTATTGGATTCTGAAATTGTTCTTGTAGACGAACATAATGTACCGCTACCGTTTGGAAGTTTAGGTATACACAAAAAGAAAGAATATAAAAACTCTAACATGTGTTTGTTCGTATTTGACTGTTTATACTTTGATGGATTCGATATGACAGACATTCCATTGTATGAACGAAGATCTTTTCTCAAAGATGTTATGGTCGAAATANCCAATAGAATAGTATTCTCAGAGTTGACGAATANTNGTNNNGAGTCTCAGTTAACTGATGTATTAGATGATGCACTAACGAGAAAATTAGAAGGATTGGTCTTAAAAGATATTAATGGCGTATACGAACCGGGAAAGAGAAGATGGTTAAAAATAAAGCGAGACTATTTGAACGAGGGTTCCATGGCAGATTCTGCCGATTTAGTAGTACTAGGTGCCTACTATGGTAAAGGAGGAAAGGGTGGTATCATGGCAGTCTTTCTAATGGGTTGTTNNGACGATGAATNCGGTAAATGGNNGACGGNNNNNNNNNNNNNNNNNNNNNNNNNNNNNNNNNNNNNNNNNNNNNNNNNNNNNNNNNNNNNNNNNNNNNNNNNNNNNNNNNNNNNNNNNNNNNNNNNNNNNNNNNNNNNNNNNNNNNNNNNNNNNNCCGATTTTGTAGTAGATGATNCGNAACAATCTCAGATATGGGAAATTTCAGGAGCAGNNNNNNNNNNNNNNNNNNNNNNNNNNNNNNNNNNNNNNNNNNNNNNNNNNNNNNNNNNNNNNNNNNNNNNNNNNNNNNNNNNNNNNNNNNNNNNNNNNNNNNNNNNNNNNNNNNNNNNNNNNNNNNNNNNNNNNNNNNNNNNNNNNNNNNNNNNNNNNNNNNNNNNNNNNNNNNNNNNNNNNNNNNNNNNNNNNNNNNNNNNNNNNNNNNNNNNNNNNNNNNNNNNNNNNNNNNNNNNANGAAATAAATCATATNNAAAATGATTTCATGATTAAACCATGTTGTGAAAGAGTTTGTNAAAAAGTCAAGAACGTTCACATCGGCGGACAATCTAAAAACAATACAGTGATTGCAGATTTGCCATATCTGGATAATGCTGTATCAGATGTATGCAAATCAATATATAAAAAGAATGTATCAAGAATATCCAGATTTGCTAATTTGATAAAAATAGATGACGATGACAAGACTCCTACCGGCGTATATAATTATTTTAAACCTAAAGATGCTATTCCTGTTATTATATCCATAGGAAAGGATAAAGATGTCTGTGAACTATTAATCTCATCTGATAAAGCGTGTGCGTGTATAAAGTTAAATTTATATAAAGTAGCCATTCTTCCCATGGATGTTTCNTTTTTTACCAAAGGAAATGCATCATTGATTATTCTCCTGTTTGATTTCTCTATCGATGCGGCACCTCTCTTAAGAAGTGTAACCGATAATAATGTTATTATATCTAGACACCAGCGCCTACATGACGAGCTTCCGAGTTCCAATTGGTTCAAGTTTTACATAAGTATAAAGTCCGACTATTGTTCTATATTATATATGGTTGTTGATGGATCTATGATGTATGCGATAGCTGATAATAGAACTCACGCAATTATTAGCAAAAATATATTAGACAATACTACGATTAACGATGAGTGTAGATGCTGTTATTCTGAACCACAGATTAGGATTCTTGATAGAGATGAGATGCTCAATGGATCATCGTGTTATATGAACAGACATTGTATTATGATGAATTTACCTGATGTAGGCGAATTTGGATCTAGTATGTTGGGGAAATATGAACCTGACATGATTAAGATTGCTCTTTCGGTGGCTGGTAATTTAATAAGAAATCNAGACTACATTNNNGGGAGACGAGGCTATAGCTACTACGTTTACGGTATAGCCTCTAGATANNTTTTTTTAAGCACGAAATNAAAACATAATTTTAAACAATCTATTTCATACTATTTTGTGTGCTCACCATGAACATAAAGATAGATATATTAGTATTTCTGGTGATAAATTTACGGCGACTGCTANGAGGGAAAATGAAGAAAGAAAAAATATCTACCTCTCCNNAAAGNAAAACTACTGATGTTATCAAACCTGATTATCTTGAGTACAATGACTTGTTAGATAGAGATGAGATGTCTACTATTCTAGAGGAATATTAGGCCTTAGAATAAAATATGGACGACTCTTAACGAAATTAGAAAATTCGATAATGATGTTGAAGAACAATTCGGTACTATAGAAGAACTCAAGCAGAAGCTTAGATTAAATTCTGAAGAGNGAGCAGATAATTTTATAGATTATATAAAGGTACAAAAACAGGATATCATCAAACTTACTGTATACGATTGCATATATCTATGATAGGATTGTGTGCGTGCGTGGTAGATGTTTGGAGAAATGAGAAACTGTTTTCTAGATGGAAATATTGGTTACAAGCGATTAAACTGTTTATTGATGATCACATGCTTGATAAGATAAAATCTATTGTAGAATAGACTAGTGTATGTGGAAATGTCATAGAAAGTTAAAAGTTAATGAGAGCAAAAATATATAAGGTTGTATTCCATATTTGTTANTTTTTTCTGTAATAGTTANAAANNACNNNCGATGGTCTATCTACCAGATTATNNNNTGTTATAAGGTACTTTNNTCNTAATAAACTAGAGTATGAGTAAGATAGTGNTTTTCAAAAACATATAAATCTAAAATTGATGGATGAGATATACAGCTATTAATTTCGAAAATATATTTTAATCTGATAACTTTAAACATGGATTTTTGATGGTGGTTTAAGTTTNAAAAAGATTTTGTTATTGTAGTATGATAATATCAAAAAGATGGATATAAAGAATTTACTGACTACATGTACTATTTTACATTACTACATTGGCTACGGCATATATACCTATTTCGTCACTTCCACACGCTCCGGTAAACGGGTGTCATGTGACGAGGGAGAATCTTGATAAGAGGCATAATCAATGTTGTAATCCGATGTCCACCTGGAGAATTTGCCAAGGTCAGATGTAGAGTTGGTAGTGATAACACAAAATGTGAACACTGCCCACCTCATACATATACCGCAATCCCCAATTATTCTAATAGATGTCATCAATGTAGAAAATGCCCAACAGGATCATTTGATAAGGTAAAGTGTACCGGAACACAGAACAAATGTTCGTGTCATCCTGGTTGGTATACGCTACTGATTCTTCACAGACTGAAGATTGTCGAGATTTGTGTACCAAAAAAGGAGATGTCCATGCGGATACTTTGGTGGAATAGATGAAGGAAATCCTATTTGTAAATCGTGTTGTGTTGGTGAATATTGCGACTACCTACGTAATTATAGACTTGATCCATTTCCTCCATGCAAACTATCTATCTAAATGTAATTAATTATGATTTTGATGATAATGTTACCATACATTATATCACTACTTGGTTAGTGTGTATTATTTAGTATGGAAGACCTATTAATAATTACTTATCTTTTGACGATCTTGTTATAATTATAATATAAAAATACTTATGACATAGTAACTCATAATTGCTGACGCGATAAATTCGTAATAATCTGTTTTGTTCAAATTTTTATAAGGAATCTACAGGCATAAAAATAAAAATATAATCTATAATATACTCTTACAACGCCATCATGAATAGCAGTGAATTAATTGCTGTTATTATGGATTTAGAAATAGTGGACGATTTTGTGATATTAATATAGTTATTAATGATGAAAGGATAAACGCGCATAGACTCATCCTATCTGGAGCCTCCGAATATTTTTTCCATTCTGTTTTCCAATAATTTTATCGATTCTAATGAATACGAAGTTAATCTAAGTCATTTAGATTATCAAAGTGTTAACTATTTGATCGATTACATTTATGGGATACCTTTGAGCCTAACTAACAATAACGTGAAATATATTCTTTCAACCGCTGATTTTTTTACAAATTGGATCTGTCATTACTGAGTGCGAAAAATACATACTTAAAAATCTTTGTTTTAGAAACTGTATCGATTTCTACATATACGCTGATAAATATAATAACAAGAAAATAGAATTAGCATCGTTTAACACAATATTACGAAATATTTTGAGACTCATCAACAATGAAAACTTTAAATACTTAACAGAGGAATCAATGATAAAAATTTTAAGCGATGATATGTTATATATAAAAAATGAGGATTTCACCCCACTGATTCTCATTAAATGGTTAGAGAGTACACCAACCATGTACCGTCGAGTTACTTAGATGCCTCAGAATATCATTTCTTTCCCCACAAGTTATAAAATCACTTTATAGTCATCGACTGGTTAGTTCAATCTACGAATGTATAACATTCTTAAACAATATAGCATTCTTGGATAAATCATTTCCTAGATACCATATCATCGAGTTGATATCTATCGGTATAAGTAATTCACATGATAAGATTTCCATAAACTGCTACAATCATAAAAAAATTCATGGGAAATGATATCTTCACGTAGATATAGGTGTAGTTTCGCAGTGACCGTCCTGGATAATATTATCTATATGATGGGTGGATATGATCAGTCCCTGTATAGAAGTTCAAAGGTTATAGCGTACAATACATGTACTAATTCTTGGATATATGATATACCAGAGCTAAAAATATCATCGTTCTAATTGCGGAGGAGTTGCCAATGACGAATACATTTATTGTATAGGCGGTATACGCGATCAGGAGTCATCGTTGATATCTAGTATCGATAGATGGAAGCCATCAAAACCATATTGGCAGAAGTATGCTAAAATGTGCGAACCAAAATGTGATATGGGGTTGCGATTTTAAACGGATTAATATATGTCATAGGTGGAGTCGTTAAAGGTGACACACATATACCAACGCACTAGAGAGTTTATCAGAAGATGGATGGATGAATCATCAACGTCTTCCAATNAAAATGTNCAATATGTCGACGATTGTTCATGCTGGAAAGATTTATATATCTAGAGGTTACAACAATAGTAGTGTAGTTAATGTAATATCGAATCTAGTCCTTAGCTATAATCCGATATATGATGAATGGACCAAATTATCATCATTAAATATTCCTAGAATTAATCCTGCTCTATGGTCAGTGTATAATAAATTATANNTAGGAGGAGTAATATCTGATGATGTTCAAACTAATACATCTGNAACATACGATAAAGNAAAAGATTGTTGGACATTNGATAATGGTCACTTGTTACCACATAATTATATAATGNNTAAATGCGAACCGTTTNAACATAGATATCCATTNNNNAAAANACAGTACACGAATGATTTTCNNNAGTATTTNNNAAGNTTTATANNNAGTTNATAGAACAAAATACATNANTTTGTNAAAATAAATCACNTTTTATACTAATATGNNACAATTANCAATACTTTTGTTACTANNNNCNNTAGTATACGCTACACCTNNTCCTCNGANATCTNAAAAAATAGGTGATGATGCAACTATATCATGTAGTCGAAATAATACAAATTACTACGTTGTTATGAGTGCTTGGTATAAGGAGCCCAATTCCATTATTCTCTTAGCTGCCAAAAGCGACGTCTTGTATTTTGATAATTATACCAAGGATNAAATATCTTACGACTCTCCATACGATGATCTAGTTACAACTATCACAATTAAATCATTGACTGCTGGAGATGCCGGTACTTATATATGTNNANNCTTTATGACATCGACTACAAATGATACTGATAAAGTAGATTATGAAGAATACTCCATAGAGTTGATTGTAAATACAGATAGTGAATCGACTATAGACATAATACTATCTGGATCTACACCAGAAACTATTTCTGAGAAACCAGAGGATATAGATAATTCTAATTGCTCGTCTGTATTCGAAATCACGACTCCGGAACCAATTACTGATAATGTAGACGACCATACAGACACCGTCACATACACTAGTGATAGCATTAATACAGTAAATGCATCATCTGGAGAATCCACAACAGACGAGATTCCGGAACCAATTACTGATAAAGAAGAAGATCATACAGTAACAGACACTGTCTCATACACTACAGTAAGTACATCATCTGGAATTGTCACTACTAAATCAACCACCGATGATGCGGATCTTTATGATACATACAATGATAATGATACAGTACCGCCAACTACTGTAGGTGGTAGTACAACCTCTATTAGCAATTATAAAACCAAGGACTTCGTAGAAATATTTGGTATTACCACATTAATTATATTGTCAGCAGTGGCGATTTTCTGTATTACGTATTATATATGTAATAAACACCCACGTAAATACAAAACAGAGAACAAAGTCTAGATTTTTGACTTACATAAATATCTGGGATAATAAAATCTATCATATTGAGAGGACCATCTGGTTCAGGAAAGACAGCCATAACCNAAAGACTGTTAAAAGACTATGGGAATATATTTGGATTTGTGGTGTCCCATACCACTAGATTNNNTCGTCCTATGGAACGAGAAGGTGTTGATTACCTTACGTTAACAGAGAGGCCATCTGGAAGGGAATAGCCGCCGGAAACTTTCTAGAACATACTGAGTTTTTAGGAAATATTTACGGAACTTNTAAAACAGCTGTAAATACAGCGGTTATTAATAATCGTATTTGCGCGATGGATTTAAACATCAACGGTGTTAGAAGTCTTAAAAATACTTACCTAATGCATTACTTGGGTATATAAGACCTACCTCTCTTAAAATGGTTGAGACCAATCTTCGTCGTAGAAACACTGAAGCGGACGACGAATCTCATCGTCGCGTGATGTTGGCAAAAAACGGATATGGATGAGGTCAACGAAGCAGGTCTATTCGACACNATTATTATTGAAGATGATGTGAATTTAGCATATAGTAAGTGTTAATTCAGATACTACAGGACCGTATTAGAATGTATTTTAACACTAATTAGAGACTTAAGATTTGACTTAAAACTTGATAATTAATAATATAACTCGTTTTTATATGTGGCTATTTCAACGTCTAATGTATTAGTTAAATATTAAAACTTACCACGTAAAACTTAAAATTTAAAATGGTATTTCATTGACAGATCATACATTATGAAGTTTCAAGGACTTGTGTTAATTGACAATTGCAAAAATCAATGGGTCGTTGGACCATTAATAGGAAAAGGTGGATTCGGTAGTATTTATACTACTAATGACAATAATTATGTAGTAAAAATAGAGCCCAAAGCTAACGGATCATTATTTACCGAACAGGCATTTTATACTAGAGTACTTAAACCATCCGTTATCGAAGAATGGAAAAAATCTCACAATATAAAGCACGTAGGTCTTATCACATGCAAGGCATTTGNTTTATACAAATCCATTAATGTNGAATATCGATTCTTNGTAATAAATAGATTAGGTGCAGATCTAGATGCGGTGATCAGAGCCAANAATAATAGACTACCAGNAAGGTCGGTGATGTTGATCGGAATCGAAATCTTAAATACCATACAATTTATGCACGAGCAAGGATATTCTCACGGAGATATTAAAGCGAGNAATATAGTCTTGGATCAAATAGATAAGAATAAANTATATCTAGTGGATTACGGATTGGTTTCTAAANTCATGTCTAACGGCGAACATGNTCCNNNNATAAGAAANCCAAATNAAATNGATNNNNGTACTCNAGAATTTACACCTATAGATTCGCATNAANNATACNNNNNNNNNAGACGTGGTGATCTAGAAACANNNNGATATTGTANGANNNNNNNNNNNNNNNNNNNNNNNNNNNNNNNNNNNNNNNNNNNNNNNNNNNNNNNNNNNNNNNNNNNNNNNNNNNNNNNNNNNNNNNNNNNNNNNNNNNNNNNNNNNNNNNNNNNNNNNNNNNNNNNNNNNNNNNNNNAATTGCNNNAATATATTANNATGGTANNNNNNNNNACATATTTTGAGGAACCNAATTACGACNANNTTNNNNNNNTNTTNNNNNNNNGANNTNNNNNAAATTTTTNTNGATNNAANNNNNNNAAAAATAACTNNGTTATTATCANNCTCGTGAGNANAATNGAAACANNNNGNNGNNNNACGNNCACGCTTTNNNNNNNNNNNNNNNNNNNNNNNNNNNNNNNNNNNNNNNNNCATCGACNGTTGCCNNNNNNGTCANGAAATATTCTGTTGTGTCAGTTTATAATNAAAAGTATAACATTGTAAAAAACAAANATATGTGGTGTAACAGTCAAGNGAACAAGAGATATATTNGAGNNNNNNNNNNNNNGTNTGAATGCAATGAATATCTACAAATTGGAGATNCAATCCATGATCTAGAAGGAAATCAAATNTNTATTGTCACATATCGCCACAAAAACNACNATGNTCNAAGTGGAANNNNNNACGAGAGTCTAGANNTGTGNTTGGAANGAGTANGGATNNNTCATCACGTACTTGAAACAGGAANNNNGGTATATGGAAAAGTNCAACATGAGTATTCTACTATCAAAGAGAAGGCCAAAGAAATGAATGCACTCAAACCAGGACCTATCATCGATTACCACGTCTGGATAGGAGATTGNGNCTGCCAAGTTACTACTGTAGACGTGCATGGAAAGGAAATTATGAGAATGAGATTCAAAAGGGGTGCGGTGCTTCCGATTCCAAATCTGGTAAAAGTTAAAGTTGGGGAGGAAAATGATACAATAAATCTTTCCACTTCCATATCAGCTCTCCTAAATTCCGGTGGCGGCACCATCGAGGTAACATCTAAGGAAGAACGTGTAGATTATGTACTCATGAAACGTTTGGAATCTATACATCATTTGTGGTCNNTAGTGTATGATCATCTTAATGTTGTGAATGGCGAAGAACGATGTTATGTACATATGCATTCATCTCATCAAAGTCCTATGCTGAGTACTGTAAAAACAAATTTGTACATGAAGACTATGGGAGCATGTCTTCAAATGGACTCCATGGAAGCTCTAGAGTATCTTAGTGAACTGAAGGAATCAGGTGGGCGGAGTCCCAGACCAGAATTGCAGAAATTTGAATATCCAGATGGAGNGAAAGACACTGAATCAATNGAGAGATTGGCAGAGGAGTTCTTCAATAGATCAGAACTTCAGGCCGGTGAATCAGTCAAATTTGGTAATTCTATTAATGTTAAACATACATCTGTTTCAGCTAAGCAACTAAGAACACGTATACGACAGCAGCTTCCTTCTATACTCTCATCTTTTGCCAACACAAAGGGTGGATATTTGTTCATTGGAGTTGATAATAATACACACAAAGTAATTGGATTCACGGTGGGTCATGACTACCTCAAACTGGTAGAGAGTGATATAGAAAAGTATATCCAAAAACTTCCTGTTGTGCATTTCTGCAAGAAAAAAGAGGACATCAAGTACGCATGTAGATTCATCAAGGTGTATAAACCTGGTGATGAGACTACCTCGACATATGTGTGCGCAATCAAAGTGGAAAGATGCTGCTGTGCTGTGTTTGCGGATTGGCCAGAATCATGGTACATGGATACTAGTGGTAGTATGAAGAAGTATTCTCCAGATGAATGGGTGTCACATATAAAATTTTAATTAGGGTAAGGTAAAACTATATATAATAACTAACAATTTGTGTATCATATAGACAATTAATTAGGTAACTGTTATCTCTTTTTAACTAACTAACTAACTAACTAACTCTTATATACTATTAATAATACATCTATTAATCATTGATTAGCTTATTGCTTTAATTGTTTTTGTAAACTAACACTGTTCATTGAAAAGGGATAACATGTTACAGAATATAAATTATATATGGATTTTTTTAAAAAGGAAATACTTGACTGGAGTATATATTTATTTCTTCATTACATAACACGTCTGTGTTCTAATTCTTCCAATTCTTCCACATCTCATATAATACAGGAATATAATCTTGTTCGAAAATATGAGAAAGTGGATAAAACAATAGTTGATTTTTTATCTAGGTGGCCAAATTTATTCCATATTTTAGAATATNGGGAAAATATTCTACATATTTATTTTATAGATGCTGCTAATACGAATATTATGANTTTTTTTCTAGATAGAGTATTAAATATTAATAAGAACCGTGGGTCATTTATACATAATCTCGGGTTATCATCCATTAATATAAAAGAATATGTATATCAATTAGTTAATAATGATCATCTAGATAATAGTATAAGACTAATGCTTGAAAATGGACGTAGAACAAGACNTTTTTTGTCTTATATATTGGATACAGTTAATATCTATATAAGTATTTTAATAAATCATAGATTTTATATAGATGCCGAAGACAGTTACGGTTGTACATTATTACATAGATGTATATATAACTATAAGAAATCAGAATCAGAATCATATAATGAATTAATTAAGATATTGTTAAATAATGGATCAGATGTAGATNNAAAAGATACGTACGGAAACACACCGTTTATCCTATTATGTAAACACGATATCGACAACGCGGAATTGTTTGAGATATGTTTAGAGAATGCTAATATAGACTCTGTAGACTTTAATGGATATACACCTCTTCATTATGTCTCATGTCGTAATAAATATGATTTTGTAAAGTTATTAATTTCTAAAGGAGCAAATGTTAATGCACGTAATAGATTCGGAACTACTCCATTTTATTGTGGAATTATACACGGTATCTCGCTTATAAAACTATATTTGGAATCAGACACAGAGTTAGAAATAGATAATGAACATATAGTTCGTCATTTAATAATTTTTGATGCTGTTGAATCTTTAGATTATCTATTGTCCAGAGGAGTTATTGATATTAACTATCGTACTATATACAACGAAACATCTATTTACGACGCTGTCAGTTATAATGCGTATAATACGTTAGTCTATCTATTAAACAGAAATGGTGATTTTGAGACGATTACTACTAGTGGATGTACATGTATTTCGGAAGCAGTCGCGAACAACAACAAAATAATAATGGATATACTATTGTCTAAACGACCATCTTTGAAAATTATGATACCATCTATGATAGCAATTACTAAACATAAACAACATAATGCAGATTTATTGAAAATGTGTATAAAATATACTGCGTGTATGACCGATTATGATACTCTTATAGATGTACAATCGCTACATCAATATAAATGGTATATTTTAAAATGTTTTGATGAAATAGATATCATGAAGAGATGTTATATAAAAAATAAAACTGTATTCCAATTAGNTTTTTGTATCAAAGACATTAATACTTTAATGAGATACNGTAGACATCCTTCTTTCGTGAAATGTAATATTCTTGACGTATACGGAAGTTGTGTACGTAATATCATAGCATCTATTAGATATCGTCAGAGATTAATTAGTCTATTATNCAAGAAGCTGGATGCTGGAGATAAATGGTCGTGTTTTCCTAACGAAATAAAATATAAAATATTGGAAAACTTTAACGATAACGAACTGACCACATATCTNNNNNNNNNNNNNNNNNNNNNNNNNNNNNNNNNNNNNNNNNNNNNNNNNNNNNNNNNNNNNNNNNNNNNNNNNNNNNNNNNNNNNNNNNNNNNNNNNNNNNNNNNNNNNNNNNNNNNNNNNNNNNNNNNNNNNNNNNNNNNNNNNNNNNNNNNNNNNNNNNNNNNNNNNNNNNNNNNNNNNNNNNNNNNNNNNNNNNNNNNNNNNNNNNNNNNNNNNNNNNNNNNNNNNNNNNNNNNNNNNNNNNNNNNNNNNNNNNNNNNNNNNNNNNNNNNNNNNNNNNNNNNNNNNNNNNNNNNNNNNNNNNNNNNNNNNNNNNNNNNNNNNNNNNNNNNNNNNNNNNNNNNNNNNNNNNNNNNNNNNNNNNNNNNNNNNNNNNNNNNNNNNNNNNNNNNNNNNNNNNNNNNNNNNNNNNNNNNNNNNNNNNNNNNNNNNNNNNNNNNNNNNNNNNNNNNNNNNNNNAAATACTTNNNNGAATGATANNNTCNCGTGTCCTANNNNNNNNNNNNNNNNNNNNCNNTTAGAACACNGATCNTNNNNNNNNNNNNNNNNNNNNNNNNNNNNNNNNNNNNNNNNNNNNNNNNNNNNNNNNNNNNNNNNNNNNNNNNNNNNNNNNNNNNNNNNNNNNNNNNNNNNNNNNNNNNNNNNNNNNNNNNNNNNNNNNNNNNNNNNNNNNNNNNNNNNNNNNNNNNNNNNNNNNNNNNNNNNNNNNNNNNNNNNNNNNNNNNNNNNNNNNNNNNNNNNNNNNNNNNNNNNNNNNNNNNNNNNNNNNNNNNNNNNNNNNNNNNNNNNNNNNNNNNNNNNNNNNNNNNNNNNNNNCCCAACANNTGTACGATCTAACGAAGAATTTGATCCAGTNGATGATGGTNCCGACGATGAGACAGATCTGAGCAAACTCTCGAAAGACGTTGTACAATATGAACAAGAAATAGAATCGTTAGAAGCAACTTATCATATAATCATAATGGCGTTGACAATTATGGGTGTCATATTTCTAATCTCCATTATAGTATTAGTTTGTTCCTGTGACAAAAATAATGACCAATATAAGTTCCATAAATTGCTACCGTGAATATAAATCCGTTAAAATAATTAATAATTAATAATTAATAACGAACAAGTATCAAAAGATTAAAGAATNAGCTAGAATCAATTAGATGTCTTCTTCAGTNGATGTTGATATCTACGATGCCGTTAGAGCANTTTTACTCAGGCACTATTATGACAAGAGATTTATTGTGTATGGAAGAAGTAACACCATATTACATAATATATACAGGCTATTTACAAGATGCACCGTTATACCGTTNGATGATATAGTACGTACTATGCCAAATGAATCACGTGTTAAACAATGGGTGATGGATACANTTAATGGTATAATGATGAATGAATTCGATACNGTATGNGTGGGTACCGGACTACGATTCATGGAAATGTTTTTCGATTACAATAAAAATAATCCCAAAAATAGCATCAACAATCAAATAATGTATGATATAATTAATAGCGTAGCCATAATTNTAGCTAATNNGAGATATNGAAGCGCGTTTAACGACGATAGAATATACATCCGTAGAACTATGATGGACAAATTGTACGAATACGCATCTCTAACTACTATTGGTACGATCACTGGAGGTGTTTGTTATTTTATCTGTTGATGCATCTAGTTAGTTTGTATAAATAATTATTTCGATATACTAGTTAAAATTTTAAGATTTTAAATGTATNAAAAACTAATAACGTTTTTATTTGTAATAGGTGCAGNTGCATCCTATTCGAATAATGAGTACANNCCGTTTAATAAACTGAGTGTNAAACTCNATATNGANNGAGTAGATAATATAGNAAATNNNNATACTGATGATAATAATNAANNNGTGTTANNNNNTNAANAGTNNACNNTTTCTATTATTACAGAGTCATGTGACGTCGGATTTGATTCCANAGATATAGATGTTATAAACGACTATAAAATTATTGATATGTATACCATTGACTCGTCTACTATTCAACGCAGAGGACATACGTGTAGAATATCTACCAAATTATCATGCCATTATGATAAGTACCCTTATATCCACAAATATGAGGGTGATGAACGACAATATTCTATTACCGCAGAGGGAAAATGCTATAAAGGAATAAAATATGAAATAAGTATGATGAACGATGATACTCTATTGAGAAAACATACTCTTAAAATTGGATTTACTTATATATTCGATCGTCATGGGCATAGTAATACATATTATTCAAAATATGATTTTTAAAAATTTAAAATATATTATCACTTCAGTGACAGTAGTCAAATAACAAACAACACCATGAGATATATTATAATTCTCGCAGTTTTGTTCATTAATAGTATACATGCTAAAATAACTAGTTATAAGTTTGAATCCGTCAATTTTGATTCCNAAATTGAATGGACTGGGGATGGTCTATACAATATATCCCTTAAAAATTATGGCATCAAGACGTGGCAAACAATGTATACAAATGTACCAGAAGGAACATACGACATATCCGGATTTCCAAAGAATGATTTCGTATCTTTCTGGGTTAAATTTGAACAAGGCGACTATAAAGTGGAAGAGTATTGTACGGGACTATGTGTCGAAGTAAAAATTGGACCACCAACTGTAATATTGACTGAATATGACGATCATATCAATTTGTTCATCGAGCATCCGTATGCTACTAGAGGTAGCAAGAAGATTCCTATTTACAAACGCGGTGACATGTGTGATATCTACTTGTTGTATACGGCTAACTTCACATTCGGAGATTCTGAAGAACCAGTAACATATGATATCGATGACTACGATTGCACGTCTACAGGTTGCAGTATAGACTTTGCCACAACAGAAAAAGTGTGTGTGACAGCACAGGGAGCCACAGAAGGGTTTCTCGAAAAAATTACTCCATGGAGTTCGGAAGTATGTCTGACACCTAAAAAGAATGTATATACGTGCGCAATTAGATCTAAAGAAGATGTTCCCAATTTCAAGGACAAAATAGCCAGAGTTATCACGAGAAAATTTAATAAACAGTCTCAATCTTATTTGACTAAATTTCTCGGTAGCACATCGAATGATGTTACAACNTTTTTTAGCATTCTTGACTAAATATTCATAACTAATTTTTATTAATGATACAAAAATGAAATAAACTGTATATTATACACTGGTTAACGCCCTTGGCTCTAACCATTTTCAAGATGAGGTCCCTGATTATAGTCCTTCTGTTCCCCTCNATCATCTACTCAATGTCTATTAGACGATGCGAGAAGACTGAAGANNNAACATNGGGNTTAAAAATANGGTTGTGTATAATTGCCAAAGATTTCTATCCCGAAAGAACTGATTGCAGTGTTCATCGCCCAACTGCAAGTGGAGGATTGATAACNGAAGGCAATGGATTCAGAGTAGTTATATATGATCAATGTACAGAACCCCATGACTTTATTATCACCGATACNCAACAAACACGTCTTGGATCATCNCATACATATATTAAATTCAGTAACATGAATACAGGTGTCCCATCTAGTATTCNAAAATGTTCCAGAACTCTCTGTATTTCTGTATATTGTGATCAAGAGGCNGGAGACATAAAATTTGAGGAGNATANNNNNNNNNNNNNNNNNNNNNNNNNNNNNNNNNNNNNNNNNNNNNNNNNNNNNNNNNNNNNNNNNNNNNNNNNNNNNNNNNNNNNNNNNNNNNNNNNNNNNNNNNNNNNNNNNNNNNNNNNNNNNNNNNNNNNNNNNNNNNNNNNNNNNNNNNNNNNNNNNNNNNNNNNNNNNNNNNNNNNNNNNNNNNNNNNNNNNNNNNNNNNNNNNNNNNNNNNNNNNNNNNNNNNNNNNNNNNNNNNNNNNNNNNNNNNNNNNNNNNNNNNNNNNNNNNNNNNNNNNNNNNNNNNNNNNNNNNNNNNNNNNNNNNNNNNNNNNNNNNNNNNNNNNNNNNNNNNNNNNNNNNNNNNNNNNNNNNNNNNNNNNNNNNNNNNNNNNNNNNNNNCTTAACAATTNGTTCTGATCTACAGACANNNNTTAATAAATCANCTTATTTTNCCAANATATTAAAANNNNNNNACNNCACNNNTNNNNNNNNATTGTGCGNCTTTCAAGATGATGNNNNATATAGGGTNNTACAGTTTTANNNNNNNNNNNNNNNNNNNNNNNNNNNNNNNNNNNNNNNNNNNNNNNNNNNNNNNNNNNNNNNNNNNNNNNNNNNNNNNNNNNNNNNNNNNNNNNNAAAATTATTTGNNTAGAATATTACGNNTATAATTGTTTNGAAATTTATAGAATAACNNNTATTAATACATTATNGTATATCTACAACGNTATNAGNNNCTTCATATTNGATAATATTACTANTAATATNNNNGGATNCAGANTTNNNNNNNTTGCCTAAATACATNNTTATAGNNTTACNNNGACAATCACCTAAATGTTTTNNACGNAGATAANNNGGTAAAGATTATNNNNACTTNTANNNCTTCCGATATCTACAAGGATATTCCATATCATCATTGTGTAAACTAAATAACGTTTTCTATGGCATTTAATAAGGACATTGGATANNTGGAAAAGTGATGTATGGAAGTTAGTACATTATCAACTTCTCCTTATTGATTGAAAATGAAAATATAAATAGTTTTTATGTATAGCGGTATCTACCCTATNGTTTTATTGCTTACTACTAACATGGATTCAGATACAGATACAGATACAGATACAGATACAGATACAGATACAGATANAGATGTAGAAGATATCATGAATGAAATAGATAGAGANAAAGAAGAAATACTNAAAAATGTAGAAATTGAAAATAATAAAAACATTAACAAGAATCATCCCAGTGAATATATTANAGAAGCACTTGTTATTAATACCAGTAGTAATAGTGATTCCATTGATAAAGAAGTTATAGAATATATCAGTCNCGATGTAGGAATATAGANCATATCTACTAATTTTTATAATCGATACAAAACATAAAAACAACTCGNTATTACATAGCAGGTATGGAATCCTTCAAGTATTGTTTTGATAACGATGGTAAGAAATGGATTATCGGAAATACTTTATATTCTGGTAATTCAATACTCTATAAGGTCAGAAAAAATTTCACTAGTTCGTTCTACAATTACGTAATGAAGATNGATCATAAATCACNCAAGCCATTGTTGTCCGAAATACGATTCTATATATCTGTATTGGATCCNTTGACTATCAACAACTGGACACGAGAANNTGGTATAAANNATTTGGCTATTNCAGATCTGNNTNNAATTGGAGAAACCGATGATTATATGTTCNTCATTATAANNAATTTGGGAAGAGNANTCGCCCCAAAGGATNGTGAATCAGNNTTCGAAGCATGTGTCACTATGATNAACACGTTNGAGTTTATACACTCTCAAGGATTTACTCATGGNAAAATNGAACCGATGAATATACTGATTAGAAATAAACGTATTTCACTAATTGACTATTCTAGAACTAACAAACTATACAAAAGTGGAACACATATAGATTACAACGAGGACATGATAACTTCAGGAAATATCAATTATATGTGTGTAGACAATCATCTTGGAGCAACAGTTTCAAGACGANGAGATTTAGAAATGTTGGGATATTGCATGATAGAATGGTTCGGTGGTAAACTTNCATGGNNAAACGAAAGTAGTATNNAAGTAATAAAACNNAAAAAAGAATATAAACAATTTATAGCTACNTTTTTTGAGGACTGTTTTCCTGAAGGAAATGAACCTCTGGAATTAGTTAGATATATAGAANTAGTATACATGTTAGATTATTCTCAAACTCCTAATTATGACAGACTACGTAGACTGTTTATACAAGATTGAAATNNTANNCNNNTTTTTATAGAGTGTGGGGGTAGTGTTACGGATATCNAATATTAATATTAGACTATCTCTANCGCGCTACACGACCAATATCGATTACTATGGATATCTTCAGGGAAATCGCANCTTCTATGAAAGGAAAGAATGTATTCATTTCTCCAGCGTCAATCTCGTCAGTATTGACAATACTGTATTATGGAGCTAATGGATCCACTGCTGAACAGCTATCAAAATATGTAGAAAAGGAGGAGAACATGGATAAGGTTAGCGCTCAGAATATCTCATTCAAATCCATGAATAAAGTATATGGGCGATATTCTGCCGTGTTTAAAGATNNCNNNNNNNNNNNNNNNNNNNNNNNNNNNNNNNNNNNNNNNNNNNNNNNNNNNNNNNNNNNNNNNNNNNNNNNNNNNNNNNNNNNNNNNNNNNNNNNNNNNNNNNNNNNNNNNNNNNNNNNNNNNNNNNNNNNNNNNNNNNNNNNNNNNNNNNNNNNNNNNNNNNTNNNCTTTAAAGCNNAATNGTTGATGCCATTCGAAANNGAATTTACCAGTGATTATCCCNNNNACGTATCTCCNNNNNAAATNGTAGNNGNNNGNATNATGTCTATTTNCGGCGAGCCANTTAATCACGCATCTNNNNNNNAATCATTCNGNAACNNNNCAATCATAGAACTGCCATATGTTGGNGATNCTAGTATGATGGNCATTCTTCCNAACAAGATTGATGGATTAGAATCCANNNNNNNNNNNNNNNNNNNNNNNNNNNNNNNNNNNNNNNNNNNNNNNNNNNNNNNNNNNNNNNNNNNNNNNNNNNNNNNNNNNNNNNNNNNNNNNNNNNNNNATAATCTTGTNNATACGCTAATAAAGTTGGGACTGACAGATGTGTTCTATTCAACTGGTGATTATATCAATATGTGTAATTCAGATGNGAGTGTTGACGCTATGATTCACAAAACGTATATAGATGTCAATGANGNGTATACAGAAGCAGCTGCAGCAACTTCNGTACTAGTGGCAGACTGTGCATCAACAGTTACAAATGAGTTCTGTGCAGATCATCCGTTCATCTATGTGATTAGACATGTCGATGGTAAAATTCNTTTCGTTGGTAGATATTGCTNTNCAACAACTAATTAAGCACATTCTTAATATTAGAATATTATATAGTTAAGATTTTTACTAACAGGTTAACANNTTTTTTTAAAAATAGNAAAAACATGTGGTATTAGTGCAGGTCGTTATTCTTCCAATTGCAATTNGTAAGATGACGGCCAACTTTAGTACCCACGTCTTTTCACCACAACACTGTNGATNTGACAGACTGACCAGTATTGATGACGTCAGACAATGTTTGACTGAATATATTTATTGGTCGTCGTATGCATACCGCAACAGGCAATGCGCTGGNNNNNNNNANNNNNNNNNNNNNNNNTTTAAAGATGATGNGGAATCAGTNNNNATCGACGTNCGTGAGCTGGNNAAAAATATGCCGTGGGATAATGTTNAGGATTGTACAGAGATCATCCGTTGTTATANANNGGATGAGCNAAAANNNATCNNNNAGATTTCGGCCATCATTGGANTTTNTNNATATGCTGCTACTTACTGGGGAGGTGAAGACCATCCCACTAGTAACAGTCTGAACGCATTGTTTGTGATGCTTGAGATGCTCAATTACATGGATTATACCATCATATTCTGGCGTATGAATTGATGAGTTACAGCTTGACATTTCTTCTTTCCTCCCTCTTCTTCTACCTTTCCCAGAAACAAACNTTTTTTACCCACTATAAAATAAAATGAGTATACTACCTGTTATATTTCTTCCTATANTTTTTTATTCTCCATTCGTTCAGACTTTTAACGTGCCTGAATGTATCGACAAAGGGCAATATTTTGCATCATTCATGGAGTTAGAAAACGAGCCAGTAATCTTACCATGTCCTCAAATAAATACGCTATCATCCGGATATAATATATTAGATATTTTATGGGAAAAACGAGGAGCGGATAATGATAGAATTATACAGATAGATAATGGTAGCAATATGCTAATTCTGAACCCGACACAATCAGACTCTGGTATTTATATATACATTACCACGAACGAAACCTACTGTGACATGATGTCGTTAAATTTGACAATCGTGTCTGTCTCAGAATCAAATATAGATCTTATCTCGTATCCACAAATAGTAAATGAGAGATCTACTGGTAAAATGGTATGTCCCAATATTAATGCATTTATTTCTAGTAACGTAAACACAGAATTATATGGAGCGGACATCGACGCCTTAGAAATAAGAGACTTAAACAACGGACACCTGGAATTATTACCATAGAAGATGTTAGAAAAAATGATGCTGGTTATTATACATGTGTTTTAGANNATATATATATATATATATATATATATATATATATATATGGGCAAAACATATAACGTAACCAGAATTATAAAATTAGAGGTACGGGATAGAATAATACCTCCTACTATGAAATTACCAGAAGGAGTAGTAACNTCAATAGGTNGTAATTTGACTATTGCATGTAGAGTATCGTTGAGACTTCCCACAACGGACACCGACGTCTTTTNGATAAGTAATGGTATATGTATTACGAAGAAGAAGACGAGGACGGAGACGGTAGAATAAGTGTAGCAAATAAAATCTATATGACCGATAAGAGACGTGTTATTACATCCTGGTTAAACATTAATCCTGTCAAGGAAGAAGATGCTACAACGTTTACGTGTATGGCGTTTACTATTCCTAGCATCAGCAAAACAGTTACTGTTAGTAAACGTGAATGTATGTTGTTACATTTCCATATCAATTGAGTTTATAAGAATTTTTTATACATTATCTTCCAACAAACAATTGACGAACGTATTGCTATGATTAACTCCCACAATACNNTATATATTATTAATCATTAACTTGCAGACTATACCTAGTAGTGCTATTTTGACATACTCATGTTCTTGTGTAATCGCAGTATCTATATTATTAAAGTACGTAAATCTAGCTATAGTTTTATTATTTAATTTTAGATAATATACTGTCTCCGTATTTTTAAAAAATTACCACATCCTTTATTAAATCATGAATGGGAATTTCTGTGTCATCGTTAGTATATTGTGAACAACAAGANCAGATATCTATAGGAAAGGGTGGAATGCGATACATTGATCTATGTAGTTTTAAAACATACGCGAACTTTGAAGAATTTATATAAATCATCTCACGAGATATTGNTCTCNGTCATATTCATACACCTGTATAAACTTTCTAGACATCTTACAATGTGTTATTTTATGATCATATTTACATATTTACTGGTATATCAAAGATGTTAGATTAGTTAATNGGAATCGTCTATAATAATGAATATTAAACAATTATAGGAGGAGTTTATACCTACAAAAACATCATAAAAATGAGTCATCGTCCGATTTATGTTTTAAATATACTAACATTACTACCTTCAGAAATTATATACGAAATATTNTACATGCTGACAATTAACGATCTTTATAATATATAGTATCCACCTACCAAAGTATAATTGTATTTTTCTCATGTGATGTGTGTNAAAAACTGATATTATATAATTATCTTAGTACCTATGATGAAGATGAAGATGAAGATGAAGATGATGGTCCGTATATATTTTGTATCATTATCGTTATTGCTATTCCATAGTTACGCCATAGACATCGAAAATGAAATCACCGAATTCTTCAATAAAATGAGAGATACTCTACCAGCTAAAGACTCTAAATGGTTGAATCCAGTATGTATGTTTGGAGGCACAATGAATGATATGGCCGCTCTAGGAGAGCCATTCAGTGCAAAGTGTCCTCCTATTGAAGACAGTCTTTTATCGCATAGATATAAAGACTATGTGGTTAAATGGGAAAGGCTAGAAAAGAATAGACGGCGACAGGTTTCTAATAAACGTGTTAAACATGGTGATTTATGGATAGCCAACTATACATCTAAATTCAGTAACCGTAGGTATTTATGTACCGTAACCACAAAGAATGGTGACTGTGTTCAGGGTGTAGTTAGATCTCATGTGTGGAAACCTTCTTCATGCATTCCAAAAACATATGAACTAGGTACTTATGATAAGTATGGCATAGACTTATACTGTGGAATTCTTTATGCGAACCATTATAATAATATAACTTGGTATAAAGATAATAAGGAAATTAATATCGACGATTTTAAGTATTCACAAGCGGGAAAGGAATTAATTATTCATAATCCAGAGTTAGAAGATAGTGGAAGATACGACTGTTACGTTCATTACGACGACGTTAGAATCAAGAATGATATCGTAGTATCAAGATGTAAAATACTTACGGTTATACCGTCACAAGACCACAGGTTTAAACTAATACTAGATCCGAAAATCAACGTAACGATAGGAGAACCTGCCAATATAACATGCAGTGCTGTGTCAACGTCATTATTTGTCGACGATGTACTGATTGAATGGGAAAATCCATCCGGATGGATTATAGGATTAGATTTTGGTGTATACTCTATTTTAACTAGTAGAGGCGGTATCACCGAGGCGACTTTGTATTTTGAAAATGTTACTGAAGAATATATAGGCAATACATATACATGTCGTGGACACAACTATTATTTTGATAAAACTCTTACAACTACAGTAGTATTGGAGTAAATACACAATGCATTTTTATATACATTACTGAATTATTATTATTAATTATATCGTATTTGTGCTATAGAATGGATGAAGATACGCGACTATCTAGGTATTTGTATCTCACCGATAGAGAACATATAAATGTAGACTCTATTAAACAGTTGTGTAAAATATCAGATCCTAATGCATGTTATAGATGTGGATGTACGGCTTTACATGAGTACTTTTATAATTATAGATCAGTCAACGGAAAATACAAGTATAGATACAACGGTTACTATCAATATTATTCATCTAGCGATTATGAAAATTATAATGAATATTATTATGATGATTATGATAGAACTGGTATGAACAGTGAGAGTGATAATATATCAATCAAAACAGAATACGAGAATGAATATGAATTCTATGATGAAACACAAGATCAAAGTACACAACTAGTAGATTACGACATTAAACTCAAAACCAATGAGGATGATTTTGTTGATGAATTCTATGGTTATGATAGATCAGTGGGTGTCCATGATTATATAGATGTATCAATTAATAAAGTAGTATATGGAAGAGAGTCTCACGTAAGATGGTGGGATATATGGCAAGAACATAATGATGGCGTATACAGTATAGGAAAGGAGTGCATAGATAATATATACGAAGACAGACATACCGTAGACGAATTCTACAAGATAGACAGCGTATCAGATGTAGATGACGCAGAACATATATCTCAGATAACTAATGATGTATCTACACAAACATGGGAAAAGAAATCAGAGTTAGATAGATACATGGAAATGTATCCTCGTCATAGATATGGTAAGCATTCTGTCTTTAAGGGATTTTCTGACAAAGTTAGAAAAAATGATTTAGACATGAACGTGGTAAAAGAATTACTTTCTAACGGTGCATCTCTAACAATCAAGGATAGCAGTAATAAGGATCCAATTGCTGTTTATTTTAGAAGAACAATAATGAATTTAGAAATGATTGATATCATTAACAAACATACAACTATCTATGAACGCAGGTATATAGTACACTCCTATCTAAAAAATTATAGAAATTTCGATTATCCATTTTTCAGAAAGTTAGTTTTGACTAATAAACATTGTCTCAACAATTATTGTAATATAAGCGACGGCAAATATGGAACACCACTACATATATTAGCATCTAATNAAAAAATAATAACTCCTAATTACATGAAGTTATTAGTGTATAACGGAAATGATATAAACGCACGAGGTGAAGATACACAAATGCGAACTCCATTACACAAATATTTGTGTAAATTTGTATATCATAATATTGAATATGGTATCCGATACTATAATGAAAAGATTATAGACGCATTTATAGAGTTAGGAGCCGATCTAACTATTCCAAATGACGATGGAATGATACCAGTAGTTTACTGTATACACTCAAATGCCGAATATGGTTATAACAATATTACTAACATAAAGATAATACGTANNNNNNNNNNNNNNNNNNNNNNNNNNNNNNNNNNNNNNNNNNNNNNNNNNNNNNNNNNNNNNNNNNNNNNNNNNNNNNNNNNNNNNNNNNNNNGTTTAGATANCNNTAGATCACTTGATGGGTANGATATTAATTGTTACTTNNNNNGACGTACACCACTTCATTGCGCTATACAATATNNNNNNACNCAGATTGCTGANTACTNNNNNNNNNNNNNNNNNNNNNNNNNNNNNNNNNNNNNNNNNNNNNNNNNNNNNNNNNNNNNNNNNNNNNNNNNNNNNNNNNNNNNNNNNNNNNNNNNNNTTTTGATTAGTCGTCTACCGCCTAAAAGNGTNATATGCTCACTGACTAACCATATAATAGATTATGTTCNTACGAACAATAGACGTATTATTTGGCAGAGTCAAATGATTANNAAGTACGTACTGTTACTGGACCCATCNNNNTATTATAGANTCAGNAATGTTATCGNNNACAAATTAGACCAATACAATAATCGTTATAATATNTTCGAACACGATANNGACGTTAANNNNNNNTATGGCAAAGTCNTACATGANCTNGATACATATATCNAGGNNGTNNAAGTATTAAAATCTACTTCCATCNNNAATAATATAACACTATACGACACNNNTATAAATAATAAGTCAGAGTTTCCTATACGTCGTGTAAACGACAAACAATTAATTAANCTNATAAAATCCAATACATATCATAATCTTATCGAAAAAGTTATTAAAAATACATTANAGAAATATACTTTAACTNATATAGTCCTNNAGNATATGATCTCATCTCNNTCTCAATCATCTTATTTGAGTCGTATTNCTAATGAGANNNTACTCGAAATATTATATAAANTCGACATGTACGATTTANNNNNNNNNNNNNNNNNNNATATNNGAGAGAATGATATCACAGAGTATCATATNNAGAATACGANGTCTGTTTCTNNANNGACATGAATAATGANNACACATNNAACNNNNTTTTTAATCTTNGATATAACACTAATTACATCNAGATTATATATTNAAATCGTAATTTGAGTTGTCTGATCNNCATGNNNNNNNNAAANNATATACNNAACATNNGCAATCTTTTAGATGATTGATATATTATTATGCGATGTAATCATAACTATCGGAGATGTAGAANNNNNNNNNNNNNNNNNNNNNNNNNNNNNNNNNNNNNNNNNNNNNNNNNNNNNNNNNNNNNNNNNNNNNNNNNNNNNNNNNNNNNNNNNNNNNNNNNNNNNNNNNNNNNNNNNNNNNNNNNNNNNNNNNNNNNNNNNNNNNNNNNNNNNNNNNNNNNNNNNNNNNNNNNNNNNNNNNNNNNNNNNNNNNNNNNNNNNNNNNNNNNNNNNNNNNNNNNNNNNNNNNNNNNNNNNNNNNNNNNNNNNNNNNNNNNNNNNNNNNNNNNNNNNNNNNNNNNNNNNNNNNNNNNNNNNNNNNNNNNNNNNNNNNNNNNNNNNNNNNNNNNNNNNNNNNNNNNNNNNNNNNNNNNNNNNNNNNNNNNNNNNNNNNNNNNNNNNNNNNNNNNNNNNNNNNNNNNNNNNNNNNNNNNNNNNNNNNNNNNNNNNNNNNNNNNNNNNNNNNNNNNNNNNNNNNNNNNNNNNNNNNNNNNNNNNNNNNNNNNNNNNNNNNNNNNNNNNNNNNNNNNNNNNNNNNNNNNNNNNNNNNNNNNNNNNNNNNNNNNNNNNNNNNNNNNNNNNNNNNNNNNNNNNNNNNNNNNNNNNNNNNNNNNNNNNNNNNNNNNNNNNNNNNNNNNNNNNNNNNNNNNNNNNNNNNNNNNNNNNNNNNNNNNNNNNNNNNNNNNNNNNNNNNNNNNNNNNNNNNNNNNNNNNNNNNNNNNNNNNNNNNNNNNNNNNNNNNNNNNNNNNNNNNNNNNNNNNNNNNNNNNNNNNNNNNNNNNNNNNNNNNNNNNNNNNNNNNNNNNNNNNNNNNNNNNNNNNNNNNNNNNNNNNNNNNNNNNNNNNNNNNNNNNNNNNNNNNNNNNNNNNNNNNNNNNNNNNNNNNNNNNNNNNNNNNNNNNNNNNNNNNNNNNNNNNNNNNNNNNNNNNNNNNNNNNNNNNNNNNNNNNNNNNNNNNNNNNNNNNNNNNNNNNNNNNNNNNNNNNNNNNNNNNNNNNNNNNNNNNNNNNNNNNNNNNNNNNNNNNNNNNNNNNNNNNNNNNNNNNNNNNNNNNNNNNNNNNNNNNNNNNNNNNNNNNNNNNNNNNNNNNNNNNNNNNNNNNNNNNNNNNNNNNNNNNNNNNNNNNNNNNNNNNNNNNNNNNNNNNNNNNNNNNNNNNNNNNNNNNNNNNNNNNNNNNNNATATAGAGAATATGAATGAGAATACANCCGATGATNAGNAGGATGACAATAATGACATGGACGTAGATATTCCGTATTGCGCGACACTAGCTACCGCAAATAAAATATACGGTAGTGATAGTATCGAGTTCCATGCCTCATTCCTACNAAAAATAAAAGACGATTTTCAAACTGTAAACTTTAATAATGCGAACCAAACAAAGGAACTAATCAACGAATGGGTTAAGACAATGACAAATGGTAAAATTAATTCCTTATTGACTAGTCCGCTATCCATTAATACTCGTATGATAGTTATTAGCGCCGTCCATTTTAAAGCAATGTGGAAATATCCATTTTCTAAACATCTTACATATACAGACAAGTTTTATATTTCTAAGAATATAGTTACCAGTGTTGATATGATGGTGGGTACCGAGAATGACTTGCAATATGTACATATTAATGAATTATTCGGAGGATTCTCTATTATCGATATTCCATACGAGGGAAACTCTAGTATGGTGATTATACTGCCGGACGACATAGAAGGTATATATAACATAGNAAAAAATATAACAGATGAAAAATTTAAAAAATGGTGTGGTATGTTATCTACTAAAAGTATAGACTTGTATATGCCAAAGTTTAAAGTGGAAATGACGGAACCGTATAATCTGGTACCGATTCTAGAAAATTTAGGACTTACTAATATATTTGGATATTATGCAGATTTTAGTAAGATGTGTAATGAAACTATCACTGTANAAAAATTTCTACATACGACGTTTATAGATGTTAATGAGGAGTATACAGAAGTATCGGCCGTTACAGGAGTATTCATGACTAACTTTTCGATGGTATATCGTATGAAGGTCTACATAAACCATCCATTCATATACATGATTAAAGATAACACCGGACATACACTTTTTATAGGGAAATACTGCTATCCGCAATAAATATAAACAATAGACTTTTATCACGTTATCTCATGTATAAAATATTACAAATAGTATAGCATAAACTAAAGTCGATACATACATTAAAACTTAAATAATAATGTAATTTACAATTAATAGTATAAACTNAAAAAATTAAAAAATTAAAAACAATATCATTATTATAAGTAATATCAAAATGACGATATACGGATTAATAGCGTATCTTATATTCGTGACTTCATCCATCGCTAGTCCACTTTACATTCCCGTTATTCCGCCCATTTCGGAAGATAAATCGTTCAATAGTGTAGAGGTATTAGTTTCTTTGTTTCCCGATGACCNAAAAGACTATACAGTAACTTCTCAGTTCAATAACTACACTATCGGTACCNAAGACTGGACTATCAACGTACTATCCACACCTGATGGTCTGGACATACCATTGACTAATATAACTTATTGGTCACGGTTTACTATAGGTCGTGCATTGTTCAAATCAGAGTCTGAGGATATTTTCCAAAAGAAAATGAGTATTCTAGGTGTTTCTATAGAATGTAAGAAGCCGTCGACATTACTTACTTTTTTAACCGTGCGTAAAATGACTCGAGTATTTAATAGATTTCCAGATATGGCTTATTATCGAGGAGACTGTCTAGAAGCCGTTTATGTAACAATGACTTATAAAAATACTAAAACTGGAGAGACTGATTACACGTACCTCTCTAATNGGGGGTTGCCTGCATACTATCGTAATGGGGTCGATGGTTGATTATTGATTAGTATATTCCTTATATTCCTTATTCTTTTTATTCACACAAAAAGAACATTTTTATAAACATGAAACCACTGTCTAAATGTAATTATGATCTTGATTTATAGATGATGATCAGCCTTCAGAGGATTTTGACCAGTATGTTTAATATGNAAAAAAACATAACTATTAAGCGCTATTGCGCTATTGTGCTTAATTATTTTGCTCTATAAACTGAATATATAGCCACAATTATTGACGGGCTTGTTTGTGACCGACAATCATGAATTTTCAGAAATTATCTCTGGCTATATATCTTACGGTGACATGTTCGTGGTGTTATGAAACATGTATGAGAAAAACTGCGTTGTATCATGACATTCAATTGGAGCATGTAGAAGACAATAAAGATAGTGTAGCATCGCTACCGTACAAGTATCTACAAGTAGTCAAACAAAGAGAACGTAGTAGATTGTTGGCTACATTTAATTGGACGGATATAGCTGAGGGTGTTAGAAATGAGTTCATTAAAATATGTGATATCAACGGAACATATCTATATAATTATACTATTGCTGTTAGTATAATTATTGATTCCACGGAAGAACTACCAACAGTTACTCCAATTACAACATATGAACCTTCTATATATAATTATACTATCGATTATAGCACTGNTATTACTACTGAAGAACTACAAGTGACTCCAACATATGCACCTGTAACAACTCCTCTTCCAACATCAGCAGTTCCTTATGATCAACGATCGAATAACAATGTAAGTACTATATCTATTCAGGTACTGAGTAAAATATTGGGAGTCAATGAAACAGAATTAACTAATTATCTTATTATNCATAAAAATGACACTGTTGACAATAACACNATGGTTGATNANGNGACANCTAACAATAACACATTACATGGTAATATAGGANTTTTNGAAANAAATAATTGTTATANNNTTTCNGTGTCAGATGCTAGTTTTAGAATAACATTAGTAAACGATACTTCTGAAGAAATTTTGCTAATGCTAACANGAACNAGTTCATCCGACACCTTCATATCTTCCACCAATATCACTGAATGTTTGAAAACATTAATCAATAATGTGTCGATTAATGATGTACTTATAACACAAAATATGAATGTAACATCTAATTGTGATAAATGCTCAATGAATTTNATGGCATCCGTTATTCCTGCAGTTAANGAANTTAACAATACGTTGATGAAAATTGGTGTNAAAGATNATGAAAACAATACGGTATATAAATATTATAATTGTAAACTAACTACAAATTNTACATGTGATGAGTTAATCAATTTAGATGAAGTCATTAACAACATAACTCTGACAAATATTANNNACAATAGTGTTTCGACNNCTAACAGCAGNAAAAGACGAGATCTGAATGATGAGTTTGAATTTTNCACTTNCNAGGAATTAGATTGTCTTTACGAATCNTATGGTGTAAANNATGATATAAGTCATTGTTTTGCATCACCTAGACNNNGACGATCNGACGACNAAANNNNGTACATGGACNNGNAATTATTCNACNNCGNNNANNNNNANTTNGGAATAGACAGTGTTATTNCTNGANGTACAACCCATTTNCAAGTANGTGCATCTGGTGCNNNTGGTGGTGTTGNAGGAGATAGNTTNNCATTTCAAAATGTTAAATCGCGTGCCAGTCTATTGGCGGNAAAAATAATGCCTAGAGTACCTATTACTGCTACCGAAGCTGATCTATATGCAACTGTAAATAGACAACCCAAGTTACCAGCAGGTGTTAAAAGTACTCCGTTTACAGAGGCGCTTGTGTCTACGATAAACCNAAAGCTTTCTAATGTTNGAGAGGTAACTTATGCTTCGCTCAATCTGCCAGGATCAAGTGGCTATGTTCATAGACCATCTGATTCTGTTATTTATAGCAGTATAAGACGGTCACGTTTACCTAGTGATAGCGATAGTGATTATGAGGATATACAAACTGTTGTTAAGGAATATAATGAAAGATATGGTAGATCAGTCAGTAGAACACAGTCATCAAGTAGTGAAAGCGATTTTGAAGATATAGATACTGTTGTTAGGGAATATAGACAAAAATATGGCAATGCAATGGCNAAAGGACGTAGTAGTTCCCCTAAACCTGATCCATTATATAGTACTGTTAAGAAAACAACTAAAAGTCTATCTACTGGTGTAGACATAGTTACAAAACAATCAGACTATTCTCTATTACCTGACGTTAATACTGGCAGTTCTATTGTGTCACCTCTCACCAGAAAAGGAGCTACTAGACGACGACCTAGACGCCCTACAAATGATGGTCTACAGAGTCCAAATCCTCCTCTCCGTAATCCACTTCCTCAACATGATGATTATTCTCCTCCACAAGTACACAGACCTCCACCACTTCCTCCTAAACCAGTCCAAAATTCGCCACAACTTNCCCCTAGACCAGTAGGTCAATTACTACCTCCTCCTATAGATCAACCAGATAAAGGATTTAGTAAGTTTGTATCACCTAGACGGTGTAGAAGAGCAAGCTCTGGAGTCATATGTGGTATGATACAATCAAAACCAAACGATGATACCTATTCACTTCTTCAACGATCAAAAATTGAACCAGAATATGTGGAGGTTGGTAATGGTATACCCAAGAACAATGTTCCTGTAATAGGTAATAAACATAGTNAAAAATATACATCGACGATGTCAAAAATATCAACAAAATTTGATAAATCTACGGCATTTGGAGCAGCAATGTTACTAACTGGTCAGCAGGCCATTAGCCAACAGACTAGATCAACTACGTTGAGTAGAAAAGATCAGATGAGTAAGGAAGAAAAGATATTCGAAGCAGTTACAATGAGTCTATCAACTATAGGTTCAACGTTGACGTCTGCAGGTATGACGGGTGGTCCAAAACTAATGATTGCAGGAATGGCTATAACGGCTATAACTGGTATAATAGATACGATAAAAGATATATATTACATGTTTTCAGGACAGGAGAGGCCAGTAGATCCTGTTATTAAATTATTTAATAAGTACACTGGCTTAATGTCCGATAATAATAAAATGGGTGTAAGAAAATGTTTGACACCCGGTGACGACACACTTATTTATATCGCATACAGAAACGATACCAGTTTTAAACAGAATACGGATGCGATGGCTTTGTATTTCTTAGATGTTATCGATTCAGAGATCCTATATCTAAACACATCAAATTTAGTTCTAGAGTATCAACTAAAGGTGGCTTGCCCCATAGGAACATTAAGATCTGTAGATGTGGACATAACTGCGTATACAATATTATATGATACAGCGGATAATATTAAGAAATACAAGTTTATCAGAATGGCAACGCTACTATCCAAACATCCAGTTATTAGATTGACATGTGGTTTAGCAGCAACATTGGTGATNAAACCGTACGAGGTACCCATCAGTGATATGCAACTACTNAAAATGGCGACGCCTGGTGAACCAGAATCCACTAAATCTATACCATCNGATGTCTGTGATAGGTATCCTCTAAAGAAATTCTATNTTTTAGCTGGTGGTTGTCCCTATGATACATCTCAAACTTTTATTGTACATACTACTTGCAGTATTCTACTAAGAACAGCTACACGGGATCAGTTTAGAAACAGATGGGTGTNACAAAATCCATTTAGACAAGAAGGGACATATAAGCNACTGTTTACCTTTAGCAAATACGATTTTAACGACACCATAATCGATCCTAATGGTGTGGTGGGTCATGCTAGCTTTTGTACCAATAGAAGCAGCAACCAATGTTTCTGGTCCGAANCTATGATATTGGAAGATGTATNATNGTGTAGTTCTAGAACTAGNAAAATATACGTAAAACTGGGAATATTTAATGCTGAAGGTTTTAATAGTTTTGTACTAAATTGTCCAACTGGGTCTACACCTACATACATCAAACATAAAAATGCGGACAGTAACAATGTTATCATAGAGCTACCTGTAGGTGATTACGGCACAGCCAAATTGTATTCAGCAACAAAACCATCGAGGATAGCTGTGTTCTGCACACATAACTATGATAAACGATTCAAATCAGATATTATAGTTCTAATGTTTAATAAAAACAGCGGTATTCCATTTTGGAGCATGTACACAGGAAGTGTAACTAGTAAAAATAGAATGTTTGCCACATTGGCTAGAGGAATGCCGTTTAGATCAACGTATTGCGATAACAGACGACGATCAGGTTGTTATTATGCAGGAATACCATTTCATGAAGATAGTGTAGAAACAGATATACATTATGGACCAGAAATAATGTTAAAGGAAACATATGACATAAACAGTATTGACCCACGAGTTATAACAAAGTCAAAGACCCATTTTCCTGCTCCATTGAGTGTAAAATTCATGGTTGACAATTTAGGAAATGGATATGACAACCCTAATTCATTTTGGGAAGATGCTAAAACTAAGAAACGGACATATAGTGCAATGACGATAAAAGTCCTACCATGTACAGTGAGAAATAAAAATATAGACTTTGGATATAACTATGGAGATATTATTTCTAATATGGTTTATCTACAATCTACTAGTCAGGATTATGGAGATGGTACCAAATATACATTTAAATCCGTAACTAGATCAGATCATGAGTGTGAATCTAGCTTAGATCTAACGTCTAAGGAAGTAACTGTGACATGTCCTGCGTTTAGTATACCAAGAAATATATCAACATATGAAGGTCTATGCTTTAGTGTTACTACATCTAAAGATCATTGTGCTACAGGTATTGGTTGGTTAAAATCTAGTGGTTATGGGAAGGAAGATGCTGATAAACCACGTGCTTGTTTTCATCATTGGAATTATTACACACTGTCGTTGGATTATTACTGTTCATACGAAGATATTTGGAGAAGCACCTGGCCTGACTATGATCCATGTAAGTCATATATCCATATAGAGTATAGAGATACATGGATAGAATCTAATGTNTTACAGCAACCTCCTTNNACATTCGAATTCATTCNTGACAATTCTAACGAATATGTGGATANAGAAATTAGTAACAAATTAAATGATCTGTACAATGAATACAAGAAGATTATGGAATATAGCGACGGATCATTGNCGGCGTCTATAAACAGATTAGCAAAGGCATTGACTTCAGAGGGTAGAGAAATAGCAAGTGTTAATATAGATGGTAATCTGTTAGATATCGCATATCAAGCAGATAAGGAAAAGATGGCCGACATACAGACAAGAATAAATGATATTATTAGAGATTTGTTTATACACACTCTATCAGACNAAGATATAAAAGACATTATAGAATCCGAAGAANGTAAGAGATNNNNTATAATAGATGTTANNAACAATCTTGTTNNAAAGTACTATTNTATTGATAATTATCNNTGTNATACTTTAGATGATTATATATACANCTCTGNAGAATATAACAANNNCTNTNTGTTAGNAAACGATACTTNTATANGCNNNNACTATCNTGAATCATCAGNNGTANNNGTTNTATCATNTTATGAAANGACTANAATCTCCTTNGATACNAAAGACGCCAAAGATGCTATAGAAGATGTGANNNTAGCNAGTGCNGTAGCCGAAGCATTGAATGACATGTTTAAGGAATTTGATNAAAACGTAAGTGCTATTATAATAAAAGAAGAAGATAATTATCTAAACAGTTCGCCCGATATCTACCATATAATATATATCATAGGTGGCACTATTCTGCTACTGTTAGTCATTATTTTAATATTGGCAATTTATATAGCGCGCAATAAATACAGAACCAGGAAATATGAAATAATGAAATATGACAATATGAGCATTAAATCTGAGCATCATGATAGTCTTGAAACAGTGTCTATGGAAATTATTGATAATCGGTACTAATAAAATAGTTTAACTCTTTTAGAACCAGTTTGGTACTGTAATTTCAGTTCATTACTCGTTGAGAATATTGATGANNTTTTTTAAATGAGTATCGGTAGTTACANNNTACCATATCATCCATTATATAATCGATGATGCATGTATTAGAATACTTTCCGAATAAGTCTTCTAAATATTGTATTAATTATGAAAAACTATGCTATGTGAGTATGATTCAAAGATGTTTAATGATACGATACTAGATTTTATCTCTANCNNGATTGTTTAGAATCATTTATCATAACTATGTTTAATAAATTCATCAACGAATATCGATAAAGACCTCTTGTAATTCGAGTATAGGAAGTAGTATTACCATATCAACTTCCGAGTTAACAATTACTCTAAAACATGAGGATTGTACTCCTGTCTTTATTGGAGATCACTATTTAGTCGTTGATAAACTAGTAACCTCAGGTTTCTTTACAAACGATAAAGTACAACATCAAGACCTCACAACACAGTGCAAGATTAATCTAGAAATCAAATGTAATTCTGGAGGAGAATCTAGACAACTAACACCCACGGCGAAGTATACTTTATGCCTCATTCAGAAACGGTAACTGTAGTAGGAGACTGTCTCTCTAATCTCGATGTATATATAATATATACCAATACGGACGCGATATATTCCGACATGAATGGCGTCGCTTATCATATGTTATATCCTAAATGTTGATCATATTCCACAAATGATTGTGAACGAGATTAAATCATCTAACAAATAATTAGNTTTTTATGACATTAACATATAATAAATAAATTAATCATTATTGACTTAACGATGACGAAAGTTATCATTATCTTAGGATTCTTGATTATTAATACAAATTCGTTGTGTCTATGAAATGTGAACAAGGTGTCTCATATTATAATGCACAAGAATTAAAGTGTTGTAAACTATCTAGCCAGGAACATATTCAGATTATCGATGTGATAAATACAGCGATACCATCTGTGGACATTGTCCAAGTGACACATTCACGTCAATATATAATCGTTCTCCTCGGTGTCATAGTTGTAGAGGTCACACCTTGNNCACCTACCACAAATAGAATATGTCATTGTGACTCGAATAGTTATCGTCTCCTTAAAGCTTCTGATGGTAACTGTGTTACATGTGCTCCTAAAACAAAATATGGTCGTGTGTACGGAAAGAAAGGNGAAAATGATATGGAATACCATTTGTAAGAAATGTCGGAANGGTACTTATNCAGATATTGTATCTGACTCTGATCAATGTAAACCTATGACAAGATAAGACTTACTCGCATCTACTGGATAGACATAAATATCCTCCTCGTNATAATGAAATATAATATACACTAATTATTAATATCAATCGAGTATTAACATATAAGTTATTTTTAAACCCCTTTTGGGTTCCGTCCTAAACGGCGTTTCGGTCTGTGTCGCCACCATGGTCACACCGAGCCTCTGCGTGCTCCTCCATCGAGGACGACTTCAACTATGACAGCTCGGTGGCGTCTGCCAGCGTGTACATACGAATGGCATTTCTAAGAAAAGTCTACGGTATCCTTTCTACAATTTCCTTTAACAACGGCAACAGCTGCAGTATTTTTATACTTTGAATGCATCGGACATTTATACAAGGGAGTCCTGTTCTAATATTGGCATCAATGTTCGGATCTATAGGCNNGATTTTCGCATTGACTTTACACAGACATGAACATNCCCTGAATCTGTACATACTTTGTGGATTTACACTGTTAGAATCTCTAACGCTGGCCTCTGTTGTTACTTTCTATGATGCACGTATCGTTATGCAAGCTTTCATGTTGACTACTGCAGTGTTTCTTGCTCTGACTACATATACTCTACAATCAANGAGAGATTTCAGTAAACTTGTAACAGGATTGTTTGCTGCTTTCTGGATTTTAATTTTGTCAGGAGTCTTGAGGATAAAGTTTAAAATAGAATTAATAAAGAACATATAGGTCATTTTTTAAACATGGATAGAAACCAAGGTTGTTAGTTAATAATATACAAGATATTTTTTCTCACTCTGATCCATGTAAACCAAGGACGAGAGACACTCTCATTCCTCATTCACGACACCATTAAAAATGGAAATTAAAGCCCTCTATTAAGCACAGACGGCTACAGGTCTACCATCAGGTTACCTTCGTCTACCTTCACAATGGCCTCTCCTTGTGCCCAGTTCAGTCCCTGTCATTGCCACGCTACTAAGGACTCCCTGAATACCGTGACTGACGTCAGACATTGTCTGACTGAATACATCCTGTGGGTTTCTCATAGATGGACCCATAGAGAAAGCGCAGGGCCTCTCTACAGGCTTCTCATCTCTTNCAGAATTGATGCAATGGAGCTATTTGGTAGCGAGTTGAAGGAGTTCTCGAATTCACTTCCGTGGGACAATATCGACAATTGCGTGGAGATCATTAAATGTTTCATCAGAAATGACTCCATGAAAACCGCCAAAGAACTTTGTGCAATAATTGGACTTTGTACTCAATCAGCTATTGTCACTGGAAGAGTCTTCAATGATAAGTATATCGACATACTACTTATGCTGCGAAAGATTCTGAACGAGAACGACTATCTCACCCTCTTGGATCATATCCTCACTGCTAAATACTAAATCTCCTTCATGCTCTCTCACTAATACTCTTACTCACTACACTTTTTATCATCTTATGATGAATGATTGCCTTCATCATTTTTTCGTGGAATATAATATAGGAATAATTAGCACCAGAATAGCTATGGATATCTCGTTAAGAATATTCTCTATAAGAGACATAATGTAGACATAGTTATTATATCCTTCTTAGATAAGTGTTACGCTACTGGAAAGTTTCCATCGTTATTATTACATGAAGATGATATAATTAAACCAACATTGAGATTGGCTCTTATGTTAGCTGGATTGAATTACTGTAATAAATGCATCGAGTATAGAGGGATATAGCAATTCTCGATAATAGTCATGCAATATTTGAATGAGACTGATAATTTAGGTAATACAGTACTACACACATATCTTTCTAGATTATATATCGTTAAAAATCTGTAAGATGTATATTTCTCATAAGTATCCACTGTGTAATATTATTAATGGATATATAGATAACGCAATAGGGACTAATAGTATTGTAAAAGATATAATCGACTATTTACGTACATATCCAGATATCTATATTCCTACTAGTTTGCTGCGTAGTTGCATCATTGATATGCATGATTTATCAGGATTCAGAGATGAATTACTAAGTAAACTACAATCCCACAATAAGTAAGAATCAAATATCNAAAACTCACTTTTGANTTTTCTAGTCTTAAGTAATACATATATTTATTAATAGACCTATGAAATNAAAAAAGGTAACAATGGATTCGCGTATAGCTATTTACGTATTAGTATCGGCATCTCTTTTGTATCTTGTTAATTGTCACAAACTAGTACATTACTTCAATCTGAAAATAAATNNAANTGATATAACNAATACAGCAGATATATTGCTNGACAATTATCCAATTATGACCTTTGATGGAAAGGATATTTATCCATCTATCTCGTTCATGGTCGGTAATAAACTTTTCCTAGATCTTTATAAAAATATCTTTGAAGAATTTTTCAGACTATTTCGAGTATCTGTAAGTAGTCAATACGAGGAATTAGAATATTATTATTCATGTGATTATACTAACAACCGTCCTACAATTAAACAACATTACTTTTATAACGGCGACGAATATACTGAAATTGATAGATCGNAAAAAGCCACTAATAAAAACAGTTGGTTAATTACTTCAGGCTTTAGACTACAAAAATGGTTCGATAGCGAAGATTGTATAATTTATCTCAGATCTTTAGTTAGAAGAATGGAAGACAGTAACAAAAACAGTAAAAAAACTTAGTACTTAGATATCGAAAAAATATATTTTTGTAGACTCTTGAGAATAGAAGGAAAACATGTACATAATTATAAAAAATGAAAATCAATGGCGAATAAGACAGTGCGATTCGCACCATGGAGTCGGTAGATTTCATGGCTGTCGATGAGCAGTTTCACGACGACCTCGATCTTTGGTCATTATCTTTGGTAGATGATTATNAAAAACATGGATTAGGTGTTGACTGTTATGTTCTAGAACCAGTTGTTGACAGGAAAATATTTGATAGATTTCTCCTTGAACCAATTTGTGATCCTGTAGATGTTCTGTATGATTATTTTAGGATTCATAGAGATAATATTGATCAGTATATAGTAGATAGACTGTTTGCATATATTACATATAAAGATATTATATCTGCATTAGTGTCAAAGAATTATATGGAAGATATTTTCTCTATAATTATTAAGAATTGTAATTCTGTGCAAGATCTCTTACTTTACTATCTATCTAATGCATATGTAGAAATAGACATTGTTGATCTTATGGTAGATCATGGGGCTGTAATATATAAAATAGAATGCTTGAATGCCTATTTTAGGGGAATATGTAAAAAGGAAAGTAGTGTTGTTGAGTTTATTTTGAATTGTGGTATCCCAGATGAAAATGATGTTAAATTAGATCTATATAAAATAATTCAGTATACTAGGGGATTCCTTGTAGATGAACCCACAGTATTAGAAATTTATAAGCTTTGTATCCCATATATTGAAGATATCAATCAACTAGATGCTGGTGGAAGGACCTTGCTTTATCGCGCTATCTATGCAGGTTATATAGATTTAGTATCATGGCTATTAGAAAATGGAGCAAATGTCAACGCAGTAATGAGTAATGGATATACATGTCTTGACGTGGCCGTGGATAGGGGATCTGTCATCGCCCGTAGGGAAGCACATCTTAAAATATTAGAAATATTGCTTAGAGAACCATTGTCTATTGACTGTATAAAATTAGCTATACTTAATAATACAATTGAAAACCATGATGTGATAAAGCTCTGTATCAAGTATTTTATGATGGTAGATTATTCACTTTGTAATGTGTATGCATCATCACTCTTTGATTATATAATTGATTGTAAACAAGAATTGGAGTACATTAGGCAGATGAAAATTCATAATACAACCATGTATGAGTTAATCTATAATAGAGACAAAAACAAGCATGCTTCCCATATTCTACATAGGTATTCTAAACATCCAGTTTTGACACAGTGTATCACTAAAGGATTCAAGATTTACACAGAAGTAACCGAGCAGGTCACTAAAGCTCTAAACAGACGTGCTCTAATAGATGAGATAATAAACAATGTATCAACTGATGACAATCTCCTATCAAAACTTCCATTAGAAATTAGGGATCTAATTGTTTCACAAGCTGTCATATAGAGTTCTATCCACCCACCTTTCTTGAAATGAGTTAATAGTCATAAGTTAGTTAAGTCATAAGNNAGTTAAGTNNTNNGTTAGNNNNNNNNNNNNNNNNNNNNNNNNNNNNNNNNNNNNNNNNNNNNNNNNNNNNNNNNNNNNNNNNNNNNNNNNNNNNNGNNNNAAGTTAGTTAAGTCATAAGTTAGTTAAGTCATAAGTTAGTTAAGTCATAAGTTAGTTAAGTCATAAGTTAGTTAAGTCATAAGTTAGTTAAGTCATAAGTTAGTTAAGTCATAAGTTAGTTAAGTCATAAGTTAGTTAAGTCATAAGTTAGTTAAGTCATAAGTTAGTTAAGTCATAAGTTAGTTAAGTCATAAGTTAGTTAAGTCATAAGTTAGTTAAGTCATAAGTTAGTTTATAGTCTAACACTTCTAATTTTTATACCTTGATCTTTTTCTCTAATTATGAAAAAGTAAATCATTATGAAGATGGATGAAATGGACGAGATTGTGCGCATCGTTAACGATAGTATGTGGTACGTACCTAACGCATTTATGGACGACGGTGATAATGAAGGTCACATTTCTGTCAATAATGTCTGTCATATGTATCTCGCATTCTTTGATGTGGATATATCATCTCATCTGTTTAAATTAGTTATTAAACACTGCGATCTGAATAAACGACTAAAATGTGGTAACTCTCCATTACATTGCTATACGATGAATACACGATTTAATCCATCTGTATTAAAGATATTGTTACGCCACGGCATGCGTAACTTTGATAGCAAGGATNAAAAAGGACATATTCCTCTACACCACTATCTGATTCATTCACTATCAATCGATAACAAGATCTTTGATATACTAACGGACCCCATTGATGACTTTAGTAAATCATCCGATCTATTGCTGTGTTATCTTAGATATAAATTCAATGGGAGCTTAAACTATTACGTTCTGTACAAATTATTGACTAAAGGATCTGACCCTAATTGCGTCGATGAGGATGGACTCACTTCTCTTCATTACTACTGTAAACACATATCCGCGTTCCACGAAAGCAATTATTACAAGTCAAAGAGTCACACTAAGATGCGAGCTGAGAAGCGATTCATCTACGCGATAATAGATCATGGAGCAAACATTAACGCGGTTACGAAAATCGGAAATACGCCGTTACACACTTACCTTCAACAGTATACCAAACATAGTCCTCGTGTGGTGTATGCTCTTTTATCTCGAGGAGCCGATACGAGGATACGTAATAATCTTAATTGTACACCCATCATGGAATACATAAAGAACGATTGTGCAACAGGTCATATTCTCATAATGTTACTCAATTGGCACGAACAAAAATACGGGAAATTACAAAAGGAAGAAGGACAACATCTACTTTATCTATTCATAAAACATAATCAAGGATATGGAAGTCGCTCTCTCAATATACTACGGTATCTACTAGATAGATTCGACATTCAGAAAGACGAATACTATAATACAATGACTCCTCTTCATACCGCCTTCCAGAATTGCAATAACAATGTTGCCTCATACCTCGTATACATTGGATACGACATCAACCTTCCGACTAAAGACGATAAGACAGTATTCGACTTGGTGTTTGAAAACAGAAACATTATATACAAGGCGGATGTCGTTAATGACATTATCCACCACAGACTGAAAGTATCTCTACCTATGATTAAATCGTTGTTCTACAAGATGTCGGAGTTCTCTCCCTACGACGATCACTACGTAAAGAAGATAATAGCCTACTGCCTATTAAGGGACGAGTCATTTGCGGNACTACATACTAAATTCTGTTTAAACGAGGACTATAAAAGTGTATTTATGAAAAATATATCATTCGATAAGATAGATTCCATCATCGAAAAATGTAGTCGTGACATAAGTCTCCTCAAAGAGATTCGAATCTCAGACACCGACTTGTATACGGTATTGAGAACAGAAGACATTCGGTATCACACATATCTCGAAGCCATACATTCAGACAAACGCATTTCATTTCCCATGTACGACGATCTCATAGAACAGTGTCATCTATCGATGGAGCATAAAAGTAAACTCGTCGACAAAGCACTCAATAAATTNGAGTCTACCATCGATAGTCAATCTAGACTATCGTATTTGCCTCCGGAAATTATGCGCAATATCATAACCAAGCTAAGCGACTACCATCTAAACAGTATGTTGTACGGAAAGAACCATTACAAATATTATCCATGATAGAAAGAAAATNTTTAAAAAATAATCTATATGATTGGAGAAGTAGGAAACAAACAGTAACAAGACGACGATTACTACATTATTAAATCATGAGGTCCGTATTATACTCGTATATATTGTTTCTCTCATGTATAATAATAAACGGAAGAGATATAGCACCACATGCACCATCCAATGGAAAGTGTAAAGACAACGAATACAGAAGCCGTAATCTATGTTGTCTATCGTGTCCTCCGGGAACTTACGCTTTCAGATTATGTGATAGCAAGACTAATACACAATGTACACCGTGTGGTTCGGATACCTTTNCATCTCACAATAATCATTTACAGGCTTGTCTAAGTTGTAACGGAAGATGTGATAGTAATCAGGTAGAGACGCGATCGTGTAACACGACTCACAATAGAATCTGTGAATGCTCTCCNGGATATTATTGTCTTCTCAAAGGAGCATCAGGGTGTAGAACATGTATTTCTAAANNAAAGTGTGGAATAGGATACGGAGTATCCGGNTACACGTCTACCGGAGACGTCATCTGTTCTCCGTGTGGTNCCGGAACATATTCTCACACCGTCTCTTCCACAGATAAATGCGAACCCGTCGTAACCAGCAATACATTTAACTATATCGATGTGGAAATTAACCTGTATCCAGTCAACGACACATCGTGTACTCGGACGACCACTACCGGTCTCAGCGAATCCATCTCAACGTCGGAACTAACTATTACCATGAATCATAAAGATTGTGATCCAGTCTTTCGTGCAGAATACTTCTCTGTCCTTAATAATGTANNAACTTCAGGATTCTTTACNGGAGAAAATAGATATCAGAATACTTCAAAGATATGTACTCTGAATTTNGNNNNNNNNNNNNNNAACAAAGATTCATCTTCCAAACAGTTAACGAAAACAAAGAATGATACTATCATGCCGCATTCAGAGACGGTAACTCTAGTGGGCGACTGTCTATCTAGCGTCGACATCTACATACTATATAGTAATACCAATACTCAAGACTACGAAAATGATACAATCTCTTATCATATGGGTAATGTTNTCGATGTCAATAGCCATATGCCCGCTAGTTGCGATATACATAAACTGATCACTAATTCCCAGAATNCCACCCACTTATAGTAAGTTTTTTTACCTATAAATAATAAATACAATAATTAATTTCTCGTAAAAGTAGAAAATATATTCTAATTTATTATATGGTAAGAAAGTAGAATCATCTAGAACAGTAATCAATCAATAGCAATCATGAAACAATATATTGTCCTGGCATGCATGTGCCTAGTGNNAGNTNCTATGCCTACTAGTCTTCAACAATCTTCATCCTCGTGTACTGAAGAAGAAAACAAACATCATATGGGAATCGATGTTATTATCAAAGTCACAAAGCAAGACCAAACACCGACCAATGATAAGATTTGTCAATCCGTAACGGAAGTTACAGAGACCGAAGATGATGANNTATCCGAAGAAGTTGTNAAAGGAGATCCCACCACTTATTACACTATCGTCNGTGCGGGTCTTNNNATGAACTTTGGATTCACCAAATGCCCAAAGATTTCATCCATCTCCGAATCCTCTGATGGAAACANTGTGAATACTAGATTGTCCAGCGTGTCACCNGGACAAGGTAAGGANNCTNCCGCGATCACGCGTGAAGAAGCTCTGGCTATGATCAAAGACTGTGAGATGTCTATCGACATCAGATGTAGCGAAGAAGAGAAAGACAGTGACATCAAGACCCATNCAGTACTTGGGTCTAACATCTCACATAAGAAAGTGAGTTACAAAGATATCATCGGTTCAACGATCGNTGATACAAAATGTGTCAAGAACCTAGAGTTTAGCGNNCGTATCGGAGACATGTGTGAGGAATCATCTGAACTTGAAGTCAAGGATGGATTCAAGTATGTCNNCGGATCGGCATCTGAAGGTGCAACCGATGATACTTNACTCATCGATTCAACAAAACTCAAAGCATGTGTCTGAATCGATAACTCTATTCATCTGNAAATGGATGAGTTNGGTTAATCGAACGATTCAGACACCGCACCACGAATTAAAAAAGACCGGGCACTATATTCCGGTTTGCAAAACNAAAATATTTAACTACATTCACAAAAAGTTANNNCTCGTTACTTCTTCNTTCTGTTTCAATATGTGATACGATATGATCACTATTCNTNTTCTCTTGGTCTCATNNNNNNNNNNNNNNNNNNNNNNNNNNNNNNNNNNNNNNNNNNNNNNNNNNNNNNNNNNNNNNNNNNNNNNNNNNNNNNNNNNNNNNNNNNNNNNNNNNNNNNNNNNNNNNNNNNNNNNNNNNNNNNNNNNNNNNNNNNNNNNNNNNNNNNNNNNNNNNNNNNNNNNNNNNNNNNNNNNNNNNNNNNNNNNNNNNNNNNNNNNNNNNNNNNNNNNNNNNNNNNNNNNNNNNNNNNNNNNNNNNNNNNNNNNNNNNNNNNNNNNNNNNNNNNNNNNNNNNNNNNNNNNNNNNNNNNNNNNNNNNNNNNNNNNNNNNNNNNNNNNNNNNNNNNNNNNNNNNNNNNNNNNNNNNNNNNNNNNNNNNNNNNNNNNNNNNNNNNNNNNNNNNNNNNNNNNNNNNNNNNNNNNNNNNNNNNNNNNN

>2022_Belgium_Monkeypox_ITM001

nnnnnnnnnnnnnnnnnnnnnnnnnnnnnnnnnnnnnnnnnnnnnnnnnnnnnnnnnnnnnnnnnnnnnnnnnnnnnnnnnnnnnnnnnnnnnnnnnnnnnnnnnnnnnnnnnnnnnnnnnnnnnnnnnnnnnnnnnnnnnnnnnnnnnnnnnnnnnnnnnnnnnnnnnnnnnnnnnnnnnnnnnnnnnnnnnnnnnnnnnnnnnnnnnnnnnnnnnnnnnnnnnnnnnnnnnnnnnnnnnnnnnnnnnnnnnnnnnnnnnnnnnnnnnnnnnnnnnnnnnnnnnnnnnnnnnnnnnnnnnnnnnnnnnnnnnnnnnnnnnnnnnnnnnnnnnnnnnnnnnnnnnnnnnnnnnnnnnnnnnnnnnnnnnnnnnnnnnnnnnnnnnnnnnnnnnnnnnnnnnnnnnnnnncaaaaatattaactatatacnnntattatatgatgttaacgtgtaaantataaacattattttatgatgcaattgtctgacaacctagattggtataaggatgttgataagctctacgagaatatattgttggacgttatcgtttacgaaatagttgagacatcagaaagaggtttaatatttttgtgagaccatcgaagagagaaagagaataaaaatatttttttttttgtaaaacttttttatgagaccaagagaatacgaatagtgatcatatcgtatcacatattgaaacagaaagaagaagtaacgagaggtaactttttgtgaatgtagttaaatatttttgttttgcaaaccggaatatagtgcccggtcttttttaattcgtggtgcggtgtctgaatcgttcgattaacccaactcatccattttcagatgaatagagttatcgattcagacacatgctttgagttttgttgaatcgatgagtgaagtatcatcggttgcaccttcagatgccgatccgtcgacatacttgaatccatccttgacttcaagttcagatgattcctcacacatgtctccgatacgtacgctaaactctaggttcttgacacattttgtatcaacgatcgttgaaccgatgatatctttgtaactcactttcttatgtgagatgttagacccaagtactggatgggtcttgatgtcactgtctttctcttcttcgctacatctgatgtcgatagacatctcacagtctttgatcatagccagagcttcttcacgcgtgatcgcgggagagtccttaccttgtcccggtgacacgctggacaatctagtattcacagtgtttccatcagaggattcggagatggataaaatctttgggcatttggtgaatccaaagttcatgttaagacccgcaccgacgatagtgtaataagtggtgggatctccttttacaacttcttcggatacctcatcatcttcggtctctgtaacttccgttacggattgacaaatcttatcattggtcggtgtttggtcttgctttgtgactttgataataacatcgattcccatatgatgtttgttttcttcttcagtacacgaggatgaagattgttgaagactagtaggcatagcagctgccactaggcacatgcatgccaggacaatatattgtttcatgattgctattgattgattactgttctagatgattctactttcttaccatataataaattagaatatattttctacttttacgagaaattaattattgtatttattatttataggtaaaaaaacttactataagtgggtgggattctgggaattagtgatcagtttatgtatatcgcaactagcgggcatatggctattgacatcgagaacattacccatatgataagagattgtatcattttcgtagtcttgagtattggtattactatatagtatgtagatgtcgacgctagatagacagtcgcccactagagttaccgtctctgaatgcggcatgatagtatcattctttgttttcgttaactgtttggaagatgaatctttgttgttacatttaatctcgaaattcagagtacatatctttgaagtattctgatatctattttctcctgtaaagaatcctgaagttgctacattattaaggacagagaagtattctgcacgaaagactggatcacaatctttatgattcatggtaatagttagttccgacgttgagatggattcgctgagaccggtagtggtcgtccgagtacacgatgtgtcgttgactggatacaggttaatttccacatcgatatagttaaatgtattgctggttacgacgggttcgcatttatctgtggaagagacggtgtgagaatatgttccgggaccacacggagaacagatgacgtctccggtagacgtgtatccggatactccgtatcctattccacactttgttttagaaatacatgttctacaccctgatgctcctttgagaagacaataatatcctggagagcattcacagattctattgtgagtcgtgttacacgatcgcgtctctacctgattactatcacatcttccgttacaacttagacaagcctgtaaatgattattgtgagatgtaaaggtatccgaaccacacggtgtacattgtgtattagtcttgctatcacataatctgaaagcgtaagttcccggaggacacgatagacaacatagattacggcttctgtattcgttgtctttacactttccattggatggtgcatgtggtgctatatctcttccgtttattattatacatgagagaaacaatatatacgagtataatacggacctcatgatttaataatgtagtaatcgtcgtcttgttactgtttgtttcctacttctccaatcatatagattattttttaaatattttctttctatcatggataatatttgtaatggttctttccgtacaacatactgtttagatggtagtcgcttagcttggttatgatattgcgcataatttccggaggcaaatacgatagtctagattgactatcgatggtagactctaatttattgagtgctttgtcgacgagtttacttttatgctccatcgatagatgacactgttctatgagatcgtcgtacatgggaaatgaaatgcgtttgtctgaatgtatggcttcgagatatgtgtgataccgaatgtcttctgttctcaataccgtatacaagtcggtgtctgagattcgaatctctttgaggagacttatgtcacgactacatttttcgatgatggaatctatcttatcgaatgatatatttttcataaatacacttttatagtcctcgtttaaacagaatttagtatgtagttccgcaaatgactcgtcccttaataggcagtaggctattatcttctttacgtagtgatcgtcgtagggagagaactccgacatcttgtagaacaacgatttaatcataggtagagatactttcagtctgtggtggataatgtcattaacgacatccgccttgtatataatgtttctgttttcaaacaccaagtcgaatactgtcttatcgtctttagtcggaaggttgatgtcgtatccaatgtatacgaggtatgaggcaacattgttattgcaattctggaaggcggtatgaagaggagtcattgtattatagtattcgtctttctgaatgtcgaatctatctagtagataccgtagtatattgagagagcgacttccatatccttgattatgttttatgaatagataaagtagatgttgtccttcttccttttgtaatttcccgtatttttgttcgtgccaattgagtaacattatgagaatatgacctgttgcacaatcgttctttatgtattccatgatgggtgtacaattaagattattacgtatcctcgtatcggctcctcgagataaaagagcatacaccacacgaggactatgtttggtatactgttgaaggtaagtgtgtaacggcgtatttccgattttcgtaaccgcgttaatgtttgctccatgatctattatcgcgtagatgaatcgcttctcagctcgcatcttagtgtgactctttgacttgtaataattgctttcgtggaacgcggatatgtgtttacagtagtaatgaagagaagtgagtccatcctcatcgacgcaattagggtcagatcctttagtcaataatttgtacagaacgtaatagtttaagctcccattgaatttatatctaagataacacagcaatagatcggatgatttactaaagtcatcaatggggtccgttagtatatcaaagatcttgttatcgattgatagtgaatgaatcagatagtggtgtagaggaatatgtccttttttatccttgctatcaaagttacgcatgccgtggcgtaacaatatctttaatacagatggattaaatcgtgtattcatcgtatagcaatgtaatggagagttaccacattttagtcgtttattcagatcgcagtgtttaataactaatttaaacagatgagatgatatatccacatcaaagaatgcgagatacatatgacagacattattgacagaaatgtgaccttcattatcaccgtcgtccataaatgcgttaggtacgtaccacatactatcgttaacgatgcgcacaatctcgtccatttcatccatcttcataatgatttactttttcataattagagaaaaagatcaaggtataaaaattagaagtgttagactataaactaacttatgacttaactaacttatgacttaactaacttatgacttaactaacttatgacttaactaacttatgacttaactaacttatgacttaactaacttatgacttaactaacttatgacttaactaacttatgacttaactaacttatgacttaactaacttatgacttaactaacttatgacttaactaacttatgacttaactaacttatgacttaactaacktatgmcttannnnnnnnnnnnnnnnnnnnnnnnnnnnnnnnnnnnnnnnnnnnnnnnnnnnnaactaacttatgacttaactaacttatgacttaactaacttatgacttaactaacttatgacttaactaacttatgacttaactaacttatgacttaactaacttatgacttaactaacttatgacttaactaacttatgacttaactaacttatgacttaactaacttatgacttaactaacttatgacttaactaacttatgacttaactaacttatgacttaactaacttatgacttaactaacttatgactattaactcatttcaagaaaggtgggtggatagaactctatatgacagcttgtgaaacaattagatccctaatttctaatggaagttttgataggagattgtcatcagttgatacattgtttattatctcatctattagagcacgtctgtttagagctttagtgacctgctcggttacttctgtgtaaatcttgaatcctttagtgatacactgtgtcaaaactggatgtttagaatacctatgtagaatatgggaagcatgcttgtttttgtctctattatagattaactcatacatggttgtattatgaattttcatctgcctaatgtactccaattcttgtttacaatcaattatataatcaaagagtgatgatgcatacacattacaaagtgaataatctaccatcataaaatacttgatacagagctttatcacatcatggttttcaattgtattattaagtatagctaattttatacagtcaatagacaatggttctctaagcaatatttctaatattttaagatgtgcttccctacgggcgatgacagatcccctatccacggccacgtcaagacatgtatatccattactcattactgcgttgacatttgctccattttctaatagccatgatactaaatctatataacctgcatagatagcgcgataaagcaaggtccttccaccagcatctagttgattgatatcttcaatatatgggatacaaagcttataaatttctaatactgtgggttcatctacaaggaatcccctagtatactgaattattttatatagatctaatttaacatcattttcatctgggataccacaattcaaaataaactcaacaacactactttcctttttacatattcccctaaaataggcattcaagcattctattttatatattacagccccatgatctaccataagatcaacaatgtctatttctacatatgcattagatagatagtaaagtaagagatcttgcacagaattacaattcttaataattatagagaaaatatcttccatataattctttgacactaatgcagatataatatctttatatgtaatatatgcaaacagtctatctactatatactgatcaatattatctctatgaatcctaaaataatcatacagaacatctacaggatcacaaattggttcaaggagaaatctatcaaatattttcctgtcaacaactggttctagaacataacagtcaacacctaatccatgttttttataatcatctaccaaagataatgaccaaagatcgaggtcgtcgtgaaactgctcatcgacagccatgaaatctaccgactccatggtgcgaatcgcactgtcttattcgccattgattttcattttttataattatgtacatgttttccttctattctcaagagtctacaaaaatatattttttcgatatctaagtactaagtttttttactgtttttgttactgtcttccattcttctaactaaagatctgagataaattatacaatcttcgctatcgaaccatttttgtagtctaaagcctgaagtaattaaccaactgtttttattagtggcttttttcgatctatcctcgtcctctcatcatccttatattattatcattattatcatagtctattaaacacaaatcatctacgtttataacaacattctcattattaattagttctgtagaatatctttaataatttggctatacatctgttcaatactatctattgatgatttcttttttaagacttaaactagttatggtaatgacgatgaaatcgagtagtaacttctaataaagacttgatatcattatcatatgtttgatcgtcatagttaatagtgtggctaaatggtactgttaataagtttatagacaatatcatagtattttctttccagaattagattattttttaaatactgatcctcacaattccgtgatgtagcagtagttggtgcatggtctatatcgttaaaatgtatcatatataatagttttctgacgtggagtacagaattttcgattaatgagttcatggtaaggaagggcaaatgcctgtatataatatacataagttaatagttttttatcatattttctaataccataataaaaattatcattattgcgtttggtagttctgccctatcatctatatcactgtcactctcgctctcactatatcttctaaaattacaaacaactggatattcgataacagcatttgtgtagtttttgtcttttacagtatatacgttattgtcaaaatctaaacaaatattagcataatacatctatctataagatcaggatccatgttcgagcatactagccatgtatatttgtaacttcgtcgtacagcgttagatcaatagaataaacaatcgtgtgacgcaacttttttacgatctagttgtatgagtttatcgtttacataagcaattaacggctttaacagatgatctgagtaataatatacctctgttatacgtttaatgttcacggtcttagtatttttagatatcaattgtgatttacaccatattcgactcccttgtgtgtaacgttagaaattctaaatctatattatctattacagcgtaaaacacattcaatattgtattgttatttttatattatttacacaattaacaatgtattattagtttatattactgaattaataatataaaattcccaatcttgtcataaacacaaaatccattaaaaatgtcgataaaatatctgatgttgttgttcgctgctatgataatcagatcattagccgatagtggtaatgctatcgaaacaacatcgccagaaattacaaacgctacaacagatattccagctatcagrttanrtggtccagagggagatggatattgtttacacggtgactgtatccacgctagagatattgacggtatgtattgtagatgctctcatggttatacaggtattagatgtcagcatgtagtattagtagactatcaacgttcaggaaaaccagacactacaacatcatatatcccatctctaggtattgtgcttgtattagtaggcattattattacgtgttgtctattatctgtttatatgttcactcgaagaactaaactacctatacaagatatggttgtgttatattttttataaaatttttttatgagtatttttacaaaaatgtatatgtatnaaaaaaatactaagtatacgatgtatcctgtattatttgtatttatctaaacaatacttctgcctctagatgggatacaaaaattttttatttcagcatattaaagtaaaatctagttaccttgaaaatgaatacagtgggtggttccgtatcaccagtaagaacataatagtcgaatacagtatccgattgagattttgcatacaatactagtctagaaagaaatttgtaatcatcctctgtgacaggagtccatatatctgtatcatcgtctagtttttatcagtgtcctatgctatattcctgttatcatcattagttaatgaaaataactctcgtgcttcagaaaagtcaaatattgtatccatacatatatctccaaaactatcacttatacgtttatctttaacgaacatatacctagatggttatttactaacagacattttttcaagatctattgacaataactcctatagtttccacatcaaccaagtaatgatcatctattgttatataacaataacataactcttttccatttttatcagtatctatatcaacgtcgttgtagtgaatagtagtcattgatctattatatgaaacggatatgtctagttaatattttctttgatttaaagtctatagtctttacaaacataatatccttatccgactttatatttcctgtagggtggcataattttattctgcctccacaatcagtgtttccaaatatattactagacaatattccatatagttattagttaagggacccaattagaacacgtacgcgcttattcatcatttggatcgtatttcataaaagttattatgttatagatgtcaacacattctacatttttttaattgtctatatagtatttttctgatattttctataatatcagaattgtcttccataggaagttgtatactattcggaatcagttacatgtttaaataattctctgatgtcattccttatacaatcaaattcattattaaacagtttaatagtctgtagacctttatcgtcgtacatatccattgtcttattagttacgcttatttttatgggttttacgttgctttattatattttataataatgattgtttgacaatgtcgtagtatagatatattattagaggaggtataattataaaaagtttctgagtacgatgttataagaggagaggacacattaacatcatacatcaattaactacattcttataacattgtaatcaaaagaattgcaattttgatgtataacaactgtcaatggaattgtatattacaaattacggtatgttgtaacaacaaataccgatcggtaattgtctctgccgctgtaatagaattaattatatatctattacaccggccttgtatcataataaagttgtggtagtatgatctccatatttataatttagtactttgtatttagtttttttggaatcataaaaaaagttttactaatttaaaatttaaaaagtatttacatttttcactgtttagtcgcggatatggaattcgatcctgtcaaaatcaatacatcatctatagatcatgtaacaatattacaatacatagatgaaccaaatgatataagactaacagtatgcattatccgaaatgttaataacattacatattatatcaatatcacaaaaataaatccacatttggctaatcgatttcgggcttggaaaaaacgtatcgccggaagggactatatgactaacttatctagagatacaggaatacaacaatcaaaacttactgaaactatacgtaactgtcaaaaaaataaaaacatatatggtctatatatacactacaatttagttattaatgtggttattgattgataaccgatgtgattgttcaatcaatattaagagggttggtaaattggtacatagctaataatacatatactccaaatacacccaataatactacaaccatttctgagttggatatcatcaaaatactggataaatacgaggacatgtatagagtaagtaaagaaaaagaatgtggaatttgctatgaagttgtttactcaaaacgattagaaaacgatagatactttggtttattggattcgtgtaatcatatattttgcataacatgtatcaatatatggcatagaacacgaagagaaaccggtgcgtcagataattgtcctatatgccgtacacgttttaaaaaaataacaatgagcaagttctataagctagttaactaataaataaaaagtttaattatcgacgatatatgtcgttatttttctctcatatgaaagattaatttgattctaatataatcttcagtattggatgaatctcaattcaaattaattccattagattagattagattagatcataaataaaaatagtagcacgcactacttcagccaaatattcttttttgaaacgccatctagcgtaatgagaacacaagtgaacctataatgagcaaatttattagtatcggttacatgaaggactttacgtagagtggtgattcctccatctgtggtacgaacggtttcatcttctttgatgccaccacccagatgttctataaacttggtatcctcgtccgatttcatatcatttgccaaccaatacatatagctaaacccaggcatacgttccacacatccggaacaatgaaattctccagaagatgttacaatggctagatttggacatttggtttcaaccgcgttaacatatgagtgaacacacccatacatgaaagcgatgagaaataggattctcatcttgccaaaatatcactataaaaaatttatttatcaattttaaaggtataaaaaaatacttattgttgctcgaatattttgtatttgatggtatacggaagattagaaatgtaggtattatcatcaactgattttatgatggttttatgaattttattatgcttcactattgcatcggaaataatatcatatgcttccacgtatattttattttgttttgactcataatacgcacgtaattctggattattggcatatcgatgaataattttagctccatgctcagtaaatattaatgagaacatagtgttgcctcctaccattatttttttcatctcattcaattcttgattgcagagatctatataatcattatagcgttgacttatggactctggaatcttagacgatgtacagtcatctataatcatggtatatttaatacattgttttatagcataggcattatctacgatattagatacttcactcaatgaatcaatcacacaatctaatgtaggtttatgacataatagcattttcagcagttcaatgtttctagattcgttgatggcaatggttatacatgtatatccgttatttgatctaatgttgacatctgaaccggattctagcagtaaagatactagagattgtttattatatctaacagccttgtgaagaagtgtttctcctcgtttgtcaatcatgttaatgtctttaagataaggtagacaaatgtttatagtactaagaattgggcaaacataagacatgtcacaaagaccctttttgtatgtataagtgtagaaattataacatccatagttggattcacataggtgtccaatcgggatctctccatcatcgagatgattgacggcatctcccccttccttttttagtagatatttcatcgtgtaagaatcaatattaatatttctaaagtatctgtgtatagcctctttatttaccacagctccatattccaacatgcattccactagagggatatcgatatcgccgaatgtcatatactcaattagtatatgttggaggacatccgagttcattgttttcaatatcaaagagatggtttccttatcatttctccatagtggtacaatactacacattattccgtacggctttccattctccaaaaacaattttaccaaatctaaatctacatctttattgtatctataatcactatttagataatcagccataattcctcgagtgcaacatgttagatcgtctatatatgaataagccgtgttatctattcctttcattaacaatttaacgatgtctatatctatatgagatgacttaatataatattgaagagctgtacaatagtttttatctatagaagacggcttgattccgtgattaattagacatttaacaacttccggacgcacatatgctctcgtatccgactctgaatacagatgagcgatgatatacagatgcaatacggtaccacaatttcgtggttgataatcatcatacacgtatccgtactcgtcatcctcataaagaacactgcatccattttctatgaacaaatcaataatttcaggaacaggatcatctgtcattacataattttctataactgaacgatggttttcacatttaacactcaagtcaaatccatgttctaccaacacctttatcaagtcaacgtctacattttttgatttcatatagctgaatatattaaagtcatttatgttgctatatccagtagcttctagtagagccatcgctatatccttattgactttaacatgtctactatttgtgtattcttctattggggtaaactgtctccaatttttatgtaatggattagtgccactatctagtagtagtttgacgacctcaacattattacaatgctcattgaaaaggtatgcgtgtaaagcattattcttgaattggttcctggtatcattaggatctctgtctctcaacatctgtttaagttcatcgagaaccacctcctcattttccagatagtcaaacattttgactgaatagaagtgaatgagctactgtgaactctatacacccgcacaactaatgtcattaaatatcatttttgaatgtatttataccatgtcaaaaacttgtacaattattaataaaaataattagtgtttaaattttaccagttccagattttacacctccgttaacacctccattaaccccactttttacaccactggacgatcctcctccccacattccactgccactagatgtataagttttagatcctttattactaccatcatgtccatggataaagacactccacatgccgccactactaccccctttagacgacatattaataagacaagtttaacaataaaattaatcacgagtaccctactaccaaccactattatatgattacagtaccttgactaaagtctctagtcacaagatcaatactaccaacctacgctattatatgattatagtttctatttttataggaacgcgtacgagaaaatcaaatgtctaagttctaacggtagtgttgataaacgattgttatccgcggatacctcatctatcatgttgtctattttcttactttgttctattaacctattagcattatatattatttgattataaaacttatattgcttattagcccaatctgtaaatatcggattattaacatatcgtttctttgtaggtttatttaacttgtacatcactgtaagcatgtccgtaccatttattttaatttgacacatatcagcaatttctttttcgcagtcggttatatattctatataagatggatacgtatcacatatgtacttatagtctactaatatgaagtacttaatacatattttcagtaacgatttagccttattacctattaataagtgcctgtcgttggataggtaatcaactgttttcttaatacattcgatggttggtaatttactcaaaataatttccaatatcttaatatatatttctgctatttctggtatacatgcatgtgccattataacacaaataccaatacatgtagacccatatgttgttgttatattaatatctgcgccattatctattaaccattctattaggtcaacactatgcgactcgatacaataataaagtatactacgtccatgtttatctattttgtttatatcatcgatatacggcttacaaatttctagtatcgataacacttctgactcgtgaataaataaggtagggaataacggcataatatttattatgttatcatcattaacaactacgtttccattttttaaaatatactctacaactttaggatccctattgtcaaatcttttaaaatatttatttatatgcttaaatctatataatatagctccttccctaatcatacatttgataacattgatgtacactgtatgataagatacatattctgacaatagatcttgtatagaatctgtatatcttttaagaattgaggatattatgacattattacgtaaactattacacaattctaaaatataaaatgtatcatgggcagataatagtttatccactatataattatctattttatgatttttcttcctatattgtttacgtaaatagatagatagaatatgcattagttcattaccgctatagttactatcgaataacacgtcaaatatttcccgtttaatatcgcatttgtcaacataataatagagtatggtacgttcacgataagtataatgacacatctcgttttcgtgcgaaattaaatagtttatcacgtccaaagatgtcacataaccatcttgtgacctagtaataatataataatagagaactgttttacccattctattatcataatcagtggtgtagtcataatctaaataatcaaactcgtcatcccaattaaaataaatataatcagtacattgaatgggtatgatattgtacccatactgtatgttgctacatgtaggtattcctttatccaataatagtttaaatacatctatattaggatttgatgttgtcgcgtatttctctacaatattaataccatttttgatactatttatttctatacctttcgaaattagtaatttcaataagtctatatcgatgttatcagaacatagatattcaaatatatcaaaatcattgatatttttatagtcgactgacgacaataacaaaatcacaacatcgtttttgatattattattttttttggtaacgtatgcctttaatggagtttcaccatcatactcatataatggatttgcaccactttctattaatgattgtgcactactggcatcgatgttaaatgttttacaactatcatagagtatcttatcgttaaccatgattggttgttgatgttatcacattttttggtttctttcatttcagttatgtatggatttagcacgtttgggaagcatgagctcatatgatttcagtactgtagtgtcagtactattagtttcgatcagatcaatgtctagatctatagaatcaaaacacaataggtcagaagataatgaatatctgtacgcttcttcttgtactgtaacttctggttttgttagatggttgcatcgtgctttaacgtcaatggtacaaattttatcctcgctttgtgtatatcatattcgtctctagtataaaattctatattcaaattatcatgcgatgtgtgtacgctaacggtatcaataaacggagcacagcatttagtcaacagtaatccaaaattttttaaagtatatcttaacgaaagaagttgtcatcgttagagtgtggtaaatcattgtctacggtactagatcctcataagtgtatatatctagagtaatatttaatttatcaaatggttgataatatggatgttgtggcaatttcctaatacggaaataagacataaacacgcaataaatctaattgcggacatgttacactccttaaaaaatacgaataaacactttggcttttagtaagtgtcatttaacactatactcatattaatccatggactcataatctctatacgggattaacggatgttctatatacggggatgagtagttctcttctttaactttatactttttactaatcatatttagactgatgtatgggtaatagtgtttgaagagctcgttctcatcatcagaataaatcaatatctctgtttttttgttatacagatgtattacagcctcatatattacgtaatagaacgtgtaatctaccttattaactttcaccgcatagttgtttgcaaatacggttaatcctttgacctcgtcgatttccgaccaatctgggcgtataatgaatcttaactttaatttcttgtaatcattcgaaataatttttagtttgcatccgtagttatctcctctatgtaactgtaaatttctcaacgcgatatctccattaataatgatgtcgaattcgtgttgtatacccatactgaattgatgaactaataccgacggtattaatagtaatttacttttcatctttacatacttggtaatagttttactatcataagtttataaattccacaagctactatggaatataccaaccatcttagtatagaacacatgtcttaaagttattaattaattacatgttgttttatatatcgctacgaatttaaacagaggaatcagttaggaagaaaaaattatctgtcatcatcatcatctattggataacatctctgtattctacgatagagtgctattttaagatgtgacagatccgtgtcatcaaatatatactccattaaaatgattattccggcagcgaacttgatattggatacatcacgacctttgttaatatccacgacaatagacagcaatcccattgttccataaacagtgagtttatctttctttgaagtgatattttgtagagatcttataaaactgtcgaatgacatcgtatttatatctttagctaaatcatatatgttaccatcataatatctaacagcatctatcttaaacgtttccatcgctgtaaagacgtttccgatagatggtctcgtttcatcagtcatactgagccaacaaatgtaatcgtgtataacatctttgatagaatcagactctaaagaaaaggaatcggctttattatacacattcatgataaacttaatgaaaaatgtttttcgttgtttaagttggatgaatagtatgtcttaataattgttattatttcactaattaatatttagtaacgagtacactctataaaaacgagaatgacataactaatcataactagttatcaaagaatgtctaggacgcgtaattttttatggtatagatcctgtaagcgttgtctgtattctggagctattttctctatcgcattagtgagttcagaatatgttataaatttaaatcgaataacgaacataactttagtaaagtcgtctatattaactcttttattttctagccatcgtaataccatgtttaagatagtatattctctagttactacgatctcatcgttgtctagaatatcgcatactgaatctacatccaattttagaaattggtctgtgctacatatctcttctatattattgttgatgtattgtcgtagaaaactattacgtagaccattttctttataaaacgaatatatagtactccaattatctttaccgatatatttgcatacataatccattctctcaatcactacatctttaagagtttggttgttaagatatttggctaaactatataattctattagatcatcaacagaatcagtatatatttttctagatccaaagatgaactctttggcatcctctataatattatcagaaaagatattttcgtgttttagtttatcaagatttaacctgttcatatccatgattaacgacgtcatataaccacataaaataaaaatccattttcatttttagcacaatactattcataattgatattgatgtaatattttgttactttgaacgtaaagacagtacacgggtccgtatctccaacaagcacgtagtaatcaaatttggtgttgttaaacttcgcaatattcatcaatttagatagaaacatatactcatcatctgttttaggaatccatgtattattaccactttccaacttatcattatcccaggctatgtttcgcccatcatcgttgcacagagtgaataattcttttgtattcggtagttcaaatatatgatccatgcatatatcgacaaagctattgtagatgtgatttttcctaaatctaatataaaactcgtttactagcaaacattttcctgatttatcgaccaagacacacatggtttctaaatctatcaagtggtggggatccatagttataacgcagtaacataaattattaccttcttgactgtcgctaatatctatatacttattgttatcgtattggattctacatatagatggcttgtatatcaaagatatagaacacataaccaatttatattctcgctttgtattttcgaatctaaagttaagagatttagaaaacattatatcatcggatgatgatatcactgtttncagagtaggatatattaaagtctttaaagattttgtccgattcaaataaatcactaaataatatcccatattatcatctgttatagtcntgtcattaaatctattatattttatgaaagatatatcactgctcannnntatatttcgtacatttttaaactgtttgtataatatctctctaatacaatcagatatatctattgtgtcggtagacgataccgttacatttgaattaatggtgttccattttacaacttttaacaagttgaccaattcatttctaatagtatcaaactctccatgattaaatattttaatagtatccattttatatcactacggacataaaccattgtatattttttatgtttattagtgtacacattttggaagtaagttcctggatcggatgtcaccgcagtaatattgttgattatttctgacatcgacgtattatatagttttttaattccatatcttttagaaaagttaaacatccttatacaatttgtggaattaatattatgaatcatggtttttacacatagatctattacaggcggaacatcaattattatggcagcaactagtatcatttctacattgtttatggtgatgtttatcttcttccagcgcatatagtctaatatcgattcaaacgcgtgatagtttataccattcaatataatcgcttcatcctttagatggtgatcctgaatgtgtttaaaaaattatacggagacgccgtaataatttccccattgatagaaaatatcacgctttccattttcttgaagtactaaaagtaattataatataatgtaaaggtttatatattcaatannnnnnnnnaaaaaaatcattttgacattaattcctttttaaatttcagtctatcatctatagaaacatattctatgaatttataaaatgcttttacgtatcctatcgtaggtgatagaaccgctaaaaaacctatcnaatttctacaaaagaatctgttatatggnntagggagagtatnaaacattaaatgtnnnnncttattnawgtattcagtagccaatcctaactctttcgaataattattaatggctcttattctgtacgaatctatttttttgaacaatggacctagtggtatatcttgttctatgtatctaaaataatgtctgactagatccgttagtttaatatccgcagtcatcttgtctagaatggcaaatctaactgcgggtttaggcgtaggcgttagtttagtttctatatctacatctatgtctttatctaacaccaaaaatataatagctaatattttattacaatcatccggatattcttctacgatctcactaactaatgtttctttggttatactagtatagtcacgatcagacaaataaagaaaatcagatgatcgatgaataatacatttaaattcatcatctgtaagatttttgagatgtctcattaaaatattattagtgtcagttctcattatcatatattgacagcagctattacacttattttatttttctgtattttattacttttcaccatatagatcagtcattagatcatcaaaatacttttcaatcatcctaaagagtatggtgaacgaatcttcccatctaatttctgaacgtctaccaatgtctctagccactttggcactaatagcgatcattcgcttaacatcttctacattattaactggttgattcaatctatctagcaatggaccgtcggatagcgtcattctcatgttcttaatcaatgtacatacatcgtcatcatctaccaattcatcaaacaatataagctttttaaaatcatcattataataggatggatcgccgtcatttctccaaagaatatatctaataagtagagtcctcatgcttagtaatttaactattttagttaacaactattttttatgttaaatcaattagtaacaccgctatgtttaatacttattcatattttagttttaggatcgagaatcaatacaaaaattaatacatcaattttggaaatacttagtttccacgtagtcaatgaaacatttgagctcatcgtaaaggacgttctcgtacaagacgtaactataaattggtttatatttgttcaagatagatacaaatccgataacttttttgacgaattctacgggattcactttaaaagtgtcataccgggttctttttattcttttaaacagatcgattgtgtgatgttgattaggtcttttacgaatttgatacagaatagcgtttacatatccaccatagtaatcaatagccatttgttcgtatgtcataaattctttaattatatgacactgtgtattatttagttcgtccttgttcatcattaggaatctatccaatatggcaattatattagaactataactgcgttgtatgcgcatgttgatgtgtctgtttatacaatcaattatactaggatccataccactacaatcgggtaaaattgtagcatcatataccatttctagtactttagattcattgttatccattgcagaggacgtcatgatcgaatccaaaaaaatatattatttttatgttattttgttaaaaataatcatcaaatacttcgtaagatactccttcatgaacataatcagttacaaaacgtttatatgaagtaaagtatctacaatttttacaaaagtcaggatgcataagtacaaagtacgcgataaacggaataataatagatttatctagtttatctttttctatctctttcatagttatatacatggtctcagaagtcggattatgtaacatcagcttcgataaaatgactgggttatttagtcttacacattcgctcatacatgtatgaccgttaactatagagtctacactaaaatgattgaataatagatagtctaccattgtttcgtattcagatagtacagcgtagtacatggcatcttcacaaattatatcattatctaatagatatttgacgcatcttatggatcccacttcaacagccatcttaaaatcggtagaatcatattgctttcctttatcgttaataatttctagaacatcatctctatcataaaagatacaaatattaactgtttgatcagtaataacattgctagtcgatatcaatttgttaataagatgcgctgggctcaatgtcttaataagaagtgtaagaggactatctccaaatttgttttgtttattaacatccgttgatggaagtaaaagatttataatgtctacatacttgactgttttagagcatacaatatggagaggcgtatttccatcatgatctggttttgagggactaattcctagtttcatcatccatgagattgtagaagcttttggattgtctgacataagatgtctatgaatatgatttttgccaaatttatccactatcctggcttcgaatccgatagacattatttttttaaacactctttctgaaggatctgtacacgccaacaacggaccacatccttcttcatcaaccgagttgttaatcttggctccatactgtaccaataaatttattctctctatgacttcatcatctgttcccgagagataatatagaggtgttttattatgtttatcacatgcgtttggatctgcgccgtgcaccagcagcatcgcgactattctattattattaattttagaagctatatgcaatggataatttccatcatcatccgtctcatttggagagtatcctctatgaagaagttcttctataaatcgttcatctagtcctttaatgccacaatacgcatgtagaatgtgataatttccagagggttcgataacttgtagcatattcctaaatacatctaaatttttactattatatttggcataaagagatagataatactcgaccgacataatgttgtgttgtccattatagtataaaaattaatatttctatttctatatatttgcaacaatttactctctataacaaatatcataacttagttcttttatgtcaagaaggcactggtttaattcatctataaatgtcactccataactaccacgcatactatactcagaattatgataaagatatttattcttggggtgtaagtaatggggattaatctttgttggatcagtctctaagttaacacatgtcacacatgatccatttatagttatatcacacgatgatgatttatgaattgattccggaagatcgctattgtattttgtagttccacaattcatttccatacatgttattgtcacactaatattatgatgaactttatctagccgctgagtggtaaacaacagaacagatagtttattatctttaccaacaccctcagccgctgccacaaatctctgatccgtatccatgatggtcatgtttacttttagtccgtatccagtcaacactatgttagcatttctgtcgatatagctttcactcatatgacactcaccaataattgtagaattaatgtcgtaatttacaccaatagtgagttcggcgacaaagtaccagtaccggtaatcttgtcgatgaggacatatagtattcttgtattctaccgaatacccgagagatgcgatacaaaagagtaagactaatttgtaaaccatcttactcaaaaatatgcgacaataggaaatctatcttatacacataattattctatcaattttaccaattagttagtgtaatgttaacaaaaatgtgggataatttaatagtttttccttacacaattgacatacatgagtctgagttcctcgtttttgctaattatttcgtccaatttattattcttgacatcgtcaagatcttttgtataggagtcagacttgtattcaacatgtttttctataatcatcttagctatttcggcatcatccaatagtacattttccagattaacagaatagatattaatgtcgtatttgaacagagcctgtaacatctcaatgtctttattatctatagccaatttgatgtccggaatgaagagaagggaattggtgtttgttgacgtcatatagtcgagcaagagaatcatcatatccacgtgtccattttttatagtggtgtgaatacaactaaggagaatagccagatcaaaaggagatggtatctctgaaagaaagtaggaaacaatacttacatcattaagcatgacagcatgataaaatgaagttttccatccagttttcccatagaacatcagtctccaatttttcttaacaaacagttttaccgtttgcatgttaccactatcaaccgcataatacaatgcggtgtttcctttgtcatcaaattgtgaatcatccattccactgaatagcaaaatctttactattttggtatcttctaatgtggctgcctgatgtaatggaaattcattatctagaagatttttcaatgctccagcgttcaacaacgtacatactagacgcacgttattatcagctattgcataatacaagacactatgaccattgatatccgccttaaatgcatctttgctagagagaaagcttttcagttgcttagacttccaagtattaattcgtgacagatccatgtctgaaacgagacgctaattagtgtatattttttcattttttataattttgtcatattgtaccagaattaataatatctctaatagatctgattagtagatatatggctatcgcaaaacaacatatacacatttaataaaaataatattcattaagaagattcagattccactgtacccatcaatataaaataaaataattattccttacatcgtacccataaacaatatattaagtatattccaccttacccataaacaatataaatccagtaatatcatgtctaatgatgaacacaaatggtgtattaaattccagttcttcaggagatgatctcgccgtagctaccatgatagtagatgcctccgctacagttccttgttcgtctacatctatctttacattctgaaacattttataaatatataatgggtccctagtcatatgtttaaacgacgccttatctggattaaacatactaggagccatcatttcggctatcgacttaatatccctcttgttttcgatagaaaatctagggagtttaagattgtacattttattccctaattgagatgaccaatattctaattttgcaggcgtgatagaatctgtgaaatgggtcatattatcacctattgccaggtacatactaatattagcatccttatacagaaggcgcaccatatcatattcttcgtcatcgattgtgattgtatttccttgcaatttagtaactacgttcatcatgggaaccgttttcgtaccgtacttattagtaaaactagcattgtgtgttttagtgatatcaaacggatattgccacgtacctttaaaatatatagtattaatgattgcccatagagtattatcgtcgagcatagtagaatcaactacattagacataccagatctacgttctactatagaattaattttattaaccgcatctcgtctaaagtttaatctatataggccgaatctatgatattgttgataatacgacggtttaatacacacagtactatcgacgaaactttgatacgttagatctgtgtacgtatatttagatgttttcatcttagctaatccggatattaattctgtaaatgctggacccagatctctttttctcaaattcatagtattcaataattctactctagtattacctgatgcagacaatagcgacataaacatagaaaacgaatacccaaacggtgagaagacaatattatcattatcatcctcatccccattttgaatatttttatacgctaatataccagcattgataaatccctgcagacgatatgcggatactgaacacgctaatgatagtatcaataacgcaatcatgatttttatggtattaataattaaccttatttttatgtttggtataaaaattattgatgtctacacatcctttataatcaactctaatcactttaacttttacagttttccctacaagtttatccctatattcaacatatctatccatatgcatctcttaacactctgccaagatagcttcaaagtgaggatagtcaaaaagataaatatatagagtataatcattctcgtatactctgccctttattacatcgcccgcattgggcaacgaataacaaaatgcaagcatcgtgttaacgggctcgtaaattgggataaaattatgtttttattgtttatctattttattcaagagaatattcaggaagttccttttctggttgtatctcgtcgcagtatatatcatttgtacattgtttcatattttttaatagtctacaccttttagtaggactagtatcgtacaattcatagctgtattttgaattccaatcacgtataaaaatatcttccaattgttgacgaagacctaatccatcatccggtgtaatattaatagatgctccacatatatccgtaaagtaatttcctgtccaatttgatgtacctatatacgccgttttatcggttaccatatattttgcatggtttaccctagaatacggaatgggaggatcagcatctggtacaataaatagctttacttctatatctatgtttttagattttagcatagctatagatcttaaaaagtttctcatgataaacgaagatcgttgccagcaactaatcaatagcttaacggatacttgtctgtctatagcggatcttcttaattcatcttctatataaggccaaaacaaaattttacccgcctttgaataaataataggaataaagttcataacagatacataaacgaatttactcgcatttccgatacatgacaataaagcggttaaatcattggttctttccatagtacataattgttgtggtgcagaagcaataaatacagagtgtggaacaccgcttacgttaatactaagaggatgatctgtattataatacgacggataaaagtttttccaattatatggtagattgttaactccaagataccagtatacctcaaaaatttgagtgagatccgctgccaagttcctattattgaagatcgcaatacccaattccttgacctgagttagtgatctccaatccatgttagcgcttcctaaataaatatgtgtattatcagatatccaaaattttgtatgaagaactcctcctaggatatttgtaatatctatgtatcgtacttcaactccggccatttgtagtctttcaacatcctttaatggtttgttggatttattgacggctactctaactcttactcctcttttgggtaattgtacaatctcgtttaatattaccgtgccgaaattcgtacccacttcatccgataaactccaataaaaagatgatatatctagtgtttttatggtattggatagaatttccctccacatgttaaatgtagtcaaatatactttatcaaattgcatacctataggaatagtctctgtaatcactgcgattgtattatccggattcattttatttgttaaaaaaataatcctatatcacttcactctattaaaaatccaagtttctatttctttcatgactgattttttaacttcatccgtttccttatgaagatgatgtttggcgccttcataaatttttatttccctattacaatttgcatgttgcatgaaataatatgcacctgaaacatcgctaatctcattgtttgttccctggagtatgagagtcggggtgttaatcttgggaattatttttctaaccttgttggtagccttcaagacctgactagcaaatccagccttaattttttcatgattgactaatggatcgtattggtatttataaacttcatccatatctctagatactgattctggacatagctttccgactgacgcatttggtgtaatggttcccataagttttgcagctagcagattcagtcttggaacagcgtctgcattaactagaggagacattagaatcattgctgtaaacaagtttggattatcgcaagcagctagtatagaaattgttgctcccatggaatgacccaataagaagactggaactcctggataagtagatttaatagtcaccacgtgctgtaccacatctctaacatacttaccaaagtcatcaatcatcattttttcaccattacttcttccatggccaatatgatcatgtgagaatactaaaattcctaacgatgatatgttttcagctagttcgtcataacgtccagaatgttcaccagctccatgacttatgaatactaatgccttaggatatgtaataggtttccaatatttacaatatatgtaatcattgtccagattgaacatacagtttgtactcatgattcactatataactatcaatattaacagttcgtttaatgatcatattatttttatgttttattgataattgtaaaaatatacaattaaatcaatatagaggaaggagacggtactgtattttgtgagatagtaatggagactaaatcagattatgaggatgctgttttttactttgtggatgatgatgaaatatgtagtcgcgactccatcattgatctaatagatgaatatatcacgtggagaaatcatgttatagtgtttaataaagatattaccagttgtggaagactgtacaaggaattgataaagttcgatgatgtcgctatacggtactatggtattgataaaattaatgagattgtcgaggctatgagtgaaggagaccactacatcaatcttacagaagtccatgatcaggaaagtctattcgctaccataggaatatgtgctaaaatcactgaacattggggatacaaaaagatttcagaatctaaattccaatcattgggaaacattacagatctgatgaccgacgataatataaacatcttgatactttttctagaaaaaaaaatgaattgatgatataagtgtcttcataacgcattattacgttagcattctattatccagtgttaaaaaaattatcctatcatgtatttgagagtcttatatgtagcaaacatgataactgcaatacccataatctttagatattcacgcgtgctatggatggcattatcccgcggtgcggaaatgtacgttatataatctacaaaataatcatcgcatatagtatgagatagtagagtaaacatttttatcgtttctactgggttcatacatcgtctacccaattcggtaatgaatgaaattgtcgccaatcttacacccaaacccttgttgttcattagtatagtattaacttcattatttatgtcataaactgtaaatgattctgtagatgccatatcacacatgatattcatgtcactattataatcattattaactttatcacaatacgtgttgataatatctacatatgatctagtttttgtgggtaattgtctatacaagtcgtctaaacgttgtttactcatatagtattgaacagccatcattacatggtcccgttccgttgatagataatcgagtatgttagtagacttgtcaaatctatataccatattttctggaagcggatatacatagtcgcgatcatcattatcactagcctcatcctctatatcatgtacatgtacataatctatgatattattatacataaacatcgacaacatactattgtctattatctaagtcctgttgatccaaacccttgatctcctctatctgtactatctagagattgtacttcttcaagttctggataatatatacgttgatagattagctgagctattctatctccagtatttacattaaacgtacattttccattattaataagaatgactcctatgcttcccctataatcttcgtctattacaccgcctcctatatcaatgccttttagggacagaccagacctaggagctattctaccatagcagaacttaggcatggacatactaatatctgtcttaattaactgtcgttctccaggagggatagtataatcgtaagcgctatacaaatcatatccggcagcacccggcgattgcctagtaggtgatttagctctgttagtttccttaacaaatctaactggtgagttaatattcatgttgaacataaaaaatatcattttatttcaaaattatttaccattccattccatcccatatattccatgaataagtgcgattattgtacacttctatagtatctatatacgatccacgataaaatcctcctatcaatagcagtttattatccactatgatcaattctggattatccctcggataaataggatcatctatcagagtccatgtattgctggattcacaataaaattccgcatttctaccaaccaagaataaccttctaccaaacactaacgcacatgatttataatgaggataataagtggatggtccaaactgccactgatcatgattgggtagcaaatattctgtagttgtatcagtttcagaatgtcctcccattacgtatataacattgtttatggatgccactgctggattacatctaggtttcagaagactcggcatattaacccaagcagcatccccgtggaaccaacgctcaacagatgtgggatttggtagacctcctactacgtataatttattgttagcgggtatcccgctagcatacagtctggggctattcatcggaggaattggaatccaattgtttgatatataatttaccgctatagcattgttatgtatttcattgttcatccatccaccgataagatatactacttctccaacatgagtacttgtacacatatggaatatatctataatttgatccatgttcataggatactctatgaatggatacttgtatgatttgcgtggttgtttatcacaatgaaatattatgttacagtctagtatccattttacattatgtatacctctgggagaaagataatttgacctgattacatttttgataagaagtagcagatttcctaatctatttcttcgcctcatataccacttaatgacaaaatcaactacataatcctcatctggaacatttagttcgtcgctttctagaataagtttcatagatagataatcaaaattgtctatgatgtcatcttccagttccaaaaagtgtttggtaataaagtctttagtatgacataagagattggatagtccgtattctatacccatcatgtaacactcgatacaatattcctttctaaaatctcgtaggataaagtttatacaagtgtagatgataaattctacagatgttaatatagaagcacgtaataaattgacgacgttatgactatctatatatacctttccagtatatgagtaaataactatagaagttagactgtgaatgtcaaggtctagacaaaccctcgtaactggatctttattttttgtgtatttttggcgtaaatgtgtgcaaaagtatggagataactttttcaatatcgtagaattgactattatattacctcctatagcttcaataattgttttgaatttcttagtcgtgtacaatgctaatatattcttacagtacacagtattgacaaatatcggcatttatgtttctttaaaagtcaacatctaaagaaaaatgattgtcttcttgagacataactcccattttttggtattcacccacacgtttttcgaaaaaattagtttttccttccaatgatatattttccatgaaatcaaacggattggtaacattgtaaatttttttaaatcccaattcagaaatcaatctatctgcgacgaattctatatatgttttcatcatttcacaattcattcctatgagtttaactggaagagccacagtaagaaattcttgttcaatggataccgcatttgttataataaatctaacggtttcttcactcggtggatgcaataaatgtttaaacatcaaacatgcgaaatcgcagtgcagaccctcgtctctactaattaattcgttagaaaacgtgagtccgggcattaggccacgctttttaagccaaaatatggaagcgaatgatccggaaaagaagattccttctactgcagcaaaggcaataagtctctctccataaccggcgctgtcatgtatccacttttgagcccaatcggccttcttttttacacaaggcatcgtttctatggcattaaagaggtagtttttttcattactatctttaacataagtatcgatcaaaagactatacatttccgaatgaatgttttcaatggccatctgaaatccgtagaaacatctagcctcggtaatctgcacttctgtacaaaatcgttctgctaaattttcattcactattccgtcactggctgcaaaaaacgccaatacatgttttataaaatatttttcgtctggtgttagtttattccagtcattgatatctttagatatatccacttcttccactgtccaaaatgatgcctctgcctttttatacatattccagatgtcatgatattggattgggaaaataacaaatctatttggatttggtgcaaggataggttccataactaaattaacaatagtagtaattttttttcagttatctgtatgacgactgtacttggatcttttgtatatcgctatcgccgcaatcactacaataattacaagtattattgatagcattgttattactactatcataattaaattatcgacattcatgggtgttgaataatcgttatcatcattttgtaattgtgacgtcatactagataaatcatttgtgagattgttgtgggaagcgggcacggaagatgcattatcattattatttaacgcctcccattcggattcacaaatgttacgcacgttcaaagttttatggaaactataattttgtgaaaacagataacaagaaaactcgtcatcgttcaaatttttaacgatagtaaaccgattaaacgtcgagctaatttctaacgctagcgactctgttggatatgggtttccagatatatatcttttcagttcccctacgtatctataatcatctgtaggaaatggaagatatttccatttatctactgttcctaatatcatatgcggtggtgtagaaccattaagcgcgaaagatgttatttcgcatcgtattttaacttcgcaataatttctggttagataacgcactctaccagtcaagtcaatgatattagcctttacagatatattcatagtagtcgtaacgatgactccatcttttagatgtgatactcctttgtatgtaccagaatcttcgtacctcaaactcgatatatttaaacaagttaatgatatattaacgcgttttatgaatgatgatatataaccagaagttttatcctctgtggctagcgctataaccttatcattataataccaactagtgtaattaatatgtgacatgacagtgtgggtacaaatatgtacattatcgtctacgtcgtatttgatacatccgcattcagccaacaaatataaaattacaaaaactctaacgacgtttgtacacatcttgatgcggtttaataaatgttttgatttcaatttattgtaaaaaaagattcggttttatactgttcgatattctcattgcttatattctcatctatcatctccacacagtcaaatccatggttaacatgtacctcatcaaccggtaaaagactatcggattcttctatcatcataactcgagaatatttaatttggtggtcattattaatcaagtcaattatcttatttttaacaaacgtaagtattttactcatttttttataaaaacttttagaaatatacagactctatcgtgtgtctatatcttctttttatatccaatgtatttatgtctgatttttcttcatttatcatatataatggtccaaattctacacgtgcttcggattcatccagatcattaaggttcttataatcgcaacatccttctcttccatcttctacatcttccttcttattcttagcgtcacagaatctaccacagcaggatcccatgacgagagtcacattaaactaattcattttcaattataatatactgattagtaatgacaattaaaataaaaatattcttcataaccggtaagaaagtaaaaagttcacattgaaactatgtcagtagttatacatcatgagatgatatactctattttggtggaggattatatgatataattcgtggataatcattcttaagacacatttcttcattcgtaaatcttttcacattaaatgagtgtccatattttgcaatttcttcatatgatggcggtgtacgtggacgaggctgctcctgttcttgtagtcgtcgactgtcgtgtttgcgtttagatccctccattatcgcgatcgcgtagtgagtactatttataccttgtaattaaatttttttattaattaaacgtataaaaacgttccgtatctgtatttaagagccagatttcgtctaatagaacaaatagctacagtaaaaataactagaataatcgctacacccactagaaaccacggatcgtaatacggcaatcggttttcgataataggtggaacgtatattttatttaaggacttaacaattgtctgtaaaccacaatttgcttccgccgatcctgtattaactatctgtaaaagcatatgttgaccgggaggagccgaacattctccgatattcaatttttgtatatctataatgttattaacctccgcatacgcattacagttcttttctagcttggatactacactaggtacatcatctaaatctattcctatttcctcagcgatagctcttctatccttttccgaaagtaatgaaatcacttcaataaatgattcaaccatgagtgtgaaactaagtcgagaattactcatgcatttgttagttattcggagcgcgcaatttttaaactgtcctataacctctcctatatgaatagcacaagtgacattagtagggatagaatgttgagctaatttttgtaaataactatctataaaaagattatacaaagttttaaactctttagtttctgccatttatccagtctgagaaaatgtctctcataataaatttttccaagaaactaattgggtgaagaatggaaacctttaatctatatttatcacagtctgttttggtacacatgatgaattcttctaatgctgtactaaattcgatatctttttcgatttctggatatgtttttaataaagtatgaacaaagaaatggaaatcgtaataccagttatgttcaactttgaaattgttttttattttcttgttaatgattccagccacttgggaaaagtcaaagtcgtttaatgccgatttaatacgttcattaaaaacaaactttttattctttagatgaattattattggttcattggaatcaaaaagtaagatattatcgggtttaagatctgcatgtaaaaagttgtcacaacagggtagttcgtagattttaatgtataacagagacatctgtaaaaagataaactttatgtattgtaccaaagatttaaatcctaatttgatagctaactcggtatctactttatctgccgaatacagtgctaggggaaaaattataatatttcctctttcgtattcgtaattagttctcttttcatgttcgaaaaagtgaaacatgcggttaaaatagtttataacattaatattactgttaataactgccggataaaagtgggatagtaatttcacgaatttgatactgtcctttctctcgttaaacgcctttagaaaaactttagaagaatatctcaatgagagttcctgaccatccatagtttgtatcaataatagcaacatatgaagaacccgcttatacagagtatgtaaaaatgttaatttatagtttaatcccatggcccacgcacacacgattaattttttttcatctccctttagattgttgtatagaaatttgggtactgtaaactccgccgtagtttccatgggactatataattttgtggcctcgaatacaaattttactacatagttatctatcttaaagactataccatatcctcctgtagatatgtgataaaaatcgtcgtttataggataaaatcgtttatctttttgttggaaaaaggatgaattaatgtaatcattctcttctatctttagtagtgtttccttattaaaattcttaaaataatttaacaatctaactgacggagcccaattttggtgtaaatctaattgggacattatgttgttaaaatataaacagtctcctaatataacagtatctgataatctatggggagacatccattgatattcaggggatgaatcattggcaacacccatttattgtacaaaaagccccaatttacaaacgaaagtccaggtttgatagagataaactattaactattttgtctctgtttttaacacctccacagtttttaatttctttggtaatgaaattattcacaatatcagtatcttctttatctaccagagattttactaacttgataaccttggctgtctcattcaatagggtagtgatatttgtatgtatgatattgatatctttttgaattgtttcttttagaagtgattctttgatggtatcagcatacgaattacaataatgcagaaactcagttaacatgcaggaattatagtaagccaattccaattgttgcctgtattgtattagagtattaatatgcgcaatgatgtccttgcgtttctctgatagaatgcgagcagcgattttggcgttatcatttgacgatatttctggaatgacgaatcctgtttctactaacttcttggtaggacaaagtgaaacaatcaagaaaatagcttctcctcctatttgtggaagaaattgaactcctctagatgatctactgacgatagtatctccttgacagatattggaccgaactacggaagtacctggaatgtaaagccctgaaaccccctcattttttaagcagattgttgccgtaaatcctgcactatgcccaagatagagagctcctttggtgaatccatcactatgtttcagtttaaccaagaaacagtcagctggtctaaaatttccatctctatctaatacagaatccaacttgatgtcagggactatgaccggtttaatgttatatgtaacattgagtaaatcnttaagttcataatcatcgttgtcatcagttatgtacgatccaaacaatgtttctaccggcatggtggatacgaagatgctatccatcagaatgtttccctgattagtattttctatatagctattcttctttaaacgattttccgaatcagtaactatgttcatttttttaggagtaggacgtctagccagtatggaagaggattttctagatactctcttcaacatctttgatctcaatggaatgcaaaaccccatggtgtaacaaccaacgataaaaataatattgttttttcactttttataattttaccatctgactcatggattcattaatatctttacaagagctactaacgtataattctttataactaaactgagatatatacaccggatctatggtttccataattgagtaaatgaatgctcggcaataactaatggcaaatgtatagaacaacgaaattatactagagttgttaaagttaatattttctatgagttgttccaataaattatttgttgtgactgcgttcaagtcataaattatcttgatactatccagtaaacagtctttaagttctggaatattatcatcccattgtaaagcccctagttcgactatcgaatatcctgctctgatagcagtttcaatatcgacggacgtcaatactgtaataaaggtggtagtattgtcatcatcgtgataaactacgggaatatggtcgttagtaggtaccgtgactttacacaacgcgatatataactttccttttgtaccatttttaacgtagttgggacgtcctgcagggtattgttttgaagaaatgatatcgagaacagatttgatacgatatttgttggattcctgattattcactataatataatctagacagatagatgattcgataaatagagaaggtatatcgttggtaggataatacatccccattccagtattctcggatactctattgatgacactagttaagaacatgtcttctattctagaaaacgaaaacatcctacatggactcattaaaacttctaacgctcctgattgtgtttcgaatgcctcgtacaaggatttcaaggatgccatagattctttgaccaacgatttagtattgcgtttagcatctgatttttttattaaatcaaatggtcggctctctggtttactaccccaatgataacaatagtcttgtaaagataaaccgcaagaaaatttatacacatccatccaaataaccctagcaccgtcggatgatattaatgtattattatagattttccatccacagttattgggccagtatactgttagcaacggtatatcgaatagattactcatgtaacctactagaatgatagttcgtgtactagtcataatatctttaatccaatctaagaaatctaaaattagatcttttacactattaaagttaacaaaggtattacccgggtacgtggatatcatatatggtattggtccattatcagtaatggctccataaactgatacggcaatggtttttatatgtgtttgatctaatgaggacgaaattcgcgcccacaattcatctctagatatgcatttaatatcgaacggtaacacatcaatctcgggacgcgtatatgtttctaaattcttaatccaaatataatgatgacctatatgccctattatcatactgtcaactatagtatacctagagaactttcgatacatctgctgtttcctgtaatcgttaaattttacaaatctataacatgctaaaccttttgacgacagccattcattaatttctgatatggaatctgtattcttaataccgtatcgttctaaagccagtgctatatctccctgttcgtgggaacgctttcgtataatatcgatcaatggataatatgaagtttttggagaataatatgattcatgatctatttcgtccataaacaatctagacataggaattggaggcgatgatcttaattttgtgcaatgggtcaatcctataacttctaatattgtaatattcatcatcgacataacactatctatgttatcatcgtatattagtataccacgaccttcttcatttcgtgccaaaatgatatacagtcttaaataattacgcaatatctcaatagtttcataattgttagctgttttcatcaaggtttgtatcctgtttaacatgatggcgttctataacgtctctattttctatttttaatttttttaaatttttaacgatttactgtggctagatacccaatctctttcaaatatttttttagccttgcttacaagctgtttatctatactattaaaactgacgaatccgtgattttggtaatgggttccgtcgaaatttgccgaagtgatatgaacatattcgtcgtcgactattaacaattttgtattattctgaatagtgaaaaccttcacagatagatcattttgaacacacaacgcatctagacttctggcggttgccatagaatatacgtcgttcttatcccaattaccaactagaagtctgatcttaactcctctattaatggctgcttctataatggagttgtaaatgtcaggccaatagtagctattaccgtcgacacgtgtagtgggaactatggccaaatgttcaatatctatactagtcttagccgacttgagtttatcaataactacatcagtgtctagatctctagaatatcccaataggtgttctggagaatcagtaaagaacactccacctataggattcttaatatgatacgcagtgctaactggcagacaacaagccgcagagcataaattcaaccatgaattttttgcgctattaaaggctttaaaagtatcaaatcttctacgaagatctgtggccagcggaggataatcagaatatacgcctaacgttttaatcgtatgtatagatcctccagtaaatgacgcgtttcctacataacatctttcatcatcagacacccaaaaacaaccgagtagtagtcccacattattttttttatctatattaacggttataaaatttatatccggggagtgactttgtagctctcccagatttcttttccctcgttcatctagcaaaactattattttaatccctttttcagatacctcttttagtttatcaaaaataagcgctcccctagtagtactcagaggattacaacaaaaagatgctatgtatatatatttcttagctagagtgataatttcgttaaaacattcaaatgttgtcaaatgatcggatctaaaatccatattttctggtagtgtttctaccagcctacattttgctcccgcaggtaccgatgcaaatggccacatttagttaacataaaaacttatatatcctgttctatcaacgattctagaatatcatcggctatatcgctaaaattttcatcaaagtcgacatcacaacctaactcagtcaatatattaagaagttccatgatgtcatcttcgtctatttctatatccgtatccattgtagattgttgaccgattatcgagtttaaatcattactaatactcaatccttcagaatacaatctgtgtttcattgtaaatttataggcggtgtatttaagttggtagattttcaattatgtatcaatatagcaacagtagttcttgctcctccttgattttagcatcctcttcattattttcttctacgtacataatcatgtctaatacgttagacaacacaccgacgatggtggccgccacagacacgaatatgactagaccgatgaccatttaaaaaatactctctagctttaacttaaactgtatcgatcattcttttagcacatgtataatataaaaacattattctatttcgaatttaggcttccaaaaatttttcatccgtaaaccgataataatatatatagacttgttaatagtcggaataaatatattaatgcttaaactatcatcatctccacgattagagatacaatatttacattctttttgctgtttcgaaactttatcaatacacgttaatacaaacccaggaaggagatattgaaactgaggctgttgaaaatgaaacggcgaatacaataattcagataatgtaaaatcatgattccgtattctgatgatattagaactgctaatggatgtcgatggtatgtatctaggagtatctattttaacaaagcatcgatttgctaatatacaattatccttttgattaattgttattttattcatattcttaaaaggtttcatatttatcaattcttctacattaaaaatttccatttttaatttatctagccccgcaatactcctcattacgtttcattttttgtctagaatacccattttgttcatcttggtacatagattatccaattgagaagcgcatttagtagttttgtacattttaagtttattaacgaatcgtcgaaaactagttatagttaacattttattatttgataccctgatattaatacccctgccgttactattatttataactgatgtaacccacgtaacattggaattaattatcgatagtaatgcatcgacacttccaaaattgtctattataaactcaccgataatttttttattgcatgttttcatattcattaggattatcaaatctttaatcttattacgattgtatgcgttgatattacaagacgtcattctaaaagacggaggatttccatcaaatgccagacaatcacgtacaaagtacatggaaataggttttgttctattacgcatcatagattcatataaaacacccgtagaaatactaatttgttttactctataaaatactattgcatctatttcatcgttttgtataacgtctttccaagtgtcaaattccaattttttttcattgatagtaccaaattcttctatctctttaactacttgcatagataggtaattacagtgatgcctacatgccgttttttgaaactgaatagatgcatctagaagcgatgctacactagtcacgatcaccactttcatatttagaatatatgtatgtaaaaatatagtagaatttcattttgtttttttctatgctataaatgaattctcattttgtatccgcacatactccgttttatatcaataccaaagaaggaagatatctggttctaaaagccgttaaagtatgcgatgttagaactgtagaatgcgaaggaagtaaagcttcctgcgtactcaaagtagataaaccctcatcacccacgtgtgagagaagaccttcgtccccgtccagatgcgagagaatgaataaccctggaaaacaagtcccgtttatgagaacggacatgttacaaaatatgtttgctgctaatcgcgacaacgtaacgtcaagacttttgaactaaaatacaattatatctttttcgatattaataaatccgtgtctcccgggttttttatctctttcagtatgtgaatagataggtattttatctctattcatcatcgaatttaagagatccgataaacattgtttgtattctccagatgtcagcatctgatacaacaatatatgtgcacataaacctctggcacttatttcatgtaccttccccttatcactaaggagaatagtatttgagaaatatgtatacatgatattatcatgtattagatatacagaatttgtaacactctcgaaatcacacgatgtgtcggcgttaagatctaatatatcactcgataacacattttcatctagatacactagacattttttaaagctaaaatagtctttagtagtaacagtaactatgcgattattttcatcgatgatacatttcatcggcatattattacgcgtaccatcaaagactataccatgtgtatatctaacgtattctagcatagttgccatacgtacattaaacttttcaggatctttggatagatcttccaatctatctatttgagaaaacatttttatcatgttcaatagttgaaacgtcggatccactatatagatattatctataaagattttaggaactatgttcatggtatcctggcgaatattaaaactatcaatgatatgattatcgttttcatcttttatcaccatatagtttctaagatatgggattttacttaatataatattatttcccgtaataaattttattagaaatgccaaatctataagaaaagtcctagaattagtctgaagaatatctatatcaccgtaccgtatatttggattaattagatatagagaatatgatccgtaacatatacaacttttattatgacgtctaagatattcttccatcaacttattaacatttttgactagggaagatacattatgacgtcccattacttttgccttgtctattacagcgacgttcatagaatttagcatatctcttgccaattcttccattgatgttacattataagaaattttagatgaaattacatttggagctttaatagtaagaactcctaatatatccgtgtatgtggtcactaatacagattgtagttctataatcgtaaataatttacctatattatatgtttgagtttgtttagaaaagtagctaagtatacgatcttttatttctgatgccgatgtatcaacatcgaaaaaaaatctttttttattcttttttactaacgatacgaatatgtctttgttaaaaacagttattttctgaatatttctagcttgtaattttaacatatgatattcgttcacactaggtactctgcctaaataggtttctataatctttaatgtaatattaggaagagtattctgatcaggattcctattcattttgaggatttaaaacnctgattattrtctaatatggtctcaacacaaactttttcacagagtgatagagtttttgataactcgtttttcttaagaaatataaaactactgtctccagagctcgctctatcttttattttatctaattcgatacaaactcctgatactggttcagaaagtaattcattaattttcagtcctttatagaagatatttaatatagataatacaaaatcttcagttcttgatatcgatctgattgatcctagaactagatatattaataacgtgctcattaggcagtttatggcagcttgataattagatatagtatattccagttcatatttattagataccgcattgcccagattttgatattctatgaattcctctgaaaataaatccaaaataactagacattctattttttgtggattagtgtactctcttccctctatcatgttcactactggtgtccacaatgataaatatctagagggaatataatatagtccatatgatgccaatctagcaatgtcgaataactgtaattttattcttcgctcttcattatgaattgaatcttgaggtataaacctaacacaaattatatcattagacttttcgtatgtaatgtctttcatgttataagtttttaatcctggaatagaatctattttaatgaggcttttaaacgcagcgttctccaacgagtcaaagcataatactctgttggttttcttatattcaatattacgattttcttctttgaatggaataggtttttgaattagtttataattacaacataatagataaggaagtgtgtaaatagtacgcggaaaaaacataatagctcccctgttttcatccatggttttaagtaaatgatcactggcttctttagtcaatggatattcgaacattaaccgtttcatcatcattggacagaatccatatttcttaatgtaaagagtgatcaaatcattgtgtttattgtaccatcttgttgtaaatgtgtattcggttatcggatctgctcctttttctattaaagtatcgatatcgatctcgtctaagaattcaactatatcgacatatttcatttgtatacacataaccattactaacgtagaatgtataggaagagatgtaacgggaacagggtttgttgattcgcaaactattctaatacataattcttctgttaatacgtcttgcacgtaatctattatagatgccaagatatctatataattattttgtaagatgatgttaacaatgtgatctatataagtagtgtaataattcatgtatttcgatatatgttccaactctgtctttgtgatgtctagtttcgtaatatctatagcgtcctcaaaaaatatattcgcatatattcccaagtcttcagttctatcttctaaaaaatcttcaacgtatggaatataataatctattttacctcttctgatgtcattaatgatatagtttttgacactattttccgtcaattgattcttattcactatgtctaaaaaccggatagcgtccctaggacgaactactgccattaatatctctattatagcttctggacataaatcatctattataccagaattaatgggaactattccgtatctatctaacatagttttaagaaagtcagaatctaagacctgatgttcatatattggttcatacatgaaatgatctctattgatgatagtgactatttcattctctgaaaattggtaactcattctatacacgccttcnttgttgataaaggatnntatatactcaatggaatttgtaccaacaaactgttctcntatgaatcgtatatcatcatctgaaatgatcatgtaangcatacatttaacaataagagacttgtntnctgttatcaatatactattcttgtgataatttatgtgtgcggcaaatttgtccacgttctttaattttgttatagtagatatcaaatccaatggagatacagttcttggcttaaacagatatagtttttctggaacgaattctacaacattattataaaggactttgggtatataagtgggatgaaatcctattttaattaatgcgatagccttgtcctcgtgcagatatccaaacgcttttgtgatagtatggcattcattgtctagaaacgctctacgaatatctgtaacagatatcatctttagagaatactagtcgcgttaatagtactaaaatttgtattttttaatctatctcaataaaaaattaatatgtatgattcaatgtataactaaactactaactgttattgataactagaatcagaatctaatgatgacataactaagaagtttatctacagccaatttagctgcattatttttagcatctcgtttagattttccatctgccttatcgaatactcttccgtcaatgtctacacaggcataaaatgtaggagagttactaggccccactgattcaatacgaaaagaccaatctctcctagttatttgacagtactcattaataacggtgacagggttaacacctttccaataaataatttttttaaccggaataacatcatcaaaagacttatgatcctctctcattgatttttcgcgggatacatcatctattatagcatcagcatcagaatctgtaggccgtgtatcagcatccattgtcgtagaccaacgaggaggagtatcgttggagctgtaaaccatagcactacgttgaagatcatacagagctttattaacttctcgcttctccatattaagttgtttagttagttgtacagcagtagctccttagtccaatgtttttaataaccgcacacaatctctgtgtcagaacgctcgtcaatatagatcttagaaatttttttagagagaactaacgcaactagcaataaaactgatcttattttatcattttttttattcatcatcctctggtggttcgtcgttcctatcgaatgtagctctgattaacccgtcatctataggtgatgctggttctggagattctggaggagatggattattatctggaagaatctctgttatttccttgttttcatgtatcgattgcgttgtaacattaagattgcgaaatgctctaaatttgggaggcttaaagtgttgtttacaatctctacacgcgtgtctaactaatggaggttcatcagcggctctagtttgaatcatcatcggtgtagtattcctacttttacagttaggacacggtgtattgtatttctcgtcgagaacgttaaaataatcgttgtaactcacatcctttattttatctatattgtattctactcctttcttaatgcattttataccgaacaagagatagcgaaggaattctttttcggtaccgctagtacccttaatcatatcacatagtgttttatattctaaatgtgtggcaatggacggtttatttctatacgatagtttgtttttggaatcctttgagtattctataccaatattattctttgattcgaatttagtttcttcgatattagattttgtattacctatattcttgatgtagtactttgatgatttttccatggcccattctattaagttttccaagttggcatcatccacatattgtgatagtaattctcggatatcagtagtgactaccgccattgatatttgttcatttgatgagtaactactaatgtatacattttccatttataacacttatgtattaactttgtttatttatattttttcattattatgttgatattaataatcgtattgtggttatatggctacaatttcataatgagttgaagtcagtgtcctatgatcaatgacgatagctttactctgaaaagaaagtatcaaatcgatagtgcagagtcaacaatgaaaatggataagacgatgacaaagtttcagaatagagtcaaaatggtaaaagaaataaatcagacgataagagcagcacaaactcattacgagacattgaaactaggatatataaaatttaagggaatgattaggactactactctagaagatatagcaccatctattccaaataatcagaaaacttataaactattctcggacatttcagtcattggcaaagcatcacagaatccgagtaagatgatatatgctcgctgctttacatgtttcccaatttgtttggagatgaccatagattcatttgttatagaatgcatccaacattgttcatgatatagttgaatcatgtatgcctgttcgtatgcctgtggctaagatactgtgtaaagaaatggtaaataaatactttgagaatctttaagagtgcattgactttgttagtgaataggcattccatctttctccaatactaattcaaattgttaaattaataatggaatagtataaatagttattagtgataggatagtaaacataattattagaatagtagtgtagtatcatagataactctcttctataaaaaatggattttattcgtagaaagtatcttatatacacagtagaaaataatatagattttttaaaggatgatacattaagtaaagtaaacaattttaccctcaatcatgtactagctctcaagtatctagttagcaattttcctcaacatgttattactaaggatgtattagctaataccaatttttttgttttcatacatatggtacgatgctgtaaagtatacgaagcggttttacgacacgcatttgatgcacccacgttgtacgttaaagcattgactaagaattatttatcgtttagtaacacaatacagtcgtacaaggaaacagtgcataaactaacacaagatgaaaaatttttagaggttgccgaatacatggacgaattaggagaacttataggcgtaaattatgacttagttcttaatccattatttcacggaggggaacccatcaaagatatggaaatcatttttttaaaactgtttaagaaaacagacttcaaagttgttaaaaaattaagtgttataagattacttatttgggcatacctaagcaagaaagatacaggcatagagtttgcggataatgatagacaagatatatatactctatttcaacaaactggtagaatagtccatagcaatctaacagaaacgtttagggattatatctttcccggagataagactagctattgggtgtggttaaacgaaagtatagctaatgatgcggatatcgttattaatagacccgccattaccatgtatgataaaattcttagttatatatactctgagataaaacagggacgcgttaataaaaacatgcttaagttagtttatatctttgagcctgaaaaagatatcagagaacttctgctagaaatcatatatgatattcctggagatatcctatctattattgatgcaaaaaacgacgattggaaaaaatattttattagtttttacaaagctaattttattaacggtaatacatttattagtgatagaacgtttaacgatgacttattcagagttgttgttcaaatagatcccgaatatttcgataatgaacgaattatgtctttattctatacgagtgctgcggacattaaacgatttgatgagttagatattaataacagttatatatctaatataatttatgaggtgaacgatatcacattagatacaatggatgatatgaagaagtgtcaaatctttaacgaggatacgttgtattatgttaaggaatacaatacatacctgtttttgcacgagtcggatcccatggtcatagagaacggaatactaaagaaactgtcatctataaaatccaagagtagacggctgaacttgtttagcaaaaacattttaaaatattatttagacggacaattggctcgtctaggtcttgtgttagatgattataaaggagacttattagttaaaatgataaaccatctcaaatctgtggaggatgtatccgcattcgttagattttctacagataaaaaccctagtattcttccatcgctaatcaaaactattttagctagttataatatttccatcatcgtcttatttcaaaggtttttaagagataatctatatcatgtagaagaattcttggataaaagcatccatctaaccaagacggataagaaatatatacttcaattgataaggcacggtagatcatagaacaaaccaaatatattattaataatttgtatatacatagatataattatcacatattaaaaaataacacatttttgataaatggaaaccgttgcaacaattcagactcccaccaaattaatgaataaagaaaatgcagaaatgattttggaaaaaattgttaatcatatagctatgtatattagtgacgaatcaatatattcagaaaataatcctgaatatattgattttcgtaacagatacggagactatagatctctcattataaaaagtgatcacgagtttgtaaagctatgtaaagatcatgcagagaaaagttctccagaaacgcaacaaatgattatcaaacacatatacgaacaatatcttattccagtatctgaagtactattaaaacctataatgtccatgggtgacatatttacatataacggatgtaaagacaatgaatggatgctagaacaactctctaccctaaactttaacaatctctacacatggaactcatgtagcataggcaatgtaacgcgtctgttttatacattttttagttatctgatgaaagataaactaaatatataagtataatcccattctaatactttaacctgatgtattattacctgcatcttattagaatattaacctaactaaaagacataaaaagcggtaggatataaatattatggccgcaaccgttccgcgttttgacgatgtgtacaaaaatgcacaaagaagaattctagatcaagaaacattttttagtagaggtctaagtagaccgttaatgaaaaacacatatctatttgataattacgcgtatggatggataccagaaactgcaatttggagtagtagatacgcaaacctagatgctagtgactattatcccatttcgttgggattacttaaaaagtttgaatttctcatgtctctatataaaggtcctattcccgtatatgaagaaaaagtaaatactgaattcattgctaatggatctttctccggtagatacgtatcatatcttagaaagttttctgcccttccaacaaacgagtttattagttttttattattgacctccatccctatctataatatcttattctggtttaaaaacacacagtttgatattactaaacacacattattcagatacgtctatacagataataccaaacaccttgcgttggctaggtatatacatcaaacaggagactataagcctttgtttagtcgtctcaaagagaattatatatttactggtcccgttccaataggtatcaaagatatagatcaccctaatcttagtagagcaagaagtccatccgattatgagacattagctaatattagtactatattgtactttaccaagtatgatccagtattaatgtttttattgttttacgtacctgggtattcaattactacaaaaattactccagccgtagaatatctaatggataaactgaatctaacaaagagcgacgtacaactgttgtaaattattttatgcttcgtaaaatgtaggtcttgaaccaaacattctttgaaaaaatgagatgcataaaactttattatccaatagattaactatttcagacgtcaatcgtttaaagtaaacttcgtaaaatattctttgattgctgccgagtttaaaacttctatcgataattgtttcatatgttttaatatttacaagttttttggtccatggtacattagctggacagatatatgcaaaataatatcgttctccaagttctatagtctctggattgtttttattatattcagtaaccaaatacatattagggttatctgcggatttataatttgagtgatgcattcgactcaacataaataattctagaggagacgatctactatcaaattcggatcgtaaatctgtttctaaagaacggagaatatctatacatacctgattagaattcatccgtccttcagacaacatctcagacagtctggtcttgtatgtcttaatcatattcttatgaaacttggaaacatctcttctagtttcactagtacctttattaattctctcaggtacagattttgaattcgacgatgccgagtatttcatcgttgtatatttcttcttcgattgcataatcaaattcttatataccgcctcaaactctattttaaaattattaaacaatactctactattaatcagtcgttctaactcctttgctatttctatggacttatctacatcttgactgtctatctctgtaaacacggagtcggtatctccatacacgctacgaaaacgaaatctataatctataggcaacgatgttttcacaatcggattaatatctctatcgtccatataaaatggattacttaatgtattggcaaaccgtaacataccgttggataactctgctccatttagtaccgattctagatacaatatcattctacgtcctatggatgtgcaactcttagccgaagcgtatgagtatagagcactatttctaaatcccatcagaccatatactgagttggctactatcttgtacgtatattgcatggaatcatagatggccttttcagttgaactggtagcctgttttaacatctttttatatctggctctctctgccaaaaatgttcttaatagtctaggaatggttccttctattgatctatcgaaaattgctatttcagagatgaggttcggtagtctaggttcacaatgaaccgtaatatatctaggaggtggatatttctgaagcaagagttgattatttatttcttcttccaatctattggtactaacaacgacaccgactaatgtttccggagatagatttccaaagatacacacattaggatacagactgttataatcaaagattaatacattattactaaacattttttgttttggagcaaataccttaccgccttcataaggaaacttttgttttgtttctgatctaactaagatagttttagtttccaacaatagctttaacagtggacccttgatgattgtactcgctctatattcgaataccatggattgaggaagcacatatgttgccgcaccagcgtctgtttttgtttctactccataatactcccacaaatactgacacaaacaagcatcatgaatacagtatctagccatatctaaagctatgtttagattataatccttatacatctgagctaaatcaatgtcatcctttccgaaagataatttatatatatcattaggtaaagtaggacatgatagtacgactttaaatccattttccaaaatatctttacgaattactttacatataatatcctcatcaacagtcacgtaattacctgtggttaaaacctttgcaaatgtatcggctttgcctttcgcgtccgtagtatcgtcaccgatgaacgtcatttctctaactcctctatttaatactttacccatgcaactgaacgcgttcttggatatagaatccaatttgtacgaatccaatttttcagatttttgaatgaatgaatatagatcgaaaaatatagttccattattgttattaacgtgaaacgtagtattggccatgccgcatactcccttatgactagactgatttctctcataaatacagagatgtacagcttcctttttgtctggagatctaaagataatcttctctcctgttaataactctagacgattagtaatatatctcagatcaaagttatgtccgttaaaggtaacgacgtagtcgaacgttagttccaacaattgtttagctattcgtaacaaaactatttcagaacatagaactagttctcgttcgtaatccatttccattagcgactgtatcctcaaacatcctctatcgacggcttcttgtatttcctgttccgttaacatctcttcattaatgagcgtaaacagtaatcgtttaccacttaaatcgatataacagtaacttgtatgcgagattgggttaataaatacagaaggaaacttcttatcgaagtgacactctatatctagaaataagtacgatcttgggatatcgaatctaggtatttctttagcgaaacagttacgtggatcgtcacaatgataacatccattgttaatctttgtcaaatattgctcgtccaacgagtaacatccgtctggagatatcccgttagaaatataaaaccaactaatattgaaaaattcatccatggtggcattttgtatgctgcgtttctttggctcttctatcaaccacatatctgcgacggagcattttctatctttaatatctagattataacttattgtctcgtcaatgtctatagttctcatctttcccatcggcctcgcattaaatggaggaggagataatgactgatatatttcgtccgtcactacgtaataaaagtaatgaggaaatcgtataaatacggtctcgccatttcgacatctggatttcagatataaaaatctgttttcaccgtgactttcaaaccaattaatacacctaacatccatttctagaatttagaaatatattttcatttaaatgaatcccaaacattggggaagagccgtatggaccattatttttatagtactttcgcaagcgggtttagacggcaacatagaagcgtgtaaacgaaaactatatactatagtcagcactcttccatgtcctgcatgtagacgacacgcgactatcgctatagagaacaataatgtcatgtctagcgatgatctgaattatatttattattttttcatcagattatttaacaatttggcatttgatcccaaatacgcaatcgatgtgtcaaaggttaaacctttataaacttaacccattataaaacttatgattagtcacgactgaaataaccgcgtgattattttttggtataattctacacggcatggtttctgtgactatgaattcaacacctgttatcttagtgaaatctttaacaaacagcaagggttcgtcaaagacataaaactcattgtttacgatcgaaatagaccccctatcacacttaaaataaaaaatatccttatcctttaccaccaaataaaattctgattggtcaatgtgaatgtattcacttaacagttccacaaatttatttattaactccgaggcacatacatcgtcggtattttttatgacaaactttactcttccagcatccgtttctaaaaaaatattaacgagttccatttatatcatccaatattattgaaatgacgttgatggacagatgatataaataagaaggtacagtacctttgtccaccatctcctccaattcatactctattttgtcattaactttaatgtgtgaaaacagtacgccacatgcttccatgacagtgtgtaacactttggatacaaaatgtttgacattagtataattgtccaagactgtcaatctataatagatagtagctataatatattctatgatggtattgaagaagatgacaaccttggcatattgatcatttaacacagacatggtatcaacaaatagcttaaatgaaagagaatcagtaattggaataagcgtcttctcgatgtagtgtccgtataccaacatgtctgatattttgatgtattccattaaattatttagttttttctttttattctcgttaaacagaatttctgtcaatggaccccaacatcgttgacctattaagttttgattgatttttccgtgtaaggcgtatctagtcagatcgtatagcctatccaataatccatcgtctgtgcgtagatcacatcgtacactttttaattttctatagaagagtgacagacatctggagcaattacagacarcaatttctttattctctacagatgtaagatacttgaagatattnctatgatgatgcagaattttggataacacggtattgatggtatctgttaccayaattgctttgactgatagtgtcaaagtacaagatttncaatcttttgcaattttcngtaccattatctttgttttgatatctatatcagacagcatggtacgtctgacaacacnnggattaagacggaaagatgaaatgattntctcaacatcttcaatagataccttgctattttttttggcattatctatatgtgagagaatatcctctagagaatcagtatcctttttgatgatagtggatctcaatgacatgggacgtctaaaccttcttattctatcaccagattgcatggtgatttgtcttctttcttttatcatgatgtaatctctaaattcatcggcaaattgtctatatctaaaatcataatatgagatgtttacctctacaaatatctgttcgtccaatgttagagtatctatatcagttttgtattccaaattaaacatggcaacggatttaattttatattcctctattaagtcctcgtcgataataacagaatgtagataatcatttaatccatcgtacatggttggaagatgctcgttgacaaaatctttaattgtcttgatgaaggtgggactatatctaacatcttgattaataaaatttataacattgtccataggatactttgtaactagttttatacacatctcttcattggtaagtttagacagaatatcgtgaacaggtggtatattatattcatcagatatacgaagaataatgtccaaatctatattgtttaatatattatatagatgtagtgtagctcctacaggaatatctttaactaagtcaatgatttcatcaacagttagatctattttaaagttaatcatataggcattgatttttaaaaggtatgtagccttgactacattctcattaattaaccattccaagtcactgtgtgtaagaagattatattctatcataagcttgactacatttggtcccgataccattaaagaattcttatgatataaggaaacagcttttaggtactcatctactctacaagaattttggagagccttaacgatatcagtgacgtttattatttcaggaggaaagaacctaacattgagaatatctgaattaatagcttccagatacagtgattttggcaatagtccgtgtaatccataatccagtaacacgagctggtgcttgctagacaccttttcaatgtttaatttttttgaaataagctttgataaagccttcctcgcaaattccggatacatgaacatgtcgccaacatgattaagtattgtttttcattatttttatattttctcaacaagttctcaataccccaatagataatagaatatcacccaatgcgtccatgttgtctatttccaacaggtcgctatatccaccaatagaagttttcccaaaaaagattctaggaacagttctaccaccagtaatttgttcaaaatagtcacgcaattcattttcgggtttaaattctttaatatctacaatttcatacgctcctcttttgaaactaaacttatttagaatatccagtgcgtttctacaaaaaggacatgtaaacttgacaaaaattgtcactttgttattggccaacctttgttgtacaaattcctcggccatttttaatatttaagtgatacaaaactatctcgacttatttaactctttagtcgagatatatggacacagatagctatatgataaccaactacagaagacaaacgctataaaaaacataattacgacgagcatatttataaatatttttattcagtattacttgatatagtaatattaggcacagtcaaacattcaaccactctagatacattaactctctcattttctttaacaaattctgcaatatcttcgtaaaaagattcttgaaactttttagaatatctatcgactctagatgaaatagcgttcgtcaacatactatgttttgtatacataaaggcgcccattttaacagtttctagtgacaaaatgctagcgatcctaggatcctttagaatcacatagattgacgattcgtctctcttagtaactctagtaaaataatcatacaatctagtacgcgaaataatattatccttgacttgaggagatctaaacaatctagttttgagaacatcgataagttcatcgggaattacatacatactatctttaatagaactcttttcatccagttgaatggattcgtccttaaccaactgattaatgagatcttctattttatcattttctagatgatatgtatgtccattaaagttaaattgtgtagcgcttctttttagcctagcagccaatactttaacatcactaatatcgatatacaaaggagatgatttatcgatggtattaagaattcgtttttcgacatccgtcaaaaccaattcctttttgcctgtatcatccagtttgccattctttgtaaagaaattattttctactagactattaataagactgataaggattcctccataattgcacaatccaaactttttcacaaaactagactttacgagatctacaggaatgcgtacttcaggtttcttagcttgtgattttttcttttgtggacattttctagtgaccaactcatctaccatttcattgattttagcagtgaaataagctttcaatgcacgggcactgatactattgaaaacgagttgatcttcaaattccgccatttaagttcaccaaacaacttttaaatacaaatatatcaatagtagtagaataagaactataaaaaaaataataattaaccaataccaaccccaacaacctgtattattagttgatgtgacagttttctcatcacttagaacagatttaacaatttctataaagtctgtcaaatcatcttcctgagaacccataaatacaccaaatatagcagcgtacaacttatccatttatacattgaatattggcttttctttatcgctatcttcatcatattcatcatcaatatcaacaagtcccagattacgaaccagatcttcttctacattttcagtcattgatacgcgttcactatctccagagagtccgataacgttagccactacttctctatcaatgattagtttcttgagcgcgaatgtaatttttgtttccgttccggatctatagaaaactacaggtgtaataattgccttggctaattgtctttctcttttactgagtgattctagttcaccttctatagatctgagaatggatgattctccagtcgaaacatattctaccatggctccgtttaatttgttgatgaagatggattcatccttaaatgttttctctgtaatagtttccgccgaaagactatgcaaagaatttggaatgcgttccttgtgtgtaatgtttccatagacagcttctagaagttgatacaacataggactagccgcggtaacttttatttttagaaagtatccatcgcttctatcttgtttagatttatttttataaagtttagtctctccttccaacataataaaagtggaagtcatctgactagataaactatcagtaagttttatagagatagatgaacaattagcgtattgagaagcatttagtgtaacgcattcgatacattttgcattagatttactaatcgattttgcatactctataacacccgcacaagtctgtagagaatcgctagatgctgtaggtcttggtgaagtttcaactctcttcttgattaccttactcatgattaaacctaaataattgtactttgtaatataatgatatatattttcactttatctcatttgagaataaaaatgtttttgttaaccactgcatgatgtacagatttcggaatcgcaaaccacttgtggttttattttatccttgtccaatgtgaattgaatgggagcggatgcgggtttcgtacgtagatagtacattcccgtttttagaccgagactccatccgtaaaaatgcatactcgttagtttggaataactcggatctgctatatggatattcatagattgactttgatcgatgaaggctcccctgtctgcagccatttttatgatcgtcttttgtggaatttcccaaatagttttataaactcgcttaatatcttctggaaggtttgtattctgaatggatccaccatctaccataatcctattcttgatctcatcattccataattttctctcggttaaaactctaaggagatgcgggttaactacttggaattctccagacaatactctccgagtgtaaatattactggtatacggttccaccgactcattatttcccaaaatttgagcagttgatgcagtcggcataggtgccaccaataaactatttctaagaccgtatgttctgattttatcttttagaggttcccaattccaaagatccgacggtacaacattccaaagatcatattgtagaataccgttactggcgtacgatcctacatatgtatcatatggtccttccttctcagctagtttacaactcgcctctaatgcaccgtaataaatggtttcaaagatcttcttatttagatcttgtgcttccaggctatcaaatggataatttaagagaataaacgcgtccgctaatccttgaacaccaataccgataggtctatgtctcttattagagatttcagcttctggaataggataataattaatatctataattttattgagatttctgacaattactttgaccacatccttcagtttgagaaaatcaaatcgcccatctattacaaacatgttcaatgcaacagatgccagattacacacggctacctcattagcatccgcatattgtattatctcagtgcaaagattactacacttgatggttcctaaattttgttgattactctttttgttacacgcatccttataaagaatgaatggagtaccagtttcaatctgagattctataatcgctttccagacgactcgagcctttattatacatttgtatctcctttctctttcgtatagtgtatacaatcgttcgaactcgtctccccaaacattgtccaatccaggacattcatccggacacatcaacgaccactctccgtcatccttcactcgtttcataaagagatcaggaatccaaagagctataaatagatctcttgttctatgttcatcgtttcctgtattctttttaagatcgaggaacgccataatatcagaatgccacggttccaagtatatggccataactccaggccgtttgtttcctccctgatctatgtatctagcggtgttattataaactctcaacattggaataataccgtttgatataccattggtaccggagatatagcttccactggcacgaatattactaattgatagacctattccccctgccattttagagattaatgcgcatcgttttaacgtgtcatagatgccttctatgctatcatcgatcatgttaagtagaaaacagctagacatttggtgacgagtagttcccgcattaaataaggtaggagaagcgtgcgtaaaccatttttcagaaagtagattgtacgtctcaatagctgagtctatatcccattgatgaattcctactgcgacacgcattaacatgtgctgaggtctttcaacaattttgttgtttattttcaacaagtaggatttttccaaagttttaaaaccaaaatagttgtatgaaaagtctcgttcgtaaataataaccgaattgagcttatccttatatttgttaactatatccatggtaatacttgaaataatcggagaatgtttcccatttttaggattaacatagttgaataaatcctccatcacttcactaaatagtttttttgtttccttgtgtagatttgatatggctattctggcggctagaatggcataatccggatgttgtgtagtacaagtggctgctatttcggctgccagagtgtccaattctaccgttgttactccattatatattccttgaataaccttcatagctattttaataggatctatatgatcagtgtttaagccatagcacaattttctaatacgagacgtgattttatcaaacatgacattttccttgtatccatttcgtttaatgacaaacatttttgttggtgtaataaaaaaaattatttaatttttcattaatagggatttgacgtatgtagcgtacaaaattatcgttcctggtatatagataaagagtcctatatatttgaaaatcgttacggttcgattaaactttaatgattgcattgtgaatatatcattaggatttaactccttgactatcatggcggtgccagaaattaccatcaaaagcattaatacagttatgcagatcgcagttagaacggttatagcatccaccatttatatctaaaaattagatcaaagaatatgtgacaacgtcctagttgtatactgagaattgacgaaacaatgtttcttacatatttttttcttattagtaaccgacttaatagtaggaactggaaaactagacttgattattctataagtatagatacccttccaaataatgttctctttgataaaagttccagaaaatgtagaattttttaaaaagttatcttttgctattactaatatcgtggttagacgcttattattaatatgagtgatgaaatccacaccgcttctagatatcgcttttatttccacattagatggtaaatccaatagtgaaactatctttttaggaatgtatggactcgcgtttagaggagtgaacgtcttcggagtagtaaaggatgattcgtcaaatgaataaacaatttcacaaatggatgttaatgtattagtaggaaattttttgacgctagtggaattgaagattctaatggatgatgttctacctatttcatccgataacatgttaatttccaataccaacggttttaatatttcgatgatatacggtagtctctctttcggacttatatagcttattccacaatacgagtcattatatactccaaaaaacaaaataactagtataaaatctgtatcgaatgggaaaaacgaaattatcgatataggtatagaatccggaacattgaacgtattaatacttaattctttttcagtggtaagaaccgataggttattgacattgtatggttttaaatattctataacttgagacttgatagatattaatgacgaattgaaaattatttttatcaccacgtgtgtttcaggatcatcgtcgacgccagttaaccaaccgaatggagtaaaataaatatcattaatatatgctctagatattagtatttttattaatcctttgattatcatcttctcgtacgcgaatgattccatgatcaagagtgatttgagaacatcctccggagtattaatgggtttagtaaacagtccatcgttgcaataataaaagttgtccaagttaaaggatattatgcattcgtttaaagatatcacctcatctaacggagacaattttttggtaggttttagagactttgaagctacttgtttaacaaagttattcatcgtcgtctactattctatttaattttgtagttaatttatcacatatcacattaattgactttttggtccacttttccatacgtttatattcttttaatcctgcgttatccgtttccgttatatacagggatagatcttgcaagttaaatagaatgctcttaaataatgtcatttttttatccgctaaaaatttaaagaatgtataaacttttttcaaagatttaaaacttttaggtggagttctggtacacaatatcataaacaaactaataaacatcccacattcagattccaacaattgattaacttccacattaatacagcctattttcgctccaaatgtacattcgaaaaatctgaataaaacatcaatatcgcaatttgtattatccaatacagaatgtctgtgattcgtgttaaaaccatcggaaaaagaatagaaataaaaattattataatggtggaattcagttggaatattgcctccggagtcataaaaggatactaaacattgttttttatcgtaaattacacatttccaatgagacaaataacaaaatccaaacattacaaatctagaggtagaacttttaattttgtctttaagtatatacgataagatatgtttattcataaacgcgtcaaatttttcatgaatagctaaggagtttaagaatctcatgtcaaattgtcctatataatccacttcggatccataagcaaactgagagactaagttcttaatacttcgattgctcatccaggctcctctctcaggctctattttcatcttgacgacctttggattttcaccagtatgtattcctttacgtgataaatcatcaattttcaaatccatttgtgagaagtctatcgccttagatactttttcccgtagttgaggtttaaagaaatacgctaacggtatactagtaggtaactcaaagacatcatatatagaatggtaacgcgtcgttaactcgtcggttaactctttcttttgatcgagttcatcgctactattgggtctgctcaggtgccccgactctactagttccaacatcataccgataggaatacaagacactttgccagcggttgtagatttatcatatttctccaccacatatccgttacaatttgttaagaatttagatacatctatattgctacataatccagctagtgaatatatatgacataataaattggtaaatcctagttctggtattttactaattactaaatctgtatatctttccatttatcatggaaaagaatttaccagatatcttcttttttccaaactgcgttaatgtattctcttacaaatattcacaagatgaattcagtaatatgagtaaaacggaacgtgataatttctcattggctgtgtttccagtgataaaacatagatggcataacgcacacgttgtaaaacataaaggaatatacaaagttagtacagaagcacgtggaaaaaaagtatctcctccatcactaggaaaacccgcacatataaacctaatgtcgaagcaatatatatatagtgagtatgcaataagctttgaatgttatagttttctaaaatgtataacaaatacagaaatcaattcgttcgatgagtatatattaagaggactattagaagctggtaatagtttacagatattttccaattccgtaggtaaacgaatagatactataggtgtactagggaataagtatccatttagcaaaattccattggcntcattaacyyctaaagcacaacgagagatatttttagcgtggatttctcatagacctgtagttttaactggaggaaccggagtgggtaagacgtcacaggtacccaagttattgctttggtttaattatttatttggtggattctcttctctagataaaatcactgactttcacgaaagaccagtcattctatctcttcctaggatagctttagttagattgcatagcaataccattttaaaatcattgggatttaaggtactagatggatctcctatctctttacggtacggatctataccggaagaattaataaacaaacaaccaaaaaaatatggaattgtattttctacccataagttatctctaacaaaactatttagttatggcactattattatagacgaagttcatgagcatgatcaaataggagatattattatagcagtagcgagaaaacatcatacgaaaatagattctatgtttttaatgactgccacgttagaggatgacagggaacgtctaaaaatatttttacctaatcccgcatttatacatattcctggagatacactgtttaaaattagcgaggtatttattcataataagataaatccatcttccagaatggcatatatagaagaagaaaagagaaatttagttactgctatacagatgtatactcctcctgatggatcatccggtatagtctttgtggcatccgttgcacagtgtcacgaatataaatcatatttagaaaaaagattaccgtatgatatgtatattattcatggtaaggtcttagatatagacgaaatattagaaaaagtgtattcatcacctaatgtatcgataattatttctactccttatttggaatccagcgttactatacgcaatgttacacacatttatgatatgggtagagtttttgtccccgctccttttggaggatcacaacaatttatttctaaatctatgagagatcaacgaaaaggaagagtaggaagagttaatcctggaacatacgtatatttctatgatctgtcttatatgaaatctatacagcgaatagattcagaatttctacataattatatattgtacgctaataagtttaatctaacactccccgaagatttgtttataatccctacaaatttggatattctatggcgtacaaaggaatatatagactcgttcgatattagtacagaaacatggaataaattattatccaattattatatgaagatgatagagtatgctaaactttatgtactaagtcctattctcgctgaggagttggataactttgagaggacgggagaattaactagtattgtacaagaagccattttatctctaaatttacaaattaagattttaaaatttaaacataaagatgatgatacgtatatacacttttgtagaatattattcggtgtctataacggaacaaacgctactatatattatcatagacctctaacgggatatatgaatatgatttcagatactatatttgttcctgtagataataactaaaaatcaaaatctaatgaccacatctttttttagagatgaaaaattttccacatctccttttgtagacacgactaaacattttgcagaaaaaagtttattattatttagataatcgtatacttcatcagtgtagatagtaaatgtgaacagataaaaggtattcttgctcaatagattggtaaattccatagaatatattaatcctttcttcttgagatcccacatcatttcaaccaaagacgttttatccaatgatttacctcgtactataccacatacaaaactagattttgcagtgatgtcgtacctggtattcctaccaaacaaaattttacttttagttcttttagaaaattctaaggtagaatctctatttgtcaatatgtcatctatggaattaccactagcaaaaaatgatagaaatatatattgatacatcgcagctggttttgatctactatactttaaaaacgaatcagattccataattgcttgtatatcatcagctgaaaaactatgttttacacgtattccttcggcatttctttttaatgatatatcttgtttagacaatgataaagttatcatgtccatgagagacgcgtctccgtatcgtataaatatttcattagatgttagacgcttcattaggggtatacttctataaggtttcttaattagtccatcattggttgcgtcaagaactactatctgatgttgttgggtatctctagtgttacacatggccttactaaagtttgggtaaataactatgatatctctattaattatagatgtatatatttcattcgtcaaggatattaatatcgacttactatcgtcattaatacgtgtaatgtaatcatataaatcatgcgatagccaaggaaaattcaaatagatgttcatcatataatcgtcgctataattcatattaatactttgacattgactaatttgtaatatagcctcgccacgaagaaagctctcgtattcagtttcatcgataaaggataccgttaaatataactggttgccgatagtctcatagtctattaagtggtaagtttcgtataaatacagaatccctaaaatattatctaatgtgggattaatccttaccataactgtataaaatggagccggagtcataactattttaccgtttgtacttactggaatagatgaaggaataatctccggacatgatggtaaagacccaaatgtctgtttgaagaaatccaatgttccaggtcctaatctcttgacaaaaattacgatattcgatcccgatatcctttgcattctatttaccagcatatcacgaactatattaagattatctatcatgtctattctcccaccgttatataaatcgcctccgctaagaaacgttagtatatccatacaatggaatacttcatttctaaaatagtattcgttttctaattctttaatgtgaaatcgtatactagaaagggaaaaattatctttgagttttccattagaaaagaaccacgaaactaatgttctgattgcgtctgactccgtcgctgaattaatagatttacaccaaaaactcatataacttctagatgtagaagcattcgctaaaaaattagtagaatcaaaggatataagtagatgttccaacaagtgagcaattcccaagatttcatctatatcattctcgaatccgaaattagaaattcccaagtagatatcctttttcatccgatcattgatgaaaatacgaactttattcggtaagacgatcatttactaaggagtaaaataggaagtaacgttcgtatatcgttatcgtcgtataaattaaaggtgtgttttttgccattaagagacattataattttaccaatattggaattataatataggtgtatttgagcactagaaacggtcgatgcatcggtaaatatagctgtatctaatgttctagtcggtatttcttcatttcgctgtctaatgatagcgttttctctatctgtttccattacagctgcctgaagtttattggtcggataatatgtaaaataataagaaatacatacgaataacaaaaataaaataagatataataaagatgccatttagagatctaattttgttcaacttgtccaaattcctacttacagaagatgaggaatcgttggagatagtatcttccttatgtagaggatttgaaatatcttacgatgacttaatatcgtactttccagataggaaataccataaatatatttctaaggtatttgaacatgtagatttatcggaggaattaagtatggaattccatgatacaactctgagagatttagtatatcttagattgtacaagtattccaagtatatacggccgtgttataaattaggagataatctaaaaggtatagttgttataaaggacagaaatatatatattagagaagcaaatgatgacttgatagaatatctcctcaaggaatacactcctcagatttatacatattctaatgagcgagttcccatagctggttcaaaattaattctttgtggattttctcaagttacatttatggcgtatacaacgtcgcatataacaacaaataaaaaggtagatgttctcgtttccaaaaaatgtatagatgaactagtcgatccaataaattatcaaatacttcaaaatttatttgataaaggaagcggaacaataaacaaaatactcaggaagatattttattcggtaacaggtggccaaactccataggtagctttttctatttcggattttagaatttccaaattcaccagcgatttatcggttttggtgaaatccaaggatttattaatgtccacaaatgccatttgttttgtctgtggattgtatttgaaaatggaaacgatgtagttagatagatgcgcggcgaagtttcctattagggttccgcgcttcacgtcacccaacatacttgaatcaccatcctttaaaaaaaatgataagatatcaacatggagtatatcatactcggattttaattcttctactgcctcactgacattttcacaaatactacaatacggtttaccgaaaataatcagtacgttcttcatttatgggtatcaaaaacttaaaatcgttactgctggaaaataaatcactgacgatattagatgataatttatacaaagtatacaatggaatatttgtggatacaatgagtatttatatagccgtcgccaattgtgtcagaaacttagaagagttaactacggtattcataaaatacgtaaacggatgggtaaaaaagggaggacatgtaaccctttttatcgatagaggaagtataaaaattaaacaagacgttagagacaagagacgtaaatattctaaattaaccaaggacagaaaaatgttagaattagaaaagtgtacatccgaaatacaaaatgttaccggatttatggaagaagaaataaaggcagaaatgcaattaaaaatcgataaactcacatttcaaatatatttatctgattatgataacataaaaatatcattgaatgagatactaacacatttcaacaataatgagaatgttacattattttattgtgatgaacgagacgcagaattcgttatgtgtctagaggctaaaacacagttctctaccacaggagaatggccgttaataataagtaccgatcaggatactatgctattcgcgtctgctgataatcatcctaagatgataaaaaacttaactcaactgtttaaatttgttccctcggcagaggataactatttagcaaaattaactgcattagtgaatggatgtgatttctttcctggactctatggggcatctataacacccaacaacttaaacaaaatacaattgtttagtgattttacaatcgataatatagtcactagtttggcaattaaaaattattatagaaagactaactctaccgtagacgtgcgtaatattgttacgtttataaacgattacgctaatttagacgatgtctactcgtatattcctccttgtcaatgcactgttcaagaatttatattctccgcattagatgaaaaatggaatgaatttaaatcatcttatttagagagcgtgccgttaccctgccaattaatgtacgcattagaaccacgtaaggagattgatgtttcagaagttaaaactttatcatcttatatagatttcgaaaatactaaatcagatatcgatgttataaaatctatatcctcgatttttggatattctaacgaaaactgtaacaccatagtgttcggcatctataaggataatttactactgagtataaataattcattttactttaacgatagtctgttaataaccaatactaaaagtgataatataataaatataggttactagattaaaaaatggtgttccagctcgtgtgttctacatgcggcaaagatatttctcacgaacgatataaattgattatacgaaaaaaatcattaaaggatgtactagtcagtgtaaagaacgaatgttgtaggttaaaattatctacacaaatagaacctcaacgtaacttaacagtgcaacctctattggatataaactaatggatccggttaattttatcaagacatatgcgcctagaggttctattatttttattaattatgccatgtcattaactagtcatttgaatccatcgatagaaaaacatgtgggtatttattatggtacgttattatcggaacacttggtagttgaatctacctatagaaaaggagttagaatagtcccattggatagattttttgaaggatatcttagtgcaaaagtatacatgttagagaatattcaagttatgaaaatagcagctgatatgtcgttaactttactaggtattccatatggatttggtcatgatagaatgtattgttttaaattggtagctgaatgttataaaaatgccggtattgatacatcgtctaaacgaatattaggtaaagatatttttctgagccaaaactttacagatgataatagatggataaagatatatgattctaataatttaacattttggcaaattgattaccttaaagggtgagttaatatgcataactactcctccgttgttttttccctcgttctttttcttaacgttgtttgccatcactctcataatgtaaagatattctaaaatggtaaacttttgcatatcggatgcagaaattggtataaatgttgtaattgtattatttcccgtcaatggactagtcacagctccatcagttttatatcctttagagtatttctcactcgtgtctagcattctagagcattccatgatctgtttatcgttgatattggccggaaagatagattttttattttttattatattactattggcaattgtagatataacttctggtaaatatttttctaccttttcaatctcttctattttcaagccggctatatattctgctatattgttactagtatcaataccttttctggctaagaagtcatatgtggtattcactatatcagttttaactggtagttccattagcctttccacttctgcagaataattagaaattggttctttaccagaaaatccagctactataataggctcaccgatgatcattggcaaaatcctatattgtaccagattaatgagagcatatttcatttccaataattctgctagttcttgagacattgatttatttgatgaatctatttggttctctagatactctaccatttctgccgcatacaataacttgttagataaaatcagggttatcaaagtgtttagtgtggctagaatagtgggcttgcacgtattaaagaatgctgtagtatgagtaaaccgttttaacgaattatatagtctccagaaatctgtggcgttgcatacatgaactgaatgacatcgaagattgtccaatatttttaatagctgctctttgtccattatttctatatttgactcgcaacaattgtagataccattaatcactgattcctttttcgatgccggacaatagcacaattgtttagctttggactctatgtattcagaattaatagatatatctctcaatacagattgcactatacattttgaaactatgtcaaaaattgtagaacgacgctgttctgtagccatttaactttaaataatttacaaaaatttaaaatgagcatccgtataaaaatcgataaattgcgccaaattgtggcatatttttcagagttcagcgaagaagtgtctataaatgtagacttgacggatgaattaatgtatatttttgccgccttgggcggatctgtaaacatttgggccattatacctctcagtgcatcagtgttctaccgcggagccgaaaatattgtgtttaaccttccagtgtccaaggtaaaatcgtgtttgtgtagttttcacaatgatgctatcataaatatagaacctgatctggaaaataatctagtaaaactttctagttatcatgtagtaagtgtcgattgtaacaaggaactgatgcctattaggacagatactactatttgtctaagtatagatcaaaagaaatcttacgtatttaattttcacaagtatgaagaaaaatgttgtggtagaaccgtcattcatctagaatggttgttgggctttatcaagtgtattagtcagcatcagcatttggctattatgtttaaagatgacaatattattatgaagactcctggtaatactgatgcgttttccagggaatattctatgactgaatgttctcaagaactacaaaagttttctttcaaaatagctatctcgtctctcaacaaactacgaggattcaaaaagagagtcaatgtttttgaaactagaatcgtaatggataatgacgataacattctaggaatgttgttttcggatagagttcaatcctttaagattaacatctttatggcgtttttagactaatactttcaatgagataaatatgggtggcggagtaagtgttgagctccctaaacgggatccacctccgggagtacccactgatgagatgttattaaacgtggataaaatgcatgacgtgatagctcccgctaagcttttagaatatgtgcatataggaccactaacaaaagataaagaggataaagtaaagaaaagatatccagagtttagattagtcaacacaggacccggtggtctttcggcattattaagacaatcatataatggaaccgcacccaattgctgtcgcacttttaatcgtactcattattggaagaaggatggaaagatatcagataagtatgaagagggtgcagtattagaatcgtgttggcccgacgtccacgacactggaaaatgcgatgttgatttattcgactggtgtcagggggatacgttcgatataaacatatgccatcagtggatcggttcagcctttaataggagtgatagaactgtagagggtcgacaatcgttaataaatctgtataataagatgcaaagattatgtagtaaagatgctagtgtaccaatatgtgaattatttttgcatcatttacgcgcacacaatacagaagatagtaaagagatgatcgattatattctaagacaacagtcggcggactttaaacagaaatatatgagatgtagttatcccactagagataagttagaagagtcattaaaatatgcggaacctcgagaatgttgggatccagagtgttcgaatgccaatgttaatttcttactaacacgtaattataataatttaggactttgcaatattgtacgatgtaatacgagcgtgaataacttacagatggataaaacttcctcattaagattatcatgtggattaagcaatagtgatagattttctactgttcccgtcaatagagcaaaagtagttcaacataatattaaacattcgttcgacctaaaattgcatttgatcagtttattatctctcttggtaatatggatactaattgtagctatttaaatgggtgccgcagcaagcatacagacgactgtgaatacactcagtgaacgtatctcgtctaaattagaacaagaagcgaacgctagtgctcaaacaaaatgtgatatagaaatcggaaatttttatatccgacaaaaccatggatgtaacatcactgttaaaaatatgtgctctgcggacgcggatgctcagttggatgctgtgttatcagccgctacagaaacatatagtggattaacaccggaacaaaaagcatacgtaccagctatgtttactgctgcgttaaacattcagacgagtgtaaacactgttgttagagattttgaaaattatgtgaaacagacttgtaattctagcgctgttgtcgataacaaattaaagatacaaaacgtaattatagatgaatgttacggagccccaggatctccaacaaatttggaatttattaatacaggatctagcaaaggaaattgtgccattaaggcgttgatgcaattgactactaaggccactactcaaatagcacctagacaagttgctggtacaggagttcagttttatatgattgttatcggtgttataatattggcagcgttgtttatgtactatgccaagcgtatgctgttcacatccaccaatgataaaatcaaacttattttagccaataaggaaaacgtccattggactacttacatggacacattctttagaacttctccgatgattattgctaccacggatatacaaaactgaaaatatattgataatattttaatagattaacatggaagttatcgctgatcgtctagacgatatagtgaaacaaaatatagcggatgaaaaatttgtagattttgttatacacggtctagagcatcaatgtcctgctatacttcgaccattaattaggttgtttattgatatactattatttgttatagtaatttatatttttacggtacgtctagtaagtagaaattatcaaatattgttggtgttggtggcgctagtcatcacattaactatttttttattactttatactataatagtactagactgacttctaacaaacatctcacctgccataaataaatgcttgatattaaagtcttctatttctaacactattccatctgtggaaaataatactctgacattatcgctaattgatacatcggtaagtgatatgcctataaagtaataatcttctttgggcacatataccagtgtaccaggttctaacaacctatttactggtgctcctgtagcatacttttttaccttgagaatatccattgtttgcttggtcaatagtgatatgtgattttttatcaaccactcaaaaaagtaattggagtgttcatatcctctacgggctattgtctcatgaccgtgtatgaaatttaagtaacacgactgtggtagatttgttctatagagccggttgccgcaaatagatagaactaccaatatgtctgtacaaatgttaaacattaattgattaacagaaaaaacaatgttcgttctgggaatagaaaccagattaaaacaaaattcattagaatatatgccacgtttatacatggaatataaaataactacagtttgaaaaataacagtatcatttaaacatttaacttgcggggttaatctcacaactttactgtttttgaactgttcaaaatatagcatagatccatgagaaatacgtttagccgcctttaatagaggaaatccaaccgcctttctggatctcaccaacgacgatagttctgaccagcaactcatttcttcatcatccacctgttttaacatataataggcaggagatagatatccatcattgcaatattccttctcgtaggcacacaatctaatattgataaaatctccattctcttctctgtatttattatcttgtctcggtggctgattaggctgtggtctatcgttgttgaatctattttggtcattaaatctttcatttcttcctggtatatttctatcacctcgtttggttggatttttgtctatattatcgtttgtaacatcggtacgggtattcatttatcacaaaaaaacttctctaaatgagtctactactagaaaacctcatcgaagaagataccatattttttgcaggaagtatatctgagtatgatgatttacaaatggttattgctggtgcaaaatccaaatttccaagatctatgctttctatttttaatatagtacctagaacgatgtcaaaatatgagttggagttgattcataacgagaatatcacaggggcaatgtttaccacaatgtataatataagaaacaatttgggtctaggcgatgataaactaactattgaagccattgaaaactatttcttggatcctaacaatgaggttatgcctcttatcattaataatacggatatgactaccgtcattcctaaaaaaagtggtaggagaaagaataagaacatggttatcttccgtcaaggatcatcacctatcttgtgtattttcgaaactcgtaaaaagattaatatttataaagaaaatatggaatccgtatcgactaagtatacacctatcggagacaacaaggctttgatatctaaatatgcgggaattaatatcctgaatgtgtattctccttccacgtccatgagattgaatgccatttacggattcaccaataaaaataaactagagaaacttagtactaataaggaactagaatcgtatagttctagccctcttcaagaacccattaggttaaatgattttctgggactattggaatgtgttaaaaagaatattcctctaacagatattccgacaaaggattgattactataaatggagaatgttcctaatgtatactttaatcctgtgtttatagagcccacgtttaaacattctttattaagtgtttataaacacagattaatagttttatttgaagtattcgttgtattcattctaatatatgtattttttagatctgaattaaatatgttcttcatgcctaaacgaaaaatacccgatcctattgatagattacgacgtgctaatctagcgtgtgaagacgataaattaatgatctatggattaccatggataacaactcaaacatctgcgttatcaataaatagtaaaccgatagtgtataaagattgtgcaaagcttttgcgatcaataaatggatcacaaccagtatctcttaacgatgttcttcgcagatgatgattcattttttaagtattttgctagtcaagatgatgaatcttcattatctgatatattgcaaatcactcaatatctagactttctgttattattattgatccaatcaaaaaataaattagaagctgtgggtcattgttatgaatctctttcagaggaatacagacaattgacaaaattcacagactctcaagattttaaaaaactgtttaacaaggtccctattgttacagatggaagggtcaaacttaataaaggatatttgttcgactttgtgattagtttgatgcgattcaaaaaagaatcagctctagctaccaccgcaatagatcctgttagatacatagatcctcgtcgtgatatcgcattttctaacgtgatggatatattaaagtcgaataaagttgaaaaataattaattctttattgttatcatgaacggcggacatattcagttgataatcggccccatgttttcaggtaaaagtacagaattaattagacgagttagacgttatcaaatagctcaatataaatgtgtgactataaaatattctaacgataatagatacggaacgggactatggacacatgataagaataattttgcagcattggaagtaactaaactatgtgatgtcttggaagcaattacagatttctccgtgataggtatagatgaaggacagttctttccagacattgttgaattctgtgagcgtatggcaaacgaaggaaaaatagttatagtagccgcgctcgatgggacatttcaacgtagaccgtttaataatattttgaatcttattccattatctgaaatggtggtaaaactaactgcagtgtgtatgaaatgctttaaggaggcttccttttctaaacgattaggtacagaaaccgagatagaaataataggaggtaatgatatgtatcaatctgtgtgtagaaagtgttacatcgactcataatattatattttttatctaaaaaactaaaaataaacattgattaaattttaatataatacttaaaaatggatgttgtgtcgttagataaaccgtttatgtattttgaggaaattgataatgagttagattacgaaccagaaagtgcaaatgaggtcgcaaaaaaactgccgtatcaaggacagttaaaactattactaggagaattattttttcttagtaagttacagcgacacggtatattagatggcgccaccgtagtgtatataggatctgctccaggtacacatatacgttatttgagagatcatttctataatttaggagtgatcatcaaatggatgctaattgacggccgccatcatgatcctattctaaatggattgcgtgatgtgactctagtgactcggtttgttgatgaggaatatctacgatccatcaaaaaacaactacatccttctaagattattttaatttctgatgtgcgatccaaacgaggaggaaatgaacctagtactgcggatttactaagtaattatgctctacaaaatgtcatgattagtattttaaaccccgtggcgtctagtcttaaatggagatgcccgtttccagatcaatggatcaaggacttttatatcccacacggtaataaaatgttacaaccttttgctccttcatattcagctgaaatgagattattaagtatttataccggtgagaatatgagactgactcgagttaccaaatcagacgctgtaaattatgaaaaaaagatgtattaccttaataagatagtccgcaacaaagtagttattaactttgattatcctaatcaggaatatgactattttcacatgtactttatgttgaggaccgtatactgcaataaaacatttcctactactaaagcaaagatactatttctacaacaatctatatttcgtttcttaaatattccaacgacatcaactgaaaaagttagtcatgaaccaatacaacgtaaaatatctagcaaagattctatgtctaaaaacagaaatagcaagagatccgtacgcggtaataaatagaaacgtactactgagatatactaccgatatagagtataatgatttagttactttaataaccgttagacataaaattgattctatgaaaactgtgtttcaggtatttaacgaatcatccataaattatactccggttgatgatgattatggagaaccaatcattataacatcgtatcttcaaaaaggtcataacaagtttcnngtaaattttctatacatagatgtggtaatatctgacttatttcctagctttgttagactagatactacagaaactaatatagttaatagtgtactacaaacaggcgatggtnaaaagactcttcgtctyyctnaaatgttagagacggaaatagttgtcaagattctctatcgtnytaatataccattaaaaattgttagatttttccgcaayaacatggtaactggagtagagatagccgatagatctgttatttcagtcgctgattaatcaattagtagagatgagataagaacattataataatcaataatatatcttatatctgtttagaaaaatgctaatattaaaatagctaacgctagtaatccaatcggaagccatttgatatctataatagggtatctaatttcctgattcagatagcgtacggctatattctcggtagctactcgtttggaatcacagacattatttacatctaatttactatctgtaatggaaacgtttcccaatgaaatggtacaatcagatacattacatcttgatatatttttttttaaagaggctggtaacaacgcatcgcttcgtttacatggctcgtaccaacaataatagggtaatcttgtatctattcctatccgtactatacttttatcaggataaatacatttacatcgtatatcgtctttgttagtatcacagaatgcataaatttgttcgtccgtcatgataaaaatttaaagtgtaaatataactattatttttatagttataataaaaagggaaatttgattgtataccttcggttctttaaaagaaactgacttgataaaaatggctgtaatctctaaggttacgtatagtctatacgatcaaaaagagattaatgccacagatattatcattagtcatattaaaaatgacgacgatatcggtaccgttaaagatggtagactaggtgctatggatggggcattatgtaaaacttgtgggaaaacggaattggaatgtttcggtcactggggtaaagtaagtatttataaaactcatatagttaagcctgaatttatttcagaaattattcgtttactgaatcatatatgtattcattgcggattattgcgttcacgagaaccgtattccgacgatattaacctaaaagagttatcggtacacgctcttaggagattaaaggataaaatattatccaagaaaaagtcatgttggaacagcgaatgtatgcaaccgtatcaaaaaattactttttcaaagaaaaaggtttgtttcgtcaacaagttggatgatattaacgttcctaattctctcatctatcaaaagttaatttctattcatgaaaagttttggccattattagaaattcatcaatatccagctaacttattttatacagactactttcccatccctccgttgattattagaccggctattagtttttggatagatagtatacccaaagagacaaatgaattaacttacttattaggtatgatcgttaagaattgtaacttgaatgctgatgaacaggttatccagaaggcggtaatagaatacgatgatattaaaattatttctaataacactaccagtatcaatttatcatatatcacatccggcaaaaataatatgattagaagttatatcgtcgctcggcgaaaagatcagaccgctagatccgtaattggtcccagtacatctatcaccgttaatgaggtaggaatgcccacatatattagaaatacacttacagaaaagatatttgttaatgcctttacagtggataaagttaaacaactattagcatcaaaccaagttaaattttactttaataaacgattaaaccaattaacaagaatacgtcaaggaaagtttatcaaaaataaaatacatttattgcctggtgattgggtagaagtagctgttcaagaatatacaagtattatttttggaagacaaccgtctctacatagatacaacgtcatcgcttcatctatcagagctaccgaaggagatactatcaaaatatctcccggaattgccaactctcaaaatgctgattttgacggagatgaagaatggatgatattggagcaaaatcctaaagccgtagttgaacaaagtattcttatgtatccgacaacgttactcaaacacgatattcatggagcccccgtttatggatctattcaagatgaaatcgtagcagcgtattcattgtttaggatacaagatctttgtttagatgaagtattgaacatcttggggaaatatggaagagagttcgatcctaaaggtaaatgtaaattcagcggtaaagatatctatacttacttgataggtgaaaagattaattatccgggtctcttaaaggatggtgaaattattgcaaacgacgtagatagtaattttgttgtagctatgaggcatctgtcattggctggactcttatccgatcataaatcgaacgtggaaggtatcaactttattatcaagtcatcttatgtttttaagagatatctatctatatacggttttggggtgacattcaaagatctgagaccaaattcgacgttcactaataaattggaggctatcaacgtagaaaaaatagaacttatcaaagaagcatacgccaaatatctcaaagatgtaagagacgggaaaatagttccattatctaaagctttagaggcggactacttggaatccatgttatccaacttgacaaatcttaatatcagagagatagaagaacatatgagacaaacgctgatagatgatccagataataacctcctgaaaatggccaaagcgggttataaagtaaatcccacagaactaatgtatattctaggtacttatggacaacagaggatagatggcgaaccagcagagactcgagtattgggtagagtcttaccttactatcttccagactctaaggatccagaaggaagaggttatattcttaattctttaacaaaaggattaacgggttctcaatattactttttgatgctggttgcaagatctcaatctactgatattgtctgtgaaacatcacgtaccggaacactggctagaaaaatcattaaaaagatggaggatatggtggtcgacggatacggacaagtagttataggtaatacgctcatcaagtacgcagccaattataccaaaattctaggctcagtatgtaaacctgtagatcttatctatccagatgagtccatgacttggtatttggaaattagtgctttgtggaataaaataaaacagggattcgtttactctcagaaacagaaacttgcaaagaagacattggcgccgtttaatttcctagtattcgtcaaacccaccactgaggataatgctattaaggttaaggatctgtacgatatgattcataacgtcattgatgatgtgagagagaaatacttctttacggtatctaatatagattttatggagtatatattcttgacgcatcttaatccttctagaattagaattacaaaagaaacggctattactatctttgaaaagttctatgaaaaactcaattatactctaggtggtggaactcctattggaattatttctgcacaggtattgtctgagaagtttacacaacaagccctgtccagttttcacactactgaaaagagtggtgctgtaaaacaaaaacttggtttcaacgagtttaataacttgactaatttgagtaagaataagaccgaaattatcactctggtatccgatgatatctctaaacttcaatctgttaagattaatttcgaatttgtatgtttgggagaattaaatccagacatcactcttcgaaaagaaacagatagatatgtagtagacataatagtcaatagattatacatcaagagagcagaaataaccgaattagtcgtcgaatatatgattgaacgatttatctcctttagcgtcattgtaaaggaatggggtatggagacattcattgaggacgaggataatattagatttactatctacctaaatttcgttgaaccggaggaattgaatcttagtaagtttatgatggttcttccaggtgccgccaacaagggcaagattagtaaattcaagattcctatctctgactatacgggatataacgacttcaatcaaacaaaaaagctcaataagatgactgtagaactcatgaatctaaaagaattgggttctttcgatttggagaacgtcaacgtgtatcctggagtatggaatacatacgatatctttggtattgaggccgctcgtggatacttgtgcgaagccatgttaaacacctatggagaaggtttcgattatctgtaccagccttgtgatcttctcgctagtttactatgtgctagttacgaaccagaatcagttaataaattcaagttcggtgcagctagtactcttaagagagctacgttcggagataataaagcattgttaaacgcggctcttcataaaaagtcagaacctattaacgataatagtagctgccacttttttagcaaggtccctaatataggaactggatattacaaatactttatcgacttgggtcttctcatgagaatggaaaggaaactatctgataagatatcttctcaaaagatcaaggagatagaagaaacagaagacttttaattcttatcaataacatatttttctatgatctgtcttttaaacgatggattttccacaaatgcgcctctcaagtccctcatagaatgatacacgtataaaaaatatagcataggtgatgactccttatttttagacattagatatgccaaaatcatagccccgcttctatttactcctgcaacacaatgaaccaacacgggctcgtttcgttgatcacatttagataagaaggcggtcacgtcgtcaaaatatttactaatatcagtagttgtatcatctaccaacggtatatgaataatattaatattagagttaggtaatgtatatttatccatcgtcaaatttaaaacatatttgaacttaacttcagatgatggtgcatccatagcatttttataatttcccaaatacacattatttgttactcttgtcattatagtgggagatttggctctgtgcatatctccagttgaacgtagtagtaagtatttatacaaacttttcttatccatttataacgtacaaatggataaaactactttatcagtaaacgcatgcaatttagaatacgttagagaaaaggctatagtaggcgtacaagcagccaagacatcaacacttatattttttgttattatattggcaattagtgcgctattactctggtttcagacgtctgataatccagtctttaatgaattaacgagatatatgcgaattaaaaatacggttaacgattggaaatcattaacggatagcaaaacaaaattagaaagcgatagaggtagacttctagccgctggtaaggatgatatattcgaattcaaatgtgtggatttcggcgcctattttatagctatgcgattggataagaaaacatatctgccgcaagctattaggcgaggtactggagacgcgtggatggttaaaaaggcggcaaaagtcgatccatctgctcaacaattttgtcagtatttgataaaacacaagtctaataatgttattacttgtggtaatgagatgttaaatgaattaggttatagcggttattttatgtcaccgcattggtgttccgatcttagtaatatggaataagtgttagataaatgcggtaacaaatgttcctgtaaggaaccataacagtttagatttaacattaaagatgagcataaacataataaacaaaattacaatcaaacctataacattaatatcaaacaatccaaaaaatgaaatcaatggagtagtaaacgtgtacataactcctggataacgtttagcagctaccgttcctattctagaccaaaaatttggtttcatggtttcgaagcggtgttctgcaacaagacgaggatcgtgttctacatatttggcagagttatccattatttgcctgttaatcttcatttcgttttcgattctggctatttcaaaataaaatcccgatgatagacctccagactttataatttcatctacgatgttcagcgccgtagtaactctaataatataggcggataagctaacatcataccctcctgtatatgtaaatatggcatgatctttgtctattacaagctcggttttaactttatttcctgtaataatttctctcatctgtaggatatctattttcttgtcatgtattgccttcaagacgggacgaagaaacgtaatatcctcaataacgttatcgttttctataataactacatattctacatttttattttctagctcggtaaaaaatttagaatcccatagggctaaatgtctagcgatatttcttttcgtttcctctgtacacatagtgttacaaaaccctgaaaagaagtgagtatacttgtcatcatctctaatatttcctccagtccattgtataaacacataatccttgtaatgatctggatcatcattgactatcacaacatctcttttttcttgcataacttcattgtccttcacatcatcgaacttctgatcattaatatgctcatgaacattaggaaatgtttctgatggaggtctatcaataactggcacaacaataacaggagttttcaccgccgccatttagttattgaaattaatcatatacaactctctaatacgagttatattttcgtctatccattgtttcacattgacatatttcgacaaaaagatataaaatgcgtattccaatgcttctctgtttaatgaattactaaaatatacaaacacgtcactgtctggtaataaataatatcttagaatattgtaacaatttattttgtattgcacatgttcgtgatctatgagttcttcttcaaatggcataggatctccgaatctgaaaacgtataaataggagttagaataataatatttgagagtattggtaatgtataaactctttagcggtataattagtttttttctctcgatttctatttttagatgtgatggaaaaatgactaattttgtagcattagtatcatgaactctaatcaaaatcttaatatcttcgtcacatgttagctctttgaagtttttaagagatgcatcagttggttttacagatggagtaggtgcaacaattttttgtttaatgcatgcatgtattggagccattgtcttaactataatggtgcttgtatcgaaaaactttaatgcggataacggaagctcttcgccgcgactttctacgtcgtaattgggttctaatgccgatctctgaatggatactagttttctaagttctaatgtaattctctgaaaatgtaaatccaattcctccggcattatagatgtgtatacatcggtaaataaaactatagtatccaacgatcccttctcgcaaattctagtcttaaccaagaaatcgtatataactacggagatggcgtatttaagagtggattcttctaccgttttgttcttggatttcatataagaaactataaagtccgcactactgttaagaatgatcactaacgcaactatatagttcaaattaagcatcttggaaacataaaataactctgtagatgatacttgactttcgaataagtttgcagacaaacgaagaaagaacagacctctcttaatttcagaagaaaactttttttcgtattcctgacgtctagagtttatatcaataagaaagttaagaattagtcggttaatgttgtatttcattacccaagtttgagatttcataatattgtcaaaagacatgataatattaaagataaagcgctgactatgaacgaaatagctatatggttcgctcaagaatatagtcttgttaaacgtggaaacgataactgtatttttaatcacgtcagcggcatctaaattaaatataggtatatttattccacacactctacaatatgccacaccatcttcataataaataaattcgttagcaaaattattaattttagtgaaatagttagcgtcaactttcatagcttccttcaatctaatttgatgctcacatggcgcgaattctactctaacatcccttttccatgcctcaggttcatcgatctctataatatctagtttcttgcgtttcacaaacacaggctcgtctctcgcgatgagatctgtatagtaactatgtaaatgataactagatagaaagatgtagctatatagatgacgatcctttaagagaggtataataactttaccccaatcagatagactgttgttatggtcttcggaaaaagaatttttataaatttttccagtattttctaaatatacgtacttgatatctaagaaatccttaataataataggaatggataatccgtctattttataaagaaatacatatcgcatattatacttttttttggaaattggaataccgatgtgtctacataaatacgcaaagtctaaatattttttagagaatcttagttggtccaaattcttttccaagtacggtaatagatttttcatattgaacggtatcttcttgatctctggttctaattccgcattaaatgatgaaactaagtcactatttttataactaacgattacatcacctctaacatcatcatttaccaggatactgatcttcttttgtcgtaaatacatgtctaatgtgttaaaaaaaagatcatacaagttatacgtcatttcatctgtagtattcttgtcattgaaggataaactcgtactaatctcttctttaacagtctgttcaaatttatatcctatatatgaaaaaatagcaaccagtgtttgatcatccgcgtcaatattctgttctatcgtagtgtataacaatcttatatcttcttctgtgatagtcgatacgttataaaggttgataacgaaaatatttttatttcgtgaaataaagtcattgtaggattttggacttatattcgtgtctagtagatatgattttatttttggaatgatctcaattaaaatagtctctttagagtccatttaaagttacaaacaactaggaaattggtttatgatgtataatttttttagtttttatagattctttattctatacttaaaaaatgaaaataaatacaaaggttcttgagggttgtgttaattgaaagcgataaataatcataaattatttcattatcgcgatatccgttaagtttgtatcgtaatggcgtggtcaattacgaataaagcggatactagtagtttcacaaagatggctgaaatcagagctcatctaagaaatagcgctgaaaataaagataaaaacgaggatattttcccggaagatgtaataattccatctactaagcccaaaaccaaacgaaccactactcctcgtaaaccagcggctactaaaagatcaaccaaaaaggataaagaaaaggaggaagtggaagaagtagttatagaggaatatcatcaaacaactgaagaaaattctccacctccgtcatcatctcctggagtcggcgacattgtagaaagcgtggccgctgtagagctcgatgatagcgacggggatgatgaacctatggtacaagttgaagctggtaaagtaaatcatagtgctagaagcgatctctctgacctaaaggtggctaccgacaatatcgttaaagatcttaagaaaattattactagaatctctgcagtatcgactgttctagaggatgttcaagcagctggtatctctagacaatttacttctatgactaaagctattacaacactatctgatctagtcaccgagggaaaatctaaagttgttcgtaaaaaagttaaaacttgtaagaagtaaatgcgtgcacttttttataaagatggtaaactgtttaccgataataattttttaaatcctgtatcagacgataatccagcgtatgaggttttgcaacatgttaaaattcctactcatttaacagatgtagtagtatatgaacaaacgtgggaagaggcattaactagattaatttttgtgggaagtgattcaaaaggacgtagacaatacttttacggaaaaatgcatatacagaatcgcaatgctaaaagagatcgtatttttgttagagtatataacgttatgaaacgaattaattgttttataaacaaaaatataaagaaatcgtccacagattccaattatcagttggcggtttttatgttaatggaaactatgttttttattagatttggtaaaatgaaatatcttaaggagaatgaaacagtagggttattaacactaaaaaataaacacatagaaataagtcccgatgaaatagttatcaagtttgtaggaaaggacaaagtttcacatgaatttgttgttcataagtctaatagactatataaaccgctattgaaactgactgatgattctagtcctgaagaatttctgttcaacaaactaagtgaacgaaaggtatatgaatgtatcaaacagtttggtattagaatcaaggatctccgaacgtatggagtcaattatacgtttttatataatttttggacaaatgtaaagtccgtatctcctcttccatcaccaaaaaagttgatagcattaactatcaaacaaactgctgaagtggtaggtcatactccatcaatttcaaaaagagcttatatggcaacgactattttagaaatggtaaaggataaaaattttttagacgtagtatctaaaactacgttcgatgaattcctatctatagtcgtagatcacgttaaatcatctacggatggatgataatagatctttacacaaataattacaagaccgataaatggaaatggataaacggatgaaatctctcgctatgacagctttcttcggagagctaaacacgttagatattatggcattgataatgtctatatttaaacaccatccaaacaataccattttttcagtggataaggatggtcaatttatgattgatttcgaatacgataattataaggcttctcaatatttggatctgaccctcactccgatatctggaaatgaatgcaagactcacgcatctagtatagccgaacaattggcgtgtgtggatattattaaagaggatattagcgaatatatcaaaactactccccgtcttaaacgatttataaaaaaataccgcaatagatcatatactcgtatcagtcgagatacagaaaagcttaaaatagctctagctaaaggcatagattacgaatatataaaagacgcttgttaataagtaaatgaaaaaaactagtcgtttataataaaacacaatatggatgccaacatagtatcatcttctactattgcgacgtatatagacgctttagcgaagaatgcttcagaattagaacagaggtctaccgcatacgaaataaataatgaattggaactagtatttattaaaccgccattgattacgttgacaaatgtagtaaatatctccacgattcaggaatcgtttattcgatttaccgttactaataaggaaggtatcaaaattagaactaagattccattatctaaggtacatggtctagatgtaaaaaatgtgcagttggtagatgctatagataacatagtttgggaaaagaaatcattagtgacggaaaatcgtcttcacaaagaatgcttgttgagactatcaacagaggaacgtcatatatttttggattacaagaaatatggatcctctatccgactagaattagtcaatcttattcaagcaaaaacaaaaaactttacgatagactttaagctaaaatattttctaggatctggcgctcaatctaaaagttctttattgcacgctattaatcatccaaagtcaaggcctaatacatctctggaaatagaatttacacctagagacaatgaaacagttccatatgatgaactaataaaggaattgacgactctctcgcgtcatatatttatggcttctccagagaatgtaattctttctccacctattaacgcacctataaagacttttatgttgcctaaacaagatatagtaggtctggatctggaaaatctatatgccgtaactaagactgacggcattcctataactatcagagttacatcaaaagggttgtattgttattttacacatcttggttatattattagatatccagttaagagaacaatagattccgaagtagtagtctttggtgaggcagttaaggataagaactggaccgtatatctcattaagctaatagagcccgtaaatgcaatcagtgatagactagaagaaagtaagtatgttgaatctaaactagtggatatttgtgatcggatagtattcaagtcaaagaaatacgaaggtccgtttactacaactagtgaagtcgtcgatatgttatctacatatttaccaaagcaaccagaaggtgttattctgttctattcaaagggacctaaatctaacattgattttaaaatcaaaaaggagaatactatagaccaaactgcaaatgtagtatttaggtacatgtccagtgaaccaattatctttggagagtcgtctatctttatagagtataagaaatttaccaacgataaaggctttcctaaagaatatggttctggtaagattgtgttatataacggcgttaattatctaaataatatctattgtttggaatatattaatacacataatgaagtgggtattaagtccgttgttgtacctattaagtttatagcagaattcttagtcaatggagaaatacttaaacctagaatcgataaaaccatgaaatatattaactcagaagactattatggaaatcaacataatatcatagtcgaacatttaagagatcaaagcatcaaaataggagatgtctttaacgaggataaactatcggatgttggacatcaatacgctgccaacaacgataaatttagattaaatccagaagttagttattttactaataaacgaactagagggccgttgggaattttatcaaactacgtcaagactcttcttatttctatgtattgttccaaaacatttttagacgattccaacaaacgaaaggtattagcgattgattttggaaacggtgctgacctggaaaaatacttttatggagagattgcgttattggtagcgacggatccggatgctgatgctatagctagaggaaatgaaagatacaacaaattaaattctggaattaaaaccaagtactacaaatttgactacattcaggaaactattcgatccgatacatttgtctctagtgtcagagaagtattctattttggaaagtttaatatcattgactggcagttcgctattcattattcttttcatccaagacattatgctacagtcatgaataacttatccgaactaactgcttctggaggcaaggtattaattactaccatggatggagacaaattatcaaaattaaccgataaaaagacttttataattcataagaatctacctagtagcgaaaactatatgtctgtagaaaaaatagctgatgatagaatagtggtatataatccatcaacaatgtctactccaatgactgaatacattatcaaaaagaacgatatagtcagagtgtttaacgaatacggatttgttcttgtagataatgttgatttcgctacaattatagaacgaagtaaaaagtttattaatggcgcatctacaatggaagatagaccgtctacaagaaactttttcgaactaaatagaggagccattaaatgtgaaggtttagatgtcgaagacttacttagttactatgttgtttatgtcttttctaagcggtaaataataatatggtatgggttctgatatccccgttctaaatgcattaaataattccaatagagcgatttttgttcctataggaccttccaactgtggatactctgtattattaatagatatattaatacttttgtagggtaacagaggttctacgtcttctaaaaataaaagttttataacatctggcctgttcataaataaaaacttggcgattctatatatactcttattatcaaatctagccattgtcttatagatgtgagctactgtaggtgtaccatttgattttctttctaatactatatatttctctcgaagaagttcttgcagatcatctgggaataaaatactactgttgagtaaatcagttattttttttatatcgatattgatggacatttttatagttaaggataataagtatcccaaagtagataacgacgataacgaagtatttatacttttaggaaatcacaatgactttatcagatcaaaattaacaaaattaaaggagcatgtatttttttctgaatatattgtgactccagatacatatggatctttatgcgtcgaattaaatgggtctagttttcagcacggtggtagatatatagaggtggaggaatttatagatgctggaagacaagttagatggtgttctacatccaatcatatatctgaagatatacacactgataaatttgtcatttatgatatttatacgtttgattcgttcaagaataaacgattggtatttgtacaggtacctccatcattaggagatgatagctatttaactaatccgttattgtctccgtattatcgtaattcagtagccagacaaatggtcaatgatatgatttttaatcaagattcatttttaaaatatttattagaacatctgattagaagccactatagagtttctaaacatataacaatagttagatacaaggataccgaagaattaaatctaacaagaatatgttataatagagataagtttaaggcgtttgtattcgcttggtttaacggcgtttcggaaaatgaaaaggtactagatacgtataaaaaggtatctgatttgatataatgaattcagtgactatatcacacgcaccatatactattacttatcacgatgattgggaaccagtaatgagtcaattggtagagttttataacgaagtagccagttggttgctacgcgacgagacgtcgcctattcctgataagttctttatacaattgaaacagccgcttagaaataaacgagtatgtgtgtgtggtatagatccgtatccaaaagatggaactggtgtaccgttcgaatcaccaaattttacaaaaaaatcaattaaggagatagcttcatctatatctagattaaccggagtaattgattataaaggttataaccttaatataatagacggggttataccctggaattattacttaagttgtaaattaggagaaacaaaaagtcacgcgatttactgggataagatttccaagttactgctacagcatataactaaacacgttagtgttctttattgtttgggtaaaacagatttctcgaatatacgggcaaagttagaatccccggtaactaccatagtgggatatcatccagcggccagagaccaccaattcgagaaagatcgatcatttgaaattatcaacgttttactggaattagacaacaagacacctataaattgggctcaagggtttatttattaatgctttagtgaaattttaacttgtgttctaaatggatgcggctattagaggtaatgatgttatctttgttcttaagactataggtgtcccgtcagcatgcagacaaaatgaagatccaagattcgtagaagcatttaaatgcgacgagttagaaagatatattgataataatccagaatgtacactattcgaaagtcttagggatgaggaagcatactctatagtcagaattttcatggatgtagatttagacgcgtgtctagacgaaatagattatttaacggctattcaagattttattatcgaggtgtcaaactgtgtagctagattcgcatttacagaatgcggtgccattcatgaaaatgtaataaaatccatgagatctaatttttcattgactaagtctacaaatagagataaaacaagttttcatattatctttttagacacgtataccactatggatacattgatagctatgaaacgaacactattagaattaagtagatcatctgaaaatccactaacaagatcgatagacactgccgtatataggagaaaaacaactcttcgggttgtaggtactaggaaaaatccaaattgcgacactattcatgtaatgcaaccacctcacgataatatagaagattacctattcacttacgtggatatgaacaacaatagttattacttttctctacaacgacgattggaggatttagttcctgataagttatgggaaccagggtttatttcgttcgaagacgctataaaaagagtttcaaaaatattcattaattctataataaactttaatgatctcgatgaaaataattttacaacggtaccactggtcatagattatgtaacaccttgtgcattatgtaaaaaacgatcgcataaacatccgcatcaactatcgttggaaaatggtgctattagaatttacaaaactggtaatccacatagttgtaaagttaaaattgttccgttggatggtaataaactgtttaatattgcacaaagaattttagacactaactctgttttattaaccgaacgaggagaccatatagtttggattaataattcatggaaatttaacagcgaagaacccttgataacaaaactaattctatcaataagacatcaactacctaaggaatattcaagcgaattactctgtccgaggaaacgaaagactgtagaagctaacatacgagacatgttagtagattcagtagagaccgatacctatccggataaacttccgtttaaaaatggtgtattggacctggtagacggaatgttttactctggagatgatgctaaaaaatatacgtgtactgtatcgaccggatttaaatttgacgatacaaaattcgtcgaagacagtccagaaatggaagagttaatgaatatcattaacgatatccaaccattaacggatgaaaataagaaaaatagagagctgtatgaaaaaacattatctagttgtttatgtggtgctaccaaaggatgtttaacattcttttttggagaaaccgcaactgggaagtcgacaaccaaacgtttgttaaagtctgctatcggtgacctgtttgtcgagacgggtcaaacaattttaacagatgtattggataaaggacctaatccatttatcgctaatatgcatttaaaaagatctgtattctgtagcgaactacctgattttgcatgtagtggatcaaagaaaattagatctgataatattaaaaagttgacagaaccttgtgtcattggaagaccgtgtttctccaataaaattaataatagaaaccatgcgacaatcattatcgatactaattacaaacctgtctttgatangatagataacgcattaatgagaagaattgccgtcgtgcgattcagaacacacttttctcaaccttctggtagagaggctgctgaaaataatgacgcgtacgataaagtcaaactattagacgagggattagatggtaaaatacagaataatagatatagattcgcatttctatacttgttggttaaatggtacaaaaaatatcatattcctattatgaaactatatcctacaccggaagagattccggactttgcattctatctcaaaataggtactctgttggtatctagctctgtaaagcatattccattaatgacggacctctccaaaaagggatatatattgtacgataatgtggttactcttccgttgactactttccaacagaaaatatccaagtattttaattctagactatttggacacgatatagagagcttcatcaatagacataagaaatttgccaatgttagtgatgaatatctgcaatatatattcatagaggatatttcatctccgtaaatatatgccatatatttatagaatatatcacatatctaaatgaataccggaatcatagatttatttgataatcatgttgatagtataccaactatattacctcatcagttagctactttagattatctagttagaactatcatagatgagaacagaagcgtgttattgttccatattatgggatcgggtaaaacaataatcgctttgttgttcgccttggtagcttccagatttaaaaaggtttacattttagtaccgaacatcaacatcttaaaattttcaattataatatggcagtgtagctatgaacttgtttaatgacgaattcatagctgagaatatctttattcattccacaacaagtttttattctcttaattataacgataacgtcattaattataacggattaagtcgctacaataactctatttttatcgttgatgaggcacataatatttttgggaataatactggagaacttatgaccgtgataaaaaataaaaacaagattccttttctactattgtctggatctcccattactaacacacctaatacgctgggtcatattatagatttaatgtccgaagagacgatagattttggtgagattattagtcgtggtaagaaagtaattcagacacttcttaacgaacgcggagtgaatgtactcaaggatttgcttaaaggaagaatatcatattacgaaatgccggacaaagatctaccaacaataagatatcacggacgtaaatttctagatactcgagtagtatattgtcacatgtctaaacttcaagagaaagattatatgattactagacggcagctatgttatcatgaaatgtttgataaaaatatgtataacgtgtcaatggcagtattgggacaacttaatctgatgaataatttagatacgttatttcaggaacaggataaggaattgtacccaaatctgaaaataaataatggagtgttatacggtgaagaattggtaacgttaaacattagttccaaatttaagtactttatcaatcggatacagacactcaagggaaaacactttatatacttctctaattctacatatggtggattggtaattaaatatatcatgctcagtaatggatattctgaatataatggttctcagggaactaatccacatatgataaacggcaaaccaaaaacatttgctatcgttactagtaaaatgaaatcgtctttagaggatctattagatgtgtataattctcctgaaaacgatgatggcaatcaattgatgtttttgttttcgtcaaacattatgtctgaatcctatactctgaaagaggtaaggcatatttggtttatgactatcccggatactttttctcaatacaaccaaattcttggacgatctattagaaaattctcttacgtcgatatttctgaacccgttaatgtatatcttttagcagccgtatattcagatttcaatgacgaagtgacgtcattaaacgattatacacaggatgaattgattaatgttttaccctttgacatcaaaaagctgttgtatctaaaatttaagactaaagaaacgaatagaatatactctattcttcaagagatgtctgaaacgtattctcttccaccacatccatcaattgtaaaagttttattgggagaattggtcagacaatttttttataataattctcgtattaagtataacgactccaagttacttaaaatggttacatcagttataaaaaataaagaagacgctaggaattacatagatgatattgtaaacggtcacttctttgtatcgaataaagtatttgataaatctcttttatacaaatacgaaaacgatattattacagtaccgtttagactttcctacgaaccatttgtttggggagttaactttcgtaaagaatataatgtggtatcttctccataaaactgatgagatatataaagaaataaatgtcgagctttgttaccaatggatatcttccagttacattggaaccacatgagttgacgttagacataaaaactaatattaggaatgccgtatataaggcgtatctccatagagaaattagtggtaaaatggccaagaaaatagaaattcgtgaagacgtggaattacctctcggtgaaatagttaataattctgtagttataaacgttccgtgtgtaataacctacgcatattatcacgttggggatatagtcagaggaacattaaacatcgaagatgaatcaaatgtaactattcaatgtggagatttaatctgtaaactaagtagagattcgggtactgtatcatttagcgattcaaagtactgcttttttcgaaatggtaatgcgtatgataacggcatcgaagtctccgccgttctaatggaggctcaacaaggtaccgaatctagttttgtttttctcgcgaatatcgttgactcataagaaagagaatagcggtgagtataaatacgaatactatggcaataattgcgaatgttttattcccttcgatatatttttgataatatgaaaaacatgcctctctcaaatcagacaaccatttcataaaatagttctctcgcactggtgaggtggttgcagctcgtataatctccccagaataatatacttgcgtgtcgtcgttcaatttatacggatttctataattctctgttatataatgaggtttaccctcatgattagacgacgacaatagtgttctgaatttagatagttgatcagaatgaatgtttattggtgttggaaaaattatccatgctgcgtctgcagagtggttgatagttgttcctagatatgtaaaataatccaacgtactaggtagcaaattgtctagataaaatactgaatcaaatggcgcagacatattagcggatctaatggaatccaattgattgactatcttttgaaaatatacatttttatgatctgatacttgtaagaatatagcaataatgataattccatcatcgtgtttttttgcctcttcataagaactatattttttcttattccaatgaaccagattaatctctccagagtatttgtatacatctatcaagtgattggatccataatcgtcttcctttccccaatatatatgtattgttgataacacatattcattggggagaaaccctccacttatatatcctcctttaaaattaatccttactagttttccagtattctggatagtggttggtttcgactcattataatgtatgtctaacgtcttcaatcgcgcgtcagaaattgcttttttagtttctatattaataggagatagttgttgaggcatagtaaaaatgaaatgataactgtctagaaatagctcttagtatgggatttacaatggatgaggaagtgatatttgaaactcctagagaattaatatctattaaacgaataaaagatattccaagatcaaaagacacgcacgtgtttgctgcgtgtataacaagtgacggatatccgttaataggagctagaagaacttcattcgcattccaggcgatattatctcaacaaaattcagattctatctttagagtatccactaaactattacggtttatgtactacaatgaactaagagaaatctttagacggttgagaaaaggttctatcaacaatatcgatcctcacttcgaagagttaatattattgggtggtaaactagataaaaaggaatctattaaagattgtttaagaagagaattaaaagaggaaagtgatgaacatataacagtaaaagaattcggaaatgtaattctaaaacttacaacgagtgataaattatttaataaagtatatataggttattgcatggcatgttttattaatcaatcgttggaggatttatcacatactagtatttacaatgtagaaattagaaagattaaatcgttaaatgattgtattaacgacgataaatacgaatatctgtcttatatttataatatactaattaatagtaaatgagcttttacagatctagtataattagtcagattattaagtataatagacgactagctaagtctattatttgcgaggatgactctcaaattattacactcacggcattcgttaaccaatgcctatggtgtcataaacgagtatccgtgtccgctattttattaactactgataacaaaatattagtatgtaacagacgagatagttttctctattctgaaataattagaactagaaacatgtatagaaagaaacgattatttctgaattattccaattatttgaacaaacaggaaagaagtatactatcgtcatttttttctctagatccagctactgctgataatgatagaataaacgctatttatccgggtggtatacccaaaaggggtgagaacgttccagagtgtttatccagggaaatcaaagaagaagttaatatagacaattcttttgtattcatagacactcgtttttttattcatggtatcatagaagataccattattaacaaattttttgaggtaattttctttgttggacgaatatctctaacgagtgatcaaattattgatacctttaaaagtaatcatgaaataaaggatctaatatttttagatccaaattcaggtaatggactccaatacgaaattgcaaaatatgctctagatactgcaaaacttaaatgttacggtcatagaggatgttattatgaatcattaaaaaaattaactgaggatgattgattagaaaatataaattaatttaccatcgtgtatttttataacgggattgtctggcatatcatgtagatagttaccgtctacatcgtatactctaccatctacgcctttaaatcctctatttattgatattaatctattagaattggaataccaaatattagtaccctcaattagtttattggtaatattttttttagacgatagatcgatggctcttgaaaccaaggttttccaaccggactcattgtctatcggtgagaagtctttttcattagcatgaatccattctaatgatgtatgtttaaacactctaaacaattgtacaaattcttttgatttgttttgaatgatttcaaataggtcttcgtctacagtaggcataccattagataatctagccattataaagtgcacgtttacatatctacgttctggaggagtaagaacgtgactattgagacgaatggctcttcctactatctgacgaagagacgcctcgttccatgtcatatctaaaatgaagatatcattgattgagaagaaactaataccctcgcctccgctagaagagaatacgcatgttttaatgcattctccgttagtgtttgattcttggttaaactcagccaccgccttgattctagtatcttttgttctagatgagaactctatattagagataccaaagactttgaaatatagtaataagatttctattcctgactgattaacaaatggttcaaagactagacatttaccatgggatgctaatattcccaaacatacatctataaatttgacgcttttctcttttaattcagtaaatagagagatatcagccgcaatagcatcccctcccaatagttctccctttttaaaggtgtctaatgcggatttagaaaattctctatctcttaatgaatttttaaaatcattatatagggttgctatctcttgtgcgtattctcccggatcacgattttgtctttcaggaaagctatcgaatgtaaacgtagtagccatacgtctcagaattctaaatgatgatataccagtttttatttctgcgagtttagccttttgataaatctcttcttgcttttttgacatattaacgtatcgcattaatactgttttcttagcgaatgatgcagacccttccacatcatcaaaaatagaaaactcgttattaactatgtacgaacataggcctcctagtttggagactaattctttttcatcgactagacgtttattctcaaatagcgattggtgttgtaaggatcctggtcgcagtaagttaaccaacatggtgaattcttgcacactattaacgataggtgtagccgataaacaaatcatcttatggttttttaacgcaatggtcttagataaaaaattatatactgaccgagtaggacggatcttaccatcttctttgattaatgatttagaaatgaagttatgacattcatcaatgatgacgcatattctactcttggaattaatagttttgatattagtaaaaaatttatttctaaaattttgatcatcgtaattaataaaaatacaatccttcgttatctctggagcgtatctgagtatagtgtttatccaaggatcttctatcaaagccttttttaccaataagataattgcccaattcgtataaatatccttaagatgtttgagaatatatacagtagtcattgttttaccgacacctgtttcatggaacaataaaagagaatgcatactgtctaatcctaagaaaactcttgctacaaaatgttgataatccttgaggcgtactacgtctgaccccatcatttcaacgggcatattagtagttctgcgtaaggcataatcgatataggccgcgtgtgatttactcatttatgagtgataagtaataactatgttttaaaaatcacagcagtagtttaactagccttctctgatgtttgttttcgatactttttgaatcagaagtcatactagaataaagcagcgagtgaacgtaatagagagcttcgtatactctattcgaaaactctaagaacttattaatgaattccgtatccactggatcgtttaaaatactaaattgaacagtgttcacatccttccaagacgaagacttagtgacggacttaacatgagacataaataaatccaaattttttttataaacatcactagccaccataatggcgctatctttcaaccaactatcgcttacgcattttaacagtctaacatttttaaagagactacaatatattctcatagtatcgattacacctctaccgaatagagtgggaagtttaataatacaatatttttcgtttacaaaatcaaataatggtcgaaacacgtcgaaggttaacatcttataatcgctaatgtatagattgttttcagtgagatgattattagatttaatagcatctcgttcacgtttgaacagtttattgcgtgcgctgaggtcggcaactacggcatccgctctagtactcctcccataatactttacgctattaatctttaaaatttcatagactttatctagatcgctttctggtaacatgatatcatgtgtaaaaagttttaacatgtcggtcggcattctatttagatcattaactctagaaatctgaagaaagtaattagctccatattccagactaggtaatgggcttttacctaaagacaagttaagttctggcaatgtttcataaaatggaagaaggacatgtgtcccctcccggatattttttacaatttcatccatttacaactctatagtttgttttcattattattagttattatctcccataatcttggtaatacttacaccttgatcataagataccttatacaggtcattacatacaactaccaattgtttttgtacataatagattggatgattgatatccatggtggaataaactactcgaacagatagtttatctttccccctagatacattggccgtaatagttgtcggcctaaagaatatctttggtgtaaagttaaaagttagggttcttgttccattattgctttttgtcagtagttcgttataaattctcgagatgggcccgttctctgaatatagaacatcatttccaaatctaacttctagtctagaaataatatcggtcttatttttaaaatctattcccttgatgaatggatcgttaataaacaaatccttggcctttgattcggctgatctattatctccgttatagacgttacgttgactagtccaaagacttacaggaatagatgtatcgatgatgttgatagtatgtgatatgtgagcaaagactgttctcttggtggcgtcgctatatgttccagtaatggcggaaaactttttagaaatgttatatataaaagaattttttcgggttccaaacattaacagattagtatgaagataaacactcatattatcaggaacattatcaatttttacataaacatcggcatcttgaatagaaacaacaccatcttctggaacctctacgatctcggcagattccggataaccagtcggtggaccatcactaacaataactagatcatccaacaatctactcacatatgcgtctatataatctttttcatcttgtgagtaccctggatacgaaataaatttgttatcagtatttccataataaggtttagtataaacagagagagatgttgctgcatgaacttcggttactgtcgccgttggttggtttatttgacctattactctcctaggtttctctataaatgatggtttaatttgtacattcttaaccatatatccaataaagctcaattcaggaacataaacaaattctttgttgaacgtttcaaagtcgaacgaagagtcacgaataacgatatcggatactggattgaaggtcaccgttacggtaatttttgaatcggatagtttaagactactgaatgtatcttccacatcaaacggagttttaatataaacgtatactgtagatggttctttaatagtgtcattaggagttaggccaatagaaatatcattaagttcactagaatatccagaatgtttcaaagcaattgtattattgatacaattattatataattcttcgccctcaatttcccaaataacaccgttacacgaagagacagatacatgattaatacatttatatccaacatatggcacgtaaccgaatcttcccatacctttaacttctggaagttccaaactcagaaccaaatgattaagcgcagtaatatactgatccctaatttcgaagctagcgatagcctgattgtctggcccatcgtttgtcataactccggatagagaaatatattgcggcatatataaagttggaatttgactatcaactgcgaagacattagaccgtttaataaagtcatccccaccgatcaaagaattaatgatagtattattcattttctatttaaaatggaaaaagcttacaataaactccgtagagaaatatctataatttgtgagttttccttaaagtaacagcttccgtaaacaccgtctttatctcttagtaagtttattgtatttatgaccttttccttatcttcatagaatactaaaggcaataaagaaatttttggttcttctctaagagctacgtgagacttaaccatagacgccaacgaatccctacatattttagaacagaaatacccaacttcaccacccttgaatgtctcaatactaataggtctaaaaaccaaatcttgattacaaaaccaacacttatcaattacactatttgtcttaatagacatatctgccatagatttataatactttggtagtatacaagcgagtgcttcttctttagcgggcttaaagactgctttaggtgctgaaataaccacatctggaagacttactcgcttagccatttaattacggaactatttttttatacttctaatgaacaagtagaaaacctctcatctacaaaaacatactcgtgtccataatcctctaccatagtaacacgttttttagatctcatatgtgctaaaaagttttcccatactaattggttactattatttttcgtataatttttaacagtttgaggttttagatttttagttacagaagtgatatcgaatattttatccaaaaagaatgagtaattaattgtcttagaaggagtgttttcttggcaaaagaataccaagtgcttaaatatttctactacttcattaatcttttctgtactcagattcagtttctcatcttttacttgattgattatttcaaagactaacttataatcctttttatttattctctcgttagccttaagaaaactagatacaaaatttgcatctacatcatccgtggatatttgatttttttccatgatatccaatagttccgagataatttctccagaacattgatgagacaataatctccgcaatacatttctcaaatgaataagtttattagacacgtggaagtttgactttttttgtacctttgtacatttttgaaatacagactcgcaaaaaatacaatattcatatccttgttcagatactataccgttatgtctacaacagctacataatcgtagattcatgttaacactctacgtatctcgtcgtccaatattttatataaaaacattttatttctagacgttgtcagaaaatcctgtaatatttttagtttttttggttgtgaataaagtatcgccctaataatattggtaccgtcttccgacaatatagtagttaaattatccgagcatgtagaagaacaccgcttaggcggattcagtacaatgttatatttttcgtaccaactcatttaaatatcataatctaaaatagttctgtaatatgtctagcgctaatatattgatcataatcctgtgcataaattaagatacaacaatgtcttgaaatcatcgacatggcttcttccatagttagaagatcatcgtcaaagttagcaacgtgattcatcaacatttgctgttttgaggcagcaaatactgaaccatcaccattcaaccattcataaaaaccatcgtctgaatccattgataatttcttgtactggtttttgagagctcgcatcaatctagcatttctagctcccggattgaaaacagaaagaggatcgtacatccaaggtccattttctgtaaatagaatcgtataatgtcccttcaagaagatatcagacgatccacaatcaaagaattggtctccgagtttgtaacagactgcggactttaacctatacatgataccgtttagcatgatttctggtgatacgtcaatcggagtatcatctattagagatctaaagccggtgtaacattctccgccaaacatattcttattctgacgtcgttctacataaaacatcattgctccattaacgataacaggtgaatgaacagcactacccatcacattagttcccaatggatcaatgtgtgtaactccagaacatcttccatagcctatgttaggaggagcgaacaccactcttccactattgccatcgaatgccatagaataaatatccttggaattgatagaaatcggactgtcggatgttgttatcatcttcataggattaacaacgatgtatggtgcagcctgaagtttcatatcgtaactgatgccgttcataggtctagccacagaaaccaacgtaggtctaaatccaactatagacaaaatagaagctaatatctgttcctcatctgtcataacttgagagcatccagtatgaataatcttcattagatggggatctaccgcatcatcatcgttacaataaaaaattcccattctaatgttcataattgcttttctaatcatggtatgaatgtttgctctctgaatctctgtggaaattagatctgatacacctgtaatcactatcggattatcctccgtaagacgattaaccaacaacatataattataagactttactcttctaaattcataaagttgctggattagactatatgtgtctccatgtacatacgcgttctcgagcgcaggaagtttaataccgaatagtgccatcagaataggatgaatgtagtaattagtttctggttttctataaataaaagacaaatcttgtgaactagacatatcggtaaaatgcatggattggaatcgtgtagtcgacagaagaatatgatgattagatggagagtatattttatctaactctttgagttggtcaccgattctaggactagctcgagaatgaataagtactaagggatgagtacatttcacagaaacactggcgttgttcaacgtactctttacatgggaaaggagttgaaatagctcgtttctatttgtcctgacaatatttagtttattcataatattaagcatatcctgaatagtaaagttagatgtgtcatacttgttagtagttagatatttagcaattgcattcccatcatttctcaatctcgtactccaatcatgtgtggatgctacttcgtcgatggaaaccatacaatcctttttgataggctgttgagattgatcatttcctgtacgtttaggtttggtacgttgatttctagcccctgctgatataaagtcatcgtctacaatttgggataatgaattacatacactacaagacaaagatttatcagaagtgtgaatatgatcttcatctaccaaagaaagagtttgattagtataactagattttagtcccgcgttagatgttaaaaaaacatcgctattgaccacggcttccattatttatattcgtagtttttactcgaaagcgtgattttaatatccaatcttattacttttggaatcgttcaaaacctttgactagttgtagaatttgatctattgccctacgcgtatactcccttgcatcatatacgttcgtcaccagatcgtttgtttcggcctgaagttgacgcatatctttttcaacactcgacatgagatccttaagggtcatatcgtctagattttgttgagatgctgctcctggatttggattttgttgtgctgttgtacatactgtaccaccagtaggtgtaggagtacatacagtggccacaataggaggttgaagaggtgtaaccgttggagtagtacaagaaatacttccatccgattgttgtgtacatgtggttgttggtaacgtctgagaaggttgggtagatggcggtgtcgtcatcttttgatctttattaaatttagagataatatcctgaacagtattgctcggcgtcaacgctggaaggagtgtactcgccggcgcatcagtatctgtagacaaccaatcaaaaagattagacatatcagatgatgtattagtttgttgacgtggttttagtacaggagcagtactactaggtagaagaataggagccggtgtaggtgtcggaaccggctgtggagttatatgaatagttggttgtagcggttgggtaggctgtctgctggcggtcatcatattatctctagctagttgttctcgcaactgtctttgataatacgactcttgagactttagtcctatttcaatcgcttcatcctttttcgtatccggatccttttcttcagaataatagattgacgactttggtgtagaggattctgccagcccctgtgagaacttgttaaagaagtccatttaaggctttaaaattgaattgcgattataagattaaatggcagacacggacgatattatcgactatgaatccgatgatctcaccgaatacgaggatgatgaagaagatggagagtcactagaaactagtgatatagatcccaaatcttcttataagattgtagaatcaacatccactcatatagaagatgcgcattccaatcttaaacatatagggaatcatatatctgctcttaaacgacgctatactagacgtataagtctatttgaaatagcgggtataatagcagaaagctataacttgcttcaacgaggaagattacctctagtttcagaattttctgacgaaacgatgaagcaaaatatgctacatgtaattatacaagagatagaggagggttcttgtcctatagtcatcgaaaagaacggagaattgttgtcggtaaacgattttgataaagatggtctaaaattccatctagactatattatcaaaatttggaaacttcaaaaacgatattagaatttatacgaatatcgttctctaaatgtcacaatcaagtctctcatattcagcagtttattgtcgtactttatatcgtgttcattaacgatattttgcaaaatagtaatgattctatcttccttcgatagatattcttcagagattattgtcttatattctttcttgttatccgatatgaatttgataagactttgaacattattaatacccgtctgtttaattttttctatagatattttagttttggtagattctatggtgtctgttaataggcatccaacatcgacattcgacgtcaattgtctataaatcagagtataaattttagaaataacattagcaaattgttgtgcgttgatgtcgttattctgaaacagtatgattttaggtagcattttcttaacaaagagaacgtatttattgttactcagttgaacagatgatatatccagattactaacgcatctgattccatataccaaactttcagaagaaatggtgtacaattgtttgtattcattcaatgtctccttttcagaaattagtttagagtcgaatactgcaataattttcaagagatagttttcatcagataagattttatttagtgtagatatgataaaactattgttttgttggagaacttgatacgccgcattctctgtagtcgacgctctcaaatgggaaacaatctctattatttttttggaatcggatactatatcttcggtatcttgacgcagtctagtatacatagagttaagagaaattagagtttgtacattaagcaacatgtctctaaatgtggctacaaacttttctttttccacatcatctagtttattatataccgatttcacaacggcaccagatttaaggaaccagaatgaaaaactctgataactacaatatttcatcatagttacgattttatcatcttctatagttggtgtgataacacatacctttttctccaagactggaaccaacgtcataaaaatgtttaaatcaaaatccatatcaacatctgatgcgctaagaccagtctcgcgttcaagattatctttactaatggtgacgaactcatcgtatagaactctaagtttgtccattatttatttacagatttagttgtttaatttatttgtgctcttccagagttgggatagtatttttctaacgtcggtattatattattaggatctacgttcatatgtatcataatattaatcatccacgttttgataaatctatctttagcttctgaaataacgtatttaaacaaaggagaaaaatatttagttacggcatcagacgcgataacattttttgtaaatgtaacgtatttagacgacagatcttcgttaaaaagttttccatctatgtagaatccatcggttgttaacaccattcccgcgtcagagtgaataggagtttgaatagtttgttttggaaatagatccttcaataacttatagttgggtgggaaaaaatcgattttatcactagactctttcttttttactatcattacctcatgaactatttcttgaatgagtatatgtattttctttcctatatcggtcgcgttcattggaaaatatatcatgtcgttaactataagaatatttttatcctcgtttacaaactgaataatatcagatatagttcgtaaacgaactatatcatcaccagcacaacatctaactatatgatatccactagtttcctttagccgtttattatcttgttccatattagcagtcattccatcatttaagaaggcgtcaaagataatagggagaaatgacattttggattctgttacgactttaccaaaattaaggatatacggacttactatctttttctcaacgtcgatttgatgaacacacgatgaaaatgtacttcgatgagattgatcatgtagaaaacaacaagggatacaatatttccgcatatcatgaaatatattaagaaatcccactttattatatttccccaaaggatcaatgcatgtaaacattataccgttatcattaataaagacttctttctcatcggatctgtaaaagttgttactgatttttttcattccaggatctagataattaataataatgggttttctattcttattctttgtattttgacatatcctagaccagtaaacagtttccactttggtaaaatcagaagacttttgaacgctattaaacatggcattaatggcaataactaaaaatgtaaaatatttttctatgttaggaatatggtttttcactttaatagatatatggttttttgccaaaatgatagatatttttttatccgatgatagtaaaatattattagtcgccgtctctataaaaatgaagctagtctcgatatccaattttattctagaattgataggagtcgccaaatgtaccttatacgttatatctcccttgatgcgttccatttgtgtatctatatcggacacaagatctgtaaatagttttacgttattaatcatcacggtatcgccatcgctagataatgctaatgtactatccaagtcccaaatggagagatttaactgttcatcgtttagaataaaatgattacctgtcatattaataaagtgttcatcgtatctagataacaacgacttataattaatgtccaagtcttgaactcgctgaatgatcttttttaacccagttagttttagattggtacgaaatatattgttaaactttgattctacagtaatgtccaaatctagttgtggaaatacttccatcaacattgtttcaaacttgataatattattatctacatcttcgtacgatccaaattccggaatagatgtatcgcacgctctggccacccagataaccaaaaagtcacacgctccagaatatacattgtataaaaagctatcgttttttagtagtgtttttttctgagtatatacgaaaggattaaaaatagtattatcaacgtaactatattccaaattattcttatgagaatagataataatatcgtccttaatatctaacaaatttcctaaatatccctttaattgagtcattcgaagcgttaataaaatatgtctcttaactatttccggccgttgtatatttaaatgacttcgtaagaaataatatataggcgacttctcatctatgtaatcatatggagtgagatatagggctcgttctacctcctgccccttacccacctgtaataccaattgcggactcactatatatcgcatatttatatcgtggggtaaagtgaaaatctactaccgatgatgtaagtcttacaatgttcgaaccagtaccagatcttaatttggaggcctccgtagaactaggggaggtaaatatagatcaaacaacacctatgataaaggagaatagcggttttatatcccgtagtagacgtctattcgcccatagatctaaggatgatgagagaaaactagcactacgattctttttacaaagactttattttttagatcatagagagattcattatttgttcagatgcgtcgacgctgtaaaagacgtcactattaccaaaaaaaataacattatcgtggcgccttatatagcacttttaactatcgcatcaaaaggatgcaaacttacagaaacaatgattgaagcattctttccagaactatataatgaacatagtaagaaattcaaattcaactctcaagtatccatcatccaagaaaaactcggataccagtctggaaactatcacgtttatgattttgaaccgtattactctacagtagctctggctattcgagatgaacattcatctggcatttttaatatccgtcaagagagttatcttgtaagttcattatctgaaataacatatagattttatctaattaatctaaaatctgatcttgttcaatggagtgctagtacgggcgctgtaattaatcaaatggtaaatactgtattgattacagtgtatgaaaaattacaactggccatagaaaatgattcacaatttacatgttcattggctgtggaatcagaacttccaataaaattacttaaagatagaaatgaattatttacaaaattcattaacgagttaaaaaagaccagttcattcaagataagcaaacgtgataaggatacgctattaaaacattttacttatgactggagttagaatttatagacgacacatttcgtttatcattgttactattactatcattattagtattcttcttgtcatcttgttcagaaatatacagcaatgctatacctaatactaaatacattatcatgcttgcaatggctctaacaacaacgaaccaaaatgaatttggtcgtagcttttgttcacaaaaatacataaagaaatgtctacataaatctatggcgccattggctacttgaaatagcgccagtcctcctacagattttaatatagctgtataacatgacatttattcatcatcaaaagagacagagtcaccatctgtcatatttagattttttttcatgtgttcaaagtatcctctactcatttcattataatagtttatcatgcttagaattttaggacggatcaatgagtaagacttgactagatcgtcagtagtaatttgtgcatcatctattctgcatccgcttcgtcgaataatgtatagcatcgctttgagattctccatagctatcaagtctttatataatgacatggaaatatctgtgaatgctttatacttctccaacatcgatgccttaacatcatcacatactttagcattgaaaatacgttctattgtgtagatggatgtagcaagatttttaaacaacaatgccatcttacatgatgattgtctcaagtctccaatcgtttgtttagaacgattagctacagagtccaatgcttggctaactagcatattattatctttagaaattgtattcttcaatgaggcgtttatcatatctgtgatttcgttagtcatattacagtctgactgggttgtaatgttatccaacatatcacctatggatacggtacacgtaccagcatttgtaataatcctatctaagatgttgtatggcattgcgcagaaaatatcttctcctgtaatatctccactctcgataaatctactcagattattcttaaatgccttattctctggagaaaagatatcagtgtccatcatttcattaatagtatacgcagaaaagataccacgagtatcaattctatccaagatacttatcggttccgagtcacagataattgtttcctctccttcgggagatcctgcatagaaatatctaggacaatagtttctatactgtctgtaactctgataatctctaaagtcactaactgataccatgaaattgagaagatcaaacgctgaagtaatcaatttttctgcctcgtttttactacaactagttttcatcaatgtagtgacgatgtattgtttagttactcttggtctaatactgatgatagagatattattgcttcccataatggatcttctagtagtcaccttaaagcccattgatgcgaatagcagatagataaagtcttggtatgactcctttctaatatagtacggactacctttgtcacccaactttatacccacataagccataacaacctctttaatagccgtttcatgaggtttatcagccatgagcctgagtagttgaaagaatcgcatgaatcccgtctcagaaagtcctatatgcatgatagatttatctttcctgggaaactctcgtatagttatagatgaaatactcttcaaagtttctgaaataagattagtaacagtcttacctccgactactctgggtaacaaacatactctaataggtgttttctctgcggagataatatcagaaaggatagagcaataagtagtattattgtgattataaagaccgaatacataacaggtagaatttataaacatcatgtcctgaagggttttagacttgtattcctcgtaatctataccgtcccaaaacatggatttggtaactttgatagccgtagatctttgttccttcgctaacaggttaaagaaattaataaagaatttgttgtttctatttatgtccacaaattgcacgtttggaagcgccacggttacattcactgcagcattttgaggatcgcgagtatgaagtacgatgttattgtttactggtatatctggaaagaaatctaccagtctaggaataagagattgatatcgcatagaaatagtaaagtttataatctcatcattgaagattactctgttaccattgtaataaattggtactctatcataatcatcgacaaagtactgttcatacatgatgagatgtttatatgttggcatagtagtgagatcgacgtttggtaatggcaatgtattaagattaactccataatgtctagcagcatctgcgatgttataagtgatgtcaaagcggggttgatcttgtgctgttatatattgtctaacacctataagattatcaaaatcttgtctgcttaatacaccgttaacaatttttgccttgaattcttttattggtgcattaataacatccttatagaggatgttaaacaaataagtattatcaaagttaagatctgggtatttcttttctgctagaacatccattgagtcggagccatctggtttaatataaccaccgataaatctagctctgtattctgtatccgtcaatctaatattaagaaggtgttgagtgaaaggtggaagatcgtaaaagctgtgagtattaataatagggttagtttccgaactaatgttaattggatgattaataatatttatatttccagcgttaagtgtaacattaaacagttttaattcacgtgacgtggtatcaattaaataattaatgcccaatttggatatagtagcctgaagctcatcttgtttagttacggatcctaatgagttattaagaaatacatcgaacggatgaacgaaggttgttttaagttggtcacatactttgtaatctagacatagatgtggaagaacggtagaaactatacgaaatagatattcagagtcctctaattgatcaagagtaactattgacttaataggcatcatttatttagtattaaatgacgaccgtaccagtgacagatatacaaaacgacttaattacagagttttcagaagataattatccatctaacaaaaattatgaaataactcttcgccaaatgtctattctaactcacgttaacaacgtggtagatagagaacataatgccgccgtagtgtcatctccagaggaaatatcatcacaacttaatgaagatctatttccagatgatgattcaccggccactattatcgaacgagtacaacctcatactactattattgacgatacgccacctcctacttttcgtagagagttattgatatcggaacaacgtcaacaacgagaaaaaagatttaatattacagtatcaaaaaattctgaagcaataatggaatctagatctatgataacttctatgccaacacaaacaccatccttgggagtagtttatgataaagataaaagaattcagatgctagaggatgaagtggttaatcttagaaatcaacgatctaatacaaaatcatctgataatttagataattttaccagaatactatttggtaagactccgtataaatcaaccgaagttaataagcgtatagccatcgttaattatgcaaatttgaacgggtcccccttatcagtcgaggacttggatgtctgttcggaggatgaaatagatagaatctataaaacgattaaacaatatcacgaaagtagaaaacgaaaaattatcgtcactaacgtgattattattgtcataaacattattgagcaggcattgctaaaactcggatttgaagaaatcaaaggactgagtaccgatatcacttcagaaattatcgatgtggagatcggagatgactgcgatgctgtagcatcaaaactaggaatcggtaacagtccggttcttaatattgtattgtttatactcaagatattcgttaaacgaattaaaattatttaatttaatacattcccatatccagacaacaatcgtctggattaatctgttcctgtcgtctcataccggacgacatattaatctttttattagtgggcatctttttagatggtttctttttcccagcattaactgattcgatacctagaagatcgtgattgatttctccgaccattccacgaacttctaattggccgtctctaacggtaccataaactattttaccagcattagtaacagcttggacaatctgaccatccattgcgttgaatgatgtagttgctgttgttctacgtctaggagcaccagaggtatttttagagctcttggatgttgatgtagaagacgaggattttgattttggtttacatgtaatacattttgaactctttgattttgtatcacatgcaccggcagtcacatctgtttgagaattaagattattgttgcctcctttgacggctgcatctccaccgatctgcgctagtagatttttaagctgtggtgtaatcttattaactgtttcaatataatcatcgtaactacttctaacggctaaattttttttatccgccatttagaagctaaaaatatttttatttatgcagaagatttaactagattatacaatgaactaatatgatccttttctagattatttacgaacttggtatttcttgtttctggaggaggagaatttaaattcggacttggattcggattttgtgggttcttgatcttattatacagcgtgtataggatggtgacggtaactgctacacaaataccgatcaacagaagaataccaatcatttattgacaataacttcactatgatcaagtatgtaataatcatcttttcactaagtaagtagtaataatgattcaacaatgacacgatatatggacgataataatttagttcatggaaatatcgctatgattggtgtgaatgactccgctaactctgtggggtgcacagtgctttccccacatagaataaattagcattccgactgtgataataataccaagtataaacgccataatactcaatactttccatgtacgagtgggactggtagacttactaaagtcaataaaggcgaagatacacgaaagaatcaaaagaatgattccagcgattagcacgccagaaaaataatttccaatcataagcatcatgtccatttaactaataaaaattttaaatcgccgaataaacaaagtggaatataaaccatataaaaacaatagtttgtactgcaaaaataatatctatttttgttttcgaagatatggtaaaattaaatagtagtacacagcatgttataactaacagcagcaacggctcgtaattacttatcatttactagacgaaaaggtggtgggatattttcttgctcaaataatacgaatatatcacccatccattttatacgatgtttatatactctaatctttaatagatctatagatgacgggtttaccaataatatagattttatcgattcatctaatttaaacccttccttaaacgtgaatgatctattatctggcataatgatgaccctacctgatgaatctgacaatgtactgggccatgtagaataaattatcaacgaattatcgtctacgaacatttatatcatttgttttaattttaggacgtgaataaatagatataaaatagaaaataacagatattacaaccagtgttatggacgcacccaaccatgtaggcagttttattttatcgtttactacaggttctcctggatgtacgtcaccaactgcagacgtagttctagtacaattagacgtaagttccgcttgggaattttttaacgctaaagagttaacgttgatcgtacacccaacgtatttacatctagttctttgaacatcttgattataatataaccattttctatctctagattcgtcagtgcactcatgtaaccaacataccctaggtcctaaatatttatctccggaattagattttggataattcgcgcaccaacaatttctatttcctttatggtcgttacaaaagacgtataatgccgtatccccaaaagtaaaataatcaggacgaataattctaataaactcagaacaatatctcgcatccatatgtttggagcaaatatcggaataagtagacatagccggtttccgttttacacgtaaccattctaaacaattggggtttccaggatcgtttctacaaaaaccagtcatgaaatcgtcacaatgttctgtcttgtaattattattaaatatttttggacagtgtttggtatttgtcttagaacaacattttgccacgctatcactatcacccaggagataatccttttttataaaatgacatcgttgcccggatgctatataatcagtagcatattttaaatccttaatatattcaggagttacctcgttctgataatagattaatgatccaggacgaaatttgaaagaactacatggttctccatgaattaatacatattgtttagcaaattcaggaactataaaactactacaatgatctatcgacataccatctatcaaacaaaatttgggtttaatttctcctggagacgtttcataataatacatataactttcttcggcaaacctaacagctctattatattcaggataattaaaatctaataccatatatttgtctcgtatatctgctattcctgtctctattttgattctattaagagtaacagctgcccccattcttaataatcatcagtatttaaactgttaaatgttggtatatcaacatctatcttatttcccgcagtataaggtttgttgcaggtatactgttcaggaatgggtacatttatacttcttttatagtcctgtctttcgatgttcatcacaaatgcaaagaacagaataaacaaaataatgtaagaaataatattaaatatctgtgaattcgtaaatacattgattgccataataattacagcagctacaatacacacaatagacattcccacagtgttgccattacctccacgatacatttgagttactaagcaataggtaataactaagctagtaagaggcaatagaaaagatgagataaatatcatcaatatagagattagaggagggctatatagagccaagacgaacaaaatcaaaccgagtaacgttctaacatcattatttttgaagattcccaaataatcattcattattcctccataatcgttttgcatcatacccccatctttaggcataaacgattgctgctgttcctctgtaaataaatctttatcaagcactccagcacccgcagagaagtcatcaagcatattgtaatatcttaaataactcatttatatattaaaaaatgtcactattaaagatggagtataatctttatgccgaactaaaaaaaatgacttgtggtcagaccataagtctttttaatgaagacggcgatttcgtagaagttgaaccaggatcatcctttaagtttctaatacctaagggattttactcctctccttgtgtaaagacgagtctagtattcaagacattaacaacgaccgataataaaattactagtatcaatccaacaaatgcgccaaagttatatcctcttcaacgcaaagtcgtatctgaagtagtttctaatatgaggaaaatgatcgaattaaaacgtcctctatacatcactcttcacttggcatgtggatttggtaagactattaccacgtgttatcttatgaccacacacggcagaaaaaccatcatttgcgtacccaataaaatgttaatacatcaatggaagacacaggtagaggcagtcggattggaacataagatatctatagatggagttagtagtctattaaaggaactaaagactcaaagtccggatgtattaatcgtagtcagtagacatctgacaaacgatgcattttgtaaatatatcaataagcattatgatttgtttatcttggatgaatcacatacgtataatctgatgaacaatacagcagttacaagatttttagcgtattatcctccgatgatgtgttattttttaactgctacacctagaccagctaaccaaatttattgtaatagtattattaatattgccaagttatccgatctaaaaaaaactatctatatagtagatagtttttttgagccatattccacagacaatattagaaatatggtaaaacgactagatggaccatctaataaatatcatatatataccgagaagttattatctgtagacgagcctagaaaccaacttattcttgataccctggtagaagaattcaagtcaggaactattaatagaattttagttattactaaactacgtgaacatatggtattcttctacaaacgattattagatcttttcggagcagaggttgtatttataggagacgcccaaaatagacgtactccagatatggtcaaatcgattaaggaactaaatagatttatattcgtatccaccttattttattccggcactggtttagatattccgagtttggattctttgttcatttgctcggcagtaatcaacaatatgcaaatagagcaattactagggagggtatgtcgagaaacagaactattagataggacggtatatgtatttcctaacacatccatcaaaaaaataaagtacatgataggaaatttcgtgcaacgaattattagtctgtctgtagataaactcggatttaaacaagaaagttatcagaaacatcaggaatctgaacccgcttccgtaccaacatcctccagagaagaacgtgtattaaatagaatatttaactcgcaaaatcgttaagaagtttaagagacgatccacatgctgagcaggccagtgtattacccctcatagtattaatataatccaatgatacttttgtgatgtcggaaatcttaaccaatttagactgacaggcagaacacgtcatacaatcatcatcgtcatcgataactgtagtcttgggcttctttttgcgactcttcattccggaacgcatattggtgctatccatttaggtagtaaaaaataagtcagaatatgccctataacacgatcgtgcaaaacctggtatatcgtctctatctttatcacaatatagtgtatcaacatctttattattgacctcgtttatcttggaacatggaatgggaacatttttgttaacggccacctttgccttaattccagatgttgtaaaattataactaaacagtctatcatcgacacaaatgaaattcttgtttagacgtttgtagtttacgtatgcggctcgttctcgtctcattttttcagatattgcaggtactataatattaaaaataagaatgaaataacataggattaaaaataaagttatcatgacttctagtgctgatttaactaacttaaaagaattacttagtctgtacaaaagtttgagattttcagattctgtggctatagagaagtataattctttggtagaatggggaacatctacttactggaaaataggcgtacaaaaggtaactaatgtcgagacgtccatatctgattattatgatgaggtaaaaaataaaccgtttaatattgatccggggtattatattttcttaccagtatattttggaagcgtctttatttattcaaagggtaaaaatatggtagaacttggatctggaaactcttttcaaataccggatgagattcgaagtgcgtgtaacaaagtattagatagtgataacggaatagactttctgagatttgttttgttaaacaatagatggataatggaagacgctatatcaaaataccagtctccagttaatatatttaaactagctagtgagtacggattaaacatacccaactatttagaaattgaaatagaggaagacacattatttgacgatgagttatactctattatggaacgctctttcgatgatacatttccaaaaatatctatatcgtatattaagttgggagaacttaagcggcaagttgtagactttttcaaattctcattcatgtatattgagtcaatcaaggtagatcgtataggagataatatttttattcctagcgttataacaaaatcaggaaaaaagatattagtaaaagatgtagaccatttaatacgatccaaggttagagaacatacatttgtaaaagtaaaaaagaaaaacacattttccattttatacgactatgatgggaacggaacagaaactagaggagaagtaataaaacgaattatagacactataggacgagactattatgttaatggaaagtatttctctaaggttggtattgcaggcttaaagcaattgactaataaattagatattaatgagtgtgcaactgtcgatgagttagttgatgagattaataaatccggaactgtaaaacgaaaaataaaaaaccaatcagtatttgatttaagcagagaatgtttgggatatccagaagcggattttataacgttagttaataacatgcggttcaaaatagaaaattgtaaggttgtaaatttcaatattgaaaatactaattgtttaaataacccgagtattgaaactatatatggaaacttcaaccagttcgtctcaatctttaataccgttaccgatgtcaaaaaaagattattcgagtgaaataatatgcgcctttgatataggtgcaaaaaatcctgccagaactgttttagaagtcaaggataactccgttagggtattggatatatcaaaattagactggagttctgattgggaaaggcgcatagctcaagatttgtcacaatatgaatacactacagttcttctagaacgtcagcctagaaggtcaccgtacgtcaaatttatctattttattaaaggctttttatatcatacatctgctgccaaagttatttgcgtctcacctgtcatgtctggtaattcatatagagatcgaaaaaagagatctgttgaagcatttcttgattggatggacacattcggattgcgagactccgttccggatagacgcaaattagacgatgtagcggatagtttcaatttggctatgagatacgtattagataaatggaatactaattatacaccttataataggtgtaaatatagaaattacataaaaaaaatgtaataacgttagtaacgccattatggataatctatttacctttctacatgaaatagaagatagatatgccagaactatttttaactttcatctaataagttgtgatgaaataggagatatatatggtcttatgaaagaacgcatttcctcagaggatatgtttgacaatatagtatataataaagatatacatcctgccattaagaaactagtttattgcgacatccaacttactaaacatattattaatcagaatacgtatccggtatttaacgattcttcacaagtgaaatgttgtcattatttcgatataaactcaaataatagcaatattagctctcgtacagtagagatatttgagagtgaaaagtcatctcttgtatcatatattaaaactaccaataagaagagaaaggtcaattacggcgaaataaagaaaactgtacatggaggcactaatgcaaattacttttccggtaaaaagtctgatgagtatctgagcactacagtcaggtccaacattaatcaaccttggatcaaaaccatttctaagagaatgagagtagatatcattaatcactctatagtaacgcgtggaaaaagctctatattacaaactatagaaattatttttactaatagaacatgtgtgaaaatattcaaggattctactatgcacattattctatccaaggacaaggatgaaaagggatgtataaacatgattgataaattattctatgtatattataatttatttctgttgttcgaggatatcatccaaaacgattactttaaagaagtagctaatgttgtaaaccatgtactcatggctacggcattagatgagaaattattcctaattaagaaaatggctgaacacgatgtttatggagttagcaatttcaaaatagggatgtttaacctgacatttattaagttgttggatcataccgttttcccctctctgttagatgaggatagcaaaataaagttttttaaggggaaaaagctcaatattgtagcattacgatctctggaggattgtacaaattacgtgactaaatccgagaatatgatagaaatgatgaaggaaagatcgactattttaaatagcatagatatagaaacggaatcggtagatcgtctaaaagaattgcttctaaaatgaaaaaaaacactgattcagaaatggatcaacgactcgggtataagtttttggtgcctgatcctaaagccggagttttttatagaccgttacatttccaatatgtatcgtattctaattttatattgcatcgattgcatgaaatcttgaccgtcaagcggccactcttatcgtttaagaataatacagaacgaattatgatagaaattagcaatgttaaagtgactcctccagattactcacctataattgcgagtattaaaggtaagagttatgacgcattagccacgttcactgtaaatatctttaaagaggtaatgaccaaagagggtatatccatcactaaaataagtagttatgagggaaaagattctcatttgataaaaattccgctactaataggatatgggaataaaaatccacttgatacagccaagtatcttgttcctaatgtcataggtggagtctttatcaataaacaatctgtcgaaaaagtaggaattaatctagtagaaaagattacaacatggccaaaatttagggttgttaagccaaactcattcactttctcgttttcctccgtatcccctcctaatgtattaccgacaagatatcgccattacaagatatctctggatatatcacaattggaagcgtcgaatatatcatcgacaaagacatttataacggtcaatattgttttgctgtctcaatatttatctagagtgagtctagaattcattagacgtagtttatcatacgatatgcctccagaagttgtctatctagtaaacgcgataatagatagtgctaaacgacttaccgaatctattactgactttaatattgatacatacattaatgacctggtggaagctgaacacattaaacaaaaatctcagttaacgattaacgagtttaaatatgaaatgctgcataactttttacctcatatgaactatacacccgatcaactaaagggattttatatgatatctttactaagaaagtttctctactgtatctaccacacttctagatatccagatagagattcgatggtttgtcatcgcatcctaacgtacggcaaatattttgagacgttagcacatgatgaattagagaattacataggtaacatccgaaacgatatcatgaacaatcacaagaacagaggcacttacgcagtaaacattcatgtactaacaactcctggacttaatcatgcattttctagtctattgagtggaaagttcaaaaagtcagacggtagttatcgaacacatcctcactattcatggatgcagaatatttctattcctagaagtgttggattttatccggatcaagtaaagatttcaaagatgttttctgtcagaaaataccatccaagccaatatctttacttttgttcatcagacgttccggaaagaggtcctcaggtaggtttagtatctcaattgtctgtcttgagttccattacaaatatactaacgtctgagtatttggatttggaaaagaaaatttgtgagtatatcagatcatattataaagatgatataagttactttgaaacaggatttccaatcactatagaaaatgctctagtcgcatctcttaatccaaatatgatatgtgattttgtaactgactttagacgtagaaaacggatgggattcttcggtaacttggaggtaggtattactttagttagggatcacatgaatgaaattcgcattaatattggagcaggaagattagtcagaccattcttggttgtggataacggagagctcatgatggatgtgtgtccggagttagaaagcagattagacgacatgacattctctgacattcagaaagagtttccacatgtcatcgaaatggtagatatagaacaatttacttttagtaacgtatgtgaatcggttcaaaaatttagaatgatgtcaaaggatgaaagaaagcaatacgatttatgtgactttcctgccgaatttagagatggatatgtagcatcttcactagtgggaatcaatcacaattctggacccagagctattcttggatgtgctcaagctaaacaagctatctcttgtctgagttcggatatacgaaataaaatagacaatggaattcatttgatgtatccagagaggccaattgtgattagtaaggctttagaaacttcaaagattgcggctaattgcttcggacaacatgttactatagcattaatgtcgtacaaaggtatcaatcaagaggatggaattatcatcaaaaaacaatttattcagagaggcggtctcgatattgttacagccaagaaacatcaagtagaaattccattggaaaactttaataacaaagaaagagataggtctaacgcctattcgaaattagaaagtaatggattagttagactgaatgctttcttggaatccggagacgctatggcaagaaatatctcatcaagaactcttgaagatgattttgctagagataatcagattagctttgatgtttccgagaaatatacagatatgtacaaatctcgcgttgaacgagtacaagtagaacttactgacaaagttaaggtgcgagtattaaccatgaaagaaagaagacccattctaggagacaaatttactactagaacgagtcaaaagggaacagtcgcgtatatcgcagatgaaacggaacttccgtacgacgaaaatggtatcacaccagatgtcattattaattctacatccatcttctctagaaaaactatatctatgttgatagaagttattttaacagccgcatattctactaagccgtacaacaataanggagnaaacmracctgtctgttttcctagtngtaacgaaacatctannnnnnnnnnnnnnnnnnnnnnnnnnnnnnnnnnnnnnnnnnnnnnnnnnnnnnnnnnnnnnnnnnnnnnntcggataaaatcttttgtgaaaagattctctatgatcctgaaacggataagccttatgaatccaaagtantttttggaccaatttattacttgcgtctgagacatttaactcaggacaaggcaaccgttagatgtagaggtaaaaagacgaagctcattagacaagcgaatgagggacgaaaacgtggaggaggtatcaagtttggagaaatggagagagactgtttaatagcacatggtgcagccaatactattacagaagttttaaaagactcagaagaggattatcaagatgtgtatatttgtgaaaattgtggagacatagcagcacaaatcaaaagtattaatacatgtcttagatgttcaaaacttaatctctctctctcttaacaaaaattgataccacgcacgtatctaaagtatttcttactcaaatgaacgccagaggcgtaaaagttaaattagatttcgaacgaaggcctccttcgttttataaaccattagataaagttgatcttaaaccgtcttttctggtataatattgtttagtagatactcatcaagataagctaattcactaaacatattatcggattcggtattgttactcgagaatagagttcgttatgctcctgatattcggaaatctgtggagtttcaggttttggtggaagtgtaactgctacttggtgggatactgaaggatatttcagagagttgtggatgttcgggttcgacatccaccgatggtgtcacgccactaatcggttcggtaacgtctgtggatggaggtgctacttctacagaacctgtagcctcagttgtcaacggagatacatattcaatgcgcggaaatgtataatttggtaatggtttctcatgtggatcttaagaagaagaggtaagatatctacgaaagataccgatcacgtttctagttctcttttgtagaactttaactttttctttctcagcatctagttgatattccgacctcttcacgtttcgcatgggttacctccgcagtttttacaagcgatttcacgttccagatcacgttcagccttcatacgtctctccctctctctatcgagtttatcagagcagtctttctgaaggcgatcgaactccataaatttctccaacgctttgattgtttccatagatttccgaagtttagcttctaggacggcgattcttttttttttttttttttttttttcgaattcacggggtacaaccgtttccattaccaccatctctatgtttcttttctagatcggcaatctttctcaacatttcatccccataccttttcattcctcgagtctattgtcgtcgaaatatcgttccagctccttttcgacctcaataactttagcacgttgtttcatcaagctctctcttgtagtactatcatttttatctgattccctgacacgtttaagatcttcatgtaattgagtcagctcttgacgcaatctcttaactaacttcctctcttgcttcttcgtcatagtacttacaatcactatgggatccattgttaccacgtctgtactcgacgagctcacgtttaagagattcaatttccagtttgtatcggtccatgtctccattgctacaccaccattagatttacaggctgctagttgtcgttcgagatcagaaatacgtgttttcttggaatggatttcgtcgatgtacttgtcatgattggcatcgaaacacttattaagttcttttttcaattctacgattttatttctttcgcgagtcaattccctcctgtagtaactatcagttttgtcagattcacgctctctacgtagactttcttgtaagttactaatttgttccctggcattaccgagttcagttttatatgccgaatagagttctgattcatcctttgagaagatctctagcgatcgttcaagatccctgattctagtctttagcctatttacctcctcagaagatgctccgttaccgtttttacaatcgttaagatgtctatcaagatccatgattctatcttttttccatatcagcattgatttcattattacgttcgcagtcgttcaactgtatttcaagatctgagattctagattgtaatctctgtagcatttccacggcattcactcagttgtctttcaagatctgagattctagattggagtctgctaatctctgtaagatttcctcctccgctctcgatgcagtcggtcaacttattctctagttctctaatacgcgaacgcagtgcatcaacttcttgtgtgtcttcttgattgcgtgtgcattcatcgagtctagattcgagatctctaacgtgacgtcgttcttcctcaagttctctgtgtactacagaaagcgtgtccctatcttgttgatatttagcaatttctgattctagagtactgattctactcacgtatgtactaatagttgtcttagccttatcaagatcctccttgtatttgtcacattccttgatatccatacgaagtctggacagttcccattcgacattacgacgtttatcgatttcagctcggagatcgtcgtcgcgttgttttagccacatacgactaagttcaagttctcgttgacaagatccatctacttttccatccctaatagtatccagttccttttctagttctgaccgcatttctcgttccatatcaagagattctctcaattctcgtatagtcttcttatcaatttctgatgaatctgaaccatcatctgtcccattttgttgcatatccctgagttctttgatctctgttgtaagtctgtcgattctttcggttttataaacagaatccctttccaaagtcctaatcttactgagtttatcattaagttcttcattcaattcagtgagttttctcttggcttcttccaagtctgttttaaactctccatcatttccgcattcttcctcgcatttatctaaccattcaattagtttattaataactagttggtaatcagcgattcctatagccgttcttgtatttgtgggaacataattaggatcttctaatggattgtatggcttgatagcatcatctttatcattattaggtgggggatggacaaccttaattggttggtcctccttatctcctccagtagcatgtggttcttcaataccagtattagtaataggcttagacaaatgcttgtcgtacgcgggcacttcctcatccatcaagtatttataatcgggttctgtttcagaatattcttttctaagagacgcgacttcaggagttagtagaagaactctgtttctgtatctatcaacgctggaatcgatactcaagttaaggatagcgaatacctcatcgtcatcatccgtatcttctgaaacgccatcatatgacatttcatgaagtctaacgtattgataaacagaatcagatttagtattaaacagatccttgacctttttagtaaatgcatatgtatattttagatctccagatttcataatatgatcgcatgccttaaatgtcaatgcttccatgatatagtctggaacactaatgggtgacgaaaaagatacagcaccatatgctacgttgataaatagatctgaaccactaagtagataatgattaatgttaaggaagaggaaatattcagtatatagatatgccttagcatcatatcttgtactaaacacgctaaacagtttattgatgtgatcaatttccaacagaacaattagagcggcaggaataccaacaaacatattaccacatccgtattttctatgaatatcacatatcatattaaaaaatcttgatagaagagcgaatatctcgtctgacttaatgagatgtagttcagcagcataagtcataactgtaaatagaacatactttcctgtagtgttgattctagactccacatcaacaccattattaaaaatagttttatatacatctttaatctgctctccgttaatcgtcgaacgttctagtatacggaaacactttgatttcttatctgtagttaatgacttagtgatatcacgaagaatattacgaattacatttcttgtttttcttgagagacctgattcagaactcaactcatcgttccatagtttttctacctcagtggcgaaatctttggagtgtttggtacattttttaataaggttcgtgacctccatttattataaaaaatttttattcaaaacttaactacaatcgggtaattataagatcgtagatctcccatgtggtggaatactaccatctatcgcatgttgatggacagtaggtaatggccatgggaacagtaatgtttgcatatttatctttcttgctagtattactgtatattgtcccaatgtttcaatgtgatgttctaacctatcaactgccactgtatcacaacaataatgtccgatggaattaagattatgatccaatgtgtttaatatatgattatcaagtcttatacgatccgcgtcttttttgacaggatcaggctcttctacaggaagaagtttcggcctcttatgatagtcatgtctgggaaatggtggtctaggatgaggatcaggtatcggagtaggttttggattataatcatcatcatcatcatcatcatcatcatcatcatcatcatctatgatatcatcatcttcgatatttattttgctatcttgataatgtcctatatcagttgcattttcagcactcgactgaatattagtacattcattgtctattattaacgtatttctaaacccaaaatgtatatgttgaacatcactactatagttgatgagtcttatagcatgaattcgcttatcgttatcgggtttatcttctgtcaccttaacaattccttttttattaaactctgcataatcataaccatttctattgtttgttctaatataaacgagtatagcatcattgctaaatttttcaatagtatcaaaaacagaatatcctaaaccatataatatatattcaggaacactcaaactaaatgtccaggattctcctaaatacgtaaactttaatagtgctaaatcattcaaaaatctaccgcttatagatagatagtacatgaatgcgtatagtagtctacctatctctttattatgaaaaccgacattacgatcatatatttcgtgatatacatgtgacccgtttacgttaaaccataaatacatgggtgatcctataaacatgaatttatttctaattctcagagccatagttaattgaccgtgtaatatttgtttacatgcatacttgatacgatcattaataagatttttatcattgctcgttatttcagaatcgtatatataaggagtaccatcatgattcttaccagatattatacaaaatactatatataaaatatattgacccacgttagtaatcatgtaaatgtttaatgttttaaattttgtatttaatgatccatcatcatatgctagcatggtcttgtgatattcattctttaaaatataatattgtgttagccattgcattggagctcctaatggagattttctattctcgtccattttaggatatgctttcataaagtccctaataacttcgtgaataatgtttctatgttttctactgatgcatgtatttgcttcgatttttttatcccatgtttcatctatcatagatttaaacgcagtaatgctcgcaacattaacatcttgaaccattggtacaattccgttccataaatttataatgttcgccatttatataactcattttttgaatatacttttaattgaacaaaagagttaagttactcatatggatgccgtccagtctgtacatcaatctttttagccagagatatcatagccgctcttagagtttcagcgtgattttccaacctaaatagaacttcatcgttgcgtttacaacacttttctatttgttcaaactttgttgttatattagtaatctttttttccaaattagttagccgttgtttgagagtttcctcattatcgtctccataggctttaacaattgcttcgcgtttagtctctggatttttagcagcctttgtagagaaaaattcagttgctggaattgcaagatcgtcatctccggggaaaagagttccgtccatttaaagtacagattttagaaactgacactctgtgttatttatatttggcgcaatacatggattataaatatcgatgttaataacatcagaaaatgtaaagtctatacattgtcgcatcgtgttaaattttctaatggatctagtattattgggtccaacttctgcctgaaatccaaatatggaagcggatacaaaaccgtttcctggataaaccacacatctccacttttgctttacatcagaaattgtgtcattgacatcttgaactctcctatctaatgccggtgttccacctatagattttgaatactcgaatgctgcatgagtagcattgaattccttaatattgccataattttcatatattgagtaactctggataaaaagtaaacacaccgcagccgtcgctactacaataaaaaaaattgatagagagttcatttataatctattagaagctgataaaatttttttacacgcgtcagacaatgctttaataaatagttcaacatctacttttgtcatatcgaaccgatggtatgattctaacctagaattacatccgaaaaagttgactatgttcatagtcattaagtcattaacgaacaacattccagactctggattataagacgatactgtttcgtcacaatcacccaccttaatcatgtgattatgaatattggctattagagtaccttctaagaaatctataatatctttgaaacacgatttaaaatcaaaccacgaatatacttctacgaagaaagttagtttacccataggagagataactataaatggagatctagatacaaaatccggatctatgatagttttaacattattatattctctattaaatacctccacatctaaaaatgttaattttgaaactatgtcttcgtttattaccgtacctgaactaaacgctataagctctattgtttgagaactctttaaacgatattcttgaaatacatgtaacaaagtttcctttaactcggtcggtttatctaccatagttacagaatttgtatccttatctataatataataatcaaaatcgtataaagttatataattatcgtgttcagattgtgatcttttcaaatagactaaaaaccccatttctctagtaagtatcttatgtatatgtttgtaaaatatcttcatggtgggaatatgctctacagcagttagccattcctcattgacagctgtagatgtattatacaaaactactccaatgtttaacaagggccattttacgagattattaaatccttgtttgataaatgtagccaatgcgggttcgagttcaacgacgattgaattctcttcccgtggatgctgcatgatgaacgacgggatgttgttgttctattgatttggaattctttttcgactttttgtttatattaaatattttaaaatttatggctgatagtaattcatgtactacggataatgtagacgtgtattgcatatcgatatctttattattagataaatttatcaataaatgtgagaagtttgcctcgttaaggtcttccatttaaatattatataaatatttgtgtttgtattttattcgtcttttatgggatagtttttaactagtaaagctgtaattacatactttgtccgtaaaacataaatataaatacccgcttttatcaaacgttccaaaaagtcggcagctgacatttttaacatgacatctattttaaatacacttaggtttttagaaaaaacatcattttataattgtaacgattcaataactaaagaaaagattaagattaaacataagggaatgttatttgtattttataagccaagcattctaccgttgttaaatacttgtctggaggaggtatatatcatgatgatttggttgtattggggaaggtaacaattaatgatctaaagatgatgctattttacatggatttatcatatcatggagtgacaagtagtggagcaatttacaaattgggatcgtctatcgatagactttctctaaataggactattgttacaaaagttaataacaattataacaattataacaattataacaattataattgttataataattataattgttataattatgatgatacattttttgacgatgatgattgatcactattacacaattttgtttttgtactttctaatatagtgtttaggttctttttcatatgagaatattgacttactaaaatatctatgtttaacttttgttctataacgtccttatcggcggtatcggtacatatacgtaattcaccttcacaaaatacggagtcttcgataataatagccaatcgattattggatctagctgtctgtatcatattcaacatgtttaatatatcctttcgtttcccctttacaggcatcgatcgtagcatattttccgcgtctgagatggaaatgttaaaactgcaaaaatgcgtaatgttagcccgtcctaatattggtacgtgtctataagtttggcatagtagaataatagacgtgtttaaatgccttccaaagtttaagaattctattagagtattacattttgatagtttatcacctacatcatcaaaaataagtaaaaagtgtgctgattttttatgattttgtgcgacagcaatacatttttctatgttacttttagttcgtatcagattatattctagagcttcctgactactaacgaaattaatatgatttggccaaatgtatccatcataatctgggttataaacgggtgtaaacaagaatatatgtttatattttttaactagtgtagaaaacagagatagtaaatagatagtttttccagatccagatcctcctgttaaaaccattctaaacggcatttttaataaattttctcttgaaaattgtttttcttgaaaacaattcataattatatttacagttactaaattaatttgataataaatcaaaatatggaaaactaaggtcgttagtagggaggagaacaacgaaggcatatcgtgatataaataacatttattatcatgatgacaccagaaaacgacgaagagcagacatctgtgttctccgctactgtttacggagacaaaattcagggaaagaataaacgcaaacgcgtgattggtctatgtattagaatatctatggttatttcactactatctatgattaccatgtccgcgtttctcatagtgcgcctaaatcaatgcatgtctgctaacaaggctgctattactgactccgctgttgccgttgctgcggcatcatctactcatagaaaggttgtgtctagcactacacaatatgatcacaaagaaagctgtaatggtttatattaccagggttcttgttatatattacattcagactataagtcattcgaggatgctaaagcaaactgcgctgcggaatcatcaacactacccaataaatccgatgtcttgactacctggctcattgattatgttgaggatacatggggatctgatggtaatccaattacaaaaactacatccgattatcaagattctgatgtatcacaagaagttagaaagtatttttgtacataaataaatgaaatcgcttaatagacaaactgtaagtaggtttaggaagttgtcggtgccggccgctataatgatgttactctcaaccattattagcggcataggaacatttctgcattacagagaagaactgatgcctagtgcttgcgccaatggatggatacaatacgataaacattgttatctggataccaacattaaaatgtctacggataatgcagtttatcagtgtcgcaaattacgagctagattgcctagacctgatactagacatctgagagtattgtttagtattttttataaagattattgggtaagtttaaaaaagaccaatgataaatggttagatattaataatgataaagatatagatattagtaaattaacaaattttaagcaactaaacagcacgacggattctgaggcgtgttatatatacaagtctggaaaactggttaaaacagtatgtaaaagtactcaatctgtactatgcgttaaaagattctacaagtgacaacaaaaaatgaattaatagtaagtcgttaacgtacgccgccatggacgccgcgtttgttattactccaatgggtgtgttgactataacagatacattgtatgatgatctcgatatctcaatcatggactttataggaccatacattataggtaacataaaaattgtccaaatagatgtacgggatataaaatattccgacatgcaaaaatgctactttagctataagggtaaaatagttcctcaggattctaatgatttggctagattcaacatttatagtatttgtacagcatacagatcaaaaaataccatcatcatagcatgcgactatgatatcatgttagatatagaaggtaaacatcaaccattttatctattcccatctattgatgtttttaacgctacaatcatagaagcgtataatctgtatacagctggagattatcatctgatcatcaatccttcagataatctgaaaatgaaattgtcgtttaattcttcattttgtatatcagacggcaatggatggattataattgatgggaaatgtaatagtaattttttatcataaaagttgtaaagtaaataataaaacaataaatattgaactagtagtatgttgtatattgagcaatcagagatgatgctggtacctcttatcacggtgaccgtagttgcgggaacaatattagtatgttatatattatatatttgtaggaaaaagatacgtactgtctataatgacaataaaattatcatgacaaaattaaaaaagataaagagtcctaattccagcaaatctagtaaatcaactgatagcgaatcagactgggaggatcactgtagtgctatggaacaaaacaatgacgtagataatatttctagaaatgagatattgaacgatgatagcttcgctggtagtttaatatgggataacgaatccaatatcatggcgcctagcacagaacacatttacgatagtgttgctggaagcacgctgctaataaataatgatcgtaatgaacagactatttatcagaatactacagtagtaattaatgatacagagactgttgaaatacttaatgaagataccaaacagattcctagctattcttccaatcctttcgtaaattataataaaaccagtatttgtagcaagtcaaatccgttcattgcagaactcaacaataaatttagtgataataatccgtttaggagagcacatagtgacgattatcttaataagcaacaagatcatgaatacgatgatatagaatcatcggttgtatcattggtctgattagtttcctttttataaaattgaagtaatatttagtattaattaccgccgatgcattatacaaatatggagatattccctgtattcggcatttctaaaattagcaattttattgctaataatgactgtagatattatatagatgtagagcatcaaaaaattatatctgatgagatcaatagacagatggatgaaacggtacttcttaccaacatcttaagcgtagaagttgtaaatgacaatgagatgtaccatcttattccccatagactatcgactattatactctgtattagttctgttggaggatgtgttatctctatagataatgacgtcaatgacaaaaatattctaacatttcccattgatcatgctgtaatcatatccccactgagtaaatgtgtcgtagttagcaagggccctacaaccatactggttgttaaagcggatatacccagcaaacgattggtaacatcgtttacaaacgacatactgtatgtaaacaatctatcactgattaattatttaccgtcgtctgtattcattattagacgagtcaccgactatttggatagacacatatgtgatcagatatttgctaataataagtggtattccattataactatcgacgataagcaatatcctattccatcaaattgtataggtatgtcttctgccaagtacataaattcgagcatcgagcaagatattttgatccatgtttgtaacctcgagcatccattcgactcagtctacaaaaaaatgcagtcgtacaattctctacctatcaaggaacaaatattgtatggtagaattgataatataaatatgagcattagtatttcggtggattaatagatttctctagtatgggatcattaatcatctctaaatacatcataaaaaagctattatcaaatactgtactgaatggattcattcttttctctttttatgaaactctgttgtatatctacggataaaactagaagcaaaaaatctgataggaagaataatgattatatggaggaacacgattattataaaataacaatagttcctggttcctcttccacgtctactagctcgtggtattatacacatgcctagtaatagtctctttgcgttgacggaaagcagactagaaataacaggccaaaatgttcagacaccataatagttcccaacccagataataacagagttccatcaacacattcctttaaactcaatcccaaacccaaaaccgttaaaatgtatccagccaattgatagtagataatgaggtgtacagcacatgataatttacacagtaaccaaaatgaaaacactttagtaattataagaaatatagacggtaatgtcatcatcaacaatccaataatatgcctgagagtaaacattgacggataaaacaaaaatgccccgcataactctatcatggcaataacgcaaccaaacacttgtaaaattcctaaattagtagaaaatacaactgatatcgatgtataagcgatttcgaggaataataagaacaaagtaattcccgtaaagataaacatcaacattgtttggtaatcattaaaccaattagtatgacgttgaattaatttcacagtatattttattccagtattatccccgcatgtatacgtacctggtaagatatctttatattccataatcaatgagacatcactatccgataacgaatgaagtctagcactagtatgccatttacttaatatggtcgtcttggaagttttattataagttaaaatatcatgattgtccaatttccatctaatatactttgtcggattatctataatacacggaataatgatggtatcattacatgctgtatattctatagtctttgtagatgttataaccacaaaagtacagaggtatatcaacaatattctaactcttaacatttttatttatttaaaatgatacctttgttatttattttattcttattttgctaacggtatcgaatggcataagtttgaaacgagtgaagaaataatttctacttacttaatagattatgtggtaacgggtgttattaatggggatgtatatacattttcaaataatgaactaaacaaaactgggttaactaataacaataattatatcacaacatctataaaagtagaggataaggatacattagtagtatgcggaaccaataacggaaatcccaaatgttggaaaatagacggttcatactacccaaaacatataggtagaggatacgatcatcaaaatagcaaagtaacgataatcagtcacaatgaatgtgtactatccgacataaacatataaaaagaaggaattaaacgatggagaagatttgacggaccatgtggttatgattaaacgagttaagttttttaagaagccttagaagagaggctattgggtatgagaatccgaaatattaaaccagacaaccccatataattttatagctaagaatgccgcgaagaatggaactaataaaaacggaaatatttgtagcacaacgaataactcccaaactgcattcatgttacactatataacactacttcggttagatgttttagaaaaaataaatatcaccgtaccgttttgttgtataaaaataacaattaacaattatcaatttttttctttaatattttacgtggttgaccattcttggtggtaaaataatctcttagtgttggaatggaatgctgtttaatgtttccacactcatcgtatattttgacgtatgcagtcacatcgtttacgcaatagtcagactgtagttctatcatgcttcctacgttagaaggaggaacagttttaaagtctcttggttttaatctattgtcattagttttcatgaaatcctttgttttatccacttcacattttaaataaatgtccactatacattcttctgttaattttactagatcatcatgagtcatagaattcataggttccgtagtccatggatccaaactagcaaacttcgcgtatacggtatcgcgattagtgtatacaccaactgtatgaaaattaagaaaacagtttaataaatctacagaaatatttaatcctccgtttgatacagatgcgccatatttatggatttcggattcacacgttgtttgtctaaggggttcgtctagtgttgcttctacatagacttcgattcccatatattctttattgccagaatcacataccgatttatcatacgctggttcacttgtttgaaaactaaatggtagtagatacatcaaaataataaataataagtacattctgcaatattgttatcgtaattggaaaattggtattcaagtgagctggattatntgannntattggattgtgrtattttattttatattttatannnnnnnnnnnnnnnnnnnnnnnnnnnnnnnnnnnnnnnnnnnnnnnnnnnnnnnnnnnnnnnnnnnnnnnnnnnnnnnnnnnnnnnnnnnnnnnnnnnnnnnnnnnnnnnnnnnnnnnnnnnnnnnnnnnnnnnnnnnnnnnnnnnnnnnnnnnnnnnnnnnnnnnnnnnnnnnnnnnnnnnnnnnnnnnnnnnnnnnnaaagaatgttctagcggctaacttnntaacagaacatttgcaaagattaatccgggtgaagttattccctcatcactaatcataatattctaaaacctcttattgtcantcatgaaattttgtattgtatatactaacttctctaatggatgagaacacgtatgctatggagttgcttactgggtacgcccctgtatctccgatcgttatagcgagaactcataccgcacttatatttttgatgggtaagccaacaacatccagacgtgatgtgtatagaacatgtagagatcacgctacccgtgtacgtgcaactggtaattaaaataaaaagtaatattcatatgtagtgtcaattttaaatgatgatgaaatggataatatccatattgacgatgtcaataatgccggtattgacatacagctcatcgatttttagatttcattcagaggatattgaattatgttatgggaatttgtattttgataggatctataataatgtagtaaatataaaatatattcctgagcatattccatatagatataattttattaatcgtacgttctccgtagatgaactagatgataatgtcttttttacacatggttattttttaaaacacaaatatggttgttcacttaatcctagtttgattgtctcattatcaggaaacttaaaatataatgatatacaatgctcagtaaatgtatcgtgtctcattaaaaatttggcaacgagtacatctactatattaacatctaaacataagacttattctctatatcggtccatgtgtattgctataataggatacgattctattatatggtataaatatataaatgacaggtataatgacatctatgattttactgcaatatgtatgctaatagcgtctacattgatagtgatcatatacgtgtttaaaaaaataaaaatgaactcttaattatgttatactattagaaatggataaaatcaaaattacgattgattcaaaaattggtaatgttgttaccatatcgtataacttggaaaagataactattgatgtcacgccaaaaaagaaaaaagaaaaggatgtattattagcgcaatcagttgctgtcgaagaggcaaaagatgtcaaggtggaagaaaaaaatattatcgatattgaagatgacgatgatatggatatagaaaacacgtaatacgatctataaaaataagtattaaatactttttatttacggtactcttgtagtggtgataccactaatcgattattttttttaaaaaaatacttattctgattcttctagccatttccgtgttcgttcgaatgccacatcgacgtcaaaaataggggagtagttgaaatctagttctgcattgttggtacgcacctcaaatgtagtgttggatatcttcaacgtatagttgttgagtattgatggttttctaaatagaattctcttcatatcattcttgcacgcgtacatttttagcatccatcttggaatcctagatccttgttctattcccaatggtttcatcaatagaagattaaacatatcgtaagaacacgatggagagtaatcgtagcaaaagtaagcatttcctttaatcgcagatcccggatactggatatattttgcagccaacacgtgcatccatgcaacatttcctacatatacccggctatgcacagcgtcatcatcgactgtacgatacataatgttaccgtgttgcttacattgctcgtaaaagactttcgtcaatttgtctccttctccgtaaattccagtgggtcttaggcaacaagtatacaattttgcgccattcatgattacggaattattggctttcataaccagttgctcggccatacgtttactttttgcgtatacatgtcctggtgatatatcataaagggtatgctcatgaccgatgaatggattaccgtgtttatttggtcctattgcttccatgctactagtatagatcaaatacttgattcctaggtccacacaagctgccaatatagtctgtgttccataatagtttactttcatgatttcattatcagtgtattttccaaatacatccactagagcagccgtatgaataatcagatttaccccatctagcgcttctctcaccttatcaaagtcgtttatatcacattgtatatagtttataaccttaactttcgaggttattggttgtggatcttctacaatatctatgactcttatttcttgaacatcatctgcgctaattaaaagttttactatatacctgcctagaaatccggcaccgccagtaaccgcgtacacggccattgctgccactcataatatcagactacttattctattttactaaataatggctgtttgtataatagaccacgataatatcagaggagttatttacgttgaacaagtccatggaaaagataaagttttaggatcagttattggattaaaatccggaacgtatagtttgataattcatcgttacggagatattagtcgaggatgtgattccataggcagtccagaaatatttatcggtaacatctttgtaaacagatatggtgtagcatatgtttatttagatacagatgtaaatatatctacaattattggaaaggcgttatctatttcaaaaaatgatcagagattagcgtgtggagttattggtatttcgtacataaatgaaaagataatacattttcttacaattaacgagaatggcgtttgatatatcagttaatgcgtctaaaacaataaatgcattagtttacttttctactcagcaagataaattagtcatacgtaatgaagttaatgatatacactacactgtcgaatttgatagggacaaagtagttgatacgtttatttcatataatagacataatgactccatagagataagaggggtgcttccagaggaaactaatattggtcgcgtggttaatacgccggttagtatgacttacttgtataataagtatagttttaaaccgattttagcagaatatataagacacagaaatactatatccggcaacatttattcggcattgatgacgctagatgatttggttattaaacagtatggagacattgatctattatttaatgagaaacttaaagtagactccgattcgggactatttgactttgtcaactttgtaaaggatatgatatgttgtgattctagaatagtagtagctctatctagtctagtatctaaacattgggaattgacaaataaaaagtataggtgtatggcattagccgaacatatagctgatagtattccaatatctgagctatctagactacgatacaatctatgtaagtatctacgcggacacactgatagcatagaggatgaatttgattattttgaagacgatgatttgtctacatgttctgccgtaaccgatagggaaacggatgtataattttttttatagtatgaaggatatgatggatatgatnatatgaaggatatgatgatatgatggatatgatggatatgatggatatgatggatatgataaaaaaatataattgttgtatccattcccattcaaatcaccttatatgattctgtaacacaatgaaggagtctcatagatatatagaggtcagatactggtttgataaactttttattccacatgagcatgtttgacttatggttagacacacatactttaacaaatcactgaaaattggagttaggtattcctctcagaatcagttgccgttctggaacattaaatgtattttttatgatatactccaacgcatttatgtgggtatacaacaagtcattaataatgagtatttccaagagttttagttgtctagtatttaacaagagaagagatttcaacagactgtttatgaactcgaataccgcctcattgtcgcttatattgatgacatgacatcgaattcccaatatcaatctcatcagtgatgagtagctcaatcttgttatcgggatccaatttctaaagatgtcattaaaccctcgatcgtgaatggatttatcatcatcgtttttatgttggacatgagcttagtccgtttgtccacatctatatacgatgatttctgaattatttcatatatctctcgttaactccaggaacttgtcagggatctaactttaatatgttctcgtctaagagatgaaaatctttggatggttgcatgtgacttttctctaaaggatgatgttacccgatcctctcttaaatgactccatcttatccttggacaagatggacagtctattttccttagatggtttaatattttttacccatgatctataaaggtagacagacctaatcgtctcggatgaccatatattattttccgttttattatacgcataaattgtaaaaaatatgttaggtttacgaaaatgtctcgtggggcattaatcgtttttgaaggattggacaaatctggaaaaacaacacaatgtatgaacatcatggaatctataccggcaaacacgataaaatatcttaactttcctcagcgatccacagtcactggaaagatgatagatgactatctaactcgtaaaaaaacctataatgatcatatagttaatctattattttgtgcaaatagatgggagtttgcatcttttatacaagaacaactagaacagggaattactttaatagttgacagatacgcgttctctggagtagcgtatgccaccgctaaaggcgcgtcaatgactctcagtaagagttatgaatctggattgcctaaacccgacttagttatattcttggaatctggtagcaaagaaattaatagaaacatcggcgaggaaatttatgaagatgttgaattccaacaaaaggtattacaagaatataaaaaaatgattgaagaaggagatattcattggcaaattatttcttctgaattcgaggaagatgtaaagaaggagttgattaagaatatagttatagaggctatacacacggttactggaccagtggggcaactgtggatgtaataaaatgaaattacatttttataaatagatgttagtacagtgttataaatggatgaagcatattactctggcaacttggaatcagtactcggggatacgtgtccgatatgcataccgaactcgcatcaatatctcaattagttattgccaagatagaaactatagataatgattattaaacaaggacattgtaaattttatcatgtgtagatcaaacttggataatccatttatctctttcctagatactgcatatactatcatagatcaagagatctatcagaacgagttgattaattcattagacgataatgaaattatcgattgtatagttaacaagtttatgagcttttataaggataacctagaaaatatggtagatgctatcattactctaaaatattataattaataatccagattttaaaactacgtatgtggaagtactcggttccagaatagctgatatagatattaaacaagtgatacgtaagaatataatacaattgtctaatgatccgcgaacgatatttgtgaaaatattaaaaaaaaatacnnnttttattaaatgacgtctcttsscgaatttagaaaattatgctgtgnnatatatcacgcatcaggatataaagaaaaatctaaattaattagagactttataacagatagagatgataccgatacatntttgatcattaagctattgcttnncggattagacgatagaatgnataacatgaacgataaacaaattataaaattatatagtataatatttaaacaatctcaggaagatatgctacaagatttaggatacggatatataggagacactattaggactttcttcaaagagaacacggaaatccgtccacgagataaaagcattttaactttagaagaagtggatagttttttaactacgttatcatcagtaactaaagaatcacatcaaataaaattattgactgatatagcatctgtttgtacatgtaatgatttaaaatgtgtagtcatgcttattgataaagatctaaaaattaaagcgggtcctcggtacgtgcttaacgctattagtcctcatgcctatgatgtttttagaaaatctaataacttgaaagagataatagaaaatgcagctaaacaaaatctagactctatatctatttctgttatgactccaattaatcccatgttagcggaatcatgtgattctgtcaataaggcgtttaaaaaatttccatcaggaatgtttgcggaagtcaaatacgatggtgaaagagtacaagttcataaaaaaaataacgagtttgcattctttagtagaaacatgaaaccagtactctctcataaagtggattatctcaaagaatacataccgaaagcatttaaaaaagctacgtctatcgtattggattctgaaattgttcttgtagacgaacataatgtaccgctaccgtttggaagtttaggtatacacaaaaagaaagaatataaaaactctaacatgtgtttgttcgtatttgactgtttatactttgatggattcgatatgacagacattccattgtatgaacgaagatcttttctcaaagatgttatggtcgaaatacccaatagaatagtattctcagagttgacgaatattagtaacgagtctcagttaactgatgtattagatgatgcactaacgagaaaattagaaggattggtcttaaaagatattaatggcgtatacgaaccgggaaagagaagatggttaaaaataaagcgagactatttgaacgagggttccatggcagattctgccgatttagtagtactaggtgcctactatggtaaaggaggaaagggtggtatcatggcagtctttctaatgggttgttacgacgatgaatccggtaaatggaagacggtaactaaatgttccggtcacgatgataatacgttaagggttttgcaagaccaattaacgatggttaaaattaacaaggatcccaaaaaaattccagagtggttggtagttaataaaatctatattcccgattttgtagtagatgatccgaaacaatctcagatatgggaaatttcaggagcagagtttacatcttccaagtcacatacagcgaatggaatatcgattagatttcctagatttactaggattagagaagataaaacgtggaaagaatctactcatctaaacgatttagtaaacttgactaaatctcttaatagttacatataaactgaaaaataaaataacactattttagttggtagtcgccatggatggtgttatcgtatactgtctaaatgcgttagtaaaacatggcgaggaaataaatcatataaaaaatgatttcatgattaaaccatgttgtgaaagagtttgtgaaaaagtcaagaacgttcacatcggcggacaatctaaaaacaatacagtgattgcagatttgccatatctggataatgctgtatcagatgtatgcaaatcaatatataaaaagaatgtatcaagaatatccagatttgctaatttgataaaaatagatgacgatgacaagactcctaccggcgtatataattattttaaacctaaagatgctattcctgttattatatccataggaaaggataaagatgtctgtgaactattaatctcatctgataaagcgtgtgcgtgtataaagttaaatttatataaagtagccattcttcccatggatgtttccttttttaccaaaggaaatgcatcattgattattctcctgtttgatttctctatcgatgcggcacctctcttaagaagtgtaaccgataataatgttattatatctagacaccagcgcctacatgacgagcttccgagttccaattggttcaagttttacataagtataaagtccgactattgttctatattatatatggttgttgatggatctatgatgtatgcgatagctgataatagaactcacgcaattattagcaaaaatatattagacaatactacgattaacgatgagtgtagatgctgttattctgaaccacagattaggattcttgatagagatgagatgctcaatggatcatcgtgttatatgaacagacattgtattatgatgaatttacctgatgtaggcgaatttggatctagtatgttggggaaatatgaacctgacatgattaagattgctctttcggtggctggtaatttaataagaaatcgagactacattcccgggagacgaggctatagctactacgtttacggtatagcctctagataatttttttaagcacgaaataaaaacataattttaaacaatctatttcatactattttgtgtgctcaccatgaacataaagatagatatattagtatttctggtgataaatttacggcgactgctaggagggaaaatgaagaaagaaaaaatatctacctctccaaaaagaaaaactactgatgttatcaaacctgattatcttgagtacaatgacttgttagatagagatgagatgtctactattctagaggaatattaggccttagaataaaatatggacgactcttaacgaaattagaaaattcgataatgatgttgaagaacaattcggtactatagaagaactcaagcagaagcttagattaaattctgaagagagagcagataattttatagattatataaaggtacaaaaacaggatatcatcaaacttactgtatacgattgcatatatctatgataggattgtgtgcgtgcgtggtagatgtttggagaaatgagaaactgttttctagatggaaatattggttacaagcgattaaactgtttattgatgatcacatgcttgataagataaaatctattgtagaatagactagtgtatgtggaaatgtcatagaaagttaaaagttaatgagagcaaaaatatataaggttgtattccatatttgttatttttttctgtaatagttagaaaatacattcgatggtctatctaccagattattatgtgttataaggtacttttctcataataaactagagtatgagtaagatagtgtttttcaaaaacatataaatctaaaattgatggatgagatatacagctattaatttcgaaaatatattttaatctgataactttaaacatggatttttgatggtggtttaagtttaaaaaagattttgttattgtagtatgataatatcaaaaagatggatataaagaatttactgactacatgtactattttacattactacattggctacggcatatatacctatttcgtcacttccacacgctccggtaaacgggtgtcatgtgacgagggagaatcttgataagaggcataatcaatgttgtaatccgatgtccacctggagaatttgccaaggtcagatgtagagttggtagtgataacacaaaatgtgaacactgcccacctcatacatataccgcaatccccaattattctaatagatgtcatcaatgtagaaaatgcccaacaggatcatttgataaggtaaagtgtaccggaacacagaacaaatgttcgtgtcatcctggttggtatacgctactgattcttcacagactgaagattgtcgagatttgtgtaccaaaaaaggagatgtccatgcggatactttggtggaatagatgaaggaaatcctatttgtaaatcgtgttgtgttggtgaatattgcgactacctacgtaattatagacttgatccatttcctccatgcaaactatctatctaaatgtaattaattatgattttgatgataatgttaccatacattatatcactacttggttagtgtgtattatttagtatggaagacctattaataattacttatcttttgacgatcttgttataattataatataaaaatacttatgacatagtaactcataattgctgacgcgataaattcgtaataatctgttttgttcaaatttttataaggaatctacaggcataaaaataaaaatataatctataatatactcttacaacgccatcatgaatagcagtgaattaattgctgttattatggatttagaaatagtggacgattttgtgatattaatatagttattaatgatgaaaggataaacgcgcatagactcatcctatctggagcctccgaatattttttccattctgttttccaataattttatcgattctaatgaatacgaagttaatctaagtcatttagattatcaaagtgttaactatttgatcgattacatttatgggatacctttgagcctaactaacaataacgtgaaatatattctttcaaccgctgatttttttacaaattggatctgtcattactgagtgcgaaaaatacatacttaaaaatctttgttttagaaactgtatcgatttctacatatacgctgataaatataataacaagaaaatagaattagcatcgtttaacacaatattacgaaatattttgagactcatcaacaatgaaaactttaaatacttaacagaggaatcaatgataaaaattttaagcgatgatatgttatatataaaaaatgaggatttcaccccactgattctcattaaatggttagagagtacaccaaccatgtaccgtcgagttacttagatgcctcagaatatcatttctttccccacaagttataaaatcactttatagtcatcgactggttagttcaatctacgaatgtataacattcttaaacaatatagcattcttggataaatcatttcctagataccatatcatcgagttgatatctatcggtataagtaattcacatgataagatttccataaactgctacaatcataaaaaaattcatgggaaatgatatcttcacgtagatataggtgtagtttcgcagtgaccgtcctggataatattatctatatgatgggtggatatgatcagtccctgtatagaagttcaaaggttatagcgtacaatacatgtactaattcttggatatatgatataccagagctaaaaatatcatcgttctaattgcggaggagttgccaatgacgaatacatttattgtataggcggtatacgcgatcaggagtcatcgttgatatctagtatcgatagatggaagccatcaaaaccatattggcagaagtatgctaaaatgtgcgaaccaaaatgtgatatggggttgcgattttaaacggattaatatatgtcataggtggagtcgttaaaggtgacacacatataccaacgcactagagagtttatcagaagatggatggatgaatcatcaacgtcttccaataaaaatgtccaatatgtcgacgattgttcatgctggaaagatttatatatctagaggttacaacaatagtagtgtagttaatgtaatatcgaatctagtccttagctataatccgatatatgatgaatggaccaaattatcatcattaaatattcctagaattaatcctgctctatggtcagtgtataataaattatatgtaggaggagtaatatctgatgatgttcaaactaatacatctgaaacatacgataaagaaaaagattgttggacattggataatggtcacttgttaccacataattatataatgtataaatgcgaaccgtttaaacatagatatccattggaaaaaacacagtacacgaatgattttctaaagtatttggaaagttttataggtagttgatagaacaaaatacataattttgtaaaaataaatcactttttatactaatatgacacaattaccaatacttttgttactaatatcattagtatacgctacaccttctcctcagacatctaaaaaataggtgatgatgcaactatatcatgtagtcgaaataatacaaattactacgttgttatgagtgcttggtataaggagcccaattccattattctcttagctgccaaaagcgacgtcttgtattttgataattataccaaggataaaatatcttacgactctccatacgatgatctagttacaactatcacaattaaatcattgactgctggagatgccggtacttatatatgtgcattctttatgacatcgactacaaatgatactgataaagtagattatgaagaatactccatagagttgattgtaaatacagatagtgaatcgactatagacataatactatctggatctacaccagaaactatttctgagaaaccagaggatatagataattctaattgctcgtctgtattcgaaatcacgactccggaaccaattactgataatgtagacgaccatacagacaccgtcacatacactagtgatagcattaatacagtaaatgcatcatctggagaatccacaacagacgagattccggaaccaattactgataaagaagaagatcatacagtaacagacactgtctcatacactacagtaagtacatcatctggaattgtcactactaaatcaaccaccgatgatgcggatctttatgatacatacaatgataatgatacagtaccgccaactactgtaggtggtagtacaacctctattagcaattataaaaccaaggacttcgtagaaatatttggtattaccacattaattatattgtcagcagtggcgattttctgtattacgtattatatatgtaataaacacccacgtaaatacaaaacagagaacaaagtctagatttttgacttacataaatatctgggataataaaatctatcatattgagaggaccatctggttcaggaaagacagccataaccaaaagactgttaaaagactatgggaatatatttggatttgtggtgtcccataccactagatttcctcgtcctatggaacgagaaggtgttgattaccttacgttaacagagaggccatctggaagggaatagccgccggaaactttctagaacatactgagtttttaggaaatatttacggaacttctaaaacagctgtaaatacagcggttattaataatcgtatttgcgcgatggatttaaacatcaacggtgttagaagtcttaaaaatacttacctaatgcattacttgggtatataagacctacctctcttaaaatggttgagaccaatcttcgtcgtagaaacactgaagcggacgacgaatctcatcgtcgcgtgatgttggcaaaaaacggatatggatgaggtcaacgaagcaggtctattcgacactattattattgaagatgatgtgaatttagcatatagtaagtgttaattcagatactacaggaccgtattagaatgtattttaacactaattagagacttaagatttgacttaaaacttgataattaataatataactcgtttttatatgtggctatttcaacgtctaatgtattagttaaatattaaaacttaccacgtaaaacttaaaatttaaaatggtatttcattgacagatcatacattatgaagtttcaaggacttgtgttaattgacaattgcaaaaatcaatgggtcgttggaccattaataggaaaaggtggattcggtagtatttatactactaatgacaataattatgtagtaaaaatagagcccaaagctaacggatcattatttaccgaacaggcattttatactagagtacttaaaccatccgttatcgaagaatggaaaaaatctcacaatataaagcacgtaggtcttatcacatgcaaggcatttggtttatacaaatccattaatgtggaatatcgattcttggtaataaatagattaggtgcagatctagatgcggtgatcagagccaataataatagactaccagaaaggtcggtgatgttgatcggaatcgaaatcttaaataccatacaatttatgcacgagcaaggatattctcacggagatattaaagcgagtaatatagtcttggatcaaatagataagaataaattatatctagtggattacggattggtttctaaattcatgtctaacggcgaacatgttccatttataagaaatccaaataaaatggataacggtactctagaatttacacctatagattcgcataaaggatacgttgtatctagacgtggtgatctagaaacacttggatattgtatgattagatggttgggaggtatcttgccatggactaagatatctgaaacaaagaattctgcattagtaagtgccgcaaaacagaaatatgttaacaatactgcgactttgttaatgaccagtttgcaatatgcacctagagaattgctgcaatatattaccatggtaaactctttgacatattttgaggaacccaattacgacgagtttcgtcgagtattaatgaatggagttatgaaaaatttttgttgataaaaaaattaaaaaaataacttagttattatcactctcgtgagtacaatagaaacatggcgatgttttacgcacacgctttcggtgggtacgacgagaaccttcatgcatttcctggaatatcatcgacggttgccaatgatgtcaggaaatattctgttgtgtcagtttataataaaaagtataacattgtaaaaaacaaatatatgtggtgtaacagtcaagtgaacaagagatatattggagcactactgcctatgtttgaatgcaatgaatatctacaaattggagatccaatccatgatctagaaggaaatcaaatctctattgtcacatatcgccacaaaaactactatgctctaagtggaattgggtacgagagtctagacttgtgtttggaaggagtagggattcatcatcacgtacttgaaacaggaaacgcggtatatggaaaagttcaacatgagtattctactatcaaagagaaggccaaagaaatgaatgcactcaaaccaggacctatcatcgattaccacgtctggataggagattgtgtctgccaagttactactgtagacgtgcatggaaaggaaattatgagaatgagattcaaaaggggtgcggtgcttccgattccaaatctggtaaaagttaaagttggggaggaaaatgatacaataaatctttccacttccatatcagctctcctaaattccggtggcggcaccatcgaggtaacatctaaggaagaacgtgtagattatgtactcatgaaacgtttggaatctatacatcatttgtggtctgtagtgtatgatcatcttaatgttgtgaatggcgaagaacgatgttatgtacatatgcattcatctcatcaaagtcctatgctgagtactgtaaaaacaaatttgtacatgaagactatgggagcatgtcttcaaatggactccatggaagctctagagtatcttagtgaactgaaggaatcaggtgggcggagtcccagaccagaattgcagaaatttgaatatccagatggagtgaaagacactgaatcaattgagagattggcagaggagttcttcaatagatcagaacttcaggccggtgaatcagtcaaatttggtaattctattaatgttaaacatacatctgtttcagctaagcaactaagaacacgtatacgacagcagcttccttctatactctcatcttttgccaacacaaagggtggatatttgttcattggagttgataataatacacacaaagtaattggattcacggtgggtcatgactacctcaaactggtagagagtgatatagaaaagtatatccaaaaacttcctgttgtgcatttctgcaagaaaaaagaggacatcaagtacgcatgtagattcatcaaggtgtataaacctggtgatgagactacctcgacatatgtgtgcgcaatcaaagtggaaagatgctgctgtgctgtgtttgcggattggccagaatcatggtacatggatactagtggtagtatgaagaagtattctccagatgaatgggtgtcacatataaaattttaattagggtaaggtaaaactatatataataactaacaatttgtgtatcatatagacaattaattaggtaactgttatctctttttaactaactaactaactaactaactcttatatactattaataatacatctattaatcattgattagcttattgctttaattgttttgtaaactaacactgttcattgaaaagggataacatgttacagaatataaattatatatggatttttttaaaaaggaaatacttgactggagtatatatttatttcttcattacataacacgtctgtgttctaattcttccaattcttccacatctcatataatacaggaatataatcttgttcgaaaatatgagaaagtggataaaacaatagttgattttttatctaggtggccaaatttattccatattttagaatatggggaaaatattctacatatttattttatagatgctgctaatacgaatattatgattttttttctagatagagtattaaatattaataagaaccgtgggtcatttatacataatctcgggttatcatccattannataaaagaatatgtatatcaattagttaataatgatcatctagataatagtataagactaatgcttgaaaatggacgtagaacaagacattttttgtcttatatattggatacagttaatatctatataagtattttaataaatcatagattttatatagatgccgaagacagttacggttgtacattattacatagatgtatatataactataagaaatcagaatcagaatcatataatgaattaattaagatattgttaaataatggatcagatgtagataaaaaagatacgtacggaaacacaccgtttatcctattatgtaaacacgatatcgacaacgcggaattgtttgagatatgtttagagaatgctaatatagactctgtagactttaatggatatacacctcttcattatgtctcatgtcgtaataaatatgattttgtaaagttattaatttctaaaggagcaaatgttaatgcacgtaatagattcggaactactccattttattgtggaattatacacggtatctcgcttataaaactatatttggaatcagacacagagttagaaatagataatgaacatatagttcgtcatttaataatttttgatgctgttgaatctttagattatctattgtccagaggagttattgatattaactatcgtactatatacaacgaaacatctatttacgacgctgtcagttataatgcgtataatacgttagtctatctattaaacagaaatggtgattttgagacgattactactagtggatgtacatgtatttcggaagcagtcgcgaacaacaacaaaataataatggatatactattgtctaaacgaccatctttgaaaattatgataccatctatgatagcaattactaaacataaacaacataatgcagatttattgaaaatgtgtataaaatatactgcgtgtatgaccgattatgatactcttatagatgtacaatcgctacatcaatataaatggtatattttaaaatgttttgatgaaatagatatcatgaagagatgttatataaaaaataaaactgtattccaattagttttttgtatcaaagacattaatactttaatgagatacggtagacatccttctttcgtgaaatgtaatattcttgacgtatacggaagttgtgtacgtaatatcatagcatctattagatatcgtcagagattaattagtctattatccaagaagctggatgctggagataaatggtcgtgttttcctaacgaaataaaatataaaatattggaaaactttaacgataacgaactgaccacatatctgaaaatcttataaacactattaaaatataaaatctaagtaggataaaatcacactacatcattgtttccttttagtgctcgacagtgtatactatttttaacactcataaataaaaatgaaaacgatttccgttgttacgttgttatgcgtactacctgctgttgtttattcaacatgtactgtacccactatgaataacgctaaattaacgtctaccgaaacatcgtttaatgataaacagaaagttacgtttacatgtgattcaggatatcattctttggatccaaatgctgtctgtgaaacagataaatggaaatacgaaaatccatgcaagaaaatgtgcacagtttctgattatgtctctgaactatatgataagccattatacgaagtgaattccaccatgacactaagttgcaacggtgaaacaaaatattttcgttgtgaagaaaaaaatggaaatacttcttggaatgatactgtcacgtgtcctaatgcggaatgtcaacctcttcaattagaacacggatcgtgtcaaccagttaaagaaaaatactcatttggggaatatatgactatcaactgtgatgttggatatgaggttattggtgtttcgtatataagttgtacggctaattcttggaatgttattccatcatgtcaacaaaaatgtgatataccgtccctatctaatggattaatttccggatctacattttctatcggtggcgttatacatcttagttgtaaaagtggttttacactaacggggtctccatcatccacatgtatcgacggtaaatggaatcccatactcccaacatgtgtacgatctaacgaagaatttgatccagtggatgatggtcccgacgatgagacagatctgagcaaactctcgaaagacgttgtacaatatgaacaagaaatagaatcgttagaagcaacttatcatataatcataatggcgttgacaattatgggtgtcatatttctaatctccattatagtattagtttgttcctgtgacaaaaataatgaccaatataagttccataaattgnnnccgtgaatataaatccgttaaarkaattaataattaatartnnrtaacgaacaagtatcaaaagattaangaannnttagctagaatcaattagatgtcttcttcagtggatgttgatatctacgatgccgttannagnnnnttttactcaggcactattatgacaagagatttattgtgtatggaagaagtaacaccatattacataatatatacaggctatttacaagatgcaccgttataccgttcgatgatatagtacgtactatgccaaatgaatcacgtgttaaacaatgggtgatggatacacttaatggtataatgatgaatgaattcgatactgtatgtgtgggtaccggactacgattcatggaaatgtttttcgattacaataaaaataatcccaaaaatagcatcaacaatcaaataatgtatgatataattaatagcgtagccataattctagctaatgagagatatagaagcgcgtttaacgacgatagaatatacatccgtagaactatgatggacaaattgtacgaatacgcatctctaactactattggtacgatcactggaggtgtttgttattttatctgttgatgcatctagttagtttgtataaataattatttcgatatactagttaaaattttaagattttaaatgtataaaaaactaataacgtttttatttgtaataggtgcagttgcatcctattcgaataatgagtacactccgtttaataaactgagtgtaaaactctatatagatggagtagataatatagaaaattcatatactgatgataataatgaattggtgttaaattttaaagagtacacaatttctattattacagagtcatgtgacgtcggatttgattccatagatatagatgttataaacgactataaaattattgatatgtataccattgactcgtctactattcaacgcagaggacatacgtgtagaatatctaccaaattatcatgccattatgataagtacccttatatccacaaatatgagggtgatgaacgacaatattctattaccgcagagggaaaatgctataaaggaataaaatatgaaataagtatgatgaacgatgatactctattgagaaaacatactcttaaaattggatttacttatatattcgatcgtcatgggcatagtaatacatattattcaaaatatgatttttaaaaatttaaaatatattatcacttcagtgacagtagtcaaataacaaacaacaccatgagatatattataattctcgcagttttgttcattaatagtatacatgctaaaataactagttataagtttgaatccgtcaattttgattccaaaattgaatggactggggatggtctatacaatatatcccttaaaaattatggcatcaagacgtggcaaacaatgtatacaaatgtaccagaaggaacatacgacatatccggatttccaaagaatgatttcgtatctttctgggttaaatttgaacaaggcgactataaagtggaagagtattgtacgggactatgtgtcgaagtaaaaattggaccaccaactgtaatattgactgaatatgacgatcatatcaatttgttcatcgagcatccgtatgctactagaggtagcaagaagattcctatttacaaacgcggtgacatgtgtgatatctacttgttgtatacggctaacttcacattcggagattctgaagaaccagtaacatatgatatcgatgactacgattgcacgtctacaggttgcagtatagactttgccacaacagaaaaagtgtgtgtgacagcacagggagccacagaagggtttctcgaaaaaattactccatggagttcggaagtatgtctgacacctaaaaagaatgtatatacgtgcgcaattagatctaaagaagatgttcccaatttcaaggacaaaatagccagagttatcacgagaaaatttaataaacagtctcaatcttatttgactaaatttctcggtagcacatcgaatgatgttacaactttttttagcattcttgactaaatattcataactaatttttattaatgatacaaaaatgaaataaactgtatattatacactggttaacgcccttggctctaaccattttcaagatgaggtccctgattatagtccttctgttcccctctatcatctactcaatgtctattagacgatgcgagaagactgaagaggaaacatggggattaaaaatagggttgtgtataattgccaaagatttctatcccgaaagaactgattgcagtgttcatcgcccaactgcaagtggaggattgataactgaaggcaatggattcagagtagttatatatgatcaatgtacagaaccccatgactttattatcaccgatactcaacaaacacgtcttggatcatctcatacatatattaaattcagtaacatgaatacaggtgtcccatctagtattccaaaatgttccagaactctctgtatttctgtatattgtgatcaagaggcgggagacataaaatttgaggagtatactcaagaatcaagtgatatcagtattagagttaagtatgattcatcatgtattgattatctgggtattaatcaaagtttcatgaatgaatgtattcgaagaattacaacatgggatagagaatcatgcgtcagaattgatacacagactataaataaatatcttaagtcttgcaccaacacaaaattcgaccgtaatgtctacaaaaggtacatactgaagagtaaagcactccatgctaaaacagagttgtaatagatataaaatactttttataataattaggctagaaaaatctcactcacatgtaatcttaaaaaaatgatatgatagttcttacaagtagcgattgagttttaaatggattctattaattaccggggaacttaacaattcgttctgatctacagacattggttaataaatcatcttattttgccaatatattaaaatgtggaaactccactaataatattacattgtgcgactttcaagatgatgtgatatatagggttatacagttttaacaattatataatagagatagaaagtacaaaagatgtagaatcaatgatatggcacgctaaacagttgggtgtggaatcattgctaaaagaatgtcaaaattatttgcttagaatattacgtatataattgtttagaaatttatagaataactaatattaatacattatcgtatatctacaacgatataagaaacttcatattggataatattactattaatatataaggatccagattttatatatttgcctaaatacattattatagatttactaggacaatcacctaaatgtttttaacgaagataatgtggtaaagattatatacacttatatatcttccgatatctacaaggatattccatatcatcattgtgtaaactaaataacgttttctatggcatttaataaggacattggatatgtggaaaagtgatgtatggaagttagtacattatcaacttctccttattgattgaaaatgaaaatataaatagtttttatgtatagcggtatctaccctatagttttattgcttactactaacatggattcagatacagatacagatacagatacagatacagatacagatacagatacagatgtagaagatatcatgaatgaaatagatagagagaaagaagaaatactaaaaaatgtagaaattgaaaataataaaaacattaacaagaatcatcccagtgaatatattagagaagcacttgttattaataccagtagtaatagtgattccattgataaagaagttatagaatatatcagtcacgatgtaggaatatagatcatatctactaatttttataatcgatacaaaacataaaaacaactcgttattacatagcaggtatggaatccttcaagtattgttttgataacgatggtaagaaatggattatcggaaatactttatattctggtaattcaatactctataaggtcagaaaaaatttcactagttcgttctacaattacgtaatgaagatagatcataaatcacacaagccattgttgtccgaaatacgattctatatatctgtattggatcctttgactatcaacaactggacacgagaacgtggtataaagtatttggctattccagatctgtatggaattggagaaaccgatgattatatgttcttcattataaagaatttgggaagagtattcgccccaaaggatagtgaatcagttttcgaagcatgtgtcactatgataaacacgttagagtttatacactctcaaggatttactcatggaaaaatagaaccgatgaatatactgattagaaataaacgtatttcactaattgactattctagaactaacaaactatacaaaagtggaacacatatagattacaacgaggacatgataacttcaggaaatatcaattatatgtgtgtagacaatcatcttggagcaacagtttcaagacgaggagatttagaaatgttgggatattgcatgatagaatggttcggtggtaaacttccatggaaaaacgaaagtagtataaaagtaataaaacaaaaaaaagaatataaacaatttatagctactttttttgaggactgttttcctgaaggaaatgaacctctggaattagttagatatatagaattagtatacatgttagattattctcaaactcctaattatgacagactacgtagactgtttatacaagattgaaattatattctttttttatagagtgtgggggtagtgttacggatatctaatattaatattagactatctctatcgcgctacacgaccaatatcgattactatggatatcttcagggaaatcgcatcttctatgaaaggaaagaatgtattcatttctccagcgtcaatctcgtcagtattgacaatactgtattatggagctaatggatccactgctgaacagctatcaaaatatgtagaaaaggaggagaacatggataaggttagcgctcagaatatctcattcaaatccatgaataaagtatatgggcgatattctgccgtgtttaaagattcctttttgggaaaaattggcgataagtttcaaactgttgacttcactgattgtcgcactatagatgcaatcaataagtgtgtagatatctttactgagggaaaaatcaatccactattggatgaaccattgtctcctgatacctgtctcctagcaattagtgccgtatactttaaagcaaaatggttgatgccattcgaaaaggaatttaccagtgattatcccttttacgtatctccaacggaaatggtagatgtaagtatgatgtctatttacggcgagccatttaatcacgcatctgtaaaagaatcattcggtaacttttcaatcatagaactgccatatgttggagatactagtatgatggtcattcttccaaacaagattgatggattagaatccatagaacaaaatctaacagatacaaattttaagaaatggtgtaactctctgaaagctacgtttatcgatgtgcacattcctaagtttaaggtaataggttcgtataatcttgtggatacgctaataaagttgggactgacagatgtgttctattcaactggtgattatatcaatatgtgtaattcagatgtgagtgttgacgctatgattcacaaaacgtatatagatgtcaatgaagagtatacagaagcagctgcagcaacttctgtactagtggcagactgtgcatcaacagttacaaatgagttctgtgcagatcatccgttcatctatgtgattagacatgtcgatggtaaaattcttttcgttggtagatattgctctccaacaactaattaagcacattcttaatattagaatattatatagttaagatttttactaacaggttaacatttttttaaaaatagaaaaaacatgtggtattagtgcaggtcgttattcttccaattgcaattggtaagatgacggccaactttagtacccacgtcttttcaccacaacactgtggatgtgacagactgaccagtattgatgacgtcagacaatgtttgactgaatatatttattggtcgtcgtatgcataccgcaacaggcaatgcgctggacaactgtatgacacactcctctcttttaaagatgatgcggaatcagtgttcatcgacgttcgtgagctggtaaaaaatatgccgtgggataatgttaaggattgtacagagatcatccgttgttatataccggatgagcaaaaaaccatcagagagatttcggccatcattggactttgtgcatatgctgctacttactggggaggtgaagaccatcccactagtaacagtctgaacgcattgtttgtgatgcttgagatgctcaattacatggattataccatcatattctggcgtatgaattgatgagttacagcttgacatttcttctttcctccctcttcttctacctttcccagaaacaaactttttttacccactataaaataaaatgagtatactacctgttatatttcttcctatatttttttattctccattcgttcagacttttaacgtgcctgaatgtatcgacaaagggcaatattttgcatcattcatggagttagaaaacgagccagtaatcttaccatgtcctcaaataaatacgctatcatccggatataatatattagatattttatgggaaaaacgaggagcggataatgatagaattatacagatagataatggtagcaatatgctaattctgaacccgacacaatcagactctggtatttatatatacattaccacgaacgaaacctactgtgacatgatgtcgttaaatttgacaatcgtgtctgtctcagaatcaaatatagatcttatctcgtatccacaaatagtaaatgagagatctactggtaaaatggtatgtcccaatattaatgcatttatttctagtaacgtaaacacagaattatatggagcggacatcgacgccttagaaataagnnagacttaaacaacggrcacnnnnnnnnnnnnnnnnnnnnnnnnnnnnnnnnnnnnnnnnnnnnnnnnnnnnnnnnnnnnnnnnnnnnnnnnnnnnnnnnnnnnnnnnnnnnnnnnnnnnnnnnnnnnnnnnnnnnnnnnnnnnnnnnnnnnnnnnnnnnnnnnnnnnnnnnnnnnnnnnnnnnnnnnnnnnnnnnnnnnnnnnnnnnnnnnnnnnnnnnnnnnnnnnnnnnnnnnnnnnnnnnnnnnnnnnncgtcttttggataagtaatggtatatgtattacgaagaagaagacgaggacggagacggtagaataagtgtagcaaataaaatctatatgaccgataagagacgtgttattacatcctggttaaacattaatcctgtcaaggaagaagatgctacaacgtttacgtgtatggcgtttactattcctagcatcagcaaaacagttactgttagtaaacgtgaatgtatgttgttacatttccatatcaattgagtttataagaattttttatacattatcttccaacaaacaattgacgaacgtattgctatgattaactcccacaatactatatatattattaatcattaacttgcagactatacctagtagtgctattttgacatactcatgttcttgtgtaatcgcagtatctatattattaaagtacgtaaatctagctatagttttattatttaattttagataatatactgtctccgtatttttaaaaaattaccacatcctttattaaatcatgaatgggaatttctgtgtcatcgttagtatattgtgaacaacaagagcagatatctataggaaagggtggaatgcgatacattgatctatgtagttttaaaacatacgcgaactttgaagaatttatataaatcatctcacgagatattgctctctgtcatattcatacacctgtataaactttctagacatcttacaatgtgttattttatgatcatatttacatatttactggtatatcaaagatgttagattagttaatgggaatcgtctataataatgaatattaaacaattataggaggagtttatacctacaaaaacatcataaaaatgagtcatcgtccgatttatgttttaaatatactaacattactaccttcagaaattatatacgaaatattatacatgctgacaattaacgatctttataatatatagtatccacctaccaaagtataattgtatttttctcatgtgatgtgtgtaaaaaactgatattatataattatcttagtacctatgatgaagatgaagatgaagatgaagatgatggtccgtatatattttgtatcattatcgttattgctattccatagttacgccatagacatcgaaaatgaaatcaccgaattcttcaataaaatgagagatactctaccagctaaagactctaaatggttgaatccagtatgtatgtttggaggcacaatgaatgatatggccgctctaggagagccattcagtgcaaagtgtcctcctattgaagacagtcttttatcgcatagatataaagactatgtggttaaatgggaaaggctagaaaagaatagacggcgacaggtttctaataaacgtgttaaacatggtgatttatggatagccaactatacatctaaattcagtaaccgtaggtatttatgtaccgtaaccacaaagaatggtgactgtgttcagggtgtagttagatctcatgtgtggaaaccttcttcatgcattccaaaaacatatgaactaggtacttatgataagtatggcatagacttatactgtggaattctttatgcgaaccattataataatataacttggtataaagataataaggaaattaatatcgacgattttaagtattcacaagcgggaaaggaattaattattcataatccagagttagaagatagtggaagatacgactgttacgttcattacgacgacgttagaatcaagaatgatatcgtagtatcaagatgtaaaatacttacggttataccgtcacaagaccacaggtttaaactaatactagatccgaaaatcaacgtaacgataggagaacctgccaatataacatgcagtgctgtgtcaacgtcattatttgtcgacgatgtactgattgaatgggaaaatccatccggatggattataggattagattttggtgtatactctattttaactagtagaggcggtatcaccgaggcgactttgtattttgaaaatgttactgaagaatatataggcaatacatatacatgtcgtggacacaactattattttgataaaactcttacaactacagtagtattggagtaaatacacaatgcatttttatatacattactgaattattattattaattatatcgtatttgtgctatagaatggatgaagatacgcgactatctaggtatttgtatctcaccgatagagaacatataaatgtagactctattaaacagttgtgtaaaatatcagatcctaatgcatgttatagatgtggatgtacggctttacatgagtacttttataattatagatcagtcaacggaaaatacaagtatagatacaacggttactatcaatattattcatctagcgattatgaaaattataatgaatattattatgatgattatgatagaactggtatgaacagtgagagtgataatatatcaatcaaaacagaatacgagaatgaatatgaattctatgatgaaacacaagatcaaagtacacaactagtagattacgacattaaactcaaaaccaatgaggatgattttgttgatgaattctatggttatgatagatcagtgggtgtccatgattatatagatgtatcaattaataaagtagtatatggaagagagtctcacgtaagatggtgggatatatggcaagaacataatgatggcgtatacagtataggaaaggagtgcatagataatatatacgaagacagacataccgtagacgaattctacaagatagacagcgtatcagatgtagatgacgcagaacatatatctcagataactaatgatgtatctacacaaacatgggaaaagaaatcagagttagatagatacatggaaatgtatcctcgtcatagatatggtaagcattctgtctttaagggattttctgacaaagttagaaaaaatgatttagacatgaacgtggtaaaagaattactttctaacggtgcatctctaacaatcaaggatagcagtaataaggatccaattgctgtttattttagaagaacaataatgaatttagaaatgattgatatcattaacaaacatacaactatctatgaacgcaggtatatagtacactcctatctaaaaaattatagaaatttcgattatccatttttcagaaagttagttttgactaataaacattgtctcaacaattattgtaatataagcgacggcaaatatggaacaccactacatatattagcatctaataaaaaaataataactcctaattacatgaagttattagtgtataacggaaatgatataaacgcacgaggtgaagatacacaaatgcgaactccattacacaaatatttgtgtaaatttgtatatcataatattgaatatggtatccgatactataatgaaaagattatagacgcatttatagagttaggagccgatctaactattccaaatgacgatggaatgataccagtagtttactgtatacactcaaatgccgaatatggttataacaatattactaacataaagataatacgtaaactacttaatcttagtagacatgcgtcacataatctatttagagatcgagtcatgcacgattatataagtaatacatatattgatcttgagtgtttagatatcattagatcacttgatgggtacgatattaattgttactttgaaggacgtacaccacttcattgcgctatacaatataacttcactcagattgctgagtacttattagatcgaggagctgatatatcattaaagacagacgatggtaaaactgtatttgatttatcgttatgtagttacattcctcttaaatggactagctttttgattagtcgtctaccgcctaaaagtgtcatatgctcactgactaaccatataatagattatgttcttacgaacaatagacgtattatttggcagagtcaaatgattaataagtacgtactgttactggacccatccttttattatagattcagaaatgttatcgaaaacaaattagaccaatacaataatcgttataatatgttcgaacacgatagggacgttaatgaaaagtatggcaaagtcttacatgacctcgatacatatatcaaggatgtacaagtattaaaatctacttccatcactaataatataacactatacgacactattataaataataagtcagagtttcctatacgtcgtgtaaacgacaaacaattaattaatctcataaaatccaatacatatcataatcttatcgaaaaagttattaaaaatacattagagaaatatactttaactaatatagtcctcgagtatatgatctcatctcgatctcaatcatcttatttgagtcgtattcctaatgagatattactcgaaatattatataaactcgacatgtacgatttacgtaatctatatacaagatatatgagagagaatgatatcacagagtatcatatagagaatacgaggtctgtttctacacagacatgaataatgaatacacatacaacgttttttttaatcttagatataacactaattacatcaagattatatattgaaatcgtaatttgagttgtctgatcatcatggatatcgaaaataatatacgtaacattagcaatcttttagatgattgatatattattatgcgatgtaatcataactatcggagatgtagaaattaaagcgcataaaactattttggttgccggatctacgtattttaaaacaatgttcacaacatctatgatagcgagagatctagcaactagagtaaatatacagatgttcgataaagatgccgtcaaaaatattgtacagtacttatacaataggtatataagttctatgaatgtgatagacatattaaaatgcaccgactaagaacgtaaaacgaactatagaatgttatacaatgggtgatgataagtagaagatgttacccgatatacccatagcattatctagttatggcatgtgtgtattagatcaatacatatacattataagcggtcgtacccaacactgattatacatcggtacatacagtaaatagcatagatatggaggaggatacaaatatttcaaataaagttatgagatacgcgctgtcaataatatatggaagacattacctaacttctgaactggaactataaatccaggctctcgcataaagatgaatatatatgttgtatgcgacatcaaagatgaaaaaatgttaagacttatatatttagatataacacgaatatgtatgacggatgggaattggtaacgatgacagaaagcagattgtcagctctgcatactattcttcatgacaataccataatgatgttacattgttatgaagcgtatatgttacaagatacatttaatgtgcttacggaacatatatttagaaacatctactaacgattttttatgnttgtattattaatggtatgtaatatggatttaattgattgtgtacacgataccaatttgnnnnnnnnnnnnnnnnnnnnnnnnnnnnnnnnnnnnnnnnnnnnnnnnnnnnnnnnnnnnnnnnnnnnnnnnnnnnnnnnnnnnnnnnnnnnnnnnnnnnnnnnnnnnnnnnnnnnnnnnnnnnnnnnnnnnnnnnnnnnnnnnnnnnnnnnnnnnnnnnnnnnnnnnnnnnnnnnnnnnnnnnnnnnnnnnnnnnnnnnnnnnnnnnnnnnnnnnnnnnnnnnnnnnnnnnnnnnnnnnnnnnnnnnnnnnnnnnnnnnnnnnnnnnnnnnnnnnnnnnnnnnnnnnnnnnnnnnnnnnnnnnnnnnnnnnnnnnnnnnnnnnnnnnnnnnnnnnnnnnnnnnnnnnnnnnnnnnnnnnnnctgaacaactatcaaaatatatnnagaatatgaatgagaatacacccgatgataagaaggatgacaataatgacatggacgtagatattccgtattgcgcgacactagctaccgcaaataaaatatacggtagtgatagtatcgagttccatgcctcattcctacaaaaaataaaagacgattttcaaactgtaaactttaataatgcgaaccaaacaaaggaactaatcaacgaatgggttaagacaatgacaaatggtaaaattaattccttattgactagtccgctatccattaatactcgtatgatagttattagcgccgtccattttaaagcaatgtggaaatatccattttctaaacatcttacatatacagacaagttttatatttctaagaatatagttaccagtgttgatatgatggtgggtaccgagaatgacttgcaatatgtacatattaatgaattattcggaggattctctattatcgatattccatacgagggaaactctagtatggtgattatactgccggacgacatagaaggtatatataacatagaaaaaaatataacagatgaaaaatttaaaaaatggtgtggtatgttatctactaaaagtatagacttgtatatgccaaagtttaaagtggaaatgacggaaccgtataatctggtaccgattctagaaaatttaggacttactaatatatttggatattatgcagattttagtaagatgtgtaatgaaactatcactgtagaaaaatttctacatacgacgtttatagatgttaatgaggagtatacagaagtatcggccgttacaggagtattcatgactaacttttcgatggtatatcgtatgaaggtctacataaaccatccattcatatacatgattaaagataacaccggacatacactttttatagggaaatactgctatccgcaataaatataaacaatagacttttatcacgttatctcatgtataaaatattacaaatagtatagcataaactaaagtcgatacatacattaaaacttaaataataatgtaatttacaattaatagtataaactaaaaaaattaaaaaattaaaaacaatatcattattataagtaatatcaaaatgacgatatacggattaatagcgtatcttatattcgtgacttcatccatcgctagtccactttacattcccgttattccgcccatttcggaagataaatcgttcaatagtgtagaggtattagtttctttgtttcccgatgaccaaaaagactatacagtaacttctcagttcaataactacactatcggtaccaaagactggactatcaacgtactatccacacctgatggtctggacataccattgactaatataacttattggtcacggtttactataggtcgtgcattgttcaaatcagagtctgaggatattttccaaaagaaaatgagtattctaggtgtttctatagaatgtaagaagccgtcgacattacttacttttttaaccgtgcgtaaaatgactcgagtatttaatagatttccagatatggcttattatcgaggagactgtctagaagccgtttatgtaacaatgacttataaaaatactaaaactggagagactgattacacgtacctctctaatggggggttgcctgcatactatcgtaatggggtcgatggttgattattgattagtatattccttatattccttattctttttattcacacaaaaagaacatttttataaacatgaaaccactgtctaaatgtaattatgatcttgatttatagatgatgatcagccttcagaggattttgaccagtatgtttaatatgaaaaaaaacataactattaagcgctattgcgctattgtgcttaattattttgctctataaactgaatatatagccacaattattgacgggcttgtttgtgaccgacaatcatgaattttcagaaattatctctggctatatatcttacggtgacatgttcgtggtgttatgaaacatgtatgagaaaaactgcgttgtatcatgacattcaattggagcatgtagaagacaataaagatagtgtagcatcgctaccgtacaagtatctacaagtagtcaaacaaagagaacgtagtagattgttggctacatttaattggacggatatagctgagggtgttagaaatgagttcattaaaatatgtgatatcaacggaacatatctatataattatactattgctgttagtataattattgattccacggaagaactaccaacagttactccaattacaacatatgaaccttctatatataattatactatcgattatagcactgttattactactgaagaactacaagtgactccaacatatgcacctgtaacaactcctcttccaacatcagcagttccttatgatcaacgatcgaataacaatgtaagtactatatctattcaggtactgagtaaaatattgggagtcaatgaaacagaattaactaattatcttattatgcataaaaatgacactgttgacaataacaccatggttgatgatgagacatctaacaataacacattacatggtaatataggatttttggaaataaataattgttataatgtttctgtgtcagatgctagttttagaataacattagtaaacgatacttctgaagaaattttgctaatgctaacaggaactagttcatccgacaccttcatatcttccaccaatatcactgaatgtttgaaaacattaatcaataatgtgtcgattaatgatgtacttataacacaaaatatgaatgtaacatctaattgtgataaatgctcaatgaatttgatggcatccgttattcctgcagttaatgaatttaacaatacgttgatgaaaattggtgtaaaagatgatgaaaacaatacggtatataaatattataattgtaaactaactacaaattctacatgtgatgagttaatcaatttagatgaagtcattaacaacataactctgacaaatattatacacaatagtgtttcgacaactaacagcagaaaaagacgagatctgaatgatgagtttgaattttccacttccaaggaattagattgtctttacgaatcatatggtgtaaacgatgatataagtcattgttttgcatcacctagacgtagacgatctgacgacaaaaaggagtacatggacatgaaattattcgaccacgcgaaaaaagatttaggaatagacagtgttattcctagaggtacaacccatttccaagtaggtgcatctggtgcaagtggtggtgttgtaggagatagtttcccatttcaaaatgttaaatcgcgtgccagtctattggcggaaaaaataatgcctagagtacctattactgctaccgaagctgatctatatgcaactgtaaatagacaacccaagttaccagcaggtgttaaaagtactccgtttacagaggcgcttgtgtctacgataaaccaaaagctttctaatgttagagaggtaacttatgcttcgctcaatctgccaggatcaagtggctatgttcatagaccatctgattctgttatttatagcagtataagacggtcacgtttacctagtgatagcgatagtgattatgaggatatacaaactgttgttaaggaatataatgaaagatatggtagatcagtcagtagaacacagtcatcaagtagtgaaagcgattttgaagatatagatactgttgttagggaatatagacaaaaatatggcaatgcaatggcaaaaggacgtagtagttcccctaaacctgatccattatatagtactgttaagaaaacaactaaaagtctatctactggtgtagacatagttacaaaacaatcagactattctctattacctgacgttaatactggcagttctattgtgtcacctctcaccagaaaaggagctactagacgacgacctagacgccctacaaatgatggtctacagagtccaaatcctcctctccgtaatccacttcctcaacatgatgattattctcctccacaagtacacagacctccaccacttcctcctaaaccagtccaaaattcgccacaacttccccctagaccagtaggtcaattactacctcctcctatagatcaaccagataaaggatttagtaagtttgtatcacctagacggtgtagaagagcaagctctggagtcatatgtggtatgatacaatcaaaaccaaacgatgatacctattcacttcttcaacgatcaaaaattgaaccagaatatgtggaggttggtaatggtatacccaagaacaatgttcctgtaataggtaataaacatagtaaaaaatatacatcgacgatgtcaaaaatatcaacaaaatttgataaatctacggcatttggagcagcaatgttactaactggtcagcaggccattagccaacagactagatcaactacgttgagtagaaaagatcagatgagtaaggaagaaaagatattcgaagcagttacaatgagtctatcaactataggttcaacgttgacgtctgcaggtatgacgggtggtccaaaactaatgattgcaggaatggctataacggctataactggtataatagatacgataaaagatatatattacatgttttcaggacaggagaggccagtagatcctgttattaaattatttaataagtacactggcttaatgtccgataataataaaatgggtgtaagaaaatgtttgacacccggtgacgacacacttatttatatcgcatacagaaacgataccagttttaaacagaatacggatgcgatggctttgtatttcttagatgttatcgattcagagatcctatatctaaacacatcaaatttagttctagagtatcaactaaaggtggcttgccccataggaacattaagatctgtagatgtggacataactgcgtatacaatattatatgatacagcggataatattaagaaatacaagtttatcagaatggcaacgctactatccaaacatccagttattagattgacatgtggtttagcagcaacattggtgattaaaccgtacgaggtacccatcagtgatatgcaactactaaaaatggcgacgcctggtgaaccagaatccactaaatctataccatccgatgtcygtgataggtatcctctaaagaaattntatcttttagctggtggttgtccctatgatacatctcaaacttttattgtacatactacttgcagtattctactaagaacagctacacgggatcagtttagaaacagatgggtgttacaaaatccatttagacaagaagggacatataagcaactgtttacctttagcaaatacgattttaacgacaccataatcgatcctaatggtgtggtgggtcatgctagcttttgtaccaatagaagcagcaaccaatgtttctggtccgaacctatgatattggaagatgtatcatcgtgtagttctagaactagaaaaatatacgtaaaactgggaatatttaatgctgaaggttttaatagttttgtactaaattgtccaactgggtctacacctacatacatcaaacataaaaatgcggacagtaacaatgttatcatagagctacctgtaggtgattacggcacagccaaattgtattcagcaacaaaaccatcgaggatagctgtgttctgcacacataactatgataaacgattcaaatcagatattatagttctaatgtttaataaaaacagcggtattccattttggagcatgtacacaggaagtgtaactagtaaaaatagaatgtttgccacattggctagaggaatgccgtttagatcaacgtattgcgataacagacgacgatcaggttgttattatgcaggaataccatttcatgaagatagtgtagaaacagatatacattatggaccagaaataatgttaaaggaaacatatgacataaacagtattgacccacgagttataacaaagtcaaagacccattttcctgctccattgagtgtaaaattcatggttgacaatttaggaaatggatatgacaaccctaattcattttgggaagatgctaaaactaagaaacggacatatagtgcaatgacgataaaagtcctaccatgtacagtgagaaataaaaatatagactttggatataactatggagatattatttctaatatggtttatctacaatctactagtcaggattatggagatggtaccaaatatacatttaaatccgtaactagatcagatcatgagtgtgaatctagcttagatctaacgtctaaggaagtaactgtgacatgtcctgcgtttagtataccaagaaatatatcaacatatgaaggtctatgctttagtgttactacatctaaagatcattgtgctacaggtattggttggttaaaatctagtggttatgggaaggaagatgctgataaaccacgtgcttgttttcatcattggaattattacacactgtcgttggattattactgttcatacgaagatatttggagaagcacctggcctgactatgatccatgtaagtcatatatccatatagagtatagagatacatggatagaatctaatgtgttacagcaacctccttacacattcgaattcattcatgacaattctaacgaatatgtggataaagaaattagtaacaaattaaatgatctgtacaatgaatacaagaagattatggaatatagcgacggatcattgccggcgtctataaacagattagcaaaggcattgacttcagagggtagagaaatagcaagtgttaatatagatggtaatctgttagatatcgcatatcaagcagataaggaaaagatggccgacatacagacaagaataaatgatattattagagatttgtttatacacactctatcagacaaagatataaaagacattatagaatccgaagaaggtaagagatgttgtataatagatgttaagaacaatcttgttaaaaagtactattctattgataattatctatgtgatactttagatgattatatatacacctctgtagaatataacaaatcctatgtgttagtaaacgatacttatataagctatgactatcttgaatcatcaggtgtagttgttctatcatgttatgaaatgactataatctccttggatacaaaagacgccaaagatgctatagaagatgtgatagtagcaagtgcggtagccgaagcattgaatgacatgtttaaggaatttgataaaaacgtaagtgctattataataaaagaagaagataattatctaaacagttcgcccgatatctaccatataatatatatcataggtggcactattctgctactgttagtcattattttaatattggcaatttatatagcgcgcaataaatacagaaccaggaaatatgaaataatgaaatatgacaatatgagcattaaatctgagcatcatgatagtcttgaaacagtgtctatggaaattattgataatcggtactaataaaatagtttaactcttttagaaccagtttggtactgtaatttcagttcattactcgttgagaatattgatgatttttttaaatgagtatcggtagttacatattaccatatcatccattatataatcgatgatgcatgtattagaatactttccgaataagtcttctaaatattgtattaattatgaaaaactatgctatgtgagtatgattcaaagatgtttaatgatacgatactagattttatctctagcgagattgtttagaatcatttatcataactatgtttaataaattcatcaacgaatatcgataaagacctcttgtaattcgagtataggaagtagtattaccatatcaacttccgagttaacaattactctaaaacatgaggattgtactcctgtctttattggagatcactatttagtcgttgataaactagtaacctcaggtttctttacaaacgataaagtacaacatcaagacctcacaacacagtgcaagattaatctagaaatcaaatgtaattctggaggagaatctagacaactaacacccacggcgaagtatactttnatgccnnnnnnnnnnnnnnnnnnnnnnnnnnnnnnnnnnnnnnnnnnnnnnnnnnnnnnnnnnnnnnnnnnnnnnnnnnnnnnnatatattccgacatgaatggcgtcgcttatcatatgttatatcctaaatgttgatcatattccacaaatgattgtgaacgagattaaatcatctaacaaataattagttttttatgacattaacatataataaataaattaatcattattgacttaacgatgacgaaagttatcattatcttaggattcttgattattaatacaaattcgttgtgtctatgaaatgtgaacaaggtgtctcatattataatgcacaagaattaaagtgttgtaaactatctagccaggaacatattcagattatcgatgtgataaatacagcgataccatctgtggacattgtccaagtgnnacattcacgtcaatatataatcgttctcctcggtgtcatagttgtagaggtcacaccttgtacacctaccacaaatagaatatgtcattgtgactcgaatagttatcgtctccttaaagcttctgatggtaactgtgttacatgtgctcctaaaacaaaatatggtcgtgtgtacggaaagaaaggagaaaatgatatggaataccatttgtaagaaatgtcggaagggtacttattcagatattgtatctgactctgatcaatgtaaacctatgacaagataagacttactngcatctactggnnnnnnnnnnnnnnnnnnnnnnnnnnnnnnnnnnnnnnnnnnnnnnnnnnnnnnnnnnnnnnnnnnnnnnnnnnnnnnnnnnnnnnnnnnnnnnnnnnnnnnnnnnnnnnnnnnnnnnntttcggtctgtgtcgccannatggycacaccgagccntctgcgtgctcctccatcgaggacgacttcaactatgacagctcggtggcgtctgccagcgtgtacanntacgaatggcatttctaagaaaagtctncggtatcctttctacaatttcctttaacaacggcaacagctgcagtatttttatactttgaatgcatcggacatttatacaagggagtcctgttctaatattggcatcaatgttcggatctataggcttgattttcgcattgactttacacagacatgaacatcccctgaatctgtacatactttgtggatttacactgttagaatctctaacgctggcctctgttgttactttctatgatgcacgtatcgttatgcaagctttcatgttgactactgcagtgtttcttgctctgactacatatactctacaatcaaagagagatttcagtaaacttgtaacaggattgtttgctgctttctggattttaattttgtcaggagtcttgaggataaagtttaaaatagaattaataaagaacatataggtcattttttaaacatggatagaaaccaaggttgttagttaataatatacaagatattttttctcactctgatccatgtaaaccaaggacgagagacactctcattcctcattcacgacaccattaaaaatggaaattaaagccctctattaagcacagacggctacaggtctaccatcaggttaccttcgtctaccttcacaatggcctctccttgtgcccagttcagtccctgtcattgccacgctactaaggactccctgaataccgtgactgacgtcagacattgtctgactgaatacatcctgtgggtttctcatagatggacccatagagaaagcgcagggcctctctacaggcttctcatctctttcagaattgatgcaatggagctatttggtagcgagttgaaggagttctcgaattcacttccgtgggacaatatcgacaattgcgtggagatcattaaatgtttcatcagaaatgactccatgaaaaccgccaaagaactttgtgcaataattggactttgtactcaatcagctattgtcactggaagagtcttcaatgataagtatatcgacatactacttatgctgcgaaagattctgaacgagaacgactatctcaccctcttggatcatatcctcactgctaaatactaaatctccttcatgctctctcactaatactcttactcactacactttttatcatcttatgatgaatgattgccttcatcattttttcgtggaatataatataggaataattagcaccagaatagctatggatatctcgttaagaatattctctataagagacataatgtagacatagttattatatccttcttagataagtgttacgctactggaaagtttccatcgttattattacatgaagatgatataattaaaccaacattgagattggctcttatgttagctggattgaattactgtaataaatgcatcgagtatagagggatatagcaattctcgataatagtcatgcaatatttgaatgagactgataatttaggtaatacagtactacacacatatctttctagattatatatcgttaaaaatctgtaagatgtatatttctcataagtatccactgtgtaatattattaatggatatatagataacgcaatagggactaatagtattgtaaaagatataatcgactatttacgtacatatccagatatctatattcctactagtttgctgcgtagttgcatcattgatatgcatgatttatcaggattcagagatgaattactaagtaaactacaatcccacaataagtaagaatcaaatatcaaaaactcacttttgatttttctagtcttaagtaatacatatatttattaatagacctatgaaataaaaaaaggtaacaatggattcgcgtatagctatttacgtattagtatcggcatctcttttgtatcttgttaattgtcacaaactagtacattacttcaatctgaaaataaatggaagtgatataactaatacagcagatatattgctggacaattatccaattatgacctttgatggaaaggatatttatccatctatctcgttcatggtcggtaataaacttttcctagatctttataaaaatatctttgaagaatttttcagactatttcgagtatctgtaagtagtcaatacgaggaattagaatattattattcatgtgattatactaacaaccgtcctacaattaaacaacattacttttataacggcgacgaatatactgaaattgatagatcgaaaaaagccactaataaaaacagttggttaattacttcaggctttagactacaaaaatggttcgatagcgaagattgtataatttatctcagatctttagttagaagaatggaagacagtaacaaaaacagtaaaaaaacttagtacttagatatcgaaaaaatatatttttgtagactcttgagaatagaaggaaaacatgtacataattataaaaaatgaaaatcaatggcgaataagacagtgcgattcgcaccatggagtcggtagatttcatggctgtcgatgagcagtttcacgacgacctcgatctttggtcattatctttggtagatgattataaaaaacatggattaggtgttgactgttatgttctagaaccagttgttgacaggaaaatatttgatagatttctccttgaaccaatttgtgatcctgtagatgttctgtatgattattttaggattcatagagataatattgatcagtatatagtagatagactgtttgcatatattacatataaagatattatatctgcattagtgtcaaagaattatatggaagatattttctctataattattaagaattgtaattctgtgcaagatctcttactttactatctatctaatgcatatgtagaaatagacattgttgatcttatggtagatcatggggctgtaatatataaaatagaatgcttgaatgcctattttaggggaatatgtaaaaaggaaagtagtgttgttgagtttattttgaattgtggtatcccagatgaaaatgatgttaaattagatctatataaaataattcagtatactagggattccttgtagatgaacccacagtattagaaatttataagctttgtatcccatatattgaagatatcaatcaactagatgctggtggaaggaccttgctttatcgcgctatctatgcaggttatatagatttagtatcatggctattagaaaatggagcaaatgtcaacgcagtaatgagtaatggatatacatgtcttgacgtggccgtggataggggatctgtcatcgcccgtagggaagcacatcttaaaatattagaaatattgcttagagaaccattgtctattgactgtataaaattagctatacttaataatacaattgaaaaccatgatgtgataaagctctgtatcaagtattttatgatggtagattattcactttgtaatgtgtatgcatcatcactctttgattatataattgattgtaaacaagaattggagtacattaggcagatgaaaattcataatacaaccatgtatgagttaatctataatagagacaaaaacaagcatgcttcccatattctacataggtattctaaacatccagttttgacacagtgtatcactaaaggattcaagatttacacagaagtaaccgagcaggtcactaaagctctaaacagacgtgctctaatagatgagataataaacaatgtatcaactgatgacaatctcctatcaaaacttccattagaaattagggatctaattgtttcacaagctgtcatatagagttctatccacccacctttcttgaaatgagttaatagtcataagttagttaagtcataagttagttaagtcatnntgtnnnnnnnnnnnnnnnnnaagtcannnnnnnnnnntanntnnnnnncataagttnnttaagtcataagttagttaagtcataagttagttaagtcataagttagttaagtcataagttagttaagtcataagttagttaagtcataagttagttaagtcataagttagttaagtcataagttagttaagtcataagttagttaagtcataagttagttaagtcataagttagttaagtcataagttagttaagtcataagttagttaagtcataagttagtttatagtctaacacttctaatttttataccttgatctttttctctaattatgaaaaagtaaatcattatgaagatggatgaaatggacgagattgtgcgcatcgttaacgatagtatgtggtacgtacctaacgcatttatggacgacggtgataatgaaggtcacatttctgtcaataatgtctgtcatatgtatctcgcattctttgatgtggatatatcatctcatctgtttaaattagttattaaacactgcgatctgaataaacgactaaaatgtggtaactctccattacattgctatacgatgaatacacgatttaatccatctgtattaaagatattgttacgccacggcatgcgtaactttgatagcaaggataaaaaaggacatattcctctacaccactatctgattcattcactatcaatcgataacaagatctttgatatactaacggaccccattgatgactttagtaaatcatccgatctattgctgtgttatcttagatataaattcaatgggagcttaaactattacgttctgtacaaattattgactaaaggatctgaccctaattgcgtcgatgaggatggactcacttctcttcattactactgtaaacacatatccgcgttccacgaaagcaattattacaagtcaaagagtcacactaagatgcgagctgagaagcgattcatctacgcgataatagatcatggagcaaacattaacgcggttacgaaaatcggaaatacgccgttacacacttaccttcaacagtataccaaacatagtcctcgtgtggtgtatgctcttttatctcgaggagccgatacgaggatacgtaataatcttaattgtacacccatcatggaatacataaagaacgattgtgcaacaggtcatattctcataatgttactcaattggcacgaacaaaaatacgggaaattacaaaaggaagaaggacaacatctactttatctattcataaaacataatcaaggatatggaagtcgctctctcaatatactacggtatctactagatagattcgacattcagaaagacgaatactataatacaatgactcctcttcataccgccttccagaattgcaataacaatgttgcctcatacctcgtatacattggatacgacatcaaccttccgactaaagacgataagacagtattcgacttggtgtttgaaaacagaaacattatatacaaggcggatgtcgttaatgacattatccaccacagactgaaagtatctctacctatgattaaatcgttgttctacaagatgtcggagttctctccctacgacgatcactacgtaaagaagataatagcctactgcctattaagggacgagtcatttgcggaactacatactaaattctgtttaaacgaggactataaaagtgtatttatgaaaaatatatcattcgataagatagattccatcatcgaaaaatgtagtcgtgacataagtctcctcaaagagattcgaatctcagacaccgacttgtatacggtattgagaacagaagacattcggtatcacacatatctcgaagccatacattcagacaaacgcatttcatttcccatgtacgacgatctcatagaacagtgtcatctatcgatggagcataaaagtaaactcgtcgacaaagcactcaataaattagagtctaccatcgatagtcaatctagactatcgtatttgcctccggaaattatgcgcaatatcataaccaagctaagcgactaccatctaaacagtatgttgtacggaaagaaccattacaaatattatccatgatagaaagaaaatatttaaaaaataatctatatgattggagaagtaggaaacaaacagtaacaagacgacgattactacattattaaatcatgaggtccgtattatactcgtatatattgtttctctcatgtataataataaacggaagagatatagcaccacatgcaccatccaatggaaagtgtaaagacaacgaatacagaagccgtaatctatgttgtctatcgtgtcctccgggaacttacgctttcagattatgtgatagcaagactaatacacaatgtacaccgtgtggttcggatacctttacatctcacaataatcatttacaggcttgtctaagttgtaacggaagatgtgatagtaatcaggtagagacgcgatcgtgtaacacgactcacaatagaatctgtgaatgctctccaggatattattgtcttctcaaaggagcatcagggtgtagaacatgtatttctaaaacaaagtgtggaataggatacggagtatccggatacacgtctaccggagacgtcatctgttctccgtgtggtcccggaacatattctcacaccgtctcttccacagataaatgcgaacccgtcgtaaccagcaatacatttaactatatcgatgtggaaattaacctgtatccagtcaacgacacatcgtgtactcggacgaccactaccggtctcagcgaatccatctcaacgtcggaactaactattaccatgaatcataaagattgtgatccagtctttcgtgcagaatacttctctgtccttaataatgtagcaacttcaggattctttacaggagaaaatagatatcagaatacttcaaagatatgtactctgaatttcgagattaaatgtaacaacaaagattcatcttccaaacagttaacgaaaacaaagaatgatactatcatgccgcattcagagacggtaactctagtgggcgactgtctatctagcgtcgacatctacatactatatagtaataccaatactcaagactacgaaaatgatacaatctcttatcatatgggtaatgttctcgatgtcaatagccatatgcccgctagttgcgatatacataaactgatcactaattcccagaatcccacccacttatagtaagtttttttacctataaataataaatacaataattaatttctcgtaaaagtagaaaatatattctaatttattatatggtaagaaagtagaatcatctagaacagtaatcaatcaatagcaatcatgaaacaatatattgtcctggcatgcatgtgcctagtggcagctgctatgcctactagtcttcaacaatcttcatcctcgtgtactgaagaagaaaacaaacatcatatgggaatcgatgttattatcaaagtcacaaagcaagaccaaacaccgaccaatgataagatttgtcaatccgtaacggaagttacagagaccgaagatgatgaggtatccgaagaagttgtaaaaggagatcccaccacttattacactatcgtcggtgcgggtcttaacatgaactttggattcaccaaatgcccaaagattttatccatctccgaatcctctgatggaaacactgtgaatactagattgtccagcgtgtcaccgggacaaggtaaggactctcccgcgatcacgcgtgaagaagctctggctatgatcaaagactgtgagatgtctatcgacatcagatgtagcgaagaagagaaagacagtgacatcaagacccatccagtacttgggtctaacatctcacataagaaagtgagttacaaagatatcatcggttcaacgatcgttgatacaaaatntgtcaagaacctagagtttagcgtacgtatcggagacatgtgtgaggaatcatctgaacttgaagtcaaggatggattcaagtatgtcgacggatcggcatctgaaggtgcaaccgatgatacttcactcatcgattcaacaaaactcaaagcatgtgtctgaatcgataactctattcatctgaaaatggatgagttgggttaatcgaacgattcagacaccgcaccacgaattnaaaaagaccgggcactatattccggtttgcaaaacaaaaatatttaactacattcacaaaaagttacctctcgttacttcttctttctgtttcaatatgtgatacgatatgatcactattcgtattctcttggtctcataaaaaagttttacaaaaaaaaaaatatttttattctctttctctcttcgatggtctcacaaaaatattaaacctctttctgatgtctcaactatttcgtaaacgataacgtccaacaatatattctcgtagagcttatcnacanccttatnnnnnnnnnnnnnnnnnnnnnnnnnnnnnnnnnnnnnnnnnnnnnnnnnnnnnnnnnnnnnnnnnnnnnnnnnnnnnnnnnnnnnnnnnnnnnnnnnnnnnnnnnnnnnnnnnnnnnnnnnnnnnnnnnnnnnnnnnnnnnnnnnnnnnnnnnnnnnnnnnnnnnnnnnnnnnnnnnnnnnnnnnnnnnnnnnnnnnnnnnnnnnnnnnnnnnnnnnnnnnnnnnnnnnnnnnnnnnnnnnnnnnnnnnnnnnnnnnnnnnnnnnnnnnnnnnnnnnnnnnnnnnnnnnnnnnnnnnnnnnnnnnnnnnnnnnnnnnnnnnnnnnnnnnnnnnnnnnnnnnnnnnnnnnnnnnnnnnnnnnnnnnnnnnnnnnnnnnnnnnnnnnnnnnnnnnnnnnnnnnnnnnnnnnnnnnnnnnnnnnnnnnnnnnnnnnnnnnnnnnnnnnnnnnnnnnnnnnnnnnnnnnnnnnnnnnnnnnnnnnnnnnnnnnnnnnnnnnnnnnnnn

>2018_Israel_Monkeypox_MN648051.1

CGAAGAGAGAAAGAGATGGTTAGTCAAGATATTTTTCTTAGTACAAAAGTCAATGTTTTAAAATATATGGACGAGAATTAATTTGTCTGTATAAAAACTTGTGTGAAATTATGTACTAGAGAAAAAACGTGAGCAGTGTCCCCTACATGGATTTTACAGATCATTTATATTCCAAAAATATTAACTATATACGTTTATTATATGATGTTAACGTGTAAATTATAAACATTATTTTATGATGCAATTGTCTGACAACCTAGATTGGTATAAGGATGTTGATAAGCTCTACGAGAATATATTGTTGGACGTTATCGTTTACGAAATAGTTGAGACATCAGAAAGAGGTTTAATATTTTTGTGAGACCATCGAAGAGAGAAAGAGAATAAAAATATTTTTTTTTTTTTTGTAAAACTTTTTTATGAGACCAAGAGAATACGAATAGTGATCATATCGTATCACATATTGAAACAGAAAGAAGAAGTAACGAGAGGTAACTTTTTGTGAATGTAGTTAAATATTTTTGTTTTGCAAACCGGAATATAGTGCCCGGTCTTTTTTAATTCGTGGTGCGGTGTCTGAATCGTTCGATTAACCCAACTCATCCATTTTCAGATGAATAGAGTTATCGATTCAGACACATGCTTTGAGTTTTGTTGAATCGATGAGTGAAGTATCATCGGTTGCACCTTCAGATGCCGATCCGTCGACATACTTGAATCCATCCTTGACTTCAAGTTCAGATGATTCCTCACACATGTCTCCGATACGTACGCTAAACTCTAGGTTCTTGACACATTTTGTATCAACGATCGTTGAACCGATGATATCTTTGTAACTCACTTTCTTATGTGAGATGTTAGACCCAAGTACTGGATGGGTCTTGATGTCACTGTCTTTCTCTTCTTCGCTACATCTGATGTCGATAGACATCTCACAGTCTTTGATCATAGCCAGAGCTTCTTCACGCGTGATCGCGGGAGAGTCCTTACCTTGTCCCGGTGACACGCTGGACAATCTAGTATTCACAGTGTTTCCATCAGAGGATTCGGAGATGGATGAAATCTTTGGGCATTTGGTGAATCCAAAGTTCATGTTAAGACCCGCACCGACGATAGTGTAATAAGTGGTGGGATCTCCTTTTACAACTTCTTCGGATACCTCATCATCTTCGGTCTCTGTAACTTCCGTTACGGATTGACAAATCTTATCATTGGTCGGTGTTTGGTCTTGCTTTGTGACTTTGATAATAACATCGATTCCCATATGATGTTTGTTTTCTTCTTCAGTACACGAGGATGAAGATTGTTGAAGACTAGTAGGCATAGCAGCTGCCACTAGGCACATGCATGCCAGGACAATATATTGTTTCATGATTGCTATTGATTGATTACTGTTCTAGATGATTCTACTTTCTTACCATATAATAAATTAGAATATATTTTCTACTTTTACGAGAAATTAATTATTGTATTTATTATTTATAGGTAAAAAAACTTACTATAAGTGGGTGGGATTCTGGGAATTAGTGATCAGTTTATGTATATCGCAACTAGCGGGCATATGGCTATTGACATCGAGAACATTACCCATATGATAAGAGATTGTATCATTTTCGTAGTCTTGAGTATTGGTATTACTATATAGTATGTAGATGTCGACGCTAGATAGACAGTCGCCCACTAGAGTTACCGTCTCTGAATGCGGCATGATAGTATCATTCTTTGTTTTCGTTAACTGTTTGGAAGATGAATCTTTGTTGTTACATTTAATCTCGAAATTCAGAGTACATATCTTTGAAGTATTCTGATATCTATTTTCTCCTGTAAAGAATCCTGAAGTTGCTACATTATTAAGGACAGAGAAGTATTCTGCACGAAAGACTGGATCACAATCTTTATGATTCATGGTAATAGTTAGTTCCGACGTTGAGATGGATTCGCTGAGACCGGTAGTGGTCGTCCGAGTACACGATGTGTCGTTGACTGGATACAGGTTAATTTCCACATCGATATAGTTAAATGTATTGCTGGTTACGACGGGTTCGCATTTATCTGTGGAAGAGACGGTGTGAGAATATGTTCCGGGACCACACGGAGAACAGATGACGTCTCCGGTAGACGTGTATCCGGATACTCCGTATCCTATTCCACACTTTGTTTTAGAAATACATGTTCTACACCCTGATGCTCCTTTGAGAAGACAATAATATCCTGGAGAGCATTCACAGATTCTATTGTGAGTCGTGTTACACGATCGCGTCTCTACCTGATTACTATCACATCTTCCGTTACAACTTAGACAAGCCTGTAAATGATTATTGTGAGATGTAAAGGTATCCGAACCACACGGTGTACATTGTGTATTAGTCTTGCTATCACATAATCTGGAAGCGTAAGTTCCCGGAGGACACGATAGACAACATAGATTACGGCTTCTGTATTCGTTGTCTTTACACTTTCCATTGGATGGTGCATGTGGTGCTATATCTCTTCCGTTTATTATTATACATGAGAGAAACAATATATACGAGTATAATACGGACCTCATGATTTAATAATGTAGTAATCGTCGTCTTGTTACTGTTTGTTTCCTACTTCTCCAATCATATAGATTATTTTTTAAATATTTTCTTTCTATCATGGATAATATTTGTAATGGTTCTTTCCGTACAACATACTGTTTAGATGGTAGTCGCTTAGCTTGGTTATGATATTGCGCATAATTTCCGGAGGCAAATACGATAGTCTAGATTGACTATCGATGGTAGACTCTAATTTATTGAGTGCTTTGTCGACGAGTTTACTTTTATGCTCCATCGATAGATGACACTGTTCTATGAGATCGTCGTACATGGGAAATGAAATGCGTTTGTCTGAATGTATGGCTTCGAGATATGTGTGATACCGGATGTCTTCTGTTCTCAATACCGTATACAAGTCGGTGTCTGAGATTCGAATCTCTTTGAGGAGACTTATGTCACGACTACATTTTTCGATGATGGAATCTATCTTATCGAATGATATATTTTTCATAAATACACTTTTATAGTCCTCGTTTAAACAGAATTTAGTATGTAGTTCCGCAAATGACTCGTCCCTTAATAGGCAGTAGGCTATTATCTTCTTTACGTAGTGATCGTCGTAGGGAGAGAACTCCGACATCTTGTAGAACAACGATTTAATCATAGGTAGAGATACTTTCAGTCTGTGGTGGATAATGTCATTAACGACATCCGCCTTGTATATAATGTTTCTGTTTTCAAACACCAAGTCGAATACTGTCTTATCGTCTTTAGTCGGAAGGTTGATGTCGTATCCGATGTATACGAGGTATGAGGCAACATTGTTATTGCAATTCTGGAAGGCGGTATGAAGAGGAGTCATTGTATTATAGTATTCGTCTTTCTGAATGTCGAATCTATCTAGTAGATACCGTAGTATATTGAGAGAGCGACTTCCATATCCTTGATTATGTTTTATGAATAGATAAAGTAGATGTTGTCCTTCTTCCTTTTGTAATTTCCCGTATTTTTGTTCGTGCCAATTGAGTAACATTATGAGAATATGACCTGTTGCACAATCGTTCTTTATGTATTCCATGATGGGTGTACAATCAAGATTATTACGTATCCTCGTATCGGCTCCTCGAGATAAAAGAGCATACACCACACGAGGACTATGTTTGGTATACTGTTGAAGGTAAGTGTGTAACGGCGTATTTCCGATTTTCGTAACCGCGTTAATGTTTGCTCCATGATCTATTATCGCGTAGATGAATCGCTTCTCAGCTCGCATCTTAGTGTGACTCTTTGACTTGTAATAATTGCTTTCGTGGAACGCGGATATGTGTTTACAGTAGTAATGAAGAGAAGTGAGTCCATCCTCATCGACGCAATTAGGGTCAGATCCTTTAGTCAATAATTTGTACAGAACGTAATAGTTTAAGCTCCCATTGAATTTATATCTAAGATAACACAGCAATAGATCGGATGATTTACTAAAGTCATCAATGGGGTCCGTTAGTATATCAAAGATCTTGTTATCGATTGATAGTGAATGAATCAGATAGTGGTGTAGAGGAATATGTCCTTTTTTATCCTTGCTATCAAAGTTACGCATGCCGTGGCGTAACAATATCTTTAATACAGATGGATTAAATCGTGTATTCATCGTATAGCAATGTAATGGAGAGTTACCACATTTTAGTCGTTTATTCAGATCGCAGTGTTTAATAACTAATTTAAACAGATGAGATGATATATCCACATCAAAGAATGCGAGATACATATGACAGACATTATTGACAGAAATGTGACCTTCATTATCACCGTCGTCCATAAATGCGTTAGGTACGTACCACATACTATCGTTAACGATGCGCACAATCTCGTCCATTTCATCCATCTTCATAATGATTTACTTTTTCATAATTAGAGAAAAAGATCAAGGTATAAAAATTAGAAGTGTTAGACTATAAACTAACTTATGACTTAACTAACTTATGACTTAACTAACTTATGACTTAACTAACTTATGACTTAACTAACTTATGACTTAACTAACTTATGACTTAACTAACTTATGACTTAACTAACTTATGACTTAACTAACTTATGACTTAACTAACTTATGACTTAACTAACTTATGACTTAACTAACTTATGACTTAACTAACTTATGACTTAACTAACTTATGACTTAACTAACTTATGACTTAACTAACTTATGACTTAACTAACTTATGACTATTAACTAACTAACTTATGACTTAACTAACTTATGACTTAACTAACTTATGACTTAACTAACTTATGACTTAACTAACTTATGACTTAACTAACTTATGACTTAACTAACTTATGACTTAACTAACTTATGACTTAACTAACTTATGACTTAACTAACTTATGACTTAACTAACTTATGACTTAACTAACTTATGACTTAACTAACTTATGACTTAACTAACTTATGACTTAACTAACTTATGACTTAACTAACTTATGACTTAACTAACTTATGACTATTAACTCATTTCAAGAAAGGTGGGTGGATAGAACTCTATATGACAGCTTGTGAAACAATTAGATCCCTAATTTCTAATGGAAGTTTTGATAGGAGATTGTCATCAGTTGATACATTGTTTATTATCTCATCTATTAGAGCACGTCTGTTTAGAGCTTTAGTGACCTGCTCGGTTACTTCTGTGTAAATCTTGAATCCTTTAGTGATACACTGTGTCAAAACTGGATGTTTAGAATACCTATGTAGAATATGGGAAGCATGCTTGTTTTTGTCTCTATTATAGATTAACTCATACATGGTTGTATTATGAATTTTCATCTGCCTAATGTACTCCAATTCTTGTTTACAATCAATTATATAATCAAAGAGTGATGATGCATACACATTACAAAGTGAATAATCTACCATCATAAAATACTTGATACAGAGCTTTATCACATCATGGTTTTCAATTGTATTATTAAGTATAGCTAATTTTATACAGTCAATAGACAATGGTTCTCTAAGCAATATTTCTAATATTTTAAGATGTGCTTCCCTACGGGCGATGACAGATCCCCTATCCACGGCCACGTCAAGACATGTATATCCATTACTCATTACTGCGTTGACATTTGCTCCATTTTCTAATAGCCATGATACTAAATCTATATAACCTGCATAGATAGCGCGATAAAGCAAGGTCCTTCCACCAGCATCTAGTTGATTGATATCTTCAATATATGGGATACAAAGCTTATAAATTTCTAATACTGTGGGTTCATCTACAAGGAATCCCCTAGTATACTGAATTATTTTATATAGATCTAATTTAACATCATTTTCATCTGGGATACCACAATTCAAAATAAACTCAACAACACTACTTTCCTTTTTACATATTCCCCTAAAATAGGCATTCAAGCATTCTATTTTATATATTACAGCCCCATGATCTACCATAAGATCAACAATGTCTATTTCTACATATGCATTAGATAGATAGTAAAGTAAGAGATCTTGCACAGAATTACAATTCTTAATAATTATAGAGAAAATATCTTCCATATAATTCTTTGACACTAATGCAGATATAATATCTTTATATGTAATATATGCAAACAGTCTATCTACTATATACTGATCAATATTATCTCTATGAATCCTAAAATAATCATACAGAACATCTACAGGATCACAAATTGGTTCAAGGAGAAATCTATCAAATATTTTCCTGTCAACAACTGGTTCTAGAACATAACAGTCAACACCTAATCCATGTTTTTTATAATCATCTACCAAAGATAATGACCAAAGATCGAGGTCGTCGTGAAACTGCTCATCGACAGCCATGAAATCTACCGACTCCATGGTGCGAATCGCACTGTCTTATTCGCCATTGATTTTCATTTTTTATAATTATGTACATGTTTTCCTTCTATTCTCAAGAGTCTACAAAAATATATTTTTTCGATATCTAAGTACTAAGTTTTTTTACTGTTTTTGTTACTGTCTTCCATTCTTCTAACTAAAGATCTGAGATAAATTATACAATCTTCGCTATCGAACCATTTTTGTAGTCTAAAGCCTGAAGTAATTAACCAACTGTTTTTATTAGTGGCTTTTTTCGATCTATCCTCGTCCTCTCATCATCCTTATATTATTATCATTATTATCATAGTCTATTAAACACAAATCATCTACGTTTATAACAACATTCTCATTATTAATTAGTTCTGTAGAATATCTTTAATAATTTGGCTATACATCTGTTCAATACTATCTATTGATGATTTCTTTTTTAAGACTTAAACTAGTTATGGTAATGACGATGAAATCGAGTAGTAACTTCTAATAAAGACTTGATATCATTATCATATGTTTGATCGTCATAGTTAATAGTGTGGCTAAATGGTACTGTTAATAAGTTTATAGACAATATCATAGTATTTTCTTTCCAGAATTAGATTATTTTTTTAAATACTGATCCTCACAATTCCGTGATGTAGCAGTAGTTGGTGCATGGTCTATATCGTTAAAATGTATCATATATAATAGTTTTCTGACGTGGAGTACAGAATTTTCGATTAATGAGTTCATGGTAAGGAAGGGCAAATGCCTGTATATAATATACATAAGTTAATAGTTTTTTATCATATTTTCTAATACCATAATAAAAATTATCATTATTGCGTTTGGTAGTTCTGCCCTATCATCTATATCACTGTCACTCTCGCTCTCACTATATCTTCTAAAATTACAAACAACTGGATATTCGATAACAGCATTTGTGTAGTTTTTGTCTTTTACAGTATATACGTTATTGTCAAAATCTAAACAAATATTAGCATAATACATCTATCTATAAGATCAGGATCCATGTTCGAGCATACTAGCCATGTATATTTGTAACTTCGTCGTACAGCGTTAGATCAATAGAATAAACAATCGTGTGACGCAACTTTTTTACGATCTAGTTGTATGAGTTTATCGTTTACATAAGCAATTAACGGCTTTAACAGATGATCTGAGTAATAATATACCTCTGTTATACGTTTAATGTTCACGGTCTTAGTATTTTTAGATATCAATTGTGATTTACACCATATTCGACTCCCTTGTGTGTAACGTTAGAAATTCTAAATCTATATTATCTATTACAGCGTAAAACACATTCAATATTGTATTGTTATTTTTATATTATTTACACAATTAACAATGTATTATTAGTTTATATTACTGAATTAATAATATAAAATTCCCAATCTTGTCATAAACACAAAATCCATTAAAAATGTCGATAAAATATCTGATGTTGTTGTTCGCTGCTATGATAATCAGATCATTAGCCGATAGTGGTAATGCTATCGAAACAACATCGCCAGAAATTACAAACGCTACAACAGATATTCCAGCTATCAGATTATGTGGTCCAGAGGGAGATGGATATTGTTTACACGGTGACTGTATCCACGCTAGAGATATCGACGGTATGTATTGTAGATGCTCTCATGGTTATACAGGTATTAGATGTCAGCATGTAGTATTAGTAGACTATCAACGTTCAGGAAAACCAGACACTACAACATCATATATCCCATCTCTAGGTATTGTGCTTGTATTAGTAGGCATTATTATTACGTGTTGTCTATTATCTGTTTATATGTTCACTCGAAGAACTAAACTACCTATACAAGATATGGTTGTGTTATATTTTTTATAAAATTTTTTTATGAGTATTTTTACAAAAATGTATATGTATAAAAAAAATACTAAGTATACGATGTATCCTGTATTATTTGTATTTATCTAAACAATACTTCTGCCTCTAGATGGGATACAAAAATTTTTTATTTCAGCATATTAAAGTAAAATCTAGTTACCTTGAAAATGAATACAGTGGGTGGTTCCGTATCACCAGTAAGAACATAATAGTCGAATACAGTATCCGATTGAGATTTTGCATACAATACTAGTCTAGAAAGAAATTTGTAATCATCCTCTGTGACAGGAGTCCATATATCTGTATCATCGTCTAGTTTTTATCAGTGTCCTATGCTATATTCCTGTTATCATCATTAGTTAATGAAAATAACTCTCGTGCTTCAGAAAAGTCAAATATTGTATCCATACATATATCTCCAAAACTATCACTTATACGTTTATCTTTAACGAACATATACCTAGATGGTTATTTACTAACAGACATTTTTTCAAGATCTATTGACAATAACTCCTATAGTTTCCACATCAACCAAGTAATGATCATCTATTGTTATATAACAATAACATAACTCTTTTCCATTTTTATCAGTATCTATATCAACGTCGTTGTAGTGAATAGTAGTCATTGATCTATTATATGAAACGGATATGTCTAGTTAATATTTTCTTTGATTTAAAGTCTATAGTCTTTACAAACATAATATCCTTATCCGACTTTATATTTCCTGTAGGGTGGCATAATTTTATTCTGCCTCCACAATTAGTGTTTCCAAATATATTACTAGACAATATTCCATATAGTTATTAGTTAAGGGACCCAATTAGAACACGTACGCGCTTATTCATCATTTGGATCGTATTTCATAAAAGTTATTATGTTATAGATGTCAACACATTCTACATTTTTTTAATTGTCTATATAGTATTTTTCTGATATTTTCTATAATATCAGAATTGTCTTCCATAGGAAGTTGTATACTATTCGGAATCAGTTACATGTTTAAATAATTCTCTGATGTCATTCCTTATACAATCAAATTCATTATTAAACAGTTTAATAGTCTGTAGACCTTTATCGTCGTACATATCCATTGTCTTATTAGTTACGCTTATTTTTATGGGTTTTACGTTGCTTTATTATATTTTATAATAATGATTGTTTGACAATGTCGTAGTATAGATATATTATTAGAGGAGGTATAATTATAAAAAGTTTCTGAGTACGATGTTATAAGAGGAGAGGACACATTAACATCATACATCAATTAACTACATTCTTATAACATTGTAATCAAAAGAATTGCAATTTTGATGTATAACAACTGTCAATGGAATTGTATATTACAAATTACGGTATGTTGTAACAACAAATACCGATCGGTAATTGTCTCTGCCGCTGTAATAGAATTAATTATATATCTATTACACCGGCCTTGTATCATAATAAAGTTGTGGTAGTATGATCTCCATATTTATAATTTAGTACTTTGTATTTAGTTTTTTTGGAATCATAAAAAAAGTTTTACTAATTTAAAATTTAAAAAGTATTTACATTTTTCACTGTTTAGTCGCGGATATGGAATTCGATCCTGTCAAAATCAATACATCATCTATAGATCATGTAACAATATTACAATACATAGATGAACCAAATGATATAAGACTAACAGTATGCATTATCCGAAATGTTAATAACATTACATATTATATCAATATCACAAAAATAAATCCACATTTGGCTAATCGATTTCGGGCTTGGAAAAAACGTATCGCCGGAAGGGACTATATGACTAACTTATCTAGAGATACAGGAATACAACAATCAAAACTTACTGAAACTATACGTAACTGTCAAAAAAATAAAAACATATATGGTCTATATATACACTACAATTTAGTTATTAATGTGGTTATTGATTGGATAACCGATGTGATTGTTCAATCAATATTAAGAGGGTTGGTAAATTGGTACATAGCTAATAATACATATACTCCAAATACACCCAATAATACTACAACCATTTCTGAGTTGGATATCATCAAAATACTGGATAAATACGAGGACATGTATAGAGTAAGTAAAGAAAAAGAATGTGGAATTTGCTATGAAGTTGTTTACTCAAAACGATTAGAAAACGATAGATACTTTGGTTTATTGGATTCGTGTAATCATATATTTTGCATAACATGTATCAATATATGGCATAGAACACGAAGAGAAACCGGTGCGTCAGATAATTGTCCTATATGCCGTACACGTTTTAAAAAAATAACAATGAGCAAGTTCTATAAGCTAGTTAACTAATAAATAAAAAGTTTAATTATCGACGATATATGTCGTTATTTTTCTCTCATATGAAAGATTAATTTGATTCTAATATAATCTTCAGTATTGGATGAATCTCAATTCAAATTAATTCCATTAGATTAGATTAGATTAGATCATAAATAAAAATAGTAGCACGCACTACTTCAGCCAAATATTCTTTTTTGAAACGCCATCTAGCGTAATGAGAACACAAGTGAACCTATAATGAGCAAATTTATTAGTATCGGTTACATGAAGGACTTTACGTAGAGTGGTGATTCCTCCATCTGTGGTACGAACGGTTTCATCTTCTTTGATGCCACCACCCAGATGTTCTATAAACTTGGTATCCTCGTCCGATTTCATATCATTTGCCAACCAATACATATAGCTAAACCCAGGCATACGTTCCACACATCCGGAACAATGAAATTCTCCAGAAGATGTTACAATGGCTAGATTTGGACATTTGGTTTCAACCGCGTTAACATATGAGTGAACACACCCATACATGAAAGCGATGAGAAATAGGATTCTCATCTTGCCAAAATATCACTATAAAAAATTTATTTATCAATTTTAAAGGTATAAAAAAATACTTATTGTTGCTCGAATATTTTGTATTTGATGGTATACGGAAGATTAGAAATGTAGGTATTATCATCAACTGATTTTATGATGGTTTTATGAATTTTATTATGCTTCACTATTGCATCGGAAATAATATCATATGCTTCCACGTATATTTTATTTTGTTTTGACTCATAATACGCACGTAATTCTGGATTATTGGCATATCGATGAATAATTTTAGCTCCATGCTCAGTAAATATTAATGAGAACATAGTGTTGCCTCCTACCATTATTTTTTTCATCTCATTCAATTCTTGATTGCAGAGATCTATATAATCATTATAGCGTTGACTTATGGACTCTGGAATCTTAGACGATGTACAGTCATCTATAATCATGGTATATTTAATACATTGTTTTATAGCATAGGCATTATCTACGATATTAGATACTTCACTCAATGAATCAATCACACAATCTAATGTAGGTTTATGACATAATAGCATTTTCAGCAGTTCAATGTTTCTAGATTCGTTGATGGCAATGGTTATACATGTATATCCGTTATTTGATCTAATGTTGACATCTGAACCGGATTCTAGCAGTAAAGATACTAGAGATTGTTTATTATATCTAACAGCCTTGTGAAGAAGTGTTTCTCCTCGTTTGTCAATCATGTTAATGTCTTTAAGATAAGGTAGACAAATGTTTATAGTACTAAGAATTGGGCAAACATAAGACATGTCACAAAGACCCTTTTTGTATGTATAAGTGTAGAAATTATAACATCCATAGTTGGATTCACATAGGTGTCCAATCGGGATCTCTCCATCATCGAGATGATTGACGGCATCTCCCCCTTCCTTTTTTAGTAGATATTTCATCGTGTAAGAATCAATATTAATATTTCTAAAGTATCTGTGTATAGCCTCTTTATTTACCACAGCTCCATATTCCAACATGCATTCCACTAGAGGGATATCGATATCGCCGAATGTCATATACTCAATTAGTATATGTTGGAGGACATCCGAGTTCATTGTTTTCAATATCAAAGAGATGGTTTCCTTATCATTTCTCCATAGTGGTACAATACTACACATTATTCCGTACGGCTTTCCATTCTCCAAAAACAATTTTACCAAATCTAAATCTACATCTTTATTGTATCTATAATCACTATTTAGATAATCAGCCATAATTCCTCGAGTGCAACATGTTAGATCGTCTATATATGAATAAGCCGTGTTATCTATTCCTTTCATTAACAATTTAACGATGTCTATATCTATATGAGATGACTTAATATAATATTGAAGAGCTGTACAATAGTTTTTATCTATAGAAGACGGCTTGATTCCGTGATTAATTAGACATTTAACAACTTCCGGACGCACATATGCTCTCGTATCCGACTCTGAATACAGATGAGCGATGATATACAGATGCAATACGGTACCACAATTTCGTGGTTGATAATCATCATACACGTATCCGTACTCGTCATCCTCATAAAGAACACTGCATCCATTTTCTATGAACAAATCAATAATTTCAGGAACAGGATCATCTGTCATTACATAATTTTCTATAACTGAACGATGGTTTTCACATTTAACACTCAAGTCAAATCCATGTTCTACCAACACCTTTATCAAGTCAACGTCTACATTTTTTGATTTCATATAGCTGAATATATTAAAGTCATTTATGTTGCTATATCCAGTAGCTTCTAGTAGAGCCATCGCTATATCCTTATTGACTTTAACATGTCTACTATTTGTGTATTCTTCTATTGGGGTAAACTGTCTCCAATTTTTATGTAATGGATTAGTGCCACTATCTAGTAGTAGTTTGACGACCTCAACATTATTACAATGCTCATTGAAAAGGTATGCGTGTAAAGCATTATTCTTGAATTGGTTCCTGGTATCATTAGGATCTCTGTCTCTCAACATCTGTTTAAGTTCATCGAGAACCACCTCCTCATTTTCCAGATAGTCAAACATTTTGACTGAATAGAAGTGAATGAGCTACTGTGAACTCTATACACCCGCACAACTAATGTCATTAAATATCATTTTTGAATGTATTTATACCATGTCAAAAACTTGTACAATTATTAATAAAAATAATTAGTGTTTAAATTTTACCAGTTCCAGATTTTACACCTCCGTTAACACCTCCATTAACCCCACTTTTTACACCACTGGACGATCCTCCTCCCCACATTCCACTGCCACTAGATGTATAAGTTTTAGATCCTTTATTACTACCATCATGTCCATGGATAAAGACACTCCACATGCCGCCACTACTACCCCCTTTAGACGACATATTAATAAGACAAGTTTAACAATAAAATTAATCACGAGTACCCTACTACCAACCACTATTATATGATTACAGTACCTTGACTAAAGTCTCTAGTCACAAGATCAATACTACCAACCTACGCTATTATATGATTATAGTTTCTATTTTTATAGGAACGCGTACGAGAAAATCAAATGTCTAAGTTCTAACGGTAGTGTTGATAAACGATTGTTATCCGCGGATACCTCATCTATCATGTTGTCTATTTTCTTACTTTGTTCTATTAACCTATTAGCATTATATATTATTTGATTATAAAACTTATATTGCTTATTAGCCCAATCTGTAAATATCGGATTATTAACATATCGTTTCTTTGTAGGTTTATTTAACTTGTACATCACTGTAAGCATGTCCGTACCATTTATTTTAATTTGACACATATCAGCAATTTCTTTTTCGCAGTCGGTTATATATTCTATATAAGATGGATACGTATCACATATGTACTTATAGTCTACTAATATGAAGTACTTAATACATATTTTCAGTAACGATTTAGCCTTATTACCTATTAATAAGTGCCTGTCGTTGGATAGGTAATCAACTGTTTTCTTAATACATTCGATGGTTGGTAATTTACTCAAAATAATTTCCAATATCTTAATATATATTTCTGCTATTTCTGGTATACATGCATGTGCCATTATAACACAAATACCAATACATGTAGACCCATATGTTGTTGTTATATTAATATCTGCGCCATTATCTATTAACCATTCTATTAGGGCAACACTATGCGACTCGATACAATAATAAAGTATACTACGTCCATGTTTATCTATTTTGTTTATATCATCGATATACGGCTTACAAATTTCTAGTATCGATAACACTTCTGACTCGTGAATAAATAAGGTAGGGAATAACGGCATAATATTTATTATGTTATCATCATTAACAACTACGTTTCCATTTTTTAAAATATACTCTACAACTTTAGGATCCCTATTGTCAAATCTTTTAAAATATTTATTTATATGCTTAAATCTATATAATATAGCTCCTTCCCTAATCATACATTTGATAACATTGATGTACACTGTATGATAAGATACATATTCTGACAATAGATCTTGTATAGAATCTGTATATCTTTTAAGAATTGAGGATATTATGACATTATTACGTAAACTATTACACAATTCTAAAATATAAAATGTATCATGGGCAGATAATAGTTTATCCACTATATAATTATCTATTTTATGATTTTTCTTCCTATATTGTTTACGTAAATAGATAGATAGAATATGCATTAGTTCATTACCGCTATAGTTACTATCGAATAACACGTCAAATATTTCCCGTTTAATATCGCATTTGTCAACATAATAATAGAGTATGGTACGTTCACGATAAGTATAATGACACATCTCGTTTTCGTGCGAAATTAAATAGTTTATCACGTCCAAAGATGTCACATAACCATCTTGTGACCTAGTAATAATATAATAATAGAGAACTGTTTTACCCATTCTATTATCATAATCAGTGGTGTAGTCATAATCTAAATAATCAAACTCGTCATCCCAATTAAAATAAATATAATCAGTACATTGAATGGGTATGATATTGTACCCATACTGTATGTTGCTACATGTAGGTATTCCTTTATCCAATAATAGTTTAAATACATCTATATTAGGATTTGATGTTGTCGCGTATTTCTCTACAATATTAATACCATTTTTGATACTATTTATTTCTATACCTTTCGAAATTAGTAATTTCAATAAGTCTATATCGATGTTATCAGAACATAGATATTCAAATATATCAAAATCATTGATATTTTTATAGTCGACTGACGACAATAACAAAATCACAACATCGTTTTTGATATTATTATTTTTTTTGGTAACGTATGCCTTTAATGGAGTTTCACCATCATACTCATATAATGGATTTGCACCACTTTCTATTAATGATTGTGCACTACTGGCATCGATGTTAAATGTTTTACAACTATCATAGAGTATCTTATCGTTAACCATGATTGGTTGTTGATGTTATCACATTTTTTTGGTTTCTTTCATTTCAGTTATGTATGGATTTAGCACGTTTGGGAAGCATGAGCTCATATGATTTCAGTACTGTAGTGTCAGTACTATTAGTTTCGATCAGATCAATGTCTAGATCTATAGAATCAAAACACGATAGGTCAGAAGATAATGAATATCTGTACGCTTCTTCTTGTACTGTAACTTCTGGTTTTGTTAGATGGTTGCATCGTGCTTTAACGTCAATGGTACAAATTTTATCCTCGCTTTGTGTATATCATATTCGTCTCTAGTATAAAATTCTATATTCAAATTATCATGCGATGTGTGTACGCTAACGGTATCAATAAACGGAGCACAGCATTTAGTCAACAGTAATCCAAAATTTTTTAAAGTATATCTTAACGAAAGAAGTTGTCATCGTTAGAGTGTGGTAAATCATTGTCTACGGTACTAGATCCTCATAAGTGTATATATCTAGAGTAATATTTAATTTATCAAATGGTTGATAATATGGATGTTGTGGCAATTTCCTAATACGGAAATAAGACATAAACACGCAATAAATCTAATTGCGGACATGTTACACTCCTTAAAAAATACGAATAAACACTTTGGCTTTTAGTAAGTGTCATTTAACACTATACTCATATTAATCCATGGACTCATAATCTCTATACGGGATTAACGGATGTTCTATATACGGGGATGAGTAGTTCTCTTCTTTAACTTTATACTTTTTACTAATCATATTTAGACTGATGTATGGGTAATAGTGTTTGAAGAGCTCGTTCTCATCATCAGAATAAATCAATATCTCTGTTTTTTTGTTATACAGATGTATTACAGCCTCATATATTACGTAATAGAACGTGTAATCTACCTTATTAACTTTCACCGCATAGTTGTTTGCAAATACGGTTAATCCTTTGACCTCGTCGATTTCCGACCAATCTGGGCGTATAATGAATCTTAACTTTAATTTCTTGTAATCATTCGAAATAATTTTTAGTTTGCATCCGTAGTTATCTCCTCTATGTAACTGTAAATTTCTCAACGCGATATCTCCATTAATAATGATGTCGAATTCGTGTTGTATACCCATACTGAATTGATGAACTAATACCGACGGTATTAATAGTAATTTACTTTTCATCTTTACATACTTGGTAATAGTTTTACTATCATAAGTTTATAAATTCCACAAGCTACTATGGAATATACCAACCATCTTAGTATAGAACACATGTCTTAAAGTTATTAATTAATTACATGTTGTTTTATATATCGCTACGAATTTAAACAGAGGAATCAGTTAGGAAGAAAAAATTATCTGTCATCATCATCATCTATTGGATAACATCTCTGTATTCTACGATAGAGTGCTATTTTAAGATGTGACAGATCCGTGTCATCAAATATATACTCCATTAAAATGATTATTCCGGCAGCGAACTTGATATTGGATACATCACGACCTTTGTTAATATCCACGACAATAGACAGCAATCCCATTGTTCCATAAACAGTGAGTTTATCTTTCTTTGAAGTGATATTTTGTAGAGATCTTATAAAACTGTCGAATGACATCGTATTTATATCTTTAGCTAAATCATATATGTTACCATCATAATATCTAACAGCATCTATCTTAAACGTTTCCATCGCTGTAAAGACGTTTCCGATAGATGGTCTCGTTTCATCAGTCATACTGAGCCAACAAATGTAATCGTGTATAACATCTTTGATAGAATCAGACTCTAAAGAAAAGGAATCGGCTTTATTATACACATTCATGATAAACTTAATGAAAAATGTTTTTCGTTGTTTAAGTTGGATGAATAGTATGTCTTAATAATTGTTATTATTTCACTAATTAATATTTAGTAACGAGTACACTCTATAAAAACGAGAATGACATAACTAATCATAACTAGTTATCAAAGAATGTCTAGGACGCGTAATTTTTTATGGTATAGATCCTGTAAGCGTTGTCTGTATTCTGGAGCTATTTTCTCTATCGCATTAGTGAGTTCAGAATATGTTATAAATTTAAATCGAATAACGAACATAACTTTAGTAAAGTCGTCTATATTAACTCTTTTATTTTCTAGCCATCGTAATACCATGTTTAAGATAGTATATTCTCTAGTTACTACGATCTCATCGTTGTCTAGAATATCGCATACTGAATCTACATCCAATTTTAGAAATTGGTCTGTGCTACATATCTCTTCTATATTATTGTTGATGTATTGTCGTAGAAAACTATTACGTAGACCATTTTCTTTATAAAACGAATATATAGTACTCCAATTATCTTTACCGATATATTTGCATACATAATCCATTCTCTCAATCACTACATCTTTAAGAGTTTGGTTGTTAAGATATTTGGCTAAACTATATAATTCTATTAGATCATCAACAGAATCAGTATATATTTTTCTAGATCCAAAGATGAACTCTTTGGCATCCTCTATAATATTATCAGAAAAGATATTTTCGTGTTTTAGTTTATCAAGATTTAACCTGTTCATATCCATGATTAACGACGTCATATAACCACATAAAATAAAAATCCATTTTCATTTTTAGCACAATACTATTCATAATTGATATTGATGTAATATTTTGTTACTTTGAACGTAAAGACAGTACACGGGTCCGTATCTCCAACAAGCACGTAGTAATCAAATTTGGTGTTGTTAAACTTCGCAATATTCATCAATTTAGATAGAAACATATACTCATCATCTGTTTTAGGAATCCATGTATTATTACCACTTTCCAACTTATCATTATCCCAGGCTATGTTTCGCCCATCATCGTTGCACAGAGTGAATAATTCTTTTGTATTCGGTAGTTCAAATATATGATCCATGCATATATCGACAAAGCTATTGTAGATGTGATTTTTCCTAAATCTAATATAAAACTCGTTTACTAGCAAACATTTTCCTGATTTATCGACCAAGACACACATGGTTTCTAAATCTATCAAGTGGTGGGGATCCATAGTTATAACGCAGTAACATAAATTATTACCTTCTTGACTGTCGCTAATATCTATATACTTATTGTTATCGTATTGGATTCTACATATAGATGGCTTGTATATCAAAGATATAGAACACATAACCAATTTATATTCTCGCTTTGTATTTTCGAATCTAAAGTTAAGAGATTTAGAAAACATTATATCATCGGATGATGATATCACTGTTTCCAGAGTAGGATATATTAAAGTCTTTAAAGATTTTGTCCGATTCAAATAAATCACTAAATAATATCCCATATTATCATCTGTTATAGTCGTGTCATTAAATCTATTATATTTTATGAAAGATATATCACTGCTCACCTCTATATTTCGTACATTTTTAAACTGTTTGTATAATATCTCTCTAATACAATCAGATATATCTATTGTGTCGGTAGACGATACCGTTACATTTGAATTAATGGTGTTCCATTTTACAACTTTTAACAAGTTGACCAATTCATTTCTAATAGTATCAAACTCTCCATGATTAAATATTTTAATAGTATCCATTTTATATCACTACGGACATAAACCATTGTATATTTTTTATGTTTATTAGTGTACACATTTTGGAAGTAAGTTCCTGGATCGGATGTCACCGCAGTAATATTGTTGATTATTTCTGACATCGACGTATTATATAGTTTTTTAATTCCATATCTTTTAGAAAAGTTAAACATCCTTATACAATTTGTGGAATTAATATTATGAATCATGGTTTTTACACATAGATCTATTACAGGCGGAACATCAATTATTATGGCAGCAACTAGTATCATTTCTACATTGTTTATGGTGATGTTTATCTTCTTCCAGCGCATATAGTCTAATATCGATTCAAACGCGTGATAGTTTATACCATTCAATATAATCGCTTCATCCTTTAGATGGTGATCCTGAATGTGTTTAAAAAATTATACGGAGACGCCGTAATAATTTCCCCATTGATAGAAAATATCACGCTTTCCATTTTCTTGAAGTACTAAAAGTAATTATAATATAATGTAAAGGTTTATATATTCAATATTTTTTATAAAAAAAATCATTTTGACATTAATTCCTTTTTAAATTTCAGTCTATCATCTATAGAAACATATTCTATGAATTTATAAAATGCTTTTACGTATCCTATCGTAGGTGATAGAACCGCTAAAAAACCTATCGAATTTCTACAAAAGAATCTGTTATATGGTATAGGGAGAGTATAAAACATTAAATGTCCGTACTTATTAAAGTATTCAGTAGCCAATCCTAACTCTTTCGAATAATTATTAATGGCTCTTATTCTGTACGAATCTATTTTTTTGAACAATGGACCTAGTGGTATATCTTGTTCTATGTATCTAAAATAATGTCTGACTAGATCCGTTAGTTTAATATCCGCAGTCATCTTGTCTAGAATGGCAAATCTAACTGCGGGTTTAGGCGTAGGCGTTAGTTTAGTTTCTATATCTACATCTATGTCTTTATCTAACACCAAAAATATAATAGCTAATATTTTATTACAATCATCCGGATATTCTTCTACGATCTCACTAACTAATGTTTCTTTGGTTATACTAGTATAGTCACGATCAGACAAATAAAGAAAATCAGATGATCGATGAATAATACATTTAAATTCATCATCTGTAAGATTTTTGAGATGTCTCATTAAAATATTATTAGTGTCAGTTCTCATTATCATATATTGACAGCAGCTATTACACTTATTTTATTTTTCTGTATTTTATTACTTTTCACCATATAGATCAGTCATTAGATCATCAAAATACTTTTCAATCATCCTAAAGAGTATGGTGAACGAATCTTCCCATCTAATTTCTGAACGTCTACCAATGTCTCTAGCCACTTTGGCACTAATAGCGATCATTCGCTTAACATCTTCTACATTATTAACTGGTTGATTCAATCTATCTAGCAATGGACCGTCGGATAGCGTCATTCTCATGTTCTTAATCAATGTACATACATCGTCATCATCTACCAATTCATCAAACAATATAAGCTTTTTAAAATCATCATTATAATAGGATGGATCGCCGTCATTTCTCCAAAGAATATATCTAATAAGTAGAGTCCTCATGCTTAGTAATTTAACTATTTTAGTTAACAACTATTTTTTATGTTAAATCAATTAGTAACACCGCTATGTTTAATACTTATTCATATTTTAGTTTTAGGATCGAGAATCAATACAAAAATTAATACATCAATTTTGGAAATACTTAGTTTCCACGTAGTCAATGAAACATTTGAGCTCATCGTAAAGGACGTTCTCGTACAAGACGTAACTATAAATTGGTTTATATTTGTTCAAGATAGATACAAATCCGATAACTTTTTTGACGAATTCTACGGGATTCACTTTAAAAGTGTCATACCGGGTTCTTTTTATTCTTTTAAACAGATCGATTGTGTGATGTTGATTAGGTCTTTTACGAATTTGATACAGAATAGCGTTTACATATCCACCATAGTAATCAATAGCCATTTGTTCGTATGTCATAAATTCTTTAATTATATGACACTGTGTATTATTTAGTTCGTCCTTGTTCATCATTAGGAATCTATCCAATATGGCAATTATATTAGAACTATAACTGCGTTGTATGCGCATGTTGATGTGTCTGTTTATACAATCAATTATACTAGGATCCATACCACTACAATCGGGTAAAATTGTAGCATCATATACCATTTCTAGTACTTTAGATTCATTGTTATCCATTGCAGAGGACGTCATGATCGAATCCAAAAAAATATATTATTTTTATGTTATTTTGTTAAAAATAATCATCAAATACTTCGTAAGATACTCCTTCATGAACATAATCAGTTACAAAACGTTTATATGAAGTAAAGTATCTACAATTTTTACAAAAGTCAGGATGCATAAGTACAAAGTACGCGATAAACGGAATAATAATAGATTTATCTAGTTTATCTTTTTCTATCTCTTTCATAGTTATATACATGGTCTCAGAAGTCGGATTATGTAACATCAGCTTCGATAAAATGACTGGGTTATTTAGTCTTACACATTCGCTCATACATGTATGACCGTTAACTATAGAGTCTACACTAAAATGATTGAATAATAGATAGTCTACCATTGTTTCGTATTCAGATAGTACAGCGTAGTACATGGCATCTTCACAAATTATATCATTATCTAATAGATATTTGACGCATCTTATGGATCCCACTTCAACAGCCATCTTAAAATCGGTAGAATCATATTGCTTTCCTTTATCGTTAATAATTTCTAGAACATCATCTCTATCATAAAAGATACAAATATTAACTGTTTGATCAGTAATAACATTGCTAGTCGATATCAATTTGTTAATAAGATGCGCTGGGCTCAATGTCTTAATAAGAAGTGTAAGAGGACTATCTCCGAATTTGTTTTGTTTATTAACATCCGTTGATGGAAGTAAAAGATTTATAATGTCTACATACTTGACTGTTTTAGAGCATACAATATGGAGAGGCGTATTTCCATCATGATCTGGTTTTGAGGGACTAATTCCTAGTTTCATCATCCATGAGATTGTAGAAGCTTTTGGATTGTCTGACATAAGATGTCTATGAATATGATTTTTGCCAAATTTATCCACTATCCTGGCTTCGAATCCGATAGACATTATTTTTTTAAACACTCTTTCTGAAGGATCTGTACACGCCAACAACGGACCACATCCTTCTTCATCAACCGAGTTGTTAATCTTGGCTCCATACTGTACCAATAAATTTATTCTCTCTATGACTTCATCATCTGTTCCCGAGAGATAATATAGAGGTGTTTTATTATGTTTATCACATGCGTTTGGATCTGCGCCGTGCACCAGCAGCATCGCGACTATTCTATTATTATTAATTTTAGAAGCTATATGCAATGGATAATTTCCATCATCATCCGTCTCATTTGGAGAGTATCCTCTATGAAGAAGTTCTTCTATAAATCGTTCATCTAGTCCTTTAATGCCACAATACGCATGTAGAATGTGATAATTTCCAGAGGGTTCGATAACTTGTAGCATATTCCTAAATACATCTAAATTTTTACTATTATATTTGGCATAAAGAGATAGATAATACTCGACCGACATAATGTTGTGTTGTCCATTATAGTATAAAAATTAATATTTCTATTTCTATATATTTGCAACAATTTACTCTCTATAACAAATATCATAACTTAGTTCTTTTATGTCAAGAAGGCACTGGTTTAATTCATCTATAAATGTCACTCCATAACTACCACGCATACTATACTCAGAATTATGATAAAGATATTTATTCTTGGGGTGTAAGTAATGGGGATTAATCTTTGTTGGATCAGTCTCTAAGTTAACACATGTCACACATGATCCATTTATAGTTATATCACACGATGATGATTTATGAATTGATTCCGGAAGATCGCTATTGTATTTTGTAGTTCCACAATTCATTTCCATACATGTTATTGTCACACTAATATTATGATGAACTTTATCTAGCCGCTGAGTGGTAAACAACAGAACAGATAGTTTATTATCTTTACCAACACCCTCAGCCGCTGCCACAAATCTCTGATCCGTATCCATGATGGTCATGTTTACTTTTAGTCCGTATCCAGTCAACACTATGTTAGCATTTCTGTCGATATAGCTTTCACTCATATGACACTCACCAATAATTGTAGAATTAATGTCGTAATTTACACCAATAGTGAGTTCGGCGACAAAGTACCAGTACCGGTAATCTTGTCGATGAGGACATATAGTATTCTTGTATTCTACCGAATACCCGAGAGATGCGATACAAAAGAGTAAGACTAATTTGTAAACCATCTTACTCAAAAATATGCGACAATAGGAAATCTATCTTATACACATAATTATTCTATCAATTTTACCAATTAGTTAGTGTAATGTTAACAAAAATGTGGGATAATTTAATAGTTTTTCCTTACACAATTGACATACATGAGTCTGAGTTCCTCGTTTTTGCTAATTATTTCGTCCAATTTATTATTCTTGACATCGTCAAGATCTTTTGTATAGGAGTCAGACTTGTATTCAACATGTTTTTCTATAATCATCTTAGCTATTTCGGCATCATCCAATAGTACATTTTCCAGATTAACAGAATAGATATTAATGTCGTATTTGAACAGAGCCTGTAACATCTCAATGTCTTTATTATCTATAGCCAATTTGATGTCCGGAATGAAGAGAAGGGAATTGGTGTTTGTTGACGTCATATAGTCGAGCAAGAGAATCATCATATCCACGTGTCCATTTTTTATAGTGGTGTGAATACAACTAAGGAGAATAGCCAGATCAAAAGGAGATGGTATCTCTGAAAGAAAGTAGGAAACAATACTTACATCATTAAGCATGACAGCATGATAAAATGAAGTTTTCCATCCAGTTTTCCCATAGAACATCAGTCTCCAATTTTTCTTAACAAACAGTTTTACCGTTTGCATGTTACCACTATCAACCGCATAATACAATGCGGTGTTTCCTTTGTCATCAAATTGTGAATCATCCATTCCACTGAATAGCAAAATCTTTACTATTTTGGTATCTTCTAATGTGGCTGCCTGATGTAATGGAAATTCATTATCTAGAAGATTTTTCAATGCTCCAGCGTTCAACAACGTACATACTAGACGCACGTTATTATCAGCTATTGCATAATACAAGACACTATGACCATTGATATCCGCCTTAAATGCATCTTTGCTAGAGAGAAAGCTTTTCAGTTGCTTAGACTTCCAAGTATTAATTCGTGACAGATCCATGTCTGAAACGAGACGCTAATTAGTGTATATTTTTTCATTTTTTATAATTTTGTCATATTGTACCAGAATTAATAATATCTCTAATAGATCTGATTAGTAGATATATGGCTATCGCAAAACAACATATACACATTTAATAAAAATAATATTCATTAAGAAGATTCAGATTCCACTGTACCCATCAATATAAAATAAAATAATTATTCCTTACATCGTACCCATAAACAATATATTAAGTATATTCCACCTTACCCATAAACAATATAAATCCAGTAATATCATGTCTAATGATGAACACAAATGGTGTATTAAATTCCAGTTCTTCAGGAGATGATCTCGCCGTAGCTACCATGATAGTAGATGCCTCCGCTACAGTTCCTTGTTCGTCTACATCTATCTTTACATTCTGAAACATTTTATAAATATATAATGGGTCCCTAGTCATATGTTTAAACGACGCCTTATCTGGATTAAACATACTAGGAGCCATCATTTCGGCTATCGACTTAATATCCCTCTTGTTTTCGATAGAAAATCTAGGGAGTTTAAGATTGTACATTTTATTCCCTAATTGAGATGACCAATATTCTAATTTTGCAGGCGTGATAGAATCTGTGAAATGGGTCATATTATCACCTATTGCCAGGTACATACTAATATTAGCATCCTTATACAGAAGGCGCACCATATCATATTCTTCGTCATCGATTGTGATTGTATTTCCTTGCAATTTAGTAACTACGTTCATCATGGGAACCGTTTTCGTACCGTACTTATTAGTAAAACTAGCATTGTGTGTTTTAGTGATATCAAACGGATATTGCCACGTACCTTTAAAATATATAGTATTAATGATTGCCCATAGAGTATTATCGTCGAGCATAGTAGAATCAACTACATTAGACATACCAGATCTACGTTCTACTATAGAATTAATTTTATTAACCGCATCTCGTCTAAAGTTTAATCTATATAGGCCGAATCTATGATATTGTTGATAATACGACGGTTTAATACACACAGTACTATCGACGAAACTTTGATACGTTAGATCTGTGTACGTATATTTAGATGTTTTCATCTTAGCTAATCCGGATATTAATTCTGTAAATGCTGGACCCAGATCTCTTTTTCTCAAATTCATAGTATTCAATAATTCTACTCTAGTATTACCTGATGCAGACAATAGCGACATAAACATAGAAAACGAATACCCAAACGGTGAGAAGACAATATTATCATTATCATCCTCATCCCCATTTTGAATATTTTTATACGCTAATATACCAGCATTGATAAATCCCTGCAGACGATATGCGGATACTGAACACGCTAATGATAGTATCAATAACGCAATCATGATTTTTATGGTATTAATAATTAACCTTATTTTTATGTTTGGTATAAAAATTATTGATGTCTACACATCCTTTATAATCAACTCTAATCACTTTAACTTTTACAGTTTTCCCTACAAGTTTATCCCTATATTCAACATATCTATCCATATGCATCTCTTAACACTCTGCCAAGATAGCTTCAGAGTGAGGATAGTCAAAAAGATAAATATATAGAGTATAATCATTCTCGTATACTCTGCCCTTTATTACATCGCCCGCATTGGGCAACGAATAACAAAATGCAAGCATCGTGTTAACGGGCTCGTAAATTGGGATAAAATTATGTTTTTATTGTTTATCTATTTTATTCAAGAGAATATTCAGGAAGTTCCTTTTCTGGTTGTATCTCGTCGCAGTATATATCATTTGTACATTGTTTCATATTTTTTAATAGTCTACACCTTTTAGTAGGACTAGTATCGTACAATTCATAGCTGTATTTTGAATTCCAATCACGTATAAAAATATCTTCCAATTGTTGACGAAGACCTAATCCATCATCCGGTGTAATATTAATAGATGCTCCACATATATCCGTAAAGTAATTTCCTGTCCAATTTGATGTACCTATATACGCCGTTTTATCGGTTACCATATATTTTGCATGGTTTACCCTAGAATACGGAATGGGAGGATCAGCATCTGGTACAATAAATAGCTTTACTTCTATATCTATGTTTTTAGATTTTAGCATAGCTATAGATCTTAAAAAGTTTCTCATGATAAACGAAGATCGTTGCCAGCAACTAATCAATAGCTTAACGGATACTTGTCTGTCTATAGCGGATCTTCTTAATTCATCTTCTATATAAGGCCAAAACAAAATTTTACCCGCCTTTGAATAAATAATAGGAATAAAGTTCATAACAGATACATAAACGAATTTACTCGCATTTCCGATACATGACAATAAAGCGGTTAAATCATTGGTTCTTTCCATAGTACATAATTGTTGTGGTGCAGAAGCAATAAATACAGAGTGTGGAACACCGCTTACGTTAATACTAAGAGGATGATCTGTATTATAATACGACGGATAAAAGTTTTTCCAATTATATGGTAGATTGTTAACTCCAAGATACCAGTATACCTCAAAAATTTGAGTGAGATCCGCTGCCAAGTTCCTATTATTGAAGATCGCAATACCCAATTCCTTGACCTGAGTTAGTGATCTCCAATCCATGTTAGCGCTTCCTAAATAAATATGTGTATTATCAGATATCCAAAATTTTGTATGAAGAACTCCTCCTAGGATATTTGTAATATCTATGTATCGTACTTCAACTCCGGCCATTTGTAGTCTTTCAACATCCTTTAATGGTTTGTTGGATTTATTGACGGCTACTCTAACTCTTACTCCTCTTTTGGGTAATTGTACAATCTCGTTTAATATTACCGTGCCGAAATTCGTACCCACTTCATCCGATAAACTCCAATAAAAAGATGATATATCTAGTGTTTTTATGGTATTGGATAGAATTTCCCTCCACATGTTAAATGTAGTCAAATATACTTTATCAAATTGCATACCTATAGGAATAGTCTCTGTAATCACTGCGATTGTATTATCCGGATTCATTTTATTTGTTAAAAAAATAATCCTATATCACTTCACTCTATTAAAAATCCAAGTTTCTATTTCTTTCATGACTGATTTTTTAACTTCATCCGTTTCCTTATGAAGATGATGTTTGGCGCCTTCATAAATTTTTATTTCCCTATTACAATTTGCATGTTGCATGAAATAATATGCACCTGAAACATCGCTAATCTCATTGTTTGTTCCCTGGAGTATGAGAGTCGGGGTGTTAATCTTGGGAATTATTTTTCTAACCTTGTTGGTAGCCTTCAAGACCTGACTAGCAAATCCAGCCTTAATTTTTTCATGATTGACTAATGGATCGTATTGGTATTTATAAACTTCATCCATATCTCTAGATACTGATTCTGGACATAGCTTTCCGACTGACGCATTTGGTGTAATGGTTCCCATAAGTTTTGCAGCTAGCAGATTCAGTCTTGGAACAGCGTCTGCATTAACTAGAGGAGACATTAGAATCATTGCTGTAAACAAGTTTGGATTATCGCAAGCAGCTAGTATAGAAATTGTTGCTCCCATGGAATGACCCAATAAGAAGACTGGAACTCCTGGATAAGTAGATTTAATAGTCACCACGTGCTGTACCACATCTCTAACATACTTACCAAAGTCATCAATCATCATTTTTTCACCATTACTTCTTCCATGGCCAATATGATCATGTGAGAATACTAAAATTCCTAACGATGATATGTTTTCAGCTAGTTCGTCATAACGTCCAGAATGTTCACCAGCTCCATGACTTATGAATACTAATGCCTTAGGATATGTAATAGGTTTCCAATATTTACAATATATGTAATCATTGTCCAGATTGAACATACAGTTTGTACTCATGATTCACTATATAACTATCAATATTAACAGTTCGTTTAATGATCATATTATTTTTATGTTTTATTGATAATTGTAAAAATATACAATTAAATCAATATAGAGGAAGGAGACGGTACTGTATTTTGTGAGATAGTAATGGAGACTAAATCAGATTATGAGGATGCTGTTTTTTACTTTGTGGATGATGATGAAATATGTAGTCGCGACTCCATCATTGATCTAATAGATGAATATATCACGTGGAGAAATCATGTTATAGTGTTTAATAAAGATATTACCAGTTGTGGAAGACTGTACAAGGAATTGATAAAGTTCGATGATGTCGCTATACGGTACTATGGTATTGATAAAATTAATGAGATTGTCGAGGCTATGAGTGAAGGAGACCACTACATCAATCTTACAGAAGTCCATGATCAGGAAAGTCTATTCGCTACCATAGGAATATGTGCTAAAATCACTGAACATTGGGGATACAAAAAGATTTCAGAATCTAAATTCCAATCATTGGGAAACATTACAGATCTGATGACCGACGATAATATAAACATCTTGATACTTTTTCTAGAAAAAAAAATGAATTGATGATATAAGTGTCTTCATAACGCATTATTACGTTAGCATTCTATTATCCAGTGTTAAAAAAATTATCCTATCATGTATTTGAGAGTCTTATATGTAGCAAACATGATAACTGCAATACCCATAATCTTTAGATATTCACGCGTGCTATGGATGGCATTATCCCGCGGTGCGGAAATGTACGTTATATAATCTACAAAATAATCATCGCATATAGTATGAGATAGTAGAGTAAACATTTTTATCGTTTCTACTGGGTTCATACATCGTCTACCCAATTCGGTAATGAATGAAATTGTCGCCAATCTTACACCCAAACCCTTGTTGTTCATTAGTATAGTATTAACTTCATTATTTATGTCATAAACTGTAAATGATTCTGTAGATGCCATATCACACATGATATTCATGTCACTATTATAATCATTATTAACTTTATCACAATACGTGTTGATAATATCTACATATGATCTAGTTTTTGTGGGTAATTGTCTATACAAGTCGTCTAAACGTTGTTTACTCATATAGTATTGAACAGCCATCATTACATGGTCCCGTTCCGTTGATAGATAATCGAGTATGTTAGTAGACTTGTCAAATCTATATACCATATTTTCTGGAAGCGGATATACATAGTCGCGATCATCATTATCACTAGCCTCATCCTCTATATCATGTACATGTACATAATCTATGATATTATTATACATAAACATCGACAACATACTATTGTCTATTATCTAAGTCCTGTTGATCCAAACCCTTGATCTCCTCTATCTGTACTATCTAGAGATTGTACTTCTTCAAGTTCTGGATAATATATACGTTGATAGATTAGCTGAGCTATTCTATCTCCAGTATTTACATTAAACGTACATTTTCCATTATTAATAAGAATGACTCCTATGCTTCCCCTATAATCTTCGTCTATTACACCGCCTCCTATATCAATGCCTTTTAGGGACAGACCAGACCTAGGAGCTATTCTACCATAGCAGAACTTAGGCATGGACATACTAATATCTGTCTTAATTAACTGTCGTTCTCCAGGAGGGATAGTATAATCGTAAGCGCTATACAAATCATATCCGGCAGCACCCGGCGATTGCCTAGTAGGTGATTTAGCTCTGTTAGTTTCCTTAACAAATCTAACTGGTGAGTTAATATTCATGTTGAACATAAAAAATATCATTTTATTTCAAAATTATTTACCATTCCATTCCATCCCATATATTCCATGAATAAGTGCGATTATTGTACACTTCTATAGTATCTATATACGATCCACGATAAAATCCTCCTATCAATAGCAGTTTATTATCCACTATGATCAATTCTGGATTATCCCTCGGATAAATAGGATCATCTATCAGAGTCCATGTATTGCTGGATTCACAATAAAATTCCGCATTTCTACCAACCAAGAATAACCTTCTACCAAACACTAACGCACATGATTTATAATGAGGATAATAAGTGGATGGTCCAAACTGCCACTGATCATGATTGGGTAGCAAATATTCTGTAGTTGTATCAGTTTCAGAATGTCCTCCCATTACGTATATAACATTGTTTATGGATGCCACTGCTGGATTACATCTAGGTTTCAGAAGACTCGGCATATTAACCCAAGCAGCATCCCCGTGGAACCAACGCTCAACAGATGTGGGATTTGGTAGACCTCCTACTACGTATAATTTATTGTTAGCGGGTATCCCGCTAGCATACAGTCTGGGGCTATTCATCGGAGGAATTGGAATCCAATTGTTTGATATATAATTTACCGCTATAGCATTGTTATGTATTTCATTGTTCATCCATCCACCGATGAGATATACTACTTCTCCAACATGAGTACTTGTACACATATGGAATATATCTATAATTTGATCCATGTTCATAGGATACTCTATGAATGGATACTTGTATGATTTGCGTGGTTGTTTATCACAATGAAATATTATGTTACAGTCTAGTATCCATTTTACATTATGTATACCTCTGGGAGAAAGATAATTTGACCTGATTACATTTTTGATAAGAAGTAGCAGATTTCCTAATCTATTTCTTCGCCTCATATACCACTTAATGACAAAATCAACTACATAATCCTCATCTGGAACATTTAGTTCGTCGCTTTCTAGAATAAGTTTCATAGATAGATAATCAAAATTGTCTATGATGTCATCTTCCAGTTCCAAAAAGTGTTTGGTAATAAAGTCTTTAGTATGACATAAGAGATTGGATAGTCCGTATTCTATACCCATCATGTAACACTCGATACAATATTCCTTTCTAAAATCTCGTAGGATAAAGTTTATACAAGTGTAGATGATAAATTCTACAGATGTTAATATAGAAGCACGTAATAAATTGACGACGTTATGACTATCTATATATACCTTTCCAGTATATGAGTAAATAACTATAGAAGTTAGACTGTGAATGTCAAGGTCTAGACAAACCCTCGTAACTGGATCTTTATTTTTTGTGTATTTTTGGCGTAAATGTGTGCGAAAGTATGGAGATAACTTTTTCAATATCGTAGAATTGACTATTATATTACCTCCTATAGCTTCAATAATTGTTTTGAATTTCTTAGTCGTGTACAATGCTAATATATTCTTACAGTACACAGTATTGACAAATATCGGCATTTATGTTTCTTTAAAAGTCAACATCTAAAGAAAAATGATTGTCTTCTTGAGACATAACTCCCATTTTTTGGTATTCACCCACACGTTTTTCGAAAAAATTAGTTTTTCCTTCCAATGATATATTTTCCATGAAATCAAACGGATTGGTAACATTGTAAATTTTTTTAAATCCCAATTCAGAAATCAATCTATCTGCGACGAATTCTATATATGTTTTCATCATTTCACAATTCATTCCTATGAGTTTAACTGGAAGAGCCACAGTAAGAAATTCTTGTTCAATGGATACCGCATTTGTTATAATAAATCTAACGGTTTCTTCACTCGGTGGATGCAATAAATGTTTAAACATCAAACATGCGAAATCGCAGTGCAGACCCTCGTCTCTACTAATTAATTCGTTAGAAAACGTGAGTCCGGGCATTAGGCCACGCTTTTTAAGCCAAAATATGGAAGCGAATGATCCGGAAAAGAAGATTCCTTCTACTGCAGCAAAGGCAATAAGTCTCTCTCCATAACCGGCGCTGTCATGTATCCACTTTTGAGCCCAATCGGCCTTCTTTTTTACACAAGGCATCGTTTCTATGGCATTAAAGAGGTAGTTTTTTTCATTACTATCTTTAACATAAGTATCGATCAAAAGACTATACATTTCCGAATGAATGTTTTCAATGGCCATCTGAAATCCGTAGAAACATCTAGCCTCGGTAATCTGCACTTCTGTACAAAATCGTTCTGCTAAATTTTCATTCACTATTCCGTCACTGGCTGCAAAAAACGCCAATACATGTTTTATAAAATATTTTTCGTCTGGTGTTAGTTTATTCCAGTCATTGATATCTTTAGATATATCCACTTCTTCCACTGTCCAAAATGATGCCTCTGCCTTTTTATACATATTCCAGATGTCATGATATTGGATTGGGAAAATAACAAATCTATTTGGATTTGGTGCAAGGATAGGTTCCATAACTAAATTAACAATAGTAGTAATTTTTTTTCAGTTATCTGTATGACGACTGTACTTGGATCTTTTGTATATCGCTATCGCCGCAATCACTACAATAATTACAAGTATTATTGATAGCATTGTTATTACTACTATCATAATTAAATTATCGACATTCATGGGTGTTGAATAATCGTTATCATCATTTTGTAATTGTGACGTCATACTAGATAAATCATTTGTGAGATTGTTGTGGGAAGCGGGCACGGAAGATGCATTATCATTATTATTTAACGCCTCCCATTCGGATTCACAAATGTTACGCACGTTCAAAGTTTTATGGAAACTATAATTTTGTGAAAACAGATAACAAGAAAACTCGTCATCGTTCAAATTTTTAACGATAGTAAACCGATTAAACGTCGAGCTAATTTCTAACGCTAGCGACTCTGTTGGATATGGGTTTCCAGATATATATCTTTTCAGTTCCCCTACGTATCTATAATCATCTGTAGGAAATGGAAGATATTTCCATTTATCTACTGTTCCTAATATCATATGCGGTGGTGTAGAACCATTAAGCGCGAAAGATGTTATTTCGCATCGTATTTTAACTTCGCAATAATTTCTGGTTAGATAACGCACTCTACCAGTCAAGTCAATGATATTAGCCTTTACAGATATATTCATAGTAGTCGTAACGATGACTCCATCTTTTAGATGTGATACTCCTTTGTATGTACCAGAATCTTCGTACCTCAAACTCGATATATTTAAACAAGTTAATGATATATTAACGCGTTTTATGAATGATGATATATAACCAGAAGTTTTATCCTCTGTGGCTAGCGCTATAACCTTATCATTATAATACCAACTAGTGTAATTAATATGTGACATGACAGTGTGGGTACAAATATGTACATTATCGTCTACGTCGTATTTGATACATCCGCATTCAGCCAACAAATATAAAATTACAAAAACTCTAACGACGTTTGTACACATCTTGATGCGGTTTAATAAATGTTTTGATTTCAATTTATTGTAAAAAAAGATTCGGTTTTATACTGTTCGATATTCTCATTGCTTATATTCTCATCTATCATCTCCACACAGTCAAATCCATGGTTAACATGTACCTCATCAACCGGTAAAAGACTATCGGATTCTTCTATCATCATAACTCGAGAATATTTAATTTGGTGGTCATTATTAATCAAGTCAATTATCTTATTTTTAACAAACGTAAGTATTTTACTCATTTTTTTATAAAAACTTTTAGAAATATACAGACTCTATCGTGTGTCTATATCTTCTTTTTATATCCAATGTATTTATGTCTGATTTTTCTTCATTTATCATATATAATGGTCCAAATTCTACACGTGCTTCGGATTCATCCAGATCATTAAGGTTCTTATAATCGCAACATCCTTCTCTTCCATCTTCTACATCTTCCTTCTTATTCTTAGCGTCACAGAATCTACCACAGCAGGATCCCATGACGAGAGTCACATTAAACTAATTCATTTTCAATTATAATATACTGATTAGTAATGACAATTAAAATAAAAATATTCTTCATAACCGGTAAGAAAGTAAAAAGTTCACATTGAAACTATGTCAGTAGTTATACATCATGAGATGATATACTCTATTTTGGTGGAGGATTATATGATATAATTCGTGGATAATCATTCTTAAGACACATTTCTTCATTCGTAAATCTTTTCACATTAAATGAGTGTCCATATTTTGCAATTTCTTCATATGATGGCGGTGTACGTGGACGAGGCTGCTCCTGTTCTTGTAGTCGTCGACTGTCGTGTTTGCGTTTAGATCCCTCCATTATCGCGATCGCGTAGTGAGTACTATTTATACCTTGTAATTAAATTTTTTTATTAATTAAACGTATAAAAACGTTCCGTATCTGTATTTAAGAGCCAGATTTCGTCTAATAGAACAAATAGCTACAGTAAAAATAACTAGAATAATCGCTACACCCACTAGAAACCACGGATCGTAATACGGCAATCGGTTTTCGATAATAGGTGGAACGTATATTTTATTTAAGGACTTAACAATTGTCTGTAAACCACAATTTGCTTCCGCCGATCCTGTATTAACTATCTGTAAAAGCATATGTTGACCGGGAGGAGCCGAACATTCTCCGATATTCAATTTTTGTATATCTATAATGTTATTAACCTCCGCATACGCATTACAGTTCTTTTCTAGCTTGGATACTACACTAGGTACATCATCTAAATCTATTCCTATTTCCTCAGCGATAGCTCTTCTATCCTTTTCCGGAAGTAATGAAATCACTTCAATAAATGATTCAACCATGAGTGTGAAACTAAGTCGAGAATTACTCATGCATTTGTTAGTTATTCGGAGCGCGCAATTTTTAAACTGTCCTATAACCTCTCCTATATGAATAGCACAAGTGACATTAGTAGGGATAGAATGTTGAGCTAATTTTTGTAAATAACTATCTATAAAAAGATTATACAAAGTTTTAAACTCTTTAGTTTCTGCCATTTATCCAGTCTGAGAAAATGTCTCTCATAATAAATTTTTCCAAGAAACTAATTGGGTGAAGAATGGAAACCTTTAATCTATATTTATCACAGTCTGTTTTGGTACACATGATGAATTCTTCTAATGCTGTACTAAATTCGATATCTTTTTCGATTTCTGGATATGTTTTTAATAAAGTATGAACAAAGAAATGGAAATCGTAATACCAGTTATGTTCAACTTTGAAATTGTTTTTTATTTTCTTGTTAATGATTCCAGCCACTTGGGAAAAGTCAAAGTCGTTTAATGCCGATTTAATACGTTCATTAAAAACAAACTTTTTATTCTTTAGATGAATTATTATTGGTTCATTGGAATCAAAAAGTAAGATATTATCGGGTTTAAGATCTGCATGTAAAAAGTTGTCACAACAGGGTAGTTCGTAGATTTTAATGTATAACAGAGACATCTGTAAAAAGATAAACTTTATGTATTGTACCAAAGATTTAAATCCTAATTTGATAGCTAACTCGGTATCTACTTTATCTGCCGAATACAGTGCTAGGGGAAAAATTATAATATTTCCTCTTTCGTATTCGTAATTAGTTCTCTTTTCATGTTCGAAAAAGTGAAACATGCGGTTAAAATAGTTTATAACATTAATATTACTGTTAATAACTGCCGGATAAAAGTGGGATAGTAATTTCACGAATTTGATACTGTCCTTTCTCTCGTTAAACGCCTTTAGAAAAACTTTAGAAGAATATCTCAATGAGAGTTCCTGACCATCCATAGTTTGTATCAATAATAGCAACATATGAAGAACCCGCTTATACAGAGTATGTAAAAATGTTAATTTATAGTTTAATCCCATGGCCCACGCACACACGATTAATTTTTTTTCATCTCCCTTTAGATTGTTGTATAGAAATTTGGGTACTGTAAACTCCGCCGTAGTTTCCATGGGACTATATAATTTTGTGGCCTCGAATACAAATTTTACTACATAGTTATCTATCTTAAAGACTATACCATATCCTCCTGTAGATATGTGATAAAAATCGTCGTTTATAGGATAAAATCGTTTATCTTTTTGTTGGAAAAAGGATGAATTAATGTAATCATTCTCTTCTATCTTTAGTAGTGTTTCCTTATTAAAATTCTTAAAATAATTTAACAATCTAACTGACGGAGCCCAATTTTGGTGTAAATCTAATTGGGACATTATGTTGTTAAAATATAAACAGTCTCCTAATATAACAGTATCTGATAATCTATGGGGAGACATCCATTGATATTCAGGGGATGAATCATTGGCAACACCCATTTATTGTACAAAAAGCCCCAATTTACAAACGAAAGTCCAGGTTTGATAGAGATAAACTATTAACTATTTTGTCTCTGTTTTTAACACCTCCACAGTTTTTAATTTCTTTGGTAATGAAATTATTCACAATATCAGTATCTTCTTTATCTACCAGAGATTTTACTAACTTGATAACCTTGGCTGTCTCATTCAATAGGGTAGTGATATTTGTATGTATGATATTGATATCTTTTTGAATTGTTTCTTTTAGAAGTGATTCTTTGATGGTATCAGCATACGAATTACAATAATGCAGAAACTCAGTTAACATGCAGGAATTATAGTAAGCCAATTCCAATTGTTGCCTGTATTGTATTAGAGTATTAATATGCGCAATGATGTCCTTGCGTTTCTCTGATAGAATGCGAGCAGCGATTTTGGCGTTATCATTTGACGATATTTCTGGAATGACGAATCCTGTTTCTACTAACTTCTTGGTAGGACAAAGTGAAACAATCAAGAAAATAGCTTCTCCTCCTATTTGTGGAAGAAATTGAACTCCTCTAGATGATCTACTGACGATAGTATCTCCTTGACAGATATTGGACCGAACTACGGAAGTACCTGGAATGTAAAGCCCTGAAACCCCCTCATTTTTTAAGCAGATTGTTGCCGTAAATCCTGCACTATGCCCAAGATAGAGAGCTCCTTTGGTGAATCCATCACTATGTTTCAGTTTAACCAAGAAACAGTCAGCTGGTCTAAAATTTCCATCTCTATCTAATACAGAATCCAACTTGATGTCAGGGACTATGACCGGTTTAATGTTATATGTAACATTGAGTAAATCCTTAAGTTCATAATCATCGTTGTCATCAGTTATGTACGATCCAAACAATGTTTCTACCGGCATGGTGGATACGAAGATGCTATCCATCAGAATGTTTCCCTGATTAGTATTTTCTATATAGCTATTCTTCTTTAAACGATTTTCCGAATCAGTAACTATGTTCATTTTTTTAGGAGTAGGACGTCTAGCCAGTATGGAAGAGGATTTTCTAGATACTCTCTTCAACATCTTTGATCTCAATGGAATGCAAAACCCCATGGTGTAACAACCAACGATAAAAATAATATTGTTTTTTCACTTTTTATAATTTTACCATCTGACTCATGGATTCATTAATATCTTTACAAGAGCTACTAACGTATAATTCTTTATAACTGAACTGAGATATATACACCGGATCTATGGTTTCCATAATTGAGTAAATGAATGCTCGGCAATAACTAATGGCAAATGTATAGAACAACGAAATTATACTAGAGTTGTTAAAGTTAATATTTTCTATGAGTTGTTCCAATAAATTATTTGTTGTGACTGCGTTCAAGTCATAAATTATCTTGATACTATCCAGTAAACAGTCTTTAAGTTCTGGAATATTATCATCCCATTGTAAAGCCCCTAGTTCGACTATCGAATATCCTGCTCTGATAGCAGTTTCAATATCGACGGACGTCAATACTGTAATAAAGGTGGTAGTATTGTCATCATCGTGATAAACTACGGGAATATGGTCGTTAGTAGGTACCGTGACTTTACACAACGCGATATATAACTTTCCTTTTGTACCATTTTTAACGTAGTTGGGACGTCCTGCAGGGTATTGTTTTGAAGAAATGATATCGAGAACAGATTTGATACGATATTTGTTGGATTCCTGATTATTCACTATAATATAATCTAGACAGATAGATGATTCGATAAATAGAGAAGGTATATCGTTGGTAGGATAATACATCCCCATTCCAGTATTCTCGGATACTCTATTGATGACACTAGTTAAGAACATGTCTTCTATTCTAGAAAACGAAAACATCCTACATGGACTCATTAAAACTTCTAACGCTCCTGATTGTGTTTCGAATGCCTCGTACAAGGATTTCAAGGATGCCATAGATTCTTTGACCAACGATTTAGTATTGCGTTTAGCATCTGATTTTTTTATTAAATCAAATGGTCGGCTCTCTGGTTTACTACCCCAATGATAACAATAGTCTTGTAAAGATAAACCGCAAGAAAATTTATACACATCCATCCAAATAACCCTAGCACCGTCGGATGATATTAATGTATTATTATAGATTTTCCATCCACAGTTATTGGGCCAGTATACTGTTAGCAACGGTATATCGAATAGATTACTCATGTAACCTACTAGAATGATAGTTCGTGTACTAGTCATAATATCTTTAATCCAATCTAAGAAATCTAAAATTAGATCTTTTACACTATTAAAGTTAACAAAGGTATTACCCGGGTACGTGGATATCATATATGGTATTGGTCCATTATCAGTAATGGCTCCATAAACTGATACGGCGATGGTTTTTATATGTGTTTGATCTAATGAGGACGAAATTCGCGCCCACAATTCATCTCTAGATATGCATTTAATATCGAACGGTAACACATCAATCTCGGGACGCGTATATGTTTCTAAATTCTTAATCCAAATATAATGATGACCTATATGCCCTATTATCATACTGTCAACTATAGTATACCTAGAGAACTTTCGATACATCTGCTGTTTCCTGTAATCGTTAAATTTTACAAATCTATAACATGCTAAACCTTTTGACGACAGCCATTCATTAATTTCTGATATGGAATCTGTATTCTCAATACCGTATCGTTCTAAAGCCAGTGCTATATCTCCCTGTTCGTGGGAACGCTTTCGTATAATATCGATCAATGGATAATATGAAGTTTTTGGAGAATAATATGATTCATGATCTATTTCGTCCATAAACAATCTAGACATAGGAATTGGAGGCGATGATCTTAATTTTGTGCAATGGGTCAATCCTATAACTTCTAATATTGTAATATTCATCATCGACATAACACTATCTATGTTATCATCGTATATTAGTATACCACGACCTTCTTCATTTCGTGCCAAAATGATATACAGTCTTAAATAATTACGCAATATCTCAATAGTTTCATAATTGTTAGCTGTTTTCATCAAGGTTTGTATCCTGTTTAACATGATGGCGTTCTATAACGTCTCTATTTTCTATTTTTAATTTTTTTAAATTTTTAACGATTTACTGTGGCTAGATACCCAATCTCTCTCAAATATTTTTTTAGCCTCGCTTACAAGCTGTTTATCTATACTATTAAAACTGACGAATCCGTGATTTTGGTAATGGGTTCCGTCGAAATTTGCCGAAGTGATATGAACATATTCGTCGTCGACTATTAACAATTTTGTATTATTCTGAATAGTGAAAACCTTCACAGATAGATCATTTTGAACACACAACGCATCTAGACTTCTGGCGGTTGCCATAGAATATACGTCGTTCTTATCCCAATTACCAACTAGAAGTCTGATCTTAACTCCTCTATTAATGGCTGCTTCTATAATGGAGTTGTAAATGTCAGGCCAATAGTAGCTATTACCGTCGACACGTGTAGTGGGAACTATGGCCAAATGTTCAATATCTATACTAGTCTTAGCCGACTTGAGTTTATCAATAACTACATCAGTGTCTAGATCTCTAGAATATCCCAATAGGTGTTCTGGAGAATCAGTAAAGAACACTCCACCTATAGGATTCTTAATATGATACGCAGTGCTAACTGGCAGACAACAAGCCGCAGAGCATAAATTCAACCATGAATTTTTTGCGCTATTAAAGGCTTTAAAAGTATCAAATCTTCTACGAAGATCTGTGGCCAGCGGAGGATAATCAGAATATACGCCTAACGTTTTAATCGTATGTATAGATCCTCCAGTAAATGACGCGTTTCCTACATAACATCTTTCATCATCAGACACCCAAAAACAACCGAGTAGTAGTCCCACATTATTTTTTTTATCTATATTAACGGTTATAAAATTTATATCCGGGGAGTGACTTTGTAGCTCTCCCAGATTTCTTTTCCCTCGTTCATCTAGCAAAACTATTATTTTAATCCCTTTTTCAGATACCTCTTTTAGTTTATCAAAAATAAGCGCTCCCCTAGTAGTACTCAGAGGATTACAACAAAAAGATGCTATGTATATATATTTCTTAGCTAGAGTGATAATTTCGTTAAAACATTCAAATGTTGTCAAATGATCGGATCTAAAATCCATATTTTCTGGTAGTGTTTCTACCAGCCTACATTTTGCTCCCGCAGGTACCGATGCAAATGGCCACATTTAGTTAACATAAAAACTTATATATCCTGTTCTATCAACGATTCTAGAATATCATCGGCTATATCGCTAAAATTTTCATCAAAGTCGACATCACAACCTAACTCAGTCAATATATTAAGAAGTTCCATGATGTCATCTTCGTCTATTTCTATATCCGTATCCATTGTAGATTGTTGACCGATTATCGAGTTTAAATCATTACTAATACTCAATCCTTCAGAATACAATCTGTGTTTCATTGTAAATTTATAGGCGGTGTATTTAAGTTGGTAGATTTTCAATTATGTATCAATATAGCAACAGTAGTTCTTGCTCCTCCTTGATTTTAGCATCCTCTTCATTATTTTCTTCTACGTACATAATCATGTCTAATACGTTAGACAACACACCGACGATGGTGGCCGCCACAGACACGAATATGACTAGACCGATGACCATTTAAAAAATACTCTCTAGCTTTAACTTAAACTGTATCGATCATTCTTTTAGCACATGTATAATATAAAAACATTATTCTATTTCGAATTTAGGCTTCCAAAAATTTTTCATCCGTAAACCGATAATAATATATATAGACTTGTTAATAGTCGGAATAAATATATTAATGCTTAAACTATCATCATCTCCACGATTAGAGATACAATATTTACATTCTTTTTGCTGTTTCGAAACTTTATCAATACACGTTAATACAAACCCAGGAAGGAGATATTGAAACTGAGGCTGTTGAAAATGAAACGGCGAATACAATAATTCAGATAATGTAAAATCATGATTCCGTATTCTGATGATATTAGAACTGCTAATGGATGTCGATGGTATGTATCTAGGAGTATCTATTTTAACAAAGCATCGATTTGCTAATATACAATTATCCTTTTGATTAATTGTTATTTTATTCATATTCTTAAAAGGTTTCATATTTATCAATTCTTCTACATTAAAAATTTCCATTTTTAATTTATCTAGCCCCGCAATACTCCTCATTACGTTTCATTTTTTGTCTAGAATACCCATTTTGTTCATCTTGGTACATAGATTATCCAATTGAGAAGCGCATTTAGTAGTTTTGTACATTTTAAGTTTATTAACGAATCGTCGAAAACTAGTTATAGTTAACATTTTATTATTTGATACCCTGATATTAATACCCCTGCCGTTACTATTATTTATAACTGATGTAACCCACGTAACATTGGAATTAATTATCGATAGTAATGCATCGACACTTCCAAAATTGTCTATTATAAACTCACCGATAATTTTTTTATTGCATGTTTTCATATTCATTAGGATTATCAAATCTTTAATCTTATTACGATTGTATGCGTTGATATTACAAGACGTCATTCTAAAAGACGGAGGATTTCCATCAAATGCCAGACAATCACGTACAAAGTACATGGAAATAGGTTTTGTTCTATTACGCATCATAGATTCATATAAAACACCCGTAGAAATACTAATTTGTTTTACTCTATAAAATACTATTGCATCTATTTCATCGTTTTGTATAACGTCTTTCCAAGTGTCAAATTCCAATTTTTTTTCATTGATAGTACCAAATTCTTCTATCTCTTTAACTACTTGCATAGATAGGTAATTACAGTGATGCCTACATGCCGTTTTTTGAAACTGAATAGATGCATCTAGAAGCGATGCTACACTAGTCACGATCACCACTTTCATATTTAGAATATATGTATGTAAAAATATAGTAGAATTTCATTTTGTTTTTTTCTATGCTATAAATGAATTCTCATTTTGTATCCGCACATACTCCGTTTTATATCAATACCAAAGAAGGAAGATATCTGGTTCTAAAAGCCGTTAAAGTATGCGATGTTAGAACTGTAGAATGCGAAGGAAGTAAAGCTTCCTGCGTACTCAAAGTAGATAAACCCTCATCACCCACGTGTGAGAGAAGACCTTCGTCCCCGTCCAGATGCGAGAGAATGAATAACCCTGGAAAACAAGTCCCGTTTATGAGAACGGACATGTTACAAAATATGTTTGCTGCTAATCGCGACAACGTAACGTCAAGACTTTTGAACTAAAATACAATTATATCTTTTTCGATATTAATAAATCCGTGTCTCCCGGGTTTTTTATCTCTTTCAGTATGTGAATAGATAGGTATTTTATCTCTATTCATCATCGAATTTAAGAGATCCGATAAACATTGTTTGTATTCTCCAGATGTCAGCATCTGATACAACAATATATGTGCACATAAACCTCTGGCACTTATTTCATGTACCTTCCCCTTATCACTAAGGAGAATAGTATTTGAGAAATATGTATACATGATATTATCATGTATTAGATATACAGAATTTGTAACACTCTCGAAATCACACGATGTGTCGGCGTTAAGATCTAATATATCACTCGATAACACATTTTCATCTAGATACACTAGACATTTTTTAAAGCTAAAATAGTCTTTAGTAGTAACAGTAACTATGCGATTATTTTCATCGATGATACATTTCATCGGCATATTATTACGCGTACCATCAAAGACTATACCATGTGTATATCTAACGTATTCTAGCATAGTTGCCATACGTACATTAAACTTTTCAGGATCTTTGGATAGATCTTCCAATCTATCTATTTGAGAAAACATTTTTATCATGTTCAATAGTTGAAACGTCGGATCCACTATATAGATATTATCTATAAAGATTTTAGGAACTATGTTCATGGTATCCTGGCGAATATTAAAACTATCAATGATATGATTATCGTTTTCATCTTTTATCACCATATAGTTTCTAAGATATGGGATTTTACTTAATATAATATTATTTCCCGTAATAAATTTTATTAGAAATGCCAAATCTATAAGAAAAGTCCTAGAATTAGTCTGAAGAATATCTATATCACCGTACCGTATATTTGGATTAATTAGATATAGAGAATATGATCCGTAACATATACAACTTTTATTATGACGTCTAAGATATTCTTCCATCAACTTATTAACATTTTTGACTAGGGAAGATACATTATGACGTCCCATTACTTTTGCCTTGTCTATTACAGCGACGTTCATAGAATTTAGCATATCTCTTGCCAATTCTTCCATTGATGTTACATTATAAGAAATTTTAGATGAAATTACATTTGGAGCTTTAATAGTAAGAACTCCTAATATATCCGTGTATGTGGTCACTAATACAGATTGTAGTTCTATAATCGTAAATAATTTACCTATATTATATGTTTGAGTTTGTTTAGAAAAGTAGCTAAGTATACGATCTTTTATTTCTGATGCCGATGTATCAACATCGAAAAAAAATCTTTTTTTATTCTTTTTTACTAACGATACGAATATGTCTTTGTTAAAAACAGTTATTTTCTGAATATTTCTAGCTTGTAATTTTAACATATGATATTCGTTCACACTAGGTACTCTGCCTAAATAGGTTTCTATAATCTTTAATGTAATATTAGGAAGAGTATTCTGATCAGGATTCCTATTCATTTTGAGGATTTAAAACTCTGATTATTGTCTAATATGGTCTCAACACAAACTTTTTCACAGAGTGATAGAGTTTTTGATAACTCGTTTTTCTTAAGAAATATAAAACTACTGTCTCCAGAGCTCGCTCTATCTTTTATTTTATCTAATTCGATACAAACTCCTGATACTGGTTCAGAAAGTAATTCATTAATTTTCAGTCCTTTATAGAAGATATTTAATATAGATAATACAAAATCTTCAGTTCTTGATATCGATCTGATTGATCCTAGAACTAGATATATTAATAACGTGCTCATTAGGCAGTTTATGGCAGCTTGATAATTAGATATAGTATATTCCAGTTCATATTTATTAGATACCGCATTGCCCAGATTTTGATATTCTATGAATTCCTCTGAAAATAAATCCAAAATAACTAGACATTCTATTTTTTGTGGATTAGTGTACTCTCTTCCCTCTATCATGTTCACTACTGGTGTCCACAATGATAAATATCTAGAGGGAATATAATATAGTCCATATGATGCCAATCTAGCAATGTCGAATAACTGTAATTTTATTCTTCGCTCTTCATTATGAATTGAATCTTGAGGTATAAACCTAACACAAATTATATCATTAGACTTTTCGTATGTAATGTCTTTCATGTTATAAGTTTTTAATCCTGGAATAGAATCTATTTTAATGAGGCTTTTAAACGCAGCGTTCTCCAACGAGTCAAAGCATAATACTCTGTTGGTTTTCTTATATTCAATATTACGATTTTCTTCTTTGAATGGAATAGGTTTTTGAATTAGTTTATAATTACAACATAATAGATAAGGAAGTGTGTAAATAGTACGCGGAAAAAACATAATAGCTCCCCTGTTTTCATCCATGGTTTTAAGTAAATGATCACTGGCTTCTTTAGTCAATGGATATTCGAACATTAACCGTTTCATCATCATTGGACAGAATCCATATTTCTTAATGTAAAGAGTGATCAAATCATTGTGTTTATTGTACCATCTTGTTGTAAATGTGTATTCGGTTATCGGATCTGCTCCTTTTTCTATTAAAGTATCGATATCGATCTCGTCTAAGAATTCAACTATATCGACATATTTCATTTGTATACACATAACCATTACTAACGTAGAATGTATAGGAAGAGATGTAACGGGAACAGGGTTTGTTGATTCGCAAACTATTCTAATACATAATTCTTCTGTTAATACGTCTTGCACGTAATCTATTATAGATGCCAAGATATCTATATAATTATTTTGTAAGATGATGTTAACAATGTGATCTATATAAGTAGTGTAATAATTCATGTATTTCGATATATGTTCCAACTCTGTCTTTGTGATGTCTAGTTTCGTAATATCTATAGCGTCCTCAAAAAATATATTCGCATATATTCCCAAGTCTTCAGTTCTATCTTCTAAAAAATCTTCAACGTATGGAATATAATAATCTATTTTACCTCTTCTGATGTCATTAATGATATAGTTTTTGACACTATTTTCCGTCAATTGATTCTTATTCACTATGTCTAAAAACCGGATAGCGTCCCTAGGACGAACTACTGCCATTAATATCTCTATTATAGCTTCTGGACATAAATCATCTATTATACCAGAATTAATGGGAACTATTCCGTATCTATCTAACATAGTTTTAAGAAAGTCAGAATCTAAGACCTGATGTTCATATATTGGTTCATACATGAAATGATCTCTATTGATGATAGTGACTATTTCATTCTCTGAAAATTGGTAACTCATTCTATACACGCTTTCCTTGTTGATAAAGGATAGTATATACTCAATGGAATTTGTACCAACAAACTGTTCTCTTATGAATCGTATATCATCATCTGAAATGATCATGTAAGGCATACATTTAACAATAAGAGACTTGTCTCCTGTTATCAATATACTATTCTTGTGATAATTTATGTGTGCGGCAAATTTGTCCACGTTCTTTAATTTTGTTATAGTAGATATCAAATCCAATGGAGATACAGTTCTTGGCTTAAACAGATATAGTTTTTCTGGAACGAATTCTACAACATTATTATAAAGGACTTTGGGTATATAAGTGGGATGAAATCCTATTTTAATTAATGCGATAGCCTTGTCCTCGTGCAGATATCCAAACGCTTTTGTGATAGTATGGCATTCATTGTCTAGAAACGCTCTACGAATATCTGTAACAGATATCATCTTTAGAGAATACTAGTCGCGTTAATAGTACTAAAATTTGTATTTTTTAATCTATCTCAATAAAAAATTAATATGTATGATTCAATGTATAACTAAACTACTAACTGTTATTGATAACTAGAATCAGAATCTAATGATGACATAACTAAGAAGTTTATCTACAGCCAATTTAGCTGCATTATTTTTAGCATCTCGTTTAGATTTTCCATCTGCCTTATCGAATACTCTTCCGTCAATGTCTACACAGGCATAAAATGTAGGAGAGTTACTAGGCCCCACTGATTCAATACGAAAAGACCAATCTCTCCTAGTTATTTGACAGTACTCATTAATAACGGTGACAGGGTTAACACCTTTCCAATAAATAATTTTTTTAACCGGAATAACATCATCAAAAGACTTATGATCCTCTCTCATTGATTTTTCGCGGGATACATCATCTATTATAGCATCAGCATCAGAATCTGTAGGCCGTGTATCAGCATCCATTGTCGTAGACCAACGAGGAGGAGTATCGTTGGAGCTGTAAACCATAGCACTACGTTGAAGATCATACAGAGCTTTATTAACTTCTCGCTTCTCCATATTAAGTTGTTTAGTTAGTTGTACAGCAGTAGCTCCTTAGTCCAATGTTTTTAATAACCGCACACAATCTCTGTGTCAGAACGCTCGTCAATATAGATCTTAGAAATTTTTTTAGAGAGAACTAACGCAACTAGCAATAAAACTGATCTTATTTTATCATTTTTTTTATTCATCATCCTCTGGTGGTTCGTCGTTCCTATCGAATGTAGCTCTGATTAACCCGTCATCTATAGGTGATGCTGGTTCTGGAGATTCTGGAGGAGATGGATTATTATCTGGAAGAATCTCTGTTATTTCCTTGTTTTCATGTATCGATTGCGTTGTAACATTAAGATTGCGAAATGCTCTAAATTTGGGAGGCTTAAAGTGTTGTTTACAATCTCTACACGCGTGTCTAACTAATGGAGGTTCATCAGCGGCTCTAGTTTGAATCATCATCGGTGTAGTATTCCTACTTTTACAGTTAGGACACGGTGTATTGTATTTCTCGTCGAGAACGTTAAAATAATCGTTGTAACTCACATCCTTTATTTTATCTATATTGTATTCTACTCCTTTCTTAATGCATTTTATACCGAACAAGAGATAGCGAAGGAATTCTTTTTCGGTACCGCTAGTACCCTTAATCATATCACATAGTGTTTTATATTCTAAATGTGTGGCAATGGACGGTTTATTTCTATACGATAGTTTGTTTTTGGAATCCTTTGAGTATTCTATACCAATATTATTCTTTGATTCGAATTTAGTTTCTTCGATATTAGATTTTGTATTACCTATATTCTTGATGTAGTACTTTGATGATTTTTCCATGGCCCATTCTATTAAGTTTTCCAAGTTGGCATCATCCACATATTGTGATAGTAATTCTCGGATATCAGTAGTGACTACCGCCATTGATATTTGTTCATTTGATGAGTAACTACTAATGTATACATTTTCCATTTATAACACTTATGTATTAACTTTGTTTATTTATATTTTTTCATTATTATGTTGATATTAATAATCGTATTGTGGTTATATGGCTACAATTTCATAATGAGTTGAAGTCAGTGTCCTATGATCAATGACGATAGCTTTACTCTGAAAAGAAAGTATCAAATCGATAGTGCAGAGTCAACAATGAAAATGGATAAGACGATGACAAAGTTTCAGAATAGAGTCAAAATGGTAAAAGAAATAAATCAGACGATAAGAGCAGCACAAACTCATTACGAGACATTGAAACTAGGATATATAAAATTTAAGGGAATGATTAGGACTACTACTCTAGAAGATATAGCACCATCTATTCCAAATAATCAGAAAACTTATAAACTATTCTCGGACATTTCAGTCATTGGCAAAGCATCACAGAATCCGAGTAAGATGATATATGCTCGCTGCTTTACATGTTTCCCAATTTGTTTGGAGATGACCATAGATTCATTTGTTATAGAATGCATCCAACATTGTTCATGATATAGTTGAATCATGTATGCCTGTTCGTATGCCTGTGGCTAAGATACTGTGTAAAGAAATGGTAAATAAATACTTTGAGAATCTTTAAGAGTGCATTGACTTTGTTAGTGAATAGGCATTCCATCTTTCTCCAATACTAATTCAAATTGTTAAATTAATAATGGAATAGTATAAATAGTTATTAGTGATAGGATAGTAAACATAATTATTAGAATAGTAGTGTAGTATCATAGATAACTCTCTTCTATAAAAAATGGATTTTATTCGTAGAAAGTATCTTATATACACAGTAGAAAATAATATAGATTTTTTAAAGGATGATACATTAAGTAAAGTAAACAATTTTACCCTCAATCATGTACTAGCTCTCAAGTATCTAGTTAGCAATTTTCCTCAACATGTTATTACTAAGGATGTATTAGCTAATACCAATTTTTTTGTTTTCATACATATGGTACGATGCTGTAAAGTATACGAAGCGGTTTTACGACACGCATTTGATGCACCCACGTTGTACGTTAAAGCATTGACTAAGAATTATTTATCGTTTAGTAACACAATACAGTCGTACAAGGAAACAGTGCATAAACTAACACAAGATGAAAAATTTTTAGAGGTTGCCGAATACATGGACGAATTAGGAGAACTTATAGGCGTAAATTATGACTTAGTTCTTAATCCATTATTTCACGGAGGGGAACCCATCAAAGATATGGAAATCATTTTTTTAAAACTGTTTAAGAAAACAGACTTCAAAGTTGTTAAAAAATTAAGTGTTATAAGATTACTTATTTGGGCATACCTAAGCAAGAAAGATACAGGCATAGAGTTTGCGGATAATGATAGACAAGATATATATACTCTATTTCAACAAACTGGTAGAATAGTCCATAGCAATCTAACAGAAACGTTTAGGGATTATATCTTTCCCGGAGATAAGACTAGCTATTGGGTGTGGTTAAACGAAAGTATAGCTAATGATGCGGATATCGTTATTAATAGACCCGCCATTACCATGTATGATAAAATTCTTAGTTATATATACTCTGAGATAAAACAGGGACGCGTTAATAAAAACATGCTTAAGTTAGTTTATATCTTTGAGCCTGAAAAAGATATCAGAGAACTTCTGCTAGAAATCATATATGATATTCCTGGAGATATCCTATCTATTATTGATGCAAAAAACGACGATTGGAAAAAATATTTTATTAGTTTTTACAAAGCTAATTTTATTAACGGTAATACATTTATTAGTGATAGAACGTTTAACGATGACTTATTCAGAGTTGTTGTTCAAATAGATCCCGAATATTTCGATAATGAACGAATTATGTCTTTATTCTATACGAGTGCTGCGGACATTAAACGATTTGATGAGTTAGATATTAATAACAGTTATATATCTAATATAATTTATGAGGTGAACGATATCACATTAGATACAATGGATGATATGAAGAAGTGTCAAATCTTTAACGAGGATACGTTGTATTATGTTAAGGAATACAATACATACCTGTTTTTGCACGAGTCGGATCCCATGGTCATAGAGAACGGAATACTAAAGAAACTGTCATCTATAAAATCCAAGAGTAGACGGCTGAACTTGTTTAGCAAAAACATTTTAAAATATTATTTAGACGGACAATTGGCTCGTCTAGGTCTTGTGTTAGATGATTATAAAGGAGACTTATTAGTTAAAATGATAAACCATCTCAAATCTGTGGAGGATGTATCCGCATTCGTTAGATTTTCTACAGATAAAAACCCTAGTATTCTTCCATCGCTAATCAAAACTATTTTAGCTAGTTATAATATTTCCATCATCGTCTTATTTCAAAGGTTTTTAAGAGATAATCTATATCATGTAGAAGAATTCTTGGATAAAAGCATCCATCTAACCAAGACGGATAAGAAATATATACTTCAATTGATAAGGCACGGTAGATCATAGAACAAACCAAATATATTATTAATAATTTGTATATACATAGATATAATTATCACATATTAAAAAATAACACATTTTTGATAAATGGAAACCGTTGCAACAATTCAGACTCCCACCAAATTAATGAATAAAGAAAATGCAGAAATGATTTTGGAAAAAATTGTTAATCATATAGCTATGTATATTAGTGACGAATCAATATATTCAGAAAATAATCCTGAATATATTGATTTTCGTAACAGATACGGAGACTATAGATCTCTCATTATAAAAAGTGATCACGAGTTTGTAAAGCTATGTAAAGATCATGCAGAGAAAAGTTCTCCAGAAACGCAACAAATGATTATCAAACACATATACGAACAATATCTTATTCCAGTATCTGAAGTACTATTAAAACCTATAATGTCCATGGGTGACATATTTACATATAACGGATGTAAAGACAATGAATGGATGCTAGAACAACTCTCTACCCTAAACTTTAACAATCTCTACACATGGAACTCATGTAGCATAGGCAATGTAACGCGTCTGTTTTATACATTTTTTAGTTATCTGATGAAAGATAAACTAAATATATAAGTATAATCCCATTCTAATACTTTAACCTGATGTATTATTACCTGCATCTTATTAGAATATTAACCTAACTAAAAGACATAAAAAGCGGTAGGATATAAATATTATGGCCGCAACCGTTCCGCGTTTTGACGATGTGTACAAAAATGCACAAAGAAGAATTCTAGATCAAGAAACATTTTTTAGTAGAGGTCTAAGTAGACCGTTAATGAAAAACACATATCTATTTGATAATTACGCGTATGGATGGATACCAGAAACTGCAATTTGGAGTAGTAGATACGCAAACCTAGATGCTAGTGACTATTATCCCATTTCGTTGGGATTACTTAAAAAGTTTGAATTTCTCATGTCTCTATATAAAGGTCCTATTCCCGTATATGAAGAAAAAGTAAATACTGAATTCATTGCTAATGGATCTTTCTCCGGTAGATACGTATCATATCTTAGAAAGTTTTCTGCCCTTCCAACAAACGAGTTTATTAGTTTTTTATTATTGACCTCCATCCCTATCTATAATATCTTATTCTGGTTTAAAAACACACAGTTTGATATTACTAAACACACATTATTCAGATACGTCTATACAGATAATACCAAACACCTTGCGTTGGCTAGGTATATACATCAAACAGGAGACTATAAGCCTTTGTTTAGTCGTCTCAAAGAGAATTATATATTTACTGGTCCCGTTCCAATAGGTATCAAAGATATAGATCACCCTAATCTTAGTAGAGCAAGAAGTCCATCCGATTATGAGACATTAGCTAATATTAGTACTATATTGTACTTTACCAAGTATGATCCAGTATTAATGTTTTTATTGTTTTACGTACCTGGGTATTCAATTACTACAAAAATTACTCCAGCCGTAGAATATCTAATGGATAAACTGAATCTAACAAAGAGCGACGTACAACTGTTGTAAATTATTTTATGCTTCGTAAAATGTAGGTCTTGAACCAAACATTCTTTGAAAAAATGAGATGCATAAAACTTTATTATCCAATAGATTAACTATTTCAGACGTCAATCGTTTAAAGTAAACTTCGTAAAATATTCTTTGATTGCTGCCGAGTTTAAAACTTCTATCGATAATTGTTTCATATGTTTTAATATTTACAAGTTTTTTGGTCCATGGTACATTAGCTGGACAGATATATGCAAAATAATATCGTTCTCCAAGTTCTATAGTCTCTGGATTGTTTTTATTATATTCAGTAACCAAATACATATTAGGGTTATCTGCGGATTTATAATTTGAGTGATGCATTCGACTCAACATAAATAATTCTAGAGGAGACGATCTACTATCAAATTCGGATCGTAAATCTGTTTCTAAAGAACGGAGAATATCTATACATACCTGATTAGAATTCATCCGTCCTTCAGACAACATCTCAGACAGTCTGGTCTTGTATGTCTTAATCATATTCTTATGAAACTTGGAAACATCTCTTCTAGTTTCACTAGTACCTTTATTAATTCTCTCAGGTACAGATTTTGAATTCGACGATGCCGAGTATTTCATCGTTGTATATTTCTTCTTCGATTGCATAATCAAATTCTTATATACCGCCTCAAACTCTATTTTAAAATTATTAAACAATACTCTACTATTAATCAGTCGTTCTAACTCCTTTGCTATTTCTATGGACTTATCTACATCTTGACTGTCTATCTCTGTAAACACGGAGTCGGTATCTCCATACACGCTACGAAAACGAAATCTATAATCTATAGGCAACGATGTTTTCACAATCGGATTAATATCTCTATCGTCCATATAAAATGGATTACTTAATGTATTGGCAAACCGTAACATACCGTTGGATAACTCTGCTCCATTTAGTACCGATTCTAGATACAATATCATTCTACGTCCTATGGATGTGCAACTCTTAGCCGAAGCGTATGAGTATAGAGCACTATTTCTAAATCCCATCAGACCATATACTGAGTTGGCTACTATCTTGTACGTATATTGCATGGAATCATAGATGGCCTTTTCAGTTGAACTGGTAGCCTGTTTTAACATCTTTTTATATCTGGCTCTCTCTGCCAAAAATGTTCTTAATAGTCTAGGAATGGTTCCTTCTATTGATCTATCGAAAATTGCTATTTCAGAGATGAGGTTCGGTAGTCTAGGTTCACAATGAACCGTAATATATCTAGGAGGTGGATATTTCTGAAGCAAGAGTTGATTATTTATTTCTTCTTCCAATCTATTGGTACTAACAACGACACCGACTAATGTTTCCGGAGATAGATTTCCAAAGATACACACATTAGGATACAGACTGTTATAATCAAAGATTAATACATTATTACTAAACATTTTTTGTTTTGGAGCAAATACCTTACCGCCTTCATAAGGAAACTTTTGTTTTGTTTCTGATCTGACTAAGATAGTTTTAGTTTCCAACAATAGCTTTAACAGTGGACCCTTGATGATTGTACTCGCTCTATATTCGAATACCATGGATTGAGGAAGCACATATGTTGCCGCACCAGCGTCTGTTTTTGTTTCTACTCCATAATACTCCCACAAATACTGACACAAACAAGCATCATGAATACAGTATCTAGCCATATCTAAAGCTATGTTTAGATTATAATCCTTATACATCTGAGCTAAATCAATGTCATCCTTTCCGAAAGATAATTTATATATATCATTAGGTAAAGTAGGACATGATAGTACGACTTTAAATCCATTTTCCAAAATATCTTTACGAATTACTTTACATATAATATCCTCATCAACAGTCACGTAATTACCTGTGGTTAAAACCTTTGCAAATGTATCGGCTTTGCCTTTCGCGTCCGTAGTATCGTCACCGATGAACGTCATTTCTCTAACTCCTCTATTTAATACTTTACCCATGCAACTGAACGCGTTCTTGGATATAGAATCCAATTTGTACGAATCCAATTTTTCAGATTTTTGAATGAATGAATATAGATCGAAAAATATAGTTCCATTATTGTTATTAACGTGAAACGTAGTATTGGCCATGCCGCATACTCCCTTATGACTAGACTGATTTCTCTCATAAATACAGAGATGTACAGCTTCCTTTTTGTCTGGAGATCTAAAGATAATCTTCTCTCCTGTTAATAACTCTAGACGATTAGTAATATATCTCAGATCAAAGTTATGTCCGTTAAAGGTAACGACGTAGTCGAACGTTAGTTCCAACAATTGTTTAGCTATTCGTAACAAAACTATTTCAGAACATAGAACTAGTTCTCGTTCGTAATCCATTTCCATTAGCGACTGTATCCTCAAACATCCTCTATCGACGGCTTCTTGTATTTCCTGTTCCGTTAACATCTCTTCATTAATGAGCGTAAACAGTAATCGTTTACCACTTAAATCGATATAACAGTAACTTGTATGCGAGATTGGGTTAATAAATACAGAAGGAAACTTCTTATCGAAGTGACACTCTATATCTAGAAATAAGTACGATCTTGGGATATCGAATCTAGGTATTTCTTTAGCGAAACAGTTACGTGGATCGTCACAATGATAACATCCATTGTTAATCTTTGTCAAATATTGCTCGTCCAACGAGTAACATCCGTCTGGAGATATCCCGTTAGAAATATAAAACCAACTAATATTGAGAAATTCATCCATGGTGGCATTTTGTATGCTGCGTTTCTTTGGCTCTTCTATCAACCACATATCTGCGACGGAGCATTTTCTATCTTTAATATCTAGATTATAACTTATTGTCTCGTCAATGTCTATAGTTCTCATCTTTCCCATCGGCCTCGCATTAAATGGAGGAGGAGATAATGACTGATATATTTCGTCCGTCACTACGTAATAAAAGTAATGAGGAAATCGTATAAATACGGTCTCGCCATTTCGACATCTGGATTTCAGATATAAAAATCTGTTTTCACCGTGACTTTCAAACCAATTAATACACCTAACATCCATTTCTAGAATTTAGAAATATATTTTCATTTAAATGAATCCCAAACATTGGGGAAGAGCCGTATGGACCATTATTTTTATAGTACTTTCGCAAGCGGGTTTAGACGGCAACATAGAAGCGTGTAAACGAAAACTATATACTATAGTCAGCACTCTTCCATGTCCTGCATGTAGACGACACGCGACTATCGCTATAGAGGACAATAATGTCATGTCTAGCGATGATCTGAATTATATTTATTATTTTTTCATCAGATTATTTAACAATTTGGCATTTGATCCCAAATACGCAATCGATGTGTCAAAGGTTAAACCTTTATAAACTTAACCCATTATAAAACTTATGATTAGTCACGACTGAAATAACCGCGTGATTATTTTTTGGTATAATTCTACACGGCATGGTTTCTGTGACTATGAATTCAACACCTGTTATCTTAGTGAAATCTTTAACAAACAGCAAGGGTTCGTCAAAGACATAAAACTCATTGTTTACGATCGAAATAGACCCCCTATCACACTTAAAATAAAAAATATCCTTATCCTTTACCACCAAATAAAATTCTGATTGGTCAATGTGAATGTATTCACTTAACAGTTCCACAAATTTATTTATTAACTCCGAGGCACATACATCGTCGGTATTTTTTATGACAAACTTTACTCTTCCAGCATCCGTTTCTAAAAAAATATTAACGAGTTCCATTTATATCATCCAATATTATTGAAATGACGTTGATGGACAGATGATATAAATAAGAAGGTACAGTACCTTTGTCCACCATCTCCTCCAATTCATACTCTATTTTGTCATTAACTTTAATGTGTGAAAACAGTACGCCACATGCTTCCATGACAGTGTGTAACACTTTGGATACAAAATGTTTGACATTAGTATAATTGTCCAAGACTGTCAATCTATAATAGATAGTAGCTATAATATATTCTATGATGGTATTGAAGAAGATGACAACCTTGGCATATTGATCATTTAACACAGACATGGTATCAACAAATAGCTTAAATGAAAGAGAATCAGTAATTGGAATAAGCGTCTTCTCGATGTAGTGTCCGTATACCAACATGTCTGATATTTTGATGTATTCCATTAAATTATTTAGTTTTTTCTTTTTATTCTCGTTAAACAGAATTTCTGTCAATGGACCCCAACATCGTTGACCTATTAAGTTTTGATTGATTTTTCCGTGTAAGGCGTATCTAGTCAGATCGTATAGCCTATCCAATAATCCATCGTCTGTGCGTAGATCACATCGTACACTTTTTAATTTTCTATAGAAGAGTGACAGACATCTGGAGCAATTACAGACAGCAATTTCTTTATTCTCTACAGATGTAAGATACTTGAAGATATTCCTATGATGATGCAGAATTTTGGATAACACGGTATTGATGGTATCTGTTACCATAATTCCTTTGACTGATAGTGTCAAAGTACAAGATTTCCAATCTTTTGCAATTTTCAGTACCATTATCTTTGTTTTGATATCTATATCAGACAGCATGGTACGTCTGACAACACAGGGATTAAGACGGAAAGATGAAATGATTCTCTCAACATCTTCAATAGATACCTTGCTATTTTTTTTGGCATTATCTATATGTGAGAGAATATCCTCTAGAGAATCAGTATCCTTTTTGATGATAGTGGATCTCAATGACATGGGACGTCTAAACCTTCTTATTCTATCACCAGATTGCATGGTGATTTGTCTTCTTTCTTTTATCATGATGTAATCTCTAAATTCATCGGCAAATTGTCTATATCTAAAATCATAATATGAGATGTTTACCTCTACAAATATCTGTTCGTCCAATGTTAGAGTATCTATATCAGTTTTGTATTCCAAATTAAACATGGCAACGGATTTAATTTTATATTCCTCTATTAAGTCCTCGTCGATAATAACAGAATGTAGATAATCATTTAATCCATCGTACATGGTTGGAAGATGCTCGTTGACAAAATCTTTAATTGTCTTGATGAAGGTGGGACTATATCTAACATCTTGATTAATAAAATTTATAACATTGTCCATAGGATACTTTGTAACTAGTTTTATACACATCTCTTCATTGGTAAGTTTAGACAGAATATCGTGAACAGGTGGTATATTATATTCATCAGATATACGAAGAATAATGTCCAAATCTATATTGTTTAATATATTATATAGATGTAGTGTAGCTCCTACAGGAATATCTTTAACTAAGTCAATGATTTCATCAACAGTTAGATCTATTTTAAAGTTAATCATATAGGCATTGATTTTTAAAAGGTATGTAGCCTTGACTACATTCTCATTAATTAACCATTCCAAGTCACTGTGTGTAAGAAGATTATATTCTATCATAAGCTTGACTACATTTGGTCCCGATACCATTAAAGAATTCTTATGATATAAGGAAACAGCTTTTAGGTACTCATCTACTCTACAAGAATTTTGGAGAGCCTTAACGATATCAGTGACGTTTATTATTTCAGGAGGAAAGAACCTAACATTGAGAATATCTGAATTAATAGCTTCCAGATACAGTGATTTTGGCAATAGTCCGTGTAATCCATAATCCAGTAACACGAGCTGGTGCTTGCTAGACACCTTTTCAATGTTTAATTTTTTTGAAATAAGCTTTGATAAAGCCTTCCTCGCAAATTCCGGATACATGAACATGTCGCCAACATGATTAAGTATTGTTTTTCATTATTTTTATATTTTCTCAACAAGTTCTCAATACCCCAATAGATAATAGAATATCACCCAATGCGTCCATGTTGTCTATTTCCAACAGGTCGCTATATCCACCAATAGAAGTTTTCCCAAAAAAGATTCTAGGAACAGTTCTACCACCAGTAATTTGTTCAAAATAGTCACGCAATTCATTTTCGGGTTTAAATTCTTTAATATCTACAATTTCATACGCTCCTCTTTTGAAACTAAACTTATTTAGAATATCCAGTGCGTTTCTACAAAAAGGACATGTAAACTTGACAAAAATTGTCACTTTGTTATTGGCCAACCTTTGTTGTACAAATTCCTCGGCCATTTTTAATATTTAAGTGATACAAAACTATCTCGACTTATTTAACTCTTTAGTCGAGATATATGGACACAGATAGCTATATGATAACCAACTACAGAAGACAAACGCTATAAAAAACATAATTACGACGAGCATATTTATAAATATTTTTATTCAGTATTACTTGATATAGTAATATTAGGCACAGTCAAACATTCAACCACTCTAGATACATTAACTCTCTCATTTTCTTTAACAAATTCTGCAATATCTTCGTAAAAAGATTCTTGAAACTTTTTAGAATATCTATCGACTCTAGATGAAATAGCGTTCGTCAACATACTATGTTTTGTATACATAAAGGCGCCCATTTTAACAGTTTCTAGTGACAAAATGCTAGCGATCCTAGGATCCTTTAGAATCACATAGATTGACGATTCGTCTCTCTTAGTAACTCTAGTAAAATAATCATACAATCTAGTACGCGAAATAATATTATCCTTGACTTGAGGAGATCTAAACAATCTAGTTTTGAGAACATCGATAAGTTCATCGGGAATTACATACATACTATCTTTAATAGAACTCTTTTCATCCAGTTGAATGGATTCGTCCTTAACCAACTGATTAATGAGATCTTCTATTTTATCATTTTCTAGATGATATGTATGTCCATTAAAGTTAAATTGTGTAGCGCTTCTTTTTAGCCTAGCAGCCAATACTTTAACATCACTAATATCGATATACAAAGGAGATGATTTATCGATGGTATTAAGAATTCGTTTTTCGACATCCGTCAAAACCAATTCCTTTTTGCCTGTATCATCCAGTTTGCCATTCTTTGTAAAGAAATTATTTTCTACTAGACTATTAATAAGACTGATAAGGATTCCTCCATAATTGCACAATCCAAACTTTTTCACAAAACTAGACTTTACGAGATCTACAGGAATGCGTACTTCAGGTTTCTTAGCTTGTGATTTTTTCTTTTGTGGACATTTTCTAGTGACCAACTCATCTACCATTTCATTGATTTTAGCAGTGAAATAAGCTTTCAATGCACGGGCACTGATACTATTGAAAACGAGTTGATCTTCAAATTCCGCCATTTAAGTTCACCAAACAACTTTTAAATACAAATATATCAATAGTAGTAGAATAAGAACTATAAAAAAAATAATAATTAACCAATACCAACCCCAACAACCTGTATTATTAGTTGATGTGACAGTTTTCTCATCACTTAGAACAGATTTAACAATTTCTATAAAGTCTGTCAAATCATCTTCCTGAGAACCCATAAATACACCAAATATAGCAGCGTACAACTTATCCATTTATACATTGAATATTGGCTTTTCTTTATCGCTATCTTCATCATATTCATCATCAATATCAACAAGTCCCAGATTACGAACCAGATCTTCTTCTACATTTTCAGTCATTGATACGCGTTCACTATCTCCAGAGAGTCCGATAACGTTAGCCACTACTTCTCTATCAATGATTAGTTTCTTGAGCGCGAATGTAATTTTTGTTTCCGTTCCGGATCTATAGAAAACTACAGGTGTAATAATTGCCTTGGCTAATTGTCTTTCTCTTTTACTGAGTGATTCTAGTTCACCTTCTATAGATCTGAGAATGGATGATTCTCCAGTCGAAACATATTCTACCATGGCTCCGTTTAATTTGTTGATGAAGATGGATTCATCCTTAAATGTTTTCTCTGTAATAGTTTCCGCCGAAAGACTATGCAAAGAATTTGGAATGCGTTCCTTGTGTGTAATGTTTCCATAGACAGCTTCTAGAAGTTGATACAACATAGGACTAGCCGCGGTAACTTTTATTTTTAGAAAGTATCCATCGCTTCTATCTTGTTTAGATTTATTTTTATAAAGTTTAGTCTCTCCTTCCAACATAATAAAAGTGGAAGTCATCTGACTAGATAAACTATCAGTAAGTTTTATAGAGATAGATGAACAATTAGCGTATTGAGAAGCATTTAGTGTAACGCATTCGATACATTTTGCATTAGATTTACTAATCGATTTTGCATACTCTATAACACCCGCACAAGTCTGTAGAGAATCGCTAGATGCTGTAGGTCTTGGTGAAGTTTCAACTCTCTTCTTGATTACCTTACTCATGATTAAACCTAAATAATTGTACTTTGTAATATAATGATATATATTTTCACTTTATCTCATTTGAGAATAAAAATGTTTTTGTTAACCACTGCATGATGTACAGATTTCGGAATCGCAAACCACTTGTGGTTTTATTTTATCCTTGTCCAATGTGAATTGAATGGGAGCGGATGCGGGTTTCGTACGTAGATAGTACATTCCCGTTTTTAGACCGAGACTCCATCCGTAAAAATGCATACTCGTTAGTTTGGAATAACTCGGATCTGCTATATGGATATTCATAGATTGACTTTGATCGATGAAGGCTCCCCTGTCTGCAGCCATTTTTATGATCGTCTTTTGTGGAATTTCCCAAATAGTTTTATAAACTCGCTTAATATCTTCTGGAAGGTTTGTATTCTGAATGGATCCACCATCTACCATAATCCTATTCTTGATCTCATCATTCCATAATTTTCTCTCGGTTAAAACTCTAAGGAGATGCGGGTTAACTACTTGGAATTCTCCAGACAATACTCTCCGAGTGTAAATATTACTGGTATACGGTTCCACCGACTCATTATTTCCCAAAATTTGAGCAGTTGATGCAGTCGGCATAGGTGCCACCAATAAACTATTTCTAAGACCGTATGTTCTGATTTTATCTTTTAGAGGTTCCCAATTCCAAAGATCCGACGGTACAACATTCCAAAGATCATATTGTAGAATACCGTTACTGGCGTACGATCCTACATATGTATCATATGGTCCTTCCTTCTCAGCTAGTTTACAACTCGCCTCTAATGCACCGTAATAAATGGTTTCAAAGATCTTCTTATTTAGATCTTGTGCTTCCAGGCTATCAAATGGATAATTTAAGAGAATAAACGCGTCCGCTAATCCTTGAACACCAATACCGATAGGTCTATGTCTCTTATTAGAGATTTCAGCTTCTGGAATAGGATAATAATTAATATCTATAATTTTATTGAGATTTCTGACAATTACTTTGACCACATCCTTCAGTTTGAGAAAATCAAATCGCCCATCTATTACAAACATGTTCAATGCAACAGATGCCAGATTACACACGGCTACCTCATTAGCATCCGCATATTGTATTATCTCAGTGCAAAGATTACTACACTTGATGGTTCCTAAATTTTGTTGATTACTCTTTTTGTTACACGCATCCTTATAAAGAATGAATGGAGTACCAGTTTCAATCTGAGATTCTATAATCGCTTTCCAGACGACTCGAGCCTTTATTATACATTTGTATCTCCTTTCTCTTTCGTATAGTGTATACAATCGTTCGAACTCGTCTCCCCAAACATTGTCCAATCCAGGACATTCATCCGGACACATCAACGACCACTCTCCGTCATCCTTCACTCGTTTCATAAAGAGATCAGGAATCCAAAGAGCTATAAATAGATCTCTTGTTCTATGTTCATCGTTTCCTGTATTCTTTTTAAGATCGAGGAACGCCATAATATCAGAATGCCACGGTTCCAAGTATATGGCCATAACTCCAGGCCGTTTGTTTCCTCCCTGATCTATGTATCTAGCGGTGTTATTATAAACTCTCAACATTGGAATAATACCGTTTGATATACCATTGGTACCGGAGATATAGCTTCCACTGGCACGAATATTACTAATTGATAGACCTATTCCCCCTGCCATTTTAGAGATTAATGCGCATCGTTTTAACGTGTCATAGATGCCTTCTATGCTATCATCGATCATGTTAAGTAGAAAACAGCTAGACATTTGGTGACGAGTAGTTCCCGCATTAAATAAGGTAGGAGAAGCGTGCGTAAACCATTTTTCAGAAAGTAGATTGTACGTCTCAATAGCTGAGTCTATATCCCATTGATGAATTCCTACTGCGACACGCATTAACATGTGCTGAGGTCTTTCAACAATTTTGTTGTTTATTTTCAACAAGTAGGATTTTTCCAAAGTTTTAAAACCAAAATAGTTGTATGAAAAGTCTCGTTCGTAAATAATAACCGAATTGAGCTTATCCTTATATTTGTTAACTATATCCATGGTAATACTTGAAATAATCGGAGAATGTTTCCCATTTTTAGGATTAACATAGTTGAATAAATCCTCCATCACTTCACTAAATAGTTTTTTTGTTTCCTTGTGTAGATTTGATATGGCTATTCTGGCGGCTAGAATGGCATAATCCGGATGTTGTGTAGTACAAGTGGCTGCTATTTCGGCTGCCAGAGTGTCCAATTCTACCGTTGTTACTCCATTATATATTCCTTGAATAACCTTCATAGCTATTTTAATAGGATCTATATGATCAGTGTTTAAGCCATAGCACAATTTTCTAATACGAGACGTGATTTTATCAAACATGACATTTTCCTTGTATCCATTTCGTTTAATGACAAACATTTTTGTTGGTGTAATAAAAAAAATTATTTAATTTTTCATTAATAGGGATTTGACGTATGTAGCGTACAAAATTATCGTTCCTGGTATATAGATAAAGAGTCCTATATATTTGAAAATCGTTACGGTTCGATTAAACTTTAATGATTGCATTGTGAATATATCATTAGGATTTAACTCCTTGACTATCATGGCGGTGCCAGAAATTACCATCAAAAGCATTAATACAGTTATGCAGATCGCAGTTAGAACGGTTATAGCATCCACCATTTATATCTAAAAATTAGATCAAAGAATATGTGACAACGTCCTAGTTGTATACTGAGAATTGACGAAACAATGTTTCTTACATATTTTTTTCTTATTAGTAACCGACTTAATAGTAGGAACTGGAAAACTAGACTTGATTATTCTATAAGTATAGATACCCTTCCAAATAATGTTCTCTTTGATAAAAGTTCCAGAAAATGTAGAATTTTTTAAAAAGTTATCTTTTGCTATTACTAATATCGTGGTTAGACGCTTATTATTAATATGAGTGATGAAATCCACACCGCTTCTAGATATCGCTTTTATTTCCACATTAGATGGTAAATCCAATAGTGAAACTATCTTTTTAGGAATGTATGGACTCGCGTTTAGAGGAGTGAACGTCTTCGGAGTAGTAAAGGATGATTCGTCAAATGAATAAACAATTTCACAAATGGATGTTAATGTATTAGTAGGAAATTTTTTGACGCTAGTGGAATTGAAGATTCTAATGGATGATGTTCTACCTATTTCATCCGATAACATGTTAATTTCCAATACCAACGGTTTTAATATTTCGATGATATACGGTAGTCTCTCTTTCGGACTTATATAGCTTATTCCACAATACGAGTCATTATATACTCCAAAAAACAAAATAACTAGTATAAAATCTGTATCGAATGGGAAAAACGAAATTATCGATATAGGTATAGAATCCGGAACATTGAACGTATTAATACTTAATTCTTTTTCAGTGGTAAGAACCGATAGGTTATTGACATTGTATGGTTTTAAATATTCTATAACTTGAGACTTGATAGATATTAATGACGAATTGAAAATTATTTTTATCACCACGTGTGTTTCAGGATCATCGTCGACGCCAGTTAACCAACCGAATGGAGTAAAATAAATATCATTAATATATGCTCTAGATATTAGTATTTTTATTAATCCTTTGATTATCATCTTCTCGTACGCGAATGATTCCATGATCAAGAGTGATTTGAGAACATCCTCCGGAGTATTAATGGGTTTAGTAAACAGTCCATCGTTGCAATAATAAAAGTTGTCCAAGTTAAAGGATATTATGCATTCGTTTAAAGATATCACCTCATCTAACGGAGACAATTTTTTGGTAGGTTTTAGAGACTTTGAAGCTACTTGTTTAACAAAGTTATTCATCGTCGTCTACTATTCTATTTAATTTTGTAGTTAATTTATCACATATCACATTAATTGACTTTTTGGTCCACTTTTCCATACGTTTATATTCTTTTAATCCTGCGTTATCCGTTTCCGTTATATACAGGGATAGATCTTGCAAGTTAAATAGAATGCTCTTAAATAATGTCATTTTTTTATCCGCTAAAAATTTAAAGAATGTATAAACTTTTTTCAAAGATTTAAAACTTTTAGGTGGAGTTCTGGTACACAATATCATAAACAAACTAATAAACATCCCACATTCAGATTCCAACAATTGATTAACTTCCACATTAATACAGCCTATTTTCGCTCCAAATGTACATTCGAAAAATCTGAATAAAACATCAATATCGCAATTTGTATTATCCAATACAGAATGTCTGTGATTCGTGTTAAAACCATCGGAAAAAGAATAGAAATAAAAATTATTATAATGGTGGAATTCAGTTGGAATATTGCCTCCGGAGTCATAAAAGGATACTAAACATTGTTTTTTATCGTAAATTACACATTTCCAATGAGACAAATAACAAAATCCAAACATTACAAATCTAGAGGTAGAACTTTTAATTTTGTCTTTAAGTATATACGATAAGATATGTTTATTCATAAACGCGTCAAATTTTTCATGAATAGCTAAGGAGTTTAAGAATCTCATGTCAAATTGTCCTATATAATCCACTTCGGATCCATAAGCAAACTGAGAGACTAAGTTCTTAATACTTCGATTGCTCATCCAGGCTCCTCTCTCAGGCTCTATTTTCATCTTGACGACCTTTGGATTTTCACCAGTATGTATTCCTTTACGTGATAAATCATCGATTTTCAAATCCATTTGTGAGAAGTCTATCGCCTTAGATACTTTTTCCCGTAGTTGAGGTTTAAAGAAATACGCTAACGGTATACTAGTAGGTAACTCAAAGACATCATATATAGAATGGTAACGCGTCGTTAACTCGTCGGTTAACTCTTTCTTTTGATCGAGTTCATCGCTACTATTGGGTCTGCTCAGGTGCCCCGACTCTACTAGTTCCAACATCATACCGATAGGAATACAAGACACTTTGCCAGCGGTTGTAGATTTATCATATTTCTCCACCACATATCCGTTACAATTTGTTAAGAATTTAGATACATCTATATTGCTACATAATCCAGCTAGTGAATATATATGACATAATAAATTGGTAAATCCTAGTTCTGGTATTTTACTAATTACTAAATCTGTATATCTTTCCATTTATCATGGAAAAGAATTTACCAGATATCTTCTTTTTTCCAAACTGCGTTAATGTATTCTCTTACAAATATTCACAAGATGAATTCAGTAATATGAGTAAAACGGAACGTGATAATTTCTCATTGGCTGTGTTTCCAGTGATAAAACATAGATGGCATAACGCACACGTTGTAAAACATAAAGGAATATACAAAGTTAGTACAGAAGCACGTGGAAAAAAAGTATCTCCTCCATCACTAGGAAAACCCGCACATATAAACCTAATGTCGAAGCAATATATATATAGTGAGTATGCAATAAGCTTTGAATGTTATAGTTTTCTAAAATGTATAACAAATACAGAAATCAATTCGTTCGATGAGTATATATTAAGAGGACTATTAGAAGCTGGTAATAGTTTACAGATATTTTCCAATTCCGTAGGTAAACGAATAGATACTATAGGTGTACTAGGGAATAAGTATCCATTTAGCAAAATTCCATTGGCCTCATTAACTCCTAAAGCACAACGAGAGATATTTTTAGCGTGGATTTCTCATAGACCTGTAGTTTTAACTGGAGGAACCGGAGTGGGTAAGACGTCACAGGTACCCAAGTTATTGCTTTGGTTTAATTATTTATTTGGTGGATTCTCTTCTCTAGATAAAATCACTGACTTTCACGAAAGACCAGTCATTCTATCTCTTCCTAGGATAGCTTTAGTTAGATTGCATAGCAATACCATTTTAAAATCATTGGGATTTAAGGTACTAGATGGATCTCCTATCTCTTTACGGTACGGATCTATACCGGAAGAATTAATAAACAAACAACCAAAAAAATATGGAATTGTATTTTCTACCCATAAGTTATCTCTAACAAAACTATTTAGTTATGGCACTATTATTATAGACGAAGTTCATGAGCATGATCAAATAGGAGATATTATTATAGCAGTAGCGAGAAAACATCATACGAAAATAGATTCTATGTTTTTAATGACTGCCACGTTAGAGGATGACAGGGAACGTCTAAAAATATTTTTACCTAATCCCGCATTTATACATATTCCTGGAGATACACTGTTTAAAATTAGCGAGGTATTTATTCATAATAAGATAAATCCATCTTCCAGAATGGCATATATAGAAGAAGAAAAGAGAAATTTAGTTACTGCTATACAGATGTATACTCCTCCTGATGGATCATCCGGTATAGTCTTTGTGGCATCCGTTGCACAGTGTCACGAATATAAATCATATTTAGAAAAAAGATTACCGTATGATATGTATATTATTCATGGTAAGGTCTTAGATATAGACGAAATATTAGAAAAAGTGTATTCATCACCTAATGTATCGATAATTATTTCTACTCCTTATTTGGAATCCAGCGTTACTATACGCAATGTTACACACATTTATGATATGGGTAGAGTTTTTGTCCCCGCTCCTTTTGGAGGATCACAACAATTTATTTCTAAATCTATGAGAGATCAACGAAAAGGAAGAGTAGGAAGAGTTAATCCTGGAACATACGTATATTTCTATGATCTGTCTTATATGAAATCTATACAGCGAATAGATTCAGAATTTCTACATAATTATATATTGTACGCTAATAAGTTTAATCTAACACTCCCCGAAGATTTGTTTATAATCCCTACAAATTTGGATATTCTATGGCGTACAAAGGAATATATAGACTCGTTCGATATTAGTACAGAAACATGGAATAAATTATTATCCAATTATTATATGAAGATGATAGAGTATGCTAAACTTTATGTACTAAGTCCTATTCTCGCTGAGGAGTTGGATAACTTTGAGAGGACGGGAGAATTAACTAGTATTGTACAAGAAGCCATTTTATCTCTAAATTTACAAATTAAGATTTTAAAATTTAAACATAAAGATGATGATACGTATATACACTTTTGTAGAATATTATTCGGTGTCTATAACGGAACAAACGCTACTATATATTATCATAGACCTCTAACGGGATATATGAATATGATTTCAGATACTATATTTGTTCCTGTAGATAATAACTAAAAATCAAAATCTAATGACCACATCTTTTTTTAGAGATGAAAAATTTTCCACATCTCCTTTTGTAGACACGACTAAACATTTTGCAGAAAAAAGTTTATTATTATTTAGATAATCGTATACTTCATCAGTGTAGATAGTAAATGTGAACAGATAAAAGGTATTCTTGCTCAATAGATTGGTAAATTCCATAGAATATATTAATCCTTTCTTCTTGAGATCCCACATCATTTCAACCAAAGACGTTTTATCCAATGATTTACCTCGTACTATACCACATACAAAACTAGATTTTGCAGTGATGTCGTACCTGGTATTCCTACCAAACAAAATTTTACTTTTAGTTCTTTTAGAAAATTCTAAGGTAGAATCTCTATTTGTCAATATGTCATCTATGGAATTACCACTAGCAAAAAATGATAGAAATATATATTGATACATCGCAGCTGGTTTTGATCTACTATACTTTAAAAACGAATCAGATTCCATAATTGCTTGTATATCATCAGCTGAAAAACTATGTTTTACACGTATTCCTTCGGCATTTCTTTTTAATGATATATCTTGTTTAGACAATGATAAAGTTATCATGTCCATGAGAGACGCGTCTCCGTATCGTATAAATATTTCATTAGATGTTAGACGCTTCATTAGGGGTATACTTCTATAAGGTTTCTTAATTAGTCCATCATTGGTTGCGTCAAGAACTACTATCTGATGTTGTTGGGTATCTCTAGTGTTACACATGGCCTTACTAAAGTTTGGGTAAATAACTATGATATCTCTATTAATTATAGATGTATATATTTCATTCGTCAAGGATATTAATATCGACTTACTATCGTCATTAATACGTGTAATGTAATCATATAAATCATGCGATAGCCAAGGAAAATTCAAATAGATGTTCATCATATAATCGTCGCTATAATTCATATTAATACTTTGACATTGACTAATTTGTAATATAGCCTCGCCACGAAGAAAGCTCTCGTATTCAGTTTCATCGATAAAGGATACCGTTAAATATAACTGGTTGCCGATAGTCTCATAGTCTATTAAGTGGTAAGTTTCGTATAAATACAGAATCCCTAAAATATTATCTAATGTGGGATTAATCCTTACCATAACTGTATAAAATGGAGCCGGAGTCATAACTATTTTACCGTTTGTACTTACTGGAATAGATGAAGGAATAATCTCCGGACATGATGGTAAAGACCCAAATGTCTGTTTGAAGAAATCCAATGTTCCAGGTCCTAATCTCTTGACAAAAATTACGATATTCGATCCCGATATCCTTTGCATTCTATTTACCAGCATATCACGAACTATATTAAGATTATCTATCATGTCTATTCTCCCACCGTTATATAAATCGCCTCCGCTAAGAAACGTTAGTATATCCATACAATGGAATACTTCATTTCTAAAATAGTATTCGTTTTCTAATTCTTTAATGTGAAATCGTATACTAGAAAGGGAAAAATTATCTTTGAGTTTTCCATTAGAAAAGAACCACGAAACTAATGTTCTGATTGCGTCTGACTCCGTCGCTGAATTAATAGATTTACACCAAAAACTCATATAACTTCTAGATGTAGAAGCATTCGCTAAAAAATTAGTAGAATCAAAGGATATAAGTAGATGTTCCAACAAGTGAGCAATTCCCAAGATTTCATCTATATCATTCTCGAATCCGAAATTAGAAATTCCCAAGTAGATATCCTTTTTCATCCGATCATTGATGAAAATACGAACTTTATTCGGTAAGACGATCATTTACTAAGGAGTAAAATAGGAAGTAACGTTCGTATATCGTTATCGTCGTATAAATTAAAGGTGTGTTTTTTGCCATTAAGAGACATTATAATTTTACCAATATTGGAATTATAATATAGGTGTATTTGAGCACTAGAAACGGTCGATGCATCGGTAAATATAGCTGTATCTAATGTTCTAGTCGGTATTTCTTCATTTCGCTGTCTAATGATAGCGTTTTCTCTATCTGTTTCCATTACAGCTGCCTGAAGTTTATTGGTCGGATAATATGTAAAATAATAAGAAATACATACGAATAACAAAAATAAAATAAGATATAATAAAGATGCCATTTAGAGATCTAATTTTGTTCAACTTGTCCAAATTCCTACTTACAGAAGATGAGGAATCGTTGGAGATAGTATCTTCCTTATGTAGAGGATTTGAAATATCTTACGATGACTTAATATCGTACTTTCCAGATAGGAAATACCATAAATATATTTCTAAGGTATTTGAACATGTAGATTTATCGGAGGAATTAAGTATGGAATTCCATGATACAACTCTGAGAGATTTAGTATATCTTAGATTGTACAAGTATTCCAAGTATATACGGCCGTGTTATAAATTAGGAGATAATCTAAAAGGTATAGTTGTTATAAAGGACAGAAATATATATATTAGAGAAGCAAATGATGACTTGATAGAATATCTCCTCAAGGAATACACTCCTCAGATTTATACATATTCTAATGAGCGAGTTCCCATAGCTGGTTCAAAATTAATTCTTTGTGGATTTTCTCAAGTTACATTTATGGCGTATACAACGTCGCATATAACAACAAATAAAAAGGTAGATGTTCTCGTTTCCAAAAAATGTATAGATGAACTAGTCGATCCAATAAATTATCAAATACTTCAAAATTTATTTGATAAAGGAAGCGGAACAATAAACAAAATACTCAGGAAGATATTTTATTCGGTAACAGGTGGCCAAACTCCATAGGTAGCTTTTTCTATTTCGGATTTTAGAATTTCCAAATTCACCAGCGATTTATCGGTTTTGGTGAAATCCAAGGATTTATTAATGTCCACAAATGCCATTTGTTTTGTCTGTGGATTGTATTTGAAAATGGAAACGATGTAGTTAGATAGATGCGCGGCGAAGTTTCCTATTAGGGTTCCGCGCTTCACGTCACCCAACATACTTGAATCACCATCCTTTAAAAAAAATGATAAGATATCAACATGGAGTATATCATACTCGGATTTTAATTCTTCTACTGCCTCACTGACATTTTCACAAATACTACAATACGGTTTACCGAAAATAATCAGTACGTTCTTCATTTATGGGTATCAAAAACTTAAAATCGTTACTGCTGGAAAATAAATCACTGACGATATTAGATGATAATTTATACAAAGTATACAATGGAATATTTGTGGATACAATGAGTATTTATATAGCCGTCGCCAATTGTGTCAGAAACTTAGAAGAGTTAACTACGGTATTCATAAAATACGTAAACGGATGGGTAAAAAAGGGAGGACATGTAACCCTTTTTATCGATAGAGGAAGTATAAAAATTAAACAAGACGTTAGAGACAAGAGACGTAAATATTCTAAATTAACCAAGGACAGAAAAATGTTAGAATTAGAAAAGTGTACATCCGAAATACAAAATGTTACCGGATTTATGGAAGAAGAAATAAAGGCAGAAATGCAATTAAAAATCGATAAACTCACATTTCAAATATATTTATCTGATTATGATAACATAAAAATATCATTGAATGAGATACTAACACATTTCAACAATAATGAGAATGTTACATTATTTTATTGTGATGAACGAGACGCAGAATTCGTTATGTGTCTAGAGGCTAAAACACAGTTCTCTACCACAGGAGAATGGCCGTTAATAATAAGTACCGATCAGGATACTATGCTATTCGCGTCTGCTGATAATCATCCTAAGATGATAAAAAACTTAACTCAACTGTTTAAATTTGTTCCCTCGGCAGAGGATAACTATTTAGCAAAATTAACTGCATTAGTGAATGGATGTGATTTCTTTCCTGGACTCTATGGGGCATCTATAACACCCAACAACTTAAACAAAATACAATTGTTTAGTGATTTTACAATCGATAATATAGTCACTAGTTTGGCAATTAAAAATTATTATAGAAAGACTAACTCTACCGTAGACGTGCGTAATATTGTTACGTTTATAAACGATTACGCTAATTTAGACGATGTCTACTCGTATATTCCTCCTTGTCAATGCACTGTTCAAGAATTTATATTCTCCGCATTAGATGAAAAATGGAATGAATTTAAATCATCTTATTTAGAGAGCGTGCCGTTACCCTGCCAATTAATGTACGCATTAGAACCACGTAAGGAGATTGATGTTTCAGAAGTTAAAACTTTATCATCTTATATAGATTTCGAAAATACTAAATCAGATATCGATGTTATAAAATCTATATCCTCGATTTTTGGATATTCTAACGAAAACTGTAACACCATAGTGTTCGGCATCTATAAGGATAATTTACTACTGAGTATAAATAATTCATTTTACTTTAACGATAGTCTGTTAATAACCAATACTAAAAGTGATAATATAATAAATATAGGTTACTAGATTAAAAAATGGTGTTCCAGCTCGTGTGTTCTACATGCGGCAAAGATATTTCTCACGAACGATATAAATTGATTATACGAAAAAAATCATTAAAGGATGTACTAGTCAGTGTAAAGAACGAATGTTGTAGGTTAAAATTATCTACACAAATAGAACCTCAACGTAACTTAACAGTGCAACCTCTATTGGATATAAACTAATGGATCCGGTTAATTTTATCAAGACATATGCGCCTAGAGGTTCTATTATTTTTATTAATTATGCCATGTCATTAACTAGTCATTTGAATCCATCGATAGAAAAACATGTGGGTATTTATTATGGTACGTTATTATCGGAACACTTGGTAGTTGAATCTACCTATAGAAAAGGAGTTAGAATAGTCCCATTGGATAGATTTTTTGAAGGATATCTTAGTGCAAAAGTATACATGTTAGAGAATATTCAAGTTATGAAAATAGCAGCTGATATGTCGTTAACTTTACTAGGTATTCCATATGGATTTGGTCATGATAGAATGTATTGTTTTAAATTGGTAGCTGAATGTTATAAAAATGCCGGTATTGATACATCGTCTAAACGAATATTAGGTAAAGATATTTTTCTGAGCCAAAACTTTACAGATGATAATAGATGGATAAAGATATATGATTCTAATAATTTAACATTTTGGCAAATTGATTACCTTAAAGGGTGAGTTAATATGCATAACTACTCCTCCGTTGTTTTTTCCCTCGTTCTTTTTCTTAACGTTGTTTGCCATCACTCTCATAATGTAAAGATATTCTAAAATGGTAAACTTTTGCATATCGGATGCAGAAATTGGTATAAATGTTGTAATTGTATTATTTCCCGTCAATGGACTAGTCACAGCTCCATCAGTTTTATATCCTTTAGAGTATTTCTCACTCGTGTCTAGCATTCTAGAGCATTCCATGATCTGTTTATCGTTGATATTGGCCGGAAAGATAGATTTTTTATTTTTTATTATATTACTATTGGCAATTGTAGATATAACTTCTGGTAAATATTTTTCTACCTTTTCAATCTCTTCTATTTTCAAGCCGGCTATATATTCTGCTATATTGTTACTAGTATCAATACCTTTTCTGGCTAAGAAGTCATATGTGGTATTCACTATATCAGTTTTAACTGGTAGTTCCATTAGCCTTTCCACTTCTGCAGAATAATTAGAAATTGGTTCTTTACCAGAAAATCCAGCTACTATAATAGGCTCACCGATGATCATTGGCAAAATCCTATATTGTACCAGATTAATGAGAGCATATTTCATTTCCAATAATTCTGCTAGTTCTTGAGACATTGATTTATTTGATGAATCTATTTGGTTCTCTAGATACTCTACCATTTCTGCCGCATACAATAACTTGTTAGATAAAATCAGGGTTATCAAAGTGTTTAGTGTGGCTAGAATAGTGGGCTTGCACGTATTAAAGAATGCTGTAGTATGAGTAAACCGTTTTAACGAATTATATAGTCTCCAGAAATCTGTGGCGTTGCATACATGAACTGAATGACATCGAAGATTGTCCAATATTTTTAATAGCTGCTCTTTGTCCATTATTTCTATATTTGACTCGCAACAATTGTAGATACCATTAATCACTGATTCCTTTTTCGATGCCGGACAATAGCACAATTGTTTAGCTTTGGACTCTATGTATTCAGAATTAATAGATATATCTCTCAATACAGATTGCACTATACATTTTGAAACTATGTCAAAAATTGTAGAACGACGCTGTTCTGTAGCCATTTAACTTTAAATAATTTACAAAAATTTAAAATGAGCATCCGTATAAAAATCGATAAATTGCGCCAAATTGTGGCATATTTTTCAGAGTTCAGCGAAGAAGTGTCTATAAATGTAGACTCGACGGATGAATTAATGTATATTTTTGCCGCCTTGGGCGGATCTGTAAACATTTGGGCCATTATACCTCTCAGTGCATCAGTGTTCTACCGCGGAGCCGAAAATATTGTGTTTAACCTTCCAGTGTCCAAGGTAAAATCGTGTTTGTGTAGTTTTCACAATGATGCTATCATAGATATAGAACCTGATCTGGAAAATAATCTAGTAAAACTTTCTAGTTATCATGTAGTAAGTGTCGATTGTAACAAGGAACTGATGCCTATTAGGACAGATACTACTATTTGTCTAAGTATAGATCAAAAGAAATCTTACGTATTTAATTTTCACAAGTATGAAGAAAAATGTTGTGGTAGAACCGTCATTCATCTAGAATGGTTGTTGGGCTTTATCAAGTGTATTAGTCAGCATCAGCATTTGGCTATTATGTTTAAAGATGACAATATTATTATGAAGACTCCTGGTAATACTGATGCGTTTTCCAGGGAATATTCTATGACTGAATGTTCTCAAGAACTACAAAAGTTTTCTTTCAAAATAGCTATCTCGTCTCTCAACAAACTACGAGGATTCAAAAAGAGAGTCAATGTTTTTGAAACTAGAATCGTAATGGATAATGACGATAACATTCTAGGAATGTTGTTTTCGGATAGAGTTCAATCCTTTAAGATTAACATCTTTATGGCGTTTTTAGACTAATACTTTCAATGAGATAAATATGGGTGGCGGAGTAAGTGTTGAGCTCCCTAAACGGGATCCACCTCCGGGAGTACCCACTGATGAGATGTTATTAAACGTGGATAAAATGCATGACGTGATAGCTCCCGCTAAGCTTTTAGAATATGTGCATATAGGACCACTAACAAAAGATAAAGAGGATAAAGTAAAGAAAAGATATCCAGAGTTTAGATTAGTCAACACAGGACCCGGTGGTCTTTCGGCATTATTAAGACAATCATATAATGGAACCGCACCCAATTGCTGTCGCACTTTTAATCGTACTCATTATTGGAAGAAGGATGGAAAGATATCAGATAAGTATGAAGAGGGTGCAGTATTAGAATCGTGTTGGCCCGACGTCCACGACACTGGAAAATGCGATGTTGATTTATTCGACTGGTGTCAGGGGGATACGTTCGATATGAACATATGCCATCAGTGGATCGGTTCAGCCTTTAATAGGAGTGATAGAACTGTAGAGGGTCGACAATCGTTAATAAATCTGTATAATAAGATGCAAAGATTATGTAGTAAAGATGCTAGTGTACCAATATGTGAATTATTTTTGCATCATTTACGCGCACACAATACAGAAGATAGTAAAGAGATGATCGATTATATTCTAAGACAACAGTCGGCGGACTTTAAACAGAAATATATGAGATGTAGTTATCCCACTAGAGATAAGTTAGAAGAGTCATTAAAATATGCGGAACCTCGAGAATGTTGGGATCCAGAGTGTTCGAATGCCAATGTTAATTTCTTACTAACACGTAATTATAATAATTTAGGACTTTGCAATATTGTACGATGTAATACGAGCGTGAATAACTTACAGATGGATAAAACTTCCTCATTAAGATTATCATGTGGATTAAGCAATAGTGATAGATTTTCTACTGTTCCCGTCAATAGAGCAAAAGTAGTTCAACATAATATTAAACATTCGTTCGACCTAAAATTGCATTTGATCAGTTTATTATCTCTCTTGGTAATATGGATACTAATTGTAGCTATTTAAATGGGTGCCGCAGCAAGCATACAGACGACTGTGAATACACTCAGTGAACGTATCTCGTCTAAATTAGAACAAGAAGCGAACGCTAGTGCTCAAACAAAATGTGATATAGAAATCGGAAATTTTTATATCCGACAAAACCATGGATGTAACATCACTGTTAAAAATATGTGCTCTGCGGACGCGGATGCTCAGTTGGATGCTGTGTTATCAGCCGCTACAGAAACATATAGTGGATTAACACCGGAACAAAAAGCATACGTACCAGCTATGTTTACTGCTGCGTTAAACATTCAGACGAGTGTAAACACTGTTGTTAGAGATTTTGAAAATTATGTGAAACAGACTTGTAATTCTAGCGCTGTTGTCGATAACAAATTAAAGATACAAAACGTAATTATAGATGAATGTTACGGAGCCCCAGGATCTCCAACAAATTTGGAATTTATTAATACAGGATCTAGCAAAGGAAATTGTGCCATTAAGGCGTTGATGCAATTGACTACTAAGGCCACTACTCAAATAGCACCTAGACAAGTTGCTGGTACAGGAGTTCAGTTTTATATGATTGTTATCGGTGTTATAATATTGGCAGCGTTGTTTATGTACTATGCCAAGCGTATGCTGTTCACATCCACCAATGATAAAATCAAACTTATTTTAGCCAATAAGGAAAACGTCCATTGGACTACTTACATGGACACATTCTTTAGAACTTCTCCGATGATTATTGCTACCACGGATATACAAAACTGAAAATATATTGATAATATTTTAATAGATTAACATGGAAGTTATCGCTGATCGTCTAGACGATATAGTGAAACAAAATATAGCGGATGAAAAATTTGTAGATTTTGTTATACACGGTCTAGAGCATCAATGTCCTGCTATACTTCGACCATTAATTAGGTTGTTTATTGATATACTATTATTTGTTATAGTAATTTATATTTTTACGGTACGTCTAGTAAGTAGAAATTATCAAATATTGTTGGTGTTGGTGGCGCTAGTCATCACATTAACTATTTTTTTATTACTTTATACTATAATAGTACTAGACTGACTTCTAACAAACATCTCACCTGCCATAAATAAATGCTTGATATTAAAGTCTTCTATTTCTAACACTATTCCATCTGTGGAAAATAATACTCTGACATTATCGCTAATTGATACATCGGTAAGTGATATGCCTATAAAGTAATAATCTTCTTTGGGCACATATACCAGTGTACCAGGTTCTAACAACCTATTTACTGGTGCTCCTGTAGCATACTTTTTTTTTACCTTGAGAATATCCATTGTTTGCTTGGTCAATAGTGATATGTGATTTTTTATCAACCACTCAAAAAAGTAATTGGAGTGTTCATATCCTCTACGGGCTATTGTCTCATGACCGTGTATGAAATTTAAGTAACACGACTGTGGTAGATTTGTTCTATAGAGCCGGTTGCCGCAAATAGATAGAACTACCAATATGTCTGTACAAATGTTAAACATTAATTGATTAACAGAAAAAACAATGTTCGTTCTGGGAATAGAAACCAGATTAAAACAAAATTCATTAGAATATATGCCACGTTTATACATGGAATATAAAATAACTACAGTTTGAAAAATAACAGTATCATTTAAACATTTAACTTGCGGGGTTAATCTCACAACTTTACTGTTTTTGAACTGTTCAAAATATAGCATAGATCCATGAGAAATACGTTTAGCCGCCTTTAATAGAGGAAATCCAACCGCCTTTCTGGATCTCACCAACGACGATAGTTCTGACCAGCAACTCATTTCTTCATCATCCACCTGTTTTAACATATAATAGGCAGGAGATAGATATCCATCATTGCAATATTCCTTCTCGTAGGCACACAATCTAATATTGATAAAATCTCCATTCTCTTCTCTGTATTTATTATCTTGTCTCGGTGGCTGATTAGGCTGTGGTCTATCGTTGTTGAATCTATTTTGGTCATTAAATCTTTCATTTCTTCCTGGTATATTTCTATCACCTCGTTTGGTTGGATTTTTGTCTATATTATCGTTTGTAACATCGGTACGGGTATTCATTTATCACAAAAAAAACTTCTCTAAATGAGTCTACTACTAGAAAACCTCATCGAAGAAGATACCATATTTTTTGCAGGAAGTATATCTGAGTATGATGATTTACAAATGGTTATTGCTGGTGCAAAATCCAAATTTCCAAGATCTATGCTTTCTATTTTTAATATAGTACCTAGAACGATGTCAAAATATGAGTTGGAGTTGATTCATAACGAGAATATCACAGGGGCAATGTTTACCACAATGTATAATATAAGAAACAATTTGGGTCTAGGCGATGATAAACTAACTATTGAAGCCATTGAAAACTATTTCTTGGATCCTAACAATGAGGTTATGCCTCTTATCATTAATAATACGGATATGACTACCGTCATTCCTAAAAAAAGTGGTAGGAGAAAGAATAAGAACATGGTTATCTTCCGTCAAGGATCATCACCTATCTTGTGTATTTTCGAAACTCGTAAAAAGATTAATATTTATAAAGAAAATATGGAATCCGTATCGACTGAGTATACACCTATCGGAGACAACAAGGCTTTGATATCTAAATATGCGGGAATTAATATCCTGAATGTGTATTCTCCTTCCACGTCCATGAGATTGAATGCCATTTACGGATTCACCAATAAAAATAAACTAGAGAAACTTAGTACTAATAAGGAACTAGAATCGTATAGTTCTAGCCCTCTTCAAGAACCCATTAGGTTAAATGATTTTCTGGGACTATTGGAATGTGTTAAAAAGAATATTCCTCTAACAGATATTCCGACAAAGGATTGATTACTATAAATGGAGAATGTTCCTAATGTATACTTTAATCCTGTGTTTATAGAGCCCACGTTTAAACATTCTTTATTAAGTGTTTATAAACACAGATTAATAGTTTTATTTGAAGTATTCGTTGTATTCATTCTAATATATGTATTTTTTAGATCTGAATTAAATATGTTCTTCATGCCTAAACGAAAAATACCCGATCCTATTGATAGATTACGACGTGCTAATCTAGCGTGTGAAGACGATAAATTAATGATCTATGGATTACCATGGATAACAACTCAAACATCTGCGTTATCAATAAATAGTAAACCGATAGTGTATAAAGATTGTGCAAAGCTTTTGCGATCAATAAATGGATCACAACCAGTATCTCTTAACGATGTTCTTCGCAGATGATGATTCATTTTTTAAGTATTTTGCTAGTCAAGATGATGAATCTTCATTATCTGATATATTGCAAATCACTCAATATCTAGACTTTCTGTTATTATTATTGATCCAATCAAAAAATAAATTAGAAGCTGTGGGTCATTGTTATGAATCTCTTTCAGAGGAATACAGACAATTGACAAAATTCACAGACTCTCAAGATTTTAAAAAACTGTTTAACAAGGTCCCTATTGTTACAGATGGAAGGGTCAAACTTAATAAAGGATATTTGTTCGACTTTGTGATTAGTTTGATGCGATTCAAAAAAGAATCAGCTCTAGCTACCACCGCAATAGATCCTGTTAGATACATAGATCCTCGTCGTGATATCGCATTTTCTAACGTGATGGATATATTAAAGTCGAATAAAGTTGAAAAATAATTAATTCTTTATTGTTATCATGAACGGCGGACATATTCAGTTGATAATCGGCCCCATGTTTTCAGGTAAAAGTACAGAATTAATTAGACGAGTTAGACGTTATCAAATAGCTCAATATAAATGTGTGACTATAAAATATTCTAACGATAATAGATACGGAACGGGACTATGGACACATGATAAGAATAATTTTGCAGCATTGGAAGTAACTAAACTATGTGATGTCTTGGAAGCAATTACAGATTTCTCCGTGATAGGTATAGATGAAGGACAGTTCTTTCCAGACATTGTTGAATTCTGTGAGCGTATGGCAAACGAAGGAAAAATAGTTATAGTAGCCGCGCTCGATGGGACATTTCAACGTAGACCGTTTAATAATATTTTGAATCTTATTCCATTATCTGAAATGGTGGTAAAACTAACTGCAGTGTGTATGAAATGCTTTAAGGAGGCTTCCTTTTCTAAACGATTAGGTACAGAAACCGAGATAGAAATAATAGGAGGTAATGATATGTATCAATCTGTGTGTAGAAAGTGTTACATCGACTCATAATATTATATTTTTTATCTAAAAAACTAAAAATAAACATTGATTAAATTTTAATATAATACTTAAAAATGGATGTTGTGTCGTTAGATAAACCGTTTATGTATTTTGAGGAAATTGATAATGAGTTAGATTACGAACCAGAAAGTGCAAATGAGGTCGCAAAAAAACTGCCGTATCAAGGACAGTTAAAACTATTACTAGGAGAATTATTTTTTCTTAGTAAGTTACAGCGACACGGTATATTAGATGGCGCCACCGTAGTGTATATAGGATCTGCTCCAGGTACACATATACGTTATTTGAGAGATCATTTCTATAATTTAGGAGTGATCATCAAATGGATGCTAATTGACGGCCGCCATCATGATCCTATTCTAAATGGATTGCGTGATGTGACTCTAGTGACTCGGTTTGTTGATGAGGAATATCTACGATCCATCAAAAAACAACTACATCCTTCTAAGATTATTTTAATTTCTGATGTGCGATCCAAACGAGGAGGAAATGAACCTAGTACTGCGGATTTACTAAGTAATTATGCTCTACAAAATGTCATGATTAGTATTTTAAACCCCGTGGCGTCTAGTCTTAAATGGAGATGCCCGTTTCCAGATCAATGGATCAAGGACTTTTATATCCCACACGGTAATAAAATGTTACAACCTTTTGCTCCTTCATATTCAGCTGAAATGAGATTATTAAGTATTTATACCGGTGAGAATATGAGACTGACTCGAGTTACCAAATCAGACGCTGTAAATTATGAAAAAAAGATGTATTACCTTAATAAGATAGTCCGCAACAAAGTAGTTATTAACTTTGATTATCCTAATCAGGAATATGACTATTTTCACATGTACTTTATGTTGAGGACCGTATACTGCAATAAAACATTTCCTACTACTAAAGCAAAGATACTATTTCTACAACAATCTATATTTCGTTTCTTAAATATTCCAACGACATCAACTGAAAAAGTTAGTCATGAACCAATACAACGTAAAATATCTAGCAAAGATTCTATGTCTAAAAACAGAAATAGCAAGAGATCCGTACGCGGTAATAAATAGAAACGTACTACTGAGATATACTACCGATATAGAGTATAATGATTTAGTTACTTTAATAACCGTTAGACATAAAATTGATTCTATGAAAACTGTGTTTCAGGTATTTAACGAATCATCCATAAATTATACTCCGGTTGATGATGATTATGGAGAACCAATCATTATAACATCGTATCTTCAAAAAGGTCATAACAAGTTTCCTGTAAATTTTCTATACATAGATGTGGTAATATCTGACTTATTTCCTAGCTTTGTTAGACTAGATACTACAGAAACTAATATAGTTAATAGTGTACTACAAACAGGCGATGGTAAAAAGACTCTTCGTCTTCCTAAAATGTTAGAGACGGAAATAGTTGTCAAGATTCTCTATCGTCCTAATATACCATTAAAAATTGTTAGATTTTTCCGCAATAACATGGTAACTGGAGTAGAGATAGCCGATAGATCTGTTATTTCAGTCGCTGATTAATCAATTAGTAGAGATGAGATAAGAACATTATAATAATCAATAATATATCTTATATCTGTTTAGAAAAATGCTAATATTAAAATAGCTAACGCTAGTAATCCAATCGGAAGCCATTTGATATCTATAATAGGGTATCTAATTTCCTGATTCAGATAGCGTACGGCTATATTCTCGGTAGCTACTCGTTTGGAATCACAGACATTATTTACATCTAATTTACTATCTGTAATGGAAACGTTTCCCAATGAAATGGTACAATCAGATACATTACATCTTGATATATTTTTTTTTAAAGAGGCTGGTAACAACGCATCGCTTCGTTTACATGGCTCGTACCAACAATAATAGGGTAATCTTGTATCTATTCCTATCCGTACTATACTTTTATCAGGATAAATACATTTACATCGTATATCGTCTTTGTTAGTATCACAGAATGCATAAATTTGTTCGTCCGTCATGATAAAAATTTAAAGTGTAAATATAACTATTATTTTTATAGTTATAATAAAAAGGGAAATTTGATTGTATACCTTCGGTTCTTTAAAAGAAACTGACTTGATAAAAATGGCTGTAATCTCTAAGGTTACGTATAGTCTATACGATCAAAAAGAGATTAATGCCACAGATATTATCATTAGTCATATTAAAAATGACGACGATATCGGTACCGTTAAAGATGGTAGACTAGGTGCTATGGATGGGGCATTATGTAAGACTTGTGGGAAAACGGAATTGGAATGTTTCGGTCACTGGGGTAAAGTAAGTATTTATAAAACTCATATAGTTAAGCCTGAATTTATTTCAGAAATTATTCGTTTACTGAATCATATATGTATTCATTGCGGATTATTGCGTTCACGAGAACCGTATTCCGACGATATTAACCTAAAAGAGTTATCGGTACACGCTCTTAGGAGATTAAAGGATAAAATATTATCCAAGAAAAAGTCATGTTGGAACAGCGAATGTATGCAACCGTATCAAAAAATTACTTTTTCAAAGAAAAAGGTTTGTTTCGTCAACAAGTTGGATGATATTAACGTTCCTAATTCTCTCATCTATCAAAAGTTAATTTCTATTCATGAAAAGTTTTGGCCATTATTAGAAATTCATCAATATCCAGCTAACTTATTTTATACAGACTACTTTCCCATCCCTCCGTTGATTATTAGACCGGCTATTAGTTTTTGGATAGATAGTATACCCAAAGAGACAAATGAATTAACTTACTTATTAGGTATGATCGTTAAGAATTGTAACTTGAATGCTGATGAACAGGTTATCCAGAAGGCGGTAATAGAATACGATGATATTAAAATTATTTCTAATAACACTACCAGTATCAATTTATCATATATCACATCCGGCAAAAATAATATGATTAGAAGTTATATCGTCGCTCGGCGAAAAGATCAGACCGCTAGATCCGTAATTGGTCCCAGTACATCTATCACCGTTAATGAGGTAGGAATGCCCACATATATTAGAAATACACTTACAGAAAAGATATTTGTTAATGCCTTTACAGTGGATAAAGTTAAACAACTATTAGCATCAAACCAAGTTAAATTTTACTTTAATAAACGATTAAACCAATTAACAAGAATACGTCAAGGAAAGTTTATCAAAAATAAAATACATTTATTGCCTGGTGATTGGGTAGAAGTAGCTGTTCAAGAATATACAAGTATTATTTTTGGAAGACAACCGTCTCTACATAGATACAACGTCATCGCTTCATCTATCAGAGCTACCGAAGGAGATACTATCAAAATATCTCCCGGAATTGCCAACTCTCAAAATGCTGATTTCGACGGAGATGAAGAATGGATGATATTGGAGCAAAATCCTAAAGCCGTAGTTGAACAAAGTATTCTTATGTATCCGACGACGTTACTCAAACACGATATTCATGGAGCCCCCGTTTATGGATCTATTCAAGATGAAATCGTAGCAGCGTATTCATTGTTTAGGATACAAGATCTTTGTTTAGATGAAGTATTGAACATCTTGGGGAAATATGGAAGAGAGTTCGATCCTAAAGGTAAATGTAAATTCAGCGGTAAAGATATCTATACTTACTTGATAGGTGAAAAGATTAATTATCCGGGTCTCTTAAAGGATGGTGAAATTATTGCAAACGACGTAGATAGTAATTTTGTTGTAGCTATGAGGCATCTGTCATTGGCTGGACTCTTATCCGATCATAAATCGAACGTGGAAGGTATCAACTTTATTATCAAGTCATCTTATGTTTTTAAGAGATATCTATCTATATACGGTTTTGGGGTGACATTCAAAGATCTGAGACCAAATTCGACGTTCACTAATAAATTGGAGGCTATCAACGTAGAAAAAATAGAACTTATCAAAGAAGCATACGCCAAATATCTCAAAGATGTAAGAGACGGGAAAATAGTTCCATTATCTAAAGCTTTAGAGGCGGACTACTTGGAATCCATGTTATCCAACTTGACAAATCTTAATATCAGAGAGATAGAAGAACATATGAGACAAACGCTGATAGATGATCCAGATAATAACCTCCTGAAAATGGCCAAAGCGGGTTATAAAGTAAATCCCACAGAACTAATGTATATTCTAGGTACTTATGGACAACAGAGGATAGATGGCGAACCAGCAGAGACTCGAGTATTGGGTAGAGTCTTACCTTACTATCTTCCAGACTCTAAGGATCCAGAAGGAAGAGGTTATATTCTTAATTCTTTAACAAAAGGATTAACGGGTTCTCAATATTACTTTTTGATGCTGGTTGCAAGATCTCAATCTACTGATATTGTCTGTGAAACATCACGTACCGGAACACTGGCTAGAAAAATCATTAAAAAGATGGAGGATATGGTGGTCGACGGATACGGACAAGTAGTTATAGGTAATACGCTCATCAAGTACGCAGCCAATTATACCAAAATTCTAGGCTCAGTATGTAAACCTGTAGATCTTATCTATCCAGATGAGTCCATGACTTGGTATTTGGAAATTAGTGCTTTGTGGAATAAAATAAAACAGGGATTCGTTTACTCTCAGAAACAGAAACTTGCAAAGAAGACATTGGCGCCGTTTAATTTCCTAGTATTCGTCAAACCCACCACTGAGGATAATGCTATTAAGGTTAAGGATCTGTACGATATGATTCATAACGTCATTGATGATGTGAGAGAGAAATACTTCTTTACGGTATCTAATATAGATTTTATGGAGTATATATTCTTGACGCATCTTAATCCTTCTAGAATTAGAATTACAAAAGAAACGGCTATTACTATCTTTGAAAAGTTCTATGAAAAACTCAATTATACTCTAGGTGGTGGAACTCCTATTGGAATTATTTCTGCACAGGTATTGTCTGAGAAGTTTACACAACAAGCCCTGTCCAGTTTTCACACTACTGAAAAGAGTGGTGCTGTAAAACAAAAACTTGGTTTCAACGAGTTTAATAACTTGACTAATTTGAGTAAGAATAAGACCGAAATTATCACTCTGGTATCCGATGATATCTCTAAACTTCAATCTGTTAAGATTAATTTCGAATTTGTATGTTTGGGAGAATTAAATCCAGACATCACTCTTCGAAAAGAAACAGATAGATATGTAGTAGACATAATAGTCAATAGATTATACATCAAGAGAGCAGAAATAACCGAATTAGTCGTCGAATATATGATTGAACGATTTATCTCCTTTAGCGTCATTGTAAAGGAATGGGGTATGGAGACATTCATTGAGGACGAGGATAATATTAGATTTACTATCTACCTAAATTTCGTTGAACCGGAGGAATTGAATCTTAGTAAGTTTATGATGGTTCTTCCAGGTGCCGCCAACAAGGGCAAGATTAGTAAATTCAAGATTCCTATCTCTGACTATACGGGATATAACGACTTCAATCAAACAAAAAAGCTCAATAAGATGACTGTAGAACTCATGAATCTAAAAGAATTGGGTTCTTTCGATTTGGAGAACGTCAACGTGTATCCTGGAGTATGGAATACATACGATATCTTCGGTATTGAGGCCGCTCGTGGATACTTGTGCGAAGCCATGTTAAACACCTATGGAGAAGGTTTCGATTATCTGTACCAGCCTTGTGATCTTCTCGCTAGTTTACTATGTGCTAGTTACGAACCAGAATCAGTTAATAAATTCAAGTTCGGTGCAGCTAGTACTCTTAAGAGAGCTACGTTCGGAGATAATAAAGCATTGTTAAACGCGGCTCTTCATAAAAAGTCAGAACCTATTAACGATAATAGTAGCTGCCACTTTTTTAGCAAGGTCCCTAATATAGGAACTGGATATTACAAATACTTTATCGACTTGGGTCTTCTCATGAGAATGGAAAGGAAACTATCTGATAAGATATCTTCTCAAAAGATCAAGGAGATAGAAGAAACAGAAGACTTTTAATTCTTATCAATAACATATTTTTCTATGATCTGTCTTTTAAACGATGGATTTTCCACAAATGCGCCTCTCAAGTCCCTCATAGAATGATACACGTATAAAAAATATAGCATAGGTGATGACTCCTTATTTTTAGACATTAGATATGCCAAAATCATAGCCCCGCTTCTATTTACTCCTGCAACACAATGAACCAACACGGGCTCGTTTCGTTGATCACATTTAGATAAGAAGGCGGTCACGTCGTCAAAATATTTACTAATATCAGTAGTTGTATCATCTACCAACGGTATATGAATAATATTAATATTAGAGTTAGGTAATGTATATTTATCCATCGTCAAATTTAAAACATATTTGAACTTAACTTCAGATGATGGTGCATCCATAGCATTTTTATAATTTCCCAAATACACATTATTTGTTACTCTTGTCATTATAGTGGGAGATTTGGCTCTGTGCATATCTCCAGTTGAACGTAGTAGTAAGTATTTATACAAACTTTTCTTATCCATTTATAACGTACAAATGGATAAAACTACTTTATCAGTAAACGCATGCAATTTAGAATACGTTAGAGAAAAGGCTATAGTAGGCGTACAAGCAGCCAAGACATCAACACTTATATTTTTTGTTATTATATTGGCAATTAGTGCGCTATTACTCTGGTTTCAGACGTCTGATAATCCAGTCTTTAATGAATTAACGAGATATATGCGAATTAAAAATACGGTTAACGATTGGAAATCATTAACGGATAGCAAAACAAAATTAGAAAGCGATAGAGGTAGACTTCTAGCCGCTGGTAAGGATGATATATTCGAATTCAAATGTGTGGATTTCGGCGCCTATTTTATAGCTATGCGATTGGATAAGAAAACATATCTGCCGCAAGCTATTAGGCGAGGTACTGGAGACGCGTGGATGGTTAAAAAGGCGGCAAAAGTCGATCCATCTGCTCAACAATTTTGTCAGTATTTGATAAAACACAAGTCTAATAATGTTATTACTTGTGGTAATGAGATGTTAAATGAATTAGGTTATAGCGGTTATTTTATGTCACCGCATTGGTGTTCCGATCTTAGTAATATGGAATAAGTGTTAGATAAATGCGGTAACAAATGTTCCTGTAAGGAACCATAACAGTTTAGATTTAACATTAAAGATGAGCATAAACATAATAAACAAAATTACAATCAAACCTATAACATTAATATCAAACAATCCAAAAAATGAAATCAATGGAGTAGTAAACGTGTACATAACTCCTGGATAACGTTTAGCAGCTACCGTTCCTATTCTAGACCAAAAATTTGGTTTCATGGTTTCGAAGCGGTGTTCTGCAACAAGACGAGGATCGTGTTCTACATATTTGGCAGAGTTATCCATTATTTGCCTGTTAATCTTCATTTCGTTTTCGATTCTGGCTATTTCAAAATAAAATCCCGATGATAGACCTCCAGACTTTATAATTTCATCTACGATGTTCAGCGCCGTAGTAACTCTAATAATATAGGCGGATAAGCTAACATCATACCCTCCTGTATATGTAAATATGGCATGATCTTTGTCTATTACAAGCTCGGTTTTAACTTTATTTCCTGTAATAATTTCTCTCATCTGTAGGATATCTATTTTCTTGTCATGTATTGCCTTCAAGACGGGACGAAGAAACGTAATATCCTCAATAACGTTATCGTTTTCTATAATAACTACATATTCTACATTTTTATTTTCTAGCTCGGTAAAAAATTTAGAATCCCATAGGGCTAAATGTCTAGCGATATTTCTTTTCGTTTCCTCTGTACACATAGTGTTACAAAACCCTGAAAAGAAGTGAGTATACTTGTCATCATCTCTAATATTTCCTCCAGTCCATTGTATAAACACATAATCCTTGTAATGATCTGGATCATCATTGACTATCACAACATCTCTTTTTTCTTGCATAACTTCATTGTCCTTCACATCATCGAACTTCTGATCATTAATATGCTCATGAACATTAGGAAATGTTTCTGATGGAGGTCTATCAATAACTGGCACAACAATAACAGGAGTTTTCACCGCCGCCATTTAGTTATTGAAATTAATCATATACAACTCTCTAATACGAGTTATATTTTCGTCTATCCATTGTTTCACATTGACATATTTCGACAAAAAGATATAAAATGCGTATTCCAATGCTTCTCTGTTTAATGAATTACTAAAATATACAAACACGTCACTGTCTGGTAATAAATAATATCTTAGAATATTGTAACAATTTATTTTGTATTGCACATGTTCGTGATCTATGAGTTCTTCTTCAAATGGCATAGGATCTCCGAATCTGAAAACGTATAAATAGGAGTTAGAATAATAATATTTGAGAGTATTGGTAATGTATAAACTCTTTAGCGGTATAATTAGTTTTTTTCTCTCGATTTCTATTTTTAGATGTGATGGAAAAATGACTAATTTTGTAGCATTAGTATCATGAACTCTAATCAAAATCTTAATATCTTCGTCACATGTTAGCTCTTTGAAGTTTTTAAGAGATGCATCAGTTGGTTTTACAGATGGAGTAGGTGCAACAATTTTTTGTTTAATGCATGCATGTATTGGAGCCATTGTCTTAACTATAATGGTGCTTGTATCGAAAAACTTTAATGCGGATAACGGAAGCTCTTCGCCGCGACTTTCTACGTCGTAATTGGGTTCTAATGCCGATCTCTGAATGGATACTAGTTTTCTAAGTTCTAATGTAATTCTCTGAAAATGTAAATCCAATTCCTCCGGCATTATAGATGTGTATACATCGGTAAATAAAACTATAGTATCCAACGATCCCTTCTCGCAAATTCTAGTCTTAACCAAGAAATCGTATATAACTACGGAGATGGCGTATTTAAGAGTGGATTCTTCTACCGTTTTGTTCTTGGATTTCATATAAGAAACTATAAAGTCCGCACTACTGTTAAGAATGATCACTAACGCAACTATATAGTTCAAATTAAGCATCTTGGAAACATAAAATAACTCTGTAGATGATACTTGACTTTCGAATAAGTTTGCAGACAAACGAAGAAAGAACAGACCTCTCTTAATTTCAGAAGAAAACTTTTTTTCGTATTCCTGACGTCTAGAGTTTATATCAATAAGAAAGTTAAGAATTAGTCGGTTAATGTTGTATTTCATTACCCAAGTTTGAGATTTCATAATATTGTCAAAAGACATGATAATATTAAAGATAAAGCGCTGACTATGAACGAAATAGCTATATGGTTCGCTCAAGAATATAGTCTTGTTAAACGTGGAAACGATAACTGTATTTTTAATCACGTCAGCGGCATCTAAATTAAATATAGGTATATTTATTCCACACACTCTACAATATGCCACACCATCTTCATAATAAATAAATTCGTTAGCAAAATTATTAATTTTAGTGAAATAGTTAGCGTCAACTTTCATAGCTTCCTTCAATCTAATTTGATGCTCACATGGCGCGAATTCTACTCTAACATCCCTTTTCCATGCCTCAGGTTCATCGATCTCTATAATATCTAGTTTCTTGCGTTTCACAAACACAGGCTCGTCTCTCGCGATGAGATCTGTATAGTAACTATGTAAATGATAACTAGATAGAAAGATGTAGCTATATAGATGACGATCCTTTAAGAGAGGTATAATAACTTTACCCCAATCAGATAGACTGTTGTTATGGTCTTCGGAAAAAGAATTTTTATAAATTTTTCCAGTATTTTCTAAATATACGTACTTGATATCTAAGAAATCCTTAATAATAATAGGAATGGATAATCCGTCTATTTTATAAAGAAATACATATCGCATATTATACTTTTTTTTGGAAATTGGAATACCGATGTGTCTACATAAATACGCAAAGTCTAAATATTTTTTAGAGAATCTTAGTTGGTCCAAATTCTTTTCCAAGTACGGTAATAGATTTTTCATATTGAACGGTATCTTCTTGATCTCTGGTTCTAATTCCGCATTAAATGATGAAACTAAGTCACTATTTTTATAACTAACGATTACATCACCTCTAACATCATCATTTACCAGGATACTGATCTTCTTTTGTCGTAAATACATGTCTAATGTGTTAAAAAAAAGATCATACAAGTTATACGTCATTTCATCTGTAGTATTCTTGTCATTGAAGGATAAACTCGTACTAATCTCTTCTTTAACAGTCTGTTCAAATTTATATCCTATATATGAAAAAATAGCAACCAGTGTTTGATCATCCGCGTCAATATTCTGTTCTATCGTAGTGTATAACAATCTTATATCTTCTTCTGTGATAGTCGATACGTTATAAAGGTTGATAACGAAAATATTTTTATTTCGTGAAATAAAGTCATTGTAGGATTTTGGACTTATATTCGTGTCTAGTAGATATGATTTTATTTTTGGAATGATCTCAATTAAAATAGTCTCTTTAGAGTCCATTTAAAGTTACAAACAACTAGGAAATTGGTTTATGATGTATAATTTTTTTAGTTTTTATAGATTCTTTATTCTATACTTAAAAAATGAAAATAAATACAAAGGTTCTTGAGGGTTGTGTTAATTGAAAGCGATAAATAATCATAAATTATTTCATTATCGCGATATCCGTTAAGTTTGTATCGTAATGGCGTGGTCAATTACGAATAAAGCGGATACTAGTAGTTTCACAAAGATGGCTGAAATCAGAGCTCATCTAAGAAATAGCGCTGAAAATAAAGATAAAAACGAGGATATTTTCCCGGAAGATGTAATAATTCCATCTACTAAGCCCAAAACCAAACGAACCACTACTCCTCGTAAACCAGCGGCTACTAAAAGATCAACCAAAAAGGATAAAGAAAAGGAGGAAGTGGAAGAAGTAGTTATAGAGGAATATCATCAAACAACTGAAGAAAATTCTCCACCTCCGTCATCATCTCCTGGAGTCGGCGACATTGTAGAAAGCGTGGCCGCTGTAGAGCTCGATGATAGCGACGGGGATGATGAACCTATGGTACAAGTTGAAGCTGGTAAAGTAAATCATAGTGCTAGAAGCGATCTCTCTGACCTAAAGGTGGCTACCGACAATATCGTTAAAGATCTTAAGAAAATTATTACTAGAATCTCTGCAGTATCGACTGTTCTAGAGGATGTTCAAGCAGCTGGTATCTCTAGACAATTTACTTCTATGACTAAAGCTATTACAACACTATCTGATCTAGTCACCGAGGGAAAATCTAAAGTTGTTCGTAAAAAAGTTAAAACTTGTAAGAAGTAAATGCGTGCACTTTTTTATAAAGATGGTAAACTGTTTACCGATAATAATTTTTTAAATCCTGTATCAGACGATAATCCAGCGTATGAGGTTTTGCAACATGTTAAAATTCCTACTCATTTAACAGATGTAGTAGTATATGAACAAACGTGGGAAGAGGCATTAACTAGATTAATTTTTGTGGGAAGTGATTCAAAAGGACGTAGACAATACTTTTACGGAAAAATGCATATACAGAATCGCAATGCTAAAAGAGATCGTATTTTTGTTAGAGTATATAACGTTATGAAACGAATTAATTGTTTTATAAACAAAAATATAAAGAAATCGTCCACAGATTCCAATTATCAGTTGGCGGTTTTTATGTTAATGGAAACTATGTTTTTTATTAGATTTGGTAAAATGAAATATCTTAAGGAGAATGAAACAGTAGGGTTATTAACACTAAAAAATAAACACATAGAAATAAGTCCCGATGAAATAGTTATCAAGTTTGTAGGAAAGGACAAAGTTTCACATGAATTTGTTGTTCATAAGTCTAATAGACTATATAAACCGCTATTGAAACTGACTGATGATTCTAGTCCCGAAGAATTTCTGTTCAACAAACTAAGTGAACGAAAGGTATATGAATGTATCAAACAGTTTGGTATTAGAATCAAGGATCTCCGAACGTATGGAGTCAATTATACGTTTTTATATAATTTTTGGACAAATGTAAAGTCCGTATCTCCTCTTCCATCACCAAAAAAGTTGATAGCATTAACTATCAAACAAACTGCTGAAGTGGTAGGTCATACTCCATCAATTTCAAAAAGAGCTTATATGGCAACGACTATTTTAGAAATGGTAAAGGATAAAAATTTTTTAGACGTAGTATCTAAAACTACGTTCGATGAATTCCTATCTATAGTCGTAGATCACGTTAAATCATCTACGGATGGATGATAATAGATCTTTACACAAATAATTACAAGACCGATAAATGGAAATGGATAAACGGATGAAATCTCTCGCTATGACAGCTTTCTTCGGAGAGCTAAACACGTTAGATATTATGGCATTGATAATGTCTATATTTAAACACCATCCAAACAATACTATTTTTTCAGTGGATAAGGATGGTCAATTTATGATTGATTTCGAATACGATAATTATAAGGCTTCTCAATATTTGGATCTGACCCTCACTCCGATATCTGGAAATGAATGCAAGACTCACGCATCTAGTATAGCCGAACAATTGGCGTGTGTGGATATTATTAAAGAGGATATTAGCGAATATATCAAAACTACTCCCCGTCTTAAACGATTTATAAAAAAATACCGCAATAGATCATATACTCGTATCAGTCGAGATACAGAAAAGCTTAAAATAGCTCTAGCTAAAGGCATAGATTACGAATATATAAAAGACGCTTGTTAATAAGTAAATGAAAAAAAACTAGTCGTTTATAATAAAACACAATATGGATGCCAACATAGTATCATCTTCTACTATTGCGACGTATATAGACGCTTTAGCGAAGAATGCTTCAGAATTAGAACAGAGGTCTACCGCATACGAAATAAATAATGAATTGGAACTAGTATTTATTAAACCGCCATTGATTACGTTGACAAATGTAGTAAATATCTCCACGATTCAGGAATCGTTTATTCGATTTACCGTTACTAATAAGGAAGGTATCAAAATTAGAACTAAGATTCCATTATCTAAGGTACATGGTCTAGATGTAAAAAATGTGCAGTTGGTAGATGCTATAGATAACATAGTTTGGGAAAAGAAATCATTAGTGACGGAAAATCGTCTTCACAAAGAATGCTTGTTGAGACTATCAACAGAGGAACGTCATATATTTTTGGATTACAAGAAATATGGATCCTCTATCCGACTAGAATTAGTCAATCTTATTCAAGCAAAAACAAAAAACTTTACGATAGACTTTAAGCTAAAATATTTTCTAGGATCTGGCGCTCAATCTAAAAGTTCTTTATTGCACGCTATTAATCATCCAAAGTCAAGGCCTAATACATCTCTGGAAATAGAATTTACACCTAGAGACAATGAAACAGTTCCATATGATGAACTAATAAAGGAATTGACGACTCTCTCGCGTCATATATTTATGGCTTCTCCAGAGAATGTAATTCTTTCTCCACCTATTAACGCACCTATAAAGACTTTTATGTTGCCTAAACAAGATATAGTAGGTCTGGATCTGGAAAATCTATATGCCGTAACTAAGACTGACGGCATTCCTATAACTATCAGAGTTACATCAAAAGGGTTGTATTGTTATTTTACACATCTTGGTTATATTATTAGATATCCAGTTAAGAGAACAATAGATTCCGAAGTAGTAGTCTTTGGTGAGGCAGTTAAGGATAAGAACTGGACCGTATATCTCATTAAGCTAATAGAGCCCGTAAATGCAATCAGTGATAGACTAGAAGAAAGTAAGTATGTTGAATCTAAACTAGTGGATATTTGTGATCGGATAGTATTCAAGTCAAAGAAATACGAAGGTCCGTTTACTACAACTAGTGAAGTCGTCGATATGTTATCTACATATTTACCAAAGCAACCAGAAGGTGTTATTCTGTTCTATTCAAAGGGACCTAAATCTAACATTGATTTTAAAATCAAAAAGGAGAATACTATAGACCAAACTGCAAATGTAGTATTTAGGTACATGTCCAGTGAACCAATTATCTTTGGAGAGTCGTCTATCTTTATAGAGTATAAGAAATTTACCAACGATAAAGGCTTTCCTAAAGAATATGGTTCTGGTAAGATTGTGTTATATAACGGCGTTAATTATCTAAATAATATCTATTGTTTGGAATATATTAATACACATAATGAAGTGGGTATTAAGTCCGTTGTTGTACCTATTAAGTTTATAGCAGAATTCTTAGTCAATGGAGAAATACTTAAACCTAGAATCGATAAAACCATGAAATATATTAACTCAGAAGACTATTATGGAAATCAACATAATATCATAGTCGAACATTTAAGAGATCAAAGCATCAAAATAGGAGATGTCTTTAACGAGGATAAACTATCGGATGTTGGACATCAATACGCTGCCAACAACGATAAATTTAGATTAAATCCAGAAGTTAGTTATTTTACTAATAAACGAACTAGAGGGCCGTTGGGAATTTTATCAAACTACGTCAAGACTCTTCTTATTTCTATGTATTGTTCCAAAACATTTTTAGACGATTCCAACAAACGAAAGGTATTAGCGATTGATTTTGGAAACGGTGCTGACCTGGAAAAATACTTTTATGGAGAGATTGCGTTATTGGTAGCGACGGATCCGGATGCTGATGCTATAGCTAGAGGAAATGAAAGATACAACAAATTAAATTCTGGAATTAAAACCAAGTACTACAAATTTGACTACATTCAGGAAACTATTCGATCCGATACATTTGTCTCTAGTGTCAGAGAAGTATTCTATTTTGGAAAGTTTAATATCATTGACTGGCAGTTCGCTATTCATTATTCTTTTCATCCAAGACATTATGCTACAGTCATGAATAACTTATCCGAACTAACTGCTTCTGGAGGCAAGGTATTAATTACTACCATGGATGGAGACAAATTATCAAAATTAACCGATAAAAAGACTTTTATAATTCATAAGAATCTACCTAGTAGCGAAAACTATATGTCTGTAGAAAAAATAGCTGATGATAGAATAGTGGTATATAATCCATCAACAATGTCTACTCCAATGACTGAATACATTATCAAAAAGAACGATATAGTCAGAGTGTTTAACGAATACGGATTTGTTCTTGTAGATAATGTTGATTTCGCTACAATTATAGAACGAAGTAAAAAGTTTATTAATGGCGCATCTACAATGGAAGATAGACCGTCTACAAGAAACTTTTTCGAACTAAATAGAGGAGCCATTAAATGTGAAGGTTTAGATGTCGAAGACTTACTTAGTTACTATGTTGTTTATGTCTTTTCTAAGCGGTAAATAATAATATGGTATGGGTTCTGATATCCCCGTTCTAAATGCATTAAATAATTCCAATAGAGCGATTTTTGTTCCTATAGGACCTTCCAACTGTGGATACTCTGTATTATTAATAGATATATTAATACTTTTGTAGGGTAACAGAGGTTCTACGTCTTCTAAAAATAAAAGTTTTATAACATCTGGCCTGTTCATAAATAAAAACTTGGCGATTCTATATATACTCTTATTATCAAATCTAGCCATTGTCTTATAGATGTGAGCTACTGTAGGTGTACCATTTGATTTTCTTTCTAATACTATATATTTCTCTCGAAGAAGTTCTTGCAGATCATCTGGGAATAAAATACTACTGTTGAGTAAATCAGTTATTTTTTTTATATCGATATTGATGGACATTTTTATAGTTAAGGATAATAAGTATCCCAAAGTAGATAACGACGATAACGAAGTATTTATACTTTTAGGAAATCACAATGACTTTATCAGATCAAAATTAACAAAATTAAAGGAGCATGTATTTTTTTCTGAATATATTGTGACTCCAGATACATATGGATCTTTATGCGTCGAATTAAATGGGTCTAGTTTTCAGCACGGTGGTAGATATATAGAGGTGGAGGAATTTATAGATGCTGGAAGACAAGTTAGATGGTGTTCTACATCCAATCATATATCTGAAGATATACACACTGATAAATTTGTCATTTATGATATTTATACGTTTGATTCGTTCAAGAATAAACGATTGGTATTTGTACAGGTGCCTCCATCATTAGGAGATGATAGCTATTTAACTAATCCGTTATTGTCTCCGTATTATCGTAATTCAGTAGCCAGACAAATGGTCAATGATATGATTTTTAATCAAGATTCATTTTTAAAATATTTATTAGAACATCTGATTAGAAGCCACTATAGAGTTTCTAAACATATAACAATAGTTAGATACAAGGATACCGAAGAATTAAATCTAACAAGAATATGTTATAATAGAGATAAGTTTAAGGCGTTTGTATTCGCTTGGTTTAACGGCGTTTCGGAAAATGAAAAGGTACTAGATACGTATAAAAAGGTATCTGATTTGATATAATGAATTCAGTGACTATATCACACGCACCATATACTATTACTTATCACGATGATTGGGAACCAGTAATGAGTCAATTGGTAGAGTTTTATAACGAAGTAGCCAGTTGGTTGCTACGCGACGAGACGTCGCCTATTCCTGATAAGTTCTTTATACAATTGAAACAGCCGCTTAGAAATAAACGAGTATGTGTGTGTGGTATAGATCCGTATCCAAAAGATGGAACTGGTGTACCGTTCGAATCACCAAATTTTACAAAAAAATCAATTAAGGAGATAGCTTCATCTATATCTAGATTAACCGGAGTAATTGATTATAAAGGTTATAACCTTAATATAATAGACGGGGTTATACCCTGGAATTATTACTTAAGTTGTAAATTAGGAGAAACAAAAAGTCACGCGATTTACTGGGATAAGATTTCCAAGTTACTGCTACAGCATATAACTAAACACGTTAGTGTTCTTTATTGTTTGGGTAAAACAGATTTCTCGAATATACGGGCAAAGTTAGAATCCCCGGTAACTACCATAGTGGGATATCATCCAGCGGCCAGAGACCACCAATTCGAGAAAGATCGATCATTTGAAATTATCAACGTTTTACTGGAATTAGACAACAAGACACCTATAAATTGGGCTCAAGGGTTTATTTATTAATGCTTTAGTGAAATTTTAACTTGTGTTCTAAATGGATGCGGCTATTAGAGGTAATGATGTTATCTTTGTTCTTAAGACTATAGGTGTCCCGTCAGCATGCAGACAAAATGAAGATCCAAGATTCGTAGAAGCATTTAAATGCGACGAGTTAGAAAGATATATTGATAATAATCCAGAATGTACACTATTCGAAAGTCTTAGGGATGAGGAAGCATACTCTATAGTCAGAATTTTCATGGATGTAGATTTAGACGCGTGTCTAGACGAAATAGATTATTTAACGGCTATTCAAGATTTTATTATCGAGGTGTCAAACTGTGTAGCTAGATTCGCATTTACAGAATGCGGTGCCATTCATGAAAATGTAATAAAATCCATGAGATCTAATTTTTCATTGACTAAGTCTACAAATAGAGATAAAACAAGTTTTCATATTATCTTTTTAGACACGTATACCACTATGGATACATTGATAGCTATGAAACGAACACTATTAGAATTAAGTAGATCATCTGAAAATCCACTAACAAGATCGATAGACACTGCCGTATATAGGAGAAAAACAACTCTTCGGGTTGTAGGTACTAGGAAAAATCCAAATTGCGACACTATTCATGTAATGCAACCACCTCACGATAATATAGAAGATTACCTATTCACTTACGTGGATATGAACAACAATAGTTATTACTTTTCTCTACAACGACGATTGGAGGATTTAGTTCCTGATAAGTTATGGGAACCAGGGTTTATTTCGTTCGAAGACGCTATAAAAAGAGTTTCAAAAATATTCATTAATTCTATAATAAACTTTAATGATCTCGATGAAAATAATTTTACAACGGTACCACTGGTCATAGATTATGTAACACCTTGTGCATTATGTAAAAAACGATCGCATAAACATCCGCATCAACTATCGTTGGAAAATGGTGCTATTAGAATTTACAAAACTGGTAATCCACATAGTTGTAAAGTTAAAATTGTTCCGTTGGATGGTAATAAACTGTTTAATATTGCACAAAGAATTTTAGACACTAACTCTGTTTTATTAACCGAACGAGGAGACCATATAGTTTGGATTAATAATTCATGGAAATTTAACAGCGAAGAACCCTTGATAACAAAACTAATTCTATCAATAAGACATCAACTACCTAAGGAATATTCAAGCGAATTACTCTGTCCGAGGAAACGAAAGACTGTAGAAGCTAACATACGAGACATGTTAGTAGATTCAGTAGAGACCGATACCTATCCGGATAAACTTCCGTTTAAAAATGGTGTATTGGACCTGGTAGACGGAATGTTTTACTCTGGAGATGATGCTAAAAAATATACGTGTACTGTATCGACCGGATTTAAATTTGACGATACAAAATTCGTCGAAGACAGTCCAGAAATGGAAGAGTTAATGAATATCATTAACGATATCCAACCATTAACGGATGAAAATAAGAAAAATAGAGAGCTGTATGAAAAAACATTATCTAGTTGTTTATGTGGTGCTACCAAAGGATGTTTAACATTCTTTTTTGGAGAAACCGCAACTGGGAAGTCGACAACCAAACGTTTGTTAAAGTCTGCTATCGGTGACCTGTTTGTCGAGACGGGTCAAACAATTTTAACAGATGTATTGGATAAAGGACCTAATCCATTTATCGCTAATATGCATTTAAAAAGATCTGTATTCTGTAGCGAACTACCTGATTTTGCATGTAGTGGATCAAAGAAAATTAGATCTGATAATATTAAAAAGTTGACAGAACCTTGTGTCATTGGAAGACCGTGTTTCTCCAATAAAATTAATAATAGAAACCATGCGACAATCATTATCGATACTAATTACAAACCTGTCTTTGATAGGATAGATAACGCATTAATGAGAAGAATTGCCGTCGTGCGATTCAGAACACACTTTTCTCAACCTTCTGGTAGAGAGGCTGCTGAAAATAATGACGCGTACGATAAAGTCAAACTATTAGACGAGGGATTAGATGGTAAAATACAGAATAATAGATATAGATTCGCATTTCTATACTTGTTGGTTAAATGGTACAAAAAATATCATATTCCTATTATGAAACTATATCCTACACCGGAAGAGATTCCGGACTTTGCATTCTATCTCAAAATAGGTACTCTGTTGGTATCTAGCTCTGTAAAGCATATTCCATTAATGACGGACCTCTCCAAAAAGGGATATATATTGTACGATAATGTGGTTACTCTTCCGTTGACTACTTTCCAACAGAAAATATCCAAGTATTTTAATTCTAGACTATTTGGACACGATATAGAGAGCTTCATCAATAGACATAAGAAATTTGCCAATGTTAGTGATGAATATCTGCAATATATATTCATAGAGGATATTTCATCTCCGTAAATATATGCCATATATTTATAGAATATATCACATATCTAAATGAATACCGGAATCATAGATTTATTTGATAATCATGTTGATAGTATACCAACTATATTACCTCATCAGTTAGCTACTTTAGATTATCTAGTTAGAACTATCATAGATGAGAACAGAAGCGTGTTATTGTTCCATATTATGGGATCGGGTAAAACAATAATCGCTTTGTTGTTCGCCTTGGTAGCTTCCAGATTTAAAAAGGTTTACATTTTAGTACCGAACATCAACATCTTAAAAATTTTCAATTATAATATGGGTGTAGCTATGAACTTGTTTAATGACGAATTCATAGCTGAGAATATCTTTATTCATTCCACAACAAGTTTTTATTCTCTTAATTATAACGATAACGTCATTAATTATAACGGATTAAGTCGCTACAATAACTCTATTTTTATCGTTGATGAGGCACATAATATTTTTGGGAATAATACTGGAGAACTTATGACCGTGATAAAAAATAAAAACAAGATTCCTTTTCTACTATTGTCTGGATCTCCCATTACTAACACACCTAATACGCTGGGTCATATTATAGATTTAATGTCCGAAGAGACGATAGATTTTGGTGAGATTATTAGTCGTGGTAAGAAAGTAATTCAGACACTTCTTAACGAACGCGGAGTGAATGTACTCAAGGATTTGCTTAAAGGAAGAATATCATATTACGAAATGCCGGACAAAGATCTACCAACAATAAGATATCACGGACGTAAATTTCTAGATACTCGAGTAGTATATTGTCACATGTCTAAACTTCAAGAGAAAGATTATATGATTACTAGACGGCAGCTATGTTATCATGAAATGTTTGATAAAAATATGTATAACGTGTCAATGGCAGTATTGGGACAACTTAATCTGATGAATAATTTAGATACGTTATTTCAGGAACAGGATAAGGAATTGTACCCAAATCTGAAAATAAATAATGGAGTGTTATACGGTGAAGAATTGGTAACGTTAAACATTAGTTCCAAATTTAAGTACTTTATCAATCGGATACAGACACTCAAGGGAAAACACTTTATATACTTCTCTAATTCTACATATGGTGGATTGGTAATTAAATATATCATGCTCAGTAATGGATATTCTGAATATAATGGTTCTCAGGGAACTAATCCACATATGATAAACGGCAAACCAAAAACATTTGCTATCGTTACTAGTAAAATGAAATCGTCTTTAGAGGATCTATTAGATGTGTATAATTCTCCTGAAAACGATGATGGCAATCAATTGATGTTTTTGTTTTCGTCAAACATTATGTCTGAATCCTATACTCTGAAAGAGGTAAGGCATATTTGGTTTATGACTATCCCGGATACTTTTTCTCAATACAACCAAATTCTTGGACGATCTATTAGAAAATTCTCTTACGTCGATATTTCTGAACCCGTTAATGTATATCTTTTAGCAGCCGTATATTCAGATTTCAATGACGAAGTGACGTCATTAAACGATTATACACAGGATGAATTGATTAATGTTTTACCCTTTGACATCAAAAAGCTGTTGTATCTAAAATTTAAGACTAAAGAAACGAATAGAATATACTCTATTCTTCAAGAGATGTCTGAAACGTATTCTCTTCCACCACATCCATCAATTGTAAAAGTTTTATTGGGAGAATTGGTCAGACAATTTTTTTATAATAATTCTCGTATTAAGTATAACGACTCCAAGTTACTTAAAATGGTTACATCAGTTATAAAAAATAAAGAAGACGCTAGGAATTACATAGATGATATTGTAAACGGTCACTTCTTTGTATCGAATAAAGTATTTGATAAATCTCTTTTATACAAATACGAAAACGATATTATTACAGTACCGTTTAGACTTTCCTACGAACCATTTGTTTGGGGAGTTAACTTTCGTAAAGAATATAATGTGGTATCTTCTCCATAAAACTGATGAGATATATAAAGAAATAAATGTCGAGCTTTGTTACCAATGGATATCTTCCAGTTACATTGGAACCACATGAGTTGACGTTAGACATAAAAACTAATATTAGGAATGCCGTATATAAGGCGTATCTCCATAGAGAAATTAGTGGTAAAATGGCCAAGAAAATAGAAATTCGTGAAGACGTGGAATTACCTCTCGGTGAAATAGTTAATAATTCTGTAGTTATAAACGTTCCGTGTGTAATAACCTACGCATATTATCACGTTGGGGATATAGTCAGAGGAACATTAAACATCGAAGATGAATCAAATGTAACTATTCAATGTGGAGATTTAATCTGTAAACTAAGTAGAGATTCGGGTACTGTATCATTTAGCGATTCAAAGTACTGCTTTTTTCGAAATGGTAATGCGTATGATAACGGCATCGAAGTCTCCGCCGTTCTAATGGAGGCTCAACAAGGTACCGAATCTAGTTTTGTTTTTCTCGCGAATATCGTTGACTCATAAGAAAGAGAATAGCGGTGAGTATAAATACGAATACTATGGCAATAATTGCGAATGTTTTATTCCCTTCGATATATTTTTGATAATATGAAAAACATGCCTCTCTCAAATCAGACAACCATTTCATAAAATAGTTCTCTCGCACTGGTGAGGTGGTTGCAGCTCGTATAATCTCCCCAGAATAATATACTTGCGTGTCGTCGTTCAATTTATACGGATTTCTATAATTCTCTGTTATATAATGAGGTTTACCCTCATGATTAGACGACGACAATAGTGTTCTGAATTTAGATAGTTGATCAGAATGAATGTTTATTGGTGTTGGAAAAATTATCCATGCTGCGTCTGCAGAGTGGTTGATAGTTGTTCCTAGATATGTAAAATAATCCAACGTACTAGGTAGCAAATTGTCTAGATAAAATACTGAATCAAATGGCGCAGACATATTAGCGGATCTAATGGAATCCAATTGATTGACTATCTTTTGAAAATATACATTTTTATGATCTGATACTTGTAAGAATATAGCAATAATGATAATTCCATCATCGTGTTTTTTTGCCTCTTCATAAGAACTATATTTTTTCTTATTCCAATGAACCAGATTAATCTCTCCAGAGTATTTGTATACATCTATCAAGTGATTGGATCCATAATCGTCTTCCTTTCCCCAATATATATGTATTGTTGATAACACATATTCATTGGGGAGAAACCCTCCACTTATATATCCTCCTTTAAAATTAATCCTTACTAGTTTTCCAGTATTCTGGATAGTGGTTGGTTTCGACTCATTATAATGTATGTCTAACGTCTTCAATCGCGCGTCAGAAATTGCTTTTTTAGTTTCTATATTAATAGGAGATAGTTGTTGAGGCATAGTAAAAATGAAATGATAACTGTCTAGAAATAGCTCTTAGTATGGGATTTACAATGGATGAGGAAGTGATATTTGAAACTCCTAGAGAATTAATATCTATTAAACGAATAAAAGATATTCCAAGATCAAAAGACACGCACGTGTTTGCTGCGTGTATAACAAGTGACGGATATCCGTTAATAGGAGCTAGAAGAACTTCATTCGCATTCCAGGCGATATTATCTCAACAAAATTCAGATTCTATCTTTAGAGTATCCACTAAACTATTACGGTTTATGTACTACAATGAACTAAGAGAAATCTTTAGACGGTTGAGAAAAGGTTCTATCAACAATATCGATCCTCACTTCGAAGAGTTAATATTATTGGGTGGTAAACTAGATAAAAAGGAATCTATTAAAGATTGTTTAAGAAGAGAATTAAAAGAGGAAAGTGATGAACATATAACAGTAAAAGAATTCGGAAATGTAATTCTAAAACTTACAACGAGTGATAAATTATTTAATAAAGTATATATAGGTTATTGCATGGCATGTTTTATTAATCAATCGTTGGAGGATTTATCACATACTAGTATTTACAATGTAGAAATTAGAAAGATTAAATCGTTAAATGATTGTATTAACGACGATAAATACGAATATCTGTCTTATATTTATAATATACTAATTAATAGTAAATGAGCTTTTACAGATCTAGTATAATTAGTCAGATTATTAAGTATAATAGACGACTAGCTAAGTCTATTATTTGCGAGGATGACTCTCAAATTATTACACTCACGGCATTCGTTAACCAATGCCTATGGTGTCATAAACGAGTATCCGTGTCCGCTATTTTATTAACTACTGATAACAAAATATTAGTATGTAACAGACGAGATAGTTTTCTCTATTCTGAAATAATTAGAACTAGAAACATGTATAGAAAGAAACGATTATTTCTGAATTATTCCAATTATTTGAACAAACAGGAAAGAAGTATACTATCGTCATTTTTTTCTCTAGATCCAGCTACTGCTGATAATGATAGAATAAACGCTATTTATCCGGGTGGTATACCCAAAAGGGGTGAGAACGTTCCAGAGTGTTTATCCAGGGAAATCAAAGAAGAAGTTAATATAGACAATTCTTTTGTATTCATAGACACTCGTTTTTTTATTCATGGTATCATAGAAGATACCATTATTAACAAATTTTTTGAGGTAATTTTCTTTGTTGGACGAATATCTCTAACGAGTGATCAAATTATTGATACCTTTAAAAGTAATCATGAAATAAAGGATCTAATATTTTTAGATCCAAATTCAGGTAATGGACTCCAATACGAAATTGCAAAATATGCTCTAGATACTGCAAAACTTAAATGTTACGGTCATAGAGGATGTTATTATGAATCATTAAAAAAATTAACTGAGGATGATTGATTAGAAAATATAAATTAATTTACCATCGTGTATTTTTATAACGGGATTGTCTGGCATATCATGTAGATAGTTACCGTCTACATCGTATACTCTACCATCTACGCCTTTAAATCCTCTATTTATTGATATTAATCTATTAGAATTGGAATACCAAATATTAGTACCCTCAATTAGTTTATTGGTAATATTTTTTTTAGACGATAGATCGATGGCTCTTGAAACCAAGGTTTTCCAACCGGACTCATTGTCTATCGGTGAGAAGTCTTTTTCATTAGCATGAATCCATTCTAATGATGTATGTTTAAACACTCTAAACAATTGTACAAATTCTTTTGATTTGTTTTGAATGATTTCAAATAGGTCTTCGTCTACAGTAGGCATACCATTAGATAATCTAGCCATTATAAAGTGCACGTTTACATATCTACGTTCTGGAGGAGTAAGAACGTGACTATTGAGACGAATGGCTCTTCCTACTATCTGACGAAGAGACGCCTCGTTCCATGTCATATCTAAAATGAAGATATCATTGATTGAGAAGAAACTAATACCCTCGCCTCCGCTAGAAGAGAATACGCATGTTTTAATGCATTCTCCGTTAGTGTTTGATTCTTGGTTAAACTCAGCCACCGCCTTGATTCTAGTATCTTTTGTTCTAGATGAGAACTCTATATTAGAGATACCAAAGACTTTGAAATATAGTAATAAGATTTCTATTCCTGACTGATTAACAAATGGTTCAAAGACTAGACATTTACCATGGGATGCTAATATTCCCAAACATACATCTATAAATTTGACGCTTTTCTCTTTTAATTCAGTAAATAGAGAGATATCAGCCGCAATAGCATCCCCTCCCAATAGTTCTCCCTTTTTAAAGGTGTCTAATGCGGATTTAGAAAATTCTCTATCTCTTAATGAATTTTTAAAATCATTATATAGGGTTGCTATCTCTTGTGCGTATTCTCCCGGATCACGATTTTGTCTTTCAGGAAAGCTATCGAATGTAAACGTAGTAGCCATACGTCTCAGAATTCTAAATGATGATATACCAGTTTTTATTTCTGCGAGTTTAGCCTTTTGATAAATCTCTTCTTGCTTTTTTGACATATTAACGTATCGCATTAATACTGTTTTCTTAGCGAATGATGCAGACCCTTCCACATCATCAAAAATAGAAAACTCGTTATTAACTATGTACGAACATAGGCCTCCTAGTTTGGAGACTAATTCTTTTTCATCGACTAGACGTTTATTCTCAAATAGCGATTGGTGTTGTAAGGATCCTGGTCGCAGTAAGTTAACCAACATGGTGAATTCTTGCACACTATTAACGATAGGTGTAGCCGATAAACAAATCATCTTATGGTTTTTTAACGCAATGGTCTTAGATAAAAAATTATATACTGACCGAGTAGGACGGATCTTACCATCTTCTTTGATTAATGATTTAGAAATGAAGTTATGACATTCATCAATGATGACGCATATTCTACTCTTGGAATTAATAGTTTTGATATTAGTAAAAAATTTATTTCTAAAATTTTGATCATCGTAATTAATAAAAATACAATCCTTCGTTATCTCTGGAGCGTATCTGAGTATAGTGTTTATCCAAGGATCTTCTATCAAAGCCTTTTTTACCAATAAGATAATTGCCCAATTCGTATAAATATCCTTAAGATGTTTGAGAATATATACAGTAGTCATTGTTTTACCGACACCTGTTTCATGGAACAATAAAAGAGAATGCATACTGTCTAATCCTAAGAAAACTCTTGCTACAAAATGTTGATAATCCTTGAGGCGTACTACGTCTGACCCCATCATTTCAACGGGCATATTAGTAGTTCTGCGTAAGGCATAATCGATATAGGCCGCGTGTGATTTACTCATTTATGAGTGATAAGTAATAACTATGTTTTAAAAATCACAGCAGTAGTTTAACTAGCCTTCTCTGATGTTTGTTTTCGATACTTTTTGAATCAGAAGTCATACTAGAATAAAGCAGCGAGTGAACGTAATAGAGAGCTTCGTATACTCTATTCGAAAACTCTAAGAACTTATTAATGAATTCCGTATCCACTGGATCGTTTAAAATACTAAATTGAACAGTGTTCACATCCTTCCAAGACGAAGACTTAGTGACGGACTTAACATGAGACATAAATAAATCCAAATTTTTTTTATAAACATCACTAGCCACCATAATGGCGCTATCTTTCAACCAACTATCGCTTACGCATTTTAACAGTCTAACATTTTTAAAGAGACTACAATATATTCTCATAGTATCGATTACACCTCTACCGAATAGAGTGGGAAGTTTAATAATACAATATTTTTCGTTTACAAAATCAAATAATGGTCGAAACACGTCGAAGGTTAACATCTTATAATCGCTAATGTATAGATTGTTTTCAGTGAGATGATTATTAGATTTAATAGCATCTCGTTCACGTTTGAACAGTTTATTGCGTGCGCTGAGGTCGGCAACTACGGCATCCGCTCTAGTACTCCTCCCATAATACTTTACGCTATTAATCTTTAAAATTTCATAGACTTTATCTAGATCGCTTTCTGGTAACATGATATCATGTGTAAAAAGTTTTAACATGTCGGTCGGCATTCTATTTAGATCATTAACTCTAGAAATCTGAAGAAAGTAATTAGCTCCATATTCCAGACTAGGTAATGGGCTTTTACCTAAAGACAAGTTAAGTTCTGGCAATGTTTCATAAAATGGAAGAAGGACATGTGTCCCCTCCCGGATATTTTTTACAATTTCATCCATTTACAACTCTATAGTTTGTTTTCATTATTATTAGTTATTATCTCCCATAATCTTGGTAATACTTACACCTTGATCATAAGATACCTTATACAGGTCATTACATACAACTACCAATTGTTTTTGTACATAATAGATTGGATGATTGATATCCATGGTGGAATAAACTACTCGAACAGATAGTTTATCTTTCCCCCTAGATACATTGGCCGTAATAGTTGTCGGCCTAAAGAATATCTTTGGTGTAAAGTTAAAAGTTAGGGTTCTTGTTCCATTATTGCTTTTTGTCAGTAGTTCGTTATAAATTCTCGAGATGGGCCCGTTCTCTGAATATAGAACATCATTTCCAAATCTAACTTCTAGTCTAGAAATAATATCGGTCTTATTTTTAAAATCTATTCCCTTGATGAATGGATCGTTAATAAACAAATCCTTGGCCTTTGATTCGGCTGATCTATTATCTCCGTTATAGACGTTACGTTGACTAGTCCAAAGACTTACAGGAATAGATGTATCGATGATGTTGATAGTATGTGATATGTGAGCAAAGACTGTTCTCTTGGTGGCGTCGCTATATGTTCCAGTAATGGCGGAAAACTTTTTAGAAATGTTATATATAAAAGAATTTTTTCGGGTTCCAAACATTAACAGATTAGTATGAAGATAAACACTCATATTATCAGGAACATTATCAATTTTTACATAAACATCGGCATCTTGAATAGAAACAACACCATCTTCTGGAACCTCTACGATCTCGGCAGATTCCGGATAACCAGTCGGTGGACCATCACTAACAATAACTAGATCATCCAACAATCTACTCACATATGCGTCTATATAATCTTTTTCATCTTGTGAGTACCCTGGATACGAAATAAATTTGTTATCAGTATTTCCATAATAAGGTTTAGTATAAACAGAGAGAGATGTTGCTGCATGAACTTCGGTTACTGTCGCCGTTGGTTGGTTTATTTGACCTATTACTCTCCTAGGTTTCTCTATAAATGATGGTTTAATTTGTACATTCTTAACCATATATCCAATAAAGCTCAATTCAGGAACATAAACAAATTCTTTGTTGAACGTTTCAAAGTCGAACGAAGAGTCACGAATAACGATATCGGATACTGGATTGAAGGTCACCGTTACGGTAATTTTTGAATCGGATAGTTTAAGACTACTGAATGTATCTTCCACATCAAACGGAGTTTTAATATAAACGTATACTGTAGATGGTTCTTTAATAGTGTCATTAGGAGTTAGGCCAATAGAAATATCATTAAGTTCACTAGAATATCCAGAATGTTTCAAAGCAATTGTATTATTGATACAATTATTATATAATTCTTCGCCCTCAATTTCCCAAATAACACCGTTACACGAAGAGACAGATACATGATTAATACATTTATATCCAACATATGGCACGTAACCGAATCTTCCCATACCTTTAACTTCTGGAAGTTCCAAACTCAGAACCAAATGATTAAGCGCAGTAATATACTGATCCCTAATTTCGAAGCTAGCGATAGCCTGATTGTCTGGCCCATCGTTTGTCATAACTCCGGATAGAGAAATATATTGCGGCATATATAAAGTTGGAATTTGACTATCAACTGCGAAGACATTAGACCGTTTAATAAAGTCATCCCCACCGATCAAAGAATTAATGATAGTATTATTCATTTTCTATTTAAAATGGAAAAAGCTTACAATAAACTCCGTAGAGAAATATCTATAATTTGTGAGTTTTCCTTAAAGTAACAGCTTCCGTAAACACCGTCTTTATCTCTTAGTAAGTTTATTGTATTTATGACCTTTTCCTTATCTTCATAGAATACTAAAGGCAATAAAGAAATTTTTGGTTCTTCTCTAAGAGCTACGTGAGACTTAACCATAGACGCCAACGAATCCCTACATATTTTAGAACAGAAATACCCAACTTCACCACCCTTGAATGTCTCAATACTAATAGGTCTAAAAACCAAATCTTGATTACAAAACCAACACTTATCAATTACACTATTTGTCTTAATAGACATATCTGCCATAGATTTATAATACTTTGGTAGTATACAAGCGAGTGCTTCTTCTTTAGCGGGCTTAAAGACTGCTTTAGGTGCTGAAATAACCACATCTGGAAGACTTACTCGCTTAGCCATTTAATTACGGAACTATTTTTTTATACTTCTAATGAACAAGTAGAAAACCTCTCATCTACAAAAACATACTCGTGTCCATAATCCTCTACCATAGTAACACGTTTTTTAGATCTCATATGTGCTAAAAAGTTTTCCCATACTAATTGGTTACTATTATTTTTCGTATAATTTTTAACAGTTTGAGGTTTTAGATTTTTAGTTACAGAAGTGATATCGAATATTTTATCCAAAAAGAATGAGTAATTAATTGTCTTAGAAGGAGTGTTTTCTTGGCAAAAGAATACCAAGTGCTTAAATATTTCTACTACTTCATTAATCTTTTCTGTACTCAGATTCAGTTTCTCATCTTTTACTTGATTGATTATTTCAAAGACTAACTTATAATCCTTTTTATTTATTCTCTCGTTAGCCTTAAGAAAACTAGATACAAAATTTGCATCTACATCATCCGTGGATATTTGATTTTTTTCCATGATATCCAATAGTTCCGAGATAATTTCTCCAGAACATTGATGAGACAATAATCTCCGCAATACATTTCTCAAATGAATAAGTTTATTAGACACGTGGAAGTTTGACTTTTTTTGTACCTTTGTACATTTTTGAAATACAGACTCGCAAAAAATACAATATTCATATCCTTGTTCAGATACTATACCGTTATGTCTACAACAGCTACATAATCGTAGATTCATGTTAACACTCTACGTATCTCGTCGTCCAATATTTTATATAAAAACATTTTATTTCTAGACGTTGTCAGAAAATCCTGTAATATTTTTAGTTTTTTTGGTTGTGAATAAAGTATCGCCCTAATAATATTGGTACCGTCTTCCGACAATATAGTAGTTAAATTATCCGAGCATGTAGAAGAACACCGCTTAGGCGGATTCAGTACAATGTTATATTTTTCGTACCAACTCATTTAAATATCATAATCTAAAATAGTTCTGTAATATGTCTAGCGCTAATATATTGATCATAATCCTGTGCATAAATTAAGATACAACAATGTCTTGAAATCATCGACATGGCTTCTTCCATAGTTAGAAGATCATCGTCAAAGTTAGCAACGTGATTCATCAACATTTGCTGTTTTGAGGCAGCAAATACTGAACCATCACCATTCAACCATTCATAAAAACCATCGTCTGAATCCATTGATAATTTCTTGTACTGGTTTTTGAGAGCTCGCATCAATCTAGCATTTCTAGCTCCCGGATTGAAAACAGAAAGAGGATCGTACATCCAAGGTCCATTTTCTGTAAATAGAATCGTATAATGTCCCTTCAAGAAGATATCAGACGATCCACAATCAAAGAATTGGTCTCCGAGTTTGTAACAGACTGCGGACTTTAACCTATACATGATACCGTTTAGCATGATTTCTGGTGATACGTCAATCGGAGTATCATCTATTAGAGATCTAAAGCCGGTGTAACATTCTCCGCCAAACATATTCTTATTCTGACGTCGTTCTACATAAAACATCATTGCTCCATTAACGATAACAGGTGAATGAACAGCACTACCCATCACATTAGTTCCCAATGGATCAATGTGTGTAACTCCAGAACATCTTCCATAGCCTATGTTAGGAGGAGCGAACACCACTCTTCCACTATTGCCATCGAATGCCATAGAATAAATATCCTTGGAATTGATAGAAATCGGACTGTCGGATGTTGTTATCATCTTCATAGGATTAACAACGATGTATGGTGCAGCCTGAAGTTTCATATCGTAACTGATGCCGTTCATAGGTCTAGCCACAGAAACCAACGTAGGTCTAAATCCAACTATAGACAAAATAGAAGCTAATATCTGTTCCTCATCTGTCATAACTTGAGAGCATCCAGTATGAATAATCTTCATTAGATGGGGATCTACCGCATCATCATCGTTACAATAAAAAATTCCCATTCTAATGTTCATAATTGCTTTTCTAATCATGGTATGAATGTTTGCTCTCTGAATCTCTGTGGAAATTAGATCTGATACACCTGTAATCACTATCGGATTATCCTCCGTAAGACGATTAACCAACAACATATAATTATAAGACTTTACTCTTCTAAATTCATAAAGTTGCTGGATTAGACTATATGTGTCTCCATGTACATACGCGTTCTCGAGCGCAGGAAGTTTAATACCGAATAGTGCCATCAGAATAGGATGAATGTAGTAATTAGTTTCTGGTTTTCTATAAATAAAAGACAAATCTTGTGAACTAGACATATCGGTAAAATGCATGGATTGGAATCGTGTAGTCGACAGAAGAATATGATGATTAGATGGAGAGTATATTTTATCTAACTCTTTGAGTTGGTCACCGATTCTAGGACTAGCTCGAGAATGAATAAGTACTAAGGGATGAGTACATTTCACAGAAACACTGGCGTTGTTCAACGTACTCTTTACATGGGAAAGGAGTTGAAATAGCTCGTTTCTATTTGTCCTGACAATATTTAGTTTATTCATAATATTAAGCATATCCTGAATAGTAAAGTTAGATGTGTCATACTTGTTAGTAGTTAGATATTTAGCAATTGCATTCCCATCATTTCTCAATCTCGTACTCCAATCATGTGTGGATGCTACTTCGTCGATGGAAACCATACAATCCTTTTTGATAGGCTGTTGAGATTGATCATTTCCTGTACGTTTAGGTTTGGTACGTTGATTTCTAGCCCCTGCTGATATAAAGTCATCGTCTACAATTTGGGATAATGAATTACATACACTACAAGACAAAGATTTATCAGAAGTGTGAATATGATCTTCATCTACCAAAGAAAGAGTTTGATTAGTATAACTAGATTTTAGTCCCGCGTTAGATGTTAAAAAAACATCGCTATTGACCACGGCTTCCATTATTTATATTCGTAGTTTTTACTCGAAAGCGTGATTTTAATATCCAATCTTATTACTTTTGGAATCGTTCAAAACCTTTGACTAGTTGTAGAATTTGATCTATTGCCCTACGCGTATACTCCCTTGCATCATATACGTTCGTCACCAGATCGTTTGTTTCGGCCTGAAGTTGACGCATATCTTTTTCAACACTCGACATGAGATCCTTAAGGGTCATATCGTCTAGATTTTGTTGAGATGCTGCTCCTGGATTTGGATTTTGTTGTGCTGTTGTACATACTGTACCACCAGTAGGTGTAGGAGTACATACAGTGGCCACAATAGGAGGTTGAAGAGGTGTAACCGTTGGAGTAGTACAAGAAATACTTCCATCCGATTGTTGTGTACATGTGGTTGTTGGTAACGTCTGAGAAGGTTGGGTAGATGGCGGTGTCGTCATCTTTTGATCTTTATTAAATTTAGAGATAATATCCTGAACAGTATTGCTCGGCGTCAACGCTGGAAGGAGTGTACTCGCCGGCGCATCAGTATCTGTAGACAACCAATCAAAAAGATTAGACATATCAGATGATGTATTAGTTTGTTGACGTGGTTTTAGTACAGGAGCAGTACTACTAGGTAGAAGAATAGGAGCCGGTGTAGGTGTCGGAACCGGCTGTGGAGTTATATGAATAGTTGGTTGTAGCGGTTGGGTAGGCTGTCTGCTGGCGGTCATCATATTATCTCTAGCTAGTTGTTCTCGCAACTGTCTTTGATAATACGACTCTTGAGACTTTAGTCCTATTTCAATCGCTTCATCCTTTTTCGTATCCGGATCCTTTTCTTCAGAATAATAGATTGACGACTTTGGTGTAGAGGATTCTGCCAGCCCCTGTGAGAACTTGTTAAAGAAGTCCATTTAAGGCTTTAAAATTGAATTGCGATTATAAGATTAAATGGCAGACACGGACGATATTATCGACTATGAATCCGATGATCTCACCGAATACGAGGATGATGAAGAAGATGGAGAGTCACTAGAAACTAGTGATATAGATCCCAAATCTTCTTATAAGATTGTAGAATCAACATCCACTCATATAGAAGATGCGCATTCCAATCTTAAACATATAGGGAATCATATATCTGCTCTTAAACGACGCTATACTAGACGTATAAGTCTATTTGAAATAGCGGGTATAATAGCAGAAAGCTATAACTTGCTTCAACGAGGAAGATTACCTCTAGTTTCAGAATTTTCTGACGAAACGATGAAGCAAAATATGCTACATGTAATTATACAAGAGATAGAGGAGGGTTCTTGTCCTATAGTCATCGAAAAGAACGGAGAATTGTTGTCGGTAAACGATTTTGATAAAGATGGTCTAAAATTCCATCTAGACTATATTATCAAAATTTGGAAACTTCAAAAACGATATTAGAATTTATACGAATATCGTTCTCTAAATGTCACAATCAAGTCTCTCATATTCAGCAGTTTATTGTCGTACTTTATATCGTGTTCATTAACGATATTTTGCAAAATAGTAATGATTCTATCTTCCTTCGATAGATATTCTTCAGAGATTATTGTCTTATATTCTTTCTTGTTATCCGATATGAATTTGATAAGACTTTGAACATTATTAATACCCGTCTGTTTAATTTTTTCTATAGATATTTTAGTTTTGGTAGATTCTATGGTGTCTGTTAATAGGCATCCAACATCGACATTCGACGTCAATTGTCTATAAATCAGAGTATAAATTTTAGAAATAACATTAGCAAATTGTTGTGCGTTGATGTCGTTATTCTGAAACAGTATGATTTTAGGTAGCATTTTCTTAACAAAGAGAACGTATTTATTGTTACTCAGTTGAACAGATGATATATCCAGATTACTAACGCATCTGATTCCATATACCAAACTTTCAGAAGAAATGGTGTACAATTGTTTGTATTCATTCAATGTCTCCTTTTCAGAAATTAGTTTAGAGTCGAATACTGCAATAATTTTCAAGAGATAGTTTTCATCAGATAAGATTTTATTTAGTGTAGATATGATAAAACTATTGTTTTGTTGGAGAACTTGATACGCCGCATTCTCTGTAGTCGACGCTCTCAAATGGGAAACAATCTCTATTATTTTTTTGGAATCGGATACTATATCTTCGGTATCTTGACGCAGTCTAGTATACATAGAGTTAAGAGAAATTAGAGTTTGTACATTAAGCAACATGTCTCTAAATGTGGCTACAAACTTTTCTTTTTCCACATCATCTAGTTTATTATATACCGATTTCACAACGGCACCAGATTTAAGGAACCAGAATGAAAAACTCTGATAACTACAATATTTCATCATAGTTACGATTTTATCATCTTCTATAGTTGGTGTGATAACACATACCTTTTTCTCCAAGACTGGAACCAACGTCATAAAAATGTTTAAATCAAAATCCATATCAACATCTGATGCGCTAAGACCAGTCTCGCGTTCAAGATTATCTTTACTAATGGTGACGAACTCATCGTATAGAACTCTAAGTTTGTCCATTATTTATTTACAGATTTAGTTGTTTAATTTATTTGTGCTCTTCCAGAGTTGGGATAGTATTTTTCTAACGTCGGTATTATATTATTAGGATCTACGTTCATATGTATCATAATATTAATCATCCACGTTTTGATAAATCTATCTTTAGCTTCTGAAATAACGTATTTAAACAAAGGAGAAAAATATTTAGTTACGGCATCAGACGCGATAACATTTTTTGTAAATGTAACGTATTTAGACGACAGATCTTCGTTAAAAAGTTTTCCATCTATGTAGAATCCATCGGTTGTTAACACCATTCCCGCGTCAGAGTGAATAGGAGTTTGAATAGTTTGTTTTGGAAATAGATCCTTCAATAACTTATAGTTGGGTGGGAAAAAATCGATTTTATCACTAGACTCTTTCTTTTTTACTATCATTACCTCATGAACTATTTCTTGAATGAGTATATGTATTTTCTTTCCTATATCGGTCGCGTTCATTGGAAAATATATCATGTCGTTAACTATAAGAATATTTTTATCCTCGTTTACAAACTGAATAATATCAGATATAGTTCGTAAACGAACTATATCATCACCAGCACAACATCTAACTATATGATATCCACTAGTTTCCTTTAGCCGTTTATTATCTTGTTCCATATTAGCAGTCATTCCATCATTTAAGAAGGCGTCAAAGATAATAGGGAGAAATGACATTTTGGATTCTGTTACGACTTTACCAAAATTAAGGATATACGGACTTACTATCTTTTTCTCAACGTCGATTTGATGAACACACGATGAAAATGTACTTCGATGAGATTGATCATGTAGAAAACAACAAGGGATACAATATTTCCGCATATCATGAAATATATTAAGAAATCCCACTTTATTATATTTCCCCAAAGGATCAATGCATGTAAACATTATACCGTTATCATTAATAAAGACTTCTTTCTCATCGGATCTGTAAAAGTTGTTACTGATTTTTTTCATTCCAGGATCTAGATAATTAATAATAATGGGTTTTCTATTCTTATTCTTTGTATTTTGACATATCCTAGACCAGTAAACAGTTTCCACTTTGGTAAAATCAGAAGACTTTTGAACGCTATTAAACATGGCATTAATGGCAATAACTAAAAATGTAAAATATTTTTCTATGTTAGGAATATGGTTTTTCACTTTAATAGATATATGGTTTTTTGCCAAAATGATAGATATTTTTTTATCCGATGATAGTAAAATATTATTAGTCGCCGTCTCTATAAAAATGAAGCTAGTCTCGATATCCAATTTTATTCTAGAATTGATAGGAGTCGCCAAATGTACCTTATACGTTATATCTCCCTTGATGCGTTCCATTTGTGTATCTATATCGGACACAAGATCTGTAAATAGTTTTACGTTATTAATCATCACGGTATCGCCATCGCTAGATAATGCTAATGTACTATCCAAGTCCCAAATGGAGAGATTTAACTGTTCATCGTTTAGAATAAAATGATTACCTGTCATATTAATAAAGTGTTCATCGTATCTAGATAACAACGACTTATAATTAATGTCCAAGTCTTGAACTCGCTGAATGATCTTTTTTAACCCAGTTAGTTTTAGATTGGTACGAAATATATTGTTAAACTTTGATTCTACAGTAATGTCCAAATCTAGTTGTGGAAATACTTCCATCAACATTGTTTCAAACTTGATAATATTATTATCTACATCTTCGTACGATCCAAATTCCGGAATAGATGTATCGCACGCTCTGGCCACCCAGATAACCAAAAAGTCACACGCTCCAGAATATACATTGTATAAAAAGCTATCGTTTTTTAGTAGTGTTTTTTTCTGAGTATATACGAAAGGATTAAAAATAGTATTATCAACGTAACTATATTCCAAATTATTCTTATGAGAATAGATAATAATATCGTCCTTAATATCTAACAAATTTCCTAAATATCCCTTTAATTGAGTCATTCGAAGCGTTAATAAAATATGTCTCTTAACTATTTCCGGCCGTTGTATATTTAAATGACTTCGTAAGAAATAATATATAGGCGACTTCTCATCTATGTAATCATATGGAGTGAGATATAGGGCTCGTTCTACCTCCTGCCCCTTACCCACCTGTAATACCAATTGCGGACTCACTATATATCGCATATTTATATCGTGGGGTAAAGTGAAAATCTACTACCGATGATGTAAGTCTTACAATGTTCGAACCAGTACCAGATCTTAATTTGGAGGCCTCCGTAGAACTAGGGGAGGTAAATATAGATCAAACAACACCTATGATAAAGGAGAATAGCGGTTTTATATCCCGTAGTAGACGTCTATTCGCCCATAGATCTAAGGATGATGAGAGAAAACTAGCACTACGATTCTTTTTACAAAGACTTTATTTTTTAGATCATAGAGAGATTCATTATTTGTTCAGATGCGTCGACGCTGTAAAAGACGTCACTATTACCAAAAAAAATAACATTATCGTGGCGCCTTATATAGCACTTTTAACTATCGCATCAAAAGGATGCAAACTTACAGAAACAATGATTGAAGCATTCTTTCCAGAACTATATAATGAACATAGTAAGAAATTCAAATTCAACTCTCAAGTATCCATCATCCAAGAAAAACTCGGATACCAGTCTGGAAACTATCACGTTTATGATTTTGAACCGTATTACTCTACAGTAGCTCTGGCTATTCGAGATGAACATTCATCTGGCATTTTTAATATCCGTCAAGAGAGTTATCTTGTAAGTTCATTATCTGAAATAACATATAGATTTTATCTAATTAATCTAAAATCTGATCTTGTTCAATGGAGTGCTAGTACGGGCGCTGTAATTAATCAAATGGTAAATACTGTATTGATTACAGTGTATGAAAAATTACAACTGGCCATAGAAAATGATTCACAATTTACATGTTCATTGGCTGTGGAATCAGAACTTCCAATAAAATTACTTAAAGATAGAAATGAATTATTTACAAAATTCATTAACGAGTTAAAAAAGACCAGTTCATTCAAGATAAGCAAACGTGATAAGGATACGCTATTAAAACATTTTACTTATGACTGGAGTTAGAATTTATAGACGACACATTTCGTTTATCATTGTTACTATTACTATCATTATTAGTATTCTTCTTGTCATCTTGTTCAGAAATATACAGCAATGCTATACCTAATACTAAATACATTATCATGCTTGCAATGGCTCTAACAACAACGAACCAAAATGAATTTGGTCGTAGCTTTTGTTCACAAAAATACATAAAGAAATGTCTACATAAATCTATGGCGCCATTGGCTACTTGAAATAGCGCCAGTCCTCCTACAGATTTTAATATAGCTGTATAACATGACATTTATTCATCATCAAAAGAGACAGAGTCACCATCTGTCATATTTAGATTTTTTTTCATGTGTTCAAAGTATCCTCTACTCATTTCATTATAATAGTTTATCATGCTTAGAATTTTAGGACGGATCAATGAGTAAGACTTGACTAGATCGTCAGTAGTAATTTGTGCATCATCTATTCTGCATCCGCTTCGTCGAATAATGTATAGCATCGCTTTGAGATTCTCCATAGCTATCAAGTCTTTATATAATGACATGGAAATATCTGTGAATGCTTTATACTTCTCCAACATCGATGCCTTAACATCATCACATACTTTAGCATTGAAAATACGTTCTATTGTGTAGATGGATGTAGCAAGATTTTTAAACAACAATGCCATCTTACATGATGATTGTCTCAAGTCTCCAATCGTTTGTTTAGAACGATTAGCTACAGAGTCCAATGCTTGGCTAACTAGCATATTATTATCTTTAGAAATTGTATTCTTCAATGAGGCGTTTATCATATCTGTGATTTCGTTAGTCATATTACAGTCTGACTGGGTTGTAATGTTATCCAACATATCACCTATGGATACGGTACACGTACCAGCATTTGTAATAATCCTATCTAAGATGTTGTATGGCATTGCGCAGAAAATATCTTCTCCTGTAATATCTCCACTCTCGATAAATCTACTCAGATTATTCTTAAATGCCTTATTCTCTGGAGAAAAGATATCAGTGTCCATCATTTCATTAATAGTATACGCAGAAAAGATACCACGAGTATCAATTCTATCCAAGATACTTATCGGTTCCGAGTCACAGATAATTGTTTCCTCTCCTTCGGGAGATCCTGCATAGAAATATCTAGGACAATAGTTTCTATACTGTCTGTAACTCTGATAATCTCTAAAGTCACTAACTGATACCATGAAATTGAGAAGATCAAACGCTGAAGTAATCAATTTTTCTGCCTCGTTTTTACTACAACTAGTTTTCATCAATGTAGTGACGATGTATTGTTTAGTTACTCTTGGTCTAATACTGATGATAGAGATATTATTGCTTCCCATAATGGATCTTCTAGTAGTCACCTTAAAGCCCATTGATGCGAATAGCAGATAGATAAAGTCTTGGTATGACTCCTTTCTAATATAGTACGGACTACCTTTGTCACCCAACTTTATACCCACATAAGCCATAACAACCTCTTTAATAGCCGTTTCATGAGGTTTATCAGCCATGAGCCTGAGTAGTTGAAAGAATCGCATGAATCCCGTCTCAGAAAGTCCTATATGCATGATAGATTTATCTTTCCTGGGAAACTCTCGTATAGTTATAGATGAAATACTCTTCAAAGTTTCTGAAATAAGATTAGTAACAGTCTTACCTCCGACTACTCTGGGTAACAAACATACTCTAATAGGTGTTTTCTCTGCGGAGATAATATCAGAAAGGATAGAGCAATAAGTAGTATTATTGTGATTATAAAGACCGAATACATAACAGGTAGAATTTATAAACATCATGTCCTGAAGGGTTTTAGACTTGTATTCCTCGTAATCTATACCGTCCCAAAACATGGATTTGGTAACTTTGATAGCCGTAGATCTTTGTTCCTTCGCTAACAGGTTAAAGAAATTAATAAAGAATTTGTTGTTTCTATTTATGTCCACAAATTGCACGTTTGGAAGCGCCACGGTTACATTCACTGCAGCATTTTGAGGATCGCGAGTATGAAGTACGATGTTATTGTTTACTGGTATATCTGGAAAGAAATCTACCAGTCTAGGAATAAGAGATTGATATCGCATAGAAATAGTAAAGTTTATAATCTCATCATTGAAGATTACTCTGTTACCATTGTAATAAATTGGTACTCTATCATAATCATCGACAAAGTACTGTTCATACATGATGAGATGTTTATATGTTGGCATAGTAGTGAGATCGACGTTTGGTAATGGCAATGTATTAAGATTAACTCCATAATGTCTAGCAGCATCTGCGATGTTATAAGTGATGTCAAAGCGGGGTTGATCTTGTGCTGTTATATATTGTCTAACACCTATAAGATTATCAAAATCTTGTCTGCTTAATACACCGTTAACAATTTTTGCCTTGAATTCTTTTATTGGTGCATTAATAACATCCTTATAGAGGATGTTAAACAAATAAGTATTATCAAAGTTAAGATCTGGGTATTTCTTTTCTGCTAGAACATCCATTGAGTCGGAGCCATCTGGTTTAATATAACCACCGATAAATCTAGCTCTGTATTCTGTATCCGTCAATCTAATATTAAGAAGGTGTTGAGTGAAAGGTGGAAGATCGTAAAAGCTGTGAGTATTAATAATAGGGTTAGTTTCCGAACTAATGTTAATTGGATGATTAATAATATTTATATTTCCAGCGTTAAGTGTAACATTAAACAGTTTTAATTCACGTGACGTGGTATCAATTAAATAATTAATGCCCAATTTGGATATAGTAGCCTGAAGCTCATCTTGTTTAGTTACGGATCCTAATGAGTTATTAAGAAATACATCGAACGGATGAACGAAGGTTGTTTTAAGTTGGTCACATACTTTGTAATCTAGACATAGATGTGGAAGAACGGTAGAAACTATACGAAATAGATATTCAGAGTCCTCTAATTGATCAAGAGTAACTATTGACTTAATAGGCATCATTTATTTAGTATTAAATGACGACCGTACCAGTGACAGATATACAAAACGACTTAATTACAGAGTTTTCAGAAGATAATTATCCATCTAACAAAAATTATGAAATAACTCTTCGCCAAATGTCTATTCTAACTCACGTTAACAACGTGGTAGATAGAGAACATAATGCCGCCGTAGTGTCATCTCCAGAGGAAATATCATCACAACTTAATGAAGATCTATTTCCAGATGATGATTCACCGGCCACTATTATCGAACGAGTACAACCTCATACTACTATTATTGACGATACGCCACCTCCTACTTTTCGTAGAGAGTTATTGATATCGGAACAACGTCAACAACGAGAAAAAAGATTTAATATTACAGTATCAAAAAATTCTGAAGCAATAATGGAATCTAGATCTATGATAACTTCTATGCCAACACAAACACCATCCTTGGGAGTAGTTTATGATAAAGATAAAAGAATTCAGATGCTAGAGGATGAAGTGGTTAATCTTAGAAATCAACGATCTAATACAAAATCATCTGATAATTTAGATAATTTTACCAGAATACTATTTGGTAAGACTCCGTATAAATCAACCGAAGTTAATAAGCGTATAGCCATCGTTAATTATGCAAATTTGAACGGGTCCCCCTTATCAGTCGAGGACTTGGATGTCTGTTCGGAGGATGAAATAGATAGAATCTATAAAACGATTAAACAATATCACGAAAGTAGAAAACGAAAAATTATCGTCACTAACGTGATTATTATTGTCATAAACATTATTGAGCAGGCATTGCTAAAACTCGGATTTGAAGAAATCAAAGGACTGAGTACCGATATCACTTCAGAAATTATCGATGTGGAGATCGGAGATGACTGCGATGCTGTAGCATCAAAACTAGGAATCGGTAACAGTCCGGTTCTTAATATTGTATTGTTTATACTCAAGATATTCGTTAAACGAATTAAAATTATTTAATTTAATACATTCCCATATCCAGACAACAATCGTCTGGATTAATCTGTTCCTGTCGTCTCATACCGGACGACATATTAATCTTTTTATTAGTGGGCATCTTTTTAGATGGTTTCTTTTTCCCAGCATTAACTGATTCGATACCTAGAAGATCGTGATTGATTTCTCCGACCATTCCACGAACTTCTAATTGGCCGTCTCTAACGGTACCATAAACTATTTTACCAGCATTAGTAACAGCTTGGACAATCTGACCATCCATTGCGTTGAATGATGTAGTTGCTGTTGTTCTACGTCTAGGAGCACCAGAGGTATTTTTAGAGCTCTTGGATGTTGATGTAGAAGACGAGGATTTTGATTTTGGTTTACATGTAATACATTTTGAACTCTTTGATTTTGTATCACATGCACCGGCAGTCACATCTGTTTGAGAATTAAGATTATTGTTGCCTCCTTTGACGGCTGCATCTCCACCGATCTGCGCTAGTAGATTTTTAAGCTGTGGTGTAATCTTATTAACTGTTTCAATATAATCATCGTAACTACTTCTAACGGCTAAATTTTTTTTATCCGCCATTTAGAAGCTAAAAATATTTTTATTTATGCAGAAGATTTAACTAGATTATACAATGAACTAATATGATCCTTTTCTAGATTATTTACGAACTTGGTATTTCTTGTTTCTGGAGGAGGAGAATTTAAATTCGGACTTGGATTCGGATTTTGTGGGTTCTTGATCTTATTATACAGCGTGTATAGGATGGTGACGGTAACTGCTACACAAATACCGATCAACAGAAGAATACCAATCATTTATTGACAATAACTTCACTATGATCAAGTATGTAATAATCATCTTTTCACTAAGTAAGTAGTAATAATGATTCAACAATGACACGATATATGGACGATAATAATTTAGTTCATGGAAATATCGCTATGATTGGTGTGAATGACTCCGCTAACTCTGTGGGGTGCACAGTGCTTTCCCCACATAGAATAAATTAGCATTCCGACTGTGATAATAATACCAAGTATAAACGCCATAATACTCAATACTTTCCATGTACGAGTGGGACTGGTAGACTTACTAAAGTCAATAAAGGCGAAGATACACGAAAGAATCAAAAGAATGATTCCAGCGATTAGCACGCCAGAAAAATAATTTCCAATCATAAGCATCATGTCCATTTAACTAATAAAAATTTTAAATCGCCGAATAAACAAAGTGGAATATAAACCATATAAAAACAATAGTTTGTACTGCAAAAATAATATCTATTTTTGTTTTCGAAGATATGGTAAAATTAAATAGTAGTACACAGCATGTTATAACTAACAGCAGCAACGGCTCGTAATTACTTATCATTTACTAGACGAAAAGGTGGTGGGATATTTTCTTGCTCAAATAATACGAATATATCACCCATCCATTTTATACGATGTTTATATACTCTAATCTTTAATAGATCTATAGATGACGGGTTTACCAATAATATAGATTTTATCGATTCATCTAATTTAAACCCTTCCTTAAACGTGAATGATCTATTATCTGGCATAATGATGACCCTACCTGATGAATCTGACAATGTACTGGGCCATGTAGAATAAATTATCAACGAATTATCGTCTACGAACATTTATATCATTTGTTTTAATTTTAGGACGTGAATAAATAGATATAAAATAGAAAATAACAGATATTACAACCAGTGTTATGGACGCACCCAACCATGTAGGCAGTTTTATTTTATCGTTTACTACAGGTTCTCCTGGATGTACGTCACCAACTGCAGACGTAGTTCTAGTACAATTAGACGTAAGTTCCGCTTGGGAATTTTTTAACGCTAAAGAGTTAACGTTGATCGTACACCCAACGTATTTACATCTAGTTCTTTGAACATCTTGATTATAATATAACCATTTTCTATCTCTAGATTCGTCAGTGCACTCATGTAACCAACATACCCTAGGTCCTAAATATTTATCTCCGGAATTAGATTTTGGATAATTCGCGCACCAACAATTTCTATTTCCTTTATGGTCGTTACAAAAGACGTATAATGCCGTATCCCCAAAAGTAAAATAATCAGGACGAATAATTCTAATAAACTCAGAACAATATCTCGCATCCATATGTTTGGAGCAAATATCGGAATAAGTAGACATAGCCGGTTTCCGTTTTACACGTAACCATTCTAAACAATTGGGGTTTCCAGGATCGTTTCTACAAAAACCAGTCATGAAATCGTCACAATGTTCTGTCTTGTAATTATTATTAAATATTTTTGGACAGTGTTTGGTATTTGTCTTAGAACAACATTTTGCCACGCTATCACTATCACCCAGGAGATAATCCTTTTTTATAAAATGACATCGTTGCCCGGATGCTATATAATCAGTAGCATATTTTAAATCCTTAATATATTCAGGAGTTACCTCGTTCTGATAATAGATTAATGATCCAGGACGAAATTTGAAAGAACTACATGGTTCTCCATGAATTAATACATATTGTTTAGCAAATTCAGGAACTATAAAACTACTACAATGATCTATCGACATACCATCTATCAAACAAAATTTGGGTTTAATTTCTCCTGGAGACGTTTCATAATAATACATATAACTTTCTTCGGCAAACCTAACAGCTCTATTATATTCAGGATAATTAAAATCTAATACCATATATTTGTCTCGTATATCTGCTATTCCTGTCTCTATTTTGATTCTATTAAGAGTAACAGCTGCCCCCATTCTTAATAATCATCAGTATTTAAACTGTTAAATGTTGGTATATCAACATCTATCTTATTTCCCGCAGTATAAGGTTTGTTGCAGGTATACTGTTCAGGAATGGGTACATTTATACTTCTTTTATAGTCCTGTCTTTCGATGTTCATCACAAATGCAAAGAACAGAATAAACAAAATAATGTAAGAAATAATATTAAATATCTGTGAATTCGTAAATACATTGATTGCCATAATAATTACAGCAGCTACAATACACACAATAGACATTCCCACAGTGTTGCCATTACCTCCACGATACATTTGAGTTACTAAGCAATAGGTAATAACTAAGCTAGTAAGAGGCAATAGAAAAGATGAGATAAATATCATCAATATAGAGATTAGAGGAGGGCTATATAGAGCCAAGACGAACAAAATCAAACCGAGTAACGTTCTAACATCATTATTTTTGAAGATTCCCAAATAATCATTCATTATTCCTCCATAATCGTTTTGCATCATACCCCCATCTTTAGGCATAAACGATTGCTGCTGTTCCTCTGTAAATAAATCTTTATCAAGCACTCCAGCACCCGCAGAGAAGTCATCAAGCATATTGTAATATCTTAAATAACTCATTTATATATTAAAAAATGTCACTATTAAAGATGGAGTATAATCTTTATGCCGAACTAAAAAAAATGACTTGTGGTCAGACCATAAGTCTTTTTAATGAAGACGGCGATTTCGTAGAAGTTGAACCAGGATCATCCTTTAAGTTTCTAATACCTAAGGGATTTTACTCCTCTCCTTGTGTAAAGACGAGTCTAGTATTCGAGACATTAACAACGACCGATAATAAAATTACTAGTATCAATCCAACAAATGCGCCAAAGTTATATCCTCTTCAACGCAAAGTCGTATCTGAAGTAGTTTCTAATATGAGGAAAATGATCGAATTAAAACGTCCTCTATACATCACTCTTCACTTGGCATGTGGATTTGGTAAGACTATTACCACGTGTTATCTTATGACCACACACGGCAGAAAAACCATCATTTGCGTACCCAATAAAATGTTAATACATCAATGGAAGACACAGGTAGAGGCAGTCGGATTGGAACATAAGATATCTATAGATGGAGTTAGTAGTCTATTAAAGGAACTAAAGACTCAAAGTCCGGATGTATTAATCGTAGTCAGTAGACATCTGACAAACGATGCATTTTGTAAATATATCAATAAGCATTATGATTTGTTTATCTTGGATGAATCACATACGTATAATCTGATGAACAATACAGCAGTTACAAGATTTTTAGCGTATTATCCTCCGATGATGTGTTATTTTTTAACTGCTACACCTAGACCAGCTAACCGAATTTATTGTAATAGTATTATTAATATTGCCAAGTTATCCGATCTAAAAAAAACTATCTATATAGTAGATAGTTTTTTTGAGCCATATTCCACAGACAATATTAGAAATATGGTAAAACGACTAGATGGACCATCTAATAAATATCATATATATACCGAGAAGTTATTATCTGTAGACGAGCCTAGAAACCAACTTATTCTTGATACCCTGGTAGAAGAATTCAAGTCAGGAACTATTAATAGAATTTTAGTTATTACTAAACTACGTGAACATATGGTATTCTTCTACAAACGATTATTAGATCTTTTCGGAGCAGAGGTTGTATTTATAGGAGACGCCCAAAATAGACGTACTCCAGATATGGTCAAATCGATTAAGGAACTAAATAGATTTATATTCGTATCCACCTTATTTTATTCCGGCACTGGTTTAGATATTCCGAGTTTGGATTCTTTGTTCATTTGCTCGGCAGTAATCAACAATATGCAAATAGAGCAATTACTAGGGAGGGTATGTCGAGAAACAGAACTATTAGATAGGACGGTATATGTATTTCCTAACACATCCATCAAAAAAATAAAGTACATGATAGGAAATTTCGTGCAACGAATTATTAGTCTGTCTGTAGATAAACTCGGATTTAAACAAGAAAGTTATCAGAAACATCAGGAATCTGAACCCGCTTCCGTACCAACATCCTCCAGAGAAGAACGTGTATTAAATAGAATATTTAACTCGCAAAATCGTTAAGAAGTTTAAGAGACGATCCACATGCTGAGCAGGCCAGTGTATTACCCCTCATAGTATTAATATAATCCAATGATACTTTTGTGATGTCGGAAATCTTAACCAATTTAGACTGACAGGCAGAACACGTCATACAATCATCATCGTCATCGATAACTGTAGTCTTGGGCTTCTTTTTGCGACTCTTCATTCCGGAACGCATATTGGTGCTATCCATTTAGGTAGTAAAAAATAAGTCAGAATATGCCCTATAACACGATCGTGCAAAACCTGGTATATCGTCTCTATCTTTATCACAATATAGTGTATCAACATCTTTATTATTGACCTCGTTTATCTTGGAACATGGAATGGGAACATTTTTGTTAACGGCCACCTTTGCCTTAATTCCAGATGTTGTAAAATTATAACTAAACAGTCTATCATCGACACAAATGAAATTCTTGTTTAGACGTTTGTAGTTTACGTATGCGGCTCGTTCTCGTCTCATTTTTTCAGATATTGCAGGTACTATAATATTAAAAATAAGAATGAAATAACATAGGATTAAAAATAAAGTTATCATGACTTCTAGTGCTGATTTAACTAACTTAAAAGAATTACTTAGTCTGTACAAAAGTTTGAGATTTTCAGATTCTGTGGCTATAGAGAAGTATAATTCTTTGGTAGAATGGGGAACATCTACTTACTGGAAAATAGGCGTACAAAAGGTAACTAATGTCGAGACGTCCATATCTGATTATTATGATGAGGTAAAAAATAAACCGTTTAATATTGATCCGGGGTATTATATTTTCTTACCAGTATATTTTGGAAGCGTCTTTATTTATTCAAAGGGTAAAAATATGGTAGAACTTGGATCTGGAAACTCTTTTCAAATACCGGATGAGATTCGAAGTGCGTGTAACAAAGTATTAGATAGTGATAACGGAATAGACTTTCTGAGATTTGTTTTGTTAAACAATAGATGGATAATGGAAGACGCTATATCAAAATACCAGTCTCCAGTTAATATATTTAAACTAGCTAGTGAGTACGGATTAAACATACCCAACTATTTAGAAATTGAAATAGAGGAAGACACATTATTTGACGATGAGTTATACTCTATTATGGAACGCTCTTTCGATGATACATTTCCAAAAATATCTATATCGTATATTAAGTTGGGAGAACTTAAGCGGCAAGTTGTAGACTTTTTCAAATTCTCATTCATGTATATTGAGTCAATCAAGGTAGATCGTATAGGAGATAATATTTTTATTCCTAGCGTTATAACAAAATCAGGAAAAAAGATATTAGTAAAAGATGTAGACCATTTAATACGATCCAAGGTTAGAGAACATACATTTGTAAAAGTAAAAAAGAAAAACACATTTTCCATTTTATACGACTATGATGGGAACGGAACAGAAACTAGAGGAGAAGTAATAAAACGAATTATAGACACTATAGGACGAGACTATTATGTTAATGGAAAGTATTTCTCTAAGGTTGGTATTGCAGGCTTAAAGCAATTGACTAATAAATTAGATATTAATGAGTGTGCAACTGTCGATGAGTTAGTTGATGAGATTAATAAATCCGGAACTGTAAAACGAAAAATAAAAAACCAATCAGTATTTGATTTAAGCAGAGAATGTTTGGGATATCCAGAAGCGGATTTTATAACGTTAGTTAATAACATGCGGTTCAAAATAGAAAATTGTAAGGTTGTAAATTTCAATATTGAAAATACTAATTGTTTAAATAACCCGAGTATTGAAACTATATATGGAAACTTCAACCAGTTCGTCTCAATCTTTAATACCGTTACCGATGTCAAAAAAAGATTATTCGAGTGAAATAATATGCGCCTTTGATATAGGTGCAAAAAATCCTGCCAGAACTGTTTTAGAAGTCAAGGATAACTCCGTTAGGGTATTGGATATATCAAAATTAGACTGGAGTTCTGATTGGGAAAGGCGCATAGCTCAAGATTTGTCACAATATGAATACACTACAGTTCTTCTAGAACGTCAGCCTAGAAGGTCACCGTACGTCAAATTTATCTATTTTATTAAAGGCTTTTTATATCATACATCTGCTGCCAAAGTTATTTGCGTCTCACCTGTCATGTCTGGTAATTCATATAGAGATCGAAAAAAGAGATCTGTTGAAGCATTTCTTGATTGGATGGACACATTCGGATTGCGAGACTCCGTTCCGGATAGACGCAAATTAGACGATGTAGCGGATAGTTTCAATTTGGCTATGAGATACGTATTAGATAAATGGAATACTAATTATACACCTTATAATAGGTGTAAATATAGAAATTACATAAAAAAAATGTAATAACGTTAGTAACGCCATTATGGATAATCTATTTACCTTTCTACATGAAATAGAAGATAGATATGCCAGAACTATTTTTAACTTTCATCTAATAAGTTGTGATGAAATAGGAGATATATATGGTCTTATGAAAGAACGCATTTCCTCAGAGGATATGTTTGACAATATAGTATATAATAAAGATATACATCCTGCCATTAAGAAACTAGTTTATTGCGACATCCAACTTACTAAACATATTATTAATCAGAATACGTATCCGGTATTTAACGATTCTTCACAAGTGAAATGTTGTCATTATTTCGATATAAACTCAAATAATAGCAATATTAGCTCTCGTACAGTAGAGATATTTGAGAGTGAAAAGTCATCTCTTGTATCATATATTAAAACTACCAATAAGAAGAGAAAGGTCAATTACGGCGAAATAAAGAAAACTGTACATGGAGGCACTAATGCAAATTACTTTTCCGGTAAAAAGTCTGATGAGTATCTGAGCACTACAGTCAGGTCCAACATTAATCAACCTTGGATCAAAACCATTTCTAAGAGAATGAGAGTAGATATCATTAATCACTCTATAGTAACGCGTGGAAAAAGCTCTATATTACAAACTATAGAAATTATTTTTACTAATAGAACATGTGTGAAAATATTCAAGGATTCTACTATGCACATTATTCTATCCAAGGACAAGGATGAAAAGGGATGTATAAACATGATTGATAAATTATTCTATGTATATTATAATTTATTTCTGTTGTTCGAGGATATCATCCAAAACGATTACTTTAAAGAAGTAGCTAATGTTGTAAACCATGTACTCATGGCTACGGCATTAGATGAGAAATTATTCCTAATTAAGAAAATGGCTGAACACGATGTTTATGGAGTTAGCAATTTCAAAATAGGGATGTTTAACCTGACATTTATTAAGTCGTTGGATCATACCGTTTTCCCCTCTCTGTTAGATGAGGATAGCAAAATAAAGTTTTTTAAGGGGAAAAAGCTCAATATTGTAGCATTACGATCTCTGGAGGATTGTACAAATTACGTGACTAAATCCGAGAATATGATAGAAATGATGAAGGAAAGATCGACTATTTTAAATAGCATAGATATAGAAACGGAATCGGTAGATCGTCTAAAAGAATTGCTTCTAAAATGAAAAAAAACACTGATTCAGAAATGGATCAACGACTCGGGTATAAGTTTTTGGTGCCTGATCCTAAAGCCGGAGTTTTTTATAGACCGTTACATTTCCAATATGTATCGTATTCTAATTTTATATTGCATCGATTGCATGAAATCTTGACCGTCAAGCGGCCACTCTTATCGTTTAAGAATAATACAGAACGAATTATGATAGAAATTAGCAATGTTAAAGTGACTCCTCCAGATTACTCACCTATAATTGCGAGTATTAAAGGTAAGAGTTATGACGCATTAGCCACGTTCACTGTAAATATCTTTAAAGAGGTAATGACCAAAGAGGGTATATCCATCACTAAAATAAGTAGTTATGAGGGAAAAGATTCTCATTTGATAAAAATTCCGCTACTAATAGGATATGGGAATAAAAATCCACTTGATACAGCCAAGTATCTTGTTCCTAATGTCATAGGTGGAGTCTTTATCAATAAACAATCTGTCGAAAAAGTAGGAATTAATCTAGTAGAAAAGATTACAACATGGCCAAAATTTAGGGTTGTTAAGCCAAACTCATTCACTTTCTCGTTTTCCTCCGTATCCCCTCCTAATGTATTACCGACAAGATATCGCCATTACAAGATATCTCTGGATATATCACAATTGGAAGCGTCGAATATATCATCGACAAAGACATTTATAACGGTCAATATTGTTTTGCTGTCTCAATATTTATCTAGAGTGAGTCTAGAATTCATTAGACGTAGTTTATCATACGATATGCCTCCAGAAGTTGTCTATCTAGTAAACGCGATAATAGATAGTGCTAAACGACTTACCGAATCTATTACTGACTTTAATATTGATACATACATTAATGACCTGGTGGAAGCTGAACACATTAAACAAAAATCTCAGTTAACGATTAACGAGTTTAAATATGAAATGCTGCATAACTTTTTACCTCATATGAACTATACACCCGATCAACTAAAGGGATTTTATATGATATCTTTACTAAGAAAGTTTCTCTACTGTATCTACCACACTTCTAGATATCCAGATAGAGATTCGATGGTTTGTCATCGCATCCTAACGTACGGCAAATATTTTGAGACGTTAGCACATGATGAATTAGAGAATTACATAGGTAACATCCGAAACGATATCATGAACAATCACAAGAACAGAGGCACTTACGCAGTAAACATTCATGTACTAACAACTCCTGGACTTAATCATGCATTTTCTAGTCTATTGAGTGGAAAGTTCAAAAAGTCAGACGGTAGTTATCGAACACATCCTCACTATTCATGGATGCAGAATATTTCTATTCCTAGAAGTGTTGGATTTTATCCGGATCAAGTAAAGATTTCAAAGATGTTTTCTGTCAGAAAATACCATCCAAGCCAATATCTTTACTTTTGTTCATCAGACGTTCCGGAAAGAGGTCCTCAGGTAGGTTTAGTATCTCAATTGTCTGTCTTGAGTTCCATTACAAATATACTAACGTCTGAGTATTTGGATTTGGAAAAGAAAATTTGTGAGTATATCAGATCATATTATAAAGATGATATAAGTTACTTTGAAACAGGATTTCCAATCACTATAGAAAATGCTCTAGTCGCATCTCTTAATCCAAATATGATATGTGATTTTGTAACTGACTTTAGACGTAGAAAACGGATGGGATTCTTCGGTAACTTGGAGGTAGGTATTACTTTAGTTAGGGATCACATGAATGAAATTCGCATTAATATTGGAGCAGGAAGATTAGTCAGACCATTCTTGGTTGTGGATAACGGAGAGCTCATGATGGATGTGTGTCCGGAGTTAGAAAGCAGATTAGACGACATGACATTCTCTGACATTCAGAAAGAGTTTCCACATGTCATCGAAATGGTAGATATAGAACAATTTACTTTTAGTAACGTATGTGAATCGGTTCAAAAATTTAGAATGATGTCAAAGGATGAAAGAAAGCAATACGATTTATGTGACTTTCCTGCCGAATTTAGAGATGGATATGTAGCATCTTCACTAGTGGGAATCAATCACAATTCTGGACCCAGAGCTATTCTTGGATGTGCTCAAGCTAAACAAGCTATCTCTTGTCTGAGTTCGGATATACGAAATAAAATAGACAATGGAATTCATTTGATGTATCCAGAGAGGCCAATTGTGATTAGTAAGGCTTTAGAAACTTCAAAGATTGCGGCTAATTGCTTCGGACAACATGTTACTATAGCATTAATGTCGTACAAAGGTATCAATCAAGAGGATGGAATTATCATCAAAAAACAATTTATTCAGAGAGGCGGTCTCGATATTGTTACAGCCAAGAAACATCAAGTAGAAATTCCATTGGAAAACTTTAATAACAAAGAAAGAGATAGGTCTAACGCCTATTCGAAATTAGAAAGTAATGGATTAGTTAGACTGAATGCTTTCTTGGAATCCGGAGACGCTATGGCAAGAAATATCTCATCAAGAACTCTTGAAGATGATTTTGCTAGAGATAATCAGATTAGCTTTGATGTTTCCGAGAAATATACAGATATGTACAAATCTCGCGTTGAACGAGTACAAGTAGAACTTACTGACAAAGTTAAGGTGCGAGTATTAACCATGAAAGAAAGAAGACCCATTCTAGGAGACAAATTTACTACTAGAACGAGTCAAAAGGGAACAGTCGCGTATATCGCAGATGAAACGGAACTTCCGTACGACGAAAATGGTATCACACCAGATGTCATTATTAATTCTACATCCATCTTCTCTAGAAAAACTATATCTATGTTGATAGAAGTTATTTTAACAGCCGCATATTCTACTAAGCCGTACAACAATAAGGGAGAAAACCGACCTGTCTGTTTTCCTAGTAGTAACGAAACATCTATCGATGCATATATGCAATTCGCTAAACAATGTTATGAGTATTCAAATCCGAAATTGTCCGAGGAAGAATTATCGGATAAAATCTTTTGTGAAAAGATTCTCTATGATCCTGAAACGGATAAGCCTTATGAATCCAAAGTATTTTTTGGACCAATTTATTACTTGCGTCTGAGACATTTAACTCAGGACAAGGCAACCGTTAGATGTAGAGGTAAAAAGACGAAGCTCATTAGACAAGCGAATGAGGGACGAAAACGTGGAGGAGGTATCAAGTTTGGAGAAATGGAGAGAGACTGTTTAATAGCACATGGTGCAGCCAATACTATTACAGAAGTTTTAAAAGACTCAGAAGAGGATTATCAAGATGTGTATATTTGTGAAAATTGTGGAGACATAGCAGCACAAATCAAAAGTATTAATACATGTCTTAGATGTTCAAAACTTAATCTCTCTCCTCTCTTAACAAAAATTGATACCACGCACGTATCTAAAGTATTTCTTACTCAAATGAACGCCAGAGGCGTAAAAGTTAAATTAGATTTCGAACGAAGGCCTCCTTCGTTTTATAAACCATTAGATAAAGTTGATCTTAAACCGTCTTTTCTGGTATAATATTGTTTAGTAGATACTCATCAAGATAAGCTAATTCACTAAACATATTATCGGATTCGGTATTGTTACTCGAGAATAGAGTTCGTTATGCTCCTGATATTCGGAAATCTGTGGAGTTTCAGGTTTTGGTGGAAGTGTAACTGCTACTTGGTGGGATACTGAAGGATATTTCAGAGAGTTGTGGATGTTCGGGTTCGACATCCACCGATGGTGTCACGCCACTAATCGGTTCGGTAACGTCTGTGGATGGAGGTGCTACTTCTACAGAACCTGTAGCCTCAGTTGTCAACGGAGATACATATTCAATGCGCGGAAATGTATAATTTGGTAATGGTTTCTCATGTGGATCTTAAGAAGAAGAGGTAAGATATCTACGAAAGATACCGATCACGTTTCTAGTTCTCTTTTGTAGAACTTTAACTTTTTCTTTCTCAGCATCTAGTTGATATTCCGACCTCTTCACGTTTCGCATGGGTTACCTCCGCAGTTTTTACAAGCGATTTCACGTTCCAGATCACGTTCAGCCTTCATACGTCTCTCCCTCTCTCTATCGAGTTTATCAGAGCAGTCTTTCTGAAGGCGATCGAACTCCATAAATTTCTCCAACGCTTTGATTGTTTCCATAGATTTCCGAAGTTTAGCTTCTAGGACGGCGATTCTTTTTTTTTTTTTTTTTTTCGAATTCACGGGGTACAACCGTTTCCATTACCACCATCTCTATGTTTCTTTTCTAGATCGGCAATCTTTCTCAATCTTTCTCAACATTTCATCCCCATACCTTTTCATTCCTCGAGTCTATTGTCGTCGAAATATCGTTCCAGCTCCTTTTCGACCTCAATAACTTTAGCACGTTGTTTCATCAAGCTCTCTCTTGTAGTACTATCATTTTTATCTGATTCCCTGACACGTTTAAGATCTTCATGTAATTGAGTCAGCTCTTGACGCAATCTCTTAACTAACTTCCTCTCTTGCTTCTTCGTCATAGTACTTACAATCACTATGGGATCCATTGTTACCACGTCTGTACTCGACGAGCTCACGTTTAAGAGATTCAATTTCCAGTTTGTATCGGTCCATGTCTCCATTGCTACACCACCATTAGATTTACAGGCTGCTAGTTGTCGTTCGAGATCAGAAATACGTGTTTTCTTGGAATGGATTTCGTCGATGTACTTGTCATGATTGGCATCGAAACACTTATTAAGTTCTTTTTTTCAATTCTACGATTTTATTTCTTTCGCGAGTCAATTCCCTCCTGTAGTAACTATCAGTTTTGTCAGATTCACGCTCTCTACGTAGACTTTCTTGTAAGTTACTAATTTGTTCCCTGGCATTACCGAGTTCAGTTTTATATGCCGAATAGAGTTCTGATTCATCCTTTGAGAAGATCTCTAGCGATCGTTCAAGATCCCTGATTCTAGTCTTTAGCCTATTTACCTCCTCAGAAGATGCTCCGTTACCGTTTTTACAATCGTTAAGATGTCTATCAAGATCCATGATTCTATCTCTTTTCCATATCAGCATTGATTTCATTATTACGTTCGCAGTCGTTCAACTGTATTTCAAGATCTGAGATTCTAGATTGTAATCTCTGTAGCATTTCCACGGCATTCACTCAGTTGTCTTTCAAGATCTGAGATTCTAGATTGGAGTCTGCTAATCTCTGTAAGATTTCCTCCTCCGCTCTCGATGCAGTCGGTCAACTTATTCTCTAGTTCTCTAATACGCGAACGCAGTGCATCAACTTCTTGTGTGTCTTCTTGATTGCGTGTGCATTCATCGAGTCTAGATTCGAGATCTCTAACGTGACGTCGTTCTTCCTCAAGTTCTCTGTGTACTACAGAAAGCGTGTCCCTATCTTGTTGATATTTAGCAATTTCTGATTCTAGAGTACTGATTCTACTCACGTATGTACTAATAGTTGTCTTAGCCTTATCAAGATCCTCCTTGTATTTGTCACATTCCTTGATATCCATACGAAGTCTGGACAGTTCCCATTCGACATTACGACGTTTATCGATTTCAGCTCGGAGATCGTCGTCGCGTTGTTTTAGCCACATACGACTAAGTTCAAGTTCTCGTTGACAAGATCCATCTACTTTTCCATCCCTAATAGTATCCAGTTCCTTTTCTAGTTCTGACCGCATTTCTCGTTCCATATCAAGAGATTCTCTCAATTCTCGTATAGTCTTCTTATCAATTTCTGATGAATCTGAACCATCATCTGTCCCATTTTGTTGCATATCCCTGAGTTCTTTGATCTCTGTTGTAAGTCTGTCGATTCTTTCGGTTTTATAAACAGAATCCCTTTCCAAAGTCCTAATCTTACTGAGTTTATCATTAAGTTCTTCATTCAATTCAGTGAGTTTTCTCTTGGCTTCTTCCAAGTCTGTTTTAAACTCTCCATCATTTCCGCATTCTTCCTCGCATTTATCTAACCATTCAATTAGTTTATTAATAACTAGTTGGTAATCAGCGATTCCTATAGCCGTTCTTGTATTTGTGGGAACATAATTAGGATCTTCTAATGGATTGTATGGCTTGATAGCATCATCTTTATCATTATTAGGTGGGGGATGGACAACCTTAATTGGTTGGTCCTCCTTATCTCCTCCAGTAGCATGTGGTTCTTCAATACCAGTATTAGTAATAGGCTTAGACAAATGCTTGTCGTACGCGGGCACTTCCTCATCCATCAAGTATTTATAATCGGGTTCTGTTTCAGAATATTCTTTTCTAAGAGACGCGACTTCAGGAGTTAGTAGAAGAACTCTGTTTCTGTATCTATCAACGCTGGAATCGATACTCAAGTTAAGGATAGCGAATACCTCATCGTCATCATCCGTATCTTCTGAAACGCCATCATATGACATTTCATGAAGTCTAACGTATTGATAAACAGAATCAGATTTAGTATTAAACAGATCCTTGACCTTTTTAGTAAATGCATATGTATATTTTAGATCTCCAGATTTCATAATATGATCGCATGCCTTAAATGTCAATGCTTCCATGATATAGTCTGGAACACTAATGGGTGACGAAAAAGATACAGCACCATATGCTACGTTGATAAATAGATCTGAACCACTAAGTAGATAATGATTAATGTTAAGGAAGAGGAAATATTCAGTATATAGATATGCCTTAGCATCATATCTTGTACTAAACACGCTAAACAGTTTATTGATGTGATCAATTTCCAACAGAACAATTAGAGCGGCAGGAATACCAACAAACATATTACCACATCCGTATTTTCTATGAATATCACATATCATATTAAAAAATCTTGATAGAAGAGCGAATATCTCGTCTGACTTAATGAGATGTAGTTCAGCAGCATAAGTCATAACTGTAAATAGAACATACTTTCCTGTAGTGTTGATTCTAGACTCCACATCAACACCATTATTAAAAATAGTTTTATATACATCTTTAATCTGCTCTCCGTTAATCGTCGAACGTTCTAGTATACGGAAACACTTTGATTTCTTATCTGTAGTTAATGACTTAGTGATATCACGAAGAATATTACGAATTACATTTCTTGTTTTTCTTGAGAGACCTGATTCAGAACTCAACTCATCGTTCCATAGTTTTTCTACCTCAGTGGCGAAATCTTTGGAGTGTTTGGTACATTTTTTAATAAGGTTCGTGACCTCCATTTATTATAAAAAATTTTTATTCAAAACTTAACTACAATCGGGTAATTATAAGATCGTAGATCTCCCATGTGGTGGAATACTACCATCTATCGCATGTTGATGGACAGTAGGTAATGGCCATGGGAACAGTAATGTTTGCATATTTATCTTTCTTGCTAGTATTACTGTATATTGTCCCAATGTTTCAATGTGATGTTCTAACCTATCAACTGCCACTGTATCACAACAATAATGTCCGATGGAATTAAGATTATGATCCAATGTGTTTAATATATGATTATCAAGTCTTATACGATCCGCGTCTTTTTTGACAGGATCAGGCTCTTCTACAGGAAGAAGTTTCGGCCTCTTATGATAGTCATGTCTGGGAAATGGTGGTCTAGGATGAGGATCAGGTATCGGAGTAGGTTTTGGATTATAATCATCATCATCATCATCATCATCATCATCATCATCATCATCTATGATATCATCATCTTCGATATTTATTTTGCTATCTTGATAATGTCCTATATCAGTTGCATTTTCAGCACTCGACTGAATATTAGTACATTCATTGTCTATTATTAACGTATTTCTAAACCCAAAATGTATATGTTGAACATCACTACTATAGTTGATGAGTCTTATAGCATGAATTCGCTTATCGTTATCGGGTTTATCTTCTGTCACCTTAACAATTCCTTTTTTATTAAACTCTGCATAATCATAACCATTTCTATTGTTTGTTCTAATATAAACGAGTATAGCATCATTGCTAAATTTTTCAATAGTATCAAAAACAGAATATCCTAAACCATATAATATATATTCAGGAACACTCAAACTAAATGTCCAGGATTCTCCTAAATACGTAAACTTTAATAGTGCTAAATCATTCAAAAATCTACCGCTTATAGATAGATAGTACATGAATGCGTATAGTAGTCTACCTATCTCTTTATTATGAAAACCGACATTACGATCATATATTTCGTGATATACATGTGACCCGTTTACGTTAAACCATAAATACATGGGTGATCCTATAAACATGAATTTATTTCTAATTCTCAGAGCCATAGTTAATTGACCGTGTAATATTTGTTTACATGCATACTTGATACGATCATTAATAAGATTTTTATCATTGCTCGTTATTTCAGAATCGTATATATAAGGAGTACCATCATGATTCTTACCAGATATTATACAAAATACTATATATAAAATATATTGACCCACGTTAGTAATCATGTAAATGTTTAATGTTTTAAATTTTGTATTTAATGATCCATCATCATATGCTAGCATGGTCTTGTGATATTCATTCTTTAAAATATAATATTGTGTTAGCCATTGCATTGGAGCTCCTAATGGAGATTTTCTATTCTCGTCCATTTTAGGATATGCTTTCATAAAGTCCCTAATAACTTCGTGAATAATGTTTCTATGTTTTCTACTGATGCATGTATTTGCTTCGATTTTTTTATCCCATGTTTCATCTATCATAGATTTAAACGCAGTAATGCTCGCAACATTAACATCTTGAACCATTGGTACAATTCCGTTCCATAAATTTATAATGTTCGCCATTTATATAACTCATTTTTTGAATATACTTTTAATTGAACAAAAGAGTTAAGTTACTCATATGGATGCCGTCCAGTCTGTACATCAATCTTTTTAGCCAGAGATATCATAGCCGCTCTTAGAGTTTCAGCGTGATTTTCCAACCTAAATAGAACTTCATCGTTGCGTTTACAACACTTTTCTATTTGTTCAAACTTTGTTGTTATATTAGTAATCTTTTTTTCCAAATTAGTTAGCCGTTGTTTGAGAGTTTCCTCATTATCGTCTCCATAGGCTTTAACAATTGCTTCGCGTTTAGTCTCTGGATTTTTAGCAGCCTTTGTAGAGAAAAATTCAGTTGCTGGAATTGCAAGATCGTCATCTCCGGGGAAAAGAGTTCCGTCCATTTAAAGTACAGATTTTAGAAACTGACACTCTGTGTTATTTATATTTGGCGCAATACATGGATTATAAATATCGATGTTAATAACATCAGAAAATGTAAAGTCTATACATTGTCGCATCGTGTTAAATTTTCTAATGGATCTAGTATTATTGGGTCCAACTTCTGCCTGAAATCCAAATATGGAAGCGGATACAAAACCGTTTCCTGGATAAACCACACATCTCCACTTTTGCTTTACATCAGAAATTGTGTCATTGACATCTTGAACTCTCCTATCTAATGCCGGTGTTCCACCTATAGATTTTGAATACTCGAATGCTGCATGAGTAGCATTGAATTCCTTAATATTGCCATAATTTTCATATATTGAGTAACTCTGGATAAAAAGTAAACACACCGCAGCCGTCGCTACTACAATAAAAAAAATTGATAGAGAGTTCATTTATAATCTATTAGAAGCTGATAAAATTTTTTTACACGCGTCAGACAATGCTTTAATAAATAGTTCAACATCTACTTTTGTCATATCGAACCGATGGTATGATTCTAACCTAGAATTACATCCGAAAAAGTTGACTATGTTCATAGTCATTAAGTCATTAACGAACAACATTCCAGACTCTGGATTATAAGACGATACTGTTTCGTCACAATCACCCACCTTAATCATGTGATTATGAATATTGGCTATTAGAGTACCTTCTAAGAAATCTATAATATCTTTGAAACACGATTTAAAATCAAACCACGAATATACTTCTACGAAGAAAGTTAGTTTACCCATAGGAGAGATAACTATAAATGGAGATCTAGATACAAAATCCGGATCTATGATAGTTTTAACATTATTATATTCTCTATTAAATACCTCCACATCTAAAAATGTTAATTTTGAAACTATGTCTTCGTTTATTACCGTACCTGAACTAAACGCTATAAGCTCTATTGTTTGAGAACTCTTTAAACGATATTCTTGAAATACATGTAACAAAGTTTCCTTTAACTCGGTCGGTTTATCTACCATAGTTACAGAATTTGTATCCTTATCTATAATATAATAATCAAAATCGTATAAAGTTATATAATTATCGTGTTCAGATTGTGATCTTTTCAAATAGACTAAAAACCCCATTTCTCTAGTAAGTATCTTATGTATATGTTTGTAAAATATCTTCATGGTGGGAATATGCTCTACAGCAGTTAGCCATTCCTCATTGACAGCTGTAGATGTATTATACAAAACTACTCCAATGTTTAACAAGGGCCATTTTACGAGATTATTAAATCCTTGTTTGATAAATGTAGCCAATGCGGGTTCGAGTTCAACGACGATTGAATTCTCTTCCCGTGGATGCTGCATGATGAACGACGGGATGTTGTTGTTCTATTGATTTGGAATTCTTTTTCGACTTTTTGTTTATATTAAATATTTTAAAATTTATGGCTGATAGTAATTCATGTACTACGGATAATGTAGACGTGTATTGCATATCGATATCTTTATTATTAGATAAATTTATCAATAAATGTGAGAAGTTTGCCTCGTTAAGGTCTTCCATTTAAATATTATATAAATATTTGTGTTTGTATTTTATTCGTCTTTTATGGGATAGTTTTTAACTAGTAAAGCTGTAATTACATACTTTGTCCGTAAAACATAAATATAAATACCCGCTTTTATCAAACGTTCCAAAAAGTCGGCAGCTGACATTTTTAACATGACATCTATTTTAAATACACTTAGGTTTTTAGAAAAAACATCATTTTATAATTGTAACGATTCAATAACTAAAGAAAAGATTAAGATTAAACATAAGGGAATGTTATTTGTATTTTATAAGCCAAAGCATTCTACCGTTGTTAAATACTTGTCTGGAGGAGGTATATATCATGATGATTTGGTTGTATTGGGGAAGGTAACAATTAATGATCTAAAGATGATGCTATTTTACATGGATTTATCATATCATGGAGTGACAAGTAGTGGAGCAATTTACAAATTGGGATCGTCTATCGATAGACTTTCTCTAAATAGGACTATTGTTACAAAAGTTAATAACAATTATAACAATTATAACAATTATAACAATTATAATTGTTATAATAATTATAATTGTTATAATTATGATGATACATTTTTTGACGATGATGATTGATCACTATTACACAATTTTGTTTTTGTACTTTCTAATATAGTGTTTAGGTTCTTTTTCATATGAGAATATTGACTTACTAAAATATCTATGTTTAACTTTTGTTCTATAACGTCCTTATCGGCGGTATCGGTACATATACGTAATTCACCTTCACAAAATACGGAGTCTTCGATAATAATAGCCAATCGATTATTGGATCTAGCTGTCTGTATCATATTCAACATGTTTAATATATCCTTTCGTTTCCCCTTTACAGGCATCGATCGTAGCATATTTTCCGCGTCTGAGATGGAAATGTTAAAACTGCAAAAATGCGTAATGTTAGCCCGTCCTAATATTGGTACGTGTCTATAAGTTTGGCATAGTAGAATAATAGACGTGTTTAAATGCCTTCCAAAGTTTAAGAATTCTATTAGAGTATTACATTTTGATAGTTTATCACCTACATCATCAAAAATAAGTAAAAAGTGTGCTGATTTTTTATGATTTTGTGCGACAGCAATACATTTTTCTATGTTACTTTTAGTTCGTATCAGATTATATTCTAGAGCTTCCTGACTACTAACGAAATTAATATGATTTGGCCAAATGTATCCATCATAATCTGGGTTATAAACGGGTGTAAACAAGAATATATGTTTATATTTTTTAACTAGTGTAGAAAACAGAGATAGTAAATAGATAGTTTTTCCAGATCCAGATCCTCCTGTTAAAACCATTCTAAACGGCATTTTTAATAAATTTTCTCTTGAAAATTGTTTTTCTTGAAAACAATTCATAATTATATTTACAGTTACTAAATTAATTTGATAATAAATCAAAATATGGAAAACTAAGGTCGTTAGTAGGGAGGAGAACAACGAAGGCATATCGTGATATAAATAACATTTATTATCATGATGACACCAGAAAACGACGAAGAGCAGACATCTGTGTTCTCCGCTACTGTTTACGGAGACAAAATTCAGGGAAAGAATAAACGCAAACGCGTGATTGGTCTATGTATTAGAATATCTATGGTTATTTCACTACTATCTATGATTACCATGTCCGCGTTTCTCATAGTGCGCCTAAATCAATGCATGTCTGCTAACAAGGCTGCTATTACTGACTCCGCTGTTGCCGTTGCTGCGGCATCATCTACTCATAGAAAGGTTGTGTCTAGCACTACACAATATGATCACAAAGAAAGCTGTAATGGTTTATATTACCAGGGTTCTTGTTATATATTACATTCAGACTATAAGTCATTCGAGGATGCTAAAGCAAACTGCGCTGCGGAATCATCAACACTACCCAATAAATCCGATGTCTTGACTACCTGGCTCATTGATTATGTTGAGGATACATGGGGATCTGATGGTAATCCAATTACAAAAACTACATCCGATTATCAAGATTCTGATGTATCACAAGAAGTTAGAAAGTATTTTTGTACATAAATAAATGAAATCGCTTAATAGACAAACTGTAAGTAGGTTTAGGAAGTTGTCGGTGCCGGCCGCTATAATGATGTTACTCTCAACCATTATTAGCGGCATAGGAACATTTCTGCATTACAGAGAAGAACTGATGCCTAGTGCTTGCGCCAATGGATGGATACAATACGATAAACATTGTTATCTGGATACCAACATTAAAATGTCTACGGATAATGCAGTTTATCAGTGTCGCAAATTACGAGCTAGATTGCCTAGACCTGATACTAGACATCTGAGAGTATTGTTTAGTATTTTTTATAAAGATTATTGGGTAAGTTTAAAAAAGACCAATGATAAATGGTTAGATATTAATAATGATAAAGATATAGATATTAGTAAATTAACAAATTTTAAGCAACTAAACAGCACGACGGATTCTGAGGCGTGTTATATATACAAGTCTGGAAAACTGGTTAAAACAGTATGTAAAAGTACTCAATCTGTACTATGCGTTAAAAGATTCTACAAGTGACAACAAAAAATGAATTAATAGTAAGTCGTTAACGTACGCCGCCATGGACGCCGCGTTTGTTATTACTCCAATGGGTGTGTTGACTATAACAGATACATTGTATGATGATCTCGATATCTCAATCATGGACTTTATAGGACCATACATTATAGGTAACATAAAAATTGTCCAAATAGATGTACGGGATATAAAATATTCCGACATGCAAAAATGCTACTTTAGCTATAAGGGTAAAATAGTTCCTCAGGATTCTAATGATTTGGCTAGATTCAACATTTATAGTATTTGTACAGCATACAGATCAAAAAATACCATCATCATAGCATGCGACTATGATATCATGTTAGATATAGAAGGTAAACATCAACCATTTTATCTATTCCCATCTATTGATGTTTTTAACGCTACAATCATAGAAGCGTATAATCTGTATACAGCTGGAGATTATCATCTGATCATCAATCCTTCAGATAATCTGAAAATGAAATTGTCGTTTAATTCTTCATTTTGTATATCAGACGGCAATGGATGGATTATAATTGATGGGAAATGTAATAGTAATTTTTTATCATAAAAGTTGTAAAGTAAATAATAAAACAATAAATATTGAACTAGTAGTATGTTGTATATTGAGCAATCAGAGATGATGCTGGTACCTCTTATCACGGTGACCGTAGTTGCGGGAACAATATTAGTATGTTATATATTATATATTTGTAGGAAAAAGATACGTACTGTCTATAATGACAATAAAATTATCATGACAAAATTAAAAAAGATAAAGAGTCCTAATTCCAGCAAATCTAGTAAATCAACTGATAGCGAATCAGACTGGGAGGATCACTGTAGTGCTATGGAACAAAACAATGACGTAGATAATATTTCTAGAAATGAGATATTGAACGATGATAGCTTCGCTGGTAGTTTAATATGGGATAACGAATCCAATATCATGGCGCCTAGCACAGAACACATTTACGATAGTGTTGCTGGAAGCACGCTGCTAATAAATAATGATCGTAATGAACAGACTATTTATCAGAATACTACAGTAGTAATTAATGATACAGAGACTGTTGAAATACTTAATGAAGATACCAAACAGATTCCTAGCTATTCTTCCAATCCTTTCGTAAATTATAATAAAACCAGTATTTGTAGCAAGTCAAATCCGTTCATTGCAGAACTCAACAATAAATTTAGTGATAATAATCCGTTTAGGAGAGCACATAGTGACGATTATCTTAATAAGCAACAAGATCATGAATACGATGATATAGAATCATCGGTTGTATCATTGGTCTGATTAGTTTCCTTTTTATAAAATTGAAGTAATATTTAGTATTAATTACCGCCGATGCATTATACAAATATGGAGATATTCCCTGTATTCGGCATTTCTAAAATTAGCAATTTTATTGCTAATAATGACTGTAGATATTATATAGATGTAGAGCATCAAAAAATTATATCTGATGAGATCAATAGACAGATGGATGAAACGGTACTTCTTACCAACATCTTAAGCGTAGAAGTTGTAAATGACAATGAGATGTACCATCTTATTCCCCATAGACTATCGACTATTATACTCTGTATTAGTTCTGTTGGAGGATGTGTTATCTCTATAGATAATGACGTCAATGACAAAAATATTCTAACATTTCCCATTGATCATGCTGTAATCATATCCCCACTGAGTAAATGTGTCGTAGTTAGCAAGGGCCCTACAACCATACTGGTTGTTAAAGCGGATATACCCAGCAAACGATTGGTAACATCGTTTACAAACGACATACTGTATGTAAACAATCTATCACTGATTAATTATTTACCGTCGTCTGTATTCATTATTAGACGAGTCACCGACTATTTGGATAGACACATATGTGATCAGATATTTGCTAATAATAAGTGGTATTCCATTATAACTATCGACGATAAGCAATATCCTATTCCATCAAATTGTATAGGTATGTCTTCTGCCAAGTACATAAATTCGAGCATCGAGCAAGATATTTTGATCCATGTTTGTAACCTCGAGCATCCATTCGACTCAGTCTACAAAAAAATGCAGTCGTACAATTCTCTACCTATCAAGGAACAAATATTGTATGGTAGAATTGATAATATAAATATGAGCATTAGTATTTCGGTGGATTAATAGATTTCTCTAGTATGGGATCATTAATCATCTCTAAATACATCATAAAAAAGCTATTATCAAATACTGTACTGAATGGATTCATTCTTTTCTCTTTTTATGAAACTCTGTTGTATATCTACGGATAAAACTAGAAGCAAAAAATCTGATAGGAAGAATAATGATTATATGGAGGAACACGATTATTATAAAATAACAATAGTTCCTGGTTCCTCTTCCACGTCTACTAGCTCGTGGTATTATACACATGCCTAGTAATAGTCTCTTTGCGTTGACGGAAAGCAGACTAGAAATAACAGGCCAAAATGTTCAGACACCATAATAGTTCCCAACCCAGATAATAACAGAGTTCCATCAACACATTCCTTTAAACTCAATCCCAAACCCAAAACCGTTAAAATGTATCCAGCCAATTGATAGTAGATAATGAGGTGTACAGCACATGATAATTTACACAGTAACCAAAATGAAAACACTTTAGTAATTATAAGAAATATAGACGGTAATGTCATCATCAACAATCCAATAATATGCCTGAGAGTAAACATTGACGGATAAAACAAAAATGCCCCGCATAACTCTATCATGGCAATAACGCAACCAAACACTTGTAAAATTCCTAAATTAGTAGAAAATACAACTGATATCGATGTATAAGCGATTTCGAGGAATAATAAGAACAAAGTAATTCCCGTAAAGATAAACATCAACATTGTTTGGTAATCATTAAACCAATTAGTATGACGTTGAATTAATTTCACAGTATATTTTATTCCAGTATTATCCCCGCATGTATACGTACCTGGTAAGATATCTTTATATTCCATAATCAATGAGACATCACTATCCGATAACGAATGAAGTCTAGCACTAGTATGCCATTTACTTAATATGGTCGTCTTGGAAGTTTTATTATAAGTTAAAATATCATGATTGTCCAATTTCCATCTAATATACTTTGTCGGATTATCTATAATACACGGAATAATGATGGTATCATTACATGCTGTATATTCTATAGTCTTTGTAGATGTTATAACCACAAAAGTACAGAGGTATATCAACAATATTCTAACTCTTAACATTTTTATTTATTTAAAATGATACCTTTGTTATTTATTTTATTCTTATTTTGCTAACGGTATCGAATGGCATAAGTTTGAAACGAGTGAAGAAATAATTTCTACTTACTTAATAGATTATGTGGTAACGGGTGTTATTAATGGGGATGTATATACATTTTCAAATAATGAACTAAACAAAACTGGGTTAACTAATAACAATAATTATATCACAACATCTATAAAAGTAGAGGATAAGGATACATTAGTAGTATGCGGAACCAATAACGGAAATCCCAAATGTTGGAAAATAGACGGTTCATACTACCCAAAACATATAGGTAGAGGATACGATCATCAAAATAGCAAAGTAACGATAATCAGTCACAATGAATGTGTACTATCCGACATAAACATATAAAAAGAAGGAATTAAACGATGGAGAAGATTTGACGGACCATGTGGTTATGATTAAACGAGTTAAGTTTTTTAAGAAGCCTTAGAAGAGAGGCTATTGGGTATGAGAATCCGAAATATTAAACCAGACAACCCCATATAATTTTATAGCTAAGAATGCCGCGAAGAATGGAACTAATAAAAACGGAAATATTTGTAGCACAACGAATAACTCCCAAACTGCATTCATGTTACACTATATAACACTACTTCGGTTAGATGTTTTAGAAAAAATAAATATCACCGTACCGTTTTGTTGTATAAAAATAACAATTAACAATTATCAATTTTTTTCTTTAATATTTTACGTGGTTGACCATTCTTGGTGGTAAAATAATCTCTTAGTGTTGGAATGGAATGCTGTTTAATGTTTCCACACTCATCGTATATTTTGACGTATGCAGTCACATCGTTTACGCAATAGTCAGACTGTAGTTCTATCATGCTTCCTACGTTAGAAGGAGGAACAGTTTTAAAGTCTCTTGGTTTTAATCTATTGTCATTAGTTTTCATGAAATCCTTTGTTTTATCCACTTCACATTTTAAATAAATGTCCACTATACATTCTTCTGTTAATTTTACTAGATCATCATGAGTCATAGAATTCATAGGTTCCGTAGTCCATGGATCCAAACTAGCAAACTTCGCGTATACGGTATCGCGATTAGTGTATACACCAACTGTATGAAAATTAAGAAAACAGTTTAATAAATCTACAGAAATATTTAATCCTCCGTTTGATACAGATGCGCCATATTTATGGATTTCGGATTCACACGTTGTTTGTCTAAGGGGTTCGTCTAGTGTTGCTTCTACATAGACTTCGATTCCCATATATTCTTTATTGCCAGAATCACATACCGATTTATCATACGCTGGTTCACTTGTTTGAAAACTAAATGGTAGTAGATACATCAAAATAATAAATAATAAGTACATTCTGCAATATTGTTATCGTAATTGGAAAATTGGTATTCAAGTGAGCTGGATTATGTGAGTATTGGATTGTATATTTTATTTTATATTTTATATTTTATATTTTATTTTATATTTTATATTTTATTTTATATTTTGTAGTAAGAATAGAATGCTAATGTCAAGTTTATTCGAATAGATGTCTTATTAAAAAACATATATAATAAATAACAATGGCTGAATGGCATAAAATTATCGAGGATATCTCAAAAAATAATAAGTTCGAGGATGCCGCCATCGTTGATTACAAGACTACAAAGAATGTTCTAGCGGCTATTCCTAACAGAACATTTGCAAAGATTAATCCGGGTGAAGTTATTCCCCTCATCACTAATCATAATATTCTAAAACCTCTTATTGGTCAGAAATTTTGTATTGTATATACTAACTCTCTAATGGATGAGAACACGTATGCTATGGAGTTGCTTACTGGGTACGCCCCTGTATCTCCGATCGTTATAGCGAGAACTCATACCGCACTTATATTTTTGATGGGTAAGCCAACAACATCCAGACGTGATGTGTATAGAACATGTAGAGATCACGCTACCCGTGTACGTGCAACTGGTAATTAAAATAAAAAGTAATATTCATATGTAGTGTCAATTTTAAATGATGATGAAATGGATAATATCCATATTGACGATGTCAATAATGCCGGTATTGACATACAGCTCATCGATTTTTAGATTTCATTCAGAGGATATTGAATTATGTTATGGGAATTTGTATTTTGATAGGATCTATAATAATGTAGTAAATATAAAATATATTCCTGTGCATATTCCATATAGATATAATTTTATTAATCGTACGTTCTCCGTAGATGAACTAGATGATAATGTCTTTTTTACACATGGTTATTTTTTAAAACACAAATATGGTTGTTCACTTAATCCTAGTTTGATTGTCTCATTATCAGGAAACTTAAAATATAATGATATACAATGCTCAGTAAATGTATCGTGTCTCATTAAAAATTTGGCAACGAGTACATCTACTATATTAACATCTAAACATAAGACTTATTCTCTATATCGGTCCATGTGTATTGCTATAATAGGATACGATTCTATTATATGGTATAAATATATAAATGACAGGTATAATGACATCTATGATTTTACTGCAATATGTATGCTAATAGCGTCTACATTGATAGTGATCATATACGTGTTTAAAAAAATAAAAATGAACTCTTAATTATGTTATACTATTAGAAATGGATAAAATCAAAATTACGATTGATTCAAAAATTGGTAATGTTGTTACCATATCGTATAACTTGGAAAAGATAACTATTGATGTCACGCCAAAAAAGAAAAAAGAAAAGGATGTATTATTAGCGCAATCAGTTGCTGTCGAAGAGGCAAAAGATGTCAAGGTGGAAGAAAAAAATATTATCGATATTGAAGATGACGATGATATGGATATAGAAAACACGTAATACGATCTATAAAAATAAGTATTAAATACTTTTTATTTACGGTACTCTTGTAGTGGTGATACCACTAATCGATTATTTTTTTTAAAAAAATACTTATTCTGATTCTTCTAGCCATTTCCGTGTTCGTTCGAATGCCACATCGACGTCAAAAATAGGGGAGTAGTTGAAATCTAGTTCTGCATTGTTGGTACGCACCTCAAATGTAGTGTTGGATATCTTCAACGTATAGTTGTTGAGTATTGATGGTTTTCTAAATAGAATTCTCTTCATATCATTCTTGCACGCGTACATTTTTAGCATCCATCTTGGAATCCTAGATCCTTGTTCTATTCCCAATGGTTTCATCAATAGAAGATTAAACATATCGTAAGAACACGATGGAGAGTAATCGTAGCAAAAGTAAGCATTTCCTTTAATCGCAGATCCCGGATACTGGATATATTTTGCAGCCAACACGTGCATCCATGCAACATTTCCTACATATACCCGGCTATGCACAGCGTCATCATCGACTGTACGATACATAATGTTACCGTGTTGCTTACATTGCTCGTAAAAGACTTTCGTCAATTTGTCTCCTTCTCCGTAAATTCCAGTGGGTCTTAGGCAACAAGTATACAATTTTGCGCCATTCATGATTACGGAATTATTGGCTTTCATAACCAGTTGCTCGGCCATACGTTTACTTTTTGCGTATACATGTCCTGGTGATATATCATAAAGGGTATGCTCATGACCGATGAATGGATTACCGTGTTTATTTGGTCCTATTGCTTCCATGCTACTAGTATAGATCAAATACTTGATTCCTAGGTCCACACAAGCTGCCAATATAGTCTGTGTTCCATAATAGTTTACTTTCATGATTTCATTATCAGTGTATTTTCCAAATACATCCACTAGAGCAGCCGTATGAATAATCAGATTTACCCCATCTAGCGCTTCTCTCACCTTATCAAAGTCGTTTATATCACATTGTATATAGTTTATAACCTTAACTTTCGAGGTTATTGGTTGTGGATCTTCTACAATATCTATGACTCTTATTTCTTGAACATCATCTGCGCTAATTAAAAGTTTTACTATATACCTGCCTAGAAATCCGGCACCGCCAGTAACCGCGTACACGGCCATTGCTGCCACTCATAATATCAGACTACTTATTCTATTTTACTAAATAATGGCTGTTTGTATAATAGACCACGATAATATCAGAGGAGTTATTTACGTTGAACAAGTCCATGGAAAAGATAAAGTTTTAGGATCAGTTATTGGATTAAAATCCGGAACGTATAGTTTGATAATTCATCGTTACGGAGATATTAGTCGAGGATGTGATTCCATAGGCAGTCCAGAAATATTTATCGGTAACATCTTTGTAAACAGATATGGTGTAGCATATGTTTATTTAGATACAGATGTAAATATATCTACAATTATTGGAAAGGCGTTATCTATTTCAAAAAATGATCAGAGATTAGCGTGTGGAGTTATTGGTATTTCGTACATAAATGAAAAGATAATACATTTTCTTACAATTAACGAGAATGGCGTTTGATATATCAGTTAATGCGTCTAAAACAATAAATGCATTAGTTTACTTTTCTACTCAGCAAGATAAATTAGTCATACGTAATGAAGTTAATGATATACACTACACTGTCGAATTTGATAGGGACAAAGTAGTTGATACGTTTATTTCATATAATAGACATAATGACTCCATAGAGATAAGAGGGGTGCTTCCAGAGGAAACTAATATTGGTCGCGTGGTTAATACGCCGGTTAGTATGACTTACTTGTATAATAAGTATAGTTTTAAACCGATTTTAGCAGAATATATAAGACACAGAAATACTATATCCGGCAACATTTATTCGGCATTGATGACGCTAGATGATTTGGTTATTAAACAGTATGGAGACATTGATCTATTATTTAATGAGAAACTTAAAGTAGACTCCGATTCGGGACTATTTGACTTTGTCAACTTTGTAAAGGATATGATATGTTGTGATTCTAGAATAGTAGTAGCTCTATCTAGTCTAGTATCTAAACATTGGGAATTGACAAATAAAAAGTATAGGTGTATGGCATTAGCCGAACATATAGCTGATAGTATTCCAATATCTGAGCTATCTAGACTACGATACAATCTATGTAAGTATCTACGCGGACACACTGATAGCATAGAGGATGAATTTGATCATTTTGAAGACGATGATTTGTCTACATGTTCTGCCGTAACCGATAGGGAAACGGATGTATAATTTTTTTTATAGTATGAAGGATATGATGGATATGATAATATGATGGATATGATGATATGATGGATATGATGGATATGATGGATATGATGGATATGATAAAAAAATATAATTGTTGTATCCATTCCCATTCAAATCACCTTATATGATTCTGTAACACAATGAAGGAGTCTCATAGATATATAGAGGTCAGATACTGGTTTGATAAACTTTTTATTCCACATGAGCATGTTTGACTTATGGTTAGACACACATACTTTAACAAATCACTGAAAATTGGAGTTAGGTATTCCTCTCAGAATCAGTTGCCGTTCTGGAACATTAAATGTATTTTTTATGATATACTCCAACGCATTTATGTGGGTATACAACAAGTCATTAATAATGAGTATTTCCAAGAGTTTTAGTTGTCTAGTATTTAACAAGAGAAGAGATTTCAACAGACTGTTTATGAACTCGAATACCGCCTCATTGTCGCTTATATTGATGACATGACATCGAATTCCCAATATCAATCTCATCAGTGATGAGTAGCTCAATCTTGTTATCGGGATCCAATTTCTAAAGATGTCATTAAACCCTCGATCGTGAATGGATTTATCATCATCGTTTTTATGTTGGACATGAGCTTAGTCCGTTTGTCCACATCTATATACGATGATTTCTGAATTATTTCATATATCTCTCGTTAACTCCAGGAACTTGTCAGGGATCTAACTTTAATATGTTCTCGTCTAAGAGATGAAAATCTTTGGATGGTTGCATGTGACTTTTCTCTAAAGGATGATGTTACCCGATCCTCTCTTAAATGACTCCATCTTATCCTTGGACAAGATGGACAGTCTATTTTCCTTAGATGGTTTAATATTTTTTACCCATGATCTATAAAGGTAGACAGACCTAATCGTCTCGGATGACCATATATTATTTTCAGTTTTATTATACGCATAAATTGTAAAAAATATGTTAGGTTTACGAAAATGTCTCGTGGGGCATTAATCGTTTTTGAAGGATTGGACAAATCTGGAAAAACAACACAATGTATGAACATCATGGAATCTATACCGGCAAACACGATAAAATATCTTAACTTTCCTCAGCGATCCACAGTCACTGGAAAGATGATAGATGACTATCTAACTCGTAAAAAAACCTATAATGATCATATAGTTAATCTATTATTTTGTGCAAATAGATGGGAGTTTGCATCTTTTATACAAGAACAACTAGAACAGGGAATTACTTTAATAGTTGACAGATACGCGTTCTCTGGAGTAGCGTATGCCACCGCTAAAGGCGCGTCAATGACTCTCAGTAAGAGTTATGAATCTGGATTGCCTAAACCCGACTTAGTTATATTCTTGGAATCTGGTAGCAAAGAAATTAATAGAAACATCGGCGAGGAAATTTATGAAGATGTTGAATTCCAACAAAAGGTATTACAAGAATATAAAAAAATGATTGAAGAAGGAGATATTCATTGGCAAATTATTTCTTCTGAATTCGAGGAAGATGTAAAGAAGGAGTTGATTAAGAATATAGTTATAGAGGCTATACACACGGTTACTGGACCAGTGGGGCAACTGTGGATGTAATAAAATGAAATTACATTTTTATAAATAGATGTTAGTACAGTGTTATAAATGGATGAAGCATATTACTCTGGCAACTTGGAATCAGTACTCGGGGATACGTGTCCGATATGCATACCGAACTCGCATCAATATCTCAATTAGTTATTGCCAAGATAGAAACTATAGATAATGATTATTAAACAAGGACATTGTAAATTTTATCATGTGTAGATCAAACTTGGATAATCCATTTATCTCTTTCCTAGATACTGCATATACTATCATAGATCAAGAGATCTATCAGAACGAGTTGATTAATTCATTAGACGATAATGAAATTATCGATTGTATAGTTAACAAGTTTATGAGCTTTTATAAGGATAACCTAGAAAATATGGTAGATGCTATCATTACTCTAAAATATTATAATTAATAATCCAGATTTTAAAACTACGTATGTGGAAGTACTCGGTTCCAGAATAGCTGATATAGATATTAAACAAGTGATACGTAAGAATATAATACAATTGTCTAATGATCCGCGAACGATATTTGTGAAAATATTAAAAAAAAATACTTTTTTTATTAAATGACGTCTCTTCGCGAATTTAGAAAATTATGCTGTGATATATATCACGCATCAGGATATAAAGAAAAATCTAAATTAATTAGAGACTTTATAACAGATAGAGATGATACCGATACATATTTGATCATTAAGCTATTGCTTCCCGGATTAGACGATAGAATGTATAACATGAACGATAAACAAATTATAAAATTATATAGTATAATATTTAAACAATCTCAGGAAGATATGCTACAAGATTTAGGATACGGATATATAGGAGACACTATTAGGACTTTCTTCAAAGAGAACACGGAAATCCGTCCACGAGATAAAAGCATTTTAACTTTAGAAGAAGTGGATAGTTTTTTAACTACGTTATCATCAGTAACTAAAGAATCACATCAAATAAAATTATTGACTGATATAGCATCTGTTTGTACATGTAATGATTTAAAATGTGTAGTCATGCTTATTGATAAAGATCTAAAAATTAAAGCGGGTCCTCGGTACGTGCTTAACGCTATTAGTCCTCATGCCTATGATGTTTTTAGAAAATCTAATAACTTGAAAGAGATAATAGAAAATGCAGCTAAACAAAATCTAGACTCTATATCTATTTCTGTTATGACTCCAATTAATCCCATGTTAGCGGAATCATGTGATTCTGTCAATAAGGCGTTTAAAAAATTTCCATCAGGAATGTTTGCGGAAGTCAAATACGATGGTGAAAGAGTACAAGTTCATAAAAAAAATAACGAGTTTGCATTCTTTAGTAGAAACATGAAACCAGTACTCTCTCATAAAGTGGATTATCTCAAAGAATACATACCGAAAGCATTTAAAAAAGCTACGTCTATCGTATTGGATTCTGAAATTGTTCTTGTAGACGAACATAATGTACCGCTACCGTTTGGAAGTTTAGGTATACACAAAAAGAAAGAATATAAAAACTCTAACATGTGTTTGTTCGTATTTGACTGTTTATACTTTGATGGATTCGATATGACAGACATTCCATTGTATGAACGAAGATCTTTTCTCAAAGATGTTATGGTCGAAATACCCAATAGAATAGTATTCTCAGAGTTGACGAATATTAGTAACGAGTCTCAGTTAACTGATGTATTAGATGATGCACTAACGAGAAAATTAGAAGGATTGGTCTTAAAAGATATTAATGGCGTATACGAACCGGGAAAGAGAAGATGGTTAAAAATAAAGCGAGACTATTTGAACGAGGGTTCCATGGCAGATTCTGCCGATTTAGTAGTACTAGGTGCCTACTATGGTAAAGGAGGAAAGGGTGGTATCATGGCAGTCTTTCTAATGGGTTGTTACGACGATGAATCCGGTAAATGGAAGACGGTAACTAAATGTTCCGGTCACGATGATAATACGTTAAGGGTTTTGCAAGACCAATTAACGATGGTTAAAATTAACAAGGATCCCAAAAAAATTCCAGAGTGGTTGGTAGTTAATAAAATCTATATTCCCGATTTTGTAGTAGATGATCCGAAACAATCTCAGATATGGGAAATTTCAGGAGCAGAGTTTACATCTTCCAAGTCACATACAGCGAATGGAATATCGATTAGATTTCCTAGATTTACTAGGATTAGAGAAGATAAAACGTGGAAAGAATCTACTCATCTAAACGATTTAGTAAACTTGACTAAATCTCTTAATAGTTACATATAAACTGAAAAATAAAATAACACTATTTTAGTTGGTAGTCGCCATGGATGGTGTTATCGTATACTGTCTAAATGCGTTAGTAAAACATGGCGAGGAAATAAATCATATAAAAAATGATTTCATGATTAAACCATGTTGTGAAAGAGTTTGTGAAAAAGTCAAGAACGTTCACATCGGCGGACAATCTAAAAACAATACAGTGATTGCAGATTTGCCATATCTGGATAATGCTGTATCAGATGTATGCAAATCAATATATAAAAAGAATGTATCAAGAATATCCAGATTTGCTAATTTGATAAAAATAGATGACGATGACAAGACTCCTACCGGCGTATATAATTATTTTAAACCTAAAGATGCTATTCCTGTTATTATATCCATAGGAAAGGATAAAGATGTCTGTGAACTATTAATCTCATCTGATAAAGCGTGTGCGTGTATAAAGTTAAATTTATATAAAGTAGCCATTCTTCCCATGGATGTTTCCTTTTTTACCAAAGGAAATGCATCATTGATTATTCTCCTGTTTGATTTCTCTATCGATGCGGCACCTCTCTTAAGAAGTGTAACCGATAATAATGTTATTATATCTAGACACCAGCGCCTACATGACGAGCTTCCGAGTTCCAATTGGTTCAAGTTTTACATAAGTATAAAGTCCGACTATTGTTCTATATTATATATGGTTGTTGATGGATCTATGATGTATGCGATAGCTGATAATAGAACTCACGCAATTATTAGCAAAAATATATTAGACAATACTACGATTAACGATGAGTGTAGATGCTGTTATTCTGAACCACAGATTAGGATTCTTGATAGAGATGAGATGCTCAATGGATCATCGTGTTATATGAACAGACATTGTATTATGATGAATTTACCTGATGTAGGCGAATTTGGATCTAGTATGTTGGGGAAATATGAACCTGACATGATTAAGATTGCTCTTTCGGTGGCTGGTAATTTAATAAGAAATCGAGACTACATTCCCGGGAGACGAGGCTATAGCTACTACGTTTACGGTATAGCCTCTAGATAATTTTTTTTAAGCACGAAATAAAAACATAATTTTAAACAATCTATTTCATACTATTTTGTGTGCTCACCATGAACATAAAGATAGATATATTAGTATTTCTGGTGATAAATTTACGGCGACTGCTAGGAGGGAAAATGAAGAAAGAAAAAATATCTACCTCTCCAAAAAGAAAAACTACTGATGTTATCAAACCTGATTATCTTGAGTACAATGACTTGTTAGATAGAGATGAGATGTCTACTATTCTAGAGGAATATTAGGCCTTAGAATAAAATATGGACGACTCTTAACGAAATTAGAAAATTCGATAATGATGTTGAAGAACAATTCGGTACTATAGAAGAACTCAAGCAGAAGCTTAGATTAAATTCTGAAGAGGGAGCAGATAATTTTATAGATTATATAAAGGTACAAAAACAGGATATCATCAAACTTACTGTATACGATTGCATATATCTATGATAGGATTGTGTGCGTGCGTGGTAGATGTTTGGAGAAATGAGAAACTGTTTTCTAGATGGAAATATTGGTTACAAGCGATTAAACTGTTTATTGATGATCACATGCTTGATAAGATAAAATCTATTGTAGAATAGACTAGTGTATGTGGAAATGTCATAGAAAGTTAAAAGTTAATGAGAGCAAAAATATATAAGGTTGTATTCCATATTTGTTATTTTTTTCTGTAATAGTTAGAAAATACATTCGATGGTCTATCTACCAGATTATTATGTGTTATAAGGTACTTTTCTCATAATAAACTAGAGTATGAGTAAGATAGTGTTTTTCAAAAACATATAAATCTAAAATTGATGGATGAGATATACAGCTATTAATTTCGAAAATATATTTTAATCTGATAACTTTAAACATGGATTTTTGATGGTGGTTTAAGTTTAAAAAAGATTTTGTTATTGTAGTATGATAATATCAAAAAGATGGATATAAAGAATTTACTGACTACATGTACTATTTTACATTACTACATTGGCTACGGCATATATACCTATTTCGTCACTTCCACACGCTCCGGTAAACGGGTGTCATGTGACGAGGGAGAATCTTGATAAGAGGCATAATCAATGTTGTAATCCGATGTCCACCTGGAGAATTTGCCAAGGTCAGATGTAGAGTTGGTAGTGATAACACAAAATGTGAACACTGCCCACCTCATACATATACCGCAATCCCCAATTATTCTAATAGATGTCATCAATGTAGAAAATGCCCAACAGGATCATTTGATAAGGTAAAGTGTACCGGAACACAGAACAAATGTTCGTGTCATCCTGGTTGGTATACGCTACTGATTCTTCACAGACTGAAGATTGTCGAGATTTGTGTACCAAAAAAGGAGATGTCCATGCGGATACTTTGGTGGAATAGATGAAGGAAATCCTATTTGTAAATCGTGTTGTGTTGGTGAATATTGCGACTACCTACGTAATTATAGACTTGATCCATTTCCTCCATGCAAACTATCTATCTAAATGTAATTAATTATGATTTTGATGATAATGTTACCATACATTATATCACTACTTGGTTAGTGTGTATTATTTAGTATGGAAGACCTATTAATAATTACTTATCTTTTGACGATCTTGTTATAATTATAATATAAAAATACTTATGACATAGTAACTCATAATTGCTGACGCGATAAATTCGTAATAATCTGTTTTGTTCAAATTTTTATAAGGAATCTACAGGCATAAAAATAAAAATATAATCTATAATATACTCTTACAACGCCATCATGAATAGCAGTGAATTAATTGCTGTTATTATGGATTTAGAAATAGTGGACGATTTTGTGATATTAATATAGTTATTAATGATGAAAGGATAAACGCGCATAGACTCATCCTATCTGGAGCCTCCGAATATTTTTTCCATTCTGTTTTCCAATAATTTTATCGATTCTAATGAATACGAAGTTAATCTAAGTCATTTAGATTATCAAAGTGTTAACTATTTGATCGATTACATTTATGGGATACCTTTGAGCCTAACTAACAATAACGTGAAATATATTCTTTCAACCGCTGATTTTTTTACAAATTGGATCTGTCATTACTGAGTGCGAAAAATACATACTTAAAAATCTTTGTTTTAGAAACTGTATCGATTTCTACATATACGCTGATAAATATAATAACAAGAAAATAGAATTAGCATCGTTTAACACAATATTACGAAATATTTTGAGACTCATCAACAATGAAAACTTTAAATACTTAACAGAGGAATCAATGATAAAAATTTTAAGCGATGATATGTTATATATAAAAAATGAGGATTTCACCCCACTGATTCTCATTAAATGGTTAGAGAGTACACCAACCATGTACCGTCGAGTTACTTAGATGCCTCAGAATATCATTTCTTTCCCCACAAGTTATAAAATCACTTTATAGTCATCGACTGGTTAGTTCAATCTACGAATGTATAACATTCTTAAACAATATAGCATTCTTGGATAAATCATTTCCTAGATACCATATCATCGAGTTGATATCTATCGGTATAAGTAATTCACATGATAAGATTTCCATAAACTGCTACAATCATAAAAAAATTCATGGGAAATGATATCTTCACGTAGATATAGGTGTAGTTTCGCAGTGACCGTCCTGGATAATATTATCTATATGATGGGTGGATATGATCAGTCCCTGTATAGAAGTTCAAAGGTTATAGCGTACAATACATGTACTAATTCTTGGATATATGATATACCAGAGCTAAAAATATCATCGTTCTAATTGCGGAGGAGTTGCCAATGACGAATACATTTATTGTATAGGCGGTATACGCGATCAGGAGTCATCGTTGATATCTAGTATCGATAGATGGAAGCCATCAAAACCATATTGGCAGAAGTATGCTAAAATGTGCGAACCAAAATGTGATATGGGGTTGCGATTTTAAACGGATTAATATATGTCATAGGTGGAGTCGTTAAAGGTGACACACATATACCAACGCACTAGAGAGTTTATCAGAAGATGGATGGATGAATCATCAACGTCTTCCAATAAAAATGTCCAATATGTCGACGATTGTTCATGCTGGAAAGATTTATATATCTAGAGGTTACAACAATAGTAGTGTAGTTAATGTAATATCGAATCTAGTCCTTAGCTATAATCCGATATATGATGAATGGACCAAATTATCATCATTAAATATTCCTAGAATTAATCCTGCTCTATGGTCAGTGTATAATAAATTATATGTAGGAGGAGTAATATCTGATGATGTTCAAACTAATACATCTGAAACATACGATAAAGAAAAAGATTGTTGGACATTGGATAATGGTCACTTGTTACCACATAATTATATAATGTATAAATGCGAACCGTTTAAACATAGATATCCATTGGAAAAAACACAGTACACGAATGATTTTCTAAAGTATTTGGAAAGTTTTATAGGTAGTTGATAGAACAAAATACATAATTTTGTAAAAATAAATCACTTTTTATACTAATATGACACAATTACCAATACTTTTGTTACTAATATCATTAGTATACGCTACACCTTCTCCTCAGACATCTAAAAAAATAGGTGATGATGCAACTATATCATGTAGTCGAAATAATACAAATTACTACGTTGTTATGAGTGCTTGGTATAAGGAGCCCAATTCCATTATTCTCTTAGCTGCCAAAAGCGACGTCTTGTATTTTGATAATTATACCAAGGATAAAATATCTTACGACTCTCCATACGATGATCTAGTTACAACTATCACAATTAAATCATTGACTGCTGGAGATGCCGGTACTTATATATGTGCATTCTTTATGACATCGACTACAAATGATACTGATAAAGTAGATTATGAAGAATACTCCATAGAGTTGATTGTAAATACAGATAGTGAATCGACTATAGACATAATACTATCTGGATCTACACCAGAAACTATTTCTGAGAAACCAGAGGATATAGATAATTCTAATTGCTCGTCTGTATTCGAAATCACGACTCCGGAACCAATTACTGATAATGTAGACGACCATACAGACACCGTCACATACACTAGTGATAGCATTAATACAGTAAATGCATCATCTGGAGAATCCACAACAGACGAGATTCCGGAACCAATTACTGATAAAGAAGAAGATCATACAGTAACAGACACTGTCTCATACACTACAGTAAGTACATCATCTGGAATTGTCACTACTAAATCAACCACCGATGATGCGGATCTTTATGATACATACAATGATAATGATACAGTACCGCCAACTACTGTAGGTGGTAGTACAACCTCTATTAGCAATTATAAAACCAAGGACTTCGTAGAAATATTTGGTATTACCACATTAATTATATTGTCAGCAGTGGCGATTTTCTGTATTACGTATTATATATGTAATAAACACCCACGTAAATACAAAACAGAGAACAAAGTCTAGATTTTTGACTTACATAAATATCTGGGATAATAAAATCTATCATATTGAGAGGACCATCTGGTTCAGGAAAGACAGCCATAACCAAAAGACTGTTAAAAGACTATGGGAATATATTTGGATTTGTGGTGTCCCATACCACTAGATTTCCTCGTCCTATGGAACGAGAAGGTGTTGATTACCTTACGTTAACAGAGAGGCCATCTGGAAGGGAATAGCCGCCGGAAACTTTCTAGAACATACTGAGTTTTTAGGAAATATTTACGGAACTTCTAAAACAGCTGTAAATACAGCGGTTATTAATAATCGTATTTGCGCGATGGATTTAAACATCAACGGTGTTAGAAGTCTTAAAAATACTTACCTAATGCATTACTTGGGTATATAAGACCTACCTCTCTTAAAATGGTTGAGACCAATCTTCGTCGTAGAAACACTGAAGCGGACGACGAATCTCATCGTCGCGTGATGTTGGCAAAAAACGGATATGGATGAGGTCAACGAAGCAGGTCTATTCGACACTATTATTATTGAAGATGATGTGAATTTAGCATATAGTAAGTGTTAATTCAGATACTACAGGACCGTATTAGAATGTATTTTAACACTAATTAGAGACTTAAGATTTGACTTAAAACTTGATAATTAATAATATAACTCGTTTTTATATGTGGCTATTTCAACGTCTAATGTATTAGTTAAATATTAAAACTTACCACGTAAAACTTAAAATTTAAAATGGTATTTCATTGACAGATCATACATTATGAAGTTTCAAGGACTTGTGTTAATTGACAATTGCAAAAATCAATGGGTCGTTGGACCATTAATAGGAAAAGGTGGATTCGGTAGTATTTATACTACTAATGACAATAATTATGTAGTAAAAATAGAGCCCAAAGCTAACGGATCATTATTTACCGAACAGGCATTTTATACTAGAGTACTTAAACCATCCGTTATCGAAGAATGGAAAAAATCTCACAATATAAAGCACGTAGGTCTTATCACATGCAAGGCATTTGGTTTATACAAATCCATTAATGTGGAATATCGATTCTTGGTAATAAATAGATTAGGTGCAGATCTAGATGCGGTGATCAGAGCCAATAATAATAGACTACCAGAAAGGTCGGTGATGTTGATCGGAATCGAAATCTTAAATACCATACAATTTATGCACGAGCAAGGATATTCTCACGGAGATATTAAAGCGAGTAATATAGTCTTGGATCAAATAGATAAGAATAAATTATATCTAGTGGATTACGGATTGGTTTCTAAATTCATGTCTAACGGCGAACATGTTCCATTTATAAGAAATCCAAATAAAATGGATAACGGTACTCTAGAATTTACACCTATAGATTCGCATAAAGGATACGTTGTATCTAGACGTGGTGATCTAGAAACACTTGGATATTGTATGATTAGATGGTTGGGAGGTATCTTGCCATGGACTAAGATATCTGAAACAAAGAATTCTGCATTAGTAAGTGCCGCAAAACAGAAATATGTTAACAATACTGCGACTTTGTTAATGACCAGTTTGCAATATGCACCTAGAGAATTGCTGCAATATATTACCATGGTAAACTCTTTGACATATTTTGAGGAACCCAATTACGACGAGTTTCGTCGAGTATTAATGAATGGAGTTATGAAAAATTTTTGTTGATAAAAAAATTAAAAAAATAACTTAGTTATTATCACTCTCGTGAGTACAATAGAAACATGGCGATGTTTTACGCACACGCTTTCGGTGGGTACGACGAGAACCTTCATGCATTTCCTGGAATATCATCGACGGTTGCCAATGATGTCAGGAAATATTCTGTTGTGTCAGTTTATAATAAAAAGTATAACATTGTAAAAAACAAATATATGTGGTGTAACAGTCAAGTGAACAAGAGATATATTGGAGCACTACTGCCTATGTTTGAATGCAATGAATATCTACAAATTGGAGATCCAATCCATGATCTAGAAGGAAATCAAATCTCTATTGTCACATATCGCCACAAAAACTACTATGCTCTAAGTGGAATTGGGTACGAGAGTCTAGACTTGTGTTTGGAAGGAGTAGGGATTCATCATCACGTACTTGAAACAGGAAACGCGGTATATGGAAAAGTTCAACATGAGTATTCTACTATCAAAGAGAAGGCCAAAGAAATGAATGCACTCAAACCAGGACCTATCATCGATTACCACGTCTGGATAGGAGATTGTGTCTGCCAAGTTACTACTGTAGACGTGCATGGAAAGGAAATTATGAGAATGAGATTCAAAAGGGGTGCGGTGCTTCCGATTCCAAATCTGGTAAAAGTTAAAGTTGGGGAGGAAAATGATACAATAAATCTTTCCACTTCCATATCAGCTCTCCTGAATTCCGGTGGCGGCACCATCGAGGTAACATCTAAGGAAGAACGTGTAGATTATGTACTCATGAAACGTTTGGAATCTATACATCATCTGTGGTCTGTAGTGTATGATCATCTTAATGTTGTGAATGGCGAAGAACGATGTTATGTACATATGCATTCATCTCATCAAAGTCCTATGCTGAGTACTGTAAAAACAAATTTGTACATGAAGACTATGGGAGCATGTCTTCAAATGGACTCCATGGAAGCTCTAGAGTATCTTAGTGAACTGAAGGAATCAGGTGGGCGGAGTCCCAGACCAGAATTGCAGAAATTTGAATATCCAGATGGAGTGAAAGACACTGAATCAATTGAGAGATTGGCAGAGGAGTTCTTCAATAGATCAGAACTTCAGGCCGGTGAATCAGTCAAATTTGGTAATTCTATTAATGTTAAACATACATCTGTTTCAGCTAAGCAACTAAGAACACGTATACGACAGCAGCTTCCTTCTATACTCTCATCTTTTGCCAACACAAAGGGTGGATATTTGTTCATTGGAGTTGATAATAATACACACAAAGTAATTGGATTCACGGTGGGTCATGACTACCTCAAACTGGTAGAGAGTGATATAGAAAAGTATATCCAAAAACTTCCTGTTGTGCATTTCTGCAAGAAAAAAGAGGACATCAAGTACGCATGTAGATTCATCAAGGTGTATAAACCTGGTGATGAGACTACCTCGACATATGTGTGCGCAATCAAAGTGGAAAGATGCTGCTGTGCTGTGTTTGCGGATTGGCCAGAATCATGGTACATGGATACTAGTGGTAGTATGAAGAAGTATTCTCCAGATGAATGGGTGTCACATATAAAATTTTAATTAGGGTAAGGTAAAACTATATATAATAACTAACAATTTGTGTATCATATAGACAATTAATTAGGTAACTGTTATCTCTTTTTAACTAACTAACTAACTAACTAACTCTTATATACTATTAATAATACATCTATTAATCATTGATTAGCTTATTGCTTTAATTGTTTTTGTAAACTAACACTGTTCATTGAAAAGGGATAACATGTTACAGAATATAAATTATATATGGATTTTTTTAAAAAGGAAATACTTGACTGGAGTATATATTTATTTCTTCATTACATAACACGTCTGTGTTCTAATTCTTCCAATTCTTCCACATCTCATATAATACAGGAATATAATCTTGTTCGAAAATATGAGAAAGTGGATAAAACAATAGTTGATTTTTTATCTAGGTGGCCAAATTTATTCCATATTTTAGAATATGGGGAAAATATTCTACATATTTATTTTATAGATGCTGCTAATACGAATATTATGATTTTTTTTCTAGATAGAGTATTAAATATTAATAAGAACCGTGGGTCATTTATACATAATCTCGGGTTATCATCCATTAATATAAAAGAATATGTATATCAATTAGTTAATAATGATCATCTAGATAATAGTATAAGACTAATGCTTGAAAATGGACGTAGAACAAGACATTTTTTGTCTTATATATTGGATACAGTTAATATCTATATAAGTATTTTAATAAATCATAGATTTTATATAGATGCCGAAGACAGTTACGGTTGTACATTATTACATAGATGTATATATAACTATAAGAAATCAGAATCAGAATCATATAATGAATTAATTAAGATATTGTTAAATAATGGATCAGATGTAGATAAAAAAGATACGTACGGAAACACACCGTTTATCCTATTATGTAAACACGATATCGACAACGCGGAATTGTTTGAGATATGTTTAGAGAATGCTAATATAGACTCTGTAGACTTTAATGGATATACACCTCTTCATTATGTCTCATGTCGTAATAAATATGATTTTGTAAAGTTATTAATTTCTAAAGGAGCAAATGTTAATGCACGTAATAGATTCGGAACTACTCCATTTTATTGTGGAATTATACACGGTATCTCGCTTATAAAACTATATTTGGAATCAGACACAGAGTTAGAAATAGATAATGAACATATAGTTCGTCATTTAATAATTTTTGATGCTGTTGAATCTTTAGATTATCTATTGTCCAGAGGAGTTATTGATATTAACTATCGTACTATATACAACGAAACATCTATTTACGACGCTGTCAGTTATAATGCGTATAATACGTTAGTCTATCTATTAAACAGAAATGGTGATTTTGAGACGATTACTACTAGTGGATGTACATGTATTTCGGAAGCAGTCGCGAACAACAACAAAATAATAATGGATATACTATTGTCTAAACGACCATCTTTGAAAATTATGATACCATCTATGATAGCAATTACTAAACATAAACAACATAATGCAGATTTATTGAAAATGTGTATAAAATATACTGCGTGTATGACCGATTATGATACTCTTATAGATGTACAATCGCTACATCAATATAAATGGTATATTTTAAAATGTTTTGATGAAATAGATATCATGAAGAGATGTTATATAAAAAATAAAACTGTATTCCAATTAGTTTTTTGTATCAAAGACATTAATACTTTAATGAGATACGGTAGACATCCTTCTTTCGTGAAATGTAATATTCTTGACGTATACGGAAGTTGTGTACGTAATATCATAGCATCTATTAGATATCGTCAGAGATTAATTAGTCTATTATCCAAGAAGCTGGATGCTGGAGATAAATGGTCGTGTTTTCCTAACGAAATAAAATATAAAATATTGGAAAACTTTAACGATAACGAACTGACCACATATCTGAAAATCTTATAAACACTATTAAAATATAAAATCTAAGTAGGATAAAATCACACTACATCATTGTTTCCTTTTAGTGCTCGACAGTGTATACTATTTTTAACACTCATAAATAAAAATGAAAACGATTTCCGTTGTTACGTTGTTATGCGTACTACCTGCTGTTGTTTATTCAACATGTACTGTACCCACTATGAATAACGCTAAATTAACGTCTACCGAAACATCGTTTAATGATAAACAGAAAGTTACGTTTACATGTGATTCAGGATATCATTCTTTGGATCCAAATGCTGTCTGTGAAACAGATAAATGGAAATACGAAAATCCATGCAAGAAAATGTGCACAGTTTCTGATTATGTCTCTGAACTATATGATAAGCCATTATACGAAGTGAATTCCACCATGACACTAAGTTGCAACGGTGAAACAAAATATTTTCGTTGTGAAGAAAAAAATGGAAATACTTCTTGGAATGATACTGTCACGTGTCCTAATGCGGAATGTCAACCTCTTCAATTAGAACACGGATCGTGTCAACCAGTTAAAGAAAAATACTCATTTGGGGAATATATGACTATCAACTGTGATGTTGGATATGAGGTTATTGGTGTTTCGTATATAAGTTGTACGGCTAATTCTTGGAATGTTATTCCATCATGTCAACAAAAATGTGATATACCGTCCCTATCTAATGGATTAATTTCCGGATCTACATTTTCTATCGGTGGCGTTATACATCTTAGTTGTAAAAGTGGTTTTACACTAACGGGGTCTCCATCATCCACATGTATCGACGGTAAATGGAATCCCATACTCCCAACATGTGTACGATCTAACGAAGAATTTGATCCAGTGGATGATGGTCCCGACGATGAGACAGATCTGAGCAAACTCTCGAAAGACGTTGTACAATATGAACAAGAAATAGAATCGTTAGAAGCAACTTATCATATAATCATAATGGCGTTGACAATTATGGGTGTCATATTTCTAATCTCCATTATAGTATTAGTTTGTTCCTGTGACAAAAATAATGACCAATATAAGTTCCATAAATTGCTACCGTGAATATAAATCCGTTAAAATAATTAATAATTAATAATTAATAACGAACAAGTATCAAAAGATTAAAGAATTAGCTAGAATCAATTAGATGTCTTCTTCAGTGGATGTTGATATCTACGATGCCGTTAGAGCATTTTTACTCAGGCACTATTATGACAAGAGATTTATTGTGTATGGAAGAAGTAACACCATATTACATAATATATACAGGCTATTTACAAGATGCACCGTTATACCGTTCGATGATATAGTACGTACTATGCCAAATGAATCACGTGTTAAACAATGGGTGATGGATACACTTAATGGTATAATGATGAATGAATTCGATACTGTATGTGTGGGTACCGGACTACGATTCATGGAAATGTTTTTCGATTACAATAAAAATAATCCCAAAAATAGCATCAACAATCAAATAATGTATGATATAATTAATAGCGTAGCCATAATTCTAGCTAATGAGAGATATAGAAGCGCGTTTAACGACGATAGAATATACATCCGTAGAACTATGATGGACAAATTGTACGAATACGCATCTCTAACTACTATTGGTACGATCACTGGAGGTGTTTGTTATTTTATCTGTTGATGCATCTAGTTAGTTTGTATAAATAATTATTTCGATATACTAGTTAAAATTTTAAGATTTTAAATGTATAAAAAACTAATAACGTTTTTATTTGTAATAGGTGCAGTTGCATCCTATTCGAATAATGAGTACACTCCGTTTAATAAACTGAGTGTAAAACTCTATATAGATGGAGTAGATAATATAGAAAATTCATATACTGATGATAATAATGAATTGGTGTTAAATTTTAAAGAGTACACAATTTCTATTATTACAGAGTCATGTGACGTCGGATTTGATTCCATAGATATAGATGTTATAAACGACTATAAAATTATTGATATGTATACCATTGACTCGTCTACTATTCAACGCAGAGGACATACGTGTAGAATATCTACCAAATTATCATGCCATTATGATAAGTACCCTTATATCCACAAATATGAGGGTGATGAACGACAATATTCTATTACCGCAGAGGGAAAATGCTATAAAGGAATAAAATATGAAATAAGTATGATGAACGATGATACTCTATTGAGAAAACATACTCTTAAAATTGGATTTACTTATATATTCGATCGTCATGGGCATAGTAATACATATTATTCAAAATATGATTTTTAAAAATTTAAAATATATTATCACTTCAGTGACAGTAGTCAAATAACAAACAACACCATGAGATATATTATAATTCTCGCAGTTTTGTTCATTAATAGTATACATGCTAAAATAACTAGTTATAAGTTTGAATCCGTCAATTTTGATTCCAAAATTGAATGGACTGGGGATGGTCTATACAATATATCCCTTAAAAATTATGGCATCAAGACGTGGCAAACAATGTATACAAATGTACCAGAAGGAACATACGACATATCCGGATTTCCAAAGAATGATTTCGTATCTTTCTGGGTTAAATTTGAACAAGGCGACTATAAAGTGGAAGAGTATTGTACGGGACTATGTGTCGAAGTAAAAATTGGACCACCAACTGTAATATTGACTGAATATGACGATCATATCAATTTGTTCATCGAGCATCCGTATGCTACTAGAGGTAGCAAGAAGATTCCTATTTACAAACGCGGTGACATGTGTGATATCTACTTGTTGTATACGGCTAACTTCACATTCGGAGATTCTGAAGAACCAGTAACATATGATATCGATGACTACGATTGCACGTCTACAGGTTGCAGTATAGACTTTGCCACAACAGAAAAAGTGTGTGTGACAGCACAGGGAGCCACAGAAGGGTTTCTCGAAAAAATTACTCCATGGAGTTCGGAAGTATGTCTGACACCTAAAAAGAATGTATATACGTGCGCAATTAGATCTAAAGAAGATGTTCCCAATTTCAAGGACAAAATAGCCAGAGTTATCACGAGAAAATTTAATAAACAGTCTCAATCTTATTTGACTAAATTTCTCGGTAGCACATCGAATGATGTTACAACTTTTTTTAGCATTCTTGACTAAATATTCATAACTAATTTTTATTAATGATACAAAAATGAAATAAACTGTATATTATACACTGGTTAACGCCCTTGGCTCTAACCATTTTCAAGATGAGGTCCCTGATTATAGTCCTTCTGTTCCCCTCTATCATCTACTCAATGTCTATTAGACGATGCGAGAAGACTGAAGAGGAAACATGGGGATTAAAAATAGGGTTGTGTATAATTGCCAAAGATTTCTATCCCGAAAGAACTGATTGCAGTGTTCATCGCCCAACTGCAAGTGGAGGATTGATAACTGAAGGCAATGGATTCAGAGTAGTTATATATGATCAATGTACAGAACCCCATGACTTTATTATCACCGATACTCAACAAACACGTCTTGGATCATCTCATACATATATTAAATTCAGTAACATGAATACAGGTGTCCCATCTAGTATTCCAAAATGTTCCAGAACTCTCTGTATTTCTGTATATTGTGATCAAGAGGCGGGAGACATAAAATTTGAGGAGTATACTCAAGAATCAAGTGATATCAGTATTAGAGTTAAGTATGATTCATCATGTATTGATTATCTGGGTATTAATCAAAGTTTCATGAATGAATGTATTCGAAGAATTACAACATGGGATAGAGAATCATGCGTCAGAATTGATACACAGACTATAAATAAATATCTTAAGTCTTGCACCAACACAAAATTCGACCGTAATGTCTACAAAAGGTACATACTGAAGAGTAAAGCACTCCATGCTAAAACAGAGTTGTAATAGATATAAAATACTTTTTATAATAATTAGGCTAGAAAAATCTCACTCACATGTAATCTTAAAAAAATGATATGATAGTTCTTACAAGTAGCGATTGAGTTTTAAATGGATTCTATTAATTACCGGGGAACTTAACAATTCGTTCTGATCTACAGACATTGGTTAATAAATCATCTTATTTTGCCAATATATTAAAATGTGGAAACTCCACTAATAATATTACATTGTGCGACTTTCAAGATGATGTGATATATAGGGTTATACAGTTTTAACAATTATATAATAGAGATAGAAAGTACAAAAGATGTAGAATCAATGATATGGCACGCTAAACAGTTGGGTGTGGAATCATTGCTAAAAGAATGTCAAAATTATTTGCTTAGAATATTACGTATATAATTGTTTAGAAATTTATAGAATAACTAATATTAATACATTATCGTATATCTACAACGATATAAGAAACTTCATATTGGATAATATTACTATTAATATATAAGGATCCAGATTTTATATATTTGCCTAAATACATTATTATAGATTTACTAGGACAATCACCTAAATGTTTTTAACGAAGATAATGTGGTAAAGATTATATACACTTATATATCTTCCGATATCTACAAGGATATTCCATATCATCATTGTGTAAACTAAATAACGTTTTCTATGGCATTTAATAAGGACATTGGATATGTGGAAAAGTGATGTATGGAAGTTAGTACATTATCAACTTCTCCTTATTGATTGAAAATGAAAATATAAATAGTTTTTATGTATAGCGGTATCTACCCTATAGTTTTATTGCTTACTACTAACATGGATTCAGATACAGATACAGATACAGATACAGATACAGATACAGATACAGATACAGATGTAGAAGATATCATGAATGAAATAGATAGAGAGAAAGAAGAAATACTAAAAAATGTAGAAATTGAAAATAATAAAAACATTAACAAGAATCATCCCAGTGAATATATTAGAGAAGCACTTGTTATTAATACCAGTAGTAATAGTGATTCCATTGATAAAGAAGTTATAGAATATATCAGTCACGATGTAGGAATATAGATCATATCTACTAATTTTTATAATCGATACAAAACATAAAAACAACTCGTTATTACATAGCAGGTATGGAATCCTTCAAGTATTGTTTTGATAACGATGGTAAGAAATGGATTATCGGAAATACTTTATATTCTGGTAATTCAATACTCTATAAGGTCAGAAAAAATTTCACTAGTTCGTTCTACAATTACGTAATGAAGATAGATCATAAATCACACAAGCCATTGTTGTCCGAAATACGATTCTATATATCTGTATTGGATCCTTTGACTATCAACAACTGGACACGGGAACGTGGTATAAAGTATTTGGCTATTCCAGATCTGTATGGAATTGGAGAAACCGATGATTATATGTTCTTCATTATAAAGAATTTGGGAAGAGTATTCGCCCCAAAGGATAGTGAATCAGTTTTCGAAGCATGTGTCACTATGATAAACACGTTAGAGTTTATACACTCTCAAGGATTTACTCATGGAAAAATAGAACCGATGAATATACTGATTAGAAATAAACGTATTTCACTAATTGACTATTCTAGAACTAACAAACTATACAAAAGTGGAACACATATAGATTACAACGAGGACATGATAACTTCAGGAAATATCAATTATATGTGTGTAGACAATCATCTTGGAGCAACAGTTTCAAGACGAGGAGATTTAGAAATGTTGGGATATTGCATGATAGAATGGTTCGGTGGTAAACTTCCATGGAAAAACGAAAGTAGTATAAAAGTAATAAAACAAAAAAAAGAATATAAACAATTTATAGCTACTTTTTTTGAGGACTGTTTTCCTGAAGGAAATGAACCTCTGGAATTAGTTAGATATATAGAATTAGTATACATGTTAGATTATTCTCAAACTCCTAATTATGACAGACTACGTAGACTGTTTATACAAGATTGAAATTATATTCTTTTTTTTATAGAGTGTGGGGGTAGTGTTACGGATATCTAATATTAATATTAGACTATCTCTATCGCGCTACACGACCAATATCGATTACTATGGATATCTTCAGGGAAATCGCATCTTCTATGAAAGGAAAGAATGTATTCATTTCTCCAGCGTCAATCTCGTCAGTATTGACAATACTGTATTATGGAGCTAATGGATCCACTGCTGAACAGCTATCAAAATATGTAGAAAAGGAGGAGAACATGGATAAGGTTAGCGCTCAGAATATCTCATTCAAATCCATGAATAAAGTATATGGGCGATATTCTGCCGTGTTTAAAGATTCCTTTTTGGGAAAAATTGGCGATAAGTTTCAAACTGTTGACTTCACTGATTGTCGCACTATAGATGCAATCAATAAGTGTGTAGATATCTTTACTGAGGGAAAAATCAATCCACTATTGGATGAACCATTGTCTCCTGATACCTGTCTCCTAGCAATTAGTGCCGTATACTTTAAAGCAAAATGGTTGATGCCATTCGAAAAGGAATTTACCAGTGATTATCCCTTTTACGTATCTCCAACGGAAATGGTAGATGTAAGTATGATGTCTATTTACGGCGAGCCATTTAATCACGCATCTGTAAAAGAATCATTCGGTAACTTTTCAATCATAGAACTGCCATATGTTGGAGATACTAGTATGATGGTCATTCTTCCAAACAAGATTGATGGATTAGAATCCATAGAACAAAATCTAACAGATACAAATTTTAAGAAATGGTGTAACTCTCTGAAAGCTACGTTTATCGATGTGCACATTCCTAAGTTTAAGGTAATAGGTTCGTATAATCTTGTGGATACGCTAATAAAGTTGGGACTGACAGATGTGTTCTATTCAACTGGTGATTATATCAATATGTGTAATTCAGATGTGAGTGTTGACGCTATGATTCACAAAACGTATATAGATGTCAATGAAGAGTATACAGAAGCAGCTGCAGCAACTTCTGTACTAGTGGCAGACTGTGCATCAACAGTTACAAATGAGTTCTGTGCAGATCATCCGTTCATCTATGTGATTAGACATGTCGATGGTAAAATTCTTTTCGTTGGTAGATATTGCTCTCCAACAACTAATTAAGCACATTCTTAATATTAGAATATTATATAGTTAAGATTTTTACTAACAGGTTAACATTTTTTTTTAAAAATAGAAAAAACATGTGGTATTAGTGCAGGTCGTTATTCTTCCAATTGCAATTGGTAAGATGACGGCCAACTTTAGTACCCACGTCTTTTCACCACAACACTGTGGATGTGACAGACTGACCAGTATTGATGACGTCAGACAATGTTTGACTGAATATATTTATTGGTCGTCGTATGCATACCGCAACAGGCAATGCGCTGGACAACTGTATGACACACTCCTCTCTTTTAAAGATGATGCGGAATCAGTGTTCATCGACGTTCGTGAGCTGGTAAAAAATATGCCGTGGGATAATGTTAAGGATTGTACAGAGATCATCCGTTGTTATATACCGGATGAGCAAAAAACCATCAGAGAGATTTCGGCCATCATTGGACTTTGTGCATATGCTGCTACTTACTGGGGAGGTGAAGACCATCCCACTAGTAACAGTCTGAACGCATTGTTTGTGATGCTTGAGATGCTCAATTACATGGATTATACCATCATATTCTGGCGTATGAATTGATGAGTTACAGCTTGACATTTCTTCTTTCCTCCCTCTTCTTCTACCTTTCCCAGAAACAAACTTTTTTTACCCACTATAAAATAAAATGAGTATACTACCTGTTATATTTCTTCCTATATTTTTTTATTCTCCATTCGTTCAGACTTTTAACGTGCCTGAATGTATCGACAAAGGGCAATATTTTGCATCATTCATGGAGTTAGAAAACGAGCCAGTAATCTTACCATGTCCTCAAATAAATACGCTATCATCCGGATATAATATATTAGATATTTTATGGGAAAAACGAGGAGCGGATAATGATAGAATTATACAGATAGATAATGGTAGCAATATGCTAATTCTGAACCCGACACAATCAGACTCTGGTATTTATATATACATTACCACGAACGAAACCTACTGTGACATGATGTCGTTAAATTTGACAATCGTGTCTGTCTCAGAATCAAATATAGATCTTATCTCGTATCCACAAATAGTAAATGAGAGATCTACTGGTAAAATGGTATGTCCCAATATTAATGCATTTATTTCTAGTAACGTAAACACAGAATTATATGGAGCGGACATCGACGCCTTAGAAATAAGAGACTTAAACAACGGACACCTGGAATTATTACCATAGAAGATGTTAGAAAAAATGATGCTGGTTATTATACATGTGTTTTAGAATATATATATATATATATATATATATATATATATATATATGGGCAAAACATATAACGTAACCAGAATTATAAAATTAGAGGTACGGGATAGAATAATACCTCCTACTATGAAATTACCAGAAGGAGTAGTAACTTCAATAGGTAGTAATTTGACTATTGCATGTAGAGTATCGTTGAGACTTCCCACAACGGACACCGACGTCTTTTGGATAAGTAATGGTATATGTATTACGAAGAAGAAGACGAGGACGGAGACGGTAGAATAAGTGTAGCAAATAAAATCTATATGACCGATAAGAGACGTGTTATTACATCCTGGTTAAACATTAATCCTGTCAAGGAAGAAGATGCTACAACGTTTACGTGTATGGCGTTTACTATTCCTAGCATCAGCAAAACAGTTACTGTTAGTAAACGTGAATGTATGTTGTTACATTTCCATATCAATTGAGTTTATAAGAATTTTTTATACATTATCTTCCAACAAACAATTGACGAACGTATTGCTATGATTAACTCCCACAATACTATATATATTATTAATCATTAACTTGCAGACTATACCTAGTAGTGCTATTTTGACATACTCATGTTCTTGTGTAATCGCAGTATCTATATTATTAAAGTACGTAAATCTAGCTATAGTTTTATTATTTAATTTTAGATAATATACTGTCTCCGTATTTTTAAAAAATTACCACATCCTTTATTAAATCATGAATGGGAATTTCTGTGTCATCGTTAGTATATTGTGAACAACAAGAGCAGATATCTATAGGAAAGGGTGGAATGCGATACATTGATCTATGTAGTTTTAAAACATACGCGAACTTTGAAGAATTTATATAAATCATCTCACGAGATATTGCTCTCTGTCATATTCATACACCTGTATAAACTTTCTAGACATCTTACAATGTGTTATTTTATGATCATATTTACATATTTACTGGTATATCAAAGATGTTAGATTAGTTAATGGGAATCGTCTATAATAATGAATATTAAACAATTATAGGAGGAGTTTATACCTACAAAAACATCATAAAAATGAGTCATCGTCCGATTTATGTTTTAAATATACTAACATTACTACCTTCAGAAATTATATACGAAATATTATACATGCTGACAATTAACGATCTTTATAATATATAGTATCCACCTACCAAAGTATAATTGTATTTTTCTCATGTGATGTGTGTAAAAAACTGATATTATATAATTATCTTAGTACCTATGATGAAGATGAAGATGAAGATGAAGATGATGGTCCGTATATATTTTGTATCATTATCGTTATTGCTATTCCATAGTTACGCCATAGACATCGAAAATGAAATCACCGAATTCTTCAATAAAATGAGAGATACTCTACCAGCTAAAGACTCTAAATGGTTGAATCCAGTATGTATGTTTGGAGGCACAATGAATGATATGGCCGCTCTAGGAGAGCCATTCAGTGCAAAGTGTCCTCCTATTGAAGACAGTCTTTTATCGCATAGATATAAAGACTATGTGGTTAAATGGGAAAGGCTAGAAAAGAATAGACGGCGACAGGTTTCTAATAAACGTGTTAAACATGGTGATTTATGGATAGCCAACTATACATCTAAATTCAGTAACCGTAGGTATTTATGTACCGTAACCACAAAGAATGGTGACTGTGTTCAGGGTGTAGTTAGATCTCATGTGTGGAAACCTTCTTCATGCATTCCAAAAACATATGAACTAGGTACTTATGATAAGTATGGCATAGACTTATACTGTGGAATTCTTTATGCGAACCATTATAATAATATAACTTGGTATAAAGATAATAAGGAAATTAATATCGACGATTTTAAGTATTCACAAGCGGGAAAGGAATTAATTATTCATAATCCAGAGTTAGAAGATAGTGGAAGATACGACTGTTACGTTCATTACGACGACGTTAGAATCAAGAATGATATCGTAGTATCAAGATGTAAAATACTTACGGTTATACCGTCACAAGACCACAGGTTTAAACTAATACTAGATCCGAAAATCAACGTAACGATAGGAGAACCTGCCAATATAACATGCAGTGCTGTGTCAACGTCATTATTTGTCGACGATGTACTGATTGAATGGGAAAATCCATCCGGATGGATTATAGGATTAGATTTTGGTGTATACTCTATTTTAACTAGTAGAGGCGGTATCACCGAGGCGACTTTGTATTTTGAAAATGTTACTGAAGAATATATAGGCAATACATATACATGTCGTGGACACAACTATTATTTTGATAAAACTCTTACAACTACAGTAGTATTGGAGTAAATACACAATGCATTTTTATATACATTACTGAATTATTATTATTAATTATATCGTATTTGTGCTATAGAATGGATGAAGATACGCGACTATCTAGGTATTTGTATCTCACCGATAGAGAACATATAAATGTAGACTCTATTAAACAGTTGTGTAAAATATCAGATCCTAATGCATGTTATAGATGTGGATGTACGGCTTTACATGAGTACTTTTATAATTATAGATCAGTCAACGGAAAATACAAGTATAGATACAACGGTTACTATCAATATTATTCATCTAGCGATTATGAAAATTATAATGAATATTATTATGATGATTATGATAGAACTGGTATGAACAGTGAGAGTGATAATATATCAATCAAAACAGAATACGAGAATGAATATGAATTCTATGATGAAACACAAGATCAAAGTACACAACTAGTAGATTACGACATTAAACTCAAAACCAATGAGGATGATTTTGTTGATGAATTCTATGGTTATGATAGATCAGTGGGTGTCCATGATTATATAGATGTATCAATTAATAAAGTAGTATATGGAAGAGAGTCTCACGTAAGATGGTGGGATATATGGCAAGAACATAATGATGGCGTATACAGTATAGGAAAGGAGTGCATAGATAATATATACGAAGACAGACATACCGTAGACGAATTCTACAAGATAGACAGCGTATCAGATGTAGATGACGCAGAACATATATCTCAGATAACTAATGATGTATCTACACAAACATGGGAAAAGAAATCAGAGTTAGATAGATACATGGAAATGTATCCTCGTCATAGATATGGTAAGCATTCTGTCTTTAAGGGATTTTCTGACAAAGTTAGAAAAAATGATTTAGACATGAACGTGGTAAAAGAATTACTTTCTAACGGTGCATCTCTAACAATCAAGGATAGCAGTAATAAGGATCCAATTGCTGTTTATTTTAGAAGAACAATAATGAATTTAGAAATGATTGATATCATTAACAAACATACAACTATCTATGAACGCAGGTATATAGTACACTCCTATCTAAAAAATTATAGAAATTTCGATTATCCATTTTTCAGAAAGTTAGTTTTGACTAATAAACATTGTCTCAACAATTATTGTAATATAAGCGACGGCAAATATGGAACACCACTACATATATTAGCATCTAATAAAAAAATAATAACTCCTAATTACATGAAGTTATTAGTGTATAACGGAAATGATATAAACGCACGAGGTGAAGATACACAAATGCGAACTCCATTACACAAATATTTGTGTAAATTTGTATATCATAATATTGAATATGGTATCCGATACTATAATGAAAAGATTATAGACGCATTTATAGAGTTAGGAGCCGATCTAACTATTCCAAATGACGATGGAATGATACCAGTAGTTTACTGTATACACTCAAATGCCGAATATGGTTATAACAATATTACTAACATAAAGATAATACGTAAACTACTTAATCTTAGTAGACATGCGTCACATAATCTATTTAGAGATCGAGTCATGCACGATTATATAAGTAATACATATATTGATCTTGAGTGTTTAGATATCATTAGATCACTTGATGGGTACGATATTAATTGTTACTTTGAAGGACGTACACCACTTCATTGCGCTATACAATATAACTTCACTCAGATTGCTGAGTACTTATTAGATCGAGGAGCTAATATATCATTAAAGACAGACGATGGTAAAACTGTATTTGATTTATCGTTATGTAGTTACATTCCTCTTAAATGGACTAGCTTTTTGATTAGTCGTCTACCGCCTAAAAGTGTCATATGCTCACTGACTAACCATATAATAGATTATGTTCTTACGAACAATAGACGTATTATTTGGCAGAGTCAAATGATTAATAAGTACGTACTGTTACTGGACCCATCCTTTTATTATAGATTCAGAAATGTTATCGAAAACAAATTAGACCAATACAATAATCGTTATAATATGTTCGAACACGATAGGGACGTTAATGAAAAGTATGGCAAAGTCTTACATGACCTCGATACATATATCAAGGATGTACAAGTATTAAAATCTACTTCCATCACTAATAATATAACACTATACGACACTATTATAAATAATAAGTCAGAGTTTCCTATACGTCGTGTAAACGACAAACAATTAATTAATCTCATAAAATCCAATACATATCATAATCTTATCGAAAAAGTTATTAAAAATACATTAGAGAAATATACTTTAACTAATATAGTCCTCGAGTATATGATCTCATCTCGATCTCAATCATCTTATTTGAGTCGTATTCCTAATGAGATATTACTCGAAATATTATATAAACTCGACATGTACGATTTACGTAATCTATATACAAGATATATGAGAGAGAATGATATCACAGAGTATCATATAGAGAATACGAGGTCTGTTTCTACACAGACATGAATAATGAATACACATACAACGTTTTTTTTAATCTTAGATATAACACTAATTACATCAAGATTATATATTGAAATCGTAATTTGAGTTGTCTGATCATCATGGATATCGAAAATGATATACGTAACATTAGCAATCTTTTAGATGATTGATATATTATTATGCGATGTAATCATAACTATCGGAGATGTAGAAATTAAAGCGCATAAAACTATTTTGGTTGCCGGATCTACGTATTTTAAAACAATGTTCACAACATCTATGATAGCGAGAGATCTAGCAACTAGAGTAAATATACAGATGTTCGATAAAGATGCCGTCAAAAATATTGTACAGTACTTATACAATAGGTATATAAGTTCTATGAATGTGATAGACATATTAAAATGCACCGACTAAGAACGTAAAACGAACTATAGAATGTTATACAATGGGTGATGATAAGTAGAAGATGTTACCCGATATACCCATAGCATTATCTAGTTATGGCATGTGTGTATTAGATCAATACATATACATTATAAGCGGTCGTACCCAACACTGATTATACATCGGTACATACAGTAAATAGCATAGATATGGAGGAGGATACAAATATTTCAAATAAAGTTATGAGATACGCGCTGTCAATAATATATGGAAGACATTACCTAACTTCTGAACTGGAACTATAAATCCAGGCTCTCGCATAAAGATGAATATATATGTTGTATGCGACATCAAAGATGAAAAAATGTTAAGACTTATATATTTAGATATAACACGAATATGTATGACGGATGGGAATTGGTAACGATGACAGAAAGCAGATTGTCAGCTCTGCATACTATTCTTCATGACAATACCATAATGATGTTACATTGTTATGAAGCGTATATGTTACAAGATACATTTAATGTGCTTACGGAACATATATTTAGAAACATCTACTAACGATTTTTTATGCTTGTATTATTAATGGTATGTAATATGATTTAATTGATTGTGTACACGATACCAATTTGTCGAGTATGAATACGGAGTACAAACATAAACTGAAGTTTAACATTATTTATTTATGATATACATTATATACATTATATACATTATATACATTATATACATTATATACATTATATACATTATATACATTATATACATACATTATATACATTATATACATTATATACATTATATACATTATATACATTATATATCGTTATTGTTTGGTCTATGCCATGGATATCTTTAAAGAACTAATCTTAAAACATACGGATGAAAATGTTTTGATTTCTCCAGTTTCCATTTTATCTACTTTATCTATTCTGAATCATGGAGCAGCTGGTTCTACAGCTGAACAACTATCAAAATATATAGAGAATATGAATGAGAATACACCCGATGATAAGAAGGATGACAATAATGACATGGACGTAGATATTCCGTATTGCGCGACACTAGCTACCGCAAATAAAATATACGGTAGTGATAGTATCGAGTTCCATGCCTCATTCCTACAAAAAATAAAAGACGATTTTCAAACTGTAAACTTTAATAATGCGAACCAAACAAAGGAACTAATCAACGAATGGGTTAAGACAATGACAAATGGTAAAATTAATTCCTTATTGACTAGTCCGCTATCCATTAATACTCGTATGATAGTTATTAGCGCCGTCCATTTTAAAGCAATGTGGAAATATCCATTTTCTAAACATCTTACATATACAGACAAGTTTTATATTTCTAAGAATATAGTTACCAGTGTTGATATGATGGTGGGTACCGAGAATGACTTGCAATATGTACATATTAATGAATTATTCGGAGGATTCTCTATTATCGATATTCCATACGAGGGAAACTCTAGTATGGTGATTATACTGCCGGACGACATAGAAGGTATATATAACATAGAAAAAAATATAACAGATGAAAAATTTAAAAAATGGTGTGGTATGTTATCTACTAAAAGTATAGACTTGTATATGCCAAAGTTTAAAGTGGAAATGACGGAACCGTATAATCTGGTACCGATTCTAGAAAATTTAGGACTTACTAATATATTTGGATATTATGCAGATTTTAGTAAGATGTGTAATGAAACTATCACTGTAGAAAAATTTCTACATACGACGTTTATAGATGTTAATGAGGAGTATACAGAAGTATCGGCCGTTACAGGAGTATTCATGACTAACTTTTCGATGGTATATCGTATGAAGGTCTACATAAACCATCCATTCATATACATGATTAAAGATAACACCGGACATACACTTTTTATAGGGAAATACTGCTATCCGCAATAAATATAAACAATAGACTTTTATCACGTTATCTCATGTATAAAATATTACAAATAGTATAGCATAAACTAAAGTCGATACATACATTAAAACTTAAATAATAATGTAATTTACAATTAATAGTATAAACTAAAAAAATTAAAAAATTAAAAACAATATCATTATTATAAGTAATATCAAAATGACGATATACGGATTAATAGCGTATCTTATATTCGTGACTTCATCCATCGCTAGTCCACTTTACATTCCCGTTATTCCGCCCATTTCGGAAGATAAATCGTTCAATAGTGTAGAGGTATTAGTTTCTTTGTTTCCCGATGACCAAAAAGACTATACAGTAACTTCTCAGTTCAATAACTACACTATCGGTACCAAAGACTGGACTATCAACGTACTATCCACACCTGATGGTCTGGACATACCATTGACTAATATAACTTATTGGTCACGGTTTACTATAGGTCGTGCATTGTTCAAATCAGAGTCTGAGGATATTTTCCAAAAGAAAATGAGTATTCTAGGTGTTTCTATAGAATGTAAGAAGCCGTCGACATTACTTACTTTTTTAACCGTGCGTAAAATGACTCGAGTATTTAATAGATTTCCAGATATGGCTTATTATCGAGGAGACTGTCTAGAAGCCGTTTATGTAACAATGACTTATAAAAATACTAAAACTGGAGAGACTGATTACACGTACCTCTCTAATGGGGGGTTGCCTGCATACTATCGTAATGGGGTCGATGGTTGATTATTGATTAGTATATTCCTTATATTCCTTATTCTTTTTATTCACACAAAAAGAACATTTTTATAAACATGAAACCACTGTCTAAATGTAATTATGATCTTGATTTATAGATGATGATCAGCCTTCAGAGGATTTTGACCAGTATGTTTAATATGAAAAAAAACATAACTATTAAGCGCTATTGCGCTATTGTGCTTAATTATTTTGCTCTATAAACTGAATATATAGCCACAATTATTGACGGGCTTGTTTGTGACCGACAATCATGAATTTTCAGAAATTATCTCTGGCTATATATCTTACGGTGACATGTTCGTGGTGTTATGAAACATGTATGAGAAAAACTGCGTTGTATCATGACATTCAATTGGAGCATGTAGAAGACAATAAAGATAGTGTAGCATCGCTACCGTACAAGTATCTACAAGTAGTCAAACAAAGAGAACGTAGTAGATTGTTGGCTACATTTAATTGGACGGATATAGCTGAGGGTGTTAGAAATGAGTTCATTAAAATATGTGATATCAACGGAACATATCTATATAATTATACTATTGCTGTTAGTATAATTATTGATTCCACGGAAGAACTACCAACAGTTACTCCAATTACAACATATGAACCTTCTATATATAATTATACTATCGATTATAGCACTGTTATTACTACTGAAGAACTACAAGTGACTCCAACATATGCACCTGTAACAACTCCTCTTCCAACATCAGCAGTTCCTTATGATCAACGATCGAATAACAATGTAAGTACTATATCTATTCAGGTACTGAGTAAAATATTGGGAGTCAATGAAACAGAATTAACTAATTATCTTATTATGCATAAAAATGACACTGTTGACAATAACACCATGGTTGATGATGAGACATCTGACAATAACACATTACATGGTAATATAGGATTTTTGGAAATAAATAATTGTTATAATGTTTCTGTGTCAGATGCTAGTTTTAGAATAACATTAGTAAACGATACTTCTGAAGAAATTTTGCTAATGCTAACAGGAACTAGTTCATCCGACACCTTCATATCTTCCACCAATATCACTGAATGTTTGAAAACATTAATCAATAATGTGTCGATTAATGATGTACTTATAACACAAAATATGAATGTAACATCTAATTGTGATAAATGCTCAATGAATTTGATGGCATCCGTTATTCCTGCAGTTAATGAATTTAACAATACGTTGATGAAAATTGGTGTAAAAGATGATGAAAACAATACGGTATATAAATATTATAATTGTAAACTAACTACAAATTCTACATGTGATGAGTTAATCAATTTAGATGAAGTCATTAACAACATAACTCTGACAAATATTATACACAATAGTGTTTCGACAACTAACAGCAGAAAAAGACGAGATCTGAATGATGAGTTTGAATTTTCCACTTCCAAGGAATTAGATTGTCTTTACGAATCATATGGTGTAAACGATGATATAAGTCATTGTTTTGCATCACCTAGACGTAGACGATCTGACGACAAAAAGGAGTACATGGACATGAAATTATTCGACCACGCGAAAAAAGATTTAGGAATAGACAGTGTTATTCCTAGAGGTACAACCCATTTCCAAGTAGGTGCATCTGGTGCAAGTGGTGGTGTTGTAGGAGATAGTTTCCCATTTCAAAATGTTAAATCGCGTGCCAGTCTATTGGCGGAAAAAATAATGCCTAGAGTACCTATTACTGCTACCGAAGCTGATCTATATGCAACTGTAAATAGACAACCCAAGTTACCAGCAGGTGTTAAAAGTACTCCGTTTACAGAGGCGCTTGTGTCTACGATAAACCAAAAGCTTTCTAATGTTAGAGAGGTAACTTATGCTTCGCTCAATCTGCCAGGATCAAGTGGCTATGTTCATAGACCATCTGATTCTGTTATTTATAGCAGTATAAGACGGTCACGTTTACCTAGTGATAGCGATAGTGATTATGAGGATATACAAACTGTTGTTAAGGAATATAATGAAAGATATGGTAGATCAGTCAGTAGAACACAGTCATCAAGTAGTGAAAGCGATTTTGAAGATATAGATACTGTTGTTAGGGAATATAGACAAAAATATGGCAATGCAATGGCAAAAGGACGTAGTAGTTCCCCTAAACCTGATCCATTATATAGTACTGTTAAGAAAACAACTAAAAGTCTATCTACTGGTGTAGACATAGTTACAAAACAATCAGACTATTCTCTATTACCTGACGTTAATACTGGCAGTTCTATTGTGTCACCTCTCACCAGAAAAGGAGCTACTAGACGACGACCTAGACGCCCTACAAATGATGGTCTACAGAGTCCAAATCCTCCTCTCCGTAATCCACTTCCTCAACATGATGATTATTCTCCTCCACAAGTACACAGACCTCCACCACTTCCTCCTAAACCAGTCCAAAATCCGCCACAACTTCCCCCTAGACCAGTAGGTCAATTACTACCTCCTCCTATAGATCAACCAGATAAAGGATTTAGTAAGTTTGTATCACCTAGACGGTGTAGAAGAGCAAGCTCTGGAGTCATATGTGGTATGATACAATCAAAACCAAACGATGATACCTATTCACTTCTTCAACGATCAAAAATTGAACCAGAATATGTGGAGGTTGGTAATGGTATACCCAAGAACAATGTTCCTGTAATAGGTAATAAACATAGTAAAAAATATACATCGACGATGTCAAAAATATCAACAAAATTTGATAAATCTACGGCATTTGGAGCAGCAATGTTACTAACTGGTCAGCAGGCCATTAGCCAACAGACTAGATCAACTACGTTGAGTAGAAAAGATCAGATGAGTAAGGAAGAAAAGATATTCGAAGCAGTTACAATGAGTCTATCAACTATAGGTTCAACGTTGACGTCTGCAGGTATGACGGGTGGTCCAAAACTAATGATTGCAGGAATGGCTATAACGGCTATAACTGGTATAATAGATACGATAAAAGATATATATTACATGTTTTCAGGACAGGAGAGGCCAGTAGATCCTGTTATTAAATTATTTAATAAGTACACTGGCTTAATGTCCGATAATAATAAAATGGGTGTAAGAAAATGTTTGACACCCGGTGACGACACACTTATTTATATCGCATACAGAAACGATACCAGTTTTAAACAGAATACGGATGCGATGGCTTTGTATTTCTTAGATGTTATCGATTCAGAGATCCTATATCTAAACACATCAAATTTAGTTCTAGAGTATCAACTAAAGGTGGCTTGCCCCATAGGAACATTAAGATCTGTAGATGTGGACATAACTGCGTATACAATATTATATGATACAGCGGATAATATTAAGAAATACAAGTTTATCAGAATGGCAACGCTACTATCCAAACATCCAGTTATTAGATTGACATGTGGTTTAGCAGCAACATTGGTGATTAAACCGTACGAGGTACCCATCAGTGATATGCAACTACTAAAAATGGCGACGCCTGGTGAACCAGAATCCACTAAATCTATACCATCCGATGTCTGTGATAGGTATCCTCTAAAGAAATTCTATCTTTTAGCTGGTGGTTGTCCCTATGATACATCTCAAACTTTTATTGTACATACTACTTGCAGTATTCTACTAAGAACAGCTACACGGGATCAGTTTAGAAACAGATGGGTGTTACAAAATCCATTTAGACAAGAAGGGACATATAAGCAACTGTTTACCTTTAGCAAATACGATTTTAACGACACCATAATCGATCCTAATGGTGTGGTGGGTCATGCTAGCTTTTGTACCAATAGAAGCAGCAACCAATGTTTCTGGTCCGAACCTATGATATTGGAAGATGTATCATCGTGTAGTTCTAGAACTAGAAAAATATACGTAAAACTGGGAATATTTAATGCTGAAGGTTTTAATAGTTTTGTACTAAATTGTCCAACTGGGTCTACACCTACATACATCAAACATAAAAATGCGGACAGTAACAATGTTATCATAGAGCTACCTGTAGGTGATTACGGCACAGCCAAATTGTATTCAGCAACAAAACCATCGAGGATAGCTGTGTTCTGCACACATAACTATGATAAACGATTCAAATCAGATATTATAGTTCTAATGTTTAATAAAAACAGCGGTATTCCATTTTGGAGCATGTACACAGGAAGTGTAACTAGTAAAAATAGAATGTTTGCCACATTGGCTAGAGGAATGCCGTTTAGATCAACGTATTGCGATAACAGACGACGATCAGGTTGTTATTATGCAGGAATACCATTTCATGAAGATAGTGTAGAAACAGATATACATTATGGACCAGAAATAATGTTAAAGGAAACATATGACATAAACAGTATTGACCCACGAGTTATAACAAAGTCAAAGACCCATTTTCCTGCTCCATTGAGTGTAAAATTCATGGTTGACAATTTAGGAAATGGATATGACAACCCTAATTCATTTTGGGAAGATGCTAAAACTAAGAAACGGACATATAGTGCAATGACGATAAAAGTCCTACCATGTACAGTGAGAAATAAAAATATAGACTTTGGATATAACTATGGAGATATTATTTCTAATATGGTTTATCTACAATCTACTAGTCAGGATTATGGAGATGGTACCAAATATACATTTAAATCCGTAACTAGATCAGATCATGAGTGTGAATCTAGCTTAGATCTAACGTCTAAGGAAGTAACTGTGACATGTCCTGCGTTTAGTATACCAAGAAATATATCAACATATGAAGGTCTATGCTTTAGTGTTACTACATCTAAAGATCATTGTGCTACAGGTATTGGTTGGTTAAAATCTAGTGGTTATGGGAAGGAAGATGCTGATAAACCACGTGCTTGTTTTCATCATTGGAATTATTACACACTGTCGTTGGATTATTACTGTTCATACGAAGATATTTGGAGAAGCACCTGGCCTGACTATGATCCATGTAAGTCATATATCCATATAGAGTATAGAGATACATGGATAGAATCTAATGTGTTACAGCAACCTCCTTACACATTCGAATTCATTCATGACAATTCTAACGAATATGTGGATAAAGAAATTAGTAACAAATTAAATGATCTGTACAATGAATACAAGAAGATTATGGAATATAGCGACGGATCATTGCCGGCGTCTATAAACAGATTAGCAAAGGCATTGACTTCAGAGGGTAGAGAAATAGCAAGTGTTAATATAGATGGTAATCTGTTAGATATCGCATATCAAGCAGATAAGGAAAAGATGGCCGACATACAGACAAGAATAAATGATATTATTAGAGATTTGTTTATACACACTCTATCAGACAAAGATATAAAAGACATTATAGAATCCGAAGAAGGTAAGAGATGTTGTATAATAGATGTTAAGAACAATCTTGTTAAAAAGTACTATTCTATTGATAATTATCTATGTGATACTTTAGATGATTATATATACACCTCTGTAGAATATAACAAATCCTATGTGTTAGTAAACGATACTTATATGAGCTATGACTATCTTGAATCATCAGGTGTAGTTGTTCTATCATGTTATGAAATGACTATAATCTCCTTGGATACAAAAGACGCCAAAGATGCTATAGAAGATGTGATAGTAGCAAGTGCGGTAGCCGAAGCATTGAATGACATGTTTAAGGAATTTGATAAAAACGTAAGTGCTATTATAATAAAAGAAGAAGATAATTATCTAAACAGTTCGCCCGATATCTACCATATAATATATATCATAGGTGGCACTATTCTGCTACTGTTAGTCATTATTTTAATATTGGCAATTTATATAGCGCGCAATAAATACAGAACCAGGAAATATGAAATAATGAAATATGACAATATGAGCATTAAATCTGAGCATCATGATAGTCTTGAAACAGTGTCTATGGAAATTATTGATAATCGGTACTAATAAAATAGTTTAACTCTTTTAGAACCAGTTTGGTACTGTAATTTCAGTTCATTACTCGTTGAGAATATTGATGATTTTTTTTAAATGAGTATCGGTAGTTACATATTACCATATCATCCATTATATAATCGATGATGCATGTATTAGAATACTTTCCGAATAAGTCTTCTAAATATTGTATTAATTATGAAAAACTATGCTATGTGAGTATGATTCAAAGATGTTTAATGATACGATACTAGATTTTATCTCTAGCGAGATTGTTTAGAATCATTTATCATAACTATGTTTAATAAATTCATCAACGAATATCGATAAAGACCTCTTGTAATTCGAGTATAGGAAGTAGTATTACCATATCAACTTCCGAGTTAACAATTACTCTAAAACATGAGGATTGTACTCCTGTCTTTATTGGAGATCACTATTTAGTCGTTGATAAACTAGTAACCTCAGGTTTCTTTACAAACGATAAAGTACAACATCAAGACCTCACAACACAGTGCAAGATTAATCTAGAAATCAAATGTAATTCTGGAGGAGAATCTAGACAACTAACACCCACGGCGAAGTATACTTTATGCCTCATTCAGAAACGGTAACTGTAGTAGGAGACTGTCTCTCTAATCTCGATGTATATATAATATATACCAATACGGACGCGATATATTCCGACATGAATGGCGTCGCTTATCATATGTTATATCCTAAATGTTGATCATATTCCACAAATGATTGTGAACGAGATTAAATCATCTAACAAATAATTAGTTTTTTATGACATTAACATATAATAAATAAATTAATCATTATTGACTTAACGATGACGAAAGTTATCATTATCTTAGGATTCTTGATTATTAATACAAATTCGTTGTGTCTATGAAATGTGAACAAGGTGTCTCATATTATAATGCACAAGAATTAAAGTGTTGTAAACTATCTAGCCAGGAACATATTCAGATTATCGATGTGATAAATACAGCGATACCATCTGTGGACATTGTCCAAGTGACACATTCACGTCAATATATAATCGTTCTCCTCGGTGTCATAGTTGTAGAGGTCACACCTTGTACACCTACCACAAATAGAATATGTCATTGTGACTCGAATAGTTATCGTCTCCTTAAAGCTTCTGATGGTAACTGTGTTACATGTGCTCCTAAAACAAAATATGGTCGTGTGTACGGAAAGAAAGGAGAAAATGATATGGAATACCATTTGTAAGAAATGTCGGAAGGGTACTTATTCAGATATTGTATCTGACTCTGATCAATGTAAACCTATGACAAGATAAGACTTACTCGCATCTACTGGATAGACATAAATATCCTCCTCGTAATAATGAAATATAATATACACTAATTATTAATATCAATCGAGTATTAACATATAAGTTATTTTTAAACCCCTTTTGGGTTCCGTCCTAAACGGCGTTTCGGTCTGTGTCGCCACCATGGTCACACCGAGCCTCTGCGTGCTCCTCCATCGAGGACGACTTCAACTATGACAGCTCGGTGGCGTCTGCCAGCGTGTACATACGAATGGCATTTCTAAGAAAAGTCTACGGTATCCTTTCTACAATTTCCTTTAACAACGGCAACAGCTGCAGTATTTTTATACTTTGAATGCATCGGACATTTATACAAGGGAGTCCTGTTCTAATATTGGCATCAATGTTCGGATCTATAGGCTTGATTTTCGCATTGACTTTACACAGACATGAACATCCCCTGAATCTGTACATACTTTGTGGATTTACACTGTTAGAATCTCTAACGCTGGCCTCTGTTGTTACTTTCTATGATGCACGTATCGTTATGCAAGCTTTCATGTTGACTACTGCAGTGTTTCTTGCTCTGACTACATATACTCTACAATCAAAGAGAGATTTCAGTAAACTTGTAACAGGATTGTTTGCTGCTTTCTGGATTTTAATTTTGTCAGGAGTCTTGAGGATAAAGTTTAAAATAGAATTAATAAAGAACATATAGGTCATTTTTTAAACATGGATAGAAACCAAGGTTGTTAGTTAATAATATACAAGATATTTTTTCTCACTCTGATCCATGTAAACCAAGGACGAGAGACACTCTCATTCCTCATTCACGACACCATTAAAAATGGAAATTAAAGCCCTCTATTAAGCACAGACGGCTACAGGTCTACCATCAGGTTACCTTCGTCTACCTTCACAATGGCCTCTCCTTGTGCCCAGTTCAGTCCCTGTCATTGCCACGCTACTAAGGACTCCCTGAATACCGTGACTGACGTCAGACATTGTCTGACTGAATACATCCTGTGGGTTTCTCATAGATGGACCCATAGAGAAAGCGCAGGGCCTCTCTACAGGCTTCTCATCTCTTTCAGAATTGATGCAATGGAGCTATTTGGTAGCGAGTTGAAGGAGTTCTCGAATTCACTTCCGTGGGACAATATCGACAATTGCGTGGAGATCATTAAATGTTTCATCAGAAATGACTCCATGAAAACCGCCAAAGAACTTTGTGCAATAATTGGACTTTGTACTCAATCAGCTATTGTCACTGGAAGAGTCTTCAATGATAAGTATATCGACATACTACTTATGCTGCGAAAGATTCTGAACGAGAACGACTATCTCACCCTCTTGGATCATATCCTCACTGCTAAATACTAAATCTCCTTCATGCTCTCTCACTAATACTCTTACTCACTACACTTTTTATCATCTTATGATGAATGATTGCCTTCATCATTTTTTCGTGGAATATAATATAGGAATAATTAGCACCAGAATAGCTATGGATATCTCGTTAAGAATATTCTCTATAAGAGACATAATGTAGACATAGTTATTATATCCTTCTTAGATAAGTGTTACGCTACTGGAAAGTTTCCATCGTTATTATTACATGAAGATGATATAATTAAACCAACATTGAGATTGGCTCTTATGTTAGCTGGATTGAATTACTGTAATAAATGCATCGAGTATAGAGGGATATAGCAATTCTCGATAATAGTCATGCAATATTTGAATGAGACTGATAATTTAGGTAATACAGTACTACACACATATCTTTCTAGATTATATATCGTTAAAAATCTGTAAGATGTATATTTCTCATAAGTATCCACTGTGTAATATTATTAATGGATATATAGATAACGCAATAGGGACTAATAGTATTGTAAAAGATATAATCGACTATTTACGTACATATCCAGATATCTATATTCCTACTAGTTTGCTGCGTAGTTGCATCATTGATATGCATGATTTATCAGGATTCAGAGATGAATTACTAAGTAAACTACAATCCCACAATAAGTAAGAATCAAATATCAAAAACTCACTTTTGATTTTTCTAGTCTTAAGTAATACATATATTTATTAATAGACCTATGAAATAAAAAAAGGTAACAATGGATTCGCGTATAGCTATTTACGTATTAGTATCGGCATCTCTTTTGTATCTTGTTAATTGTCACAAACTAGTACATTACTTCAATCTGAAAATAAATGGAAGTGATATAACTAATACAGCAGATATATTGCTGGACAATTATCCAATTATGACCTTTGATGGAAAGGATATTTATCCATCTATCTCGTTCATGGTCGGTAATAAACTTTTCCTAGATCTTTATAAAAATATCTTTGAAGAATTTTTCAGACTATTTCGAGTATCTGTAAGTAGTCAATACGAGGAATTAGAATATTATTATTCATGTGATTATACTAACAACCGTCCTACAATTAAACAACATTACTTTTATAACGGCGACGAATATACTGAAATTGATAGATCGAAAAAAGCCACTAATAAAAACAGTTGGTTAATTACTTCAGGCTTTAGACTACAAAAATGGTTCGATAGCGAAGATTGTATAATTTATCTCAGATCTTTAGTTAGAAGAATGGAAGACAGTAACAAAAACAGTAAAAAAACTTAGTACTTAGATATCGAAAAAATATATTTTTGTAGACTCTTGAGAATAGAAGGAAAACATGTACATAATTATAAAAAATGAAAATCAATGGCGAATAAGACAGTGCGATTCGCACCATGGAGTCGGTAGATTTCATGGCTGTCGATGAGCAGTTTCACGACGACCTCGATCTTTGGTCATTATCTTTGGTAGATGATTATAAAAAACATGGATTAGGTGTTGACTGTTATGTTCTAGAACCAGTTGTTGACAGGAAAATATTTGATAGATTTCTCCTTGAACCAATTTGTGATCCTGTAGATGTTCTGTATGATTATTTTAGGATTCATAGAGATAATATTGATCAGTATATAGTAGATAGACTGTTTGCATATATTACATATAAAGATATTATATCTGCATTAGTGTCAAAGAATTATATGGAAGATATTTTCTCTATAATTATTAAGAATTGTAATTCTGTGCAAGATCTCTTACTTTACTATCTATCTAATGCATATGTAGAAATAGACATTGTTGATCTTATGGTAGATCATGGGGCTGTAATATATAAAATAGAATGCTTGAATGCCTATTTTAGGGGAATATGTAAAAAGGAAAGTAGTGTTGTTGAGTTTATTTTGAATTGTGGTATCCCAGATGAAAATGATGTTAAATTAGATCTATATAAAATAATTCAGTATACTAGGGGATTCCTTGTAGATGAACCCACAGTATTAGAAATTTATAAGCTTTGTATCCCATATATTGAAGATATCAATCAACTAGATGCTGGTGGAAGGACCTTGCTTTATCGCGCTATCTATGCAGGTTATATAGATTTAGTATCATGGCTATTAGAAAATGGAGCAAATGTCAACGCAGTAATGAGTAATGGATATACATGTCTTGACGTGGCCGTGGATAGGGGATCTGTCATCGCCCGTAGGGAAGCACATCTTAAAATATTAGAAATATTGCTTAGAGAACCATTGTCTATTGACTGTATAAAATTAGCTATACTTAATAATACAATTGAAAACCATGATGTGATAAAGCTCTGTATCAAGTATTTTATGATGGTAGATTATTCACTTTGTAATGTGTATGCATCATCACTCTTTGATTATATAATTGATTGTAAACAAGAATTGGAGTACATTAGGCAGATGAAAATTCATAATACAACCATGTATGAGTTAATCTATAATAGAGACAAAAACAAGCATGCTTCCCATATTCTACATAGGTATTCTAAACATCCAGTTTTGACACAGTGTATCACTAAAGGATTCAAGATTTACACAGAAGTAACCGAGCAGGTCACTAAAGCTCTAAACAGACGTGCTCTAATAGATGAGATAATAAACAATGTATCAACTGATGACAATCTCCTATCAAAACTTCCATTAGAAATTAGGGATCTAATTGTTTCACAAGCTGTCATATAGAGTTCTATCCACCCACCTTTCTTGAAATGAGTTAATAGTCATAAGTTAGTTAAGTCATAAGTTAGTTAAGTCATAAGTTAGTTAAGTCATAAGTTAGTTAAGTCATAAGTTAGTTAAGAGTTAATAGTCATAAGTTAGTTAAGTCATAAGTTAGTTAAGTCATAAGTTAGTTAAGTCATAAGTTAGTTAAGTCATAAGTTAGTTAAGTCATAAGTTAGTTAAGTCATAAGTTAGTTAAGTCATAAGTTAGTTAAGTCATAAGTTAGTTAAGTCATAAGTTAGTTAAGTCATAAGTTAGTTAAGTCATAAGTTAGTTAAGTCATAAGTTAGTTAAGTCATAAGTTAGTTAAGTCATAAGTTAGTTAAGTCATAAGTTAGTTAAGTCATAAGTTAGTTTATAGTCTAACACTTCTAATTTTTATACCTTGATCTTTTTCTCTAATTATGAAAAAGTAAATCATTATGAAGATGGATGAAATGGACGAGATTGTGCGCATCGTTAACGATAGTATGTGGTACGTACCTAACGCATTTATGGACGACGGTGATAATGAAGGTCACATTTCTGTCAATAATGTCTGTCATATGTATCTCGCATTCTTTGATGTGGATATATCATCTCATCTGTTTAAATTAGTTATTAAACACTGCGATCTGAATAAACGACTAAAATGTGGTAACTCTCCATTACATTGCTATACGATGAATACACGATTTAATCCATCTGTATTAAAGATATTGTTACGCCACGGCATGCGTAACTTTGATAGCAAGGATAAAAAAGGACATATTCCTCTACACCACTATCTGATTCATTCACTATCAATCGATAACAAGATCTTTGATATACTAACGGACCCCATTGATGACTTTAGTAAATCATCCGATCTATTGCTGTGTTATCTTAGATATAAATTCAATGGGAGCTTAAACTATTACGTTCTGTACAAATTATTGACTAAAGGATCTGACCCTAATTGCGTCGATGAGGATGGACTCACTTCTCTTCATTACTACTGTAAACACATATCCGCGTTCCACGAAAGCAATTATTACAAGTCAAAGAGTCACACTAAGATGCGAGCTGAGAAGCGATTCATCTACGCGATAATAGATCATGGAGCAAACATTAACGCGGTTACGAAAATCGGAAATACGCCGTTACACACTTACCTTCAACAGTATACCAAACATAGTCCTCGTGTGGTGTATGCTCTTTTATCTCGAGGAGCCGATACGAGGATACGTAATAATCTTGATTGTACACCCATCATGGAATACATAAAGAACGATTGTGCAACAGGTCATATTCTCATAATGTTACTCAATTGGCACGAACAAAAATACGGGAAATTACAAAAGGAAGAAGGACAACATCTACTTTATCTATTCATAAAACATAATCAAGGATATGGAAGTCGCTCTCTCAATATACTACGGTATCTACTAGATAGATTCGACATTCAGAAAGACGAATACTATAATACAATGACTCCTCTTCATACCGCCTTCCAGAATTGCAATAACAATGTTGCCTCATACCTCGTATACATCGGATACGACATCAACCTTCCGACTAAAGACGATAAGACAGTATTCGACTTGGTGTTTGAAAACAGAAACATTATATACAAGGCGGATGTCGTTAATGACATTATCCACCACAGACTGAAAGTATCTCTACCTATGATTAAATCGTTGTTCTACAAGATGTCGGAGTTCTCTCCCTACGACGATCACTACGTAAAGAAGATAATAGCCTACTGCCTATTAAGGGACGAGTCATTTGCGGAACTACATACTAAATTCTGTTTAAACGAGGACTATAAAAGTGTATTTATGAAAAATATATCATTCGATAAGATAGATTCCATCATCGAAAAATGTAGTCGTGACATAAGTCTCCTCAAAGAGATTCGAATCTCAGACACCGACTTGTATACGGTATTGAGAACAGAAGACATCCGGTATCACACATATCTCGAAGCCATACATTCAGACAAACGCATTTCATTTCCCATGTACGACGATCTCATAGAACAGTGTCATCTATCGATGGAGCATAAAAGTAAACTCGTCGACAAAGCACTCAATAAATTAGAGTCTACCATCGATAGTCAATCTAGACTATCGTATTTGCCTCCGGAAATTATGCGCAATATCATAACCAAGCTAAGCGACTACCATCTAAACAGTATGTTGTACGGAAAGAACCATTACAAATATTATCCATGATAGAAAGAAAATATTTAAAAAATAATCTATATGATTGGAGAAGTAGGAAACAAACAGTAACAAGACGACGATTACTACATTATTAAATCATGAGGTCCGTATTATACTCGTATATATTGTTTCTCTCATGTATAATAATAAACGGAAGAGATATAGCACCACATGCACCATCCAATGGAAAGTGTAAAGACAACGAATACAGAAGCCGTAATCTATGTTGTCTATCGTGTCCTCCGGGAACTTACGCTTCCAGATTATGTGATAGCAAGACTAATACACAATGTACACCGTGTGGTTCGGATACCTTTACATCTCACAATAATCATTTACAGGCTTGTCTAAGTTGTAACGGAAGATGTGATAGTAATCAGGTAGAGACGCGATCGTGTAACACGACTCACAATAGAATCTGTGAATGCTCTCCAGGATATTATTGTCTTCTCAAAGGAGCATCAGGGTGTAGAACATGTATTTCTAAAACAAAGTGTGGAATAGGATACGGAGTATCCGGATACACGTCTACCGGAGACGTCATCTGTTCTCCGTGTGGTCCCGGAACATATTCTCACACCGTCTCTTCCACAGATAAATGCGAACCCGTCGTAACCAGCAATACATTTAACTATATCGATGTGGAAATTAACCTGTATCCAGTCAACGACACATCGTGTACTCGGACGACCACTACCGGTCTCAGCGAATCCATCTCAACGTCGGAACTAACTATTACCATGAATCATAAAGATTGTGATCCAGTCTTTCGTGCAGAATACTTCTCTGTCCTTAATAATGTAGCAACTTCAGGATTCTTTACAGGAGAAAATAGATATCAGAATACTTCAAAGATATGTACTCTGAATTTCGAGATTAAATGTAACAACAAAGATTCATCTTCCAAACAGTTAACGAAAACAAAGAATGATACTATCATGCCGCATTCAGAGACGGTAACTCTAGTGGGCGACTGTCTATCTAGCGTCGACATCTACATACTATATAGTAATACCAATACTCAAGACTACGAAAATGATACAATCTCTTATCATATGGGTAATGTTCTCGATGTCAATAGCCATATGCCCGCTAGTTGCGATATACATAAACTGATCACTAATTCCCAGAATCCCACCCACTTATAGTAAGTTTTTTTACCTATAAATAATAAATACAATAATTAATTTCTCGTAAAAGTAGAAAATATATTCTAATTTATTATATGGTAAGAAAGTAGAATCATCTAGAACAGTAATCAATCAATAGCAATCATGAAACAATATATTGTCCTGGCATGCATGTGCCTAGTGGCAGCTGCTATGCCTACTAGTCTTCAACAATCTTCATCCTCGTGTACTGAAGAAGAAAACAAACATCATATGGGAATCGATGTTATTATCAAAGTCACAAAGCAAGACCAAACACCGACCAATGATAAGATTTGTCAATCCGTAACGGAAGTTACAGAGACCGAAGATGATGAGGTATCCGAAGAAGTTGTAAAAGGAGATCCCACCACTTATTACACTATCGTCGGTGCGGGTCTTAACATGAACTTTGGATTCACCAAATGCCCAAAGATTTCATCCATCTCCGAATCCTCTGATGGAAACACTGTGAATACTAGATTGTCCAGCGTGTCACCGGGACAAGGTAAGGACTCTCCCGCGATCACGCGTGAAGAAGCTCTGGCTATGATCAAAGACTGTGAGATGTCTATCGACATCAGATGTAGCGAAGAAGAGAAAGACAGTGACATCAAGACCCATCCAGTACTTGGGTCTAACATCTCACATAAGAAAGTGAGTTACAAAGATATCATCGGTTCAACGATCGTTGATACAAAATGTGTCAAGAACCTAGAGTTTAGCGTACGTATCGGAGACATGTGTGAGGAATCATCTGAACTTGAAGTCAAGGATGGATTCAAGTATGTCGACGGATCGGCATCTGAAGGTGCAACCGATGATACTTCACTCATCGATTCAACAAAACTCAAAGCATGTGTCTGAATCGATAACTCTATTCATCTGAAAATGGATGAGTTGGGTTAATCGAACGATTCAGACACCGCACCACGAATTAAAAAAGACCGGGCACTATATTCCGGTTTGCAAAACAAAAATATTTAACTACATTCACAAAAAGTTACCTCTCGTTACTTCTTCTTTCTGTTTCAATATGTGATACGATATGATCACTATTCGTATTCTCTTGGTCTCATAAAAAAGTTTTACAAAAAAAAAAAAAATATTTTTATTCTCTTTCTCTCTTCGATGGTCTCACAAAAATATTAAACCTCTTTCTGATGTCTCAACTATTTCGTAAACGATAACGTCCAACAATATATTCTCGTAGAGCTTATCAACATCCTTATACCAATCTAGGTTGTCAGACAATTGCATCATAAAATAATGTTTATAATTTACACGTTAACATCATATAATAAACGTATATAGTTAATATTTTTGGAATATAAATGATCTGTAAAATCCATGTAGGGGACACTGCTCACGTTTTTTCTCTAGTACATAATTTCACACAAGTTTTTATACAGACAAATTAATTCTCGTCCATATATTTTAAAACATTGACTTTTGTACTAAGAAAAATATCTTGACTAACCATCTCTTTCTCTCTTCG

>2017_UK_Monkeypox_MT903343.1

TATTATATTTTACTATTTTATTTAGTGTCTAGAAAAAAATGTGTGACCCACGACCGTAGGAAACTCTAGAGGGTAAGAAAAATCAATCGTTTATAGAGACCATCAGAAAGAGGTTTAATATTTTTGTGAGACCTATCGAAGAGAGAAAGGATAAAAACTTTTTACGACTCCATCAGAAAGAGGTTTAATATTTTTGTGAGACCCATCGAAGAGAGAAAGAGATGGTTAGTCAAGATATTTTTCTTAGTACAAAAGTCAATGTTTTAAAATATATGGACGAGAATTAATTTGTCTGTATAAAAACTTGTGTGAAATTATGTACTAGAGAAAAAACGTGAGCAGTGTCCCCTACATGGATTTTACAGATCATTTATATTCCAAAAATATTAACTATATACGTTTATTATATGATGTTAACGTGTAAATTATAAACATTATTTTATGATGCAATTGTCTGACAACCTAGATTGGTATAAGGATGTTGATAAGCTCTACGAGAATATATTGTTGGACGTTATCGTTTACGAAATAGTTGAGACATCAGAAAGAGGTTTAATATTTTTGTGAGACCATCGAAGAGAGAAAGAGAATAAAAATATTTTTTTTTTTTTTTTGTAAAACTTTTTTATGAGACCAAGAGAATACGAATAGTGATCATATCGTATCACATATTGAAACAGAAAGAAGAAGTAACGAGAGGTAACTTTTTGTGAATGTAGTTAAATATTTTTGTTTTGCAAACCGGAATATAGTGCCCGGTCTTTTTTAATTCGTGGTGCGGTGTCTGAATCGTTCGATTAACCCAACTCATCCATTTTCAGATGAATAGAGTTATCGATTCAGACACATGCTTTGAGTTTTGTTGAATCGATGAGTGAAGTATCATCGGTTGCACCTTCAGATGCCGATCCGTCGACATACTTGAATCCATCCTTGACTTCAAGTTCAGATGATTCCTCACACATGTCTCCGATACGTACGCTAAACTCTAGGTTCTTGACACATTTTGTATCAACGATCGTTGAACCGATGATATCTTTGTAACTCACTTTCTTATGTGAGATGTTAGACCCAAGTACTGGATGGGTCTTGATGTCACTGTCTTTCTCTTCTTCGCTACATCTGATGTCGATAGACATCTCACAGTCTTTGATCATAGCCAGAGCTTCTTCACGCGTGATCGCGGGAGAGTCCTTACCTTGTCCCGGTGACACGCTGGACAATCTAGTATTCACAGTGTTTCCATCAGAGGATTCGGAGATGGATGAAATCTTTGGGCATTTGGTGAATCCAAAGTTCATGTTAAGACCCGCACCGACGATAGTGTAATAAGTGGTGGGATCTCCTTTTACAACTTCTTCGGATACCTCATCATCTTCGGTCTCTGTAACTTCCGTTACGGATTGACAAATCTTATCATTGGTCGGTGTTTGGTCTTGCTTTGTGACTTTGATAATAACATCGATTCCCATATGATGTTTGTTTTCTTCTTCAGTACACGAGGATGAAGATTGTTGAAGACTAGTAGGCATAGCAGCTGCCACTAGGCACATGCATGCCAGGACAATATATTGTTTCATGATTGCTATTGATTGATTACTGTTCTAGATGATTCTACTTTCTTACCATATAATAAATTAGAATATATTTTCTACTTTTACGAGAAATTAATTATTGTATTTATTATTTATAGGTAAAAAAACTTACTATAAGTGGGTGGGATTCTGGGAATTAGTGATCAGTTTATGTATATCGCAACTAGCGGGCATATGGCTATTGACATCGAGAACATTACCCATATGATAAGAGATTGTATCATTTTCGTAGTCTTGAGTATTGGTATTACTATATAGTATGTAGATGTCGACGCTAGATAGACAGTCGCCCACTAGAGTTACCGTCTCTGAATGCGGCATGATAGTATCATTCTTTGTTTTCGTTAACTGTTTGGAAGATGAATCTTTGTTGTTACATTTAATCTCGAAATTCAGAGTACATATCTTTGAAGTATTCTGATATCTATTTTCTCCTGTAAAGAATCCTGAAGTTGCTACATTATTAAGGACAGAGAAGTATTCTGCACGAAAGACTGGATCACAATCTTTATGATTCATGGTAATAGTTAGTTCCGACGTTGAGATGGATTCGCTGAGACCGGTAGTGGTCGTCCGAGTACACGATGTGTCGTTGACTGGATACAGGTTAATTTCCACATCGATATAGTTAAATGTATTGCTGGTTACGACGGGTTCGCATTTATCTGTGGAAGAGACGGTGTGAGAATATGTTCCGGGACCACACGGAGAACAGATGACGTCTCCGGTAGACGTGTATCCGGATACTCCGTATCCTATTCCACACTTTGTTTTAGAAATACATGTTCTACACCCTGATGCTCCTTTGAGAAGACAATAATATCCTGGAGAGCATTCACAGATTCTATTGTGAGTCGTGTTACACGATCGCGTCTCTACCTGATTACTATCACATCTTCCGTTACAACTTAGACAAGCCTGTAAATGATTATTGTGAGATGTAAAGGTATCCGAACCACACGGTGTACATTGTGTATTAGTCTTGCTATCACATAATCTGGAAGCGTAAGTTCCCGGAGGACACGATAGACAACATAGATTACGGCTTCTGTATTCGTTGTCTTTACACTTTCCATTGGATGGTGCATGTGGTGCTATATCTCTTCCGTTTATTATTATACATGAGAGAAACAATATATACGAGTATAATACGGACCTCATGATTTAATAATGTAGTAATCGTCGTCTTGTTACTGTTTGTTTCCTACTTCTCCAATCATATAGATTATTTTTTAAATATTTTCTTTCTATCATGGATAATATTTGTAATGGTTCTTTCCGTACAACATACTGTTTAGATGGTAGTCGCTTAGCTTGGTTATGATATTGCGCATAATTTCCGGAGGCAAATACGATAGTCTAGATTGACTATCGATGGTAGACTCTAATTTATTGAGTGCTTTGTCGACGAGTTTACTTTTATGCTCCATCGATAGATGACACTGTTCTATGAGATCGTCGTACATGGGAAATGAAATGCGTTTGTCTGAATGTATGGCTTCGAGATATGTGTGATACCGGATGTCTTCTGTTCTCAATACCGTATACAAGTCGGTGTCTGAGATTCGAATCTCTTTGAGGAGACTTATGTCACGACTACATTTTTCGATGATGGAATCTATCTTATCGAATGATATATTTTTCATAAATACACTTTTATAGTCCTCGTTTAAACAGAATTTAGTATGTAGTTCCGCAAATGACTCGTCCCTTAATAGGCAGTAGGCTATTATCTTCTTTACGTAGTGATCGTCGTAGGGAGAGAACTCCGACATCTTGTAGAACAACGATTTAATCATAGGTAGAGATACTTTCAGTCTGTGGTGGATAATGTCATTAACGACATCCGCCTTGTATATAATGTTTCTGTTTTCAAACACCAAGTCGAATACTGTCTTATCGTCTTTAGTCGGAAGGTTGATGTCGTATCCGATGTATACGAGGTATGAGGCAACATTGTTATTGCAATTCTGGAAGGCGGTATGAAGAGGAGTCATTGTATTATAGTATTCGTCTTTCTGAATGTCGAATCTATCTAGTAGATACCGTAGTATATTGAGAGAGCGACTTCCATATCCTTGATTATGTTTTATGAATAGATAAAGTAGATGTTGTCCTTCTTCCTTTTGTAATTTCCCGTATTTTTGTTCGTGCCAATTGAGTAACATTATGAGAATATGACCTGTTGCACAATCGTTCTTTATGTATTCCATGATGGGTGTACAATCAAGATTATTACGTATCCTCGTATCGGCTCCTCGAGATAAAAGAGCATACACCACACGAGGACTATGTTTGGTATACTGTTGAAGGTAAGTGTGTAACGGCGTATTTCCGATTTTCGTAACCGCGTTAATGTTTGCTCCATGATCTATTATCGCGTAGATGAATCGCTTCTCAGCTCGCATCTTAGTGTGACTCTTTGACTTGTAATAATTGCTTTCGTGGAACGCGGATATGTGTTTACAGTAGTAATGAAGAGAAGTGAGTCCATCCTCATCGACGCAATTAGGGTCAGATCCTTTAGTCAATAATTTGTACAGAACGTAATAGTTTAAGCTCCCATTGAATTTATATCTAAGATAACACAGCAATAGATCGGATGATTTACTAAAGTCATCAATGGGGTCCGTTAGTATATCAAAGATCTTGTTATCGATTGATAGTGAATGAATCAGATAGTGGTGTAGAGGAATATGTCCTTTTTTATCCTTGCTATCAAAGTTACGCATGCCGTGGCGTAACAATATCTTTAATACAGATGGATTAAATCGTGTATTCATCGTATAGCAATGTAATGGAGAGTTACCACATTTTAGTCGTTTATTCAGATCGCAGTGTTTAATAACTAATTTAAACAGATGAGATGATATATCCACATCAAAGAATGCGAGATACATATGACAGACATTATTGACAGAAATGTGACCTTCATTATCACCGTCGTCCATAAATGCGTTAGGTACGTACCACATACTATCGTTAACGATGCGCACAATCTCGTCCATTTCATCCATCTTCATAATGATTTACTTTTTCATAATTAGAGAAAAAGATCAAGGTATAAAAATTAGAAGTGTTAGACTATAAACTAACTTATGACTTAACTAACTTATGACTTAACTAACTTATGACTTAACTAACTTATGACTTAACTAACTTATGACTTAACTAACTTATGACTTAACTAACTTATGACTTAACTAACTTATGACTATTAACTCATTTCAAGAAAGGTGGGTGGATAGAACTCTATATGACAGCTTGTGAAACAATTAGATCCCTAATTTCTAATGGAAGTTTTGATAGGAGATTGTCATCAGTTGATACATTGTTTATTATCTCATCTATTAGAGCACGTCTGTTTAGAGCTTTAGTGACCTGCTCGGTTACTTCTGTGTAAATCTTGAATCCTTTAGTGATACACTGTGTCAAAACTGGATGTTTAGAATACCTATGTAGAATATGGGAAGCATGCTTGTTTTTGTCTCTATTATAGATTAACTCATACATGGTTGTATTATGAATTTTCATCTGCCTAATGTACTCCAATTCTTGTTTACAATCAATTATATAATCAAAGAGTGATGATGCATACACATTACAAAGTGAATAATCTACCATCATAAAATACTTGATACAGAGCTTTATCACATCATGGTTTTCAATTGTATTATTAAGTATAGCTAATTTTATACAGTCAATAGACAATGGTTCTCTAAGCAATATTTCTAATATTTTAAGATGTGCTTCCCTACGGGCGATGACAGATCCCCTATCCACGGCCACGTCAAGACATGTATATCCATTACTCATTACTGCGTTGACATTTGCTCCATTTTCTAATAGCCATGATACTAAATCTATATAACCTGCATAGATAGCGCGATAAAGCAAGGTCCTTCCACCAGCATCTAGTTGATTGATATCTTCAATATATGGGATACAAAGCTTATAAATTTCTAATACTGTGGGTTCATCTACAAGGAATCCCCTAGTATACTGAATTATTTTATATAGATCTAATTTAACATCATTTTCATCTGGGATACCACAATTCAAAATAAACTCAACAACACTACTTTCCTTTTTACATATTCCCCTAAAATAGGCATTCAAGCATTCTATTTTATATATTACAGCCCCATGATCTACCATAAGATCAACAATGTCTATTTCTACATATGCATTAGATAGATAGTAAAGTAAGAGATCTTGCACAGAATTACAATTCTTAATAATTATAGAGAAAATATCTTCCATATAATTCTTTGACACTAATGCAGATATAATATCTTTATATGTAATATATGCAAACAGTCTATCTACTATATACTGATCAATATTATCTCTATGAATCCTAAAATAATCATACAGAACATCTACAGGATCACAAATTGGTTCAAGGAGAAATCTATCAAATATTTTCCTGTCAACAACTGGTTCTAGAACATAACAGTCAACACCTAATCCATGTTTTTTATAATCATCTACCAAAGATAATGACCAAAGATCGAGGTCGTCGTGAAACTGCTCATCGACAGCCATGAAATCTACCGACTCCATGGTGCGAATCGCACTGTCTTATTCGCCATTGATTTTCATTTTTTATAATTATGTACATGTTTTCCTTCTATTCTCAAGAGTCTACAAAAATATATTTTTTCGATATCTAAGTACTAAGTTTTTTTACTGTTTTTGTTACTGTCTTCCATTCTTCTAACTAAAGATCTGAGATAAATTATACAATCTTCGCTATCGAACCATTTTTGTAGTCTAAAGCCTGAAGTAATTAACCAACTGTTTTTATTAGTGGCTTTTTTCGATCTATCCTCGTCCTCTCATCATCCTTATATTATTATCATTATTATCATAGTCTATTAAACACAAATCATCTACGTTTATAACAACATTCTCATTATTAATTAGTTCTGTAGAATATCTTTAATAATTTGGCTATACATCTGTTCAATACTATCTATTGATGATTTCTTTTTTAAGACTTAAACTAGTTATGGTAATGACGATGAAATCGAGTAGTAACTTCTAATAAAGACTTGATATCATTATCATATGTTTGATCGTCATAGTTAATAGTGTGGCTAAATGGTACTGTTAATAAGTTTATAGACAATATCATAGTATTTTCTTTCCAGAATTAGATTATTTTTTTAAATACTGATCCTCACAATTCCGTGATGTAGCAGTAGTTGGTGCATGGTCTATATCGTTAAAATGTATCATATATAATAGTTTTCTGACGTGGAGTACAGAATTTTCGATTAATGAGTTCATGGTAAGGAAGGGCAAATGCCTGTATATAATATACATAAGTTAATAGTTTTTTATCATATTTTCTAATACCATAATAAAAATTATCATTATTGCGTTTGGTAGTTCTGCCCTATCATCTATATCACTGTCACTCTCGCTCTCACTATATCTTCTAAAATTACAAACAACTGGATATTCGATAACAGCATTTGTGTAGTTTTTGTCTTTTACAGTATATACGTTATTGTCAAAATCTAAACAAATATTAGCATAATACATCTATCTATAAGATCAGGATCCATGTTCGAGCATACTAGCCATGTATATTTGTAACTTCGTCGTACAGCGTTAGATCAATAGAATAAACAATCGTGTGACGCAACTTTTTTACGATCTAGTTGTATGAGTTTATCGTTTACATAAGCAATTAACGGCTTTAACAGATGATCTGAGTAATAATATACCTCTGTTATACGTTTAATGTTCACGGTCTTAGTATTTTTAGATATCAATTGTGATTTACACCATATTCGACTCCCTTGTGTGTAACGTTAGAAATTCTAAATCTATATTATCTATTACAGCGTAAAACACATTCAATATTGTATTGTTATTTTTATATTATTTACACAATTAACAATGTATTATTAGTTTATATTACTGAATTAATAATATAAAATTCCCAATCTTGTCATAAACACAAAATCCATTAAAAATGTCGATAAAATATCTGATGTTGTTGTTCGCTGCTATGATAATCAGATCATTAGCCGATAGTGGTAATGCTATCGAAACAACATCGCCAGAAATTACAAACGCTACAACAGATATTCCAGCTATCAGATTATGTGGTCCAGAGGGAGATGGATATTGTTTACACGGTGACTGTATCCACGCTAGAGATATCGACGGTATGTATTGTAGATGCTCTCATGGTTATACAGGTATTAGATGTCAGCATGTAGTATTAGTAGACTATCAACGTTCAGGAAAACCAGACACTACAACATCATATATCCCATCTCTAGGTATTGTGCTTGTATTAGTAGGCATTATTATTACGTGTTGTCTATTATCTGTTTATATGTTCACTCGAAGAACTAAACTACCTATACAAGATATGGTTGTGTTATATTTTTTATAAAATTTTTTTATGAGTATTTTTACAAAAATGTATATGTATAAAAAAAATACTAAGTATACGATGTATCCTGTATTATTTGTATTTATCTAAACAATACTTCTGCCTCTAGATGGGATACAAAAATTTTTTATTTCAGCATATTAAAGTAAAATCTAGTTACCTTGAAAATGAATACAGTGGGTGGTTCCGTATCACCAGTAAGAACATAATAGTCGAATACAGTATCCGATTGAGATTTTGCATACAATACTAGTCTAGAAAGAAATTTGTAATCATCCTCTGTGACAGGAGTCCATATATCTGTATCATCGTCTAGTTTTTATCAGTGTCCTATGCTATATTCCTGTTATCATCATTAGTTAATGAAAATAACTCTCGTGCTTCAGAAAAGTCAAATATTGTATCCATACATATATCTCCAAAACTATCACTTATACGTTTATCTTTAACGAACATATACCTAGATGGTTATTTACTAACAGACATTTTTTCAAGATCTATTGACAATAACTCCTATAGTTTCCACATCAACCAAGTAATGATCATCTATTGTTATATAACAATAACATAACTCTTTTCCATTTTTATCAGTATCTATATCAACGTCGTTGTAGTGAATAGTAGTCATTGATCTATTATATGAAACGGATATGTCTAGTTAATATTTTCTTTGATTTAAAGTCTATAGTCTTTACAAACATAATATCCTTATCCGACTTTATATTTCCTGTAGGGTGGCATAATTTTATTCTGCCTCCACAATCAGTGTTTCCAAATATATTACTAGACAATATTCCATATAGTTATTAGTTAAGGGACCCAATTAGAACACGTACGCGCTTATTCATCATTTGGATCGTATTTCATAAAAGTTATTATGTTATAGATGTCAACACATTCTACATTTTTTTAATTGTCTATATAGTATTTTTCTGATATTTTCTATAATATCAGAATTGTCTTCCATAGGAAGTTGTATACTATTCGGAATCAGTTACATGTTTAAATAATTCTCTGATGTCATTCCTTATACAATCAAATTCATTATTAAACAGTTTAATAGTCTGTAGACCTTTATCGTCGTACATATCCATTGTCTTATTAGTTACGCTTATTTTTATGGGTTTTACGTTGCTTTATTATATTTTATAATAATGATTGTTTGACAATGTCGTAGTATAGATATATTATTAGAGGAGGTATAATTATAAAAAGTTTCTGAGTACGATGTTATAAGAGGAGAGGACACATTAACATCATACATCAATTAACTACATTCTTATAACATTGTAATCAAAAGAATTGCAATTTTGATGTATAACAACTGTCAATGGAATTGTATATTACAAATTACGGTATGTTGTAACAACAAATACCGATCGGTAATTGTCTCTGCCGCTGTAATAGAATTAATTATATATCTATTACACCGGCCTTGTATCATAATAAAGTTGTGGTAGTATGATCTCCATATTTATAATTTAGTACTTTGTATTTAGTTTTTTTGGAATCATAAAAAAAGTTTTACTAATTTAAAATTTAAAAAGTATTTACATTTTTCACTGTTTAGTCGCGGATATGGAATTCGATCCTGTCAAAATCAATACATCATCTATAGATCATGTAACAATATTACAATACATAGATGAACCAAATGATATAAGACTAACAGTATGCATTATCCGAAATGTTAATAACATTACATATTATATCAATATCACAAAAATAAATCCACATTTGGCTAATCGATTTCGGGCTTGGAAAAAACGTATCGCCGGAAGGGACTATATGACTAACTTATCTAGAGATACAGGAATACAACAATCAAAACTTACTGAAACTATACGTAACTGTCAAAAAAATAAAAACATATATGGTCTATATATACACTACAATTTAGTTATTAATGTGGTTATTGATTGGATAACCGATGTGATTGTTCAATCAATATTAAGAGGGTTGGTAAATTGGTACATAGCTAATAATACATATACTCCAAATACACCCAATAATACTACAACCATTTCTGAGTTGGATATCATCAAAATACTGGATAAATACGAGGACATGTATAGAGTAAGTAAAGAAAAAGAATGTGGAATTTGCTATGAAGTTGTTTACTCAAAACGATTAGAAAACGATAGATACTTTGGTTTATTGGATTCGTGTAATCATATATTTTGCATAACATGTATCAATATATGGCATAGAACACGAAGAGAAACCGGTGCGTCAGATAATTGTCCTATATGCCGTACACGTTTTAAAAAAATAACAATGAGCAAGTTCTATAAGCTAGTTAACTAATAAATAAAAAGTTTAATTATCGACGATATATGTCGTTATTTTTCTCTCATATGAAAGATTAATTTGATTCTAATATAATCTTCAGTATTGGATGAATCTCAATTCAAATTAATTCCATTAGATTAGATTAGATTAGATCATAAATAAAAATAGTAGCACGCACTACTTCAGCCAAATATTCTTTTTTGAAACGCCATCTAGCGTAATGAGAACACAAGTGAACCTATAATGAGCAAATTTATTAGTATCGGTTACATGAAGGACTTTACGTAGAGTGGTGATTCCTCCATCTGTGGTACGAACGGTTTCATCTTCTTTGATGCCACCACCCAGATGTTCTATAAACTTGGTATCCTCGTCCGATTTCATATCATTTGCCAACCAATACATATAGCTAAACCCAGGCATACGTTCCACACATCCGGAACAATGAAATTCTCCAGAAGATGTTACAATGGCTAGATTTGGACATTTGGTTTCAACCGCGTTAACATATGAGTGAACACACCCATACATGAAAGCGATGAGAAATAGGATTCTCATCTTGCCAAAATATCACTATAAAAAATTTATTTATCAATTTTAAAGGTATAAAAAAATACTTATTGTTGCTCGAATATTTTGTATTTGATGGTATACGGAAGATTAGAAATGTAGGTATTATCATCAACTGATTTTATGATGGTTTTATGAATTTTATTATGCTTCACTATTGCATCGGAAATAATATCATATGCTTCCACGTATATTTTATTTTGTTTTGACTCATAATACGCACGTAATTCTGGATTATTGGCATATCGATGAATAATTTTAGCTCCATGCTCAGTAAATATTAATGAGAACATAGTGTTGCCTCCTACCATTATTTTTTTCATCTCATTCAATTCTTGATTGCAGAGATCTATATAATCATTATAGCGTTGACTTATGGACTCTGGAATCTTAGACGATGTACAGTCATCTATAATCATGGTATATTTAATACATTGTTTTATAGCATAGGCATTATCTACGATATTAGATACTTCACTCAATGAATCAATCACACAATCTAATGTAGGTTTATGACATAATAGCATTTTCAGCAGTTCAATGTTTCTAGATTCGTTGATGGCAATGGTTATACATGTATATCCGTTATTTGATCTAATGTTGACATCTGAACCGGATTCTAGCAGTAAAGATACTAGAGATTGTTTATTATATCTAACAGCCTTGTGAAGAAGTGTTTCTCCTCGTTTGTCAATCATGTTAATGTCTTTAAGATAAGGTAGACAAATGTTTATAGTACTAAGAATTGGGCAAACATAAGACATGTCACAAAGACCCTTTTTGTATGTATAAGTGTAGAAATTATAACATCCATAGTTGGATTCACATAGGTGTCCAATCGGGATCTCTCCATCATCGAGATGATTGACGGCATCTCCCCCTTCCTTTTTTAGTAGATATTTCATCGTGTAAGAATCAATATTAATATTTCTAAAGTATCTGTGTATAGCCTCTTTATTTACCACAGCTCCATATTCCAACATGCATTCCACTAGAGGGATATCGATATCGCCGAATGTCATATACTCAATTAGTATATGTTGGAGGACATCCGAGTTCATTGTTTTCAATATCAAAGAGATGGTTTCCTTATCATTTCTCCATAGTGGTACAATACTACACATTATTCCGTACGGCTTTCCATTCTCCAAAAACAATTTTACCAAATCTAAATCTACATCTTTATTGTATCTATAATCACTATTTAGATAATCAGCCATAATTCCTCGAGTGCAACATGTTAGATCGTCTATATATGAATAAGCCGTGTTATCTATTCCTTTCATTAACAATTTAACGATGTCTATATCTATATGAGATGACTTAATATAATATTGAAGAGCTGTACAATAGTTTTTATCTATAGAAGACGGCTTGATTCCGTGATTAATTAGACATTTAACAACTTCCGGACGCACATATGCTCTCGTATCCGACTCTGAATACAGATGAGCGATGATATACAGATGCAATACGGTACCACAATTTCGTGGTTGATAATCATCATACACGTATCCGTACTCGTCATCCTCATAAAGAACACTGCATCCATTTTCTATGAACAAATCAATAATTTCAGGAACAGGATCATCTGTCATTACATAATTTTCTATAACTGAACGATGGTTTTCACATTTAACACTCAAGTCAAATCCATGTTCTACCAACACCTTTATCAAGTCAACGTCTACATTTTTTGATTTCATATAGCTGAATATATTAAAGTCATTTATGTTGCTATATCCAGTAGCTTCTAGTAGAGCCATCGCTATATCCTTATTGACTTTAACATGTCTACTATTTGTGTATTCTTCTATTGGGGTAAACTGTCTCCAATTTTTATGTAATGGATTAGTGCCACTATCTAGTAGTAGTTTGACGACCTCAACATTATTACAATGCTCATTGAAAAGGTATGCGTGTAAAGCATTATTCTTGAATTGGTTCCTGGTATCATTAGGATCTCTGTCTCTCAACATCTGTTTAAGTTCATCGAGAACCACCTCCTCATTTTCCAGATAGTCAAACATTTTGACTGAATAGAAGTGAATGAGCTACTGTGAACTCTATACACCCGCACAACTAATGTCATTAAATATCATTTTTGAATGTATTTATACCATGTCAAAAACTTGTACAATTATTAATAAAAATAATTAGTGTTTAAATTTTACCAGTTCCAGATTTTACACCTCCGTTAACACCTCCATTAACCCCACTTTTTACACCACTGGACGATCCTCCTCCCCACATTCCACTGCCACTAGATGTATAAGTTTTAGATCCTTTATTACTACCATCATGTCCATGGATAAAGACACTCCACATGCCGCCACTACTACCCCCTTTAGACGACATATTAATAAGACAAGTTTAACAATAAAATTAATCACGAGTACCCTACTACCAACCACTATTATATGATTACAGTACCTTGACTAAAGTCTCTAGTCACAAGATCAATACTACCAACCTACGCTATTATATGATTATAGTTTCTATTTTTATAGGAACGCGTACGAGAAAATCAAATGTCTAAGTTCTAACGGTAGTGTTGATAAACGATTGTTATCCGCGGATACCTCATCTATCATGTTGTCTATTTTCTTACTTTGTTCTATTAACCTATTAGCATTATATATTATTTGATTATAAAACTTATATTGCTTATTAGCCCAATCTGTAAATATCGGATTATTAACATATCGTTTCTTTGTAGGTTTATTTAACTTGTACATCACTGTAAGCATGTCCGTACCATTTATTTTAATTTGACACATATCAGCAATTTCTTTTTCGCAGTCGGTTATATATTCTATATAAGATGGATACGTATCACATATGTACTTATAGTCTACTAATATGAAGTACTTAATACATATTTTCAGTAACGATTTAGCCTTATTACCTATTAATAAGTGCCTGTCGTTGGATAGGTAATCAACTGTTTTCTTAATACATTCGATGGTTGGTAATTTACTCAAAATAATTTCCAATATCTTAATATATATTTCTGCTATTTCTGGTATACATGCATGTGCCATTATAACACAAATACCAATACATGTAGACCCATATGTTGTTGTTATATTAATATCTGCGCCATTATCTATTAACCATTCTATTAGGGCAACACTATGCGACTCGATACAATAATAAAGTATACTACGTCCATGTTTATCTATTTTGTTTATATCATCGATATACGGCTTACAAATTTCTAGTATCGATAACACTTCTGACTCGTGAATAAATAAGGTAGGGAATAACGGCATAATATTTATTATGTTATCATCATTAACAACTACGTTTCCATTTTTTAAAATATACTCTACAACTTTAGGATCCCTATTGTCAAATCTTTTAAAATATTTATTTATATGCTTAAATCTATATAATATAGCTCCTTCCCTAATCATACATTTGATAACATTGATGTACACTGTATGATAAGATACATATTCTGACAATAGATCTTGTATAGAATCTGTATATCTTTTAAGAATTGAGGATATTATGACATTATTACGTAAACTATTACACAATTCTAAAATATAAAATGTATCATGGGCAGATAATAGTTTATCCACTATATAATTATCTATTTTATGATTTTTCTTCCTATATTGTTTACGTAAATAGATAGATAGAATATGCATTAGTTCATTACCGCTATAGTTACTATCGAATAACACGTCAAATATTTCCCGTTTAATATCGCATTTGTCAACATAATAATAGAGTATGGTACGTTCACGATAAGTATAATGACACATCTCGTTTTCGTGCGAAATTAAATAGTTTATCACGTCCAAAGATGTCACATAACCATCTTGTGACCTAGTAATAATATAATAATAGAGAACTGTTTTACCCATTCTATTATCATAATCAGTGGTGTAGTCATAATCTAAATAATCAAACTCGTCATCCCAATTAAAATAAATATAATCAGTACATTGAATGGGTATGATATTGTACCCATACTGTATGTTGCTACATGTAGGTATTCCTTTATCCAATAATAGTTTAAATACATCTATATTAGGATTTGATGTTGTCGCGTATTTCTCTACAATATTAATACCATTTTTGATACTATTTATTTCTATACCTTTCGAAATTAGTAATTTCAATAAGTCTATATCGATGTTATCAGAACATAGATATTCAAATATATCAAAATCATTGATATTTTTATAGTCGACTGACGACAATAACAAAATCACAACATCGTTTTTGATATTATTATTTTTTTTGGTAACGTATGCCTTTAATGGAGTTTCACCATCATACTCATATAATGGATTTGCACCACTTTCTATTAATGATTGTGCACTACTGGCATCGATGTTAAATGTTTTACAACTATCATAGAGTATCTTATCGTTAACCATGATTGGTTGTTGATGTTATCACATTTTTTGGTTTCTTTCATTTCAGTTATGTATGGATTTAGCACGTTTGGGAAGCATGAGCTCATATGATTTCAGTACTGTAGTGTCAGTACTATTAGTTTCGATCAGATCAATGTCTAGATCTATAGAATCAAAACACGATAGGTCAGAAGATAATGAATATCTGTACGCTTCTTCTTGTACTGTAACTTCTGGTTTTGTTAGATGGTTGCATCGTGCTTTAACGTCAATGGTACAAATTTTATCCTCGCTTTGTGTATATCATATTCGTCTCTAGTATAAAATTCTATATTCAAATTATCATGCGATGTGTGTACGCTAACGGTATCAATAAACGGAGCACAGCATTTAGTCAACAGTAATCCAAAATTTTTTAAAGTATATCTTAACGAAAGAAGTTGTCATCGTTAGAGTGTGGTAAATCATTGTCTACGGTACTAGATCCTCATAAGTGTATATATCTAGAGTAATATTTAATTTATCAAATGGTTGATAATATGGATGTTGTGGCAATTTCCTAATACGGAAATAAGACATAAACACGCAATAAATCTAATTGCGGACATGTTACACTCCTTAAAAAATACGAATAAACACTTTGGCTTTTAGTAAGTGTCATTTAACACTATACTCATATTAATCCATGGACTCATAATCTCTATACGGGATTAACGGATGTTCTATATACGGGGATGAGTAGTTCTCTTCTTTAACTTTATACTTTTTACTAATCATATTTAGACTGATGTATGGGTAATAGTGTTTGAAGAGCTCGTTCTCATCATCAGAATAAATCAATATCTCTGTTTTTTTGTTATACAGATGTATTACAGCCTCATATATTACGTAATAGAACGTGTAATCTACCTTATTAACTTTCACCGCATAGTTGTTTGCAAATACGGTTAATCCTTTGACCTCGTCGATTTCCGACCAATCTGGGCGTATAATGAATCTTAACTTTAATTTCTTGTAATCATTCGAAATAATTTTTAGTTTGCATCCGTAGTTATCTCCTCTATGTAACTGTAAATTTCTCAACGCGATATCTCCATTAATAATGATGTCGAATTCGTGTTGTATACCCATACTGAATTGATGAACTAATACCGACGGTATTAATAGTAATTTACTTTTCATCTTTACATACTTGGTAATAGTTTTACTATCATAAGTTTATAAATTCCACAAGCTACTATGGAATATACCAACCATCTTAGTATAGAACACATGTCTTAAAGTTATTAATTAATTACATGTTGTTTTATATATCGCTACGAATTTAAACAGAGGAATCAGTTAGGAAGAAAAAATTATCTGTCATCATCATCATCTATTGGATAACATCTCTGTATTCTACGATAGAGTGCTATTTTAAGATGTGACAGATCCGTGTCATCAAATATATACTCCATTAAAATGATTATTCCGGCAGCGAACTTGATATTGGATACATCACGACCTTTGTTAATATCCACGACAATAGACAGCAATCCCATTGTTCCATAAACAGTGAGTTTATCTTTCTTTGAAGTGATATTTTGTAGAGATCTTATAAAACTGTCGAATGACATCGTATTTATATCTTTAGCTAAATCATATATGTTACCATCATAATATCTAACAGCATCTATCTTAAACGTTTCCATCGCTGTAAAGACGTTTCCGATAGATGGTCTCGTTTCATCAGTCATACTGAGCCAACAAATGTAATCGTGTATAACATCTTTGATAGAATCAGACTCTAAAGAAAAGGAATCGGCTTTATTATACACATTCATGATAAACTTAATGAAAAATGTTTTTCGTTGTTTAAGTTGGATGAATAGTATGTCTTAATAATTGTTATTATTTCACTAATTAATATTTAGTAACGAGTACACTCTATAAAAACGAGAATGACATAACTAATCATAACTAGTTATCAAAGAATGTCTAGGACGCGTAATTTTTTATGGTATAGATCCTGTAAGCGTTGTCTGTATTCTGGAGCTATTTTCTCTATCGCATTAGTGAGTTCAGAATATGTTATAAATTTAAATCGAATAACGAACATAACTTTAGTAAAGTCGTCTATATTAACTCTTTTATTTTCTAGCCATCGTAATACCATGTTTAAGATAGTATATTCTCTAGTTACTACGATCTCATCGTTGTCTAGAATATCGCATACTGAATCTACATCCAATTTTAGAAATTGGTCTGTGCTACATATCTCTTCTATATTATTGTTGATGTATTGTCGTAGAAAACTATTACGTAGACCATTTTCTTTATAAAACGAATATATAGTACTCCAATTATCTTTACCGATATATTTGCATACATAATCCATTCTCTCAATCACTACATCTTTAAGAGTTTGGTTGTTAAGATATTTGGCTAAACTATATAATTCTATTAGATCATCAACAGAATCAGTATATATTTTTCTAGATCCAAAGATGAACTCTTTGGCATCCTCTATAATATTATCAGAAAAGATATTTTCGTGTTTTAGTTTATCAAGATTTAACCTGTTCATATCCATGATTAACGACGTCATATAACCACATAAAATAAAAATCCATTTTCATTTTTAGCACAATACTATTCATAATTGATATTGATGTAATATTTTGTTACTTTGAACGTAAAGACAGTACACGGGTCCGTATCTCCAACAAGCACGTAGTAATCAAATTTGGTGTTGTTAAACTTCGCAATATTCATCAATTTAGATAGAAACATATACTCATCATCTGTTTTAGGAATCCATGTATTATTACCACTTTCCAACTTATCATTATCCCAGGCTATGTTTCGCCCATCATCGTTGCACAGAGTGAATAATTCTTTTGTATTCGGTAGTTCAAATATATGATCCATGCATATATCGACAAAGCTATTGTAGATGTGATTTTTCCTAAATCTAATATAAAACTCGTTTACTAGCAAACATTTTCCTGATTTATCGACCAAGACACACATGGTTTCTAAATCTATCAAGTGGTGGGGATCCATAGTTATAACGCAGTAACATAAATTATTACCTTCTTGACTGTCGCTAATATCTATATACTTATTGTTATCGTATTGGATTCTACATATAGATGGCTTGTATATCAAAGATATAGAACACATAACCAATTTATATTCTCGCTTTGTATTTTCGAATCTAAAGTTAAGAGATTTAGAAAACATTATATCATCGGATGATGATATCACTGTTTCCAGAGTAGGATATATTAAAGTCTTTAAAGATTTTGTCCGATTCAAATAAATCACTAAATAATATCCCATATTATCATCTGTTATAGTCGTGTCATTAAATCTATTATATTTTATGAAAGATATATCACTGCTCACCTCTATATTTCGTACATTTTTAAACTGTTTGTATAATATCTCTCTAATACAATCAGATATATCTATTGTGTCGGTAGACGATACCGTTACATTTGAATTAATGGTGTTCCATTTTACAACTTTTAACAAGTTGACCAATTCATTTCTAATAGTATCAAACTCTCCATGATTAAATATTTTAATAGTATCCATTTTATATCACTACGGACATAAACCATTGTATATTTTTTATGTTTATTAGTGTACACATTTTGGAAGTAAGTTCCTGGATCGGATGTCACCGCAGTAATATTGTTGATTATTTCTGACATCGACGTATTATATAGTTTTTTAATTCCATATCTTTTAGAAAAGTTAAACATCCTTATACAATTTGTGGAATTAATATTATGAATCATGGTTTTTACACATAGATCTATTACAGGCGGAACATCAATTATTATGGCAGCAACTAGTATCATTTCTACATTGTTTATGGTGATGTTTATCTTCTTCCAGCGCATATAGTCTAATATCGATTCAAACGCGTGATAGTTTATACCATTCAATATAATCGCTTCATCCTTTAGATGGTGATCCTGAATGTGTTTAAAAAATTATACGGAGACGCCGTAATAATTTCCCCATTGATAGAAAATATCACGCTTTCCATTTTCTTGAAGTACTAAAAGTAATTATAATATAATGTAAAGGTTTATATATTCAATATTTTTTATAAAAAAAATCATTTTGACATTAATTCCTTTTTAAATTTCAGTCTATCATCTATAGAAACATATTCTATGAATTTATAAAATGCTTTTACGTATCCTATCGTAGGTGATAGAACCGCTAAAAAACCTATCGAATTTCTACAAAAGAATCTGTTATATGGTATAGGGAGAGTATAAAACATTAAATGTCCGTACTTATTAAAGTATTCAGTAGCCAATCCTAACTCTTTCGAATAATTATTAATGGCTCTTATTCTGTACGAATCTATTTTTTTGAACAATGGACCTAGTGGTATATCTTGTTCTATGTATCTAAAATAATGTCTGACTAGATCCGTTAGTTTAATATCCGCAGTCATCTTGTCTAGAATGGCAAATCTAACTGCGGGTTTAGGCGTAGGCGTTAGTTTAGTTTCTATATCTACATCTATGTCTTTATCTAACACCAAAAATATAATAGCTAATATTTTATTACAATCATCCGGATATTCTTCTACGATCTCACTAACTAATGTTTCTTTGGTTATACTAGTATAGTCACGATCAGACAAATAAAGAAAATCAGATGATCGATGAATAATACATTTAAATTCATCATCTGTAAGATTTTTGAGATGTCTCATTAAAATATTATTAGTGTCAGTTCTCATTATCATATATTGACAGCAGCTATTACACTTATTTTATTTTTCTGTATTTTATTACTTTTCACCATATAGATCAGTCATTAGATCATCAAAATACTTTTCAATCATCCTAAAGAGTATGGTGAACGAATCTTCCCATCTAATTTCTGAACGTCTACCAATGTCTCTAGCCACTTTGGCACTAATAGCGATCATTCGCTTAACATCTTCTACATTATTAACTGGTTGATTCAATCTATCTAGCAATGGACCGTCGGATAGCGTCATTCTCATGTTCTTAATCAATGTACATACATCGTCATCATCTACCAATTCATCAAACAATATAAGCTTTTTAAAATCATCATTATAATAGGATGGATCGCCGTCATTTCTCCAAAGAATATATCTAATAAGTAGAGTCCTCATGCTTAGTAATTTAACTATTTTAGTTAACAACTATTTTTTATGTTAAATCAATTAGTAACACCGCTATGTTTAATACTTATTCATATTTTAGTTTTAGGATCGAGAATCAATACAAAAATTAATACATCAATTTTGGAAATACTTAGTTTCCACGTAGTCAATGAAACATTTGAGCTCATCGTAAAGGACGTTCTCGTACAAGACGTAACTATAAATTGGTTTATATTTGTTCAAGATAGATACAAATCCGATAACTTTTTTGACGAATTCTACGGGATTCACTTTAAAAGTGTCATACCGGGTTCTTTTTATTCTTTTAAACAGATCGATTGTGTGATGTTGATTAGGTCTTTTACGAATTTGATACAGAATAGCGTTTACATATCCACCATAGTAATCAATAGCCATTTGTTCGTATGTCATAAATTCTTTAATTATATGACACTGTGTATTATTTAGTTCGTCCTTGTTCATCATTAGGAATCTATCCAATATGGCAATTATATTAGAACTATAACTGCGTTGTATGCGCATGTTGATGTGTCTGTTTATACAATCAATTATACTAGGATCCATACCACTACAATCGGGTAAAATTGTAGCATCATATACCATTTCTAGTACTTTAGATTCATTGTTATCCATTGCAGAGGACGTCATGATCGAATCCAAAAAAATATATTATTTTTATGTTATTTTGTTAAAAATAATCATCAAATACTTCGTAAGATACTCCTTCATGAACATAATCAGTTACAAAACGTTTATATGAAGTAAAGTATCTACAATTTTTACAAAAGTCAGGATGCATAAGTACAAAGTACGCGATAAACGGAATAATAATAGATTTATCTAGTTTATCTTTTTCTATCTCTTTCATAGTTATATACATGGTCTCAGAAGTCGGATTATGTAACATCAGCTTCGATAAAATGACTGGGTTATTTAGTCTTACACATTCGCTCATACATGTATGACCGTTAACTATAGAGTCTACACTAAAATGATTGAATAATAGATAGTCTACCATTGTTTCGTATTCAGATAGTACAGCGTAGTACATGGCATCTTCACAAATTATATCATTATCTAATAGATATTTGACGCATCTTATGGATCCCACTTCAACAGCCATCTTAAAATCGGTAGAATCATATTGCTTTCCTTTATCGTTAATAATTTCTAGAACATCATCTCTATCATAAAAGATACAAATATTAACTGTTTGATCAGTAATAACATTGCTAGTCGATATCAATTTGTTAATAAGATGCGCTGGGCTCAATGTCTTAATAAGAAGTGTAAGAGGACTATCTCCGAATTTGTTTTGTTTATTAACATCCGTTGATGGAAGTAAAAGATTTATAATGTCTACATACTTGACTGTTTTAGAGCATACAATATGGAGAGGCGTATTTCCATCATGATCTGGTTTTGAGGGACTAATTCCTAGTTTCATCATCCATGAGATTGTAGAAGCTTTTGGATTGTCTGACATAAGATGTCTATGAATATGATTTTTGCCAAATTTATCCACTATCCTGGCTTCGAATCCGATAGACATTATTTTTTTAAACACTCTTTCTGAAGGATCTGTACACGCCAACAACGGACCACATCCTTCTTCATCAACCGAGTTGTTAATCTTGGCTCCATACTGTACCAATAAATTTATTCTCTCTATGACTTCATCATCTGTTCCCGAGAGATAATATAGAGGTGTTTTATTATGTTTATCACATGCGTTTGGATCTGCGCCGTGCACCAGCAGCATCGCGACTATTCTATTATTATTAATTTTAGAAGCTATATGCAATGGATAATTTCCATCATCATCCGTCTCATTTGGAGAGTATCCTCTATGAAGAAGTTCTTCTATAAATCGTTCATCTAGTCCTTTAATGCCACAATACGCATGTAGAATGTGATAATTTCCAGAGGGTTCGATAACTTGTAGCATATTCCTAAATACATCTAAATTTTTACTATTATATTTGGCATAAAGAGATAGATAATACTCGACCGACATAATGTTGTGTTGTCCATTATAGTATAAAAATTAATATTTCTATTTCTATATATTTGCAACAATTTACTCTCTATAACAAATATCATAACTTAGTTCTTTTATGTCAAGAAGGCACTGGTTTAATTCATCTATAAATGTCACTCCATAACTACCACGCATACTATACTCAGAATTATGATAAAGATATTTATTCTTGGGGTGTAAGTAATGGGGATTAATCTTTGTTGGATCAGTCTCTAAGTTAACACATGTCACACATGATCCATTTATAGTTATATCACACGATGATGATTTATGAATTGATTCCGGAAGATCGCTATTGTATTTTGTAGTTCCACAATTCATTTCCATACATGTTATTGTCACACTAATATTATGATGAACTTTATCTAGCCGCTGAGTGGTAAACAACAGAACAGATAGTTTATTATCTTTACCAACACCCTCAGCCGCTGCCACAAATCTCTGATCCGTATCCATGATGGTCATGTTTACTTTTAGTCCGTATCCAGTCAACACTATGTTAGCATTTCTGTCGATATAGCTTTCACTCATATGACACTCACCAATAATTGTAGAATTAATGTCGTAATTTACACCAATAGTGAGTTCGGCGACAAAGTACCAGTACCGGTAATCTTGTCGATGAGGACATATAGTATTCTTGTATTCTACCGAATACCCGAGAGATGCGATACAAAAGAGTAAGACTAATTTGTAAACCATCTTACTCAAAAATATGCGACAATAGGAAATCTATCTTATACACATAATTATTCTATCAATTTTACCAATTAGTTAGTGTAATGTTAACAAAAATGTGGGATAATTTAATAGTTTTTCCTTACACAATTGACATACATGAGTCTGAGTTCCTCGTTTTTGCTAATTATTTCGTCCAATTTATTATTCTTGACATCGTCAAGATCTTTTGTATAGGAGTCAGACTTGTATTCAACATGTTTTTCTATAATCATCTTAGCTATTTCGGCATCATCCAATAGTACATTTTCCAGATTAACAGAATAGATATTAATGTCGTATTTGAACAGAGCCTGTAACATCTCAATGTCTTTATTATCTATAGCCAATTTGATGTCCGGAATGAAGAGAAGGGAATTGGTGTTTGTTGACGTCATATAGTCGAGCAAGAGAATCATCATATCCACGTGTCCATTTTTTATAGTGGTGTGAATACAACTAAGGAGAATAGCCAGATCAAAAGGAGATGGTATCTCTGAAAGAAAGTAGGAAACAATACTTACATCATTAAGCATGACAGCATGATAAAATGAAGTTTTCCATCCAGTTTTCCCATAGAACATCAGTCTCCAATTTTTCTTAACAAACAGTTTTACCGTTTGCATGTTACCACTATCAACCGCATAATACAATGCGGTGTTTCCTTTGTCATCAAATTGTGAATCATCCATTCCACTGAATAGCAAAATCTTTACTATTTTGGTATCTTCTAATGTGGCTGCCTGATGTAATGGAAATTCATTATCTAGAAGATTTTTCAATGCTCCAGCGTTCAACAACGTACATACTAGACGCACGTTATTATCAGCTATTGCATAATACAAGACACTATGACCATTGATATCCGCCTTAAATGCATCTTTGCTAGAGAGAAAGCTTTTCAGTTGCTTAGACTTCCAAGTATTAATTCGTGACAGATCCATGTCTGAAACGAGACGCTAATTAGTGTATATTTTTTCATTTTTTATAATTTTGTCATATTGTACCAGAATTAATAATATCTCTAATAGATCTGATTAGTAGATATATGGCTATCGCAAAACAACATATACACATTTAATAAAAATAATATTCATTAAGAAGATTCAGATTCCACTGTACCCATCAATATAAAATAAAATAATTATTCCTTACATCGTACCCATAAACAATATATTAAGTATATTCCACCTTACCCATAAACAATATAAATCCAGTAATATCATGTCTAATGATGAACACAAATGGTGTATTAAATTCCAGTTCTTCAGGAGATGATCTCGCCGTAGCTACCATGATAGTAGATGCCTCCGCTACAGTTCCTTGTTCGTCTACATCTATCTTTACATTCTGAAACATTTTATAAATATATAATGGGTCCCTAGTCATATGTTTAAACGACGCCTTATCTGGATTAAACATACTAGGAGCCATCATTTCGGCTATCGACTTAATATCCCTCTTGTTTTCGATAGAAAATCTAGGGAGTTTAAGATTGTACATTTTATTCCCTAATTGAGATGACCAATATTCTAATTTTGCAGGCGTGATAGAATCTGTGAAATGGGTCATATTATCACCTATTGCCAGGTACATACTAATATTAGCATCCTTATACAGAAGGCGCACCATATCATATTCTTCGTCATCGATTGTGATTGTATTTCCTTGCAATTTAGTAACTACGTTCATCATGGGAACCGTTTTCGTACCGTACTTATTAGTAAAACTAGCATTGTGTGTTTTAGTGATATCAAACGGATATTGCCACGTACCTTTAAAATATATAGTATTAATGATTGCCCATAGAGTATTATCGTCGAGCATAGTAGAATCAACTACATTAGACATACCAGATCTACGTTCTACTATAGAATTAATTTTATTAACCGCATCTCGTCTAAAGTTTAATCTATATAGGCCGAATCTATGATATTGTTGATAATACGACGGTTTAATACACACAGTACTATCGACGAAACTTTGATACGTTAGATCTGTGTACGTATATTTAGATGTTTTCATCTTAGCTAATCCGGATATTAATTCTGTAAATGCTGGACCCAGATCTCTTTTTCTCAAATTCATAGTATTCAATAATTCTACTCTAGTATTACCTGATGCAGACAATAGCGACATAAACATAGAAAACGAATACCCAAACGGTGAGAAGACAATATTATCATTATCATCCTCATCCCCATTTTGAATATTTTTATACGCTAATATACCAGCATTGATAAATCCCTGCAGACGATATGCGGATACTGAACACGCTAATGATAGTATCAATAACGCAATCATGATTTTTATGGTATTAATAATTAACCTTATTTTTATGTTTGGTATAAAAATTATTGATGTCTACACATCCTTTATAATCAACTCTAATCACTTTAACTTTTACAGTTTTCCCTACAAGTTTATCCCTATATTCAACATATCTATCCATATGCATCTCTTAACACTCTGCCAAGATAGCTTCAGAGTGAGGATAGTCAAAAAGATAAATATATAGAGTATAATCATTCTCGTATACTCTGCCCTTTATTACATCGCCCGCATTGGGCAACGAATAACAAAATGCAAGCATCGTGTTAACGGGCTCGTAAATTGGGATAAAATTATGTTTTTATTGTTTATCTATTTTATTCAAGAGAATATTCAGGAAGTTCCTTTTCTGGTTGTATCTCGTCGCAGTATATATCATTTGTACATTGTTTCATATTTTTTAATAGTCTACACCTTTTAGTAGGACTAGTATCGTACAATTCATAGCTGTATTTTGAATTCCAATCACGTATAAAAATATCTTCCAATTGTTGACGAAGACCTAATCCATCATCCGGTGTAATATTAATAGATGCTCCACATATATCCGTAAAGTAATTTCCTGTCCAATTTGATGTACCTATATACGCCGTTTTATCGGTTACCATATATTTTGCATGGTTTACCCTAGAATACGGAATGGGAGGATCAGCATCTGGTACAATAAATAGCTTTACTTCTATATCTATGTTTTTAGATTTTAGCATAGCTATAGATCTTAAAAAGTTTCTCATGATAAACGAAGATCGTTGCCAGCAACTAATCAATAGCTTAACGGATACTTGTCTGTCTATAGCGGATCTTCTTAATTCATCTTCTATATAAGGCCAAAACAAAATTTTACCCGCCTTTGAATAAATAATAGGAATAAAGTTCATAACAGATACATAAACGAATTTACTCGCATTTCCGATACATGACAATAAAGCGGTTAAATCATTGGTTCTTTCCATAGTACATAATTGTTGTGGTGCAGAAGCAATAAATACAGAGTGTGGAACACCGCTTACGTTAATACTAAGAGGATGATCTGTATTATAATACGACGGATAAAAGTTTTTCCAATTATATGGTAGATTGTTAACTCCAAGATACCAGTATACCTCAAAAATTTGAGTGAGATCCGCTGCCAAGTTCCTATTATTGAAGATCGCAATACCCAATTCCTTGACCTGAGTTAGTGATCTCCAATCCATGTTAGCGCTTCCTAAATAAATATGTGTATTATCAGATATCCAAAATTTTGTATGAAGAACTCCTCCTAGGATATTTGTAATATCTATGTATCGTACTTCAACTCCGGCCATTTGTAGTCTTTCAACATCCTTTAATGGTTTGTTGGATTTATTGACGGCTACTCTAACTCTTACTCCTCTTTTGGGTAATTGTACAATCTCGTTTAATATTACCGTGCCGAAATTCGTACCCACTTCATCCGATAAACTCCAATAAAAAGATGATATATCTAGTGTTTTTATGGTATTGGATAGAATTTCCCTCCACATGTTAAATGTAGTCAAATATACTTTATCAAATTGCATACCTATAGGAATAGTCTCTGTAATCACTGCGATTGTATTATCCGGATTCATTTTATTTGTTAAAAAAATAATCCTATATCACTTCACTCTATTAAAAATCCAAGTTTCTATTTCTTTCATGACTGATTTTTTAACTTCATCCGTTTCCTTATGAAGATGATGTTTGGCGCCTTCATAAATTTTTATTTCCCTATTACAATTTGCATGTTGCATGAAATAATATGCACCTGAAACATCGCTAATCTCATTGTTTGTTCCCTGGAGTATGAGAGTCGGGGTGTTAATCTTGGGAATTATTTTTCTAACCTTGTTGGTAGCCTTCAAGACCTGACTAGCAAATCCAGCCTTAATTTTTTCATGATTGACTAATGGATCGTATTGGTATTTATAAACTTCATCCATATCTCTAGATACTGATTCTGGACATAGCTTTCCGACTGACGCATTTGGTGTAATGGTTCCCATAAGTTTTGCAGCTAGCAGATTCAGTCTTGGAACAGCGTCTGCATTAACTAGAGGAGACATTAGAATCATTGCTGTAAACAAGTTTGGATTATCGCAAGCAGCTAGTATAGAAATTGTTGCTCCCATGGAATGACCCAATAAGAAGACTGGAACTCCTGGATAAGTAGATTTAATAGTCACCACGTGCTGTACCACATCTCTAACATACTTACCAAAGTCATCAATCATCATTTTTTCACCATTACTTCTTCCATGGCCAATATGATCATGTGAGAATACTAAAATTCCTAACGATGATATGTTTTCAGCTAGTTCGTCATAACGTCCAGAATGTTCACCAGCTCCATGACTTATGAATACTAATGCCTTAGGATATGTAATAGGTTTCCAATATTTACAATATATGTAATCATTGTCCAGATTGAACATACAGTTTGTACTCATGATTCACTATATAACTATCAATATTAACAGTTCGTTTAATGATCATATTATTTTTATGTTTTATTGATAATTGTAAAAATATACAATTAAATCAATATAGAGGAAGGAGACGGTACTGTATTTTGTGAGATAGTAATGGAGACTAAATCAGATTATGAGGATGCTGTTTTTTACTTTGTGGATGATGATGAAATATGTAGTCGCGACTCCATCATTGATCTAATAGATGAATATATCACGTGGAGAAATCATGTTATAGTGTTTAATAAAGATATTACCAGTTGTGGAAGACTGTACAAGGAATTGATAAAGTTCGATGATGTCGCTATACGGTACTATGGTATTGATAAAATTAATGAGATTGTCGAGGCTATGAGTGAAGGAGACCACTACATCAATCTTACAGAAGTCCATGATCAGGAAAGTCTATTCGCTACCATAGGAATATGTGCTAAAATCACTGAACATTGGGGATACAAAAAGATTTCAGAATCTAAATTCCAATCATTGGGAAACATTACAGATCTGATGACCGACGATAATATAAACATCTTGATACTTTTTCTAGAAAAAAAAATGAATTGATGATATAAGTGTCTTCATAACGCATTATTACGTTAGCATTCTATTATCCAGTGTTAAAAAAATTATCCTATCATGTATTTGAGAGTCTTATATGTAGCAAACATGATAACTGCAATACCCATAATCTTTAGATATTCACGCGTGCTATGGATGGCATTATCCCGCGGTGCGGAAATGTACGTTATATAATCTACAAAATAATCATCGCATATAGTATGAGATAGTAGAGTAAACATTTTTATCGTTTCTACTGGGTTCATACATCGTCTACCCAATTCGGTAATGAATGAAATTGTCACCAATCTTACACCCAAACCCTTGTTGTTCATTAGTATAGTATTAACTTCATTATTTATGTCATAAACTGTAAATGATTCTGTAGATGCCATATCACACATGATATTCATGTCACTATTATAATCATTATTAACTTTATCACAATACGTGTTGATAATATCTACATATGATCTAGTTTTTGTGGGTAATTGTCTATACAAGTCGTCTAAACGTTGTTTACTCATATAGTATTGAACAGCCATCATTACATGGTCCCGTTCCGTTGATAGATAATCGAGTATGTTAGTAGACTTGTCAAATCTATATACCATATTTTCTGGAAGCGGATATACATAGTCGCGATCATCATTATCACTAGCCTCATCCTCTATATCATGTACATGTACATAATCTATGATATTATTATACATAAACATCGACAACATACTATTGTCTATTATCTAAGTCCTGTTGATCCAAACCCTTGATCTCCTCTATCTGTACTATCTAGAGATTGTACTTCTTCAAGTTCTGGATAATATATACGTTGATAGATTAGCTGAGCTATTCTATCTCCAGTATTTACATTAAACGTACATTTTCCATTATTAATAAGAATGACTCCTATGCTTCCCCTATAATCTTCGTCTATTACACCGCCTCCTATATCAATGCCTTTTAGGGACAGACCAGACCTAGGAGCTATTCTACCATAGCAGAACTTAGGCATGGACATACTAATATCTGTCTTAATTAACTGTCGTTCTCCAGGAGGGATAGTATAATCGTAAGCGCTATACAAATCATATCCGGCAGCACCCGGCGATTGCCTAGTAGGTGATTTAGCTCTGTTAGTTTCCTTAACAAATCTAACTGGTGAGTTAATATTCATGTTGAACATAAAAAATATCATTTTATTTCAAAATTATTTACCATTCCATTCCATCCCATATATTCCATGAATAAGTGCGATTATTGTACACTTCTATAGTATCTATATACGATCCACGATAAAATCCTCCTATCAATAGCAGTTTATTATCCACTATGATCAATTCTGGATTATCCCTCGGATAAATAGGATCATCTATCAGAGTCCATGTATTGCTGGATTCACAATAAAATTCCGCATTTCTACCAACCAAGAATAACCTTCTACCAAACACTAACGCACATGATTTATAATGAGGATAATAAGTGGATGGTCCAAACTGCCACTGATCATGATTGGGTAGCAAATATTCTGTAGTTGTATCAGTTTCAGAATGTCCTCCCATTACGTATATAACATTGTTTATGGATGCCACTGCTGGATTACATCTAGGTTTCAGAAGACTCGGCATATTAACCCAAGCAGCATCCCCGTGGAACCAACGCTCAACAGATGTGGGATTTGGTAGACCTCCTACTACGTATAATTTATTGTTAGCGGGTATCCCGCTAGCATACAGTCTGGGGCTATTCATCGGAGGAATTGGAATCCAATTGTTTGATATATAATTTACCGCTATAGCATTGTTATGTATTTCATTGTTCATCCATCCACCGATGAGATATACTACTTCTCCAACATGAGTACTTGTACACATATGGAATATATCTATAATTTGATCCATGTTCATAGGATACTCTATGAATGGATACTTGTATGATTTGCGTGGTTGTTTATCACAATGAAATATTATGTTACAGTCTAGTATCCATTTTACATTATGTATACCTCTGGGAGAAAGATAATTTGACCTGATTACATTTTTGATAAGAAGTAGCAGATTTCCTAATCTATTTCTTCGCCTCATATACCACTTAATGACAAAATCAACTACATAATCCTCATCTGGAACATTTAGTTCGTCGCTTTCTAGAATAAGTTTCATAGATAGATAATCAAAATTGTCTATGATGTCATCTTCCAGTTCCAAAAAGTGTTTGGTAATAAAGTCTTTAGTATGACATAAGAGATTGGATAGTCCGTATTCTATACCCATCATGTAACACTCGATACAATATTCCTTTCTAAAATCTCGTAGGATAAAGTTTATACAAGTGTAGATGATAAATTCTACAGATGTTAATATAGAAGCACGTAATAAATTGACGACGTTATGACTATCTATATATACCTTTCCAGTATATGAGTAAATAACTATAGAAGTTAGACTGTGAATGTCAAGGTCTAGACAAACCCTCGTAACTGGATCTTTATTTTTTGTGTATTTTTGGCGTAAATGTGTGCGAAAGTATGGAGATAACTTTTTCAATATCGTAGAATTGACTATTATATTACCTCCTATAGCTTCAATAATTGTTTTGAATTTCTTAGTCGTGTACAATGCTAATATATTCTTACAGTACACAGTATTGACAAATATCGGCATTTATGTTTCTTTAAAAGTCAACATCTAAAGAAAAATGATTGTCTTCTTGAGACATAACTCCCATTTTTTGGTATTCACCCACACGTTTTTCGAAAAAATTAGTTTTTCCTTCCAATGATATATTTTCCATGAAATCAAACGGATTGGTAACATTGTAAATTTTTTTAAATCCCAATTCAGAAATCAATCTATCTGCGACGAATTCTATATATGTTTTCATCATTTCACAATTCATTCCTATGAGTTTAACTGGAAGAGCCACAGTAAGAAATTCTTGTTCAATGGATACCGCATTTGTTATAATAAATCTAACGGTTTCTTCACTCGGTGGATGCAATAAATGTTTAAACATCAAACATGCGAAATCGCAGTGCAGACCCTCGTCTCTACTAATTAATTCGTTAGAAAACGTGAGTCCGGGCATTAGGCCACGCTTTTTAAGCCAAAATATGGAAGCGAATGATCCGGAAAAGAAGATTCCTTCTACTGCAGCAAAGGCAATAAGTCTCTCTCCATAACCGGCGCTGTCATGTATCCACTTTTGAGCCCAATCGGCCTTCTTTTTTACACAAGGCATCGTTTCTATGGCATTAAAGAGGTAGTTTTTTTCATTACTATCTTTAACATAAGTATCGATCAAAAGACTATACATTTCCGAATGAATGTTTTCAATGGCCATCTGAAATCCGTAGAAACATCTAGCCTCGGTAATCTGCACTTCTGTACAAAATCGTTCTGCTAAATTTTCATTCACTATTCCGTCACTGGCTGCAAAAAACGCCAATACATGTTTTATAAAATATTTTTCGTCTGGTGTTAGTTTATTCCAGTCATTGATATCTTTAGATATATCCACTTCTTCCACTGTCCAAAATGATGCCTCTGCCTTTTTATACATATTCCAGATGTCATGATATTGGATTGGGAAAATAACAAATCTATTTGGATTTGGTGCAAGGATAGGTTCCATAACTAAATTAACAATAGTAGTAATTTTTTTTCAGTTATCTGTATGACGACTGTACTTGGATCTTTTGTATATCGCTATCGCCGCAATCACTACAATAATTACAAGTATTATTGATAGCATTGTTATTACTACTATCATAATTAAATTATCGACATTCATGGGTGTTGAATAATCGTTATCATCATTTTGTAATTGTGACGTCATACTAGATAAATCATTTGTGAGATTGTTGTGGGAAGCGGGCACGGAAGATGCATTATCATTATTATTTAACGCCTCCCATTCGGATTCACAAATGTTACGCACGTTCAAAGTTTTATGGAAACTATAATTTTGTGAAAACAGATAACAAGAAAACTCGTCATCGTTCAAATTTTTAACGATAGTAAACCGATTAAACGTCGAGCTAATTTCTAACGCTAGCGACTCTGTTGGATATGGGTTTCCAGATATATATCTTTTCAGTTCCCCTACGTATCTATAATCATCTGTAGGAAATGGAAGATATTTCCATTTATCTACTGTTCCTAATATCATATGCGGTGGTGTAGAACCATTAAGCGCGAAAGATGTTATTTCGCATCGTATTTTAACTTCGCAATAATTTCTGGTTAGATAACGCACTCTACCAGTCAAGTCAATGATATTAGCCTTTACAGATATATTCATAGTAGTCGTAACGATGACTCCATCTTTTAGATGTGATACTCCTTTGTATGTACCAGAATCTTCGTACCTCAAACTCGATATATTTAAACAAGTTAATGATATATTAACGCGTTTTATGAATGATGATATATAACCAGAAGTTTTATCCTCTGTGGCTAGCGCTATAACCTTATCATTATAATACCAACTAGTGTAATTAATATGTGACATGACAGTGTGGGTACAAATATGTACATTATCGTCTACGTCGTATTTGATACATCCGCATTCAGCCAACAAATATAAAATTACAAAAACTCTAACGACGTTTGTACACATCTTGATGCGGTTTAATAAATGTTTTGATTTCAATTTATTGTAAAAAAAGATTCGGTTTTATACTGTTCGATATTCTCATTGCTTATATTCTCATCTATCATCTCCACACAGTCAAATCCATGGTTAACATGTACCTCATCAACCGGTAAAAGACTATCGGATTCTTCTATCATCATAACTCGAGAATATTTAATTTGGTGGTCATTATTAATCAAGTCAATTATCTTATTTTTAACAAACGTAAGTATTTTACTCATTTTTTTATAAAAACTTTTAGAAATATACAGACTCTATCGTGTGTCTATATCTTCTTTTTATATCCAATGTATTTATGTCTGATTTTTCTTCATTTATCATATATAATGGTCCAAATTCTACACGTGCTTCGGATTCATCCAGATCATTAAGGTTCTTATAATCGCAACATCCTTCTCTTCCATCTTCTACATCTTCCTTCTTATTCTTAGCGTCACAGAATCTACCACAGCAGGATCCCATGACGAGAGTCACATTAAACTAATTCATTTTCAATTATAATATACTGATTAGTAATGACAATTAAAATAAAAATATTCTTCATAACCGGTAAGAAAGTAAAAAGTTCACATTGAAACTATGTCAGTAGTTATACATCATGAGATGATATACTCTATTTTGGTGGAGGATTATATGATATAATTCGTGGATAATCATTCTTAAGACACATTTCTTCATTCGTAAATCTTTTCACATTAAATGAGTGTCCATATTTTGCAATTTCTTCATATGATGGCGGTGTACGTGGACGAGGCTGCTCCTGTTCTTGTAGTCGTCGACTGTCGTGTTTGCGTTTAGATCCCTCCATTATCGCGATCGCGTAGTGAGTACTATTTATACCTTGTAATTAAATTTTTTTATTAATTAAACGTATAAAAACGTTCCGTATCTGTATTTAAGAGCCAGATTTCGTCTAATAGAACAAATAGCTACAGTAAAAATAACTAGAATAATCGCTACACCCACTAGAAACCACGGATCGTAATACGGCAATCGGTTTTCGATAATAGGTGGAACGTATATTTTATTTAAGGACTTAACAATTGTCTGTAAACCACAATTTGCTTCCGCCGATCCTGTATTAACTATCTGTAAAAGCATATGTTGACCGGGAGGAGCCGAACATTCTCCGATATTCAATTTTTGTATATCTATAATGTTATTAACCTCCGCATACGCATTACAGTTCTTTTCTAGCTTGGATACTACACTAGGTACATCATCTAAATCTATTCCTATTTCCTCAGCGATAGCTCTTCTATCCTTTTCCGGAAGTAATGAAATCACTTCAATAAATGATTCAACCATGAGTGTGAAACTAAGTCGAGAATTACTCATGCATTTGTTAGTTATTCGGAGCGCGCAATTTTTAAACTGTCCTATAACCTCTCCTATATGAATAGCACAAGTGACATTAGTAGGGATAGAATGTTGAGCTAATTTTTGTAAATAACTATCTATAAAAAGATTATACAAAGTTTTAAACTCTTTAGTTTCTGCCATTTATCCAGTCTGAGAAAATGTCTCTCATAATAAATTTTTCCAAGAAACTAATTGGGTGAAGAATGGAAACCTTTAATCTATATTTATCACAGTCTGTTTTGGTACACATGATGAATTCTTCTAATGCTGTACTAAATTCGATATCTTTTTCGATTTCTGGATATGTTTTTAATAAAGTATGAACAAAGAAATGGAAATCGTAATACCAGTTATGTTCAACTTTGAAATTGTTTTTTATTTTCTTGTTAATGATTCCAGCCACTTGGGAAAAGTCAAAGTCGTTTAATGCCGATTTAATACGTTCATTAAAAACAAACTTTTTATTCTTTAGATGAATTATTATTGGTTCATTGGAATCAAAAAGTAAGATATTATCGGGTTTAAGATCTGCATGTAAAAAGTTGTCACAACAGGGTAGTTCGTAGATTTTAATGTATAACAGAGACATCTGTAAAAAGATAAACTTTATGTATTGTACCAAAGATTTAAATCCTAATTTGATAGCTAACTCGGTATCTACTTTATCTGCCGAATACAGTGCTAGGGGAAAAATTATAATATTTCCTCTTTCGTATTCGTAATTAGTTCTCTTTTCATGTTCGAAAAAGTGAAACATGCGGTTAAAATAGTTTATAACATTAATATTACTGTTAATAACTGCCGGATAAAAGTGGGATAGTAATTTCACGAATTTGATACTGTCCTTTCTCTCGTTAAACGCCTTTAGAAAAACTTTAGAAGAATATCTCAATGAGAGTTCCTGACCATCCATAGTTTGTATCAATAATAGCAACATATGAAGAACCCGCTTATACAGAGTATGTAAAAATGTTAATTTATAGTTTAATCCCATGGCCCACGCACACACGATTAATTTTTTTTCATCTCCCTTTAGATTGTTGTATAGAAATTTGGGTACTGTAAACTCCGCCGTAGTTTCCATGGGACTATATAATTTTGTGGCCTCGAATACAAATTTTACTACATAGTTATCTATCTTAAAGACTATACCATATCCTCCTGTAGATATGTGATAAAAATCGTCGTTTATAGGATAAAATCGTTTATCTTTTTGTTGGAAAAAGGATGAATTAATGTAATCATTCTCTTCTATCTTTAGTAGTGTTTCCTTATTAAAATTCTTAAAATAATTTAACAATCTAACTGACGGAGCCCAATTTTGGTGTAAATCTAATTGGGACATTATGTTGTTAAAATATAAACAGTCTCCTAATATAACAGTATCTGATAATCTATGGGGAGACATCCATTGATATTCAGGGGATGAATCATTGGCAACACCCATTTATTGTACAAAAAGCCCCAATTTACAAACGAAAGTCCAGGTTTGATAGAGATAAACTATTAACTATTTTGTCTCTGTTTTTAACACCTCCACAGTTTTTAATTTCTTTGGTAATGAAATTATTCACAATATCAGTATCTTCTTTATCTACCAGAGATTTTACTAACTTGATAACCTTGGCTGTCTCATTCAATAGGGTAGTGATATTTGTATGTATGATATTGATATCTTTTTGAATTGTTTCTTTTAGAAGTGATTCTTTGATGGTATCAGCATACGAATTACAATAATGCAGAAACTCAGTTAACATGCAGGAATTATAGTAAGCCAATTCCAATTGTTGCCTGTATTGTATTAGAGTATTAATATGCGCAATGATGTCCTTGCGTTTCTCTGATAGAATGCGAGCAGCGATTTTGGCGTTATCATTTGACGATATTTCTGGAATGACGAATCCTGTTTCTACTAACTTCTTGGTAGGACAAAGTGAAACAATCAAGAAAATAGCTTCTCCTCCTATTTGTGGAAGAAATTGAACTCCTCTAGATGATCTACTGACGATAGTATCTCCTTGACAGATATTGGACCGAACTACGGAAGTACCTGGAATGTAAAGCCCTGAAACCCCCTCATTTTTTAAGCAGATTGTTGCCGTAAATCCTGCACTATGCCCAAGATAGAGAGCTCCTTTGGTGAATCCATCACTATGTTTCAGTTTAACCAAGAAACAGTCAGCTGGTCTAAAATTTCCATCTCTATCTAATACAGAATCCAACTTGATGTCAGGGACTATGACCGGTTTAATGTTATATGTAACATTGAGTAAATCCTTAAGTTCATAATCATCGTTGTCATCAGTTATGTACGATCCAAACAATGTTTCTACCGGCATGGTGGATACGAAGATGCTATCCATCAGAATGTTTCCCTGATTAGTATTTTCTATATAGCTATTCTTCTTTAAACGATTTTCCGAATCAGTAACTATGTTCATTTTTTTAGGAGTAGGACGTCTAGCCAGTATGGAAGAGGATTTTCTAGATACTCTCTTCAACATCTTTGATCTCAATGGAATGCAAAACCCCATGGTGTAACAACCAACGATAAAAATAATATTGTTTTTTCACTTTTTATAATTTTACCATCTGACTCATGGATTCATTAATATCTTTACAAGAGCTACTAACGTATAATTCTTTATAACTGAACTGAGATATATACACCGGATCTATGGTTTCCATAATTGAGTAAATGAATGCTCGGCAATAACTAATGGCAAATGTATAGAACAACGAAATTATACTAGAGTTGTTAAAGTTAATATTTTCTATGAGTTGTTCCAATAAATTATTTGTTGTGACTGCGTTCAAGTCATAAATTATCTTGATACTATCCAGTAAACAGTCTTTAAGTTCTGGAATATTATCATCCCATTGTAAAGCCCCTAGTTCGACTATCGAATATCCTGCTCTGATAGCAGTTTCAATATCGACGGACGTCAATACTGTAATAAAGGTGGTAGTATTGTCATCATCGTGATAAACTACGGGAATATGGTCGTTAGTAGGTACCGTGACTTTACACAACGCGATATATAACTTTCCTTTTGTACCATTTTTAACGTAGTTGGGACGTCCTGCAGGGTATTGTTTTGAAGAAATGATATCGAGAACAGATTTGATACGATATTTGTTGGATTCCTGATTATTCACTATAATATAATCTAGACAGATAGATGATTCGATAAATAGAGAAGGTATATCGTTGGTAGGATAATACATCCCCATTCCAGTATTCTCGGATACTCTATTGATGACACTAGTTAAGAACATGTCTTCTATTCTAGAAAACGAAAACATCCTACATGGACTCATTAAAACTTCTAACGCTCCTGATTGTGTTTCGAATGCCTCGTACAAGGATTTCAAGGATGCCATAGATTCTTTGACCAACGATTTAGTATTGCGTTTAGCATCTGATTTTTTTATTAAATCAAATGGTCGGCTCTCTGGTTTACTACCCCAATGATAACAATAGTCTTGTAAAGATAAACCGCAAGAAAATTTATACACATCCATCCAAATAACCCTAGCACCGTCGGATGATATTAATGTATTATTATAGATTTTCCATCCACAGTTATTGGGCCAGTATACTGTTAGCAACGGTATATCGAATAGATTACTCATGTAACCTACTAGAATGATAGTTCGTGTACTAGTCATAATATCTTTAATCCAATCTAAGAAATCTAAAATTAGATCTTTTACACTATTAAAGTTAACAAAGGTATTACCCGGGTACGTGGATATCATATATGGTATTGGTCCATTATCAGTAATGGCTCCATAAACTGATACGGCGATGGTTTTTATATGTGTTTGATCTAATGAGGACGAAATTCGCGCCCACAATTCATCTCTAGATATGCATTTAATATCGAACGGTAACACATCAATCTCGGGACGCGTATATGTTTCTAAATTCTTAATCCAAATATAATGATGACCTATATGCCCTATTATCATACTGTCAACTATAGTATACCTAGAGAACTTTCGATACATCTGCTGTTTCCTGTAATCGTTAAATTTTACAAATCTATAACATGCTAAACCTTTTGACGACAGCCATTCATTAATTTCTGATATGGAATCTGTATTCTCAATACCGTATCGTTCTAAAGCCAGTGCTATATCTCCCTGTTCGTGGGAACGCTTTCGTATAATATCGATCAATGGATAATATGAAGTTTTTGGAGAATAATATGATTCATGATCTATTTCGTCCATAAACAATCTAGACATAGGAATTGGAGGCGATGATCTTAATTTTGTGCAATGGGTCAATCCTATAACTTCTAATATTGTAATATTCATCATCGACATAACACTATCTATGTTATCATCGTATATTAGTATACCACGACCTTCTTCATTTCGTGCCAAAATGATATACAGTCTTAAATAATTACGCAATATCTCAATAGTTTCATAATTGTTAGCTGTTTTCATCAAGGTTTGTATCCTGTTTAACATGATGGCGTTCTATAACGTCTCTATTTTCTATTTTTAATTTTTTTAAATTTTTAACGATTTACTGTGGCTAGATACCCAATCTCTCTCAAATATTTTTTTAGCCTCGCTTACAAGCTGTTTATCTATACTATTAAAACTGACGAATCCGTGATTTTGGTAATGGGTTCCGTCGAAATTTGCCGAAGTGATATGAACATATTCGTCGTCGACTATTAACAATTTTGTATTATTCTGAATAGTGAAAACCTTCACAGATAGATCATTTTGAACACACAACGCATCTAGACTTCTGGCGGTTGCCATAGAATATACGTCGTTCTTATCCCAATTACCAACTAGAAGTCTGATCTTAACTCCTCTATTAATGGCTGCTTCTATAATGGAGTTGTAAATGTCAGGCCAATAGTAGCTATTACCGTCGACACGTGTAGTGGGAACTATGGCCAAATGTTCAATATCTATACTAGTCTTAGCCGACTTGAGTTTATCAATAACTACATCAGTGTCTAGATCTCTAGAATATCCCAATAGGTGTTCTGGAGAATCAGTAAAGAACACTCCACCTATAGGATTCTTAATATGATACGCAGTGCTAACTGGCAGACAACAAGCCGCAGAGCATAAATTCAACCATGAATTTTTTGCGCTATTAAAGGCTTTAAAAGTATCAAATCTTCTACGAAGATCTGTGGCCAGCGGAGGATAATCAGAATATACGCCTAACGTTTTAATCGTATGTATAGATCCTCCAGTAAATGACGCGTTTCCTACATAACATCTTTCATCATCAGACACCCAAAAACAACCGAGTAGTAGTCCCACATTATTTTTTTTATCTATATTAACGGTTATAAAATTTATATCCGGGGAGTGACTTTGTAGCTCTCCCAGATTTCTTTTCCCTCGTTCATCTAGCAAAACTATTATTTTAATCCCTTTTTCAGATACCTCTTTTAGTTTATCAAAAATAAGCGCTCCCCTAGTAGTACTCAGAGGATTACAACAAAAAGATGCTATGTATATATATTTCTTAGCTAGAGTGATAATTTCGTTAAAACATTCAAATGTTGTCAAATGATCGGATCTAAAATCCATATTTTCTGGTAGTGTTTCTACCAGCCTACATTTTGCTCCCGCAGGTACCGATGCAAATGGCCACATTTAGTTAACATAAAAACTTATATATCCTGTTCTATCAACGATTCTAGAATATCATCGGCTATATCGCTAAAATTTTCATCAAAGTCGACATCACAACCTAACTCAGTCAATATATTAAGAAGTTCCATGATGTCATCTTCGTCTATTTCTATATCCGTATCCATTGTAGATTGTTGACCGATTATCGAGTTTAAATCATTACTAATACTCAATCCTTCAGAATACAATCTGTGTTTCATTGTAAATTTATAGGCGGTGTATTTAAGTTGGTAGATTTTCAATTATGTATCAATATAGCAACAGTAGTTCTTGCTCCTCCTTGATTTTAGCATCCTCTTCATTATTTTCTTCTACGTACATAATCATGTCTAATACGTTAGACAACACACCGACGATGGTGGCCGCCACAGACACGAATATGACTAGACCGATGACCATTTAAAAAATACTCTCTAGCTTTAACTTAAACTGTATCGATCATTCTTTTAGCACATGTATAATATAAAAACATTATTCTATTTCGAATTTAGGCTTCCAAAAATTTTTCATCCGTAAACCGATAATAATATATATAGACTTGTTAATAGTCGGAATAAATATATTAATGCTTAAACTATCATCATCTCCACGATTAGAGATACAATATTTACATTCTTTTTGCTGTTTCGAAACTTTATCAATACACGTTAATACAAACCCAGGAAGGAGATATTGAAACTGAGGCTGTTGAAAATGAAACGGCGAATACAATAATTCAGATAATGTAAAATCATGATTCCGTATTCTGATGATATTAGAACTGCTAATGGATGTCGATGGTATGTATCTAGGAGTATCTATTTTAACAAAGCATCGATTTGCTAATATACAATTATCCTTTTGATTAATTGTTATTTTATTCATATTCTTAAAAGGTTTCATATTTATCAATTCTTCTACATTAAAAATTTCCATTTTTAATTTATCTAGCCCCGCAATACTCCTCATTACGTTTCATTTTTTGTCTAGAATACCCATTTTGTTCATCTTGGTACATAGATTATCCAATTGAGAAGCGCATTTAGTAGTTTTGTACATTTTAAGTTTATTAACGAATCGTCGAAAACTAGTTATAGTTAACATTTTATTATTTGATACCCTGATATTAATACCCCTGCCGTTACTATTATTTATAACTGATGTAACCCACGTAACATTGGAATTAATTATCGATAGTAATGCATCGACACTTCCAAAATTGTCTATTATAAACTCACCGATAATTTTTTTATTGCATGTTTTCATATTCATTAGGATTATCAAATCTTTAATCTTATTACGATTGTATGCGTTGATATTACAAGACGTCATTCTAAAAGACGGAGGATTTCCATCAAATGCCAGACAATCACGTACAAAGTACATGGAAATAGGTTTTGTTCTATTACGCATCATAGATTCATATAAAACACCCGTAGAAATACTAATTTGTTTTACTCTATAAAATACTATTGCATCTATTTCATCGTTTTGTATAACGTCTTTCCAAGTGTCAAATTCCAATTTTTTTTCATTGATAGTACCAAATTCTTCTATCTCTTTAACTACTTGCATAGATAGGTAATTACAGTGATGCCTACATGCCGTTTTTTGAAACTGAATAGATGCATCTAGAAGCGATGCTACACTAGTCACGATCACCACTTTCATATTTAGAATATATGTATGTAAAAATATAGTAGAATTTCATTTTGTTTTTTTCTATGCTATAAATGAATTCTCATTTTGTATCCGCACATACTCCGTTTTATATCAATACCAAAGAAGGAAGATATCTGGTTCTAAAAGCCGTTAAAGTATGCGATGTTAGAACTGTAGAATGCGAAGGAAGTAAAGCTTCCTGCGTACTCAAAGTAGATAAACCCTCATCACCCACGTGTGAGAGAAGACCTTCGTCCCCGTCCAGATGCGAGAGAATGAATAACCCTGGAAAACAAGTCCCGTTTATGAGAACGGACATGTTACAAAATATGTTTGCTGCTAATCGCGACAACGTAACGTCAAGACTTTTGAACTAAAATACAATTATATCTTTTTCGATATTAATAAATCCGTGTCTCCCGGGTTTTTTATCTCTTTCAGTATGTGAATAGATAGGTATTTTATCTCTATTCATCATCGAATTTAAGAGATCCGATAAACATTGTTTGTATTCTCCAGATGTCAGCATCTGATACAACAATATATGTGCACATAAACCTCTGGCACTTATTTCATGTACCTTCCCCTTATCACTAAGGAGAATAGTATTTGAGAAATATGTATACATGATATTATCATGTATTAGATATACAGAATTTGTAACACTCTCGAAATCACACGATGTGTCGGCGTTAAGATCTAATATATCACTCGATAACACATTTTCATCTAGATACACTAGACATTTTTTAAAGCTAAAATAGTCTTTAGTAGTAACAGTAACTATGCGATTATTTTCATCGATGATACATTTCATCGGCATATTATTACGCGTACCATCAAAGACTATACCATGTGTATATCTAACGTATTCTAGCATAGTTGCCATACGTACATTAAACTTTTCAGGATCTTTGGATAGATCTTCCAATCTATCTATTTGAGAAAACATTTTTATCATGTTCAATAGTTGAAACGTCGGATCCACTATATAGATATTATCTATAAAGATTTTAGGAACTATGTTCATGGTATCCTGGCGAATATTAAAACTATCAATGATATGATTATCGTTTTCATCTTTTATCACCATATAGTTTCTAAGATATGGGATTTTACTTAATATAATATTATTTCCCGTAATAAATTTTATTAGAAATGCCAAATCTATAAGAAAAGTCCTAGAATTAGTCTGAAGAATATCTATATCACCGTACCGTATATTTGGATTAATTAGATATAGAGAATATGATCCGTAACATATACAACTTTTATTATGACGTCTAAGATATTCTTCCATCAACTTATTAACATTTTTGACTAGGGAAGATACATTATGACGTCCCATTACTTTTGCCTTGTCTATTACAGCGACGTTCATAGAATTTAGCATATCTCTTGCCAATTCTTCCATTGATGTTACATTATAAGAAATTTTAGATGAAATTACATTTGGAGCTTTAATAGTAAGAACTCCTAATATATCCGTGTATGTGGTCACTAATACAGATTGTAGTTCTATAATCGTAAATAATTTACCTATATTATATGTTTGAGTTTGTTTAGAAAAGTAGCTAAGTATACGATCTTTTATTTCTGATGCCGATGTATCAACATCGAAAAAAAATCTTTTTTTATTCTTTTTTACTAACGATACGAATATGTCTTTGTTAAAAACAGTTATTTTCTGAATATTTCTAGCTTGTAATTTTAACATATGATATTCGTTCACACTAGGTACTCTGCCTAAATAGGTTTCTATAATCTTTAATGTAATATTAGGAAGAGTATTCTGATCAGGATTCCTATTCATTTTGAGGATTTAAAACTCTGATTATTGTCTAATATGGTCTCAACACAAACTTTTTCACAGAGTGATAGAGTTTTTGATAACTCGTTTTTCTTAAGAAATATAAAACTACTGTCTCCAGAGCTCGCTCTATCTTTTATTTTATCTAATTCGATACAAACTCCTGATACTGGTTCAGAAAGTAATTCATTAATTTTCAGTCCTTTATAGAAGATATTTAATATAGATAATACAAAATCTTCAGTTCTTGATATCGATCTGATTGATCCTAGAACTAGATATATTAATAACGTGCTCATTAGGCAGTTTATGGCAGCTTGATAATTAGATATAGTATATTCCAGTTCATATTTATTAGATACCGCATTGCCCAGATTTTGATATTCTATGAATTCCTCTGAAAATAAATCCAAAATAACTAGACATTCTATTTTTTGTGGATTAGTGTACTCTCTTCCCTCTATCATGTTCACTACTGGTGTCCACAATGATAAATATCTAGAGGGAATATAATATAGTCCATATGATGCCAATCTAGCAATGTCGAATAACTGTAATTTTATTCTTCGCTCTTCATTATGAATTGAATCTTGAGGTATAAACCTAACACAAATTATATCATTAGACTTTTCGTATGTAATGTCTTTCATGTTATAAGTTTTTAATCCTGGAATAGAATCTATTTTAATGAGGCTTTTAAACGCAGCGTTCTCCAACGAGTCAAAGCATAATACTCTGTTGGTTTTCTTATATTCAATATTACGATTTTCTTCTTTGAATGGAATAGGTTTTTGAATTAGTTTATAATTACAACATAATAGATAAGGAAGTGTGTAAATAGTACGCGGAAAAAACATAATAGCTCCCCTGTTTTCATCCATGGTTTTAAGTAAATGATCACTGGCTTCTTTAGTCAATGGATATTCGAACATTAACCGTTTCATCATCATTGGACAGAATCCATATTTCTTAATGTAAAGAGTGATCAAATCATTGTGTTTATTGTACCATCTTGTTGTAAATGTGTATTCGGTTATCGGATCTGCTCCTTTTTCTATTAAAGTATCGATATCGATCTCGTCTAAGAATTCAACTATATCGACATATTTCATTTGTATACACATAACCATTACTAACGTAGAATGTATAGGAAGAGATGTAACGGGAACAGGGTTTGTTGATTCGCAAACTATTCTAATACATAATTCTTCTGTTAATACGTCTTGCACGTAATCTATTATAGATGCCAAGATATCTATATAATTATTTTGTAAGATGATGTTAACAATGTGATCTATATAAGTAGTGTAATAATTCATGTATTTCGATATATGTTCCAACTCTGTCTTTGTGATGTCTAGTTTCGTAATATCTATAGCGTCCTCAAAAAATATATTCGCATATATTCCCAAGTCTTCAGTTCTATCTTCTAAAAAATCTTCAACGTATGGAATATAATAATCTATTTTACCTCTTCTGATGTCATTAATGATATAGTTTTTGACACTATTTTCCGTCAATTGATTCTTATTCACTATGTCTAAAAACCGGATAGCGTCCCTAGGACGAACTACTGCCATTAATATCTCTATTATAGCTTCTGGACATAAATCATCTATTATACCAGAATTAATGGGAACTATTCCGTATCTATCTAACATAGTTTTAAGAAAGTCAGAATCTAAGACCTGATGTTCATATATTGGTTCATACATGAAATGATCTCTATTGATGATAGTGACTATTTCATTCTCTGAAAATTGGTAACTCATTCTATACACGCTTTCCTTGTTGATAAAGGATAGTATATACTCAATGGAATTTGTACCAACAAACTGTTCTCTTATGAATCGTATATCATCATCTGAAATGATCATGTAAGGCATACATTTAACAATAAGAGACTTGTCTCCTGTTATCAATATACTATTCTTGTGATAATTTATGTGTGCGGCAAATTTGTCCACGTTCTTTAATTTTGTTATAGTAGATATCAAATCCAATGGAGATACAGTTCTTGGCTTAAACAGATATAGTTTTTCTGGAACGAATTCTACAACATTATTATAAAGGACTTTGGGTATATAAGTGGGATGAAATCCTATTTTAATTAATGCGATAGCCTTGTCCTCGTGCAGATATCCAAACGCTTTTGTGATAGTATGGCATTCATTGTCTAGAAACGCTCTACGAATATCTGTAACAGATATCATCTTTAGAGAATACTAGTCGCGTTAATAGTACTAAAATTTGTATTTTTTAATCTATCTCAATAAAAAATTAATATGTATGATTCAATGTATAACTAAACTACTAACTGTTATTGATAACTAGAATCAGAATCTAATGATGACATAACTAAGAAGTTTATCTACAGCCAATTTAGCTGCATTATTTTTAGCATCTCGTTTAGATTTTCCATCTGCCTTATCGAATACTCTTCCGTCAATGTCTACACAGGCATAAAATGTAGGAGAGTTACTAGGCCCCACTGATTCAATACGAAAAGACCAATCTCTCCTAGTTATTTGACAGTACTCATTAATAACGGTGACAGGGTTAACACCTTTCCAATAAATAATTTTTTTAACCGGAATAACATCATCAAAAGACTTATGATCCTCTCTCATTGATTTTTCGCGGGATACATCATCTATTATAGCATCAGCATCAGAATCTGTAGGCCGTGTATCAGCATCCATTGTCGTAGACCAACGAGGAGGAGTATCGTTGGAGCTGTAAACCATAGCACTACGTTGAAGATCATACAGAGCTTTATTAACTTCTCGCTTCTCCATATTAAGTTGTTTAGTTAGTTGTACAGCAGTAGCTCCTTAGTCCAATGTTTTTAATAACCGCACACAATCTCTGTGTCAGAACGCTCGTCAATATAGATCTTAGAAATTTTTTTAGAGAGAACTAACGCAACTAGCAATAAAACTGATCTTATTTTATCATTTTTTTTATTCATCATCCTCTGGTGGTTCGTCGTTCCTATCGAATGTAGCTCTGATTAACCCGTCATCTATAGGTGATGCTGGTTCTGGAGATTCTGGAGGAGATGGATTATTATCTGGAAGAATCTCTGTTATTTCCTTGTTTTCATGTATCGATTGCGTTGTAACATTAAGATTGCGAAATGCTCTAAATTTGGGAGGCTTAAAGTGTTGTTTACAATCTCTACACGCGTGTCTAACTAATGGAGGTTCATCAGCGGCTCTAGTTTGAATCATCATCGGTGTAGTATTCCTACTTTTACAGTTAGGACACGGTGTATTGTATTTCTCGTCGAGAACGTTAAAATAATCGTTGTAACTCACATCCTTTATTTTATCTATATTGTATTCTACTCCTTTCTTAATGCATTTTATACCGAACAAGAGATAGCGAAGGAATTCTTTTTCGGTACCGCTAGTACCCTTAATCATATCACATAGTGTTTTATATTCTAAATGTGTGGCAATGGACGGTTTATTTCTATACGATAGTTTGTTTTTGGAATCCTTTGAGTATTCTATACCAATATTATTCTTTGATTCGAATTTAGTTTCTTCGATATTAGATTTTGTATTACCTATATTCTTGATGTAGTACTTTGATGATTTTTCCATGGCCCATTCTATTAAGTTTTCCAAGTTGGCATCATCCACATATTGTGATAGTAATTCTCGGATATCAGTAGTGACTACCGCCATTGATATTTGTTCATTTGATGAGTAACTACTAATGTATACATTTTCCATTTATAACACTTATGTATTAACTTTGTTTATTTATATTTTTTCATTATTATGTTGATATTAATAATCGTATTGTGGTTATATGGCTACAATTTCATAATGAGTTGAAGTCAGTGTCCTATGATCAATGACGATAGCTTTACTCTGAAAAGAAAGTATCAAATCGATAGTGCAGAGTCAACAATGAAAATGGATAAGACGATGACAAAGTTTCAGAATAGAGTCAAAATGGTAAAAGAAATAAATCAGACGATAAGAGCAGCACAAACTCATTACGAGACATTGAAACTAGGATATATAAAATTTAAGGGAATGATTAGGACTACTACTCTAGAAGATATAGCACCATCTATTCCAAATAATCAGAAAACTTATAAACTATTCTCGGACATTTCAGTCATTGGCAAAGCATCACAGAATCCGAGTAAGATGATATATGCTCGCTGCTTTACATGTTTCCCAATTTGTTTGGAGATGACCATAGATTCATTTGTTATAGAATGCATCCAACATTGTTCATGATATAGTTGAATCATGTATGCCTGTTCGTATGCCTGTGGCTAAGATACTGTGTAAAGAAATGGTAAATAAATACTTTGAGAATCTTTAAGAGTGCATTGACTTTGTTAGTGAATAGGCATTCCATCTTTCTCCAATACTAATTCAAATTGTTAAATTAATAATGGAATAGTATAAATAGTTATTAGTGATAGGATAGTAAACATAATTATTAGAATAGTAGTGTAGTATCATAGATAACTCTCTTCTATAAAAAATGGATTTTATTCGTAGAAAGTATCTTATATACACAGTAGAAAATAATATAGATTTTTTAAAGGATGATACATTAAGTAAAGTAAACAATTTTACCCTCAATCATGTACTAGCTCTCAAGTATCTAGTTAGCAATTTTCCTCAACATGTTATTACTAAGGATGTATTAGCTAATACCAATTTTTTTGTTTTCATACATATGGTACGATGCTGTAAAGTATACGAAGCGGTTTTACGACACGCATTTGATGCACCCACGTTGTACGTTAAAGCATTGACTAAGAATTATTTATCGTTTAGTAACACAATACAGTCGTACAAGGAAACAGTGCATAAACTAACACAAGATGAAAAATTTTTAGAGGTTGCCGAATACATGGACGAATTAGGAGAACTTATAGGCGTAAATTATGACTTAGTTCTTAATCCATTATTTCACGGAGGGGAACCCATCAAAGATATGGAAATCATTTTTTTAAAACTGTTTAAGAAAACAGACTTCAAAGTTGTTAAAAAATTAAGTGTTATAAGATTACTTATTTGGGCATACCTAAGCAAGAAAGATACAGGCATAGAGTTTGCGGATAATGATAGACAAGATATATATACTCTATTTCAACAAACTGGTAGAATAGTCCATAGCAATCTAACAGAAACGTTTAGGGATTATATCTTTCCCGGAGATAAGACTAGCTATTGGGTGTGGTTAAACGAAAGTATAGCTAATGATGCGGATATCGTTATTAATAGACCCGCCATTACCATGTATGATAAAATTCTTAGTTATATATACTCTGAGATAAAACAGGGACGCGTTAATAAAAACATGCTTAAGTTAGTTTATATCTTTGAGCCTGAAAAAGATATCAGAGAACTTCTGCTAGAAATCATATATGATATTCCTGGAGATATCCTATCTATTATTGATGCAAAAAACGACGATTGGAAAAAATATTTTATTAGTTTTTACAAAGCTAATTTTATTAACGGTAATACATTTATTAGTGATAGAACGTTTAACGATGACTTATTCAGAGTTGTTGTTCAAATAGATCCCGAATATTTCGATAATGAACGAATTATGTCTTTATTCTATACGAGTGCTGCGGACATTAAACGATTTGATGAGTTAGATATTAATAACAGTTATATATCTAATATAATTTATGAGGTGAACGATATCACATTAGATACAATGGATGATATGAAGAAGTGTCAAATCTTTAACGAGGATACGTTGTATTATGTTAAGGAATACAATACATACCTGTTTTTGCACGAGTCGGATCCCATGGTCATAGAGAACGGAATACTAAAGAAACTGTCATCTATAAAATCCAAGAGTAGACGGCTGAACTTGTTTAGCAAAAACATTTTAAAATATTATTTAGACGGACAATTGGCTCGTCTAGGTCTTGTGTTAGATGATTATAAAGGAGACTTATTAGTTAAAATGATAAACCATCTCAAATCTGTGGAGGATGTATCCGCATTCGTTAGATTTTCTACAGATAAAAACCCTAGTATTCTTCCATCGCTAATCAAAACTATTTTAGCTAGTTATAATATTTCCATCATCGTCTTATTTCAAAGGTTTTTAAGAGATAATCTATATCATGTAGAAGAATTCTTGGATAAAAGCATCCATCTAACCAAGACGGATAAGAAATATATACTTCAATTGATAAGGCACGGTAGATCATAGAACAAACCAAATATATTATTAATAATTTGTATATACATAGATATAATTATCACATATTAAAAAATAACACATTTTTGATAAATGGAAACCGTTGCAACAATTCAGACTCCCACCAAATTAATGAATAAAGAAAATGCAGAAATGATTTTGGAAAAAATTGTTAATCATATAGCTATGTATATTAGTGACGAATCAATATATTCAGAAAATAATCCTGAATATATTGATTTTCGTAACAGATACGGAGACTATAGATCTCTCATTATAAAAAGTGATCACGAGTTTGTAAAGCTATGTAAAGATCATGCAGAGAAAAGTTCTCCAGAAACGCAACAAATGATTATCAAACACATATACGAACAATATCTTATTCCAGTATCTGAAGTACTATTAAAACCTATAATGTCCATGGGTGACATATTTACATATAACGGATGTAAAGACAATGAATGGATGCTAGAACAACTCTCTACCCTAAACTTTAACAATCTCTACACATGGAACTCATGTAGCATAGGCAATGTAACGCGTCTGTTTTATACATTTTTTAGTTATCTGATGAAAGATAAACTAAATATATAAGTATAATCCCATTCTAATACTTTAACCTGATGTATTATTACCTGCATCTTATTAGAATATTAACCTAACTAAAAGACATAAAAAGCGGTAGGATATAAATATTATGGCCGCAACCGTTCCGCGTTTTGACGATGTGTACAAAAATGCACAAAGAAGAATTCTAGATCAAGAAACATTTTTTAGTAGAGGTCTAAGTAGACCGTTAATGAAAAACACATATCTATTTGATAATTACGCGTATGGATGGATACCAGAAACTGCAATTTGGAGTAGTAGATACGCAAACCTAGATGCTAGTGACTATTATCCCATTTCGTTGGGATTACTTAAAAAGTTTGAATTTCTCATGTCTCTATATAAAGGTCCTATTCCCGTATATGAAGAAAAAGTAAATACTGAATTCATTGCTAATGGATCTTTCTCCGGTAGATACGTATCATATCTTAGAAAGTTTTCTGCCCTTCCAACAAACGAGTTTATTAGTTTTTTATTATTGACCTCCATCCCTATCTATAATATCTTATTCTGGTTTAAAAACACACAGTTTGATATTACTAAACACACATTATTCAGATACGTCTATACAGATAATACCAAACACCTTGCGTTGGCTAGGTATATACATCAAACAGGAGACTATAAGCCTTTGTTTAGTCGTCTCAAAGAGAATTATATATTTACTGGTCCCGTTCCAATAGGTATCAAAGATATAGATCACCCTAATCTTAGTAGAGCAAGAAGTCCATCCGATTATGAGACATTAGCTAATATTAGTACTATATTGTACTTTACCAAGTATGATCCAGTATTAATGTTTTTATTGTTTTACGTACCTGGGTATTCAATTACTACAAAAATTACTCCAGCCGTAGAATATCTAATGGATAAACTGAATCTAACAAAGAGCGACGTACAACTGTTGTAAATTATTTTATGCTTCGTAAAATGTAGGTCTTGAACCAAACATTCTTTGAAAAAATGAGATGCATAAAACTTTATTATCCAATAGATTAACTATTTCAGACGTCAATCGTTTAAAGTAAACTTCGTAAAATATTCTTTGATTGCTGCCGAGTTTAAAACTTCTATCGATAATTGTTTCATATGTTTTAATATTTACAAGTTTTTTGGTCCATGGTACATTAGCTGGACAGATATATGCAAAATAATATCGTTCTCCAAGTTCTATAGTCTCTGGATTGTTTTTATTATATTCAGTAACCAAATACATATTAGGGTTATCTGCGGATTTATAATTTGAGTGATGCATTCGACTCAACATAAATAATTCTAGAGGAGACGATCTACTATCAAATTCGGATCGTAAATCTGTTTCTAAAGAACGGAGAATATCTATACATACCTGATTAGAATTCATCCGTCCTTCAGACAACATCTCAGACAGTCTGGTCTTGTATGTCTTAATCATATTCTTATGAAACTTGGAAACATCTCTTCTAGTTTCACTAGTACCTTTATTAATTCTCTCAGGTACAGATTTTGAATTCGACGATGCCGAGTATTTCATCGTTGTATATTTCTTCTTCGATTGCATAATCAAATTCTTATATACCGCCTCAAACTCTATTTTAAAATTATTAAACAATACTCTACTATTAATCAGTCGTTCTAACTCCTTTGCTATTTCTATGGACTTATCTACATCTTGACTGTCTATCTCTGTAAACACGGAGTCGGTATCTCCATACACGCTACGAAAACGAAATCTATAATCTATAGGCAACGATGTTTTCACAATCGGATTAATATCTCTATCGTCCATATAAAATGGATTACTTAATGTATTGGCAAACCGTAACATACCGTTGGATAACTCTGCTCCATTTAGTACCGATTCTAGATACAATATCATTCTACGTCCTATGGATGTGCAACTCTTAGCCGAAGCGTATGAGTATAGAGCACTATTTCTAAATCCCATCAGACCATATACTGAGTTGGCTACTATCTTGTACGTATATTGCATGGAATCATAGATGGCCTTTTCAGTTGAACTGGTAGCCTGTTTTAACATCTTTTTATATCTGGCTCTCTCTGCCAAAAATGTTCTTAATAGTCTAGGAATGGTTCCTTCTATTGATCTATCGAAAATTGCTATTTCAGAGATGAGGTTCGGTAGTCTAGGTTCACAATGAACCGTAATATATCTAGGAGGTGGATATTTCTGAAGCAAGAGTTGATTATTTATTTCTTCTTCCAATCTATTGGTACTAACAACGACACCGACTAATGTTTCCGGAGATAGATTTCCAAAGATACACACATTAGGATACAGACTGTTATAATCAAAGATTAATACATTATTACTAAACATTTTTTGTTTTGGAGCAAATACCTTACCGCCTTCATAAGGAAACTTTTGTTTTGTTTCTGATCTGACTAAGATAGTTTTAGTTTCCAACAATAGCTTTAACAGTGGACCCTTGATGATTGTACTCGCTCTATATTCGAATACCATGGATTGAGGAAGCACATATGTTGCCGCACCAGCGTCTGTTTTTGTTTCTACTCCATAATACTCCCACAAATACTGACACAAACAAGCATCATGAATACAGTATCTAGCCATATCTAAAGCTATGTTTAGATTATAATCCTTATACATCTGAGCTAAATCAATGTCATCCTTTCCGAAAGATAATTTATATATATCATTAGGTAAAGTAGGACATGATAGTACGACTTTAAATCCATTTTCCAAAATATCTTTACGAATTACTTTACATATAATATCCTCATCAACAGTCACGTAATTACCTGTGGTTAAAACCTTTGCAAATGTATCGGCTTTGCCTTTCGCGTCCGTAGTATCGTCACCGATGAACGTCATTTCTCTAACTCCTCTATTTAATACTTTACCCATGCAACTGAACGCGTTCTTGGATATAGAATCCAATTTGTACGAATCCAATTTTTCAGATTTTTGAATGAATGAATATAGATCGAAAAATATAGTTCCATTATTGTTATTAACGTGAAACGTAGTATTGGCCATGCCGCATACTCCCTTATGACTAGACTGATTTCTCTCATAAATACAGAGATGTACAGCTTCCTTTTTGTCTGGAGATCTAAAGATAATCTTCTCTCCTGTTAATAACTCTAGACGATTAGTAATATATCTCAGATCAAAGTTATGTCCGTTAAAGGTAACGACGTAGTCGAACGTTAGTTCCAACAATTGTTTAGCTATTCGTAACAAAACTATTTCAGAACATAGAACTAGTTCTCGTTCGTAATCCATTTCCATTAGCGACTGTATCCTCAAACATCCTCTATCGACGGCTTCTTGTATTTCCTGTTCCGTTAACATCTCTTCATTAATGAGCGTAAACAGTAATCGTTTACCACTTAAATCGATATAACAGTAACTTGTATGCGAGATTGGGTTAATAAATACAGAAGGAAACTTCTTATCGAAGTGACACTCTATATCTAGAAATAAGTACGATCTTGGGATATCGAATCTAGGTATTTCTTTAGCGAAACAGTTACGTGGATCGTCACAATGATAACATCCATTGTTAATCTTTGTCAAATATTGCTCGTCCAACGAGTAACATCCGTCTGGAGATATCCCGTTAGAAATATAAAACCAACTAATATTGAGAAATTCATCCATGGTGGCATTTTGTATGCTGCGTTTCTTTGGCTCTTCTATCAACCACATATCTGCGACGGAGCATTTTCTATCTTTAATATCTAGATTATAACTTATTGTCTCGTCAATGTCTATAGTTCTCATCTTTCCCATCGGCCTCGCATTAAATGGAGGAGGAGATAATGACTGATATATTTCGTCCGTCACTACGTAATAAAAGTAATGAGGAAATCGTATAAATACGGTCTCGCCATTTCGACATCTGGATTTCAGATATAAAAATCTGTTTTCACCGTGACTTTCAAACCAATTAATACACCTAACATCCATTTCTAGAATTTAGAAATATATTTTCATTTAAATGAATCCCAAACATTGGGGAAGAGCCGTATGGACCATTATTTTTATAGTACTTTCGCAAGCGGGTTTAGACGGCAACATAGAAGCGTGTAAACGAAAACTATATACTATAGTCAGCACTCTTCCATGTCCTGCATGTAGACGACACGCGACTATCGCTATAGAGGACAATAATGTCATGTCTAGCGATGATCTGAATTATATTTATTATTTTTTCATCAGATTATTTAACAATTTGGCATTTGATCCCAAATACGCAATCGATGTGTCAAAGGTTAAACCTTTATAAACTTAACCCATTATAAAACTTATGATTAGTCACGACTGAAATAACCGCGTGATTATTTTTTGGTATAATTCTACACGGCATGGTTTCTGTGACTATGAATTCAACACCTGTTATCTTAGTGAAATCTTTAACAAACAGCAAGGGTTCGTCAAAGACATAAAACTCATTGTTTACGATCGAAATAGACCCCCTATCACACTTAAAATAAAAAATATCCTTATCCTTTACCACCAAATAAAATTCTGATTGGTCAATGTGAATGTATTCACTTAACAGTTCCACAAATTTATTTATTAACTCCGAGGCACATACATCGTCGGTATTTTTTATGACAAACTTTACTCTTCCAGCATCCGTTTCTAAAAAAATATTAACGAGTTCCATTTATATCATCCAATATTATTGAAATGACGTTGATGGACAGATGATATAAATAAGAAGGTACAGTACCTTTGTCCACCATCTCCTCCAATTCATACTCTATTTTGTCATTAACTTTAATGTGTGAAAACAGTACGCCACATGCTTCCATGACAGTGTGTAACACTTTGGATACAAAATGTTTGACATTAGTATAATTGTCCAAGACTGTCAATCTATAATAGATAGTAGCTATAATATATTCTATGATGGTATTGAAGAAGATGACAACCTTGGCATATTGATCATTTAACACAGACATGGTATCAACAAATAGCTTAAATGAAAGAGAATCAGTAATTGGAATAAGCGTCTTCTCGATGTAGTGTCCGTATACCAACATGTCTGATATTTTGATGTATTCCATTAAATTATTTAGTTTTTTCTTTTTATTCTCGTTAAACAGAATTTCTGTCAATGGACCCCAACATCGTTGACCTATTAAGTTTTGATTGATTTTTCCGTGTAAGGCGTATCTAGTCAGATCGTATAGCCTATCCAATAATCCATCGTCTGTGCGTAGATCACATCGTACACTTTTTAATTTTCTATAGAAGAGTGACAGACATCTGGAGCAATTACAGACAGCAATTTCTTTATTCTCTACAGATGTAAGATACTTGAAGATATTCCTATGATGATGCAGAATTTTGGATAACACGGTATTGATGGTATCTGTTACCATAATTCCTTTGACTGATAGTGTCAAAGTACAAGATTTCCAATCTTTTGCAATTTTCAGTACCATTATCTTTGTTTTGATATCTATATCAGACAGCATGGTACGTCTGACAACACAGGGATTAAGACGGAAAGATGAAATGATTCTCTCAACATCTTCAATAGATACCTTGCTATTTTTTTTGGCATTATCTATATGTGAGAGAATATCCTCTAGAGAATCAGTATCCTTTTTGATGATAGTGGATCTCAATGACATGGGACGTCTAAACCTTCTTATTCTATCACCAGATTGCATGGTGATTTGTCTTCTTTCTTTTATCATGATGTAATCTCTAAATTCATCGGCAAATTGTCTATATCTAAAATCATAATATGAGATGTTTACCTCTACAAATATCTGTTCGTCCAATGTTAGAGTATCTATATCAGTTTTGTATTCCAAATTAAACATGGCAACGGATTTAATTTTATATTCCTCTATTAAGTCCTCGTCGATAATAACAGAATGTAGATAATCATTTAATCCATCGTACATGGTTGGAAGATGCTCGTTGACAAAATCTTTAATTGTCTTGATGAAGGTGGGACTATATCTAACATCTTGATTAATAAAATTTATAACATTGTCCATAGGATACTTTGTAACTAGTTTTATACACATCTCTTCATTGGTAAGTTTAGACAGAATATCGTGAACAGGTGGTATATTATATTCATCAGATATACGAAGAATAATGTCCAAATCTATATTGTTTAATATATTATATAGATGTAGTGTAGCTCCTACAGGAATATCTTTAACTAAGTCAATGATTTCATCAACAGTTAGATCTATTTTAAAGTTAATCATATAGGCATTGATTTTTAAAAGGTATGTAGCCTTGACTACATTCTCATTAATTAACCATTCCAAGTCACTGTGTGTAAGAAGATTATATTCTATCATAAGCTTGACTACATTTGGTCCCGATACCATTAAAGAATTCTTATGATATAAGGAAACAGCTTTTAGGTACTCATCTACTCTACAAGAATTTTGGAGAGCCTTAACGATATCAGTGACGTTTATTATTTCAGGAGGAAAGAACCTAACATTGAGAATATCTGAATTAATAGCTTCCAGATACAGTGATTTTGGCAATAGTCCGTGTAATCCATAATCCAGTAACACGAGCTGGTGCTTGCTAGACACCTTTTCAATGTTTAATTTTTTTGAAATAAGCTTTGATAAAGCCTTCCTCGCAAATTCCGGATACATGAACATGTCGCCAACATGATTAAGTATTGTTTTTCATTATTTTTATATTTTCTCAACAAGTTCTCAATACCCCAATAGATAATAGAATATCACCCAATGCGTCCATGTTGTCTATTTCCAACAGGTCGCTATATCCACCAATAGAAGTTTTCCCAAAAAAGATTCTAGGAACAGTTCTACCACCAGTAATTTGTTCAAAATAGTCACGCAATTCATTTTCGGGTTTAAATTCTTTAATATCTACAATTTCATACGCTCCTCTTTTGAAACTAAACTTATTTAGAATATCCAGTGCGTTTCTACAAAAAGGACATGTAAACTTGACAAAAATTGTCACTTTGTTATTGGCCAACCTTTGTTGTACAAATTCCTCGGCCATTTTTAATATTTAAGTGATACAAAACTATCTCGACTTATTTAACTCTTTAGTCGAGATATATGGACACAGATAGCTATATGATAACCAACTACAGAAGACAAACGCTATAAAAAACATAATTACGACGAGCATATTTATAAATATTTTTATTCAGTATTACTTGATATAGTAATATTAGGCACAGTCAAACATTCAACCACTCTAGATACATTAACTCTCTCATTTTCTTTAACAAATTCTGCAATATCTTCGTAAAAAGATTCTTGAAACTTTTTAGAATATCTATCGACTCTAGATGAAATAGCGTTCGTCAACATACTATGTTTTGTATACATAAAGGCGCCCATTTTAACAGTTTCTAGTGACAAAATGCTAGCGATCCTAGGATCCTTTAGAATCACATAGATTGACGATTCGTCTCTCTTAGTAACTCTAGTAAAATAATCATACAATCTAGTACGCGAAATAATATTATCCTTGACTTGAGGAGATCTAAACAATCTAGTTTTGAGAACATCGATAAGTTCATCGGGAATTACATACATACTATCTTTAATAGAACTCTTTTCATCCAGTTGAATGGATTCGTCCTTAACCAACTGATTAATGAGATCTTCTATTTTATCATTTTCTAGATGATATGTATGTCCATTAAAGTTAAATTGTGTAGCGCTTCTTTTTAGCCTAGCAGCCAATACTTTAACATCACTAATATCGATATACAAAGGAGATGATTTATCGATGGTATTAAGAATTCGTTTTTCGACATCCGTCAAAACCAATTCCTTTTTGCCTGTATCATCCAGTTTGCCATTCTTTGTAAAGAAATTATTTTCTACTAGACTATTAATAAGACTGATAAGGATTCCTCCATAATTGCACAATCCAAACTTTTTCACAAAACTAGACTTTACGAGATCTACAGGAATGCGTACTTCAGGTTTCTTAGCTTGTGATTTTTTCTTTTGTGGACATTTTCTAGTGACCAACTCATCTACCATTTCATTGATTTTAGCAGTGAAATAAGCTTTCAATGCACGGGCACTGATACTATTGAAAACGAGTTGATCTTCAAATTCCGCCATTTAAGTTCACCAAACAACTTTTAAATACAAATATATCAATAGTAGTAGAATAAGAACTATAAAAAAAATAATAATTAACCAATACCAACCCCAACAACCTGTATTATTAGTTGATGTGACAGTTTTCTCATCACTTAGAACAGATTTAACAATTTCTATAAAGTCTGTCAAATCATCTTCCTGAGAACCCATAAATACACCAAATATAGCAGCGTACAACTTATCCATTTATACATTGAATATTGGCTTTTCTTTATCGCTATCTTCATCATATTCATCATCAATATCAACAAGTCCCAGATTACGAACCAGATCTTCTTCTACATTTTCAGTCATTGATACGCGTTCACTATCTCCAGAGAGTCCGATAACGTTAGCCACTACTTCTCTATCAATGATTAGTTTCTTGAGCGCGAATGTAATTTTTGTTTCCGTTCCGGATCTATAGAAAACTACAGGTGTAATAATTGCCTTGGCTAATTGTCTTTCTCTTTTACTGAGTGATTCTAGTTCACCTTCTATAGATCTGAGAATGGATGATTCTCCAGTCGAAACATATTCTACCATGGCTCCGTTTAATTTGTTGATGAAGATGGATTCATCCTTAAATGTTTTCTCTGTAATAGTTTCCGCCGAAAGACTATGCAAAGAATTTGGAATGCGTTCCTTGTGTGTAATGTTTCCATAGACAGCTTCTAGAAGTTGATACAACATAGGACTAGCCGCGGTAACTTTTATTTTTAGAAAGTATCCATCGCTTCTATCTTGTTTAGATTTATTTTTATAAAGTTTAGTCTCTCCTTCCAACATAATAAAAGTGGAAGTCATCTGACTAGATAAACTATCAGTAAGTTTTATAGAGATAGATGAACAATTAGCGTATTGAGAAGCATTTAGTGTAACGCATTCGATACATTTTGCATTAGATTTACTAATCGATTTTGCATACTCTATAACACCCGCACAAGTCTGTAGAGAATCGCTAGATGCTGTAGGTCTTGGTGAAGTTTCAACTCTCTTCTTGATTACCTTACTCATGATTAAACCTAAATAATTGTACTTTGTAATATAATGATATATATTTTCACTTTATCTCATTTGAGAATAAAAATGTTTTTGTTAACCACTGCATGATGTACAGATTTCGGAATCGCAAACCACTTGTGGTTTTATTTTATCCTTGTCCAATGTGAATTGAATGGGAGCGGATGCGGGTTTCGTACGTAGATAGTACATTCCCGTTTTTAGACCGAGACTCCATCCGTAAAAATGCATACTCGTTAGTTTGGAATAACTCGGATCTGCTATATGGATATTCATAGATTGACTTTGATCGATGAAGGCTCCCCTGTCTGCAGCCATTTTTATGATCGTCTTTTGTGGAATTTCCCAAATAGTTTTATAAACTCGCTTAATATCTTCTGGAAGGTTTGTATTCTGAATGGATCCACCATCTACCATAATCCTATTCTTGATCTCATCATTCCATAATTTTCTCTCGGTTAAAACTCTAAGGAGATGCGGGTTAACTACTTGGAATTCTCCAGACAATACTCTCCGAGTGTAAATATTACTGGTATACGGTTCCACCGACTCATTATTTCCCAAAATTTGAGCAGTTGATGCAGTCGGCATAGGTGCCACCAATAAACTATTTCTAAGACCGTATGTTCTGATTTTATCTTTTAGAGGTTCCCAATTCCAAAGATCCGACGGTACAACATTCCAAAGATCATATTGTAGAATACCGTTACTGGCGTACGATCCTACATATGTATCATATGGTCCTTCCTTCTCAGCTAGTTTACAACTCGCCTCTAATGCACCGTAATAAATGGTTTCAAAGATCTTCTTATTTAGATCTTGTGCTTCCAGGCTATCAAATGGATAATTTAAGAGAATAAACGCGTCCGCTAATCCTTGAACACCAATACCGATAGGTCTATGTCTCTTATTAGAGATTTCAGCTTCTGGAATAGGATAATAATTAATATCTATAATTTTATTGAGATTTCTGACAATTACTTTGACCACATCCTTCAGTTTGAGAAAATCAAATCGCCCATCTATTACAAACATGTTCAATGCAACAGATGCCAGATTACACACGGCTACCTCATTAGCATCCGCATATTGTATTATCTCAGTGCAAAGATTACTACACTTGATGGTTCCTAAATTTTGTTGATTACTCTTTTTGTTACACGCATCCTTATAAAGAATGAATGGAGTACCAGTTTCAATCTGAGATTCTATAATCGCTTTCCAGACGACTCGAGCCTTTATTATACATTTGTATCTCCTTTCTCTTTCGTATAGTGTATACAATCGTTCGAACTCGTCTCCCCAAACATTGTCCAATCCAGGACATTCATCCGGACACATCAACGACCACTCTCCGTCATCCTTCACTCGTTTCATAAAGAGATCAGGAATCCAAAGAGCTATAAATAGATCTCTTGTTCTATGTTCATCGTTTCCTGTATTCTTTTTAAGATCGAGGAACGCCATAATATCAGAATGCCACGGTTCCAAGTATATGGCCATAACTCCAGGCCGTTTGTTTCCTCCCTGATCTATGTATCTAGCGGTGTTATTATAAACTCTCAACATTGGAATAATACCGTTTGATATACCATTGGTACCGGAGATATAGCTTCCACTGGCACGAATATTACTAATTGATAGACCTATTCCCCCTGCCATTTTAGAGATTAATGCGCATCGTTTTAACGTGTCATAGATGCCTTCTATGCTATCATCGATCATGTTAAGTAGAAAACAGCTAGACATTTGGTGACGAGTAGTTCCCGCATTAAATAAGGTAGGAGAAGCGTGCGTAAACCATTTTTCAGAAAGTAGATTGTACGTCTCAATAGCTGAGTCTATATCCCATTGATGAATTCCTACTGCGACACGCATTAACATGTGCTGAGGTCTTTCAACAATTTTGTTGTTTATTTTCAACAAGTAGGATTTTTCCAAAGTTTTAAAACCAAAATAGTTGTATGAAAAGTCTCGTTCGTAAATAATAACCGAATTGAGCTTATCCTTATATTTGTTAACTATATCCATGGTAATACTTGAAATAATCGGAGAATGTTTCCCATTTTTAGGATTAACATAGTTGAATAAATCCTCCATCACTTCACTAAATAGTTTTTTTGTTTCCTTGTGTAGATTTGATATGGCTATTCTGGCGGCTAGAATGGCATAATCCGGATGTTGTGTAGTACAAGTGGCTGCTATTTCGGCTGCCAGAGTGTCCAATTCTACCGTTGTTACTCCATTATATATTCCTTGAATAACCTTCATAGCTATTTTAATAGGATCTATATGATCAGTGTTTAAGCCATAGCACAATTTTCTAATACGAGACGTGATTTTATCAAACATGACATTTTCCTTGTATCCATTTCGTTTAATGACAAACATTTTTGTTGGTGTAATAAAAAAAATTATTTAATTTTTCATTAATAGGGATTTGACGTATGTAGCGTACAAAATTATCGTTCCTGGTATATAGATAAAGAGTCCTATATATTTGAAAATCGTTACGGTTCGATTAAACTTTAATGATTGCATTGTGAATATATCATTAGGATTTAACTCCTTGACTATCATGGCGGTGCCAGAAATTACCATCAAAAGCATTAATACAGTTATGCAGATCGCAGTTAGAACGGTTATAGCATCCACCATTTATATCTAAAAATTAGATCAAAGAATATGTGACAACGTCCTAGTTGTATACTGAGAATTGACGAAACAATGTTTCTTACATATTTTTTTCTTATTAGTAACCGACTTAATAGTAGGAACTGGAAAACTAGACTTGATTATTCTATAAGTATAGATACCCTTCCAAATAATGTTCTCTTTGATAAAAGTTCCAGAAAATGTAGAATTTTTTAAAAAGTTATCTTTTGCTATTACTAATATCGTGGTTAGACGCTTATTATTAATATGAGTGATGAAATCCACACCGCTTCTAGATATCGCTTTTATTTCCACATTAGATGGTAAATCCAATAGTGAAACTATCTTTTTAGGAATGTATGGACTCGCGTTTAGAGGAGTGAACGTCTTCGGAGTAGTAAAGGATGATTCGTCAAATGAATAAACAATTTCACAAATGGATGTTAATGTATTAGTAGGAAATTTTTTGACGCTAGTGGAATTGAAGATTCTAATGGATGATGTTCTACCTATTTCATCCGATAACATGTTAATTTCCAATACCAACGGTTTTAATATTTCGATGATATACGGTAGTCTCTCTTTCGGACTTATATAGCTTATTCCACAATACGAGTCATTATATACTCCAAAAAACAAAATAACTAGTATAAAATCTGTATCGAATGGGAAAAACGAAATTATCGATATAGGTATAGAATCCGGAACATTGAACGTATTAATACTTAATTCTTTTTCAGTGGTAAGAACCGATAGGTTATTGACATTGTATGGTTTTAAATATTCTATAACTTGAGACTTGATAGATATTAATGACGAATTGAAAATTATTTTTATCACCACGTGTGTTTCAGGATCATCGTCGACGCCAGTTAACCAACCGAATGGAGTAAAATAAATATCATTAATATATGCTCTAGATATTAGTATTTTTATTAATCCTTTGATTATCATCTTCTCGTACGCGAATGATTCCATGATCAAGAGTGATTTGAGAACATCCTCCGGAGTATTAATGGGTTTAGTAAACAGTCCATCGTTGCAATAATAAAAGTTGTCCAAGTTAAAGGATATTATGCATTCGTTTAAAGATATCACCTCATCTAACGGAGACAATTTTTTGGTAGGTTTTAGAGACTTTGAAGCTACTTGTTTAACAAAGTTATTCATCGTCGTCTACTATTCTATTTAATTTTGTAGTTAATTTATCACATATCACATTAATTGACTTTTTGGTCCACTTTTCCATACGTTTATATTCTTTTAATCCTGCGTTATCCGTTTCCGTTATATACAGGGATAGATCTTGCAAGTTAAATAGAATGCTCTTAAATAATGTCATTTTTTTATCCGCTAAAAATTTAAAGAATGTATAAACTTTTTTCAAAGATTTAAAACTTTTAGGTGGAGTTCTGGTACACAATATCATAAACAAACTAATAAACATCCCACATTCAGATTCCAACAATTGATTAACTTCCACATTAATACAGCCTATTTTCGCTCCAAATGTACATTCGAAAAATCTGAATAAAACATCAATATCGCAATTTGTATTATCCAATACAGAATGTCTGTGATTCGTGTTAAAACCATCGGAAAAAGAATAGAAATAAAAATTATTATAATGGTGGAATTCAGTTGGAATATTGCCTCCGGAGTCATAAAAGGATACTAAACATTGTTTTTTATCGTAAATTACACATTTCCAATGAGACAAATAACAAAATCCAAACATTACAAATCTAGAGGTAGAACTTTTAATTTTGTCTTTAAGTATATACGATAAGATATGTTTATTCATAAACGCGTCAAATTTTTCATGAATAGCTAAGGAGTTTAAGAATCTCATGTCAAATTGTCCTATATAATCCACTTCGGATCCATAAGCAAACTGAGAGACTAAGTTCTTAATACTTCGATTGCTCATCCAGGCTCCTCTCTCAGGCTCTATTTTCATCTTGACGACCTTTGGATTTTCACCAGTATGTATTCCTTTACGTGATAAATCATCGATTTTCAAATCCATTTGTGAGAAGTCTATCGCCTTAGATACTTTTTCCCGTAGTTGAGGTTTAAAGAAATACGCTAACGGTATACTAGTAGGTAACTCAAAGACATCATATATAGAATGGTAACGCGTCGTTAACTCGTCGGTTAACTCTTTCTTTTGATCGAGTTCATCGCTACTATTGGGTCTGCTCAGGTGCCCCGACTCTACTAGTTCCAACATCATACCGATAGGAATACAAGACACTTTGCCAGCGGTTGTAGATTTATCATATTTCTCCACCACATATCCGTTACAATTTGTTAAGAATTTAGATACATCTATATTGCTACATAATCCAGCTAGTGAATATATATGACATAATAAATTGGTAAATCCTAGTTCTGGTATTTTACTAATTACTAAATCTGTATATCTTTCCATTTATCATGGAAAAGAATTTACCAGATATCTTCTTTTTTCCAAACTGCGTTAATGTATTCTCTTACAAATATTCACAAGATGAATTCAGTAATATGAGTAAAACGGAACGTGATAATTTCTCATTGGCTGTGTTTCCAGTGATAAAACATAGATGGCATAACGCACACGTTGTAAAACATAAAGGAATATACAAAGTTAGTACAGAAGCACGTGGAAAAAAAGTATCTCCTCCATCACTAGGAAAACCCGCACATATAAACCTAATGTCGAAGCAATATATATATAGTGAGTATGCAATAAGCTTTGAATGTTATAGTTTTCTAAAATGTATAACAAATACAGAAATCAATTCGTTCGATGAGTATATATTAAGAGGACTATTAGAAGCTGGTAATAGTTTACAGATATTTTCCAATTCCGTAGGTAAACGAATAGATACTATAGGTGTACTAGGGAATAAGTATCCATTTAGCAAAATTCCATTGGCCTCATTAACTCCTAAAGCACAACGAGAGATATTTTTAGCGTGGATTTCTCATAGACCTGTAGTTTTAACTGGAGGAACCGGAGTGGGTAAGACGTCACAGGTACCCAAGTTATTGCTTTGGTTTAATTATTTATTTGGTGGATTCTCTTCTCTAGATAAAATCACTGACTTTCACGAAAGACCAGTCATTCTATCTCTTCCTAGGATAGCTTTAGTTAGATTGCATAGCAATACCATTTTAAAATCATTGGGATTTAAGGTACTAGATGGATCTCCTATCTCTTTACGGTACGGATCTATACCGGAAGAATTAATAAACAAACAACCAAAAAAATATGGAATTGTATTTTCTACCCATAAGTTATCTCTAACAAAACTATTTAGTTATGGCACTATTATTATAGACGAAGTTCATGAGCATGATCAAATAGGAGATATTATTATAGCAGTAGCGAGAAAACATCATACGAAAATAGATTCTATGTTTTTAATGACTGCCACGTTAGAGGATGACAGGGAACGTCTAAAAATATTTTTACCTAATCCCGCATTTATACATATTCCTGGAGATACACTGTTTAAAATTAGCGAGGTATTTATTCATAATAAGATAAATCCATCTTCCAGAATGGCATATATAGAAGAAGAAAAGAGAAATTTAGTTACTGCTATACAGATGTATACTCCTCCTGATGGATCATCCGGTATAGTCTTTGTGGCATCCGTTGCACAGTGTCACGAATATAAATCATATTTAGAAAAAAGATTACCGTATGATATGTATATTATTCATGGTAAGGTCTTAGATATAGACGAAATATTAGAAAAAGTGTATTCATCACCTAATGTATCGATAATTATTTCTACTCCTTATTTGGAATCCAGCGTTACTATACGCAATGTTACACACATTTATGATATGGGTAGAGTTTTTGTCCCCGCTCCTTTTGGAGGATCACAACAATTTATTTCTAAATCTATGAGAGATCAACGAAAAGGAAGAGTAGGAAGAGTTAATCCTGGAACATACGTATATTTCTATGATCTGTCTTATATGAAATCTATACAGCGAATAGATTCAGAATTTCTACATAATTATATATTGTACGCTAATAAGTTTAATCTAACACTCCCCGAAGATTTGTTTATAATCCCTACAAATTTGGATATTCTATGGCGTACAAAGGAATATATAGACTCGTTCGATATTAGTACAGAAACATGGAATAAATTATTATCCAATTATTATATGAAGATGATAGAGTATGCTAAACTTTATGTACTAAGTCCTATTCTCGCTGAGGAGTTGGATAACTTTGAGAGGACGGGAGAATTAACTAGTATTGTACAAGAAGCCATTTTATCTCTAAATTTACAAATTAAGATTTTAAAATTTAAACATAAAGATGATGATACGTATATACACTTTTGTAGAATATTATTCGGTGTCTATAACGGAACAAACGCTACTATATATTATCATAGACCTCTAACGGGATATATGAATATGATTTCAGATACTATATTTGTTCCTGTAGATAATAACTAAAAATCAAAATCTAATGACCACATCTTTTTTTAGAGATGAAAAATTTTCCACATCTCCTTTTGTAGACACGACTAAACATTTTGCAGAAAAAAGTTTATTATTATTTAGATAATCGTATACTTCATCAGTGTAGATAGTAAATGTGAACAGATAAAAGGTATTCTTGCTCAATAGATTGGTAAATTCCATAGAATATATTAATCCTTTCTTCTTGAGATCCCACATCATTTCAACCAAAGACGTTTTATCCAATGATTTACCTCGTACTATACCACATACAAAACTAGATTTTGCAGTGATGTCGTACCTGGTATTCCTACCAAACAAAATTTTACTTTTAGTTCTTTTAGAAAATTCTAAGGTAGAATCTCTATTTGTCAATATGTCATCTATGGAATTACCACTAGCAAAAAATGATAGAAATATATATTGATACATCGCAGCTGGTTTTGATCTACTATACTTTAAAAACGAATCAGATTCCATAATTGCTTGTATATCATCAGCTGAAAAACTATGTTTTACACGTATTCCTTCGGCATTTCTTTTTAATGATATATCTTGTTTAGACAATGATAAAGTTATCATGTCCATGAGAGACGCGTCTCCGTATCGTATAAATATTTCATTAGATGTTAGACGCTTCATTAGGGGTATACTTCTATAAGGTTTCTTAATTAGTCCATCATTGGTTGCGTCAAGAACTACTATCTGATGTTGTTGGGTATCTCTAGTGTTACACATGGCCTTACTAAAGTTTGGGTAAATAACTATGATATCTCTATTAATTATAGATGTATATATTTCATTCGTCAAGGATATTAATATCGACTTACTATCGTCATTAATACGTGTAATGTAATCATATAAATCATGCGATAGCCAAGGAAAATTCAAATAGATGTTCATCATATAATCGTCGCTATAATTCATATTAATACTTTGACATTGACTAATTTGTAATATAGCCTCGCCACGAAGAAAGCTCTCGTATTCAGTTTCATCGATAAAGGATACCGTTAAATATAACTGGTTGCCGATAGTCTCATAGTCTATTAAGTGGTAAGTTTCGTATAAATACAGAATCCCTAAAATATTATCTAATGTGGGATTAATCCTTACCATAACTGTATAAAATGGAGCCGGAGTCATAACTATTTTACCGTTTGTACTTACTGGAATAGATGAAGGAATAATCTCCGGACATGATGGTAAAGACCCAAATGTCTGTTTGAAGAAATCCAATGTTCCAGGTCCTAATCTCTTGACAAAAATTACGATATTCGATCCCGATATCCTTTGCATTCTATTTACCAGCATATCACGAACTATATTAAGATTATCTATCATGTCTATTCTCCCACCGTTATATAAATCGCCTCCGCTAAGAAACGTTAGTATATCCATACAATGGAATACTTCATTTCTAAAATAGTATTCGTTTTCTAATTCTTTAATGTGAAATCGTATACTAGAAAGGGAAAAATTATCTTTGAGTTTTCCATTAGAAAAGAACCACGAAACTAATGTTCTGATTGCGTCTGACTCCGTCGCTGAATTAATAGATTTACACCAAAAACTCATATAACTTCTAGATGTAGAAGCATTCGCTAAAAAATTAGTAGAATCAAAGGATATAAGTAGATGTTCCAACAAGTGAGCAATTCCCAAGATTTCATCTATATCATTCTCGAATCCGAAATTAGAAATTCCCAAGTAGATATCCTTTTTCATCCGATCATTGATGAAAATACGAACTTTATTCGGTAAGACGATCATTTACTAAGGAGTAAAATAGGAAGTAACGTTCGTATATCGTTATCGTCGTATAAATTAAAGGTGTGTTTTTTGCCATTAAGAGACATTATAATTTTACCAATATTGGAATTATAATATAGGTGTATTTGAGCACTAGAAACGGTCGATGCATCGGTAAATATAGCTGTATCTAATGTTCTAGTCGGTATTTCTTCATTTCGCTGTCTAATGATAGCGTTTTCTCTATCTGTTTCCATTACAGCTGCCTGAAGTTTATTGGTCGGATAATATGTAAAATAATAAGAAATACATACGAATAACAAAAATAAAATAAGATATAATAAAGATGCCATTTAGAGATCTAATTTTGTTCAACTTGTCCAAATTCCTACTTACAGAAGATGAGGAATCGTTGGAGATAGTATCTTCCTTATGTAGAGGATTTGAAATATCTTACGATGACTTAATATCGTACTTTCCAGATAGGAAATACCATAAATATATTTCTAAGGTATTTGAACATGTAGATTTATCGGAGGAATTAAGTATGGAATTCCATGATACAACTCTGAGAGATTTAGTATATCTTAGATTGTACAAGTATTCCAAGTATATACGGCCGTGTTATAAATTAGGAGATAATCTAAAAGGTATAGTTGTTATAAAGGACAGAAATATATATATTAGAGAAGCAAATGATGACTTGATAGAATATCTCCTCAAGGAATACACTCCTCAGATTTATACATATTCTAATGAGCGAGTTCCCATAGCTGGTTCAAAATTAATTCTTTGTGGATTTTCTCAAGTTACATTTATGGCGTATACAACGTCGCATATAACAACAAATAAAAAGGTAGATGTTCTCGTTTCCAAAAAATGTATAGATGAACTAGTCGATCCAATAAATTATCAAATACTTCAAAATTTATTTGATAAAGGAAGCGGAACAATAAACAAAATACTCAGGAAGATATTTTATTCGGTAACAGGTGGCCAAACTCCATAGGTAGCTTTTTCTATTTCGGATTTTAGAATTTCCAAATTCACCAGCGATTTATCGGTTTTGGTGAAATCCAAGGATTTATTAATGTCCACAAATGCCATTTGTTTTGTCTGTGGATTGTATTTGAAAATGGAAACGATGTAGTTAGATAGATGCGCGGCGAAGTTTCCTATTAGGGTTCCGCGCTTCACGTCACCCAACATACTTGAATCACCATCCTTTAAAAAAAATGATAAGATATCAACATGGAGTATATCATACTCGGATTTTAATTCTTCTACTGCCTCACTGACATTTTCACAAATACTACAATACGGTTTACCGAAAATAATCAGTACGTTCTTCATTTATGGGTATCAAAAACTTAAAATCGTTACTGCTGGAAAATAAATCACTGACGATATTAGATGATAATTTATACAAAGTATACAATGGAATATTTGTGGATACAATGAGTATTTATATAGCCGTCGCCAATTGTGTCAGAAACTTAGAAGAGTTAACTACGGTATTCATAAAATACGTAAACGGATGGGTAAAAAAGGGAGGACATGTAACCCTTTTTATCGATAGAGGAAGTATAAAAATTAAACAAGACGTTAGAGACAAGAGACGTAAATATTCTAAATTAACCAAGGACAGAAAAATGTTAGAATTAGAAAAGTGTACATCCGAAATACAAAATGTTACCGGATTTATGGAAGAAGAAATAAAGGCAGAAATGCAATTAAAAATCGATAAACTCACATTTCAAATATATTTATCTGATTATGATAACATAAAAATATCATTGAATGAGATACTAACACATTTCAACAATAATGAGAATGTTACATTATTTTATTGTGATGAACGAGACGCAGAATTCGTTATGTGTCTAGAGGCTAAAACACAGTTCTCTACCACAGGAGAATGGCCGTTAATAATAAGTACCGATCAGGATACTATGCTATTCGCGTCTGCTGATAATCATCCTAAGATGATAAAAAACTTAACTCAACTGTTTAAATTTGTTCCCTCGGCAGAGGATAACTATTTAGCAAAATTAACTGCATTAGTGAATGGATGTGATTTCTTTCCTGGACTCTATGGGGCATCTATAACACCCAACAACTTAAACAAAATACAATTGTTTAGTGATTTTACAATCGATAATATAGTCACTAGTTTGGCAATTAAAAATTATTATAGAAAGACTAACTCTACCGTAGACGTGCGTAATATTGTTACGTTTATAAACGATTACGCTAATTTAGACGATGTCTACTCGTATATTCCTCCTTGTCAATGCACTGTTCAAGAATTTATATTCTCCGCATTAGATGAAAAATGGAATGAATTTAAATCATCTTATTTAGAGAGCGTGCCGTTACCCTGCCAATTAATGTACGCATTAGAACCACGTAAGGAGATTGATGTTTCAGAAGTTAAAACTTTATCATCTTATATAGATTTCGAAAATACTAAATCAGATATCGATGTTATAAAATCTATATCCTCGATTTTTGGATATTCTAACGAAAACTGTAACACCATAGTGTTCGGCATCTATAAGGATAATTTACTACTGAGTATAAATAATTCATTTTACTTTAACGATAGTCTGTTAATAACCAATACTAAAAGTGATAATATAATAAATATAGGTTACTAGATTAAAAAATGGTGTTCCAGCTCGTGTGTTCTACATGCGGCAAAGATATTTCTCACGAACGATATAAATTGATTATACGAAAAAAATCATTAAAGGATGTACTAGTCAGTGTAAAGAACGAATGTTGTAGGTTAAAATTATCTACACAAATAGAACCTCAACGTAACTTAACAGTGCAACCTCTATTGGATATAAACTAATGGATCCGGTTAATTTTATCAAGACATATGCGCCTAGAGGTTCTATTATTTTTATTAATTATGCCATGTCATTAACTAGTCATTTGAATCCATCGATAGAAAAACATGTGGGTATTTATTATGGTACGTTATTATCGGAACACTTGGTAGTTGAATCTACCTATAGAAAAGGAGTTAGAATAGTCCCATTGGATAGATTTTTTGAAGGATATCTTAGTGCAAAAGTATACATGTTAGAGAATATTCAAGTTATGAAAATAGCAGCTGATATGTCGTTAACTTTACTAGGTATTCCATATGGATTTGGTCATGATAGAATGTATTGTTTTAAATTGGTAGCTGAATGTTATAAAAATGCCGGTATTGATACATCGTCTAAACGAATATTAGGTAAAGATATTTTTCTGAGCCAAAACTTTACAGATGATAATAGATGGATAAAGATATATGATTCTAATAATTTAACATTTTGGCAAATTGATTACCTTAAAGGGTGAGTTAATATGCATAACTACTCCTCCGTTGTTTTTTCCCTCGTTCTTTTTCTTAACGTTGTTTGCCATCACTCTCATAATGTAAAGATATTCTAAAATGGTAAACTTTTGCATATCGGATGCAGAAATTGGTATAAATGTTGTAATTGTATTATTTCCCGTCAATGGACTAGTCACAGCTCCATCAGTTTTATATCCTTTAGAGTATTTCTCACTCGTGTCTAGCATTCTAGAGCATTCCATGATCTGTTTATCGTTGATATTGGCCGGAAAGATAGATTTTTTATTTTTTATTATATTACTATTGGCAATTGTAGATATAACTTCTGGTAAATATTTTTCTACCTTTTCAATCTCTTCTATTTTCAAGCCGGCTATATATTCTGCTATATTGTTACTAGTATCAATACCTTTTCTGGCTAAGAAGTCATATGTGGTATTCACTATATCAGTTTTAACTGGTAGTTCCATTAGCCTTTCCACTTCTGCAGAATAATTAGAAATTGGTTCTTTACCAGAAAATCCAGCTACTATAATAGGCTCACCGATGATCATTGGCAAAATCCTATATTGTACCAGATTAATGAGAGCATATTTCATTTCCAATAATTCTGCTAGTTCTTGAGACATTGATTTATTTGATGAATCTATTTGGTTCTCTAGATACTCTACCATTTCTGCCGCATACAATAACTTGTTAGATAAAATCAGGGTTATCAAAGTGTTTAGTGTGGCTAGAATAGTGGGCTTGCACGTATTAAAGAATGCTGTAGTATGAGTAAACCGTTTTAACGAATTATATAGTCTCCAGAAATCTGTGGCGTTGCATACATGAACTGAATGACATCGAAGATTGTCCAATATTTTTAATAGCTGCTCTTTGTCCATTATTTCTATATTTGACTCGCAACAATTGTAGATACCATTAATCACTGATTCCTTTTTCGATGCCGGACAATAGCACAATTGTTTAGCTTTGGACTCTATGTATTCAGAATTAATAGATATATCTCTCAATACAGATTGCACTATACATTTTGAAACTATGTCAAAAATTGTAGAACGACGCTGTTCTGTAGCCATTTAACTTTAAATAATTTACAAAAATTTAAAATGAGCATCCGTATAAAAATCGATAAATTGCGCCAAATTGTGGCATATTTTTCAGAGTTCAGCGAAGAAGTGTCTATAAATGTAGACTCGACGGATGAATTAATGTATATTTTTGCCGCCTTGGGCGGATCTGTAAACATTTGGGCCATTATACCTCTCAGTGCATCAGTGTTCTACCGCGGAGCCGAAAATATTGTGTTTAACCTTCCAGTGTCCAAGGTAAAATCGTGTTTGTGTAGTTTTCACAATGATGCTATCATAGATATAGAACCTGATCTGGAAAATAATCTAGTAAAACTTTCTAGTTATCATGTAGTAAGTGTCGATTGTAACAAGGAACTGATGCCTATTAGGACAGATACTACTATTTGTCTAAGTATAGATCAAAAGAAATCTTACGTATTTAATTTTCACAAGTATGAAGAAAAATGTTGTGGTAGAACCGTCATTCATCTAGAATGGTTGTTGGGCTTTATCAAGTGTATTAGTCAGCATCAGCATTTGGCTATTATGTTTAAAGATGACAATATTATTATGAAGACTCCTGGTAATACTGATGCGTTTTCCAGGGAATATTCTATGACTGAATGTTCTCAAGAACTACAAAAGTTTTCTTTCAAAATAGCTATCTCGTCTCTCAACAAACTACGAGGATTCAAAAAGAGAGTCAATGTTTTTGAAACTAGAATCGTAATGGATAATGACGATAACATTCTAGGAATGTTGTTTTCGGATAGAGTTCAATCCTTTAAGATTAACATCTTTATGGCGTTTTTAGACTAATACTTTCAATGAGATAAATATGGGTGGCGGAGTAAGTGTTGAGCTCCCTAAACGGGATCCACCTCCGGGAGTACCCACTGATGAGATGTTATTAAACGTGGATAAAATGCATGACGTGATAGCTCCCGCTAAGCTTTTAGAATATGTGCATATAGGACCACTAACAAAAGATAAAGAGGATAAAGTAAAGAAAAGATATCCAGAGTTTAGATTAGTCAACACAGGACCCGGTGGTCTTTCGGCATTATTAAGACAATCATATAATGGAACCGCACCCAATTGCTGTCGCACTTTTAATCGTACTCATTATTGGAAGAAGGATGGAAAGATATCAGATAAGTATGAAGAGGGTGCAGTATTAGAATCGTGTTGGCCCGACGTCCACGACACTGGAAAATGCGATGTTGATTTATTCGACTGGTGTCAGGGGGATACGTTCGATATGAACATATGCCATCAGTGGATCGGTTCAGCCTTTAATAGGAGTGATAGAACTGTAGAGGGTCGACAATCGTTAATAAATCTGTATAATAAGATGCAAAGATTATGTAGTAAAGATGCTAGTGTACCAATATGTGAATTATTTTTGCATCATTTACGCGCACACAATACAGAAGATAGTAAAGAGATGATCGATTATATTCTAAGACAACAGTCGGCGGACTTTAAACAGAAATATATGAGATGTAGTTATCCCACTAGAGATAAGTTAGAAGAGTCATTAAAATATGCGGAACCTCGAGAATGTTGGGATCCAGAGTGTTCGAATGCCAATGTTAATTTCTTACTAACACGTAATTATAATAATTTAGGACTTTGCAATATTGTACGATGTAATACGAGCGTGAATAACTTACAGATGGATAAAACTTCCTCATTAAGATTATCATGTGGATTAAGCAATAGTGATAGATTTTCTACTGTTCCCGTCAATAGAGCAAAAGTAGTTCAACATAATATTAAACATTCGTTCGACCTAAAATTGCATTTGATCAGTTTATTATCTCTCTTGGTAATATGGATACTAATTGTAGCTATTTAAATGGGTGCCGCAGCAAGCATACAGACGACTGTGAATACACTCAGTGAACGTATCTCGTCTAAATTAGAACAAGAAGCGAACGCTAGTGCTCAAACAAAATGTGATATAGAAATCGGAAATTTTTATATCCGACAAAACCATGGATGTAACATCACTGTTAAAAATATGTGCTCTGCGGACGCGGATGCTCAGTTGGATGCTGTGTTATCAGCCGCTACAGAAACATATAGTGGATTAACACCGGAACAAAAAGCATACGTACCAGCTATGTTTACTGCTGCGTTAAACATTCAGACGAGTGTAAACACTGTTGTTAGAGATTTTGAAAATTATGTGAAACAGACTTGTAATTCTAGCGCTGTTGTCGATAACAAATTAAAGATACAAAACGTAATTATAGATGAATGTTACGGAGCCCCAGGATCTCCAACAAATTTGGAATTTATTAATACAGGATCTAGCAAAGGAAATTGTGCCATTAAGGCGTTGATGCAATTGACTACTAAGGCCACTACTCAAATAGCACCTAGACAAGTTGCTGGTACAGGAGTTCAGTTTTATATGATTGTTATCGGTGTTATAATATTGGCAGCGTTGTTTATGTACTATGCCAAGCGTATGCTGTTCACATCCACCAATGATAAAATCAAACTTATTTTAGCCAATAAGGAAAACGTCCATTGGACTACTTACATGGACACATTCTTTAGAACTTCTCCGATGATTATTGCTACCACGGATATACAAAACTGAAAATATATTGATAATATTTTAATAGATTAACATGGAAGTTATCGCTGATCGTCTAGACGATATAGTGAAACAAAATATAGCGGATGAAAAATTTGTAGATTTTGTTATACACGGTCTAGAGCATCAATGTCCTGCTATACTTCGACCATTAATTAGGTTGTTTATTGATATACTATTATTTGTTATAGTAATTTATATTTTTACGGTACGTCTAGTAAGTAGAAATTATCAAATATTGTTGGTGTTGGTGGCGCTAGTCATCACATTAACTATTTTTTTATTACTTTATACTATAATAGTACTAGACTGACTTCTAACAAACATCTCACCTGCCATAAATAAATGCTTGATATTAAAGTCTTCTATTTCTAACACTATTCCATCTGTGGAAAATAATACTCTGACATTATCGCTAATTGATACATCGGTAAGTGATATGCCTATAAAGTAATAATCTTCTTTGGGCACATATACCAGTGTACCAGGTTCTAACAACCTATTTACTGGTGCTCCTGTAGCATACTTTTTTTTTACCTTGAGAATATCCATTGTTTGCTTGGTCAATAGTGATATGTGATTTTTTATCAACCACTCAAAAAAGTAATTGGAGTGTTCATATCCTCTACGGGCTATTGTCTCATGACCGTGTATGAAATTTAAGTAACACGACTGTGGTAGATTTGTTCTATAGAGCCGGTTGCCGCAAATAGATAGAACTACCAATATGTCTGTACAAATGTTAAACATTAATTGATTAACAGAAAAAACAATGTTCGTTCTGGGAATAGAAACCAGATTAAAACAAAATTCATTAGAATATATGCCACGTTTATACATGGAATATAAAATAACTACAGTTTGAAAAATAACAGTATCATTTAAACATTTAACTTGCGGGGTTAATCTCACAACTTTACTGTTTTTGAACTGTTCAAAATATAGCATAGATCCATGAGAAATACGTTTAGCCGCCTTTAATAGAGGAAATCCAACCGCCTTTCTGGATCTCACCAACGACGATAGTTCTGACCAGCAACTCATTTCTTCATCATCCACCTGTTTTAACATATAATAGGCAGGAGATAGATATCCATCATTGCAATATTCCTTCTCGTAGGCACACAATCTAATATTGATAAAATCTCCATTCTCTTCTCTGTATTTATTATCTTGTCTCGGTGGCTGATTAGGCTGTGGTCTATCGTTGTTGAATCTATTTTGGTCATTAAATCTTTCATTTCTTCCTGGTATATTTCTATCACCTCGTTTGGTTGGATTTTTGTCTATATTATCGTTTGTAACATCGGTACGGGTATTCATTTATCACAAAAAAAACTTCTCTAAATGAGTCTACTACTAGAAAACCTCATCGAAGAAGATACCATATTTTTTGCAGGAAGTATATCTGAGTATGATGATTTACAAATGGTTATTGCTGGTGCAAAATCCAAATTTCCAAGATCTATGCTTTCTATTTTTAATATAGTACCTAGAACGATGTCAAAATATGAGTTGGAGTTGATTCATAACGAGAATATCACAGGGGCAATGTTTACCACAATGTATAATATAAGAAACAATTTGGGTCTAGGCGATGATAAACTAACTATTGAAGCCATTGAAAACTATTTCTTGGATCCTAACAATGAGGTTATGCCTCTTATCATTAATAATACGGATATGACTACCGTCATTCCTAAAAAAAGTGGTAGGAGAAAGAATAAGAACATGGTTATCTTCCGTCAAGGATCATCACCTATCTTGTGTATTTTCGAAACTCGTAAAAAGATTAATATTTATAAAGAAAATATGGAATCCGTATCGACTGAGTATACACCTATCGGAGACAACAAGGCTTTGATATCTAAATATGCGGGAATTAATATCCTGAATGTGTATTCTCCTTCCACGTCCATGAGATTGAATGCCATTTACGGATTCACCAATAAAAATAAACTAGAGAAACTTAGTACTAATAAGGAACTAGAATCGTATAGTTCTAGCCCTCTTCAAGAACCCATTAGGTTAAATGATTTTCTGGGACTATTGGAATGTGTTAAAAAGAATATTCCTCTAACAGATATTCCGACAAAGGATTGATTACTATAAATGGAGAATGTTCCTAATGTATACTTTAATCCTGTGTTTATAGAGCCCACGTTTAAACATTCTTTATTAAGTGTTTATAAACACAGATTAATAGTTTTATTTGAAGTATTCGTTGTATTCATTCTAATATATGTATTTTTTAGATCTGAATTAAATATGTTCTTCATGCCTAAACGAAAAATACCCGATCCTATTGATAGATTACGACGTGCTAATCTAGCGTGTGAAGACGATAAATTAATGATCTATGGATTACCATGGATAACAACTCAAACATCTGCGTTATCAATAAATAGTAAACCGATAGTGTATAAAGATTGTGCAAAGCTTTTGCGATCAATAAATGGATCACAACCAGTATCTCTTAACGATGTTCTTCGCAGATGATGATTCATTTTTTAAGTATTTTGCTAGTCAAGATGATGAATCTTCATTATCTGATATATTGCAAATCACTCAATATCTAGACTTTCTGTTATTATTATTGATCCAATCAAAAAATAAATTAGAAGCTGTGGGTCATTGTTATGAATCTCTTTCAGAGGAATACAGACAATTGACAAAATTCACAGACTCTCAAGATTTTAAAAAACTGTTTAACAAGGTCCCTATTGTTACAGATGGAAGGGTCAAACTTAATAAAGGATATTTGTTCGACTTTGTGATTAGTTTGATGCGATTCAAAAAAGAATCAGCTCTAGCTACCACCGCAATAGATCCTGTTAGATACATAGATCCTCGTCGTGATATCGCATTTTCTAACGTGATGGATATATTAAAGTCGAATAAAGTTGAAAAATAATTAATTCTTTATTGTTATCATGAACGGCGGACATATTCAGTTGATAATCGGCCCCATGTTTTCAGGTAAAAGTACAGAATTAATTAGACGAGTTAGACGTTATCAAATAGCTCAATATAAATGTGTGACTATAAAATATTCTAACGATAATAGATACGGAACGGGACTATGGACACATGATAAGAATAATTTTGCAGCATTGGAAGTAACTAAACTATGTGATGTCTTGGAAGCAATTACAGATTTCTCCGTGATAGGTATAGATGAAGGACAGTTCTTTCCAGACATTGTTGAATTCTGTGAGCGTATGGCAAACGAAGGAAAAATAGTTATAGTAGCCGCGCTCGATGGGACATTTCAACGTAGACCGTTTAATAATATTTTGAATCTTATTCCATTATCTGAAATGGTGGTAAAACTAACTGCAGTGTGTATGAAATGCTTTAAGGAGGCTTCCTTTTCTAAACGATTAGGTACAGAAACCGAGATAGAAATAATAGGAGGTAATGATATGTATCAATCTGTGTGTAGAAAGTGTTACATCGACTCATAATATTATATTTTTTATCTAAAAAACTAAAAATAAACATTGATTAAATTTTAATATAATACTTAAAAATGGATGTTGTGTCGTTAGATAAACCGTTTATGTATTTTGAGGAAATTGATAATGAGTTAGATTACGAACCAGAAAGTGCAAATGAGGTCGCAAAAAAACTGCCGTATCAAGGACAGTTAAAACTATTACTAGGAGAATTATTTTTTCTTAGTAAGTTACAGCGACACGGTATATTAGATGGCGCCACCGTAGTGTATATAGGATCTGCTCCAGGTACACATATACGTTATTTGAGAGATCATTTCTATAATTTAGGAGTGATCATCAAATGGATGCTAATTGACGGCCGCCATCATGATCCTATTCTAAATGGATTGCGTGATGTGACTCTAGTGACTCGGTTTGTTGATGAGGAATATCTACGATCCATCAAAAAACAACTACATCCTTCTAAGATTATTTTAATTTCTGATGTGCGATCCAAACGAGGAGGAAATGAACCTAGTACTGCGGATTTACTAAGTAATTATGCTCTACAAAATGTCATGATTAGTATTTTAAACCCCGTGGCGTCTAGTCTTAAATGGAGATGCCCGTTTCCAGATCAATGGATCAAGGACTTTTATATCCCACACGGTAATAAAATGTTACAACCTTTTGCTCCTTCATATTCAGCTGAAATGAGATTATTAAGTATTTATACCGGTGAGAATATGAGACTGACTCGAGTTACCAAATCAGACGCTGTAAATTATGAAAAAAAGATGTATTACCTTAATAAGATAGTCCGCAACAAAGTAGTTATTAACTTTGATTATCCTAATCAGGAATATGACTATTTTCACATGTACTTTATGTTGAGGACCGTATACTGCAATAAAACATTTCCTACTACTAAAGCAAAGATACTATTTCTACAACAATCTATATTTCGTTTCTTAAATATTCCAACGACATCAACTGAAAAAGTTAGTCATGAACCAATACAACGTAAAATATCTAGCAAAGATTCTATGTCTAAAAACAGAAATAGCAAGAGATCCGTACGCGGTAATAAATAGAAACGTACTACTGAGATATACTACCGATATAGAGTATAATGATTTAGTTACTTTAATAACCGTTAGACATAAAATTGATTCTATGAAAACTGTGTTTCAGGTATTTAACGAATCATCCATAAATTATACTCCGGTTGATGATGATTATGGAGAACCAATCATTATAACATCGTATCTTCAAAAAGGTCATAACAAGTTTCCTGTAAATTTTCTATACATAGATGTGGTAATATCTGACTTATTTCCTAGCTTTGTTAGACTAGATACTACAGAAACTAATATAGTTAATAGTGTACTACAAACAGGCGATGGTAAAAAGACTCTTCGTCTTCCTAAAATGTTAGAGACGGAAATAGTTGTCAAGATTCTCTATCGTCCTAATATACCATTAAAAATTGTTAGATTTTTCCGCAATAACATGGTAACTGGAGTAGAGATAGCCGATAGATCTGTTATTTCAGTCGCTGATTAATCAATTAGTAGAGATGAGATAAGAACATTATAATAATCAATAATATATCTTATATCTGTTTAGAAAAATGCTAATATTAAAATAGCTAACGCTAGTAATCCAATCGGAAGCCATTTGATATCTATAATAGGGTATCTAATTTCCTGATTCAGATAGCGTACGGCTATATTCTCGGTAGCTACTCGTTTGGAATCACAGACATTATTTACATCTAATTTACTATCTGTAATGGAAACGTTTCCCAATGAAATGGTACAATCAGATACATTACATCTTGATATATTTTTTTTTAAAGAGGCTGGTAACAACGCATCGCTTCGTTTACATGGCTCGTACCAACAATAATAGGGTAATCTTGTATCTATTCCTATCCGTACTATACTTTTATCAGGATAAATACATTTACATCGTATATCGTCTTTGTTAGTATCACAGAATGCATAAATTTGTTCGTCCGTCATGATAAAAATTTAAAGTGTAAATATAACTATTATTTTTATAGTTATAATAAAAAGGGAAATTTGATTGTATACCTTCGGTTCTTTAAAAGAAACTGACTTGATAAAAATGGCTGTAATCTCTAAGGTTACGTATAGTCTATACGATCAAAAAGAGATTAATGCCACAGATATTATCATTAGTCATATTAAAAATGACGACGATATCGGTACCGTTAAAGATGGTAGACTAGGTGCTATGGATGGGGCATTATGTAAGACTTGTGGGAAAACGGAATTGGAATGTTTCGGTCACTGGGGTAAAGTAAGTATTTATAAAACTCATATAGTTAAGCCTGAATTTATTTCAGAAATTATTCGTTTACTGAATCATATATGTATTCATTGCGGATTATTGCGTTCACGAGAACCGTATTCCGACGATATTAACCTAAAAGAGTTATCGGTACACGCTCTTAGGAGATTAAAGGATAAAATATTATCCAAGAAAAAGTCATGTTGGAACAGCGAATGTATGCAACCGTATCAAAAAATTACTTTTTCAAAGAAAAAGGTTTGTTTCGTCAACAAGTTGGATGATATTAACGTTCCTAATTCTCTCATCTATCAAAAGTTAATTTCTATTCATGAAAAGTTTTGGCCATTATTAGAAATTCATCAATATCCAGCTAACTTATTTTATACAGACTACTTTCCCATCCCTCCGTTGATTATTAGACCGGCTATTAGTTTTTGGATAGATAGTATACCCAAAGAGACAAATGAATTAACTTACTTATTAGGTATGATCGTTAAGAATTGTAACTTGAATGCTGATGAACAGGTTATCCAGAAGGCGGTAATAGAATACGATGATATTAAAATTATTTCTAATAACACTACCAGTATCAATTTATCATATATCACATCCGGCAAAAATAATATGATTAGAAGTTATATCGTCGCTCGGCGAAAAGATCAGACCGCTAGATCCGTAATTGGTCCCAGTACATCTATCACCGTTAATGAGGTAGGAATGCCCACATATATTAGAAATACACTTACAGAAAAGATATTTGTTAATGCCTTTACAGTGGATAAAGTTAAACAACTATTAGCATCAAACCAAGTTAAATTTTACTTTAATAAACGATTAAACCAATTAACAAGAATACGTCAAGGAAAGTTTATCAAAAATAAAATACATTTATTGCCTGGTGATTGGGTAGAAGTAGCTGTTCAAGAATATACAAGTATTATTTTTGGAAGACAACCGTCTCTACATAGATACAACGTCATCGCTTCATCTATCAGAGCTACCGAAGGAGATACTATCAAAATATCTCCCGGAATTGCCAACTCTCAAAATGCTGATTTCGACGGAGATGAAGAATGGATGATATTGGAGCAAAATCCTAAAGCCGTAGTTGAACAAAGTATTCTTATGTATCCGACGACGTTACTCAAACACGATATTCATGGAGCCCCCGTTTATGGATCTATTCAAGATGAAATCGTAGCAGCGTATTCATTGTTTAGGATACAAGATCTTTGTTTAGATGAAGTATTGAACATCTTGGGGAAATATGGAAGAGAGTTCGATCCTAAAGGTAAATGTAAATTCAGCGGTAAAGATATCTATACTTACTTGATAGGTGAAAAGATTAATTATCCGGGTCTCTTAAAGGATGGTGAAATTATTGCAAACGACGTAGATAGTAATTTTGTTGTAGCTATGAGGCATCTGTCATTGGCTGGACTCTTATCCGATCATAAATCGAACGTGGAAGGTATCAACTTTATTATCAAGTCATCTTATGTTTTTAAGAGATATCTATCTATATACGGTTTTGGGGTGACATTCAAAGATCTGAGACCAAATTCGACGTTCACTAATAAATTGGAGGCTATCAACGTAGAAAAAATAGAACTTATCAAAGAAGCATACGCCAAATATCTCAAAGATGTAAGAGACGGGAAAATAGTTCCATTATCTAAAGCTTTAGAGGCGGACTACTTGGAATCCATGTTATCCAACTTGACAAATCTTAATATCAGAGAGATAGAAGAACATATGAGACAAACGCTGATAGATGATCCAGATAATAACCTCCTGAAAATGGCCAAAGCGGGTTATAAAGTAAATCCCACAGAACTAATGTATATTCTAGGTACTTATGGACAACAGAGGATAGATGGCGAACCAGCAGAGACTCGAGTATTGGGTAGAGTCTTACCTTACTATCTTCCAGACTCTAAGGATCCAGAAGGAAGAGGTTATATTCTTAATTCTTTAACAAAAGGATTAACGGGTTCTCAATATTACTTTTTGATGCTGGTTGCAAGATCTCAATCTACTGATATTGTCTGTGAAACATCACGTACCGGAACACTGGCTAGAAAAATCATTAAAAAGATGGAGGATATGGTGGTCGACGGATACGGACAAGTAGTTATAGGTAATACGCTCATCAAGTACGCAGCCAATTATACCAAAATTCTAGGCTCAGTATGTAAACCTGTAGATCTTATCTATCCAGATGAGTCCATGACTTGGTATTTGGAAATTAGTGCTTTGTGGAATAAAATAAAACAGGGATTCGTTTACTCTCAGAAACAGAAACTTGCAAAGAAGACATTGGCGCCGTTTAATTTCCTAGTATTCGTCAAACCCACCACTGAGGATAATGCTATTAAGGTTAAGGATCTGTACGATATGATTCATAACGTCATTGATGATGTGAGAGAGAAATACTTCTTTACGGTATCTAATATAGATTTTATGGAGTATATATTCTTGACGCATCTTAATCCTTCTAGAATTAGAATTACAAAAGAAACGGCTATTACTATCTTTGAAAAGTTCTATGAAAAACTCAATTATACTCTAGGTGGTGGAACTCCTATTGGAATTATTTCTGCACAGGTATTGTCTGAGAAGTTTACACAACAAGCCCTGTCCAGTTTTCACACTACTGAAAAGAGTGGTGCTGTAAAACAAAAACTTGGTTTCAACGAGTTTAATAACTTGACTAATTTGAGTAAGAATAAGACCGAAATTATCACTCTGGTATCCGATGATATCTCTAAACTTCAATCTGTTAAGATTAATTTCGAATTTGTATGTTTGGGAGAATTAAATCCAGACATCACTCTTCGAAAAGAAACAGATAGATATGTAGTAGACATAATAGTCAATAGATTATACATCAAGAGAGCAGAAATAACCGAATTAGTCGTCGAATATATGATTGAACGATTTATCTCCTTTAGCGTCATTGTAAAGGAATGGGGTATGGAGACATTCATTGAGGACGAGGATAATATTAGATTTACTATCTACCTAAATTTCGTTGAACCGGAGGAATTGAATCTTAGTAAGTTTATGATGGTTCTTCCAGGTGCCGCCAACAAGGGCAAGATTAGTAAATTCAAGATTCCTATCTCTGACTATACGGGATATAACGACTTCAATCAAACAAAAAAGCTCAATAAGATGACTGTAGAACTCATGAATCTAAAAGAATTGGGTTCTTTCGATTTGGAGAACGTCAACGTGTATCCTGGAGTATGGAATACATACGATATCTTCGGTATTGAGGCCGCTCGTGGATACTTGTGCGAAGCCATGTTAAACACCTATGGAGAAGGTTTCGATTATCTGTACCAGCCTTGTGATCTTCTCGCTAGTTTACTATGTGCTAGTTACGAACCAGAATCAGTTAATAAATTCAAGTTCGGTGCAGCTAGTACTCTTAAGAGAGCTACGTTCGGAGATAATAAAGCATTGTTAAACGCGGCTCTTCATAAAAAGTCAGAACCTATTAACGATAATAGTAGCTGCCACTTTTTTAGCAAGGTCCCTAATATAGGAACTGGATATTACAAATACTTTATCGACTTGGGTCTTCTCATGAGAATGGAAAGGAAACTATCTGATAAGATATCTTCTCAAAAGATCAAGGAGATAGAAGAAACAGAAGACTTTTAATTCTTATCAATAACATATTTTTCTATGATCTGTCTTTTAAACGATGGATTTTCCACAAATGCGCCTCTCAAGTCCCTCATAGAATGATACACGTATAAAAAATATAGCATAGGTGATGACTCCTTATTTTTAGACATTAGATATGCCAAAATCATAGCCCCGCTTCTATTTACTCCTGCAACACAATGAACCAACACGGGCTCGTTTCGTTGATCACATTTAGATAAGAAGGCGGTCACGTCGTCAAAATATTTACTAATATCAGTAGTTGTATCATCTACCAACGGTATATGAATAATATTAATATTAGAGTTAGGTAATGTATATTTATCCATCGTCAAATTTAAAACATATTTGAACTTAACTTCAGATGATGGTGCATCCATAGCATTTTTATAATTTCCCAAATACACATTATTTGTTACTCTTGTCATTATAGTGGGAGATTTGGCTCTGTGCATATCTCCAGTTGAACGTAGTAGTAAGTATTTATACAAACTTTTCTTATCCATTTATAACGTACAAATGGATAAAACTACTTTATCAGTAAACGCATGCAATTTAGAATACGTTAGAGAAAAGGCTATAGTAGGCGTACAAGCAGCCAAGACATCAACACTTATATTTTTTGTTATTATATTGGCAATTAGTGCGCTATTACTCTGGTTTCAGACGTCTGATAATCCAGTCTTTAATGAATTAACGAGATATATGCGAATTAAAAATACGGTTAACGATTGGAAATCATTAACGGATAGCAAAACAAAATTAGAAAGCGATAGAGGTAGACTTCTAGCCGCTGGTAAGGATGATATATTCGAATTCAAATGTGTGGATTTCGGCGCCTATTTTATAGCTATGCGATTGGATAAGAAAACATATCTGCCGCAAGCTATTAGGCGAGGTACTGGAGACGCGTGGATGGTTAAAAAGGCGGCAAAAGTCGATCCATCTGCTCAACAATTTTGTCAGTATTTGATAAAACACAAGTCTAATAATGTTATTACTTGTGGTAATGAGATGTTAAATGAATTAGGTTATAGCGGTTATTTTATGTCACCGCATTGGTGTTCCGATCTTAGTAATATGGAATAAGTGTTAGATAAATGCGGTAACAAATGTTCCTGTAAGGAACCATAACAGTTTAGATTTAACATTAAAGATGAGCATAAACATAATAAACAAAATTACAATCAAACCTATAACATTAATATCAAACAATCCAAAAAATGAAATCAATGGAGTAGTAAACGTGTACATAACTCCTGGATAACGTTTAGCAGCTACCGTTCCTATTCTAGACCAAAAATTTGGTTTCATGGTTTCGAAGCGGTGTTCTGCAACAAGACGAGGATCGTGTTCTACATATTTGGCAGAGTTATCCATTATTTGCCTGTTAATCTTCATTTCGTTTTCGATTCTGGCTATTTCAAAATAAAATCCCGATGATAGACCTCCAGACTTTATAATTTCATCTACGATGTTCAGCGCCGTAGTAACTCTAATAATATAGGCGGATAAGCTAACATCATACCCTCCTGTATATGTAAATATGGCATGATCTTTGTCTATTACAAGCTCGGTTTTAACTTTATTTCCTGTAATAATTTCTCTCATCTGTAGGATATCTATTTTCTTGTCATGTATTGCCTTCAAGACGGGACGAAGAAACGTAATATCCTCAATAACGTTATCGTTTTCTATAATAACTACATATTCTACATTTTTATTTTCTAGCTCGGTAAAAAATTTAGAATCCCATAGGGCTAAATGTCTAGCGATATTTCTTTTCGTTTCCTCTGTACACATAGTGTTACAAAACCCTGAAAAGAAGTGAGTATACTTGTCATCATCTCTAATATTTCCTCCAGTCCATTGTATAAACACATAATCCTTGTAATGATCTGGATCATCATTGACTATCACAACATCTCTTTTTTCTTGCATAACTTCATTGTCCTTCACATCATCGAACTTCTGATCATTAATATGCTCATGAACATTAGGAAATGTTTCTGATGGAGGTCTATCAATAACTGGCACAACAATAACAGGAGTTTTCACCGCCGCCATTTAGTTATTGAAATTAATCATATACAACTCTCTAATACGAGTTATATTTTCGTCTATCCATTGTTTCACATTGACATATTTCGACAAAAAGATATAAAATGCGTATTCCAATGCTTCTCTGTTTAATGAATTACTAAAATATACAAACACGTCACTGTCTGGTAATAAATAATATCTTAGAATATTGTAACAATTTATTTTGTATTGCACATGTTCGTGATCTATGAGTTCTTCTTCAAATGGCATAGGATCTCCGAATCTGAAAACGTATAAATAGGAGTTAGAATAATAATATTTGAGAGTATTGGTAATGTATAAACTCTTTAGCGGTATAATTAGTTTTTTTCTCTCGATTTCTATTTTTAGATGTGATGGAAAAATGACTAATTTTGTAGCATTAGTATCATGAACTCTAATCAAAATCTTAATATCTTCGTCACATGTTAGCTCTTTGAAGTTTTTAAGAGATGCATCAGTTGGTTTTACAGATGGAGTAGGTGCAACAATTTTTTGTTTAATGCATGCATGTATTGGAGCCATTGTCTTAACTATAATGGTGCTTGTATCGAAAAACTTTAATGCGGATAACGGAAGCTCTTCGCCGCGACTTTCTACGTCGTAATTGGGTTCTAATGCCGATCTCTGAATGGATACTAGTTTTCTAAGTTCTAATGTAATTCTCTGAAAATGTAAATCCAATTCCTCCGGCATTATAGATGTGTATACATCGGTAAATAAAACTATAGTATCCAACGATCCCTTCTCGCAAATTCTAGTCTTAACCAAGAAATCGTATATAACTACGGAGATGGCGTATTTAAGAGTGGATTCTTCTACCGTTTTGTTCTTGGATTTCATATAAGAAACTATAAAGTCCGCACTACTGTTAAGAATGATCACTAACGCAACTATATAGTTCAAATTAAGCATCTTGGAAACATAAAATAACTCTGTAGATGATACTTGACTTTCGAATAAGTTTGCAGACAAACGAAGAAAGAACAGACCTCTCTTAATTTCAGAAGAAAACTTTTTTTCGTATTCCTGACGTCTAGAGTTTATATCAATAAGAAAGTTAAGAATTAGTCGGTTAATGTTGTATTTCATTACCCAAGTTTGAGATTTCATAATATTGTCAAAAGACATGATAATATTAAAGATAAAGCGCTGACTATGAACGAAATAGCTATATGGTTCGCTCAAGAATATAGTCTTGTTAAACGTGGAAACGATAACTGTATTTTTAATCACGTCAGCGGCATCTAAATTAAATATAGGTATATTTATTCCACACACTCTACAATATGCCACACCATCTTCATAATAAATAAATTCGTTAGCAAAATTATTAATTTTAGTGAAATAGTTAGCGTCAACTTTCATAGCTTCCTTCAATCTAATTTGATGCTCACATGGCGCGAATTCTACTCTAACATCCCTTTTCCATGCCTCAGGTTCATCGATCTCTATAATATCTAGTTTCTTGCGTTTCACAAACACAGGCTCGTCTCTCGCGATGAGATCTGTATAGTAACTATGTAAATGATAACTAGATAGAAAGATGTAGCTATATAGATGACGATCCTTTAAGAGAGGTATAATAACTTTACCCCAATCAGATAGACTGTTGTTATGGTCTTCGGAAAAAGAATTTTTATAAATTTTTCCAGTATTTTCTAAATATACGTACTTGATATCTAAGAAATCCTTAATAATAATAGGAATGGATAATCCGTCTATTTTATAAAGAAATACATATCGCATATTATACTTTTTTTTGGAAATTGGAATACCGATGTGTCTACATAAATACGCAAAGTCTAAATATTTTTTAGAGAATCTTAGTTGGTCCAAATTCTTTTCCAAGTACGGTAATAGATTTTTCATATTGAACGGTATCTTCTTGATCTCTGGTTCTAATTCCGCATTAAATGATGAAACTAAGTCACTATTTTTATAACTAACGATTACATCACCTCTAACATCATCATTTACCAGGATACTGATCTTCTTTTGTCGTAAATACATGTCTAATGTGTTAAAAAAAAGATCATACAAGTTATACGTCATTTCATCTGTAGTATTCTTGTCATTGAAGGATAAACTCGTACTAATCTCTTCTTTAACAGTCTGTTCAAATTTATATCCTATATATGAAAAAATAGCAACCAGTGTTTGATCATCCGCGTCAATATTCTGTTCTATCGTAGTGTATAACAATCTTATATCTTCTTCTGTGATAGTCGATACGTTATAAAGGTTGATAACGAAAATATTTTTATTTCGTGAAATAAAGTCATTGTAGGATTTTGGACTTATATTCGTGTCTAGTAGATATGATTTTATTTTTGGAATGATCTCAATTAAAATAGTCTCTTTAGAGTCCATTTAAAGTTACAAACAACTAGGAAATTGGTTTATGATGTATAATTTTTTTAGTTTTTATAGATTCTTTATTCTATACTTAAAAAATGAAAATAAATACAAAGGTTCTTGAGGGTTGTGTTAATTGAAAGCGATAAATAATCATAAATTATTTCATTATCGCGATATCCGTTAAGTTTGTATCGTAATGGCGTGGTCAATTACGAATAAAGCGGATACTAGTAGTTTCACAAAGATGGCTGAAATCAGAGCTCATCTAAGAAATAGCGCTGAAAATAAAGATAAAAACGAGGATATTTTCCCGGAAGATGTAATAATTCCATCTACTAAGCCCAAAACCAAACGAACCACTACTCCTCGTAAACCAGCGGCTACTAAAAGATCAACCAAAAAGGATAAAGAAAAGGAGGAAGTGGAAGAAGTAGTTATAGAGGAATATCATCAAACAACTGAAGAAAATTCTCCACCTCCGTCATCATCTCCTGGAGTCGGCGACATTGTAGAAAGCGTGGCCGCTGTAGAGCTCGATGATAGCGACGGGGATGATGAACCTATGGTACAAGTTGAAGCTGGTAAAGTAAATCATAGTGCTAGAAGCGATCTCTCTGACCTAAAGGTGGCTACCGACAATATCGTTAAAGATCTTAAGAAAATTATTACTAGAATCTCTGCAGTATCGACTGTTCTAGAGGATGTTCAAGCAGCTGGTATCTCTAGACAATTTACTTCTATGACTAAAGCTATTACAACACTATCTGATCTAGTCACCGAGGGAAAATCTAAAGTTGTTCGTAAAAAAGTTAAAACTTGTAAGAAGTAAATGCGTGCACTTTTTTATAAAGATGGTAAACTGTTTACCGATAATAATTTTTTAAATCCTGTATCAGACGATAATCCAGCGTATGAGGTTTTGCAACATGTTAAAATTCCTACTCATTTAACAGATGTAGTAGTATATGAACAAACGTGGGAAGAGGCATTAACTAGATTAATTTTTGTGGGAAGTGATTCAAAAGGACGTAGACAATACTTTTACGGAAAAATGCATATACAGAATCGCAATGCTAAAAGAGATCGTATTTTTGTTAGAGTATATAACGTTATGAAACGAATTAATTGTTTTATAAACAAAAATATAAAGAAATCGTCCACAGATTCCAATTATCAGTTGGCGGTTTTTATGTTAATGGAAACTATGTTTTTTATTAGATTTGGTAAAATGAAATATCTTAAGGAGAATGAAACAGTAGGGTTATTAACACTAAAAAATAAACACATAGAAATAAGTCCCGATGAAATAGTTATCAAGTTTGTAGGAAAGGACAAAGTTTCACATGAATTTGTTGTTCATAAGTCTAATAGACTATATAAACCGCTATTGAAACTGACTGATGATTCTAGTCCCGAAGAATTTCTGTTCAACAAACTAAGTGAACGAAAGGTATATGAATGTATCAAACAGTTTGGTATTAGAATCAAGGATCTCCGAACGTATGGAGTCAATTATACGTTTTTATATAATTTTTGGACAAATGTAAAGTCCGTATCTCCTCTTCCATCACCAAAAAAGTTGATAGCATTAACTATCAAACAAACTGCTGAAGTGGTAGGTCATACTCCATCAATTTCAAAAAGAGCTTATATGGCAACGACTATTTTAGAAATGGTAAAGGATAAAAATTTTTTAGACGTAGTATCTAAAACTACGTTCGATGAATTCCTATCTATAGTCGTAGATCACGTTAAATCATCTACGGATGGATGATAATAGATCTTTACACAAATAATTACAAGACCGATAAATGGAAATGGATAAACGGATGAAATCTCTCGCTATGACAGCTTTCTTCGGAGAGCTAAACACGTTAGATATTATGGCATTGATAATGTCTATATTTAAACACCATCCAAACAATACCATTTTTTCAGTGGATAAGGATGGTCAATTTATGATTGATTTCGAATACGATAATTATAAGGCTTCTCAATATTTGGATCTGACCCTCACTCCGATATCTGGAAATGAATGCAAGACTCACGCATCTAGTATAGCCGAACAATTGGCGTGTGTGGATATTATTAAAGAGGATATTAGCGAATATATCAAAACTACTCCCCGTCTTAAACGATTTATAAAAAAATACCGCAATAGATCATATACTCGTATCAGTCGAGATACAGAAAAGCTTAAAATAGCTCTAGCTAAAGGCATAGATTACGAATATATAAAAGACGCTTGTTAATAAGTAAATGAAAAAAAACTAGTCGTTTATAATAAAACACAATATGGATGCCAACATAGTATCATCTTCTACTATTGCGACGTATATAGACGCTTTAGCGAAGAATGCTTCAGAATTAGAACAGAGGTCTACCGCATACGAAATAAATAATGAATTGGAACTAGTATTTATTAAACCGCCATTGATTACGTTGACAAATGTAGTAAATATCTCCACGATTCAGGAATCGTTTATTCGATTTACCGTTACTAATAAGGAAGGTATCAAAATTAGAACTAAGATTCCATTATCTAAGGTACATGGTCTAGATGTAAAAAATGTGCAGTTGGTAGATGCTATAGATAACATAGTTTGGGAAAAGAAATCATTAGTGACGGAAAATCGTCTTCACAAAGAATGCTTGTTGAGACTATCAACAGAGGAACGTCATATATTTTTGGATTACAAGAAATATGGATCCTCTATCCGACTAGAATTAGTCAATCTTATTCAAGCAAAAACAAAAAACTTTACGATAGACTTTAAGCTAAAATATTTTCTAGGATCTGGCGCTCAATCTAAAAGTTCTTTATTGCACGCTATTAATCATCCAAAGTCAAGGCCTAATACATCTCTGGAAATAGAATTTACACCTAGAGACAATGAAACAGTTCCATATGATAAACTAATAAAGGAATTGACGACTCTCTCGCGTCATATATTTATGGCTTCTCCAGAGAATGTAATTCTTTCTCCACCTATTAACGCACCTATAAAGACTTTTATGTTGCCTAAACAAGATATAGTAGGTCTGGATCTGGAAAATCTATATGCCGTAACTAAGACTGACGGCATTCCTATAACTATCAGAGTTACATCAAAAGGGTTGTATTGTTATTTTACACATCTTGGTTATATTATTAGATATCCAGTTAAGAGAACAATAGATTCCGAAGTAGTAGTCTTTGGTGAGGCAGTTAAGGATAAGAACTGGACCGTATATCTCATTAAGCTAATAGAGCCCGTAAATGCAATCAGTGATAGACTAGAAGAAAGTAAGTATGTTGAATCTAAACTAGTGGATATTTGTGATCGGATAGTATTCAAGTCAAAGAAATACGAAGGTCCGTTTACTACAACTAGTGAAGTCGTCGATATGTTATCTACATATTTACCAAAGCAACCAGAAGGTGTTATTCTGTTCTATTCAAAGGGACCTAAATCTAACATTGATTTTAAAATCAAAAAGGAGAATACTATAGACCAAACTGCAAATGTAGTATTTAGGTACATGTCCAGTGAACCAATTATCTTTGGAGAGTCGTCTATCTTTATAGAGTATAAGAAATTTACCAACGATAAAGGCTTTCCTAAAGAATATGGTTCTGGTAAGATTGTGTTATATAACGGCGTTAATTATCTAAATAATATCTATTGTTTGGAATATATTAATACACATAATGAAGTGGGTATTAAGTCCGTTGTTGTACCTATTAAGTTTATAGCAGAATTCTTAGTCAATGGAGAAATACTTAAACCTAGAATCGATAAAACCATGAAATATATTAACTCAGAAGACTATTATGGAAATCAACATAATATCATAGTCGAACATTTAAGAGATCAAAGCATCAAAATAGGAGATGTCTTTAACGAGGATAAACTATCGGATGTTGGACATCAATACGCTGCCAACAACGATAAATTTAGATTAAATCCAGAAGTTAGTTATTTTACTAATAAACGAACTAGAGGGCCGTTGGGAATTTTATCAAACTACGTCAAGACTCTTCTTATTTCTATGTATTGTTCCAAAACATTTTTAGACGATTCCAACAAACGAAAGGTATTAGCGATTGATTTTGGAAACGGTGCTGACCTGGAAAAATACTTTTATGGAGAGATTGCGTTATTGGTAGCGACGGATCCGGATGCTGATGCTATAGCTAGAGGAAATGAAAGATACAACAAATTAAATTCTGGAATTAAAACCAAGTACTACAAATTTGACTACATTCAGGAAACTATTCGATCCGATACATTTGTCTCTAGTGTCAGAGAAGTATTCTATTTTGGAAAGTTTAATATCATTGACTGGCAGTTCGCTATTCATTATTCTTTTCATCCAAGACATTATGCTACAGTCATGAATAACTTATCCGAACTAACTGCTTCTGGAGGCAAGGTATTAATTACTACCATGGATGGAGACAAATTATCAAAATTAACCGATAAAAAGACTTTTATAATTCATAAGAATCTACCTAGTAGCGAAAACTATATGTCTGTAGAAAAAATAGCTGATGATAGAATAGTGGTATATAATCCATCAACAATGTCTACTCCAATGACTGAATACATTATCAAAAAGAACGATATAGTCAGAGTGTTTAACGAATACGGATTTGTTCTTGTAGATAATGTTGATTTCGCTACAATTATAGAACGAAGTAAAAAGTTTATTAATGGCGCATCTACAATGGAAGATAGACCGTCTACAAGAAACTTTTTCGAACTAAATAGAGGAGCCATTAAATGTGAAGGTTTAGATGTCGAAGACTTACTTAGTTACTATGTTGTTTATGTCTTTTCTAAGCGGTAAATAATAATATGGTATGGGTTCTGATATCCCCGTTCTAAATGCATTAAATAATTCCAATAGAGCGATTTTTGTTCCTATAGGACCTTCCAACTGTGGATACTCTGTATTATTAATAGATATATTAATACTTTTGTAGGGTAACAGAGGTTCTACGTCTTCTAAAAATAAAAGTTTTATAACATCTGGCCTGTTCATAAATAAAAACTTGGCGATTCTATATATACTCTTATTATCAAATCTAGCCATTGTCTTATAGATGTGAGCTACTGTAGGTGTACCATTTGATTTTCTTTCTAATACTATATATTTCTCTCGAAGAAGTTCTTGCAGATCATCTGGGAATAAAATACTACTGTTGAGTAAATCAGTTATTTTTTTTATATCGATATTGATGGACATTTTTATAGTTAAGGATAATAAGTATCCCAAAGTAGATAACGACGATAACGAAGTATTTATACTTTTAGGAAATCACAATGACTTTATCAGATCAAAATTAACAAAATTAAAGGAGCATGTATTTTTTTCTGAATATATTGTGACTCCAGATACATATGGATCTTTATGCGTCGAATTAAATGGGTCTAGTTTTCAGCACGGTGGTAGATATATAGAGGTGGAGGAATTTATAGATGCTGGAAGACAAGTTAGATGGTGTTCTACATCCAATCATATATCTGAAGATATACACACTGATAAATTTGTCATTTATGATATTTATACGTTTGATTCGTTCAAGAATAAACGATTGGTATTTGTACAGGTGCCTCCATCATTAGGAGATGATAGCTATTTAACTAATCCGTTATTGTCTCCGTATTATCGTAATTCAGTAGCCAGACAAATGGTCAATGATATGATTTTTAATCAAGATTCATTTTTAAAATATTTATTAGAACATCTGATTAGAAGCCACTATAGAGTTTCTAAACATATAACAATAGTTAGATACAAGGATACCGAAGAATTAAATCTAACAAGAATATGTTATAATAGAGATAAGTTTAAGGCGTTTGTATTCGCTTGGTTTAACGGCGTTTCGGAAAATGAAAAGGTACTAGATACGTATAAAAAGGTATCTGATTTGATATAATGAATTCAGTGACTATATCACACGCACCATATACTATTACTTATCACGATGATTGGGAACCAGTAATGAGTCAATTGGTAGAGTTTTATAACGAAGTAGCCAGTTGGTTGCTACGCGACGAGACGTCGCCTATTCCTGATAAGTTCTTTATACAATTGAAACAGCCGCTTAGAAATAAACGAGTATGTGTGTGTGGTATAGATCCGTATCCAAAAGATGGAACTGGTGTACCGTTCGAATCACCAAATTTTACAAAAAAATCAATTAAGGAGATAGCTTCATCTATATCTAGATTAACCGGAGTAATTGATTATAAAGGTTATAACCTTAATATAATAGACGGGGTTATACCCTGGAATTATTACTTAAGTTGTAAATTAGGAGAAACAAAAAGTCACGCGATTTACTGGGATAAGATTTCCAAGTTACTGCTACAGCATATAACTAAACACGTTAGTGTTCTTTATTGTTTGGGTAAAACAGATTTCTCGAATATACGGGCAAAGTTAGAATCCCCGGTAACTACCATAGTGGGATATCATCCAGCGGCCAGAGACCACCAATTCGAGAAAGATCGATCATTTGAAATTATCAACGTTTTACTGGAATTAGACAACAAGACACCTATAAATTGGGCTCAAGGGTTTATTTATTAATGCTTTAGTGAAATTTTAACTTGTGTTCTAAATGGATGCGGCTATTAGAGGTAATGATGTTATCTTTGTTCTTAAGACTATAGGTGTCCCGTCAGCATGCAGACAAAATGAAGATCCAAGATTCGTAGAAGCATTTAAATGCGACGAGTTAGAAAGATATATTGATAATAATCCAGAATGTACACTATTCGAAAGTCTTAGGGATGAGGAAGCATACTCTATAGTCAGAATTTTCATGGATGTAGATTTAGACGCGTGTCTAGACGAAATAGATTATTTAACGGCTATTCAAGATTTTATTATCGAGGTGTCAAACTGTGTAGCTAGATTCGCATTTACAGAATGCGGTGCCATTCATGAAAATGTAATAAAATCCATGAGATCTAATTTTTCATTGACTAAGTCTACAAATAGAGATAAAACAAGTTTTCATATTATCTTTTTAGACACGTATACCACTATGGATACATTGATAGCTATGAAACGAACACTATTAGAATTAAGTAGATCATCTGAAAATCCACTAACAAGATCGATAGACACTGCCGTATATAGGAGAAAAACAACTCTTCGGGTTGTAGGTACTAGGAAAAATCCAAATTGCGACACTATTCATGTAATGCAACCACCTCACGATAATATAGAAGATTACCTATTCACTTACGTGGATATGAACAACAATAGTTATTACTTTTCTCTACAACGACGATTGGAGGATTTAGTTCCTGATAAGTTATGGGAACCAGGGTTTATTTCGTTCGAAGACGCTATAAAAAGAGTTTCAAAAATATTCATTAATTCTATAATAAACTTTAATGATCTCGATGAAAATAATTTTACAACGGTACCACTGGTCATAGATTATGTAACACCTTGTGCATTATGTAAAAAACGATCGCATAAACATCCGCATCAACTATCGTTGGAAAATGGTGCTATTAGAATTTACAAAACTGGTAATCCACATAGTTGTAAAGTTAAAATTGTTCCGTTGGATGGTAATAAACTGTTTAATATTGCACAAAGAATTTTAGACACTAACTCTGTTTTATTAACCGAACGAGGAGACCATATAGTTTGGATTAATAATTCATGGAAATTTAACAGCGAAGAACCCTTGATAACAAAACTAATTCTATCAATAAGACATCAACTACCTAAGGAATATTCAAGCGAATTACTCTGTCCGAGGAAACGAAAGACTGTAGAAGCTAACATACGAGACATGTTAGTAGATTCAGTAGAGACCGATACCTATCCGGATAAACTTCCGTTTAAAAATGGTGTATTGGACCTGGTAGACGGAATGTTTTACTCTGGAGATGATGCTAAAAAATATACGTGTACTGTATCGACCGGATTTAAATTTGACGATACAAAATTCGTCGAAGACAGTCCAGAAATGGAAGAGTTAATGAATATCATTAACGATATCCAACCATTAACGGATGAAAATAAGAAAAATAGAGAGCTGTATGAAAAAACATTATCTAGTTGTTTATGTGGTGCTACCAAAGGATGTTTAACATTCTTTTTTGGAGAAACCGCAACTGGGAAGTCGACAACCAAACGTTTGTTAAAGTCTGCTATCGGTGACCTGTTTGTCGAGACGGGTCAAACAATTTTAACAGATGTATTGGATAAAGGACCTAATCCATTTATCGCTAATATGCATTTAAAAAGATCTGTATTCTGTAGCGAACTACCTGATTTTGCATGTAGTGGATCAAAGAAAATTAGATCTGATAATATTAAAAAGTTGACAGAACCTTGTGTCATTGGAAGACCGTGTTTCTCCAATAAAATTAATAATAGAAACCATGCGACAATCATTATCGATACTAATTACAAACCTGTCTTTGATAGGATAGATAACGCATTAATGAGAAGAATTGCCGTCGTGCGATTCAGAACACACTTTTCTCAACCTTCTGGTAGAGAGGCTGCTGAAAATAATGACGCGTACGATAAAGTCAAACTATTAGACGAGGGATTAGATGGTAAAATACAGAATAATAGATATAGATTCGCATTTCTATACTTGTTGGTTAAATGGTACAAAAAATATCATATTCCTATTATGAAACTATATCCTACACCGGAAGAGATTCCGGACTTTGCATTCTATCTCAAAATAGGTACTCTGTTGGTATCTAGCTCTGTAAAGCATATTCCATTAATGACGGACCTCTCCAAAAAGGGATATATATTGTACGATAATGTGGTTACTCTTCCGTTGACTACTTTCCAACAGAAAATATCCAAGTATTTTAATTCTAGACTATTTGGACACGATATAGAGAGCTTCATCAATAGACATAAGAAATTTGCCAATGTTAGTGATGAATATCTGCAATATATATTCATAGAGGATATTTCATCTCCGTAAATATATGCCATATATTTATAGAATATATCACATATCTAAATGAATACCGGAATCATAGATTTATTTGATAATCATGTTGATAGTATACCAACTATATTACCTCATCAGTTAGCTACTTTAGATTATCTAGTTAGAACTATCATAGATGAGAACAGAAGCGTGTTATTGTTCCATATTATGGGATCGGGTAAAACAATAATCGCTTTGTTGTTCGCCTTGGTAGCTTCCAGATTTAAAAAGGTTTACATTTTAGTACCGAACATCAACATCTTAAAAATTTTCAATTATAATATGGGTGTAGCTATGAACTTGTTTAATGACGAATTCATAGCTGAGAATATCTTTATTCATTCCACAACAAGTTTTTATTCTCTTAATTATAACGATAACGTCATTAATTATAACGGATTAAGTCGCTACAATAACTCTATTTTTATCGTTGATGAGGCACATAATATTTTTGGGAATAATACTGGAGAACTTATGACCGTGATAAAAAATAAAAACAAGATTCCTTTTCTACTATTGTCTGGATCTCCCATTACTAACACACCTAATACGCTGGGTCATATTATAGATTTAATGTCCGAAGAGACGATAGATTTTGGTGAGATTATTAGTCGTGGTAAGAAAGTAATTCAGACACTTCTTAACGAACGCGGAGTGAATGTACTCAAGGATTTGCTTAAAGGAAGAATATCATATTACGAAATGCCGGACAAAGATCTACCAACAATAAGATATCACGGACGTAAATTTCTAGATACTCGAGTAGTATATTGTCACATGTCTAAACTTCAAGAGAAAGATTATATGATTACTAGACGGCAGCTATGTTATCATGAAATGTTTGATAAAAATATGTATAACGTGTCAATGGCAGTATTGGGACAACTTAATCTGATGAATAATTTAGATACGTTATTTCAGGAACAGGATAAGGAATTGTACCCAAATCTGAAAATAAATAATGGAGTGTTATACGGTGAAGAATTGGTAACGTTAAACATTAGTTCCAAATTTAAGTACTTTATCAATCGGATACAGACACTCAAGGGAAAACACTTTATATACTTCTCTAATTCTACATATGGTGGATTGGTAATTAAATATATCATGCTCAGTAATGGATATTCTGAATATAATGGTTCTCAGGGAACTAATCCACATATGATAAACGGCAAACCAAAAACATTTGCTATCGTTACTAGTAAAATGAAATCGTCTTTAGAGGATCTATTAGATGTGTATAATTCTCCTGAAAACGATGATGGCAATCAATTGATGTTTTTGTTTTCGTCAAACATTATGTCTGAATCCTATACTCTGAAAGAGGTAAGGCATATTTGGTTTATGACTATCCCGGATACTTTTTCTCAATACAACCAAATTCTTGGACGATCTATTAGAAAATTCTCTTACGTCGATATTTCTGAACCCGTTAATGTATATCTTTTAGCAGCCGTATATTCAGATTTCAATGACGAAGTGACGTCATTAAACGATTATACACAGGATGAATTGATTAATGTTTTACCCTTTGACATCAAAAAGCTGTTGTATCTAAAATTTAAGACTAAAGAAACGAATAGAATATACTCTATTCTTCAAGAGATGTCTGAAACGTATTCTCTTCCACCACATCCATCAATTGTAAAAGTTTTATTGGGAGAATTGGTCAGACAATTTTTTTATAATAATTCTCGTATTAAGTATAACGACTCCAAGTTACTTAAAATGGTTACATCAGTTATAAAAAATAAAGAAGACGCTAGGAATTACATAGATGATATTGTAAACGGTCACTTCTTTGTATCGAATAAAGTATTTGATAAATCTCTTTTATACAAATACGAAAACGATATTATTACAGTACCGTTTAGACTTTCCTACGAACCATTTGTTTGGGGAGTTAACTTTCGTAAAGAATATAATGTGGTATCTTCTCCATAAAACTGATGAGATATATAAAGAAATAAATGTCGAGCTTTGTTACCAATGGATATCTTCCAGTTACATTGGAACCACATGAGTTGACGTTAGACATAAAAACTAATATTAGGAATGCCGTATATAAGGCGTATCTCCATAGAGAAATTAGTGGTAAAATGGCCAAGAAAATAGAAATTCGTGAAGACGTGGAATTACCTCTCGGTGAAATAGTTAATAATTCTGTAGTTATAAACGTTCCGTGTGTAATAACCTACGCATATTATCACGTTGGGGATATAGTCAGAGGAACATTAAACATCGAAGATGAATCAAATGTAACTATTCAATGTGGAGATTTAATCTGTAAACTAAGTAGAGATTCGGGTACTGTATCATTTAGCGATTCAAAGTACTGCTTTTTTCGAAATGGTAATGCGTATGATAACGGCATCGAAGTCTCCGCCGTTCTAATGGAGGCTCAACAAGGTACCGAATCTAGTTTTGTTTTTCTCGCGAATATCGTTGACTCATAAGAAAGAGAATAGCGGTGAGTATAAATACGAATACTATGGCAATAATTGCGAATGTTTTATTCCCTTCGATATATTTTTGATAATATGAAAAACATGCCTCTCTCAAATCAGACAACCATTTCATAAAATAGTTCTCTCGCACTGGTGAGGTGGTTGCAGCTCGTATAATCTCCCCAGAATAATATACTTGCGTGTCGTCGTTCAATTTATACGGATTTCTATAATTCTCTGTTATATAATGAGGTTTACCCTCATGATTAGACGACGACAATAGTGTTCTGAATTTAGATAGTTGATCAGAATGAATGTTTATTGGTGTTGGAAAAATTATCCATGCTGCGTCTGCAGAGTGGTTGATAGTTGTTCCTAGATATGTAAAATAATCCAACGTACTAGGTAGCAAATTGTCTAGATAAAATACTGAATCAAATGGCGCAGACATATTAGCGGATCTAATGGAATCCAATTGATTGACTATCTTTTGAAAATATACATTTTTATGATCTGATACTTGTAAGAATATAGCAATAATGATAATTCCATCATCGTGTTTTTTTGCCTCTTCATAAGAACTATATTTTTTCTTATTCCAATGAACCAGATTAATCTCTCCAGAGTATTTGTATACATCTATCAAGTGATTGGATCCATAATCGTCTTCCTTTCCCCAATATATATGTATTGTTGATAACACATATTCATTGGGGAGAAACCCTCCACTTATATATCCTCCTTTAAAATTAATCCTTACTAGTTTTCCAGTATTCTGGATAGTGGTTGGTTTCGACTCATTATAATGTATGTCTAACGTCTTCAATCGCGCGTCAGAAATTGCTTTTTTAGTTTCTATATTAATAGGAGATAGTTGTTGAGGCATAGTAAAAATGAAATGATAACTGTCTAGAAATAGCTCTTAGTATGGGATTTACAATGGATGAGGAAGTGATATTTGAAACTCCTAGAGAATTAATATCTATTAAACGAATAAAAGATATTCCAAGATCAAAAGACACGCACGTGTTTGCTGCGTGTATAACAAGTGACGGATATCCGTTAATAGGAGCTAGAAGAACTTCATTCGCATTCCAGGCGATATTATCTCAACAAAATTCAGATTCTATCTTTAGAGTATCCACTAAACTATTACGGTTTATGTACTACAATGAACTAAGAGAAATCTTTAGACGGTTGAGAAAAGGTTCTATCAACAATATCGATCCTCACTTCGAAGAGTTAATATTATTGGGTGGTAAACTAGATAAAAAGGAATCTATTAAAGATTGTTTAAGAAGAGAATTAAAAGAGGAAAGTGATGAACATATAACAGTAAAAGAATTCGGAAATGTAATTCTAAAACTTACAACGAGTGATAAATTATTTAATAAAGTATATATAGGTTATTGCATGGCATGTTTTATTAATCAATCGTTGGAGGATTTATCACATACTAGTATTTACAATGTAGAAATTAGAAAGATTAAATCGTTAAATGATTGTATTAACGACGATAAATACGAATATCTGTCTTATATTTATAATATACTAATTAATAGTAAATGAGCTTTTACAGATCTAGTATAATTAGTCAGATTATTAAGTATAATAGACGACTAGCTAAGTCTATTATTTGCGAGGATGACTCTCAAATTATTACACTCACGGCATTCGTTAACCAATGCCTATGGTGTCATAAACGAGTATCCGTGTCCGCTATTTTATTAACTACTGATAACAAAATATTAGTATGTAACAGACGAGATAGTTTTCTCTATTCTGAAATAATTAGAACTAGAAACATGTATAGAAAGAAACGATTATTTCTGAATTATTCCAATTATTTGAACAAACAGGAAAGAAGTATACTATCGTCATTTTTTTCTCTAGATCCAGCTACTGCTGATAATGATAGAATAAACGCTATTTATCCGGGTGGTATACCCAAAAGGGGTGAGAACGTTCCAGAGTGTTTATCCAGGGAAATCAAAGAAGAAGTTAATATAGACAATTCTTTTGTATTCATAGACACTCGTTTTTTTATTCATGGTATCATAGAAGATACCATTATTAACAAATTTTTTGAGGTAATTTTCTTTGTTGGACGAATATCTCTAACGAGTGATCAAATTATTGATACCTTTAAAAGTAATCATGAAATAAAGGATCTAATATTTTTAGATCCAAATTCAGGTAATGGACTCCAATACGAAATTGCAAAATATGCTCTAGATACTGCAAAACTTAAATGTTACGGTCATAGAGGATGTTATTATGAATCATTAAAAAAATTAACTGAGGATGATTGATTAGAAAATATAAATTAATTTACCATCGTGTATTTTTATAACGGGATTGTCTGGCATATCATGTAGATAGTTACCGTCTACATCGTATACTCTACCATCTACGCCTTTAAATCCTCTATTTATTGATATTAATCTATTAGAATTGGAATACCAAATATTAGTACCCTCAATTAGTTTATTGGTAATATTTTTTTTAGACGATAGATCGATGGCTCTTGAAACCAAGGTTTTCCAACCGGACTCATTGTCTATCGGTGAGAAGTCTTTTTCATTAGCATGAATCCATTCTAATGATGTATGTTTAAACACTCTAAACAATTGTACAAATTCTTTTGATTTGTTTTGAATGATTTCAAATAGGTCTTCGTCTACAGTAGGCATACCATTAGATAATCTAGCCATTATAAAGTGCACGTTTACATATCTACGTTCTGGAGGAGTAAGAACGTGACTATTGAGACGAATGGCTCTTCCTACTATCTGACGAAGAGACGCCTCGTTCCATGTCATATCTAAAATGAAGATATCATTGATTGAGAAGAAACTAATACCCTCGCCTCCGCTAGAAGAGAATACGCATGTTTTAATGCATTCTCCGTTAGTGTTTGATTCTTGGTTAAACTCAGCCACCGCCTTGATTCTAGTATCTTTTGTTCTAGATGAGAACTCTATATTAGAGATACCAAAGACTTTGAAATATAGTAATAAGATTTCTATTCCTGACTGATTAACAAATGGTTCAAAGACTAGACATTTACCATGGGATGCTAATATTCCCAAACATACATCTATAAATTTGACGCTTTTCTCTTTTAATTCAGTAAATAGAGAGATATCAGCCGCAATAGCATCCCCTCCCAATAGTTCTCCCTTTTTAAAGGTGTCTAATGCGGATTTAGAAAATTCTCTATCTCTTAATGAATTTTTAAAATCATTATATAGGGTTGCTATCTCTTGTGCGTATTCTCCCGGATCACGATTTTGTCTTTCAGGAAAGCTATCGAATGTAAACGTAGTAGCCATACGTCTCAGAATTCTAAATGATGATATACCAGTTTTTATTTCTGCGAGTTTAGCCTTTTGATAAATCTCTTCTTGCTTTTTTGACATATTAACGTATCGCATTAATACTGTTTTCTTAGCGAATGATGCAGACCCTTCCACATCATCAAAAATAGAAAACTCGTTATTAACTATGTACGAACATAGGCCTCCTAGTTTGGAGACTAATTCTTTTTCATCGACTAGACGTTTATTCTCAAATAGCGATTGGTGTTGTAAGGATCCTGGTCGCAGTAAGTTAACCAACATGGTGAATTCTTGCACACTATTAACGATAGGTGTAGCCGATAAACAAATCATCTTATGGTTTTTTAACGCAATGGTCTTAGATAAAAAATTATATACTGACCGAGTAGGACGGATCTTACCATCTTCTTTGATTAATGATTTAGAAATGAAGTTATGACATTCATCAATGATGACGCATATTCTACTCTTGGAATTAATAGTTTTGATATTAGTAAAAAATTTATTTCTAAAATTTTGATCATCGTAATTAATAAAAATACAATCCTTCGTTATCTCTGGAGCGTATCTGAGTATAGTGTTTATCCAAGGATCTTCTATCAAAGCCTTTTTTACCAATAAGATAATTGCCCAATTCGTATAAATATCCTTAAGATGTTTGAGAATATATACAGTAGTCATTGTTTTACCGACACCTGTTTCATGGAACAATAAAAGAGAATGCATACTGTCTAATCCTAAGAAAACTCTTGCTACAAAATGTTGATAATCCTTGAGGCGTACTACGTCTGACCCCATCATTTCAACGGGCATATTAGTAGTTCTGCGTAAGGCATAATCGATATAGGCCGCGTGTGATTTACTCATTTATGAGTGATAAGTAATAACTATGTTTTAAAAATCACAGCAGTAGTTTAACTAGCCTTCTCTGATGTTTGTTTTCGATACTTTTTGAATCAGAAGTCATACTAGAATAAAGCAGCGAGTGAACGTAATAGAGAGCTTCGTATACTCTATTCGAAAACTCTAAGAACTTATTAATGAATTCCGTATCCACTGGATCGTTTAAAATACTAAATTGAACAGTGTTCACATCCTTCCAAGACGAAGACTTAGTGACGGACTTAACATGAGACATAAATAAATCCAAATTTTTTTTATAAACATCACTAGCCACCATAATGGCGCTATCTTTCAACCAACTATCGCTTACGCATTTTAACAGTCTAACATTTTTAAAGAGACTACAATATATTCTCATAGTATCGATTACACCTCTACCGAATAGAGTGGGAAGTTTAATAATACAATATTTTTCGTTTACAAAATCAAATAATGGTCGAAACACGTCGAAGGTTAACATCTTATAATCGCTAATGTATAGATTGTTTTCAGTGAGATGATTATTAGATTTAATAGCATCTCGTTCACGTTTGAACAGTTTATTGCGTGCGCTGAGGTCGGCAACTACGGCATCCGCTCTAGTACTCCTCCCATAATACTTTACGCTATTAATCTTTAAAATTTCATAGACTTTATCTAGATCGCTTTCTGGTAACATGATATCATGTGTAAAAAGTTTTAACATGTCGGTCGGCATTCTATTTAGATCATTAACTCTAGAAATCTGAAGAAAGTAATTAGCTCCATATTCCAGACTAGGTAATGGGCTTTTACCTAAAGACAAGTTAAGTTCTGGCAATGTTTCATAAAATGGAAGAAGGACATGTGTCCCCTCCCGGATATTTTTTACAATTTCATCCATTTACAACTCTATAGTTTGTTTTCATTATTATTAGTTATTATCTCCCATAATCTTGGTAATACTTACACCTTGATCATAAGATACCTTATACAGGTCATTACATACAACTACCAATTGTTTTTGTACATAATAGATTGGATGATTGATATCCATGGTGGAATAAACTACTCGAACAGATAGTTTATCTTTCCCCCTAGATACATTGGCCGTAATAGTTGTCGGCCTAAAGAATATCTTTGGTGTAAAGTTAAAAGTTAGGGTTCTTGTTCCATTATTGCTTTTTGTCAGTAGTTCGTTATAAATTCTCGAGATGGGCCCGTTCTCTGAATATAGAACATCATTTCCAAATCTAACTTCTAGTCTAGAAATAATATCGGTCTTATTTTTAAAATCTATTCCCTTGATGAATGGATCGTTAATAAACAAATCCTTGGCCTTTGATTCGGCTGATCTATTATCTCCGTTATAGACGTTACGTTGACTAGTCCAAAGACTTACAGGAATAGATGTATCGATGATGTTGATAGTATGTGATATGTGAGCAAAGACTGTTCTCTTGGTGGCGTCGCTATATGTTCCAGTAATGGCGGAAAACTTTTTAGAAATGTTATATATAAAAGAATTTTTTCGGGTTCCAAACATTAACAGATTAGTATGAAGATAAACACTCATATTATCAGGAACATTATCAATTTTTACATAAACATCGGCATCTTGAATAGAAACAACACCATCTTCTGGAACCTCTACGATCTCGGCAGATTCCGGATAACCAGTCGGTGGACCATCACTAACAATAACTAGATCATCCAACAATCTACTCACATATGCGTCTATATAATCTTTTTCATCTTGTGAGTACCCTGGATACGAAATAAATTTGTTATCAGTATTTCCATAATAAGGTTTAGTATAAACAGAGAGAGATGTTGCTGCATGAACTTCGGTTACTGTCGCCGTTGGTTGGTTTATTTGACCTATTACTCTCCTAGGTTTCTCTATAAATGATGGTTTAATTTGTACATTCTTAACCATATATCCAATAAAGCTCAATTCAGGAACATAAACAAATTCTTTGTTGAACGTTTCAAAGTCGAACGAAGAGTCACGAATAACGATATCGGATACTGGATTGAAGGTCACCGTTACGGTAATTTTTGAATCGGATAGTTTAAGACTACTGAATGTATCTTCCACATCAAACGGAGTTTTAATATAAACGTATACTGTAGATGGTTCTTTAATAGTGTCATTAGGAGTTAGGCCAATAGAAATATCATTAAGTTCACTAGAATATCCAGAATGTTTCAAAGCAATTGTATTATTGATACAATTATTATATAATTCTTCGCCCTCAATTTCCCAAATAACACCGTTACACGAAGAGACAGATACATGATTAATACATTTATATCCAACATATGGCACGTAACCGAATCTTCCCATACCTTTAACTTCTGGAAGTTCCAAACTCAGAACCAAATGATTAAGCGCAGTAATATACTGATCCCTAATTTCGAAGCTAGCGATAGCCTGATTGTCTGGCCCATCGTTTGTCATAACTCCGGATAGAGAAATATATTGCGGCATATATAAAGTTGGAATTTGACTATCAACTGCGAAGACATTAGACCGTTTAATAAAGTCATCCCCACCGATCAAAGAATTAATGATAGTATTATTCATTTTCTATTTAAAATGGAAAAAGCTTACAATAAACTCCGTAGAGAAATATCTATAATTTGTGAGTTTTCCTTAAAGTAACAGCTTCCGTAAACACCGTCTTTATCTCTTAGTAAGTTTATTGTATTTATGACCTTTTCCTTATCTTCATAGAATACTAAAGGCAATAAAGAAATTTTTGGTTCTTCTCTAAGAGCTACGTGAGACTTAACCATAGACGCCAACGAATCCCTACATATTTTAGAACAGAAATACCCAACTTCACCACCCTTGAATGTCTCAATACTAATAGGTCTAAAAACCAAATCTTGATTACAAAACCAACACTTATCAATTACACTATTTGTCTTAATAGACATATCTGCCATAGATTTATAATACTTTGGTAGTATACAAGCGAGTGCTTCTTCTTTAGCGGGCTTAAAGACTGCTTTAGGTGCTGAAATAACCACATCTGGAAGACTTACTCGCTTAGCCATTTAATTACGGAACTATTTTTTTATACTTCTAATGAACAAGTAGAAAACCTCTCATCTACAAAAACATACTCGTGTCCATAATCCTCTACCATAGTAACACGTTTTTTAGATCTCATATGTGCTAAAAAGTTTTCCCATACTAATTGGTTACTATTATTTTTCGTATAATTTTTAACAGTTTGAGGTTTTAGATTTTTAGTTACAGAAGTGATATCGAATATTTTATCCAAAAAGAATGAGTAATTAATTGTCTTAGAAGGAGTGTTTTCTTGGCAAAAGAATACCAAGTGCTTAAATATTTCTACTACTTCATTAATCTTTTCTGTACTCAGATTCAGTTTCTCATCTTTTACTTGATTGATTATTTCAAAGACTAACTTATAATCCTTTTTATTTATTCTCTCGTTAGCCTTAAGAAAACTAGATACAAAATTTGCATCTACATCATCCGTGGATATTTGATTTTTTTCCATGATATCCAATAGTTCCGAGATAATTTCTCCAGAACATTGATGAGACAATAATCTCCGCAATACATTTCTCAAATGAATAAGTTTATTAGACACGTGGAAGTTTGACTTTTTTTGTACCTTTGTACATTTTTGAAATACAGACTCGCAAAAAATACAATATTCATATCCTTGTTCAGATACTATACCGTTATGTCTACAACAGCTACATAATCGTAGATTCATGTTAACACTCTACGTATCTCGTCGTCCAATATTTTATATAAAAACATTTTATTTCTAGACGTTGTCAGAAAATCCTGTAATATTTTTAGTTTTTTTGGTTGTGAATAAAGTATCGCCCTAATAATATTGGTACCGTCTTCCGACAATATAGTAGTTAAATTATCCGAGCATGTAGAAGAACACCGCTTAGGCGGATTCAGTACAATGTTATATTTTTCGTACCAACTCATTTAAATATCATAATCTAAAATAGTTCTGTAATATGTCTAGCGCTAATATATTGATCATAATCCTGTGCATAAATTAAGATACAACAATGTCTTGAAATCATCGACATGGCTTCTTCCATAGTTAGAAGATCATCGTCAAAGTTAGCAACGTGATTCATCAACATTTGCTGTTTTGAGGCAGCAAATACTGAACCATCACCATTCAACCATTCATAAAAACCATCGTCTGAATCCATTGATAATTTCTTGTACTGGTTTTTGAGAGCTCGCATCAATCTAGCATTTCTAGCTCCCGGATTGAAAACAGAAAGAGGATCGTACATCCAAGGTCCATTTTCTGTAAATAGAATCGTATAATGTCCCTTCAAGAAGATATCAGACGATCCACAATCAAAGAATTGGTCTCCGAGTTTGTAACAGACTGCGGACTTTAACCTATACATGATACCGTTTAGCATGATTTCTGGTGATACGTCAATCGGAGTATCATCTATTAGAGATCTAAAGCCGGTGTAACATTCTCCGCCAAACATATTCTTATTCTGACGTCGTTCTACATAAAACATCATTGCTCCATTAACGATAACAGGTGAATGAACAGCACTACCCATCACATTAGTTCCCAATGGATCAATGTGTGTAACTCCAGAACATCTTCCATAGCCTATGTTAGGAGGAGCGAACACCACTCTTCCACTATTGCCATCGAATGCCATAGAATAAATATCCTTGGAATTGATAGAAATCGGACTGTCGGATGTTGTTATCATCTTCATAGGATTAACAACGATGTATGGTGCAGCCTGAAGTTTCATATCGTAACTGATGCCGTTCATAGGTCTAGCCACAGAAACCAACGTAGGTCTAAATCCAACTATAGACAAAATAGAAGCTAATATCTGTTCCTCATCTGTCATAACTTGAGAGCATCCAGTATGAATAATCTTCATTAGATGGGGATCTACCGCATCATCATCGTTACAATAAAAAATTCCCATTCTAATGTTCATAATTGCTTTTCTAATCATGGTATGAATGTTTGCTCTCTGAATCTCTGTGGAAATTAGATCTGATACACCTGTAATCACTATCGGATTATCCTCCGTAAGACGATTAACCAACAACATATAATTATAAGACTTTACTCTTCTAAATTCATAAAGTTGCTGGATTAGACTATATGTGTCTCCATGTACATACGCGTTCTCGAGCGCAGGAAGTTTAATACCGAATAGTGCCATCAGAATAGGATGAATGTAGTAATTAGTTTCTGGTTTTCTATAAATAAAAGACAAATCTTGTGAACTAGACATATCGGTAAAATGCATGGATTGGAATCGTGTAGTCGACAGAAGAATATGATGATTAGATGGAGAGTATATTTTATCTAACTCTTTGAGTTGGTCACCGATTCTAGGACTAGCTCGAGAATGAATAAGTACTAAGGGATGAGTACATTTCACAGAAACACTGGCGTTGTTCAACGTACTCTTTACATGGGAAAGGAGTTGAAATAGCTCGTTTCTATTTGTCCTGACAATATTTAGTTTATTCATAATATTAAGCATATCCTGAATAGTAAAGTTAGATGTGTCATACTTGTTAGTAGTTAGATATTTAGCAATTGCATTCCCATCATTTCTCAATCTCGTACTCCAATCATGTGTGGATGCTACTTCGTCGATGGAAACCATACAATCCTTTTTGATAGGCTGTTGAGATTGATCATTTCCTGTACGTTTAGGTTTGGTACGTTGATTTCTAGCCCCTGCTGATATAAAGTCATCGTCTACAATTTGGGATAATGAATTACATACACTACAAGACAAAGATTTATCAGAAGTGTGAATATGATCTTCATCTACCAAAGAAAGAGTTTGATTAGTATAACTAGATTTTAGTCCCGCGTTAGATGTTAAAAAAACATCGCTATTGACCACGGCTTCCATTATTTATATTCGTAGTTTTTACTCGAAAGCGTGATTTTAATATCCAATCTTATTACTTTTGGAATCGTTCAAAACCTTTGACTAGTTGTAGAATTTGATCTATTGCCCTACGCGTATACTCCCTTGCATCATATACGTTCGTCACCAGATCGTTTGTTTCGGCCTGAAGTTGACGCATATCTTTTTCAACACTCGACATGAGATCCTTAAGGGTCATATCGTCTAGATTTTGTTGAGATGCTGCTCCTGGATTTGGATTTTGTTGTGCTGTTGTACATACTGTACCACCAGTAGGTGTAGGAGTACATACAGTGGCCACAATAGGAGGTTGAAGAGGTGTAACCGTTGGAGTAGTACAAGAAATACTTCCATCCGATTGTTGTGTACATGTGGTTGTTGGTAACGTCTGAGAAGGTTGGGTAGATGGCGGTGTCGTCATCTTTTGATCTTTATTAAATTTAGAGATAATATCCTGAACAGTATTGCTCGGCGTCAACGCTGGAAGGAGTGTACTCGCCGGCGCATCAGTATCTGTAGACAACCAATCAAAAAGATTAGACATATCAGATGATGTATTAGTTTGTTGACGTGGTTTTAGTACAGGAGCAGTACTACTAGGTAGAAGAATAGGAGCCGGTGTAGGTGTCGGAACCGGCTGTGGAGTTATATGAATAGTTGGTTGTAGCGGTTGGGTAGGCTGTCTGCTGGCGGTCATCATATTATCTCTAGCTAGTTGTTCTCGCAACTGTCTTTGATAATACGACTCTTGAGACTTTAGTCCTATTTCAATCGCTTCATCCTTTTTCGTATCCGGATCCTTTTCTTCAGAATAATAGATTGACGACTTTGGTGTAGAGGATTCTGCCAGCCCCTGTGAGAACTTGTTAAAGAAGTCCATTTAAGGCTTTAAAATTGAATTGCGATTATAAGATTAAATGGCAGACACGGACGATATTATCGACTATGAATCCGATGATCTCACCGAATACGAGGATGATGAAGAAGATGGAGAGTCACTAGAAACTAGTGATATAGATCCCAAATCTTCTTATAAGATTGTAGAATCAACATCCACTCATATAGAAGATGCGCATTCCAATCTTAAACATATAGGGAATCATATATCTGCTCTTAAACGACGCTATACTAGACGTATAAGTCTATTTGAAATAGCGGGTATAATAGCAGAAAGCTATAACTTGCTTCAACGAGGAAGATTACCTCTAGTTTCAGAATTTTCTGACGAAACGATGAAGCAAAATATGCTACATGTAATTATACAAGAGATAGAGGAGGGTTCTTGTCCTATAGTCATCGAAAAGAACGGAGAATTGTTGTCGGTAAACGATTTTGATAAAGATGGTCTAAAATTCCATCTAGACTATATTATCAAAATTTGGAAACTTCAAAAACGATATTAGAATTTATACGAATATCGTTCTCTAAATGTCACAATCAAGTCTCTCATATTCAGCAGTTTATTGTCGTACTTTATATCGTGTTCATTAACGATATTTTGCAAAATAGTAATGATTCTATCTTCCTTCGATAGATATTCTTCAGAGATTATTGTCTTATATTCTTTCTTGTTATCCGATATGAATTTGATAAGACTTTGAACATTATTAATACCCGTCTGTTTAATTTTTTCTATAGATATTTTAGTTTTGGTAGATTCTATGGTGTCTGTTAATAGGCATCCAACATCGACATTCGACGTCAATTGTCTATAAATCAGAGTATAAATTTTAGAAATAACATTAGCAAATTGTTGTGCGTTGATGTCGTTATTCTGAAACAGTATGATTTTAGGTAGCATTTTCTTAACAAAGAGAACGTATTTATTGTTACTCAGTTGAACAGATGATATATCCAGATTACTAACGCATCTGATTCCATATACCAAACTTTCAGAAGAAATGGTGTACAATTGTTTGTATTCATTCAATGTCTCCTTTTCAGAAATTAGTTTAGAGTCGAATACTGCAATAATTTTCAAGAGATAGTTTTCATCAGATAAGATTTTATTTAGTGTAGATATGATAAAACTATTGTTTTGTTGGAGAACTTGATACGCCGCATTCTCTGTAGTCGACGCTCTCAAATGGGAAACAATCTCTATTATTTTTTTGGAATCGGATACTATATCTTCGGTATCTTGACGCAGTCTAGTATACATAGAGTTAAGAGAAATTAGAGTTTGTACATTAAGCAACATGTCTCTAAATGTGGCTACAAACTTTTCTTTTTCCACATCATCTAGTTTATTATATACCGATTTCACAACGGCACCAGATTTAAGGAACCAGAATGAAAAACTCTGATAACTACAATATTTCATCATAGTTACGATTTTATCATCTTCTATAGTTGGTGTGATAACACATACCTTTTTCTCCAAGACTGGAACCAACGTCATAAAAATGTTTAAATCAAAATCCATATCAACATCTGATGCGCTAAGACCAGTCTCGCGTTCAAGATTATCTTTACTAATGGTGACGAACTCATCGTATAGAACTCTAAGTTTGTCCATTATTTATTTACAGATTTAGTTGTTTAATTTATTTGTGCTCTTCCAGAGTTGGGATAGTATTTTTCTAACGTCGGTATTATATTATTAGGATCTACGTTCATATGTATCATAATATTAATCATCCACGTTTTGATAAATCTATCTTTAGCTTCTGAAATAACGTATTTAAACAAAGGAGAAAAATATTTAGTTACGGCATCAGACGCGATAACATTTTTTGTAAATGTAACGTATTTAGACGACAGATCTTCGTTAAAAAGTTTTCCATCTATGTAGAATCCATCGGTTGTTAACACCATTCCCGCGTCAGAGTGAATAGGAGTTTGAATAGTTTGTTTTGGAAATAGATCCTTCAATAACTTATAGTTGGGTGGGAAAAAATCGATTTTATCACTAGACTCTTTCTTTTTTACTATCATTACCTCATGAACTATTTCTTGAATGAGTATATGTATTTTCTTTCCTATATCGGTCGCGTTCATTGGAAAATATATCATGTCGTTAACTATAAGAATATTTTTATCCTCGTTTACAAACTGAATAATATCAGATATAGTTCGTAAACGAACTATATCATCACCAGCACAACATCTAACTATATGATATCCACTAGTTTCCTTTAGCCGTTTATTATCTTGTTCCATATTAGCAGTCATTCCATCATTTAAGAAGGCGTCAAAGATAATAGGGAGAAATGACATTTTGGATTCTGTTACGACTTTACCAAAATTAAGGATATACGGACTTACTATCTTTTTCTCAACGTCGATTTGATGAACACACGATGAAAATGTACTTCGATGAGATTGATCATGTAGAAAACAACAAGGGATACAATATTTCCGCATATCATGAAATATATTAAGAAATCCCACTTTATTATATTTCCCCAAAGGATCAATGCATGTAAACATTATACCGTTATCATTAATAAAGACTTCTTTCTCATCGGATCTGTAAAAGTTGTTACTGATTTTTTTCATTCCAGGATCTAGATAATTAATAATAATGGGTTTTCTATTCTTATTCTTTGTATTTTGACATATCCTAGACCAGTAAACAGTTTCCACTTTGGTAAAATCAGAAGACTTTTGAACGCTATTAAACATGGCATTAATGGCAATAACTAAAAATGTAAAATATTTTTCTATGTTAGGAATATGGTTTTTCACTTTAATAGATATATGGTTTTTTGCCAAAATGATAGATATTTTTTTATCCGATGATAGTAAAATATTATTAGTCGCCGTCTCTATAAAAATGAAGCTAGTCTCGATATCCAATTTTATTCTAGAATTGATAGGAGTCGCCAAATGTACCTTATACGTTATATCTCCCTTGATGCGTTCCATTTGTGTATCTATATCGGACACAAGATCTGTAAATAGTTTTACGTTATTAATCATCACGGTATCGCCATCGCTAGATAATGCTAATGTACTATCCAAGTCCCAAATGGAGAGATTTAACTGTTCATCGTTTAGAATAAAATGATTACCTGTCATATTAATAAAGTGTTCATCGTATCTAGATAACAACGACTTATAATTAATGTCCAAGTCTTGAACTCGCTGAATGATCTTTTTTAACCCAGTTAGTTTTAGATTGGTACGAAATATATTGTTAAACTTTGATTCTACAGTAATGTCCAAATCTAGTTGTGGAAATACTTCCATCAACATTGTTTCAAACTTGATAATATTATTATCTACATCTTCGTACGATCCAAATTCCGGAATAGATGTATCGCACGCTCTGGCCACCCAGATAACCAAAAAGTCACACGCTCCAGAATATACATTGTATAAAAAGCTATCGTTTTTTAGTAGTGTTTTTTTCTGAGTATATACGAAAGGATTAAAAATAGTATTATCAACGTAACTATATTCCAAATTATTCTTATGAGAATAGATAATAATATCGTCCTTAATATCTAACAAATTTCCTAAATATCCCTTTAATTGAGTCATTCGAAGCGTTAATAAAATATGTCTCTTAACTATTTCCGGCCGTTGTATATTTAAATGACTTCGTAAGAAATAATATATAGGCGACTTCTCATCTATGTAATCATATGGAGTGAGATATAGGGCTCGTTCTACCTCCTGCCCCTTACCCACCTGTAATACCAATTGCGGACTCACTATATATCGCATATTTATATCGTGGGGTAAAGTGAAAATCTACTACCGATGATGTAAGTCTTACAATGTTCGAACCAGTACCAGATCTTAATTTGGAGGCCTCCGTAGAACTAGGGGAGGTAAATATAGATCAAACAACACCTATGATAAAGGAGAATAGCGGTTTTATATCCCGTAGTAGACGTCTATTCGCCCATAGATCTAAGGATGATGAGAGAAAACTAGCACTACGATTCTTTTTACAAAGACTTTATTTTTTAGATCATAGAGAGATTCATTATTTGTTCAGATGCGTCGACGCTGTAAAAGACGTCACTATTACCAAAAAAAATAACATTATCGTGGCGCCTTATATAGCACTTTTAACTATCGCATCAAAAGGATGCAAACTTACAGAAACAATGATTGAAGCATTCTTTCCAGAACTATATAATGAACATAGTAAGAAATTCAAATTCAACTCTCAAGTATCCATCATCCAAGAAAAACTCGGATACCAGTCTGGAAACTATCACGTTTATGATTTTGAACCGTATTACTCTACAGTAGCTCTGGCTATTCGAGATGAACATTCATCTGGCATTTTTAATATCCGTCAAGAGAGTTATCTTGTAAGTTCATTATCTGAAATAACATATAGATTTTATCTAATTAATCTAAAATCTGATCTTGTTCAATGGAGTGCTAGTACGGGCGCTGTAATTAATCAAATGGTAAATACTGTATTGATTACAGTGTATGAAAAATTACAACTGGCCATAGAAAATGATTCACAATTTACATGTTCATTGGCTGTGGAATCAGAACTTCCAATAAAATTACTTAAAGATAGAAATGAATTATTTACAAAATTCATTAACGAGTTAAAAAAGACCAGTTCATTCAAGATAAGCAAACGTGATAAGGATACGCTATTAAAACATTTTACTTATGACTGGAGTTAGAATTTATAGACGACACATTTCGTTTATCATTGTTACTATTACTATCATTATTAGTATTCTTCTTGTCATCTTGTTCAGAAATATACAGCAATGCTATACCTAATACTAAATACATTATCATGCTTGCAATGGCTCTAACAACAACGAACCAAAATGAATTTGGTCGTAGCTTTTGTTCACAAAAATACATAAAGAAATGTCTACATAAATCTATGGCGCCATTGGCTACTTGAAATAGCGCCAGTCCTCCTACAGATTTTAATATAGCTGTATAACATGACATTTATTCATCATCAAAAGAGACAGAGTCACCATCTGTCATATTTAGATTTTTTTTCATGTGTTCAAAGTATCCTCTACTCATTTCATTATAATAGTTTATCATGCTTAGAATTTTAGGACGGATCAATGAGTAAGACTTGACTAGATCGTCAGTAGTAATTTGTGCATCATCTATTCTGCATCCGCTTCGTCGAATAATGTATAGCATCGCTTTGAGATTCTCCATAGCTATCAAGTCTTTATATAATGACATGGAAATATCTGTGAATGCTTTATACTTCTCCAACATCGATGCCTTAACATCATCACATACTTTAGCATTGAAAATACGTTCTATTGTGTAGATGGATGTAGCAAGATTTTTAAACAACAATGCCATCTTACATGATGATTGTCTCAAGTCTCCAATCGTTTGTTTAGAACGATTAGCTACAGAGTCCAATGCTTGGCTAACTAGCATATTATTATCTTTAGAAATTGTATTCTTCAATGAGGCGTTTATCATATCTGTGATTTCGTTAGTCATATTACAGTCTGACTGGGTTGTAATGTTATCCAACATATCACCTATGGATACGGTACACGTACCAGCATTTGTAATAATCCTATCTAAGATGTTGTATGGCATTGCGCAGAAAATATCTTCTCCTGTAATATCTCCACTCTCGATAAATCTACTCAGATTATTCTTAAATGCCTTATTCTCTGGAGAAAAGATATCAGTGTCCATCATTTCATTAATAGTATACGCAGAAAAGATACCACGAGTATCAATTCTATCCAAGATACTTATCGGTTCCGAGTCACAGATAATTGTTTCCTCTCCTTCGGGAGATCCTGCATAGAAATATCTAGGACAATAGTTTCTATACTGTCTGTAACTCTGATAATCTCTAAAGTCACTAACTGATACCATGAAATTGAGAAGATCAAACGCTGAAGTAATCAATTTTTCTGCCTCGTTTTTACTACAACTAGTTTTCATCAATGTAGTGACGATGTATTGTTTAGTTACTCTTGGTCTAATACTGATGATAGAGATATTATTGCTTCCCATAATGGATCTTCTAGTAGTCACCTTAAAGCCCATTGATGCGAATAGCAGATAGATAAAGTCTTGGTATGACTCCTTTCTAATATAGTACGGACTACCTTTGTCACCCAACTTTATACCCACATAAGCCATAACAACCTCTTTAATAGCCGTTTCATGAGGTTTATCAGCCATGAGCCTGAGTAGTTGAAAGAATCGCATGAATCCCGTCTCAGAAAGTCCTATATGCATGATAGATTTATCTTTCCTGGGAAACTCTCGTATAGTTATAGATGAAATACTCTTCAAAGTTTCTGAAATAAGATTAGTAACAGTCTTACCTCCGACTACTCTGGGTAACAAACATACTCTAATAGGTGTTTTCTCTGCGGAGATAATATCAGAAAGGATAGAGCAATAAGTAGTATTATTGTGATTATAAAGACCGAATACATAACAGGTAGAATTTATAAACATCATGTCCTGAAGGGTTTTAGACTTGTATTCCTCGTAATCTATACCGTCCCAAAACATGGATTTGGTAACTTTGATAGCCGTAGATCTTTGTTCCTTCGCTAACAGGTTAAAGAAATTAATAAAGAATTTGTTGTTTCTATTTATGTCCACAAATTGCACGTTTGGAAGCGCCACGGTTACATTCACTGCAGCATTTTGAGGATCGCGAGTATGAAGTACGATGTTATTGTTTACTGGTATATCTGGAAAGAAATCTACCAGTCTAGGAATAAGAGATTGATATCGCATAGAAATAGTAAAGTTTATAATCTCATCATTGAAGATTACTCTGTTACCATTGTAATAAATTGGTACTCTATCATAATCATCGACAAAGTACTGTTCATACATGATGAGATGTTTATATGTTGGCATAGTAGTGAGATCGACGTTTGGTAATGGCAATGTATTAAGATTAACTCCATAATGTCTAGCAGCATCTGCGATGTTATAAGTGATGTCAAAGCGGGGTTGATCTTGTGCTGTTATATATTGTCTAACACCTATAAGATTATCAAAATCTTGTCTGCTTAATACACCGTTAACAATTTTTGCCTTGAATTCTTTTATTGGTGCATTAATAACATCCTTATAGAGGATGTTAAACAAATAAGTATTATCAAAGTTAAGATCTGGGTATTTCTTTTCTGCTAGAACATCCATTGAGTCGGAGCCATCTGGTTTAATATAACCACCGATAAATCTAGCTCTGTATTCTGTATCCGTCAATCTAATATTAAGAAGGTGTTGAGTGAAAGGTGGAAGATCGTAAAAGCTGTGAGTATTAATAATAGGGTTAGTTTCCGAACTAATGTTAATTGGATGATTAATAATATTTATATTTCCAGCGTTAAGTGTAACATTAAACAGTTTTAATTCACGTGACGTGGTATCAATTAAATAATTAATGCCCAATTTGGATATAGTAGCCTGAAGCTCATCTTGTTTAGTTACGGATCCTAATGAGTTATTAAGAAATACATCGAACGGATGAACGAAGGTTGTTTTAAGTTGGTCACATACTTTGTAATCTAGACATAGATGTGGAAGAACGGTAGAAACTATACGAAATAGATATTCAGAGTCCTCTAATTGATCAAGAGTAACTATTGACTTAATAGGCATCATTTATTTAGTATTAAATGACGACCGTACCAGTGACAGATATACAAAACGACTTAATTACAGAGTTTTCAGAAGATAATTATCCATCTAACAAAAATTATGAAATAACTCTTCGCCAAATGTCTATTCTAACTCACGTTAACAACGTGGTAGATAGAGAACATAATGCCGCCGTAGTGTCATCTCCAGAGGAAATATCATCACAACTTAATGAAGATCTATTTCCAGATGATGATTCACCGGCCACTATTATCGAACGAGTACAACCTCATACTACTATTATTGACGATACGCCACCTCCTACTTTTCGTAGAGAGTTATTGATATCGGAACAACGTCAACAACGAGAAAAAAGATTTAATATTACAGTATCAAAAAATTCTGAAGCAATAATGGAATCTAGATCTATGATAACTTCTATGCCAACACAAACACCATCCTTGGGAGTAGTTTATGATAAAGATAAAAGAATTCAGATGCTAGAGGATGAAGTGGTTAATCTTAGAAATCAACGATCTAATACAAAATCATCTGATAATTTAGATAATTTTACCAGAATACTATTTGGTAAGACTCCGTATAAATCAACCGAAGTTAATAAGCGTATAGCCATCGTTAATTATGCAAATTTGAACGGGTCCCCCTTATCAGTCGAGGACTTGGATGTCTGTTCGGAGGATGAAATAGATAGAATCTATAAAACGATTAAACAATATCACGAAAGTAGAAAACGAAAAATTATCGTCACTAACGTGATTATTATTGTCATAAACATTATTGAGCAGGCATTGCTAAAACTCGGATTTGAAGAAATCAAAGGACTGAGTACCGATATCACTTCAGAAATTATCGATGTGGAGATCGGAGATGACTGCGATGCTGTAGCATCAAAACTAGGAATCGGTAACAGTCCGGTTCTTAATATTGTATTGTTTATACTCAAGATATTCGTTAAACGAATTAAAATTATTTAATTTAATACATTCCCATATCCAGACAACAATCGTCTGGATTAATCTGTTCCTGTCGTCTCATACCGGACGACATATTAATCTTTTTATTAGTGGGCATCTTTTTAGATGGTTTCTTTTTCCCAGCATTAACTGATTCGATACCTAGAAGATCGTGATTGATTTCTCCGACCATTCCACGAACTTCTAATTGGCCGTCTCTAACGGTACCATAAACTATTTTACCAGCATTAGTAACAGCTTGGACAATCTGACCATCCATTGCGTTGAATGATGTAGTTGCTGTTGTTCTACGTCTAGGAGCACCAGAGGTATTTTTAGAGCTCTTGGATGTTGATGTAGAAGACGAGGATTTTGATTTTGGTTTACATGTAATACATTTTGAACTCTTTGATTTTGTATCACATGCACCGGCAGTCACATCTGTTTGAGAATTAAGATTATTGTTGCCTCCTTTGACGGCTGCATCTCCACCGATCTGCGCTAGTAGATTTTTAAGCTGTGGTGTAATCTTATTAACTGTTTCAATATAATCATCGTAACTACTTCTAACGGCTAAATTTTTTTTATCCGCCATTTAGAAGCTAAAAATATTTTTATTTATGCAGAAGATTTAACTAGATTATACAATGAACTAATATGATCCTTTTCTAGATTATTTACGAACTTGGTATTTCTTGTTTCTGGAGGAGGAGAATTTAAATTCGGACTTGGATTCGGATTTTGTGGGTTCTTGATCTTATTATACAGCGTGTATAGGATGGTGACGGTAACTGCTACACAAATACCGATCAACAGAAGAATACCAATCATTTATTGACAATAACTTCACTATGATCAAGTATGTAATAATCATCTTTTCACTAAGTAAGTAGTAATAATGATTCAACAATGACACGATATATGGACGATAATAATTTAGTTCATGGAAATATCGCTATGATTGGTGTGAATGACTCCGCTAACTCTGTGGGGTGCACAGTGCTTTCCCCACATAGAATAAATTAGCATTCCGACTGTGATAATAATACCAAGTATAAACGCCATAATACTCAATACTTTCCATGTACGAGTGGGACTGGTAGACTTACTAAAGTCAATAAAGGCGAAGATACACGAAAGAATCAAAAGAATGATTCCAGCGATTAGCACGCCAGAAAAATAATTTCCAATCATAAGCATCATGTCCATTTAACTAATAAAAATTTTAAATCGCCGAATAAACAAAGTGGAATATAAACCATATAAAAACAATAGTTTGTACTGCAAAAATAATATCTATTTTTGTTTTCGAAGATATGGTAAAATTAAATAGTAGTACACAGCATGTTATAACTAACAGCAGCAACGGCTCGTAATTACTTATCATTTACTAGACGAAAAGGTGGTGGGATATTTTCTTGCTCAAATAATACGAATATATCACCCATCCATTTTATACGATGTTTATATACTCTAATCTTTAATAGATCTATAGATGACGGGTTTACCAATAATATAGATTTTATCGATTCATCTAATTTAAACCCTTCCTTAAACGTGAATGATCTATTATCTGGCATAATGATGACCCTACCTGATGAATCTGACAATGTACTGGGCCATGTAGAATAAATTATCAACGAATTATCGTCTACGAACATTTATATCATTTGTTTTAATTTTAGGACGTGAATAAATAGATATAAAATAGAAAATAACAGATATTACAACCAGTGTTATGGACGCACCCAACCATGTAGGCAGTTTTATTTTATCGTTTACTACAGGTTCTCCTGGATGTACGTCACCAACTGCAGACGTAGTTCTAGTACAATTAGACGTAAGTTCCGCTTGGGAATTTTTTAACGCTAAAGAGTTAACGTTGATCGTACACCCAACGTATTTACATCTAGTTCTTTGAACATCTTGATTATAATATAACCATTTTCTATCTCTAGATTCGTCAGTGCACTCATGTAACCAACATACCCTAGGTCCTAAATATTTATCTCCGGAATTAGATTTTGGATAATTCGCGCACCAACAATTTCTATTTCCTTTATGGTCGTTACAAAAGACGTATAATGCCGTATCCCCAAAAGTAAAATAATCAGGACGAATAATTCTAATAAACTCAGAACAATATCTCGCATCCATATGTTTGGAGCAAATATCGGAATAAGTAGACATAGCCGGTTTCCGTTTTACACGTAACCATTCTAAACAATTGGGGTTTCCAGGATCGTTTCTACAAAAACCAGTCATGAAATCGTCACAATGTTCTGTCTTGTAATTATTATTAAATATTTTTGGACAGTGTTTGGTATTTGTCTTAGAACAACATTTTGCCACGCTATCACTATCACCCAGGAGATAATCCTTTTTTATAAAATGACATCGTTGCCCGGATGCTATATAATCAGTAGCATATTTTAAATCCTTAATATATTCAGGAGTTACCTCGTTCTGATAATAGATTAATGATCCAGGACGAAATTTGAAAGAACTACATGGTTCTCCATGAATTAATACATATTGTTTAGCAAATTCAGGAACTATAAAACTACTACAATGATCTATCGACATACCATCTATCAAACAAAATTTGGGTTTAATTTCTCCTGGAGACGTTTCATAATAATACATATAACTTTCTTCGGCAAACCTAACAGCTCTATTATATTCAGGATAATTAAAATCTAATACCATATATTTGTCTCGTATATCTGCTATTCCTGTCTCTATTTTGATTCTATTAAGAGTAACAGCTGCCCCCATTCTTAATAATCATCAGTATTTAAACTGTTAAATGTTGGTATATCAACATCTATCTTATTTCCCGCAGTATAAGGTTTGTTGCAGGTATACTGTTCAGGAATGGGTACATTTATACTTCTTTTATAGTCCTGTCTTTCGATGTTCATCACAAATGCAAAGAACAGAATAAACAAAATAATGTAAGAAATAATATTAAATATCTGTGAATTCGTAAATACATTGATTGCCATAATAATTACAGCAGCTACAATACACACAATAGACATTCCCACAGTGTTGCCATTACCTCCACGATACATTTGAGTTACTAAGCAATAGGTAATAACTAAGCTAGTAAGAGGCAATAGAAAAGATGAGATAAATATCATCAATATAGAGATTAGAGGAGGGCTATATAGAGCCAAGACGAACAAAATCAAACCGAGTAACGTTCTAACATCATTATTTTTGAAGATTCCCAAATAATCATTCATTATTCCTCCATAATCGTTTTGCATCATACCCCCATCTTTAGGCATAAACGATTGCTGCTGTTCCTCTGTAAATAAATCTTTATCAAGCACTCCAGCACCCGCAGAGAAGTCATCAAGCATATTGTAATATCTTAAATAACTCATTTATATATTAAAAAATGTCACTATTAAAGATGGAGTATAATCTTTATGCCGAACTAAAAAAAATGACTTGTGGTCAGACCATAAGTCTTTTTAATGAAGACGGCGATTTCGTAGAAGTTGAACCAGGATCATCCTTTAAGTTTCTAATACCTAAGGGATTTTACTCCTCTCCTTGTGTAAAGACGAGTCTAGTATTCGAGACATTAACAACGACCGATAATAAAATTACTAGTATCAATCCAACAAATGCGCCAAAGTTATATCCTCTTCAACGCAAAGTCGTATCTGAAGTAGTTTCTAATATGAGGAAAATGATCGAATTAAAACGTCCTCTATACATCACTCTTCACTTGGCATGTGGATTTGGTAAGACTATTACCACGTGTTATCTTATGACCACACACGGCAGAAAAACCATCATTTGCGTACCCAATAAAATGTTAATACATCAATGGAAGACACAGGTAGAGGCAGTCGGATTGGAACATAAGATATCTATAGATGGAGTTAGTAGTCTATTAAAGGAACTAAAGACTCAAAGTCCGGATGTATTAATCGTAGTCAGTAGACATCTGACAAACGATGCATTTTGTAAATATATCAATAAGCATTATGATTTGTTTATCTTGGATGAATCACATACGTATAATCTGATGAACAATACAGCAGTTACAAGATTTTTAGCGTATTATCCTCCGATGATGTGTTATTTTTTAACTGCTACACCTAGACCAGCTAACCGAATTTATTGTAATAGTATTATTAATATTGCCAAGTTATCCGATCTAAAAAAAACTATCTATATAGTAGATAGTTTTTTTGAGCCATATTCCACAGACAATATTAGAAATATGGTAAAACGACTAGATGGACCATCTAATAAATATCATATATATACCGAGAAGTTATTATCTGTAGACGAGCCTAGAAACCAACTTATTCTTGATACCCTGGTAGAAGAATTCAAGTCAGGAACTATTAATAGAATTTTAGTTATTACTAAACTACGTGAACATATGGTATTCTTCTACAAACGATTATTAGATCTTTTCGGAGCAGAGGTTGTATTTATAGGAGACGCCCAAAATAGACGTACTCCAGATATGGTCAAATCGATTAAGGAACTAAATAGATTTATATTCGTATCCACCTTATTTTATTCCGGCACTGGTTTAGATATTCCGAGTTTGGATTCTTTGTTCATTTGCTCGGCAGTAATCAACAATATGCAAATAGAGCAATTACTAGGGAGGGTATGTCGAGAAACAGAACTATTAGATAGGACGGTATATGTATTTCCTAACACATCCATCAAAAAAATAAAGTACATGATAGGAAATTTCGTGCAACGAATTATTAGTCTGTCTGTAGATAAACTCGGATTTAAACAAGAAAGTTATCAGAAACATCAGGAATCTGAACCCGCTTCCGTACCAACATCCTCCAGAGAAGAACGTGTATTAAATAGAATATTTAACTCGCAAAATCGTTAAGAAGTTTAAGAGACGATCCACATGCTGAGCAGGCCAGTGTATTACCCCTCATAGTATTAATATAATCCAATGATACTTTTGTGATGTCGGAAATCTTAACCAATTTAGACTGACAGGCAGAACACGTCATACAATCATCATCGTCATCGATAACTGTAGTCTTGGGCTTCTTTTTGCGACTCTTCATTCCGGAACGCATATTGGTGCTATCCATTTAGGTAGTAAAAAATAAGTCAGAATATGCCCTATAACACGATCGTGCAAAACCTGGTATATCGTCTCTATCTTTATCACAATATAGTGTATCAACATCTTTATTATTGACCTCGTTTATCTTGGAACATGGAATGGGAACATTTTTGTTAACGGCCACCTTTGCCTTAATTCCAGATGTTGTAAAATTATAACTAAACAGTCTATCATCGACACAAATGAAATTCTTGTTTAGACGTTTGTAGTTTACGTATGCGGCTCGTTCTCGTCTCATTTTTTCAGATATTGCAGGTACTATAATATTAAAAATAAGAATGAAATAACATAGGATTAAAAATAAAGTTATCATGACTTCTAGTGCTGATTTAACTAACTTAAAAGAATTACTTAGTCTGTACAAAAGTTTGAGATTTTCAGATTCTGTGGCTATAGAGAAGTATAATTCTTTGGTAGAATGGGGAACATCTACTTACTGGAAAATAGGCGTACAAAAGGTAACTAATGTCGAGACGTCCATATCTGATTATTATGATGAGGTAAAAAATAAACCGTTTAATATTGATCCGGGGTATTATATTTTCTTACCAGTATATTTTGGAAGCGTCTTTATTTATTCAAAGGGTAAAAATATGGTAGAACTTGGATCTGGAAACTCTTTTCAAATACCGGATGAGATTCGAAGTGCGTGTAACAAAGTATTAGATAGTGATAACGGAATAGACTTTCTGAGATTTGTTTTGTTAAACAATAGATGGATAATGGAAGACGCTATATCAAAATACCAGTCTCCAGTTAATATATTTAAACTAGCTAGTGAGTACGGATTAAACATACCCAACTATTTAGAAATTGAAATAGAGGAAGACACATTATTTGACGATGAGTTATACTCTATTATGGAACGCTCTTTCGATGATACATTTCCAAAAATATCTATATCGTATATTAAGTTGGGAGAACTTAAGCGGCAAGTTGTAGACTTTTTCAAATTCTCATTCATGTATATTGAGTCAATCAAGGTAGATCGTATAGGAGATAATATTTTTATTCCTAGCGTTATAACAAAATCAGGAAAAAAGATATTAGTAAAAGATGTAGACCATTTAATACGATCCAAGGTTAGAGAACATACATTTGTAAAAGTAAAAAAGAAAAACACATTTTCCATTTTATACGACTATGATGGGAACGGAACAGAAACTAGAGGAGAAGTAATAAAACGAATTATAGACACTATAGGACGAGACTATTATGTTAATGGAAAGTATTTCTCTAAGGTTGGTATTGCAGGCTTAAAGCAATTGACTAATAAATTAGATATTAATGAGTGTGCAACTGTCGATGAGTTAGTTGATGAGATTAATAAATCCGGAACTGTAAAACGAAAAATAAAAAACCAATCAGTATTTGATTTAAGCAGAGAATGTTTGGGATATCCAGAAGCGGATTTTATAACGTTAGTTAATAACATGCGGTTCAAAATAGAAAATTGTAAGGTTGTAAATTTCAATATTGAAAATACTAATTGTTTAAATAACCCGAGTATTGAAACTATATATGGAAACTTCAACCAGTTCGTCTCAATCTTTAATACCGTTACCGATGTCAAAAAAAGATTATTCGAGTGAAATAATATGCGCCTTTGATATAGGTGCAAAAAATCCTGCCAGAACTGTTTTAGAAGTCAAGGATAACTCCGTTAGGGTATTGGATATATCAAAATTAGACTGGAGTTCTGATTGGGAAAGGCGCATAGCTCAAGATTTGTCACAATATGAATACACTACAGTTCTTCTAGAACGTCAGCCTAGAAGGTCACCGTACGTCAAATTTATCTATTTTATTAAAGGCTTTTTATATCATACATCTGCTGCCAAAGTTATTTGCGTCTCACCTGTCATGTCTGGTAATTCATATAGAGATCGAAAAAAGAGATCTGTTGAAGCATTTCTTGATTGGATGGACACATTCGGATTGCGAGACTCCGTTCCGGATAGACGCAAATTAGACGATGTAGCGGATAGTTTCAATTTGGCTATGAGATACGTATTAGATAAATGGAATACTAATTATACACCTTATAATAGGTGTAAATATAGAAATTACATAAAAAAAATGTAATAACGTTAGTAACGCCATTATGGATAATCTATTTACCTTTCTACATGAAATAGAAGATAGATATGCCAGAACTATTTTTAACTTTCATCTAATAAGTTGTGATGAAATAGGAGATATATATGGTCTTATGAAAGAACGCATTTCCTCAGAGGATATGTTTGACAATATAGTATATAATAAAGATATACATCCTGCCATTAAGAAACTAGTTTATTGCGACATCCAACTTACTAAACATATTATTAATCAGAATACGTATCCGGTATTTAACGATTCTTCACAAGTGAAATGTTGTCATTATTTCGATATAAACTCAAATAATAGCAATATTAGCTCTCGTACAGTAGAGATATTTGAGAGTGAAAAGTCATCTCTTGTATCATATATTAAAACTACCAATAAGAAGAGAAAGGTCAATTACGGCGAAATAAAGAAAACTGTACATGGAGGCACTAATGCAAATTACTTTTCCGGTAAAAAGTCTGATGAGTATCTGAGCACTACAGTCAGGTCCAACATTAATCAACCTTGGATCAAAACCATTTCTAAGAGAATGAGAGTAGATATCATTAATCACTCTATAGTAACGCGTGGAAAAAGCTCTATATTACAAACTATAGAAATTATTTTTACTAATAGAACATGTGTGAAAATATTCAAGGATTCTACTATGCACATTATTCTATCCAAGGACAAGGATGAAAAGGGATGTATAAACATGATTGATAAATTATTCTATGTATATTATAATTTATTTCTGTTGTTCGAGGATATCATCCAAAACGATTACTTTAAAGAAGTAGCTAATGTTGTAAACCATGTACTCATGGCTACGGCATTAGATGAGAAATTATTCCTAATTAAGAAAATGGCTGAACACGATGTTTATGGAGTTAGCAATTTCAAAATAGGGATGTTTAACCTGACATTTATTAAGTCGTTGGATCATACCGTTTTCCCCTCTCTGTTAGATGAGGATAGCAAAATAAAGTTTTTTAAGGGGAAAAAGCTCAATATTGTAGCATTACGATCTCTGGAGGATTGTACAAATTACGTGACTAAATCCGAGAATATGATAGAAATGATGAAGGAAAGATCGACTATTTTAAATAGCATAGATATAGAAACGGAATCGGTAGATCGTCTAAAAGAATTGCTTCTAAAATGAAAAAAAACACTGATTCAGAAATGGATCAACGACTCGGGTATAAGTTTTTGGTGCCTGATCCTAAAGCCGGAGTTTTTTATAGACCGTTACATTTCCAATATGTATCGTATTCTAATTTTATATTGCATCGATTGCATGAAATCTTGACCGTCAAGCGGCCACTCTTATCGTTTAAGAATAATACAGAACGAATTATGATAGAAATTAGCAATGTTAAAGTGACTCCTCCAGATTACTCACCTATAATTGCGAGTATTAAAGGTAAGAGTTATGACGCATTAGCCACGTTCACTGTAAATATCTTTAAAGAGGTAATGACCAAAGAGGGTATATCCATCACTAAAATAAGTAGTTATGAGGGAAAAGATTCTCATTTGATAAAAATTCCGCTACTAATAGGATATGGGAATAAAAATCCACTTGATACAGCCAAGTATCTTGTTCCTAATGTCATAGGTGGAGTCTTTATCAATAAACAATCTGTCGAAAAAGTAGGAATTAATCTAGTAGAAAAGATTACAACATGGCCAAAATTTAGGGTTGTTAAGCCAAACTCATTCACTTTCTCGTTTTCCTCCGTATCCCCTCCTAATGTATTACCGACAAGATATCGCCATTACAAGATATCTCTGGATATATCACAATTGGAAGCGTCGAATATATCATCGACAAAGACATTTATAACGGTCAATATTGTTTTGCTGTCTCAATATTTATCTAGAGTGAGTCTAGAATTCATTAGACGTAGTTTATCATACGATATGCCTCCAGAAGTTGTCTATCTAGTAAACGCGATAATAGATAGTGCTAAACGACTTACCGAATCTATTACTGACTTTAATATTGATACATACATTAATGACCTGGTGGAAGCTGAACACATTAAACAAAAATCTCAGTTAACGATTAACGAGTTTAAATATGAAATGCTGCATAACTTTTTACCTCATATGAACTATACACCCGATCAACTAAAGGGATTTTATATGATATCTTTACTAAGAAAGTTTCTCTACTGTATCTACCACACTTCTAGATATCCAGATAGAGATTCGATGGTTTGTCATCGCATCCTAACGTACGGCAAATATTTTGAGACGTTAGCACATGATGAATTAGAGAATTACATAGGTAACATCCGAAACGATATCATGAACAATCACAAGAACAGAGGCACTTACGCAGTAAACATTCATGTACTAACAACTCCTGGACTTAATCATGCATTTTCTAGTCTATTGAGTGGAAAGTTCAAAAAGTCAGACGGTAGTTATCGAACACATCCTCACTATTCATGGATGCAGAATATTTCTATTCCTAGAAGTGTTGGATTTTATCCGGATCAAGTAAAGATTTCAAAGATGTTTTCTGTCAGAAAATACCATCCAAGCCAATATCTTTACTTTTGTTCATCAGACGTTCCGGAAAGAGGTCCTCAGGTAGGTTTAGTATCTCAATTGTCTGTCTTGAGTTCCATTACAAATATACTAACGTCTGAGTATTTGGATTTGGAAAAGAAAATTTGTGAGTATATCAGATCATATTATAAAGATGATATAAGTTACTTTGAAACAGGATTTCCAATCACTATAGAAAATGCTCTAGTCGCATCTCTTAATCCAAATATGATATGTGATTTTGTAACTGACTTTAGACGTAGAAAACGGATGGGATTCTTCGGTAACTTGGAGGTAGGTATTACTTTAGTTAGGGATCACATGAATGAAATTCGCATTAATATTGGAGCAGGAAGATTAGTCAGACCATTCTTGGTTGTGGATAACGGAGAGCTCATGATGGATGTGTGTCCGGAGTTAGAAAGCAGATTAGACGACATGACATTCTCTGACATTCAGAAAGAGTTTCCACATGTCATCGAAATGGTAGATATAGAACAATTTACTTTTAGTAACGTATGTGAATCGGTTCAAAAATTTAGAATGATGTCAAAGGATGAAAGAAAGCAATACGATTTATGTGACTTTCCTGCCGAATTTAGAGATGGATATGTAGCATCTTCACTAGTGGGAATCAATCACAATTCTGGACCCAGAGCTATTCTTGGATGTGCTCAAGCTAAACAAGCTATCTCTTGTCTGAGTTCGGATATACGAAATAAAATAGACAATGGAATTCATTTGATGTATCCAGAGAGGCCAATTGTGATTAGTAAGGCTTTAGAAACTTCAAAGATTGCGGCTAATTGCTTCGGACAACATGTTACTATAGCATTAATGTCGTACAAAGGTATCAATCAAGAGGATGGAATTATCATCAAAAAACAATTTATTCAGAGAGGCGGTCTCGATATTGTTACAGCCAAGAAACATCAAGTAGAAATTCCATTGGAAAACTTTAATAACAAAGAAAGAGATAGGTCTAACGCCTATTCGAAATTAGAAAGTAATGGATTAGTTAGACTGAATGCTTTCTTGGAATCCGGAGACGCTATGGCAAGAAATATCTCATCAAGAACTCTTGAAGATGATTTTGCTAGAGATAATCAGATTAGCTTTGATGTTTCCGAGAAATATACAGATATGTACAAATCTCGCGTTGAACGAGTACAAGTAGAACTTACTGACAAAGTTAAGGTGCGAGTATTAACCATGAAAGAAAGAAGACCCATTCTAGGAGACAAATTTACTACTAGAACGAGTCAAAAGGGAACAGTCGCGTATATCGCAGATGAAACGGAACTTCCGTACGACGAAAATGGTATCACACCAGATGTCATTATTAATTCTACATCCATCTTCTCTAGAAAAACTATATCTATGTTGATAGAAGTTATTTTAACAGCCGCATATTCTACTAAGCCGTACAACAATAAGGGAGAAAACCGACCTGTCTGTTTTCCTAGTAGTAACGAAACATCTATCGATGCATATATGCAATTCGCTAAACAATGTTATGAGTATTCAAATCCGAAATTGTCCGAGGAAGAATTATCGGATAAAATCTTTTGTGAAAAGATTCTCTATGATCCTGAAACGGATAAGCCTTATGAATCCAAAGTATTTTTTGGACCAATTTATTACTTGCGTCTGAGACATTTAACTCAGGACAAGGCAACCGTTAGATGTAGAGGTAAAAAGACGAAGCTCATTAGACAAGCGAATGAGGGACGAAAACGTGGAGGAGGTATCAAGTTTGGAGAAATGGAGAGAGACTGTTTAATAGCACATGGTGCAGCCAATACTATTACAGAAGTTTTAAAAGACTCAGAAGAGGATTATCAAGATGTGTATATTTGTGAAAATTGTGGAGACATAGCAGCACAAATCAAAAGTATTAATACATGTCTTAGATGTTCAAAACTTAATCTCTCTCCTCTCTTAACAAAAATTGATACCACGCACGTATCTAAAGTATTTCTTACTCAAATGAACGCCAGAGGCGTAAAAGTTAAATTAGATTTCGAACGAAGGCCTCCTTCGTTTTATAAACCATTAGATAAAGTTGATCTTAAACCGTCTTTTCTGGTATAATATTGTTTAGTAGATACTCATCAAGATAAGCTAATTCACTAAACATATTATCGGATTCGGTATTGTTACTCGAGAATAGAGTTCGTTATGCTCCTGATATTCGGAAATCTGTGGAGTTTCAGGTTTTGGTGGAAGTGTAACTGCTACTTGGTGGGATACTGAAGGATATTTCAGAGAGTTGTGGATGTTCGGGTTCGACATCCACCGATGGTGTCACGCCACTAATCGGTTCGGTAACGTCTGTGGATGGAGGTGCTACTTCTACAGAACCTGTAGCCTCAGTTGTCAACGGAGATACATATTCAATGCGCGGAAATGTATAATTTGGTAATGGTTTCTCATGTGGATCTTAAGAAGAAGAGGTAAGATATCTACGAAAGATACCGATCACGTTTCTAGTTCTCTTTTGTAGAACTTTAACTTTTTCTTTCTCAGCATCTAGTTGATATTCCGACCTCTTCACGTTTCGCATGGGTTACCTCCGCAGTTTTTACAAGCGATTTCACGTTCCAGATCACGTTCAGCCTTCATACGTCTCTCCCTCTCTCTATCGAGTTTATCAGAGCAGTCTTTCTGAAGGCGATCGAACTCCATAAATTTCTCCAACGCTTTGATTGTTTCCATAGATTTCCGAAGTTTAGCTTCTAGGACGGCGATTCTTTTTTTTTTTTTTTTTTTTTTTTTCGAATTCACGGGGTACAACCGTTTCCATTACCACCATCTCTATGTTTCTTTTCTAGATCGGCAATCTTTCTCAATCTTTCTCAACATTTCATCCCCATACCTTTTCATTCCTCGAGTCTATTGTCGTCGAAATATCGTTCCAGCTCCTTTTCGACCTCAATAACTTTAGCACGTTGTTTCATCAAGCTCTCTCTTGTAGTACTATCATTTTTATCTGATTCCCTGACACGTTTAAGATCTTCATGTAATTGAGTCAGCTCTTGACGCAATCTCTTAACTAACTTCCTCTCTTGCTTCTTCGTCATAGTACTTACAATCACTATGGGATCCATTGTTACCACGTCTGTACTCGACGAGCTCACGTTTAAGAGATTCAATTTCCAGTTTGTATCGGTCCATGTCTCCATTGCTACACCACCATTAGATTTACAGGCTGCTAGTTGTCGTTCGAGATCAGAAATACGTGTTTTCTTGGAATGGATTTCGTCGATGTACTTGTCATGATTGGCATCGAAACACTTATTAAGTTCTTTTTTTCAATTCTACGATTTTATTTCTTTCGCGAGTCAATTCCCTCCTGTAGTAACTATCAGTTTTGTCAGATTCACGCTCTCTACGTAGACTTTCTTGTAAGTTACTAATTTGTTCCCTGGCATTACCGAGTTCAGTTTTATATGCCGAATAGAGTTCTGATTCATCCTTTGAGAAGATCTCTAGCGATCGTTCAAGATCCCTGATTCTAGTCTTTAGCCTATTTACCTCCTCAGAAGATGCTCCGTTACCGTTTTTACAATCGTTAAGATGTCTATCAAGATCCATGATTCTATCTCTTTTCCATATCAGCATTGATTTCATTATTACGTTCGCAGTCGTTCAACTGTATTTCAAGATCTGAGATTCTAGATTGTAATCTCTGTAGCATTTCCACGGCATTCACTCAGTTGTCTTTCAAGATCTGAGATTCTAGATTGGAGTCTGCTAATCTCTGTAAGATTTCCTCCTCCGCTCTCGATGCAGTCGGTCAACTTATTCTCTAGTTCTCTAATACGCGAACGCAGTGCATCAACTTCTTGTGTGTCTTCTTGATTGCGTGTGCATTCATCGAGTCTAGATTCGAGATCTCTAACGTGACGTCGTTCTTCCTCAAGTTCTCTGTGTACTACAGAAAGCGTGTCCCTATCTTGTTGATATTTAGCAATTTCTGATTCTAGAGTACTGATTCTACTCACGTATGTACTAATAGTTGTCTTAGCCTTATCAAGATCCTCCTTGTATTTGTCACATTCCTTGATATCCATACGAAGTCTGGACAGTTCCCATTCGACATTACGACGTTTATCGATTTCAGCTCGGAGATCGTCGTCGCGTTGTTTTAGCCACATACGACTAAGTTCAAGTTCTCGTTGACAAGATCCATCTACTTTTCCATCCCTAATAGTATCCAGTTCCTTTTCTAGTTCTGACCGCATTTCTCGTTCCATATCAAGAGATTCTCTCAATTCTCGTATAGTCTTCTTATCAATTTCTGATGAATCTGAACCATCATCTGTCCCATTTTGTTGCATATCCCTGAGTTCTTTGATCTCTGTTGTAAGTCTGTCGATTCTTTCGGTTTTATAAACAGAATCCCTTTCCAAAGTCCTAATCTTACTGAGTTTATCATTAAGTTCTTCATTCAATTCAGTGAGTTTTCTCTTGGCTTCTTCCAAGTCTGTTTTAAACTCTCCATCATTTCCGCATTCTTCCTCGCATTTATCTAACCATTCAATTAGTTTATTAATAACTAGTTGGTAATCAGCGATTCCTATAGCCGTTCTTGTATTTGTGGGAACATAATTAGGATCTTCTAATGGATTGTATGGCTTGATAGCATCATCTTTATCATTATTAGGTGGGGGATGGACAACCTTAATTGGTTGGTCCTCCTTATCTCCTCCAGTAGCATGTGGTTCTTCAATACCAGTATTAGTAATAGGCTTAGACAAATGCTTGTCGTACGCGGGCACTTCCTCATCCATCAAGTATTTATAATCGGGTTCTGTTTCAGAATATTCTTTTCTAAGAGACGCGACTTCAGGAGTTAGTAGAAGAACTCTGTTTCTGTATCTATCAACGCTGGAATCGATACTCAAGTTAAGGATAGCGAATACCTCATCGTCATCATCCGTATCTTCTGAAACGCCATCATATGACATTTCATGAAGTCTAACGTATTGATAAACAGAATCAGATTTAGTATTAAACAGATCCTTGACCTTTTTAGTAAATGCATATGTATATTTTAGATCTCCAGATTTCATAATATGATCGCATGCCTTAAATGTCAATGCTTCCATGATATAGTCTGGAACACTAATGGGTGACGAAAAAGATACAGCACCATATGCTACGTTGATAAATAGATCTGAACCACTAAGTAGATAATGATTAATGTTAAGGAAGAGGAAATATTCAGTATATAGATATGCCTTAGCATCATATCTTGTACTAAACACGCTAAACAGTTTATTGATGTGATCAATTTCCAACAGAACAATTAGAGCGGCAGGAATACCAACAAACATATTACCACATCCGTATTTTCTATGAATATCACATATCATATTAAAAAATCTTGATAGAAGAGCGAATATCTCGTCTGACTTAATGAGATGTAGTTCAGCAGCATAAGTCATAACTGTAAATAGAACATACTTTCCTGTAGTGTTGATTCTAGACTCCACATCAACACCATTATTAAAAATAGTTTTATATACATCTTTAATCTGCTCTCCGTTAATCGTCGAACGTTCTAGTATACGGAAACACTTTGATTTCTTATCTGTAGTTAATGACTTAGTGATATCACGAAGAATATTACGAATTACATTTCTTGTTTTTCTTGAGAGACCTGATTCAGAACTCAACTCATCGTTCCATAGTTTTTCTACCTCAGTGGCGAAATCTTTGGAGTGTTTGGTACATTTTTTAATAAGGTTCGTGACCTCCATTTATTATAAAAAATTTTTATTCAAAACTTAACTACAATCGGGTAATTATAAGATCGTAGATCTCCCATGTGGTGGAATACTACCATCTATCGCATGTTGATGGACAGTAGGTAATGGCCATGGGAACAGTAATGTTTGCATATTTATCTTTCTTGCTAGTATTACTGTATATTGTCCCAATGTTTCAATGTGATGTTCTAACCTATCAACTGCCACTGTATCACAACAATAATGTCCGATGGAATTAAGATTATGATCCAATGTGTTTAATATATGATTATCAAGTCTTATACGATCCGCGTCTTTTTTGACAGGATCAGGCTCTTCTACAGGAAGAAGTTTCGGCCTCTTATGATAGTCATGTCTGGGAAATGGTGGTCTAGGATGAGGATCAGGTATCGGAGTAGGTTTTGGATTATAATCATCATCATCATCATCATCATCATCATCATCATCATCATCTATGATATCATCATCTTCGATATTTATTTTGCTATCTTGATAATGTCCTATATCAGTTGCATTTTCAGCACTCGACTGAATATTAGTACATTCATTGTCTATTATTAACGTATTTCTAAACCCAAAATGTATATGTTGAACATCACTACTATAGTTGATGAGTCTTATAGCATGAATTCGCTTATCGTTATCGGGTTTATCTTCTGTCACCTTAACAATTCCTTTTTTATTAAACTCTGCATAATCATAACCATTTCTATTGTTTGTTCTAATATAAACGAGTATAGCATCATTGCTAAATTTTTCAATAGTATCAAAAACAGAATATCCTAAACCATATAATATATATTCAGGAACACTCAAACTAAATGTCCAGGATTCTCCTAAATACGTAAACTTTAATAGTGCTAAATCATTCAAAAATCTACCGCTTATAGATAGATAGTACATGAATGCGTATAGTAGTCTACCTATCTCTTTATTATGAAAACCGACATTACGATCATATATTTCGTGATATACATGTGACCCGTTTACGTTAAACCATAAATACATGGGTGATCCTATAAACATGAATTTATTTCTAATTCTCAGAGCCATAGTTAATTGACCGTGTAATATTTGTTTACATGCATACTTGATACGATCATTAATAAGATTTTTATCATTGCTCGTTATTTCAGAATCGTATATATAAGGAGTACCATCATGATTCTTACCAGATATTATACAAAATACTATATATAAAATATATTGACCCACGTTAGTAATCATGTAAATGTTTAATGTTTTAAATTTTGTATTTAATGATCCATCATCATATGCTAGCATGGTCTTGTGATATTCATTCTTTAAAATATAATATTGTGTTAGCCATTGCATTGGAGCTCCTAATGGAGATTTTCTATTCTCGTCCATTTTAGGATATGCTTTCATAAAGTCCCTAATAACTTCGTGAATAATGTTTCTATGTTTTCTACTGATGCATGTATTTGCTTCGATTTTTTTATCCCATGTTTCATCTATCATAGATTTAAACGCAGTAATGCTCGCAACATTAACATCTTGAACCATTGGTACAATTCCGTTCCATAAATTTATAATGTTCGCCATTTATATAACTCATTTTTTGAATATACTTTTAATTGAACAAAAGAGTTAAGTTACTCATATGGATGCCGTCCAGTCTGTACATCAATCTTTTTAGCCAGAGATATCATAGCCGCTCTTAGAGTTTCAGCGTGATTTTCCAACCTAAATAGAACTTCATCGTTGCGTTTACAACACTTTTCTATTTGTTCAAACTTTGTTGTTATATTAGTAATCTTTTTTTCCAAATTAGTTAGCCGTTGTTTGAGAGTTTCCTCATTATCGTCTCCATAGGCTTTAACAATTGCTTCGCGTTTAGTCTCTGGATTTTTAGCAGCCTTTGTAGAGAAAAATTCAGTTGCTGGAATTGCAAGATCGTCATCTCCGGGGAAAAGAGTTCCGTCCATTTAAAGTACAGATTTTAGAAACTGACACTCTGTGTTATTTATATTTGGCGCAATACATGGATTATAAATATCGATGTTAATAACATCAGAAAATGTAAAGTCTATACATTGTCGCATCGTGTTAAATTTTCTAATGGATCTAGTATTATTGGGTCCAACTTCTGCCTGAAATCCAAATATGGAAGCGGATACAAAACCGTTTCCTGGATAAACCACACATCTCCACTTTTGCTTTACATCAGAAATTGTGTCATTGACATCTTGAACTCTCCTATCTAATGCCGGTGTTCCACCTATAGATTTTGAATACTCGAATGCTGCATGAGTAGCATTGAATTCCTTAATATTGCCATAATTTTCATATATTGAGTAACTCTGGATAAAAAGTAAACACACCGCAGCCGTCGCTACTACAATAAAAAAAATTGATAGAGAGTTCATTTATAATCTATTAGAAGCTGATAAAATTTTTTTACACGCGTCAGACAATGCTTTAATAAATAGTTCAACATCTACTTTTGTCATATCGAACCGATGGTATGATTCTAACCTAGAATTACATCCGAAAAAGTTGACTATGTTCATAGTCATTAAGTCATTAACGAACAACATTCCAGACTCTGGATTATAAGACGATACTGTTTCGTCACAATCACCCACCTTAATCATGTGATTATGAATATTGGCTATTAGAGTACCTTCTAAGAAATCTATAATATCTTTGAAACACGATTTAAAATCAAACCACGAATATACTTCTACGAAGAAAGTTAGTTTACCCATAGGAGAGATAACTATAAATGGAGATCTAGATACAAAATCCGGATCTATGATAGTTTTAACATTATTATATTCTCTATTAAATACCTCCACATCTAAAAATGTTAATTTTGAAACTATGTCTTCGTTTATTACCGTACCTGAACTAAACGCTATAAGCTCTATTGTTTGAGAACTCTTTAAACGATATTCTTGAAATACATGTAACAAAGTTTCCTTTAACTCGGTCGGTTTATCTACCATAGTTACAGAATTTGTATCCTTATCTATAATATAATAATCAAAATCGTATAAAGTTATATAATTATCGTGTTCAGATTGTGATCTTTTCAAATAGACTAAAAACCCCATTTCTCTAGTAAGTATCTTATGTATATGTTTGTAAAATATCTTCATGGTGGGAATATGCTCTACAGCAGTTAGCCATTCCTCATTGACAGCTGTAGATGTATTATACAAAACTACTCCAATGTTTAACAAGGGCCATTTTACGAGATTATTAAATCCTTGTTTGATAAATGTAGCCAATGCGGGTTCGAGTTCAACGACGATTGAATTCTCTTCCCGTGGATGCTGCATGATGAACGACGGGATGTTGTTGTTCTATTGATTTGGAATTCTTTTTCGACTTTTTGTTTATATTAAATATTTTAAAATTTATGGCTGATAGTAATTCATGTACTACGGATAATGTAGACGTGTATTGCATATCGATATCTTTATTATTAGATAAATTTATCAATAAATGTGAGAAGTTTGCCTCGTTAAGGTCTTCCATTTAAATATTATATAAATATTTGTGTTTGTATTTTATTCGTCTTTTATGGGATAGTTTTTAACTAGTAAAGCTGTAATTACATACTTTGTCCGTAAAACATAAATATAAATACCCGCTTTTATCAAACGTTCCAAAAAGTCGGCAGCTGACATTTTTAACATGACATCTATTTTAAATACACTTAGGTTTTTAGAAAAAACATCATTTTATAATTGTAACGATTCAATAACTAAAGAAAAGATTAAGATTAAACATAAGGGAATGTTATTTGTATTTTATAAGCCAAAGCATTCTACCGTTGTTAAATACTTGTCTGGAGGAGGTATATATCATGATGATTTGGTTGTATTGGGGAAGGTAACAATTAATGATCTAAAGATGATGCTATTTTACATGGATTTATCATATCATGGAGTGACAAGTAGTGGAGCAATTTACAAATTGGGATCGTCTATCGATAGACTTTCTCTAAATAGGACTATTGTTACAAAAGTTAATAACAATTATAACAATTATAACAATTATAACAATTATAATTGTTATAATAATTATAATTGTTATAATTATGATGATACATTTTTTGACGATGATGATTGATCACTATTACACAATTTTGTTTTTGTACTTTCTAATATAGTGTTTAGGTTCTTTTTCATATGAGAATATTGACTTACTAAAATATCTATGTTTAACTTTTGTTCTATAACGTCCTTATCGGCGGTATCGGTACATATACGTAATTCACCTTCACAAAATACGGAGTCTTCGATAATAATAGCCAATCGATTATTGGATCTAGCTGTCTGTATCATATTCAACATGTTTAATATATCCTTTCGTTTCCCCTTTACAGGCATCGATCGTAGCATATTTTCCGCGTCTGAGATGGAAATGTTAAAACTGCAAAAATGCGTAATGTTAGCCCGTCCTAATATTGGTACGTGTCTATAAGTTTGGCATAGTAGAATAATAGACGTGTTTAAATGCCTTCCAAAGTTTAAGAATTCTATTAGAGTATTACATTTTGATAGTTTATCACCTACATCATCAAAAATAAGTAAAAAGTGTGCTGATTTTTTATGATTTTGTGCGACAGCAATACATTTTTCTATGTTACTTTTAGTTCGTATCAGATTATATTCTAGAGCTTCCTGACTACTAACGAAATTAATATGATTTGGCCAAATGTATCCATCATAATCTGGGTTATAAACGGGTGTAAACAAGAATATATGTTTATATTTTTTAACTAGTGTAGAAAACAGAGATAGTAAATAGATAGTTTTTCCAGATCCAGATCCTCCTGTTAAAACCATTCTAAACGGCATTTTTAATAAATTTTCTCTTGAAAATTGTTTTTCTTGAAAACAATTCATAATTATATTTACAGTTACTAAATTAATTTGATAATAAATCAAAATATGGAAAACTAAGGTCGTTAGTAGGGAGGAGAACAACGAAGGCATATCGTGATATAAATAACATTTATTATCATGATGACACCAGAAAACGACGAAGAGCAGACATCTGTGTTCTCCGCTACTGTTTACGGAGACAAAATTCAGGGAAAGAATAAACGCAAACGCGTGATTGGTCTATGTATTAGAATATCTATGGTTATTTCACTACTATCTATGATTACCATGTCCGCGTTTCTCATAGTGCGCCTAAATCAATGCATGTCTGCTAACAAGGCTGCTATTACTGACTCCGCTGTTGCCGTTGCTGCGGCATCATCTACTCATAGAAAGGTTGTGTCTAGCACTACACAATATGATCACAAAGAAAGCTGTAATGGTTTATATTACCAGGGTTCTTGTTATATATTACATTCAGACTATAAGTCATTCGAGGATGCTAAAGCAAACTGCGCTGCGGAATCATCAACACTACCCAATAAATCCGATGTCTTGACTACCTGGCTCATTGATTATGTTGAGGATACATGGGGATCTGATGGTAATCCAATTACAAAAACTACATCCGATTATCAAGATTCTGATGTATCACAAGAAGTTAGAAAGTATTTTTGTACATAAATAAATGAAATCGCTTAATAGACAAACTGTAAGTAGGTTTAGGAAGTTGTCGGTGCCGGCCGCTATAATGATGTTACTCTCAACCATTATTAGCGGCATAGGAACATTTCTGCATTACAGAGAAGAACTGATGCCTAGTGCTTGCGCCAATGGATGGATACAATACGATAAACATTGTTATCTGGATACCAACATTAAAATGTCTACGGATAATGCAGTTTATCAGTGTCGCAAATTACGAGCTAGATTGCCTAGACCTGATACTAGACATCTGAGAGTATTGTTTAGTATTTTTTATAAAGATTATTGGGTAAGTTTAAAAAAGACCAATGATAAATGGTTAGATATTAATAATGATAAAGATATAGATATTAGTAAATTAACAAATTTTAAGCAACTAAACAGCACGACGGATTCTGAGGCGTGTTATATATACAAGTCTGGAAAACTGGTTAAAACAGTATGTAAAAGTACTCAATCTGTACTATGCGTTAAAAGATTCTACAAGTGACAACAAAAAATGAATTAATAGTAAGTCGTTAACGTACGCCGCCATGGACGCCGCGTTTGTTATTACTCCAATGGGTGTGTTGACTATAACAGATACATTGTATGATGATCTCGATATCTCAATCATGGACTTTATAGGACCATACATTATAGGTAACATAAAAATTGTCCAAATAGATGTACGGGATATAAAATATTCCGACATGCAAAAATGCTACTTTAGCTATAAGGGTAAAATAGTTCCTCAGGATTCTAATGATTTGGCTAGATTCAACATTTATAGTATTTGTACAGCATACAGATCAAAAAATACCATCATCATAGCATGCGACTATGATATCATGTTAGATATAGAAGGTAAACATCAACCATTTTATCTATTCCCATCTATTGATGTTTTTAACGCTACAATCATAGAAGCGTATAATCTGTATACAGCTGGAGATTATCATCTGATCATCAATCCTTCAGATAATCTGAAAATGAAATTGTCGTTTAATTCTTCATTTTGTATATCAGACGGCAATGGATGGATTATAATTGATGGGAAATGTAATAGTAATTTTTTATCATAAAAGTTGTAAAGTAAATAATAAAACAATAAATATTGAACTAGTAGTATGTTGTATATTGAGCAATCAGAGATGATGCTGGTACCTCTTATCACGGTGACCGTAGTTGCGGGAACAATATTAGTATGTTATATATTATATATTTGTAGGAAAAAGATACGTACTGTCTATAATGACAATAAAATTATCATGACAAAATTAAAAAAGATAAAGAGTCCTAATTCCAGCAAATCTAGTAAATCAACTGATAGCGAATCAGACTGGGAGGATCACTGTAGTGCTATGGAACAAAACAATGACGTAGATAATATTTCTAGAAATGAGATATTGAACGATGATAGCTTCGCTGGTAGTTTAATATGGGATAACGAATCCAATATCATGGCGCCTAGCACAGAACACATTTACGATAGTGTTGCTGGAAGCACGCTGCTAATAAATAATGATCGTAATGAACAGACTATTTATCAGAATACTACAGTAGTAATTAATGATACAGAGACTGTTGAAATACTTAATGAAGATACCAAACAGATTCCTAGCTATTCTTCCAATCCTTTCGTAAATTATAATAAAACCAGTATTTGTAGCAAGTCAAATCCGTTCATTGCAGAACTCAACAATAAATTTAGTGATAATAATCCGTTTAGGAGAGCACATAGTGACGATTATCTTAATAAGCAACAAGATCATGAATACGATGATATAGAATCATCGGTTGTATCATTGGTCTGATTAGTTTCCTTTTTATAAAATTGAAGTAATATTTAGTATTAATTACCGCCGATGCATTATACAAATATGGAGATATTCCCTGTATTCGGCATTTCTAAAATTAGCAATTTTATTGCTAATAATGACTGTAGATATTATATAGATGTAGAGCATCAAAAAATTATATCTGATGAGATCAATAGACAGATGGATGAAACGGTACTTCTTACCAACATCTTAAGCGTAGAAGTTGTAAATGACAATGAGATGTACCATCTTATTCCCCATAGACTATCGACTATTATACTCTGTATTAGTTCTGTTGGAGGATGTGTTATCTCTATAGATAATGACGTCAATGACAAAAATATTCTAACATTTCCCATTGATCATGCTGTAATCATATCCCCACTGAGTAAATGTGTCGTAGTTAGCAAGGGCCCTACAACCATACTGGTTGTTAAAGCGGATATACCCAGCAAACGATTGGTAACATCGTTTACAAACGACATACTGTATGTAAACAATCTATCACTGATTAATTATTTACCGTCGTCTGTATTCATTATTAGACGAGTCACCGACTATTTGGATAGACACATATGTGATCAGATATTTGCTAATAATAAGTGGTATTCCATTATAACTATCGACGATAAGCAATATCCTATTCCATCAAATTGTATAGGTATGTCTTCTGCCAAGTACATAAATTCGAGCATCGAGCAAGATATTTTGATCCATGTTTGTAACCTCGAGCATCCATTCGACTCAGTCTACAAAAAAATGCAGTCGTACAATTCTCTACCTATCAAGGAACAAATATTGTATGGTAGAATTGATAATATAAATATGAGCATTAGTATTTCGGTGGATTAATAGATTTCTCTAGTATGGGATCATTAATCATCTCTAAATACATCATAAAAAAGCTATTATCAAATACTGTACTGAATGGATTCATTCTTTTCTCTTTTTATGAAACTCTGTTGTATATCTACGGATAAAACTAGAAGCAAAAAATCTGATAGGAAGAATAATGATTATATGGAGGAACACGATTATTATAAAATAACAATAGTTCCTGGTTCCTCTTCCACGTCTACTAGCTCGTGGTATTATACACATGCCTAGTAATAGTCTCTTTGCGTTGACGGAAAGCAGACTAGAAATAACAGGCCAAAATGTTCAGACACCATAATAGTTCCCAACCCAGATAATAACAGAGTTCCATCAACACATTCCTTTAAACTCAATCCCAAACCCAAAACCGTTAAAATGTATCCAGCCAATTGATAGTAGATAATGAGGTGTACAGCACATGATAATTTACACAGTAACCAAAATGAAAACACTTTAGTAATTATAAGAAATATAGACGGTAATGTCATCATCAACAATCCAATAATATGCCTGAGAGTAAACATTGACGGATAAAACAAAAATGCCCCGCATAACTCTATCATGGCAATAACGCAACCAAACACTTGTAAAATTCCTAAATTAGTAGAAAATACAACTGATATCGATGTATAAGCGATTTCGAGGAATAATAAGAACAAAGTAATTCCCGTAAAGATAAACATCAACATTGTTTGGTAATCATTAAACCAATTAGTATGACGTTGAATTAATTTCACAGTATATTTTATTCCAGTATTATCCCCGCATGTATACGTACCTGGTAAGATATCTTTATATTCCATAATCAATGAGACATCACTATCCGATAACGAATGAAGTCTAGCACTAGTATGCCATTTACTTAATATGGTCGTCTTGGAAGTTTTATTATAAGTTAAAATATCATGATTGTCCAATTTCCATCTAATATACTTTGTCGGATTATCTATAATACACGGAATAATGATGGTATCATTACATGCTGTATATTCTATAGTCTTTGTAGATGTTATAACCACAAAAGTACAGAGGTATATCAACAATATTCTAACTCTTAACATTTTTATTTATTTAAAATGATACCTTTGTTATTTATTTTATTCTTATTTTGCTAACGGTATCGAATGGCATAAGTTTGAAACGAGTGAAGAAATAATTTCTACTTACTTAATAGATTATGTGGTAACGGGTGTTATTAATGGGGATGTATATACATTTTCAAATAATGAACTAAACAAAACTGGGTTAACTAATAACAATAATTATATCACAACATCTATAAAAGTAGAGGATAAGGATACATTAGTAGTATGCGGAACCAATAACGGAAATCCCAAATGTTGGAAAATAGACGGTTCATACTACCCAAAACATATAGGTAGAGGATACGATCATCAAAATAGCAAAGTAACGATAATCAGTCACAATGAATGTGTACTATCCGACATAAACATATAAAAAGAAGGAATTAAACGATGGAGAAGATTTGACGGACCATGTGGTTATGATTAAACGAGTTAAGTTTTTTAAGAAGCCTTAGAAGAGAGGCTATTGGGTATGAGAATCCGAAATATTAAACCAGACAACCCCATATAATTTTATAGCTAAGAATGCCGCGAAGAATGGAACTAATAAAAACGGAAATATTTGTAGCACAACGAATAACTCCCAAACTGCATTCATGTTACACTATATAACACTACTTCGGTTAGATGTTTTAGAAAAAATAAATATCACCGTACCGTTTTGTTGTATAAAAATAACAATTAACAATTATCAATTTTTTTCTTTAATATTTTACGTGGTTGACCATTCTTGGTGGTAAAATAATCTCTTAGTGTTGGAATGGAATGCTGTTTAATGTTTCCACACTCATCGTATATTTTGACGTATGCAGTCACATCGTTTACGCAATAGTCAGACTGTAGTTCTATCATGCTTCCTACGTTAGAAGGAGGAACAGTTTTAAAGTCTCTTGGTTTTAATCTATTGTCATTAGTTTTCATGAAATCCTTTGTTTTATCCACTTCACATTTTAAATAAATGTCCACTATACATTCTTCTGTTAATTTTACTAGATCATCATGAGTCATAGAATTCATAGGTTCCGTAGTCCATGGATCCAAACTAGCAAACTTCGCGTATACGGTATCGCGATTAGTGTATACACCAACTGTATGAAAATTAAGAAAACAGTTTAATAAATCTACAGAAATATTTAATCCTCCGTTTGATACAGATGCGCCATATTTATGGATTTCGGATTCACACGTTGTTTGTCTAAGGGGTTCGTCTAGTGTTGCTTCTACATAGACTTCGATTCCCATATATTCTTTATTGCCAGAATCACATACCGATTTATCATACGCTGGTTCACTTGTTTGAAAACTAAATGGTAGTAGATACATCAAAATAATAAATAATAAGTACATTCTGCAATATTGTTATCGTAATTGGAAAATTGGTATTCAAGTGAGCTGGATTATGTGAGTATTGGATTGTATATTTTATTTTATATTTTATATTTTATATTTTATTTTATATTTTATATTTTATTTTATATTTTGTAGTAAGAATAGAATGCTAATGTCAAGTTTATTCGAATAGATGTCTTATTAAAAAACATATATAATAAATAACAATGGCTGAATGGCATAAAATTATCGAGGATATCTCAAAAAATAATAAGTTCGAGGATGCCGCCATCGTTGATTACAAGACTACAAAGAATGTTCTAGCGGCTATTCCTAACAGAACATTTGCAAAGATTAATCCGGGTGAAGTTATTCCCCTCATCACTAATCATAATATTCTAAAACCTCTTATTGGTCAGAAATTTTGTATTGTATATACTAACTCTCTAATGGATGAGAACACGTATGCTATGGAGTTGCTTACTGGGTACGCCCCTGTATCTCCGATCGTTATAGCGAGAACTCATACCGCACTTATATTTTTGATGGGTAAGCCAACAACATCCAGACGTGATGTGTATAGAACATGTAGAGATCACGCTACCCGTGTACGTGCAACTGGTAATTAAAATAAAAAGTAATATTCATATGTAGTGTCAATTTTAAATGATGATGAAATGGATAATATCCATATTGACGATGTCAATAATGCCGGTATTGACATACAGCTCATCGATTTTTAGATTTCATTCAGAGGATATTGAATTATGTTATGGGAATTTGTATTTTGATAGGATCTATAATAATGTAGTAAATATAAAATATATTCCTGAGCATATTCCATATAGATATAATTTTATTAATCGTACGTTCTCCGTAGATGAACTAGATGATAATGTCTTTTTTACACATGGTTATTTTTTAAAACACAAATATGGTTGTTCACTTAATCCTAGTTTGATTGTCTCATTATCAGGAAACTTAAAATATAATGATATACAATGCTCAGTAAATGTATCGTGTCTCATTAAAAATTTGGCAACGAGTACATCTACTATATTAACATCTAAACATAAGACTTATTCTCTATATCGGTCCATGTGTATTGCTATAATAGGATACGATTCTATTATATGGTATAAATATATAAATGACAGGTATAATGACATCTATGATTTTACTGCAATATGTATGCTAATAGCGTCTACATTGATAGTGATCATATACGTGTTTAAAAAAATAAAAATGAACTCTTAATTATGTTATACTATTAGAAATGGATAAAATCAAAATTACGATTGATTCAAAAATTGGTAATGTTGTTACCATATCGTATAACTTGGAAAAGATAACTATTGATGTCACGCCAAAAAAGAAAAAAGAAAAGGATGTATTATTAGCGCAATCAGTTGCTGTCGAAGAGGCAAAAGATGTCAAGGTGGAAGAAAAAAATATTATCGATATTGAAGATGACGATGATATGGATATAGAAAACACGTAATACGATCTATAAAAATAAGTATTAAATACTTTTTATTTACGGTACTCTTGTAGTGGTGATACCACTAATCGATTATTTTTTTTAAAAAAATACTTATTCTGATTCTTCTAGCCATTTCCGTGTTCGTTCGAATGCCACATCGACGTCAAAAATAGGGGAGTAGTTGAAATCTAGTTCTGCATTGTTGGTACGCACCTCAAATGTAGTGTTGGATATCTTCAACGTATAGTTGTTGAGTATTGATGGTTTTCTAAATAGAATTCTCTTCATATCATTCTTGCACGCGTACATTTTTAGCATCCATCTTGGAATCCTAGATCCTTGTTCTATTCCCAATGGTTTCATCAATAGAAGATTAAACATATCGTAAGAACACGATGGAGAGTAATCGTAGCAAAAGTAAGCATTTCCTTTAATCGCAGATCCCGGATACTGGATATATTTTGCAGCCAACACGTGCATCCATGCAACATTTCCTACATATACCCGGCTATGCACAGCGTCATCATCGACTGTACGATACATAATGTTACCGTGTTGCTTACATTGCTCGTAAAAGACTTTCGTCAATTTGTCTCCTTCTCCGTAAATTCCAGTGGGTCTTAGGCAACAAGTATACAATTTTGCGCCATTCATGATTACGGAATTATTGGCTTTCATAACCAGTTGCTCGGCCATACGTTTACTTTTTGCGTATACATGTCCTGGTGATATATCATAAAGGGTATGCTCATGACCGATGAATGGATTACCGTGTTTATTTGGTCCTATTGCTTCCATGCTACTAGTATAGATCAAATACTTGATTCCTAGGTCCACACAAGCTGCCAATATAGTCTGTGTTCCATAATAGTTTACTTTCATGATTTCATTATCAGTGTATTTTCCAAATACATCCACTAGAGCAGCCGTATGAATAATCAGATTTACCCCATCTAGCGCTTCTCTCACCTTATCAAAGTCGTTTATATCACATTGTATATAGTTTATAACCTTAACTTTCGAGGTTATTGGTTGTGGATCTTCTACAATATCTATGACTCTTATTTCTTGAACATCATCTGCGCTAATTAAAAGTTTTACTATATACCTGCCTAGAAATCCGGCACCGCCAGTAACCGCGTACACGGCCATTGCTGCCACTCATAATATCAGACTACTTATTCTATTTTACTAAATAATGGCTGTTTGTATAATAGACCACGATAATATCAGAGGAGTTATTTACGTTGAACAAGTCCATGGAAAAGATAAAGTTTTAGGATCAGTTATTGGATTAAAATCCGGAACGTATAGTTTGATAATTCATCGTTACGGAGATATTAGTCGAGGATGTGATTCCATAGGCAGTCCAGAAATATTTATCGGTAACATCTTTGTAAACAGATATGGTGTAGCATATGTTTATTTAGATACAGATGTAAATATATCTACAATTATTGGAAAGGCGTTATCTATTTCAAAAAATGATCAGAGATTAGCGTGTGGAGTTATTGGTATTTCGTACATAAATGAAAAGATAATACATTTTCTTACAATTAACGAGAATGGCGTTTGATATATCAGTTAATGCGTCTAAAACAATAAATGCATTAGTTTACTTTTCTACTCAGCAAGATAAATTAGTCATACGTAATGAAGTTAATGATATACACTACACTGTCGAATTTGATAGGGACAAAGTAGTTGATACGTTTATTTCATATAATAGACATAATGACTCCATAGAGATAAGAGGGGTGCTTCCAGAGGAAACTAATATTGGTCGCGTGGTTAATACGCCGGTTAGTATGACTTACTTGTATAATAAGTATAGTTTTAAACCGATTTTAGCAGAATATATAAGACACAGAAATACTATATCCGGCAACATTTATTCGGCATTGATGACGCTAGATGATTTGGTTATTAAACAGTATGGAGACATTGATCTATTATTTAATGAGAAACTTAAAGTAGACTCCGATTCGGGACTATTTGACTTTGTCAACTTTGTAAAGGATATGATATGTTGTGATTCTAGAATAGTAGTAGCTCTATCTAGTCTAGTATCTAAACATTGGGAATTGACAAATAAAAAGTATAGGTGTATGGCATTAGCCGAACATATAGCTGATAGTATTCCAATATCTGAGCTATCTAGACTACGATACAATCTATGTAAGTATCTACGCGGACACACTGATAGCATAGAGGATGAATTTGATCATTTTGAAGACGATGATTTGTCTACATGTTCTGCCGTAACCGATAGGGAAACGGATGTATAATTTTTTTTATAGTATGAAGGATATGATGGATATGATGATATGATGGATATGATGATATGATGGATATGATGGATATGATGGATATGATGGATATGATAAAAAAATATAATTGTTGTATCCATTCCCATTCAAATCACCTTATATGATTCTGTAACACAATGAAGGAGTCTCATAGATATATAGAGGTCAGATACTGGTTTGATAAACTTTTTATTCCACATGAGCATGTTTGACTTATGGTTAGACACACATACTTTAACAAATCACTGAAAATTGGAGTTAGGTATTCCTCTCAGAATCAGTTGCCGTTCTGGAACATTAAATGTATTTTTTATGATATACTCCAACGCATTTATGTGGGTATACAACAAGTCATTAATAATGAGTATTTCCAAGAGTTTTAGTTGTCTAGTATTTAACAAGAGAAGAGATTTCAACAGACTGTTTATGAACTCGAATACCGCCTCATTGTCGCTTATATTGATGACATGACATCGAATTCCCAATATCAATCTCATCAGTGATGAGTAGCTCAATCTTGTTATCGGGATCCAATTTCTAAAGATGTCATTAAACCCTCGATCGTGAATGGATTTATCATCATCGTTTTTATGTTGGACATGAGCTTAGTCCGTTTGTCCACATCTATATACGATGATTTCTGAATTATTTCATATATCTCTCGTTAACTCCAGGAACTTGTCAGGGATCTAACTTTAATATGTTCTCGTCTAAGAGATGAAAATCTTTGGATGGTTGCATGTGACTTTTCTCTAAAGGATGATGTTACCCGATCCTCTCTTAAATGACTCCATCTTATCCTTGGACAAGATGGACAGTCTATTTTCCTTAGATGGTTTAATATTTTTTACCCATGATCTATAAAGGTAGACAGACCTAATCGTCTCGGATGACCATATATTATTTTCAGTTTTATTATACGCATAAATTGTAAAAAATATGTTAGGTTTACGAAAATGTCTCGTGGGGCATTAATCGTTTTTGAAGGATTGGACAAATCTGGAAAAACAACACAATGTATGAACATCATGGAATCTATACCGGCAAACACGATAAAATATCTTAACTTTCCTCAGCGATCCACAGTCACTGGAAAGATGATAGATGACTATCTAACTCGTAAAAAAACCTATAATGATCATATAGTTAATCTATTATTTTGTGCAAATAGATGGGAGTTTGCATCTTTTATACAAGAACAACTAGAACAGGGAATTACTTTAATAGTTGACAGATACGCGTTCTCTGGAGTAGCGTATGCCACCGCTAAAGGCGCGTCAATGACTCTCAGTAAGAGTTATGAATCTGGATTGCCTAAACCCGACTTAGTTATATTCTTGGAATCTGGTAGCAAAGAAATTAATAGAAACATCGGCGAGGAAATTTATGAAGATGTTGAATTCCAACAAAAGGTATTACAAGAATATAAAAAAATGATTGAAGAAGGAGATATTCATTGGCAAATTATTTCTTCTGAATTCGAGGAAGATGTAAAGAAGGAGTTGATTAAGAATATAGTTATAGAGGCTATACACACGGTTACTGGACCAGTGGGGCAACTGTGGATGTAATAAAATGAAATTACATTTTTATAAATAGATGTTAGTACAGTGTTATAAATGGATGAAGCATATTACTCTGGCAACTTGGAATCAGTACTCGGGGATACGTGTCCGATATGCATACCGAACTCGCATCAATATCTCAATTAGTTATTGCCAAGATAGAAACTATAGATAATGATTATTAAACAAGGACATTGTAAATTTTATCATGTGTAGATCAAACTTGGATAATCCATTTATCTCTTTCCTAGATACTGCATATACTATCATAGATCAAGAGATCTATCAGAACGAGTTGATTAATTCATTAGACGATAATGAAATTATCGATTGTATAGTTAACAAGTTTATGAGCTTTTATAAGGATAACCTAGAAAATATGGTAGATGCTATCATTACTCTAAAATATTATAATTAATAATCCAGATTTTAAAACTACGTATGTGGAAGTACTCGGTTCCAGAATAGCTGATATAGATATTAAACAAGTGATACGTAAGAATATAATACAATTGTCTAATGATCCGCGAACGATATTTGTGAAAATATTAAAAAAAAATACTTTTTTTATTAAATGACGTCTCTTCGCGAATTTAGAAAATTATGCTGTGATATATATCACGCATCAGGATATAAAGAAAAATCTAAATTAATTAGAGACTTTATAACAGATAGAGATGATACCGATACATATTTGATCATTAAGCTATTGCTTCCCGGATTAGACGATAGAATGTATAACATGAACGATAAACAAATTATAAAATTATATAGTATAATATTTAAACAATCTCAGGAAGATATGCTACAAGATTTAGGATACGGATATATAGGAGACACTATTAGGACTTTCTTCAAAGAGAACACGGAAATCCGTCCACGAGATAAAAGCATTTTAACTTTAGAAGAAGTGGATAGTTTTTTAACTACGTTATCATCAGTAACTAAAGAATCACATCAAATAAAATTATTGACTGATATAGCATCTGTTTGTACATGTAATGATTTAAAATGTGTAGTCATGCTTATTGATAAAGATCTAAAAATTAAAGCGGGTCCTCGGTACGTGCTTAACGCTATTAGTCCTCATGCCTATGATGTTTTTAGAAAATCTAATAACTTGAAAGAGATAATAGAAAATGCAGCTAAACAAAATCTAGACTCTATATCTATTTCTGTTATGACTCCAATTAATCCCATGTTAGCGGAATCATGTGATTCTGTCAATAAGGCGTTTAAAAAATTTCCATCAGGAATGTTTGCGGAAGTCAAATACGATGGTGAAAGAGTACAAGTTCATAAAAAAAATAACGAGTTTGCATTCTTTAGTAGAAACATGAAACCAGTACTCTCTCATAAAGTGGATTATCTCAAAGAATACATACCGAAAGCATTTAAAAAAGCTACGTCTATCGTATTGGATTCTGAAATTGTTCTTGTAGACGAACATAATGTACCGCTACCGTTTGGAAGTTTAGGTATACACAAAAAGAAAGAATATAAAAACTCTAACATGTGTTTGTTCGTATTTGACTGTTTATACTTTGATGGATTCGATATGACAGACATTCCATTGTATGAACGAAGATCTTTTCTCAAAGATGTTATGGTCGAAATACCCAATAGAATAGTATTCTCAGAGTTGACGAATATTAGTAACGAGTCTCAGTTAACTGATGTATTAGATGATGCACTAACGAGAAAATTAGAAGGATTGGTCTTAAAAGATATTAATGGCGTATACGAACCGGGAAAGAGAAGATGGTTAAAAATAAAGCGAGACTATTTGAACGAGGGTTCCATGGCAGATTCTGCCGATTTAGTAGTACTAGGTGCCTACTATGGTAAAGGAGGAAAGGGTGGTATCATGGCAGTCTTTCTAATGGGTTGTTACGACGATGAATCCGGTAAATGGAAGACGGTAACTAAATGTTCCGGTCACGATGATAATACGTTAAGGGTTTTGCAAGACCAATTAACGATGGTTAAAATTAACAAGGATCCCAAAAAAATTCCAGAGTGGTTGGTAGTTAATAAAATCTATATTCCCGATTTTGTAGTAGATGATCCGAAACAATCTCAGATATGGGAAATTTCAGGAGCAGAGTTTACATCTTCCAAGTCACATACAGCGAATGGAATATCGATTAGATTTCCTAGATTTACTAGGATTAGAGAAGATAAAACGTGGAAAGAATCTACTCATCTAAACGATTTAGTAAACTTGACTAAATCTCTTAATAGTTACATATAAACTGAAAAATAAAATAACACTATTTTAGTTGGTAGTCGCCATGGATGGTGTTATCGTATACTGTCTAAATGCGTTAGTAAAACATGGCGAGGAAATAAATCATATAAAAAATGATTTCATGATTAAACCATGTTGTGAAAGAGTTTGTGAAAAAGTCAAGAACGTTCACATCGGCGGACAATCTAAAAACAATACAGTGATTGCAGATTTGCCATATCTGGATAATGCTGTATCAGATGTATGCAAATCAATATATAAAAAGAATGTATCAAGAATATCCAGATTTGCTAATTTGATAAAAATAGATGACGATGACAAGACTCCTACCGGCGTATATAATTATTTTAAACCTAAAGATGCTATTCCTGTTATTATATCCATAGGAAAGGATAAAGATGTCTGTGAACTATTAATCTCATCTGATAAAGCGTGTGCGTGTATAAAGTTAAATTTATATAAAGTAGCCATTCTTCCCATGGATGTTTCCTTTTTTACCAAAGGAAATGCATCATTGATTATTCTCCTGTTTGATTTCTCTATCGATGCGGCACCTCTCTTAAGAAGTGTAACCGATAATAATGTTATTATATCTAGACACCAGCGCCTACATGACGAGCTTCCGAGTTCCAATTGGTTCAAGTTTTACATAAGTATAAAGTCCGACTATTGTTCTATATTATATATGGTTGTTGATGGATCTATGATGTATGCGATAGCTGATAATAGAACTCACGCAATTATTAGCAAAAATATATTAGACAATACTACGATTAACGATGAGTGTAGATGCTGTTATTCTGAACCACAGATTAGGATTCTTGATAGAGATGAGATGCTCAATGGATCATCGTGTTATATGAACAGACATTGTATTATGATGAATTTACCTGATGTAGGCGAATTTGGATCTAGTATGTTGGGGAAATATGAACCTGACATGATTAAGATTGCTCTTTCGGTGGCTGGTAATTTAATAAGAAATCGAGACTACATTCCCGGGAGACGAGGCTATAGCTACTACGTTTACGGTATAGCCTCTAGATAATTTTTTTTAAGCACGAAATAAAAACATAATTTTAAACAATCTATTTCATACTATTTTGTGTGCTCACCATGAACATAAAGATAGATATATTAGTATTTCTGGTGATAAATTTACGGCGACTGCTAGGAGGGAAAATGAAGAAAGAAAAAATATCTACCTCTCCAAAAAGAAAAACTACTGATGTTATCAAACCTGATTATCTTGAGTACAATGACTTGTTAGATAGAGATGAGATGTCTACTATTCTAGAGGAATATTAGGCCTTAGAATAAAATATGGACGACTCTTAACGAAATTAGAAAATTCGATAATGATGTTGAAGAACAATTCGGTACTATAGAAGAACTCAAGCAGAAGCTTAGATTAAATTCTGAAGAGGGAGCAGATAATTTTATAGATTATATAAAGGTACAAAAACAGGATATCATCAAACTTACTGTATACGATTGCATATATCTATGATAGGATTGTGTGCGTGCGTGGTAGATGTTTGGAGAAATGAGAAACTGTTTTCTAGATGGAAATATTGGTTACAAGCGATTAAACTGTTTATTGATGATCACATGCTTGATAAGATAAAATCTATTGTAGAATAGACTAGTGTATGTGGAAATGTCATAGAAAGTTAAAAGTTAATGAGAGCAAAAATATATAAGGTTGTATTCCATATTTGTTATTTTTTTCTGTAATAGTTAGAAAATACATTCGATGGTCTATCTACCAGATTATTATGTGTTATAAGGTACTTTTCTCATAATAAACTAGAGTATGAGTAAGATAGTGTTTTTCAAAAACATATAAATCTAAAATTGATGGATGAGATATACAGCTATTAATTTCGAAAATATATTTTAATCTGATAACTTTAAACATGGATTTTTGATGGTGGTTTAAGTTTAAAAAAGATTTTGTTATTGTAGTATGATAATATCAAAAAGATGGATATAAAGAATTTACTGACTACATGTACTATTTTACATTACTACATTGGCTACGGCATATATACCTATTTCGTCACTTCCACACGCTCCGGTAAACGGGTGTCATGTGACGAGGGAGAATCTTGATAAGAGGCATAATCAATGTTGTAATCCGATGTCCACCTGGAGAATTTGCCAAGGTCAGATGTAGAGTTGGTAGTGATAACACAAAATGTGAACACTGCCCACCTCATACATATACCGCAATCCCCAATTATTCTAATAGATGTCATCAATGTAGAAAATGCCCAACAGGATCATTTGATAAGGTAAAGTGTACCGGAACACAGAACAAATGTTCGTGTCATCCTGGTTGGTATACGCTACTGATTCTTCACAGACTGAAGATTGTCGAGATTTGTGTACCAAAAAAGGAGATGTCCATGCGGATACTTTGGTGGAATAGATGAAGGAAATCCTATTTGTAAATCGTGTTGTGTTGGTGAATATTGCGACTACCTACGTAATTATAGACTTGATCCATTTCCTCCATGCAAACTATCTATCTAAATGTAATTAATTATGATTTTGATGATAATGTTACCATACATTATATCACTACTTGGTTAGTGTGTATTATTTAGTATGGAAGACCTATTAATAATTACTTATCTTTTGACGATCTTGTTATAATTATAATATAAAAATACTTATGACATAGTAACTCATAATTGCTGACGCGATAAATTCGTAATAATCTGTTTTGTTCAAATTTTTATAAGGAATCTACAGGCATAAAAATAAAAATATAATCTATAATATACTCTTACAACGCCATCATGAATAGCAGTGAATTAATTGCTGTTATTATGGATTTAGAAATAGTGGACGATTTTGTGATATTAATATAGTTATTAATGATGAAAGGATAAACGCGCATAGACTCATCCTATCTGGAGCCTCCGAATATTTTTTCCATTCTGTTTTCCAATAATTTTATCGATTCTAATGAATACGAAGTTAATCTAAGTCATTTAGATTATCAAAGTGTTAACTATTTGATCGATTACATTTATGGGATACCTTTGAGCCTAACTAACAATAACGTGAAATATATTCTTTCAACCGCTGATTTTTTTACAAATTGGATCTGTCATTACTGAGTGCGAAAAATACATACTTAAAAATCTTTGTTTTAGAAACTGTATCGATTTCTACATATACGCTGATAAATATAATAACAAGAAAATAGAATTAGCATCGTTTAACACAATATTACGAAATATTTTGAGACTCATCAACAATGAAAACTTTAAATACTTAACAGAGGAATCAATGATAAAAATTTTAAGCGATGATATGTTATATATAAAAAATGAGGATTTCACCCCACTGATTCTCATTAAATGGTTAGAGAGTACACCAACCATGTACCGTCGAGTTACTTAGATGCCTCAGAATATCATTTCTTTCCCCACAAGTTATAAAATCACTTTATAGTCATCGACTGGTTAGTTCAATCTACGAATGTATAACATTCTTAAACAATATAGCATTCTTGGATAAATCATTTCCTAGATACCATATCATCGAGTTGATATCTATCGGTATAAGTAATTCACATGATAAGATTTCCATAAACTGCTACAATCATAAAAAAATTCATGGGAAATGATATCTTCACGTAGATATAGGTGTAGTTTCGCAGTGACCGTCCTGGATAATATTATCTATATGATGGGTGGATATGATCAGTCCCTGTATAGAAGTTCAAAGGTTATAGCGTACAATACATGTACTAATTCTTGGATATATGATATACCAGAGCTAAAAATATCATCGTTCTAATTGCGGAGGAGTTGCCAATGACGAATACATTTATTGTATAGGCGGTATACGCGATCAGGAGTCATCGTTGATATCTAGTATCGATAGATGGAAGCCATCAAAACCATATTGGCAGAAGTATGCTAAAATGTGCGAACCAAAATGTGATATGGGGTTGCGATTTTAAACGGATTAATATATGTCATAGGTGGAGTCGTTAAAGGTGACACACATATACCAACGCACTAGAGAGTTTATCAGAAGATGGATGGATGAATCATCAACGTCTTCCAATAAAAATGTCCAATATGTCGACGATTGTTCATGCTGGAAAGATTTATATATCTAGAGGTTACAACAATAGTAGTGTAGTTAATGTAATATCGAATCTAGTCCTTAGCTATAATCCGATATATGATGAATGGACCAAATTATCATCATTAAATATTCCTAGAATTAATCCTGCTCTATGGTCAGTGTATAATAAATTATATGTAGGAGGAGTAATATCTGATGATGTTCAAACTAATACATCTGAAACATACGATAAAGAAAAAGATTGTTGGACATTGGATAATGGTCACTTGTTACCACATAATTATATAATGTATAAATGCGAACCGTTTAAACATAGATATCCATTGGAAAAAACACAGTACACGAATGATTTTCTAAAGTATTTGGAAAGTTTTATAGGTAGTTGATAGAACAAAATACATAATTTTGTAAAAATAAATCACTTTTTATACTAATATGACACAATTACCAATACTTTTGTTACTAATATCATTAGTATACGCTACACCTTCTCCTCAGACATCTAAAAAAATAGGTGATGATGCAACTATATCATGTAGTCGAAATAATACAAATTACTACGTTGTTATGAGTGCTTGGTATAAGGAGCCCAATTCCATTATTCTCTTAGCTGCCAAAAGCGACGTCTTGTATTTTGATAATTATACCAAGGATAAAATATCTTACGACTCTCCATACGATGATCTAGTTACAACTATCACAATTAAATCATTGACTGCTGGAGATGCCGGTACTTATATATGTGCATTCTTTATGACATCGACTACAAATGATACTGATAAAGTAGATTATGAAGAATACTCCATAGAGTTGATTGTAAATACAGATAGTGAATCGACTATAGACATAATACTATCTGGATCTACACCAGAAACTATTTCTGAGAAACCAGAGGATATAGATAATTCTAATTGCTCGTCTGTATTCGAAATCACGACTCCGGAACCAATTACTGATAATGTAGACGACCATACAGACACCGTCACATACACTAGTGATAGCATTAATACAGTAAATGCATCATCTGGAGAATCCACAACAGACGAGATTCCGGAACCAATTACTGATAAAGAAGAAGATCATACAGTAACAGACACTGTCTCATACACTACAGTAAGTACATCATCTGGAATTGTCACTACTAAATCAACCACCGATGATGCGGATCTTTATGATACATACAATGATAATGATACAGTACCGCCAACTACTGTAGGTGGTAGTACAACCTCTATTAGCAATTATAAAACCAAGGACTTCGTAGAAATATTTGGTATTACCACATTAATTATATTGTCAGCAGTGGCGATTTTCTGTATTACGTATTATATATGTAATAAACACCCACGTAAATACAAAACAGAGAACAAAGTCTAGATTTTTGACTTACATAAATATCTGGGATAATAAAATCTATCATATTGAGAGGACCATCTGGTTCAGGAAAGACAGCCATAACCAAAAGACTGTTAAAAGACTATGGGAATATATTTGGATTTGTGGTGTCCCATACCACTAGATTTCCTCGTCCTATGGAACGAGAAGGTGTTGATTACCTTACGTTAACAGAGAGGCCATCTGGAAGGGAATAGCCGCCGGAAACTTTCTAGAACATACTGAGTTTTTAGGAAATATTTACGGAACTTCTAAAACAGCTGTAAATACAGCGGTTATTAATAATCGTATTTGCGCGATGGATTTAAACATCAACGGTGTTAGAAGTCTTAAAAATACTTACCTAATGCATTACTTGGGTATATAAGACCTACCTCTCTTAAAATGGTTGAGACCAATCTTCGTCGTAGAAACACTGAAGCGGACGACGAATCTCATCGTCGCGTGATGTTGGCAAAAAACGGATATGGATGAGGTCAACGAAGCAGGTCTATTCGACACTATTATTATTGAAGATGATGTGAATTTAGCATATAGTAAGTGTTAATTCAGATACTACAGGACCGTATTAGAATGTATTTTAACACTAATTAGAGACTTAAGATTTGACTTAAAACTTGATAATTAATAATATAACTCGTTTTTATATGTGGCTATTTCAACGTCTAATGTATTAGTTAAATATTAAAACTTACCACGTAAAACTTAAAATTTAAAATGGTATTTCATTGACAGATCATACATTATGAAGTTTCAAGGACTTGTGTTAATTGACAATTGCAAAAATCAATGGGTCGTTGGACCATTAATAGGAAAAGGTGGATTCGGTAGTATTTATACTACTAATGACAATAATTATGTAGTAAAAATAGAGCCCAAAGCTAACGGATCATTATTTACCGAACAGGCATTTTATACTAGAGTACTTAAACCATCCGTTATCGAAGAATGGAAAAAATCTCACAATATAAAGCACGTAGGTCTTATCACATGCAAGGCATTTGGTTTATACAAATCCATTAATGTGGAATATCGATTCTTGGTAATAAATAGATTAGGTGCAGATCTAGATGCGGTGATCAGAGCCAATAATAATAGACTACCAGAAAGGTCGGTGATGTTGATCGGAATCGAAATCTTAAATACCATACAATTTATGCACGAGCAAGGATATTCTCACGGAGATATTAAAGCGAGTAATATAGTCTTGGATCAAATAGATAAGAATAAATTATATCTAGTGGATTACGGATTGGTTTCTAAATTCATGTCTAACGGCGAACATGTTCCATTTATAAGAAATCCAAATAAAATGGATAACGGTACTCTAGAATTTACACCTATAGATTCGCATAAAGGATACGTTGTATCTAGACGTGGTGATCTAGAAACACTTGGATATTGTATGATTAGATGGTTGGGAGGTATCTTGCCATGGACTAAGATATCTGAAACAAAGAATTCTGCATTAGTAAGTGCCGCAAAACAGAAATATGTTAACAATACTGCGACTTTGTTAATGACCAGTTTGCAATATGCACCTAGAGAATTGCTGCAATATATTACCATGGTAAACTCTTTGACATATTTTGAGGAACCCAATTACGACGAGTTTCGTCGAGTATTAATGAATGGAGTTATGAAAAATTTTTGTTGATAAAAAAATTAAAAAAATAACTTAGTTATTATCACTCTCGTGAGTACAATAGAAACATGGCGATGTTTTACGCACACGCTTTCGGTGGGTACGACGAGAACCTTCATGCATTTCCTGGAATATCATCGACGGTTGCCAATGATGTCAGGAAATATTCTGTTGTGTCAGTTTATAATAAAAAGTATAACATTGTAAAAAACAAATATATGTGGTGTAACAGTCAAGTGAACAAGAGATATATTGGAGCACTACTGCCTATGTTTGAATGCAATGAATATCTACAAATTGGAGATCCAATCCATGATCTAGAAGGAAATCAAATCTCTATTGTCACATATCGCCACAAAAACTACTATGCTCTAAGTGGAATTGGGTACGAGAGTCTAGACTTGTGTTTGGAAGGAGTAGGGATTCATCATCACGTACTTGAAACAGGAAACGCGGTATATGGAAAAGTTCAACATGAGTATTCTACTATCAAAGAGAAGGCCAAAGAAATGAATGCACTCAAACCAGGACCTATCATCGATTACCACGTCTGGATAGGAGATTGTGTCTGCCAAGTTACTACTGTAGACGTGCATGGAAAGGAAATTATGAGAATGAGATTCAAAAGGGGTGCGGTGCTTCCGATTCCAAATCTGGTAAAAGTTAAAGTTGGGGAGGAAAATGATACAATAAATCTTTCCACTTCCATATCAGCTCTCCTGAATTCCGGTGGCGGCACCATCGAGGTAACATCTAAGGAAGAACGTGTAGATTATGTACTCATGAAACGTTTGGAATCTATACATCATCTGTGGTCTGTAGTGTATGATCATCTTAATGTTGTGAATGGCGAAGAACGATGTTATGTACATATGCATTCATCTCATCAAAGTCCTATGCTGAGTACTGTAAAAACAAATTTGTACATGAAGACTATGGGAGCATGTCTTCAAATGGACTCCATGGAAGCTCTAGAGTATCTTAGTGAACTGAAGGAATCAGGTGGGCGGAGTCCCAGACCAGAATTGCAGAAATTTGAATATCCAGATGGAGTGAAAGACACTGAATCAATTGAGAGATTGGCAGAGGAGTTCTTCAATAGATCAGAACTTCAGGCCGGTGAATCAGTCAAATTTGGTAATTCTATTAATGTTAAACATACATCTGTTTCAGCTAAGCAACTAAGAACACGTATACGACAGCAGCTTCCTTCTATACTCTCATCTTTTGCCAACACAAAGGGTGGATATTTGTTCATTGGAGTTGATAATAATACACACAAAGTAATTGGATTCACGGTGGGTCATGACTACCTCAAACTGGTAGAGAGTGATATAGAAAAGTATATCCAAAAACTTCCTGTTGTGCATTTCTGCAAGAAAAAAGAGGACATCAAGTACGCATGTAGATTCATCAAGGTGTATAAACCTGGTGATGAGACTACCTCGACATATGTGTGCGCAATCAAAGTGGAAAGATGCTGCTGTGCTGTGTTTGCGGATTGGCCAGAATCATGGTACATGGATACTAGTGGTAGTATGAAGAAGTATTCTCCAGATGAATGGGTGTCACATATAAAATTTTAATTAGGGTAAGGTAAAACTATATATAATAACTAACAATTTGTGTATCATATAGACAATTAATTAGGTAACTGTTATCTCTTTTTAACTAACTAACTAACTAACTAACTCTTATATACTATTAATAATACATCTATTAATCATTGATTAGCTTATTGCTTTAATTGTTTTTGTAAACTAACACTGTTCATTGAAAAGGGATAACATGTTACAGAATATAAATTATATATGGATTTTTTTAAAAAGGAAATACTTGACTGGAGTATATATTTATTTCTTCATTACATAACACGTCTGTGTTCTAATTCTTCCAATTCTTCCACATCTCATATAATACAGGAATATAATCTTGTTCGAAAATATGAGAAAGTGGATAAAACAATAGTTGATTTTTTATCTAGGTGGCCAAATTTATTCCATATTTTAGAATATGGGGAAAATATTCTACATATTTATTTTATAGATGCTGCTAATACGAATATTATGATTTTTTTTCTAGATAGAGTATTAAATATTAATAAGAACCGTGGGTCATTTATACATAATCTCGGGTTATCATCCATTAATATAAAAGAATATGTATATCAATTAGTTAATAATGATCATCTAGATAATAGTATAAGACTAATGCTTGAAAATGGACGTAGAACAAGACATTTTTTGTCTTATATATTGGATACAGTTAATATCTATATAAGTATTTTAATAAATCATAGATTTTATATAGATGCCGAAGACAGTTACGGTTGTACATTATTACATAGATGTATATATAACTATAAGAAATCAGAATCAGAATCATATAATGAATTAATTAAGATATTGTTAAATAATGGATCAGATGTAGATAAAAAAGATACGTACGGAAACACACCGTTTATCCTATTATGTAAACACGATATCGACAACGCGGAATTGTTTGAGATATGTTTAGAGAATGCTAATATAGACTCTGTAGACTTTAATGGATATACACCTCTTCATTATGTCTCATGTCGTAATAAATATGATTTTGTAAAGTTATTAATTTCTAAAGGAGCAAATGTTAATGCACGTAATAGATTCGGAACTACTCCATTTTATTGTGGAATTATACACGGTATCTCGCTTATAAAACTATATTTGGAATCAGACACAGAGTTAGAAATAGATAATGAACATATAGTTCGTCATTTAATAATTTTTGATGCTGTTGAATCTTTAGATTATCTATTGTCCAGAGGAGTTATTGATATTAACTATCGTACTATATACAACGAAACATCTATTTACGACGCTGTCAGTTATAATGCGTATAATACGTTAGTCTATCTATTAAACAGAAATGGTGATTTTGAGACGATTACTACTAGTGGATGTACATGTATTTCGGAAGCAGTCGCGAACAACAACAAAATAATAATGGATATACTATTGTCTAAACGACCATCTTTGAAAATTATGATACCATCTATGATAGCAATTACTAAACATAAACAACATAATGCAGATTTATTGAAAATGTGTATAAAATATACTGCGTGTATGACCGATTATGATACTCTTATAGATGTACAATCGCTACATCAATATAAATGGTATATTTTAAAATGTTTTGATGAAATAGATATCATGAAGAGATGTTATATAAAAAATAAAACTGTATTCCAATTAGTTTTTTGTATCAAAGACATTAATACTTTAATGAGATACGGTAGACATCCTTCTTTCGTGAAATGTAATATTCTTGACGTATACGGAAGTTGTGTACGTAATATCATAGCATCTATTAGATATCGTCAGAGATTAATTAGTCTATTATCCAAGAAGCTGGATGCTGGAGATAAATGGTCGTGTTTTCCTAACGAAATAAAATATAAAATATTGGAAAACTTTAACGATAACGAACTGACCACATATCTGAAAATCTTATAAACACTATTAAAATATAAAATCTAAGTAGGATAAAATCACACTACATCATTGTTTCCTTTTAGTGCTCGACAGTGTATACTATTTTTAACACTCATAAATAAAAATGAAAACGATTTCCGTTGTTACGTTGTTATGCGTACTACCTGCTGTTGTTTATTCAACATGTACTGTACCCACTATGAATAACGCTAAATTAACGTCTACCGAAACATCGTTTAATGATAAACAGAAAGTTACGTTTACATGTGATTCAGGATATCATTCTTTGGATCCAAATGCTGTCTGTGAAACAGATAAATGGAAATACGAAAATCCATGCAAGAAAATGTGCACAGTTTCTGATTATGTCTCTGAACTATATGATAAGCCATTATACGAAGTGAATTCCACCATGACACTAAGTTGCAACGGTGAAACAAAATATTTTCGTTGTGAAGAAAAAAATGGAAATACTTCTTGGAATGATACTGTCACGTGTCCTAATGCGGAATGTCAACCTCTTCAATTAGAACACGGATCGTGTCAACCAGTTAAAGAAAAATACTCATTTGGGGAATATATGACTATCAACTGTGATGTTGGATATGAGGTTATTGGTGTTTCGTATATAAGTTGTACGGCTAATTCTTGGAATGTTATTCCATCATGTCAACAAAAATGTGATATACCGTCCCTATCTAATGGATTAATTTCCGGATCTACATTTTCTATCGGTGGCGTTATACATCTTAGTTGTAAAAGTGGTTTTACACTAACGGGGTCTCCATCATCCACATGTATCGACGGTAAATGGAATCCCATACTCCCAACATGTGTACGATCTAACGAAGAATTTGATCCAGTGGATGATGGTCCCGACGATGAGACAGATCTGAGCAAACTCTCGAAAGACGTTGTACAATATGAACAAGAAATAGAATCGTTAGAAGCAACTTATCATATAATCATAATGGCGTTGACAATTATGGGTGTCATATTTCTAATCTCCATTATAGTATTAGTTTGTTCCTGTGACAAAAATAATGACCAATATAAGTTCCATAAATTGCTACCGTGAATATAAATCCGTTAAAATAATTAATAATTAATAATTAATAACGAACAAGTATCAAAAGATTAAAGAATTAGCTAGAATCAATTAGATGTCTTCTTCAGTGGATGTTGATATCTACGATGCCGTTAGAGCATTTTTACTCAGGCACTATTATGACAAGAGATTTATTGTGTATGGAAGAAGTAACACCATATTACATAATATATACAGGCTATTTACAAGATGCACCGTTATACCGTTCGATGATATAGTACGTACTATGCCAAATGAATCACGTGTTAAACAATGGGTGATGGATACACTTAATGGTATAATGATGAATGAATTCGATACTGTATGTGTGGGTACCGGACTACGATTCATGGAAATGTTTTTCGATTACAATAAAAATAATCCCAAAAATAGCATCAACAATCAAATAATGTATGATATAATTAATAGCGTAGCCATAATTCTAGCTAATGAGAGATATAGAAGCGCGTTTAACGACGATAGAATATACATCCGTAGAACTATGATGGACAAATTGTACGAATACGCATCTCTAACTACTATTGGTACGATCACTGGAGGTGTTTGTTATTTTATCTGTTGATGCATCTAGTTAGTTTGTATAAATAATTATTTCGATATACTAGTTAAAATTTTAAGATTTTAAATGTATAAAAAACTAATAACGTTTTTATTTGTAATAGGTGCAGTTGCATCCTATTCGAATAATGAGTACACTCCGTTTAATAAACTGAGTGTAAAACTCTATATAGATGGAGTAGATAATATAGAAAATTCATATACTGATGATAATAATGAATTGGTGTTAAATTTTAAAGAGTACACAATTTCTATTATTACAGAGTCATGTGACGTCGGATTTGATTCCATAGATATAGATGTTATAAACGACTATAAAATTATTGATATGTATACCATTGACTCGTCTACTATTCAACGCAGAGGACATACGTGTAGAATATCTACCAAATTATCATGCCATTATGATAAGTACCCTTATATCCACAAATATGAGGGTGATGAACGACAATATTCTATTACCGCAGAGGGAAAATGCTATAAAGGAATAAAATATGAAATAAGTATGATGAACGATGATACTCTATTGAGAAAACATACTCTTAAAATTGGATTTACTTATATATTCGATCGTCATGGGCATAGTAATACATATTATTCAAAATATGATTTTTAAAAATTTAAAATATATTATCACTTCAGTGACAGTAGTCAAATAACAAACAACACCATGAGATATATTATAATTCTCGCAGTTTTGTTCATTAATAGTATACATGCTAAAATAACTAGTTATAAGTTTGAATCCGTCAATTTTGATTCCAAAATTGAATGGACTGGGGATGGTCTATACAATATATCCCTTAAAAATTATGGCATCAAGACGTGGCAAACAATGTATACAAATGTACCAGAAGGAACATACGACATATCCGGATTTCCAAAGAATGATTTCGTATCTTTCTGGGTTAAATTTGAACAAGGCGACTATAAAGTGGAAGAGTATTGTACGGGACTATGTGTCGAAGTAAAAATTGGACCACCAACTGTAATATTGACTGAATATGACGATCATATCAATTTGTTCATCGAGCATCCGTATGCTACTAGAGGTAGCAAGAAGATTCCTATTTACAAACGCGGTGACATGTGTGATATCTACTTGTTGTATACGGCTAACTTCACATTCGGAGATTCTGAAGAACCAGTAACATATGATATCGATGACTACGATTGCACGTCTACAGGTTGCAGTATAGACTTTGCCACAACAGAAAAAGTGTGTGTGACAGCACAGGGAGCCACAGAAGGGTTTCTCGAAAAAATTACTCCATGGAGTTCGGAAGTATGTCTGACACCTAAAAAGAATGTATATACGTGCGCAATTAGATCTAAAGAAGATGTTCCCAATTTCAAGGACAAAATAGCCAGAGTTATCACGAGAAAATTTAATAAACAGTCTCAATCTTATTTGACTAAATTTCTCGGTAGCACATCGAATGATGTTACAACTTTTTTTAGCATTCTTGACTAAATATTCATAACTAATTTTTATTAATGATACAAAAATGAAATAAACTGTATATTATACACTGGTTAACGCCCTTGGCTCTAACCATTTTCAAGATGAGGTCCCTGATTATAGTCCTTCTGTTCCCCTCTATCATCTACTCAATGTCTATTAGACGATGCGAGAAGACTGAAGAGGAAACATGGGGATTAAAAATAGGGTTGTGTATAATTGCCAAAGATTTCTATCCCGAAAGAACTGATTGCAGTGTTCATCGCCCAACTGCAAGTGGAGGATTGATAACTGAAGGCAATGGATTCAGAGTAGTTATATATGATCAATGTACAGAACCCCATGACTTTATTATCACCGATACTCAACAAACACGTCTTGGATCATCTCATACATATATTAAATTCAGTAACATGAATACAGGTGTCCCATCTAGTATTCCAAAATGTTCCAGAACTCTCTGTATTTCTGTATATTGTGATCAAGAGGCGGGAGACATAAAATTTGAGGAGTATACTCAAGAATCAAGTGATATCAGTATTAGAGTTAAGTATGATTCATCATGTATTGATTATCTGGGTATTAATCAAAGTTTCATGAATGAATGTATTCGAAGAATTACAACATGGGATAGAGAATCATGCGTCAGAATTGATACACAGACTATAAATAAATATCTTAAGTCTTGCACCAACACAAAATTCGACCGTAATGTCTACAAAAGGTACATACTGAAGAGTAAAGCACTCCATGCTAAAACAGAGTTGTAATAGATATAAAATACTTTTTATAATAATTAGGCTAGAAAAATCTCACTCACATGTAATCTTAAAAAAATGATATGATAGTTCTTACAAGTAGCGATTGAGTTTTAAATGGATTCTATTAATTACCGGGGAACTTAACAATTCGTTCTGATCTACAGACATTGGTTAATAAATCATCTTATTTTGCCAATATATTAAAATGTGGAAACTCCACTAATAATATTACATTGTGCGACTTTCAAGATGATGTGATATATAGGGTTATACAGTTTTAACAATTATATAATAGAGATAGAAAGTACAAAAGATGTAGAATCAATGATATGGCACGCTAAACAGTTGGGTGTGGAATCATTGCTAAAAGAATGTCAAAATTATTTGCTTAGAATATTACGTATATAATTGTTTAGAAATTTATAGAATAACTAATATTAATACATTATCGTATATCTACAACGATATAAGAAACTTCATATTGGATAATATTACTATTAATATATAAGGATCCAGATTTTATATATTTGCCTAAATACATTATTATAGATTTACTAGGACAATCACCTAAATGTTTTTAACGAAGATAATGTGGTAAAGATTATATACACTTATATATCTTCCGATATCTACAAGGATATTCCATATCATCATTGTGTAAACTAAATAACGTTTTCTATGGCATTTAATAAGGACATTGGATATGTGGAAAAGTGATGTATGGAAGTTAGTACATTATCAACTTCTCCTTATTGATTGAAAATGAAAATATAAATAGTTTTTATGTATAGCGGTATCTACCCTATAGTTTTATTGCTTACTACTAACATGGATTCAGATACAGATACAGATACAGATACAGATACAGATACAGATACAGATACAGATGTAGAAGATATCATGAATGAAATAGATAGAGAGAAAGAAGAAATACTAAAAAATGTAGAAATTGAAAATAATAAAAACATTAACAAGAATCATCCCAGTGAATATATTAGAGAAGCACTTGTTATTAATACCAGTAGTAATAGTGATTCCATTGATAAAGAAGTTATAGAATATATCAGTCACGATGTAGGAATATAGATCATATCTACTAATTTTTATAATCGATACAAAACATAAAAACAACTCGTTATTACATAGCAGGTATGGAATCCTTCAAGTATTGTTTTGATAACGATGGTAAGAAATGGATTATCGGAAATACTTTATATTCTGGTAATTCAATACTCTATAAGGTCAGAAAAAATTTCACTAGTTCGTTCTACAATTACGTAATGAAGATAGATCATAAATCACACAAGCCATTGTTGTCCGAAATACGATTCTATATATCTGTATTGGATCCTTTGACTATCAACAACTGGACACGGGAACGTGGTATAAAGTATTTGGCTATTCCAGATCTGTATGGAATTGGAGAAACCGATGATTATATGTTCTTCATTATAAAGAATTTGGGAAGAGTATTCGCCCCAAAGGATAGTGAATCAGTTTTCGAAGCATGTGTCACTATGATAAACACGTTAGAGTTTATACACTCTCAAGGATTTACTCATGGAAAAATAGAACCGATGAATATACTGATTAGAAATAAACGTATTTCACTAATTGACTATTCTAGAACTAACAAACTATACAAAAGTGGAACACATATAGATTACAACGAGGACATGATAACTTCAGGAAATATCAATTATATGTGTGTAGACAATCATCTTGGAGCAACAGTTTCAAGACGAGGAGATTTAGAAATGTTGGGATATTGCATGATAGAATGGTTCGGTGGTAAACTTCCATGGAAAAACGAAAGTAGTATAAAAGTAATAAAACAAAAAAAAGAATATAAACAATTTATAGCTACTTTTTTTGAGGACTGTTTTCCTGAAGGAAATGAACCTCTGGAATTAGTTAGATATATAGAATTAGTATACATGTTAGATTATTCTCAAACTCCTAATTATGACAGACTACGTAGACTGTTTATACAAGATTGAAATTATATTCTTTTTTTTATAGAGTGTGGGGGTAGTGTTACGGATATCTAATATTAATATTAGACTATCTCTATCGCGCTACACGACCAATATCGATTACTATGGATATCTTCAGGGAAATCGCATCTTCTATGAAAGGAAAGAATGTATTCATTTCTCCAGCGTCAATCTCGTCAGTATTGACAATACTGTATTATGGAGCTAATGGATCCACTGCTGAACAGCTATCAAAATATGTAGAAAAGGAGGAGAACATGGATAAGGTTAGCGCTCAGAATATCTCATTCAAATCCATGAATAAAGTATATGGGCGATATTCTGCCGTGTTTAAAGATTCCTTTTTGGGAAAAATTGGCGATAAGTTTCAAACTGTTGACTTCACTGATTGTCGCACTATAGATGCAATCAATAAGTGTGTAGATATCTTTACTGAGGGAAAAATCAATCCACTATTGGATGAACCATTGTCTCCTGATACCTGTCTCCTAGCAATTAGTGCCGTATACTTTAAAGCAAAATGGTTGATGCCATTCGAAAAGGAATTTACCAGTGATTATCCCTTTTACGTATCTCCAACGGAAATGGTAGATGTAAGTATGATGTCTATTTACGGCGAGCCATTTAATCACGCATCTGTAAAAGAATCATTCGGTAACTTTTCAATCATAGAACTGCCATATGTTGGAGATACTAGTATGATGGTCATTCTTCCAAACAAGATTGATGGATTAGAATCCATAGAACAAAATCTAACAGATACAAATTTTAAGAAATGGTGTAACTCTCTGAAAGCTACGTTTATCGATGTGCACATTCCTAAGTTTAAGGTAATAGGTTCGTATAATCTTGTGGATACGCTAATAAAGTTGGGACTGACAGATGTGTTCTATTCAACTGGTGATTATATCAATATGTGTAATTCAGATGTGAGTGTTGACGCTATGATTCACAAAACGTATATAGATGTCAATGAAGAGTATACAGAAGCAGCTGCAGCAACTTCTGTACTAGTGGCAGACTGTGCATCAACAGTTACAAATGAGTTCTGTGCAGATCATCCGTTCATCTATGTGATTAGACATGTCGATGGTAAAATTCTTTTCGTTGGTAGATATTGCTCTCCAACAACTAATTAAGCACATTCTTAATATTAGAATATTATATAGTTAAGATTTTTACTAACAGGTTAACATTTTTTTTTAAAAATAGAAAAAACATGTGGTATTAGTGCAGGTCGTTATTCTTCCAATTGCAATTGGTAAGATGACGGCCAACTTTAGTACCCACGTCTTTTCACCACAACACTGTGGATGTGACAGACTGACCAGTATTGATGACGTCAGACAATGTTTGACTGAATATATTTATTGGTCGTCGTATGCATACCGCAACAGGCAATGCGCTGGACAACTGTATGACACACTCCTCTCTTTTAAAGATGATGCGGAATCAGTGTTCATCGACGTTCGTGAGCTGGTAAAAAATATGCCGTGGGATAATGTTAAGGATTGTACAGAGATCATCCGTTGTTATATACCGGATGAGCAAAAAACCATCAGAGAGATTTCGGCCATCATTGGACTTTGTGCATATGCTGCTACTTACTGGGGAGGTGAAGACCATCCCACTAGTAACAGTCTGAACGCATTGTTTGTGATGCTTGAGATGCTCAATTACATGGATTATACCATCATATTCTGGCGTATGAATTGATGAGTTACAGCTTGACATTTCTTCTTTCCTCCCTCTTCTTCTACCTTTCCCAGAAACAAACTTTTTTTACCCACTATAAAATAAAATGAGTATACTACCTGTTATATTTCTTCCTATATTTTTTTATTCTCCATTCGTTCAGACTTTTAACGTGCCTGAATGTATCGACAAAGGGCAATATTTTGCATCATTCATGGAGTTAGAAAACGAGCCAGTAATCTTACCATGTCCTCAAATAAATACGCTATCATCCGGATATAATATATTAGATATTTTATGGGAAAAACGAGGAGCGGATAATGATAGAATTATACAGATAGATAATGGTAGCAATATGCTAATTCTGAACCCGACACAATCAGACTCTGGTATTTATATATACATTACCACGAACGAAACCTACTGTGACATGATGTCGTTAAATTTGACAATCGTGTCTGTCTCAGAATCAAATATAGATCTTATCTCGTATCCACAAATAGTAAATGAGAGATCTACTGGTAAAATGGTATGTCCCAATATTAATGCATTTATTTCTAGTAACGTAAACACAGAATTATATGGAGCGGACATCGACGCCTTAGAAATAAGAGACTTAAACAACGGACACCTGGAATTATTACCATAGAAGATGTTAGAAAAAATGATGCTGGTTATTATACATGTGTTTTAGAATATATATATATATATATATATATATATATATATATATATATATATATATATATATGGGCAAAACATATAACGTAACCAGAATTATAAAATTAGAGGTACGGGATAGAATAATACCTCCTACTATGAAATTACCAGAAGGAGTAGTAACTTCAATAGGTAGTAATTTGACTATTGCATGTAGAGTATCGTTGAGACTTCCCACAACGGACACCGACGTCTTTTGGATAAGTAATGGTATATGTATTACGAAGAAGAAGACGAGGACGGAGACGGTAGAATAAGTGTAGCAAATAAAATCTATATGACCGATAAGAGACGTGTTATTACATCCTGGTTAAACATTAATCCTGTCAAGGAAGAAGATGCTACAACGTTTACGTGTATGGCGTTTACTATTCCTAGCATCAGCAAAACAGTTACTGTTAGTAAACGTGAATGTATGTTGTTACATTTCCATATCAATTGAGTTTATAAGAATTTTTTATACATTATCTTCCAACAAACAATTGACGAACGTATTGCTATGATTAACTCCCACAATACTATATATATTATTAATCATTAACTTGCAGACTATACCTAGTAGTGCTATTTTGACATACTCATGTTCTTGTGTAATCGCAGTATCTATATTATTAAAGTACGTAAATCTAGCTATAGTTTTATTATTTAATTTTAGATAATATACTGTCTCCGTATTTTTAAAAAATTACCACATCCTTTATTAAATCATGAATGGGAATTTCTGTGTCATCGTTAGTATATTGTGAACAACAAGAGCAGATATCTATAGGAAAGGGTGGAATGCGATACATTGATCTATGTAGTTTTAAAACATACGCGAACTTTGAAGAATTTATATAAATCATCTCACGAGATATTGCTCTCTGTCATATTCATACACCTGTATAAACTTTCTAGACATCTTACAATGTGTTATTTTATGATCATATTTACATATTTACTGGTATATCAAAGATGTTAGATTAGTTAATGGGAATCGTCTATAATAATGAATATTAAACAATTATAGGAGGAGTTTATACCTACAAAAACATCATAAAAATGAGTCATCGTCCGATTTATGTTTTAAATATACTAACATTACTACCTTCAGAAATTATATACGAAATATTATACATGCTGACAATTAACGATCTTTATAATATATAGTATCCACCTACCAAAGTATAATTGTATTTTTCTCATGTGATGTGTGTAAAAAACTGATATTATATAATTATCTTAGTACCTATGATGAAGATGAAGATGAAGATGAAGATGATGGTCCGTATATATTTTGTATCATTATCGTTATTGCTATTCCATAGTTACGCCATAGACATCGAAAATGAAATCACCGAATTCTTCAATAAAATGAGAGATACTCTACCAGCTAAAGACTCTAAATGGTTGAATCCAGTATGTATGTTTGGAGGCACAATGAATGATATGGCCGCTCTAGGAGAGCCATTCAGTGCAAAGTGTCCTCCTATTGAAGACAGTCTTTTATCGCATAGATATAAAGACTATGTGGTTAAATGGGAAAGGCTAGAAAAGAATAGACGGCGACAGGTTTCTAATAAACGTGTTAAACATGGTGATTTATGGATAGCCAACTATACATCTAAATTCAGTAACCGTAGGTATTTATGTACCGTAACCACAAAGAATGGTGACTGTGTTCAGGGTGTAGTTAGATCTCATGTGTGGAAACCTTCTTCATGCATTCCAAAAACATATGAACTAGGTACTTATGATAAGTATGGCATAGACTTATACTGTGGAATTCTTTATGCGAACCATTATAATAATATAACTTGGTATAAAGATAATAAGGAAATTAATATCGACGATTTTAAGTATTCACAAGCGGGAAAGGAATTAATTATTCATAATCCAGAGTTAGAAGATAGTGGAAGATACGACTGTTACGTTCATTACGACGACGTTAGAATCAAGAATGATATCGTAGTATCAAGATGTAAAATACTTACGGTTATACCGTCACAAGACCACAGGTTTAAACTAATACTAGATCCGAAAATCAACGTAACGATAGGAGAACCTGCCAATATAACATGCAGTGCTGTGTCAACGTCATTATTTGTCGACGATGTACTGATTGAATGGGAAAATCCATCCGGATGGATTATAGGATTAGATTTTGGTGTATACTCTATTTTAACTAGTAGAGGCGGTATCACCGAGGCGACTTTGTATTTTGAAAATGTTACTGAAGAATATATAGGCAATACATATACATGTCGTGGACACAACTATTATTTTGATAAAACTCTTACAACTACAGTAGTATTGGAGTAAATACACAATGCATTTTTATATACATTACTGAATTATTATTATTAATTATATCGTATTTGTGCTATAGAATGGATGAAGATACGCGACTATCTAGGTATTTGTATCTCACCGATAGAGAACATATAAATGTAGACTCTATTAAACAGTTGTGTAAAATATCAGATCCTAATGCATGTTATAGATGTGGATGTACGGCTTTACATGAGTACTTTTATAATTATAGATCAGTCAACGGAAAATACAAGTATAGATACAACGGTTACTATCAATATTATTCATCTAGCGATTATGAAAATTATAATGAATATTATTATGATGATTATGATAGAACTGGTATGAACAGTGAGAGTGATAATATATCAATCAAAACAGAATACGAGAATGAATATGAATTCTATGATGAAACACAAGATCAAAGTACACAACTAGTAGATTACGACATTAAACTCAAAACCAATGAGGATGATTTTGTTGATGAATTCTATGGTTATGATAGATCAGTGGGTGTCCATGATTATATAGATGTATCAATTAATAAAGTAGTATATGGAAGAGAGTCTCACGTAAGATGGTGGGATATATGGCAAGAACATAATGATGGCGTATACAGTATAGGAAAGGAGTGCATAGATAATATATACGAAGACAGACATACCGTAGACGAATTCTACAAGATAGACAGCGTATCAGATGTAGATGACGCAGAACATATATCTCAGATAACTAATGATGTATCTACACAAACATGGGAAAAGAAATCAGAGTTAGATAGATACATGGAAATGTATCCTCGTCATAGATATGGTAAGCATTCTGTCTTTAAGGGATTTTCTGACAAAGTTAGAAAAAATGATTTAGACATGAACGTGGTAAAAGAATTACTTTCTAACGGTGCATCTCTAACAATCAAGGATAGCAGTAATAAGGATCCAATTGCTGTTTATTTTAGAAGAACAATAATGAATTTAGAAATGATTGATATCATTAACAAACATACAACTATCTATGAACGCAGGTATATAGTACACTCCTATCTAAAAAATTATAGAAATTTCGATTATCCATTTTTCAGAAAGTTAGTTTTGACTAATAAACATTGTCTCAACAATTATTGTAATATAAGCGACGGCAAATATGGAACACCACTACATATATTAGCATCTAATAAAAAAATAATAACTCCTAATTACATGAAGTTATTAGTGTATAACGGAAATGATATAAACGCACGAGGTGAAGATACACAAATGCGAACTCCATTACACAAATATTTGTGTAAATTTGTATATCATAATATTGAATATGGTATCCGATACTATAATGAAAAGATTATAGACGCATTTATAGAGTTAGGAGCCGATCTAACTATTCCAAATGACGATGGAATGATACCAGTAGTTTACTGTATACACTCAAATGCCGAATATGGTTATAACAATATTACTAACATAAAGATAATACGTAAACTACTTAATCTTAGTAGACATGCGTCACATAATCTATTTAGAGATCGAGTCATGCACGATTATATAAGTAATACATATATTGATCTTGAGTGTTTAGATATCATTAGATCACTTGATGGGTACGATATTAATTGTTACTTTGAAGGACGTACACCACTTCATTGCGCTATACAATATAACTTCACTCAGATTGCTGAGTACTTATTAGATCGAGGAGCTGATATATCATTAAAGACAGACGATGGTAAAACTGTATTTGATTTATCGTTATGTAGTTACATTCCTCTTAAATGGACTAGCTTTTTGATTAGTCGTCTACCGCCTAAAAGTGTCATATGCTCACTGACTAACCATATAATAGATTATGTTCTTACGAACAATAGACGTATTATTTGGCAGAGTCAAATGATTAATAAGTACGTACTGTTACTGGACCCATCCTTTTATTATAGATTCAGAAATGTTATCGAAAACAAATTAGACCAATACAATAATCGTTATAATATGTTCGAACACGATAGGGACGTTAATGAAAAGTATGGCAAAGTCTTACATGACCTCGATACATATATCAAGGATGTACAAGTATTAAAATCTACTTCCATCACTAATAATATAACACTATACGACACTATTATAAATAATAAGTCAGAGTTTCCTATACGTCGTGTAAACGACAAACAATTAATTAATCTCATAAAATCCAATACATATCATAATCTTATCGAAAAAGTTATTAAAAATACATTAGAGAAATATACTTTAACTAATATAGTCCTCGAGTATATGATCTCATCTCGATCTCAATCATCTTATTTGAGTCGTATTCCTAATGAGATATTACTCGAAATATTATATAAACTCGACATGTACGATTTACGTAATCTATATACAAGATATATGAGAGAGAATGATATCACAGAGTATCATATAGAGAATACGAGGTCTGTTTCTACACAGACATGAATAATGAATACACATACAACGTTTTTTTTAATCTTAGATATAACACTAATTACATCAAGATTATATATTGAAATCGTAATTTGAGTTGTCTGATCATCATGGATATCGAAAATGATATACGTAACATTAGCAATCTTTTAGATGATTGATATATTATTATGCGATGTAATCATAACTATCGGAGATGTAGAAATTAAAGCGCATAAAACTATTTTGGTTGCCGGATCTACGTATTTTAAAACAATGTTCACAACATCTATGATAGCGAGAGATCTAGCAACTAGAGTAAATATACAGATGTTCGATAAAGATGCCGTCAAAAATATTGTACAGTACTTATACAATAGGTATATAAGTTCTATGAATGTGATAGACATATTAAAATGCACCGACTAAGAACGTAAAACGAACTATAGAATGTTATACAATGGGTGATGATAAGTAGAAGATGTTACCCGATATACCCATAGCATTATCTAGTTATGGCATGTGTGTATTAGATCAATACATATACATTATAAGCGGTCGTACCCAACACTGATTATACATCGGTACATACAGTAAATAGCATAGATATGGAGGAGGATACAAATATTTCAAATAAAGTTATGAGATACGCGCTGTCAATAATATATGGAAGACATTACCTAACTTCTGAACTGGAACTATAAATCCAGGCTCTCGCATAAAGATGAATATATATGTTGTATGCGACATCAAAGATGAAAAAATGTTAAGACTTATATATTTAGATATAACACGAATATGTATGACGGATGGGAATTGGTAACGATGACAGAAAGCAGATTGTCAGCTCTGCATACTATTCTTCATGACAATACCATAATGATGTTACATTGTTATGAAGCGTATATGTTACAAGATACATTTAATGTGCTTACGGAACATATATTTAGAAACATCTACTAACGATTTTTTATGCTTGTATTATTAATGGTATGTAATATGATTTAATTGATTGTGTACACGATACCAATTTGTCGAGTATGAATACGGAGTACAAACATAAACTGAAGTTTAACATTATTTATTTATGNNATATACATTATATACATTATATACATTATATACATTATATACATTATATACATTATATACATTATATATCGTTATTGTTTGGTCTATGCCATGGATATCTTTAAAGAACTAATCTTAAAACATACGGATGAAAATGTTTTGATTTCTCCAGTTTCCATTTTATCTACTTTATCTATTCTGAATCATGGAGCAGCTGGTTCTACAGCTGAACAACTATCAAAATATATAGAGAATATGAATGAGAATACACCCGATGATAAGAAGGATGACAATAATGACATGGACGTAGATATTCCGTATTGCGCGACACTAGCTACCGCAAATAAAATATACGGTAGTGATAGTATCGAGTTCCATGCCTCATTCCTACAAAAAATAAAAGACGATTTTCAAACTGTAAACTTTAATAATGCGAACCAAACAAAGGAACTAATCAACGAATGGGTTAAGACAATGACAAATGGTAAAATTAATTCCTTATTGACTAGTCCGCTATCCATTAATACTCGTATGATAGTTATTAGCGCCGTCCATTTTAAAGCAATGTGGAAATATCCATTTTCTAAACATCTTACATATACAGACAAGTTTTATATTTCTAAGAATATAGTTACCAGTGTTGATATGATGGTGGGTACCGAGAATGACTTGCAATATGTACATATTAATGAATTATTCGGAGGATTCTCTATTATCGATATTCCATACGAGGGAAACTCTAGTATGGTGATTATACTGCCGGACGACATAGAAGGTATATATAACATAGAAAAAAATATAACAGATGAAAAATTTAAAAAATGGTGTGGTATGTTATCTACTAAAAGTATAGACTTGTATATGCCAAAGTTTAAAGTGGAAATGACGGAACCGTATAATCTGGTACCGATTCTAGAAAATTTAGGACTTACTAATATATTTGGATATTATGCAGATTTTAGTAAGATGTGTAATGAAACTATCACTGTAGAAAAATTTCTACATACGACGTTTATAGATGTTAATGAGGAGTATACAGAAGTATCGGCCGTTACAGGAGTATTCATGACTAACTTTTCGATGGTATATCGTATGAAGGTCTACATAAACCATCCATTCATATACATGATTAAAGATAACACCGGACATACACTTTTTATAGGGAAATACTGCTATCCGCAATAAATATAAACAATAGACTTTTATCACGTTATCTCATGTATAAAATATTACAAATAGTATAGCATAAACTAAAGTCGATACATACATTAAAACTTAAATAATAATGTAATTTACAATTAATAGTATAAACTAAAAAAATTAAAAAATTAAAAACAATATCATTATTATAAGTAATATCAAAATGACGATATACGGATTAATAGCGTATCTTATATTCGTGACTTCATCCATCGCTAGTCCACTTTACATTCCCGTTATTCCGCCCATTTCGGAAGATAAATCGTTCAATAGTGTAGAGGTATTAGTTTCTTTGTTTCCCGATGACCAAAAAGACTATACAGTAACTTCTCAGTTCAATAACTACACTATCGGTACCAAAGACTGGACTATCAACGTACTATCCACACCTGATGGTCTGGACATACCATTGACTAATATAACTTATTGGTCACGGTTTACTATAGGTCGTGCATTGTTCAAATCAGAGTCTGAGGATATTTTCCAAAAGAAAATGAGTATTCTAGGTGTTTCTATAGAATGTAAGAAGCCGTCGACATTACTTACTTTTTTAACCGTGCGTAAAATGACTCGAGTATTTAATAGATTTCCAGATATGGCTTATTATCGAGGAGACTGTCTAGAAGCCGTTTATGTAACAATGACTTATAAAAATACTAAAACTGGAGAGACTGATTACACGTACCTCTCTAATGGGGGGTTGCCTGCATACTATCGTAATGGGGTCGATGGTTGATTATTGATTAGTATATTCCTTATATTCCTTATTCTTTTTATTCACACAAAAAGAACATTTTTATAAACATGAAACCACTGTCTAAATGTAATTATGATCTTGATTTATAGATGATGATCAGCCTTCAGAGGATTTTGACCAGTATGTTTAATATGAAAAAAAACATAACTATTAAGCGCTATTGCGCTATTGTGCTTAATTATTTTGCTCTATAAACTGAATATATAGCCACAATTATTGACGGGCTTGTTTGTGACCGACAATCATGAATTTTCAGAAATTATCTCTGGCTATATATCTTACGGTGACATGTTCGTGGTGTTATGAAACATGTATGAGAAAAACTGCGTTGTATCATGACATTCAATTGGAGCATGTAGAAGACAATAAAGATAGTGTAGCATCGCTACCGTACAAGTATCTACAAGTAGTCAAACAAAGAGAACGTAGTAGATTGTTGGCTACATTTAATTGGACGGATATAGCTGAGGGTGTTAGAAATGAGTTCATTAAAATATGTGATATCAACGGAACATATCTATATAATTATACTATTGCTGTTAGTATAATTATTGATTCCACGGAAGAACTACCAACAGTTACTCCAATTACAACATATGAACCTTCTATATATAATTATACTATCGATTATAGCACTGTTATTACTACTGAAGAACTACAAGTGACTCCAACATATGCACCTGTAACAACTCCTCTTCCAACATCAGCAGTTCCTTATGATCAACGATCGAATAACAATGTAAGTACTATATCTATTCAGGTACTGAGTAAAATATTGGGAGTCAATGAAACAGAATTAACTAATTATCTTATTATGCATAAAAATGACACTGTTGACAATAACACCATGGTTGATGATGAGACATCTGACAATAACACATTACATGGTAATATAGGATTTTTGGAAATAAATAATTGTTATAATGTTTCTGTGTCAGATGCTAGTTTTAGAATAACATTAGTAAACGATACTTCTGAAGAAATTTTGCTAATGCTAACAGGAACTAGTTCATCCGACACCTTCATATCTTCCACCAATATCACTGAATGTTTGAAAACATTAATCAATAATGTGTCGATTAATGATGTACTTATAACACAAAATATGAATGTAACATCTAATTGTGATAAATGCTCAATGAATTTGATGGCATCCGTTATTCCTGCAGTTAATGAATTTAACAATACGTTGATGAAAATTGGTGTAAAAGATGATGAAAACAATACGGTATATAAATATTATAATTGTAAACTAACTACAAATTCTACATGTGATGAGTTAATCAATTTAGATGAAGTCATTAACAACATAACTCTGACAAATATTATACACAATAGTGTTTCGACAACTAACAGCAGAAAAAGACGAGATCTGAATGATGAGTTTGAATTTTCCACTTCCAAGGAATTAGATTGTCTTTACGAATCATATGGTGTAAACGATGATATAAGTCATTGTTTTGCATCACCTAGACGTAGACGATCTGACGACAAAAAGGAGTACATGGACATGAAATTATTCGACCACGCGAAAAAAGATTTAGGAATAGACAGTGTTATTCCTAGAGGTACAACCCATTTCCAAGTAGGTGCATCTGGTGCAAGTGGTGGTGTTGTAGGAGATAGTTTCCCATTTCAAAATGTTAAATCGCGTGCCAGTCTATTGGCGGAAAAAATAATGCCTAGAGTACCTATTACTGCTACCGAAGCTGATCTATATGCAACTGTAAATAGACAACCCAAGTTACCAGCAGGTGTTAAAAGTACTCCGTTTACAGAGGCGCTTGTGTCTACGATAAACCAAAAGCTTTCTAATGTTAGAGAGGTAACTTATGCTTCGCTCAATCTGCCAGGATCAAGTGGCTATGTTCATAGACCATCTGATTCTGTTATTTATAGCAGTATAAGACGGTCACGTTTACCTAGTGATAGCGATAGTGATTATGAGGATATACAAACTGTTGTTAAGGAATATAATGAAAGATATGGTAGATCAGTCAGTAGAACACAGTCATCAAGTAGTGAAAGCGATTTTGAAGATATAGATACTGTTGTTAGGGAATATAGACAAAAATATGGCAATGCAATGGCAAAAGGACGTAGTAGTTCCCCTAAACCTGATCCATTATATAGTACTGTTAAGAAAACAACTAAAAGTCTATCTACTGGTGTAGACATAGTTACAAAACAATCAGACTATTCTCTATTACCTGACGTTAATACTGGCAGTTCTATTGTGTCACCTCTCACCAGAAAAGGAGCTACTAGACGACGACCTAGACGCCCTACAAATGATGGTCTACAGAGTCCAAATCCTCCTCTCCGTAATCCACTTCCTCAACATGATGATTATTCTCCTCCACAAGTACACAGACCTCCACCACTTCCTCCTAAACCAGTCCAAAATCCGCCACAACTTCCCCCTAGACCAGTAGGTCAATTACTACCTCCTCCTATAGATCAACCAGATAAAGGATTTAGTAAGTTTGTATCACCTAGACGGTGTAGAAGAGCAAGCTCTGGAGTCATATGTGGTATGATACAATCAAAACCAAACGATGATACCTATTCACTTCTTCAACGATCAAAAATTGAACCAGAATATGTGGAGGTTGGTAATGGTATACCCAAGAACAATGTTCCTGTAATAGGTAATAAACATAGTAAAAAATATACATCGACGATGTCAAAAATATCAACAAAATTTGATAAATCTACGGCATTTGGAGCAGCAATGTTACTAACTGGTCAGCAGGCCATTAGCCAACAGACTAGATCAACTACGTTGAGTAGAAAAGATCAGATGAGTAAGGAAGAAAAGATATTCGAAGCAGTTACAATGAGTCTATCAACTATAGGTTCAACGTTGACGTCTGCAGGTATGACGGGTGGTCCAAAACTAATGATTGCAGGAATGGCTATAACGGCTATAACTGGTATAATAGATACGATAAAAGATATATATTACATGTTTTCAGGACAGGAGAGGCCAGTAGATCCTGTTATTAAATTATTTAATAAGTACACTGGCTTAATGTCCGATAATAATAAAATGGGTGTAAGAAAATGTTTGACACCCGGTGACGACACACTTATTTATATCGCATACAGAAACGATACCAGTTTTAAACAGAATACGGATGCGATGGCTTTGTATTTCTTAGATGTTATCGATTCAGAGATCCTATATCTAAACACATCAAATTTAGTTCTAGAGTATCAACTAAAGGTGGCTTGCCCCATAGGAACATTAAGATCTGTAGATGTGGACATAACTGCGTATACAATATTATATGATACAGCGGATAATATTAAGAAATACAAGTTTATCAGAATGGCAACGCTACTATCCAAACATCCAGTTATTAGATTGACATGTGGTTTAGCAGCAACATTGGTGATTAAACCGTACGAGGTACCCATCAGTGATATGCAACTACTAAAAATGGCGACGCCTGGTGAACCAGAATCCACTAAATCTATACCATCCGATGTCTGTGATAGGTATCCTCTAAAGAAATTCTATCTTTTAGCTGGTGGTTGTCCCTATGATACATCTCAAACTTTTATTGTACATACTACTTGCAGTATTCTACTAAGAACAGCTACACGGGATCAGTTTAGAAACAGATGGGTGTTACAAAATCCATTTAGACAAGAAGGGACATATAAGCAACTGTTTACCTTTAGCAAATACGATTTTAACGACACCATAATCGATCCTAATGGTGTGGTGGGTCATGCTAGCTTTTGTACCAATAGAAGCAGCAACCAATGTTTCTGGTCCGAACCTATGATATTGGAAGATGTATCATCGTGTAGTTCTAGAACTAGAAAAATATACGTAAAACTGGGAATATTTAATGCTGAAGGTTTTAATAGTTTTGTACTAAATTGTCCAACTGGGTCTACACCTACATACATCAAACATAAAAATGCGGACAGTAACAATGTTATCATAGAGCTACCTGTAGGTGATTACGGCACAGCCAAATTGTATTCAGCAACAAAACCATCGAGGATAGCTGTGTTCTGCACACATAACTATGATAAACGATTCAAATCAGATATTATAGTTCTAATGTTTAATAAAAACAGCGGTATTCCATTTTGGAGCATGTACACAGGAAGTGTAACTAGTAAAAATAGAATGTTTGCCACATTGGCTAGAGGAATGCCGTTTAGATCAACGTATTGCGATAACAGACGACGATCAGGTTGTTATTATGCAGGAATACCATTTCATGAAGATAGTGTAGAAACAGATATACATTATGGACCAGAAATAATGTTAAAGGAAACATATGACATAAACAGTATTGACCCACGAGTTATAACAAAGTCAAAGACCCATTTTCCTGCTCCATTGAGTGTAAAATTCATGGTTGACAATTTAGGAAATGGATATGACAACCCTAATTCATTTTGGGAAGATGCTAAAACTAAGAAACGGACATATAGTGCAATGACGATAAAAGTCCTACCATGTACAGTGAGAAATAAAAATATAGACTTTGGATATAACTATGGAGATATTATTTCTAATATGGTTTATCTACAATCTACTAGTCAGGATTATGGAGATGGTACCAAATATACATTTAAATCCGTAACTAGATCAGATCATGAGTGTGAATCTAGCTTAGATCTAACGTCTAAGGAAGTAACTGTGACATGTCCTGCGTTTAGTATACCAAGAAATATATCAACATATGAAGGTCTATGCTTTAGTGTTACTACATCTAAAGATCATTGTGCTACAGGTATTGGTTGGTTAAAATCTAGTGGTTATGGGAAGGAAGATGCTGATAAACCACGTGCTTGTTTTCATCATTGGAATTATTACACACTGTCGTTGGATTATTACTGTTCATACGAAGATATTTGGAGAAGCACCTGGCCTGACTATGATCCATGTAAGTCATATATCCATATAGAGTATAGAGATACATGGATAGAATCTAATGTGTTACAGCAACCTCCTTACACATTCGAATTCATTCATGACAATTCTAACGAATATGTGGATAAAGAAATTAGTAACAAATTAAATGATCTGTACAATGAATACAAGAAGATTATGGAATATAGCGACGGATCATTGCCGGCGTCTATAAACAGATTAGCAAAGGCATTGACTTCAGAGGGTAGAGAAATAGCAAGTGTTAATATAGATGGTAATCTGTTAGATATCGCATATCAAGCAGATAAGGAAAAGATGGCCGACATACAGACAAGAATAAATGATATTATTAGAGATTTGTTTATACACACTCTATCAGACAAAGATATAAAAGACATTATAGAATCCGAAGAAGGTAAGAGATGTTGTATAATAGATGTTAAGAACAATCTTGTTAAAAAGTACTATTCTATTGATAATTATCTATGTGATACTTTAGATGATTATATATACACCTCTGTAGAATATAACAAATCCTATGTGTTAGTAAACGATACTTATATGAGCTATGACTATCTTGAATCATCAGGTGTAGTTGTTCTATCATGTTATGAAATGACTATAATCTCCTTGGATACAAAAGACGCCAAAGATGCTATAGAAGATGTGATAGTAGCAAGTGCGGTAGCCGAAGCATTGAATGACATGTTTAAGGAATTTGATAAAAACGTAAGTGCTATTATAATAAAAGAAGAAGATAATTATCTAAACAGTTCGCCCGATATCTACCATATAATATATATCATAGGTGGCACTATTCTGCTACTGTTAGTCATTATTTTAATATTGGCAATTTATATAGCGCGCAATAAATACAGAACCAGGAAATATGAAATAATGAAATATGACAATATGAGCATTAAATCTGAGCATCATGATAGTCTTGAAACAGTGTCTATGGAAATTATTGATAATCGGTACTAATAAAATAGTTTAACTCTTTTAGAACCAGTTTGGTACTGTAATTTCAGTTCATTACTCGTTGAGAATATTGATGATTTTTTTTAAATGAGTATCGGTAGTTACATATTACCATATCATCCATTATATAATCGATGATGCATGTATTAGAATACTTTCCGAATAAGTCTTCTAAATATTGTATTAATTATGAAAAACTATGCTATGTGAGTATGATTCAAAGATGTTTAATGATACGATACTAGATTTTATCTCTAGCGAGATTGTTTAGAATCATTTATCATAACTATGTTTAATAAATTCATCAACGAATATCGATAAAGACCTCTTGTAATTCGAGTATAGGAAGTAGTATTACCATATCAACTTCCGAGTTAACAATTACTCTAAAACATGAGGATTGTACTCCTGTCTTTATTGGAGATCACTATTTAGTCGTTGATAAACTAGTAACCTCAGGTTTCTTTACAAACGATAAAGTACAACATCAAGACCTCACAACACAGTGCAAGATTAATCTAGAAATCAAATGTAATTCTGGAGGAGAATCTAGACAACTAACACCCACGGCGAAGTATACTTTATGCCTCATTCAGAAACGGTAACTGTAGTAGGAGACTGTCTCTCTAATCTCGATGTATATATAATATATACCAATACGGACGCGATATATTCCGACATGAATGGCGTCGCTTATCATATGTTATATCCTAAATGTTGATCATATTCCACAAATGATTGTGAACGAGATTAAATCATCTAACAAATAATTAGTTTTTTATGACATTAACATATAATAAATAAATTAATCATTATTGACTTAACGATGACGAAAGTTATCATTATCTTAGGATTCTTGATTATTAATACAAATTCGTTGTGTCTATGAAATGTGAACAAGGTGTCTCATATTATAATGCACAAGAATTAAAGTGTTGTAAACTATCTAGCCAGGAACATATTCAGATTATCGATGTGATAAATACAGCGATACCATCTGTGGACATTGTCCAAGTGACACATTCACGTCAATATATAATCGTTCTCCTCGGTGTCATAGTTGTAGAGGTCACACCTTGTACACCTACCACAAATAGAATATGTCATTGTGACTCGAATAGTTATCGTCTCCTTAAAGCTTCTGATGGTAACTGTGTTACATGTGCTCCTAAAACAAAATATGGTCGTGTGTACGGAAAGAAAGGAGAAAATGATATGGAATACCATTTGTAAGAAATGTCGGAAGGGTACTTATTCAGATATTGTATCTGACTCTGATCAATGTAAACCTATGACAAGATAAGACTTACTCGCATCTACTGGATAGACATAAATATCCTCCTCGTAATAATGAAATATAATATACACTAATTATTAATATCAATCGAGTATTAACATATAAGTTATTTTTAAACCCCTTTTGGGTTCCGTCCTAAACGGCGTTTCGGTCTGTGTCGCCACCATGGTCACACCGAGCCTCTGCGTGCTCCTCCATCGAAGACGACTTCAACTATGACAGCTCGGTGGCGTCTGCCAGCGTGTACATACGAATGGCATTTCTAAGAAAAGTCTACGGTATCCTTTCTACAATTTCCTTTAACAACGGCAACAGCTGCAGTATTTTTATACTTTGAATGCATCGGACATTTATACAAGGGAGTCCTGTTCTAATATTGGCATCAATGTTCGGATCTATAGGCTTGATTTTCGCATTGACTTTACACAGACATGAACATCCCCTGAATCTGTACATACTTTGTGGATTTACACTGTTAGAATCTCTAACGCTGGCCTCTGTTGTTACTTTCTATGATGCACGTATCGTTATGCAAGCTTTCATGTTGACTACTGCAGTGTTTCTTGCTCTGACTACATATACTCTACAATCAAAGAGAGATTTCAGTAAACTTGTAACAGGATTGTTTGCTGCTTTCTGGATTTTAATTTTGTCAGGAGTCTTGAGGATAAAGTTTAAAATAGAATTAATAAAGAACATATAGGTCATTTTTTAAACATGGATAGAAACCAAGGTTGTTAGTTAATAATATACAAGATATTTTTTCTCACTCTGATCCATGTAAACCAAGGACGAGAGACACTCTCATTCCTCATTCACGACACCATTAAAAATGGAAATTAAAGCCCTCTATTAAGCACAGACGGCTACAGGTCTACCATCAGGTTACCTTCGTCTACCTTCACAATGGCCTCTCCTTGTGCCCAGTTCAGTCCCTGTCATTGCCACGCTACTAAGGACTCCCTGAATACCGTGACTGACGTCAGACATTGTCTGACTGAATACATCCTGTGGGTTTCTCATAGATGGACCCATAGAGAAAGCGCAGGGCCTCTCTACAGGCTTCTCATCTCTTTCAGAATTGATGCAATGGAGCTATTTGGTAGCGAGTTGAAGGAGTTCTCGAATTCACTTCCGTGGGACAATATCGACAATTGCGTGGAGATCATTAAATGTTTCATCAGAAATGACTCCATGAAAACCGCCAAAGAACTTTGTGCAATAATTGGACTTTGTACTCAATCAGCTATTGTCACTGGAAGAGTCTTCAATGATAAGTATATCGACATACTACTTATGCTGCGAAAGATTCTGAACGAGAACGACTATCTCACCCTCTTGGATCATATCCTCACTGCTAAATACTAAATCTCCTTCATGCTCTCTCACTAATACTCTTACTCACTACACTTTTTATCATCTTATGATGAATGATTGCCTTCATCATTTTTTCGTGGAATATAATATAGGAATAATTAGCACCAGAATAGCTATGGATATCTCGTTAAGAATATTCTCTATAAGAGACATAATGTAGACATAGTTATTATATCCTTCTTAGATAAGTGTTACGCTACTGGAAAGTTTCCATCGTTATTATTACATGAAGATGATATAATTAAACCAACATTGAGATTGGCTCTTATGTTAGCTGGATTGAATTACTGTAATAAATGCATCGAGTATAGAGGGATATAGCAATTCTCGATAATAGTCATGCAATATTTGAATGAGACTGATAATTTAGGTAATACAGTACTACACACATATCTTTCTAGATTATATATCGTTAAAAATCTGTAAGATGTATATTTCTCATAAGTATCCACTGTGTAATATTATTAATGGATATATAGATAACGCAATAGGGACTAATAGTATTGTAAAAGATATAATCGACTATTTACGTACATATCCAGATATCTATATTCCTACTAGTTTGCTGCGTAGTTGCATCATTGATATGCATGATTTATCAGGATTCAGAGATGAATTACTAAGTAAACTACAATCCCACAATAAGTAAGAATCAAATATCAAAAACTCACTTTTGATTTTTCTAGTCTTAAGTAATACATATATTTATTAATAGACCTATGAAATAAAAAAAGGTAACAATGGATTCGCGTATAGCTATTTACGTATTAGTATCGGCATCTCTTTTGTATCTTGTTAATTGTCACAAACTAGTACATTACTTCAATCTGAAAATAAATGGAAGTGATATAACTAATACAGCAGATATATTGCTGGACAATTATCCAATTATGACCTTTGATGGAAAGGATATTTATCCATCTATCTCGTTCATGGTCGGTAATAAACTTTTCCTAGATCTTTATAAAAATATCTTTGAAGAATTTTTCAGACTATTTCGAGTATCTGTAAGTAGTCAATACGAGGAATTAGAATATTATTATTCATGTGATTATACTAACAACCGTCCTACAATTAAACAACATTACTTTTATAACGGCGACGAATATACTGAAATTGATAGATCGAAAAAAGCCACTAATAAAAACAGTTGGTTAATTACTTCAGGCTTTAGACTACAAAAATGGTTCGATAGCGAAGATTGTATAATTTATCTCAGATCTTTAGTTAGAAGAATGGAAGACAGTAACAAAAACAGTAAAAAAACTTAGTACTTAGATATCGAAAAAATATATTTTTGTAGACTCTTGAGAATAGAAGGAAAACATGTACATAATTATAAAAAATGAAAATCAATGGCGAATAAGACAGTGCGATTCGCACCATGGAGTCGGTAGATTTCATGGCTGTCGATGAGCAGTTTCACGACGACCTCGATCTTTGGTCATTATCTTTGGTAGATGATTATAAAAAACATGGATTAGGTGTTGACTGTTATGTTCTAGAACCAGTTGTTGACAGGAAAATATTTGATAGATTTCTCCTTGAACCAATTTGTGATCCTGTAGATGTTCTGTATGATTATTTTAGGATTCATAGAGATAATATTGATCAGTATATAGTAGATAGACTGTTTGCATATATTACATATAAAGATATTATATCTGCATTAGTGTCAAAGAATTATATGGAAGATATTTTCTCTATAATTATTAAGAATTGTAATTCTGTGCAAGATCTCTTACTTTACTATCTATCTAATGCATATGTAGAAATAGACATTGTTGATCTTATGGTAGATCATGGGGCTGTAATATATAAAATAGAATGCTTGAATGCCTATTTTAGGGGAATATGTAAAAAGGAAAGTAGTGTTGTTGAGTTTATTTTGAATTGTGGTATCCCAGATGAAAATGATGTTAAATTAGATCTATATAAAATAATTCAGTATACTAGGGGATTCCTTGTAGATGAACCCACAGTATTAGAAATTTATAAGCTTTGTATCCCATATATTGAAGATATCAATCAACTAGATGCTGGTGGAAGGACCTTGCTTTATCGCGCTATCTATGCAGGTTATATAGATTTAGTATCATGGCTATTAGAAAATGGAGCAAATGTCAACGCAGTAATGAGTAATGGATATACATGTCTTGACGTGGCCGTGGATAGGGGATCTGTCATCGCCCGTAGGGAAGCACATCTTAAAATATTAGAAATATTGCTTAGAGAACCATTGTCTATTGACTGTATAAAATTAGCTATACTTAATAATACAATTGAAAACCATGATGTGATAAAGCTCTGTATCAAGTATTTTATGATGGTAGATTATTCACTTTGTAATGTGTATGCATCATCACTCTTTGATTATATAATTGATTGTAAACAAGAATTGGAGTACATTAGGCAGATGAAAATTCATAATACAACCATGTATGAGTTAATCTATAATAGAGACAAAAACAAGCATGCTTCCCATATTCTACATAGGTATTCTAAACATCCAGTTTTGACACAGTGTATCACTAAAGGATTCAAGATTTACACAGAAGTAACCGAGCAGGTCACTAAAGCTCTAAACAGACGTGCTCTAATAGATGAGATAATAAACAATGTATCAACTGATGACAATCTCCTATCAAAACTTCCATTAGAAATTAGGGATCTAATTGTTTCACAAGCTGTCATATAGAGTTCTATCCACCCACCTTTCTTGAAATGAGTTAATAGTCATAAGTTAGTTAAGTCATAAGTTAGTTAAGTCATAAGTTAGTTAAGTCATAAGTTAGTTAAGTCATAAGTTAGTTAAGTCATAAGTTAGTTAAGTCATAAGTTAGTTAAGTCATAAGTTAGTTTATAGTCTAACACTTCTAATTTTTATACCTTGATCTTTTTCTCTAATTATGAAAAAGTAAATCATTATGAAGATGGATGAAATGGACGAGATTGTGCGCATCGTTAACGATAGTATGTGGTACGTACCTAACGCATTTATGGACGACGGTGATAATGAAGGTCACATTTCTGTCAATAATGTCTGTCATATGTATCTCGCATTCTTTGATGTGGATATATCATCTCATCTGTTTAAATTAGTTATTAAACACTGCGATCTGAATAAACGACTAAAATGTGGTAACTCTCCATTACATTGCTATACGATGAATACACGATTTAATCCATCTGTATTAAAGATATTGTTACGCCACGGCATGCGTAACTTTGATAGCAAGGATAAAAAAGGACATATTCCTCTACACCACTATCTGATTCATTCACTATCAATCGATAACAAGATCTTTGATATACTAACGGACCCCATTGATGACTTTAGTAAATCATCCGATCTATTGCTGTGTTATCTTAGATATAAATTCAATGGGAGCTTAAACTATTACGTTCTGTACAAATTATTGACTAAAGGATCTGACCCTAATTGCGTCGATGAGGATGGACTCACTTCTCTTCATTACTACTGTAAACACATATCCGCGTTCCACGAAAGCAATTATTACAAGTCAAAGAGTCACACTAAGATGCGAGCTGAGAAGCGATTCATCTACGCGATAATAGATCATGGAGCAAACATTAACGCGGTTACGAAAATCGGAAATACGCCGTTACACACTTACCTTCAACAGTATACCAAACATAGTCCTCGTGTGGTGTATGCTCTTTTATCTCGAGGAGCCGATACGAGGATACGTAATAATCTTGATTGTACACCCATCATGGAATACATAAAGAACGATTGTGCAACAGGTCATATTCTCATAATGTTACTCAATTGGCACGAACAAAAATACGGGAAATTACAAAAGGAAGAAGGACAACATCTACTTTATCTATTCATAAAACATAATCAAGGATATGGAAGTCGCTCTCTCAATATACTACGGTATCTACTAGATAGATTCGACATTCAGAAAGACGAATACTATAATACAATGACTCCTCTTCATACCGCCTTCCAGAATTGCAATAACAATGTTGCCTCATACCTCGTATACATCGGATACGACATCAACCTTCCGACTAAAGACGATAAGACAGTATTCGACTTGGTGTTTGAAAACAGAAACATTATATACAAGGCGGATGTCGTTAATGACATTATCCACCACAGACTGAAAGTATCTCTACCTATGATTAAATCGTTGTTCTACAAGATGTCGGAGTTCTCTCCCTACGACGATCACTACGTAAAGAAGATAATAGCCTACTGCCTATTAAGGGACGAGTCATTTGCGGAACTACATACTAAATTCTGTTTAAACGAGGACTATAAAAGTGTATTTATGAAAAATATATCATTCGATAAGATAGATTCCATCATCGAAAAATGTAGTCGTGACATAAGTCTCCTCAAAGAGATTCGAATCTCAGACACCGACTTGTATACGGTATTGAGAACAGAAGACATCCGGTATCACACATATCTCGAAGCCATACATTCAGACAAACGCATTTCATTTCCCATGTACGACGATCTCATAGAACAGTGTCATCTATCGATGGAGCATAAAAGTAAACTCGTCGACAAAGCACTCAATAAATTAGAGTCTACCATCGATAGTCAATCTAGACTATCGTATTTGCCTCCGGAAATTATGCGCAATATCATAACCAAGCTAAGCGACTACCATCTAAACAGTATGTTGTACGGAAAGAACCATTACAAATATTATCCATGATAGAAAGAAAATATTTAAAAAATAATCTATATGATTGGAGAAGTAGGAAACAAACAGTAACAAGACGACGATTACTACATTATTAAATCATGAGGTCCGTATTATACTCGTATATATTGTTTCTCTCATGTATAATAATAAACGGAAGAGATATAGCACCACATGCACCATCCAATGGAAAGTGTAAAGACAACGAATACAGAAGCCGTAATCTATGTTGTCTATCGTGTCCTCCGGGAACTTACGCTTCCAGATTATGTGATAGCAAGACTAATACACAATGTACACCGTGTGGTTCGGATACCTTTACATCTCACAATAATCATTTACAGGCTTGTCTAAGTTGTAACGGAAGATGTGATAGTAATCAGGTAGAGACGCGATCGTGTAACACGACTCACAATAGAATCTGTGAATGCTCTCCAGGATATTATTGTCTTCTCAAAGGAGCATCAGGGTGTAGAACATGTATTTCTAAAACAAAGTGTGGAATAGGATACGGAGTATCCGGATACACGTCTACCGGAGACGTCATCTGTTCTCCGTGTGGTCCCGGAACATATTCTCACACCGTCTCTTCCACAGATAAATGCGAACCCGTCGTAACCAGCAATACATTTAACTATATCGATGTGGAAATTAACCTGTATCCAGTCAACGACACATCGTGTACTCGGACGACCACTACCGGTCTCAGCGAATCCATCTCAACGTCGGAACTAACTATTACCATGAATCATAAAGATTGTGATCCAGTCTTTCGTGCAGAATACTTCTCTGTCCTTAATAATGTAGCAACTTCAGGATTCTTTACAGGAGAAAATAGATATCAGAATACTTCAAAGATATGTACTCTGAATTTCGAGATTAAATGTAACAACAAAGATTCATCTTCCAAACAGTTAACGAAAACAAAGAATGATACTATCATGCCGCATTCAGAGACGGTAACTCTAGTGGGCGACTGTCTATCTAGCGTCGACATCTACATACTATATAGTAATACCAATACTCAAGACTACGAAAATGATACAATCTCTTATCATATGGGTAATGTTCTCGATGTCAATAGCCATATGCCCGCTAGTTGCGATATACATAAACTGATCACTAATTCCCAGAATCCCACCCACTTATAGTAAGTTTTTTTACCTATAAATAATAAATACAATAATTAATTTCTCGTAAAAGTAGAAAATATATTCTAATTTATTATATGGTAAGAAAGTAGAATCATCTAGAACAGTAATCAATCAATAGCAATCATGAAACAATATATTGTCCTGGCATGCATGTGCCTAGTGGCAGCTGCTATGCCTACTAGTCTTCAACAATCTTCATCCTCGTGTACTGAAGAAGAAAACAAACATCATATGGGAATCGATGTTATTATCAAAGTCACAAAGCAAGACCAAACACCGACCAATGATAAGATTTGTCAATCCGTAACGGAAGTTACAGAGACCGAAGATGATGAGGTATCCGAAGAAGTTGTAAAAGGAGATCCCACCACTTATTACACTATCGTCGGTGCGGGTCTTAACATGAACTTTGGATTCACCAAATGCCCAAAGATTTCATCCATCTCCGAATCCTCTGATGGAAACACTGTGAATACTAGATTGTCCAGCGTGTCACCGGGACAAGGTAAGGACTCTCCCGCGATCACGCGTGAAGAAGCTCTGGCTATGATCAAAGACTGTGAGATGTCTATCGACATCAGATGTAGCGAAGAAGAGAAAGACAGTGACATCAAGACCCATCCAGTACTTGGGTCTAACATCTCACATAAGAAAGTGAGTTACAAAGATATCATCGGTTCAACGATCGTTGATACAAAATGTGTCAAGAACCTAGAGTTTAGCGTACGTATCGGAGACATGTGTGAGGAATCATCTGAACTTGAAGTCAAGGATGGATTCAAGTATGTCGACGGATCGGCATCTGAAGGTGCAACCGATGATACTTCACTCATCGATTCAACAAAACTCAAAGCATGTGTCTGAATCGATAACTCTATTCATCTGAAAATGGATGAGTTGGGTTAATCGAACGATTCAGACACCGCACCACGAATTAAAAAAGACCGGGCACTATATTCCGGTTTGCAAAACAAAAATATTTAACTACATTCACAAAAAGTTACCTCTCGTTACTTCTTCTTTCTGTTTCAATATGTGATACGATATGATCACTATTCGTATTCTCTTGGTCTCATAAAAAAGTTTTACAAAAAAAAAAAAAAAATATTTTTATTCTCTTTCTCTCTTCGATGGTCTCACAAAAATATTAAACCTCTTTCTGATGTCTCAACTATTTCGTAAACGATAACGTCCAACAATATATTCTCGTAGAGCTTATCAACATCCTTATACCAATCTAGGTTGTCAGACAATTGCATCATAAAATAATGTTTATAATTTACACGTTAACATCATATAATAAACGTATATAGTTAATATTTTTGGAATATAAATGATCTGTAAAATCCATGTAGGGGACACTGCTCACGTTTTTTCTCTAGTACATAATTTCACACAAGTTTTTATACAGACAAATTAATTCTCGTCCATATATTTTAAAACATTGACTTTTGTACTAAGAAAAATATCTTGACTAACCATCTCTTTCTCTCTTCGATGGGTCTCACAAAAATATTAAACCTCTTTCTGATGGAGTCGTAAAAAGTTTTTATCCTTTCTCTCTTCGATAGGTCTCACAAAAATATTAAACCTCTTTCTGATGGTCTCTATAAACGATTGATTTTTCTTACCCTCTAGAGTTTCCTACGGTCGTGGGTCACACATTTTTTTCTAGACACTAAATAAAATAGTAAAATATAATA

>2017_Nigeria_Monkeypox_MT903340.1

ATTTTACTATTTTATTTAGTGTCTAGAAAAAAATGTGTGACCCACGACCGTAGGAAACTCTAGAGGGTAAGAAAAATCAATCGTTTATAGAGACCATCAGAAAGAGGTTTAATATTTTTGTGAGACCTATCGAAGAGAGAAAGGATAAAAACTTTTTACGACTCCATCAGAAAGAGGTTTAATATTTTTGTGAGACCCATCGAAGAGAGAAAGAGATGGTTAGTCAAGATATTTTTCTTAGTACAAAAGTCAATGTTTTAAAATATATGGACGAGAATTAATTTGTCTGTATAAAAACTTGTGTGAAATTATGTACTAGAGAAAAAACGTGAGCAGTGTCCCCTACATGGATTTTACAGATCATTTATATTCCAAAAATATTAACTATATACGTTTATTATATGATGTTAACGTGTAAATTATAAACATTATTTTATGATGCAATTGTCTGACAACCTAGATTGGTATAAGGATGTTGATAAGCTCTACGAGAATATATTGTTGGACGTTATCGTTTACGAAATAGTTGAGACATCAGAAAGAGGTTTAATATTTTTGTGAGACCATCGAAGAGAGAAAGAGAATAAAAATATTTTTTTTTTTTTTTTTGTAAAACTTTTTTATGAGACCAAGAGAATACGAATAGTGATCATATCGTATCACATATTGAAACAGAAAGAAGAAGTAACGAGAGGTAACTTTTTGTGAATGTAGTTAAATATTTTTGTTTTGCAAACCGGAATATAGTGCCCGGTCTTTTTTAATTCGTGGTGCGGTGTCTGAATCGTTCGATTAACCCAACTCATCCATTTTCAGATGAATAGAGTTATCGATTCAGACACATGCTTTGAGTTTTGTTGAATCGATGAGTGAAGTATCATCGGTTGCACCTTCAGATGCCGATCCGTCGACATACTTGAATCCATCCTTGACTTCAAGTTCAGATGATTCCTCACACATGTCTCCGATACGTACGCTAAACTCTAGGTTCTTGACACATTTTGTATCAACGATCGTTGAACCGATGATATCTTTGTAACTCACTTTCTTATGTGAGATGTTAGACCCAAGTACTGGATGGGTCTTGATGTCACTGTCTTTCTCTTCTTCGCTACATCTGATGTCGATAGACATCTCACAGTCTTTGATCATAGCCAGAGCTTCTTCACGCGTGATCGCGGGAGAGTCCTTACCTTGTCCCGGTGACACGCTGGACAATCTAGTATTCACAGTGTTTCCATCAGAGGATTCGGAGATGGATGAAATCTTTGGGCATTTGGTGAATCCAAAGTTCATGTTAAGACCCGCACCGACGATAGTGTAATAAGTGGTGGGATCTCCTTTTACAACTTCTTCGGATACCTCATCATCTTCGGTCTCTGTAACTTCCGTTACGGATTGACAAATCTTATCATTGGTCGGTGTTTGGTCTTGCTTTGTGACTTTGATAATAACATCGATTCCCATATGATGTTTGTTTTCTTCTTCAGTACACGAGGATGAAGATTGTTGAAGACTAGTAGGCATAGCAGCTGCCACTAGGCACATGCATGCCAGGACAATATATTGTTTCATGATTGCTATTGATTGATTACTGTTCTAGATGATTCTACTTTCTTACCATATAATAAATTAGAATATATTTTCTACTTTTACGAGAAATTAATTATTGTATTTATTATTTATAGGTAAAAAAACTTACTATAAGTGGGTGGGATTCTGGGAATTAGTGATCAGTTTATGTATATCGCAACTAGCGGGCATATGGCTATTGACATCGAGAACATTACCCATATGATAAGAGATTGTATCATTTTCGTAGTCTTGAGTATTGGTATTACTATATAGTATGTAGATGTCGACGCTAGATAGACAGTCGCCCACTAGAGTTACCGTCTCTGAATGCGGCATGATAGTATCATTCTTTGTTTTCGTTAACTGTTTGGAAGATGAATCTTTGTTGTTACATTTAATCTCGAAATTCAGAGTACATATCTTTGAAGTATTCTGATATCTATTTTCTCCTGTAAAGAATCCTGAAGTTGCTACATTATTAAGGACAGAGAAGTATTCTGCACGAAAGACTGGATCACAATCTTTATGATTCATGGTAATAGTTAGTTCCGACGTTGAGATGGATTCGCTGAGACCGGTAGTGGTCGTCCGAGTACACGATGTGTCGTTGACTGGATACAGGTTAATTTCCACATCGATATAGTTAAATGTATTGCTGGTTACGACGGGTTCGCATTTATCTGTGGAAGAGACGGTGTGAGAATATGTTCCGGGACCACACGGAGAACAGATGACGTCTCCGGTAGACGTGTATCCGGATACTCCGTATCCTATTCCACACTTTGTTTTAGAAATACATGTTCTACACCCTGATGCTCCTTTGAGAAGACAATAATATCCTGGAGAGCATTCACAGATTCTATTGTGAGTCGTGTTACACGATCGCGTCTCTACCTGATTACTATCACATCTTCCGTTACAACTTAGACAAGCCTGTAAATGATTATTGTGAGATGTAAAGGTATCCGAACCACACGGTGTACATTGTGTATTAGTCTTGCTATCACATAATCTGGAAGCGTAAGTTCCCGGAGGACACGATAGACAACATAGATTACGGCTTCTGTATTCGTTGTCTTTACACTTTCCATTGGATGGTGCATGTGGTGCTATATCTCTTCCGTTTATTATTATACATGAGAGAAACAATATATACGAGTATAATACGGACCTCATGATTTAATAATGTAGTAATCGTCGTCTTGTTACTGTTTGTTTCCTACTTCTCCAATCATATAGATTATTTTTTAAATATTTTCTTTCTATCATGGATAATATTTGTAATGGTTCTTTCCGTACAACATACTGTTTAGATGGTAGTCGCTTAGCTTGGTTATGATATTGCGCATAATTTCCGGAGGCAAATACGATAGTCTAGATTGACTATCGATGGTAGACTCTAATTTATTGAGTGCTTTGTCGACGAGTTTACTTTTATGCTCCATCGATAGATGACACTGTTCTATGAGATCGTCGTACATGGGAAATGAAATGCGTTTGTCTGAATGTATGGCTTCGAGATATGTGTGATACCGGATGTCTTCTGTTCTCAATACCGTATACAAGTCGGTGTCTGAGATTCGAATCTCTTTGAGGAGACTTATGTCACGACTACATTTTTCGATGATGGAATCTATCTTATCGAATGATATATTTTTCATAAATACACTTTTATAGTCCTCGTTTAAACAGAATTTAGTATGTAGTTCCGCAAATGACTCGTCCCTTAATAGGCAGTAGGCTATTATCTTCTTTACGTAGTGATCGTCGTAGGGAGAGAACTCCGACATCTTGTAGAACAACGATTTAATCATAGGTAGAGATACTTTCAGTCTGTGGTGGATAATGTCATTAACGACATCCGCCTTGTATATAATGTTTCTGTTTTCAAACACCAAGTCGAATACTGTCTTATCGTCTTTAGTCGGAAGGTTGATGTCGTATCCGATGTATACGAGGTATGAGGCAACATTGTTATTGCAATTCTGGAAGGCGGTATGAAGAGGAGTCATTGTATTATAGTATTCGTCTTTCTGAATGTCGAATCTATCTAGTAGATACCGTAGTATATTGAGAGAGCGACTTCCATATCCTTGATTATGTTTTATGAATAGATAAAGTAGATGTTGTCCTTCTTCCTTTTGTAATTTCCCGTATTTTTGTTCGTGCCAATTGAGTAACATTATGAGAATATGACCTGTTGCACAATCGTTCTTTATGTATTCCATGATGGGTGTACAATCAAGATTATTACGTATCCTCGTATCGGCTCCTCGAGATAAAAGAGCATACACCACACGAGGACTATGTTTGGTATACTGTTGAAGGTAAGTGTGTAACGGCGTATTTCCGATTTTCGTAACCGCGTTAATGTTTGCTCCATGATCTATTATCGCGTAGATGAATCGCTTCTCAGCTCGCATCTTAGTGTGACTCTTTGACTTGTAATAATTGCTTTCGTGGAACGCGGATATGTGTTTACAGTAGTAATGAAGAGAAGTGAGTCCATCCTCATCGACGCAATTAGGGTCAGATCCTTTAGTCAATAATTTGTACAGAACGTAATAGTTTAAGCTCCCATTGAATTTATATCTAAGATAACACAGCAATAGATCGGATGATTTACTAAAGTCATCAATGGGGTCCGTTAGTATATCAAAGATCTTGTTATCGATTGATAGTGAATGAATCAGATAGTGGTGTAGAGGAATATGTCCTTTTTTATCCTTGCTATCAAAGTTACGCATGCCGTGGCGTAACAATATCTTTAATACAGATGGATTAAATCGTGTATTCATCGTATAGCAATGTAATGGAGAGTTACCACATTTTAGTCGTTTATTCAGATCGCAGTGTTTAATAACTAATTTAAACAGATGAGATGATATATCCACATCAAAGAATGCGAGATACATATGACAGACATTATTGACAGAAATGTGACCTTCATTATCACCGTCGTCCATAAATGCGTTAGGTACGTACCACATACTATCGTTAACGATGCGCACAATCTCGTCCATTTCATCCATCTTCATAATGATTTACTTTTTCATAATTAGAGAAAAAGATCAAGGTATAAAAATTAGAAGTGTTAGACTATAAACTAACTTATGACTTAACTAACTTATGACTTAACTAACTTATGACTTAACTAACTTATGACTTAACTAACTTATGACTTAACTAACTTATGACTTAACTAACTTATGACTTAACTAACTTATGACTATTAACTCATTTCAAGAAAGGTGGGTGGATAGAACTCTATATGACAGCTTGTGAAACAATTAGATCCCTAATTTCTAATGGAAGTTTTGATAGGAGATTGTCATCAGTTGATACATTGTTTATTATCTCATCTATTAGAGCACGTCTGTTTAGAGCTTTAGTGACCTGCTCGGTTACTTCTGTGTAAATCTTGAATCCTTTAGTGATACACTGTGTCAAAACTGGATGTTTAGAATACCTATGTAGAATATGGGAAGCATGCTTGTTTTTGTCTCTATTATAGATTAACTCATACATGGTTGTATTATGAATTTTCATCTGCCTAATGTACTCCAATTCTTGTTTACAATCAATTATATAATCAAAGAGTGATGATGCATACACATTACAAAGTGAATAATCTACCATCATAAAATACTTGATACAGAGCTTTATCACATCATGGTTTTCAATTGTATTATTAAGTATAGCTAATTTTATACAGTCAATAGACAATGGTTCTCTAAGCAATATTTCTAATATTTTAAGATGTGCTTCCCTACGGGCGATGACAGATCCCCTATCCACGGCCACGTCAAGACATGTATATCCATTACTCATTACTGCGTTGACATTTGCTCCATTTTCTAATAGCCATGATACTAAATCTATATAACCTGCATAGATAGCGCGATAAAGCAAGGTCCTTCCACCAGCATCTAGTTGATTGATATCTTCAATATATGGGATACAAAGCTTATAAATTTCTAATACTGTGGGTTCATCTACAAGGAATCCCCTAGTATACTGAATTATTTTATATAGATCTAATTTAACATCATTTTCATCTGGGATACCACAATTCAAAATAAACTCAACAACACTACTTTCCTTTTTACATATTCCCCTAAAATAGGCATTCAAGCATTCTATTTTATATATTACAGCCCCATGATCTACCATAAGATCAACAATGTCTATTTCTACATATGCATTAGATAGATAGTAAAGTAAGAGATCTTGCACAGAATTACAATTCTTAATAATTATAGAGAAAATATCTTCCATATAATTCTTTGACACTAATGCAGATATAATATCTTTATATGTAATATATGCAAACAGTCTATCTACTATATACTGATCAATATTATCTCTATGAATCCTAAAATAATCATACAGAACATCTACAGGATCACAAATTGGTTCAAGGAGAAATCTATCAAATATTTTCCTGTCAACAACTGGTTCTAGAACATAACAGTCAACACCTAATCCATGTTTTTTATAATCATCTACCAAAGATAATGACCAAAGATCGAGGTCGTCGTGAAACTGCTCATCGACAGCCATGAAATCTACCGACTCCATGGTGCGAATCGCACTGTCTTATTCGCCATTGATTTTCATTTTTTATAATTATGTACATGTTTTCCTTCTATTCTCAAGAGTCTACAAAAATATATTTTTTCGATATCTAAGTACTAAGTTTTTTTACTGTTTTTGTTACTGTCTTCCATTCTTCTAACTAAAGATCTGAGATAAATTATACAATCTTCGCTATCGAACCATTTTTGTAGTCTAAAGCCTGAAGTAATTAACCAACTGTTTTTATTAGTGGCTTTTTTCGATCTATCCTCGTCCTCTCATCATCCTTATATTATTATCATTATTATCATAGTCTATTAAACACAAATCATCTACGTTTATAACAACATTCTCATTATTAATTAGTTCTGTAGAATATCTTTAATAATTTGGCTATACATCTGTTCAATACTATCTATTGATGATTTCTTTTTTAAGACTTAAACTAGTTATGGTAATGACGATGAAATCGAGTAGTAACTTCTAATAAAGACTTGATATCATTATCATATGTTTGATCGTCATAGTTAATAGTGTGGCTAAATGGTACTGTTAATAAGTTTATAGACAATATCATAGTATTTTCTTTCCAGAATTAGATTATTTTTTTAAATACTGATCCTCACAATTCCGTGATGTAGCAGTAGTTGGTGCATGGTCTATATCGTTAAAATGTATCATATATAATAGTTTTCTGACGTGGAGTACAGAATTTTCGATTAATGAGTTCATGGTAAGGAAGGGCAAATGCCTGTATATAATATACATAAGTTAATAGTTTTTTATCATATTTTCTAATACCATAATAAAAATTATCATTATTGCGTTTGGTAGTTCTGCCCTATCATCTATATCACTGTCACTCTCGCTCTCACTATATCTTCTAAAATTACAAACAACTGGATATTCGATAACAGCATTTGTGTAGTTTTTGTCTTTTACAGTATATACGTTATTGTCAAAATCTAAACAAATATTAGCATAATACATCTATCTATAAGATCAGGATCCATGTTCGAGCATACTAGCCATGTATATTTGTAACTTCGTCGTACAGCGTTAGATCAATAGAATAAACAATCGTGTGACGCAACTTTTTTACGATCTAGTTGTATGAGTTTATCGTTTACATAAGCAATTAACGGCTTTAACAGATGATCTGAGTAATAATATACCTCTGTTATACGTTTAATGTTCACGGTCTTAGTATTTTTAGATATCAATTGTGATTTACACCATATTCGACTCCCTTGTGTGTAACGTTAGAAATTCTAAATCTATATTATCTATTACAGCGTAAAACACATTCAATATTGTATTGTTATTTTTATATTATTTACACAATTAACAATGTATTATTAGTTTATATTACTGAATTAATAATATAAAATTCCCAATCTTGTCATAAACACAAAATCCATTAAAAATGTCGATAAAATATCTGATGTTGTTGTTCGCTGCTATGATAATCAGATCATTAGCCGATAGTGGTAATGCTATCGAAACAACATCGCCAGAAATTACAAACGCTACAACAGATATTCCAGCTATCAGATTATGTGGTCCAGAGGGAGATGGATATTGTTTACACGGTGACTGTATCCACGCTAGAGATATCGACGGTATGTATTGTAGATGCTCTCATGGTTATACAGGTATTAGATGTCAGCATGTAGTATTAGTAGACTATCAACGTTCAGGAAAACCAGACACTACAACATCATATATCCCATCTCTAGGTATTGTGCTTGTATTAGTAGGCATTATTATTACGTGTTGTCTATTATCTGTTTATATGTTCACTCGAAGAACTAAACTACCTATACAAGATATGGTTGTGTTATATTTTTTATAAAATTTTTTTATGAGTATTTTTACAAAAATGTATATGTATAAAAAAAATACTAAGTATACGATGTATCCTGTATTATTTGTATTTATCTAAACAATACTTCTGCCTCTAGATGGGATACAAAAATTTTTTATTTCAGCATATTAAAGTAAAATCTAGTTACCTTGAAAATGAATACAGTGGGTGGTTCCGTATCACCAGTAAGAACATAATAGTCGAATACAGTATCCGATTGAGATTTTGCATACAATACTAGTCTAGAAAGAAATTTGTAATCATCCTCTGTGACAGGAGTCCATATATCTGTATCATCGTCTAGTTTTTATCAGTGTCCTATGCTATATTCCTGTTATCATCATTAGTTAATGAAAATAACTCTCGTGCTTCAGAAAAGTCAAATATTGTATCCATACATATATCTCCAAAACTATCACTTATACGTTTATCTTTAACGAACATATACCTAGATGGTTATTTACTAACAGACATTTTTTCAAGATCTATTGACAATAACTCCTATAGTTTCCACATCAACCAAGTAATGATCATCTATTGTTATATAACAATAACATAACTCTTTTCCATTTTTATCAGTATCTATATCAACGTCGTTGTAGTGAATAGTAGTCATTGATCTATTATATGAAACGGATATGTCTAGTTAATATTTTCTTTGATTTAAAGTCTATAGTCTTTACAAACATAATATCCTTATCCGACTTTATATTTCCTGTAGGGTGGCATAATTTTATTCTGCCTCCACAATCAGTGTTTCCAAATATATTACTAGACAATATTCCATATAGTTATTAGTTAAGGGACCCAATTAGAACACGTACGCGCTTATTCATCATTTGGATCGTATTTCATAAAAGTTATTATGTTATAGATGTCAACACATTCTACATTTTTTTAATTGTCTATATAGTATTTTTCTGATATTTTCTATAATATCAGAATTGTCTTCCATAGGAAGTTGTATACTATTCGGAATCAGTTACATGTTTAAATAATTCTCTGATGTCATTCCTTATACAATCAAATTCATTATTAAACAGTTTAATAGTCTGTAGACCTTTATCGTCGTACATATCCATTGTCTTATTAGTTACGCTTATTTTTATGGGTTTTACGTTGCTTTATTATATTTTATAATAATGATTGTTTGACAATGTCGTAGTATAGATATATTATTAGAGGAGGTATAATTATAAAAAGTTTCTGAGTACGATGTTATAAGAGGAGAGGACACATTAACATCATACATCAATTAACTACATTCTTATAACATTGTAATCAAAAGAATTGCAATTTTGATGTATAACAACTGTCAATGGAATTGTATATTACAAATTACGGTATGTTGTAACAACAAATACCGATCGGTAATTGTCTCTGCCGCTGTAATAGAATTAATTATATATCTATTACACCGGCCTTGTATCATAATAAAGTTGTGGTAGTATGATCTCCATATTTATAATTTAGTACTTTGTATTTAGTTTTTTTGGAATCATAAAAAAAGTTTTACTAATTTAAAATTTAAAAAGTATTTACATTTTTCACTGTTTAGTCGCGGATATGGAATTCGATCCTGTCAAAATCAATACATCATCTATAGATCATGTAACAATATTACAATACATAGATGAACCAAATGATATAAGACTAACAGTATGCATTATCCGAAATGTTAATAACATTACATATTATATCAATATCACAAAAATAAATCCACATTTGGCTAATCGATTTCGGGCTTGGAAAAAACGTATCGCCGGAAGGGACTATATGACTAACTTATCTAGAGATACAGGAATACAACAATCAAAACTTACTGAAACTATACGTAACTGTCAAAAAAATAAAAACATATATGGTCTATATATACACTACAATTTAGTTATTAATGTGGTTATTGATTGGATAACCGATGTGATTGTTCAATCAATATTAAGAGGGTTGGTAAATTGGTACATAGCTAATAATACATATACTCCAAATACACCCAATAATACTACAACCATTTCTGAGTTGGATATCATCAAAATACTGGATAAATACGAGGACATGTATAGAGTAAGTAAAGAAAAAGAATGTGGAATTTGCTATGAAGTTGTTTACTCAAAACGATTAGAAAACGATAGATACTTTGGTTTATTGGATTCGTGTAATCATATATTTTGCATAACATGTATCAATATATGGCATAGAACACGAAGAGAAACCGGTGCGTCAGATAATTGTCCTATATGCCGTACACGTTTTAAAAAAATAACAATGAGCAAGTTCTATAAGCTAGTTAACTAATAAATAAAAAGTTTAATTATCGACGATATATGTCGTTATTTTTCTCTCATATGAAAGATTAATTTGATTCTAATATAATCTTCAGTATTGGATGAATCTCAATTCAAATTAATTCCATTAGATTAGATTAGATTAGATCATAAATAAAAATAGTAGCACGCACTACTTCAGCCAAATATTCTTTTTTGAAACGCCATCTAGCGTAATGAGAACACAAGTGAACCTATAATGAGCAAATTTATTAGTATCGGTTACATGAAGGACTTTACGTAGAGTGGTGATTCCTCCATCTGTGGTACGAACGGTTTCATCTTCTTTGATGCCACCACCCAGATGTTCTATAAACTTGGTATCCTCGTCCGATTTCATATCATTTGCCAACCAATACATATAGCTAAACCCAGGCATACGTTCCACACATCCGGAACAATGAAATTCTCCAGAAGATGTTACAATGGCTAGATTTGGACATTTGGTTTCAACCGCGTTAACATATGAGTGAACACACCCATACATGAAAGCGATGAGAAATAGGATTCTCATCTTGCCAAAATATCACTATAAAAAATTTATTTATCAATTTTAAAGGTATAAAAAAATACTTATTGTTGCTCGAATATTTTGTATTTGATGGTATACGGAAGATTAGAAATGTAGGTATTATCATCAACTGATTTTATGATGGTTTTATGAATTTTATTATGCTTCACTATTGCATCGGAAATAATATCATATGCTTCCACGTATATTTTATTTTGTTTTGACTCATAATACGCACGTAATTCTGGATTATTGGCATATCGATGAATAATTTTAGCTCCATGCTCAGTAAATATTAATGAGAACATAGTGTTGCCTCCTACCATTATTTTTTTCATCTCATTCAATTCTTGATTGCAGAGATCTATATAATCATTATAGCGTTGACTTATGGACTCTGGAATCTTAGACGATGTACAGTCATCTATAATCATGGTATATTTAATACATTGTTTTATAGCATAGGCATTATCTACGATATTAGATACTTCACTCAATGAATCAATCACACAATCTAATGTAGGTTTATGACATAATAGCATTTTCAGCAGTTCAATGTTTCTAGATTCGTTGATGGCAATGGTTATACATGTATATCCGTTATTTGATCTAATGTTGACATCTGAACCGGATTCTAGCAGTAAAGATACTAGAGATTGTTTATTATATCTAACAGCCTTGTGAAGAAGTGTTTCTCCTCGTTTGTCAATCATGTTAATGTCTTTAAGATAAGGTAGACAAATGTTTATAGTACTAAGAATTGGGCAAACATAAGACATGTCACAAAGACCCTTTTTGTATGTATAAGTGTAGAAATTATAACATCCATAGTTGGATTCACATAGGTGTCCAATCGGGATCTCTCCATCATCGAGATGATTGACGGCATCTCCCCCTTCCTTTTTTAGTAGATATTTCATCGTGTAAGAATCAATATTAATATTTCTAAAGTATCTGTGTATAGCCTCTTTATTTACCACAGCTCCATATTCCAACATGCATTCCACTAGAGGGATATCGATATCGCCGAATGTCATATACTCAATTAGTATATGTTGGAGGACATCCGAGTTCATTGTTTTCAATATCAAAGAGATGGTTTCCTTATCATTTCTCCATAGTGGTACAATACTACACATTATTCCGTACGGCTTTCCATTCTCCAAAAACAATTTTACCAAATCTAAATCTACATCTTTATTGTATCTATAATCACTATTTAGATAATCAGCCATAATTCCTCGAGTGCAACATGTTAGATCGTCTATATATGAATAAGCCGTGTTATCTATTCCTTTCATTAACAATTTAACGATGTCTATATCTATATGAGATGACTTAATATAATATTGAAGAGCTGTACAATAGTTTTTATCTATAGAAGACGGCTTGATTCCGTGATTAATTAGACATTTAACAACTTCCGGACGCACATATGCTCTCGTATCCGACTCTGAATACAGATGAGCGATGATATACAGATGCAATACGGTACCACAATTTCGTGGTTGATAATCATCATACACGTATCCGTACTCGTCATCCTCATAAAGAACACTGCATCCATTTTCTATGAACAAATCAATAATTTCAGGAACAGGATCATCTGTCATTACATAATTTTCTATAACTGAACGATGGTTTTCACATTTAACACTCAAGTCAAATCCATGTTCTACCAACACCTTTATCAAGTCAACGTCTACATTTTTTGATTTCATATAGCTGAATATATTAAAGTCATTTATGTTGCTATATCCAGTAGCTTCTAGTAGAGCCATCGCTATATCCTTATTGACTTTAACATGTCTACTATTTGTGTATTCTTCTATTGGGGTAAACTGTCTCCAATTTTTATGTAATGGATTAGTGCCACTATCTAGTAGTAGTTTGACGACCTCAACATTATTACAATGCTCATTGAAAAGGTATGCGTGTAAAGCATTATTCTTGAATTGGTTCCTGGTATCATTAGGATCTCTGTCTCTCAACATCTGTTTAAGTTCATCGAGAACCACCTCCTCATTTTCCAGATAGTCAAACATTTTGACTGAATAGAAGTGAATGAGCTACTGTGAACTCTATACACCCGCACAACTAATGTCATTAAATATCATTTTTGAATGTATTTATACCATGTCAAAAACTTGTACAATTATTAATAAAAATAATTAGTGTTTAAATTTTACCAGTTCCAGATTTTACACCTCCGTTAACACCTCCATTAACCCCACTTTTTACACCACTGGACGATCCTCCTCCCCACATTCCACTGCCACTAGATGTATAAGTTTTAGATCCTTTATTACTACCATCATGTCCATGGATAAAGACACTCCACATGCCGCCACTACTACCCCCTTTAGACGACATATTAATAAGACAAGTTTAACAATAAAATTAATCACGAGTACCCTACTACCAACCACTATTATATGATTACAGTACCTTGACTAAAGTCTCTAGTCACAAGATCAATACTACCAACCTACGCTATTATATGATTATAGTTTCTATTTTTATAGGAACGCGTACGAGAAAATCAAATGTCTAAGTTCTAACGGTAGTGTTGATAAACGATTGTTATCCGCGGATACCTCATCTATCATGTTGTCTATTTTCTTACTTTGTTCTATTAACCTATTAGCATTATATATTATTTGATTATAAAACTTATATTGCTTATTAGCCCAATCTGTAAATATCGGATTATTAACATATCGTTTCTTTGTAGGTTTATTTAACTTGTACATCACTGTAAGCATGTCCGTACCATTTATTTTAATTTGACACATATCAGCAATTTCTTTTTCGCAGTCGGTTATATATTCTATATAAGATGGATACGTATCACATATGTACTTATAGTCTACTAATATGAAGTACTTAATACATATTTTCAGTAACGATTTAGCCTTATTACCTATTAATAAGTGCCTGTCGTTGGATAGGTAATCAACTGTTTTCTTAATACATTCGATGGTTGGTAATTTACTCAAAATAATTTCCAATATCTTAATATATATTTCTGCTATTTCTGGTATACATGCATGTGCCATTATAACACAAATACCAATACATGTAGACCCATATGTTGTTGTTATATTAATATCTGCGCCATTATCTATTAACCATTCTATTAGGGCAACACTATGCGACTCGATACAATAATAAAGTATACTACGTCCATGTTTATCTATTTTGTTTATATCATCGATATACGGCTTACAAATTTCTAGTATCGATAACACTTCTGACTCGTGAATAAATAAGGTAGGGAATAACGGCATAATATTTATTATGTTATCATCATTAACAACTACGTTTCCATTTTTTAAAATATACTCTACAACTTTAGGATCCCTATTGTCAAATCTTTTAAAATATTTATTTATATGCTTAAATCTATATAATATAGCTCCTTCCCTAATCATACATTTGATAACATTGATGTACACTGTATGATAAGATACATATTCTGACAATAGATCTTGTATAGAATCTGTATATCTTTTAAGAATTGAGGATATTATGACATTATTACGTAAACTATTACACAATTCTAAAATATAAAATGTATCATGGGCAGATAATAGTTTATCCACTATATAATTATCTATTTTATGATTTTTCTTCCTATATTGTTTACGTAAATAGATAGATAGAATATGCATTAGTTCATTACCGCTATAGTTACTATCGAATAACACGTCAAATATTTCCCGTTTAATATCGCATTTGTCAACATAATAATAGAGTATGGTACGTTCACGATAAGTATAATGACACATCTCGTTTTCGTGCGAAATTAAATAGTTTATCACGTCCAAAGATGTCACATAACCATCTTGTGACCTAGTAATAATATAATAATAGAGAACTGTTTTACCCATTCTATTATCATAATCAGTGGTGTAGTCATAATCTAAATAATCAAACTCGTCATCCCAATTAAAATAAATATAATCAGTACATTGAATGGGTATGATATTGTACCCATACTGTATGTTGCTACATGTAGGTATTCCTTTATCCAATAATAGTTTAAATACATCTATATTAGGATTTGATGTTGTCGCGTATTTCTCTACAATATTAATACCATTTTTGATACTATTTATTTCTATACCTTTCGAAATTAGTAATTTCAATAAGTCTATATCGATGTTATCAGAACATAGATATTCAAATATATCAAAATCATTGATATTTTTATAGTCGACTGACGACAATAACAAAATCACAACATCGTTTTTGATATTATTATTTTTTTTGGTAACGTATGCCTTTAATGGAGTTTCACCATCATACTCATATAATGGATTTGCACCACTTTCTATTAATGATTGTGCACTACTGGCATCGATGTTAAATGTTTTACAACTATCATAGAGTATCTTATCGTTAACCATGATTGGTTGTTGATGTTATCACATTTTTTGGTTTCTTTCATTTCAGTTATGTATGGATTTAGCACGTTTGGGAAGCATGAGCTCATATGATTTCAGTACTGTAGTGTCAGTACTATTAGTTTCGATCAGATCAATGTCTAGATCTATAGAATCAAAACACGATAGGTCAGAAGATAATGAATATCTGTACGCTTCTTCTTGTACTGTAACTTCTGGTTTTGTTAGATGGTTGCATCGTGCTTTAACGTCAATGGTACAAATTTTATCCTCGCTTTGTGTATATCATATTCGTCTCTAGTATAAAATTCTATATTCAAATTATCATGCGATGTGTGTACGCTAACGGTATCAATAAACGGAGCACAGCATTTAGTCAACAGTAATCCAAAATTTTTTAAAGTATATCTTAACGAAAGAAGTTGTCATCGTTAGAGTGTGGTAAATCATTGTCTACGGTACTAGATCCTCATAAGTGTATATATCTAGAGTAATATTTAATTTATCAAATGGTTGATAATATGGATGTTGTGGCAATTTCCTAATACGGAAATAAGACATAAACACGCAATAAATCTAATTGCGGACATGTTACACTCCTTAAAAAATACGAATAAACACTTTGGCTTTTAGTAAGTGTCATTTAACACTATACTCATATTAATCCATGGACTCATAATCTCTATACGGGATTAACGGATGTTCTATATACGGGGATGAGTAGTTCTCTTCTTTAACTTTATACTTTTTACTAATCATATTTAGACTGATGTATGGGTAATAGTGTTTGAAGAGCTCGTTCTCATCATCAGAATAAATCAATATCTCTGTTTTTTTGTTATACAGATGTATTACAGCCTCATATATTACGTAATAGAACGTGTAATCTACCTTATTAACTTTCACCGCATAGTTGTTTGCAAATACGGTTAATCCTTTGACCTCGTCGATTTCCGACCAATCTGGGCGTATAATGAATCTTAACTTTAATTTCTTGTAATCATTCGAAATAATTTTTAGTTTGCATCCGTAGTTATCTCCTCTATGTAACTGTAAATTTCTCAACGCGATATCTCCATTAATAATGATGTCGAATTCGTGTTGTATACCCATACTGAATTGATGAACTAATACCGACGGTATTAATAGTAATTTACTTTTCATCTTTACATACTTGGTAATAGTTTTACTATCATAAGTTTATAAATTCCACAAGCTACTATGGAATATACCAACCATCTTAGTATAGAACACATGTCTTAAAGTTATTAATTAATTACATGTTGTTTTATATATCGCTACGAATTTAAACAGAGGAATCAGTTAGGAAGAAAAAATTATCTGTCATCATCATCATCTATTGGATAACATCTCTGTATTCTACGATAGAGTGCTATTTTAAGATGTGACAGATCCGTGTCATCAAATATATACTCCATTAAAATGATTATTCCGGCAGCGAACTTGATATTGGATACATCACGACCTTTGTTAATATCCACGACAATAGACAGCAATCCCATTGTTCCATAAACAGTGAGTTTATCTTTCTTTGAAGTGATATTTTGTAGAGATCTTATAAAACTGTCGAATGACATCGTATTTATATCTTTAGCTAAATCATATATGTTACCATCATAATATCTAACAGCATCTATCTTAAACGTTTCCATCGCTGTAAAGACGTTTCCGATAGATGGTCTCGTTTCATCAGTCATACTGAGCCAACAAATGTAATCGTGTATAACATCTTTGATAGAATCAGACTCTAAAGAAAAGGAATCGGCTTTATTATACACATTCATGATAAACTTAATGAAAAATGTTTTTCGTTGTTTAAGTTGGATGAATAGTATGTCTTAATAATTGTTATTATTTCACTAATTAATATTTAGTAACGAGTACACTCTATAAAAACGAGAATGACATAACTAATCATAACTAGTTATCAAAGAATGTCTAGGACGCGTAATTTTTTATGGTATAGATCCTGTAAGCGTTGTCTGTATTCTGGAGCTATTTTCTCTATCGCATTAGTGAGTTCAGAATATGTTATAAATTTAAATCGAATAACGAACATAACTTTAGTAAAGTCGTCTATATTAACTCTTTTATTTTCTAGCCATCGTAATACCATGTTTAAGATAGTATATTCTCTAGTTACTACGATCTCATCGTTGTCTAGAATATCGCATACTGAATCTACATCCAATTTTAGAAATTGGTCTGTGCTACATATCTCTTCTATATTATTGTTGATGTATTGTCGTAGAAAACTATTACGTAGACCATTTTCTTTATAAAACGAATATATAGTACTCCAATTATCTTTACCGATATATTTGCATACATAATCCATTCTCTCAATCACTACATCTTTAAGAGTTTGGTTGTTAAGATATTTGGCTAAACTATATAATTCTATTAGATCATCAACAGAATCAGTATATATTTTTCTAGATCCAAAGATGAACTCTTTGGCATCCTCTATAATATTATCAGAAAAGATATTTTCGTGTTTTAGTTTATCAAGATTTAACCTGTTCATATCCATGATTAACGACGTCATATAACCACATAAAATAAAAATCCATTTTCATTTTTAGCACAATACTATTCATAATTGATATTGATGTAATATTTTGTTACTTTGAACGTAAAGACAGTACACGGGTCCGTATCTCCAACAAGCACGTAGTAATCAAATTTGGTGTTGTTAAACTTCGCAATATTCATCAATTTAGATAGAAACATATACTCATCATCTGTTTTAGGAATCCATGTATTATTACCACTTTCCAACTTATCATTATCCCAGGCTATGTTTCGCCCATCATCGTTGCACAGAGTGAATAATTCTTTTGTATTCGGTAGTTCAAATATATGATCCATGCATATATCGACAAAGCTATTGTAGATGTGATTTTTCCTAAATCTAATATAAAACTCGTTTACTAGCAAACATTTTCCTGATTTATCGACCAAGACACACATGGTTTCTAAATCTATCAAGTGGTGGGGATCCATAGTTATAACGCAGTAACATAAATTATTACCTTCTTGACTGTCGCTAATATCTATATACTTATTGTTATCGTATTGGATTCTACATATAGATGGCTTGTATATCAAAGATATAGAACACATAACCAATTTATATTCTCGCTTTGTATTTTCGAATCTAAAGTTAAGAGATTTAGAAAACATTATATCATCGGATGATGATATCACTGTTTCCAGAGTAGGATATATTAAAGTCTTTAAAGATTTTGTCCGATTCAAATAAATCACTAAATAATATCCCATATTATCATCTGTTATAGTCGTGTCATTAAATCTATTATATTTTATGAAAGATATATCACTGCTCACCTCTATATTTCGTACATTTTTAAACTGTTTGTATAATATCTCTCTAATACAATCAGATATATCTATTGTGTCGGTAGACGATACCGTTACATTTGAATTAATGGTGTTCCATTTTACAACTTTTAACAAGTTAACCAATTCATTTCTAATAGTATCAAACTCTCCATGATTAAATATTTTAATAGTATCCATTTTATATCACTACGGACATAAACCATTGTATATTTTTTATGTTTATTAGTGTACACATTTTGGAAGTAAGTTCCTGGATCGGATGTCACCGCAGTAATATTGTTGATTATTTCTGACATCGACGTATTATATAGTTTTTTAATTCCATATCTTTTAGAAAAGTTAAACATCCTTATACAATTTGTGGAATTAATATTATGAATCATGGTTTTTACACATAGATCTATTACAGGCGGAACATCAATTATTATGGCAGCAACTAGTATCATTTCTACATTGTTTATGGTGATGTTTATCTTCTTCCAGCGCATATAGTCTAATATCGATTCAAACGCGTGATAGTTTATACCATTCAATATAATCGCTTCATCCTTTAGATGGTGATCCTGAATGTGTTTAAAAAATTATACGGAGACGCCGTAATAATTTCCCCATTGATAGAAAATATCACGCTTTCCATTTTCTTGAAGTACTAAAAGTAATTATAATATAATGTAAAGGTTTATATATTCAATATTTTTTATAAAAAAAATCATTTTGACATTAATTCCTTTTTAAATTTCAGTCTATCATCTATAGAAACATATTCTATGAATTTATAAAATGCTTTTACGTATCCTATCGTAGGTGATAGAACCGCTAAAAAACCTATCGAATTTCTACAAAAGAATCTGTTATATGGTATAGGGAGAGTATAAAACATTAAATGTCCGTACTTATTAAAGTATTCAGTAGCCAATCCTAACTCTTTCGAATAATTATTAATGGCTCTTATTCTGTACGAATCTATTTTTTTGAACAATGGACCTAGTGGTATATCTTGTTCTATGTATCTAAAATAATGTCTGACTAGATCCGTTAGTTTAATATCCGCAGTCATCTTGTCTAGAATGGCAAATCTAACTGCGGGTTTAGGCGTAGGCGTTAGTTTAGTTTCTATATCTACATCTATGTCTTTATCTAACACCAAAAATATAATAGCTAATATTTTATTACAATCATCCGGATATTCTTCTACGATCTCACTAACTAATGTTTCTTTGGTTATACTAGTATAGTCACGATCAGACAAATAAAGAAAATCAGATGATCGATGAATAATACATTTAAATTCATCATCTGTAAGATTTTTGAGATGTCTCATTAAAATATTATTAGTGTCAGTTCTCATTATCATATATTGACAGCAGCTATTACACTTATTTTATTTTTCTGTATTTTATTACTTTTCACCATATAGATCAGTCATTAGATCATCAAAATACTTTTCAATCATCCTAAAGAGTATGGTGAACGAATCTTCCCATCTAATTTCTGAACGTCTACCAATGTCTCTAGCCACTTTGGCACTAATAGCGATCATTCGCTTAACATCTTCTACATTATTAACTGGTTGATTCAATCTATCTAGCAATGGACCGTCGGATAGCGTCATTCTCATGTTCTTAATCAATGTACATACATCGTCATCATCTACCAATTCATCAAACAATATAAGCTTTTTAAAATCATCATTATAATAGGATGGATCGCCGTCATTTCTCCAAAGAATATATCTAATAAGTAGAGTCCTCATGCTTAGTAATTTAACTATTTTAGTTAACAACTATTTTTTATGTTAAATCAATTAGTAACACCGCTATGTTTAATACTTATTCATATTTTAGTTTTAGGATCGAGAATCAATACAAAAATTAATACATCAATTTTGGAAATACTTAGTTTCCACGTAGTCAATGAAACATTTGAGCTCATCGTAAAGGACGTTCTCGTACAAGACGTAACTATAAATTGGTTTATATTTGTTCAAGATAGATACAAATCCGATAACTTTTTTGACGAATTCTACGGGATTCACTTTAAAAGTGTCATACCGGGTTCTTTTTATTCTTTTAAACAGATCGATTGTGTGATGTTGATTAGGTCTTTTACGAATTTGATACAGAATAGCGTTTACATATCCACCATAGTAATCAATAGCCATTTGTTCGTATGTCATAAATTCTTTAATTATATGACACTGTGTATTATTTAGTTCGTCCTTGTTCATCATTAGGAATCTATCCAATATGGCAATTATATTAGAACTATAACTGCGTTGTATGCGCATGTTGATGTGTCTGTTTATACAATCAATTATACTAGGATCCATACCACTACAATCGGGTAAAATTGTAGCATCATATACCATTTCTAGTACTTTAGATTCATTGTTATCCATTGCAGAGGACGTCATGATCGAATCCAAAAAAATATATTATTTTTATGTTATTTTGTTAAAAATAATCATCAAATACTTCGTAAGATACTCCTTCATGAACATAATCAGTTACAAAACGTTTATATGAAGTAAAGTATCTACAATTTTTACAAAAGTCAGGATGCATAAGTACAAAGTACGCGATAAACGGAATAATAATAGATTTATCTAGTTTATCTTTTTCTATCTCTTTCATAGTTATATACATGGTCTCAGAAGTCGGATTATGTAACATCAGCTTCGATAAAATGACTGGGTTATTTAGTCTTACACATTCGCTCATACATGTATGACCGTTAACTATAGAGTCTACACTAAAATGATTGAATAATAGATAGTCTACCATTGTTTCGTATTCAGATAGTACAGCGTAGTACATGGCATCTTCACAAATTATATCATTATCTAATAGATATTTGACGCATCTTATGGATCCCACTTCAACAGCCATCTTAAAATCGGTAGAATCATATTGCTTTCCTTTATCGTTAATAATTTCTAGAACATCATCTCTATCATAAAAGATACAAATATTAACTGTTTGATCAGTAATAACATTGCTAGTCGATATCAATTTGTTAATAAGATGCGCTGGGCTCAATGTCTTAATAAGAAGTGTAAGAGGACTATCTCCGAATTTGTTTTGTTTATTAACATCCGTTGATGGAAGTAAAAGATTTATAATGTCTACATACTTGACTGTTTTAGAGCATACAATATGGAGAGGCGTATTTCCATCATGATCTGGTTTTGAGGGACTAATTCCTAGTTTCATCATCCATGAGATTGTAGAAGCTTTTGGATTGTCTGACATAAGATGTCTATGAATATGATTTTTGCCAAATTTATCCACTATCCTGGCTTCGAATCCGATAGACATTATTTTTTTAAACACTCTTTCTGAAGGATCTGTACACGCCAACAACGGACCACATCCTTCTTCATCAACCGAGTTGTTAATCTTGGCTCCATACTGTACCAATAAATTTATTCTCTCTATGACTTCATCATCTGTTCCCGAGAGATAATATAGAGGTGTTTTATTATGTTTATCACATGCGTTTGGATCTGCGCCGTGCACCAGCAGCATCGCGACTATTCTATTATTATTAATTTTAGAAGCTATATGCAATGGATAATTTCCATCATCATCCGTCTCATTTGGAGAGTATCCTCTATGAAGAAGTTCTTCTATAAATCGTTCATCTAGTCCTTTAATGCCACAATACGCATGTAGAATGTGATAATTTCCAGAGGGTTCGATAACTTGTAGCATATTCCTAAATACATCTAAATTTTTACTATTATATTTGGCATAAAGAGATAGATAATACTCGACCGACATAATGTTGTGTTGTCCATTATAGTATAAAAATTAATATTTCTATTTCTATATATTTGCAACAATTTACTCTCTATAACAAATATCATAACTTAGTTCTTTTATGTCAAGAAGGCACTGGTTTAATTCATCTATAAATGTCACTCCATAACTACCACGCATACTATACTCAGAATTATGATAAAGATATTTATTCTTGGGGTGTAAGTAATGGGGATTAATCTTTGTTGGATCAGTCTCTAAGTTAACACATGTCACACATGATCCATTTATAGTTATATCACACGATGATGATTTATGAATTGATTCCGGAAGATCGCTATTGTATTTTGTAGTTCCACAATTCATTTCCATACATGTTATTGTCACACTAATATTATGATGAACTTTATCTAGCCGCTGAGTGGTAAACAACAGAACAGATAGTTTATTATCTTTACCAACACCCTCAGCCGCTGCCACAAATCTCTGATCCGTATCCATGATGGTCATGTTTACTTTTAGTCCGTATCCAGTCAACACTATGTTAGCATTTCTGTCGATATAGCTTTCACTCATATGACACTCACCAATAATTGTAGAATTAATGTCGTAATTTACACCAATAGTGAGTTCGGCGACAAAGTACCAGTACCGGTAATCTTGTCGATGAGGACATATAGTATTCTTGTATTCTACCGAATACCCGAGAGATGCGATACAAAAGAGTAAGACTAATTTGTAAACCATCTTACTCAAAAATATGCGACAATAGGAAATCTATCTTATACACATAATTATTCTATCAATTTTACCAATTAGTTAGTGTAATGTTAACAAAAATGTGGGATAATTTAATAGTTTTTCCTTACACAATTGACATACATGAGTCTGAGTTCCTCGTTTTTGCTAATTATTTCGTCCAATTTATTATTCTTGACATCGTCAAGATCTTTTGTATAGGAGTCAGACTTGTATTCAACATGTTTTTCTATAATCATCTTAGCTATTTCGGCATCATCCAATAGTACATTTTCCAGATTAACAGAATAGATATTAATGTCGTATTTGAACAGAGCCTGTAACATCTCAATGTCTTTATTATCTATAGCCAATTTGATGTCCGGAATGAAGAGAAGGGAATTGGTGTTTGTCGACGTCATATAGTCGAGCAAGAGAATCATCATATCCACGTGTCCATTTTTTATAGTGGTGTGAATACAACTAAGGAGAATAGCCAGATCAAAAGGAGATGGTATCTCTGAAAGAAAGTAGGAAACAATACTTACATCATTAAGCATGACAGCATGATAAAATGAAGTTTTCCATCCAGTTTTCCCATAGAACATCAGTCTCCAATTTTTCTTAACAAACAGTTTTACCGTTTGCATGTTACCACTATCAACCGCATAATACAATGCGGTGTTTCCTTTGTCATCAAATTGTGAATCATCCATTCCACTGAATAGCAAAATCTTTACTATTTTGGTATCTTCTAATGTGGCTGCCTGATGTAATGGAAATTCATTATCTAGAAGATTTTTCAATGCTCCAGCGTTCAACAACGTACATACTAGACGCACGTTATTATCAGCTATTGCATAATACAAGACACTATGACCATTGATATCCGCCTTAAATGCATCTTTGCTAGAGAGAAAGCTTTTCAGTTGCTTAGACTTCCAAGTATTAATTCGTGACAGATCCATGTCTGAAACGAGACGCTAATTAGTGTATATTTTTTCATTTTTTATAATTTTGTCATATTGTACCAGAATTAATAATATCTCTAATAGATCTGATTAGTAGATATATGGCTATCGCAAAACAACATATACACATTTAATAAAAATAATATTCATTAAGAAGATTCAGATTCCACTGTACCCATCAATATAAAATAAAATAATTATTCCTTACATCGTACCCATAAACAATATATTAAGTATATTCCACCTTACCCATAAACAATATAAATCCAGTAATATCATGTCTAATGATGAACACAAATGGTGTATTAAATTCCAGTTCTTCAGGAGATGATCTCGCCGTAGCTACCATGATAGTAGATGCCTCCGCTACAGTTCCTTGTTCGTCTACATCTATCTTTACATTCTGAAACATTTTATAAATATATAATGGGTCCCTAGTCATATGTTTAAACGACGCCTTATCTGGATTAAACATACTAGGAGCCATCATTTCGGCTATCGACTTAATATCCCTCTTGTTTTCGATAGAAAATCTAGGGAGTTTAAGATTGTACATTTTATTCCCTAATTGAGATGACCAATATTCTAATTTTGCAGGCGTGATAGAATCTGTGAAATGGGTCATATTATCACCTATTGCCAGGTACATACTAATATTAGCATCCTTATACAGAAGGCGCACCATATCATATTCTTCGTCATCGATTGTGATTGTATTTCCTTGCAATTTAGTAACTACGTTCATCATGGGAACCGTTTTCGTACCGTACTTATTAGTAAAACTAGCATTGTGTGTTTTAGTGATATCAAACGGATATTGCCACGTACCTTTAAAATATATAGTATTAATGATTGCCCATAGAGTATTATCGTCGAGCATAGTAGAATCAACTACATTAGACATACCAGATCTACGTTCTACTATAGAATTAATTTTATTAACCGCATCTCGTCTAAAGTTTAATCTATATAGGCCGAATCTATGATATTGTTGATAATACGACGGTTTAATACACACAGTACTATCGACGAAACTTTGATACGTTAGATCTGTGTACGTATATTTAGATGTTTTCATCTTAGCTAATCCGGATATTAATTCTGTAAATGCTGGACCCAGATCTCTTTTTCTCAAATTCATAGTATTCAATAATTCTACTCTAGTATTACCTGATGCAGACAATAGCGACATAAACATAGAAAACGAATACCCAAACGGTGAGAAGACAATATTATCATTATCATCCTCATCCCCATTTTGAATATTTTTATACGCTAATATACCAGCATTGATAAATCCCTGCAGACGATATGCGGATACTGAACACGCTAATGATAGTATCAATAACGCAATCATGATTTTTATGGTATTAATAATTAACCTTATTTTTATGTTTGGTATAAAAATTATTGATGTCTACACATCCTTTATAATCAACTCTAATCACTTTAACTTTTACAGTTTTCCCTACAAGTTTATCCCTATATTCAACATATCTATCCATATGCATCTCTTAACACTCTGCCAAGATAGCTTCAGAGTGAGGATAGTCAAAAAGATAAATATATAGAGTATAATCATTCTCGTATACTCTGCCCTTTATTACATCGCCCGCATTGGGCAACGAATAACAAAATGCAAGCATCGTGTTAACGGGCTCGTAAATTGGGATAAAATTATGTTTTTATTGTTTATCTATTTTATTCAAGAGAATATTCAGGAAGTTCCTTTTCTGGTTGTATCTCGTCGCAGTATATATCATTTGTACATTGTTTCATATTTTTTAATAGTCTACACCTTTTAGTAGGACTAGTATCGTACAATTCATAGCTGTATTTTGAATTCCAATCACGTATAAAAATATCTTCCAATTGTTGACGAAGACCTAATCCATCATCCGGTGTAATATTAATAGATGCTCCACATATATCCGTAAAGTAATTTCCTGTCCAATTTGATGTACCTATATACGCCGTTTTATCGGTTACCATATATTTTGCATGGTTTACCCTAGAATACGGAATGGGAGGATCAGCATCTGGTACAATAAATAGCTTTACTTCTATATCTATGTTTTTAGATTTTAGCATAGCTATAGATCTTAAAAAGTTTCTCATGATAAACGAAGATCGTTGCCAGCAACTAATCAATAGCTTAACGGATACTTGTCTGTCTATAGCGGATCTTCTTAATTCATCTTCTATATAAGGCCAAAACAAAATTTTACCCGCCTTTGAATAAATAATAGGAATAAAGTTCATAACAGATACATAAACGAATTTACTCGCATTTCCGATACATGACAATAAAGCGGTTAAATCATTGGTTCTTTCCATAGTACATAATTGTTGTGGTGCAGAAGCAATAAATACAGAGTGTGGAACACCGCTTACGTTAATACTAAGAGGATGATCTGTATTATAATACGACGGATAAAAGTTTTTCCAATTATATGGTAGATTGTTAACTCCAAGATACCAGTATACCTCAAAAATTTGAGTGAGATCCGCTGCCAAGTTCCTATTATTGAAGATCGCAATACCCAATTCCTTGACCTGAGTTAGTGATCTCCAATCCATGTTAGCGCTTCCTAAATAAATATGTGTATTATCAGATATCCAAAATTTTGTATGAAGAACTCCTCCTAGGATATTTGTAATATCTATGTATCGTACTTCAACTCCGGCCATTTGTAGTCTTTCAACATCCTTTAATGGTTTGTTGGATTTATTGACGGCTACTCTAACTCTTACTCCTCTTTTGGGTAATTGTACAATCTCGTTTAATATTACCGTGCCGAAATTCGTACCCACTTCATCCGATAAACTCCAATAAAAAGATGATATATCTAGTGTTTTTATGGTATTGGATAGAATTTCCCTCCACATGTTAAATGTAGTCAAATATACTTTATCAAATTGCATACCTATAGGAATAGTCTCTGTAATCACTGCGATTGTATTATCCGGATTCATTTTATTTGTTAAAAAAATAATCCTATATCACTTCACTCTATTAAAAATCCAAGTTTCTATTTCTTTCATGACTGATTTTTTAACTTCATCCGTTTCCTTATGAAGATGATGTTTGGCGCCTTCATAAATTTTTATTTCCCTATTACAATTTGCATGTTGCATGAAATAATATGCACCTGAAACATCGCTAATCTCATTGTTTGTTCCCTGGAGTATGAGAGTCGGGGTGTTAATCTTGGGAATTATTTTTCTAACCTTGTTGGTAGCCTTCAAGACCTGACTAGCAAATCCAGCCTTAATTTTTTCATGATTGACTAATGGATCGTATTGGTATTTATAAACTTCATCCATATCTCTAGATACTGATTCTGGACATAGCTTTCCGACTGACGCATTTGGTGTAATGGTTCCCATAAGTTTTGCAGCTAGCAGATTCAGTCTTGGAACAGCGTCTGCATTAACTAGAGGAGACATTAGAATCATTGCTGTAAACAAGTTTGGATTATCGCAAGCAGCTAGTATAGAAATTGTTGCTCCCATGGAATGACCCAATAAGAAGACTGGAACTCCTGGATAAGTAGATTTAATAGTCACCACGTGCTGTACCACATCTCTAACATACTTACCAAAGTCATCAATCATCATTTTTTCACCATTACTTCTTCCATGGCCAATATGATCATGTGAGAATACTAAAATTCCTAACGATGATATGTTTTCAGCTAGTTCGTCATAACGTCCAGAATGTTCACCAGCTCCATGACTTATGAATACTAATGCCTTAGGATATGTAATAGGTTTCCAATATTTACAATATATGTAATCATTGTCCAGATTGAACATACAGTTTGTACTCATGATTCACTATATAACTATCAATATTAACAGTTCGTTTAATGATCATATTATTTTTATGTTTTATTGATAATTGTAAAAATATACAATTAAATCAATATAGAGGAAGGAGACGGTACTGTATTTTGTGAGATAGTAATGGAGACTAAATCAGATTATGAGGATGCTGTTTTTTACTTTGTGGATGATGATGAAATATGTAGTCGCGACTCCATCATTGATTTAATAGATGAATATATCACGTGGAGAAATCATGTTATAGTGTTTAATAAAGATATTACCAGTTGTGGAAGACTGTACAAGGAATTGATAAAGTTCGATGATGTCGCTATACGGTACTATGGTATTGATAAAATTAATGAGATTGTCGAGGCTATGAGTGAAGGAGACCACTACATCAATCTTACAGAAGTCCATGATCAGGAAAGTCTATTCGCTACCATAGGAATATGTGCTAAAATCACTGAACATTGGGGATACAAAAAGATTTCAGAATCTAAATTCCAATCATTGGGAAACATTACAGATCTGATGACCGACGATAATATAAACATCTTGATACTTTTTCTAGAAAAAAAAATGAATTGATGATATAAGTGTCTTCATAACGCATTATTACGTTAGCATTCTATTATCCAGTGTTAAAAAAATTATCCTATCATGTATTTGAGAGTCTTATATGTAGCAAACATGATAACTGCAATACCCATAATCTTTAGATATTCACGCGTGCTATGGATGGCATTATCCCGCGGTGCGGAAATGTACGTTATATAATCTACAAAATAATCATCGCATATAGTATGAGATAGTAGAGTAAACATTTTTATCGTTTCTACTGGGTTCATACATCGTCTACCCAATTCGGTAATGAATGAAATTGTCGCCAATCTTACACCCAAACCCTTGTTGTTCATTAGTATAGTATTAACTTCATTATTTATGTCATAAACTGTAAATGATTCTGTAGATGCCATATCACACATGATATTCATGTCACTATTATAATCATTATTAACTTTATCACAATACGTGTTGATAATATCTACATATGATCTAGTTTTTGTGGGTAATTGTCTATACAAGTCGTCTAAACGTTGTTTACTCATATAGTATTGAACAGCCATCATTACATGGTCCCGTTCCGTTGATAGATAATCGAGTATGTTAGTAGACTTGTCAAATCTATATACCATATTTTCTGGAAGCGGATATACATAGTCGCGATCATCATTATCACTAGCCTCATCCTCTATATCATGTACATGTACATAATCTATGATATTATTATACATAAACATCGACAACATACTATTGTCTATTATCTAAGTCCTGTTGATCCAAACCCTTGATCTCCTCTATCTGTACTATCTAGAGATTGTACTTCTTCAAGTTCTGGATAATATATACGTTGATAGATTAGCTGAGCTATTCTATCTCCAGTATTTACATTAAACGTACATTTTCCATTATTAATAAGAATGACTCCTATGCTTCCCCTATAATCTTCGTCTATTACACCGCCTCCTATATCAATGCCTTTTAGGGACAGACCAGACCTAGGAGCTATTCTACCATAGCAGAACTTAGGCATGGACATACTAATATCTGTCTTAATTAACTGTCGTTCTCCAGGAGGGATAGTATAATCGTAAGCGCTATACAAATCATATCCGGCAGCACCCGGCGATTGCCTAGTAGGTGATTTAGCTCTGTTAGTTTCCTTAACAAATCTAACTGGTGAGTTAATATTCATGTTGAACATAAAAAATATCATTTTATTTCAAAATTATTTACCATTCCATTCCATCCCATATATTCCATGAATAAGTGCGATTATTGTACACTTCTATAGTATCTATATACGATCCACGATAAAATCCTCCTATCAATAGCAGTTTATTATCCACTATGATCAATTCTGGATTATCCCTCGGATAAATAGGATCATCTATCAGAGTCCATGTATTGCTGGATTCACAATAAAATTCCGCATTTCTACCAACCAAGAATAACCTTCTACCAAACACTAACGCACATGATTTATAATGAGGATAATAAGTGGATGGTCCAAACTGCCACTGATCATGATTGGGTAGCAAATATTCTGTAGTTGTATCAGTTTCAGAATGTCCTCCCATTACGTATATAACATTGTTTATGGATGCCACTGCTGGATTACATCTAGGTTTCAGAAGACTCGGCATATTAACCCAAGCAGCATCCCCGTGGAACCAACGCTCAACAGATGTGGGATTTGGTAGACCTCCTACTACGTATAATTTATTGTTAGCGGGTATCCCGCTAGCATACAGTCTGGGGCTATTCATCGGAGGAATTGGAATCCAATTGTTTGATATATAATTTACCGCTATAGCATTGTTATGTATTTCATTGTTCATCCATCCACCGATGAGATATACTACTTCTCCAACATGAGTACTTGTACACATATGGAATATATCTATAATTTGATCCATGTTCATAGGATACTCTATGAATGGATACTTGTATGATTTGCGTGGTTGTTTATCACAATGAAATATTATGTTACAGTCTAGTATCCATTTTACATTATGTATACCTCTGGGAGAAAGATAATTTGACCTGATTACATTTTTGATAAGAAGTAGCAGATTTCCTAATCTATTTCTTCGCCTCATATACCACTTAATGACAAAATCAACTACATAATCCTCATCTGGAACATTTAGTTCGTCGCTTTCTAGAATAAGTTTCATAGATAGATAATCAAAATTGTCTATGATGTCATCTTCCAGTTCCAAAAAGTGTTTGGTAATAAAGTCTTTAGTATGACATAAGAGATTGGATAGTCCGTATTCTATACCCATCATGTAACACTCGATACAATATTCCTTTCTAAAATCTCGTAGGATAAAGTTTATACAAGTGTAGATGATAAATTCTACAGATGTTAATATAGAAGCACGTAATAAATTGACGACGTTATGACTATCTATATATACCTTTCCAGTATATGAGTAAATAACTATAGAAGTTAGACTGTGAATGTCAAGGTCTAGACAAACCCTCGTAACTGGATCTTTATTTTTTGTGTATTTTTGGCGTAAATGTGTGCGAAAGTATGGAGATAACTTTTTCAATATCGTAGAATTGACTATTATATTACCTCCTATAGCTTCAATAATTGTTTTGAATTTCTTAGTCGTGTACAATGCTAATATATTCTTACAGTACACAGTATTGACAAATATCGGCATTTATGTTTCTTTAAAAGTCAACATCTAAAGAAAAATGATTGTCTTCTTGAGACATAACTCCCATTTTTTGGTATTCACCCACACGTTTTTCGAAAAAATTAGTTTTTCCTTCCAATGATATATTTTCCATGAAATCAAACGGATTGGTAACATTGTAAATTTTTTTAAATCCCAATTCAGAAATCAATCTATCTGCGACGAATTCTATATATGTTTTCATCATTTCACAATTCATTCCTATGAGTTTAACTGGAAGAGCCACAGTAAGAAATTCTTGTTCAATGGATACCGCATTTGTTATAATAAATCTAACGGTTTCTTCACTCGGTGGATGCAATAAATGTTTAAACATCAAACATGCGAAATCGCAGTGCAGACCCTCGTCTCTACTAATTAATTCGTTAGAAAACGTGAGTCCGGGCATTAGGCCACGCTTTTTAAGCCAAAATATGGAAGCGAATGATCCGGAAAAGAAGATTCCTTCTACTGCAGCAAAGGCAATAAGTCTCTCTCCATAACCGGCGCTGTCATGTATCCACTTTTGAGCCCAATCGGCCTTCTTTTTTACACAAGGCATCGTTTCTATGGCATTAAAGAGGTAGTTTTTTTCATTACTATCTTTAACATAAGTATCGATCAAAAGACTATACATTTCCGAATGAATGTTTTCAATGGCCATCTGAAATCCGTAGAAACATCTAGCCTCGGTAATCTGCACTTCTGTACAAAATCGTTCTGCTAAATTTTCATTCACTATTCCGTCACTGGCTGCAAAAAACGCCAATACATGTTTTATAAAATATTTTTCGTCTGGTGTTAGTTTATTCCAGTCATTGATATCTTTAGATATATCCACTTCTTCCACTGTCCAAAATGATGCCTCTGCCTTTTTATACATATTCCAGATGTCATGATATTGGATTGGGAAAATAACAAATCTATTTGGATTTGGTGCAAGGATAGGTTCCATAACTAAATTAACAATAGTAGTAATTTTTTTTCAGTTATCTGTATGACGACTGTACTTGGATCTTTTGTATATCGCTATCGCCGCAATCACTACAATAATTACAAGTATTATTGATAGCATTGTTATTACTACTATCATAATTAAATTATCGACATTCATGGGTGTTGAATAATCGTTATCATCATTTTGTAATTGTGACGTCATACTAGATAAATCATTTGTGAGATTGTTGTGGGAAGCGGGCACGGAAGATGCATTATCATTATTATTTAACGCCTCCCATTCGGATTCACAAATGTTACGCACGTTCAAAGTTTTATGGAAACTATAATTTTGTGAAAACAGATAACAAGAAAACTCGTCATCGTTCAAATTTTTAACGATAGTAAACCGATTAAACGTCGAGCTAATTTCTAACGCTAGCGACTCTGTTGGATATGGGTTTCCAGATATATATCTTTTCAGTTCCCCTACGTATCTATAATCATCTGTAGGAAATGGAAGATATTTCCATTTATCTACTGTTCCTAATATCATATGCGGTGGTGTAGAACCATTAAGCGCGAAAGATGTTATTTCGCATCGTATTTTAACTTCGCAATAATTTCTGGTTAGATAACGCACTCTACCAGTCAAGTCAATGATATTAGCCTTTACAGATATATTCATAGTAGTCGTAACGATGACTCCATCTTTTAGATGTGATACTCCTTTGTATGTACCAGAATCTTCGTACCTCAAACTCGATATATTTAAACAAGTTAATGATATATTAACGCGTTTTATGAATGATGATATATAACCAGAAGTTTTATCCTCTGTGGCTAGCGCTATAACCTTATCATTATAATACCAACTAGTGTAATTAATATGTGACATGACAGTGTGGGTACAAATATGTACATTATCGTCTACGTCGTATTTGATACATCCGCATTCAGCCAACAAATATAAAATTACAAAAACTCTAACGACGTTTGTACACATCTTGATGCGGTTTAATAAATGTTTTGATTTCAATTTATTGTAAAAAAAGATTCGGTTTTATACTGTTCGATATTCTCATTGCTTATATTCTCATCTATCATCTCCACACAGTCAAATCCATGGTTAACATGTACCTCATCAACCGGTAAAAGACTATCGGATTCTTCTATCATCATAACTCGAGAATATTTAATTTGGTGGTCATTATTAATCAAGTCAATTATCTTATTTTTAACAAACGTAAGTATTTTACTCATTTTTTTATAAAAACTTTTAGAAATATACAGACTCTATCGTGTGTCTATATCTTCTTTTTATATCCAATGTATTTATGTCTGATTTTTCTTCATTTATCATATATAATGGTCCAAATTCTACACGTGCTTCGGATTCATCCAGATCATTAAGGTTCTTATAATCGCAACATCCTTCTCTTCCATCTTCTACATCTTCCTTCTTATTCTTAGCGTCACAGAATCTACCACAGCAGGATCCCATGACGAGAGTCACATTAAACTAATTCATTTTCAATTATAATATACTGATTAGTAATGACAATTAAAATAAAAATATTCTTCATAACCGGTAAGAAAGTAAAAAGTTCACATTGAAACTATGTCAGTAGTTATACATCATGAGATGATATACTCTATTTTGGTGGAGGATTATATGATATAATTCGTGGATAATCATTCTTAAGACACATTTCTTCATTCGTAAATCTTTTCACATTAAATGAGTGTCCATATTTTGCAATTTCTTCATATGATGGCGGTGTACGTGGACGAGGCTGCTCCTGTTCTTGTAGTCGTCGACTGTCGTGTTTGCGTTTAGATCCCTCCATTATCGCGATCGCGTAGTGAGTACTATTTATACCTTGTAATTAAATTTTTTTATTAATTAAACGTATAAAAACGTTCCGTATCTGTATTTAAGAGCCAGATTTCGTCTAATAGAACAAATAGCTACAGTAAAAATAACTAGAATAATCGCTACACCCACTAGAAACCACGGATCGTAATACGGCAATCGGTTTTCGATAATAGGTGGAACGTATATTTTATTTAAGGACTTAACAATTGTCTGTAAACCACAATTTGCTTCCGCCGATCCTGTATTAACTATCTGTAAAAGCATATGTTGACCGGGAGGAGCCGAACATTCTCCGATATTCAATTTTTGTATATCTATAATGTTATTAACCTCCGCATACGCATTACAGTTCTTTTCTAGCTTGGATACTACACTAGGTACATCATCTAAATCTATTCCTATTTCCTCAGCGATAGCTCTTCTATCCTTTTCCGGAAGTAATGAAATCACTTCAATAAATGATTCAACCATGAGTGTGAAACTAAGTCGAGAATTACTCATGCATTTGTTAGTTATTCGGAGCGCGCAATTTTTAAACTGTCCTATAACCTCTCCTATATGAATAGCACAAGTGACATTAGTAGGGATAGAATGTTGAGCTAATTTTTGTAAATAACTATCTATAAAAAGATTATACAAAGTTTTAAACTCTTTAGTTTCTGCCATTTATCCAGTCTGAGAAAATGTCTCTCATAATAAATTTTTCCAAGAAACTAATTGGGTGAAGAATGGAAACCTTTAATCTATATTTATCACAGTCTGTTTTGGTACACATGATGAATTCTTCTAATGCTGTACTAAATTCGATATCTTTTTCGATTTCTGGATATGTTTTTAATAAAGTATGAACAAAGAAATGGAAATCGTAATACCAGTTATGTTCAACTTTGAAATTGTTTTTTATTTTCTTGTTAATGATTCCAGCCACTTGGGAAAAGTCAAAGTCGTTTAATGCCGATTTAATACGTTCATTAAAAACAAACTTTTTATTCTTTAGATGAATTATTATTGGTTCATTGGAATCAAAAAGTAAGATATTATCGGGTTTAAGATCTGCATGTAAAAAGTTGTCACAACAGGGTAGTTCGTAGATTTTAATGTATAACAGAGACATCTGTAAAAAGATAAACTTTATGTATTGTACCAAAGATTTAAATCCTAATTTGATAGCTAACTCGGTATCTACTTTATCTGCCGAATACAGTGCTAGGGGAAAAATTATAATATTTCCTCTTTCGTATTCGTAATTAGTTCTCTTTTCATGTTCGAAAAAGTGAAACATGCGGTTAAAATAGTTTATAACATTAATATTACTGTTAATAACTGCCGGATAAAAGTGGGATAGTAATTTCACGAATTTGATACTGTCCTTTCTCTCGTTAAACGCCTTTAGAAAAACTTTAGAAGAATATCTCAATGAGAGTTCCTGACCATCCATAGTTTGTATCAATAATAGCAACATATGAAGAACCCGCTTATACAGAGTATGTAAAAATGTTAATTTATAGTTTAATCCCATGGCCCACGCACACACGATTAATTTTTTTTCATCTCCCTTTAGATTGTTGTATAGAAATTTGGGTACTGTAAACTCCGCCGTAGTTTCCATGGGACTATATAATTTTGTGGCCTCGAATACAAATTTTACTACATAGTTATCTATCTTAAAGACTATACCATATCCTCCTGTAGATATGTGATAAAAATCGTCGTTTATAGGATAAAATCGTTTATCTTTTTGTTGGAAAAAGGATGAATTAATGTAATCATTCTCTTCTATCTTTAGTAGTGTTTCCTTATTAAAATTCTTAAAATAATTTAACAATCTAACTGACGGAGCCCAATTTTGGTGTAAATCTAATTGGGACATTATGTTGTTAAAATATAAACAGTCTCCTAATATAACAGTATCTGATAATCTATGGGGAGACATCCATTGATATTCAGGGGATGAATCATTGGCAACACCCATTTATTGTACAAAAAGCCCCAATTTACAAACGAAAGTCCAGGTTTGATAGAGATAAACTATTAACTATTTTGTCTCTGTTTTTAACACCTCCACAGTTTTTAATTTCTTTGGTAATGAAATTATTCACAATATCAGTATCTTCTTTATCTACCAGAGATTTTACTAACTTGATAACCTTGGCTGTCTCATTCAATAGGGTAGTGATATTTGTATGTATGATATTGATATCTTTTTGAATTGTTTCTTTTAGAAGTGATTCTTTGATGGTATCAGCATACGAATTACAATAATGCAGAAACTCAGTTAACATGCAGGAATTATAGTAAGCCAATTCCAATTGTTGCCTGTATTGTATTAGAGTATTAATATGCGCAATGATGTCCTTGCGTTTCTCTGATAGAATGCGAGCAGCGATTTTGGCGTTATCATTTGACGATATTTCTGGAATGACGAATCCTGTTTCTACTAACTTCTTGGTAGGACAAAGTGAAACAATCAAGAAAATAGCTTCTCCTCCTATTTGTGGAAGAAATTGAACTCCTCTAGATGATCTACTGACGATAGTATCTCCTTGACAGATATTGGACCGAACTACGGAAGTACCTGGAATGTAAAGCCCTGAAACCCCCTCATTTTTTAAGCAGATTGTTGCCGTAAATCCTGCACTATGCCCAAGATAGAGAGCTCCTTTGGTGAATCCATCACTATGTTTCAGTTTAACCAAGAAACAGTCAGCTGGTCTAAAATTTCCATCTCTATCTAATACAGAATCCAACTTGATGTCAGGGACTATGACCGGTTTAATGTTATATGTAACATTGAGTAAATCCTTAAGTTCATAATCATCGTTGTCATCAGTTATGTACGATCCAAACAATGTTTCTACCGGCATGGTGGATACGAAGATGCTATCCATCAGAATGTTTCCCTGATTAGTATTTTCTATATAGCTATTCTTCTTTAAACGATTTTCCGAATCAGTAACTATGTTCATTTTTTTAGGAGTAGGACGTCTAGCCAGTATGGAAGAGGATTTTCTAGATACTCTCTTCAACATCTTTGATCTCAATGGAATGCAAAACCCCATGGTGTAACAACCAACGATAAAAATAATATTGTTTTTTCACTTTTTATAATTTTACCATCTGACTCATGGATTCATTAATATCTTTACAAGAGCTACTAACGTATAATTCTTTATAACTGAACTGAGATATATACACCGGATCTATGGTTTCCATAATTGAGTAAATGAATGCTCGGCAATAACTAATGGCAAATGTATAGAACAACGAAATTATACTAGAGTTGTTAAAGTTAATATTTTCTATGAGTTGTTCCAATAAATTATTTGTTGTGACTGCGTTCAAGTCATAAATTATCTTGATACTATCCAGTAAACAGTCTTTAAGTTCTGGAATATTATCATCCCATTGTAAAGCCCCTAGTTCGACTATCGAATATCCTGCTCTGATAGCAGTTTCAATATCGACGGACGTCAATACTGTAATAAAGGTGGTAGTATTGTCATCATCGTGATAAACTACGGGAATATGGTCGTTAGTAGGTACCGTGACTTTACACAACGCGATATATAACTTTCCTTTTGTACCATTTTTAACGTAGTTGGGACGTCCTGCAGGGTATTGTTTTGAAGAAATGATATCGAGAACAGATTTGATACGATATTTGTTGGATTCCTGATTATTCACTATAATATAATCTAGACAGATAGATGATTCGATAAATAGAGAAGGTATATCGTTGGTAGGATAATACATCCCCATTCCAGTATTCTCGGATACTCTATTGATGACACTAGTTAAGAACATGTCTTCTATTCTAGAAAACGAAAACATCCTACATGGACTCATTAAAACTTCTAACGCTCCTGATTGTGTTTCGAATGCCTCGTACAAGGATTTCAAGGATGCCATAGATTCTTTGACCAACGATTTAGTATTGCGTTTAGCATCTGATTTTTTTATTAAATCAAATGGTCGGCTCTCTGGTTTACTACCCCAATGATAACAATAGTCTTGTAAAGATAAACCGCAAGAAAATTTATACACATCCATCCAAATAACCCTAGCACCGTCGGATGATATTAATGTATTATTATAGATTTTCCATCCACAGTTATTGGGCCAGTATACTGTTAGCAACGGTATATCGAATAGATTACTCATGTAACCTACTAGAATGATAGTTCGTGTACTAGTCATAATATCTTTAATCCAATCTAAGAAATCTAAAATTAGATCTTTTACACTATTAAAGTTAACAAAGGTATTACCCGGGTACGTGGATATCATATATGGTATTGGTCCATTATCAGTAATGGCTCCATAAACTGATACGGCGATGGTTTTTATATGTGTTTGATCTAATGAGGACGAAATTCGCGCCCACAATTCATCTCTAGATATGCATTTAATATCGAACGGTAACACATCAATCTCGGGACGCGTATATGTTTCTAAATTCTTAATCCAAATATAATGATGACCTATATGCCCTATTATCATACTGTCAACTATAGTATACCTAGAGAACTTTCGATACATCTGCTGTTTCCTGTAATCGTTAAATTTTACAAATCTATAACATGCTAAACCTTTTGACGACAGCCATTCATTAATTTCTGATATGGAATCTGTATTCTCAATACCGTATCGTTCTAAAGCCAGTGCTATATCTCCCTGTTCGTGGGAACGCTTTCGTATAATATCGATCAATGGATAATATGAAGTTTTTGGAGAATAATATGATTCATGATCTATTTCGTCCATAAACAATCTAGACATAGGAATTGGAGGCGATGATCTTAATTTTGTGCAATGGGTCAATCCTATAACTTCTAATATTGTAATATTCATCATCGACATAACACTATCTATGTTATCATCGTATATTAGTATACCACGACCTTCTTCATTTCGTGCCAAAATGATATACAGTCTTAAATAATTACGCAATATCTCAATAGTTTCATAATTGTTAGCTGTTTTCATCAAGGTTTGTATCCTGTTTAACATGATGGCGTTCTATAACGTCTCTATTTTCTATTTTTAATTTTTTTAAATTTTTAACGATTTACTGTGGCTAGATACCCAATCTCTCTCAAATATTTTTTTAGCCTCGCTTACAAGCTGTTTATCTATACTATTAAAACTGACGAATCCGTGATTTTGGTAATGGGTTCCGTCGAAATTTGCCGAAGTGATATGAACATATTCGTCGTCGACTATTAACAATTTTGTATTATTCTGAATAGTGAAAACCTTCACAGATAGATCATTTTGAACACACAACGCATCTAGACTTCTGGCGGTTGCCATAGAATATACGTCGTTCTTATCCCAATTACCAACTAGAAGTCTGATCTTAACTCCTCTATTAATGGCTGCTTCTATAATGGAGTTGTAAATGTCAGGCCAATAGTAGCTATTACCGTCGACACGTGTAGTGGGAACTATGGCCAAATGTTCAATATCTATACTAGTCTTAGCCGACTTGAGTTTATCAATAACTACATCAGTGTCTAGATCTCTAGAATATCCCAATAGGTGTTCTGGAGAATCAGTAAAGAACACTCCACCTATAGGATTCTTAATATGATACGCAGTGCTAACTGGCAGACAACAAGCCGCAGAGCATAAATTCAACCATGAATTTTTTGCGCTATTAAAGGCTTTAAAAGTATCAAATCTTCTACGAAGATCTGTGGCCAGCGGAGGATAATCAGAATATACGCCTAACGTTTTAATCGTATGTATAGATCCTCCAGTAAATGACGCGTTTCCTACATAACATCTTTCATCATCAGACACCCAAAAACAACCGAGTAGTAGTCCCACATTATTTTTTTTATCTATATTAACGGTTATAAAATTTATATCCGGGGAGTGACTTTGTAGCTCTCCCAGATTTCTTTTCCCTCGTTCATCTAGCAAAACTATTATTTTAATCCCTTTTTCAGATACCTCTTTTAGTTTATCAAAAATAAGCGCTCCCCTAGTAGTACTCAGAGGATTACAACAAAAAGATGCTATGTATATATATTTCTTAGCTAGAGTGATAATTTCGTTAAAACATTCAAATGTTGTCAAATGATCGGATCTAAAATCCATATTTTCTGGTAGTGTTTCTACCAGCCTACATTTTGCTCCCGCAGGTACCGATGCAAATGGCCACATTTAGTTAACATAAAAACTTATATATCCTGTTCTATCAACGATTCTAGAATATCATCGGCTATATCGCTAAAATTTTCATCAAAGTCGACATCACAACCTAACTCAGTCAATATATTAAGAAGTTCCATGATGTCATCTTCGTCTATTTCTATATCCGTATCCATTGTAGATTGTTGACCGATTATCGAGTTTAAATCATTACTAATACTCAATCCTTCAGAATACAATCTGTGTTTCATTGTAAATTTATAGGCGGTGTATTTAAGTTGGTAGATTTTCAATTATGTATCAATATAGCAACAGTAGTTCTTGCTCCTCCTTGATTTTAGCATCCTCTTCATTATTTTCTTCTACGTACATAATCATGTCTAATACGTTAGACAACACACCGACGATGGTGGCCGCCACAGACACGAATATGACTAGACCGATGACCATTTAAAAAATACTCTCTAGCTTTAACTTAAACTGTATCGATCATTCTTTTAGCACATGTATAATATAAAAACATTATTCTATTTCGAATTTAGGCTTCCAAAAATTTTTCATCCGTAAACCGATAATAATATATATAGACTTGTTAATAGTCGGAATAAATATATTAATGCTTAAACTATCATCATCTCCACGATTAGAGATACAATATTTACATTCTTTTTGCTGTTTCGAAACTTTATCAATACACGTTAATACAAACCCAGGAAGGAGATATTGAAACTGAGGCTGTTGAAAATGAAACGGCGAATACAATAATTCAGATAATGTAAAATCATGATTCCGTATTCTGATGATATTAGAACTGCTAATGGATGTCGATGGTATGTATCTAGGAGTATCTATTTTAACAAAGCATCGATTTGCTAATATACAATTATCCTTTTGATTAATTGTTATTTTATTCATATTCTTAAAAGGTTTCATATTTATCAATTCTTCTACATTAAAAATTTCCATTTTTAATTTATCTAGCCCCGCAATACTCCTCATTACGTTTCATTTTTTGTCTAGAATACCCATTTTGTTCATCTTGGTACATAGATTATCCAATTGAGAAGCGCATTTAGTAGTTTTGTACATTTTAAGTTTATTAACGAATCGTCGAAAACTAGTTATAGTTAACATTTTATTATTTGATACCCTGATATTAATACCCCTGCCGTTACTATTATTTATAACTGATGTAACCCACGTAACATTGGAATTAATTATCGATAGTAATGCATCGACACTTCCAAAATTGTCTATTATAAACTCACCGATAATTTTTTTATTGCATGTTTTCATATTCATTAGGATTATCAAATCTTTAATCTTATTACGATTGTATGCGTTGATATTACAAGACGTCATTCTAAAAGACGGAGGATTTCCATCAAATGCCAGACAATCACGTACAAAGTACATGGAAATAGGTTTTGTTCTATTACGCATCATAGATTCATATAAAACACCCGTAGAAATACTAATTTGTTTTACTCTATAAAATACTATTGCATCTATTTCATCGTTTTGTATAACGTCTTTCCAAGTGTCAAATTCCAATTTTTTTTCATTGATAGTACCAAATTCTTCTATCTCTTTAACTACTTGCATAGATAGGTAATTACAGTGATGCCTACATGCCGTTTTTTGAAACTGAATAGATGCATCTAGAAGCGATGCTACACTAGTCACGATCACCACTTTCATATTTAGAATATATGTATGTAAAAATATAGTAGAATTTCATTTTGTTTTTTTCTATGCTATAAATGAATTCTCATTTTGTATCCGCACATACTCCGTTTTATATCAATACCAAAGAAGGAAGATATCTGGTTCTAAAAGCCGTTAAAGTATGCGATGTTAGAACTGTAGAATGCGAAGGAAGTAAAGCTTCCTGCGTACTCAAAGTAGATAAACCCTCATCACCCACGTGTGAGAGAAGACCTTCGTCCCCGTCCAGATGCGAGAGAATGAATAACCCTGGAAAACAAGTCCCGTTTATGAGAACGGACATGTTACAAAATATGTTTGCTGCTAATCGCGACAACGTAACGTCAAGACTTTTGAACTAAAATACAATTATATCTTTTTCGATATTAATAAATCCGTGTCTCCCGGGTTTTTTATCTCTTTCAGTATGTGAATAGATAGGTATTTTATCTCTATTCATCATCGAATTTAAGAGATCCGATAAACATTGTTTGTATTCTCCAGATGTCAGCATCTGATACAACAATATATGTGCACATAAACCTCTGGCACTTATTTCATGTACCTTCCCCTTATCACTAAGGAGAATAGTATTTGAGAAATATGTATACATGATATTATCATGTATTAGATATACAGAATTTGTAACACTCTCGAAATCACACGATGTGTCGGCGTTAAGATCTAATATATCACTCGATAACACATTTTCATCTAGATACACTAGACATTTTTTAAAGCTAAAATAGTCTTTAGTAGTAACAGTAACTATGCGATTATTTTCATCGATGATACATTTCATCGGCATATTATTACGCGTACCATCAAAGACTATACCATGTGTATATCTAACGTATTCTAGCATAGTTGCCATACGTACATTAAACTTTTCAGGATCTTTGGATAGATCTTCCAATCTATCTATTTGAGAAAACATTTTTATCATGTTCAATAGTTGAAACGTCGGATCCACTATATAGATATTATCTATAAAGATTTTAGGAACTATGTTCATGGTATCCTGGCGAATATTAAAACTATCAATGATATGATTATCGTTTTCATCTTTTATCACCATATAGTTTCTAAGATATGGGATTTTACTTAATATAATATTATTTCCCGTAATAAATTTTATTAGAAATGCCAAATCTATAAGAAAAGTCCTAGAATTAGTCTGAAGAATATCTATATCACCGTACCGTATATTTGGATTAATTAGATATAGAGAATATGATCCGTAACATATACAACTTTTATTATGACGTCTAAGATATTCTTCCATCAACTTATTAACATTTTTGACTAGGGAAGATACATTATGACGTCCCATTACTTTTGCCTTGTCTATTACAGCGACGTTCATAGAATTTAGCATATCTCTTGCCAATTCTTCCATTGATGTTACATTATAAGAAATTTTAGATGAAATTACATTTGGAGCTTTAATAGTAAGAACTCCTAATATATCCGTGTATGTGGTCACTAATACAGATTGTAGTTCTATAATCGTAAATAATTTACCTATATTATATGTTTGAGTTTGTTTAGAAAAGTAGCTAAGTATACGATCTTTTATTTCTGATGCCGATGTATCAACATCGAAAAAAAATCTTTTTTTATTCTTTTTTACTAACGATACGAATATGTCTTTGTTAAAAACAGTTATTTTCTGAATATTTCTAGCTTGTAATTTTAACATATGATATTCGTTCACACTAGGTACTCTGCCTAAATAGGTTTCTATAATCTTTAATGTAATATTAGGAAGAGTATTCTGATCAGGATTCCTATTCATTTTGAGGATTTAAAACTCTGATTATTGTCTAATATGGTCTCAACACAAACTTTTTCACAGAGTGATAGAGTTTTTGATAACTCGTTTTTCTTAAGAAATATAAAACTACTGTCTCCAGAGCTCGCTCTATCTTTTATTTTATCTAATTCGATACAAACTCCTGATACTGGTTCAGAAAGTAATTCATTAATTTTCAGTCCTTTATAGAAGATATTTAATATAGATAATACAAAATCTTCAGTTCTTGATATCGATCTGATTGATCCTAGAACTAGATATATTAATAACGTGCTCATTAGGCAGTTTATGGCAGCTTGATAATTAGATATAGTATATTCCAGTTCATATTTATTAGATACCGCATTGCCCAGATTTTGATATTCTATGAATTCCTCTGAAAATAAATCCAAAATAACTAGACATTCTATTTTTTGTGGATTAGTGTACTCTCTTCCCTCTATCATGTTCACTACTGGTGTCCACAATGATAAATATCTAGAGGGAATATAATATAGTCCATATGATGCCAATCTAGCAATGTCGAATAACTGTAATTTTATTCTTCGCTCTTCATTATGAATTGAATCTTGAGGTATAAACCTAACACAAATTATATCATTAGACTTTTCGTATGTAATGTCTTTCATGTTATAAGTTTTTAATCCTGGAATAGAATCTATTTTAATGAGGCTTTTAAACGCAGCGTTCTCCAACGAGTCAAAGCATAATACTCTGTTGGTTTTCTTATATTCAATATTACGATTTTCTTCTTTGAATGGAATAGGTTTTTGAATTAGTTTATAATTACAACATAATAGATAAGGAAGTGTGTAAATAGTACGCGGAAAAAACATAATAGCTCCCCTGTTTTCATCCATGGTTTTAAGTAAATGATCACTGGCTTCTTTAGTCAATGGATATTCGAACATTAACCGTTTCATCATCATTGGACAGAATCCATATTTCTTAATGTAAAGAGTGATCAAATCATTGTGTTTATTGTACCATCTTGTTGTAAATGTGTATTCGGTTATCGGATCTGCTCCTTTTTCTATTAAAGTATCGATATCGATCTCGTCTAAGAATTCAACTATATCGACATATTTCATTTGTATACACATAACCATTACTAACGTAGAATGTATAGGAAGAGATGTAACGGGAACAGGGTTTGTTGATTCGCAAACTATTCTAATACATAATTCTTCTGTTAATACGTCTTGCACGTAATCTATTATAGATGCCAAGATATCTATATAATTATTTTGTAAGATGATGTTAACAATGTGATCTATATAAGTAGTGTAATAATTCATGTATTTCGATATATGTTCCAACTCTGTCTTTGTGATGTCTAGTTTCGTAATATCTATAGCGTCCTCAAAAAATATATTCGCATATATTCCCAAGTCTTCAGTTCTATCTTCTAAAAAATCTTCAACGTATGGAATATAATAATCTATTTTACCTCTTCTGATGTCATTAATGATATAGTTTTTGACACTATTTTCCGTCAATTGATTCTTATTCACTATGTCTAAAAACCGGATAGCGTCCCTAGGACGAACTACTGCCATTAATATCTCTATTATAGCTTCTGGACATAAATCATCTATTATACCAGAATTAATGGGAACTATTCCGTATCTATCTAACATAGTTTTAAGAAAGTCAGAATCTAAGACCTGATGTTCATATATTGGTTCATACATGAAATGATCTCTATTGATGATAGTGACTATTTCATTCTCTGAAAATTGGTAACTCATTCTATACACGCTTTCCTTGTTGATAAAGGATAGTATATACTCAATGGAATTTGTACCAACAAACTGTTCTCTTATGAATCGTATATCATCATCTGAAATGATCATGTAAGGCATACATTTAACAATAAGAGACTTGTCTCCTGTTATCAATATACTATTCTTGTGATAATTTATGTGTGCGGCAAATTTGTCCACGTTCTTTAATTTTGTTATAGTAGATATCAAATCCAATGGAGATACAGTTCTTGGCTTAAACAGATATAGTTTTTCTGGAACGAATTCTACAACATTATTATAAAGGACTTTGGGTATATAAGTGGGATGAAATCCTATTTTAATTAATGCGATAGCCTTGTCCTCGTGCAGATATCCAAACGCTTTTGTGATAGTATGGCATTCATTGTCTAGAAACGCTCTACGAATATCTGTAACAGATATCATCTTTAGAGAATACTAGTCGCGTTAATAGTACTAAAATTTGTATTTTTTAATCTATCTCAATAAAAAATTAATATGTATGATTCAATGTATAACTAAACTACTAACTGTTATTGATAACTAGAATCAGAATCTAATGATGACATAACTAAGAAGTTTATCTACAGCCAATTTAGCTGCATTATTTTTAGCATCTCGTTTAGATTTTCCATCTGCCTTATCGAATACTCTTCCGTCAATGTCTACACAGGCATAAAATGTAGGAGAGTTACTAGGCCCCACTGATTCAATACGAAAAGACCAATCTCTCCTAGTTATTTGACAGTACTCATTAATAACGGTGACAGGGTTAACACCTTTCCAATAAATAATTTTTTTAACCGGAATAACATCATCAAAAGACTTATGATCCTCTCTCATTGATTTTTCGCGGGATACATCATCTATTATAGCATCAGCATCAGAATCTGTAGGCCGTGTATCAGCATCCATTGTCGTAGACCAACGAGGAGGAGTATCGTTGGAGCTGTAAACCATAGCACTACGTTGAAGATCATACAGAGCTTTATTAACTTCTCGCTTCTCCATATTAAGTTGTTTAGTTAGTTGTACAGCAGTAGCTCCTTAGTCCAATGTTTTTAATAACCGCACACAATCTCTGTGTCAGAACGCTCGTCAATATAGATCTTAGAAATTTTTTTAGAGAGAACTAACGCAACTAGCAATAAAACTGATCTTATTTTATCATTTTTTTTATTCATCATCCTCTGGTGGTTCGTCGTTCCTATCGAATGTAGCTCTGATTAACCCGTCATCTATAGGTGATGCTGGTTCTGGAGATTCTGGAGGAGATGGATTATTATCTGGAAGAATCTCTGTTATTTCCTTGTTTTCATGTATCGATTGCGTTGTAACATTAAGATTGCGAAATGCTCTAAATTTGGGAGGCTTAAAGTGTTGTTTACAATCTCTACACGCGTGTCTAACTAATGGAGGTTCATCAGCGGCTCTAGTTTGAATCATCATCGGTGTAGTATTCCTACTTTTACAGTTAGGACACGGTGTATTGTATTTCTCGTCGAGAACGTTAAAATAATCGTTGTAACTCACATCCTTTATTTTATCTATATTGTATTCTACTCCTTTCTTAATGCATTTTATACCGAACAAGAGATAGCGAAGGAATTCTTTTTCGGTACCGCTAGTACCCTTAATCATATCACATAGTGTTTTATATTCTAAATGTGTGGCAATGGACGGTTTATTTCTATACGATAGTTTGTTTTTGGAATCCTTTGAGTATTCTATACCAATATTATTCTTTGATTCGAATTTAGTTTCTTCGATATTAGATTTTGTATTACCTATATTCTTGATGTAGTACTTTGATGATTTTTCCATGGCCCATTCTATTAAGTTTTCCAAGTTGGCATCATCCACATATTGTGATAGTAATTCTCGGATATCAGTAGTGACTACCGCCATTGATATTTGTTCATTTGATGAGTAACTACTAATGTATACATTTTCCATTTATAACACTTATGTATTAACTTTGTTTATTTATATTTTTTCATTATTATGTTGATATTAATAATCGTATTGTGGTTATATGGCTACAATTTCATAATGAGTTGAAGTCAGTGTCCTATGATCAATGACGATAGCTTTACTCTGAAAAGAAAGTATCAAATCGATAGTGCAGAGTCAACAATGAAAATGGATAAGACGATGACAAAGTTTCAGAATAGAGTCAAAATGGTAAAAGAAATAAATCAGACGATAAGAGCAGCACAAACTCATTACGAGACATTGAAACTAGGATATATAAAATTTAAGGGAATGATTAGGACTACTACTCTAGAAGATATAGCACCATCTATTCCAAATAATCAGAAAACTTATAAACTATTCTCGGACATTTCAGTCATTGGCAAAGCATCACAGAATCCGAGTAAGATGATATATGCTCGCTGCTTTACATGTTTCCCAATTTGTTTGGAGATGACCATAGATTCATTTGTTATAGAATGCATCCAACATTGTTCATGATATAGTTGAATCATGTATGCCTGTTCGTATGCCTGTGGCTAAGATACTGTGTAAAGAAATGGTAAATAAATACTTTGAGAATCTTTAAGAGTGCATTGACTTTGTTAGTGAATAGGCATTCCATCTTTCTCCAATACTAATTCAAATTGTTAAATTAATAATGGAATAGTATAAATAGTTATTAGTGATAGGATAGTAAACATAATTATTAGAATAGTAGTGTAGTATCATAGATAACTCTCTTCTATAAAAAATGGATTTTATTCGTAGAAAGTATCTTATATACACAGTAGAAAATAATATAGATTTTTTAAAGGATGATACATTAAGTAAAGTAAACAATTTTACCCTCAATCATGTACTAGCTCTCAAGTATCTAGTTAGCAATTTTCCTCAACATGTTATTACTAAGGATGTATTAGCTAATACCAATTTTTTTGTTTTCATACATATGGTACGATGCTGTAAAGTATACGAAGCGGTTTTACGACACGCATTTGATGCACCCACGTTGTACGTTAAAGCATTGACTAAGAATTATTTATCGTTTAGTAACACAATACAGTCGTACAAGGAAACAGTGCATAAACTAACACAAGATGAAAAATTTTTAGAGGTTGCCGAATACATGGACGAATTAGGAGAACTTATAGGCGTAAATTATGACTTAGTTCTTAATCCATTATTTCACGGAGGGGAACCCATCAAAGATATGGAAATCATTTTTTTAAAACTGTTTAAGAAAACAGACTTCAAAGTTGTTAAAAAATTAAGTGTTATAAGATTACTTATTTGGGCATACCTAAGCAAGAAAGATACAGGCATAGAGTTTGCGGATAATGATAGACAAGATATATATACTCTATTTCAACAAACTGGTAGAATAGTCCATAGCAATCTAACAGAAACGTTTAGGGATTATATCTTTCCCGGAGATAAGACTAGCTATTGGGTGTGGTTAAACGAAAGTATAGCTAATGATGCGGATATCGTTATTAATAGACCCGCCATTACCATGTATGATAAAATTCTTAGTTATATATACTCTGAGATAAAACAGGGACGCGTTAATAAAAACATGCTTAAGTTAGTTTATATCTTTGAGCCTGAAAAAGATATCAGAGAACTTCTGCTAGAAATCATATATGATATTCCTGGAGATATCCTATCTATTATTGATGCAAAAAACGACGATTGGAAAAAATATTTTATTAGTTTTTACAAAGCTAATTTTATTAACGGTAATACATTTATTAGTGATAGAACGTTTAACGATGACTTATTCAGAGTTGTTGTTCAAATAGATCCCGAATATTTCGATAATGAACGAATTATGTCTTTATTCTATACGAGTGCTGCGGACATTAAACGATTTGATGAGTTAGATATTAATAACAGTTATATATCTAATATAATTTATGAGGTGAACGATATCACATTAGATACAATGGATGATATGAAGAAGTGTCAAATCTTTAACGAGGATACGTTGTATTATGTTAAGGAATACAATACATACCTGTTTTTGCACGAGTCGGATCCCATGGTCATAGAGAACGGAATACTAAAGAAACTGTCATCTATAAAATCCAAGAGTAGACGGCTGAACTTGTTTAGCAAAAACATTTTAAAATATTATTTAGACGGACAATTGGCTCGTCTAGGTCTTGTGTTAGATGATTATAAAGGAGACTTATTAGTTAAAATGATAAACCATCTCAAATCTGTGGAGGATGTATCCGCATTCGTTAGATTTTCTACAGATAAAAACCCTAGTATTCTTCCATCGCTAATCAAAACTATTTTAGCTAGTTATAATATTTCCATCATCGTCTTATTTCAAAGGTTTTTAAGAGATAATCTATATCATGTAGAAGAATTCTTGGATAAAAGCATCCATCTAACCAAGACGGATAAGAAATATATACTTCAATTGATAAGGCACGGTAGATCATAGAACAAACCAAATATATTATTAATAATTTGTATATACATAGATATAATTATCACATATTAAAAAATAACACATTTTTGATAAATGGAAACCGTTGCAACAATTCAGACTCCCACCAAATTAATGAATAAAGAAAATGCAGAAATGATTTTGGAAAAAATTGTTAATCATATAGCTATGTATATTAGTGACGAATCAATATATTCAGAAAATAATCCTGAATATATTGATTTTCGTAACAGATACGGAGACTATAGATCTCTCATTATAAAAAGTGATCACGAGTTTGTAAAGCTATGTAAAGATCATGCAGAGAAAAGTTCTCCAGAAACGCAACAAATGATTATCAAACACATATACGAACAATATCTTATTCCAGTATCTGAAGTACTATTAAAACCTATAATGTCCATGGGTGACATATTTACATATAACGGATGTAAAGACAATGAATGGATGCTAGAACAACTCTCTACCCTAAACTTTAACAATCTCTACACATGGAACTCATGTAGCATAGGCAATGTAACGCGTCTGTTTTATACATTTTTTAGTTATCTGATGAAAGATAAACTAAATATATAAGTATAATCCCATTCTAATACTTTAACCTGATGTATTATTACCTGCATCTTATTAGAATATTAACCTAACTAAAAGACATAAAAAGCGGTAGGATATAAATATTATGGCCGCAACCGTTCCGCGTTTTGACGATGTGTACAAAAATGCACAAAGAAGAATTCTAGATCAAGAAACATTTTTTAGTAGAGGTCTAAGTAGACCGTTAATGAAAAACACATATCTATTTGATAATTACGCGTATGGATGGATACCAGAAACTGCAATTTGGAGTAGTAGATACGCAAACCTAGATGCTAGTGACTATTATCCCATTTCGTTGGGATTACTTAAAAAGTTTGAATTTCTCATGTCTCTATATAAAGGTCCTATTCCCGTATATGAAGAAAAAGTAAATACTGAATTCATTGCTAATGGATCTTTCTCCGGTAGATACGTATCATATCTTAGAAAGTTTTCTGCCCTTCCAACAAACGAGTTTATTAGTTTTTTATTATTGACCTCCATCCCTATCTATAATATCTTATTCTGGTTTAAAAACACACAGTTTGATATTACTAAACACACATTATTCAGATACGTCTATACAGATAATACCAAACACCTTGCGTTGGCTAGGTATATACATCAAACAGGAGACTATAAGCCTTTGTTTAGTCGTCTCAAAGAGAATTATATATTTACTGGTCCCGTTCCAATAGGTATCAAAGATATAGATCACCCTAATCTTAGTAGAGCAAGAAGTCCATCCGATTATGAGACATTAGCTAATATTAGTACTATATTGTACTTTACCAAGTATGATCCAGTATTAATGTTTTTATTGTTTTACGTACCTGGGTATTCAATTACTACAAAAATTACTCCAGCCGTAGAATATCTAATGGATAAACTGAATCTAACAAAGAGCGACGTACAACTGTTGTAAATTATTTTATGCTTCGTAAAATGTAGGTCTTGAACCAAACATTCTTTGAAAAAATGAGATGCATAAAACTTTATTATCCAATAGATTAACTATTTCAGACGTCAATCGTTTAAAGTAAACTTCGTAAAATATTCTTTGATTGCTGCCGAGTTTAAAACTTCTATCGATAATTGTTTCATATGTTTTAATATTTACAAGTTTTTTGGTCCATGGTACATTAGCTGGACAGATATATGCAAAATAATATCGTTCTCCAAGTTCTATAGTCTCTGGATTGTTTTTATTATATTCAGTAACCAAATACATATTAGGGTTATCTGCGGATTTATAATTTGAGTGATGCATTCGACTCAACATAAATAATTCTAGAGGAGACGATCTACTATCAAATTCGGATCGTAAATCTGTTTCTAAAGAACGGAGAATATCTATACATACCTGATTAGAATTCATCCGTCCTTCAGACAACATCTCAGACAGTCTGGTCTTGTATGTCTTAATCATATTCTTATGAAACTTGGAAACATCTCTTCTAGTTTCACTAGTACCTTTATTAATTCTCTCAGGTACAGATTTTGAATTCGACGATGCCGAGTATTTCATCGTTGTATATTTCTTCTTCGATTGCATAATCAAATTCTTATATACCGCCTCAAACTCTATTTTAAAATTATTAAACAATACTCTACTATTAATCAGTCGTTCTAACTCCTTTGCTATTTCTATGGACTTATCTACATCTTGACTGTCTATCTCTGTAAACACGGAGTCGGTATCTCCATACACGCTACGAAAACGAAATCTATAATCTATAGGCAACGATGTTTTCACAATCGGATTAATATCTCTATCGTCCATATAAAATGGATTACTTAATGTATTGGCAAACCGTAACATACCGTTGGATAACTCTGCTCCATTTAGTACCGATTCTAGATACAATATCATTCTACGTCCTATGGATGTGCAACTCTTAGCCGAAGCGTATGAGTATAGAGCACTATTTCTAAATCCCATCAGACCATATACTGAGTTGGCTACTATCTTGTACGTATATTGCATGGAATCATAGATGGCCTTTTCAGTTGAACTGGTAGCCTGTTTTAACATCTTTTTATATCTGGCTCTCTCTGCCAAAAATGTTCTTAATAGTCTAGGAATGGTTCCTTCTATTGATCTATCGAAAATTGCTATTTCAGAGATGAGGTTCGGTAGTCTAGGTTCACAATGAACCGTAATATATCTAGGAGGTGGATATTTCTGAAGCAAGAGTTGATTATTTATTTCTTCTTCCAATCTATTGGTACTAACAACGACACCGACTAATGTTTCCGGAGATAGATTTCCAAAGATACACACATTAGGATACAGACTGTTATAATCAAAGATTAATACATTATTACTAAACATTTTTTGTTTTGGAGCAAATACCTTACCGCCTTCATAAGGAAACTTTTGTTTTGTTTCTGATCTGACTAAGATAGTTTTAGTTTCCAACAATAGCTTTAACAGTGGACCCTTGATGATTGTACTCGCTCTATATTCGAATACCATGGATTGAGGAAGCACATATGTTGCCGCACCAGCGTCTGTTTTTGTTTCTACTCCATAATACTCCCACAAATACTGACACAAACAAGCATCATGAATACAGTATCTAGCCATATCTAAAGCTATGTTTAGATTATAATCCTTATACATCTGAGCTAAATCAATGTCATCCTTTCCGAAAGATAATTTATATATATCATTAGGTAAAGTAGGACATGATAGTACGACTTTAAATCCATTTTCCAAAATATCTTTACGAATTACTTTACATATAATATCCTCATCAACAGTCACGTAATTACCTGTGGTTAAAACCTTTGCAAATGTATCGGCTTTGCCTTTCGCGTCCGTAGTATCGTCACCGATGAACGTCATTTCTCTAACTCCTCTATTTAATACTTTACCCATGCAACTGAACGCGTTCTTGGATATAGAATCCAATTTGTACGAATCCAATTTTTCAGATTTTTGAATGAATGAATATAGATCGAAAAATATAGTTCCATTATTGTTATTAACGTGAAACGTAGTATTGGCCATGCCGCATACTCCCTTATGACTAGACTGATTTCTCTCATAAATACAGAGATGTACAGCTTCCTTTTTGTCTGGAGATCTAAAGATAATCTTCTCTCCTGTTAATAACTCTAGACGATTAGTAATATATCTCAGATCAAAGTTATGTCCGTTAAAGGTAACGACGTAGTCGAACGTTAGTTCCAACAATTGTTTAGCTATTCGTAACAAAACTATTTCAGAACATAGAACTAGTTCTCGTTCGTAATCCATTTCCATTAGCGACTGTATCCTCAAACATCCTCTATCGACGGCTTCTTGTATTTCCTGTTCCGTTAACATCTCTTCATTAATGAGCGTAAACAGTAATCGTTTACCACTTAAATCGATATAACAGTAACTTGTATGCGAGATTGGGTTAATAAATACAGAAGGAAACTTCTTATCGAAGTGACACTCTATATCTAGAAATAAGTACGATCTTGGGATATCGAATCTAGGTATTTCTTTAGCGAAACAGTTACGTGGATCGTCACAATGATAACATCCATTGTTAATCTTTGTCAAATATTGCTCGTCCAACGAGTAACATCCGTCTGGAGATATCCCGTTAGAAATATAAAACCAACTAATATTGAGAAATTCATCCATGGTGGCATTTTGTATGCTGCGTTTCTTTGGCTCTTCTATCAACCACATATCTGCGACGGAGCATTTTCTATCTTTAATATCTAGATTATAACTTATTGTCTCGTCAATGTCTATAGTTCTCATCTTTCCCATCGGCCTCGCATTAAATGGAGGAGGAGATAATGACTGATATATTTCGTCCGTCACTACGTAATAAAAGTAATGAGGAAATCGTATAAATACGGTCTCGCCATTTCGACATCTGGATTTCAGATATAAAAATCTGTTTTCACCGTGACTTTCAAACCAATTAATACACCTAACATCCATTTCTAGAATTTAGAAATATATTTTCATTTAAATGAATCCCAAACATTGGGGAAGAGCCGTATGGACCATTATTTTTATAGTACTTTCGCAAGCGGGTTTAGACGGCAACATAGAAGCGTGTAAACGAAAACTATATACTATAGTCAGCACTCTTCCATGTCCTGCATGTAGACGACACGCGACTATCGCTATAGAGGACAATAATGTCATGTCTAGCGATGATCTGAATTATATTTATTATTTTTTCATCAGATTATTTAACAATTTGGCATTTGATCCCAAATACGCAATCGATGTGTCAAAGGTTAAACCTTTATAAACTTAACCCATTATAAAACTTATGATTAGTCACGACTGAAATAACCGCGTGATTATTTTTTGGTATAATTCTACACGGCATGGTTTCTGTGACTATGAATTCAACACCTGTTATCTTAGTGAAATCTTTAACAAACAGCAAGGGTTCGTCAAAGACATAAAACTCATTGTTTACGATCGAAATAGACCCCCTATCACACTTAAAATAAAAAATATCCTTATCCTTTACCACCAAATAAAATTCTGATTGGTCAATGTGAATGTATTCACTTAACAGTTCCACAAATTTATTTATTAACTCCGAGGCACATACATCGTCGGTATTTTTTATGACAAACTTTACTCTTCCAGCATCCGTTTCTAAAAAAATATTAACGAGTTCCATTTATATCATCCAATATTATTGAAATGACGTTGATGGACAGATGATATAAATAAGAAGGTACAGTACCTTTGTCCACCATCTCCTCCAATTCATACTCTATTTTGTCATTAACTTTAATGTGTGAAAACAGTACGCCACATGCTTCCATGACAGTGTGTAACACTTTGGATACAAAATGTTTGACATTAGTATAATTGTCCAAGACTGTCAATCTATAATAGATAGTAGCTATAATATATTCTATGATGGTATTGAAGAAGATGACAACCTTGGCATATTGATCATTTAACACAGACATGGTATCAACAAATAGCTTAAATGAAAGAGAATCAGTAATTGGAATAAGCGTCTTCTCGATGTAGTGTCCGTATACCAACATGTCTGATATTTTGATGTATTCCATTAAATTATTTAGTTTTTTCTTTTTATTCTCGTTAAACAGAATTTCTGTCAATGGACCCCAACATCGTTGACCTATTAAGTTTTGATTGATTTTTCCGTGTAAGGCGTATCTAGTCAGATCGTATAGCCTATCCAATAATCCATCGTCTGTGCGTAGATCACATCGTACACTTTTTAATTTTCTATAGAAGAGTGACAGACATCTGGAGCAATTACAGACAGCAATTTCTTTATTCTCTACAGATGTAAGATACTTGAAGATATTCCTATGATGATGCAGAATTTTGGATAACACGGTATTGATGGTATCTGTTACCATAATTCCTTTGACTGATAGTGTCAAAGTACAAGATTTCCAATCTTTTGCAATTTTCAGTACCATTATCTTTGTTTTGATATCTATATCAGACAGCATGGTACGTCTGACAACACAGGGATTAAGACGGAAAGATGAAATGATTCTCTCAACATCTTCAATAGATACCTTGCTATTTTTTTTGGCATTATCTATATGTGAGAGAATATCCTCTAGAGAATCAGTATCCTTTTTGATGATAGTGGATCTCAATGACATGGGACGTCTAAACCTTCTTATTCTATCACCAGATTGCATGGTGATTTGTCTTCTTTCTTTTATCATGATGTAATCTCTAAATTCATCGGCAAATTGTCTATATCTAAAATCATAATATGAGATGTTTACCTCTACAAATATCTGTTCGTCCAATGTTAGAGTATCTATATCAGTTTTGTATTCCAAATTAAACATGGCAACGGATTTAATTTTATATTCCTCTATTAAGTCCTCGTCGATAATAACAGAATGTAGATAATCATTTAATCCATCGTACATGGTTGGAAGATGCTCGTTGACAAAATCTTTAATTGTCTTGATGAAGGTGGGACTATATCTAACATCTTGATTAATAAAATTTATAACATTGTCCATAGGATACTTTGTAACTAGTTTTATACACATCTCTTCATTGGTAAGTTTAGACAGAATATCGTGAACAGGTGGTATATTATATTCATCAGATATACGAAGAATAATGTCCAAATCTATATTGTTTAATATATTATATAGATGTAGTGTAGCTCCTACAGGAATATCTTTAACTAAGTCAATGATTTCATCAACAGTTAGATCTATTTTAAAGTTAATCATATAGGCATTGATTTTTAAAAGGTATGTAGCCTTGACTACATTCTCATTAATTAACCATTCCAAGTCACTGTGTGTAAGAAGATTATATTCTATCATAAGCTTGACTACATTTGGTCCCGATACCATTAAAGAATTCTTATGATATAAGGAAACAGCTTTTAGGTACTCATCTACTCTACAAGAATTTTGGAGAGCCTTAACGATATCAGTGACGTTTATTATTTCAGGAGGAAAGAACCTAACATTGAGAATATCTGAATTAATAGCTTCCAGATACAGTGATTTTGGCAATAGTCCGTGTAATCCATAATCCAGTAACACGAGCTGGTGCTTGCTAGACACCTTTTCAATGTTTAATTTTTTTGAAATAAGCTTTGATAAAGCCTTCCTCGCAAATTCCGGATACATGAACATGTCGCCAACATGATTAAGTATTGTTTTTCATTATTTTTATATTTTCTCAACAAGTTCTCAATACCCCAATAGATAATAGAATATCACCCAATGCGTCCATGTTGTCTATTTCCAACAGGTCGCTATATCCACCAATAGAAGTTTTCCCAAAAAAGATTCTAGGAACAGTTCTACCACCAGTAATTTGTTCAAAATAGTCACGCAATTCATTTTCGGGTTTAAATTCTTTAATATCTACAATTTCATACGCTCCTCTTTTGAAACTAAACTTATTTAGAATATCCAGTGCGTTTCTACAAAAAGGACATGTAAACTTGACAAAAATTGTCACTTTGTTATTGGCCAACCTTTGTTGTACAAATTCCTCGGCCATTTTTAATATTTAAGTGATACAAAACTATCTCGACTTATTTAACTCTTTAGTCGAGATATATGGACACAGATAGCTATATGATAACCAACTACAGAAGACAAACGCTATAAAAAACATAATTACGACGAGCATATTTATAAATATTTTTATTCAGTATTACTTGATATAGTAATATTAGGCACAGTCAAACATTCAACCACTCTAGATACATTAACTCTCTCATTTTCTTTAACAAATTCTGCAATATCTTCGTAAAAAGATTCTTGAAACTTTTTAGAATATCTATCGACTCTAGATGAAATAGCGTTCGTCAACATACTATGTTTTGTATACATAAAGGCGCCCATTTTAACAGTTTCTAGTGACAAAATGCTAGCGATCCTAGGATCCTTTAGAATCACATAGATTGACGATTCGTCTCTCTTAGTAACTCTAGTAAAATAATCATACAATCTAGTACGCGAAATAATATTATCCTTGACTTGAGGAGATCTAAACAATCTAGTTTTGAGAACATCGATAAGTTCATCGGGAATTACATACATACTATCTTTAATAGAACTCTTTTCATCCAGTTGAATGGATTCGTCCTTAACCAACTGATTAATGAGATCTTCTATTTTATCATTTTCTAGATGATATGTATGTCCATTAAAGTTAAATTGTGTAGCGCTTCTTTTTAGCCTAGCAGCCAATACTTTAACATCACTAATATCGATATACAAAGGAGATGATTTATCGATGGTATTAAGAATTCGTTTTTCGACATCCGTCAAAACCAATTCCTTTTTGCCTGTATCATCCAGTTTGCCATTCTTTGTAAAGAAATTATTTTCTACTAGACTATTAATAAGACTGATAAGGATTCCTCCATAATTGCACAATCCAAACTTTTTCACAAAACTAGACTTTACGAGATCTACAGGAATGCGTACTTCAGGTTTCTTAGCTTGTGATTTTTTCTTTTGTGGACATTTTCTAGTGACCAACTCATCTACCATTTCATTGATTTTAGCAGTGAAATAAGCTTTCAATGCACGGGCACTGATACTATTGAAAACGAGTTGATCTTCAAATTCCGCCATTTAAGTTCACCAAACAACTTTTAAATACAAATATATCAATAGTAGTAGAATAAGAACTATAAAAAAAATAATAATTAACCAATACCAACCCCAACAACCTGTATTATTAGTTGATGTGACAGTTTTCTCATCACTTAGAACAGATTTAACAATTTCTATAAAGTCTGTCAAATCATCTTCCTGAGAACCCATAAATACACCAAATATAGCAGCGTACAACTTATCCATTTATACATTGAATATTGGCTTTTCTTTATCGCTATCTTCATCATATTCATCATCAATATCAACAAGTCCCAGATTACGAACCAGATCTTCTTCTACATTTTCAGTCATTGATACGCGTTCACTATCTCCAGAGAGTCCGATAACGTTAGCCACTACTTCTCTATCAATGATTAGTTTCTTGAGCGCGAATGTAATTTTTGTTTCCGTTCCGGATCTATAGAAAACTACAGGTGTAATAATTGCCTTGGCTAATTGTCTTTCTCTTTTACTGAGTGATTCTAGTTCACCTTCTATAGATCTGAGAATGGATGATTCTCCAGTCGAAACATATTCTACCATGGCTCCGTTTAATTTGTTGATGAAGATGGATTCATCCTTAAATGTTTTCTCTGTAATAGTTTCCGCCGAAAGACTATGCAAAGAATTTGGAATGCGTTCCTTGTGTGTAATGTTTCCATAGACAGCTTCTAGAAGTTGATACAACATAGGACTAGCCGCGGTAACTTTTATTTTTAGAAAGTATCCATCGCTTCTATCTTGTTTAGATTTATTTTTATAAAGTTTAGTCTCTCCTTCCAACATAATAAAAGTGGAAGTCATCTGACTAGATAAACTATCAGTAAGTTTTATAGAGATAGATGAACAATTAGCGTATTGAGAAGCATTTAGTGTAACGCATTCGATACATTTTGCATTAGATTTACTAATCGATTTTGCATACTCTATAACACCCGCACAAGTCTGTAGAGAATCGCTAGATGCTGTAGGTCTTGGTGAAGTTTCAACTCTCTTCTTGATTACCTTACTCATGATTAAACCTAAATAATTGTACTTTGTAATATAATGATATATATTTTCACTTTATCTCATTTGAGAATAAAAATGTTTTTGTTAACCACTGCATGATGTACAGATTTCGGAATCGCAAACCACTTGTGGTTTTATTTTATCCTTGTCCAATGTGAATTGAATGGGAGCGGATGCGGGTTTCGTACGTAGATAGTACATTCCCGTTTTTAGACCGAGACTCCATCCGTAAAAATGCATACTCGTTAGTTTGGAATAACTCGGATCTGCTATATGGATATTCATAGATTGACTTTGATCGATGAAGGCTCCCCTGTCTGCAGCCATTTTTATGATCGTCTTTTGTGGAATTTCCCAAATAGTTTTATAAACTCGCTTAATATCTTCTGGAAGGTTTGTATTCTGAATGGATCCACCATCTACCATAATCCTATTCTTGATCTCATCATTCCATAATTTTCTCTCGGTTAAAACTCTAAGGAGATGCGGGTTAACTACTTGGAATTCTCCAGACAATACTCTCCGAGTGTAAATATTACTGGTATACGGTTCCACCGACTCATTATTTCCCAAAATTTGAGCAGTTGATGCAGTCGGCATAGGTGCCACCAATAAACTATTTCTAAGACCGTATGTTCTGATTTTATCTTTTAGAGGTTCCCAATTCCAAAGATCCGACGGTACAACATTCCAAAGATCATATTGTAGAATACCGTTACTGGCGTACGATCCTACATATGTATCATATGGTCCTTCCTTCTCAGCTAGTTTACAACTCGCCTCTAATGCACCGTAATAAATGGTTTCAAAGATCTTCTTATTTAGATCTTGTGCTTCCAGGCTATCAAATGGATAATTTAAGAGAATAAACGCGTCCGCTAATCCTTGAACACCAATACCGATAGGTCTATGTCTCTTATTAGAGATTTCAGCTTCTGGAATAGGATAATAATTAATATCTATAATTTTATTGAGATTTCTGACAATTACTTTGACCACATCCTTCAGTTTGAGAAAATCAAATCGCCCATCTATTACAAACATGTTCAATGCAACAGATGCCAGATTACACACGGCTACCTCATTAGCATCCGCATATTGTATTATCTCAGTGCAAAGATTACTACACTTGATGGTTCCTAAATTTTGTTGATTACTCTTTTTGTTACACGCATCCTTATAAAGAATGAATGGAGTACCAGTTTCAATCTGAGATTCTATAATCGCTTTCCAGACGACTCGAGCCTTTATTATACATTTGTATCTCCTTTCTCTTTCGTATAGTGTATACAATCGTTCGAACTCGTCTCCCCAAACATTGTCCAATCCAGGACATTCATCCGGACACATCAACGACCACTCTCCGTCATCCTTCACTCGTTTCATAAAGAGATCAGGAATCCAAAGAGCTATAAATAGATCTCTTGTTCTATGTTCATCGTTTCCTGTATTCTTTTTAAGATCGAGGAACGCCATAATATCAGAATGCCACGGTTCCAAGTATATGGCCATAACTCCAGGCCGTTTGTTTCCTCCCTGATCTATGTATCTAGCGGTGTTATTATAAACTCTCAACATTGGAATAATACCGTTTGATATACCATTGGTACCGGAGATATAGCTTCCACTGGCACGAATATTACTAATTGATAGACCTATTCCCCCTGCCATTTTAGAGATTAATGCGCATCGTTTTAACGTGTCATAGATGCCTTCTATGCTATCATCGATCATGTTAAGTAGAAAACAGCTAGACATTTGGTGACGAGTAGTTCCCGCATTAAATAAGGTAGGAGAAGCGTGCGTAAACCATTTTTCAGAAAGTAGATTGTACGTCTCAATAGCTGAGTCTATATCCCATTGATGAATTCCTACTGCGACACGCATTAACATGTGCTGAGGTCTTTCAACAATTTTGTTGTTTATTTTCAACAAGTAGGATTTTTCCAAAGTTTTAAAACCAAAATAGTTGTATGAAAAGTCTCGTTCGTAAATAATAACCGAATTGAGCTTATCCTTATATTTGTTAACTATATCCATGGTAATACTTGAAATAATCGGAGAATGTTTCCCATTTTTAGGATTAACATAGTTGAATAAATCCTCCATCACTTCACTAAATAGTTTTTTTGTTTCCTTGTGTAGATTTGATATGGCTATTCTGGCGGCTAGAATGGCATAATCCGGATGTTGTGTAGTACAAGTGGCTGCTATTTCGGCTGCCAGAGTGTCCAATTCTACCGTTGTTACTCCATTATATATTCCTTGAATAACCTTCATAGCTATTTTAATAGGATCTATATGATCAGTGTTTAAGCCATAGCACAATTTTCTAATACGAGACGTGATTTTATCAAACATGACATTTTCCTTGTATCCATTTCGTTTAATGACAAACATTTTTGTTGGTGTAATAAAAAAAATTATTTAATTTTTCATTAATAGGGATTTGACGTATGTAGCGTACAAAATTATCGTTCCTGGTATATAGATAAAGAGTCCTATATATTTGAAAATCGTTACGGTTCGATTAAACTTTAATGATTGCATTGTGAATATATCATTAGGATTTAACTCCTTGACTATCATGGCGGTGCCAGAAATTACCATCAAAAGCATTAATACAGTTATGCAGATCGCAGTTAGAACGGTTATAGCATCCACCATTTATATCTAAAAATTAGATCAAAGAATATGTGACAACGTCCTAGTTGTATACTGAGAATTGACGAAACAATGTTTCTTACATATTTTTTTCTTATTAGTAACCGACTTAATAGTAGGAACTGGAAAACTAGACTTGATTATTCTATAAGTATAGATACCCTTCCAAATAATGTTCTCTTTGATAAAAGTTCCAGAAAATGTAGAATTTTTTAAAAAGTTATCTTTTGCTATTACTAATATCGTGGTTAGACGCTTATTATTAATATGAGTGATGAAATCCACACCGCTTCTAGATATCGCTTTTATTTCCACATTAGATGGTAAATCCAATAGTGAAACTATCTTTTTAGGAATGTATGGACTCGCGTTTAGAGGAGTGAACGTCTTCGGAGTAGTAAAGGATGATTCGTCAAATGAATAAACAATTTCACAAATGGATGTTAATGTATTAGTAGGAAATTTTTTGACGCTAGTGGAATTGAAGATTCTAATGGATGATGTTCTACCTATTTCATCCGATAACATGTTAATTTCCAATACCAACGGTTTTAATATTTCGATGATATACGGTAGTCTCTCTTTCGGACTTATATAGCTTATTCCACAATACGAGTCATTATATACTCCAAAAAACAAAATAACTAGTATAAAATCTGTATCGAATGGGAAAAACGAAATTATCGATATAGGTATAGAATCCGGAACATTGAACGTATTAATACTTAATTCTTTTTCAGTGGTAAGAACCGATAGGTTATTGACATTGTATGGTTTTAAATATTCTATAACTTGAGACTTGATAGATATTAATGACGAATTGAAAATTATTTTTATCACCACGTGTGTTTCAGGATCATCGTCGACGCCAGTTAACCAACCGAATGGAGTAAAATAAATATCATTAATATATGCTCTAGATATTAGTATTTTTATTAATCCTTTGATTATCATCTTCTCGTACGCGAATGATTCCATGATCAAGAGTGATTTGAGAACATCCTCCGGAGTATTAATGGGTTTAGTAAACAGTCCATCGTTGCAATAATAAAAGTTGTCCAAGTTAAAGGATATTATGCATTCGTTTAAAGATATCACCTCATCTAACGGAGACAATTTTTTGGTAGGTTTTAGAGACTTTGAAGCTACTTGTTTAACAAAGTTATTCATCGTCGTCTACTATTCTATTTAATTTTGTAGTTAATTTATCACATATCACATTAATTGACTTTTTGGTCCACTTTTCCATACGTTTATATTCTTTTAATCCTGCGTTATCCGTTTCCGTTATATACAGGGATAGATCTTGCAAGTTAAATAGAATGCTCTTAAATAATGTCATTTTTTTATCCGCTAAAAATTTAAAGAATGTATAAACTTTTTTCAAAGATTTAAAACTTTTAGGTGGAGTTCTGGTACACAATATCATAAACAAACTAATAAACATCCCACATTCAGATTCCAACAATTGATTAACTTCCACATTAATACAGCCTATTTTCGCTCCAAATGTACATTCGAAAAATCTGAATAAAACATCAATATCGCAATTTGTATTATCCAATACAGAATGTCTGTGATTCGTGTTAAAACCATCGGAAAAAGAATAGAAATAAAAATTATTATAATGGTGGAATTCAGTTGGAATATTGCCTCCGGAGTCATAAAAGGATACTAAACATTGTTTTTTATCGTAAATTACACATTTCCAATGAGACAAATAACAAAATCCAAACATTACAAATCTAGAGGTAGAACTTTTAATTTTGTCTTTAAGTATATACGATAAGATATGTTTATTCATAAACGCGTCAAATTTTTCATGAATAGCTAAGGAGTTTAAGAATCTCATGTCAAATTGTCCTATATAATCCACTTCGGATCCATAAGCAAACTGAGAGACTAAGTTCTTAATACTTCGATTGCTCATCCAGGCTCCTCTCTCAGGCTCTATTTTCATCTTGACGACCTTTGGATTTTCACCAGTATGTATTCCTTTACGTGATAAATCATCGATTTTCAAATCCATTTGTGAGAAGTCTATCGCCTTAGATACTTTTTCCCGTAGTTGAGGTTTAAAGAAATACGCTAACGGTATACTAGTAGGTAACTCAAAGACATCATATATAGAATGGTAACGCGTCGTTAACTCGTCGGTTAACTCTTTCTTTTGATCGAGTTCATCGCTACTATTGGGTCTGCTCAGGTGCCCCGACTCTACTAGTTCCAACATCATACCGATAGGAATACAAGACACTTTGCCAGCGGTTGTAGATTTATCATATTTCTCCACCACATATCCGTTACAATTTGTTAAGAATTTAGATACATCTATATTGCTACATAATCCAGCTAGTGAATATATATGACATAATAAATTGGTAAATCCTAGTTCTGGTATTTTACTAATTACTAAATCTGTATATCTTTCCATTTATCATGGAAAAGAATTTACCAGATATCTTCTTTTTTCCAAACTGCGTTAATGTATTCTCTTACAAATATTCACAAGATGAATTCAGTAATATGAGTAAAACGGAACGTGATAATTTCTCATTGGCTGTGTTTCCAGTGATAAAACATAGATGGCATAACGCACACGTTGTAAAACATAAAGGAATATACAAAGTTAGTACAGAAGCACGTGGAAAAAAAGTATCTCCTCCATCACTAGGAAAACCCGCACATATAAACCTAATGTCGAAGCAATATATATATAGTGAGTATGCAATAAGCTTTGAATGTTATAGTTTTCTAAAATGTATAACAAATACAGAAATCAATTCGTTCGATGAGTATATATTAAGAGGACTATTAGAAGCTGGTAATAGTTTACAGATATTTTCCAATTCCGTAGGTAAACGAATAGATACTATAGGTGTACTAGGGAATAAGTATCCATTTAGCAAAATTCCATTGGCCTCATTAACTCCTAAAGCACAACGAGAGATATTTTTAGCGTGGATTTCTCATAGACCTGTAGTTTTAACTGGAGGAACCGGAGTGGGTAAGACGTCACAGGTACCCAAGTTATTGCTTTGGTTTAATTATTTATTTGGTGGATTCTCTTCTCTAGATAAAATCACTGACTTTCACGAAAGACCAGTCATTCTATCTCTTCCTAGGATAGCTTTAGTTAGATTGCATAGCAATACCATTTTAAAATCATTGGGATTTAAGGTACTAGATGGATCTCCTATCTCTTTACGGTACGGATCTATACCGGAAGAATTAATAAACAAACAACCAAAAAAATATGGAATTGTATTTTCTACCCATAAGTTATCTCTAACAAAACTATTTAGTTATGGCACTATTATTATAGACGAAGTTCATGAGCATGATCAAATAGGAGATATTATTATAGCAGTAGCGAGAAAACATCATACGAAAATAGATTCTATGTTTTTAATGACTGCCACGTTAGAGGATGACAGGGAACGTCTAAAAATATTTTTACCTAATCCCGCATTTATACATATTCCTGGAGATACACTGTTTAAAATTAGCGAGGTATTTATTCATAATAAGATAAATCCATCTTCCAGAATGGCATATATAGAAGAAGAAAAGAGAAATTTAGTTACTGCTATACAGATGTATACTCCTCCTGATGGATCATCCGGTATAGTCTTTGTGGCATCCGTTGCACAGTGTCACGAATATAAATCATATTTAGAAAAAAGATTACCGTATGATATGTATATTATTCATGGTAAGGTCTTAGATATAGACGAAATATTAGAAAAAGTGTATTCATCACCTAATGTATCGATAATTATTTCTACTCCTTATTTGGAATCCAGCGTTACTATACGCAATGTTACACACATTTATGATATGGGTAGAGTTTTTGTCCCCGCTCCTTTTGGAGGATCACAACAATTTATTTCTAAATCTATGAGAGATCAACGAAAAGGAAGAGTAGGAAGAGTTAATCCTGGAACATACGTATATTTCTATGATCTGTCTTATATGAAATCTATACAGCGAATAGATTCAGAATTTCTACATAATTATATATTGTACGCTAATAAGTTTAATCTAACACTCCCCGAAGATTTGTTTATAATCCCTACAAATTTGGATATTCTATGGCGTACAAAGGAATATATAGACTCGTTCGATATTAGTACAGAAACATGGAATAAATTATTATCCAATTATTATATGAAGATGATAGAGTATGCTAAACTTTATGTACTAAGTCCTATTCTCGCTGAGGAGTTGGATAACTTTGAGAGGACGGGAGAATTAACTAGTATTGTACAAGAAGCCATTTTATCTCTAAATTTACAAATTAAGATTTTAAAATTTAAACATAAAGATGATGATACGTATATACACTTTTGTAGAATATTATTCGGTGTCTATAACGGAACAAACGCTACTATATATTATCATAGACCTCTAACGGGATATATGAATATGATTTCAGATACTATATTTGTTCCTGTAGATAATAACTAAAAATCAAAATCTAATGACCACATCTTTTTTTAGAGATGAAAAATTTTCCACATCTCCTTTTGTAGACACGACTAAACATTTTGCAGAAAAAAGTTTATTATTATTTAGATAATCGTATACTTCATCAGTGTAGATAGTAAATGTGAACAGATAAAAGGTATTCTTGCTCAATAGATTGGTAAATTCCATAGAATATATTAATCCTTTCTTCTTGAGATCCCACATCATTTCAACCAAAGACGTTTTATCCAATGATTTACCTCGTACTATACCACATACAAAACTAGATTTTGCAGTGATGTCGTACCTGGTATTCCTACCAAACAAAATTTTACTTTTAGTTCTTTTAGAAAATTCTAAGGTAGAATCTCTATTTGTCAATATGTCATCTATGGAATTACCACTAGCAAAAAATGATAGAAATATATATTGATACATCGCAGCTGGTTTTGATCTACTATACTTTAAAAACGAATCAGATTCCATAATTGCTTGTATATCATCAGCTGAAAAACTATGTTTTACACGTATTCCTTCGGCATTTCTTTTTAATGATATATCTTGTTTAGACAATGATAAAGTTATCATGTCCATGAGAGACGCGTCTCCGTATCGTATAAATATTTCATTAGATGTTAGACGCTTCATTAGGGGTATACTTCTATAAGGTTTCTTAATTAGTCCATCATTGGTTGCGTCAAGAACTACTATCTGATGTTGTTGGGTATCTCTAGTGTTACACATGGCCTTACTAAAGTTTGGGTAAATAACTATGATATCTCTATTAATTATAGATGTATATATTTCATTCGTCAAGGATATTAATATCGACTTACTATCGTCATTAATACGTGTAATGTAATCATATAAATCATGCGATAGCCAAGGAAAATTCAAATAGATGTTCATCATATAATCGTCGCTATAATTCATATTAATACTTTGACATTGACTAATTTGTAATATAGCCTCGCCACGAAGAAAGCTCTCGTATTCAGTTTCATCGATAAAGGATACCGTTAAATATAACTGGTTGCCGATAGTCTCATAGTCTATTAAGTGGTAAGTTTCGTATAAATACAGAATCCCTAAAATATTATCTAATGTGGGATTAATCCTTACCATAACTGTATAAAATGGAGCCGGAGTCATAACTATTTTACCGTTTGTACTTACTGGAATAGATGAAGGAATAATCTCCGGACATGATGGTAAAGACCCAAATGTCTGTTTGAAGAAATCCAATGTTCCAGGTCCTAATCTCTTGACAAAAATTACGATATTCGATCCCGATATCCTTTGCATTCTATTTACCAGCATATCACGAACTATATTAAGATTATCTATCATGTCTATTCTCCCACCGTTATATAAATCGCCTCCGCTAAGAAACGTTAGTATATCCATACAATGGAATACTTCATTTCTAAAATAGTATTCGTTTTCTAATTCTTTAATGTGAAATCGTATACTAGAAAGGGAAAAATTATCTTTGAGTTTTCCATTAGAAAAGAACCACGAAACTAATGTTCTGATTGCGTCTGACTCCGTCGCTGAATTAATAGATTTACACCAAAAACTCATATAACTTCTAGATGTAGAAGCATTCGCTAAAAAATTAGTAGAATCAAAGGATATAAGTAGATGTTCCAACAAGTGAGCAATTCCCAAGATTTCATCTATATCATTCTCGAATCCGAAATTAGAAATTCCCAAGTAGATATCCTTTTTCATCCGATCATTGATGAAAATACGAACTTTATTCGGTAAGACGATCATTTACTAAGGAGTAAAATAGGAAGTAACGTTCGTATATCGTTATCGTCGTATAAATTAAAGGTGTGTTTTTTGCCATTAAGAGACATTATAATTTTACCAATATTGGAATTATAATATAGGTGTATTTGAGCACTAGAAACGGTCGATGCATCGGTAAATATAGCTGTATCTAATGTTCTAGTCGGTATTTCTTCATTTCGCTGTCTAATGATAGCGTTTTCTCTATCTGTTTCCATTACAGCTGCCTGAAGTTTATTGGTCGGATAATATGTAAAATAATAAGAAATACATACGAATAACAAAAATAAAATAAGATATAATAAAGATGCCATTTAGAGATCTAATTTTGTTCAACTTGTCCAAATTCCTACTTACAGAAGATGAGGAATCGTTGGAGATAGTATCTTCCTTATGTAGAGGATTTGAAATATCTTACGATGACTTAATATCGTACTTTCCAGATAGGAAATACCATAAATATATTTCTAAGGTATTTGAACATGTAGATTTATCGGAGGAATTAAGTATGGAATTCCATGATACAACTCTGAGAGATTTAGTATATCTTAGATTGTACAAGTATTCCAAGTATATACGGCCGTGTTATAAATTAGGAGATAATCTAAAAGGTATAGTTGTTATAAAGGACAGAAATATATATATTAGAGAAGCAAATGATGACTTGATAGAATATCTCCTCAAGGAATACACTCCTCAGATTTATACATATTCTAATGAGCGAGTTCCCATAGCTGGTTCAAAATTAATTCTTTGTGGATTTTCTCAAGTTACATTTATGGCGTATACAACGTCGCATATAACAACAAATAAAAAGGTAGATGTTCTCGTTTCCAAAAAATGTATAGATGAACTAGTCGATCCAATAAATTATCAAATACTTCAAAATTTATTTGATAAAGGAAGCGGAACAATAAACAAAATACTCAGGAAGATATTTTATTCGGTAACAGGTGGCCAAACTCCATAGGTAGCTTTTTCTATTTCGGATTTTAGAATTTCCAAATTCACCAGCGATTTATCGGTTTTGGTGAAATCCAAGGATTTATTAATGTCCACAAATGCCATTTGTTTTGTCTGTGGATTGTATTTGAAAATGGAAACGATGTAGTTAGATAGATGCGCGGCGAAGTTTCCTATTAGGGTTCCGCGCTTCACGTCACCCAACATACTTGAATCACCATCCTTTAAAAAAAATGATAAGATATCAACATGGAGTATATCATACTCGGATTTTAATTCTTCTACTGCCTCACTGACATTTTCACAAATACTACAATACGGTTTACCGAAAATAATCAGTACGTTCTTCATTTATGGGTATCAAAAACTTAAAATCGTTACTGCTGGAAAATAAATCACTGACGATATTAGATGATAATTTATACAAAGTATACAATGGAATATTTGTGGATACAATGAGTATTTATATAGCCGTCGCCAATTGTGTCAGAAACTTAGAAGAGTTAACTACGGTATTCATAAAATACGTAAACGGATGGGTAAAAAAGGGAGGACATGTAACCCTTTTTATCGATAGAGGAAGTATAAAAATTAAACAAGACGTTAGAGACAAGAGACGTAAATATTCTAAATTAACCAAGGACAGAAAAATGTTAGAATTAGAAAAGTGTACATCCGAAATACAAAATGTTACCGGATTTATGGAAGAAGAAATAAAGGCAGAAATGCAATTAAAAATCGATAAACTCACATTTCAAATATATTTATCTGATTATGATAACATAAAAATATCATTGAATGAGATACTAACACATTTCAACAATAATGAGAATGTTACATTATTTTATTGTGATGAACGAGACGCAGAATTCGTTATGTGTCTAGAGGCTAAAACACAGTTCTCTACCACAGGAGAATGGCCGTTAATAATAAGTACCGATCAGGATACTATGCTATTCGCGTCTGCTGATAATCATCCTAAGATGATAAAAAACTTAACTCAACTGTTTAAATTTGTTCCCTCGGCAGAGGATAACTATTTAGCAAAATTAACTGCATTAGTGAATGGATGTGATTTCTTTCCTGGACTCTATGGGGCATCTATAACACCCAACAACTTAAACAAAATACAATTGTTTAGTGATTTTACAATCGATAATATAGTCACTAGTTTGGCAATTAAAAATTATTATAGAAAGACTAACTCTACCGTAGACGTGCGTAATATTGTTACGTTTATAAACGATTACGCTAATTTAGACGATGTCTACTCGTATATTCCTCCTTGTCAATGCACTGTTCAAGAATTTATATTCTCCGCATTAGATGAAAAATGGAATGAATTTAAATCATCTTATTTAGAGAGCGTGCCGTTACCCTGCCAATTAATGTACGCATTAGAACCACGTAAGGAGATTGATGTTTCAGAAGTTAAAACTTTATCATCTTATATAGATTTCGAAAATACTAAATCAGATATCGATGTTATAAAATCTATATCCTCGATTTTTGGATATTCTAACGAAAACTGTAACACCATAGTGTTCGGCATCTATAAGGATAATTTACTACTGAGTATAAATAATTCATTTTACTTTAACGATAGTCTGTTAATAACCAATACTAAAAGTGATAATATAATAAATATAGGTTACTAGATTAAAAAATGGTGTTCCAGCTCGTGTGTTCTACATGCGGCAAAGATATTTCTCACGAACGATATAAATTGATTATACGAAAAAAATCATTAAAGGATGTACTAGTCAGTGTAAAGAACGAATGTTGTAGGTTAAAATTATCTACACAAATAGAACCTCAACGTAACTTAACAGTGCAACCTCTATTGGATATAAACTAATGGATCCGGTTAATTTTATCAAGACATATGCGCCTAGAGGTTCTATTATTTTTATTAATTATGCCATGTCATTAACTAGTCATTTGAATCCATCGATAGAAAAACATGTGGGTATTTATTATGGTACGTTATTATCGGAACACTTGGTAGTTGAATCTACCTATAGAAAAGGAGTTAGAATAGTCCCATTGGATAGATTTTTTGAAGGATATCTTAGTGCAAAAGTATACATGTTAGAGAATATTCAAGTTATGAAAATAGCAGCTGATATGTCGTTAACTTTACTAGGTATTCCATATGGATTTGGTCATGATAGAATGTATTGTTTTAAATTGGTAGCTGAATGTTATAAAAATGCCGGTATTGATACATCGTCTAAACGAATATTAGGTAAAGATATTTTTCTGAGCCAAAACTTTACAGATGATAATAGATGGATAAAGATATATGATTCTAATAATTTAACATTTTGGCAAATTGATTACCTTAAAGGGTGAGTTAATATGCATAACTACTCCTCCGTTGTTTTTTCCCTCGTTCTTTTTCTTAACGTTGTTTGCCATCACTCTCATAATGTAAAGATATTCTAAAATGGTAAACTTTTGCATATCGGATGCAGAAATTGGTATAAATGTTGTAATTGTATTATTTCCCGTCAATGGACTAGTCACAGCTCCATCAGTTTTATATCCTTTAGAGTATTTCTCACTCGTGTCTAGCATTCTAGAGCATTCCATGATCTGTTTATCGTTGATATTGGCCGGAAAGATAGATTTTTTATTTTTTATTATATTACTATTGGCAATTGTAGATATAACTTCTGGTAAATATTTTTCTACCTTTTCAATCTCTTCTATTTTCAAGCCGGCTATATATTCTGCTATATTGTTACTAGTATCAATACCTTTTCTGGCTAAGAAGTCATATGTGGTATTCACTATATCAGTTTTAACTGGTAGTTCCATTAGCCTTTCCACTTCTGCAGAATAATCAGAAATTGGTTCTTTACCAGAAAATCCAGCTACTATAATAGGCTCACCGATGATCATTGGCAAAATCCTATATTGTACCAGATTAATGAGAGCATATTTCATTTCCAATAATTCTGCTAGTTCTTGAGACATTGATTTATTTGATGAATCTATTTGGTTCTCTAGATACTCTACCATTTCTGCCGCATACAATAACTTGTTAGATAAAATCAGGGTTATCAAAGTGTTTAGTGTGGCTAGAATAGTGGGCTTGCACGTATTAAAGAATGCTGTAGTATGAGTAAACCGTTTTAACGAATTATATAGTCTCCAGAAATCTGTGGCGTTGCATACATGAACTGAATGACATCGAAGATTGTCCAATATTTTTAATAGCTGCTCTTTGTCCATTATTTCTATATTTGACTCGCAACAATTGTAGATACCATTAATCACTGATTCCTTTTTCGATGCCGGACAATAGCACAATTGTTTAGCTTTGGACTCTATGTATTCAGAATTAATAGATATATCTCTCAATACAGATTGCACTATACATTTTGAAACTATGTCAAAAATTGTAGAACGACGCTGTTCTGTAGCCATTTAACTTTAAATAATTTACAAAAATTTAAAATGAGCATCCGTATAAAAATCGATAAATTGCGCCAAATTGTGGCATATTTTTCAGAGTTCAGCGAAGAAGTGTCTATAAATGTAGACTCGACGGATGAATTAATGTATATTTTTGCCGCCTTGGGCGGATCTGTAAACATTTGGGCCATTATACCTCTCAGTGCATCAGTGTTCTACCGCGGAGCCGAAAATATTGTGTTTAACCTTCCAGTGTCCAAGGTAAAATCGTGTTTGTGTAGTTTTCACAATGATGCTATCATAGATATAGAACCTGATCTGGAAAATAATCTAGTAAAACTTTCTAGTTATCATGTAGTAAGTGTCGATTGTAACAAGGAACTGATGCCTATTAGGACAGATACTACTATTTGTCTAAGTATAGATCAAAAGAAATCTTACGTATTTAATTTTCACAAGTATGAAGAAAAATGTTGTGGTAGAACCGTCATTCATCTAGAATGGTTGTTGGGCTTTATCAAGTGTATTAGTCAGCATCAGCATTTGGCTATTATGTTTAAAGATGACAATATTATTATGAAGACTCCTGGTAATACTGATGCGTTTTCCAGGGAATATTCTATGACTGAATGTTCTCAAGAACTACAAAAGTTTTCTTTCAAAATAGCTATCTCGTCTCTCAACAAACTACGAGGATTCAAAAAGAGAGTCAATGTTTTTGAAACTAGAATCGTAATGGATAATGACGATAACATTCTAGGAATGTTGTTTTCGGATAGAGTTCAATCCTTTAAGATTAACATCTTTATGGCGTTTTTAGACTAATACTTTCAATGAGATAAATATGGGTGGCGGAGTAAGTGTTGAGCTCCCTAAACGGGATCCACCTCCGGGAGTACCCACTGATGAGATGTTATTAAACGTGGATAAAATGCATGACGTGATAGCTCCCGCTAAGCTTTTAGAATATGTGCATATAGGACCACTAACAAAAGATAAAGAGGATAAAGTAAAGAAAAGATATCCAGAGTTTAGATTAGTCAACACAGGACCCGGTGGTCTTTCGGCATTATTAAGACAATCATATAATGGAACCGCACCCAATTGCTGTCGCACTTTTAATCGTACTCATTATTGGAAGAAGGATGGAAAGATATCAGATAAGTATGAAGAGGGTGCAGTATTAGAATCGTGTTGGCCCGACGTCCACGACACTGGAAAATGCGATGTTGATTTATTCGACTGGTGTCAGGGGGATACGTTCGATATGAACATATGCCATCAGTGGATCGGTTCAGCCTTTAATAGGAGTGATAGAACTGTAGAGGGTCGACAATCGTTAATAAATCTGTATAATAAGATGCAAAGATTATGTAGTAAAGATGCTAGTGTACCAATATGTGAATTATTTTTGCATCATTTACGCGCACACAATACAGAAGATAGTAAAGAGATGATCGATTATATTCTAAGACAACAGTCGGCGGACTTTAAACAGAAATATATGAGATGTAGTTATCCCACTAGAGATAAGTTAGAAGAGTCATTAAAATATGCGGAACCTCGAGAATGTTGGGATCCAGAGTGTTCGAATGCCAATGTTAATTTCTTACTAACACGTAATTATAATAATTTAGGACTTTGCAATATTGTACGATGTAATACGAGCGTGAATAACTTACAGATGGATAAAACTTCCTCATTAAGATTATCATGTGGATTAAGCAATAGTGATAGATTTTCTACTGTTCCCGTCAATAGAGCAAAAGTAGTTCAACATAATATTAAACATTCGTTCGACCTAAAATTGCATTTGATCAGTTTATTATCTCTCTTGGTAATATGGATACTAATTGTAGCTATTTAAATGGGTGCCGCAGCAAGCATACAGACGACTGTGAATACACTCAGTGAACGTATCTCGTCTAAATTAGAACAAGAAGCGAACGCTAGTGCTCAAACAAAATGTGATATAGAAATCGGAAATTTTTATATCCGACAAAACCATGGATGTAACATCACTGTTAAAAATATGTGCTCTGCGGACGCGGATGCTCAGTTGGATGCTGTGTTATCAGCCGCTACAGAAACATATAGTGGATTAACACCGGAACAAAAAGCATACGTACCAGCTATGTTTACTGCTGCGTTAAACATTCAGACGAGTGTAAACACTGTTGTTAGAGATTTTGAAAATTATGTGAAACAGACTTGTAATTCTAGCGCTGTTGTCGATAACAAATTAAAGATACAAAACGTAATTATAGATGAATGTTACGGAGCCCCAGGATCTCCAACAAATTTGGAATTTATTAATACAGGATCTAGCAAAGGAAATTGTGCCATTAAGGCGTTGATGCAATTGACTACTAAGGCCACTACTCAAATAGCACCTAGACAAGTTGCTGGTACAGGAGTTCAGTTTTATATGATTGTTATCGGTGTTATAATATTGGCAGCGTTGTTTATGTACTATGCCAAGCGTATGCTGTTCACATCCACCAATGATAAAATCAAACTTATTTTAGCCAATAAGGAAAACGTCCATTGGACTACTTACATGGACACATTCTTTAGAACTTCTCCGATGATTATTGCTACCACGGATATACAAAACTGAAAATATATTGATAATATTTTAATAGATTAACATGGAAGTTATCGCTGATCGTCTAGACGATATAGTGAAACAAAATATAGCGGATGAAAAATTTGTAGATTTTGTTATACACGGTCTAGAGCATCAATGTCCTGCTATACTTCGACCATTAATTAGGTTGTTTATTGATATACTATTATTTGTTATAGTAATTTATATTTTTACGGTACGTCTAGTAAGTAGAAATTATCAAATATTGTTGGTGTTGGTGGCGCTAGTCATCACATTAACTATTTTTTTATTACTTTATACTATAATAGTACTAGACTGACTTCTAACAAACATCTCACCTGCCATAAATAAATGCTTGATATTAAAGTCTTCTATTTCTAACACTATTCCATCTGTGGAAAATAATACTCTGACATTATCGCTAATTGATACATCGGTAAGTGATATGCCTATAAAGTAATAATCTTCTTTGGGCACATATACCAGTGTACCAGGTTCTAACAACCTATTTACTGGTGCTCCTGTAGCATACTTTTTTTTTACCTTGAGAATATCCATTGTTTGCTTGGTCAATAGTGATATGTGATTTTTTATCAACCACTCAAAAAAGTAATTGGAGTGTTCATATCCTCTACGGGCTATTGTCTCATGACCGTGTATGAAATTTAAGTAACACGACTGTGGTAGATTTGTTCTATAGAGCCGGTTGCCGCAAATAGATAGAACTACCAATATGTCTGTACAAATGTTAAACATTAATTGATTAACAGAAAAAACAATGTTCGTTCTGGGAATAGAAACCAGATTAAAACAAAATTCATTAGAATATATGCCACGTTTATACATGGAATATAAAATAACTACAGTTTGAAAAATAACAGTATCATTTAAACATTTAACTTGCGGGGTTAATCTCACAACTTTACTGTTTTTGAACTGTTCAAAATATAGCATAGATCCATGAGAAATACGTTTAGCCGCCTTTAATAGAGGAAATCCAACCGCCTTTCTGGATCTCACCAACGACGATAGTTCTGACCAGCAACTCATTTCTTCATCATCCACCTGTTTTAACATATAATAGGCAGGAGATAGATATCCATCATTGCAATATTCCTTCTCGTAGGCACACAATCTAATATTGATAAAATCTCCATTCTCTTCTCTGTATTTATTATCTTGTCTCGGTGGCTGATTAGGCTGTGGTCTATCGTTGTTGAATCTATTTTGGTCATTAAATCTTTCATTTCTTCCTGGTATATTTCTATCACCTCGTTTGGTTGGATTTTTGTCTATATTATCGTTTGTAACATCGGTACGGGTATTCATTTATCACAAAAAAAACTTCTCTAAATGAGTCTACTACTAGAAAACCTCATCGAAGAAGATACCATATTTTTTGCAGGAAGTATATCTGAGTATGATGATTTACAAATGGTTATTGCTGGTGCAAAATCCAAATTTCCAAGATCTATGCTTTCTATTTTTAATATAGTACCTAGAACGATGTCAAAATATGAGTTGGAGTTGATTCATAACGAGAATATCACAGGGGCAATGTTTACCACAATGTATAATATAAGAAACAATTTGGGTCTAGGCGATGATAAACTAACTATTGAAGCCATTGAAAACTATTTCTTGGATCCTAACAATGAGGTTATGCCTCTTATCATTAATAATACGGATATGACTACCGTCATTCCTAAAAAAAGTGGTAGGAGAAAGAATAAGAACATGGTTATCTTCCGTCAAGGATCATCACCTATCTTGTGTATTTTCGAAACTCGTAAAAAGATTAATATTTATAAAGAAAATATGGAATCCGTATCGACTGAGTATACACCTATCGGAGACAACAAGGCTTTGATATCTAAATATGCGGGAATTAATATCCTGAATGTGTATTCTCCTTCCACGTCCATGAGATTGAATGCCATTTACGGATTCACCAATAAAAATAAACTAGAGAAACTTAGTACTAATAAGGAACTAGAATCGTATAGTTCTAGCCCTCTTCAAGAACCCATTAGGTTAAATGATTTTCTGGGACTATTGGAATGTGTTAAAAAGAATATTCCTCTAACAGATATTCCGACAAAGGATTGATTACTATAAATGGAGAATGTTCCTAATGTATACTTTAATCCTGTGTTTATAGAGCCCACGTTTAAACATTCTTTATTAAGTGTTTATAAACACAGATTAATAGTTTTATTTGAAGTATTCGTTGTATTCATTCTAATATATGTATTTTTTAGATCTGAATTAAATATGTTCTTCATGCCTAAACGAAAAATACCCGATCCTATTGATAGATTACGACGTGCTAATCTAGCGTGTGAAGACGATAAATTAATGATCTATGGATTACCATGGATAACAACTCAAACATCTGCGTTATCAATAAATAGTAAACCGATAGTGTATAAAGATTGTGCAAAGCTTTTGCGATCAATAAATGGATCACAACCAGTATCTCTTAACGATGTTCTTCGCAGATGATGATTCATTTTTTAAGTATTTTGCTAGTCAAGATGATGAATCTTCATTATCTGATATATTGCAAATCACTCAATATCTAGACTTTCTGTTATTATTATTGATCCAATCAAAAAATAAATTAGAAGCTGTGGGTCATTGTTATGAATCTCTTTCAGAGGAATACAGACAATTGACAAAATTCACAGACTCTCAAGATTTTAAAAAACTGTTTAACAAGGTCCCTATTGTTACAGATGGAAGGGTCAAACTTAATAAAGGATATTTGTTCGACTTTGTGATTAGTTTGATGCGATTCAAAAAAGAATCAGCTCTAGCTACCACCGCAATAGATCCTGTTAGATACATAGATCCTCGTCGTGATATCGCATTTTCTAACGTGATGGATATATTAAAGTCGAATAAAGTTGAAAAATAATTAATTCTTTATTGTTATCATGAACGGCGGACATATTCAGTTGATAATCGGCCCCATGTTTTCAGGTAAAAGTACAGAATTAATTAGACGAGTTAGACGTTATCAAATAGCTCAATATAAATGTGTGACTATAAAATATTCTAACGATAATAGATACGGAACGGGACTATGGACACATGATAAGAATAATTTTGCAGCATTGGAAGTAACTAAACTATGTGATGTCTTGGAAGCAATTACAGATTTCTCCGTGATAGGTATAGATGAAGGACAGTTCTTTCCAGACATTGTTGAATTCTGTGAGCGTATGGCAAACGAAGGAAAAATAGTTATAGTAGCCGCGCTCGATGGGACATTTCAACGTAGACCGTTTAATAATATTTTGAATCTTATTCCATTATCTGAAATGGTGGTAAAACTAACTGCAGTGTGTATGAAATGCTTTAAGGAGGCTTCCTTTTCTAAACGATTAGGTACAGAAACCGAGATAGAAATAATAGGAGGTAATGATATGTATCAATCTGTGTGTAGAAAGTGTTACATCGACTCATAATATTATATTTTTTATCTAAAAAACTAAAAATAAACATTGATTAAATTTTAATATAATACTTAAAAATGGATGTTGTGTCGTTAGATAAACCGTTTATGTATTTTGAGGAAATTGATAATGAGTTAGATTACGAACCAGAAAGTGCAAATGAGGTCGCAAAAAAACTGCCGTATCAAGGACAGTTAAAACTATTACTAGGAGAATTATTTTTTCTTAGTAAGTTACAGCGACACGGTATATTAGATGGCGCCACCGTAGTGTATATAGGATCTGCTCCAGGTACACATATACGTTATTTGAGAGATCATTTCTATAATTTAGGAGTGATCATCAAATGGATGCTAATTGACGGCCGCCATCATGATCCTATTCTAAATGGATTGCGTGATGTGACTCTAGTGACTCGGTTTGTTGATGAGGAATATCTACGATCCATCAAAAAACAACTACATCCTTCTAAGATTATTTTAATTTCTGATGTGCGATCCAAACGAGGAGGAAATGAACCTAGTACTGCGGATTTACTAAGTAATTATGCTCTACAAAATGTCATGATTAGTATTTTAAACCCCGTGGCGTCTAGTCTTAAATGGAGATGCCCGTTTCCAGATCAATGGATCAAGGACTTTTATATCCCACACGGTAATAAAATGTTACAACCTTTTGCTCCTTCATATTCAGCTGAAATGAGATTATTAAGTATTTATACCGGTGAGAATATGAGACTGACTCGAGTTACCAAATCAGACGCTGTAAATTATGAAAAAAAGATGTATTACCTTAATAAGATAGTCCGCAACAAAGTAGTTATTAACTTTGATTATCCTAATCAGGAATATGACTATTTTCACATGTACTTTATGTTGAGGACCGTATACTGCAATAAAACATTTCCTACTACTAAAGCAAAGATACTATTTCTACAACAATCTATATTTCGTTTCTTAAATATTCCAACGACATCAACTGAAAAAGTTAGTCATGAACCAATACAACGTAAAATATCTAGCAAAGATTCTATGTCTAAAAACAGAAATAGCAAGAGATCCGTACGCGGTAATAAATAGAAACGTACTACTGAGATATACTACCGATATAGAGTATAATGATTTAGTTACTTTAATAACCGTTAGACATAAAATTGATTCTATGAAAACTGTGTTTCAGGTATTTAACGAATCATCCATAAATTATACTCCGGTTGATGATGATTATGGAGAACCAATCATTATAACATCGTATCTTCAAAAAGGTCATAACAAGTTTCCTGTAAATTTTCTATACATAGATGTGGTAATATCTGACTTATTTCCTAGCTTTGTTAGACTAGATACTACAGAAACTAATATAGTTAATAGTGTACTACAAACAGGCGATGGTAAAAAGACTCTTCGTCTTCCTAAAATGTTAGAGACGGAAATAGTTGTCAAGATTCTCTATCGTCCTAATATACCATTAAAAATTGTTAGATTTTTCCGCAATAACATGGTAACTGGAGTAGAGATAGCCGATAGATCTGTTATTTCAGTCGCTGATTAATCAATTAGTAGAGATGAGATAAGAACATTATAATAATCAATAATATATCTTATATCTGTTTAGAAAAATGCTAATATTAAAATAGCTAACGCTAGTAATCCAATCGGAAGCCATTTGATATCTATAATAGGGTATCTAATTTCCTGATTCAGATAGCGTACGGCTATATTCTCGGTAGCTACTCGTTTGGAATCACAGACATTATTTACATCTAATTTACTATCTGTAATGGAAACGTTTCCCAATGAAATGGTACAATCAGATACATTACATCTTGATATATTTTTTTTTAAAGAGGCTGGTAACAACGCATCGCTTCGTTTACATGGCTCGTACCAACAATAATAGGGTAATCTTGTATCTATTCCTATCCGTACTATACTTTTATCAGGATAAATACATTTACATCGTATATCGTCTTTGTTAGTATCACAGAATGCATAAATTTGTTCGTCCGTCATGATAAAAATTTAAAGTGTAAATATAACTATTATTTTTATAGTTATAATAAAAAGGGAAATTTGATTGTATACCTTCGGTTCTTTAAAAGAAACTGACTTGATAAAAATGGCTGTAATCTCTAAGGTTACGTATAGTCTATACGATCAAAAAGAGATTAATGCCACAGATATTATCATTAGTCATATTAAAAATGACGACGATATCGGTACCGTTAAAGATGGTAGACTAGGTGCTATGGATGGGGCATTATGTAAGACTTGTGGGAAAACGGAATTGGAATGTTTCGGTCACTGGGGTAAAGTAAGTATTTATAAAACTCATATAGTTAAGCCTGAATTTATTTCAGAAATTATTCGTTTACTGAATCATATATGTATTCATTGCGGATTATTGCGTTCACGAGAACCGTATTCCGACGATATTAACCTAAAAGAGTTATCGGTACACGCTCTTAGGAGATTAAAGGATAAAATATTATCCAAGAAAAAGTCATGTTGGAACAGCGAATGTATGCAACCGTATCAAAAAATTACTTTTTCAAAGAAAAAGGTTTGTTTCGTCAACAAGTTGGATGATATTAACGTTCCTAATTCTCTCATCTATCAAAAGTTAATTTCTATTCATGAAAAGTTTTGGCCATTATTAGAAATTCATCAATATCCAGCTAACTTATTTTATACAGACTACTTTCCCATCCCTCCGTTGATTATTAGACCGGCTATTAGTTTTTGGATAGATAGTATACCCAAAGAGACAAATGAATTAACTTACTTATTAGGTATGATCGTTAAGAATTGTAACTTGAATGCTGATGAACAGGTTATCCAGAAGGCGGTAATAGAATACGATGATATTAAAATTATTTCTAATAACACTACCAGTATCAATTTATCATATATCACATCCGGCAAAAATAATATGATTAGAAGTTATATCGTCGCTCGGCGAAAAGATCAAACCGCTAGATCCGTAATTGGTCCCAGTACATCTATCACCGTTAATGAGGTAGGAATGCCCACATATATTAGAAATACACTTACAGAAAAGATATTTGTTAATGCCTTTACAGTGGATAAAGTTAAACAACTATTAGCATCAAACCAAGTTAAATTTTACTTTAATAAACGATTAAACCAATTAACAAGAATACGTCAAGGAAAGTTTATCAAAAATAAAATACATTTATTGCCTGGTGATTGGGTAGAAGTAGCTGTTCAAGAATATACAAGTATTATTTTTGGAAGACAACCGTCTCTACATAGATACAACGTCATCGCTTCATCTATCAGAGCTACCGAAGGAGATACTATCAAAATATCTCCCGGAATTGCCAACTCTCAAAATGCTGATTTCGACGGAGATGAAGAATGGATGATATTGGAGCAAAATCCTAAAGCCGTAGTTGAACAAAGTATTCTTATGTATCCGACGACGTTACTCAAACACGATATTCATGGAGCCCCCGTTTATGGATCTATTCAAGATGAAATCGTAGCAGCGTATTCATTGTTTAGGATACAAGATCTTTGTTTAGATGAAGTATTGAACATCTTGGGGAAATATGGAAGAGAGTTCGATCCTAAAGGTAAATGTAAATTCAGCGGTAAAGATATCTATACTTACTTGATAGGTGAAAAGATTAATTATCCGGGTCTCTTAAAGGATGGTGAAATTATTGCAAACGACGTAGATAGTAATTTTGTTGTAGCTATGAGGCATCTGTCATTGGCTGGACTCTTATCCGATCATAAATCGAACGTGGAAGGTATCAACTTTATTATCAAGTCATCTTATGTTTTTAAGAGATATCTATCTATATACGGTTTTGGGGTGACATTCAAAGATCTGAGACCAAATTCGACGTTCACTAATAAATTGGAGGCTATCAACGTAGAAAAAATAGAACTTATCAAAGAAGCATACGCCAAATATCTCAAAGATGTAAGAGACGGGAAAATAGTTCCATTATCTAAAGCTTTAGAGGCGGACTACTTGGAATCCATGTTATCCAACTTGACAAATCTTAATATCAGAGAGATAGAAGAACATATGAGACAAACGCTGATAGATGATCCAGATAATAACCTCCTGAAAATGGCCAAAGCGGGTTATAAAGTAAATCCCACAGAACTAATGTATATTCTAGGTACTTATGGACAACAGAGGATAGATGGCGAACCAGCAGAGACTCGAGTATTGGGTAGAGTCTTACCTTACTATCTTCCAGACTCTAAGGATCCAGAAGGAAGAGGTTATATTCTTAATTCTTTAACAAAAGGATTAACGGGTTCTCAATATTACTTTTCGATGCTGGTTGCAAGATCTCAATCTACTGATATTGTCTGTGAAACATCACGTACCGGAACACTGGCTAGAAAAATCATTAAAAAGATGGAGGATATGGTGGTCGACGGATACGGACAAGTAGTTATAGGTAATACGCTCATCAAGTACGCAGCCAATTATACCAAAATTCTAGGCTCAGTATGTAAACCTGTAGATCTTATCTATCCAGATGAGTCCATGACTTGGTATTTGGAAATTAGTGCTTTGTGGAATAAAATAAAACAGGGATTCGTTTACTCTCAGAAACAGAAACTTGCAAAGAAGACATTGGCGCCGTTTAATTTCCTAGTATTCGTCAAACCCACCACTGAGGATAATGCTATTAAGGTTAAGGATCTGTACGATATGATTCATAACGTCATTGATGATGTGAGAGAGAAATACTTCTTTACGGTATCTAATATAGATTTTATGGAGTATATATTCTTGACGCATCTTAATCCTTCTAGAATTAGAATTACAAAAGAAACGGCTATTACTATCTTTGAAAAGTTCTATGAAAAACTCAATTATACTCTAGGTGGTGGAACTCCTATTGGAATTATTTCTGCACAGGTATTGTCTGAGAAGTTTACACAACAAGCCCTGTCCAGTTTTCACACTACTGAAAAGAGTGGTGCTGTAAAACAAAAACTTGGTTTCAACGAGTTTAATAACTTGACTAATTTGAGTAAGAATAAGACCGAAATTATCACTCTGGTATCCGATGATATCTCTAAACTTCAATCTGTTAAGATTAATTTCGAATTTGTATGTTTGGGAGAATTAAATCCAGACATCACTCTTCGAAAAGAAACAGATAGATATGTAGTAGACATAATAGTCAATAGATTATACATCAAGAGAGCAGAAATAACCGAATTAGTCGTCGAATATATGATTGAACGATTTATCTCCTTTAGCGTCATTGTAAAGGAATGGGGTATGGAGACATTCATTGAGGACGAGGATAATATTAGATTTACTATCTACCTAAATTTCGTTGAACCGGAGGAATTGAATCTTAGTAAGTTTATGATGGTTCTTCCAGGTGCCGCCAACAAGGGCAAGATTAGTAAATTCAAGATTCCTATCTCTGACTATACGGGATATAACGACTTCAATCAAACAAAAAAGCTCAATAAGATGACTGTAGAACTCATGAATCTAAAAGAATTGGGTTCTTTCGATTTGGAGAACGTCAACGTGTATCCTGGAGTATGGAATACATACGATATCTTCGGTATTGAGGCCGCTCGTGGATACTTGTGCGAAGCCATGTTAAACACCTATGGAGAAGGTTTCGATTATCTGTACCAGCCTTGTGATCTTCTCGCTAGTTTACTATGTGCTAGTTACGAACCAGAATCAGTTAATAAATTCAAGTTCGGTGCAGCTAGTACTCTTAAGAGAGCTACGTTCGGAGATAATAAAGCATTGTTAAACGCGGCTCTTCATAAAAAGTCAGAACCTATTAACGATAATAGTAGCTGCCACTTTTTTAGCAAGGTCCCTAATATAGGAACTGGATATTACAAATACTTTATCGACTTGGGTCTTCTCATGAGAATGGAAAGGAAACTATCTGATAAGATATCTTCTCAAAAGATCAAGGAGATAGAAGAAACAGAAGACTTTTAATTCTTATCAATAACATATTTTTCTATGATCTGTCTTTTAAACGATGGATTTTCCACAAATGCGCCTCTCAAGTCCCTCATAGAATGATACACGTATAAAAAATATAGCATAGGTGATGACTCCTTATTTTTAGACATTAGATATGCCAAAATCATAGCCCCGCTTCTATTTACTCCTGCAACACAATGAACCAACACGGGCTCGTTTCGTTGATCACATTTAGATAAGAAGGCGGTCACGTCGTCAAAATATTTACTAATATCAGTAGTTGTATCATCTACCAACGGTATATGAATAATATTAATATTAGAGTTAGGTAATGTATATTTATCCATCGTCAAATTTAAAACATATTTGAACTTAACTTCAGATGATGGTGCATCCATAGCATTTTTATAATTTCCCAAATACACATTATTTGTTACTCTTGTCATTATAGTGGGAGATTTGGCTCTGTGCATATCTCCAGTTGAACGTAGTAGTAAGTATTTATACAAACTTTTCTTATCCATTTATAACGTACAAATGGATAAAACTACTTTATCAGTAAACGCATGCAATTTAGAATACGTTAGAGAAAAGGCTATAGTAGGCGTACAAGCAGCCAAGACATCAACACTTATATTTTTTGTTATTATATTGGCAATTAGTGCGCTATTACTCTGGTTTCAGACGTCTGATAATCCAGTCTTTAATGAATTAACGAGATATATGCGAATTAAAAATACGGTTAACGATTGGAAATCATTAACGGATAGCAAAACAAAATTAGAAAGCGATAGAGGTAGACTTCTAGCCGCTGGTAAGGATGATATATTCGAATTCAAATGTGTGGATTTCGGCGCCTATTTTATAGCTATGCGATTGGATAAGAAAACATATCTGCCGCAAGCTATTAGGCGAGGTACTGGAGACGCGTGGATGGTTAAAAAGGCGGCAAAAGTCGATCCATCTGCTCAACAATTTTGTCAGTATTTGATAAAACACAAGTCTAATAATGTTATTACTTGTGGTAATGAGATGTTAAATGAATTAGGTTATAGCGGTTATTTTATGTCACCGCATTGGTGTTCCGATCTTAGTAATATGGAATAAGTGTTAGATAAATGCGGTAACAAATGTTCCTGTAAGGAACCATAACAGTTTAGATTTAACATTAAAGATGAGCATAAACATAATAAACAAAATTACAATCAAACCTATAACATTAATATCAAACAATCCAAAAAATGAAATCAATGGAGTAGTAAACGTGTACATAACTCCTGGATAACGTTTAGCAGCTACCGTTCCTATTCTAGACCAAAAATTTGGTTTCATGGTTTCGAAGCGGTGTTCTGCAACAAGACGAGGATCGTGTTCTACATATTTGGCAGAGTTATCCATTATTTGCCTGTTAATCTTCATTTCGTTTTCGATTCTGGCTATTTCAAAATAAAATCCCGATGATAGACCTCCAGACTTTATAATTTCATCTACGATGTTCAGCGCCGTAGTAACTCTAATAATATAGGCGGATAAGCTAACATCATACCCTCCTGTATATGTAAATATGGCATGATCTTTGTCTATTACAAGCTCGGTTTTAACTTTATTTCCTGTAATAATTTCTCTCATCTGTAGGATATCTATTTTCTTGTCATGTATTGCCTTCAAGACGGGACGAAGAAACGTAATATCCTCAATAACGTTATCGTTTTCTATAATAACTACATATTCTACATTTTTATTTTCTAGCTCGGTAAAAAATTTAGAATCCCATAGGGCTAAATGTCTAGCGATATTTCTTTTCGTTTCCTCTGTACACATAGTGTTACAAAACCCTGAAAAGAAGTGAGTATACTTGTCATCATCTCTAATATTTCCTCCAGTCCATTGTATAAACACATAATCCTTGTAATGATCTGGATCATCATTGACTATCACAACATCTCTTTTTTCTTGCATAACTTCATTGTCCTTCACATCATCGAACTTCTGATCATTAATATGCTCATGAACATTAGGAAATGTTTCTGATGGAGGTCTATCAATAACTGGCACAACAATAACAGGAGTTTTCACCGCCGCCATTTAGTTATTGAAATTAATCATATACAACTCTCTAATACGAGTTATATTTTCGTCTATCCATTGTTTCACATTGACATATTTCGACAAAAAGATATAAAATGCGTATTCCAATGCTTCTCTGTTTAATGAATTACTAAAATATACAAACACGTCACTGTCTGGTAATAAATGATATCTTAGAATATTGTAACAATTTATTTTGTATTGCACATGTTCGTGATCTATGAGTTCTTCTTCGAATGGCATAGGATCTCCGAATCTGAAAACGTATAAATAGGAGTTAGAATAATAATATTTGAGAGTATTGGTAATGTATAAACTCTTTAGCGGTATAATTAGTTTTTTTCTCTCGATTTCTATTTTTAGATGTGATGGAAAAATGACTAATTTTGTAGCATTAGTATCATGAACTCTAATCAAAATCTTAATATCTTCGTCACATGTTAGCTCTTTGAAGTTTTTAAGAGATGCATCAGTTGGTTTTACAGATGGAGTAGGTGCAACAATTTTTTGTTTAATGCATGCATGTATTGGAGCCATTGTCTTAACTATAATGGTGCTTGTATCGAAAAACTTTAATGCGGATAACGGAAGCTCTTCGCCGCGACTTTCTACGTCGTAATTGGGTTCTAATGCCGATCTCTGAATGGATACTAGTTTTCTAAGTTCTAATGTAATTCTCTGAAAATGTAAATCCAATTCCTCCGGCATTATAGATGTGTATACATCGGTAAATAAAACTATAGTATCCAACGATCCCTTCTCGCAAATTCTAGTCTTAACCAAGAAATCGTATATAACTACGGAGATGGCGTATTTAAGAGTGGATTCTTCTACCGTTTTGTTCTTGGATTTCATATAAGAAACTATAAAGTCCGCACTACTGTTAAGAATGATCACTAACGCAACTATATAGTTCAAATTAAGCATCTTGGAAACATAAAATAACTCTGTAGATGATACTTGACTTTCGAATAAGTTTGCAGACAAACGAAGAAAGAACAGACCTCTCTTAATTTCAGAAGAAAACTTTTTTTCGTATTCCTGACGTCTAGAGTTTATATCAATAAGAAAGTTAAGAATTAGTCGGTTAATGTTGTATTTCATTACCCAAGTTTGAGATTTCATAATATTGTCAAAAGACATGATAATATTAAAGATAAAGCGCTGACTATGAACGAAATAGCTATATGGTTCGCTCAAGAATATAGTCTTGTTAAACGTGGAAACGATAACTGTATTTTTAATCACGTCAGCGGCATCTAAATTAAATATAGGTATATTTATTCCACACACTCTACAATATGCCACACCATCTTCATAATAAATAAATTCGTTAGCAAAATTATTAATTTTAGTGAAATAGTTAGCGTCAACTTTCATAGCTTCCTTCAATCTAATTTGATGCTCACATGGCGCGAATTCTACTCTAACATCCCTTTTCCATGCCTCAGGTTCATCGATCTCTATAATATCTAGTTTCTTGCGTTTCACAAACACAGGCTCGTCTCTCGCGATGAGATCTGTATAGTAACTATGTAAATGATAACTAGATAGAAAGATGTAGCTATATAGATGACGATCCTTTAAGAGAGGTATAATAACTTTACCCCAATCAGATAGACTGTTGTTATGGTCTTCGGAAAAAGAATTTTTATAAATTTTTCCAGTATTTTCTAAATATACGTACTTGATATCTAAGAAATCCTTAATAATAATAGGAATGGATAATCCGTCTATTTTATAAAGAAATACATATCGCATATTATACTTTTTTTTGGAAATTGGAATACCGATGTGTCTACATAAATACGCAAAGTCTAAATATTTTTTAGAGAATCTTAGTTGGTCCAAATTCTTTTCCAAGTACGGTAATAGATTTTTCATATTGAACGGTATCTTCTTGATCTCTGGTTCTAATTCCGCATTAAATGATGAAACTAAGTCACTATTTTTATAACTAACGATTACATCACCTCTAACATCATCATTTACCAGGATACTGATCTTCTTTTGTCGTAAATACATGTCTAATGTGTTAAAAAAAAGATCATACAAGTTATACGTCATTTCATCTGTAGTATTCTTGTCATTGAAGGATAAACTCGTACTAATCTCTTCTTTAACAGTCTGTTCAAATTTATATCCTATATATGAAAAAATAGCAACCAGTGTTTGATCATCCGCGTCAATATTCTGTTCTATCGTAGTGTATAACAATCTTATATCTTCTTCTGTGATAGTCGATACGTTATAAAGGTTGATAACGAAAATATTTTTATTTCGTGAAATAAAGTCATTGTAGGATTTTGGACTTATATTCGTGTCTAGTAGATATGATTTTATTTTTGGAATGATCTCAATTAAAATAGTCTCTTTAGAGTCCATTTAAAGTTACAAACAACTAGGAAATTGGTTTATGATGTATAATTTTTTTAGTTTTTATAGATTCTTTATTCTATACTTAAAAAATGAAAATAAATACAAAGGTTCTTGAGGGTTGTGTTAATTGAAAGCGATAAATAATCATAAATTATTTCATTATCGCGATATCCGTTAAGTTTGTATCGTAATGGCGTGGTCAATTACGAATAAAGCGGATACTAGTAGTTTCACAAAGATGGCTGAAATCAGAGCTCATCTAAGAAATAGCGCTGAAAATAAAGATAAAAACGAGGATATTTTCCCGGAAGATGTAATAATTCCATCTACTAAGCCCAAAACCAAACGAACCACTACTCCTCGTAAACCAGCGGCTACTAAAAGATCAACCAAAAAGGATAAAGAAAAGGAGGAAGTGGAAGAAGTAGTTATAGAGGAATATCATCAAACAACTGAAGAAAATTCTCCACCTCCGTCATCATCTCCTGGAGTCGGCGACATTGTAGAAAGCGTGGCCGCTGTAGAGCTCGATGATAGCGACGGGGATGATGAACCTATGGTACAAGTTGAAGCTGGTAAAGTAAATCATAGTGCTAGAAGCGATCTCTCTGACCTAAAGGTGGCTACCGACAATATCGTTAAAGATCTTAAGAAAATTATTACTAGAATCTCTGCAGTATCGACTGTTCTAGAGGATGTTCAAGCAGCTGGTATCTCTAGACAATTTACTTCTATGACTAAAGCTATTACAACACTATCTGATCTAGTCACCGAGGGAAAATCTAAAGTTGTTCGTAAAAAAGTTAAAACTTGTAAGAAGTAAATGCGTGCACTTTTTTATAAAGATGGTAAACTGTTTACCGATAATAATTTTTTAAATCCTGTATCAGACGATAATCCAGCGTATGAGGTTTTGCAACATGTTAAAATTCCTACTCATTTAACAGATGTAGTAGTATATGAACAAACGTGGGAAGAGGCATTAACTAGATTAATTTTTGTGGGAAGTGATTCAAAAGGACGTAGACAATACTTTTACGGAAAAATGCATATACAGAATCGCAATGCTAAAAGAGATCGTATTTTTGTTAGAGTATATAACGTTATGAAACGAATTAATTGTTTTATAAACAAAAATATAAAGAAATCGTCCACAGATTCCAATTATCAGTTGGCGGTTTTTATGTTAATGGAAACTATGTTTTTTATTAGATTTGGTAAAATGAAATATCTTAAGGAGAATGAAACAGTAGGGTTATTAACACTAAAAAATAAACACATAGAAATAAGTCCCGATGAAATAGTTATCAAGTTTGTAGGAAAGGACAAAGTTTCACATGAATTTGTTGTTCATAAGTCTAATAGACTATATAAACCGCTATTGAAACTGACTGATGATTCTAGTCCCGAAGAATTTCTGTTCAACAAACTAAGTGAACGAAAGGTATATGAATGTATCAAACAGTTTGGTATTAGAATCAAGGATCTCCGAACGTATGGAGTCAATTATACGTTTTTATATAATTTTTGGACAAATGTAAAGTCCGTATCTCCTCTTCCATCACCAAAAAAGTTGATAGCATTAACTATCAAACAAACTGCTGAAGTGGTAGGTCATACTCCATCAATTTCAAAAAGAGCTTATATGGCAACGACTATTTTAGAAATGGTAAAGGATAAAAATTTTTTAGACGTAGTATCTAAAACTACGTTCGATGAATTCCTATCTATAGTCGTAGATCACGTTAAATCATCTACGGATGGATGATAATAGATCTTTACACAAATAATTACAAGACCGATAAATGGAAATGGATAAACGGATGAAATCTCTCGCTATGACAGCTTTCTTCGGAGAGCTAAACACGTTAGATATTATGGCATTGATAATGTCTATATTTAAACACCATCCAAACAATACCATTTTTTCAGTGGATAAGGATGGTCAATTTATGATTGATTTCGAATACGATAATTATAAGGCTTCTCAATATTTGGATCTGACCCTCACTCCGATATCTGGAAATGAATGCAAGACTCACGCATCTAGTATAGCCGAACAATTGGCGTGTGTGGATATTATTAAAGAGGATATTAGCGAATATATCAAAACTACTCCCCGTCTTAAACGATTTATAAAAAAATACCGCAATAGATCATATACTCGTATCAGTCGAGATACAGAAAAGCTTAAAATAGCTCTAGCTAAAGGCATAGATTACGAATATATAAAAGACGCTTGTTAATAAGTAAATGAAAAAAAACTAGTCGTTTATAATAAAACACGATATGGATGCCAACATAGTATCATCTTCTACTATTGCGACGTATATAGACGCTTTAGCGAAGAATGCTTCAGAATTAGAACAGAGGTCTACCGCATACGAAATAAATAATGAATTGGAACTAGTATTTATTAAACCGCCATTGATTACGTTGACAAATGTAGTAAATATCTCCACGATTCAGGAATCGTTTATTCGATTTACCGTTACTAATAAGGAAGGTATCAAAATTAGAACTAAGATTCCATTATCTAAGGTACATGGTCTAGATGTAAAAAATGTGCAGTTGGTAGATGCTATAGATAACATAGTTTGGGAAAAGAAATCATTAGTGACGGAAAATCGTCTTCACAAAGAATGCTTGTTGAGACTATCAACAGAGGAACGTCATATATTTTTGGATTACAAGAAATATGGATCCTCTATCCGACTAGAATTAGTCAATCTTATTCAAGCAAAAACAAAAAACTTTACGATAGACTTTAAGCTAAAATATTTTCTAGGATCTGGCGCTCAATCTAAAAGTTCTTTATTGCACGCTATTAATCATCCAAAGTCAAGGCCTAATACATCTCTGGAAATAGAATTTACACCTAGAGACAATGAAACAGTTCCATATGATGAACTAATAAAGGAATTGACGACTCTCTCGCGTCATATATTTATGGCTTCTCCAGAGAATGTAATTCTTTCTCCACCTATTAACGCACCTATAAAGACTTTTATGTTGCCTAAACAAGATATAGTAGGTCTGGATCTGGAAAATCTATATGCCGTAACTAAGACTGACGGCATTCCTATAACTATCAGAGTTACATCAAAAGGGTTGTATTGTTATTTTACACATCTTGGTTATATTATTAGATATCCAGTTAAGAGAACAATAGATTCCGAAGTAGTAGTCTTTGGTGAGGCAGTTAAGGATAAGAACTGGACCGTATATCTCATTAAGCTAATAGAGCCCGTAAATGCAATCAGTGATAGACTAGAAGAAAGTAAGTATGTTGAATCTAAACTAGTGGATATTTGTGATCGGATAGTATTCAAGTCAAAGAAATACGAAGGTCCGTTTACTACAACTAGTGAAGTCGTCGATATGTTATCTACATATTTACCAAAGCAACCAGAAGGTGTTATTCTGTTCTATTCAAAGGGACCTAAATCTAACATTGATTTTAAAATCAAAAAGGAGAATACTATAGACCAAACTGCAAATGTAGTATTTAGGTACATGTCCAGTGAACCAATTATCTTTGGAGAGTCGTCTATCTTTATAGAGTATAAGAAATTTACCAACGATAAAGGCTTTCCTAAAGAATATGGTTCTGGTAAGATTGTGTTATATAACGGCGTTAATTATCTAAATAATATCTATTGTTTGGAATATATTAATACACATAATGAAGTGGGTATTAAGTCCGTTGTTGTACCTATTAAGTTTATAGCAGAATTCTTAGTCAATGGAGAAATACTTAAACCTAGAATCGATAAAACCATGAAATATATTAACTCAGAAGACTATTATGGAAATCAACATAATATCATAGTCGAACATTTAAGAGATCAAAGCATCAAAATAGGAGATGTCTTTAACGAGGATAAACTATCGGATGTTGGACATCAATACGCTGCCAACAACGATAAATTTAGATTAAATCCAGAAGTTAGTTATTTTACTAATAAACGAACTAGAGGGCCGTTGGGAATTTTATCAAACTACGTCAAGACTCTTCTTATTTCTATGTATTGTTCCAAAACATTTTTAGACGATTCCAACAAACGAAAGGTATTAGCGATTGATTTTGGAAACGGTGCTGACCTGGAAAAATACTTTTATGGAGAGATTGCGTTATTGGTAGCGACGGATCCGGATGCTGATGCTATAGCTAGAGGAAATGAAAGATACAACAAATTAAATTCTGGAATTAAAACCAAGTACTACAAATTTGACTACATTCAGGAAACTATTCGATCCGATACATTTGTCTCTAGTGTCAGAGAAGTATTCTATTTTGGAAAGTTTAATATCATTGACTGGCAGTTCGCTATTCATTATTCTTTTCATCCAAGACATTATGCTACAGTCATGAATAACTTATCCGAACTAACTGCTTCTGGAGGCAAGGTATTAATTACTACCATGGATGGAGACAAATTATCAAAATTAACCGATAAAAAGACTTTTATAATTCATAAGAATCTACCTAGTAGCGAAAACTATATGTCTGTAGAAAAAATAGCTGATGATAGAATAGTGGTATATAATCCATCAACAATGTCTACTCCAATGACTGAATACATTATCAAAAAGAACGATATAGTCAGAGTGTTTAACGAATACGGATTTGTTCTTGTAGATAATGTTGATTTCGCTACAATTATAGAACGAAGTAAAAAGTTTATTAATGGCGCATCTACAATGGAAGATAGACCGTCTACAAGAAACTTTTTCGAACTAAATAGAGGAGCCATTAAATGTGAAGGTTTAGATGTCGAAGACTTACTTAGTTACTATGTTGTTTATGTCTTTTCTAAGCGGTAAATAATAATATGGTATGGGTTCTGATATCCCCGTTCTAAATGCATTAAATAATTCCAATAGAGCGATTTTTGTTCCTATAGGACCTTCCAACTGTGGATACTCTGTATTATTAATAGATATATTAATACTTTTGTAGGGTAACAGAGGTTCTACGTCTTCTAAAAATAAAAGTTTTATAACATCTGGCCTGTTCATAAATAAAAACTTGGCGATTCTATATATACTCTTATTATCAAATCTAGCCATTGTCTTATAGATGTGAGCTACTGTAGGTGTACCATTTGATTTTCTTTCTAATACTATATATTTCTCTCGAAGAAGTTCTTGCAGATCATCTGGGAATAAAATACTACTGTTGAGTAAATCAGTTATTTTTTTTATATCGATATTGATGGACATTTTTATAGTTAAGGATAATAAGTATCCCAAAGTAGATAACGACGATAACGAAGTATTTATACTTTTAGGAAATCACAATGACTTTATCAGATCAAAATTAACAAAATTAAAGGAGCATGTATTTTTTTCTGAATATATTGTGACTCCAGATACATATGGATCTTTATGCGTCGAATTAAATGGGTCTAGTTTTCAGCACGGTGGTAGATATATAGAGGTGGAGGAATTTATAGATGCTGGAAGACAAGTTAGATGGTGTTCTACATCCAATCATATATCTGAAGATATACACACTGATAAATTTGTCATTTATGATATTTATACGTTTGATTCGTTCAAGAATAAACGATTGGTATTTGTACAGGTGCCTCCATCATTAGGAGATGATAGCTATTTAACTAATCCGTTATTGTCTCCGTATTATCGTAATTCAGTAGCCAGACAAATGGTCAATGATATGATTTTTAATCAAGATTCATTTTTAAAATATTTATTAGAACATCTGATTAGAAGCCACTATAGAGTTTCTAAACATATAACAATAGTTAGATACAAGGATACCGAAGAATTAAATCTAACAAGAATATGTTATAATAGAGATAAGTTTAAGGCGTTTGTATTCGCTTGGTTTAACGGCGTTTCGGAAAATGAAAAGGTACTAGATACGTATAAAAAGGTATCTGATTTGATATAATGAATTCAGTGACTATATCACACGCACCATATACTATTACTTATCACGATGATTGGGAACCAGTAATGAGTCAATTGGTAGAGTTTTATAACGAAGTAGCCAGTTGGTTGCTACGCGACGAGACGTCGCCTATTCCTGATAAGTTCTTTATACAATTGAAACAGCCGCTTAGAAATAAACGAGTATGTGTGTGTGGTATAGATCCGTATCCAAAAGATGGAACTGGTGTACCGTTCGAATCACCAAATTTTACAAAAAAATCAATTAAGGAGATAGCTTCATCTATATCTAGATTAACCGGAGTAATTGATTATAAAGGTTATAACCTTAATATAATAGACGGGGTTATACCCTGGAATTATTACTTAAGTTGTAAATTAGGAGAAACAAAAAGTCACGCGATTTACTGGGATAAGATTTCCAAGTTACTGCTACAGCATATAACTAAACACGTTAGTGTTCTTTATTGTTTGGGTAAAACAGATTTCTCGAATATACGGGCAAAGTTAGAATCCCCGGTAACTACCATAGTGGGATATCATCCAGCGGCCAGAGACCACCAATTCGAGAAAGATCGATCATTTGAAATTATCAACGTTTTACTGGAATTAGACAACAAGACACCTATAAATTGGGCTCAAGGGTTTATTTATTAATGCTTTAGTGAAATTTTAACTTGTGTTCTAAATGGATGCGGCTATTAGAGGTAATGATGTTATCTTTGTTCTTAAGACTATAGGTGTCCCGTCAGCATGCAGACAAAATGAAGATCCAAGATTCGTAGAAGCATTTAAATGCGACGAGTTAGAAAGATATATTGATAATAATCCAGAATGTACACTATTCGAAAGTCTTAGGGATGAGGAAGCATACTCTATAGTCAGAATTTTCATGGATGTAGATTTAGACGCGTGTCTAGACGAAATAGATTATTTAACGGCTATTCAAGATTTTATTATCGAGGTGTCAAACTGTGTAGCTAGATTCGCATTTACAGAATGCGGTGCCATTCATGAAAATGTAATAAAATCCATGAGATCTAATTTTTCATTGACTAAGTCTACAAATAGAGATAAAACAAGTTTTCATATTATCTTTTTAGACACGTATACCACTATGGATACATTGATAGCTATGAAACGAACACTATTAGAATTAAGTAGATCATCTGAAAATCCACTAACAAGATCGATAGACACTGCCGTATATAGGAGAAAAACAACTCTTCGGGTTGTAGGTACTAGGAAAAATCCAAATTGCGACACTATTCATGTAATGCAACCACCTCACGATAATATAGAAGATTACCTATTCACTTACGTGGATATGAACAACAATAGTTATTACTTTTCTCTACAACGACGATTGGAGGATTTAGTTCCTGATAAGTTATGGGAACCAGGGTTTATTTCGTTCGAAGACGCTATAAAAAGAGTTTCAAAAATATTCATTAATTCTATAATAAACTTTAATGATCTCGATGAAAATAATTTTACAACGGTACCACTGGTCATAGATTATGTAACACCTTGTGCATTATGTAAAAAACGATCGCATAAACATCCGCATCAACTATCGTTGGAAAATGGTGCTATTAGAATTTACAAAACTGGTAATCCACATAGTTGTAAAGTTAAAATTGTTCCGTTGGATGGTAATAAACTGTTTAATATTGCACAAAGAATTTTAGACACTAACTCTGTTTTATTAACCGAACGAGGAGACCATATAGTTTGGATTAATAATTCATGGAAATTTAACAGCGAAGAACCCTTGATAACAAAACTAATTCTATCAATAAGACATCAACTACCTAAGGAATATTCAAGCGAATTACTCTGTCCGAGGAAACGAAAGACTGTAGAAGCTAACATACGAGACATGTTAGTAGATTCAGTAGAGACCGATACCTATCCGGATAAACTTCCGTTTAAAAATGGTGTATTGGACCTGGTAGACGGAATGTTTTACTCTGGAGATGATGCTAAAAAATATACGTGTACTGTATCGACCGGATTTAAATTTGACGATACAAAATTCGTCGAAGACAGTCCAGAAATGGAAGAGTTAATGAATATCATTAACGATATCCAACCATTAACGGATGAAAATAAGAAAAATAGAGAGCTGTATGAAAAAACATTATCTAGTTGTTTATGTGGTGCTACCAAAGGATGTTTAACATTCTTTTTTGGAGAAACCGCAACTGGGAAGTCGACAACCAAACGTTTGTTAAAGTCTGCTATCGGTGACCTGTTTGTCGAGACGGGTCAAACAATTTTAACAGATGTATTGGATAAAGGACCTAATCCATTTATCGCTAATATGCATTTAAAAAGATCTGTATTCTGTAGCGAACTACCTGATTTTGCATGTAGTGGATCAAAGAAAATTAGATCTGATAATATTAAAAAGTTGACAGAACCTTGTGTCATTGGAAGACCGTGTTTCTCCAATAAAATTAATAATAGAAACCATGCGACAATCATTATCGATACTAATTACAAACCTGTCTTTGATAGGATAGATAACGCATTAATGAGAAGAATTGCCGTCGTGCGATTCAGAACACACTTTTCTCAACCTTCTGGTAGAGAGGCTGCTGAAAATAATGACGCGTACGATAAAGTCAAACTATTAGACGAGGGATTAGATGGTAAAATACAGAATAATAGATATAGATTCGCATTTCTATACTTGTTGGTTAAATGGTACAAAAAATATCATATTCCTATTATGAAACTATATCCTACACCGGAAGAGATTCCGGACTTTGCATTCTATCTCAAAATAGGTACTCTGTTGGTATCTAGCTCTGTAAAGCATATTCCATTAATGACGGACCTCTCCAAAAAGGGATATATATTGTACGATAATGTGGTTACTCTTCCGTTGACTACTTTCCAACAGAAAATATCCAAGTATTTTAATTCTAGACTATTTGGACACGATATAGAGAGCTTCATCAATAGACATAAGAAATTTGCCAATGTTAGTGATGAATATCTGCAATATATATTCATAGAGGATATTTCATCTCCGTAAATATATGCCATATATTTATAGAATATATCACATATCTAAATGAATACCGGAATCATAGATTTATTTGATAATCATGTTGATAGTATACCAACTATATTACCTCATCAGTTAGCTACTTTAGATTATCTAGTTAGAACTATCATAGATGAGAACAGAAGCGTGTTATTGTTCCATATTATGGGATCGGGTAAAACAATAATCGCTTTGTTGTTCGCCTTGGTAGCTTCCAGATTTAAAAAGGTTTACATTTTAGTACCGAACATCAACATCTTAAAAATTTTCAATTATAATATGGGTGTAGCTATGAACTTGTTTAATGACGAATTCATAGCTGAGAATATCTTTATTCATTCCACAACAAGTTTTTATTCTCTTAATTATAACGATAACGTCATTAATTATAACGGATTAAGTCGCTACAATAACTCTATTTTTATCGTTGATGAGGCACATAATATTTTTGGGAATAATACTGGAGAACTTATGACCGTGATAAAAAATAAAAACAAGATTCCTTTTCTACTATTGTCTGGATCTCCCATTACTAACACACCTAATACGCTGGGTCATATTATAGATTTAATGTCCGAAGAGACGATAGATTTTGGTGAGATTATTAGTCGTGGTAAGAAAGTAATTCAGACACTTCTTAACGAACGCGGAGTGAATGTACTCAAGGATTTGCTTAAAGGAAGAATATCATATTACGAAATGCCGGACAAAGATCTACCAACAATAAGATATCACGGACGTAAATTTCTAGATACTCGAGTAGTATATTGTCACATGTCTAAACTTCAAGAGAAAGATTATATGATTACTAGACGGCAGCTATGTTATCATGAAATGTTTGATAAAAATATGTATAACGTGTCAATGGCAGTATTGGGACAACTTAATCTGATGAATAATTTAGATACGTTATTTCAGGAACAGGATAAGGAATTGTACCCAAATCTGAAAATAAATAATGGAGTGTTATACGGTGAAGAATTGGTAACGTTAAACATTAGTTCCAAATTTAAGTACTTTATCAATCGGATACAGACACTCAAGGGAAAACACTTTATATACTTCTCTAATTCTACATATGGTGGATTGGTAATTAAATATATCATGCTCAGTAATGGATATTCTGAATATAATGGTTCTCAGGGAACTAATCCACATATGATAAACGGCAAACCAAAAACATTTGCTATCGTTACTAGTAAAATGAAATCGTCTTTAGAGGATCTATTAGATGTGTATAATTCTCCTGAAAACGATGATGGCAATCAATTGATGTTTTTGTTTTCGTCAAACATTATGTCTGAATCCTATACTCTGAAAGAGGTAAGGCATATTTGGTTTATGACTATCCCGGATACTTTTTCTCAATACAACCAAATTCTTGGACGATCTATTAGAAAATTCTCTTACGTCGATATTTCTGAACCCGTTAATGTATATCTTTTAGCAGCCGTATATTCAGATTTCAATGACGAAGTGACGTCATTAAACGATTATACACAGGATGAATTGATTAATGTTTTACCCTTTGACATCAAAAAGCTGTTGTATCTAAAATTTAAGACTAAAGAAACGAATAGAATATACTCTATTCTTCAAGAGATGTCTGAAACGTATTCTCTTCCACCACATCCATCAATTGTAAAAGTTTTATTGGGAGAATTGGTCAGACAATTTTTTTATAATAATTCTCGTATTAAGTATAACGACTCCAAGTTACTTAAAATGGTTACATCAGTTATAAAAAATAAAGAAGACGCTAGGAATTACATAGATGATATTGTAAACGGTCACTTCTTTGTATCGAATAAAGTATTTGATAAATCTCTTTTATACAAATACAAAAACGATATTATTACAGTACCGTTTAGACTTTCCTACGAACCATTTGTTTGGGGAGTTAACTTTCGTAAAGAATATAATGTGGTATCTTCTCCATAAAACTGATGAGATATATAAAGAAATAAATGTCGAGCTTTGTTACCAATGGATATCTTCCAGTTACATTGGAACCACATGAGTTGACGTTAGACATAAAAACTAATATTAGGAATGCCGTATATAAGGCGTATCTCCATAGAGAAATTAGTGGTAAAATGGCCAAGAAAATAGAAATTCGTGAAGACGTGGAATTACCTCTCGGTGAAATAGTTAATAATTCTGTAGTTATAAACGTTCCGTGTGTAATAACCTACGCATATTATCACGTTGGGGATATAGTCAGAGGAACATTAAACATCGAAGATGAATCAAATGTAACTATTCAATGTGGAGATTTAATCTGTAAACTAAGTAGAGATTCGGGTACTGTATCATTTAGCGATTCAAAGTACTGCTTTTTTCGAAATGGTAATGCGTATGATAACGGCATCGAAGTCTCCGCCGTTCTAATGGAGGCTCAACAAGGTACCGAATCTAGTTTTGTTTTTCTCGCGAATATCGTTGACTCATAAGAAAGAGAATAGCGGTGAGTATAAATACGAATACTATGGCAATAATTGCGAATGTTTTATTCCCTTCGATATATTTTTGATAATATGAAAAACATGCCTCTCTCAAATCAGACAACCATTTCATAAAATAGTTCTCTCGCACTGGTGAGGTGGTTGCAGCTCGTATAATCTCCCCAGAATAATATACTTGCGTGTCGTCGTTCAATTTATACGGATTTCTATAATTCTCTGTTATATAATGAGGTTTACCCTCATGATTAGACGACGACAATAGTGTTCTGAATTTAGATAGTTGATCAGAATGAATGTTTATTGGTGTTGGAAAAATTATCCATGCTGCGTCTGCAGAGTGGTTGATAGTTGTTCCTAGATATGTAAAATAATCCAACGTACTAGGTAGCAAATTGTCTAGATAAAATACTGAATCAAATGGCGCAGACATATTAGCGGATCTAATGGAATCCAATTGATTGACTATCTTTTGAAAATATACATTTTTATGATCTGATACTTGTAAGAATATAGCAATAATGATAATTCCATCATCGTGTTTTTTTGCCTCTTCATAAGAACTATATTTTTTCTTATTCCAATGAACCAGATTAATCTCTCCAGAGTATTTGTATACATCTATCAAGTGATTGGATCCATAATCGTCTTCCTTTCCCCAATATATATGTATTGTTGATAACACATATTCATTGGGGAGAAACCCTCCACTTATATATCCTCCTTTAAAATTAATCCTTACTAGTTTTCCAGTATTCTGGATAGTGGTTGGTTTCGACTCATTATAATGTATGTCTAACGTCTTCAATCGCGCGTCAGAAATTGCTTTTTTAGTTTCTATATTAATAGGAGATAGTTGTTGAGGCATAGTAAAAATGAAATGATAACTGTCTAGAAATAGCTCTTAGTATGGGATTTACAATGGATGAGGAAGTGATATTTGAAACTCCTAGAGAATTAATATCTATTAAACGAATAAAAGATATTCCAAGATCAAAAGACACGCACGTGTTTGCTGCGTGTATAACAAGTGACGGATATCCGTTAATAGGAGCTAGAAGAACTTCATTCGCATTCCAGGCGATATTATCTCAACAAAATTCAGATTCTATCTTTAGAGTATCCACTAAACTATTACGGTTTATGTACTACAATGAACTAAGAGAAATCTTTAGACGGTTGAGAAAAGGTTCTATCAACAATATCGATCCTCACTTCGAAGAGTTAATATTATTGGGTGGTAAACTAGATAAAAAGGAATCTATTAAAGATTGTTTAAGAAGAGAATTAAAAGAGGAAAGTGATGAACATATAACAGTAAAAGAATTCGGAAATGTAATTCTAAAACTTACAACGAGTGATAAATTATTTAATAAAGTATATATAGGTTATTGCATGGCATGTTTTATTAATCAATCGTTGGAGGATTTATCACATACTAGTATTTACAATGTAGAAATTAGAAAGATTAAATCGTTAAATGATTGTATTAACGACGATAAATACGAATATCTGTCTTATATTTATAATATACTAATTAATAGTAAATGAGCTTTTACAGATCTAGTATAATTAGTCAGATTATTAAGTATAATAGACGACTAGCTAAGTCTATTATTTGCGAGGATGACTCTCAAATTATTACACTCACGGCATTCGTTAACCAATGCCTATGGTGTCATAAACGAGTATCCGTGTCCGCTATTTTATTAACTACTGATAACAAAATATTAGTATGTAACAGACGAGATAGTTTTCTCTATTCTGAAATAATTAGAACTAGAAACATGTATAGAAAGAAACGATTATTTCTGAATTATTCCAATTATTTGAACAAACAGGAAAGAAGTATACTATCGTCATTTTTTTCTCTAGATCCAGCTACTGCTGATAATGATAGAATAAACGCTATTTATCCGGGTGGTATACCCAAAAGGGGTGAGAACGTTCCAGAGTGTTTATCCAGGGAAATCAAAGAAGAAGTTAATATAGACAATTCTTTTGTATTCATAGACACTCGTTTTTTTATTCATGGTATCATAGAAGATACCATTATTAACAAATTTTTTGAGGTAATTTTCTTTGTTGGACGAATATCTCTAACGAGTGATCAAATTATTGATACCTTTAAAAGTAATCATGAAATAAAGGATCTAATATTTTTAGATCCAAATTCAGGTAATGGACTCCAATACGAAATTGCAAAATATGCTCTAGATACTGCAAAACTTAAATGTTACGGTCATAGAGGATGTTATTATGAATCATTAAAAAAATTAACTGAGGATGATTGATTAGAAAATATAAATTAATTTACCATCGTGTATTTTTATAACGGGATTGTCTGGCATATCATGTAGATAGTTACCGTCTACATCGTATACTCTACCATCTACGCCTTTAAATCCTCTATTTATTGATATTAATCTATTAGAATTGGAATACCAAATATTAGTACCCTCAATTAGTTTATTGGTAATATTTTTTTTAGACGATAGATCGATGGCTCTTGAAACCAAGGTTTTCCAACCGGACTCATTGTCTATCGGTGAGAAGTCTTTTTCATTAGCATGAATCCATTCTAATGATGTATGTTTAAACACTCTAAACAATTGTACAAATTCTTTTGATTTGTTTTGAATGATTTCAAATAGGTCTTCGTCTACAGTAGGCATACCATTAGATAATCTAGCCATTATAAAGTGCACGTTTACATATCTACGTTCTGGAGGAGTAAGAACGTGACTATTGAGACGAATGGCTCTTCCTACTATCTGACGAAGAGACGCCTCGTTCCATGTCATATCTAAAATGAAGATATCATTGATTGAGAAGAAACTAATACCCTCGCCTCCGCTAGAAGAGAATACGCATGTTTTAATGCATTCTCCGTTAGTGTTTGATTCTTGGTTAAACTCAGCCACCGCCTTGATTCTAGTATCTTTTGTTCTAGATGAGAACTCTATATTAGAGATACCAAAGACTTTGAAATATAGTAATAAGATTTCTATTCCTGACTGATTAACAAATGGTTCAAAGACTAGACATTTACCATGGGATGCTAATATTCCCAAACATACATCTATAAATTTGACGCTTTTCTCTTTTAATTCAGTAAATAGAGAGATATCAGCCGCAATAGCATCCCCTCCCAATAGTTCTCCCTTTTTAAAGGTGTCTAATGCGGATTTAGAAAATTCTCTATCTCTTAATGAATTTTTAAAATCATTATATAGGGTTGCTATCTCTTGTGCGTATTCTCCCGGATCACGATTTTGTCTTTCAGGAAAGCTATCGAATGTAAACGTAGTAGCCATACGTCTCAGAATTCTAAATGATGATATACCAGTTTTTATTTCTGCGAGTTTAGCCTTTTGATAAATCTCTTCTTGCTTTTTTGACATATTAACGTATCGCATTAATACTGTTTTCTTAGCGAATGATGCAGACCCTTCCACATCATCAAAAATAGAAAACTCGTTATTAACTATGTACGAACATAGGCCTCCTAGTTTGGAGACTAATTCTTTTTCATCGACTAGACGTTTATTCTCAAATAGCGATTGGTGTTGTAAGGATCCTGGTCGCAGTAAGTTAACCAACATGGTGAATTCTTGCACACTATTAACGATAGGTGTAGCCGATAAACAAATCATCTTATGGTTTTTTAACGCAATGGTCTTAGATAAAAAATTATATACTGACCGAGTAGGACGGATCTTACCATCTTCTTTGATTAATGATTTAGAAATGAAGTTATGACATTCATCAATGATGACGCATATTCTACTCTTGGAATTAATAGTTTTGATATTAGTAAAAAATTTATTTCTAAAATTTTGATCATCGTAATTAATAAAAATACAATCCTTCGTTATCTCTGGAGCGTATCTGAGTATAGTGTTTATCCAAGGATCTTCTATCAAAGCCTTTTTTACCAATAAGATAATTGCCCAATTCGTATAAATATCCTTAAGATGTTTGAGAATATATACAGTAGTCATTGTTTTACCGACACCTGTTTCATGGAACAATAAAAGAGAATGCATACTGTCTAATCCTAAGAAAACTCTTGCTACAAAATGTTGATAATCCTTGAGGCGTACTACGTCTGACCCCATCATTTCAACGGGCATATTAGTAGTTCTGCGTAAGGCATAATCGATATAGGCCGCGTGTGATTTACTCATTTATGAGTGATAAGTAATAACTATGTTTTAAAAATCACAGCAGTAGTTTAACTAGCCTTCTCTGATGTTTGTTTTCGATACTTTTTGAATCAGAAGTCATACTAGAATAAAGCAGCGAGTGAACGTAATAGAGAGCTTCGTATACTCTATTCGAAAACTCTAAGAACTTATTAATGAATTCCGTATCCACTGGATCGTTTAAAATACTAAATTGAACAGTGTTCACATCCTTCCAAGACGAAGACTTAGTGACGGACTTAACATGAGACATAAATAAATCCAAATTTTTTTTATAAACATCACTAGCCACCATAATGGCGCTATCTTTCAACCAACTATCGCTTACGCATTTTAACAGTCTAACATTTTTAAAGAGACTACAATATATTCTCATAGTATCGATTACACCTCTACCGAATAGAGTGGGAAGTTTAATAATACAATATTTTTCGTTTACAAAATCAAATAATGGTCGAAACACGTCGAAGGTTAACATCTTATAATCGCTAATGTATAGATTGTTTTCAGTGAGATGATTATTAGATTTAATAGCATCTCGTTCACGTTTGAACAGTTTATTGCGTGCGCTGAGGTCGGCAACTACGGCATCCGCTCTAGTACTCCTCCCATAATACTTTACGCTATTAATCTTTAAAATTTCATAGACTTTATCTAGATCGCTTTCTGGTAACATGATATCATGTGTAAAAAGTTTTAACATGTCGGTCGGCATTCTATTTAGATCATTAACTCTAGAAATCTGAAGAAAGTAATTAGCTCCATATTCCAGACTAGGTAATGGGCTTTTACCTAAAGACAAGTTAAGTTCTGGCAATGTTTCATAAAATGGAAGAAGGACATGTGTCCCCTCCCGGATATTTTTTACAATTTCATCCATTTACAACTCTATAGTTTGTTTTCATTATTATTAGTTATTATCTCCCATAATCTTGGTAATACTTACACCTTGATCATAAGATACCTTATACAGGTCATTACATACAACTACCAATTGTTTTTGTACATAATAGATTGGATGATTGATATCCATGGTGGAATAAACTACTCGAACAGATAGTTTATCTTTCCCCCTAGATACATTGGCCGTAATAGTTGTCGGCCTAAAGAATATCTTTGGTGTAAAGTTAAAAGTTAGGGTTCTTGTTCCATTATTGCTTTTTGTCAGTAGTTCGTTATAAATTCTCGAGATGGGCCCGTTCTCTGAATATAGAACATCATTTCCAAATCTAACTTCTAGTCTAGAAATAATATCGGTCTTATTTTTAAAATCTATTCCCTTGATGAATGGATCGTTAATAAACAAATCCTTGGCCTTTGATTCGGCTGATCTATTATCTCCGTTATAGACGTTACGTTGACTAGTCCAAAGACTTACAGGAATAGATGTATCGATGATGTTGATAGTATGTGATATGTGAGCAAAGACTGTTCTCTTGGTGGCGTCGCTATATGTTCCAGTAATGGCGGAAAACTTTTTAGAAATGTTATATATAAAAGAATTTTTTCGGGTTCCAAACATTAACAGATTAGTATGAAGATAAACACTCATATTATCAGGAACATTATCAATTTTTACATAAACATCGGCATCTTGAATAGAAACAACACCATCTTCTGGAACCTCTACGATCTCGGCAGATTCCGGATAACCAGTCGGTGGACCATCACTAACAATAACTAGATCATCCAACAATCTACTCACATATGCGTCTATATAATCTTTTTCATCTTGTGAGTACCCTGGATACGAAATAAATTTGTTATCAGTATTTCCATAATAAGGTTTAGTATAAACAGAGAGAGATGTTGCTGCATGAACTTCGGTTACTGTCGCCGTTGGTTGGTTTATTTGACCTATTACTCTCCTAGGTTTCTCTATAAATGATGGTTTAATTTGTACATTCTTAACCATATATCCAATAAAGCTCAATTCAGGAACATAAACAAATTCTTTGTTGAACGTTTCAAAGTCGAACGAAGAGTCACGAATAACGATATCGGATACTGGATTGAAGGTCACCGTTACGGTAATTTTTGAATCGGATAGTTTAAGACTACTGAATGTATCTTCCACATCAAACGGAGTTTTAATATAAACGTATACTGTAGATGGTTCTTTAATAGTGTCATTAGGAGTTAGGCCAATAGAAATATCATTAAGTTCACTAGAATATCCAGAATGTTTCAAAGCAATTGTATTATTGATACAATTATTATATAATTCTTCGCCCTCAATTTCCCAAATAACACCGTTACACGAAGAGACAGATACATGATTAATACATTTATATCCAACATATGGCACGTAACCGAATCTTCCCATACCTTTAACTTCTGGAAGTTCCAAACTCAGAACCAAATGATTAAGCGCAGTAATATACTGATCCCTAATTTCGAAGCTAGCGATAGCCTGATTGTCTGGCCCATCGTTTGTCATAACTCCGGATAGAGAAATATATTGCGGCATATATAAAGTTGGAATTTGACTATCAACTGCGAAGACATTAGACCGTTTAATAAAGTCATCCCCACCGATCAAAGAATTAATGATAGTATTATTCATTTTCTATTTAAAATGGAAAAAGCTTACAATAAACTCCGTAGAGAAATATCTATAATTTGTGAGTTTTCCTTAAAGTAACAGCTTCCGTAAACACCGTCTTTATCTCTTAGTAAGTTTATTGTATTTATGACCTTTTCCTTATCTTCATAGAATACTAAAGGCAATAAAGAAATTTTTGGTTCTTCTCTAAGAGCTACGTGAGACTTAACCATAGACGCCAACGAATCCCTACATATTTTAGAACAGAAATACCCAACTTCACCACCCTTGAATGTCTCAATACTAATAGGTCTAAAAACCAAATCTTGATTACAAAACCAACACTTATCAATTACACTATTTGTCTTAATAGACATATCTGCCATAGATTTATAATACTTTGGTAGTATACAAGCGAGTGCTTCTTCTTTAGCGGGCTTAAAGACTGCTTTAGGTGCTGAAATAACCACATCTGGAAGACTTACTCGCTTAGCCATTTAATTACGGAACTATTTTTTTATACTTCTAATGAACAAGTAGAAAACCTCTCATCTACAAAAACATACTCGTGTCCATAATCCTCTACCATAGTAACACGTTTTTTAGATCTCATATGTGCTAAAAAGTTTTCCCATACTAATTGGTTACTATTATTTTTCGTATAATTTTTAACAGTTTGAGGTTTTAGATTTTTAGTTACAGAAGTGATATCGAATATTTTATCCAAAAAGAATGAGTAATTAATTGTCTTAGAAGGAGTGTTTTCTTGGCAAAAGAATACCAAGTGCTTAAATATTTCTACTACTTCATTAATCTTTTCTGTACTCAGATTCAGTTTCTCATCTTTTACTTGATTGATTATTTCAAAGACTAACTTATAATCCTTTTTATTTATTCTCTCGTTAGCCTTAAGAAAACTAGATACAAAATTTGCATCTACATCATCCGTGGATATTTGATTTTTTTCCATGATATCCAATAGTTCCGAGATAATTTCTCCAGAACATTGATGAGACAATAATCTCCGCAATACATTTCTCAAATGAATAAGTTTATTAGACACGTGGAAGTTTGACTTTTTTTGTACCTTTGTACATTTTTGAAATACAGACTCGCAAAAAATACAATATTCATATCCTTGTTCAGATACTATACCGTTATGTCTACAACAGCTACATAATCGTAGATTCATGTTAACACTCTACGTATCTCGTCGTCCAATATTTTATATAAAAACATTTTATTTCTAGACGTTGTCAGAAAATCCTGTAATATTTTTAGTTTTTTTGGTTGTGAATAAAGTATCGCCCTAATAATATTGGTACCGTCTTCCGACAATATAGTAGTTAAATTATCCGAGCATGTAGAAGAACACCGCTTAGGCGGATTCAGTACAATGTTATATTTTTCGTACCAACTCATTTAAATATCATAATCTAAAATAGTTCTGTAATATGTCTAGCGCTAATATATTGATCATAATCCTGTGCATAAATTAAGATACAACAATGTCTTGAAATCATCGACATGGCTTCTTCCATAGTTAGAAGATCATCGTCAAAGTTAGCAACGTGATTCATCAACATTTGCTGTTTTGAGGCAGCAAATACTGAACCATCACCATTCAACCATTCATAAAAACCATCGTCTGAATCCATTGATAATTTCTTGTACTGGTTTTTGAGAGCTCGCATCAATCTAGCATTTCTAGCTCCCGGATTGAAAACAGAAAGAGGATCGTACATCCAAGGTCCATTTTCTGTAAATAGAATCGTATAATGTCCCTTCAAGAAGATATCAGACGATCCACAATCAAAGAATTGGTCTCCGAGTTTGTAACAGACTGCGGACTTTAACCTATACATGATACCGTTTAGCATGATTTCTGGTGATACGTCAATCGGAGTATCATCTATTAGAGATCTAAAGCCGGTGTAACATTCTCCGCCAAACATATTCTTATTCTGACGTCGTTCTACATAAAACATCATTGCTCCATTAACGATAACAGGTGAATGAACAGCACTACCCATCACATTAGTTCCCAATGGATCAATGTGTGTAACTCCAGAACATCTTCCATAGCCTATGTTAGGAGGAGCGAACACCACTCTTCCACTATTGCCATCGAATGCCATAGAATAAATATCCTTGGAATTGATAGAAATCGGACTGTCGGATGTTGTTATCATCTTCATAGGATTAACAACGATGTATGGTGCAGCCTGAAGTTTCATATCGTAACTGATGCCGTTCATAGGTCTAGCCACAGAAACCAACGTAGGTCTAAATCCAACTATAGACAAAATAGAAGCTAATATCTGTTCCTCATCTGTCATAACTTGAGAGCATCCAGTATGAATAATCTTCATTAGATGGGGATCTACCGCATCATCATCGTTACAATAAAAAATTCCCATTCTAATGTTCATAATTGCTTTTCTAATCATGGTATGAATGTTTGCTCTCTGAATCTCTGTGGAAATTAGATCTGATACACCTGTAATCACTATCGGATTATCCTCCGTAAGACGATTAACCAACAACATATAATTATAAGACTTTACTCTTCTAAATTCATAAAGTTGCTGGATTAGACTATATGTGTCTCCATGTACATACGCGTTCTCGAGCGCAGGAAGTTTAATACCGAATAGTGCCATCAGAATAGGATGAATGTAGTAATTAGTTTCTGGTTTTCTATAAATAAAAGACAAATCTTGTGAACTAGACATATCGGTAAAATGCATGGATTGGAATCGTGTAGTCGACAGAAGAATATGATGATTAGATGGAGAGTATATTTTATCTAACTCTTTGAGTTGGTCACCGATTCTAGGACTAGCTCGAGAATGAATAAGTACTAAGGGATGAGTACATTTCACAGAAACACTGGCGTTGTTCAACGTACTCTTTACATGGGAAAGGAGTTGAAATAGCTCGTTTCTATTTGTCCTGACAATATTTAGTTTATTCATAATATTAAGCATATCCTGAATAGTAAAGTTAGATGTGTCATACTTGTTAGTAGTTAGATATTTAGCAATTGCATTCCCATCATTTCTCAATCTCGTACTCCAATCATGTGTGGATGCTACTTCGTCGATGGAAACCATACAATCCTTTTTGATAGGCTGTTGAGATTGATCATTTCCTGTACGTTTAGGTTTGGTACGTTGATTTCTAGCCCCTGCTGATATAAAGTCATCGTCTACAATTTGGGATAATGAATTACATACACTACAAGACAAAGATTTATCAGAAGTGTGAATATGATCTTCATCTACCAAAGAAAGAGTTTGATTAGTATAACTAGATTTTAGTCCCGCGTTAGATGTTAAAAAAACATCGCTATTGACCACGGCTTCCATTATTTATATTCGTAGTTTTTACTCGAAAGCGTGATTTTAATATCCAATCTTATTACTTTTGGAATCGTTCAAAACCTTTGACTAGTTGTAGAATTTGATCTATTGCCCTACGCGTATACTCCCTTGCATCATATACGTTCGTCACCAGATCGTTTGTTTCGGCCTGAAGTTGACGCATATCTTTTTCAACACTCGACATGAGATCCTTAAGGGTCATATCGTCTAGATTTTGTTGAGATGCTGCTCCTGGATTTGGATTTTGTTGTGCTGTTGTACATACTGTACCACCAGTAGGTGTAGGAGTACATACAGTGGCCACAATAGGAGGTTGAAGAGGTGTAACCGTTGGAGTAGTACAAGAAATACTTCCATCCGATTGTTGTGTACATGTGGTTGTTGGTAACGTCTGAGAAGGTTGGGTAGATGGCGGTGTCGTCATCTTTTGATCTTTATTAAATTTAGAGATAATATCCTGAACAGTATTGCTCGGCGTCAACGCTGGAAGGAGTGTACTCGCCGGCGCATCAGTATCTGTAGACAACCAATCAAAAAGATTAGACATATCAGATGATGTATTAGTTTGTTGACGTGGTTTTAGTACAGGAGCAGTACTACTAGGTAGAAGAATAGGAGCCGGTGTAGGTGTCGGAACCGGCTGTGGAGTTATATGAATAGTTGGTTGTAGCGGTTGGGTAGGCTGTCTGCTGGCGGTCATCATATTATCTCTAGCTAGTTGTTCTCGCAACTGTCTTTGATAATACGACTCTTGAGACTTTAGTCCTATTTCAATCGCTTCATCCTTTTTCGTATCCGGATCCTTTTCTTCAGAATAATAGATTGACGACTTTGGTGTAGAGGATTCTGCCAGCCCCTGTGAGAACTTGTTAAAGAAGTCCATTTAAGGCTTTAAAATTGAATTGCGATTATAAGATTAAATGGCAGACACGGACGATATTATCGACTATGAATCCGATGATCTCACCGAATACGAGGATGATGAAGAAGATGGAGAGTCACTAGAAACTAGTGATATAGATCCCAAATCTTCTTATAAGATTGTAGAATCAACATCCACTCATATAGAAGATGCGCATTCCAATCTTAAACATATAGGGAATCATATATCTGCTCTTAAACGACGCTATACTAGACGTATAAGTCTATTTGAAATAGCGGGTATAATAGCAGAAAGCTATAACTTGCTTCAACGAGGAAGATTACCTCTAGTTTCAGAATTTTCTGACGAAACGATGAAGCAAAATATGCTACATGTAATTATACAAGAGATAGAGGAGGGTTCTTGTCCTATAGTCATCGAAAAGAACGGAGAATTGTTGTCGGTAAACGATTTTGATAAAGATGGTCTAAAATTCCATCTAGACTATATTATCAAAATTTGGAAACTTCAAAAACGATATTAGAATTTATACGAATATCGTTCTCTAAATGTCACAATCAAGTCTCTCATATTCAGCAGTTTATTGTCGTACTTTATATCGTGTTCATTAACGATATTTTGCAAAATAGTAATGATTCTATCTTCCTTCGATAGATATTCTTCAGAGATTATTGTCTTATATTCTTTCTTGTTATCCGATATGAATTTGATAAGACTTTGAACATTATTAATACCCGTCTGTTTAATTTTTTCTATAGATATTTTAGTTTTGGTAGATTCTATGGTGTCTGTTAATAGGCATCCAACATCGACATTCGACGTCAATTGTCTATAAATCAGAGTATAAATTTTAGAAATAACATTAGCAAATTGTTGTGCGTTGATGTCGTTATTCTGAAACAGTATGATTTTAGGTAGCATTTTCTTAACAAAGAGAACGTATTTATTGTTACTCAGTTGAACAGATGATATATCCAGATTACTAACGCATCTGATTCCATATACCAAACTTTCAGAAGAAATGGTGTACAATTGTTTGTATTCATTCAATGTCTCCTTTTCAGAAATTAGTTTAGAGTCGAATACTGCAATAATTTTCAAGAGATAGTTTTCATCAGATAAGATTTTATTTAGTGTAGATATGATAAAACTATTGTTTTGTTGGAGAACTTGATACGCCGCATTCTCTGTAGTCGACGCTCTCAAATGGGAAACAATCTCTATTATTTTTTTGGAATCGGATACTATATCTTCGGTATCTTGACGCAGTCTAGTATACATAGAGTTAAGAGAAATTAGAGTTTGTACATTAAGCAACATGTCTCTAAATGTGGCTACAAACTTTTCTTTTTCCACATCATCTAGTTTATTATATACCGATTTCACAACGGCACCAGATTTAAGGAACCAGAATGAAAAACTCTGATAACTACAATATTTCATCATAGTTACGATTTTATCATCTTCTATAGTTGGTGTGATAACACATACCTTTTTCTCCAAGACTGGAACCAACGTCATAAAAATGTTTAAATCAAAATCCATATCAACATCTGATGCGCTAAGACCAGTCTCGCGTTCAAGATTATCTTTACTAATGGTGACGAACTCATCGTATAGAACTCTAAGTTTGTCCATTATTTATTTACAGATTTAGTTGTTTAATTTATTTGTGCTCTTCCAGAGTTGGGATAGTATTTTTCTAACGTCGGTATTATATTATTAGGATCTACGTTCATATGTATCATAATATTAATCATCCACGTTTTGATAAATCTATCTTTAGCTTCTGAAATAACGTATTTAAACAAAGGAGAAAAATATTTAGTTACGGCATCAGACGCGATAACATTTTTTGTAAATGTAACGTATTTAGACGACAGATCTTCGTTAAAAAGTTTTCCATCTATGTAGAATCCATCGGTTGTTAACACCATTCCCGCGTCAGAGTGAATAGGAGTTTGAATAGTTTGTTTTGGAAATAGATCCTTCAATAACTTATAGTTGGGTGGGAAAAAATCGATTTTATCACTAGACTCTTTCTTTTTTACTATCATTACCTCATGAACTATTTCTTGAATGAGTATATGTATTTTCTTTCCTATATCGGTCGCGTTCATTGGAAAATATATCATGTCGTTAACTATAAGAATATTTTTATCCTCGTTTACAAACTGAATAATATCAGATATAGTTCGTAAACGAACTATATCATCACCAGCACAACATCTAACTATATGATATCCACTAGTTTCCTTTAGCCGTTTATTATCTTGTTCCATATTAGCAGTCATTCCATCATTTAAGAAGGCGTCAAAGATAATAGGGAGAAATGACATTTTGGATTCTGTTACGACTTTACCAAAATTAAGGATATACGGACTTACTATCTTTTTCTCAACGTCGATTTGATGAACACACGATGAAAATGTACTTCGATGAGATTGATCATGTAGAAAACAACAAGGGATACAATATTTCCGCATATCATGAAATATATTAAGAAATCCCACTTTATTATATTTCCCCAAAGGATCAATGCATGTAAACATTATACCGTTATCATTAATAAAGACTTCTTTCTCATCGGATCTGTAAAAGTTGTTACTGATTTTTTTCATTCCAGGATCTAGATAATTAATAATAATGGGTTTTCTATTCTTATTCTTTGTATTTTGACATATCCTAGACCAGTAAACAGTTTCCACTTTGGTAAAATCAGAAGACTTTTGAACGCTATTAAACATGGCATTAATGGCAATAACTAAAAATGTAAAATATTTTTCTATGTTAGGAATATGGTTTTTCACTTTAATAGATATATGGTTTTTTGCCAAAATGATAGATATTTTTTTATCCGATGATAGTAAAATATTATTAGTCGCCGTCTCTATAAAAATGAAGCTAGTCTCGATATCCAATTTTATTCTAGAATTGATAGGAGTCGCCAAATGTACCTTATACGTTATATCTCCCTTGATGCGTTCCATTTGTGTATCTATATCGGACACAAGATCTGTAAATAGTTTTACGTTATTAATCATCACGGTATCGCCATCGCTAGATAATGCTAATGTACTATCCAAGTCCCAAATGGAGAGATTTAACTGTTCATCGTTTAGAATAAAATGATTACCTGTCATATTAATAAAGTGTTCATCGTATCTAGATAACAACGACTTATAATTAATGTCCAAGTCTTGAACTCGCTGAATGATCTTTTTTAACCCAGTTAGTTTTAGATTGGTACGAAATATATTGTTAAACTTTGATTCTACAGTAATGTCCAAATCTAGTTGTGGAAATACTTCCATCAACATTGTTTCAAACTTGATAATATTATTATCTACATCTTCGTACGATCCAAATTCCGGAATAGATGTATCGCACGCTCTGGCCACCCAGATAACCAAAAAGTCACACGCTCCAGAATATACATTGTATAAAAAGCTATCGTTTTTTAGTAGTGTTTTTTTCTGAGTATATACGAAAGGATTAAAAATAGTATTATCAACGTAACTATATTCCAAATTATTCTTATGAGAATAGATAATAATATCGTCCTTAATATCTAACAAATTTCCTAAATATCCCTTTAATTGAGTCATTCGAAGCGTTAATAAAATATGTCTCTTAACTATTTCCGGCCGTTGTATATTTAAATGACTTCGTAAGAAATAATATATAGGCGACTTCTCATCTATGTAATCATATGGAGTGAGATATAGGGCTCGTTCTACCTCCTGCCCCTTACCCACCTGTAATACCAATTGCGGACTCACTATATATCGCATATTTATATCGTGGGGTAAAGTGAAAATCTACTACCGATGATGTAAGTCTTACAATGTTCGAACCAGTACCAGATCTTAATTTGGAGGCCTCCGTAGAACTAGGGGAGGTAAATATAGATCAAACAACACCTATGATAAAGGAGAATAGCGGTTTTATATCCCGTAGTAGACGTCTATTCGCCCATAGATCTAAGGATGATGAGAGAAAACTAGCACTACGATTCTTTTTACAAAGACTTTATTTTTTAGATCATAGAGAGATTCATTATTTGTTCAGATGCGTCGACGCTGTAAAAGACGTCACTATTACCAAAAAAAATAACATTATCGTGGCGCCTTATATAGCACTTTTAACTATCGCATCAAAAGGATGCAAACTTACAGAAACAATGATTGAAGCATTCTTTCCAGAACTATATAATGAACATAGTAAGAAATTCAAATTCAACTCTCAAGTATCCATCATCCAAGAAAAACTCGGATACCAGTCTGGAAACTATCACGTTTATGATTTTGAACCGTATTACTCTACAGTAGCTCTGGCTATTCGAGATGAACATTCATCTGGCATTTTTAATATCCGTCAAGAGAGTTATCTTGTAAGTTCATTATCTGAAATAACATATAGATTTTATCTAATTAATCTAAAATCTGATCTTGTTCAATGGAGTGCTAGTACGGGCGCTGTAATTAATCAAATGGTAAATACTGTATTGATTACAGTGTATGAAAAATTACAACTGGCCATAGAAAATGATTCACAATTTACATGTTCATTGGCTGTGGAATCAGAACTTCCAATAAAATTACTTAAAGATAGAAATGAATTATTTACAAAATTCATTAACGAGTTAAAAAAGACCAGTTCATTCAAGATAAGCAAACGTGATAAGGATACGCTATTAAAACATTTTACTTATGACTGGAGTTAGAATTTATAGACGACACATTTCGTTTATCATTGTTACTATTACTATCATTATTAGTATTCTTCTTGTCATCTTGTTCAGAAATATACAGCAATGCTATACCTAATACTAAATACATTATCATGCTTGCAATGGCTCTAACAACAACGAACCAAAATGAATTTGGTCGTAGCTTTTGTTCACAAAAATACATAAAGAAATGTCTACATAAATCTATGGCGCCATTGGCTACTTGAAATAGCGCCAGTCCTCCTACAGATTTTAATATAGCTGTATAACATGACATTTATTCATCATCAAAAGAGACAGAGTCACCATCTGTCATATTTAGATTTTTTTTCATGTGTTCAAAGTATCCTCTACTCATTTCATTATAATAGTTTATCATGCTTAGAATTTTAGGACGGATCAATGAGTAAGACTTGACTAGATCGTCAGTAGTAATTTGTGCATCATCTATTCTGCATCCGCTTCGTCGAATAATGTATAGCATCGCTTTGAGATTCTCCATAGCTATCAAGTCTTTATATAATGACATGGAAATATCTGTGAATGCTTTATACTTCTCCAACATCGATGCCTTAACATCATCACATACTTTAGCATTGAAAATACGTTCTATTGTGTAGATGGATGTAGCAAGATTTTTAAACAACAATGCCATCTTACATGATGATTGTCTCAAGTCTCCAATCGTTTGTTTAGAACGATTAGCTACAGAGTCCAATGCTTGGCTAACTAGCATATTATTATCTTTAGAAATTGTATTCTTCAATGAGGCGTTTATCATATCTGTGATTTCGTTAGTCATATTACAGTCTGACTGGGTTGTAATGTTATCCAACATATCACCTATGGATACGGTACACGTACCAGCATTTGTAATAATCCTATCTAAGATGTTGTATGGCATTGCGCAGAAAATATCTTCTCCTGTAATATCTCCACTCTCGATAAATCTACTCAGATTATTCTTAAATGCCTTATTCTCTGGAGAAAAGATATCAGTGTCCATCATTTCATTAATAGTATACGCAGAAAAGATACCACGAGTATCAATTCTATCCAAGATACTTATCGGTTCCGAGTCACAGATAATTGTTTCCTCTCCTTCGGGAGATCCTGCATAGAAATATCTAGGACAATAGTTTCTATACTGTCTGTAACTCTGATAATCTCTAAAGTCACTAACTGATACCATGAAATTGAGAAGATCAAACGCTGAAGTAATCAATTTTTCTGCCTCGTTTTTACTACAACTAGTTTTCATCAATGTAGTGACGATGTATTGTTTAGTTACTCTTGGTCTAATACTGATGATAGAGATATTATTGCTTCCCATAATGGATCTTCTAGTAGTCACCTTAAAGCCCATTGATGCGAATAGCAGATAGATAAAGTCTTGGTATGACTCCTTTCTAATATAGTACGGACTACCTTTGTCACCCAACTTTATACCCACATAAGCCATAACAACCTCTTTAATAGCCGTTTCATGAGGTTTATCAGCCATGAGCCTGAGTAGTTGAAAGAATCGCATGAATCCCGTCTCAGAAAGTCCTATATGCATGATAGATTTATCTTTCCTGGGAAACTCTCGTATAGTTATAGATGAAATACTCTTCAAAGTTTCTGAAATAAGATTAGTAACAGTCTTACCTCCGACTACTCTGGGTAACAAACATACTCTAATAGGTGTTTTCTCTGCGGAGATAATATCAGAAAGGATAGAGCAATAAGTAGTATTATTGTGATTATAAAGACCGAATACATAACAGGTAGAATTTATAAACATCATGTCCTGAAGGGTTTTAGACTTGTATTCCTCGTAATCTATACCGTCCCAAAACATGGATTTGGTAACTTTGATAGCCGTAGATCTTTGTTCCTTCGCTAACAGGTTAAAGAAATTAATAAAGAATTTGTTGTTTCTATTTATGTCCACAAATTGCACGTTTGGAAGCGCCACGGTTACATTCACTGCAGCATTTTGAGGATCGCGAGTATGAAGTACGATGTTATTGTTTACTGGTATATCTGGAAAGAAATCTACCAGTCTAGGAATAAGAGATTGATATCGCATAGAAATAGTAAAGTTTATAATCTCATCATTGAAGATTACTCTGTTACCATTGTAATAAATTGGTACTCTATCATAATCATCGACAAAGTACTGTTCATACATGATGAGATGTTTATATGTTGGCATAGTAGTGAGATCGACGTTTGGTAATGGCAATGTATTAAGATTAACTCCATAATGTCTAGCAGCATCTGCGATGTTATAAGTGATGTCAAAGCGGGGTTGATCTTGTGCTGTTATATATTGTCTAACACCTATAAGATTATCAAAATCTTGTCTGCTTAATACACCGTTAACAATTTTTGCCTTGAATTCTTTTATTGGTGCATTAATAACATCCTTATAGAGGATGTTAAACAAATAAGTATTATCAAAGTTAAGATCTGGGTATTTCTTTTCTGCTAGAACATCCATTGAGTCGGAGCCATCTGGTTTAATATAACCACCGATAAATCTAGCTCTGTATTCTGTATCCGTCAATCTAATATTAAGAAGGTGTTGAGTGAAAGGTGGAAGATCGTAAAAGCTGTGAGTATTAATAATAGGGTTAGTTTCCGAACTAATGTTAATTGGATGATTAATAATATCTATATTTCCAGCGTTAAGTGTAACATTAAACAGTTTTAATTCACGTGACGTGGTATCAATTAAATAATTAATGCCCAATTTGGATATAGTAGCCTGAAGCTCATCTTGTTTAGTTACGGATCCTAATGAGTTATTAAGAAATACATCGAACGGATGAACGAAGGTTGTTTTAAGTTGGTCACATACTTTGTAATCTAGACATAGATGTGGAAGAACGGTAGAAACTATACGAAATAGATATTCAGAGTCCTCTAATTGATCAAGAGTAACTATTGACTTAATAGGCATCATTTATTTAGTATTAAATGACGACCGTACCAGTGACAGATATACAAAACGACTTAATTACAGAGTTTTCAGAAGATAATTATCCATCTAACAAAAATTATGAAATAACTCTTCGCCAAATGTCTATTCTAACTCACGTTAACAACGTGGTAGATAGAGAACATAATGCCGCCGTAGTGTCATCTCCAGAGGAAATATCATCACAACTTAATGAAGATCTATTTCCAGATGATGATTCACCGGCCACTATTATCGAACGAGTACAACCTCATACTACTATTATTGACGATACGCCACCTCCTACTTTTCGTAGAGAGTTATTGATATCGGAACAACGTCAACAACGAGAAAAAAGATTTAATATTACAGTATCAAAAAATTCTGAAGCAATAATGGAATCTAGATCTATGATAACTTCTATGCCAACACAAACACCATCCTTGGGAGTAGTTTATGATAAAGATAAAAGAATTCAGATGCTAGAGGATGAAGTGGTTAATCTTAGAAATCAACGATCTAATACAAAATCATCTGATAATTTAGATAATTTTACCAGAATACTATTTGGTAAGACTCCGTATAAATCAACCGAAGTTAATAAGCGTATAGCCATCGTTAATTATGCAAATTTGAACGGGTCCCCCTTATCAGTCGAGGACTTGGATGTCTGTTCGGAGGATGAAATAGATAGAATCTATAAAACGATTAAACAATATCACGAAAGTAGAAAACGAAAAATTATCGTCACTAACGTGATTATTATTGTCATAAACATTATTGAGCAGGCATTGCTAAAACTCGGATTTGAAGAAATCAAAGGACTGAGTACCGATATCACTTCAGAAATTATCGATGTGGAGATCGGAGATGACTGCGATGCTGTAGCATCAAAACTAGGAATCGGTAACAGTCCGGTTCTTAATATTGTATTGTTTATACTCAAGATATTCGTTAAACGAATTAAAATTATTTAATTTAATACATTCCCATATCCAGACAACAATCGTCTGGATTAATCTGTTCCTGTCGTCTCATACCGGACGACATATTAATCTTTTTATTAGTGGGCATCTTTTTAGATGGTTTCTTTTTCCCAGCATTAACTGATTCGATACCTAGAAGATCGTGATTGATTTCTCCGACCATTCCACGAACTTCTAATTGGCCGTCTCTAACGGTACCATAAACTATTTTACCAGCATTAGTAACAGCTTGGACAATCTGACCATCCATTGCGTTGAATGATGTAGTTGCTGTTGTTCTACGTCTAGGAGCACCAGAGGTATTTTTAGAGCTCTTGGATGTTGATGTAGAAGACGAGGATTTTGATTTTGGTTTACATGTAATACATTTTGAACTCTTTGATTTTGTATCACATGCACCGGCAGTCACATCTGTTTGAGAATTAAGATTATTGTTGCCTCCTTTGACGGCTGCATCTCCACCGATCTGCGCTAGTAGATTTTTAAGCTGTGGTGTAATCTTATTAACTGTTTCAATATAATCATCGTAACTACTTCTAACGGCTAAATTTTTTTTATCCGCCATTTAGAAGCTAAAAATATTTTTATTTATGCAGAAGATTTAACTAGATTATACAATGAACTAATATGATCCTTTTCTAGATTATTTACGAACTTGGTATTTCTTGTTTCTGGAGGAGGAGAATTTAAATTCGGACTTGGATTCGGATTTTGTGGGTTCTTGATCTTATTATACAGCGTGTATAGGATGGCGACGGTAACTGCTACACAAATACCGATCAACAGAAGAATACCAATCATTTATTGACAATAACTTCACTATGATCAAGTATGTAATAATCATCTTTTCACTAAGTAAGTAGTAATAATGATTCAACAATGACACGATATATGGACGATAATAATTTAGTTCATGGAAATATCGCTATGATTGGTGTGAATGACTCCGCTAACTCTGTGGGGTGCACAGTGCTTTCCCCACATAGAATAAATTAGCATTCCGACTGTGATAATAATACCAAGTATAAACGCCATAATACTCAATACTTTCCATGTACGAGTGGGACTGGTAGACTTACTAAAGTCAATAAAGGCGAAGATACACGAAAGAATCAAAAGAATGATTCCAGCGATTAGCACGCCAGAAAAATAATTTCCAATCATAAGCATCATGTCCATTTAACTAATAAAAATTTTAAATCGCCGAATAAACAAAGTGGAATATAAACCATATAAAAACAATAGTTTGTACTGCAAAAATAATATCTATTTTTGTTTTCGAAGATATGGTAAAATTAAATAGTAGTACACAGCATGTTATAACTAACAGCAGCAACGGCTCGTAATTACTTATCATTTACTAGACGAAAAGGTGGTGGGATATTTTCTTGCTCAAATAATACGAATATATCACCCATCCATTTTATACGATGTTTATATACTCTAATCTTTAATAGATCTATAGATGACGGGTTTACCAATAATATAGATTTTATCGATTCATCTAATTTAAACCCTTCCTTAAACGTGAATGATCTATTATCTGGCATAATGATGACCCTACCTGATGAATCTGACAATGTACTGGGCCATGTAGAATAAATTATCAACGAATTATCGTCTACGAACATTTATATCATTTGTTTTAATTTTAGGACGTGAATAAATAGATATAAAATAGAAAATAACAGATATTACAACCAGTGTTATGGACGCACCCAACCATGTAGGCAGTTTTATTTTATCGTTTACTACAGGTTCTCCTGGATGTACGTCACCAACTGCAGACGTAGTTCTAGTACAATTAGACGTAAGTTCCGCTTGGGAATTTTTTAACGCTAAAGAGTTAACGTTGATCGTACACCCAACGTATTTACATCTAGTTCTTTGAACATCTTGATTATAATATAACCATTTTCTATCTCTAGATTCGTCAGTGCACTCATGTAACCAACATACCCTAGGTCCTAAATATTTATCTCCGGAATTAGATTTTGGATAATTCGCGCACCAACAATTTCTATTTCCTTTATGGTCGTTACAAAAGACGTATAATGCCGTATCCCCAAAAGTAAAATAATCAGGACGAATAATTCTAATAAACTCAGAACAATATCTCGCATCCATATGTTTGGAGCAAATATCGGAATAAGTAGACATAGCCGGTTTCCGTTTTACACGTAACCATTCTAAACAATTGGGGTTTCCAGGATCGTTTCTACAAAAACCAGTCATGAAATCGTCACAATGTTCTGTCTTGTAATTATTATTAAATATTTTTGGACAGTGTTTGGTATTTGTCTTAGAACAACATTTTGCCACGCTATCACTATCACCCAGGAGATAATCCTTTTTTATAAAATGACATCGTTGCCCGGATGCTATATAATCAGTAGCATATTTTAAATCCTTAATATATTCAGGAGTTACCTCGTTCTGATAATAGATTAATGATCCAGGACGAAATTTGAAAGAACTACATGGTTCTCCATGAATTAATACATATTGTTTAGCAAATTCAGGAACTATAAAACTACTACAATGATCTATCGACATACCATCTATCAAACAAAATTTGGGTTTAATTTCTCCTGGAGACGTTTCATAATAATACATATAACTTTCTTCGGCAAACCTAACAGCTCTATTATATTCAGGATAATTAAAATCTAATACCATATATTTGTCTCGTATATCTGCTATTCCTGTCTCTATTTTGATTCTATTAAGAGTAACAGCTGCCCCCATTCTTAATAATCATCAGTATTTAAACTGTTAAATGTTGGTATATCAACATCTATCTTATTTCCCGCAGTATAAGGTTTGTTGCAGGTATACTGTTCAGGAATGGGTACATTTATACTTCTTTTATAGTCCTGTCTTTCGATGTTCATCACAAATGCAAAGAACAGAATAAACAAAATAATGTAAGAAATAATATTAAATATCTGTGAATTCGTAAATACATTGATTGCCATAATAATTACAGCAGCTACAATACACACAATAGACATTCCCACAGTGTTGCCATTACCTCCACGATACATTTGAGTTACTAAGCAATAGGTAATAACTAAGCTAGTAAGAGGCAATAGAAAAGATGAGATAAATATCATCAATATAGAGATTAGAGGAGGGCTATATAGAGCCAAGACGAACAAAATCAAACCGAGTAACGTTCTAACATCATTATTTTTGAAGATTCCCAAATAATCATTCATTATTCCTCCATAATCGTTTTGCATCATACCCCCATCTTTAGGCATAAACGATTGCTGCTGTTCCTCTGTAAATAAATCTTTATCAAGCACTCCAGCACCCGCAGAGAAGTCATCAAGCATATTGTAATATCTTAAATAACTCATTTATATATTAAAAAATGTCACTATTAAAGATGGAGTATAATCTTTATGCCGAACTAAAAAAAATGACTTGTGGTCAGACCATAAGTCTTTTTAATGAAGACGGCGATTTCGTAGAAGTTGAACCAGGATCATCCTTTAAGTTTCTAATACCTAAGGGATTTTACTCCTCTCCTTGTGTAAAGACGAGTCTAGTATTCGAGACATTAACAACGACCGATAATAAAATTACTAGTATCAATCCAACAAATGCGCCAAAGTTATATCCTCTTCAACGCAAAGTCGTATCTGAAGTAGTTTCTAATATGAGGAAAATGATCGAATTAAAACGTCCTCTATACATCACTCTTCACTTGGCATGTGGATTTGGTAAGACTATTACCACGTGTTATCTTATGACCACACACGGCAGAAAAACCATCATTTGCGTACCCAATAAAATGTTAATACATCAATGGAAGACACAGGTAGAGGCAGTCGGATTGGAACATAAGATATCTATAGATGGAGTTAGTAGTCTATTAAAGGAACTAAAGACTCAAAGTCCGGATGTATTAATCGTAGTCAGTAGACATCTGACAAACGATGCATTTTGTAAATATATCAATAAGCATTATGATTTGTTTATCTTGGATGAATCACATACGTATAATCTGATGAACAATACAGCAGTTACAAGATTTTTAGCGTATTATCCTCCGATGATGTGTTATTTTTTAACTGCTACACCTAGACCAGCTAACCGAATTTATTGTAATAGTATTATTAATATTGCCAAGTTATCCGATCTAAAAAAAACTATCTATATAGTAGATAGTTTTTTTGAGCCATATTCCACAGACAATATTAGAAATATGGTAAAACGACTAGATGGACCATCTAATAAATATCATATATATACCGAGAAGTTATTATCTGTAGACGAGCCTAGAAACCAACTTATTCTTGATACCCTGGTAGAAGAATTCAAGTCAGGAACTATTAATAGAATTTTAGTTATTACTAAACTACGTGAACATATGGTATTCTTCTACAAACGATTATTAGATCTTTTCGGAGCAGAGGTTGTATTTATAGGAGACGCCCAAAATAGACGTACTCCAGATATGGTCAAATCGATTAAGGAACTAAATAGATTTATATTCGTATCCACCTTATTTTATTCCGGCACTGGTTTAGATATTCCGAGTTTGGATTCTTTGTTCATTTGCTCGGCAGTAATCAACAATATGCAAATAGAGCAATTACTAGGGAGGGTATGTCGAGAAACAGAACTATTAGATAGGACGGTATATGTATTTCCTAACACATCCATCAAAGAAATAAAGTACATGATAGGAAATTTCGTGCAACGAATTATTAGTCTGTCTGTAGATAAACTCGGATTTAAACAAGAAAGTTATCAGAAACATCAGGAATCTGAACCCGCTTCCGTACCAACATCCTCCAGAGAAGAACGTGTATTAAATAGAATATTTAACTCGCAAAATCGTTAAGAAGTTTAAGAGACGATCCACATGCTGAGCAGGCCAGTGTATTACCCCTCATAGTATTAATATAATCCAATGATACTTTTGTGATGTCGGAAATCTTAACCAATTTAGACTGACAGGCAGAACACGTCATACAATCATCATCGTCATCGATAACTGTAGTCTTGGGCTTCTTTTTGCGACTCTTCATTCCGGAACGCATATTGGTGCTATCCATTTAGGTAGTAAAAAATAAGTCAGAATATGCCCTATAACACGATCGTGCAAAACCTGGTATATCGTCTCTATCTTTATCACAATATAGTGTATCAACATCTTTATTATTGACCTCGTTTATCTTGGAACATGGAATGGGAACATTTTTGTTAACGGCCACCTTTGCCTTAATTCCAGATGTTGTAAAATTATAACTAAACAGTCTATCATCGACACAAATGAAATTCTTGTTTAGACGTTTGTAGTTTACGTATGCGGCTCGTTCTCGTCTCATTTTTTCAGATATTGCAGGTACTATAATATTAAAAATAAGAATGAAATAACATAGGATTAAAAATAAAGTTATCATGACTTCTAGTGCTGATTTAACTAACTTAAAAGAATTACTTAGTCTGTACAAAAGTTTGAGATTTTCAGATTCTGTGGCTATAGAGAAGTATAATTCTTTGGTAGAATGGGGAACATCTACTTACTGGAAAATAGGCGTACAAAAGGTAACTAATGTCGAGACGTCCATATCTGATTATTATGATGAGGTAAAAAATAAACCGTTTAATATTGATCCGGGGTATTATATTTTCTTACCAGTATATTTTGGAAGCGTCTTTATTTATTCAAAGGGTAAAAATATGGTAGAACTTGGATCTGGAAACTCTTTTCAAATACCGGATGAGATTCGAAGTGCGTGTAACAAAGTATTAGATAGTGATAACGGAATAGACTTTCTGAGATTTGTTTTGTTAAACAATAGATGGATAATGGAAGACGCTATATCAAAATACCAGTCTCCAGTTAATATATTTAAACTAGCTAGTGAGTACGGATTAAACATACCCAACTATTTAGAAATTGAAATAGAGGAAGACACATTATTTGACGATGAGTTATACTCTATTATGGAACGCTCTTTCGATGATACATTTCCAAAAATATCTATATCGTATATTAAGTTGGGAGAACTTAAGCGGCAAGTTGTAGACTTTTTCAAATTCTCATTCATGTATATTGAGTCAATCAAGGTAGATCGTATAGGAGATAATATTTTTATTCCTAGCGTTATAACAAAATCAGGAAAAAAGATATTAGTAAAAGATGTAGACCATTTAATACGATCCAAGGTTAGAGAACATACATTTGTAAAAGTAAAAAAGAAAAACACATTTTCCATTTTATACGACTATGATGGGAACGGAACAGAAACTAGAGGAGAAGTAATAAAACGAATTATAGACACTATAGGACGAGACTATTATGTTAATGGAAAGTATTTCTCTAAGGTTGGTATTGCAGGCTTAAAGCAATTGACTAATAAATTAGATATTAATGAGTGTGCAACTGTCGATGAGTTAGTTGATGAGATTAATAAATCCGGAACTGTAAAACGAAAAATAAAAAACCAATCAGTATTTGATTTAAGCAGAGAATGTTTGGGATATCCAGAAGCGGATTTTATAACGTTAGTTAATAACATGCGGTTCAAAATAGAAAATTGTAAGGTTGTAAATTTCAATATTGAAAATACTAATTGTTTAAATAACCCGAGTATTGAAACTATATATGGAAACTTCAACCAGTTCGTCTCAATCTTTAATACCGTTACCGATGTCAAAAAAAGATTATTCGAGTGAAATAATATGCGCCTTTGATATAGGTGCAAAAAATCCTGCCAGAACTGTTTTAGAAGTCAAGGATAACTCCGTTAGGGTATTGGATATATCAAAATTAGACTGGAGTTCTGATTGGGAAAGGCGCATAGCTCAAGATTTGTCACAATATGAATACACTACAGTTCTTCTAGAACGTCAGCCTAGAAGGTCACCGTACGTCAAATTTATCTATTTTATTAAAGGCTTTTTATATCATACATCTGCTGCCAAAGTTATTTGCGTCTCACCTGTCATGTCTGGTAATTCATATAGAGATCGAAAAAAGAGATCTGTTGAAGCATTTCTTGATTGGATGGACACATTCGGATTGCGAGACTCCGTTCCGGATAGACGCAAATTAGACGATGTAGCGGATAGTTTCAATTTGGCTATGAGATACGTATTAGATAAATGGAATACTAATTATACACCTTATAATAGGTGTAAATATAGAAATTACATAAAAAAAATGTAATAACGTTAGTAACGCCATTATGGATAATCTATTTACCTTTCTACATGAAATAGAAGATAGATATGCCAGAACTATTTTTAACTTTCATCTAATAAGTTGTGATGAAATAGGAGATATATATGGTCTTATGAAAGAACGCATTTCCTCAGAGGATATGTTTGACAATATAGTATATAATAAAGATATACATCCTGCCATTAAGAAACTAGTTTATTGCGACATCCAACTTACTAAACATATTATTAATCAGAATACGTATCCGGTATTTAACGATTCTTCACAAGTGAAATGTTGTCATTATTTCGATATAAACTCAAATAATAGCAATATTAGCTCTCGTACAGTAGAGATATTTGAGAGTGAAAAGTCATCTCTTGTATCATATATTAAAACTACCAATAAGAAGAGAAAGGTCAATTACGGCGAAATAAAGAAAACTGTACATGGAGGCACTAATGCAAATTACTTTTCCGGTAAAAAGTCTGATGAGTATCTGAGCACTACAGTCAGGTCCAACATTAATCAACCTTGGATCAAAACCATTTCTAAGAGAATGAGAGTAGATATCATTAATCACTCTATAGTAACGCGTGGAAAAAGCTCTATATTACAAACTATAGAAATTATTTTTACTAATAGAACATGTGTGAAAATATTCAAGGATTCTACTATGCACATTATTCTATCCAAGGACAAGGATGAAAAGGGATGTATAAACATGATTGATAAATTATTCTATGTATATTATAATTTATTTCTGTTGTTCGAGGATATCATCCAAAACGATTACTTTAAAGAAGTAGCTAATGTTGTAAACCATGTACTCATGGCTACGGCATTAGATGAGAAATTATTCCTAATTAAGAAAATGGCTGAACACGATGTTTATGGAGTTAGCAATTTCAAAATAGGGATGTTTAACCTGACATTTATTAAGTCGTTGGATCATACCGTTTTCCCCTCTCTGTTAGATGAGGATAGCAAAATAAAGTTTTTTAAGGGGAAAAAGCTCAATATTGTAGCATTACGATCTCTGGAGGATTGTACAAATTACGTGACTAAATCCGAGAATATGATAGAAATGATGAAGGAAAGATCGACTATTTTAAATAGCATAGATATAGAAACGGAATCGGTAGATCGTCTAAAAGAATTGCTTCTAAAATGAAAAAAAACACTGATTCAGAAATGGATCAACGACTCGGGTATAAGTTTTTGGTGCCTGATCCTAAAGCCGGAGTTTTTTATAGACCGTTACATTTCCAATATGTATCGTATTCTAATTTTATATTGCATCGATTGCATGAAATCTTGACCGTCAAGCGGCCACTCTTATCGTTTAAGAATAATACAGAACGAATTATGATAGAAATTAGCAATGTTAAAGTGACTCCTCCAGATTACTCACCTATAATTGCGAGTATTAAAGGTAAGAGTTATGACGCATTAGCCACGTTCACTGTAAATATCTTTAAAGAGGTAATGACCAAAGAGGGTATATCCATCACTAAAATAAGTAGTTATGAGGGAAAAGATTCTCATTTGATAAAAATTCCGCTACTAATAGGATATGGGAATAAAAATCCACTTGATACAGCCAAGTATCTTGTTCCTAATGTCATAGGTGGAGTCTTTATCAATAAACAATCTGTCGAAAAAGTAGGAATTAATCTAGTAGAAAAGATTACAACATGGCCAAAATTTAGGGTTGTTAAGCCAAACTCATTCACTTTCTCGTTTTCCTCCGTATCCCCTCCTAATGTATTACCGACAAGATATCGCCATTACAAGATATCTCTGGATATATCACAATTGGAAGCGTCGAATATATCATCGACAAAGACATTTATAACGGTCAATATTGTTTTGCTGTCTCAATATTTATCTAGAGTGAGTCTAGAATTCATTAGACGTAGTTTATCATACGATATGCCTCCAGAAGTTGTCTATCTAGTAAACGCGATAATAGATAGTGCTAAACGACTTACCGAATCTATTACTGACTTTAATATTGATACATACATTAATGACCTGGTGGAAGCTGAACACATTAAACAAAAATCTCAGTTAACGATTAACGAGTTTAAATATGAAATGCTGCATAACTTTTTACCTCATATGAACTATACACCCGATCAACTAAAGGGATTTTATATGATATCTTTACTAAGAAAGTTTCTCTACTGTATCTACCACACTTCTAGATATCCAGATAGAGATTCGATGGTTTGTCATCGCATCCTAACGTACGGCAAATATTTTGAGACGTTAGCACATGATGAATTAGAGAATTACATAGGTAACATCCGAAACGATATCATGAACAATCACAAGAACAGAGGCACTTACGCAGTAAACATTCATGTACTAACAACTCCTGGACTTAATCATGCATTTTCTAGTCTATTGAGTGGAAAGTTCAAAAAGTCAGACGGTAGTTATCGAACACATCCTCACTATTCATGGATGCAGAATATTTCTATTCCTAGAAGTGTTGGATTTTATCCGGATCAAGTAAAGATTTCAAAGATGTTTTCTGTCAGAAAATACCATCCAAGCCAATATCTTTACTTTTGTTCATCAGACGTTCCGGAAAGAGGTCCTCAGGTAGGTTTAGTATCTCAATTGTCTGTCTTGAGTTCCATTACAAATATACTAACGTCTGAGTATTTGGATTTGGAAAAGAAAATTTGTGAGTATATCAGATCATATTATAAAGATGATATAAGTTACTTTGAAACAGGATTTCCAATCACTATAGAAAATGCTCTAGTCGCATCTCTTAATCCAAATATGATATGTGATTTTGTAACTGACTTTAGACGTAGAAAACGGATGGGATTCTTCGGTAACTTGGAGGTAGGTATTACTTTAGTTAGGGATCACATGAATGAAATTCGCATTAATATTGGAGCAGGAAGATTAGTCAGACCATTCTTGGTTGTGGATAACGGAGAGCTCATGATGGATGTGTGTCCGGAGTTAGAAAGCAGATTAGACGACATGACATTCTCTGACATTCAGAAAGAGTTTCCACATGTCATCGAAATGGTAGATATAGAACAATTTACTTTTAGTAACGTATGTGAATCGGTTCAAAAATTTAGAATGATGTCAAAGGATGAAAGAAAGCAATACGATTTATGTGACTTTCCTGCCGAATTTAGAGATGGATATGTAGCATCTTCACTAGTGGGAATCAATCACAATTCTGGACCCAGAGCTATTCTTGGATGTGCTCAAGCTAAACAAGCTATCTCTTGTCTGAGTTCGGATATACGAAATAAAATAGACAATGGAATTCATTTGATGTATCCAGAGAGGCCAATTGTGATTAGTAAGGCTTTAGAAACTTCAAAGATTGCGGCTAATTGCTTCGGACAACATGTTACTATAGCATTAATGTCGTACAAAGGTATCAATCAAGAGGATGGAATTATCATCAAAAAACAATTTATTCAGAGAGGCGGTCTCGATATTGTTACAGCCAAGAAACATCAAGTAGAAATTCCATTGGAAAACTTTAATAACAAAGAAAGAGATAGGTCTAACGCCTATTCGAAATTAGAAAGTAATGGATTAGTTAGACTGAATGCTTTCTTGGAATCCGGAGACGCTATGGCAAGAAATATCTCATCAAGAACTCTTGAAGATGATTTTGCTAGAGATAATCAGATTAGCTTTGATGTTTCCGAGAAATATACAGATATGTACAAATCTCGCGTTGAACGAGTACAAGTAGAACTTACTGACAAAGTTAAGGTGCGAGTATTAACCATGAAAGAAAGAAGACCCATTCTAGGAGACAAATTTACTACTAGAACGAGTCAAAAGGGAACAGTCGCGTATATCGCAGATGAAACGGAACTTCCGTACGACGAAAATGGTATCACACCAGATGTCATTATTAATTCTACATCCATCTTCTCTAGAAAAACTATATCTATGTTGATAGAAGTTATTTTAACAGCCGCATATTCTACTAAGCCGTACAACAATAAGGGAGAAAACCGACCTGTCTGTTTTCCTAGTAGTAACGAAACATCTATCGATGCATATATGCAATTCGCTAAACAATGTTATGAGTATTCAAATCCGAAATTGTCCGAGGAAGAATTATCGGATAAAATCTTTTGTGAAAAGATTCTCTATGATCCTGAAACGGATAAGCCTTATGAATCCAAAGTATTTTTTGGACCAATTTATTACTTGCGTCTGAGACATTTAACTCAGGACAAGGCAACCGTTAGATGTAGAGGTAAAAAGACGAAGCTCATTAGACAAGCGAATGAGGGACGAAAACGTGGAGGAGGTATCAAGTTTGGAGAAATGGAGAGAGACTGTTTAATAGCACATGGTGCAGCCAATACTATTACAGAAGTTTTAAAAGACTCAGAAGAGGATTATCAAGATGTGTATATTTGTGAAAATTGTGGAGACATAGCAGCACAAATCAAAAGTATTAATACATGTCTTAGATGTTCAAAACTTAATCTCTCTCCTCTCTTAACAAAAATTGATACCACGCACGTATCTAAAGTATTTCTTACTCAAATGAACGCCAGAGGCGTAAAAGTTAAATTAGATTTCGAACGAAGGCCTCCTTCGTTTTATAAACCATTAGATAAAGTTGATCTTAAACCGTCTTTTCTGGTATAATATTGTTTAGTAGATACTCATCAAGATAAGCTAATTCACTAAACATATTATCGGATTCGGTATTGTTACTCGAGAATAGAGTTCGTTATGCTCCTGATATTCGGAAATCTGTGGAGTTTCAGGTTTTGGTGGAAGTGTAACTGCTACTTGGTGGGATACTGAAGGATATTTCAGAGAGTTGTGGATGTTCGGGTTCGACATCCACCGATGGTGTCACGCCACTAATCGGTTCGGTAACGTCTGTGGATGGAGGTGCTACTTCTACAGAACCTGTAGCCTCAGTTGTCAACGGAGATACATATTCAATGCGCGGAAATGTATAATTTGGTAATGGTTTCTCATGTGGATCTTAAGAAGAAGAGGTAAGATATCTACGAAAGATACCGATCACGTTTCTAGTTCTCTTTTGTAGAACTTTAACTTTTTCTTTCTCAGCATCTAGTTGATATTCCGACCTCTTCACGTTTCGCATGGGTTACCTCCGCAGTTTTTACAAGCGATTTCACGTTCCAGATCACGTTCAGCCTTCATACGTCTCTCCCTCTCTCTATCGAGTTTATCAGAGCAGTCTTTCTGAAGGCGATCGAACTCCATAAATTTCTCCAACGCTTTGATTGTTTCCATAGATTTCCGAAGTTTAGCTTCTAGGACGGCGATTCTTTTTTTTTTTTTTTTTTTTTTTTTCGAATTCACGGGGTACAACCGTTTCCATTACCACCATCTCTATGTTTCTTTTCTAGATCGGCAATCTTTCTCAATCTTTCTCAACATTTCATCCCCATACCTTTTCATTCCTCGAGTCTATTGTCGTCGAAATATCGTTCCAGCTCCTTTTCGACCTCAATAACTTTAGCACGTTGTTTCATCAAGCTCTCTCTTGTAGTACTATCATTTTTATCTGATTCCCTGACACGTTTAAGATCTTCATGTAATTGAGTCAGCTCTTGACGCAATCTCTTAACTAACTTCCTCTCTTGCTTCTTCGTCATAGTACTTACAATCACTATGGGATCCATTGTTACCACGTCTGTACTCGACGAGCTCACGTTTAAGAGATTCAATTTCCAGTTTGTATCGGTCCATGTCTCCATTGCTACACCACCATTAGATTTACAGGCTGCTAGTTGTCGTTCGAGATCAGAAATACGTGTTTTCTTGGAATGGATTTCGTCGATGTACTTGTCATGATTGGCATCGAAACACTTATTAAGTTCTTTTTTTCAATTCTACGATTTTATTTCTTTCGCGAGTCAATTCCCTCCTGTAGTAACTATCAGTTTTGTCAGATTCACGCTCTCTACGTAGACTTTCTTGTAAGTTACTAATTTGTTCCCTGGCATTACCGAGTTCAGTTTTATATGCCGAATAGAGTTCTGATTCATCCTTTGAGAAGATCTCTAGCGATCGTTCAAGATCCCTGATTCTAGTCTTTAGCCTATTTACCTCCTCAGAAGATGCTCCGTTACCGTTTTTACAATCGTTAAGATGTCTATCAAGATCCATGATTCTATCTCTTTTCCATATCAGCATTGATTTCATTATTACGTTCGCAGTCGTTCAACTGTATTTCAAGATCTGAGATTCTAGATTGTAATCTCTGTAGCATTTCCACGGCATTCACTCAGTTGTCTTTCAAGATCTGAGATTCTAGATTGGAGTCTGCTAATCTCTGTAAGATTTCCTCCTCCGCTCTCGATGCAGTCGGTCAACTTATTCTCTAGTTCTCTAATACGCGAACGCAGTGCATCAACTTCTTGTGTGTCTTCTTGATTGCGTGTGCATTCATCGAGTCTAGATTCGAGATCTCTAACGTGACGTCGTTCTTCCTCAAGTTCTCTGTGTACTACAGAAAGCGTGTCCCTATCTTGTTGATATTTAGCAATTTCTGATTCTAGAGTACTGATTCTACTCACGTATGTACTAATAGTTGTCTTAGCCTTATCAAGATCCTCCTTGTATTTGTCACATTCCTTGATATCCATACGAAGTCTGGACAGTTCCCATTCGACATTACGACGTTTATCGATTTCAGCTCGGAGATCGTCGTCGCGTTGTTTTAGCCACATACGACTAAGTTCAAGTTCTCGTTGACAAGATCCATCTACTTTTCCATCCCTAATAGTATCCAGTTCCTTTTCTAGTTCTGACCGCATTTCTCGTTCCATATCAAGAGATTCTCTCAATTCTCGTATAGTCTTCTTATCAATTTCTGATGAATCTGAACCATCATCTGTCCCATTTTGTTGCATATCCCTGAGTTCTTTGATCTCTGTTGTAAGTCTGTCGATTCTTTCGGTTTTATAAACAGAATCCCTTTCCAAAGTCCTAATCTTACTGAGTTTATCATTAAGTTCTTCATTCAATTCAGTGAGTTTTCTCTTGGCTTCTTCCAAGTCTGTTTTAAACTCTCCATCATTTCCGCATTCTTCCTCGCATTTATCTAACCATTCAATTAGTTTATTAATAACTAGTTGGTAATCAGCGATTCCTATAGCCGTTCTTGTATTTGTGGGAACATAATTAGGATCTTCTAATGGATTGTATGGCTTGATAGCATCATCTTTATCATTATTAGGTGGGGGATGGACAACCTTAATTGGTTGGTCCTCCTTATCTCCTCCAGTAGCATGTGGTTCTTCAATACCAGTATTAGTAATAGGCTTAGACAAATGCTTGTCGTACGCGGGCACTTCCTCATCCATCAAGTATTTATAATCGGGTTCTGTTTCAGAATATTCTTTTCTAAGAGACGCGACTTCAGGAGTTAGTAGAAGAACTCTGTTTCTGTATCTATCAACGCTGGAATCGATACTCAAGTTAAGGATAGCGAATACCTCATCGTCATCATCCGTATCTTCTGAAACGCCATCATATGACATTTCATGAAGTCTAACGTATTGATAAACAGAATCAGATTTAGTATTAAACAGATCCTTGACCTTTTTAGTAAATGCATATGTATATTTTAGATCTCCAGATTTCATAATATGATCGCATGCCTTAAATGTCAATGCTTCCATGATATAGTCTGGAACACTAATGGGTGACGAAAAAGATACAGCACCATATGCTACGTTGATAAATAGATCTGAACCACTAAGTAGATAATGATTAATGTTAAGGAAGAGGAAATATTCAGTATATAGATATGCCTTAGCATCATATCTTGTACTAAACACGCTAAACAGTTTATTGATGTGATCAATTTCCAACAGAACAATTAGAGCGGCAGGAATACCAACAAACATATTACCACATCCGTATTTTCTATGAATATCACATATCATATTAAAAAATCTTGATAGAAGAGCGAATATCTCGTCTGACTTAATGAGATGTAGTTCAGCAGCATAAGTCATAACTGTAAATAGAACATACTTTCCTGTAGTGTTGATTCTAGACTCCACATCAACACCATTATTAAAAATAGTTTTATATACATCTTTAATCTGCTCTCCGTTAATCGTCGAACGTTCTAGTATACGGAAACACTTTGATTTCTTATCTGTAGTTAATGACTTAGTGATATCACGAAGAATATTACGAATTACATTTCTTGTTTTTCTTGAGAGACCTGATTCAGAACTCAACTCATCGTTCCATAGTTTTTCTACCTCAGTGGCGAAATCTTTGGAGTGTTTGGTACATTTTTTAATAAGGTTCGTGACCTCCATTTATTATAAAAAATTTTTATTCAAAACTTAACTACAATCGGGTAATTATAAGATCGTAGATCTCCCATGTGGTGGAATACTACCATCTATCGCATGTTGATGGACAGTAGGTAATGGCCATGGGAACAGTAATGTTTGCATATTTATCTTTCTTGCTAGTATTACTGTATATTGTCCCAATGTTTCAATGTGATGTTCTAACCTATCAACTGCCACTGTATCACAACAATAATGTCCGATGGAATTAAGATTATGATCCAATGTGTTTAATATATGATTATCAAGTCTTATACGATCCGCGTCTTTTTTGACAGGATCAGGCTCTTCTACAGGAAGAAGTTTCGGCCTCTTATGATAGTCATGTCTGGGAAATGGTGGTCTAGGATGAGGATCAGGTATCGGAGTAGGTTTTGGATTATAATCATCATCATCATCATCATCATCATCATCATCATCATCATCTATGATATCATCATCTTCGATATTTATTTTGCTATCTTGATAATGTCCTATATCAGTTGCATTTTCAGCACTCGACTGAATATTAGTACATTCATTGTCTATTATTAACGTATTTCTAAACCCAAAATGTATATGTTGAACATCACTACTATAGTTGATGAGTCTTATAGCATGAATTCGCTTATCGTTATCGGGTTTATCTTCTGTCACCTTAACAATTCCTTTTTTATTAAACTCTGCATAATCATAACCATTTCTATTGTTTGTTCTAATATAAACGAGTATAGCATCATTGCTAAATTTTTCAATAGTATCAAAAACAGAATATCCTAAACCATATAATATATATTCAGGAACACTCAAACTAAATGTCCAGGATTCTCCTAAATACGTAAACTTTAATAGTGCTAAATCATTCAAAAATCTACCGCTTATAGATAGATAGTACATGAATGCGTATAGTAGTCTACCTATCTCTTTATTATGAAAACCGACATTACGATCATATATTTCGTGATATACATGTGACCCGTTTACGTTAAACCATAAATACATGGGTGATCCTATAAACATGAATTTATTTCTAATTCTCAGAGCCATAGTTAATTGACCGTGTAATATTTGTTTACATGCATACTTGATACGATCATTAATAAGATTTTTATCATTGCTCGTTATTTCAGAATCGTATATATAAGGAGTACCATCATGATTCTTACCAGATATTATACAAAATACTATATATAAAATATATTGACCCACGTTAGTAATCATGTAAATGTTTAATGTTTTAAATTTTGTATTTAATGATCCATCATCATATGCTAGCATGGTCTTGTGATATTCATTCTTTAAAATATAATATTGTGTTAGCCATTGCATTGGAGCTCCTAATGGAGATTTTCTATTCTCGTCCATTTTAGGATATGCTTTCATAAAGTCCCTAATAACTTCGTGAATAATGTTTCTATGTTTTCTACTGATGCATGTATTTGCTTCGATTTTTTTATCCCATGTTTCATCTATCATAGATTTAAACGCAGTAATGCTCGCAACATTAACATCTTGAACCATTGGTACAATTCCGTTCCATAAATTTATAATGTTCGCCATTTATATAACTCATTTTTTGAATATACTTTTAATTGAACAAAAGAGTTAAGTTACTCATATGGATGCCGTCCAGTCTGTACATCAATCTTTTTAGCCAGAGATATCATAGCCGCTCTTAGAGTTTCAGCGTGATTTTCCAACCTAAATAGAACTTCATCGTTGCGTTTACAACACTTTTCTATTTGTTCAAACTTTGTTGTTATATTAGTAATCTTTTTTTCCAAATTAGTTAGCCGTTGTTTGAGAGTTTCCTCATTATCGTCTCCATAGGCTTTAACAATTGCTTCGCGTTTAGTCTCTGGATTTTTAGCAGCCTTTGTAGAGAAAAATTCAGTTGCTGGAATTGCAAGATCGTCATCTCCGGGGAAAAGAGTTCCGTCCATTTAAAGTACAGATTTTAGAAACTGACACTCTGTGTTATTTATATTTGGCGCAATACATGGATTATAAATATCGATGTTAATAACATCAGAAAATGTAAAGTCTATACATTGTCGCATCGTGTTAAATTTTCTAATGGATCTAGTATTATTGGGTCCAACTTCTGCCTGAAATCCAAATATGGAAGCGGATACAAAACCGTTTCCTGGATAAACCACACATCTCCACTTTTGCTTTACATCAGAAATTGTGTCATTGACATCTTGAACTCTCCTATCTAATGCCGGTGTTCCACCTATAGATTTTGAATACTCGAATGCTGCATGAGTAGCATTGAATTCCTTAATATTGCCATAATTTTCATATATTGAGTAACTCTGGATAAAAAGTAAACACACCGCAGCCGTCGCTACTACAATAAAAAAAATTGATAGAGAGTTCATTTATAATCTATTAGAAGCTGATAAAATTTTTTTACACGCGTCAGACAATGCTTTAATAAATAGTTCAACATCTACTTTTGTCATATCGAACCGATGGTATGATTCTAACCTAGAATTACATCCGAAAAAGTTGACTATGTTCATAGTCATTAAGTCATTAACGAACAACATTCCAGACTCTGGATTATAAGACGATACTGTTTCGTCACAATCACCCACCTTAATCATGTGATTATGAATATTGGCTATTAGAGTACCTTCTAAGAAATCTATAATATCTTTGAAACACGATTTAAAATCAAACCACGAATATACTTCTACGAAGAAAGTTAGTTTACCCATAGGAGAGATAACTATAAATGGAGATCTAGATACAAAATCCGGATCTATGATAGTTTTAACATTATTATATTCTCTATTAAATACCTCCACATCTAAAAATGTTAATTTTGAAACTATGTCTTCGTTTATTACCGTACCTGAACTAAACGCTATAAGCTCTATTGTTTGAGAACTCTTTAAACGATATTCTTGAAATACATGTAACAAAGTTTCCTTTAACTCGGTCGGTTTATCTACCATAGTTACAGAATTTGTATCCTTATCTATAATATAATAATCAAAATCGTATAAAGTTATATAATTATCGTGTTCAGATTGTGATCTTTTCAAATAGACTAAAAACCCCATTTCTCTAGTAAGTATCTTATGTATATGTTTGTAAAATATCTTCATGGTGGGAATATGCTCTACAGCAGTTAGCCATTCCTCATTGACAGCTGTAGATGTATTATACAAAACTACTCCAATGTTTAACAAGGGCCATTTTACGAGATTATTAAATCCTTGTTTGATAAATGTAGCCAATGCGGGTTCGAGTTCAACGACGATTGAATTCTCTTCCCGTGGATGCTGCATGATGAACGACGGGATGTTGTTGTTCTATTGATTTGGAATTCTTTTTCGACTTTTTGTTTATATTAAATATTTTAAAATTTATGGCTGATAGTAATTCATGTACTACGGATAATGTAGACGTGTATTGCATATCGATATCTTTATTATTAGATAAATTTATCAATAAATGTGAGAAGTTTGCCTCGTTAAGGTCTTCCATTTAAATATTATATAAATATTTGTGTTTGTATTTTATTCGTCTTTTATGGGATAGTTTTTAACTAGTAAAGCTGTAATTACATACTTTGTCCGTAAAACATAAATATAAATACCCGCTTTTATCAAACGTTCCAAAAAGTCGGCAGCTGACATTTTTAACATGACATCTATTTTAAATACACTTAGGTTTTTAGAAAAAACATCATTTTATAATTGTAACGATTCAATAACTAAAGAAAAGATTAAGATTAAACATAAGGGAATGTTATTTGTATTTTATAAGCCAAAGCATTCTACCGTTGTTAAATACTTGTCTGGAGGAGGTATATATCATGATGATTTGGTTGTATTGGGGAAGGTAACAATTAATGATCTAAAGATGATGCTATTTTACATGGATTTATCATATCATGGAGTGACAAGTAGTGGAGCAATTTACAAATTGGGATCGTCTATCGATAGACTTTCTCTAAATAGGACTATTGTTACAAAAGTTAATAACAATTATAACAATTATAACAATTATAACAATTATAATTGTTATAATAATTATAATTGTTATAATTATGATGATACATTTTTTGACGATGATGATTGATCACTATTACACAATTTTGTTTTTGTACTTTCTAATATAGTGTTTAGGTTCTTTTTCATATGAGAATATTGACTTACTAAAATATCTATGTTTAACTTTTGTTCTATAACGTCCTTATCGGCGGTATCGGTACATATACGTAATTCACCTTCACAAAATACGGAGTCTTCGATAATAATAGCCAATCGATTATTGGATCTAGCTGTCTGTATCATATTCAACATGTTTAATATATCCTTTCGTTTCCCCTTTACAGGCATCGATCGTAGCATATTTTCCGCGTCTGAGATGGAAATGTTAAAACTGCAAAAATGCGTAATGTTAGCCCGTCCTAATATTGGTACGTGTCTATAAGTTTGGCATAGTAGAATAATAGACGTGTTTAAATGCCTTCCAAAGTTTAAGAATTCTATTAGAGTATTACATTTTGATAGTTTATCACCTACATCATCAAAAATAAGTAAAAAGTGTGCTGATTTTTTATGATTTTGTGCGACAGCAATACATTTTTCTATGTTACTTTTAGTTCGTATCAGATTATATTCTAGAGCTTCCTGACTACTAACGAAATTAATATGATTTGGCCAAATGTATCCATCATAATCTGGGTTATAAACGGGTGTAAACAAGAATATATGTTTATATTTTTTAACTAGTGTAGAAAACAGAGATAGTAAATAGATAGTTTTTCCAGATCCAGATCCTCCTGTTAAAACCATTCTAAACGGCATTTTTAATAAATTTTCTCTTGAAAATTGTTTTTCTTGAAAACAATTCATAATTATATTTACAGTTACTAAATTAATTTGATAATAAATCAAAATATGGAAAACTAAGGTCGTTAGTAGGGAGGAGAACAACGAAGGCATATCGTGATATAAATAACATTTATTATCATGATGACACCAGAAAACGACGAAGAGCAGACATCTGTGTTCTCCGCTACTGTTTACGGAGACAAAATTCAGGGAAAGAATAAACGCAAACGCGTGATTGGTCTATGTATTAGAATATCTATGGTTATTTCACTACTATCTATGATTACCATGTCCGCGTTTCTCATAGTGCGCCTAAATCAATGCATGTCTGCTAACAAGGCTGCTATTACTGACTCCGCTGTTGCCGTTGCTGCGGCATCATCTACTCATAGAAAGGTTGTGTCTAGCACTACACAATATGATCACAAAGAAAGCTGTAATGGTTTATATTACCAGGGTTCTTGTTATATATTACATTCAGACTATAAGTCATTCGAGGATGCTAAAGCAAACTGCGCTGCGGAATCATCAACACTACCCAATAAATCCGATGTCTTGACTACCTGGCTCATTGATTATGTTGAGGATACATGGGGATCTGATGGTAATCCAATTACAAAAACTACATCCGATTATCAAGATTCTGATGTATCACAAGAAGTTAGAAAGTATTTTTGTACATAAATAAATGAAATCGCTTAATAGACAAACTGTAAGTAGGTTTAGGAAGTTGTCGGTGCCGGCCGCTATAATGATGTTACTCTCAACCATTATTAGCGGCATAGGAACATTTCTGCATTACAGAGAAGAACTGATGCCTAGTGCTTGCGCCAATGGATGGATACAATACGATAAACATTGTTATCTGGATACCAACATTAAAATGTCTACGGATAATGCAGTTTATCAGTGTCGCAAATTACGAGCTAGATTGCCTAGACCTGATACTAGACATCTGAGAGTATTGTTTAGTATTTTTTATAAAGATTATTGGGTAAGTTTAAAAAAGACCAATGATAAATGGTTAGATATTAATAATGATAAAGATATAGATATTAGTAAATTAACAAATTTTAAGCAACTAAACAGCACGACGGATTCTGAGGCGTGTTATATATACAAGTCTGGAAAACTGGTTAAAACAGTATGTAAAAGTACTCAATCTGTACTATGCGTTAAAAGATTCTACAAGTGACAACAAAAAATGAATTAATAGTAAGTCGTTAACGTACGCCGCCATGGACGCCGCGTTTGTTATTACTCCAATGGGTGTGTTGACTATAACAGATACATTGTATGATGATCTCGATATCTCAATCATGGACTTTATAGGACCATACATTATAGGTAACATAAAAATTGTCCAAATAGATGTACGGGATATAAAATATTCCGACATGCAAAAATGCTACTTTAGCTATAAGGGTAAAATAGTTCCTCAGGATTCTAATGATTTGGCTAGATTCAACATTTATAGTATTTGTACAGCATACAGATCAAAAAATACCATCATCATAGCATGCGACTATGATATCATGTTAGATATAGAAGGTAAACATCAACCATTTTATCTATTCCCATCTATTGATGTTTTTAACGCTACAATCATAGAAGCGTATAATCTGTATACAGCTGGAGATTATCATCTGATCATCAATCCTTCAGATAATCTGAAAATGAAATTGTCGTTTAATTCTTCATTTTGTATATCAGACGGCAATGGATGGATTATAATTGATGGGAAATGTAATAGTAATTTTTTATCATAAAAGTTGTAAAGTAAATAATAAAACAATAAATATTGAACTAGTAGTATGTTGTATATTGAGCAATCAGAGATGATGCTGGTACCTCTTATCACGGTGACCGTAGTTGCGGGAACAATATTAGTATGTTATATATTATATATTTGTAGGAAAAAGATACGTACTGTCTATAATGACAATAAAATTATCATGACAAAATTAAAAAAGATAAAGAGTCCTAATTCCAGCAAATCTAGTAAATCAACTGATAGCGAATCAGACTGGGAGGATCACTGTAGTGCTATGGAACAAAACAATGACGTAGATAATATTTCTAGAAATGAGATATTGAACGATGATAGCTTCGCTGGTAGTTTAATATGGGATAACGAATCCAATATCATGGCGCCTAGCACAGAACACATTTACGATAGTGTTGCTGGAAGCACGCTGCTAATAAATAATGATCGTAATGAACAGACTATTTATCAGAATACTACAGTAGTAATTAATGATACAGAGACTGTTGAAATACTTAATGAAGATACCAAACAGATTCCTAGCTATTCTTCCAATCCTTTCGTAAATTATAATAAAACCAGTATTTGTAGCAAGTCAAATCCGTTCATTGCAGAACTCAACAATAAATTTAGTGATAATAATCCGTTTAGGAGAGCACATAGTGACGATTATCTTAATAAGCAACAAGATCATGAATACGATGATATAGAATCATCGGTTGTATCATTGGTCTGATTAGTTTCCTTTTTATAAAATTGAAGTAATATTTAGTATTAATTACCGCCGATGCATTATACAAATATGGAGATATTCCCTGTATTCGGCATTTCTAAAATTAGCAATTTTATTGCTAATAATGACTGTAGATATTATATAGATGTAGAGCATCAAAAAATTATATCTGATGAGATCAATAGACAGATGGATGAAACGGTACTTCTTACCAACATCTTAAGCGTAGAAGTTGTAAATGACAATGAGATGTACCATCTTATTCCCCATAGACTATCGACTATTATACTCTGTATTAGTTCTGTTGGAGGATGTGTTATCTCTATAGATAATGACGTCAATGACAAAAATATTCTAACATTTCCCATTGATCATGCTGTAATCATATCCCCACTGAGTAAATGTGTCGTAGTTAGCAAGGGCCCTACAACCATACTGGTTGTTAAAGCGGATATACCCAGCAAACGATTGGTAACATCGTTTACAAACGACATACTGTATGTAAACAATCTATCACTGATTAATTATTTACCGTCGTCTGTATTCATTATTAGACGAGTCACCGACTATTTGGATAGACACATATGTGATCAGATATTTGCTAATAATAAGTGGTATTCCATTATAACTATCGACGATAAGCAATATCCTATTCCATCAAATTGTATAGGTATGTCTTCTGCCAAGTACATAAATTCGAGCATCGAGCAAGATATTTTGATCCATGTTTGTAACCTCGAGCATCCATTCGACTCAGTCTACAAAAAAATGCAGTCGTACAATTCTCTACCTATCAAGGAACAAATATTGTATGGTAGAATTGATAATATAAATATGAGCATTAGTATTTCGGTGGATTAATAGATTTCTCTAGTATGGGATCATTAATCATCTCTAAATACATCATAAAAAAGCTATTATCAAATACTGTACTGAATGGATTCATTCTTTTCTCTTTTTATGAAACTCTGTTGTATATCTACGGATAAAACTAGAAGCAAAAAATCTGATAGGAAGAATAATGATTATATGGAGGAACACGATTATTATAAAATAACAATAGTTCCTGGTTCCTCTTCCACGTCTACTAGCTCGTGGTATTATACACATGCCTAGTAATAGTCTCTTTGCGTTGACGGAAAGCAGACTAGAAATAACAGGCCAAAATGTTCAGACACCATAATAGTTCCCAACCCAGATAATAACAGAGTTCCATCAACACATTCCTTTAAACTCAATCCCAAACCCAAAACCGTTAAAATGTATCCAGCCAATTGATAGTAGATAATGAGGTGTACAGCACATGATAATTTACACAGTAACCAAAATGAAAACACTTTAGTAATTATAAGAAATATAGACGGTAATGTCATCATCAACAATCCAATAATATGCCTGAGAGTAAACATTGACGGATAAAACAAAAATGCCCCGCATAACTCTATCATGGCAATAACGCAACCAAACACTTGTAAAATTCCTAAATTAGTAGAAAATACAACTGATATCGATGTATAAGCGATTTCGAGGAATAATAAGAACAAAGTAATTCCCGTAAAGATAAACATCAACATTGTTTGGTAATCATTAAACCAATTAGTATGACGTTGAATTAATTTCACAGTATATTTTATTCCAGTATTATCCCCGCATGTATACGTACCTGGTAAGATATCTTTATATTCCATAATCAATGAGACATCACTATCCGATAACGAATGAAGTCTAGCACTAGTATGCCATTTACTTAATATGGTCGTCTTGGAAGTTTTATTATAAGTTAAAATATCATGATTGTCCAATTTCCATCTAATATACTTTGTCGGATTATCTATAATACACGGAATAATGATGGTATCATTACATGCTGTATATTCTATAGTCTTTGTAGATGTTATAACCACAAAAGTACAGAGGTATATCAACAATATTCTAACTCTTAACATTTTTATTTATTTAAAATGATACCTTTGTTATTTATTTTATTCTTATTTTGCTAACGGTATCGAATGGCATAAGTTTGAAACGAGTGAAGAAATAATTTCTACTTACTTAATAGATTATGTGGTAACGGGTGTTATTAATGGGGATGTATATACATTTTCAAATAATGAACTAAACAAAACTGGGTTAACTAATAACAATAATTATATCACAACATCTATAAAAGTAGAGGATAAGGATACATTAGTAGTATGCGGAACCAATAACGGAAATCCCAAATGTTGGAAAATAGACGGTTCATACTACCCAAAACATATAGGTAGAGGATACGATCATCAAAATAGCAAAGTAACGATAATCAGTCACAATGAATGTGTACTATCCGACATAAACATATAAAAAGAAGGAATTAAACGATGGAGAAGATTTGACGGACCATGTGGTTATGATTAAACGAGTTAAGTTTTTTAAGAAGCCTTAGAAGAGAGGCTATTGGGTATGAGAATCCGAAATATTAAACCAGACAACCCCATATAATTTTATAGCTAAGAATGCCGCGAAGAATGGAACTAATAAAAACGGAAATATTTGTAGCACAACGAATAACTCCCAAACTGCATTCATGTTACACTATATAACACTACTTCGGTTAGATGTTTTAGAAAAAATAAATATCACCGTACCGTTTTGTTGTATAAAAATAACAATTAACAATTATCAATTTTTTTCTTTAATATTTTACGTGGTTGACCATTCTTGGTGGTAAAATAATCTCTTAGTGTTGGAATGGAATGCTGTTTAATGTTTCCACACTCATCGTATATTTTGACGTATGCAGTCACATCGTTTACGCAATAGTCAGACTGTAGTTCTATCATGCTTCCTACGTTAGAAGGAGGAACAGTTTTAAAGTCTCTTGGTTTTAATCTATTGTCATTAGTTTTCATGAAATCCTTTGTTTTATCCACTTCACATTTTAAATAAATGTCCACTATACATTCTTCTGTTAATTTTACTAGATCATCATGAGTCATAGAATTCATAGGTTCCGTAGTCCATGGATCCAAACTAGCAAACTTCGCGTATACGGTATCGCGATTAGTGTATACACCAACTGTATGAAAATTAAGAAAACAGTTTAATAAATCTACAGAAATATTTAATCCTCCGTTTGATACAGATGCGCCATATTTATGGATTTCGGATTCACACGTTGTTTGTCTAAGGGGTTCGTCTAGTGTTGCTTCTACATAGACTTCGATTCCCATATATTCTTTATTGCCAGAATCACATACCGATTTATCATACGCTGGTTCACTTGTTTGAAAACTAAATGGTAGTAGATACATCAAAATAATAAATAATAAGTACATTCTGCAATATTGTTATCGTAATTGGAAAATTGGTATTCAAGTGAGCTGGATTATGTGAGTATTGGATTGTATATTTTATTTTATATTTTATATTTTATATTTTATTTTATATTTTATATTTTATTTTATATTTTGTAGTAAGAATAGAATGCTAATGTCAAGTTTATTCGAATAGATGTCTTATTAAAAAACATATATAATAAATAACAATGGCTGAATGGCATAAAATTATCGAGGATATCTCAAAAAATAATAAGTTCGAGGATGCCGCCATCGTTGATTACAAGACTACAAAGAATGTTCTAGCGGCTATTCCTAACAGAACATTTGCAAAGATTAATCCGGGTGAAGTTATTCCCCTCATCACTAATCATAATATTCTAAAACCTCTTATTGGTCAGAAATTTTGTATTGTATATACTAACTCTCTAATGGATGAGAACACGTATGCTATGGAGTTGCTTACTGGGTACGCCCCTGTATCTCCGATCGTTATAGCGAGAACTCATACCGCACTTATATTTTTGATGGGTAAGCCAACAACATCCAGACGTGATGTGTATAGAACATGTAGAGATCACGCTACCCGTGTACGTGCAACTGGTAATTAAAATAAAAAGTAATATTCATATGTAGTGTCAATTTTAAATGATGATGAAATGGATAATATCCATATTGACGATGTCAATAATGCCGGTATTGACATACAGCTCATCGATTTTTAGATTTCATTCAGAGGATATTGAATTATGTTATGGGAATTTGTATTTTGATAGGATCTATAATAATGTAGTAAATATAAAATATATTCCTGAGCATATTCCATATAGATATAATTTTATTAATCGTACGTTCTCCGTAGATGAACTAGATGATAATGTCTTTTTTACACATGGTTATTTTTTAAAACACAAATATGGTTGTTCACTTAATCCTAGTTTGATTGTCTCATTATCAGGAAACTTAAAATATAATGATATACAATGCTCAGTAAATGTATCGTGTCTCATTAAAAATTTGGCAACGAGTACATCTACTATATTAACATCTAAACATAAGACTTATTCTCTATATCGGTCCATGTGTATTGCTATAATAGGATACGATTCTATTATATGGTATAAATATATAAATGACAGGTATAATGACATCTATGATTTTACTGCAATATGTATGCTAATAGCGTCTACATTGATAGTGATCATATACGTGTTTAAAAAAATAAAAATGAACTCTTAATTATGTTATACTATTAGAAATGGATAAAATCAAAATTACGATTGATTCAAAAATTGGTAATGTTGTTACCATATCGTATAACTTGGAAAAGATAACTATTGATGTCACGCCAAAAAAGAAAAAAGAAAAGGATGTATTATTAGCGCAATCAGTTGCTGTCGAAGAGGCAAAAGATGTCAAGGTGGAAGAAAAAAATATTATCGATATTGAAGATGACGATGATATGGATATAGAAAACACGTAATACGATCTATAAAAATAAGTATTAAATACTTTTTATTTACGGTACTCTTGTAGTGGTGATACCACTAATCGATTATTTTTTTTAAAAAAATACTTATTCTGATTCTTCTAGCCATTTCCGTGTTCGTTCGAATGCCACATCGACGTCAAAGATAGGGGAGTAGTTGAAATCTAGTTCTGCATTGTTGGTACGCACCTCAAATGTAGTGTTGGATATCTTCAACGTATAGTTGTTGAGTATTGATGGTTTTCTAAATAGAATTCTCTTCATATCATTCTTGCACGCGTACATTTTTAGCATCCATCTTGGAATCCTAGATCCTTGTTCTATTCCCAATGGTTTCATCAATAGAAGATTAAACATATCGTAAGAACACGATGGAGAGTAATCGTAGCAAAAGTAAGCATTTCCTTTAATCGCAGATCCCGGATACTGGATATATTTTGCAGCCAACACGTGCATCCATGCAACATTTCCTACATATACCCGGCTATGCACAGCGTCATCATCGACTGTACGATACATAATGTTACCGTGTTGCTTACATTGCTCGTAAAAGACTTTCGTCAATTTGTCTCCTTCTCCGTAAATTCCAGTGGGTCTTAGGCAACAAGTATACAATTTTGCGCCATTCATGATTACGGAATTATTGGCTTTCATAACCAGTTGCTCGGCCATACGTTTACTTTTTGCGTATACATGTCCTGGTGATATATCATAAAGGGTATGCTCATGACCGATGAATGGATTACCGTGTTTATTTGGTCCTATTGCTTCCATGCTACTAGTATAGATCAAATACTTGATTCCTAGGTCCACACAAGCTGCCAATATAGTCTGTGTTCCATAATAGTTTACTTTCATGATTTCATTATCAGTGTATTTTCCAAATACATCCACTAGAGCAGCCGTATGAATAATCAGATTTACCCCATCTAGCGCTTCTCTCACCTTATCAAAGTCGTTTATATCACATTGTATATAGTTTATAACCTTAACTTTCGAGGTTATTGGTTGTGGATCTTCTACAATATCTATGACTCTTATTTCTTGAACATCATCTGCGCTAATTAAAAGTTTTACTATATACCTGCCTAGAAATCCGGCACCGCCAGTAACCGCGTACACGGCCATTGCTGCCACTCATAATATCAGACTACTTATTCTATTTTACTAAATAATGGCTGTTTGTATAATAGACCACGATAATATCAGAGGAGTTATTTACGTTGAACAAGTCCATGGAAAAGATAAAGTTTTAGGATCAGTTATTGGATTAAAATCCGGAACGTATAGTTTGATAATTCATCGTTACGGAGATATTAGTCGAGGATGTGATTCCATAGGCAGTCCAGAAATATTTATCGGTAACATCTTTGTAAACAGATATGGTGTAGCATATGTTTATTTAGATACAGATGTAAATATATCTACAATTATTGGAAAGGCGTTATCTATTTCAAAAAATGATCAGAGATTAGCGTGTGGAGTTATTGGTATTTCGTACATAAATGAAAAGATAATACATTTTCTTACAATTAACGAGAATGGCGTTTGATATATCAGTTAATGCGTCTAAAACAATAAATGCATTAGTTTACTTTTCTACTCAGCAAGATAAATTAGTCATACGTAATGAAGTTAATGATATACACTACACTGTCGAATTTGATAGGGACAAAGTAGTTGATACGTTTATTTCATATAATAGACATAATGACTCCATAGAGATAAGAGGGGTGCTTCCAGAGGAAACTAATATTGGTCGCGTGGTTAATACGCCGGTTAGTATGACTTACTTGTATAATAAGTATAGTTTTAAACCGATTTTAGCAGAATATATAAGACACAGAAATACTATATCCGGCAACATTTATTCGGCATTGATGACGCTAGATGATTTGGTTATTAAACAGTATGGAGACATTGATCTATTATTTAATGAGAAACTTAAAGTAGACTCCGATTCGGGACTATTTGACTTTGTCAACTTTGTAAAGGATATGATATGTTGTGATTCTAGAATAGTAGTAGCTCTATCTAGTCTAGTATCTAAACATTGGGAATTGACAAATAAAAAGTATAGGTGTATGGCATTAGCCGAACATATAGCTGATAGTATTCCAATATCTGAGCTATCTAGACTACGATACAATCTATGTAAGTATCTACGCGGACACACTGATAGCATAGAGGATGAATTTGATCATTTTGAAGACGATGATTTGTCTACATGTTCTGCCGTAACCGATAGGGAAACGGATGTATAATTTTTTTTATAGTATGAAGGATATGATGGATATGATGATATGATGGATATGATGATATGATGGATATGATGGATATGATGGATATGATGGATATGATAAAAAAATATAATTGTTGTATCCATTCCCATTCAAATCACCTTATATGATTCTGTAACACAATGAAGGAGTCTCATAGATATATAGAGGTCAGATACTGGTTTGATAAACTTTTTATTCCACATGAGCATGTTTGACTTATGGTTAGACACACATACTTTAACAAATCACTGAAAATTGGAGTTAGGTATTCCTCTCAGAATCAGTTGCCGTTCTGGAACATTAAATGTATTTTTTATGATATACTCCAACGCATTTATGTGGGTATACAACAAGTCATTAATAATGAGTATTTCCAAGAGTTTTAGTTGTCTAGTATTTAACAAGAGAAGAGATTTCAACAGACTGTTTATGAACTCGAATACCGCCTCATTGTCGCTTATATTGATGACATGACATCGAATTCCCAATATCAATCTCATCAGTGATGAGTAGCTCAATCTTGTTATCGGGATCCAATTTCTAAAGATGTCATTAAACCCTCGATCGTGAATGGATTTATCATCATCGTTTTTATGTTGGACATGAGCTTAGTCCGTTTGTCCACATCTATATACGATGATTTCTGAATTATTTCATATATCTCTCGTTAACTCCAGGAACTTGTCAGGGATCTAACTTTAATATGTTCTCGTCTAAGAGATGAAAATCTTTGGATGGTTGCATGTGACTTTTCTCTAAAGGATGATGTTACCCGATCCTCTCTTAAATGACTCCATCTTATCCTTGGACAAGATGGACAGTCTATTTTCCTTAGATGGTTTAATATTTTTTACCCATGATCTATAAAGGTAGACAGACCTAATCGTCTCGGATGACCATATATTATTTTCAGTTTTATTATACGCATAAATTGTAAAAAATATGTTAGGTTTACGAAAATGTCTCGTGGGGCATTAATCGTTTTTGAAGGATTGGACAAATCTGGAAAAACAACACAATGTATGAACATCATGGAATCTATACCGGCAAACACGATAAAATATCTTAACTTTCCTCAGCGATCCACAGTCACTGGAAAGATGATAGATGACTATCTAACTCGTAAAAAAACCTATAATGATCATATAGTTAATCTATTATTTTGTGCAAATAGATGGGAGTTTGCATCTTTTATACAAGAACAACTAGAACAGGGAATTACTTTAATAGTTGACAGATACGCGTTCTCTGGAGTAGCGTATGCCACCGCTAAAGGCGCGTCAATGACTCTCAGTAAGAGTTATGAATCTGGATTGCCTAAACCCGACTTAGTTATATTCTTGGAATCTGGTAGCAAAGAAATTAATAGAAACATCGGCGAGGAAATTTATGAAGATGTTGAATTCCAACAAAAGGTATTACAAGAATATAAAAAAATGATTGAAGAAGGAGATATTCATTGGCAAATTATTTCTTCTGAATTCGAGGAAGATGTAAAGAAGGAGTTGATTAAGAATATAGTTATAGAGGCTATACACACGGTTACTGGACCAGTGGGGCAACTGTGGATGTAATAAAATGAAATTACATTTTTATAAATAGATGTTAGTACAGTGTTATAAATGGATGAAGCATATTACTCTGGCAACTTGGAATCAGTACTCGGGGATACGTGTCCGATATGCATACCGAACTCGCATCAATATCTCAATTAGTTATTGCCAAGATAGAAACTATAGATAATGATTATTAAACAAGGACATTGTAAATTTTATCATGTGTAGATCAAACTTGGATAATCCATTTATCTCTTTCCTAGATACTGCATATACTATCATAGATCAAGAGATCTATCAGAACGAGTTGATTAATTCATTAGACGATAATGAAATTATCGATTGTATAGTTAACAAGTTTATGAGCTTTTATAAGGATAACCTAGAAAATATGGTAGATGCTATCATTACTCTAAAATATTATAATTAATAATCCAGATTTTAAAACTACGTATGTGGAAGTACTCGGTTCCAGAATAGCTGATATAGATATTAAACAAGTGATACGTAAGAATATAATACAATTGTCTAATGATCCGCGAACGATATTTGTGAAAATATTAAAAAAAAATACTTTTTTTATTAAATGACGTCTCTTCGCGAATTTAGAAAATTATGCTGTGATATATATCACGCATCAGGATATAAAGAAAAATCTAAATTAATTAGAGACTTTATAACAGATAGAGATGATACCGATACATATTTGATCATTAAGCTATTGCTTCCCGGATTAGACGATAGAATGTATAACATGAACGATAAACAAATTATAAAATTATATAGTATAATATTTAAACAATCTCAGGAAGATATGCTACAAGATTTAGGATACGGATATATAGGAGACACTATTAGGACTTTCTTCAAAGAGAACACGGAAATCCGTCCACGAGATAAAAGCATTTTAACTTTAGAAGAAGTGGATAGTTTTTTAACTACGTTATCATCAGTAACTAAAGAATCACATCAAATAAAATTATTGACTGATATAGCATCTGTTTGTACATGTAATGATTTAAAATGTGTAGTCATGCTTATTGATAAAGATCTAAAAATTAAAGCGGGTCCTCGGTACGTGCTTAACGCTATTAGTCCTCATGCCTATGATGTTTTTAGAAAATCTAATAACTTGAAAGAGATAATAGAAAATGCAGCTAAACAAAATCTAGACTCTATATCTATTTCTGTTATGACTCCAATTAATCCCATGTTAGCGGAATCATGTGATTCTGTCAATAAGGCGTTTAAAAAATTTCCATCAGGAATGTTTGCGGAAGTCAAATACGATGGTGAAAGAGTACAAGTTCATAAAAAAAATAACGAGTTTGCATTCTTTAGTAGAAACATGAAACCAGTACTCTCTCATAAAGTGGATTATCTCAAAGAATACATACCGAAAGCATTTAAAAAAGCTACGTCTATCGTATTGGATTCTGAAATTGTTCTTGTAGACGAACATAATGTACCGCTACCGTTTGGAAGTTTAGGTATACACAAAAAGAAAGAATATAAAAACTCTAACATGTGTTTGTTCGTATTTGACTGTTTATACTTTGATGGATTCGATATGACAGACATTCCATTGTATGAACGAAGATCTTTTCTCAAAGATGTTATGGTCGAAATACCCAATAGAATAGTATTCTCAGAGTTGACGAATATTAGTAACGAGTCTCAGTTAACTGATGTATTAGATGATGCACTAACGAGAAAATTAGAAGGATTGGTCTTAAAAGATATTAATGGCGTATACGAACCGGGAAAGAGAAGATGGTTAAAAATAAAGCGAGACTATTTGAACGAGGGTTCCATGGCAGATTCTGCCGATTTAGTAGTACTAGGTGCCTACTATGGTAAAGGAGGAAAGGGTGGTATCATGGCAGTCTTTCTAATGGGTTGTTACGACGATGAATCCGGTAAATGGAAGACGGTAACTAAATGTTCCGGTCACGATGATAATACGTTAAGGGTTTTGCAAGACCAATTAACGATGGTTAAAATTAACAAGGATCCCAAAAAAATTCCAGAGTGGTTGGTAGTTAATAAAATCTATATTCCCGATTTTGTAGTAGATGATCCGAAACAATCTCAGATATGGGAAATTTCAGGAGCAGAGTTTACATCTTCCAAGTCACATACAGCGAATGGAATATCGATTAGATTTCCTAGATTTACTAGGATTAGAGAAGATAAAACGTGGAAAGAATCTACTCATCTAAACGATTTAGTAAACTTGACTAAATCTCTTAATAGTTACATATAAACTGAAAAATAAAATAACACTATTTTAGTTGGTAGTTGCCATGGATGGTGTTATCGTATACTGTCTAAATGCGTTAGTAAAACATGGCGAGGAAATAAATCATATAAAAAATGATTTCATGATTAAACCATGTTGTGAAAGAGTTTGTGAAAAAGTCAAGAACGTTCACATCGGCGGACAATCTAAAAACAATACAGTGATTGCAGATTTGCCATATCTGGATAATGCTGTATCAGATGTATGCAAATCAATATATAAAAAGAATGTATCAAGAATATCCAGATTTGCTAATTTGATAAAAATAGATGACGATGACAAGACTCCTACCGGCGTATATAATTATTTTAAACCTAAAGATGCTATTCCTGTTATTATATCCATAGGAAAGGATAAAGATGTCTGTGAACTATTAATCTCATCTGATAAAGCGTGTGCGTGTATAAAGTTAAATTTATATAAAGTAGCCATTCTTCCCATGGATGTTTCCTTTTTTACCAAAGGAAATGCATCATTGATTATTCTCCTGTTTGATTTCTCTATCGATGCGGCACCTCTCTTAAGAAGTGTAACCGATAATAATGTTATTATATCTAGACACCAGCGCCTACATGACGAGCTTCCGAGTTCCAATTGGTTCAAGTTTTACATAAGTATAAAGTCCGACTATTGTTCTATATTATATATGGTTGTTGATGGATCTATGATGTATGCGATAGCTGATAATAGAACTCACGCAATTATTAGCAAAAATATATTAGACAATACTACGATTAACGATGAGTGTAGATGCTGTTATTCTGAACCACAGATTAGGATTCTTGATAGAGATGAGATGCTCAATGGATCATCGTGTTATATGAACAGACATTGTATTATGATGAATTTACCTGATGTAGGCGAATTTGGATCTAGTATGTTGGGGAAATATGAACCTGACATGATTAAGATTGCTCTTTCGGTGGCTGGTAATTTAATAAGAAATCGAGACTACATTCCCGGGAGACGAGGCTATAGCTACTACGTTTACGGTATAGCCTCTAGATAATTTTTTTTAAGCACGAAATAAAAACATAATTTTAAACAATCTATTTCATACTATTTTGTGTGCTCACCATGAACATAAAGATAGATATATTAGTATTTCTGGTGATAAATTTACGGCGACTGCTAGGAGGGAAAATGAAGAAAGAAAAAATATCTACCTCTCCAAAAAGAAAAACTACTGATGTTATCAAACCTGATTATCTTGAGTACAATGACTTGTTAGATAGAGATGAGATGTCTACTATTCTAGAGGAATATTAGGCCTTAGAATAAAATATGGACGACTCTTAACGAAATTAGAAAATTCGATAATGATGTTGAAGAACAATTCGGTACTATAGAAGAACTCAAGCAGAAGCTTAGATTAAATTCTGAAGAGGGAGCAGATAATTTTATAGATTATATAAAGGTACAAAAACAGGATATCATCAAACTTACTGTATACGATTGCATATATCTATGATAGGATTGTGTGCGTGCGTGGTAGATGTTTGGAGAAATGAGAAACTGTTTTCTAGATGGAAATATTGGTTACAAGCGATTAAACTGTTTATTGATGATCACATGCTTGATAAGATAAAATCTATTGTAGAATAGACTAGTGTATGTGGAAATGTCATAGAAAGTTAAAAGTTAATGAGAGCAAAAATATATAAGGTTGTATTCCATATTTGTTATTTTTTTCTGTAATAGTTAGAAAATACATTCGATGGTCTATCTACCAGATTATTATGTGTTATAAGGTACTTTTCTCATAATAAACTAGAGTATGAGTAAGATAGTGTTTTTCAAAAACATATAAATCTAAAATTGATGGATGAGATATACAGCTATTAATTTCGAAAATATATTTTAATCTGATAACTTTAAACATGGATTTTTGATGGTGGTTTAAGTTTAAAAAAGATTTTGTTATTGTAGTATGATAATATCAAAAAGATGGATATAAAGAATTTACTGACTACATGTACTATTTTACATTACTACATTGGCTACGGCATATATACCTATTTCGTCACTTCCACACGCTCCGGTAAACGGGTGTCATGTGACGAGGGAGAATCTTGATAAGAGGCATAATCAATGTTGTAATCCGATGTCCACCTGGAGAATTTGCCAAGGTCAGATGTAGAGTTGGTAGTGATAACACAAAATGTGAACACTGCCCACCTCATACATATACCGCAATCCCCAATTATTCTAATAGATGTCATCAATGTAGAAAATGCCCAACAGGATCATTTGATAAGGTAAAGTGTACCGGAACACAGAACAAATGTTCGTGTCATCCTGGTTGGTATACGCTACTGATTCTTCACAGACTGAAGATTGTCGAGATTTGTGTACCAAAAAAGGAGATGTCCATGCGGATACTTTGGTGGAATAGATGAAGGAAATCCTATTTGTAAATCGTGTTGTGTTGGTGAATATTGCGACTACCTACGTAATTATAGACTTGATCCATTTCCTCCATGCAAACTATCTATCTAAATGTAATTAATTATGATTTTGATGATAATGTTACCATACATTATATCACTACTTGGTTAGTGTGTATTATTTAGTATGGAAGACCTATTAATAATTACTTATCTTTTGACGATCTTGTTATAATTATAATATAAAAATACTTATGACATAGTAACTCATAATTGCTGACGCGATAAATTCGTAATAATCTGTTTTGTTCAAATTTTTATAAGGAATCTACAGGCATAAAAATAAAAATATAATCTATAATATACTCTTACAACGCCATCATGAATAGCAGTGAATTAATTGCTGTTATTATGGATTTAGAAATAGTGGACGATTTTGTGATATTAATATAGTTATTAATGATGAAAGGATAAACGCGCATAGACTCATCCTATCTGGAGCCTCCGAATATTTTTTCCATTCTGTTTTCCAATAATTTTATCGATTCTAATGAATACGAAGTTAATCTAAGTCATTTAGATTATCAAAGTGTTAACTATTTGATCGATTACATTTATGGGATACCTTTGAGCCTAACTAACAATAACGTGAAATATATTCTTTCAACCGCTGATTTTTTTACAAATTGGATCTGTCATTACTGAGTGCGAAAAATACATACTTAAAAATCTTTGTTTTAGAAACTGTATCGATTTCTACATATACGCTGATAAATATAATAACAAGAAAATAGAATTAGCATCGTTTAACACAATATTACGAAATATTTTGAGACTCATCAACAATGAAAACTTTAAATACTTAACAGAGGAATCAATGATAAAAATTTTAAGCGATGATATGTTATATATAAAAAATGAGGATTTCACCCCACTGATTCTCATTAAATGGTTAGAGAGTACACCAACCATGTACCGTCGAGTTACTTAGATGCCTCAGAATATCATTTCTTTCCCCACAAGTTATAAAATCACTTTATAGTCATCGACTGGTTAGTTCAATCTACGAATGTATAACATTCTTAAACAATATAGCATTCTTGGATAAATCATTTCCTAGATACCATATCATCGAGTTGATATCTATCGGTATAAGTAATTCACATGATAAGATTTCCATAAACTGCTACAATCATAAAAAAATTCATGGGAAATGATATCTTCACGTAGATATAGGTGTAGTTTCGCAGTGACCGTCCTGGATAATATTATCTATATGATGGGTGGATATGATCAGTCCCTGTATAGAAGTTCAAAGGTTATAGCGTACAATACATGTACTAATTCTTGGATATATGATATACCAGAGCTAAAAATATCATCGTTCTAATTGCGGAGGAGTTGCCAATGACGAATACATTTATTGTATAGGCGGTATACGCGATCAGGAGTCATCGTTGATATCTAGTATCGATAGATGGAAGCCATCAAAACCATATTGGCAGAAGTATGCTAAAATGTGCGAACCAAAATGTGATATGGGGTTGCGATTTTAAACGGATTAATATATGTCATAGGTGGAGTCGTTAAAGGTGACACACATATACCAACGCACTAGAGAGTTTATCAGAAGATGGATGGATGAATCATCAACGTCTTCCAATAAAAATGTCCAATATGTCGACGATTGTTCATGCTGGAAAGATTTATATATCTAGAGGTTACAACAATAGTAGTGTAGTTAATGTAATATCGAATCTAGTCCTTAGCTATAATCCGATATATGATGAATGGACCAAATTATCATCATTAAATATTCCTAGAATTAATCCTGCTCTATGGTCAGTGTATAATAAATTATATGTAGGAGGAGTAATATCTGATGATGTTCAAACTAATACATCTGAAACATACGATAAAGAAAAAGATTGTTGGACATTGGATAATGGTCACTTGTTACCACATAATTATATAATGTATAAATGCGAACCGTTTAAACATAGATATCCATTGGAAAAAACACAGTACACGAATGATTTTCTAAAGTATTTGGAAAGTTTTATAGGTAGTTGATAGAACAAAATACATAATTTTGTAAAAATAAATCACTTTTTATACTAATATGACACAATTACCAATACTTTTGTTACTAATATCATTAGTATACGCTACACCTTCTCCTCAGACATCTAAAAAAATAGGTGATGATGCAACTATATCATGTAGTCGAAATAATACAAATTACTACGTTGTTATGAGTGCTTGGTATAAGGAGCCCAATTCCATTATTCTCTTAGCTGCCAAAAGCGACGTCTTGTATTTTGATAATTATACCAAGGATAAAATATCTTACGACTCTCCATACGATGATCTAGTTACAACTATCACAATTAAATCATTGACTGCTGGAGATGCCGGTACTTATATATGTGCATTCTTTATGACATCGACTACAAATGATACTGATAAAGTAGATTATGAAGAATACTCCATAGAGTTGATTGTAAATACAGATAGTGAATCGACTATAGACATAATACTATCTGGATCTACACCAGAAACTATTTCTGAGAAACCAGAGGATATAGATAATTCTAATTGCTCGTCTGTATTCGAAATCACGACTCCGGAACCAATTACTGATAATGTAGACGACCATACAGACACCGTCACATACACTAGTGATAGCATTAATACAGTAAATGCATCATCTGGAGAATCCACAACAGACGAGATTCCGGAACCAATTACTGATAAAGAAGAAGATCATACAGTAACAGACACTGTCTCATACACTACAGTAAGTACATCATCTGGAATTGTCACTACTAAATCAACCACCGATGATGCGGATCTTTATGATACATACAATGATAATGATACAGTACCGCCAACTACTGTAGGTGGTAGTACAACCTCTATTAGCAATTATAAAACCAAGGACTTCGTAGAAATATTTGGTATTACCACATTAATTATATTGTCAGCAGTGGCGATTTTCTGTATTACGTATTATATATGTAATAAACACCCACGTAAATACAAAACAGAGAACAAAGTCTAGATTTTTGACTTACATAAATATCTGGGATAATAAAATCTATCATATTGAGAGGACCATCTGGTTCAGGAAAGACAGCCATAACCAAAAGACTGTTAAAAGACTATGGGAATATATTTGGATTTGTGGTGTCCCATACCACTAGATTTCCTCGTCCTATGGAACGAGAAGGTGTTGATTACCTTACGTTAACAGAGAGGCCATCTGGAAGGGAATAGCCGCCGGAAACTTTCTAGAACATACTGAGTTTTTAGGAAATATTTACGGAACTTCTAAAACAGCTGTAAATACAGCGGTTATTAATAATCGTATTTGCGCGATGGATTTAAACATCAACGGTGTTAGAAGTCTTAAAAATACTTACCTAATGCATTACTTGGGTATATAAGACCTACCTCTCTTAAAATGGTTGAGACCAATCTTCGTCGTAGAAACACTGAAGCGGACGACGAATCTCATCGTCGCGTGATGTTGGCAAAAAACGGATATGGATGAGGTCAACGAAGCAGGTCTATTCGACACTATTATTATTGAAGATGATGTGAATTTAGCATATAGTAAGTGTTAATTCAGATACTACAGGACCGTATTAGAATGTATTTTAACACTAATTAGAGACTTAAGATTTGACTTAAAACTTGATAATTAATAATATAACTCGTTTTTATATGTGGCTATTTCAACGTCTAATGTATTAGTTAAATATTAAAACTTACCACGTAAAACTTAAAATTTAAAATGGTATTTCATTGACAGATCATACATTATGAAGTTTCAAGGACTTGTGTTAATTGACAATTGCAAAAATCAATGGGTCGTTGGACCATTAATAGGAAAAGGTGGATTCGGTAGTATTTATACTACTAATGACAATAATTATGTAGTAAAAATAGAGCCCAAAGCTAACGGATCATTATTTACCGAACAGGCATTTTATACTAGAGTACTTAAACCATCCGTTATCGAAGAATGGAAAAAATCTCACAATATAAAGCACGTAGGTCTTATCACATGCAAGGCATTTGGTTTATACAAATCCATTAATGTGGAATATCGATTCTTGGTAATAAATAGATTAGGTGCAGATCTAGATGCGGTGATCAGAGCCAATAATAATAGACTACCAGAAAGGTCGGTGATGTTGATCGGAATCGAAATCTTAAATACCATACAATTTATGCACGAGCAAGGATATTCTCACGGAGATATTAAAGCGAGTAATATAGTCTTGGATCAAATAGATAAGAATAAATTATATCTAGTGGATTACGGATTGGTTTCTAAATTCATGTCTAACGGCGAACATGTTCCATTTATAAGAAATCCAAATAAAATGGATAACGGTACTCTAGAATTTACACCTATAGATTCGCATAAAGGATACGTTGTATCTAGACGTGGTGATCTAGAAACACTTGGATATTGTATGATTAGATGGTTGGGAGGTATCTTGCCATGGACTAAGATATCTGAAACAAAGAATTCTGCATTAGTAAGTGCCGCAAAACAGAAATATGTTAACAATACTGCGACTTTGTTAATGACCAGTTTGCAATATGCACCTAGAGAATTGCTGCAATATATTACCATGGTAAACTCTTTGACATATTTTGAGGAACCCAATTACGACGAGTTTCGTCGAGTATTAATGAATGGAGTTATGAAAAATTTTTGTTGATAAAAAAATTAAAAAAATAACTTAGTTATTATCACTCTCGTGAGTACAATAGAAACATGGCGATGTTTTACGCACACGCTTTCGGTGGGTACGACGAGAACCTTCATGCATTTCCTGGAATATCATCGACGGTTGCCAATGATGTCAGGAAATATTCTGTTGTGTCAGTTTATAATAAAAAGTATAACATTGTAAAAAACAAATATATGTGGTGTAACAGTCAAGTGAACAAGAGATATATTGGAGCACTACTGCCTATGTTTGAATGCAATGAATATCTACAAATTGGAGATCCAATCCATGATCTAGAAGGAAATCAAATCTCTATTGTCACATATCGCCACAAAAACTACTATGCTCTAAGTGGAATTGGGTACGAGAGTCTAGACTTGTGTTTGGAAGGAGTAGGGATTCATCATCACGTACTTGAAACAGGAAACGCGGTATATGGAAAAGTTCAACATGAGTATTCTACTATCAAAGAGAAGGCCAAAGAAATGAATGCACTCAAACCAGGACCTATCATCGATTACCACGTCTGGATAGGAGATTGTGTCTGCCAAGTTACTACTGTAGACGTGCATGGAAAGGAAATTATGAGAATGAGATTCAAAAGGGGTGCGGTGCTTCCGATTCCAAATCTGGTAAAAGTTAAAGTTGGGGAGGAAAATGATACAATAAATCTTTCCACTTCCATATCAGCTCTCCTGAATTCCGGTGGCGGCACCATCGAGGTAACATCTAAGGAAGAACGTGTAGATTATGTACTCATGAAACGTTTGGAATCTATACATCATCTGTGGTCTGTAGTGTATGATCATCTTAATGTTGTGAATGGCGAAGAACGATGTTATGTACATATGCATTCATCTCATCAAAGTCCTATGCTGAGTACTGTAAAAACAAATTTGTACATGAAGACTATGGGAGCATGTCTTCAAATGGACTCCATGGAAGCTCTAGAGTATCTTAGTGAACTGAAGGAATCAGGTGGGCGGAGTCCCAGACCAGAATTGCAGAAATTTGAATATCCAGATGGAGTGAAAGACACTGAATCAATTGAGAGATTGGCAGAGGAGTTCTTCAATAGATCAGAACTTCAGGCCGGTGAATCAGTCAAATTTGGTAATTCTATTAATGTTAAACATACATCTGTTTCAGCTAAGCAACTAAGAACACGTATACGACAGCAGCTTCCTTCTATACTCTCATCTTTTGCCAACACAAAGGGTGGATATTTGTTCATTGGAGTTGATAATAATACACACAAAGTAATTGGATTCACGGTGGGTCATGACTACCTCAAACTGGTAGAGAGTGATATAGAAAAGTATATCCAAAAACTTCCTGTTGTGCATTTCTGCAAGAAAAAAGAGGACATCAAGTACGCATGTAGATTCATCAAGGTGTATAAACCTGGTGATGAGACTACCTCGACATATGTGTGCGCAATCAAAGTGGAAAGATGCTGCTGTGCTGTGTTTGCGGATTGGCCAGAATCATGGTACATGGATACTAGTGGTAGTATGAAGAAGTATTCTCCAGATGAATGGGTGTCACATATAAAATTTTAATTAGGGTAAGGTAAAACTATATATAATAACTAACAATTTGTGTATCATATAGACAATTAATTAGGTAACTGTTATCTCTTTTTAACTAACTAACTAACTAACTAACTCTTATATACTATTAATAATACATCTATTAATCATTGATTAGCTTATTGCTTTAATTGTTTTTGTAAACTAACACTGTTCATTGAAAAGGGATAACATGTTACAGAATATAAATTATATATGGATTTTTTTAAAAAGGAAATACTTGACTGGAGTATATATTTATTTCTTCATTACATAACACGTCTGTGTTCTAATTCTTCCAATTCTTCCACATCTCATATAATACAGGAATATAATCTTGTTCGAAAATATGAGAAAGTGGATAAAACAATAGTTGATTTTTTATCTAGGTGGCCAAATTTATTCCATATTTTAGAATATGGGGAAAATATTCTACATATTTATTTTATAGATGCTGCTAATACGAATATTATGATTTTTTTTCTAGATAGAGTATTAAATATTAATAAGAACCGTGGGTCATTTATACATAATCTCGGGTTATCATCCATTAATATAAAAGAATATGTATATCAATTAGTTAATAATGATCATCTAGATAATAGTATAAGACTAATGCTTGAAAATGGACGTAGAACAAGACATTTTTTGTCTTATATATTGGATACAGTTAATATCTATATAAGTATTTTAATAAATCATAGATTTTATATAGATGCCGAAGACAGTTACGGTTGTACATTATTACATAGATGTATATATAACTATAAGAAATCAGAATCAGAATCATATAATGAATTAATTAAGATATTGTTAAATAATGGATCAGATGTAGATAAAAAAGATACGTACGGAAACACACCGTTTATCCTATTATGTAAACACGATATCGACAACGCGGAATTGTTTGAGATATGTTTAGAGAATGCTAATATAGACTCTGTAGACTTTAATGGATATACACCTCTTCATTATGTCTCATGTCGTAATAAATATGATTTTGTAAAGTTATTAATTTCTAAAGGAGCAAATGTTAATGCACGTAATAGATTCGGAACTACTCCATTTTATTGTGGAATTATACACGGTATCTCGCTTATAAAACTATATTTGGAATCAGACACAGAGTTAGAAATAGATAATGAACATATAGTTCGTCATTTAATAATTTTTGATGCTGTTGAATCTTTAGATTATCTATTGTCCAGAGGAGTTATTGATATTAACTATCGTACTATATACAACGAAACATCTATTTACGACGCTGTCAGTTATAATGCGTATAATACGTTAGTCTATCTATTAAACAGAAATGGTGATTTTGAGACGATTACTACTAGTGGATGTACATGTATTTCGGAAGCAGTCGCGAACAACAACAAAATAATAATGGATATACTATTGTCTAAACGACCATCTTTGAAAATTATGATACCATCTATGATAGCAATTACTAAACATAAACAACATAATGCAGATTTATTGAAAATGTGTATAAAATATACTGCGTGTATGACCGATTATGATACTCTTATAGATGTACAATCGCTACATCAATATAAATGGTATATTTTAAAATGTTTTGATGAAATAGATATCATGAAGAGATGTTATATAAAAAATAAAACTGTATTCCAATTAGTTTTTTGTATCAAAGACATTAATACTTTAATGAGATACGGTAGACATCCTTCTTTCGTGAAATGTAATATTCTCGACGTATACGGAAGTTGTGTACGTAATATCATAGCATCTATTAGATATCGTCAGAGATTAATTAGTCTATTATCCAAGAAGCTGGATGCTGGAGATAAATGGTCGTGTTTTCCTAACGAAATAAAATATAAAATATTGGAAAACTTTAACGATAACGAACTGACCACATATCTGAAAATCTTATAAACACTATTAAAATATAAAATCTAAGTAGGATAAAATCACACTACATCATTGTTTCCTTTTAGTGCTCGACAGTGTATACTATTTTTAACACTCATAAATAAAAATGAAAACGATTTCCGTTGTTACGTTGTTATGCGTACTACCTGCTGTTGTTTATTCAACATGTACTGTACCCACTATGAATAACGCTAAATTAACGTCTACCGAAACATCGTTTAATGATAAACAGAAAGTTACGTTTACATGTGATTCAGGATATCATTCTTTGGATCCAAATGCTGTCTGTGAAACAGATAAATGGAAATACGAAAATCCATGCAAGAAAATGTGCACAGTTTCTGATTATGTCTCTGAACTATATGATAAGCCATTATACGAAGTGAATTCCACCATGACACTAAGTTGCAACGGTGAAACAAAATATTTTCGTTGTGAAGAAAAAAATGGAAATACTTCTTGGAATGATACTGTCACGTGTCCTAATGCGGAATGTCAACCTCTTCAATTAGAACACGGATCGTGTCAACCAGTTAAAGAAAAATACTCATTTGGGGAATATATGACTATCAACTGTGATGTTGGATATGAGGTTATTGGTGTTTCGTATATAAGTTGTACGGCTAATTCTTGGAATGTTATTCCATCATGTCAACAAAAATGTGATATACCGTCCCTATCTAATGGATTAATTTCCGGATCTACATTTTCTATCGGTGGCGTTATACATCTTAGTTGTAAAAGTGGTTTTACACTAACGGGGTCTCCATCATCCACATGTATCGACGGTAAATGGAATCCCATACTCCCAACATGTGTACGATCTAACGAAGAATTTGATCCAGTGGATGATGGTCCCGACGATGAGACAGATCTGAGCAAACTCTCGAAAGACGTTGTACAATATGAACAAGAAATAGAATCGTTAGAAGCAACTTATCATATAATCATAATGGCGTTGACAATTATGGGTGTCATATTTCTAATCTCCATTATAGTATTAGTTTGTTCCTGTGACAAAAATAATGACCAATATAAGTTCCATAAATTGCTACCGTGAATATAAATCCGTTAAAATAATTAATAATTAATAATTAATAACGAACAAGTATCAAAAGATTAAAGAATTAGCTAGAATCAATTAGATGTCTTCTTCAGTGGATGTTGATATCTACGATGCCGTTAGAGCATTTTTACTCAGGCACTATTATGACAAGAGATTTATTGTGTATGGAAGAAGTAACACCATATTACATAATATATACAGGCTATTTACAAGATGCACCGTTATACCGTTCGATGATATAGTACGTACTATGCCAAATGAATCACGTGTTAAACAATGGGTGATGGATACACTTAATGGTATAATGATGAATGAATTCGATACTGTATGTGTGGGTACCGGACTACGATTCATGGAAATGTTTTTCGATTACAATAAAAATAATCCCAAAAATAGCATCAACAATCAAATAATGTATGATATAATTAATAGCGTAGCCATAATTCTAGCTAATGAGAGATATAGAAGCGCGTTTAACGACGATAGAATATACATCCGTAGAACTATGATGGACAAATTGTACGAATACGCATCTCTAACTACTATTGGTACGATCACTGGAGGTGTTTGTTATTTTATCTGTTGATGCATCTAGTTAGTTTGTATAAATAATTATTTCGATATACTAGTTAAAATTTTAAGATTTTAAATGTATAAAAAACTAATAACGTTTTTATTTGTAATAGGTGCAGTTGCATCCTATTCGAATAATGAGTACACTCCGTTTAATAAACTGAGTGTAAAACTCTATATAGATGGAGTAGATAATATAGAAAATTCATATACTGATGATAATAATGAATTGGTGTTAAATTTTAAAGAGTACACAATTTCTATTATTACAGAGTCATGTGACGTCGGATTTGATTCCATAGATATAGATGTTATAAACGACTATAAAATTATTGATATGTATACCATTGACTCGTCTACTATTCAACGCAGAGGACATACGTGTAGAATATCTACCAAATTATCATGCCATTATGATAAGTACCCTTATATCCACAAATATGAGGGTGATGAACGACAATATTCTATTACCGCAGAGGGAAAATGCTATAAAGGAATAAAATATGAAATAAGTATGATGAACGATGATACTCTATTGAGAAAACATACTCTTAAAATTGGATTTACTTATATATTCGATCGTCATGGGCATAGTAATACATATTATTCAAAATATGATTTTTAAAAATTTAAAATATATTATCACTTCAGTGACAGTAGTCAAATAACAAACAACACCATGAGATATATTATAATTCTCGCAGTTTTGTTCATTAATAGTATACATGCTAAAATAACTAGTTATAAGTTTGAATCCGTCAATTTTGATTCCAAAATTGAATGGACTGGGGATGGTCTATACAATATATCCCTTAAAAATTATGGCATCAAGACGTGGCAAACAATGTATACAAATGTACCAGAAGGAACATACGACATATCCGGATTTCCAAAGAATGATTTCGTATCTTTCTGGGTTAAATTTGAACAAGGCGACTATAAAGTGGAAGAGTATTGTACGGGACTATGTGTCGAAGTAAAAATTGGACCACCAACTGTAATATTGACTGAATATGACGATCATATCAATTTGTTCATCGAGCATCCGTATGCTACTAGAGGTAGCAAGAAGATTCCTATTTACAAACGCGGTGACATGTGTGATATCTACTTGTTGTATACGGCTAACTTCACATTCGGAGATTCTGAAGAACCAGTAACATATGATATCGATGACTACGATTGCACGTCTACAGGTTGCAGTATAGACTTTGCCACAACAGAAAAAGTGTGTGTGACAGCACAGGGAGCCACAGAAGGGTTTCTCGAAAAAATTACTCCATGGAGTTCGGAAGTATGTCTGACACCTAAAAAGAATGTATATACGTGCGCAATTAGATCTAAAGAAGATGTTCCCAATTTCAAGGACAAAATAGCCAGAGTTATCACGAGAAAATTTAATAAACAGTCTCAATCTTATTTGACTAAATTTCTCGGTAGCACATCGAATGATGTTACAACTTTTCTTAGCATTCTTGACTAAATATTCATAACTAATTTTTATTAATGATACAAAAATGAAATAAACTGTATATTATACACTGGTTAACGCCCTTGGCTCTAACCATTTTCAAGATGAGGTCCCTGATTATAGTCCTTCTGTTCCCCTCTATCATCTACTCAATGTCTATTAGACGATGCGAGAAGACTGAAGAGGAAACATGGGGATTAAAAATAGGGTTGTGTATAATTGCCAAAGATTTCTATCCCGAAAGAACTGATTGCAGTGTTCATCGCCCAACTGCAAGTGGAGGATTGATAACTGAAGGCAATGGATTCAGAGTAGTTATATATGATCAATGTACAGAACCCCATGACTTTATTATCACCGATACTCAACAAACACGTCTTGGATCATCTCATACATATATTAAATTCAGTAACATGAATACAGGTGTCCCATCTAGTATTCCAAAATGTTCCAGAACTCTCTGTATTTCTGTATATTGTGATCAAGAGGCGGGAGACATAAAATTTGAGGAGTATACTCAAGAATCAAGTGATATCAGTATTAGAGTTAAGTATGATTCATCATGTATTGATTATCTGGGTATTAATCAAAGTTTCATGAATGAATGTATTCGAAGAATTACAACATGGGATAGAGAATCATGCGTCAGAATTGATACACAGACTATAAATAAATATCTTAAGTCTTGCACCAACACAAAATTCGACCGTAATGTCTACAAAAGGTACATACTGAAGAGTAAAGCACTCCATGCTAAAACAGAGTTGTAATAGATATAAAATACTTTTTATAATAATTAGGCTAGAAAAATCTCACTCACATGTAATCTTAAAAAAATGATATGATAGTTCTTACAAGTAGCGATTGAGTTTTAAATGGATTCTATTAATTACCGGGGAACTTAACAATTCGTTCTGATCTACAGACATTGGTTAATAAATCATCTTATTTTGCCAATATATTAAAATGTGGAAACTCCACTAATAATATTACATTGTGCGACTTTCAAGATGATGTGATATATAGGGTTATACAGTTTTAACAATTATATAATAGAGATAGAAAGTACAAAAGATGTAGAATCAATGATATGGCACGCTAAACAGTTGGGTGTGGAATCATTGCTAAAAGAATGTCAAAATTATTTGCTTAGAATATTACGTATATAATTGTTTAGAAATTTATAGAATAACTAATATTAATACATTATCGTATATCTACAACGATATAAGAAACTTCATATTGGATAATATTACTATTAATATATAAGGATCCAGATTTTATATATTTGCCTAAATACATTATTATAGATTTACTAGGACAATCACCTAAATGTTTTTAACGAAGATAATGTGGTAAAGATTATATACACTTATATATCTTCCGATATCTACAAGGATATTCCATATCATCATTGTGTAAACTAAATAACGTTTTCTATGGCATTTAATAAGGACATTGGATATGTGGAAAAGTGATGTATGGAAGTTAGTACATTATCAACTTCTCCTTATTGATTGAAAATGAAAATATAAATAGTTTTTATGTATAGCGGTATCTACCCTATAGTTTTATTGCTTACTACTAACATGGATTCAGATACAGATACAGATACAGATACAGATACAGATACAGATACAGATACAGATGTAGAAGATATCATGAATGAAATAGATAGAGAGAAAGAAGAAATACTAAAAAATGTAGAAATTGAAAATAATAAAAACATTAACAAGAATCATCCCAGTGAATATATTAGAGAAGCACTTGTTATTAATACCAGTAGTAATAGTGATTCCATTGATAAAGAAGTTATAGAATATATCAGTCACGATGTAGGAATATAGATCATATCTACTAATTTTTATAATCGATACAAAACATAAAAACAACTCGTTATTACATAGCAGGTATGGAATCCTTCAAGTATTGTTTTGATAACGATGGTAAGAAATGGATTATCGGAAATACTTTATATTCTGGTAATTCAATACTCTATAAGGTCAGAAAAAATTTCACTAGTTCGTTCTACAATTACGTAATGAAGATAGATCATAAATCACACAAGCCATTGTTGTCCGAAATACGATTCTATATATCTGTATTGGATCCTTTGACTATCAACAACTGGACACGGGAACGTGGTATAAAGTATTTGGCTATTCCAGATCTGTATGGAATTGGAGAAACCGATGATTATATGTTCTTCATTATAAAGAATTTGGGAAGAGTATTCGCCCCAAAGGATAGTGAATCAGTTTTCGAAGCATGTGTCACTATGATAAACACGTTAGAGTTTATACACTCTCAAGGATTTACTCATGGAAAAATAGAACCGATGAATATACTGATTAGAAATAAACGTATTTCACTAATTGACTATTCTAGAACTAACAAACTATACAAAAGTGGAACACATATAGATTACAACGAGGACATGATAACTTCAGGAAATATCAATTATATGTGTGTAGACAATCATCTTGGAGCAACAGTTTCAAGACGAGGAGATTTAGAAATGTTGGGATATTGCATGATAGAATGGTTCGGTGGTAAACTTCCATGGAAAAACGAAAGTAGTATAAAAGTAATAAAACAAAAAAAAGAATATAAACAATTTATAGCTACTTTTTTTGAGGACTGTTTTCCTGAAGGAAATGAACCTCTGGAATTAGTTAGATATATAGAATTAGTATACATGTTAGATTATTCTCAAACTCCTAATTATGACAGACTACGTAGACTGTTTATACAAGATTGAAATTATATTCTTTTTTTTATAGAGTGTGGGGGTAGTGTTACGGATATCTAATATTAATATTAGACTATCTCTATCGCGCTACACGACCAATATCGATTACTATGGATATCTTCAGGGAAATCGCATCTTCTATGAAAGGAAAGAATGTATTCATTTCTCCAGCGTCAATCTCGTCAGTATTGACAATACTGTATTATGGAGCTAATGGATCCACTGCTGAACAGCTATCAAAATATGTAGAAAAGGAGGAGAACATGGATAAGGTTAGCGCTCAGAATATCTCATTCAAATCCATGAATAAAGTATATGGGCGATATTCTGCCGTGTTTAAAGATTCCTTTTTGGGAAAAATTGGCGATAAGTTTCAAACTGTTGACTTCACTGATTGTCGCACTATAGATGCAATCAATAAGTGTGTAGATATCTTTACTGAGGGAAAAATCAATCCACTATTGGATGAACCATTGTCTCCTGATACCTGTCTCCTAGCAATTAGTGCCGTATACTTTAAAGCAAAATGGTTGATGCCATTCGAAAAGGAATTTACCAGTGATTATCCCTTTTACGTATCTCCAACGGAAATGGTAGATGTAAGTATGATGTCTATTTACGGCGAGCCATTTAATCACGCATCTGTAAAAGAATCATTCGGTAACTTTTCAATCATAGAACTGCCATATGTTGGAGATACTAGTATGATGGTCATTCTTCCAAACAAGATTGATGGATTAGAATCCATAGAACAAAATCTAACAGATACAAATTTTAAGAAATGGTGTAACTCTCTGAAAGCTACGTTTATCGATGTGCACATTCCTAAGTTTAAGGTAATAGGTTCGTATAATCTTGTGGATACGCTAATAAAGTTGGGACTGACAGATGTGTTCTATTCAACTGGTGATTATATCAATATGTGTAATTCAGATGTGAGTGTTGACGCTATGATTCACAAAACGTATATAGATGTCAATGAAGAGTATACAGAAGCAGCTGCAGCAACTTCTGTACTAGTGGCAGACTGTGCATCAACAGTTACAAATGAGTTCTGTGCAGATCATCCGTTCATCTATGTGATTAGACATGTCGATGGTAAAATTCTTTTCGTTGGTAGATATTGCTCTCCAACAACTAATTAAGCACATTCTTAATATTAGAATATTATATAGTTAAGATTTTTACTAACAGGTTAACATTTTTTTTTAAAAATAGAAAAAACATGTGGTATTAGTGCAGGTCGTTATTCTTCCAATTGCAATTGGTAAGATGACGGCCAACTTTAGTACCCACGTCTTTTCACCACAACACTGTGGATGTGACAGACTGACCAGTATTGATGACGTCAGACAATGTTTGACTGAATATATTTATTGGTCGTCGTATGCATACCGCAACAGGCAATGCGCTGGACAACTGTATGACACACTCCTCTCTTTTAAAGATGATGCGGAATCAGTGTTCATCGACGTTCGTGAGCTGGTAAAAAATATGCCGTGGGATAATGTTAAGGATTGTACAGAGATCATCCGTTGTTATATACCGGATGAGCAAAAAACCATCAGAGAGATTTCGGCCATCATTGGACTTTGTGCATATGCTGCTACTTACTGGGGAGGTGAAGACCATCCCACTAGTAACAGTCTGAACGCATTGTTTGTGATGCTTGAGATGCTCAATTACATGGATTATACCATCATATTCTGGCGTATGAATTGATGAGTTACAGCTTGACATTTCTTCTTTCCTCCCTCTTCTTCTACCTTTCCCAGAAACAAACTTTTTTTACCCACTATAAAATAAAATGAGTATACTACCTGTTATATTTCTTCCTATATTTTTTTATTCTCCATTCGTTCAGACTTTTAACGTGCCTGAATGTATCGACAAAGGGCAATATTTTGCATCATTCATGGAGTTAGAAAACGAGCCAGTAATCTTACCATGTCCTCAAATAAATACGCTATCATCCGGATATAATATATTAGATATTTTATGGGAAAAACGAGGAGCGGATAATGATAGAATTATACAGATAGATAATGGTAGCAATATGCTAATTCTGAACCCGACACAATCAGACTCTGGTATTTATATATACATTACCACGAACGAAACCTACTGTGACATGATGTCGTTAAATTTGACAATCGTGTCTGTCTCAGAATCAAATATAGATCTTATCTCGTATCCACAAATAGTAAATGAGAGATCTACTGGTAAAATGGTATGTCCCAATATTAATGCATTTATTTCTAGTAACGTAAACACAGAATTATATGGAGCGGACATCGACGCCTTAGAAATAAGAGACTTAAACAACGGACACCTGGAATTATTACCATAGAAGATGTTAGAAAAAATGATGCTGGTTATTATACATGTGTTTTAGAATATATATATATATATATATATATATATATATATATATATATATGGGCAAAACATATAACGTAACCAGAATTATAAAATTAGAGGTACGGGATAGAATAATACCTCCTACTATGAAATTACCAGAAGGAGTAGTAACTTCAATAGGTAGTAATTTGACTATTGCATGTAGAGTATCGTTGAGACTTCCCACAACGGACACCGACGTCTTTTGGATAAGTAATGGTATATGTATTACGAAGAAGAAGACGAGGACGGAGACGGTAGAATAAGTGTAGCAAATAAAATCTATATGACCGATAAGAGACGTGTTATTACATCCTGGTTAAACATTAATCCTGTCAAGGAAGAAGATGCTACAACGTTTACGTGTATGGCGTTTACTATTCCTAGCATCAGCAAAACAGTTACTGTTAGTAAACGTGAATGTATGTTGTTACATTTCCATATCAATTGAGTTTATAAGAATTTTTTATACATTATCTTCCAACAAACAATTGACGAACGTATTGCTATGATTAACTCCCACAATACTATATATATTATTAATCATTAACTTGCAGACTATACCTAGTAGTGCTATTTTGACATACTCATGTTCTTGTGTAATCGCAGTATCTATATTATTAAAGTACGTAAATCTAGCTATAGTTTTATTATTTAATTTTAGATAATATACTGTCTCCGTATTTTTAAAAAATTACCACATCCTTTATTAAATCATGAATGGGAATTTCTGTGTCATCGTTAGTATATTGTGAACAACAAGAGCAGATATCTATAGGAAAGGGTGGAATGCGATACATTGATCTATGTAGTTTTAAAACATACGCGAACTTTGAAGAATTTATATAAATCATCTCACGAGATATTGCTCTCTGTCATATTCATACACCTGTATAAACTTTCTAGACATCTTACAATGTGTTATTTTATGATCATATTTACATATTTACTGGTATATCAAAGATGTTAGATTAGTTAATGGGAATCGTCTATAATAATGAATATTAAACAATTATAGGAGGAGTTTATACCTACAAAAACATCATAAAAATGAGTCATCGTCCGATTTATGTTTTAAATATACTAACATTACTACCTTCAGAAATTATATACGAAATATTATACATGCTGACAATTAACGATCTTTATAATATATAGTATCCACCTACCAAAGTATAATTGTATTTTTCTCATGTGATGTGTGTAAAAAACTGATATTATATAATTATCTTAGTACCTATGATGAAGATGAAGATGAAGATGAAGATGATGGTCCGTATATATTTTGTATCATTATCGTTATTGCTATTCCATAGTTACGCCATAGACATCGAAAATGAAATCACCGAATTCTTCAATAAAATGAGAGATACTCTACCAGCTAAAGACTCTAAATGGTTGAATCCAGTATGTATGTTTGGAGGCACAATGAATGATATGGCCGCTCTAGGAGAGCCATTCAGTGCAAAGTGTCCTCCTATTGAAGACAGTCTTTTATCGCATAGATATAAAGACTATGTGGTTAAATGGGAAAGGCTAGAAAAGAATAGACGGCGACAGGTTTCTAATAAACGTGTTAAACATGGTGATTTATGGATAGCCAACTATACATCTAAATTCAGTAACCGTAGGTATTTATGTACCGTAACCACAAAGAATGGTGACTGTGTTCAGGGTGTAGTTAGATCTCATGTGTGGAAACCTTCTTCATGCATTCCAAAAACATATGAACTAGGTACTTATGATAAGTATGGCATAGACTTATACTGTGGAATTCTTTATGCGAACCATTATAATAATATAACTTGGTATAAAGATAATAAGGAAATTAATATCGACGATTTTAAGTATTCACAAGCGGGAAAGGAATTAATTATTCATAATCCAGAGTTAGAAGATAGTGGAAGATACGACTGTTACGTTCATTACGACGACGTTAGAATCAAGAATGATATCGTAGTATCAAGATGTAAAATACTTACGGTTATACCGTCACAAGACCACAGGTTTAAACTAATACTAGATCCGAAAATCAACGTAACGATAGGAGAACCTGCCAATATAACATGCAGTGCTGTGTCAACGTCATTATTTGTCGACGATGTACTGATTGAATGGGAAAATCCATCCGGATGGATTATAGGATTAGATTTTGGTGTATACTCTATTTTAACTAGTAGAGGCGGTATCACCGAGGCGACTTTGTATTTTGAAAATGTTACTGAAGAATATATAGGCAATACATATACATGTCGTGGACACAACTATTATTTTGATAAAACTCTTACAACTACAGTAGTATTGGAGTAAATACACAATGCATTTTTATATACATTACTGAATTATTATTATTAATTATATCGTATTTGTGCTATAGAATGGATGAAGATACGCGACTATCTAGGTATTTGTATCTCACCGATAGAGAACATATAAATGTAGACTCTATTAAACAGTTGTGTAAAATATCAGATCCTAATGCATGTTATAGATGTGGATGTACGGCTTTACATGAGTACTTTTATAATTATAGATCAGTCAACGGAAAATACAAGTATAGATACAACGGTTACTATCAATATTATTCATCTAGCGATTATGAAAATTATAATGAATATTATTATGATGATTATGATAGAACTGGTATGAACAGTGAGAGTGATAATATATCAATCAAAACAGAATACGAGAATGAATATGAATTCTATGATGAAACACAAGATCAAAGTACACAACTAGTAGATTACGACATTAAACTCAAAACCAATGAGGATGATTTTGTTGATGAATTCTATGGTTATGATAGATCAGTGGGTGTCCATGATTATATAGATGTATCAATTAATAAAGTAGTATATGGAAGAGAGTCTCACGTAAGATGGTGGGATATATGGCAAGAACATAATGATGGCGTATACAGTATAGGAAAGGAGTGCATAGATAATATATACGAAGACAGACATACCGTAGACGAATTCTACAAGATAGACAGCGTATCAGATGTAGATGACGCAGAACATATATCTCAGATAACTAATGATGTATCTACACAAACATGGGAAAAGAAATCAGAGTTAGATAGATACATGGAAATGTATCCTCGTCATAGATATGGTAAGCATTCTGTCTTTAAGGGATTTTCTGACAAAGTTAGAAAAAATGATTTAGACATGAACGTGGTAAAAGAATTACTTTCTAACGGTGCATCTCTAACAATCAAGGATAGCAGTAATAAGGATCCAATTGCTGTTTATTTTAGAAGAACAATAATGAATTTAGAAATGATTGATATCATTAACAAACATACAACTATCTATGAACGCAGGTATATAGTACACTCCTATCTAAAAAATTATAGAAATTTCGATTATCCATTTTTCAGAAAGTTAGTTTTGACTAATAAACATTGTCTCAACAATTATTGTAATATAAGCGACGGCAAATATGGAACACCACTACATATATTAGCATCTAATAAAAAAATAATAACTCCTAATTACATGAAGTTATTAGTGTATAACGGAAATGATATAAACGCACGAGGTGAAGATACACAAATGCGAACTCCATTACACAAATATTTGTGTAAATTTGTATATCATAATATTGAATATGGTATCCGATACTATAATGAAAAGATTATAGACGCATTTATAGAGTTAGGAGCCGATCTAACTATTCCAAATGACGATGGAATGATACCAGTAGTTTACTGTATACACTCAAATGCCGAATATGGTTATAACAATATTACTAACATAAAGATAATACGTAAACTACTTAATCTTAGTAGACATGCGTCACATAATCTATTTAGAGATCGAGTCATGCACGATTATATAAGTAATACATATATTGATCTTGAGTGTTTAGATATCATTAGATCACTTGATGGGTACGATATTAATTGTTACTTTGAAGGACGTACACCACTTCATTGCGCTATACAATATAACTTCACTCAGATTGCTGAGTACTTATTAGATCGAGGAGCTGATATATCATTAAAGACAGACGATGGTAAAACTGTATTTGATTTATCGTTATGTAGTTACATTCCTCTTAAATGGACTAGCTTTTTGATTAGTCGTCTACCGCCTAAAAGTGTCATATGCTCACTGACTAACCATATAATAGATTATGTTCTTACGAACAATAGACGTATTATTTGGCAGAGTCAAATGATTAATAAGTACGTACTGTTACTGGACCCATCCTTTTATTATAGATTCAGAAATGTTATCGAAAACAAATTAGACCAATACAATAATCGTTATAATATGTTCGAACACGATAGGGACGTTAATGAAAAGTATGGCAAAGTCTTACATGACCTCGATACATATATCAAGGATGTACAAGTATTAAAATCTACTTCCATCACTAATAATATAACACTATACGACACTATTATAAATAATAAGTCAGAGTTTCCTATACGTCGTGTAAACGACAAACAATTAATTAATCTCATAAAATCCAATACATATCATAATCTTATCGAAAAAGTTATTAAAAATACATTAGAGAAATATACTTTAACTAATATAGTCCTCGAGTATATGATCTCATCTCGATCTCAATCATCTTATTTGAGTCGTATTCCTAATGAGATATTACTCGAAATATTATATAAACTCGACATGTACGATTTACGTAATCTATATACAAGATATATGAGAGAGAATGATATCACAGAGTATCATATAGAGAATACGAGGTCTGTTTCTACACAGACATGAATAATGAATACACATACAACGTTTTTTTTAATCTTAGATATAACACTAATTACATCAAGATTATATATTGAAATCGTAATTTGAGTTGTCTGATCATCATGGATATCGAAAATGATATACGTAACATTAGCAATCTTTTAGATGATTGATATATTATTATGCGATGTAATCATAACTATCGGAGATGTAGAAATTAAAGCGCATAAAACTATTTTGGTTGCCGGATCTACGTATTTTAAAACAATGTTCACAACATCTATGATAGCGAGAGATCTAGCAACTAGAGTAAATATACAGATGTTCGATAAAGATGCCGTCAAAAATATTGTACAGTACTTATACAATAGGTATATAAGTTCTATGAATGTGATAGACATATTAAAATGCACCGACTAAGAACGTAAAACGAACTATAGAATGTTATACAATGGGTGATGATAAGTAGAAGATGTTACCCGATATACCCATAGCATTATCTAGTTATGGCATGTGTGTATTAGATCAATACATATACATTATAAGCGGTCGTACCCAACACTGATTATACATCGGTACATACAGTAAATAGCATAGATATGGAGGAGGATACAAATATTTCAAATAAAGTTATGAGATACGCGCTGTCAATAATATATGGAAGACATTACCTAACTTCTGAACTGGAACTATAAATCCAGGCTCTCGCATAAAGATGAATATATATGTTGTATGCGACATCAAAGATGAAAAAATGTTAAGACTTATATATTTAGATATAACACGAATATGTATGACGGATGGGAATTGGTAACGATGACAGAAAGCAGATTGTCAGCTCTGCATACTATTCTTCATGACAATACCATAATGATGTTACATTGTTATGAAGCGTATATGTTACAAGATACATTTAATGTGCTTACGGAACATATATTTAGAAACATCTACTAACGATTTTTTATGCTTGTATTATTAATGGTATGTAATATGATTTAATTGATTGTGTACACGATACCAATTTGTCGAGTATGAATACGGAGTACAAACATAAACTGAAGTTTAACATTATTTATTTATGATATACATTATATACATTATATACATTATATACATTATATACATTATATACATTATATACATTATATACATTATATACATTATATACATTATATACATTATATACATTATATACATTATATACATTATATACATTATATACATTATATACATTATATACATTATATATCGTTATTGTTTGGTCTATGCCATGGATATCTTTAAAGAACTAATCTTAAAACATACGGATGAAAATGTTTTGATTTCTCCAGTTTCCATTTTATCTACTTTATCTATTCTGAATCATGGAGCAGCTGGTTCTACAGCTGAACAACTATCAAAATATATAGAGAATATGAATGAGAATACACCCGATGATAAGAAGGATGACAATAATGACATGGACGTAGATATTCCGTATTGCGCGACACTAGCTACCGCAAATAAAATATACGGTAGTGATAGTATCGAGTTCCATGCCTCATTCCTACAAAAAATAAAAGACGATTTTCAAACTGTAAACTTTAATAATGCGAACCAAACAAAGGAACTAATCAACGAATGGGTTAAGACAATGACAAATGGTAAAATTAATTCCTTATTGACTAGTCCGCTATCCATTAATACTCGTATGATAGTTATTAGCGCCGTCCATTTTAAAGCAATGTGGAAATATCCATTTTCTAAACATCTTACATATACAGACAAGTTTTATATTTCTAAGAATATAGTTACCAGTGTTGATATGATGGTGGGTACCGAGAATGACTTGCAATATGTACATATTAATGAATTATTCGGAGGATTCTCTATTATCGATATTCCATACGAGGGAAACTCTAGTATGGTGATTATACTGCCGGACGACATAGAAGGTATATATAACATAGAAAAAAATATAACAGATGAAAAATTTAAAAAATGGTGTGGTATGTTATCTACTAAAAGTATAGACTTGTATATGCCAAAGTTTAAAGTGGAAATGACGGAACCGTATAATCTGGTACCGATTCTAGAAAATTTAGGACTTACTAATATATTTGGATATTATGCAGATTTTAGTAAGATGTGTAATGAAACTATCACTGTAGAAAAATTTCTACATACGACGTTTATAGATGTTAATGAGGAGTATACAGAAGTATCGGCCGTTACAGGAGTATTCATGACTAACTTTTCGATGGTATATCGTATGAAGGTCTACATAAACCATCCATTCATATACATGATTAAAGATAACACCGGACATACACTTTTTATAGGGAAATACTGCTATCCGCAATAAATATAAACAATAGACTTTTATCACGTTATCTCATGTATAAAATATTACAAATAGTATAGCATAAACTAAAGTCGATACATACATTAAAACTTAAATAATAATGTAATTTACAATTAATAGTATAAACTAAAAAAATTAAAAAATTAAAAACAATATCATTATTATAAGTAATATCAAAATGACGATATACGGATTAATAGCGTATCTTATATTCGTGACTTCATCCATCGCTAGTCCACTTTACATTCCCGTTATTCCGCCCATTTCGGAAGATAAATCGTTCAATAGTGTAGAGGTATTAGTTTCTTTGTTTCCCGATGACCAAAAAGACTATACAGTAACTTCTCAGTTCAATAACTACACTATCGGTACCAAAGACTGGACTATCAACGTACTATCCACACCTGATGGTCTGGACATACCATTGACTAATATAACTTATTGGTCACGGTTTACTATAGGTCGTGCATTGTTCAAATCAGAGTCTGAGGATATTTTCCAAAAGAAAATGAGTATTCTAGGTGTTTCTATAGAATGTAAGAAGCCGTCGACATTACTTACTTTTTTAACCGTGCGTAAAATGACTCGAGTATTTAATAGATTTCCAGATATGGCTTATTATCGAGGAGACTGTCTAGAAGCCGTTTATGTAACAATGACTTATAAAAATACTAAAACTGGAGAGACTGATTACACGTACCTCTCTAATGGGGGGTTGCCTGCATACTATCGTAATGGGGTCGATGGTTGATTATTGATTAGTATATTCCTTATATTCCTTATTCTTTTTATTCACACAAAAAGAACATTTTTATAAACATGAAACCACTGTCTAAATGTAATTATGATCTTGATTTATAGATGATGATCAGCCTTCAGAGGATTTTGACCAGTATGTTTAATATGAAAAAAAACATAACTATTAAGCGCTATTGCGCTATTGTGCTTAATTATTTTGCTCTATAAACTGAATATATAGCCACAATTATTGACGGGCTTGTTTGTGACCGACAATCATGAATTTTCAGAAATTATCTCTGGCTATATATCTTACGGTGACATGTTCGTGGTGTTATGAAACATGTATGAGAAAAACTGCGTTGTATCATGACATTCAATTGGAGCATGTAGAAGACAATAAAGATAGTGTAGCATCGCTACCGTACAAGTATCTACAAGTAGTCAAACAAAGAGAACGTAGTAGATTGTTGGCTACATTTAATTGGACGGATATAGCTGAGGGTGTTAGAAATGAGTTCATTAAAATATGTGATATCAACGGAACATATCTATATAATTATACTATTGCTGTTAGTATAATTATTGATTCCACGGAAGAACTACCAACAGTTACTCCAATTACAACATATGAACCTTCTATATATAATTATACTATCGATTATAGCACTGTTATTACTACTGAAGAACTACAAGTGACTCCAACATATGCACCTGTAACAACTCCTCTTCCAACATCAGCAGTTCCTTATGATCAACGATCGAATAACAATGTAAGTACTATATCTATTCAGGTACTGAGTAAAATATTGGGAGTCAATGAAACAGAATTAACTAATTATCTTATTATGCATAAAAATGACACTGTTGACAATAACACCATGGTTGATGATGAGACATCTGACAATAACACATTACATGGTAATATAGGATTTTTGGAAATAAATAATTGTTATAATGTTTCTGTGTCAGATGCTAGTTTTAGAATAACATTAGTAAACGATACTTCTGAAGAAATTTTGCTAATGCTAACAGGAACTAGTTCATCCGACACCTTCATATCTTCCACCAATATCACTGAATGTTTGAAAACATTAATCAATAATGTGTCGATTAATGATGTACTTATAACACAAAATATGAATGTAACATCTAATTGTGATAAATGCTCAATGAATTTGATGGCATCCGTTATTCCTGCAGTTAATGAATTTAACAATACGTTGATGAAAATTGGTGTAAAAGATGATGAAAACAATACGGTATATAAATATTATAATTGTAAACTAACTACAAATTCTACATGTGATGAGTTAATCAATTTAGATGAAGTCATTAACAACATAACTCTGACAAATATTATACACAATAGTGTTTCGACAACTAACAGCAGAAAAAGACGAGATCTGAATGATGAGTTTGAATTTTCCACTTCCAAGGAATTAGATTGTCTTTACGAATCATATGGTGTAAACGATGATATAAGTCATTGTTTTGCATCACCTAGACGTAGACGATCTGACGACAAAAAGGAGTACATGGACATGAAATTATTCGACCACGCGAAAAAAGATTTAGGAATAGACAGTGTTATTCCTAGAGGTACAACCCATTTCCAAGTAGGTGCATCTGGTGCAAGTGGTGGTGTTGTAGGAGATAGTTTCCCATTTCAAAATGTTAAATCGCGTGCCAGTCTATTGGCGGAAAAAATAATGCCTAGAGTACCTATTACTGCTACCGAAGCTGATCTATATGCAACTGTAAATAGACAACCCAAGTTACCAGCAGGTGTTAAAAGTACTCCGTTTACAGAGGCGCTTGTGTCTACGATAAACCAAAAGCTTTCTAATGTTAGAGAGGTAACTTATGCTTCGCTCAATCTGCCAGGATCAAGTGGCTATGTTCATAGACCATCTGATTCTGTTATTTATAGCAGTATAAGACGGTCACGTTTACCTAGTGATAGCGATAGTGATTATGAGGATATACAAACTGTTGTTAAGGAATATAATGAAAGATATGGTAGATCAGTCAGTAGAACACAGTCATCAAGTAGTGAAAGCGATTTTGAAGATATAGATACTGTTGTTAGGGAATATAGACAAAAATATGGCAATGCAATGGCAAAAGGACGTAGTAGTTCCCCTAAACCTGATCCATTATATAGTACTGTTAAGAAAACAACTAAAAGTCTATCTACTGGTGTAGACATAGTTACAAAACAATCAGACTATTCTCTATTACCTGACGTTAATACTGGCAGTTCTATTGTGTCACCTCTCACCAGAAAAGGAGCTACTAGACGACGACCTAGACGCCCTACAAATGATGGTCTACAGAGTCCAAATCCTCCTCTCCGTAATCCACTTCCTCAACATGATGATTATTCTCCTCCACAAGTACACAGACCTCCACCACTTCCTCCTAAACCAGTCCAAAATCCGCCACAACTTCCCCCTAGACCAGTAGGTCAATTACTACCTCCTCCTATAGATCAACCAGATAAAGGATTTAGTAAGTTTGTATCACCTAGACGGTGTAGAAGAGCAAGCTCTGGAGTCATATGTGGTATGATACAATCAAAACCAAACGATGATACCTATTCACTTCTTCAACGATCAAAAATTGAACCAGAATATGTGGAGGTTGGTAATGGTATACCCAAGAACAATGTTCCTGTAATAGGTAATAAACATAGTAAAAAATATACATCGACGATGTCAAAAATATCAACAAAATTTGATAAATCTACGGCATTTGGAGCAGCAATGTTACTAACTGGTCAGCAGGCCATTAGCCAACAGACTAGATCAACTACGTTGAGTAGAAAAGATCAGATGAGTAAGGAAGAAAAGATATTCGAAGCAGTTACAATGAGTCTATCAACTATAGGTTCAACGTTGACGTCTGCAGGTATGACGGGTGGTCCAAAACTAATGATTGCAGGAATGGCTATAACGGCTATAACTGGTATAATAGATACGATAAAAGATATATATTACATGTTTTCAGGACAGGAGAGGCCAGTAGATCCTGTTATTAAATTATTTAATAAGTACACTGGCTTAATGTCCGATAATAATAAAATGGGTGTAAGAAAATGTTTGACACCCGGTGACGACACACTTATTTATATCGCATACAGAAACGATACCAGTTTTAAACAGAATACGGATGCGATGGCTTTGTATTTCTTAGATGTTATCGATTCAGAGATCCTATATCTAAACACATCAAATTTAGTTCTAGAGTATCAACTAAAGGTGGCTTGCCCCATAGGAACATTAAGATCTGTAGATGTGGACATAACTGCGTATACAATATTATATGATACAGCGGATAATATTAAGAAATACAAGTTTATCAGAATGGCAACGCTACTATCCAAACATCCAGTTATTAGATTGACATGTGGTTTAGCAGCAACATTGGTGATTAAACCGTACGAGGTACCCATCAGTGATATGCAACTACTAAAAATGGCGACGCCTGGTGAACCAGAATCCACTAAATCTATACCATCCGATGTCTGTGATAGGTATCCTCTAAAGAAATTCTATCTTTTAGCTGGTGGTTGTCCCTATGATACATCTCAAACTTTTATTGTACATACTACTTGCAGTATTCTACTAAGAACAGCTACACGGGATCAGTTTAGAAACAGATGGGTGTTACAAAATCCATTTAGACAAGAAGGGACATATAAGCAACTGTTTACCTTTAGCAAATACGATTTTAACGACACCATAATCGATCCTAATGGTGTGGTGGGTCATGCTAGCTTTTGTACCAATAGAAGCAGCAACCAATGTTTCTGGTCCGAACCTATGATATTGGAAGATGTATCATCGTGTAGTTCTAGAACTAGAAAAATATACGTAAAACTGGGAATATTTAATGCTGAAGGTTTTAATAGTTTTGTACTAAATTGTCCAACTGGGTCTACACCTACATACATCAAACATAAAAATGCGGACAGTAACAATGTTATCATAGAGCTACCTGTAGGTGATTACGGCACAGCCAAATTGTATTCAGCAACAAAACCATCGAGGATAGCTGTGTTCTGCACACATAACTATGATAAACGATTCAAATCAGATATTATAGTTCTAATGTTTAATAAAAACAGCGGTATTCCATTTTGGAGCATGTACACAGGAAGTGTAACTAGTAAAAATAGAATGTTTGCCACATTGGCTAGAGGAATGCCGTTTAGATCAACGTATTGCGATAACAGACGACGATCAGGTTGTTATTATGCAGGAATACCATTTCATGAAGATAGTGTAGAAACAGATATACATTATGGACCAGAAATAATGTTAAAGGAAACATATGACATAAACAGTATTGACCCACGAGTTATAACAAAGTCAAAGACCCATTTTCCTGCTCCATTGAGTGTAAAATTCATGGTTGACAATTTAGGAAATGGATATGACAACCCTAATTCATTTTGGGAAGATGCTAAAACTAAGAAACGGACATATAGTGCAATGACGATAAAAGTCCTACCATGTACAGTGAGAAATAAAAATATAGACTTTGGATATAACTATGGAGATATTATTTCTAATATGGTTTATCTACAATCTACTAGTCAGGATTATGGAGATGGTACCAAATATACATTTAAATCCGTAACTAGATCAGATCATGAGTGTGAATCTAGCTTAGATCTAACGTCTAAGGAAGTAACTGTGACATGTCCTGCGTTTAGTATACCAAGAAATATATCAACATATGAAGGTCTATGCTTTAGTGTTACTACATCTAAAGATCATTGTGCTACAGGTATTGGTTGGTTAAAATCTAGTGGTTATGGGAAGGAAGATGCTGATAAACCACGTGCTTGTTTTCATCATTGGAATTATTACACACTGTCGTTGGATTATTACTGTTCATACGAAGATATTTGGAGAAGCACCTGGCCTGACTATGATCCATGTAAGTCATATATCCATATAGAGTATAGAGATACATGGATAGAATCTAATGTGTTACAGCAACCTCCTTACACATTCGAATTCATTCATGACAATTCTAACGAATATGTGGATAAAGAAATTAGTAACAAATTAAATGATCTGTACAATGAATACAAGAAGATTATGGAATATAGCGACGGATCATTGCCGGCGTCTATAAACAGATTAGCAAAGGCATTGACTTCAGAGGGTAGAGAAATAGCAAGTGTTAATATAGATGGTAATCTGTTAGATATCGCATATCAAGCAGATAAGGAAAAGATGGCCGACATACAGACAAGAATAAATGATATTATTAGAGATTTGTTTATACACACTCTATCAGACAAAGATATAAAAGACATTATAGAATCCGAAGAAGGTAAGAGATGTTGTATAATAGATGTTAAGAACAATCTTGTTAAAAAGTACTATTCTATTGATAATTATCTATGTGATACTTTAGATGATTATATATACACCTCTGTAGAATATAACAAATCCTATGTGTTAGTAAACGATACTTATATGAGCTATGACTATCTTGAATCATCAGGTGTAGTTGTTCTATCATGTTATGAAATGACTATAATCTCCTTGGATACAAAAGACGCCAAAGATGCTATAGAAGATGTGATAGTAGCAAGTGCGGTAGCCGAAGCATTGAATGACATGTTTAAGGAATTTGATAAAAACGTAAGTGCTATTATAATAAAAGAAGAAGATAATTATCTAAACAGTTCGCCCGATATCTACCATATAATATATATCATAGGTGGCACTATTCTGCTACTGTTAGTCATTATTTTAATATTGGCAATTTATATAGCGCGCAATAAATACAGAACCAGGAAATATGAAATAATGAAATATGACAATATGAGCATTAAATCTGAGCATCATGATAGTCTTGAAACAGTGTCTATGGAAATTATTGATAATCGGTACTAATAAAATAGTTTAACTCTTTTAGAACCAGTTTGGTACTGTAATTTCAGTTCATTACTCGTTGAGAATATTGATGATTTTTTTTAAATGAGTATCGGTAGTTACATATTACCATATCATCCATTATATAATCGATGATGCATGTATTAGAATACTTTCCGAATAAGTCTTCTAAATATTGTATTAATTATGAAAAACTATGCTATGTGAGTATGATTCAAAGATGTTTAATGATACGATACTAGATTTTATCTCTAGCGAGATTGTTTAGAATCATTTATCATAACTATGTTTAATAAATTCATCAACGAATATCGATAAAGACCTCTTGTAATTCGAGTATAGGAAGTAGTATTACCATATCAACTTCCGAGTTAACAATTACTCTAAAACATGAGGATTGTACTCCTGTCTTTATTGGAGATCACTATTCAGTCGTTGATAAACTAGTAACCTCAGGTTTCTTTACAAACGATAAAGTACAACATCAAGACCTCACAACACAGTGCAAGATTAATCTAGAAATCAAATGTAATTCTGGAGGAGAATCTAGACAACTAACACCCACGGCGAAGTATACTTTATGCCTCATTCAGAAACGGTAACTGTAGTAGGAGACTGTCTCTCTAATCTCGATGTATATATAATATATACCAATACGGACGCGATATATTCCGACATGAATGGCGTCGCTTATCATATGTTATATCCTAAATGTTGATCATATTCCACAAATGATTGTGAACGAGATTAAATCATCTAACAAATAATTAGTTTTTTATGACATTAACATATAATAAATAAATTAATCATTATTGACTTAACGATGACGAAAGTTATCATTATCTTAGGATTCTTGATTATTAATACAAATTCGTTGTGTCTATGAAATGTGAACAAGGTGTCTCATATTATAATGCACAAGAATTAAAGTGTTGTAAACTATCTAGCCAGGAACATATTCAGATTATCGATGTGATAAATACAGCGATACCATCTGTGGACATTGTCCAAGTGACACATTCACGTCAATATATAATCGTTCTCCTCGGTGTCATAGTTGTAGAGGTCACACCTTGTACACCTACCACAAATAGAATATGTCATTGTGACTCGAATAGTTATCGTCTCCTTAAAGCTTCTGATGGTAACTGTGTTACATGTGCTCCTAAAACAAAATATGGTCGTGTGTACGGAAAGAAAGGAGAAAATGATATGGAATACCATTTGTAAGAAATGTCGGAAGGGTACTTATTCAGATATTGTATCTGACTCTGATCAATGTAAACCTATGACAAGATAAGACTTACTCGCATCTACTGGATAGACATAAATATCCTCCTCGTAATAATGAAATATAATATACACTAATTATTAATATCAATCGAGTATTAACATATAAGTTATTTTTAAACCCCTTTTGGGTTCCGTCCTAAACGGCGTTTCGGTCTGTGTCGCCACCATGGTCACACCGAGCCTCTGCGTGCTCCTCCATCGAGGACGACTTCAACTATGACAGCTCGGTGGCGTCTGCCAGCGTGTACATACGAATGGCATTTCTAAGAAAAGTCTACGGTATCCTTTCTACAATTTCCTTTAACAACGGCAACAGCTGCAGTATTTTTATACTTTGAATGCATCGGACATTTATACAAGGGAGTCCTGTTCTAATATTGGCATCAATGTTCGGATCTATAGGCTTGATTTTCGCATTGACTTTACACAGACATGAACATCCCCTGAATCTGTACATACTTTGTGGATTTACACTGTTAGAATCTCTAACGCTGGCCTCTGTTGTTACTTTCTATGATGCACGTATCGTTATGCAAGCTTTCATGTTGACTACTGCAGTGTTTCTTGCTCTGACTACATATACTCTACAATCAAAGAGAGATTTCAGTAAACTTGTAACAGGATTGTTTGCTGCTTTCTGGATTTTAATTTTGTCAGGAGTCTTGAGGATAAAGTTTAAAATAGAATTAATAAAGAACATATAGGTCATTTTTTAAACATGGATAGAAACCAAGGTTGTTAGTTAATAATATACAAGATATTTTTTCTCACTCTGATCCATGTAAACCAAGGACGAGAGACACTCTCATTCCTCATTCACGACACCATTAAAAATGGAAATTAAAGCCCTCTATTAAGCACAGACGGCTACAGGTCTACCATCAGGTTACCTTCGTCTACCTTCACAATGGCCTCTCCTTGTGCCCAGTTCAGTCCCTGTCATTGCCACGCTACTAAGGACTCCCTGAATACCGTGACTGACGTCAGACATTGTCTGACTGAATACATCCTGTGGGTTTCTCATAGATGGACCCATAGAGAAAGCGCAGGGCCTCTCTACAGGCTTCTCATCTCTTTCAGAATTGATGCAATGGAGCTATTTGGTAGCGAGTTGAAGGAGTTCTCGAATTCACTTCCGTGGGACAATATCGACAATTGCGTGGAGATCATTAAATGTTTCATCAGAAATGACTCCATGAAAACCGCCAAAGAACTTTGTGCAATAATTGGACTTTGTACTCAATCAGCTATTGTCACTGGAAGAGTCTTCAATGATAAGTATATCGACATACTACTTATGCTGCGAAAGATTCTGAACGAGAACGACTATCTCACCCTCTTGGATCATATCCTCACTGCTAAATACTAAATCTCCTTCATGCTCTCTCACTAATACTCTTACTCACTACACTTTTTATCATCTTATGATGAATGATTGCCTTCATCATTTTTTCGTGGAATATAATATAGGAATAATTAGCACCAGAATAGCTATGGATATCTCGTTAAGAATATTCTCTATAAGAGACATAATGTAGACATAGTTATTATATCCTTCTTAGATAAGTGTTACGCTACTGGAAAGTTTCCATCGTTATTATTACATGAAGATGATATAATTAAACCAACATTGAGATTGGCTCTTATGTTAGCTGGATTGAATTACTGTAATAAATGCATCGAGTATAGAGGGATATAGCAATTCTCGATAATAGTCATGCAATATTTGAATGAGACTGATAATTTAGGTAATACAGTACTACACACATATCTTTCTAGATTATATATCGTTAAAAATCTGTAAGATGTATATTTCTCATAAGTATCCACTGTGTAATATTATTAATGGATATATAGATAACGCAATAGGGACTAATAGTATTGTAAAAGATATAATCGACTATTTACGTACATATCCAGATATCTATATTCCTACTAGTTTGCTGCGTAGTTGCATCATTGATATGCATGATTTATCAGGATTCAGAGATGAATTACTAAGTAAACTACAATCCCACAATAAGTAAGAATCAAATATCAAAAACTCACTTTTGATTTTTCTAGTCTTAAGTAATACATATATTTATTAATAGACCTATGAAATAAAAAAAGGTAACAATGGATTCGCGTATAGCTATTTACGTATTAGTATCGGCATCTCTTTTGTATCTTGTTAATTGTCACAAACTAGTACATTACTTCAATCTGAAAATAAATGGAAGTGATATAACTAATACAGCAGATATATTGCTGGACAATTATCCAATTATGACCTTTGATGGAAAGGATATTTATCCATCTATCTCGTTCATGGTCGGTAATAAACTTTTCCTAGATCTTTATAAAAATATCTTTGAAGAATTTTTCAGACTATTTCGAGTATCTGTAAGTAGTCAATACGAGGAATTAGAATATTATTATTCATGTGATTATACTAACAACCGTCCTACAATTAAACAACATTACTTTTATAACGGCGACGAATATACTGAAATTGATAGATCGAAAAAAGCCACTAATAAAAACAGTTGGTTAATTACTTCAGGCTTTAGACTACAAAAATGGTTCGATAGCGAAGATTGTATAATTTATCTCAGATCTTTAGTTAGAAGAATGGAAGACAGTAACAAAAACAGTAAAAAAACTTAGTACTTAGATATCGAAAAAATATATTTTTGTAGACTCTTGAGAATAGAAGGAAAACATGTACATAATTATAAAAAATGAAAATCAATGGCGAATAAGACAGTGCGATTCGCACCATGGAGTCGGTAGATTTCATGGCTGTCGATGAGCAGTTTCACGACGACCTCGATCTTTGGTCATTATCTTTGGTAGATGATTATAAAAAACATGGATTAGGTGTTGACTGTTATGTTCTAGAACCAGTTGTTGACAGGAAAATATTTGATAGATTTCTCCTTGAACCAATTTGTGATCCTGTAGATGTTCTGTATGATTATTTTAGGATTCATAGAGATAATATTGATCAGTATATAGTAGATAGACTGTTTGCATATATTACATATAAAGATATTATATCTGCATTAGTGTCAAAGAATTATATGGAAGATATTTTCTCTATAATTATTAAGAATTGTAATTCTGTGCAAGATCTCTTACTTTACTATCTATCTAATGCATATGTAGAAATAGACATTGTTGATCTTATGGTAGATCATGGGGCTGTAATATATAAAATAGAATGCTTGAATGCCTATTTTAGGGGAATATGTAAAAAGGAAAGTAGTGTTGTTGAGTTTATTTTGAATTGTGGTATCCCAGATGAAAATGATGTTAAATTAGATCTATATAAAATAATTCAGTATACTAGGGGATTCCTTGTAGATGAACCCACAGTATTAGAAATTTATAAGCTTTGTATCCCATATATTGAAGATATCAATCAACTAGATGCTGGTGGAAGGACCTTGCTTTATCGCGCTATCTATGCAGGTTATATAGATTTAGTATCATGGCTATTAGAAAATGGAGCAAATGTCAACGCAGTAATGAGTAATGGATATACATGTCTTGACGTGGCCGTGGATAGGGGATCTGTCATCGCCCGTAGGGAAGCACATCTTAAAATATTAGAAATATTGCTTAGAGAACCATTGTCTATTGACTGTATAAAATTAGCTATACTTAATAATACAATTGAAAACCATGATGTGATAAAGCTCTGTATCAAGTATTTTATGATGGTAGATTATTCACTTTGTAATGTGTATGCATCATCACTCTTTGATTATATAATTGATTGTAAACAAGAATTGGAGTACATTAGGCAGATGAAAATTCATAATACAACCATGTATGAGTTAATCTATAATAGAGACAAAAACAAGCATGCTTCCCATATTCTACATAGGTATTCTAAACATCCAGTTTTGACACAGTGTATCACTAAAGGATTCAAGATTTACACAGAAGTAACCGAGCAGGTCACTAAAGCTCTAAACAGACGTGCTCTAATAGATGAGATAATAAACAATGTATCAACTGATGACAATCTCCTATCAAAACTTCCATTAGAAATTAGGGATCTAATTGTTTCACAAGCTGTCATATAGAGTTCTATCCACCCACCTTTCTTGAAATGAGTTAATAGTCATAAGTTAGTTAAGTCATAAGTTAGTTAAGTCATAAGTTAGTTAAGTCATAAGTTAGTTAAGTCATAAGTTAGTTAAGTCATAAGTTAGTTAAGTCATAAGTTAGTTAAGTCATAAGTTAGTTTATAGTCTAACACTTCTAATTTTTATACCTTGATCTTTTTCTCTAATTATGAAAAAGTAAATCATTATGAAGATGGATGAAATGGACGAGATTGTGCGCATCGTTAACGATAGTATGTGGTACGTACCTAACGCATTTATGGACGACGGTGATAATGAAGGTCACATTTCTGTCAATAATGTCTGTCATATGTATCTCGCATTCTTTGATGTGGATATATCATCTCATCTGTTTAAATTAGTTATTAAACACTGCGATCTGAATAAACGACTAAAATGTGGTAACTCTCCATTACATTGCTATACGATGAATACACGATTTAATCCATCTGTATTAAAGATATTGTTACGCCACGGCATGCGTAACTTTGATAGCAAGGATAAAAAAGGACATATTCCTCTACACCACTATCTGATTCATTCACTATCAATCGATAACAAGATCTTTGATATACTAACGGACCCCATTGATGACTTTAGTAAATCATCCGATCTATTGCTGTGTTATCTTAGATATAAATTCAATGGGAGCTTAAACTATTACGTTCTGTACAAATTATTGACTAAAGGATCTGACCCTAATTGCGTCGATGAGGATGGACTCACTTCTCTTCATTACTACTGTAAACACATATCCGCGTTCCACGAAAGCAATTATTACAAGTCAAAGAGTCACACTAAGATGCGAGCTGAGAAGCGATTCATCTACGCGATAATAGATCATGGAGCAAACATTAACGCGGTTACGAAAATCGGAAATACGCCGTTACACACTTACCTTCAACAGTATACCAAACATAGTCCTCGTGTGGTGTATGCTCTTTTATCTCGAGGAGCCGATACGAGGATACGTAATAATCTTGATTGTACACCCATCATGGAATACATAAAGAACGATTGTGCAACAGGTCATATTCTCATAATGTTACTCAATTGGCACGAACAAAAATACGGGAAATTACAAAAGGAAGAAGGACAACATCTACTTTATCTATTCATAAAACATAATCAAGGATATGGAAGTCGCTCTCTCAATATACTACGGTATCTACTAGATAGATTCGACATTCAGAAAGACGAATACTATAATACAATGACTCCTCTTCATACCGCCTTCCAGAATTGCAATAACAATGTTGCCTCATACCTCGTATACATCGGATACGACATCAACCTTCCGACTAAAGACGATAAGACAGTATTCGACTTGGTGTTTGAAAACAGAAACATTATATACAAGGCGGATGTCGTTAATGACATTATCCACCACAGACTGAAAGTATCTCTACCTATGATTAAATCGTTGTTCTACAAGATGTCGGAGTTCTCTCCCTACGACGATCACTACGTAAAGAAGATAATAGCCTACTGCCTATTAAGGGACGAGTCATTTGCGGAACTACATACTAAATTCTGTTTAAACGAGGACTATAAAAGTGTATTTATGAAAAATATATCATTCGATAAGATAGATTCCATCATCGAAAAATGTAGTCGTGACATAAGTCTCCTCAAAGAGATTCGAATCTCAGACACCGACTTGTATACGGTATTGAGAACAGAAGACATCCGGTATCACACATATCTCGAAGCCATACATTCAGACAAACGCATTTCATTTCCCATGTACGACGATCTCATAGAACAGTGTCATCTATCGATGGAGCATAAAAGTAAACTCGTCGACAAAGCACTCAATAAATTAGAGTCTACCATCGATAGTCAATCTAGACTATCGTATTTGCCTCCGGAAATTATGCGCAATATCATAACCAAGCTAAGCGACTACCATCTAAACAGTATGTTGTACGGAAAGAACCATTACAAATATTATCCATGATAGAAAGAAAATATTTAAAAAATAATCTATATGATTGGAGAAGTAGGAAACAAACAGTAACAAGACGACGATTACTACATTATTAAATCATGAGGTCCGTATTATACTCGTATATATTGTTTCTCTCATGTATAATAATAAACGGAAGAGATATAGCACCACATGCACCATCCAATGGAAAGTGTAAAGACAACGAATACAGAAGCCGTAATCTATGTTGTCTATCGTGTCCTCCGGGAACTTACGCTTCCAGATTATGTGATAGCAAGACTAATACACAATGTACACCGTGTGGTTCGGATACCTTTACATCTCACAATAATCATTTACAGGCTTGTCTAAGTTGTAACGGAAGATGTGATAGTAATCAGGTAGAGACGCGATCGTGTAACACGACTCACAATAGAATCTGTGAATGCTCTCCAGGATATTATTGTCTTCTCAAAGGAGCATCAGGGTGTAGAACATGTATTTCTAAAACAAAGTGTGGAATAGGATACGGAGTATCCGGATACACGTCTACCGGAGACGTCATCTGTTCTCCGTGTGGTCCCGGAACATATTCTCACACCGTCTCTTCCACAGATAAATGCGAACCCGTCGTAACCAGCAATACATTTAACTATATCGATGTGGAAATTAACCTGTATCCAGTCAACGACACATCGTGTACTCGGACGACCACTACCGGTCTCAGCGAATCCATCTCAACGTCGGAACTAACTATTACCATGAATCATAAAGATTGTGATCCAGTCTTTCGTGCAGAATACTTCTCTGTCCTTAATAATGTAGCAACTTCAGGATTCTTTACAGGAGAAAATAGATATCAGAATACTTCAAAGATATGTACTCTGAATTTCGAGATTAAATGTAACAACAAAGATTCATCTTCCAAACAGTTAACGAAAACAAAGAATGATACTATCATGCCGCATTCAGAGACGGTAACTCTAGTGGGCGACTGTCTATCTAGCGTCGACATCTACATACTATATAGTAATACCAATACTCAAGACTACGAAAATGATACAATCTCTTATCATATGGGTAATGTTCTCGATGTCAATAGCCATATGCCCGCTAGTTGCGATATACATAAACTGATCACTAATTCCCAGAATCCCACCCACTTATAGTAAGTTTTTTTACCTATAAATAATAAATACAATAATTAATTTCTCGTAAAAGTAGAAAATATATTCTAATTTATTATATGGTAAGAAAGTAGAATCATCTAGAACAGTAATCAATCAATAGCAATCATGAAACAATATATTGTCCTGGCATGCATGTGCCTAGTGGCAGCTGCTATGCCTACTAGTCTTCAACAATCTTCATCCTCGTGTACTGAAGAAGAAAACAAACATCATATGGGAATCGATGTTATTATCAAAGTCACAAAGCAAGACCAAACACCGACCAATGATAAGATTTGTCAATCCGTAACGGAAGTTACAGAGACCGAAGATGATGAGGTATCCGAAGAAGTTGTAAAAGGAGATCCCACCACTTATTACACTATCGTCGGTGCGGGTCTTAACATGAACTTTGGATTCACCAAATGCCCAAAGATTTCATCCATCTCCGAATCCTCTGATGGAAACACTGTGAATACTAGATTGTCCAGCGTGTCACCGGGACAAGGTAAGGACTCTCCCGCGATCACGCGTGAAGAAGCTCTGGCTATGATCAAAGACTGTGAGATGTCTATCGACATCAGATGTAGCGAAGAAGAGAAAGACAGTGACATCAAGACCCATCCAGTACTTGGGTCTAACATCTCACATAAGAAAGTGAGTTACAAAGATATCATCGGTTCAACGATCGTTGATACAAAATGTGTCAAGAACCTAGAGTTTAGCGTACGTATCGGAGACATGTGTGAGGAATCATCTGAACTTGAAGTCAAGGATGGATTCAAGTATGTCGACGGATCGGCATCTGAAGGTGCAACCGATGATACTTCACTCATCGATTCAACAAAACTCAAAGCATGTGTCTGAATCGATAACTCTATTCATCTGAAAATGGATGAGTTGGGTTAATCGAACGATTCAGACACCGCACCACGAATTAAAAAAGACCGGGCACTATATTCCGGTTTGCAAAACAAAAATATTTAACTACATTCACAAAAAGTTACCTCTCGTTACTTCTTCTTTCTGTTTCAATATGTGATACGATATGATCACTATTCGTATTCTCTTGGTCTCATAAAAAAGTTTTACAAAAAAAAAAAAAAAAATATTTTTATTCTCTTTCTCTCTTCGATGGTCTCACAAAAATATTAAACCTCTTTCTGATGTCTCAACTATTTCGTAAACGATAACGTCCAACAATATATTCTCGTAGAGCTTATCAACATCCTTATACCAATCTAGGTTGTCAGACAATTGCATCATAAAATAATGTTTATAATTTACACGTTAACATCATATAATAAACGTATATAGTTAATATTTTTGGAATATAAATGATCTGTAAAATCCATGTAGGGGACACTGCTCACGTTTTTTCTCTAGTACATAATTTCACACAAGTTTTTATACAGACAAATTAATTCTCGTCCATATATTTTAAAACATTGACTTTTGTACTAAGAAAAATATCTTGACTAACCATCTCTTTCTCTCTTCGATGGGTCTCACAAAAATATTAAACCTCTTTCTGATGGAGTCGTAAAAAGTTTTTATCCTTTCTCTCTTCGATAGGTCTCACAAAAATATTAAACCTCTTTCTGATGGTCTCTATAAACGATTGATTTTTCTTACCCTCTAGAGTTTCCTACGGTCGTGGGTCACACATTTTTTTCTAGACACTAAATAAAATAGTAAAAT

>2017_UK_Monkeypox_MT903344.1

TAATTAATTTAATTTTACTATTTTATTTAGTGTCTAGAAAAAAATGTGTGACCCACGACCGTAGGAAACTCTAGAGGGTAAGAAAAATCAATCGTTTATAGAGACCATCAGAAAGAGGTTTAATATTTTTGTGAGACCTATCGAAGAGAGAAAGGATAAAAACTTTTTACGACTCCATCAGAAAGAGGTTTAATATTTTTGTGAGACCCATCGAAGAGAGAAAGAGATGGTTAGTCAAGATATTTTTCTTAGTACAAAAGTCAATGTTTTAAAATATATGGACGAGAATTAATTTGTCTGTATAAAAACTTGTGTGAAATTATGTACTAGAGAAAAAACGTGAGCAGTGTCCCCTACATGGATTTTACAGATCATTTATATTCCAAAAATATTAACTATATACGTTTATTATATGATGTTAACGTGTAAATTATAAACATTATTTTATGATGCAATTGTCTGACAACCTAGATTGGTATAAGGATGTTGATAAGCTCTACGAGAATATATTGTTGGACGTTATCGTTTACGAAATAGTTGAGACATCAGAAAGAGGTTTAATATTTTTGTGAGACCATCGAAGAGAGAAAGAGAATAAAAATATTTTTTTTTTTTTTTGTAAAACTTTTTTATGAGACCAAGAGAATACGAATAGTGATCATATCGTATCACATATTGAAACAGAAAGAAGAAGTAACGAGAGGTAACTTTTTGTGAATGTAGTTAAATATTTTTGTTTTGCAAACCGGAATATAGTGCCCGGTCTTTTTTAATTCGTGGTGCGGTGTCTGAATCGTTCGATTAACCCAACTCATCCATTTTCAGATGAATAGAGTTATCGATTCAGACACATGCTTTGAGTTTTGTTGAATCGATGAGTGAAGTATCATCGGTTGCACCTTCAGATGCCGATCCGTCGACATACTTGAATCCATCCTTGACTTCAAGTTCAGATGATTCCTCACACATGTCTCCGATACGTACGCTAAACTCTAGGTTCTTGACACATTTTGTATCAACGATCGTTGAACCGATGATATCTTTGTAACTCACTTTCTTATGTGAGATGTTAGACCCAAGTACTGGATGGGTCTTGATGTCACTGTCTTTCTCTTCTTCGCTACATCTGATGTCGATAGACATCTCACAGTCTTTGATCATAGCCAGAGCTTCTTCACGCGTGATCGCGGGAGAGTCCTTACCTTGTCCCGGTGACACGCTGGACAATCTAGTATTCACAGTGTTTCCATCAGAGGATTCGGAGATGGATGAAATCTTTGGGCATTTGGTGAATCCAAAGTTCATGTTAAGACCCGCACCGACGATAGTGTAATAAGTGGTGGGATCTCCTTTTACAACTTCTTCGGATACCTCATCATCTTCGGTCTCTGTAACTTCCGTTACGGATTGACAAATCTTATCATTGGTCGGTGTTTGGTCTTGCTTTGTGACTTTGATAATAACATCGATTCCCATATGATGTTTGTTTTCTTCTTCAGTACACGAGGATGAAGATTGTTGAAGACTAGTAGGCATAGCAGCTGCCACTAGGCACATGCATGCCAGGACAATATATTGTTTCATGATTGCTATTGATTGATTACTGTTCTAGATGATTCTACTTTCTTACCATATAATAAATTAGAATATATTTTCTACTTTTACGAGAAATTAATTATTGTATTTATTATTTATAGGTAAAAAAACTTACTATAAGTGGGTGGGATTCTGGGAATTAGTGATCAGTTTATGTATATCGCAACTAGCGGGCATATGGCTATTGACATCGAGAACATTACCCATATGATAAGAGATTGTATCATTTTCGTAGTCTTGAGTATTGGTATTACTATATAGTATGTAGATGTCGACGCTAGATAGACAGTCGCCCACTAGAGTTACCGTCTCTGAATGCGGCATGATAGTATCATTCTTTGTTTTCGTTAACTGTTTGGAAGATGAATCTTTGTTGTTACATTTAATCTCGAAATTCAGAGTACATATCTTTGAAGTATTCTGATATCTATTTTCTCCTGTAAAGAATCCTGAAGTTGCTACATTATTAAGGACAGAGAAGTATTCTGCACGAAAGACTGGATCACAATCTTTATGATTCATGGTAATAGTTAGTTCCGACGTTGAGATGGATTCGCTGAGACCGGTAGTGGTCGTCCGAGTACACGATGTGTCGTTGACTGGATACAGGTTAATTTCCACATCGATATAGTTAAATGTATTGCTGGTTACGACGGGTTCGCATTTATCTGTGGAAGAGACGGTGTGAGAATATGTTCCGGGACCACACGGAGAACAGATGACGTCTCCGGTAGACGTGTATCCGGATACTCCGTATCCTATTCCACACTTTGTTTTAGAAATACATGTTCTACACCCTGATGCTCCTTTGAGAAGACAATAATATCCTGGAGAGCATTCACAGATTCTATTGTGAGTCGTGTTACACGATCGCGTCTCTACCTGATTACTATCACATCTTCCGTTACAACTTAGACAAGCCTGTAAATGATTATTGTGAGATGTAAAGGTATCCGAACCACACGGTGTACATTGTGTATTAGTCTTGCTATCACATAATCTGGAAGCGTAAGTTCCCGGAGGACACGATAGACAACATAGATTACGGCTTCTGTATTCGTTGTCTTTACACTTTCCATTGGATGGTGCATGTGGTGCTATATCTCTTCCGTTTATTATTATACATGAGAGAAACAATATATACGAGTATAATACGGACCTCATGATTTAATAATGTAGTAATCGTCGTCTTGTTACTGTTTGTTTCCTACTTCTCCAATCATATAGATTATTTTTTAAATATTTTCTTTCTATCATGGATAATATTTGTAATGGTTCTTTCCGTACAACATACTGTTTAGATGGTAGTCGCTTAGCTTGGTTATGATATTGCGCATAATTTCCGGAGGCAAATACGATAGTCTAGATTGACTATCGATGGTAGACTCTAATTTATTGAGTGCTTTGTCGACGAGTTTACTTTTATGCTCCATCGATAGATGACACTGTTCTATGAGATCGTCGTACATGGGAAATGAAATGCGTTTGTCTGAATGTATGGCTTCGAGATATGTGTGATACCGGATGTCTTCTGTTCTCAATACCGTATACAAGTCGGTGTCTGAGATTCGAATCTCTTTGAGGAGACTTATGTCACGACTACATTTTTCGATGATGGAATCTATCTTATCGAATGATATATTTTTCATAAATACACTTTTATAGTCCTCGTTTAAACAGAATTTAGTATGTAGTTCCGCAAATGACTCGTCCCTTAATAGGCAGTAGGCTATTATCTTCTTTACGTAGTGATCGTCGTAGGGAGAGAACTCCGACATCTTGTAGAACAACGATTTAATCATAGGTAGAGATACTTTCAGTCTGTGGTGGATAATGTCATTAACGACATCCGCCTTGTATATAATGTTTCTGTTTTCAAACACCAAGTCGAATACTGTCTTATCGTCTTTAGTCGGAAGGTTGATGTCGTATCCGATGTATACGAGGTATGAGGCAACATTGTTATTGCAATTCTGGAAGGCGGTATGAAGAGGAGTCATTGTATTATAGTATTCGTCTTTCTGAATGTCGAATCTATCTAGTAGATACCGTAGTATATTGAGAGAGCGACTTCCATATCCTTGATTATGTTTTATGAATAGATAAAGTAGATGTTGTCCTTCTTCCTTTTGTAATTTCCCGTATTTTTGTTCGTGCCAATTGAGTAACATTATGAGAATATGACCTGTTGCACAATCGTTCTTTATGTATTCCATGATGGGTGTACAATCAAGATTATTACGTATCCTCGTATCGGCTCCTCGAGATAAAAGAGCATACACCACACGAGGACTATGTTTGGTATACTGTTGAAGGTAAGTGTGTAACGGCGTATTTCCGATTTTCGTAACCGCGTTAATGTTTGCTCCATGATCTATTATCGCGTAGATGAATCGCTTCTCAGCTCGCATCTTAGTGTGACTCTTTGACTTGTAATAATTGCTTTCGTGGAACGCGGATATGTGTTTACAGTAGTAATGAAGAGAAGTGAGTCCATCCTCATCGACGCAATTAGGGTCAGATCCTTTAGTCAATAATTTGTACAGAACGTAATAGTTTAAGCTCCCATTGAATTTATATCTAAGATAACACAGCAATAGATCGGATGATTTACTAAAGTCATCAATGGGGTCCGTTAGTATATCAAAGATCTTGTTATCGATTGATAGTGAATGAATCAGATAGTGGTGTAGAGGAATATGTCCTTTTTTATCCTTGCTATCAAAGTTACGCATGCCGTGGCGTAACAATATCTTTAATACAGATGGATTAAATCGTGTATTCATCGTATAGCAATGTAATGGAGAGTTACCACATTTTAGTCGTTTATTCAGATCGCAGTGTTTAATAACTAATTTAAACAGATGAGATGATATATCCACATCAAAGAATGCGAGATACATATGACAGACATTATTGACAGAAATGTGACCTTCATTATCACCGTCGTCCATAAATGCGTTAGGTACGTACCACATACTATCGTTAACGATGCGCACAATCTCGTCCATTTCATCCATCTTCATAATGATTTACTTTTTCATAATTAGAGAAAAAGATCAAGGTATAAAAATTAGAAGTGTTAGACTATAAACTAACTTATGACTTAACTAACTTATGACTTAACTAACTTATGACTTAACTAACTTATGACTTAACTAACTTATGACTTAACTAACTTATGACTTAACTAACTTATGACTTAACTAACTTATGACTATTAACTCATTTCAAGAAAGGTGGGTGGATAGAACTCTATATGACAGCTTGTGAAACAATTAGATCCCTAATTTCTAATGGAAGTTTTGATAGGAGATTGTCATCAGTTGATACATTGTTTATTATCTCATCTATTAGAGCACGTCTGTTTAGAGCTTTAGTGACCTGCTCGGTTACTTCTGTGTAAATCTTGAATCCTTTAGTGATACACTGTGTCAAAACTGGATGTTTAGAATACCTATGTAGAATATGGGAAGCATGCTTGTTTTTGTCTCTATTATAGATTAACTCATACATGGTTGTATTATGAATTTTCATCTGCCTAATGTACTCCAATTCTTGTTTACAATCAATTATATAATCAAAGAGTGATGATGCATACACATTACAAAGTGAATAATCTACCATCATAAAATACTTGATACAGAGCTTTATCACATCATGGTTTTCAATTGTATTATTAAGTATAGCTAATTTTATACAGTCAATAGACAATGGTTCTCTAAGCAATATTTCTAATATTTTAAGATGTGCTTCCCTACGGGCGATGACAGATCCCCTATCCACGGCCACGTCAAGACATGTATATCCATTACTCATTACTGCGTTGACATTTGCTCCATTTTCTAATAGCCATGATACTAAATCTATATAACCTGCATAGATAGCGCGATAAAGCAAGGTCCTTCCACCAGCATCTAGTTGATTGATATCTTCAATATATGGGATACAAAGCTTATAAATTTCTAATACTGTGGGTTCATCTACAAGGAATCCCCTAGTATACTGAATTATTTTATATAGATCTAATTTAACATCATTTTCATCTGGGATACCACAATTCAAAATAAACTCAACAACACTACTTTCCTTTTTACATATTCCCCTAAAATAGGCATTCAAGCATTCTATTTTATATATTACAGCCCCATGATCTACCATAAGATCAACAATGTCTATTTCTACATATGCATTAGATAGATAGTAAAGTAAGAGATCTTGCACAGAATTACAATTCTTAATAATTATAGAGAAAATATCTTCCATATAATTCTTTGACACTAATGCAGATATAATATCTTTATATGTAATATATGCAAACAGTCTATCTACTATATACTGATCAATATTATCTCTATGAATCCTAAAATAATCATACAGAACATCTACAGGATCACAAATTGGTTCAAGGAGAAATCTATCAAATATTTTCCTGTCAACAACTGGTTCTAGAACATAACAGTCAACACCTAATCCATGTTTTTTATAATCATCTACCAAAGATAATGACCAAAGATCGAGGTCGTCGTGAAACTGCTCATCGACAGCCATGAAATCTACCGACTCCATGGTGCGAATCGCACTGTCTTATTCGCCATTGATTTTCATTTTTTATAATTATGTACATGTTTTCCTTCTATTCTCAAGAGTCTACAAAAATATATTTTTTCGATATCTAAGTACTAAGTTTTTTTACTGTTTTTGTTACTGTCTTCCATTCTTCTAACTAAAGATCTGAGATAAATTATACAATCTTCGCTATCGAACCATTTTTGTAGTCTAAAGCCTGAAGTAATTAACCAACTGTTTTTATTAGTGGCTTTTTTCGATCTATCCTCGTCCTCTCATCATCCTTATATTATTATCATTATTATCATAGTCTATTAAACACAAATCATCTACGTTTATAACAACATTCTCATTATTAATTAGTTCTGTAGAATATCTTTAATAATTTGGCTATACATCTGTTCAATACTATCTATTGATGATTTCTTTTTTAAGACTTAAACTAGTTATGGTAATGACGATGAAATCGAGTAGTAACTTCTAATAAAGACTTGATATCATTATCATATGTTTGATCGTCATAGTTAATAGTGTGGCTAAATGGTACTGTTAATAAGTTTATAGACAATATCATAGTATTTTCTTTCCAGAATTAGATTATTTTTTTAAATACTGATCCTCACAATTCCGTGATGTAGCAGTAGTTGGTGCATGGTCTATATCGTTAAAATGTATCATATATAATAGTTTTCTGACGTGGAGTACAGAATTTTCGATTAATGAGTTCATGGTAAGGAAGGGCAAATGCCTGTATATAATATACATAAGTTAATAGTTTTTTATCATATTTTCTAATACCATAATAAAAATTATCATTATTGCGTTTGGTAGTTCTGCCCTATCATCTATATCACTGTCACTCTCGCTCTCACTATATCTTCTAAAATTACAAACAACTGGATATTCGATAACAGCATTTGTGTAGTTTTTGTCTTTTACAGTATATACGTTATTGTCAAAATCTAAACAAATATTAGCATAATACATCTATCTATAAGATCAGGATCCATGTTCGAGCATACTAGCCATGTATATTTGTAACTTCGTCGTACAGCGTTAGATCAATAGAATAAACAATCGTGTGACGCAACTTTTTTACGATCTAGTTGTATGAGTTTATCGTTTACATAAGCAATTAACGGCTTTAACAGATGATCTGAGTAATAATATACCTCTGTTATACGTTTAATGTTCACGGTCTTAGTATTTTTAGATATCAATTGTGATTTACACCATATTCGACTCCCTTGTGTGTAACGTTAGAAATTCTAAATCTATATTATCTATTACAGCGTAAAACACATTCAATATTGTATTGTTATTTTTATATTATTTACACAATTAACAATGTATTATTAGTTTATATTACTGAATTAATAATATAAAATTCCCAATCTTGTCATAAACACAAAATCCATTAAAAATGTCGATAAAATATCTGATGTTGTTGTTCGCTGCTATGATAATCAGATCATTAGCCGATAGTGGTAATGCTATCGAAACAACATCGCCAGAAATTACAAACGCTACAACAGATATTCCAGCTATCAGATTATGTGGTCCAGAGGGAGATGGATATTGTTTACACGGTGACTGTATCCACGCTAGAGATATCGACGGTATGTATTGTAGATGCTCTCATGGTTATACAGGTATTAGATGTCAGCATGTAGTATTAGTAGACTATCAACGTTCAGGAAAACCAGACACTACAACATCATATATCCCATCTCTAGGTATTGTGCTTGTATTAGTAGGCATTATTATTACGTGTTGTCTATTATCTGTTTATATGTTCACTCGAAGAACTAAACTACCTATACAAGATATGGTTGTGTTATATTTTTTATAAAATTTTTTTATGAGTATTTTTACAAAAATGTATATGTATAAAAAAAATACTAAGTATACGATGTATCCTGTATTATTTGTATTTATCTAAACAATACTTCTGCCTCTAGATGGGATACAAAAATTTTTTATTTCAGCATATTAAAGTAAAATCTAGTTACCTTGAAAATGAATACAGTGGGTGGTTCCGTATCACCAGTAAGAACATAATAGTCGAATACAGTATCCGATTGAGATTTTGCATACAATACTAGTCTAGAAAGAAATTTGTAATCATCCTCTGTGACAGGAGTCCATATATCTGTATCATCGTCTAGTTTTTATCAGTGTCCTATGCTATATTCCTGTTATCATCATTAGTTAATGAAAATAACTCTCGTGCTTCAGAAAAGTCAAATATTGTATCCATACATATATCTCCAAAACTATCACTTATACGTTTATCTTTAACGAACATATACCTAGATGGTTATTTACTAACAGACATTTTTTCAAGATCTATTGACAATAACTCCTATAGTTTCCACATCAACCAAGTAATGATCATCTATTGTTATATAACAATAACATAACTCTTTTCCATTTTTATCAGTATCTATATCAACGTCGTTGTAGTGAATAGTAGTCATTGATCTATTATATGAAACGGATATGTCTAGTTAATATTTTCTTTGATTTAAAGTCTATAGTCTTTACAAACATAATATCCTTATCCGACTTTATATTTCCTGTAGGGTGGCATAATTTTATTCTGCCTCCACAATCAGTGTTTCCAAATATATTACTAGACAATATTCCATATAGTTATTAGTTAAGGGACCCAATTAGAACACGTACGCGCTTATTCATCATTTGGATCGTATTTCATAAAAGTTATTATGTTATAGATGTCAACACATTCTACATTTTTTTAATTGTCTATATAGTATTTTTCTGATATTTTCTATAATATCAGAATTGTCTTCCATAGGAAGTTGTATACTATTCGGAATCAGTTACATGTTTAAATAATTCTCTGATGTCATTCCTTATACAATCAAATTCATTATTAAACAGTTTAATAGTCTGTAGACCTTTATCGTCGTACATATCCATTGTCTTATTAGTTACGCTTATTTTTATGGGTTTTACGTTGCTTTATTATATTTTATAATAATGATTGTTTGACAATGTCGTAGTATAGATATATTATTAGAGGAGGTATAATTATAAAAAGTTTCTGAGTACGATGTTATAAGAGGAGAGGACACATTAACATCATACATCAATTAACTACATTCTTATAACATTGTAATCAAAAGAATTGCAATTTTGATGTATAACAACTGTCAATGGAATTGTATATTACAAATTACGGTATGTTGTAACAACAAATACCGATCGGTAATTGTCTCTGCCGCTGTAATAGAATTAATTATATATCTATTACACCGGCCTTGTATCATAATAAAGTTGTGGTAGTATGATCTCCATATTTATAATTTAGTACTTTGTATTTAGTTTTTTTGGAATCATAAAAAAAGTTTTACTAATTTAAAATTTAAAAAGTATTTACATTTTTCACTGTTTAGTCGCGGATATGGAATTCGATCCTGTCAAAATCAATACATCATCTATAGATCATGTAACAATATTACAATACATAGATGAACCAAATGATATAAGACTAACAGTATGCATTATCCGAAATGTTAATAACATTACATATTATATCAATATCACAAAAATAAATCCACATTTGGCTAATCGATTTCGGGCTTGGAAAAAACGTATCGCCGGAAGGGACTATATGACTAACTTATCTAGAGATACAGGAATACAACAATCAAAACTTACTGAAACTATACGTAACTGTCAAAAAAATAAAAACATATATGGTCTATATATACACTACAATTTAGTTATTAATGTGGTTATTGATTGGATAACCGATGTGATTGTTCAATCAATATTAAGAGGGTTGGTAAATTGGTACATAGCTAATAATACATATACTCCAAATACACCCAATAATACTACAACCATTTCTGAGTTGGATATCATCAAAATACTGGATAAATACGAGGACATGTATAGAGTAAGTAAAGAAAAAGAATGTGGAATTTGCTATGAAGTTGTTTACTCAAAACGATTAGAAAACGATAGATACTTTGGTTTATTGGATTCGTGTAATCATATATTTTGCATAACATGTATCAATATATGGCATAGAACACGAAGAGAAACCGGTGCGTCAGATAATTGTCCTATATGCCGTACACGTTTTAAAAAAATAACAATGAGCAAGTTCTATAAGCTAGTTAACTAATAAATAAAAAGTTTAATTATCGACGATATATGTCGTTATTTTTCTCTCATATGAAAGATTAATTTGATTCTAATATAATCTTCAGTATTGGATGAATCTCAATTCAAATTAATTCCATTAGATTAGATTAGATTAGATCATAAATAAAAATAGTAGCACGCACTACTTCAGCCAAATATTCTTTTTTGAAACGCCATCTAGCGTAATGAGAACACAAGTGAACCTATAATGAGCAAATTTATTAGTATCGGTTACATGAAGGACTTTACGTAGAGTGGTGATTCCTCCATCTGTGGTACGAACGGTTTCATCTTCTTTGATGCCACCACCCAGATGTTCTATAAACTTGGTATCCTCGTCCGATTTCATATCATTTGCCAACCAATACATATAGCTAAACCCAGGCATACGTTCCACACATCCGGAACAATGAAATTCTCCAGAAGATGTTACAATGGCTAGATTTGGACATTTGGTTTCAACCGCGTTAACATATGAGTGAACACACCCATACATGAAAGCGATGAGAAATAGGATTCTCATCTTGCCAAAATATCACTATAAAAAATTTATTTATCAATTTTAAAGGTATAAAAAAATACTTATTGTTGCTCGAATATTTTGTATTTGATGGTATACGGAAGATTAGAAATGTAGGTATTATCATCAACTGATTTTATGATGGTTTTATGAATTTTATTATGCTTCACTATTGCATCGGAAATAATATCATATGCTTCCACGTATATTTTATTTTGTTTTGACTCATAATACGCACGTAATTCTGGATTATTGGCATATCGATGAATAATTTTAGCTCCATGCTCAGTAAATATTAATGAGAACATAGTGTTGCCTCCTACCATTATTTTTTTCATCTCATTCAATTCTTGATTGCAGAGATCTATATAATCATTATAGCGTTGACTTATGGACTCTGGAATCTTAGACGATGTACAGTCATCTATAATCATGGTATATTTAATACATTGTTTTATAGCATAGGCATTATCTACGATATTAGATACTTCACTCAATGAATCAATCACACAATCTAATGTAGGTTTATGACATAATAGCATTTTCAGCAGTTCAATGTTTCTAGATTCGTTGATGGCAATGGTTATACATGTATATCCGTTATTTGATCTAATGTTGACATCTGAACCGGATTCTAGCAGTAAAGATACTAGAGATTGTTTATTATATCTAACAGCCTTGTGAAGAAGTGTTTCTCCTCGTTTGTCAATCATGTTAATGTCTTTAAGATAAGGTAGACAAATGTTTATAGTACTAAGAATTGGGCAAACATAAGACATGTCACAAAGACCCTTTTTGTATGTATAAGTGTAGAAATTATAACATCCATAGTTGGATTCACATAGGTGTCCAATCGGGATCTCTCCATCATCGAGATGATTGACGGCATCTCCCCCTTCCTTTTTTAGTAGATATTTCATCGTGTAAGAATCAATATTAATATTTCTAAAGTATCTGTGTATAGCCTCTTTATTTACCACAGCTCCATATTCCAACATGCATTCCACTAGAGGGATATCGATATCGCCGAATGTCATATACTCAATTAGTATATGTTGGAGGACATCCGAGTTCATTGTTTTCAATATCAAAGAGATGGTTTCCTTATCATTTCTCCATAGTGGTACAATACTACACATTATTCCGTACGGCTTTCCATTCTCCAAAAACAATTTTACCAAATCTAAATCTACATCTTTATTGTATCTATAATCACTATTTAGATAATCAGCCATAATTCCTCGAGTGCAACATGTTAGATCGTCTATATATGAATAAGCCGTGTTATCTATTCCTTTCATTAACAATTTAACGATGTCTATATCTATATGAGATGACTTAATATAATATTGAAGAGCTGTACAATAGTTTTTATCTATAGAAGACGGCTTGATTCCGTGATTAATTAGACATTTAACAACTTCCGGACGCACATATGCTCTCGTATCCGACTCTGAATACAGATGAGCGATGATATACAGATGCAATACGGTACCACAATTTCGTGGTTGATAATCATCATACACGTATCCGTACTCGTCATCCTCATAAAGAACACTGCATCCATTTTCTATGAACAAATCAATAATTTCAGGAACAGGATCATCTGTCATTACATAATTTTCTATAACTGAACGATGGTTTTCACATTTAACACTCAAGTCAAATCCATGTTCTACCAACACCTTTATCAAGTCAACGTCTACATTTTTTGATTTCATATAGCTGAATATATTAAAGTCATTTATGTTGCTATATCCAGTAGCTTCTAGTAGAGCCATCGCTATATCCTTATTGACTTTAACATGTCTACTATTTGTGTATTCTTCTATTGGGGTAAACTGTCTCCAATTTTTATGTAATGGATTAGTGCCACTATCTAGTAGTAGTTTGACGACCTCAACATTATTACAATGCTCATTGAAAAGGTATGCGTGTAAAGCATTATTCTTGAATTGGTTCCTGGTATCATTAGGATCTCTGTCTCTCAACATCTGTTTAAGTTCATCGAGAACCACCTCCTCATTTTCCAGATAGTCAAACATTTTGACTGAATAGAAGTGAATGAGCTACTGTGAACTCTATACACCCGCACAACTAATGTCATTAAATATCATTTTTGAATGTATTTATACCATGTCAAAAACTTGTACAATTATTAATAAAAATAATTAGTGTTTAAATTTTACCAGTTCCAGATTTTACACCTCCGTTAACACCTCCATTAACCCCACTTTTTACACCACTGGACGATCCTCCTCCCCACATTCCACTGCCACTAGATGTATAAGTTTTAGATCCTTTATTACTACCATCATGTCCATGGATAAAGACACTCCACATGCCGCCACTACTACCCCCTTTAGACGACATATTAATAAGACAAGTTTAACAATAAAATTAATCACGAGTACCCTACTACCAACCACTATTATATGATTACAGTACCTTGACTAAAGTCTCTAGTCACAAGATCAATACTACCAACCTACGCTATTATATGATTATAGTTTCTATTTTTATAGGAACGCGTACGAGAAAATCAAATGTCTAAGTTCTAACGGTAGTGTTGATAAACGATTGTTATCCGCGGATACCTCATCTATCATGTTGTCTATTTTCTTACTTTGTTCTATTAACCTATTAGCATTATATATTATTTGATTATAAAACTTATATTGCTTATTAGCCCAATCTGTAAATATCGGATTATTAACATATCGTTTCTTTGTAGGTTTATTTAACTTGTACATCACTGTAAGCATGTCCGTACCATTTATTTTAATTTGACACATATCAGCAATTTCTTTTTCGCAGTCGGTTATATATTCTATATAAGATGGATACGTATCACATATGTACTTATAGTCTACTAATATGAAGTACTTAATACATATTTTCAGTAACGATTTAGCCTTATTACCTATTAATAAGTGCCTGTCGTTGGATAGGTAATCAACTGTTTTCTTAATACATTCGATGGTTGGTAATTTACTCAAAATAATTTCCAATATCTTAATATATATTTCTGCTATTTCTGGTATACATGCATGTGCCATTATAACACAAATACCAATACATGTAGACCCATATGTTGTTGTTATATTAATATCTGCGCCATTATCTATTAACCATTCTATTAGGGCAACACTATGCGACTCGATACAATAATAAAGTATACTACGTCCATGTTTATCTATTTTGTTTATATCATCGATATACGGCTTACAAATTTCTAGTATCGATAACACTTCTGACTCGTGAATAAATAAGGTAGGGAATAACGGCATAATATTTATTATGTTATCATCATTAACAACTACGTTTCCATTTTTTAAAATATACTCTACAACTTTAGGATCCCTATTGTCAAATCTTTTAAAATATTTATTTATATGCTTAAATCTATATAATATAGCTCCTTCCCTAATCATACATTTGATAACATTGATGTACACTGTATGATAAGATACATATTCTGACAATAGATCTTGTATAGAATCTGTATATCTTTTAAGAATTGAGGATATTATGACATTATTACGTAAACTATTACACAATTCTAAAATATAAAATGTATCATGGGCAGATAATAGTTTATCCACTATATAATTATCTATTTTATGATTTTTCTTCCTATATTGTTTACGTAAATAGATAGATAGAATATGCATTAGTTCATTACCGCTATAGTTACTATCGAATAACACGTCAAATATTTCCCGTTTAATATCGCATTTGTCAACATAATAATAGAGTATGGTACGTTCACGATAAGTATAATGACACATCTCGTTTTCGTGCGAAATTAAATAGTTTATCACGTCCAAAGATGTCACATAACCATCTTGTGACCTAGTAATAATATAATAATAGAGAACTGTTTTACCCATTCTATTATCATAATCAGTGGTGTAGTCATAATCTAAATAATCAAACTCGTCATCCCAATTAAAATAAATATAATCAGTACATTGAATGGGTATGATATTGTACCCATACTGTATGTTGCTACATGTAGGTATTCCTTTATCCAATAATAGTTTAAATACATCTATATTAGGATTTGATGTTGTCGCGTATTTCTCTACAATATTAATACCATTTTTGATACTATTTATTTCTATACCTTTCGAAATTAGTAATTTCAATAAGTCTATATCGATGTTATCAGAACATAGATATTCAAATATATCAAAATCATTGATATTTTTATAGTCGACTGACGACAATAACAAAATCACAACATCGTTTTTGATATTATTATTTTTTTTGGTAACGTATGCCTTTAATGGAGTTTCACCATCATACTCATATAATGGATTTGCACCACTTTCTATTAATGATTGTGCACTACTGGCATCGATGTTAAATGTTTTACAACTATCATAGAGTATCTTATCGTTAACCATGATTGGTTGTTGATGTTATCACATTTTTTGGTTTCTTTCATTTCAGTTATGTATGGATTTAGCACGTTTGGGAAGCATGAGCTCATATGATTTCAGTACTGTAGTGTCAGTACTATTAGTTTCGATCAGATCAATGTCTAGATCTATAGAATCAAAACACGATAGGTCAGAAGATAATGAATATCTGTACGCTTCTTCTTGTACTGTAACTTCTGGTTTTGTTAGATGGTTGCATCGTGCTTTAACGTCAATGGTACAAATTTTATCCTCGCTTTGTGTATATCATATTCGTCTCTAGTATAAAATTCTATATTCAAATTATCATGCGATGTGTGTACGCTAACGGTATCAATAAACGGAGCACAGCATTTAGTCAACAGTAATCCAAAATTTTTTAAAGTATATCTTAACGAAAGAAGTTGTCATCGTTAGAGTGTGGTAAATCATTGTCTACGGTACTAGATCCTCATAAGTGTATATATCTAGAGTAATATTTAATTTATCAAATGGTTGATAATATGGATGTTGTGGCAATTTCCTAATACGGAAATAAGACATAAACACGCAATAAATCTAATTGCGGACATGTTACACTCCTTAAAAAATACGAATAAACACTTTGGCTTTTAGTAAGTGTCATTTAACACTATACTCATATTAATCCATGGACTCATAATCTCTATACGGGATTAACGGATGTTCTATATACGGGGATGAGTAGTTCTCTTCTTTAACTTTATACTTTTTACTAATCATATTTAGACTGATGTATGGGTAATAGTGTTTGAAGAGCTCGTTCTCATCATCAGAATAAATCAATATCTCTGTTTTTTTGTTATACAGATGTATTACAGCCTCATATATTACGTAATAGAACGTGTAATCTACCTTATTAACTTTCACCGCATAGTTGTTTGCAAATACGGTTAATCCTTTGACCTCGTCGATTTCCGACCAATCTGGGCGTATAATGAATCTTAACTTTAATTTCTTGTAATCATTCGAAATAATTTTTAGTTTGCATCCGTAGTTATCTCCTCTATGTAACTGTAAATTTCTCAACGCGATATCTCCATTAATAATGATGTCGAATTCGTGTTGTATACCCATACTGAATTGATGAACTAATACCGACGGTATTAATAGTAATTTACTTTTCATCTTTACATACTTGGTAATAGTTTTACTATCATAAGTTTATAAATTCCACAAGCTACTATGGAATATACCAACCATCTTAGTATAGAACACATGTCTTAAAGTTATTAATTAATTACATGTTGTTTTATATATCGCTACGAATTTAAACAGAGGAATCAGTTAGGAAGAAAAAATTATCTGTCATCATCATCATCTATTGGATAACATCTCTGTATTCTACGATAGAGTGCTATTTTAAGATGTGACAGATCCGTGTCATCAAATATATACTCCATTAAAATGATTATTCCGGCAGCGAACTTGATATTGGATACATCACGACCTTTGTTAATATCCACGACAATAGACAGCAATCCCATTGTTCCATAAACAGTGAGTTTATCTTTCTTTGAAGTGATATTTTGTAGAGATCTTATAAAACTGTCGAATGACATCGTATTTATATCTTTAGCTAAATCATATATGTTACCATCATAATATCTAACAGCATCTATCTTAAACGTTTCCATCGCTGTAAAGACGTTTCCGATAGATGGTCTCGTTTCATCAGTCATACTGAGCCAACAAATGTAATCGTGTATAACATCTTTGATAGAATCAGACTCTAAAGAAAAGGAATCGGCTTTATTATACACATTCATGATAAACTTAATGAAAAATGTTTTTCGTTGTTTAAGTTGGATGAATAGTATGTCTTAATAATTGTTATTATTTCACTAATTAATATTTAGTAACGAGTACACTCTATAAAAACGAGAATGACATAACTAATCATAACTAGTTATCAAAGAATGTCTAGGACGCGTAATTTTTTATGGTATAGATCCTGTAAGCGTTGTCTGTATTCTGGAGCTATTTTCTCTATCGCATTAGTGAGTTCAGAATATGTTATAAATTTAAATCGAATAACGAACATAACTTTAGTAAAGTCGTCTATATTAACTCTTTTATTTTCTAGCCATCGTAATACCATGTTTAAGATAGTATATTCTCTAGTTACTACGATCTCATCGTTGTCTAGAATATCGCATACTGAATCTACATCCAATTTTAGAAATTGGTCTGTGCTACATATCTCTTCTATATTATTGTTGATGTATTGTCGTAGAAAACTATTACGTAGACCATTTTCTTTATAAAACGAATATATAGTACTCCAATTATCTTTACCGATATATTTGCATACATAATCCATTCTCTCAATCACTACATCTTTAAGAGTTTGGTTGTTAAGATATTTGGCTAAACTATATAATTCTATTAGATCATCAACAGAATCAGTATATATTTTTCTAGATCCAAAGATGAACTCTTTGGCATCCTCTATAATATTATCAGAAAAGATATTTTCGTGTTTTAGTTTATCAAGATTTAACCTGTTCATATCCATGATTAACGACGTCATATAACCACATAAAATAAAAATCCATTTTCATTTTTAGCACAATACTATTCATAATTGATATTGATGTAATATTTTGTTACTTTGAACGTAAAGACAGTACACGGGTCCGTATCTCCAACAAGCACGTAGTAATCAAATTTGGTGTTGTTAAACTTCGCAATATTCATCAATTTAGATAGAAACATATACTCATCATCTGTTTTAGGAATCCATGTATTATTACCACTTTCCAACTTATCATTATCCCAGGCTATGTTTCGCCCATCATCGTTGCACAGAGTGAATAATTCTTTTGTATTCGGTAGTTCAAATATATGATCCATGCATATATCGACAAAGCTATTGTAGATGTGATTTTTCCTAAATCTAATATAAAACTCGTTTACTAGCAAACATTTTCCTGATTTATCGACCAAGACACACATGGTTTCTAAATCTATCAAGTGGTGGGGATCCATAGTTATAACGCAGTAACATAAATTATTACCTTCTTGACTGTCGCTAATATCTATATACTTATTGTTATCGTATTGGATTCTACATATAGATGGCTTGTATATCAAAGATATAGAACACATAACCAATTTATATTCTCGCTTTGTATTTTCGAATCTAAAGTTAAGAGATTTAGAAAACATTATATCATCGGATGATGATATCACTGTTTCCAGAGTAGGATATATTAAAGTCTTTAAAGATTTTGTCCGATTCAAATAAATCACTAAATAATATCCCATATTATCATCTGTTATAGTCGTGTCATTAAATCTATTATATTTTATGAAAGATATATCACTGCTCACCTCTATATTTCGTACATTTTTAAACTGTTTGTATAATATCTCTCTAATACAATCAGATATATCTATTGTGTCGGTAGACGATACCGTTACATTTGAATTAATGGTGTTCCATTTTACAACTTTTAACAAGTTGACCAATTCATTTCTAATAGTATCAAACTCTCCATGATTAAATATTTTAATAGTATCCATTTTATATCACTACGGACATAAACCATTGTATATTTTTTATGTTTATTAGTGTACACATTTTGGAAGTAAGTTCCTGGATCGGATGTCACCGCAGTAATATTGTTGATTATTTCTGACATCGACGTATTATATAGTTTTTTAATTCCATATCTTTTAGAAAAGTTAAACATCCTTATACAATTTGTGGAATTAATATTATGAATCATGGTTTTTACACATAGATCTATTACAGGCGGAACATCAATTATTATGGCAGCAACTAGTATCATTTCTACATTGTTTATGGTGATGTTTATCTTCTTCCAGCGCATATAGTCTAATATCGATTCAAACGCGTGATAGTTTATACCATTCAATATAATCGCTTCATCCTTTAGATGGTGATCCTGAATGTGTTTAAAAAATTATACGGAGACGCCGTAATAATTTCCCCATTGATAGAAAATATCACGCTTTCCATTTTCTTGAAGTACTAAAAGTAATTATAATATAATGTAAAGGTTTATATATTCAATATTTTTTATAAAAAAAATCATTTTGACATTAATTCCTTTTTAAATTTCAGTCTATCATCTATAGAAACATATTCTATGAATTTATAAAATGCTTTTACGTATCCTATCGTAGGTGATAGAACCGCTAAAAAACCTATCGAATTTCTACAAAAGAATCTGTTATATGGTATAGGGAGAGTATAAAACATTAAATGTCCGTACTTATTAAAGTATTCAGTAGCCAATCCTAACTCTTTCGAATAATTATTAATGGCTCTTATTCTGTACGAATCTATTTTTTTGAACAATGGACCTAGTGGTATATCTTGTTCTATGTATCTAAAATAATGTCTGACTAGATCCGTTAGTTTAATATCCGCAGTCATCTTGTCTAGAATGGCAAATCTAACTGCGGGTTTAGGCGTAGGCGTTAGTTTAGTTTCTATATCTACATCTATGTCTTTATCTAACACCAAAAATATAATAGCTAATATTTTATTACAATCATCCGGATATTCTTCTACGATCTCACTAACTAATGTTTCTTTGGTTATACTAGTATAGTCACGATCAGACAAATAAAGAAAATCAGATGATCGATGAATAATACATTTAAATTCATCATCTGTAAGATTTTTGAGATGTCTCATTAAAATATTATTAGTGTCAGTTCTCATTATCATATATTGACAGCAGCTATTACACTTATTTTATTTTTCTGTATTTTATTACTTTTCACCATATAGATCAGTCATTAGATCATCAAAATACTTTTCAATCATCCTAAAGAGTATGGTGAACGAATCTTCCCATCTAATTTCTGAACGTCTACCAATGTCTCTAGCCACTTTGGCACTAATAGCGATCATTCGCTTAACATCTTCTACATTATTAACTGGTTGATTCAATCTATCTAGCAATGGACCGTCGGATAGCGTCATTCTCATGTTCTTAATCAATGTACATACATCGTCATCATCTACCAATTCATCAAACAATATAAGCTTTTTAAAATCATCATTATAATAGGATGGATCGCCGTCATTTCTCCAAAGAATATATCTAATAAGTAGAGTCCTCATGCTTAGTAATTTAACTATTTTAGTTAACAACTATTTTTTATGTTAAATCAATTAGTAACACCGCTATGTTTAATACTTATTCATATTTTAGTTTTAGGATCGAGAATCAATACAAAAATTAATACATCAATTTTGGAAATACTTAGTTTCCACGTAGTCAATGAAACATTTGAGCTCATCGTAAAGGACGTTCTCGTACAAGACGTAACTATAAATTGGTTTATATTTGTTCAAGATAGATACAAATCCGATAACTTTTTTGACGAATTCTACGGGATTCACTTTAAAAGTGTCATACCGGGTTCTTTTTATTCTTTTAAACAGATCGATTGTGTGATGTTGATTAGGTCTTTTACGAATTTGATACAGAATAGCGTTTACATATCCACCATAGTAATCAATAGCCATTTGTTCGTATGTCATAAATTCTTTAATTATATGACACTGTGTATTATTTAGTTCGTCCTTGTTCATCATTAGGAATCTATCCAATATGGCAATTATATTAGAACTATAACTGCGTTGTATGCGCATGTTGATGTGTCTGTTTATACAATCAATTATACTAGGATCCATACCACTACAATCGGGTAAAATTGTAGCATCATATACCATTTCTAGTACTTTAGATTCATTGTTATCCATTGCAGAGGACGTCATGATCGAATCCAAAAAAATATATTATTTTTATGTTATTTTGTTAAAAATAATCATCAAATACTTCGTAAGATACTCCTTCATGAACATAATCAGTTACAAAACGTTTATATGAAGTAAAGTATCTACAATTTTTACAAAAGTCAGGATGCATAAGTACAAAGTACGCGATAAACGGAATAATAATAGATTTATCTAGTTTATCTTTTTCTATCTCTTTCATAGTTATATACATGGTCTCAGAAGTCGGATTATGTAACATCAGCTTCGATAAAATGACTGGGTTATTTAGTCTTACACATTCGCTCATACATGTATGACCGTTAACTATAGAGTCTACACTAAAATGATTGAATAATAGATAGTCTACCATTGTTTCGTATTCAGATAGTACAGCGTAGTACATGGCATCTTCACAAATTATATCATTATCTAATAGATATTTGACGCATCTTATGGATCCCACTTCAACAGCCATCTTAAAATCGGTAGAATCATATTGCTTTCCTTTATCGTTAATAATTTCTAGAACATCATCTCTATCATAAAAGATACAAATATTAACTGTTTGATCAGTAATAACATTGCTAGTCGATATCAATTTGTTAATAAGATGCGCTGGGCTCAATGTCTTAATAAGAAGTGTAAGAGGACTATCTCCGAATTTGTTTTGTTTATTAACATCCGTTGATGGAAGTAAAAGATTTATAATGTCTACATACTTGACTGTTTTAGAGCATACAATATGGAGAGGCGTATTTCCATCATGATCTGGTTTTGAGGGACTAATTCCTAGTTTCATCATCCATGAGATTGTAGAAGCTTTTGGATTGTCTGACATAAGATGTCTATGAATATGATTTTTGCCAAATTTATCCACTATCCTGGCTTCGAATCCGATAGACATTATTTTTTTAAACACTCTTTCTGAAGGATCTGTACACGCCAACAACGGACCACATCCTTCTTCATCAACCGAGTTGTTAATCTTGGCTCCATACTGTACCAATAAATTTATTCTCTCTATGACTTCATCATCTGTTCCCGAGAGATAATATAGAGGTGTTTTATTATGTTTATCACATGCGTTTGGATCTGCGCCGTGCACCAGCAGCATCGCGACTATTCTATTATTATTAATTTTAGAAGCTATATGCAATGGATAATTTCCATCATCATCCGTCTCATTTGGAGAGTATCCTCTATGAAGAAGTTCTTCTATAAATCGTTCATCTAGTCCTTTAATGCCACAATACGCATGTAGAATGTGATAATTTCCAGAGGGTTCGATAACTTGTAGCATATTCCTAAATACATCTAAATTTTTACTATTATATTTGGCATAAAGAGATAGATAATACTCGACCGACATAATGTTGTGTTGTCCATTATAGTATAAAAATTAATATTTCTATTTCTATATATTTGCAACAATTTACTCTCTATAACAAATATCATAACTTAGTTCTTTTATGTCAAGAAGGCACTGGTTTAATTCATCTATAAATGTCACTCCATAACTACCACGCATACTATACTCAGAATTATGATAAAGATATTTATTCTTGGGGTGTAAGTAATGGGGATTAATCTTTGTTGGATCAGTCTCTAAGTTAACACATGTCACACATGATCCATTTATAGTTATATCACACGATGATGATTTATGAATTGATTCCGGAAGATCGCTATTGTATTTTGTAGTTCCACAATTCATTTCCATACATGTTATTGTCACACTAATATTATGATGAACTTTATCTAGCCGCTGAGTGGTAAACAACAGAACAGATAGTTTATTATCTTTACCAACACCCTCAGCCGCTGCCACAAATCTCTGATCCGTATCCATGATGGTCATGTTTACTTTTAGTCCGTATCCAGTCAACACTATGTTAGCATTTCTGTCGATATAGCTTTCACTCATATGACACTCACCAATAATTGTAGAATTAATGTCGTAATTTACACCAATAGTGAGTTCGGCGACAAAGTACCAGTACCGGTAATCTTGTCGATGAGGACATATAGTATTCTTGTATTCTACCGAATACCCGAGAGATGCGATACAAAAGAGTAAGACTAATTTGTAAACCATCTTACTCAAAAATATGCGACAATAGGAAATCTATCTTATACACATAATTATTCTATCAATTTTACCAATTAGTTAGTGTAATGTTAACAAAAATGTGGGATAATTTAATAGTTTTTCCTTACACAATTGACATACATGAGTCTGAGTTCCTCGTTTTTGCTAATTATTTCGTCCAATTTATTATTCTTGACATCGTCAAGATCTTTTGTATAGGAGTCAGACTTGTATTCAACATGTTTTTCTATAATCATCTTAGCTATTTCGGCATCATCCAATAGTACATTTTCCAGATTAACAGAATAGATATTAATGTCGTATTTGAACAGAGCCTGTAACATCTCAATGTCTTTATTATCTATAGCCAATTTGATGTCCGGAATGAAGAGAAGGGAATTGGTGTTTGTTGACGTCATATAGTCGAGCAAGAGAATCATCATATCCACGTGTCCATTTTTTATAGTGGTGTGAATACAACTAAGGAGAATAGCCAGATCAAAAGGAGATGGTATCTCTGAAAGAAAGTAGGAAACAATACTTACATCATTAAGCATGACAGCATGATAAAATGAAGTTTTCCATCCAGTTTTCCCATAGAACATCAGTCTCCAATTTTTCTTAACAAACAGTTTTACCGTTTGCATGTTACCACTATCAACCGCATAATACAATGCGGTGTTTCCTTTGTCATCAAATTGTGAATCATCCATTCCACTGAATAGCAAAATCTTTACTATTTTGGTATCTTCTAATGTGGCTGCCTGATGTAATGGAAATTCATTATCTAGAAGATTTTTCAATGCTCCAGCGTTCAACAACGTACATACTAGACGCACGTTATTATCAGCTATTGCATAATACAAGACACTATGACCATTGATATCCGCCTTAAATGCATCTTTGCTAGAGAGAAAGCTTTTCAGTTGCTTAGACTTCCAAGTATTAATTCGTGACAGATCCATGTCTGAAACGAGACGCTAATTAGTGTATATTTTTTCATTTTTTATAATTTTGTCATATTGTACCAGAATTAATAATATCTCTAATAGATCTGATTAGTAGATATATGGCTATCGCAAAACAACATATACACATTTAATAAAAATAATATTCATTAAGAAGATTCAGATTCCACTGTACCCATCAATATAAAATAAAATAATTATTCCTTACATCGTACCCATAAACAATATATTAAGTATATTCCACCTTACCCATAAACAATATAAATCCAGTAATATCATGTCTAATGATGAACACAAATGGTGTATTAAATTCCAGTTCTTCAGGAGATGATCTCGCCGTAGCTACCATGATAGTAGATGCCTCCGCTACAGTTCCTTGTTCGTCTACATCTATCTTTACATTCTGAAACATTTTATAAATATATAATGGGTCCCTAGTCATATGTTTAAACGACGCCTTATCTGGATTAAACATACTAGGAGCCATCATTTCGGCTATCGACTTAATATCCCTCTTGTTTTCGATAGAAAATCTAGGGAGTTTAAGATTGTACATTTTATTCCCTAATTGAGATGACCAATATTCTAATTTTGCAGGCGTGATAGAATCTGTGAAATGGGTCATATTATCACCTATTGCCAGGTACATACTAATATTAGCATCCTTATACAGAAGGCGCACCATATCATATTCTTCGTCATCGATTGTGATTGTATTTCCTTGCAATTTAGTAACTACGTTCATCATGGGAACCGTTTTCGTACCGTACTTATTAGTAAAACTAGCATTGTGTGTTTTAGTGATATCAAACGGATATTGCCACGTACCTTTAAAATATATAGTATTAATGATTGCCCATAGAGTATTATCGTCGAGCATAGTAGAATCAACTACATTAGACATACCAGATCTACGTTCTACTATAGAATTAATTTTATTAACCGCATCTCGTCTAAAGTTTAATCTATATAGGCCGAATCTATGATATTGTTGATAATACGACGGTTTAATACACACAGTACTATCGACGAAACTTTGATACGTTAGATCTGTGTACGTATATTTAGATGTTTTCATCTTAGCTAATCCGGATATTAATTCTGTAAATGCTGGACCCAGATCTCTTTTTCTCAAATTCATAGTATTCAATAATTCTACTCTAGTATTACCTGATGCAGACAATAGCGACATAAACATAGAAAACGAATACCCAAACGGTGAGAAGACAATATTATCATTATCATCCTCATCCCCATTTTGAATATTTTTATACGCTAATATACCAGCATTGATAAATCCCTGCAGACGATATGCGGATACTGAACACGCTAATGATAGTATCAATAACGCAATCATGATTTTTATGGTATTAATAATTAACCTTATTTTTATGTTTGGTATAAAAATTATTGATGTCTACACATCCTTTATAATCAACTCTAATCACTTTAACTTTTACAGTTTTCCCTACAAGTTTATCCCTATATTCAACATATCTATCCATATGCATCTCTTAACACTCTGCCAAGATAGCTTCAAAGTGAGGATAGTCAAAAAGATAAATATATAGAGTATAATCATTCTCGTATACTCTGCCCTTTATTACATCGCCCGCATTGGGCAACGAATAACAAAATGCAAGCATCGTGTTAACGGGCTCGTAAATTGGGATAAAATTATGTTTTTATTGTTTATCTATTTTATTCAAGAGAATATTCAGGAAGTTCCTTTTCTGGTTGTATCTCGTCGCAGTATATATCATTTGTACATTGTTTCATATTTTTTAATAGTCTACACCTTTTAGTAGGACTAGTATCGTACAATTCATAGCTGTATTTTGAATTCCAATCACGTATAAAAATATCTTCCAATTGTTGACGAAGACCTAATCCATCATCCGGTGTAATATTAATAGATGCTCCACATATATCCGTAAAGTAATTTCCTGTCCAATTTGATGTACCTATATACGCCGTTTTATCGGTTACCATATATTTTGCATGGTTTACCCTAGAATACGGAATGGGAGGATCAGCATCTGGTACAATAAATAGCTTTACTTCTATATCTATGTTTTTAGATTTTAGCATAGCTATAGATCTTAAAAAGTTTCTCATGATAAACGAAGATCGTTGCCAGCAACTAATCAATAGCTTAACGGATACTTGTCTGTCTATAGCGGATCTTCTTAATTCATCTTCTATATAAGGCCAAAACAAAATTTTACCCGCCTTTGAATAAATAATAGGAATAAAGTTCATAACAGATACATAAACGAATTTACTCGCATTTCCGATACATGACAATAAAGCGGTTAAATCATTGGTTCTTTCCATAGTACATAATTGTTGTGGTGCAGAAGCAATAAATACAGAGTGTGGAACACCGCTTACGTTAATACTAAGAGGATGATCTGTATTATAATACGACGGATAAAAGTTTTTCCAATTATATGGTAGATTGTTAACTCCAAGATACCAGTATACCTCAAAAATTTGAGTGAGATCCGCTGCCAAGTTCCTATTATTGAAGATCGCAATACCCAATTCCTTGACCTGAGTTAGTGATCTCCAATCCATGTTAGCGCTTCCTAAATAAATATGTGTATTATCAGATATCCAAAATTTTGTATGAAGAACTCCTCCTAGGATATTTGTAATATCTATGTATCGTACTTCAACTCCGGCCATTTGTAGTCTTTCAACATCCTTTAATGGTTTGTTGGATTTATTGACGGCTACTCTAACTCTTACTCCTCTTTTGGGTAATTGTACAATCTCGTTTAATATTACCGTGCCGAAATTCGTACCCACTTCATCCGATAAACTCCAATAAAAAGATGATATATCTAGTGTTTTTATGGTATTGGATAGAATTTCCCTCCACATGTTAAATGTAGTCAAATATACTTTATCAAATTGCATACCTATAGGAATAGTCTCTGTAATCACTGCGATTGTATTATCCGGATTCATTTTATTTGTTAAAAAAATAATCCTATATCACTTCACTCTATTAAAAATCCAAGTTTCTATTTCTTTCATGACTGATTTTTTAACTTCATCCGTTTCCTTATGAAGATGATGTTTGGCGCCTTCATAAATTTTTATTTCCCTATTACAATTTGCATGTTGCATGAAATAATATGCACCTGAAACATCGCTAATCTCATTGTTTGTTCCCTGGAGTATGAGAGTCGGGGTGTTAATCTTGGGAATTATTTTTCTAACCTTGTTGGTAGCCTTCAAGACCTGACTAGCAAATCCAGCCTTAATTTTTTCATGATTGACTAATGGATCGTATTGGTATTTATAAACTTCATCCATATCTCTAGATACTGATTCTGGACATAGCTTTCCGACTGACGCATTTGGTGTAATGGTTCCCATAAGTTTTGCAGCTAGCAGATTCAGTCTTGGAACAGCGTCTGCATTAACTAGAGGAGACATTAGAATCATTGCTGTAAACAAGTTTGGATTATCGCAAGCAGCTAGTATAGAAATTGTTGCTCCCATGGAATGACCCAATAAGAAGACTGGAACTCCTGGATAAGTAGATTTAATAGTCACCACGTGCTGTACCACATCTCTAACATACTTACCAAAGTCATCAATCATCATTTTTTCACCATTACTTCTTCCATGGCCAATATGATCATGTGAGAATACTAAAATTCCTAACGATGATATGTTTTCAGCTAGTTCGTCATAACGTCCAGAATGTTCACCAGCTCCATGACTTATGAATACTAATGCCTTAGGATATGTAATAGGTTTCCAATATTTACAATATATGTAATCATTGTCCAGATTGAACATACAGTTTGTACTCATGATTCACTATATAACTATCAATATTAACAGTTCGTTTAATGATCATATTATTTTTATGTTTTATTGATAATTGTAAAAATATACAATTAAATCAATATAGAGGAAGGAGACGGTACTGTATTTTGTGAGATAGTAATGGAGACTAAATCAGATTATGAGGATGCTGTTTTTTACTTTGTGGATGATGATGAAATATGTAGTCGCGACTCCATCATTGATCTAATAGATGAATATATCACGTGGAGAAATCATGTTATAGTGTTTAATAAAGATATTACCAGTTGTGGAAGACTGTACAAGGAATTGATAAAGTTCGATGATGTCGCTATACGGTACTATGGTATTGATAAAATTAATGAGATTGTCGAGGCTATGAGTGAAGGAGACCACTACATCAATCTTACAGAAGTCCATGATCAGGAAAGTCTATTCGCTACCATAGGAATATGTGCTAAAATCACTGAACATTGGGGATACAAAAAGATTTCAGAATCTAAATTCCAATCATTGGGAAACATTACAGATCTGATGACCGACGATAATATAAACATCTTGATACTTTTTCTAGAAAAAAAAATGAATTGATGATATAAGTGTCTTCATAACGCATTATTACGTTAGCATTCTATTATCCAGTGTTAAAAAAATTATCCTATCATGTATTTGAGAGTCTTATATGTAGCAAACATGATAACTGCAATACCCATAATCTTTAGATATTCACGCGTGCTATGGATGGCATTATCCCGCGGTGCGGAAATGTACGTTATATAATCTACAAAATAATCATCGCATATAGTATGAGATAGTAGAGTAAACATTTTTATCGTTTCTACTGGGTTCATACATCGTCTACCCAATTCGGTAATGAATGAAATTGTCGCCAATCTTACACCCAAACCCTTGTTGTTCATTAGTATAGTATTAACTTCATTATTTATGTCATAAACTGTAAATGATTCTGTAGATGCCATATCACACATGATATTCATGTCACTATTATAATCATTATTAACTTTATCACAATACGTGTTGATAATATCTACATATGATCTAGTTTTTGTGGGTAATTGTCTATACAAGTCGTCTAAACGTTGTTTACTCATATAGTATTGAACAGCCATCATTACATGGTCCCGTTCCGTTGATAGATAATCGAGTATGTTAGTAGACTTGTCAAATCTATATACCATATTTTCTGGAAGCGGATATACATAGTCGCGATCATCATTATCACTAGCCTCATCCTCTATATCATGTACATGTACATAATCTATGATATTATTATACATAAACATCGACAACATACTATTGTCTATTATCTAAGTCCTGTTGATCCAAACCCTTGATCTCCTCTATCTGTACTATCTAGAGATTGTACTTCTTCAAGTTCTGGATAATATATACGTTGATAGATTAGCTGAGCTATTCTATCTCCAGTATTTACATTAAACGTACATTTTCCATTATTAATAAGAATGACTCCTATGCTTCCCCTATAATCTTCGTCTATTACACCGCCTCCTATATCAATGCCTTTTAGGGACAGACCAGACCTAGGAGCTATTCTACCATAGCAGAACTTAGGCATGGACATACTAATATCTGTCTTAATTAACTGTCGTTCTCCAGGAGGGATAGTATAATCGTAAGCGCTATACAAATCATATCCGGCAGCACCCGGCGATTGCCTAGTAGGTGATTTAGCTCTGTTAGTTTCCTTAACAAATCTAACTGGTGAGTTAATATTCATGTTGAACATAAAAAATATCATTTTATTTCAAAATTATTTACCATTCCATTCCATCCCATATATTCCATGAATAAGTGCGATTATTGTACACTTCTATAGTATCTATATACGATCCACGATAAAATCCTCCTATCAATAGCAGTTTATTATCCACTATGATCAATTCTGGATTATCCCTCGGATAAATAGGATCATCTATCAGAGTCCATGTATTGCTGGATTCACAATAAAATTCCGCATTTCTACCAACCAAGAATAACCTTCTACCAAACACTAACGCACATGATTTATAATGAGGATAATAAGTGGATGGTCCAAACTGCCACTGATCATGATTGGGTAGCAAATATTCTGTAGTTGTATCAGTTTCAGAATGTCCTCCCATTACGTATATAACATTGTTTATGGATGCCACTGCTGGATTACATCTAGGTTTCAGAAGACTCGGCATATTAACCCAAGCAGCATCCCCGTGGAACCAACGCTCAACAGATGTGGGATTTGGTAGACCTCCTACTACGTATAATTTATTGTTAGCGGGTATCCCGCTAGCATACAGTCTGGGGCTATTCATCGGAGGAATTGGAATCCAATTGTTTGATATATAATTTACCGCTATAGCATTGTTATGTATTTCATTGTTCATCCATCCACCGATGAGATATACTACTTCTCCAACATGAGTACTTGTACACATATGGAATATATCTATAATTTGATCCATGTTCATAGGATACTCTATGAATGGATACTTGTATGATTTGCGTGGTTGTTTATCACAATGAAATATTATGTTACAGTCTAGTATCCATTTTACATTATGTATACCTCTGGGAGAAAGATAATTTGACCTGATTACATTTTTGATAAGAAGTAGCAGATTTCCTAATCTATTTCTTCGCCTCATATACCACTTAATGACAAAATCAACTACATAATCCTCATCTGGAACATTTAGTTCGTCGCTTTCTAGAATAAGTTTCATAGATAGATAATCAAAATTGTCTATGATGTCATCTTCCAGTTCCAAAAAGTGTTTGGTAATAAAGTCTTTAGTATGACATAAGAGATTGGATAGTCCGTATTCTATACCCATCATGTAACACTCGATACAATATTCCTTTCTAAAATCTCGTAGGATAAAGTTTATACAAGTGTAGATGATAAATTCTACAGATGTTAATATAGAAGCACGTAATAAATTGACGACGTTATGACTATCTATATATACCTTTCCAGTATATGAGTAAATAACTATAGAAGTTAGACTGTGAATGTCAAGGTCTAGACAAACCCTCGTAACTGGATCTTTATTTTTTGTGTATTTTTGGCGTAAATGTGTGCGAAAGTATGGAGATAACTTTTTCAATATCGTAGAATTGACTATTATATTACCTCCTATAGCTTCAATAATTGTTTTGAATTTCTTAGTCGTGTACAATGCTAATATATTCTTACAGTACACAGTATTGACAAATATCGGCATTTATGTTTCTTTAAAAGTCAACATCTAAAGAAAAATGATTGTCTTCTTGAGACATAACTCCCATTTTTTGGTATTCACCCACACGTTTTTCGAAAAAATTAGTTTTTCCTTCCAATGATATATTTTCCATGAAATCAAACGGATTGGTAACATTGTAAATTTTTTTAAATCCCAATTCAGAAATCAATCTATCTGCGACGAATTCTATATATGTTTTCATCATTTCACAATTCATTCCTATGAGTTTAACTGGAAGAGCCACAGTAAGAAATTCTTGTTCAATGGATACCGCATTTGTTATAATAAATCTAACGGTTTCTTCACTCGGTGGATGCAATAAATGTTTAAACATCAAACATGCGAAATCGCAGTGCAGACCCTCGTCTCTACTAATTAATTCGTTAGAAAACGTGAGTCCGGGCATTAGGCCACGCTTTTTAAGCCAAAATATGGAAGCGAATGATCCGGAAAAGAAGATTCCTTCTACTGCAGCAAAGGCAATAAGTCTCTCTCCATAACCGGCGCTGTCATGTATCCACTTTTGAGCCCAATCGGCCTTCTTTTTTACACAAGGCATCGTTTCTATGGCATTAAAGAGGTAGTTTTTTTCATTACTATCTTTAACATAAGTATCGATCAAAAGACTATACATTTCCGAATGAATGTTTTCAATGGCCATCTGAAATCCGTAGAAACATCTAGCCTCGGTAATCTGCACTTCTGTACAAAATCGTTCTGCTAAATTTTCATTCACTATTCCGTCACTGGCTGCAAAAAACGCCAATACATGTTTTATAAAATATTTTTCGTCTGGTGTTAGTTTATTCCAGTCATTGATATCTTTAGATATATCCACTTCTTCCACTGTCCAAAATGATGCCTCTGCCTTTTTATACATATTCCAGATGTCATGATATTGGATTGGGAAAATAACAAATCTATTTGGATTTGGTGCAAGGATAGGTTCCATAACTAAATTAACAATAGTAGTAATTTTTTTTCAGTTATCTGTATGACGACTGTACTTGGATCTTTTGTATATCGCTATCGCCGCAATCACTACAATAATTACAAGTATTATTGATAGCATTGTTATTACTACTATCATAATTAAATTATCGACATTCATGGGTGTTGAATAATCGTTATCATCATTTTGTAATTGTGACGTCATACTAGATAAATCATTTGTGAGATTGTTGTGGGAAGCGGGCACGGAAGATGCATTATCATTATTATTTAACGCCTCCCATTCGGATTCACAAATGTTACGCACGTTCAAAGTTTTATGGAAACTATAATTTTGTGAAAACAGATAACAAGAAAACTCGTCATCGTTCAAATTTTTAACGATAGTAAACCGATTAAACGTCGAGCTAATTTCTAACGCTAGCGACTCTGTTGGATATGGGTTTCCAGATATATATCTTTTCAGTTCCCCTACGTATCTATAATCATCTGTAGGAAATGGAAGATATTTCCATTTATCTACTGTTCCTAATATCATATGCGGTGGTGTAGAACCATTAAGCGCGAAAGATGTTATTTCGCATCGTATTTTAACTTCGCAATAATTTCTGGTTAGATAACGCACTCTACCAGTCAAGTCAATGATATTAGCCTTTACAGATATATTCATAGTAGTCGTAACGATGACTCCATCTTTTAGATGTGATACTCCTTTGTATGTACCAGAATCTTCGTACCTCAAACTCGATATATTTAAACAAGTTAATGATATATTAACGCGTTTTATGAATGATGATATATAACCAGAAGTTTTATCCTCTGTGGCTAGCGCTATAACCTTATCATTATAATACCAACTAGTGTAATTAATATGTGACATGACAGTGTGGGTACAAATATGTACATTATCGTCTACGTCGTATTTGATACATCCGCATTCAGCCAACAAATATAAAATTACAAAAACTCTAACGACGTTTGTACACATCTTGATGCGGTTTAATAAATGTTTTGATTTCAATTTATTGTAAAAAAAGATTCGGTTTTATACTGTTCGATATTCTCATTGCTTATATTCTCATCTATCATCTCCACACAGTCAAATCCATGGTTAACATGTACCTCATCAACCGGTAAAAGACTATCGGATTCTTCTATCATCATAACTCGAGAATATTTAATTTGGTGGTCATTATTAATCAAGTCAATTATCTTATTTTTAACAAACGTAAGTATTTTACTCATTTTTTTATAAAAACTTTTAGAAATATACAGACTCTATCGTGTGTCTATATCTTCTTTTTATATCCAATGTATTTATGTCTGATTTTTCTTCATTTATCATATATAATGGTCCAAATTCTACACGTGCTTCGGATTCATCCAGATCATTAAGGTTCTTATAATCGCAACATCCTTCTCTTCCATCTTCTACATCTTCCTTCTTATTCTTAGCGTCACAGAATCTACCACAGCAGGATCCCATGACGAGAGTCACATTAAACTAATTCATTTTCAATTATAATATACTGATTAGTAATGACAATTAAAATAAAAATATTCTTCATAACCGGTAAGAAAGTAAAAAGTTCACATTGAAACTATGTCAGTAGTTATACATCATGAGATGATATACTCTATTTTGGTGGAGGATTATATGATATAATTCGTGGATAATCATTCTTAAGACACATTTCTTCATTCGTAAATCTTTTCACATTAAATGAGTGTCCATATTTTGCAATTTCTTCATATGATGGCGGTGTACGTGGACGAGGCTGCTCCTGTTCTTGTAGTCGTCGACTGTCGTGTTTGCGTTTAGATCCCTCCATTATCGCGATCGCGTAGTGAGTACTATTTATACCTTGTAATTAAATTTTTTTATTAATTAAACGTATAAAAACGTTCCGTATCTGTATTTAAGAGCCAGATTTCGTCTAATAGAACAAATAGCTACAGTAAAAATAACTAGAATAATCGCTACACCCACTAGAAACCACGGATCGTAATACGGCAATCGGTTTTCGATAATAGGTGGAACGTATATTTTATTTAAGGACTTAACAATTGTCTGTAAACCACAATTTGCTTCCGCCGATCCTGTATTAACTATCTGTAAAAGCATATGTTGACCGGGAGGAGCCGAACATTCTCCGATATTCAATTTTTGTATATCTATAATGTTATTAACCTCCGCATACGCATTACAGTTCTTTTCTAGCTTGGATACTACACTAGGTACATCATCTAAATCTATTCCTATTTCCTCAGCGATAGCTCTTCTATCCTTTTCCGGAAGTAATGAAATCACTTCAATAAATGATTCAACCATGAGTGTGAAACTAAGTCGAGAATTACTCATGCATTTGTTAGTTATTCGGAGCGCGCAATTTTTAAACTGTCCTATAACCTCTCCTATATGAATAGCACAAGTGACATTAGTAGGGATAGAATGTTGAGCTAATTTTTGTAAATAACTATCTATAAAAAGATTATACAAAGTTTTAAACTCTTTAGTTTCTGCCATTTATCCAGTCTGAGAAAATGTCTCTCATAATAAATTTTTCCAAGAAACTAATTGGGTGAAGAATGGAAACCTTTAATCTATATTTATCACAGTCTGTTTTGGTACACATGATGAATTCTTCTAATGCTGTACTAAATTCGATATCTTTTTCGATTTCTGGATATGTTTTTAATAAAGTATGAACAAAGAAATGGAAATCGTAATACCAGTTATGTTCAACTTTGAAATTGTTTTTTATTTTCTTGTTAATGATTCCAGCCACTTGGGAAAAGTCAAAGTCGTTTAATGCCGATTTAATACGTTCATTAAAAACAAACTTTTTATTCTTTAGATGAATTATTATTGGTTCATTGGAATCAAAAAGTAAGATATTATCGGGTTTAAGATCTGCATGTAAAAAGTTGTCACAACAGGGTAGTTCGTAGATTTTAATGTATAACAGAGACATCTGTAAAAAGATAAACTTTATGTATTGTACCAAAGATTTAAATCCTAATTTGATAGCTAACTCGGTATCTACTTTATCTGCCGAATACAGTGCTAGGGGAAAAATTATAATATTTCCTCTTTCGTATTCGTAATTAGTTCTCTTTTCATGTTCGAAAAAGTGAAACATGCGGTTAAAATAGTTTATAACATTAATATTACTGTTAATAACTGCCGGATAAAAGTGGGATAGTAATTTCACGAATTTGATACTGTCCTTTCTCTCGTTAAACGCCTTTAGAAAAACTTTAGAAGAATATCTCAATGAGAGTTCCTGACCATCCATAGTTTGTATCAATAATAGCAACATATGAAGAACCCGCTTATACAGAGTATGTAAAAATGTTAATTTATAGTTTAATCCCATGGCCCACGCACACACGATTAATTTTTTTTCATCTCCCTTTAGATTGTTGTATAGAAATTTGGGTACTGTAAACTCCGCCGTAGTTTCCATGGGACTATATAATTTTGTGGCCTCGAATACAAATTTTACTACATAGTTATCTATCTTAAAGACTATACCATATCCTCCTGTAGATATGTGATAAAAATCGTCGTTTATAGGATAAAATCGTTTATCTTTTTGTTGGAAAAAGGATGAATTAATGTAATCATTCTCTTCTATCTTTAGTAGTGTTTCCTTATTAAAATTCTTAAAATAATTTAACAATCTAACTGACGGAGCCCAATTTTGGTGTAAATCTAATTGGGACATTATGTTGTTAAAATATAAACAGTCTCCTAATATAACAGTATCTGATAATCTATGGGGAGACATCCATTGATATTCAGGGGATGAATCATTGGCAACACCCATTTATTGTACAAAAAGCCCCAATTTACAAACGAAAGTCCAGGTTTGATAGAGATAAACTATTAACTATTTTGTCTCTGTTTTTAACACCTCCACAGTTTTTAATTTCTTTGGTAATGAAATTATTCACAATATCAGTATCTTCTTTATCTACCAGAGATTTTACTAACTTGATAACCTTGGCTGTCTCATTCAATAGGGTAGTGATATTTGTATGTATGATATTGATATCTTTTTGAATTGTTTCTTTTAGAAGTGATTCTTTGATGGTATCAGCATACGAATTACAATAATGCAGAAACTCAGTTAACATGCAGGAATTATAGTAAGCCAATTCCAATTGTTGCCTGTATTGTATTAGAGTATTAATATGCGCAATGATGTCCTTGCGTTTCTCTGATAGAATGCGAGCAGCGATTTTGGCGTTATCATTTGACGATATTTCTGGAATGACGAATCCTGTTTCTACTAACTTCTTGGTAGGACAAAGTGAAACAATCAAGAAAATAGCTTCTCCTCCTATTTGTGGAAGAAATTGAACTCCTCTAGATGATCTACTGACGATAGTATCTCCTTGACAGATATTGGACCGAACTACGGAAGTACCTGGAATGTAAAGCCCTGAAACCCCCTCATTTTTTAAGCAGATTGTTGCCGTAAATCCTGCACTATGCCCAAGATAGAGAGCTCCTTTGGTGAATCCATCACTATGTTTCAGTTTAACCAAGAAACAGTCAGCTGGTCTAAAATTTCCATCTCTATCTAATACAGAATCCAACTTGATGTCAGGGACTATGACCGGTTTAATGTTATATGTAACATTGAGTAAATCCTTAAGTTCATAATCATCGTTGTCATCAGTTATGTACGATCCAAACAATGTTTCTACCGGCATGGTGGATACGAAGATGCTATCCATCAGAATGTTTCCCTGATTAGTATTTTCTATATAGCTATTCTTCTTTAAACGATTTTCCGAATCAGTAACTATGTTCATTTTTTTAGGAGTAGGACGTCTAGCCAGTATGGAAGAGGATTTTCTAGATACTCTCTTCAACATCTTTGATCTCAATGGAATGCAAAACCCCATGGTGTAACAACCAACGATAAAAATAATATTGTTTTTTCACTTTTTATAATTTTACCATCTGACTCATGGATTCATTAATATCTTTACAAGAGCTACTAACGTATAATTCTTTATAACTGAACTGAGATATATACACCGGATCTATGGTTTCCATAATTGAGTAAATGAATGCTCGGCAATAACTAATGGCAAATGTATAGAACAACGAAATTATACTAGAGTTGTTAAAGTTAATATTTTCTATGAGTTGTTCCAATAAATTATTTGTTGTGACTGCGTTCAAGTCATAAATTATCTTGATACTATCCAGTAAACAGTCTTTAAGTTCTGGAATATTATCATCCCATTGTAAAGCCCCTAGTTCGACTATCGAATATCCTGCTCTGATAGCAGTTTCAATATCGACGGACGTCAATACTGTAATAAAGGTGGTAGTATTGTCATCATCGTGATAAACTACGGGAATATGGTCGTTAGTAGGTACCGTGACTTTACACAACGCGATATATAACTTTCCTTTTGTACCATTTTTAACGTAGTTGGGACGTCCTGCAGGGTATTGTTTTGAAGAAATGATATCGAGAACAGATTTGATACGATATTTGTTGGATTCCTGATTATTCACTATAATATAATCTAGACAGATAGATGATTCGATAAATAGAGAAGGTATATCGTTGGTAGGATAATACATCCCCATTCCAGTATTCTCGGATACTCTATTGATGACACTAGTTAAGAACATGTCTTCTATTCTAGAAAACGAAAACATCCTACATGGACTCATTAAAACTTCTAACGCTCCTGATTGTGTTTCGAATGCCTCGTACAAGGATTTCAAGGATGCCATAGATTCTTTGACCAACGATTTAGTATTGCGTTTAGCATCTGATTTTTTTATTAAATCAAATGGTCGGCTCTCTGGTTTACTACCCCAATGATAACAATAGTCTTGTAAAGATAAACCGCAAGAAAATTTATACACATCCATCCAAATAACCCTAGCACCGTCGGATGATATTAATGTATTATTATAGATTTTCCATCCACAGTTATTGGGCCAGTATACTGTTAGCAACGGTATATCGAATAGATTACTCATGTAACCTACTAGAATGATAGTTCGTGTACTAGTCATAATATCTTTAATCCAATCTAAGAAATCTAAAATTAGATCTTTTACACTATTAAAGTTAACAAAGGTATTACCCGGGTACGTGGATATCATATATGGTATTGGTCCATTATCAGTAATGGCTCCATAAACTGATACGGCGATGGTTTTTATATGTGTTTGATCTAATGAGGACGAAATTCGCGCCCACAATTCATCTCTAGATATGCATTTAATATCGAACGGTAACACATCAATCTCGGGACGCGTATATGTTTCTAAATTCTTAATCCAAATATAATGATGACCTATATGCCCTATTATCATACTGTCAACTATAGTATACCTAGAGAACTTTCGATACATCTGCTGTTTCCTGTAATCGTTAAATTTTACAAATCTATAACATGCTAAACCTTTTGACGACAGCCATTCATTAATTTCTGATATGGAATCTGTATTCTCAATACCGTATCGTTCTAAAGCCAGTGCTATATCTCCCTGTTCGTGGGAACGCTTTCGTATAATATCGATCAATGGATAATATGAAGTTTTTGGAGAATAATATGATTCATGATCTATTTCGTCCATAAACAATCTAGACATAGGAATTGGAGGCGATGATCTTAATTTTGTGCAATGGGTCAATCCTATAACTTCTAATATTGTAATATTCATCATCGACATAACACTATCTATGTTATCATCGTATATTAGTATACCACGACCTTCTTCATTTCGTGCCAAAATGATATACAGTCTTAAATAATTACGCAATATCTCAATAGTTTCATAATTGTTAGCTGTTTTCATCAAGGTTTGTATCCTGTTTAACATGATGGCGTTCTATAACGTCTCTATTTTCTATTTTTAATTTTTTTAAATTTTTAACGATTTACTGTGGCTAGATACCCAATCTCTTTCAAATATTTTTTTAGCCTCGCTTACAAGCTGTTTATCTATACTATTAAAACTGACGAATCCGTGATTTTGGTAATGGGTTCCGTCGAAATTTGCCGAAGTGATATGAACATATTCGTCGTCGACTATTAACAATTTTGTATTATTCTGAATAGTGAAAACCTTCACAGATAGATCATTTTGAACACACAACGCATCTAGACTTCTGGCGGTTGCCATAGAATATACGTCGTTCTTATCCCAATTACCAACTAGAAGTCTGATCTTAACTCCTCTATTAATGGCTGCTTCTATAATGGAGTTGTAAATGTCAGGCCAATAGTAGCTATTACCGTCGACACGTGTAGTGGGAACTATGGCCAAATGTTCAATATCTATACTAGTCTTAGCCGACTTGAGTTTATCAATAACTACATCAGTGTCTAGATCTCTAGAATATCCCAATAGGTGTTCTGGAGAATCAGTAAAGAACACTCCACCTATAGGATTCTTAATATGATACGCAGTGCTAACTGGCAGACAACAAGCCGCAGAGCATAAATTCAACCATGAATTTTTTGCGCTATTAAAGGCTTTAAAAGTATCAAATCTTCTACGAAGATCTGTGGCCAGCGGAGGATAATCAGAATATACGCCTAACGTTTTAATCGTATGTATAGATCCTCCAGTAAATGACGCGTTTCCTACATAACATCTTTCATCATCAGACACCCAAAAACAACCGAGTAGTAGTCCCACATTATTTTTTTTATCTATATTAACGGTTATAAAATTTATATCCGGGGAGTGACTTTGTAGCTCTCCCAGATTTCTTTTCCCTCGTTCATCTAGCAAAACTATTATTTTAATCCCTTTTTCAGATACCTCTTTTAGTTTATCAAAAATAAGCGCTCCCCTAGTAGTACTCAGAGGATTACAACAAAAAGATGCTATGTATATATATTTCTTAGCTAGAGTGATAATTTCGTTAAAACATTCAAATGTTGTCAAATGATCGGATCTAAAATCCATATTTTCTGGTAGTGTTTCTACCAGCCTACATTTTGCTCCCGCAGGTACCGATGCAAATGGCCACATTTAGTTAACATAAAAACTTATATATCCTGTTCTATCAACGATTCTAGAATATCATCGGCTATATCGCTAAAATTTTCATCAAAGTCGACATCACAACCTAACTCAGTCAATATATTAAGAAGTTCCATGATGTCATCTTCGTCTATTTCTATATCCGTATCCATTGTAGATTGTTGACCGATTATCGAGTTTAAATCATTACTAATACTCAATCCTTCAGAATACAATCTGTGTTTCATTGTAAATTTATAGGCGGTGTATTTAAGTTGGTAGATTTTCAATTATGTATCAATATAGCAACAGTAGTTCTTGCTCCTCCTTGATTTTAGCATCCTCTTCATTATTTTCTTCTACGTACATAATCATGTCTAATACGTTAGACAACACACCGACGATGGTGGCCGCCACAGACACGAATATGACTAGACCGATGACCATTTAAAAAATACTCTCTAGCTTTAACTTAAACTGTATCGATCATTCTTTTAGCACATGTATAATATAAAAACATTATTCTATTTCGAATTTAGGCTTCCAAAAATTTTTCATCCGTAAACCGATAATAATATATATAGACTTGTTAATAGTCGGAATAAATATATTAATGCTTAAACTATCATCATCTCCACGATTAGAGATACAATATTTACATTCTTTTTGCTGTTTCGAAACTTTATCAATACACGTTAATACAAACCCAGGAAGGAGATATTGAAACTGAGGCTGTTGAAAATGAAACGGCGAATACAATAATTCAGATAATGTAAAATCATGATTCCGTATTCTGATGATATTAGAACTGCTAATGGATGTCGATGGTATGTATCTAGGAGTATCTATTTTAACAAAGCATCGATTTGCTAATATACAATTATCCTTTTGATTAATTGTTATTTTATTCATATTCTTAAAAGGTTTCATATTTATCAATTCTTCTACATTAAAAATTTCCATTTTTAATTTATCTAGCCCCGCAATACTCCTCATTACGTTTCATTTTTTGTCTAGAATACCCATTTTGTTCATCTTGGTACATAGATTATCCAATTGAGAAGCGCATTTAGTAGTTTTGTACATTTTAAGTTTATTAACGAATCGTCGAAAACTAGTTATAGTTAACATTTTATTATTTGATACCCTGATATTAATACCCCTGCCGTTACTATTATTTATAACTGATGTAACCCACGTAACATTGGAATTAATTATCGATAGTAATGCATCGACACTTCCAAAATTGTCTATTATAAACTCACCGATAATTTTTTTATTGCATGTTTTCATATTCATTAGGATTATCAAATCTTTAATCTTATTACGATTGTATGCGTTGATATTACAAGACGTCATTCTAAAAGACGGAGGATTTCCATCAAATGCCAGACAATCACGTACAAAGTACATGGAAATAGGTTTTGTTCTATTACGCATCATAGATTCATATAAAACACCCGTAGAAATACTAATTTGTTTTACTCTATAAAATACTATTGCATCTATTTCATCGTTTTGTATAACGTCTTTCCAAGTGTCAAATTCCAATTTTTTTTCATTGATAGTACCAAATTCTTCTATCTCTTTAACTACTTGCATAGATAGGTAATTACAGTGATGCCTACATGCCGTTTTTTGAAACTGAATAGATGCATCTAGAAGCGATGCTACACTAGTCACGATCACCACTTTCATATTTAGAATATATGTATGTAAAAATATAGTAGAATTTCATTTTGTTTTTTTCTATGCTATAAATGAATTCTCATTTTGTATCCGCACATACTCCGTTTTATATCAATACCAAAGAAGGAAGATATCTGGTTCTAAAAGCCGTTAAAGTATGCGATGTTAGAACTGTAGAATGCGAAGGAAGTAAAGCTTCCTGCGTACTCAAAGTAGATAAACCCTCATCACCCACGTGTGAGAGAAGACCTTCGTCCCCGTCCAGATGCGAGAGAATGAATAACCCTGGAAAACAAGTCCCGTTTATGAGAACGGACATGTTACAAAATATGTTTGCTGCTAATCGCGACAACGTAACGTCAAGACTTTTGAACTAAAATACAATTATATCTTTTTCGATATTAATAAATCCGTGTCTCCCGGGTTTTTTATCTCTTTCAGTATGTGAATAGATAGGTATTTTATCTCTATTCATCATCGAATTTAAGAGATCCGATAAACATTGTTTGTATTCTCCAGATGTCAGCATCTGATACAACAATATATGTGCACATAAACCTCTGGCACTTATTTCATGTACCTTCCCCTTATCACTAAGGAGAATAGTATTTGAGAAATATGTATACATGATATTATCATGTATTAGATATACAGAATTTGTAACACTCTCGAAATCACACGATGTGTCGGCGTTAAGATCTAATATATCACTCGATAACACATTTTCATCTAGATACACTAGACATTTTTTAAAGCTAAAATAGTCTTTAGTAGTAACAGTAACTATGCGATTATTTTCATCGATGATACATTTCATCGGCATATTATTACGCGTACCATCAAAGACTATACCATGTGTATATCTAACGTATTCTAGCATAGTTGCCATACGTACATTAAACTTTTCAGGATCTTTGGATAGATCTTCCAATCTATCTATTTGAGAAAACATTTTTATCATGTTCAATAGTTGAAACGTCGGATCCACTATATAGATATTATCTATAAAGATTTTAGGAACTATGTTCATGGTATCCTGGCGAATATTAAAACTATCAATGATATGATTATCGTTTTCATCTTTTATCACCATATAGTTTCTAAGATATGGGATTTTACTTAATATAATATTATTTCCCGTAATAAATTTTATTAGAAATGCCAAATCTATAAGAAAAGTCCTAGAATTAGTCTGAAGAATATCTATATCACCGTACCGTATATTTGGATTAATTAGATATAGAGAATATGATCCGTAACATATACAACTTTTATTATGACGTCTAAGATATTCTTCCATCAACTTATTAACATTTTTGACTAGGGAAGATACATTATGACGTCCCATTACTTTTGCCTTGTCTATTACAGCGACGTTCATAGAATTTAGCATATCTCTTGCCAATTCTTCCATTGATGTTACATTATAAGAAATTTTAGATGAAATTACATTTGGAGCTTTAATAGTAAGAACTCCTAATATATCCGTGTATGTGGTCACTAATACAGATTGTAGTTCTATAATCGTAAATAATTTACCTATATTATATGTTTGAGTTTGTTTAGAAAAGTAGCTAAGTATACGATCTTTTATTTCTGATGCCGATGTATCAACATCGAAAAAAAATCTTTTTTTATTCTTTTTTACTAACGATACGAATATGTCTTTGTTAAAAACAGTTATTTTCTGAATATTTCTAGCTTGTAATTTTAACATATGATATTCGTTCACACTAGGTACTCTGCCTAAATAGGTTTCTATAATCTTTAATGTAATATTAGGAAGAGTATTCTGATCAGGATTCCTATTCATTTTGAGGATTTAAAACTCTGATTATTGTCTAATATGGTCTCAACACAAACTTTTTCACAGAGTGATAGAGTTTTTGATAACTCGTTTTTCTTAAGAAATATAAAACTACTGTCTCCAGAGCTCGCTCTATCTTTTATTTTATCTAATTCGATACAAACTCCTGATACTGGTTCAGAAAGTAATTCATTAATTTTCAGTCCTTTATAGAAGATATTTAATATAGATAATACAAAATCTTCAGTTCTTGATATCGATCTGATTGATCCTAGAACTAGATATATTAATAACGTGCTCATTAGGCAGTTTATGGCAGCTTGATAATTAGATATAGTATATTCCAGTTCATATTTATTAGATACCGCATTGCCCAGATTTTGATATTCTATGAATTCCTCTGAAAATAAATCCAAAATAACTAGACATTCTATTTTTTGTGGATTAGTGTACTCTCTTCCCTCTATCATGTTCACTACTGGTGTCCACAATGATAAATATCTAGAGGGAATATAATATAGTCCATATGATGCCAATCTAGCAATGTCGAATAACTGTAATTTTATTCTTCGCTCTTCATTATGAATTGAATCTTGAGGTATAAACCTAACACAAATTATATCATTAGACTTTTCGTATGTAATGTCTTTCATGTTATAAGTTTTTAATCCTGGAATAGAATCTATTTTAATGAGGCTTTTAAACGCAGCGTTCTCCAACGAGTCAAAGCATAATACTCTGTTGGTTTTCTTATATTCAATATTACGATTTTCTTCTTTGAATGGAATAGGTTTTTGAATTAGTTTATAATTACAACATAATAGATAAGGAAGTGTGTAAATAGTACGCGGAAAAAACATAATAGCTCCCCTGTTTTCATCCATGGTTTTAAGTAAATGATCACTGGCTTCTTTAGTCAATGGATATTCGAACATTAACCGTTTCATCATCATTGGACAGAATCCATATTTCTTAATGTAAAGAGTGATCAAATCATTGTGTTTATTGTACCATCTTGTTGTAAATGTGTATTCGGTTATCGGATCTGCTCCTTTTTCTATTAAAGTATCGATATCGATCTCGTCTAAGAATTCAACTATATCGACATATTTCATTTGTATACACATAACCATTACTAACGTAGAATGTATAGGAAGAGATGTAACGGGAACAGGGTTTGTTGATTCGCAAACTATTCTAATACATAATTCTTCTGTTAATACGTCTTGCACGTAATCTATTATAGATGCCAAGATATCTATATAATTATTTTGTAAGATGATGTTAACAATGTGATCTATATAAGTAGTGTAATAATTCATGTATTTCGATATATGTTCCAACTCTGTCTTTGTGATGTCTAGTTTCGTAATATCTATAGCGTCCTCAAAAAATATATTCGCATATATTCCCAAGTCTTCAGTTCTATCTTCTAAAAAATCTTCAACGTATGGAATATAATAATCTATTTTACCTCTTCTGATGTCATTAATGATATAGTTTTTGACACTATTTTCCGTCAATTGATTCTTATTCACTATGTCTAAAAACCGGATAGCGTCCCTAGGACGAACTACTGCCATTAATATCTCTATTATAGCTTCTGGACATAAATCATCTATTATACCAGAATTAATGGGAACTATTCCGTATCTATCTAACATAGTTTTAAGAAAGTCAGAATCTAAGACCTGATGTTCATATATTGGTTCATACATGAAATGATCTCTATTGATGATAGTGACTATTTCATTCTCTGAAAATTGGTAACTCATTCTATACACGCTTTCCTTGTTGATAAAGGATAGTATATACTCAATGGAATTTGTACCAACAAACTGTTCTCTTATGAATCGTATATCATCATCTGAAATGATCATGTAAGGCATACATTTAACAATAAGAGACTTGTCTCCTGTTATCAATATACTATTCTTGTGATAATTTATGTGTGCGGCAAATTTGTCCACGTTCTTTAATTTTGTTATAGTAGATATCAAATCCAATGGAGATACAGTTCTTGGCTTAAACAGATATAGTTTTTCTGGAACGAATTCTACAACATTATTATAAAGGACTTTGGGTATATAAGTGGGATGAAATCCTATTTTAATTAATGCGATAGCCTTGTCCTCGTGCAGATATCCAAACGCTTTTGTGATAGTATGGCATTCATTGTCTAGAAACGCTCTACGAATATCTGTAACAGATATCATCTTTAGAGAATACTAGTCGCGTTAATAGTACTAAAATTTGTATTTTTTAATCTATCTCAATAAAAAATTAATATGTATGATTCAATGTATAACTAAACTACTAACTGTTATTGATAACTAGAATCAGAATCTAATGATGACATAACTAAGAAGTTTATCTACAGCCAATTTAGCTGCATTATTTTTAGCATCTCGTTTAGATTTTCCATCTGCCTTATCGAATACTCTTCCGTCAATGTCTACACAGGCATAAAATGTAGGAGAGTTACTAGGCCCCACTGATTCAATACGAAAAGACCAATCTCTCCTAGTTATTTGACAGTACTCATTAATAACGGTGACAGGGTTAACACCTTTCCAATAAATAATTTTTTTAACCGGAATAACATCATCAAAAGACTTATGATCCTCTCTCATTGATTTTTCGCGGGATACATCATCTATTATAGCATCAGCATCAGAATCTGTAGGCCGTGTATCAGCATCCATTGTCGTAGACCAACGAGGAGGAGTATCGTTGGAGCTGTAAACCATAGCACTACGTTGAAGATCATACAGAGCTTTATTAACTTCTCGCTTCTCCATATTAAGTTGTTTAGTTAGTTGTACAGCAGTAGCTCCTTAGTCCAATGTTTTTAATAACCGCACACAATCTCTGTGTCAGAACGCTCGTCAATATAGATCTTAGAAATTTTTTTAGAGAGAACTAACGCAACTAGCAATAAAACTGATCTTATTTTATCATTTTTTTTATTCATCATCCTCTGGTGGTTCGTCGTTCCTATCGAATGTAGCTCTGATTAACCCGTCATCTATAGGTGATGCTGGTTCTGGAGATTCTGGAGGAGATGGATTATTATCTGGAAGAATCTCTGTTATTTCCTTGTTTTCATGTATCGATTGCGTTGTAACATTAAGATTGCGAAATGCTCTAAATTTGGGAGGCTTAAAGTGTTGTTTACAATCTCTACACGCGTGTCTAACTAATGGAGGTTCATCAGCGGCTCTAGTTTGAATCATCATCGGTGTAGTATTCCTACTTTTACAGTTAGGACACGGTGTATTGTATTTCTCGTCGAGAACGTTAAAATAATCGTTGTAACTCACATCCTTTATTTTATCTATATTGTATTCTACTCCTTTCTTAATGCATTTTATACCGAACAAGAGATAGCGAAGGAATTCTTTTTCGGTACCGCTAGTACCCTTAATCATATCACATAGTGTTTTATATTCTAAATGTGTGGCAATGGACGGTTTATTTCTATACGATAGTTTGTTTTTGGAATCCTTTGAGTATTCTATACCAATATTATTCTTTGATTCGAATTTAGTTTCTTCGATATTAGATTTTGTATTACCTATATTCTTGATGTAGTACTTTGATGATTTTTCCATGGCCCATTCTATTAAGTTTTCCAAGTTGGCATCATCCACATATTGTGATAGTAATTCTCGGATATCAGTAGTGACTACCGCCATTGATATTTGTTCATTTGATGAGTAACTACTAATGTATACATTTTCCATTTATAACACTTATGTATTAACTTTGTTTATTTATATTTTTTCATTATTATGTTGATATTAATAATCGTATTGTGGTTATATGGCTACAATTTCATAATGAGTTGAAGTCAGTGTCCTATGATCAATGACGATAGCTTTACTCTGAAAAGAAAGTATCAAATCGATAGTGCAGAGTCAACAATGAAAATGGATAAGACGATGACAAAGTTTCAGAATAGAGTCAAAATGGTAAAAGAAATAAATCAGACGATAAGAGCAGCACAAACTCATTACGAGACATTGAAACTAGGATATATAAAATTTAAGGGAATGATTAGGACTACTACTCTAGAAGATATAGCACCATCTATTCCAAATAATCAGAAAACTTATAAACTATTCTCGGACATTTCAGTCATTGGCAAAGCATCACAGAATCCGAGTAAGATGATATATGCTCGCTGCTTTACATGTTTCCCAATTTGTTTGGAGATGACCATAGATTCATTTGTTATAGAATGCATCCAACATTGTTCATGATATAGTTGAATCATGTATGCCTGTTCGTATGCCTGTGGCTAAGATACTGTGTAAAGAAATGGTAAATAAATACTTTGAGAATCTTTAAGAGTGCATTGACTTTGTTAGTGAATAGGCATTCCATCTTTCTCCAATACTAATTCAAATTGTTAAATTAATAATGGAATAGTATAAATAGTTATTAGTGATAGGATAGTAAACATAATTATTAGAATAGTAGTGTAGTATCATAGATAACTCTCTTCTATAAAAAATGGATTTTATTCGTAGAAAGTATCTTATATACACAGTAGAAAATAATATAGATTTTTTAAAGGATGATACATTAAGTAAAGTAAACAATTTTACCCTCAATCATGTACTAGCTCTCAAGTATCTAGTTAGCAATTTTCCTCAACATGTTATTACTAAGGATGTATTAGCTAATACCAATTTTTTTGTTTTCATACATATGGTACGATGCTGTAAAGTATACGAAGCGGTTTTACGACACGCATTTGATGCACCCACGTTGTACGTTAAAGCATTGACTAAGAATTATTTATCGTTTAGTAACACAATACAGTCGTACAAGGAAACAGTGCATAAACTAACACAAGATGAAAAATTTTTAGAGGTTGCCGAATACATGGACGAATTAGGAGAACTTATAGGCGTAAATTATGACTTAGTTCTTAATCCATTATTTCACGGAGGGGAACCCATCAAAGATATGGAAATCATTTTTTTAAAACTGTTTAAGAAAACAGACTTCAAAGTTGTTAAAAAATTAAGTGTTATAAGATTACTTATTTGGGCATACCTAAGCAAGAAAGATACAGGCATAGAGTTTGCGGATAATGATAGACAAGATATATATACTCTATTTCAACAAACTGGTAGAATAGTCCATAGCAATCTAACAGAAACGTTTAGGGATTATATCTTTCCCGGAGATAAGACTAGCTATTGGGTGTGGTTAAACGAAAGTATAGCTAATGATGCGGATATCGTTATTAATAGACCCGCCATTACCATGTATGATAAAATTCTTAGTTATATATACTCTGAGATAAAACAGGGACGCGTTAATAAAAACATGCTTAAGTTAGTTTATATCTTTGAGCCTGAAAAAGATATCAGAGAACTTCTGCTAGAAATCATATATGATATTCCTGGAGATATCCTATCTATTATTGATGCAAAAAACGACGATTGGAAAAAATATTTTATTAGTTTTTACAAAGCTAATTTTATTAACGGTAATACATTTATTAGTGATAGAACGTTTAACGATGACTTATTCAGAGTTGTTGTTCAAATAGATCCCGAATATTTCGATAATGAACGAATTATGTCTTTATTCTATACGAGTGCTGCGGACATTAAACGATTTGATGAGTTAGATATTAATAACAGTTATATATCTAATATAATTTATGAGGTGAACGATATCACATTAGATACAATGGATGATATGAAGAAGTGTCAAATCTTTAACGAGGATACGTTGTATTATGTTAAGGAATACAATACATACCTGTTTTTGCACGAGTCGGATCCCATGGTCATAGAGAACGGAATACTAAAGAAACTGTCATCTATAAAATCCAAGAGTAGACGGCTGAACTTGTTTAGCAAAAACATTTTAAAATATTATTTAGACGGACAATTGGCTCGTCTAGGTCTTGTGTTAGATGATTATAAAGGAGACTTATTAGTTAAAATGATAAACCATCTCAAATCTGTGGAGGATGTATCCGCATTCGTTAGATTTTCTACAGATAAAAACCCTAGTATTCTTCCATCGCTAATCAAAACTATTTTAGCTAGTTATAATATTTCCATCATCGTCTTATTTCAAAGGTTTTTAAGAGATAATCTATATCATGTAGAAGAATTCTTGGATAAAAGCATCCATCTAACCAAGACGGATAAGAAATATATACTTCAATTGATAAGGCACGGTAGATCATAGAACAAACCAAATATATTATTAATAATTTGTATATACATAGATATAATTATCACATATTAAAAAATAACACATTTTTGATAAATGGAAACCGTTGCAACAATTCAGACTCCCACCAAATTAATGAATAAAGAAAATGCAGAAATGATTTTGGAAAAAATTGTTAATCATATAGCTATGTATATTAGTGACGAATCAATATATTCAGAAAATAATCCTGAATATATTGATTTTCGTAACAGATACGGAGACTATAGATCTCTCATTATAAAAAGTGATCACGAGTTTGTAAAGCTATGTAAAGATCATGCAGAGAAAAGTTCTCCAGAAACGCAACAAATGATTATCAAACACATATACGAACAATATCTTATTCCAGTATCTGAAGTACTATTAAAACCTATAATGTCCATGGGTGACATATTTACATATAACGGATGTAAAGACAATGAATGGATGCTAGAACAACTCTCTACCCTAAACTTTAACAATCTCTACACATGGAACTCATGTAGCATAGGCAATGTAACGCGTCTGTTTTATACATTTTTTAGTTATCTGATGAAAGATAAACTAAATATATAAGTATAATCCCATTCTAATACTTTAACCTGATGTATTATTACCTGCATCTTATTAGAATATTAACCTAACTAAAAGACATAAAAAGCGGTAGGATATAAATATTATGGCCGCAACCGTTCCGCGTTTTGACGATGTGTACAAAAATGCACAAAGAAGAATTCTAGATCAAGAAACATTTTTTAGTAGAGGTCTAAGTAGACCGTTAATGAAAAACACATATCTATTTGATAATTACGCGTATGGATGGATACCAGAAACTGCAATTTGGAGTAGTAGATACGCAAACCTAGATGCTAGTGACTATTATCCCATTTCGTTGGGATTACTTAAAAAGTTTGAATTTCTCATGTCTCTATATAAAGGTCCTATTCCCGTATATGAAGAAAAAGTAAATACTGAATTCATTGCTAATGGATCTTTCTCCGGTAGATACGTATCATATCTTAGAAAGTTTTCTGCCCTTCCAACAAACGAGTTTATTAGTTTTTTATTATTGACCTCCATCCCTATCTATAATATCTTATTCTGGTTTAAAAACACACAGTTTGATATTACTAAACACACATTATTCAGATACGTCTATACAGATAATACCAAACACCTTGCGTTGGCTAGGTATATACATCAAACAGGAGACTATAAGCCTTTGTTTAGTCGTCTCAAAGAGAATTATATATTTACTGGTCCCGTTCCAATAGGTATCAAAGATATAGATCACCCTAATCTTAGTAGAGCAAGAAGTCCATCCGATTATGAGACATTAGCTAATATTAGTACTATATTGTACTTTACCAAGTATGATCCAGTATTAATGTTTTTATTGTTTTACGTACCTGGGTATTCAATTACTACAAAAATTACTCCAGCCGTAGAATATCTAATGGATAAACTGAATCTAACAAAGAGCGACGTACAACTGTTGTAAATTATTTTATGCTTCGTAAAATGTAGGTCTTGAACCAAACATTCTTTGAAAAAATGAGATGCATAAAACTTTATTATCCAATAGATTAACTATTTCAGACGTCAATCGTTTAAAGTAAACTTCGTAAAATATTCTTTGATTGCTGCCGAGTTTAAAACTTCTATCGATAATTGTTTCATATGTTTTAATATTTACAAGTTTTTTGGTCCATGGTACATTAGCTGGACAGATATATGCAAAATAATATCGTTCTCCAAGTTCTATAGTCTCTGGATTGTTTTTATTATATTCAGTAACCAAATACATATTAGGGTTATCTGCGGATTTATAATTTGAGTGATGCATTCGACTCAACATAAATAATTCTAGAGGAGACGATCTACTATCAAATTCGGATCGTAAATCTGTTTCTAAAGAACGGAGAATATCTATACATACCTGATTAGAATTCATCCGTCCTTCAGACAACATCTCAGACAGTCTGGTCTTGTATGTCTTAATCATATTCTTATGAAACTTGGAAACATCTCTTCTAGTTTCACTAGTACCTTTATTAATTCTCTCAGGTACAGATTTTGAATTCGACGATGCCGAGTATTTCATCGTTGTATATTTCTTCTTCGATTGCATAATCAAATTCTTATATACCGCCTCAAACTCTATTTTAAAATTATTAAACAATACTCTACTATTAATCAGTCGTTCTAACTCCTTTGCTATTTCTATGGACTTATCTACATCTTGACTGTCTATCTCTGTAAACACGGAGTCGGTATCTCCATACACGCTACGAAAACGAAATCTATAATCTATAGGCAACGATGTTTTCACAATCGGATTAATATCTCTATCGTCCATATAAAATGGATTACTTAATGTATTGGCAAACCGTAACATACCGTTGGATAACTCTGCTCCATTTAGTACCGATTCTAGATACAATATCATTCTACGTCCTATGGATGTGCAACTCTTAGCCGAAGCGTATGAGTATAGAGCACTATTTCTAAATCCCATCAGACCATATACTGAGTTGGCTACTATCTTGTACGTATATTGCATGGAATCATAGATGGCCTTTTCAGTTGAACTGGTAGCCTGTTTTAACATCTTTTTATATCTGGCTCTCTCTGCCAAAAATGTTCTTAATAGTCTAGGAATGGTTCCTTCTATTGATCTATCGAAAATTGCTATTTCAGAGATGAGGTTCGGTAGTCTAGGTTCACAATGAACCGTAATATATCTAGGAGGTGGATATTTCTGAAGCAAGAGTTGATTATTTATTTCTTCTTCCAATCTATTGGTACTAACAACGACACCGACTAATGTTTCCGGAGATAGATTTCCAAAGATACACACATTAGGATACAGACTGTTATAATCAAAGATTAATACATTATTACTAAACATTTTTTGTTTTGGAGCAAATACCTTACCGCCTTCATAAGGAAACTTTTGTTTTGTTTCTGATCTGACTAAGATAGTTTTAGTTTCCAACAATAGCTTTAACAGTGGACCCTTGATGATTGTACTCGCTCTATATTCGAATACCATGGATTGAGGAAGCACATATGTTGCCGCACCAGCGTCTGTTTTTGTTTCTACTCCATAATACTCCCACAAATACTGACACAAACAAGCATCATGAATACAGTATCTAGCCATATCTAAAGCTATGTTTAGATTATAATCCTTATACATCTGAGCTAAATCAATGTCATCCTTTCCGAAAGATAATTTATATATATCATTAGGTAAAGTAGGACATGATAGTACGACTTTAAATCCATTTTCCAAAATATCTTTACGAATTACTTTACATATAATATCCTCATCAACAGTCACGTAATTACCTGTGGTTAAAACCTTTGCAAATGTATCGGCTTTGCCTTTCGCGTCCGTAGTATCGTCACCGATGAACGTCATTTCTCTAACTCCTCTATTTAATACTTTACCCATGCAACTGAACGCGTTCTTGGATATAGAATCCAATTTGTACGAATCCAATTTTTCAGATTTTTGAATGAATGAATATAGATCGAAAAATATAGTTCCATTATTGTTATTAACGTGAAACGTAGTATTGGCCATGCCGCATACTCCCTTATGACTAGACTGATTTCTCTCATAAATACAGAGATGTACAGCTTCCTTTTTGTCTGGAGATCTAAAGATAATCTTCTCTCCTGTTAATAACTCTAGACGATTAGTAATATATCTCAGATCAAAGTTATGTCCGTTAAAGGTAACGACGTAGTCGAACGTTAGTTCCAACAATTGTTTAGCTATTCGTAACAAAACTATTTCAGAACATAGAACTAGTTCTCGTTCGTAATCCATTTCCATTAGCGACTGTATCCTCAAACATCCTCTATCGACGGCTTCTTGTATTTCCTGTTCCGTTAACATCTCTTCATTAATGAGCGTAAACAGTAATCGTTTACCACTTAAATCGATATAACAGTAACTTGTATGCGAGATTGGGTTAATAAATACAGAAGGAAACTTCTTATCGAAGTGACACTCTATATCTAGAAATAAGTACGATCTTGGGATATCGAATCTAGGTATTTCTTTAGCGAAACAGTTACGTGGATCGTCACAATGATAACATCCATTGTTAATCTTTGTCAAATATTGCTCGTCCAACGAGTAACATCCGTCTGGAGATATCCCGTTAGAAATATAAAACCAACTAATATTGAGAAATTCATCCATGGTGGCATTTTGTATGCTGCGTTTCTTTGGCTCTTCTATCAACCACATATCTGCGACGGAGCATTTTCTATCTTTAATATCTAGATTATAACTTATTGTCTCGTCAATGTCTATAGTTCTCATCTTTCCCATCGGCCTCGCATTAAATGGAGGAGGAGATAATGACTGATATATTTCGTCCGTCACTACGTAATAAAAGTAATGAGGAAATCGTATAAATACGGTCTCGCCATTTCGACATCTGGATTTCAGATATAAAAATCTGTTTTCACCGTGACTTTCAAACCAATTAATACACCTAACATCCATTTCTAGAATTTAGAAATATATTTTCATTTAAATGAATCCCAAACATTGGGGAAGAGCCGTATGGACCATTATTTTTATAGTACTTTCGCAAGCGGGTTTAGACGGCAACATAGAAGCGTGTAAACGAAAACTATATACTATAGTCAGCACTCTTCCATGTCCTGCATGTAGACGACACGCGACTATCGCTATAGAGGACAATAATGTCATGTCTAGCGATGATCTGAATTATATTTATTATTTTTTCATCAGATTATTTAACAATTTGGCATTTGATCCCAAATACGCAATCGATGTGTCAAAGGTTAAACCTTTATAAACTTAACCCATTATAAAACTTATGATTAGTCACGACTGAAATAACCGCGTGATTATTTTTTGGTATAATTCTACACGGCATGGTTTCTGTGACTATGAATTCAACACCTGTTATCTTAGTGAAATCTTTAACAAACAGCAAGGGTTCGTCAAAGACATAAAACTCATTGTTTACGATCGAAATAGACCCCCTATCACACTTAAAATAAAAAATATCCTTATCCTTTACCACCAAATAAAATTCTGATTGGTCAATGTGAATGTATTCACTTAACAGTTCCACAAATTTATTTATTAACTCCGAGGCACATACATCGTCGGTATTTTTTATGACAAACTTTACTCTTCCAGCATCCGTTTCTAAAAAAATATTAACGAGTTCCATTTATATCATCCAATATTATTGAAATGACGTTGATGGACAGATGATATAAATAAGAAGGTACAGTACCTTTGTCCACCATCTCCTCCAATTCATACTCTATTTTGTCATTAACTTTAATGTGTGAAAACAGTACGCCACATGCTTCCATGACAGTGTGTAACACTTTGGATACAAAATGTTTGACATTAGTATAATTGTCCAAGACTGTCAATCTATAATAGATAGTAGCTATAATATATTCTATGATGGTATTGAAGAAGATGACAACCTTGGCATATTGATCATTTAACACAGACATGGTATCAACAAATAGCTTAAATGAAAGAGAATCAGTAATTGGAATAAGCGTCTTCTCGATGTAGTGTCCGTATACCAACATGTCTGATATTTTGATGTATTCCATTAAATTATTTAGTTTTTTCTTTTTATTCTCGTTAAACAGAATTTCTGTCAATGGACCCCAACATCGTTGACCTATTAAGTTTTGATTGATTTTTCCGTGTAAGGCGTATCTAGTCAGATCGTATAGCCTATCCAATAATCCATCGTCTGTGCGTAGATCACATCGTACACTTTTTAATTTTCTATAGAAGAGTGACAGACATCTGGAGCAATTACAGACAGCAATTTCTTTATTCTCTACAGATGTAAGATACTTGAAGATATTCCTATGATGATGCAGAATTTTGGATAACACGGTATTGATGGTATCTGTTACCATAATTCCTTTGACTGATAGTGTCAAAGTACAAGATTTCCAATCTTTTGCAATTTTCAGTACCATTATCTTTGTTTTGATATCTATATCAGACAGCATGGTACGTCTGACAACACAGGGATTAAGACGGAAAGATGAAATGATTCTCTCAACATCTTCAATAGATACCTTGCTATTTTTTTTGGCATTATCTATATGTGAGAGAATATCCTCTAGAGAATCAGTATCCTTTTTGATGATAGTGGATCTCAATGACATGGGACGTCTAAACCTTCTTATTCTATCACCAGATTGCATGGTGATTTGTCTTCTTTCTTTTATCATGATGTAATCTCTAAATTCATCGGCAAATTGTCTATATCTAAAATCATAATATGAGATGTTTACCTCTACAAATATCTGTTCGTCCAATGTTAGAGTATCTATATCAGTTTTGTATTCCAAATTAAACATGGCAACGGATTTAATTTTATATTCCTCTATTAAGTCCTCGTCGATAATAACAGAATGTAGATAATCATTTAATCCATCGTACATGGTTGGAAGATGCTCGTTGACAAAATCTTTAATTGTCTTGATGAAGGTGGGACTATATCTAACATCTTGATTAATAAAATTTATAACATTGTCCATAGGATACTTTGTAACTAGTTTTATACACATCTCTTCATTGGTAAGTTTAGACAGAATATCGTGAACAGGTGGTATATTATATTCATCAGATATACGAAGAATAATGTCCAAATCTATATTGTTTAATATATTATATAGATGTAGTGTAGCTCCTACAGGAATATCTTTAACTAAGTCAATGATTTCATCAACAGTTAGATCTATTTTAAAGTTAATCATATAGGCATTGATTTTTAAAAGGTATGTAGCCTTGACTACATTCTCATTAATTAACCATTCCAAGTCACTGTGTGTAAGAAGATTATATTCTATCATAAGCTTGACTACATTTGGTCCCGATACCATTAAAGAATTCTTATGATATAAGGAAACAGCTTTTAGGTACTCATCTACTCTACAAGAATTTTGGAGAGCCTTAACGATATCAGTGACGTTTATTATTTCAGGAGGAAAGAACCTAACATTGAGAATATCTGAATTAATAGCTTCCAGATACAGTGATTTTGGCAATAGTCCGTGTAATCCATAATCCAGTAACACGAGCTGGTGCTTGCTAGACACCTTTTCAATGTTTAATTTTTTTGAAATAAGCTTTGATAAAGCCTTCCTCGCAAATTCCGGATACATGAACATGTCGCCAACATGATTAAGTATTGTTTTTCATTATTTTTATATTTTCTCAACAAGTTCTCAATACCCCAATAGATAATAGAATATCACCCAATGCGTCCATGTTGTCTATTTCCAACAGGTCGCTATATCCACCAATAGAAGTTTTCCCAAAAAAGATTCTAGGAACAGTTCTACCACCAGTAATTTGTTCAAAATAGTCACGCAATTCATTTTCGGGTTTAAATTCTTTAATATCTACAATTTCATACGCTCCTCTTTTGAAACTAAACTTATTTAGAATATCCAGTGCGTTTCTACAAAAAGGACATGTAAACTTGACAAAAATTGTCACTTTGTTATTGGCCAACCTTTGTTGTACAAATTCCTCGGCCATTTTTAATATTTAAGTGATACAAAACTATCTCGACTTATTTAACTCTTTAGTCGAGATATATGGACACAGATAGCTATATGATAACCAACTACAGAAGACAAACGCTATAAAAAACATAATTACGACGAGCATATTTATAAATATTTTTATTCAGTATTACTTGATATAGTAATATTAGGCACAGTCAAACATTCAACCACTCTAGATACATTAACTCTCTCATTTTCTTTAACAAATTCTGCAATATCTTCGTAAAAAGATTCTTGAAACTTTTTAGAATATCTATCGACTCTAGATGAAATAGCGTTCGTCAACATACTATGTTTTGTATACATAAAGGCGCCCATTTTAACAGTTTCTAGTGACAAAATGCTAGCGATCCTAGGATCCTTTAGAATCACATAGATTGACGATTCGTCTCTCTTAGTAACTCTAGTAAAATAATCATACAATCTAGTACGCGAAATAATATTATCCTTGACTTGAGGAGATCTAAACAATCTAGTTTTGAGAACATCGATAAGTTCATCGGGAATTACATACATACTATCTTTAATAGAACTCTTTTCATCCAGTTGAATGGATTCGTCCTTAACCAACTGATTAATGAGATCTTCTATTTTATCATTTTCTAGATGATATGTATGTCCATTAAAGTTAAATTGTGTAGCGCTTCTTTTTAGCCTAGCAGCCAATACTTTAACATCACTAATATCGATATACAAAGGAGATGATTTATCGATGGTATTAAGAATTCGTTTTTCGACATCCGTCAAAACCAATTCCTTTTTGCCTGTATCATCCAGTTTGCCATTCTTTGTAAAGAAATTATTTTCTACTAGACTATTAATAAGACTGATAAGGATTCCTCCATAATTGCACAATCCAAACTTTTTCACAAAACTAGACTTTACGAGATCTACAGGAATGCGTACTTCAGGTTTCTTAGCTTGTGATTTTTTCTTTTGTGGACATTTTCTAGTGACCAACTCATCTACCATTTCATTGATTTTAGCAGTGAAATAAGCTTTCAATGCACGGGCACTGATACTATTGAAAACGAGTTGATCTTCAAATTCCGCCATTTAAGTTCACCAAACAACTTTTAAATACAAATATATCAATAGTAGTAGAATAAGAACTATAAAAAAAATAATAATTAACCAATACCAACCCCAACAACCTGTATTATTAGTTGATGTGACAGTTTTCTCATCACTTAGAACAGATTTAACAATTTCTATAAAGTCTGTCAAATCATCTTCCTGAGAACCCATAAATACACCAAATATAGCAGCGTACAACTTATCCATTTATACATTGAATATTGGCTTTTCTTTATCGCTATCTTCATCATATTCATCATCAATATCAACAAGTCCCAGATTACGAACCAGATCTTCTTCTACATTTTCAGTCATTGATACGCGTTCACTATCTCCAGAGAGTCCGATAACGTTAGCCACTACTTCTCTATCAATGATTAGTTTCTTGAGCGCGAATGTAATTTTTGTTTCCGTTCCGGATCTATAGAAAACTACAGGTGTAATAATTGCCTTGGCTAATTGTCTTTCTCTTTTACTGAGTGATTCTAGTTCACCTTCTATAGATCTGAGAATGGATGATTCTCCAGTCGAAACATATTCTACCATGGCTCCGTTTAATTTGTTGATGAAGATGGATTCATCCTTAAATGTTTTCTCTGTAATAGTTTCCGCCGAAAGACTATGCAAAGAATTTGGAATGCGTTCCTTGTGTGTAATGTTTCCATAGACAGCTTCTAGAAGTTGATACAACATAGGACTAGCCGCGGTAACTTTTATTTTTAGAAAGTATCCATCGCTTCTATCTTGTTTAGATTTATTTTTATAAAGTTTAGTCTCTCCTTCCAACATAATAAAAGTGGAAGTCATCTGACTAGATAAACTATCAGTAAGTTTTATAGAGATAGATGAACAATTAGCGTATTGAGAAGCATTTAGTGTAACGCATTCGATACATTTTGCATTAGATTTACTAATCGATTTTGCATACTCTATAACACCCGCACAAGTCTGTAGAGAATCGCTAGATGCTGTAGGTCTTGGTGAAGTTTCAACTCTCTTCTTGATTACCTTACTCATGATTAAACCTAAATAATTGTACTTTGTAATATAATGATATATATTTTCACTTTATCTCATTTGAGAATAAAAATGTTTTTGTTAACCACTGCATGATGTACAGATTTCGGAATCGCAAACCACTTGTGGTTTTATTTTATCCTTGTCCAATGTGAATTGAATGGGAGCGGATGCGGGTTTCGTACGTAGATAGTACATTCCCGTTTTTAGACCGAGACTCCATCCGTAAAAATGCATACTCGTTAGTTTGGAATAACTCGGATCTGCTATATGGATATTCATAGATTGACTTTGATCGATGAAGGCTCCCCTGTCTGCAGCCATTTTTATGATCGTCTTTTGTGGAATTTCCCAAATAGTTTTATAAACTCGCTTAATATCTTCTGGAAGGTTTGTATTCTGAATGGATCCACCATCTACCATAATCCTATTCTTGATCTCATCATTCCATAATTTTCTCTCGGTTAAAACTCTAAGGAGATGCGGGTTAACTACTTGGAATTCTCCAGACAATACTCTCCGAGTGTAAATATTACTGGTATACGGTTCCACCGACTCATTATTTCCCAAAATTTGAGCAGTTGATGCAGTCGGCATAGGTGCCACCAATAAACTATTTCTAAGACCGTATGTTCTGATTTTATCTTTTAGAGGTTCCCAATTCCAAAGATCCGACGGTACAACATTCCAAAGATCATATTGTAGAATACCGTTACTGGCGTACGATCCTACATATGTATCATATGGTCCTTCCTTCTCAGCTAGTTTACAACTCGCCTCTAATGCACCGTAATAAATGGTTTCAAAGATCTTCTTATTTAGATCTTGTGCTTCCAGGCTATCAAATGGATAATTTAAGAGAATAAACGCGTCCGCTAATCCTTGAACACCAATACCGATAGGTCTATGTCTCTTATTAGAGATTTCAGCTTCTGGAATAGGATAATAATTAATATCTATAATTTTATTGAGATTTCTGACAATTACTTTGACCACATCCTTCAGTTTGAGAAAATCAAATCGCCCATCTATTACAAACATGTTCAATGCAACAGATGCCAGATTACACACGGCTACCTCATTAGCATCCGCATATTGTATTATCTCAGTGCAAAGATTACTACACTTGATGGTTCCTAAATTTTGTTGATTACTCTTTTTGTTACACGCATCCTTATAAAGAATGAATGGAGTACCAGTTTCAATCTGAGATTCTATAATCGCTTTCCAGACGACTCGAGCCTTTATTATACATTTGTATCTCCTTTCTCTTTCGTATAGTGTATACAATCGTTCGAACTCGTCTCCCCAAACATTGTCCAATCCAGGACATTCATCCGGACACATCAACGACCACTCTCCGTCATCCTTCACTCGTTTCATAAAGAGATCAGGAATCCAAAGAGCTATAAATAGATCTCTTGTTCTATGTTCATCGTTTCCTGTATTCTTTTTAAGATCGAGGAACGCCATAATATCAGAATGCCACGGTTCCAAGTATATGGCCATAACTCCAGGCCGTTTGTTTCCTCCCTGATCTATGTATCTAGCGGTGTTATTATAAACTCTCAACATTGGAATAATACCGTTTGATATACCATTGGTACCGGAGATATAGCTTCCACTGGCACGAATATTACTAATTGATAGACCTATTCCCCCTGCCATTTTAGAGATTAATGCGCATCGTTTTAACGTGTCATAGATGCCTTCTATGCTATCATCGATCATGTTAAGTAGAAAACAGCTAGACATTTGGTGACGAGTAGTTCCCGCATTAAATAAGGTAGGAGAAGCGTGCGTAAACCATTTTTCAGAAAGTAGATTGTACGTCTCAATAGCTGAGTCTATATCCCATTGATGAATTCCTACTGCGACACGCATTAACATGTGCTGAGGTCTTTCAACAATTTTGTTGTTTATTTTCAACAAGTAGGATTTTTCCAAAGTTTTAAAACCAAAATAGTTGTATGAAAAGTCTCGTTCGTAAATAATAACCGAATTGAGCTTATCCTTATATTTGTTAACTATATCCATGGTAATACTTGAAATAATCGGAGAATGTTTCCCATTTTTAGGATTAACATAGTTGAATAAATCCTCCATCACTTCACTAAATAGTTTTTTTGTTTCCTTGTGTAGATTTGATATGGCTATTCTGGCGGCTAGAATGGCATAATCCGGATGTTGTGTAGTACAAGTGGCTGCTATTTCGGCTGCCAGAGTGTCCAATTCTACCGTTGTTACTCCATTATATATTCCTTGAATAACCTTCATAGCTATTTTAATAGGATCTATATGATCAGTGTTTAAGCCATAGCACAATTTTCTAATACGAGACGTGATTTTATCAAACATGACATTTTCCTTGTATCCATTTCGTTTAATGACAAACATTTTTGTTGGTGTAATAAAAAAAATTATTTAATTTTTCATTAATAGGGATTTGACGTATGTAGCGTACAAAATTATCGTTCCTGGTATATAGATAAAGAGTCCTATATATTTGAAAATCGTTACGGTTCGATTAAACTTTAATGATTGCATTGTGAATATATCATTAGGATTTAACTCCTTGACTATCATGGCGGTGCCAGAAATTACCATCAAAAGCATTAATACAGTTATGCAGATCGCAGTTAGAACGGTTATAGCATCCACCATTTATATCTAAAAATTAGATCAAAGAATATGTGACAACGTCCTAGTTGTATACTGAGAATTGACGAAACAATGTTTCTTACATATTTTTTTCTTATTAGTAACCGACTTAATAGTAGGAACTGGAAAACTAGACTTGATTATTCTATAAGTATAGATACCCTTCCAAATAATGTTCTCTTTGATAAAAGTTCCAGAAAATGTAGAATTTTTTAAAAAGTTATCTTTTGCTATTACTAATATCGTGGTTAGACGCTTATTATTAATATGAGTGATGAAATCCACACCGCTTCTAGATATCGCTTTTATTTCCACATTAGATGGTAAATCCAATAGTGAAACTATCTTTTTAGGAATGTATGGACTCGCGTTTAGAGGAGTGAACGTCTTCGGAGTAGTAAAGGATGATTCGTCAAATGAATAAACAATTTCACAAATGGATGTTAATGTATTAGTAGGAAATTTTTTGACGCTAGTGGAATTGAAGATTCTAATGGATGATGTTCTACCTATTTCATCCGATAACATGTTAATTTCCAATACCAACGGTTTTAATATTTCGATGATATACGGTAGTCTCTCTTTCGGACTTATATAGCTTATTCCACAATACGAGTCATTATATACTCCAAAAAACAAAATAACTAGTATAAAATCTGTATCGAATGGGAAAAACGAAATTATCGATATAGGTATAGAATCCGGAACATTGAACGTATTAATACTTAATTCTTTTTCAGTGGTAAGAACCGATAGGTTATTGACATTGTATGGTTTTAAATATTCTATAACTTGAGACTTGATAGATATTAATGACGAATTGAAAATTATTTTTATCACCACGTGTGTTTCAGGATCATCGTCGACGCCAGTTAACCAACCGAATGGAGTAAAATAAATATCATTAATATATGCTCTAGATATTAGTATTTTTATTAATCCTTTGATTATCATCTTCTCGTACGCGAATGATTCCATGATCAAGAGTGATTTGAGAACATCCTCCGGAGTATTAATGGGTTTAGTAAACAGTCCATCGTTGCAATAATAAAAGTTGTCCAAGTTAAAGGATATTATGCATTCGTTTAAAGATATCACCTCATCTAACGGAGACAATTTTTTGGTAGGTTTTAGAGACTTTGAAGCTACTTGTTTAACAAAGTTATTCATCGTCGTCTACTATTCTATTTAATTTTGTAGTTAATTTATCACATATCACATTAATTGACTTTTTGGTCCACTTTTCCATACGTTTATATTCTTTTAATCCTGCGTTATCCGTTTCCGTTATATACAGGGATAGATCTTGCAAGTTAAATAGAATGCTCTTAAATAATGTCATTTTTTTATCCGCTAAAAATTTAAAGAATGTATAAACTTTTTTCAAAGATTTAAAACTTTTAGGTGGAGTTCTGGTACACAATATCATAAACAAACTAATAAACATCCCACATTCAGATTCCAACAATTGATTAACTTCCACATTAATACAGCCTATTTTCGCTCCAAATGTACATTCGAAAAATCTGAATAAAACATCAATATCGCAATTTGTATTATCCAATACAGAATGTCTGTGATTCGTGTTAAAACCATCGGAAAAAGAATAGAAATAAAAATTATTATAATGGTGGAATTCAGTTGGAATATTGCCTCCGGAGTCATAAAAGGATACTAAACATTGTTTTTTATCGTAAATTACACATTTCCAATGAGACAAATAACAAAATCCAAACATTACAAATCTAGAGGTAGAACTTTTAATTTTGTCTTTAAGTATATACGATAAGATATGTTTATTCATAAACGCGTCAAATTTTTCATGAATAGCTAAGGAGTTTAAGAATCTCATGTCAAATTGTCCTATATAATCCACTTCGGATCCATAAGCAAACTGAGAGACTAAGTTCTTAATACTTCGATTGCTCATCCAGGCTCCTCTCTCAGGCTCTATTTTCATCTTGACGACCTTTGGATTTTCACCAGTATGTATTCCTTTACGTGATAAATCATCGATTTTCAAATCCATTTGTGAGAAGTCTATCGCCTTAGATACTTTTTCCCGTAGTTGAGGTTTAAAGAAATACGCTAACGGTATACTAGTAGGTAACTCAAAGACATCATATATAGAATGGTAACGCGTCGTTAACTCGTCGGTTAACTCTTTCTTTTGATCGAGTTCATCGCTACTATTGGGTCTGCTCAGGTGCCCCGACTCTACTAGTTCCAACATCATACCGATAGGAATACAAGACACTTTGCCAGCGGTTGTAGATTTATCATATTTCTCCACCACATATCCGTTACAATTTGTTAAGAATTTAGATACATCTATATTGCTACATAATCCAGCTAGTGAATATATATGACATAATAAATTGGTAAATCCTAGTTCTGGTATTTTACTAATTACTAAATCTGTATATCTTTCCATTTATCATGGAAAAGAATTTACCAGATATCTTCTTTTTTCCAAACTGCGTTAATGTATTCTCTTACAAATATTCACAAGATGAATTCAGTAATATGAGTAAAACGGAACGTGATAATTTCTCATTGGCTGTGTTTCCAGTGATAAAACATAGATGGCATAACGCACACGTTGTAAAACATAAAGGAATATACAAAGTTAGTACAGAAGCACGTGGAAAAAAAGTATCTCCTCCATCACTAGGAAAACCCGCACATATAAACCTAATGTCGAAGCAATATATATATAGTGAGTATGCAATAAGCTTTGAATGTTATAGTTTTCTAAAATGTATAACAAATACAGAAATCAATTCGTTCGATGAGTATATATTAAGAGGACTATTAGAAGCTGGTAATAGTTTACAGATATTTTCCAATTCCGTAGGTAAACGAATAGATACTATAGGTGTACTAGGGAATAAGTATCCATTTAGCAAAATTCCATTGGCCTCATTAACTCCTAAAGCACAACGAGAGATATTTTTAGCGTGGATTTCTCATAGACCTGTAGTTTTAACTGGAGGAACCGGAGTGGGTAAGACGTCACAGGTACCCAAGTTATTGCTTTGGTTTAATTATTTATTTGGTGGATTCTCTTCTCTAGATAAAATCACTGACTTTCACGAAAGACCAGTCATTCTATCTCTTCCTAGGATAGCTTTAGTTAGATTGCATAGCAATACCATTTTAAAATCATTGGGATTTAAGGTACTAGATGGATCTCCTATCTCTTTACGGTACGGATCTATACCGGAAGAATTAATAAACAAACAACCAAAAAAATATGGAATTGTATTTTCTACCCATAAGTTATCTCTAACAAAACTATTTAGTTATGGCACTATTATTATAGACGAAGTTCATGAGCATGATCAAATAGGAGATATTATTATAGCAGTAGCGAGAAAACATCATACGAAAATAGATTCTATGTTTTTAATGACTGCCACGTTAGAGGATGACAGGGAACGTCTAAAAATATTTTTACCTAATCCCGCATTTATACATATTCCTGGAGATACACTGTTTAAAATTAGCGAGGTATTTATTCATAATAAGATAAATCCATCTTCCAGAATGGCATATATAGAAGAAGAAAAGAGAAATTTAGTTACTGCTATACAGATGTATACTCCTCCTGATGGATCATCCGGTATAGTCTTTGTGGCATCCGTTGCACAGTGTCACGAATATAAATCATATTTAGAAAAAAGATTACCGTATGATATGTATATTATTCATGGTAAGGTCTTAGATATAGACGAAATATTAGAAAAAGTGTATTCATCACCTAATGTATCGATAATTATTTCTACTCCTTATTTGGAATCCAGCGTTACTATACGCAATGTTACACACATTTATGATATGGGTAGAGTTTTTGTCCCCGCTCCTTTTGGAGGATCACAACAATTTATTTCTAAATCTATGAGAGATCAACGAAAAGGAAGAGTAGGAAGAGTTAATCCTGGAACATACGTATATTTCTATGATCTGTCTTATATGAAATCTATACAGCGAATAGATTCAGAATTTCTACATAATTATATATTGTACGCTAATAAGTTTAATCTAACACTCCCCGAAGATTTGTTTATAATCCCTACAAATTTGGATATTCTATGGCGTACAAAGGAATATATAGACTCGTTCGATATTAGTACAGAAACATGGAATAAATTATTATCCAATTATTATATGAAGATGATAGAGTATGCTAAACTTTATGTACTAAGTCCTATTCTCGCTGAGGAGTTGGATAACTTTGAGAGGACGGGAGAATTAACTAGTATTGTACAAGAAGCCATTTTATCTCTAAATTTACAAATTAAGATTTTAAAATTTAAACATAAAGATGATGATACGTATATACACTTTTGTAGAATATTATTCGGTGTCTATAACGGAACAAACGCTACTATATATTATCATAGACCTCTAACGGGATATATGAATATGATTTCAGATACTATATTTGTTCCTGTAGATAATAACTAAAAATCAAAATCTAATGACCACATCTTTTTTTAGAGATGAAAAATTTTCCACATCTCCTTTTGTAGACACGACTAAACATTTTGCAGAAAAAAGTTTATTATTATTTAGATAATCGTATACTTCATCAGTGTAGATAGTAAATGTGAACAGATAAAAGGTATTCTTGCTCAATAGATTGGTAAATTCCATAGAATATATTAATCCTTTCTTCTTGAGATCCCACATCATTTCAACCAAAGACGTTTTATCCAATGATTTACCTCGTACTATACCACATACAAAACTAGATTTTGCAGTGATGTCGTACCTGGTATTCCTACCAAACAAAATTTTACTTTTAGTTCTTTTAGAAAATTCTAAGGTAGAATCTCTATTTGTCAATATGTCATCTATGGAATTACCACTAGCAAAAAATGATAGAAATATATATTGATACATCGCAGCTGGTTTTGATCTACTATACTTTAAAAACGAATCAGATTCCATAATTGCTTGTATATCATCAGCTGAAAAACTATGTTTTACACGTATTCCTTCGGCATTTCTTTTTAATGATATATCTTGTTTAGACAATGATAAAGTTATCATGTCCATGAGAGACGCGTCTCCGTATCGTATAAATATTTCATTAGATGTTAGACGCTTCATTAGGGGTATACTTCTATAAGGTTTCTTAATTAGTCCATCATTGGTTGCGTCAAGAACTACTATCTGATGTTGTTGGGTATCTCTAGTGTTACACATGGCCTTACTAAAGTTTGGGTAAATAACTATGATATCTCTATTAATTATAGATGTATATATTTCATTCGTCAAGGATATTAATATCGACTTACTATCGTCATTAATACGTGTAATGTAATCATATAAATCATGCGATAGCCAAGGAAAATTCAAATAGATGTTCATCATATAATCGTCGCTATAATTCATATTAATACTTTGACATTGACTAATTTGTAATATAGCCTCGCCACGAAGAAAGCTCTCGTATTCAGTTTCATCGATAAAGGATACCGTTAAATATAACTGGTTGCCGATAGTCTCATAGTCTATTAAGTGGTAAGTTTCGTATAAATACAGAATCCCTAAAATATTATCTAATGTGGGATTAATCCTTACCATAACTGTATAAAATGGAGCCGGAGTCATAACTATTTTACCGTTTGTACTTACTGGAATAGATGAAGGAATAATCTCCGGACATGATGGTAAAGACCCAAATGTCTGTTTGAAGAAATCCAATGTTCCAGGTCCTAATCTCTTGACAAAAATTACGATATTCGATCCCGATATCCTTTGCATTCTATTTACCAGCATATCACGAACTATATTAAGATTATCTATCATGTCTATTCTCCCACCGTTATATAAATCGCCTCCGCTAAGAAACGTTAGTATATCCATACAATGGAATACTTCATTTCTAAAATAGTATTCGTTTTCTAATTCTTTAATGTGAAATCGTATACTAGAAAGGGAAAAATTATCTTTGAGTTTTCCATTAGAAAAGAACCACGAAACTAATGTTCTGATTGCGTCTGACTCCGTCGCTGAATTAATAGATTTACACCAAAAACTCATATAACTTCTAGATGTAGAAGCATTCGCTAAAAAATTAGTAGAATCAAAGGATATAAGTAGATGTTCCAACAAGTGAGCAATTCCCAAGATTTCATCTATATCATTCTCGAATCCGAAATTAGAAATTCCCAAGTAGATATCCTTTTTCATCCGATCATTGATGAAAATACGAACTTTATTCGGTAAGACGATCATTTACTAAGGAGTAAAATAGGAAGTAACGTTCGTATATCGTTATCGTCGTATAAATTAAAGGTGTGTTTTTTGCCATTAAGAGACATTATAATTTTACCAATATTGGAATTATAATATAGGTGTATTTGAGCACTAGAAACGGTCGATGCATCGGTAAATATAGCTGTATCTAATGTTCTAGTCGGTATTTCTTCATTTCGCTGTCTAATGATAGCGTTTTCTCTATCTGTTTCCATTACAGCTGCCTGAAGTTTATTGGTCGGATAATATGTAAAATAATAAGAAATACATACGAATAACAAAAATAAAATAAGATATAATAAAGATGCCATTTAGAGATCTAATTTTGTTCAACTTGTCCAAATTCCTACTTACAGAAGATGAGGAATCGTTGGAGATAGTATCTTCCTTATGTAGAGGATTTGAAATATCTTACGATGACTTAATATCGTACTTTCCAGATAGGAAATACCATAAATATATTTCTAAGGTATTTGAACATGTAGATTTATCGGAGGAATTAAGTATGGAATTCCATGATACAACTCTGAGAGATTTAGTATATCTTAGATTGTACAAGTATTCCAAGTATATACGGCCGTGTTATAAATTAGGAGATAATCTAAAAGGTATAGTTGTTATAAAGGACAGAAATATATATATTAGAGAAGCAAATGATGACTTGATAGAATATCTCCTCAAGGAATACACTCCTCAGATTTATACATATTCTAATGAGCGAGTTCCCATAGCTGGTTCAAAATTAATTCTTTGTGGATTTTCTCAAGTTACATTTATGGCGTATACAACGTCGCATATAACAACAAATAAAAAGGTAGATGTTCTCGTTTCCAAAAAATGTATAGATGAACTAGTCGATCCAATAAATTATCAAATACTTCAAAATTTATTTGATAAAGGAAGCGGAACAATAAACAAAATACTCAGGAAGATATTTTATTCGGTAACAGGTGGCCAAACTCCATAGGTAGCTTTTTCTATTTCGGATTTTAGAATTTCCAAATTCACCAGCGATTTATCGGTTTTGGTGAAATCCAAGGATTTATTAATGTCCACAAATGCCATTTGTTTTGTCTGTGGATTGTATTTGAAAATGGAAACGATGTAGTTAGATAGATGCGCGGCGAAGTTTCCTATTAGGGTTCCGCGCTTCACGTCACCCAACATACTTGAATCACCATCCTTTAAAAAAAATGATAAGATATCAACATGGAGTATATCATACTCGGATTTTAATTCTTCTACTGCCTCACTGACATTTTCACAAATACTACAATACGGTTTACCGAAAATAATCAGTACGTTCTTCATTTATGGGTATCAAAAACTTAAAATCGTTACTGCTGGAAAATAAATCACTGACGATATTAGATGATAATTTATACAAAGTATACAATGGAATATTTGTGGATACAATGAGTATTTATATAGCCGTCGCCAATTGTGTCAGAAACTTAGAAGAGTTAACTACGGTATTCATAAAATACGTAAACGGATGGGTAAAAAAGGGAGGACATGTAACCCTTTTTATCGATAGAGGAAGTATAAAAATTAAACAAGACGTTAGAGACAAGAGACGTAAATATTCTAAATTAACCAAGGACAGAAAAATGTTAGAATTAGAAAAGTGTACATCCGAAATACAAAATGTTACCGGATTTATGGAAGAAGAAATAAAGGCAGAAATGCAATTAAAAATCGATAAACTCACATTTCAAATATATTTATCTGATTATGATAACATAAAAATATCATTGAATGAGATACTAACACATTTCAACAATAATGAGAATGTTACATTATTTTATTGTGATGAACGAGACGCAGAATTCGTTATGTGTCTAGAGGCTAAAACACAGTTCTCTACCACAGGAGAATGGCCGTTAATAATAAGTACCGATCAGGATACTATGCTATTCGCGTCTGCTGATAATCATCCTAAGATGATAAAAAACTTAACTCAACTGTTTAAATTTGTTCCCTCGGCAGAGGATAACTATTTAGCAAAATTAACTGCATTAGTGAATGGATGTGATTTCTTTCCTGGACTCTATGGGGCATCTATAACACCCAACAACTTAAACAAAATACAATTGTTTAGTGATTTTACAATCGATAATATAGTCACTAGTTTGGCAATTAAAAATTATTATAGAAAGACTAACTCTACCGTAGACGTGCGTAATATTGTTACGTTTATAAACGATTACGCTAATTTAGACGATGTCTACTCGTATATTCCTCCTTGTCAATGCACTGTTCAAGAATTTATATTCTCCGCATTAGATGAAAAATGGAATGAATTTAAATCATCTTATTTAGAGAGCGTGCCGTTACCCTGCCAATTAATGTACGCATTAGAACCACGTAAGGAGATTGATGTTTCAGAAGTTAAAACTTTATCATCTTATATAGATTTCGAAAATACTAAATCAGATATCGATGTTATAAAATCTATATCCTCGATTTTTGGATATTCTAACGAAAACTGTAACACCATAGTGTTCGGCATCTATAAGGATAATTTACTACTGAGTATAAATAATTCATTTTACTTTAACGATAGTCTGTTAATAACCAATACTAAAAGTGATAATATAATAAATATAGGTTACTAGATTAAAAAATGGTGTTCCAGCTCGTGTGTTCTACATGCGGCAAAGATATTTCTCACGAACGATATAAATTGATTATACGAAAAAAATCATTAAAGGATGTACTAGTCAGTGTAAAGAACGAATGTTGTAGGTTAAAATTATCTACACAAATAGAACCTCAACGTAACTTAACAGTGCAACCTCTATTGGATATAAACTAATGGATCCGGTTAATTTTATCAAGACATATGCGCCTAGAGGTTCTATTATTTTTATTAATTATGCCATGTCATTAACTAGTCATTTGAATCCATCGATAGAAAAACATGTGGGTATTTATTATGGTACGTTATTATCGGAACACTTGGTAGTTGAATCTACCTATAGAAAAGGAGTTAGAATAGTCCCATTGGATAGATTTTTTGAAGGATATCTTAGTGCAAAAGTATACATGTTAGAGAATATTCAAGTTATGAAAATAGCAGCTGATATGTCGTTAACTTTACTAGGTATTCCATATGGATTTGGTCATGATAGAATGTATTGTTTTAAATTGGTAGCTGAATGTTATAAAAATGCCGGTATTGATACATCGTCTAAACGAATATTAGGTAAAGATATTTTTCTGAGCCAAAACTTTACAGATGATAATAGATGGATAAAGATATATGATTCTAATAATTTAACATTTTGGCAAATTGATTACCTTAAAGGGTGAGTTAATATGCATAACTACTCCTCCGTTGTTTTTTCCCTCGTTCTTTTTCTTAACGTTGTTTGCCATCACTCTCATAATGTAAAGATATTCTAAAATGGTAAACTTTTGCATATCGGATGCAGAAATTGGTATAAATGTTGTAATTGTATTATTTCCCGTCAATGGACTAGTCACAGCTCCATCAGTTTTATATCCTTTAGAGTATTTCTCACTCGTGTCTAGCATTCTAGAGCATTCCATGATCTGTTTATCGTTGATATTGGCCGGAAAGATAGATTTTTTATTTTTTATTATATTACTATTGGCAATTGTAGATATAACTTCTGGTAAATATTTTTCTACCTTTTCAATCTCTTCTATTTTCAAGCCGGCTATATATTCTGCTATATTGTTACTAGTATCAATACCTTTTCTGGCTAAGAAGTCATATGTGGTATTCACTATATCAGTTTTAACTGGTAGTTCCATTAGCCTTTCCACTTCTGCAGAATAATTAGAAATTGGTTCTTTACCAGAAAATCCAGCTACTATAATAGGCTCACCGATGATCATTGGCAAAATCCTATATTGTACCAGATTAATGAGAGCATATTTCATTTCCAATAATTCTGCTAGTTCTTGAGACATTGATTTATTTGATGAATCTATTTGGTTCTCTAGATACTCTACCATTTCTGCCGCATACAATAACTTGTTAGATAAAATCAGGGTTATCAAAGTGTTTAGTGTGGCTAGAATAGTGGGCTTGCACGTATTAAAGAATGCTGTAGTATGAGTAAACCGTTTTAACGAATTATATAGTCTCCAGAAATCTGTGGCGTTGCATACATGAACTGAATGACATCGAAGATTGTCCAATATTTTTAATAGCTGCTCTTTGTCCATTATTTCTATATTTGACTCGCAACAATTGTAGATACCATTAATCACTGATTCCTTTTTCGATGCCGGACAATAGCACAATTGTTTAGCTTTGGACTCTATGTATTCAGAATTAATAGATATATCTCTCAATACAGATTGCACTATACATTTTGAAACTATGTCAAAAATTGTAGAACGACGCTGTTCTGTAGCCATTTAACTTTAAATAATTTACAAAAATTTAAAATGAGCATCCGTATAAAAATCGATAAATTGCGCCAAATTGTGGCATATTTTTCAGAGTTCAGCGAAGAAGTGTCTATAAATGTAGACTCGACGGATGAATTAATGTATATTTTTGCCGCCTTGGGCGGATCTGTAAACATTTGGGCCATTATACCTCTCAGTGCATCAGTGTTCTACCGCGGAGCCGAAAATATTGTGTTTAACCTTCCAGTGTCCAAGGTAAAATCGTGTTTGTGTAGTTTTCACAATGATGCTATCATAGATATAGAACCTGATCTGGAAAATAATCTAGTAAAACTTTCTAGTTATCATGTAGTAAGTGTCGATTGTAACAAGGAACTGATGCCTATTAGGACAGATACTACTATTTGTCTAAGTATAGATCAAAAGAAATCTTACGTATTTAATTTTCACAAGTATGAAGAAAAATGTTGTGGTAGAACCGTCATTCATCTAGAATGGTTGTTGGGCTTTATCAAGTGTATTAGTCAGCATCAGCATTTGGCTATTATGTTTAAAGATGACAATATTATTATGAAGACTCCTGGTAATACTGATGCGTTTTCCAGGGAATATTCTATGACTGAATGTTCTCAAGAACTACAAAAGTTTTCTTTCAAAATAGCTATCTCGTCTCTCAACAAACTACGAGGATTCAAAAAGAGAGTCAATGTTTTTGAAACTAGAATCGTAATGGATAATGACGATAACATTCTAGGAATGTTGTTTTCGGATAGAGTTCAATCCTTTAAGATTAACATCTTTATGGCGTTTTTAGACTAATACTTTCAATGAGATAAATATGGGTGGCGGAGTAAGTGTTGAGCTCCCTAAACGGGATCCACCTCCGGGAGTACCCACTGATGAGATGTTATTAAACGTGGATAAAATGCATGACGTGATAGCTCCCGCTAAGCTTTTAGAATATGTGCATATAGGACCACTAACAAAAGATAAAGAGGATAAAGTAAAGAAAAGATATCCAGAGTTTAGATTAGTCAACACAGGACCCGGTGGTCTTTCGGCATTATTAAGACAATCATATAATGGAACCGCACCCAATTGCTGTCGCACTTTTAATCGTACTCATTATTGGAAGAAGGATGGAAAGATATCAGATAAGTATGAAGAGGGTGCAGTATTAGAATCGTGTTGGCCCGACGTCCACGACACTGGAAAATGCGATGTTGATTTATTCGACTGGTGTCAGGGGGATACGTTCGATATGAACATATGCCATCAGTGGATCGGTTCAGCCTTTAATAGGAGTGATAGAACTGTAGAGGGTCGACAATCGTTAATAAATCTGTATAATAAGATGCAAAGATTATGTAGTAAAGATGCTAGTGTACCAATATGTGAATTATTTTTGCATCATTTACGCGCACACAATACAGAAGATAGTAAAGAGATGATCGATTATATTCTAAGACAACAGTCGGCGGACTTTAAACAGAAATATATGAGATGTAGTTATCCCACTAGAGATAAGTTAGAAGAGTCATTAAAATATGCGGAACCTCGAGAATGTTGGGATCCAGAGTGTTCGAATGCCAATGTTAATTTCTTACTAACACGTAATTATAATAATTTAGGACTTTGCAATATTGTACGATGTAATACGAGCGTGAATAACTTACAGATGGATAAAACTTCCTCATTAAGATTATCATGTGGATTAAGCAATAGTGATAGATTTTCTACTGTTCCCGTCAATAGAGCAAAAGTAGTTCAACATAATATTAAACATTCGTTCGACCTAAAATTGCATTTGATCAGTTTATTATCTCTCTTGGTAATATGGATACTAATTGTAGCTATTTAAATGGGTGCCGCAGCAAGCATACAGACGACTGTGAATACACTCAGTGAACGTATCTCGTCTAAATTAGAACAAGAAGCGAACGCTAGTGCTCAAACAAAATGTGATATAGAAATCGGAAATTTTTATATCCGACAAAACCATGGATGTAACATCACTGTTAAAAATATGTGCTCTGCGGACGCGGATGCTCAGTTGGATGCTGTGTTATCAGCCGCTACAGAAACATATAGTGGATTAACACCGGAACAAAAAGCATACGTACCAGCTATGTTTACTGCTGCGTTAAACATTCAGACGAGTGTAAACACTGTTGTTAGAGATTTTGAAAATTATGTGAAACAGACTTGTAATTCTAGCGCTGTTGTCGATAACAAATTAAAGATACAAAACGTAATTATAGATGAATGTTACGGAGCCCCAGGATCTCCAACAAATTTGGAATTTATTAATACAGGATCTAGCAAAGGAAATTGTGCCATTAAGGCGTTGATGCAATTGACTACTAAGGCCACTACTCAAATAGCACCTAGACAAGTTGCTGGTACAGGAGTTCAGTTTTATATGATTGTTATCGGTGTTATAATATTGGCAGCGTTGTTTATGTACTATGCCAAGCGTATGCTGTTCACATCCACCAATGATAAAATCAAACTTATTTTAGCCAATAAGGAAAACGTCCATTGGACTACTTACATGGACACATTCTTTAGAACTTCTCCGATGATTATTGCTACCACGGATATACAAAACTGAAAATATATTGATAATATTTTAATAGATTAACATGGAAGTTATCGCTGATCGTCTAGACGATATAGTGAAACAAAATATAGCGGATGAAAAATTTGTAGATTTTGTTATACACGGTCTAGAGCATCAATGTCCTGCTATACTTCGACCATTAATTAGGTTGTTTATTGATATACTATTATTTGTTATAGTAATTTATATTTTTACGGTACGTCTAGTAAGTAGAAATTATCAAATATTGTTGGTGTTGGTGGCGCTAGTCATCACATTAACTATTTTTTTATTACTTTATACTATAATAGTACTAGACTGACTTCTAACAAACATCTCACCTGCCATAAATAAATGCTTGATATTAAAGTCTTCTATTTCTAACACTATTCCATCTGTGGAAAATAATACTCTGACATTATCGCTAATTGATACATCGGTAAGTGATATGCCTATAAAGTAATAATCTTCTTTGGGCACATATACCAGTGTACCAGGTTCTAACAACCTATTTACTGGTGCTCCTGTAGCATACTTTTTTTTTACCTTGAGAATATCCATTGTTTGCTTGGTCAATAGTGATATGTGATTTTTTATCAACCACTCAAAAAAGTAATTGGAGTGTTCATATCCTCTACGGGCTATTGTCTCATGACCGTGTATGAAATTTAAGTAACACGACTGTGGTAGATTTGTTCTATAGAGCCGGTTGCCGCAAATAGATAGAACTACCAATATGTCTGTACAAATGTTAAACATTAATTGATTAACAGAAAAAACAATGTTCGTTCTGGGAATAGAAACCAGATTAAAACAAAATTCATTAGAATATATGCCACGTTTATACATGGAATATAAAATAACTACAGTTTGAAAAATAACAGTATCATTTAAACATTTAACTTGCGGGGTTAATCTCACAACTTTACTGTTTTTGAACTGTTCAAAATATAGCATAGATCCATGAGAAATACGTTTAGCCGCCTTTAATAGAGGAAATCCAACCGCCTTTCTGGATCTCACCAACGACGATAGTTCTGACCAGCAACTCATTTCTTCATCATCCACCTGTTTTAACATATAATAGGCAGGAGATAGATATCCATCATTGCAATATTCCTTCTCGTAGGCACACAATCTAATATTGATAAAATCTCCATTCTCTTCTCTGTATTTATTATCTTGTCTCGGTGGCTGATTAGGCTGTGGTCTATCGTTGTTGAATCTATTTTGGTCATTAAATCTTTCATTTCTTCCTGGTATATTTCTATCACCTCGTTTGGTTGGATTTTTGTCTATATTATCGTTTGTAACATCGGTACGGGTATTCATTTATCACAAAAAAAACTTCTCTAAATGAGTCTACTACTAGAAAACCTCATCGAAGAAGATACCATATTTTTTGCAGGAAGTATATCTGAGTATGATGATTTACAAATGGTTATTGCTGGTGCAAAATCCAAATTTCCAAGATCTATGCTTTCTATTTTTAATATAGTACCTAGAACGATGTCAAAATATGAGTTGGAGTTGATTCATAACGAGAATATCACAGGGGCAATGTTTACCACAATGTATAATATAAGAAACAATTTGGGTCTAGGCGATGATAAACTAACTATTGAAGCCATTGAAAACTATTTCTTGGATCCTAACAATGAGGTTATGCCTCTTATCATTAATAATACGGATATGACTACCGTCATTCCTAAAAAAAGTGGTAGGAGAAAGAATAAGAACATGGTTATCTTCCGTCAAGGATCATCACCTATCTTGTGTATTTTCGAAACTCGTAAAAAGATTAATATTTATAAAGAAAATATGGAATCCGTATCGACTGAGTATACACCTATCGGAGACAACAAGGCTTTGATATCTAAATATGCGGGAATTAATATCCTGAATGTGTATTCTCCTTCCACGTCCATGAGATTGAATGCCATTTACGGATTCACCAATAAAAATAAACTAGAGAAACTTAGTACTAATAAGGAACTAGAATCGTATAGTTCTAGCCCTCTTCAAGAACCCATTAGGTTAAATGATTTTCTGGGACTATTGGAATGTGTTAAAAAGAATATTCCTCTAACAGATATTCCGACAAAGGATTGATTACTATAAATGGAGAATGTTCCTAATGTATACTTTAATCCTGTGTTTATAGAGCCCACGTTTAAACATTCTTTATTAAGTGTTTATAAACACAGATTAATAGTTTTATTTGAAGTATTCGTTGTATTCATTCTAATATATGTATTTTTTAGATCTGAATTAAATATGTTCTTCATGCCTAAACGAAAAATACCCGATCCTATTGATAGATTACGACGTGCTAATCTAGCGTGTGAAGACGATAAATTAATGATCTATGGATTACCATGGATAACAACTCAAACATCTGCGTTATCAATAAATAGTAAACCGATAGTGTATAAAGATTGTGCAAAGCTTTTGCGATCAATAAATGGATCACAACCAGTATCTCTTAACGATGTTCTTCGCAGATGATGATTCATTTTTTAAGTATTTTGCTAGTCAAGATGATGAATCTTCATTATCTGATATATTGCAAATCACTCAATATCTAGACTTTCTGTTATTATTATTGATCCAATCAAAAAATAAATTAGAAGCTGTGGGTCATTGTTATGAATCTCTTTCAGAGGAATACAGACAATTGACAAAATTCACAGACTCTCAAGATTTTAAAAAACTGTTTAACAAGGTCCCTATTGTTACAGATGGAAGGGTCAAACTTAATAAAGGATATTTGTTCGACTTTGTGATTAGTTTGATGCGATTCAAAAAAGAATCAGCTCTAGCTACCACCGCAATAGATCCTGTTAGATACATAGATCCTCGTCGTGATATCGCATTTTCTAACGTGATGGATATATTAAAGTCGAATAAAGTTGAAAAATAATTAATTCTTTATTGTTATCATGAACGGCGGACATATTCAGTTGATAATCGGCCCCATGTTTTCAGGTAAAAGTACAGAATTAATTAGACGAGTTAGACGTTATCAAATAGCTCAATATAAATGTGTGACTATAAAATATTCTAACGATAATAGATACGGAACGGGACTATGGACACATGATAAGAATAATTTTGCAGCATTGGAAGTAACTAAACTATGTGATGTCTTGGAAGCAATTACAGATTTCTCCGTGATAGGTATAGATGAAGGACAGTTCTTTCCAGACATTGTTGAATTCTGTGAGCGTATGGCAAACGAAGGAAAAATAGTTATAGTAGCCGCGCTCGATGGGACATTTCAACGTAGACCGTTTAATAATATTTTGAATCTTATTCCATTATCTGAAATGGTGGTAAAACTAACTGCAGTGTGTATGAAATGCTTTAAGGAGGCTTCCTTTTCTAAACGATTAGGTACAGAAACCGAGATAGAAATAATAGGAGGTAATGATATGTATCAATCTGTGTGTAGAAAGTGTTACATCGACTCATAATATTATATTTTTTATCTAAAAAACTAAAAATAAACATTGATTAAATTTTAATATAATACTTAAAAATGGATGTTGTGTCGTTAGATAAACCGTTTATGTATTTTGAGGAAATTGATAATGAGTTAGATTACGAACCAGAAAGTGCAAATGAGGTCGCAAAAAAACTGCCGTATCAAGGACAGTTAAAACTATTACTAGGAGAATTATTTTTTCTTAGTAAGTTACAGCGACACGGTATATTAGATGGCGCCACCGTAGTGTATATAGGATCTGCTCCAGGTACACATATACGTTATTTGAGAGATCATTTCTATAATTTAGGAGTGATCATCAAATGGATGCTAATTGACGGCCGCCATCATGATCCTATTCTAAATGGATTGCGTGATGTGACTCTAGTGACTCGGTTTGTTGATGAGGAATATCTACGATCCATCAAAAAACAACTACATCCTTCTAAGATTATTTTAATTTCTGATGTGCGATCCAAACGAGGAGGAAATGAACCTAGTACTGCGGATTTACTAAGTAATTATGCTCTACAAAATGTCATGATTAGTATTTTAAACCCCGTGGCGTCTAGTCTTAAATGGAGATGCCCGTTTCCAGATCAATGGATCAAGGACTTTTATATCCCACACGGTAATAAAATGTTACAACCTTTTGCTCCTTCATATTCAGCTGAAATGAGATTATTAAGTATTTATACCGGTGAGAATATGAGACTGACTCGAGTTACCAAATCAGACGCTGTAAATTATGAAAAAAAGATGTATTACCTTAATAAGATAGTCCGCAACAAAGTAGTTATTAACTTTGATTATCCTAATCAGGAATATGACTATTTTCACATGTACTTTATGTTGAGGACCGTATACTGCAATAAAACATTTCCTACTACTAAAGCAAAGATACTATTTCTACAACAATCTATATTTCGTTTCTTAAATATTCCAACGACATCAACTGAAAAAGTTAGTCATGAACCAATACAACGTAAAATATCTAGCAAAGATTCTATGTCTAAAAACAGAAATAGCAAGAGATCCGTACGCGGTAATAAATAGAAACGTACTACTGAGATATACTACCGATATAGAGTATAATGATTTAGTTACTTTAATAACCGTTAGACATAAAATTGATTCTATGAAAACTGTGTTTCAGGTATTTAACGAATCATCCATAAATTATACTCCGGTTGATGATGATTATGGAGAACCAATCATTATAACATCGTATCTTCAAAAAGGTCATAACAAGTTTCCTGTAAATTTTCTATACATAGATGTGGTAATATCTGACTTATTTCCTAGCTTTGTTAGACTAGATACTACAGAAACTAATATAGTTAATAGTGTACTACAAACAGGCGATGGTAAAAAGACTCTTCGTCTTCCTAAAATGTTAGAGACGGAAATAGTTGTCAAGATTCTCTATCGTCCTAATATACCATTAAAAATTGTTAGATTTTTCCGCAATAACATGGTAACTGGAGTAGAGATAGCCGATAGATCTGTTATTTCAGTCGCTGATTAATCAATTAGTAGAGATGAGATAAGAACATTATAATAATCAATAATATATCTTATATCTGTTTAGAAAAATGCTAATATTAAAATAGCTAACGCTAGTAATCCAATCGGAAGCCATTTGATATCTATAATAGGGTATCTAATTTCCTGATTCAGATAGCGTACGGCTATATTCTCGGTAGCTACTCGTTTGGAATCACAGACATTATTTACATCTAATTTACTATCTGTAATGGAAACGTTTCCCAATGAAATGGTACAATCAGATACATTACATCTTGATATATTTTTTTTTAAAGAGGCTGGTAACAACGCATCGCTTCGTTTACATGGCTCGTACCAACAATAATAGGGTAATCTTGTATCTATTCCTATCCGTACTATACTTTTATCAGGATAAATACATTTACATCGTATATCGTCTTTGTTAGTATCACAGAATGCATAAATTTGTTCGTCCGTCATGATAAAAATTTAAAGTGTAAATATAACTATTATTTTTATAGTTATAATAAAAAGGGAAATTTGATTGTATACCTTCGGTTCTTTAAAAGAAACTGACTTGATAAAAATGGCTGTAATCTCTAAGGTTACGTATAGTCTATACGATCAAAAAGAGATTAATGCCACAGATATTATCATTAGTCATATTAAAAATGACGACGATATCGGTACCGTTAAAGATGGTAGACTAGGTGCTATGGATGGGGCATTATGTAAGACTTGTGGGAAAACGGAATTGGAATGTTTCGGTCACTGGGGTAAAGTAAGTATTTATAAAACTCATATAGTTAAGCCTGAATTTATTTCAGAAATTATTCGTTTACTGAATCATATATGTATTCATTGCGGATTATTGCGTTCACGAGAACCGTATTCCGACGATATTAACCTAAAAGAGTTATCGGTACACGCTCTTAGGAGATTAAAGGATAAAATATTATCCAAGAAAAAGTCATGTTGGAACAGCGAATGTATGCAACCGTATCAAAAAATTACTTTTTCAAAGAAAAAGGTTTGTTTCGTCAACAAGTTGGATGATATTAACGTTCCTAATTCTCTCATCTATCAAAAGTTAATTTCTATTCATGAAAAGTTTTGGCCATTATTAGAAATTCATCAATATCCAGCTAACTTATTTTATACAGACTACTTTCCCATCCCTCCGTTGATTATTAGACCGGCTATTAGTTTTTGGATAGATAGTATACCCAAAGAGACAAATGAATTAACTTACTTATTAGGTATGATCGTTAAGAATTGTAACTTGAATGCTGATGAACAGGTTATCCAGAAGGCGGTAATAGAATACGATGATATTAAAATTATTTCTAATAACACTACCAGTATCAATTTATCATATATCACATCCGGCAAAAATAATATGATTAGAAGTTATATCGTCGCTCGGCGAAAAGATCAGACCGCTAGATCCGTAATTGGTCCCAGTACATCTATCACCGTTAATGAGGTAGGAATGCCCACATATATTAGAAATACACTTACAGAAAAGATATTTGTTAATGCCTTTACAGTGGATAAAGTTAAACAACTATTAGCATCAAACCAAGTTAAATTTTACTTTAATAAACGATTAAACCAATTAACAAGAATACGTCAAGGAAAGTTTATCAAAAATAAAATACATTTATTGCCTGGTGATTGGGTAGAAGTAGCTGTTCAAGAATATACAAGTATTATTTTTGGAAGACAACCGTCTCTACATAGATACAACGTCATCGCTTCATCTATCAGAGCTACCGAAGGAGATACTATCAAAATATCTCCCGGAATTGCCAACTCTCAAAATGCTGATTTCGACGGAGATGAAGAATGGATGATATTGGAGCAAAATCCTAAAGCCGTAGTTGAACAAAGTATTCTTATGTATCCGACGACGTTACTCAAACACGATATTCATGGAGCCCCCGTTTATGGATCTATTCAAGATGAAATCGTAGCAGCGTATTCATTGTTTAGGATACAAGATCTTTGTTTAGATGAAGTATTGAACATCTTGGGGAAATATGGAAGAGAGTTCGATCCTAAAGGTAAATGTAAATTCAGCGGTAAAGATATCTATACTTACTTGATAGGTGAAAAGATTAATTATCCGGGTCTCTTAAAGGATGGTGAAATTATTGCAAACGACGTAGATAGTAATTTTGTTGTAGCTATGAGGCATCTGTCATTGGCTGGACTCTTATCCGATCATAAATCGAACGTGGAAGGTATCAACTTTATTATCAAGTCATCTTATGTTTTTAAGAGATATCTATCTATATACGGTTTTGGGGTGACATTCAAAGATCTGAGACCAAATTCGACGTTCACTAATAAATTGGAGGCTATCAACGTAGAAAAAATAGAACTTATCAAAGAAGCATACGCCAAATATCTCAAAGATGTAAGAGACGGGAAAATAGTTCCATTATCTAAAGCTTTAGAGGCGGACTACTTGGAATCCATGTTATCCAACTTGACAAATCTTAATATCAGAGAGATAGAAGAACATATGAGACAAACGCTGATAGATGATCCAGATAATAACCTCCTGAAAATGGCCAAAGCGGGTTATAAAGTAAATCCCACAGAACTAATGTATATTCTAGGTACTTATGGACAACAGAGGATAGATGGCGAACCAGCAGAGACTCGAGTATTGGGTAGAGTCTTACCTTACTATCTTCCAGACTCTAAGGATCCAGAAGGAAGAGGTTATATTCTTAATTCTTTAACAAAAGGATTAACGGGTTCTCAATATTACTTTTTGATGCTGGTTGCAAGATCTCAATCTACTGATATTGTCTGTGAAACATCACGTACCGGAACACTGGCTAGAAAAATCATTAAAAAGATGGAGGATATGGTGGTCGACGGATACGGACAAGTAGTTATAGGTAATACGCTCATCAAGTACGCAGCCAATTATACCAAAATTCTAGGCTCAGTATGTAAACCTGTAGATCTTATCTATCCAGATGAGTCCATGACTTGGTATTTGGAAATTAGTGCTTTGTGGAATAAAATAAAACAGGGATTCGTTTACTCTCAGAAACAGAAACTTGCAAAGAAGACATTGGCGCCGTTTAATTTCCTAGTATTCGTCAAACCCACCACTGAGGATAATGCTATTAAGGTTAAGGATCTGTACGATATGATTCATAACGTCATTGATGATGTGAGAGAGAAATACTTCTTTACGGTATCTAATATAGATTTTATGGAGTATATATTCTTGACGCATCTTAATCCTTCTAGAATTAGAATTACAAAAGAAACGGCTATTACTATCTTTGAAAAGTTCTATGAAAAACTCAATTATACTCTAGGTGGTGGAACTCCTATTGGAATTATTTCTGCACAGGTATTGTCTGAGAAGTTTACACAACAAGCCCTGTCCAGTTTTCACACTACTGAAAAGAGTGGTGCTGTAAAACAAAAACTTGGTTTCAACGAGTTTAATAACTTGACTAATTTGAGTAAGAATAAGACCGAAATTATCACTCTGGTATCCGATGATATCTCTAAACTTCAATCTGTTAAGATTAATTTCGAATTTGTATGTTTGGGAGAATTAAATCCAGACATCACTCTTCGAAAAGAAACAGATAGATATGTAGTAGACATAATAGTCAATAGATTATACATCAAGAGAGCAGAAATAACCGAATTAGTCGTCGAATATATGATTGAACGATTTATCTCCTTTAGCGTCATTGTAAAGGAATGGGGTATGGAGACATTCATTGAGGACGAGGATAATATTAGATTTACTATCTACCTAAATTTCGTTGAACCGGAGGAATTGAATCTTAGTAAGTTTATGATGGTTCTTCCAGGTGCCGCCAACAAGGGCAAGATTAGTAAATTCAAGATTCCTATCTCTGACTATACGGGATATAACGACTTCAATCAAACAAAAAAGCTCAATAAGATGACTGTAGAACTCATGAATCTAAAAGAATTGGGTTCTTTCGATTTGGAGAACGTCAACGTGTATCCTGGAGTATGGAATACATACGATATCTTCGGTATTGAGGCCGCTCGTGGATACTTGTGCGAAGCCATGTTAAACACCTATGGAGAAGGTTTCGATTATCTGTACCAGCCTTGTGATCTTCTCGCTAGTTTACTATGTGCTAGTTACGAACCAGAATCAGTTAATAAATTCAAGTTCGGTGCAGCTAGTACTCTTAAGAGAGCTACGTTCGGAGATAATAAAGCATTGTTAAACGCGGCTCTTCATAAAAAGTCAGAACCTATTAACGATAATAGTAGCTGCCACTTTTTTAGCAAGGTCCCTAATATAGGAACTGGATATTACAAATACTTTATCGACTTGGGTCTTCTCATGAGAATGGAAAGGAAACTATCTGATAAGATATCTTCTCAAAAGATCAAGGAGATAGAAGAAACAGAAGACTTTTAATTCTTATCAATAACATATTTTTCTATGATCTGTCTTTTAAACGATGGATTTTCCACAAATGCGCCTCTCAAGTCCCTCATAGAATGATACACGTATAAAAAATATAGCATAGGTGATGACTCCTTATTTTTAGACATTAGATATGCCAAAATCATAGCCCCGCTTCTATTTACTCCTGCAACACAATGAACCAACACGGGCTCGTTTCGTTGATCACATTTAGATAAGAAGGCGGTCACGTCGTCAAAATATTTACTAATATCAGTAGTTGTATCATCTACCAACGGTATATGAATAATATTAATATTAGAGTTAGGTAATGTATATTTATCCATCGTCAAATTTAAAACATATTTGAACTTAACTTCAGATGATGGTGCATCCATAGCATTTTTATAATTTCCCAAATACACATTATTTGTTACTCTTGTCATTATAGTGGGAGATTTGGCTCTGTGCATATCTCCAGTTGAACGTAGTAGTAAGTATTTATACAAACTTTTCTTATCCATTTATAACGTACAAATGGATAAAACTACTTTATCAGTAAACGCATGCAATTTAGAATACGTTAGAGAAAAGGCTATAGTAGGCGTACAAGCAGCCAAGACATCAACACTTATATTTTTTGTTATTATATTGGCAATTAGTGCGCTATTACTCTGGTTTCAGACGTCTGATAATCCAGTCTTTAATGAATTAACGAGATATATGCGAATTAAAAATACGGTTAACGATTGGAAATCATTAACGGATAGCAAAACAAAATTAGAAAGCGATAGAGGTAGACTTCTAGCCGCTGGTAAGGATGATATATTCGAATTCAAATGTGTGGATTTCGGCGCCTATTTTATAGCTATGCGATTGGATAAGAAAACATATCTGCCGCAAGCTATTAGGCGAGGTACTGGAGACGCGTGGATGGTTAAAAAGGCGGCAAAAGTCGATCCATCTGCTCAACAATTTTGTCAGTATTTGATAAAACACAAGTCTAATAATGTTATTACTTGTGGTAATGAGATGTTAAATGAATTAGGTTATAGCGGTTATTTTATGTCACCGCATTGGTGTTCCGATCTTAGTAATATGGAATAAGTGTTAGATAAATGCGGTAACAAATGTTCCTGTAAGGAACCATAACAGTTTAGATTTAACATTAAAGATGAGCATAAACATAATAAACAAAATTACAATCAAACCTATAACATTAATATCAAACAATCCAAAAAATGAAATCAATGGAGTAGTAAACGTGTACATAACTCCTGGATAACGTTTAGCAGCTACCGTTCCTATTCTAGACCAAAAATTTGGTTTCATGGTTTCGAAGCGGTGTTCTGCAACAAGACGAGGATCGTGTTCTACATATTTGGCAGAGTTATCCATTATTTGCCTGTTAATCTTCATTTCGTTTTCGATTCTGGCTATTTCAAAATAAAATCCCGATGATAGACCTCCAGACTTTATAATTTCATCTACGATGTTCAGCGCCGTAGTAACTCTAATAATATAGGCGGATAAGCTAACATCATACCCTCCTGTATATGTAAATATGGCATGATCTTTGTCTATTACAAGCTCGGTTTTAACTTTATTTCCTGTAATAATTTCTCTCATCTGTAGGATATCTATTTTCTTGTCATGTATTGCCTTCAAGACGGGACGAAGAAACGTAATATCCTCAATAACGTTATCGTTTTCTATAATAACTACATATTCTACATTTTTATTTTCTAGCTCGGTAAAAAATTTAGAATCCCATAGGGCTAAATGTCTAGCGATATTTCTTTTCGTTTCCTCTGTACACATAGTGTTACAAAACCCTGAAAAGAAGTGAGTATACTTGTCATCATCTCTAATATTTCCTCCAGTCCATTGTATAAACACATAATCCTTGTAATGATCTGGATCATCATTGACTATCACAACATCTCTTTTTTCTTGCATAACTTCATTGTCCTTCACATCATCGAACTTCTGATCATTAATATGCTCATGAACATTAGGAAATGTTTCTGATGGAGGTCTATCAATAACTGGCACAACAATAACAGGAGTTTTCACCGCCGCCATTTAGTTATTGAAATTAATCATATACAACTCTCTAATACGAGTTATATTTTCGTCTATCCATTGTTTCACATTGACATATTTCGACAAAAAGATATAAAATGCGTATTCCAATGCTTCTCTGTTTAATGAATTACTAAAATATACAAACACGTCACTGTCTGGTAATAAATAATATCTTAGAATATTGTAACAATTTATTTTGTATTGCACATGTTCGTGATCTATGAGTTCTTCTTCAAATGGCATAGGATCTCCGAATCTGAAAACGTATAAATAGGAGTTAGAATAATAATATTTGAGAGTATTGGTAATGTATAAACTCTTTAGCGGTATAATTAGTTTTTTTCTCTCGATTTCTATTTTTAGATGTGATGGAAAAATGACTAATTTTGTAGCATTAGTATCATGAACTCTAATCAAAATCTTAATATCTTCGTCACATGTTAGCTCTTTGAAGTTTTTAAGAGATGCATCAGTTGGTTTTACAGATGGAGTAGGTGCAACAATTTTTTGTTTAATGCATGCATGTATTGGAGCCATTGTCTTAACTATAATGGTGCTTGTATCGAAAAACTTTAATGCGGATAACGGAAGCTCTTCGCCGCGACTTTCTACGTCGTAATTGGGTTCTAATGCCGATCTCTGAATGGATACTAGTTTTCTAAGTTCTAATGTAATTCTCTGAAAATGTAAATCCAATTCCTCCGGCATTATAGATGTGTATACATCGGTAAATAAAACTATAGTATCCAACGATCCCTTCTCGCAAATTCTAGTCTTAACCAAGAAATCGTATATAACTACGGAGATGGCGTATTTAAGAGTGGATTCTTCTACCGTTTTGTTCTTGGATTTCATATAAGAAACTATAAAGTCCGCACTACTGTTAAGAATGATCACTAACGCAACTATATAGTTCAAATTAAGCATCTTGGAAACATAAAATAACTCTGTAGATGATACTTGACTTTCGAATAAGTTTGCAGACAAACGAAGAAAGAACAGACCTCTCTTAATTTCAGAAGAAAACTTTTTTTCGTATTCCTGACGTCTAGAGTTTATATCAATAAGAAAGTTAAGAATTAGTCGGTTAATGTTGTATTTCATTACCCAAGTTTGAGATTTCATAATATTGTCAAAAGACATGATAATATTAAAGATAAAGCGCTGACTATGAACGAAATAGCTATATGGTTCGCTCAAGAATATAGTCTTGTTAAACGTGGAAACGATAACTGTATTTTTAATCACGTCAGCGGCATCTAAATTAAATATAGGTATATTTATTCCACACACTCTACAATATGCCACACCATCTTCATAATAAATAAATTCGTTAGCAAAATTATTAATTTTAGTGAAATAGTTAGCGTCAACTTTCATAGCTTCCTTCAATCTAATTTGATGCTCACATGGCGCGAATTCTACTCTAACATCCCTTTTCCATGCCTCAGGTTCATCGATCTCTATAATATCTAGTTTCTTGCGTTTCACAAACACAGGCTCGTCTCTCGCGATGAGATCTGTATAGTAACTATGTAAATGATAACTAGATAGAAAGATGTAGCTATATAGATGACGATCCTTTAAGAGAGGTATAATAACTTTACCCCAATCAGATAGACTGTTGTTATGGTCTTCGGAAAAAGAATTTTTATAAATTTTTCCAGTATTTTCTAAATATACGTACTTGATATCTAAGAAATCCTTAATAATAATAGGAATGGATAATCCGTCTATTTTATAAAGAAATACATATCGCATATTATACTTTTTTTTGGAAATTGGAATACCGATGTGTCTACATAAATACGCAAAGTCTAAATATTTTTTAGAGAATCTTAGTTGGTCCAAATTCTTTTCCAAGTACGGTAATAGATTTTTCATATTGAACGGTATCTTCTTGATCTCTGGTTCTAATTCCGCATTAAATGATGAAACTAAGTCACTATTTTTATAACTAACGATTACATCACCTCTAACATCATCATTTACCAGGATACTGATCTTCTTTTGTCGTAAATACATGTCTAATGTGTTAAAAAAAAGATCATACAAGTTATACGTCATTTCATCTGTAGTATTCTTGTCATTGAAGGATAAACTCGTACTAATCTCTTCTTTAACAGTCTGTTCAAATTTATATCCTATATATGAAAAAATAGCAACCAGTGTTTGATCATCCGCGTCAATATTCTGTTCTATCGTAGTGTATAACAATCTTATATCTTCTTCTGTGATAGTCGATACGTTATAAAGGTTGATAACGAAAATATTTTTATTTCGTGAAATAAAGTCATTGTAGGATTTTGGACTTATATTCGTGTCTAGTAGATATGATTTTATTTTTGGAATGATCTCAATTAAAATAGTCTCTTTAGAGTCCATTTAAAGTTACAAACAACTAGGAAATTGGTTTATGATGTATAATTTTTTTAGTTTTTATAGATTCTTTATTCTATACTTAAAAAATGAAAATAAATACAAAGGTTCTTGAGGGTTGTGTTAATTGAAAGCGATAAATAATCATAAATTATTTCATTATCGCGATATCCGTTAAGTTTGTATCGTAATGGCGTGGTCAATTACGAATAAAGCGGATACTAGTAGTTTCACAAAGATGGCTGAAATCAGAGCTCATCTAAGAAATAGCGCTGAAAATAAAGATAAAAACGAGGATATTTTCCCGGAAGATGTAATAATTCCATCTACTAAGCCCAAAACCAAACGAACCACTACTCCTCGTAAACCAGCGGCTACTAAAAGATCAACCAAAAAGGATAAAGAAAAGGAGGAAGTGGAAGAAGTAGTTATAGAGGAATATCATCAAACAACTGAAGAAAATTCTCCACCTCCGTCATCATCTCCTGGAGTCGGCGACATTGTAGAAAGCGTGGCCGCTGTAGAGCTCGATGATAGCGACGGGGATGATGAACCTATGGTACAAGTTGAAGCTGGTAAAGTAAATCATAGTGCTAGAAGCGATCTCTCTGACCTAAAGGTGGCTACCGACAATATCGTTAAAGATCTTAAGAAAATTATTACTAGAATCTCTGCAGTATCGACTGTTCTAGAGGATGTTCAAGCAGCTGGTATCTCTAGACAATTTACTTCTATGACTAAAGCTATTACAACACTATCTGATCTAGTCACCGAGGGAAAATCTAAAGTTGTTCGTAAAAAAGTTAAAACTTGTAAGAAGTAAATGCGTGCACTTTTTTATAAAGATGGTAAACTGTTTACCGATAATAATTTTTTAAATCCTGTATCAGACGATAATCCAGCGTATGAGGTTTTGCAACATGTTAAAATTCCTACTCATTTAACAGATGTAGTAGTATATGAACAAACGTGGGAAGAGGCATTAACTAGATTAATTTTTGTGGGAAGTGATTCAAAAGGACGTAGACAATACTTTTACGGAAAAATGCATATACAGAATCGCAATGCTAAAAGAGATCGTATTTTTGTTAGAGTATATAACGTTATGAAACGAATTAATTGTTTTATAAACAAAAATATAAAGAAATCGTCCACAGATTCCAATTATCAGTTGGCGGTTTTTATGTTAATGGAAACTATGTTTTTTATTAGATTTGGTAAAATGAAATATCTTAAGGAGAATGAAACAGTAGGGTTATTAACACTAAAAAATAAACACATAGAAATAAGTCCCGATGAAATAGTTATCAAGTTTGTAGGAAAGGACAAAGTTTCACATGAATTTGTTGTTCATAAGTCTAATAGACTATATAAACCGCTATTGAAACTGACTGATGATTCTAGTCCCGAAGAATTTCTGTTCAACAAACTAAGTGAACGAAAGGTATATGAATGTATCAAACAGTTTGGTATTAGAATCAAGGATCTCCGAACGTATGGAGTCAATTATACGTTTTTATATAATTTTTGGACAAATGTAAAGTCCGTATCTCCTCTTCCATCACCAAAAAAGTTGATAGCATTAACTATCAAACAAACTGCTGAAGTGGTAGGTCATACTCCATCAATTTCAAAAAGAGCTTATATGGCAACGACTATTTTAGAAATGGTAAAGGATAAAAATTTTTTAGACGTAGTATCTAAAACTACGTTCGATGAATTCCTATCTATAGTCGTAGATCACGTTAAATCATCTACGGATGGATGATAATAGATCTTTACACAAATAATTACAAGACCGATAAATGGAAATGGATAAACGGATGAAATCTCTCGCTATGACAGCTTTCTTCGGAGAGCTAAACACGTTAGATATTATGGCATTGATAATGTCTATATTTAAACACCATCCAAACAATACCATTTTTTCAGTGGATAAGGATGGTCAATTTATGATTGATTTCGAATACGATAATTATAAGGCTTCTCAATATTTGGATCTGACCCTCACTCCGATATCTGGAAATGAATGCAAGACTCACGCATCTAGTATAGCCGAACAATTGGCGTGTGTGGATATTATTAAAGAGGATATTAGCGAATATATCAAAACTACTCCCCGTCTTAAACGATTTATAAAAAAATACCGCAATAGATCATATACTCGTATCAGTCGAGATACAGAAAAGCTTAAAATAGCTCTAGCTAAAGGCATAGATTACGAATATATAAAAGACGCTTGTTAATAAGTAAATGAAAAAAAACTAGTCGTTTATAATAAAACACAATATGGATGCCAACATAGTATCATCTTCTACTATTGCGACGTATATAGACGCTTTAGCGAAGAATGCTTCAGAATTAGAACAGAGGTCTACCGCATACGAAATAAATAATGAATTGGAACTAGTATTTATTAAACCGCCATTGATTACGTTGACAAATGTAGTAAATATCTCCACGATTCAGGAATCGTTTATTCGATTTACCGTTACTAATAAGGAAGGTATCAAAATTAGAACTAAGATTCCATTATCTAAGGTACATGGTCTAGATGTAAAAAATGTGCAGTTGGTAGATGCTATAGATAACATAGTTTGGGAAAAGAAATCATTAGTGACGGAAAATCGTCTTCACAAAGAATGCTTGTTGAGACTATCAACAGAGGAACGTCATATATTTTTGGATTACAAGAAATATGGATCCTCTATCCGACTAGAATTAGTCAATCTTATTCAAGCAAAAACAAAAAACTTTACGATAGACTTTAAGCTAAAATATTTTCTAGGATCTGGCGCTCAATCTAAAAGTTCTTTATTGCACGCTATTAATCATCCAAAGTCAAGGCCTAATACATCTCTGGAAATAGAATTTACACCTAGAGACAATGAAACAGTTCCATATGATGAACTAATAAAGGAATTGACGACTCTCTCGCGTCATATATTTATGGCTTCTCCAGAGAATGTAATTCTTTCTCCACCTATTAACGCACCTATAAAGACTTTTATGTTGCCTAAACAAGATATAGTAGGTCTGGATCTGGAAAATCTATATGCCGTAACTAAGACTGACGGCATTCCTATAACTATCAGAGTTACATCAAAAGGGTTGTATTGTTATTTTACACATCTTGGTTATATTATTAGATATCCAGTTAAGAGAACAATAGATTCCGAAGTAGTAGTCTTTGGTGAGGCAGTTAAGGATAAGAACTGGACCGTATATCTCATTAAGCTAATAGAGCCCGTAAATGCAATCAGTGATAGACTAGAAGAAAGTAAGTATGTTGAATCTAAACTAGTGGATATTTGTGATCGGATAGTATTCAAGTCAAAGAAATACGAAGGTCCGTTTACTACAACTAGTGAAGTCGTCGATATGTTATCTACATATTTACCAAAGCAACCAGAAGGTGTTATTCTGTTCTATTCAAAGGGACCTAAATCTAACATTGATTTTAAAATCAAAAAGGAGAATACTATAGACCAAACTGCAAATGTAGTATTTAGGTACATGTCCAGTGAACCAATTATCTTTGGAGAGTCGTCTATCTTTATAGAGTATAAGAAATTTACCAACGATAAAGGCTTTCCTAAAGAATATGGTTCTGGTAAGATTGTGTTATATAACGGCGTTAATTATCTAAATAATATCTATTGTTTGGAATATATTAATACACATAATGAAGTGGGTATTAAGTCCGTTGTTGTACCTATTAAGTTTATAGCAGAATTCTTAGTCAATGGAGAAATACTTAAACCTAGAATCGATAAAACCATGAAATATATTAACTCAGAAGACTATTATGGAAATCAACATAATATCATAGTCGAACATTTAAGAGATCAAAGCATCAAAATAGGAGATGTCTTTAACGAGGATAAACTATCGGATGTTGGACATCAATACGCTGCCAACAACGATAAATTTAGATTAAATCCAGAAGTTAGTTATTTTACTAATAAACGAACTAGAGGGCCGTTGGGAATTTTATCAAACTACGTCAAGACTCTTCTTATTTCTATGTATTGTTCCAAAACATTTTTAGACGATTCCAACAAACGAAAGGTATTAGCGATTGATTTTGGAAACGGTGCTGACCTGGAAAAATACTTTTATGGAGAGATTGCGTTATTGGTAGCGACGGATCCGGATGCTGATGCTATAGCTAGAGGAAATGAAAGATACAACAAATTAAATTCTGGAATTAAAACCAAGTACTACAAATTTGACTACATTCAGGAAACTATTCGATCCGATACATTTGTCTCTAGTGTCAGAGAAGTATTCTATTTTGGAAAGTTTAATATCATTGACTGGCAGTTCGCTATTCATTATTCTTTTCATCCAAGACATTATGCTACAGTCATGAATAACTTATCCGAACTAACTGCTTCTGGAGGCAAGGTATTAATTACTACCATGGATGGAGACAAATTATCAAAATTAACCGATAAAAAGACTTTTATAATTCATAAGAATCTACCTAGTAGCGAAAACTATATGTCTGTAGAAAAAATAGCTGATGATAGAATAGTGGTATATAATCCATCAACAATGTCTACTCCAATGACTGAATACATTATCAAAAAGAACGATATAGTCAGAGTGTTTAACGAATACGGATTTGTTCTTGTAGATAATGTTGATTTCGCTACAATTATAGAACGAAGTAAAAAGTTTATTAATGGCGCATCTACAATGGAAGATAGACCGTCTACAAGAAACTTTTTCGAACTAAATAGAGGAGCCATTAAATGTGAAGGTTTAGATGTCGAAGACTTACTTAGTTACTATGTTGTTTATGTCTTTTCTAAGCGGTAAATAATAATATGGTATGGGTTCTGATATCCCCGTTCTAAATGCATTAAATAATTCCAATAGAGCGATTTTTGTTCCTATAGGACCTTCCAACTGTGGATACTCTGTATTATTAATAGATATATTAATACTTTTGTAGGGTAACAGAGGTTCTACGTCTTCTAAAAATAAAAGTTTTATAACATCTGGCCTGTTCATAAATAAAAACTTGGCGATTCTATATATACTCTTATTATCAAATCTAGCCATTGTCTTATAGATGTGAGCTACTGTAGGTGTACCATTTGATTTTCTTTCTAATACTATATATTTCTCTCGAAGAAGTTCTTGCAGATCATCTGGGAATAAAATACTACTGTTGAGTAAATCAGTTATTTTTTTTATATCGATATTGATGGACATTTTTATAGTTAAGGATAATAAGTATCCCAAAGTAGATAACGACGATAACGAAGTATTTATACTTTTAGGAAATCACAATGACTTTATCAGATCAAAATTAACAAAATTAAAGGAGCATGTATTTTTTTCTGAATATATTGTGACTCCAGATACATATGGATCTTTATGCGTCGAATTAAATGGGTCTAGTTTTCAGCACGGTGGTAGATATATAGAGGTGGAGGAATTTATAGATGCTGGAAGACAAGTTAGATGGTGTTCTACATCCAATCATATATCTGAAGATATACACACTGATAAATTTGTCATTTATGATATTTATACGTTTGATTCGTTCAAGAATAAACGATTGGTATTTGTACAGGTGCCTCCATCATTAGGAGATGATAGCTATTTAACTAATCCGTTATTGTCTCCGTATTATCGTAATTCAGTAGCCAGACAAATGGTCAATGATATGATTTTTAATCAAGATTCATTTTTAAAATATTTATTAGAACATCTGATTAGAAGCCACTATAGAGTTTCTAAACATATAACAATAGTTAGATACAAGGATACCGAAGAATTAAATCTAACAAGAATATGTTATAATAGAGATAAGTTTAAGGCGTTTGTATTCGCTTGGTTTAACGGCGTTTCGGAAAATGAAAAGGTACTAGATACGTATAAAAAGGTATCTGATTTGATATAATGAATTCAGTGACTATATCACACGCACCATATACTATTACTTATCACGATGATTGGGAACCAGTAATGAGTCAATTGGTAGAGTTTTATAACGAAGTAGCCAGTTGGTTGCTACGCGACGAGACGTCGCCTATTCCTGATAAGTTCTTTATACAATTGAAACAGCCGCTTAGAAATAAACGAGTATGTGTGTGTGGTATAGATCCGTATCCAAAAGATGGAACTGGTGTACCGTTCGAATCACCAAATTTTACAAAAAAATCAATTAAGGAGATAGCTTCATCTATATCTAGATTAACCGGAGTAATTGATTATAAAGGTTATAACCTTAATATAATAGACGGGGTTATACCCTGGAATTATTACTTAAGTTGTAAATTAGGAGAAACAAAAAGTCACGCGATTTACTGGGATAAGATTTCCAAGTTACTGCTACAGCATATAACTAAACACGTTAGTGTTCTTTATTGTTTGGGTAAAACAGATTTCTCGAATATACGGGCAAAGTTAGAATCCCCGGTAACTACCATAGTGGGATATCATCCAGCGGCCAGAGACCACCAATTCGAGAAAGATCGATCATTTGAAATTATCAACGTTTTACTGGAATTAGACAACAAGACACCTATAAATTGGGCTCAAGGGTTTATTTATTAATGCTTTAGTGAAATTTTAACTTGTGTTCTAAATGGATGCGGCTATTAGAGGTAATGATGTTATCTTTGTTCTTAAGACTATAGGTGTCCCGTCAGCATGCAGACAAAATGAAGATCCAAGATTCGTAGAAGCATTTAAATGCGACGAGTTAGAAAGATATATTGATAATAATCCAGAATGTACACTATTCGAAAGTCTTAGGGATGAGGAAGCATACTCTATAGTCAGAATTTTCATGGATGTAGATTTAGACGCGTGTCTAGACGAAATAGATTATTTAACGGCTATTCAAGATTTTATTATCGAGGTGTCAAACTGTGTAGCTAGATTCGCATTTACAGAATGCGGTGCCATTCATGAAAATGTAATAAAATCCATGAGATCTAATTTTTCATTGACTAAGTCTACAAATAGAGATAAAACAAGTTTTCATATTATCTTTTTAGACACGTATACCACTATGGATACATTGATAGCTATGAAACGAACACTATTAGAATTAAGTAGATCATCTGAAAATCCACTAACAAGATCGATAGACACTGCCGTATATAGGAGAAAAACAACTCTTCGGGTTGTAGGTACTAGGAAAAATCCAAATTGCGACACTATTCATGTAATGCAACCACCTCACGATAATATAGAAGATTACCTATTCACTTACGTGGATATGAACAACAATAGTTATTACTTTTCTCTACAACGACGATTGGAGGATTTAGTTCCTGATAAGTTATGGGAACCAGGGTTTATTTCGTTCGAAGACGCTATAAAAAGAGTTTCAAAAATATTCATTAATTCTATAATAAACTTTAATGATCTCGATGAAAATAATTTTACAACGGTACCACTGGTCATAGATTATGTAACACCTTGTGCATTATGTAAAAAACGATCGCATAAACATCCGCATCAACTATCGTTGGAAAATGGTGCTATTAGAATTTACAAAACTGGTAATCCACATAGTTGTAAAGTTAAAATTGTTCCGTTGGATGGTAATAAACTGTTTAATATTGCACAAAGAATTTTAGACACTAACTCTGTTTTATTAACCGAACGAGGAGACCATATAGTTTGGATTAATAATTCATGGAAATTTAACAGCGAAGAACCCTTGATAACAAAACTAATTCTATCAATAAGACATCAACTACCTAAGGAATATTCAAGCGAATTACTCTGTCCGAGGAAACGAAAGACTGTAGAAGCTAACATACGAGACATGTTAGTAGATTCAGTAGAGACCGATACCTATCCGGATAAACTTCCGTTTAAAAATGGTGTATTGGACCTGGTAGACGGAATGTTTTACTCTGGAGATGATGCTAAAAAATATACGTGTACTGTATCGACCGGATTTAAATTTGACGATACAAAATTCGTCGAAGACAGTCCAGAAATGGAAGAGTTAATGAATATCATTAACGATATCCAACCATTAACGGATGAAAATAAGAAAAATAGAGAGCTGTATGAAAAAACATTATCTAGTTGTTTATGTGGTGCTACCAAAGGATGTTTAACATTCTTTTTTGGAGAAACCGCAACTGGGAAGTCGACAACCAAACGTTTGTTAAAGTCTGCTATCGGTGACCTGTTTGTCGAGACGGGTCAAACAATTTTAACAGATGTATTGGATAAAGGACCTAATCCATTTATCGCTAATATGCATTTAAAAAGATCTGTATTCTGTAGCGAACTACCTGATTTTGCATGTAGTGGATCAAAGAAAATTAGATCTGATAATATTAAAAAGTTGACAGAACCTTGTGTCATTGGAAGACCGTGTTTCTCCAATAAAATTAATAATAGAAACCATGCGACAATCATTATCGATACTAATTACAAACCTGTCTTTGATAGGATAGATAACGCATTAATGAGAAGAATTGCCGTCGTGCGATTCAGAACACACTTTTCTCAACCTTCTGGTAGAGAGGCTGCTGAAAATAATGACGCGTACGATAAAGTCAAACTATTAGACGAGGGATTAGATGGTAAAATACAGAATAATAGATATAGATTCGCATTTCTATACTTGTTGGTTAAATGGTACAAAAAATATCATATTCCTATTATGAAACTATATCCTACACCGGAAGAGATTCCGGACTTTGCATTCTATCTCAAAATAGGTACTCTGTTGGTATCTAGCTCTGTAAAGCATATTCCATTAATGACGGACCTCTCCAAAAAGGGATATATATTGTACGATAATGTGGTTACTCTTCCGTTGACTACTTTCCAACAGAAAATATCCAAGTATTTTAATTCTAGACTATTTGGACACGATATAGAGAGCTTCATCAATAGACATAAGAAATTTGCCAATGTTAGTGATGAATATCTGCAATATATATTCATAGAGGATATTTCATCTCCGTAAATATATGCCATATATTTATAGAATATATCACATATCTAAATGAATACCGGAATCATAGATTTATTTGATAATCATGTTGATAGTATACCAACTATATTACCTCATCAGTTAGCTACTTTAGATTATCTAGTTAGAACTATCATAGATGAGAACAGAAGCGTGTTATTGTTCCATATTATGGGATCGGGTAAAACAATAATCGCTTTGTTGTTCGCCTTGGTAGCTTCCAGATTTAAAAAGGTTTACATTTTAGTACCGAACATCAACATCTTAAAAATTTTCAATTATAATATGGGTGTAGCTATGAACTTGTTTAATGACGAATTCATAGCTGAGAATATCTTTATTCATTCCACAACAAGTTTTTATTCTCTTAATTATAACGATAACGTCATTAATTATAACGGATTAAGTCGCTACAATAACTCTATTTTTATCGTTGATGAGGCACATAATATTTTTGGGAATAATACTGGAGAACTTATGACCGTGATAAAAAATAAAAACAAGATTCCTTTTCTACTATTGTCTGGATCTCCCATTACTAACACACCTAATACGCTGGGTCATATTATAGATTTAATGTCCGAAGAGACGATAGATTTTGGTGAGATTATTAGTCGTGGTAAGAAAGTAATTCAGACACTTCTTAACGAACGCGGAGTGAATGTACTCAAGGATTTGCTTAAAGGAAGAATATCATATTACGAAATGCCGGACAAAGATCTACCAACAATAAGATATCACGGACGTAAATTTCTAGATACTCGAGTAGTATATTGTCACATGTCTAAACTTCAAGAGAAAGATTATATGATTACTAGACGGCAGCTATGTTATCATGAAATGTTTGATAAAAATATGTATAACGTGTCAATGGCAGTATTGGGACAACTTAATCTGATGAATAATTTAGATACGTTATTTCAGGAACAGGATAAGGAATTGTACCCAAATCTGAAAATAAATAATGGAGTGTTATACGGTGAAGAATTGGTAACGTTAAACATTAGTTCCAAATTTAAGTACTTTATCAATCGGATACAGACACTCAAGGGAAAACACTTTATATACTTCTCTAATTCTACATATGGTGGATTGGTAATTAAATATATCATGCTCAGTAATGGATATTCTGAATATAATGGTTCTCAGGGAACTAATCCACATATGATAAACGGCAAACCAAAAACATTTGCTATCGTTACTAGTAAAATGAAATCGTCTTTAGAGGATCTATTAGATGTGTATAATTCTCCTGAAAACGATGATGGCAATCAATTGATGTTTTTGTTTTCGTCAAACATTATGTCTGAATCCTATACTCTGAAAGAGGTAAGGCATATTTGGTTTATGACTATCCCGGATACTTTTTCTCAATACAACCAAATTCTTGGACGATCTATTAGAAAATTCTCTTACGTCGATATTTCTGAACCCGTTAATGTATATCTTTTAGCAGCCGTATATTCAGATTTCAATGACGAAGTGACGTCATTAAACGATTATACACAGGATGAATTGATTAATGTTTTACCCTTTGACATCAAAAAGCTGTTGTATCTAAAATTTAAGACTAAAGAAACGAATAGAATATACTCTATTCTTCAAGAGATGTCTGAAACGTATTCTCTTCCACCACATCCATCAATTGTAAAAGTTTTATTGGGAGAATTGGTCAGACAATTTTTTTATAATAATTCTCGTATTAAGTATAACGACTCCAAGTTACTTAAAATGGTTACATCAGTTATAAAAAATAAAGAAGACGCTAGGAATTACATAGATGATATTGTAAACGGTCACTTCTTTGTATCGAATAAAGTATTTGATAAATCTCTTTTATACAAATACGAAAACGATATTATTACAGTACCGTTTAGACTTTCCTACGAACCATTTGTTTGGGGAGTTAACTTTCGTAAAGAATATAATGTGGTATCTTCTCCATAAAACTGATGAGATATATAAAGAAATAAATGTCGAGCTTTGTTACCAATGGATATCTTCCAGTTACATTGGAACCACATGAGTTGACGTTAGACATAAAAACTAATATTAGGAATGCCGTATATAAGGCGTATCTCCATAGAGAAATTAGTGGTAAAATGGCCAAGAAAATAGAAATTCGTGAAGACGTGGAATTACCTCTCGGTGAAATAGTTAATAATTCTGTAGTTATAAACGTTCCGTGTGTAATAACCTACGCATATTATCACGTTGGGGATATAGTCAGAGGAACATTAAACATCGAAGATGAATCAAATGTAACTATTCAATGTGGAGATTTAATCTGTAAACTAAGTAGAGATTCGGGTACTGTATCATTTAGCGATTCAAAGTACTGCTTTTTTCGAAATGGTAATGCGTATGATAACGGCATCGAAGTCTCCGCCGTTCTAATGGAGGCTCAACAAGGTACCGAATCTAGTTTTGTTTTTCTCGCGAATATCGTTGACTCATAAGAAAGAGAATAGCGGTGAGTATAAATACGAATACTATGGCAATAATTGCGAATGTTTTATTCCCTTCGATATATTTTTGATAATATGAAAAACATGCCTCTCTCAAATCAGACAACCATTTCATAAAATAGTTCTCTCGCACTGGTGAGGTGGTTGCAGCTCGTATAATCTCCCCAGAATAATATACTTGCGTGTCGTCGTTCAATTTATACGGATTTCTATAATTCTCTGTTATATAATGAGGTTTACCCTCATGATTAGACGACGACAATAGTGTTCTGAATTTAGATAGTTGATCAGAATGAATGTTTATTGGTGTTGGAAAAATTATCCATGCTGCGTCTGCAGAGTGGTTGATAGTTGTTCCTAGATATGTAAAATAATCCAACGTACTAGGTAGCAAATTGTCTAGATAAAATACTGAATCAAATGGCGCAGACATATTAGCGGATCTAATGGAATCCAATTGATTGACTATCTTTTGAAAATATACATTTTTATGATCTGATACTTGTAAGAATATAGCAATAATGATAATTCCATCATCGTGTTTTTTTGCCTCTTCATAAGAACTATATTTTTTCTTATTCCAATGAACCAGATTAATCTCTCCAGAGTATTTGTATACATCTATCAAGTGATTGGATCCATAATCGTCTTCCTTTCCCCAATATATATGTATTGTTGATAACACATATTCATTGGGGAGAAACCCTCCACTTATATATCCTCCTTTAAAATTAATCCTTACTAGTTTTCCAGTATTCTGGATAGTGGTTGGTTTCGACTCATTATAATGTATGTCTAACGTCTTCAATCGCGCGTCAGAAATTGCTTTTTTAGTTTCTATATTAATAGGAGATAGTTGTTGAGGCATAGTAAAAATGAAATGATAACTGTCTAGAAATAGCTCTTAGTATGGGATTTACAATGGATGAGGAAGTGATATTTGAAACTCCTAGAGAATTAATATCTATTAAACGAATAAAAGATATTCCAAGATCAAAAGACACGCACGTGTTTGCTGCGTGTATAACAAGTGACGGATATCCGTTAATAGGAGCTAGAAGAACTTCATTCGCATTCCAGGCGATATTATCTCAACAAAATTCAGATTCTATCTTTAGAGTATCCACTAAACTATTACGGTTTATGTACTACAATGAACTAAGAGAAATCTTTAGACGGTTGAGAAAAGGTTCTATCAACAATATCGATCCTCACTTCGAAGAGTTAATATTATTGGGTGGTAAACTAGATAAAAAGGAATCTATTAAAGATTGTTTAAGAAGAGAATTAAAAGAGGAAAGTGATGAACATATAACAGTAAAAGAATTCGGAAATGTAATTCTAAAACTTACAACGAGTGATAAATTATTTAATAAAGTATATATAGGTTATTGCATGGCATGTTTTATTAATCAATCGTTGGAGGATTTATCACATACTAGTATTTACAATGTAGAAATTAGAAAGATTAAATCGTTAAATGATTGTATTAACGACGATAAATACGAATATCTGTCTTATATTTATAATATACTAATTAATAGTAAATGAGCTTTTACAGATCTAGTATAATTAGTCAGATTATTAAGTATAATAGACGACTAGCTAAGTCTATTATTTGCGAGGATGACTCTCAAATTATTACACTCACGGCATTCGTTAACCAATGCCTATGGTGTCATAAACGAGTATCCGTGTCCGCTATTTTATTAACTACTGATAACAAAATATTAGTATGTAACAGACGAGATAGTTTTCTCTATTCTGAAATAATTAGAACTAGAAACATGTATAGAAAGAAACGATTATTTCTGAATTATTCCAATTATTTGAACAAACAGGAAAGAAGTATACTATCGTCATTTTTTTCTCTAGATCCAGCTACTGCTGATAATGATAGAATAAACGCTATTTATCCGGGTGGTATACCCAAAAGGGGTGAGAACGTTCCAGAGTGTTTATCCAGGGAAATCAAAGAAGAAGTTAATATAGACAATTCTTTTGTATTCATAGACACTCGTTTTTTTATTCATGGTATCATAGAAGATACCATTATTAACAAATTTTTTGAGGTAATTTTCTTTGTTGGACGAATATCTCTAACGAGTGATCAAATTATTGATACCTTTAAAAGTAATCATGAAATAAAGGATCTAATATTTTTAGATCCAAATTCAGGTAATGGACTCCAATACGAAATTGCAAAATATGCTCTAGATACTGCAAAACTTAAATGTTACGGTCATAGAGGATGTTATTATGAATCATTAAAAAAATTAACTGAGGATGATTGATTAGAAAATATAAATTAATTTACCATCGTGTATTTTTATAACGGGATTGTCTGGCATATCATGTAGATAGTTACCGTCTACATCGTATACTCTACCATCTACGCCTTTAAATCCTCTATTTATTGATATTAATCTATTAGAATTGGAATACCAAATATTAGTACCCTCAATTAGTTTATTGGTAATATTTTTTTTAGACGATAGATCGATGGCTCTTGAAACCAAGGTTTTCCAACCGGACTCATTGTCTATCGGTGAGAAGTCTTTTTCATTAGCATGAATCCATTCTAATGATGTATGTTTAAACACTCTAAACAATTGTACAAATTCTTTTGATTTGTTTTGAATGATTTCAAATAGGTCTTCGTCTACAGTAGGCATACCATTAGATAATCTAGCCATTATAAAGTGCACGTTTACATATCTACGTTCTGGAGGAGTAAGAACGTGACTATTGAGACGAATGGCTCTTCCTACTATCTGACGAAGAGACGCCTCGTTCCATGTCATATCTAAAATGAAGATATCATTGATTGAGAAGAAACTAATACCCTCGCCTCCGCTAGAAGAGAATACGCATGTTTTAATGCATTCTCCGTTAGTGTTTGATTCTTGGTTAAACTCAGCCACCGCCTTGATTCTAGTATCTTTTGTTCTAGATGAGAACTCTATATTAGAGATACCAAAGACTTTGAAATATAGTAATAAGATTTCTATTCCTGACTGATTAACAAATGGTTCAAAGACTAGACATTTACCATGGGATGCTAATATTCCCAAACATACATCTATAAATTTGACGCTTTTCTCTTTTAATTCAGTAAATAGAGAGATATCAGCCGCAATAGCATCCCCTCCCAATAGTTCTCCCTTTTTAAAGGTGTCTAATGCGGATTTAGAAAATTCTCTATCTCTTAATGAATTTTTAAAATCATTATATAGGGTTGCTATCTCTTGTGCGTATTCTCCCGGATCACGATTTTGTCTTTCAGGAAAGCTATCGAATGTAAACGTAGTAGCCATACGTCTCAGAATTCTAAATGATGATATACCAGTTTTTATTTCTGCGAGTTTAGCCTTTTGATAAATCTCTTCTTGCTTTTTTGACATATTAACGTATCGCATTAATACTGTTTTCTTAGCGAATGATGCAGACCCTTCCACATCATCAAAAATAGAAAACTCGTTATTAACTATGTACGAACATAGGCCTCCTAGTTTGGAGACTAATTCTTTTTCATCGACTAGACGTTTATTCTCAAATAGCGATTGGTGTTGTAAGGATCCTGGTCGCAGTAAGTTAACCAACATGGTGAATTCTTGCACACTATTAACGATAGGTGTAGCCGATAAACAAATCATCTTATGGTTTTTTAACGCAATGGTCTTAGATAAAAAATTATATACTGACCGAGTAGGACGGATCTTACCATCTTCTTTGATTAATGATTTAGAAATGAAGTTATGACATTCATCAATGATGACGCATATTCTACTCTTGGAATTAATAGTTTTGATATTAGTAAAAAATTTATTTCTAAAATTTTGATCATCGTAATTAATAAAAATACAATCCTTCGTTATCTCTGGAGCGTATCTGAGTATAGTGTTTATCCAAGGATCTTCTATCAAAGCCTTTTTTACCAATAAGATAATTGCCCAATTCGTATAAATATCCTTAAGATGTTTGAGAATATATACAGTAGTCATTGTTTTACCGACACCTGTTTCATGGAACAATAAAAGAGAATGCATACTGTCTAATCCTAAGAAAACTCTTGCTACAAAATGTTGATAATCCTTGAGGCGTACTACGTCTGACCCCATCATTTCAACGGGCATATTAGTAGTTCTGCGTAAGGCATAATCGATATAGGCCGCGTGTGATTTACTCATTTATGAGTGATAAGTAATAACTATGTTTTAAAAATCACAGCAGTAGTTTAACTAGCCTTCTCTGATGTTTGTTTTCGATACTTTTTGAATCAGAAGTCATACTAGAATAAAGCAGCGAGTGAACGTAATAGAGAGCTTCGTATACTCTATTCGAAAACTCTAAGAACTTATTAATGAATTCCGTATCCACTGGATCGTTTAAAATACTAAATTGAACAGTGTTCACATCCTTCCAAGACGAAGACTTAGTGACGGACTTAACATGAGACATAAATAAATCCAAATTTTTTTTATAAACATCACTAGCCACCATAATGGCGCTATCTTTCAACCAACTATCGCTTACGCATTTTAACAGTCTAACATTTTTAAAGAGACTACAATATATTCTCATAGTATCGATTACACCTCTACCGAATAGAGTGGGAAGTTTAATAATACAATATTTTTCGTTTACAAAATCAAATAATGGTCGAAACACGTCGAAGGTTAACATCTTATAATCGCTAATGTATAGATTGTTTTCAGTGAGATGATTATTAGATTTAATAGCATCTCGTTCACGTTTGAACAGTTTATTGCGTGCGCTGAGGTCGGCAACTACGGCATCCGCTCTAGTACTCCTCCCATAATACTTTACGCTATTAATCTTTAAAATTTCATAGACTTTATCTAGATCGCTTTCTGGTAACATGATATCATGTGTAAAAAGTTTTAACATGTCGGTCGGCATTCTATTTAGATCATTAACTCTAGAAATCTGAAGAAAGTAATTAGCTCCATATTCCAGACTAGGTAATGGGCTTTTACCTAAAGACAAGTTAAGTTCTGGCAATGTTTCATAAAATGGAAGAAGGACATGTGTCCCCTCCCGGATATTTTTTACAATTTCATCCATTTACAACTCTATAGTTTGTTTTCATTATTATTAGTTATTATCTCCCATAATCTTGGTAATACTTACACCTTGATCATAAGATACCTTATACAGGTCATTACATACAACTACCAATTGTTTTTGTACATAATAGATTGGATGATTGATATCCATGGTGGAATAAACTACTCGAACAGATAGTTTATCTTTCCCCCTAGATACATTGGCCGTAATAGTTGTCGGCCTAAAGAATATCTTTGGTGTAAAGTTAAAAGTTAGGGTTCTTGTTCCATTATTGCTTTTTGTCAGTAGTTCGTTATAAATTCTCGAGATGGGCCCGTTCTCTGAATATAGAACATCATTTCCAAATCTAACTTCTAGTCTAGAAATAATATCGGTCTTATTTTTAAAATCTATTCCCTTGATGAATGGATCGTTAATAAACAAATCCTTGGCCTTTGATTCGGCTGATCTATTATCTCCGTTATAGACGTTACGTTGACTAGTCCAAAGACTTACAGGAATAGATGTATCGATGATGTTGATAGTATGTGATATGTGAGCAAAGACTGTTCTCTTGGTGGCGTCGCTATATGTTCCAGTAATGGCGGAAAACTTTTTAGAAATGTTATATATAAAAGAATTTTTTCGGGTTCCAAACATTAACAGATTAGTATGAAGATAAACACTCATATTATCAGGAACATTATCAATTTTTACATAAACATCGGCATCTTGAATAGAAACAACACCATCTTCTGGAACCTCTACGATCTCGGCAGATTCCGGATAACCAGTCGGTGGACCATCACTAACAATAACTAGATCATCCAACAATCTACTCACATATGCGTCTATATAATCTTTTTCATCTTGTGAGTACCCTGGATACGAAATAAATTTGTTATCAGTATTTCCATAATAAGGTTTAGTATAAACAGAGAGAGATGTTGCTGCATGAACTTCGGTTACTGTCGCCGTTGGTTGGTTTATTTGACCTATTACTCTCCTAGGTTTCTCTATAAATGATGGTTTAATTTGTACATTCTTAACCATATATCCAATAAAGCTCAATTCAGGAACATAAACAAATTCTTTGTTGAACGTTTCAAAGTCGAACGAAGAGTCACGAATAACGATATCGGATACTGGATTGAAGGTCACCGTTACGGTAATTTTTGAATCGGATAGTTTAAGACTACTGAATGTATCTTCCACATCAAACGGAGTTTTAATATAAACGTATACTGTAGATGGTTCTTTAATAGTGTCATTAGGAGTTAGGCCAATAGAAATATCATTAAGTTCACTAGAATATCCAGAATGTTTCAAAGCAATTGTATTATTGATACAATTATTATATAATTCTTCGCCCTCAATTTCCCAAATAACACCGTTACACGAAGAGACAGATACATGATTAATACATTTATATCCAACATATGGCACGTAACCGAATCTTCCCATACCTTTAACTTCTGGAAGTTCCAAACTCAGAACCAAATGATTAAGCGCAGTAATATACTGATCCCTAATTTCGAAGCTAGCGATAGCCTGATTGTCTGGCCCATCGTTTGTCATAACTCCGGATAGAGAAATATATTGCGGCATATATAAAGTTGGAATTTGACTATCAACTGCGAAGACATTAGACCGTTTAATAAAGTCATCCCCACCGATCAAAGAATTAATGATAGTATTATTCATTTTCTATTTAAAATGGAAAAAGCTTACAATAAACTCCGTAGAGAAATATCTATAATTTGTGAGTTTTCCTTAAAGTAACAGCTTCCGTAAACACCGTCTTTATCTCTTAGTAAGTTTATTGTATTTATGACCTTTTCCTTATCTTCATAGAATACTAAAGGCAATAAAGAAATTTTTGGTTCTTCTCTAAGAGCTACGTGAGACTTAACCATAGACGCCAACGAATCCCTACATATTTTAGAACAGAAATACCCAACTTCACCACCCTTGAATGTCTCAATACTAATAGGTCTAAAAACCAAATCTTGATTACAAAACCAACACTTATCAATTACACTATTTGTCTTAATAGACATATCTGCCATAGATTTATAATACTTTGGTAGTATACAAGCGAGTGCTTCTTCTTTAGCGGGCTTAAAGACTGCTTTAGGTGCTGAAATAACCACATCTGGAAGACTTACTCGCTTAGCCATTTAATTACGGAACTATTTTTTTATACTTCTAATGAACAAGTAGAAAACCTCTCATCTACAAAAACATACTCGTGTCCATAATCCTCTACCATAGTAACACGTTTTTTAGATCTCATATGTGCTAAAAAGTTTTCCCATACTAATTGGTTACTATTATTTTTCGTATAATTTTTAACAGTTTGAGGTTTTAGATTTTTAGTTACAGAAGTGATATCGAATATTTTATCCAAAAAGAATGAGTAATTAATTGTCTTAGAAGGAGTGTTTTCTTGGCAAAAGAATACCAAGTGCTTAAATATTTCTACTACTTCATTAATCTTTTCTGTACTCAGATTCAGTTTCTCATCTTTTACTTGATTGATTATTTCAAAGACTAACTTATAATCCTTTTTATTTATTCTCTCGTTAGCCTTAAGAAAACTAGATACAAAATTTGCATCTACATCATCCGTGGATATTTGATTTTTTTCCATGATATCCAATAGTTCCGAGATAATTTCTCCAGAACATTGATGAGACAATAATCTCCGCAATACATTTCTCAAATGAATAAGTTTATTAGACACGTGGAAGTTTGACTTTTTTTGTACCTTTGTACATTTTTGAAATACAGACTCGCAAAAAATACAATATTCATATCCTTGTTCAGATACTATACCGTTATGTCTACAACAGCTACATAATCGTAGATTCATGTTAACACTCTACGTATCTCGTCGTCCAATATTTTATATAAAAACATTTTATTTCTAGACGTTGTCAGAAAATCCTGTAATATTTTTAGTTTTTTTGGTTGTGAATAAAGTATCGCCCTAATAATATTGGTACCGTCTTCCGACAATATAGTAGTTAAATTATCCGAGCATGTAGAAGAACACCGCTTAGGCGGATTCAGTACAATGTTATATTTTTCGTACCAACTCATTTAAATATCATAATCTAAAATAGTTCTGTAATATGTCTAGCGCTAATATATTGATCATAATCCTGTGCATAAATTAAGATACAACAATGTCTTGAAATCATCGACATGGCTTCTTCCATAGTTAGAAGATCATCGTCAAAGTTAGCAACGTGATTCATCAACATTTGCTGTTTTGAGGCAGCAAATACTGAACCATCACCATTCAACCATTCATAAAAACCATCGTCTGAATCCATTGATAATTTCTTGTACTGGTTTTTGAGAGCTCGCATCAATCTAGCATTTCTAGCTCCCGGATTGAAAACAGAAAGAGGATCGTACATCCAAGGTCCATTTTCTGTAAATAGAATCGTATAATGTCCCTTCAAGAAGATATCAGACGATCCACAATCAAAGAATTGGTCTCCGAGTTTGTAACAGACTGCGGACTTTAACCTATACATGATACCGTTTAGCATGATTTCTGGTGATACGTCAATCGGAGTATCATCTATTAGAGATCTAAAGCCGGTGTAACATTCTCCGCCAAACATATTCTTATTCTGACGTCGTTCTACATAAAACATCATTGCTCCATTAACGATAACAGGTGAATGAACAGCACTACCCATCACATTAGTTCCCAATGGATCAATGTGTGTAACTCCAGAACATCTTCCATAGCCTATGTTAGGAGGAGCGAACACCACTCTTCCACTATTGCCATCGAATGCCATAGAATAAATATCCTTGGAATTGATAGAAATCGGACTGTCGGATGTTGTTATCATCTTCATAGGATTAACAACGATGTATGGTGCAGCCTGAAGTTTCATATCGTAACTGATGCCGTTCATAGGTCTAGCCACAGAAACCAACGTAGGTCTAAATCCAACTATAGACAAAATAGAAGCTAATATCTGTTCCTCATCTGTCATAACTTGAGAGCATCCAGTATGAATAATCTTCATTAGATGGGGATCTACCGCATCATCATCGTTACAATAAAAAATTCCCATTCTAATGTTCATAATTGCTTTTCTAATCATGGTATGAATGTTTGCTCTCTGAATCTCTGTGGAAATTAGATCTGATACACCTGTAATCACTATCGGATTATCCTCCGTAAGACGATTAACCAACAACATATAATTATAAGACTTTACTCTTCTAAATTCATAAAGTTGCTGGATTAGACTATATGTGTCTCCATGTACATACGCGTTCTCGAGCGCAGGAAGTTTAATACCGAATAGTGCCATCAGAATAGGATGAATGTAGTAATTAGTTTCTGGTTTTCTATAAATAAAAGACAAATCTTGTGAACTAGACATATCGGTAAAATGCATGGATTGGAATCGTGTAGTCGACAGAAGAATATGATGATTAGATGGAGAGTATATTTTATCTAACTCTTTGAGTTGGTCACCGATTCTAGGACTAGCTCGAGAATGAATAAGTACTAAGGGATGAGTACATTTCACAGAAACACTGGCGTTGTTCAACGTACTCTTTACATGGGAAAGGAGTTGAAATAGCTCGTTTCTATTTGTCCTGACAATATTTAGTTTATTCATAATATTAAGCATATCCTGAATAGTAAAGTTAGATGTGTCATACTTGTTAGTAGTTAGATATTTAGCAATTGCATTCCCATCATTTCTCAATCTCGTACTCCAATCATGTGTGGATGCTACTTCGTCGATGGAAACCATACAATCCTTTTTGATAGGCTGTTGAGATTGATCATTTCCTGTACGTTTAGGTTTGGTACGTTGATTTCTAGCCCCTGCTGATATAAAGTCATCGTCTACAATTTGGGATAATGAATTACATACACTACAAGACAAAGATTTATCAGAAGTGTGAATATGATCTTCATCTACCAAAGAAAGAGTTTGATTAGTATAACTAGATTTTAGTCCCGCGTTAGATGTTAAAAAAACATCGCTATTGACCACGGCTTCCATTATTTATATTCGTAGTTTTTACTCGAAAGCGTGATTTTAATATCCAATCTTATTACTTTTGGAATCGTTCAAAACCTTTGACTAGTTGTAGAATTTGATCTATTGCCCTACGCGTATACTCCCTTGCATCATATACGTTCGTCACCAGATCGTTTGTTTCGGCCTGAAGTTGACGCATATCTTTTTCAACACTCGACATGAGATCCTTAAGGGTCATATCGTCTAGATTTTGTTGAGATGCTGCTCCTGGATTTGGATTTTGTTGTGCTGTTGTACATACTGTACCACCAGTAGGTGTAGGAGTACATACAGTGGCCACAATAGGAGGTTGAAGAGGTGTAACCGTTGGAGTAGTACAAGAAATACTTCCATCCGATTGTTGTGTACATGTGGTTGTTGGTAACGTCTGAGAAGGTTGGGTAGATGGCGGTGTCGTCATCTTTTGATCTTTATTAAATTTAGAGATAATATCCTGAACAGTATTGCTCGGCGTCAACGCTGGAAGGAGTGTACTCGCCGGCGCATCAGTATCTGTAGACAACCAATCAAAAAGATTAGACATATCAGATGATGTATTAGTTTGTTGACGTGGTTTTAGTACAGGAGCAGTACTACTAGGTAGAAGAATAGGAGCCGGTGTAGGTGTCGGAACCGGCTGTGGAGTTATATGAATAGTTGGTTGTAGCGGTTGGGTAGGCTGTCTGCTGGCGGTCATCATATTATCTCTAGCTAGTTGTTCTCGCAACTGTCTTTGATAATACGACTCTTGAGACTTTAGTCCTATTTCAATCGCTTCATCCTTTTTCGTATCCGGATCCTTTTCTTCAGAATAATAGATTGACGACTTTGGTGTAGAGGATTCTGCCAGCCCCTGTGAGAACTTGTTAAAGAAGTCCATTTAAGGCTTTAAAATTGAATTGCGATTATAAGATTAAATGGCAGACACGGACGATATTATCGACTATGAATCCGATGATCTCACCGAATACGAGGATGATGAAGAAGATGGAGAGTCACTAGAAACTAGTGATATAGATCCCAAATCTTCTTATAAGATTGTAGAATCAACATCCACTCATATAGAAGATGCGCATTCCAATCTTAAACATATAGGGAATCATATATCTGCTCTTAAACGACGCTATACTAGACGTATAAGTCTATTTGAAATAGCGGGTATAATAGCAGAAAGCTATAACTTGCTTCAACGAGGAAGATTACCTCTAGTTTCAGAATTTTCTGACGAAACGATGAAGCAAAATATGCTACATGTAATTATACAAGAGATAGAGGAGGGTTCTTGTCCTATAGTCATCGAAAAGAACGGAGAATTGTTGTCGGTAAACGATTTTGATAAAGATGGTCTAAAATTCCATCTAGACTATATTATCAAAATTTGGAAACTTCAAAAACGATATTAGAATTTATACGAATATCGTTCTCTAAATGTCACAATCAAGTCTCTCATATTCAGCAGTTTATTGTCGTACTTTATATCGTGTTCATTAACGATATTTTGCAAAATAGTAATGATTCTATCTTCCTTCGATAGATATTCTTCAGAGATTATTGTCTTATATTCTTTCTTGTTATCCGATATGAATTTGATAAGACTTTGAACATTATTAATACCCGTCTGTTTAATTTTTTCTATAGATATTTTAGTTTTGGTAGATTCTATGGTGTCTGTTAATAGGCATCCAACATCGACATTCGACGTCAATTGTCTATAAATCAGAGTATAAATTTTAGAAATAACATTAGCAAATTGTTGTGCGTTGATGTCGTTATTCTGAAACAGTATGATTTTAGGTAGCATTTTCTTAACAAAGAGAACGTATTTATTGTTACTCAGTTGAACAGATGATATATCCAGATTACTAACGCATCTGATTCCATATACCAAACTTTCAGAAGAAATGGTGTACAATTGTTTGTATTCATTCAATGTCTCCTTTTCAGAAATTAGTTTAGAGTCGAATACTGCAATAATTTTCAAGAGATAGTTTTCATCAGATAAGATTTTATTTAGTGTAGATATGATAAAACTATTGTTTTGTTGGAGAACTTGATACGCCGCATTCTCTGTAGTCGACGCTCTCAAATGGGAAACAATCTCTATTATTTTTTTGGAATCGGATACTATATCTTCGGTATCTTGACGCAGTCTAGTATACATAGAGTTAAGAGAAATTAGAGTTTGTACATTAAGCAACATGTCTCTAAATGTGGCTACAAACTTTTCTTTTTCCACATCATCTAGTTTATTATATACCGATTTCACAACGGCACCAGATTTAAGGAACCAGAATGAAAAACTCTGATAACTACAATATTTCATCATAGTTACGATTTTATCATCTTCTATAGTTGGTGTGATAACACATACCTTTTTCTCCAAGACTGGAACCAACGTCATAAAAATGTTTAAATCAAAATCCATATCAACATCTGATGCGCTAAGACCAGTCTCGCGTTCAAGATTATCTTTACTAATGGTGACGAACTCATCGTATAGAACTCTAAGTTTGTCCATTATTTATTTACAGATTTAGTTGTTTAATTTATTTGTGCTCTTCCAGAGTTGGGATAGTATTTTTCTAACGTCGGTATTATATTATTAGGATCTACGTTCATATGTATCATAATATTAATCATCCACGTTTTGATAAATCTATCTTTAGCTTCTGAAATAACGTATTTAAACAAAGGAGAAAAATATTTAGTTACGGCATCAGACGCGATAACATTTTTTGTAAATGTAACGTATTTAGACGACAGATCTTCGTTAAAAAGTTTTCCATCTATGTAGAATCCATCGGTTGTTAACACCATTCCCGCGTCAGAGTGAATAGGAGTTTGAATAGTTTGTTTTGGAAATAGATCCTTCAATAACTTATAGTTGGGTGGGAAAAAATCGATTTTATCACTAGACTCTTTCTTTTTTACTATCATTACCTCATGAACTATTTCTTGAATGAGTATATGTATTTTCTTTCCTATATCGGTCGCGTTCATTGGAAAATATATCATGTCGTTAACTATAAGAATATTTTTATCCTCGTTTACAAACTGAATAATATCAGATATAGTTCGTAAACGAACTATATCATCACCAGCACAACATCTAACTATATGATATCCACTAGTTTCCTTTAGCCGTTTATTATCTTGTTCCATATTAGCAGTCATTCCATCATTTAAGAAGGCGTCAAAGATAATAGGGAGAAATGACATTTTGGATTCTGTTACGACTTTACCAAAATTAAGGATATACGGACTTACTATCTTTTTCTCAACGTCGATTTGATGAACACACGATGAAAATGTACTTCGATGAGATTGATCATGTAGAAAACAACAAGGGATACAATATTTCCGCATATCATGAAATATATTAAGAAATCCCACTTTATTATATTTCCCCAAAGGATCAATGCATGTAAACATTATACCGTTATCATTAATAAAGACTTCTTTCTCATCGGATCTGTAAAAGTTGTTACTGATTTTTTTCATTCCAGGATCTAGATAATTAATAATAATGGGTTTTCTATTCTTATTCTTTGTATTTTGACATATCCTAGACCAGTAAACAGTTTCCACTTTGGTAAAATCAGAAGACTTTTGAACGCTATTAAACATGGCATTAATGGCAATAACTAAAAATGTAAAATATTTTTCTATGTTAGGAATATGGTTTTTCACTTTAATAGATATATGGTTTTTTGCCAAAATGATAGATATTTTTTTATCCGATGATAGTAAAATATTATTAGTCGCCGTCTCTATAAAAATGAAGCTAGTCTCGATATCCAATTTTATTCTAGAATTGATAGGAGTCGCCAAATGTACCTTATACGTTATATCTCCCTTGATGCGTTCCATTTGTGTATCTATATCGGACACAAGATCTGTAAATAGTTTTACGTTATTAATCATCACGGTATCGCCATCGCTAGATAATGCTAATGTACTATCCAAGTCCCAAATGGAGAGATTTAACTGTTCATCGTTTAGAATAAAATGATTACCTGTCATATTAATAAAGTGTTCATCGTATCTAGATAACAACGACTTATAATTAATGTCCAAGTCTTGAACTCGCTGAATGATCTTTTTTAACCCAGTTAGTTTTAGATTGGTACGAAATATATTGTTAAACTTTGATTCTACAGTAATGTCCAAATCTAGTTGTGGAAATACTTCCATCAACATTGTTTCAAACTTGATAATATTATTATCTACATCTTCGTACGATCCAAATTCCGGAATAGATGTATCGCACGCTCTGGCCACCCAGATAACCAAAAAGTCACACGCTCCAGAATATACATTGTATAAAAAGCTATCGTTTTTTAGTAGTGTTTTTTTCTGAGTATATACGAAAGGATTAAAAATAGTATTATCAACGTAACTATATTCCAAATTATTCTTATGAGAATAGATAATAATATCGTCCTTAATATCTAACAAATTTCCTAAATATCCCTTTAATTGAGTCATTCGAAGCGTTAATAAAATATGTCTCTTAACTATTTCCGGCCGTTGTATATTTAAATGACTTCGTAAGAAATAATATATAGGCGACTTCTCATCTATGTAATCATATGGAGTGAGATATAGGGCTCGTTCTACCTCCTGCCCCTTACCCACCTGTAATACCAATTGCGGACTCACTATATATCGCATATTTATATCGTGGGGTAAAGTGAAAATCTACTACCGATGATGTAAGTCTTACAATGTTCGAACCAGTACCAGATCTTAATTTGGAGGCCTCCGTAGAACTAGGGGAGGTAAATATAGATCAAACAACACCTATGATAAAGGAGAATAGCGGTTTTATATCCCGTAGTAGACGTCTATTCGCCCATAGATCTAAGGATGATGAGAGAAAACTAGCACTACGATTCTTTTTACAAAGACTTTATTTTTTAGATCATAGAGAGATTCATTATTTGTTCAGATGCGTCGACGCTGTAAAAGACGTCACTATTACCAAAAAAAATAACATTATCGTGGCGCCTTATATAGCACTTTTAACTATCGCATCAAAAGGATGCAAACTTACAGAAACAATGATTGAAGCATTCTTTCCAGAACTATATAATGAACATAGTAAGAAATTCAAATTCAACTCTCAAGTATCCATCATCCAAGAAAAACTCGGATACCAGTCTGGAAACTATCACGTTTATGATTTTGAACCGTATTACTCTACAGTAGCTCTGGCTATTCGAGATGAACATTCATCTGGCATTTTTAATATCCGTCAAGAGAGTTATCTTGTAAGTTCATTATCTGAAATAACATATAGATTTTATCTAATTAATCTAAAATCTGATCTTGTTCAATGGAGTGCTAGTACGGGCGCTGTAATTAATCAAATGGTAAATACTGTATTGATTACAGTGTATGAAAAATTACAACTGGCCATAGAAAATGATTCACAATTTACATGTTCATTGGCTGTGGAATCAGAACTTCCAATAAAATTACTTAAAGATAGAAATGAATTATTTACAAAATTCATTAACGAGTTAAAAAAGACCAGTTCATTCAAGATAAGCAAACGTGATAAGGATACGCTATTAAAACATTTTACTTATGACTGGAGTTAGAATTTATAGACGACACATTTCGTTTATCATTGTTACTATTACTATCATTATTAGTATTCTTCTTGTCATCTTGTTCAGAAATATACAGCAATGCTATACCTAATACTAAATACATTATCATGCTTGCAATGGCTCTAACAACAACGAACCAAAATGAATTTGGTCGTAGCTTTTGTTCACAAAAATACATAAAGAAATGTCTACATAAATCTATGGCGCCATTGGCTACTTGAAATAGCGCCAGTCCTCCTACAGATTTTAATATAGCTGTATAACATGACATTTATTCATCATCAAAAGAGACAGAGTCACCATCTGTCATATTTAGATTTTTTTTCATGTGTTCAAAGTATCCTCTACTCATTTCATTATAATAGTTTATCATGCTTAGAATTTTAGGACGGATCAATGAGTAAGACTTGACTAGATCGTCAGTAGTAATTTGTGCATCATCTATTCTGCATCCGCTTCGTCGAATAATGTATAGCATCGCTTTGAGATTCTCCATAGCTATCAAGTCTTTATATAATGACATGGAAATATCTGTGAATGCTTTATACTTCTCCAACATCGATGCCTTAACATCATCACATACTTTAGCATTGAAAATACGTTCTATTGTGTAGATGGATGTAGCAAGATTTTTAAACAACAATGCCATCTTACATGATGATTGTCTCAAGTCTCCAATCGTTTGTTTAGAACGATTAGCTACAGAGTCCAATGCTTGGCTAACTAGCATATTATTATCTTTAGAAATTGTATTCTTCAATGAGGCGTTTATCATATCTGTGATTTCGTTAGTCATATTACAGTCTGACTGGGTTGTAATGTTATCCAACATATCACCTATGGATACGGTACACGTACCAGCATTTGTAATAATCCTATCTAAGATGTTGTATGGCATTGCGCAGAAAATATCTTCTCCTGTAATATCTCCACTCTCGATAAATCTACTCAGATTATTCTTAAATGCCTTATTCTCTGGAGAAAAGATATCAGTGTCCATCATTTCATTAATAGTATACGCAGAAAAGATACCACGAGTATCAATTCTATCCAAGATACTTATCGGTTCCGAGTCACAGATAATTGTTTCCTCTCCTTCGGGAGATCCTGCATAGAAATATCTAGGACAATAGTTTCTATACTGTCTGTAACTCTGATAATCTCTAAAGTCACTAACTGATACCATGAAATTGAGAAGATCAAACGCTGAAGTAATCAATTTTTCTGCCTCGTTTTTACTACAACTAGTTTTCATCAATGTAGTGACGATGTATTGTTTAGTTACTCTTGGTCTAATACTGATGATAGAGATATTATTGCTTCCCATAATGGATCTTCTAGTAGTCACCTTAAAGCCCATTGATGCGAATAGCAGATAGATAAAGTCTTGGTATGACTCCTTTCTAATATAGTACGGACTACCTTTGTCACCCAACTTTATACCCACATAAGCCATAACAACCTCTTTAATAGCCGTTTCATGAGGTTTATCAGCCATGAGCCTGAGTAGTTGAAAGAATCGCATGAATCCCGTCTCAGAAAGTCCTATATGCATGATAGATTTATCTTTCCTGGGAAACTCTCGTATAGTTATAGATGAAATACTCTTCAAAGTTTCTGAAATAAGATTAGTAACAGTCTTACCTCCGACTACTCTGGGTAACAAACATACTCTAATAGGTGTTTTCTCTGCGGAGATAATATCAGAAAGGATAGAGCAATAAGTAGTATTATTGTGATTATAAAGACCGAATACATAACAGGTAGAATTTATAAACATCATGTCCTGAAGGGTTTTAGACTTGTATTCCTCGTAATCTATACCGTCCCAAAACATGGATTTGGTAACTTTGATAGCCGTAGATCTTTGTTCCTTCGCTAACAGGTTAAAGAAATTAATAAAGAATTTGTTGTTTCTATTTATGTCCACAAATTGCACGTTTGGAAGCGCCACGGTTACATTCACTGCAGCATTTTGAGGATCGCGAGTATGAAGTACGATGTTATTGTTTACTGGTATATCTGGAAAGAAATCTACCAGTCTAGGAATAAGAGATTGATATCGCATAGAAATAGTAAAGTTTATAATCTCATCATTGAAGATTACTCTGTTACCATTGTAATAAATTGGTACTCTATCATAATCATCGACAAAGTACTGTTCATACATGATGAGATGTTTATATGTTGGCATAGTAGTGAGATCGACGTTTGGTAATGGCAATGTATTAAGATTAACTCCATAATGTCTAGCAGCATCTGCGATGTTATAAGTGATGTCAAAGCGGGGTTGATCTTGTGCTGTTATATATTGTCTAACACCTATAAGATTATCAAAATCTTGTCTGCTTAATACACCGTTAACAATTTTTGCCTTGAATTCTTTTATTGGTGCATTAATAACATCCTTATAGAGGATGTTAAACAAATAAGTATTATCAAAGTTAAGATCTGGGTATTTCTTTTCTGCTAGAACATCCATTGAGTCGGAGCCATCTGGTTTAATATAACCACCGATAAATCTAGCTCTGTATTCTGTATCCGTCAATCTAATATTAAGAAGGTGTTGAGTGAAAGGTGGAAGATCGTAAAAGCTGTGAGTATTAATAATAGGGTTAGTTTCCGAACTAATGTTAATTGGATGATTAATAATATTTATATTTCCAGCGTTAAGTGTAACATTAAACAGTTTTAATTCACGTGACGTGGTATCAATTAAATAATTAATGCCCAATTTGGATATAGTAGCCTGAAGCTCATCTTGTTTAGTTACGGATCCTAATGAGTTATTAAGAAATACATCGAACGGATGAACGAAGGTTGTTTTAAGTTGGTCACATACTTTGTAATCTAGACATAGATGTGGAAGAACGGTAGAAACTATACGAAATAGATATTCAGAGTCCTCTAATTGATCAAGAGTAACTATTGACTTAATAGGCATCATTTATTTAGTATTAAATGACGACCGTACCAGTGACAGATATACAAAACGACTTAATTACAGAGTTTTCAGAAGATAATTATCCATCTAACAAAAATTATGAAATAACTCTTCGCCAAATGTCTATTCTAACTCACGTTAACAACGTGGTAGATAGAGAACATAATGCCGCCGTAGTGTCATCTCCAGAGGAAATATCATCACAACTTAATGAAGATCTATTTCCAGATGATGATTCACCGGCCACTATTATCGAACGAGTACAACCTCATACTACTATTATTGACGATACGCCACCTCCTACTTTTCGTAGAGAGTTATTGATATCGGAACAACGTCAACAACGAGAAAAAAGATTTAATATTACAGTATCAAAAAATTCTGAAGCAATAATGGAATCTAGATCTATGATAACTTCTATGCCAACACAAACACCATCCTTGGGAGTAGTTTATGATAAAGATAAAAGAATTCAGATGCTAGAGGATGAAGTGGTTAATCTTAGAAATCAACGATCTAATACAAAATCATCTGATAATTTAGATAATTTTACCAGAATACTATTTGGTAAGACTCCGTATAAATCAACCGAAGTTAATAAGCGTATAGCCATCGTTAATTATGCAAATTTGAACGGGTCCCCCTTATCAGTCGAGGACTTGGATGTCTGTTCGGAGGATGAAATAGATAGAATCTATAAAACGATTAAACAATATCACGAAAGTAGAAAACGAAAAATTATCGTCACTAACGTGATTATTATTGTCATAAACATTATTGAGCAGGCATTGCTAAAACTCGGATTTGAAGAAATCAAAGGACTGAGTACCGATATCACTTCAGAAATTATCGATGTGGAGATCGGAGATGACTGCGATGCTGTAGCATCAAAACTAGGAATCGGTAACAGTCCGGTTCTTAATATTGTATTGTTTATACTCAAGATATTCGTTAAACGAATTAAAATTATTTAATTTAATACATTCCCATATCCAGACAACAATCGTCTGGATTAATCTGTTCCTGTCGTCTCATACCGGACGACATATTAATCTTTTTATTAGTGGGCATCTTTTTAGATGGTTTCTTTTTCCCAGCATTAACTGATTCGATACCTAGAAGATCGTGATTGATTTCTCCGACCATTCCACGAACTTCTAATTGGCCGTCTCTAACGGTACCATAAACTATTTTACCAGCATTAGTAACAGCTTGGACAATCTGACCATCCATTGCGTTGAATGATGTAGTTGCTGTTGTTCTACGTCTAGGAGCACCAGAGGTATTTTTAGAGCTCTTGGATGTTGATGTAGAAGACGAGGATTTTGATTTTGGTTTACATGTAATACATTTTGAACTCTTTGATTTTGTATCACATGCACCGGCAGTCACATCTGTTTGAGAATTAAGATTATTGTTGCCTCCTTTGACGGCTGCATCTCCACCGATCTGCGCTAGTAGATTTTTAAGCTGTGGTGTAATCTTATTAACTGTTTCAATATAATCATCGTAACTACTTCTAACGGCTAAATTTTTTTTATCCGCCATTTAGAAGCTAAAAATATTTTTATTTATGCAGAAGATTTAACTAGATTATACAATGAACTAATATGATCCTTTTCTAGATTATTTACGAACTTGGTATTTCTTGTTTCTGGAGGAGGAGAATTTAAATTCGGACTTGGATTCGGATTTTGTGGGTTCTTGATCTTATTATACAGCGTGTATAGGATGGTGACGGTAACTGCTACACAAATACCGATCAACAGAAGAATACCAATCATTTATTGACAATAACTTCACTATGATCAAGTATGTAATAATCATCTTTTCACTAAGTAAGTAGTAATAATGATTCAACAATGACACGATATATGGACGATAATAATTTAGTTCATGGAAATATCGCTATGATTGGTGTGAATGACTCCGCTAACTCTGTGGGGTGCACAGTGCTTTCCCCACATAGAATAAATTAGCATTCCGACTGTGATAATAATACCAAGTATAAACGCCATAATACTCAATACTTTCCATGTACGAGTGGGACTGGTAGACTTACTAAAGTCAATAAAGGCGAAGATACACGAAAGAATCAAAAGAATGATTCCAGCGATTAGCACGCCAGAAAAATAATTTCCAATCATAAGCATCATGTCCATTTAACTAATAAAAATTTTAAATCGCCGAATAAACAAAGTGGAATATAAACCATATAAAAACAATAGTTTGTACTGCAAAAATAATATCTATTTTTGTTTTCGAAGATATGGTAAAATTAAATAGTAGTACACAGCATGTTATAACTAACAGCAGCAACGGCTCGTAATTACTTATCATTTACTAGACGAAAAGGTGGTGGGATATTTTCTTGCTCAAATAATACGAATATATCACCCATCCATTTTATACGATGTTTATATACTCTAATCTTTAATAGATCTATAGATGACGGGTTTACCAATAATATAGATTTTATCGATTCATCTAATTTAAACCCTTCCTTAAACGTGAATGATCTATTATCTGGCATAATGATGACCCTACCTGATGAATCTGACAATGTACTGGGCCATGTAGAATAAATTATCAACGAATTATCGTCTACGAACATTTATATCATTTGTTTTAATTTTAGGACGTGAATAAATAGATATAAAATAGAAAATAACAGATATTACAACCAGTGTTATGGACGCACCCAACCATGTAGGCAGTTTTATTTTATCGTTTACTACAGGTTCTCCTGGATGTACGTCACCAACTGCAGACGTAGTTCTAGTACAATTAGACGTAAGTTCCGCTTGGGAATTTTTTAACGCTAAAGAGTTAACGTTGATCGTACACCCAACGTATTTACATCTAGTTCTTTGAACATCTTGATTATAATATAACCATTTTCTATCTCTAGATTCGTCAGTGCACTCATGTAACCAACATACCCTAGGTCCTAAATATTTATCTCCGGAATTAGATTTTGGATAATTCGCGCACCAACAATTTCTATTTCCTTTATGGTCGTTACAAAAGACGTATAATGCCGTATCCCCAAAAGTAAAATAATCAGGACGAATAATTCTAATAAACTCAGAACAATATCTCGCATCCATATGTTTGGAGCAAATATCGGAATAAGTAGACATAGCCGGTTTCCGTTTTACACGTAACCATTCTAAACAATTGGGGTTTCCAGGATCGTTTCTACAAAAACCAGTCATGAAATCGTCACAATGTTCTGTCTTGTAATTATTATTAAATATTTTTGGACAGTGTTTGGTATTTGTCTTAGAACAACATTTTGCCACGCTATCACTATCACCCAGGAGATAATCCTTTTTTATAAAATGACATCGTTGCCCGGATGCTATATAATCAGTAGCATATTTTAAATCCTTAATATATTCAGGAGTTACCTCGTTCTGATAATAGATTAATGATCCAGGACGAAATTTGAAAGAACTACATGGTTCTCCATGAATTAATACATATTGTTTAGCAAATTCAGGAACTATAAAACTACTACAATGATCTATCGACATACCATCTATCAAACAAAATTTGGGTTTAATTTCTCCTGGAGACGTTTCATAATAATACATATAACTTTCTTCGGCAAACCTAACAGCTCTATTATATTCAGGATAATTAAAATCTAATACCATATATTTGTCTCGTATATCTGCTATTCCTGTCTCTATTTTGATTCTATTAAGAGTAACAGCTGCCCCCATTCTTAATAATCATCAGTATTTAAACTGTTAAATGTTGGTATATCAACATCTATCTTATTTCCCGCAGTATAAGGTTTGTTGCAGGTATACTGTTCAGGAATGGGTACATTTATACTTCTTTTATAGTCCTGTCTTTCGATGTTCATCACAAATGCAAAGAACAGAATAAACAAAATAATGTAAGAAATAATATTAAATATCTGTGAATTCGTAAATACATTGATTGCCATAATAATTACAGCAGCTACAATACACACAATAGACATTCCCACAGTGTTGCCATTACCTCCACGATACATTTGAGTTACTAAGCAATAGGTAATAACTAAGCTAGTAAGAGGCAATAGAAAAGATGAGATAAATATCATCAATATAGAGATTAGAGGAGGGCTATATAGAGCCAAGACGAACAAAATCAAACCGAGTAACGTTCTAACATCATTATTTTTGAAGATTCCCAAATAATCATTCATTATTCCTCCATAATCGTTTTGCATCATACCCCCATCTTTAGGCATAAACGATTGCTGCTGTTCCTCTGTAAATAAATCTTTATCAAGCACTCCAGCACCCGCAGAGAAGTCATCAAGCATATTGTAATATCTTAAATAACTCATTTATATATTAAAAAATGTCACTATTAAAGATGGAGTATAATCTTTATGCCGAACTAAAAAAAATGACTTGTGGTCAGACCATAAGTCTTTTTAATGAAGACGGCGATTTCGTAGAAGTTGAACCAGGATCATCCTTTAAGTTTCTAATACCTAAGGGATTTTACTCCTCTCCTTGTGTAAAGACGAGTCTAGTATTCGAGACATTAACAACGACCGATAATAAAATTACTAGTATCAATCCAACAAATGCGCCAAAGTTATATCCTCTTCAACGCAAAGTCGTATCTGAAGTAGTTTCTAATATGAGGAAAATGATCGAATTAAAACGTCCTCTATACATCACTCTTCACTTGGCATGTGGATTTGGTAAGACTATTACCACGTGTTATCTTATGACCACACACGGCAGAAAAACCATCATTTGCGTACCCAATAAAATGTTAATACATCAATGGAAGACACAGGTAGAGGCAGTCGGATTGGAACATAAGATATCTATAGATGGAGTTAGTAGTCTATTAAAGGAACTAAAGACTCAAAGTCCGGATGTATTAATCGTAGTCAGTAGACATCTGACAAACGATGCATTTTGTAAATATATCAATAAGCATTATGATTTGTTTATCTTGGATGAATCACATACGTATAATCTGATGAACAATACAGCAGTTACAAGATTTTTAGCGTATTATCCTCCGATGATGTGTTATTTTTTAACTGCTACACCTAGACCAGCTAACCGAATTTATTGTAATAGTATTATTAATATTGCCAAGTTATCCGATCTAAAAAAAACTATCTATATAGTAGATAGTTTTTTTGAGCCATATTCCACAGACAATATTAGAAATATGGTAAAACGACTAGATGGACCATCTAATAAATATCATATATATACCGAGAAGTTATTATCTGTAGACGAGCCTAGAAACCAACTTATTCTTGATACCCTGGTAGAAGAATTCAAGTCAGGAACTATTAATAGAATTTTAGTTATTACTAAACTACGTGAACATATGGTATTCTTCTACAAACGATTATTAGATCTTTTCGGAGCAGAGGTTGTATTTATAGGAGACGCCCAAAATAGACGTACTCCAGATATGGTCAAATCGATTAAGGAACTAAATAGATTTATATTCGTATCCACCTTATTTTATTCCGGCACTGGTTTAGATATTCCGAGTTTGGATTCTTTGTTCATTTGCTCGGCAGTAATCAACAATATGCAAATAGAGCAATTACTAGGGAGGGTATGTCGAGAAACAGAACTATTAGATAGGACGGTATATGTATTTCCTAACACATCCATCAAAAAAATAAAGTACATGATAGGAAATTTCGTGCAACGAATTATTAGTCTGTCTGTAGATAAACTCGGATTTAAACAAGAAAGTTATCAGAAACATCAGGAATCTGAACCCGCTTCCGTACCAACATCCTCCAGAGAAGAACGTGTATTAAATAGAATATTTAACTCGCAAAATCGTTAAGAAGTTTAAGAGACGATCCACATGCTGAGCAGGCCAGTGTATTACCCCTCATAGTATTAATATAATCCAATGATACTTTTGTGATGTCGGAAATCTTAACCAATTTAGACTGACAGGCAGAACACGTCATACAATCATCATCGTCATCGATAACTGTAGTCTTGGGCTTCTTTTTGCGACTCTTCATTCCGGAACGCATATTGGTGCTATCCATTTAGGTAGTAAAAAATAAGTCAGAATATGCCCTATAACACGATCGTGCAAAACCTGGTATATCGTCTCTATCTTTATCACAATATAGTGTATCAACATCTTTATTATTGACCTCGTTTATCTTGGAACATGGAATGGGAACATTTTTGTTAACGGCCACCTTTGCCTTAATTCCAGATGTTGTAAAATTATAACTAAACAGTCTATCATCGACACAAATGAAATTCTTGTTTAGACGTTTGTAGTTTACGTATGCGGCTCGTTCTCGTCTCATTTTTTCAGATATTGCAGGTACTATAATATTAAAAATAAGAATGAAATAACATAGGATTAAAAATAAAGTTATCATGACTTCTAGTGCTGATTTAACTAACTTAAAAGAATTACTTAGTCTGTACAAAAGTTTGAGATTTTCAGATTCTGTGGCTATAGAGAAGTATAATTCTTTGGTAGAATGGGGAACATCTACTTACTGGAAAATAGGCGTACAAAAGGTAACTAATGTCGAGACGTCCATATCTGATTATTATGATGAGGTAAAAAATAAACCGTTTAATATTGATCCGGGGTATTATATTTTCTTACCAGTATATTTTGGAAGCGTCTTTATTTATTCAAAGGGTAAAAATATGGTAGAACTTGGATCTGGAAACTCTTTTCAAATACCGGATGAGATTCGAAGTGCGTGTAACAAAGTATTAGATAGTGATAACGGAATAGACTTTCTGAGATTTGTTTTGTTAAACAATAGATGGATAATGGAAGACGCTATATCAAAATACCAGTCTCCAGTTAATATATTTAAACTAGCTAGTGAGTACGGATTAAACATACCCAACTATTTAGAAATTGAAATAGAGGAAGACACATTATTTGACGATGAGTTATACTCTATTATGGAACGCTCTTTCGATGATACATTTCCAAAAATATCTATATCGTATATTAAGTTGGGAGAACTTAAGCGGCAAGTTGTAGACTTTTTCAAATTCTCATTCATGTATATTGAGTCAATCAAGGTAGATCGTATAGGAGATAATATTTTTATTCCTAGCGTTATAACAAAATCAGGAAAAAAGATATTAGTAAAAGATGTAGACCATTTAATACGATCCAAGGTTAGAGAACATACATTTGTAAAAGTAAAAAAGAAAAACACATTTTCCATTTTATACGACTATGATGGGAACGGAACAGAAACTAGAGGAGAAGTAATAAAACGAATTATAGACACTATAGGACGAGACTATTATGTTAATGGAAAGTATTTCTCTAAGGTTGGTATTGCAGGCTTAAAGCAATTGACTAATAAATTAGATATTAATGAGTGTGCAACTGTCGATGAGTTAGTTGATGAGATTAATAAATCCGGAACTGTAAAACGAAAAATAAAAAACCAATCAGTATTTGATTTAAGCAGAGAATGTTTGGGATATCCAGAAGCGGATTTTATAACGTTAGTTAATAACATGCGGTTCAAAATAGAAAATTGTAAGGTTGTAAATTTCAATATTGAAAATACTAATTGTTTAAATAACCCGAGTATTGAAACTATATATGGAAACTTCAACCAGTTCGTCTCAATCTTTAATACCGTTACCGATGTCAAAAAAAGATTATTCGAGTGAAATAATATGCGCCTTTGATATAGGTGCAAAAAATCCTGCCAGAACTGTTTTAGAAGTCAAGGATAACTCCGTTAGGGTATTGGATATATCAAAATTAGACTGGAGTTCTGATTGGGAAAGGCGCATAGCTCAAGATTTGTCACAATATGAATACACTACAGTTCTTCTAGAACGTCAGCCTAGAAGGTCACCGTACGTCAAATTTATCTATTTTATTAAAGGCTTTTTATATCATACATCTGCTGCCAAAGTTATTTGCGTCTCACCTGTCATGTCTGGTAATTCATATAGAGATCGAAAAAAGAGATCTGTTGAAGCATTTCTTGATTGGATGGACACATTCGGATTGCGAGACTCCGTTCCGGATAGACGCAAATTAGACGATGTAGCGGATAGTTTCAATTTGGCTATGAGATACGTATTAGATAAATGGAATACTAATTATACACCTTATAATAGGTGTAAATATAGAAATTACATAAAAAAAATGTAATAACGTTAGTAACGCCATTATGGATAATCTATTTACCTTTCTACATGAAATAGAAGATAGATATGCCAGAACTATTTTTAACTTTCATCTAATAAGTTGTGATGAAATAGGAGATATATATGGTCTTATGAAAGAACGCATTTCCTCAGAGGATATGTTTGACAATATAGTATATAATAAAGATATACATCCTGCCATTAAGAAACTAGTTTATTGCGACATCCAACTTACTAAACATATTATTAATCAGAATACGTATCCGGTATTTAACGATTCTTCACAAGTGAAATGTTGTCATTATTTCGATATAAACTCAAATAATAGCAATATTAGCTCTCGTACAGTAGAGATATTTGAGAGTGAAAAGTCATCTCTTGTATCATATATTAAAACTACCAATAAGAAGAGAAAGGTCAATTACGGCGAAATAAAGAAAACTGTACATGGAGGCACTAATGCAAATTACTTTTCCGGTAAAAAGTCTGATGAGTATCTGAGCACTACAGTCAGGTCCAACATTAATCAACCTTGGATCAAAACCATTTCTAAGAGAATGAGAGTAGATATCATTAATCACTCTATAGTAACGCGTGGAAAAAGCTCTATATTACAAACTATAGAAATTATTTTTACTAATAGAACATGTGTGAAAATATTCAAGGATTCTACTATGCACATTATTCTATCCAAGGACAAGGATGAAAAGGGATGTATAAACATGATTGATAAATTATTCTATGTATATTATAATTTATTTCTGTTGTTCGAGGATATCATCCAAAACGATTACTTTAAAGAAGTAGCTAATGTTGTAAACCATGTACTCATGGCTACGGCATTAGATGAGAAATTATTCCTAATTAAGAAAATGGCTGAACACGATGTTTATGGAGTTAGCAATTTCAAAATAGGGATGTTTAACCTGACATTTATTAAGTCGTTGGATCATACCGTTTTCCCCTCTCTGTTAGATGAGGATAGCAAAATAAAGTTTTTTAAGGGGAAAAAGCTCAATATTGTAGCATTACGATCTCTGGAGGATTGTACAAATTACGTGACTAAATCCGAGAATATGATAGAAATGATGAAGGAAAGATCGACTATTTTAAATAGCATAGATATAGAAACGGAATCGGTAGATCGTCTAAAAGAATTGCTTCTAAAATGAAAAAAAACACTGATTCAGAAATGGATCAACGACTCGGGTATAAGTTTTTGGTGCCTGATCCTAAAGCCGGAGTTTTTTATAGACCGTTACATTTCCAATATGTATCGTATTCTAATTTTATATTGCATCGATTGCATGAAATCTTGACCGTCAAGCGGCCACTCTTATCGTTTAAGAATAATACAGAACGAATTATGATAGAAATTAGCAATGTTAAAGTGACTCCTCCAGATTACTCACCTATAATTGCGAGTATTAAAGGTAAGAGTTATGACGCATTAGCCACGTTCACTGTAAATATCTTTAAAGAGGTAATGACCAAAGAGGGTATATCCATCACTAAAATAAGTAGTTATGAGGGAAAAGATTCTCATTTGATAAAAATTCCGCTACTAATAGGATATGGGAATAAAAATCCACTTGATACAGCCAAGTATCTTGTTCCTAATGTCATAGGTGGAGTCTTTATCAATAAACAATCTGTCGAAAAAGTAGGAATTAATCTAGTAGAAAAGATTACAACATGGCCAAAATTTAGGGTTGTTAAGCCAAACTCATTCACTTTCTCGTTTTCCTCCGTATCCCCTCCTAATGTATTACCGACAAGATATCGCCATTACAAGATATCTCTGGATATATCACAATTGGAAGCGTCGAATATATCATCGACAAAGACATTTATAACGGTCAATATTGTTTTGCTGTCTCAATATTTATCTAGAGTGAGTCTAGAATTCATTAGACGTAGTTTATCATACGATATGCCTCCAGAAGTTGTCTATCTAGTAAACGCGATAATAGATAGTGCTAAACGACTTACCGAATCTATTACTGACTTTAATATTGATACATACATTAATGACCTGGTGGAAGCTGAACACATTAAACAAAAATCTCAGTTAACGATTAACGAGTTTAAATATGAAATGCTGCATAACTTTTTACCTCATATGAACTATACACCCGATCAACTAAAGGGATTTTATATGATATCTTTACTAAGAAAGTTTCTCTACTGTATCTACCACACTTCTAGATATCCAGATAGAGATTCGATGGTTTGTCATCGCATCCTAACGTACGGCAAATATTTTGAGACGTTAGCACATGATGAATTAGAGAATTACATAGGTAACATCCGAAACGATATCATGAACAATCACAAGAACAGAGGCACTTACGCAGTAAACATTCATGTACTAACAACTCCTGGACTTAATCATGCATTTTCTAGTCTATTGAGTGGAAAGTTCAAAAAGTCAGACGGTAGTTATCGAACACATCCTCACTATTCATGGATGCAGAATATTTCTATTCCTAGAAGTGTTGGATTTTATCCGGATCAAGTAAAGATTTCAAAGATGTTTTCTGTCAGAAAATACCATCCAAGCCAATATCTTTACTTTTGTTCATCAGACGTTCCGGAAAGAGGTCCTCAGGTAGGTTTAGTATCTCAATTGTCTGTCTTGAGTTCCATTACAAATATACTAACGTCTGAGTATTTGGATTTGGAAAAGAAAATTTGTGAGTATATCAGATCATATTATAAAGATGATATAAGTTACTTTGAAACAGGATTTCCAATCACTATAGAAAATGCTCTAGTCGCATCTCTTAATCCAAATATGATATGTGATTTTGTAACTGACTTTAGACGTAGAAAACGGATGGGATTCTTCGGTAACTTGGAGGTAGGTATTACTTTAGTTAGGGATCACATGAATGAAATTCGCATTAATATTGGAGCAGGAAGATTAGTCAGACCATTCTTGGTTGTGGATAACGGAGAGCTCATGATGGATGTGTGTCCGGAGTTAGAAAGCAGATTAGACGACATGACATTCTCTGACATTCAGAAAGAGTTTCCACATGTCATCGAAATGGTAGATATAGAACAATTTACTTTTAGTAACGTATGTGAATCGGTTCAAAAATTTAGAATGATGTCAAAGGATGAAAGAAAGCAATACGATTTATGTGACTTTCCTGCCGAATTTAGAGATGGATATGTAGCATCTTCACTAGTGGGAATCAATCACAATTCTGGACCCAGAGCTATTCTTGGATGTGCTCAAGCTAAACAAGCTATCTCTTGTCTGAGTTCGGATATACGAAATAAAATAGACAATGGAATTCATTTGATGTATCCAGAGAGGCCAATTGTGATTAGTAAGGCTTTAGAAACTTCAAAGATTGCGGCTAATTGCTTCGGACAACATGTTACTATAGCATTAATGTCGTACAAAGGTATCAATCAAGAGGATGGAATTATCATCAAAAAACAATTTATTCAGAGAGGCGGTCTCGATATTGTTACAGCCAAGAAACATCAAGTAGAAATTCCATTGGAAAACTTTAATAACAAAGAAAGAGATAGGTCTAACGCCTATTCGAAATTAGAAAGTAATGGATTAGTTAGACTGAATGCTTTCTTGGAATCCGGAGACGCTATGGCAAGAAATATCTCATCAAGAACTCTTGAAGATGATTTTGCTAGAGATAATCAGATTAGCTTTGATGTTTCCGAGAAATATACAGATATGTACAAATCTCGCGTTGAACGAGTACAAGTAGAACTTACTGACAAAGTTAAGGTGCGAGTATTAACCATGAAAGAAAGAAGACCCATTCTAGGAGACAAATTTACTACTAGAACGAGTCAAAAGGGAACAGTCGCGTATATCGCAGATGAAACGGAACTTCCGTACGACGAAAATGGTATCACACCAGATGTCATTATTAATTCTACATCCATCTTCTCTAGAAAAACTATATCTATGTTGATAGAAGTTATTTTAACAGCCGCATATTCTACTAAGCCGTACAACAATAAGGGAGAAAACCGACCTGTCTGTTTTCCTAGTAGTAACGAAACATCTATCGATGCATATATGCAATTCGCTAAACAATGTTATGAGTATTCAAATCCGAAATTGTCCGAGGAAGAATTATCGGATAAAATCTTTTGTGAAAAGATTCTCTATGATCCTGAAACGGATAAGCCTTATGAATCCAAAGTATTTTTTGGACCAATTTATTACTTGCGTCTGAGACATTTAACTCAGGACAAGGCAACCGTTAGATGTAGAGGTAAAAAGACGAAGCTCATTAGACAAGCGAATGAGGGACGAAAACGTGGAGGAGGTATCAAGTTTGGAGAAATGGAGAGAGACTGTTTAATAGCACATGGTGCAGCCAATACTATTACAGAAGTTTTAAAAGACTCAGAAGAGGATTATCAAGATGTGTATATTTGTGAAAATTGTGGAGACATAGCAGCACAAATCAAAAGTATTAATACATGTCTTAGATGTTCAAAACTTAATCTCTCTCCTCTCTTAACAAAAATTGATACCACGCACGTATCTAAAGTATTTCTTACTCAAATGAACGCCAGAGGCGTAAAAGTTAAATTAGATTTCGAACGAAGGCCTCCTTCGTTTTATAAACCATTAGATAAAGTTGATCTTAAACCGTCTTTTCTGGTATAATATTGTTTAGTAGATACTCATCAAGATAAGCTAATTCACTAAACATATTATCGGATTCGGTATTGTTACTCGAGAATAGAGTTCGTTATGCTCCTGATATTCGGAAATCTGTGGAGTTTCAGGTTTTGGTGGAAGTGTAACTGCTACTTGGTGGGATACTGAAGGATATTTCAGAGAGTTGTGGATGTTCGGGTTCGACATCCACCGATGGTGTCACGCCACTAATCGGTTCGGTAACGTCTGTGGATGGAGGTGCTACTTCTACAGAACCTGTAGCCTCAGTTGTCAACGGAGATACATATTCAATGCGCGGAAATGTATAATTTGGTAATGGTTTCTCATGTGGATCTTAAGAAGAAGAGGTAAGATATCTACGAAAGATACCGATCACGTTTCTAGTTCTCTTTTGTAGAACTTTAACTTTTTCTTTCTCAGCATCTAGTTGATATTCCGACCTCTTCACGTTTCGCATGGGTTACCTCCGCAGTTTTTACAAGCGATTTCACGTTCCAGATCACGTTCAGCCTTCATACGTCTCTCCCTCTCTCTATCGAGTTTATCAGAGCAGTCTTTCTGAAGGCGATCGAACTCCATAAATTTCTCCAACGCTTTGATTGTTTCCATAGATTTCCGAAGTTTAGCTTCTAGGACGGCGATTCTTTTTTTTTTTTTTTTTTTTTTTTTTTCGAATTCACGGGGTACAACCGTTTCCATTACCACCATCTCTATGTTTCTTTTCTAGATCGGCAATCTTTCTCAATCTTTCTCAACATTTCATCCCCATACCTTTTCATTCCTCGAGTCTATTGTCGTCGAAATATCGTTCCAGCTCCTTTTCGACCTCAATAACTTTAGCACGTTGTTTCATCAAGCTCTCTCTTGTAGTACTATCATTTTTATCTGATTCCCTGACACGTTTAAGATCTTCATGTAATTGAGTCAGCTCTTGACGCAATCTCTTAACTAACTTCCTCTCTTGCTTCTTCGTCATAGTACTTACAATCACTATGGGATCCATTGTTACCACGTCTGTACTCGACGAGCTCACGTTTAAGAGATTCAATTTCCAGTTTGTATCGGTCCATGTCTCCATTGCTACACCACCATTAGATTTACAGGCTGCTAGTTGTCGTTCGAGATCAGAAATACGTGTTTTCTTGGAATGGATTTCGTCGATGTACTTGTCATGATTGGCATCGAAACACTTATTAAGTTCTTTTTTTCAATTCTACGATTTTATTTCTTTCGCGAGTCAATTCCCTCCTGTAGTAACTATCAGTTTTGTCAGATTCACGCTCTCTACGTAGACTTTCTTGTAAGTTACTAATTTGTTCCCTGGCATTACCGAGTTCAGTTTTATATGCCGAATAGAGTTCTGATTCATCCTTTGAGAAGATCTCTAGCGATCGTTCAAGATCCCTGATTCTAGTCTTTAGCCTATTTACCTCCTCAGAAGATGCTCCGTTACCGTTTTTACAATCGTTAAGATGTCTATCAAGATCCATGATTCTATCTCTTTTCCATATCAGCATTGATTTCATTATTACGTTCGCAGTCGTTCAACTGTATTTCAAGATCTGAGATTCTAGATTGTAATCTCTGTAGCATTTCCACGGCATTCACTCAGTTGTCTTTCAAGATCTGAGATTCTAGATTGGAGTCTGCTAATCTCTGTAAGATTTCCTCCTCCGCTCTCGATGCAGTCGGTCAACTTATTCTCTAGTTCTCTAATACGCGAACGCAGTGCATCAACTTCTTGTGTGTCTTCTTGATTGCGTGTGCATTCATCGAGTCTAGATTCGAGATCTCTAACGTGACGTCGTTCTTCCTCAAGTTCTCTGTGTACTACAGAAAGCGTGTCCCTATCTTGTTGATATTTAGCAATTTCTGATTCTAGAGTACTGATTCTACTCACGTATGTACTAATAGTTGTCTTAGCCTTATCAAGATCCTCCTTGTATTTGTCACATTCCTTGATATCCATACGAAGTCTGGACAGTTCCCATTCGACATTACGACGTTTATCGATTTCAGCTCGGAGATCGTCGTCGCGTTGTTTTAGCCACATACGACTAAGTTCAAGTTCTCGTTGACAAGATCCATCTACTTTTCCATCCCTAATAGTATCCAGTTCCTTTTCTAGTTCTGACCGCATTTCTCGTTCCATATCAAGAGATTCTCTCAATTCTCGTATAGTCTTCTTATCAATTTCTGATGAATCTGAACCATCATCTGTCCCATTTTGTTGCATATCCCTGAGTTCTTTGATCTCTGTTGTAAGTCTGTCGATTCTTTCGGTTTTATAAACAGAATCCCTTTCCAAAGTCCTAATCTTACTGAGTTTATCATTAAGTTCTTCATTCAATTCAGTGAGTTTTCTCTTGGCTTCTTCCAAGTCTGTTTTAAACTCTCCATCATTTCCGCATTCTTCCTCGCATTTATCTAACCATTCAATTAGTTTATTAATAACTAGTTGGTAATCAGCGATTCCTATAGCCGTTCTTGTATTTGTGGGAACATAATTAGGATCTTCTAATGGATTGTATGGCTTGATAGCATCATCTTTATCATTATTAGGTGGGGGATGGACAACCTTAATTGGTTGGTCCTCCTTATCTCCTCCAGTAGCATGTGGTTCTTCAATACCAGTATTAGTAATAGGCTTAGACAAATGCTTGTCGTACGCGGGCACTTCCTCATCCATCAAGTATTTATAATCGGGTTCTGTTTCAGAATATTCTTTTCTAAGAGACGCGACTTCAGGAGTTAGTAGAAGAACTCTGTTTCTGTATCTATCAACGCTGGAATCGATACTCAAGTTAAGGATAGCGAATACCTCATCGTCATCATCCGTATCTTCTGAAACGCCATCATATGACATTTCATGAAGTCTAACGTATTGATAAACAGAATCAGATTTAGTATTAAACAGATCCTTGACCTTTTTAGTAAATGCATATGTATATTTTAGATCTCCAGATTTCATAATATGATCGCATGCCTTAAATGTCAATGCTTCCATGATATAGTCTGGAACACTAATGGGTGACGAAAAAGATACAGCACCATATGCTACGTTGATAAATAGATCTGAACCACTAAGTAGATAATGATTAATGTTAAGGAAGAGGAAATATTCAGTATATAGATATGCCTTAGCATCATATCTTGTACTAAACACGCTAAACAGTTTATTGATGTGATCAATTTCCAACAGAACAATTAGAGCGGCAGGAATACCAACAAACATATTACCACATCCGTATTTTCTATGAATATCACATATCATATTAAAAAATCTTGATAGAAGAGCGAATATCTCGTCTGACTTAATGAGATGTAGTTCAGCAGCATAAGTCATAACTGTAAATAGAACATACTTTCCTGTAGTGTTGATTCTAGACTCCACATCAACACCATTATTAAAAATAGTTTTATATACATCTTTAATCTGCTCTCCGTTAATCGTCGAACGTTCTAGTATACGGAAACACTTTGATTTCTTATCTGTAGTTAATGACTTAGTGATATCACGAAGAATATTACGAATTACATTTCTTGTTTTTCTTGAGAGACCTGATTCAGAACTCAACTCATCGTTCCATAGTTTTTCTACCTCAGTGGCGAAATCTTTGGAGTGTTTGGTACATTTTTTAATAAGGTTCGTGACCTCCATTTATTATAAAAAATTTTTATTCAAAACTTAACTACAATCGGGTAATTATAAGATCGTAGATCTCCCATGTGGTGGAATACTACCATCTATCGCATGTTGATGGACAGTAGGTAATGGCCATGGGAACAGTAATGTTTGCATATTTATCTTTCTTGCTAGTATTACTGTATATTGTCCCAATGTTTCAATGTGATGTTCTAACCTATCAACTGCCACTGTATCACAACAATAATGTCCGATGGAATTAAGATTATGATCCAATGTGTTTAATATATGATTATCAAGTCTTATACGATCCGCGTCTTTTTTGACAGGATCAGGCTCTTCTACAGGAAGAAGTTTCGGCCTCTTATGATAGTCATGTCTGGGAAATGGTGGTCTAGGATGAGGATCAGGTATCGGAGTAGGTTTTGGATTATAATCATCATCATCATCATCATCATCATCATCATCATCATCATCTATGATATCATCATCTTCGATATTTATTTTGCTATCTTGATAATGTCCTATATCAGTTGCATTTTCAGCACTCGACTGAATATTAGTACATTCATTGTCTATTATTAACGTATTTCTAAACCCAAAATGTATATGTTGAACATCACTACTATAGTTGATGAGTCTTATAGCATGAATTCGCTTATCGTTATCGGGTTTATCTTCTGTCACCTTAACAATTCCTTTTTTATTAAACTCTGCATAATCATAACCATTTCTATTGTTTGTTCTAATATAAACGAGTATAGCATCATTGCTAAATTTTTCAATAGTATCAAAAACAGAATATCCTAAACCATATAATATATATTCAGGAACACTCAAACTAAATGTCCAGGATTCTCCTAAATACGTAAACTTTAATAGTGCTAAATCATTCAAAAATCTACCGCTTATAGATAGATAGTACATGAATGCGTATAGTAGTCTACCTATCTCTTTATTATGAAAACCGACATTACGATCATATATTTCGTGATATACATGTGACCCGTTTACGTTAAACCATAAATACATGGGTGATCCTATAAACATGAATTTATTTCTAATTCTCAGAGCCATAGTTAATTGACCGTGTAATATTTGTTTACATGCATACTTGATACGATCATTAATAAGATTTTTATCATTGCTCGTTATTTCAGAATCGTATATATAAGGAGTACCATCATGATTCTTACCAGATATTATACAAAATACTATATATAAAATATATTGACCCACGTTAGTAATCATGTAAATGTTTAATGTTTTAAATTTTGTATTTAATGATCCATCATCATATGCTAGCATGGTCTTGTGATATTCATTCTTTAAAATATAATATTGTGTTAGCCATTGCATTGGAGCTCCTAATGGAGATTTTCTATTCTCGTCCATTTTAGGATATGCTTTCATAAAGTCCCTAATAACTTCGTGAATAATGTTTCTATGTTTTCTACTGATGCATGTATTTGCTTCGATTTTTTTATCCCATGTTTCATCTATCATAGATTTAAACGCAGTAATGCTCGCAACATTAACATCTTGAACCATTGGTACAATTCCGTTCCATAAATTTATAATGTTCGCCATTTATATAACTCATTTTTTGAATATACTTTTAATTGAACAAAAGAGTTAAGTTACTCATATGGATGCCGTCCAGTCTGTACATCAATCTTTTTAGCCAGAGATATCATAGCCGCTCTTAGAGTTTCAGCGTGATTTTCCAACCTAAATAGAACTTCATCGTTGCGTTTACAACACTTTTCTATTTGTTCAAACTTTGTTGTTATATTAGTAATCTTTTTTTCCAAATTAGTTAGCCGTTGTTTGAGAGTTTCCTCATTATCGTCTCCATAGGCTTTAACAATTGCTTCGCGTTTAGTCTCTGGATTTTTAGCAGCCTTTGTAGAGAAAAATTCAGTTGCTGGAATTGCAAGATCGTCATCTCCGGGGAAAAGAGTTCCGTCCATTTAAAGTACAGATTTTAGAAACTGACACTCTGTGTTATTTATATTTGGCGCAATACATGGATTATAAATATCGATGTTAATAACATCAGAAAATGTAAAGTCTATACATTGTCGCATCGTGTTAAATTTTCTAATGGATCTAGTATTATTGGGTCCAACTTCTGCCTGAAATCCAAATATGGAAGCGGATACAAAACCGTTTCCTGGATAAACCACACATCTCCACTTTTGCTTTACATCAGAAATTGTGTCATTGACATCTTGAACTCTCCTATCTAATGCCGGTGTTCCACCTATAGATTTTGAATACTCGAATGCTGCATGAGTAGCATTGAATTCCTTAATATTGCCATAATTTTCATATATTGAGTAACTCTGGATAAAAAGTAAACACACCGCAGCCGTCGCTACTACAATAAAAAAAATTGATAGAGAGTTCATTTATAATCTATTAGAAGCTGATAAAATTTTTTTACACGCGTCAGACAATGCTTTAATAAATAGTTCAACATCTACTTTTGTCATATCGAACCGATGGTATGATTCTAACCTAGAATTACATCCGAAAAAGTTGACTATGTTCATAGTCATTAAGTCATTAACGAACAACATTCCAGACTCTGGATTATAAGACGATACTGTTTCGTCACAATCACCCACCTTAATCATGTGATTATGAATATTGGCTATTAGAGTACCTTCTAAGAAATCTATAATATCTTTGAAACACGATTTAAAATCAAACCACGAATATACTTCTACGAAGAAAGTTAGTTTACCCATAGGAGAGATAACTATAAATGGAGATCTAGATACAAAATCCGGATCTATGATAGTTTTAACATTATTATATTCTCTATTAAATACCTCCACATCTAAAAATGTTAATTTTGAAACTATGTCTTCGTTTATTACCGTACCTGAACTAAACGCTATAAGCTCTATTGTTTGAGAACTCTTTAAACGATATTCTTGAAATACATGTAACAAAGTTTCCTTTAACTCGGTCGGTTTATCTACCATAGTTACAGAATTTGTATCCTTATCTATAATATAATAATCAAAATCGTATAAAGTTATATAATTATCGTGTTCAGATTGTGATCTTTTCAAATAGACTAAAAACCCCATTTCTCTAGTAAGTATCTTATGTATATGTTTGTAAAATATCTTCATGGTGGGAATATGCTCTACAGCAGTTAGCCATTCCTCATTGACAGCTGTAGATGTATTATACAAAACTACTCCAATGTTTAACAAGGGCCATTTTACGAGATTATTAAATCCTTGTTTGATAAATGTAGCCAATGCGGGTTCGAGTTCAACGACGATTGAATTCTCTTCCCGTGGATGCTGCATGATGAACGACGGGATGTTGTTGTTCTATTGATTTGGAATTCTTTTTCGACTTTTTGTTTATATTAAATATTTTAAAATTTATGGCTGATAGTAATTCATGTACTACGGATAATGTAGACGTGTATTGCATATCGATATCTTTATTATTAGATAAATTTATCAATAAATGTGAGAAGTTTGCCTCGTTAAGGTCTTCCATTTAAATATTATATAAATATTTGTGTTTGTATTTTATTCGTCTTTTATGGGATAGTTTTTAACTAGTAAAGCTGTAATTACATACTTTGTCCGTAAAACATAAATATAAATACCCGCTTTTATCAAACGTTCCAAAAAGTCGGCAGCTGACATTTTTAACATGACATCTATTTTAAATACACTTAGGTTTTTAGAAAAAACATCATTTTATAATTGTAACGATTCAATAACTAAAGAAAAGATTAAGATTAAACATAAGGGAATGTTATTTGTATTTTATAAGCCAAAGCATTCTACCGTTGTTAAATACTTGTCTGGAGGAGGTATATATCATGATGATTTGGTTGTATTGGGGAAGGTAACAATTAATGATCTAAAGATGATGCTATTTTACATGGATTTATCATATCATGGAGTGACAAGTAGTGGAGCAATTTACAAATTGGGATCGTCTATCGATAGACTTTCTCTAAATAGGACTATTGTTACAAAAGTTAATAACAATTATAACAATTATAACAATTATAACAATTATAATTGTTATAATAATTATAATTGTTATAATTATGATGATACATTTTTTGACGATGATGATTGATCACTATTACACAATTTTGTTTTTGTACTTTCTAATATAGTGTTTAGGTTCTTTTTCATATGAGAATATTGACTTACTAAAATATCTATGTTTAACTTTTGTTCTATAACGTCCTTATCGGCGGTATCGGTACATATACGTAATTCACCTTCACAAAATACGGAGTCTTCGATAATAATAGCCAATCGATTATTGGATCTAGCTGTCTGTATCATATTCAACATGTTTAATATATCCTTTCGTTTCCCCTTTACAGGCATCGATCGTAGCATATTTTCCGCGTCTGAGATGGAAATGTTAAAACTGCAAAAATGCGTAATGTTAGCCCGTCCTAATATTGGTACGTGTCTATAAGTTTGGCATAGTAGAATAATAGACGTGTTTAAATGCCTTCCAAAGTTTAAGAATTCTATTAGAGTATTACATTTTGATAGTTTATCACCTACATCATCAAAAATAAGTAAAAAGTGTGCTGATTTTTTATGATTTTGTGCGACAGCAATACATTTTTCTATGTTACTTTTAGTTCGTATCAGATTATATTCTAGAGCTTCCTGACTACTAACGAAATTAATATGATTTGGCCAAATGTATCCATCATAATCTGGGTTATAAACGGGTGTAAACAAGAATATATGTTTATATTTTTTAACTAGTGTAGAAAACAGAGATAGTAAATAGATAGTTTTTCCAGATCCAGATCCTCCTGTTAAAACCATTCTAAACGGCATTTTTAATAAATTTTCTCTTGAAAATTGTTTTTCTTGAAAACAATTCATAATTATATTTACAGTTACTAAATTAATTTGATAATAAATCAAAATATGGAAAACTAAGGTCGTTAGTAGGGAGGAGAACAACGAAGGCATATCGTGATATAAATAACATTTATTATCATGATGACACCAGAAAACGACGAAGAGCAGACATCTGTGTTCTCCGCTACTGTTTACGGAGACAAAATTCAGGGAAAGAATAAACGCAAACGCGTGATTGGTCTATGTATTAGAATATCTATGGTTATTTCACTACTATCTATGATTACCATGTCCGCGTTTCTCATAGTGCGCCTAAATCAATGCATGTCTGCTAACAAGGCTGCTATTACTGACTCCGCTGTTGCCGTTGCTGCGGCATCATCTACTCATAGAAAGGTTGTGTCTAGCACTACACAATATGATCACAAAGAAAGCTGTAATGGTTTATATTACCAGGGTTCTTGTTATATATTACATTCAGACTATAAGTCATTCGAGGATGCTAAAGCAAACTGCGCTGCGGAATCATCAACACTACCCAATAAATCCGATGTCTTGACTACCTGGCTCATTGATTATGTTGAGGATACATGGGGATCTGATGGTAATCCAATTACAAAAACTACATCCGATTATCAAGATTCTGATGTATCACAAGAAGTTAGAAAGTATTTTTGTACATAAATAAATGAAATCGCTTAATAGACAAACTGTAAGTAGGTTTAGGAAGTTGTCGGTGCCGGCCGCTATAATGATGTTACTCTCAACCATTATTAGCGGCATAGGAACATTTCTGCATTACAGAGAAGAACTGATGCCTAGTGCTTGCGCCAATGGATGGATACAATACGATAAACATTGTTATCTGGATACCAACATTAAAATGTCTACGGATAATGCAGTTTATCAGTGTCGCAAATTACGAGCTAGATTGCCTAGACCTGATACTAGACATCTGAGAGTATTGTTTAGTATTTTTTATAAAGATTATTGGGTAAGTTTAAAAAAGACCAATGATAAATGGTTAGATATTAATAATGATAAAGATATAGATATTAGTAAATTAACAAATTTTAAGCAACTAAACAGCACGACGGATTCTGAGGCGTGTTATATATACAAGTCTGGAAAACTGGTTAAAACAGTATGTAAAAGTACTCAATCTGTACTATGCGTTAAAAGATTCTACAAGTGACAACAAAAAATGAATTAATAGTAAGTCGTTAACGTACGCCGCCATGGACGCCGCGTTTGTTATTACTCCAATGGGTGTGTTGACTATAACAGATACATTGTATGATGATCTCGATATCTCAATCATGGACTTTATAGGACCATACATTATAGGTAACATAAAAATTGTCCAAATAGATGTACGGGATATAAAATATTCCGACATGCAAAAATGCTACTTTAGCTATAAGGGTAAAATAGTTCCTCAGGATTCTAATGATTTGGCTAGATTCAACATTTATAGTATTTGTACAGCATACAGATCAAAAAATACCATCATCATAGCATGCGACTATGATATCATGTTAGATATAGAAGGTAAACATCAACCATTTTATCTATTCCCATCTATTGATGTTTTTAACGCTACAATCATAGAAGCGTATAATCTGTATACAGCTGGAGATTATCATCTGATCATCAATCCTTCAGATAATCTGAAAATGAAATTGTCGTTTAATTCTTCATTTTGTATATCAGACGGCAATGGATGGATTATAATTGATGGGAAATGTAATAGTAATTTTTTATCATAAAAGTTGTAAAGTAAATAATAAAACAATAAATATTGAACTAGTAGTATGTTGTATATTGAGCAATCAGAGATGATGCTGGTACCTCTTATCACGGTGACCGTAGTTGCGGGAACAATATTAGTATGTTATATATTATATATTTGTAGGAAAAAGATACGTACTGTCTATAATGACAATAAAATTATCATGACAAAATTAAAAAAGATAAAGAGTCCTAATTCCAGCAAATCTAGTAAATCAACTGATAGCGAATCAGACTGGGAGGATCACTGTAGTGCTATGGAACAAAACAATGACGTAGATAATATTTCTAGAAATGAGATATTGAACGATGATAGCTTCGCTGGTAGTTTAATATGGGATAACGAATCCAATATCATGGCGCCTAGCACAGAACACATTTACGATAGTGTTGCTGGAAGCACGCTGCTAATAAATAATGATCGTAATGAACAGACTATTTATCAGAATACTACAGTAGTAATTAATGATACAGAGACTGTTGAAATACTTAATGAAGATACCAAACAGATTCCTAGCTATTCTTCCAATCCTTTCGTAAATTATAATAAAACCAGTATTTGTAGCAAGTCAAATCCGTTCATTGCAGAACTCAACAATAAATTTAGTGATAATAATCCGTTTAGGAGAGCACATAGTGACGATTATCTTAATAAGCAACAAGATCATGAATACGATGATATAGAATCATCGGTTGTATCATTGGTCTGATTAGTTTCCTTTTTATAAAATTGAAGTAATATTTAGTATTAATTACCGCCGATGCATTATACAAATATGGAGATATTCCCTGTATTCGGCATTTCTAAAATTAGCAATTTTATTGCTAATAATGACTGTAGATATTATATAGATGTAGAGCATCAAAAAATTATATCTGATGAGATCAATAGACAGATGGATGAAACGGTACTTCTTACCAACATCTTAAGCGTAGAAGTTGTAAATGACAATGAGATGTACCATCTTATTCCCCATAGACTATCGACTATTATACTCTGTATTAGTTCTGTTGGAGGATGTGTTATCTCTATAGATAATGACGTCAATGACAAAAATATTCTAACATTTCCCATTGATCATGCTGTAATCATATCCCCACTGAGTAAATGTGTCGTAGTTAGCAAGGGCCCTACAACCATACTGGTTGTTAAAGCGGATATACCCAGCAAACGATTGGTAACATCGTTTACAAACGACATACTGTATGTAAACAATCTATCACTGATTAATTATTTACCGTCGTCTGTATTCATTATTAGACGAGTCACCGACTATTTGGATAGACACATATGTGATCAGATATTTGCTAATAATAAGTGGTATTCCATTATAACTATCGACGATAAGCAATATCCTATTCCATCAAATTGTATAGGTATGTCTTCTGCCAAGTACATAAATTCGAGCATCGAGCAAGATATTTTGATCCATGTTTGTAACCTCGAGCATCCATTCGACTCAGTCTACAAAAAAATGCAGTCGTACAATTCTCTACCTATCAAGGAACAAATATTGTATGGTAGAATTGATAATATAAATATGAGCATTAGTATTTCGGTGGATTAATAGATTTCTCTAGTATGGGATCATTAATCATCTCTAAATACATCATAAAAAAGCTATTATCAAATACTGTACTGAATGGATTCATTCTTTTCTCTTTTTATGAAACTCTGTTGTATATCTACGGATAAAACTAGAAGCAAAAAATCTGATAGGAAGAATAATGATTATATGGAGGAACACGATTATTATAAAATAACAATAGTTCCTGGTTCCTCTTCCACGTCTACTAGCTCGTGGTATTATACACATGCCTAGTAATAGTCTCTTTGCGTTGACGGAAAGCAGACTAGAAATAACAGGCCAAAATGTTCAGACACCATAATAGTTCCCAACCCAGATAATAACAGAGTTCCATCAACACATTCCTTTAAACTCAATCCCAAACCCAAAACCGTTAAAATGTATCCAGCCAATTGATAGTAGATAATGAGGTGTACAGCACATGATAATTTACACAGTAACCAAAATGAAAACACTTTAGTAATTATAAGAAATATAGACGGTAATGTCATCATCAACAATCCAATAATATGCCTGAGAGTAAACATTGACGGATAAAACAAAAATGCCCCGCATAACTCTATCATGGCAATAACGCAACCAAACACTTGTAAAATTCCTAAATTAGTAGAAAATACAACTGATATCGATGTATAAGCGATTTCGAGGAATAATAAGAACAAAGTAATTCCCGTAAAGATAAACATCAACATTGTTTGGTAATCATTAAACCAATTAGTATGACGTTGAATTAATTTCACAGTATATTTTATTCCAGTATTATCCCCGCATGTATACGTACCTGGTAAGATATCTTTATATTCCATAATCAATGAGACATCACTATCCGATAACGAATGAAGTCTAGCACTAGTATGCCATTTACTTAATATGGTCGTCTTGGAAGTTTTATTATAAGTTAAAATATCATGATTGTCCAATTTCCATCTAATATACTTTGTCGGATTATCTATAATACACGGAATAATGATGGTATCATTACATGCTGTATATTCTATAGTCTTTGTAGATGTTATAACCACAAAAGTACAGAGGTATATCAACAATATTCTAACTCTTAACATTTTTATTTATTTAAAATGATACCTTTGTTATTTATTTTATTCTTATTTTGCTAACGGTATCGAATGGCATAAGTTTGAAACGAGTGAAGAAATAATTTCTACTTACTTAATAGATTATGTGGTAACGGGTGTTATTAATGGGGATGTATATACATTTTCAAATAATGAACTAAACAAAACTGGGTTAACTAATAACAATAATTATATCACAACATCTATAAAAGTAGAGGATAAGGATACATTAGTAGTATGCGGAACCAATAACGGAAATCCCAAATGTTGGAAAATAGACGGTTCATACTACCCAAAACATATAGGTAGAGGATACGATCATCAAAATAGCAAAGTAACGATAATCAGTCACAATGAATGTGTACTATCCGACATAAACATATAAAAAGAAGGAATTAAACGATGGAGAAGATTTGACGGACCATGTGGTTATGATTAAACGAGTTAAGTTTTTTAAGAAGCCTTAGAAGAGAGGCTATTGGGTATGAGAATCCGAAATATTAAACCAGACAACCCCATATAATTTTATAGCTAAGAATGCCGCGAAGAATGGAACTAATAAAAACGGAAATATTTGTAGCACAACGAATAACTCCCAAACTGCATTCATGTTACACTATATAACACTACTTCGGTTAGATGTTTTAGAAAAAATAAATATCACCGTACCGTTTTGTTGTATAAAAATAACAATTAACAATTATCAATTTTTTTCTTTAATATTTTACGTGGTTGACCATTCTTGGTGGTAAAATAATCTCTTAGTGTTGGAATGGAATGCTGTTTAATGTTTCCACACTCATCGTATATTTTGACGTATGCAGTCACATCGTTTACGCAATAGTCAGACTGTAGTTCTATCATGCTTCCTACGTTAGAAGGAGGAACAGTTTTAAAGTCTCTTGGTTTTAATCTATTGTCATTAGTTTTCATGAAATCCTTTGTTTTATCCACTTCACATTTTAAATAAATGTCCACTATACATTCTTCTGTTAATTTTACTAGATCATCATGAGTCATAGAATTCATAGGTTCCGTAGTCCATGGATCCAAACTAGCAAACTTCGCGTATACGGTATCGCGATTAGTGTATACACCAACTGTATGAAAATTAAGAAAACAGTTTAATAAATCTACAGAAATATTTAATCCTCCGTTTGATACAGATGCGCCATATTTATGGATTTCGGATTCACACGTTGTTTGTCTAAGGGGTTCGTCTAGTGTTGCTTCTACATAGACTTCGATTCCCATATATTCTTTATTGCCAGAATCACATACCGATTTATCATACGCTGGTTCACTTGTTTGAAAACTAAATGGTAGTAGATACATCAAAATAATAAATAATAAGTACATTCTGCAATATTGTTATCGTAATTGGAAAATTGGTATTCAAGTGAGCTGGATTATGTGAGTATTGGATTGTATATTTTATTTTATATTTTATATTTTATATTTTATTTTATATTTTATATTTTATTTTATATTTTGTAGTAAGAATAGAATGCTAATGTCAAGTTTATTCGAATAGATGTCTTATTAAAAAACATATATAATAAATAACAATGGCTGAATGGCATAAAATTATCGAGGATATCTCAAAAAATAATAAGTTCGAGGATGCCGCCATCGTTGATTACAAGACTACAAAGAATGTTCTAGCGGCTATTCCTAACAGAACATTTGCAAAGATTAATCCGGGTGAAGTTATTCCCCTCATCACTAATCATAATATTCTAAAACCTCTTATTGGTCAGAAATTTTGTATTGTATATACTAACTCTCTAATGGATGAGAACACGTATGCTATGGAGTTGCTTACTGGGTACGCCCCTGTATCTCCGATCGTTATAGCGAGAACTCATACCGCACTTATATTTTTGATGGGTAAGCCAACAACATCCAGACGTGATGTGTATAGAACATGTAGAGATCACGCTACCCGTGTACGTGCAACTGGTAATTAAAATAAAAAGTAATATTCATATGTAGTGTCAATTTTAAATGATGATGAAATGGATAATATCCATATTGACGATGTCAATAATGCCGGTATTGACATACAGCTCATCGATTTTTAGATTTCATTCAGAGGATATTGAATTATGTTATGGGAATTTGTATTTTGATAGGATCTATAATAATGTAGTAAATATAAAATATATTCCTGAGCATATTCCATATAGATATAATTTTATTAATCGTACGTTCTCCGTAGATGAACTAGATGATAATGTCTTTTTTACACATGGTTATTTTTTAAAACACAAATATGGTTGTTCACTTAATCCTAGTTTGATTGTCTCATTATCAGGAAACTTAAAATATAATGATATACAATGCTCAGTAAATGTATCGTGTCTCATTAAAAATTTGGCAACGAGTACATCTACTATATTAACATCTAAACATAAGACTTATTCTCTATATCGGTCCATGTGTATTGCTATAATAGGATACGATTCTATTATATGGTATAAATATATAAATGACAGGTATAATGACATCTATGATTTTACTGCAATATGTATGCTAATAGCGTCTACATTGATAGTGATCATATACGTGTTTAAAAAAATAAAAATGAACTCTTAATTATGTTATACTATTAGAAATGGATAAAATCAAAATTACGATTGATTCAAAAATTGGTAATGTTGTTACCATATCGTATAACTTGGAAAAGATAACTATTGATGTCACGCCAAAAAAGAAAAAAGAAAAGGATGTATTATTAGCGCAATCAGTTGCTGTCGAAGAGGCAAAAGATGTCAAGGTGGAAGAAAAAAATATTATCGATATTGAAGATGACGATGATATGGATATAGAAAACACGTAATACGATCTATAAAAATAAGTATTAAATACTTTTTATTTACGGTACTCTTGTAGTGGTGATACCACTAATCGATTATTTTTTTTAAAAAAATACTTATTCTGATTCTTCTAGCCATTTCCGTGTTCGTTCGAATGCCACATCGACGTCAAAAATAGGGGAGTAGTTGAAATCTAGTTCTGCATTGTTGGTACGCACCTCAAATGTAGTGTTGGATATCTTCAACGTATAGTTGTTGAGTATTGATGGTTTTCTAAATAGAATTCTCTTCATATCATTCTTGCACGCGTACATTTTTAGCATCCATCTTGGAATCCTAGATCCTTGTTCTATTCCCAATGGTTTCATCAATAGAAGATTAAACATATCGTAAGAACACGATGGAGAGTAATCGTAGCAAAAGTAAGCATTTCCTTTAATCGCAGATCCCGGATACTGGATATATTTTGCAGCCAACACGTGCATCCATGCAACATTTCCTACATATACCCGGCTATGCACAGCGTCATCATCGACTGTACGATACATAATGTTACCGTGTTGCTTACATTGCTCGTAAAAGACTTTCGTCAATTTGTCTCCTTCTCCGTAAATTCCAGTGGGTCTTAGGCAACAAGTATACAATTTTGCGCCATTCATGATTACGGAATTATTGGCTTTCATAACCAGTTGCTCGGCCATACGTTTACTTTTTGCGTATACATGTCCTGGTGATATATCATAAAGGGTATGCTCATGACCGATGAATGGATTACCGTGTTTATTTGGTCCTATTGCTTCCATGCTACTAGTATAGATCAAATACTTGATTCCTAGGTCCACACAAGCTGCCAATATAGTCTGTGTTCCATAATAGTTTACTTTCATGATTTCATTATCAGTGTATTTTCCAAATACATCCACTAGAGCAGCCGTATGAATAATCAGATTTACCCCATCTAGCGCTTCTCTCACCTTATCAAAGTCGTTTATATCACATTGTATATAGTTTATAACCTTAACTTTCGAGGTTATTGGTTGTGGATCTTCTACAATATCTATGACTCTTATTTCTTGAACATCATCTGCGCTAATTAAAAGTTTTACTATATACCTGCCTAGAAATCCGGCACCGCCAGTAACCGCGTACACGGCCATTGCTGCCACTCATAATATCAGACTACTTATTCTATTTTACTAAATAATGGCTGTTTGTATAATAGACCACGATAATATCAGAGGAGTTATTTACGTTGAACAAGTCCATGGAAAAGATAAAGTTTTAGGATCAGTTATTGGATTAAAATCCGGAACGTATAGTTTGATAATTCATCGTTACGGAGATATTAGTCGAGGATGTGATTCCATAGGCAGTCCAGAAATATTTATCGGTAACATCTTTGTAAACAGATATGGTGTAGCATATGTTTATTTAGATACAGATGTAAATATATCTACAATTATTGGAAAGGCGTTATCTATTTCAAAAAATGATCAGAGATTAGCGTGTGGAGTTATTGGTATTTCGTACATAAATGAAAAGATAATACATTTTCTTACAATTAACGAGAATGGCGTTTGATATATCAGTTAATGCGTCTAAAACAATAAATGCATTAGTTTACTTTTCTACTCAGCAAGATAAATTAGTCATACGTAATGAAGTTAATGATATACACTACACTGTCGAATTTGATAGGGACAAAGTAGTTGATACGTTTATTTCATATAATAGACATAATGACTCCATAGAGATAAGAGGGGTGCTTCCAGAGGAAACTAATATTGGTCGCGTGGTTAATACGCCGGTTAGTATGACTTACTTGTATAATAAGTATAGTTTTAAACCGATTTTAGCAGAATATATAAGACACAGAAATACTATATCCGGCAACATTTATTCGGCATTGATGACGCTAGATGATTTGGTTATTAAACAGTATGGAGACATTGATCTATTATTTAATGAGAAACTTAAAGTAGACTCCGATTCGGGACTATTTGACTTTGTCAACTTTGTAAAGGATATGATATGTTGTGATTCTAGAATAGTAGTAGCTCTATCTAGTCTAGTATCTAAACATTGGGAATTGACAAATAAAAAGTATAGGTGTATGGCATTAGCCGAACATATAGCTGATAGTATTCCAATATCTGAGCTATCTAGACTACGATACAATCTATGTAAGTATCTACGCGGACACACTGATAGCATAGAGGATGAATTTGATCATTTTGAAGACGATGATTTGTCTACATGTTCTGCCGTAACCGATAGGGAAACGGATGTATAATTTTTTTTATAGTATGAAGGATATGATGGATATGATGATATGATGGATATGATGATATGATGGATATGATGGATATGATGGATATGATGGATATGATAAAAAAATATAATTGTTGTATCCATTCCCATTCAAATCACCTTATATGATTCTGTAACACAATGAAGGAGTCTCATAGATATATAGAGGTCAGATACTGGTTTGATAAACTTTTTATTCCACATGAGCATGTTTGACTTATGGTTAGACACACATACTTTAACAAATCACTGAAAATTGGAGTTAGGTATTCCTCTCAGAATCAGTTGCCGTTCTGGAACATTAAATGTATTTTTTATGATATACTCCAACGCATTTATGTGGGTATACAACAAGTCATTAATAATGAGTATTTCCAAGAGTTTTAGTTGTCTAGTATTTAACAAGAGAAGAGATTTCAACAGACTGTTTATGAACTCGAATACCGCCTCATTGTCGCTTATATTGATGACATGACATCGAATTCCCAATATCAATCTCATCAGTGATGAGTAGCTCAATCTTGTTATCGGGATCCAATTTCTAAAGATGTCATTAAACCCTCGATCGTGAATGGATTTATCATCATCGTTTTTATGTTGGACATGAGCTTAGTCCGTTTGTCCACATCTATATACGATGATTTCTGAATTATTTCATATATCTCTCGTTAACTCCAGGAACTTGTCAGGGATCTAACTTTAATATGTTCTCGTCTAAGAGATGAAAATCTTTGGATGGTTGCATGTGACTTTTCTCTAAAGGATGATGTTACCCGATCCTCTCTTAAATGACTCCATCTTATCCTTGGACAAGATGGACAGTCTATTTTCCTTAGATGGTTTAATATTTTTTACCCATGATCTATAAAGGTAGACAGACCTAATCGTCTCGGATGACCATATATTATTTTCAGTTTTATTATACGCATAAATTGTAAAAAATATGTTAGGTTTACGAAAATGTCTCGTGGGGCATTAATCGTTTTTGAAGGATTGGACAAATCTGGAAAAACAACACAATGTATGAACATCATGGAATCTATACCGGCAAACACGATAAAATATCTTAACTTTCCTCAGCGATCCACAGTCACTGGAAAGATGATAGATGACTATCTAACTCGTAAAAAAACCTATAATGATCATATAGTTAATCTATTATTTTGTGCAAATAGATGGGAGTTTGCATCTTTTATACAAGAACAACTAGAACAGGGAATTACTTTAATAGTTGACAGATACGCGTTCTCTGGAGTAGCGTATGCCACCGCTAAAGGCGCGTCAATGACTCTCAGTAAGAGTTATGAATCTGGATTGCCTAAACCCGACTTAGTTATATTCTTGGAATCTGGTAGCAAAGAAATTAATAGAAACATCGGCGAGGAAATTTATGAAGATGTTGAATTCCAACAAAAGGTATTACAAGAATATAAAAAAATGATTGAAGAAGGAGATATTCATTGGCAAATTATTTCTTCTGAATTCGAGGAAGATGTAAAGAAGGAGTTGATTAAGAATATAGTTATAGAGGCTATACACACGGTTACTGGACCAGTGGGGCAACTGTGGATGTAATAAAATGAAATTACATTTTTATAAATAGATGTTAGTACAGTGTTATAAATGGATGAAGCATATTACTCTGGCAACTTGGAATCAGTACTCGGGGATACGTGTCCGATATGCATACCGAACTCGCATCAATATCTCAATTAGTTATTGCCAAGATAGAAACTATAGATAATGATTATTAAACAAGGACATTGTAAATTTTATCATGTGTAGATCAAACTTGGATAATCCATTTATCTCTTTCCTAGATACTGCATATACTATCATAGATCAAGAGATCTATCAGAACGAGTTGATTAATTCATTAGACGATAATGAAATTATCGATTGTATAGTTAACAAGTTTATGAGCTTTTATAAGGATAACCTAGAAAATATGGTAGATGCTATCATTACTCTAAAATATTATAATTAATAATCCAGATTTTAAAACTACGTATGTGGAAGTACTCGGTTCCAGAATAGCTGATATAGATATTAAACAAGTGATACGTAAGAATATAATACAATTGTCTAATGATCCGCGAACGATATTTGTGAAAATATTAAAAAAAAATACTTTTTTTATTAAATGACGTCTCTTCGCGAATTTAGAAAATTATGCTGTGATATATATCACGCATCAGGATATAAAGAAAAATCTAAATTAATTAGAGACTTTATAACAGATAGAGATGATACCGATACATATTTGATCATTAAGCTATTGCTTCCCGGATTAGACGATAGAATGTATAACATGAACGATAAACAAATTATAAAATTATATAGTATAATATTTAAACAATCTCAGGAAGATATGCTACAAGATTTAGGATACGGATATATAGGAGACACTATTAGGACTTTCTTCAAAGAGAACACGGAAATCCGTCCACGAGATAAAAGCATTTTAACTTTAGAAGAAGTGGATAGTTTTTTAACTACGTTATCATCAGTAACTAAAGAATCACATCAAATAAAATTATTGACTGATATAGCATCTGTTTGTACATGTAATGATTTAAAATGTGTAGTCATGCTTATTGATAAAGATCTAAAAATTAAAGCGGGTCCTCGGTACGTGCTTAACGCTATTAGTCCTCATGCCTATGATGTTTTTAGAAAATCTAATAACTTGAAAGAGATAATAGAAAATGCAGCTAAACAAAATCTAGACTCTATATCTATTTCTGTTATGACTCCAATTAATCCCATGTTAGCGGAATCATGTGATTCTGTCAATAAGGCGTTTAAAAAATTTCCATCAGGAATGTTTGCGGAAGTCAAATACGATGGTGAAAGAGTACAAGTTCATAAAAAAAATAACGAGTTTGCATTCTTTAGTAGAAACATGAAACCAGTACTCTCTCATAAAGTGGATTATCTCAAAGAATACATACCGAAAGCATTTAAAAAAGCTACGTCTATCGTATTGGATTCTGAAATTGTTCTTGTAGACGAACATAATGTACCGCTACCGTTTGGAAGTTTAGGTATACACAAAAAGAAAGAATATAAAAACTCTAACATGTGTTTGTTCGTATTTGACTGTTTATACTTTGATGGATTCGATATGACAGACATTCCATTGTATGAACGAAGATCTTTTCTCAAAGATGTTATGGTCGAAATACCCAATAGAATAGTATTCTCAGAGTTGACGAATATTAGTAACGAGTCTCAGTTAACTGATGTATTAGATGATGCACTAACGAGAAAATTAGAAGGATTGGTCTTAAAAGATATTAATGGCGTATACGAACCGGGAAAGAGAAGATGGTTAAAAATAAAGCGAGACTATTTGAACGAGGGTTCCATGGCAGATTCTGCCGATTTAGTAGTACTAGGTGCCTACTATGGTAAAGGAGGAAAGGGTGGTATCATGGCAGTCTTTCTAATGGGTTGTTACGACGATGAATCCGGTAAATGGAAGACGGTAACTAAATGTTCCGGTCACGATGATAATACGTTAAGGGTTTTGCAAGACCAATTAACGATGGTTAAAATTAACAAGGATCCCAAAAAAATTCCAGAGTGGTTGGTAGTTAATAAAATCTATATTCCCGATTTTGTAGTAGATGATCCGAAACAATCTCAGATATGGGAAATTTCAGGAGCAGAGTTTACATCTTCCAAGTCACATACAGCGAATGGAATATCGATTAGATTTCCTAGATTTACTAGGATTAGAGAAGATAAAACGTGGAAAGAATCTACTCATCTAAACGATTTAGTAAACTTGACTAAATCTCTTAATAGTTACATATAAACTGAAAAATAAAATAACACTATTTTAGTTGGTAGTCGCCATGGATGGTGTTATCGTATACTGTCTAAATGCGTTAGTAAAACATGGCGAGGAAATAAATCATATAAAAAATGATTTCATGATTAAACCATGTTGTGAAAGAGTTTGTGAAAAAGTCAAGAACGTTCACATCGGCGGACAATCTAAAAACAATACAGTGATTGCAGATTTGCCATATCTGGATAATGCTGTATCAGATGTATGCAAATCAATATATAAAAAGAATGTATCAAGAATATCCAGATTTGCTAATTTGATAAAAATAGATGACGATGACAAGACTCCTACCGGCGTATATAATTATTTTAAACCTAAAGATGCTATTCCTGTTATTATATCCATAGGAAAGGATAAAGATGTCTGTGAACTATTAATCTCATCTGATAAAGCGTGTGCGTGTATAAAGTTAAATTTATATAAAGTAGCCATTCTTCCCATGGATGTTTCCTTTTTTACCAAAGGAAATGCATCATTGATTATTCTCCTGTTTGATTTCTCTATCGATGCGGCACCTCTCTTAAGAAGTGTAACCGATAATAATGTTATTATATCTAGACACCAGCGCCTACATGACGAGCTTCCGAGTTCCAATTGGTTCAAGTTTTACATAAGTATAAAGTCCGACTATTGTTCTATATTATATATGGTTGTTGATGGATCTATGATGTATGCGATAGCTGATAATAGAACTCACGCAATTATTAGCAAAAATATATTAGACAATACTACGATTAACGATGAGTGTAGATGCTGTTATTCTGAACCACAGATTAGGATTCTTGATAGAGATGAGATGCTCAATGGATCATCGTGTTATATGAACAGACATTGTATTATGATGAATTTACCTGATGTAGGCGAATTTGGATCTAGTATGTTGGGGAAATATGAACCTGACATGATTAAGATTGCTCTTTCGGTGGCTGGTAATTTAATAAGAAATCGAGACTACATTCCCGGGAGACGAGGCTATAGCTACTACGTTTACGGTATAGCCTCTAGATAATTTTTTTTAAGCACGAAATAAAAACATAATTTTAAACAATCTATTTCATACTATTTTGTGTGCTCACCATGAACATAAAGATAGATATATTAGTATTTCTGGTGATAAATTTACGGCGACTGCTAGGAGGGAAAATGAAGAAAGAAAAAATATCTACCTCTCCAAAAAGAAAAACTACTGATGTTATCAAACCTGATTATCTTGAGTACAATGACTTGTTAGATAGAGATGAGATGTCTACTATTCTAGAGGAATATTAGGCCTTAGAATAAAATATGGACGACTCTTAACGAAATTAGAAAATTCGATAATGATGTTGAAGAACAATTCGGTACTATAGAAGAACTCAAGCAGAAGCTTAGATTAAATTCTGAAGAGGGAGCAGATAATTTTATAGATTATATAAAGGTACAAAAACAGGATATCATCAAACTTACTGTATACGATTGCATATATCTATGATAGGATTGTGTGCGTGCGTGGTAGATGTTTGGAGAAATGAGAAACTGTTTTCTAGATGGAAATATTGGTTACAAGCGATTAAACTGTTTATTGATGATCACATGCTTGATAAGATAAAATCTATTGTAGAATAGACTAGTGTATGTGGAAATGTCATAGAAAGTTAAAAGTTAATGAGAGCAAAAATATATAAGGTTGTATTCCATATTTGTTATTTTTTTCTGTAATAGTTAGAAAATACATTCGATGGTCTATCTACCAGATTATTATGTGTTATAAGGTACTTTTCTCATAATAAACTAGAGTATGAGTAAGATAGTGTTTTTCAAAAACATATAAATCTAAAATTGATGGATGAGATATACAGCTATTAATTTCGAAAATATATTTTAATCTGATAACTTTAAACATGGATTTTTGATGGTGGTTTAAGTTTAAAAAAGATTTTGTTATTGTAGTATGATAATATCAAAAAGATGGATATAAAGAATTTACTGACTACATGTACTATTTTACATTACTACATTGGCTACGGCATATATACCTATTTCGTCACTTCCACACGCTCCGGTAAACGGGTGTCATGTGACGAGGGAGAATCTTGATAAGAGGCATAATCAATGTTGTAATCCGATGTCCACCTGGAGAATTTGCCAAGGTCAGATGTAGAGTTGGTAGTGATAACACAAAATGTGAACACTGCCCACCTCATACATATACCGCAATCCCCAATTATTCTAATAGATGTCATCAATGTAGAAAATGCCCAACAGGATCATTTGATAAGGTAAAGTGTACCGGAACACAGAACAAATGTTCGTGTCATCCTGGTTGGTATACGCTACTGATTCTTCACAGACTGAAGATTGTCGAGATTTGTGTACCAAAAAAGGAGATGTCCATGCGGATACTTTGGTGGAATAGATGAAGGAAATCCTATTTGTAAATCGTGTTGTGTTGGTGAATATTGCGACTACCTACGTAATTATAGACTTGATCCATTTCCTCCATGCAAACTATCTATCTAAATGTAATTAATTATGATTTTGATGATAATGTTACCATACATTATATCACTACTTGGTTAGTGTGTATTATTTAGTATGGAAGACCTATTAATAATTACTTATCTTTTGACGATCTTGTTATAATTATAATATAAAAATACTTATGACATAGTAACTCATAATTGCTGACGCGATAAATTCGTAATAATCTGTTTTGTTCAAATTTTTATAAGGAATCTACAGGCATAAAAATAAAAATATAATCTATAATATACTCTTACAACGCCATCATGAATAGCAGTGAATTAATTGCTGTTATTATGGATTTAGAAATAGTGGACGATTTTGTGATATTAATATAGTTATTAATGATGAAAGGATAAACGCGCATAGACTCATCCTATCTGGAGCCTCCGAATATTTTTTCCATTCTGTTTTCCAATAATTTTATCGATTCTAATGAATACGAAGTTAATCTAAGTCATTTAGATTATCAAAGTGTTAACTATTTGATCGATTACATTTATGGGATACCTTTGAGCCTAACTAACAATAACGTGAAATATATTCTTTCAACCGCTGATTTTTTTACAAATTGGATCTGTCATTACTGAGTGCGAAAAATACATACTTAAAAATCTTTGTTTTAGAAACTGTATCGATTTCTACATATACGCTGATAAATATAATAACAAGAAAATAGAATTAGCATCGTTTAACACAATATTACGAAATATTTTGAGACTCATCAACAATGAAAACTTTAAATACTTAACAGAGGAATCAATGATAAAAATTTTAAGCGATGATATGTTATATATAAAAAATGAGGATTTCACCCCACTGATTCTCATTAAATGGTTAGAGAGTACACCAACCATGTACCGTCGAGTTACTTAGATGCCTCAGAATATCATTTCTTTCCCCACAAGTTATAAAATCACTTTATAGTCATCGACTGGTTAGTTCAATCTACGAATGTATAACATTCTTAAACAATATAGCATTCTTGGATAAATCATTTCCTAGATACCATATCATCGAGTTGATATCTATCGGTATAAGTAATTCACATGATAAGATTTCCATAAACTGCTACAATCATAAAAAAATTCATGGGAAATGATATCTTCACGTAGATATAGGTGTAGTTTCGCAGTGACCGTCCTGGATAATATTATCTATATGATGGGTGGATATGATCAGTCCCTGTATAGAAGTTCAAAGGTTATAGCGTACAATACATGTACTAATTCTTGGATATATGATATACCAGAGCTAAAAATATCATCGTTCTAATTGCGGAGGAGTTGCCAATGACGAATACATTTATTGTATAGGCGGTATACGCGATCAGGAGTCATCGTTGATATCTAGTATCGATAGATGGAAGCCATCAAAACCATATTGGCAGAAGTATGCTAAAATGTGCGAACCAAAATGTGATATGGGGTTGCGATTTTAAACGGATTAATATATGTCATAGGTGGAGTCGTTAAAGGTGACACACATATACCAACGCACTAGAGAGTTTATCAGAAGATGGATGGATGAATCATCAACGTCTTCCAATAAAAATGTCCAATATGTCGACGATTGTTCATGCTGGAAAGATTTATATATCTAGAGGTTACAACAATAGTAGTGTAGTTAATGTAATATCGAATCTAGTCCTTAGCTATAATCCGATATATGATGAATGGACCAAATTATCATCATTAAATATTCCTAGAATTAATCCTGCTCTATGGTCAGTGTATAATAAATTATATGTAGGAGGAGTAATATCTGATGATGTTCAAACTAATACATCTGAAACATACGATAAAGAAAAAGATTGTTGGACATTGGATAATGGTCACTTGTTACCACATAATTATATAATGTATAAATGCGAACCGTTTAAACATAGATATCCATTGGAAAAAACACAGTACACGAATGATTTTCTAAAGTATTTGGAAAGTTTTATAGGTAGTTGATAGAACAAAATACATAATTTTGTAAAAATAAATCACTTTTTATACTAATATGACACAATTACCAATACTTTTGTTACTAATATCATTAGTATACGCTACACCTTCTCCTCAGACATCTAAAAAAATAGGTGATGATGCAACTATATCATGTAGTCGAAATAATACAAATTACTACGTTGTTATGAGTGCTTGGTATAAGGAGCCCAATTCCATTATTCTCTTAGCTGCCAAAAGCGACGTCTTGTATTTTGATAATTATACCAAGGATAAAATATCTTACGACTCTCCATACGATGATCTAGTTACAACTATCACAATTAAATCATTGACTGCTGGAGATGCCGGTACTTATATATGTGCATTCTTTATGACATCGACTACAAATGATACTGATAAAGTAGATTATGAAGAATACTCCATAGAGTTGATTGTAAATACAGATAGTGAATCGACTATAGACATAATACTATCTGGATCTACACCAGAAACTATTTCTGAGAAACCAGAGGATATAGATAATTCTAATTGCTCGTCTGTATTCGAAATCACGACTCCGGAACCAATTACTGATAATGTAGACGACCATACAGACACCGTCACATACACTAGTGATAGCATTAATACAGTAAATGCATCATCTGGAGAATCCACAACAGACGAGATTCCGGAACCAATTACTGATAAAGAAGAAGATCATACAGTAACAGACACTGTCTCATACACTACAGTAAGTACATCATCTGGAATTGTCACTACTAAATCAACCACCGATGATGCGGATCTTTATGATACATACAATGATAATGATACAGTACCGCCAACTACTGTAGGTGGTAGTACAACCTCTATTAGCAATTATAAAACCAAGGACTTCGTAGAAATATTTGGTATTACCACATTAATTATATTGTCAGCAGTGGCGATTTTCTGTATTACGTATTATATATGTAATAAACACCCACGTAAATACAAAACAGAGAACAAAGTCTAGATTTTTGACTTACATAAATATCTGGGATAATAAAATCTATCATATTGAGAGGACCATCTGGTTCAGGAAAGACAGCCATAACCAAAAGACTGTTAAAAGACTATGGGAATATATTTGGATTTGTGGTGTCCCATACCACTAGATTTCCTCGTCCTATGGAACGAGAAGGTGTTGATTACCTTACGTTAACAGAGAGGCCATCTGGAAGGGAATAGCCGCCGGAAACTTTCTAGAACATACTGAGTTTTTAGGAAATATTTACGGAACTTCTAAAACAGCTGTAAATACAGCGGTTATTAATAATCGTATTTGCGCGATGGATTTAAACATCAACGGTGTTAGAAGTCTTAAAAATACTTACCTAATGCATTACTTGGGTATATAAGACCTACCTCTCTTAAAATGGTTGAGACCAATCTTCGTCGTAGAAACACTGAAGCGGACGACGAATCTCATCGTCGCGTGATGTTGGCAAAAAACGGATATGGATGAGGTCAACGAAGCAGGTCTATTCGACACTATTATTATTGAAGATGATGTGAATTTAGCATATAGTAAGTGTTAATTCAGATACTACAGGACCGTATTAGAATGTATTTTAACACTAATTAGAGACTTAAGATTTGACTTAAAACTTGATAATTAATAATATAACTCGTTTTTATATGTGGCTATTTCAACGTCTAATGTATTAGTTAAATATTAAAACTTACCACGTAAAACTTAAAATTTAAAATGGTATTTCATTGACAGATCATACATTATGAAGTTTCAAGGACTTGTGTTAATTGACAATTGCAAAAATCAATGGGTCGTTGGACCATTAATAGGAAAAGGTGGATTCGGTAGTATTTATACTACTAATGACAATAATTATGTAGTAAAAATAGAGCCCAAAGCTAACGGATCATTATTTACCGAACAGGCATTTTATACTAGAGTACTTAAACCATCCGTTATCGAAGAATGGAAAAAATCTCACAATATAAAGCACGTAGGTCTTATCACATGCAAGGCATTTGGTTTATACAAATCCATTAATGTGGAATATCGATTCTTGGTAATAAATAGATTAGGTGCAGATCTAGATGCGGTGATCAGAGCCAATAATAATAGACTACCAGAAAGGTCGGTGATGTTGATCGGAATCGAAATCTTAAATACCATACAATTTATGCACGAGCAAGGATATTCTCACGGAGATATTAAAGCGAGTAATATAGTCTTGGATCAAATAGATAAGAATAAATTATATCTAGTGGATTACGGATTGGTTTCTAAATTCATGTCTAACGGCGAACATGTTCCATTTATAAGAAATCCAAATAAAATGGATAACGGTACTCTAGAATTTACACCTATAGATTCGCATAAAGGATACGTTGTATCTAGACGTGGTGATCTAGAAACACTTGGATATTGTATGATTAGATGGTTGGGAGGTATCTTGCCATGGACTAAGATATCTGAAACAAAGAATTCTGCATTAGTAAGTGCCGCAAAACAGAAATATGTTAACAATACTGCGACTTTGTTAATGACCAGTTTGCAATATGCACCTAGAGAATTGCTGCAATATATTACCATGGTAAACTCTTTGACATATTTTGAGGAACCCAATTACGACGAGTTTCGTCGAGTATTAATGAATGGAGTTATGAAAAATTTTTGTTGATAAAAAAATTAAAAAAATAACTTAGTTATTATCACTCTCGTGAGTACAATAGAAACATGGCGATGTTTTACGCACACGCTTTCGGTGGGTACGACGAGAACCTTCATGCATTTCCTGGAATATCATCGACGGTTGCCAATGATGTCAGGAAATATTCTGTTGTGTCAGTTTATAATAAAAAGTATAACATTGTAAAAAACAAATATATGTGGTGTAACAGTCAAGTGAACAAGAGATATATTGGAGCACTACTGCCTATGTTTGAATGCAATGAATATCTACAAATTGGAGATCCAATCCATGATCTAGAAGGAAATCAAATCTCTATTGTCACATATCGCCACAAAAACTACTATGCTCTAAGTGGAATTGGGTACGAGAGTCTAGACTTGTGTTTGGAAGGAGTAGGGATTCATCATCACGTACTTGAAACAGGAAACGCGGTATATGGAAAAGTTCAACATGAGTATTCTACTATCAAAGAGAAGGCCAAAGAAATGAATGCACTCAAACCAGGACCTATCATCGATTACCACGTCTGGATAGGAGATTGTGTCTGCCAAGTTACTACTGTAGACGTGCATGGAAAGGAAATTATGAGAATGAGATTCAAAAGGGGTGCGGTGCTTCCGATTCCAAATCTGGTAAAAGTTAAAGTTGGGGAGGAAAATGATACAATAAATCTTTCCACTTCCATATCAGCTCTCCTAAATTCCGGTGGCGGCACCATCGAGGTAACATCTAAGGAAGAACGTGTAGATTATGTACTCATGAAACGTTTGGAATCTATACATCATCTGTGGTCTGTAGTGTATGATCATCTTAATGTTGTGAATGGCGAAGAACGATGTTATGTACATATGCATTCATCTCATCAAAGTCCTATGCTGAGTACTGTAAAAACAAATTTGTACATGAAGACTATGGGAGCATGTCTTCAAATGGACTCCATGGAAGCTCTAGAGTATCTTAGTGAACTGAAGGAATCAGGTGGGCGGAGTCCCAGACCAGAATTGCAGAAATTTGAATATCCAGATGGAGTGAAAGACACTGAATCAATTGAGAGATTGGCAGAGGAGTTCTTCAATAGATCAGAACTTCAGGCCGGTGAATCAGTCAAATTTGGTAATTCTATTAATGTTAAACATACATCTGTTTCAGCTAAGCAACTAAGAACACGTATACGACAGCAGCTTCCTTCTATACTCTCATCTTTTGCCAACACAAAGGGTGGATATTTGTTCATTGGAGTTGATAATAATACACACAAAGTAATTGGATTCACGGTGGGTCATGACTACCTCAAACTGGTAGAGAGTGATATAGAAAAGTATATCCAAAAACTTCCTGTTGTGCATTTCTGCAAGAAAAAAGAGGACATCAAGTACGCATGTAGATTCATCAAGGTGTATAAACCTGGTGATGAGACTACCTCGACATATGTGTGCGCAATCAAAGTGGAAAGATGCTGCTGTGCTGTGTTTGCGGATTGGCCAGAATCATGGTACATGGATACTAGTGGTAGTATGAAGAAGTATTCTCCAGATGAATGGGTGTCACATATAAAATTTTAATTAGGGTAAGGTAAAACTATATATAATAACTAACAATTTGTGTATCATATAGACAATTAATTAGGTAACTGTTATCTCTTTTTAACTAACTAACTAACTAACTAACTCTTATATACTATTAATAATACATCTATTAATCATTGATTAGCTTATTGCTTTAATTGTTTTTGTAAACTAACACTGTTCATTGAAAAGGGATAACATGTTACAGAATATAAATTATATATGGATTTTTTTAAAAAGGAAATACTTGACTGGAGTATATATTTATTTCTTCATTACATAACACGTCTGTGTTCTAATTCTTCCAATTCTTCCACATCTCATATAATACAGGAATATAATCTTGTTCGAAAATATGAGAAAGTGGATAAAACAATAGTTGATTTTTTATCTAGGTGGCCAAATTTATTCCATATTTTAGAATATGGGGAAAATATTCTACATATTTATTTTATAGATGCTGCTAATACGAATATTATGATTTTTTTTCTAGATAGAGTATTAAATATTAATAAGAACCGTGGGTCATTTATACATAATCTCGGGTTATCATCCATTAATATAAAAGAATATGTATATCAATTAGTTAATAATGATCATCTAGATAATAGTATAAGACTAATGCTTGAAAATGGACGTAGAACAAGACATTTTTTGTCTTATATATTGGATACAGTTAATATCTATATAAGTATTTTAATAAATCATAGATTTTATATAGATGCCGAAGACAGTTACGGTTGTACATTATTACATAGATGTATATATAACTATAAGAAATCAGAATCAGAATCATATAATGAATTAATTAAGATATTGTTAAATAATGGATCAGATGTAGATAAAAAAGATACGTACGGAAACACACCGTTTATCCTATTATGTAAACACGATATCGACAACGCGGAATTGTTTGAGATATGTTTAGAGAATGCTAATATAGACTCTGTAGACTTTAATGGATATACACCTCTTCATTATGTCTCATGTCGTAATAAATATGATTTTGTAAAGTTATTAATTTCTAAAGGAGCAAATGTTAATGCACGTAATAGATTCGGAACTACTCCATTTTATTGTGGAATTATACACGGTATCTCGCTTATAAAACTATATTTGGAATCAGACACAGAGTTAGAAATAGATAATGAACATATAGTTCGTCATTTAATAATTTTTGATGCTGTTGAATCTTTAGATTATCTATTGTCCAGAGGAGTTATTGATATTAACTATCGTACTATATACAACGAAACATCTATTTACGACGCTGTCAGTTATAATGCGTATAATACGTTAGTCTATCTATTAAACAGAAATGGTGATTTTGAGACGATTACTACTAGTGGATGTACATGTATTTCGGAAGCAGTCGCGAACAACAACAAAATAATAATGGATATACTATTGTCTAAACGACCATCTTTGAAAATTATGATACCATCTATGATAGCAATTACTAAACATAAACAACATAATGCAGATTTATTGAAAATGTGTATAAAATATACTGCGTGTATGACCGATTATGATACTCTTATAGATGTACAATCGCTACATCAATATAAATGGTATATTTTAAAATGTTTTGATGAAATAGATATCATGAAGAGATGTTATATAAAAAATAAAACTGTATTCCAATTAGTTTTTTGTATCAAAGACATTAATACTTTAATGAGATACGGTAGACATCCTTCTTTCGTGAAATGTAATATTCTTGACGTATACGGAAGTTGTGTACGTAATATCATAGCATCTATTAGATATCGTCAGAGATTAATTAGTCTATTATCCAAGAAGCTGGATGCTGGAGATAAATGGTCGTGTTTTCCTAACGAAATAAAATATAAAATATTGGAAAACTTTAACGATAACGAACTGACCACATATCTGAAAATCTTATAAACACTATTAAAATATAAAATCTAAGTAGGATAAAATCACACTACATCATTGTTTCCTTTTAGTGCTCGACAGTGTATACTATTTTTAACACTCATAAATAAAAATGAAAACGATTTCCGTTGTTACGTTGTTATGCGTACTACCTGCTGTTGTTTATTCAACATGTACTGTACCCACTATGAATAACGCTAAATTAACGTCTACCGAAACATCGTTTAATGATAAACAGAAAGTTACGTTTACATGTGATTCAGGATATCATTCTTTGGATCCAAATGCTGTCTGTGAAACAGATAAATGGAAATACGAAAATCCATGCAAGAAAATGTGCACAGTTTCTGATTATGTCTCTGAACTATATGATAAGCCATTATACGAAGTGAATTCCACCATGACACTAAGTTGCAACGGTGAAACAAAATATTTTCGTTGTGAAGAAAAAAATGGAAATACTTCTTGGAATGATACTGTCACGTGTCCTAATGCGGAATGTCAACCTCTTCAATTAGAACACGGATCGTGTCAACCAGTTAAAGAAAAATACTCATTTGGGGAATATATGACTATCAACTGTGATGTTGGATATGAGGTTATTGGTGTTTCGTATATAAGTTGTACGGCTAATTCTTGGAATGTTATTCCATCATGTCAACAAAAATGTGATATACCGTCCCTATCTAATGGATTAATTTCCGGATCTACATTTTCTATCGGTGGCGTTATACATCTTAGTTGTAAAAGTGGTTTTACACTAACGGGGTCTCCATCATCCACATGTATCGACGGTAAATGGAATCCCATACTCCCAACATGTGTACGATCTAACGAAGAATTTGATCCAGTGGATGATGGTCCCGACGATGAGACAGATCTGAGCAAACTCTCGAAAGACGTTGTACAATATGAACAAGAAATAGAATCGTTAGAAGCAACTTATCATATAATCATAATGGCGTTGACAATTATGGGTGTCATATTTCTAATCTCCATTATAGTATTAGTTTGTTCCTGTGACAAAAATAATGACCAATATAAGTTCCATAAATTGCTACCGTGAATATAAATCCGTTAAAATAATTAATAATTAATAATTAATAACGAACAAGTATCAAAAGATTAAAGAATTAGCTAGAATCAATTAGATGTCTTCTTCAGTGGATGTTGATATCTACGATGCCGTTAGAGCATTTTTACTCAGGCACTATTATGACAAGAGATTTATTGTGTATGGAAGAAGTAACACCATATTACATAATATATACAGGCTATTTACAAGATGCACCGTTATACCGTTCGATGATATAGTACGTACTATGCCAAATGAATCACGTGTTAAACAATGGGTGATGGATACACTTAATGGTATAATGATGAATGAATTCGATACTGTATGTGTGGGTACCGGACTACGATTCATGGAAATGTTTTTCGATTACAATAAAAATAATCCCAAAAATAGCATCAACAATCAAATAATGTATGATATAATTAATAGCGTAGCCATAATTCTAGCTAATGAGAGATATAGAAGCGCGTTTAACGACGATAGAATATACATCCGTAGAACTATGATGGACAAATTGTACGAATACGCATCTCTAACTACTATTGGTACGATCACTGGAGGTGTTTGTTATTTTATCTGTTGATGCATCTAGTTAGTTTGTATAAATAATTATTTCGATATACTAGTTAAAATTTTAAGATTTTAAATGTATAAAAAACTAATAACGTTTTTATTTGTAATAGGTGCAGTTGCATCCTATTCGAATAATGAGTACACTCCGTTTAATAAACTGAGTGTAAAACTCTATATAGATGGAGTAGATAATATAGAAAATTCATATACTGATGATAATAATGAATTGGTGTTAAATTTTAAAGAGTACACAATTTCTATTATTACAGAGTCATGTGACGTCGGATTTGATTCCATAGATATAGATGTTATAAACGACTATAAAATTATTGATATGTATACCATTGACTCGTCTACTATTCAACGCAGAGGACATACGTGTAGAATATCTACCAAATTATCATGCCATTATGATAAGTACCCTTATATCCACAAATATGAGGGTGATGAACGACAATATTCTATTACCGCAGAGGGAAAATGCTATAAAGGAATAAAATATGAAATAAGTATGATGAACGATGATACTCTATTGAGAAAACATACTCTTAAAATTGGATTTACTTATATATTCGATCGTCATGGGCATAGTAATACATATTATTCAAAATATGATTTTTAAAAATTTAAAATATATTATCACTTCAGTGACAGTAGTCAAATAACAAACAACACCATGAGATATATTATAATTCTCGCAGTTTTGTTCATTAATAGTATACATGCTAAAATAACTAGTTATAAGTTTGAATCCGTCAATTTTGATTCCAAAATTGAATGGACTGGGGATGGTCTATACAATATATCCCTTAAAAATTATGGCATCAAGACGTGGCAAACAATGTATACAAATGTACCAGAAGGAACATACGACATATCCGGATTTCCAAAGAATGATTTCGTATCTTTCTGGGTTAAATTTGAACAAGGCGACTATAAAGTGGAAGAGTATTGTACGGGACTATGTGTCGAAGTAAAAATTGGACCACCAACTGTAATATTGACTGAATATGACGATCATATCAATTTGTTCATCGAGCATCCGTATGCTACTAGAGGTAGCAAGAAGATTCCTATTTACAAACGCGGTGACATGTGTGATATCTACTTGTTGTATACGGCTAACTTCACATTCGGAGATTCTGAAGAACCAGTAACATATGATATCGATGACTACGATTGCACGTCTACAGGTTGCAGTATAGACTTTGCCACAACAGAAAAAGTGTGTGTGACAGCACAGGGAGCCACAGAAGGGTTTCTCGAAAAAATTACTCCATGGAGTTCGGAAGTATGTCTGACACCTAAAAAGAATGTATATACGTGCGCAATTAGATCTAAAGAAGATGTTCCCAATTTCAAGGACAAAATAGCCAGAGTTATCACGAGAAAATTTAATAAACAGTCTCAATCTTATTTGACTAAATTTCTCGGTAGCACATCGAATGATGTTACAACTTTTTTTAGCATTCTTGACTAAATATTCATAACTAATTTTTATTAATGATACAAAAATGAAATAAACTGTATATTATACACTGGTTAACGCCCTTGGCTCTAACCATTTTCAAGATGAGGTCCCTGATTATAGTCCTTCTGTTCCCCTCTATCATCTACTCAATGTCTATTAGACGATGCGAGAAGACTGAAGAGGAAACATGGGGATTAAAAATAGGGTTGTGTATAATTGCCAAAGATTTCTATCCCGAAAGAACTGATTGCAGTGTTCATCGCCCAACTGCAAGTGGAGGATTGATAACTGAAGGCAATGGATTCAGAGTAGTTATATATGATCAATGTACAGAACCCCATGACTTTATTATCACCGATACTCAACAAACACGTCTTGGATCATCTCATACATATATTAAATTCAGTAACATGAATACAGGTGTCCCATCTAGTATTCCAAAATGTTCCAGAACTCTCTGTATTTCTGTATATTGTGATCAAGAGGCGGGAGACATAAAATTTGAGGAGTATACTCAAGAATCAAGTGATATCAGTATTAGAGTTAAGTATGATTCATCATGTATTGATTATCTGGGTATTAATCAAAGTTTCATGAATGAATGTATTCGAAGAATTACAACATGGGATAGAGAATCATGCGTCAGAATTGATACACAGACTATAAATAAATATCTTAAGTCTTGCACCAACACAAAATTCGACCGTAATGTCTACAAAAGGTACATACTGAAGAGTAAAGCACTCCATGCTAAAACAGAGTTGTAATAGATATAAAATACTTTTTATAATAATTAGGCTAGAAAAATCTCACTCACATGTAATCTTAAAAAAATGATATGATAGTTCTTACAAGTAGCGATTGAGTTTTAAATGGATTCTATTAATTACCGGGGAACTTAACAATTCGTTCTGATCTACAGACATTGGTTAATAAATCATCTTATTTTGCCAATATATTAAAATGTGGAAACTCCACTAATAATATTACATTGTGCGACTTTCAAGATGATGTGATATATAGGGTTATACAGTTTTAACAATTATATAATAGAGATAGAAAGTACAAAAGATGTAGAATCAATGATATGGCACGCTAAACAGTTGGGTGTGGAATCATTGCTAAAAGAATGTCAAAATTATTTGCTTAGAATATTACGTATATAATTGTTTAGAAATTTATAGAATAACTAATATTAATACATTATCGTATATCTACAACGATATAAGAAACTTCATATTGGATAATATTACTATTAATATATAAGGATCCAGATTTTATATATTTGCCTAAATACATTATTATAGATTTACTAGGACAATCACCTAAATGTTTTTAACGAAGATAATGTGGTAAAGATTATATACACTTATATATCTTCCGATATCTACAAGGATATTCCATATCATCATTGTGTAAACTAAATAACGTTTTCTATGGCATTTAATAAGGACATTGGATATGTGGAAAAGTGATGTATGGAAGTTAGTACATTATCAACTTCTCCTTATTGATTGAAAATGAAAATATAAATAGTTTTTATGTATAGCGGTATCTACCCTATAGTTTTATTGCTTACTACTAACATGGATTCAGATACAGATACAGATACAGATACAGATACAGATACAGATACAGATACAGATGTAGAAGATATCATGAATGAAATAGATAGAGAGAAAGAAGAAATACTAAAAAATGTAGAAATTGAAAATAATAAAAACATTAACAAGAATCATCCCAGTGAATATATTAGAGAAGCACTTGTTATTAATACCAGTAGTAATAGTGATTCCATTGATAAAGAAGTTATAGAATATATCAGTCACGATGTAGGAATATAGATCATATCTACTAATTTTTATAATCGATACAAAACATAAAAACAACTCGTTATTACATAGCAGGTATGGAATCCTTCAAGTATTGTTTTGATAACGATGGTAAGAAATGGATTATCGGAAATACTTTATATTCTGGTAATTCAATACTCTATAAGGTCAGAAAAAATTTCACTAGTTCGTTCTACAATTACGTAATGAAGATAGATCATAAATCACACAAGCCATTGTTGTCCGAAATACGATTCTATATATCTGTATTGGATCCTTTGACTATCAACAACTGGACACGGGAACGTGGTATAAAGTATTTGGCTATTCCAGATCTGTATGGAATTGGAGAAACCGATGATTATATGTTCTTCATTATAAAGAATTTGGGAAGAGTATTCGCCCCAAAGGATAGTGAATCAGTTTTCGAAGCATGTGTCACTATGATAAACACGTTAGAGTTTATACACTCTCAAGGATTTACTCATGGAAAAATAGAACCGATGAATATACTGATTAGAAATAAACGTATTTCACTAATTGACTATTCTAGAACTAACAAACTATACAAAAGTGGAACACATATAGATTACAACGAGGACATGATAACTTCAGGAAATATCAATTATATGTGTGTAGACAATCATCTTGGAGCAACAGTTTCAAGACGAGGAGATTTAGAAATGTTGGGATATTGCATGATAGAATGGTTCGGTGGTAAACTTCCATGGAAAAACGAAAGTAGTATAAAAGTAATAAAACAAAAAAAAGAATATAAACAATTTATAGCTACTTTTTTTGAGGACTGTTTTCCTGAAGGAAATGAACCTCTGGAATTAGTTAGATATATAGAATTAGTATACATGTTAGATTATTCTCAAACTCCTAATTATGACAGACTACGTAGACTGTTTATACAAGATTGAAATTATATTCTTTTTTTTATAGAGTGTGGGGGTAGTGTTACGGATATCTAATATTAATATTAGACTATCTCTATCGCGCTACACGACCAATATCGATTACTATGGATATCTTCAGGGAAATCGCATCTTCTATGAAAGGAAAGAATGTATTCATTTCTCCAGCGTCAATCTCGTCAGTATTGACAATACTGTATTATGGAGCTAATGGATCCACTGCTGAACAGCTATCAAAATATGTAGAAAAGGAGGAGAACATGGATAAGGTTAGCGCTCAGAATATCTCATTCAAATCCATGAATAAAGTATATGGGCGATATTCTGCCGTGTTTAAAGATTCCTTTTTGGGAAAAATTGGCGATAAGTTTCAAACTGTTGACTTCACTGATTGTCGCACTATAGATGCAATCAATAAGTGTGTAGATATCTTTACTGAGGGAAAAATCAATCCACTATTGGATGAACCATTGTCTCCTGATACCTGTCTCCTAGCAATTAGTGCCGTATACTTTAAAGCAAAATGGTTGATGCCATTCGAAAAGGAATTTACCAGTGATTATCCCTTTTACGTATCTCCAACGGAAATGGTAGATGTAAGTATGATGTCTATTTACGGCGAGCCATTTAATCACGCATCTGTAAAAGAATCATTCGGTAACTTTTCAATCATAGAACTGCCATATGTTGGAGATACTAGTATGATGGTCATTCTTCCAAACAAGATTGATGGATTAGAATCCATAGAACAAAATCTAACAGATACAAATTTTAAGAAATGGTGTAACTCTCTGAAAGCTACGTTTATCGATGTGCACATTCCTAAGTTTAAGGTAATAGGTTCGTATAATCTTGTGGATACGCTAATAAAGTTGGGACTGACAGATGTGTTCTATTCAACTGGTGATTATATCAATATGTGTAATTCAGATGTGAGTGTTGACGCTATGATTCACAAAACGTATATAGATGTCAATGAAGAGTATACAGAAGCAGCTGCAGCAACTTCTGTACTAGTGGCAGACTGTGCATCAACAGTTACAAATGAGTTCTGTGCAGATCATCCGTTCATCTATGTGATTAGACATGTCGATGGTAAAATTCTTTTCGTTGGTAGATATTGCTCTCCAACAACTAATTAAGCACATTCTTAATATTAGAATATTATATAGTTAAGATTTTTACTAACAGGTTAACATTTTTTTTTAAAAATAGAAAAAACATGTGGTATTAGTGCAGGTCGTTATTCTTCCAATTGCAATTGGTAAGATGACGGCCAACTTTAGTACCCACGTCTTTTCACCACAACACTGTGGATGTGACAGACTGACCAGTATTGATGACGTCAGACAATGTTTGACTGAATATATTTATTGGTCGTCGTATGCATACCGCAACAGGCAATGCGCTGGACAACTGTATGACACACTCCTCTCTTTTAAAGATGATGCGGAATCAGTGTTCATCGACGTTCGTGAGCTGGTAAAAAATATGCCGTGGGATAATGTTAAGGATTGTACAGAGATCATCCGTTGTTATATACCGGATGAGCAAAAAACCATCAGAGAGATTTCGGCCATCATTGGACTTTGTGCATATGCTGCTACTTACTGGGGAGGTGAAGACCATCCCACTAGTAACAGTCTGAACGCATTGTTTGTGATGCTTGAGATGCTCAATTACATGGATTATACCATCATATTCTGGCGTATGAATTGATGAGTTACAGCTTGACATTTCTTCTTTCCTCCCTCTTCTTCTACCTTTCCCAGAAACAAACTTTTTTTACCCACTATAAAATAAAATGAGTATACTACCTGTTATATTTCTTCCTATATTTTTTTATTCTCCATTCGTTCAGACTTTTAACGTGCCTGAATGTATCGACAAAGGGCAATATTTTGCATCATTCATGGAGTTAGAAAACGAGCCAGTAATCTTACCATGTCCTCAAATAAATACGCTATCATCCGGATATAATATATTAGATATTTTATGGGAAAAACGAGGAGCGGATAATGATAGAATTATACAGATAGATAATGGTAGCAATATGCTAATTCTGAACCCGACACAATCAGACTCTGGTATTTATATATACATTACCACGAACGAAACCTACTGTGACATGATGTCGTTAAATTTGACAATCGTGTCTGTCTCAGAATCAAATATAGATCTTATCTCGTATCCACAAATAGTAAATGAGAGATCTACTGGTAAAATGGTATGTCCCAATATTAATGCATTTATTTCTAGTAACGTAAACACAGAATTATATGGAGCGGACATCGACGCCTTAGAAATAAGAGACTTAAACAACGGACACCTGGAATTATTACCATAGAAGATGTTAGAAAAAATGATGCTGGTTATTATACATGTGTTTTAGAANNNNNNNNNNTATATATATATATATATATATATATATATATATATATATATATATATATATATATATATATATATATATATATATATATATATATATATATATATATATATATATATATGGGCAAAACATATAACGTAACCAGAATTATAAAATTAGAGGTACGGGATAGAATAATACCTCCTACTATGAAATTACCAGAAGGAGTAGTAACTTCAATAGGTAGTAATTTGACTATTGCATGTAGAGTATCGTTGAGACTTCCCACAACGGACACCGACGTCTTTTGGATAAGTAATGGTATATGTATTACGAAGAAGAAGACGAGGACGGAGACGGTAGAATAAGTGTAGCAAATAAAATCTATATGACCGATAAGAGACGTGTTATTACATCCTGGTTAAACATTAATCCTGTCAAGGAAGAAGATGCTACAACGTTTACGTGTATGGCGTTTACTATTCCTAGCATCAGCAAAACAGTTACTGTTAGTAAACGTGAATGTATGTTGTTACATTTCCATATCAATTGAGTTTATAAGAATTTTTTATACATTATCTTCCAACAAACAATTGACGAACGTATTGCTATGATTAACTCCCACAATACTATATATATTATTAATCATTAACTTGCAGACTATACCTAGTAGTGCTATTTTGACATACTCATGTTCTTGTGTAATCGCAGTATCTATATTATTAAAGTACGTAAATCTAGCTATAGTTTTATTATTTAATTTTAGATAATATACTGTCTCCGTATTTTTAAAAAATTACCACATCCTTTATTAAATCATGAATGGGAATTTCTGTGTCATCGTTAGTATATTGTGAACAACAAGAGCAGATATCTATAGGAAAGGGTGGAATGCGATACATTGATCTATGTAGTTTTAAAACATACGCGAACTTTGAAGAATTTATATAAATCATCTCACGAGATATTGCTCTCTGTCATATTCATACACCTGTATAAACTTTCTAGACATCTTACAATGTGTTATTTTATGATCATATTTACATATTTACTGGTATATCAAAGATGTTAGATTAGTTAATGGGAATCGTCTATAATAATGAATATTAAACAATTATAGGAGGAGTTTATACCTACAAAAACATCATAAAAATGAGTCATCGTCCGATTTATGTTTTAAATATACTAACATTACTACCTTCAGAAATTATATACGAAATATTATACATGCTGACAATTAACGATCTTTATAATATATAGTATCCACCTACCAAAGTATAATTGTATTTTTCTCATGTGATGTGTGTAAAAAACTGATATTATATAATTATCTTAGTACCTATGATGAAGATGAAGATGAAGATGAAGATGATGGTCCGTATATATTTTGTATCATTATCGTTATTGCTATTCCATAGTTACGCCATAGACATCGAAAATGAAATCACCGAATTCTTCAATAAAATGAGAGATACTCTACCAGCTAAAGACTCTAAATGGTTGAATCCAGTATGTATGTTTGGAGGCACAATGAATGATATGGCCGCTCTAGGAGAGCCATTCAGTGCAAAGTGTCCTCCTATTGAAGACAGTCTTTTATCGCATAGATATAAAGACTATGTGGTTAAATGGGAAAGGCTAGAAAAGAATAGACGGCGACAGGTTTCTAATAAACGTGTTAAACATGGTGATTTATGGATAGCCAACTATACATCTAAATTCAGTAACCGTAGGTATTTATGTACCGTAACCACAAAGAATGGTGACTGTGTTCAGGGTGTAGTTAGATCTCATGTGTGGAAACCTTCTTCATGCATTCCAAAAACATATGAACTAGGTACTTATGATAAGTATGGCATAGACTTATACTGTGGAATTCTTTATGCGAACCATTATAATAATATAACTTGGTATAAAGATAATAAGGAAATTAATATCGACGATTTTAAGTATTCACAAGCGGGAAAGGAATTAATTATTCATAATCCAGAGTTAGAAGATAGTGGAAGATACGACTGTTACGTTCATTACGACGACGTTAGAATCAAGAATGATATCGTAGTATCAAGATGTAAAATACTTACGGTTATACCGTCACAAGACCACAGGTTTAAACTAATACTAGATCCGAAAATCAACGTAACGATAGGAGAACCTGCCAATATAACATGCAGTGCTGTGTCAACGTCATTATTTGTCGACGATGTACTGATTGAATGGGAAAATCCATCCGGATGGATTATAGGATTAGATTTTGGTGTATACTCTATTTTAACTAGTAGAGGCGGTATCACCGAGGCGACTTTGTATTTTGAAAATGTTACTGAAGAATATATAGGCAATACATATACATGTCGTGGACACAACTATTATTTTGATAAAACTCTTACAACTACAGTAGTATTGGAGTAAATACACAATGCATTTTTATATACATTACTGAATTATTATTATTAATTATATCGTATTTGTGCTATAGAATGGATGAAGATACGCGACTATCTAGGTATTTGTATCTCACCGATAGAGAACATATAAATGTAGACTCTATTAAACAGTTGTGTAAAATATCAGATCCTAATGCATGTTATAGATGTGGATGTACGGCTTTACATGAGTACTTTTATAATTATAGATCAGTCAACGGAAAATACAAGTATAGATACAACGGTTACTATCAATATTATTCATCTAGCGATTATGAAAATTATAATGAATATTATTATGATGATTATGATAGAACTGGTATGAACAGTGAGAGTGATAATATATCAATCAAAACAGAATACGAGAATGAATATGAATTCTATGATGAAACACAAGATCAAAGTACACAACTAGTAGATTACGACATTAAACTCAAAACCAATGAGGATGATTTTGTTGATGAATTCTATGGTTATGATAGATCAGTGGGTGTCCATGATTATATAGATGTATCAATTAATAAAGTAGTATATGGAAGAGAGTCTCACGTAAGATGGTGGGATATATGGCAAGAACATAATGATGGCGTATACAGTATAGGAAAGGAGTGCATAGATAATATATACGAAGACAGACATACCGTAGACGAATTCTACAAGATAGACAGCGTATCAGATGTAGATGACGCAGAACATATATCTCAGATAACTAATGATGTATCTACACAAACATGGGAAAAGAAATCAGAGTTAGATAGATACATGGAAATGTATCCTCGTCATAGATATGGTAAGCATTCTGTCTTTAAGGGATTTTCTGACAAAGTTAGAAAAAATGATTTAGACATGAACGTGGTAAAAGAATTACTTTCTAACGGTGCATCTCTAACAATCAAGGATAGCAGTAATAAGGATCCAATTGCTGTTTATTTTAGAAGAACAATAATGAATTTAGAAATGATTGATATCATTAACAAACATACAACTATCTATGAACGCAGGTATATAGTACACTCCTATCTAAAAAATTATAGAAATTTCGATTATCCATTTTTCAGAAAGTTAGTTTTGACTAATAAACATTGTCTCAACAATTATTGTAATATAAGCGACGGCAAATATGGAACACCACTACATATATTAGCATCTAATAAAAAAATAATAACTCCTAATTACATGAAGTTATTAGTGTATAACGGAAATGATATAAACGCACGAGGTGAAGATACACAAATGCGAACTCCATTACACAAATATTTGTGTAAATTTGTATATCATAATATTGAATATGGTATCCGATACTATAATGAAAAGATTATAGACGCATTTATAGAGTTAGGAGCCGATCTAACTATTCCAAATGACGATGGAATGATACCAGTAGTTTACTGTATACACTCAAATGCCGAATATGGTTATAACAATATTACTAACATAAAGATAATACGTAAACTACTTAATCTTAGTAGACATGCGTCACATAATCTATTTAGAGATCGAGTCATGCACGATTATATAAGTAATACATATATTGATCTTGAGTGTTTAGATATCATTAGATCACTTGATGGGTACGATATTAATTGTTACTTTGAAGGACGTACACCACTTCATTGCGCTATACAATATAACTTCACTCAGATTGCTGAGTACTTATTAGATCGAGGAGCTGATATATCATTAAAGACAGACGATGGTAAAACTGTATTTGATTTATCGTTATGTAGTTACATTCCTCTTAAATGGACTAGCTTTTTGATTAGTCGTCTACCGCCTAAAAGTGTCATATGCTCACTGACTAACCATATAATAGATTATGTTCTTACGAACAATAGACGTATTATTTGGCAGAGTCAAATGATTAATAAGTACGTACTGTTACTGGACCCATCCTTTTATTATAGATTCAGAAATGTTATCGAAAACAAATTAGACCAATACAATAATCGTTATAATATGTTCGAACACGATAGGGACGTTAATGAAAAGTATGGCAAAGTCTTACATGACCTCGATACATATATCAAGGATGTACAAGTATTAAAATCTACTTCCATCACTAATAATATAACACTATACGACACTATTATAAATAATAAGTCAGAGTTTCCTATACGTCGTGTAAACGACAAACAATTAATTAATCTCATAAAATCCAATACATATCATAATCTTATCGAAAAAGTTATTAAAAATACATTAGAGAAATATACTTTAACTAATATAGTCCTCGAGTATATGATCTCATCTCGATCTCAATCATCTTATTTGAGTCGTATTCCTAATGAGATATTACTCGAAATATTATATAAACTCGACATGTACGATTTACGTAATCTATATACAAGATATATGAGAGAGAATGATATCACAGAGTATCATATAGAGAATACGAGGTCTGTTTCTACACAGACATGAATAATGAATACACATACAACGTTTTTTTTAATCTTAGATATAACACTAATTACATCAAGATTATATATTGAAATCGTAATTTGAGTTGTCTGATCATCATGGATATCGAAAATGATATACGTAACATTAGCAATCTTTTAGATGATTGATATATTATTATGCGATGTAATCATAACTATCGGAGATGTAGAAATTAAAGCGCATAAAACTATTTTGGTTGCCGGATCTACGTATTTTAAAACAATGTTCACAACATCTATGATAGCGAGAGATCTAGCAACTAGAGTAAATATACAGATGTTCGATAAAGATGCCGTCAAAAATATTGTACAGTACTTATACAATAGGTATATAAGTTCTATGAATGTGATAGACATATTAAAATGCACCGACTAAGAACGTAAAACGAACTATAGAATGTTATACAATGGGTGATGATAAGTAGAAGATGTTACCCGATATACCCATAGCATTATCTAGTTATGGCATGTGTGTATTAGATCAATACATATACATTATAAGCGGTCGTACCCAACACTGATTATACATCGGTACATACAGTAAATAGCATAGATATGGAGGAGGATACAAATATTTCAAATAAAGTTATGAGATACGCGCTGTCAATAATATATGGAAGACATTACCTAACTTCTGAACTGGAACTATAAATCCAGGCTCTCGCATAAAGATGAATATATATGTTGTATGCGACATCAAAGATGAAAAAATGTTAAGACTTATATATTTAGATATAACACGAATATGTATGACGGATGGGAATTGGTAACGATGACAGAAAGCAGATTGTCAGCTCTGCATACTATTCTTCATGACAATACCATAATGATGTTACATTGTTATGAAGCGTATATGTTACAAGATACATTTAATGTGCTTACGGAACATATATTTAGAAACATCTACTAACGATTTTTTATGCTTGTATTATTAATGGTATGTAATATGATTTAATTGATTGTGTACACGATACCAATTTGTCGAGTATGAATACGGAGTACAAACATAAACTGAAGTTTAACATTATTTATTTATGATATACATTATATACATTATATACATTATATACATTATATACATTATATACATTATATACATTATATACATTATATACATTATATACATTATATATCGTTATTGTTTGGTCTATGCCATGGATATCTTTAAAGAACTAATCTTAAAACATACGGATGAAAATGTTTTGATTTCTCCAGTTTCCATTTTATCTACTTTATCTATTCTGAATCATGGAGCAGCTGGTTCTACAGCTGAACAACTATCAAAATATATAGAGAATATGAATGAGAATACACCCGATGATAAGAAGGATGACAATAATGACATGGACGTAGATATTCCGTATTGCGCGACACTAGCTACCGCAAATAAAATATACGGTAGTGATAGTATCGAGTTCCATGCCTCATTCCTACAAAAAATAAAAGACGATTTTCAAACTGTAAACTTTAATAATGCGAACCAAACAAAGGAACTAATCAACGAATGGGTTAAGACAATGACAAATGGTAAAATTAATTCCTTATTGACTAGTCCGCTATCCATTAATACTCGTATGATAGTTATTAGCGCCGTCCATTTTAAAGCAATGTGGAAATATCCATTTTCTAAACATCTTACATATACAGACAAGTTTTATATTTCTAAGAATATAGTTACCAGTGTTGATATGATGGTGGGTACCGAGAATGACTTGCAATATGTACATATTAATGAATTATTCGGAGGATTCTCTATTATCGATATTCCATACGAGGGAAACTCTAGTATGGTGATTATACTGCCGGACGACATAGAAGGTATATATAACATAGAAAAAAATATAACAGATGAAAAATTTAAAAAATGGTGTGGTATGTTATCTACTAAAAGTATAGACTTGTATATGCCAAAGTTTAAAGTGGAAATGACGGAACCGTATAATCTGGTACCGATTCTAGAAAATTTAGGACTTACTAATATATTTGGATATTATGCAGATTTTAGTAAGATGTGTAATGAAACTATCACTGTAGAAAAATTTCTACATACGACGTTTATAGATGTTAATGAGGAGTATACAGAAGTATCGGCCGTTACAGGAGTATTCATGACTAACTTTTCGATGGTATATCGTATGAAGGTCTACATAAACCATCCATTCATATACATGATTAAAGATAACACCGGACATACACTTTTTATAGGGAAATACTGCTATCCGCAATAAATATAAACAATAGACTTTTATCACGTTATCTCATGTATAAAATATTACAAATAGTATAGCATAAACTAAAGTCGATACATACATTAAAACTTAAATAATAATGTAATTTACAATTAATAGTATAAACTAAAAAAATTAAAAAATTAAAAACAATATCATTATTATAAGTAATATCAAAATGACGATATACGGATTAATAGCGTATCTTATATTCGTGACTTCATCCATCGCTAGTCCACTTTACATTCCCGTTATTCCGCCCATTTCGGAAGATAAATCGTTCAATAGTGTAGAGGTATTAGTTTCTTTGTTTCCCGATGACCAAAAAGACTATACAGTAACTTCTCAGTTCAATAACTACACTATCGGTACCAAAGACTGGACTATCAACGTACTATCCACACCTGATGGTCTGGACATACCATTGACTAATATAACTTATTGGTCACGGTTTACTATAGGTCGTGCATTGTTCAAATCAGAGTCTGAGGATATTTTCCAAAAGAAAATGAGTATTCTAGGTGTTTCTATAGAATGTAAGAAGCCGTCGACATTACTTACTTTTTTAACCGTGCGTAAAATGACTCGAGTATTTAATAGATTTCCAGATATGGCTTATTATCGAGGAGACTGTCTAGAAGCCGTTTATGTAACAATGACTTATAAAAATACTAAAACTGGAGAGACTGATTACACGTACCTCTCTAATGGGGGGTTGCCTGCATACTATCGTAATGGGGTCGATGGTTGATTATTGATTAGTATATTCCTTATATTCCTTATTCTTTTTATTCACACAAAAAGAACATTTTTATAAACATGAAACCACTGTCTAAATGTAATTATGATCTTGATTTATAGATGATGATCAGCCTTCAGAGGATTTTGACCAGTATGTTTAATATGAAAAAAAACATAACTATTAAGCGCTATTGCGCTATTGTGCTTAATTATTTTGCTCTATAAACTGAATATATAGCCACAATTATTGACGGGCTTGTTTGTGACCGACAATCATGAATTTTCAGAAATTATCTCTGGCTATATATCTTACGGTGACATGTTCGTGGTGTTATGAAACATGTATGAGAAAAACTGCGTTGTATCATGACATTCAATTGGAGCATGTAGAAGACAATAAAGATAGTGTAGCATCGCTACCGTACAAGTATCTACAAGTAGTCAAACAAAGAGAACGTAGTAGATTGTTGGCTACATTTAATTGGACGGATATAGCTGAGGGTGTTAGAAATGAGTTCATTAAAATATGTGATATCAACGGAACATATCTATATAATTATACTATTGCTGTTAGTATAATTATTGATTCCACGGAAGAACTACCAACAGTTACTCCAATTACAACATATGAACCTTCTATATATAATTATACTATCGATTATAGCACTGTTATTACTACTGAAGAACTACAAGTGACTCCAACATATGCACCTGTAACAACTCCTCTTCCAACATCAGCAGTTCCTTATGATCAACGATCGAATAACAATGTAAGTACTATATCTATTCAGGTACTGAGTAAAATATTGGGAGTCAATGAAACAGAATTAACTAATTATCTTATTATGCATAAAAATGACACTGTTGACAATAACACCATGGTTGATGATGAGACATCTGACAATAACACATTACATGGTAATATAGGATTTTTGGAAATAAATAATTGTTATAATGTTTCTGTGTCAGATGCTAGTTTTAGAATAACATTAGTAAACGATACTTCTGAAGAAATTTTGCTAATGCTAACAGGAACTAGTTCATCCGACACCTTCATATCTTCCACCAATATCACTGAATGTTTGAAAACATTAATCAATAATGTGTCGATTAATGATGTACTTATAACACAAAATATGAATGTAACATCTAATTGTGATAAATGCTCAATGAATTTGATGGCATCCGTTATTCCTGCAGTTAATGAATTTAACAATACGTTGATGAAAATTGGTGTAAAAGATGATGAAAACAATACGGTATATAAATATTATAATTGTAAACTAACTACAAATTCTACATGTGATGAGTTAATCAATTTAGATGAAGTCATTAACAACATAACTCTGACAAATATTATACACAATAGTGTTTCGACAACTAACAGCAGAAAAAGACGAGATCTGAATGATGAGTTTGAATTTTCCACTTCCAAGGAATTAGATTGTCTTTACGAATCATATGGTGTAAACGATGATATAAGTCATTGTTTTGCATCACCTAGACGTAGACGATCTGACGACAAAAAGGAGTACATGGACATGAAATTATTCGACCACGCGAAAAAAGATTTAGGAATAGACAGTGTTATTCCTAGAGGTACAACCCATTTCCAAGTAGGTGCATCTGGTGCAAGTGGTGGTGTTGTAGGAGATAGTTTCCCATTTCAAAATGTTAAATCGCGTGCCAGTCTATTGGCGGAAAAAATAATGCCTAGAGTACCTATTACTGCTACCGAAGCTGATCTATATGCAACTGTAAATAGACAACCCAAGTTACCAGCAGGTGTTAAAAGTACTCCGTTTACAGAGGCGCTTGTGTCTACGATAAACCAAAAGCTTTCTAATGTTAGAGAGGTAACTTATGCTTCGCTCAATCTGCCAGGATCAAGTGGCTATGTTCATAGACCATCTGATTCTGTTATTTATAGCAGTATAAGACGGTCACGTTTACCTAGTGATAGCGATAGTGATTATGAGGATATACAAACTGTTGTTAAGGAATATAATGAAAGATATGGTAGATCAGTCAGTAGAACACAGTCATCAAGTAGTGAAAGCGATTTTGAAGATATAGATACTGTTGTTAGGGAATATAGACAAAAATATGGCAATGCAATGGCAAAAGGACGTAGTAGTTCCCCTAAACCTGATCCATTATATAGTACTGTTAAGAAAACAACTAAAAGTCTATCTACTGGTGTAGACATAGTTACAAAACAATCAGACTATTCTCTATTACCTGACGTTAATACTGGCAGTTCTATTGTGTCACCTCTCACCAGAAAAGGAGCTACTAGACGACGACCTAGACGCCCTACAAATGATGGTCTACAGAGTCCAAATCCTCCTCTCCGTAATCCACTTCCTCAACATGATGATTATTCTCCTCCACAAGTACACAGACCTCCACCACTTCCTCCTAAACCAGTCCAAAATCCGCCACAACTTCCCCCTAGACCAGTAGGTCAATTACTACCTCCTCCTATAGATCAACCAGATAAAGGATTTAGTAAGTTTGTATCACCTAGACGGTGTAGAAGAGCAAGCTCTGGAGTCATATGTGGTATGATACAATCAAAACCAAACGATGATACCTATTCACTTCTTCAACGATCAAAAATTGAACCAGAATATGTGGAGGTTGGTAATGGTATACCCAAGAACAATGTTCCTGTAATAGGTAATAAACATAGTAAAAAATATACATCGACGATGTCAAAAATATCAACAAAATTTGATAAATCTACGGCATTTGGAGCAGCAATGTTACTAACTGGTCAGCAGGCCATTAGCCAACAGACTAGATCAACTACGTTGAGTAGAAAAGATCAGATGAGTAAGGAAGAAAAGATATTCGAAGCAGTTACAATGAGTCTATCAACTATAGGTTCAACGTTGACGTCTGCAGGTATGACGGGTGGTCCAAAACTAATGATTGCAGGAATGGCTATAACGGCTATAACTGGTATAATAGATACGATAAAAGATATATATTACATGTTTTCAGGACAGGAGAGGCCAGTAGATCCTGTTATTAAATTATTTAATAAGTACACTGGCTTAATGTCCGATAATAATAAAATGGGTGTAAGAAAATGTTTGACACCCGGTGACGACACACTTATTTATATCGCATACAGAAACGATACCAGTTTTAAACAGAATACGGATGCGATGGCTTTGTATTTCTTAGATGTTATCGATTCAGAGATCCTATATCTAAACACATCAAATTTAGTTCTAGAGTATCAACTAAAGGTGGCTTGCCCCATAGGAACATTAAGATCTGTAGATGTGGACATAACTGCGTATACAATATTATATGATACAGCGGATAATATTAAGAAATACAAGTTTATCAGAATGGCAACGCTACTATCCAAACATCCAGTTATTAGATTGACATGTGGTTTAGCAGCAACATTGGTGATTAAACCGTACGAGGTACCCATCAGTGATATGCAACTACTAAAAATGGCGACGCCTGGTGAACCAGAATCCACTAAATCTATACCATCCGATGTCTGTGATAGGTATCCTCTAAAGAAATTCTATCTTTTAGCTGGTGGTTGTCCCTATGATACATCTCAAACTTTTATTGTACATACTACTTGCAGTATTCTACTAAGAACAGCTACACGGGATCAGTTTAGAAACAGATGGGTGTTACAAAATCCATTTAGACAAGAAGGGACATATAAGCAACTGTTTACCTTTAGCAAATACGATTTTAACGACACCATAATCGATCCTAATGGTGTGGTGGGTCATGCTAGCTTTTGTACCAATAGAAGCAGCAACCAATGTTTCTGGTCCGAACCTATGATATTGGAAGATGTATCATCGTGTAGTTCTAGAACTAGAAAAATATACGTAAAACTGGGAATATTTAATGCTGAAGGTTTTAATAGTTTTGTACTAAATTGTCCAACTGGGTCTACACCTACATACATCAAACATAAAAATGCGGACAGTAACAATGTTATCATAGAGCTACCTGTAGGTGATTACGGCACAGCCAAATTGTATTCAGCAACAAAACCATCGAGGATAGCTGTGTTCTGCACACATAACTATGATAAACGATTCAAATCAGATATTATAGTTCTAATGTTTAATAAAAACAGCGGTATTCCATTTTGGAGCATGTACACAGGAAGTGTAACTAGTAAAAATAGAATGTTTGCCACATTGGCTAGAGGAATGCCGTTTAGATCAACGTATTGCGATAACAGACGACGATCAGGTTGTTATTATGCAGGAATACCATTTCATGAAGATAGTGTAGAAACAGATATACATTATGGACCAGAAATAATGTTAAAGGAAACATATGACATAAACAGTATTGACCCACGAGTTATAACAAAGTCAAAGACCCATTTTCCTGCTCCATTGAGTGTAAAATTCATGGTTGACAATTTAGGAAATGGATATGACAACCCTAATTCATTTTGGGAAGATGCTAAAACTAAGAAACGGACATATAGTGCAATGACGATAAAAGTCCTACCATGTACAGTGAGAAATAAAAATATAGACTTTGGATATAACTATGGAGATATTATTTCTAATATGGTTTATCTACAATCTACTAGTCAGGATTATGGAGATGGTACCAAATATACATTTAAATCCGTAACTAGATCAGATCATGAGTGTGAATCTAGCTTAGATCTAACGTCTAAGGAAGTAACTGTGACATGTCCTGCGTTTAGTATACCAAGAAATATATCAACATATGAAGGTCTATGCTTTAGTGTTACTACATCTAAAGATCATTGTGCTACAGGTATTGGTTGGTTAAAATCTAGTGGTTATGGGAAGGAAGATGCTGATAAACCACGTGCTTGTTTTCATCATTGGAATTATTACACACTGTCGTTGGATTATTACTGTTCATACGAAGATATTTGGAGAAGCACCTGGCCTGACTATGATCCATGTAAGTCATATATCCATATAGAGTATAGAGATACATGGATAGAATCTAATGTGTTACAGCAACCTCCTTACACATTCGAATTCATTCATGACAATTCTAACGAATATGTGGATAAAGAAATTAGTAACAAATTAAATGATCTGTACAATGAATACAAGAAGATTATGGAATATAGCGACGGATCATTGCCGGCGTCTATAAACAGATTAGCAAAGGCATTGACTTCAGAGGGTAGAGAAATAGCAAGTGTTAATATAGATGGTAATCTGTTAGATATCGCATATCAAGCAGATAAGGAAAAGATGGCCGACATACAGACAAGAATAAATGATATTATTAGAGATTTGTTTATACACACTCTATCAGACAAAGATATAAAAGACATTATAGAATCCGAAGAAGGTAAGAGATGTTGTATAATAGATGTTAAGAACAATCTTGTTAAAAAGTACTATTCTATTGATAATTATCTATGTGATACTTTAGATGATTATATATACACCTCTGTAGAATATAACAAATCCTATGTGTTAGTAAACGATACTTATATGAGCTATGACTATCTTGAATCATCAGGTGTAGTTGTTCTATCATGTTATGAAATGACTATAATCTCCTTGGATACAAAAGACGCCAAAGATGCTATAGAAGATGTGATAGTAGCAAGTGCGGTAGCCGAAGCATTGAATGACATGTTTAAGGAATTTGATAAAAACGTAAGTGCTATTATAATAAAAGAAGAAGATAATTATCTAAACAGTTCGCCCGATATCTACCATATAATATATATCATAGGTGGCACTATTCTGCTACTGTTAGTCATTATTTTAATATTGGCAATTTATATAGCGCGCAATAAATACAGAACCAGGAAATATGAAATAATGAAATATGACAATATGAGCATTAAATCTGAGCATCATGATAGTCTTGAAACAGTGTCTATGGAAATTATTGATAATCGGTACTAATAAAATAGTTTAACTCTTTTAGAACCAGTTTGGTACTGTAATTTCAGTTCATTACTCGTTGAGAATATTGATGATTTTTTTTAAATGAGTATCGGTAGTTACATATTACCATATCATCCATTATATAATCGATGATGCATGTATTAGAATACTTTCCGAATAAGTCTTCTAAATATTGTATTAATTATGAAAAACTATGCTATGTGAGTATGATTCAAAGATGTTTAATGATACGATACTAGATTTTATCTCTAGCGAGATTGTTTAGAATCATTTATCATAACTATGTTTAATAAATTCATCAACGAATATCGATAAAGACCTCTTGTAATTCGAGTATAGGAAGTAGTATTACCATATCAACTTCCGAGTTAACAATTACTCTAAAACATGAGGATTGTACTCCTGTCTTTATTGGAGATCACTATTTAGTCGTTGATAAACTAGTAACCTCAGGTTTCTTTACAAACGATAAAGTACAACATCAAGACCTCACAACACAGTGCAAGATTAATCTAGAAATCAAATGTAATTCTGGAGGAGAATCTAGACAACTAACACCCACGGCGAAGTATACTTTATGCCTCATTCAGAAACGGTAACTGTAGTAGGAGACTGTCTCTCTAATCTCGATGTATATATAATATATACCAATACGGACGCGATATATTCCGACATGAATGGCGTCGCTTATCATATGTTATATCCTAAATGTTGATCATATTCCACAAATGATTGTGAACGAGATTAAATCATCTAACAAATAATTAGTTTTTTATGACATTAACATATAATAAATAAATTAATCATTATTGACTTAACGATGACGAAAGTTATCATTATCTTAGGATTCTTGATTATTAATACAAATTCGTTGTGTCTATGAAATGTGAACAAGGTGTCTCATATTATAATGCACAAGAATTAAAGTGTTGTAAACTATCTAGCCAGGAACATATTCAGATTATCGATGTGATAAATACAGCGATACCATCTGTGGACATTGTCCAAGTGACACATTCACGTCAATATATAATCGTTCTCCTCGGTGTCATAGTTGTAGAGGTCACACCTTGTACACCTACCACAAATAGAATATGTCATTGTGACTCGAATAGTTATCGTCTCCTTAAAGCTTCTGATGGTAACTGTGTTACATGTGCTCCTAAAACAAAATATGGTCGTGTGTACGGAAAGAAAGGAGAAAATGATATGGAATACCATTTGTAAGAAATGTCGGAAGGGTACTTATTCAGATATTGTATCTGACTCTGATCAATGTAAACCTATGACAAGATAAGACTTACTCGCATCTACTGGATAGACATAAATATCCTCCTCGTAATAATGAAATATAATATACACTAATTATTAATATCAATCGAGTATTAACATATAAGTTATTTTTAAACCCCTTTTGGGTTCCGTCCTAAACGGCGTTTCGGTCTGTGTCGCCACCATGGTCACACCGAGCCTCTGCGTGCTCCTCCATCGAGGACGACTTCAACTATGACAGCTCGGTGGCGTCTGCCAGCGTGTACATACGAATGGCATTTCTAAGAAAAGTCTACGGTATCCTTTCTACAATTTCCTTTAACAACGGCAACAGCTGCAGTATTTTTATACTTTGAATGCATCGGACATTTATACAAGGGAGTCCTGTTCTAATATTGGCATCAATGTTCGGATCTATAGGCTTGATTTTCGCATTGACTTTACACAGACATGAACATCCCCTGAATCTGTACATACTTTGTGGATTTACACTGTTAGAATCTCTAACGCTGGCCTCTGTTGTTACTTTCTATGATGCACGTATCGTTATGCAAGCTTTCATGTTGACTACTGCAGTGTTTCTTGCTCTGACTACATATACTCTACAATCAAAGAGAGATTTCAGTAAACTTGTAACAGGATTGTTTGCTGCTTTCTGGATTTTAATTTTGTCAGGAGTCTTGAGGATAAAGTTTAAAATAGAATTAATAAAGAACATATAGGTCATTTTTTAAACATGGATAGAAACCAAGGTTGTTAGTTAATAATATACAAGATATTTTTTCTCACTCTGATCCATGTAAACCAAGGACGAGAGACACTCTCATTCCTCATTCACGACACCATTAAAAATGGAAATTAAAGCCCTCTATTAAGCACAGACGGCTACAGGTCTACCATCAGGTTACCTTCGTCTACCTTCACAATGGCCTCTCCTTGTGCCCAGTTCAGTCCCTGTCATTGCCACGCTACTAAGGACTCCCTGAATACCGTGACTGACGTCAGACATTGTCTGACTGAATACATCCTGTGGGTTTCTCATAGATGGACCCATAGAGAAAGCGCAGGGCCTCTCTACAGGCTTCTCATCTCTTTCAGAATTGATGCAATGGAGCTATTTGGTAGCGAGTTGAAGGAGTTCTCGAATTCACTTCCGTGGGACAATATCGACAATTGCGTGGAGATCATTAAATGTTTCATCAGAAATGACTCCATGAAAACCGCCAAAGAACTTTGTGCAATAATTGGACTTTGTACTCAATCAGCTATTGTCACTGGAAGAGTCTTCAATGATAAGTATATCGACATACTACTTATGCTGCGAAAGATTCTGAACGAGAACGACTATCTCACCCTCTTGGATCATATCCTCACTGCTAAATACTAAATCTCCTTCATGCTCTCTCACTAATACTCTTACTCACTACACTTTTTATCATCTTATGATGAATGATTGCCTTCATCATTTTTTCGTGGAATATAATATAGGAATAATTAGCACCAGAATAGCTATGGATATCTCGTTAAGAATATTCTCTATAAGAGACATAATGTAGACATAGTTATTATATCCTTCTTAGATAAGTGTTACGCTACTGGAAAGTTTCCATCGTTATTATTACATGAAGATGATATAATTAAACCAACATTGAGATTGGCTCTTATGTTAGCTGGATTGAATTACTGTAATAAATGCATCGAGTATAGAGGGATATAGCAATTCTCGATAATAGTCATGCAATATTTGAATGAGACTGATAATTTAGGTAATACAGTACTACACACATATCTTTCTAGATTATATATCGTTAAAAATCTGTAAGATGTATATTTCTCATAAGTATCCACTGTGTAATATTATTAATGGATATATAGATAACGCAATAGGGACTAATAGTATTGTAAAAGATATAATCGACTATTTACGTACATATCCAGATATCTATATTCCTACTAGTTTGCTGCGTAGTTGCATCATTGATATGCATGATTTATCAGGATTCAGAGATGAATTACTAAGTAAACTACAATCCCACAATAAGTAAGAATCAAATATCAAAAACTCACTTTTGATTTTTCTAGTCTTAAGTAATACATATATTTATTAATAGACCTATGAAATAAAAAAAGGTAACAATGGATTCGCGTATAGCTATTTACGTATTAGTATCGGCATCTCTTTTGTATCTTGTTAATTGTCACAAACTAGTACATTACTTCAATCTGAAAATAAATGGAAGTGATATAACTAATACAGCAGATATATTGCTGGACAATTATCCAATTATGACCTTTGATGGAAAGGATATTTATCCATCTATCTCGTTCATGGTCGGTAATAAACTTTTCCTAGATCTTTATAAAAATATCTTTGAAGAATTTTTCAGACTATTTCGAGTATCTGTAAGTAGTCAATACGAGGAATTAGAATATTATTATTCATGTGATTATACTAACAACCGTCCTACAATTAAACAACATTACTTTTATAACGGCGACGAATATACTGAAATTGATAGATCGAAAAAAGCCACTAATAAAAACAGTTGGTTAATTACTTCAGGCTTTAGACTACAAAAATGGTTCGATAGCGAAGATTGTATAATTTATCTCAGATCTTTAGTTAGAAGAATGGAAGACAGTAACAAAAACAGTAAAAAAACTTAGTACTTAGATATCGAAAAAATATATTTTTGTAGACTCTTGAGAATAGAAGGAAAACATGTACATAATTATAAAAAATGAAAATCAATGGCGAATAAGACAGTGCGATTCGCACCATGGAGTCGGTAGATTTCATGGCTGTCGATGAGCAGTTTCACGACGACCTCGATCTTTGGTCATTATCTTTGGTAGATGATTATAAAAAACATGGATTAGGTGTTGACTGTTATGTTCTAGAACCAGTTGTTGACAGGAAAATATTTGATAGATTTCTCCTTGAACCAATTTGTGATCCTGTAGATGTTCTGTATGATTATTTTAGGATTCATAGAGATAATATTGATCAGTATATAGTAGATAGACTGTTTGCATATATTACATATAAAGATATTATATCTGCATTAGTGTCAAAGAATTATATGGAAGATATTTTCTCTATAATTATTAAGAATTGTAATTCTGTGCAAGATCTCTTACTTTACTATCTATCTAATGCATATGTAGAAATAGACATTGTTGATCTTATGGTAGATCATGGGGCTGTAATATATAAAATAGAATGCTTGAATGCCTATTTTAGGGGAATATGTAAAAAGGAAAGTAGTGTTGTTGAGTTTATTTTGAATTGTGGTATCCCAGATGAAAATGATGTTAAATTAGATCTATATAAAATAATTCAGTATACTAGGGGATTCCTTGTAGATGAACCCACAGTATTAGAAATTTATAAGCTTTGTATCCCATATATTGAAGATATCAATCAACTAGATGCTGGTGGAAGGACCTTGCTTTATCGCGCTATCTATGCAGGTTATATAGATTTAGTATCATGGCTATTAGAAAATGGAGCAAATGTCAACGCAGTAATGAGTAATGGATATACATGTCTTGACGTGGCCGTGGATAGGGGATCTGTCATCGCCCGTAGGGAAGCACATCTTAAAATATTAGAAATATTGCTTAGAGAACCATTGTCTATTGACTGTATAAAATTAGCTATACTTAATAATACAATTGAAAACCATGATGTGATAAAGCTCTGTATCAAGTATTTTATGATGGTAGATTATTCACTTTGTAATGTGTATGCATCATCACTCTTTGATTATATAATTGATTGTAAACAAGAATTGGAGTACATTAGGCAGATGAAAATTCATAATACAACCATGTATGAGTTAATCTATAATAGAGACAAAAACAAGCATGCTTCCCATATTCTACATAGGTATTCTAAACATCCAGTTTTGACACAGTGTATCACTAAAGGATTCAAGATTTACACAGAAGTAACCGAGCAGGTCACTAAAGCTCTAAACAGACGTGCTCTAATAGATGAGATAATAAACAATGTATCAACTGATGACAATCTCCTATCAAAACTTCCATTAGAAATTAGGGATCTAATTGTTTCACAAGCTGTCATATAGAGTTCTATCCACCCACCTTTCTTGAAATGAGTTAATAGTCATAAGTTAGTTAAGTCATAAGTTAGTTAAGTCATAAGTTAGTTAAGTCATAAGTTAGTTAAGTCATAAGTTAGTTAAGTCATAAGTTAGTTAAGTCATAAGTTAGTTAAGTCATAAGTTAGTTTATAGTCTAACACTTCTAATTTTTATACCTTGATCTTTTTCTCTAATTATGAAAAAGTAAATCATTATGAAGATGGATGAAATGGACGAGATTGTGCGCATCGTTAACGATAGTATGTGGTACGTACCTAACGCATTTATGGACGACGGTGATAATGAAGGTCACATTTCTGTCAATAATGTCTGTCATATGTATCTCGCATTCTTTGATGTGGATATATCATCTCATCTGTTTAAATTAGTTATTAAACACTGCGATCTGAATAAACGACTAAAATGTGGTAACTCTCCATTACATTGCTATACGATGAATACACGATTTAATCCATCTGTATTAAAGATATTGTTACGCCACGGCATGCGTAACTTTGATAGCAAGGATAAAAAAGGACATATTCCTCTACACCACTATCTGATTCATTCACTATCAATCGATAACAAGATCTTTGATATACTAACGGACCCCATTGATGACTTTAGTAAATCATCCGATCTATTGCTGTGTTATCTTAGATATAAATTCAATGGGAGCTTAAACTATTACGTTCTGTACAAATTATTGACTAAAGGATCTGACCCTAATTGCGTCGATGAGGATGGACTCACTTCTCTTCATTACTACTGTAAACACATATCCGCGTTCCACGAAAGCAATTATTACAAGTCAAAGAGTCACACTAAGATGCGAGCTGAGAAGCGATTCATCTACGCGATAATAGATCATGGAGCAAACATTAACGCGGTTACGAAAATCGGAAATACGCCGTTACACACTTACCTTCAACAGTATACCAAACATAGTCCTCGTGTGGTGTATGCTCTTTTATCTCGAGGAGCCGATACGAGGATACGTAATAATCTTGATTGTACACCCATCATGGAATACATAAAGAACGATTGTGCAACAGGTCATATTCTCATAATGTTACTCAATTGGCACGAACAAAAATACGGGAAATTACAAAAGGAAGAAGGACAACATCTACTTTATCTATTCATAAAACATAATCAAGGATATGGAAGTCGCTCTCTCAATATACTACGGTATCTACTAGATAGATTCGACATTCAGAAAGACGAATACTATAATACAATGACTCCTCTTCATACCGCCTTCCAGAATTGCAATAACAATGTTGCCTCATACCTCGTATACATCGGATACGACATCAACCTTCCGACTAAAGACGATAAGACAGTATTCGACTTGGTGTTTGAAAACAGAAACATTATATACAAGGCGGATGTCGTTAATGACATTATCCACCACAGACTGAAAGTATCTCTACCTATGATTAAATCGTTGTTCTACAAGATGTCGGAGTTCTCTCCCTACGACGATCACTACGTAAAGAAGATAATAGCCTACTGCCTATTAAGGGACGAGTCATTTGCGGAACTACATACTAAATTCTGTTTAAACGAGGACTATAAAAGTGTATTTATGAAAAATATATCATTCGATAAGATAGATTCCATCATCGAAAAATGTAGTCGTGACATAAGTCTCCTCAAAGAGATTCGAATCTCAGACACCGACTTGTATACGGTATTGAGAACAGAAGACATCCGGTATCACACATATCTCGAAGCCATACATTCAGACAAACGCATTTCATTTCCCATGTACGACGATCTCATAGAACAGTGTCATCTATCGATGGAGCATAAAAGTAAACTCGTCGACAAAGCACTCAATAAATTAGAGTCTACCATCGATAGTCAATCTAGACTATCGTATTTGCCTCCGGAAATTATGCGCAATATCATAACCAAGCTAAGCGACTACCATCTAAACAGTATGTTGTACGGAAAGAACCATTACAAATATTATCCATGATAGAAAGAAAATATTTAAAAAATAATCTATATGATTGGAGAAGTAGGAAACAAACAGTAACAAGACGACGATTACTACATTATTAAATCATGAGGTCCGTATTATACTCGTATATATTGTTTCTCTCATGTATAATAATAAACGGAAGAGATATAGCACCACATGCACCATCCAATGGAAAGTGTAAAGACAACGAATACAGAAGCCGTAATCTATGTTGTCTATCGTGTCCTCCGGGAACTTACGCTTCCAGATTATGTGATAGCAAGACTAATACACAATGTACACCGTGTGGTTCGGATACCTTTACATCTCACAATAATCATTTACAGGCTTGTCTAAGTTGTAACGGAAGATGTGATAGTAATCAGGTAGAGACGCGATCGTGTAACACGACTCACAATAGAATCTGTGAATGCTCTCCAGGATATTATTGTCTTCTCAAAGGAGCATCAGGGTGTAGAACATGTATTTCTAAAACAAAGTGTGGAATAGGATACGGAGTATCCGGATACACGTCTACCGGAGACGTCATCTGTTCTCCGTGTGGTCCCGGAACATATTCTCACACCGTCTCTTCCACAGATAAATGCGAACCCGTCGTAACCAGCAATACATTTAACTATATCGATGTGGAAATTAACCTGTATCCAGTCAACGACACATCGTGTACTCGGACGACCACTACCGGTCTCAGCGAATCCATCTCAACGTCGGAACTAACTATTACCATGAATCATAAAGATTGTGATCCAGTCTTTCGTGCAGAATACTTCTCTGTCCTTAATAATGTAGCAACTTCAGGATTCTTTACAGGAGAAAATAGATATCAGAATACTTCAAAGATATGTACTCTGAATTTCGAGATTAAATGTAACAACAAAGATTCATCTTCCAAACAGTTAACGAAAACAAAGAATGATACTATCATGCCGCATTCAGAGACGGTAACTCTAGTGGGCGACTGTCTATCTAGCGTCGACATCTACATACTATATAGTAATACCAATACTCAAGACTACGAAAATGATACAATCTCTTATCATATGGGTAATGTTCTCGATGTCAATAGCCATATGCCCGCTAGTTGCGATATACATAAACTGATCACTAATTCCCAGAATCCCACCCACTTATAGTAAGTTTTTTTACCTATAAATAATAAATACAATAATTAATTTCTCGTAAAAGTAGAAAATATATTCTAATTTATTATATGGTAAGAAAGTAGAATCATCTAGAACAGTAATCAATCAATAGCAATCATGAAACAATATATTGTCCTGGCATGCATGTGCCTAGTGGCAGCTGCTATGCCTACTAGTCTTCAACAATCTTCATCCTCGTGTACTGAAGAAGAAAACAAACATCATATGGGAATCGATGTTATTATCAAAGTCACAAAGCAAGACCAAACACCGACCAATGATAAGATTTGTCAATCCGTAACGGAAGTTACAGAGACCGAAGATGATGAGGTATCCGAAGAAGTTGTAAAAGGAGATCCCACCACTTATTACACTATCGTCGGTGCGGGTCTTAACATGAACTTTGGATTCACCAAATGCCCAAAGATTTCATCCATCTCCGAATCCTCTGATGGAAACACTGTGAATACTAGATTGTCCAGCGTGTCACCGGGACAAGGTAAGGACTCTCCCGCGATCACGCGTGAAGAAGCTCTGGCTATGATCAAAGACTGTGAGATGTCTATCGACATCAGATGTAGCGAAGAAGAGAAAGACAGTGACATCAAGACCCATCCAGTACTTGGGTCTAACATCTCACATAAGAAAGTGAGTTACAAAGATATCATCGGTTCAACGATCGTTGATACAAAATGTGTCAAGAACCTAGAGTTTAGCGTACGTATCGGAGACATGTGTGAGGAATCATCTGAACTTGAAGTCAAGGATGGATTCAAGTATGTCGACGGATCGGCATCTGAAGGTGCAACCGATGATACTTCACTCATCGATTCAACAAAACTCAAAGCATGTGTCTGAATCGATAACTCTATTCATCTGAAAATGGATGAGTTGGGTTAATCGAACGATTCAGACACCGCACCACGAATTAAAAAAGACCGGGCACTATATTCCGGTTTGCAAAACAAAAATATTTAACTACATTCACAAAAAGTTACCTCTCGTTACTTCTTCTTTCTGTTTCAATATGTGATACGATATGATCACTATTCGTATTCTCTTGGTCTCATAAAAAAGTTTTACAAAAAAAAAAAAAAATATTTTTATTCTCTTTCTCTCTTCGATGGTCTCACAAAAATATTAAACCTCTTTCTGATGTCTCAACTATTTCGTAAACGATAACGTCCAACAATATATTCTCGTAGAGCTTATCAACATCCTTATACCAATCTAGGTTGTCAGACAATTGCATCATAAAATAATGTTTATAATTTACACGTTAACATCATATAATAAACGTATATAGTTAATATTTTTGGAATATAAATGATCTGTAAAATCCATGTAGGGGACACTGCTCACGTTTTTTCTCTAGTACATAATTTCACACAAGTTTTTATACAGACAAATTAATTCTCGTCCATATATTTTAAAACATTGACTTTTGTACTAAGAAAAATATCTTGACTAACCATCTCTTTCTCTCTTCGATGGGTCTCACAAAAATATTAAACCTCTTTCTGATGGAGTCGTAAAAAGTTTTTATCCTTTCTCTCTTCGATAGGTCTCACAAAAATATTAAACCTCTTTCTGATGGTCTCTATAAACGATTGATTTTTCTTACCCTCTAGAGTTTCCTACGGTCGTGGGTCACACATTTTTTTCTAGACACTAAATAAAATAGTAAAATTAAATTAATTA

>2017_UK_Monkeypox_MT903345.1

ATAAGTTTTAGTACATTAATATTATATTTTACTATTTTATTTAGTGTCTAGAAAAAAATGTGTGACCCACGACCGTAGGAAACTCTAGAGGGTAAGAAAAATCAATCGTTTATAGAGACCATCAGAAAGAGGTTTAATATTTTTGTGAGACCTATCGAAGAGAGAAAGGATAAAAACTTTTTACGACTCCATCAGAAAGAGGTTTAATATTTTTGTGAGACCCATCGAAGAGAGAAAGAGATGGTTAGTCAAGATATTTTTCTTAGTACAAAAGTCAATGTTTTAAAATATATGGACGAGAATTAATTTGTCTGTATAAAAACTTGTGTGAAATTATGTACTAGAGAAAAAACGTGAGCAGTGTCCCCTACATGGATTTTACAGATCATTTATATTCCAAAAATATTAACTATATACGTTTATTATATGATGTTAACGTGTAAATTATAAACATTATTTTATGATGCAATTGTCTGACAACCTAGATTGGTATAAGGATGTTGATAAGCTCTACGAGAATATATTGTTGGACGTTATCGTTTACGAAATAGTTGAGACATCAGAAAGAGGTTTAATATTTTTGTGAGACCATCGAAGAGAGAAAGAGAATAAAAATATTTTTTTTTTTTTTGTAAAACTTTTTTATGAGACCAAGAGAATACGAATAGTGATCATATCGTATCACATATTGAAACAGAAAGAAGAAGTAACGAGAGGTAACTTTTTGTGAATGTAGTTAAATATTTTTGTTTTGCAAACCGGAATATAGTGCCCGGTCTTTTTTAATTCGTGGTGCGGTGTCTGAATCGTTCGATTAACCCAACTCATCCATTTTCAGATGAATAGAGTTATCGATTCAGACACATGCTTTGAGTTTTGTTGAATCGATGAGTGAAGTATCATCGGTTGCACCTTCAGATGCCGATCCGTCGACATACTTGAATCCATCCTTGACTTCAAGTTCAGATGATTCCTCACACATGTCTCCGATACGTACGCTAAACTCTAGGTTCTTGACACATTTTGTATCAACGATCGTTGAACCGATGATATCTTTGTAACTCACTTTCTTATGTGAGATGTTAGACCCAAGTACTGGATGGGTCTTGATGTCACTGTCTTTCTCTTCTTCGCTACATCTGATGTCGATAGACATCTCACAGTCTTTGATCATAGCCAGAGCTTCTTCACGCGTGATCGCGGGAGAGTCCTTACCTTGTCCCGGTGACACGCTGGACAATCTAGTATTCACAGTGTTTCCATCAGAGGATTCGGAGATGGATGAAATCTTTGGGCATTTGGTGAATCCAAAGTTCATGTTAAGACCCGCACCGACGATAGTGTAATAAGTGGTGGGATCTCCTTTTACAACTTCTTCGGATACCTCATCATCTTCGGTCTCTGTAACTTCCGTTACGGATTGACAAATCTTATCATTGGTCGGTGTTTGGTCTTGCTTTGTGACTTTGATAATAACATCGATTCCCATATGATGTTTGTTTTCTTCTTCAGTACACGAGGATGAAGATTGTTGAAGACTAGTAGGCATAGCAGCTGCCACTAGGCACATGCATGCCAGGACAATATATTGTTTCATGATTGCTATTGATTGATTACTGTTCTAGATGATTCTACTTTCTTACCATATAATAAATTAGAATATATTTTCTACTTTTACGAGAAATTAATTATTGTATTTATTATTTATAGGTAAAAAAACTTACTATAAGTGGGTGGGATTCTGGGAATTAGTGATCAGTTTATGTATATCGCAACTAGCGGGCATATGGCTATTGACATCGAGAACATTACCCATATGATAAGAGATTGTATCATTTTCGTAGTCTTGAGTATTGGTATTACTATATAGTATGTAGATGTCGACGCTAGATAGACAGTCGCCCACTAGAGTTACCGTCTCTGAATGCGGCATGATAGTATCATTCTTTGTTTTCGTTAACTGTTTGGAAGATGAATCTTTGTTGTTACATTTAATCTCGAAATTCAGAGTACATATCTTTGAAGTATTCTGATATCTATTTTCTCCTGTAAAGAATCCTGAAGTTGCTACATTATTAAGGACAGAGAAGTATTCTGCACGAAAGACTGGATCACAATCTTTATGATTCATGGTAATAGTTAGTTCCGACGTTGAGATGGATTCGCTGAGACCGGTAGTGGTCGTCCGAGTACACGATGTGTCGTTGACTGGATACAGGTTAATTTCCACATCGATATAGTTAAATGTATTGCTGGTTACGACGGGTTCGCATTTATCTGTGGAAGAGACGGTGTGAGAATATGTTCCGGGACCACACGGAGAACAGATGACGTCTCCGGTAGACGTGTATCCGGATACTCCGTATCCTATTCCACACTTTGTTTTAGAAATACATGTTCTACACCCTGATGCTCCTTTGAGAAGACAATAATATCCTGGAGAGCATTCACAGATTCTATTGTGAGTCGTGTTACACGATCGCGTCTCTACCTGATTACTATCACATCTTCCGTTACAACTTAGACAAGCCTGTAAATGATTATTGTGAGATGTAAAGGTATCCGAACCACACGGTGTACATTGTGTATTAGTCTTGCTATCACATAATCTGGAAGCGTAAGTTCCCGGAGGACACGATAGACAACATAGATTACGGCTTCTGTATTCGTTGTCTTTACACTTTCCATTGGATGGTGCATGTGGTGCTATATCTCTTCCGTTTATTATTATACATGAGAGAAACAATATATACGAGTATAATACGGACCTCATGATTTAATAATGTAGTAATCGTCGTCTTGTTACTGTTTGTTTCCTACTTCTCCAATCATATAGATTATTTTTTAAATATTTTCTTTCTATCATGGATAATATTTGTAATGGTTCTTTCCGTACAACATACTGTTTAGATGGTAGTCGCTTAGCTTGGTTATGATATTGCGCATAATTTCCGGAGGCAAATACGATAGTCTAGATTGACTATCGATGGTAGACTCTAATTTATTGAGTGCTTTGTCGACGAGTTTACTTTTATGCTCCATCGATAGATGACACTGTTCTATGAGATCGTCGTACATGGGAAATGAAATGCGTTTGTCTGAATGTATGGCTTCGAGATATGTGTGATACCGGATGTCTTCTGTTCTCAATACCGTATACAAGTCGGTGTCTGAGATTCGAATCTCTTTGAGGAGACTTATGTCACGACTACATTTTTCGATGATGGAATCTATCTTATCGAATGATATATTTTTCATAAATACACTTTTATAGTCCTCGTTTAAACAGAATTTAGTATGTAGTTCCGCAAATGACTCGTCCCTTAATAGGCAGTAGGCTATTATCTTCTTTACGTAGTGATCGTCGTAGGGAGAGAACTCCGACATCTTGTAGAACAACGATTTAATCATAGGTAGAGATACTTTCAGTCTGTGGTGGATAATGTCATTAACGACATCCGCCTTGTATATAATGTTTCTGTTTTCAAACACCAAGTCGAATACTGTCTTATCGTCTTTAGTCGGAAGGTTGATGTCGTATCCGATGTATACGAGGTATGAGGCAACATTGTTATTGCAATTCTGGAAGGCGGTATGAAGAGGAGTCATTGTATTATAGTATTCGTCTTTCTGAATGTCGAATCTATCTAGTAGATACCGTAGTATATTGAGAGAGCGACTTCCATATCCTTGATTATGTTTTATGAATAGATAAAGTAGATGTTGTCCTTCTTCCTTTTGTAATTTCCCGTATTTTTGTTCGTGCCAATTGAGTAACATTATGAGAATATGACCTGTTGCACAATCGTTCTTTATGTATTCCATGATGGGTGTACAATCAAGATTATTACGTATCCTCGTATCGGCTCCTCGAGATAAAAGAGCATACACCACACGAGGACTATGTTTGGTATACTGTTGAAGGTAAGTGTGTAACGGCGTATTTCCGATTTTCGTAACCGCGTTAATGTTTGCTCCATGATCTATTATCGCGTAGATGAATCGCTTCTCAGCTCGCATCTTAGTGTGACTCTTTGACTTGTAATAATTGCTTTCGTGGAACGCGGATATGTGTTTACAGTAGTAATGAAGAGAAGTGAGTCCATCCTCATCGACGCAATTAGGGTCAGATCCTTTAGTCAATAATTTGTACAGAACGTAATAGTTTAAGCTCCCATTGAATTTATATCTAAGATAACACAGCAATAGATCGGATGATTTACTAAAGTCATCAATGGGGTCCGTTAGTATATCAAAGATCTTGTTATCGATTGATAGTGAATGAATCAGATAGTGGTGTAGAGGAATATGTCCTTTTTTATCCTTGCTATCAAAGTTACGCATGCCGTGGCGTAACAATATCTTTAATACAGATGGATTAAATCGTGTATTCATCGTATAGCAATGTAATGGAGAGTTACCACATTTTAGTCGTTTATTCAGATCGCAGTGTTTAATAACTAATTTAAACAGATGAGATGATATATCCACATCAAAGAATGCGAGATACATATGACAGACATTATTGACAGAAATGTGACCTTCATTATCACCGTCGTCCATAAATGCGTTAGGTACGTACCACATACTATCGTTAACGATGCGCACAATCTCGTCCATTTCATCCATCTTCATAATGATTTACTTTTTCATAATTAGAGAAAAAGATCAAGGTATAAAAATTAGAAGTGTTAGACTATAAACTAACTTATGACTTAACTAACTTATGACTTAACTAACTTATGACTTAACTAACTTATGACTTAACTAACTTATGACTTAACTAACTTATGACTTAACTAACTTATGACTTAACTAACTTATGACTATTAACTCATTTCAAGAAAGGTGGGTGGATAGAACTCTATATGACAGCTTGTGAAACAATTAGATCCCTAATTTCTAATGGAAGTTTTGATAGGAGATTGTCATCAGTTGATACATTGTTTATTATCTCATCTATTAGAGCACGTCTGTTTAGAGCTTTAGTGACCTGCTCGGTTACTTCTGTGTAAATCTTGAATCCTTTAGTGATACACTGTGTCAAAACTGGATGTTTAGAATACCTATGTAGAATATGGGAAGCATGCTTGTTTTTGTCTCTATTATAGATTAACTCATACATGGTTGTATTATGAATTTTCATCTGCCTAATGTACTCCAATTCTTGTTTACAATCAATTATATAATCAAAGAGTGATGATGCATACACATTACAAAGTGAATAATCTACCATCATAAAATACTTGATACAGAGCTTTATCACATCATGGTTTTCAATTGTATTATTAAGTATAGCTAATTTTATACAGTCAATAGACAATGGTTCTCTAAGCAATATTTCTAATATTTTAAGATGTGCTTCCCTACGGGCGATGACAGATCCCCTATCCACGGCCACGTCAAGACATGTATATCCATTACTCATTACTGCGTTGACATTTGCTCCATTTTCTAATAGCCATGATACTAAATCTATATAACCTGCATAGATAGCGCGATAAAGCAAGGTCCTTCCACCAGCATCTAGTTGATTGATATCTTCAATATATGGGATACAAAGCTTATAAATTTCTAATACTGTGGGTTCATCTACAAGGAATCCCCTAGTATACTGAATTATTTTATATAGATCTAATTTAACATCATTTTCATCTGGGATACCACAATTCAAAATAAACTCAACAACACTACTTTCCTTTTTACATATTCCCCTAAAATAGGCATTCAAGCATTCTATTTTATATATTACAGCCCCATGATCTACCATAAGATCAACAATGTCTATTTCTACATATGCATTAGATAGATAGTAAAGTAAGAGATCTTGCACAGAATTACAATTCTTAATAATTATAGAGAAAATATCTTCCATATAATTCTTTGACACTAATGCAGATATAATATCTTTATATGTAATATATGCAAACAGTCTATCTACTATATACTGATCAATATTATCTCTATGAATCCTAAAATAATCATACAGAACATCTACAGGATCACAAATTGGTTCAAGGAGAAATCTATCAAATATTTTCCTGTCAACAACTGGTTCTAGAACATAACAGTCAACACCTAATCCATGTTTTTTATAATCATCTACCAAAGATAATGACCAAAGATCGAGGTCGTCGTGAAACTGCTCATCGACAGCCATGAAATCTACCGACTCCATGGTGCGAATCGCACTGTCTTATTCGCCATTGATTTTCATTTTTTATAATTATGTACATGTTTTCCTTCTATTCTCAAGAGTCTACAAAAATATATTTTTTCGATATCTAAGTACTAAGTTTTTTTACTGTTTTTGTTACTGTCTTCCATTCTTCTAACTAAAGATCTGAGATAAATTATACAATCTTCGCTATCGAACCATTTTTGTAGTCTAAAGCCTGAAGTAATTAACCAACTGTTTTTATTAGTGGCTTTTTTCGATCTATCCTCGTCCTCTCATCATCCTTATATTATTATCATTATTATCATAGTCTATTAAACACAAATCATCTACGTTTATAACAACATTCTCATTATTAATTAGTTCTGTAGAATATCTTTAATAATTTGGCTATACATCTGTTCAATACTATCTATTGATGATTTCTTTTTTAAGACTTAAACTAGTTATGGTAATGACGATGAAATCGAGTAGTAACTTCTAATAAAGACTTGATATCATTATCATATGTTTGATCGTCATAGTTAATAGTGTGGCTAAATGGTACTGTTAATAAGTTTATAGACAATATCATAGTATTTTCTTTCCAGAATTAGATTATTTTTTTAAATACTGATCCTCACAATTCCGTGATGTAGCAGTAGTTGGTGCATGGTCTATATCGTTAAAATGTATCATATATAATAGTTTTCTGACGTGGAGTACAGAATTTTCGATTAATGAGTTCATGGTAAGGAAGGGCAAATGCCTGTATATAATATACATAAGTTAATAGTTTTTTATCATATTTTCTAATACCATAATAAAAATTATCATTATTGCGTTTGGTAGTTCTGCCCTATCATCTATATCACTGTCACTCTCGCTCTCACTATATCTTCTAAAATTACAAACAACTGGATATTCGATAACAGCATTTGTGTAGTTTTTGTCTTTTACAGTATATACGTTATTGTCAAAATCTAAACAAATATTAGCATAATACATCTATCTATAAGATCAGGATCCATGTTCGAGCATACTAGCCATGTATATTTGTAACTTCGTCGTACAGCGTTAGATCAATAGAATAAACAATCGTGTGACGCAACTTTTTTACGATCTAGTTGTATGAGTTTATCGTTTACATAAGCAATTAACGGCTTTAACAGATGATCTGAGTAATAATATACCTCTGTTATACGTTTAATGTTCACGGTCTTAGTATTTTTAGATATCAATTGTGATTTACACCATATTCGACTCCCTTGTGTGTAACGTTAGAAATTCTAAATCTATATTATCTATTACAGCGTAAAACACATTCAATATTGTATTGTTATTTTTATATTATTTACACAATTAACAATGTATTATTAGTTTATATTACTGAATTAATAATATAAAATTCCCAATCTTGTCATAAACACAAAATCCATTAAAAATGTCGATAAAATATCTGATGTTGTTGTTCGCTGCTATGATAATCAGATCATTAGCCGATAGTGGTAATGCTATCGAAACAACATCGCCAGAAATTACAAACGCTACAACAGATATTCCAGCTATCAGATTATGTGGTCCAGAGGGAGATGGATATTGTTTACACGGTGACTGTATCCACGCTAGAGATATCGACGGTATGTATTGTAGATGCTCTCATGGTTATACAGGTATTAGATGTCAGCATGTAGTATTAGTAGACTATCAACGTTCAGGAAAACCAGACACTACAACATCATATATCCCATCTCTAGGTATTGTGCTTGTATTAGTAGGCATTATTATTACGTGTTGTCTATTATCTGTTTATATGTTCACTCGAAGAACTAAACTACCTATACAAGATATGGTTGTGTTATATTTTTTATAAAATTTTTTTATGAGTATTTTTACAAAAATGTATATGTATAAAAAAAATACTAAGTATACGATGTATCCTGTATTATTTGTATTTATCTAAACAATACTTCTGCCTCTAGATGGGATACAAAAATTTTTTATTTCAGCATATTAAAGTAAAATCTAGTTACCTTGAAAATGAATACAGTGGGTGGTTCCGTATCACCAGTAAGAACATAATAGTCGAATACAGTATCCGATTGAGATTTTGCATACAATACTAGTCTAGAAAGAAATTTGTAATCATCCTCTGTGACAGGAGTCCATATATCTGTATCATCGTCTAGTTTTTATCAGTGTCCTATGCTATATTCCTGTTATCATCATTAGTTAATGAAAATAACTCTCGTGCTTCAGAAAAGTCAAATATTGTATCCATACATATATCTCCAAAACTATCACTTATACGTTTATCTTTAACGAACATATACCTAGATGGTTATTTACTAACAGACATTTTTTCAAGATCTATTGACAATAACTCCTATAGTTTCCACATCAACCAAGTAATGATCATCTATTGTTATATAACAATAACATAACTCTTTTCCATTTTTATCAGTATCTATATCAACGTCGTTGTAGTGAATAGTAGTCATTGATCTATTATATGAAACGGATATGTCTAGTTAATATTTTCTTTGATTTAAAGTCTATAGTCTTTACAAACATAATATCCTTATCCGACTTTATATTTCCTGTAGGGTGGCATAATTTTATTCTGCCTCCACAATCAGTGTTTCCAAATATATTACTAGACAATATTCCATATAGTTATTAGTTAAGGGACCCAATTAGAACACGTACGCGCTTATTCATCATTTGGATCGTATTTCATAAAAGTTATTATGTTATAGATGTCAACACATTCTACATTTTTTTAATTGTCTATATAGTATTTTTCTGATATTTTCTATAATATCAGAATTGTCTTCCATAGGAAGTTGTATACTATTCGGAATCAGTTACATGTTTAAATAATTCTCTGATGTCATTCCTTATACAATCAAATTCATTATTAAACAGTTTAATAGTCTGTAGACCTTTATCGTCGTACATATCCATTGTCTTATTAGTTACGCTTATTTTTATGGGTTTTACGTTGCTTTATTATATTTTATAATAATGATTGTTTGACAATGTCGTAGTATAGATATATTATTAGAGGAGGTATAATTATAAAAAGTTTCTGAGTACGATGTTATAAGAGGAGAGGACACATTAACATCATACATCAATTAACTACATTCTTATAACATTGTAATCAAAAGAATTGCAATTTTGATGTATAACAACTGTCAATGGAATTGTATATTACAAATTACGGTATGTTGTAACAACAAATACCGATCGGTAATTGTCTCTGCCGCTGTAATAGAATTAATTATATATCTATTACACCGGCCTTGTATCATAATAAAGTTGTGGTAGTATGATCTCCATATTTATAATTTAGTACTTTGTATTTAGTTTTTTTGGAATCATAAAAAAAGTTTTACTAATTTAAAATTTAAAAAGTATTTACATTTTTCACTGTTTAGTCGCGGATATGGAATTCGATCCTGTCAAAATCAATACATCATCTATAGATCATGTAACAATATTACAATACATAGATGAACCAAATGATATAAGACTAACAGTATGCATTATCCGAAATGTTAATAACATTACATATTATATCAATATCACAAAAATAAATCCACATTTGGCTAATCGATTTCGGGCTTGGAAAAAACGTATCGCCGGAAGGGACTATATGACTAACTTATCTAGAGATACAGGAATACAACAATCAAAACTTACTGAAACTATACGTAACTGTCAAAAAAATAAAAACATATATGGTCTATATATACACTACAATTTAGTTATTAATGTGGTTATTGATTGGATAACCGATGTGATTGTTCAATCAATATTAAGAGGGTTGGTAAATTGGTACATAGCTAATAATACATATACTCCAAATACACCCAATAATACTACAACCATTTCTGAGTTGGATATCATCAAAATACTGGATAAATACGAGGACATGTATAGAGTAAGTAAAGAAAAAGAATGTGGAATTTGCTATGAAGTTGTTTACTCAAAACGATTAGAAAACGATAGATACTTTGGTTTATTGGATTCGTGTAATCATATATTTTGCATAACATGTATCAATATATGGCATAGAACACGAAGAGAAACCGGTGCGTCAGATAATTGTCCTATATGCCGTACACGTTTTAAAAAAATAACAATGAGCAAGTTCTATAAGCTAGTTAACTAATAAATAAAAAGTTTAATTATCGACGATATATGTCGTTATTTTTCTCTCATATGAAAGATTAATTTGATTCTAATATAATCTTCAGTATTGGATGAATCTCAATTCAAATTAATTCCATTAGATTAGATTAGATTAGATCATAAATAAAAATAGTAGCACGCACTACTTCAGCCAAATATTCTTTTTTGAAACGCCATCTAGCGTAATGAGAACACAAGTGAACCTATAATGAGCAAATTTATTAGTATCGGTTACATGAAGGACTTTACGTAGAGTGGTGATTCCTCCATCTGTGGTACGAACGGTTTCATCTTCTTTGATGCCACCACCCAGATGTTCTATAAACTTGGTATCCTCGTCCGATTTCATATCATTTGCCAACCAATACATATAGCTAAACCCAGGCATACGTTCCACACATCCGGAACAATGAAATTCTCCAGAAGATGTTACAATGGCTAGATTTGGACATTTGGTTTCAACCGCGTTAACATATGAGTGAACACACCCATACATGAAAGCGATGAGAAATAGGATTCTCATCTTGCCAAAATATCACTATAAAAAATTTATTTATCAATTTTAAAGGTATAAAAAAATACTTATTGTTGCTCGAATATTTTGTATTTGATGGTATACGGAAGATTAGAAATGTAGGTATTATCATCAACTGATTTTATGATGGTTTTATGAATTTTATTATGCTTCACTATTGCATCGGAAATAATATCATATGCTTCCACGTATATTTTATTTTGTTTTGACTCATAATACGCACGTAATTCTGGATTATTGGCATATCGATGAATAATTTTAGCTCCATGCTCAGTAAATATTAATGAGAACATAGTGTTGCCTCCTACCATTATTTTTTTCATCTCATTCAATTCTTGATTGCAGAGATCTATATAATCATTATAGCGTTGACTTATGGACTCTGGAATCTTAGACGATGTACAGTCATCTATAATCATGGTATATTTAATACATTGTTTTATAGCATAGGCATTATCTACGATATTAGATACTTCACTCAATGAATCAATCACACAATCTAATGTAGGTTTATGACATAATAGCATTTTCAGCAGTTCAATGTTTCTAGATTCGTTGATGGCAATGGTTATACATGTATATCCGTTATTTGATCTAATGTTGACATCTGAACCGGATTCTAGCAGTAAAGATACTAGAGATTGTTTATTATATCTAACAGCCTTGTGAAGAAGTGTTTCTCCTCGTTTGTCAATCATGTTAATGTCTTTAAGATAAGGTAGACAAATGTTTATAGTACTAAGAATTGGGCAAACATAAGACATGTCACAAAGACCCTTTTTGTATGTATAAGTGTAGAAATTATAACATCCATAGTTGGATTCACATAGGTGTCCAATCGGGATCTCTCCATCATCGAGATGATTGACGGCATCTCCCCCTTCCTTTTTTAGTAGATATTTCATCGTGTAAGAATCAATATTAATATTTCTAAAGTATCTGTGTATAGCCTCTTTATTTACCACAGCTCCATATTCCAACATGCATTCCACTAGAGGGATATCGATATCGCCGAATGTCATATACTCAATTAGTATATGTTGGAGGACATCCGAGTTCATTGTTTTCAATATCAAAGAGATGGTTTCCTTATCATTTCTCCATAGTGGTACAATACTACACATTATTCCGTACGGCTTTCCATTCTCCAAAAACAATTTTACCAAATCTAAATCTACATCTTTATTGTATCTATAATCACTATTTAGATAATCAGCCATAATTCCTCGAGTGCAACATGTTAGATCGTCTATATATGAATAAGCCGTGTTATCTATTCCTTTCATTAACAATTTAACGATGTCTATATCTATATGAGATGACTTAATATAATATTGAAGAGCTGTACAATAGTTTTTATCTATAGAAGACGGCTTGATTCCGTGATTAATTAGACATTTAACAACTTCCGGACGCACATATGCTCTCGTATCCGACTCTGAATACAGATGAGCGATGATATACAGATGCAATACGGTACCACAATTTCGTGGTTGATAATCATCATACACGTATCCGTACTCGTCATCCTCATAAAGAACACTGCATCCATTTTCTATGAACAAATCAATAATTTCAGGAACAGGATCATCTGTCATTACATAATTTTCTATAACTGAACGATGGTTTTCACATTTAACACTCAAGTCAAATCCATGTTCTACCAACACCTTTATCAAGTCAACGTCTACATTTTTTGATTTCATATAGCTGAATATATTAAAGTCATTTATGTTGCTATATCCAGTAGCTTCTAGTAGAGCCATCGCTATATCCTTATTGACTTTAACATGTCTACTATTTGTGTATTCTTCTATTGGGGTAAACTGTCTCCAATTTTTATGTAATGGATTAGTGCCACTATCTAGTAGTAGTTTGACGACCTCAACATTATTACAATGCTCATTGAAAAGGTATGCGTGTAAAGCATTATTCTTGAATTGGTTCCTGGTATCATTAGGATCTCTGTCTCTCAACATCTGTTTAAGTTCATCGAGAACCACCTCCTCATTTTCCAGATAGTCAAACATTTTGACTGAATAGAAGTGAATGAGCTACTGTGAACTCTATACACCCGCACAACTAATGTCATTAAATATCATTTTTGAATGTATTTATACCATGTCAAAAACTTGTACAATTATTAATAAAAATAATTAGTGTTTAAATTTTACCAGTTCCAGATTTTACACCTCCGTTAACACCTCCATTAACCCCACTTTTTACACCACTGGACGATCCTCCTCCCCACATTCCACTGCCACTAGATGTATAAGTTTTAGATCCTTTATTACTACCATCATGTCCATGGATAAAGACACTCCACATGCCGCCACTACTACCCCCTTTAGACGACATATTAATAAGACAAGTTTAACAATAAAATTAATCACGAGTACCCTACTACCAACCACTATTATATGATTACAGTACCTTGACTAAAGTCTCTAGTCACAAGATCAATACTACCAACCTACGCTATTATATGATTATAGTTTCTATTTTTATAGGAACGCGTACGAGAAAATCAAATGTCTAAGTTCTAACGGTAGTGTTGATAAACGATTGTTATCCGCGGATACCTCATCTATCATGTTGTCTATTTTCTTACTTTGTTCTATTAACCTATTAGCATTATATATTATTTGATTATAAAACTTATATTGCTTATTAGCCCAATCTGTAAATATCGGATTATTAACATATCGTTTCTTTGTAGGTTTATTTAACTTGTACATCACTGTAAGCATGTCCGTACCATTTATTTTAATTTGACACATATCAGCAATTTCTTTTTCGCAGTCGGTTATATATTCTATATAAGATGGATACGTATCACATATGTACTTATAGTCTACTAATATGAAGTACTTAATACATATTTTCAGTAACGATTTAGCCTTATTACCTATTAATAAGTGCCTGTCGTTGGATAGGTAATCAACTGTTTTCTTAATACATTCGATGGTTGGTAATTTACTCAAAATAATTTCCAATATCTTAATATATATTTCTGCTATTTCTGGTATACATGCATGTGCCATTATAACACAAATACCAATACATGTAGACCCATATGTTGTTGTTATATTAATATCTGCGCCATTATCTATTAACCATTCTATTAGGGCAACACTATGCGACTCGATACAATAATAAAGTATACTACGTCCATGTTTATCTATTTTGTTTATATCATCGATATACGGCTTACAAATTTCTAGTATCGATAACACTTCTGACTCGTGAATAAATAAGGTAGGGAATAACGGCATAATATTTATTATGTTATCATCATTAACAACTACGTTTCCATTTTTTAAAATATACTCTACAACTTTAGGATCCCTATTGTCAAATCTTTTAAAATATTTATTTATATGCTTAAATCTATATAATATAGCTCCTTCCCTAATCATACATTTGATAACATTGATGTACACTGTATGATAAGATACATATTCTGACAATAGATCTTGTATAGAATCTGTATATCTTTTAAGAATTGAGGATATTATGACATTATTACGTAAACTATTACACAATTCTAAAATATAAAATGTATCATGGGCAGATAATAGTTTATCCACTATATAATTATCTATTTTATGATTTTTCTTCCTATATTGTTTACGTAAATAGATAGATAGAATATGCATTAGTTCATTACCGCTATAGTTACTATCGAATAACACGTCAAATATTTCCCGTTTAATATCGCATTTGTCAACATAATAATAGAGTATGGTACGTTCACGATAAGTATAATGACACATCTCGTTTTCGTGCGAAATTAAATAGTTTATCACGTCCAAAGATGTCACATAACCATCTTGTGACCTAGTAATAATATAATAATAGAGAACTGTTTTACCCATTCTATTATCATAATCAGTGGTGTAGTCATAATCTAAATAATCAAACTCGTCATCCCAATTAAAATAAATATAATCAGTACATTGAATGGGTATGATATTGTACCCATACTGTATGTTGCTACATGTAGGTATTCCTTTATCCAATAATAGTTTAAATACATCTATATTAGGATTTGATGTTGTCGCGTATTTCTCTACAATATTAATACCATTTTTGATACTATTTATTTCTATACCTTTCGAAATTAGTAATTTCAATAAGTCTATATCGATGTTATCAGAACATAGATATTCAAATATATCAAAATCATTGATATTTTTATAGTCGACTGACGACAATAACAAAATCACAACATCGTTTTTGATATTATTATTTTTTTTGGTAACGTATGCCTTTAATGGAGTTTCACCATCATACTCATATAATGGATTTGCACCACTTTCTATTAATGATTGTGCACTACTGGCATCGATGTTAAATGTTTTACAACTATCATAGAGTATCTTATCGTTAACCATGATTGGTTGTTGATGTTATCACATTTTTTGGTTTCTTTCATTTCAGTTATGTATGGATTTAGCACGTTTGGGAAGCATGAGCTCATATGATTTCAGTACTGTAGTGTCAGTACTATTAGTTTCGATCAGATCAATGTCTAGATCTATAGAATCAAAACACGATAGGTCAGAAGATAATGAATATCTGTACGCTTCTTCTTGTACTGTAACTTCTGGTTTTGTTAGATGGTTGCATCGTGCTTTAACGTCAATGGTACAAATTTTATCCTCGCTTTGTGTATATCATATTCGTCTCTAGTATAAAATTCTATATTCAAATTATCATGCGATGTGTGTACGCTAACGGTATCAATAAACGGAGCACAGCATTTAGTCAACAGTAATCCAAAATTTTTTAAAGTATATCTTAACGAAAGAAGTTGTCATCGTTAGAGTGTGGTAAATCATTGTCTACGGTACTAGATCCTCATAAGTGTATATATCTAGAGTAATATTTAATTTATCAAATGGTTGATAATATGGATGTTGTGGCAATTTCCTAATACGGAAATAAGACATAAACACGCAATAAATCTAATTGCGGACATGTTACACTCCTTAAAAAATACGAATAAACACTTTGGCTTTTAGTAAGTGTCATTTAACACTATACTCATATTAATCCATGGACTCATAATCTCTATACGGGATTAACGGATGTTCTATATACGGGGATGAGTAGTTCTCTTCTTTAACTTTATACTTTTTACTAATCATATTTAGACTGATGTATGGGTAATAGTGTTTGAAGAGCTCGTTCTCATCATCAGAATAAATCAATATCTCTGTTTTTTTGTTATACAGATGTATTACAGCCTCATATATTACGTAATAGAACGTGTAATCTACCTTATTAACTTTCACCGCATAGTTGTTTGCAAATACGGTTAATCCTTTGACCTCGTCGATTTCCGACCAATCTGGGCGTATAATGAATCTTAACTTTAATTTCTTGTAATCATTCGAAATAATTTTTAGTTTGCATCCGTAGTTATCTCCTCTATGTAACTGTAAATTTCTCAACGCGATATCTCCATTAATAATGATGTCGAATTCGTGTTGTATACCCATACTGAATTGATGAACTAATACCGACGGTATTAATAGTAATTTACTTTTCATCTTTACATACTTGGTAATAGTTTTACTATCATAAGTTTATAAATTCCACAAGCTACTATGGAATATACCAACCATCTTAGTATAGAACACATGTCTTAAAGTTATTAATTAATTACATGTTGTTTTATATATCGCTACGAATTTAAACAGAGGAATCAGTTAGGAAGAAAAAATTATCTGTCATCATCATCATCTATTGGATAACATCTCTGTATTCTACGATAGAGTGCTATTTTAAGATGTGACAGATCCGTGTCATCAAATATATACTCCATTAAAATGATTATTCCGGCAGCGAACTTGATATTGGATACATCACGACCTTTGTTAATATCCACGACAATAGACAGCAATCCCATTGTTCCATAAACAGTGAGTTTATCTTTCTTTGAAGTGATATTTTGTAGAGATCTTATAAAACTGTCGAATGACATCGTATTTATATCTTTAGCTAAATCATATATGTTACCATCATAATATCTAACAGCATCTATCTTAAACGTTTCCATCGCTGTAAAGACGTTTCCGATAGATGGTCTCGTTTCATCAGTCATACTGAGCCAACAAATGTAATCGTGTATAACATCTTTGATAGAATCAGACTCTAAAGAAAAGGAATCGGCTTTATTATACACATTCATGATAAACTTAATGAAAAATGTTTTTCGTTGTTTAAGTTGGATGAATAGTATGTCTTAATAATTGTTATTATTTCACTAATTAATATTTAGTAACGAGTACACTCTATAAAAACGAGAATGACATAACTAATCATAACTAGTTATCAAAGAATGTCTAGGACGCGTAATTTTTTATGGTATAGATCCTGTAAGCGTTGTCTGTATTCTGGAGCTATTTTCTCTATCGCATTAGTGAGTTCAGAATATGTTATAAATTTAAATCGAATAACGAACATAACTTTAGTAAAGTCGTCTATATTAACTCTTTTATTTTCTAGCCATCGTAATACCATGTTTAAGATAGTATATTCTCTAGTTACTACGATCTCATCGTTGTCTAGAATATCGCATACTGAATCTACATCCAATTTTAGAAATTGGTCTGTGCTACATATCTCTTCTATATTATTGTTGATGTATTGTCGTAGAAAACTATTACGTAGACCATTTTCTTTATAAAACGAATATATAGTACTCCAATTATCTTTACCGATATATTTGCATACATAATCCATTCTCTCAATCACTACATCTTTAAGAGTTTGGTTGTTAAGATATTTGGCTAAACTATATAATTCTATTAGATCATCAACAGAATCAGTATATATTTTTCTAGATCCAAAGATGAACTCTTTGGCATCCTCTATAATATTATCAGAAAAGATATTTTCGTGTTTTAGTTTATCAAGATTTAACCTGTTCATATCCATGATTAACGACGTCATATAACCACATAAAATAAAAATCCATTTTCATTTTTAGCACAATACTATTCATAATTGATATTGATGTAATATTTTGTTACTTTGAACGTAAAGACAGTACACGGGTCCGTATCTCCAACAAGCACGTAGTAATCAAATTTGGTGTTGTTAAACTTCGCAATATTCATCAATTTAGATAGAAACATATACTCATCATCTGTTTTAGGAATCCATGTATTATTACCACTTTCCAACTTATCATTATCCCAGGCTATGTTTCGCCCATCATCGTTGCACAGAGTGAATAATTCTTTTGTATTCGGTAGTTCAAATATATGATCCATGCATATATCGACAAAGCTATTGTAGATGTGATTTTTCCTAAATCTAATATAAAACTCGTTTACTAGCAAACATTTTCCTGATTTATCGACCAAGACACACATGGTTTCTAAATCTATCAAGTGGTGGGGATCCATAGTTATAACGCAGTAACATAAATTATTACCTTCTTGACTGTCGCTAATATCTATATACTTATTGTTATCGTATTGGATTCTACATATAGATGGCTTGTATATCAAAGATATAGAACACATAACCAATTTATATTCTCGCTTTGTATTTTCGAATCTAAAGTTAAGAGATTTAGAAAACATTATATCATCGGATGATGATATCACTGTTTCCAGAGTAGGATATATTAAAGTCTTTAAAGATTTTGTCCGATTCAAATAAATCACTAAATAATATCCCATATTATCATCTGTTATAGTCGTGTCATTAAATCTATTATATTTTATGAAAGATATATCACTGCTCACCTCTATATTTCGTACATTTTTAAACTGTTTGTATAATATCTCTCTAATACAATCAGATATATCTATTGTGTCGGTAGACGATACCGTTACATTTGAATTAATGGTGTTCCATTTTACAACTTTTAACAAGTTGACCAATTCATTTCTAATAGTATCAAACTCTCCATGATTAAATATTTTAATAGTATCCATTTTATATCACTACGGACATAAACCATTGTATATTTTTTATGTTTATTAGTGTACACATTTTGGAAGTAAGTTCCTGGATCGGATGTCACCGCAGTAATATTGTTGATTATTTCTGACATCGACGTATTATATAGTTTTTTAATTCCATATCTTTTAGAAAAGTTAAACATCCTTATACAATTTGTGGAATTAATATTATGAATCATGGTTTTTACACATAGATCTATTACAGGCGGAACATCAATTATTATGGCAGCAACTAGTATCATTTCTACATTGTTTATGGTGATGTTTATCTTCTTCCAGCGCATATAGTCTAATATCGATTCAAACGCGTGATAGTTTATACCATTCAATATAATCGCTTCATCCTTTAGATGGTGATCCTGAATGTGTTTAAAAAATTATACGGAGACGCCGTAATAATTTCCCCATTGATAGAAAATATCACGCTTTCCATTTTCTTGAAGTACTAAAAGTAATTATAATATAATGTAAAGGTTTATATATTCAATATTTTTTATAAAAAAAATCATTTTGACATTAATTCCTTTTTAAATTTCAGTCTATCATCTATAGAAACATATTCTATGAATTTATAAAATGCTTTTACGTATCCTATCGTAGGTGATAGAACCGCTAAAAAACCTATCGAATTTCTACAAAAGAATCTGTTATATGGTATAGGGAGAGTATAAAACATTAAATGTCCGTACTTATTAAAGTATTCAGTAGCCAATCCTAACTCTTTCGAATAATTATTAATGGCTCTTATTCTGTACGAATCTATTTTTTTGAACAATGGACCTAGTGGTATATCTTGTTCTATGTATCTAAAATAATGTCTGACTAGATCCGTTAGTTTAATATCCGCAGTCATCTTGTCTAGAATGGCAAATCTAACTGCGGGTTTAGGCGTAGGCGTTAGTTTAGTTTCTATATCTACATCTATGTCTTTATCTAACACCAAAAATATAATAGCTAATATTTTATTACAATCATCCGGATATTCTTCTACGATCTCACTAACTAATGTTTCTTTGGTTATACTAGTATAGTCACGATCAGACAAATAAAGAAAATCAGATGATCGATGAATAATACATTTAAATTCATCATCTGTAAGATTTTTGAGATGTCTCATTAAAATATTATTAGTGTCAGTTCTCATTATCATATATTGACAGCAGCTATTACACTTATTTTATTTTTCTGTATTTTATTACTTTTCACCATATAGATCAGTCATTAGATCATCAAAATACTTTTCAATCATCCTAAAGAGTATGGTGAACGAATCTTCCCATCTAATTTCTGAACGTCTACCAATGTCTCTAGCCACTTTGGCACTAATAGCGATCATTCGCTTAACATCTTCTACATTATTAACTGGTTGATTCAATCTATCTAGCAATGGACCGTCGGATAGCGTCATTCTCATGTTCTTAATCAATGTACATACATCGTCATCATCTACCAATTCATCAAACAATATAAGCTTTTTAAAATCATCATTATAATAGGATGGATCGCCGTCATTTCTCCAAAGAATATATCTAATAAGTAGAGTCCTCATGCTTAGTAATTTAACTATTTTAGTTAACAACTATTTTTTATGTTAAATCAATTAGTAACACCGCTATGTTTAATACTTATTCATATTTTAGTTTTAGGATCGAGAATCAATACAAAAATTAATACATCAATTTTGGAAATACTTAGTTTCCACGTAGTCAATGAAACATTTGAGCTCATCGTAAAGGACGTTCTCGTACAAGACGTAACTATAAATTGGTTTATATTTGTTCAAGATAGATACAAATCCGATAACTTTTTTGACGAATTCTACGGGATTCACTTTAAAAGTGTCATACCGGGTTCTTTTTATTCTTTTAAACAGATCGATTGTGTGATGTTGATTAGGTCTTTTACGAATTTGATACAGAATAGCGTTTACATATCCACCATAGTAATCAATAGCCATTTGTTCGTATGTCATAAATTCTTTAATTATATGACACTGTGTATTATTTAGTTCGTCCTTGTTCATCATTAGGAATCTATCCAATATGGCAATTATATTAGAACTATAACTGCGTTGTATGCGCATGTTGATGTGTCTGTTTATACAATCAATTATACTAGGATCCATACCACTACAATCGGGTAAAATTGTAGCATCATATACCATTTCTAGTACTTTAGATTCATTGTTATCCATTGCAGAGGACGTCATGATCGAATCCAAAAAAATATATTATTTTTATGTTATTTTGTTAAAAATAATCATCAAATACTTCGTAAGATACTCCTTCATGAACATAATCAGTTACAAAACGTTTATATGAAGTAAAGTATCTACAATTTTTACAAAAGTCAGGATGCATAAGTACAAAGTACGCGATAAACGGAATAATAATAGATTTATCTAGTTTATCTTTTTCTATCTCTTTCATAGTTATATACATGGTCTCAGAAGTCGGATTATGTAACATCAGCTTCGATAAAATGACTGGGTTATTTAGTCTTACACATTCGCTCATACATGTATGACCGTTAACTATAGAGTCTACACTAAAATGATTGAATAATAGATAGTCTACCATTGTTTCGTATTCAGATAGTACAGCGTAGTACATGGCATCTTCACAAATTATATCATTATCTAATAGATATTTGACGCATCTTATGGATCCCACTTCAACAGCCATCTTAAAATCGGTAGAATCATATTGCTTTCCTTTATCGTTAATAATTTCTAGAACATCATCTCTATCATAAAAGATACAAATATTAACTGTTTGATCAGTAATAACATTGCTAGTCGATATCAATTTGTTAATAAGATGCGCTGGGCTCAATGTCTTAATAAGAAGTGTAAGAGGACTATCTCCGAATTTGTTTTGTTTATTAACATCCGTTGATGGAAGTAAAAGATTTATAATGTCTACATACTTGACTGTTTTAGAGCATACAATATGGAGAGGCGTATTTCCATCATGATCTGGTTTTGAGGGACTAATTCCTAGTTTCATCATCCATGAGATTGTAGAAGCTTTTGGATTGTCTGACATAAGATGTCTATGAATATGATTTTTGCCAAATTTATCCACTATCCTGGCTTCGAATCCGATAGACATTATTTTTTTAAACACTCTTTCTGAAGGATCTGTACACGCCAACAACGGACCACATCCTTCTTCATCAACCGAGTTGTTAATCTTGGCTCCATACTGTACCAATAAATTTATTCTCTCTATGACTTCATCATCTGTTCCCGAGAGATAATATAGAGGTGTTTTATTATGTTTATCACATGCGTTTGGATCTGCGCCGTGCACCAGCAGCATCGCGACTATTCTATTATTATTAATTTTAGAAGCTATATGCAATGGATAATTTCCATCATCATCCGTCTCATTTGGAGAGTATCCTCTATGAAGAAGTTCTTCTATAAATCGTTCATCTAGTCCTTTAATGCCACAATACGCATGTAGAATGTGATAATTTCCAGAGGGTTCGATAACTTGTAGCATATTCCTAAATACATCTAAATTTTTACTATTATATTTGGCATAAAGAGATAGATAATACTCGACCGACATAATGTTGTGTTGTCCATTATAGTATAAAAATTAATATTTCTATTTCTATATATTTGCAACAATTTACTCTCTATAACAAATATCATAACTTAGTTCTTTTATGTCAAGAAGGCACTGGTTTAATTCATCTATAAATGTCACTCCATAACTACCACGCATACTATACTCAGAATTATGATAAAGATATTTATTCTTGGGGTGTAAGTAATGGGGATTAATCTTTGTTGGATCAGTCTCTAAGTTAACACATGTCACACATGATCCATTTATAGTTATATCACACGATGATGATTTATGAATTGATTCCGGAAGATCGCTATTGTATTTTGTAGTTCCACAATTCATTTCCATACATGTTATTGTCACACTAATATTATGATGAACTTTATCTAGCCGCTGAGTGGTAAACAACAGAACAGATAGTTTATTATCTTTACCAACACCCTCAGCCGCTGCCACAAATCTCTGATCCGTATCCATGATGGTCATGTTTACTTTTAGTCCGTATCCAGTCAACACTATGTTAGCATTTCTGTCGATATAGCTTTCACTCATATGACACTCACCAATAATTGTAGAATTAATGTCGTAATTTACACCAATAGTGAGTTCGGCGACAAAGTACCAGTACCGGTAATCTTGTCGATGAGGACATATAGTATTCTTGTATTCTACCGAATACCCGAGAGATGCGATACAAAAGAGTAAGACTAATTTGTAAACCATCTTACTCAAAAATATGCGACAATAGGAAATCTATCTTATACACATAATTATTCTATCAATTTTACCAATTAGTTAGTGTAATGTTAACAAAAATGTGGGATAATTTAATAGTTTTTCCTTACACAATTGACATACATGAGTCTGAGTTCCTCGTTTTTGCTAATTATTTCGTCCAATTTATTATTCTTGACATCGTCAAGATCTTTTGTATAGGAGTCAGACTTGTATTCAACATGTTTTTCTATAATCATCTTAGCTATTTCGGCATCATCCAATAGTACATTTTCCAGATTAACAGAATAGATATTAATGTCGTATTTGAACAGAGCCTGTAACATCTCAATGTCTTTATTATCTATAGCCAATTTGATGTCCGGAATGAAGAGAAGGGAATTGGTGTTTGTTGACGTCATATAGTCGAGCAAGAGAATCATCATATCCACGTGTCCATTTTTTATAGTGGTGTGAATACAACTAAGGAGAATAGCCAGATCAAAAGGAGATGGTATCTCTGAAAGAAAGTAGGAAACAATACTTACATCATTAAGCATGACAGCATGATAAAATGAAGTTTTCCATCCAGTTTTCCCATAGAACATCAGTCTCCAATTTTTCTTAACAAACAGTTTTACCGTTTGCATGTTACCACTATCAACCGCATAATACAATGCGGTGTTTCCTTTGTCATCAAATTGTGAATCATCCATTCCACTGAATAGCAAAATCTTTACTATTTTGGTATCTTCTAATGTGGCTGCCTGATGTAATGGAAATTCATTATCTAGAAGATTTTTCAATGCTCCAGCGTTCAACAACGTACATACTAGACGCACGTTATTATCAGCTATTGCATAATACAAGACACTATGACCATTGATATCCGCCTTAAATGCATCTTTGCTAGAGAGAAAGCTTTTCAGTTGCTTAGACTTCCAAGTATTAATTCGTGACAGATCCATGTCTGAAACGAGACGCTAATTAGTGTATATTTTTTCATTTTTTATAATTTTGTCATATTGTACCAGAATTAATAATATCTCTAATAGATCTGATTAGTAGATATATGGCTATCGCAAAACAACATATACACATTTAATAAAAATAATATTCATTAAGAAGATTCAGATTCCACTGTACCCATCAATATAAAATAAAATAATTATTCCTTACATCGTACCCATAAACAATATATTAAGTATATTCCACCTTACCCATAAACAATATAAATCCAGTAATATCATGTCTAATGATGAACACAAATGGTGTATTAAATTCCAGTTCTTCAGGAGATGATCTCGCCGTAGCTACCATGATAGTAGATGCCTCCGCTACAGTTCCTTGTTCGTCTACATCTATCTTTACATTCTGAAACATTTTATAAATATATAATGGGTCCCTAGTCATATGTTTAAACGACGCCTTATCTGGATTAAACATACTAGGAGCCATCATTTCGGCTATCGACTTAATATCCCTCTTGTTTTCGATAGAAAATCTAGGGAGTTTAAGATTGTACATTTTATTCCCTAATTGAGATGACCAATATTCTAATTTTGCAGGCGTGATAGAATCTGTGAAATGGGTCATATTATCACCTATTGCCAGGTACATACTAATATTAGCATCCTTATACAGAAGGCGCACCATATCATATTCTTCGTCATCGATTGTGATTGTATTTCCTTGCAATTTAGTAACTACGTTCATCATGGGAACCGTTTTCGTACCGTACTTATTAGTAAAACTAGCATTGTGTGTTTTAGTGATATCAAACGGATATTGCCACGTACCTTTAAAATATATAGTATTAATGATTGCCCATAGAGTATTATCGTCGAGCATAGTAGAATCAACTACATTAGACATACCAGATCTACGTTCTACTATAGAATTAATTTTATTAACCGCATCTCGTCTAAAGTTTAATCTATATAGGCCGAATCTATGATATTGTTGATAATACGACGGTTTAATACACACAGTACTATCGACGAAACTTTGATACGTTAGATCTGTGTACGTATATTTAGATGTTTTCATCTTAGCTAATCCGGATATTAATTCTGTAAATGCTGGACCCAGATCTCTTTTTCTCAAATTCATAGTATTCAATAATTCTACTCTAGTATTACCTGATGCAGACAATAGCGACATAAACATAGAAAACGAATACCCAAACGGTGAGAAGACAATATTATCATTATCATCCTCATCCCCATTTTGAATATTTTTATACGCTAATATACCAGCATTGATAAATCCCTGCAGACGATATGCGGATACTGAACACGCTAATGATAGTATCAATAACGCAATCATGATTTTTATGGTATTAATAATTAACCTTATTTTTATGTTTGGTATAAAAATTATTGATGTCTACACATCCTTTATAATCAACTCTAATCACTTTAACTTTTACAGTTTTCCCTACAAGTTTATCCCTATATTCAACATATCTATCCATATGCATCTCTTAACACTCTGCCAAGATAGCTTCAAAGTGAGGATAGTCAAAAAGATAAATATATAGAGTATAATCATTCTCGTATACTCTGCCCTTTATTACATCGCCCGCATTGGGCAACGAATAACAAAATGCAAGCATCGTGTTAACGGGCTCGTAAATTGGGATAAAATTATGTTTTTATTGTTTATCTATTTTATTCAAGAGAATATTCAGGAAGTTCCTTTTCTGGTTGTATCTCGTCGCAGTATATATCATTTGTACATTGTTTCATATTTTTTAATAGTCTACACCTTTTAGTAGGACTAGTATCGTACAATTCATAGCTGTATTTTGAATTCCAATCACGTATAAAAATATCTTCCAATTGTTGACGAAGACCTAATCCATCATCCGGTGTAATATTAATAGATGCTCCACATATATCCGTAAAGTAATTTCCTGTCCAATTTGATGTACCTATATACGCCGTTTTATCGGTTACCATATATTTTGCATGGTTTACCCTAGAATACGGAATGGGAGGATCAGCATCTGGTACAATAAATAGCTTTACTTCTATATCTATGTTTTTAGATTTTAGCATAGCTATAGATCTTAAAAAGTTTCTCATGATAAACGAAGATCGTTGCCAGCAACTAATCAATAGCTTAACGGATACTTGTCTGTCTATAGCGGATCTTCTTAATTCATCTTCTATATAAGGCCAAAACAAAATTTTACCCGCCTTTGAATAAATAATAGGAATAAAGTTCATAACAGATACATAAACGAATTTACTCGCATTTCCGATACATGACAATAAAGCGGTTAAATCATTGGTTCTTTCCATAGTACATAATTGTTGTGGTGCAGAAGCAATAAATACAGAGTGTGGAACACCGCTTACGTTAATACTAAGAGGATGATCTGTATTATAATACGACGGATAAAAGTTTTTCCAATTATATGGTAGATTGTTAACTCCAAGATACCAGTATACCTCAAAAATTTGAGTGAGATCCGCTGCCAAGTTCCTATTATTGAAGATCGCAATACCCAATTCCTTGACCTGAGTTAGTGATCTCCAATCCATGTTAGCGCTTCCTAAATAAATATGTGTATTATCAGATATCCAAAATTTTGTATGAAGAACTCCTCCTAGGATATTTGTAATATCTATGTATCGTACTTCAACTCCGGCCATTTGTAGTCTTTCAACATCCTTTAATGGTTTGTTGGATTTATTGACGGCTACTCTAACTCTTACTCCTCTTTTGGGTAATTGTACAATCTCGTTTAATATTACCGTGCCGAAATTCGTACCCACTTCATCCGATAAACTCCAATAAAAAGATGATATATCTAGTGTTTTTATGGTATTGGATAGAATTTCCCTCCACATGTTAAATGTAGTCAAATATACTTTATCAAATTGCATACCTATAGGAATAGTCTCTGTAATCACTGCGATTGTATTATCCGGATTCATTTTATTTGTTAAAAAAATAATCCTATATCACTTCACTCTATTAAAAATCCAAGTTTCTATTTCTTTCATGACTGATTTTTTAACTTCATCCGTTTCCTTATGAAGATGATGTTTGGCGCCTTCATAAATTTTTATTTCCCTATTACAATTTGCATGTTGCATGAAATAATATGCACCTGAAACATCGCTAATCTCATTGTTTGTTCCCTGGAGTATGAGAGTCGGGGTGTTAATCTTGGGAATTATTTTTCTAACCTTGTTGGTAGCCTTCAAGACCTGACTAGCAAATCCAGCCTTAATTTTTTCATGATTGACTAATGGATCGTATTGGTATTTATAAACTTCATCCATATCTCTAGATACTGATTCTGGACATAGCTTTCCGACTGACGCATTTGGTGTAATGGTTCCCATAAGTTTTGCAGCTAGCAGATTCAGTCTTGGAACAGCGTCTGCATTAACTAGAGGAGACATTAGAATCATTGCTGTAAACAAGTTTGGATTATCGCAAGCAGCTAGTATAGAAATTGTTGCTCCCATGGAATGACCCAATAAGAAGACTGGAACTCCTGGATAAGTAGATTTAATAGTCACCACGTGCTGTACCACATCTCTAACATACTTACCAAAGTCATCAATCATCATTTTTTCACCATTACTTCTTCCATGGCCAATATGATCATGTGAGAATACTAAAATTCCTAACGATGATATGTTTTCAGCTAGTTCGTCATAACGTCCAGAATGTTCACCAGCTCCATGACTTATGAATACTAATGCCTTAGGATATGTAATAGGTTTCCAATATTTACAATATATGTAATCATTGTCCAGATTGAACATACAGTTTGTACTCATGATTCACTATATAACTATCAATATTAACAGTTCGTTTAATGATCATATTATTTTTATGTTTTATTGATAATTGTAAAAATATACAATTAAATCAATATAGAGGAAGGAGACGGTACTGTATTTTGTGAGATAGTAATGGAGACTAAATCAGATTATGAGGATGCTGTTTTTTACTTTGTGGATGATGATGAAATATGTAGTCGCGACTCCATCATTGATCTAATAGATGAATATATCACGTGGAGAAATCATGTTATAGTGTTTAATAAAGATATTACCAGTTGTGGAAGACTGTACAAGGAATTGATAAAGTTCGATGATGTCGCTATACGGTACTATGGTATTGATAAAATTAATGAGATTGTCGAGGCTATGAGTGAAGGAGACCACTACATCAATCTTACAGAAGTCCATGATCAGGAAAGTCTATTCGCTACCATAGGAATATGTGCTAAAATCACTGAACATTGGGGATACAAAAAGATTTCAGAATCTAAATTCCAATCATTGGGAAACATTACAGATCTGATGACCGACGATAATATAAACATCTTGATACTTTTTCTAGAAAAAAAAATGAATTGATGATATAAGTGTCTTCATAACGCATTATTACGTTAGCATTCTATTATCCAGTGTTAAAAAAATTATCCTATCATGTATTTGAGAGTCTTATATGTAGCAAACATGATAACTGCAATACCCATAATCTTTAGATATTCACGCGTGCTATGGATGGCATTATCCCGCGGTGCGGAAATGTACGTTATATAATCTACAAAATAATCATCGCATATAGTATGAGATAGTAGAGTAAACATTTTTATCGTTTCTACTGGGTTCATACATCGTCTACCCAATTCGGTAATGAATGAAATTGTCGCCAATCTTACACCCAAACCCTTGTTGTTCATTAGTATAGTATTAACTTCATTATTTATGTCATAAACTGTAAATGATTCTGTAGATGCCATATCACACATGATATTCATGTCACTATTATAATCATTATTAACTTTATCACAATACGTGTTGATAATATCTACATATGATCTAGTTTTTGTGGGTAATTGTCTATACAAGTCGTCTAAACGTTGTTTACTCATATAGTATTGAACAGCCATCATTACATGGTCCCGTTCCGTTGATAGATAATCGAGTATGTTAGTAGACTTGTCAAATCTATATACCATATTTTCTGGAAGCGGATATACATAGTCGCGATCATCATTATCACTAGCCTCATCCTCTATATCATGTACATGTACATAATCTATGATATTATTATACATAAACATCGACAACATACTATTGTCTATTATCTAAGTCCTGTTGATCCAAACCCTTGATCTCCTCTATCTGTACTATCTAGAGATTGTACTTCTTCAAGTTCTGGATAATATATACGTTGATAGATTAGCTGAGCTATTCTATCTCCAGTATTTACATTAAACGTACATTTTCCATTATTAATAAGAATGACTCCTATGCTTCCCCTATAATCTTCGTCTATTACACCGCCTCCTATATCAATGCCTTTTAGGGACAGACCAGACCTAGGAGCTATTCTACCATAGCAGAACTTAGGCATGGACATACTAATATCTGTCTTAATTAACTGTCGTTCTCCAGGAGGGATAGTATAATCGTAAGCGCTATACAAATCATATCCGGCAGCACCCGGCGATTGCCTAGTAGGTGATTTAGCTCTGTTAGTTTCCTTAACAAATCTAACTGGTGAGTTAATATTCATGTTGAACATAAAAAATATCATTTTATTTCAAAATTATTTACCATTCCATTCCATCCCATATATTCCATGAATAAGTGCGATTATTGTACACTTCTATAGTATCTATATACGATCCACGATAAAATCCTCCTATCAATAGCAGTTTATTATCCACTATGATCAATTCTGGATTATCCCTCGGATAAATAGGATCATCTATCAGAGTCCATGTATTGCTGGATTCACAATAAAATTCCGCATTTCTACCAACCAAGAATAACCTTCTACCAAACACTAACGCACATGATTTATAATGAGGATAATAAGTGGATGGTCCAAACTGCCACTGATCATGATTGGGTAGCAAATATTCTGTAGTTGTATCAGTTTCAGAATGTCCTCCCATTACGTATATAACATTGTTTATGGATGCCACTGCTGGATTACATCTAGGTTTCAGAAGACTCGGCATATTAACCCAAGCAGCATCCCCGTGGAACCAACGCTCAACAGATGTGGGATTTGGTAGACCTCCTACTACGTATAATTTATTGTTAGCGGGTATCCCGCTAGCATACAGTCTGGGGCTATTCATCGGAGGAATTGGAATCCAATTGTTTGATATATAATTTACCGCTATAGCATTGTTATGTATTTCATTGTTCATCCATCCACCGATGAGATATACTACTTCTCCAACATGAGTACTTGTACACATATGGAATATATCTATAATTTGATCCATGTTCATAGGATACTCTATGAATGGATACTTGTATGATTTGCGTGGTTGTTTATCACAATGAAATATTATGTTACAGTCTAGTATCCATTTTACATTATGTATACCTCTGGGAGAAAGATAATTTGACCTGATTACATTTTTGATAAGAAGTAGCAGATTTCCTAATCTATTTCTTCGCCTCATATACCACTTAATGACAAAATCAACTACATAATCCTCATCTGGAACATTTAGTTCGTCGCTTTCTAGAATAAGTTTCATAGATAGATAATCAAAATTGTCTATGATGTCATCTTCCAGTTCCAAAAAGTGTTTGGTAATAAAGTCTTTAGTATGACATAAGAGATTGGATAGTCCGTATTCTATACCCATCATGTAACACTCGATACAATATTCCTTTCTAAAATCTCGTAGGATAAAGTTTATACAAGTGTAGATGATAAATTCTACAGATGTTAATATAGAAGCACGTAATAAATTGACGACGTTATGACTATCTATATATACCTTTCCAGTATATGAGTAAATAACTATAGAAGTTAGACTGTGAATGTCAAGGTCTAGACAAACCCTCGTAACTGGATCTTTATTTTTTGTGTATTTTTGGCGTAAATGTGTGCGAAAGTATGGAGATAACTTTTTCAATATCGTAGAATTGACTATTATATTACCTCCTATAGCTTCAATAATTGTTTTGAATTTCTTAGTCGTGTACAATGCTAATATATTCTTACAGTACACAGTATTGACAAATATCGGCATTTATGTTTCTTTAAAAGTCAACATCTAAAGAAAAATGATTGTCTTCTTGAGACATAACTCCCATTTTTTGGTATTCACCCACACGTTTTTCGAAAAAATTAGTTTTTCCTTCCAATGATATATTTTCCATGAAATCAAACGGATTGGTAACATTGTAAATTTTTTTAAATCCCAATTCAGAAATCAATCTATCTGCGACGAATTCTATATATGTTTTCATCATTTCACAATTCATTCCTATGAGTTTAACTGGAAGAGCCACAGTAAGAAATTCTTGTTCAATGGATACCGCATTTGTTATAATAAATCTAACGGTTTCTTCACTCGGTGGATGCAATAAATGTTTAAACATCAAACATGCGAAATCGCAGTGCAGACCCTCGTCTCTACTAATTAATTCGTTAGAAAACGTGAGTCCGGGCATTAGGCCACGCTTTTTAAGCCAAAATATGGAAGCGAATGATCCGGAAAAGAAGATTCCTTCTACTGCAGCAAAGGCAATAAGTCTCTCTCCATAACCGGCGCTGTCATGTATCCACTTTTGAGCCCAATCGGCCTTCTTTTTTACACAAGGCATCGTTTCTATGGCATTAAAGAGGTAGTTTTTTTCATTACTATCTTTAACATAAGTATCGATCAAAAGACTATACATTTCCGAATGAATGTTTTCAATGGCCATCTGAAATCCGTAGAAACATCTAGCCTCGGTAATCTGCACTTCTGTACAAAATCGTTCTGCTAAATTTTCATTCACTATTCCGTCACTGGCTGCAAAAAACGCCAATACATGTTTTATAAAATATTTTTCGTCTGGTGTTAGTTTATTCCAGTCATTGATATCTTTAGATATATCCACTTCTTCCACTGTCCAAAATGATGCCTCTGCCTTTTTATACATATTCCAGATGTCATGATATTGGATTGGGAAAATAACAAATCTATTTGGATTTGGTGCAAGGATAGGTTCCATAACTAAATTAACAATAGTAGTAATTTTTTTTCAGTTATCTGTATGACGACTGTACTTGGATCTTTTGTATATCGCTATCGCCGCAATCACTACAATAATTACAAGTATTATTGATAGCATTGTTATTACTACTATCATAATTAAATTATCGACATTCATGGGTGTTGAATAATCGTTATCATCATTTTGTAATTGTGACGTCATACTAGATAAATCATTTGTGAGATTGTTGTGGGAAGCGGGCACGGAAGATGCATTATCATTATTATTTAACGCCTCCCATTCGGATTCACAAATGTTACGCACGTTCAAAGTTTTATGGAAACTATAATTTTGTGAAAACAGATAACAAGAAAACTCGTCATCGTTCAAATTTTTAACGATAGTAAACCGATTAAACGTCGAGCTAATTTCTAACGCTAGCGACTCTGTTGGATATGGGTTTCCAGATATATATCTTTTCAGTTCCCCTACGTATCTATAATCATCTGTAGGAAATGGAAGATATTTCCATTTATCTACTGTTCCTAATATCATATGCGGTGGTGTAGAACCATTAAGCGCGAAAGATGTTATTTCGCATCGTATTTTAACTTCGCAATAATTTCTGGTTAGATAACGCACTCTACCAGTCAAGTCAATGATATTAGCCTTTACAGATATATTCATAGTAGTCGTAACGATGACTCCATCTTTTAGATGTGATACTCCTTTGTATGTACCAGAATCTTCGTACCTCAAACTCGATATATTTAAACAAGTTAATGATATATTAACGCGTTTTATGAATGATGATATATAACCAGAAGTTTTATCCTCTGTGGCTAGCGCTATAACCTTATCATTATAATACCAACTAGTGTAATTAATATGTGACATGACAGTGTGGGTACAAATATGTACATTATCGTCTACGTCGTATTTGATACATCCGCATTCAGCCAACAAATATAAAATTACAAAAACTCTAACGACGTTTGTACACATCTTGATGCGGTTTAATAAATGTTTTGATTTCAATTTATTGTAAAAAAAGATTCGGTTTTATACTGTTCGATATTCTCATTGCTTATATTCTCATCTATCATCTCCACACAGTCAAATCCATGGTTAACATGTACCTCATCAACCGGTAAAAGACTATCGGATTCTTCTATCATCATAACTCGAGAATATTTAATTTGGTGGTCATTATTAATCAAGTCAATTATCTTATTTTTAACAAACGTAAGTATTTTACTCATTTTTTTATAAAAACTTTTAGAAATATACAGACTCTATCGTGTGTCTATATCTTCTTTTTATATCCAATGTATTTATGTCTGATTTTTCTTCATTTATCATATATAATGGTCCAAATTCTACACGTGCTTCGGATTCATCCAGATCATTAAGGTTCTTATAATCGCAACATCCTTCTCTTCCATCTTCTACATCTTCCTTCTTATTCTTAGCGTCACAGAATCTACCACAGCAGGATCCCATGACGAGAGTCACATTAAACTAATTCATTTTCAATTATAATATACTGATTAGTAATGACAATTAAAATAAAAATATTCTTCATAACCGGTAAGAAAGTAAAAAGTTCACATTGAAACTATGTCAGTAGTTATACATCATGAGATGATATACTCTATTTTGGTGGAGGATTATATGATATAATTCGTGGATAATCATTCTTAAGACACATTTCTTCATTCGTAAATCTTTTCACATTAAATGAGTGTCCATATTTTGCAATTTCTTCATATGATGGCGGTGTACGTGGACGAGGCTGCTCCTGTTCTTGTAGTCGTCGACTGTCGTGTTTGCGTTTAGATCCCTCCATTATCGCGATCGCGTAGTGAGTACTATTTATACCTTGTAATTAAATTTTTTTATTAATTAAACGTATAAAAACGTTCCGTATCTGTATTTAAGAGCCAGATTTCGTCTAATAGAACAAATAGCTACAGTAAAAATAACTAGAATAATCGCTACACCCACTAGAAACCACGGATCGTAATACGGCAATCGGTTTTCGATAATAGGTGGAACGTATATTTTATTTAAGGACTTAACAATTGTCTGTAAACCACAATTTGCTTCCGCCGATCCTGTATTAACTATCTGTAAAAGCATATGTTGACCGGGAGGAGCCGAACATTCTCCGATATTCAATTTTTGTATATCTATAATGTTATTAACCTCCGCATACGCATTACAGTTCTTTTCTAGCTTGGATACTACACTAGGTACATCATCTAAATCTATTCCTATTTCCTCAGCGATAGCTCTTCTATCCTTTTCCGGAAGTAATGAAATCACTTCAATAAATGATTCAACCATGAGTGTGAAACTAAGTCGAGAATTACTCATGCATTTGTTAGTTATTCGGAGCGCGCAATTTTTAAACTGTCCTATAACCTCTCCTATATGAATAGCACAAGTGACATTAGTAGGGATAGAATGTTGAGCTAATTTTTGTAAATAACTATCTATAAAAAGATTATACAAAGTTTTAAACTCTTTAGTTTCTGCCATTTATCCAGTCTGAGAAAATGTCTCTCATAATAAATTTTTCCAAGAAACTAATTGGGTGAAGAATGGAAACCTTTAATCTATATTTATCACAGTCTGTTTTGGTACACATGATGAATTCTTCTAATGCTGTACTAAATTCGATATCTTTTTCGATTTCTGGATATGTTTTTAATAAAGTATGAACAAAGAAATGGAAATCGTAATACCAGTTATGTTCAACTTTGAAATTGTTTTTTATTTTCTTGTTAATGATTCCAGCCACTTGGGAAAAGTCAAAGTCGTTTAATGCCGATTTAATACGTTCATTAAAAACAAACTTTTTATTCTTTAGATGAATTATTATTGGTTCATTGGAATCAAAAAGTAAGATATTATCGGGTTTAAGATCTGCATGTAAAAAGTTGTCACAACAGGGTAGTTCGTAGATTTTAATGTATAACAGAGACATCTGTAAAAAGATAAACTTTATGTATTGTACCAAAGATTTAAATCCTAATTTGATAGCTAACTCGGTATCTACTTTATCTGCCGAATACAGTGCTAGGGGAAAAATTATAATATTTCCTCTTTCGTATTCGTAATTAGTTCTCTTTTCATGTTCGAAAAAGTGAAACATGCGGTTAAAATAGTTTATAACATTAATATTACTGTTAATAACTGCCGGATAAAAGTGGGATAGTAATTTCACGAATTTGATACTGTCCTTTCTCTCGTTAAACGCCTTTAGAAAAACTTTAGAAGAATATCTCAATGAGAGTTCCTGACCATCCATAGTTTGTATCAATAATAGCAACATATGAAGAACCCGCTTATACAGAGTATGTAAAAATGTTAATTTATAGTTTAATCCCATGGCCCACGCACACACGATTAATTTTTTTTCATCTCCCTTTAGATTGTTGTATAGAAATTTGGGTACTGTAAACTCCGCCGTAGTTTCCATGGGACTATATAATTTTGTGGCCTCGAATACAAATTTTACTACATAGTTATCTATCTTAAAGACTATACCATATCCTCCTGTAGATATGTGATAAAAATCGTCGTTTATAGGATAAAATCGTTTATCTTTTTGTTGGAAAAAGGATGAATTAATGTAATCATTCTCTTCTATCTTTAGTAGTGTTTCCTTATTAAAATTCTTAAAATAATTTAACAATCTAACTGACGGAGCCCAATTTTGGTGTAAATCTAATTGGGACATTATGTTGTTAAAATATAAACAGTCTCCTAATATAACAGTATCTGATAATCTATGGGGAGACATCCATTGATATTCAGGGGATGAATCATTGGCAACACCCATTTATTGTACAAAAAGCCCCAATTTACAAACGAAAGTCCAGGTTTGATAGAGATAAACTATTAACTATTTTGTCTCTGTTTTTAACACCTCCACAGTTTTTAATTTCTTTGGTAATGAAATTATTCACAATATCAGTATCTTCTTTATCTACCAGAGATTTTACTAACTTGATAACCTTGGCTGTCTCATTCAATAGGGTAGTGATATTTGTATGTATGATATTGATATCTTTTTGAATTGTTTCTTTTAGAAGTGATTCTTTGATGGTATCAGCATACGAATTACAATAATGCAGAAACTCAGTTAACATGCAGGAATTATAGTAAGCCAATTCCAATTGTTGCCTGTATTGTATTAGAGTATTAATATGCGCAATGATGTCCTTGCGTTTCTCTGATAGAATGCGAGCAGCGATTTTGGCGTTATCATTTGACGATATTTCTGGAATGACGAATCCTGTTTCTACTAACTTCTTGGTAGGACAAAGTGAAACAATCAAGAAAATAGCTTCTCCTCCTATTTGTGGAAGAAATTGAACTCCTCTAGATGATCTACTGACGATAGTATCTCCTTGACAGATATTGGACCGAACTACGGAAGTACCTGGAATGTAAAGCCCTGAAACCCCCTCATTTTTTAAGCAGATTGTTGCCGTAAATCCTGCACTATGCCCAAGATAGAGAGCTCCTTTGGTGAATCCATCACTATGTTTCAGTTTAACCAAGAAACAGTCAGCTGGTCTAAAATTTCCATCTCTATCTAATACAGAATCCAACTTGATGTCAGGGACTATGACCGGTTTAATGTTATATGTAACATTGAGTAAATCCTTAAGTTCATAATCATCGTTGTCATCAGTTATGTACGATCCAAACAATGTTTCTACCGGCATGGTGGATACGAAGATGCTATCCATCAGAATGTTTCCCTGATTAGTATTTTCTATATAGCTATTCTTCTTTAAACGATTTTCCGAATCAGTAACTATGTTCATTTTTTTAGGAGTAGGACGTCTAGCCAGTATGGAAGAGGATTTTCTAGATACTCTCTTCAACATCTTTGATCTCAATGGAATGCAAAACCCCATGGTGTAACAACCAACGATAAAAATAATATTGTTTTTTCACTTTTTATAATTTTACCATCTGACTCATGGATTCATTAATATCTTTACAAGAGCTACTAACGTATAATTCTTTATAACTGAACTGAGATATATACACCGGATCTATGGTTTCCATAATTGAGTAAATGAATGCTCGGCAATAACTAATGGCAAATGTATAGAACAACGAAATTATACTAGAGTTGTTAAAGTTAATATTTTCTATGAGTTGTTCCAATAAATTATTTGTTGTGACTGCGTTCAAGTCATAAATTATCTTGATACTATCCAGTAAACAGTCTTTAAGTTCTGGAATATTATCATCCCATTGTAAAGCCCCTAGTTCGACTATCGAATATCCTGCTCTGATAGCAGTTTCAATATCGACGGACGTCAATACTGTAATAAAGGTGGTAGTATTGTCATCATCGTGATAAACTACGGGAATATGGTCGTTAGTAGGTACCGTGACTTTACACAACGCGATATATAACTTTCCTTTTGTACCATTTTTAACGTAGTTGGGACGTCCTGCAGGGTATTGTTTTGAAGAAATGATATCGAGAACAGATTTGATACGATATTTGTTGGATTCCTGATTATTCACTATAATATAATCTAGACAGATAGATGATTCGATAAATAGAGAAGGTATATCGTTGGTAGGATAATACATCCCCATTCCAGTATTCTCGGATACTCTATTGATGACACTAGTTAAGAACATGTCTTCTATTCTAGAAAACGAAAACATCCTACATGGACTCATTAAAACTTCTAACGCTCCTGATTGTGTTTCGAATGCCTCGTACAAGGATTTCAAGGATGCCATAGATTCTTTGACCAACGATTTAGTATTGCGTTTAGCATCTGATTTTTTTATTAAATCAAATGGTCGGCTCTCTGGTTTACTACCCCAATGATAACAATAGTCTTGTAAAGATAAACCGCAAGAAAATTTATACACATCCATCCAAATAACCCTAGCACCGTCGGATGATATTAATGTATTATTATAGATTTTCCATCCACAGTTATTGGGCCAGTATACTGTTAGCAACGGTATATCGAATAGATTACTCATGTAACCTACTAGAATGATAGTTCGTGTACTAGTCATAATATCTTTAATCCAATCTAAGAAATCTAAAATTAGATCTTTTACACTATTAAAGTTAACAAAGGTATTACCCGGGTACGTGGATATCATATATGGTATTGGTCCATTATCAGTAATGGCTCCATAAACTGATACGGCGATGGTTTTTATATGTGTTTGATCTAATGAGGACGAAATTCGCGCCCACAATTCATCTCTAGATATGCATTTAATATCGAACGGTAACACATCAATCTCGGGACGCGTATATGTTTCTAAATTCTTAATCCAAATATAATGATGACCTATATGCCCTATTATCATACTGTCAACTATAGTATACCTAGAGAACTTTCGATACATCTGCTGTTTCCTGTAATCGTTAAATTTTACAAATCTATAACATGCTAAACCTTTTGACGACAGCCATTCATTAATTTCTGATATGGAATCTGTATTCTCAATACCGTATCGTTCTAAAGCCAGTGCTATATCTCCCTGTTCGTGGGAACGCTTTCGTATAATATCGATCAATGGATAATATGAAGTTTTTGGAGAATAATATGATTCATGATCTATTTCGTCCATAAACAATCTAGACATAGGAATTGGAGGCGATGATCTTAATTTTGTGCAATGGGTCAATCCTATAACTTCTAATATTGTAATATTCATCATCGACATAACACTATCTATGTTATCATCGTATATTAGTATACCACGACCTTCTTCATTTCGTGCCAAAATGATATACAGTCTTAAATAATTACGCAATATCTCAATAGTTTCATAATTGTTAGCTGTTTTCATCAAGGTTTGTATCCTGTTTAACATGATGGCGTTCTATAACGTCTCTATTTTCTATTTTTAATTTTTTTAAATTTTTAACGATTTACTGTGGCTAGATACCCAATCTCTTTCAAATATTTTTTTAGCCTCGCTTACAAGCTGTTTATCTATACTATTAAAACTGACGAATCCGTGATTTTGGTAATGGGTTCCGTCGAAATTTGCCGAAGTGATATGAACATATTCGTCGTCGACTATTAACAATTTTGTATTATTCTGAATAGTGAAAACCTTCACAGATAGATCATTTTGAACACACAACGCATCTAGACTTCTGGCGGTTGCCATAGAATATACGTCGTTCTTATCCCAATTACCAACTAGAAGTCTGATCTTAACTCCTCTATTAATGGCTGCTTCTATAATGGAGTTGTAAATGTCAGGCCAATAGTAGCTATTACCGTCGACACGTGTAGTGGGAACTATGGCCAAATGTTCAATATCTATACTAGTCTTAGCCGACTTGAGTTTATCAATAACTACATCAGTGTCTAGATCTCTAGAATATCCCAATAGGTGTTCTGGAGAATCAGTAAAGAACACTCCACCTATAGGATTCTTAATATGATACGCAGTGCTAACTGGCAGACAACAAGCCGCAGAGCATAAATTCAACCATGAATTTTTTGCGCTATTAAAGGCTTTAAAAGTATCAAATCTTCTACGAAGATCTGTGGCCAGCGGAGGATAATCAGAATATACGCCTAACGTTTTAATCGTATGTATAGATCCTCCAGTAAATGACGCGTTTCCTACATAACATCTTTCATCATCAGACACCCAAAAACAACCGAGTAGTAGTCCCACATTATTTTTTTTATCTATATTAACGGTTATAAAATTTATATCCGGGGAGTGACTTTGTAGCTCTCCCAGATTTCTTTTCCCTCGTTCATCTAGCAAAACTATTATTTTAATCCCTTTTTCAGATACCTCTTTTAGTTTATCAAAAATAAGCGCTCCCCTAGTAGTACTCAGAGGATTACAACAAAAAGATGCTATGTATATATATTTCTTAGCTAGAGTGATAATTTCGTTAAAACATTCAAATGTTGTCAAATGATCGGATCTAAAATCCATATTTTCTGGTAGTGTTTCTACCAGCCTACATTTTGCTCCCGCAGGTACCGATGCAAATGGCCACATTTAGTTAACATAAAAACTTATATATCCTGTTCTATCAACGATTCTAGAATATCATCGGCTATATCGCTAAAATTTTCATCAAAGTCGACATCACAACCTAACTCAGTCAATATATTAAGAAGTTCCATGATGTCATCTTCGTCTATTTCTATATCCGTATCCATTGTAGATTGTTGACCGATTATCGAGTTTAAATCATTACTAATACTCAATCCTTCAGAATACAATCTGTGTTTCATTGTAAATTTATAGGCGGTGTATTTAAGTTGGTAGATTTTCAATTATGTATCAATATAGCAACAGTAGTTCTTGCTCCTCCTTGATTTTAGCATCCTCTTCATTATTTTCTTCTACGTACATAATCATGTCTAATACGTTAGACAACACACCGACGATGGTGGCCGCCACAGACACGAATATGACTAGACCGATGACCATTTAAAAAATACTCTCTAGCTTTAACTTAAACTGTATCGATCATTCTTTTAGCACATGTATAATATAAAAACATTATTCTATTTCGAATTTAGGCTTCCAAAAATTTTTCATCCGTAAACCGATAATAATATATATAGACTTGTTAATAGTCGGAATAAATATATTAATGCTTAAACTATCATCATCTCCACGATTAGAGATACAATATTTACATTCTTTTTGCTGTTTCGAAACTTTATCAATACACGTTAATACAAACCCAGGAAGGAGATATTGAAACTGAGGCTGTTGAAAATGAAACGGCGAATACAATAATTCAGATAATGTAAAATCATGATTCCGTATTCTGATGATATTAGAACTGCTAATGGATGTCGATGGTATGTATCTAGGAGTATCTATTTTAACAAAGCATCGATTTGCTAATATACAATTATCCTTTTGATTAATTGTTATTTTATTCATATTCTTAAAAGGTTTCATATTTATCAATTCTTCTACATTAAAAATTTCCATTTTTAATTTATCTAGCCCCGCAATACTCCTCATTACGTTTCATTTTTTGTCTAGAATACCCATTTTGTTCATCTTGGTACATAGATTATCCAATTGAGAAGCGCATTTAGTAGTTTTGTACATTTTAAGTTTATTAACGAATCGTCGAAAACTAGTTATAGTTAACATTTTATTATTTGATACCCTGATATTAATACCCCTGCCGTTACTATTATTTATAACTGATGTAACCCACGTAACATTGGAATTAATTATCGATAGTAATGCATCGACACTTCCAAAATTGTCTATTATAAACTCACCGATAATTTTTTTATTGCATGTTTTCATATTCATTAGGATTATCAAATCTTTAATCTTATTACGATTGTATGCGTTGATATTACAAGACGTCATTCTAAAAGACGGAGGATTTCCATCAAATGCCAGACAATCACGTACAAAGTACATGGAAATAGGTTTTGTTCTATTACGCATCATAGATTCATATAAAACACCCGTAGAAATACTAATTTGTTTTACTCTATAAAATACTATTGCATCTATTTCATCGTTTTGTATAACGTCTTTCCAAGTGTCAAATTCCAATTTTTTTTCATTGATAGTACCAAATTCTTCTATCTCTTTAACTACTTGCATAGATAGGTAATTACAGTGATGCCTACATGCCGTTTTTTGAAACTGAATAGATGCATCTAGAAGCGATGCTACACTAGTCACGATCACCACTTTCATATTTAGAATATATGTATGTAAAAATATAGTAGAATTTCATTTTGTTTTTTTCTATGCTATAAATGAATTCTCATTTTGTATCCGCACATACTCCGTTTTATATCAATACCAAAGAAGGAAGATATCTGGTTCTAAAAGCCGTTAAAGTATGCGATGTTAGAACTGTAGAATGCGAAGGAAGTAAAGCTTCCTGCGTACTCAAAGTAGATAAACCCTCATCACCCACGTGTGAGAGAAGACCTTCGTCCCCGTCCAGATGCGAGAGAATGAATAACCCTGGAAAACAAGTCCCGTTTATGAGAACGGACATGTTACAAAATATGTTTGCTGCTAATCGCGACAACGTAACGTCAAGACTTTTGAACTAAAATACAATTATATCTTTTTCGATATTAATAAATCCGTGTCTCCCGGGTTTTTTATCTCTTTCAGTATGTGAATAGATAGGTATTTTATCTCTATTCATCATCGAATTTAAGAGATCCGATAAACATTGTTTGTATTCTCCAGATGTCAGCATCTGATACAACAATATATGTGCACATAAACCTCTGGCACTTATTTCATGTACCTTCCCCTTATCACTAAGGAGAATAGTATTTGAGAAATATGTATACATGATATTATCATGTATTAGATATACAGAATTTGTAACACTCTCGAAATCACACGATGTGTCGGCGTTAAGATCTAATATATCACTCGATAACACATTTTCATCTAGATACACTAGACATTTTTTAAAGCTAAAATAGTCTTTAGTAGTAACAGTAACTATGCGATTATTTTCATCGATGATACATTTCATCGGCATATTATTACGCGTACCATCAAAGACTATACCATGTGTATATCTAACGTATTCTAGCATAGTTGCCATACGTACATTAAACTTTTCAGGATCTTTGGATAGATCTTCCAATCTATCTATTTGAGAAAACATTTTTATCATGTTCAATAGTTGAAACGTCGGATCCACTATATAGATATTATCTATAAAGATTTTAGGAACTATGTTCATGGTATCCTGGCGAATATTAAAACTATCAATGATATGATTATCGTTTTCATCTTTTATCACCATATAGTTTCTAAGATATGGGATTTTACTTAATATAATATTATTTCCCGTAATAAATTTTATTAGAAATGCCAAATCTATAAGAAAAGTCCTAGAATTAGTCTGAAGAATATCTATATCACCGTACCGTATATTTGGATTAATTAGATATAGAGAATATGATCCGTAACATATACAACTTTTATTATGACGTCTAAGATATTCTTCCATCAACTTATTAACATTTTTGACTAGGGAAGATACATTATGACGTCCCATTACTTTTGCCTTGTCTATTACAGCGACGTTCATAGAATTTAGCATATCTCTTGCCAATTCTTCCATTGATGTTACATTATAAGAAATTTTAGATGAAATTACATTTGGAGCTTTAATAGTAAGAACTCCTAATATATCCGTGTATGTGGTCACTAATACAGATTGTAGTTCTATAATCGTAAATAATTTACCTATATTATATGTTTGAGTTTGTTTAGAAAAGTAGCTAAGTATACGATCTTTTATTTCTGATGCCGATGTATCAACATCGAAAAAAAATCTTTTTTTATTCTTTTTTACTAACGATACGAATATGTCTTTGTTAAAAACAGTTATTTTCTGAATATTTCTAGCTTGTAATTTTAACATATGATATTCGTTCACACTAGGTACTCTGCCTAAATAGGTTTCTATAATCTTTAATGTAATATTAGGAAGAGTATTCTGATCAGGATTCCTATTCATTTTGAGGATTTAAAACTCTGATTATTGTCTAATATGGTCTCAACACAAACTTTTTCACAGAGTGATAGAGTTTTTGATAACTCGTTTTTCTTAAGAAATATAAAACTACTGTCTCCAGAGCTCGCTCTATCTTTTATTTTATCTAATTCGATACAAACTCCTGATACTGGTTCAGAAAGTAATTCATTAATTTTCAGTCCTTTATAGAAGATATTTAATATAGATAATACAAAATCTTCAGTTCTTGATATCGATCTGATTGATCCTAGAACTAGATATATTAATAACGTGCTCATTAGGCAGTTTATGGCAGCTTGATAATTAGATATAGTATATTCCAGTTCATATTTATTAGATACCGCATTGCCCAGATTTTGATATTCTATGAATTCCTCTGAAAATAAATCCAAAATAACTAGACATTCTATTTTTTGTGGATTAGTGTACTCTCTTCCCTCTATCATGTTCACTACTGGTGTCCACAATGATAAATATCTAGAGGGAATATAATATAGTCCATATGATGCCAATCTAGCAATGTCGAATAACTGTAATTTTATTCTTCGCTCTTCATTATGAATTGAATCTTGAGGTATAAACCTAACACAAATTATATCATTAGACTTTTCGTATGTAATGTCTTTCATGTTATAAGTTTTTAATCCTGGAATAGAATCTATTTTAATGAGGCTTTTAAACGCAGCGTTCTCCAACGAGTCAAAGCATAATACTCTGTTGGTTTTCTTATATTCAATATTACGATTTTCTTCTTTGAATGGAATAGGTTTTTGAATTAGTTTATAATTACAACATAATAGATAAGGAAGTGTGTAAATAGTACGCGGAAAAAACATAATAGCTCCCCTGTTTTCATCCATGGTTTTAAGTAAATGATCACTGGCTTCTTTAGTCAATGGATATTCGAACATTAACCGTTTCATCATCATTGGACAGAATCCATATTTCTTAATGTAAAGAGTGATCAAATCATTGTGTTTATTGTACCATCTTGTTGTAAATGTGTATTCGGTTATCGGATCTGCTCCTTTTTCTATTAAAGTATCGATATCGATCTCGTCTAAGAATTCAACTATATCGACATATTTCATTTGTATACACATAACCATTACTAACGTAGAATGTATAGGAAGAGATGTAACGGGAACAGGGTTTGTTGATTCGCAAACTATTCTAATACATAATTCTTCTGTTAATACGTCTTGCACGTAATCTATTATAGATGCCAAGATATCTATATAATTATTTTGTAAGATGATGTTAACAATGTGATCTATATAAGTAGTGTAATAATTCATGTATTTCGATATATGTTCCAACTCTGTCTTTGTGATGTCTAGTTTCGTAATATCTATAGCGTCCTCAAAAAATATATTCGCATATATTCCCAAGTCTTCAGTTCTATCTTCTAAAAAATCTTCAACGTATGGAATATAATAATCTATTTTACCTCTTCTGATGTCATTAATGATATAGTTTTTGACACTATTTTCCGTCAATTGATTCTTATTCACTATGTCTAAAAACCGGATAGCGTCCCTAGGACGAACTACTGCCATTAATATCTCTATTATAGCTTCTGGACATAAATCATCTATTATACCAGAATTAATGGGAACTATTCCGTATCTATCTAACATAGTTTTAAGAAAGTCAGAATCTAAGACCTGATGTTCATATATTGGTTCATACATGAAATGATCTCTATTGATGATAGTGACTATTTCATTCTCTGAAAATTGGTAACTCATTCTATACACGCTTTCCTTGTTGATAAAGGATAGTATATACTCAATGGAATTTGTACCAACAAACTGTTCTCTTATGAATCGTATATCATCATCTGAAATGATCATGTAAGGCATACATTTAACAATAAGAGACTTGTCTCCTGTTATCAATATACTATTCTTGTGATAATTTATGTGTGCGGCAAATTTGTCCACGTTCTTTAATTTTGTTATAGTAGATATCAAATCCAATGGAGATACAGTTCTTGGCTTAAACAGATATAGTTTTTCTGGAACGAATTCTACAACATTATTATAAAGGACTTTGGGTATATAAGTGGGATGAAATCCTATTTTAATTAATGCGATAGCCTTGTCCTCGTGCAGATATCCAAACGCTTTTGTGATAGTATGGCATTCATTGTCTAGAAACGCTCTACGAATATCTGTAACAGATATCATCTTTAGAGAATACTAGTCGCGTTAATAGTACTAAAATTTGTATTTTTTAATCTATCTCAATAAAAAATTAATATGTATGATTCAATGTATAACTAAACTACTAACTGTTATTGATAACTAGAATCAGAATCTAATGATGACATAACTAAGAAGTTTATCTACAGCCAATTTAGCTGCATTATTTTTAGCATCTCGTTTAGATTTTCCATCTGCCTTATCGAATACTCTTCCGTCAATGTCTACACAGGCATAAAATGTAGGAGAGTTACTAGGCCCCACTGATTCAATACGAAAAGACCAATCTCTCCTAGTTATTTGACAGTACTCATTAATAACGGTGACAGGGTTAACACCTTTCCAATAAATAATTTTTTTAACCGGAATAACATCATCAAAAGACTTATGATCCTCTCTCATTGATTTTTCGCGGGATACATCATCTATTATAGCATCAGCATCAGAATCTGTAGGCCGTGTATCAGCATCCATTGTCGTAGACCAACGAGGAGGAGTATCGTTGGAGCTGTAAACCATAGCACTACGTTGAAGATCATACAGAGCTTTATTAACTTCTCGCTTCTCCATATTAAGTTGTTTAGTTAGTTGTACAGCAGTAGCTCCTTAGTCCAATGTTTTTAATAACCGCACACAATCTCTGTGTCAGAACGCTCGTCAATATAGATCTTAGAAATTTTTTTAGAGAGAACTAACGCAACTAGCAATAAAACTGATCTTATTTTATCATTTTTTTTATTCATCATCCTCTGGTGGTTCGTCGTTCCTATCGAATGTAGCTCTGATTAACCCGTCATCTATAGGTGATGCTGGTTCTGGAGATTCTGGAGGAGATGGATTATTATCTGGAAGAATCTCTGTTATTTCCTTGTTTTCATGTATCGATTGCGTTGTAACATTAAGATTGCGAAATGCTCTAAATTTGGGAGGCTTAAAGTGTTGTTTACAATCTCTACACGCGTGTCTAACTAATGGAGGTTCATCAGCGGCTCTAGTTTGAATCATCATCGGTGTAGTATTCCTACTTTTACAGTTAGGACACGGTGTATTGTATTTCTCGTCGAGAACGTTAAAATAATCGTTGTAACTCACATCCTTTATTTTATCTATATTGTATTCTACTCCTTTCTTAATGCATTTTATACCGAACAAGAGATAGCGAAGGAATTCTTTTTCGGTACCGCTAGTACCCTTAATCATATCACATAGTGTTTTATATTCTAAATGTGTGGCAATGGACGGTTTATTTCTATACGATAGTTTGTTTTTGGAATCCTTTGAGTATTCTATACCAATATTATTCTTTGATTCGAATTTAGTTTCTTCGATATTAGATTTTGTATTACCTATATTCTTGATGTAGTACTTTGATGATTTTTCCATGGCCCATTCTATTAAGTTTTCCAAGTTGGCATCATCCACATATTGTGATAGTAATTCTCGGATATCAGTAGTGACTACCGCCATTGATATTTGTTCATTTGATGAGTAACTACTAATGTATACATTTTCCATTTATAACACTTATGTATTAACTTTGTTTATTTATATTTTTTCATTATTATGTTGATATTAATAATCGTATTGTGGTTATATGGCTACAATTTCATAATGAGTTGAAGTCAGTGTCCTATGATCAATGACGATAGCTTTACTCTGAAAAGAAAGTATCAAATCGATAGTGCAGAGTCAACAATGAAAATGGATAAGACGATGACAAAGTTTCAGAATAGAGTCAAAATGGTAAAAGAAATAAATCAGACGATAAGAGCAGCACAAACTCATTACGAGACATTGAAACTAGGATATATAAAATTTAAGGGAATGATTAGGACTACTACTCTAGAAGATATAGCACCATCTATTCCAAATAATCAGAAAACTTATAAACTATTCTCGGACATTTCAGTCATTGGCAAAGCATCACAGAATCCGAGTAAGATGATATATGCTCGCTGCTTTACATGTTTCCCAATTTGTTTGGAGATGACCATAGATTCATTTGTTATAGAATGCATCCAACATTGTTCATGATATAGTTGAATCATGTATGCCTGTTCGTATGCCTGTGGCTAAGATACTGTGTAAAGAAATGGTAAATAAATACTTTGAGAATCTTTAAGAGTGCATTGACTTTGTTAGTGAATAGGCATTCCATCTTTCTCCAATACTAATTCAAATTGTTAAATTAATAATGGAATAGTATAAATAGTTATTAGTGATAGGATAGTAAACATAATTATTAGAATAGTAGTGTAGTATCATAGATAACTCTCTTCTATAAAAAATGGATTTTATTCGTAGAAAGTATCTTATATACACAGTAGAAAATAATATAGATTTTTTAAAGGATGATACATTAAGTAAAGTAAACAATTTTACCCTCAATCATGTACTAGCTCTCAAGTATCTAGTTAGCAATTTTCCTCAACATGTTATTACTAAGGATGTATTAGCTAATACCAATTTTTTTGTTTTCATACATATGGTACGATGCTGTAAAGTATACGAAGCGGTTTTACGACACGCATTTGATGCACCCACGTTGTACGTTAAAGCATTGACTAAGAATTATTTATCGTTTAGTAACACAATACAGTCGTACAAGGAAACAGTGCATAAACTAACACAAGATGAAAAATTTTTAGAGGTTGCCGAATACATGGACGAATTAGGAGAACTTATAGGCGTAAATTATGACTTAGTTCTTAATCCATTATTTCACGGAGGGGAACCCATCAAAGATATGGAAATCATTTTTTTAAAACTGTTTAAGAAAACAGACTTCAAAGTTGTTAAAAAATTAAGTGTTATAAGATTACTTATTTGGGCATACCTAAGCAAGAAAGATACAGGCATAGAGTTTGCGGATAATGATAGACAAGATATATATACTCTATTTCAACAAACTGGTAGAATAGTCCATAGCAATCTAACAGAAACGTTTAGGGATTATATCTTTCCCGGAGATAAGACTAGCTATTGGGTGTGGTTAAACGAAAGTATAGCTAATGATGCGGATATCGTTATTAATAGACCCGCCATTACCATGTATGATAAAATTCTTAGTTATATATACTCTGAGATAAAACAGGGACGCGTTAATAAAAACATGCTTAAGTTAGTTTATATCTTTGAGCCTGAAAAAGATATCAGAGAACTTCTGCTAGAAATCATATATGATATTCCTGGAGATATCCTATCTATTATTGATGCAAAAAACGACGATTGGAAAAAATATTTTATTAGTTTTTACAAAGCTAATTTTATTAACGGTAATACATTTATTAGTGATAGAACGTTTAACGATGACTTATTCAGAGTTGTTGTTCAAATAGATCCCGAATATTTCGATAATGAACGAATTATGTCTTTATTCTATACGAGTGCTGCGGACATTAAACGATTTGATGAGTTAGATATTAATAACAGTTATATATCTAATATAATTTATGAGGTGAACGATATCACATTAGATACAATGGATGATATGAAGAAGTGTCAAATCTTTAACGAGGATACGTTGTATTATGTTAAGGAATACAATACATACCTGTTTTTGCACGAGTCGGATCCCATGGTCATAGAGAACGGAATACTAAAGAAACTGTCATCTATAAAATCCAAGAGTAGACGGCTGAACTTGTTTAGCAAAAACATTTTAAAATATTATTTAGACGGACAATTGGCTCGTCTAGGTCTTGTGTTAGATGATTATAAAGGAGACTTATTAGTTAAAATGATAAACCATCTCAAATCTGTGGAGGATGTATCCGCATTCGTTAGATTTTCTACAGATAAAAACCCTAGTATTCTTCCATCGCTAATCAAAACTATTTTAGCTAGTTATAATATTTCCATCATCGTCTTATTTCAAAGGTTTTTAAGAGATAATCTATATCATGTAGAAGAATTCTTGGATAAAAGCATCCATCTAACCAAGACGGATAAGAAATATATACTTCAATTGATAAGGCACGGTAGATCATAGAACAAACCAAATATATTATTAATAATTTGTATATACATAGATATAATTATCACATATTAAAAAATAACACATTTTTGATAAATGGAAACCGTTGCAACAATTCAGACTCCCACCAAATTAATGAATAAAGAAAATGCAGAAATGATTTTGGAAAAAATTGTTAATCATATAGCTATGTATATTAGTGACGAATCAATATATTCAGAAAATAATCCTGAATATATTGATTTTCGTAACAGATACGGAGACTATAGATCTCTCATTATAAAAAGTGATCACGAGTTTGTAAAGCTATGTAAAGATCATGCAGAGAAAAGTTCTCCAGAAACGCAACAAATGATTATCAAACACATATACGAACAATATCTTATTCCAGTATCTGAAGTACTATTAAAACCTATAATGTCCATGGGTGACATATTTACATATAACGGATGTAAAGACAATGAATGGATGCTAGAACAACTCTCTACCCTAAACTTTAACAATCTCTACACATGGAACTCATGTAGCATAGGCAATGTAACGCGTCTGTTTTATACATTTTTTAGTTATCTGATGAAAGATAAACTAAATATATAAGTATAATCCCATTCTAATACTTTAACCTGATGTATTATTACCTGCATCTTATTAGAATATTAACCTAACTAAAAGACATAAAAAGCGGTAGGATATAAATATTATGGCCGCAACCGTTCCGCGTTTTGACGATGTGTACAAAAATGCACAAAGAAGAATTCTAGATCAAGAAACATTTTTTAGTAGAGGTCTAAGTAGACCGTTAATGAAAAACACATATCTATTTGATAATTACGCGTATGGATGGATACCAGAAACTGCAATTTGGAGTAGTAGATACGCAAACCTAGATGCTAGTGACTATTATCCCATTTCGTTGGGATTACTTAAAAAGTTTGAATTTCTCATGTCTCTATATAAAGGTCCTATTCCCGTATATGAAGAAAAAGTAAATACTGAATTCATTGCTAATGGATCTTTCTCCGGTAGATACGTATCATATCTTAGAAAGTTTTCTGCCCTTCCAACAAACGAGTTTATTAGTTTTTTATTATTGACCTCCATCCCTATCTATAATATCTTATTCTGGTTTAAAAACACACAGTTTGATATTACTAAACACACATTATTCAGATACGTCTATACAGATAATACCAAACACCTTGCGTTGGCTAGGTATATACATCAAACAGGAGACTATAAGCCTTTGTTTAGTCGTCTCAAAGAGAATTATATATTTACTGGTCCCGTTCCAATAGGTATCAAAGATATAGATCACCCTAATCTTAGTAGAGCAAGAAGTCCATCCGATTATGAGACATTAGCTAATATTAGTACTATATTGTACTTTACCAAGTATGATCCAGTATTAATGTTTTTATTGTTTTACGTACCTGGGTATTCAATTACTACAAAAATTACTCCAGCCGTAGAATATCTAATGGATAAACTGAATCTAACAAAGAGCGACGTACAACTGTTGTAAATTATTTTATGCTTCGTAAAATGTAGGTCTTGAACCAAACATTCTTTGAAAAAATGAGATGCATAAAACTTTATTATCCAATAGATTAACTATTTCAGACGTCAATCGTTTAAAGTAAACTTCGTAAAATATTCTTTGATTGCTGCCGAGTTTAAAACTTCTATCGATAATTGTTTCATATGTTTTAATATTTACAAGTTTTTTGGTCCATGGTACATTAGCTGGACAGATATATGCAAAATAATATCGTTCTCCAAGTTCTATAGTCTCTGGATTGTTTTTATTATATTCAGTAACCAAATACATATTAGGGTTATCTGCGGATTTATAATTTGAGTGATGCATTCGACTCAACATAAATAATTCTAGAGGAGACGATCTACTATCAAATTCGGATCGTAAATCTGTTTCTAAAGAACGGAGAATATCTATACATACCTGATTAGAATTCATCCGTCCTTCAGACAACATCTCAGACAGTCTGGTCTTGTATGTCTTAATCATATTCTTATGAAACTTGGAAACATCTCTTCTAGTTTCACTAGTACCTTTATTAATTCTCTCAGGTACAGATTTTGAATTCGACGATGCCGAGTATTTCATCGTTGTATATTTCTTCTTCGATTGCATAATCAAATTCTTATATACCGCCTCAAACTCTATTTTAAAATTATTAAACAATACTCTACTATTAATCAGTCGTTCTAACTCCTTTGCTATTTCTATGGACTTATCTACATCTTGACTGTCTATCTCTGTAAACACGGAGTCGGTATCTCCATACACGCTACGAAAACGAAATCTATAATCTATAGGCAACGATGTTTTCACAATCGGATTAATATCTCTATCGTCCATATAAAATGGATTACTTAATGTATTGGCAAACCGTAACATACCGTTGGATAACTCTGCTCCATTTAGTACCGATTCTAGATACAATATCATTCTACGTCCTATGGATGTGCAACTCTTAGCCGAAGCGTATGAGTATAGAGCACTATTTCTAAATCCCATCAGACCATATACTGAGTTGGCTACTATCTTGTACGTATATTGCATGGAATCATAGATGGCCTTTTCAGTTGAACTGGTAGCCTGTTTTAACATCTTTTTATATCTGGCTCTCTCTGCCAAAAATGTTCTTAATAGTCTAGGAATGGTTCCTTCTATTGATCTATCGAAAATTGCTATTTCAGAGATGAGGTTCGGTAGTCTAGGTTCACAATGAACCGTAATATATCTAGGAGGTGGATATTTCTGAAGCAAGAGTTGATTATTTATTTCTTCTTCCAATCTATTGGTACTAACAACGACACCGACTAATGTTTCCGGAGATAGATTTCCAAAGATACACACATTAGGATACAGACTGTTATAATCAAAGATTAATACATTATTACTAAACATTTTTTGTTTTGGAGCAAATACCTTACCGCCTTCATAAGGAAACTTTTGTTTTGTTTCTGATCTGACTAAGATAGTTTTAGTTTCCAACAATAGCTTTAACAGTGGACCCTTGATGATTGTACTCGCTCTATATTCGAATACCATGGATTGAGGAAGCACATATGTTGCCGCACCAGCGTCTGTTTTTGTTTCTACTCCATAATACTCCCACAAATACTGACACAAACAAGCATCATGAATACAGTATCTAGCCATATCTAAAGCTATGTTTAGATTATAATCCTTATACATCTGAGCTAAATCAATGTCATCCTTTCCGAAAGATAATTTATATATATCATTAGGTAAAGTAGGACATGATAGTACGACTTTAAATCCATTTTCCAAAATATCTTTACGAATTACTTTACATATAATATCCTCATCAACAGTCACGTAATTACCTGTGGTTAAAACCTTTGCAAATGTATCGGCTTTGCCTTTCGCGTCCGTAGTATCGTCACCGATGAACGTCATTTCTCTAACTCCTCTATTTAATACTTTACCCATGCAACTGAACGCGTTCTTGGATATAGAATCCAATTTGTACGAATCCAATTTTTCAGATTTTTGAATGAATGAATATAGATCGAAAAATATAGTTCCATTATTGTTATTAACGTGAAACGTAGTATTGGCCATGCCGCATACTCCCTTATGACTAGACTGATTTCTCTCATAAATACAGAGATGTACAGCTTCCTTTTTGTCTGGAGATCTAAAGATAATCTTCTCTCCTGTTAATAACTCTAGACGATTAGTAATATATCTCAGATCAAAGTTATGTCCGTTAAAGGTAACGACGTAGTCGAACGTTAGTTCCAACAATTGTTTAGCTATTCGTAACAAAACTATTTCAGAACATAGAACTAGTTCTCGTTCGTAATCCATTTCCATTAGCGACTGTATCCTCAAACATCCTCTATCGACGGCTTCTTGTATTTCCTGTTCCGTTAACATCTCTTCATTAATGAGCGTAAACAGTAATCGTTTACCACTTAAATCGATATAACAGTAACTTGTATGCGAGATTGGGTTAATAAATACAGAAGGAAACTTCTTATCGAAGTGACACTCTATATCTAGAAATAAGTACGATCTTGGGATATCGAATCTAGGTATTTCTTTAGCGAAACAGTTACGTGGATCGTCACAATGATAACATCCATTGTTAATCTTTGTCAAATATTGCTCGTCCAACGAGTAACATCCGTCTGGAGATATCCCGTTAGAAATATAAAACCAACTAATATTGAGAAATTCATCCATGGTGGCATTTTGTATGCTGCGTTTCTTTGGCTCTTCTATCAACCACATATCTGCGACGGAGCATTTTCTATCTTTAATATCTAGATTATAACTTATTGTCTCGTCAATGTCTATAGTTCTCATCTTTCCCATCGGCCTCGCATTAAATGGAGGAGGAGATAATGACTGATATATTTCGTCCGTCACTACGTAATAAAAGTAATGAGGAAATCGTATAAATACGGTCTCGCCATTTCGACATCTGGATTTCAGATATAAAAATCTGTTTTCACCGTGACTTTCAAACCAATTAATACACCTAACATCCATTTCTAGAATTTAGAAATATATTTTCATTTAAATGAATCCCAAACATTGGGGAAGAGCCGTATGGACCATTATTTTTATAGTACTTTCGCAAGCGGGTTTAGACGGCAACATAGAAGCGTGTAAACGAAAACTATATACTATAGTCAGCACTCTTCCATGTCCTGCATGTAGACGACACGCGACTATCGCTATAGAGGACAATAATGTCATGTCTAGCGATGATCTGAATTATATTTATTATTTTTTCATCAGATTATTTAACAATTTGGCATTTGATCCCAAATACGCAATCGATGTGTCAAAGGTTAAACCTTTATAAACTTAACCCATTATAAAACTTATGATTAGTCACGACTGAAATAACCGCGTGATTATTTTTTGGTATAATTCTACACGGCATGGTTTCTGTGACTATGAATTCAACACCTGTTATCTTAGTGAAATCTTTAACAAACAGCAAGGGTTCGTCAAAGACATAAAACTCATTGTTTACGATCGAAATAGACCCCCTATCACACTTAAAATAAAAAATATCCTTATCCTTTACCACCAAATAAAATTCTGATTGGTCAATGTGAATGTATTCACTTAACAGTTCCACAAATTTATTTATTAACTCCGAGGCACATACATCGTCGGTATTTTTTATGACAAACTTTACTCTTCCAGCATCCGTTTCTAAAAAAATATTAACGAGTTCCATTTATATCATCCAATATTATTGAAATGACGTTGATGGACAGATGATATAAATAAGAAGGTACAGTACCTTTGTCCACCATCTCCTCCAATTCATACTCTATTTTGTCATTAACTTTAATGTGTGAAAACAGTACGCCACATGCTTCCATGACAGTGTGTAACACTTTGGATACAAAATGTTTGACATTAGTATAATTGTCCAAGACTGTCAATCTATAATAGATAGTAGCTATAATATATTCTATGATGGTATTGAAGAAGATGACAACCTTGGCATATTGATCATTTAACACAGACATGGTATCAACAAATAGCTTAAATGAAAGAGAATCAGTAATTGGAATAAGCGTCTTCTCGATGTAGTGTCCGTATACCAACATGTCTGATATTTTGATGTATTCCATTAAATTATTTAGTTTTTTCTTTTTATTCTCGTTAAACAGAATTTCTGTCAATGGACCCCAACATCGTTGACCTATTAAGTTTTGATTGATTTTTCCGTGTAAGGCGTATCTAGTCAGATCGTATAGCCTATCCAATAATCCATCGTCTGTGCGTAGATCACATCGTACACTTTTTAATTTTCTATAGAAGAGTGACAGACATCTGGAGCAATTACAGACAGCAATTTCTTTATTCTCTACAGATGTAAGATACTTGAAGATATTCCTATGATGATGCAGAATTTTGGATAACACGGTATTGATGGTATCTGTTACCATAATTCCTTTGACTGATAGTGTCAAAGTACAAGATTTCCAATCTTTTGCAATTTTCAGTACCATTATCTTTGTTTTGATATCTATATCAGACAGCATGGTACGTCTGACAACACAGGGATTAAGACGGAAAGATGAAATGATTCTCTCAACATCTTCAATAGATACCTTGCTATTTTTTTTGGCATTATCTATATGTGAGAGAATATCCTCTAGAGAATCAGTATCCTTTTTGATGATAGTGGATCTCAATGACATGGGACGTCTAAACCTTCTTATTCTATCACCAGATTGCATGGTGATTTGTCTTCTTTCTTTTATCATGATGTAATCTCTAAATTCATCGGCAAATTGTCTATATCTAAAATCATAATATGAGATGTTTACCTCTACAAATATCTGTTCGTCCAATGTTAGAGTATCTATATCAGTTTTGTATTCCAAATTAAACATGGCAACGGATTTAATTTTATATTCCTCTATTAAGTCCTCGTCGATAATAACAGAATGTAGATAATCATTTAATCCATCGTACATGGTTGGAAGATGCTCGTTGACAAAATCTTTAATTGTCTTGATGAAGGTGGGACTATATCTAACATCTTGATTAATAAAATTTATAACATTGTCCATAGGATACTTTGTAACTAGTTTTATACACATCTCTTCATTGGTAAGTTTAGACAGAATATCGTGAACAGGTGGTATATTATATTCATCAGATATACGAAGAATAATGTCCAAATCTATATTGTTTAATATATTATATAGATGTAGTGTAGCTCCTACAGGAATATCTTTAACTAAGTCAATGATTTCATCAACAGTTAGATCTATTTTAAAGTTAATCATATAGGCATTGATTTTTAAAAGGTATGTAGCCTTGACTACATTCTCATTAATTAACCATTCCAAGTCACTGTGTGTAAGAAGATTATATTCTATCATAAGCTTGACTACATTTGGTCCCGATACCATTAAAGAATTCTTATGATATAAGGAAACAGCTTTTAGGTACTCATCTACTCTACAAGAATTTTGGAGAGCCTTAACGATATCAGTGACGTTTATTATTTCAGGAGGAAAGAACCTAACATTGAGAATATCTGAATTAATAGCTTCCAGATACAGTGATTTTGGCAATAGTCCGTGTAATCCATAATCCAGTAACACGAGCTGGTGCTTGCTAGACACCTTTTCAATGTTTAATTTTTTTGAAATAAGCTTTGATAAAGCCTTCCTCGCAAATTCCGGATACATGAACATGTCGCCAACATGATTAAGTATTGTTTTTCATTATTTTTATATTTTCTCAACAAGTTCTCAATACCCCAATAGATAATAGAATATCACCCAATGCGTCCATGTTGTCTATTTCCAACAGGTCGCTATATCCACCAATAGAAGTTTTCCCAAAAAAGATTCTAGGAACAGTTCTACCACCAGTAATTTGTTCAAAATAGTCACGCAATTCATTTTCGGGTTTAAATTCTTTAATATCTACAATTTCATACGCTCCTCTTTTGAAACTAAACTTATTTAGAATATCCAGTGCGTTTCTACAAAAAGGACATGTAAACTTGACAAAAATTGTCACTTTGTTATTGGCCAACCTTTGTTGTACAAATTCCTCGGCCATTTTTAATATTTAAGTGATACAAAACTATCTCGACTTATTTAACTCTTTAGTCGAGATATATGGACACAGATAGCTATATGATAACCAACTACAGAAGACAAACGCTATAAAAAACATAATTACGACGAGCATATTTATAAATATTTTTATTCAGTATTACTTGATATAGTAATATTAGGCACAGTCAAACATTCAACCACTCTAGATACATTAACTCTCTCATTTTCTTTAACAAATTCTGCAATATCTTCGTAAAAAGATTCTTGAAACTTTTTAGAATATCTATCGACTCTAGATGAAATAGCGTTCGTCAACATACTATGTTTTGTATACATAAAGGCGCCCATTTTAACAGTTTCTAGTGACAAAATGCTAGCGATCCTAGGATCCTTTAGAATCACATAGATTGACGATTCGTCTCTCTTAGTAACTCTAGTAAAATAATCATACAATCTAGTACGCGAAATAATATTATCCTTGACTTGAGGAGATCTAAACAATCTAGTTTTGAGAACATCGATAAGTTCATCGGGAATTACATACATACTATCTTTAATAGAACTCTTTTCATCCAGTTGAATGGATTCGTCCTTAACCAACTGATTAATGAGATCTTCTATTTTATCATTTTCTAGATGATATGTATGTCCATTAAAGTTAAATTGTGTAGCGCTTCTTTTTAGCCTAGCAGCCAATACTTTAACATCACTAATATCGATATACAAAGGAGATGATTTATCGATGGTATTAAGAATTCGTTTTTCGACATCCGTCAAAACCAATTCCTTTTTGCCTGTATCATCCAGTTTGCCATTCTTTGTAAAGAAATTATTTTCTACTAGACTATTAATAAGACTGATAAGGATTCCTCCATAATTGCACAATCCAAACTTTTTCACAAAACTAGACTTTACGAGATCTACAGGAATGCGTACTTCAGGTTTCTTAGCTTGTGATTTTTTCTTTTGTGGACATTTTCTAGTGACCAACTCATCTACCATTTCATTGATTTTAGCAGTGAAATAAGCTTTCAATGCACGGGCACTGATACTATTGAAAACGAGTTGATCTTCAAATTCCGCCATTTAAGTTCACCAAACAACTTTTAAATACAAATATATCAATAGTAGTAGAATAAGAACTATAAAAAAAATAATAATTAACCAATACCAACCCCAACAACCTGTATTATTAGTTGATGTGACAGTTTTCTCATCACTTAGAACAGATTTAACAATTTCTATAAAGTCTGTCAAATCATCTTCCTGAGAACCCATAAATACACCAAATATAGCAGCGTACAACTTATCCATTTATACATTGAATATTGGCTTTTCTTTATCGCTATCTTCATCATATTCATCATCAATATCAACAAGTCCCAGATTACGAACCAGATCTTCTTCTACATTTTCAGTCATTGATACGCGTTCACTATCTCCAGAGAGTCCGATAACGTTAGCCACTACTTCTCTATCAATGATTAGTTTCTTGAGCGCGAATGTAATTTTTGTTTCCGTTCCGGATCTATAGAAAACTACAGGTGTAATAATTGCCTTGGCTAATTGTCTTTCTCTTTTACTGAGTGATTCTAGTTCACCTTCTATAGATCTGAGAATGGATGATTCTCCAGTCGAAACATATTCTACCATGGCTCCGTTTAATTTGTTGATGAAGATGGATTCATCCTTAAATGTTTTCTCTGTAATAGTTTCCGCCGAAAGACTATGCAAAGAATTTGGAATGCGTTCCTTGTGTGTAATGTTTCCATAGACAGCTTCTAGAAGTTGATACAACATAGGACTAGCCGCGGTAACTTTTATTTTTAGAAAGTATCCATCGCTTCTATCTTGTTTAGATTTATTTTTATAAAGTTTAGTCTCTCCTTCCAACATAATAAAAGTGGAAGTCATCTGACTAGATAAACTATCAGTAAGTTTTATAGAGATAGATGAACAATTAGCGTATTGAGAAGCATTTAGTGTAACGCATTCGATACATTTTGCATTAGATTTACTAATCGATTTTGCATACTCTATAACACCCGCACAAGTCTGTAGAGAATCGCTAGATGCTGTAGGTCTTGGTGAAGTTTCAACTCTCTTCTTGATTACCTTACTCATGATTAAACCTAAATAATTGTACTTTGTAATATAATGATATATATTTTCACTTTATCTCATTTGAGAATAAAAATGTTTTTGTTAACCACTGCATGATGTACAGATTTCGGAATCGCAAACCACTTGTGGTTTTATTTTATCCTTGTCCAATGTGAATTGAATGGGAGCGGATGCGGGTTTCGTACGTAGATAGTACATTCCCGTTTTTAGACCGAGACTCCATCCGTAAAAATGCATACTCGTTAGTTTGGAATAACTCGGATCTGCTATATGGATATTCATAGATTGACTTTGATCGATGAAGGCTCCCCTGTCTGCAGCCATTTTTATGATCGTCTTTTGTGGAATTTCCCAAATAGTTTTATAAACTCGCTTAATATCTTCTGGAAGGTTTGTATTCTGAATGGATCCACCATCTACCATAATCCTATTCTTGATCTCATCATTCCATAATTTTCTCTCGGTTAAAACTCTAAGGAGATGCGGGTTAACTACTTGGAATTCTCCAGACAATACTCTCCGAGTGTAAATATTACTGGTATACGGTTCCACCGACTCATTATTTCCCAAAATTTGAGCAGTTGATGCAGTCGGCATAGGTGCCACCAATAAACTATTTCTAAGACCGTATGTTCTGATTTTATCTTTTAGAGGTTCCCAATTCCAAAGATCCGACGGTACAACATTCCAAAGATCATATTGTAGAATACCGTTACTGGCGTACGATCCTACATATGTATCATATGGTCCTTCCTTCTCAGCTAGTTTACAACTCGCCTCTAATGCACCGTAATAAATGGTTTCAAAGATCTTCTTATTTAGATCTTGTGCTTCCAGGCTATCAAATGGATAATTTAAGAGAATAAACGCGTCCGCTAATCCTTGAACACCAATACCGATAGGTCTATGTCTCTTATTAGAGATTTCAGCTTCTGGAATAGGATAATAATTAATATCTATAATTTTATTGAGATTTCTGACAATTACTTTGACCACATCCTTCAGTTTGAGAAAATCAAATCGCCCATCTATTACAAACATGTTCAATGCAACAGATGCCAGATTACACACGGCTACCTCATTAGCATCCGCATATTGTATTATCTCAGTGCAAAGATTACTACACTTGATGGTTCCTAAATTTTGTTGATTACTCTTTTTGTTACACGCATCCTTATAAAGAATGAATGGAGTACCAGTTTCAATCTGAGATTCTATAATCGCTTTCCAGACGACTCGAGCCTTTATTATACATTTGTATCTCCTTTCTCTTTCGTATAGTGTATACAATCGTTCGAACTCGTCTCCCCAAACATTGTCCAATCCAGGACATTCATCCGGACACATCAACGACCACTCTCCGTCATCCTTCACTCGTTTCATAAAGAGATCAGGAATCCAAAGAGCTATAAATAGATCTCTTGTTCTATGTTCATCGTTTCCTGTATTCTTTTTAAGATCGAGGAACGCCATAATATCAGAATGCCACGGTTCCAAGTATATGGCCATAACTCCAGGCCGTTTGTTTCCTCCCTGATCTATGTATCTAGCGGTGTTATTATAAACTCTCAACATTGGAATAATACCGTTTGATATACCATTGGTACCGGAGATATAGCTTCCACTGGCACGAATATTACTAATTGATAGACCTATTCCCCCTGCCATTTTAGAGATTAATGCGCATCGTTTTAACGTGTCATAGATGCCTTCTATGCTATCATCGATCATGTTAAGTAGAAAACAGCTAGACATTTGGTGACGAGTAGTTCCCGCATTAAATAAGGTAGGAGAAGCGTGCGTAAACCATTTTTCAGAAAGTAGATTGTACGTCTCAATAGCTGAGTCTATATCCCATTGATGAATTCCTACTGCGACACGCATTAACATGTGCTGAGGTCTTTCAACAATTTTGTTGTTTATTTTCAACAAGTAGGATTTTTCCAAAGTTTTAAAACCAAAATAGTTGTATGAAAAGTCTCGTTCGTAAATAATAACCGAATTGAGCTTATCCTTATATTTGTTAACTATATCCATGGTAATACTTGAAATAATCGGAGAATGTTTCCCATTTTTAGGATTAACATAGTTGAATAAATCCTCCATCACTTCACTAAATAGTTTTTTTGTTTCCTTGTGTAGATTTGATATGGCTATTCTGGCGGCTAGAATGGCATAATCCGGATGTTGTGTAGTACAAGTGGCTGCTATTTCGGCTGCCAGAGTGTCCAATTCTACCGTTGTTACTCCATTATATATTCCTTGAATAACCTTCATAGCTATTTTAATAGGATCTATATGATCAGTGTTTAAGCCATAGCACAATTTTCTAATACGAGACGTGATTTTATCAAACATGACATTTTCCTTGTATCCATTTCGTTTAATGACAAACATTTTTGTTGGTGTAATAAAAAAAATTATTTAATTTTTCATTAATAGGGATTTGACGTATGTAGCGTACAAAATTATCGTTCCTGGTATATAGATAAAGAGTCCTATATATTTGAAAATCGTTACGGTTCGATTAAACTTTAATGATTGCATTGTGAATATATCATTAGGATTTAACTCCTTGACTATCATGGCGGTGCCAGAAATTACCATCAAAAGCATTAATACAGTTATGCAGATCGCAGTTAGAACGGTTATAGCATCCACCATTTATATCTAAAAATTAGATCAAAGAATATGTGACAACGTCCTAGTTGTATACTGAGAATTGACGAAACAATGTTTCTTACATATTTTTTTCTTATTAGTAACCGACTTAATAGTAGGAACTGGAAAACTAGACTTGATTATTCTATAAGTATAGATACCCTTCCAAATAATGTTCTCTTTGATAAAAGTTCCAGAAAATGTAGAATTTTTTAAAAAGTTATCTTTTGCTATTACTAATATCGTGGTTAGACGCTTATTATTAATATGAGTGATGAAATCCACACCGCTTCTAGATATCGCTTTTATTTCCACATTAGATGGTAAATCCAATAGTGAAACTATCTTTTTAGGAATGTATGGACTCGCGTTTAGAGGAGTGAACGTCTTCGGAGTAGTAAAGGATGATTCGTCAAATGAATAAACAATTTCACAAATGGATGTTAATGTATTAGTAGGAAATTTTTTGACGCTAGTGGAATTGAAGATTCTAATGGATGATGTTCTACCTATTTCATCCGATAACATGTTAATTTCCAATACCAACGGTTTTAATATTTCGATGATATACGGTAGTCTCTCTTTCGGACTTATATAGCTTATTCCACAATACGAGTCATTATATACTCCAAAAAACAAAATAACTAGTATAAAATCTGTATCGAATGGGAAAAACGAAATTATCGATATAGGTATAGAATCCGGAACATTGAACGTATTAATACTTAATTCTTTTTCAGTGGTAAGAACCGATAGGTTATTGACATTGTATGGTTTTAAATATTCTATAACTTGAGACTTGATAGATATTAATGACGAATTGAAAATTATTTTTATCACCACGTGTGTTTCAGGATCATCGTCGACGCCAGTTAACCAACCGAATGGAGTAAAATAAATATCATTAATATATGCTCTAGATATTAGTATTTTTATTAATCCTTTGATTATCATCTTCTCGTACGCGAATGATTCCATGATCAAGAGTGATTTGAGAACATCCTCCGGAGTATTAATGGGTTTAGTAAACAGTCCATCGTTGCAATAATAAAAGTTGTCCAAGTTAAAGGATATTATGCATTCGTTTAAAGATATCACCTCATCTAACGGAGACAATTTTTTGGTAGGTTTTAGAGACTTTGAAGCTACTTGTTTAACAAAGTTATTCATCGTCGTCTACTATTCTATTTAATTTTGTAGTTAATTTATCACATATCACATTAATTGACTTTTTGGTCCACTTTTCCATACGTTTATATTCTTTTAATCCTGCGTTATCCGTTTCCGTTATATACAGGGATAGATCTTGCAAGTTAAATAGAATGCTCTTAAATAATGTCATTTTTTTATCCGCTAAAAATTTAAAGAATGTATAAACTTTTTTCAAAGATTTAAAACTTTTAGGTGGAGTTCTGGTACACAATATCATAAACAAACTAATAAACATCCCACATTCAGATTCCAACAATTGATTAACTTCCACATTAATACAGCCTATTTTCGCTCCAAATGTACATTCGAAAAATCTGAATAAAACATCAATATCGCAATTTGTATTATCCAATACAGAATGTCTGTGATTCGTGTTAAAACCATCGGAAAAAGAATAGAAATAAAAATTATTATAATGGTGGAATTCAGTTGGAATATTGCCTCCGGAGTCATAAAAGGATACTAAACATTGTTTTTTATCGTAAATTACACATTTCCAATGAGACAAATAACAAAATCCAAACATTACAAATCTAGAGGTAGAACTTTTAATTTTGTCTTTAAGTATATACGATAAGATATGTTTATTCATAAACGCGTCAAATTTTTCATGAATAGCTAAGGAGTTTAAGAATCTCATGTCAAATTGTCCTATATAATCCACTTCGGATCCATAAGCAAACTGAGAGACTAAGTTCTTAATACTTCGATTGCTCATCCAGGCTCCTCTCTCAGGCTCTATTTTCATCTTGACGACCTTTGGATTTTCACCAGTATGTATTCCTTTACGTGATAAATCATCGATTTTCAAATCCATTTGTGAGAAGTCTATCGCCTTAGATACTTTTTCCCGTAGTTGAGGTTTAAAGAAATACGCTAACGGTATACTAGTAGGTAACTCAAAGACATCATATATAGAATGGTAACGCGTCGTTAACTCGTCGGTTAACTCTTTCTTTTGATCGAGTTCATCGCTACTATTGGGTCTGCTCAGGTGCCCCGACTCTACTAGTTCCAACATCATACCGATAGGAATACAAGACACTTTGCCAGCGGTTGTAGATTTATCATATTTCTCCACCACATATCCGTTACAATTTGTTAAGAATTTAGATACATCTATATTGCTACATAATCCAGCTAGTGAATATATATGACATAATAAATTGGTAAATCCTAGTTCTGGTATTTTACTAATTACTAAATCTGTATATCTTTCCATTTATCATGGAAAAGAATTTACCAGATATCTTCTTTTTTCCAAACTGCGTTAATGTATTCTCTTACAAATATTCACAAGATGAATTCAGTAATATGAGTAAAACGGAACGTGATAATTTCTCATTGGCTGTGTTTCCAGTGATAAAACATAGATGGCATAACGCACACGTTGTAAAACATAAAGGAATATACAAAGTTAGTACAGAAGCACGTGGAAAAAAAGTATCTCCTCCATCACTAGGAAAACCCGCACATATAAACCTAATGTCGAAGCAATATATATATAGTGAGTATGCAATAAGCTTTGAATGTTATAGTTTTCTAAAATGTATAACAAATACAGAAATCAATTCGTTCGATGAGTATATATTAAGAGGACTATTAGAAGCTGGTAATAGTTTACAGATATTTTCCAATTCCGTAGGTAAACGAATAGATACTATAGGTGTACTAGGGAATAAGTATCCATTTAGCAAAATTCCATTGGCCTCATTAACTCCTAAAGCACAACGAGAGATATTTTTAGCGTGGATTTCTCATAGACCTGTAGTTTTAACTGGAGGAACCGGAGTGGGTAAGACGTCACAGGTACCCAAGTTATTGCTTTGGTTTAATTATTTATTTGGTGGATTCTCTTCTCTAGATAAAATCACTGACTTTCACGAAAGACCAGTCATTCTATCTCTTCCTAGGATAGCTTTAGTTAGATTGCATAGCAATACCATTTTAAAATCATTGGGATTTAAGGTACTAGATGGATCTCCTATCTCTTTACGGTACGGATCTATACCGGAAGAATTAATAAACAAACAACCAAAAAAATATGGAATTGTATTTTCTACCCATAAGTTATCTCTAACAAAACTATTTAGTTATGGCACTATTATTATAGACGAAGTTCATGAGCATGATCAAATAGGAGATATTATTATAGCAGTAGCGAGAAAACATCATACGAAAATAGATTCTATGTTTTTAATGACTGCCACGTTAGAGGATGACAGGGAACGTCTAAAAATATTTTTACCTAATCCCGCATTTATACATATTCCTGGAGATACACTGTTTAAAATTAGCGAGGTATTTATTCATAATAAGATAAATCCATCTTCCAGAATGGCATATATAGAAGAAGAAAAGAGAAATTTAGTTACTGCTATACAGATGTATACTCCTCCTGATGGATCATCCGGTATAGTCTTTGTGGCATCCGTTGCACAGTGTCACGAATATAAATCATATTTAGAAAAAAGATTACCGTATGATATGTATATTATTCATGGTAAGGTCTTAGATATAGACGAAATATTAGAAAAAGTGTATTCATCACCTAATGTATCGATAATTATTTCTACTCCTTATTTGGAATCCAGCGTTACTATACGCAATGTTACACACATTTATGATATGGGTAGAGTTTTTGTCCCCGCTCCTTTTGGAGGATCACAACAATTTATTTCTAAATCTATGAGAGATCAACGAAAAGGAAGAGTAGGAAGAGTTAATCCTGGAACATACGTATATTTCTATGATCTGTCTTATATGAAATCTATACAGCGAATAGATTCAGAATTTCTACATAATTATATATTGTACGCTAATAAGTTTAATCTAACACTCCCCGAAGATTTGTTTATAATCCCTACAAATTTGGATATTCTATGGCGTACAAAGGAATATATAGACTCGTTCGATATTAGTACAGAAACATGGAATAAATTATTATCCAATTATTATATGAAGATGATAGAGTATGCTAAACTTTATGTACTAAGTCCTATTCTCGCTGAGGAGTTGGATAACTTTGAGAGGACGGGAGAATTAACTAGTATTGTACAAGAAGCCATTTTATCTCTAAATTTACAAATTAAGATTTTAAAATTTAAACATAAAGATGATGATACGTATATACACTTTTGTAGAATATTATTCGGTGTCTATAACGGAACAAACGCTACTATATATTATCATAGACCTCTAACGGGATATATGAATATGATTTCAGATACTATATTTGTTCCTGTAGATAATAACTAAAAATCAAAATCTAATGACCACATCTTTTTTTAGAGATGAAAAATTTTCCACATCTCCTTTTGTAGACACGACTAAACATTTTGCAGAAAAAAGTTTATTATTATTTAGATAATCGTATACTTCATCAGTGTAGATAGTAAATGTGAACAGATAAAAGGTATTCTTGCTCAATAGATTGGTAAATTCCATAGAATATATTAATCCTTTCTTCTTGAGATCCCACATCATTTCAACCAAAGACGTTTTATCCAATGATTTACCTCGTACTATACCACATACAAAACTAGATTTTGCAGTGATGTCGTACCTGGTATTCCTACCAAACAAAATTTTACTTTTAGTTCTTTTAGAAAATTCTAAGGTAGAATCTCTATTTGTCAATATGTCATCTATGGAATTACCACTAGCAAAAAATGATAGAAATATATATTGATACATCGCAGCTGGTTTTGATCTACTATACTTTAAAAACGAATCAGATTCCATAATTGCTTGTATATCATCAGCTGAAAAACTATGTTTTACACGTATTCCTTCGGCATTTCTTTTTAATGATATATCTTGTTTAGACAATGATAAAGTTATCATGTCCATGAGAGACGCGTCTCCGTATCGTATAAATATTTCATTAGATGTTAGACGCTTCATTAGGGGTATACTTCTATAAGGTTTCTTAATTAGTCCATCATTGGTTGCGTCAAGAACTACTATCTGATGTTGTTGGGTATCTCTAGTGTTACACATGGCCTTACTAAAGTTTGGGTAAATAACTATGATATCTCTATTAATTATAGATGTATATATTTCATTCGTCAAGGATATTAATATCGACTTACTATCGTCATTAATACGTGTAATGTAATCATATAAATCATGCGATAGCCAAGGAAAATTCAAATAGATGTTCATCATATAATCGTCGCTATAATTCATATTAATACTTTGACATTGACTAATTTGTAATATAGCCTCGCCACGAAGAAAGCTCTCGTATTCAGTTTCATCGATAAAGGATACCGTTAAATATAACTGGTTGCCGATAGTCTCATAGTCTATTAAGTGGTAAGTTTCGTATAAATACAGAATCCCTAAAATATTATCTAATGTGGGATTAATCCTTACCATAACTGTATAAAATGGAGCCGGAGTCATAACTATTTTACCGTTTGTACTTACTGGAATAGATGAAGGAATAATCTCCGGACATGATGGTAAAGACCCAAATGTCTGTTTGAAGAAATCCAATGTTCCAGGTCCTAATCTCTTGACAAAAATTACGATATTCGATCCCGATATCCTTTGCATTCTATTTACCAGCATATCACGAACTATATTAAGATTATCTATCATGTCTATTCTCCCACCGTTATATAAATCGCCTCCGCTAAGAAACGTTAGTATATCCATACAATGGAATACTTCATTTCTAAAATAGTATTCGTTTTCTAATTCTTTAATGTGAAATCGTATACTAGAAAGGGAAAAATTATCTTTGAGTTTTCCATTAGAAAAGAACCACGAAACTAATGTTCTGATTGCGTCTGACTCCGTCGCTGAATTAATAGATTTACACCAAAAACTCATATAACTTCTAGATGTAGAAGCATTCGCTAAAAAATTAGTAGAATCAAAGGATATAAGTAGATGTTCCAACAAGTGAGCAATTCCCAAGATTTCATCTATATCATTCTCGAATCCGAAATTAGAAATTCCCAAGTAGATATCCTTTTTCATCCGATCATTGATGAAAATACGAACTTTATTCGGTAAGACGATCATTTACTAAGGAGTAAAATAGGAAGTAACGTTCGTATATCGTTATCGTCGTATAAATTAAAGGTGTGTTTTTTGCCATTAAGAGACATTATAATTTTACCAATATTGGAATTATAATATAGGTGTATTTGAGCACTAGAAACGGTCGATGCATCGGTAAATATAGCTGTATCTAATGTTCTAGTCGGTATTTCTTCATTTCGCTGTCTAATGATAGCGTTTTCTCTATCTGTTTCCATTACAGCTGCCTGAAGTTTATTGGTCGGATAATATGTAAAATAATAAGAAATACATACGAATAACAAAAATAAAATAAGATATAATAAAGATGCCATTTAGAGATCTAATTTTGTTCAACTTGTCCAAATTCCTACTTACAGAAGATGAGGAATCGTTGGAGATAGTATCTTCCTTATGTAGAGGATTTGAAATATCTTACGATGACTTAATATCGTACTTTCCAGATAGGAAATACCATAAATATATTTCTAAGGTATTTGAACATGTAGATTTATCGGAGGAATTAAGTATGGAATTCCATGATACAACTCTGAGAGATTTAGTATATCTTAGATTGTACAAGTATTCCAAGTATATACGGCCGTGTTATAAATTAGGAGATAATCTAAAAGGTATAGTTGTTATAAAGGACAGAAATATATATATTAGAGAAGCAAATGATGACTTGATAGAATATCTCCTCAAGGAATACACTCCTCAGATTTATACATATTCTAATGAGCGAGTTCCCATAGCTGGTTCAAAATTAATTCTTTGTGGATTTTCTCAAGTTACATTTATGGCGTATACAACGTCGCATATAACAACAAATAAAAAGGTAGATGTTCTCGTTTCCAAAAAATGTATAGATGAACTAGTCGATCCAATAAATTATCAAATACTTCAAAATTTATTTGATAAAGGAAGCGGAACAATAAACAAAATACTCAGGAAGATATTTTATTCGGTAACAGGTGGCCAAACTCCATAGGTAGCTTTTTCTATTTCGGATTTTAGAATTTCCAAATTCACCAGCGATTTATCGGTTTTGGTGAAATCCAAGGATTTATTAATGTCCACAAATGCCATTTGTTTTGTCTGTGGATTGTATTTGAAAATGGAAACGATGTAGTTAGATAGATGCGCGGCGAAGTTTCCTATTAGGGTTCCGCGCTTCACGTCACCCAACATACTTGAATCACCATCCTTTAAAAAAAATGATAAGATATCAACATGGAGTATATCATACTCGGATTTTAATTCTTCTACTGCCTCACTGACATTTTCACAAATACTACAATACGGTTTACCGAAAATAATCAGTACGTTCTTCATTTATGGGTATCAAAAACTTAAAATCGTTACTGCTGGAAAATAAATCACTGACGATATTAGATGATAATTTATACAAAGTATACAATGGAATATTTGTGGATACAATGAGTATTTATATAGCCGTCGCCAATTGTGTCAGAAACTTAGAAGAGTTAACTACGGTATTCATAAAATACGTAAACGGATGGGTAAAAAAGGGAGGACATGTAACCCTTTTTATCGATAGAGGAAGTATAAAAATTAAACAAGACGTTAGAGACAAGAGACGTAAATATTCTAAATTAACCAAGGACAGAAAAATGTTAGAATTAGAAAAGTGTACATCCGAAATACAAAATGTTACCGGATTTATGGAAGAAGAAATAAAGGCAGAAATGCAATTAAAAATCGATAAACTCACATTTCAAATATATTTATCTGATTATGATAACATAAAAATATCATTGAATGAGATACTAACACATTTCAACAATAATGAGAATGTTACATTATTTTATTGTGATGAACGAGACGCAGAATTCGTTATGTGTCTAGAGGCTAAAACACAGTTCTCTACCACAGGAGAATGGCCGTTAATAATAAGTACCGATCAGGATACTATGCTATTCGCGTCTGCTGATAATCATCCTAAGATGATAAAAAACTTAACTCAACTGTTTAAATTTGTTCCCTCGGCAGAGGATAACTATTTAGCAAAATTAACTGCATTAGTGAATGGATGTGATTTCTTTCCTGGACTCTATGGGGCATCTATAACACCCAACAACTTAAACAAAATACAATTGTTTAGTGATTTTACAATCGATAATATAGTCACTAGTTTGGCAATTAAAAATTATTATAGAAAGACTAACTCTACCGTAGACGTGCGTAATATTGTTACGTTTATAAACGATTACGCTAATTTAGACGATGTCTACTCGTATATTCCTCCTTGTCAATGCACTGTTCAAGAATTTATATTCTCCGCATTAGATGAAAAATGGAATGAATTTAAATCATCTTATTTAGAGAGCGTGCCGTTACCCTGCCAATTAATGTACGCATTAGAACCACGTAAGGAGATTGATGTTTCAGAAGTTAAAACTTTATCATCTTATATAGATTTCGAAAATACTAAATCAGATATCGATGTTATAAAATCTATATCCTCGATTTTTGGATATTCTAACGAAAACTGTAACACCATAGTGTTCGGCATCTATAAGGATAATTTACTACTGAGTATAAATAATTCATTTTACTTTAACGATAGTCTGTTAATAACCAATACTAAAAGTGATAATATAATAAATATAGGTTACTAGATTAAAAAATGGTGTTCCAGCTCGTGTGTTCTACATGCGGCAAAGATATTTCTCACGAACGATATAAATTGATTATACGAAAAAAATCATTAAAGGATGTACTAGTCAGTGTAAAGAACGAATGTTGTAGGTTAAAATTATCTACACAAATAGAACCTCAACGTAACTTAACAGTGCAACCTCTATTGGATATAAACTAATGGATCCGGTTAATTTTATCAAGACATATGCGCCTAGAGGTTCTATTATTTTTATTAATTATGCCATGTCATTAACTAGTCATTTGAATCCATCGATAGAAAAACATGTGGGTATTTATTATGGTACGTTATTATCGGAACACTTGGTAGTTGAATCTACCTATAGAAAAGGAGTTAGAATAGTCCCATTGGATAGATTTTTTGAAGGATATCTTAGTGCAAAAGTATACATGTTAGAGAATATTCAAGTTATGAAAATAGCAGCTGATATGTCGTTAACTTTACTAGGTATTCCATATGGATTTGGTCATGATAGAATGTATTGTTTTAAATTGGTAGCTGAATGTTATAAAAATGCCGGTATTGATACATCGTCTAAACGAATATTAGGTAAAGATATTTTTCTGAGCCAAAACTTTACAGATGATAATAGATGGATAAAGATATATGATTCTAATAATTTAACATTTTGGCAAATTGATTACCTTAAAGGGTGAGTTAATATGCATAACTACTCCTCCGTTGTTTTTTCCCTCGTTCTTTTTCTTAACGTTGTTTGCCATCACTCTCATAATGTAAAGATATTCTAAAATGGTAAACTTTTGCATATCGGATGCAGAAATTGGTATAAATGTTGTAATTGTATTATTTCCCGTCAATGGACTAGTCACAGCTCCATCAGTTTTATATCCTTTAGAGTATTTCTCACTCGTGTCTAGCATTCTAGAGCATTCCATGATCTGTTTATCGTTGATATTGGCCGGAAAGATAGATTTTTTATTTTTTATTATATTACTATTGGCAATTGTAGATATAACTTCTGGTAAATATTTTTCTACCTTTTCAATCTCTTCTATTTTCAAGCCGGCTATATATTCTGCTATATTGTTACTAGTATCAATACCTTTTCTGGCTAAGAAGTCATATGTGGTATTCACTATATCAGTTTTAACTGGTAGTTCCATTAGCCTTTCCACTTCTGCAGAATAATTAGAAATTGGTTCTTTACCAGAAAATCCAGCTACTATAATAGGCTCACCGATGATCATTGGCAAAATCCTATATTGTACCAGATTAATGAGAGCATATTTCATTTCCAATAATTCTGCTAGTTCTTGAGACATTGATTTATTTGATGAATCTATTTGGTTCTCTAGATACTCTACCATTTCTGCCGCATACAATAACTTGTTAGATAAAATCAGGGTTATCAAAGTGTTTAGTGTGGCTAGAATAGTGGGCTTGCACGTATTAAAGAATGCTGTAGTATGAGTAAACCGTTTTAACGAATTATATAGTCTCCAGAAATCTGTGGCGTTGCATACATGAACTGAATGACATCGAAGATTGTCCAATATTTTTAATAGCTGCTCTTTGTCCATTATTTCTATATTTGACTCGCAACAATTGTAGATACCATTAATCACTGATTCCTTTTTCGATGCCGGACAATAGCACAATTGTTTAGCTTTGGACTCTATGTATTCAGAATTAATAGATATATCTCTCAATACAGATTGCACTATACATTTTGAAACTATGTCAAAAATTGTAGAACGACGCTGTTCTGTAGCCATTTAACTTTAAATAATTTACAAAAATTTAAAATGAGCATCCGTATAAAAATCGATAAATTGCGCCAAATTGTGGCATATTTTTCAGAGTTCAGCGAAGAAGTGTCTATAAATGTAGACTCGACGGATGAATTAATGTATATTTTTGCCGCCTTGGGCGGATCTGTAAACATTTGGGCCATTATACCTCTCAGTGCATCAGTGTTCTACCGCGGAGCCGAAAATATTGTGTTTAACCTTCCAGTGTCCAAGGTAAAATCGTGTTTGTGTAGTTTTCACAATGATGCTATCATAGATATAGAACCTGATCTGGAAAATAATCTAGTAAAACTTTCTAGTTATCATGTAGTAAGTGTCGATTGTAACAAGGAACTGATGCCTATTAGGACAGATACTACTATTTGTCTAAGTATAGATCAAAAGAAATCTTACGTATTTAATTTTCACAAGTATGAAGAAAAATGTTGTGGTAGAACCGTCATTCATCTAGAATGGTTGTTGGGCTTTATCAAGTGTATTAGTCAGCATCAGCATTTGGCTATTATGTTTAAAGATGACAATATTATTATGAAGACTCCTGGTAATACTGATGCGTTTTCCAGGGAATATTCTATGACTGAATGTTCTCAAGAACTACAAAAGTTTTCTTTCAAAATAGCTATCTCGTCTCTCAACAAACTACGAGGATTCAAAAAGAGAGTCAATGTTTTTGAAACTAGAATCGTAATGGATAATGACGATAACATTCTAGGAATGTTGTTTTCGGATAGAGTTCAATCCTTTAAGATTAACATCTTTATGGCGTTTTTAGACTAATACTTTCAATGAGATAAATATGGGTGGCGGAGTAAGTGTTGAGCTCCCTAAACGGGATCCACCTCCGGGAGTACCCACTGATGAGATGTTATTAAACGTGGATAAAATGCATGACGTGATAGCTCCCGCTAAGCTTTTAGAATATGTGCATATAGGACCACTAACAAAAGATAAAGAGGATAAAGTAAAGAAAAGATATCCAGAGTTTAGATTAGTCAACACAGGACCCGGTGGTCTTTCGGCATTATTAAGACAATCATATAATGGAACCGCACCCAATTGCTGTCGCACTTTTAATCGTACTCATTATTGGAAGAAGGATGGAAAGATATCAGATAAGTATGAAGAGGGTGCAGTATTAGAATCGTGTTGGCCCGACGTCCACGACACTGGAAAATGCGATGTTGATTTATTCGACTGGTGTCAGGGGGATACGTTCGATATGAACATATGCCATCAGTGGATCGGTTCAGCCTTTAATAGGAGTGATAGAACTGTAGAGGGTCGACAATCGTTAATAAATCTGTATAATAAGATGCAAAGATTATGTAGTAAAGATGCTAGTGTACCAATATGTGAATTATTTTTGCATCATTTACGCGCACACAATACAGAAGATAGTAAAGAGATGATCGATTATATTCTAAGACAACAGTCGGCGGACTTTAAACAGAAATATATGAGATGTAGTTATCCCACTAGAGATAAGTTAGAAGAGTCATTAAAATATGCGGAACCTCGAGAATGTTGGGATCCAGAGTGTTCGAATGCCAATGTTAATTTCTTACTAACACGTAATTATAATAATTTAGGACTTTGCAATATTGTACGATGTAATACGAGCGTGAATAACTTACAGATGGATAAAACTTCCTCATTAAGATTATCATGTGGATTAAGCAATAGTGATAGATTTTCTACTGTTCCCGTCAATAGAGCAAAAGTAGTTCAACATAATATTAAACATTCGTTCGACCTAAAATTGCATTTGATCAGTTTATTATCTCTCTTGGTAATATGGATACTAATTGTAGCTATTTAAATGGGTGCCGCAGCAAGCATACAGACGACTGTGAATACACTCAGTGAACGTATCTCGTCTAAATTAGAACAAGAAGCGAACGCTAGTGCTCAAACAAAATGTGATATAGAAATCGGAAATTTTTATATCCGACAAAACCATGGATGTAACATCACTGTTAAAAATATGTGCTCTGCGGACGCGGATGCTCAGTTGGATGCTGTGTTATCAGCCGCTACAGAAACATATAGTGGATTAACACCGGAACAAAAAGCATACGTACCAGCTATGTTTACTGCTGCGTTAAACATTCAGACGAGTGTAAACACTGTTGTTAGAGATTTTGAAAATTATGTGAAACAGACTTGTAATTCTAGCGCTGTTGTCGATAACAAATTAAAGATACAAAACGTAATTATAGATGAATGTTACGGAGCCCCAGGATCTCCAACAAATTTGGAATTTATTAATACAGGATCTAGCAAAGGAAATTGTGCCATTAAGGCGTTGATGCAATTGACTACTAAGGCCACTACTCAAATAGCACCTAGACAAGTTGCTGGTACAGGAGTTCAGTTTTATATGATTGTTATCGGTGTTATAATATTGGCAGCGTTGTTTATGTACTATGCCAAGCGTATGCTGTTCACATCCACCAATGATAAAATCAAACTTATTTTAGCCAATAAGGAAAACGTCCATTGGACTACTTACATGGACACATTCTTTAGAACTTCTCCGATGATTATTGCTACCACGGATATACAAAACTGAAAATATATTGATAATATTTTAATAGATTAACATGGAAGTTATCGCTGATCGTCTAGACGATATAGTGAAACAAAATATAGCGGATGAAAAATTTGTAGATTTTGTTATACACGGTCTAGAGCATCAATGTCCTGCTATACTTCGACCATTAATTAGGTTGTTTATTGATATACTATTATTTGTTATAGTAATTTATATTTTTACGGTACGTCTAGTAAGTAGAAATTATCAAATATTGTTGGTGTTGGTGGCGCTAGTCATCACATTAACTATTTTTTTATTACTTTATACTATAATAGTACTAGACTGACTTCTAACAAACATCTCACCTGCCATAAATAAATGCTTGATATTAAAGTCTTCTATTTCTAACACTATTCCATCTGTGGAAAATAATACTCTGACATTATCGCTAATTGATACATCGGTAAGTGATATGCCTATAAAGTAATAATCTTCTTTGGGCACATATACCAGTGTACCAGGTTCTAACAACCTATTTACTGGTGCTCCTGTAGCATACTTTTTTTTTACCTTGAGAATATCCATTGTTTGCTTGGTCAATAGTGATATGTGATTTTTTATCAACCACTCAAAAAAGTAATTGGAGTGTTCATATCCTCTACGGGCTATTGTCTCATGACCGTGTATGAAATTTAAGTAACACGACTGTGGTAGATTTGTTCTATAGAGCCGGTTGCCGCAAATAGATAGAACTACCAATATGTCTGTACAAATGTTAAACATTAATTGATTAACAGAAAAAACAATGTTCGTTCTGGGAATAGAAACCAGATTAAAACAAAATTCATTAGAATATATGCCACGTTTATACATGGAATATAAAATAACTACAGTTTGAAAAATAACAGTATCATTTAAACATTTAACTTGCGGGGTTAATCTCACAACTTTACTGTTTTTGAACTGTTCAAAATATAGCATAGATCCATGAGAAATACGTTTAGCCGCCTTTAATAGAGGAAATCCAACCGCCTTTCTGGATCTCACCAACGACGATAGTTCTGACCAGCAACTCATTTCTTCATCATCCACCTGTTTTAACATATAATAGGCAGGAGATAGATATCCATCATTGCAATATTCCTTCTCGTAGGCACACAATCTAATATTGATAAAATCTCCATTCTCTTCTCTGTATTTATTATCTTGTCTCGGTGGCTGATTAGGCTGTGGTCTATCGTTGTTGAATCTATTTTGGTCATTAAATCTTTCATTTCTTCCTGGTATATTTCTATCACCTCGTTTGGTTGGATTTTTGTCTATATTATCGTTTGTAACATCGGTACGGGTATTCATTTATCACAAAAAAAACTTCTCTAAATGAGTCTACTACTAGAAAACCTCATCGAAGAAGATACCATATTTTTTGCAGGAAGTATATCTGAGTATGATGATTTACAAATGGTTATTGCTGGTGCAAAATCCAAATTTCCAAGATCTATGCTTTCTATTTTTAATATAGTACCTAGAACGATGTCAAAATATGAGTTGGAGTTGATTCATAACGAGAATATCACAGGGGCAATGTTTACCACAATGTATAATATAAGAAACAATTTGGGTCTAGGCGATGATAAACTAACTATTGAAGCCATTGAAAACTATTTCTTGGATCCTAACAATGAGGTTATGCCTCTTATCATTAATAATACGGATATGACTACCGTCATTCCTAAAAAAAGTGGTAGGAGAAAGAATAAGAACATGGTTATCTTCCGTCAAGGATCATCACCTATCTTGTGTATTTTCGAAACTCGTAAAAAGATTAATATTTATAAAGAAAATATGGAATCCGTATCGACTGAGTATACACCTATCGGAGACAACAAGGCTTTGATATCTAAATATGCGGGAATTAATATCCTGAATGTGTATTCTCCTTCCACGTCCATGAGATTGAATGCCATTTACGGATTCACCAATAAAAATAAACTAGAGAAACTTAGTACTAATAAGGAACTAGAATCGTATAGTTCTAGCCCTCTTCAAGAACCCATTAGGTTAAATGATTTTCTGGGACTATTGGAATGTGTTAAAAAGAATATTCCTCTAACAGATATTCCGACAAAGGATTGATTACTATAAATGGAGAATGTTCCTAATGTATACTTTAATCCTGTGTTTATAGAGCCCACGTTTAAACATTCTTTATTAAGTGTTTATAAACACAGATTAATAGTTTTATTTGAAGTATTCGTTGTATTCATTCTAATATATGTATTTTTTAGATCTGAATTAAATATGTTCTTCATGCCTAAACGAAAAATACCCGATCCTATTGATAGATTACGACGTGCTAATCTAGCGTGTGAAGACGATAAATTAATGATCTATGGATTACCATGGATAACAACTCAAACATCTGCGTTATCAATAAATAGTAAACCGATAGTGTATAAAGATTGTGCAAAGCTTTTGCGATCAATAAATGGATCACAACCAGTATCTCTTAACGATGTTCTTCGCAGATGATGATTCATTTTTTAAGTATTTTGCTAGTCAAGATGATGAATCTTCATTATCTGATATATTGCAAATCACTCAATATCTAGACTTTCTGTTATTATTATTGATCCAATCAAAAAATAAATTAGAAGCTGTGGGTCATTGTTATGAATCTCTTTCAGAGGAATACAGACAATTGACAAAATTCACAGACTCTCAAGATTTTAAAAAACTGTTTAACAAGGTCCCTATTGTTACAGATGGAAGGGTCAAACTTAATAAAGGATATTTGTTCGACTTTGTGATTAGTTTGATGCGATTCAAAAAAGAATCAGCTCTAGCTACCACCGCAATAGATCCTGTTAGATACATAGATCCTCGTCGTGATATCGCATTTTCTAACGTGATGGATATATTAAAGTCGAATAAAGTTGAAAAATAATTAATTCTTTATTGTTATCATGAACGGCGGACATATTCAGTTGATAATCGGCCCCATGTTTTCAGGTAAAAGTACAGAATTAATTAGACGAGTTAGACGTTATCAAATAGCTCAATATAAATGTGTGACTATAAAATATTCTAACGATAATAGATACGGAACGGGACTATGGACACATGATAAGAATAATTTTGCAGCATTGGAAGTAACTAAACTATGTGATGTCTTGGAAGCAATTACAGATTTCTCCGTGATAGGTATAGATGAAGGACAGTTCTTTCCAGACATTGTTGAATTCTGTGAGCGTATGGCAAACGAAGGAAAAATAGTTATAGTAGCCGCGCTCGATGGGACATTTCAACGTAGACCGTTTAATAATATTTTGAATCTTATTCCATTATCTGAAATGGTGGTAAAACTAACTGCAGTGTGTATGAAATGCTTTAAGGAGGCTTCCTTTTCTAAACGATTAGGTACAGAAACCGAGATAGAAATAATAGGAGGTAATGATATGTATCAATCTGTGTGTAGAAAGTGTTACATCGACTCATAATATTATATTTTTTATCTAAAAAACTAAAAATAAACATTGATTAAATTTTAATATAATACTTAAAAATGGATGTTGTGTCGTTAGATAAACCGTTTATGTATTTTGAGGAAATTGATAATGAGTTAGATTACGAACCAGAAAGTGCAAATGAGGTCGCAAAAAAACTGCCGTATCAAGGACAGTTAAAACTATTACTAGGAGAATTATTTTTTCTTAGTAAGTTACAGCGACACGGTATATTAGATGGCGCCACCGTAGTGTATATAGGATCTGCTCCAGGTACACATATACGTTATTTGAGAGATCATTTCTATAATTTAGGAGTGATCATCAAATGGATGCTAATTGACGGCCGCCATCATGATCCTATTCTAAATGGATTGCGTGATGTGACTCTAGTGACTCGGTTTGTTGATGAGGAATATCTACGATCCATCAAAAAACAACTACATCCTTCTAAGATTATTTTAATTTCTGATGTGCGATCCAAACGAGGAGGAAATGAACCTAGTACTGCGGATTTACTAAGTAATTATGCTCTACAAAATGTCATGATTAGTATTTTAAACCCCGTGGCGTCTAGTCTTAAATGGAGATGCCCGTTTCCAGATCAATGGATCAAGGACTTTTATATCCCACACGGTAATAAAATGTTACAACCTTTTGCTCCTTCATATTCAGCTGAAATGAGATTATTAAGTATTTATACCGGTGAGAATATGAGACTGACTCGAGTTACCAAATCAGACGCTGTAAATTATGAAAAAAAGATGTATTACCTTAATAAGATAGTCCGCAACAAAGTAGTTATTAACTTTGATTATCCTAATCAGGAATATGACTATTTTCACATGTACTTTATGTTGAGGACCGTATACTGCAATAAAACATTTCCTACTACTAAAGCAAAGATACTATTTCTACAACAATCTATATTTCGTTTCTTAAATATTCCAACGACATCAACTGAAAAAGTTAGTCATGAACCAATACAACGTAAAATATCTAGCAAAGATTCTATGTCTAAAAACAGAAATAGCAAGAGATCCGTACGCGGTAATAAATAGAAACGTACTACTGAGATATACTACCGATATAGAGTATAATGATTTAGTTACTTTAATAACCGTTAGACATAAAATTGATTCTATGAAAACTGTGTTTCAGGTATTTAACGAATCATCCATAAATTATACTCCGGTTGATGATGATTATGGAGAACCAATCATTATAACATCGTATCTTCAAAAAGGTCATAACAAGTTTCCTGTAAATTTTCTATACATAGATGTGGTAATATCTGACTTATTTCCTAGCTTTGTTAGACTAGATACTACAGAAACTAATATAGTTAATAGTGTACTACAAACAGGCGATGGTAAAAAGACTCTTCGTCTTCCTAAAATGTTAGAGACGGAAATAGTTGTCAAGATTCTCTATCGTCCTAATATACCATTAAAAATTGTTAGATTTTTCCGCAATAACATGGTAACTGGAGTAGAGATAGCCGATAGATCTGTTATTTCAGTCGCTGATTAATCAATTAGTAGAGATGAGATAAGAACATTATAATAATCAATAATATATCTTATATCTGTTTAGAAAAATGCTAATATTAAAATAGCTAACGCTAGTAATCCAATCGGAAGCCATTTGATATCTATAATAGGGTATCTAATTTCCTGATTCAGATAGCGTACGGCTATATTCTCGGTAGCTACTCGTTTGGAATCACAGACATTATTTACATCTAATTTACTATCTGTAATGGAAACGTTTCCCAATGAAATGGTACAATCAGATACATTACATCTTGATATATTTTTTTTTAAAGAGGCTGGTAACAACGCATCGCTTCGTTTACATGGCTCGTACCAACAATAATAGGGTAATCTTGTATCTATTCCTATCCGTACTATACTTTTATCAGGATAAATACATTTACATCGTATATCGTCTTTGTTAGTATCACAGAATGCATAAATTTGTTCGTCCGTCATGATAAAAATTTAAAGTGTAAATATAACTATTATTTTTATAGTTATAATAAAAAGGGAAATTTGATTGTATACCTTCGGTTCTTTAAAAGAAACTGACTTGATAAAAATGGCTGTAATCTCTAAGGTTACGTATAGTCTATACGATCAAAAAGAGATTAATGCCACAGATATTATCATTAGTCATATTAAAAATGACGACGATATCGGTACCGTTAAAGATGGTAGACTAGGTGCTATGGATGGGGCATTATGTAAGACTTGTGGGAAAACGGAATTGGAATGTTTCGGTCACTGGGGTAAAGTAAGTATTTATAAAACTCATATAGTTAAGCCTGAATTTATTTCAGAAATTATTCGTTTACTGAATCATATATGTATTCATTGCGGATTATTGCGTTCACGAGAACCGTATTCCGACGATATTAACCTAAAAGAGTTATCGGTACACGCTCTTAGGAGATTAAAGGATAAAATATTATCCAAGAAAAAGTCATGTTGGAACAGCGAATGTATGCAACCGTATCAAAAAATTACTTTTTCAAAGAAAAAGGTTTGTTTCGTCAACAAGTTGGATGATATTAACGTTCCTAATTCTCTCATCTATCAAAAGTTAATTTCTATTCATGAAAAGTTTTGGCCATTATTAGAAATTCATCAATATCCAGCTAACTTATTTTATACAGACTACTTTCCCATCCCTCCGTTGATTATTAGACCGGCTATTAGTTTTTGGATAGATAGTATACCCAAAGAGACAAATGAATTAACTTACTTATTAGGTATGATCGTTAAGAATTGTAACTTGAATGCTGATGAACAGGTTATCCAGAAGGCGGTAATAGAATACGATGATATTAAAATTATTTCTAATAACACTACCAGTATCAATTTATCATATATCACATCCGGCAAAAATAATATGATTAGAAGTTATATCGTCGCTCGGCGAAAAGATCAGACCGCTAGATCCGTAATTGGTCCCAGTACATCTATCACCGTTAATGAGGTAGGAATGCCCACATATATTAGAAATACACTTACAGAAAAGATATTTGTTAATGCCTTTACAGTGGATAAAGTTAAACAACTATTAGCATCAAACCAAGTTAAATTTTACTTTAATAAACGATTAAACCAATTAACAAGAATACGTCAAGGAAAGTTTATCAAAAATAAAATACATTTATTGCCTGGTGATTGGGTAGAAGTAGCTGTTCAAGAATATACAAGTATTATTTTTGGAAGACAACCGTCTCTACATAGATACAACGTCATCGCTTCATCTATCAGAGCTACCGAAGGAGATACTATCAAAATATCTCCCGGAATTGCCAACTCTCAAAATGCTGATTTCGACGGAGATGAAGAATGGATGATATTGGAGCAAAATCCTAAAGCCGTAGTTGAACAAAGTATTCTTATGTATCCGACGACGTTACTCAAACACGATATTCATGGAGCCCCCGTTTATGGATCTATTCAAGATGAAATCGTAGCAGCGTATTCATTGTTTAGGATACAAGATCTTTGTTTAGATGAAGTATTGAACATCTTGGGGAAATATGGAAGAGAGTTCGATCCTAAAGGTAAATGTAAATTCAGCGGTAAAGATATCTATACTTACTTGATAGGTGAAAAGATTAATTATCCGGGTCTCTTAAAGGATGGTGAAATTATTGCAAACGACGTAGATAGTAATTTTGTTGTAGCTATGAGGCATCTGTCATTGGCTGGACTCTTATCCGATCATAAATCGAACGTGGAAGGTATCAACTTTATTATCAAGTCATCTTATGTTTTTAAGAGATATCTATCTATATACGGTTTTGGGGTGACATTCAAAGATCTGAGACCAAATTCGACGTTCACTAATAAATTGGAGGCTATCAACGTAGAAAAAATAGAACTTATCAAAGAAGCATACGCCAAATATCTCAAAGATGTAAGAGACGGGAAAATAGTTCCATTATCTAAAGCTTTAGAGGCGGACTACTTGGAATCCATGTTATCCAACTTGACAAATCTTAATATCAGAGAGATAGAAGAACATATGAGACAAACGCTGATAGATGATCCAGATAATAACCTCCTGAAAATGGCCAAAGCGGGTTATAAAGTAAATCCCACAGAACTAATGTATATTCTAGGTACTTATGGACAACAGAGGATAGATGGCGAACCAGCAGAGACTCGAGTATTGGGTAGAGTCTTACCTTACTATCTTCCAGACTCTAAGGATCCAGAAGGAAGAGGTTATATTCTTAATTCTTTAACAAAAGGATTAACGGGTTCTCAATATTACTTTTTGATGCTGGTTGCAAGATCTCAATCTACTGATATTGTCTGTGAAACATCACGTACCGGAACACTGGCTAGAAAAATCATTAAAAAGATGGAGGATATGGTGGTCGACGGATACGGACAAGTAGTTATAGGTAATACGCTCATCAAGTACGCAGCCAATTATACCAAAATTCTAGGCTCAGTATGTAAACCTGTAGATCTTATCTATCCAGATGAGTCCATGACTTGGTATTTGGAAATTAGTGCTTTGTGGAATAAAATAAAACAGGGATTCGTTTACTCTCAGAAACAGAAACTTGCAAAGAAGACATTGGCGCCGTTTAATTTCCTAGTATTCGTCAAACCCACCACTGAGGATAATGCTATTAAGGTTAAGGATCTGTACGATATGATTCATAACGTCATTGATGATGTGAGAGAGAAATACTTCTTTACGGTATCTAATATAGATTTTATGGAGTATATATTCTTGACGCATCTTAATCCTTCTAGAATTAGAATTACAAAAGAAACGGCTATTACTATCTTTGAAAAGTTCTATGAAAAACTCAATTATACTCTAGGTGGTGGAACTCCTATTGGAATTATTTCTGCACAGGTATTGTCTGAGAAGTTTACACAACAAGCCCTGTCCAGTTTTCACACTACTGAAAAGAGTGGTGCTGTAAAACAAAAACTTGGTTTCAACGAGTTTAATAACTTGACTAATTTGAGTAAGAATAAGACCGAAATTATCACTCTGGTATCCGATGATATCTCTAAACTTCAATCTGTTAAGATTAATTTCGAATTTGTATGTTTGGGAGAATTAAATCCAGACATCACTCTTCGAAAAGAAACAGATAGATATGTAGTAGACATAATAGTCAATAGATTATACATCAAGAGAGCAGAAATAACCGAATTAGTCGTCGAATATATGATTGAACGATTTATCTCCTTTAGCGTCATTGTAAAGGAATGGGGTATGGAGACATTCATTGAGGACGAGGATAATATTAGATTTACTATCTACCTAAATTTCGTTGAACCGGAGGAATTGAATCTTAGTAAGTTTATGATGGTTCTTCCAGGTGCCGCCAACAAGGGCAAGATTAGTAAATTCAAGATTCCTATCTCTGACTATACGGGATATAACGACTTCAATCAAACAAAAAAGCTCAATAAGATGACTGTAGAACTCATGAATCTAAAAGAATTGGGTTCTTTCGATTTGGAGAACGTCAACGTGTATCCTGGAGTATGGAATACATACGATATCTTCGGTATTGAGGCCGCTCGTGGATACTTGTGCGAAGCCATGTTAAACACCTATGGAGAAGGTTTCGATTATCTGTACCAGCCTTGTGATCTTCTCGCTAGTTTACTATGTGCTAGTTACGAACCAGAATCAGTTAATAAATTCAAGTTCGGTGCAGCTAGTACTCTTAAGAGAGCTACGTTCGGAGATAATAAAGCATTGTTAAACGCGGCTCTTCATAAAAAGTCAGAACCTATTAACGATAATAGTAGCTGCCACTTTTTTAGCAAGGTCCCTAATATAGGAACTGGATATTACAAATACTTTATCGACTTGGGTCTTCTCATGAGAATGGAAAGGAAACTATCTGATAAGATATCTTCTCAAAAGATCAAGGAGATAGAAGAAACAGAAGACTTTTAATTCTTATCAATAACATATTTTTCTATGATCTGTCTTTTAAACGATGGATTTTCCACAAATGCGCCTCTCAAGTCCCTCATAGAATGATACACGTATAAAAAATATAGCATAGGTGATGACTCCTTATTTTTAGACATTAGATATGCCAAAATCATAGCCCCGCTTCTATTTACTCCTGCAACACAATGAACCAACACGGGCTCGTTTCGTTGATCACATTTAGATAAGAAGGCGGTCACGTCGTCAAAATATTTACTAATATCAGTAGTTGTATCATCTACCAACGGTATATGAATAATATTAATATTAGAGTTAGGTAATGTATATTTATCCATCGTCAAATTTAAAACATATTTGAACTTAACTTCAGATGATGGTGCATCCATAGCATTTTTATAATTTCCCAAATACACATTATTTGTTACTCTTGTCATTATAGTGGGAGATTTGGCTCTGTGCATATCTCCAGTTGAACGTAGTAGTAAGTATTTATACAAACTTTTCTTATCCATTTATAACGTACAAATGGATAAAACTACTTTATCAGTAAACGCATGCAATTTAGAATACGTTAGAGAAAAGGCTATAGTAGGCGTACAAGCAGCCAAGACATCAACACTTATATTTTTTGTTATTATATTGGCAATTAGTGCGCTATTACTCTGGTTTCAGACGTCTGATAATCCAGTCTTTAATGAATTAACGAGATATATGCGAATTAAAAATACGGTTAACGATTGGAAATCATTAACGGATAGCAAAACAAAATTAGAAAGCGATAGAGGTAGACTTCTAGCCGCTGGTAAGGATGATATATTCGAATTCAAATGTGTGGATTTCGGCGCCTATTTTATAGCTATGCGATTGGATAAGAAAACATATCTGCCGCAAGCTATTAGGCGAGGTACTGGAGACGCGTGGATGGTTAAAAAGGCGGCAAAAGTCGATCCATCTGCTCAACAATTTTGTCAGTATTTGATAAAACACAAGTCTAATAATGTTATTACTTGTGGTAATGAGATGTTAAATGAATTAGGTTATAGCGGTTATTTTATGTCACCGCATTGGTGTTCCGATCTTAGTAATATGGAATAAGTGTTAGATAAATGCGGTAACAAATGTTCCTGTAAGGAACCATAACAGTTTAGATTTAACATTAAAGATGAGCATAAACATAATAAACAAAATTACAATCAAACCTATAACATTAATATCAAACAATCCAAAAAATGAAATCAATGGAGTAGTAAACGTGTACATAACTCCTGGATAACGTTTAGCAGCTACCGTTCCTATTCTAGACCAAAAATTTGGTTTCATGGTTTCGAAGCGGTGTTCTGCAACAAGACGAGGATCGTGTTCTACATATTTGGCAGAGTTATCCATTATTTGCCTGTTAATCTTCATTTCGTTTTCGATTCTGGCTATTTCAAAATAAAATCCCGATGATAGACCTCCAGACTTTATAATTTCATCTACGATGTTCAGCGCCGTAGTAACTCTAATAATATAGGCGGATAAGCTAACATCATACCCTCCTGTATATGTAAATATGGCATGATCTTTGTCTATTACAAGCTCGGTTTTAACTTTATTTCCTGTAATAATTTCTCTCATCTGTAGGATATCTATTTTCTTGTCATGTATTGCCTTCAAGACGGGACGAAGAAACGTAATATCCTCAATAACGTTATCGTTTTCTATAATAACTACATATTCTACATTTTTATTTTCTAGCTCGGTAAAAAATTTAGAATCCCATAGGGCTAAATGTCTAGCGATATTTCTTTTCGTTTCCTCTGTACACATAGTGTTACAAAACCCTGAAAAGAAGTGAGTATACTTGTCATCATCTCTAATATTTCCTCCAGTCCATTGTATAAACACATAATCCTTGTAATGATCTGGATCATCATTGACTATCACAACATCTCTTTTTTCTTGCATAACTTCATTGTCCTTCACATCATCGAACTTCTGATCATTAATATGCTCATGAACATTAGGAAATGTTTCTGATGGAGGTCTATCAATAACTGGCACAACAATAACAGGAGTTTTCACCGCCGCCATTTAGTTATTGAAATTAATCATATACAACTCTCTAATACGAGTTATATTTTCGTCTATCCATTGTTTCACATTGACATATTTCGACAAAAAGATATAAAATGCGTATTCCAATGCTTCTCTGTTTAATGAATTACTAAAATATACAAACACGTCACTGTCTGGTAATAAATAATATCTTAGAATATTGTAACAATTTATTTTGTATTGCACATGTTCGTGATCTATGAGTTCTTCTTCAAATGGCATAGGATCTCCGAATCTGAAAACGTATAAATAGGAGTTAGAATAATAATATTTGAGAGTATTGGTAATGTATAAACTCTTTAGCGGTATAATTAGTTTTTTTCTCTCGATTTCTATTTTTAGATGTGATGGAAAAATGACTAATTTTGTAGCATTAGTATCATGAACTCTAATCAAAATCTTAATATCTTCGTCACATGTTAGCTCTTTGAAGTTTTTAAGAGATGCATCAGTTGGTTTTACAGATGGAGTAGGTGCAACAATTTTTTGTTTAATGCATGCATGTATTGGAGCCATTGTCTTAACTATAATGGTGCTTGTATCGAAAAACTTTAATGCGGATAACGGAAGCTCTTCGCCGCGACTTTCTACGTCGTAATTGGGTTCTAATGCCGATCTCTGAATGGATACTAGTTTTCTAAGTTCTAATGTAATTCTCTGAAAATGTAAATCCAATTCCTCCGGCATTATAGATGTGTATACATCGGTAAATAAAACTATAGTATCCAACGATCCCTTCTCGCAAATTCTAGTCTTAACCAAGAAATCGTATATAACTACGGAGATGGCGTATTTAAGAGTGGATTCTTCTACCGTTTTGTTCTTGGATTTCATATAAGAAACTATAAAGTCCGCACTACTGTTAAGAATGATCACTAACGCAACTATATAGTTCAAATTAAGCATCTTGGAAACATAAAATAACTCTGTAGATGATACTTGACTTTCGAATAAGTTTGCAGACAAACGAAGAAAGAACAGACCTCTCTTAATTTCAGAAGAAAACTTTTTTTCGTATTCCTGACGTCTAGAGTTTATATCAATAAGAAAGTTAAGAATTAGTCGGTTAATGTTGTATTTCATTACCCAAGTTTGAGATTTCATAATATTGTCAAAAGACATGATAATATTAAAGATAAAGCGCTGACTATGAACGAAATAGCTATATGGTTCGCTCAAGAATATAGTCTTGTTAAACGTGGAAACGATAACTGTATTTTTAATCACGTCAGCGGCATCTAAATTAAATATAGGTATATTTATTCCACACACTCTACAATATGCCACACCATCTTCATAATAAATAAATTCGTTAGCAAAATTATTAATTTTAGTGAAATAGTTAGCGTCAACTTTCATAGCTTCCTTCAATCTAATTTGATGCTCACATGGCGCGAATTCTACTCTAACATCCCTTTTCCATGCCTCAGGTTCATCGATCTCTATAATATCTAGTTTCTTGCGTTTCACAAACACAGGCTCGTCTCTCGCGATGAGATCTGTATAGTAACTATGTAAATGATAACTAGATAGAAAGATGTAGCTATATAGATGACGATCCTTTAAGAGAGGTATAATAACTTTACCCCAATCAGATAGACTGTTGTTATGGTCTTCGGAAAAAGAATTTTTATAAATTTTTCCAGTATTTTCTAAATATACGTACTTGATATCTAAGAAATCCTTAATAATAATAGGAATGGATAATCCGTCTATTTTATAAAGAAATACATATCGCATATTATACTTTTTTTTGGAAATTGGAATACCGATGTGTCTACATAAATACGCAAAGTCTAAATATTTTTTAGAGAATCTTAGTTGGTCCAAATTCTTTTCCAAGTACGGTAATAGATTTTTCATATTGAACGGTATCTTCTTGATCTCTGGTTCTAATTCCGCATTAAATGATGAAACTAAGTCACTATTTTTATAACTAACGATTACATCACCTCTAACATCATCATTTACCAGGATACTGATCTTCTTTTGTCGTAAATACATGTCTAATGTGTTAAAAAAAAGATCATACAAGTTATACGTCATTTCATCTGTAGTATTCTTGTCATTGAAGGATAAACTCGTACTAATCTCTTCTTTAACAGTCTGTTCAAATTTATATCCTATATATGAAAAAATAGCAACCAGTGTTTGATCATCCGCGTCAATATTCTGTTCTATCGTAGTGTATAACAATCTTATATCTTCTTCTGTGATAGTCGATACGTTATAAAGGTTGATAACGAAAATATTTTTATTTCGTGAAATAAAGTCATTGTAGGATTTTGGACTTATATTCGTGTCTAGTAGATATGATTTTATTTTTGGAATGATCTCAATTAAAATAGTCTCTTTAGAGTCCATTTAAAGTTACAAACAACTAGGAAATTGGTTTATGATGTATAATTTTTTTAGTTTTTATAGATTCTTTATTCTATACTTAAAAAATGAAAATAAATACAAAGGTTCTTGAGGGTTGTGTTAATTGAAAGCGATAAATAATCATAAATTATTTCATTATCGCGATATCCGTTAAGTTTGTATCGTAATGGCGTGGTCAATTACGAATAAAGCGGATACTAGTAGTTTCACAAAGATGGCTGAAATCAGAGCTCATCTAAGAAATAGCGCTGAAAATAAAGATAAAAACGAGGATATTTTCCCGGAAGATGTAATAATTCCATCTACTAAGCCCAAAACCAAACGAACCACTACTCCTCGTAAACCAGCGGCTACTAAAAGATCAACCAAAAAGGATAAAGAAAAGGAGGAAGTGGAAGAAGTAGTTATAGAGGAATATCATCAAACAACTGAAGAAAATTCTCCACCTCCGTCATCATCTCCTGGAGTCGGCGACATTGTAGAAAGCGTGGCCGCTGTAGAGCTCGATGATAGCGACGGGGATGATGAACCTATGGTACAAGTTGAAGCTGGTAAAGTAAATCATAGTGCTAGAAGCGATCTCTCTGACCTAAAGGTGGCTACCGACAATATCGTTAAAGATCTTAAGAAAATTATTACTAGAATCTCTGCAGTATCGACTGTTCTAGAGGATGTTCAAGCAGCTGGTATCTCTAGACAATTTACTTCTATGACTAAAGCTATTACAACACTATCTGATCTAGTCACCGAGGGAAAATCTAAAGTTGTTCGTAAAAAAGTTAAAACTTGTAAGAAGTAAATGCGTGCACTTTTTTATAAAGATGGTAAACTGTTTACCGATAATAATTTTTTAAATCCTGTATCAGACGATAATCCAGCGTATGAGGTTTTGCAACATGTTAAAATTCCTACTCATTTAACAGATGTAGTAGTATATGAACAAACGTGGGAAGAGGCATTAACTAGATTAATTTTTGTGGGAAGTGATTCAAAAGGACGTAGACAATACTTTTACGGAAAAATGCATATACAGAATCGCAATGCTAAAAGAGATCGTATTTTTGTTAGAGTATATAACGTTATGAAACGAATTAATTGTTTTATAAACAAAAATATAAAGAAATCGTCCACAGATTCCAATTATCAGTTGGCGGTTTTTATGTTAATGGAAACTATGTTTTTTATTAGATTTGGTAAAATGAAATATCTTAAGGAGAATGAAACAGTAGGGTTATTAACACTAAAAAATAAACACATAGAAATAAGTCCCGATGAAATAGTTATCAAGTTTGTAGGAAAGGACAAAGTTTCACATGAATTTGTTGTTCATAAGTCTAATAGACTATATAAACCGCTATTGAAACTGACTGATGATTCTAGTCCCGAAGAATTTCTGTTCAACAAACTAAGTGAACGAAAGGTATATGAATGTATCAAACAGTTTGGTATTAGAATCAAGGATCTCCGAACGTATGGAGTCAATTATACGTTTTTATATAATTTTTGGACAAATGTAAAGTCCGTATCTCCTCTTCCATCACCAAAAAAGTTGATAGCATTAACTATCAAACAAACTGCTGAAGTGGTAGGTCATACTCCATCAATTTCAAAAAGAGCTTATATGGCAACGACTATTTTAGAAATGGTAAAGGATAAAAATTTTTTAGACGTAGTATCTAAAACTACGTTCGATGAATTCCTATCTATAGTCGTAGATCACGTTAAATCATCTACGGATGGATGATAATAGATCTTTACACAAATAATTACAAGACCGATAAATGGAAATGGATAAACGGATGAAATCTCTCGCTATGACAGCTTTCTTCGGAGAGCTAAACACGTTAGATATTATGGCATTGATAATGTCTATATTTAAACACCATCCAAACAATACCATTTTTTCAGTGGATAAGGATGGTCAATTTATGATTGATTTCGAATACGATAATTATAAGGCTTCTCAATATTTGGATCTGACCCTCACTCCGATATCTGGAAATGAATGCAAGACTCACGCATCTAGTATAGCCGAACAATTGGCGTGTGTGGATATTATTAAAGAGGATATTAGCGAATATATCAAAACTACTCCCCGTCTTAAACGATTTATAAAAAAATACCGCAATAGATCATATACTCGTATCAGTCGAGATACAGAAAAGCTTAAAATAGCTCTAGCTAAAGGCATAGATTACGAATATATAAAAGACGCTTGTTAATAAGTAAATGAAAAAAAACTAGTCGTTTATAATAAAACACAATATGGATGCCAACATAGTATCATCTTCTACTATTGCGACGTATATAGACGCTTTAGCGAAGAATGCTTCAGAATTAGAACAGAGGTCTACCGCATACGAAATAAATAATGAATTGGAACTAGTATTTATTAAACCGCCATTGATTACGTTGACAAATGTAGTAAATATCTCCACGATTCAGGAATCGTTTATTCGATTTACCGTTACTAATAAGGAAGGTATCAAAATTAGAACTAAGATTCCATTATCTAAGGTACATGGTCTAGATGTAAAAAATGTGCAGTTGGTAGATGCTATAGATAACATAGTTTGGGAAAAGAAATCATTAGTGACGGAAAATCGTCTTCACAAAGAATGCTTGTTGAGACTATCAACAGAGGAACGTCATATATTTTTGGATTACAAGAAATATGGATCCTCTATCCGACTAGAATTAGTCAATCTTATTCAAGCAAAAACAAAAAACTTTACGATAGACTTTAAGCTAAAATATTTTCTAGGATCTGGCGCTCAATCTAAAAGTTCTTTATTGCACGCTATTAATCATCCAAAGTCAAGGCCTAATACATCTCTGGAAATAGAATTTACACCTAGAGACAATGAAACAGTTCCATATGATGAACTAATAAAGGAATTGACGACTCTCTCGCGTCATATATTTATGGCTTCTCCAGAGAATGTAATTCTTTCTCCACCTATTAACGCACCTATAAAGACTTTTATGTTGCCTAAACAAGATATAGTAGGTCTGGATCTGGAAAATCTATATGCCGTAACTAAGACTGACGGCATTCCTATAACTATCAGAGTTACATCAAAAGGGTTGTATTGTTATTTTACACATCTTGGTTATATTATTAGATATCCAGTTAAGAGAACAATAGATTCCGAAGTAGTAGTCTTTGGTGAGGCAGTTAAGGATAAGAACTGGACCGTATATCTCATTAAGCTAATAGAGCCCGTAAATGCAATCAGTGATAGACTAGAAGAAAGTAAGTATGTTGAATCTAAACTAGTGGATATTTGTGATCGGATAGTATTCAAGTCAAAGAAATACGAAGGTCCGTTTACTACAACTAGTGAAGTCGTCGATATGTTATCTACATATTTACCAAAGCAACCAGAAGGTGTTATTCTGTTCTATTCAAAGGGACCTAAATCTAACATTGATTTTAAAATCAAAAAGGAGAATACTATAGACCAAACTGCAAATGTAGTATTTAGGTACATGTCCAGTGAACCAATTATCTTTGGAGAGTCGTCTATCTTTATAGAGTATAAGAAATTTACCAACGATAAAGGCTTTCCTAAAGAATATGGTTCTGGTAAGATTGTGTTATATAACGGCGTTAATTATCTAAATAATATCTATTGTTTGGAATATATTAATACACATAATGAAGTGGGTATTAAGTCCGTTGTTGTACCTATTAAGTTTATAGCAGAATTCTTAGTCAATGGAGAAATACTTAAACCTAGAATCGATAAAACCATGAAATATATTAACTCAGAAGACTATTATGGAAATCAACATAATATCATAGTCGAACATTTAAGAGATCAAAGCATCAAAATAGGAGATGTCTTTAACGAGGATAAACTATCGGATGTTGGACATCAATACGCTGCCAACAACGATAAATTTAGATTAAATCCAGAAGTTAGTTATTTTACTAATAAACGAACTAGAGGGCCGTTGGGAATTTTATCAAACTACGTCAAGACTCTTCTTATTTCTATGTATTGTTCCAAAACATTTTTAGACGATTCCAACAAACGAAAGGTATTAGCGATTGATTTTGGAAACGGTGCTGACCTGGAAAAATACTTTTATGGAGAGATTGCGTTATTGGTAGCGACGGATCCGGATGCTGATGCTATAGCTAGAGGAAATGAAAGATACAACAAATTAAATTCTGGAATTAAAACCAAGTACTACAAATTTGACTACATTCAGGAAACTATTCGATCCGATACATTTGTCTCTAGTGTCAGAGAAGTATTCTATTTTGGAAAGTTTAATATCATTGACTGGCAGTTCGCTATTCATTATTCTTTTCATCCAAGACATTATGCTACAGTCATGAATAACTTATCCGAACTAACTGCTTCTGGAGGCAAGGTATTAATTACTACCATGGATGGAGACAAATTATCAAAATTAACCGATAAAAAGACTTTTATAATTCATAAGAATCTACCTAGTAGCGAAAACTATATGTCTGTAGAAAAAATAGCTGATGATAGAATAGTGGTATATAATCCATCAACAATGTCTACTCCAATGACTGAATACATTATCAAAAAGAACGATATAGTCAGAGTGTTTAACGAATACGGATTTGTTCTTGTAGATAATGTTGATTTCGCTACAATTATAGAACGAAGTAAAAAGTTTATTAATGGCGCATCTACAATGGAAGATAGACCGTCTACAAGAAACTTTTTCGAACTAAATAGAGGAGCCATTAAATGTGAAGGTTTAGATGTCGAAGACTTACTTAGTTACTATGTTGTTTATGTCTTTTCTAAGCGGTAAATAATAATATGGTATGGGTTCTGATATCCCCGTTCTAAATGCATTAAATAATTCCAATAGAGCGATTTTTGTTCCTATAGGACCTTCCAACTGTGGATACTCTGTATTATTAATAGATATATTAATACTTTTGTAGGGTAACAGAGGTTCTACGTCTTCTAAAAATAAAAGTTTTATAACATCTGGCCTGTTCATAAATAAAAACTTGGCGATTCTATATATACTCTTATTATCAAATCTAGCCATTGTCTTATAGATGTGAGCTACTGTAGGTGTACCATTTGATTTTCTTTCTAATACTATATATTTCTCTCGAAGAAGTTCTTGCAGATCATCTGGGAATAAAATACTACTGTTGAGTAAATCAGTTATTTTTTTTATATCGATATTGATGGACATTTTTATAGTTAAGGATAATAAGTATCCCAAAGTAGATAACGACGATAACGAAGTATTTATACTTTTAGGAAATCACAATGACTTTATCAGATCAAAATTAACAAAATTAAAGGAGCATGTATTTTTTTCTGAATATATTGTGACTCCAGATACATATGGATCTTTATGCGTCGAATTAAATGGGTCTAGTTTTCAGCACGGTGGTAGATATATAGAGGTGGAGGAATTTATAGATGCTGGAAGACAAGTTAGATGGTGTTCTACATCCAATCATATATCTGAAGATATACACACTGATAAATTTGTCATTTATGATATTTATACGTTTGATTCGTTCAAGAATAAACGATTGGTATTTGTACAGGTGCCTCCATCATTAGGAGATGATAGCTATTTAACTAATCCGTTATTGTCTCCGTATTATCGTAATTCAGTAGCCAGACAAATGGTCAATGATATGATTTTTAATCAAGATTCATTTTTAAAATATTTATTAGAACATCTGATTAGAAGCCACTATAGAGTTTCTAAACATATAACAATAGTTAGATACAAGGATACCGAAGAATTAAATCTAACAAGAATATGTTATAATAGAGATAAGTTTAAGGCGTTTGTATTCGCTTGGTTTAACGGCGTTTCGGAAAATGAAAAGGTACTAGATACGTATAAAAAGGTATCTGATTTGATATAATGAATTCAGTGACTATATCACACGCACCATATACTATTACTTATCACGATGATTGGGAACCAGTAATGAGTCAATTGGTAGAGTTTTATAACGAAGTAGCCAGTTGGTTGCTACGCGACGAGACGTCGCCTATTCCTGATAAGTTCTTTATACAATTGAAACAGCCGCTTAGAAATAAACGAGTATGTGTGTGTGGTATAGATCCGTATCCAAAAGATGGAACTGGTGTACCGTTCGAATCACCAAATTTTACAAAAAAATCAATTAAGGAGATAGCTTCATCTATATCTAGATTAACCGGAGTAATTGATTATAAAGGTTATAACCTTAATATAATAGACGGGGTTATACCCTGGAATTATTACTTAAGTTGTAAATTAGGAGAAACAAAAAGTCACGCGATTTACTGGGATAAGATTTCCAAGTTACTGCTACAGCATATAACTAAACACGTTAGTGTTCTTTATTGTTTGGGTAAAACAGATTTCTCGAATATACGGGCAAAGTTAGAATCCCCGGTAACTACCATAGTGGGATATCATCCAGCGGCCAGAGACCACCAATTCGAGAAAGATCGATCATTTGAAATTATCAACGTTTTACTGGAATTAGACAACAAGACACCTATAAATTGGGCTCAAGGGTTTATTTATTAATGCTTTAGTGAAATTTTAACTTGTGTTCTAAATGGATGCGGCTATTAGAGGTAATGATGTTATCTTTGTTCTTAAGACTATAGGTGTCCCGTCAGCATGCAGACAAAATGAAGATCCAAGATTCGTAGAAGCATTTAAATGCGACGAGTTAGAAAGATATATTGATAATAATCCAGAATGTACACTATTCGAAAGTCTTAGGGATGAGGAAGCATACTCTATAGTCAGAATTTTCATGGATGTAGATTTAGACGCGTGTCTAGACGAAATAGATTATTTAACGGCTATTCAAGATTTTATTATCGAGGTGTCAAACTGTGTAGCTAGATTCGCATTTACAGAATGCGGTGCCATTCATGAAAATGTAATAAAATCCATGAGATCTAATTTTTCATTGACTAAGTCTACAAATAGAGATAAAACAAGTTTTCATATTATCTTTTTAGACACGTATACCACTATGGATACATTGATAGCTATGAAACGAACACTATTAGAATTAAGTAGATCATCTGAAAATCCACTAACAAGATCGATAGACACTGCCGTATATAGGAGAAAAACAACTCTTCGGGTTGTAGGTACTAGGAAAAATCCAAATTGCGACACTATTCATGTAATGCAACCACCTCACGATAATATAGAAGATTACCTATTCACTTACGTGGATATGAACAACAATAGTTATTACTTTTCTCTACAACGACGATTGGAGGATTTAGTTCCTGATAAGTTATGGGAACCAGGGTTTATTTCGTTCGAAGACGCTATAAAAAGAGTTTCAAAAATATTCATTAATTCTATAATAAACTTTAATGATCTCGATGAAAATAATTTTACAACGGTACCACTGGTCATAGATTATGTAACACCTTGTGCATTATGTAAAAAACGATCGCATAAACATCCGCATCAACTATCGTTGGAAAATGGTGCTATTAGAATTTACAAAACTGGTAATCCACATAGTTGTAAAGTTAAAATTGTTCCGTTGGATGGTAATAAACTGTTTAATATTGCACAAAGAATTTTAGACACTAACTCTGTTTTATTAACCGAACGAGGAGACCATATAGTTTGGATTAATAATTCATGGAAATTTAACAGCGAAGAACCCTTGATAACAAAACTAATTCTATCAATAAGACATCAACTACCTAAGGAATATTCAAGCGAATTACTCTGTCCGAGGAAACGAAAGACTGTAGAAGCTAACATACGAGACATGTTAGTAGATTCAGTAGAGACCGATACCTATCCGGATAAACTTCCGTTTAAAAATGGTGTATTGGACCTGGTAGACGGAATGTTTTACTCTGGAGATGATGCTAAAAAATATACGTGTACTGTATCGACCGGATTTAAATTTGACGATACAAAATTCGTCGAAGACAGTCCAGAAATGGAAGAGTTAATGAATATCATTAACGATATCCAACCATTAACGGATGAAAATAAGAAAAATAGAGAGCTGTATGAAAAAACATTATCTAGTTGTTTATGTGGTGCTACCAAAGGATGTTTAACATTCTTTTTTGGAGAAACCGCAACTGGGAAGTCGACAACCAAACGTTTGTTAAAGTCTGCTATCGGTGACCTGTTTGTCGAGACGGGTCAAACAATTTTAACAGATGTATTGGATAAAGGACCTAATCCATTTATCGCTAATATGCATTTAAAAAGATCTGTATTCTGTAGCGAACTACCTGATTTTGCATGTAGTGGATCAAAGAAAATTAGATCTGATAATATTAAAAAGTTGACAGAACCTTGTGTCATTGGAAGACCGTGTTTCTCCAATAAAATTAATAATAGAAACCATGCGACAATCATTATCGATACTAATTACAAACCTGTCTTTGATAGGATAGATAACGCATTAATGAGAAGAATTGCCGTCGTGCGATTCAGAACACACTTTTCTCAACCTTCTGGTAGAGAGGCTGCTGAAAATAATGACGCGTACGATAAAGTCAAACTATTAGACGAGGGATTAGATGGTAAAATACAGAATAATAGATATAGATTCGCATTTCTATACTTGTTGGTTAAATGGTACAAAAAATATCATATTCCTATTATGAAACTATATCCTACACCGGAAGAGATTCCGGACTTTGCATTCTATCTCAAAATAGGTACTCTGTTGGTATCTAGCTCTGTAAAGCATATTCCATTAATGACGGACCTCTCCAAAAAGGGATATATATTGTACGATAATGTGGTTACTCTTCCGTTGACTACTTTCCAACAGAAAATATCCAAGTATTTTAATTCTAGACTATTTGGACACGATATAGAGAGCTTCATCAATAGACATAAGAAATTTGCCAATGTTAGTGATGAATATCTGCAATATATATTCATAGAGGATATTTCATCTCCGTAAATATATGCCATATATTTATAGAATATATCACATATCTAAATGAATACCGGAATCATAGATTTATTTGATAATCATGTTGATAGTATACCAACTATATTACCTCATCAGTTAGCTACTTTAGATTATCTAGTTAGAACTATCATAGATGAGAACAGAAGCGTGTTATTGTTCCATATTATGGGATCGGGTAAAACAATAATCGCTTTGTTGTTCGCCTTGGTAGCTTCCAGATTTAAAAAGGTTTACATTTTAGTACCGAACATCAACATCTTAAAAATTTTCAATTATAATATGGGTGTAGCTATGAACTTGTTTAATGACGAATTCATAGCTGAGAATATCTTTATTCATTCCACAACAAGTTTTTATTCTCTTAATTATAACGATAACGTCATTAATTATAACGGATTAAGTCGCTACAATAACTCTATTTTTATCGTTGATGAGGCACATAATATTTTTGGGAATAATACTGGAGAACTTATGACCGTGATAAAAAATAAAAACAAGATTCCTTTTCTACTATTGTCTGGATCTCCCATTACTAACACACCTAATACGCTGGGTCATATTATAGATTTAATGTCCGAAGAGACGATAGATTTTGGTGAGATTATTAGTCGTGGTAAGAAAGTAATTCAGACACTTCTTAACGAACGCGGAGTGAATGTACTCAAGGATTTGCTTAAAGGAAGAATATCATATTACGAAATGCCGGACAAAGATCTACCAACAATAAGATATCACGGACGTAAATTTCTAGATACTCGAGTAGTATATTGTCACATGTCTAAACTTCAAGAGAAAGATTATATGATTACTAGACGGCAGCTATGTTATCATGAAATGTTTGATAAAAATATGTATAACGTGTCAATGGCAGTATTGGGACAACTTAATCTGATGAATAATTTAGATACGTTATTTCAGGAACAGGATAAGGAATTGTACCCAAATCTGAAAATAAATAATGGAGTGTTATACGGTGAAGAATTGGTAACGTTAAACATTAGTTCCAAATTTAAGTACTTTATCAATCGGATACAGACACTCAAGGGAAAACACTTTATATACTTCTCTAATTCTACATATGGTGGATTGGTAATTAAATATATCATGCTCAGTAATGGATATTCTGAATATAATGGTTCTCAGGGAACTAATCCACATATGATAAACGGCAAACCAAAAACATTTGCTATCGTTACTAGTAAAATGAAATCGTCTTTAGAGGATCTATTAGATGTGTATAATTCTCCTGAAAACGATGATGGCAATCAATTGATGTTTTTGTTTTCGTCAAACATTATGTCTGAATCCTATACTCTGAAAGAGGTAAGGCATATTTGGTTTATGACTATCCCGGATACTTTTTCTCAATACAACCAAATTCTTGGACGATCTATTAGAAAATTCTCTTACGTCGATATTTCTGAACCCGTTAATGTATATCTTTTAGCAGCCGTATATTCAGATTTCAATGACGAAGTGACGTCATTAAACGATTATACACAGGATGAATTGATTAATGTTTTACCCTTTGACATCAAAAAGCTGTTGTATCTAAAATTTAAGACTAAAGAAACGAATAGAATATACTCTATTCTTCAAGAGATGTCTGAAACGTATTCTCTTCCACCACATCCATCAATTGTAAAAGTTTTATTGGGAGAATTGGTCAGACAATTTTTTTATAATAATTCTCGTATTAAGTATAACGACTCCAAGTTACTTAAAATGGTTACATCAGTTATAAAAAATAAAGAAGACGCTAGGAATTACATAGATGATATTGTAAACGGTCACTTCTTTGTATCGAATAAAGTATTTGATAAATCTCTTTTATACAAATACGAAAACGATATTATTACAGTACCGTTTAGACTTTCCTACGAACCATTTGTTTGGGGAGTTAACTTTCGTAAAGAATATAATGTGGTATCTTCTCCATAAAACTGATGAGATATATAAAGAAATAAATGTCGAGCTTTGTTACCAATGGATATCTTCCAGTTACATTGGAACCACATGAGTTGACGTTAGACATAAAAACTAATATTAGGAATGCCGTATATAAGGCGTATCTCCATAGAGAAATTAGTGGTAAAATGGCCAAGAAAATAGAAATTCGTGAAGACGTGGAATTACCTCTCGGTGAAATAGTTAATAATTCTGTAGTTATAAACGTTCCGTGTGTAATAACCTACGCATATTATCACGTTGGGGATATAGTCAGAGGAACATTAAACATCGAAGATGAATCAAATGTAACTATTCAATGTGGAGATTTAATCTGTAAACTAAGTAGAGATTCGGGTACTGTATCATTTAGCGATTCAAAGTACTGCTTTTTTCGAAATGGTAATGCGTATGATAACGGCATCGAAGTCTCCGCCGTTCTAATGGAGGCTCAACAAGGTACCGAATCTAGTTTTGTTTTTCTCGCGAATATCGTTGACTCATAAGAAAGAGAATAGCGGTGAGTATAAATACGAATACTATGGCAATAATTGCGAATGTTTTATTCCCTTCGATATATTTTTGATAATATGAAAAACATGCCTCTCTCAAATCAGACAACCATTTCATAAAATAGTTCTCTCGCACTGGTGAGGTGGTTGCAGCTCGTATAATCTCCCCAGAATAATATACTTGCGTGTCGTCGTTCAATTTATACGGATTTCTATAATTCTCTGTTATATAATGAGGTTTACCCTCATGATTAGACGACGACAATAGTGTTCTGAATTTAGATAGTTGATCAGAATGAATGTTTATTGGTGTTGGAAAAATTATCCATGCTGCGTCTGCAGAGTGGTTGATAGTTGTTCCTAGATATGTAAAATAATCCAACGTACTAGGTAGCAAATTGTCTAGATAAAATACTGAATCAAATGGCGCAGACATATTAGCGGATCTAATGGAATCCAATTGATTGACTATCTTTTGAAAATATACATTTTTATGATCTGATACTTGTAAGAATATAGCAATAATGATAATTCCATCATCGTGTTTTTTTGCCTCTTCATAAGAACTATATTTTTTCTTATTCCAATGAACCAGATTAATCTCTCCAGAGTATTTGTATACATCTATCAAGTGATTGGATCCATAATCGTCTTCCTTTCCCCAATATATATGTATTGTTGATAACACATATTCATTGGGGAGAAACCCTCCACTTATATATCCTCCTTTAAAATTAATCCTTACTAGTTTTCCAGTATTCTGGATAGTGGTTGGTTTCGACTCATTATAATGTATGTCTAACGTCTTCAATCGCGCGTCAGAAATTGCTTTTTTAGTTTCTATATTAATAGGAGATAGTTGTTGAGGCATAGTAAAAATGAAATGATAACTGTCTAGAAATAGCTCTTAGTATGGGATTTACAATGGATGAGGAAGTGATATTTGAAACTCCTAGAGAATTAATATCTATTAAACGAATAAAAGATATTCCAAGATCAAAAGACACGCACGTGTTTGCTGCGTGTATAACAAGTGACGGATATCCGTTAATAGGAGCTAGAAGAACTTCATTCGCATTCCAGGCGATATTATCTCAACAAAATTCAGATTCTATCTTTAGAGTATCCACTAAACTATTACGGTTTATGTACTACAATGAACTAAGAGAAATCTTTAGACGGTTGAGAAAAGGTTCTATCAACAATATCGATCCTCACTTCGAAGAGTTAATATTATTGGGTGGTAAACTAGATAAAAAGGAATCTATTAAAGATTGTTTAAGAAGAGAATTAAAAGAGGAAAGTGATGAACATATAACAGTAAAAGAATTCGGAAATGTAATTCTAAAACTTACAACGAGTGATAAATTATTTAATAAAGTATATATAGGTTATTGCATGGCATGTTTTATTAATCAATCGTTGGAGGATTTATCACATACTAGTATTTACAATGTAGAAATTAGAAAGATTAAATCGTTAAATGATTGTATTAACGACGATAAATACGAATATCTGTCTTATATTTATAATATACTAATTAATAGTAAATGAGCTTTTACAGATCTAGTATAATTAGTCAGATTATTAAGTATAATAGACGACTAGCTAAGTCTATTATTTGCGAGGATGACTCTCAAATTATTACACTCACGGCATTCGTTAACCAATGCCTATGGTGTCATAAACGAGTATCCGTGTCCGCTATTTTATTAACTACTGATAACAAAATATTAGTATGTAACAGACGAGATAGTTTTCTCTATTCTGAAATAATTAGAACTAGAAACATGTATAGAAAGAAACGATTATTTCTGAATTATTCCAATTATTTGAACAAACAGGAAAGAAGTATACTATCGTCATTTTTTTCTCTAGATCCAGCTACTGCTGATAATGATAGAATAAACGCTATTTATCCGGGTGGTATACCCAAAAGGGGTGAGAACGTTCCAGAGTGTTTATCCAGGGAAATCAAAGAAGAAGTTAATATAGACAATTCTTTTGTATTCATAGACACTCGTTTTTTTATTCATGGTATCATAGAAGATACCATTATTAACAAATTTTTTGAGGTAATTTTCTTTGTTGGACGAATATCTCTAACGAGTGATCAAATTATTGATACCTTTAAAAGTAATCATGAAATAAAGGATCTAATATTTTTAGATCCAAATTCAGGTAATGGACTCCAATACGAAATTGCAAAATATGCTCTAGATACTGCAAAACTTAAATGTTACGGTCATAGAGGATGTTATTATGAATCATTAAAAAAATTAACTGAGGATGATTGATTAGAAAATATAAATTAATTTACCATCGTGTATTTTTATAACGGGATTGTCTGGCATATCATGTAGATAGTTACCGTCTACATCGTATACTCTACCATCTACGCCTTTAAATCCTCTATTTATTGATATTAATCTATTAGAATTGGAATACCAAATATTAGTACCCTCAATTAGTTTATTGGTAATATTTTTTTTAGACGATAGATCGATGGCTCTTGAAACCAAGGTTTTCCAACCGGACTCATTGTCTATCGGTGAGAAGTCTTTTTCATTAGCATGAATCCATTCTAATGATGTATGTTTAAACACTCTAAACAATTGTACAAATTCTTTTGATTTGTTTTGAATGATTTCAAATAGGTCTTCGTCTACAGTAGGCATACCATTAGATAATCTAGCCATTATAAAGTGCACGTTTACATATCTACGTTCTGGAGGAGTAAGAACGTGACTATTGAGACGAATGGCTCTTCCTACTATCTGACGAAGAGACGCCTCGTTCCATGTCATATCTAAAATGAAGATATCATTGATTGAGAAGAAACTAATACCCTCGCCTCCGCTAGAAGAGAATACGCATGTTTTAATGCATTCTCCGTTAGTGTTTGATTCTTGGTTAAACTCAGCCACCGCCTTGATTCTAGTATCTTTTGTTCTAGATGAGAACTCTATATTAGAGATACCAAAGACTTTGAAATATAGTAATAAGATTTCTATTCCTGACTGATTAACAAATGGTTCAAAGACTAGACATTTACCATGGGATGCTAATATTCCCAAACATACATCTATAAATTTGACGCTTTTCTCTTTTAATTCAGTAAATAGAGAGATATCAGCCGCAATAGCATCCCCTCCCAATAGTTCTCCCTTTTTAAAGGTGTCTAATGCGGATTTAGAAAATTCTCTATCTCTTAATGAATTTTTAAAATCATTATATAGGGTTGCTATCTCTTGTGCGTATTCTCCCGGATCACGATTTTGTCTTTCAGGAAAGCTATCGAATGTAAACGTAGTAGCCATACGTCTCAGAATTCTAAATGATGATATACCAGTTTTTATTTCTGCGAGTTTAGCCTTTTGATAAATCTCTTCTTGCTTTTTTGACATATTAACGTATCGCATTAATACTGTTTTCTTAGCGAATGATGCAGACCCTTCCACATCATCAAAAATAGAAAACTCGTTATTAACTATGTACGAACATAGGCCTCCTAGTTTGGAGACTAATTCTTTTTCATCGACTAGACGTTTATTCTCAAATAGCGATTGGTGTTGTAAGGATCCTGGTCGCAGTAAGTTAACCAACATGGTGAATTCTTGCACACTATTAACGATAGGTGTAGCCGATAAACAAATCATCTTATGGTTTTTTAACGCAATGGTCTTAGATAAAAAATTATATACTGACCGAGTAGGACGGATCTTACCATCTTCTTTGATTAATGATTTAGAAATGAAGTTATGACATTCATCAATGATGACGCATATTCTACTCTTGGAATTAATAGTTTTGATATTAGTAAAAAATTTATTTCTAAAATTTTGATCATCGTAATTAATAAAAATACAATCCTTCGTTATCTCTGGAGCGTATCTGAGTATAGTGTTTATCCAAGGATCTTCTATCAAAGCCTTTTTTACCAATAAGATAATTGCCCAATTCGTATAAATATCCTTAAGATGTTTGAGAATATATACAGTAGTCATTGTTTTACCGACACCTGTTTCATGGAACAATAAAAGAGAATGCATACTGTCTAATCCTAAGAAAACTCTTGCTACAAAATGTTGATAATCCTTGAGGCGTACTACGTCTGACCCCATCATTTCAACGGGCATATTAGTAGTTCTGCGTAAGGCATAATCGATATAGGCCGCGTGTGATTTACTCATTTATGAGTGATAAGTAATAACTATGTTTTAAAAATCACAGCAGTAGTTTAACTAGCCTTCTCTGATGTTTGTTTTCGATACTTTTTGAATCAGAAGTCATACTAGAATAAAGCAGCGAGTGAACGTAATAGAGAGCTTCGTATACTCTATTCGAAAACTCTAAGAACTTATTAATGAATTCCGTATCCACTGGATCGTTTAAAATACTAAATTGAACAGTGTTCACATCCTTCCAAGACGAAGACTTAGTGACGGACTTAACATGAGACATAAATAAATCCAAATTTTTTTTATAAACATCACTAGCCACCATAATGGCGCTATCTTTCAACCAACTATCGCTTACGCATTTTAACAGTCTAACATTTTTAAAGAGACTACAATATATTCTCATAGTATCGATTACACCTCTACCGAATAGAGTGGGAAGTTTAATAATACAATATTTTTCGTTTACAAAATCAAATAATGGTCGAAACACGTCGAAGGTTAACATCTTATAATCGCTAATGTATAGATTGTTTTCAGTGAGATGATTATTAGATTTAATAGCATCTCGTTCACGTTTGAACAGTTTATTGCGTGCGCTGAGGTCGGCAACTACGGCATCCGCTCTAGTACTCCTCCCATAATACTTTACGCTATTAATCTTTAAAATTTCATAGACTTTATCTAGATCGCTTTCTGGTAACATGATATCATGTGTAAAAAGTTTTAACATGTCGGTCGGCATTCTATTTAGATCATTAACTCTAGAAATCTGAAGAAAGTAATTAGCTCCATATTCCAGACTAGGTAATGGGCTTTTACCTAAAGACAAGTTAAGTTCTGGCAATGTTTCATAAAATGGAAGAAGGACATGTGTCCCCTCCCGGATATTTTTTACAATTTCATCCATTTACAACTCTATAGTTTGTTTTCATTATTATTAGTTATTATCTCCCATAATCTTGGTAATACTTACACCTTGATCATAAGATACCTTATACAGGTCATTACATACAACTACCAATTGTTTTTGTACATAATAGATTGGATGATTGATATCCATGGTGGAATAAACTACTCGAACAGATAGTTTATCTTTCCCCCTAGATACATTGGCCGTAATAGTTGTCGGCCTAAAGAATATCTTTGGTGTAAAGTTAAAAGTTAGGGTTCTTGTTCCATTATTGCTTTTTGTCAGTAGTTCGTTATAAATTCTCGAGATGGGCCCGTTCTCTGAATATAGAACATCATTTCCAAATCTAACTTCTAGTCTAGAAATAATATCGGTCTTATTTTTAAAATCTATTCCCTTGATGAATGGATCGTTAATAAACAAATCCTTGGCCTTTGATTCGGCTGATCTATTATCTCCGTTATAGACGTTACGTTGACTAGTCCAAAGACTTACAGGAATAGATGTATCGATGATGTTGATAGTATGTGATATGTGAGCAAAGACTGTTCTCTTGGTGGCGTCGCTATATGTTCCAGTAATGGCGGAAAACTTTTTAGAAATGTTATATATAAAAGAATTTTTTCGGGTTCCAAACATTAACAGATTAGTATGAAGATAAACACTCATATTATCAGGAACATTATCAATTTTTACATAAACATCGGCATCTTGAATAGAAACAACACCATCTTCTGGAACCTCTACGATCTCGGCAGATTCCGGATAACCAGTCGGTGGACCATCACTAACAATAACTAGATCATCCAACAATCTACTCACATATGCGTCTATATAATCTTTTTCATCTTGTGAGTACCCTGGATACGAAATAAATTTGTTATCAGTATTTCCATAATAAGGTTTAGTATAAACAGAGAGAGATGTTGCTGCATGAACTTCGGTTACTGTCGCCGTTGGTTGGTTTATTTGACCTATTACTCTCCTAGGTTTCTCTATAAATGATGGTTTAATTTGTACATTCTTAACCATATATCCAATAAAGCTCAATTCAGGAACATAAACAAATTCTTTGTTGAACGTTTCAAAGTCGAACGAAGAGTCACGAATAACGATATCGGATACTGGATTGAAGGTCACCGTTACGGTAATTTTTGAATCGGATAGTTTAAGACTACTGAATGTATCTTCCACATCAAACGGAGTTTTAATATAAACGTATACTGTAGATGGTTCTTTAATAGTGTCATTAGGAGTTAGGCCAATAGAAATATCATTAAGTTCACTAGAATATCCAGAATGTTTCAAAGCAATTGTATTATTGATACAATTATTATATAATTCTTCGCCCTCAATTTCCCAAATAACACCGTTACACGAAGAGACAGATACATGATTAATACATTTATATCCAACATATGGCACGTAACCGAATCTTCCCATACCTTTAACTTCTGGAAGTTCCAAACTCAGAACCAAATGATTAAGCGCAGTAATATACTGATCCCTAATTTCGAAGCTAGCGATAGCCTGATTGTCTGGCCCATCGTTTGTCATAACTCCGGATAGAGAAATATATTGCGGCATATATAAAGTTGGAATTTGACTATCAACTGCGAAGACATTAGACCGTTTAATAAAGTCATCCCCACCGATCAAAGAATTAATGATAGTATTATTCATTTTCTATTTAAAATGGAAAAAGCTTACAATAAACTCCGTAGAGAAATATCTATAATTTGTGAGTTTTCCTTAAAGTAACAGCTTCCGTAAACACCGTCTTTATCTCTTAGTAAGTTTATTGTATTTATGACCTTTTCCTTATCTTCATAGAATACTAAAGGCAATAAAGAAATTTTTGGTTCTTCTCTAAGAGCTACGTGAGACTTAACCATAGACGCCAACGAATCCCTACATATTTTAGAACAGAAATACCCAACTTCACCACCCTTGAATGTCTCAATACTAATAGGTCTAAAAACCAAATCTTGATTACAAAACCAACACTTATCAATTACACTATTTGTCTTAATAGACATATCTGCCATAGATTTATAATACTTTGGTAGTATACAAGCGAGTGCTTCTTCTTTAGCGGGCTTAAAGACTGCTTTAGGTGCTGAAATAACCACATCTGGAAGACTTACTCGCTTAGCCATTTAATTACGGAACTATTTTTTTATACTTCTAATGAACAAGTAGAAAACCTCTCATCTACAAAAACATACTCGTGTCCATAATCCTCTACCATAGTAACACGTTTTTTAGATCTCATATGTGCTAAAAAGTTTTCCCATACTAATTGGTTACTATTATTTTTCGTATAATTTTTAACAGTTTGAGGTTTTAGATTTTTAGTTACAGAAGTGATATCGAATATTTTATCCAAAAAGAATGAGTAATTAATTGTCTTAGAAGGAGTGTTTTCTTGGCAAAAGAATACCAAGTGCTTAAATATTTCTACTACTTCATTAATCTTTTCTGTACTCAGATTCAGTTTCTCATCTTTTACTTGATTGATTATTTCAAAGACTAACTTATAATCCTTTTTATTTATTCTCTCGTTAGCCTTAAGAAAACTAGATACAAAATTTGCATCTACATCATCCGTGGATATTTGATTTTTTTCCATGATATCCAATAGTTCCGAGATAATTTCTCCAGAACATTGATGAGACAATAATCTCCGCAATACATTTCTCAAATGAATAAGTTTATTAGACACGTGGAAGTTTGACTTTTTTTGTACCTTTGTACATTTTTGAAATACAGACTCGCAAAAAATACAATATTCATATCCTTGTTCAGATACTATACCGTTATGTCTACAACAGCTACATAATCGTAGATTCATGTTAACACTCTACGTATCTCGTCGTCCAATATTTTATATAAAAACATTTTATTTCTAGACGTTGTCAGAAAATCCTGTAATATTTTTAGTTTTTTTGGTTGTGAATAAAGTATCGCCCTAATAATATTGGTACCGTCTTCCGACAATATAGTAGTTAAATTATCCGAGCATGTAGAAGAACACCGCTTAGGCGGATTCAGTACAATGTTATATTTTTCGTACCAACTCATTTAAATATCATAATCTAAAATAGTTCTGTAATATGTCTAGCGCTAATATATTGATCATAATCCTGTGCATAAATTAAGATACAACAATGTCTTGAAATCATCGACATGGCTTCTTCCATAGTTAGAAGATCATCGTCAAAGTTAGCAACGTGATTCATCAACATTTGCTGTTTTGAGGCAGCAAATACTGAACCATCACCATTCAACCATTCATAAAAACCATCGTCTGAATCCATTGATAATTTCTTGTACTGGTTTTTGAGAGCTCGCATCAATCTAGCATTTCTAGCTCCCGGATTGAAAACAGAAAGAGGATCGTACATCCAAGGTCCATTTTCTGTAAATAGAATCGTATAATGTCCCTTCAAGAAGATATCAGACGATCCACAATCAAAGAATTGGTCTCCGAGTTTGTAACAGACTGCGGACTTTAACCTATACATGATACCGTTTAGCATGATTTCTGGTGATACGTCAATCGGAGTATCATCTATTAGAGATCTAAAGCCGGTGTAACATTCTCCGCCAAACATATTCTTATTCTGACGTCGTTCTACATAAAACATCATTGCTCCATTAACGATAACAGGTGAATGAACAGCACTACCCATCACATTAGTTCCCAATGGATCAATGTGTGTAACTCCAGAACATCTTCCATAGCCTATGTTAGGAGGAGCGAACACCACTCTTCCACTATTGCCATCGAATGCCATAGAATAAATATCCTTGGAATTGATAGAAATCGGACTGTCGGATGTTGTTATCATCTTCATAGGATTAACAACGATGTATGGTGCAGCCTGAAGTTTCATATCGTAACTGATGCCGTTCATAGGTCTAGCCACAGAAACCAACGTAGGTCTAAATCCAACTATAGACAAAATAGAAGCTAATATCTGTTCCTCATCTGTCATAACTTGAGAGCATCCAGTATGAATAATCTTCATTAGATGGGGATCTACCGCATCATCATCGTTACAATAAAAAATTCCCATTCTAATGTTCATAATTGCTTTTCTAATCATGGTATGAATGTTTGCTCTCTGAATCTCTGTGGAAATTAGATCTGATACACCTGTAATCACTATCGGATTATCCTCCGTAAGACGATTAACCAACAACATATAATTATAAGACTTTACTCTTCTAAATTCATAAAGTTGCTGGATTAGACTATATGTGTCTCCATGTACATACGCGTTCTCGAGCGCAGGAAGTTTAATACCGAATAGTGCCATCAGAATAGGATGAATGTAGTAATTAGTTTCTGGTTTTCTATAAATAAAAGACAAATCTTGTGAACTAGACATATCGGTAAAATGCATGGATTGGAATCGTGTAGTCGACAGAAGAATATGATGATTAGATGGAGAGTATATTTTATCTAACTCTTTGAGTTGGTCACCGATTCTAGGACTAGCTCGAGAATGAATAAGTACTAAGGGATGAGTACATTTCACAGAAACACTGGCGTTGTTCAACGTACTCTTTACATGGGAAAGGAGTTGAAATAGCTCGTTTCTATTTGTCCTGACAATATTTAGTTTATTCATAATATTAAGCATATCCTGAATAGTAAAGTTAGATGTGTCATACTTGTTAGTAGTTAGATATTTAGCAATTGCATTCCCATCATTTCTCAATCTCGTACTCCAATCATGTGTGGATGCTACTTCGTCGATGGAAACCATACAATCCTTTTTGATAGGCTGTTGAGATTGATCATTTCCTGTACGTTTAGGTTTGGTACGTTGATTTCTAGCCCCTGCTGATATAAAGTCATCGTCTACAATTTGGGATAATGAATTACATACACTACAAGACAAAGATTTATCAGAAGTGTGAATATGATCTTCATCTACCAAAGAAAGAGTTTGATTAGTATAACTAGATTTTAGTCCCGCGTTAGATGTTAAAAAAACATCGCTATTGACCACGGCTTCCATTATTTATATTCGTAGTTTTTACTCGAAAGCGTGATTTTAATATCCAATCTTATTACTTTTGGAATCGTTCAAAACCTTTGACTAGTTGTAGAATTTGATCTATTGCCCTACGCGTATACTCCCTTGCATCATATACGTTCGTCACCAGATCGTTTGTTTCGGCCTGAAGTTGACGCATATCTTTTTCAACACTCGACATGAGATCCTTAAGGGTCATATCGTCTAGATTTTGTTGAGATGCTGCTCCTGGATTTGGATTTTGTTGTGCTGTTGTACATACTGTACCACCAGTAGGTGTAGGAGTACATACAGTGGCCACAATAGGAGGTTGAAGAGGTGTAACCGTTGGAGTAGTACAAGAAATACTTCCATCCGATTGTTGTGTACATGTGGTTGTTGGTAACGTCTGAGAAGGTTGGGTAGATGGCGGTGTCGTCATCTTTTGATCTTTATTAAATTTAGAGATAATATCCTGAACAGTATTGCTCGGCGTCAACGCTGGAAGGAGTGTACTCGCCGGCGCATCAGTATCTGTAGACAACCAATCAAAAAGATTAGACATATCAGATGATGTATTAGTTTGTTGACGTGGTTTTAGTACAGGAGCAGTACTACTAGGTAGAAGAATAGGAGCCGGTGTAGGTGTCGGAACCGGCTGTGGAGTTATATGAATAGTTGGTTGTAGCGGTTGGGTAGGCTGTCTGCTGGCGGTCATCATATTATCTCTAGCTAGTTGTTCTCGCAACTGTCTTTGATAATACGACTCTTGAGACTTTAGTCCTATTTCAATCGCTTCATCCTTTTTCGTATCCGGATCCTTTTCTTCAGAATAATAGATTGACGACTTTGGTGTAGAGGATTCTGCCAGCCCCTGTGAGAACTTGTTAAAGAAGTCCATTTAAGGCTTTAAAATTGAATTGCGATTATAAGATTAAATGGCAGACACGGACGATATTATCGACTATGAATCCGATGATCTCACCGAATACGAGGATGATGAAGAAGATGGAGAGTCACTAGAAACTAGTGATATAGATCCCAAATCTTCTTATAAGATTGTAGAATCAACATCCACTCATATAGAAGATGCGCATTCCAATCTTAAACATATAGGGAATCATATATCTGCTCTTAAACGACGCTATACTAGACGTATAAGTCTATTTGAAATAGCGGGTATAATAGCAGAAAGCTATAACTTGCTTCAACGAGGAAGATTACCTCTAGTTTCAGAATTTTCTGACGAAACGATGAAGCAAAATATGCTACATGTAATTATACAAGAGATAGAGGAGGGTTCTTGTCCTATAGTCATCGAAAAGAACGGAGAATTGTTGTCGGTAAACGATTTTGATAAAGATGGTCTAAAATTCCATCTAGACTATATTATCAAAATTTGGAAACTTCAAAAACGATATTAGAATTTATACGAATATCGTTCTCTAAATGTCACAATCAAGTCTCTCATATTCAGCAGTTTATTGTCGTACTTTATATCGTGTTCATTAACGATATTTTGCAAAATAGTAATGATTCTATCTTCCTTCGATAGATATTCTTCAGAGATTATTGTCTTATATTCTTTCTTGTTATCCGATATGAATTTGATAAGACTTTGAACATTATTAATACCCGTCTGTTTAATTTTTTCTATAGATATTTTAGTTTTGGTAGATTCTATGGTGTCTGTTAATAGGCATCCAACATCGACATTCGACGTCAATTGTCTATAAATCAGAGTATAAATTTTAGAAATAACATTAGCAAATTGTTGTGCGTTGATGTCGTTATTCTGAAACAGTATGATTTTAGGTAGCATTTTCTTAACAAAGAGAACGTATTTATTGTTACTCAGTTGAACAGATGATATATCCAGATTACTAACGCATCTGATTCCATATACCAAACTTTCAGAAGAAATGGTGTACAATTGTTTGTATTCATTCAATGTCTCCTTTTCAGAAATTAGTTTAGAGTCGAATACTGCAATAATTTTCAAGAGATAGTTTTCATCAGATAAGATTTTATTTAGTGTAGATATGATAAAACTATTGTTTTGTTGGAGAACTTGATACGCCGCATTCTCTGTAGTCGACGCTCTCAAATGGGAAACAATCTCTATTATTTTTTTGGAATCGGATACTATATCTTCGGTATCTTGACGCAGTCTAGTATACATAGAGTTAAGAGAAATTAGAGTTTGTACATTAAGCAACATGTCTCTAAATGTGGCTACAAACTTTTCTTTTTCCACATCATCTAGTTTATTATATACCGATTTCACAACGGCACCAGATTTAAGGAACCAGAATGAAAAACTCTGATAACTACAATATTTCATCATAGTTACGATTTTATCATCTTCTATAGTTGGTGTGATAACACATACCTTTTTCTCCAAGACTGGAACCAACGTCATAAAAATGTTTAAATCAAAATCCATATCAACATCTGATGCGCTAAGACCAGTCTCGCGTTCAAGATTATCTTTACTAATGGTGACGAACTCATCGTATAGAACTCTAAGTTTGTCCATTATTTATTTACAGATTTAGTTGTTTAATTTATTTGTGCTCTTCCAGAGTTGGGATAGTATTTTTCTAACGTCGGTATTATATTATTAGGATCTACGTTCATATGTATCATAATATTAATCATCCACGTTTTGATAAATCTATCTTTAGCTTCTGAAATAACGTATTTAAACAAAGGAGAAAAATATTTAGTTACGGCATCAGACGCGATAACATTTTTTGTAAATGTAACGTATTTAGACGACAGATCTTCGTTAAAAAGTTTTCCATCTATGTAGAATCCATCGGTTGTTAACACCATTCCCGCGTCAGAGTGAATAGGAGTTTGAATAGTTTGTTTTGGAAATAGATCCTTCAATAACTTATAGTTGGGTGGGAAAAAATCGATTTTATCACTAGACTCTTTCTTTTTTACTATCATTACCTCATGAACTATTTCTTGAATGAGTATATGTATTTTCTTTCCTATATCGGTCGCGTTCATTGGAAAATATATCATGTCGTTAACTATAAGAATATTTTTATCCTCGTTTACAAACTGAATAATATCAGATATAGTTCGTAAACGAACTATATCATCACCAGCACAACATCTAACTATATGATATCCACTAGTTTCCTTTAGCCGTTTATTATCTTGTTCCATATTAGCAGTCATTCCATCATTTAAGAAGGCGTCAAAGATAATAGGGAGAAATGACATTTTGGATTCTGTTACGACTTTACCAAAATTAAGGATATACGGACTTACTATCTTTTTCTCAACGTCGATTTGATGAACACACGATGAAAATGTACTTCGATGAGATTGATCATGTAGAAAACAACAAGGGATACAATATTTCCGCATATCATGAAATATATTAAGAAATCCCACTTTATTATATTTCCCCAAAGGATCAATGCATGTAAACATTATACCGTTATCATTAATAAAGACTTCTTTCTCATCGGATCTGTAAAAGTTGTTACTGATTTTTTTCATTCCAGGATCTAGATAATTAATAATAATGGGTTTTCTATTCTTATTCTTTGTATTTTGACATATCCTAGACCAGTAAACAGTTTCCACTTTGGTAAAATCAGAAGACTTTTGAACGCTATTAAACATGGCATTAATGGCAATAACTAAAAATGTAAAATATTTTTCTATGTTAGGAATATGGTTTTTCACTTTAATAGATATATGGTTTTTTGCCAAAATGATAGATATTTTTTTATCCGATGATAGTAAAATATTATTAGTCGCCGTCTCTATAAAAATGAAGCTAGTCTCGATATCCAATTTTATTCTAGAATTGATAGGAGTCGCCAAATGTACCTTATACGTTATATCTCCCTTGATGCGTTCCATTTGTGTATCTATATCGGACACAAGATCTGTAAATAGTTTTACGTTATTAATCATCACGGTATCGCCATCGCTAGATAATGCTAATGTACTATCCAAGTCCCAAATGGAGAGATTTAACTGTTCATCGTTTAGAATAAAATGATTACCTGTCATATTAATAAAGTGTTCATCGTATCTAGATAACAACGACTTATAATTAATGTCCAAGTCTTGAACTCGCTGAATGATCTTTTTTAACCCAGTTAGTTTTAGATTGGTACGAAATATATTGTTAAACTTTGATTCTACAGTAATGTCCAAATCTAGTTGTGGAAATACTTCCATCAACATTGTTTCAAACTTGATAATATTATTATCTACATCTTCGTACGATCCAAATTCCGGAATAGATGTATCGCACGCTCTGGCCACCCAGATAACCAAAAAGTCACACGCTCCAGAATATACATTGTATAAAAAGCTATCGTTTTTTAGTAGTGTTTTTTTCTGAGTATATACGAAAGGATTAAAAATAGTATTATCAACGTAACTATATTCCAAATTATTCTTATGAGAATAGATAATAATATCGTCCTTAATATCTAACAAATTTCCTAAATATCCCTTTAATTGAGTCATTCGAAGCGTTAATAAAATATGTCTCTTAACTATTTCCGGCCGTTGTATATTTAAATGACTTCGTAAGAAATAATATATAGGCGACTTCTCATCTATGTAATCATATGGAGTGAGATATAGGGCTCGTTCTACCTCCTGCCCCTTACCCACCTGTAATACCAATTGCGGACTCACTATATATCGCATATTTATATCGTGGGGTAAAGTGAAAATCTACTACCGATGATGTAAGTCTTACAATGTTCGAACCAGTACCAGATCTTAATTTGGAGGCCTCCGTAGAACTAGGGGAGGTAAATATAGATCAAACAACACCTATGATAAAGGAGAATAGCGGTTTTATATCCCGTAGTAGACGTCTATTCGCCCATAGATCTAAGGATGATGAGAGAAAACTAGCACTACGATTCTTTTTACAAAGACTTTATTTTTTAGATCATAGAGAGATTCATTATTTGTTCAGATGCGTCGACGCTGTAAAAGACGTCACTATTACCAAAAAAAATAACATTATCGTGGCGCCTTATATAGCACTTTTAACTATCGCATCAAAAGGATGCAAACTTACAGAAACAATGATTGAAGCATTCTTTCCAGAACTATATAATGAACATAGTAAGAAATTCAAATTCAACTCTCAAGTATCCATCATCCAAGAAAAACTCGGATACCAGTCTGGAAACTATCACGTTTATGATTTTGAACCGTATTACTCTACAGTAGCTCTGGCTATTCGAGATGAACATTCATCTGGCATTTTTAATATCCGTCAAGAGAGTTATCTTGTAAGTTCATTATCTGAAATAACATATAGATTTTATCTAATTAATCTAAAATCTGATCTTGTTCAATGGAGTGCTAGTACGGGCGCTGTAATTAATCAAATGGTAAATACTGTATTGATTACAGTGTATGAAAAATTACAACTGGCCATAGAAAATGATTCACAATTTACATGTTCATTGGCTGTGGAATCAGAACTTCCAATAAAATTACTTAAAGATAGAAATGAATTATTTACAAAATTCATTAACGAGTTAAAAAAGACCAGTTCATTCAAGATAAGCAAACGTGATAAGGATACGCTATTAAAACATTTTACTTATGACTGGAGTTAGAATTTATAGACGACACATTTCGTTTATCATTGTTACTATTACTATCATTATTAGTATTCTTCTTGTCATCTTGTTCAGAAATATACAGCAATGCTATACCTAATACTAAATACATTATCATGCTTGCAATGGCTCTAACAACAACGAACCAAAATGAATTTGGTCGTAGCTTTTGTTCACAAAAATACATAAAGAAATGTCTACATAAATCTATGGCGCCATTGGCTACTTGAAATAGCGCCAGTCCTCCTACAGATTTTAATATAGCTGTATAACATGACATTTATTCATCATCAAAAGAGACAGAGTCACCATCTGTCATATTTAGATTTTTTTTCATGTGTTCAAAGTATCCTCTACTCATTTCATTATAATAGTTTATCATGCTTAGAATTTTAGGACGGATCAATGAGTAAGACTTGACTAGATCGTCAGTAGTAATTTGTGCATCATCTATTCTGCATCCGCTTCGTCGAATAATGTATAGCATCGCTTTGAGATTCTCCATAGCTATCAAGTCTTTATATAATGACATGGAAATATCTGTGAATGCTTTATACTTCTCCAACATCGATGCCTTAACATCATCACATACTTTAGCATTGAAAATACGTTCTATTGTGTAGATGGATGTAGCAAGATTTTTAAACAACAATGCCATCTTACATGATGATTGTCTCAAGTCTCCAATCGTTTGTTTAGAACGATTAGCTACAGAGTCCAATGCTTGGCTAACTAGCATATTATTATCTTTAGAAATTGTATTCTTCAATGAGGCGTTTATCATATCTGTGATTTCGTTAGTCATATTACAGTCTGACTGGGTTGTAATGTTATCCAACATATCACCTATGGATACGGTACACGTACCAGCATTTGTAATAATCCTATCTAAGATGTTGTATGGCATTGCGCAGAAAATATCTTCTCCTGTAATATCTCCACTCTCGATAAATCTACTCAGATTATTCTTAAATGCCTTATTCTCTGGAGAAAAGATATCAGTGTCCATCATTTCATTAATAGTATACGCAGAAAAGATACCACGAGTATCAATTCTATCCAAGATACTTATCGGTTCCGAGTCACAGATAATTGTTTCCTCTCCTTCGGGAGATCCTGCATAGAAATATCTAGGACAATAGTTTCTATACTGTCTGTAACTCTGATAATCTCTAAAGTCACTAACTGATACCATGAAATTGAGAAGATCAAACGCTGAAGTAATCAATTTTTCTGCCTCGTTTTTACTACAACTAGTTTTCATCAATGTAGTGACGATGTATTGTTTAGTTACTCTTGGTCTAATACTGATGATAGAGATATTATTGCTTCCCATAATGGATCTTCTAGTAGTCACCTTAAAGCCCATTGATGCGAATAGCAGATAGATAAAGTCTTGGTATGACTCCTTTCTAATATAGTACGGACTACCTTTGTCACCCAACTTTATACCCACATAAGCCATAACAACCTCTTTAATAGCCGTTTCATGAGGTTTATCAGCCATGAGCCTGAGTAGTTGAAAGAATCGCATGAATCCCGTCTCAGAAAGTCCTATATGCATGATAGATTTATCTTTCCTGGGAAACTCTCGTATAGTTATAGATGAAATACTCTTCAAAGTTTCTGAAATAAGATTAGTAACAGTCTTACCTCCGACTACTCTGGGTAACAAACATACTCTAATAGGTGTTTTCTCTGCGGAGATAATATCAGAAAGGATAGAGCAATAAGTAGTATTATTGTGATTATAAAGACCGAATACATAACAGGTAGAATTTATAAACATCATGTCCTGAAGGGTTTTAGACTTGTATTCCTCGTAATCTATACCGTCCCAAAACATGGATTTGGTAACTTTGATAGCCGTAGATCTTTGTTCCTTCGCTAACAGGTTAAAGAAATTAATAAAGAATTTGTTGTTTCTATTTATGTCCACAAATTGCACGTTTGGAAGCGCCACGGTTACATTCACTGCAGCATTTTGAGGATCGCGAGTATGAAGTACGATGTTATTGTTTACTGGTATATCTGGAAAGAAATCTACCAGTCTAGGAATAAGAGATTGATATCGCATAGAAATAGTAAAGTTTATAATCTCATCATTGAAGATTACTCTGTTACCATTGTAATAAATTGGTACTCTATCATAATCATCGACAAAGTACTGTTCATACATGATGAGATGTTTATATGTTGGCATAGTAGTGAGATCGACGTTTGGTAATGGCAATGTATTAAGATTAACTCCATAATGTCTAGCAGCATCTGCGATGTTATAAGTGATGTCAAAGCGGGGTTGATCTTGTGCTGTTATATATTGTCTAACACCTATAAGATTATCAAAATCTTGTCTGCTTAATACACCGTTAACAATTTTTGCCTTGAATTCTTTTATTGGTGCATTAATAACATCCTTATAGAGGATGTTAAACAAATAAGTATTATCAAAGTTAAGATCTGGGTATTTCTTTTCTGCTAGAACATCCATTGAGTCGGAGCCATCTGGTTTAATATAACCACCGATAAATCTAGCTCTGTATTCTGTATCCGTCAATCTAATATTAAGAAGGTGTTGAGTGAAAGGTGGAAGATCGTAAAAGCTGTGAGTATTAATAATAGGGTTAGTTTCCGAACTAATGTTAATTGGATGATTAATAATATTTATATTTCCAGCGTTAAGTGTAACATTAAACAGTTTTAATTCACGTGACGTGGTATCAATTAAATAATTAATGCCCAATTTGGATATAGTAGCCTGAAGCTCATCTTGTTTAGTTACGGATCCTAATGAGTTATTAAGAAATACATCGAACGGATGAACGAAGGTTGTTTTAAGTTGGTCACATACTTTGTAATCTAGACATAGATGTGGAAGAACGGTAGAAACTATACGAAATAGATATTCAGAGTCCTCTAATTGATCAAGAGTAACTATTGACTTAATAGGCATCATTTATTTAGTATTAAATGACGACCGTACCAGTGACAGATATACAAAACGACTTAATTACAGAGTTTTCAGAAGATAATTATCCATCTAACAAAAATTATGAAATAACTCTTCGCCAAATGTCTATTCTAACTCACGTTAACAACGTGGTAGATAGAGAACATAATGCCGCCGTAGTGTCATCTCCAGAGGAAATATCATCACAACTTAATGAAGATCTATTTCCAGATGATGATTCACCGGCCACTATTATCGAACGAGTACAACCTCATACTACTATTATTGACGATACGCCACCTCCTACTTTTCGTAGAGAGTTATTGATATCGGAACAACGTCAACAACGAGAAAAAAGATTTAATATTACAGTATCAAAAAATTCTGAAGCAATAATGGAATCTAGATCTATGATAACTTCTATGCCAACACAAACACCATCCTTGGGAGTAGTTTATGATAAAGATAAAAGAATTCAGATGCTAGAGGATGAAGTGGTTAATCTTAGAAATCAACGATCTAATACAAAATCATCTGATAATTTAGATAATTTTACCAGAATACTATTTGGTAAGACTCCGTATAAATCAACCGAAGTTAATAAGCGTATAGCCATCGTTAATTATGCAAATTTGAACGGGTCCCCCTTATCAGTCGAGGACTTGGATGTCTGTTCGGAGGATGAAATAGATAGAATCTATAAAACGATTAAACAATATCACGAAAGTAGAAAACGAAAAATTATCGTCACTAACGTGATTATTATTGTCATAAACATTATTGAGCAGGCATTGCTAAAACTCGGATTTGAAGAAATCAAAGGACTGAGTACCGATATCACTTCAGAAATTATCGATGTGGAGATCGGAGATGACTGCGATGCTGTAGCATCAAAACTAGGAATCGGTAACAGTCCGGTTCTTAATATTGTATTGTTTATACTCAAGATATTCGTTAAACGAATTAAAATTATTTAATTTAATACATTCCCATATCCAGACAACAATCGTCTGGATTAATCTGTTCCTGTCGTCTCATACCGGACGACATATTAATCTTTTTATTAGTGGGCATCTTTTTAGATGGTTTCTTTTTCCCAGCATTAACTGATTCGATACCTAGAAGATCGTGATTGATTTCTCCGACCATTCCACGAACTTCTAATTGGCCGTCTCTAACGGTACCATAAACTATTTTACCAGCATTAGTAACAGCTTGGACAATCTGACCATCCATTGCGTTGAATGATGTAGTTGCTGTTGTTCTACGTCTAGGAGCACCAGAGGTATTTTTAGAGCTCTTGGATGTTGATGTAGAAGACGAGGATTTTGATTTTGGTTTACATGTAATACATTTTGAACTCTTTGATTTTGTATCACATGCACCGGCAGTCACATCTGTTTGAGAATTAAGATTATTGTTGCCTCCTTTGACGGCTGCATCTCCACCGATCTGCGCTAGTAGATTTTTAAGCTGTGGTGTAATCTTATTAACTGTTTCAATATAATCATCGTAACTACTTCTAACGGCTAAATTTTTTTTATCCGCCATTTAGAAGCTAAAAATATTTTTATTTATGCAGAAGATTTAACTAGATTATACAATGAACTAATATGATCCTTTTCTAGATTATTTACGAACTTGGTATTTCTTGTTTCTGGAGGAGGAGAATTTAAATTCGGACTTGGATTCGGATTTTGTGGGTTCTTGATCTTATTATACAGCGTGTATAGGATGGTGACGGTAACTGCTACACAAATACCGATCAACAGAAGAATACCAATCATTTATTGACAATAACTTCACTATGATCAAGTATGTAATAATCATCTTTTCACTAAGTAAGTAGTAATAATGATTCAACAATGACACGATATATGGACGATAATAATTTAGTTCATGGAAATATCGCTATGATTGGTGTGAATGACTCCGCTAACTCTGTGGGGTGCACAGTGCTTTCCCCACATAGAATAAATTAGCATTCCGACTGTGATAATAATACCAAGTATAAACGCCATAATACTCAATACTTTCCATGTACGAGTGGGACTGGTAGACTTACTAAAGTCAATAAAGGCGAAGATACACGAAAGAATCAAAAGAATGATTCCAGCGATTAGCACGCCAGAAAAATAATTTCCAATCATAAGCATCATGTCCATTTAACTAATAAAAATTTTAAATCGCCGAATAAACAAAGTGGAATATAAACCATATAAAAACAATAGTTTGTACTGCAAAAATAATATCTATTTTTGTTTTCGAAGATATGGTAAAATTAAATAGTAGTACACAGCATGTTATAACTAACAGCAGCAACGGCTCGTAATTACTTATCATTTACTAGACGAAAAGGTGGTGGGATATTTTCTTGCTCAAATAATACGAATATATCACCCATCCATTTTATACGATGTTTATATACTCTAATCTTTAATAGATCTATAGATGACGGGTTTACCAATAATATAGATTTTATCGATTCATCTAATTTAAACCCTTCCTTAAACGTGAATGATCTATTATCTGGCATAATGATGACCCTACCTGATGAATCTGACAATGTACTGGGCCATGTAGAATAAATTATCAACGAATTATCGTCTACGAACATTTATATCATTTGTTTTAATTTTAGGACGTGAATAAATAGATATAAAATAGAAAATAACAGATATTACAACCAGTGTTATGGACGCACCCAACCATGTAGGCAGTTTTATTTTATCGTTTACTACAGGTTCTCCTGGATGTACGTCACCAACTGCAGACGTAGTTCTAGTACAATTAGACGTAAGTTCCGCTTGGGAATTTTTTAACGCTAAAGAGTTAACGTTGATCGTACACCCAACGTATTTACATCTAGTTCTTTGAACATCTTGATTATAATATAACCATTTTCTATCTCTAGATTCGTCAGTGCACTCATGTAACCAACATACCCTAGGTCCTAAATATTTATCTCCGGAATTAGATTTTGGATAATTCGCGCACCAACAATTTCTATTTCCTTTATGGTCGTTACAAAAGACGTATAATGCCGTATCCCCAAAAGTAAAATAATCAGGACGAATAATTCTAATAAACTCAGAACAATATCTCGCATCCATATGTTTGGAGCAAATATCGGAATAAGTAGACATAGCCGGTTTCCGTTTTACACGTAACCATTCTAAACAATTGGGGTTTCCAGGATCGTTTCTACAAAAACCAGTCATGAAATCGTCACAATGTTCTGTCTTGTAATTATTATTAAATATTTTTGGACAGTGTTTGGTATTTGTCTTAGAACAACATTTTGCCACGCTATCACTATCACCCAGGAGATAATCCTTTTTTATAAAATGACATCGTTGCCCGGATGCTATATAATCAGTAGCATATTTTAAATCCTTAATATATTCAGGAGTTACCTCGTTCTGATAATAGATTAATGATCCAGGACGAAATTTGAAAGAACTACATGGTTCTCCATGAATTAATACATATTGTTTAGCAAATTCAGGAACTATAAAACTACTACAATGATCTATCGACATACCATCTATCAAACAAAATTTGGGTTTAATTTCTCCTGGAGACGTTTCATAATAATACATATAACTTTCTTCGGCAAACCTAACAGCTCTATTATATTCAGGATAATTAAAATCTAATACCATATATTTGTCTCGTATATCTGCTATTCCTGTCTCTATTTTGATTCTATTAAGAGTAACAGCTGCCCCCATTCTTAATAATCATCAGTATTTAAACTGTTAAATGTTGGTATATCAACATCTATCTTATTTCCCGCAGTATAAGGTTTGTTGCAGGTATACTGTTCAGGAATGGGTACATTTATACTTCTTTTATAGTCCTGTCTTTCGATGTTCATCACAAATGCAAAGAACAGAATAAACAAAATAATGTAAGAAATAATATTAAATATCTGTGAATTCGTAAATACATTGATTGCCATAATAATTACAGCAGCTACAATACACACAATAGACATTCCCACAGTGTTGCCATTACCTCCACGATACATTTGAGTTACTAAGCAATAGGTAATAACTAAGCTAGTAAGAGGCAATAGAAAAGATGAGATAAATATCATCAATATAGAGATTAGAGGAGGGCTATATAGAGCCAAGACGAACAAAATCAAACCGAGTAACGTTCTAACATCATTATTTTTGAAGATTCCCAAATAATCATTCATTATTCCTCCATAATCGTTTTGCATCATACCCCCATCTTTAGGCATAAACGATTGCTGCTGTTCCTCTGTAAATAAATCTTTATCAAGCACTCCAGCACCCGCAGAGAAGTCATCAAGCATATTGTAATATCTTAAATAACTCATTTATATATTAAAAAATGTCACTATTAAAGATGGAGTATAATCTTTATGCCGAACTAAAAAAAATGACTTGTGGTCAGACCATAAGTCTTTTTAATGAAGACGGCGATTTCGTAGAAGTTGAACCAGGATCATCCTTTAAGTTTCTAATACCTAAGGGATTTTACTCCTCTCCTTGTGTAAAGACGAGTCTAGTATTCGAGACATTAACAACGACCGATAATAAAATTACTAGTATCAATCCAACAAATGCGCCAAAGTTATATCCTCTTCAACGCAAAGTCGTATCTGAAGTAGTTTCTAATATGAGGAAAATGATCGAATTAAAACGTCCTCTATACATCACTCTTCACTTGGCATGTGGATTTGGTAAGACTATTACCACGTGTTATCTTATGACCACACACGGCAGAAAAACCATCATTTGCGTACCCAATAAAATGTTAATACATCAATGGAAGACACAGGTAGAGGCAGTCGGATTGGAACATAAGATATCTATAGATGGAGTTAGTAGTCTATTAAAGGAACTAAAGACTCAAAGTCCGGATGTATTAATCGTAGTCAGTAGACATCTGACAAACGATGCATTTTGTAAATATATCAATAAGCATTATGATTTGTTTATCTTGGATGAATCACATACGTATAATCTGATGAACAATACAGCAGTTACAAGATTTTTAGCGTATTATCCTCCGATGATGTGTTATTTTTTAACTGCTACACCTAGACCAGCTAACCGAATTTATTGTAATAGTATTATTAATATTGCCAAGTTATCCGATCTAAAAAAAACTATCTATATAGTAGATAGTTTTTTTGAGCCATATTCCACAGACAATATTAGAAATATGGTAAAACGACTAGATGGACCATCTAATAAATATCATATATATACCGAGAAGTTATTATCTGTAGACGAGCCTAGAAACCAACTTATTCTTGATACCCTGGTAGAAGAATTCAAGTCAGGAACTATTAATAGAATTTTAGTTATTACTAAACTACGTGAACATATGGTATTCTTCTACAAACGATTATTAGATCTTTTCGGAGCAGAGGTTGTATTTATAGGAGACGCCCAAAATAGACGTACTCCAGATATGGTCAAATCGATTAAGGAACTAAATAGATTTATATTCGTATCCACCTTATTTTATTCCGGCACTGGTTTAGATATTCCGAGTTTGGATTCTTTGTTCATTTGCTCGGCAGTAATCAACAATATGCAAATAGAGCAATTACTAGGGAGGGTATGTCGAGAAACAGAACTATTAGATAGGACGGTATATGTATTTCCTAACACATCCATCAAAAAAATAAAGTACATGATAGGAAATTTCGTGCAACGAATTATTAGTCTGTCTGTAGATAAACTCGGATTTAAACAAGAAAGTTATCAGAAACATCAGGAATCTGAACCCGCTTCCGTACCAACATCCTCCAGAGAAGAACGTGTATTAAATAGAATATTTAACTCGCAAAATCGTTAAGAAGTTTAAGAGACGATCCACATGCTGAGCAGGCCAGTGTATTACCCCTCATAGTATTAATATAATCCAATGATACTTTTGTGATGTCGGAAATCTTAACCAATTTAGACTGACAGGCAGAACACGTCATACAATCATCATCGTCATCGATAACTGTAGTCTTGGGCTTCTTTTTGCGACTCTTCATTCCGGAACGCATATTGGTGCTATCCATTTAGGTAGTAAAAAATAAGTCAGAATATGCCCTATAACACGATCGTGCAAAACCTGGTATATCGTCTCTATCTTTATCACAATATAGTGTATCAACATCTTTATTATTGACCTCGTTTATCTTGGAACATGGAATGGGAACATTTTTGTTAACGGCCACCTTTGCCTTAATTCCAGATGTTGTAAAATTATAACTAAACAGTCTATCATCGACACAAATGAAATTCTTGTTTAGACGTTTGTAGTTTACGTATGCGGCTCGTTCTCGTCTCATTTTTTCAGATATTGCAGGTACTATAATATTAAAAATAAGAATGAAATAACATAGGATTAAAAATAAAGTTATCATGACTTCTAGTGCTGATTTAACTAACTTAAAAGAATTACTTAGTCTGTACAAAAGTTTGAGATTTTCAGATTCTGTGGCTATAGAGAAGTATAATTCTTTGGTAGAATGGGGAACATCTACTTACTGGAAAATAGGCGTACAAAAGGTAACTAATGTCGAGACGTCCATATCTGATTATTATGATGAGGTAAAAAATAAACCGTTTAATATTGATCCGGGGTATTATATTTTCTTACCAGTATATTTTGGAAGCGTCTTTATTTATTCAAAGGGTAAAAATATGGTAGAACTTGGATCTGGAAACTCTTTTCAAATACCGGATGAGATTCGAAGTGCGTGTAACAAAGTATTAGATAGTGATAACGGAATAGACTTTCTGAGATTTGTTTTGTTAAACAATAGATGGATAATGGAAGACGCTATATCAAAATACCAGTCTCCAGTTAATATATTTAAACTAGCTAGTGAGTACGGATTAAACATACCCAACTATTTAGAAATTGAAATAGAGGAAGACACATTATTTGACGATGAGTTATACTCTATTATGGAACGCTCTTTCGATGATACATTTCCAAAAATATCTATATCGTATATTAAGTTGGGAGAACTTAAGCGGCAAGTTGTAGACTTTTTCAAATTCTCATTCATGTATATTGAGTCAATCAAGGTAGATCGTATAGGAGATAATATTTTTATTCCTAGCGTTATAACAAAATCAGGAAAAAAGATATTAGTAAAAGATGTAGACCATTTAATACGATCCAAGGTTAGAGAACATACATTTGTAAAAGTAAAAAAGAAAAACACATTTTCCATTTTATACGACTATGATGGGAACGGAACAGAAACTAGAGGAGAAGTAATAAAACGAATTATAGACACTATAGGACGAGACTATTATGTTAATGGAAAGTATTTCTCTAAGGTTGGTATTGCAGGCTTAAAGCAATTGACTAATAAATTAGATATTAATGAGTGTGCAACTGTCGATGAGTTAGTTGATGAGATTAATAAATCCGGAACTGTAAAACGAAAAATAAAAAACCAATCAGTATTTGATTTAAGCAGAGAATGTTTGGGATATCCAGAAGCGGATTTTATAACGTTAGTTAATAACATGCGGTTCAAAATAGAAAATTGTAAGGTTGTAAATTTCAATATTGAAAATACTAATTGTTTAAATAACCCGAGTATTGAAACTATATATGGAAACTTCAACCAGTTCGTCTCAATCTTTAATACCGTTACCGATGTCAAAAAAAGATTATTCGAGTGAAATAATATGCGCCTTTGATATAGGTGCAAAAAATCCTGCCAGAACTGTTTTAGAAGTCAAGGATAACTCCGTTAGGGTATTGGATATATCAAAATTAGACTGGAGTTCTGATTGGGAAAGGCGCATAGCTCAAGATTTGTCACAATATGAATACACTACAGTTCTTCTAGAACGTCAGCCTAGAAGGTCACCGTACGTCAAATTTATCTATTTTATTAAAGGCTTTTTATATCATACATCTGCTGCCAAAGTTATTTGCGTCTCACCTGTCATGTCTGGTAATTCATATAGAGATCGAAAAAAGAGATCTGTTGAAGCATTTCTTGATTGGATGGACACATTCGGATTGCGAGACTCCGTTCCGGATAGACGCAAATTAGACGATGTAGCGGATAGTTTCAATTTGGCTATGAGATACGTATTAGATAAATGGAATACTAATTATACACCTTATAATAGGTGTAAATATAGAAATTACATAAAAAAAATGTAATAACGTTAGTAACGCCATTATGGATAATCTATTTACCTTTCTACATGAAATAGAAGATAGATATGCCAGAACTATTTTTAACTTTCATCTAATAAGTTGTGATGAAATAGGAGATATATATGGTCTTATGAAAGAACGCATTTCCTCAGAGGATATGTTTGACAATATAGTATATAATAAAGATATACATCCTGCCATTAAGAAACTAGTTTATTGCGACATCCAACTTACTAAACATATTATTAATCAGAATACGTATCCGGTATTTAACGATTCTTCACAAGTGAAATGTTGTCATTATTTCGATATAAACTCAAATAATAGCAATATTAGCTCTCGTACAGTAGAGATATTTGAGAGTGAAAAGTCATCTCTTGTATCATATATTAAAACTACCAATAAGAAGAGAAAGGTCAATTACGGCGAAATAAAGAAAACTGTACATGGAGGCACTAATGCAAATTACTTTTCCGGTAAAAAGTCTGATGAGTATCTGAGCACTACAGTCAGGTCCAACATTAATCAACCTTGGATCAAAACCATTTCTAAGAGAATGAGAGTAGATATCATTAATCACTCTATAGTAACGCGTGGAAAAAGCTCTATATTACAAACTATAGAAATTATTTTTACTAATAGAACATGTGTGAAAATATTCAAGGATTCTACTATGCACATTATTCTATCCAAGGACAAGGATGAAAAGGGATGTATAAACATGATTGATAAATTATTCTATGTATATTATAATTTATTTCTGTTGTTCGAGGATATCATCCAAAACGATTACTTTAAAGAAGTAGCTAATGTTGTAAACCATGTACTCATGGCTACGGCATTAGATGAGAAATTATTCCTAATTAAGAAAATGGCTGAACACGATGTTTATGGAGTTAGCAATTTCAAAATAGGGATGTTTAACCTGACATTTATTAAGTCGTTGGATCATACCGTTTTCCCCTCTCTGTTAGATGAGGATAGCAAAATAAAGTTTTTTAAGGGGAAAAAGCTCAATATTGTAGCATTACGATCTCTGGAGGATTGTACAAATTACGTGACTAAATCCGAGAATATGATAGAAATGATGAAGGAAAGATCGACTATTTTAAATAGCATAGATATAGAAACGGAATCGGTAGATCGTCTAAAAGAATTGCTTCTAAAATGAAAAAAAACACTGATTCAGAAATGGATCAACGACTCGGGTATAAGTTTTTGGTGCCTGATCCTAAAGCCGGAGTTTTTTATAGACCGTTACATTTCCAATATGTATCGTATTCTAATTTTATATTGCATCGATTGCATGAAATCTTGACCGTCAAGCGGCCACTCTTATCGTTTAAGAATAATACAGAACGAATTATGATAGAAATTAGCAATGTTAAAGTGACTCCTCCAGATTACTCACCTATAATTGCGAGTATTAAAGGTAAGAGTTATGACGCATTAGCCACGTTCACTGTAAATATCTTTAAAGAGGTAATGACCAAAGAGGGTATATCCATCACTAAAATAAGTAGTTATGAGGGAAAAGATTCTCATTTGATAAAAATTCCGCTACTAATAGGATATGGGAATAAAAATCCACTTGATACAGCCAAGTATCTTGTTCCTAATGTCATAGGTGGAGTCTTTATCAATAAACAATCTGTCGAAAAAGTAGGAATTAATCTAGTAGAAAAGATTACAACATGGCCAAAATTTAGGGTTGTTAAGCCAAACTCATTCACTTTCTCGTTTTCCTCCGTATCCCCTCCTAATGTATTACCGACAAGATATCGCCATTACAAGATATCTCTGGATATATCACAATTGGAAGCGTCGAATATATCATCGACAAAGACATTTATAACGGTCAATATTGTTTTGCTGTCTCAATATTTATCTAGAGTGAGTCTAGAATTCATTAGACGTAGTTTATCATACGATATGCCTCCAGAAGTTGTCTATCTAGTAAACGCGATAATAGATAGTGCTAAACGACTTACCGAATCTATTACTGACTTTAATATTGATACATACATTAATGACCTGGTGGAAGCTGAACACATTAAACAAAAATCTCAGTTAACGATTAACGAGTTTAAATATGAAATGCTGCATAACTTTTTACCTCATATGAACTATACACCCGATCAACTAAAGGGATTTTATATGATATCTTTACTAAGAAAGTTTCTCTACTGTATCTACCACACTTCTAGATATCCAGATAGAGATTCGATGGTTTGTCATCGCATCCTAACGTACGGCAAATATTTTGAGACGTTAGCACATGATGAATTAGAGAATTACATAGGTAACATCCGAAACGATATCATGAACAATCACAAGAACAGAGGCACTTACGCAGTAAACATTCATGTACTAACAACTCCTGGACTTAATCATGCATTTTCTAGTCTATTGAGTGGAAAGTTCAAAAAGTCAGACGGTAGTTATCGAACACATCCTCACTATTCATGGATGCAGAATATTTCTATTCCTAGAAGTGTTGGATTTTATCCGGATCAAGTAAAGATTTCAAAGATGTTTTCTGTCAGAAAATACCATCCAAGCCAATATCTTTACTTTTGTTCATCAGACGTTCCGGAAAGAGGTCCTCAGGTAGGTTTAGTATCTCAATTGTCTGTCTTGAGTTCCATTACAAATATACTAACGTCTGAGTATTTGGATTTGGAAAAGAAAATTTGTGAGTATATCAGATCATATTATAAAGATGATATAAGTTACTTTGAAACAGGATTTCCAATCACTATAGAAAATGCTCTAGTCGCATCTCTTAATCCAAATATGATATGTGATTTTGTAACTGACTTTAGACGTAGAAAACGGATGGGATTCTTCGGTAACTTGGAGGTAGGTATTACTTTAGTTAGGGATCACATGAATGAAATTCGCATTAATATTGGAGCAGGAAGATTAGTCAGACCATTCTTGGTTGTGGATAACGGAGAGCTCATGATGGATGTGTGTCCGGAGTTAGAAAGCAGATTAGACGACATGACATTCTCTGACATTCAGAAAGAGTTTCCACATGTCATCGAAATGGTAGATATAGAACAATTTACTTTTAGTAACGTATGTGAATCGGTTCAAAAATTTAGAATGATGTCAAAGGATGAAAGAAAGCAATACGATTTATGTGACTTTCCTGCCGAATTTAGAGATGGATATGTAGCATCTTCACTAGTGGGAATCAATCACAATTCTGGACCCAGAGCTATTCTTGGATGTGCTCAAGCTAAACAAGCTATCTCTTGTCTGAGTTCGGATATACGAAATAAAATAGACAATGGAATTCATTTGATGTATCCAGAGAGGCCAATTGTGATTAGTAAGGCTTTAGAAACTTCAAAGATTGCGGCTAATTGCTTCGGACAACATGTTACTATAGCATTAATGTCGTACAAAGGTATCAATCAAGAGGATGGAATTATCATCAAAAAACAATTTATTCAGAGAGGCGGTCTCGATATTGTTACAGCCAAGAAACATCAAGTAGAAATTCCATTGGAAAACTTTAATAACAAAGAAAGAGATAGGTCTAACGCCTATTCGAAATTAGAAAGTAATGGATTAGTTAGACTGAATGCTTTCTTGGAATCCGGAGACGCTATGGCAAGAAATATCTCATCAAGAACTCTTGAAGATGATTTTGCTAGAGATAATCAGATTAGCTTTGATGTTTCCGAGAAATATACAGATATGTACAAATCTCGCGTTGAACGAGTACAAGTAGAACTTACTGACAAAGTTAAGGTGCGAGTATTAACCATGAAAGAAAGAAGACCCATTCTAGGAGACAAATTTACTACTAGAACGAGTCAAAAGGGAACAGTCGCGTATATCGCAGATGAAACGGAACTTCCGTACGACGAAAATGGTATCACACCAGATGTCATTATTAATTCTACATCCATCTTCTCTAGAAAAACTATATCTATGTTGATAGAAGTTATTTTAACAGCCGCATATTCTACTAAGCCGTACAACAATAAGGGAGAAAACCGACCTGTCTGTTTTCCTAGTAGTAACGAAACATCTATCGATGCATATATGCAATTCGCTAAACAATGTTATGAGTATTCAAATCCGAAATTGTCCGAGGAAGAATTATCGGATAAAATCTTTTGTGAAAAGATTCTCTATGATCCTGAAACGGATAAGCCTTATGAATCCAAAGTATTTTTTGGACCAATTTATTACTTGCGTCTGAGACATTTAACTCAGGACAAGGCAACCGTTAGATGTAGAGGTAAAAAGACGAAGCTCATTAGACAAGCGAATGAGGGACGAAAACGTGGAGGAGGTATCAAGTTTGGAGAAATGGAGAGAGACTGTTTAATAGCACATGGTGCAGCCAATACTATTACAGAAGTTTTAAAAGACTCAGAAGAGGATTATCAAGATGTGTATATTTGTGAAAATTGTGGAGACATAGCAGCACAAATCAAAAGTATTAATACATGTCTTAGATGTTCAAAACTTAATCTCTCTCCTCTCTTAACAAAAATTGATACCACGCACGTATCTAAAGTATTTCTTACTCAAATGAACGCCAGAGGCGTAAAAGTTAAATTAGATTTCGAACGAAGGCCTCCTTCGTTTTATAAACCATTAGATAAAGTTGATCTTAAACCGTCTTTTCTGGTATAATATTGTTTAGTAGATACTCATCAAGATAAGCTAATTCACTAAACATATTATCGGATTCGGTATTGTTACTCGAGAATAGAGTTCGTTATGCTCCTGATATTCGGAAATCTGTGGAGTTTCAGGTTTTGGTGGAAGTGTAACTGCTACTTGGTGGGATACTGAAGGATATTTCAGAGAGTTGTGGATGTTCGGGTTCGACATCCACCGATGGTGTCACGCCACTAATCGGTTCGGTAACGTCTGTGGATGGAGGTGCTACTTCTACAGAACCTGTAGCCTCAGTTGTCAACGGAGATACATATTCAATGCGCGGAAATGTATAATTTGGTAATGGTTTCTCATGTGGATCTTAAGAAGAAGAGGTAAGATATCTACGAAAGATACCGATCACGTTTCTAGTTCTCTTTTGTAGAACTTTAACTTTTTCTTTCTCAGCATCTAGTTGATATTCCGACCTCTTCACGTTTCGCATGGGTTACCTCCGCAGTTTTTACAAGCGATTTCACGTTCCAGATCACGTTCAGCCTTCATACGTCTCTCCCTCTCTCTATCGAGTTTATCAGAGCAGTCTTTCTGAAGGCGATCGAACTCCATAAATTTCTCCAACGCTTTGATTGTTTCCATAGATTTCCGAAGTTTAGCTTCTAGGACGGCGATTCTTTTTTTTTTTTTTTTTTTTTTTTTTTTCGAATTCACGGGGTACAACCGTTTCCATTACCACCATCTCTATGTTTCTTTTCTAGATCGGCAATCTTTCTCAATCTTTCTCAACATTTCATCCCCATACCTTTTCATTCCTCGAGTCTATTGTCGTCGAAATATCGTTCCAGCTCCTTTTCGACCTCAATAACTTTAGCACGTTGTTTCATCAAGCTCTCTCTTGTAGTACTATCATTTTTATCTGATTCCCTGACACGTTTAAGATCTTCATGTAATTGAGTCAGCTCTTGACGCAATCTCTTAACTAACTTCCTCTCTTGCTTCTTCGTCATAGTACTTACAATCACTATGGGATCCATTGTTACCACGTCTGTACTCGACGAGCTCACGTTTAAGAGATTCAATTTCCAGTTTGTATCGGTCCATGTCTCCATTGCTACACCACCATTAGATTTACAGGCTGCTAGTTGTCGTTCGAGATCAGAAATACGTGTTTTCTTGGAATGGATTTCGTCGATGTACTTGTCATGATTGGCATCGAAACACTTATTAAGTTCTTTTTTTCAATTCTACGATTTTATTTCTTTCGCGAGTCAATTCCCTCCTGTAGTAACTATCAGTTTTGTCAGATTCACGCTCTCTACGTAGACTTTCTTGTAAGTTACTAATTTGTTCCCTGGCATTACCGAGTTCAGTTTTATATGCCGAATAGAGTTCTGATTCATCCTTTGAGAAGATCTCTAGCGATCGTTCAAGATCCCTGATTCTAGTCTTTAGCCTATTTACCTCCTCAGAAGATGCTCCGTTACCGTTTTTACAATCGTTAAGATGTCTATCAAGATCCATGATTCTATCTCTTTTCCATATCAGCATTGATTTCATTATTACGTTCGCAGTCGTTCAACTGTATTTCAAGATCTGAGATTCTAGATTGTAATCTCTGTAGCATTTCCACGGCATTCACTCAGTTGTCTTTCAAGATCTGAGATTCTAGATTGGAGTCTGCTAATCTCTGTAAGATTTCCTCCTCCGCTCTCGATGCAGTCGGTCAACTTATTCTCTAGTTCTCTAATACGCGAACGCAGTGCATCAACTTCTTGTGTGTCTTCTTGATTGCGTGTGCATTCATCGAGTCTAGATTCGAGATCTCTAACGTGACGTCGTTCTTCCTCAAGTTCTCTGTGTACTACAGAAAGCGTGTCCCTATCTTGTTGATATTTAGCAATTTCTGATTCTAGAGTACTGATTCTACTCACGTATGTACTAATAGTTGTCTTAGCCTTATCAAGATCCTCCTTGTATTTGTCACATTCCTTGATATCCATACGAAGTCTGGACAGTTCCCATTCGACATTACGACGTTTATCGATTTCAGCTCGGAGATCGTCGTCGCGTTGTTTTAGCCACATACGACTAAGTTCAAGTTCTCGTTGACAAGATCCATCTACTTTTCCATCCCTAATAGTATCCAGTTCCTTTTCTAGTTCTGACCGCATTTCTCGTTCCATATCAAGAGATTCTCTCAATTCTCGTATAGTCTTCTTATCAATTTCTGATGAATCTGAACCATCATCTGTCCCATTTTGTTGCATATCCCTGAGTTCTTTGATCTCTGTTGTAAGTCTGTCGATTCTTTCGGTTTTATAAACAGAATCCCTTTCCAAAGTCCTAATCTTACTGAGTTTATCATTAAGTTCTTCATTCAATTCAGTGAGTTTTCTCTTGGCTTCTTCCAAGTCTGTTTTAAACTCTCCATCATTTCCGCATTCTTCCTCGCATTTATCTAACCATTCAATTAGTTTATTAATAACTAGTTGGTAATCAGCGATTCCTATAGCCGTTCTTGTATTTGTGGGAACATAATTAGGATCTTCTAATGGATTGTATGGCTTGATAGCATCATCTTTATCATTATTAGGTGGGGGATGGACAACCTTAATTGGTTGGTCCTCCTTATCTCCTCCAGTAGCATGTGGTTCTTCAATACCAGTATTAGTAATAGGCTTAGACAAATGCTTGTCGTACGCGGGCACTTCCTCATCCATCAAGTATTTATAATCGGGTTCTGTTTCAGAATATTCTTTTCTAAGAGACGCGACTTCAGGAGTTAGTAGAAGAACTCTGTTTCTGTATCTATCAACGCTGGAATCGATACTCAAGTTAAGGATAGCGAATACCTCATCGTCATCATCCGTATCTTCTGAAACGCCATCATATGACATTTCATGAAGTCTAACGTATTGATAAACAGAATCAGATTTAGTATTAAACAGATCCTTGACCTTTTTAGTAAATGCATATGTATATTTTAGATCTCCAGATTTCATAATATGATCGCATGCCTTAAATGTCAATGCTTCCATGATATAGTCTGGAACACTAATGGGTGACGAAAAAGATACAGCACCATATGCTACGTTGATAAATAGATCTGAACCACTAAGTAGATAATGATTAATGTTAAGGAAGAGGAAATATTCAGTATATAGATATGCCTTAGCATCATATCTTGTACTAAACACGCTAAACAGTTTATTGATGTGATCAATTTCCAACAGAACAATTAGAGCGGCAGGAATACCAACAAACATATTACCACATCCGTATTTTCTATGAATATCACATATCATATTAAAAAATCTTGATAGAAGAGCGAATATCTCGTCTGACTTAATGAGATGTAGTTCAGCAGCATAAGTCATAACTGTAAATAGAACATACTTTCCTGTAGTGTTGATTCTAGACTCCACATCAACACCATTATTAAAAATAGTTTTATATACATCTTTAATCTGCTCTCCGTTAATCGTCGAACGTTCTAGTATACGGAAACACTTTGATTTCTTATCTGTAGTTAATGACTTAGTGATATCACGAAGAATATTACGAATTACATTTCTTGTTTTTCTTGAGAGACCTGATTCAGAACTCAACTCATCGTTCCATAGTTTTTCTACCTCAGTGGCGAAATCTTTGGAGTGTTTGGTACATTTTTTAATAAGGTTCGTGACCTCCATTTATTATAAAAAATTTTTATTCAAAACTTAACTACAATCGGGTAATTATAAGATCGTAGATCTCCCATGTGGTGGAATACTACCATCTATCGCATGTTGATGGACAGTAGGTAATGGCCATGGGAACAGTAATGTTTGCATATTTATCTTTCTTGCTAGTATTACTGTATATTGTCCCAATGTTTCAATGTGATGTTCTAACCTATCAACTGCCACTGTATCACAACAATAATGTCCGATGGAATTAAGATTATGATCCAATGTGTTTAATATATGATTATCAAGTCTTATACGATCCGCGTCTTTTTTGACAGGATCAGGCTCTTCTACAGGAAGAAGTTTCGGCCTCTTATGATAGTCATGTCTGGGAAATGGTGGTCTAGGATGAGGATCAGGTATCGGAGTAGGTTTTGGATTATAATCATCATCATCATCATCATCATCATCATCATCATCATCATCTATGATATCATCATCTTCGATATTTATTTTGCTATCTTGATAATGTCCTATATCAGTTGCATTTTCAGCACTCGACTGAATATTAGTACATTCATTGTCTATTATTAACGTATTTCTAAACCCAAAATGTATATGTTGAACATCACTACTATAGTTGATGAGTCTTATAGCATGAATTCGCTTATCGTTATCGGGTTTATCTTCTGTCACCTTAACAATTCCTTTTTTATTAAACTCTGCATAATCATAACCATTTCTATTGTTTGTTCTAATATAAACGAGTATAGCATCATTGCTAAATTTTTCAATAGTATCAAAAACAGAATATCCTAAACCATATAATATATATTCAGGAACACTCAAACTAAATGTCCAGGATTCTCCTAAATACGTAAACTTTAATAGTGCTAAATCATTCAAAAATCTACCGCTTATAGATAGATAGTACATGAATGCGTATAGTAGTCTACCTATCTCTTTATTATGAAAACCGACATTACGATCATATATTTCGTGATATACATGTGACCCGTTTACGTTAAACCATAAATACATGGGTGATCCTATAAACATGAATTTATTTCTAATTCTCAGAGCCATAGTTAATTGACCGTGTAATATTTGTTTACATGCATACTTGATACGATCATTAATAAGATTTTTATCATTGCTCGTTATTTCAGAATCGTATATATAAGGAGTACCATCATGATTCTTACCAGATATTATACAAAATACTATATATAAAATATATTGACCCACGTTAGTAATCATGTAAATGTTTAATGTTTTAAATTTTGTATTTAATGATCCATCATCATATGCTAGCATGGTCTTGTGATATTCATTCTTTAAAATATAATATTGTGTTAGCCATTGCATTGGAGCTCCTAATGGAGATTTTCTATTCTCGTCCATTTTAGGATATGCTTTCATAAAGTCCCTAATAACTTCGTGAATAATGTTTCTATGTTTTCTACTGATGCATGTATTTGCTTCGATTTTTTTATCCCATGTTTCATCTATCATAGATTTAAACGCAGTAATGCTCGCAACATTAACATCTTGAACCATTGGTACAATTCCGTTCCATAAATTTATAATGTTCGCCATTTATATAACTCATTTTTTGAATATACTTTTAATTGAACAAAAGAGTTAAGTTACTCATATGGATGCCGTCCAGTCTGTACATCAATCTTTTTAGCCAGAGATATCATAGCCGCTCTTAGAGTTTCAGCGTGATTTTCCAACCTAAATAGAACTTCATCGTTGCGTTTACAACACTTTTCTATTTGTTCAAACTTTGTTGTTATATTAGTAATCTTTTTTTCCAAATTAGTTAGCCGTTGTTTGAGAGTTTCCTCATTATCGTCTCCATAGGCTTTAACAATTGCTTCGCGTTTAGTCTCTGGATTTTTAGCAGCCTTTGTAGAGAAAAATTCAGTTGCTGGAATTGCAAGATCGTCATCTCCGGGGAAAAGAGTTCCGTCCATTTAAAGTACAGATTTTAGAAACTGACACTCTGTGTTATTTATATTTGGCGCAATACATGGATTATAAATATCGATGTTAATAACATCAGAAAATGTAAAGTCTATACATTGTCGCATCGTGTTAAATTTTCTAATGGATCTAGTATTATTGGGTCCAACTTCTGCCTGAAATCCAAATATGGAAGCGGATACAAAACCGTTTCCTGGATAAACCACACATCTCCACTTTTGCTTTACATCAGAAATTGTGTCATTGACATCTTGAACTCTCCTATCTAATGCCGGTGTTCCACCTATAGATTTTGAATACTCGAATGCTGCATGAGTAGCATTGAATTCCTTAATATTGCCATAATTTTCATATATTGAGTAACTCTGGATAAAAAGTAAACACACCGCAGCCGTCGCTACTACAATAAAAAAAATTGATAGAGAGTTCATTTATAATCTATTAGAAGCTGATAAAATTTTTTTACACGCGTCAGACAATGCTTTAATAAATAGTTCAACATCTACTTTTGTCATATCGAACCGATGGTATGATTCTAACCTAGAATTACATCCGAAAAAGTTGACTATGTTCATAGTCATTAAGTCATTAACGAACAACATTCCAGACTCTGGATTATAAGACGATACTGTTTCGTCACAATCACCCACCTTAATCATGTGATTATGAATATTGGCTATTAGAGTACCTTCTAAGAAATCTATAATATCTTTGAAACACGATTTAAAATCAAACCACGAATATACTTCTACGAAGAAAGTTAGTTTACCCATAGGAGAGATAACTATAAATGGAGATCTAGATACAAAATCCGGATCTATGATAGTTTTAACATTATTATATTCTCTATTAAATACCTCCACATCTAAAAATGTTAATTTTGAAACTATGTCTTCGTTTATTACCGTACCTGAACTAAACGCTATAAGCTCTATTGTTTGAGAACTCTTTAAACGATATTCTTGAAATACATGTAACAAAGTTTCCTTTAACTCGGTCGGTTTATCTACCATAGTTACAGAATTTGTATCCTTATCTATAATATAATAATCAAAATCGTATAAAGTTATATAATTATCGTGTTCAGATTGTGATCTTTTCAAATAGACTAAAAACCCCATTTCTCTAGTAAGTATCTTATGTATATGTTTGTAAAATATCTTCATGGTGGGAATATGCTCTACAGCAGTTAGCCATTCCTCATTGACAGCTGTAGATGTATTATACAAAACTACTCCAATGTTTAACAAGGGCCATTTTACGAGATTATTAAATCCTTGTTTGATAAATGTAGCCAATGCGGGTTCGAGTTCAACGACGATTGAATTCTCTTCCCGTGGATGCTGCATGATGAACGACGGGATGTTGTTGTTCTATTGATTTGGAATTCTTTTTCGACTTTTTGTTTATATTAAATATTTTAAAATTTATGGCTGATAGTAATTCATGTACTACGGATAATGTAGACGTGTATTGCATATCGATATCTTTATTATTAGATAAATTTATCAATAAATGTGAGAAGTTTGCCTCGTTAAGGTCTTCCATTTAAATATTATATAAATATTTGTGTTTGTATTTTATTCGTCTTTTATGGGATAGTTTTTAACTAGTAAAGCTGTAATTACATACTTTGTCCGTAAAACATAAATATAAATACCCGCTTTTATCAAACGTTCCAAAAAGTCGGCAGCTGACATTTTTAACATGACATCTATTTTAAATACACTTAGGTTTTTAGAAAAAACATCATTTTATAATTGTAACGATTCAATAACTAAAGAAAAGATTAAGATTAAACATAAGGGAATGTTATTTGTATTTTATAAGCCAAAGCATTCTACCGTTGTTAAATACTTGTCTGGAGGAGGTATATATCATGATGATTTGGTTGTATTGGGGAAGGTAACAATTAATGATCTAAAGATGATGCTATTTTACATGGATTTATCATATCATGGAGTGACAAGTAGTGGAGCAATTTACAAATTGGGATCGTCTATCGATAGACTTTCTCTAAATAGGACTATTGTTACAAAAGTTAATAACAATTATAACAATTATAACAATTATAACAATTATAATTGTTATAATAATTATAATTGTTATAATTATGATGATACATTTTTTGACGATGATGATTGATCACTATTACACAATTTTGTTTTTGTACTTTCTAATATAGTGTTTAGGTTCTTTTTCATATGAGAATATTGACTTACTAAAATATCTATGTTTAACTTTTGTTCTATAACGTCCTTATCGGCGGTATCGGTACATATACGTAATTCACCTTCACAAAATACGGAGTCTTCGATAATAATAGCCAATCGATTATTGGATCTAGCTGTCTGTATCATATTCAACATGTTTAATATATCCTTTCGTTTCCCCTTTACAGGCATCGATCGTAGCATATTTTCCGCGTCTGAGATGGAAATGTTAAAACTGCAAAAATGCGTAATGTTAGCCCGTCCTAATATTGGTACGTGTCTATAAGTTTGGCATAGTAGAATAATAGACGTGTTTAAATGCCTTCCAAAGTTTAAGAATTCTATTAGAGTATTACATTTTGATAGTTTATCACCTACATCATCAAAAATAAGTAAAAAGTGTGCTGATTTTTTATGATTTTGTGCGACAGCAATACATTTTTCTATGTTACTTTTAGTTCGTATCAGATTATATTCTAGAGCTTCCTGACTACTAACGAAATTAATATGATTTGGCCAAATGTATCCATCATAATCTGGGTTATAAACGGGTGTAAACAAGAATATATGTTTATATTTTTTAACTAGTGTAGAAAACAGAGATAGTAAATAGATAGTTTTTCCAGATCCAGATCCTCCTGTTAAAACCATTCTAAACGGCATTTTTAATAAATTTTCTCTTGAAAATTGTTTTTCTTGAAAACAATTCATAATTATATTTACAGTTACTAAATTAATTTGATAATAAATCAAAATATGGAAAACTAAGGTCGTTAGTAGGGAGGAGAACAACGAAGGCATATCGTGATATAAATAACATTTATTATCATGATGACACCAGAAAACGACGAAGAGCAGACATCTGTGTTCTCCGCTACTGTTTACGGAGACAAAATTCAGGGAAAGAATAAACGCAAACGCGTGATTGGTCTATGTATTAGAATATCTATGGTTATTTCACTACTATCTATGATTACCATGTCCGCGTTTCTCATAGTGCGCCTAAATCAATGCATGTCTGCTAACAAGGCTGCTATTACTGACTCCGCTGTTGCCGTTGCTGCGGCATCATCTACTCATAGAAAGGTTGTGTCTAGCACTACACAATATGATCACAAAGAAAGCTGTAATGGTTTATATTACCAGGGTTCTTGTTATATATTACATTCAGACTATAAGTCATTCGAGGATGCTAAAGCAAACTGCGCTGCGGAATCATCAACACTACCCAATAAATCCGATGTCTTGACTACCTGGCTCATTGATTATGTTGAGGATACATGGGGATCTGATGGTAATCCAATTACAAAAACTACATCCGATTATCAAGATTCTGATGTATCACAAGAAGTTAGAAAGTATTTTTGTACATAAATAAATGAAATCGCTTAATAGACAAACTGTAAGTAGGTTTAGGAAGTTGTCGGTGCCGGCCGCTATAATGATGTTACTCTCAACCATTATTAGCGGCATAGGAACATTTCTGCATTACAGAGAAGAACTGATGCCTAGTGCTTGCGCCAATGGATGGATACAATACGATAAACATTGTTATCTGGATACCAACATTAAAATGTCTACGGATAATGCAGTTTATCAGTGTCGCAAATTACGAGCTAGATTGCCTAGACCTGATACTAGACATCTGAGAGTATTGTTTAGTATTTTTTATAAAGATTATTGGGTAAGTTTAAAAAAGACCAATGATAAATGGTTAGATATTAATAATGATAAAGATATAGATATTAGTAAATTAACAAATTTTAAGCAACTAAACAGCACGACGGATTCTGAGGCGTGTTATATATACAAGTCTGGAAAACTGGTTAAAACAGTATGTAAAAGTACTCAATCTGTACTATGCGTTAAAAGATTCTACAAGTGACAACAAAAAATGAATTAATAGTAAGTCGTTAACGTACGCCGCCATGGACGCCGCGTTTGTTATTACTCCAATGGGTGTGTTGACTATAACAGATACATTGTATGATGATCTCGATATCTCAATCATGGACTTTATAGGACCATACATTATAGGTAACATAAAAATTGTCCAAATAGATGTACGGGATATAAAATATTCCGACATGCAAAAATGCTACTTTAGCTATAAGGGTAAAATAGTTCCTCAGGATTCTAATGATTTGGCTAGATTCAACATTTATAGTATTTGTACAGCATACAGATCAAAAAATACCATCATCATAGCATGCGACTATGATATCATGTTAGATATAGAAGGTAAACATCAACCATTTTATCTATTCCCATCTATTGATGTTTTTAACGCTACAATCATAGAAGCGTATAATCTGTATACAGCTGGAGATTATCATCTGATCATCAATCCTTCAGATAATCTGAAAATGAAATTGTCGTTTAATTCTTCATTTTGTATATCAGACGGCAATGGATGGATTATAATTGATGGGAAATGTAATAGTAATTTTTTATCATAAAAGTTGTAAAGTAAATAATAAAACAATAAATATTGAACTAGTAGTATGTTGTATATTGAGCAATCAGAGATGATGCTGGTACCTCTTATCACGGTGACCGTAGTTGCGGGAACAATATTAGTATGTTATATATTATATATTTGTAGGAAAAAGATACGTACTGTCTATAATGACAATAAAATTATCATGACAAAATTAAAAAAGATAAAGAGTCCTAATTCCAGCAAATCTAGTAAATCAACTGATAGCGAATCAGACTGGGAGGATCACTGTAGTGCTATGGAACAAAACAATGACGTAGATAATATTTCTAGAAATGAGATATTGAACGATGATAGCTTCGCTGGTAGTTTAATATGGGATAACGAATCCAATATCATGGCGCCTAGCACAGAACACATTTACGATAGTGTTGCTGGAAGCACGCTGCTAATAAATAATGATCGTAATGAACAGACTATTTATCAGAATACTACAGTAGTAATTAATGATACAGAGACTGTTGAAATACTTAATGAAGATACCAAACAGATTCCTAGCTATTCTTCCAATCCTTTCGTAAATTATAATAAAACCAGTATTTGTAGCAAGTCAAATCCGTTCATTGCAGAACTCAACAATAAATTTAGTGATAATAATCCGTTTAGGAGAGCACATAGTGACGATTATCTTAATAAGCAACAAGATCATGAATACGATGATATAGAATCATCGGTTGTATCATTGGTCTGATTAGTTTCCTTTTTATAAAATTGAAGTAATATTTAGTATTAATTACCGCCGATGCATTATACAAATATGGAGATATTCCCTGTATTCGGCATTTCTAAAATTAGCAATTTTATTGCTAATAATGACTGTAGATATTATATAGATGTAGAGCATCAAAAAATTATATCTGATGAGATCAATAGACAGATGGATGAAACGGTACTTCTTACCAACATCTTAAGCGTAGAAGTTGTAAATGACAATGAGATGTACCATCTTATTCCCCATAGACTATCGACTATTATACTCTGTATTAGTTCTGTTGGAGGATGTGTTATCTCTATAGATAATGACGTCAATGACAAAAATATTCTAACATTTCCCATTGATCATGCTGTAATCATATCCCCACTGAGTAAATGTGTCGTAGTTAGCAAGGGCCCTACAACCATACTGGTTGTTAAAGCGGATATACCCAGCAAACGATTGGTAACATCGTTTACAAACGACATACTGTATGTAAACAATCTATCACTGATTAATTATTTACCGTCGTCTGTATTCATTATTAGACGAGTCACCGACTATTTGGATAGACACATATGTGATCAGATATTTGCTAATAATAAGTGGTATTCCATTATAACTATCGACGATAAGCAATATCCTATTCCATCAAATTGTATAGGTATGTCTTCTGCCAAGTACATAAATTCGAGCATCGAGCAAGATATTTTGATCCATGTTTGTAACCTCGAGCATCCATTCGACTCAGTCTACAAAAAAATGCAGTCGTACAATTCTCTACCTATCAAGGAACAAATATTGTATGGTAGAATTGATAATATAAATATGAGCATTAGTATTTCGGTGGATTAATAGATTTCTCTAGTATGGGATCATTAATCATCTCTAAATACATCATAAAAAAGCTATTATCAAATACTGTACTGAATGGATTCATTCTTTTCTCTTTTTATGAAACTCTGTTGTATATCTACGGATAAAACTAGAAGCAAAAAATCTGATAGGAAGAATAATGATTATATGGAGGAACACGATTATTATAAAATAACAATAGTTCCTGGTTCCTCTTCCACGTCTACTAGCTCGTGGTATTATACACATGCCTAGTAATAGTCTCTTTGCGTTGACGGAAAGCAGACTAGAAATAACAGGCCAAAATGTTCAGACACCATAATAGTTCCCAACCCAGATAATAACAGAGTTCCATCAACACATTCCTTTAAACTCAATCCCAAACCCAAAACCGTTAAAATGTATCCAGCCAATTGATAGTAGATAATGAGGTGTACAGCACATGATAATTTACACAGTAACCAAAATGAAAACACTTTAGTAATTATAAGAAATATAGACGGTAATGTCATCATCAACAATCCAATAATATGCCTGAGAGTAAACATTGACGGATAAAACAAAAATGCCCCGCATAACTCTATCATGGCAATAACGCAACCAAACACTTGTAAAATTCCTAAATTAGTAGAAAATACAACTGATATCGATGTATAAGCGATTTCGAGGAATAATAAGAACAAAGTAATTCCCGTAAAGATAAACATCAACATTGTTTGGTAATCATTAAACCAATTAGTATGACGTTGAATTAATTTCACAGTATATTTTATTCCAGTATTATCCCCGCATGTATACGTACCTGGTAAGATATCTTTATATTCCATAATCAATGAGACATCACTATCCGATAACGAATGAAGTCTAGCACTAGTATGCCATTTACTTAATATGGTCGTCTTGGAAGTTTTATTATAAGTTAAAATATCATGATTGTCCAATTTCCATCTAATATACTTTGTCGGATTATCTATAATACACGGAATAATGATGGTATCATTACATGCTGTATATTCTATAGTCTTTGTAGATGTTATAACCACAAAAGTACAGAGGTATATCAACAATATTCTAACTCTTAACATTTTTATTTATTTAAAATGATACCTTTGTTATTTATTTTATTCTTATTTTGCTAACGGTATCGAATGGCATAAGTTTGAAACGAGTGAAGAAATAATTTCTACTTACTTAATAGATTATGTGGTAACGGGTGTTATTAATGGGGATGTATATACATTTTCAAATAATGAACTAAACAAAACTGGGTTAACTAATAACAATAATTATATCACAACATCTATAAAAGTAGAGGATAAGGATACATTAGTAGTATGCGGAACCAATAACGGAAATCCCAAATGTTGGAAAATAGACGGTTCATACTACCCAAAACATATAGGTAGAGGATACGATCATCAAAATAGCAAAGTAACGATAATCAGTCACAATGAATGTGTACTATCCGACATAAACATATAAAAAGAAGGAATTAAACGATGGAGAAGATTTGACGGACCATGTGGTTATGATTAAACGAGTTAAGTTTTTTAAGAAGCCTTAGAAGAGAGGCTATTGGGTATGAGAATCCGAAATATTAAACCAGACAACCCCATATAATTTTATAGCTAAGAATGCCGCGAAGAATGGAACTAATAAAAACGGAAATATTTGTAGCACAACGAATAACTCCCAAACTGCATTCATGTTACACTATATAACACTACTTCGGTTAGATGTTTTAGAAAAAATAAATATCACCGTACCGTTTTGTTGTATAAAAATAACAATTAACAATTATCAATTTTTTTCTTTAATATTTTACGTGGTTGACCATTCTTGGTGGTAAAATAATCTCTTAGTGTTGGAATGGAATGCTGTTTAATGTTTCCACACTCATCGTATATTTTGACGTATGCAGTCACATCGTTTACGCAATAGTCAGACTGTAGTTCTATCATGCTTCCTACGTTAGAAGGAGGAACAGTTTTAAAGTCTCTTGGTTTTAATCTATTGTCATTAGTTTTCATGAAATCCTTTGTTTTATCCACTTCACATTTTAAATAAATGTCCACTATACATTCTTCTGTTAATTTTACTAGATCATCATGAGTCATAGAATTCATAGGTTCCGTAGTCCATGGATCCAAACTAGCAAACTTCGCGTATACGGTATCGCGATTAGTGTATACACCAACTGTATGAAAATTAAGAAAACAGTTTAATAAATCTACAGAAATATTTAATCCTCCGTTTGATACAGATGCGCCATATTTATGGATTTCGGATTCACACGTTGTTTGTCTAAGGGGTTCGTCTAGTGTTGCTTCTACATAGACTTCGATTCCCATATATTCTTTATTGCCAGAATCACATACCGATTTATCATACGCTGGTTCACTTGTTTGAAAACTAAATGGTAGTAGATACATCAAAATAATAAATAATAAGTACATTCTGCAATATTGTTATCGTAATTGGAAAATTGGTATTCAAGTGAGCTGGATTATGTGAGTATTGGATTGTATATTTTATTTTATATTTTATATTTTATATTTTATTTTATATTTTATATTTTATTTTATATTTTGTAGTAAGAATAGAATGCTAATGTCAAGTTTATTCGAATAGATGTCTTATTAAAAAACATATATAATAAATAACAATGGCTGAATGGCATAAAATTATCGAGGATATCTCAAAAAATAATAAGTTCGAGGATGCCGCCATCGTTGATTACAAGACTACAAAGAATGTTCTAGCGGCTATTCCTAACAGAACATTTGCAAAGATTAATCCGGGTGAAGTTATTCCCCTCATCACTAATCATAATATTCTAAAACCTCTTATTGGTCAGAAATTTTGTATTGTATATACTAACTCTCTAATGGATGAGAACACGTATGCTATGGAGTTGCTTACTGGGTACGCCCCTGTATCTCCGATCGTTATAGCGAGAACTCATACCGCACTTATATTTTTGATGGGTAAGCCAACAACATCCAGACGTGATGTGTATAGAACATGTAGAGATCACGCTACCCGTGTACGTGCAACTGGTAATTAAAATAAAAAGTAATATTCATATGTAGTGTCAATTTTAAATGATGATGAAATGGATAATATCCATATTGACGATGTCAATAATGCCGGTATTGACATACAGCTCATCGATTTTTAGATTTCATTCAGAGGATATTGAATTATGTTATGGGAATTTGTATTTTGATAGGATCTATAATAATGTAGTAAATATAAAATATATTCCTGAGCATATTCCATATAGATATAATTTTATTAATCGTACGTTCTCCGTAGATGAACTAGATGATAATGTCTTTTTTACACATGGTTATTTTTTAAAACACAAATATGGTTGTTCACTTAATCCTAGTTTGATTGTCTCATTATCAGGAAACTTAAAATATAATGATATACAATGCTCAGTAAATGTATCGTGTCTCATTAAAAATTTGGCAACGAGTACATCTACTATATTAACATCTAAACATAAGACTTATTCTCTATATCGGTCCATGTGTATTGCTATAATAGGATACGATTCTATTATATGGTATAAATATATAAATGACAGGTATAATGACATCTATGATTTTACTGCAATATGTATGCTAATAGCGTCTACATTGATAGTGATCATATACGTGTTTAAAAAAATAAAAATGAACTCTTAATTATGTTATACTATTAGAAATGGATAAAATCAAAATTACGATTGATTCAAAAATTGGTAATGTTGTTACCATATCGTATAACTTGGAAAAGATAACTATTGATGTCACGCCAAAAAAGAAAAAAGAAAAGGATGTATTATTAGCGCAATCAGTTGCTGTCGAAGAGGCAAAAGATGTCAAGGTGGAAGAAAAAAATATTATCGATATTGAAGATGACGATGATATGGATATAGAAAACACGTAATACGATCTATAAAAATAAGTATTAAATACTTTTTATTTACGGTACTCTTGTAGTGGTGATACCACTAATCGATTATTTTTTTTAAAAAAATACTTATTCTGATTCTTCTAGCCATTTCCGTGTTCGTTCGAATGCCACATCGACGTCAAAAATAGGGGAGTAGTTGAAATCTAGTTCTGCATTGTTGGTACGCACCTCAAATGTAGTGTTGGATATCTTCAACGTATAGTTGTTGAGTATTGATGGTTTTCTAAATAGAATTCTCTTCATATCATTCTTGCACGCGTACATTTTTAGCATCCATCTTGGAATCCTAGATCCTTGTTCTATTCCCAATGGTTTCATCAATAGAAGATTAAACATATCGTAAGAACACGATGGAGAGTAATCGTAGCAAAAGTAAGCATTTCCTTTAATCGCAGATCCCGGATACTGGATATATTTTGCAGCCAACACGTGCATCCATGCAACATTTCCTACATATACCCGGCTATGCACAGCGTCATCATCGACTGTACGATACATAATGTTACCGTGTTGCTTACATTGCTCGTAAAAGACTTTCGTCAATTTGTCTCCTTCTCCGTAAATTCCAGTGGGTCTTAGGCAACAAGTATACAATTTTGCGCCATTCATGATTACGGAATTATTGGCTTTCATAACCAGTTGCTCGGCCATACGTTTACTTTTTGCGTATACATGTCCTGGTGATATATCATAAAGGGTATGCTCATGACCGATGAATGGATTACCGTGTTTATTTGGTCCTATTGCTTCCATGCTACTAGTATAGATCAAATACTTGATTCCTAGGTCCACACAAGCTGCCAATATAGTCTGTGTTCCATAATAGTTTACTTTCATGATTTCATTATCAGTGTATTTTCCAAATACATCCACTAGAGCAGCCGTATGAATAATCAGATTTACCCCATCTAGCGCTTCTCTCACCTTATCAAAGTCGTTTATATCACATTGTATATAGTTTATAACCTTAACTTTCGAGGTTATTGGTTGTGGATCTTCTACAATATCTATGACTCTTATTTCTTGAACATCATCTGCGCTAATTAAAAGTTTTACTATATACCTGCCTAGAAATCCGGCACCGCCAGTAACCGCGTACACGGCCATTGCTGCCACTCATAATATCAGACTACTTATTCTATTTTACTAAATAATGGCTGTTTGTATAATAGACCACGATAATATCAGAGGAGTTATTTACGTTGAACAAGTCCATGGAAAAGATAAAGTTTTAGGATCAGTTATTGGATTAAAATCCGGAACGTATAGTTTGATAATTCATCGTTACGGAGATATTAGTCGAGGATGTGATTCCATAGGCAGTCCAGAAATATTTATCGGTAACATCTTTGTAAACAGATATGGTGTAGCATATGTTTATTTAGATACAGATGTAAATATATCTACAATTATTGGAAAGGCGTTATCTATTTCAAAAAATGATCAGAGATTAGCGTGTGGAGTTATTGGTATTTCGTACATAAATGAAAAGATAATACATTTTCTTACAATTAACGAGAATGGCGTTTGATATATCAGTTAATGCGTCTAAAACAATAAATGCATTAGTTTACTTTTCTACTCAGCAAGATAAATTAGTCATACGTAATGAAGTTAATGATATACACTACACTGTCGAATTTGATAGGGACAAAGTAGTTGATACGTTTATTTCATATAATAGACATAATGACTCCATAGAGATAAGAGGGGTGCTTCCAGAGGAAACTAATATTGGTCGCGTGGTTAATACGCCGGTTAGTATGACTTACTTGTATAATAAGTATAGTTTTAAACCGATTTTAGCAGAATATATAAGACACAGAAATACTATATCCGGCAACATTTATTCGGCATTGATGACGCTAGATGATTTGGTTATTAAACAGTATGGAGACATTGATCTATTATTTAATGAGAAACTTAAAGTAGACTCCGATTCGGGACTATTTGACTTTGTCAACTTTGTAAAGGATATGATATGTTGTGATTCTAGAATAGTAGTAGCTCTATCTAGTCTAGTATCTAAACATTGGGAATTGACAAATAAAAAGTATAGGTGTATGGCATTAGCCGAACATATAGCTGATAGTATTCCAATATCTGAGCTATCTAGACTACGATACAATCTATGTAAGTATCTACGCGGACACACTGATAGCATAGAGGATGAATTTGATCATTTTGAAGACGATGATTTGTCTACATGTTCTGCCGTAACCGATAGGGAAACGGATGTATAATTTTTTTTATAGTATGAAGGATATGATGGATATGATGATATGATGGATATGATGATATGATGGATATGATGGATATGATGGATATGATGGATATGATAAAAAAATATAATTGTTGTATCCATTCCCATTCAAATCACCTTATATGATTCTGTAACACAATGAAGGAGTCTCATAGATATATAGAGGTCAGATACTGGTTTGATAAACTTTTTATTCCACATGAGCATGTTTGACTTATGGTTAGACACACATACTTTAACAAATCACTGAAAATTGGAGTTAGGTATTCCTCTCAGAATCAGTTGCCGTTCTGGAACATTAAATGTATTTTTTATGATATACTCCAACGCATTTATGTGGGTATACAACAAGTCATTAATAATGAGTATTTCCAAGAGTTTTAGTTGTCTAGTATTTAACAAGAGAAGAGATTTCAACAGACTGTTTATGAACTCGAATACCGCCTCATTGTCGCTTATATTGATGACATGACATCGAATTCCCAATATCAATCTCATCAGTGATGAGTAGCTCAATCTTGTTATCGGGATCCAATTTCTAAAGATGTCATTAAACCCTCGATCGTGAATGGATTTATCATCATCGTTTTTATGTTGGACATGAGCTTAGTCCGTTTGTCCACATCTATATACGATGATTTCTGAATTATTTCATATATCTCTCGTTAACTCCAGGAACTTGTCAGGGATCTAACTTTAATATGTTCTCGTCTAAGAGATGAAAATCTTTGGATGGTTGCATGTGACTTTTCTCTAAAGGATGATGTTACCCGATCCTCTCTTAAATGACTCCATCTTATCCTTGGACAAGATGGACAGTCTATTTTCCTTAGATGGTTTAATATTTTTTACCCATGATCTATAAAGGTAGACAGACCTAATCGTCTCGGATGACCATATATTATTTTCAGTTTTATTATACGCATAAATTGTAAAAAATATGTTAGGTTTACGAAAATGTCTCGTGGGGCATTAATCGTTTTTGAAGGATTGGACAAATCTGGAAAAACAACACAATGTATGAACATCATGGAATCTATACCGGCAAACACGATAAAATATCTTAACTTTCCTCAGCGATCCACAGTCACTGGAAAGATGATAGATGACTATCTAACTCGTAAAAAAACCTATAATGATCATATAGTTAATCTATTATTTTGTGCAAATAGATGGGAGTTTGCATCTTTTATACAAGAACAACTAGAACAGGGAATTACTTTAATAGTTGACAGATACGCGTTCTCTGGAGTAGCGTATGCCACCGCTAAAGGCGCGTCAATGACTCTCAGTAAGAGTTATGAATCTGGATTGCCTAAACCCGACTTAGTTATATTCTTGGAATCTGGTAGCAAAGAAATTAATAGAAACATCGGCGAGGAAATTTATGAAGATGTTGAATTCCAACAAAAGGTATTACAAGAATATAAAAAAATGATTGAAGAAGGAGATATTCATTGGCAAATTATTTCTTCTGAATTCGAGGAAGATGTAAAGAAGGAGTTGATTAAGAATATAGTTATAGAGGCTATACACACGGTTACTGGACCAGTGGGGCAACTGTGGATGTAATAAAATGAAATTACATTTTTATAAATAGATGTTAGTACAGTGTTATAAATGGATGAAGCATATTACTCTGGCAACTTGGAATCAGTACTCGGGGATACGTGTCCGATATGCATACCGAACTCGCATCAATATCTCAATTAGTTATTGCCAAGATAGAAACTATAGATAATGATTATTAAACAAGGACATTGTAAATTTTATCATGTGTAGATCAAACTTGGATAATCCATTTATCTCTTTCCTAGATACTGCATATACTATCATAGATCAAGAGATCTATCAGAACGAGTTGATTAATTCATTAGACGATAATGAAATTATCGATTGTATAGTTAACAAGTTTATGAGCTTTTATAAGGATAACCTAGAAAATATGGTAGATGCTATCATTACTCTAAAATATTATAATTAATAATCCAGATTTTAAAACTACGTATGTGGAAGTACTCGGTTCCAGAATAGCTGATATAGATATTAAACAAGTGATACGTAAGAATATAATACAATTGTCTAATGATCCGCGAACGATATTTGTGAAAATATTAAAAAAAAATACTTTTTTTATTAAATGACGTCTCTTCGCGAATTTAGAAAATTATGCTGTGATATATATCACGCATCAGGATATAAAGAAAAATCTAAATTAATTAGAGACTTTATAACAGATAGAGATGATACCGATACATATTTGATCATTAAGCTATTGCTTCCCGGATTAGACGATAGAATGTATAACATGAACGATAAACAAATTATAAAATTATATAGTATAATATTTAAACAATCTCAGGAAGATATGCTACAAGATTTAGGATACGGATATATAGGAGACACTATTAGGACTTTCTTCAAAGAGAACACGGAAATCCGTCCACGAGATAAAAGCATTTTAACTTTAGAAGAAGTGGATAGTTTTTTAACTACGTTATCATCAGTAACTAAAGAATCACATCAAATAAAATTATTGACTGATATAGCATCTGTTTGTACATGTAATGATTTAAAATGTGTAGTCATGCTTATTGATAAAGATCTAAAAATTAAAGCGGGTCCTCGGTACGTGCTTAACGCTATTAGTCCTCATGCCTATGATGTTTTTAGAAAATCTAATAACTTGAAAGAGATAATAGAAAATGCAGCTAAACAAAATCTAGACTCTATATCTATTTCTGTTATGACTCCAATTAATCCCATGTTAGCGGAATCATGTGATTCTGTCAATAAGGCGTTTAAAAAATTTCCATCAGGAATGTTTGCGGAAGTCAAATACGATGGTGAAAGAGTACAAGTTCATAAAAAAAATAACGAGTTTGCATTCTTTAGTAGAAACATGAAACCAGTACTCTCTCATAAAGTGGATTATCTCAAAGAATACATACCGAAAGCATTTAAAAAAGCTACGTCTATCGTATTGGATTCTGAAATTGTTCTTGTAGACGAACATAATGTACCGCTACCGTTTGGAAGTTTAGGTATACACAAAAAGAAAGAATATAAAAACTCTAACATGTGTTTGTTCGTATTTGACTGTTTATACTTTGATGGATTCGATATGACAGACATTCCATTGTATGAACGAAGATCTTTTCTCAAAGATGTTATGGTCGAAATACCCAATAGAATAGTATTCTCAGAGTTGACGAATATTAGTAACGAGTCTCAGTTAACTGATGTATTAGATGATGCACTAACGAGAAAATTAGAAGGATTGGTCTTAAAAGATATTAATGGCGTATACGAACCGGGAAAGAGAAGATGGTTAAAAATAAAGCGAGACTATTTGAACGAGGGTTCCATGGCAGATTCTGCCGATTTAGTAGTACTAGGTGCCTACTATGGTAAAGGAGGAAAGGGTGGTATCATGGCAGTCTTTCTAATGGGTTGTTACGACGATGAATCCGGTAAATGGAAGACGGTAACTAAATGTTCCGGTCACGATGATAATACGTTAAGGGTTTTGCAAGACCAATTAACGATGGTTAAAATTAACAAGGATCCCAAAAAAATTCCAGAGTGGTTGGTAGTTAATAAAATCTATATTCCCGATTTTGTAGTAGATGATCCGAAACAATCTCAGATATGGGAAATTTCAGGAGCAGAGTTTACATCTTCCAAGTCACATACAGCGAATGGAATATCGATTAGATTTCCTAGATTTACTAGGATTAGAGAAGATAAAACGTGGAAAGAATCTACTCATCTAAACGATTTAGTAAACTTGACTAAATCTCTTAATAGTTACATATAAACTGAAAAATAAAATAACACTATTTTAGTTGGTAGTCGCCATGGATGGTGTTATCGTATACTGTCTAAATGCGTTAGTAAAACATGGCGAGGAAATAAATCATATAAAAAATGATTTCATGATTAAACCATGTTGTGAAAGAGTTTGTGAAAAAGTCAAGAACGTTCACATCGGCGGACAATCTAAAAACAATACAGTGATTGCAGATTTGCCATATCTGGATAATGCTGTATCAGATGTATGCAAATCAATATATAAAAAGAATGTATCAAGAATATCCAGATTTGCTAATTTGATAAAAATAGATGACGATGACAAGACTCCTACCGGCGTATATAATTATTTTAAACCTAAAGATGCTATTCCTGTTATTATATCCATAGGAAAGGATAAAGATGTCTGTGAACTATTAATCTCATCTGATAAAGCGTGTGCGTGTATAAAGTTAAATTTATATAAAGTAGCCATTCTTCCCATGGATGTTTCCTTTTTTACCAAAGGAAATGCATCATTGATTATTCTCCTGTTTGATTTCTCTATCGATGCGGCACCTCTCTTAAGAAGTGTAACCGATAATAATGTTATTATATCTAGACACCAGCGCCTACATGACGAGCTTCCGAGTTCCAATTGGTTCAAGTTTTACATAAGTATAAAGTCCGACTATTGTTCTATATTATATATGGTTGTTGATGGATCTATGATGTATGCGATAGCTGATAATAGAACTCACGCAATTATTAGCAAAAATATATTAGACAATACTACGATTAACGATGAGTGTAGATGCTGTTATTCTGAACCACAGATTAGGATTCTTGATAGAGATGAGATGCTCAATGGATCATCGTGTTATATGAACAGACATTGTATTATGATGAATTTACCTGATGTAGGCGAATTTGGATCTAGTATGTTGGGGAAATATGAACCTGACATGATTAAGATTGCTCTTTCGGTGGCTGGTAATTTAATAAGAAATCGAGACTACATTCCCGGGAGACGAGGCTATAGCTACTACGTTTACGGTATAGCCTCTAGATAATTTTTTTTAAGCACGAAATAAAAACATAATTTTAAACAATCTATTTCATACTATTTTGTGTGCTCACCATGAACATAAAGATAGATATATTAGTATTTCTGGTGATAAATTTACGGCGACTGCTAGGAGGGAAAATGAAGAAAGAAAAAATATCTACCTCTCCAAAAAGAAAAACTACTGATGTTATCAAACCTGATTATCTTGAGTACAATGACTTGTTAGATAGAGATGAGATGTCTACTATTCTAGAGGAATATTAGGCCTTAGAATAAAATATGGACGACTCTTAACGAAATTAGAAAATTCGATAATGATGTTGAAGAACAATTCGGTACTATAGAAGAACTCAAGCAGAAGCTTAGATTAAATTCTGAAGAGGGAGCAGATAATTTTATAGATTATATAAAGGTACAAAAACAGGATATCATCAAACTTACTGTATACGATTGCATATATCTATGATAGGATTGTGTGCGTGCGTGGTAGATGTTTGGAGAAATGAGAAACTGTTTTCTAGATGGAAATATTGGTTACAAGCGATTAAACTGTTTATTGATGATCACATGCTTGATAAGATAAAATCTATTGTAGAATAGACTAGTGTATGTGGAAATGTCATAGAAAGTTAAAAGTTAATGAGAGCAAAAATATATAAGGTTGTATTCCATATTTGTTATTTTTTTCTGTAATAGTTAGAAAATACATTCGATGGTCTATCTACCAGATTATTATGTGTTATAAGGTACTTTTCTCATAATAAACTAGAGTATGAGTAAGATAGTGTTTTTCAAAAACATATAAATCTAAAATTGATGGATGAGATATACAGCTATTAATTTCGAAAATATATTTTAATCTGATAACTTTAAACATGGATTTTTGATGGTGGTTTAAGTTTAAAAAAGATTTTGTTATTGTAGTATGATAATATCAAAAAGATGGATATAAAGAATTTACTGACTACATGTACTATTTTACATTACTACATTGGCTACGGCATATATACCTATTTCGTCACTTCCACACGCTCCGGTAAACGGGTGTCATGTGACGAGGGAGAATCTTGATAAGAGGCATAATCAATGTTGTAATCCGATGTCCACCTGGAGAATTTGCCAAGGTCAGATGTAGAGTTGGTAGTGATAACACAAAATGTGAACACTGCCCACCTCATACATATACCGCAATCCCCAATTATTCTAATAGATGTCATCAATGTAGAAAATGCCCAACAGGATCATTTGATAAGGTAAAGTGTACCGGAACACAGAACAAATGTTCGTGTCATCCTGGTTGGTATACGCTACTGATTCTTCACAGACTGAAGATTGTCGAGATTTGTGTACCAAAAAAGGAGATGTCCATGCGGATACTTTGGTGGAATAGATGAAGGAAATCCTATTTGTAAATCGTGTTGTGTTGGTGAATATTGCGACTACCTACGTAATTATAGACTTGATCCATTTCCTCCATGCAAACTATCTATCTAAATGTAATTAATTATGATTTTGATGATAATGTTACCATACATTATATCACTACTTGGTTAGTGTGTATTATTTAGTATGGAAGACCTATTAATAATTACTTATCTTTTGACGATCTTGTTATAATTATAATATAAAAATACTTATGACATAGTAACTCATAATTGCTGACGCGATAAATTCGTAATAATCTGTTTTGTTCAAATTTTTATAAGGAATCTACAGGCATAAAAATAAAAATATAATCTATAATATACTCTTACAACGCCATCATGAATAGCAGTGAATTAATTGCTGTTATTATGGATTTAGAAATAGTGGACGATTTTGTGATATTAATATAGTTATTAATGATGAAAGGATAAACGCGCATAGACTCATCCTATCTGGAGCCTCCGAATATTTTTTCCATTCTGTTTTCCAATAATTTTATCGATTCTAATGAATACGAAGTTAATCTAAGTCATTTAGATTATCAAAGTGTTAACTATTTGATCGATTACATTTATGGGATACCTTTGAGCCTAACTAACAATAACGTGAAATATATTCTTTCAACCGCTGATTTTTTTACAAATTGGATCTGTCATTACTGAGTGCGAAAAATACATACTTAAAAATCTTTGTTTTAGAAACTGTATCGATTTCTACATATACGCTGATAAATATAATAACAAGAAAATAGAATTAGCATCGTTTAACACAATATTACGAAATATTTTGAGACTCATCAACAATGAAAACTTTAAATACTTAACAGAGGAATCAATGATAAAAATTTTAAGCGATGATATGTTATATATAAAAAATGAGGATTTCACCCCACTGATTCTCATTAAATGGTTAGAGAGTACACCAACCATGTACCGTCGAGTTACTTAGATGCCTCAGAATATCATTTCTTTCCCCACAAGTTATAAAATCACTTTATAGTCATCGACTGGTTAGTTCAATCTACGAATGTATAACATTCTTAAACAATATAGCATTCTTGGATAAATCATTTCCTAGATACCATATCATCGAGTTGATATCTATCGGTATAAGTAATTCACATGATAAGATTTCCATAAACTGCTACAATCATAAAAAAATTCATGGGAAATGATATCTTCACGTAGATATAGGTGTAGTTTCGCAGTGACCGTCCTGGATAATATTATCTATATGATGGGTGGATATGATCAGTCCCTGTATAGAAGTTCAAAGGTTATAGCGTACAATACATGTACTAATTCTTGGATATATGATATACCAGAGCTAAAAATATCATCGTTCTAATTGCGGAGGAGTTGCCAATGACGAATACATTTATTGTATAGGCGGTATACGCGATCAGGAGTCATCGTTGATATCTAGTATCGATAGATGGAAGCCATCAAAACCATATTGGCAGAAGTATGCTAAAATGTGCGAACCAAAATGTGATATGGGGTTGCGATTTTAAACGGATTAATATATGTCATAGGTGGAGTCGTTAAAGGTGACACACATATACCAACGCACTAGAGAGTTTATCAGAAGATGGATGGATGAATCATCAACGTCTTCCAATAAAAATGTCCAATATGTCGACGATTGTTCATGCTGGAAAGATTTATATATCTAGAGGTTACAACAATAGTAGTGTAGTTAATGTAATATCGAATCTAGTCCTTAGCTATAATCCGATATATGATGAATGGACCAAATTATCATCATTAAATATTCCTAGAATTAATCCTGCTCTATGGTCAGTGTATAATAAATTATATGTAGGAGGAGTAATATCTGATGATGTTCAAACTAATACATCTGAAACATACGATAAAGAAAAAGATTGTTGGACATTGGATAATGGTCACTTGTTACCACATAATTATATAATGTATAAATGCGAACCGTTTAAACATAGATATCCATTGGAAAAAACACAGTACACGAATGATTTTCTAAAGTATTTGGAAAGTTTTATAGGTAGTTGATAGAACAAAATACATAATTTTGTAAAAATAAATCACTTTTTATACTAATATGACACAATTACCAATACTTTTGTTACTAATATCATTAGTATACGCTACACCTTCTCCTCAGACATCTAAAAAAATAGGTGATGATGCAACTATATCATGTAGTCGAAATAATACAAATTACTACGTTGTTATGAGTGCTTGGTATAAGGAGCCCAATTCCATTATTCTCTTAGCTGCCAAAAGCGACGTCTTGTATTTTGATAATTATACCAAGGATAAAATATCTTACGACTCTCCATACGATGATCTAGTTACAACTATCACAATTAAATCATTGACTGCTGGAGATGCCGGTACTTATATATGTGCATTCTTTATGACATCGACTACAAATGATACTGATAAAGTAGATTATGAAGAATACTCCATAGAGTTGATTGTAAATACAGATAGTGAATCGACTATAGACATAATACTATCTGGATCTACACCAGAAACTATTTCTGAGAAACCAGAGGATATAGATAATTCTAATTGCTCGTCTGTATTCGAAATCACGACTCCGGAACCAATTACTGATAATGTAGACGACCATACAGACACCGTCACATACACTAGTGATAGCATTAATACAGTAAATGCATCATCTGGAGAATCCACAACAGACGAGATTCCGGAACCAATTACTGATAAAGAAGAAGATCATACAGTAACAGACACTGTCTCATACACTACAGTAAGTACATCATCTGGAATTGTCACTACTAAATCAACCACCGATGATGCGGATCTTTATGATACATACAATGATAATGATACAGTACCGCCAACTACTGTAGGTGGTAGTACAACCTCTATTAGCAATTATAAAACCAAGGACTTCGTAGAAATATTTGGTATTACCACATTAATTATATTGTCAGCAGTGGCGATTTTCTGTATTACGTATTATATATGTAATAAACACCCACGTAAATACAAAACAGAGAACAAAGTCTAGATTTTTGACTTACATAAATATCTGGGATAATAAAATCTATCATATTGAGAGGACCATCTGGTTCAGGAAAGACAGCCATAACCAAAAGACTGTTAAAAGACTATGGGAATATATTTGGATTTGTGGTGTCCCATACCACTAGATTTCCTCGTCCTATGGAACGAGAAGGTGTTGATTACCTTACGTTAACAGAGAGGCCATCTGGAAGGGAATAGCCGCCGGAAACTTTCTAGAACATACTGAGTTTTTAGGAAATATTTACGGAACTTCTAAAACAGCTGTAAATACAGCGGTTATTAATAATCGTATTTGCGCGATGGATTTAAACATCAACGGTGTTAGAAGTCTTAAAAATACTTACCTAATGCATTACTTGGGTATATAAGACCTACCTCTCTTAAAATGGTTGAGACCAATCTTCGTCGTAGAAACACTGAAGCGGACGACGAATCTCATCGTCGCGTGATGTTGGCAAAAAACGGATATGGATGAGGTCAACGAAGCAGGTCTATTCGACACTATTATTATTGAAGATGATGTGAATTTAGCATATAGTAAGTGTTAATTCAGATACTACAGGACCGTATTAGAATGTATTTTAACACTAATTAGAGACTTAAGATTTGACTTAAAACTTGATAATTAATAATATAACTCGTTTTTATATGTGGCTATTTCAACGTCTAATGTATTAGTTAAATATTAAAACTTACCACGTAAAACTTAAAATTTAAAATGGTATTTCATTGACAGATCATACATTATGAAGTTTCAAGGACTTGTGTTAATTGACAATTGCAAAAATCAATGGGTCGTTGGACCATTAATAGGAAAAGGTGGATTCGGTAGTATTTATACTACTAATGACAATAATTATGTAGTAAAAATAGAGCCCAAAGCTAACGGATCATTATTTACCGAACAGGCATTTTATACTAGAGTACTTAAACCATCCGTTATCGAAGAATGGAAAAAATCTCACAATATAAAGCACGTAGGTCTTATCACATGCAAGGCATTTGGTTTATACAAATCCATTAATGTGGAATATCGATTCTTGGTAATAAATAGATTAGGTGCAGATCTAGATGCGGTGATCAGAGCCAATAATAATAGACTACCAGAAAGGTCGGTGATGTTGATCGGAATCGAAATCTTAAATACCATACAATTTATGCACGAGCAAGGATATTCTCACGGAGATATTAAAGCGAGTAATATAGTCTTGGATCAAATAGATAAGAATAAATTATATCTAGTGGATTACGGATTGGTTTCTAAATTCATGTCTAACGGCGAACATGTTCCATTTATAAGAAATCCAAATAAAATGGATAACGGTACTCTAGAATTTACACCTATAGATTCGCATAAAGGATACGTTGTATCTAGACGTGGTGATCTAGAAACACTTGGATATTGTATGATTAGATGGTTGGGAGGTATCTTGCCATGGACTAAGATATCTGAAACAAAGAATTCTGCATTAGTAAGTGCCGCAAAACAGAAATATGTTAACAATACTGCGACTTTGTTAATGACCAGTTTGCAATATGCACCTAGAGAATTGCTGCAATATATTACCATGGTAAACTCTTTGACATATTTTGAGGAACCCAATTACGACGAGTTTCGTCGAGTATTAATGAATGGAGTTATGAAAAATTTTTGTTGATAAAAAAATTAAAAAAATAACTTAGTTATTATCACTCTCGTGAGTACAATAGAAACATGGCGATGTTTTACGCACACGCTTTCGGTGGGTACGACGAGAACCTTCATGCATTTCCTGGAATATCATCGACGGTTGCCAATGATGTCAGGAAATATTCTGTTGTGTCAGTTTATAATAAAAAGTATAACATTGTAAAAAACAAATATATGTGGTGTAACAGTCAAGTGAACAAGAGATATATTGGAGCACTACTGCCTATGTTTGAATGCAATGAATATCTACAAATTGGAGATCCAATCCATGATCTAGAAGGAAATCAAATCTCTATTGTCACATATCGCCACAAAAACTACTATGCTCTAAGTGGAATTGGGTACGAGAGTCTAGACTTGTGTTTGGAAGGAGTAGGGATTCATCATCACGTACTTGAAACAGGAAACGCGGTATATGGAAAAGTTCAACATGAGTATTCTACTATCAAAGAGAAGGCCAAAGAAATGAATGCACTCAAACCAGGACCTATCATCGATTACCACGTCTGGATAGGAGATTGTGTCTGCCAAGTTACTACTGTAGACGTGCATGGAAAGGAAATTATGAGAATGAGATTCAAAAGGGGTGCGGTGCTTCCGATTCCAAATCTGGTAAAAGTTAAAGTTGGGGAGGAAAATGATACAATAAATCTTTCCACTTCCATATCAGCTCTCCTAAATTCCGGTGGCGGCACCATCGAGGTAACATCTAAGGAAGAACGTGTAGATTATGTACTCATGAAACGTTTGGAATCTATACATCATCTGTGGTCTGTAGTGTATGATCATCTTAATGTTGTGAATGGCGAAGAACGATGTTATGTACATATGCATTCATCTCATCAAAGTCCTATGCTGAGTACTGTAAAAACAAATTTGTACATGAAGACTATGGGAGCATGTCTTCAAATGGACTCCATGGAAGCTCTAGAGTATCTTAGTGAACTGAAGGAATCAGGTGGGCGGAGTCCCAGACCAGAATTGCAGAAATTTGAATATCCAGATGGAGTGAAAGACACTGAATCAATTGAGAGATTGGCAGAGGAGTTCTTCAATAGATCAGAACTTCAGGCCGGTGAATCAGTCAAATTTGGTAATTCTATTAATGTTAAACATACATCTGTTTCAGCTAAGCAACTAAGAACACGTATACGACAGCAGCTTCCTTCTATACTCTCATCTTTTGCCAACACAAAGGGTGGATATTTGTTCATTGGAGTTGATAATAATACACACAAAGTAATTGGATTCACGGTGGGTCATGACTACCTCAAACTGGTAGAGAGTGATATAGAAAAGTATATCCAAAAACTTCCTGTTGTGCATTTCTGCAAGAAAAAAGAGGACATCAAGTACGCATGTAGATTCATCAAGGTGTATAAACCTGGTGATGAGACTACCTCGACATATGTGTGCGCAATCAAAGTGGAAAGATGCTGCTGTGCTGTGTTTGCGGATTGGCCAGAATCATGGTACATGGATACTAGTGGTAGTATGAAGAAGTATTCTCCAGATGAATGGGTGTCACATATAAAATTTTAATTAGGGTAAGGTAAAACTATATATAATAACTAACAATTTGTGTATCATATAGACAATTAATTAGGTAACTGTTATCTCTTTTTAACTAACTAACTAACTAACTAACTCTTATATACTATTAATAATACATCTATTAATCATTGATTAGCTTATTGCTTTAATTGTTTTTGTAAACTAACACTGTTCATTGAAAAGGGATAACATGTTACAGAATATAAATTATATATGGATTTTTTTAAAAAGGAAATACTTGACTGGAGTATATATTTATTTCTTCATTACATAACACGTCTGTGTTCTAATTCTTCCAATTCTTCCACATCTCATATAATACAGGAATATAATCTTGTTCGAAAATATGAGAAAGTGGATAAAACAATAGTTGATTTTTTATCTAGGTGGCCAAATTTATTCCATATTTTAGAATATGGGGAAAATATTCTACATATTTATTTTATAGATGCTGCTAATACGAATATTATGATTTTTTTTCTAGATAGAGTATTAAATATTAATAAGAACCGTGGGTCATTTATACATAATCTCGGGTTATCATCCATTAATATAAAAGAATATGTATATCAATTAGTTAATAATGATCATCTAGATAATAGTATAAGACTAATGCTTGAAAATGGACGTAGAACAAGACATTTTTTGTCTTATATATTGGATACAGTTAATATCTATATAAGTATTTTAATAAATCATAGATTTTATATAGATGCCGAAGACAGTTACGGTTGTACATTATTACATAGATGTATATATAACTATAAGAAATCAGAATCAGAATCATATAATGAATTAATTAAGATATTGTTAAATAATGGATCAGATGTAGATAAAAAAGATACGTACGGAAACACACCGTTTATCCTATTATGTAAACACGATATCGACAACGCGGAATTGTTTGAGATATGTTTAGAGAATGCTAATATAGACTCTGTAGACTTTAATGGATATACACCTCTTCATTATGTCTCATGTCGTAATAAATATGATTTTGTAAAGTTATTAATTTCTAAAGGAGCAAATGTTAATGCACGTAATAGATTCGGAACTACTCCATTTTATTGTGGAATTATACACGGTATCTCGCTTATAAAACTATATTTGGAATCAGACACAGAGTTAGAAATAGATAATGAACATATAGTTCGTCATTTAATAATTTTTGATGCTGTTGAATCTTTAGATTATCTATTGTCCAGAGGAGTTATTGATATTAACTATCGTACTATATACAACGAAACATCTATTTACGACGCTGTCAGTTATAATGCGTATAATACGTTAGTCTATCTATTAAACAGAAATGGTGATTTTGAGACGATTACTACTAGTGGATGTACATGTATTTCGGAAGCAGTCGCGAACAACAACAAAATAATAATGGATATACTATTGTCTAAACGACCATCTTTGAAAATTATGATACCATCTATGATAGCAATTACTAAACATAAACAACATAATGCAGATTTATTGAAAATGTGTATAAAATATACTGCGTGTATGACCGATTATGATACTCTTATAGATGTACAATCGCTACATCAATATAAATGGTATATTTTAAAATGTTTTGATGAAATAGATATCATGAAGAGATGTTATATAAAAAATAAAACTGTATTCCAATTAGTTTTTTGTATCAAAGACATTAATACTTTAATGAGATACGGTAGACATCCTTCTTTCGTGAAATGTAATATTCTTGACGTATACGGAAGTTGTGTACGTAATATCATAGCATCTATTAGATATCGTCAGAGATTAATTAGTCTATTATCCAAGAAGCTGGATGCTGGAGATAAATGGTCGTGTTTTCCTAACGAAATAAAATATAAAATATTGGAAAACTTTAACGATAACGAACTGACCACATATCTGAAAATCTTATAAACACTATTAAAATATAAAATCTAAGTAGGATAAAATCACACTACATCATTGTTTCCTTTTAGTGCTCGACAGTGTATACTATTTTTAACACTCATAAATAAAAATGAAAACGATTTCCGTTGTTACGTTGTTATGCGTACTACCTGCTGTTGTTTATTCAACATGTACTGTACCCACTATGAATAACGCTAAATTAACGTCTACCGAAACATCGTTTAATGATAAACAGAAAGTTACGTTTACATGTGATTCAGGATATCATTCTTTGGATCCAAATGCTGTCTGTGAAACAGATAAATGGAAATACGAAAATCCATGCAAGAAAATGTGCACAGTTTCTGATTATGTCTCTGAACTATATGATAAGCCATTATACGAAGTGAATTCCACCATGACACTAAGTTGCAACGGTGAAACAAAATATTTTCGTTGTGAAGAAAAAAATGGAAATACTTCTTGGAATGATACTGTCACGTGTCCTAATGCGGAATGTCAACCTCTTCAATTAGAACACGGATCGTGTCAACCAGTTAAAGAAAAATACTCATTTGGGGAATATATGACTATCAACTGTGATGTTGGATATGAGGTTATTGGTGTTTCGTATATAAGTTGTACGGCTAATTCTTGGAATGTTATTCCATCATGTCAACAAAAATGTGATATACCGTCCCTATCTAATGGATTAATTTCCGGATCTACATTTTCTATCGGTGGCGTTATACATCTTAGTTGTAAAAGTGGTTTTACACTAACGGGGTCTCCATCATCCACATGTATCGACGGTAAATGGAATCCCATACTCCCAACATGTGTACGATCTAACGAAGAATTTGATCCAGTGGATGATGGTCCCGACGATGAGACAGATCTGAGCAAACTCTCGAAAGACGTTGTACAATATGAACAAGAAATAGAATCGTTAGAAGCAACTTATCATATAATCATAATGGCGTTGACAATTATGGGTGTCATATTTCTAATCTCCATTATAGTATTAGTTTGTTCCTGTGACAAAAATAATGACCAATATAAGTTCCATAAATTGCTACCGTGAATATAAATCCGTTAAAATAATTAATAATTAATAATTAATAACGAACAAGTATCAAAAGATTAAAGAATTAGCTAGAATCAATTAGATGTCTTCTTCAGTGGATGTTGATATCTACGATGCCGTTAGAGCATTTTTACTCAGGCACTATTATGACAAGAGATTTATTGTGTATGGAAGAAGTAACACCATATTACATAATATATACAGGCTATTTACAAGATGCACCGTTATACCGTTCGATGATATAGTACGTACTATGCCAAATGAATCACGTGTTAAACAATGGGTGATGGATACACTTAATGGTATAATGATGAATGAATTCGATACTGTATGTGTGGGTACCGGACTACGATTCATGGAAATGTTTTTCGATTACAATAAAAATAATCCCAAAAATAGCATCAACAATCAAATAATGTATGATATAATTAATAGCGTAGCCATAATTCTAGCTAATGAGAGATATAGAAGCGCGTTTAACGACGATAGAATATACATCCGTAGAACTATGATGGACAAATTGTACGAATACGCATCTCTAACTACTATTGGTACGATCACTGGAGGTGTTTGTTATTTTATCTGTTGATGCATCTAGTTAGTTTGTATAAATAATTATTTCGATATACTAGTTAAAATTTTAAGATTTTAAATGTATAAAAAACTAATAACGTTTTTATTTGTAATAGGTGCAGTTGCATCCTATTCGAATAATGAGTACACTCCGTTTAATAAACTGAGTGTAAAACTCTATATAGATGGAGTAGATAATATAGAAAATTCATATACTGATGATAATAATGAATTGGTGTTAAATTTTAAAGAGTACACAATTTCTATTATTACAGAGTCATGTGACGTCGGATTTGATTCCATAGATATAGATGTTATAAACGACTATAAAATTATTGATATGTATACCATTGACTCGTCTACTATTCAACGCAGAGGACATACGTGTAGAATATCTACCAAATTATCATGCCATTATGATAAGTACCCTTATATCCACAAATATGAGGGTGATGAACGACAATATTCTATTACCGCAGAGGGAAAATGCTATAAAGGAATAAAATATGAAATAAGTATGATGAACGATGATACTCTATTGAGAAAACATACTCTTAAAATTGGATTTACTTATATATTCGATCGTCATGGGCATAGTAATACATATTATTCAAAATATGATTTTTAAAAATTTAAAATATATTATCACTTCAGTGACAGTAGTCAAATAACAAACAACACCATGAGATATATTATAATTCTCGCAGTTTTGTTCATTAATAGTATACATGCTAAAATAACTAGTTATAAGTTTGAATCCGTCAATTTTGATTCCAAAATTGAATGGACTGGGGATGGTCTATACAATATATCCCTTAAAAATTATGGCATCAAGACGTGGCAAACAATGTATACAAATGTACCAGAAGGAACATACGACATATCCGGATTTCCAAAGAATGATTTCGTATCTTTCTGGGTTAAATTTGAACAAGGCGACTATAAAGTGGAAGAGTATTGTACGGGACTATGTGTCGAAGTAAAAATTGGACCACCAACTGTAATATTGACTGAATATGACGATCATATCAATTTGTTCATCGAGCATCCGTATGCTACTAGAGGTAGCAAGAAGATTCCTATTTACAAACGCGGTGACATGTGTGATATCTACTTGTTGTATACGGCTAACTTCACATTCGGAGATTCTGAAGAACCAGTAACATATGATATCGATGACTACGATTGCACGTCTACAGGTTGCAGTATAGACTTTGCCACAACAGAAAAAGTGTGTGTGACAGCACAGGGAGCCACAGAAGGGTTTCTCGAAAAAATTACTCCATGGAGTTCGGAAGTATGTCTGACACCTAAAAAGAATGTATATACGTGCGCAATTAGATCTAAAGAAGATGTTCCCAATTTCAAGGACAAAATAGCCAGAGTTATCACGAGAAAATTTAATAAACAGTCTCAATCTTATTTGACTAAATTTCTCGGTAGCACATCGAATGATGTTACAACTTTTTTTAGCATTCTTGACTAAATATTCATAACTAATTTTTATTAATGATACAAAAATGAAATAAACTGTATATTATACACTGGTTAACGCCCTTGGCTCTAACCATTTTCAAGATGAGGTCCCTGATTATAGTCCTTCTGTTCCCCTCTATCATCTACTCAATGTCTATTAGACGATGCGAGAAGACTGAAGAGGAAACATGGGGATTAAAAATAGGGTTGTGTATAATTGCCAAAGATTTCTATCCCGAAAGAACTGATTGCAGTGTTCATCGCCCAACTGCAAGTGGAGGATTGATAACTGAAGGCAATGGATTCAGAGTAGTTATATATGATCAATGTACAGAACCCCATGACTTTATTATCACCGATACTCAACAAACACGTCTTGGATCATCTCATACATATATTAAATTCAGTAACATGAATACAGGTGTCCCATCTAGTATTCCAAAATGTTCCAGAACTCTCTGTATTTCTGTATATTGTGATCAAGAGGCGGGAGACATAAAATTTGAGGAGTATACTCAAGAATCAAGTGATATCAGTATTAGAGTTAAGTATGATTCATCATGTATTGATTATCTGGGTATTAATCAAAGTTTCATGAATGAATGTATTCGAAGAATTACAACATGGGATAGAGAATCATGCGTCAGAATTGATACACAGACTATAAATAAATATCTTAAGTCTTGCACCAACACAAAATTCGACCGTAATGTCTACAAAAGGTACATACTGAAGAGTAAAGCACTCCATGCTAAAACAGAGTTGTAATAGATATAAAATACTTTTTATAATAATTAGGCTAGAAAAATCTCACTCACATGTAATCTTAAAAAAATGATATGATAGTTCTTACAAGTAGCGATTGAGTTTTAAATGGATTCTATTAATTACCGGGGAACTTAACAATTCGTTCTGATCTACAGACATTGGTTAATAAATCATCTTATTTTGCCAATATATTAAAATGTGGAAACTCCACTAATAATATTACATTGTGCGACTTTCAAGATGATGTGATATATAGGGTTATACAGTTTTAACAATTATATAATAGAGATAGAAAGTACAAAAGATGTAGAATCAATGATATGGCACGCTAAACAGTTGGGTGTGGAATCATTGCTAAAAGAATGTCAAAATTATTTGCTTAGAATATTACGTATATAATTGTTTAGAAATTTATAGAATAACTAATATTAATACATTATCGTATATCTACAACGATATAAGAAACTTCATATTGGATAATATTACTATTAATATATAAGGATCCAGATTTTATATATTTGCCTAAATACATTATTATAGATTTACTAGGACAATCACCTAAATGTTTTTAACGAAGATAATGTGGTAAAGATTATATACACTTATATATCTTCCGATATCTACAAGGATATTCCATATCATCATTGTGTAAACTAAATAACGTTTTCTATGGCATTTAATAAGGACATTGGATATGTGGAAAAGTGATGTATGGAAGTTAGTACATTATCAACTTCTCCTTATTGATTGAAAATGAAAATATAAATAGTTTTTATGTATAGCGGTATCTACCCTATAGTTTTATTGCTTACTACTAACATGGATTCAGATACAGATACAGATACAGATACAGATACAGATACAGATACAGATACAGATGTAGAAGATATCATGAATGAAATAGATAGAGAGAAAGAAGAAATACTAAAAAATGTAGAAATTGAAAATAATAAAAACATTAACAAGAATCATCCCAGTGAATATATTAGAGAAGCACTTGTTATTAATACCAGTAGTAATAGTGATTCCATTGATAAAGAAGTTATAGAATATATCAGTCACGATGTAGGAATATAGATCATATCTACTAATTTTTATAATCGATACAAAACATAAAAACAACTCGTTATTACATAGCAGGTATGGAATCCTTCAAGTATTGTTTTGATAACGATGGTAAGAAATGGATTATCGGAAATACTTTATATTCTGGTAATTCAATACTCTATAAGGTCAGAAAAAATTTCACTAGTTCGTTCTACAATTACGTAATGAAGATAGATCATAAATCACACAAGCCATTGTTGTCCGAAATACGATTCTATATATCTGTATTGGATCCTTTGACTATCAACAACTGGACACGGGAACGTGGTATAAAGTATTTGGCTATTCCAGATCTGTATGGAATTGGAGAAACCGATGATTATATGTTCTTCATTATAAAGAATTTGGGAAGAGTATTCGCCCCAAAGGATAGTGAATCAGTTTTCGAAGCATGTGTCACTATGATAAACACGTTAGAGTTTATACACTCTCAAGGATTTACTCATGGAAAAATAGAACCGATGAATATACTGATTAGAAATAAACGTATTTCACTAATTGACTATTCTAGAACTAACAAACTATACAAAAGTGGAACACATATAGATTACAACGAGGACATGATAACTTCAGGAAATATCAATTATATGTGTGTAGACAATCATCTTGGAGCAACAGTTTCAAGACGAGGAGATTTAGAAATGTTGGGATATTGCATGATAGAATGGTTCGGTGGTAAACTTCCATGGAAAAACGAAAGTAGTATAAAAGTAATAAAACAAAAAAAAGAATATAAACAATTTATAGCTACTTTTTTTGAGGACTGTTTTCCTGAAGGAAATGAACCTCTGGAATTAGTTAGATATATAGAATTAGTATACATGTTAGATTATTCTCAAACTCCTAATTATGACAGACTACGTAGACTGTTTATACAAGATTGAAATTATATTCTTTTTTTTATAGAGTGTGGGGGTAGTGTTACGGATATCTAATATTAATATTAGACTATCTCTATCGCGCTACACGACCAATATCGATTACTATGGATATCTTCAGGGAAATCGCATCTTCTATGAAAGGAAAGAATGTATTCATTTCTCCAGCGTCAATCTCGTCAGTATTGACAATACTGTATTATGGAGCTAATGGATCCACTGCTGAACAGCTATCAAAATATGTAGAAAAGGAGGAGAACATGGATAAGGTTAGCGCTCAGAATATCTCATTCAAATCCATGAATAAAGTATATGGGCGATATTCTGCCGTGTTTAAAGATTCCTTTTTGGGAAAAATTGGCGATAAGTTTCAAACTGTTGACTTCACTGATTGTCGCACTATAGATGCAATCAATAAGTGTGTAGATATCTTTACTGAGGGAAAAATCAATCCACTATTGGATGAACCATTGTCTCCTGATACCTGTCTCCTAGCAATTAGTGCCGTATACTTTAAAGCAAAATGGTTGATGCCATTCGAAAAGGAATTTACCAGTGATTATCCCTTTTACGTATCTCCAACGGAAATGGTAGATGTAAGTATGATGTCTATTTACGGCGAGCCATTTAATCACGCATCTGTAAAAGAATCATTCGGTAACTTTTCAATCATAGAACTGCCATATGTTGGAGATACTAGTATGATGGTCATTCTTCCAAACAAGATTGATGGATTAGAATCCATAGAACAAAATCTAACAGATACAAATTTTAAGAAATGGTGTAACTCTCTGAAAGCTACGTTTATCGATGTGCACATTCCTAAGTTTAAGGTAATAGGTTCGTATAATCTTGTGGATACGCTAATAAAGTTGGGACTGACAGATGTGTTCTATTCAACTGGTGATTATATCAATATGTGTAATTCAGATGTGAGTGTTGACGCTATGATTCACAAAACGTATATAGATGTCAATGAAGAGTATACAGAAGCAGCTGCAGCAACTTCTGTACTAGTGGCAGACTGTGCATCAACAGTTACAAATGAGTTCTGTGCAGATCATCCGTTCATCTATGTGATTAGACATGTCGATGGTAAAATTCTTTTCGTTGGTAGATATTGCTCTCCAACAACTAATTAAGCACATTCTTAATATTAGAATATTATATAGTTAAGATTTTTACTAACAGGTTAACATTTTTTTTTAAAAATAGAAAAAACATGTGGTATTAGTGCAGGTCGTTATTCTTCCAATTGCAATTGGTAAGATGACGGCCAACTTTAGTACCCACGTCTTTTCACCACAACACTGTGGATGTGACAGACTGACCAGTATTGATGACGTCAGACAATGTTTGACTGAATATATTTATTGGTCGTCGTATGCATACCGCAACAGGCAATGCGCTGGACAACTGTATGACACACTCCTCTCTTTTAAAGATGATGCGGAATCAGTGTTCATCGACGTTCGTGAGCTGGTAAAAAATATGCCGTGGGATAATGTTAAGGATTGTACAGAGATCATCCGTTGTTATATACCGGATGAGCAAAAAACCATCAGAGAGATTTCGGCCATCATTGGACTTTGTGCATATGCTGCTACTTACTGGGGAGGTGAAGACCATCCCACTAGTAACAGTCTGAACGCATTGTTTGTGATGCTTGAGATGCTCAATTACATGGATTATACCATCATATTCTGGCGTATGAATTGATGAGTTACAGCTTGACATTTCTTCTTTCCTCCCTCTTCTTCTACCTTTCCCAGAAACAAACTTTTTTTACCCACTATAAAATAAAATGAGTATACTACCTGTTATATTTCTTCCTATATTTTTTTATTCTCCATTCGTTCAGACTTTTAACGTGCCTGAATGTATCGACAAAGGGCAATATTTTGCATCATTCATGGAGTTAGAAAACGAGCCAGTAATCTTACCATGTCCTCAAATAAATACGCTATCATCCGGATATAATATATTAGATATTTTATGGGAAAAACGAGGAGCGGATAATGATAGAATTATACAGATAGATAATGGTAGCAATATGCTAATTCTGAACCCGACACAATCAGACTCTGGTATTTATATATACATTACCACGAACGAAACCTACTGTGACATGATGTCGTTAAATTTGACAATCGTGTCTGTCTCAGAATCAAATATAGATCTTATCTCGTATCCACAAATAGTAAATGAGAGATCTACTGGTAAAATGGTATGTCCCAATATTAATGCATTTATTTCTAGTAACGTAAACACAGAATTATATGGAGCGGACATCGACGCCTTAGAAATAAGAGACTTAAACAACGGACACCTGGAATTATTACCATAGAAGATGTTAGAAAAAATGATGCTGGTTATTATACATGTGTTTTAGANNNNNNNNNNATATATATATATATATATATATATATATATATATATATATATATATATATATATATATATATATATATATATATATATATGGGCAAAACATATAACGTAACCAGAATTATAAAATTAGAGGTACGGGATAGAATAATACCTCCTACTATGAAATTACCAGAAGGAGTAGTAACTTCAATAGGTAGTAATTTGACTATTGCATGTAGAGTATCGTTGAGACTTCCCACAACGGACACCGACGTCTTTTGGATAAGTAATGGTATATGTATTACGAAGAAGAAGACGAGGACGGAGACGGTAGAATAAGTGTAGCAAATAAAATCTATATGACCGATAAGAGACGTGTTATTACATCCTGGTTAAACATTAATCCTGTCAAGGAAGAAGATGCTACAACGTTTACGTGTATGGCGTTTACTATTCCTAGCATCAGCAAAACAGTTACTGTTAGTAAACGTGAATGTATGTTGTTACATTTCCATATCAATTGAGTTTATAAGAATTTTTTATACATTATCTTCCAACAAACAATTGACGAACGTATTGCTATGATTAACTCCCACAATACTATATATATTATTAATCATTAACTTGCAGACTATACCTAGTAGTGCTATTTTGACATACTCATGTTCTTGTGTAATCGCAGTATCTATATTATTAAAGTACGTAAATCTAGCTATAGTTTTATTATTTAATTTTAGATAATATACTGTCTCCGTATTTTTAAAAAATTACCACATCCTTTATTAAATCATGAATGGGAATTTCTGTGTCATCGTTAGTATATTGTGAACAACAAGAGCAGATATCTATAGGAAAGGGTGGAATGCGATACATTGATCTATGTAGTTTTAAAACATACGCGAACTTTGAAGAATTTATATAAATCATCTCACGAGATATTGCTCTCTGTCATATTCATACACCTGTATAAACTTTCTAGACATCTTACAATGTGTTATTTTATGATCATATTTACATATTTACTGGTATATCAAAGATGTTAGATTAGTTAATGGGAATCGTCTATAATAATGAATATTAAACAATTATAGGAGGAGTTTATACCTACAAAAACATCATAAAAATGAGTCATCGTCCGATTTATGTTTTAAATATACTAACATTACTACCTTCAGAAATTATATACGAAATATTATACATGCTGACAATTAACGATCTTTATAATATATAGTATCCACCTACCAAAGTATAATTGTATTTTTCTCATGTGATGTGTGTAAAAAACTGATATTATATAATTATCTTAGTACCTATGATGAAGATGAAGATGAAGATGAAGATGATGGTCCGTATATATTTTGTATCATTATCGTTATTGCTATTCCATAGTTACGCCATAGACATCGAAAATGAAATCACCGAATTCTTCAATAAAATGAGAGATACTCTACCAGCTAAAGACTCTAAATGGTTGAATCCAGTATGTATGTTTGGAGGCACAATGAATGATATGGCCGCTCTAGGAGAGCCATTCAGTGCAAAGTGTCCTCCTATTGAAGACAGTCTTTTATCGCATAGATATAAAGACTATGTGGTTAAATGGGAAAGGCTAGAAAAGAATAGACGGCGACAGGTTTCTAATAAACGTGTTAAACATGGTGATTTATGGATAGCCAACTATACATCTAAATTCAGTAACCGTAGGTATTTATGTACCGTAACCACAAAGAATGGTGACTGTGTTCAGGGTGTAGTTAGATCTCATGTGTGGAAACCTTCTTCATGCATTCCAAAAACATATGAACTAGGTACTTATGATAAGTATGGCATAGACTTATACTGTGGAATTCTTTATGCGAACCATTATAATAATATAACTTGGTATAAAGATAATAAGGAAATTAATATCGACGATTTTAAGTATTCACAAGCGGGAAAGGAATTAATTATTCATAATCCAGAGTTAGAAGATAGTGGAAGATACGACTGTTACGTTCATTACGACGACGTTAGAATCAAGAATGATATCGTAGTATCAAGATGTAAAATACTTACGGTTATACCGTCACAAGACCACAGGTTTAAACTAATACTAGATCCGAAAATCAACGTAACGATAGGAGAACCTGCCAATATAACATGCAGTGCTGTGTCAACGTCATTATTTGTCGACGATGTACTGATTGAATGGGAAAATCCATCCGGATGGATTATAGGATTAGATTTTGGTGTATACTCTATTTTAACTAGTAGAGGCGGTATCACCGAGGCGACTTTGTATTTTGAAAATGTTACTGAAGAATATATAGGCAATACATATACATGTCGTGGACACAACTATTATTTTGATAAAACTCTTACAACTACAGTAGTATTGGAGTAAATACACAATGCATTTTTATATACATTACTGAATTATTATTATTAATTATATCGTATTTGTGCTATAGAATGGATGAAGATACGCGACTATCTAGGTATTTGTATCTCACCGATAGAGAACATATAAATGTAGACTCTATTAAACAGTTGTGTAAAATATCAGATCCTAATGCATGTTATAGATGTGGATGTACGGCTTTACATGAGTACTTTTATAATTATAGATCAGTCAACGGAAAATACAAGTATAGATACAACGGTTACTATCAATATTATTCATCTAGCGATTATGAAAATTATAATGAATATTATTATGATGATTATGATAGAACTGGTATGAACAGTGAGAGTGATAATATATCAATCAAAACAGAATACGAGAATGAATATGAATTCTATGATGAAACACAAGATCAAAGTACACAACTAGTAGATTACGACATTAAACTCAAAACCAATGAGGATGATTTTGTTGATGAATTCTATGGTTATGATAGATCAGTGGGTGTCCATGATTATATAGATGTATCAATTAATAAAGTAGTATATGGAAGAGAGTCTCACGTAAGATGGTGGGATATATGGCAAGAACATAATGATGGCGTATACAGTATAGGAAAGGAGTGCATAGATAATATATACGAAGACAGACATACCGTAGACGAATTCTACAAGATAGACAGCGTATCAGATGTAGATGACGCAGAACATATATCTCAGATAACTAATGATGTATCTACACAAACATGGGAAAAGAAATCAGAGTTAGATAGATACATGGAAATGTATCCTCGTCATAGATATGGTAAGCATTCTGTCTTTAAGGGATTTTCTGACAAAGTTAGAAAAAATGATTTAGACATGAACGTGGTAAAAGAATTACTTTCTAACGGTGCATCTCTAACAATCAAGGATAGCAGTAATAAGGATCCAATTGCTGTTTATTTTAGAAGAACAATAATGAATTTAGAAATGATTGATATCATTAACAAACATACAACTATCTATGAACGCAGGTATATAGTACACTCCTATCTAAAAAATTATAGAAATTTCGATTATCCATTTTTCAGAAAGTTAGTTTTGACTAATAAACATTGTCTCAACAATTATTGTAATATAAGCGACGGCAAATATGGAACACCACTACATATATTAGCATCTAATAAAAAAATAATAACTCCTAATTACATGAAGTTATTAGTGTATAACGGAAATGATATAAACGCACGAGGTGAAGATACACAAATGCGAACTCCATTACACAAATATTTGTGTAAATTTGTATATCATAATATTGAATATGGTATCCGATACTATAATGAAAAGATTATAGACGCATTTATAGAGTTAGGAGCCGATCTAACTATTCCAAATGACGATGGAATGATACCAGTAGTTTACTGTATACACTCAAATGCCGAATATGGTTATAACAATATTACTAACATAAAGATAATACGTAAACTACTTAATCTTAGTAGACATGCGTCACATAATCTATTTAGAGATCGAGTCATGCACGATTATATAAGTAATACATATATTGATCTTGAGTGTTTAGATATCATTAGATCACTTGATGGGTACGATATTAATTGTTACTTTGAAGGACGTACACCACTTCATTGCGCTATACAATATAACTTCACTCAGATTGCTGAGTACTTATTAGATCGAGGAGCTGATATATCATTAAAGACAGACGATGGTAAAACTGTATTTGATTTATCGTTATGTAGTTACATTCCTCTTAAATGGACTAGCTTTTTGATTAGTCGTCTACCGCCTAAAAGTGTCATATGCTCACTGACTAACCATATAATAGATTATGTTCTTACGAACAATAGACGTATTATTTGGCAGAGTCAAATGATTAATAAGTACGTACTGTTACTGGACCCATCCTTTTATTATAGATTCAGAAATGTTATCGAAAACAAATTAGACCAATACAATAATCGTTATAATATGTTCGAACACGATAGGGACGTTAATGAAAAGTATGGCAAAGTCTTACATGACCTCGATACATATATCAAGGATGTACAAGTATTAAAATCTACTTCCATCACTAATAATATAACACTATACGACACTATTATAAATAATAAGTCAGAGTTTCCTATACGTCGTGTAAACGACAAACAATTAATTAATCTCATAAAATCCAATACATATCATAATCTTATCGAAAAAGTTATTAAAAATACATTAGAGAAATATACTTTAACTAATATAGTCCTCGAGTATATGATCTCATCTCGATCTCAATCATCTTATTTGAGTCGTATTCCTAATGAGATATTACTCGAAATATTATATAAACTCGACATGTACGATTTACGTAATCTATATACAAGATATATGAGAGAGAATGATATCACAGAGTATCATATAGAGAATACGAGGTCTGTTTCTACACAGACATGAATAATGAATACACATACAACGTTTTTTTTAATCTTAGATATAACACTAATTACATCAAGATTATATATTGAAATCGTAATTTGAGTTGTCTGATCATCATGGATATCGAAAATGATATACGTAACATTAGCAATCTTTTAGATGATTGATATATTATTATGCGATGTAATCATAACTATCGGAGATGTAGAAATTAAAGCGCATAAAACTATTTTGGTTGCCGGATCTACGTATTTTAAAACAATGTTCACAACATCTATGATAGCGAGAGATCTAGCAACTAGAGTAAATATACAGATGTTCGATAAAGATGCCGTCAAAAATATTGTACAGTACTTATACAATAGGTATATAAGTTCTATGAATGTGATAGACATATTAAAATGCACCGACTAAGAACGTAAAACGAACTATAGAATGTTATACAATGGGTGATGATAAGTAGAAGATGTTACCCGATATACCCATAGCATTATCTAGTTATGGCATGTGTGTATTAGATCAATACATATACATTATAAGCGGTCGTACCCAACACTGATTATACATCGGTACATACAGTAAATAGCATAGATATGGAGGAGGATACAAATATTTCAAATAAAGTTATGAGATACGCGCTGTCAATAATATATGGAAGACATTACCTAACTTCTGAACTGGAACTATAAATCCAGGCTCTCGCATAAAGATGAATATATATGTTGTATGCGACATCAAAGATGAAAAAATGTTAAGACTTATATATTTAGATATAACACGAATATGTATGACGGATGGGAATTGGTAACGATGACAGAAAGCAGATTGTCAGCTCTGCATACTATTCTTCATGACAATACCATAATGATGTTACATTGTTATGAAGCGTATATGTTACAAGATACATTTAATGTGCTTACGGAACATATATTTAGAAACATCTACTAACGATTTTTTATGCTTGTATTATTAATGGTATGTAATATGATTTAATTGATTGTGTACACGATACCAATTTGTCGAGTATGAATACGGAGTACAAACATAAACTGAAGTTTAACATTATTTATTTATGNNATATACATTATATACATTATATACATTATATACATTATATACATTATATACATTATATACATTATATATCGTTATTGTTTGGTCTATGCCATGGATATCTTTAAAGAACTAATCTTAAAACATACGGATGAAAATGTTTTGATTTCTCCAGTTTCCATTTTATCTACTTTATCTATTCTGAATCATGGAGCAGCTGGTTCTACAGCTGAACAACTATCAAAATATATAGAGAATATGAATGAGAATACACCCGATGATAAGAAGGATGACAATAATGACATGGACGTAGATATTCCGTATTGCGCGACACTAGCTACCGCAAATAAAATATACGGTAGTGATAGTATCGAGTTCCATGCCTCATTCCTACAAAAAATAAAAGACGATTTTCAAACTGTAAACTTTAATAATGCGAACCAAACAAAGGAACTAATCAACGAATGGGTTAAGACAATGACAAATGGTAAAATTAATTCCTTATTGACTAGTCCGCTATCCATTAATACTCGTATGATAGTTATTAGCGCCGTCCATTTTAAAGCAATGTGGAAATATCCATTTTCTAAACATCTTACATATACAGACAAGTTTTATATTTCTAAGAATATAGTTACCAGTGTTGATATGATGGTGGGTACCGAGAATGACTTGCAATATGTACATATTAATGAATTATTCGGAGGATTCTCTATTATCGATATTCCATACGAGGGAAACTCTAGTATGGTGATTATACTGCCGGACGACATAGAAGGTATATATAACATAGAAAAAAATATAACAGATGAAAAATTTAAAAAATGGTGTGGTATGTTATCTACTAAAAGTATAGACTTGTATATGCCAAAGTTTAAAGTGGAAATGACGGAACCGTATAATCTGGTACCGATTCTAGAAAATTTAGGACTTACTAATATATTTGGATATTATGCAGATTTTAGTAAGATGTGTAATGAAACTATCACTGTAGAAAAATTTCTACATACGACGTTTATAGATGTTAATGAGGAGTATACAGAAGTATCGGCCGTTACAGGAGTATTCATGACTAACTTTTCGATGGTATATCGTATGAAGGTCTACATAAACCATCCATTCATATACATGATTAAAGATAACACCGGACATACACTTTTTATAGGGAAATACTGCTATCCGCAATAAATATAAACAATAGACTTTTATCACGTTATCTCATGTATAAAATATTACAAATAGTATAGCATAAACTAAAGTCGATACATACATTAAAACTTAAATAATAATGTAATTTACAATTAATAGTATAAACTAAAAAAATTAAAAAATTAAAAACAATATCATTATTATAAGTAATATCAAAATGACGATATACGGATTAATAGCGTATCTTATATTCGTGACTTCATCCATCGCTAGTCCACTTTACATTCCCGTTATTCCGCCCATTTCGGAAGATAAATCGTTCAATAGTGTAGAGGTATTAGTTTCTTTGTTTCCCGATGACCAAAAAGACTATACAGTAACTTCTCAGTTCAATAACTACACTATCGGTACCAAAGACTGGACTATCAACGTACTATCCACACCTGATGGTCTGGACATACCATTGACTAATATAACTTATTGGTCACGGTTTACTATAGGTCGTGCATTGTTCAAATCAGAGTCTGAGGATATTTTCCAAAAGAAAATGAGTATTCTAGGTGTTTCTATAGAATGTAAGAAGCCGTCGACATTACTTACTTTTTTAACCGTGCGTAAAATGACTCGAGTATTTAATAGATTTCCAGATATGGCTTATTATCGAGGAGACTGTCTAGAAGCCGTTTATGTAACAATGACTTATAAAAATACTAAAACTGGAGAGACTGATTACACGTACCTCTCTAATGGGGGGTTGCCTGCATACTATCGTAATGGGGTCGATGGTTGATTATTGATTAGTATATTCCTTATATTCCTTATTCTTTTTATTCACACAAAAAGAACATTTTTATAAACATGAAACCACTGTCTAAATGTAATTATGATCTTGATTTATAGATGATGATCAGCCTTCAGAGGATTTTGACCAGTATGTTTAATATGAAAAAAAACATAACTATTAAGCGCTATTGCGCTATTGTGCTTAATTATTTTGCTCTATAAACTGAATATATAGCCACAATTATTGACGGGCTTGTTTGTGACCGACAATCATGAATTTTCAGAAATTATCTCTGGCTATATATCTTACGGTGACATGTTCGTGGTGTTATGAAACATGTATGAGAAAAACTGCGTTGTATCATGACATTCAATTGGAGCATGTAGAAGACAATAAAGATAGTGTAGCATCGCTACCGTACAAGTATCTACAAGTAGTCAAACAAAGAGAACGTAGTAGATTGTTGGCTACATTTAATTGGACGGATATAGCTGAGGGTGTTAGAAATGAGTTCATTAAAATATGTGATATCAACGGAACATATCTATATAATTATACTATTGCTGTTAGTATAATTATTGATTCCACGGAAGAACTACCAACAGTTACTCCAATTACAACATATGAACCTTCTATATATAATTATACTATCGATTATAGCACTGTTATTACTACTGAAGAACTACAAGTGACTCCAACATATGCACCTGTAACAACTCCTCTTCCAACATCAGCAGTTCCTTATGATCAACGATCGAATAACAATGTAAGTACTATATCTATTCAGGTACTGAGTAAAATATTGGGAGTCAATGAAACAGAATTAACTAATTATCTTATTATGCATAAAAATGACACTGTTGACAATAACACCATGGTTGATGATGAGACATCTGACAATAACACATTACATGGTAATATAGGATTTTTGGAAATAAATAATTGTTATAATGTTTCTGTGTCAGATGCTAGTTTTAGAATAACATTAGTAAACGATACTTCTGAAGAAATTTTGCTAATGCTAACAGGAACTAGTTCATCCGACACCTTCATATCTTCCACCAATATCACTGAATGTTTGAAAACATTAATCAATAATGTGTCGATTAATGATGTACTTATAACACAAAATATGAATGTAACATCTAATTGTGATAAATGCTCAATGAATTTGATGGCATCCGTTATTCCTGCAGTTAATGAATTTAACAATACGTTGATGAAAATTGGTGTAAAAGATGATGAAAACAATACGGTATATAAATATTATAATTGTAAACTAACTACAAATTCTACATGTGATGAGTTAATCAATTTAGATGAAGTCATTAACAACATAACTCTGACAAATATTATACACAATAGTGTTTCGACAACTAACAGCAGAAAAAGACGAGATCTGAATGATGAGTTTGAATTTTCCACTTCCAAGGAATTAGATTGTCTTTACGAATCATATGGTGTAAACGATGATATAAGTCATTGTTTTGCATCACCTAGACGTAGACGATCTGACGACAAAAAGGAGTACATGGACATGAAATTATTCGACCACGCGAAAAAAGATTTAGGAATAGACAGTGTTATTCCTAGAGGTACAACCCATTTCCAAGTAGGTGCATCTGGTGCAAGTGGTGGTGTTGTAGGAGATAGTTTCCCATTTCAAAATGTTAAATCGCGTGCCAGTCTATTGGCGGAAAAAATAATGCCTAGAGTACCTATTACTGCTACCGAAGCTGATCTATATGCAACTGTAAATAGACAACCCAAGTTACCAGCAGGTGTTAAAAGTACTCCGTTTACAGAGGCGCTTGTGTCTACGATAAACCAAAAGCTTTCTAATGTTAGAGAGGTAACTTATGCTTCGCTCAATCTGCCAGGATCAAGTGGCTATGTTCATAGACCATCTGATTCTGTTATTTATAGCAGTATAAGACGGTCACGTTTACCTAGTGATAGCGATAGTGATTATGAGGATATACAAACTGTTGTTAAGGAATATAATGAAAGATATGGTAGATCAGTCAGTAGAACACAGTCATCAAGTAGTGAAAGCGATTTTGAAGATATAGATACTGTTGTTAGGGAATATAGACAAAAATATGGCAATGCAATGGCAAAAGGACGTAGTAGTTCCCCTAAACCTGATCCATTATATAGTACTGTTAAGAAAACAACTAAAAGTCTATCTACTGGTGTAGACATAGTTACAAAACAATCAGACTATTCTCTATTACCTGACGTTAATACTGGCAGTTCTATTGTGTCACCTCTCACCAGAAAAGGAGCTACTAGACGACGACCTAGACGCCCTACAAATGATGGTCTACAGAGTCCAAATCCTCCTCTCCGTAATCCACTTCCTCAACATGATGATTATTCTCCTCCACAAGTACACAGACCTCCACCACTTCCTCCTAAACCAGTCCAAAATCCGCCACAACTTCCCCCTAGACCAGTAGGTCAATTACTACCTCCTCCTATAGATCAACCAGATAAAGGATTTAGTAAGTTTGTATCACCTAGACGGTGTAGAAGAGCAAGCTCTGGAGTCATATGTGGTATGATACAATCAAAACCAAACGATGATACCTATTCACTTCTTCAACGATCAAAAATTGAACCAGAATATGTGGAGGTTGGTAATGGTATACCCAAGAACAATGTTCCTGTAATAGGTAATAAACATAGTAAAAAATATACATCGACGATGTCAAAAATATCAACAAAATTTGATAAATCTACGGCATTTGGAGCAGCAATGTTACTAACTGGTCAGCAGGCCATTAGCCAACAGACTAGATCAACTACGTTGAGTAGAAAAGATCAGATGAGTAAGGAAGAAAAGATATTCGAAGCAGTTACAATGAGTCTATCAACTATAGGTTCAACGTTGACGTCTGCAGGTATGACGGGTGGTCCAAAACTAATGATTGCAGGAATGGCTATAACGGCTATAACTGGTATAATAGATACGATAAAAGATATATATTACATGTTTTCAGGACAGGAGAGGCCAGTAGATCCTGTTATTAAATTATTTAATAAGTACACTGGCTTAATGTCCGATAATAATAAAATGGGTGTAAGAAAATGTTTGACACCCGGTGACGACACACTTATTTATATCGCATACAGAAACGATACCAGTTTTAAACAGAATACGGATGCGATGGCTTTGTATTTCTTAGATGTTATCGATTCAGAGATCCTATATCTAAACACATCAAATTTAGTTCTAGAGTATCAACTAAAGGTGGCTTGCCCCATAGGAACATTAAGATCTGTAGATGTGGACATAACTGCGTATACAATATTATATGATACAGCGGATAATATTAAGAAATACAAGTTTATCAGAATGGCAACGCTACTATCCAAACATCCAGTTATTAGATTGACATGTGGTTTAGCAGCAACATTGGTGATTAAACCGTACGAGGTACCCATCAGTGATATGCAACTACTAAAAATGGCGACGCCTGGTGAACCAGAATCCACTAAATCTATACCATCCGATGTCTGTGATAGGTATCCTCTAAAGAAATTCTATCTTTTAGCTGGTGGTTGTCCCTATGATACATCTCAAACTTTTATTGTACATACTACTTGCAGTATTCTACTAAGAACAGCTACACGGGATCAGTTTAGAAACAGATGGGTGTTACAAAATCCATTTAGACAAGAAGGGACATATAAGCAACTGTTTACCTTTAGCAAATACGATTTTAACGACACCATAATCGATCCTAATGGTGTGGTGGGTCATGCTAGCTTTTGTACCAATAGAAGCAGCAACCAATGTTTCTGGTCCGAACCTATGATATTGGAAGATGTATCATCGTGTAGTTCTAGAACTAGAAAAATATACGTAAAACTGGGAATATTTAATGCTGAAGGTTTTAATAGTTTTGTACTAAATTGTCCAACTGGGTCTACACCTACATACATCAAACATAAAAATGCGGACAGTAACAATGTTATCATAGAGCTACCTGTAGGTGATTACGGCACAGCCAAATTGTATTCAGCAACAAAACCATCGAGGATAGCTGTGTTCTGCACACATAACTATGATAAACGATTCAAATCAGATATTATAGTTCTAATGTTTAATAAAAACAGCGGTATTCCATTTTGGAGCATGTACACAGGAAGTGTAACTAGTAAAAATAGAATGTTTGCCACATTGGCTAGAGGAATGCCGTTTAGATCAACGTATTGCGATAACAGACGACGATCAGGTTGTTATTATGCAGGAATACCATTTCATGAAGATAGTGTAGAAACAGATATACATTATGGACCAGAAATAATGTTAAAGGAAACATATGACATAAACAGTATTGACCCACGAGTTATAACAAAGTCAAAGACCCATTTTCCTGCTCCATTGAGTGTAAAATTCATGGTTGACAATTTAGGAAATGGATATGACAACCCTAATTCATTTTGGGAAGATGCTAAAACTAAGAAACGGACATATAGTGCAATGACGATAAAAGTCCTACCATGTACAGTGAGAAATAAAAATATAGACTTTGGATATAACTATGGAGATATTATTTCTAATATGGTTTATCTACAATCTACTAGTCAGGATTATGGAGATGGTACCAAATATACATTTAAATCCGTAACTAGATCAGATCATGAGTGTGAATCTAGCTTAGATCTAACGTCTAAGGAAGTAACTGTGACATGTCCTGCGTTTAGTATACCAAGAAATATATCAACATATGAAGGTCTATGCTTTAGTGTTACTACATCTAAAGATCATTGTGCTACAGGTATTGGTTGGTTAAAATCTAGTGGTTATGGGAAGGAAGATGCTGATAAACCACGTGCTTGTTTTCATCATTGGAATTATTACACACTGTCGTTGGATTATTACTGTTCATACGAAGATATTTGGAGAAGCACCTGGCCTGACTATGATCCATGTAAGTCATATATCCATATAGAGTATAGAGATACATGGATAGAATCTAATGTGTTACAGCAACCTCCTTACACATTCGAATTCATTCATGACAATTCTAACGAATATGTGGATAAAGAAATTAGTAACAAATTAAATGATCTGTACAATGAATACAAGAAGATTATGGAATATAGCGACGGATCATTGCCGGCGTCTATAAACAGATTAGCAAAGGCATTGACTTCAGAGGGTAGAGAAATAGCAAGTGTTAATATAGATGGTAATCTGTTAGATATCGCATATCAAGCAGATAAGGAAAAGATGGCCGACATACAGACAAGAATAAATGATATTATTAGAGATTTGTTTATACACACTCTATCAGACAAAGATATAAAAGACATTATAGAATCCGAAGAAGGTAAGAGATGTTGTATAATAGATGTTAAGAACAATCTTGTTAAAAAGTACTATTCTATTGATAATTATCTATGTGATACTTTAGATGATTATATATACACCTCTGTAGAATATAACAAATCCTATGTGTTAGTAAACGATACTTATATGAGCTATGACTATCTTGAATCATCAGGTGTAGTTGTTCTATCATGTTATGAAATGACTATAATCTCCTTGGATACAAAAGACGCCAAAGATGCTATAGAAGATGTGATAGTAGCAAGTGCGGTAGCCGAAGCATTGAATGACATGTTTAAGGAATTTGATAAAAACGTAAGTGCTATTATAATAAAAGAAGAAGATAATTATCTAAACAGTTCGCCCGATATCTACCATATAATATATATCATAGGTGGCACTATTCTGCTACTGTTAGTCATTATTTTAATATTGGCAATTTATATAGCGCGCAATAAATACAGAACCAGGAAATATGAAATAATGAAATATGACAATATGAGCATTAAATCTGAGCATCATGATAGTCTTGAAACAGTGTCTATGGAAATTATTGATAATCGGTACTAATAAAATAGTTTAACTCTTTTAGAACCAGTTTGGTACTGTAATTTCAGTTCATTACTCGTTGAGAATATTGATGATTTTTTTTAAATGAGTATCGGTAGTTACATATTACCATATCATCCATTATATAATCGATGATGCATGTATTAGAATACTTTCCGAATAAGTCTTCTAAATATTGTATTAATTATGAAAAACTATGCTATGTGAGTATGATTCAAAGATGTTTAATGATACGATACTAGATTTTATCTCTAGCGAGATTGTTTAGAATCATTTATCATAACTATGTTTAATAAATTCATCAACGAATATCGATAAAGACCTCTTGTAATTCGAGTATAGGAAGTAGTATTACCATATCAACTTCCGAGTTAACAATTACTCTAAAACATGAGGATTGTACTCCTGTCTTTATTGGAGATCACTATTTAGTCGTTGATAAACTAGTAACCTCAGGTTTCTTTACAAACGATAAAGTACAACATCAAGACCTCACAACACAGTGCAAGATTAATCTAGAAATCAAATGTAATTCTGGAGGAGAATCTAGACAACTAACACCCACGGCGAAGTATACTTTATGCCTCATTCAGAAACGGTAACTGTAGTAGGAGACTGTCTCTCTAATCTCGATGTATATATAATATATACCAATACGGACGCGATATATTCCGACATGAATGGCGTCGCTTATCATATGTTATATCCTAAATGTTGATCATATTCCACAAATGATTGTGAACGAGATTAAATCATCTAACAAATAATTAGTTTTTTATGACATTAACATATAATAAATAAATTAATCATTATTGACTTAACGATGACGAAAGTTATCATTATCTTAGGATTCTTGATTATTAATACAAATTCGTTGTGTCTATGAAATGTGAACAAGGTGTCTCATATTATAATGCACAAGAATTAAAGTGTTGTAAACTATCTAGCCAGGAACATATTCAGATTATCGATGTGATAAATACAGCGATACCATCTGTGGACATTGTCCAAGTGACACATTCACGTCAATATATAATCGTTCTCCTCGGTGTCATAGTTGTAGAGGTCACACCTTGTACACCTACCACAAATAGAATATGTCATTGTGACTCGAATAGTTATCGTCTCCTTAAAGCTTCTGATGGTAACTGTGTTACATGTGCTCCTAAAACAAAATATGGTCGTGTGTACGGAAAGAAAGGAGAAAATGATATGGAATACCATTTGTAAGAAATGTCGGAAGGGTACTTATTCAGATATTGTATCTGACTCTGATCAATGTAAACCTATGACAAGATAAGACTTACTCGCATCTACTGGATAGACATAAATATCCTCCTCGTAATAATGAAATATAATATACACTAATTATTAATATCAATCGAGTATTAACATATAAGTTATTTTTAAACCCCTTTTGGGTTCCGTCCTAAACGGCGTTTCGGTCTGTGTCGCCACCATGGTCACACCGAGCCTCTGCGTGCTCCTCCATCGAGGACGACTTCAACTATGACAGCTCGGTGGCGTCTGCCAGCGTGTACATACGAATGGCATTTCTAAGAAAAGTCTACGGTATCCTTTCTACAATTTCCTTTAACAACGGCAACAGCTGCAGTATTTTTATACTTTGAATGCATCGGACATTTATACAAGGGAGTCCTGTTCTAATATTGGCATCAATGTTCGGATCTATAGGCTTGATTTTCGCATTGACTTTACACAGACATGAACATCCCCTGAATCTGTACATACTTTGTGGATTTACACTGTTAGAATCTCTAACGCTGGCCTCTGTTGTTACTTTCTATGATGCACGTATCGTTATGCAAGCTTTCATGTTGACTACTGCAGTGTTTCTTGCTCTGACTACATATACTCTACAATCAAAGAGAGATTTCAGTAAACTTGTAACAGGATTGTTTGCTGCTTTCTGGATTTTAATTTTGTCAGGAGTCTTGAGGATAAAGTTTAAAATAGAATTAATAAAGAACATATAGGTCATTTTTTAAACATGGATAGAAACCAAGGTTGTTAGTTAATAATATACAAGATATTTTTTCTCACTCTGATCCATGTAAACCAAGGACGAGAGACACTCTCATTCCTCATTCACGACACCATTAAAAATGGAAATTAAAGCCCTCTATTAAGCACAGACGGCTACAGGTCTACCATCAGGTTACCTTCGTCTACCTTCACAATGGCCTCTCCTTGTGCCCAGTTCAGTCCCTGTCATTGCCACGCTACTAAGGACTCCCTGAATACCGTGACTGACGTCAGACATTGTCTGACTGAATACATCCTGTGGGTTTCTCATAGATGGACCCATAGAGAAAGCGCAGGGCCTCTCTACAGGCTTCTCATCTCTTTCAGAATTGATGCAATGGAGCTATTTGGTAGCGAGTTGAAGGAGTTCTCGAATTCACTTCCGTGGGACAATATCGACAATTGCGTGGAGATCATTAAATGTTTCATCAGAAATGACTCCATGAAAACCGCCAAAGAACTTTGTGCAATAATTGGACTTTGTACTCAATCAGCTATTGTCACTGGAAGAGTCTTCAATGATAAGTATATCGACATACTACTTATGCTGCGAAAGATTCTGAACGAGAACGACTATCTCACCCTCTTGGATCATATCCTCACTGCTAAATACTAAATCTCCTTCATGCTCTCTCACTAATACTCTTACTCACTACACTTTTTATCATCTTATGATGAATGATTGCCTTCATCATTTTTTCGTGGAATATAATATAGGAATAATTAGCACCAGAATAGCTATGGATATCTCGTTAAGAATATTCTCTATAAGAGACATAATGTAGACATAGTTATTATATCCTTCTTAGATAAGTGTTACGCTACTGGAAAGTTTCCATCGTTATTATTACATGAAGATGATATAATTAAACCAACATTGAGATTGGCTCTTATGTTAGCTGGATTGAATTACTGTAATAAATGCATCGAGTATAGAGGGATATAGCAATTCTCGATAATAGTCATGCAATATTTGAATGAGACTGATAATTTAGGTAATACAGTACTACACACATATCTTTCTAGATTATATATCGTTAAAAATCTGTAAGATGTATATTTCTCATAAGTATCCACTGTGTAATATTATTAATGGATATATAGATAACGCAATAGGGACTAATAGTATTGTAAAAGATATAATCGACTATTTACGTACATATCCAGATATCTATATTCCTACTAGTTTGCTGCGTAGTTGCATCATTGATATGCATGATTTATCAGGATTCAGAGATGAATTACTAAGTAAACTACAATCCCACAATAAGTAAGAATCAAATATCAAAAACTCACTTTTGATTTTTCTAGTCTTAAGTAATACATATATTTATTAATAGACCTATGAAATAAAAAAAGGTAACAATGGATTCGCGTATAGCTATTTACGTATTAGTATCGGCATCTCTTTTGTATCTTGTTAATTGTCACAAACTAGTACATTACTTCAATCTGAAAATAAATGGAAGTGATATAACTAATACAGCAGATATATTGCTGGACAATTATCCAATTATGACCTTTGATGGAAAGGATATTTATCCATCTATCTCGTTCATGGTCGGTAATAAACTTTTCCTAGATCTTTATAAAAATATCTTTGAAGAATTTTTCAGACTATTTCGAGTATCTGTAAGTAGTCAATACGAGGAATTAGAATATTATTATTCATGTGATTATACTAACAACCGTCCTACAATTAAACAACATTACTTTTATAACGGCGACGAATATACTGAAATTGATAGATCGAAAAAAGCCACTAATAAAAACAGTTGGTTAATTACTTCAGGCTTTAGACTACAAAAATGGTTCGATAGCGAAGATTGTATAATTTATCTCAGATCTTTAGTTAGAAGAATGGAAGACAGTAACAAAAACAGTAAAAAAACTTAGTACTTAGATATCGAAAAAATATATTTTTGTAGACTCTTGAGAATAGAAGGAAAACATGTACATAATTATAAAAAATGAAAATCAATGGCGAATAAGACAGTGCGATTCGCACCATGGAGTCGGTAGATTTCATGGCTGTCGATGAGCAGTTTCACGACGACCTCGATCTTTGGTCATTATCTTTGGTAGATGATTATAAAAAACATGGATTAGGTGTTGACTGTTATGTTCTAGAACCAGTTGTTGACAGGAAAATATTTGATAGATTTCTCCTTGAACCAATTTGTGATCCTGTAGATGTTCTGTATGATTATTTTAGGATTCATAGAGATAATATTGATCAGTATATAGTAGATAGACTGTTTGCATATATTACATATAAAGATATTATATCTGCATTAGTGTCAAAGAATTATATGGAAGATATTTTCTCTATAATTATTAAGAATTGTAATTCTGTGCAAGATCTCTTACTTTACTATCTATCTAATGCATATGTAGAAATAGACATTGTTGATCTTATGGTAGATCATGGGGCTGTAATATATAAAATAGAATGCTTGAATGCCTATTTTAGGGGAATATGTAAAAAGGAAAGTAGTGTTGTTGAGTTTATTTTGAATTGTGGTATCCCAGATGAAAATGATGTTAAATTAGATCTATATAAAATAATTCAGTATACTAGGGGATTCCTTGTAGATGAACCCACAGTATTAGAAATTTATAAGCTTTGTATCCCATATATTGAAGATATCAATCAACTAGATGCTGGTGGAAGGACCTTGCTTTATCGCGCTATCTATGCAGGTTATATAGATTTAGTATCATGGCTATTAGAAAATGGAGCAAATGTCAACGCAGTAATGAGTAATGGATATACATGTCTTGACGTGGCCGTGGATAGGGGATCTGTCATCGCCCGTAGGGAAGCACATCTTAAAATATTAGAAATATTGCTTAGAGAACCATTGTCTATTGACTGTATAAAATTAGCTATACTTAATAATACAATTGAAAACCATGATGTGATAAAGCTCTGTATCAAGTATTTTATGATGGTAGATTATTCACTTTGTAATGTGTATGCATCATCACTCTTTGATTATATAATTGATTGTAAACAAGAATTGGAGTACATTAGGCAGATGAAAATTCATAATACAACCATGTATGAGTTAATCTATAATAGAGACAAAAACAAGCATGCTTCCCATATTCTACATAGGTATTCTAAACATCCAGTTTTGACACAGTGTATCACTAAAGGATTCAAGATTTACACAGAAGTAACCGAGCAGGTCACTAAAGCTCTAAACAGACGTGCTCTAATAGATGAGATAATAAACAATGTATCAACTGATGACAATCTCCTATCAAAACTTCCATTAGAAATTAGGGATCTAATTGTTTCACAAGCTGTCATATAGAGTTCTATCCACCCACCTTTCTTGAAATGAGTTAATAGTCATAAGTTAGTTAAGTCATAAGTTAGTTAAGTCATAAGTTAGTTAAGTCATAAGTTAGTTAAGTCATAAGTTAGTTAAGTCATAAGTTAGTTAAGTCATAAGTTAGTTAAGTCATAAGTTAGTTTATAGTCTAACACTTCTAATTTTTATACCTTGATCTTTTTCTCTAATTATGAAAAAGTAAATCATTATGAAGATGGATGAAATGGACGAGATTGTGCGCATCGTTAACGATAGTATGTGGTACGTACCTAACGCATTTATGGACGACGGTGATAATGAAGGTCACATTTCTGTCAATAATGTCTGTCATATGTATCTCGCATTCTTTGATGTGGATATATCATCTCATCTGTTTAAATTAGTTATTAAACACTGCGATCTGAATAAACGACTAAAATGTGGTAACTCTCCATTACATTGCTATACGATGAATACACGATTTAATCCATCTGTATTAAAGATATTGTTACGCCACGGCATGCGTAACTTTGATAGCAAGGATAAAAAAGGACATATTCCTCTACACCACTATCTGATTCATTCACTATCAATCGATAACAAGATCTTTGATATACTAACGGACCCCATTGATGACTTTAGTAAATCATCCGATCTATTGCTGTGTTATCTTAGATATAAATTCAATGGGAGCTTAAACTATTACGTTCTGTACAAATTATTGACTAAAGGATCTGACCCTAATTGCGTCGATGAGGATGGACTCACTTCTCTTCATTACTACTGTAAACACATATCCGCGTTCCACGAAAGCAATTATTACAAGTCAAAGAGTCACACTAAGATGCGAGCTGAGAAGCGATTCATCTACGCGATAATAGATCATGGAGCAAACATTAACGCGGTTACGAAAATCGGAAATACGCCGTTACACACTTACCTTCAACAGTATACCAAACATAGTCCTCGTGTGGTGTATGCTCTTTTATCTCGAGGAGCCGATACGAGGATACGTAATAATCTTGATTGTACACCCATCATGGAATACATAAAGAACGATTGTGCAACAGGTCATATTCTCATAATGTTACTCAATTGGCACGAACAAAAATACGGGAAATTACAAAAGGAAGAAGGACAACATCTACTTTATCTATTCATAAAACATAATCAAGGATATGGAAGTCGCTCTCTCAATATACTACGGTATCTACTAGATAGATTCGACATTCAGAAAGACGAATACTATAATACAATGACTCCTCTTCATACCGCCTTCCAGAATTGCAATAACAATGTTGCCTCATACCTCGTATACATCGGATACGACATCAACCTTCCGACTAAAGACGATAAGACAGTATTCGACTTGGTGTTTGAAAACAGAAACATTATATACAAGGCGGATGTCGTTAATGACATTATCCACCACAGACTGAAAGTATCTCTACCTATGATTAAATCGTTGTTCTACAAGATGTCGGAGTTCTCTCCCTACGACGATCACTACGTAAAGAAGATAATAGCCTACTGCCTATTAAGGGACGAGTCATTTGCGGAACTACATACTAAATTCTGTTTAAACGAGGACTATAAAAGTGTATTTATGAAAAATATATCATTCGATAAGATAGATTCCATCATCGAAAAATGTAGTCGTGACATAAGTCTCCTCAAAGAGATTCGAATCTCAGACACCGACTTGTATACGGTATTGAGAACAGAAGACATCCGGTATCACACATATCTCGAAGCCATACATTCAGACAAACGCATTTCATTTCCCATGTACGACGATCTCATAGAACAGTGTCATCTATCGATGGAGCATAAAAGTAAACTCGTCGACAAAGCACTCAATAAATTAGAGTCTACCATCGATAGTCAATCTAGACTATCGTATTTGCCTCCGGAAATTATGCGCAATATCATAACCAAGCTAAGCGACTACCATCTAAACAGTATGTTGTACGGAAAGAACCATTACAAATATTATCCATGATAGAAAGAAAATATTTAAAAAATAATCTATATGATTGGAGAAGTAGGAAACAAACAGTAACAAGACGACGATTACTACATTATTAAATCATGAGGTCCGTATTATACTCGTATATATTGTTTCTCTCATGTATAATAATAAACGGAAGAGATATAGCACCACATGCACCATCCAATGGAAAGTGTAAAGACAACGAATACAGAAGCCGTAATCTATGTTGTCTATCGTGTCCTCCGGGAACTTACGCTTCCAGATTATGTGATAGCAAGACTAATACACAATGTACACCGTGTGGTTCGGATACCTTTACATCTCACAATAATCATTTACAGGCTTGTCTAAGTTGTAACGGAAGATGTGATAGTAATCAGGTAGAGACGCGATCGTGTAACACGACTCACAATAGAATCTGTGAATGCTCTCCAGGATATTATTGTCTTCTCAAAGGAGCATCAGGGTGTAGAACATGTATTTCTAAAACAAAGTGTGGAATAGGATACGGAGTATCCGGATACACGTCTACCGGAGACGTCATCTGTTCTCCGTGTGGTCCCGGAACATATTCTCACACCGTCTCTTCCACAGATAAATGCGAACCCGTCGTAACCAGCAATACATTTAACTATATCGATGTGGAAATTAACCTGTATCCAGTCAACGACACATCGTGTACTCGGACGACCACTACCGGTCTCAGCGAATCCATCTCAACGTCGGAACTAACTATTACCATGAATCATAAAGATTGTGATCCAGTCTTTCGTGCAGAATACTTCTCTGTCCTTAATAATGTAGCAACTTCAGGATTCTTTACAGGAGAAAATAGATATCAGAATACTTCAAAGATATGTACTCTGAATTTCGAGATTAAATGTAACAACAAAGATTCATCTTCCAAACAGTTAACGAAAACAAAGAATGATACTATCATGCCGCATTCAGAGACGGTAACTCTAGTGGGCGACTGTCTATCTAGCGTCGACATCTACATACTATATAGTAATACCAATACTCAAGACTACGAAAATGATACAATCTCTTATCATATGGGTAATGTTCTCGATGTCAATAGCCATATGCCCGCTAGTTGCGATATACATAAACTGATCACTAATTCCCAGAATCCCACCCACTTATAGTAAGTTTTTTTACCTATAAATAATAAATACAATAATTAATTTCTCGTAAAAGTAGAAAATATATTCTAATTTATTATATGGTAAGAAAGTAGAATCATCTAGAACAGTAATCAATCAATAGCAATCATGAAACAATATATTGTCCTGGCATGCATGTGCCTAGTGGCAGCTGCTATGCCTACTAGTCTTCAACAATCTTCATCCTCGTGTACTGAAGAAGAAAACAAACATCATATGGGAATCGATGTTATTATCAAAGTCACAAAGCAAGACCAAACACCGACCAATGATAAGATTTGTCAATCCGTAACGGAAGTTACAGAGACCGAAGATGATGAGGTATCCGAAGAAGTTGTAAAAGGAGATCCCACCACTTATTACACTATCGTCGGTGCGGGTCTTAACATGAACTTTGGATTCACCAAATGCCCAAAGATTTCATCCATCTCCGAATCCTCTGATGGAAACACTGTGAATACTAGATTGTCCAGCGTGTCACCGGGACAAGGTAAGGACTCTCCCGCGATCACGCGTGAAGAAGCTCTGGCTATGATCAAAGACTGTGAGATGTCTATCGACATCAGATGTAGCGAAGAAGAGAAAGACAGTGACATCAAGACCCATCCAGTACTTGGGTCTAACATCTCACATAAGAAAGTGAGTTACAAAGATATCATCGGTTCAACGATCGTTGATACAAAATGTGTCAAGAACCTAGAGTTTAGCGTACGTATCGGAGACATGTGTGAGGAATCATCTGAACTTGAAGTCAAGGATGGATTCAAGTATGTCGACGGATCGGCATCTGAAGGTGCAACCGATGATACTTCACTCATCGATTCAACAAAACTCAAAGCATGTGTCTGAATCGATAACTCTATTCATCTGAAAATGGATGAGTTGGGTTAATCGAACGATTCAGACACCGCACCACGAATTAAAAAAGACCGGGCACTATATTCCGGTTTGCAAAACAAAAATATTTAACTACATTCACAAAAAGTTACCTCTCGTTACTTCTTCTTTCTGTTTCAATATGTGATACGATATGATCACTATTCGTATTCTCTTGGTCTCATAAAAAAGTTTTACAAAAAAAAAAAAAATATTTTTATTCTCTTTCTCTCTTCGATGGTCTCACAAAAATATTAAACCTCTTTCTGATGTCTCAACTATTTCGTAAACGATAACGTCCAACAATATATTCTCGTAGAGCTTATCAACATCCTTATACCAATCTAGGTTGTCAGACAATTGCATCATAAAATAATGTTTATAATTTACACGTTAACATCATATAATAAACGTATATAGTTAATATTTTTGGAATATAAATGATCTGTAAAATCCATGTAGGGGACACTGCTCACGTTTTTTCTCTAGTACATAATTTCACACAAGTTTTTATACAGACAAATTAATTCTCGTCCATATATTTTAAAACATTGACTTTTGTACTAAGAAAAATATCTTGACTAACCATCTCTTTCTCTCTTCGATGGGTCTCACAAAAATATTAAACCTCTTTCTGATGGAGTCGTAAAAAGTTTTTATCCTTTCTCTCTTCGATAGGTCTCACAAAAATATTAAACCTCTTTCTGATGGTCTCTATAAACGATTGATTTTTCTTACCCTCTAGAGTTTCCTACGGTCGTGGGTCACACATTTTTTTCTAGACACTAAATAAAATAGTAAAATATAATATTAATGTACTAAAACTTAT

>2017_Nigeria_Monkeypox_MT903341.1

CTAAATAAATTAGTATAATTAATATTAATGTACTAAAACTTATGTATTATTAATTTATCTAACTAAAGTTAGTAAATTATATACATAATTTTATAATTAATTTAATTTTACTATTTTATTTAGTGTCTAGAAAAAAATGTGTGACCCACGACCGTAGGAAACTCTAGAGGGTAAGAAAAATCAATCGTTTATAGAGACCATCAGAAAGAGGTTTAATATTTTTGTGAGACCTATCGAAGAGAGAAAGGATAAAAACTTTTTACGACTCCATCAGAAAGAGGTTTAATATTTTTGTGAGACCCATCGAAGAGAGAAAGAGATGGTTAGTCAAGATATTTTTCTTAGTACAAAAGTCAATGTTTTAAAATATATGGACGAGAATTAATTTGTCTGTATAAAAACTTGTGTGAAATTATGTACTAGAGAAAAAACGTGAGCAGTGTCCCCTACATGGATTTTACAGATCATTTATATTCCAAAAATATTAACTATATACGTTTATTATATGATGTTAACGTGTAAATTATAAACATTATTTTATGATGCAATTGTCTGACAACCTAGATTGGTATAAGGATGTTGATAAGCTCTACGAGAATATATTGTTGGACGTTATCGTTTACGAAATAGTTGAGACATCAGAAAGAGGTTTAATATTTTTGTGAGACCATCGAAGAGAGAAAGAGAATAAAAATATTTTTTTTTTTTTTTTGTAAAACTTTTTTATGAGACCAAGAGAATACGAATAGTGATCATATCGTATCACATATTGAAACAGAAAGAAGAAGTAACGAGAGGTAACTTTTTGTGAATGTAGTTAAATATTTTTGTTTTGCAAACCGGAATATAGTGCCCGGTCTTTTTTAATTCGTGGTGCGGTGTCTGAATCGTTCGATTAACCCAACTCATCCATTTTCAGATGAATAGAGTTATCGATTCAGACACATGCTTTGAGTTTTGTTGAATCGATGAGTGAAGTATCATCGGTTGCACCTTCAGATGCCGATCCGTCGACATACTTGAATCCATCCTTGACTTCAAGTTCAGATGATTCCTCACACATGTCTCCGATACGTACGCTAAACTCTAGGTTCTTGACACATTTTGTATCAACGATCGTTGAACCGATGATATCTTTGTAACTCACTTTCTTATGTGAGATGTTAGACCCAAGTACTGGATGGGTCTTGATGTCACTGTCTTTCTCTTCTTCGCTACATCTGATGTCGATAGACATCTCACAGTCTTTGATCATAGCCAGAGCTTCTTCACGCGTGATCGCGGGAGAGTCCTTACCTTGTCCCGGTGACACGCTGGACAATCTAGTATTCACAGTGTTTCCATCAGAGGATTCGGAGATGGATGAAATCTTTGGGCATTTGGTGAATCCAAAGTTCATGTTAAGACCCGCACCGACGATAGTGTAATAAGTGGTGGGATCTCCTTTTACAACTTCTTCGGATACCTCATCATCTTCGGTCTCTGTAACTTCCGTTACGGATTGACAAATCTTATCATTGGTCGGTGTTTGGTCTTGCTTTGTGACTTTGATAATAACATCGATTCCCATATGATGTTTGTTTTCTTCTTCAGTACACGAGGATGAAGATTGTTGAAGACTAGTAGGCATAGCAGCTGCCACTAGGCACATGCATGCCAGGACAATATATTGTTTCATGATTGCTATTGATTGATTACTGTTCTAGATGATTCTACTTTCTTACCATATAATAAATTAGAATATATTTTCTACTTTTACGAGAAATTAATTATTGTATTTATTATTTATAGGTAAAAAAACTTACTATAAGTGGGTGGGATTCTGGGAATTAGTGATCAGTTTATGTATATCGCAACTAGCGGGCATATGGCTATTGACATCGAGAACATTACCCATATGATAAGAGATTGTATCATTTTCGTAGTCTTGAGTATTGGTATTACTATATAGTATGTAGATGTCGACGCTAGATAGACAGTCGCCCACTAGAGTTACCGTCTCTGAATGCGGCATGATAGTATCATTCTTTGTTTTCGTTAACTGTTTGGAAGATGAATCTTTGTTGTTACATTTAATCTCGAAATTCAGAGTACATATCTTTGAAGTATTCTGATATCTATTTTCTCCTGTAAAGAATCCTGAAGTTGCTACATTATTAAGGACAGAGAAGTATTCTGCACGAAAGACTGGATCACAATCTTTATGATTCATGGTAATAGTTAGTTCCGACGTTGAGATGGATTCGCTGAGACCGGTAGTGGTCGTCCGAGTACACGATGTGTCGTTGACTGGATACAGGTTAATTTCCACATCGATATAGTTAAATGTATTGCTGGTTACGACGGGTTCGCATTTATCTGTGGAAGAGACGGTGTGAGAATATGTTCCGGGACCACACGGAGAACAGATGACGTCTCCGGTAGACGTGTATCCGGATACTCCGTATCCTATTCCACACTTTGTTTTAGAAATACATGTTCTACACCCTGATGCTCCTTTGAGAAGACAATAATATCCTGGAGAGCATTCACAGATTCTATTGTGAGTCGTGTTACACGATCGCGTCTCTACCTGATTACTATCACATCTTCCGTTACAACTTAGACAAGCCTGTAAATGATTATTGTGAGATGTAAAGGTATCCGAACCACACGGTGTACATTGTGTATTAGTCTTGCTATCACATAATCTGGAAGCGTAAGTTCCCGGAGGACACGATAGACAACATAGATTACGGCTTCTGTATTCGTTGTCTTTACACTTTCCATTGGATGGTGCATGTGGTGCTATATCTCTTCCGTTTATTATTATACATGAGAGAAACAATATATACGAGTATAATACGGACCTCATGATTTAATAATGTAGTAATCGTCGTCTTGTTACTGTTTGTTTCCTACTTCTCCAATCATATAGATTATTTTTTAAATATTTTCTTTCTATCATGGATAATATTTGTAATGGTTCTTTCCGTACAACATACTGTTTAGATGGTAGTCGCTTAGCTTGGTTATGATATTGCGCATAATTTCCGGAGGCAAATACGATAGTCTAGATTGACTATCGATGGTAGACTCTAATTTATTGAGTGCTTTGTCGACGAGTTTACTTTTATGCTCCATCGATAGATGACACTGTTCTATGAGATCGTCGTACATGGGAAATGAAATGCGTTTGTCTGAATGTATGGCTTCGAGATATGTGTGATACCGGATGTCTTCTGTTCTCAATACCGTATACAAGTCGGTGTCTGATCATTGAGATGATTGACGGCATCTCCCCCTTCCTTTTTTAGTAGATATTTCATCGTGTAAGAATCAATATTAATATTTCTAAAGTATCTGTGTATAGCCTCTTTATTTACCACAGCTCCATATTCCAACATGCATTCCACTAGAGGGATATCGATATCGCCGAATGTCATATACTCAATTAGTATATGTTGGAGGACATCCGAGTTCATTGTTTTCAATATCAAAGAGATGGTTTCCTTATCATTTCTCCATAGTGGTACAATACTACACATTATTCCGTACGGCTTTCCATTCTCCAAAAACAATTTTACCAAATCTAAATCTACATCTTTATTGTATCTATAATCACTATTTAGATAATCAGCCATAATTCCTCGAGTGCAACATGTTAGATCGTCTATATATGAATAAGCCGTGTTATCTATTCCTTTCATTAACAATTTAACGATGTCTATATCTATATGAGATGACTTAATATAATATTGAAGAGCTGTACAATAGTTTTTATCTATAGAAGACGGCTTGATTCCGTGATTAATTAGACATTTAACAACTTCCGGACGCACATATGCTCTCGTATCCGACTCTGAATACAGATGAGCGATGATATACAGATGCAATACGGTACCACAATTTCGTGGTTGATAATCATCATACACGTATCCGTACTCGTCATCCTCATAAAGAACACTGCATCCATTTTCTATGAACAAATCAATAATTTCAGGAACAGGATCATCTGTCATTACATAATTTTCTATAACTGAACGATGGTTTTCACATTTAACACTCAAGTCAAATCCATGTTCTACCAACACCTTTATCAAGTCAACGTCTACATTTTTTGATTTCATATAGCTGAATATATTAAAGTCATTTATGTTGCTATATCCAGTAGCTTCTAGTAGAGCCATCGCTATATCCTTATTGACTTTAACATGTCTACTATTTGTGTATTCTTCTATTGGGGTAAACTGTCTCCAATTTTTATGTAATGGATTAGTGCCACTATCTAGTAGTAGTTTGACGACCTCAACATTATTACAATGCTCATTGAAAAGGTATGCGTGTAAAGCATTATTCTTGAATTGGTTCCTGGTATCATTAGGATCTCTGTCTCTCAACATCTGTTTAAGTTCATCGAGAACCACCTCCTCATTTTCCAGATAGTCAAACATTTTGACTGAATAGAAGTGAATGAGCTACTGTGAACTCTATACACCCGCACAACTAATGTCATTAAATATCATTTTTGAATGTATTTATACCATGTCAAAAACTTGTACAATTATTAATAAAAATAATTAGTGTTTAAATTTTACCAGTTCCAGATTTTACACCTCCGTTAACACCTCCATTAACCCCACTTTTTACACCACTGGACGATCCTCCTCCCCACATTCCACTGCCACTAGATGTATAAGTTTTAGATCCTTTATTACTACCATCATGTCCATGGATAAAGACACTCCACATGCCGCCACTACTACCCCCTTTAGACGACATATTAATAAGACAAGTTTAACAATAAAATTAATCACGAGTACCCTACTACCAACCACTATTATATGATTACAGTACCTTGACTAAAGTCTCTAGTCACAAGATCAATACTACCAACCTACGCTATTATATGATTATAGTTTCTATTTTTATAGGAACGCGTACGAGAAAATCAAATGTCTAAGTTCTAACGGTAGTGTTGATAAACGATTGTTATCCGCGGATACCTCATCTATCATGTTGTCTATTTTCTTACTTTGTTCTATTAACCTATTAGCATTATATATTATTTGATTATAAAACTTATATTGCTTATTAGCCCAATCTGTAAATATCGGATTATTAACATATCGTTTCTTTGTAGGTTTATTTAACTTGTACATCACTGTAAGCATGTCCGTACCATTTATTTTAATTTGACACATATCAGCAATTTCTTTTTCGCAGTCGGTTATATATTCTATATAAGATGGATACGTATCACATATGTACTTATAGTCTACTAATATGAAGTACTTAATACATATTTTCAGTAACGATTTAGCCTTATTACCTATTAATAAGTGCCTGTCGTTGGATAGGTAATCAACTGTTTTCTTAATACATTCGATGGTTGGTAATTTACTCAAAATAATTTCCAATATCTTAATATATATTTCTGCTATTTCTGGTATACATGCATGTGCCATTATAACACAAATACCAATACATGTAGACCCATATGTTGTTGTTATATTAATATCTGCGCCATTATCTATTAACCATTCTATTAGGGCAACACTATGCGACTCGATACAATAATAAAGTATACTACGTCCATGTTTATCTATTTTGTTTATATCATCGATATACGGCTTACAAATTTCTAGTATCGATAACACTTCTGACTCGTGAATAAATAAGGTAGGGAATAACGGCATAATATTTATTATGTTATCATCATTAACAACTACGTTTCCATTTTTTAAAATATACTCTACAACTTTAGGATCCCTATTGTCAAATCTTTTAAAATATTTATTTATATGCTTAAATCTATATAATATAGCTCCTTCCCTAATCATACATTTGATAACATTGATGTACACTGTATGATAAGATACATATTCTGACAATAGATCTTGTATAGAATCTGTATATCTTTTAAGAATTGAGGATATTATGACATTATTACGTAAACTATTACACAATTCTAAAATATAAAATGTATCATGGGCAGATAATAGTTTATCCACTATATAATTATCTATTTTATGATTTTTCTTCCTATATTGTTTACGTAAATAGATAGATAGAATATGCATTAGTTCATTACCGCTATAGTTACTATCGAATAACACGTCAAATATTTCCCGTTTAATATCGCATTTGTCAACATAATAATAGAGTATGGTACGTTCACGATAAGTATAATGACACATCTCGTTTTCGTGCGAAATTAAATAGTTTATCACGTCCAAAGATGTCACATAACCATCTTGTGACCTAGTAATAATATAATAATAGAGAACTGTTTTACCCATTCTATTATCATAATCAGTGGTGTAGTCATAATCTAAATAATCAAACTCGTCATCCCAATTAAAATAAATATAATCAGTACATTGAATGGGTATGATATTGTACCCATACTGTATGTTGCTACATGTAGGTATTCCTTTATCCAATAATAGTTTAAATACATCTATATTAGGATTTGATGTTGTCGCGTATTTCTCTACAATATTAATACCATTTTTGATACTATTTATTTCTATACCTTTCGAAATTAGTAATTTCAATAAGTCTATATCGATGTTATCAGAACATAGATATTCAAATATATCAAAATCATTGATATTTTTATAGTCGACTGACGACAATAACAAAATCACAACATCGTTTTTGATATTATTATTTTTTTTGGTAACGTATGCCTTTAATGGAGTTTCACCATCATACTCATATAATGGATTTGCACCACTTTCTATTAATGATTGTGCACTACTGGCATCGATGTTAAATGTTTTACAACTATCATAGAGTATCTTATCGTTAACCATGATTGGTTGTTGATGTTATCACATTTTTTGGTTTCTTTCATTTCAGTTATGTATGGATTTAGCACGTTTGGGAAGCATGAGCTCATATGATTTCAGTACTGTAGTGTCAGTACTATTAGTTTCGATCAGATCAATGTCTAGATCTATAGAATCAAAACACGATAGGTCAGAAGATAATGAATATCTGTACGCTTCTTCTTGTACTGTAACTTCTGGTTTTGTTAGATGGTTGCATCGTGCTTTAACGTCAATGGTACAAATTTTATCCTCGCTTTGTGTATATCATATTCGTCTCTAGTATAAAATTCTATATTCAAATTATCATGCGATGTGTGTACGCTAACGGTATCAATAAACGGAGCACAGCATTTAGTCAACAGTAATCCAAAATTTTTTAAAGTATATCTTAACGAAAGAAGTTGTCATCGTTAGAGTGTGGTAAATCATTGTCTACGGTACTAGATCCTCATAAGTGTATATATCTAGAGTAATATTTAATTTATCAAATGGTTGATAATATGGATGTTGTGGCAATTTCCTAATACGGAAATAAGACATAAACACGCAATAAATCTAATTGCGGACATGTTACACTCCTTAAAAAATACGAATAAACACTTTGGCTTTTAGTAAGTGTCATTTAACACTATACTCATATTAATCCATGGACTCATAATCTCTATACGGGATTAACGGATGTTCTATATACGGGGATGAGTAGTTCTCTTCTTTAACTTTATACTTTTTACTAATCATATTTAGACTGATGTATGGGTAATAGTGTTTGAAGAGCTCGTTCTCATCATCAGAATAAATCAATATCTCTGTTTTTTTGTTATACAGATGTATTACAGCCTCATATATTACGTAATAGAACGTGTAATCTACCTTATTAACTTTCACCGCATAGTTGTTTGCAAATACGGTTAATCCTTTGACCTCGTCGATTTCCGACCAATCTGGGCGTATAATGAATCTTAACTTTAATTTCTTGTAATCATTCGAAATAATTTTTAGTTTGCATCCGTAGTTATCTCCTCTATGTAACTGTAAATTTCTCAACGCGATATCTCCATTAATAATGATGTCGAATTCGTGTTGTATACCCATACTGAATTGATGAACTAATACCGACGGTATTAATAGTAATTTACTTTTCATCTTTACATACTTGGTAATAGTTTTACTATCATAAGTTTATAAATTCCACAAGCTACTATGGAATATACCAACCATCTTAGTATAGAACACATGTCTTAAAGTTATTAATTAATTACATGTTGTTTTATATATCGCTACGAATTTAAACAGAGGAATCAGTTAGGAAGAAAAAATTATCTGTCATCATCATCATCTATTGGATAACATCTCTGTATTCTACGATAGAGTGCTATTTTAAGATGTGACAGATCCGTGTCATCAAATATATACTCCATTAAAATGATTATTCCGGCAGCGAACTTGATATTGGATACATCACGACCTTTGTTAATATCCACGACAATAGACAGCAATCCCATTGTTCCATAAACAGTGAGTTTATCTTTCTTTGAAGTGATATTTTGTAGAGATCTTATAAAACTGTCGAATGACATCGTATTTATATCTTTAGCTAAATCATATATGTTACCATCATAATATCTAACAGCATCTATCTTAAACGTTTCCATCGCTGTAAAGACGTTTCCGATAGATGGTCTCGTTTCATCAGTCATACTGAGCCAACAAATGTAATCGTGTATAACATCTTTGATAGAATCAGACTCTAAAGAAAAGGAATCGGCTTTATTATACACATTCATGATAAACTTAATGAAAAATGTTTTTCGTTGTTTAAGTTGGATGAATAGTATGTCTTAATAATTGTTATTATTTCACTAATTAATATTTAGTAACGAGTACACTCTATAAAAACGAGAATGACATAACTAATCATAACTAGTTATCAAAGAATGTCTAGGACGCGTAATTTTTTATGGTATAGATCCTGTAAGCGTTGTCTGTATTCTGGAGCTATTTTCTCTATCGCATTAGTGAGTTCAGAATATGTTATAAATTTAAATCGAATAACGAACATAACTTTAGTAAAGTCGTCTATATTAACTCTTTTATTTTCTAGCCATCGTAATACCATGTTTAAGATAGTATATTCTCTAGTTACTACGATCTCATCGTTGTCTAGAATATCGCATACTGAATCTACATCCAATTTTAGAAATTGGTCTGTGCTACATATCTCTTCTATATTATTGTTGATGTATTGTCGTAGAAAACTATTACGTAGACCATTTTCTTTATAAAACGAATATATAGTACTCCAATTATCTTTACCGATATATTTGCATACATAATCCATTCTCTCAATCACTACATCTTTAAGAGTTTGGTTGTTAAGATATTTGGCTAAACTATATAATTCTATTAGATCATCAACAGAATCAGTATATATTTTTCTAGATCCAAAGATGAACTCTTTGGCATCCTCTATAATATTATCAGAAAAGATATTTTCGTGTTTTAGTTTATCAAGATTTAACCTGTTCATATCCATGATTAACGACGTCATATAACCACATAAAATAAAAATCCATTTTCATTTTTAGCACAATACTATTCATAATTGATATTGATGTAATATTTTGTTACTTTGAACGTAAAGACAGTACACGGGTCCGTATCTCCAACAAGCACGTAGTAATCAAATTTGGTGTTGTTAAACTTCGCAATATTCATCAATTTAGATAGAAACATATACTCATCATCTGTTTTAGGAATCCATGTATTATTACCACTTTCCAACTTATCATTATCCCAGGCTATGTTTCGCCCATCATCGTTGCACAGAGTGAATAATTCTTTTGTATTCGGTAGTTCAAATATATGATCCATGCATATATCGACAAAGCTATTGTAGATGTGATTTTTCCTAAATCTAATATAAAACTCGTTTACTAGCAAACATTTTCCTGATTTATCGACCAAGACACACATGGTTTCTAAATCTATCAAGTGGTGGGGATCCATAGTTATAACGCAGTAACATAAATTATTACCTTCTTGACTGTCGCTAATATCTATATACTTATTGTTATCGTATTGGATTCTACATATAGATGGCTTGTATATCAAAGATATAGAACACATAACCAATTTATATTCTCGCTTTGTATTTTCGAATCTAAAGTTAAGAGATTTAGAAAACATTATATCATCGGATGATGATATCACTGTTTCCAGAGTAGGATATATTAAAGTCTTTAAAGATTTTGTCCGATTCAAATAAATCACTAAATAATATCCCATATTATCATCTGTTATAGTCGTGTCATTAAATCTATTATATTTTATGAAAGATATATCACTGCTCACCTCTATATTTCGTACATTTTTAAACTGTTTGTATAATATCTCTCTAATACAATCAGATATATCTATTGTGTCGGTAGACGATACCGTTACATTTGAATTAATGGTGTTCCATTTTACAACTTTTAACAAGTTGACCAATTCATTTCTAATAGTATCAAACTCTCCATGATTAAATATTTTAATAGTATCCATTTTATATCACTACGGACATAAACCATTGTATATTTTTTATGTTTATTAGTGTACACATTTTGGAAGTAAGTTCCTGGATCGGATGTCACCGCAGTAATATTGTTGATTATTTCTGACATCGACGTATTATATAGTTTTTTAATTCCATATCTTTTAGAAAAGTTAAACATCCTTATACAATTTGTGGAATTAATATTATGAATCATGGTTTTTACACATAGATCTATTACAGGCGGAACATCAATTATTATGGCAGCAACTAGTATCATTTCTACATTGTTTATGGTGATGTTTATCTTCTTCCAGCGCATATAGTCTAATATCGATTCAAACGCGTGATAGTTTATACCATTCAATATAATCGCTTCATCCTTTAGATGGTGATCCTGAATGTGTTTAAAAAATTATACGGAGACGCCGTAATAATTTCCCCATTGATAGAAAATATCACGCTTTCCATTTTCTTGAAGTACTAAAAGTAATTATAATATAATGTAAAGGTTTATATATTCAATATTTTTTATAAAAAAAATCATTTTGACATTAATTCCTTTTTAAATTTCAGTCTATCATCTATAGAAACATATTCTATGAATTTATAAAATGCTTTTACGTATCCTATCGTAGGTGATAGAACCGCTAAAAAACCTATCGAATTTCTACAAAAGAATCTGTTATATGGTATAGGGAGAGTATAAAACATTAAATGTCCGTACTTATTAAAGTATTCAGTAGCCAATCCTAACTCTTTCGAATAATTATTAATGGCTCTTATTCTGTACGAATCTATTTTTTTGAACAATGGACCTAGTGGTATATCTTGTTCTATGTATCTAAAATAATGTCTGACTAGATCCGTTAGTTTAATATCCGCAGTCATCTTGTCTAGAATGGCAAATCTAACTGCGGGTTTAGGCGTAGGCGTTAGTTTAGTTTCTATATCTACATCTATGTCTTTATCTAACACCAAAAATATAATAGCTAATATTTTATTACAATCATCCGGATATTCTTCTACGATCTCACTAACTAATGTTTCTTTGGTTATACTAGTATAGTCACGATCAGACAAATAAAGAAAATCAGATGATCGATGAATAATACATTTAAATTCATCATCTGTAAGATTTTTGAGATGTCTCATTAAAATATTATTAGTGTCAGTTCTCATTATCATATATTGACAGCAGCTATTACACTTATTTTATTTTTCTGTATTTTATTACTTTTCACCATATAGATCAGTCATTAGATCATCAAAATACTTTTCAATCATCCTAAAGAGTATGGTGAACGAATCTTCCCATCTAATTTCTGAACGTCTACCAATGTCTCTAGCCACTTTGGCACTAATAGCGATCATTCGCTTAACATCTTCTACATTATTAACTGGTTGATTCAATCTATCTAGCAATGGACCGTCGGATAGCGTCATTCTCATGTTCTTAATCAATGTACATACATCGTCATCATCTACCAATTCATCAAACAATATAAGCTTTTTAAAATCATCATTATAATAGGATGGATCGCCGTCATTTCTCCAAAGAATATATCTAATAAGTAGAGTCCTCATGCTTAGTAATTTAACTATTTTAGTTAACAACTATTTTTTATGTTAAATCAATTAGTAACACCGCTATGTTTAATACTTATTCATATTTTAGTTTTAGGATCGAGAATCAATACAAAAATTAATACATCAATTTTGGAAATACTTAGTTTCCACGTAGTCAATGAAACATTTGAGCTCATCGTAAAGGACGTTCTCGTACAAGACGTAACTATAAATTGGTTTATATTTGTTCAAGATAGATACAAATCCGATAACTTTTTTGACGAATTCTACGGGATTCACTTTAAAAGTGTCATACCGGGTTCTTTTTATTCTTTTAAACAGATCGATTGTGTGATGTTGATTAGGTCTTTTACGAATTTGATACAGAATAGCGTTTACATATCCACCATAGTAATCAATAGCCATTTGTTCGTATGTCATAAATTCTTTAATTATATGACACTGTGTATTATTTAGTTCGTCCTTGTTCATCATTAGGAATCTATCCAATATGGCAATTATATTAGAACTATAACTGCGTTGTATGCGCATGTTGATGTGTCTGTTTATACAATCAATTATACTAGGATCCATACCACTACAATCGGGTAAAATTGTAGCATCATATACCATTTCTAGTACTTTAGATTCATTGTTATCCATTGCAGAGGACGTCATGATCGAATCCAAAAAAATATATTATTTTTATGTTATTTTGTTAAAAATAATCATCAAATACTTCGTAAGATACTCCTTCATGAACATAATCAGTTACAAAACGTTTATATGAAGTAAAGTATCTACAATTTTTACAAAAGTCAGGATGCATAAGTACAAAGTACGCGATAAACGGAATAATAATAGATTTATCTAGTTTATCTTTTTCTATCTCTTTCATAGTTATATACATGGTCTCAGAAGTCGGATTATGTAACATCAGCTTCGATAAAATGACTGGGTTATTTAGTCTTACACATTCGCTCATACATGTATGACCGTTAACTATAGAGTCTACACTAAAATGATTGAATAATAGATAGTCTACCATTGTTTCGTATTCAGATAGTACAGCGTAGTACATGGCATCTTCACAAATTATATCATTATCTAATAGATATTTGACGCATCTTATGGATCCCACTTCAACAGCCATCTTAAAATCGGTAGAATCATATTGCTTTCCTTTATCGTTAATAATTTCTAGAACATCATCTCTATCATAAAAGATACAAATATTAACTGTTTGATCAGTAATAACATTGCTAGTCGATATCAATTTGTTAATAAGATGCGCTGGGCTCAATGTCTTAATAAGAAGTGTAAGAGGACTATCTCCGAATTTGTTTTGTTTATTAACATCCGTTGATGGAAGTAAAAGATTTATAATGTCTACATACTTGACTGTTTTAGAGCATACAATATGGAGAGGCGTATTTCCATCATGATCTGGTTTTGAGGGACTAATTCCTAGTTTCATCATCCATGAGATTGTAGAAGCTTTTGGATTGTCTGACATAAGATGTCTATGAATATGATTTTTGCCAAATTTATCCACTATCCTGGCTTCGAATCCGATAGACATTATTTTTTTAAACACTCTTTCTGAAGGATCTGTACACGCCAACAACGGACCACATCCTTCTTCATCAACCGAGTTGTTAATCTTGGCTCCATACTGTACCAATAAATTTATTCTCTCTATGACTTCATCATCTGTTCCCGAGAGATAATATAGAGGTGTTTTATTATGTTTATCACATGCGTTTGGATCTGCGCCGTGCACCAGCAGCATCGCGACTATTCTATTATTATTAATTTTAGAAGCTATATGCAATGGATAATTTCCATCATCATCCGTCTCATTTGGAGAGTATCCTCTATGAAGAAGTTCTTCTATAAATCGTTCATCTAGTCCTTTAATGCCACAATACGCATGTAGAATGTGATAATTTCCAGAGGGTTCGATAACTTGTAGCATATTCCTAAATACATCTAAATTTTTACTATTATATTTGGCATAAAGAGATAGATAATACTCGACCGACATAATGTTGTGTTGTCCATTATAGTATAAAAATTAATATTTCTATTTCTATATATTTGCAACAATTTACTCTCTATAACAAATATCATAACTTAGTTCTTTTATGTCAAGAAGGCACTGGTTTAATTCATCTATAAATGTCACTCCATAACTACCACGCATACTATACTCAGAATTATGATAAAGATATTTATTCTTGGGGTGTAAGTAATGGGGATTAATCTTTGTTGGATCAGTCTCTAAGTTAACACATGTCACACATGATCCATTTATAGTTATATCACACGATGATGATTTATGAATTGATTCCGGAAGATCGCTATTGTATTTTGTAGTTCCACAATTCATTTCCATACATGTTATTGTCACACTAATATTATGATGAACTTTATCTAGCCGCTGAGTGGTAAACAACAGAACAGATAGTTTATTATCTTTACCAACACCCTCAGCCGCTGCCACAAATCTCTGATCCGTATCCATGATGGTCATGTTTACTTTTAGTCCGTATCCAGTCAACACTATGTTAGCATTTCTGTCGATATAGCTTTCACTCATATGACACTCACCAATAATTGTAGAATTAATGTCGTAATTTACACCAATAGTGAGTTCGGCGACAAAGTACCAGTACCGGTAATCTTGTCGATGAGGACATATAGTATTCTTGTATTCTACCGAATACCCGAGAGATGCGATACAAAAGAGTAAGACTAATTTGTAAACCATCTTACTCAAAAATATGCGACAATAGGAAATCTATCTTATACACATAATTATTCTATCAATTTTACCAATTAGTTAGTGTAATGTTAACAAAAATGTGGGATAATTTAATAGTTTTTCCTTACACAATTGACATACATGAGTCTGAGTTCCTCGTTTTTGCTAATTATTTCGTCCAATTTATTATTCTTGACATCGTCAAGATCTTTTGTATAGGAGTCAGACTTGTATTCAACATGTTTTTCTATAATCATCTTAGCTATTTCGGCATCATCCAATAGTACATTTTCCAGATTAACAGAATAGATATTAATGTCGTATTTGAACAGAGCCTGTAACATCTCAATGTCTTTATTATCTATAGCCAATTTGATGTCCGGAATGAAGAGAAGGGAATTGGTGTTTGTTGACGTCATATAGTCGAGCAAGAGAATCATCATATCCACGTGTCCATTTTTTATAGTGGTGTGAATACAACTAAGGAGAATAGCCAGATCAAAAGGAGATGGTATCTCTGAAAGAAAGTAGGAAACAATACTTACATCATTAAGCATGACAGCATGATAAAATGAAGTTTTCCATCCAGTTTTCCCATAGAACATCAGTCTCCAATTTTTCTTAACAAACAGTTTTACCGTTTGCATGTTACCACTATCAACCGCATAATACAATGCGGTGTTTCCTTTGTCATCAAATTGTGAATCATCCATTCCACTGAATAGCAAAATCTTTACTATTTTGGTATCTTCTAATGTGGCTGCCTGATGTAATGGAAATTCATTATCTAGAAGATTTTTCAATGCTCCAGCGTTCAACAACGTACATACTAGACGCACGTTATTATCAGCTATTGCATAATACAAGACACTATGACCATTGATATCCGCCTTAAATGCATCTTTGCTAGAGAGAAAGCTTTTCAGTTGCTTAGACTTCCAAGTATTAATTCGTGACAGATCCATGTCTGAAACGAGACGCTAATTAGTGTATATTTTTTCATTTTTTATAATTTTGTCATATTGTACCAGAATTAATAATATCTCTAATAGATCTGATTAGTAGATATATGGCTATCGCAAAACAACATATACACATTTAATAAAAATAATATTCATTAAGAAGATTCAGATTCCACTGTACCCATCAATATAAAATAAAATAATTATTCCTTACATCGTACCCATAAACAATATATTAAGTATATTCCACCTTACCCATAAACAATATAAATCCAGTAATATCATGTCTAATGATGAACACAAATGGTGTATTAAATTCCAGTTCTTCAGGAGATGATCTCGCCGTAGCTACCATGATAGTAGATGCCTCCGCTACAGTTCCTTGTTCGTCTACATCTATCTTTACATTCTGAAACATTTTATAAATATATAATGGGTCCCTAGTCATATGTTTAAACGACGCCTTATCTGGATTAAACATACTAGGAGCCATCATTTCGGCTATCGACTTAATATCCCTCTTGTTTTCGATAGAAAATCTAGGGAGTTTAAGATTGTACATTTTATTCCCTAATTGAGATGACCAATATTCTAATTTTGCAGGCGTGATAGAATCTGTGAAATGGGTCATATTATCACCTATTGCCAGGTACATACTAATATTAGCATCCTTATACAGAAGGCGCACCATATCATATTCTTCGTCATCGATTGTGATTGTATTTCCTTGCAATTTAGTAACTACGTTCATCATGGGAACCGTTTTCGTACCGTACTTATTAGTAAAACTAGCATTGTGTGTTTTAGTGATATCAAACGGATATTGCCACGTACCTTTAAAATATATAGTATTAATGATTGCCCATAGAGTATTATCGTCGAGCATAGTAGAATCAACTACATTAGACATACCAGATCTACGTTCTACTATAGAATTAATTTTATTAACCGCATCTCGTCTAAAGTTTAATCTATATAGGCCGAATCTATGATATTGTTGATAATACGACGGTTTAATACACACAGTACTATCGACGAAACTTTGATACGTTAGATCTGTGTACGTATATTTAGATGTTTTCATCTTAGCTAATCCGGATATTAATTCTGTAAATGCTGGACCCAGATCTCTTTTTCTCAAATTCATAGTATTCAATAATTCTACTCTAGTATTACCTGATGCAGACAATAGCGACATAAACATAGAAAACGAATACCCAAACGGTGAGAAGACAATATTATCATTATCATCCTCATCCCCATTTTGAATATTTTTATACGCTAATATACCAGCATTGATAAATCCCTGCAGACGATATGCGGATACTGAACACGCTAATGATAGTATCAATAACGCAATCATGATTTTTATGGTATTAATAATTAACCTTATTTTTATGTTTGGTATAAAAATTATTGATGTCTACACATCCTTTATAATCAACTCTAATCACTTTAACTTTTACAGTTTTCCCTACAAGTTTATCCCTATATTCAACATATCTATCCATATGCATCTCTTAACACTCTGCCAAGATAGCTTCAGAGTGAGGATAGTCAAAAAGATAAATATATAGAGTATAATCATTCTCGTATACTCTGCCCTTTATTACATCGCCCGCATTGGGCAACGAATAACAAAATGCAAGCATCGTGTTAACGGGCTCGTAAATTGGGATAAAATTATGTTTTTATTGTTTATCTATTTTATTCAAGAGAATATTCAGGAAGTTCCTTTTCTGGTTGTATCTCGTCGCAGTATATATCATTTGTACATTGTTTCATATTTTTTAATAGTCTACACCTTTTAGTAGGACTAGTATCGTACAATTCATAGCTGTATTTTGAATTCCAATCACGTATAAAAATATCTTCCAATTGTTGACGAAGACCTAATCCATCATCCGGTGTAATATTAATAGATGCTCCACATATATCCGTAAAGTAATTTCCTGTCCAATTTGATGTACCTATATACGCCGTTTTATCGGTTACCATATATTTTGCATGGTTTACCCTAGAATACGGAATGGGAGGATCAGCATCTGGTACAATAAATAGCTTTACTTCTATATCTATGTTTTTAGATTTTAGCATAGCTATAGATCTTAAAAAGTTTCTCATGATAAACGAAGATCGTTGCCAGCAACTAATCAATAGCTTAACGGATACTTGTCTGTCTATAGCGGATCTTCTTAATTCATCTTCTATATAAGGCCAAAACAAAATTTTACCCGCCTTTGAATAAATAATAGGAATAAAGTTCATAACAGATACATAAACGAATTTACTCGCATTTCCGATACATGACAATAAAGCGGTTAAATCATTGGTTCTTTCCATAGTACATAATTGTTGTGGTGCAGAAGCAATAAATACAGAGTGTGGAACACCGCTTACGTTAATACTAAGAGGATGATCTGTATTATAATACGACGGATAAAAGTTTTTCCAATTATATGGTAGATTGTTAACTCCAAGATACCAGTATACCTCAAAAATTTGAGTGAGATCCGCTGCCAAGTTCCTATTATTGAAGATCGCAATACCCAATTCCTTGACCTGAGTTAGTGATCTCCAATCCATGTTAGCGCTTCCTAAATAAATATGTGTATTATCAGATATCCAAAATTTTGTATGAAGAACTCCTCCTAGGATATTTGTAATATCTATGTATCGTACTTCAACTCCGGCCATTTGTAGTCTTTCAACATCCTTTAATGGTTTGTTGGATTTATTGACGGCTACTCTAACTCTTACTCCTCTTTTGGGTAATTGTACAATCTCGTTTAATATTACCGTGCCGAAATTCGTACCCACTTCATCCGATAAACTCCAATAAAAAGATGATATATCTAGTGTTTTTATGGTATTGGATAGAATTTCCCTCCACATGTTAAATGTAGTCAAATATACTTTATCAAATTGCATACCTATAGGAATAGTCTCTGTAATCACTGCGATTGTATTATCCGGATTCATTTTATTTGTTAAAAAAATAATCCTATATCACTTCACTCTATTAAAAATCCAAGTTTCTATTTCTTTCATGACTGATTTTTTAACTTCATCCGTTTCCTTATGAAGATGATGTTTGGCGCCTTCATAAATTTTTATTTCCCTATTACAATTTGCATGTTGCATGAAATAATATGCACCTGAAACATCGCTAATCTCATTGTTTGTTCCCTGGAGTATGAGAGTCGGGGTGTTAATCTTGGGAATTATTTTTCTAACCTTGTTGGTAGCCTTCAAGACCTGACTAGCAAATCCAGCCTTAATTTTTTCATGATTGACTAATGGATCGTATTGGTATTTATAAACTTCATCCATATCTCTAGATACTGATTCTGGACATAGCTTTCCGACTGACGCATTTGGTGTAATGGTTCCCATAAGTTTTGCAGCTAGCAGATTCAGTCTTGGAACAGCGTCTGCATTAACTAGAGGAGACATTAGAATCATTGCTGTAAACAAGTTTGGATTATCGCAAGCAGCTAGTATAGAAATTGTTGCTCCCATGGAATGACCCAATAAGAAGACTGGAACTCCTGGATAAGTAGATTTAATAGTCACCACGTGCTGTACCACATCTCTAACATACTTACCAAAGTCATCAATCATCATTTTTTCACCATTACTTCTTCCATGGCCAATATGATCATGTGAGAATACTAAAATTCCTAACGATGATATGTTTTCAGCTAGTTCGTCATAACGTCCAGAATGTTCACCAGCTCCATGACTTATGAATACTAATGCCTTAGGATATGTAATAGGTTTCCAATATTTACAATATATGTAATCATTGTCCAGATTGAACATACAGTTTGTACTCATGATTCACTATATAACTATCAATATTAACAGTTCGTTTAATGATCATATTATTTTTATGTTTTATTGATAATTGTAAAAATATACAATTAAATCAATATAGAGGAAGGAGACGGTACTGTATTTTGTGAGATAGTAATGGAGACTAAATCAGATTATGAGGATGCTGTTTTTTACTTTGTGGATGATGATGAAATATGTAGTCGCGACTCCATCATTGATCTAATAGATGAATATATCACGTGGAGAAATCATGTTATAGTGTTTAATAAAGATATTACCAGTTGTGGAAGACTGTACAAGGAATTGATAAAGTTCGATGATGTCGCTATACGGTACTATGGTATTGATAAAATTAATGAGATTGTCGAGGCTATGAGTGAAGGAGACCACTACATCAATCTTACAGAAGTCCATGATCAGGAAAGTCTATTCGCTACCATAGGAATATGTGCTAAAATCACTGAACATTGGGGATACAAAAAGATTTCAGAATCTAAATTCCAATCATTGGGAAACATTACAGATCTGATGACCGACGATAATATAAACATCTTGATACTTTTTCTAGAAAAAAAAATGAATTGATGATATAAGTGTCTTCATAACGCATTATTACGTTAGCATTCTATTATCCAGTGTTAAAAAAATTATCCTATCATGTATTTGAGAGTCTTATATGTAGCAAACATGATAACTGCAATACCCATAATCTTTAGATATTCACGCGTGCTATGGATGGCATTATCCCGCGGTGCGGAAATGTACGTTATATAATCTACAAAATAATCATCGCATATAGTATGAGATAGTAGAGTAAACATTTTTATCGTTTCTACTGGGTTCATACATCGTCTACCCAATTCGGTAATGAATGAAATTGTCGCCAATCTTACACCCAAACCCTTGTTGTTCATTAGTATAGTATTAACTTCATTATTTATGTCATAAACTGTAAATGATTCTGTAGATGCCATATCACACATGATATTCATGTCACTATTATAATCATTATTAACTTTATCACAATACGTGTTGATAATATCTACATATGATCTAGTTTTTGTGGGTAATTGTCTATACAAGTCGTCTAAACGTTGTTTACTCATATAGTATTGAACAGCCATCATTACATGGTCCCGTTCCGTTGATAGATAATCGAGTATGTTAGTAGACTTGTCAAATCTATATACCATATTTTCTGGAAGCGGATATACATAGTCGCGATCATCATTATCACTAGCCTCATCCTCTATATCATGTACATGTACATAATCTATGATATTATTATACATAAACATCGACAACATACTATTGTCTATTATCTAAGTCCTGTTGATCCAAACCCTTGATCTCCTCTATCTGTACTATCTAGAGATTGTACTTCTTCAAGTTCTGGATAATATATACGTTGATAGATTAGCTGAGCTATTCTATCTCCAGTATTTACATTAAACGTACATTTTCCATTATTAATAAGAATGACTCCTATGCTTCCCCTATAATCTTCGTCTATTACACCGCCTCCTATATCAATGCCTTTTAGGGACAGACCAGACCTAGGAGCTATTCTACCATAGCAGAACTTAGGCATGGACATACTAATATCTGTCTTAATTAACTGTCGTTCTCCAGGAGGGATAGTATAATCGTAAGCGCTATACAAATCATATCCGGCAGCACCCGGCGATTGCCTAGTAGGTGATTTAGCTCTGTTAGTTTCCTTAACAAATCTAACTGGTGAGTTAATATTCATGTTGAACATAAAAAATATCATTTTATTTCAAAATTATTTACCATTCCATTCCATCCCATATATTCCATGAATAAGTGCGATTATTGTACACTTCTATAGTATCTATATACGATCCACGATAAAATCCTCCTATCAATAGCAGTTTATTATCCACTATGATCAATTCTGGATTATCCCTCGGATAAATAGGATCATCTATCAGAGTCCATGTATTGCTGGATTCACAATAAAATTCCGCATTTCTACCAACCAAGAATAACCTTCTACCAAACACTAACGCACATGATTTATAATGAGGATAATAAGTGGATGGTCCAAACTGCCACTGATCATGATTGGGTAGCAAATATTCTGTAGTTGTATCAGTTTCAGAATGTCCTCCCATTACGTATATAACATTGTTTATGGATGCCACTGCTGGATTACATCTAGGTTTCAGAAGACTCGGCATATTAACCCAAGCAGCATCCCCGTGGAACCAACGCTCAACAGATGTGGGATTTGGTAGACCTCCTACTACGTATAATTTATTGTTAGCGGGTATCCCGCTAGCATACAGTCTGGGGCTATTCATCGGAGGAATTGGAATCCAATTGTTTGATATATAATTTACCGCTATAGCATTGTTATGTATTTCATTGTTCATCCATCCACCGATGAGATATACTACTTCTCCAACATGAGTACTTGTACACATATGGAATATATCTATAATTTGATCCATGTTCATAGGATACTCTATGAATGGATACTTGTATGATTTGCGTGGTTGTTTATCACAATGAAATATTATGTTACAGTCTAGTATCCATTTTACATTATGTATACCTCTGGGAGAAAGATAATTTGACCTGATTACATTTTTGATAAGAAGTAGCAGATTTCCTAATCTATTTCTTCGCCTCATATACCACTTAATGACAAAATCAACTACATAATCCTCATCTGGAACATTTAGTTCGTCGCTTTCTAGAATAAGTTTCATAGATAGATAATCAAAATTGTCTATGATGTCATCTTCCAGTTCCAAAAAGTGTTTGGTAATAAAGTCTTTAGTATGACATAAGAGATTGGATAGTCCGTATTCTATACCCATCATGTAACACTCGATACAATATTCCTTTCTAAAATCTCGTAGGATAAAGTTTATACAAGTGTAGATGATAAATTCTACAGATGTTAATATAGAAGCACGTAATAAATTGACGACGTTATGACTATCTATATATACCTTTCCAGTATATGAGTAAATAACTATAGAAGTTAGACTGTGAATGTCAAGGTCTAGACAAACCCTCGTAACTGGATCTTTATTTTTTGTGTATTTTTGGCGTAAATGTGTGCGAAAGTATGGAGATAACTTTTTCAATATCGTAGAATTGACTATTATATTACCTCCTATAGCTTCAATAATTGTTTTGAATTTCTTAGTCGTGTACAATGCTAATATATTCTTACAGTACACAGTATTGACAAATATCGGCATTTATGTTTCTTTAAAAGTCAACATCTAAAGAAAAATGATTGTCTTCTTGAGACATAACTCCCATTTTTTGGTATTCACCCACACGTTTTTCGAAAAAATTAGTTTTTCCTTCCAATGATATATTTTCCATGAAATCAAACGGATTGGTAACATTGTAAATTTTTTTAAATCCCAATTCAGAAATCAATCTATCTGCGACGAATTCTATATATGTTTTCATCATTTCACAATTCATTCCTATGAGTTTAACTGGAAGAGCCACAGTAAGAAATTCTTGTTCAATGGATACCGCATTTGTTATAATAAATCTAACGGTTTCTTCACTCGGTGGATGCAATAAATGTTTAAACATCAAACATGCGAAATCGCAGTGCAGACCCTCGTCTCTACTAATTAATTCGTTAGAAAACGTGAGTCCGGGCATTAGGCCACGCTTTTTAAGCCAAAATATGGAAGCGAATGATCCGGAAAAGAAGATTCCTTCTACTGCAGCAAAGGCAATAAGTCTCTCTCCATAACCGGCGCTGTCATGTATCCACTTTTGAGCCCAATCGGCCTTCTTTTTTACACAAGGCATCGTTTCTATGGCATTAAAGAGGTAGTTTTTTTCATTACTATCTTTAACATAAGTATCGATCAAAAGACTATACATTTCCGAATGAATGTTTTCAATGGCCATCTGAAATCCGTAGAAACATCTAGCCTCGGTAATCTGCACTTCTGTACAAAATCGTTCTGCTAAATTTTCATTCACTATTCCGTCACTGGCTGCAAAAAACGCCAATACATGTTTTATAAAATATTTTTCGTCTGGTGTTAGTTTATTCCAGTCATTGATATCTTTAGATATATCCACTTCTTCCACTGTCCAAAATGATGCCTCTGCCTTTTTATACATATTCCAGATGTCATGATATTGGATTGGGAAAATAACAAATCTATTTGGATTTGGTGCAAGGATAGGTTCCATAACTAAATTAACAATAGTAGTAATTTTTTTTCAGTTATCTGTATGACGACTGTACTTGGATCTTTTGTATATCGCTATCGCCGCAATCACTACAATAATTACAAGTATTATTGATAGCATTGTTATTACTACTATCATAATTAAATTATCGACATTCATGGGTGTTGAATAATCGTTATCATCATTTTGTAATTGTGACGTCATACTAGATAAATCATTTGTGAGATTGTTGTGGGAAGCGGGCACGGAAGATGCATTATCATTATTATTTAACGCCTCCCATTCGGATTCACAAATGTTACGCACGTTCAAAGTTTTATGGAAACTATAATTTTGTGAAAACAGATAACAAGAAAACTCGTCATCGTTCAAATTTTTAACGATAGTAAACCGATTAAACGTCGAGCTAATTTCTAACGCTAGCGACTCTGTTGGATATGGGTTTCCAGATATATATCTTTTCAGTTCCCCTACGTATCTATAATCATCTGTAGGAAATGGAAGATATTTCCATTTATCTACTGTTCCTAATATCATATGCGGTGGTGTAGAACCATTAAGCGCGAAAGATGTTATTTCGCATCGTATTTTAACTTCGCAATAATTTCTGGTTAGATAACGCACTCTACCAGTCAAGTCAATGATATTAGCCTTTACAGATATATTCATAGTAGTCGTAACGATGACTCCATCTTTTAGATGTGATACTCCTTTGTATGTACCAGAATCTTCGTACCTCAAACTCGATATATTTAAACAAGTTAATGATATATTAACGCGTTTTATGAATGATGATATATAACCAGAAGTTTTATCCTCTGTGGCTAGCGCTATAACCTTATCATTATAATACCAACTAGTGTAATTAATATGTGACATGACAGTGTGGGTACAAATATGTACATTATCGTCTACGTCGTATTTGATACATCCGCATTCAGCCAACAAATATAAAATTACAAAAACTCTAACGACGTTTGTACACATCTTGATGCGGTTTAATAAATGTTTTGATTTCAATTTATTGTAAAAAAAGATTCGGTTTTATACTGTTCGATATTCTCATTGCTTATATTCTCATCTATCATCTCCACACAGTCAAATCCATGGTTAACATGTACCTCATCAACCGGTAAAAGACTATCGGATTCTTCTATCATCATAACTCGAGAATATTTAATTTGGTGGTCATTATTAATCAAGTCAATTATCTTATTTTTAACAAACGTAAGTATTTTACTCATTTTTTTATAAAAACTTTTAGAAATATACAGACTCTATCGTGTGTCTATATCTTCTTTTTATATCCAATGTATTTATGTCTGATTTTTCTTCATTTATCATATATAATGGTCCAAATTCTACACGTGCTTCGGATTCATCCAGATCATTAAGGTTCTTATAATCGCAACATCCTTCTCTTCCATCTTCTACATCTTCCTTCTTATTCTTAGCGTCACAGAATCTACCACAGCAGGATCCCATGACGAGAGTCACATTAAACTAATTCATTTTCAATTATAATATACTGATTAGTAATGACAATTAAAATAAAAATATTCTTCATAACCGGTAAGAAAGTAAAAAGTTCACATTGAAACTATGTCAGTAGTTATACATCATGAGATGATATACTCTATTTTGGTGGAGGATTATATGATATAATTCGTGGATAATCATTCTTAAGACACATTTCTTCATTCGTAAATCTTTTCACATTAAATGAGTGTCCATATTTTGCAATTTCTTCATATGATGGCGGTGTACGTGGACGAGGCTGCTCCTGTTCTTGTAGTCGTCGACTGTCGTGTTTGCGTTTAGATCCCTCCATTATCGCGATCGCGTAGTGAGTACTATTTATACCTTGTAATTAAATTTTTTTATTAATTAAACGTATAAAAACGTTCCGTATCTGTATTTAAGAGCCAGATTTCGTCTAATAGAACAAATAGCTACAGTAAAAATAACTAGAATAATCGCTACACCCACTAGAAACCACGGATCGTAATACGGCAATCGGTTTTCGATAATAGGTGGAACGTATATTTTATTTAAGGACTTAACAATTGTCTGTAAACCACAATTTGCTTCCGCCGATCCTGTATTAACTATCTGTAAAAGCATATGTTGACCGGGAGGAGCCGAACATTCTCCGATATTCAATTTTTGTATATCTATAATGTTATTAACCTCCGCATACGCATTACAGTTCTTTTCTAGCTTGGATACTACACTAGGTACATCATCTAAATCTATTCCTATTTCCTCAGCGATAGCTCTTCTATCCTTTTCCGGAAGTAATGAAATCACTTCAATAAATGATTCAACCATGAGTGTGAAACTAAGTCGAGAATTACTCATGCATTTGTTAGTTATTCGGAGCGCGCAATTTTTAAACTGTCCTATAACCTCTCCTATATGAATAGCACAAGTGACATTAGTAGGGATAGAATGTTGAGCTAATTTTTGTAAATAACTATCTATAAAAAGATTATACAAAGTTTTAAACTCTTTAGTTTCTGCCATTTATCCAGTCTGAGAAAATGTCTCTCATAATAAATTTTTCCAAGAAACTAATTGGGTGAAGAATGGAAACCTTTAATCTATATTTATCACAGTCTGTTTTGGTACACATGATGAATTCTTCTAATGCTGTACTAAATTCGATATCTTTTTCGATTTCTGGATATGTTTTTAATAAAGTATGAACAAAGAAATGGAAATCGTAATACCAGTTATGTTCAACTTTGAAATTGTTTTTTATTTTCTTGTTAATGATTCCAGCCACTTGGGAAAAGTCAAAGTCGTTTAATGCCGATTTAATACGTTCATTAAAAACAAACTTTTTATTCTTTAGATGAATTATTATTGGTTCATTGGAATCAAAAAGTAAGATATTATCGGGTTTAAGATCTGCATGTAAAAAGTTGTCACAACAGGGTAGTTCGTAGATTTTAATGTATAACAGAGACATCTGTAAAAAGATAAACTTTATGTATTGTACCAAAGATTTAAATCCTAATTTGATAGCTAACTCGGTATCTACTTTATCTGCCGAATACAGTGCTAGGGGAAAAATTATAATATTTCCTCTTTCGTATTCGTAATTAGTTCTCTTTTCATGTTCGAAAAAGTGAAACATGCGGTTAAAATAGTTTATAACATTAATATTACTGTTAATAACTGCCGGATAAAAGTGGGATAGTAATTTCACGAATTTGATACTGTCCTTTCTCTCGTTAAACGCCTTTAGAAAAACTTTAGAAGAATATCTCAATGAGAGTTCCTGACCATCCATAGTTTGTATCAATAATAGCAACATATGAAGAACCCGCTTATACAGAGTATGTAAAAATGTTAATTTATAGTTTAATCCCATGGCCCACGCACACACGATTAATTTTTTTTCATCTCCCTTTAGATTGTTGTATAGAAATTTGGGTACTGTAAACTCCGCCGTAGTTTCCATGGGACTATATAATTTTGTGGCCTCGAATACAAATTTTACTACATAGTTATCTATCTTAAAGACTATACCATATCCTCCTGTAGATATGTGATAAAAATCGTCGTTTATAGGATAAAATCGTTTATCTTTTTGTTGGAAAAAGGATGAATTAATGTAATCATTCTCTTCTATCTTTAGTAGTGTTTCCTTATTAAAATTCTTAAAATAATTTAACAATCTAACTGACGGAGCCCAATTTTGGTGTAAATCTAATTGGGACATTATGTTGTTAAAATATAAACAGTCTCCTAATATAACAGTATCTGATAATCTATGGGGAGACATCCATTGATATTCAGGGGATGAATCATTGGCAACACCCATTTATTGTACAAAAAGCCCCAATTTACAAACGAAAGTCCAGGTTTGATAGAGATAAACTATTAACTATTTTGTCTCTGTTTTTAACACCTCCACAGTTTTTAATTTCTTTGGTAATGAAATTATTCACAATATCAGTATCTTCTTTATCTACCAGAGATTTTACTAACTTGATAACCTTGGCTGTCTCATTCAATAGGGTAGTGATATTTGTATGTATGATATTGATATCTTTTTGAATTGTTTCTTTTAGAAGTGATTCTTTGATGGTATCAGCATACGAATTACAATAATGCAGAAACTCAGTTAACATGCAGGAATTATAGTAAGCCAATTCCAATTGTTGCCTGTATTGTATTAGAGTATTAATATGCGCAATGATGTCCTTGCGTTTCTCTGATAGAATGCGAGCAGCGATTTTGGCGTTATCATTTGACGATATTTCTGGAATGACGAATCCTGTTTCTACTAACTTCTTGGTAGGACAAAGTGAAACAATCAAGAAAATAGCTTCTCCTCCTATTTGTGGAAGAAATTGAACTCCTCTAGATGATCTACTGACGATAGTATCTCCTTGACAGATATTGGACCGAACTACGGAAGTACCTGGAATGTAAAGCCCTGAAACCCCCTCATTTTTTAAGCAGATTGTTGCCGTAAATCCTGCACTATGCCCAAGATAGAGAGCTCCTTTGGTGAATCCATCACTATGTTTCAGTTTAACCAAGAAACAGTCAGCTGGTCTAAAATTTCCATCTCTATCTAATACAGAATCCAACTTGATGTCAGGGACTATGACCGGTTTAATGTTATATGTAACATTGAGTAAATCCTTAAGTTCATAATCATCGTTGTCATCAGTTATGTACGATCCAAACAATGTTTCTACCGGCATGGTGGATACGAAGATGCTATCCATCAGAATGTTTCCCTGATTAGTATTTTCTATATAGCTATTCTTCTTTAAACGATTTTCCGAATCAGTAACTATGTTCATTTTTTTAGGAGTAGGACGTCTAGCCAGTATGGAAGAGGATTTTCTAGATACTCTCTTCAACATCTTTGATCTCAATGGAATGCAAAACCCCATGGTGTAACAACCAACGATAAAAATAATATTGTTTTTTCACTTTTTATAATTTTACCATCTGACTCATGGATTCATTAATATCTTTACAAGAGCTACTAACGTATAATTCTTTATAACTGAACTGAGATATATACACCGGATCTATGGTTTCCATAATTGAGTAAATGAATGCTCGGCAATAACTAATGGCAAATGTATAGAACAACGAAATTATACTAGAGTTGTTAAAGTTAATATTTTCTATGAGTTGTTCCAATAAATTATTTGTTGTGACTGCGTTCAAGTCATAAATTATCTTGATACTATCCAGTAAACAGTCTTTAAGTTCTGGAATATTATCATCCCATTGTAAAGCCCCTAGTTCGACTATCGAATATCCTGCTCTGATAGCAGTTTCAATATCGACGGACGTCAATACTGTAATAAAGGTGGTAGTATTGTCATCATCGTGATAAACTACGGGAATATGGTCGTTAGTAGGTACCGTGACTTTACACAACGCGATATATAACTTTCCTTTTGTACCATTTTTAACGTAGTTGGGACGTCCTGCAGGGTATTGTTTTGAAGAAATGATATCGAGAACAGATTTGATACGATATTTGTTGGATTCCTGATTATTCACTATAATATAATCTAGACAGATAGATGATTCGATAAATAGAGAAGGTATATCGTTGGTAGGATAATACATCCCCATTCCAGTATTCTCGGATACTCTATTGATGACACTAGTTAAGAACATGTCTTCTATTCTAGAAAACGAAAACATCCTACATGGACTCATTAAAACTTCTAACGCTCCTGATTGTGTTTCGAATGCCTCGTACAAGGATTTCAAGGATGCCATAGATTCTTTGACCAACGATTTAGTATTGCGTTTAGCATCTGATTTTTTTATTAAATCAAATGGTCGGCTCTCTGGTTTACTACCCCAATGATAACAATAGTCTTGTAAAGATAAACCGCAAGAAAATTTATACACATCCATCCAAATAACCCTAGCACCGTCGGATGATATTAATGTATTATTATAGATTTTCCATCCACAGTTATTGGGCCAGTATACTGTTAGCAACGGTATATCGAATAGATTACTCATGTAACCTACTAGAATGATAGTTCGTGTACTAGTCATAATATCTTTAATCCAATCTAAGAAATCTAAAATTAGATCTTTTACACTATTAAAGTTAACAAAGGTATTACCCGGGTACGTGGATATCATATATGGTATTGGTCCATTATCAGTAATGGCTCCATAAACTGATACGGCGATGGTTTTTATATGTGTTTGATCTAATGAGGACGAAATTCGCGCCCACAATTCATCTCTAGATATGCATTTAATATCGAACGGTAACACATCAATCTCGGGACGCGTATATGTTTCTAAATTCTTAATCCAAATATAATGATGACCTATATGCCCTATTATCATACTGTCAACTATAGTATACCTAGAGAACTTTCGATACATCTGCTGTTTCCTGTAATCGTTAAATTTTACAAATCTATAACATGCTAAACCTTTTGACGACAGCCATTCATTAATTTCTGATATGGAATCTGTATTCTCAATACCGTATCGTTCTAAAGCCAGTGCTATATCTCCCTGTTCGTGGGAACGCTTTCGTATAATATCGATCAATGGATAATATGAAGTTTTTGGAGAATAATATGATTCATGATCTATTTCGTCCATAAACAATCTAGACATAGGAATTGGAGGCGATGATCTTAATTTTGTGCAATGGGTCAATCCTATAACTTCTAATATTGTAATATTCATCATCGACATAACACTATCTATGTTATCATCGTATATTAGTATACCACGACCTTCTTCATTTCGTGCCAAAATGATATACAGTCTTAAATAATTACGCAATATCTCAATAGTTTCATAATTGTTAGCTGTTTTCATCAAGGTTTGTATCCTGTTTAACATGATGGCGTTCTATAACGTCTCTATTTTCTATTTTTAATTTTTTTAAATTTTTAACGATTTACTGTGGCTAGATACCCAATCTCTCTCAAATATTTTTTTAGCCTTGCTTACAAGCTGTTTATCTATACTATTAAAACTGACGAATCCGTGATTTTGGTAATGGGTTCCGTCGAAATTTGCCGAAGTGATATGAACATATTCGTCGTCGACTATTAACAATTTTGTATTATTCTGAATAGTGAAAACCTTCACAGATAGATCATTTTGAACACACAACGCATCTAGACTTCTGGCGGTTGCCATAGAATATACGTCGTTCTTATCCCAATTACCAACTAGAAGTCTGATCTTAACTCCTCTATTAATGGCTGCTTCTATAATGGAGTTGTAAATGTCAGGCCAATAGTAGCTATTACCGTCGACACGTGTAGTGGGAACTATGGCCAAATGTTCAATATCTATACTAGTCTTAGCCGACTTGAGTTTATCAATAACTACATCAGTGTCTAGATCTCTAGAATATCCCAATAGGTGTTCTGGAGAATCAGTAAAGAACACTCCACCTATAGGATTCTTAATATGATACGCAGTGCTAACTGGCAGACAACAAGCCGCAGAGCATAAATTCAACCATGAATTTTTTGCGCTATTAAAGGCTTTAAAAGTATCAAATCTTCTACGAAGATCTGTGGCCAGCGGAGGATAATCAGAATATACGCCTAACGTTTTAATCGTATGTATAGATCCTCCAGTAAATGACGCGTTTCCTACATAACATCTTTCATCATCAGACACCCAAAAACAACCGAGTAGTAGTCCCACATTATTTTTTTTATCTATATTAACGGTTATAAAATTTATATCCGGGGAGTGACTTTGTAGCTCTCCCAGATTTCTTTTCCCTCGTTCATCTAGCAAAACTATTATTTTAATCCCTTTTTCAGATACCTCTTTTAGTTTATCAAAAATAAGCGCTCCCCTAGTAGTACTCAGAGGATTACAACAAAAAGATGCTATGTATATATATTTCTTAGCTAGAGTGATAATTTCGTTAAAACATTCAAATGTTGTCAAATGATCGGATCTAAAATCCATATTTTCTGGTAGTGTTTCTACCAGCCTACATTTTGCTCCCGCAGGTACCGATGCAAATGGCCACATTTAGTTAACATAAAAACTTATATATCCTGTTCTATCAACGATTCTAGAATATCATCGGCTATATCGCTAAAATTTTCATCAAAGTCGACATCACAACCTAACTCAGTCAATATATTAAGAAGTTCCATGATGTCATCTTCGTCTATTTCTATATCCGTATCCATTGTAGATTGTTGACCGATTATCGAGTTTAAATCATTACTAATACTCAATCCTTCAGAATACAATCTGTGTTTCATTGTAAATTTATAGGCGGTGTATTTAAGTTGGTAGATTTTCAATTATGTATCAATATAGCAACAGTAGTTCTTGCTCCTCCTTGATTTTAGCATCCTCTTCATTATTTTCTTCTACGTACATAATCATGTCTAATACGTTAGACAACACACCGACGATGGTGGCCGCCACAGACACGAATATGACTAGACCGATGACCATTTAAAAAATACTCTCTAGCTTTAACTTAAACTGTATCGATCATTCTTTTAGCACATGTATAATATAAAAACATTATTCTATTTCGAATTTAGGCTTCCAAAAATTTTTCATCCGTAAACCGATAATAATATATATAGACTTGTTAATAGTCGGAATAAATATATTAATGCTTAAACTATCATCATCTCCACGATTAGAGATACAATATTTACATTCTTTTTGCTGTTTCGAAACTTTATCAATACACGTTAATACAAACCCAGGAAGGAGATATTGAAACTGAGGCTGTTGAAAATGAAACGGCGAATACAATAATTCAGATAATGTAAAATCATGATTCCGTATTCTGATGATATTAGAACTGCTAATGGATGTCGATGGTATGTATCTAGGAGTATCTATTTTAACAAAGCATCGATTTGCTAATATACAATTATCCTTTTGATTAATTGTTATTTTATTCATATTCTTAAAAGGTTTCATATTTATCAATTCTTCTACATTAAAAATTTCCATTTTTAATTTATCTAGCCCCGCAATACTCCTCATTACGTTTCATTTTTTGTCTAGAATACCCATTTTGTTCATCTTGGTACATAGATTATCCAATTGAGAAGCGCATTTAGTAGTTTTGTACATTTTAAGTTTATTAACGAATCGTCGAAAACTAGTTATAGTTAACATTTTATTATTTGATACCCTGATATTAATACCCCTGCCGTTACTATTATTTATAACTGATGTAACCCACGTAACATTGGAATTAATTATCGATAGTAATGCATCGACACTTCCAAAATTGTCTATTATAAACTCACCGATAATTTTTTTATTGCATGTTTTCATATTCATTAGGATTATCAAATCTTTAATCTTATTACGATTGTATGCGTTGATATTACAAGACGTCATTCTAAAAGACGGAGGATTTCCATCAAATGCCAGACAATCACGTACAAAGTACATGGAAATAGGTTTTGTTCTATTACGCATCATAGATTCATATAAAACACCCGTAGAAATACTAATTTGTTTTACTCTATAAAATACTATTGCATCTATTTCATCGTTTTGTATAACGTCTTTCCAAGTGTCAAATTCCAATTTTTTTTCATTGATAGTACCAAATTCTTCTATCTCTTTAACTACTTGCATAGATAGGTAATTACAGTGATGCCTACATGCCGTTTTTTGAAACTGAATAGATGCATCTAGAAGCGATGCTACACTAGTCACGATCACCACTTTCATATTTAGAATATATGTATGTAAAAATATAGTAGAATTTCATTTTGTTTTTTTCTATGCTATAAATGAATTCTCATTTTGTATCCGCACATACTCCGTTTTATATCAATACCAAAGAAGGAAGATATCTGGTTCTAAAAGCCGTTAAAGTATGCGATGTTAGAACTGTAGAATGCGAAGGAAGTAAAGCTTCCTGCGTACTCAAAGTAGATAAACCCTCATCACCCACGTGTGAGAGAAGACCTTCGTCCCCGTCCAGATGCGAGAGAATGAATAACCCTGGAAAACAAGTCCCGTTTATGAGAACGGACATGTTACAAAATATGTTTGCTGCTAATCGCGACAACGTAACGTCAAGACTTTTGAACTAAAATACAATTATATCTTTTTCGATATTAATAAATCCGTGTCTCCCGGGTTTTTTATCTCTTTCAGTATGTGAATAGATAGGTATTTTATCTCTATTCATCATCGAATTTAAGAGATCCGATAAACATTGTTTGTATTCTCCAGATGTCAGCATCTGATACAACAATATATGTGCACATAAACCTCTGGCACTTATTTCATGTACCTTCCCCTTATCACTAAGGAGAATAGTATTTGAGAAATATGTATACATGATATTATCATGTATTAGATATACAGAATTTGTAACACTCTCGAAATCACACGATGTGTCGGCGTTAAGATCTAATATATCACTCGATAACACATTTTCATCTAGATACACTAGACATTTTTTAAAGCTAAAATAGTCTTTAGTAGTAACAGTAACTATGCGATTATTTTCATCGATGATACATTTCATCGGCATATTATTACGCGTACCATCAAAGACTATACCATGTGTATATCTAACGTATTCTAGCATAGTTGCCATACGTACATTAAACTTTTCAGGATCTTTGGATAGATCTTCCAATCTATCTATTTGAGAAAACATTTTTATCATGTTCAATAGTTGAAACGTCGGATCCACTATATAGATATTATCTATAAAGATTTTAGGAACTATGTTCATGGTATCCTGGCGAATATTAAAACTATCAATGATATGATTATCGTTTTCATCTTTTATCACCATATAGTTTCTAAGATATGGGATTTTACTTAATATAATATTATTTCCCGTAATAAATTTTATTAGAAATGCCAAATCTATAAGAAAAGTCCTAGAATTAGTCTGAAGAATATCTATATCACCGTACCGTATATTTGGATTAATTAGATATAGAGAATATGATCCGTAACATATACAACTTTTATTATGACGTCTAAGATATTCTTCCATCAACTTATTAACATTTTTGACTAGGGAAGATACATTATGACGTCCCATTACTTTTGCCTTGTCTATTACAGCGACGTTCATAGAATTTAGCATATCTCTTGCCAATTCTTCCATTGATGTTACATTATAAGAAATTTTAGATGAAATTACATTTGGAGCTTTAATAGTAAGAACTCCTAATATATCCGTGTATGTGGTCACTAATACAGATTGTAGTTCTATAATCGTAAATAATTTACCTATATTATATGTTTGAGTTTGTTTAGAAAAGTAGCTAAGTATACGATCTTTTATTTCTGATGCCGATGTATCAACATCGAAAAAAAATCTTTTTTTATTCTTTTTTACTAACGATACGAATATGTCTTTGTTAAAAACAGTTATTTTCTGAATATTTCTAGCTTGTAATTTTAACATATGATATTCGTTCACACTAGGTACTCTGCCTAAATAGGTTTCTATAATCTTTAATGTAATATTAGGAAGAGTATTCTGATCAGGATTCCTATTCATTTTGAGGATTTAAAACTCTGATTATTGTCTAATATGGTCTCAACACAAACTTTTTCACAGAGTGATAGAGTTTTTGATAACTCGTTTTTCTTAAGAAATATAAAACTACTGTCTCCAGAGCTCGCTCTATCTTTTATTTTATCTAATTCGATACAAACTCCTGATACTGGTTCAGAAAGTAATTCATTAATTTTCAGTCCTTTATAGAAGATATTTAATATAGATAATACAAAATCTTCAGTTCTTGATATCGATCTGATTGATCCTAGAACTAGATATATTAATAACGTGCTCATTAGGCAGTTTATGGCAGCTTGATAATTAGATATAGTATATTCCAGTTCATATTTATTAGATACCGCATTGCCCAGATTTTGATATTCTATGAATTCCTCTGAAAATAAATCCAAAATAACTAGACATTCTATTTTTTGTGGATTAGTGTACTCTCTTCCCTCTATCATGTTCACTACTGGTGTCCACAATGATAAATATCTAGAGGGAATATAATATAGTCCATATGATGCCAATCTAGCAATGTCGAATAACTGTAATTTTATTCTTCGCTCTTCATTATGAATTGAATCTTGAGGTATAAACCTAACACAAATTATATCATTAGACTTTTCGTATGTAATGTCTTTCATGTTATAAGTTTTTAATCCTGGAATAGAATCTATTTTAATGAGGCTTTTAAACGCAGCGTTCTCCAACGAGTCAAAGCATAATACTCTGTTGGTTTTCTTATATTCAATATTACGATTTTCTTCTTTGAATGGAATAGGTTTTTGAATTAGTTTATAATTACAACATAATAGATAAGGAAGTGTGTAAATAGTACGCGGAAAAAACATAATAGCTCCCCTGTTTTCATCCATGGTTTTAAGTAAATGATCACTGGCTTCTTTAGTCAATGGATATTCGAACATTAACCGTTTCATCATCATTGGACAGAATCCATATTTCTTAATGTAAAGAGTGATCAAATCATTGTGTTTATTGTACCATCTTGTTGTAAATGTGTATTCGGTTATCGGATCTGCTCCTTTTTCTATTAAAGTATCGATATCGATCTCGTCTAAGAATTCAACTATATCGACATATTTCATTTGTATACACATAACCATTACTAACGTAGAATGTATAGGAAGAGATGTAACGGGAACAGGGTTTGTTGATTCGCAAACTATTCTAATACATAATTCTTCTGTTAATACGTCTTGCACGTAATCTATTATAGATGCCAAGATATCTATATAATTATTTTGTAAGATGATGTTAACAATGTGATCTATATAAGTAGTGTAATAATTCATGTATTTCGATATATGTTCCAACTCTGTCTTTGTGATGTCTAGTTTCGTAATATCTATAGCGTCCTCAAAAAATATATTCGCATATATTCCCAAGTCTTCAGTTCTATCTTCTAAAAAATCTTCAACGTATGGAATATAATAATCTATTTTACCTCTTCTGATGTCATTAATGATATAGTTTTTGACACTATTTTCCGTCAATTGATTCTTATTCACTATGTCTAAAAACCGGATAGCGTCCCTAGGACGAACTACTGCCATTAATATCTCTATTATAGCTTCTGGACATAAATCATCTATTATACCAGAATTAATGGGAACTATTCCGTATCTATCTAACATAGTTTTAAGAAAGTCAGAATCTAAGACCTGATGTTCATATATTGGTTCATACATGAAATGATCTCTATTGATGATAGTGACTATTTCATTCTCTGAAAATTGGTAACTCATTCTATACACGCTTTCCTTGTTGATAAAGGATAGTATATACTCAATGGAATTTGTACCAACAAACTGTTCTCTTATGAATCGTATATCATCATCTGAAATGATCATGTAAGGCATACATTTAACAATAAGAGACTTGTCTCCTGTTATCAATATACTATTCTTGTGATAATTTATGTGTGCGGCAAATTTGTCCACGTTCTTTAATTTTGTTATAGTAGATATCAAATCCAATGGAGATACAGTTCTTGGCTTAAACAGATATAGTTTTTCTGGAACGAATTCTACAACATTATTATAAAGGACTTTGGGTATATAAGTGGGATGAAATCCTATTTTAATTAATGCGATAGCCTTGTCCTCGTGCAGATATCCAAACGCTTTTGTGATAGTATGGCATTCATTGTCTAGAAACGCTCTACGAATATCTGTAACAGATATCATCTTTAGAGAATACTAGTCGCGTTAATAGTACTAAAATTTGTATTTTTTAATCTATCTCAATAAAAAATTAATATGTATGATTCAATGTATAACTAAACTACTAACTGTTATTGATAACTAGAATCAGAATCTAATGATGACATAACTAAGAAGTTTATCTACAGCCAATTTAGCTGCATTATTTTTAGCATCTCGTTTAGATTTTCCATCTGCCTTATCGAATACTCTTCCGTCAATGTCTACACAGGCATAAAATGTAGGAGAGTTACTAGGCCCCACTGATTCAATACGAAAAGACCAATCTCTCCTAGTTATTTGACAGTACTCATTAATAACGGTGACAGGGTTAACACCTTTCCAATAAATAATTTTTTTAACCGGAATAACATCATCAAAAGACTTATGATCCTCTCTCATTGATTTTTCGCGGGATACATCATCTATTATAGCATCAGCATCAGAATCTGTAGGCCGTGTATCAGCATCCATTGTCGTAGACCAACGAGGAGGAGTATCGTTGGAGCTGTAAACCATAGCACTACGTTGAAGATCATACAGAGCTTTATTAACTTCTCGCTTCTCCATATTAAGTTGTTTAGTTAGTTGTACAGCAGTAGCTCCTTAGTCCAATGTTTTTAATAACCGCACACAATCTCTGTGTCAGAACGCTCGTCAATATAGATCTTAGAAATTTTTTTAGAGAGAACTAACGCAACTAGCAATAAAACTGATCTTATTTTATCATTTTTTTTATTCATCATCCTCTGGTGGTTCGTCGTTCCTATCGAATGTAGCTCTGATTAACCCGTCATCTATAGGTGATGCTGGTTCTGGAGATTCTGGAGGAGATGGATTATTATCTGGAAGAATCTCTGTTATTTCCTTGTTTTCATGTATCGATTGCGTTGTAACATTAAGATTGCGAAATGCTCTAAATTTGGGAGGCTTAAAGTGTTGTTTACAATCTCTACACGCGTGTCTAACTAATGGAGGTTCATCAGCGGCTCTAGTTTGAATCATCATCGGTGTAGTATTCCTACTTTTACAGTTAGGACACGGTGTATTGTATTTCTCGTCGAGAACGTTAAAATAATCGTTGTAACTCACATCCTTTATTTTATCTATATTGTATTCTACTCCTTTCTTAATGCATTTTATACCGAACAAGAGATAGCGAAGGAATTCTTTTTCGGTACCGCTAGTACCCTTAATCATATCACATAGTGTTTTATATTCTAAATGTGTGGCAATGGACGGTTTATTTCTATACGATAGTTTGTTTTTGGAATCCTTTGAGTATTCTATACCAATATTATTCTTTGATTCGAATTTAGTTTCTTCGATATTAGATTTTGTATTACCTATATTCTTGATGTAGTACTTTGATGATTTTTCCATGGCCCATTCTATTAAGTTTTCCAAGTTGGCATCATCCACATATTGTGATAGTAATTCTCGGATATCAGTAGTGACTACCGCCATTGATATTTGTTCATTTGATGAGTAACTACTAATGTATACATTTTCCATTTATAACACTTATGTATTAACTTTGTTTATTTATATTTTTTCATTATTATGTTGATATTAATAATCGTATTGTGGTTATATGGCTACAATTTCATAATGAGTTGAAGTCAGTGTCCTATGATCAATGACGATAGCTTTACTCTGAAAAGAAAGTATCAAATCGATAGTGCAGAGTCAACAATGAAAATGGATAAGACGATGACAAAGTTTCAGAATAGAGTCAAAATGGTAAAAGAAATAAATCAGACGATAAGAGCAGCACAAACTCATTACGAGACATTGAAACTAGGATATATAAAATTTAAGGGAATGATTAGGACTACTACTCTAGAAGATATAGCACCATCTATTCCAAATAATCAGAAAACTTATAAACTATTCTCGGACATTTCAGTCATTGGCAAAGCATCACAGAATCCGAGTAAGATGATATATGCTCGCTGCTTTACATGTTTCCCAATTTGTTTGGAGATGACCATAGATTCATTTGTTATAGAATGCATCCAACATTGTTCATGATATAGTTGAATCATGTATGCCTGTTCGTATGCCTGTGGCTAAGATACTGTGTAAAGAAATGGTAAATAAATACTTTGAGAATCTTTAAGAGTGCATTGACTTTGTTAGTGAATAGGCATTCCATCTTTCTCCAATACTAATTCAAATTGTTAAATTAATAATGGAATAGTATAAATAGTTATTAGTGATAGGATAGTAAACATAATTATTAGAATAGTAGTGTAGTATCATAGATAACTCTCTTCTATAAAAAATGGATTTTATTCGTAGAAAGTATCTTATATACACAGTAGAAAATAATATAGATTTTTTAAAGGATGATACATTAAGTAAAGTAAACAATTTTACCCTCAATCATGTACTAGCTCTCAAGTATCTAGTTAGCAATTTTCCTCAACATGTTATTACTAAGGATGTATTAGCTAATACCAATTTTTTTGTTTTCATACATATGGTACGATGCTGTAAAGTATACGAAGCGGTTTTACGACACGCATTTGATGCACCCACGTTGTACGTTAAAGCATTGACTAAGAATTATTTATCGTTTAGTAACACAATACAGTCGTACAAGGAAACAGTGCATAAACTAACACAAGATGAAAAATTTTTAGAGGTTGCCGAATACATGGACGAATTAGGAGAACTTATAGGCGTAAATTATGACTTAGTTCTTAATCCATTATTTCACGGAGGGGAACCCATCAAAGATATGGAAATCATTTTTTTAAAACTGTTTAAGAAAACAGACTTCAAAGTTGTTAAAAAATTAAGTGTTATAAGATTACTTATTTGGGCATACCTAAGCAAGAAAGATACAGGCATAGAGTTTGCGGATAATGATAGACAAGATATATATACTCTATTTCAACAAACTGGTAGAATAGTCCATAGCAATCTAACAGAAACGTTTAGGGATTATATCTTTCCCGGAGATAAGACTAGCTATTGGGTGTGGTTAAACGAAAGTATAGCTAATGATGCGGATATCGTTATTAATAGACCCGCCATTACCATGTATGATAAAATTCTTAGTTATATATACTCTGAGATAAAACAGGGACGCGTTAATAAAAACATGCTTAAGTTAGTTTATATCTTTGAGCCTGAAAAAGATATCAGAGAACTTCTGCTAGAAATCATATATGATATTCCTGGAGATATCCTATCTATTATTGATGCAAAAAACGACGATTGGAAAAAATATTTTATTAGTTTTTACAAAGCTAATTTTATTAACGGTAATACATTTATTAGTGATAGAACGTTTAACGATGACTTATTCAGAGTTGTTGTTCAAATAGATCCCGAATATTTCGATAATGAACGAATTATGTCTTTATTCTATACGAGTGCTGCGGACATTAAACGATTTGATGAGTTAGATATTAATAACAGTTATATATCTAATATAATTTATGAGGTGAACGATATCACATTAGATACAATGGATGATATGAAGAAGTGTCAAATCTTTAACGAGGATACGTTGTATTATGTTAAGGAATACAATACATACCTGTTTTTGCACGAGTCGGATCCCATGGTCATAGAGAACGGAATACTAAAGAAACTGTCATCTATAAAATCCAAGAGTAGACGGCTGAACTTGTTTAGCAAAAACATTTTAAAATATTATTTAGACGGACAATTGGCTCGTCTAGGTCTTGTGTTAGATGATTATAAAGGAGACTTATTAGTTAAAATGATAAACCATCTCAAATCTGTGGAGGATGTATCCGCATTCGTTAGATTTTCTACAGATAAAAACCCTAGTATTCTTCCATCGCTAATCAAAACTATTTTAGCTAGTTATAATATTTCCATCATCGTCTTATTTCAAAGGTTTTTAAGAGATAATCTATATCATGTAGAAGAATTCTTGGATAAAAGCATCCATCTAACCAAGACGGATAAGAAATATATACTTCAATTGATAAGGCACGGTAGATCATAGAACAAACCAAATATATTATTAATAATTTGTATATACATAGATATAATTATCACATATTAAAAAATAACACATTTTTGATAAATGGAAACCGTTGCAACAATTCAGACTCCCACCAAATTAATGAATAAAGAAAATGCAGAAATGATTTTGGAAAAAATTGTTAATCATATAGCTATGTATATTAGTGACGAATCAATATATTCAGAAAATAATCCTGAATATATTGATTTTCGTAACAGATACGGAGACTATAGATCTCTCATTATAAAAAGTGATCACGAGTTTGTAAAGCTATGTAAAGATCATGCAGAGAAAAGTTCTCCAGAAACGCAACAAATGATTATCAAACACATATACGAACAATATCTTATTCCAGTATCTGAAGTACTATTAAAACCTATAATGTCCATGGGTGACATATTTACATATAACGGATGTAAAGACAATGAATGGATGCTAGAACAACTCTCTACCCTAAACTTTAACAATCTCTACACATGGAACTCATGTAGCATAGGCAATGTAACGCGTCTGTTTTATACATTTTTTAGTTATCTGATGAAAGATAAACTAAATATATAAGTATAATCCCATTCTAATACTTTAACCTGATGTATTATTACCTGCATCTTATTAGAATATTAACCTAACTAAAAGACATAAAAAGCGGTAGGATATAAATATTATGGCCGCAACCGTTCCGCGTTTTGACGATGTGTACAAAAATGCACAAAGAAGAATTCTAGATCAAGAAACATTTTTTAGTAGAGGTCTAAGTAGACCGTTAATGAAAAACACATATCTATTTGATAATTACGCGTATGGATGGATACCAGAAACTGCAATTTGGAGTAGTAGATACGCAAACCTAGATGCTAGTGACTATTATCCCATTTCGTTGGGATTACTTAAAAAGTTTGAATTTCTCATGTCTCTATATAAAGGTCCTATTCCCGTATATGAAGAAAAAGTAAATACTGAATTCATTGCTAATGGATCTTTCTCCGGTAGATACGTATCATATCTTAGAAAGTTTTCTGCCCTTCCAACAAACGAGTTTATTAGTTTTTTATTATTGACCTCCATCCCTATCTATAATATCTTATTCTGGTTTAAAAACACACAGTTTGATATTACTAAACACACATTATTCAGATACGTCTATACAGATAATACCAAACACCTTGCGTTGGCTAGGTATATACATCAAACAGGAGACTATAAGCCTTTGTTTAGTCGTCTCAAAGAGAATTATATATTTACTGGTCCCGTTCCAATAGGTATCAAAGATATAGATCACCCTAATCTTAGTAGAGCAAGAAGTCCATCCGATTATGAGACATTAGCTAATATTAGTACTATATTGTACTTTACCAAGTATGATCCAGTATTAATGTTTTTATTGTTTTACGTACCTGGGTATTCAATTACTACAAAAATTACTCCAGCCGTAGAATATCTAATGGATAAACTGAATCTAACAAAGAGCGACGTACAACTGTTGTAAATTATTTTATGCTTCGTAAAATGTAGGTCTTGAACCAAACATTCTTTGAAAAAATGAGATGCATAAAACTTTATTATCCAATAGATTAACTATTTCAGACGTCAATCGTTTAAAGTAAACTTCGTAAAATATTCTTTGATTGCTGCCGAGTTTAAAACTTCTATCGATAATTGTTTCATATGTTTTAATATTTACAAGTTTTTTGGTCCATGGTACATTAGCTGGACAGATATATGCAAAATAATATCGTTCTCCAAGTTCTATAGTCTCTGGATTGTTTTTATTATATTCAGTAACCAAATACATATTAGGGTTATCTGCGGATTTATAATTTGAGTGATGCATTCGACTCAACATAAATAATTCTAGAGGAGACGATCTACTATCAAATTCGGATCGTAAATCTGTTTCTAAAGAACGGAGAATATCTATACATACCTGATTAGAATTCATCCGTCCTTCAGACAACATCTCAGACAGTCTGGTCTTGTATGTCTTAATCATATTCTTATGAAACTTGGAAACATCTCTTCTAGTTTCACTAGTACCTTTATTAATTCTCTCAGGTACAGATTTTGAATTCGACGATGCCGAGTATTTCATCGTTGTATATTTCTTCTTCGATTGCATAATCAAATTCTTATATACCGCCTCAAACTCTATTTTAAAATTATTAAACAATACTCTACTATTAATCAGTCGTTCTAACTCCTTTGCTATTTCTATGGACTTATCTACATCTTGACTGTCTATCTCTGTAAACACGGAGTCGGTATCTCCATACACGCTACGAAAACGAAATCTATAATCTATAGGCAACGATGTTTTCACAATCGGATTAATATCTCTATCGTCCATATAAAATGGATTACTTAATGTATTGGCAAACCGTAACATACCGTTGGATAACTCTGCTCCATTTAGTACCGATTCTAGATACAATATCATTCTACGTCCTATGGATGTGCAACTCTTAGCCGAAGCGTATGAGTATAGAGCACTATTTCTAAATCCCATCAGACCATATACTGAGTTGGCTACTATCTTGTACGTATATTGCATGGAATCATAGATGGCCTTTTCAGTTGAACTGGTAGCCTGTTTTAACATCTTTTTATATCTGGCTCTCTCTGCCAAAAATGTTCTTAATAGTCTAGGAATGGTTCCTTCTATTGATCTATCGAAAATTGCTATTTCAGAGATGAGGTTCGGTAGTCTAGGTTCACAATGAACCGTAATATATCTAGGAGGTGGATATTTCTGAAGCAAGAGTTGATTATTTATTTCTTCTTCCAATCTATTGGTACTAACAACGACACCGACTAATGTTTCCGGAGATAGATTTCCAAAGATACACACATTAGGATACAGACTGTTATAATCAAAGATTAATACATTATTACTAAACATTTTTTGTTTTGGAGCAAATACCTTACCGCCTTCATAAGGAAACTTTTGTTTTGTTTCTGATCTGACTAAGATAGTTTTAGTTTCCAACAATAGCTTTAACAGTGGACCCTTGATGATTGTACTCGCTCTATATTCGAATACCATGGATTGAGGAAGCACATATGTTGCCGCACCAGCGTCTGTTTTTGTTTCTACTCCATAATACTCCCACAAATACTGACACAAACAAGCATCATGAATACAGTATCTAGCCATATCTAAAGCTATGTTTAGATTATAATCCTTATACATCTGAGCTAAATCAATGTCATCCTTTCCGAAAGATAATTTATATATATCATTAGGTAAAGTAGGACATGATAGTACGACTTTAAATCCATTTTCCAAAATATCTTTACGAATTACTTTACATATAATATCCTCATCAACAGTCACGTAATTACCTGTGGTTAAAACCTTTGCAAATGTATCGGCTTTGCCTTTCGCGTCCGTAGTATCGTCACCGATGAACGTCATTTCTCTAACTCCTCTATTTAATACTTTACCCATGCAACTGAACGCGTTCTTGGATATAGAATCCAATTTGTACGAATCCAATTTTTCAGATTTTTGAATGAATGAATATAGATCGAAAAATATAGTTCCATTATTGTTATTAACGTGAAACGTAGTATTGGCCATGCCGCATACTCCCTTATGACTAGACTGATTTCTCTCATAAATACAGAGATGTACAGCTTCCTTTTTGTCTGGAGATCTAAAGATAATCTTCTCTCCTGTTAATAACTCTAGACGATTAGTAATATATCTCAGATCAAAGTTATGTCCGTTAAAGGTAACGACGTAGTCGAACGTTAGTTCCAACAATTGTTTAGCTATTCGTAACAAAACTATTTCAGAACATAGAACTAGTTCTCGTTCGTAATCCATTTCCATTAGCGACTGTATCCTCAAACATCCTCTATCGACGGCTTCTTGTATTTCCTGTTCCGTTAACATCTCTTCATTAATGAGCGTAAACAGTAATCGTTTACCACTTAAATCGATATAACAGTAACTTGTATGCGAGATTGGGTTAATAAATACAGAAGGAAACTTCTTATCGAAGTGACACTCTATATCTAGAAATAAGTACGATCTTGGGATATCGAATCTAGGTATTTCTTTAGCGAAACAGTTACGTGGATCGTCACAATGATAACATCCATTGTTAATCTTTGTCAAATATTGCTCGTCCAACGAGTAACATCCGTCTGGAGATATCCCGTTAGAAATATAAAACCAACTAATATTGAGAAATTCATCCATGGTGGCATTTTGTATGCTGCGTTTCTTTGGCTCTTCTATCAACCACATATCTGCGACGGAGCATTTTCTATCTTTAATATCTAGATTATAACTTATTGTCTCGTCAATGTCTATAGTTCTCATCTTTCCCATCGGCCTCGCATTAAATGGAGGAGGAGATAATGACTGATATATTTCGTCCGTCACTACGTAATAAAAGTAATGAGGAAATCGTATAAATACGGTCTCGCCATTTCGACATCTGGATTTCAGATATAAAAATCTGTTTTCACCGTGACTTTCAAACCAATTAATACACCTAACATCCATTTCTAGAATTTAGAAATATATTTTCATTTAAATGAATCCCAAACATTGGGGAAGAGCCGTATGGACCATTATTTTTATAGTACTTTCGCAAGCGGGTTTAGACGGCAACATAGAAGCGTGTAAACGAAAACTATATACTATAGTCAGCACTCTTCCATGTCCTGCATGTAGACGACACGCGACTATCGCTATAGAGGACAATAATGTCATGTCTAGCGATGATCTGAATTATATTTATTATTTTTTCATCAGATTATTTAACAATTTGGCATTTGATCCCAAATACGCAATCGATGTGTCAAAGGTTAAACCTTTATAAACTTAACCCATTATAAAACTTATGATTAGTCACGACTGAAATAACCGCGTGATTATTTTTTGGTATAATTCTACACGGCATGGTTTCTGTGACTATGAATTCAACACCTGTTATCTTAGTGAAATCTTTAACAAACAGCAAGGGTTCGTCAAAGACATAAAACTCATTGTTTACGATCGAAATAGACCCCCTATCACACTTAAAATAAAAAATATCCTTATCCTTTACCACCAAATAAAATTCTGATTGGTCAATGTGAATGTATTCACTTAACAGTTCCACAAATTTATTTATTAACTCCGAGGCACATACATCGTCGGTATTTTTTATGACAAACTTTACTCTTCCAGCATCCGTTTCTAAAAAAATATTAACGAGTTCCATTTATATCATCCAATATTATTGAAATGACGTTGATGGACAGATGATATAAATAAGAAGGTACAGTACCTTTGTCCACCATCTCCTCCAATTCATACTCTATTTTGTCATTAACTTTAATGTGTGAAAACAGTACGCCACATGCTTCCATGACAGTGTGTAACACTTTGGATACAAAATGTTTGACATTAGTATAATTGTCCAAGACTGTCAATCTATAATAGATAGTAGCTATAATATATTCTATGATGGTATTGAAGAAGATGACAACCTTGGCATATTGATCATTTAACACAGACATGGTATCAACAAATAGCTTAAATGAAAGAGAATCAGTAATTGGAATAAGCGTCTTCTCGATGTAGTGTCCGTATACCAACATGTCTGATATTTTGATGTATTCCATTAAATTATTTAGTTTTTTCTTTTTATTCTCGTTAAACAGAATTTCTGTCAATGGACCCCAACATCGTTGACCTATTAAGTTTTGATTGATTTTTCCGTGTAAGGCGTATCTAGTCAGATCGTATAGCCTATCCAATAATCCATCGTCTGTGCGTAGATCACATCGTACACTTTTTAATTTTCTATAGAAGAGTGACAGACATCTGGAGCAATTACAGACAGCAATTTCTTTATTCTCTACAGATGTAAGATACTTGAAGATATTCCTATGATGATGCAGAATTTTGGATAACACGGTATTGATGGTATCTGTTACCATAATTCCTTTGACTGATAGTGTCAAAGTACAAGATTTCCAATCTTTTGCAATTTTCAGTACCATTATCTTTGTTTTGATATCTATATCAGACAGCATGGTACGTCTGACAACACAGGGATTAAGACGGAAAGATGAAATGATTCTCTCAACATCTTCAATAGATACCTTGCTATTTTTTTTGGCATTATCTATATGTGAGAGAATATCCTCTAGAGAATCAGTATCCTTTTTGATGATAGTGGATCTCAATGACATGGGACGTCTAAACCTTCTTATTCTATCACCAGATTGCATGGTGATTTGTCTTCTTTCTTTTATCATGATGTAATCTCTAAATTCATCGGCAAATTGTCTATATCTAAAATCATAATATGAGATGTTTACCTCTACAAATATCTGTTCGTCCAATGTTAGAGTATCTATATCAGTTTTGTATTCCAAATTAAACATGGCAACGGATTTAATTTTATATTCCTCTATTAAGTCCTCGTCGATAATAACAGAATGTAGATAATCATTTAATCCATCGTACATGGTTGGAAGATGCTCGTTGACAAAATCTTTAATTGTCTTGATGAAGGTGGGACTATATCTAACATCTTGATTAATAAAATTTATAACATTGTCCATAGGATACTTTGTAACTAGTTTTATACACATCTCTTCATTGGTAAGTTTAGACAGAATATCGTGAACAGGTGGTATATTATATTCATCAGATATACGAAGAATAATGTCCAAATCTATATTGTTTAATATATTATATAGATGTAGTGTAGCTCCTACAGGAATATCTTTAACTAAGTCAATGATTTCATCAACAGTTAGATCTATTTTAAAGTTAATCATATAGGCATTGATTTTTAAAAGGTATGTAGCCTTGACTACATTCTCATTAATTAACCATTCCAAGTCACTGTGTGTAAGAAGATTATATTCTATCATAAGCTTGACTACATTTGGTCCCGATACCATTAAAGAATTCTTATGATATAAGGAAACAGCTTTTAGGTACTCATCTACTCTACAAGAATTTTGGAGAGCCTTAACGATATCAGTGACGTTTATTATTTCAGGAGGAAAGAACCTAACATTGAGAATATCTGAATTAATAGCTTCCAGATACAGTGATTTTGGCAATAGTCCGTGTAATCCATAATCCAGTAACACGAGCTGGTGCTTGCTAGACACCTTTTCAATGTTTAATTTTTTTGAAATAAGCTTTGATAAAGCCTTCCTCGCAAATTCCGGATACATGAACATGTCGCCAACATGATTAAGTATTGTTTTTCATTATTTTTATATTTTCTCAACAAGTTCTCAATACCCCAATAGATAATAGAATATCACCCAATGCGTCCATGTTGTCTATTTCCAACAGGTCGCTATATCCACCAATAGAAGTTTTCCCAAAAAAGATTCTAGGAACAGTTCTACCACCAGTAATTTGTTCAAAATAGTCACGCAATTCATTTTCGGGTTTAAATTCTTTAATATCTACAATTTCATACGCTCCTCTTTTGAAACTAAACTTATTTAGAATATCCAGTGCGTTTCTACAAAAAGGACATGTAAACTTGACAAAAATTGTCACTTTGTTATTGGCCAACCTTTGTTGTACAAATTCCTCGGCCATTTTTAATATTTAAGTGATACAAAACTATCTCGACTTATTTAACTCTTTAGTCGAGATATATGGACACAGATAGCTATATGATAACCAACTACAGAAGACAAACGCTATAAAAAACATAATTACGACGAGCATATTTATAAATATTTTTATTCAGTATTACTTGATATAGTAATATTAGGCACAGTCAAACATTCAACCACTCTAGATACATTAACTCTCTCATTTTCTTTAACAAATTCTGCAATATCTTCGTAAAAAGATTCTTGAAACTTTTTAGAATATCTATCGACTCTAGATGAAATAGCGTTCGTCAACATACTATGTTTTGTATACATAAAGGCGCCCATTTTAACAGTTTCTAGTGACAAAATGCTAGCGATCCTAGGATCCTTTAGAATCACATAGATTGACGATTCGTCTCTCTTAGTAACTCTAGTAAAATAATCATACAATCTAGTACGCGAAATAATATTATCCTTGACTTGAGGAGATCTAAACAATCTAGTTTTGAGAACATCGATAAGTTCATCGGGAATTACATACATACTATCTTTAATAGAACTCTTTTCATCCAGTTGAATGGATTCGTCCTTAACCAACTGATTAATGAGATCTTCTATTTTATCATTTTCTAGATGATATGTATGTCCATTAAAGTTAAATTGTGTAGCGCTTCTTTTTAGCCTAGCAGCCAATACTTTAACATCACTAATATCGATATACAAAGGAGATGATTTATCGATGGTATTAAGAATTCGTTTTTCGACATCCGTCAAAACCAATTCCTTTTTGCCTGTATCATCCAGTTTGCCATTCTTTGTAAAGAAATTATTTTCTACTAGACTATTAATAAGACTGATAAGGATTCCTCCATAATTGCACAATCCAAACTTTTTCACAAAACTAGACTTTACGAGATCTACAGGAATGCGTACTTCAGGTTTCTTAGCTTGTGATTTTTTCTTTTGTGGACATTTTCTAGTGACCAACTCATCTACCATTTCATTGATTTTAGCAGTGAAATAAGCTTTCAATGCACGGGCACTGATACTATTGAAAACGAGTTGATCTTCAAATTCCGCCATTTAAGTTCACCAAACAACTTTTAAATACAAATATATCAATAGTAGTAGAATAAGAACTATAAAAAAAATAATAATTAACCAATACCAACCCCAACAACCTGTATTATTAGTTGATGTGACAGTTTTCTCATCACTTAGAACAGATTTAACAATTTCTATAAAGTCTGTCAAATCATCTTCCTGAGAACCCATAAATACACCAAATATAGCAGCGTACAACTTATCCATTTATACATTGAATATTGGCTTTTCTTTATCGCTATCTTCATCATATTCATCATCAATATCAACAAGTCCCAGATTACGAACCAGATCTTCTTCTACATTTTCAGTCATTGATACGCGTTCACTATCTCCAGAGAGTCCGATAACGTTAGCCACTACTTCTCTATCAATGATTAGTTTCTTGAGCGCGAATGTAATTTTTGTTTCCGTTCCGGATCTATAGAAAACTACAGGTGTAATAATTGCCTTGGCTAATTGTCTTTCTCTTTTACTGAGTGATTCTAGTTCACCTTCTATAGATCTGAGAATGGATGATTCTCCAGTCGAAACATATTCTACCATGGCTCCGTTTAATTTGTTGATGAAGATGGATTCATCCTTAAATGTTTTCTCTGTAATAGTTTCCGCCGAAAGACTATGCAAAGAATTTGGAATGCGTTCCTTGTGTGTAATGTTTCCATAGACAGCTTCTAGAAGTTGATACAACATAGGACTAGCCGCGGTAACTTTTATTTTTAGAAAGTATCCATCGCTTCTATCTTGTTTAGATTTATTTTTATAAAGTTTAGTCTCTCCTTCCAACATAATAAAAGTGGAAGTCATCTGACTAGATAAACTATCAGTAAGTTTTATAGAGATAGATGAACAATTAGCGTATTGAGAAGCATTTAGTGTAACGCATTCGATACATTTTGCATTAGATTTACTAATCGATTTTGCATACTCTATAACACCCGCACAAGTCTGTAGAGAATCGCTAGATGCTGTAGGTCTTGGTGAAGTTTCAACTCTCTTCTTGATTACCTTACTCATGATTAAACCTAAATAATTGTACTTTGTAATATAATGATATATATTTTCACTTTATCTCATTTGAGAATAAAAATGTTTTTGTTAACCACTGCATGATGTACAGATTTCGGAATCGCAAACCACTTGTGGTTTTATTTTATCCTTGTCCAATGTGAATTGAATGGGAGCGGATGCGGGTTTCGTACGTAGATAGTACATTCCCGTTTTTAGACCGAGACTCCATCCGTAAAAATGCATACTCGTTAGTTTGGAATAACTCGGATCTGCTATATGGATATTCATAGATTGACTTTGATCGATGAAGGCTCCCCTGTCTGCAGCCATTTTTATGATCGTCTTTTGTGGAATTTCCCAAATAGTTTTATAAACTCGCTTAATATCTTCTGGAAGGTTTGTATTCTGAATGGATCCACCATCTACCATAATCCTATTCTTGATCTCATCATTCCATAATTTTCTCTCGGTTAAAACTCTAAGGAGATGCGGGTTAACTACTTGGAATTCTCCAGACAATACTCTCCGAGTGTAAATATTACTGGTATACGGTTCCACCGACTCATTATTTCCCAAAATTTGAGCAGTTGATGCAGTCGGCATAGGTGCCACCAATAAACTATTTCTAAGACCGTATGTTCTGATTTTATCTTTTAGAGGTTCCCAATTCCAAAGATCCGACGGTACAACATTCCAAAGATCATATTGTAGAATACCGTTACTGGCGTACGATCCTACATATGTATCATATGGTCCTTCCTTCTCAGCTAGTTTACAACTCGCCTCTAATGCACCGTAATAAATGGTTTCAAAGATCTTCTTATTTAGATCTTGTGCTTCCAGGCTATCAAATGGATAATTTAAGAGAATAAACGCGTCCGCTAATCCTTGAACACCAATACCGATAGGTCTATGTCTCTTATTAGAGATTTCAGCTTCTGGAATAGGATAATAATTAATATCTATAATTTTATTGAGATTTCTGACAATTACTTTGACCACATCCTTCAGTTTGAGAAAATCAAATCGCCCATCTATTACAAACATGTTCAATGCAACAGATGCCAGATTACACACGGCTACCTCATTAGCATCCGCATATTGTATTATCTCAGTGCAAAGATTACTACACTTGATGGTTCCTAAATTTTGTTGATTACTCTTTTTGTTACACGCATCCTTATAAAGAATGAATGGAGTACCAGTTTCAATCTGAGATTCTATAATCGCTTTCCAGACGACTCGAGCCTTTATTATACATTTGTATCTCCTTTCTCTTTCGTATAGTGTATACAATCGTTCGAACTCGTCTCCCCAAACATTGTCCAATCCAGGACATTCATCCGGACACATCAACGACCACTCTCCGTCATCCTTCACTCGTTTCATAAAGAGATCAGGAATCCAAAGAGCTATAAATAGATCTCTTGTTCTATGTTCATCGTTTCCTGTATTCTTTTTAAGATCGAGGAACGCCATAATATCAGAATGCCACGGTTCCAAGTATATGGCCATAACTCCAGGCCGTTTGTTTCCTCCCTGATCTATGTATCTAGCGGTGTTATTATAAACTCTCAACATTGGAATAATACCGTTTGATATACCATTGGTACCGGAGATATAGCTTCCACTGGCACGAATATTACTAATTGATAGACCTATTCCCCCTGCCATTTTAGAGATTAATGCGCATCGTTTTAACGTGTCATAGATGCCTTCTATGCTATCATCGATCATGTTAAGTAGAAAACAGCTAGACATTTGGTGACGAGTAGTTCCCGCATTAAATAAGGTAGGAGAAGCGTGCGTAAACCATTTTTCAGAAAGTAGATTGTACGTCTCAATAGCTGAGTCTATATCCCATTGATGAATTCCTACTGCGACACGCATTAACATGTGCTGAGGTCTTTCAACAATTTTGTTGTTTATTTTCAACAAGTAGGATTTTTCCAAAGTTTTAAAACCAAAATAGTTGTATGAAAAGTCTCGTTCGTAAATAATAACCGAATTGAGCTTATCCTTATATTTGTTAACTATATCCATGGTAATACTTGAAATAATCGGAGAATGTTTCCCATTTTTAGGATTAACATAGTTGAATAAATCCTCCATCACTTCACTAAATAGTTTTTTTGTTTCCTTGTGTAGATTTGATATGGCTATTCTGGCGGCTAGAATGGCATAATCCGGATGTTGTGTAGTACAAGTGGCTGCTATTTCGGCTGCCAGAGTGTCCAATTCTACCGTTGTTACTCCATTATATATTCCTTGAATAACCTTCATAGCTATTTTAATAGGATCTATATGATCAGTGTTTAAGCCATAGCACAATTTTCTAATACGAGACGTGATTTTATCAAACATGACATTTTCCTTGTATCCATTTCGTTTAATGACAAACATTTTTGTTGGTGTAATAAAAAAAATTATTTAATTTTTCATTAATAGGGATTTGACGTATGTAGCGTACAAAATTATCGTTCCTGGTATATAGATAAAGAGTCCTATATATTTGAAAATCGTTACGGTTCGATTAAACTTTAATGATTGCATTGTGAATATATCATTAGGATTTAACTCCTTGACTATCATGGCGGTGCCAGAAATTACCATCAAAAGCATTAATACAGTTATGCAGATCGCAGTTAGAACGGTTATAGCATCCACCATTTATATCTAAAAATTAGATCAAAGAATATGTGACAACGTCCTAGTTGTATACTGAGAATTGACGAAACAATGTTTCTTACATATTTTTTTCTTATTAGTAACCGACTTAATAGTAGGAACTGGAAAACTAGACTTGATTATTCTATAAGTATAGATACCCTTCCAAATAATGTTCTCTTTGATAAAAGTTCCAGAAAATGTAGAATTTTTTAAAAAGTTATCTTTTGCTATTACTAATATCGTGGTTAGACGCTTATTATTAATATGAGTGATGAAATCCACACCGCTTCTAGATATCGCTTTTATTTCCACATTAGATGGTAAATCCAATAGTGAAACTATCTTTTTAGGAATGTATGGACTCGCGTTTAGAGGAGTGAACGTCTTCGGAGTAGTAAAGGATGATTCGTCAAATGAATAAACAATTTCACAAATGGATGTTAATGTATTAGTAGGAAATTTTTTGACGCTAGTGGAATTGAAGATTCTAATGGATGATGTTCTACCTATTTCATCCGATAACATGTTAATTTCCAATACCAACGGTTTTAATATTTCGATGATATACGGTAGTCTCTCTTTCGGACTTATATAGCTTATTCCACAATACGAGTCATTATATACTCCAAAAAACAAAATAACTAGTATAAAATCTGTATCGAATGGGAAAAACGAAATTATCGATATAGGTATAGAATCCGGAACATTGAACGTATTAATACTTAATTCTTTTTCAGTGGTAAGAACCGATAGGTTATTGACATTGTATGGTTTTAAATATTCTATAACTTGAGACTTGATAGATATTAATGACGAATTGAAAATTATTTTTATCACCACGTGTGTTTCAGGATCATCGTCGACGCCAGTTAACCAACCGAATGGAGTAAAATAAATATCATTAATATATGCTCTAGATATTAGTATTTTTATTAATCCTTTGATTATCATCTTCTCGTACGCGAATGATTCCATGATCAAGAGTGATTTGAGAACATCCTCCGGAGTATTAATGGGTTTAGTAAACAGTCCATCGTTGCAATAATAAAAGTTGTCCAAGTTAAAGGATATTATGCATTCGTTTAAAGATATCACCTCATCTAACGGAGACAATTTTTTGGTAGGTTTTAGAGACTTTGAAGCTACTTGTTTAACAAAGTTATTCATCGTCGTCTACTATTCTATTTAATTTTGTAGTTAATTTATCACATATCACATTAATTGACTTTTTGGTCCACTTTTCCATACGTTTATATTCTTTTAATCCTGCGTTATCCGTTTCCGTTATATACAGGGATAGATCTTGCAAGTTAAATAGAATGCTCTTAAATAATGTCATTTTTTTATCCGCTAAAAATTTAAAGAATGTATAAACTTTTTTCAAAGATTTAAAACTTTTAGGTGGAGTTCTGGTACACAATATCATAAACAAACTAATAAACATCCCACATTCAGATTCCAACAATTGATTAACTTCCACATTAATACAGCCTATTTTCGCTCCAAATGTACATTCGAAAAATCTGAATAAAACATCAATATCGCAATTTGTATTATCCAATACAGAATGTCTGTGATTCGTGTTAAAACCATCGGAAAAAGAATAGAAATAAAAATTATTATAATGGTGGAATTCAGTTGGAATATTGCCTCCGGAGTCATAAAAGGATACTAAACATTGTTTTTTATCGTAAATTACACATTTCCAATGAGACAAATAACAAAATCCAAACATTACAAATCTAGAGGTAGAACTTTTAATTTTGTCTTTAAGTATATACGATAAGATATGTTTATTCATAAACGCGTCAAATTTTTCATGAATAGCTAAGGAGTTTAAGAATCTCATGTCAAATTGTCCTATATAATCCACTTCGGATCCATAAGCAAACTGAGAGACTAAGTTCTTAATACTTCGATTGCTCATCCAGGCTCCTCTCTCAGGCTCTATTTTCATCTTGACGACCTTTGGATTTTCACCAGTATGTATTCCTTTACGTGATAAATCATCGATTTTCAAATCCATTTGTGAGAAGTCTATCGCCTTAGATACTTTTTCCCGTAGTTGAGGTTTAAAGAAATACGCTAACGGTATACTAGTAGGTAACTCAAAGACATCATATATAGAATGGTAACGCGTCGTTAACTCGTCGGTTAACTCTTTCTTTTGATCGAGTTCATCGCTACTATTGGGTCTGCTCAGGTGCCCCGACTCTACTAGTTCCAACATCATACCGATAGGAATACAAGACACTTTGCCAGCGGTTGTAGATTTATCATATTTCTCCACCACATATCCGTTACAATTTGTTAAGAATTTAGATACATCTATATTGCTACATAATCCAGCTAGTGAATATATATGACATAATAAATTGGTAAATCCTAGTTCTGGTATTTTACTAATTACTAAATCTGTATATCTTTCCATTTATCATGGAAAAGAATTTACCAGATATCTTCTTTTTTCCAAACTGCGTTAATGTATTCTCTTACAAATATTCACAAGATGAATTCAGTAATATGAGTAAAACGGAACGTGATAATTTCTCATTGGCTGTGTTTCCAGTGATAAAACATAGATGGCATAACGCACACGTTGTAAAACATAAAGGAATATACAAAGTTAGTACAGAAGCACGTGGAAAAAAAGTATCTCCTCCATCACTAGGAAAACCCGCACATATAAACCTAATGTCGAAGCAATATATATATAGTGAGTATGCAATAAGCTTTGAATGTTATAGTTTTCTAAAATGTATAACAAATACAGAAATCAATTCGTTCGATGAGTATATATTAAGAGGACTATTAGAAGCTGGTAATAGTTTACAGATATTTTCCAATTCCGTAGGTAAACGAATAGATACTATAGGTGTACTAGGGAATAAGTATCCATTTAGCAAAATTCCATTGGCCTCATTAACTCCTAAAGCACAACGAGAGATATTTTTAGCGTGGATTTCTCATAGACCTGTAGTTTTAACTGGAGGAACCGGAGTGGGTAAGACGTCACAGGTACCCAAGTTATTGCTTTGGTTTAATTATTTATTTGGTGGATTCTCTTCTCTAGATAAAATCACTGACTTTCACGAAAGACCAGTCATTCTATCTCTTCCTAGGATAGCTTTAGTTAGATTGCATAGCAATACCATTTTAAAATCATTGGGATTTAAGGTACTAGATGGATCTCCTATCTCTTTACGGTACGGATCTATACCGGAAGAATTAATAAACAAACAACCAAAAAAATATGGAATTGTATTTTCTACCCATAAGTTATCTCTAACAAAACTATTTAGTTATGGCACTATTATTATAGACGAAGTTCATGAGCATGATCAAATAGGAGATATTATTATAGCAGTAGCGAGAAAACATCATACGAAAATAGATTCTATGTTTTTAATGACTGCCACGTTAGAGGATGACAGGGAACGTCTAAAAATATTTTTACCTAATCCCGCATTTATACATATTCCTGGAGATACACTGTTTAAAATTAGCGAGGTATTTATTCATAATAAGATAAATCCATCTTCCAGAATGGCATATATAGAAGAAGAAAAGAGAAATTTAGTTACTGCTATACAGATGTATACTCCTCCTGATGGATCATCCGGTATAGTCTTTGTGGCATCCGTTGCACAGTGTCACGAATATAAATCATATTTAGAAAAAAGATTACCGTATGATATGTATATTATTCATGGTAAGGTCTTAGATATAGACGAAATATTAGAAAAAGTGTATTCATCACCTAATGTATCGATAATTATTTCTACTCCTTATTTGGAATCCAGCGTTACTATACGCAATGTTACACACATTTATGATATGGGTAGAGTTTTTGTCCCCGCTCCTTTTGGAGGATCACAACAATTTATTTCTAAATCTATGAGAGATCAACGAAAAGGAAGAGTAGGAAGAGTTAATCCTGGAACATACGTATATTTCTATGATCTGTCTTATATGAAATCTATACAGCGAATAGATTCAGAATTTCTACATAATTATATATTGTACGCTAATAAGTTTAATCTAACACTCCCCGAAGATTTGTTTATAATCCCTACAAATTTGGATATTCTATGGCGTACAAAGGAATATATAGACTCGTTCGATATTAGTACAGAAACATGGAATAAATTATTATCCAATTATTATATGAAGATGATAGAGTATGCTAAACTTTATGTACTAAGTCCTATTCTCGCTGAGGAGTTGGATAACTTTGAGAGGACGGGAGAATTAACTAGTATTGTACAAGAAGCCATTTTATCTCTAAATTTACAAATTAAGATTTTAAAATTTAAACATAAAGATGATGATACGTATATACACTTTTGTAGAATATTATTCGGTGTCTATAACGGAACAAACGCTACTATATATTATCATAGACCTCTAACGGGATATATGAATATGATTTCAGATACTATATTTGTTCCTGTAGATAATAACTAAAAATCAAAATCTAATGACCACATCTTTTTTTAGAGATGAAAAATTTTCCACATCTCCTTTTGTAGACACGACTAAACATTTTGCAGAAAAAAGTTTATTATTATTTAGATAATCGTATACTTCATCAGTGTAGATAGTAAATGTGAACAGATAAAAGGTATTCTTGCTCAATAGATTGGTAAATTCCATAGAATATATTAATCCTTTCTTCTTGAGATCCCACATCATTTCAACCAAAGACGTTTTATCCAATGATTTACCTCGTACTATACCACATACAAAACTAGATTTTGCAGTGATGTCGTACCTGGTATTCCTACCAAACAAAATTTTACTTTTAGTTCTTTTAGAAAATTCTAAGGTAGAATCTCTATTTGTCAATATGTCATCTATGGAATTACCACTAGCAAAAAATGATAGAAATATATATTGATACATCGCAGCTGGTTTTGATCTACTATACTTTAAAAACGAATCAGATTCCATAATTGCTTGTATATCATCAGCTGAAAAACTATGTTTTACACGTATTCCTTCGGCATTTCTTTTTAATGATATATCTTGTTTAGACAATGATAAAGTTATCATGTCCATGAGAGACGCGTCTCCGTATCGTATAAATATTTCATTAGATGTTAGACGCTTCATTAGGGGTATACTTCTATAAGGTTTCTTAATTAGTCCATCATTGGTTGCGTCAAGAACTACTATCTGATGTTGTTGGGTATCTCTAGTGTTACACATGGCCTTACTAAAGTTTGGGTAAATAACTATGATATCTCTATTAATTATAGATGTATATATTTCATTCGTCAAGGATATTAATATCGACTTACTATCGTCATTAATACGTGTAATGTAATCATATAAATCATGCGATAGCCAAGGAAAATTCAAATAGATGTTCATCATATAATCGTCGCTATAATTCATATTAATACTTTGACATTGACTAATTTGTAATATAGCCTCGCCACGAAGAAAGCTCTCGTATTCAGTTTCATCGATAAAGGATACCGTTAAATATAACTGGTTGCCGATAGTCTCATAGTCTATTAAGTGGTAAGTTTCGTATAAATACAGAATCCCTAAAATATTATCTAATGTGGGATTAATCCTTACCATAACTGTATAAAATGGAGCCGGAGTCATAACTATTTTACCGTTTGTACTTACTGGAATAGATGAAGGAATAATCTCCGGACATGATGGTAAAGACCCAAATGTCTGTTTGAAGAAATCCAATGTTCCAGGTCCTAATCTCTTGACAAAAATTACGATATTCGATCCCGATATCCTTTGCATTCTATTTACCAGCATATCACGAACTATATTAAGATTATCTATCATGTCTATTCTCCCACCGTTATATAAATCGCCTCCGCTAAGAAACGTTAGTATATCCATACAATGGAATACTTCATTTCTAAAATAGTATTCGTTTTCTAATTCTTTAATGTGAAATCGTATACTAGAAAGGGAAAAATTATCTTTGAGTTTTCCATTAGAAAAGAACCACGAAACTAATGTTCTGATTGCGTCTGACTCCGTCGCTGAATTAATAGATTTACACCAAAAACTCATATAACTTCTAGATGTAGAAGCATTCGCTAAAAAATTAGTAGAATCAAAGGATATAAGTAGATGTTCCAACAAGTGAGCAATTCCCAAGATTTCATCTATATCATTCTCGAATCCGAAATTAGAAATTCCCAAGTAGATATCCTTTTTCATCCGATCATTGATGAAAATACGAACTTTATTCGGTAAGACGATCATTTACTAAGGAGTAAAATAGGAAGTAACGTTCGTATATCGTTATCGTCGTATAAATTAAAGGTGTGTTTTTTGCCATTAAGAGACATTATAATTTTACCAATATTGGAATTATAATATAGGTGTATTTGAGCACTAGAAACGGTCGATGCATCGGTAAATATAGCTGTATCTAATGTTCTAGTCGGTATTTCTTCATTTCGCTGTCTAATGATAGCGTTTTCTCTATCTGTTTCCATTACAGCTGCCTGAAGTTTATTGGTCGGATAATATGTAAAATAATAAGAAATACATACGAATAACAAAAATAAAATAAGATATAATAAAGATGCCATTTAGAGATCTAATTTTGTTCAACTTGTCCAAATTCCTACTTACAGAAGATGAGGAATCGTTGGAGATAGTATCTTCCTTATGTAGAGGATTTGAAATATCTTACGATGACTTAATATCGTACTTTCCAGATAGGAAATACCATAAATATATTTCTAAGGTATTTGAACATGTAGATTTATCGGAGGAATTAAGTATGGAATTCCATGATACAACTCTGAGAGATTTAGTATATCTTAGATTGTACAAGTATTCCAAGTATATACGGCCGTGTTATAAATTAGGAGATAATCTAAAAGGTATAGTTGTTATAAAGGACAGAAATATATATATTAGAGAAGCAAATGATGACTTGATAGAATATCTCCTCAAGGAATACACTCCTCAGATTTATACATATTCTAATGAGCGAGTTCCCATAGCTGGTTCAAAATTAATTCTTTGTGGATTTTCTCAAGTTACATTTATGGCGTATACAACGTCGCATATAACAACAAATAAAAAGGTAGATGTTCTCGTTTCCAAAAAATGTATAGATGAACTAGTCGATCCAATAAATTATCAAATACTTCAAAATTTATTTGATAAAGGAAGCGGAACAATAAACAAAATACTCAGGAAGATATTTTATTCGGTAACAGGTGGCCAAACTCCATAGGTAGCTTTTTCTATTTCGGATTTTAGAATTTCCAAATTCACCAGCGATTTATCGGTTTTGGTGAAATCCAAGGATTTATTAATGTCCACAAATGCCATTTGTTTTGTCTGTGGATTGTATTTGAAAATGGAAACGATGTAGTTAGATAGATGCGCGGCGAAGTTTCCTATTAGGGTTCCGCGCTTCACGTCACCCAACATACTTGAATCACCATCCTTTAAAAAAAATGATAAGATATCAACATGGAGTATATCATACTCGGATTTTAATTCTTCTACTGCCTCACTGACATTTTCACAAATACTACAATACGGTTTACCGAAAATAATCAGTACGTTCTTCATTTATGGGTATCAAAAACTTAAAATCGTTACTGCTGGAAAATAAATCACTGACGATATTAGATGATAATTTATACAAAGTATACAATGGAATATTTGTGGATACAATGAGTATTTATATAGCCGTCGCCAATTGTGTCAGAAACTTAGAAGAGTTAACTACGGTATTCATAAAATACGTAAACGGATGGGTAAAAAAGGGAGGACATGTAACCCTTTTTATCGATAGAGGAAGTATAAAAATTAAACAAGACGTTAGAGACAAGAGACGTAAATATTCTAAATTAACCAAGGACAGAAAAATGTTAGAATTAGAAAAGTGTACATCCGAAATACAAAATGTTACCGGATTTATGGAAGAAGAAATAAAGGCAGAAATGCAATTAAAAATCGATAAACTCACATTTCAAATATATTTATCTGATTATGATAACATAAAAATATCATTGAATGAGATACTAACACATTTCAACAATAATGAGAATGTTACATTATTTTATTGTGATGAACGAGACGCAGAATTCGTTATGTGTCTAGAGGCTAAAACACAGTTCTCTACCACAGGAGAATGGCCGTTAATAATAAGTACCGATCAGGATACTATGCTATTCGCGTCTGCTGATAATCATCCTAAGATGATAAAAAACTTAACTCAACTGTTTAAATTTGTTCCCTCGGCAGAGGATAACTATTTAGCAAAATTAACTGCATTAGTGAATGGATGTGATTTCTTTCCTGGACTCTATGGGGCATCTATAACACCCAACAACTTAAACAAAATACAATTGTTTAGTGATTTTACAATCGATAATATAGTCACTAGTTTGGCAATTAAAAATTATTATAGAAAGACTAACTCTACCGTAGACGTGCGTAATATTGTTACGTTTATAAACGATTACGCTAATTTAGACGATGTCTACTCGTATATTCCTCCTTGTCAATGCACTGTTCAAGAATTTATATTCTCCGCATTAGATGAAAAATGGAATGAATTTAAATCATCTTATTTAGAGAGCGTGCCGTTACCCTGCCAATTAATGTACGCATTAGAACCACGTAAGGAGATTGATGTTTCAGAAGTTAAAACTTTATCATCTTATATAGATTTCGAAAATACTAAATCAGATATCGATGTTATAAAATCTATATCCTCGATTTTTGGATATTCTAACGAAAACTGTAACACCATAGTGTTCGGCATCTATAAGGATAATTTACTACTGAGTATAAATAATTCATTTTACTTTAACGATAGTCTGTTAATAACCAATACTAAAAGTGATAATATAATAAATATAGGTTACTAGATTAAAAAATGGTGTTCCAGCTCGTGTGTTCTACATGCGGCAAAGATATTTCTCACGAACGATATAAATTGATTATACGAAAAAAATCATTAAAGGATGTACTAGTCAGTGTAAAGAACGAATGTTGTAGGTTAAAATTATCTACACAAATAGAACCTCAACGTAACTTAACAGTGCAACCTCTATTGGATATAAACTAATGGATCCGGTTAATTTTATCAAGACATATGCGCCTAGAGGTTCTATTATTTTTATTAATTATGCCATGTCATTAACTAGTCATTTGAATCCATCGATAGAAAAACATGTGGGTATTTATTATGGTACGTTATTATCGGAACACTTGGTAGTTGAATCTACCTATAGAAAAGGAGTTAGAATAGTCCCATTGGATAGATTTTTTGAAGGATATCTTAGTGCAAAAGTATACATGTTAGAGAATATTCAAGTTATGAAAATAGCAGCTGATATGTCGTTAACTTTACTAGGTATTCCATATGGATTTGGTCATGATAGAATGTATTGTTTTAAATTGGTAGCTGAATGTTATAAAAATGCCGGTATTGATACATCGTCTAAACGAATATTAGGTAAAGATATTTTTCTGAGCCAAAACTTTACAGATGATAATAGATGGATAAAGATATATGATTCTAATAATTTAACATTTTGGCAAATTGATTACCTTAAAGGGTGAGTTAATATGCATAACTACTCCTCCGTTGTTTTTTCCCTCGTTCTTTTTCTTAACGTTGTTTGCCATCACTCTCATAATGTAAAGATATTCTAAAATGGTAAACTTTTGCATATCGGATGCAGAAATTGGTATAAATGTTGTAATTGTATTATTTCCCGTCAATGGACTAGTCACAGCTCCATCAGTTTTATATCCTTTAGAGTATTTCTCACTCGTGTCTAGCATTCTAGAGCATTCCATGATCTGTTTATCGTTGATATTGGCCGGAAAGATAGATTTTTTATTTTTTATTATATTACTATTGGCAATTGTAGATATAACTTCTGGTAAATATTTTTCTACCTTTTCAATCTCTTCTATTTTCAAGCCGGCTATATATTCTGCTATATTGTTACTAGTATCAATACCTTTTCTGGCTAAGAAGTCATATGTGGTATTCACTATATCAGTTTTAACTGGTAGTTCCATTAGCCTTTCCACTTCTGCAGAATAATTAGAAATTGGTTCTTTACCAGAAAATCCAGCTACTATAATAGGCTCACCGATGATCATTGGCAAAATCCTATATTGTACCAGATTAATGAGAGCATATTTCATTTCCAATAATTCTGCTAGTTCTTGAGACATTGATTTATTTGATGAATCTATTTGGTTCTCTAGATACTCTACCATTTCTGCCGCATACAATAACTTGTTAGATAAAATCAGGGTTATCAAAGTGTTTAGTGTGGCTAGAATAGTGGGCTTGCACGTATTAAAGAATGCTGTAGTATGAGTAAACCGTTTTAACGAATTATATAGTCTCCAGAAATCTGTGGCGTTGCATACATGAACTGAATGACATCGAAGATTGTCCAATATTTTTAATAGCTGCTCTTTGTCCATTATTTCTATATTTGACTCGCAACAATTGTAGATACCATTAATCACTGATTCCTTTTTCGATGCCGGACAATAGCACAATTGTTTAGCTTTGGACTCTATGTATTCAGAATTAATAGATATATCTCTCAATACAGATTGCACTATACATTTTGAAACTATGTCAAAAATTGTAGAACGACGCTGTTCTGTAGCCATTTAACTTTAAATAATTTACAAAAATTTAAAATGAGCATCCGTATAAAAATCGATAAATTGCGCCAAATTGTGGCATATTTTTCAGAGTTCAGCGAAGAAGTGTCTATAAATGTAGACTCGACGGATGAATTAATGTATATTTTTGCCGCCTTGGGCGGATCTGTAAACATTTGGGCCATTATACCTCTCAGTGCATCAGTGTTCTACCGCGGAGCCGAAAATATTGTGTTTAACCTTCCAGTGTCCAAGGTAAAATCGTGTTTGTGTAGTTTTCACAATGATGCTATCATAGATATAGAACCTGATCTGGAAAATAATCTAGTAAAACTTTCTAGTTATCATGTAGTAAGTGTCGATTGTAACAAGGAACTGATGCCTATTAGGACAGATACTACTATTTGTCTAAGTATAGATCAAAAGAAATCTTACGTATTTAATTTTCACAAGTATGAAGAAAAATGTTGTGGTAGAACCGTCATTCATCTAGAATGGTTGTTGGGCTTTATCAAGTGTATTAGTCAGCATCAGCATTTGGCTATTATGTTTAAAGATGACAATATTATTATGAAGACTCCTGGTAATACTGATGCGTTTTCCAGGGAATATTCTATGACTGAATGTTCTCAAGAACTACAAAAGTTTTCTTTCAAAATAGCTATCTCGTCTCTCAACAAACTACGAGGATTCAAAAAGAGAGTCAATGTTTTTGAAACTAGAATCGTAATGGATAATGACGATAACATTCTAGGAATGTTGTTTTCGGATAGAGTTCAATCCTTTAAGATTAACATCTTTATGGCGTTTTTAGACTAATACTTTCAATGAGATAAATATGGGTGGCGGAGTAAGTGTTGAGCTCCCTAAACGGGATCCACCTCCGGGAGTACCCACTGATGAGATGTTATTAAACGTGGATAAAATGCATGACGTGATAGCTCCCGCTAAGCTTTTAGAATATGTGCATATAGGACCACTAACAAAAGATAAAGAGGATAAAGTAAAGAAAAGATATCCAGAGTTTAGATTAGTCAACACAGGACCCGGTGGTCTTTCGGCATTATTAAGACAATCATATAATGGAACCGCACCCAATTGCTGTCGCACTTTTAATCGTACTCATTATTGGAAGAAGGATGGAAAGATATCAGATAAGTATGAAGAGGGTGCAGTATTAGAATCGTGTTGGCCCGACGTCCACGACACTGGAAAATGCGATGTTGATTTATTCGACTGGTGTCAGGGGGATACGTTCGATATGAACATATGCCATCAGTGGATCGGTTCAGCCTTTAATAGGAGTGATAGAACTGTAGAGGGTCGACAATCGTTAATAAATCTGTATAATAAGATGCAAAGATTATGTAGTAAAGATGCTAGTGTACCAATATGTGAATTATTTTTGCATCATTTACGCGCACACAATACAGAAGATAGTAAAGAGATGATCGATTATATTCTAAGACAACAGTCGGCGGACTTTAAACAGAAATATATGAGATGTAGTTATCCCACTAGAGATAAGTTAGAAGAGTCATTAAAATATGCGGAACCTCGAGAATGTTGGGATCCAGAGTGTTCGAATGCCAATGTTAATTTCTTACTAACACGTAATTATAATAATTTAGGACTTTGCAATATTGTACGATGTAATACGAGCGTGAATAACTTACAGATGGATAAAACTTCCTCATTAAGATTATCATGTGGATTAAGCAATAGTGATAGATTTTCTACTGTTCCCGTCAATAGAGCAAAAGTAGTTCAACATAATATTAAACATTCGTTCGACCTAAAATTGCATTTGATCAGTTTATTATCTCTCTTGGTAATATGGATACTAATTGTAGCTATTTAAATGGGTGCCGCAGCAAGCATACAGACGACTGTGAATACACTCAGTGAACGTATCTCGTCTAAATTAGAACAAGAAGCGAACGCTAGTGCTCAAACAAAATGTGATATAGAAATCGGAAATTTTTATATCCGACAAAACCATGGATGTAACATCACTGTTAAAAATATGTGCTCTGCGGACGCGGATGCTCAGTTGGATGCTGTGTTATCAGCCGCTACAGAAACATATAGTGGATTAACACCGGAACAAAAAGCATACGTACCAGCTATGTTTACTGCTGCGTTAAACATTCAGACGAGTGTAAACACTGTTGTTAGAGATTTTGAAAATTATGTGAAACAGACTTGTAATTCTAGCGCTGTTGTCGATAACAAATTAAAGATACAAAACGTAATTATAGATGAATGTTACGGAGCCCCAGGATCTCCAACAAATTTGGAATTTATTAATACAGGATCTAGCAAAGGAAATTGTGCCATTAAGGCGTTGATGCAATTGACTACTAAGGCCACTACTCAAATAGCACCTAGACAAGTTGCTGGTACAGGAGTTCAGTTTTATATGATTGTTATCGGTGTTATAATATTGGCAGCGTTGTTTATGTACTATGCCAAGCGTATGCTGTTCACATCCACCAATGATAAAATCAAACTTATTTTAGCCAATAAGGAAAACGTCCATTGGACTACTTACATGGACACATTCTTTAGAACTTCTCCGATGATTATTGCTACCACGGATATACAAAACTGAAAATATATTGATAATATTTTAATAGATTAACATGGAAGTTATCGCTGATCGTCTAGACGATATAGTGAAACAAAATATAGCGGATGAAAAATTTGTAGATTTTGTTATACACGGTCTAGAGCATCAATGTCCTGCTATACTTCGACCATTAATTAGGTTGTTTATTGATATACTATTATTTGTTATAGTAATTTATATTTTTACGGTACGTCTAGTAAGTAGAAATTATCAAATATTGTTGGTGTTGGTGGCGCTAGTCATCACATTAACTATTTTTTTATTACTTTATACTATAATAGTACTAGACTGACTTCTAACAAACATCTCACCTGCCATAAATAAATGCTTGATATTAAAGTCTTCTATTTCTAACACTATTCCATCTGTGGAAAATAATACTCTGACATTATCGCTAATTGATACATCGGTAAGTGATATGCCTATAAAGTAATAATCTTCTTTGGGCACATATACCAGTGTACCAGGTTCTAACAACCTATTTACTGGTGCTCCTGTAGCATACTTTTTTTTTACCTTGAGAATATCCATTGTTTGCTTGGTCAATAGTGATATGTGATTTTTTATCAACCACTCAAAAAAGTAATTGGAGTGTTCATATCCTCTACGGGCTATTGTCTCATGACCGTGTATGAAATTTAAGTAACACGACTGTGGTAGATTTGTTCTATAGAGCCGGTTGCCGCAAATAGATAGAACTACCAATATGTCTGTACAAATGTTAAACATTAATTGATTAACAGAAAAAACAATGTTCGTTCTGGGAATAGAAACCAGATTAAAACAAAATTCATTAGAATATATGCCACGTTTATACATGGAATATAAAATAACTACAGTTTGAAAAATAACAGTATCATTTAAACATTTAACTTGCGGGGTTAATCTCACAACTTTACTGTTTTTGAACTGTTCAAAATATAGCATAGATCCATGAGAAATACGTTTAGCCGCCTTTAATAGAGGAAATCCAACCGCCTTTCTGGATCTCACCAACGACGATAGTTCTGACCAGCAACTCATTTCTTCATCATCCACCTGTTTTAACATATAATAGGCAGGAGATAGATATCCATCATTGCAATATTCCTTCTCGTAGGCACACAATCTAATATTGATAAAATCTCCATTCTCTTCTCTGTATTTATTATCTTGTCTCGGTGGCTGATTAGGCTGTGGTCTATCGTTGTTGAATCTATTTTGGTCATTAAATCTTTCATTTCTTCCTGGTATATTTCTATCACCTCGTTTGGTTGGATTTTTGTCTATATTATCGTTTGTAACATCGGTACGGGTATTCATTTATCACAAAAAAAACTTCTCTAAATGAGTCTACTACTAGAAAACCTCATCGAAGAAGATACCATATTTTTTGCAGGAAGTATATCTGAGTATGATGATTTACAAATGGTTATTGCTGGTGCAAAATCCAAATTTCCAAGATCTATGCTTTCTATTTTTAATATAGTACCTAGAACGATGTCAAAATATGAGTTGGAGTTGATTCATAACGAGAATATCACAGGGGCAATGTTTACCACAATGTATAATATAAGAAACAATTTGGGTCTAGGCGATGATAAACTAACTATTGAAGCCATTGAAAACTATTTCTTGGATCCTAACAATGAGGTTATGCCTCTTATCATTAATAATACGGATATGACTACCGTCATTCCTAAAAAAAGTGGTAGGAGAAAGAATAAGAACATGGTTATCTTCCGTCAAGGATCATCACCTATCTTGTGTATTTTCGAAACTCGTAAAAAGATTAATATTTATAAAGAAAATATGGAATCCGTATCGACTGAGTATACACCTATCGGAGACAACAAGGCTTTGATATCTAAATATGCGGGAATTAATATCCTGAATGTGTATTCTCCTTCCACGTCCATGAGATTGAATGCCATTTACGGATTCACCAATAAAAATAAACTAGAGAAACTTAGTACTAATAAGGAACTAGAATCGTATAGTTCTAGCCCTCTTCAAGAACCCATTAGGTTAAATGATTTTCTGGGACTATTGGAATGTGTTAAAAAGAATATTCCTCTAACAGATATTCCGACAAAGGATTGATTACTATAAATGGAGAATGTTCCTAATGTATACTTTAATCCTGTGTTTATAGAGCCCACGTTTAAACATTCTTTATTAAGTGTTTATAAACACAGATTAATAGTTTTATTTGAAGTATTCGTTGTATTCATTCTAATATATGTATTTTTTAGATCTGAATTAAATATGTTCTTCATGCCTAAACGAAAAATACCCGATCCTATTGATAGATTACGACGTGCTAATCTAGCGTGTGAAGACGATAAATTAATGATCTATGGATTACCATGGATAACAACTCAAACATCTGCGTTATCAATAAATAGTAAACCGATAGTGTATAAAGATTGTGCAAAGCTTTTGCGATCAATAAATGGATCACAACCAGTATCTCTTAACGATGTTCTTCGCAGATGATGATTCATTTTTTAAGTATTTTGCTAGTCAAGATGATGAATCTTCATTATCTGATATATTGCAAATCACTCAATATCTAGACTTTCTGTTATTATTATTGATCCAATCAAAAAATAAATTAGAAGCTGTGGGTCATTGTTATGAATCTCTTTCAGAGGAATACAGACAATTGACAAAATTCACAGACTCTCAAGATTTTAAAAAACTGTTTAACAAGGTCCCTATTGTTACAGATGGAAGGGTCAAACTTAATAAAGGATATTTGTTCGACTTTGTGATTAGTTTGATGCGATTCAAAAAAGAATCAGCTCTAGCTACCACCGCAATAGATCCTGTTAGATACATAGATCCTCGTCGTGATATCGCATTTTCTAACGTGATGGATATATTAAAGTCGAATAAAGTTGAAAAATAATTAATTCTTTATTGTTATCATGAACGGCGGACATATTCAGTTGATAATCGGCCCCATGTTTTCAGGTAAAAGTACAGAATTAATTAGACGAGTTAGACGTTATCAAATAGCTCAATATAAATGTGTGACTATAAAATATTCTAACGATAATAGATACGGAACGGGACTATGGACACATGATAAGAATAATTTTGCAGCATTGGAAGTAACTAAACTATGTGATGTCTTGGAAGCAATTACAGATTTCTCCGTGATAGGTATAGATGAAGGACAGTTCTTTCCAGACATTGTTGAATTCTGTGAGCGTATGGCAAACGAAGGAAAAATAGTTATAGTAGCCGCGCTCGATGGGACATTTCAACGTAGACCGTTTAATAATATTTTGAATCTTATTCCATTATCTGAAATGGTGGTAAAACTAACTGCAGTGTGTATGAAATGCTTTAAGGAGGCTTCCTTTTCTAAACGATTAGGTACAGAAACCGAGATAGAAATAATAGGAGGTAATGATATGTATCAATCTGTGTGTAGAAAGTGTTACATCGACTCATAATATTATATTTTTTATCTAAAAAACTAAAAATAAACATTGATTAAATTTTAATATAATACTTAAAAATGGATGTTGTGTCGTTAGATAAACCGTTTATGTATTTTGAGGAAATTGATAATGAGTTAGATTACGAACCAGAAAGTGCAAATGAGGTCGCAAAAAAACTGCCGTATCAAGGACAGTTAAAACTATTACTAGGAGAATTATTTTTTCTTAGTAAGTTACAGCGACACGGTATATTAGATGGCGCCACCGTAGTGTATATAGGATCTGCTCCAGGTACACATATACGTTATTTGAGAGATCATTTCTATAATTTAGGAGTGATCATCAAATGGATGCTAATTGACGGCCGCCATCATGATCCTATTCTAAATGGATTGCGTGATGTGACTCTAGTGACTCGGTTTGTTGATGAGGAATATCTACGATCCATCAAAAAACAACTACATCCTTCTAAGATTATTTTAATTTCTGATGTGCGATCCAAACGAGGAGGAAATGAACCTAGTACTGCGGATTTACTAAGTAATTATGCTCTACAAAATGTCATGATTAGTATTTTAAACCCCGTGGCGTCTAGTCTTAAATGGAGATGCCCGTTTCCAGATCAATGGATCAAGGACTTTTATATCCCACACGGTAATAAAATGTTACAACCTTTTGCTCCTTCATATTCAGCTGAAATGAGATTATTAAGTATTTATACCGGTGAGAATATGAGACTGACTCGAGTTACCAAATCAGACGCTGTAAATTATGAAAAAAAGATGTATTACCTTAATAAGATAGTCCGCAACAAAGTAGTTATTAACTTTGATTATCCTAATCAGGAATATGACTATTTTCACATGTACTTTATGTTGAGGACCGTATACTGCAATAAAACATTTCCTACTACTAAAGCAAAGATACTATTTCTACAACAATCTATATTTCGTTTCTTAAATATTCCAACGACATCAACTGAAAAAGTTAGTCATGAACCAATACAACGTAAAATATCTAGCAAAGATTCTATGTCTAAAAACAGAAATAGCAAGAGATCCGTACGCGGTAATAAATAGAAACGTACTACTGAGATATACTACCGATATAGAGTATAATGATTTAGTTACTTTAATAACCGTTAGACATAAAATTGATTCTATGAAAACTGTGTTTCAGGTATTTAACGAATCATCCATAAATTATACTCCGGTTGATGATGATTATGGAGAACCAATCATTATAACATCGTATCTTCAAAAAGGTCATAACAAGTTTCCTGTAAATTTTCTATACATAGATGTGGTAATATCTGACTTATTTCCTAGCTTTGTTAGACTAGATACTACAGAAACTAATATAGTTAATAGTGTACTACAAACAGGCGATGGTAAAAAGACTCTTCGTCTTCCTAAAATGTTAGAGACGGAAATAGTTGTCAAGATTCTCTATCGTCCTAATATACCATTAAAAATTGTTAGATTTTTCCGCAATAACATGGTAACTGGAGTAGAGATAGCCGATAGATCTGTTATTTCAGTCGCTGATTAATCAATTAGTAGAGATGAGATAAGAACATTATAATAATCAATAATATATCTTATATCTGTTTAGAAAAATGCTAATATTAAAATAGCTAACGCTAGTAATCCAATCGGAAGCCATTTGATATCTATAATAGGGTATCTAATTTCCTGATTCAGATAGCGTACGGCTATATTCTCGGTAGCTACTCGTTTGGAATCACAGACATTATTTACATCTAATTTACTATCTGTAATGGAAACGTTTCCCAATGAAATGGTACAATCAGATACATTACATCTTGATATATTTTTTTTTAAAGAGGCTGGTAACAACGCATCGCTTCGTTTACATGGCTCGTACCAACAATAATAGGGTAATCTTGTATCTATTCCTATCCGTACTATACTTTTATCAGGATAAATACATTTACATCGTATATCGTCTTTGTTAGTATCACAGAATGCATAAATTTGTTCGTCCGTCATGATAAAAATTTAAAGTGTAAATATAACTATTATTTTTATAGTTATAATAAAAAGGGAAATTTGATTGTATACCTTCGGTTCTTTAAAAGAAACTGACTTGATAAAAATGGCTGTAATCTCTAAGGTTACGTATAGTCTATACGATCAAAAAGAGATTAATGCCACAGATATTATCATTAGTCATATTAAAAATGACGACGATATCGGTACCGTTAAAGATGGTAGACTAGGTGCTATGGATGGGGCATTATGTAAGACTTGTGGGAAAACGGAATTGGAATGTTTCGGTCACTGGGGTAAAGTAAGTATTTATAAAACTCATATAGTTAAGCCTGAATTTATTTCAGAAATTATTCGTTTACTGAATCATATATGTATTCATTGCGGATTATTGCGTTCACGAGAACCGTATTCCGACGATATTAACCTAAAAGAGTTATCGGTACACGCTCTTAGGAGATTAAAGGATAAAATATTATCCAAGAAAAAGTCATGTTGGAACAGCGAATGTATGCAACCGTATCAAAAAATTACTTTTTCAAAGAAAAAGGTTTGTTTCGTCAACAAGTTGGATGATATTAACGTTCCTAATTCTCTCATCTATCAAAAGTTAATTTCTATTCATGAAAAGTTTTGGCCATTATTAGAAATTCATCAATATCCAGCTAACTTATTTTATACAGACTACTTTCCCATCCCTCCGTTGATTATTAGACCGGCTATTAGTTTTTGGATAGATAGTATACCCAAAGAGACAAATGAATTAACTTACTTATTAGGTATGATCGTTAAGAATTGTAACTTGAATGCTGATGAACAGGTTATCCAGAAGGCGGTAATAGAATACGATGATATTAAAATTATTTCTAATAACACTACCAGTATCAATTTATCATATATCACATCCGGCAAAAATAATATGATTAGAAGTTATATCGTCGCTCGGCGAAAAGATCAGACCGCTAGATCCGTAATTGGTCCCAGTACATCTATCACCGTTAATGAGGTAGGAATGCCCACATATATTAGAAATACACTTACAGAAAAGATATTTGTTAATGCCTTTACAGTGGATAAAGTTAAACAACTATTAGCATCAAACCAAGTTAAATTTTACTTTAATAAACGATTAAACCAATTAACAAGAATACGTCAAGGAAAGTTTATCAAAAATAAAATACATTTATTGCCTGGTGATTGGGTAGAAGTAGCTGTTCAAGAATATACAAGTATTATTTTTGGAAGACAACCGTCTCTACATAGATACAACGTCATCGCTTCATCTATCAGAGCTACCGAAGGAGATACTATCAAAATATCTCCCGGAATTGCCAACTCTCAAAATGCTGATTTCGACGGAGATGAAGAATGGATGATATTGGAGCAAAATCCTAAAGCCGTAGTTGAACAAAGTATTCTTATGTATCCGACGACGTTACTCAAACACGATATTCATGGAGCCCCCGTTTATGGATCTATTCAAGATGAAATCGTAGCAGCGTATTCATTGTTTAGGATACAAGATCTTTGTTTAGATGAAGTATTGAACATCTTGGGGAAATATGGAAGAGAGTTCGATCCTAAAGGTAAATGTAAATTCAGCGGTAAAGATATCTATACTTACTTGATAGGTGAAAAGATTAATTATCCGGGTCTCTTAAAGGATGGTGAAATTATTGCAAACGACGTAGATAGTAATTTTGTTGTAGCTATGAGGCATCTGTCATTGGCTGGACTCTTATCCGATCATAAATCGAACGTGGAAGGTATCAACTTTATTATCAAGTCATCTTATGTTTTTAAGAGATATCTATCTATATACGGTTTTGGGGTGACATTCAAAGATCTGAGACCAAATTCGACGTTCACTAATAAATTGGAGGCTATCAACGTAGAAAAAATAGAACTTATCAAAGAAGCATACGCCAAATATCTCAAAGATGTAAGAGACGGGAAAATAGTTCCATTATCTAAAGCTTTAGAGGCGGACTACTTGGAATCCATGTTATCCAACTTGACAAATCTTAATATCAGAGAGATAGAAGAACATATGAGACAAACGCTGATAGATGATCCAGATAATAACCTCCTGAAAATGGCCAAAGCGGGTTATAAAGTAAATCCCACAGAACTAATGTATATTCTAGGTACTTATGGACAACAGAGGATAGATGGCGAACCAGCAGAGACTCGAGTATTGGGTAGAGTCTTACCTTACTATCTTCCAGACTCTAAGGATCCAGAAGGAAGAGGTTATATTCTTAATTCTTTAACAAAAGGATTAACGGGTTCTCAATATTACTTTTTGATGCTGGTTGCAAGATCTCAATCTACTGATATTGTCTGTGAAACATCACGTACCGGAACACTGGCTAGAAAAATCATTAAAAAGATGGAGGATATGGTGGTCGACGGATACGGACAAGTAGTTATAGGTAATACGCTCATCAAGTACGCAGCCAATTATACCAAAATTCTAGGCTCAGTATGTAAACCTGTAGATCTTATCTATCCAGATGAGTCCATGACTTGGTATTTGGAAATTAGTGCTTTGTGGAATAAAATAAAACAGGGATTCGTTTACTCTCAGAAACAGAAACTTGCAAAGAAGACATTGGCGCCGTTTAATTTCCTAGTATTCGTCAAACCCACCACTGAGGATAATGCTATTAAGGTTAAGGATCTGTACGATATGATTCATAACGTCATTGATGATGTGAGAGAGAAATACTTCTTTACGGTATCTAATATAGATTTTATGGAGTATATATTCTTGACGCATCTTAATCCTTCTAGAATTAGAATTACAAAAGAAACGGCTATTACTATCTTTGAAAAGTTCTATGAAAAACTCAATTATACTCTAGGTGGTGGAACTCCTATTGGAATTATTTCTGCACAGGTATTGTCTGAGAAGTTTACACAACAAGCCCTGTCCAGTTTTCACACTACTGAAAAGAGTGGTGCTGTAAAACAAAAACTTGGTTTCAACGAGTTTAATAACTTGACTAATTTGAGTAAGAATAAGACCGAAATTATCACTCTGGTATCCGATGATATCTCTAAACTTCAATCTGTTAAGATTAATTTCGAATTTGTATGTTTGGGAGAATTAAATCCAGACATCACTCTTCGAAAAGAAACAGATAGATATGTAGTAGACATAATAGTCAATAGATTATACATCAAGAGAGCAGAAATAACCGAATTAGTCGTCGAATATATGATTGAACGATTTATCTCCTTTAGCGTCATTGTAAAGGAATGGGGTATGGAGACATTCATTGAGGACGAGGATAATATTAGATTTACTATCTACCTAAATTTCGTTGAACCGGAGGAATTGAATCTTAGTAAGTTTATGATGGTTCTTCCAGGTGCCGCCAACAAGGGCAAGATTAGTAAATTCAAGATTCCTATCTCTGACTATACGGGATATAACGACTTCAATCAAACAAAAAAGCTCAATAAGATGACTGTAGAACTCATGAATCTAAAAGAATTGGGTTCTTTCGATTTGGAGAACGTCAACGTGTATCCTGGAGTATGGAATACATACGATATCTTCGGTATTGAGGCCGCTCGTGGATACTTGTGCGAAGCCATGTTAAACACCTATGGAGAAGGTTTCGATTATCTGTACCAGCCTTGTGATCTTCTCGCTAGTTTACTATGTGCTAGTTACGAACCAGAATCAGTTAATAAATTCAAGTTCGGTGCAGCTAGTACTCTTAAGAGAGCTACGTTCGGAGATAATAAAGCATTGTTAAACGCGGCTCTTCATAAAAAGTCAGAACCTATTAACGATAATAGTAGCTGCCACTTTTTTAGCAAGGTCCCTAATATAGGAACTGGATATTACAAATACTTTATCGACTTGGGTCTTCTCATGAGAATGGAAAGGAAACTATCTGATAAGATATCTTCTCAAAAGATCAAGGAGATAGAAGAAACAGAAGACTTTTAATTCTTATCAATAACATATTTTTCTATGATCTGTCTTTTAAACGATGGATTTTCCACAAATGCGCCTCTCAAGTCCCTCATAGAATGATACACGTATAAAAAATATAGCATAGGTGATGACTCCTTATTTTTAGACATTAGATATGCCAAAATCATAGCCCCGCTTCTATTTACTCCTGCAACACAATGAACCAACACGGGCTCGTTTCGTTGATCACATTTAGATAAGAAGGCGGTCACGTCGTCAAAATATTTACTAATATCAGTAGTTGTATCATCTACCAACGGTATATGAATAATATTAATATTAGAGTTAGGTAATGTATATTTATCCATCGTCAAATTTAAAACATATTTGAACTTAACTTCAGATGATGGTGCATCCATAGCATTTTTATAATTTCCCAAATACACATTATTTGTTACTCTTGTCATTATAGTGGGAGATTTGGCTCTGTGCATATCTCCAGTTGAACGTAGTAGTAAGTATTTATACAAACTTTTCTTATCCATTTATAACGTACAAATGGATAAAACTACTTTATCAGTAAACGCATGCAATTTAGAATACGTTAGAGAAAAGGCTATAGTAGGCGTACAAGCAGCCAAGACATCAACACTTATATTTTTTGTTATTATATTGGCAATTAGTGCGCTATTACTCTGGTTTCAGACGTCTGATAATCCAGTCTTTAATGAATTAACGAGATATATGCGAATTAAAAATACGGTTAACGATTGGAAATCATTAACGGATAGCAAAACAAAATTAGAAAGCGATAGAGGTAGACTTCTAGCCGCTGGTAAGGATGATATATTCGAATTCAAATGTGTGGATTTCGGCGCCTATTTTATAGCTATGCGATTGGATAAGAAAACATATCTGCCGCAAGCTATTAGGCGAGGTACTGGAGACGCGTGGATGGTTAAAAAGGCGGCAAAAGTCGATCCATCTGCTCAACAATTTTGTCAGTATTTGATAAAACACAAGTCTAATAATGTTATTACTTGTGGTAATGAGATGTTAAATGAATTAGGTTATAGCGGTTATTTTATGTCACCGCATTGGTGTTCCGATCTTAGTAATATGGAATAAGTGTTAGATAAATGCGGTAACAAATGTTCCTGTAAGGAACCATAACAGTTTAGATTTAACATTAAAGATGAGCATAAACATAATAAACAAAATTACAATCAAACCTATAACATTAATATCAAACAATCCAAAAAATGAAATCAATGGAGTAGTAAACGTGTACATAACTCCTGGATAACGTTTAGCAGCTACCGTTCCTATTCTAGACCAAAAATTTGGTTTCATGGTTTCGAAGCGGTGTTCTGCAACAAGACGAGGATCGTGTTCTACATATTTGGCAGAGTTATCCATTATTTGCCTGTTAATCTTCATTTCGTTTTCGATTCTGGCTATTTCAAAATAAAATCCCGATGATAGACCTCCAGACTTTATAATTTCATCTACGATGTTCAGCGCCGTAGTAACTCTAATAATATAGGCGGATAAGCTAACATCATACCCTCCTGTATATGTAAATATGGCATGATCTTTGTCTATTACAAGCTCGGTTTTAACTTTATTTCCTGTAATAATTTCTCTCATCTGTAGGATATCTATTTTCTTGTCATGTATTGCCTTCAAGACGGGACGAAGAAACGTAATATCCTCAATAACGTTATCGTTTTCTATAATAACTACATATTCTACATTTTTATTTTCTAGCTCGGTAAAAAATTTAGAATCCCATAGGGCTAAATGTCTAGCGATATTTCTTTTCGTTTCCTCTGTACACATAGTGTTACAAAACCCTGAAAAGAAGTGAGTATACTTGTCATCATCTCTAATATTTCCTCCAGTCCATTGTATAAACACATAATCCTTGTAATGATCTGGATCATCATTGACTATCACAACATCTCTTTTTTCTTGCATAACTTCATTGTCCTTCACATCATCGAACTTCTGATCATTAATATGCTCATGAACATTAGGAAATGTTTCTGATGGAGGTCTATCAATAACTGGCACAACAATAACAGGAGTTTTCACCGCCGCCATTTAGTTATTGAAATTAATCATATACAACTCTCTAATACGAGTTATATTTTCGTCTATCCATTGTTTCACATTGACATATTTCGACAAAAAGATATAAAATGCGTATTCCAATGCTTCTCTGTTTAATGAATTACTAAAATATACAAACACGTCACTGTCTGGTAATAAATAATATCTTAGAATATTGTAACAATTTATTTTGTATTGCACATGTTCGTGATCTATGAGTTCTTCTTCAAATGGCATAGGATCTCCGAATCTGAAAACGTATAAATAGGAGTTAGAATAATAATATTTGAGAGTATTGGTAATGTATAAACTCTTTAGCGGTATAATTAGTTTTTTTCTCTCGATTTCTATTTTTAGATGTGATGGAAAAATGACTAATTTTGTAGCATTAGTATCATGAACTCTAATCAAAATCTTAATATCTTCGTCACATGTTAGCTCTTTGAAGTTTTTAAGAGATGCATCAGTTGGTTTTACAGATGGAGTAGGTGCAACAATTTTTTGTTTAATGCATGCATGTATTGGAGCCATTGTCTTAACTATAATGGTGCTTGTATCGAAAAACTTTAATGCGGATAACGGAAGCTCTTCGCCGCGACTTTCTACGTCGTAATTGGGTTCTAATGCCGATCTCTGAATGGATACTAGTTTTCTAAGTTCTAATGTAATTCTCTGAAAATGTAAATCCAATTCCTCCGGCATTATAGATGTGTATACATCGGTAAATAAAACTATAGTATCCAACGATCCCTTCTCGCAAATTCTAGTCTTAACCAAGAAATCGTATATAACTACGGAGATGGCGTATTTAAGAGTGGATTCTTCTACCGTTTTGTTCTTGGATTTCATATAAGAAACTATAAAGTCCGCACTACTGTTAAGAATGATCACTAACGCAACTATATAGTTCAAATTAAGCATCTTGGAAACATAAAATAACTCTGTAGATGATACTTGACTTTCGAATAAGTTTGCAGACAAACGAAGAAAGAACAGACCTCTCTTAATTTCAGAAGAAAACTTTTTTTCGTATTCCTGACGTCTAGAGTTTATATCAATAAGAAAGTTAAGAATTAGTCGGTTAATGTTGTATTTCATTACCCAAGTTTGAGATTTCATAATATTGTCAAAAGACATGATAATATTAAAGATAAAGCGCTGACTATGAACGAAATAGCTATATGGTTCGCTCAAGAATATAGTCTTGTTAAACGTGGAAACGATAACTGTATTTTTAATCACGTCAGCGGCATCTAAATTAAATATAGGTATATTTATTCCACACACTCTACAATATGCCACACCATCTTCATAATAAATAAATTCGTTAGCAAAATTATTAATTTTAGTGAAATAGTTAGCGTCAACTTTCATAGCTTCCTTCAATCTAATTTGATGCTCACATGGCGCGAATTCTACTCTAACATCCCTTTTCCATGCCTCAGGTTCATCGATCTCTATAATATCTAGTTTCTTGCGTTTCACAAACACAGGCTCGTCTCTCGCGATGAGATCTGTATAGTAACTATGTAAATGATAACTAGATAGAAAGATGTAGCTATATAGATGACGATCCTTTAAGAGAGGTATAATAACTTTACCCCAATCAGATAGACTGTTGTTATGGTCTTCGGAAAAAGAATTTTTATAAATTTTTCCAGTATTTTCTAAATATACGTACTTGATATCTAAGAAATCCTTAATAATAATAGGAATGGATAATCCGTCTATTTTATAAAGAAATACATATCGCATATTATACTTTTTTTTGGAAATTGGAATACCGATGTGTCTACATAAATACGCAAAGTCTAAATATTTTTTAGAGAATCTTAGTTGGTCCAAATTCTTTTCCAAGTACGGTAATAGATTTTTCATATTGAACGGTATCTTCTTGATCTCTGGTTCTAATTCCGCATTAAATGATGAAACTAAGTCACTATTTTTATAACTAACGATTACATCACCTCTAACATCATCATTTACCAGGATACTGATCTTCTTTTGTCGTAAATACATGTCTAATGTGTTAAAAAAAAGATCATACAAGTTATACGTCATTTCATCTGTAGTATTCTTGTCATTGAAGGATAAACTCGTACTAATCTCTTCTTTAACAGTCTGTTCAAATTTATATCCTATATATGAAAAAATAGCAACCAGTGTTTGATCATCCGCGTCAATATTCTGTTCTATCGTAGTGTATAACAATCTTATATCTTCTTCTGTGATAGTCGATACGTTATAAAGGTTGATAACGAAAATATTTTTATTTCGTGAAATAAAGTCATTGTAGGATTTTGGACTTATATTCGTGTCTAGTAGATATGATTTTATTTTTGGAATGATCTCAATTAAAATAGTCTCTTTAGAGTCCATTTAAAGTTACAAACAACTAGGAAATTGGTTTATGATGTATAATTTTTTTAGTTTTTATAGATTCTTTATTCTATACTTAAAAAATGAAAATAAATACAAAGGTTCTTGAGGGTTGTGTTAATTGAAAGCGATAAATAATCATAAATTATTTCATTATCGCGATATCCGTTAAGTTTGTATCGTAATGGCGTGGTCAATTACGAATAAAGCGGATACTAGTAGTTTCACAAAGATGGCTGAAATCAGAGCTCATCTAAGAAATAGCGCTGAAAATAAAGATAAAAACGAGGATATTTTCCCGGAAGATGTAATAATTCCATCTACTAAGCCCAAAACCAAACGAACCACTACTCCTCGTAAACCAGCGGCTACTAAAAGATCAACCAAAAAGGATAAAGAAAAGGAGGAAGTGGAAGAAGTAGTTATAGAGGAATATCATCAAACAACTGAAGAAAATTCTCCACCTCCGTCATCATCTCCTGGAGTCGGCGACATTGTAGAAAGCGTGGCCGCTGTAGAGCTCGATGATAGCGACGGGGATGATGAACCTATGGTACAAGTTGAAGCTGGTAAAGTAAATCATAGTGCTAGAAGCGATCTCTCTGACCTAAAGGTGGCTACCGACAATATCGTTAAAGATCTTAAGAAAATTATTACTAGAATCTCTGCAGTATCGACTGTTCTAGAGGATGTTCAAGCAGCTGGTATCTCTAGACAATTTACTTCTATGACTAAAGCTATTACAACACTATCTGATCTAGTCACCGAGGGAAAATCTAAAGTTGTTCGTAAAAAAGTTAAAACTTGTAAGAAGTAAATGCGTGCACTTTTTTATAAAGATGGTAAACTGTTTACCGATAATAATTTTTTAAATCCTGTATCAGACGATAATCCAGCGTATGAGGTTTTGCAACATGTTAAAATTCCTACTCATTTAACAGATGTAGTAGTATATGAACAAACGTGGGAAGAGGCATTAACTAGATTAATTTTTGTGGGAAGTGATTCAAAAGGACGTAGACAATACTTTTACGGAAAAATGCATATACAGAATCGCAATGCTAAAAGAGATCGTATTTTTGTTAGAGTATATAACGTTATGAAACGAATTAATTGTTTTATAAACAAAAATATAAAGAAATCGTCCACAGATTCCAATTATCAGTTGGCGGTTTTTATGTTAATGGAAACTATGTTTTTTATTAGATTTGGTAAAATGAAATATCTTAAGGAGAATGAAACAGTAGGGTTATTAACACTAAAAAATAAACACATAGAAATAAGTCCCGATGAAATAGTTATCAAGTTTGTAGGAAAGGACAAAGTTTCACATGAATTTGTTGTTCATAAGTCTAATAGACTATATAAACCGCTATTGAAACTGACTGATGATTCTAGTCCCGAAGAATTTCTGTTCAACAAACTAAGTGAACGAAAGGTATATGAATGTATCAAACAGTTTGGTATTAGAATCAAGGATCTCCGAACGTATGGAGTCAATTATACGTTTTTATATAATTTTTGGACAAATGTAAAGTCCGTATCTCCTCTTCCATCACCAAAAAAGTTGATAGCATTAACTATCAAACAAACTGCTGAAGTGGTAGGTCATACTCCATCAATTTCAAAAAGAGCTTATATGGCAACGACTATTTTAGAAATGGTAAAGGATAAAAATTTTTTAGACGTAGTATCTAAAACTACGTTCGATGAATTCCTATCTATAGTCGTAGATCACGTTAAATCATCTACGGATGGATGATAATAGATCTTTACACAAATAATTACAAGACCGATAAATGGAAATGGATAAACGGATGAAATCTCTCGCTATGACAGCTTTCTTCGGAGAGCTAAACACGTTAGATATTATGGCATTGATAATGTCTATATTTAAACACCATCCAAACAATACCATTTTTTCAGTGGATAAGGATGGTCAATTTATGATTGATTTCGAATACGATAATTATAAGGCTTCTCAATATTTGGATCTGACCCTCACTCCGATATCTGGAAATGAATGCAAGACTCACGCATCTAGTATAGCCGAACAATTGGCGTGTGTGGATATTATTAAAGAGGATATTAGCGAATATATCAAAACTACTCCCCGTCTTAAACGATTTATAAAAAAATACCGCAATAGATCATATACTCGTATCAGTCGAGATACAGAAAAGCTTAAAATAGCTCTAGCTAAAGGCATAGATTACGAATATATAAAAGACGCTTGTTAATAAGTAAATGAAAAAAAACTAGTCGTTTATAATAAAACACAATATGGATGCCAACATAGTATCATCTTCTACTATTGCGACGTATATAGACGCTTTAGCGAAGAATGCTTCAGAATTAGAACAGAGGTCTACCGCATACGAAATAAATAATGAATTGGAACTAGTATTTATTAAACCGCCATTGATTACGTTGACAAATGTAGTAAATATCTCCACGATTCAGGAATCGTTTATTCGATTTACCGTTACTAATAAGGAAGGTATCAAAATTAGAACTAAGATTCCATTATCTAAGGTACATGGTCTAGATGTAAAAAATGTGCAGTTGGTAGATGCTATAGATAACATAGTTTGGGAAAAGAAATCATTAGTGACGGAAAATCGTCTTCACAAAGAATGCTTGTTGAGACTATCAACAGAGGAACGTCATATATTTTTGGATTACAAGAAATATGGATCCTCTATCCGACTAGAATTAGTCAATCTTATTCAAGCAAAAACAAAAAACTTTACGATAGACTTTAAGCTAAAATATTTTCTAGGATCTGGCGCTCAATCTAAAAGTTCTTTATTGCACGCTATTAATCATCCAAAGTCAAGGCCTAATACATCTCTGGAAATAGAATTTACACCTAGAGACAATGAAACAGTTCCATATGATGAACTAATAAAGGAATTGACGACTCTCTCGCGTCATATATTTATGGCTTCTCCAGAGAATGTAATTCTTTCTCCACCTATTAACGCACCTATAAAGACTTTTATGTTGCCTAAACAAGATATAGTAGGTCTGGATCTGGAAAATCTATATGCCGTAACTAAGACTGACGGCATTCCTATAACTATCAGAGTTACATCAAAAGGGTTGTATTGTTATTTTACACATCTTGGTTATATTATTAGATATCCAGTTAAGAGAACAATAGATTCCGAAGTAGTAGTCTTTGGTGAGGCAGTTAAGGATAAGAACTGGACCGTATATCTCATTAAGCTAATAGAGCCCGTAAATGCAATCAGTGATAGACTAGAAGAAAGTAAGTATGTTGAATCTAAACTAGTGGATATTTGTGATCGGATAGTATTCAAGTCAAAGAAATACGAAGGTCCGTTTACTACAACTAGTGAAGTCGTCGATATGTTATCTACATATTTACCAAAGCAACCAGAAGGTGTTATTCTGTTCTATTCAAAGGGACCTAAATCTAACATTGATTTTAAAATCAAAAAGGAGAATACTATAGACCAAACTGCAAATGTAGTATTTAGGTACATGTCCAGTGAACCAATTATCTTTGGAGAGTCGTCTATCTTTATAGAGTATAAGAAATTTACCAACGATAAAGGCTTTCCTAAAGAATATGGTTCTGGTAAGATTGTGTTATATAACGGCGTTAATTATCTAAATAATATCTATTGTTTGGAATATATTAATACACATAATGAAGTGGGTATTAAGTCCGTTGTTGTACCTATTAAGTTTATAGCAGAATTCTTAGTCAATGGAGAAATACTTAAACCTAGAATCGATAAAACCATGAAATATATTAACTCAGAAGACTATTATGGAAATCAACATAATATCATAGTCGAACATTTAAGAGATCAAAGCATCAAAATAGGAGATGTCTTTAACGAGGATAAACTATCGGATGTTGGACATCAATACGCTGCCAACAACGATAAATTTAGATTAAATCCAGAAGTTAGTTATTTTACTAATAAACGAACTAGAGGGCCGTTGGGAATTTTATCAAACTACGTCAAGACTCTTCTTATTTCTATGTATTGTTCCAAAACATTTTTAGACGATTCCAACAAACGAAAGGTATTAGCGATTGATTTTGGAAACGGTGCTGACCTGGAAAAATACTTTTATGGAGAGATTGCGTTATTGGTAGCGACGGATCCGGATGCTGATGCTATAGCTAGAGGAAATGAAAGATACAACAAATTAAATTCTGGAATTAAAACCAAGTACTACAAATTTGACTACATTCAGGAAACTATTCGATCCGATACATTTGTCTCTAGTGTCAGAGAAGTATTCTATTTTGGAAAGTTTAATATCATTGACTGGCAGTTCGCTATTCATTATTCTTTTCATCCAAGACATTATGCTACAGTCATGAATAACTTATCCGAACTAACTGCTTCTGGAGGCAAGGTATTAATTACTACCATGGATGGAGACAAATTATCAAAATTAACCGATAAAAAGACTTTTATAATTCATAAGAATCTACCTAGTAGCGAAAACTATATGTCTGTAGAAAAAATAGCTGATGATAGAATAGTGGTATATAATCCATCAACAATGTCTACTCCAATGACTGAATACATTATCAAAAAGAACGATATAGTCAGAGTGTTTAACGAATACGGATTTGTTCTTGTAGATAATGTTGATTTCGCTACAATTATAGAACGAAGTAAAAAGTTTATTAATGGCGCATCTACAATGGAAGATAGACCGTCTACAAGAAACTTTTTCGAACTAAATAGAGGAGCCATTAAATGTGAAGGTTTAGATGTCGAAGACTTACTTAGTTACTATGTTGTTTATGTCTTTTCTAAGCGGTAAATAATAATATGGTATGGGTTCTGATATCCCCGTTCTAAATGCATTAAATAATTCCAATAGAGCGATTTTTGTTCCTATAGGACCTTCCAACTGTGGATACTCTGTATTATTAATAGATATATTAATACTTTTGTAGGGTAACAGAGGTTCTACGTCTTCTAAAAATAAAAGTTTTATAACATCTGGCCTGTTCATAAATAAAAACTTGGCGATTCTATATATACTCTTATTATCAAATCTAGCCATTGTCTTATAGATGTGAGCTACTGTAGGTGTACCATTTGATTTTCTTTCTAATACTATATATTTCTCTCGAAGAAGTTCTTGCAGATCATCTGGGAATAAAATACTACTGTTGAGTAAATCAGTTATTTTTTTTATATCGATATTGATGGACATTTTTATAGTTAAGGATAATAAGTATCCCAAAGTAGATAACGACGATAACGAAGTATTTATACTTTTAGGAAATCACAATGACTTTATCAGATCAAAATTAACAAAATTAAAGGAGCATGTATTTTTTTCTGAATATATTGTGACTCCAGATACATATGGATCTTTATGCGTCGAATTAAATGGGTCTAGTTTTCAGCACGGTGGTAGATATATAGAGGTGGAGGAATTTATAGATGCTGGAAGACAAGTTAGATGGTGTTCTACATCCAATCATATATCTGAAGATATACACACTGATAAATTTGTCATTTATGATATTTATACGTTTGATTCGTTCAAGAATAAACGATTGGTATTTGTACAGGTGCCTCCATCATTAGGAGATGATAGCTATTTAACTAATCCGTTATTGTCTCCGTATTATCGTAATTCAGTAGCCAGACAAATGGTCAATGATATGATTTTTAATCAAGATTCATTTTTAAAATATTTATTAGAACATCTGATTAGAAGCCACTATAGAGTTTCTAAACATATAACAATAGTTAGATACAAGGATACCGAAGAATTAAATCTAACAAGAATATGTTATAATAGAGATAAGTTTAAGGCGTTTGTATTCGCTTGGTTTAACGGCGTTTCGGAAAATGAAAAGGTACTAGATACGTATAAAAAGGTATCTGATTTGATATAATGAATTCAGTGACTATATCACACGCACCATATACTATTACTTATCACGATGATTGGGAACCAGTAATGAGTCAATTGGTAGAGTTTTATAACGAAGTAGCCAGTTGGTTGCTACGCGACGAGACGTCGCCTATTCCTGATAAGTTCTTTATACAATTGAAACAGCCGCTTAGAAATAAACGAGTATGTGTGTGTGGTATAGATCCGTATCCAAAAGATGGAACTGGTGTACCGTTCGAATCACCAAATTTTACAAAAAAATCAATTAAGGAGATAGCTTCATCTATATCTAGATTAACCGGAGTAATTGATTATAAAGGTTATAACCTTAATATAATAGACGGGGTTATACCCTGGAATTATTACTTAAGTTGTAAATTAGGAGAAACAAAAAGTCACGCGATTTACTGGGATAAGATTTCCAAGTTACTGCTACAGCATATAACTAAACACGTTAGTGTTCTTTATTGTTTGGGTAAAACAGATTTCTCGAATATACGGGCAAAGTTAGAATCCCCGGTAACTACCATAGTGGGATATCATCCAGCGGCCAGAGACCACCAATTCGAGAAAGATCGATCATTTGAAATTATCAACGTTTTACTGGAATTAGACAACAAGACACCTATAAATTGGGCTCAAGGGTTTATTTATTAATGCTTTAGTGAAATTTTAACTTGTGTTCTAAATGGATGCGGCTATTAGAGGTAATGATGTTATCTTTGTTCTTAAGACTATAGGTGTCCCGTCAGCATGCAGACAAAATGAAGATCCAAGATTCGTAGAAGCATTTAAATGCGACGAGTTAGAAAGATATATTGATAATAATCCAGAATGTACACTATTCGAAAGTCTTAGGGATGAGGAAGCATACTCTATAGTCAGAATTTTCATGGATGTAGATTTAGACGCGTGTCTAGACGAAATAGATTATTTAACGGCTATTCAAGATTTTATTATCGAGGTGTCAAACTGTGTAGCTAGATTCGCATTTACAGAATGCGGTGCCATTCATGAAAATGTAATAAAATCCATGAGATCTAATTTTTCATTGACTAAGTCTACAAATAGAGATAAAACAAGTTTTCATATTATCTTTTTAGACACGTATACCACTATGGATACATTGATAGCTATGAAACGAACACTATTAGAATTAAGTAGATCATCTGAAAATCCACTAACAAGATCGATAGACACTGCCGTATATAGGAGAAAAACAACTCTTCGGGTTGTAGGTACTAGGAAAAATCCAAATTGCGACACTATTCATGTAATGCAACCACCTCACGATAATATAGAAGATTACCTATTCACTTACGTGGATATGAACAACAATAGTTATTACTTTTCTCTACAACGACGATTGGAGGATTTAGTTCCTGATAAGTTATGGGAACCAGGGTTTATTTCGTTCGAAGACGCTATAAAAAGAGTTTCAAAAATATTCATTAATTCTATAATAAACTTTAATGATCTCGATGAAAATAATTTTACAACGGTACCACTGGTCATAGATTATGTAACACCTTGTGCATTATGTAAAAAACGATCGCATAAACATCCGCATCAACTATCGTTGGAAAATGGTGCTATTAGAATTTACAAAACTGGTAATCCACATAGTTGTAAAGTTAAAATTGTTCCGTTGGATGGTAATAAACTGTTTAATATTGCACAAAGAATTTTAGACACTAACTCTGTTTTATTAACCGAACGAGGAGACCATATAGTTTGGATTAATAATTCATGGAAATTTAACAGCGAAGAACCCTTGATAACAAAACTAATTCTATCAATAAGACATCAACTACCTAAGGAATATTCAAGCGAATTACTCTGTCCGAGGAAACGAAAGACTGTAGAAGCTAACATACGAGACATGTTAGTAGATTCAGTAGAGACCGATACCTATCCGGATAAACTTCCGTTTAAAAATGGTGTATTGGACCTGGTAGACGGAATGTTTTACTCTGGAGATGATGCTAAAAAATATACGTGTACTGTATCGACCGGATTTAAATTTGACGATACAAAATTCGTCGAAGACAGTCCAGAAATGGAAGAGTTAATGAATATCATTAACGATATCCAACCATTAACGGATGAAAATAAGAAAAATAGAGAGCTGTATGAAAAAACATTATCTAGTTGTTTATGTGGTGCTACCAAAGGATGTTTAACATTCTTTTTTGGAGAAACCGCAACTGGGAAGTCGACAACCAAACGTTTGTTAAAGTCTGCTATCGGTGACCTGTTTGTCGAGACGGGTCAAACAATTTTAACAGATGTATTGGATAAAGGACCTAATCCATTTATCGCTAATATGCATTTAAAAAGATCTGTATTCTGTAGCGAACTACCTGATTTTGCATGTAGTGGATCAAAGAAAATTAGATCTGATAATATTAAAAAGTTGACAGAACCTTGTGTCATTGGAAGACCGTGTTTCTCCAATAAAATTAATAATAGAAACCATGCGACAATCATTATCGATACTAATTACAAACCTGTCTTTGATAGGATAGATAACGCATTAATGAGAAGAATTGCCGTCGTGCGATTCAGAACACACTTTTCTCAACCTTCTGGTAGAGAGGCTGCTGAAAATAATGACGCGTACGATAAAGTCAAACTATTAGACGAGGGATTAGATGGTAAAATACAGAATAATAGATATAGATTCGCATTTCTATACTTGTTGGTTAAATGGTACAAAAAATATCATATTCCTATTATGAAACTATATCCTACACCGGAAGAGATTCCGGACTTTGCATTCTATCTCAAAATAGGTACTCTGTTGGTATCTAGCTCTGTAAAGCATATTCCATTAATGACGGACCTCTCCAAAAAGGGATATATATTGTACGATAATGTGGTTACTCTTCCGTTGACTACTTTCCAACAGAAAATATCCAAGTATTTTAATTCTAGACTATTTGGACACGATATAGAGAGCTTCATCAATAGACATAAGAAATTTGCCAATGTTAGTGATGAATATCTGCAATATATATTCATAGAGGATATTTCATCTCCGTAAATATATGCCATATATTTATAGAATATATCACATATCTAAATGAATACCGGAATCATAGATTTATTTGATAATCATGTTGATAGTATACCAACTATATTACCTCATCAGTTAGCTACTTTAGATTATCTAGTTAGAACTATCATAGATGAGAACAGAAGCGTGTTATTGTTCCATATTATGGGATCGGGTAAAACAATAATCGCTTTGTTGTTCGCCTTGGTAGCTTCCAGATTTAAAAAGGTTTACATTTTAGTACCGAACATCAACATCTTAAAAATTTTCAATTATAATATGGGTGTAGCTATGAACTTGTTTAATGACGAATTCATAGCTGAGAATATCTTTATTCATTCCACAACAAGTTTTTATTCTCTTAATTATAACGATAACGTCATTAATTATAACGGATTAAGTCGCTACAATAACTCTATTTTTATCGTTGATGAGGCACATAATATTTTTGGGAATAATACTGGAGAACTTATGACCGTGATAAAAAATAAAAACAAGATTCCTTTTCTACTATTGTCTGGATCTCCCATTACTAACACACCTAATACGCTGGGTCATATTATAGATTTAATGTCCGAAGAGACGATAGATTTTGGTGAGATTATTAGTCGTGGTAAGAAAGTAATTCAGACACTTCTTAACGAACGCGGAGTGAATGTACTCAAGGATTTGCTTAAAGGAAGAATATCATATTACGAAATGCCGGACAAAGATCTACCAACAATAAGATATCACGGACGTAAATTTCTAGATACTCGAGTAGTATATTGTCACATGTCTAAACTTCAAGAGAAAGATTATATGATTACTAGACGGCAGCTATGTTATCATGAAATGTTTGATAAAAATATGTATAACGTGTCAATGGCAGTATTGGGACAACTTAATCTGATGAATAATTTAGATACGTTATTTCAGGAACAGGATAAGGAATTGTACCCAAATCTGAAAATAAATAATGGAGTGTTATACGGTGAAGAATTGGTAACGTTAAACATTAGTTCCAAATTTAAGTACTTTATCAATCGGATACAGACACTCAAGGGAAAACACTTTATATACTTCTCTAATTCTACATATGGTGGATTGGTAATTAAATATATCATGCTCAGTAATGGATATTCTGAATATAATGGTTCTCAGGGAACTAATCCACATATGATAAACGGCAAACCAAAAACATTTGCTATCGTTACTAGTAAAATGAAATCGTCTTTAGAGGATCTATTAGATGTGTATAATTCTCCTGAAAACGATGATGGCAATCAATTGATGTTTTTGTTTTCGTCAAACATTATGTCTGAATCCTATACTCTGAAAGAGGTAAGGCATATTTGGTTTATGACTATCCCGGATACTTTTTCTCAATACAACCAAATTCTTGGACGATCTATTAGAAAATTCTCTTACGTCGATATTTCTGAACCCGTTAATGTATATCTTTTAGCAGCCGTATATTCAGATTTCAATGACGAAGTGACGTCATTAAACGATTATACACAGGATGAATTGATTAATGTTTTACCCTTTGACATCAAAAAGCTGTTGTATCTAAAATTTAAGACTAAAGAAACGAATAGAATATACTCTATTCTTCAAGAGATGTCTGAAACGTATTCTCTTCCACCACATCCATCAATTGTAAAAGTTTTATTGGGAGAATTGGTCAGACAATTTTTTTATAATAATTCTCGTATTAAGTATAACGACTCCAAGTTACTTAAAATGGTTACATCAGTTATAAAAAATAAAGAAGACGCTAGGAATTACATAGATGATATTGTAAACGGTCACTTCTTTGTATCGAATAAAGTATTTGATAAATCTCTTTTATACAAATACGAAAACGATATTATTACAGTACCGTTTAGACTTTCCTACGAACCATTTGTTTGGGGAGTTAACTTTCGTAAAGAATATAATGTGGTATCTTCTCCATAAAACTGATGAGATATATAAAGAAATAAATGTCGAGCTTTGTTACCAATGGATATCTTCCAGTTACATTGGAACCACATGAGTTGACGTTAGACATAAAAACTAATATTAGGAATGCCGTATATAAGGCGTATCTCCATAGAGAAATTAGTGGTAAAATGGCCAAGAAAATAGAAATTCGTGAAGACGTGGAATTACCTCTCGGTGAAATAGTTAATAATTCTGTAGTTATAAACGTTCCGTGTGTAATAACCTACGCATATTATCACGTTGGGGATATAGTCAGAGGAACATTAAACATCGAAGATGAATCAAATGTAACTATTCAATGTGGAGATTTAATCTGTAAACTAAGTAGAGATTCGGGTACTGTATCATTTAGCGATTCAAAGTACTGCTTTTTTCGAAATGGTAATGCGTATGATAACGGCATCGAAGTCTCCGCCGTTCTAATGGAGGCTCAACAAGGTACCGAATCTAGTTTTGTTTTTCTCGCGAATATCGTTGACTCATAAGAAAGAGAATAGCGGTGAGTATAAATACGAATACTATGGCAATAATTGCGAATGTTTTATTCCCTTCGATATATTTTTGATAATATGAAAAACATGCCTCTCTCAAATCAGACAACCATTTCATAAAATAGTTCTCTCGCACTGGTGAGGTGGTTGCAGCTCGTATAATCTCCCCAGAATAATATACTTGCGTGTCGTCGTTCAATTTATACGGATTTCTATAATTCTCTGTTATATAATGAGGTTTACCCTCATGATTAGACGACGACAATAGTGTTCTGAATTTAGATAGTTGATCAGAATGAATGTTTATTGGTGTTGGAAAAATTATCCATGCTGCGTCTGCAGAGTGGTTGATAGTTGTTCCTAGATATGTAAAATAATCCAACGTACTAGGTAGCAAATTGTCTAGATAAAATACTGAATCAAATGGCGCAGACATATTAGCGGATCTAATGGAATCCAATTGATTGACTATCTTTTGAAAATATACATTTTTATGATCTGATACTTGTAAGAATATAGCAATAATGATAATTCCATCATCGTGTTTTTTTGCCTCTTCATAAGAACTATATTTTTTCTTATTCCAATGAACCAGATTAATCTCTCCAGAGTATTTGTATACATCTATCAAGTGATTGGATCCATAATCGTCTTCCTTTCCCCAATATATATGTATTGTTGATAACACATATTCATTGGGGAGAAACCCTCCACTTATATATCCTCCTTTAAAATTAATCCTTACTAGTTTTCCAGTATTCTGGATAGTGGTTGGTTTCGACTCATTATAATGTATGTCTAACGTCTTCAATCGCGCGTCAGAAATTGCTTTTTTAGTTTCTATATTAATAGGAGATAGTTGTTGAGGCATAGTAAAAATGAAATGATAACTGTCTAGAAATAGCTCTTAGTATGGGATTTACAATGGATGAGGAAGTGATATTTGAAACTCCTAGAGAATTAATATCTATTAAACGAATAAAAGATATTCCAAGATCAAAAGACACGCACGTGTTTGCTGCGTGTATAACAAGTGACGGATATCCGTTAATAGGAGCTAGAAGAACTTCATTCGCATTCCAGGCGATATTATCTCAACAAAATTCAGATTCTATCTTTAGAGTATCCACTAAACTATTACGGTTTATGTACTACAATGAACTAAGAGAAATCTTTAGACGGTTGAGAAAAGGTTCTATCAACAATATCGATCCTCACTTCGAAGAGTTAATATTATTGGGTGGTAAACTAGATAAAAAGGAATCTATTAAAGATTGTTTAAGAAGAGAATTAAAAGAGGAAAGTGATGAACATATAACAGTAAAAGAATTCGGAAATGTAATTCTAAAACTTACAACGAGTGATAAATTATTTAATAAAGTATATATAGGTTATTGCATGGCATGTTTTATTAATCAATCGTTGGAGGATTTATCACATACTAGTATTTACAATGTAGAAATTAGAAAGATTAAATCGTTAAATGATTGTATTAACGACGATAAATACGAATATCTGTCTTATATTTATAATATACTAATTAATAGTAAATGAGCTTTTACAGATCTAGTATAATTAGTCAGATTATTAAGTATAATAGACGACTAGCTAAGTCTATTATTTGCGAGGATGACTCTCAAATTATTACACTCACGGCATTCGTTAACCAATGCCTATGGTGTCATAAACGAGTATCCGTGTCCGCTATTTTATTAACTACTGATAACAAAATATTAGTATGTAACAGACGAGATAGTTTTCTCTATTCTGAAATAATTAGAACTAGAAACATGTATAGAAAGAAACGATTATTTCTGAATTATTCCAATTATTTGAACAAACAGGAAAGAAGTATACTATCGTCATTTTTTTCTCTAGATCCAGCTACTGCTGATAATGATAGAATAAACGCTATTTATCCGGGTGGTATACCCAAAAGGGGTGAGAACGTTCCAGAGTGTTTATCCAGGGAAATCAAAGAAGAAGTTAATATAGACAATTCTTTTGTATTCATAGACACTCGTTTTTTTATTCATGGTATCATAGAAGATACCATTATTAACAAATTTTTTGAGGTAATTTTCTTTGTTGGACGAATATCTCTAACGAGTGATCAAATTATTGATACCTTTAAAAGTAATCATGAAATAAAGGATCTAATATTTTTAGATCCAAATTCAGGTAATGGACTCCAATACGAAATTGCAAAATATGCTCTAGATACTGCAAAACTTAAATGTTACGGTCATAGAGGATGTTATTATGAATCATTAAAAAAATTAACTGAGGATGATTGATTAGAAAATATAAATTAATTTACCATCGTGTATTTTTATAACGGGATTGTCTGGCATATCATGTAGATAGTTACCGTCTACATCGTATACTCTACCATCTACGCCTTTAAATCCTCTATTTATTGATATTAATCTATTAGAATTGGAATACCAAATATTAGTACCCTCAATTAGTTTATTGGTAATATTTTTTTTAGACGATAGATCGATGGCTCTTGAAACCAAGGTTTTCCAACCGGACTCATTGTCTATCGGTGAGAAGTCTTTTTCATTAGCATGAATCCATTCTAATGATGTATGTTTAAACACTCTAAACAATTGTACAAATTCTTTTGATTTGTTTTGAATGATTTCAAATAGGTCTTCGTCTACAGTAGGCATACCATTAGATAATCTAGCCATTATAAAGTGCACGTTTACATATCTACGTTCTGGAGGAGTAAGAACGTGACTATTGAGACGAATGGCTCTTCCTACTATCTGACGAAGAGACGCCTCGTTCCATGTCATATCTAAAATGAAGATATCATTGATTGAGAAGAAACTAATACCCTCGCCTCCGCTAGAAGAGAATACGCATGTTTTAATGCATTCTCCGTTAGTGTTTGATTCTTGGTTAAACTCAGCCACCGCCTTGATTCTAGTATCTTTTGTTCTAGATGAGAACTCTATATTAGAGATACCAAAGACTTTGAAATATAGTAATAAGATTTCTATTCCTGACTGATTAACAAATGGTTCAAAGACTAGACATTTACCATGGGATGCTAATATTCCCAAACATACATCTATAAATTTGACGCTTTTCTCTTTTAATTCAGTAAATAGAGAGATATCAGCCGCAATAGCATCCCCTCCCAATAGTTCTCCCTTTTTAAAGGTGTCTAATGCGGATTTAGAAAATTCTCTATCTCTTAATGAATTTTTAAAATCATTATATAGGGTTGCTATCTCTTGTGCGTATTCTCCCGGATCACGATTTTGTCTTTCAGGAAAGCTATCGAATGTAAACGTAGTAGCCATACGTCTCAGAATTCTAAATGATGATATACCAGTTTTTATTTCTGCGAGTTTAGCCTTTTGATAAATCTCTTCTTGCTTTTTTGACATATTAACGTATCGCATTAATACTGTTTTCTTAGCGAATGATGCAGACCCTTCCACATCATCAAAAATAGAAAACTCGTTATTAACTATGTACGAACATAGGCCTCCTAGTTTGGAGACTAATTCTTTTTCATCGACTAGACGTTTATTCTCAAATAGCGATTGGTGTTGTAAGGATCCTGGTCGCAGTAAGTTAACCAACATGGTGAATTCTTGCACACTATTAACGATAGGTGTAGCCGATAAACAAATCATCTTATGGTTTTTTAACGCAATGGTCTTAGATAAAAAATTATATACTGACCGAGTAGGACGGATCTTACCATCTTCTTTGATTAATGATTTAGAAATGAAGTTATGACATTCATCAATGATGACGCATATTCTACTCTTGGAATTAATAGTTTTGATATTAGTAAAAAATTTATTTCTAAAATTTTGATCATCGTAATTAATAAAAATACAATCCTTCGTTATCTCTGGAGCGTATCTGAGTATAGTGTTTATCCAAGGATCTTCTATCAAAGCCTTTTTTACCAATAAGATAATTGCCCAATTCGTATAAATATCCTTAAGATGTTTGAGAATATATACAGTAGTCATTGTTTTACCGACACCTGTTTCATGGAACAATAAAAGAGAATGCATACTGTCTAATCCTAAGAAAACTCTTGCTACAAAATGTTGATAATCCTTGAGGCGTACTACGTCTGACCCCATCATTTCAACGGGCATATTAGTAGTTCTGCGTAAGGCATAATCGATATAGGCCGCGTGTGATTTACTCATTTATGAGTGATAAGTAATAACTATGTTTTAAAAATCACAGCAGTAGTTTAACTAGCCTTCTCTGATGTTTGTTTTCGATACTTTTTGAATCAGAAGTCATACTAGAATAAAGCAGCGAGTGAACGTAATAGAGAGCTTCGTATACTCTATTCGAAAACTCTAAGAACTTATTAATGAATTCCGTATCCACTGGATCGTTTAAAATACTAAATTGAACAGTGTTCACATCCTTCCAAGACGAAGACTTAGTGACGGACTTAACATGAGACATAAATAAATCCAAATTTTTTTTATAAACATCACTAGCCACCATAATGGCGCTATCTTTCAACCAACTATCGCTTACGCATTTTAACAGTCTAACATTTTTAAAGAGACTACAATATATTCTCATAGTATCGATTACACCTCTACCGAATAGAGTGGGAAGTTTAATAATACAATATTTTTCGTTTACAAAATCAAATAATGGTCGAAACACGTCGAAGGTTAACATCTTATAATCGCTAATGTATAGATTGTTTTCAGTGAGATGATTATTAGATTTAATAGCATCTCGTTCACGTTTGAACAGTTTATTGCGTGCGCTGAGGTCGGCAACTACGGCATCCGCTCTAGTACTCCTCCCATAATACTTTACGCTATTAATCTTTAAAATTTCATAGACTTTATCTAGATCGCTTTCTGGTAACATGATATCATGTGTAAAAAGTTTTAACATGTCGGTCGGCATTCTATTTAGATCATTAACTCTAGAAATCTGAAGAAAGTAATTAGCTCCATATTCCAGACTAGGTAATGGGCTTTTACCTAAAGACAAGTTAAGTTCTGGCAATGTTTCATAAAATGGAAGAAGGACATGTGTCCCCTCCCGGATATTTTTTACAATTTCATCCATTTACAACTCTATAGTTTGTTTTCATTATTATTAGTTATTATCTCCCATAATCTTGGTAATACTTACACCTTGATCATAAGATACCTTATACAGGTCATTACATACAACTACCAATTGTTTTTGTACATAATAGATTGGATGATTGATATCCATGGTGGAATAAACTACTCGAACAGATAGTTTATCTTTCCCCCTAGATACATTGGCCGTAATAGTTGTCGGCCTAAAGAATATCTTTGGTGTAAAGTTAAAAGTTAGGGTTCTTGTTCCATTATTGCTTTTTGTCAGTAGTTCGTTATAAATTCTCGAGATGGGCCCGTTCTCTGAATATAGAACATCATTTCCAAATCTAACTTCTAGTCTAGAAATAATATCGGTCTTATTTTTAAAATCTATTCCCTTGATGAATGGATCGTTAATAAACAAATCCTTGGCCTTTGATTCGGCTGATCTATTATCTCCGTTATAGACGTTACGTTGACTAGTCCAAAGACTTACAGGAATAGATGTATCGATGATGTTGATAGTATGTGATATGTGAGCAAAGACTGTTCTCTTGGTGGCGTCGCTATATGTTCCAGTAATGGCGGAAAACTTTTTAGAAATGTTATATATAAAAGAATTTTTTCGGGTTCCAAACATTAACAGATTAGTATGAAGATAAACACTCATATTATCAGGAACATTATCAATTTTTACATAAACATCGGCATCTTGAATAGAAACAACACCATCTTCTGGAACCTCTACGATCTCGGCAGATTCCGGATAACCAGTCGGTGGACCATCACTAACAATAACTAGATCATCCAACAATCTACTCACATATGCGTCTATATAATCTTTTTCATCTTGTGAGTACCCTGGATACGAAATAAATTTGTTATCAGTATTTCCATAATAAGGTTTAGTATAAACAGAGAGAGATGTTGCTGCATGAACTTCGGTTACTGTCGCCGTTGGTTGGTTTATTTGACCTATTACTCTCCTAGGTTTCTCTATAAATGATGGTTTAATTTGTACATTCTTAACCATATATCCAATAAAGCTCAATTCAGGAACATAAACAAATTCTTTGTTGAACGTTTCAAAGTCGAACGAAGAGTCACGAATAACGATATCGGATACTGGATTGAAGGTCACCGTTACGGTAATTTTTGAATCGGATAGTTTAAGACTACTGAATGTATCTTCCACATCAAACGGAGTTTTAATATAAACGTATACTGTAGATGGTTCTTTAATAGTGTCATTAGGAGTTAGGCCAATAGAAATATCATTAAGTTCACTAGAATATCCAGAATGTTTCAAAGCAATTGTATTATTGATACAATTATTATATAATTCTTCGCCCTCAATTTCCCAAATAACACCGTTACACGAAGAGACAGATACATGATTAATACATTTATATCCAACATATGGCACGTAACCGAATCTTCCCATACCTTTAACTTCTGGAAGTTCCAAACTCAGAACCAAATGATTAAGCGCAGTAATATACTGATCCCTAATTTCGAAGCTAGCGATAGCCTGATTGTCTGGCCCATCGTTTGTCATAACTCCGGATAGAGAAATATATTGCGGCATATATAAAGTTGGAATTTGACTATCAACTGCGAAGACATTAGACCGTTTAATAAAGTCATCCCCACCGATCAAAGAATTAATGATAGTATTATTCATTTTCTATTTAAAATGGAAAAAGCTTACAATAAACTCCGTAGAGAAATATCTATAATTTGTGAGTTTTCCTTAAAGTAACAGCTTCCGTAAACACCGTCTTTATCTCTTAGTAAGTTTATTGTATTTATGACCTTTTCCTTATCTTCATAGAATACTAAAGGCAATAAAGAAATTTTTGGTTCTTCTCTAAGAGCTACGTGAGACTTAACCATAGACGCCAACGAATCCCTACATATTTTAGAACAGAAATACCCAACTTCACCACCCTTGAATGTCTCAATACTAATAGGTCTAAAAACCAAATCTTGATTACAAAACCAACACTTATCAATTACACTATTTGTCTTAATAGACATATCTGCCATAGATTTATAATACTTTGGTAGTATACAAGCGAGTGCTTCTTCTTTAGCGGGCTTAAAGACTGCTTTAGGTGCTGAAATAACCACATCTGGAAGACTTACTCGCTTAGCCATTTAATTACGGAACTATTTTTTTATACTTCTAATGAACAAGTAGAAAACCTCTCATCTACAAAAACATACTCGTGTCCATAATCCTCTACCATAGTAACACGTTTTTTAGATCTCATATGTGCTAAAAAGTTTTCCCATACTAATTGGTTACTATTATTTTTCGTATAATTTTTAACAGTTTGAGGTTTTAGATTTTTAGTTACAGAAGTGATATCGAATATTTTATCCAAAAAGAATGAGTAATTAATTGTCTTAGAAGGAGTGTTTTCTTGGCAAAAGAATACCAAGTGCTTAAATATTTCTACTACTTCATTAATCTTTTCTGTACTCAGATTCAGTTTCTCATCTTTTACTTGATTGATTATTTCAAAGACTAACTTATAATCCTTTTTATTTATTCTCTCGTTAGCCTTAAGAAAACTAGATACAAAATTTGCATCTACATCATCCGTGGATATTTGATTTTTTTCCATGATATCCAATAGTTCCGAGATAATTTCTCCAGAACATTGATGAGACAATAATCTCCGCAATACATTTCTCAAATGAATAAGTTTATTAGACACGTGGAAGTTTGACTTTTTTTGTACCTTTGTACATTTTTGAAATACAGACTCGCAAAAAATACAATATTCATATCCTTGTTCAGATACTATACCGTTATGTCTACAACAGCTACATAATCGTAGATTCATGTTAACACTCTACGTATCTCGTCGTCCAATATTTTATATAAAAACATTTTATTTCTAGACGTTGTCAGAAAATCCTGTAATATTTTTAGTTTTTTTGGTTGTGAATAAAGTATCGCCCTAATAATATTGGTACCGTCTTCCGACAATATAGTAGTTAAATTATCCGAGCATGTAGAAGAACACCGCTTAGGCGGATTCAGTACAATGTTATATTTTTCGTACCAACTCATTTAAATATCATAATCTAAAATAGTTCTGTAATATGTCTAGCGCTAATATATTGATCATAATCCTGTGCATAAATTAAGATACAACAATGTCTTGAAATCATCGACATGGCTTCTTCCATAGTTAGAAGATCATCGTCAAAGTTAGCAACGTGATTCATCAACATTTGCTGTTTTGAGGCAGCAAATACTGAACCATCACCATTCAACCATTCATAAAAACCATCGTCTGAATCCATTGATAATTTCTTGTACTGGTTTTTGAGAGCTCGCATCAATCTAGCATTTCTAGCTCCCGGATTGAAAACAGAAAGAGGATCGTACATCCAAGGTCCATTTTCTGTAAATAGAATCGTATAATGTCCCTTCAAGAAGATATCAGACGATCCACAATCAAAGAATTGGTCTCCGAGTTTGTAACAGACTGCGGACTTTAACCTATACATGATACCGTTTAGCATGATTTCTGGTGATACGTCAATCGGAGTATCATCTATTAGAGATCTAAAGCCGGTGTAACATTCTCCGCCAAACATATTCTTATTCTGACGTCGTTCTACATAAAACATCATTGCTCCATTAACGATAACAGGTGAATGAACAGCACTACCCATCACATTAGTTCCCAATGGATCAATGTGTGTAACTCCAGAACATCTTCCATAGCCTATGTTAGGAGGAGCGAACACCACTCTTCCACTATTGCCATCGAATGCCATAGAATAAATATCCTTGGAATTGATAGAAATCGGACTGTCGGATGTTGTTATCATCTTCATAGGATTAACAACGATGTATGGTGCAGCCTGAAGTTTCATATCGTAACTGATGCCGTTCATAGGTCTAGCCACAGAAACCAACGTAGGTCTAAATCCAACTATAGACAAAATAGAAGCTAATATCTGTTCCTCATCTGTCATAACTTGAGAGCATCCAGTATGAATAATCTTCATTAGATGGGGATCTACCGCATCATCATCGTTACAATAAAAAATTCCCATTCTAATGTTCATAATTGCTTTTCTAATCATGGTATGAATGTTTGCTCTCTGAATCTCTGTGGAAATTAGATCTGATACACCTGTAATCACTATCGGATTATCCTCCGTAAGACGATTAACCAACAACATATAATTATAAGACTTTACTCTTCTAAATTCATAAAGTTGCTGGATTAGACTATATGTGTCTCCATGTACATACGCGTTCTCGAGCGCAGGAAGTTTAATACCGAATAGTGCCATCAGAATAGGATGAATGTAGTAATTAGTTTCTGGTTTTCTATAAATAAAAGACAAATCTTGTGAACTAGACATATCGGTAAAATGCATGGATTGGAATCGTGTAGTCGACAGAAGAATATGATGATTAGATGGAGAGTATATTTTATCTAACTCTTTGAGTTGGTCACCGATTCTAGGACTAGCTCGAGAATGAATAAGTACTAAGGGATGAGTACATTTCACAGAAACACTGGCGTTGTTCAACGTACTCTTTACATGGGAAAGGAGTTGAAATAGCTCGTTTCTATTTGTCCTGACAATATTTAGTTTATTCATAATATTAAGCATATCCTGAATAGTAAAGTTAGATGTGTCATACTTGTTAGTAGTTAGATATTTAGCAATTGCATTCCCATCATTTCTCAATCTCGTACTCCAATCATGTGTGGATGCTACTTCGTCGATGGAAACCATACAATCCTTTTTGATAGGCTGTTGAGATTGATCATTTCCTGTACGTTTAGGTTTGGTACGTTGATTTCTAGCCCCTGCTGATATAAAGTCATCGTCTACAATTTGGGATAATGAATTACATACACTACAAGACAAAGATTTATCAGAAGTGTGAATATGATCTTCATCTACCAAAGAAAGAGTTTGATTAGTATAACTAGATTTTAGTCCCGCGTTAGATGTTAAAAAAACATCGCTATTGACCACGGCTTCCATTATTTATATTCGTAGTTTTTACTCGAAAGCGTGATTTTAATATCCAATCTTATTACTTTTGGAATCGTTCAAAACCTTTGACTAGTTGTAGAATTTGATCTATTGCCCTACGCGTATACTCCCTTGCATCATATACGTTCGTCACCAGATCGTTTGTTTCGGCCTGAAGTTGACGCATATCTTTTTCAACACTCGACATGAGATCCTTAAGGGTCATATCGTCTAGATTTTGTTGAGATGCTGCTCCTGGATTTGGATTTTGTTGTGCTGTTGTACATACTGTACCACCAGTAGGTGTAGGAGTACATACAGTGGCCACAATAGGAGGTTGAAGAGGTGTAACCGTTGGAGTAGTACAAGAAATACTTCCATCCGATTGTTGTGTACATGTGGTTGTTGGTAACGTCTGAGAAGGTTGGGTAGATGGCGGTGTCGTCATCTTTTGATCTTTATTAAATTTAGAGATAATATCCTGAACAGTATTGCTCGGCGTCAACGCTGGAAGGAGTGTACTCGCCGGCGCATCAGTATCTGTAGACAACCAATCAAAAAGATTAGACATATCAGATGATGTATTAGTTTGTTGACGTGGTTTTAGTACAGGAGCAGTACTACTAGGTAGAAGAATAGGAGCCGGTGTAGGTGTCGGAACCGGCTGTGGAGTTATATGAATAGTTGGTTGTAGCGGTTGGGTAGGCTGTCTGCTGGCGGTCATCATATTATCTCTAGCTAGTTGTTCTCGCAACTGTCTTTGATAATACGACTCTTGAGACTTTAGTCCTATTTCAATCGCTTCATCCTTTTTCGTATCCGGATCCTTTTCTTCAGAATAATAGATTGACGACTTTGGTGTAGAGGATTCTGCCAGCCCCTGTGAGAACTTGTTAAAGAAGTCCATTTAAGGCTTTAAAATTGAATTGCGATTATAAGATTAAATGGCAGACACGGACGATATTATCGACTATGAATCCGATGATCTCACCGAATACGAGGATGATGAAGAAGATGGAGAGTCACTAGAAACTAGTGATATAGATCCCAAATCTTCTTATAAGATTGTAGAATCAACATCCACTCATATAGAAGATGCGCATTCCAATCTTAAACATATAGGGAATCATATATCTGCTCTTAAACGACGCTATACTAGACGTATAAGTCTATTTGAAATAGCGGGTATAATAGCAGAAAGCTATAACTTGCTTCAACGAGGAAGATTACCTCTAGTTTCAGAATTTTCTGACGAAACGATGAAGCAAAATATGCTACATGTAATTATACAAGAGATAGAGGAGGGTTCTTGTCCTATAGTCATCGAAAAGAACGGAGAATTGTTGTCGGTAAACGATTTTGATAAAGATGGTCTAAAATTCCATCTAGACTATATTATCAAAATTTGGAAACTTCAAAAACGATATTAGAATTTATACGAATATCGTTCTCTAAATGTCACAATCAAGTCTCTCATATTCAGCAGTTTATTGTCGTACTTTATATCGTGTTCATTAACGATATTTTGCAAAATAGTAATGATTCTATCTTCCTTCGATAGATATTCTTCAGAGATTATTGTCTTATATTCTTTCTTGTTATCCGATATGAATTTGATAAGACTTTGAACATTATTAATACCCGTCTGTTTAATTTTTTCTATAGATATTTTAGTTTTGGTAGATTCTATGGTGTCTGTTAATAGGCATCCAACATCGACATTCGACGTCAATTGTCTATAAATCAGAGTATAAATTTTAGAAATAACATTAGCAAATTGTTGTGCGTTGATGTCGTTATTCTGAAACAGTATGATTTTAGGTAGCATTTTCTTAACAAAGAGAACGTATTTATTGTTACTCAGTTGAACAGATGATATATCCAGATTACTAACGCATCTGATTCCATATACCAAACTTTCAGAAGAAATGGTGTACAATTGTTTGTATTCATTCAATGTCTCCTTTTCAGAAATTAGTTTAGAGTCGAATACTGCAATAATTTTCAAGAGATAGTTTTCATCAGATAAGATTTTATTTAGTGTAGATATGATAAAACTATTGTTTTGTTGGAGAACTTGATACGCCGCATTCTCTGTAGTCGACGCTCTCAAATGGGAAACAATCTCTATTATTTTTTTGGAATCGGATACTATATCTTCGGTATCTTGACGCAGTCTAGTATACATAGAGTTAAGAGAAATTAGAGTTTGTACATTAAGCAACATGTCTCTAAATGTGGCTACAAACTTTTCTTTTTCCACATCATCTAGTTTATTATATACCGATTTCACAACGGCACCAGATTTAAGGAACCAGAATGAAAAACTCTGATAACTACAATATTTCATCATAGTTACGATTTTATCATCTTCTATAGTTGGTGTGATAACACATACCTTTTTCTCCAAGACTGGAACCAACGTCATAAAAATGTTTAAATCAAAATCCATATCAACATCTGATGCGCTAAGACCAGTCTCGCGTTCAAGATTATCTTTACTAATGGTGACGAACTCATCGTATAGAACTCTAAGTTTGTCCATTATTTATTTACAGATTTAGTTGTTTAATTTATTTGTGCTCTTCCAGAGTTGGGATAGTATTTTTCTAACGTCGGTATTATATTATTAGGATCTACGTTCATATGTATCATAATATTAATCATCCACGTTTTGATAAATCTATCTTTAGCTTCTGAAATAACGTATTTAAACAAAGGAGAAAAATATTTAGTTACGGCATCAGACGCGATAACATTTTTTGTAAATGTAACGTATTTAGACGACAGATCTTCGTTAAAAAGTTTTCCATCTATGTAGAATCCATCGGTTGTTAACACCATTCCCGCGTCAGAGTGAATAGGAGTTTGAATAGTTTGTTTTGGAAATAGATCCTTCAATAACTTATAGTTGGGTGGGAAAAAATCGATTTTATCACTAGACTCTTTCTTTTTTACTATCATTACCTCATGAACTATTTCTTGAATGAGTATATGTATTTTCTTTCCTATATCGGTCGCGTTCATTGGAAAATATATCATGTCGTTAACTATAAGAATATTTTTATCCTCGTTTACAAACTGAATAATATCAGATATAGTTCGTAAACGAACTATATCATCACCAGCACAACATCTAACTATATGATATCCACTAGTTTCCTTTAGCCGTTTATTATCTTGTTCCATATTAGCAGTCATTCCATCATTTAAGAAGGCGTCAAAGATAATAGGGAGAAATGACATTTTGGATTCTGTTACGACTTTACCAAAATTAAGGATATACGGACTTACTATCTTTTTCTCAACGTCGATTTGATGAACACACGATGAAAATGTACTTCGATGAGATTGATCATGTAGAAAACAACAAGGGATACAATATTTCCGCATATCATGAAATATATTAAGAAATCCCACTTTATTATATTTCCCCAAAGGATCAATGCATGTAAACATTATACCGTTATCATTAATAAAGACTTCTTTCTCATCGGATCTGTAAAAGTTGTTACTGATTTTTTTCATTCCAGGATCTAGATAATTAATAATAATGGGTTTTCTATTCTTATTCTTTGTATTTTGACATATCCTAGACCAGTAAACAGTTTCCACTTTGGTAAAATCAGAAGACTTTTGAACGCTATTAAACATGGCATTAATGGCAATAACTAAAAATGTAAAATATTTTTCTATGTTAGGAATATGGTTTTTCACTTTAATAGATATATGGTTTTTTGCCAAAATGATAGATATTTTTTTATCCGATGATAGTAAAATATTATTAGTCGCCGTCTCTATAAAAATGAAGCTAGTCTCGATATCCAATTTTATTCTAGAATTGATAGGAGTCGCCAAATGTACCTTATACGTTATATCTCCCTTGATGCGTTCCATTTGTGTATCTATATCGGACACAAGATCTGTAAATAGTTTTACGTTATTAATCATCACGGTATCGCCATCGCTAGATAATGCTAATGTACTATCCAAGTCCCAAATGGAGAGATTTAACTGTTCATCGTTTAGAATAAAATGATTACCTGTCATATTAATAAAGTGTTCATCGTATCTAGATAACAACGACTTATAATTAATGTCCAAGTCTTGAACTCGCTGAATGATCTTTTTTAACCCAGTTAGTTTTAGATTGGTACGAAATATATTGTTAAACTTTGATTCTACAGTAATGTCCAAATCTAGTTGTGGAAATACTTCCATCAACATTGTTTCAAACTTGATAATATTATTATCTACATCTTCGTACGATCCAAATTCCGGAATAGATGTATCGCACGCTCTGGCCACCCAGATAACCAAAAAGTCACACGCTCCAGAATATACATTGTATAAAAAGCTATCGTTTTTTAGTAGTGTTTTTTTCTGAGTATATACGAAAGGATTAAAAATAGTATTATCAACGTAACTATATTCCAAATTATTCTTATGAGAATAGATAATAATATCGTCCTTAATATCTAACAAATTTCCTAAATATCCCTTTAATTGAGTCATTCGAAGCGTTAATAAAATATGTCTCTTAACTATTTCCGGCCGTTGTATATTTAAATGACTTCGTAAGAAATAATATATAGGCGACTTCTCATCTATGTAATCATATGGAGTGAGATATAGGGCTCGTTCTACCTCCTGCCCCTTACCCACCTGTAATACCAATTGCGGACTCACTATATATCGCATATTTATATCGTGGGGTAAAGTGAAAATCTACTACCGATGATGTAAGTCTTACAATGTTCGAACCAGTACCAGATCTTAATTTGGAGGCCTCCGTAGAACTAGGGGAGGTAAATATAGATCAAACAACACCTATGATAAAGGAGAATAGCGGTTTTATATCCCGTAGTAGACGTCTATTCGCCCATAGATCTAAGGATGATGAGAGAAAACTAGCACTACGATTCTTTTTACAAAGACTTTATTTTTTAGATCATAGAGAGATTCATTATTTGTTCAGATGCGTCGACGCTGTAAAAGACGTCACTATTACCAAAAAAAATAACATTATCGTGGCGCCTTATATAGCACTTTTAACTATCGCATCAAAAGGATGCAAACTTACAGAAACAATGATTGAAGCATTCTTTCCAGAACTATATAATGAACATAGTAAGAAATTCAAATTCAACTCTCAAGTATCCATCATCCAAGAAAAACTCGGATACCAGTCTGGAAACTATCACGTTTATGATTTTGAACCGTATTACTCTACAGTAGCTCTGGCTATTCGAGATGAACATTCATCTGGCATTTTTAATATCCGTCAAGAGAGTTATCTTGTAAGTTCATTATCTGAAATAACATATAGATTTTATCTAATTAATCTAAAATCTGATCTTGTTCAATGGAGTGCTAGTACGGGCGCTGTAATTAATCAAATGGTAAATACTGTATTGATTACAGTGTATGAAAAATTACAACTGGCCATAGAAAATGATTCACAATTTACATGTTCATTGGCTGTGGAATCAGAACTTCCAATAAAATTACTTAAAGATAGAAATGAATTATTTACAAAATTCATTAACGAGTTAAAAAAGACCAGTTCATTCAAGATAAGCAAACGTGATAAGGATACGCTATTAAAACATTTTACTTATGACTGGAGTTAGAATTTATAGACGACACATTTCGTTTATCATTGTTACTATTACTATCATTATTAGTATTCTTCTTGTCATCTTGTTCAGAAATATACAGCAATGCTATACCTAATACTAAATACATTATCATGCTTGCAATGGCTCTAACAACAACGAACCAAAATGAATTTGGTCGTAGCTTTTGTTCACAAAAATACATAAAGAAATGTCTACATAAATCTATGGCGCCATTGGCTACTTGAAATAGCGCCAGTCCTCCTACAGATTTTAATATAGCTGTATAACATGACATTTATTCATCATCAAAAGAGACAGAGTCACCATCTGTCATATTTAGATTTTTTTTCATGTGTTCAAAGTATCCTCTACTCATTTCATTATAATAGTTTATCATGCTTAGAATTTTAGGACGGATCAATGAGTAAGACTTGACTAGATCGTCAGTAGTAATTTGTGCATCATCTATTCTGCATCCGCTTCGTCGAATAATGTATAGCATCGCTTTGAGATTCTCCATAGCTATCAAGTCTTTATATAATGACATGGAAATATCTGTGAATGCTTTATACTTCTCCAACATCGATGCCTTAACATCATCACATACTTTAGCATTGAAAATACGTTCTATTGTGTAGATGGATGTAGCAAGATTTTTAAACAACAATGCCATCTTACATGATGATTGTCTCAAGTCTCCAATCGTTTGTTTAGAACGATTAGCTACAGAGTCCAATGCTTGGCTAACTAGCATATTATTATCTTTAGAAATTGTATTCTTCAATGAGGCGTTTATCATATCTGTGATTTCGTTAGTCATATTACAGTCTGACTGGGTTGTAATGTTATCCAACATATCACCTATGGATACGGTACACGTACCAGCATTTGTAATAATCCTATCTAAGATGTTGTATGGCATTGCGCAGAAAATATCTTCTCCTGTAATATCTCCACTCTCGATAAATCTACTCAGATTATTCTTAAATGCCTTATTCTCTGGAGAAAAGATATCAGTGTCCATCATTTCATTAATAGTATACGCAGAAAAGATACCACGAGTATCAATTCTATCCAAGATACTTATCGGTTCCGAGTCACAGATAATTGTTTCCTCTCCTTCGGGAGATCCTGCATAGAAATATCTAGGACAATAGTTTCTATACTGTCTGTAACTCTGATAATCTCTAAAGTCACTAACTGATACCATGAAATTGAGAAGATCAAACGCTGAAGTAATCAATTTTTCTGCCTCGTTTTTACTACAACTAGTTTTCATCAATGTAGTGACGATGTATTGTTTAGTTACTCTTGGTCTAATACTGATGATAGAGATATTATTGCTTCCCATAATGGATCTTCTAGTAGTCACCTTAAAGCCCATTGATGCGAATAGCAGATAGATAAAGTCTTGGTATGACTCCTTTCTAATATAGTACGGACTACCTTTGTCACCCAACTTTATACCCACATAAGCCATAACAACCTCTTTAATAGCCGTTTCATGAGGTTTATCAGCCATGAGCCTGAGTAGTTGAAAGAATCGCATGAATCCCGTCTCAGAAAGTCCTATATGCATGATAGATTTATCTTTCCTGGGAAACTCTCGTATAGTTATAGATGAAATACTCTTCAAAGTTTCTGAAATAAGATTAGTAACAGTCTTACCTCCGACTACTCTGGGTAACAAACATACTCTAATAGGTGTTTTCTCTGCGGAGATAATATCAGAAAGGATAGAGCAATAAGTAGTATTATTGTGATTATAAAGACCGAATACATAACAGGTAGAATTTATAAACATCATGTCCTGAAGGGTTTTAGACTTGTATTCCTCGTAATCTATACCGTCCCAAAACATGGATTTGGTAACTTTGATAGCCGTAGATCTTTGTTCCTTCGCTAACAGGTTAAAGAAATTAATAAAGAATTTGTTGTTTCTATTTATGTCCACAAATTGCACGTTTGGAAGCGCCACGGTTACATTCACTGCAGCATTTTGAGGATCGCGAGTATGAAGTACGATGTTATTGTTTACTGGTATATCTGGAAAGAAATCTACCAGTCTAGGAATAAGAGATTGATATCGCATAGAAATAGTAAAGTTTATAATCTCATCATTGAAGATTACTCTGTTACCATTGTAATAAATTGGTACTCTATCATAATCATCGACAAAGTACTGTTCATACATGATGAGATGTTTATATGTTGGCATAGTAGTGAGATCGACGTTTGGTAATGGCAATGTATTAAGATTAACTCCATAATGTCTAGCAGCATCTGCGATGTTATAAGTGATGTCAAAGCGGGGTTGATCTTGTGCTGTTATATATTGTCTAACACCTATAAGATTATCAAAATCTTGTCTGCTTAATACACCGTTAACAATTTTTGCCTTGAATTCTTTTATTGGTGCATTAATAACATCCTTATAGAGGATGTTAAACAAATAAGTATTATCAAAGTTAAGATCTGGGTATTTCTTTTCTGCTAGAACATCCATTGAGTCGGAGCCATCTGGTTTAATATAACCACCGATAAATCTAGCTCTGTATTCTGTATCCGTCAATCTAATATTAAGAAGGTGTTGAGTGAAAGGTGGAAGATCGTAAAAGCTGTGAGTATTAATAATAGGGTTAGTTTCCGAACTAATGTTAATTGGATGATTAATAATATTTATATTTCCAGCGTTAAGTGTAACATTAAACAGTTTTAATTCACGTGACGTGGTATCAATTAAATAATTAATGCCCAATTTGGATATAGTAGCCTGAAGCTCATCTTGTTTAGTTACGGATCCTAATGAGTTATTAAGAAATACATCGAACGGATGAACGAAGGTTGTTTTAAGTTGGTCACATACTTTGTAATCTAGACATAGATGTGGAAGAACGGTAGAAACTATACGAAATAGATATTCAGAGTCCTCTAATTGATCAAGAGTAACTATTGACTTAATAGGCATCATTTATTTAGTATTAAATGACGACCGTACCAGTGACAGATATACAAAACGACTTAATTACAGAGTTTTCAGAAGATAATTATCCATCTAACAAAAATTATGAAATAACTCTTCGCCAAATGTCTATTCTAACTCACGTTAACAACGTGGTAGATAGAGAACATAATGCCGCCGTAGTGTCATCTCCAGAGGAAATATCATCACAACTTAATGAAGATCTATTTCCAGATGATGATTCACCGGCCACTATTATCGAACGAGTACAACCTCATACTACTATTATTGACGATACGCCACCTCCTACTTTTCGTAGAGAGTTATTGATATCGGAACAACGTCAACAACGAGAAAAAAGATTTAATATTACAGTATCAAAAAATTCTGAAGCAATAATGGAATCTAGATCTATGATAACTTCTATGCCAACACAAACACCATCCTTGGGAGTAGTTTATGATAAAGATAAAAGAATTCAGATGCTAGAGGATGAAGTGGTTAATCTTAGAAATCAACGATCTAATACAAAATCATCTGATAATTTAGATAATTTTACCAGAATACTATTTGGTAAGACTCCGTATAAATCAACCGAAGTTAATAAGCGTATAGCCATCGTTAATTATGCAAATTTGAACGGGTCCCCCTTATCAGTCGAGGACTTGGATGTCTGTTCGGAGGATGAAATAGATAGAATCTATAAAACGATTAAACAATATCACGAAAGTAGAAAACGAAAAATTATCGTCACTAACGTGATTATTATTGTCATAAACATTATTGAGCAGGCATTGCTAAAACTCGGATTTGAAGAAATCAAAGGACTGAGTACCGATATCACTTCAGAAATTATCGATGTGGAGATCGGAGATGACTGCGATGCTGTAGCATCAAAACTAGGAATCGGTAACAGTCCGGTTCTTAATATTGTATTGTTTATACTCAAGATATTCGTTAAACGAATTAAAATTATTTAATTTAATACATTCCCATATCCAGACAACAATCGTCTGGATTAATCTGTTCCTGTCGTCTCATACCGGACGACATATTAATCTTTTTATTAGTGGGCATCTTTTTAGATGGTTTCTTTTTCCCAGCATTAACTGATTCGATACCTAGAAGATCGTGATTGATTTCTCCGACCATTCCACGAACTTCTAATTGGCCGTCTCTAACGGTACCATAAACTATTTTACCAGCATTAGTAACAGCTTGGACAATCTGACCATCCATTGCGTTGAATGATGTAGTTGCTGTTGTTCTACGTCTAGGAGCACCAGAGGTATTTTTAGAGCTCTTGGATGTTGATGTAGAAGACGAGGATTTTGATTTTGGTTTACATGTAATACATTTTGAACTCTTTGATTTTGTATCACATGCACCGGCAGTCACATCTGTTTGAGAATTAAGATTATTGTTGCCTCCTTTGACGGCTGCATCTCCACCGATCTGCGCTAGTAGATTTTTAAGCTGTGGTGTAATCTTATTAACTGTTTCAATATAATCATCGTAACTACTTCTAACGGCTAAATTTTTTTTATCCGCCATTTAGAAGCTAAAAATATTTTTATTTATGCAGAAGATTTAACTAGATTATACAATGAACTAATATGATCCTTTTCTAGATTATTTACGAACTTGGTATTTCTTGTTTCTGGAGGAGGAGAATTTAAATTCGGACTTGGATTCGGATTTTGTGGGTTCTTGATCTTATTATACAGCGTGTATAGGATGGTGACGGTAACTGCTACACAAATACCGATCAACAGAAGAATACCAATCATTTATTGACAATAACTTCACTATGATCAAGTATGTAATAATCATCTTTTCACTAAGTAAGTAGTAATAATGATTCAACAATGACACGATATATGGACGATAATAATTTAGTTCATGGAAATATCGCTATGATTGGTGTGAATGACTCCGCTAACTCTGTGGGGTGCACAGTGCTTTCCCCACATAGAATAAATTAGCATTCCGACTGTGATAATAATACCAAGTATAAACGCCATAATACTCAATACTTTCCATGTACGAGTGGGACTGGTAGACTTACTAAAGTCAATAAAGGCGAAGATACACGAAAGAATCAAAAGAATGATTCCAGCGATTAGCACGCCAGAAAAATAATTTCCAATCATAAGCATCATGTCCATTTAACTAATAAAAATTTTAAATCGCCGAATAAACAAAGTGGAATATAAACCATATAAAAACAATAGTTTGTACTGCAAAAATAATATCTATTTTTGTTTTCGAAGATATGGTAAAATTAAATAGTAGTACACAGCATGTTATAACTAACAGCAGCAACGGCTCGTAATTACTTATCATTTACTAGACGAAAAGGTGGTGGGATATTTTCTTGCTCAAATAATACGAATATATCACCCATCCATTTTATACGATGTTTATATACTCTAATCTTTAATAGATCTATAGATGACGGGTTTACCAATAATATAGATTTTATCGATTCATCTAATTTAAACCCTTCCTTAAACGTGAATGATCTATTATCTGGCATAATGATGACCCTACCTGATGAATCTGACAATGTACTGGGCCATGTAGAATAAATTATCAACGAATTATCGTCTACGAACATTTATATCATTTGTTTTAATTTTAGGACGTGAATAAATAGATATAAAATAGAAAATAACAGATATTACAACCAGTGTTATGGACGCACCCAACCATGTAGGCAGTTTTATTTTATCGTTTACTACAGGTTCTCCTGGATGTACGTCACCAACTGCAGACGTAGTTCTAGTACAATTAGACGTAAGTTCCGCTTGGGAATTTTTTAACGCTAAAGAGTTAACGTTGATCGTACACCCAACGTATTTACATCTAGTTCTTTGAACATCTTGATTATAATATAACCATTTTCTATCTCTAGATTCGTCAGTGCACTCATGTAACCAACATACCCTAGGTCCTAAATATTTATCTCCGGAATTAGATTTTGGATAATTCGCGCACCAACAATTTCTATTTCCTTTATGGTCGTTACAAAAGACGTATAATGCCGTATCCCCAAAAGTAAAATAATCAGGACGAATAATTCTAATAAACTCAGAACAATATCTCGCATCCATATGTTTGGAGCAAATATCGGAATAAGTAGACATAGCCGGTTTCCGTTTTACACGTAACCATTCTAAACAATTGGGGTTTCCAGGATCGTTTCTACAAAAACCAGTCATGAAATCGTCACAATGTTCTGTCTTGTAATTATTATTAAATATTTTTGGACAGTGTTTGGTATTTGTCTTAGAACAACATTTTGCCACGCTATCACTATCACCCAGGAGATAATCCTTTTTTATAAAATGACATCGTTGCCCGGATGCTATATAATCAGTAGCATATTTTAAATCCTTAATATATTCAGGAGTTACCTCGTTCTGATAATAGATTAATGATCCAGGACGAAATTTGAAAGAACTACATGGTTCTCCATGAATTAATACATATTGTTTAGCAAATTCAGGAACTATAAAACTACTACAATGATCTATCGACATACCATCTATCAAACAAAATTTGGGTTTAATTTCTCCTGGAGACGTTTCATAATAATACATATAACTTTCTTCGGCAAACCTAACAGCTCTATTATATTCAGGATAATTAAAATCTAATACCATATATTTGTCTCGTATATCTGCTATTCCTGTCTCTATTTTGATTCTATTAAGAGTAACAGCTGCCCCCATTCTTAATAATCATCAGTATTTAAACTGTTAAATGTTGGTATATCAACATCTATCTTATTTCCCGCAGTATAAGGTTTGTTGCAGGTATACTGTTCAGGAATGGGTACATTTATACTTCTTTTATAGTCCTGTCTTTCGATGTTCATCACAAATGCAAAGAACAGAATAAACAAAATAATGTAAGAAATAATATTAAATATCTGTGAATTCGTAAATACATTGATTGCCATAATAATTACAGCAGCTACAATACACACAATAGACATTCCCACAGTGTTGCCATTACCTCCACGATACATTTGAGTTACTAAGCAATAGGTAATAACTAAGCTAGTAAGAGGCAATAGAAAAGATGAGATAAATATCATCAATATAGAGATTAGAGGAGGGCTATATAGAGCCAAGACGAACAAAATCAAACCGAGTAACGTTCTAACATCATTATTTTTGAAGATTCCCAAATAATCATTCATTATTCCTCCATAATCGTTTTGCATCATACCCCCATCTTTAGGCATAAACGATTGCTGCTGTTCCTCTGTAAATAAATCTTTATCAAGCACTCCAGCACCCGCAGAGAAGTCATCAAGCATATTGTAATATCTTAAATAACTCATTTATATATTAAAAAATGTCACTATTAAAGATGGAGTATAATCTTTATGCCGAACTAAAAAAAATGACTTGTGGTCAGACCATAAGTCTTTTTAATGAAGACGGCGATTTCGTAGAAGTTGAACCAGGATCATCCTTTAAGTTTCTAATACCTAAGGGATTTTACTCCTCTCCTTGTGTAAAGACGAGTCTAGTATTCGAGACATTAACAACGACCGATAATAAAATTACTAGTATCAATCCAACAAATGCGCCAAAGTTATATCCTCTTCAACGCAAAGTCGTATCTGAAGTAGTTTCTAATATGAGGAAAATGATCGAATTAAAACGTCCTCTATACATCACTCTTCACTTGGCATGTGGATTTGGTAAGACTATTACCACGTGTTATCTTATGACCACACACGGCAGAAAAACCATCATTTGCGTACCCAATAAAATGTTAATACATCAATGGAAGACACAGGTAGAGGCAGTCGGATTGGAACATAAGATATCTATAGATGGAGTTAGTAGTCTATTAAAGGAACTAAAGACTCAAAGTCCGGATGTATTAATCGTAGTCAGTAGACATCTGACAAACGATGCATTTTGTAAATATATCAATAAGCATTATGATTTGTTTATCTTGGATGAATCACATACGTATAATCTGATGAACAATACAGCAGTTACAAGATTTTTAGCGTATTATCCTCCGATGATGTGTTATTTTTTAACTGCTACACCTAGACCAGCTAACCGAATTTATTGTAATAGTATTATTAATATTGCCAAGTTATCCGATCTAAAAAAAACTATCTATATAGTAGATAGTTTTTTTGAGCCATATTCCACAGACAATATTAGAAATATGGTAAAACGACTAGATGGACCATCTAATAAATATCATATATATACCGAGAAGTTATTATCTGTAGACGAGCCTAGAAACCAACTTATTCTTGATACCCTGGTAGAAGAATTCAAGTCAGGAACTATTAATAGAATTTTAGTTATTACTAAACTACGTGAACATATGGTATTCTTCTACAAACGATTATTAGATCTTTTCGGAGCAGAGGTTGTATTTATAGGAGACGCCCAAAATAGACGTACTCCAGATATGGTCAAATCGATTAAGGAACTAAATAGATTTATATTCGTATCCACCTTATTTTATTCCGGCACTGGTTTAGATATTCCGAGTTTGGATTCTTTGTTCATTTGCTCGGCAGTAATCAACAATATGCAAATAGAGCAATTACTAGGGAGGGTATGTCGAGAAACAGAACTATTAGATAGGACGGTATATGTATTTCCTAACACATCCATCAAAAAAATAAAGTACATGATAGGAAATTTCGTGCAACGAATTATTAGTCTGTCTGTAGATAAACTCGGATTTAAACAAGAAAGTTATCAGAAACATCAGGAATCTGAACCCGCTTCCGTACCAACATCCTCCAGAGAAGAACGTGTATTAAATAGAATATTTAACTCGCAAAATCGTTAAGAAGTTTAAGAGACGATCCACATGCTGAGCAGGCCAGTGTATTACCCCTCATAGTATTAATATAATCCAATGATACTTTTGTGATGTCGGAAATCTTAACCAATTTAGACTGACAGGCAGAACACGTCATACAATCATCATCGTCATCGATAACTGTAGTCTTGGGCTTCTTTTTGCGACTCTTCATTCCGGAACGCATATTGGTGCTATCCATTTAGGTAGTAAAAAATAAGTCAGAATATGCCCTATAACACGATCGTGCAAAACCTGGTATATCGTCTCTATCTTTATCACAATATAGTGTATCAACATCTTTATTATTGACCTCGTTTATCTTGGAACATGGAATGGGAACATTTTTGTTAACGGCCACCTTTGCCTTAATTCCAGATGTTGTAAAATTATAACTAAACAGTCTATCATCGACACAAATGAAATTCTTGTTTAGACGTTTGTAGTTTACGTATGCGGCTCGTTCTCGTCTCATTTTTTCAGATATTGCAGGTACTATAATATTAAAAATAAGAATGAAATAACATAGGATTAAAAATAAAGTTATCATGACTTCTAGTGCTGATTTAACTAACTTAAAAGAATTACTTAGTCTGTACAAAAGTTTGAGATTTTCAGATTCTGTGGCTATAGAGAAGTATAATTCTTTGGTAGAATGGGGAACATCTACTTACTGGAAAATAGGCGTACAAAAGGTAACTAATGTCGAGACGTCCATATCTGATTATTATGATGAGGTAAAAAATAAACCGTTTAATATTGATCCGGGGTATTATATTTTCTTACCAGTATATTTTGGAAGCGTCTTTATTTATTCAAAGGGTAAAAATATGGTAGAACTTGGATCTGGAAACTCTTTTCAAATACCGGATGAGATTCGAAGTGCGTGTAACAAAGTATTAGATAGTGATAACGGAATAGACTTTCTGAGATTTGTTTTGTTAAACAATAGATGGATAATGGAAGACGCTATATCAAAATACCAGTCTCCAGTTAATATATTTAAACTAGCTAGTGAGTACGGATTAAACATACCCAACTATTTAGAAATTGAAATAGAGGAAGACACATTATTTGACGATGAGTTATACTCTATTATGGAACGCTCTTTCGATGATACATTTCCAAAAATATCTATATCGTATATTAAGTTGGGAGAACTTAAGCGGCAAGTTGTAGACTTTTTCAAATTCTCATTCATGTATATTGAGTCAATCAAGGTAGATCGTATAGGAGATAATATTTTTATTCCTAGCGTTATAACAAAATCAGGAAAAAAGATATTAGTAAAAGATGTAGACCATTTAATACGATCCAAGGTTAGAGAACATACATTTGTAAAAGTAAAAAAGAAAAACACATTTTCCATTTTATACGACTATGATGGGAACGGAACAGAAACTAGAGGAGAAGTAATAAAACGAATTATAGACACTATAGGACGAGACTATTATGTTAATGGAAAGTATTTCTCTAAGGTTGGTATTGCAGGCTTAAAGCAATTGACTAATAAATTAGATATTAATGAGTGTGCAACTGTCGATGAGTTAGTTGATGAGATTAATAAATCCGGAACTGTAAAACGAAAAATAAAAAACCAATCAGTATTTGATTTAAGCAGAGAATGTTTGGGATATCCAGAAGCGGATTTTATAACGTTAGTTAATAACATGCGGTTCAAAATAGAAAATTGTAAGGTTGTAAATTTCAATATTGAAAATACTAATTGTTTAAATAACCCGAGTATTGAAACTATATATGGAAACTTCAACCAGTTCGTCTCAATCTTTAATACCGTTACCGATGTCAAAAAAAGATTATTCGAGTGAAATAATATGCGCCTTTGATATAGGTGCAAAAAATCCTGCCAGAACTGTTTTAGAAGTCAAGGATAACTCCGTTAGGGTATTGGATATATCAAAATTAGACTGGAGTTCTGATTGGGAAAGGCGCATAGCTCAAGATTTGTCACAATATGAATACACTACAGTTCTTCTAGAACGTCAGCCTAGAAGGTCACCGTACGTCAAATTTATCTATTTTATTAAAGGCTTTTTATATCATACATCTGCTGCCAAAGTTATTTGCGTCTCACCTGTCATGTCTGGTAATTCATATAGAGATCGAAAAAAGAGATCTGTTGAAGCATTTCTTGATTGGATGGACACATTCGGATTGCGAGACTCCGTTCCGGATAGACGCAAATTAGACGATGTAGCGGATAGTTTCAATTTGGCTATGAGATACGTATTAGATAAATGGAATACTAATTATACACCTTATAATAGGTGTAAATATAGAAATTACATAAAAAAAATGTAATAACGTTAGTAACGCCATTATGGATAATCTATTTACCTTTCTACATGAAATAGAAGATAGATATGCCAGAACTATTTTTAACTTTCATCTAATAAGTTGTGATGAAATAGGAGATATATATGGTCTTATGAAAGAACGCATTTCCTCAGAGGATATGTTTGACAATATAGTATATAATAAAGATATACATCCTGCCATTAAGAAACTAGTTTATTGCGACATCCAACTTACTAAACATATTATTAATCAGAATACGTATCCGGTATTTAACGATTCTTCACAAGTGAAATGTTGTCATTATTTCGATATAAACTCAAATAATAGCAATATTAGCTCTCGTACAGTAGAGATATTTGAGAGTGAAAAGTCATCTCTTGTATCATATATTAAAACTACCAATAAGAAGAGAAAGGTCAATTACGGCGAAATAAAGAAAACTGTACATGGAGGCACTAATGCAAATTACTTTTCCGGTAAAAAGTCTGATGAGTATCTGAGCACTACAGTCAGGTCCAACATTAATCAACCTTGGATCAAAACCATTTCTAAGAGAATGAGAGTAGATATCATTAATCACTCTATAGTAACGCGTGGAAAAAGCTCTATATTACAAACTATAGAAATTATTTTTACTAATAGAACATGTGTGAAAATATTCAAGGATTCTACTATGCACATTATTCTATCCAAGGACAAGGATGAAAAGGGATGTATAAACATGATTGATAAATTATTCTATGTATATTATAATTTATTTCTGTTGTTCGAGGATATCATCCAAAACGATTACTTTAAAGAAGTAGCTAATGTTGTAAACCATGTACTCATGGCTACGGCATTAGATGAGAAATTATTCCTAATTAAGAAAATGGCTGAACACGATGTTTATGGAGTTAGCAATTTCAAAATAGGGATGTTTAACCTGACATTTATTAAGTCGTTGGATCATACCGTTTTCCCCTCTCTGTTAGATGAGGATAGCAAAATAAAGTTTTTTAAGGGGAAAAAGCTCAATATTGTAGCATTACGATCTCTGGAGGATTGTACAAATTACGTGACTAAATCCGAGAATATGATAGAAATGATGAAGGAAAGATCGACTATTTTAAATAGCATAGATATAGAAACGGAATCGGTAGATCGTCTAAAAGAATTGCTTCTAAAATGAAAAAAAACACTGATTCAGAAATGGATCAACGACTCGGGTATAAGTTTTTGGTGCCTGATCCTAAAGCCGGAGTTTTTTATAGACCGTTACATTTCCAATATGTATCGTATTCTAATTTTATATTGCATCGATTGCATGAAATCTTGACCGTCAAGCGGCCACTCTTATCGTTTAAGAATAATACAGAACGAATTATGATAGAAATTAGCAATGTTAAAGTGACTCCTCCAGATTACTCACCTATAATTGCGAGTATTAAAGGTAAGAGTTATGACGCATTAGCCACGTTCACTGTAAATATCTTTAAAGAGGTAATGACCAAAGAGGGTATATCCATCACTAAAATAAGTAGTTATGAGGGAAAAGATTCTCATTTGATAAAAATTCCGCTACTAATAGGATATGGGAATAAAAATCCACTTGATACAGCCAAGTATCTTGTTCCTAATGTCATAGGTGGAGTCTTTATCAATAAACAATCTGTCGAAAAAGTAGGAATTAATCTAGTAGAAAAGATTACAACATGGCCAAAATTTAGGGTTGTTAAGCCAAACTCATTCACTTTCTCGTTTTCCTCCGTATCCCCTCCTAATGTATTACCGACAAGATATCGCCATTACAAGATATCTCTGGATATATCACAATTGGAAGCGTCGAATATATCATCGACAAAGACATTTATAACGGTCAATATTGTTTTGCTGTCTCAATATTTATCTAGAGTGAGTCTAGAATTCATTAGACGTAGTTTATCATACGATATGCCTCCAGAAGTTGTCTATCTAGTAAACGCGATAATAGATAGTGCTAAACGACTTACCGAATCTATTACTGACTTTAATATTGATACATACATTAATGACCTGGTGGAAGCTGAACACATTAAACAAAAATCTCAGTTAACGATTAACGAGTTTAAATATGAAATGCTGCATAACTTTTTACCTCATATGAACTATACACCCGATCAACTAAAGGGATTTTATATGATATCTTTACTAAGAAAGTTTCTCTACTGTATCTACCACACTTCTAGATATCCAGATAGAGATTCGATGGTTTGTCATCGCATCCTAACGTACGGCAAATATTTTGAGACGTTAGCACATGATGAATTAGAGAATTACATAGGTAACATCCGAAACGATATCATGAACAATCACAAGAACAGAGGCACTTACGCAGTAAACATTCATGTACTAACAACTCCTGGACTTAATCATGCATTTTCTAGTCTATTGAGTGGAAAGTTCAAAAAGTCAGACGGTAGTTATCGAACACATCCTCACTATTCATGGATGCAGAATATTTCTATTCCTAGAAGTGTTGGATTTTATCCGGATCAAGTAAAGATTTCAAAGATGTTTTCTGTCAGAAAATACCATCCAAGCCAATATCTTTACTTTTGTTCATCAGACGTTCCGGAAAGAGGTCCTCAGGTAGGTTTAGTATCTCAATTGTCTGTCTTGAGTTCCATTACAAATATACTAACGTCTGAGTATTTGGATTTGGAAAAGAAAATTTGTGAGTATATCAGATCATATTATAAAGATGATATAAGTTACTTTGAAACAGGATTTCCAATCACTATAGAAAATGCTCTAGTCGCATCTCTTAATCCAAATATGATATGTGATTTTGTAACTGACTTTAGACGTAGAAAACGGATGGGATTCTTCGGTAACTTGGAGGTAGGTATTACTTTAGTTAGGGATCACATGAATGAAATTCGCATTAATATTGGAGCAGGAAGATTAGTCAGACCATTCTTGGTTGTGGATAACGGAGAGCTCATGATGGATGTGTGTCCGGAGTTAGAAAGCAGATTAGACGACATGACATTCTCTGACATTCAGAAAGAGTTTCCACATGTCATCGAAATGGTAGATATAGAACAATTTACTTTTAGTAACGTATGTGAATCGGTTCAAAAATTTAGAATGATGTCAAAGGATGAAAGAAAGCAATACGATTTATGTGACTTTCCTGCCGAATTTAGAGATGGATATGTAGCATCTTCACTAGTGGGAATCAATCACAATTCTGGACCCAGAGCTATTCTTGGATGTGCTCAAGCTAAACAAGCTATCTCTTGTCTGAGTTCGGATATACGAAATAAAATAGACAATGGAATTCATTTGATGTATCCAGAGAGGCCAATTGTGATTAGTAAGGCTTTAGAAACTTCAAAGATTGCGGCTAATTGCTTCGGACAACATGTTACTATAGCATTAATGTCGTACAAAGGTATCAATCAAGAGGATGGAATTATCATCAAAAAACAATTTATTCAGAGAGGCGGTCTCGATATTGTTACAGCCAAGAAACATCAAGTAGAAATTCCATTGGAAAACTTTAATAACAAAGAAAGAGATAGGTCTAACGCCTATTCGAAATTAGAAAGTAATGGATTAGTTAGACTGAATGCTTTCTTGGAATCCGGAGACGCTATGGCAAGAAATATCTCATCAAGAACTCTTGAAGATGATTTTGCTAGAGATAATCAGATTAGCTTTGATGTTTCCGAGAAATATACAGATATGTACAAATCTCGCGTTGAACGAGTACAAGTAGAACTTACTGACAAAGTTAAGGTGCGAGTATTAACCATGAAAGAAAGAAGACCCATTCTAGGAGACAAATTTACTACTAGAACGAGTCAAAAGGGAACAGTCGCGTATATCGCAGATGAAACGGAACTTCCGTACGACGAAAATGGTATCACACCAGATGTCATTATTAATTCTACATCCATCTTCTCTAGAAAAACTATATCTATGTTGATAGAAGTTATTTTAACAGCCGCATATTCTACTAAGCCGTACAACAATAAGGGAGAAAACCGACCTGTCTGTTTTCCTAGTAGTAACGAAACATCTATCGATGCATATATGCAATTCGCTAAACAATGTTATGAGTATTCAAATCCGAAATTGTCCGAGGAAGAATTATCGGATAAAATCTTTTGTGAAAAGATTCTCTATGATCCTGAAACGGATAAGCCTTATGAATCCAAAGTATTTTTTGGACCAATTTATTACTTGCGTCTGAGACATTTAACTCAGGACAAGGCAACCGTTAGATGTAGAGGTAAAAAGACGAAGCTCATTAGACAAGCGAATGAGGGACGAAAACGTGGAGGAGGTATCAAGTTTGGAGAAATGGAGAGAGACTGTTTAATAGCACATGGTGCAGCCAATACTATTACAGAAGTTTTAAAAGACTCAGAAGAGGATTATCAAGATGTGTATATTTGTGAAAATTGTGGAGACATAGCAGCACAAATCAAAAGTATTAATACATGTCTTAGATGTTCAAAACTTAATCTCTCTCCTCTCTTAACAAAAATTGATACCACGCACGTATCTAAAGTATTTCTTACTCAAATGAACGCCAGAGGCGTAAAAGTTAAATTAGATTTCGAACGAAGGCCTCCTTCGTTTTATAAACCATTAGATAAAGTTGATCTTAAACCGTCTTTTCTGGTATAATATTGTTTAGTAGATACTCATCAAGATAAGCTAATTCACTAAACATATTATCGGATTCGGTATTGTTACTCGAGAATAGAGTTCGTTATGCTCCTGATATTCGGAAATCTGTGGAGTTTCAGGTTTTGGTGGAAGTGTAACTGCTACTTGGTGGGATACTGAAGGATATTTCAGAGAGTTGTGGATGTTCGGGTTCGACATCCACCGATGGTGTCACGCCACTAATCGGTTCGGTAACGTCTGTGGATGGAGGTGCTACTTCTACAGAACCTGTAGCCTCAGTTGTCAACGGAGATACATATTCAATGCGCGGAAATGTATAATTTGGTAATGGTTTCTCATGTGGATCTTAAGAAGAAGAGGTAAGATATCTACGAAAGATACCGATCACGTTTCTAGTTCTCTTTTGTAGAACTTTAACTTTTTCTTTCTCAGCATCTAGTTGATATTCCGACCTCTTCACGTTTCGCATGGGTTACCTCCGCAGTTTTTACAAGCGATTTCACGTTCCAGATCACGTTCAGCCTTCATACGTCTCTCCCTCTCTCTATCGAGTTTATCAGAGCAGTCTTTCTGAAGGCGATCGAACTCCATAAATTTCTCCAACGCTTTGATTGTTTCCATAGATTTCCGAAGTTTAGCTTCTAGGACGGCGATTCTTTTTTTTTTTTTTTTTTTTTTTTTTCGAATTCACGGGGTACAACCGTTTCCATTACCACCATCTCTATGTTTCTTTTCTAGATCGGCAATCTTTCTCAATCTTTCTCAACATTTCATCCCCATACCTTTTCATTCCTCGAGTCTATTGTCGTCGAAATATCGTTCCAGCTCCTTTTCGACCTCAATAACTTTAGCACGTTGTTTCATCAAGCTCTCTCTTGTAGTACTATCATTTTTATCTGATTCCCTGACACGTTTAAGATCTTCATGTAATTGAGTCAGCTCTTGACGCAATCTCTTAACTAACTTCCTCTCTTGCTTCTTCGTCATAGTACTTACAATCACTATGGGATCCATTGTTACCACGTCTGTACTCGACGAGCTCACGTTTAAGAGATTCAATTTCCAGTTTGTATCGGTCCATGTCTCCATTGCTACACCACCATTAGATTTACAGGCTGCTAGTTGTCGTTCGAGATCAGAAATACGTGTTTTCTTGGAATGGATTTCGTCGATGTACTTGTCATGATTGGCATCGAAACACTTATTAAGTTCTTTTTTTCAATTCTACGATTTTATTTCTTTCGCGAGTCAATTCCCTCCTGTAGTAACTATCAGTTTTGTCAGATTCACGCTCTCTACGTAGACTTTCTTGTAAGTTACTAATTTGTTCCCTGGCATTACCGAGTTCAGTTTTATATGCCGAATAGAGTTCTGATTCATCCTTTGAGAAGATCTCTAGCGATCGTTCAAGATCCCTGATTCTAGTCTTTAGCCTATTTACCTCCTCAGAAGATGCTCCGTTACCGTTTTTACAATCGTTAAGATGTCTATCAAGATCCATGATTCTATCTCTTTTCCATATCAGCATTGATTTCATTATTACGTTCGCAGTCGTTCAACTGTATTTCAAGATCTGAGATTCTAGATTGTAATCTCTGTAGCATTTCCACGGCATTCACTCAGTTGTCTTTCAAGATCTGAGATTCTAGATTGGAGTCTGCTAATCTCTGTAAGATTTCCTCCTCCGCTCTCGATGCAGTCGGTCAACTTATTCTCTAGTTCTCTAATACGCGAACGCAGTGCATCAACTTCTTGTGTGTCTTCTTGATTGCGTGTGCATTCATCGAGTCTAGATTCGAGATCTCTAACGTGACGTCGTTCTTCCTCAAGTTCTCTGTGTACTACAGAAAGCGTGTCCCTATCTTGTTGATATTTAGCAATTTCTGATTCTAGAGTACTGATTCTACTCACGTATGTACTAATAGTTGTCTTAGCCTTATCAAGATCCTCCTTGTATTTGTCACATTCCTTGATATCCATACGAAGTCTGGACAGTTCCCATTCGACATTACGACGTTTATCGATTTCAGCTCGGAGATCGTCGTCGCGTTGTTTTAGCCACATACGACTAAGTTCAAGTTCTCGTTGACAAGATCCATCTACTTTTCCATCCCTAATAGTATCCAGTTCCTTTTCTAGTTCTGACCGCATTTCTCGTTCCATATCAAGAGATTCTCTCAATTCTCGTATAGTCTTCTTATCAATTTCTGATGAATCTGAACCATCATCTGTCCCATTTTGTTGCATATCCCTGAGTTCTTTGATCTCTGTTGTAAGTCTGTCGATTCTTTCGGTTTTATAAACAGAATCCCTTTCCAAAGTCCTAATCTTACTGAGTTTATCATTAAGTTCTTCATTCAATTCAGTGAGTTTTCTCTTGGCTTCTTCCAAGTCTGTTTTAAACTCTCCATCATTTCCGCATTCTTCCTCGCATTTATCTAACCATTCAATTAGTTTATTAATAACTAGTTGGTAATCAGCGATTCCTATAGCCGTTCTTGTATTTGTGGGAACATAATTAGGATCTTCTAATGGATTGTATGGCTTGATAGCATCATCTTTATCATTATTAGGTGGGGGATGGACAACCTTAATTGGTTGGTCCTCCTTATCTCCTCCAGTAGCATGTGGTTCTTCAATACCAGTATTAGTAATAGGCTTAGACAAATGCTTGTCGTACGCGGGCACTTCCTCATCCATCAAGTATTTATAATCGGGTTCTGTTTCAGAATATTCTTTTCTAAGAGACGCGACTTCAGGAGTTAGTAGAAGAACTCTGTTTCTGTATCTATCAACGCTGGAATCGATACTCAAGTTAAGGATAGCGAATACCTCATCGTCATCATCCGTATCTTCTGAAACGCCATCATATGACATTTCATGAAGTCTAACGTATTGATAAACAGAATCAGATTTAGTATTAAACAGATCCTTGACCTTTTTAGTAAATGCATATGTATATTTTAGATCTCCAGATTTCATAATATGATCGCATGCCTTAAATGTCAATGCTTCCATGATATAGTCTGGAACACTAATGGGTGACGAAAAAGATACAGCACCATATGCTACGTTGATAAATAGATCTGAACCACTAAGTAGATAATGATTAATGTTAAGGAAGAGGAAATATTCAGTATATAGATATGCCTTAGCATCATATCTTGTACTAAACACGCTAAACAGTTTATTGATGTGATCAATTTCCAACAGAACAATTAGAGCGGCAGGAATACCAACAAACATATTACCACATCCGTATTTTCTATGAATATCACATATCATATTAAAAAATCTTGATAGAAGAGCGAATATCTCGTCTGACTTAATGAGATGTAGTTCAGCAGCATAAGTCATAACTGTAAATAGAACATACTTTCCTGTAGTGTTGATTCTAGACTCCACATCAACACCATTATTAAAAATAGTTTTATATACATCTTTAATCTGCTCTCCGTTAATCGTCGAACGTTCTAGTATACGGAAACACTTTGATTTCTTATCTGTAGTTAATGACTTAGTGATATCACGAAGAATATTACGAATTACATTTCTTGTTTTTCTTGAGAGACCTGATTCAGAACTCAACTCATCGTTCCATAGTTTTTCTACCTCAGTGGCGAAATCTTTGGAGTGTTTGGTACATTTTTTAATAAGGTTCGTGACCTCCATTTATTATAAAAAATTTTTATTCAAAACTTAACTACAATCGGGTAATTATAAGATCGTAGATCTCCCATGTGGTGGAATACTACCATCTATCGCATGTTGATGGACAGTAGGTAATGGCCATGGGAACAGTAATGTTTGCATATTTATCTTTCTTGCTAGTATTACTGTATATTGTCCCAATGTTTCAATGTGATGTTCTAACCTATCAACTGCCACTGTATCACAACAATAATGTCCGATGGAATTAAGATTATGATCCAATGTGTTTAATATATGATTATCAAGTCTTATACGATCCGCGTCTTTTTTGACAGGATCAGGCTCTTCTACAGGAAGAAGTTTCGGCCTCTTATGATAGTCATGTCTGGGAAATGGTGGTCTAGGATGAGGATCAGGTATCGGAGTAGGTTTTGGATTATAATCATCATCATCATCATCATCATCATCATCATCATCATCATCTATGATATCATCATCTTCGATATTTATTTTGCTATCTTGATAATGTCCTATATCAGTTGCATTTTCAGCACTCGACTGAATATTAGTACATTCATTGTCTATTATTAACGTATTTCTAAACCCAAAATGTATATGTTGAACATCACTACTATAGTTGATGAGTCTTATAGCATGAATTCGCTTATCGTTATCGGGTTTATCTTCTGTCACCTTAACAATTCCTTTTTTATTAAACTCTGCATAATCATAACCATTTCTATTGTTTGTTCTAATATAAACGAGTATAGCATCATTGCTAAATTTTTCAATAGTATCAAAAACAGAATATCCTAAACCATATAATATATATTCAGGAACACTCAAACTAAATGTCCAGGATTCTCCTAAATACGTAAACTTTAATAGTGCTAAATCATTCAAAAATCTACCGCTTATAGATAGATAGTACATGAATGCGTATAGTAGTCTACCTATCTCTTTATTATGAAAACCGACATTACGATCATATATTTCGTGATATACATGTGACCCGTTTACGTTAAACCATAAATACATGGGTGATCCTATAAACATGAATTTATTTCTAATTCTCAGAGCCATAGTTAATTGACCGTGTAATATTTGTTTACATGCATACTTGATACGATCATTAATAAGATTTTTATCATTGCTCGTTATTTCAGAATCGTATATATAAGGAGTACCATCATGATTCTTACCAGATATTATACAAAATACTATATATAAAATATATTGACCCACGTTAGTAATCATGTAAATGTTTAATGTTTTAAATTTTGTATTTAATGATCCATCATCATATGCTAGCATGGTCTTGTGATATTCATTCTTTAAAATATAATATTGTGTTAGCCATTGCATTGGAGCTCCTAATGGAGATTTTCTATTCTCGTCCATTTTAGGATATGCTTTCATAAAGTCCCTAATAACTTCGTGAATAATGTTTCTATGTTTTCTACTGATGCATGTATTTGCTTCGATTTTTTTATCCCATGTTTCATCTATCATAGATTTAAACGCAGTAATGCTCGCAACATTAACATCTTGAACCATTGGTACAATTCCGTTCCATAAATTTATAATGTTCGCCATTTATATAACTCATTTTTTGAATATACTTTTAATTGAACAAAAGAGTTAAGTTACTCATATGGATGCCGTCCAGTCTGTACATCAATCTTTTTAGCCAGAGATATCATAGCCGCTCTTAGAGTTTCAGCGTGATTTTCCAACCTAAATAGAACTTCATCGTTGCGTTTACAACACTTTTCTATTTGTTCAAACTTTGTTGTTATATTAGTAATCTTTTTTTCCAAATTAGTTAGCCGTTGTTTGAGAGTTTCCTCATTATCGTCTCCATAGGCTTTAACAATTGCTTCGCGTTTAGTCTCTGGATTTTTAGCAGCCTTTGTAGAGAAAAATTCAGTTGCTGGAATTGCAAGATCGTCATCTCCGGGGAAAAGAGTTCCGTCCATTTAAAGTACAGATTTTAGAAACTGACACTCTGTGTTATTTATATTTGGCGCAATACATGGATTATAAATATCGATGTTAATAACATCAGAAAATGTAAAGTCTATACATTGTCGCATCGTGTTAAATTTTCTAATGGATCTAGTATTATTGGGTCCAACTTCTGCCTGAAATCCAAATATGGAAGCGGATACAAAACCGTTTCCTGGATAAACCACACATCTCCACTTTTGCTTTACATCAGAAATTGTGTCATTGACATCTTGAACTCTCCTATCTAATGCCGGTGTTCCACCTATAGATTTTGAATACTCGAATGCTGCATGAGTAGCATTGAATTCCTTAATATTGCCATAATTTTCATATATTGAGTAACTCTGGATAAAAAGTAAACACACCGCAGCCGTCGCTACTACAATAAAAAAAATTGATAGAGAGTTCATTTATAATCTATTAGAAGCTGATAAAATTTTTTTACACGCGTCAGACAATGCTTTAATAAATAGTTCAACATCTACTTTTGTCATATCGAACCGATGGTATGATTCTAACCTAGAATTACATCCGAAAAAGTTGACTATGTTCATAGTCATTAAGTCATTAACGAACAACATTCCAGACTCTGGATTATAAGACGATACTGTTTCGTCACAATCACCCACCTTAATCATGTGATTATGAATATTGGCTATTAGAGTACCTTCTAAGAAATCTATAATATCTTTGAAACACGATTTAAAATCAAACCACGAATATACTTCTACGAAGAAAGTTAGTTTACCCATAGGAGAGATAACTATAAATGGAGATCTAGATACAAAATCCGGATCTATGATAGTTTTAACATTATTATATTCTCTATTAAATACCTCCACATCTAAAAATGTTAATTTTGAAACTATGTCTTCGTTTATTACCGTACCTGAACTAAACGCTATAAGCTCTATTGTTTGAGAACTCTTTAAACGATATTCTTGAAATACATGTAACAAAGTTTCCTTTAACTCGGTCGGTTTATCTACCATAGTTACAGAATTTGTATCCTTATCTATAATATAATAATCAAAATCGTATAAAGTTATATAATTATCGTGTTCAGATTGTGATCTTTTCAAATAGACTAAAAACCCCATTTCTCTAGTAAGTATCTTATGTATATGTTTGTAAAATATCTTCATGGTGGGAATATGCTCTACAGCAGTTAGCCATTCCTCATTGACAGCTGTAGATGTATTATACAAAACTACTCCAATGTTTAACAAGGGCCATTTTACGAGATTATTAAATCCTTGTTTGATAAATGTAGCCAATGCGGGTTCGAGTTCAACGACGATTGAATTCTCTTCCCGTGGATGCTGCATGATGAACGACGGGATGTTGTTGTTCTATTGATTTGGAATTCTTTTTCGACTTTTTGTTTATATTAAATATTTTAAAATTTATGGCTGATAGTAATTCATGTACTACGGATAATGTAGACGTGTATTGCATATCGATATCTTTATTATTAGATAAATTTATCAATAAATGTGAGAAGTTTGCCTCGTTAAGGTCTTCCATTTAAATATTATATAAATATTTGTGTTTGTATTTTATTCGTCTTTTATGGGATAGTTTTTAACTAGTAAAGCTGTAATTACATACTTTGTCCGTAAAACATAAATATAAATACCCGCTTTTATCAAACGTTCCAAAAAGTCGGCAGCTGACATTTTTAACATGACATCTATTTTAAATACACTTAGGTTTTTAGAAAAAACATCATTTTATAATTGTAACGATTCAATAACTAAAGAAAAGATTAAGATTAAACATAAGGGAATGTTATTTGTATTTTATAAGCCAAAGCATTCTACCGTTGTTAAATACTTGTCTGGAGGAGGTATATATCATGATGATTTGGTTGTATTGGGGAAGGTAACAATTAATGATCTAAAGATGATGCTATTTTACATGGATTTATCATATCATGGAGTGACAAGTAGTGGAGCAATTTACAAATTGGGATCGTCTATCGATAGACTTTCTCTAAATAGGACTATTGTTACAAAAGTTAATAACAATTATAACAATTATAACAATTATAACAATTATAATTGTTATAATAATTATAATTGTTATAATTATGATGATACATTTTTTGACGATGATGATTGATCACTATTACACAATTTTGTTTTTGTACTTTCTAATATAGTGTTTAGGTTCTTTTTCATATGAGAATATTGACTTACTAAAATATCTATGTTTAACTTTTGTTCTATAACGTCCTTATCGGCGGTATCGGTACATATACGTAATTCACCTTCACAAAATACGGAGTCTTCGATAATAATAGCCAATCGATTATTGGATCTAGCTGTCTGTATCATATTCAACATGTTTAATATATCCTTTCGTTTCCCCTTTACAGGCATCGATCGTAGCATATTTTCCGCGTCTGAGATGGAAATGTTAAAACTGCAAAAATGCGTAATGTTAGCCCGTCCTAATATTGGTACGTGTCTATAAGTTTGGCATAGTAGAATAATAGACGTGTTTAAATGCCTTCCAAAGTTTAAGAATTCTATTAGAGTATTACATTTTGATAGTTTATCACCTACATCATCAAAAATAAGTAAAAAGTGTGCTGATTTTTTATGATTTTGTGCGACAGCAATACATTTTTCTATGTTACTTTTAGTTCGTATCAGATTATATTCTAGAGCTTCCTGACTACTAACGAAATTAATATGATTTGGCCAAATGTATCCATCATAATCTGGGTTATAAACGGGTGTAAACAAGAATATATGTTTATATTTTTTAACTAGTGTAGAAAACAGAGATAGTAAATAGATAGTTTTTCCAGATCCAGATCCTCCTGTTAAAACCATTCTAAACGGCATTTTTAATAAATTTTCTCTTGAAAATTGTTTTTCTTGAAAACAATTCATAATTATATTTACAGTTACTAAATTAATTTGATAATAAATCAAAATATGGAAAACTAAGGTCGTTAGTAGGGAGGAGAACAACGAAGGCATATCGTGATATAAATAACATTTATTATCATGATGACACCAGAAAACGACGAAGAGCAGACATCTGTGTTCTCCGCTACTGTTTACGGAGACAAAATTCAGGGAAAGAATAAACGCAAACGCGTGATTGGTCTATGTATTAGAATATCTATGGTTATTTCACTACTATCTATGATTACCATGTCCGCGTTTCTCATAGTGCGCCTAAATCAATGCATGTCTGCTAACAAGGCTGCTATTACTGACTCCGCTGTTGCCGTTGCTGCGGCATCATCTACTCATAGAAAGGTTGTGTCTAGCACTACACAATATGATCACAAAGAAAGCTGTAATGGTTTATATTACCAGGGTTCTTGTTATATATTACATTCAGACTATAAGTCATTCGAGGATGCTAAAGCAAACTGCGCTGCGGAATCATCAACACTACCCAATAAATCCGATGTCTTGACTACCTGGCTCATTGATTATGTTGAGGATACATGGGGATCTGATGGTAATCCAATTACAAAAACTACATCCGATTATCAAGATTCTGATGTATCACAAGAAGTTAGAAAGTATTTTTGTACATAAATAAATGAAATCGCTTAATAGACAAACTGTAAGTAGGTTTAGGAAGTTGTCGGTGCCGGCCGCTATAATGATGTTACTCTCAACCATTATTAGCGGCATAGGAACATTTCTGCATTACAGAGAAGAACTGATGCCTAGTGCTTGCGCCAATGGATGGATACAATACGATAAACATTGTTATCTGGATACCAACATTAAAATGTCTACGGATAATGCAGTTTATCAGTGTCGCAAATTACGAGCTAGATTGCCTAGACCTGATACTAGACATCTGAGAGTATTGTTTAGTATTTTTTATAAAGATTATTGGGTAAGTTTAAAAAAGACCAATGATAAATGGTTAGATATTAATAATGATAAAGATATAGATATTAGTAAATTAACAAATTTTAAGCAACTAAACAGCACGACGGATTCTGAGGCGTGTTATATATACAAGTCTGGAAAACTGGTTAAAACAGTATGTAAAAGTACTCAATCTGTACTATGCGTTAAAAGATTCTACAAGTGACAACAAAAAATGAATTAATAGTAAGTCGTTAACGTACGCCGCCATGGACGCCGCGTTTGTTATTACTCCAATGGGTGTGTTGACTATAACAGATACATTGTATGATGATCTCGATATCTCAATCATGGACTTTATAGGACCATACATTATAGGTAACATAAAAATTGTCCAAATAGATGTACGGGATATAAAATATTCCGACATGCAAAAATGCTACTTTAGCTATAAGGGTAAAATAGTTCCTCAGGATTCTAATGATTTGGCTAGATTCAACATTTATAGTATTTGTACAGCATACAGATCAAAAAATACCATCATCATAGCATGCGACTATGATATCATGTTAGATATAGAAGGTAAACATCAACCATTTTATCTATTCCCATCTATTGATGTTTTTAACGCTACAATCATAGAAGCGTATAATCTGTATACAGCTGGAGATTATCATCTGATCATCAATCCTTCAGATAATCTGAAAATGAAATTGTCGTTTAATTCTTCATTTTGTATATCAGACGGCAATGGATGGATTATAATTGATGGGAAATGTAATAGTAATTTTTTATCATAAAAGTTGTAAAGTAAATAATAAAACAATAAATATTGAACTAGTAGTATGTTGTATATTGAGCAATCAGAGATGATGCTGGTACCTCTTATCACGGTGACCGTAGTTGCGGGAACAATATTAGTATGTTATATATTATATATTTGTAGGAAAAAGATACGTACTGTCTATAATGACAATAAAATTATCATGACAAAATTAAAAAAGATAAAGAGTCCTAATTCCAGCAAATCTAGTAAATCAACTGATAGCGAATCAGACTGGGAGGATCACTGTAGTGCTATGGAACAAAACAATGACGTAGATAATATTTCTAGAAATGAGATATTGAACGATGATAGCTTCGCTGGTAGTTTAATATGGGATAACGAATCCAATATCATGGCGCCTAGCACAGAACACATTTACGATAGTGTTGCTGGAAGCACGCTGCTAATAAATAATGATCGTAATGAACAGACTATTTATCAGAATACTACAGTAGTAATTAATGATACAGAGACTGTTGAAATACTTAATGAAGATACCAAACAGATTCCTAGCTATTCTTCCAATCCTTTCGTAAATTATAATAAAACCAGTATTTGTAGCAAGTCAAATCCGTTCATTGCAGAACTCAACAATAAATTTAGTGATAATAATCCGTTTAGGAGAGCACATAGTGACGATTATCTTAATAAGCAACAAGATCATGAATACGATGATATAGAATCATCGGTTGTATCATTGGTCTGATTAGTTTCCTTTTTATAAAATTGAAGTAATATTTAGTATTAATTACCGCCGATGCATTATACAAATATGGAGATATTCCCTGTATTCGGCATTTCTAAAATTAGCAATTTTATTGCTAATAATGACTGTAGATATTATATAGATGTAGAGCATCAAAAAATTATATCTGATGAGATCAATAGACAGATGGATGAAACGGTACTTCTTACCAACATCTTAAGCGTAGAAGTTGTAAATGACAATGAGATGTACCATCTTATTCCCCATAGACTATCGACTATTATACTCTGTATTAGTTCTGTTGGAGGATGTGTTATCTCTATAGATAATGACGTCAATGACAAAAATATTCTAACATTTCCCATTGATCATGCTGTAATCATATCCCCACTGAGTAAATGTGTCGTAGTTAGCAAGGGCCCTACAACCATACTGGTTGTTAAAGCGGATATACCCAGCAAACGATTGGTAACATCGTTTACAAACGACATACTGTATGTAAACAATCTATCACTGATTAATTATTTACCGTCGTCTGTATTCATTATTAGACGAGTCACCGACTATTTGGATAGACACATATGTGATCAGATATTTGCTAATAATAAGTGGTATTCCATTATAACTATCGACGATAAGCAATATCCTATTCCATCAAATTGTATAGGTATGTCTTCTGCCAAGTACATAAATTCGAGCATCGAGCAAGATATTTTGATCCATGTTTGTAACCTCGAGCATCCATTCGACTCAGTCTACAAAAAAATGCAGTCGTACAATTCTCTACCTATCAAGGAACAAATATTGTATGGTAGAATTGATAATATAAATATGAGCATTAGTATTTCGGTGGATTAATAGATTTCTCTAGTATGGGATCATTAATCATCTCTAAATACATCATAAAAAAGCTATTATCAAATACTGTACTGAATGGATTCATTCTTTTCTCTTTTTATGAAACTCTGTTGTATATCTACGGATAAAACTAGAAGCAAAAAATCTGATAGGAAGAATAATGATTATATGGAGGAACACGATTATTATAAAATAACAATAGTTCCTGGTTCCTCTTCCACGTCTACTAGCTCGTGGTATTATACACATGCCTAGTAATAGTCTCTTTGCGTTGACGGAAAGCAGACTAGAAATAACAGGCCAAAATGTTCAGACACCATAATAGTTCCCAACCCAGATAATAACAGAGTTCCATCAACACATTCCTTTAAACTCAATCCCAAACCCAAAACCGTTAAAATGTATCCAGCCAATTGATAGTAGATAATGAGGTGTACAGCACATGATAATTTACACAGTAACCAAAATGAAAACACTTTAGTAATTATAAGAAATATAGACGGTAATGTCATCATCAACAATCCAATAATATGCCTGAGAGTAAACATTGACGGATAAAACAAAAATGCCCCGCATAACTCTATCATGGCAATAACGCAACCAAACACTTGTAAAATTCCTAAATTAGTAGAAAATACAACTGATATCGATGTATAAGCGATTTCGAGGAATAATAAGAACAAAGTAATTCCCGTAAAGATAAACATCAACATTGTTTGGTAATCATTAAACCAATTAGTATGACGTTGAATTAATTTCACAGTATATTTTATTCCAGTATTATCCCCGCATGTATACGTACCTGGTAAGATATCTTTATATTCCATAATCAATGAGACATCACTATCCGATAACGAATGAAGTCTAGCACTAGTATGCCATTTACTTAATATGGTCGTCTTGGAAGTTTTATTATAAGTTAAAATATCATGATTGTCCAATTTCCATCTAATATACTTTGTCGGATTATCTATAATACACGGAATAATGATGGTATCATTACATGCTGTATATTCTATAGTCTTTGTAGATGTTATAACCACAAAAGTACAGAGGTATATCAACAATATTCTAACTCTTAACATTTTTATTTATTTAAAATGATACCTTTGTTATTTATTTTATTCTTATTTTGCTAACGGTATCGAATGGCATAAGTTTGAAACGAGTGAAGAAATAATTTCTACTTACTTAATAGATTATGTGGTAACGGGTGTTATTAATGGGGATGTATATACATTTTCAAATAATGAACTAAACAAAACTGGGTTAACTAATAACAATAATTATATCACAACATCTATAAAAGTAGAGGATAAGGATACATTAGTAGTATGCGGAACCAATAACGGAAATCCCAAATGTTGGAAAATAGACGGTTCATACTACCCAAAACATATAGGTAGAGGATACGATCATCAAAATAGCAAAGTAACGATAATCAGTCACAATGAATGTGTACTATCCGACATAAACATATAAAAAGAAGGAATTAAACGATGGAGAAGATTTGACGGACCATGTGGTTATGATTAAACGAGTTAAGTTTTTTAAGAAGCCTTAGAAGAGAGGCTATTGGGTATGAGAATCCGAAATATTAAACCAGACAACCCCATATAATTTTATAGCTAAGAATGCCGCGAAGAATGGAACTAATAAAAACGGAAATATTTGTAGCACAACGAATAACTCCCAAACTGCATTCATGTTACACTATATAACACTACTTCGGTTAGATGTTTTAGAAAAAATAAATATCACCGTACCGTTTTGTTGTATAAAAATAACAATTAACAATTATCAATTTTTTTCTTTAATATTTTACGTGGTTGACCATTCTTGGTGGTAAAATAATCTCTTAGTGTTGGAATGGAATGCTGTTTAATGTTTCCACACTCATCGTATATTTTGACGTATGCAGTCACATCGTTTACGCAATAGTCAGACTGTAGTTCTATCATGCTTCCTACGTTAGAAGGAGGAACAGTTTTAAAGTCTCTTGGTTTTAATCTATTGTCATTAGTTTTCATGAAATCCTTTGTTTTATCCACTTCACATTTTAAATAAATGTCCACTATACATTCTTCTGTTAATTTTACTAGATCATCATGAGTCATAGAATTCATAGGTTCCGTAGTCCATGGATCCAAACTAGCAAACTTCGCGTATACGGTATCGCGATTAGTGTATACACCAACTGTATGAAAATTAAGAAAACAGTTTAATAAATCTACAGAAATATTTAATCCTCCGTTTGATACAGATGCGCCATATTTATGGATTTCGGATTCACACGTTGTTTGTCTAAGGGGTTCGTCTAGTGTTGCTTCTACATAGACTTCGATTCCCATATATTCTTTATTGCCAGAATCACATACCGATTTATCATACGCTGGTTCACTTGTTTGAAAACTAAATGGTAGTAGATACATCAAAATAATAAATAATAAGTACATTCTGCAATATTGTTATCGTAATTGGAAAATTGGTATTCAAGTGAGCTGGATTATGTGAGTATTGGATTGTATATTTTATTTTATATTTTATATTTTATATTTTATATTTTATTTTATATTTTATATTTTATTTTATATTTTGTAGTAAGAATAGAATGCTAATGTCAAGTTTATTCGAATAGATGTCTTATTAAAAAACATATATAATAAATAACAATGGCTGAATGGCATAAAATTATCGAGGATATCTCAAAAAATAATAAGTTCGAGGATGCCGCCATCGTTGATTACAAGACTACAAAGAATGTTCTAGCGGCTATTCCTAACAGAACATTTGCAAAGATTAATCCGGGTGAAGTTATTCCCCTCATCACTAATCATAATATTCTAAAACCTCTTATTGGTCAGAAATTTTGTATTGTATATACTAACTCTCTAATGGATGAGAACACGTATGCTATGGAGTTGCTTACTGGGTACGCCCCTGTATCTCCGATCGTTATAGCGAGAACTCATACCGCACTTATATTTTTGATGGGTAAGCCAACAACATCCAGACGTGATGTGTATAGAACATGTAGAGATCACGCTACCCGTGTACGTGCAACTGGTAATTAAAATAAAAAGTAATATTCATATGTAGTGTCAATTTTAAATGATGATGAAATGGATAATATCCATATTGACGATGTCAATAATGCCGGTATTGACATACAGCTCATCGATTTTTAGATTTCATTCAGAGGATATTGAATTATGTTATGGGAATTTGTATTTTGATAGGATCTATAATAATGTAGTAAATATAAAATATATTCCTGAGCATATTCCATATAGATATAATTTTATTAATCGTACGTTCTCCGTAGATGAACTAGATGATAATGTCTTTTTTACACATGGTTATTTTTTAAAACACAAATATGGTTGTTCACTTAATCCTAGTTTGATTGTCTCATTATCAGGAAACTTAAAATATAATGATATACAATGCTCAGTAAATGTATCGTGTCTCATTAAAAATTTGGCAACGAGTACATCTACTATATTAACATCTAAACATAAGACTTATTCTCTATATCGGTCCATGTGTATTGCTATAATAGGATACGATTCTATTATATGGTATAAATATATAAATGACAGGTATAATGACATCTATGATTTTACTGCAATATGTATGCTAATAGCGTCTACATTGATAGTGATCATATACGTGTTTAAAAAAATAAAAATGAACTCTTAATTATGTTATACTATTAGAAATGGATAAAATCAAAATTACGATTGATTCAAAAATTGGTAATGTTGTTACCATATCGTATAACTTGGAAAAGATAACTATTGATGTCACGCCAAAAAAGAAAAAAGAAAAGGATGTATTATTAGCGCAATCAGTTGCTGTCGAAGAGGCAAAAGATGTCAAGGTGGAAGAAAAAAATATTATCGATATTGAAGATGACGATGATATGGATATAGAAAACACGTAATACGATCTATAAAAATAAGTATTAAATACTTTTTATTTACGGTACTCTTGTAGTGGTGATACCACTAATCGATTATTTTTTTTAAAAAAATACTTATTCTGATTCTTCTAGCCATTTCCGTGTTCGTTCGAATGCCACATCGACGTCAAAAATAGGGGAGTAGTTGAAATCTAGTTCTGCATTGTTGGTACGCACCTCAAATGTAGTGTTGGATATCTTCAACGTATAGTTGTTGAGTATTGATGGTTTTCTAAATAGAATTCTCTTCATATCATTCTTGCACGCGTACATTTTTAGCATCCATCTTGGAATCCTAGATCCTTGTTCTATTCCCAATGGTTTCATCAATAGAAGATTAAACATATCGTAAGAACACGATGGAGAGTAATCGTAGCAAAAGTAAGCATTTCCTTTAATCGCAGATCCCGGATACTGGATATATTTTGCAGCCAACACGTGCATCCATGCAACATTTCCTACATATACCCGGCTATGCACAGCGTCATCATCGACTGTACGATACATAATGTTACCGTGTTGCTTACATTGCTCGTAAAAGACTTTCGTCAATTTGTCTCCTTCTCCGTAAATTCCAGTGGGTCTTAGGCAACAAGTATACAATTTTGCGCCATTCATGATTACGGAATTATTGGCTTTCATAACCAGTTGCTCGGCCATACGTTTACTTTTTGCGTATACATGTCCTGGTGATATATCATAAAGGGTATGCTCATGACCGATGAATGGATTACCGTGTTTATTTGGTCCTATTGCTTCCATGCTACTAGTATAGATCAAATACTTGATTCCTAGGTCCACACAAGCTGCCAATATAGTCTGTGTTCCATAATAGTTTACTTTCATGATTTCATTATCAGTGTATTTTCCAAATACATCCACTAGAGCAGCCGTATGAATAATCAGATTTACCCCATCTAGCGCTTCTCTCACCTTATCAAAGTCGTTTATATCACATTGTATATAGTTTATAACCTTAACTTTCGAGGTTATTGGTTGTGGATCTTCTACAATATCTATGACTCTTATTTCTTGAACATCATCTGCGCTAATTAAAAGTTTTACTATATACCTGCCTAGAAATCCGGCACCGCCAGTAACCGCGTACACGGCCATTGCTGCCACTCATAATATCAGACTACTTATTCTATTTTACTAAATAATGGCTGTTTGTATAATAGACCACGATAATATCAGAGGAGTTATTTACGTTGAACAAGTCCATGGAAAAGATAAAGTTTTAGGATCAGTTATTGGATTAAAATCCGGAACGTATAGTTTGATAATTCATCGTTACGGAGATATTAGTCGAGGATGTGATTCCATAGGCAGTCCAGAAATATTTATCGGTAACATCTTTGTAAACAGATATGGTGTAGCATATGTTTATTTAGATACAGATGTAAATATATCTACAATTATTGGAAAGGCGTTATCTATTTCAAAAAATGATCAGAGATTAGCGTGTGGAGTTATTGGTATTTCGTACATAAATGAAAAGATAATACATTTTCTTACAATTAACGAGAATGGCGTTTGATATATCAGTTAATGCGTCTAAAACAATAAATGCATTAGTTTACTTTTCTACTCAGCAAGATAAATTAGTCATACGTAATGAAGTTAATGATATACACTACACTGTCGAATTTGATAGGGACAAAGTAGTTGATACGTTTATTTCATATAATAGACATAATGACTCCATAGAGATAAGAGGGGTGCTTCCAGAGGAAACTAATATTGGTCGCGTGGTTAATACGCCGGTTAGTATGACTTACTTGTATAATAAGTATAGTTTTAAACCGATTTTAGCAGAATATATAAGACACAGAAATACTATATCCGGCAACATTTATTCGGCATTGATGACGCTAGATGATTTGGTTATTAAACAGTATGGAGACATTGATCTATTATTTAATGAGAAACTTAAAGTAGACTCCGATTCGGGACTATTTGACTTTGTCAACTTTGTAAAGGATATGATATGTTGTGATTCTAGAATAGTAGTAGCTCTATCTAGTCTAGTATCTAAACATTGGGAATTGACAAATAAAAAGTATAGGTGTATGGCATTAGCCGAACATATAGCTGATAGTATTCCAATATCTGAGCTATCTAGACTACGATACAATCTATGTAAGTATCTACGCGGACACACTGATAGCATAGAGGATGAATTTGATCATTTTGAAGACGATGATTTGTCTACATGTTCTGCCGTAACCGATAGGGAAACGGATGTATAATTTTTTTTATAGTATGAAGGATATGATGGATATGATGATATGATGGATATGATGATATGATGGATATGATGGATATGATGGATATGATGGATATGATAAAAAAATATAATTGTTGTATCCATTCCCATTCAAATCACCTTATATGATTCTGTAACACAATGAAGGAGTCTCATAGATATATAGAGGTCAGATACTGGTTTGATAAACTTTTTATTCCACATGAGCATGTTTGACTTATGGTTAGACACACATACTTTAACAAATCACTGAAAATTGGAGTTAGGTATTCCTCTCAGAATCAGTTGCCGTTCTGGAACATTAAATGTATTTTTTATGATATACTCCAACGCATTTATGTGGGTATACAACAAGTCATTAATAATGAGTATTTCCAAGAGTTTTAGTTGTCTAGTATTTAACAAGAGAAGAGATTTCAACAGACTGTTTATGAACTCGAATACCGCCTCATTGTCGCTTATATTGATGACATGACATCGAATTCCCAATATCAATCTCATCAGTGATGAGTAGCTCAATCTTGTTATCGGGATCCAATTTCTAAAGATGTCATTAAACCCTCGATCGTGAATGGATTTATCATCATCGTTTTTATGTTGGACATGAGCTTAGTCCGTTTGTCCACATCTATATACGATGATTTCTGAATTATTTCATATATCTCTCGTTAACTCCAGGAACTTGTCAGGGATCTAACTTTAATATGTTCTCGTCTAAGAGATGAAAATCTTTGGATGGTTGCATGTGACTTTTCTCTAAAGGATGATGTTACCCGATCCTCTCTTAAATGACTCCATCTTATCCTTGGACAAGATGGACAGTCTATTTTCCTTAGATGGTTTAATATTTTTTACCCATGATCTATAAAGGTAGACAGACCTAATCGTCTCGGATGACCATATATTATTTTCAGTTTTATTATACGCATAAATTGTAAAAAATATGTTAGGTTTACGAAAATGTCTCGTGGGGCATTAATCGTTTTTGAAGGATTGGACAAATCTGGAAAAACAACACAATGTATGAACATCATGGAATCTATACCGGCAAACACGATAAAATATCTTAACTTTCCTCAGCGATCCACAGTCACTGGAAAGATGATAGATGACTATCTAACTCGTAAAAAAACCTATAATGATCATATAGTTAATCTATTATTTTGTGCAAATAGATGGGAGTTTGCATCTTTTATACAAGAACAACTAGAACAGGGAATTACTTTAATAGTTGACAGATACGCGTTCTCTGGAGTAGCGTATGCCACCGCTAAAGGCGCGTCAATGACTCTCAGTAAGAGTTATGAATCTGGATTGCCTAAACCCGACTTAGTTATATTCTTGGAATCTGGTAGCAAAGAAATTAATAGAAACATCGGCGAGGAAATTTATGAAGATGTTGAATTCCAACAAAAGGTATTACAAGAATATAAAAAAATGATTGAAGAAGGAGATATTCATTGGCAAATTATTTCTTCTGAATTCGAGGAAGATGTAAAGAAGGAGTTGATTAAGAATATAGTTATAGAGGCTATACACACGGTTACTGGACCAGTGGGGCAACTGTGGATGTAATAAAATGAAATTACATTTTTATAAATAGATGTTAGTACAGTGTTATAAATGGATGAAGCATATTACTCTGGCAACTTGGAATCAGTACTCGGGGATACGTGTCCGATATGCATACCGAACTCGCATCAATATCTCAATTAGTTATTGCCAAGATAGAAACTATAGATAATGATTATTAAACAAGGACATTGTAAATTTTATCATGTGTAGATCAAACTTGGATAATCCATTTATCTCTTTCCTAGATACTGCATATACTATCATAGATCAAGAGATCTATCAGAACGAGTTGATTAATTCATTAGACGATAATGAAATTATCGATTGTATAGTTAACAAGTTTATGAGCTTTTATAAGGATAACCTAGAAAATATGGTAGATGCTATCATTACTCTAAAATATTATAATTAATAATCCAGATTTTAAAACTACGTATGTGGAAGTACTCGGTTCCAGAATAGCTGATATAGATATTAAACAAGTGATACGTAAGAATATAATACAATTGTCTAATGATCCGCGAACGATATTTGTGAAAATATTAAAAAAAAATACTTTTTTTATTAAATGACGTCTCTTCGCGAATTTAGAAAATTATGCTGTGATATATATCACGCATCAGGATATAAAGAAAAATCTAAATTAATTAGAGACTTTATAACAGATAGAGATGATACCGATACATATTTGATCATTAAGCTATTGCTTCCCGGATTAGACGATAGAATGTATAACATGAACGATAAACAAATTATAAAATTATATAGTATAATATTTAAACAATCTCAGGAAGATATGCTACAAGATTTAGGATACGGATATATAGGAGACACTATTAGGACTTTCTTCAAAGAGAACACGGAAATCCGTCCACGAGATAAAAGCATTTTAACTTTAGAAGAAGTGGATAGTTTTTTAACTACGTTATCATCAGTAACTAAAGAATCACATCAAATAAAATTATTGACTGATATAGCATCTGTTTGTACATGTAATGATTTAAAATGTGTAGTCATGCTTATTGATAAAGATCTAAAAATTAAAGCGGGTCCTCGGTACGTGCTTAACGCTATTAGTCCTCATGCCTATGATGTTTTTAGAAAATCTAATAACTTGAAAGAGATAATAGAAAATGCAGCTAAACAAAATCTAGACTCTATATCTATTTCTGTTATGACTCCAATTAATCCCATGTTAGCGGAATCATGTGATTCTGTCAATAAGGCGTTTAAAAAATTTCCATCAGGAATGTTTGCGGAAGTCAAATACGATGGTGAAAGAGTACAAGTTCATAAAAAAAATAACGAGTTTGCATTCTTTAGTAGAAACATGAAACCAGTACTCTCTCATAAAGTGGATTATCTCAAAGAATACATACCGAAAGCATTTAAAAAAGCTACGTCTATCGTATTGGATTCTGAAATTGTTCTTGTAGACGAACATAATGTACCGCTACCGTTTGGAAGTTTAGGTATACACAAAAAGAAAGAATATAAAAACTCTAACATGTGTTTGTTCGTATTTGACTGTTTATACTTTGATGGATTCGATATGACAGACATTCCATTGTATGAACGAAGATCTTTTCTCAAAGATGTTATGGTCGAAATACCCAATAGAATAGTATTCTCAGAGTTGACGAATATTAGTAACGAGTCTCAGTTAACTGATGTATTAGATGATGCACTAACGAGAAAATTAGAAGGATTGGTCTTAAAAGATATTAATGGCGTATACGAACCGGGAAAGAGAAGATGGTTAAAAATAAAGCGAGACTATTTGAACGAGGGTTCCATGGCAGATTCTGCCGATTTAGTAGTACTAGGTGCCTACTATGGTAAAGGAGGAAAGGGTGGTATCATGGCAGTCTTTCTAATGGGTTGTTACGACGATGAATCCGGTAAATGGAAGACGGTAACTAAATGTTCCGGTCACGATGATAATACGTTAAGGGTTTTGCAAGACCAATTAACGATGGTTAAAATTAACAAGGATCCCAAAAAAATTCCAGAGTGGTTGGTAGTTAATAAAATCTATATTCCCGATTTTGTAGTAGATGATCCGAAACAATCTCAGATATGGGAAATTTCAGGAGCAGAGTTTACATCTTCCAAGTCACATACAGCGAATGGAATATCGATTAGATTTCCTAGATTTACTAGGATTAGAGAAGATAAAACGTGGAAAGAATCTACTCATCTAAACGATTTAGTAAACTTGACTAAATCTCTTAATAGTTACATATAAACTGAAAAATAAAATAACACTATTTTAGTTGGTAGTCGCCATGGATGGTGTTATCGTATACTGTCTAAATGCGTTAGTAAAACATGGCGAGGAAATAAATCATATAAAAAATGATTTCATGATTAAACCATGTTGTGAAAGAGTTTGTGAAAAAGTCAAGAACGTTCACATCGGCGGACAATCTAAAAACAATACAGTGATTGCAGATTTGCCATATCTGGATAATGCTGTATCAGATGTATGCAAATCAATATATAAAAAGAATGTATCAAGAATATCCAGATTTGCTAATTTGATAAAAATAGATGACGATGACAAGACTCCTACCGGCGTATATAATTATTTTAAACCTAAAGATGCTATTCCTGTTATTATATCCATAGGAAAGGATAAAGATGTCTGTGAACTATTAATCTCATCTGATAAAGCGTGTGCGTGTATAAAGTTAAATTTATATAAAGTAGCCATTCTTCCCATGGATGTTTCCTTTTTTACCAAAGGAAATGCATCATTGATTATTCTCCTGTTTGATTTCTCTATCGATGCGGCACCTCTCTTAAGAAGTGTAACCGATAATAATGTTATTATATCTAGACACCAGCGCCTACATGACGAGCTTCCGAGTTCCAATTGGTTCAAGTTTTACATAAGTATAAAGTCCGACTATTGTTCTATATTATATATGGTTGTTGATGGATCTATGATGTATGCGATAGCTGATAATAGAACTCACGCAATTATTAGCAAAAATATATTAGACAATACTACGATTAACGATGAGTGTAGATGCTGTTATTCTGAACCACAGATTAGGATTCTTGATAGAGATGAGATGCTCAATGGATCATCGTGTTATATGAACAGACATTGTATTATGATGAATTTACCTGATGTAGGCGAATTTGGATCTAGTATGTTGGGGAAATATGAACCTGACATGATTAAGATTGCTCTTTCGGTGGCTGGTAATTTAATAAGAAATCGAGACTACATTCCCGGGAGACGAGGCTATAGCTACTACGTTTACGGTATAGCCTCTAGATAATTTTTTTTAAGCACGAAATAAAAACATAATTTTAAACAATCTATTTCATACTATTTTGTGTGCTCACCATGAACATAAAGATAGATATATTAGTATTTCTGGTGATAAATTTACGGCGACTGCTAGGAGGGAAAATGAAGAAAGAAAAAATATCTACCTCTCCAAAAAGAAAAACTACTGATGTTATCAAACCTGATTATCTTGAGTACAATGACTTGTTAGATAGAGATGAGATGTCTACTATTCTAGAGGAATATTAGGCCTTAGAATAAAATATGGACGACTCTTAACGAAATTAGAAAATTCGATAATGATGTTGAAGAACAATTCGGTACTATAGAAGAACTCAAGCAGAAGCTTAGATTAAATTCTGAAGAGGGAGCAGATAATTTTATAGATTATATAAAGGTACAAAAACAGGATATCATCAAACTTACTGTATACGATTGCATATATCTATGATAGGATTGTGTGCGTGCGTGGTAGATGTTTGGAGAAATGAGAAACTGTTTTCTAGATGGAAATATTGGTTACAAGCGATTAAACTGTTTATTGATGATCACATGCTTGATAAGATAAAATCTATTGTAGAATAGACTAGTGTATGTGGAAATGTCATAGAAAGTTAAAAGTTAATGAGAGCAAAAATATATAAGGTTGTATTCCATATTTGTTATTTTTTTCTGTAATAGTTAGAAAATACATTCGATGGTCTATCTACCAGATTATTATGTGTTATAAGGTACTTTTCTCATAATAAACTAGAGTATGAGTAAGATAGTGTTTTTCAAAAACATATAAATCTAAAATTGATGGATGAGATATACAGCTATTAATTTCGAAAATATATTTTAATCTGATAACTTTAAACATGGATTTTTGATGGTGGTTTAAGTTTAAAAAAGATTTTGTTATTGTAGTATGATAATATCAAAAAGATGGATATAAAGAATTTACTGACTACATGTACTATTTTACATTACTACATTGGCTACGGCATATATACCTATTTCGTCACTTCCACACGCTCCGGTAAACGGGTGTCATGTGACGAGGGAGAATCTTGATAAGAGGCATAATCAATGTTGTAATCCGATGTCCACCTGGAGAATTTGCCAAGGTCAGATGTAGAGTTGGTAGTGATAACACAAAATGTGAACACTGCCCACCTCATACATATACCGCAATCCCCAATTATTCTAATAGATGTCATCAATGTAGAAAATGCCCAACAGGATCATTTGATAAGGTAAAGTGTACCGGAACACAGAACAAATGTTCGTGTCATCCTGGTTGGTATACGCTACTGATTCTTCACAGACTGAAGATTGTCGAGATTTGTGTACCAAAAAAGGAGATGTCCATGCGGATACTTTGGTGGAATAGATGAAGGAAATCCTATTTGTAAATCGTGTTGTGTTGGTGAATATTGCGACTACCTACGTAATTATAGACTTGATCCATTTCCTCCATGCAAACTATCTATCTAAATGTAATTAATTATGATTTTGATGATAATGTTACCATACATTATATCACTACTTGGTTAGTGTGTATTATTTAGTATGGAAGACCTATTAATAATTACTTATCTTTTGACGATCTTGTTATAATTATAATATAAAAATACTTATGACATAGTAACTCATAATTGCTGACGCGATAAATTCGTAATAATCTGTTTTGTTCAAATTTTTATAAGGAATCTACAGGCATAAAAATAAAAATATAATCTATAATATACTCTTACAACGCCATCATGAATAGCAGTGAATTAATTGCTGTTATTATGGATTTAGAAATAGTGGACGATTTTGTGATATTAATATAGTTATTAATGATGAAAGGATAAACGCGCATAGACTCATCCTATCTGGAGCCTCCGAATATTTTTTCCATTCTGTTTTCCAATAATTTTATCGATTCTAATGAATACGAAGTTAATCTAAGTCATTTAGATTATCAAAGTGTTAACTATTTGATCGATTACATTTATGGGATACCTTTGAGCCTAACTAACAATAACGTGAAATATATTCTTTCAACCGCTGATTTTTTTACAAATTGGATCTGTCATTACTGAGTGCGAAAAATACATACTTAAAAATCTTTGTTTTAGAAACTGTATCGATTTCTACATATACGCTGATAAATATAATAACAAGAAAATAGAATTAGCATCGTTTAACACAATATTACGAAATATTTTGAGACTCATCAACAATGAAAACTTTAAATACTTAACAGAGGAATCAATGATAAAAATTTTAAGCGATGATATGTTATATATAAAAAATGAGGATTTCACCCCACTGATTCTCATTAAATGGTTAGAGAGTACACCAACCATGTACCGTCGAGTTACTTAGATGCCTCAGAATATCATTTCTTTCCCCACAAGTTATAAAATCACTTTATAGTCATCGACTGGTTAGTTCAATCTACGAATGTATAACATTCTTAAACAATATAGCATTCTTGGATAAATCATTTCCTAGATACCATATCATCGAGTTGATATCTATCGGTATAAGTAATTCACATGATAAGATTTCCATAAACTGCTACAATCATAAAAAAATTCATGGGAAATGATATCTTCACGTAGATATAGGTGTAGTTTCGCAGTGACCGTCCTGGATAATATTATCTATATGATGGGTGGATATGATCAGTCCCTGTATAGAAGTTCAAAGGTTATAGCGTACAATACATGTACTAATTCTTGGATATATGATATACCAGAGCTAAAAATATCATCGTTCTAATTGCGGAGGAGTTGCCAATGACGAATACATTTATTGTATAGGCGGTATACGCGATCAGGAGTCATCGTTGATATCTAGTATCGATAGATGGAAGCCATCAAAACCATATTGGCAGAAGTATGCTAAAATGTGCGAACCAAAATGTGATATGGGGTTGCGATTTTAAACGGATTAATATATGTCATAGGTGGAGTCGTTAAAGGTGACACACATATACCAACGCACTAGAGAGTTTATCAGAAGATGGATGGATGAATCATCAACGTCTTCCAATAAAAATGTCCAATATGTCGACGATTGTTCATGCTGGAAAGATTTATATATCTAGAGGTTACAACAATAGTAGTGTAGTTAATGTAATATCGAATCTAGTCCTTAGCTATAATCCGATATATGATGAATGGACCAAATTATCATCATTAAATATTCCTAGAATTAATCCTGCTCTATGGTCAGTGTATAATAAATTATATGTAGGAGGAGTAATATCTGATGATGTTCAAACTAATACATCTGAAACATACGATAAAGAAAAAGATTGTTGGACATTGGATAATGGTCACTTGTTACCACATAATTATATAATGTATAAATGCGAACCGTTTAAACATAGATATCCATTGGAAAAAACACAGTACACGAATGATTTTCTAAAGTATTTGGAAAGTTTTATAGGTAGTTGATAGAACAAAATACATAATTTTGTAAAAATAAATCACTTTTTATACTAATATGACACAATTACCAATACTTTTGTTACTAATATCATTAGTATACGCTACACCTTCTCCTCAGACATCTAAAAAAATAGGTGATGATGCAACTATATCATGTAGTCGAAATAATACAAATTACTACGTTGTTATGAGTGCTTGGTATAAGGAGCCCAATTCCATTATTCTCTTAGCTGCCAAAAGCGACGTCTTGTATTTTGATAATTATACCAAGGATAAAATATCTTACGACTCTCCATACGATGATCTAGTTACAACTATCACAATTAAATCATTGACTGCTGGAGATGCCGGTACTTATATATGTGCATTCTTTATGACATCGACTACAAATGATACTGATAAAGTAGATTATGAAGAATACTCCATAGAGTTGATTGTAAATACAGATAGTGAATCGACTATAGACATAATACTATCTGGATCTACACCAGAAACTATTTCTGAGAAACCAGAGGATATAGATAATTCTAATTGCTCGTCTGTATTCGAAATCACGACTCCGGAACCAATTACTGATAATGTAGACGACCATACAGACACCGTCACATACACTAGTGATAGCATTAATACAGTAAATGCATCATCTGGAGAATCCACAACAGACGAGATTCCGGAACCAATTACTGATAAAGAAGAAGATCATACAGTAACAGACACTGTCTCATACACTACAGTAAGTACATCATCTGGAATTGTCACTACTAAATCAACCACCGATGATGCGGATCTTTATGATACATACAATGATAATGATACAGTACCGCCAACTACTGTAGGTGGTAGTACAACCTCTATTAGCAATTATAAAACCAAGGACTTCGTAGAAATATTTGGTATTACCACATTAATTATATTGTCAGCAGTGGCGATTTTCTGTATTACGTATTATATATGTAATAAACACCCACGTAAATACAAAACAGAGAACAAAGTCTAGATTTTTGACTTACATAAATATCTGGGATAATAAAATCTATCATATTGAGAGGACCATCTGGTTCAGGAAAGACAGCCATAACCAAAAGACTGTTAAAAGACTATGGGAATATATTTGGATTTGTGGTGTCCCATACCACTAGATTTCCTCGTCCTATGGAACGAGAAGGTGTTGATTACCTTACGTTAACAGAGAGGCCATCTGGAAGGGAATAGCCGCCGGAAACTTTCTAGAACATACTGAGTTTTTAGGAAATATTTACGGAACTTCTAAAACAGCTGTAAATACAGCGGTTATTAATAATCGTATTTGCGCGATGGATTTAAACATCAACGGTGTTAGAAGTCTTAAAAATACTTACCTAATGCATTACTTGGGTATATAAGACCTACCTCTCTTAAAATGGTTGAGACCAATCTTCGTCGTAGAAACACTGAAGCGGACGACGAATCTCATCGTCGCGTGATGTTGGCAAAAAACGGATATGGATGAGGTCAACGAAGCAGGTCTATTCGACACTATTATTATTGAAGATGATGTGAATTTAGCATATAGTAAGTGTTAATTCAGATACTACAGGACCGTATTAGAATGTATTTTAACACTAATTAGAGACTTAAGATTTGACTTAAAACTTGATAATTAATAATATAACTCGTTTTTATATGTGGCTATTTCAACGTCTAATGTATTAGTTAAATATTAAAACTTACCACGTAAAACTTAAAATTTAAAATGGTATTTCATTGACAGATCATACATTATGAAGTTTCAAGGACTTGTGTTAATTGACAATTGCAAAAATCAATGGGTCGTTGGACCATTAATAGGAAAAGGTGGATTCGGTAGTATTTATACTACTAATGACAATAATTATGTAGTAAAAATAGAGCCCAAAGCTAACGGATCATTATTTACCGAACAGGCATTTTATACTAGAGTACTTAAACCATCCGTTATCGAAGAATGGAAAAAATCTCACAATATAAAGCACGTAGGTCTTATCACATGCAAGGCATTTGGTTTATACAAATCCATTAATGTGGAATATCGATTCTTGGTAATAAATAGATTAGGTGCAGATCTAGATGCGGTGATCAGAGCCAATAATAATAGACTACCAGAAAGGTCGGTGATGTTGATCGGAATCGAAATCTTAAATACCATACAATTTATGCACGAGCAAGGATATTCTCACGGAGATATTAAAGCGAGTAATATAGTCTTGGATCAAATAGATAAGAATAAATTATATCTAGTGGATTACGGATTGGTTTCTAAATTCATGTCTAACGGCGAACATGTTCCATTTATAAGAAATCCAAATAAAATGGATAACGGTACTCTAGAATTTACACCTATAGATTCGCATAAAGGATACGTTGTATCTAGACGTGGTGATCTAGAAACACTTGGATATTGTATGATTAGATGGTTGGGAGGTATCTTGCCATGGACTAAGATATCTGAAACAAAGAATTCTGCATTAGTAAGTGCCGCAAAACAGAAATATGTTAACAATACTGCGACTTTGTTAATGACCAGTTTGCAATATGCACCTAGAGAATTGCTGCAATATATTACCATGGTAAACTCTTTGACATATTTTGAGGAACCCAATTACGACGAGTTTCGTCGAGTATTAATGAATGGAGTTATGAAAAATTTTTGTTGATAAAAAAATTAAAAAAATAACTTAGTTATTATCACTCTCGTGAGTACAATAGAAACATGGCGATGTTTTACGCACACGCTTTCGGTGGGTACGACGAGAACCTTCATGCATTTCCTGGAATATCATCGACGGTTGCCAATGATGTCAGGAAATATTCTGTTGTGTCAGTTTATAATAAAAAGTATAACATTGTAAAAAACAAATATATGTGGTGTAACAGTCAAGTGAACAAGAGATATATTGGAGCACTACTGCCTATGTTTGAATGCAATGAATATCTACAAATTGGAGATCCAATCCATGATCTAGAAGGAAATCAAATCTCTATTGTCACATATCGCCACAAAAACTACTATGCTCTAAGTGGAATTGGGTACGAGAGTCTAGACTTGTGTTTGGAAGGAGTAGGGATTCATCATCACGTACTTGAAACAGGAAACGCGGTATATGGAAAAGTTCAACATGAGTATTCTACTATCAAAGAGAAGGCCAAAGAAATGAATGCACTCAAACCAGGACCTATCATCGATTACCACGTCTGGATAGGAGATTGTGTCTGCCAAGTTACTACTGTAGACGTGCATGGAAAGGAAATTATGAGAATGAGATTCAAAAGGGGTGCGGTGCTTCCGATTCCAAATCTGGTAAAAGTTAAAGTTGGGGAGGAAAATGATACAATAAATCTTTCCACTTCCATATCAGCTCTCCTGAATTCCGGTGGCGGCACCATCGAGGTAACATCTAAGGAAGAACGTGTAGATTATGTACTCATGAAACGTTTGGAATCTATACATCATCTGTGGTCTGTAGTGTATGATCATCTTAATGTTGTGAATGGCGAAGAACGATGTTATGTACATATGCATTCATCTCATCAAAGTCCTATGCTGAGTACTGTAAAAACAAATTTGTACATGAAGACTATGGGAGCATGTCTTCAAATGGACTCCATGGAAGCTCTAGAGTATCTTAGTGAACTGAAGGAATCAGGTGGGCGGAGTCCCAGACCAGAATTGCAGAAATTTGAATATCCAGATGGAGTGAAAGACACTGAATCAATTGAGAGATTGGCAGAGGAGTTCTTCAATAGATCAGAACTTCAGGCCGGTGAATCAGTCAAATTTGGTAATTCTATTAATGTTAAACATACATCTGTTTCAGCTAAGCAACTAAGAACACGTATACGACAGCAGCTTCCTTCTATACTCTCATCTTTTGCCAACACAAAGGGTGGATATTTGTTCATTGGAGTTGATAATAATACACACAAAGTAATTGGATTCACGGTGGGTCATGACTACCTCAAACTGGTAGAGAGTGATATAGAAAAGTATATCCAAAAACTTCCTGTTGTGCATTTCTGCAAGAAAAAAGAGGACATCAAGTACGCATGTAGATTCATCAAGGTGTATAAACCTGGTGATGAGACTACCTCGACATATGTGTGCGCAATCAAAGTGGAAAGATGCTGCTGTGCTGTGTTTGCGGATTGGCCAGAATCATGGTACATGGATACTAGTGGTAGTATGAAGAAGTATTCTCCAGATGAATGGGTGTCACATATAAAATTTTAATTAGGGTAAGGTAAAACTATATATAATAACTAACAATTTGTGTATCATATAGACAATTAATTAGGTAACTGTTATCTCTTTTTAACTAACTAACTAACTAACTAACTCTTATATACTATTAATAATACATCTATTAATCATTGATTAGCTTATTGCTTTAATTGTTTTTGTAAACTAACACTGTTCATTGAAAAGGGATAACATGTTACAGAATATAAATTATATATGGATTTTTTTAAAAAGGAAATACTTGACTGGAGTATATATTTATTTCTTCATTACATAACACGTCTGTGTTCTAATTCTTCCAATTCTTCCACATCTCATATAATACAGGAATATAATCTTGTTCGAAAATATGAGAAAGTGGATAAAACAATAGTTGATTTTTTATCTAGGTGGCCAAATTTATTCCATATTTTAGAATATGGGGAAAATATTCTACATATTTATTTTATAGATGCTGCTAATACGAATATTATGATTTTTTTTCTAGATAGAGTATTAAATATTAATAAGAACCGTGGGTCATTTATACATAATCTCGGGTTATCATCCATTAATATAAAAGAATATGTATATCAATTAGTTAATAATGATCATCTAGATAATAGTATAAGACTAATGCTTGAAAATGGACGTAGAACAAGACATTTTTTGTCTTATATATTGGATACAGTTAATATCTATATAAGTATTTTAATAAATCATAGATTTTATATAGATGCCGAAGACAGTTACGGTTGTACATTATTACATAGATGTATATATAACTATAAGAAATCAGAATCAGAATCATATAATGAATTAATTAAGATATTGTTAAATAATGGATCAGATGTAGATAAAAAAGATACGTACGGAAACACACCGTTTATCCTATTATGTAAACACGATATCGACAACGCGGAATTGTTTGAGATATGTTTAGAGAATGCTAATATAGACTCTGTAGACTTTAATGGATATACACCTCTTCATTATGTCTCATGTCGTAATAAATATGATTTTGTAAAGTTATTAATTTCTAAAGGAGCAAATGTTAATGCACGTAATAGATTCGGAACTACTCCATTTTATTGTGGAATTATACACGGTATCTCGCTTATAAAACTATATTTGGAATCAGACACAGAGTTAGAAATAGATAATGAACATATAGTTCGTCATTTAATAATTTTTGATGCTGTTGAATCTTTAGATTATCTATTGTCCAGAGGAGTTATTGATATTAACTATCGTACTATATACAACGAAACATCTATTTACGACGCTGTCAGTTATAATGCGTATAATACGTTAGTCTATCTATTAAACAGAAATGGTGATTTTGAGACGATTACTACTAGTGGATGTACATGTATTTCGGAAGCAGTCGCGAACAACAACAAAATAATAATGGATATACTATTGTCTAAACGACCATCTTTGAAAATTATGATACCATCTATGATAGCAATTACTAAACATAAACAACATAATGCAGATTTATTGAAAATGTGTATAAAATATACTGCGTGTATGACCGATTATGATACTCTTATAGATGTACAATCGCTACATCAATATAAATGGTATATTTTAAAATGTTTTGATGAAATAGATATCATGAAGAGATGTTATATAAAAAATAAAACTGTATTCCAATTAGTTTTTTGTATCAAAGACATTAATACTTTAATGAGATACGGTAGACATCCTTCTTTCGTGAAATGTAATATTCTTGACGTATACGGAAGTTGTGTACGTAATATCATAGCATCTATTAGATATCGTCAGAGATTAATTAGTCTATTATCCAAGAAGCTGGATGCTGGAGATAAATGGTCGTGTTTTCCTAACGAAATAAAATATAAAATATTGGAAAACTTTAACGATAACGAACTGACCACATATCTGAAAATCTTATAAACACTATTAAAATATAAAATCTAAGTAGGATAAAATCACACTACATCATTGTTTCCTTTTAGTGCTCGACAGTGTATACTATTTTTAACACTCATAAATAAAAATGAAAACGATTTCCGTTGTTACGTTGTTATGCGTACTACCTGCTGTTGTTTATTCAACATGTACTGTACCCACTATGAATAACGCTAAATTAACGTCTACCGAAACATCGTTTAATGATAAACAGAAAGTTACGTTTACATGTGATTCAGGATATCATTCTTTGGATCCAAATGCTGTCTGTGAAACAGATAAATGGAAATACGAAAATCCATGCAAGAAAATGTGCACAGTTTCTGATTATGTCTCTGAACTATATGATAAGCCATTATACGAAGTGAATTCCACCATGACACTAAGTTGCAACGGTGAAACAAAATATTTTCGTTGTGAAGAAAAAAATGGAAATACTTCTTGGAATGATACTGTCACGTGTCCTAATGCGGAATGTCAACCTCTTCAATTAGAACACGGATCGTGTCAACCAGTTAAAGAAAAATACTCATTTGGGGAATATATGACTATCAACTGTGATGTTGGATATGAGGTTATTGGTGTTTCGTATATAAGTTGTACGGCTAATTCTTGGAATGTTATTCCATCATGTCAACAAAAATGTGATATACCGTCCCTATCTAATGGATTAATTTCCGGATCTACATTTTCTATCGGTGGCGTTATACATCTTAGTTGTAAAAGTGGTTTTACACTAACGGGGTCTCCATCATCCACATGTATCGACGGTAAATGGAATCCCATACTCCCAACATGTGTACGATCTAACGAAGAATTTGATCCAGTGGATGATGGTCCCGACGATGAGACAGATCTGAGCAAACTCTCGAAAGACGTTGTACAATATGAACAAGAAATAGAATCGTTAGAAGCAACTTATCATATAATCATAATGGCGTTGACAATTATGGGTGTCATATTTCTAATCTCCATTATAGTATTAGTTTGTTCCTGTGACAAAAATAATGACCAATATAAGTTCCATAAATTGCTACCGTGAATATAAATCCGTTAAAATAATTAATAATTAATAATTAATAACGAACAAGTATCAAAAGATTAAAGAATTAGCTAGAATCAATTAGATGTCTTCTTCAGTGGATGTTGATATCTACGATGCCGTTAGAGCATTTTTACTCAGGCACTATTATGACAAGAGATTTATTGTGTATGGAAGAAGTAACACCATATTACATAATATATACAGGCTATTTACAAGATGCACCGTTATACCGTTCGATGATATAGTACGTACTATGCCAAATGAATCACGTGTTAAACAATGGGTGATGGATACACTTAATGGTATAATGATGAATGAATTCGATACTGTATGTGTGGGTACCGGACTACGATTCATGGAAATGTTTTTCGATTACAATAAAAATAATCCCAAAAATAGCATCAACAATCAAATAATGTATGATATAATTAATAGCGTAGCCATAATTCTAGCTAATGAGAGATATAGAAGCGCGTTTAACGACGATAGAATATACATCCGTAGAACTATGATGGACAAATTGTACGAATACGCATCTCTAACTACTATTGGTACGATCACTGGAGGTGTTTGTTATTTTATCTGTTGATGCATCTAGTTAGTTTGTATAAATAATTATTTCGATATACTAGTTAAAATTTTAAGATTTTAAATGTATAAAAAACTAATAACGTTTTTATTTGTAATAGGTGCAGTTGCATCCTATTCGAATAATGAGTACACTCCGTTTAATAAACTGAGTGTAAAACTCTATATAGATGGAGTAGATAATATAGAAAATTCATATACTGATGATAATAATGAATTGGTGTTAAATTTTAAAGAGTACACAATTTCTATTATTACAGAGTCATGTGACGTCGGATTTGATTCCATAGATATAGATGTTATAAACGACTATAAAATTATTGATATGTATACCATTGACTCGTCTACTATTCAACGCAGAGGACATACGTGTAGAATATCTACCAAATTATCATGCCATTATGATAAGTACCCTTATATCCACAAATATGAGGGTGATGAACGACAATATTCTATTACCGCAGAGGGAAAATGCTATAAAGGAATAAAATATGAAATAAGTATGATGAACGATGATACTCTATTGAGAAAACATACTCTTAAAATTGGATTTACTTATATATTCGATCGTCATGGGCATAGTAATACATATTATTCAAAATATGATTTTTAAAAATTTAAAATATATTATCACTTCAGTGACAGTAGTCAAATAACAAACAACACCATGAGATATATTATAATTCTCGCAGTTTTGTTCATTAATAGTATACATGCTAAAATAACTAGTTATAAGTTTGAATCCGTCAATTTTGATTCCAAAATTGAATGGACTGGGGATGGTCTATACAATATATCCCTTAAAAATTATGGCATCAAGACGTGGCAAACAATGTATACAAATGTACCAGAAGGAACATACGACATATCCGGATTTCCAAAGAATGATTTCGTATCTTTCTGGGTTAAATTTGAACAAGGCGACTATAAAGTGGAAGAGTATTGTACGGGACTATGTGTCGAAGTAAAAATTGGACCACCAACTGTAATATTGACTGAATATGACGATCATATCAATTTGTTCATCGAGCATCCGTATGCTACTAGAGGTAGCAAGAAGATTCCTATTTACAAACGCGGTGACATGTGTGATATCTACTTGTTGTATACGGCTAACTTCACATTCGGAGATTCTGAAGAACCAGTAACATATGATATCGATGACTACGATTGCACGTCTACAGGTTGCAGTATAGACTTTGCCACAACAGAAAAAGTGTGTGTGACAGCACAGGGAGCCACAGAAGGGTTTCTCGAAAAAATTACTCCATGGAGTTCGGAAGTATGTCTGACACCTAAAAAGAATGTATATACGTGCGCAATTAGATCTAAAGAAGATGTTCCCAATTTCAAGGACAAAATAGCCAGAGTTATCACGAGAAAATTTAATAAACAGTCTCAATCTTATTTGACTAAATTTCTCGGTAGCACATCGAATGATGTTACAACTTTTTTTAGCATTCTTGACTAAATATTCATAACTAATTTTTATTAATGATACAAAAATGAAATAAACTGTATATTATACACTGGTTAACGCCCTTGGCTCTAACCATTTTCAAGATGAGGTCCCTGATTATAGTCCTTCTGTTCCCCTCTATCATCTACTCAATGTCTATTAGACGATGCGAGAAGACTGAAGAGGAAACATGGGGATTAAAAATAGGGTTGTGTATAATTGCCAAAGATTTCTATCCCGAAAGAACTGATTGCAGTGTTCATCGCCCAACTGCAAGTGGAGGATTGATAACTGAAGGCAATGGATTCAGAGTAGTTATATATGATCAATGTACAGAACCCCATGACTTTATTATCACCGATACTCAACAAACACGTCTTGGATCATCTCATACATATATTAAATTCAGTAACATGAATACAGGTGTCCCATCTAGTATTCCAAAATGTTCCAGAACTCTCTGTATTTCTGTATATTGTGATCAAGAGGCGGGAGACATAAAATTTGAGGAGTATACTCAAGAATCAAGTGATATCAGTATTAGAGTTAAGTATGATTCATCATGTATTGATTATCTGGGTATTAATCAAAGTTTCATGAATGAATGTATTCGAAGAATTACAACATGGGATAGAGAATCATGCGTCAGAATTGATACACAGACTATAAATAAATATCTTAAGTCTTGCACCAACACAAAATTCGACCGTAATGTCTACAAAAGGTACATACTGAAGAGTAAAGCACTCCATGCTAAAACAGAGTTGTAATAGATATAAAATACTTTTTATAATAATTAGGCTAGAAAAATCTCACTCACATGTAATCTTAAAAAAATGATATGATAGTTCTTACAAGTAGCGATTGAGTTTTAAATGGATTCTATTAATTACCGGGGAACTTAACAATTCGTTCTGATCTACAGACATTGGTTAATAAATCATCTTATTTTGCCAATATATTAAAATGTGGAAACTCCACTAATAATATTACATTGTGCGACTTTCAAGATGATGTGATATATAGGGTTATACAGTTTTAACAATTATATAATAGAGATAGAAAGTACAAAAGATGTAGAATCAATGATATGGCACGCTAAACAGTTGGGTGTGGAATCATTGCTAAAAGAATGTCAAAATTATTTGCTTAGAATATTACGTATATAATTGTTTAGAAATTTATAGAATAACTAATATTAATACATTATCGTATATCTACAACGATATAAGAAACTTCATATTGGATAATATTACTATTAATATATAAGGATCCAGATTTTATATATTTGCCTAAATACATTATTATAGATTTACTAGGACAATCACCTAAATGTTTTTAACGAAGATAATGTGGTAAAGATTATATACACTTATATATCTTCCGATATCTACAAGGATATTCCATATCATCATTGTGTAAACTAAATAACGTTTTCTATGGCATTTAATAAGGACATTGGATATGTGGAAAAGTGATGTATGGAAGTTAGTACATTATCAACTTCTCCTTATTGATTGAAAATGAAAATATAAATAGTTTTTATGTATAGCGGTATCTACCCTATAGTTTTATTGCTTACTACTAACATGGATTCAGATACAGATACAGATACAGATACAGATACAGATACAGATACAGATACAGATGTAGAAGATATCATGAATGAAATAGATAGAGAGAAAGAAGAAATACTAAAAAATGTAGAAATTGAAAATAATAAAAACATTAACAAGAATCATCCCAGTGAATATATTAGAGAAGCACTTGTTATTAATACCAGTAGTAATAGTGATTCCATTGATAAAGAAGTTATAGAATATATCAGTCACGATGTAGGAATATAGATCATATCTACTAATTTTTATAATCGATACAAAACATAAAAACAACTCGTTATTACATAGCAGGTATGGAATCCTTCAAGTATTGTTTTGATAACGATGGTAAGAAATGGATTATCGGAAATACTTTATATTCTGGTAATTCAATACTCTATAAGGTCAGAAAAAATTTCACTAGTTCGTTCTACAATTACGTAATGAAGATAGATCATAAATCACACAAGCCATTGTTGTCCGAAATACGATTCTATATATCTGTATTGGATCCTTTGACTATCAACAACTGGACACGGGAACGTGGTATAAAGTATTTGGCTATTCCAGATCTGTATGGAATTGGAGAAACCGATGATTATATGTTCTTCATTATAAAGAATTTGGGAAGAGTATTCGCCCCAAAGGATAGTGAATCAGTTTTCGAAGCATGTGTCACTATGATAAACACGTTAGAGTTTATACACTCTCAAGGATTTACTCATGGAAAAATAGAACCGATGAATATACTGATTAGAAATAAACGTATTTCACTAATTGACTATTCTAGAACTAACAAACTATACAAAAGTGGAACACATATAGATTACAACGAGGACATGATAACTTCAGGAAATATCAATTATATGTGTGTAGACAATCATCTTGGAGCAACAGTTTCAAGACGAGGAGATTTAGAAATGTTGGGATATTGCATGATAGAATGGTTCGGTGGTAAACTTCCATGGAAAAACGAAAGTAGTATAAAAGTAATAAAACAAAAAAAAGAATATAAACAATTTATAGCTACTTTTTTTGAGGACTGTTTTCCTGAAGGAAATGAACCTCTGGAATTAGTTAGATATATAGAATTAGTATACATGTTAGATTATTCTCAAACTCCTAATTATGACAGACTACGTAGACTGTTTATACAAGATTGAAATTATATTCTTTTTTTTATAGAGTGTGGGGGTAGTGTTACGGATATCTAATATTAATATTAGACTATCTCTATCGCGCTACACGACCAATATCGATTACTATGGATATCTTCAGGGAAATCGCATCTTCTATGAAAGGAAAGAATGTATTCATTTCTCCAGCGTCAATCTCGTCAGTATTGACAATACTGTATTATGGAGCTAATGGATCCACTGCTGAACAGCTATCAAAATATGTAGAAAAGGAGGAGAACATGGATAAGGTTAGCGCTCAGAATATCTCATTCAAATCCATGAATAAAGTATATGGGCGATATTCTGCCGTGTTTAAAGATTCCTTTTTGGGAAAAATTGGCGATAAGTTTCAAACTGTTGACTTCACTGATTGTCGCACTATAGATGCAATCAATAAGTGTGTAGATATCTTTACTGAGGGAAAAATCAATCCACTATTGGATGAACCATTGTCTCCTGATACCTGTCTCCTAGCAATTAGTGCCGTATACTTTAAAGCAAAATGGTTGATGCCATTCGAAAAGGAATTTACCAGTGATTATCCCTTTTACGTATCTCCAACGGAAATGGTAGATGTAAGTATGATGTCTATTTACGGCGAGCCATTTAATCACGCATCTGTAAAAGAATCATTCGGTAACTTTTCAATCATAGAACTGCCATATGTTGGAGATACTAGTATGATGGTCATTCTTCCAAACAAGATTGATGGATTAGAATCCATAGAACAAAATCTAACAGATACAAATTTTAAGAAATGGTGTAACTCTCTGAAAGCTACGTTTATCGATGTGCACATTCCTAAGTTTAAGGTAATAGGTTCGTATAATCTTGTGGATACGCTAATAAAGTTGGGACTGACAGATGTGTTCTATTCAACTGGTGATTATATCAATATGTGTAATTCAGATGTGAGTGTTGACGCTATGATTCACAAAACGTATATAGATGTCAATGAAGAGTATACAGAAGCAGCTGCAGCAACTTCTGTACTAGTGGCAGACTGTGCATCAACAGTTACAAATGAGTTCTGTGCAGATCATCCGTTCATCTATGTGATTAGACATGTCGATGGTAAAATTCTTTTCGTTGGTAGATATTGCTCTCCAACAACTAATTAAGCACATTCTTAATATTAGAATATTATATAGTTAAGATTTTTACTAACAGGTTAACATTTTTTTTTAAAAATAGAAAAAACATGTGGTATTAGTGCAGGTCGTTATTCTTCCAATTGCAATTGGTAAGATGACGGCCAACTTTAGTACCCACGTCTTTTCACCACAACACTGTGGATGTGACAGACTGACCAGTATTGATGACGTCAGACAATGTTTGACTGAATATATTTATTGGTCGTCGTATGCATACCGCAACAGGCAATGCGCTGGACAACTGTATGACACACTCCTCTCTTTTAAAGATGATGCGGAATCAGTGTTCATCGACGTTCGTGAGCTGGTAAAAAATATGCCGTGGGATAATGTTAAGGATTGTACAGAGATCATCCGTTGTTATATACCGGATGAGCAAAAAACCATCAGAGAGATTTCGGCCATCATTGGACTTTGTGCATATGCTGCTACTTACTGGGGAGGTGAAGACCATCCCACTAGTAACAGTCTGAACGCATTGTTTGTGATGCTTGAGATGCTCAATTACATGGATTATACCATCATATTCTGGCGTATGAATTGATGAGTTACAGCTTGACATTTCTTCTTTCCTCCCTCTTCTTCTACCTTTCCCAGAAACAAACTTTTTTTACCCACTATAAAATAAAATGAGTATACTACCTGTTATATTTCTTCCTATATTTTTTTATTCTCCATTCGTTCAGACTTTTAACGTGCCTGAATGTATCGACAAAGGGCAATATTTTGCATCATTCATGGAGTTAGAAAACGAGCCAGTAATCTTACCATGTCCTCAAATAAATACGCTATCATCCGGATATAATATATTAGATATTTTATGGGAAAAACGAGGAGCGGATAATGATAGAATTATACAGATAGATAATGGTAGCAATATGCTAATTCTGAACCCGACACAATCAGACTCTGGTATTTATATATACATTACCACGAACGAAACCTACTGTGACATGATGTCGTTAAATTTGACAATCGTGTCTGTCTCAGAATCAAATATAGATCTTATCTCGTATCCACAAATAGTAAATGAGAGATCTACTGGTAAAATGGTATGTCCCAATATTAATGCATTTATTTCTAGTAACGTAAACACAGAATTATATGGAGCGGACATCGACGCCTTAGAAATAAGAGACTTAAACAACGGACACCTGGAATTATTACCATAGAAGATGTTAGAAAAAATGATGCTGGTTATTATACATGTGTTTTAGAATATATATATATATATATATATATATATATATATATATATATATATATATATATATATATATATATGGGCAAAACATATAACGTAACCAGAATTATAAAATTAGAGGTACGGGATAGAATAATACCTCCTACTATGAAATTACCAGAAGGAGTAGTAACTTCAATAGGTAGTAATTTGACTATTGCATGTAGAGTATCGTTGAGACTTCCCACAACGGACACCGACGTCTTTTGGATAAGTAATGGTATATGTATTACGAAGAAGAAGACGAGGACGGAGACGGTAGAATAAGTGTAGCAAATAAAATCTATATGACCGATAAGAGACGTGTTATTACATCCTGGTTAAACATTAATCCTGTCAAGGAAGAAGATGCTACAACGTTTACGTGTATGGCGTTTACTATTCCTAGCATCAGCAAAACAGTTACTGTTAGTAAACGTGAATGTATGTTGTTACATTTCCATATCAATTGAGTTTATAAGAATTTTTTATACATTATCTTCCAACAAACAATTGACGAACGTATTGCTATGATTAACTCCCACAATACTATATATATTATTAATCATTAACTTGCAGACTATACCTAGTAGTGCTATTTTGACATACTCATGTTCTTGTGTAATCGCAGTATCTATATTATTAAAGTACGTAAATCTAGCTATAGTTTTATTATTTAATTTTAGATAATATACTGTCTCCGTATTTTTAAAAAATTACCACATCCTTTATTAAATCATGAATGGGAATTTCTGTGTCATCGTTAGTATATTGTGAACAACAAGAGCAGATATCTATAGGAAAGGGTGGAATGCGATACATTGATCTATGTAGTTTTAAAACATACGCGAACTTTGAAGAATTTATATAAATCATCTCACGAGATATTGCTCTCTGTCATATTCATACACCTGTATAAACTTTCTAGACATCTTACAATGTGTTATTTTATGATCATATTTACATATTTACTGGTATATCAAAGATGTTAGATTAGTTAATGGGAATCGTCTATAATAATGAATATTAAACAATTATAGGAGGAGTTTATACCTACAAAAACATCATAAAAATGAGTCATCGTCCGATTTATGTTTTAAATATACTAACATTACTACCTTCAGAAATTATATACGAAATATTATACATGCTGACAATTAACGATCTTTATAATATATAGTATCCACCTACCAAAGTATAATTGTATTTTTCTCATGTGATGTGTGTAAAAAACTGATATTATATAATTATCTTAGTACCTATGATGAAGATGAAGATGAAGATGAAGATGATGGTCCGTATATATTTTGTATCATTATCGTTATTGCTATTCCATAGTTACGCCATAGACATCGAAAATGAAATCACCGAATTCTTCAATAAAATGAGAGATACTCTACCAGCTAAAGACTCTAAATGGTTGAATCCAGTATGTATGTTTGGAGGCACAATGAATGATATGGCCGCTCTAGGAGAGCCATTCAGTGCAAAGTGTCCTCCTATTGAAGACAGTCTTTTATCGCATAGATATAAAGACTATGTGGTTAAATGGGAAAGGCTAGAAAAGAATAGACGGCGACAGGTTTCTAATAAACGTGTTAAACATGGTGATTTATGGATAGCCAACTATACATCTAAATTCAGTAACCGTAGGTATTTATGTACCGTAACCACAAAGAATGGTGACTGTGTTCAGGGTGTAGTTAGATCTCATGTGTGGAAACCTTCTTCATGCATTCCAAAAACATATGAACTAGGTACTTATGATAAGTATGGCATAGACTTATACTGTGGAATTCTTTATGCGAACCATTATAATAATATAACTTGGTATAAAGATAATAAGGAAATTAATATCGACGATTTTAAGTATTCACAAGCGGGAAAGGAATTAATTATTCATAATCCAGAGTTAGAAGATAGTGGAAGATACGACTGTTACGTTCATTACGACGACGTTAGAATCAAGAATGATATCGTAGTATCAAGATGTAAAATACTTACGGTTATACCGTCACAAGACCACAGGTTTAAACTAATACTAGATCCGAAAATCAACGTAACGATAGGAGAACCTGCCAATATAACATGCAGTGCTGTGTCAACGTCATTATTTGTCGACGATGTACTGATTGAATGGGAAAATCCATCCGGATGGATTATAGGATTAGATTTTGGTGTATACTCTATTTTAACTAGTAGAGGCGGTATCACCGAGGCGACTTTGTATTTTGAAAATGTTACTGAAGAATATATAGGCAATACATATACATGTCGTGGACACAACTATTATTTTGATAAAACTCTTACAACTACAGTAGTATTGGAGTAAATACACAATGCATTTTTATATACATTACTGAATTATTATTATTAATTATATCGTATTTGTGCTATAGAATGGATGAAGATACGCGACTATCTAGGTATTTGTATCTCACCGATAGAGAACATATAAATGTAGACTCTATTAAACAGTTGTGTAAAATATCAGATCCTAATGCATGTTATAGATGTGGATGTACGGCTTTACATGAGTACTTTTATAATTATAGATCAGTCAACGGAAAATACAAGTATAGATACAACGGTTACTATCAATATTATTCATCTAGCGATTATGAAAATTATAATGAATATTATTATGATGATTATGATAGAACTGGTATGAACAGTGAGAGTGATAATATATCAATCAAAACAGAATACGAGAATGAATATGAATTCTATGATGAAACACAAGATCAAAGTACACAACTAGTAGATTACGACATTAAACTCAAAACCAATGAGGATGATTTTGTTGATGAATTCTATGGTTATGATAGATCAGTGGGTGTCCATGATTATATAGATGTATCAATTAATAAAGTAGTATATGGAAGAGAGTCTCACGTAAGATGGTGGGATATATGGCAAGAACATAATGATGGCGTATACAGTATAGGAAAGGAGTGCATAGATAATATATACGAAGACAGACATACCGTAGACGAATTCTACAAGATAGACAGCGTATCAGATGTAGATGACGCAGAACATATATCTCAGATAACTAATGATGTATCTACACAAACATGGGAAAAGAAATCAGAGTTAGATAGATACATGGAAATGTATCCTCGTCATAGATATGGTAAGCATTCTGTCTTTAAGGGATTTTCTGACAAAGTTAGAAAAAATGATTTAGACATGAACGTGGTAAAAGAATTACTTTCTAACGGTGCATCTCTAACAATCAAGGATAGCAGTAATAAGGATCCAATTGCTGTTTATTTTAGAAGAACAATAATGAATTTAGAAATGATTGATATCATTAACAAACATACAACTATCTATGAACGCAGGTATATAGTACACTCCTATCTAAAAAATTATAGAAATTTCGATTATCCATTTTTCAGAAAGTTAGTTTTGACTAATAAACATTGTCTCAACAATTATTGTAATATAAGCGACGGCAAATATGGAACACCACTACATATATTAGCATCTAATAAAAAAATAATAACTCCTAATTACATGAAGTTATTAGTGTATAACGGAAATGATATAAACGCACGAGGTGAAGATACACAAATGCGAACTCCATTACACAAATATTTGTGTAAATTTGTATATCATAATATTGAATATGGTATCCGATACTATAATGAAAAGATTATAGACGCATTTATAGAGTTAGGAGCCGATCTAACTATTCCAAATGACGATGGAATGATACCAGTAGTTTACTGTATACACTCAAATGCCGAATATGGTTATAACAATATTACTAACATAAAGATAATACGTAAACTACTTAATCTTAGTAGACATGCGTCACATAATCTATTTAGAGATCGAGTCATGCACGATTATATAAGTAATACATATATTGATCTTGAGTGTTTAGATATCATTAGATCACTTGATGGGTACGATATTAATTGTTACTTTGAAGGACGTACACCACTTCATTGCGCTATACAATATAACTTCACTCAGATTGCTGAGTACTTATTAGATCGAGGAGCTGATATATCATTAAAGACAGACGATGGTAAAACTGTATTTGATTTATCGTTATGTAGTTACATTCCTCTTAAATGGACTAGCTTTTTGATTAGTCGTCTACCGCCTAAAAGTGTCATATGCTCACTGACTAACCATATAATAGATTATGTTCTTACGAACAATAGACGTATTATTTGGCAGAGTCAAATGATTAATAAGTACGTACTGTTACTGGACCCATCCTTTTATTATAGATTCAGAAATGTTATCGAAAACAAATTAGACCAATACAATAATCGTTATAATATGTTCGAACACGATAGGGACGTTAATGAAAAGTATGGCAAAGTCTTACATGACCTCGATACATATATCAAGGATGTACAAGTATTAAAATCTACTTCCATCACTAATAATATAACACTATACGACACTATTATAAATAATAAGTCAGAGTTTCCTATACGTCGTGTAAACGACAAACAATTAATTAATCTCATAAAATCCAATACATATCATAATCTTATCGAAAAAGTTATTAAAAATACATTAGAGAAATATACTTTAACTAATATAGTCCTCGAGTATATGATCTCATCTCGATCTCAATCATCTTATTTGAGTCGTATTCCTAATGAGATATTACTCGAAATATTATATAAACTCGACATGTACGATTTACGTAATCTATATACAAGATATATGAGAGAGAATGATATCACAGAGTATCATATAGAGAATACGAGGTCTGTTTCTACACAGACATGAATAATGAATACACATACAACGTTTTTTTTAATCTTAGATATAACACTAATTACATCAAGATTATATATTGAAATCGTAATTTGAGTTGTCTGATCATCATGGATATCGAAAATGATATACGTAACATTAGCAATCTTTTAGATGATTGATATATTATTATGCGATGTAATCATAACTATCGGAGATGTAGAAATTAAAGCGCATAAAACTATTTTGGTTGCCGGATCTACGTATTTTAAAACAATGTTCACAACATCTATGATAGCGAGAGATCTAGCAACTAGAGTAAATATACAGATGTTCGATAAAGATGCCGTCAAAAATATTGTACAGTACTTATACAATAGGTATATAAGTTCTATGAATGTGATAGACATATTAAAATGCACCGACTAAGAACGTAAAACGAACTATAGAATGTTATACAATGGGTGATGATAAGTAGAAGATGTTACCCGATATACCCATAGCATTATCTAGTTATGGCATGTGTGTATTAGATCAATACATATACATTATAAGCGGTCGTACCCAACACTGATTATACATCGGTACATACAGTAAATAGCATAGATATGGAGGAGGATACAAATATTTCAAATAAAGTTATGAGATACGCGCTGTCAATAATATATGGAAGACATTACCTAACTTCTGAACTGGAACTATAAATCCAGGCTCTCGCATAAAGATGAATATATATGTTGTATGCGACATCAAAGATGAAAAAATGTTAAGACTTATATATTTAGATATAACACGAATATGTATGACGGATGGGAATTGGTAACGATGACAGAAAGCAGATTGTCAGCTCTGCATACTATTCTTCATGACAATACCATAATGATGTTACATTGTTATGAAGCGTATATGTTACAAGATACATTTAATGTGCTTACGGAACATATATTTAGAAACATCTACTAACGATTTTTTATGCTTGTATTATTAATGGTATGTAATATGATTTAATTGATTGTGTACACGATACCAATTTGTCGAGTATGAATACGGAGTACAAACATAAACTGAAGTTTAACATTATTTATTTATGATATACATTATATACATTATATACATTATATACATTATATACATTATATACATTATATACATTATATACATTATATACATTATATACATTATATACATTATATACATTATATACATTATATACATTATATACATTATATACATTATATATCGTTATTGTTTGGTCTATGCCATGGATATCTTTAAAGAACTAATCTTAAAACATACGGATGAAAATGTTTTGATTTCTCCAGTTTCCATTTTATCTACTTTATCTATTCTGAATCATGGAGCAGCTGGTTCTACAGCTGAACAACTATCAAAATATATAGAGAATATGAATGAGAATACACCCGATGATAAGAAGGATGACAATAATGACATGGACGTAGATATTCCGTATTGCGCGACACTAGCTACCGCAAATAAAATATACGGTAGTGATAGTATCGAGTTCCATGCCTCATTCCTACAAAAAATAAAAGACGATTTTCAAACTGTAAACTTTAATAATGCGAACCAAACAAAGGAACTAATCAACGAATGGGTTAAGACAATGACAAATGGTAAAATTAATTCCTTATTGACTAGTCCGCTATCCATTAATACTCGTATGATAGTTATTAGCGCCGTCCATTTTAAAGCAATGTGGAAATATCCATTTTCTAAACATCTTACATATACAGACAAGTTTTATATTTCTAAGAATATAGTTACCAGTGTTGATATGATGGTGGGTACCGAGAATGACTTGCAATATGTACATATTAATGAATTATTCGGAGGATTCTCTATTATCGATATTCCATACGAGGGAAACTCTAGTATGGTGATTATACTGCCGGACGACATAGAAGGTATATATAACATAGAAAAAAATATAACAGATGAAAAATTTAAAAAATGGTGTGGTATGTTATCTACTAAAAGTATAGACTTGTATATGCCAAAGTTTAAAGTGGAAATGACGGAACCGTATAATCTGGTACCGATTCTAGAAAATTTAGGACTTACTAATATATTTGGATATTATGCAGATTTTAGTAAGATGTGTAATGAAACTATCACTGTAGAAAAATTTCTACATACGACGTTTATAGATGTTAATGAGGAGTATACAGAAGTATCGGCCGTTACAGGAGTATTCATGACTAACTTTTCGATGGTATATCGTATGAAGGTCTACATAAACCATCCATTCATATACATGATTAAAGATAACACCGGACATACACTTTTTATAGGGAAATACTGCTATCCGCAATAAATATAAACAATAGACTTTTATCACGTTATCTCATGTATAAAATATTACAAATAGTATAGCATAAACTAAAGTCGATACATACATTAAAACTTAAATAATAATGTAATTTACAATTAATAGTATAAACTAAAAAAATTAAAAAATTAAAAACAATATCATTATTATAAGTAATATCAAAATGACGATATACGGATTAATAGCGTATCTTATATTCGTGACTTCATCCATCGCTAGTCCACTTTACATTCCCGTTATTCCGCCCATTTCGGAAGATAAATCGTTCAATAGTGTAGAGGTATTAGTTTCTTTGTTTCCCGATGACCAAAAAGACTATACAGTAACTTCTCAGTTCAATAACTACACTATCGGTACCAAAGACTGGACTATCAACGTACTATCCACACCTGATGGTCTGGACATACCATTGACTAATATAACTTATTGGTCACGGTTTACTATAGGTCGTGCATTGTTCAAATCAGAGTCTGAGGATATTTTCCAAAAGAAAATGAGTATTCTAGGTGTTTCTATAGAATGTAAGAAGCCGTCGACATTACTTACTTTTTTAACCGTGCGTAAAATGACTCGAGTATTTAATAGATTTCCAGATATGGCTTATTATCGAGGAGACTGTCTAGAAGCCGTTTATGTAACAATGACTTATAAAAATACTAAAACTGGAGAGACTGATTACACGTACCTCTCTAATGGGGGGTTGCCTGCATACTATCGTAATGGGGTCGATGGTTGATTATTGATTAGTATATTCCTTATATTCCTTATTCTTTTTATTCACACAAAAAGAACATTTTTATAAACATGAAACCACTGTCTAAATGTAATTATGATCTTGATTTATAGATGATGATCAGCCTTCAGAGGATTTTGACCAGTATGTTTAATATGAAAAAAAACATAACTATTAAGCGCTATTGCGCTATTGTGCTTAATTATTTTGCTCTATAAACTGAATATATAGCCACAATTATTGACGGGCTTGTTTGTGACCGACAATCATGAATTTTCAGAAATTATCTCTGGCTATATATCTTACGGTGACATGTTCGTGGTGTTATGAAACATGTATGAGAAAAACTGCGTTGTATCATGACATTCAATTGGAGCATGTAGAAGACAATAAAGATAGTGTAGCATCGCTACCGTACAAGTATCTACAAGTAGTCAAACAAAGAGAACGTAGTAGATTGTTGGCTACATTTAATTGGACGGATATAGCTGAGGGTGTTAGAAATGAGTTCATTAAAATATGTGATATCAACGGAACATATCTATATAATTATACTATTGCTGTTAGTATAATTATTGATTCCACGGAAGAACTACCAACAGTTACTCCAATTACAACATATGAACCTTCTATATATAATTATACTATCGATTATAGCACTGTTATTACTACTGAAGAACTACAAGTGACTCCAACATATGCACCTGTAACAACTCCTCTTCCAACATCAGCAGTTCCTTATGATCAACGATCGAATAACAATGTAAGTACTATATCTATTCAGGTACTGAGTAAAATATTGGGAGTCAATGAAACAGAATTAACTAATTATCTTATTATGCATAAAAATGACACTGTTGACAATAACACCATGGTTGATGATGAGACATCTGACAATAACACATTACATGGTAATATAGGATTTTTGGAAATAAATAATTGTTATAATGTTTCTGTGTCAGATGCTAGTTTTAGAATAACATTAGTAAACGATACTTCTGAAGAAATTTTGCTAATGCTAACAGGAACTAGTTCATCCGACACCTTCATATCTTCCACCAATATCACTGAATGTTTGAAAACATTAATCAATAATGTGTCGATTAATGATGTACTTATAACACAAAATATGAATGTAACATCTAATTGTGATAAATGCTCAATGAATTTGATGGCATCCGTTATTCCTGCAGTTAATGAATTTAACAATACGTTGATGAAAATTGGTGTAAAAGATGATGAAAACAATACGGTATATAAATATTATAATTGTAAACTAACTACAAATTCTACATGTGATGAGTTAATCAATTTAGATGAAGTCATTAACAACATAACTCTGACAAATATTATACACAATAGTGTTTCGACAACTAACAGCAGAAAAAGACGAGATCTGAATGATGAGTTTGAATTTTCCACTTCCAAGGAATTAGATTGTCTTTACGAATCATATGGTGTAAACGATGATATAAGTCATTGTTTTGCATCACCTAGACGTAGACGATCTGACGACAAAAAGGAGTACATGGACATGAAATTATTCGACCACGCGAAAAAAGATTTAGGAATAGACAGTGTTATTCCTAGAGGTACAACCCATTTCCAAGTAGGTGCATCTGGTGCAAGTGGTGGTGTTGTAGGAGATAGTTTCCCATTTCAAAATGTTAAATCGCGTGCCAGTCTATTGGCGGAAAAAATAATGCCTAGAGTACCTATTACTGCTACCGAAGCTGATCTATATGCAACTGTAAATAGACAACCCAAGTTACCAGCAGGTGTTAAAAGTACTCCGTTTACAGAGGCGCTTGTGTCTACGATAAACCAAAAGCTTTCTAATGTTAGAGAGGTAACTTATGCTTCGCTCAATCTGCCAGGATCAAGTGGCTATGTTCATAGACCATCTGATTCTGTTATTTATAGCAGTATAAGACGGTCACGTTTACCTAGTGATAGCGATAGTGATTATGAGGATATACAAACTGTTGTTAAGGAATATAATGAAAGATATGGTAGATCAGTCAGTAGAACACAGTCATCAAGTAGTGAAAGCGATTTTGAAGATATAGATACTGTTGTTAGGGAATATAGACAAAAATATGGCAATGCAATGGCAAAAGGACGTAGTAGTTCCCCTAAACCTGATCCATTATATAGTACTGTTAAGAAAACAACTAAAAGTCTATCTACTGGTGTAGACATAGTTACAAAACAATCAGACTATTCTCTATTACCTGACGTTAATACTGGCAGTTCTATTGTGTCACCTCTCACCAGAAAAGGAGCTACTAGACGACGACCTAGACGCCCTACAAATGATGGTCTACAGAGTCCAAATCCTCCTCTCCGTAATCCACTTCCTCAACATGATGATTATTCTCCTCCACAAGTACACAGACCTCCACCACTTCCTCCTAAACCAGTCCAAAATCCGCCACAACTTCCCCCTAGACCAGTAGGTCAATTACTACCTCCTCCTATAGATCAACCAGATAAAGGATTTAGTAAGTTTGTATCACCTAGACGGTGTAGAAGAGCAAGCTCTGGAGTCATATGTGGTATGATACAATCAAAACCAAACGATGATACCTATTCACTTCTTCAACGATCAAAAATTGAACCAGAATATGTGGAGGTTGGTAATGGTATACCCAAGAACAATGTTCCTGTAATAGGTAATAAACATAGTAAAAAATATACATCGACGATGTCAAAAATATCAACAAAATTTGATAAATCTACGGCATTTGGAGCAGCAATGTTACTAACTGGTCAGCAGGCCATTAGCCAACAGACTAGATCAACTACGTTGAGTAGAAAAGATCAGATGAGTAAGGAAGAAAAGATATTCGAAGCAGTTACAATGAGTCTATCAACTATAGGTTCAACGTTGACGTCTGCAGGTATGACGGGTGGTCCAAAACTAATGATTGCAGGAATGGCTATAACGGCTATAACTGGTATAATAGATACGATAAAAGATATATATTACATGTTTTCAGGACAGGAGAGGCCAGTAGATCCTGTTATTAAATTATTTAATAAGTACACTGGCTTAATGTCCGATAATAATAAAATGGGTGTAAGAAAATGTTTGACACCCGGTGACGACACACTTATTTATATCGCATACAGAAACGATACCAGTTTTAAACAGAATACGGATGCGATGGCTTTGTATTTCTTAGATGTTATCGATTCAGAGATCCTATATCTAAACACATCAAATTTAGTTCTAGAGTATCAACTAAAGGTGGCTTGCCCCATAGGAACATTAAGATCTGTAGATGTGGACATAACTGCGTATACAATATTATATGATACAGCGGATAATATTAAGAAATACAAGTTTATCAGAATGGCAACGCTACTATCCAAACATCCAGTTATTAGATTGACATGTGGTTTAGCAGCAACATTGGTGATTAAACCGTACGAGGTACCCATCAGTGATATGCAACTACTAAAAATGGCGACGCCTGGTGAACCAGAATCCACTAAATCTATACCATCCGATGTCTGTGATAGGTATCCTCTAAAGAAATTCTATCTTTTAGCTGGTGGTTGTCCCTATGATACATCTCAAACTTTTATTGTACATACTACTTGCAGTATTCTACTAAGAACAGCTACACGGGATCAGTTTAGAAACAGATGGGTGTTACAAAATCCATTTAGACAAGAAGGGACATATAAGCAACTGTTTACCTTTAGCAAATACGATTTTAACGACACCATAATCGATCCTAATGGTGTGGTGGGTCATGCTAGCTTTTGTACCAATAGAAGCAGCAACCAATGTTTCTGGTCCGAACCTATGATATTGGAAGATGTATCATCGTGTAGTTCTAGAACTAGAAAAATATACGTAAAACTGGGAATATTTAATGCTGAAGGTTTTAATAGTTTTGTACTAAATTGTCCAACTGGGTCTACACCTACATACATCAAACATAAAAATGCGGACAGTAACAATGTTATCATAGAGCTACCTGTAGGTGATTACGGCACAGCCAAATTGTATTCAGCAACAAAACCATCGAGGATAGCTGTGTTCTGCACACATAACTATGATAAACGATTCAAATCAGATATTATAGTTCTAATGTTTAATAAAAACAGCGGTATTCCATTTTGGAGCATGTACACAGGAAGTGTAACTAGTAAAAATTTTAATTTTGTCAGGAGTCTTGAGGATAAAGTTTAAAATAGAATTAATAAAGAACATATAGGTCATTTTTTAAACATGGATAGAAACCAAGGTTGTTAGTTAATAATATACAAGATATTTTTTCTCACTCTGATCCATGTAAACCAAGGACGAGAGACACTCTCATTCCTCATTCACGACACCATTAAAAATGGAAATTAAAGCCCTCTATTAAGCACAGACGGCTACAGGTCTACCATCAGGTTACCTTCGTCTACCTTCACAATGGCCTCTCCTTGTGCCCAGTTCAGTCCCTGTCATTGCCACGCTACTAAGGACTCCCTGAATACCGTGACTGACGTCAGACATTGTCTGACTGAATACATCCTGTGGGTTTCTCATAGATGGACCCATAGAGAAAGCGCAGGGCCTCTCTACAGGCTTCTCATCTCTTTCAGAATTGATGCAATGGAGCTATTTGGTAGCGAGTTGAAGGAGTTCTCGAATTCACTTCCGTGGGACAATATCGACAATTGCGTGGAGATCATTAAATGTTTCATCAGAAATGACTCCATGAAAACCGCCAAAGAACTTTGTGCAATAATTGGACTTTGTACTCAATCAGCTATTGTCACTGGAAGAGTCTTCAATGATAAGTATATCGACATACTACTTATGCTGCGAAAGATTCTGAACGAGAACGACTATCTCACCCTCTTGGATCATATCCTCACTGCTAAATACTAAATCTCCTTCATGCTCTCTCACTAATACTCTTACTCACTACACTTTTTATCATCTTATGATGAATGATTGCCTTCATCATTTTTTCGTGGAATATAATATAGGAATAATTAGCACCAGAATAGCTATGGATATCTCGTTAAGAATATTCTCTATAAGAGACATAATGTAGACATAGTTATTATATCCTTCTTAGATAAGTGTTACGCTACTGGAAAGTTTCCATCGTTATTATTACATGAAGATGATATAATTAAACCAACATTGAGATTGGCTCTTATGTTAGCTGGATTGAATTACTGTAATAAATGCATCGAGTATAGAGGGATATAGCAATTCTCGATAATAGTCATGCAATATTTGAATGAGACTGATAATTTAGGTAATACAGTACTACACACATATCTTTCTAGATTATATATCGTTAAAAATCTGTAAGATGTATATTTCTCATAAGTATCCACTGTGTAATATTATTAATGGATATATAGATAACGCAATAGGGACTAATAGTATTGTAAAAGATATAATCGACTATTTACGTACATATCCAGATATCTATATTCCTACTAGTTTGCTGCGTAGTTGCATCATTGATATGCATGATTTATCAGGATTCAGAGATGAATTACTAAGTAAACTACAATCCCACAATAAGTAAGAATCAAATATCAAAAACTCACTTTTGATTTTTCTAGTCTTAAGTAATACATATATTTATTAATAGACCTATGAAATAAAAAAAGGTAACAATGGATTCGCGTATAGCTATTTACGTATTAGTATCGGCATCTCTTTTGTATCTTGTTAATTGTCACAAACTAGTACATTACTTCAATCTGAAAATAAATGGAAGTGATATAACTAATACAGCAGATATATTGCTGGACAATTATCCAATTATGACCTTTGATGGAAAGGATATTTATCCATCTATCTCGTTCATGGTCGGTAATAAACTTTTCCTAGATCTTTATAAAAATATCTTTGAAGAATTTTTCAGACTATTTCGAGTATCTGTAAGTAGTCAATACGAGGAATTAGAATATTATTATTCATGTGATTATACTAACAACCGTCCTACAATTAAACAACATTACTTTTATAACGGCGACGAATATACTGAAATTGATAGATCGAAAAAAGCCACTAATAAAAACAGTTGGTTAATTACTTCAGGCTTTAGACTACAAAAATGGTTCGATAGCGAAGATTGTATAATTTATCTCAGATCTTTAGTTAGAAGAATGGAAGACAGTAACAAAAACAGTAAAAAAACTTAGTACTTAGATATCGAAAAAATATATTTTTGTAGACTCTTGAGAATAGAAGGAAAACATGTACATAATTATAAAAAATGAAAATCAATGGCGAATAAGACAGTGCGATTCGCACCATGGAGTCGGTAGATTTCATGGCTGTCGATGAGCAGTTTCACGACGACCTCGATCTTTGGTCATTATCTTTGGTAGATGATTATAAAAAACATGGATTAGGTGTTGACTGTTATGTTCTAGAACCAGTTGTTGACAGGAAAATATTTGATAGATTTCTCCTTGAACCAATTTGTGATCCTGTAGATGTTCTGTATGATTATTTTAGGATTCATAGAGATAATATTGATCAGTATATAGTAGATAGACTGTTTGCATATATTACATATAAAGATATTATATCTGCATTAGTGTCAAAGAATTATATGGAAGATATTTTCTCTATAATTATTAAGAATTGTAATTCTGTGCAAGATCTCTTACTTTACTATCTATCTAATGCATATGTAGAAATAGACATTGTTGATCTTATGGTAGATCATGGGGCTGTAATATATAAAATAGAATGCTTGAATGCCTATTTTAGGGGAATATGTAAAAAGGAAAGTAGTGTTGTTGAGTTTATTTTGAATTGTGGTATCCCAGATGAAAATGATGTTAAATTAGATCTATATAAAATAATTCAGTATACTAGGGGATTCCTTGTAGATGAACCCACAGTATTAGAAATTTATAAGCTTTGTATCCCATATATTGAAGATATCAATCAACTAGATGCTGGTGGAAGGACCTTGCTTTATCGCGCTATCTATGCAGGTTATATAGATTTAGTATCATGGCTATTAGAAAATGGAGCAAATGTCAACGCAGTAATGAGTAATGGATATACATGTCTTGACGTGGCCGTGGATAGGGGATCTGTCATCGCCCGTAGGGAAGCACATCTTAAAATATTAGAAATATTGCTTAGAGAACCATTGTCTATTGACTGTATAAAATTAGCTATACTTAATAATACAATTGAAAACCATGATGTGATAAAGCTCTGTATCAAGTATTTTATGATGGTAGATTATTCACTTTGTAATGTGTATGCATCATCACTCTTTGATTATATAATTGATTGTAAACAAGAATTGGAGTACATTAGGCAGATGAAAATTCATAATACAACCATGTATGAGTTAATCTATAATAGAGACAAAAACAAGCATGCTTCCCATATTCTACATAGGTATTCTAAACATCCAGTTTTGACACAGTGTATCACTAAAGGATTCAAGATTTACACAGAAGTAACCGAGCAGGTCACTAAAGCTCTAAACAGACGTGCTCTAATAGATGAGATAATAAACAATGTATCAACTGATGACAATCTCCTATCAAAACTTCCATTAGAAATTAGGGATCTAATTGTTTCACAAGCTGTCATATAGAGTTCTATCCACCCACCTTTCTTGAAATGAGTTAATAGTCATAAGTTAGTTAAGTCATAAGTTAGTTAAGTCATAAGTTAGTTAAGTCATAAGTTAGTTAAGTCATAAGTTAGTTAAGTCATAAGTTAGTTAAGTCATAAGTTAGTTAAGTCATAAGTTAGTTAAGTCATAAGTTAGTTAAGTCATAAGTTAGTTAAGTCATAAGTTAGTTAAGTCATAAGTTAGTTAAGTCATAAGTTAGTTAAGTCATAAGTTAGTTAAGTCATAAGTTAGTTAAGTCATAAGTTAGTTTATAGTCTAACACTTCTAATTTTTATACCTTGATCTTTTTCTCTAATTATGAAAAAGTAAATCATTATGAAGATGGATGAAATGGACGAGATTGTGCGCATCGTTAACGATAGTATGTGGTACGTACCTAACGCATTTATGGACGACGGTGATAATGAAGGTCACATTTCTGTCAATAATGTCTGTCATATGTATCTCGCATTCTTTGATGTGGATATATCATCTCATCTGTTTAAATTAGTTATTAAACACTGCGATCTGAATAAACGACTAAAATGTGGTAACTCTCCATTACATTGCTATACGATGAATACACGATTTAATCCATCTGTATTAAAGATATTGTTACGCCACGGCATGCGTAACTTTGATAGCAAGGATAAAAAAGGACATATTCCTCTACACCACTATCTGATTCATTCACTATCAATCGATAACAAGATCTTTGATATACTAACGGACCCCATTGATGACTTTAGTAAATCATCCGATCTATTGCTGTGTTATCTTAGATATAAATTCAATGGGAGCTTAAACTATTACGTTCTGTACAAATTATTGACTAAAGGATCTGACCCTAATTGCGTCGATGAGGATGGACTCACTTCTCTTCATTACTACTGTAAACACATATCCGCGTTCCACGAAAGCAATTATTACAAGTCAAAGAGTCACACTAAGATGCGAGCTGAGAAGCGATTCATCTACGCGATAATAGATCATGGAGCAAACATTAACGCGGTTACGAAAATCGGAAATACGCCGTTACACACTTACCTTCAACAGTATACCAAACATAGTCCTCGTGTGGTGTATGCTCTTTTATCTCGAGGAGCCGATACGAGGATACGTAATAATCTTGATTGTACACCCATCATGGAATACATAAAGAACGATTGTGCAACAGGTCATATTCTCATAATGTTACTCAATTGGCACGAACAAAAATACGGGAAATTACAAAAGGAAGAAGGACAACATCTACTTTATCTATTCATAAAACATAATCAAGGATATGGAAGTCGCTCTCTCAATATACTACGGTATCTACTAGATAGATTCGACATTCAGAAAGACGAATACTATAATACAATGACTCCTCTTCATACCGCCTTCCAGAATTGCAATAACAATGTTGCCTCATACCTCGTATACATCGGATACGACATCAACCTTCCGACTAAAGACGATAAGACAGTATTCGACTTGGTGTTTGAAAACAGAAACATTATATACAAGGCGGATGTCGTTAATGACATTATCCACCACAGACTGAAAGTATCTCTACCTATGATTAAATCGTTGTTCTACAAGATGTCGGAGTTCTCTCCCTACGACGATCACTACGTAAAGAAGATAATAGCCTACTGCCTATTAAGGGACGAGTCATTTGCGGAACTACATACTAAATTCTGTTTAAACGAGGACTATAAAAGTGTATTTATGAAAAATATATCATTCGATAAGATAGATTCCATCATCGAAAAATGTAGTCGTGACATAAGTCTCCTCAAAGAGATTCGAATCTCAGACACCGACTTGTATACGGTATTGAGAACAGAAGACATCCGGTATCACACATATCTCGAAGCCATACATTCAGACAAACGCATTTCATTTCCCATGTACGACGATCTCATAGAACAGTGTCATCTATCGATGGAGCATAAAAGTAAACTCGTCGACAAAGCACTCAATAAATTAGAGTCTACCATCGATAGTCAATCTAGACTATCGTATTTGCCTCCGGAAATTATGCGCAATATCATAACCAAGCTAAGCGACTACCATCTAAACAGTATGTTGTACGGAAAGAACCATTACAAATATTATCCATGATAGAAAGAAAATATTTAAAAAATAATCTATATGATTGGAGAAGTAGGAAACAAACAGTAACAAGACGACGATTACTACATTATTAAATCATGAGGTCCGTATTATACTCGTATATATTGTTTCTCTCATGTATAATAATAAACGGAAGAGATATAGCACCACATGCACCATCCAATGGAAAGTGTAAAGACAACGAATACAGAAGCCGTAATCTATGTTGTCTATCGTGTCCTCCGGGAACTTACGCTTCCAGATTATGTGATAGCAAGACTAATACACAATGTACACCGTGTGGTTCGGATACCTTTACATCTCACAATAATCATTTACAGGCTTGTCTAAGTTGTAACGGAAGATGTGATAGTAATCAGGTAGAGACGCGATCGTGTAACACGACTCACAATAGAATCTGTGAATGCTCTCCAGGATATTATTGTCTTCTCAAAGGAGCATCAGGGTGTAGAACATGTATTTCTAAAACAAAGTGTGGAATAGGATACGGAGTATCCGGATACACGTCTACCGGAGACGTCATCTGTTCTCCGTGTGGTCCCGGAACATATTCTCACACCGTCTCTTCCACAGATAAATGCGAACCCGTCGTAACCAGCAATACATTTAACTATATCGATGTGGAAATTAACCTGTATCCAGTCAACGACACATCGTGTACTCGGACGACCACTACCGGTCTCAGCGAATCCATCTCAACGTCGGAACTAACTATTACCATGAATCATAAAGATTGTGATCCAGTCTTTCGTGCAGAATACTTCTCTGTCCTTAATAATGTAGCAACTTCAGGATTCTTTACAGGAGAAAATAGATATCAGAATACTTCAAAGATATGTACTCTGAATTTCGAGATTAAATGTAACAACAAAGATTCATCTTCCAAACAGTTAACGAAAACAAAGAATGATACTATCATGCCGCATTCAGAGACGGTAACTCTAGTGGGCGACTGTCTATCTAGCGTCGACATCTACATACTATATAGTAATACCAATACTCAAGACTACGAAAATGATACAATCTCTTATCATATGGGTAATGTTCTCGATGTCAATAGCCATATGCCCGCTAGTTGCGATATACATAAACTGATCACTAATTCCCAGAATCCCACCCACTTATAGTAAGTTTTTTTACCTATAAATAATAAATACAATAATTAATTTCTCGTAAAAGTAGAAAATATATTCTAATTTATTATATGGTAAGAAAGTAGAATCATCTAGAACAGTAATCAATCAATAGCAATCATGAAACAATATATTGTCCTGGCATGCATGTGCCTAGTGGCAGCTGCTATGCCTACTAGTCTTCAACAATCTTCATCCTCGTGTACTGAAGAAGAAAACAAACATCATATGGGAATCGATGTTATTATCAAAGTCACAAAGCAAGACCAAACACCGACCAATGATAAGATTTGTCAATCCGTAACGGAAGTTACAGAGACCGAAGATGATGAGGTATCCGAAGAAGTTGTAAAAGGAGATCCCACCACTTATTACACTATCGTCGGTGCGGGTCTTAACATGAACTTTGGATTCACCAAATGCCCAAAGATTTCATCCATCTCCGAATCCTCTGATGGAAACACTGTGAATACTAGATTGTCCAGCGTGTCACCGGGACAAGGTAAGGACTCTCCCGCGATCACGCGTGAAGAAGCTCTGGCTATGATCAAAGACTGTGAGATGTCTATCGACATCAGATGTAGCGAAGAAGAGAAAGACAGTGACATCAAGACCCATCCAGTACTTGGGTCTAACATCTCACATAAGAAAGTGAGTTACAAAGATATCATCGGTTCAACGATCGTTGATACAAAATGTGTCAAGAACCTAGAGTTTAGCGTACGTATCGGAGACATGTGTGAGGAATCATCTGAACTTGAAGTCAAGGATGGATTCAAGTATGTCGACGGATCGGCATCTGAAGGTGCAACCGATGATACTTCACTCATCGATTCAACAAAACTCAAAGCATGTGTCTGAATCGATAACTCTATTCATCTGAAAATGGATGAGTTGGGTTAATCGAACGATTCAGACACCGCACCACGAATTAAAAAAGACCGGGCACTATATTCCGGTTTGCAAAACAAAAATATTTAACTACATTCACAAAAAGTTACCTCTCGTTACTTCTTCTTTCTGTTTCAATATGTGATACGATATGATCACTATTCGTATTCTCTTGGTCTCATAAAAAAGTTTTACAAAAAAAAAAAAAAAATATTTTTATTCTCTTTCTCTCTTCGATGGTCTCACAAAAATATTAAACCTCTTTCTGATGTCTCAACTATTTCGTAAACGATAACGTCCAACAATATATTCTCGTAGAGCTTATCAACATCCTTATACCAATCTAGGTTGTCAGACAATTGCATCATAAAATAATGTTTATAATTTACACGTTAACATCATATAATAAACGTATATAGTTAATATTTTTGGAATATAAATGATCTGTAAAATCCATGTAGGGGACACTGCTCACGTTTTTTCTCTAGTACATAATTTCACACAAGTTTTTATACAGACAAATTAATTCTCGTCCATATATTTTAAAACATTGACTTTTGTACTAAGAAAAATATCTTGACTAACCATCTCTTTCTCTCTTCGATGGGTCTCACAAAAATATTAAACCTCTTTCTGATGGAGTCGTAAAAAGTTTTTATCCTTTCTCTCTTCGATAGGTCTCACAAAAATATTAAACCTCTTTCTGATGGTCTCTATAAACGATTGATTTTTCTTACCCTCTAGAGTTTCCTACGGTCGTGGGTCACACATTTTTTTCTAGACACTAAATAAAATAGTAAAATTAAATTAATTATAAAATTATGTATATAATTTACTAACTTTAGTTAGATAAATTAATAATACATAAGTTTTAGTACATTAATATTAATTATACTAATTTATTTAG

>2017_Nigeria_Monkeypox_MG693723.1
