## Supplementary figures and images for "Genomic analysis of the recent monkeypox outbreak"

### Supplementary File S2

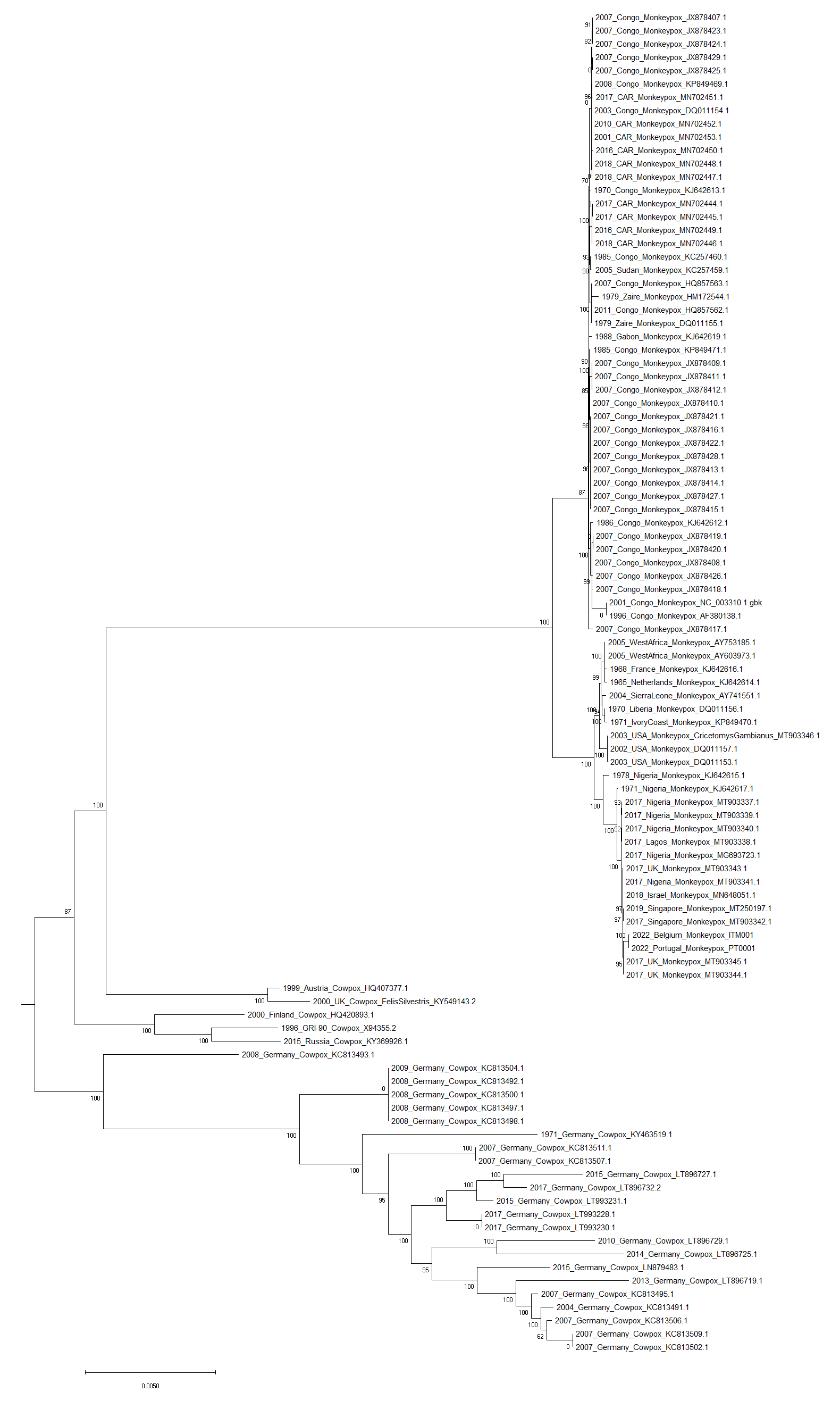
